# Atom-level Machine Learning of Protein-Glycan Interactions and Cross-chiral Recognition in Glycobiology

**DOI:** 10.1101/2025.01.21.633632

**Authors:** Eric J. Carpenter, Chuanhao Peng, Sheng-Kai Wang, Russell Greiner, Ratmir Derda

## Abstract

Cross-chiral recognition in glycobiology is the interactions between biologically conventional proteins and the enantiomers of biological glycans (e.g., L-proteins binding with L-hexoses) from organisms across all kingdoms of life. By symmetry, it also describes the interactions of chirally mirrored proteins with normal D-glycans. Knowledge of cross-chiral recognition is critical to understanding the potential interactions of existing life forms with artificial mirror-life forms, but currently known rules of protein-glycan interaction are insufficient. To build a methodology for learning such interactions, we used machine learned models that can predict binding strength between a set of proteins and glycans represented as graphs of atoms, rather than monosaccharides. Atomic *q*-gram and Morgan fingerprint (MF) based representation of glycans made it possible to train ML models that predict lectin binding properties of glycans, glycomimetic compounds, and enantiomers of all natural glycans. Critical to this training was merging disparate data—some with relative fluorescence units (RFU) from glycan microarrays and others with *K*_d_ values from ITC—using a universal “fraction bound” parameter *f* at a specific lectin concentration. A fully-connected neural network architecture, MCNet takes a MF and concentration (C) as inputs and returns *f* for 147 lectins. Performance of MCNet is comparable to the GlyNet models, and by proxy to other state-of-the art models that predict strength of protein-glycan interactions. MCNet effectively predicts binding of glycomimetic compounds to Galectins 1, 3, and 7. Breaking from a monosaccharide-based description makes it possible for MCNet to predict cross-chiral recognition. We employed a Liquid Glycan Array to validate some predictions, such as the lack of interactions of L-mannose with D-mannose binding lectins, purified ConA, and DC-SIGN displayed on cells, and weak binding of L-Man to galactose-binding lectins and L-Glc binding by canonical fucose binding lectins. MCNet’s atom-level input makes it possible to agglomerate protein-glycan data from diverse glycans across all kingdoms of life and non-glycan structures (e.g. glycomimetic compounds). The universal fraction bound parameter makes it possible to unify disparate quantitative observations (*K*_d_/*IC*_50_, RFU, chromatographic retention times, etc.). We believe that such an approach will facilitate a merger of knowledge from diverse glycobiology datasets and predict protein interactions with uncommon/unnatural glycans not attainable from current ML models.

## Introduction

Glycans are a major building block of life and major constituent of the glycocalyx that coats cells in all kingdoms of life.^1^ Recognition of glycans by glycan-binding proteins (GBPs or lectins) is critical for cell-to-cell communication, immune responses, self vs. non-self recognition,^2^ pathological responses like tumorigenesis,^2^ and normal physiological processes like fertilization.^3,4^ Unlike DNA, RNA or proteins, glycans form linear and branched oligomers with diverse stereochemistry of the monomers and linkages giving rise to a vast “feasible complexity” of glycans.^5^ Number of feasible glycan structures is orders of magnitude higher than nucleotides or proteins, even in a short sequence.^6,7^ However, the estimate of “biologically relevant complexity”—the number of distinct glycan structures synthesized by a given organism—is smaller by many orders of magnitude. For example, the human glycome is estimated to be limited to ∼7,000 structures created by known glycosynthetic enzymes.^8^ Building on this dichotomy, inquiries in glycobiology are skewed to glycans of “biologically relevant complexity”. Such glycans dominate glycan microarrays, like those produced by Consortium for Functional Glycomics (CFG).^9^ Data from CFG and related arrays is used for training of machine learning (ML) that predict protein-glycan interaction.^10^ The resulting ML models, again, are focused on predicting interactions between the biologically relevant glycan motifs and well-understood lectins in their training data. Some researchers may questions the need to study “biologically irrelevant” protein-glycan recognition, while others (e.g. Thomas Kuhn^11^) teach the need to break away from a dogmatic paradigm. For example, the elegant report of Canner and Gray used ML models to ask the fundamental question, what is a glycan binding protein,^12-14^ and uncovered the intriguing possibility that many organisms contain GBPs that have not been yet acknowledged as such. An interesting example are C-type lectins, it is not known how many of the C-type lectins actually bind to traditional or non-traditional glycans^15,16^. In this manuscript, we train ML models that predict quantitative glycan-GBP interactions; however, our primary goal is to break away from the “working complexity” and instead examine binding in i) the feasible complexity of glycans and ii) glycomimetic structures.

An emerging open-ended question regarding the feasible glycan complexity type pertains to enantiomers of known glycans. Such glycans would be produced by mirror-image life, for which the plausibility of laboratory construction has been discussed for the last few decades^17^. The feasibility of creating mirror-life was boosted by the recent synthesis of critical D-enzymes such as D-DNA-ligase, D-RNA polymerase and enantiomeric ribozymes,^18-20^ as well as enzymes for ligation of D-proteins.^21^ The pioneering work of Kent and co-workers experimentally proved that enzymes made of D-amino acids do not recognize peptide and peptidomimetic substrates of the natural enzyme, but they effectively process the enantiomers of natural substrates^22^. Mirror-image phage display^23^ showed that L-peptides evolved to recognize mirror-image D-proteins do not bind to L-proteins, however, D-peptides binds to L-protein. The resistance of mirror-image proteins^24^ and enantiomeric aptamers^18-20^ to degradation by natural peptidases and nucleases *in vivo* is well established. These combined observations lead to a general notion of an absence of cross-chiral recognition between peptides, nucleotides and metabolites. This limited cross-chiral recognition is an important consideration raised in a 2024 report in Science, cautioning that if mirror image lifeforms were constructed, they may evade many aspects of immune recognition, response, and ecological control that are present in conventional-chiral organisms^25^. A critical element that received attention in a followup commentary^26^ is the overwhelming abundance of glycans and critical role of glycan-GBP interactions in self vs. non-self determination. Mirror-image glycans will be one of the first entities encountered by our immune systems^26^; however, our understanding of cross-chiral recognition between proteins and glycans is very limited. For example, D-mannose is abundant in the glycocalyx of most organisms; the interaction of its 1,2-bis-equatorial diol with Ca^2+^ in C-type lectins is at the heart of the immune response elicited by these lectins^15,16^. However, the L-mannose enantiomer is rare in biology^27^ and its protein binding profile is rarely studied^28^. The core recognition motif of D- and L-mannose is mirror-symmetric, 1,2-bis-equatorial diol. Reflection of this diol produces the same 1,2-bis-equatorial diol (**Figure 1A-B**). Antibodies to L-rhamnose (6-deoxy-L-mannose) are abundant in human serum^25^ but recognition of L-mannose by L-rhamnose-binding proteins is poorly understood. The spatial arrangement of three adjacent hydroxyls in L-mannose mimics that in D-galactose (**Figure 1C**). Indeed, there are scarce reports of L-mannose serving as a ligand for galactose-binding proteins^28^, and complementary reports that L-galactose is an efficient mimic of D-mannose and is recognized by the D-mannose binding immune lectin DC-SIGN^26^. *Osmerus eperlanus mordax* lectins recognize the monosaccharides L-Rha, L-Man, and D-Gal^27^, while RSL from *Ralstonia solanacearum* recognises another set, L-Fuc, L-Gal, D-Ara, D-Man^29^ (Figure **1D**). In turn, L-galactose, the enantiomer of common galactose, and a close analog of natural L-fucose was reported as a viable substrate of α1,2-L-fucosyltransferase^30^. Further, the discovery of L-glucosidases points to possible use of L-glucose in glyco-conjugates, and the existence of lectins for their recognition^27,31^. This critical difference in symmetry inversion between glycans and other biological oligomers highlights that cross-chiral glycan-GBP interaction is more plastic than cross-chiral recognition between polypeptides, oligonucleotides and metabolites^25^. We propose that ML models that effectively embed atomic composition and chirality of such atoms can be serve as tools for the first evaluation of cross-chiral recognition between glycans and GBPs. Such ML models can also unify our understanding of traditional glycans and the “rare glycans” observed in the microbial world.

**Fig. 1.**
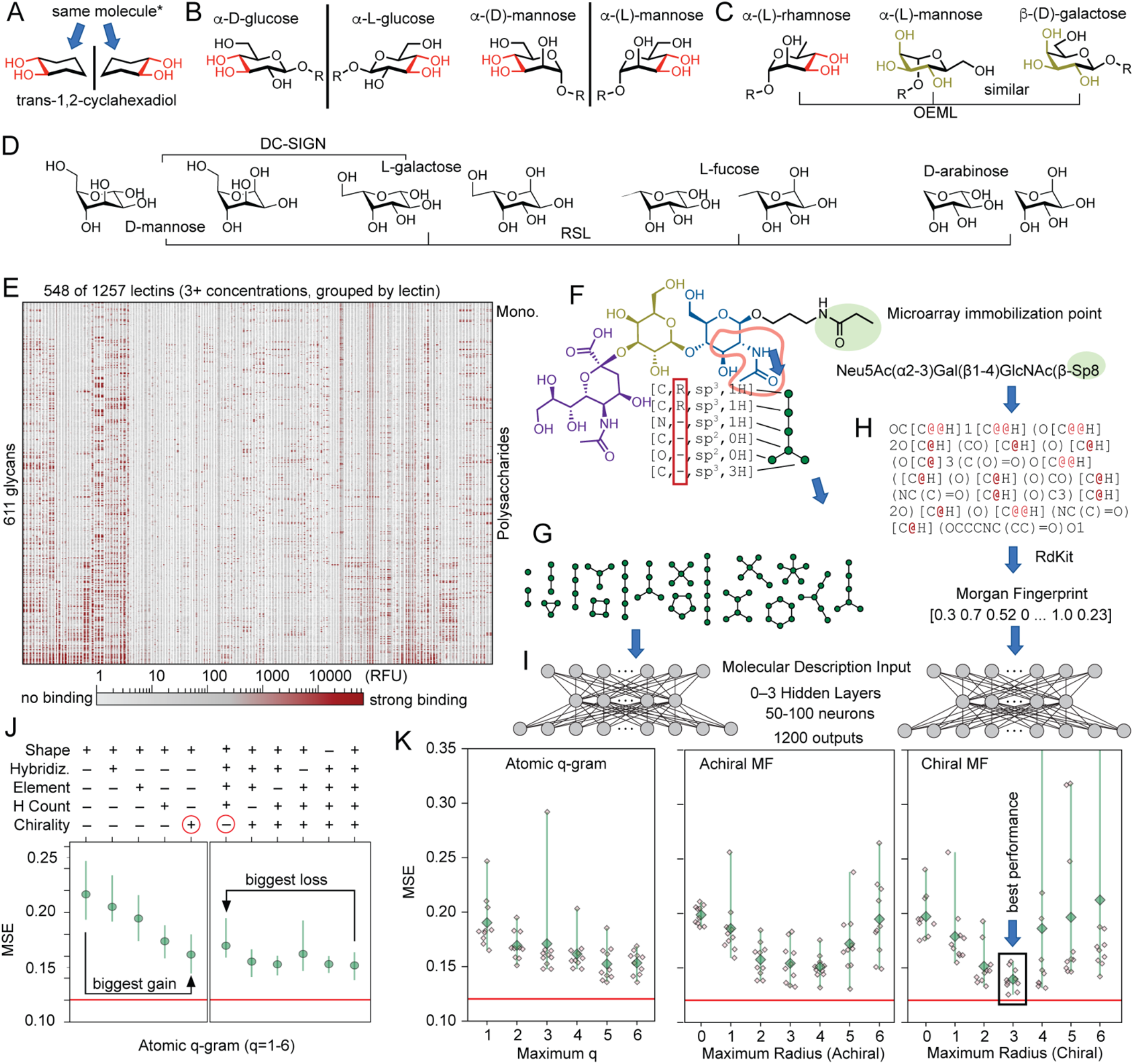
Machine learning (ML) of glycan:GBP interactions from all-atomic description. (A-D) Examples of preservation of stereochemical motifs between enantiomers, unnatural L-glycans and natural glycans. (E) Glycan:GBP binding data from CFG used for ML. (F-H) Each glycan is converted to counts of (G) atom *q*-grams (AQG) or (H) Morgan fingerprint (MF). Features of nodes in AQG, (F) include element, chiral state, hybridization, and attached hydrogen count, and overall shape. (I) Either MQG or MF counts are used as input into a fully-connected neural network which is trained to produce binding strength estimates at its outputs. (J) Mean square error (MSE) measure the performance of the models that employ different features in AQG nodes. Models trained using AQG with only a minimal set of features rank importance of chirality > H-count > element > hybridization. Similarly, systematic removal of features from AQG show that removal of chirality diminishes performance, whereas removal of hybridization, element identity or H-count is inconsequential. (K) Comparison of AQG of size 1=6 and MF or r=1-6 show that AQG with 6 atoms achieve best performance in AQG, chiral MF3-representation as best across all representations. There is a marked improvement in model trained using chiral vs. achiral MF3. Each value represents MSE measured in separate training fold; green scatter plot shows mean and standard deviation of MSE; the red line denotes the performance of GlyNet with monosaccharide embedding.

Outside of the traditional domain of mammalian N- and O-linked glycans, data for glycan-GBP recognition is not systematic. Even in a human glycome, recognition of heparin sulfate proteoglycans and mucins is only starting to emerge^32,33^. Rules for recognition by GBPs for plant glycans^34^ or the eubacterial glycans are built progressively using complex organic synthesis of glycan arrays^35-38^ and specialized lectin arrays^39,40^. N-linked glycans in Archaea contain exotic building blocks with different architectures and subunits^41^. Intriguingly, many orthologous functions, including speciation in sexually reproducing Archaea were proposed to be driven by interactions of these exotic glycans with as yet uncharacterized GBPs^41^. One barrier towards integrating all such information is the ever-increasing, but still incomplete, alphabet of all glycan building blocks across all kingdoms of life. A mirror inversion of the building blocks exacerbates the problem by effectively doubling the complexity. An ML model, like SweetTalk^42^ and GlycanML^43^ which represents glycans as collections of carbohydrate monomers can predict whether bacterial glycans are likely to be immunogenic in humans as long as such glycans contain only “natural” building blocks. However, these ML models cannot interpret L-Mannose and other “unnatural” building blocks although a recent preprint^44^ and publication^45^ that construct monosaccharide-level descriptions from atom-level ones may be able to bypass this issue. As a step towards bypassing all such problems, we examined the possibility of completely foregoing the “monosaccharide alphabet” and predicting GBP-binding properties of glycans from their atom-level structures.

Atom-level structural insights such as crystallography and molecular dynamics are workhorses in understanding glycan-protein interactions^46^. These investigations make is clear that a specific conformation of glycan and specific conformation of protein define glycan:GBP recognition^47^. This realization led to development of methods that estimate binding properties by computing conformations of proteins and glycans to find an optimal match (e.g., by docking)^46,48^. Calculation of thermodynamic values by exhaustive computation of the ensemble of all conformers is contrasted by Anfinsen’s thermodynamic hypothesis “the folded structure is determined [solely] by the amino acid sequence”.^49^ It is reinforced by the 2024 Nobel prize for AlphaFold^50^ that predicts a folded structure and subsequently a binding pose in protein-ligand complexes^51^ from primary sequences of the protein and ligand. From these observations, we infer the existence of a lightweight ML model that predicts the interaction between two molecules using only atomic connectivity of these two partners. Here we examine the plausibility of lightweight ML models that predict binding of protein to glycan from the mere atomic connectivity of glycan. A caveat in the Anfinsen hypothesis is the role of the environment: Anfinsen’s RNAse adopted two distinct conformations in two different buffers (folding and denaturing buffer)^49^. Similarly, all molecular interactions depend on the environment. A common shortcut^52-55^ is to focus on binding events measured in nearly uniform binding environments (similar temperature, buffer compositions, pH). Models from such measurements are not universal, but they should achieve reasonable performance in predicting binding strengths in a standard environment.

Models predicting quantitative parameters (e.g., *K*_d_ or IC_50_) for interactions of proteins with other proteins, antibodies or small molecules are common due to availability of curated datasets of *K*_d_ and IC_50_ values. Historically, many glycoinformatic ML models focused on classification—binder vs. non-binder^42,55,56^—and similar binary decisions (e.g., has motif / does not have motif).^54,57^ The reasons for the prevalence of such binary models is rooted in semi-quantitative measurements^58^. More recently, regression-type models have been developed for the same problems^42,44,53,59,60^. We find this binary view of binding unsatisfactory because the binding tends to zero as the concentration tends to zero. At low enough concentration of a binding partner, the answer is always “no”. Previous reports avoided this problem by training a regression ML model that predicts continuous variables, the Relative Fluorescent Units (RFU) measured in glycan microarray experiments^42,44,53,59,60^. Arthur and coworkers further generalized this problem by extrapolating *K*_d_ surrogates from the microarray data^61^. The CarboGrove tool developed by Brian Haab and coworkers elegantly merged data from disparate experimental sources and further associated the binding at a specific concentration with the presence or absence of specific motifs^57,62^. We envisioned that the problem of merging glycobiology datasets for ML^10^ can benefit from merger with broader chemical datasets like BindingDB^63^ or ChEMBL^64^. However, this merger is not possible if glycans are described as oligomers of monosaccharides^10^. Rather than cataloguing an ever-growing catalogue of monomers, we focus on more productive atomic compositions. We paraphrase the dream of “*I want to know if this oligosaccharide motif binds to the GBP*” to the more fine-detail request, “*Given a GBP and a glycan of a definitive atomic composition, I want to know the fraction of glycan* ***f*** *bound to this GBP*.” Our manuscript shows the plausibility of training a model that takes a glycan, GBP, and the circumstances as input and returns ***f***. A model that forgoes monosaccharide composition should also be able to handle any glycans across all kingdoms of life. It can account for well-documented effects of spacers, linkers, underlying scaffolds, (**Figure S1** and **S2**) in diverse arrays. It can bridge glycan structure with composition of protein sequences on which such glycans are displayed and it can give the first important glimpse into cross-chiral recognition between glycans and GBPs.

## Results

### Embedding of all-atomic representation of glycans

As in previous reports^9,53,59,60^, the core dataset for the training of ML models was binding of GBP to glycans on microarrays produced by the consortium of Functional Glycomics (CFG, Mammalian Printed Array version 5). In short, this data describes performance of 611 glyco-conjugates printed in discrete spots on the glass plate exposed to 1257 fluorescently-labelled GBP compositions and imaged by fluorescent scanner. The amount of protein bound to each glycan is reported as Relative Fluorescent Units (RFU, **Figure 1E**). We previously trained GlyNet neural network to predicted 1257 RFU values for CFG glycan structures represented as monosaccharides q-grams (MQG)^59^. To test whether atom-level representation of glycans can be used for prediction of qualitative interactions with GBP, we converted all CFG glycan structures to SMILES strings and tested two types of fixed-length representations of SMILE strings: Morgan fingerprints (MF) and atom q-grams (AQG) using RDKit (**Figure 1F-H**). We included atom-q-gram inspired by genomic k-mer statistical methods^65,66^ and to be able to draw an explainable parallel between our prior monosaccharide-tree embedding^59^ and the new atom-level embedding. Replacing the MQG inputs of the GlyNet architecture with MF or AQG inputs gave rise to several neural network architectures with 2-3 hidden layers and 50-100 neurons per layer (**Figure 1I**); each architecture was optimized using Adam^67^ and tested in 10-fold cross validation for their ability to predict 1257 RFU values for lectins given a MF or AQG representation of CGF glycans.

**Figure 1J-K** describes optimal performance for each neural network model after selecting an optimal number of hidden layers, nodes in hidden layers, and weight decay parameters. The distribution of glycans to 10 folds was identical for all models and identical to previously published GlyNet^59^; hence distribution of mean square errors (MSE) across the folds made it possible to compare all models with MSE of the GlyNet model (red line in **Figure 1K**), and by proxy many other state-of-the art models^52-55^. A simple, explainable representation of glycans using AQG of size 6 yielded MSE=0.15, which was only modestly worse than state-of-the-art GlyNet with glycan monomer embedding (MSE=0.12). Systematic subtraction or addition of graph topology, element, hybridization, chirality, number of attached hydrogens (H-count), and chirality of the node (R, S or none) rank chirality as the most important feature and H-count as the most important (**Figure 1J**). Deletion of identity atoms or their hybridization (proxy for bond orders) was inconsequential to predictions. Describing glycans by featureless graphs with 6 nodes yielded MSE=0.22 but adding only chirality information yielded MSE=0.16 (**Figure 1J**). Both observations show an expected critical role of chirality in predicting binding of glycans to lectins. An intriguing observation that was that reasonable prediction of glycan:protein interactions can be achieved by representing glycans as graphs that know only chirality of the node. This observation shows how different glycan:GBP recognition is from a interaction of proteins with classical small molecules that contain few to no chiral centers.

MF of radius 3 with chirality (MF3_c_) yielded the most optimal performance, which was comparable to performance of GlyNet (**Figure 1K**). This observation clearly demonstrated it is possible to predict glycan-binding performance by replacing a domain-specific monomer composition with universal all-atom-representation. A simple learning architecture like fully connected neural networks can be trained even in relatively small dataset of 600 glycans despite relatively large number of atoms in some glycans (500 non-hydrogen atoms). Elimination of chirality in MF led to notable decrease in performance (**Figure 1K**). Decreasing the maximum radius below 3 was clearly detrimental whereas increasing the radius above three (from 1385-dimensional to 3142-dimensional vectors at *r* ≤ 4) decreased median performance and increased fluctuations between the folds. Both observations may be the result of small size of the dataset. A mere 600 glycans is likely not sufficient to learn from high-dimensional MF4-vectors, but it possible that datasets with many more than 600 glycans and MF4 or higher may yield even better performance than current model.

### Expansion of training beyond microarray datasets

All atom representation in principle makes it possible to combine learning for natural glycans and “unnatural” glycomimetic (GM) compounds. To test this hypothesis, we selected 274 GM structures with reported binding strength (association constants, K_a_) to Galectin-1, -3, or -7 manually curated from 42 publications (see **Supplementary Table 1**). We embedded GM structures using MF3_c_ and attempted to train a multi-output neural network that predicted RFU values in some outputs and K_a_ values for other outputs. The training failed likely due to the disparate structure of the datasets: RFU values were available for CFG glycans, but not GMs while Ka data was available only for the GMs but not the CFG glycans (see supplementary discussion and **Figures S3-S5**). We were able to resolve this problem inspired by the work of Haab and coworkers (CarboGrove^52^), who unified disparate datasets by examining dose response curves. Previous reports employed RFU values to extrapolate K_a_ values. However, we elected to convert both RFU and K_a_ to a series of values describing the fraction, *f*, of the glycans bound to GBP at the specific concentrations, C (**Figure 2A-D** and **S6-S8**). K_a_ is trivially converted to *f*(C) via the Henderson-Hasselbalch equation (**Figure S7**). For the CFG glycans *f* was interpolated between the available concentrations using an array specific minimum and a global maximum RFU (**Figure 2A-D**, detailed in **Figure S6, Supplementary Data 2**). The *f* value is uniquely suited for learning because it is always bounded to the (0;1) region. It allows the qualitative dichotomy where f∼0 and f∼1 are equivalent to “no binding” and “yes binding”: it also brings an important reminder that such assessment can be made only when concentration is specified. Finally, *f* has an intuitive interpretation (e.g., *f* = 0.5 at a concentration nearing K_a_ , IC_50_ or EC_50_) although such generalization needs to be done with caution as highlighted in numerous publications^70-74.^

**Fig. 2.**
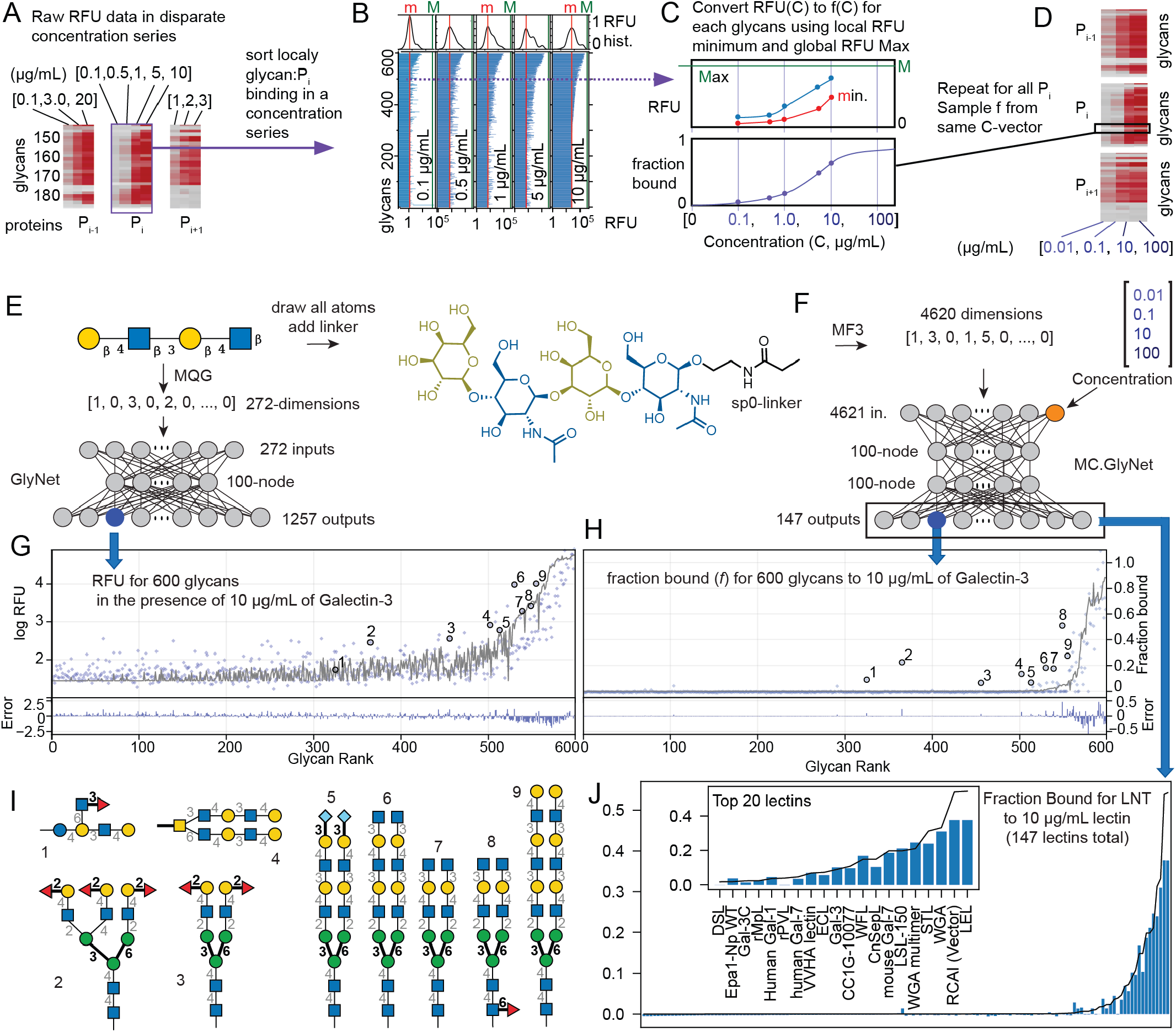
Workflow of data in training of MCNet. A) 147 lectins tested at multiple concentrations in the CFG v5 data were identified. B) Microarray RFU distributions were processed to extract a background and maximum which were then C) used to rescale into a fraction bound and D) resampled at multiple concentrations. We compared E) our prior monosachharide-tree based RFU model to F) an atom-graph (includes spacer structures) model using Morgan fingerprints and protein concentrations. The results (G & H respectively) show similar trends. In particular, I) a group of glycans predicted by the MF Net model to bind more strongly to Gal-3 than the CFG data, overlaps significantly (6 of 10) with over-predictions in the prior GlyNet model, and are of structures reported^68,69^ to have binding. J) Model predictions for binding of LNT to the 147 lectins.

To test *f* as a predictive output, we trained a fully connected neural networks which has the form, ***f*** = *Net*(**M**,C) where inputs **M** and C are MF3 description vector and concentration, *Net* is the neural network function, and ***f*** the resultant fractions bound. This architecture, referred to as MCNet, has 3931 inputs, one for protein concentration (C) and the 3930-dimensional **M**-vector MF3_c_ and 147-dimensional output ***f*** vector for 147 lectins from CFG dataset (**Figure 2F**). Models were trained on CFG glycans using the same ten-fold cross-validation structure as above with one clarification: all concentrations were in the same hold-out fold. Due to novelty of the *f*-value, MSE for MCNet cannot be compared directly to MSE of GlyNet or MSE of any state-of-the-art models; instead, we examined individual prediction of glycan binding profiles for lectins with well-known binding properties, with a focus on Galectins (**Figure 2G** vs. **2H**). Supplementary information contains analogous alignment of predicted f-values for other lectins juxtaposed to predictions made by monosaccharide-based models (e.g., GlyNet).

From 599 glycans, MCNet predicted ∼10 with modest upward deviation from the predicted trend (i.e., “false positive binders” **Figure 2H**). Examination of the individual structures, we noted that 7 of the 10 glycans contain definitive galectin-binding motifs (LacNAc repeats, **Figure 2I**) and 6 of the 10 of these had the same over-predicted property in the prior GlyNet model, many more than random chance would predict. Atom-level models are learning that similar features of the molecules as GlyNet (**Figure 2E-H**), even though one is looking at smaller groups of atoms and the other at the groups of monosaccharides. Glycans 2 and 3 contained terminal LacNAc with a α-2 fucosylation on terminal Gal (group-H, **Figure 2I**); however α–2 fucosylated lactose have been reported to bind more strongly than lactose by Klassen and coworkers^68^ and the α–2 fucosylated LacNAc was confirmed as ligand for Galectin-1 by Surolia and coworkers (**Table S3**)^69^. A 2022 re-run of Galectin-3 binding to CFG arrays by Arthur, Stowel and co-workers (data not used in MCNet training) confirmed group-H glycans on CFG arrays are modest but detectable binders to Galectin-3^61^. The same glycans 2 and 3 in GlyNet model, were predicted to have an order of magnitude higher RFU than the “ground truth” (**Figure 2G**). We conclude that these glycans are experimental false negatives of glycan microarray measurement rather than “false positives” of the ML model. Atom-level MCNet that predicts fraction bound, not only achieved desired performance but also highlighted experimental false negative observations in CFG dataset. This is not a criticism of CFG dataset but merely a reminder of well-established notion that the experimental uncertainty defines a natural upper limit to the predictive performance of machine learning models^74^.

We tested the ability of MCNet architecture to make out-of-domain predictions. MCNet1 trained only on CFG data was unable to predict any binding for any glycomimetic molecules (**Figure 3C**). Training MCNet2 only on the GM dataset made it possible to predict the binding of GM compounds in held-out datasets, but MCNet2 was completely wrong in predicting the binding of CFG glycans to galectins (**Figure 3D**). In contrast, the **MCNet3** model trained on all GM and all CFG data effectively predicted GM binding to Gal-1, 3, and 7 for held out GM datasets and all lectin binding for all held out CFG datasets (**Figure 3E**, and supporting file **Per Glycan Prediction Plots.pdf**). Glycan and glycomimetic recognitions are non-overlapping and knowledge of one does not guarantee to knowledge of the other dataset. And conversely, blending of datasets is positively synergistic: accuracy of the prediction of binding of GM to galectins and even binding prediction for CFG glycans to 144 other lectins increased when the MCNet3 model was trained using both CFG and GM molecules. The number of compounds increased from 600 in CFG to 800 in CFG+GM dataset, but the mere 30% increase in the size of the dataset cannot account for an exponential increase in accuracy.

**Fig. 3.**
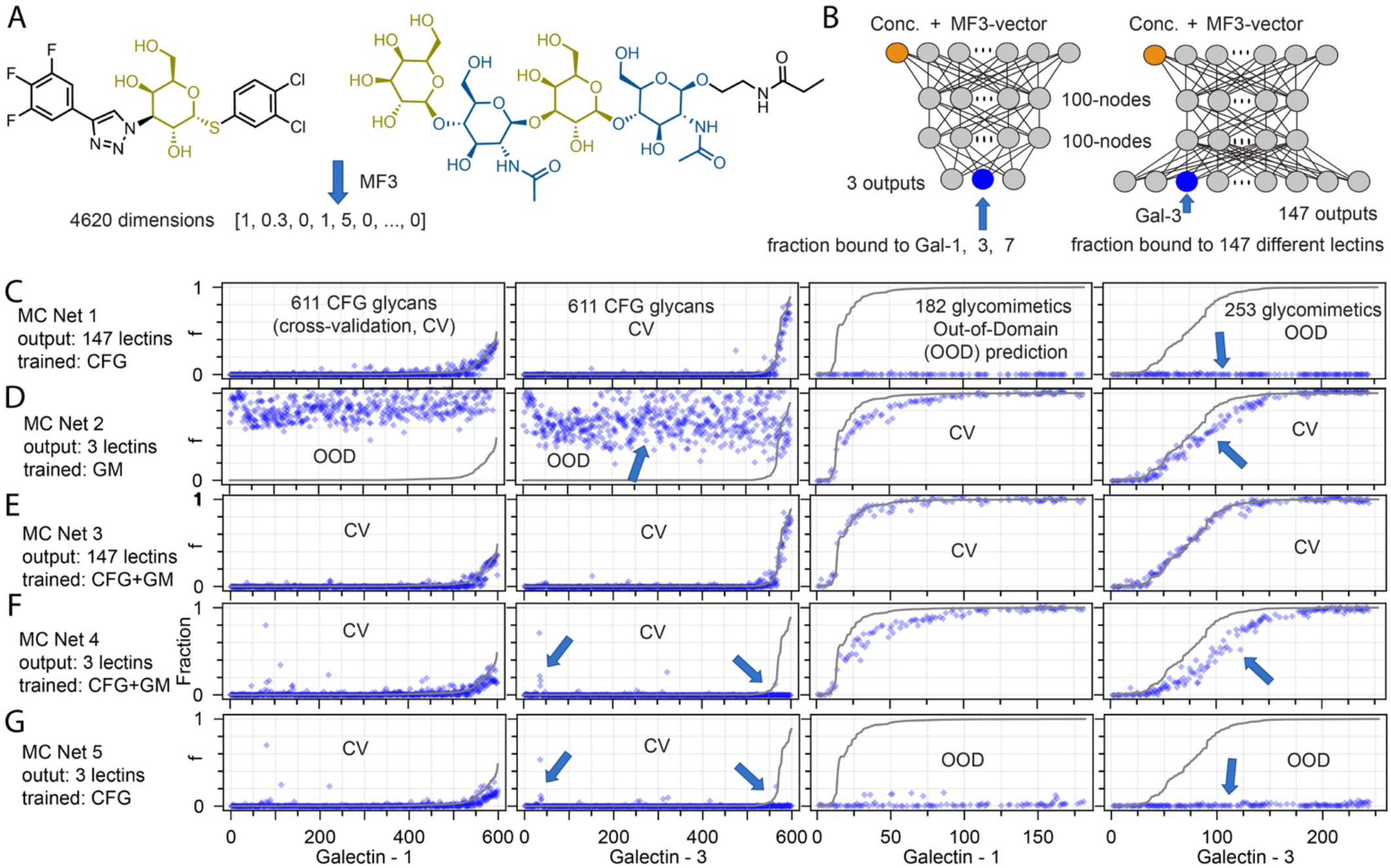
Predicted binding for the CFG glycans and the glycomimetics (GM) across different model classes. All plots are at ∼10 µg/mL. (A) Both glycans and GM compounds can be embedded as MF3 vector. (B) MCNet architectures with either 3 or 147 outputs were used in this test. (C) MCNet1 trained using CFG glycan data (analogous to **Fig. 2F, H**) predict binding of CFG glycans to Galectins in held-out datasets but failed to predict binding of GM to Galectins. (D) MCNet2 trained on GM data exhibits reasonable prediction of GM:Galectin binding but fails in predicting binding properties of CFG glycans. (E) MCNet3 trained using both datasets exhibits desired predictions and exhibits improvement in predictions of GM:Galectin interactions when compared to MCNet2 trained only on GM data or (F) MCNet4 trained only on galectin binding data of GM and CFG compounds. (G) MCNet5 trained only on Galectin:CFG data makes inferior predictions for binding of CFG glycans to Galectin-3 and it can learn only some trends in Galectin-1:CFG dataset.

Control MCNet4, trained on all GM data and only on a subset of CFG data describing binding to Galectin-1, -3, and -7 was also able to predict binding for held out GM and CFG datasets, albeit its accuracy dropped when compared to MCNet3 (**Figure 3G**). This observation illustrates how knowledge from seemingly unrelated glycan:GBP interactions, used in training of MCNet3, enhanced the accuracy of prediction of glycan:Galectin interactions. Another control MCNet5 trained only on a subset of CFG data describing binding to Galectins 1, 3, and 7 to CFG glycans, failed to predict GM binding (**Figure 3G**). We noticed that prediction of Galectin-3 binding deteriorated in MCNet4 and 5 but Galectin-1 predictions remained similar to MCNet1 and 3 (**Figure 3F-G**). It is tempting to blame the ML architecture for these failures, but the reasons are likely more fundamental. These failure of MCNet to extrapolate between CFG and GM datasets indicate that the rules of protein recognition by chiral hydrophilic sp^3^-centres in CFG dataset and achiral hydrophobic sp^2^-centres in GM dataset cannot be extrapolated from one another.

We anticipated that the learning the dose response should not be different from learning parameters of the curves that describes them. We made a curious observation, however MCNet did not always learn the monotonously increasing sigmoidal nature of the binding curves (**Fig. S9**). Models were trained using 50 f values at 50 input concentrations between 10^−5^ and 100 µM for each molecule (**Fig. S9**). Still, for many GM compounds, models failed to predict an obvious boundary condition: f must become zero at infinitely low concentrations. Instead for many GM compounds f plateaued at arbitrary values. Predictions for some molecules were satisfyingly monotonous, whereas for other we observed unexpected bumps and dips reminiscent of overfitting the monotonous “ground truth” with complex-shaped high-degree polynomial. We anticipate these problems to be resolved in future algorithms that learn f from discrete binding values.

### Out-of-domain extrapolation of binding properties to other datasets

MCNet architectures are capable of extrapolating lectin binding not only to glycan structures from GlyTouCan but also to any molecules (e.g., BindingDB, see **Fig. S10**). High-level observations mirror the observations already made in **Figure 3**. Specifically, MCNet1, trained only on CFG glycans, predicts zero binding performance for all molecules that are not *bona fide* glycans. Despite these deficiencies, MCNet1 effectively identifies glycans that are reasonable predictions as ligands for specific lectins. As there are currently no systematic ground truths for these predictions only some of these predictions can be evaluated. An interesting case is the bacterial glycans, which are not part of the training set. MCNet predicts that Galectin-1 and -3 interact with Gal-GalNAc oligomers, structures reminiscent of the E. coli O127 outer polysaccharide (OPS) and the Klebsiella pneumoniae O-antigens and other bacterial polysaccharides reported by Drickamer and co-workers^39^, Kiessling, Imperiali and co-workers^40^ and Cummings, Arthur, Stowell and co-workers^75,76^. Such data provides a baseline benchmark for all future predictions or experimental validations (see Supplementary file Models.zip).

### Mirror image glycan extrapolation

Inversion of all stereocenters in every molecule yields enantiomeric glycans and GM compounds. We tested the ability of MCNet to predict recognition between lectins and enantiomers of the molecules from CFG and GM datasets. Such observations have no “ground truth” only an “expert opinion”. For example, with 100% certainty, we can assert that every GM compound should bind to Galectin 1, 3 and 7 significantly better than the enantiomer of the same compound. The absence of cross-chiral recognition arises from the well-understood structural role of the chiral centers around the galactose ring. Interestingly, not all MCNet models predicted a drop in cross-chiral recognition. MCNet2 trained only on GM compounds placed the activity of enantiomers at random, either above or below the performance of the parental compounds (**Figure 4A**). This observation is recurrent in the ML literature and other reports have observed that chirality is one of the most difficult concepts for ML models to understand^77^. In contrast, an MCNet3 model trained on GM and CFG data—the latter contains numerous examples of epimers and diastereomers—was able to predict that nearly of enantiomers of GM compounds are less active than parental compounds (**Figure 4B**). It was important to include data for binding to all 147 lectins: MCNet4 trained using data for 300 GM and 600 CFG glycans binding only to Galectins 1,3,7 has diminished understanding of cross-chiral recognition (**Figure 4C**) when compared to MCNet3 (**Figure 4B**). MCNet appears to agglomerate the knowledge from various proteins effectively. Indirect knowledge of how epimerization of individual atoms influences glycan:protein recognition helps MCNet3 to produce the right answers for out-of-domain cross-chiral recognition.

**Figure 4.**
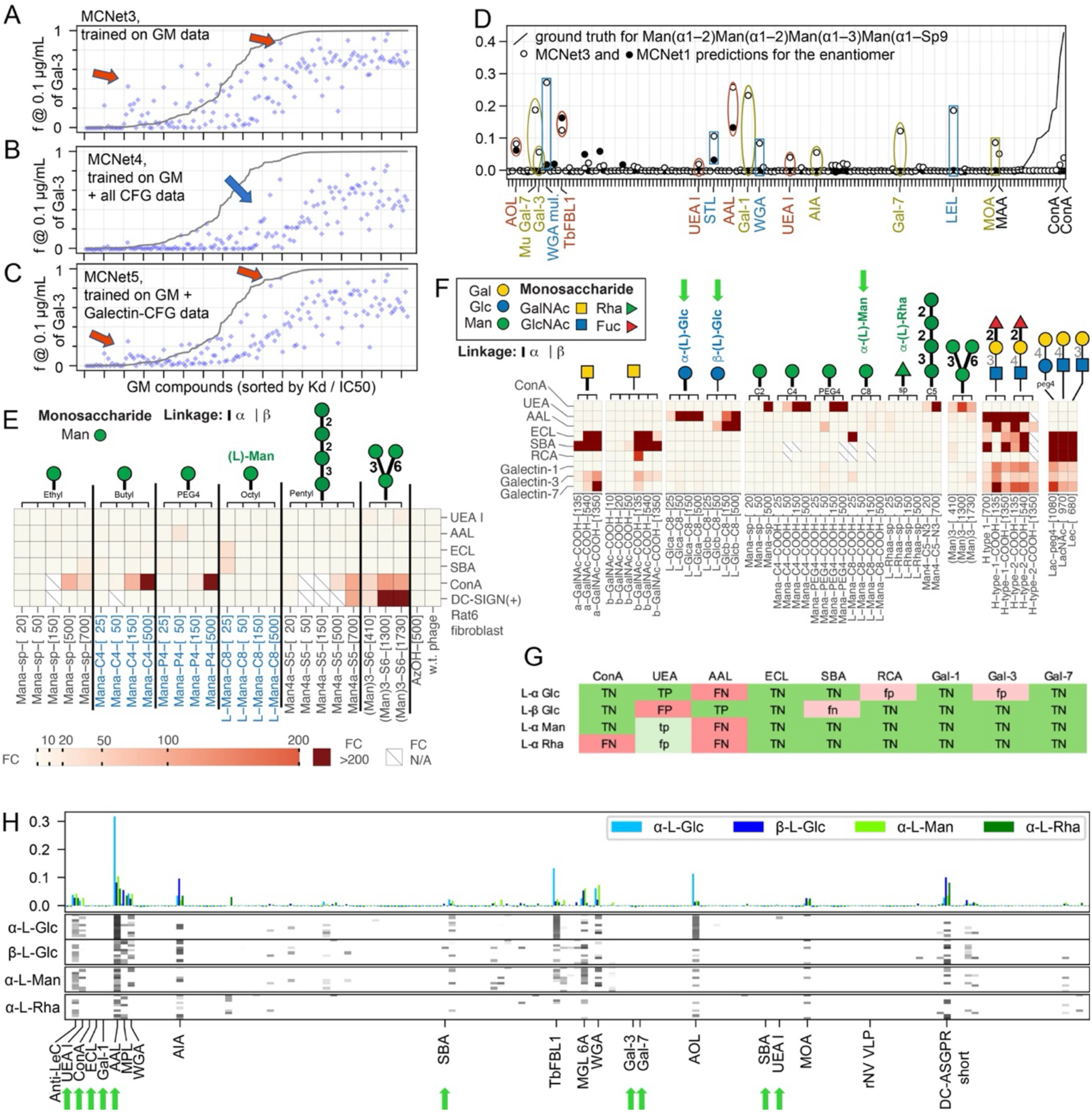
Prediction of Cross-chiral recognition by MCNet. (**A-C**) Solid line describes the ground truth: binding of glycomimetic (GM) compounds to Galectin-3. Blue dots are predicted binding strength of the GM-enantiomer. Expert knowledge of GM:Galectin-3 interactions dictates that all predictions should be below the solid line (i.e., activity of enantiomer must be lower than the activity of the parental compound). Only MCNet3 (**B**) produced answers aligned with “expert knowledge” of cross-chiral recognition. Galectins-1 and 7 exhibit similar trend (Figure **S14**). (**D**) MCNet3 predictions of binding of 147 Lectins to *α*-L-Glu, *β*-L-Glu, *α*-L-Man, *α*-L-Rha monomers displayed on a glycan array by a two-carbon linker (a.k.a., “sp0” in CFG notation). Predictions for lectins denoted with green arrows (ConA, UEA, AAL, ECL, SBA, Galectin-1, 3, 7) have been validated using monomers displayed with Liquid Glycan Array (**E-F**) confirming positive binding of L-Glu to AAL and UEA to and lack of binding of all other lectins. (**E**) Experimental confirmation of the cross-chiral recognition. L-Mannose, displayed at 25 to 500 copies on M13 virion fails to bind to ConA; however, glycans with D-Mannose exhibit an expected performance in the same liquid array. (**F**) Among nine lectins, the two canonical fucose binders bind to L-Glc. The rightmost seven columns are D-sugars as positive controls. More details are shown in Figure **S13**. (**G**) Numeric values (fraction bound-ML model predictions) and fold change (LiGA) were thresholded to classify each as binding or non-binding for comparison. Mean predictions were taken over the models from the cross-validation hold-out fold. Notably, 28 of the 36 predictions agreed between the two. (**H**) Details of fraction bound predictions for 4 L-sugars. Mean values are shown in coloured bars (top) and values for individual cross-validation folds in heatmaps (bottom).

Epimerization of a glycan may yield another common glycan (e.g. changing a glucose into mannose), but it may also give rise to an atypical glycoform (e.g. β-Glc to α-L-Ido). We examined the ability of MCNet to predict binding-strength for epimers of existing glycans (i.e. molecules that differ from the original by the inversion of one chiral centre, see Supplementary Epimer Heatmaps.pdf). MCNet 1 and 3 generally agree on their predictions with patterns of strongly bound lectins common to both models, but some predictions diverge where binding of the parent glycan in weak. Unsurprisingly, for many glycans both MCNet models identified that epimerizations in small glycans that destroy recognition by lectins of the original glycan, while often having no effect in larger glycans, see Supplementary Epimer Heatmaps.pdf. Further summary of observations is available in the supplementary materials.

We then tested the ability of MCNet to predict properties of enantiomers of the CFG glycans. A ground truth for these do not exist because it is unreasonable to propose to manufacture the CFG arrays with 600 enantiomers of existing glycans. However, Peter Kim’s mirror-image paradigm^78^ postulates that the binding of L-lectins to the enantiomers of CFG glycans can be measured by probing D-lectins against the existing CFG array. Unfortunately, synthesizing C-type lectins (DC-SIGN, DC-SIGNR, Langerin) from all D-amino acids is not trivial, and our attempts are still ongoing. Nevertheless, we established the experimental ground truth for these observations first by measuring binding of several lectins to various valences of L-Mannose and D-Mannose in the same Liquid Glycan Array^79^. Unlike D-Mannose, L-mannose was completely inactive in binding to every calcium-dependent lectin tested (ConA, DC-SIGN on the surface of live cells). Both MCNet1 and MCNet3 were able to corroborate these observations. For Man(α1-6)Man(β-Sp10 and Man(α1-2)Man(α1-2)Man(α1-3)Man(α-Sp9 both MCNet1 and MCNet3 predict that the mirror forms of the glycans have a significant decrease in binding to ConA (**Figure 4D**). MCNet predicted that a tetra-mannose enantiomer is recognized by lectins that traditionally bind galactose (Galectin, MOA, AIA), fucose (AOL, TbFBL-1, UEA-I) and GlcNAc, GlcNAc oligomers and LacDiNAc structures (Stl, WGA, LEL). Recognition by galactose-binding lectins is not entirely surprising given the resemblance of a triol configuration in L-Man and D-galactose and a preliminary LiGA experiment suggested weak binding between L-Man and prototypical D-Gal binding lectins (**Figure 4E-F**). Recognition of fucose and GalNac/LacNAc-binding lectins is less clear; we tested one of them and observed no binding of L-Man monomers to UEA. Further experiments are needed to confirm whether larger oligomers of L-Man possess UEA binding. Intrigued by the initial observations, we performed follow-up measurements of *α*- and *β*-L-Glu, *α*-L-Man, and *α*-L-Rha monomers displayed with Liquid Glycan Array binding to eight lectins: ConA, UEA, AAL, ECL, SBA, Galectin-1, 3, 7 (**Figures 4F, S13**). We observed an unexpected binding of L-Glu to the classical fucose binding lectins UEA and AAL, and we were pleased to observe that this binding was anticipated by MCNet3 (**Figure 4G-H**). We also observed no binding of ConA, ECL, SBA, Galectin-1, 3, 7 to any of the four L-monomers, the lack of such binding as was also anticipated by MCNet3 (**Figure 4G-H**).

Learning from the glass-based CFG glycan array and testing on the M13-bacteriophage based LiGA^79^ is not ideal, even though measurements made with these two arrays have been shown to correlate^79^. MCNet3 is trained on glass-surface display which is largely silent for all tested monosaccharides (**Figure 1E**). In contrast, the bacteriophage display of glycans, can detect monosaccharide binding after appropriate tuning of surface density. Curiously, the details of the L-Glu anomeric configurations are not matched between the observed AAL and UEA binding and the predictions (compare **Figure 4F-H** and **Figure S13**); however, the mere observation that MCNet3 anticipates binding to L-glucose by a series of fucose-binding lectins was very rewarding.

These predictions highlight the power of complete datasets, such as CFG array datasets for training of ML models that are capable of interpreting outside-of-domain prediction of cross-chiral recognition. The sheer number of chiral centres, diastereomers and near-stereoisomers provides rich training ground for training models focused on predictions of the effect of chirality rather than the effect of addition and substation of atoms and functional groups.

## Discussion

In this manuscript, we put contemporary understanding of protein:glycan interaction to a test by aiming to solve “biologically irrelevant” questions. We investigate whether it is possible to predict the binding ability of naturally non-existant or rare glycans, such as enantiomers of natural carbohydrates. Such extrapolation is not without its benefits. Glycans across all kingdoms of life appear to be tightly linked to speciation. Species recognize each other and make decision such as “friend” and “danger” by integrating data from glycan:protein interactions; glycan based pathogen-associated molecular patterns (PAMPs) and damage-associated molecular patterns (DAMPs) are central to our immunity. The prediction of strength of glycan:GBP interaction offers a universally comparable property and give rise to taxonomy and speciation orthogonal to the traditional 16S RNA and DNA taxonomy^80^. Synthesis of the mirror-image life forms is not the question of “if” but “when”^17,81^ These new species will not use “unknown” glycans but rather a very well-defined enantiomer of known glycans. It is important to understand how this new class of species coated with mirror image glycans will be interpreted using canonical protein receptors; will they be recognized by self/danger recognition receptors? Our manuscript does not resolve this question but builds the ML framework for addressing such question with higher and higher degree of confidence.

The atoms and their connections within a glycan appears to be necessary and sufficient for lightweight ML models to predict the recognition of that glycan by protein receptors. More generally, this atomic connectivity of any molecule is a minimal description of the molecule, and is sufficient to describe its properties. Identity and connectivity of atoms relates to Kolmogorov complexity of a molecule: it is sufficient to compute the properties of a molecule. For example: quantum mechanics (QM) can estimate electronic structure and Boltzmann ensemble of conformations from mere atomic connectivity. ML models can be trained using atomic coordinates; but such models produce properties of specific conformers. This ensemble needs to be computed and integrated over a Boltzmann ensemble to yield a property of such ensemble. We believe that thermodynamic scalar such as strength of binding between glycan and protein at a given concentration can be predicted without the knowledge of the Boltzmann ensemble of all conformations. We show that binding properties are possible to predict from mere atomic connectivity without any knowledge of any conformations. This observation forms a generalization of well-accepted Anfinsen hypothesis across the entire domain of molecular interactions. It can be phrased simply as “*atomic connectivity of the interacting partners are necessary and sufficient to determine the strength of the binding between these partners*”.

The goal of this manuscript is not to achieve a best-in-class performance of an ML model but rather to raise an awareness of the problems for future ML models to focus on. Growing advances in conversion of SMILES to fixed-length representation should be tested alongside MF and AQG in the subsequent manuscripts. This exploration will find the most effective representation of a chemical structure that drives MSE or other loss function to an optimum^43^. We leave such benchmarking and competition for future reports. We find it productive to re-examine what are perhaps the most bare-bone, and interpretable representations of molecules, the atomic *q*-gram (AQG) counts. Predictions of glycan properties from molecules represented with *q*-grams or MF of one atom were surprising. These models are only have counts of atom identities as inputs, with no information about any connections between the atoms. In retrospect, the marginal drop in MSE from 0.12 to 0.20 is not surprising because the 611 CFG glycans are rather diverse. There are only a few isomeric glycans and zero enantiomeric glycans in CFG collection. The binding of large glycans can be correlated with “expanded elemental composition” of such glycan without any consideration for topology. Mere memorization of composition may be a fallacy of models learning from datasets that contain molecules with diverse sets of atoms. The true challenge for molecular interaction is prediction of binding for *bona fide* isomers and stereoisomers. MCNet3 can predict binding to a diverse set of diastereomeric glycans enabling first exploration of mirror-recognition glycobiology. We noticed, however that MCNet struggles to predict obvious expert knowledge such as “binding of Galectin-3 to LacNAc must be ablated when carbons 2, 3, 4 or 5 are epimerized in any ring,” (see pages 50, 51 in Epimer Heatmaps.pdf). This is “obvious knowledge” for every glycobiologist. Yet it is not clear how to integrate such obvious expert knowledge into the ML training without exhaustive synthesis and measurement of all epimers of LacNAc.

We proposed that a “simple” total chemical synthesis of a lectin made of D-amino acids and testing of such a lectin on a CFG microarray can test many predictions made in this manuscript. Docking^82-87^ is another fast way to test some of the predicted interactions between enantiomeric glycans and common lectins. Such techniques could rapidly advance exploration of cross-chiral recognition in glycobiology. The lack of L-mannose binding to all lectins that bind D-mannose (**Figure 4**) was anticipated by Goldstein et al.^88^ The dichotomy between L- and D-mannose may rationalized based on the relation of L-mannose and L-rhamnose. L-rhamnose binding lectins do not recognize D-mannose^89-91^ and D-mannose-binding proteins rarely bind to L-rhamnose. But counterexamples are known, some lectins bind mannose and rhamnose equally.^92^ A report of Tatibouët and co-workers measured the binding of a multivalent BSA-L-Rha_17_ conjugate to ConA and DC-SIGN with IC_50_ of 3 and 0.6 µM while BSA-D-Man_16_ achieved similar activity (IC_50_ of 0.2 and 0.6 µM). Hence, even extrapolating between L-mannose, L-rhamnose and D-mannose may not be obvious to human experts and needs to be supported by experimental validation. Binding of L-glucose to lectins that recognize L-fucose is another experimental observation that was confirmed by MCNet. A minor discrepancy in alpha vs. beta anomers of L-glucose (Figure **4H** vs. **S13**) may be a result of a mismatch in predictions (extrapolated from glass-array data) and testing (made in liquid array). Part of the problem may be inability of glass-based CFG array to detect binding of glycans to monosaccharides (Figure **1E**). It is possible that signals from monosacharides are overshadowed by stronger binding to a larger oligosaccharide on the same array. Suboptimal presentation of glycans on the glass surface has been shown to be improved in LiGA, where density of the glycan can be varied systematically.^79^ Integration of LiGA data and CFG data *via* a universal “fraction bound” will reintroduce critical information about monosaccharide:lectin interactions. An agglomeration of such measurements would form an effective ML training dataset for learning about the role of stereochemistry in glycan recognition and in overall molecular recognition. Applying an ML model trained on aggregated datasets to problems of cross-chiral recognition in glycobiology may uncover further unexpected observations, which eventually will reinforce the overall protein-glycan “rules of recognition”.

Integration of microarray data from multiple protein concentrations into a surrogate of binding strength (e.g. EC_50_)^61^ is one approach, but not free of caveats. For example, line shape in dose responses are not always trivial to convert to definitive EC_50_ values and steep, cooperative responses should be delineated from non-cooperative sigmoidal curves (i.e., curves need Hill coefficients). We do not use the experimental errors in our training process, but in models that do, propagation of errors from responses to errors in fits adds another layer of complexity. Both problems are avoided by learning fraction bound instead of Kd. The atom-level glycan representation overcomes the deficiencies observed in monosaccharide-based representations of glycans. For example, the CFG dataset does not contain xylose or L-mannose, hence, a GlyNet-type model or LectinOracle^48^ that embeds monosaccharides simply cannot predict binding of oligosaccharides that contain xylose or L-mannose. Embedding an atom-level description of glycans solves this problem and gives the first glimpse into the unseen mirror-universe of glycans: such bizarre mirror-image monomers have never been considered by any ML model due to lack of any training data. These observations will help to improve understanding of the risks associated with the synthesis of living mirror-image cells.^25^

The ability to represent a structure does not imply the ability to predict its binding accurately. The atomic q-gram approach has an obvious out-of-domain deficiency: If the CFG and GM dataset does not contain an element (e.g., Br), how can a model even know what a bromine atom is? Any molecule with Br atom appears to be a “cold start problem”. An analogous rationale can be made for groups of atoms (functional groups) that are absent from the training dataset. GlyTouCan and BindingDB contains cyclopropane rings, which is completely absent from CFG and GM training dataset, **Table S2**. Interestingly, the chirality of individual atoms presents another “cold start problem” and in the absence of any examples of chiral differences, the ML model (e.g., MCNet2) cannot make any meaningful predictions for inversion of stereochemistry. Chemical intuition can explain the properties of Br from trends in periodic table (e.g., H, F, Cl, Br, I, etc) from simple features like count of electrons and trends in Periodic Table of Elements. Chemical intuition suggests that ML models could take into the account electronic structure of atoms and functional groups (orbital structure, polarizability, Milliken charge) instead of their linguistic description (Bromine, Chlorine, cyclopropane, etc.). Such models might be able to overcome some “cold start” problems. Indeed Jeff Kelly and co-workers profiled 357 quantum mechanical descriptors to predict those that contribute the most to protein-glycan interactions^93^. However, *simple* electronic structure theory cannot explain long-range effects caused by chirality. The effect of chirality on molecular recognition is easier to teach “by example”. This observation highlights critical importance of datasets emanating from glycans, glycobiology and protein:glycan interactions for global efforts in design of “chirality aware” ML models of molecular recognition. Integration of datasets that describe protein:glycan interactions into any ML models introduces the most complete description of the effect of local changes in chirality on the overall molecular recognition; no other fields offer such exhaustive coverage of the properties of diastereomers. Our work helps such integration by proposing strategies for integration of disparate quantitative data via universal fraction bound and encouraging all-atomic molecular representation in the field forged on a monomer-centric alphabet.

## Supporting information

Raw RFU to Fraction Bound Conversions

Per Glycan Prediction Plots

Predictions Across Concentrations

Mannose Enantiomer Prediction Plots

All Enantiomer Prediction Plots

Epimer Heatmaps

SMILES-GlyTouCan

Models

## Acknowledgements

We thank Natalia Baranova and Jared Zhang for their efforts in searching for small-molecule galectin inhibitors and reported *K*_d_ values in the published literature. We thank Todd Lowary, Matt Macauley and Chris Cairo for their comments and feedback. We thank James Smith and co-workers at the Mirror Biology Dialogues Fund (https://www.mbdialogues.org/) for providing comments of the mirror-image life discussion in this manuscript. This work was supported by Canadian Institutes of Health Research (CIHR) (no. 180445), Natural Sciences and Engineering Research Council of Canada (NSERC) Discovery Grant (RGPIN-2016-402511) and GlycoNet (CR-29 and TP−22).

