## Supplement Information for "Atom-level Machine Learning of Protein-Glycan Interactions and Cross-chiral Recognition in Glycobiology"

#### Methods

##### Comparison of Morgan fingerprint and atomic *q*-gram encoding

Two types of embedding were used to trained models that take molecular embedding of glycans as input and predict RFU values for 1257 lectin compositions as output.
we decided that our atom-graph models would not directly include the commonly mis-used "bond order" and replaced it with hybridization of atoms (nodes) as sp, sp2, sp3. To differentiate between q-grams with and without edges to other neighbouring atoms we included a count of attached hydrogen atoms.
Training was performed using 10-fold cross-validation and hold out sets were selected as in the previous GlyNet paper. Indistinguishable molecules were either all used for training, or all excluded from training even though the ability to encode the linker/spacer structures means that some previously indistinguishable glycans are now different. We compared MSE (loss) of models with different levels of details in the atoms/nodes, different maximum subgraph sizes $q = \leq 1 - 6$, numbers of hidden layers, nodes in hidden layers, and choices of the weight decay parameter, and compare atom-level structure results to our prior monosaccharide-level work, GlyNet.

##### Preprocessing and merger of disparate datasets via fraction bound

We identified 147 cases were the multiple CFG arrays were run on the same protein, but with at least three different input concentrations. We did not use the clamping of the bottom third to zero as in our prior report[28]. Instead, we identified an average minimum RFU signal (see Suppl. Fig. 6) from abundance peaks that we take to correspond to the non-binding condition. We then estimate the fraction bound of the glycan by rescaling each glycoconjugate's mean RFU (across flowcell replicates). The per experiment RFU minimum becomes 0, and the maximum RFU experimentally observed across all data becomes 1. Values above and below this range were clamped to 1 and 0 respectively. For these 147 cases, we calculated fractions bound from CFG array results.

For each glycoconjugate, we took the fractions bound at different concentrations and extrapolated these to a set of fifty concentrations between 0.01 μM and 100 μM.

Note that each molecule was provided as input fifty times, once for each concentration. Concentrations are from the set $(1, 1.5, 2, 3, 4, 5, 7) \times 10^{(-5, -4, -3, -2, -1, 0, 1)}$ and 100 μM.

In cases where we had a published Ka or Kd value, such as the glycomimetic compounds, fractions bound were calculated from the published Ka/Kd values at the same set of concentrations using the Henderson-Hasselbalch equation. We then trained models with these concentration-molecule inputs with the fractions bound as the desired outputs.

##### Training and evaluation of the models

We explored variations in the maximum sizes of the graphlets and in the per atom features. Adding more features, and larger graphlets gave better results (Fig. 3), except for the largest radii (4 and greater) of Morgan fingerprints. The atom-level models are comparable to monosaccharide ones on the same CFG dataset in our prior work. Especially for the Morgan fingerprint-based ones, MSE performance is comparable. Not all hyperparameters are equally good, but we can choose acceptable ones. In particular, the models are sensitive to number of hidden layers, the size of the hidden layers, choice of atom labels, and the types of graphlets used.

**LiGA**

LiGA experiments were performed as described in prior reports [1,2]. The most substantial changes was the addition of the L-Man decorated phage to the array mixture. Fuller details will be provided in subsequent reports.

1. Sojitra, M. et al (2021). Genetically encoded multivalent liquid glycan array displayed on M13 bacteriophage. Nature Chemical Biology, 17(7), 806–816. https://doi.org/10.1038/s41589-021-00788-5
2. Sojitra, M et al. (2024). Measuring carbohydrate recognition profile of lectins on live cells using liquid glycan array (LiGA). Nature Protocols, https://doi.org/10.1038/s41596-024-01070-3

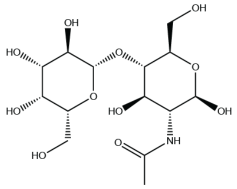

unmodified glycan (LAcNAc)
Used for ITC, ESI-MS

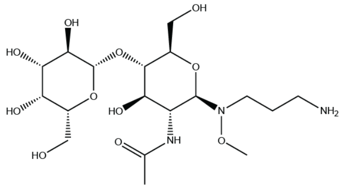

N-alkyl-methoxyamine bi-functional linkers
For microarrays
dot:10.1007/s10719-017-9785-4 M.S.M. Timmer and coworkers

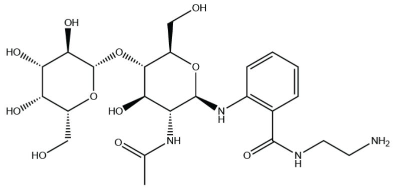

2-amino-N-(2-amino-ethyl)-benzamide (AEAB)
For microarrays
doi: 10.1016/j.chembiol.2008.11.004 R. Cummings and coworkers

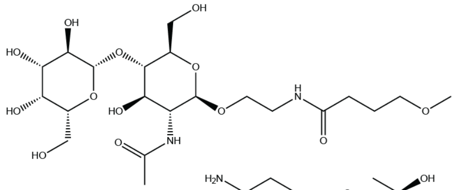

CFG Glass Array Sp0

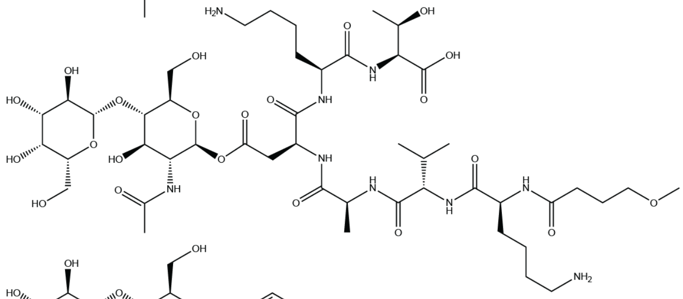

CFG Glass Array Sp24

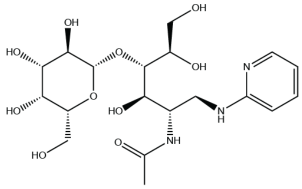

frontal affinity chromatography with
fluorescence detection (FAC-FD) pyridylaminated (PA)

**Fig. S1.** A diversity of chemical structures are used to represent a glycan.
Here the same glycan (LacNAc) is shown with different linkage structures employed in various measurement assays. Notice that the linker may be compatible in size to (or even larger than) the disaccharide.

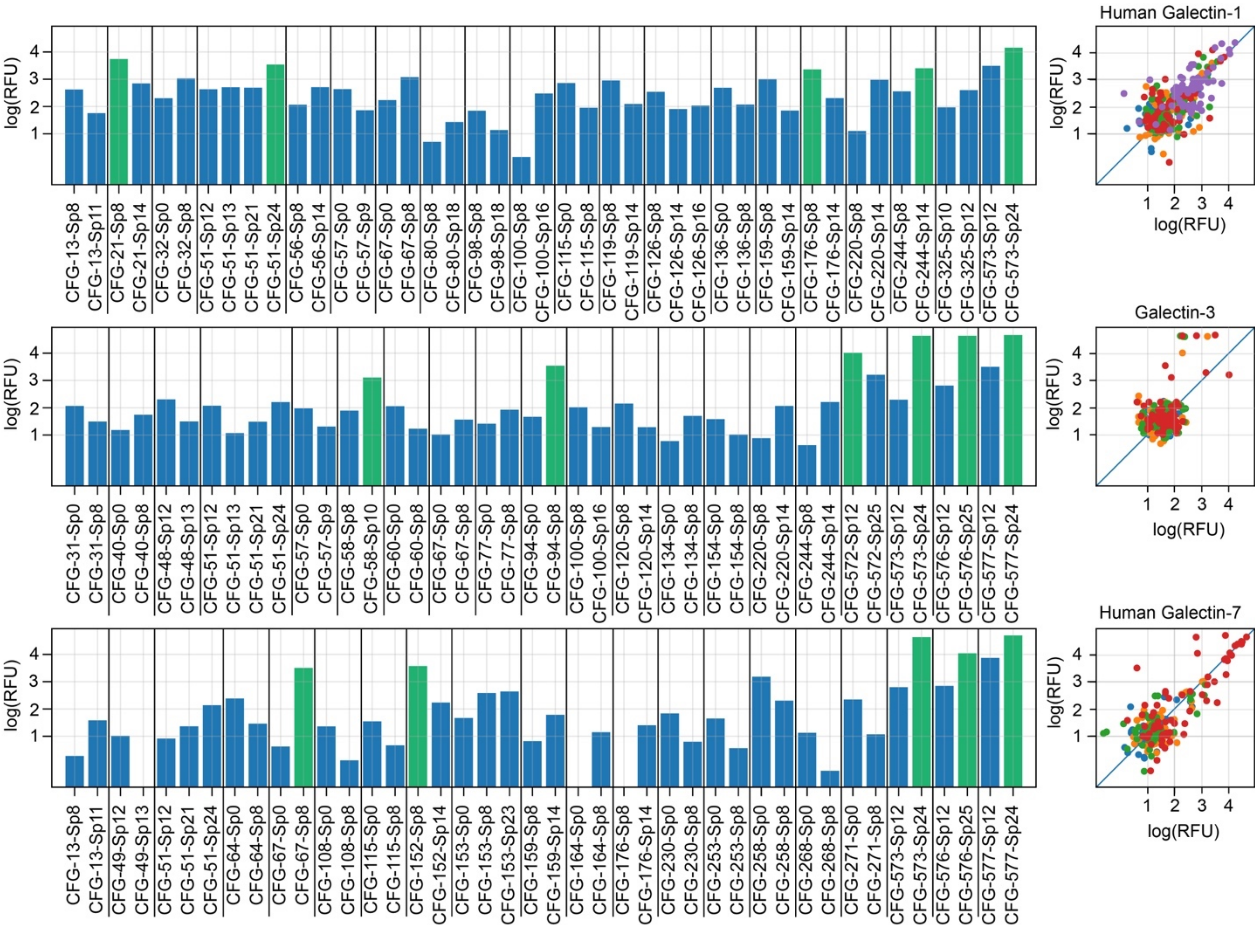

**Fig. S2.** Effect of different linkages on the binding among glycans in the CFG data.
At left blue bars show the reported binding (in RFU) for twenty glycans with multiple spacers.
At right the scatterplots plot the RFUs of pairs with identical glycans but differing spacers.
Different colours represent different concentrations of the galectins.

A

| | No Confident Zeros | | Assymetric Pseudo-zeros | | Symmetric Pseudo-zeros | |
|---|---|---|---|---|---|---|
| alpha-Gal - Sp8 | n/a | 1.4 | **3.0** | 1.4 | 3.0 | 1.4 |
| beta-Glc - Sp8 | n/a | 1.3 | **3.0** | 1.3 | 3.0 | 1.3 |
| beta-Man - Sp8 | n/a | 2.2 | n/a | 2.2 | n/a | 2.2 |
| 13-saccharide | n/a | 4.0 | n/a | 4.0 | n/a | 4.0 |
| Neu5Gc + above | n/a | 1.5 | **3.0** | 1.5 | 3.0 | 1.5 |
| 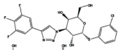 | 7.3 | n/a | 7.3 | n/a | 7.3 | n/a |
| 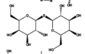 | 4.3 | n/a | 4.3 | n/a | 4.3 | n/a |
| 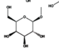 | 2.4 | n/a | 2.4 | n/a | 2.4 | **2.0** |

B

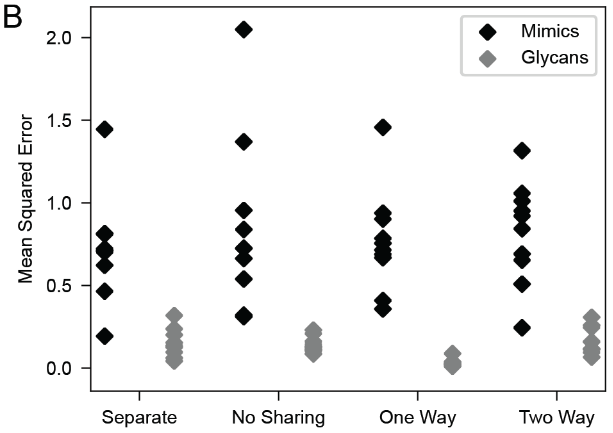

**Fig. S3.** Pseudo-labeling RFU and Kd datasets with "confident zeros".

(A) Example of data that combines RFU values for CFG glycans (small value = no binding) and log(Ka) values for glycomimetic (GM) compounds (small log(Ka) = strong binding). In asymmetric pseudo-labeling, any compound that has low log(RFU) value was also assigned a low log(Ka) value (i.e., compounds that show no binding in CFG microarrays are very likely to possess a very low Ka). In symmetric pseudo-labeling, any compound that has low log(Ka) value was also assigned a low log(RFU) value. Labeling is made under assumption that compounds that show weak binding should not show any binding on glycan arrays. We hypothesized

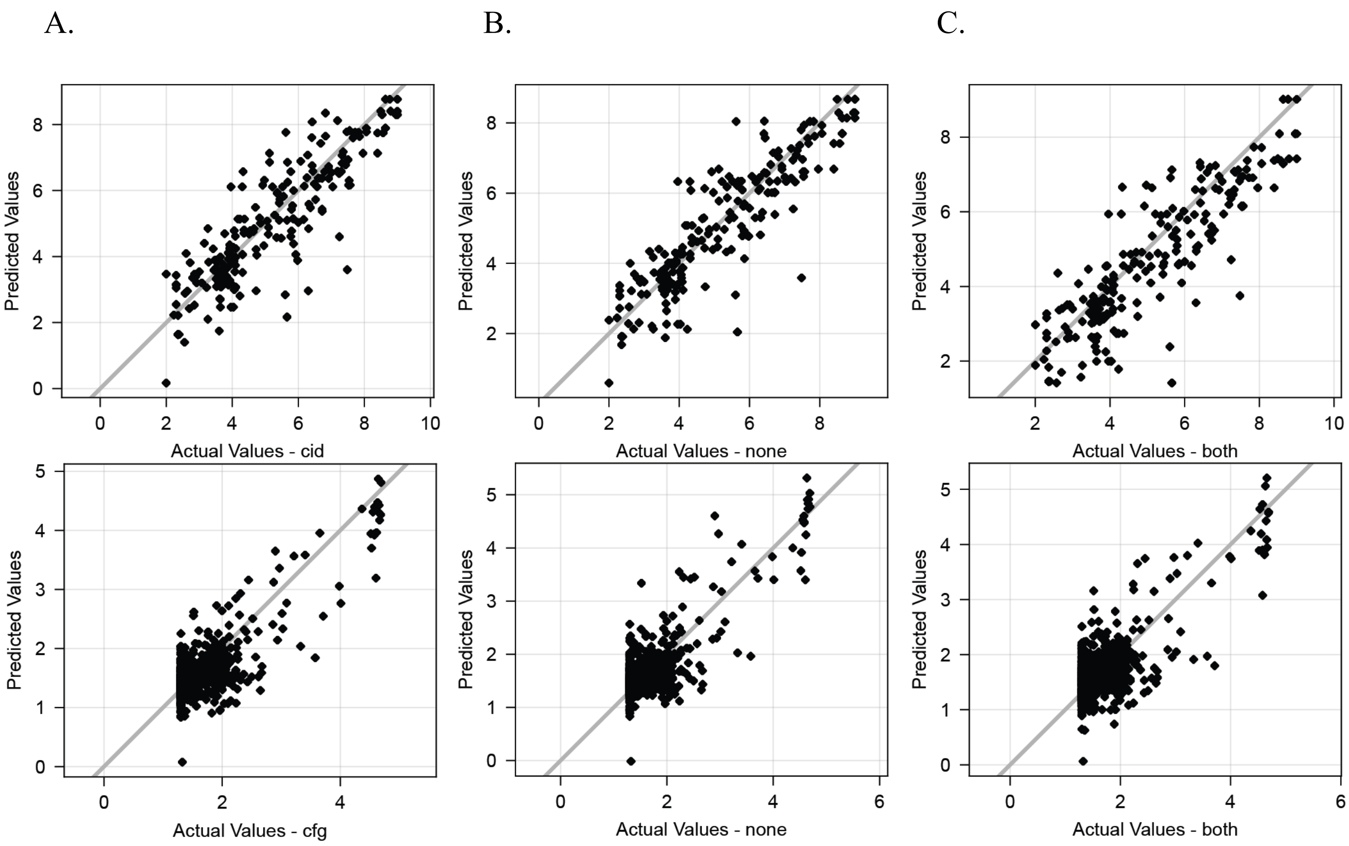

**Fig. S4.** Plots of predicted vs. actual values (on hold-out sets) of CFG glycans binding to Galectin-3 sample (upper panels) and the GM compounds binding to Galectin-3 (lower panels). (A) Model1 trained on original RFU-Kd data; (B) Model2 trained on data with asymmetric pseudo-labels; (C) Model3 trained on data with symmetric pseudo-labels. All three models are reproducing the data but show no sensitivity to any additional pseudo-labels.

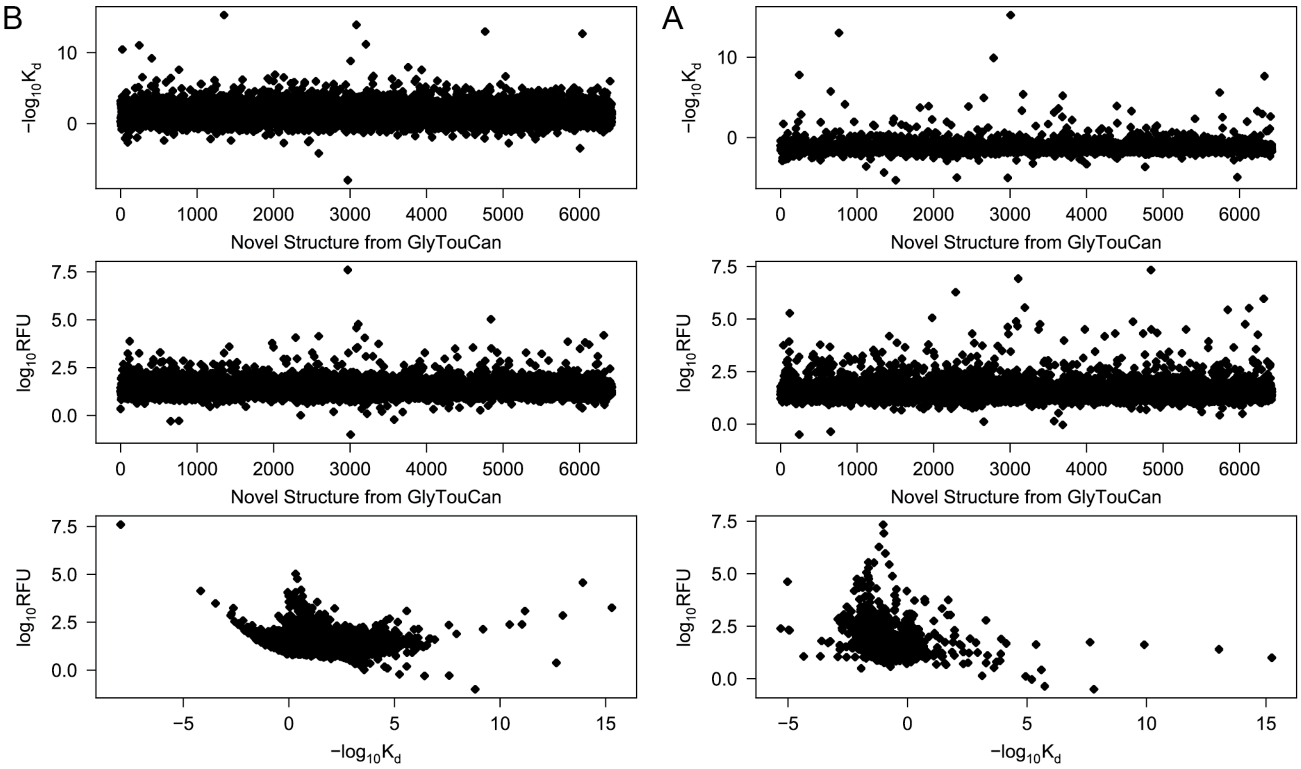

**Fig. S5.** Two model trained on combined RFU+Kd data

(A) with pseudolabels (symmetric) and (B) without pseudolabels. Models were tested for their ability to predict binding of glycans from GlyTouCan to Galectin-3. Although models can make predictions, there exists no anticipated direct correlation between predicted log(RFU) and predicted log(Ka). The trend is exactly the opposite: compounds with high log(RFU) (i.e., strong binders in microarray) are predicted to have low Ka (not a binders) and compounds with high Ka (strong binders) are predicted to have no detectable RFU signal on microarray.

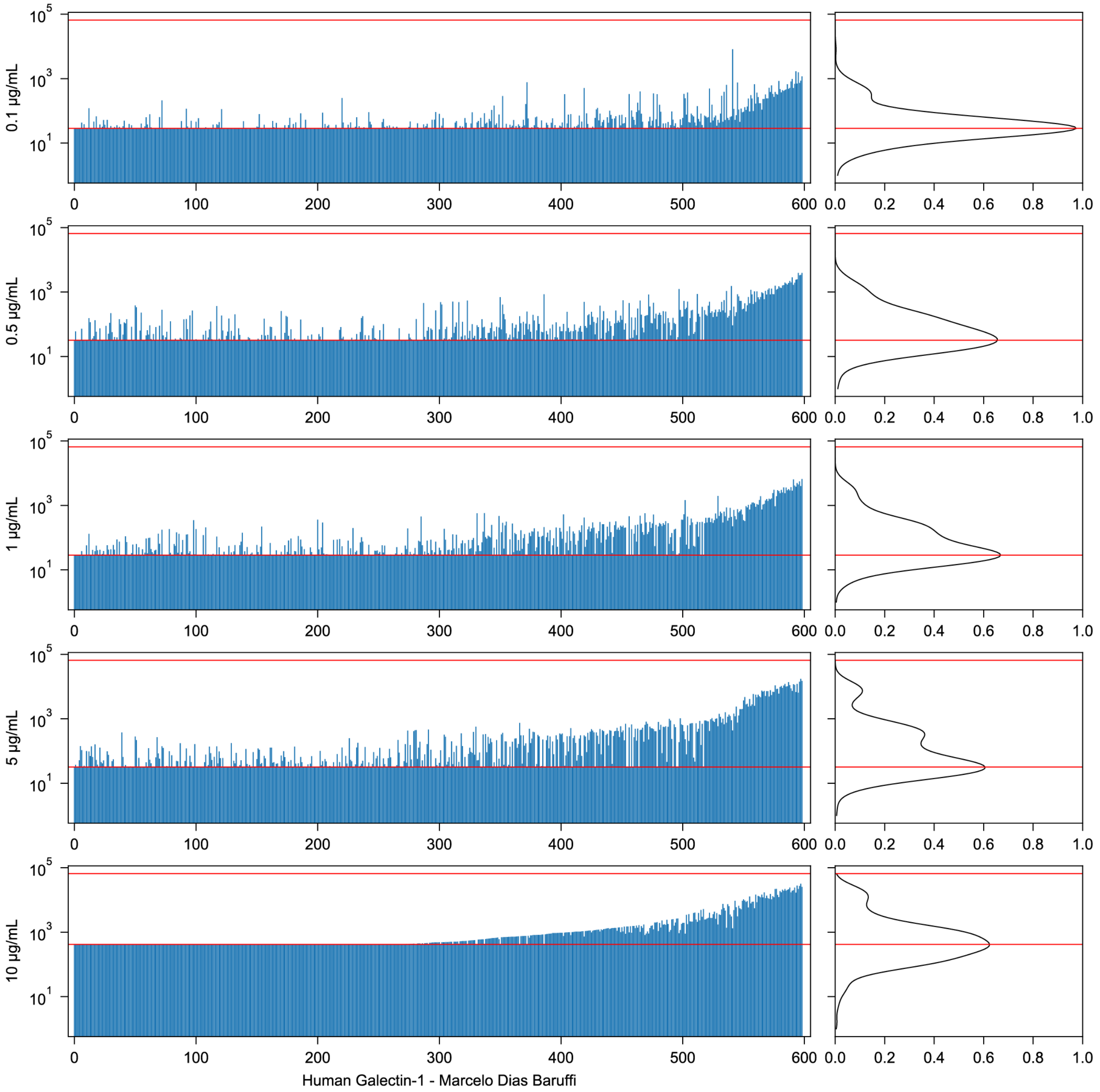

**Fig. S6.** Conversion of glycan microarray RFU values to fractions bound.

Shown are RFU data from five arrays with different concentrations of Galectin-1. In each of the five, many of the RFUs are clustered about a low value. This is shown as peaks in the kernel density estimation plots (right of figure). The levels of these peaks (lower red lines) are taken as background RFUs, and assumed to be non-binding ($f = 0$). The maximum RFU across all samples (upper red lines) is taken as full binding ($f = 1$).

### Some Henderson–Hasselbalch Mathematics

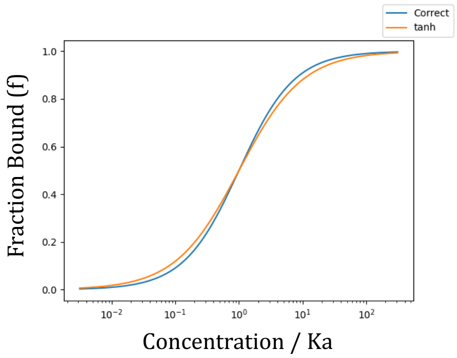

At Concentration = Ka:
$f$ = 1 / (1 + Ka/Ka) = 0.5

At C = 10 Ka: f = 0.909,
tanh sigmoid = 0.881

| | |
|---|---|
| $X + Y \rightleftharpoons XY$ | reaction |
| $K_a = \frac{1}{K_d} = \frac{[X][Y]}{[XY]}$ | kinetic equation |
| $[Y_0] = [Y] + [XY]$ | conservation of mass |
| $f = \frac{[XY]}{[Y_0]}$ | fraction of $Y$ bound |
| $1 = \frac{[Y]}{[Y_0]} + f$ | fraction unbound |
| $\frac{K_a}{[X]} = \frac{[Y][Y_0]}{[XY][Y_0]} = \frac{1-f}{f}$ | rearranged with substitutions |
| $f = \frac{1}{1 + K_a/[X]} = \left(1 + 10^{-\log K_d}/[X]\right)^{-1}$ | solved for $f$ using Ka/Kd and [X] |

**Fig. S7.** Some mathematics of the Henderson-Hasselbalch equation.

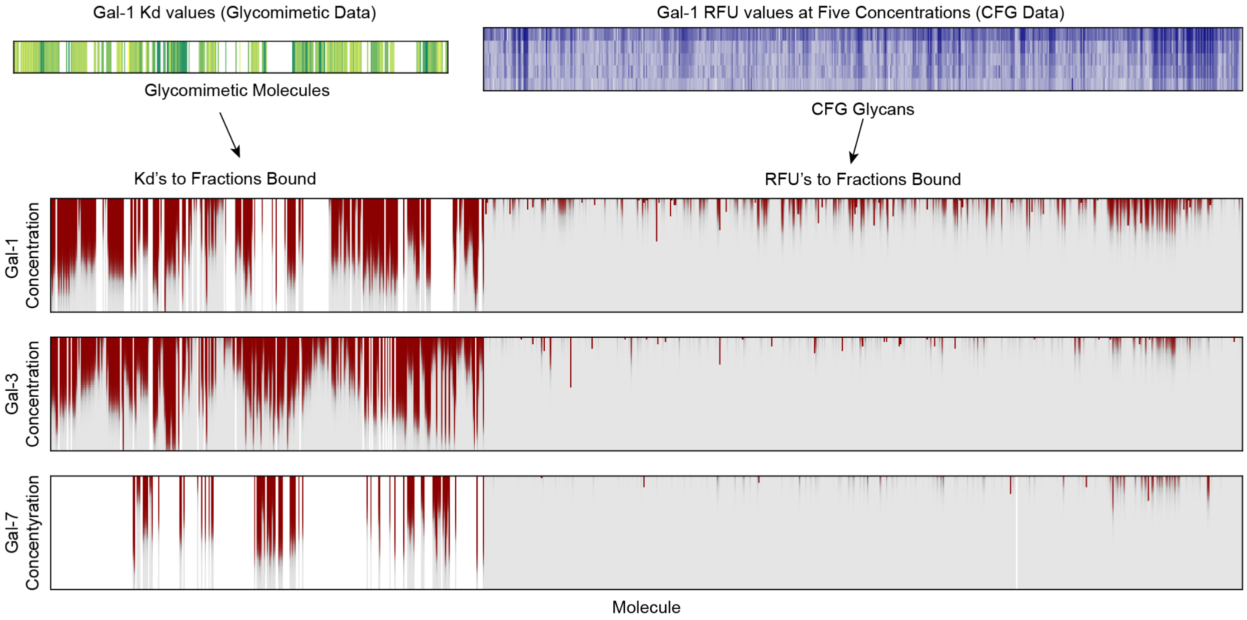

**Fig. S8.** Overview of the merger of data from multiple sources to unified fractions bound. Here Kd values for Galectin-1 binding to the glycomimetic molecules and RFU values for five Galectin-1 CFG arrays are converted into set of fractions bound, *f*s, across the 50 concentrations (vertical axes). These are then combined into a single set across all the molecules. Unknown (missing) values propagate through the processing and produce holes in the results (white columns). Also shown are the results for Galectins-3 and 7.

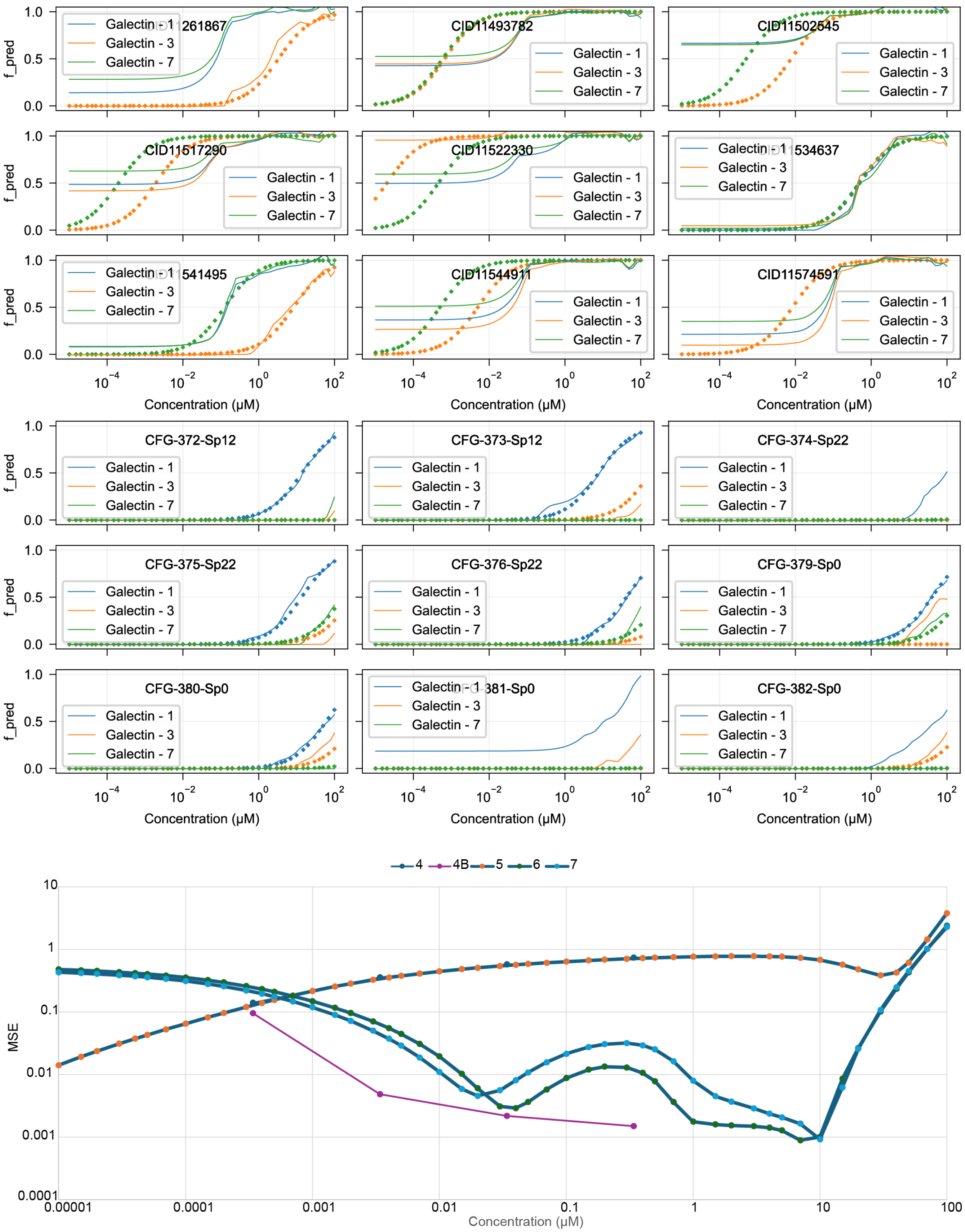

**Fig. S9.** The prediction performance of the MCNet3 model as a function of concentration. Shown at top are predicted fractions bound to Galectin-1, -3, and -7 (solid curves) against the data points used to train the model (diamonds). While the CFG glycans (top 3 rows) in general reproduce the curves, the glycomimetics (middle 3 rows) have significant mismatches, although there are some exceptions. The MSE loss over the hold-out molecules varies with the concentration.

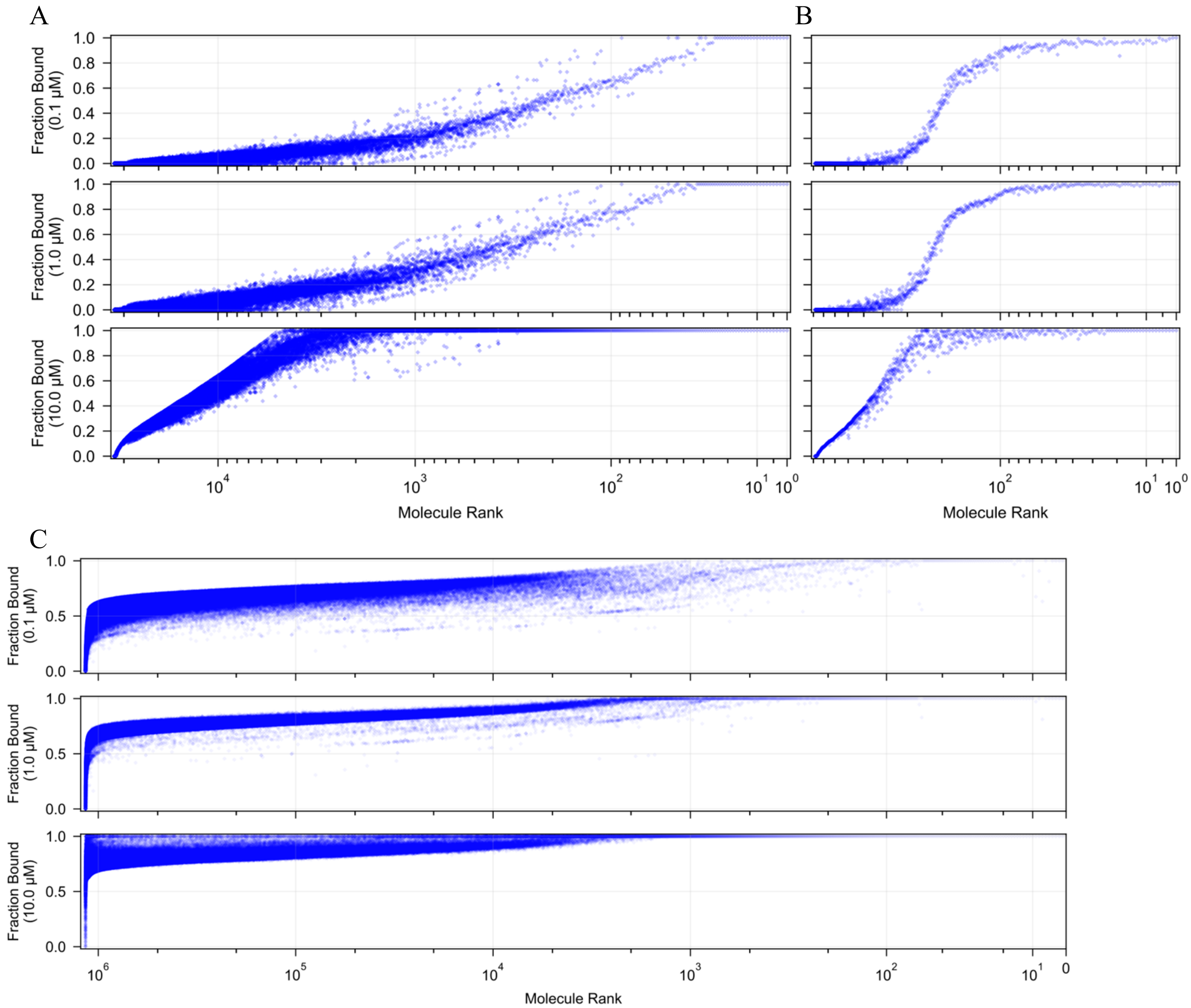

**Fig. S10.** Predictions of binding of GlyTouCan and CFG glycans to Galectin-1.
A) GlyToucan and B) CFG. The molecules have been ranked by the sums of the fractions bound across the three concentrations (0.1 μM, 1.0 μM, and 10 μM). A non-linear axis is used to emphasize the strongest binders. C). Predicted binding over the BindingDB set, by Model MCNet at 0.1 μM concentration. See SMILES-GlyTouCan.txt for corresponding molecule identifiers. In contrast to panels A and B, nearly all one million small molecules are predicted to have $f > 0.5$ at 0.1 μM concentration.

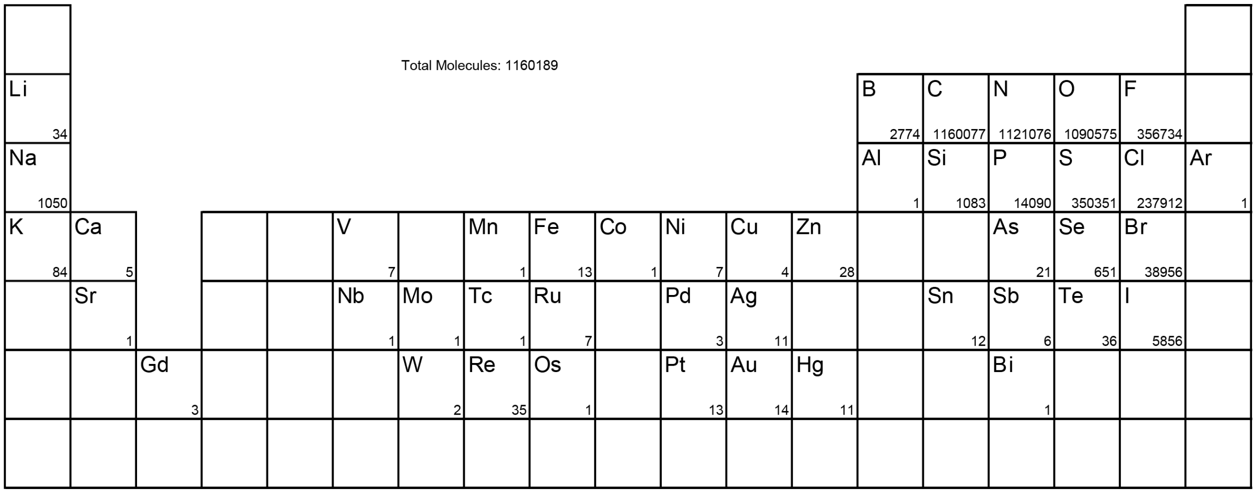

**Fig. S11.** The elements in the BindingDB molecules.

Numbers in the boxes are the count of molecules containing at least one atom of that element.

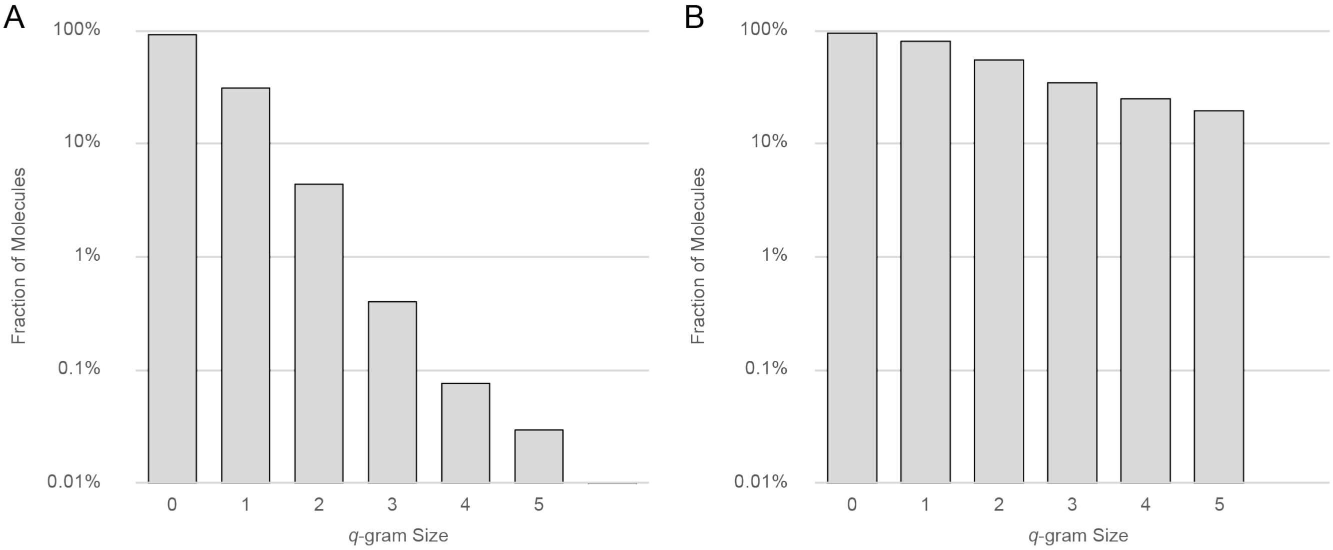

**Fig. S12.** Effect of *q*-gram size on the representability of molecules.

The fraction of molecules for A) the BindingDB dataset and B) the GlyTouCan dataset that can be represented using only q-grams present in a training set comprised of the combined CFG glycan and glycomimetic molecule sets. Notice that the glycans in GlyTouCan have greater similarity to the training set than the assorted small molecules of BindDB as evidenced by slower drop in the fraction of representable molecules as the maximal q-gram size increases.

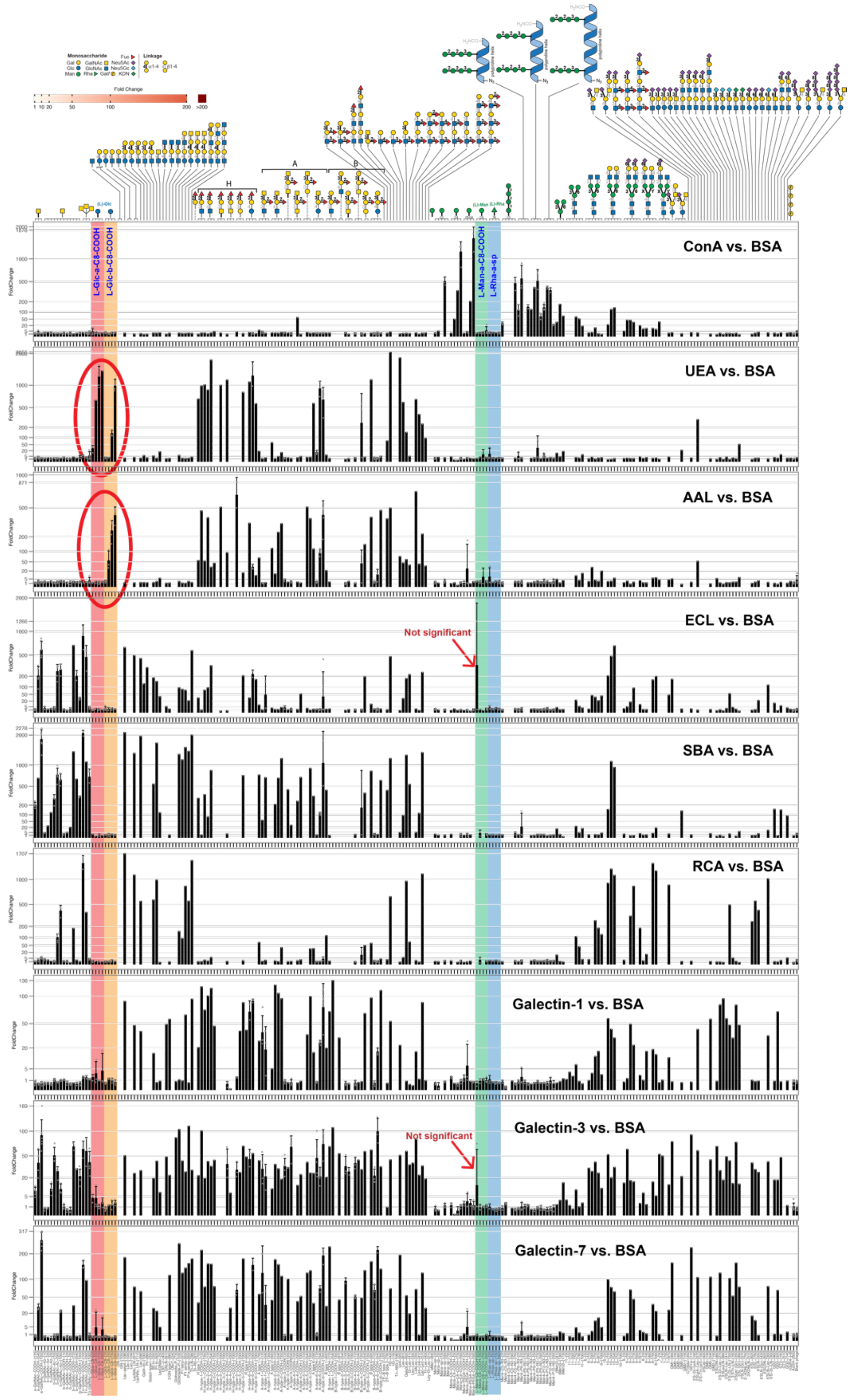

**Fig. S13.** Experimental data showing binding of a Liquid Glycan Array to eight lectins. Displays of four L-sugars (α-L-Glc, β-L-Glc, α-L-Man, α-L-Rha) in the coloured columns show substantial binding (large fold-changes) for the L-glucoses when presented to the lectins UEA-I and AAL but not in other cases.

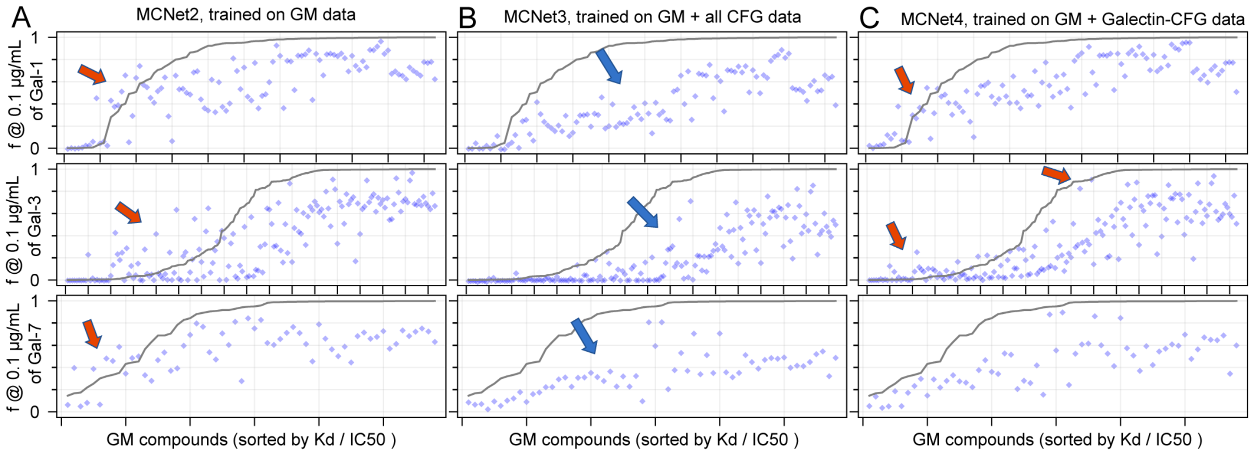

**Fig. S14.** Prediction of cross-chiral recognition by MCNet.
Solid lines describe the ground truth: binding of glycomimetic compounds to Galectins-1, 3 and 7. Blue dots are predicted binding strength of the GM-enantiomer. Expert knowledge of GM:Galectin interactions dictates that all predictions should be below the solid line (i.e., activity of enantiomer must be lower than the activity of the parental compound). Only MCNet3 (B) produced answers aligned with the "expert knowledge" of cross-chiral recognition.

**Table S1.** Thermodynamic Galectin binding data from Brewer's and Surolia's groups. Column #2 lists RFU values of the glycans in CFG array binding at 5.0 µg/mL of Galectins 1, 3, and 7. Columns 4-5-6 list Ka values ($M^{-1}$) measured by ITC. Yellow highlights values are from Surolia and co-workers.

| CFG code | 1 | 3 | 7 | Name | Gal-1 300 | Gal-1 287 | Gal-3 300 | Gal-3 287 | Gal-7 300 | Gal-7 287 | IUPAC |
|---|---|---|---|---|---|---|---|---|---|---|---|
| 154-Sp0 | 51 | 28 | 4 | Lactose | | 6100 | 802 | 1160 | 2200 | 4900 | Gal(β1-4)Glc(β |
| 154-Sp8 | 48 | 35 | 16 | | | | | | | | |
| 134-Sp0 | 246 | 16 | 8 | LacNAc-I<br>LNB | 9000 | | 8200<br>1500 | 2880 | | | Gal(β1-3)GlcNAc(β |
| 134-Sp8 | 397 | 38 | 12 | | | | | | | | |
| 153-Sp0 | 200 | 42 | 4 | LacNAc-II<br>LacNAc | 11000 | 24000 | 19000<br>5238 | 37000<br>8645 | 1700 | 6300 | Gal(β1-4)GlcNAc(β |
| 153-Sp8 | 281 | 49 | 8 | | | | | | | | |
| 153-Sp23 | 276 | 8 | 0 | | | | | | | | |
| | | | | ThiodiGal<br>TDG | 18000 | | 9800<br>1780 | 1930 | | 4000 | Gal(β1-1β)Gal |
| 147-Sp0 | 823 | 43 | 10 | DiLacNAc | 7400 | | 35000 | | | 7400 | Gal(β1-4)GlcNAc(β1-3)Gal(β1-4)GlcNAc(β |
| 146-Sp0 | 1002 | 8999 | 18 | TriLacNAc | 22000 | | 63000 | | | 6500 | Gal(β1-4)GlcNAc(β1-3)Gal(β1-4)GlcNAc(β1-3)Gal(β1-4)GlcNAc(β |
| 133-Sp10 | 305 | 31 | 48 | Lacto-N-tetraose<br>LNTet | 10000 | | 26000<br>8040 | 18200 | | | Gal(β1-3)GlcNAc(β1-3)Gal(β1-4)Glc(β |
| 148-Sp0 | 838 | 61 | 23 | Lacto-N-neo-tetraose | | | | | | | Gal(β1-4)GlcNAc(β1-3)Gal(β1-4)Glc(β |
| 148-Sp8 | 960 | 25 | 49 | | | | | | | | |
| 258-Sp0 | 177 | 44 | 7 | 2,3-Sialyl LacNAc | | 23000 | | 33000 | | | Neu5Ac(α2-3)Gal(β1-4)GlcNAc(β |
| 258-Sp8 | 328 | 35 | 5 | | | | | | | | |
| 268-Sp0 | 025 | 26 | 13 | 2,6-Sialyl LacNAc | | | | | | | Neu5Ac(α2-6)Gal(β1-4)GlcNAc(β |
| 268-Sp8 | 13 | 18 | -5 | | | | | | | | |
| 270-Sp0 | 55 | 28 | 31 | 2,6-Sialyl DiLacNAc | | | 32000 | 48000 | | | Neu5Ac(α2-6)Gal(β1-4)GlcNAc(β1-3)Gal(β1-4)GlcNAc(β |
| | | | | 2,6-Sialyl Lacto-N-neo-tetraose | | | | 21000 | | | Neu5Ac(α2-6)Gal(β1-4)GlcNAc(β1-3)Gal(β1-4)Glc(β |
| 166-Sp0 | 37 | 390 | -5 | Trisaccharide | | | | 8800 | | | GlcNAc(β1-3)Gal(β1-4)Glc(β |
| | | | | C2-DiLacNAc | 7400 | | 21000 | | | | Gal(β1-4)GlcNAc(β1-)[Gal(β1-4)GlcNAc(β1-)](CH2)2 |
| | | | | C3-DiLacNAc | 15000 | | 23000 | | | | Gal(β1-4)GlcNAc(β1-)[Gal(β1-4)GlcNAc(β1-)](CH2)3 |
| | | | | C4-DiLacNAc | 12000 | | 22000 | | | | Gal(β1-4)GlcNAc(β1-)[Gal(β1-4)GlcNAc(β1-)](CH2)4 |
| | | | | Lacto-N-hexaose<br>LNhex | | 18000 | 65000<br>23277 | 147000<br>34309 | | | Gal(β1-4)GlcNAc(β1-6)[Gal(β1-4)GlcNAc(β1-3)]Gal(β1-4)Glc(β |
| | | | | 2,4-Pentasaccharide | | 31000 | 26000 | 61000 | | | Gal(β1-4)GlcNAc(β1-4)[Gal(β1-4)GlcNAc(β1-2)]Gal(β |
| | | | | 2,6-Pentasaccharide | | 12000 | 22000 | 61000 | | | Gal(β1-4)GlcNAc(β1-6)[Gal(β1-4)GlcNAc(β1-2)]Gal(β |
| | | | | 3,6-Pentasaccharide | | 25000 | 27000 | 53000 | | | Gal(β1-4)GlcNAc(β1-6)[Gal(β1-4)GlcNAc(β1-3)]Gal(β |
| | | | | 4,6-Pentasaccharide | | 51000 | 25000 | 59000 | | | Gal(β1-4)GlcNAc(β1-6)[Gal(β1-4)GlcNAc(β1-4)]Gal(β |
| | | | | 2'FL | | | 2420 | 2310 | | | Fuc(a1-4)Gal(β1-4)Glc(β |
| | | | | A-Tri | | | 18672 | 22928 | | | Gal(a1-4)Gal(β1-4)GlcNAc(β |
| | | | | B-tri | | | 13391 | 28200 | | | GalNAc(a1-3)[Fuca1-2)]Gal(β |
| | | | | A-tetra | | | 13830 | 43100 | | | GalNAc(a1-3)[Fuca1-2)]Gal(β1-4)Glc(β |

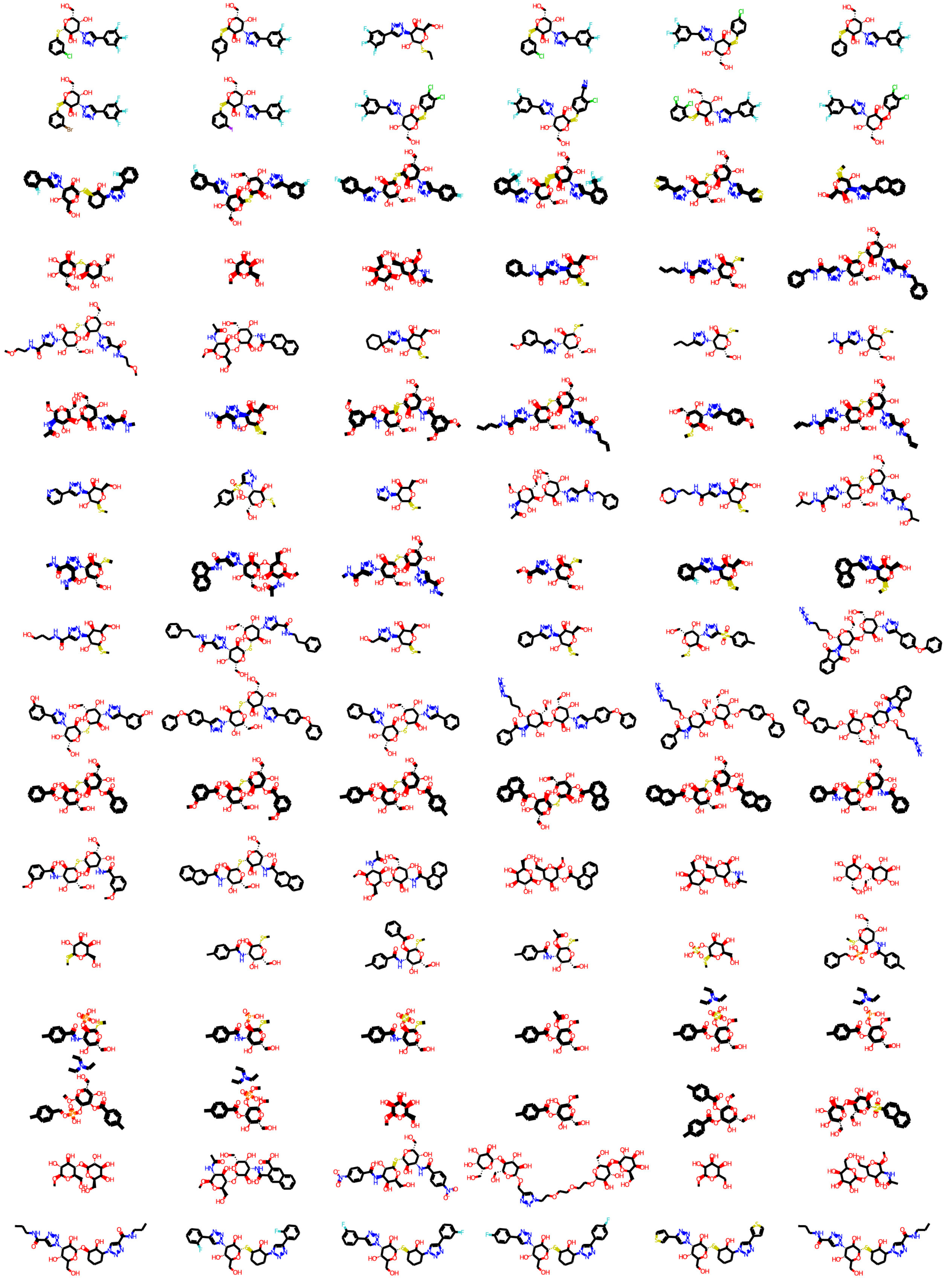

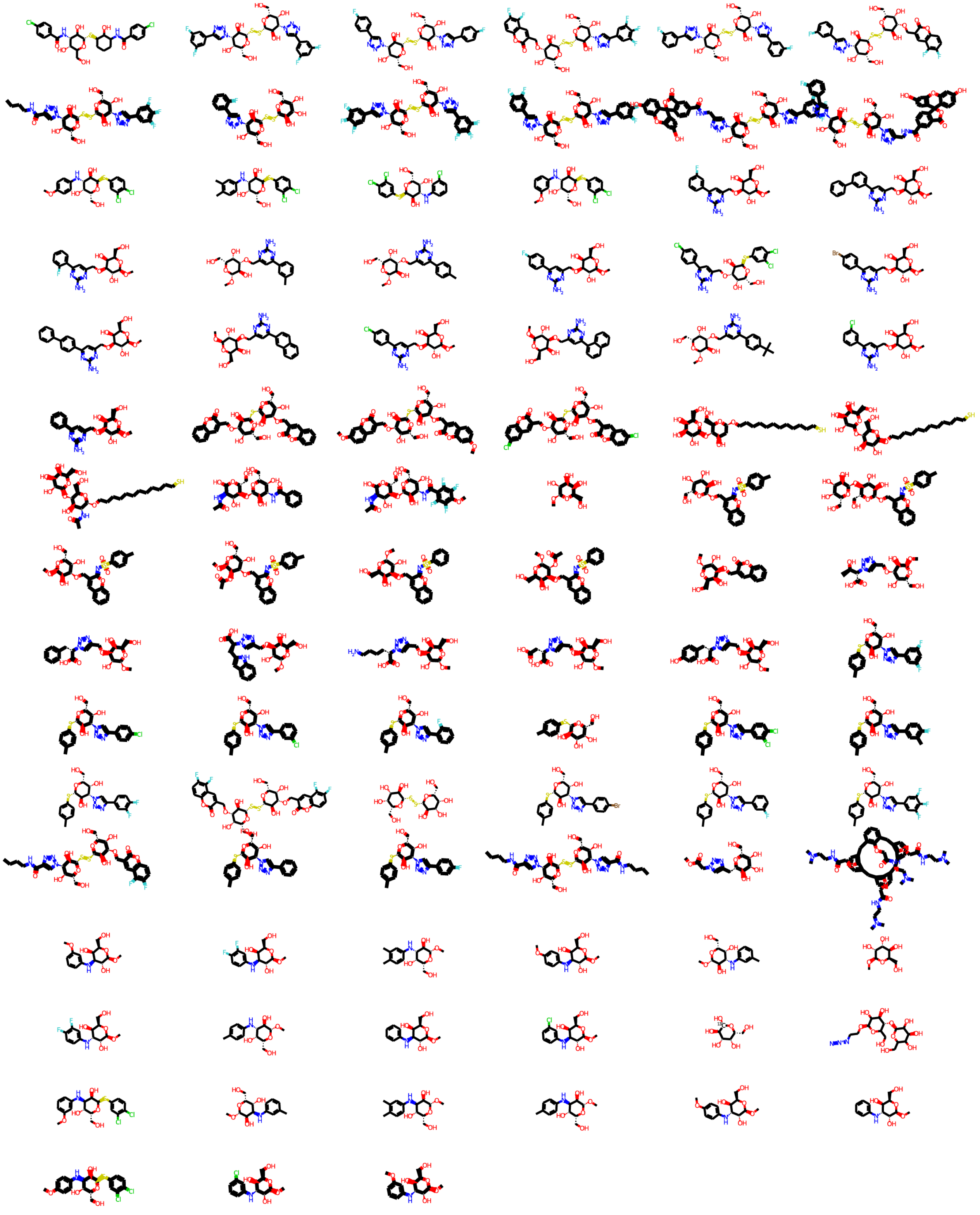

**Table S2**. Structures of Glycomimetic Molecules.

**Table S3**. List of lectin multi-concentration groups (N = number of concentrations)

| # | First Sample | N | cbpId | Investigator | Protein Name |
|---|---|---|---|---|---|
| 1 | 1 | 9 | cbp_2281 | Lara K. Mahal | SNA |
| 2 | 10 | 7 | | Nicolai Bovin | Anti-LeC |
| 3 | 17 | 7 | cbp_2787 | Lara K. Mahal | UEA I |
| 4 | 24 | 6 | cbp_2554 | Lara K. Mahal | ConA |
| 5 | 30 | 6 | cbp_2456 | Lara K. Mahal | ECL |
| 6 | 36 | 5 | cbp_hum_Stlect_00116 | Marcelo Dias Baruffi | Human Galectin1 |
| 7 | 41 | 5 | cbp_1304 | Marcelo Dias Baruffi | Mouse Galectin1 |
| 8 | 46 | 5 | cbp_2450 | Lara K. Mahal | AAL |
| 9 | 51 | 5 | cbp_2469 | Lara K. Mahal | MPL |
| 10 | 56 | 5 | cbp_2390 | Lara K. Mahal | WGA |
| 11 | 61 | 5 | cbp_2553 | Lara K. Mahal | WGA |
| 12 | 66 | 5 | cbp_2685 | Lara K. Mahal | CTB |
| 13 | 71 | 5 | cbp_hum_Stlect_00118 | Richard D. Cummings | Gal-3C |
| 14 | 76 | 4 | cbp_1608 | Elizabeth M. Meiering | Threefoil |
| 15 | 80 | 4 | cbp_2440 | Karel Bezouska | Mouse NKR-P1C |
| 16 | 84 | 4 | cbp_2452 | Lara K. Mahal | ACL |
| 17 | 88 | 4 | cbp_2561 | Lara K. Mahal | AIA |
| 18 | 92 | 4 | cbp_1478 | Janko Kos | Mono1-CNL |
| 19 | 96 | 4 | cbp_1478 | Janko Kos | Mono2R-CNL |
| 20 | 100 | 4 | cbp_1622 | Bidadi Venkataram Prasad | P11 VP8 |
| 21 | 104 | 4 | cbp_2689 | Kinjiro Morimoto | PFL |
| 22 | 108 | 4 | cbp_2493 | Lara K. Mahal | CA |
| 23 | 112 | 4 | cbp_2541 | Lara K. Mahal | CAA |
| 24 | 116 | 4 | cbp_2453 | Lara K. Mahal | ConA |
| 25 | 120 | 4 | cbp_2460 | Lara K. Mahal | GSL-2 |
| 26 | 124 | 4 | cbp_2459 | Lara K. Mahal | GSL-1 |
| 27 | 128 | 4 | cbp_2458 | Lara K. Mahal | GNL |
| 28 | 132 | 4 | cbp_2457 | Lara K. Mahal | EEL |
| 29 | 136 | 4 | cbp_2455 | Lara K. Mahal | DSL |
| 30 | 140 | 4 | cbp_2454 | Lara K. Mahal | DBA |
| 31 | 144 | 4 | cbp_2462 | Lara K. Mahal | HHL |
| 32 | 148 | 4 | cbp_2544 | Lara K. Mahal | HAA |
| 33 | 152 | 4 | cbp_2770 | Lara K. Mahal | HPA |
| 34 | 156 | 4 | cbp_2542 | Lara K. Mahal | CSA(EY Lab) |
| 35 | 160 | 4 | cbp_2461 | Lara K. Mahal | GSL-I-B4 |
| 36 | 164 | 4 | cbp_2464 | Lara K. Mahal | LCA |
| 37 | 168 | 4 | cbp_2511 | Lara K. Mahal | LcH |
| 38 | 172 | 4 | cbp_2465 | Lara K. Mahal | LEL |
| 39 | 176 | 4 | cbp_2470 | Lara K. Mahal | NPA |
| 40 | 180 | 4 | cbp_2466 | Lara K. Mahal | LTL |
| 41 | 184 | 4 | cbp_2467 | Lara K. Mahal | MAL I |
| 42 | 188 | 4 | cbp_2555 | Lara K. Mahal | MAA |
| 43 | 192 | 4 | cbp_2471 | Lara K. Mahal | PHA-E |
| 44 | 196 | 4 | cbp_2774 | Lara K. Mahal | MAA(Seikagaku) |

| | | | | | |
|---|---|---|---|---|---|
| 45 | 200 | 4 | cbp_2468 | Lara K. Mahal | MAL II |
| 46 | 204 | 4 | cbp_2558 | Lara K. Mahal | PHA-E |
| 47 | 208 | 4 | cbp_2522 | Lara K. Mahal | PSL |
| 48 | 212 | 4 | cbp_2472 | Lara K. Mahal | PHA-L |
| 49 | 216 | 4 | cbp_2547 | Lara K. Mahal | PTA-Gal |
| 50 | 220 | 4 | cbp_2475 | Lara K. Mahal | PTL-I |
| 51 | 224 | 4 | cbp_2476 | Lara K. Mahal | PTL-II |
| 52 | 228 | 4 | cbp_2276 | Lara K. Mahal | RCAI (Vector) |
| 53 | 232 | 4 | cbp_2473 | Lara K. Mahal | PNA |
| 54 | 236 | 4 | cbp_2474 | Lara K. Mahal | PSA |
| 55 | 240 | 4 | cbp_2478 | Lara K. Mahal | SBA |
| 56 | 244 | 4 | cbp_2479 | Lara K. Mahal | STL |
| 57 | 248 | 4 | cbp_2550 | Lara K. Mahal | TJA-I |
| 58 | 252 | 4 | cbp_2551 | Lara K. Mahal | TJA-II |
| 59 | 256 | 4 | cbp_2531 | Lara K. Mahal | TL(EY Lab) |
| 60 | 260 | 4 | cbp_2548 | Lara K. Mahal | UDA |
| 61 | 264 | 4 | cbp_2533 | Lara K. Mahal | UEAII |
| 62 | 268 | 4 | cbp_2482 | Lara K. Mahal | VVL |
| 63 | 272 | 4 | cbp_2483 | Lara K. Mahal | WFL |
| 64 | 276 | 4 | cbp_2784 | Lara K. Mahal | PA-IL |
| 65 | 280 | 4 | cbp_2788 | Lara K. Mahal | WGA(Seikagaku Bio) |
| 66 | 284 | 4 | cbp_2690 | Miriam Dwek | Native HPA |
| 67 | 288 | 4 | cbp_2703 | Jatinder Singh | SGA |
| 68 | 292 | 4 | cbp_2602 | Jatinder Singh | AHL |
| 69 | 296 | 4 | cbp_2751 | Shashikala Inamdar | SSR1 |
| 70 | 300 | 4 | cbp_1606 | Janko Kos | rMpL |
| 71 | 304 | 4 | cbp_2905 | Anne Imberty | TbFBL1 |
| 72 | 308 | 4 | cbp_2937 | Markus Kuenzler | CC1G-10077 |
| 73 | 312 | 4 | cbp_2916 | Yvette Van Kooyk | MGL 6B |
| 74 | 316 | 4 | cbp_2917 | Yvette Van Kooyk | MGL 6A-H262T |
| 75 | 320 | 4 | cbp_hum_Ctlect_292 | Yvette Van Kooyk | MGL 6A |
| 76 | 324 | 4 | cbp_2951 | Rongsheng Jin | HA70 complex-Alexa |
| 77 | 328 | 4 | cbp_2390 | Brian Haab | WGA multimer |
| 78 | 332 | 4 | cbp_3020 | Anne Imberty | rPVL |
| 79 | 336 | 4 | cbp_3061 | Rich Olson | VVHA lectin |
| 80 | 340 | 4 | cbp_pla_other_334 | Carole A. Bewley | BanLec |
| 81 | 344 | 4 | cbp_3129 | Pavel Lukyanov | CGL |
| 82 | 348 | 4 | cbp_hum_Stlect_00118 | Richard D. Cummings | Gal-3 |
| 83 | 352 | 4 | cbp_3484 | Richard D. Cummings | human Gal-7 |
| 84 | 356 | 4 | cbp_3486 | Richard D. Cummings | mouse Gal-7 |
| 85 | 360 | 3 | | Brian Haab | CA19.9-1B844 |
| 86 | 363 | 3 | | Brian Haab | CA19.9-9L426 |
| 87 | 366 | 3 | | Brian Haab | CA19.9-121SLE |
| 88 | 369 | 3 | | Hans-Jörg Büring | IPS-K-4A2B8 |
| 89 | 372 | 3 | cbp_1672 | Marie Hanigan | Microvirin |
| 90 | 375 | 3 | | Hiroto Kawashima | ab5 |
| 91 | 378 | 3 | cbp_2552 | Lara K. Mahal | AOL (Mahal Lab) |
| 92 | 381 | 3 | cbp_2487 | Lara K. Mahal | ABA |
| 93 | 384 | 3 | cbp_1478 | Janko Kos | Mono2W-CNL |

| | | | | | |
|---|---|---|---|---|---|
| 94 | 387 | 3 | cbp_2451 | Lara K. Mahal | BPL |
| 95 | 390 | 3 | cbp_2506 | Lara K. Mahal | GS-II (EY Lab) |
| 96 | 393 | 3 | cbp_2505 | Lara K. Mahal | EEA(EY Lab) |
| 97 | 396 | 3 | cbp_2507 | Lara K. Mahal | GNA (EY Lab) |
| 98 | 399 | 3 | cbp_2509 | Lara K. Mahal | HHA(EY Lab) |
| 99 | 402 | 3 | cbp_2560 | Lara K. Mahal | PHA-L |
| 100 | 405 | 3 | cbp_2563 | Lara K. Mahal | PNA |
| 101 | 408 | 3 | cbp_2562 | Lara K. Mahal | SBA |
| 102 | 411 | 3 | cbp_2480 | Lara K. Mahal | SJA |
| 103 | 414 | 3 | cbp_2481 | Lara K. Mahal | UEA I |
| 104 | 417 | 3 | cbp_2549 | Lara K. Mahal | VGA(EY Lab) |
| 105 | 420 | 3 | cbp_2687 | Lara K. Mahal | WGA |
| 106 | 423 | 3 | cbp_2789 | Lara K. Mahal | MNA-M |
| 107 | 426 | 3 | cbp_2790 | Lara K. Mahal | MOA |
| 108 | 429 | 3 | cbp_1716 | Rich Olson | WT-VCC |
| 109 | 432 | 3 | cbp_hum_Ctlect_228 | Irma Van Die | ME5(detected with a-HUD) |
| 110 | 435 | 3 | cbp_1477 | Hui Sun | AAL |
| 111 | 438 | 3 | cbp_2445 | José M. Mancheño | LSL-150 |
| 112 | 441 | 3 | cbp_2446 | José M. Mancheño | LBL-152 |
| 113 | 444 | 3 | cbp_hum_Ctlect_228 | Irma Van Die | ME7(detected with a-HUD) |
| 114 | 447 | 3 | cbp_2733 | John Fraser | SSL0-488 |
| 115 | 450 | 3 | cbp_2744 | Zeev Pancer | VLRB.aGPA.23 |
| 116 | 453 | 3 | cbp_1714 | Robert L. Atmar | rNV VLP-Cy3 |
| 117 | 456 | 3 | cbp_2904 | Anne Imberty | AFCNVH |
| 118 | 459 | 3 | cbp_2903 | Anne Imberty | HML-Native |
| 119 | 462 | 3 | | Lara K. Mahal | #8 Lewis B (T218) (Abcam) |
| 120 | 465 | 3 | | Lara K. Mahal | #15 Sialyl Lewis X (BD) |
| 121 | 468 | 3 | | Lara K. Mahal | #14 Sialyl Lewis X (Calbio) |
| 122 | 471 | 3 | cbp_hum_Stlect_00116 | Francoise Guerlesquin | hGal-1-488+BCR |
| 123 | 474 | 3 | cbp_2938 | Anne Imberty | TbFBL2 |
| 124 | 477 | 3 | cbp_hum_Ctlect_217 | Kurt Drickamer | MGL 416 |
| 125 | 480 | 3 | cbp_2948 | Anne Imberty | BTL native |
| 126 | 483 | 3 | | John Kearney | GAC78 |
| 127 | 486 | 3 | cbp_2984 | Gerard Zurawski | DC-ASGPR short |
| 128 | 489 | 3 | | Luc Teyton | Sp Serotype 14.2 |
| 129 | 492 | 3 | | Luc Teyton | Sp Serotype 14.15 |
| 130 | 495 | 3 | cbp_2750 | Shashikala Inamdar | rSRL |
| 131 | 498 | 3 | cbp_oth_Other_933 | Shashikala Inamdar | SRL |
| 132 | 501 | 3 | cbp_hum_Stlect_00116 | Linda Baum | Galectin-1 |
| 133 | 504 | 3 | cbp_3021 | Horacio Heras | PsSC |
| 134 | 507 | 3 | cbp_3074 | Nobuyuki Matoba | Avaren-Fc |
| 135 | 510 | 3 | cbp_3134 | Janko Kos | CnSepL |
| 136 | 513 | 3 | | Brian Haab | CA19-9 Ab (DRG) |
| 137 | 516 | 3 | cbp_mou_Stlect_203 | Oscar Campetella | WT Galectin-8 |

| | | | | | |
|---|---|---|---|---|---|
| 138 | 519 | 3 | cbp_3151 | Markus Kuenzler | CML1 |
| 139 | 522 | 3 | cbp_3239 | Almudena Fernández Briera | PhoSPL001 |
| 140 | 525 | 3 | cbp_oth_other_339 | Ronnie Willaert | Epa1-Np mutant Y228W |
| 141 | 528 | 3 | cbp_oth_other_339 | Ronnie Willaert | Epa1-Np mutant Y228A |
| 142 | 531 | 3 | cbp_oth_other_339 | Ronnie Willaert | Epa1-Np mutant E227G |
| 143 | 534 | 3 | cbp_oth_other_339 | Ronnie Willaert | Epa1-Np mutant E227A |
| 144 | 537 | 3 | cbp_oth_other_339 | Ronnie Willaert | Epa1-Np WT |
| 145 | 540 | 3 | cbp_3345 | Janko Kos | CnMBL |
| 146 | 543 | 3 | cbp_3245 | Shashikala Inamdar | DBL-1 |
| 147 | 546 | 3 | | Kate Rittenhouse-Olson | hJAA-F11-B11 |

**Table S3.** Galectin Inhibitor Data from Literature

| | | |
|---|---|---|
| 3 | 134 | https://doi.org/10.1016/j.bmc.2010.05.040 |
| 4 | 87 | https://doi.org/10.1021/acsmedchemlett.9b00396 |
| 5 | 72 | https://doi.org/10.1021/acs.jmedchem.7b01626 |
| 6 | 62 | https://doi.org/10.1021/jm801077j |
| 7 | 57 | https://doi.org/10.1016/j.bmcl.2014.05.063 |
| 8 | 39 | https://doi.org/10.1039/C9MD00183B |
| 9 | 36 | https://doi.org/10.1021/acs.jmedchem.6b00957 |
| 10 | 28 | https://doi.org/10.1016/j.bmcl.2010.12.049 |
| 11 | 25 | https://doi.org/10.1016/j.bmcl.2008.05.066 |
| 12 | 17 | https://doi.org/10.1021/jm301677r |
| 13 | 17 | https://pubmed.ncbi.nlm.nih.gov/11921396/ (doi not found) |
| 14 | 12 | https://doi.org/10.1002/cmdc.201700744 |
| 15 | 10 | https://doi.org/10.1016/j.bmc.2008.06.044 |
| 16 | 9 | https://doi.org/10.1016/j.bmc.2015.04.044 |
| 17 | 9 | https://doi.org/10.1021/jm701266y |
| 18 | 5 | https://doi.org/10.1002/cbic.201600285 |
| 19 | 3 | https://pubs.acs.org/doi/10.1021/jm500325k |
| 20 | 2 | https://doi.org/10.1093/glycob/cwm026 |
| 21 | 1 | https://doi.org/10.1021/jm300014q |

**Table S4.** Galectin Inhibitor References
