## Supplementary material for "Atom-level Machine Learning of Protein-Glycan Interactions and Cross-chiral Recognition in Glycobiology": Raw RFU to Fraction Bound Conversions

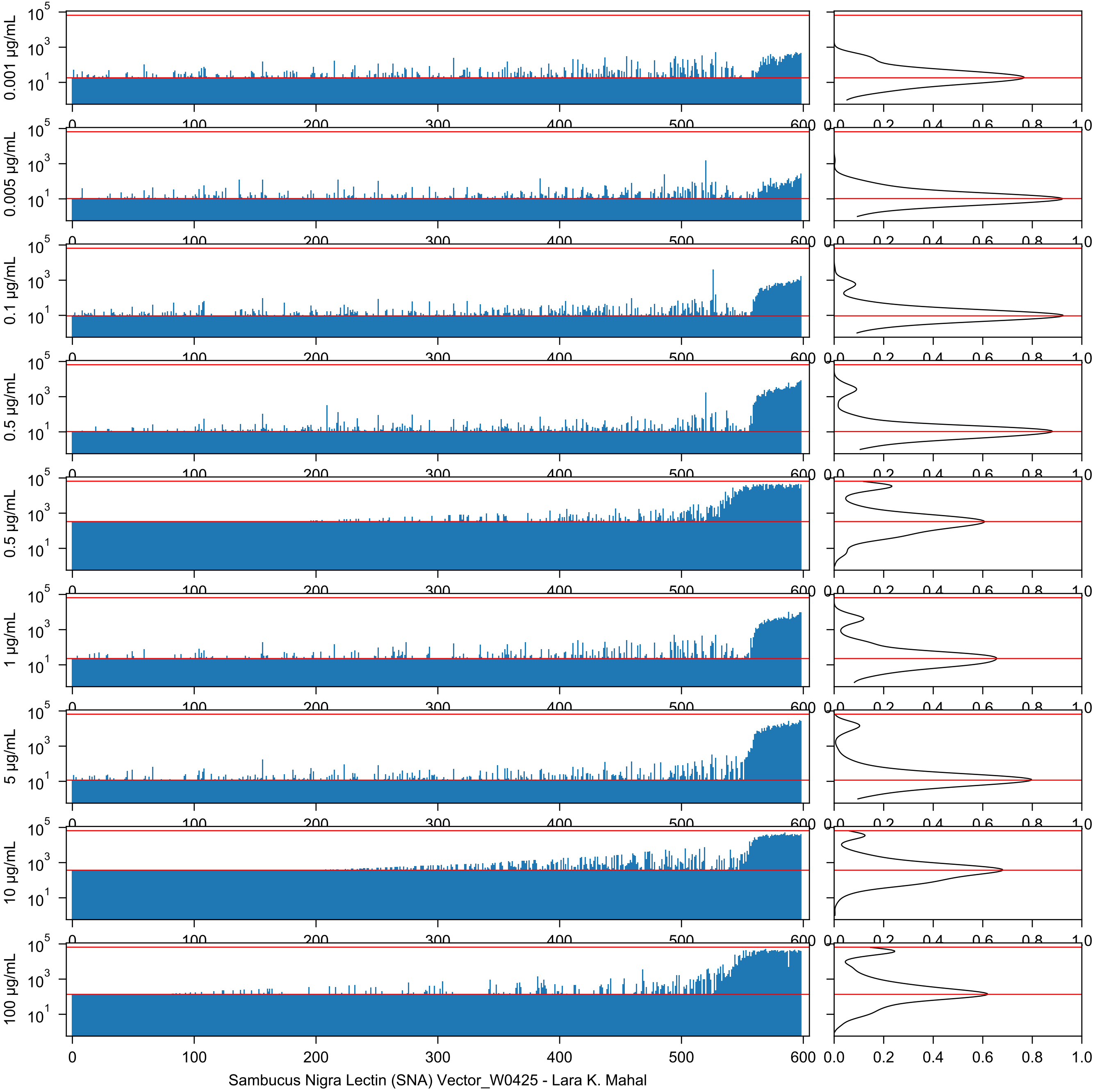
0.001 µg/mL
0.005 µg/mL
0.1 µg/mL
0.5 µg/mL
0.5 µg/mL
1 µg/mL
5 µg/mL
10 µg/mL
100 µg/mL
Sambucus Nigra Lectin (SNA) Vector_W0425 - Lara K. Mahal

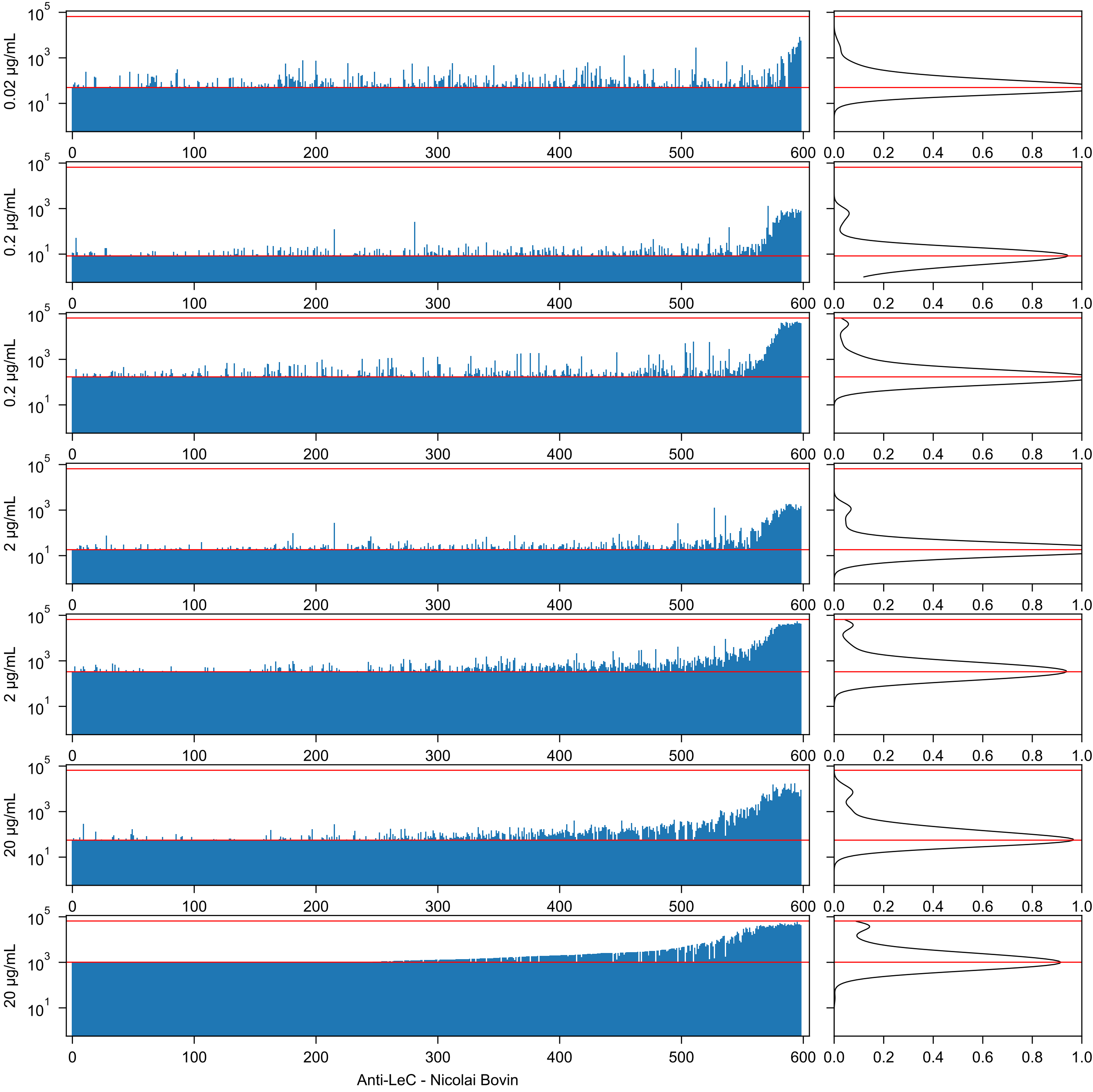
0.02 μg/mL
0.2 μg/mL
0.2 μg/mL
2 μg/mL
2 μg/mL
20 μg/mL
20 μg/mL
Anti-LeC - Nicolai Bovin

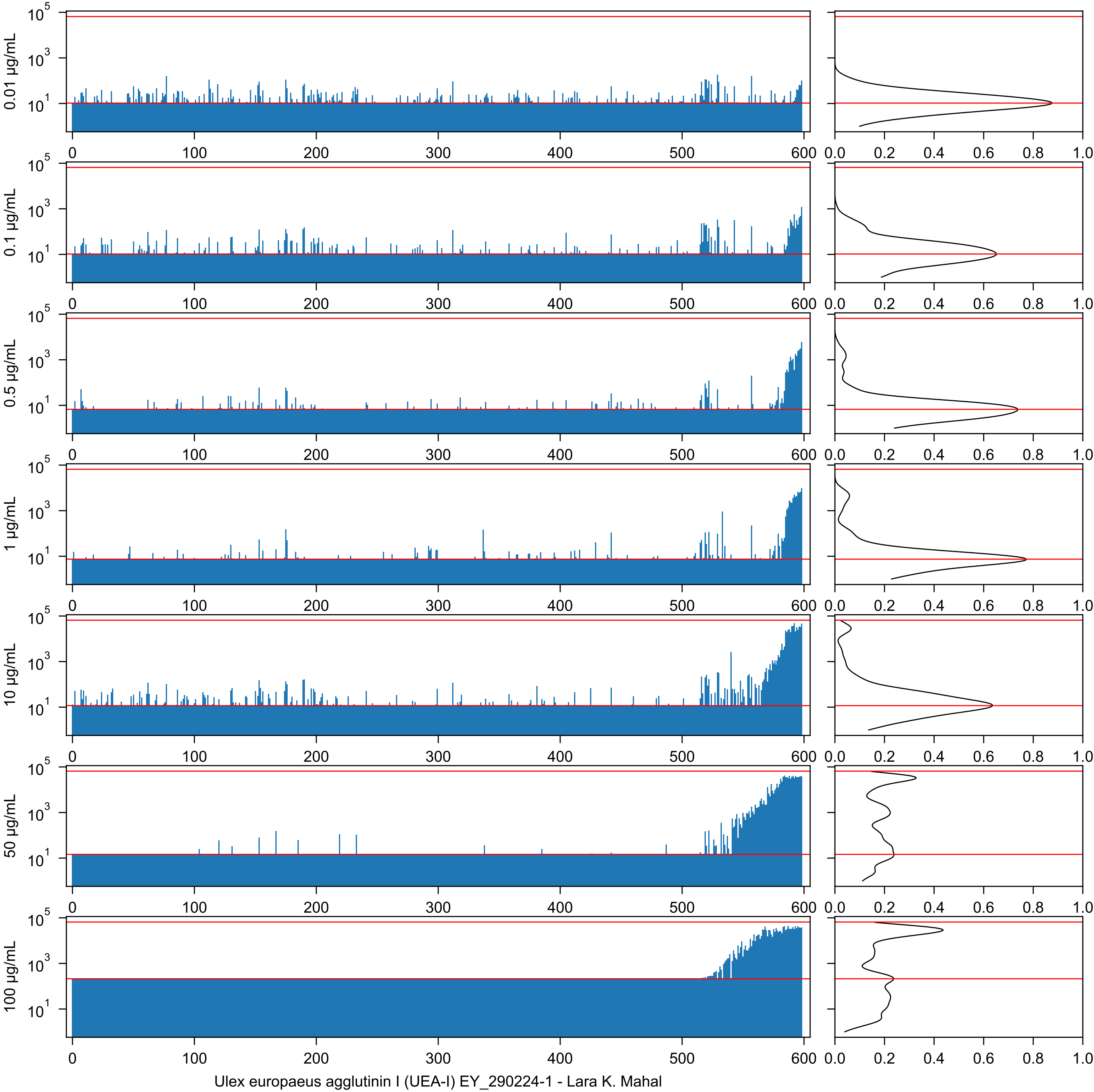


Ulex europaeus agglutinin I (UEA-I) EY_290224-1 - Lara K. Mahal

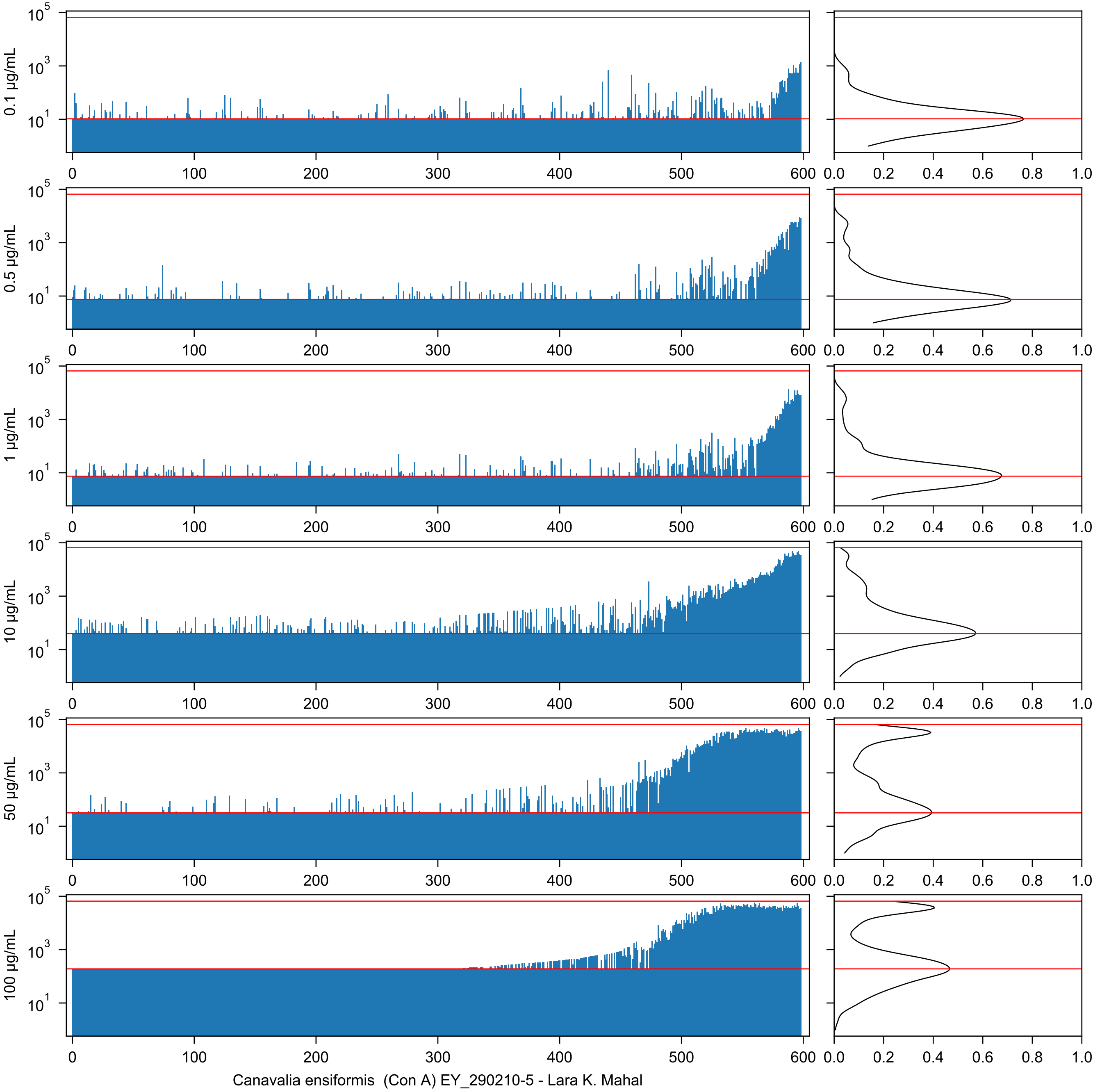
0.1 μg/mL
0.5 μg/mL
1 μg/mL
10 μg/mL
50 μg/mL
100 μg/mL
Canavalia ensiformis (Con A) EY_290210-5 - Lara K. Mahal

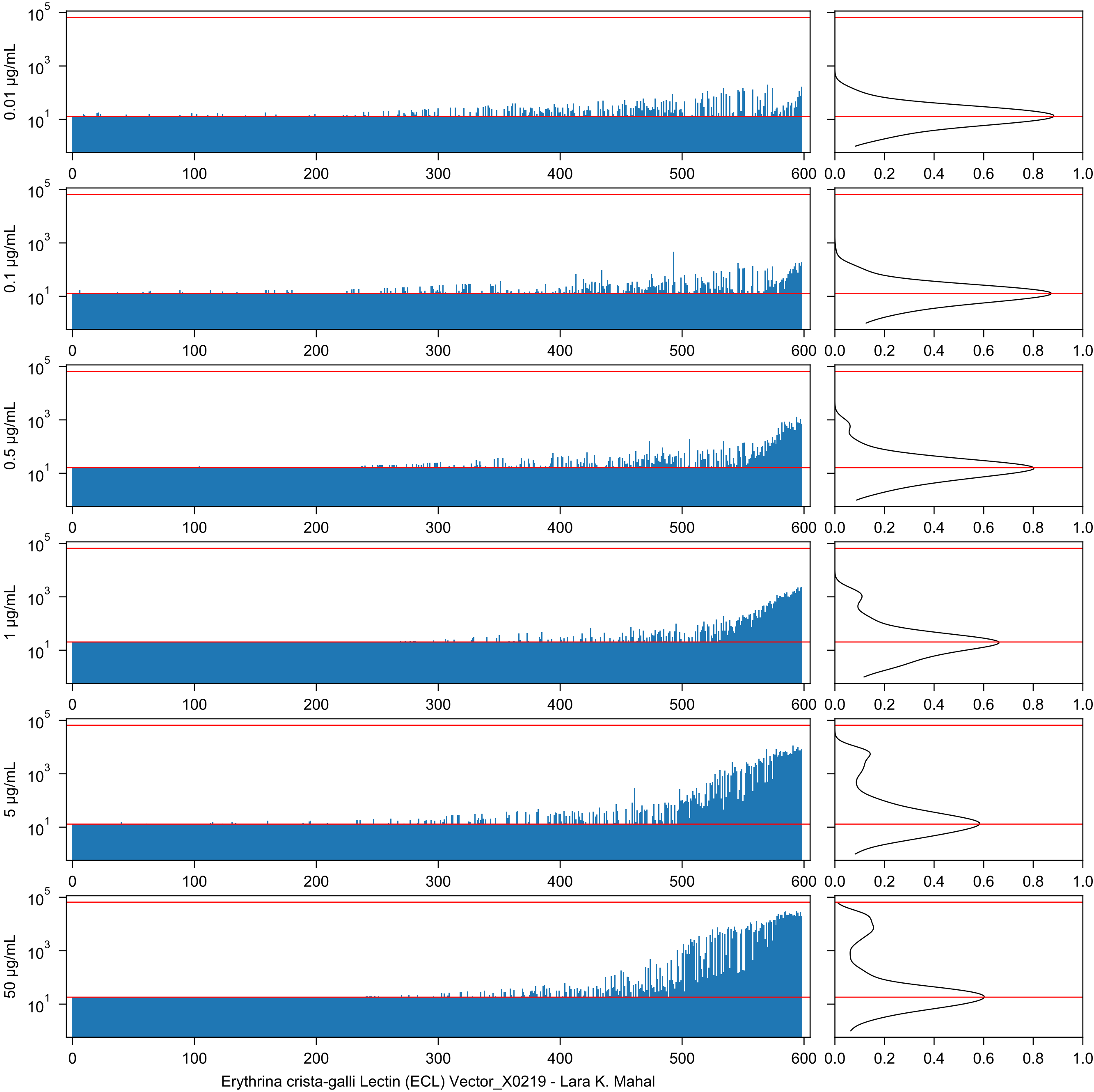


Erythrina crista-galli Lectin (ECL) Vector_X0219 - Lara K. Mahal

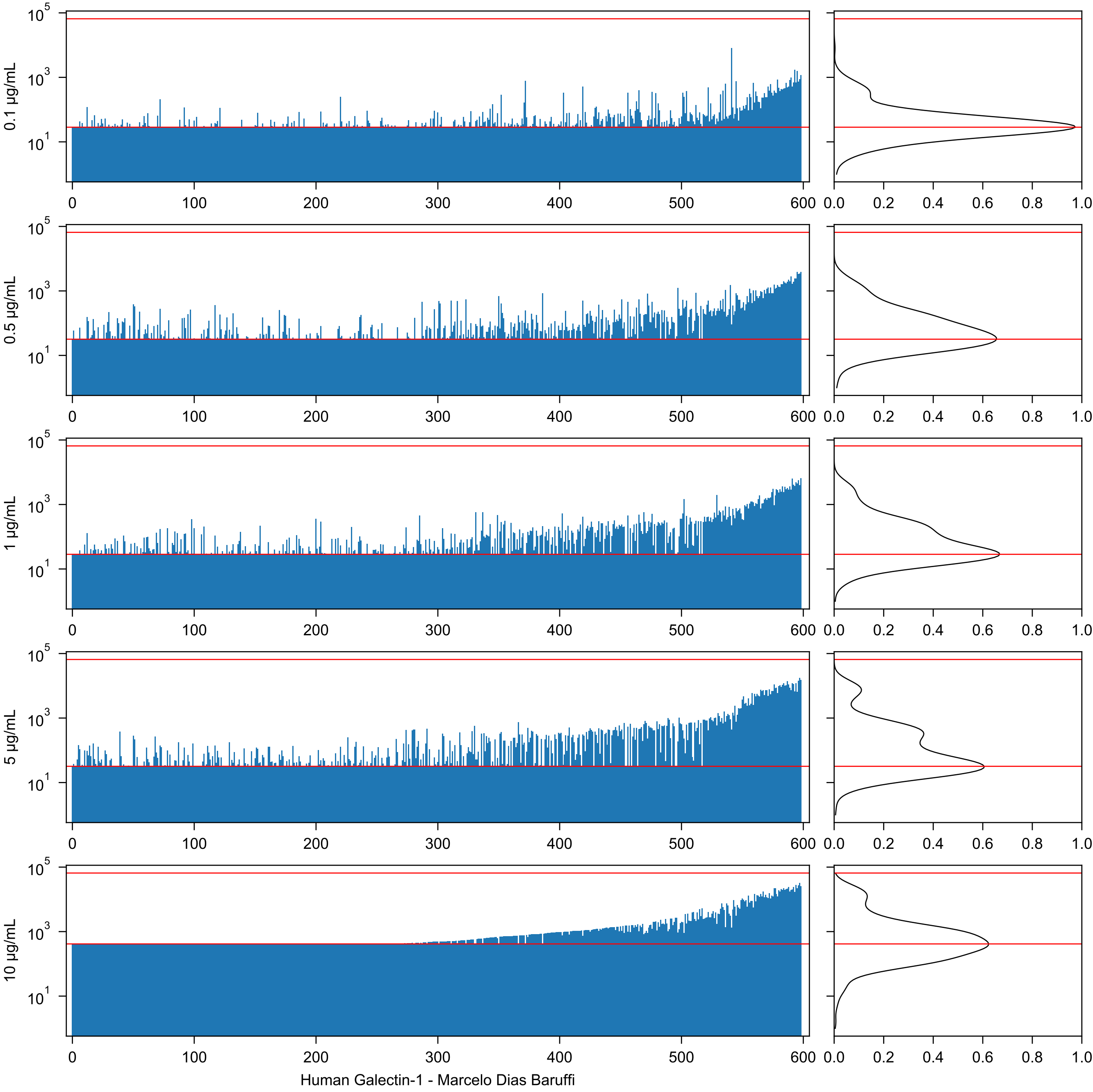
0.1 µg/mL
0.5 µg/mL
1 µg/mL
5 µg/mL
10 µg/mL
$10^5$
$10^3$
$10^1$
0
100
200
300
400
500
600
0.0
0.2
0.4
0.6
0.8
1.0
Human Galectin-1 - Marcelo Dias Baruffi

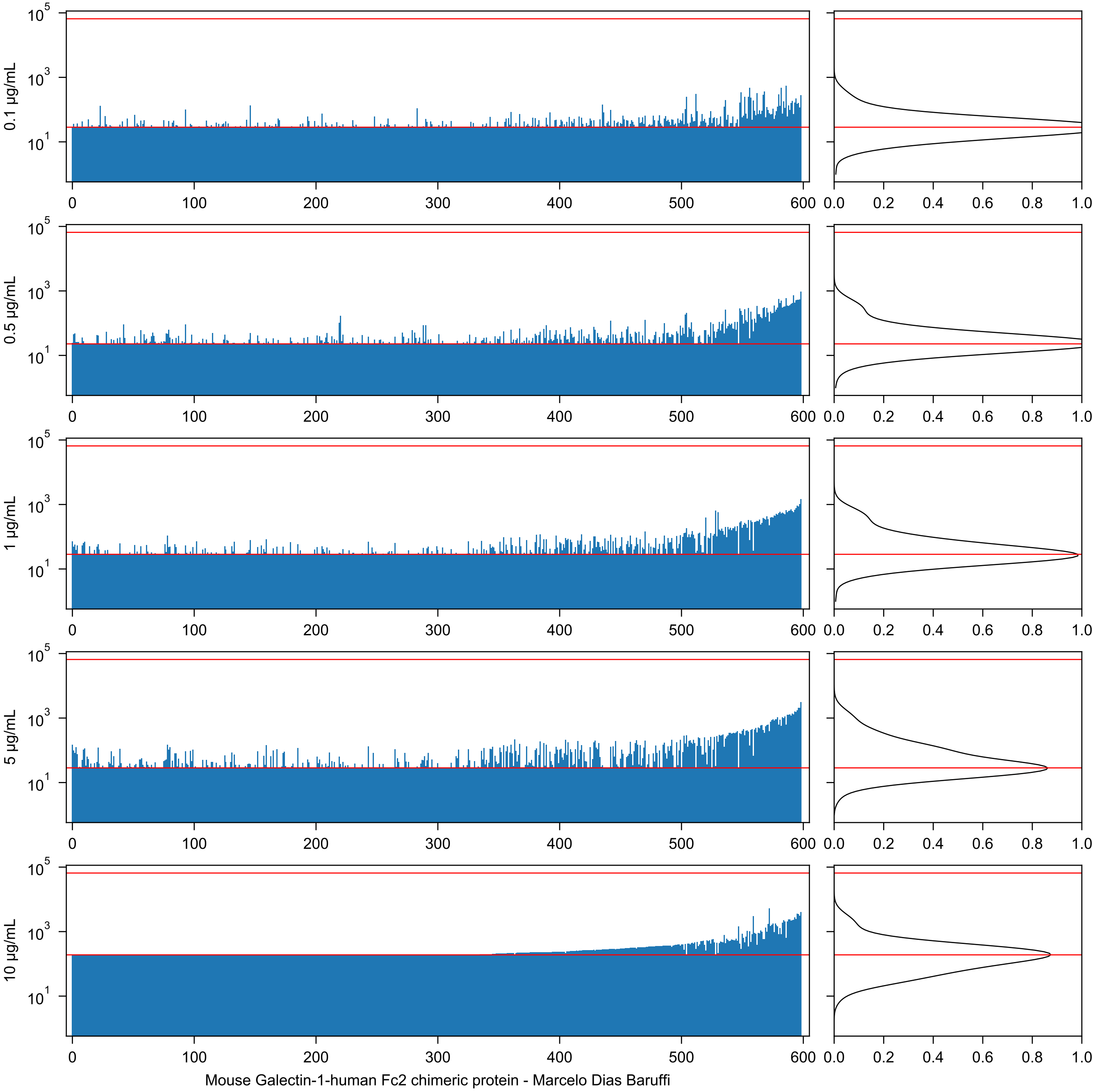

0.1 µg/mL
0.5 µg/mL
1 µg/mL
5 µg/mL
10 µg/mL
$10^5$
$10^3$
$10^1$
0
100
200
300
400
500
600
0.0
0.2
0.4
0.6
0.8
1.0
Mouse Galectin-1-human Fc2 chimeric protein - Marcelo Dias Baruffi

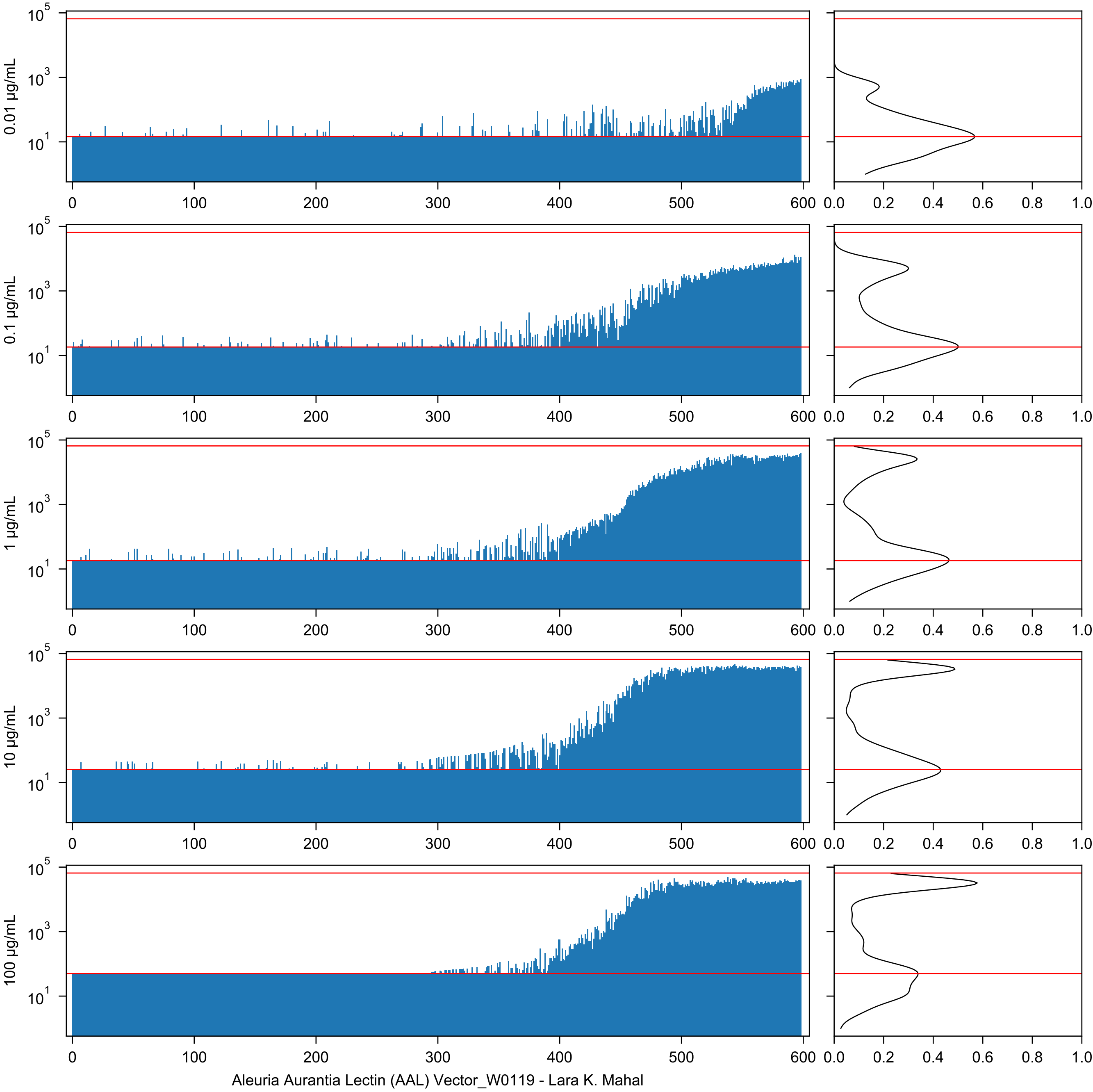

0.01 µg/mL
0.1 µg/mL
1 µg/mL
10 µg/mL
100 µg/mL
Aleuria Aurantia Lectin (AAL) Vector_W0119 - Lara K. Mahal

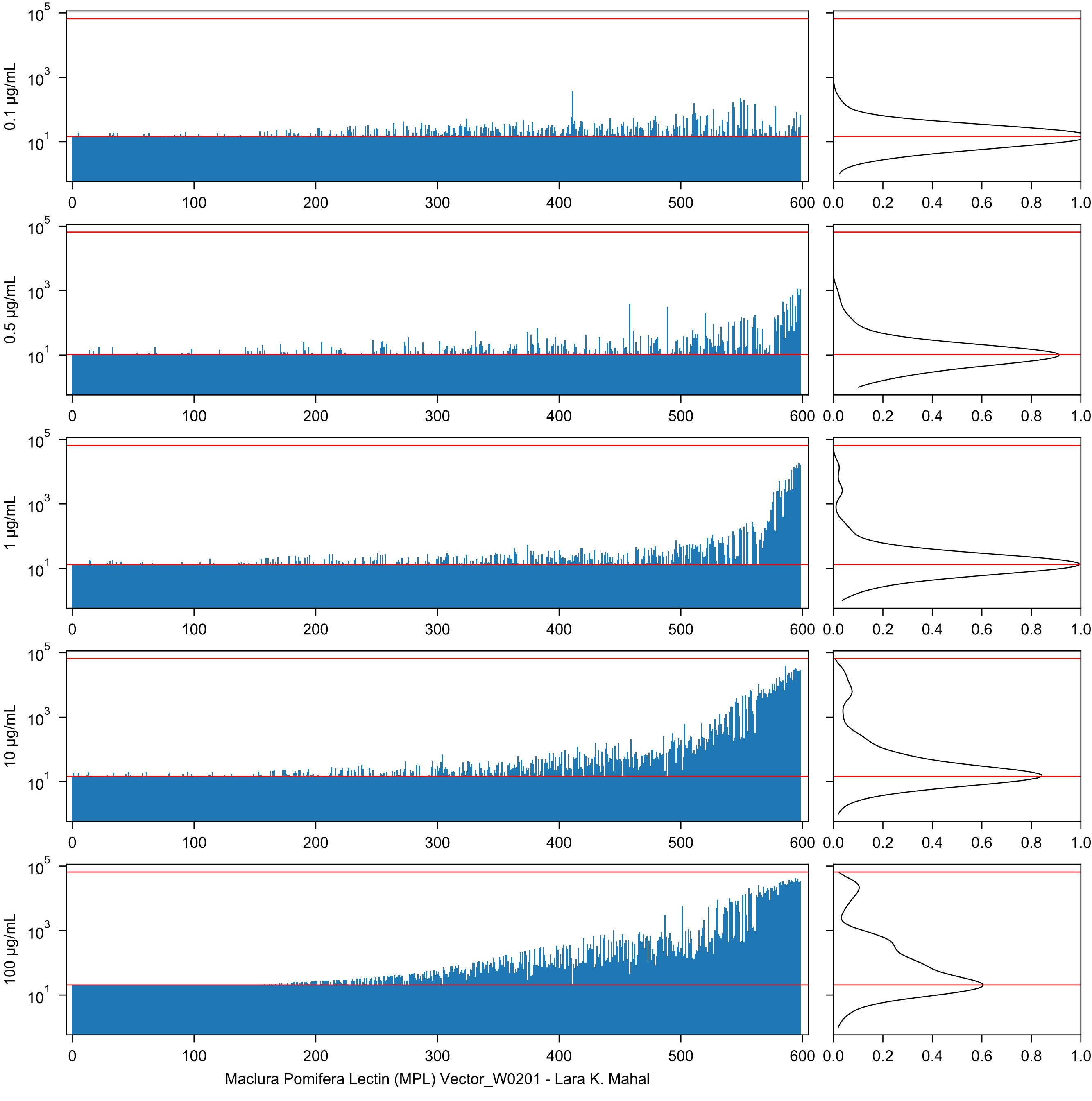
0.1 µg/mL
0.5 µg/mL
1 µg/mL
10 µg/mL
100 µg/mL
Maclura Pomifera Lectin (MPL) Vector_W0201 - Lara K. Mahal

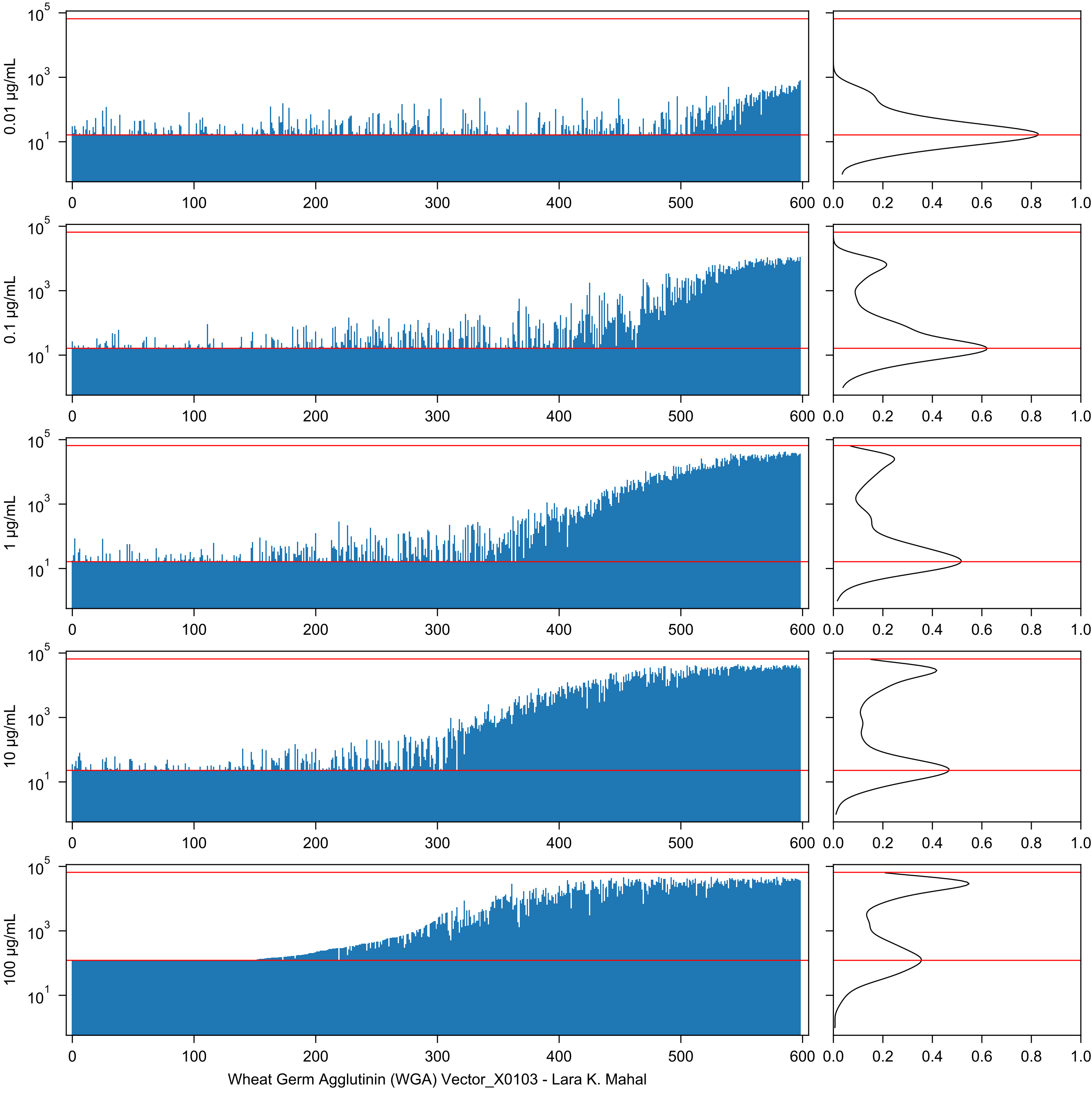
0.01 µg/mL
0.1 µg/mL
1 µg/mL
10 µg/mL
100 µg/mL
Wheat Germ Agglutinin (WGA) Vector_X0103 - Lara K. Mahal

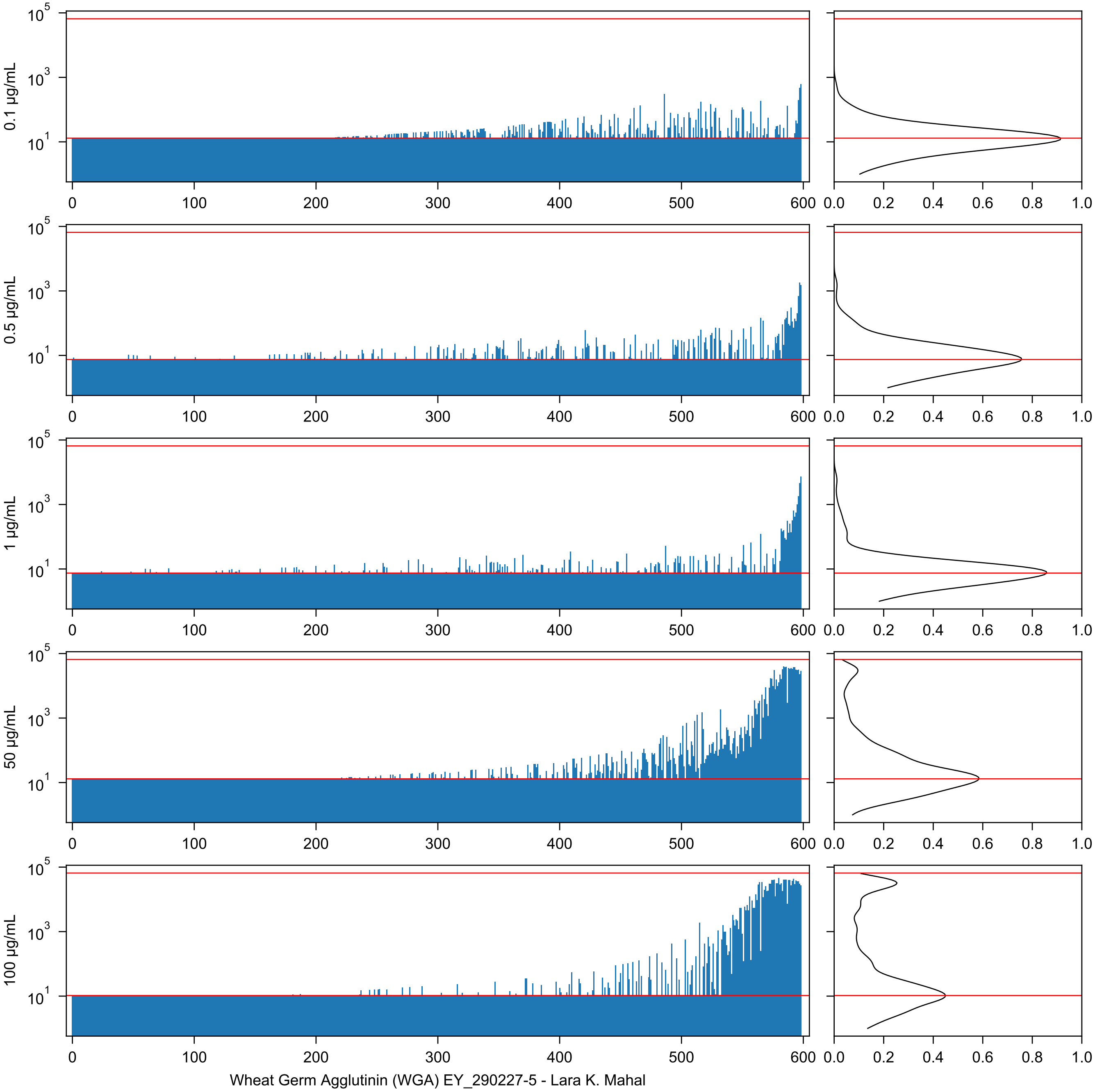
0.1 µg/mL
0.5 µg/mL
1 µg/mL
50 µg/mL
100 µg/mL
Wheat Germ Agglutinin (WGA) EY_290227-5 - Lara K. Mahal

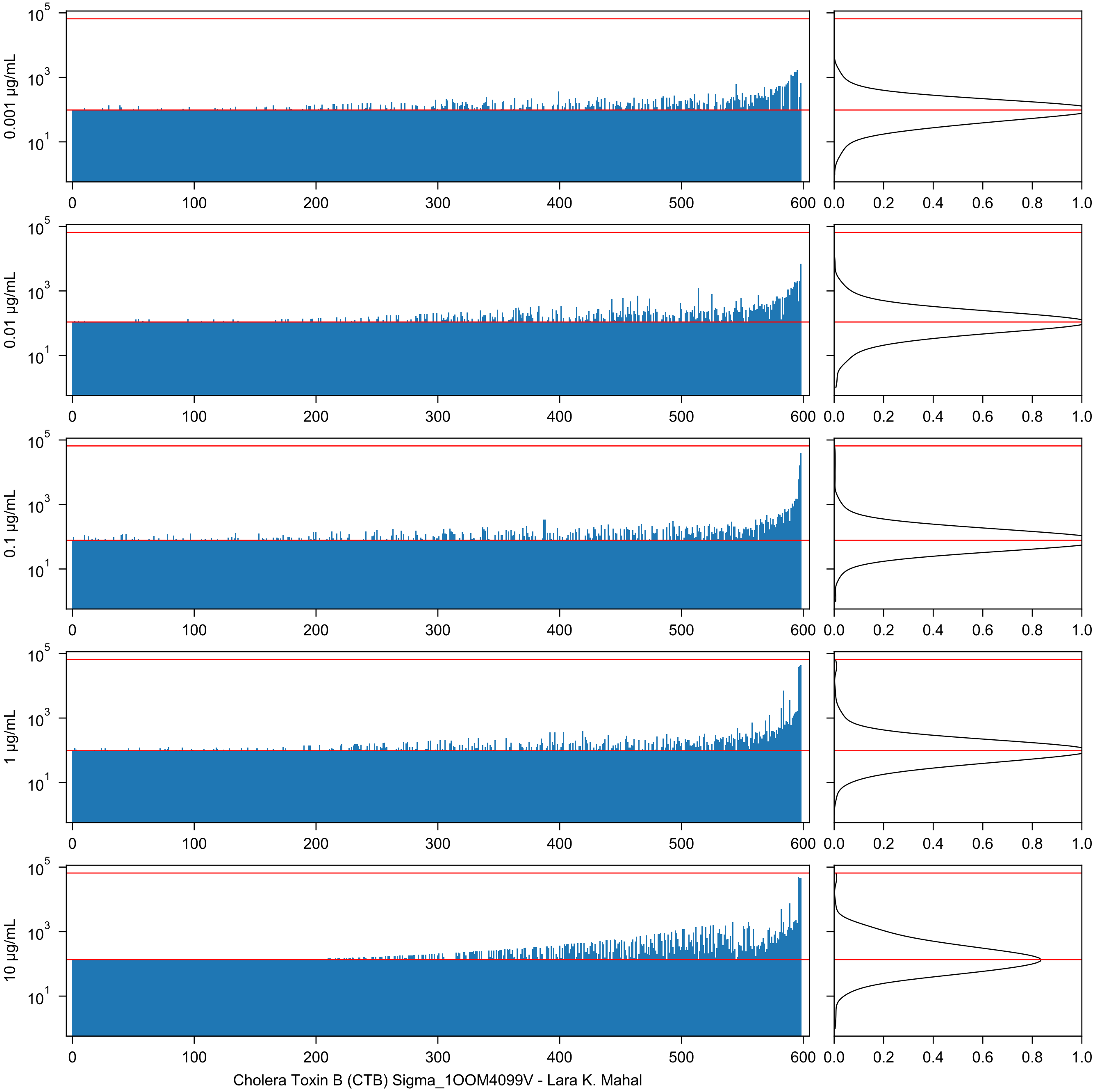
0.001 μg/mL
0.01 μg/mL
0.1 μg/mL
1 μg/mL
10 μg/mL
10^5
10^3
10^1
0
100
200
300
400
500
600
0.0
0.2
0.4
0.6
0.8
1.0
Cholera Toxin B (CTB) Sigma_1OOM4099V - Lara K. Mahal

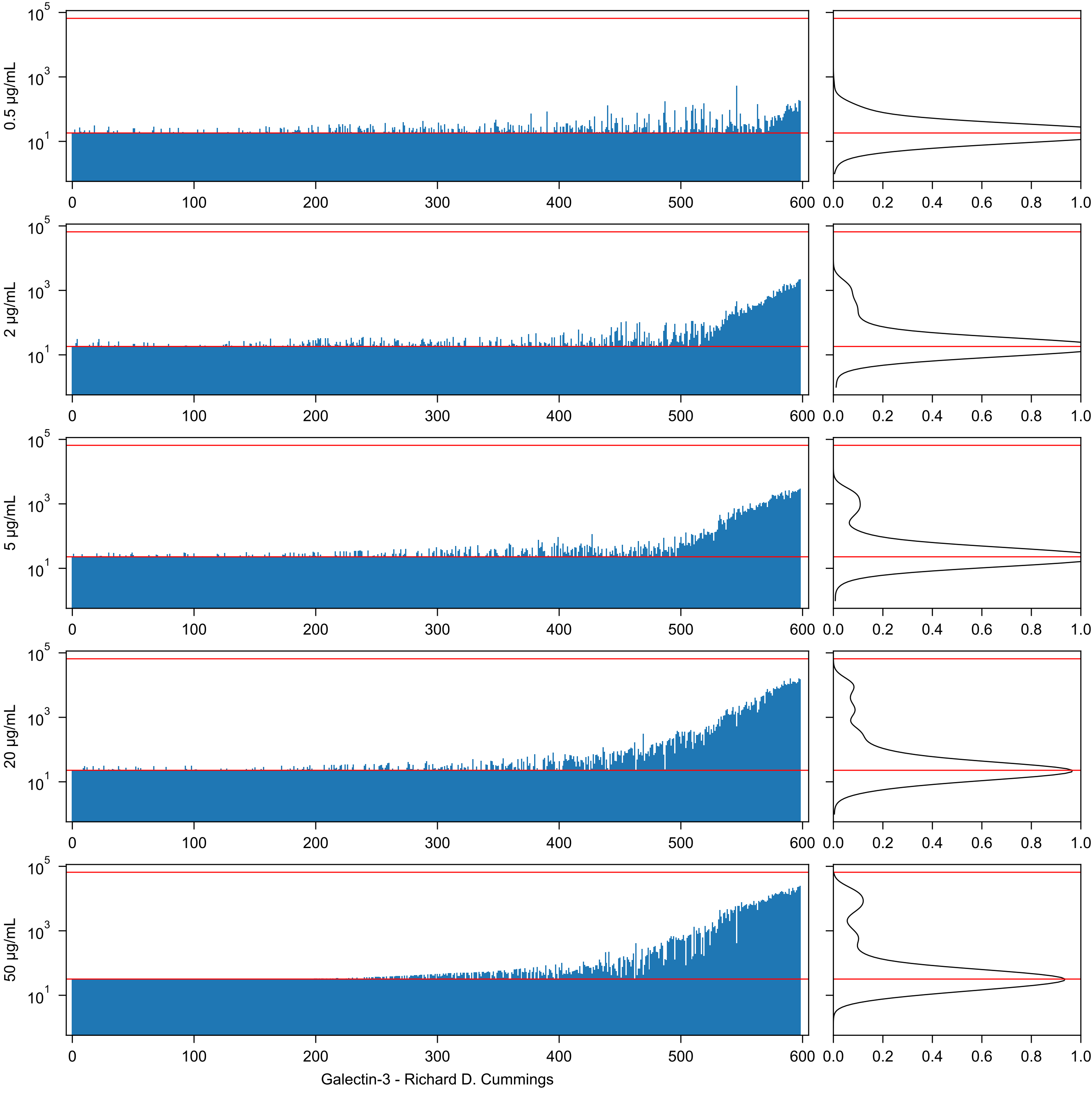
0.5 µg/mL
2 µg/mL
5 µg/mL
20 µg/mL
50 µg/mL
$10^5$
$10^3$
$10^1$
0
100
200
300
400
500
600
0.0
0.2
0.4
0.6
0.8
1.0
Galectin-3 - Richard D. Cummings

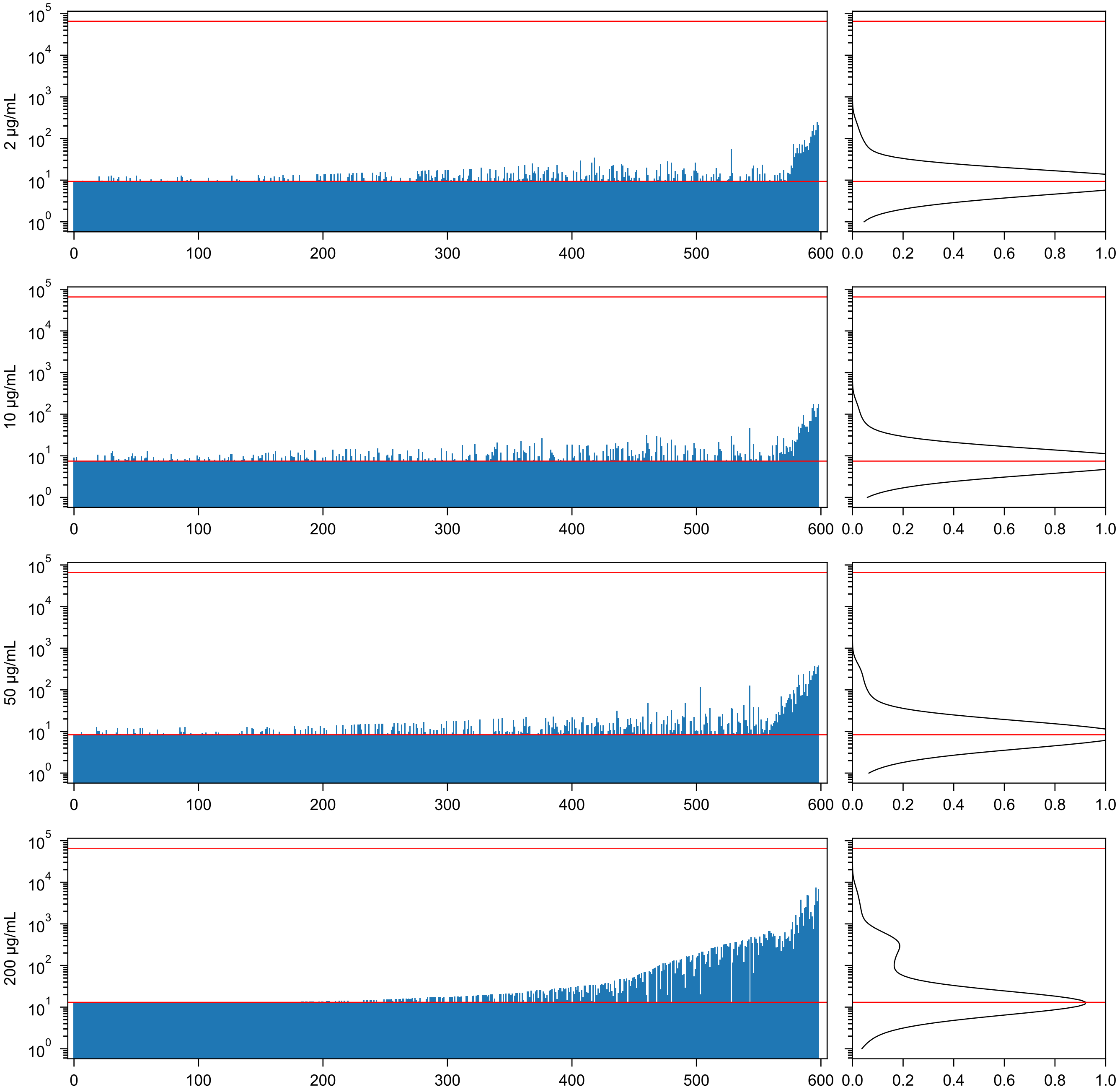


ThreeFoil - Elizabeth M. Meiering

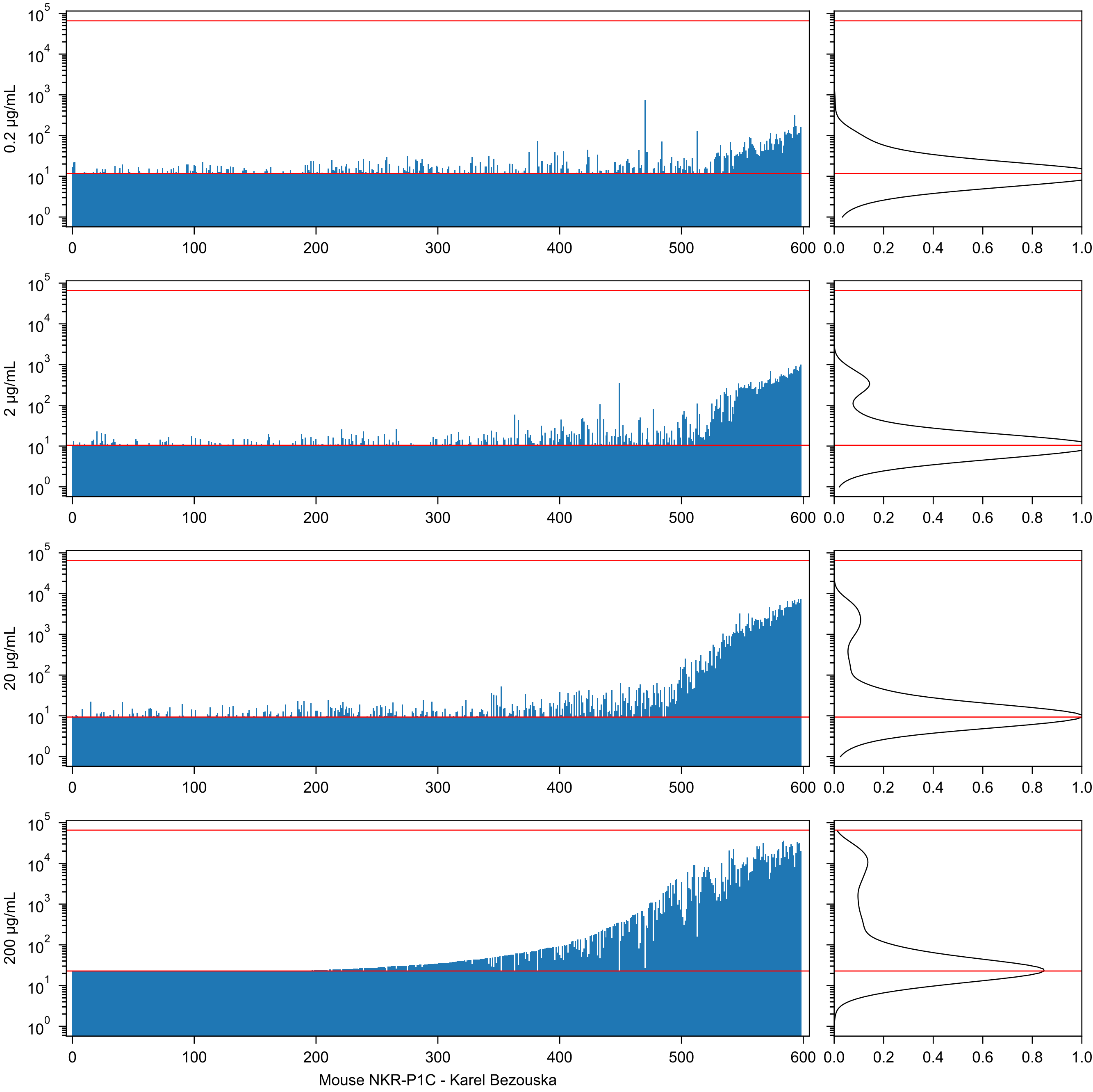
0.2 µg/mL
2 µg/mL
20 µg/mL
200 µg/mL
Mouse NKR-P1C - Karel Bezouska

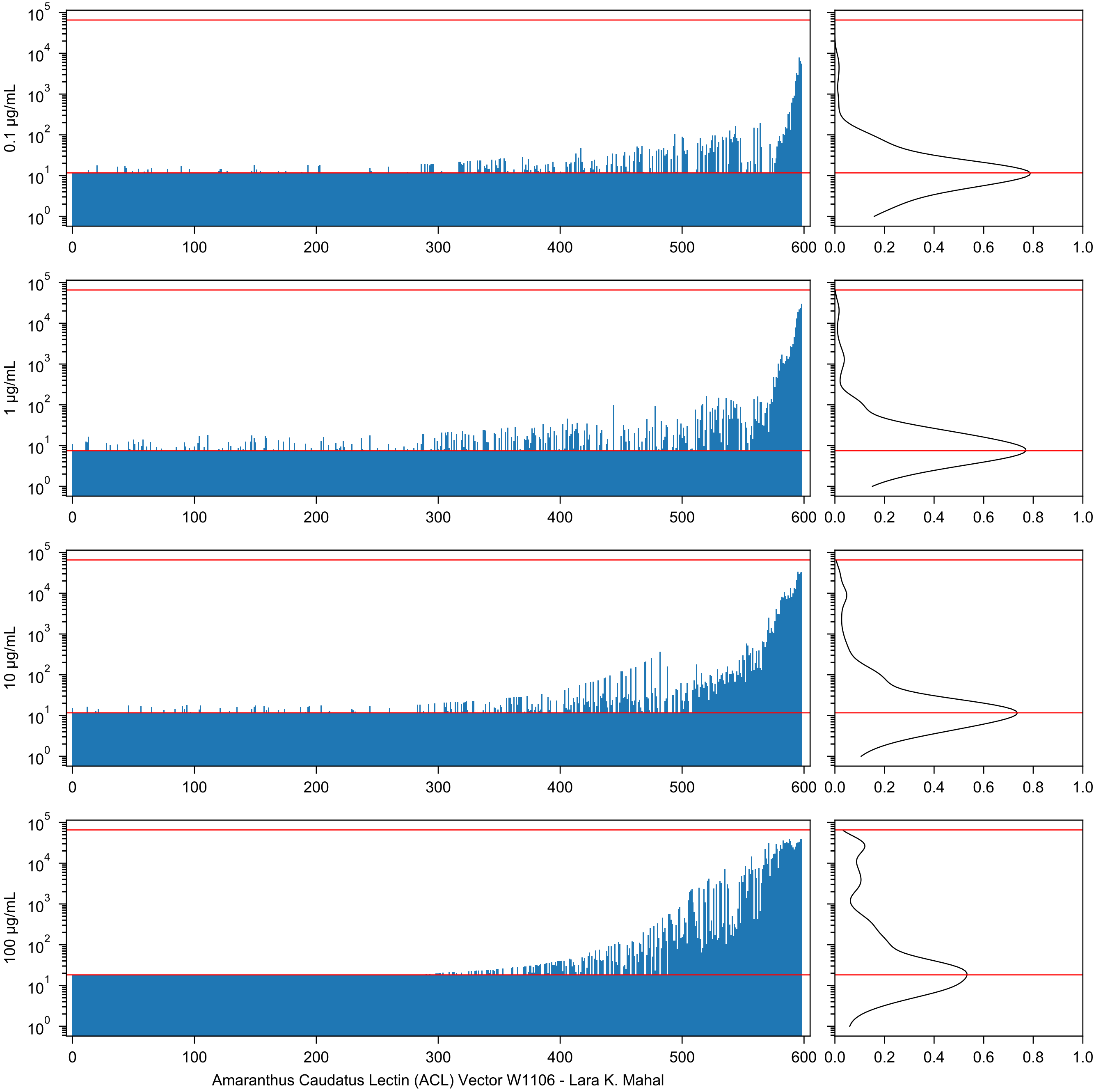
0.1 µg/mL
1 µg/mL
10 µg/mL
100 µg/mL
Amaranthus Caudatus Lectin (ACL) Vector W1106 - Lara K. Mahal

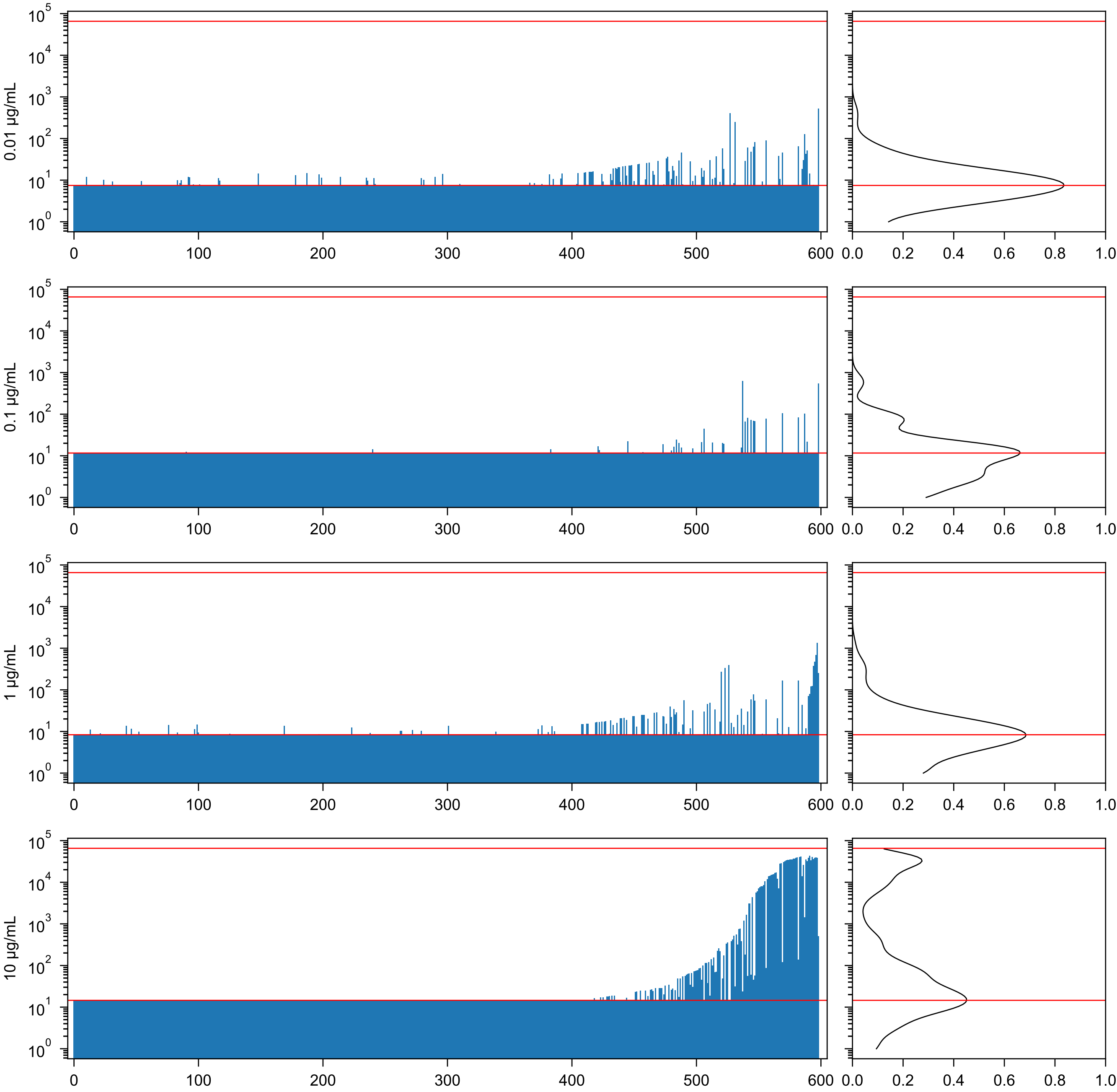

Artocarpus integrifolia Agglutinin (AIA) EY_281128-1 - Lara K. Mahal

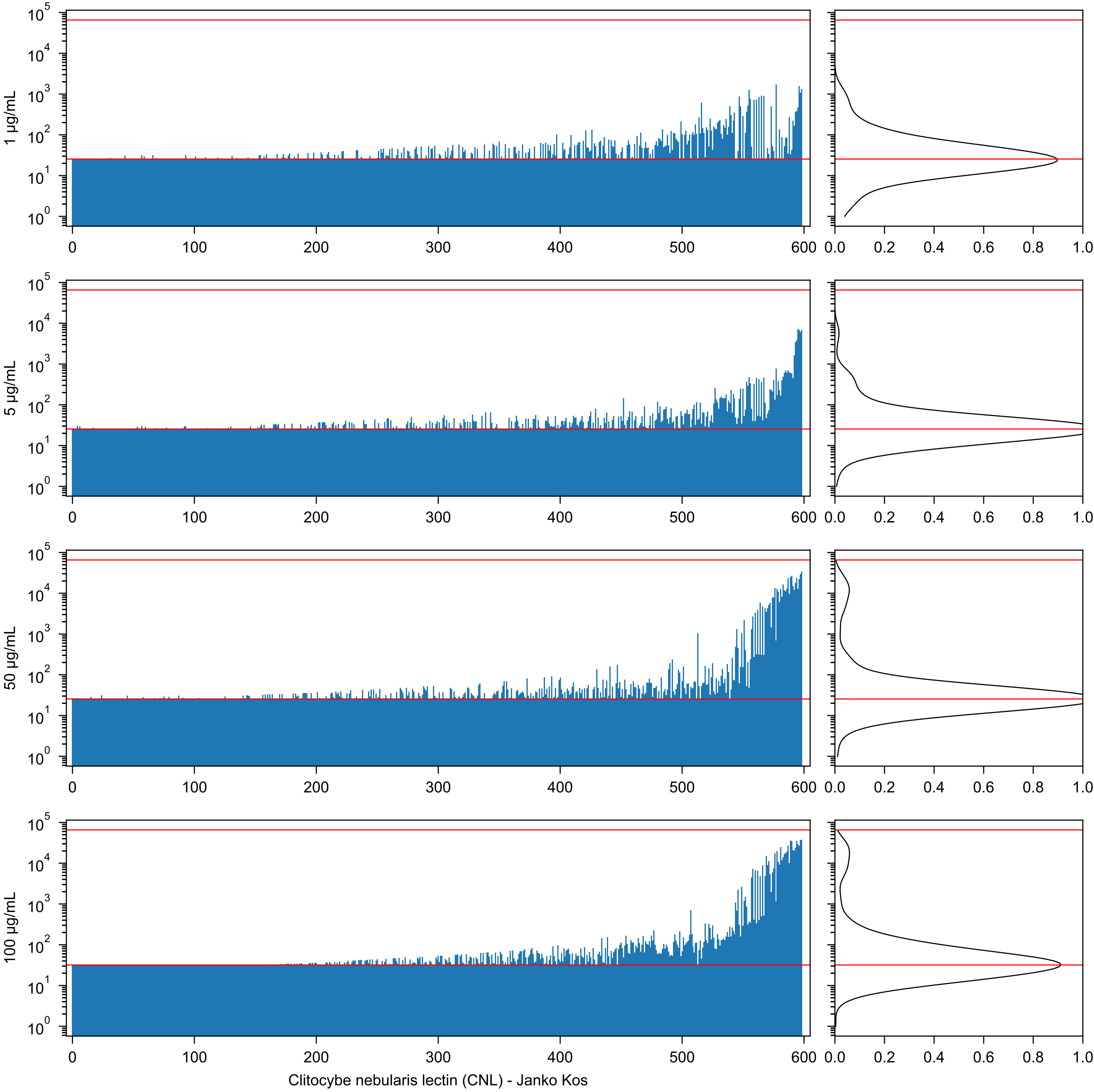
1 µg/mL
5 µg/mL
50 µg/mL
100 µg/mL
Clitocybe nebularis lectin (CNL) - Janko Kos

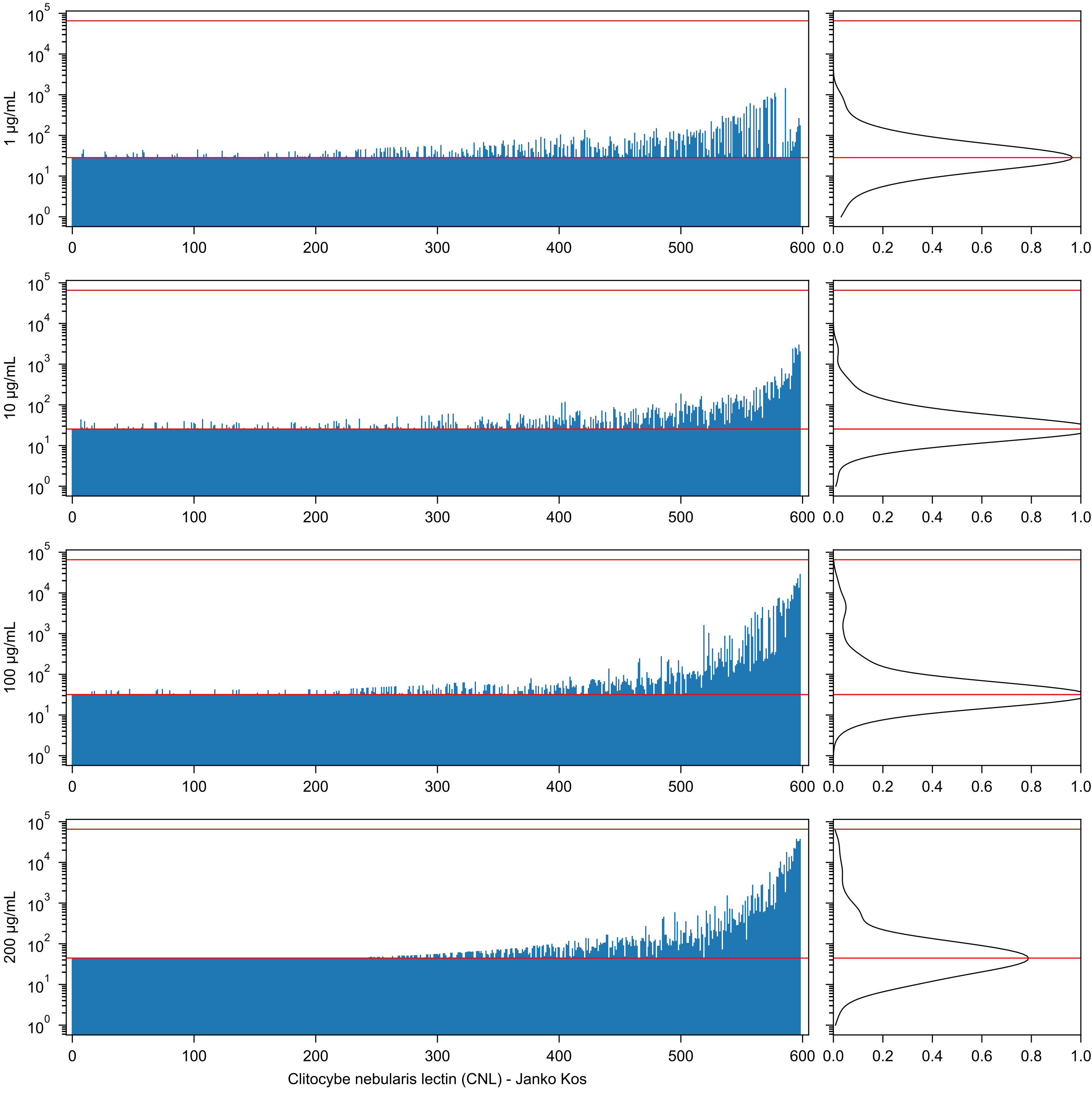

1 µg/mL
10 µg/mL
100 µg/mL
200 µg/mL
10^0 10^1 10^2 10^3 10^4 10^5
0 100 200 300 400 500 600
0.0 0.2 0.4 0.6 0.8 1.0
Clitocybe nebularis lectin (CNL) - Janko Kos

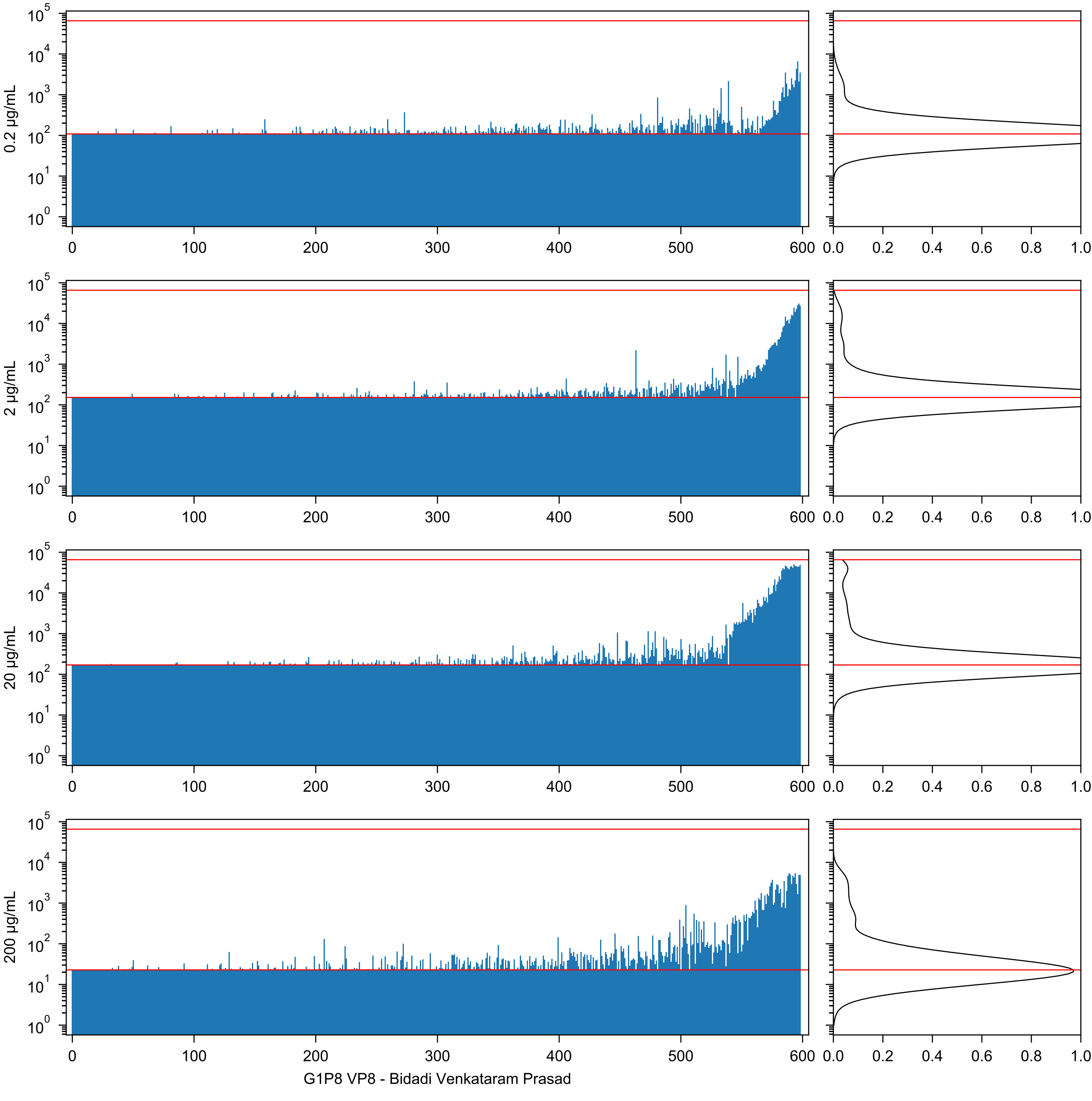
0.2 µg/mL
2 µg/mL
20 µg/mL
200 µg/mL
G1P8 VP8 - Bidadi Venkataram Prasad

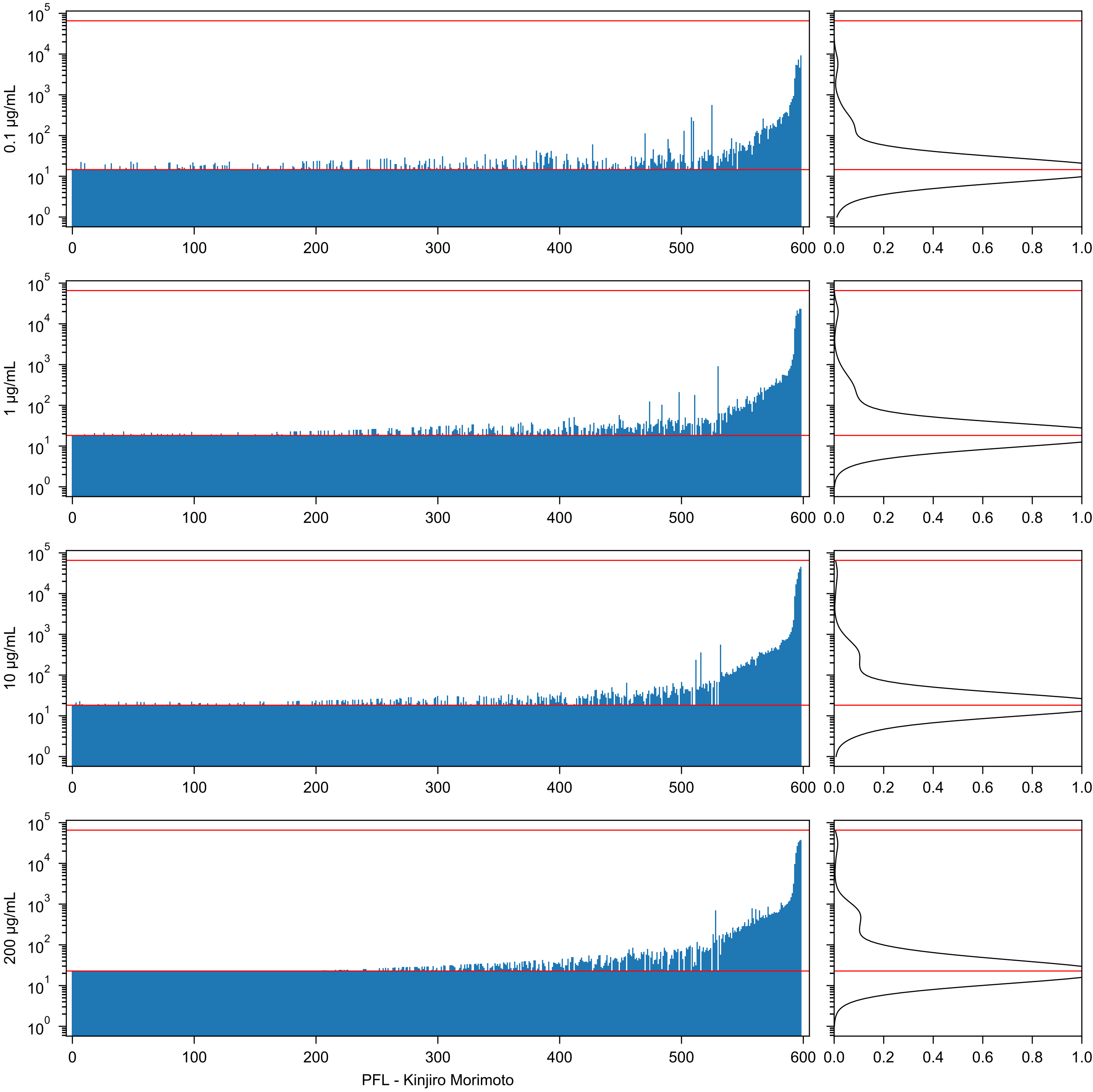
0.1 µg/mL
1 µg/mL
10 µg/mL
200 µg/mL
PFL - Kinjiro Morimoto

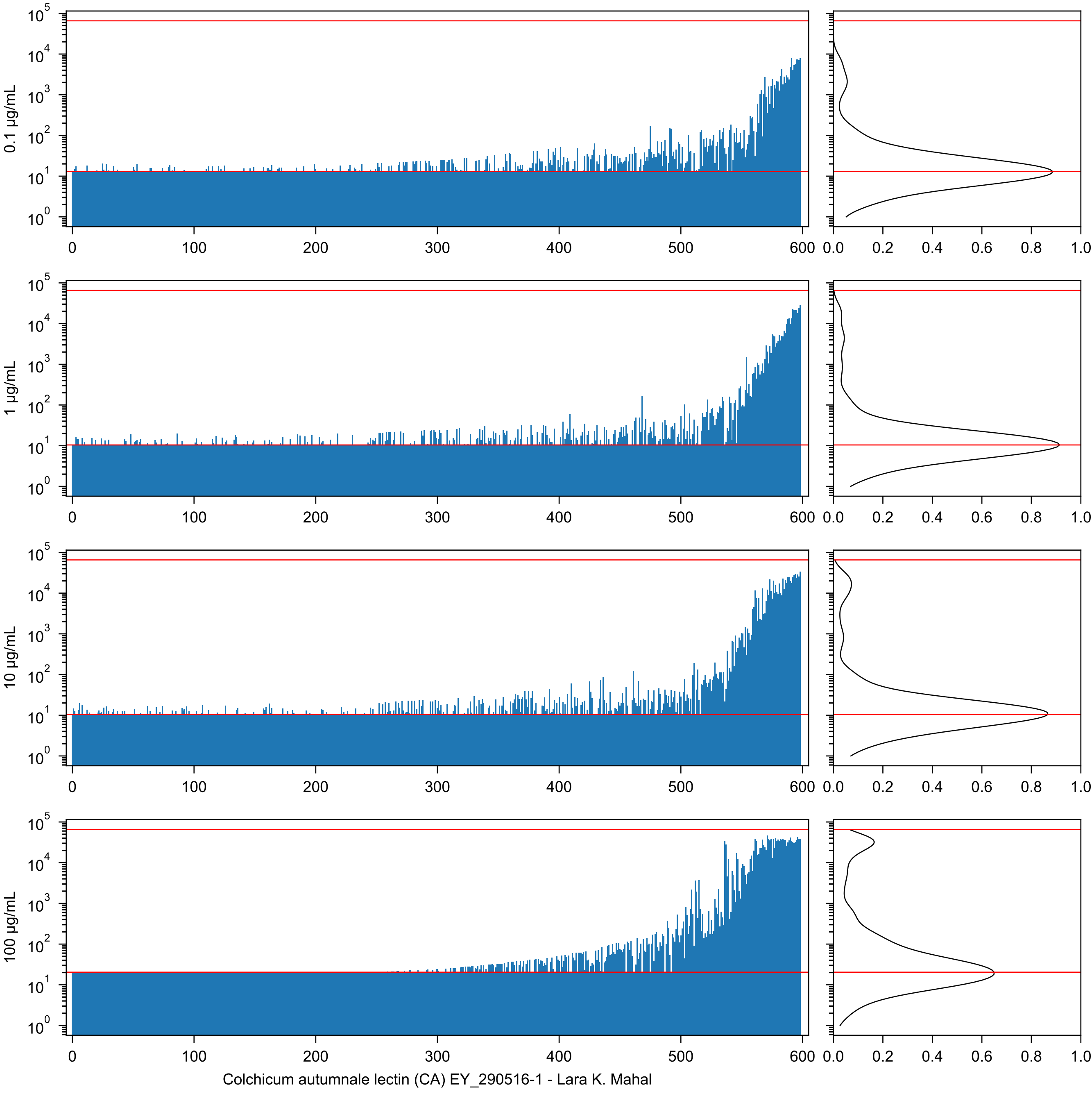

0.1 µg/mL
1 µg/mL
10 µg/mL
100 µg/mL
Colchicum autumnale lectin (CA) EY_290516-1 - Lara K. Mahal

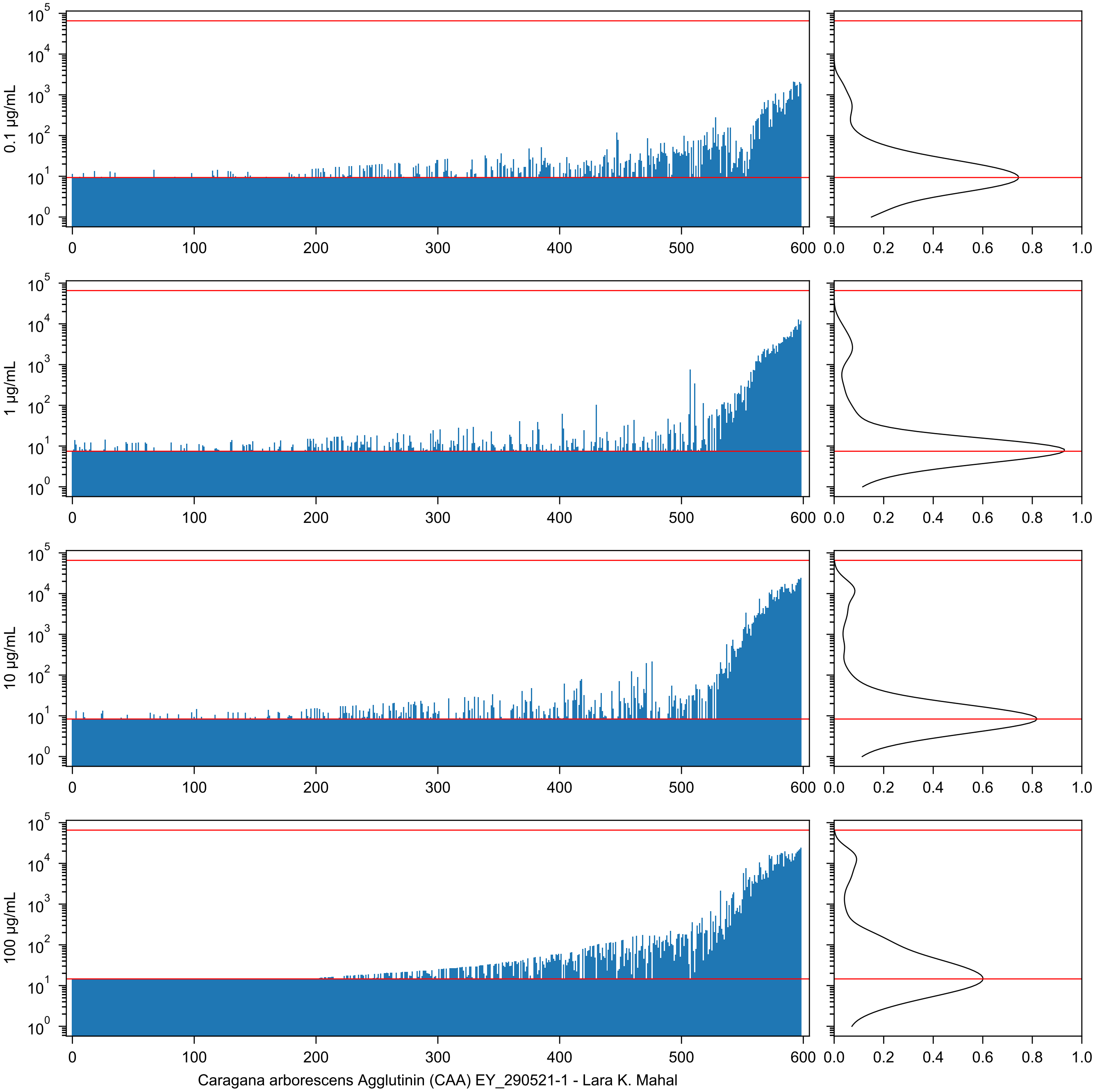
0.1 µg/mL
1 µg/mL
10 µg/mL
100 µg/mL
Caragana arborescens Agglutinin (CAA) EY_290521-1 - Lara K. Mahal

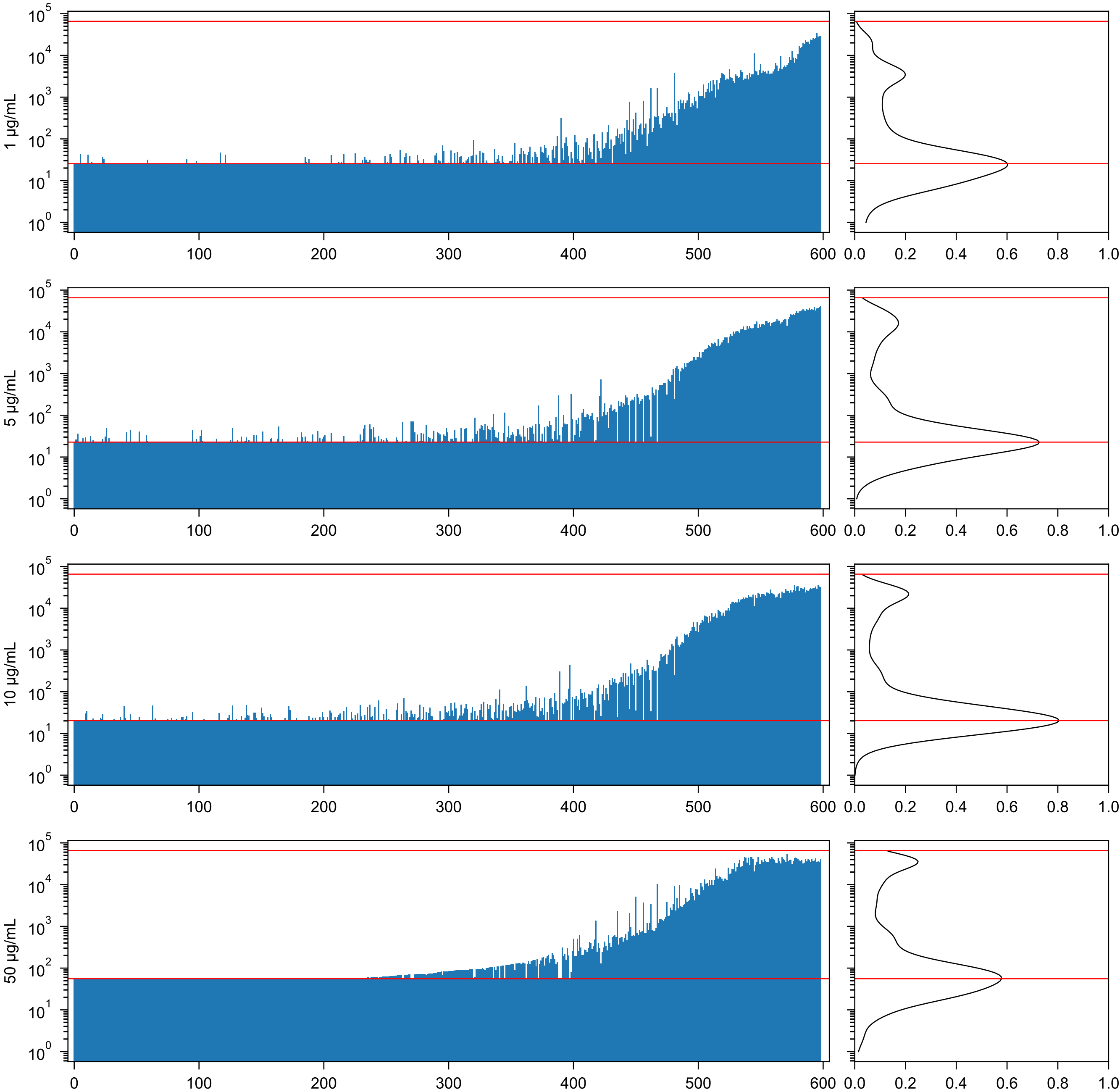


Canavalia A (Con A) Vector_W0828 - Lara K. Mahal

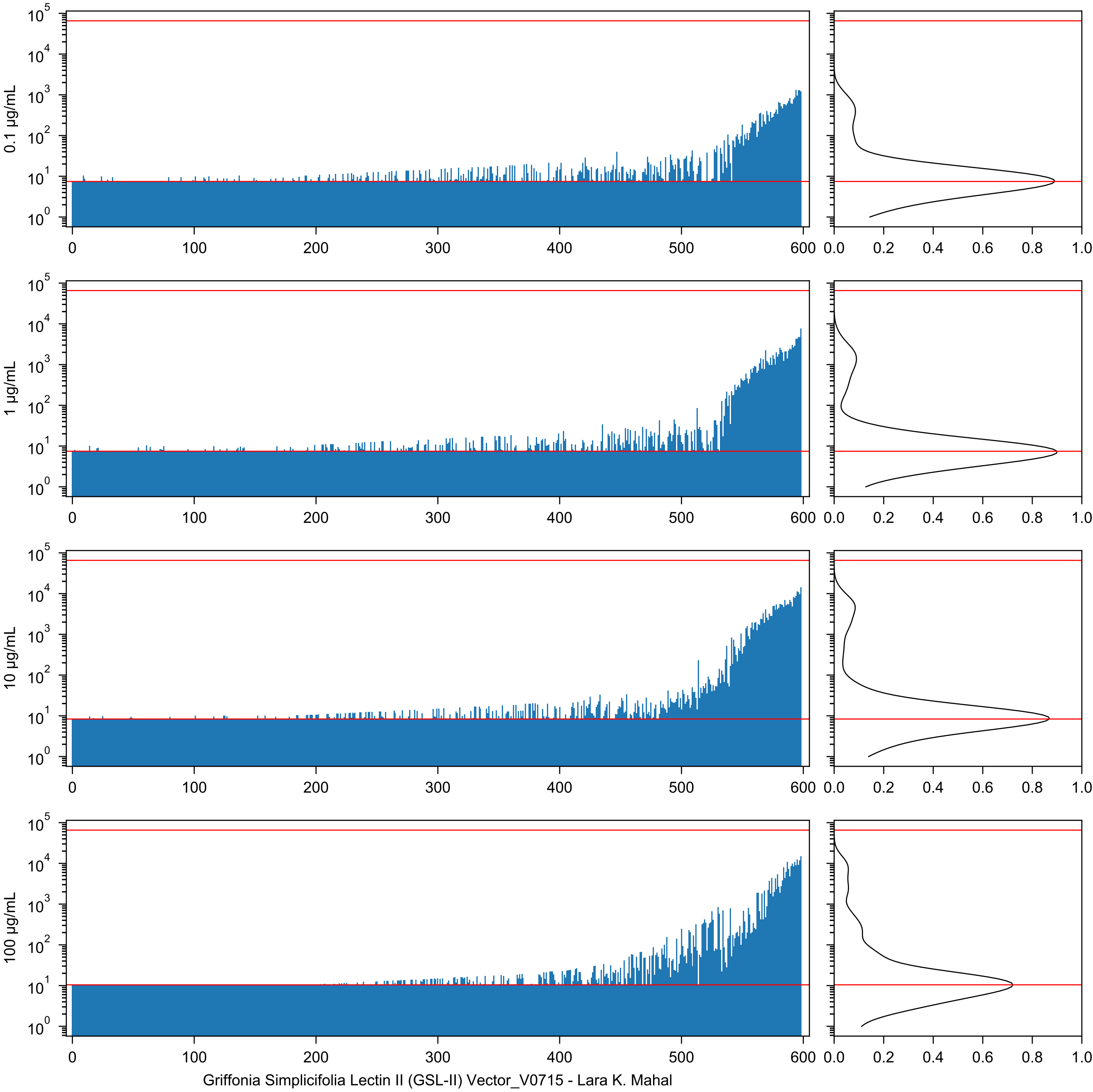

0.1 µg/mL
1 µg/mL
10 µg/mL
100 µg/mL
Griffonia Simplicifolia Lectin II (GSL-II) Vector_V0715 - Lara K. Mahal

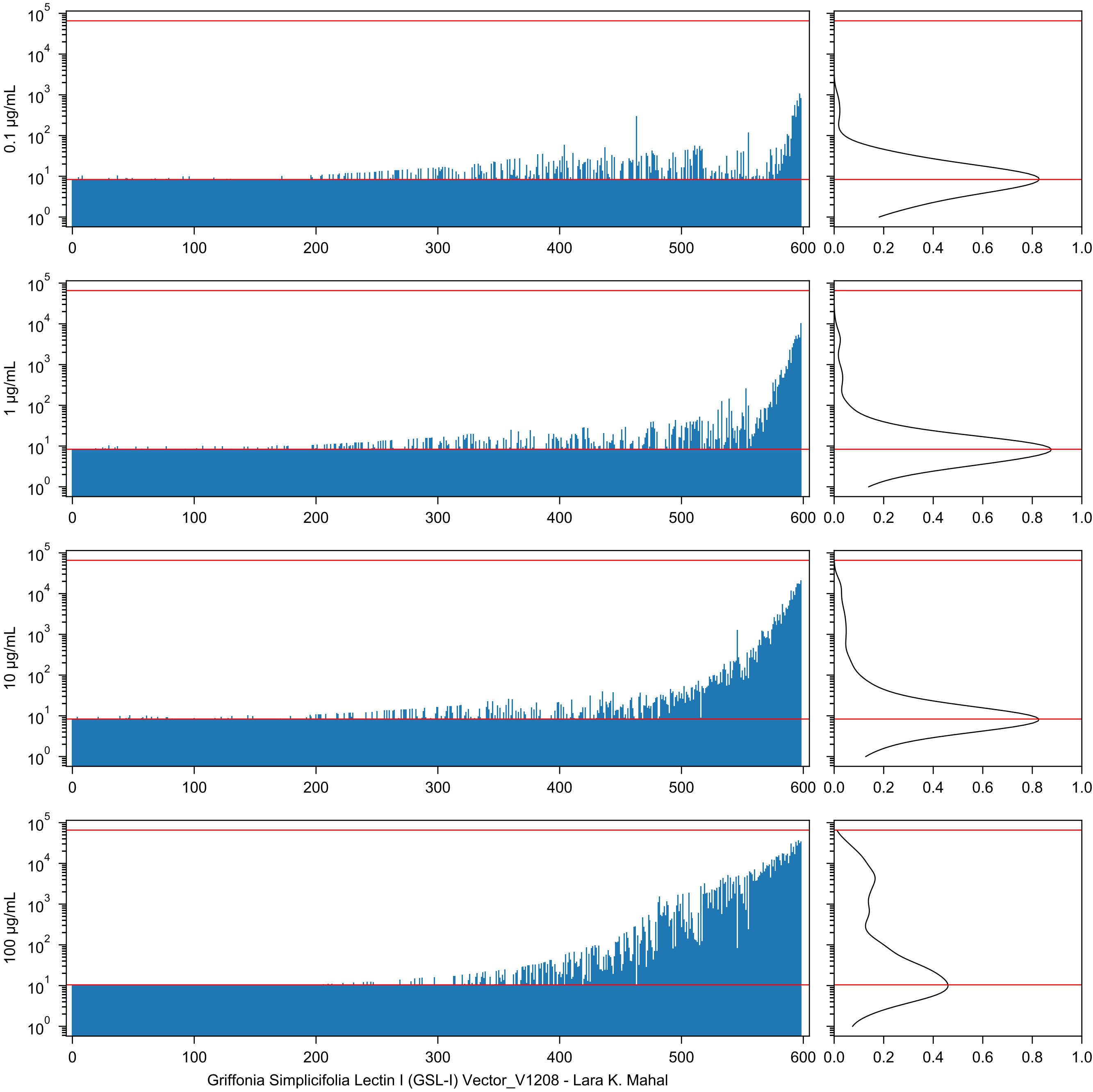
0.1 μg/mL
1 μg/mL
10 μg/mL
100 μg/mL
Griffonia Simplicifolia Lectin I (GSL-I) Vector_V1208 - Lara K. Mahal

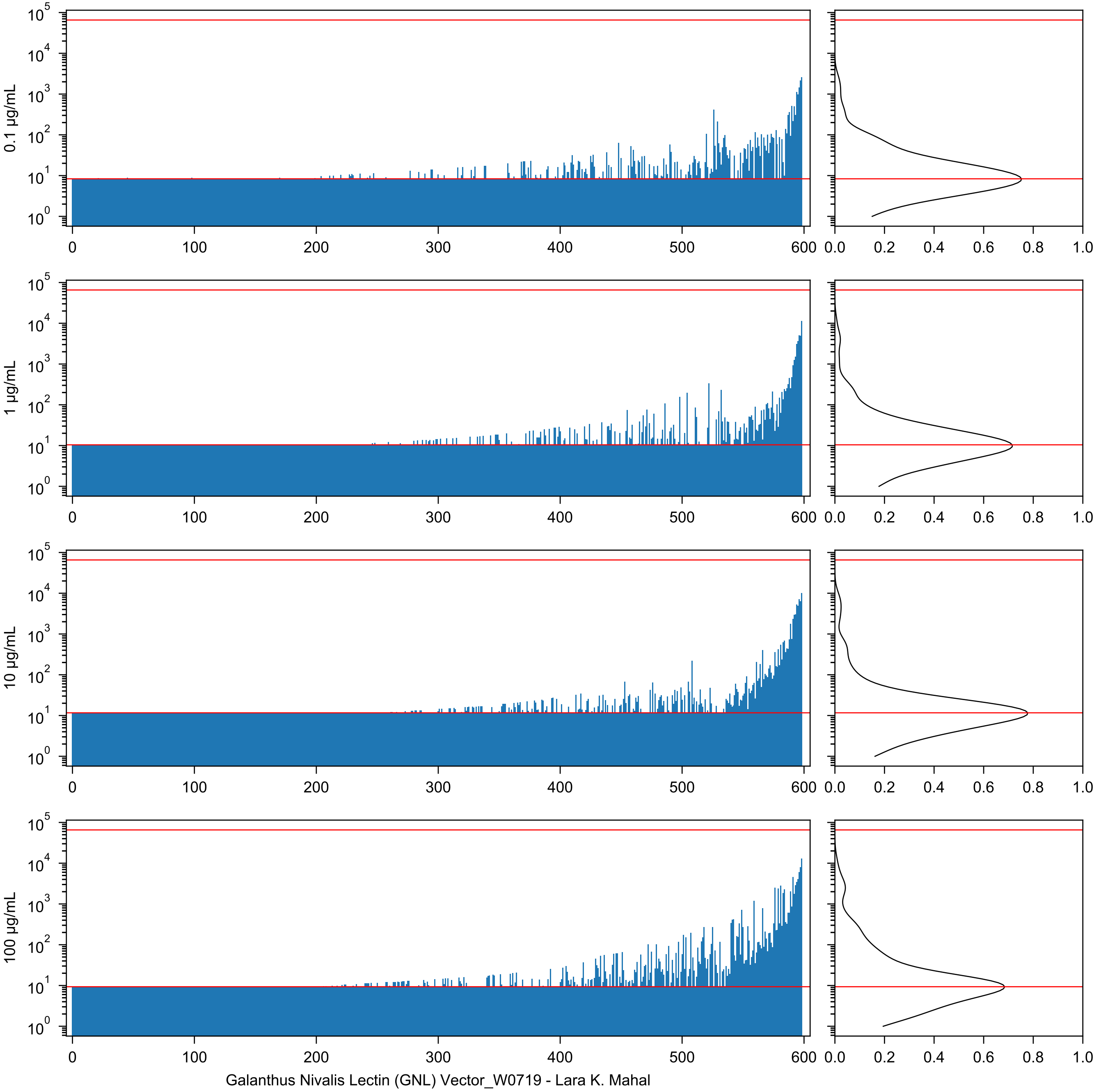

0.1 µg/mL
1 µg/mL
10 µg/mL
100 µg/mL
Galanthus Nivalis Lectin (GNL) Vector_W0719 - Lara K. Mahal

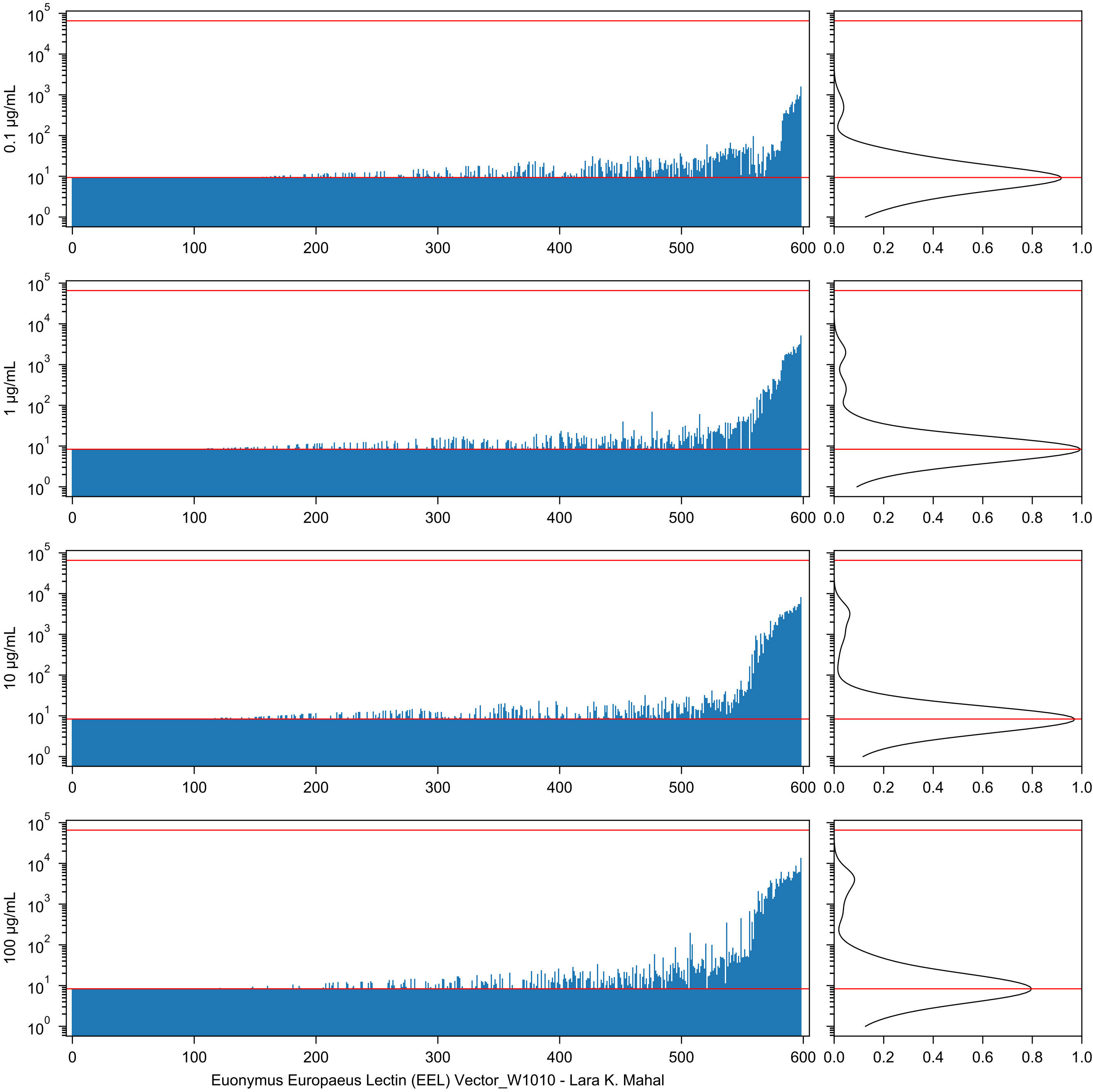
0.1 µg/mL
1 µg/mL
10 µg/mL
100 µg/mL
Euonymus Europaeus Lectin (EEL) Vector_W1010 - Lara K. Mahal

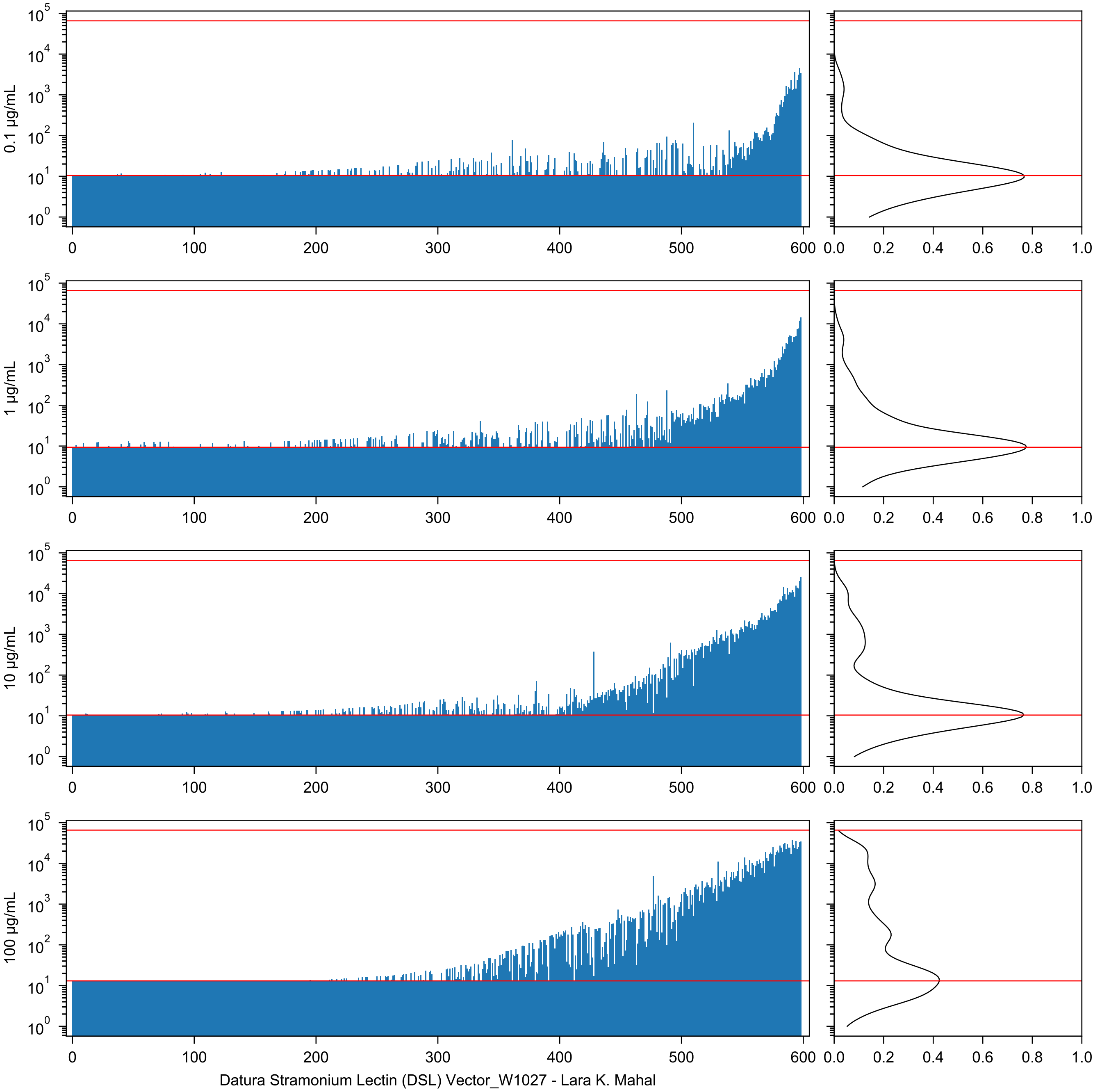
0.1 µg/mL
1 µg/mL
10 µg/mL
100 µg/mL
Datura Stramonium Lectin (DSL) Vector_W1027 - Lara K. Mahal

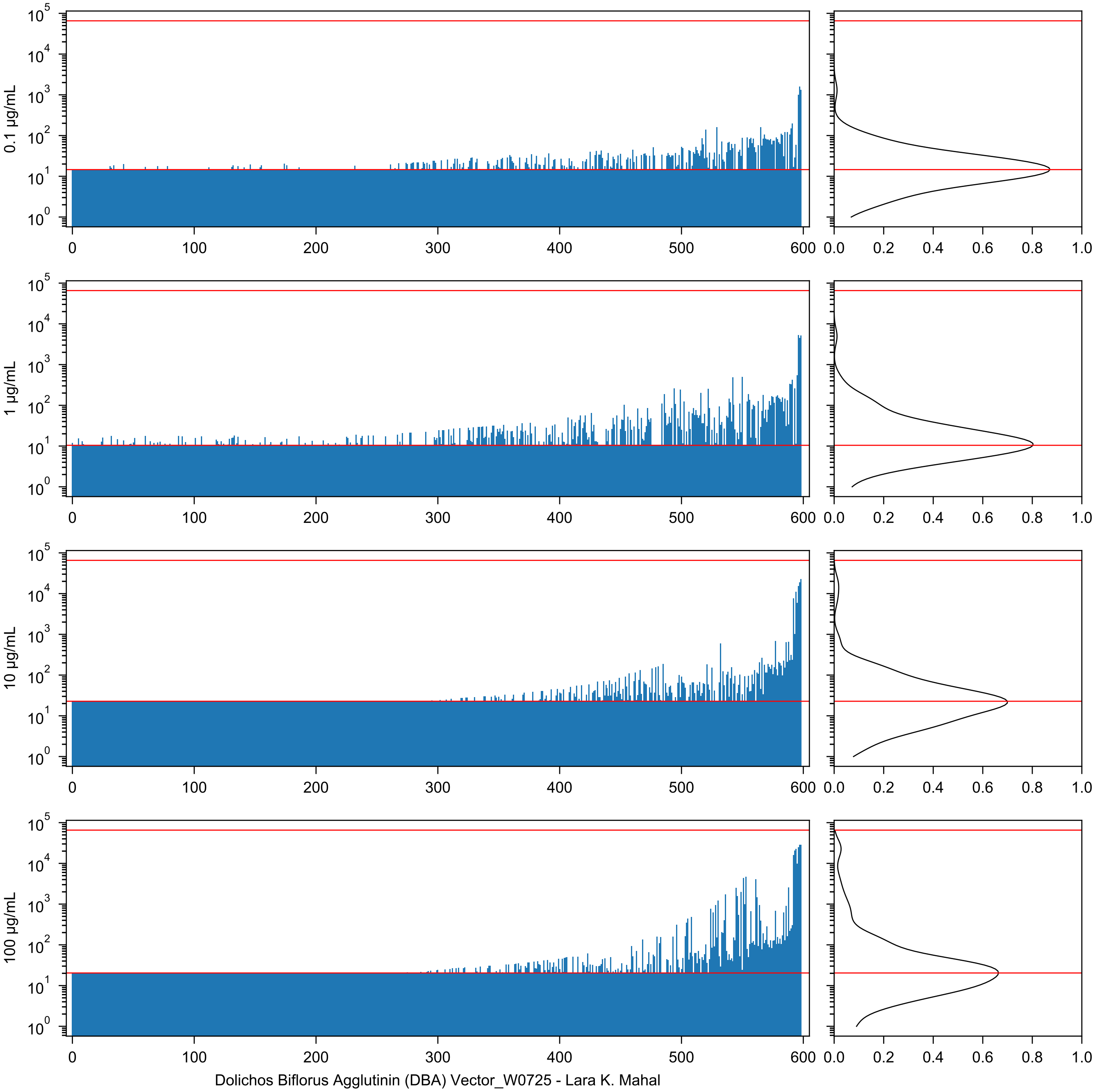

0.1 µg/mL
1 µg/mL
10 µg/mL
100 µg/mL
Dolichos Biflorus Agglutinin (DBA) Vector_W0725 - Lara K. Mahal

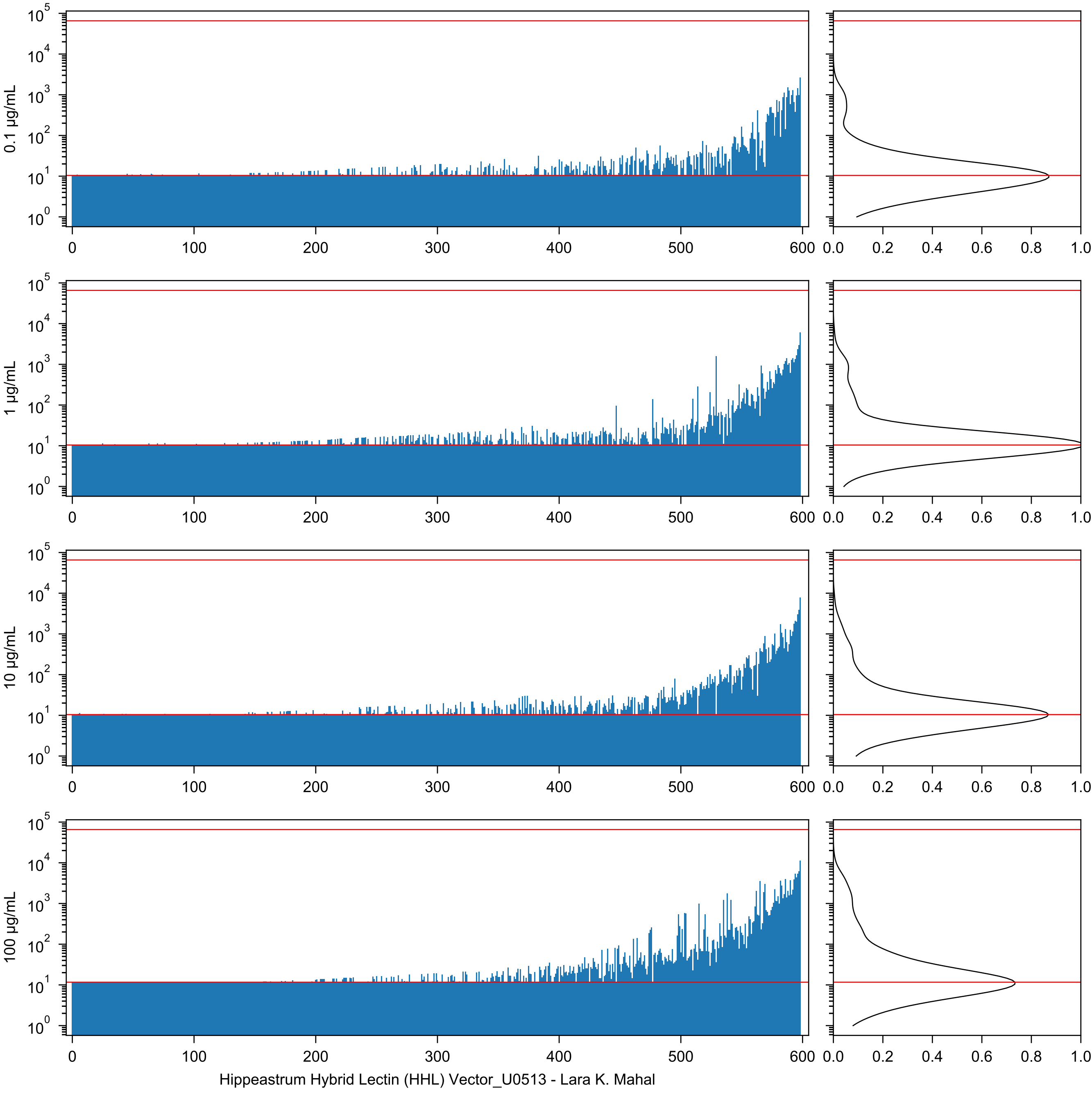

0.1 µg/mL
1 µg/mL
10 µg/mL
100 µg/mL
Hippeastrum Hybrid Lectin (HHL) Vector_U0513 - Lara K. Mahal

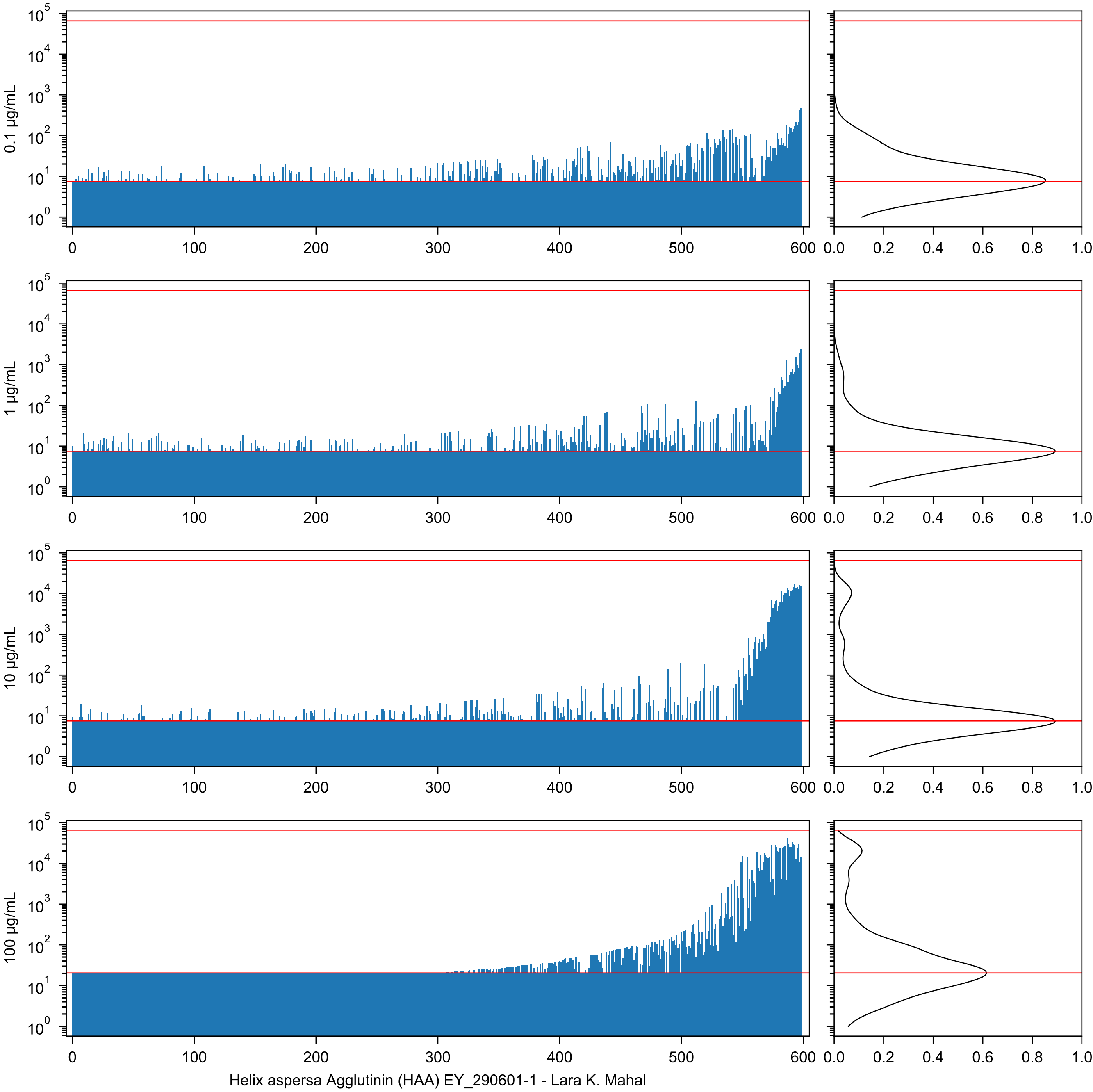

0.1 µg/mL
1 µg/mL
10 µg/mL
100 µg/mL
Helix aspersa Agglutinin (HAA) EY_290601-1 - Lara K. Mahal

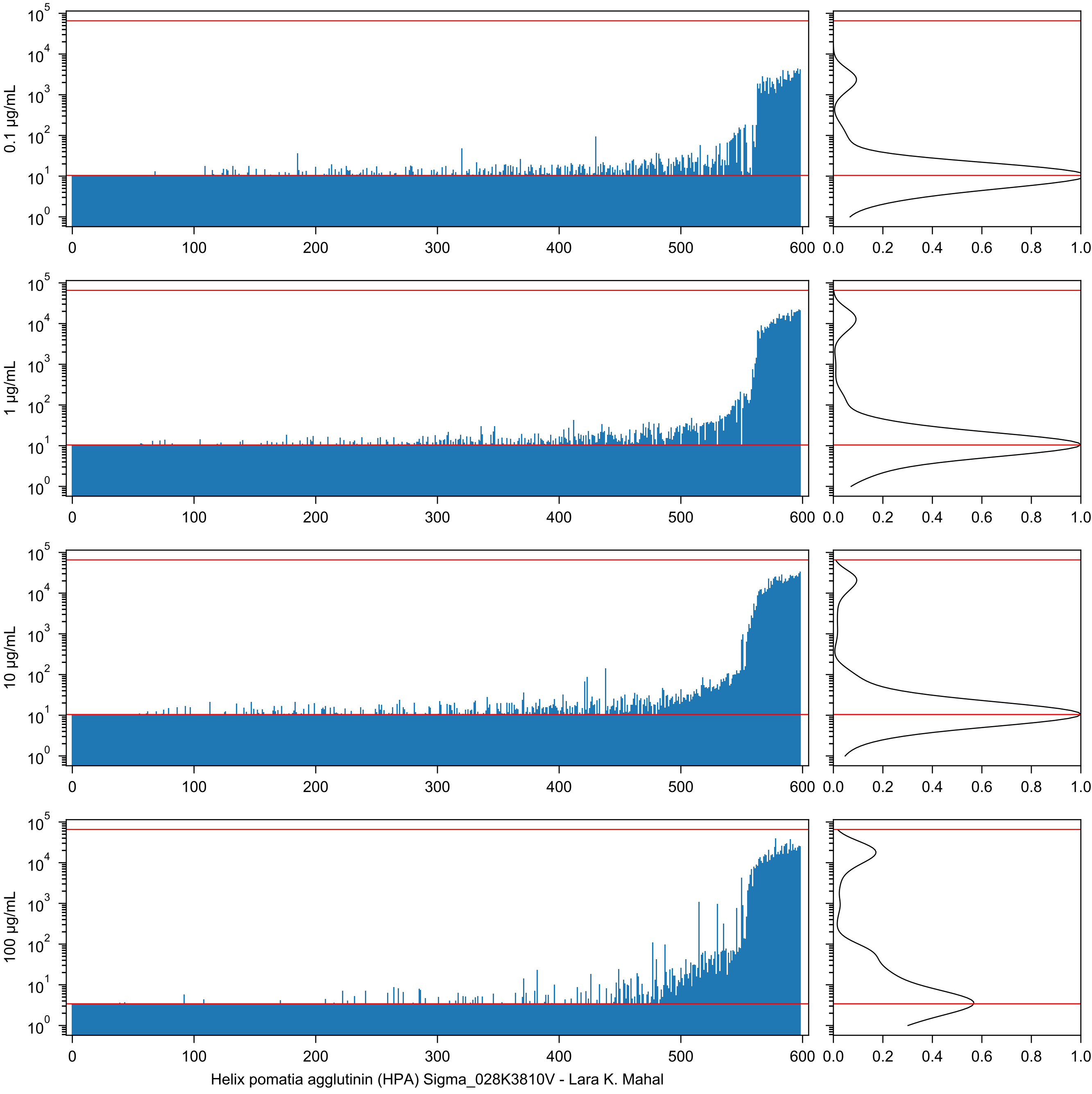
0.1 µg/mL
1 µg/mL
10 µg/mL
100 µg/mL
Helix pomatia agglutinin (HPA) Sigma_028K3810V - Lara K. Mahal

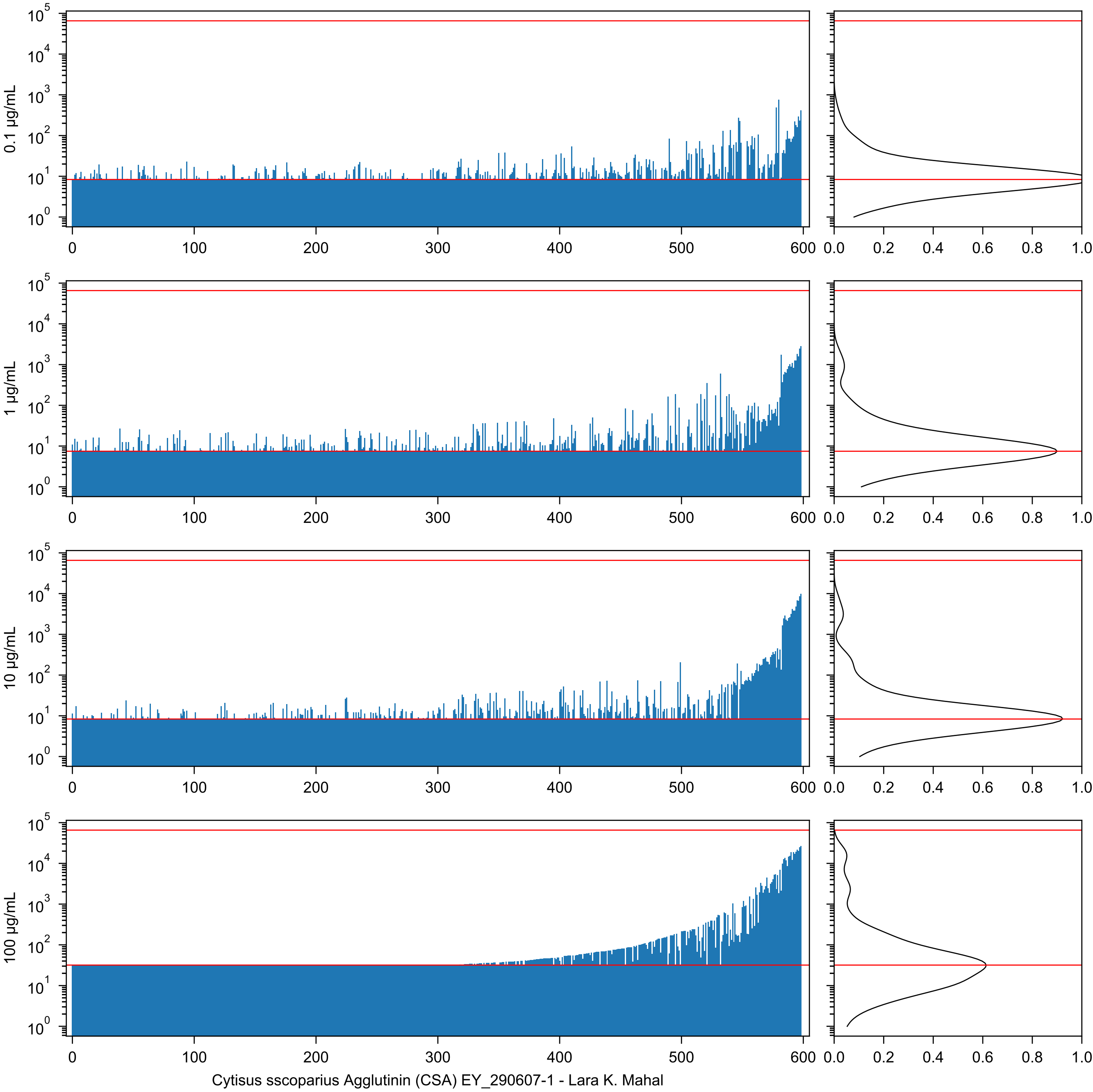
0.1 µg/mL
1 µg/mL
10 µg/mL
100 µg/mL
0
100
200
300
400
500
600
0.0
0.2
0.4
0.6
0.8
1.0
Cytisus sscoparius Agglutinin (CSA) EY_290607-1 - Lara K. Mahal

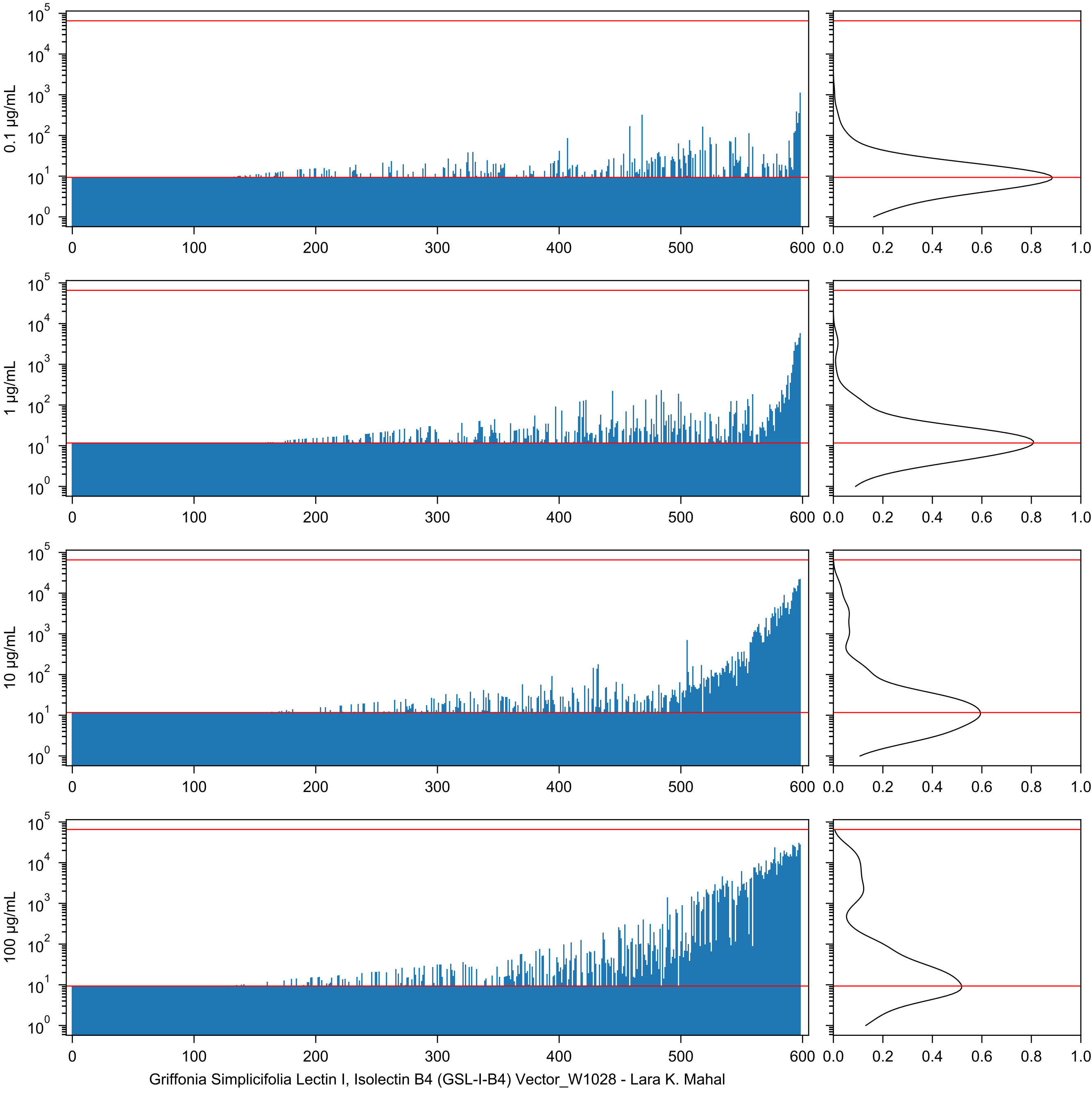
0.1 µg/mL
1 µg/mL
10 µg/mL
100 µg/mL
Griffonia Simplicifolia Lectin I, Isolectin B4 (GSL-I-B4) Vector_W1028 - Lara K. Mahal

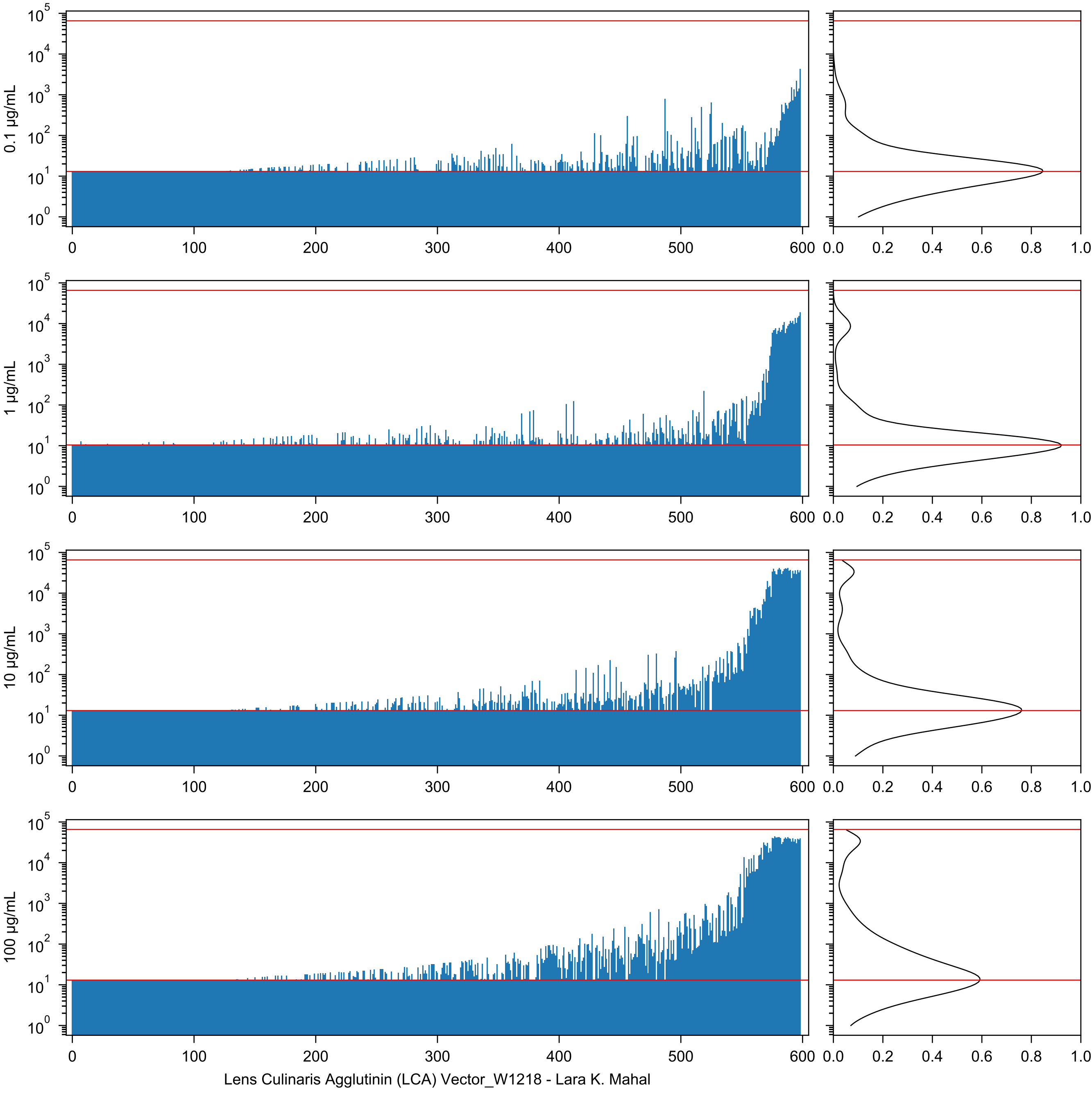
0.1 µg/mL
1 µg/mL
10 µg/mL
100 µg/mL
0
100
200
300
400
500
600
0.0
0.2
0.4
0.6
0.8
1.0
Lens Culinaris Agglutinin (LCA) Vector_W1218 - Lara K. Mahal

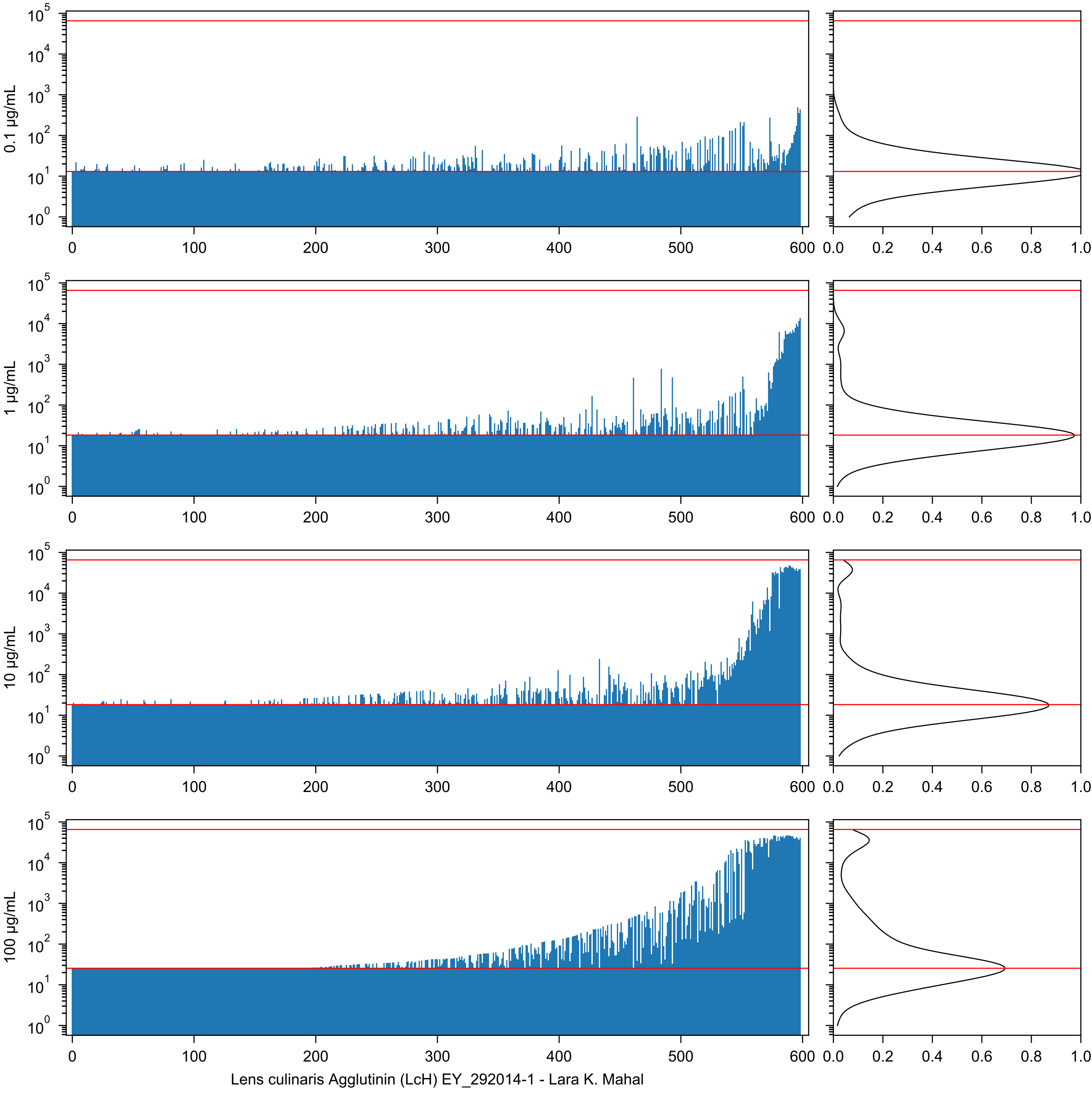

0.1 µg/mL
1 µg/mL
10 µg/mL
100 µg/mL
0
100
200
300
400
500
600
0.0
0.2
0.4
0.6
0.8
1.0
Lens culinaris Agglutinin (LcH) EY_292014-1 - Lara K. Mahal

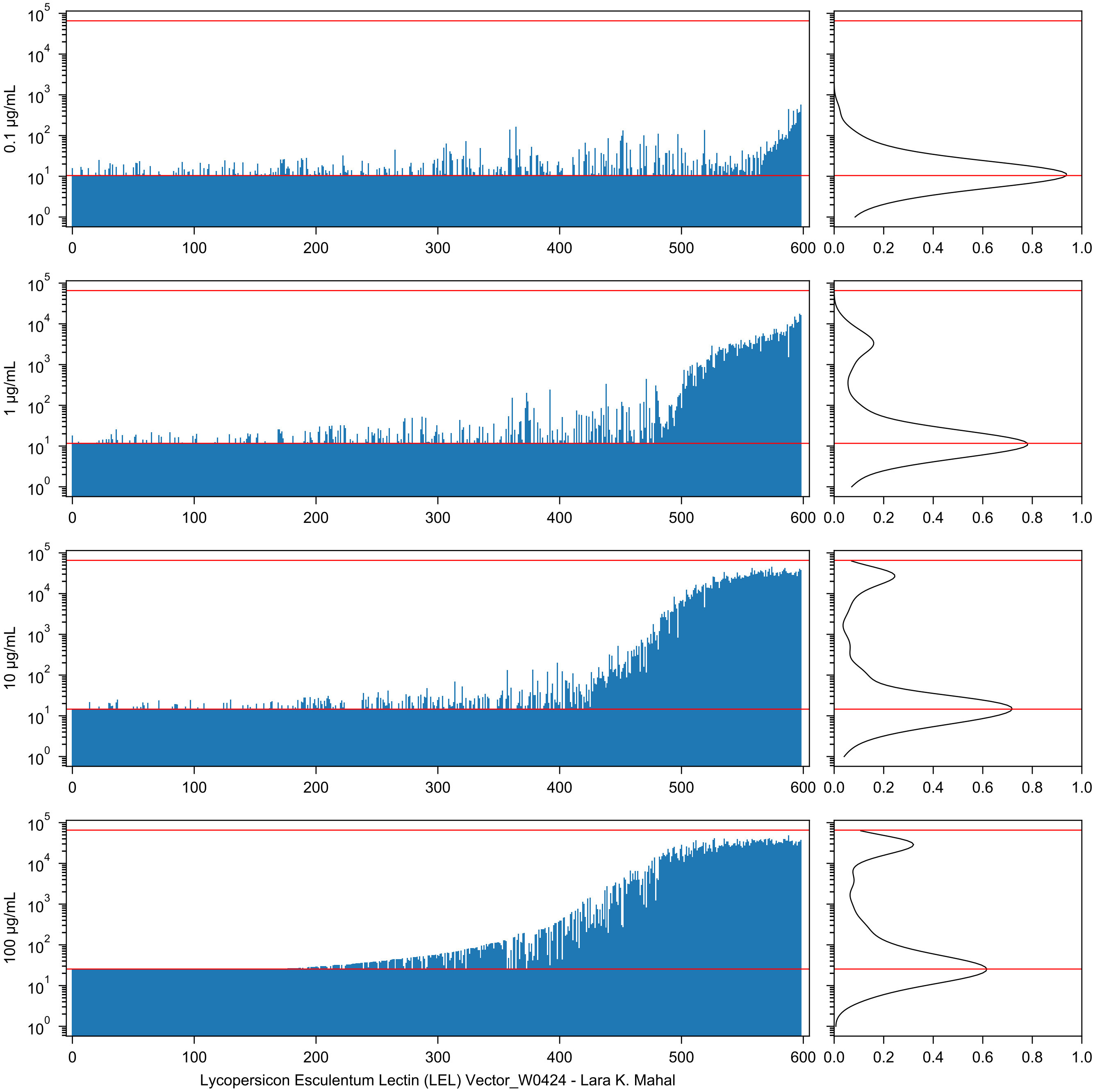
0.1 µg/mL
1 µg/mL
10 µg/mL
100 µg/mL
Lycopersicon Esculentum Lectin (LEL) Vector_W0424 - Lara K. Mahal

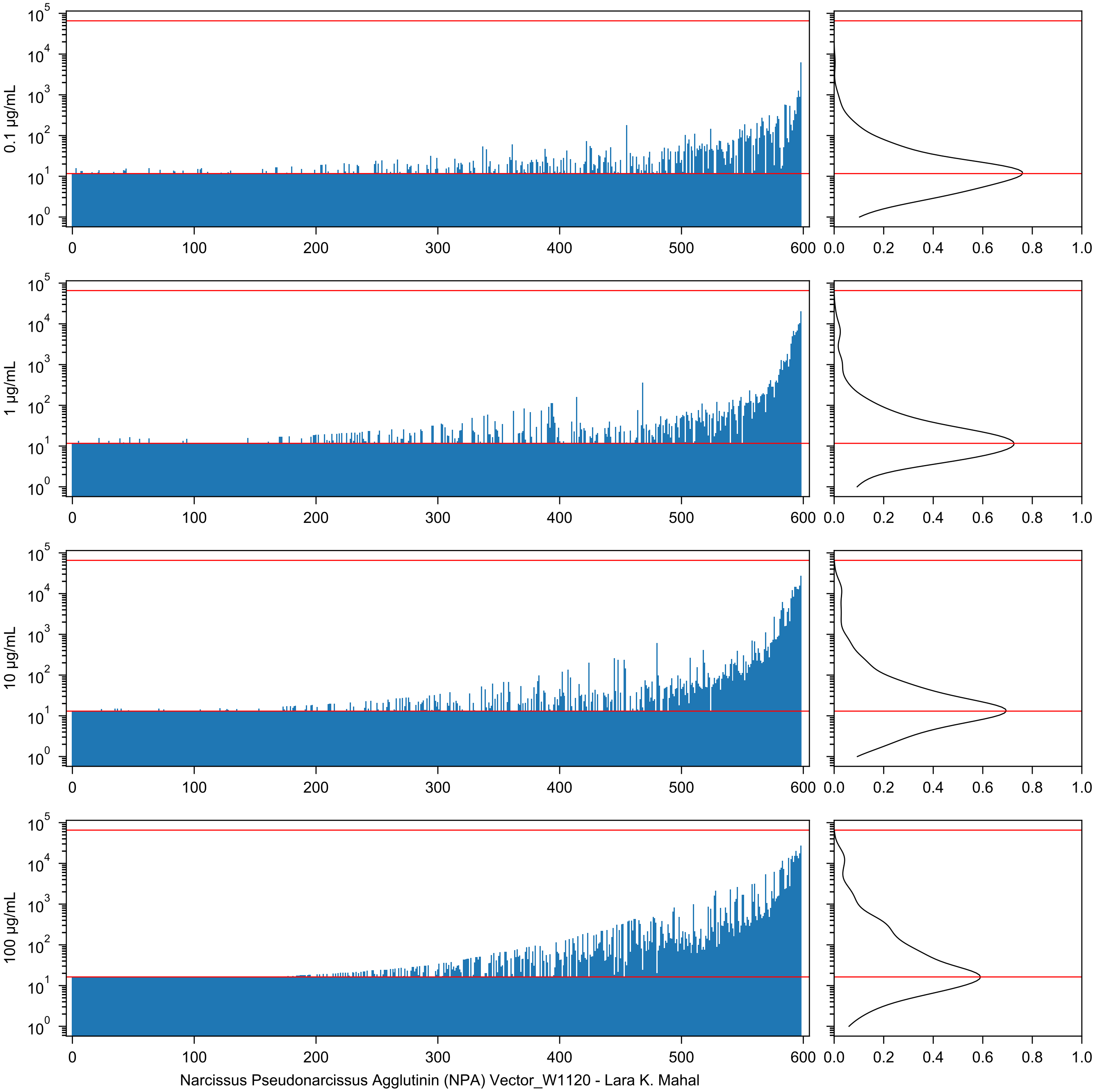
0.1 µg/mL
1 µg/mL
10 µg/mL
100 µg/mL
Narcissus Pseudonarcissus Agglutinin (NPA) Vector_W1120 - Lara K. Mahal

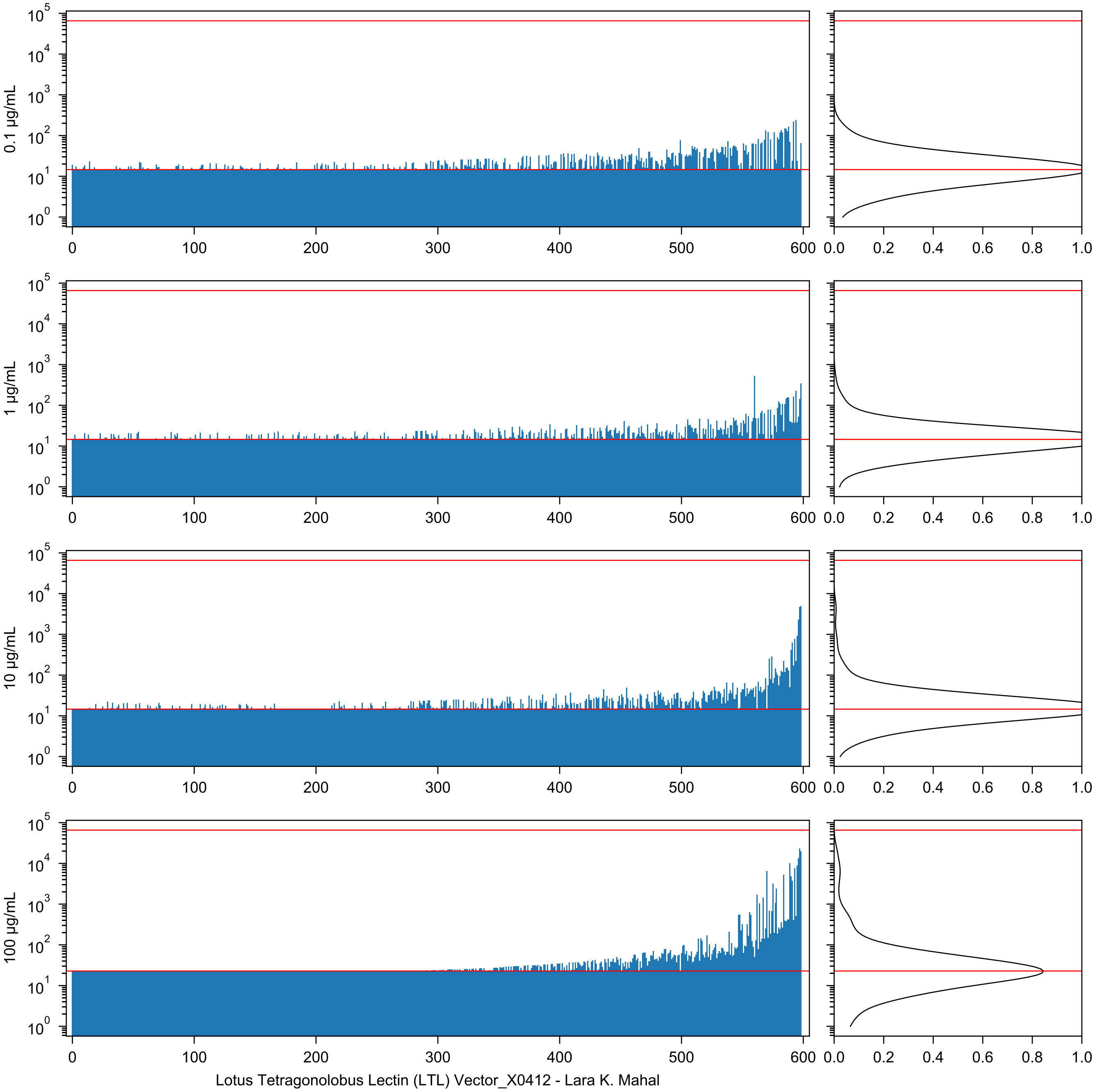

0.1 µg/mL
1 µg/mL
10 µg/mL
100 µg/mL
Lotus Tetragonolobus Lectin (LTL) Vector_X0412 - Lara K. Mahal

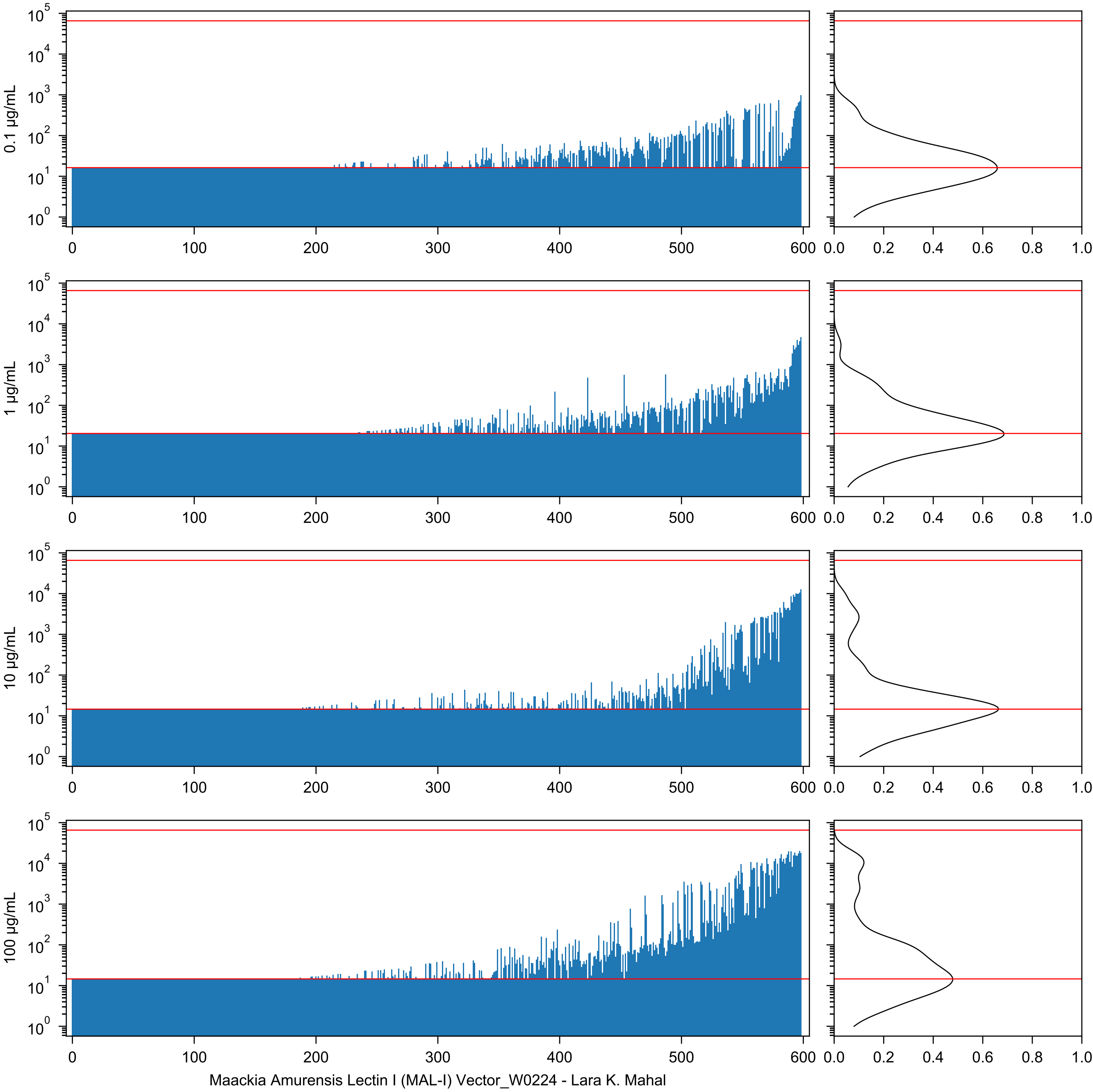
0.1 µg/mL
1 µg/mL
10 µg/mL
100 µg/mL
Maackia Amurensis Lectin I (MAL-I) Vector_W0224 - Lara K. Mahal

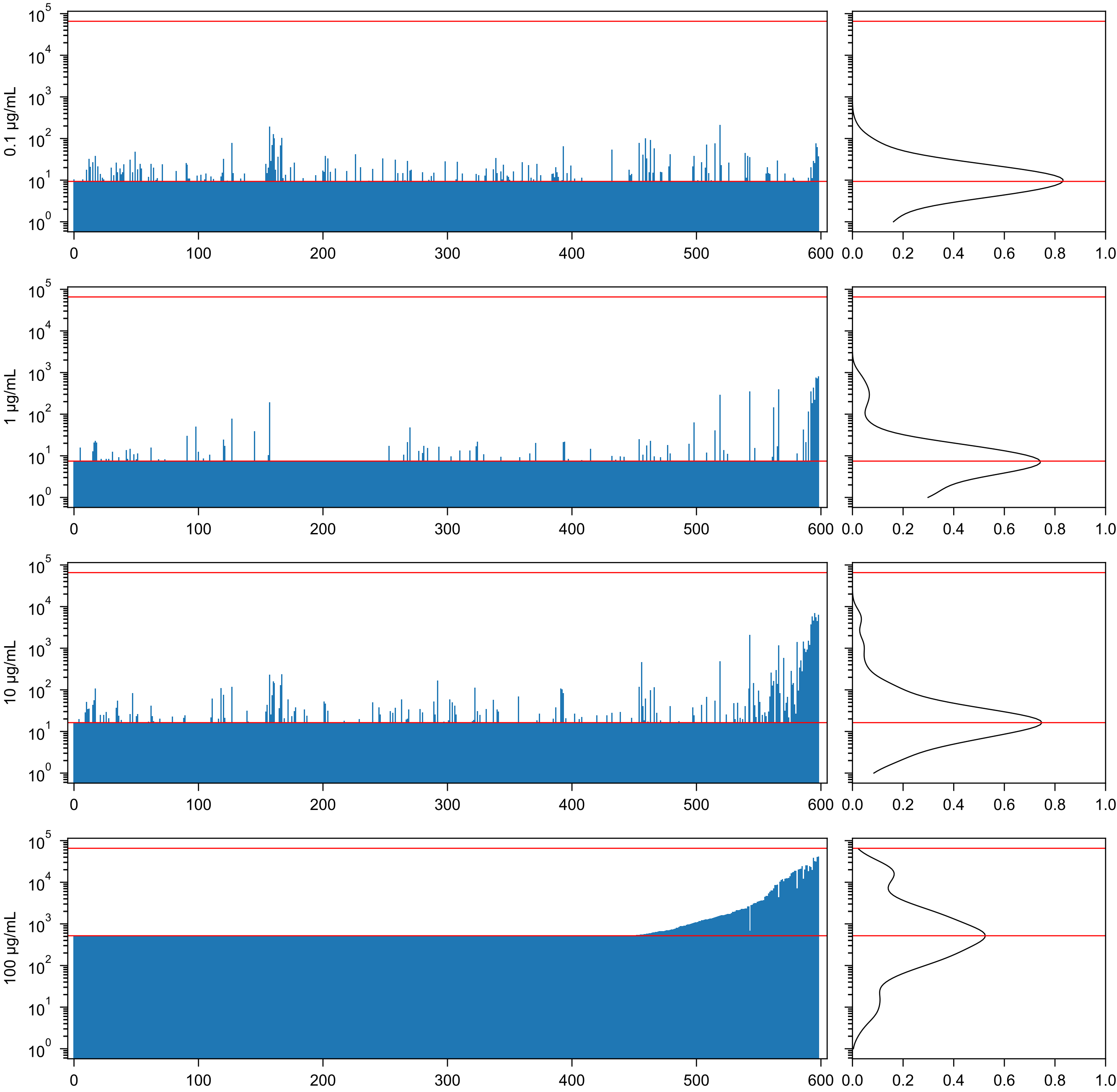

Maackia amurensis Agglutinin (MAA) EY_290224-5 - Lara K. Mahal

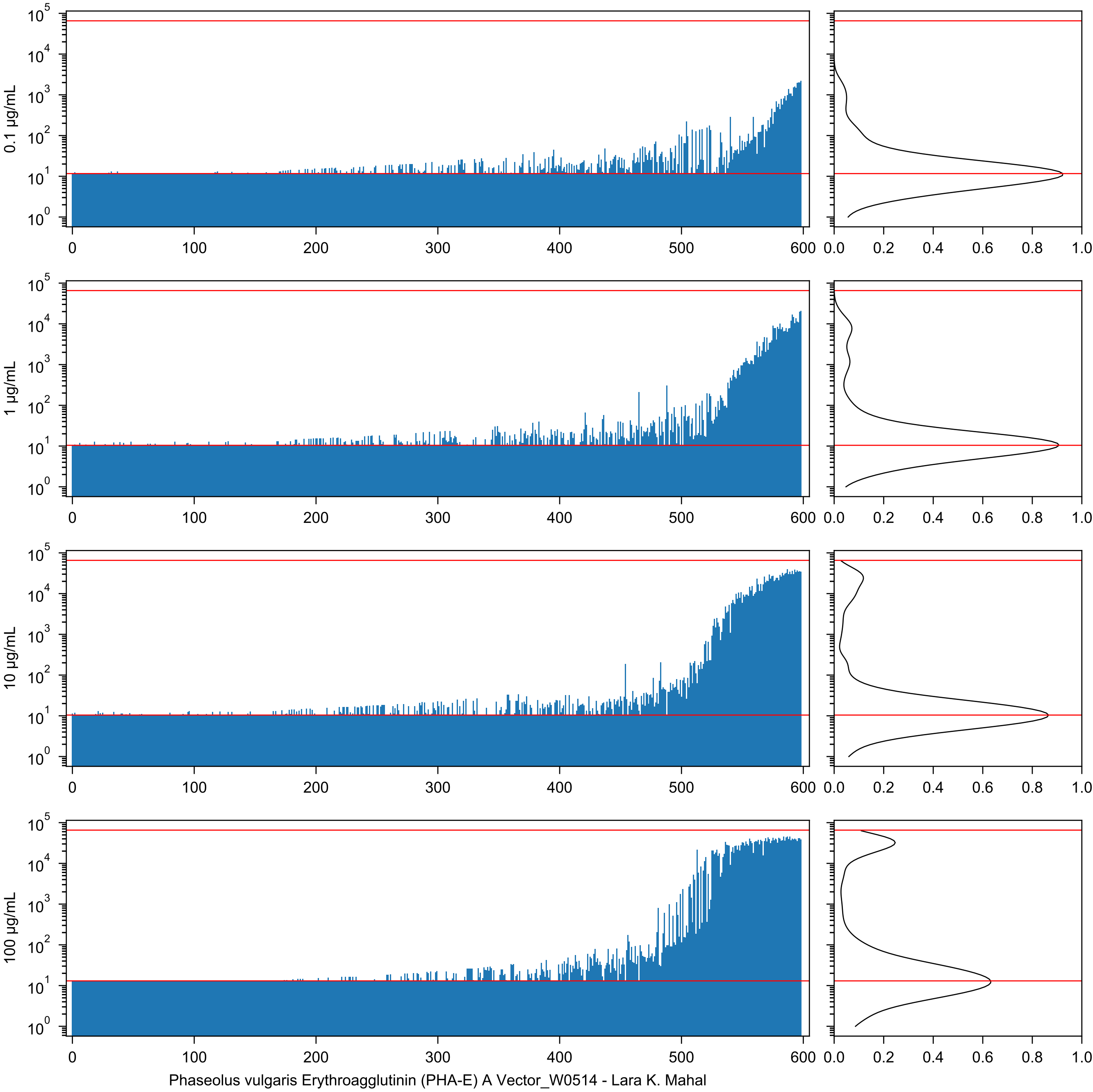

0.1 µg/mL
1 µg/mL
10 µg/mL
100 µg/mL
Phaseolus vulgaris Erythroagglutinin (PHA-E) A Vector_W0514 - Lara K. Mahal

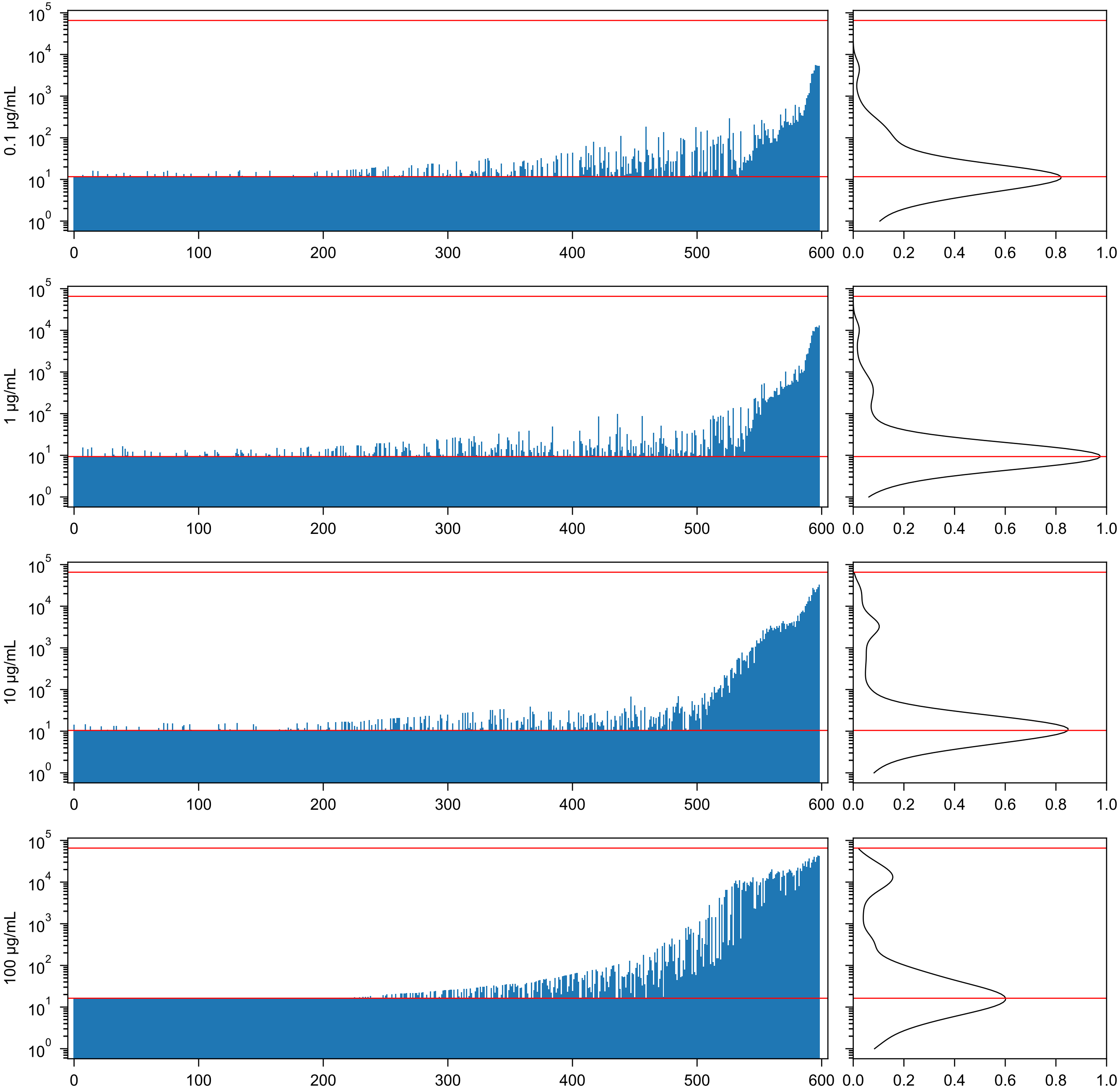


Maackia amurensis agglutinin (MAA) Seikagaku_22101C - Lara K. Mahal

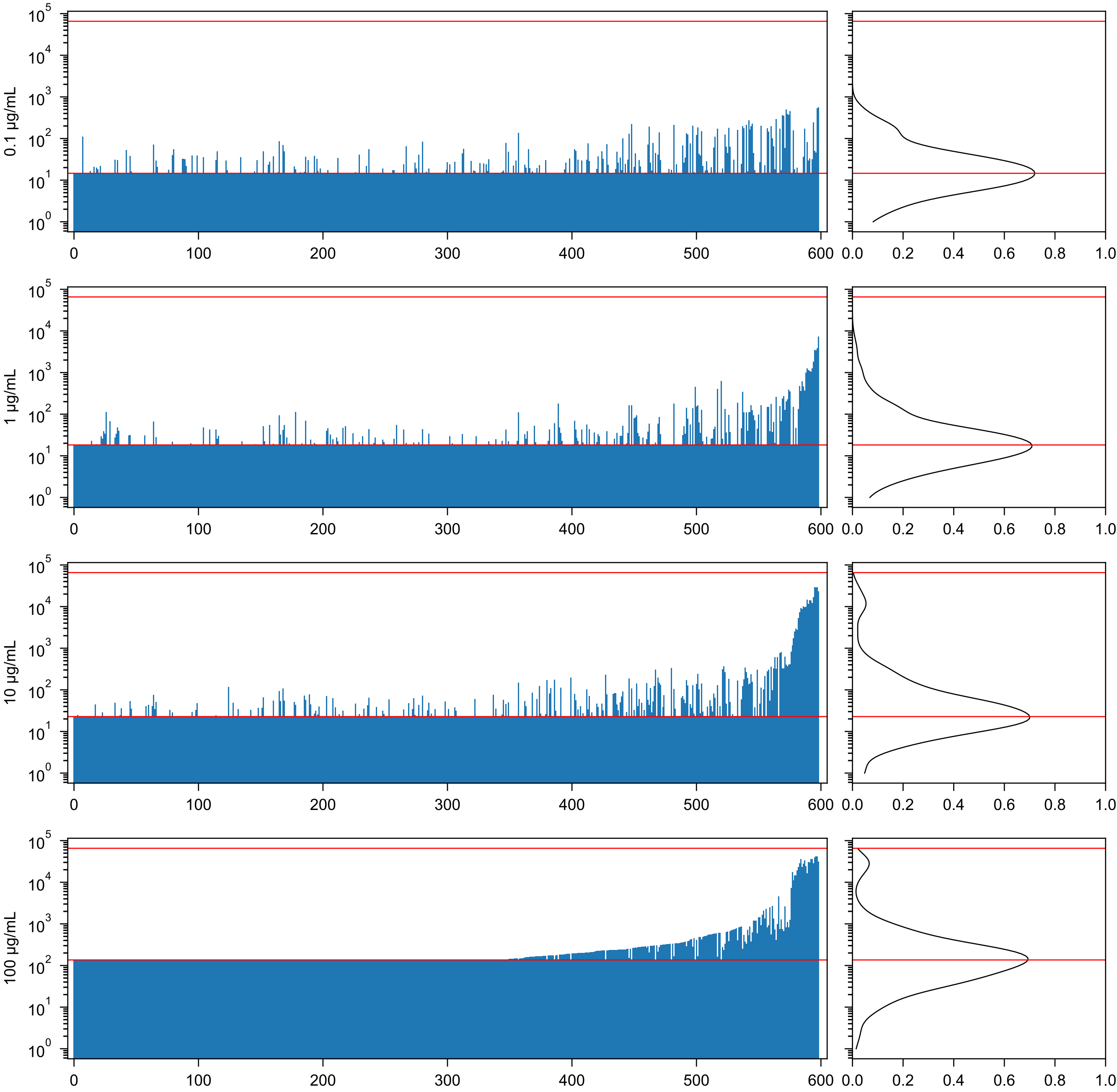

Maackia Amurensis Lectin II (MAL-II) Vector_W1014 - Lara K. Mahal

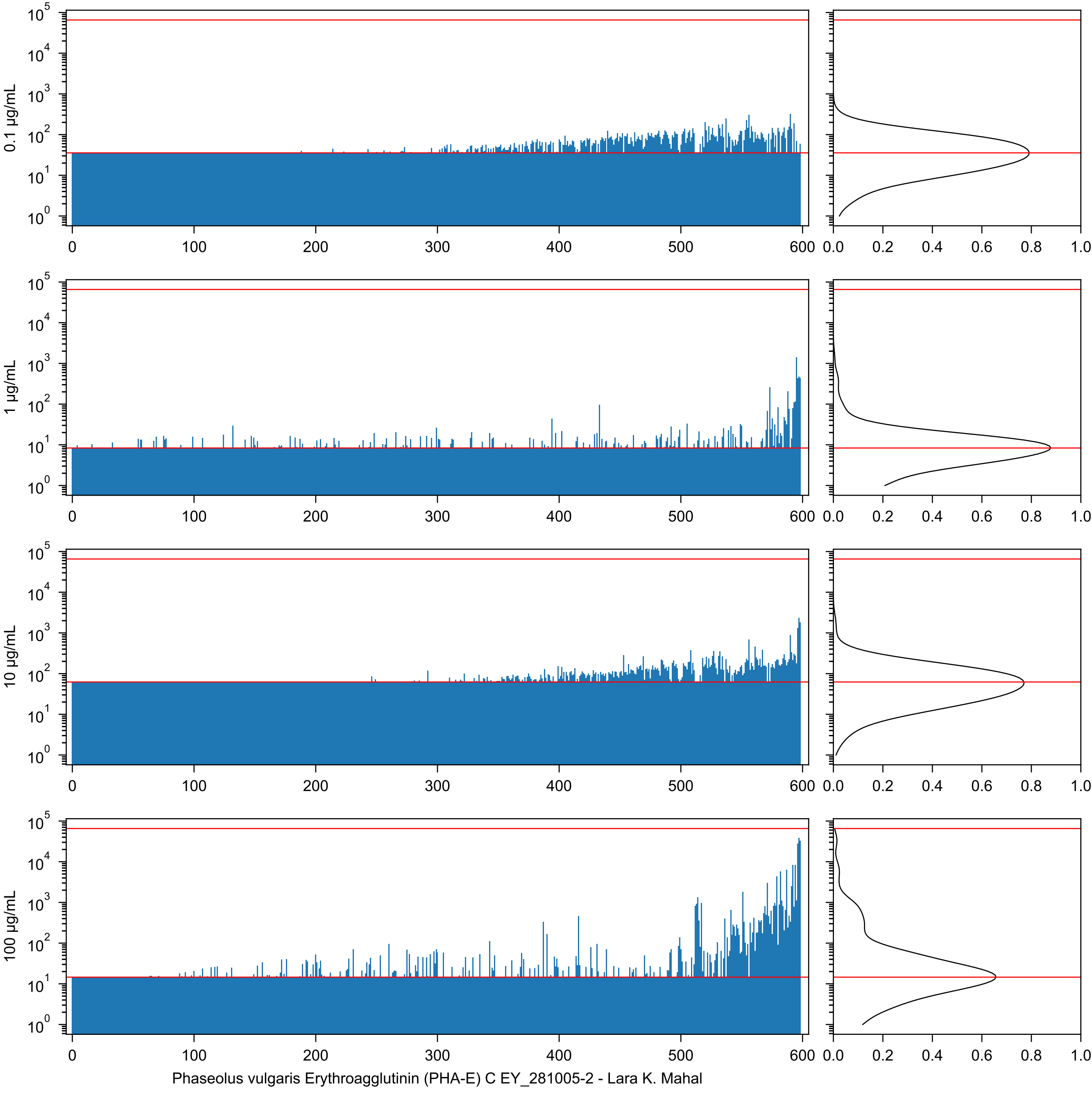
0.1 µg/mL
1 µg/mL
10 µg/mL
100 µg/mL
Phaseolus vulgaris Erythroagglutinin (PHA-E) C EY_281005-2 - Lara K. Mahal

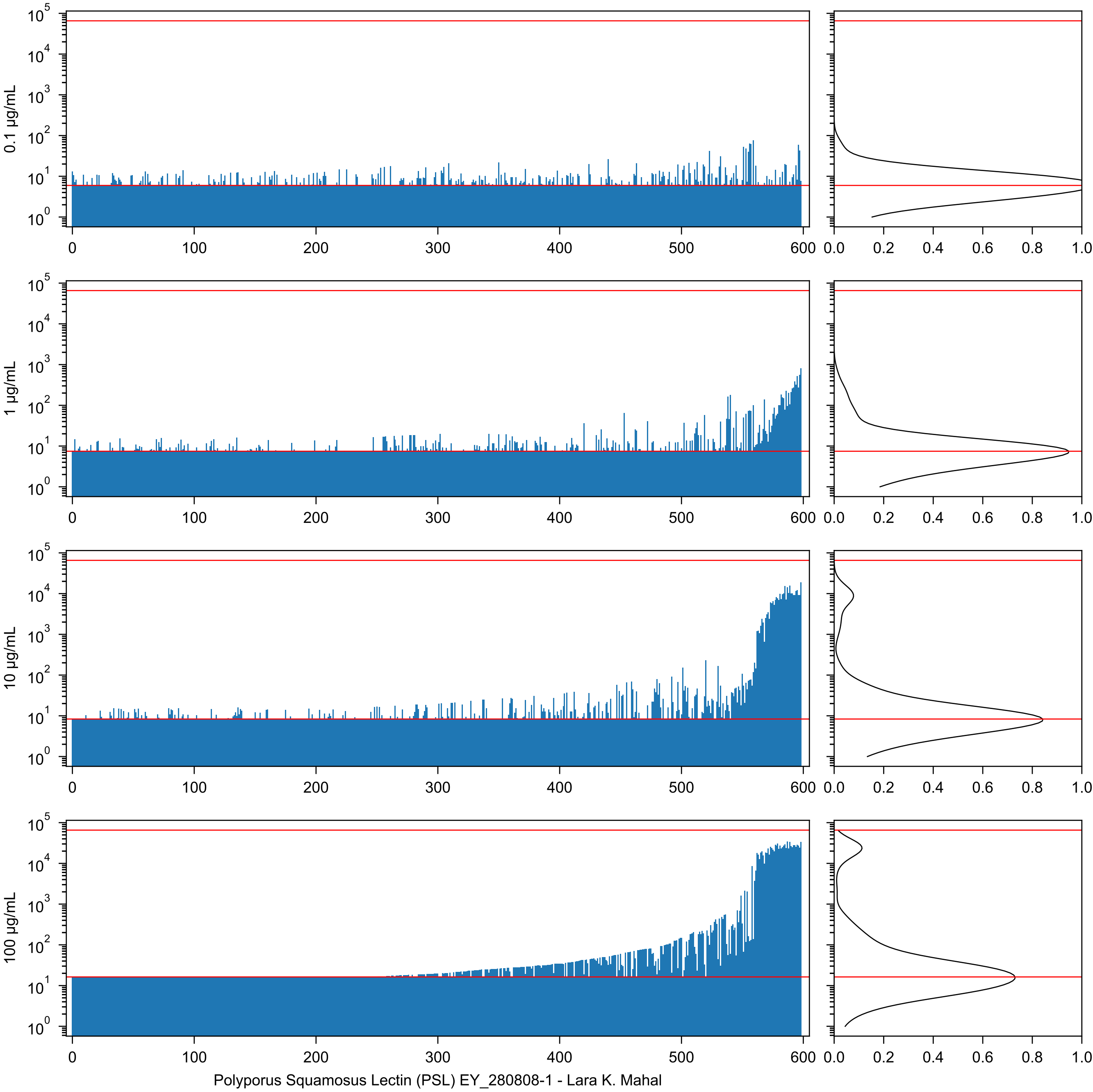

0.1 µg/mL
1 µg/mL
10 µg/mL
100 µg/mL
Polyporus Squamosus Lectin (PSL) EY_280808-1 - Lara K. Mahal

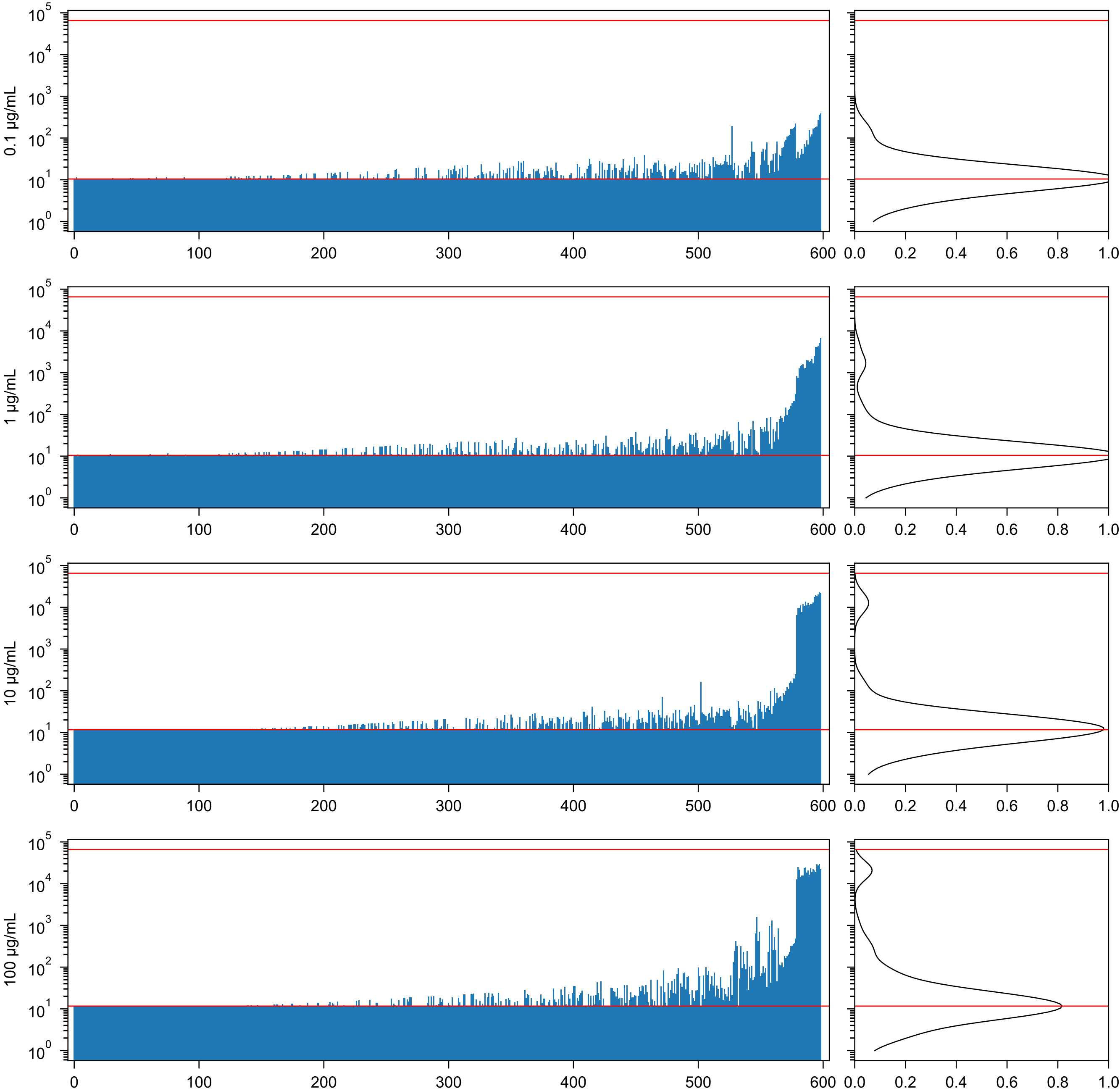

Phaseolus vulgaris Leucoagglutinin (PHA-L) A Vector_W1031 - Lara K. Mahal

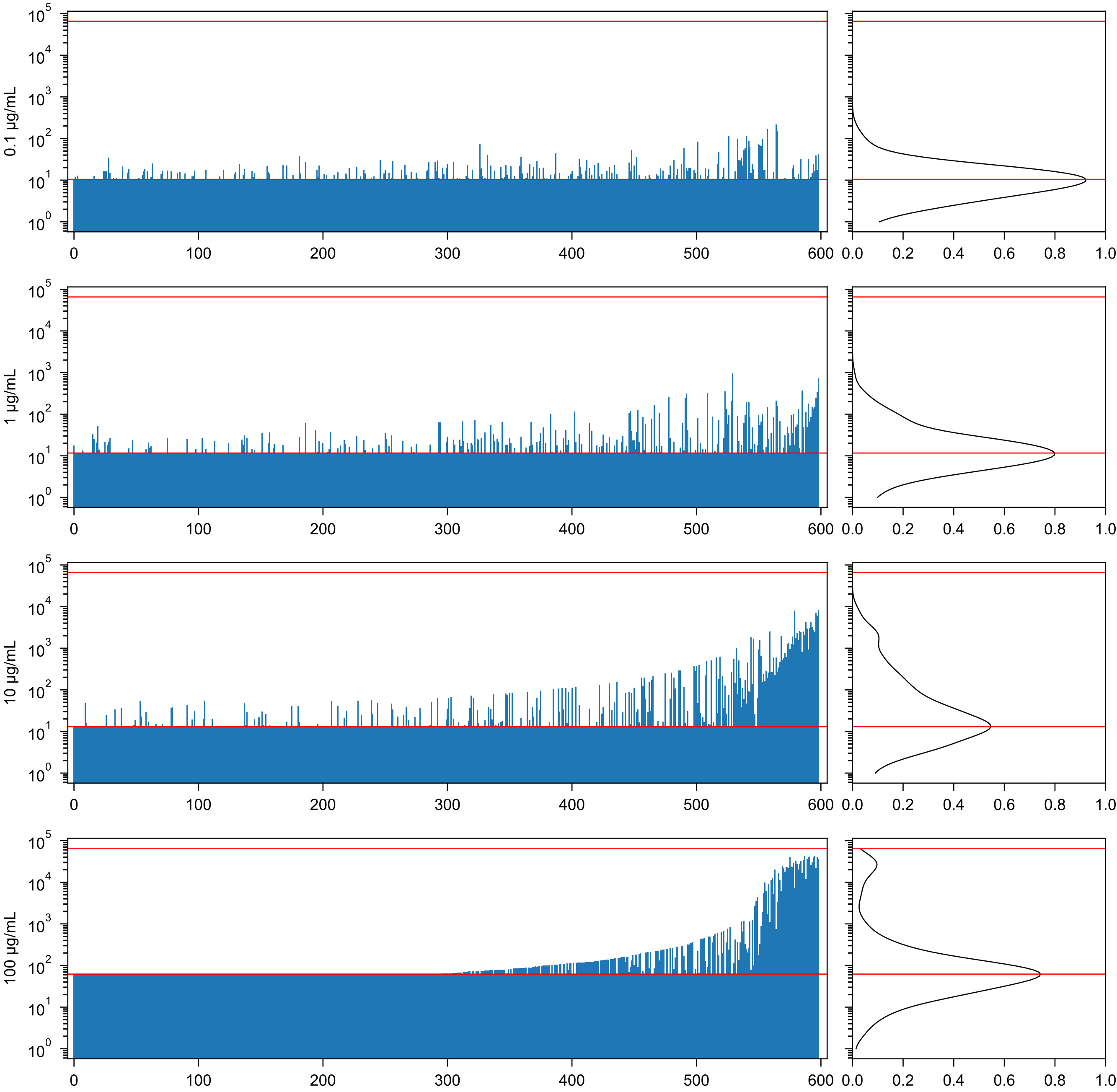


Psophocarpus tetragonolobus Agglutinin, Galactose specific (PTA-Gal) EY_290606-1 - Lara K. Mahal

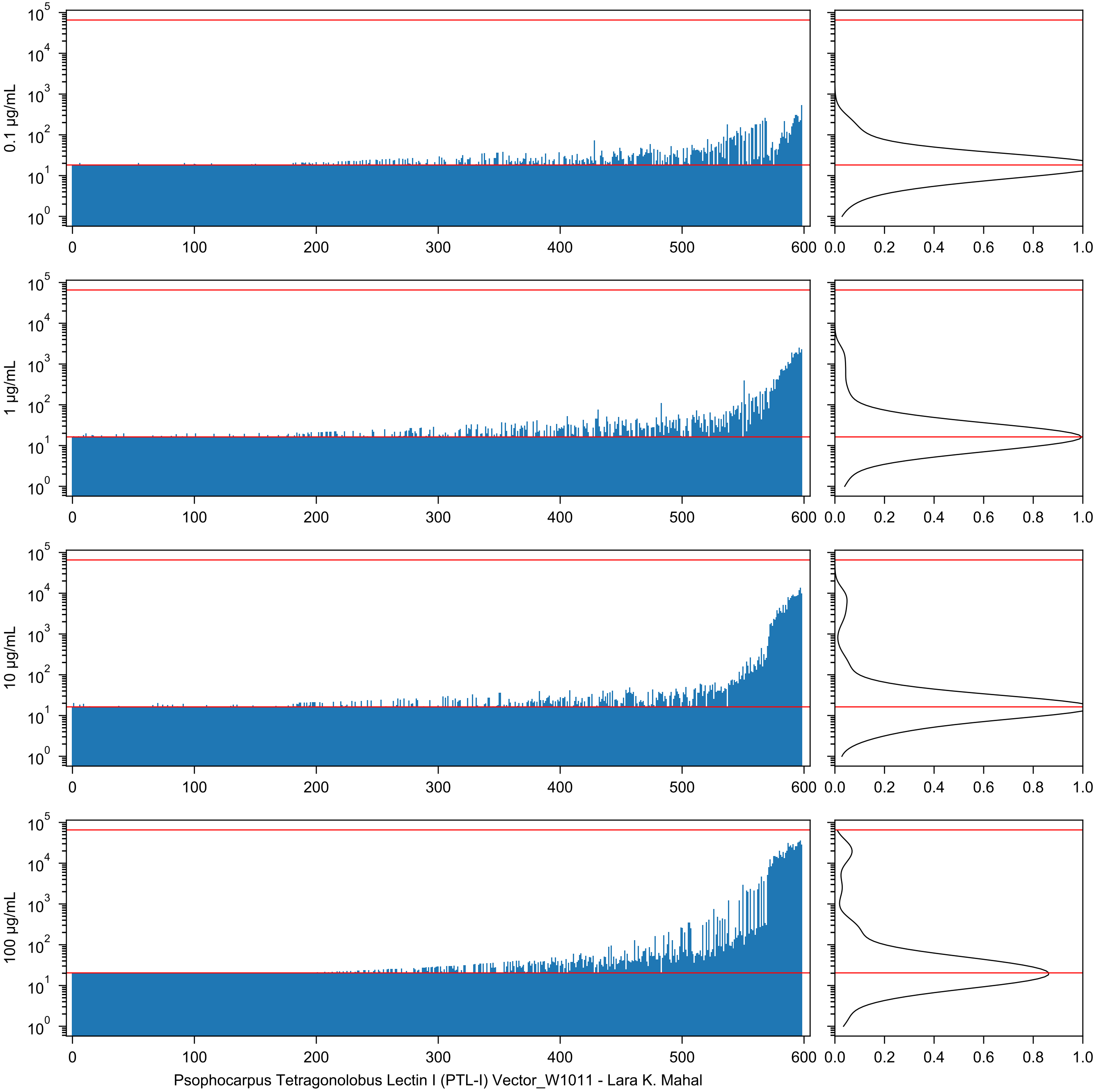
0.1 µg/mL
1 µg/mL
10 µg/mL
100 µg/mL
Psophocarpus Tetragonolobus Lectin I (PTL-I) Vector_W1011 - Lara K. Mahal

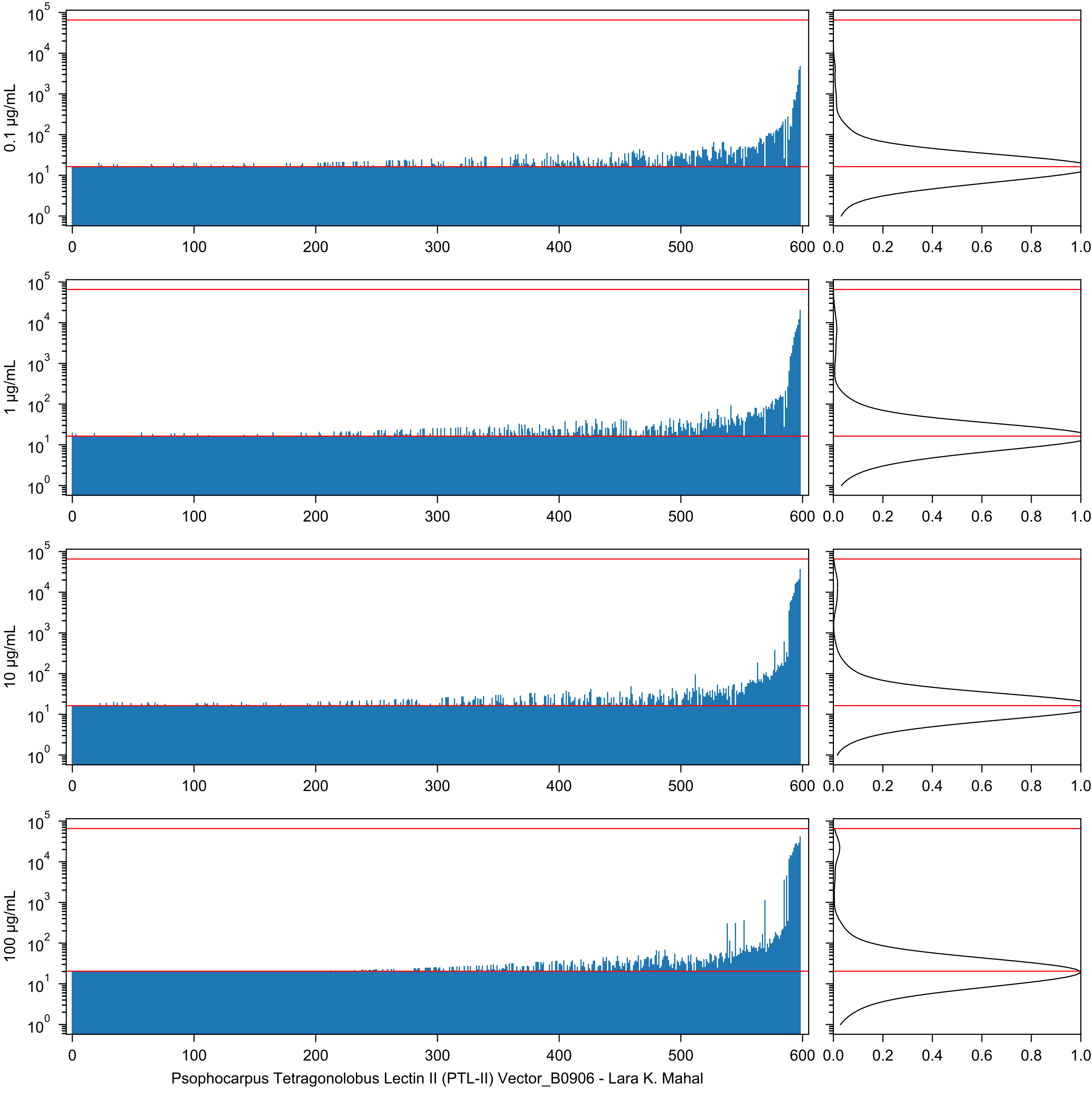

0.1 µg/mL
1 µg/mL
10 µg/mL
100 µg/mL
Psophocarpus Tetragonolobus Lectin II (PTL-II) Vector_B0906 - Lara K. Mahal

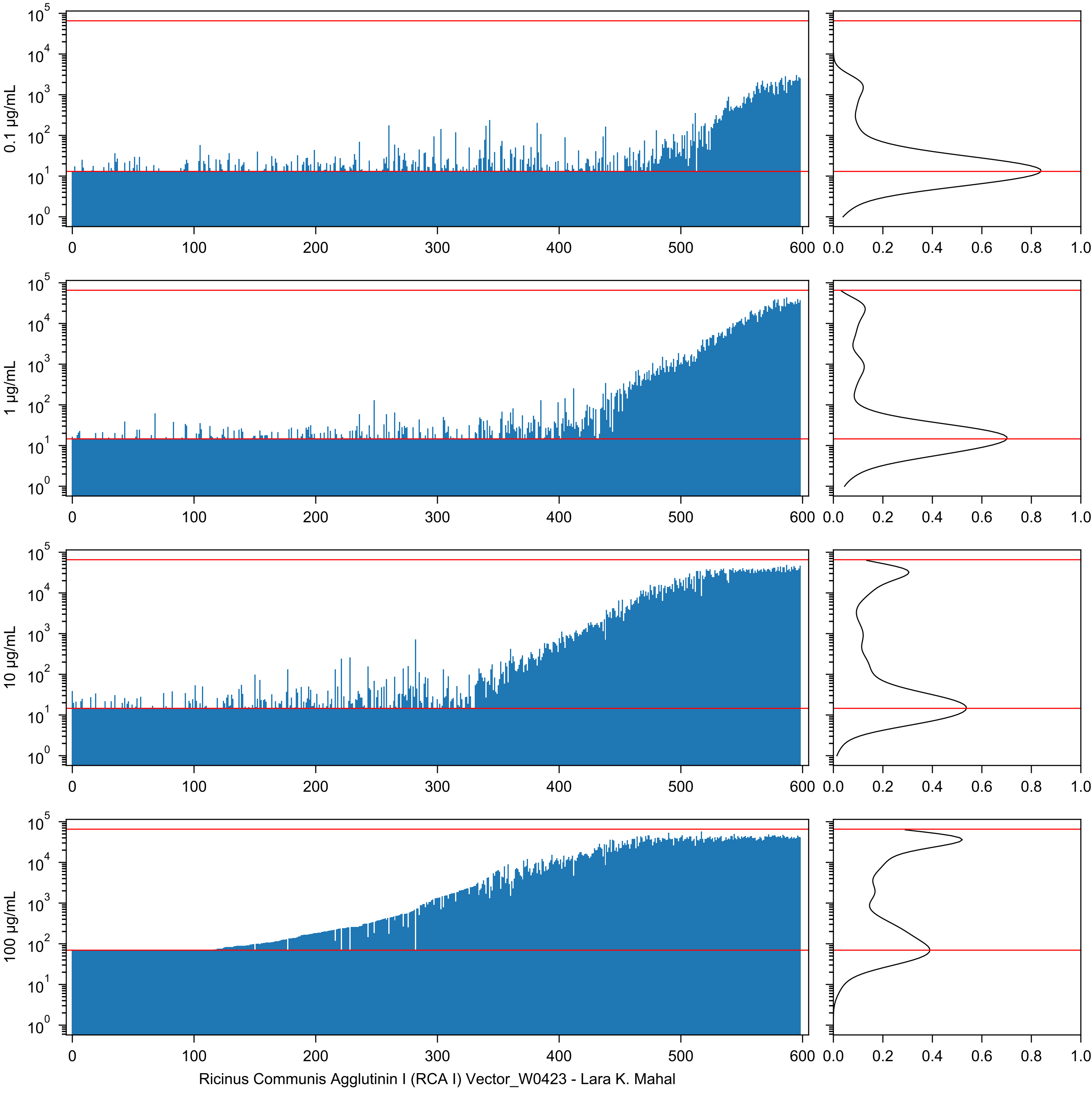

0.1 µg/mL
1 µg/mL
10 µg/mL
100 µg/mL
Ricinus Communis Agglutinin I (RCA I) Vector_W0423 - Lara K. Mahal

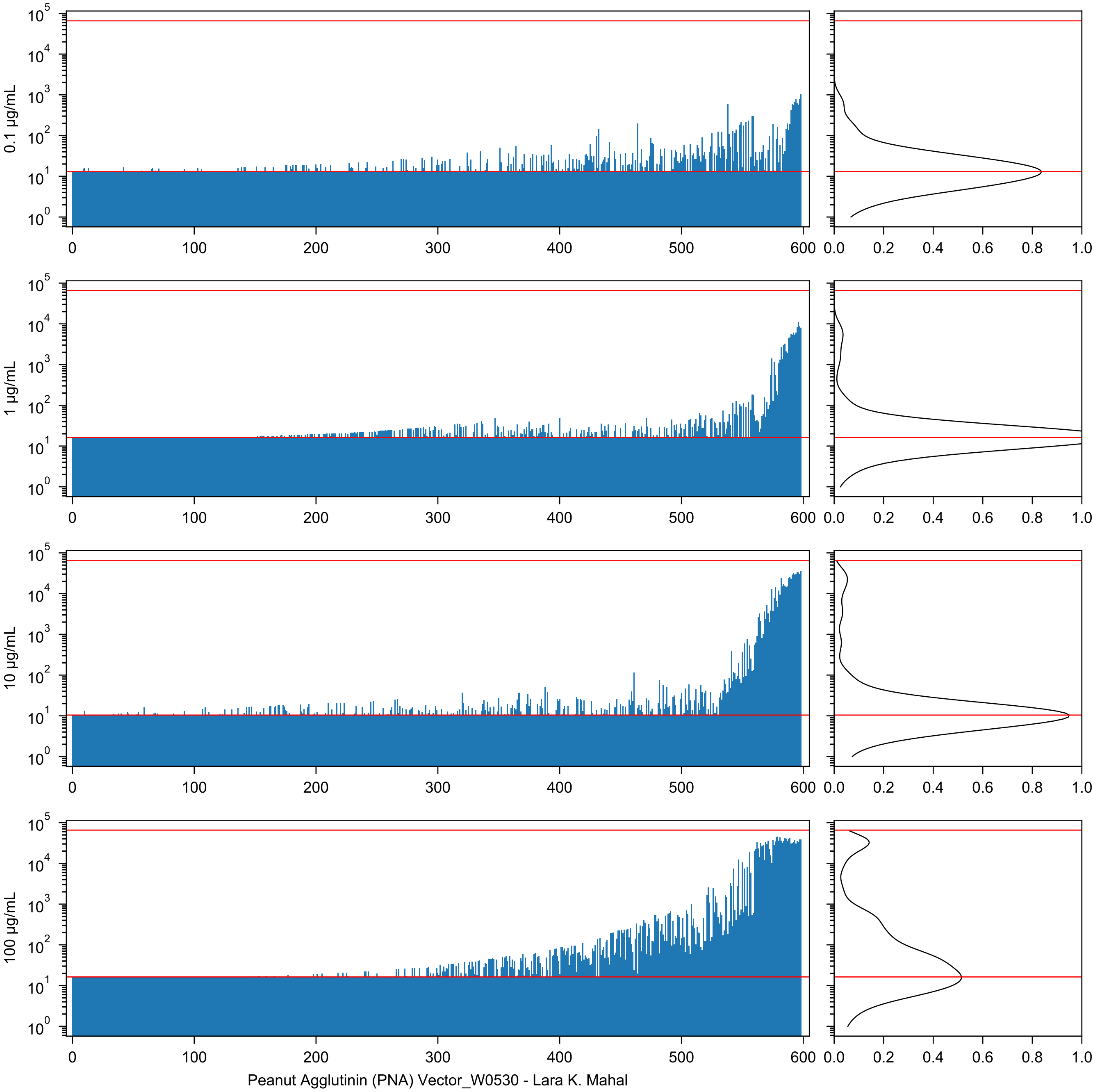
0.1 µg/mL
1 µg/mL
10 µg/mL
100 µg/mL
Peanut Agglutinin (PNA) Vector_W0530 - Lara K. Mahal

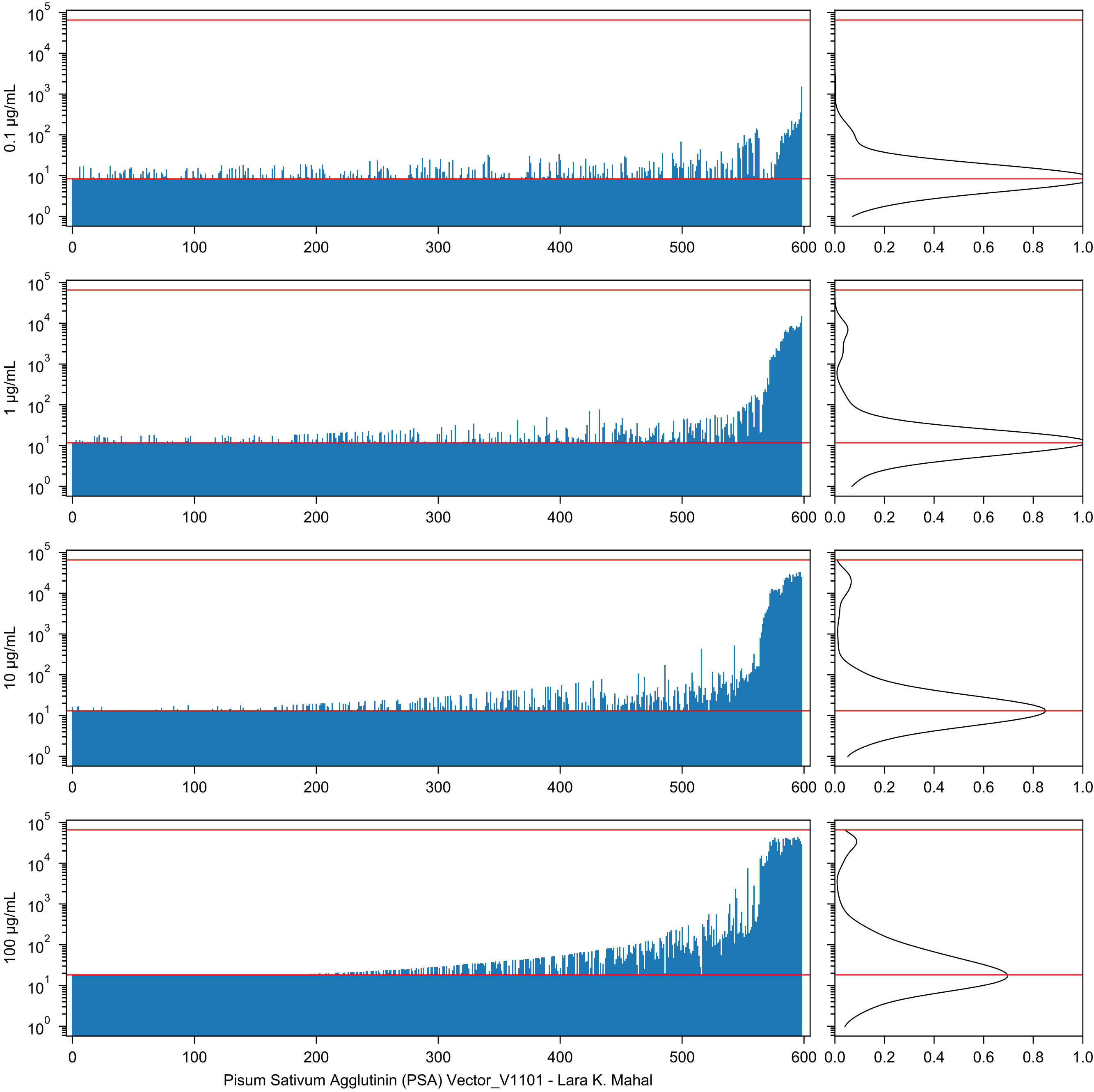

0.1 µg/mL
1 µg/mL
10 µg/mL
100 µg/mL
Pisum Sativum Agglutinin (PSA) Vector_V1101 - Lara K. Mahal

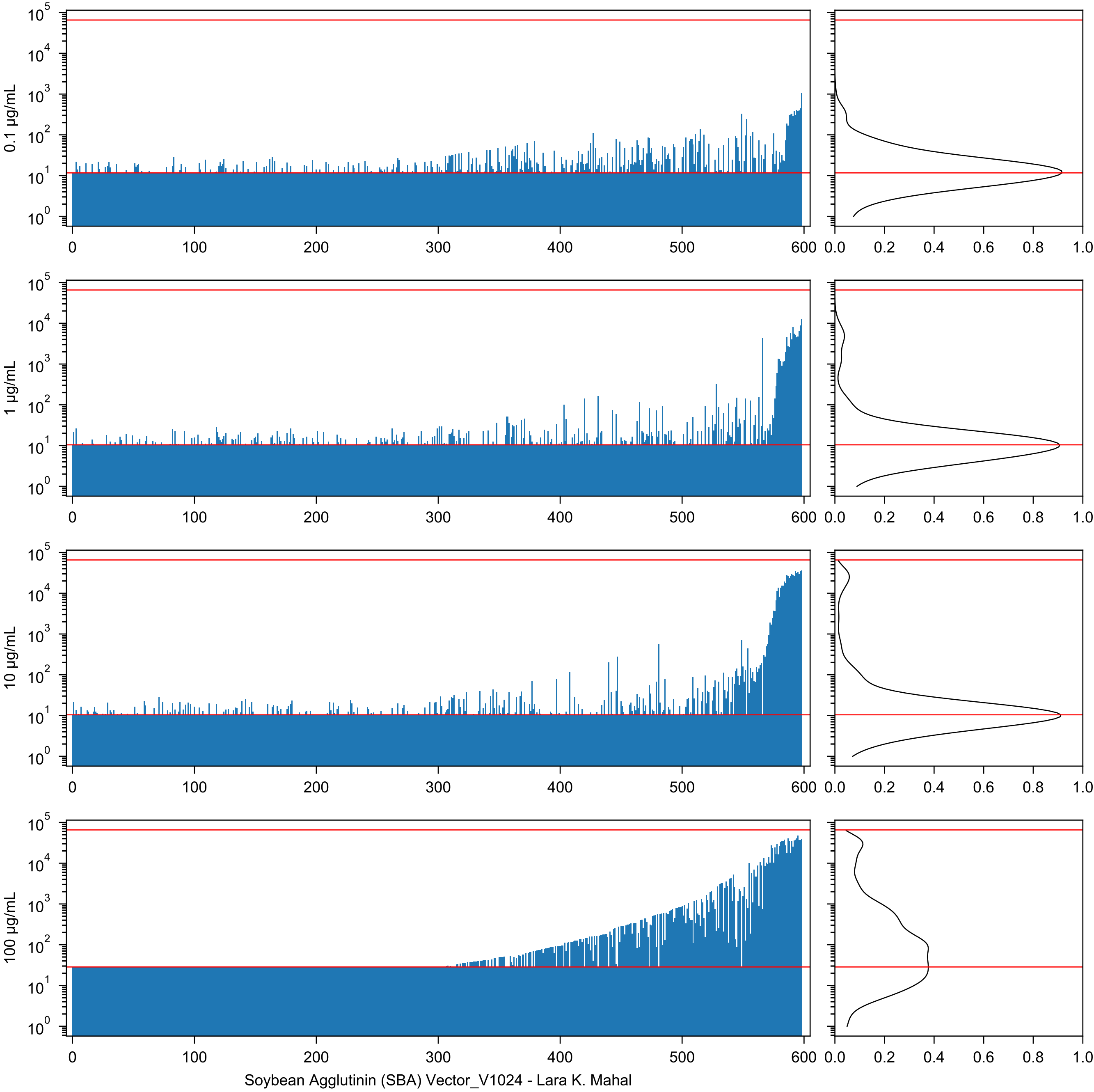
0.1 µg/mL
1 µg/mL
10 µg/mL
100 µg/mL
Soybean Agglutinin (SBA) Vector_V1024 - Lara K. Mahal

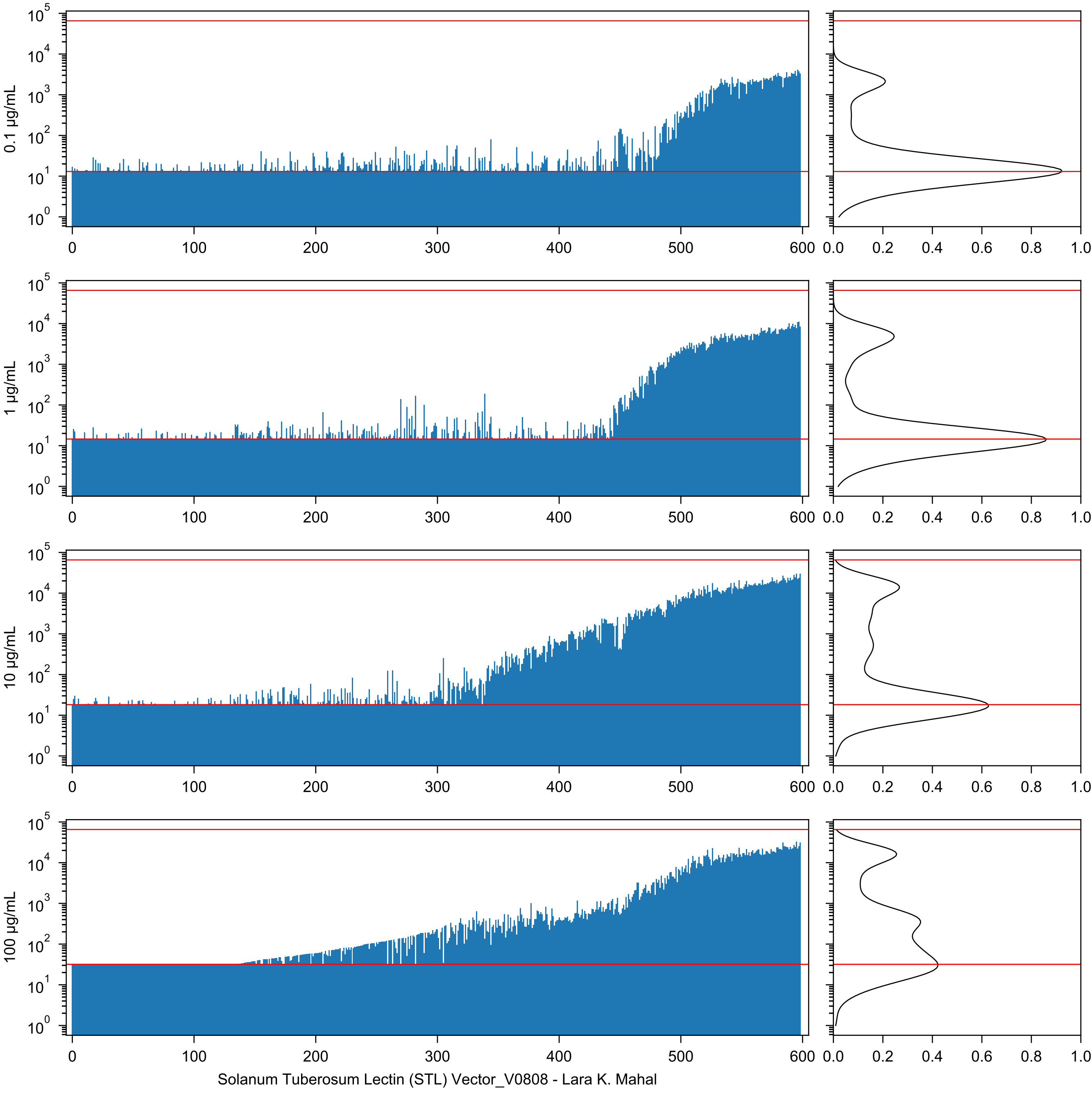
0.1 µg/mL
1 µg/mL
10 µg/mL
100 µg/mL
Solanum Tuberosum Lectin (STL) Vector_V0808 - Lara K. Mahal

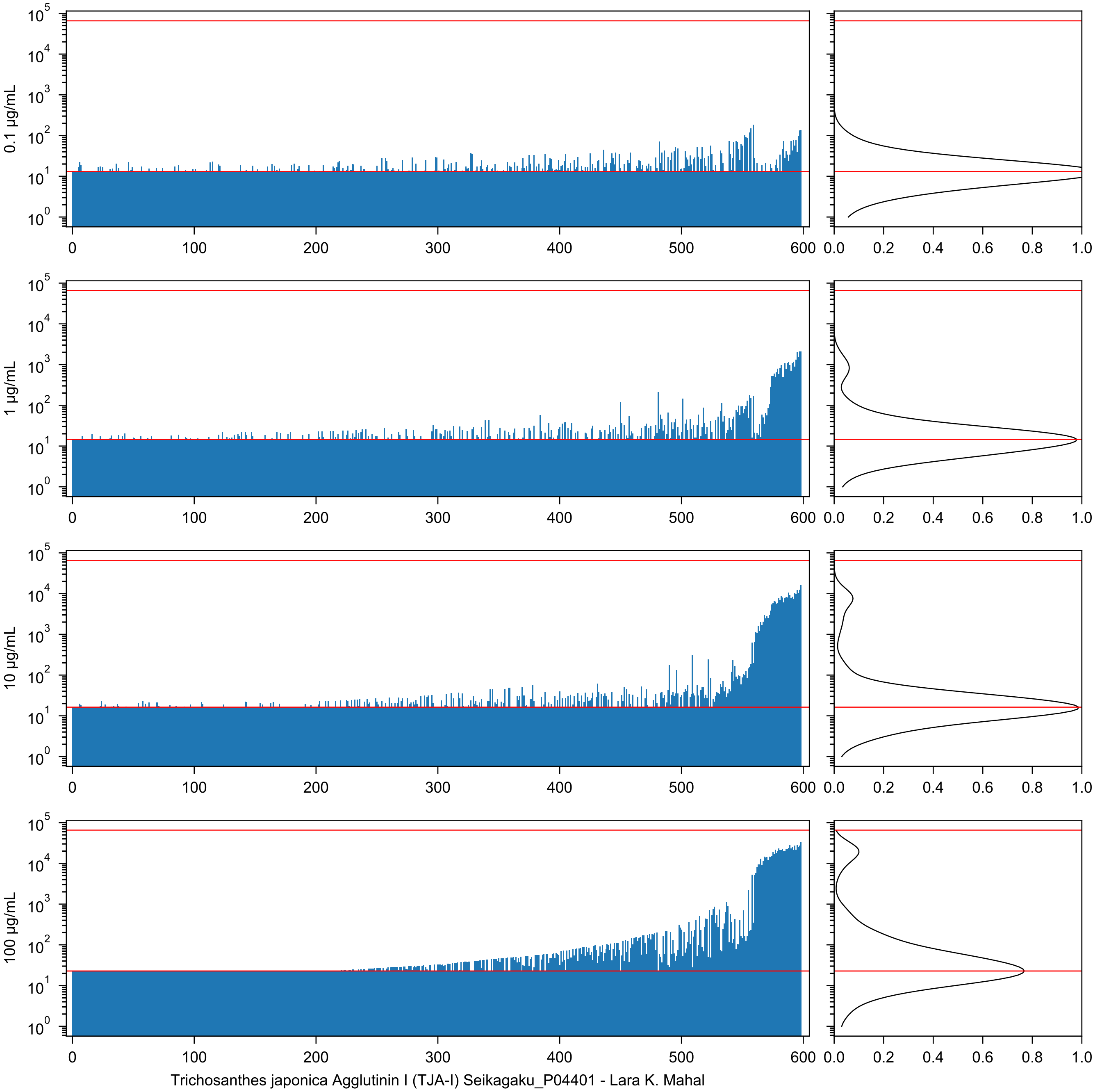
0.1 µg/mL
1 µg/mL
10 µg/mL
100 µg/mL
Trichosanthes japonica Agglutinin I (TJA-I) Seikagaku_P04401 - Lara K. Mahal

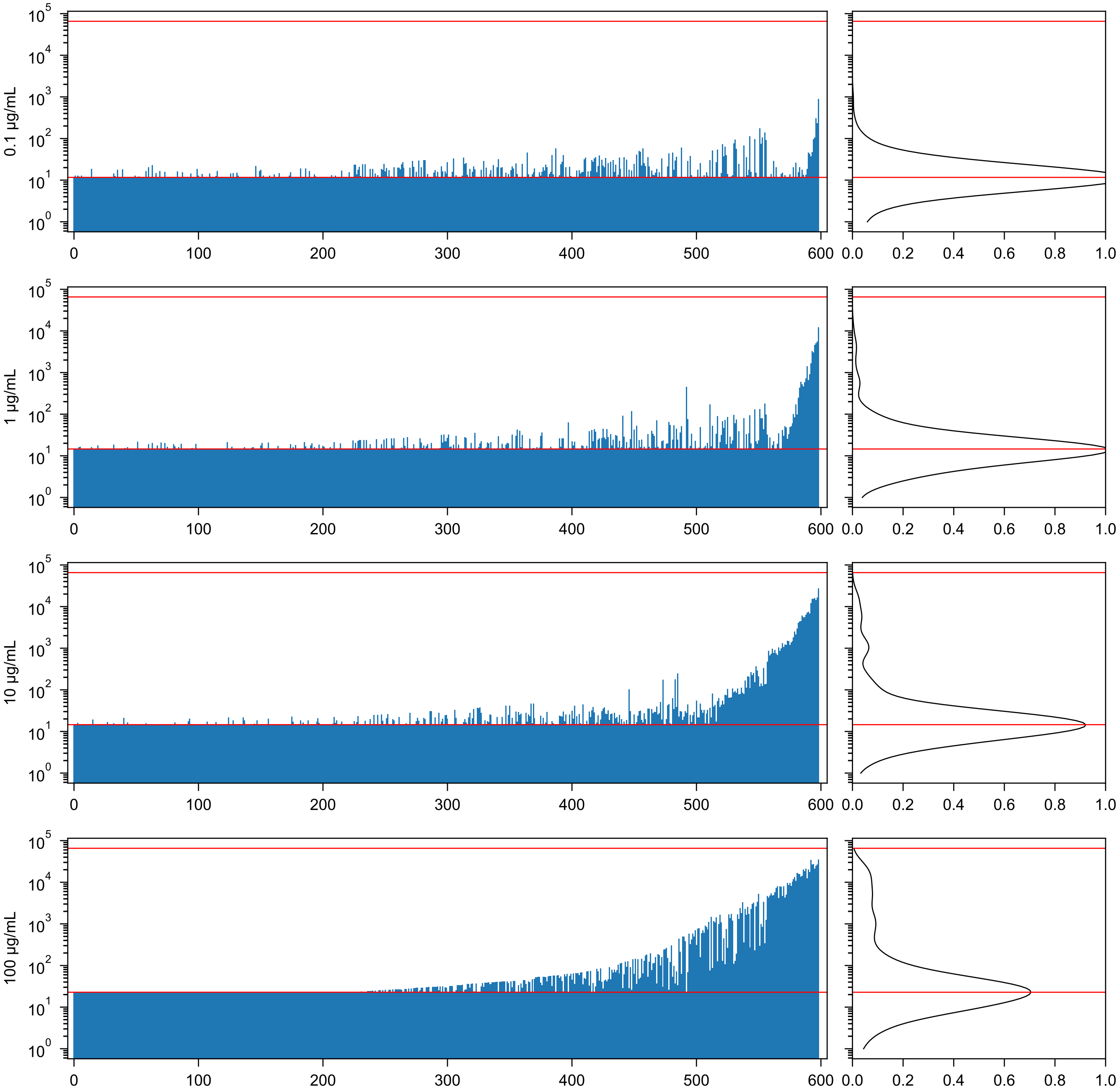

Trichosanthes japonica Agglutinin II (TJA-II) Seikagaku_P04Y01 - Lara K. Mahal

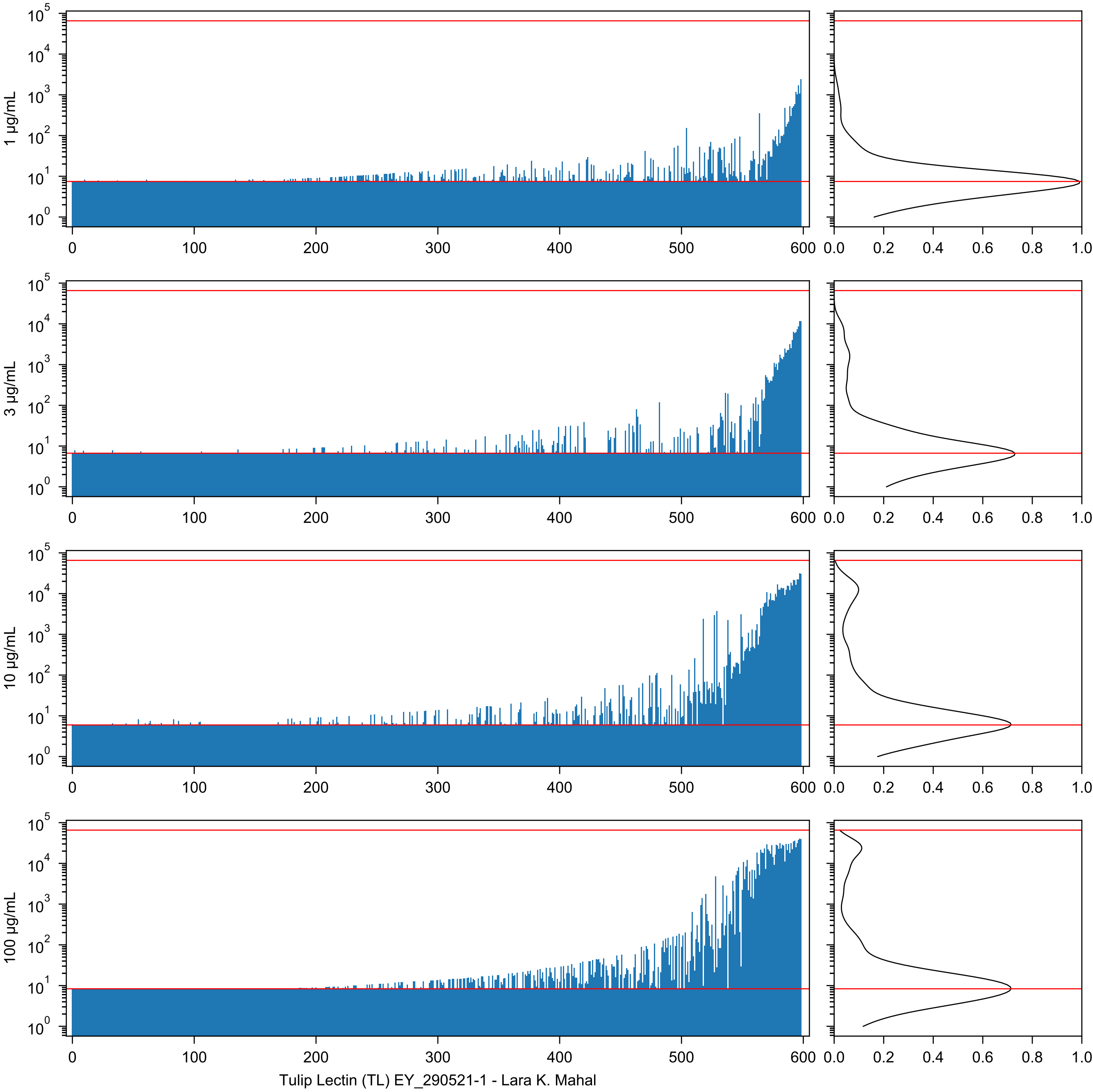
1 µg/mL
3 µg/mL
10 µg/mL
100 µg/mL
Tulip Lectin (TL) EY_290521-1 - Lara K. Mahal

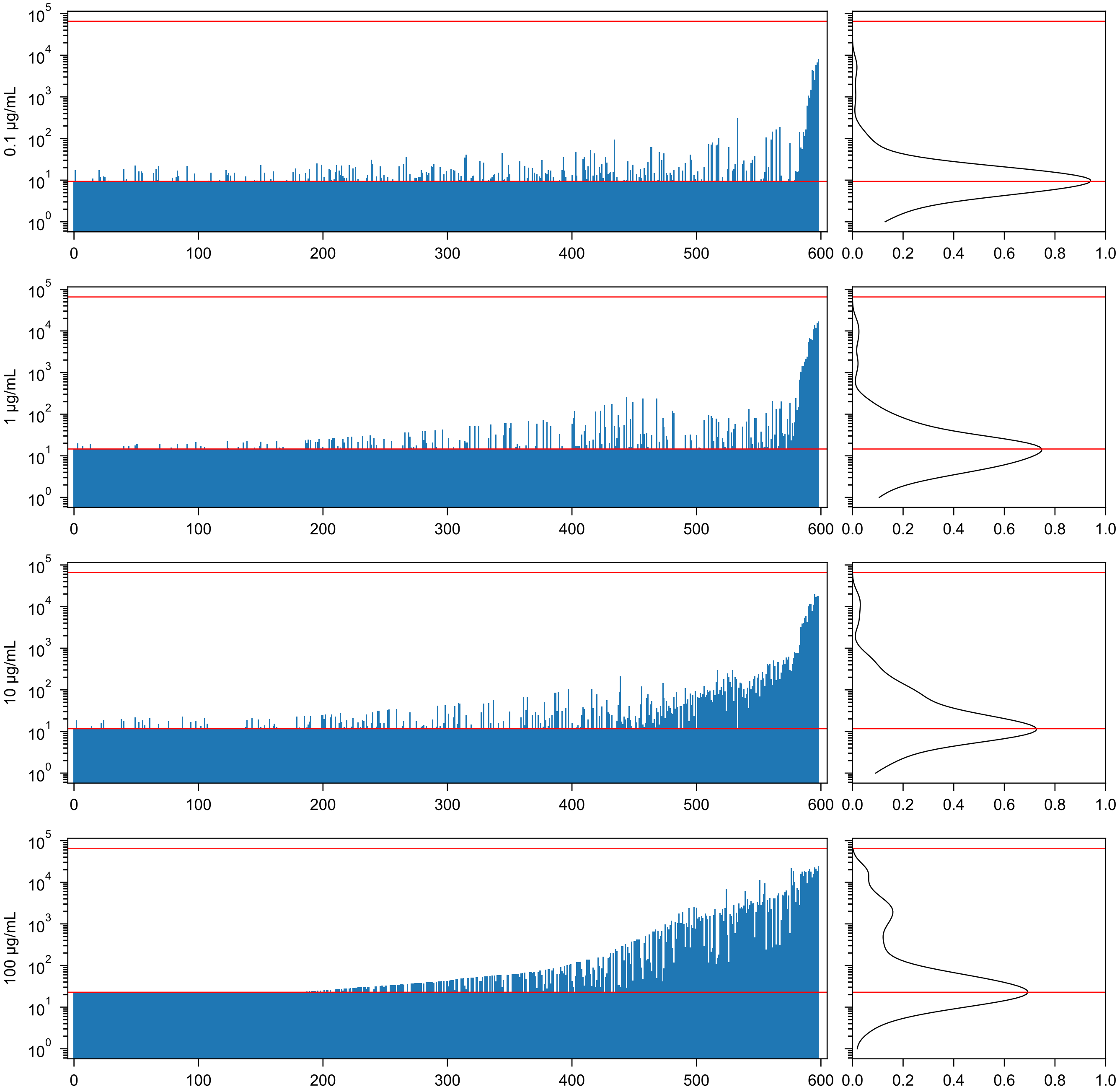


Urtica dioica Agglutinin (UDA) EY_29021-1 - Lara K. Mahal

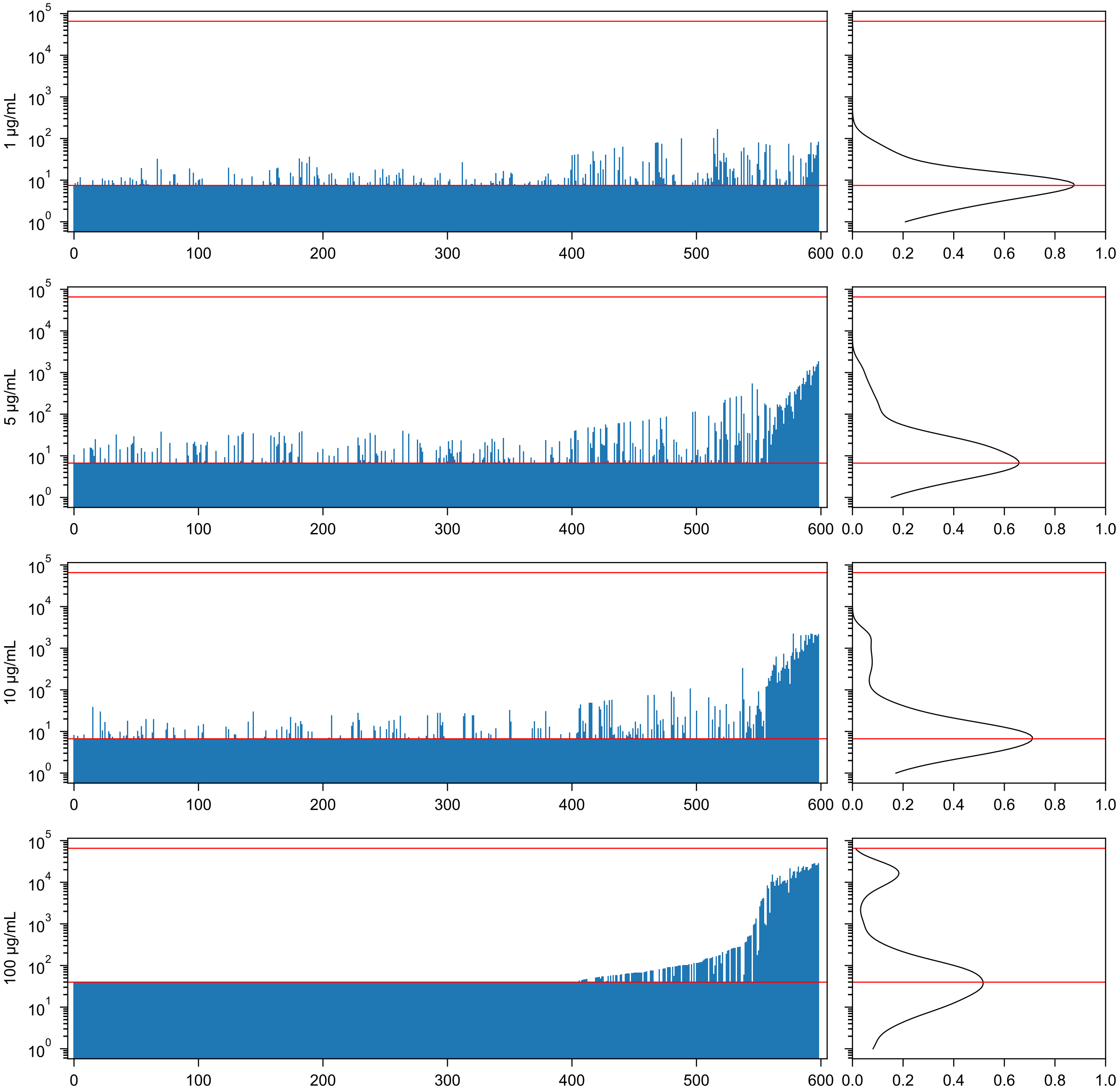

Ulex europaeus Agglutinin II (UEA-II) EY_290521-2 - Lara K. Mahal

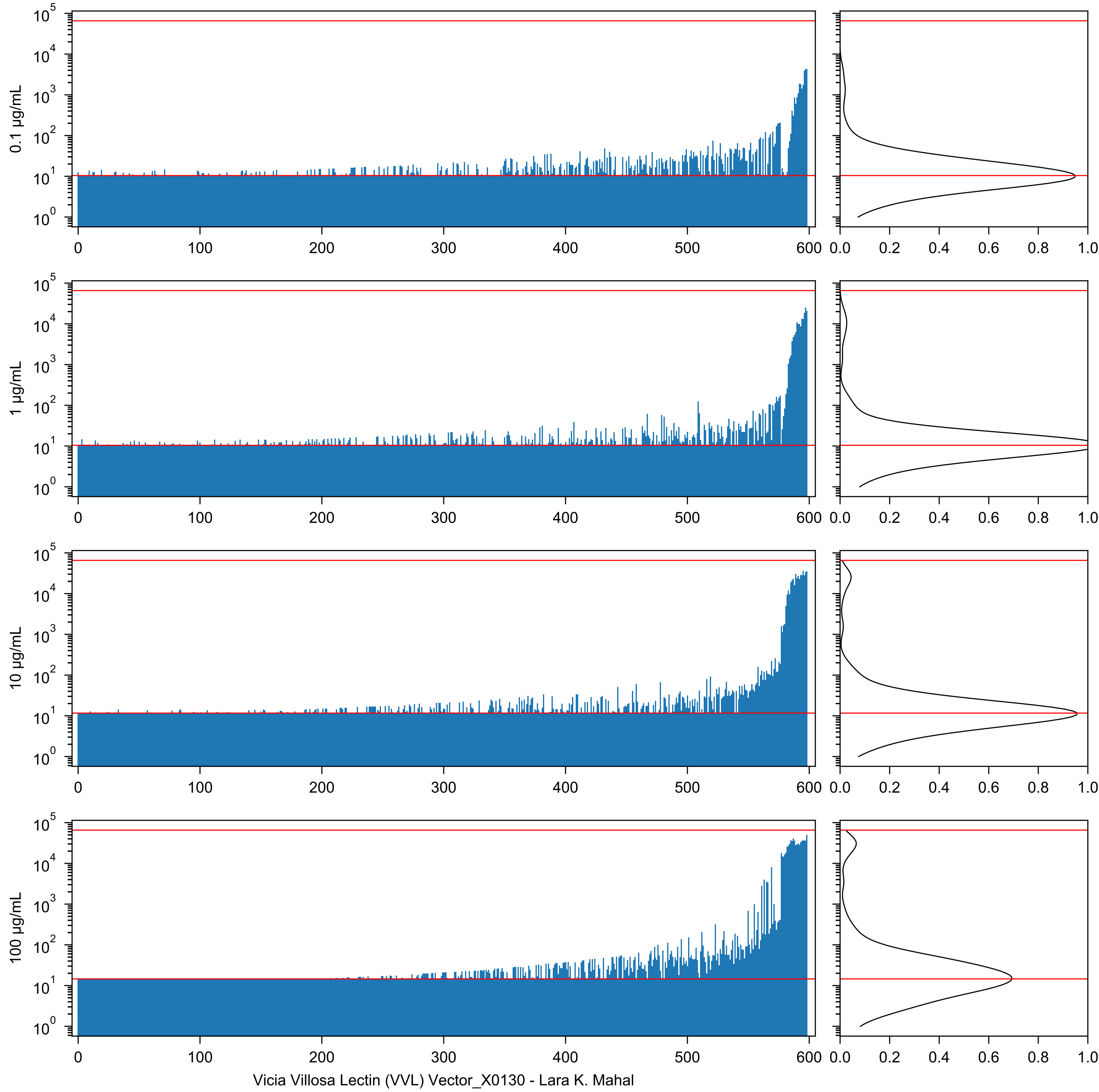
0.1 µg/mL
1 µg/mL
10 µg/mL
100 µg/mL
Vicia Villosa Lectin (VVL) Vector_X0130 - Lara K. Mahal

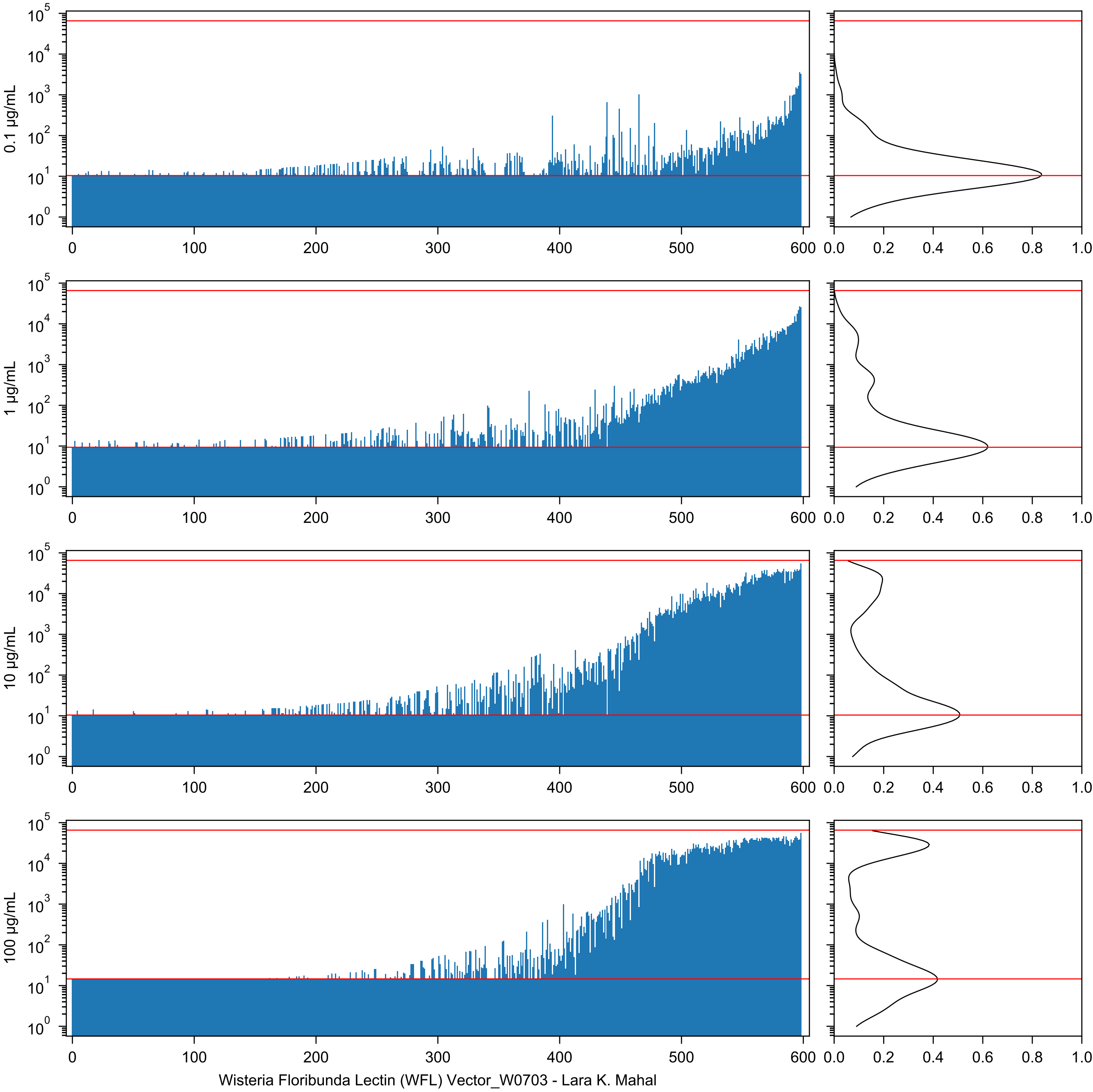
0.1 µg/mL
1 µg/mL
10 µg/mL
100 µg/mL
Wisteria Floribunda Lectin (WFL) Vector_W0703 - Lara K. Mahal

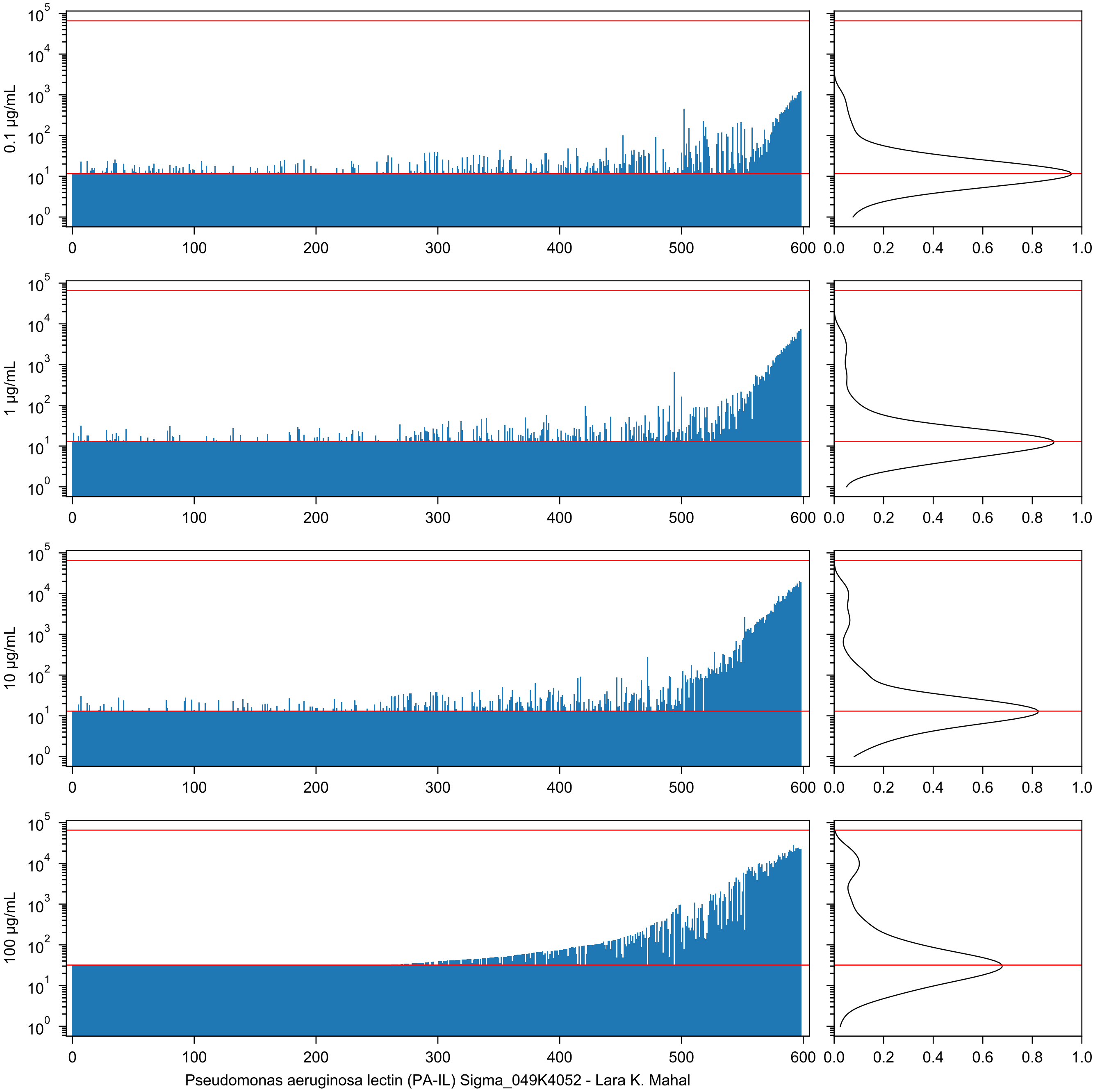

0.1 µg/mL
1 µg/mL
10 µg/mL
100 µg/mL
Pseudomonas aeruginosa lectin (PA-IL) Sigma_049K4052 - Lara K. Mahal

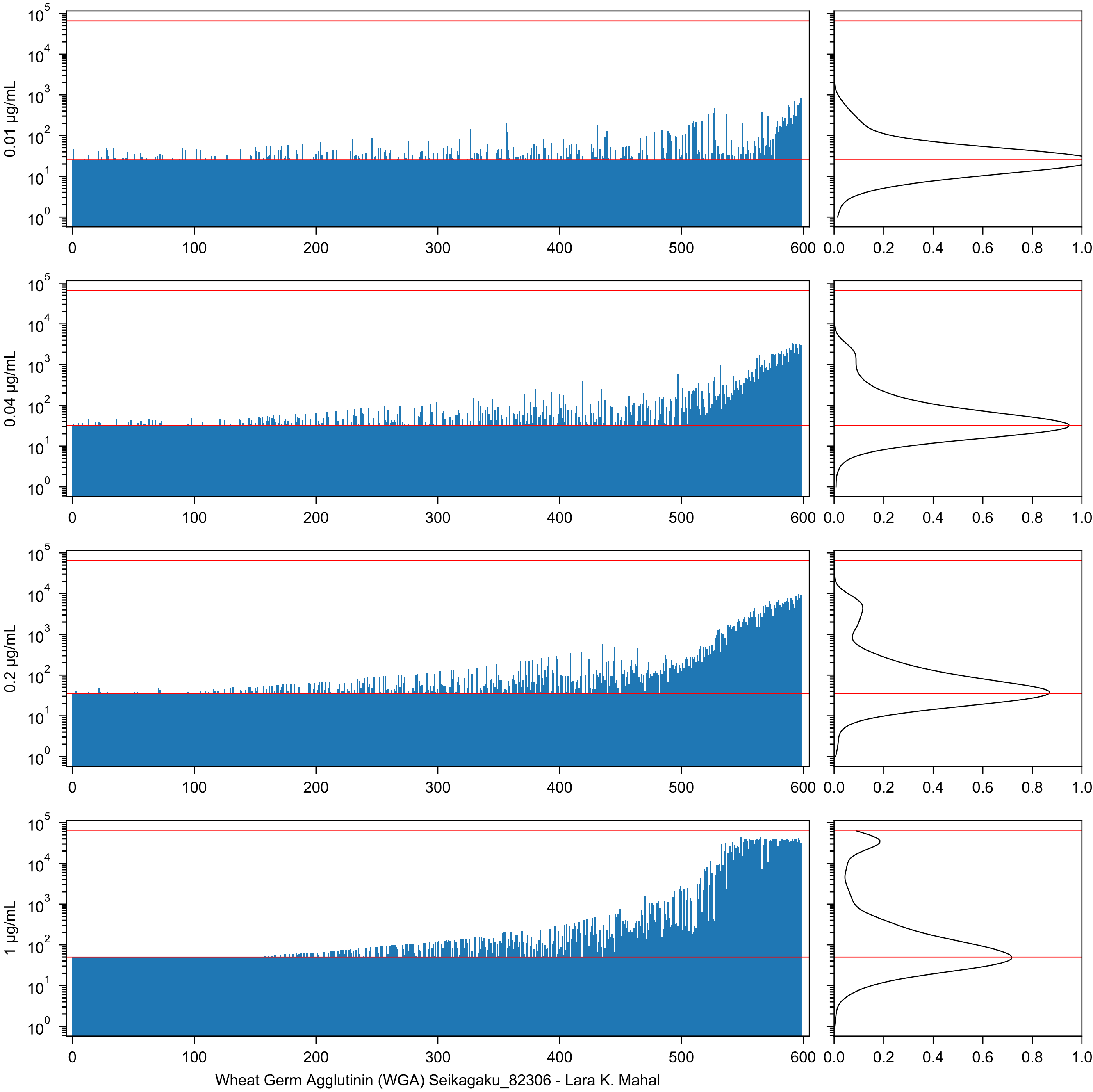
0.01 µg/mL
0.04 µg/mL
0.2 µg/mL
1 µg/mL
Wheat Germ Agglutinin (WGA) Seikagaku_82306 - Lara K. Mahal

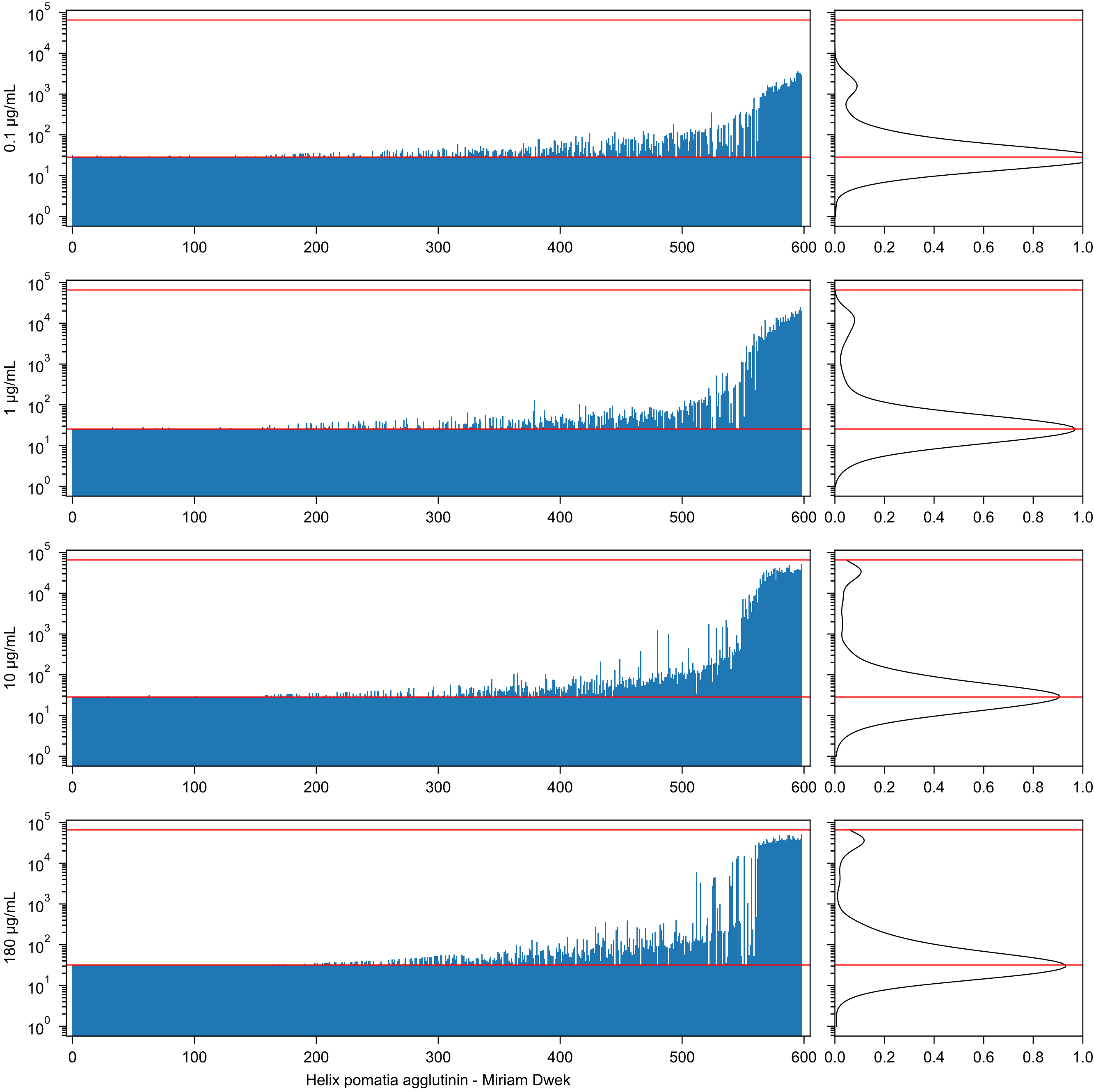

0.1 µg/mL
1 µg/mL
10 µg/mL
180 µg/mL
Helix pomatia agglutinin - Miriam Dwek

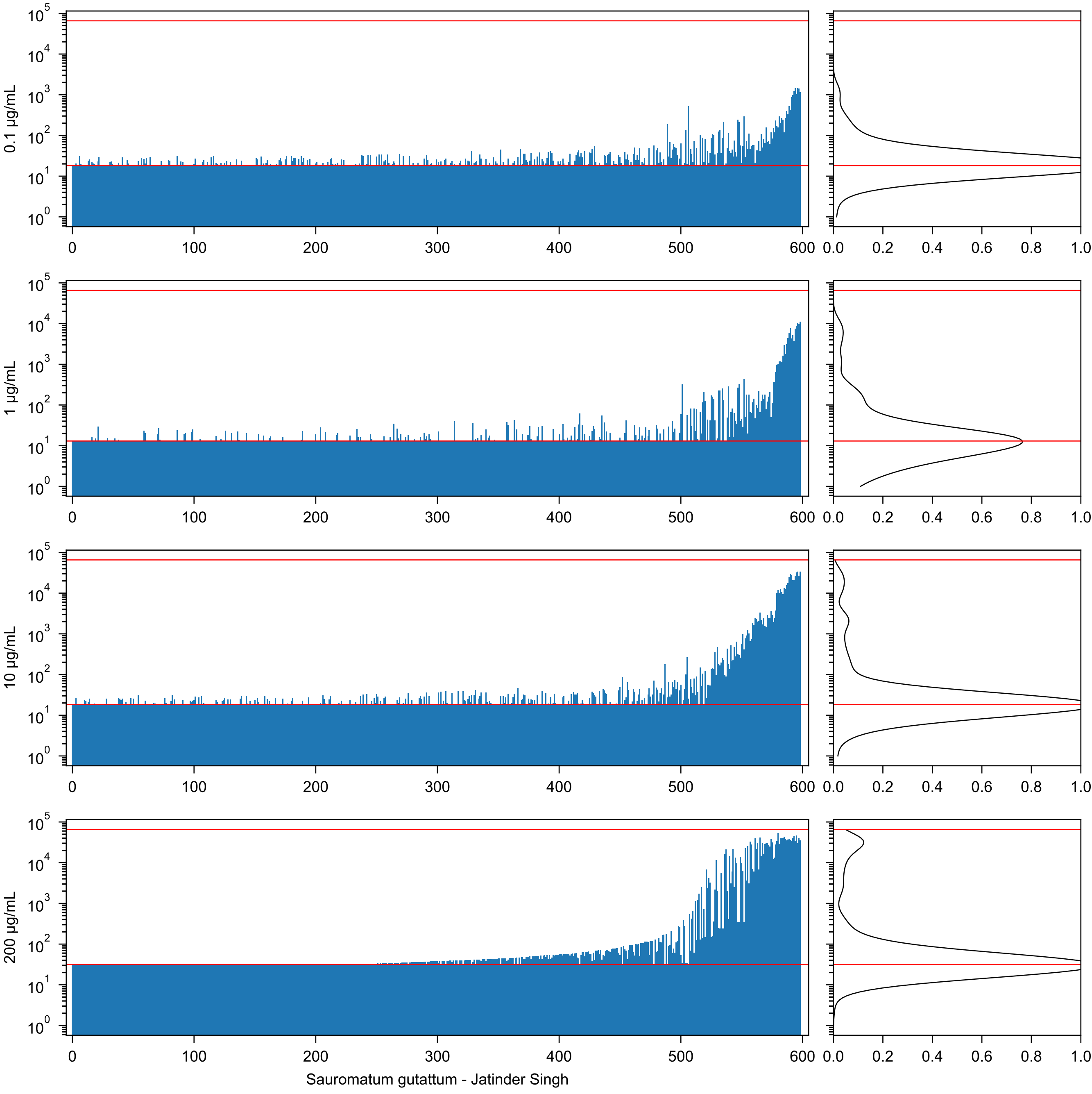
0.1 µg/mL
1 µg/mL
10 µg/mL
200 µg/mL
Sauromatum gutattum - Jatinder Singh

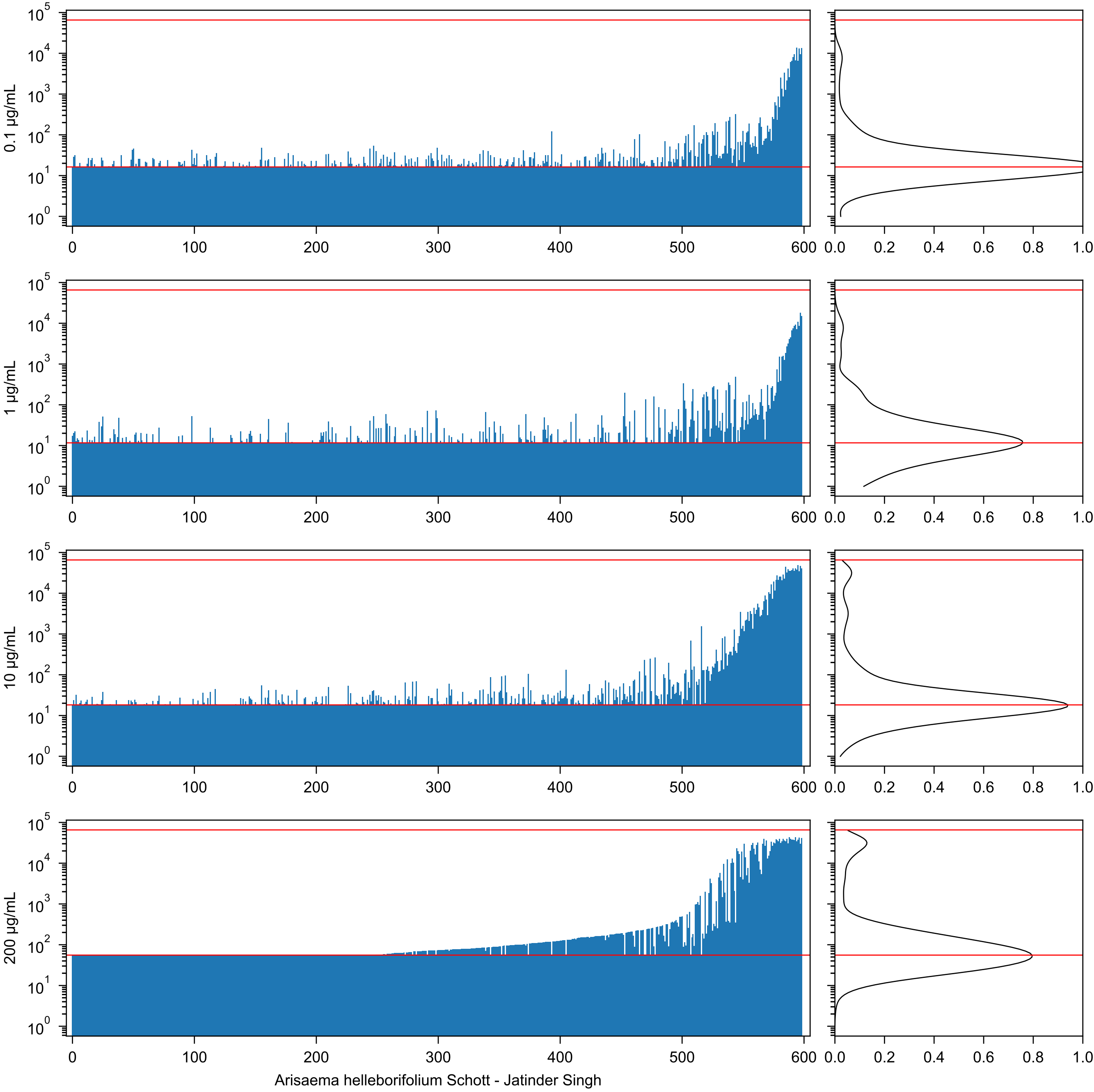

0.1 µg/mL
1 µg/mL
10 µg/mL
200 µg/mL
Arisaema helleborifolium Schott - Jatinder Singh

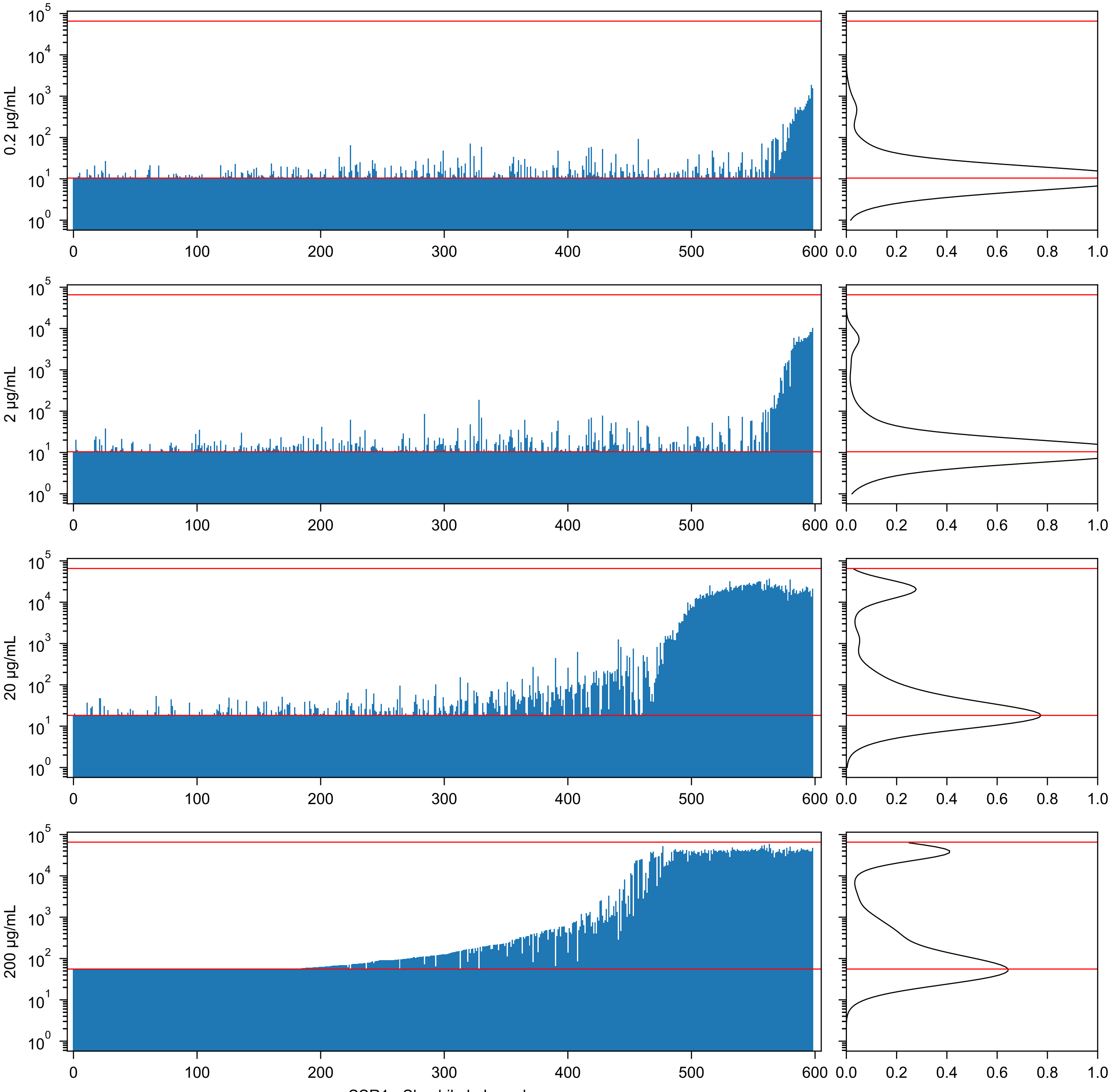


SSR1 - Shashikala Inamdar

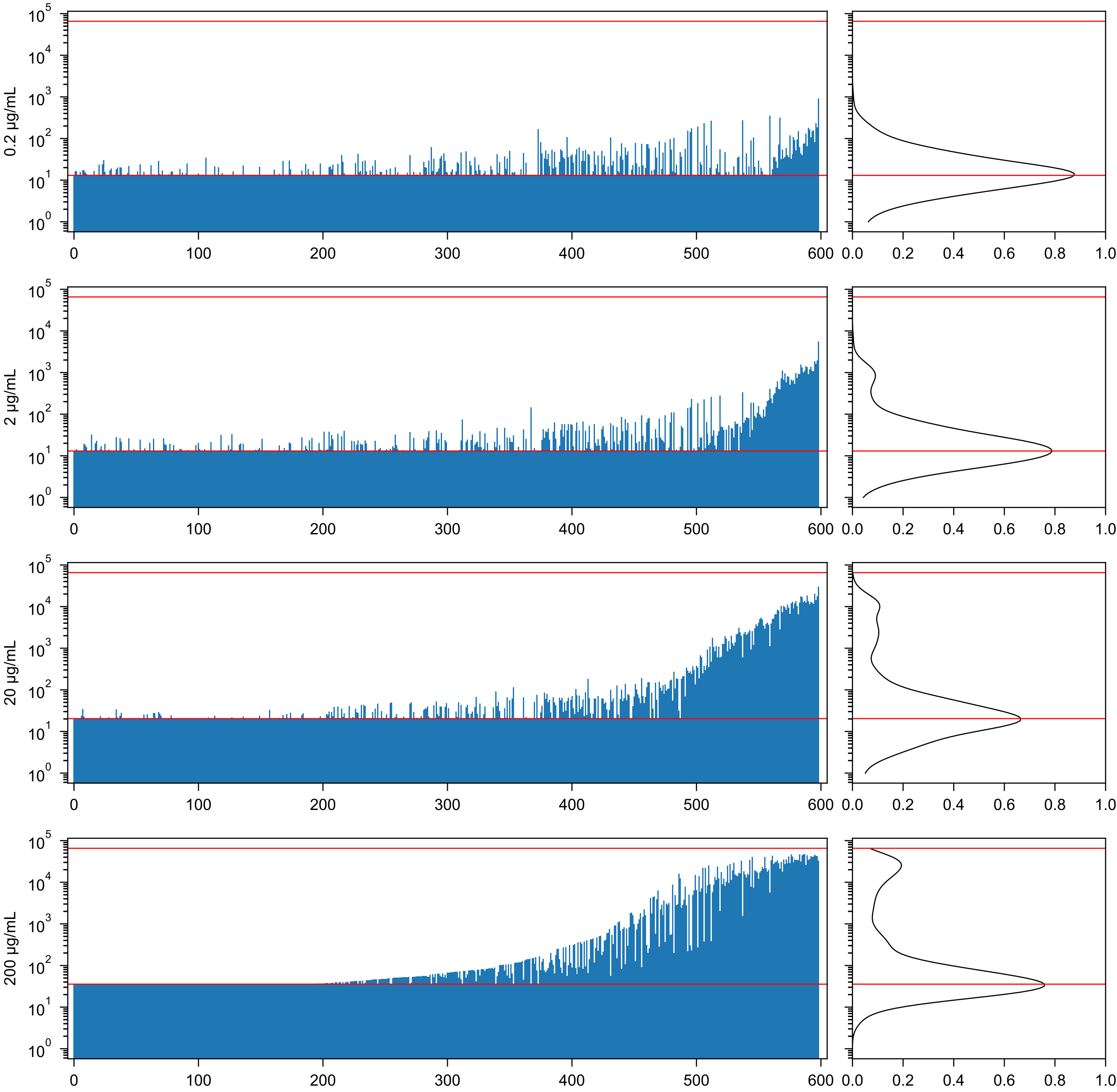


MpL-1 (Macrolepiota procera lectin isolectin 1) - Janko Kos

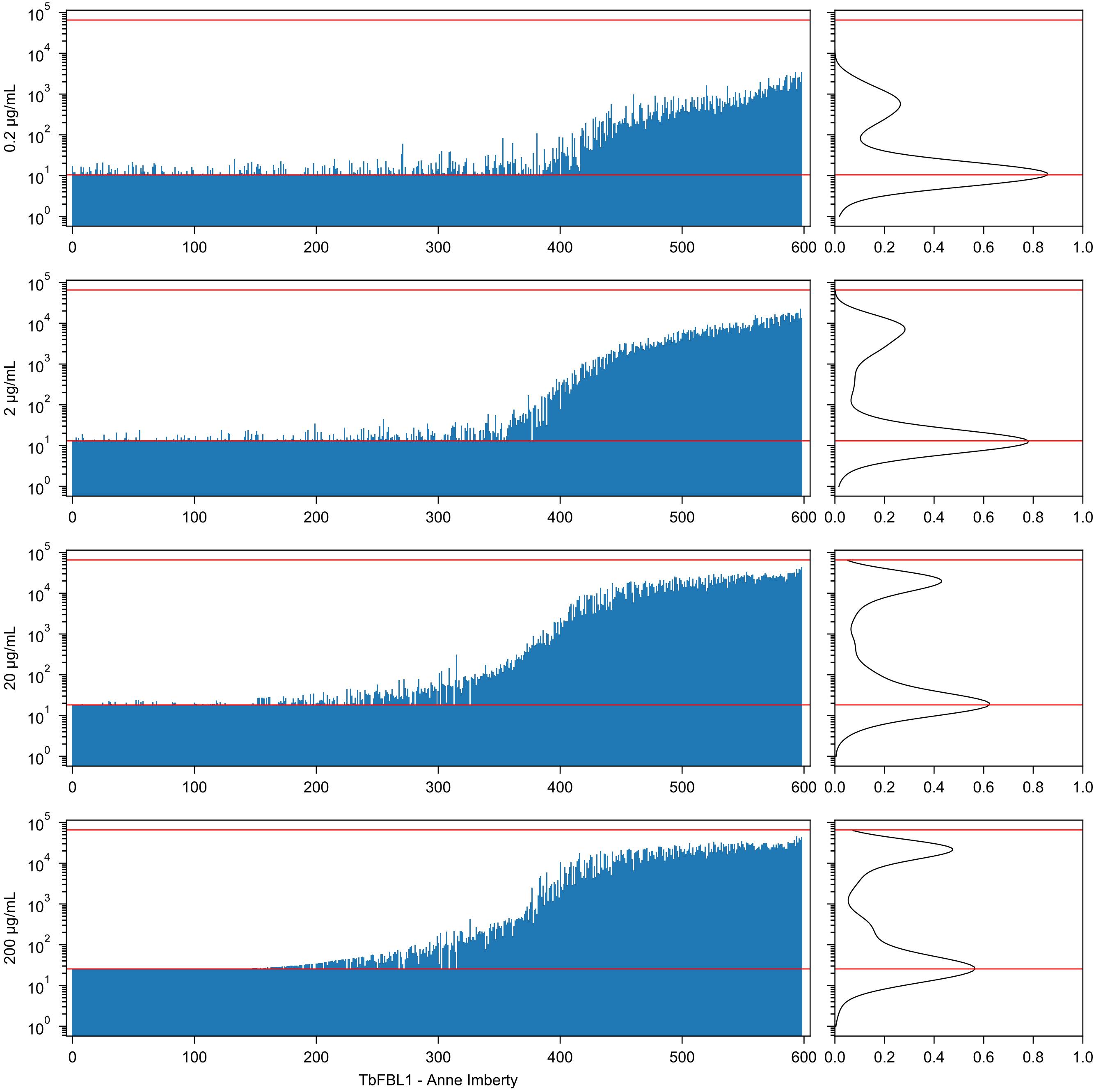
0.2 µg/mL
2 µg/mL
20 µg/mL
200 µg/mL
TbFBL1 - Anne Imberty

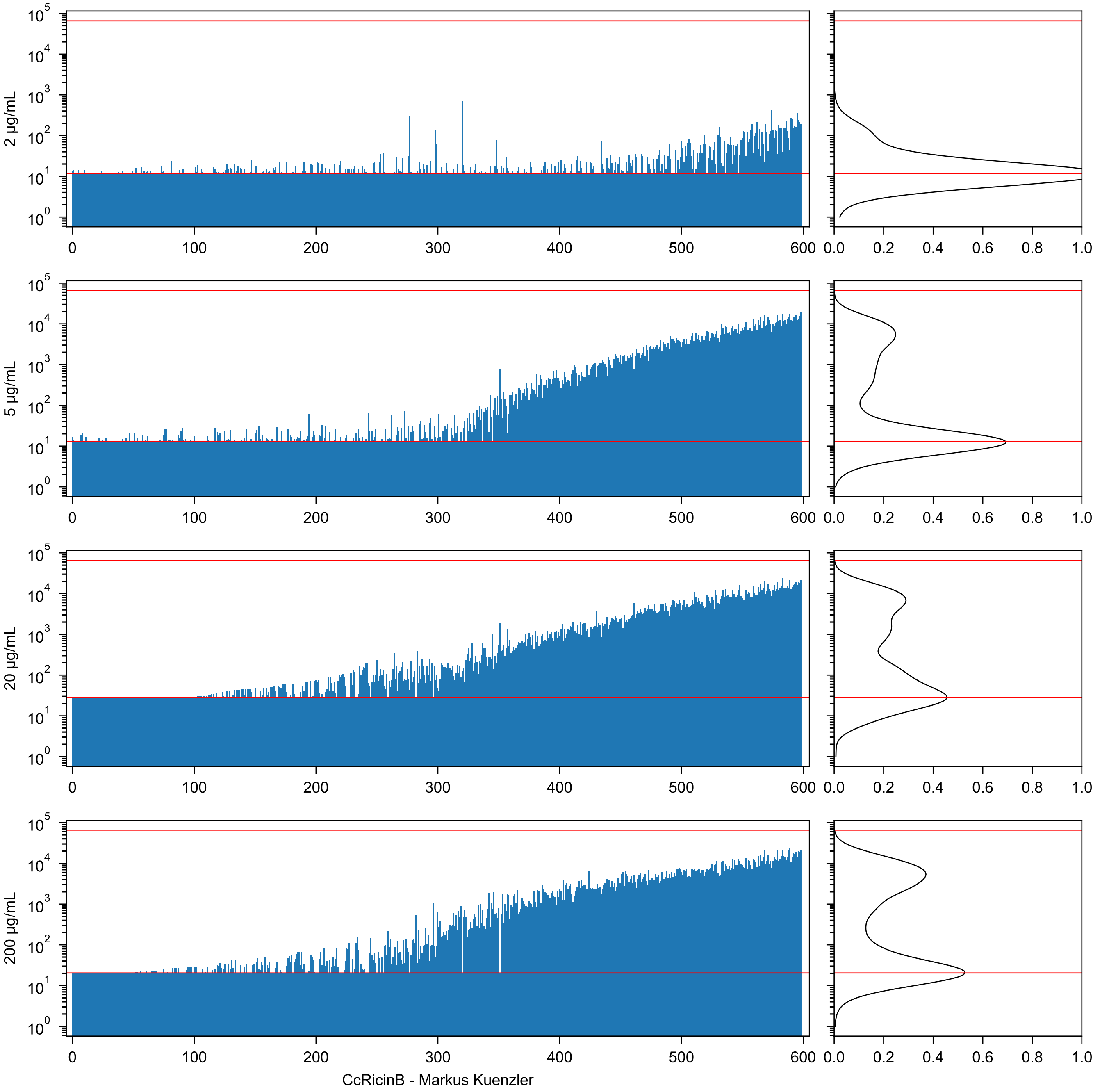
2 μg/mL
5 μg/mL
20 μg/mL
200 μg/mL
CcRicinB - Markus Kuenzler

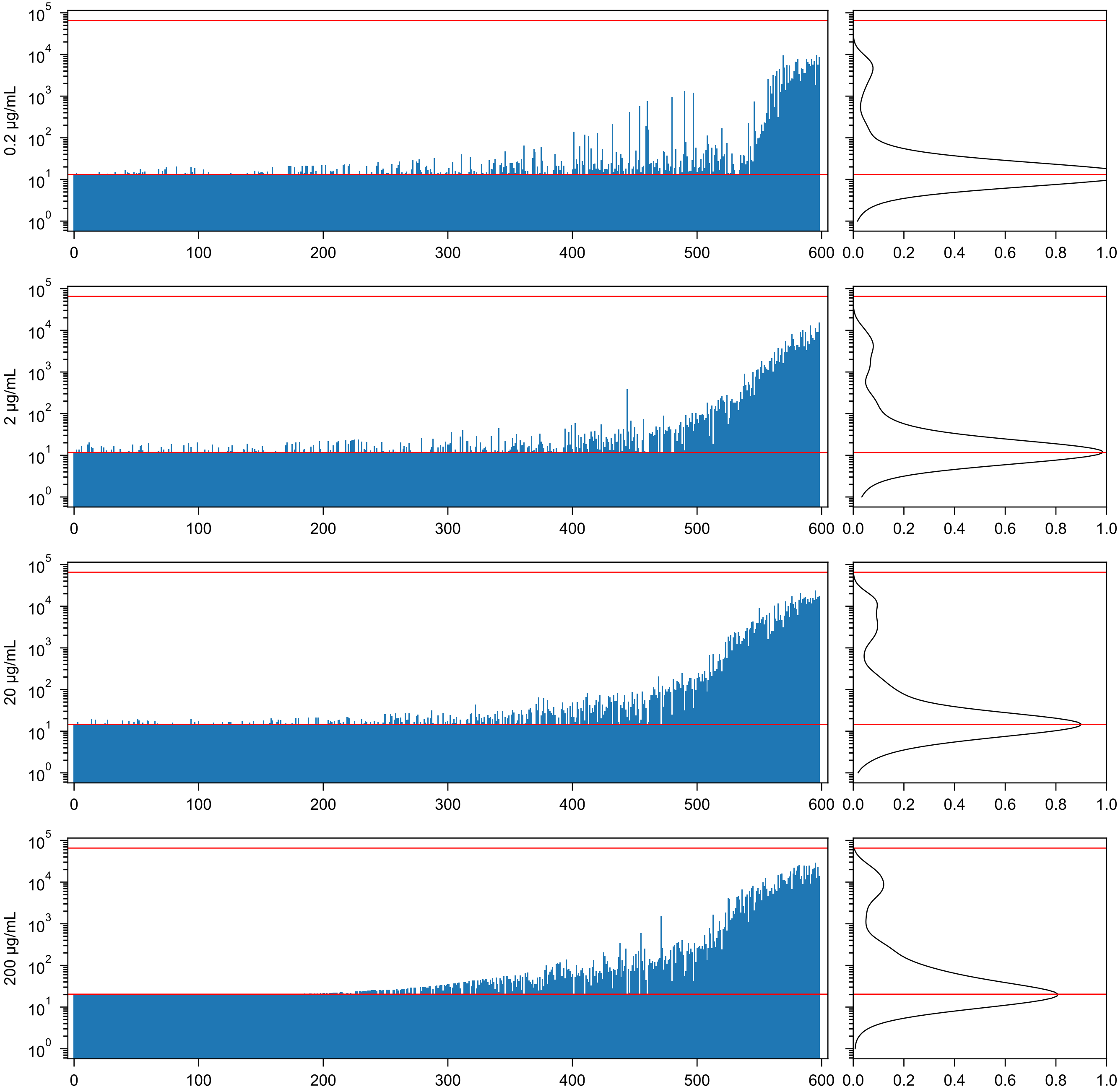

MGL 6B - Yvette Van Kooyk

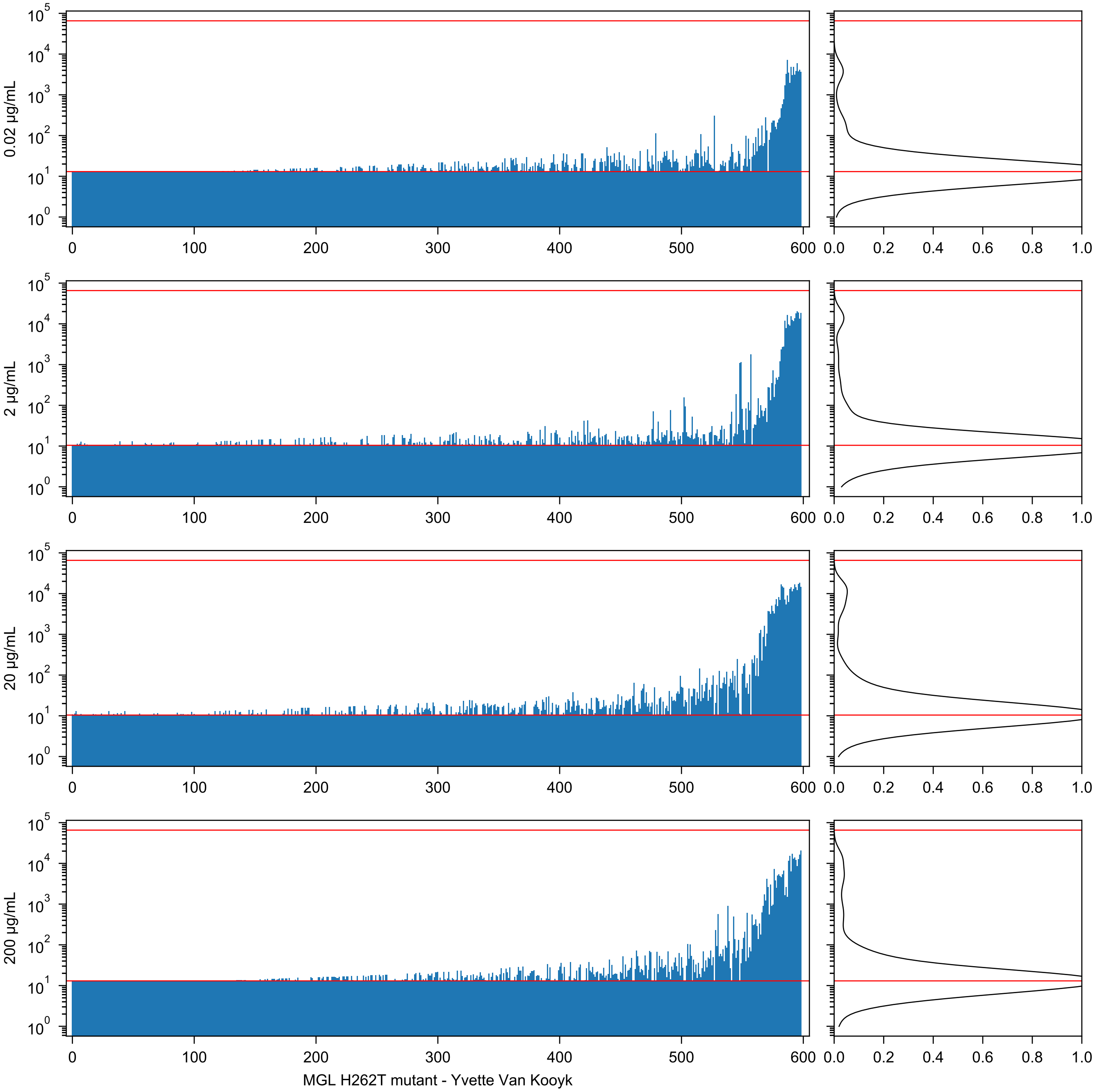

0.02 µg/mL
2 µg/mL
20 µg/mL
200 µg/mL
MGL H262T mutant - Yvette Van Kooyk

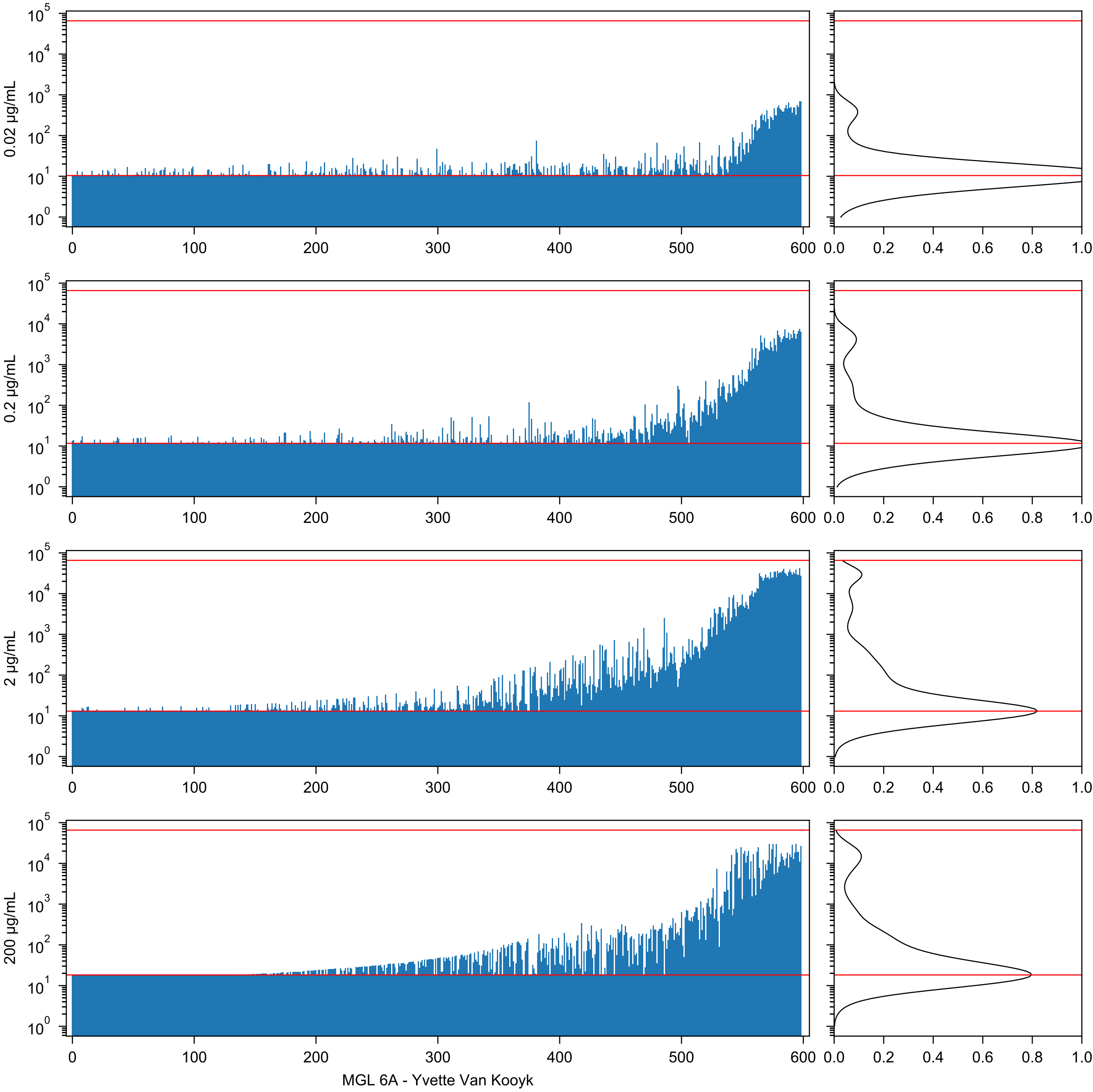

0.02 µg/mL
0.2 µg/mL
2 µg/mL
200 µg/mL
MGL 6A - Yvette Van Kooyk

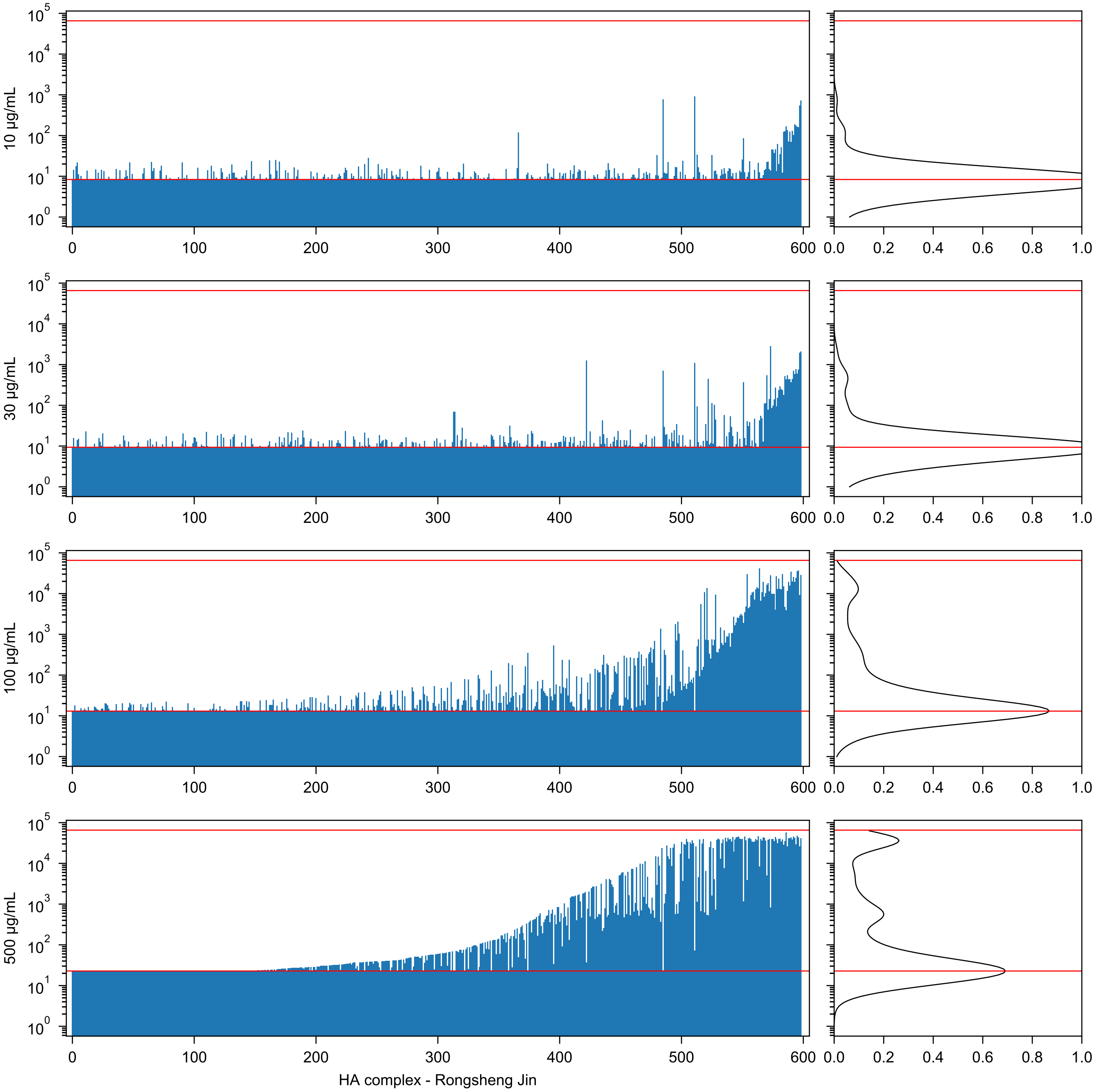

10 µg/mL
30 µg/mL
100 µg/mL
500 µg/mL
HA complex - Rongsheng Jin

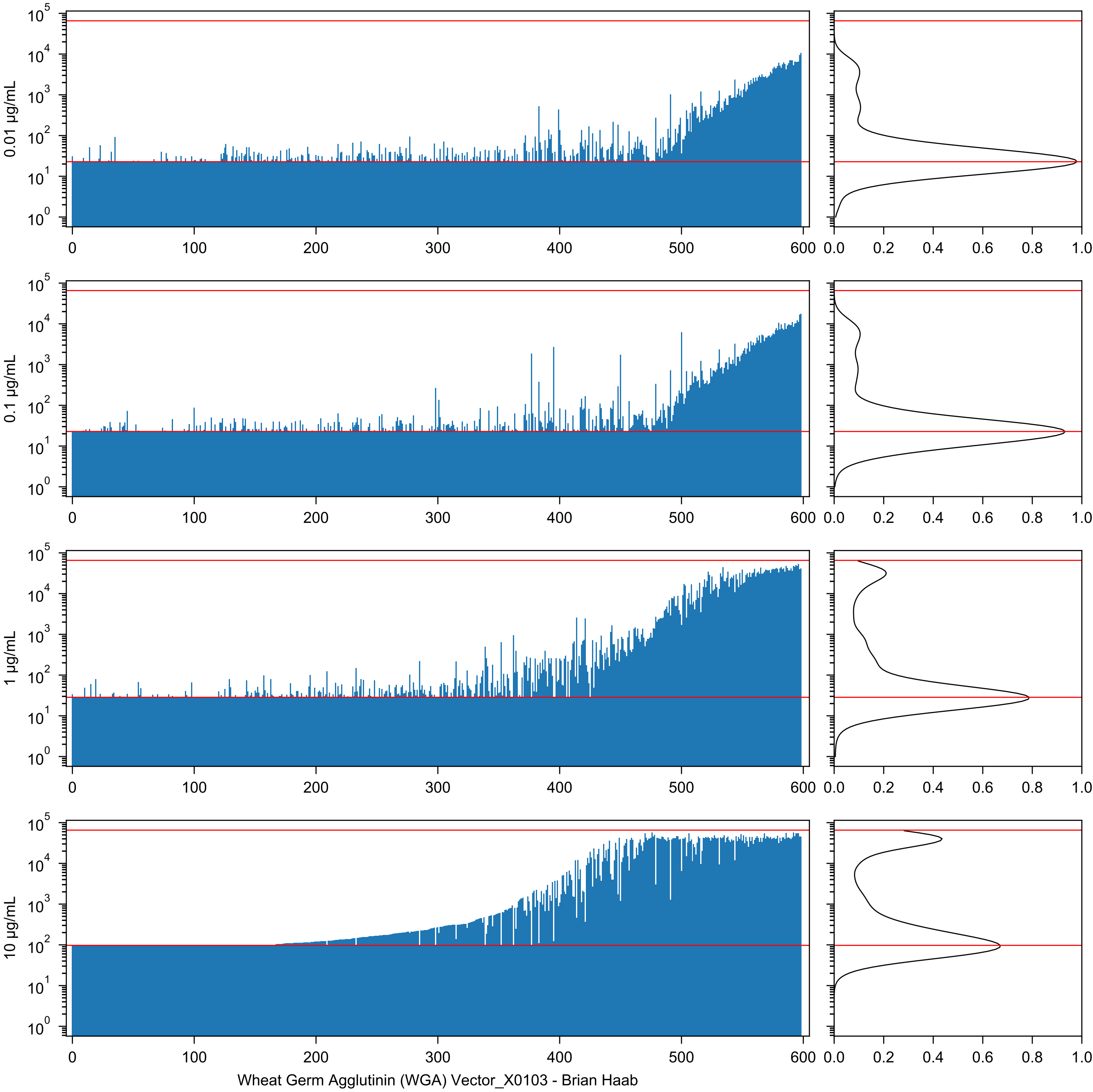

0.01 µg/mL
0.1 µg/mL
1 µg/mL
10 µg/mL
$10^0$
$10^1$
$10^2$
$10^3$
$10^4$
$10^5$
0
100
200
300
400
500
600
0.0
0.2
0.4
0.6
0.8
1.0
Wheat Germ Agglutinin (WGA) Vector_X0103 - Brian Haab

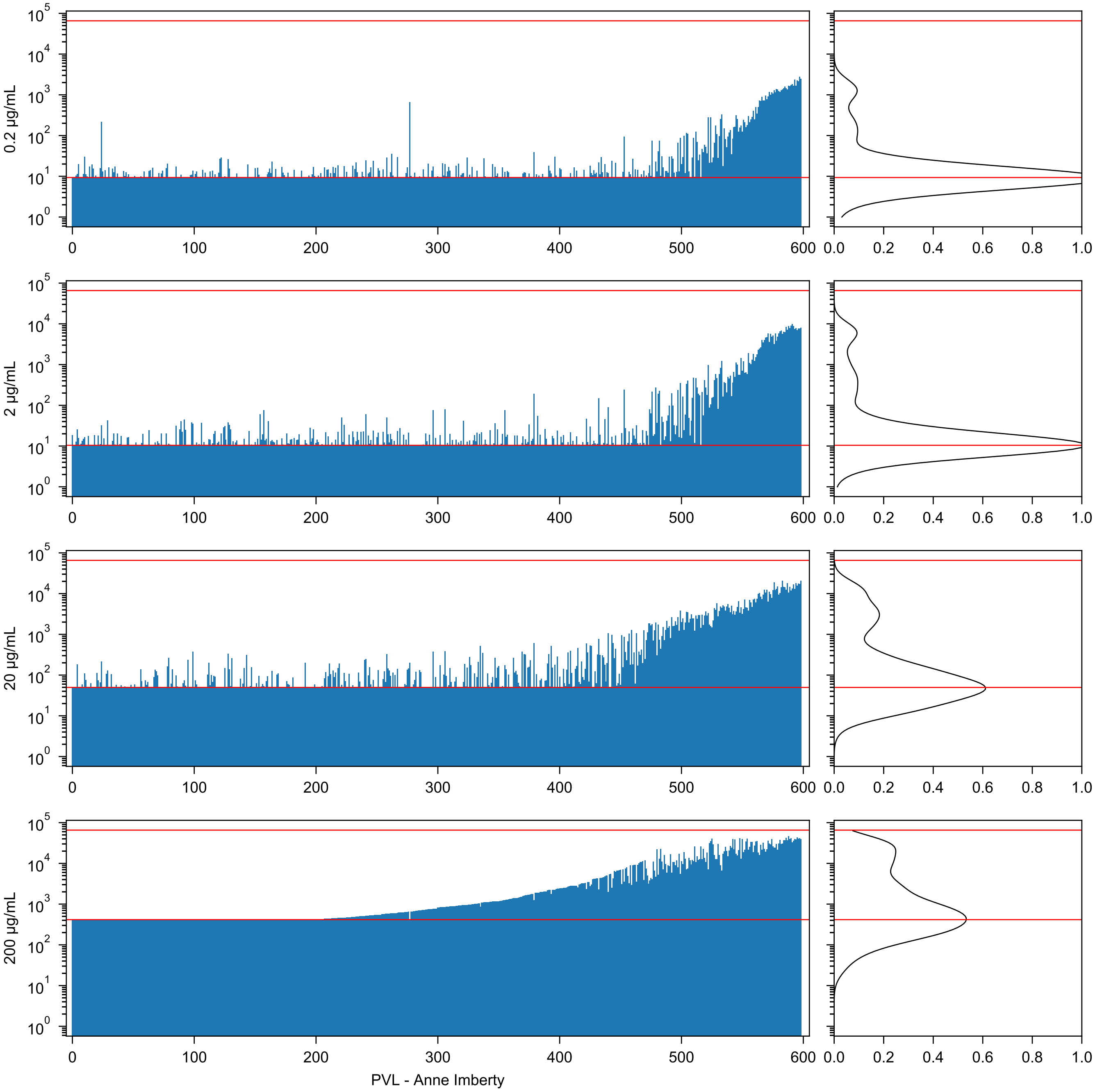
0.2 µg/mL
2 µg/mL
20 µg/mL
200 µg/mL
PVL - Anne Imberty

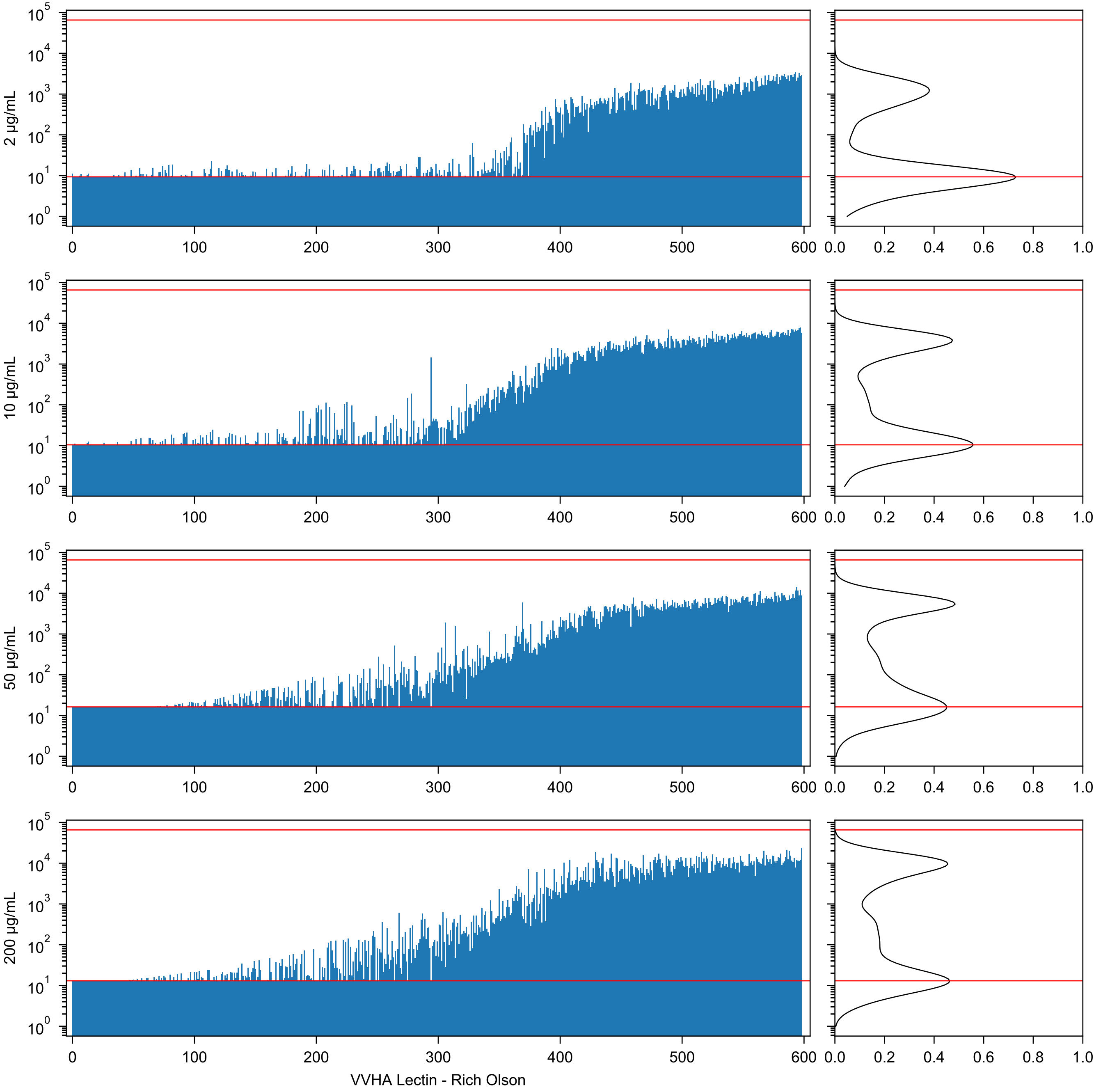
2 µg/mL
10 µg/mL
50 µg/mL
200 µg/mL
VVHA Lectin - Rich Olson

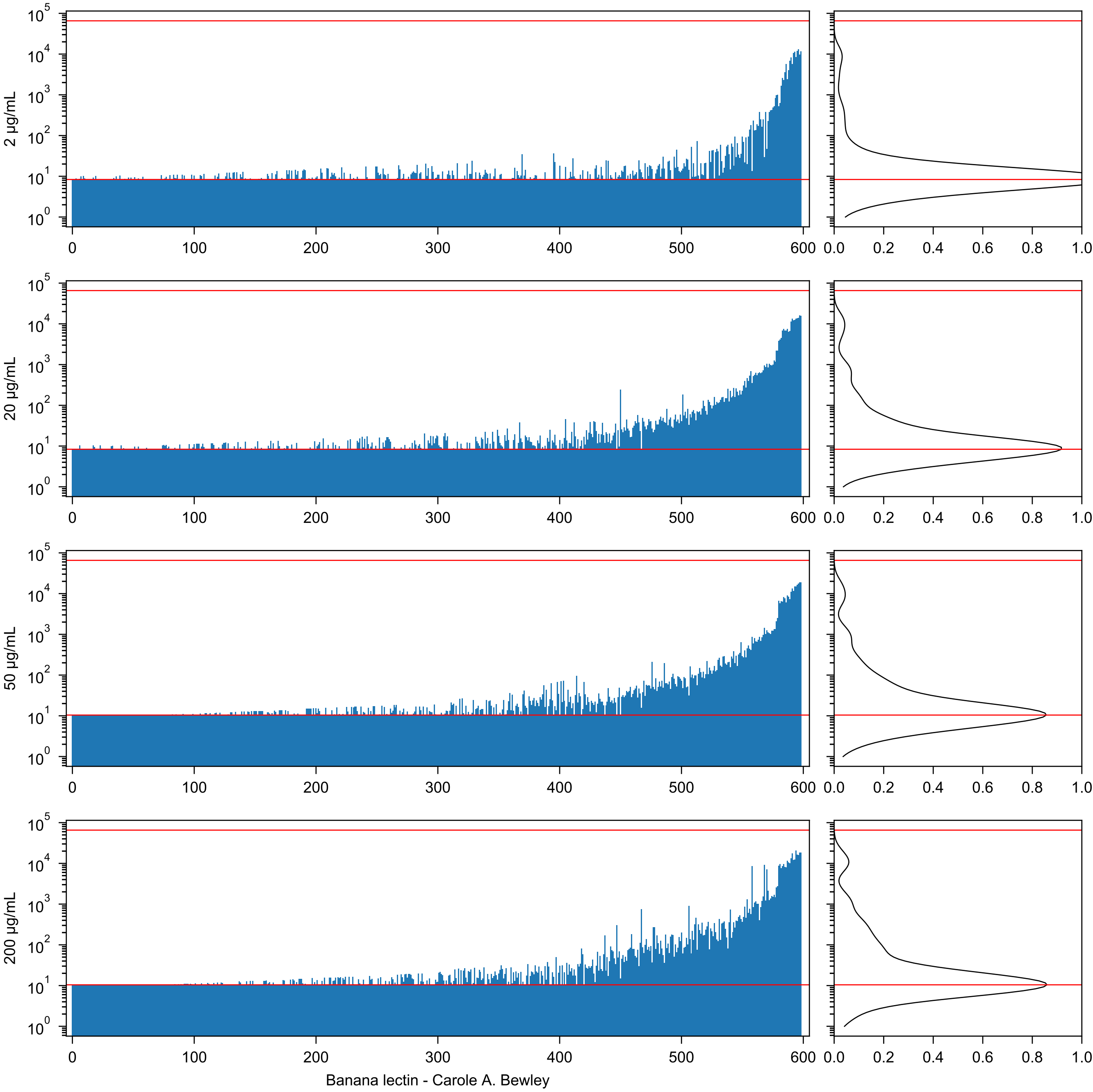

2 μg/mL
20 μg/mL
50 μg/mL
200 μg/mL
Banana lectin - Carole A. Bewley

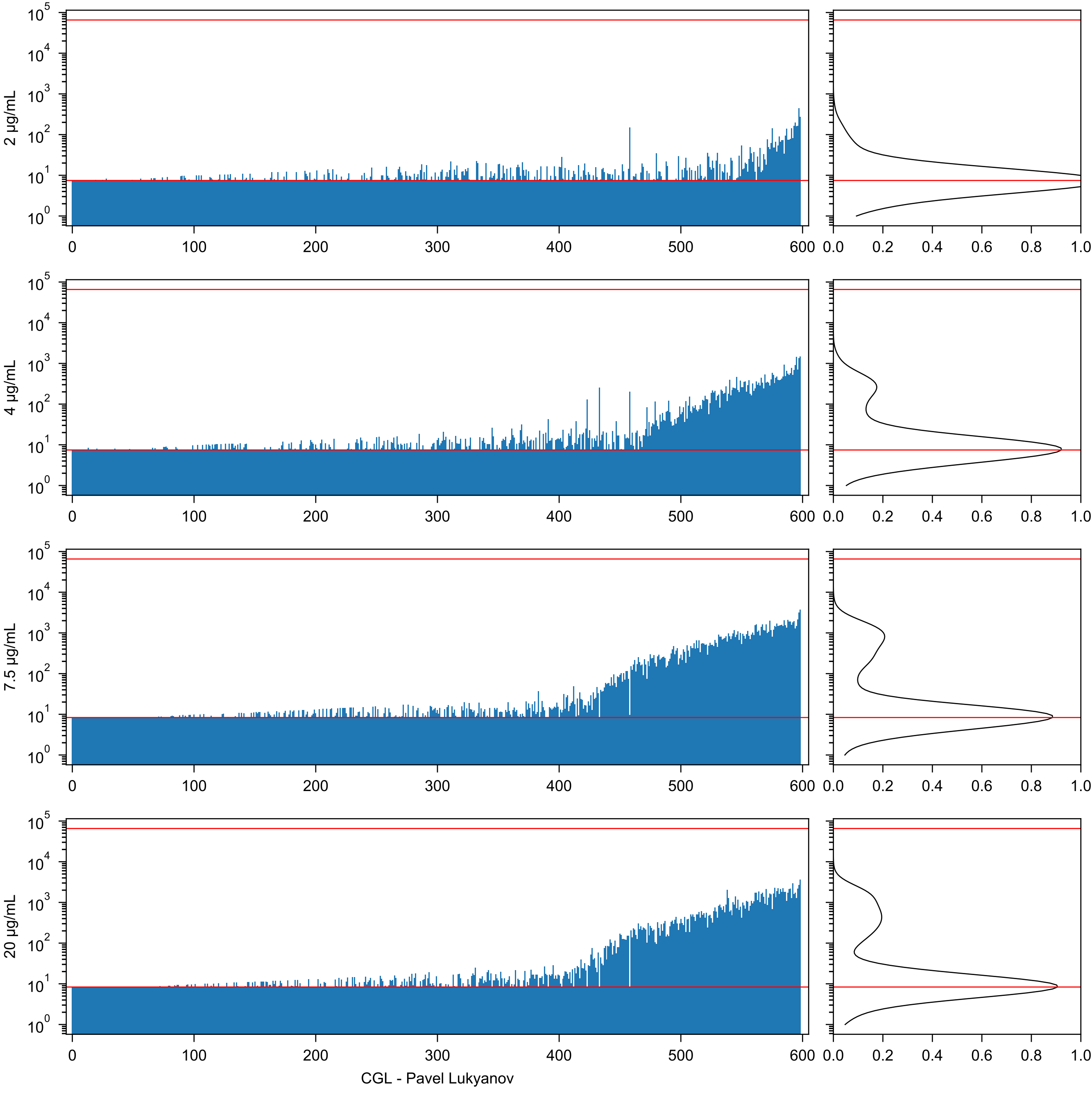

2 μg/mL
4 μg/mL
7.5 μg/mL
20 μg/mL
CGL - Pavel Lukyanov

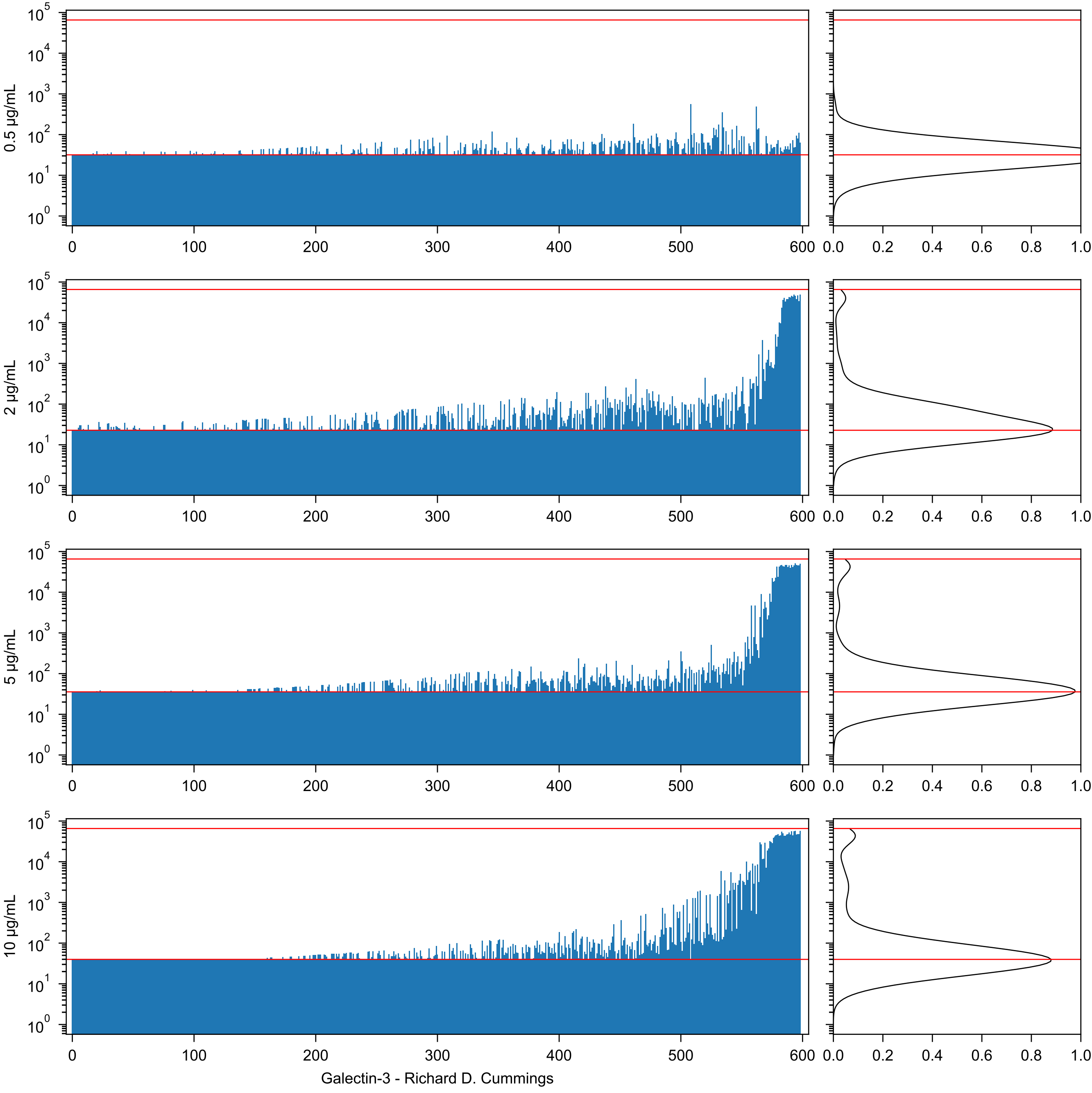
0.5 µg/mL
2 µg/mL
5 µg/mL
10 µg/mL
Galectin-3 - Richard D. Cummings

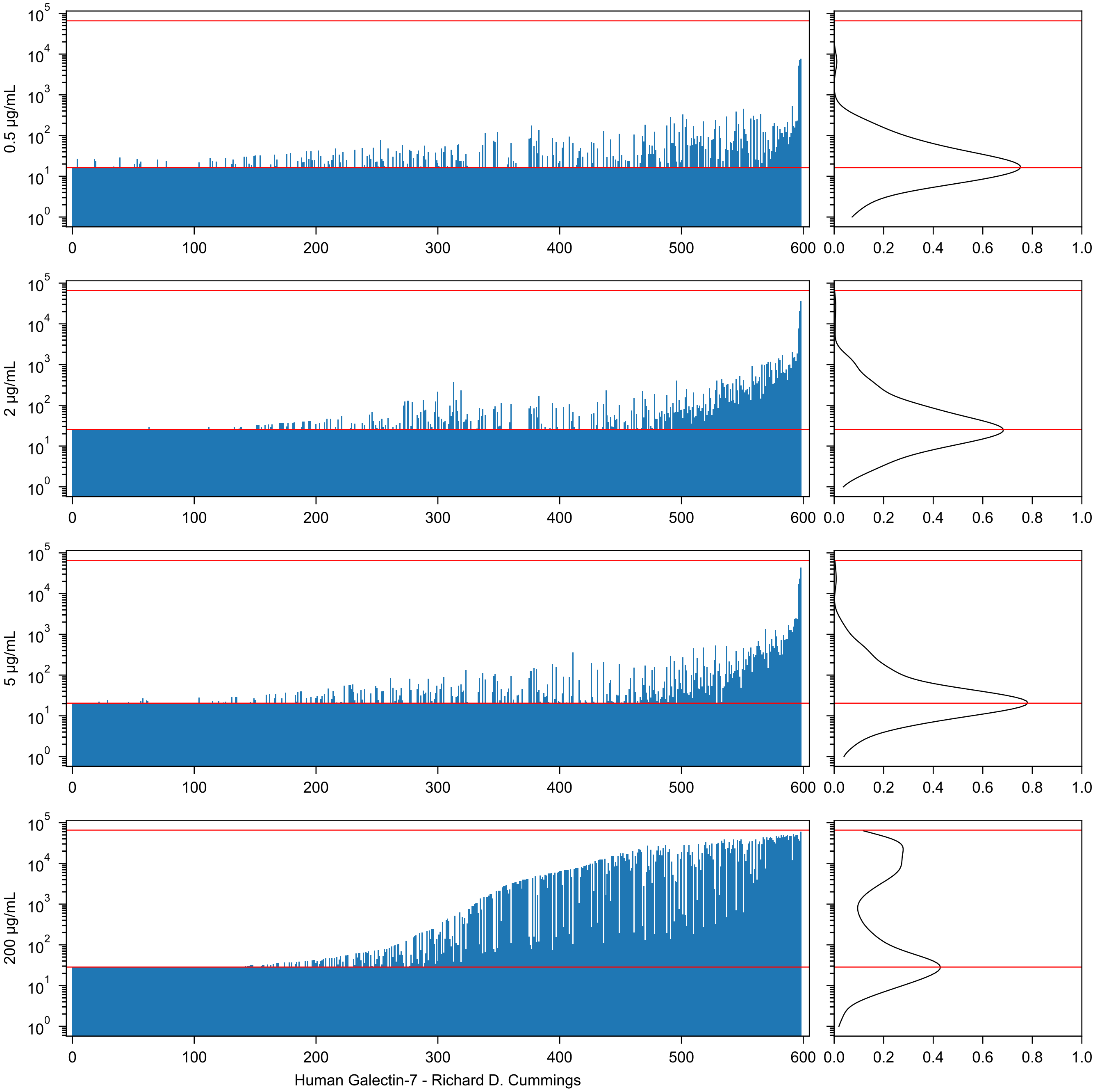
0.5 µg/mL
2 µg/mL
5 µg/mL
200 µg/mL
Human Galectin-7 - Richard D. Cummings

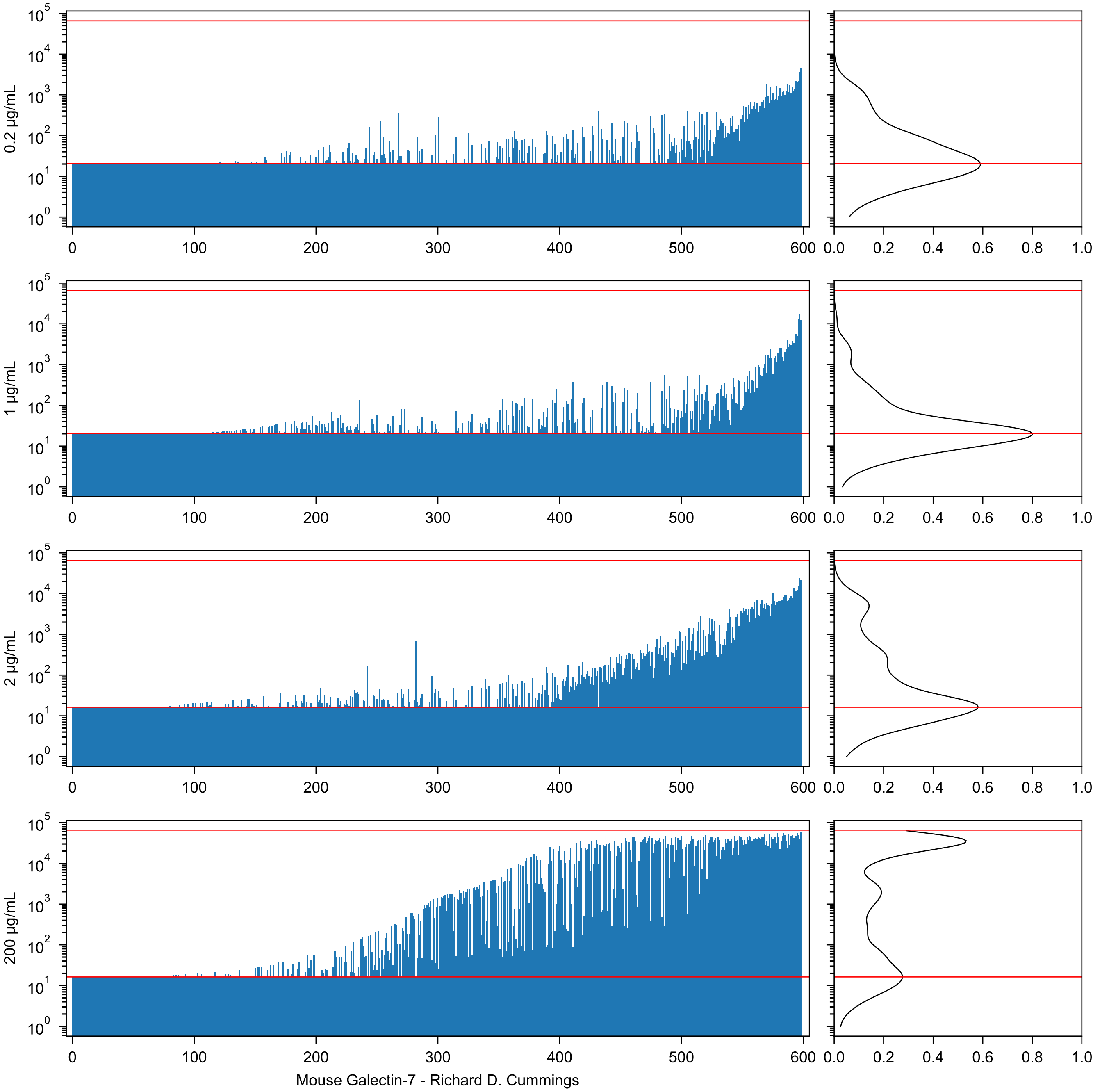
0.2 µg/mL
1 µg/mL
2 µg/mL
200 µg/mL
Mouse Galectin-7 - Richard D. Cummings

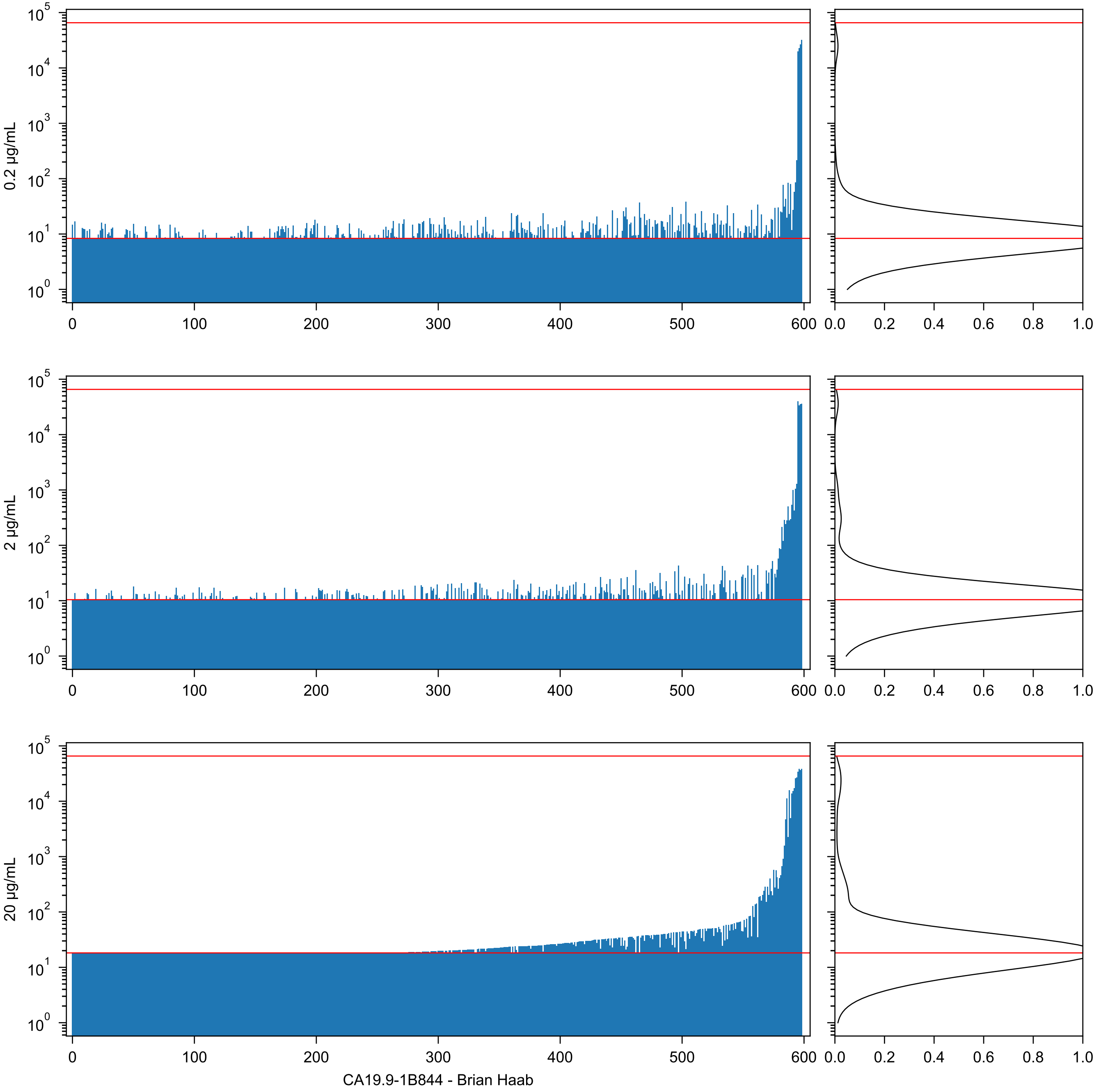
0.2 µg/mL
2 µg/mL
20 µg/mL
CA19.9-1B844 - Brian Haab

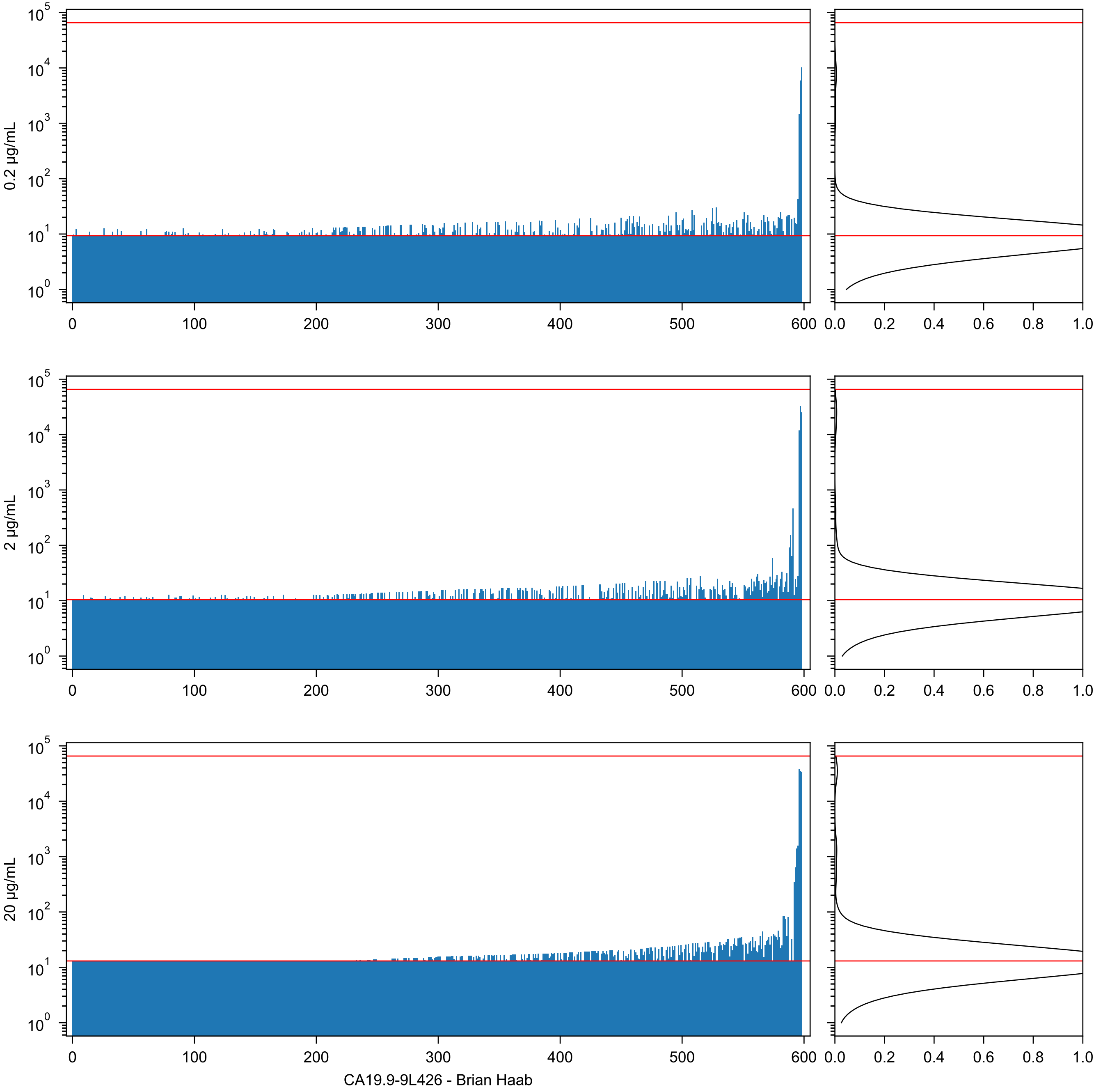

0.2 µg/mL
2 µg/mL
20 µg/mL
CA19.9-9L426 - Brian Haab

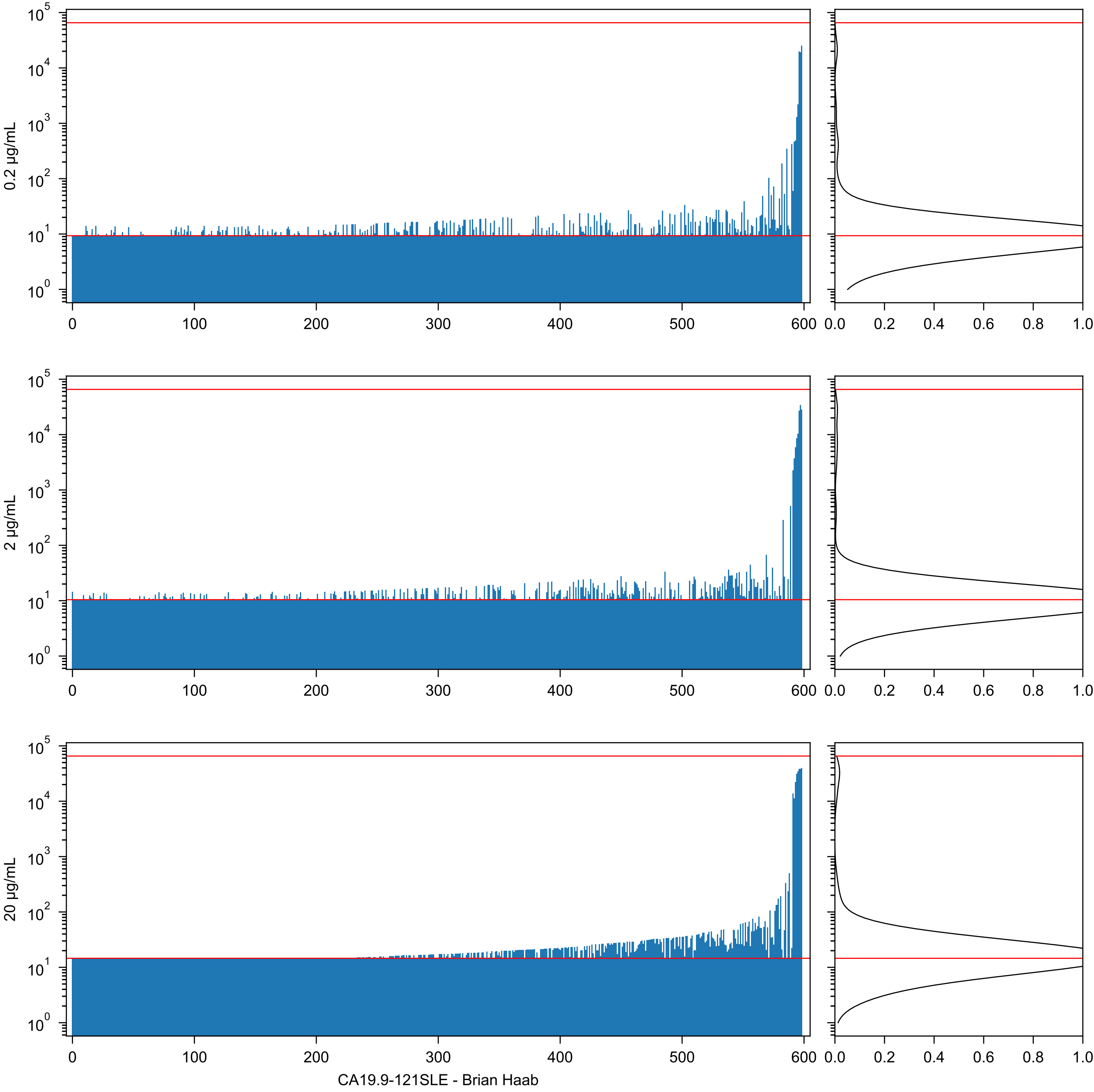

0.2 μg/mL
2 μg/mL
20 μg/mL
CA19.9-121SLE - Brian Haab

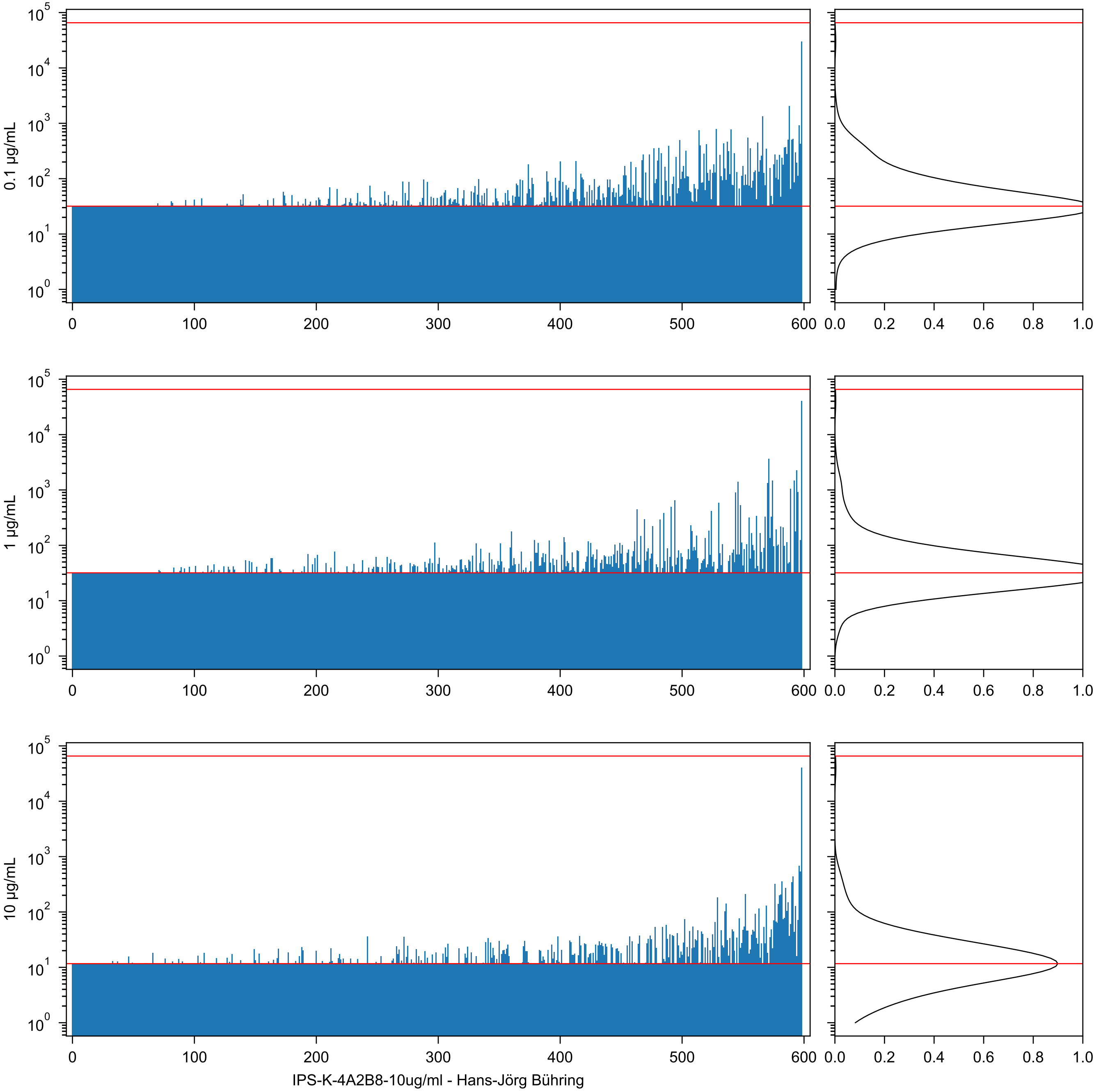
0.1 µg/mL
1 µg/mL
10 µg/mL
IPS-K-4A2B8-10ug/ml - Hans-Jörg Bühring

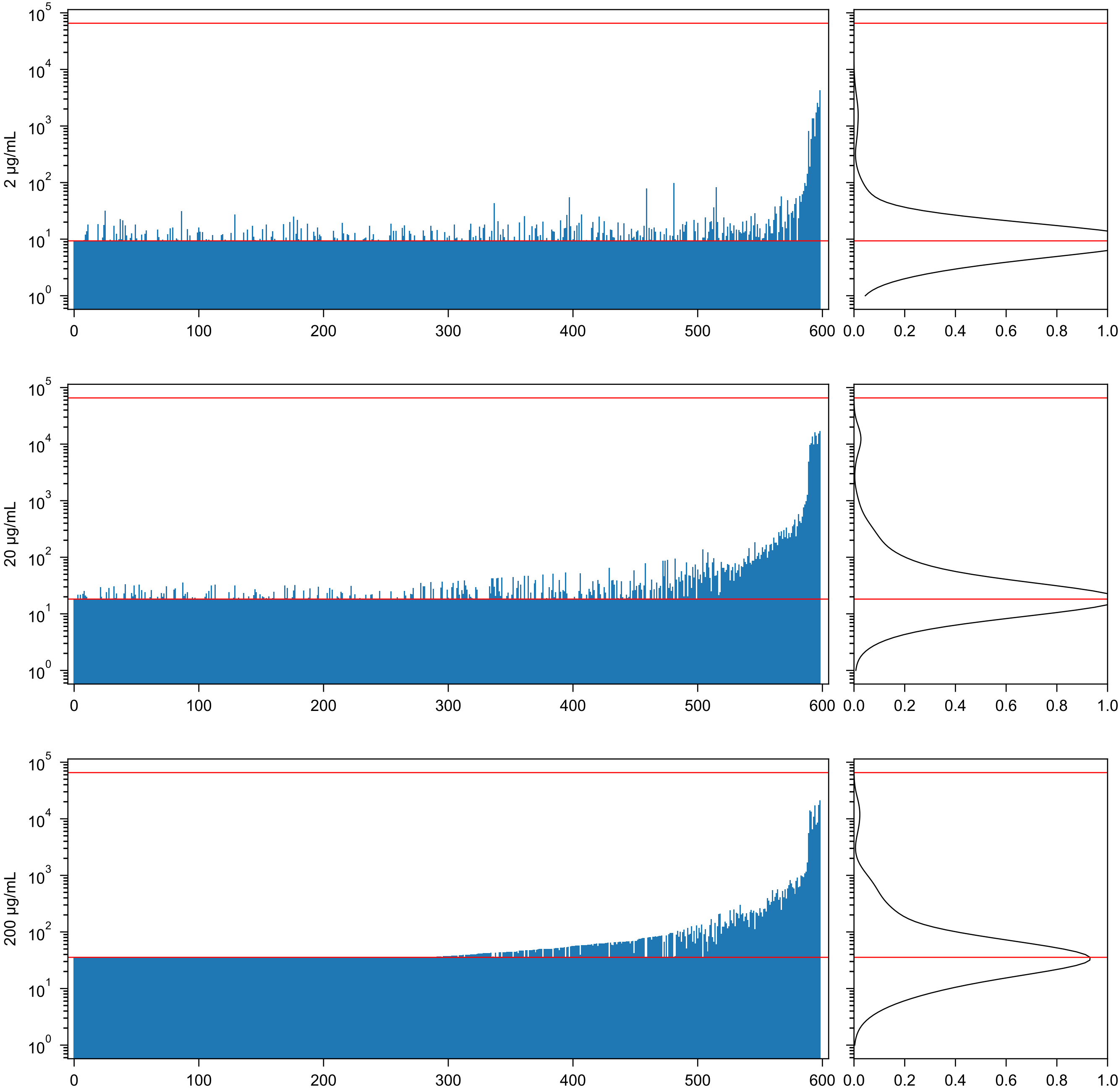


MVN22 Microvirin dimer with N-terminal hexahistidine tag - Marie Hanigan

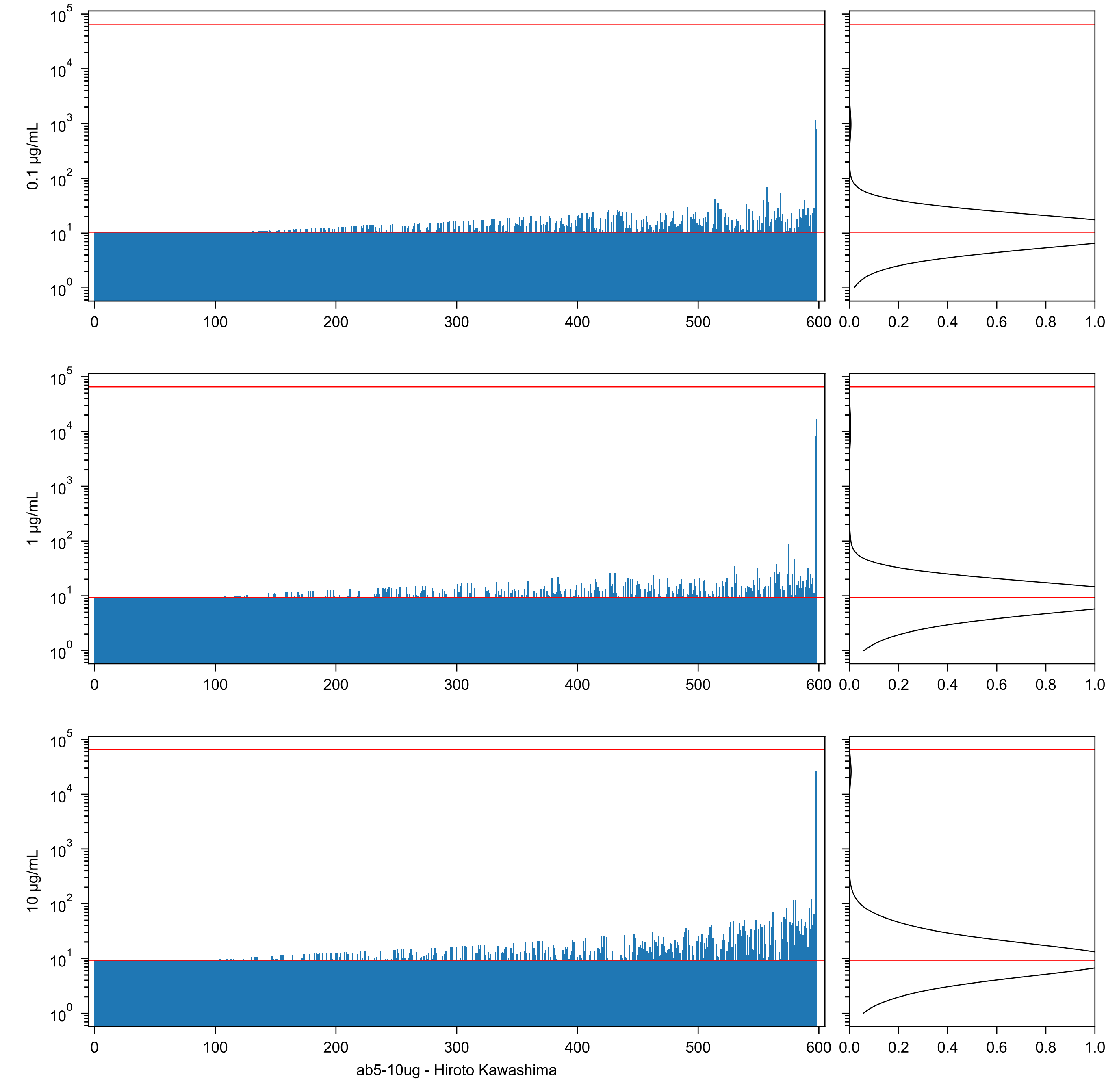
0.1 µg/mL
1 µg/mL
10 µg/mL
ab5-10ug - Hiroto Kawashima

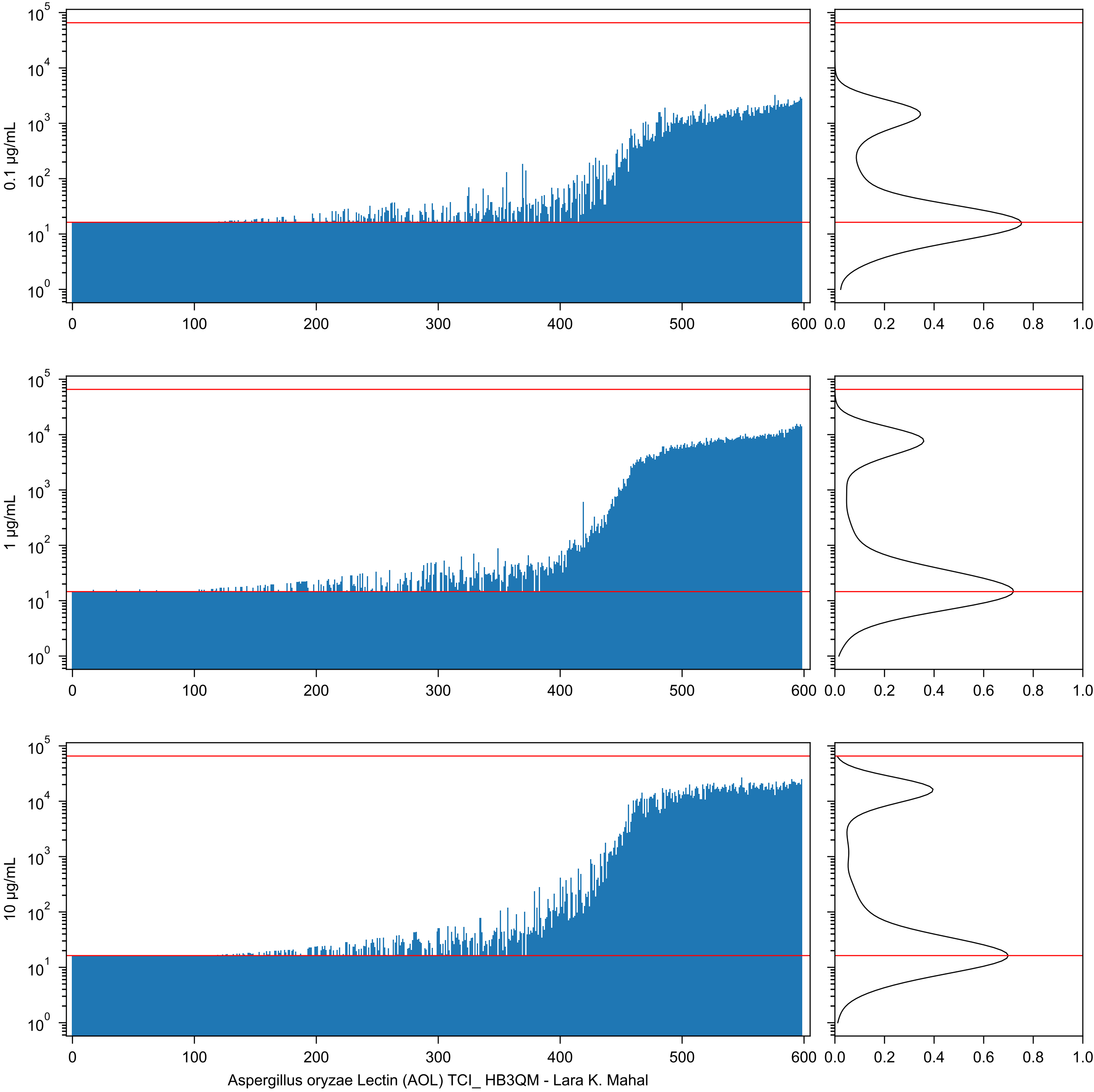

0.1 µg/mL
1 µg/mL
10 µg/mL
Aspergillus oryzae Lectin (AOL) TCI_ HB3QM - Lara K. Mahal

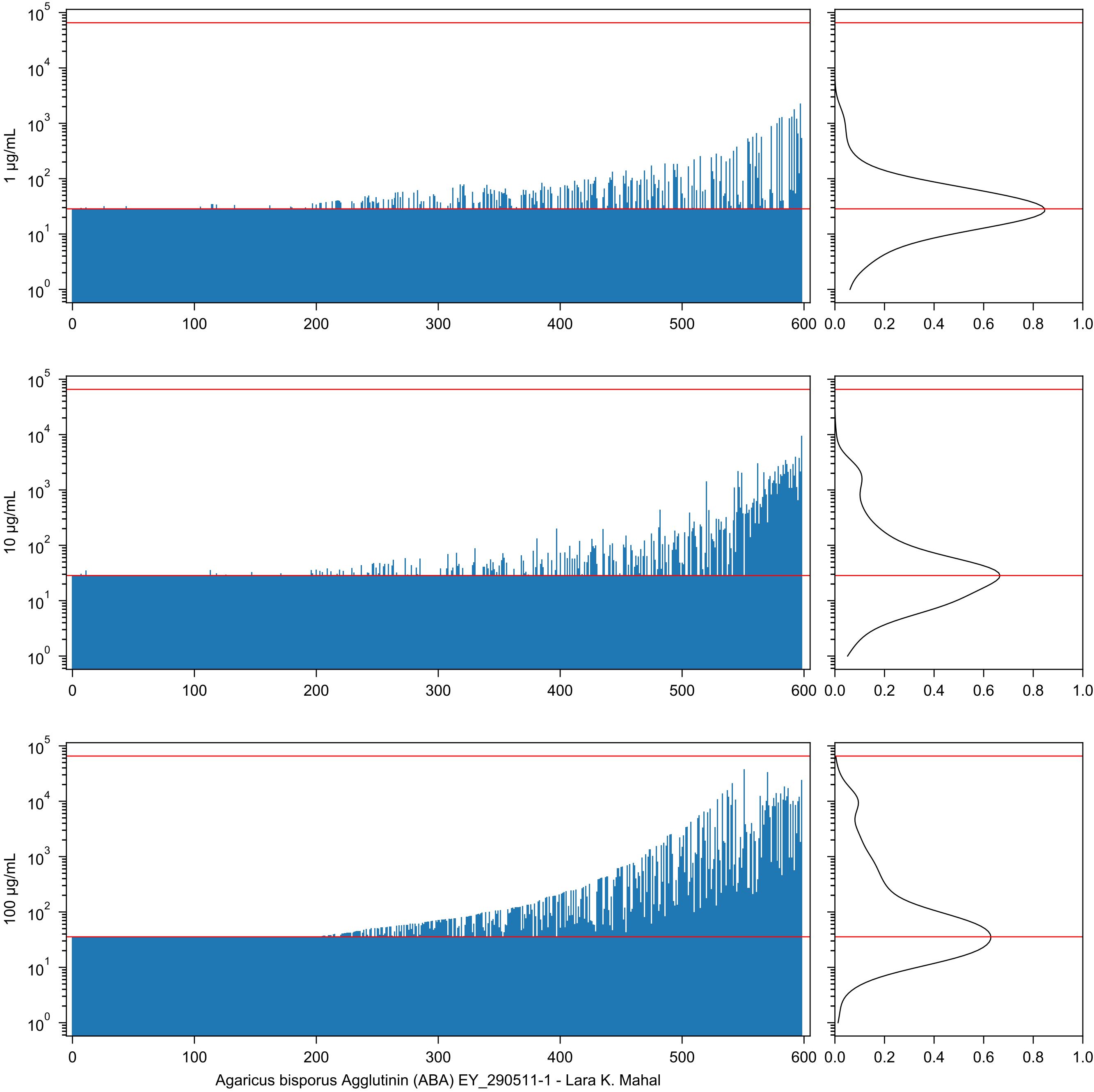
1 µg/mL
10 µg/mL
100 µg/mL
0 100 200 300 400 500 600
0.0 0.2 0.4 0.6 0.8 1.0
Agaricus bisporus Agglutinin (ABA) EY_290511-1 - Lara K. Mahal

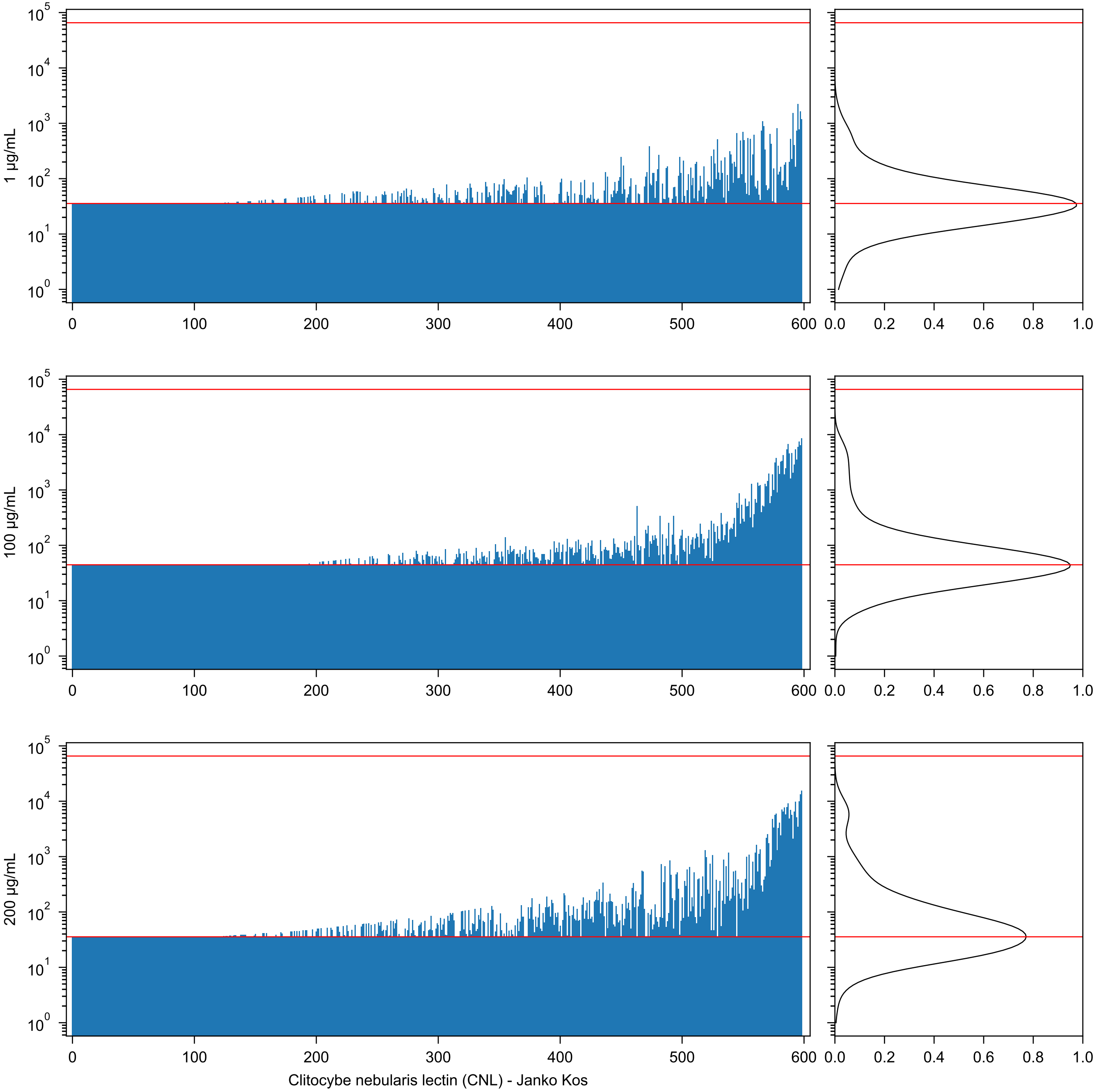
1 μg/mL
100 μg/mL
200 μg/mL
Clitocybe nebularis lectin (CNL) - Janko Kos

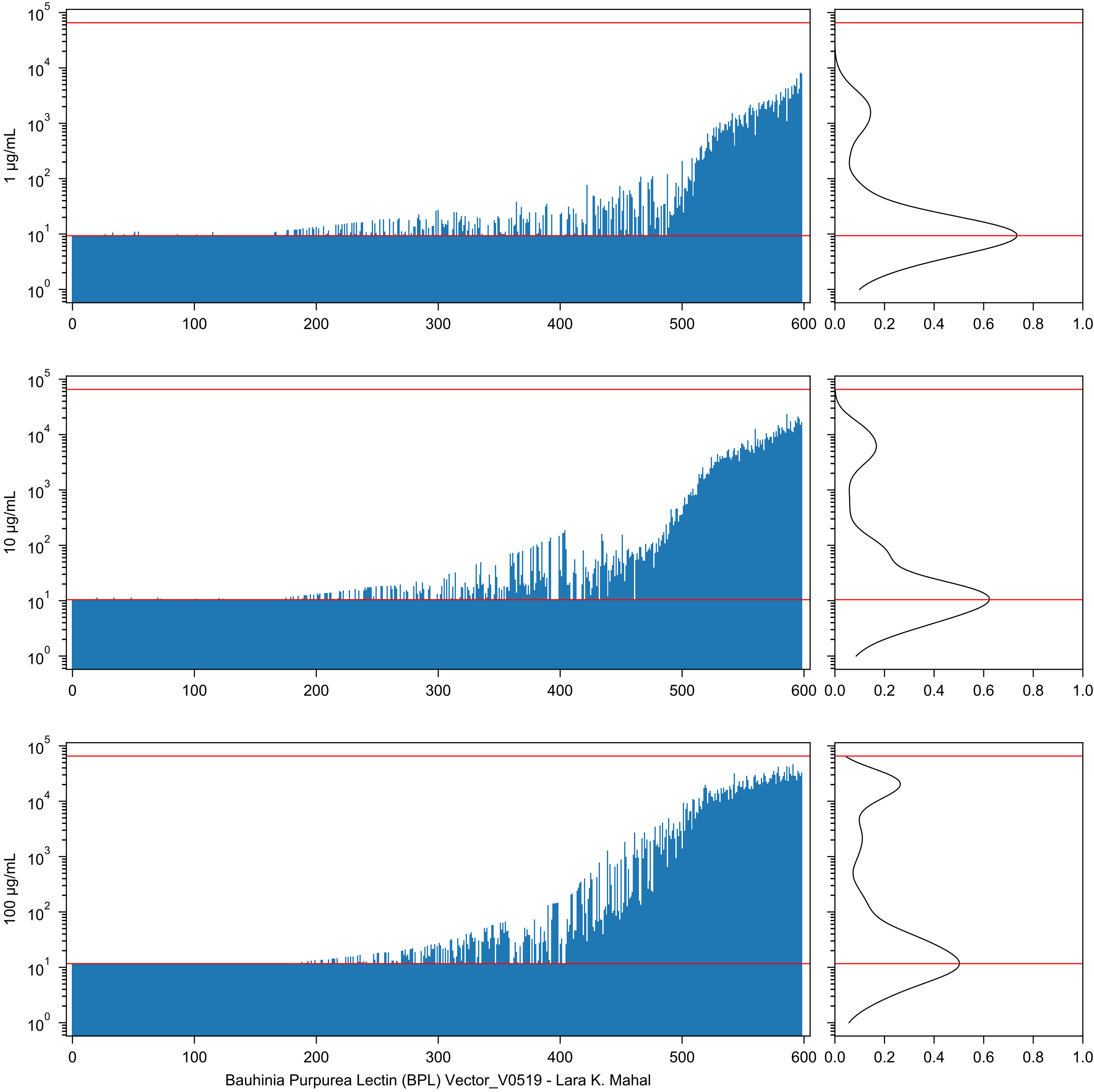
1 µg/mL
10 µg/mL
100 µg/mL
Bauhinia Purpurea Lectin (BPL) Vector_V0519 - Lara K. Mahal

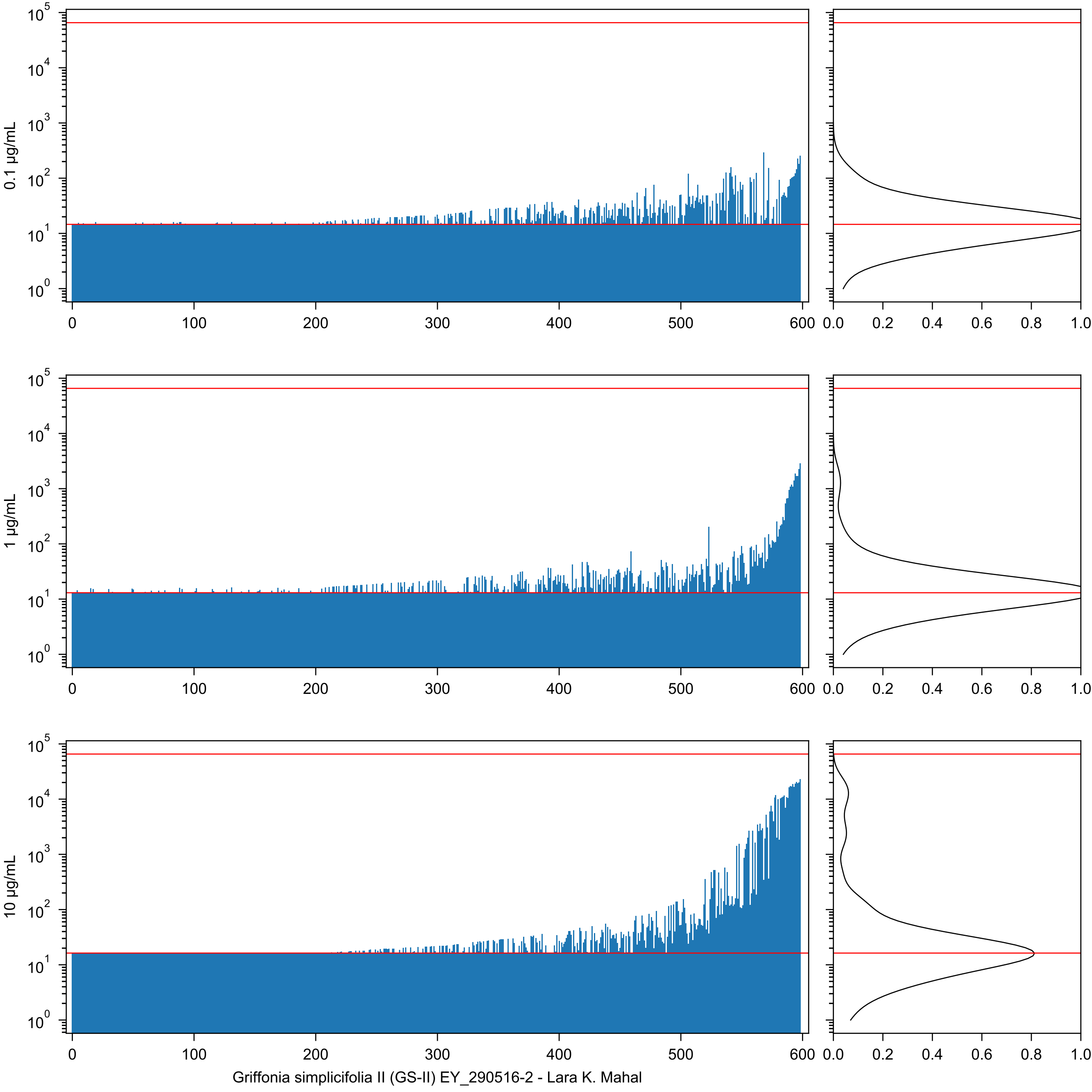
0.1 μg/mL
1 μg/mL
10 μg/mL
Griffonia simplicifolia II (GS-II) EY_290516-2 - Lara K. Mahal

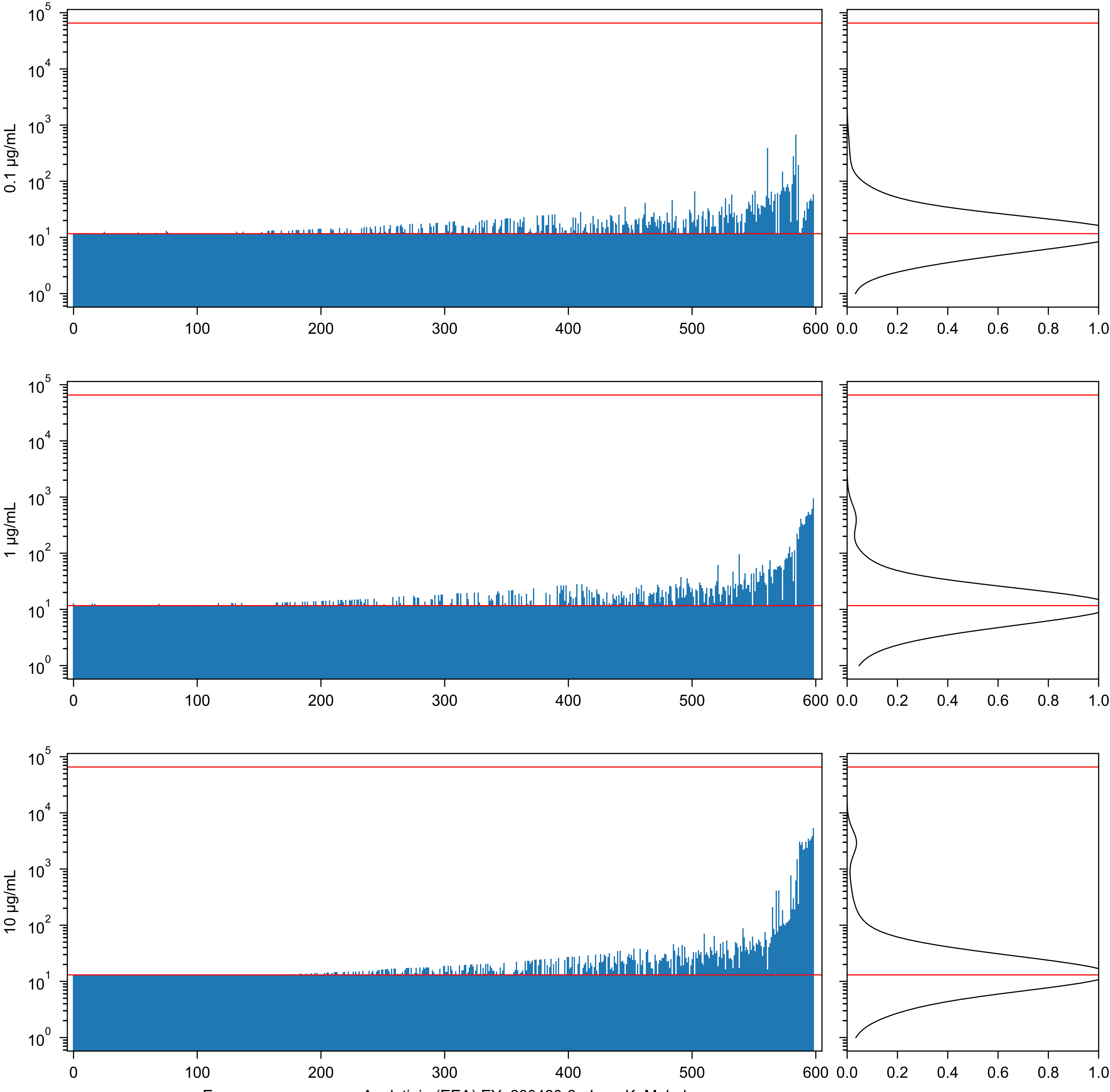

0.1 µg/mL
1 µg/mL
10 µg/mL
Euonymus europaeus Agglutinin (EEA) EY_280420-2 - Lara K. Mahal

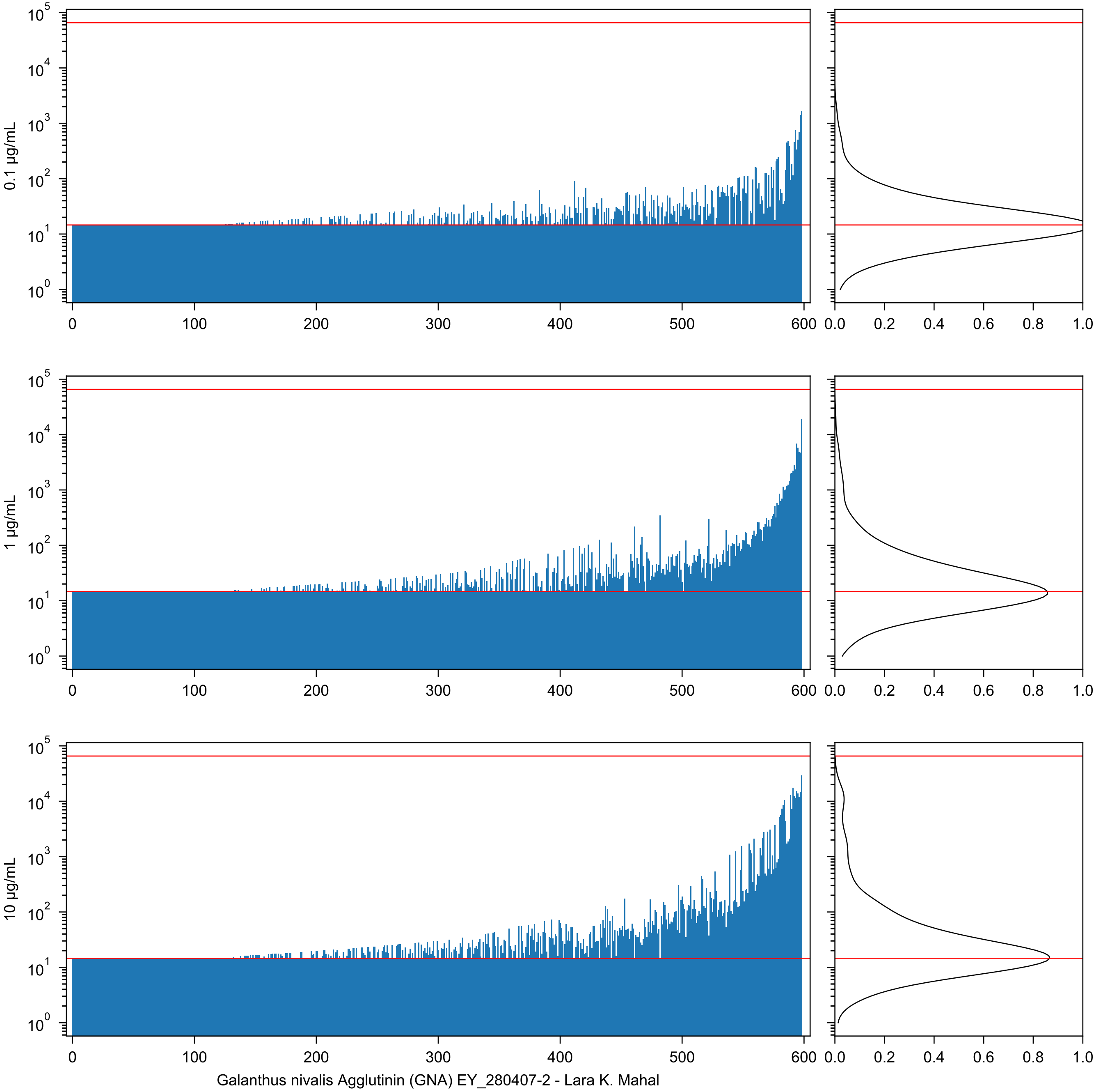
0.1 µg/mL
1 µg/mL
10 µg/mL
Galanthus nivalis Agglutinin (GNA) EY_280407-2 - Lara K. Mahal

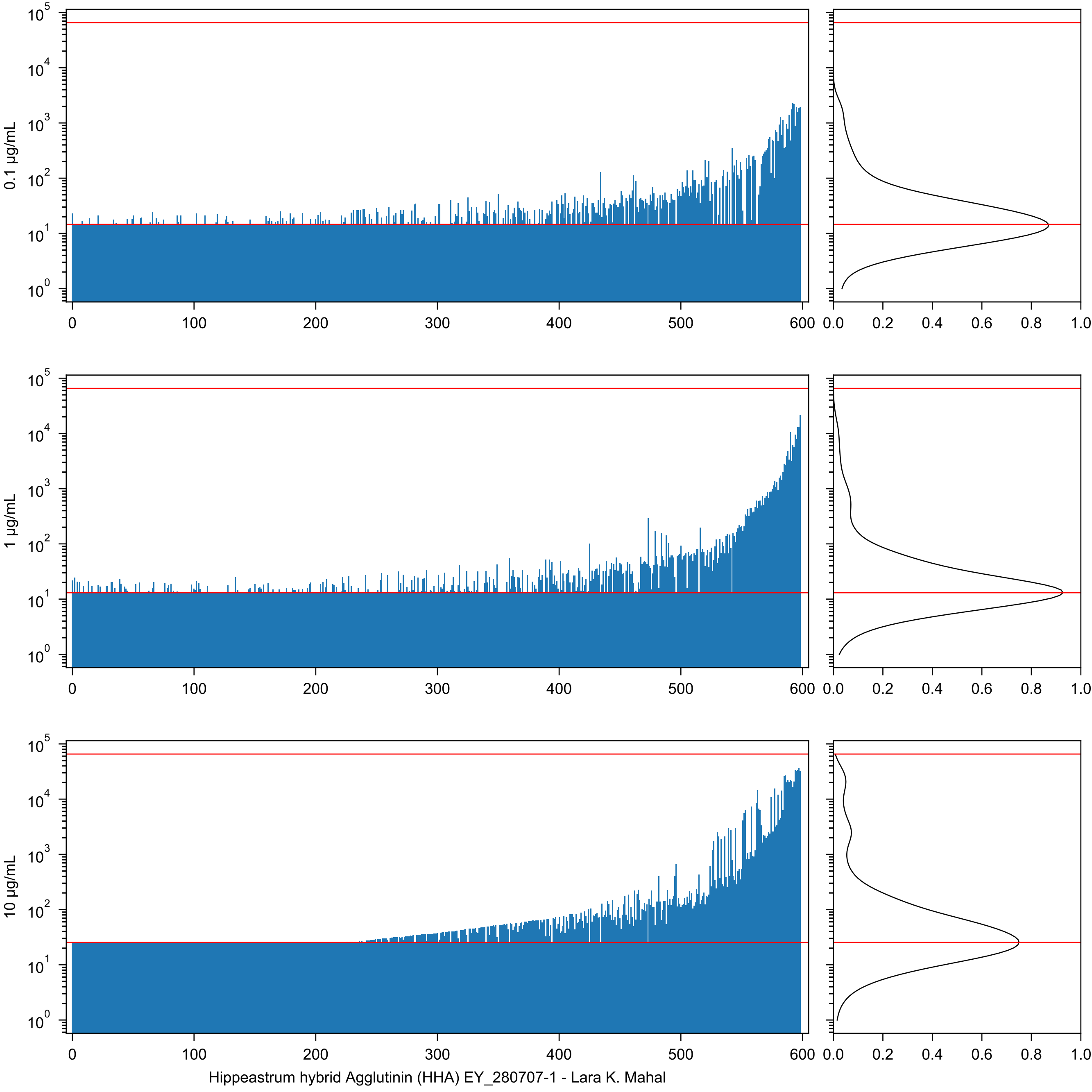
0.1 µg/mL
1 µg/mL
10 µg/mL
Hippeastrum hybrid Agglutinin (HHA) EY_280707-1 - Lara K. Mahal

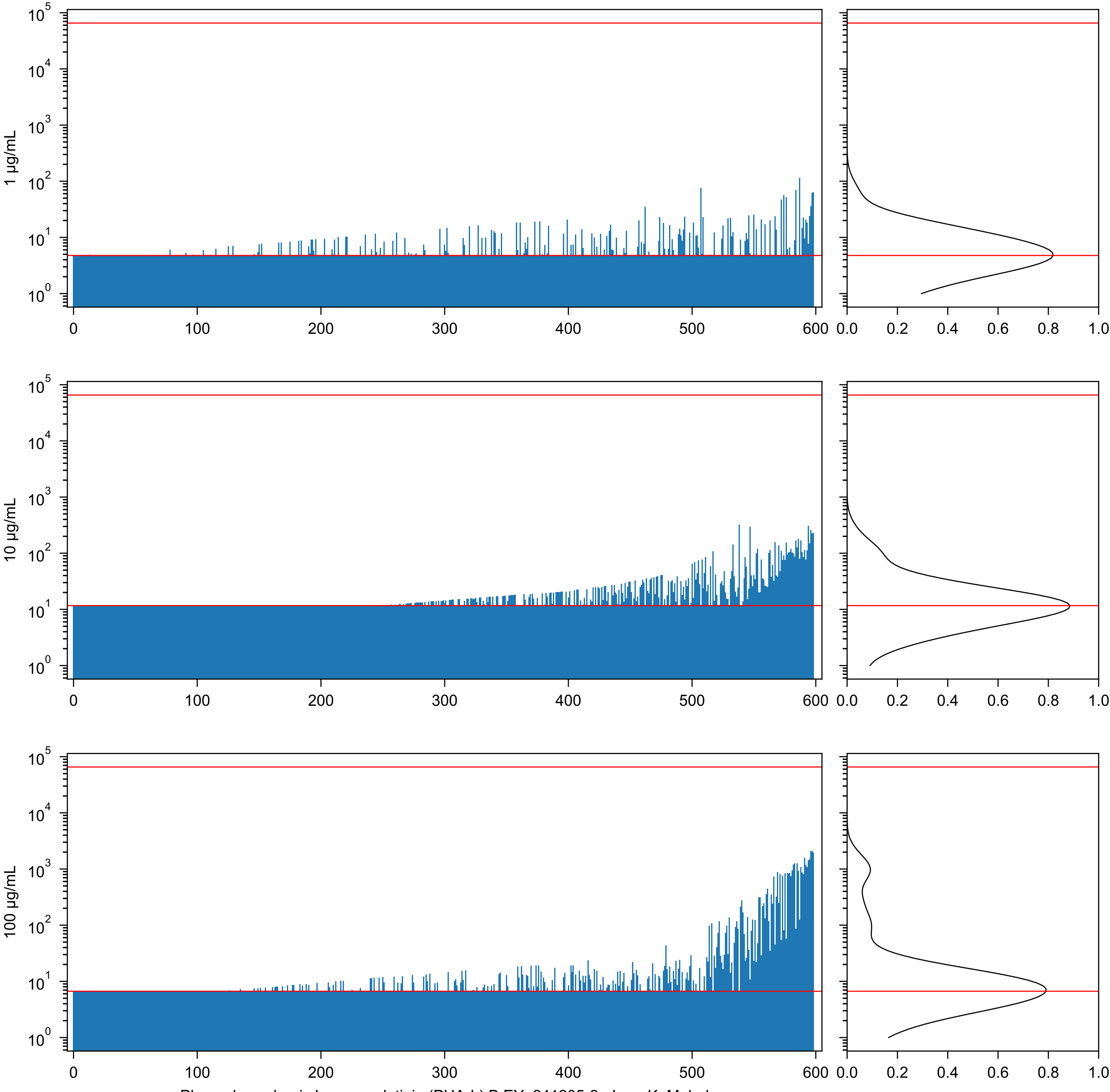


Phaseolus vulgaris Leucoagglutinin (PHA-L) B EY_241205-2 - Lara K. Mahal

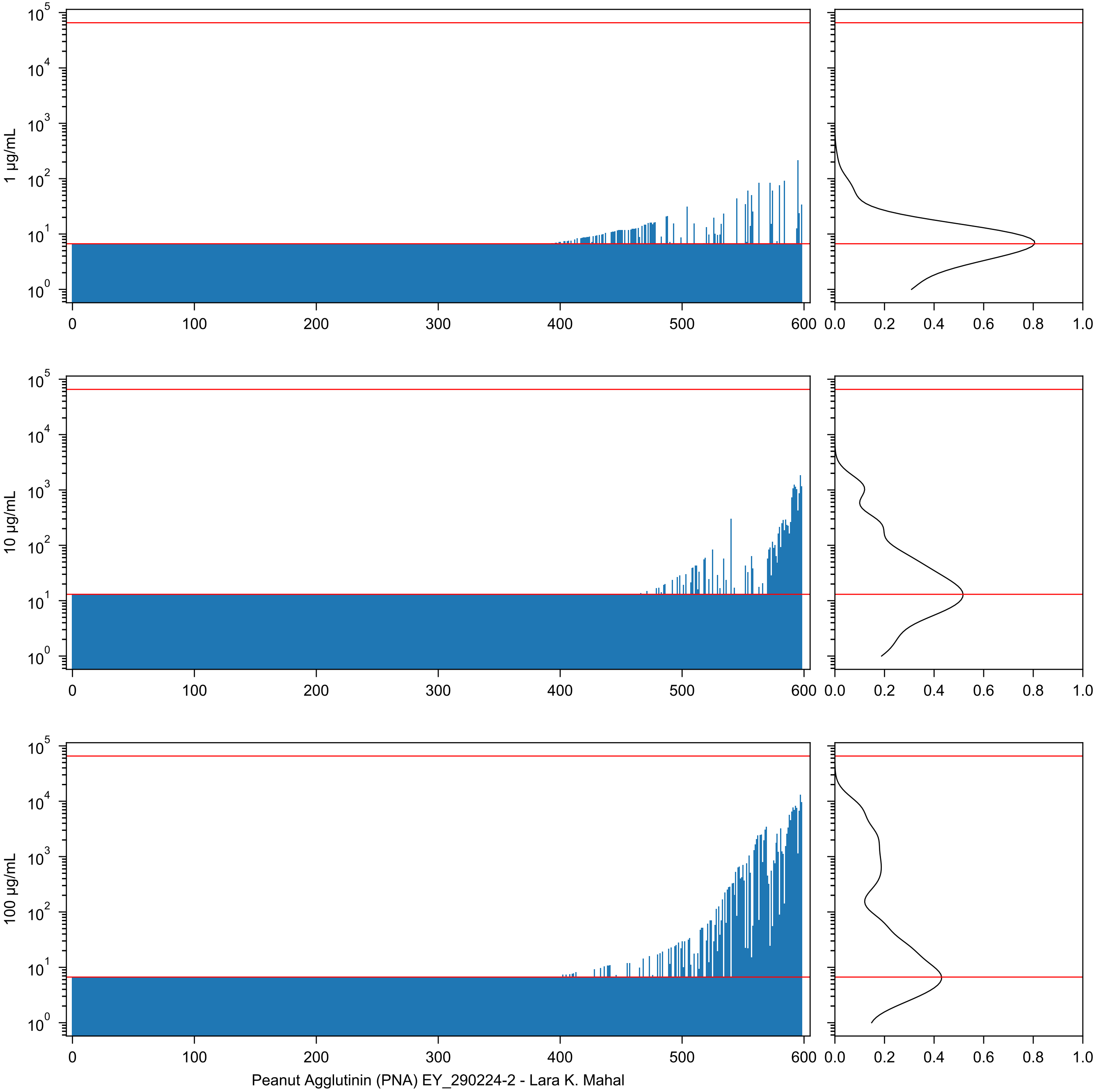

1 µg/mL
10 µg/mL
100 µg/mL
Peanut Agglutinin (PNA) EY_290224-2 - Lara K. Mahal

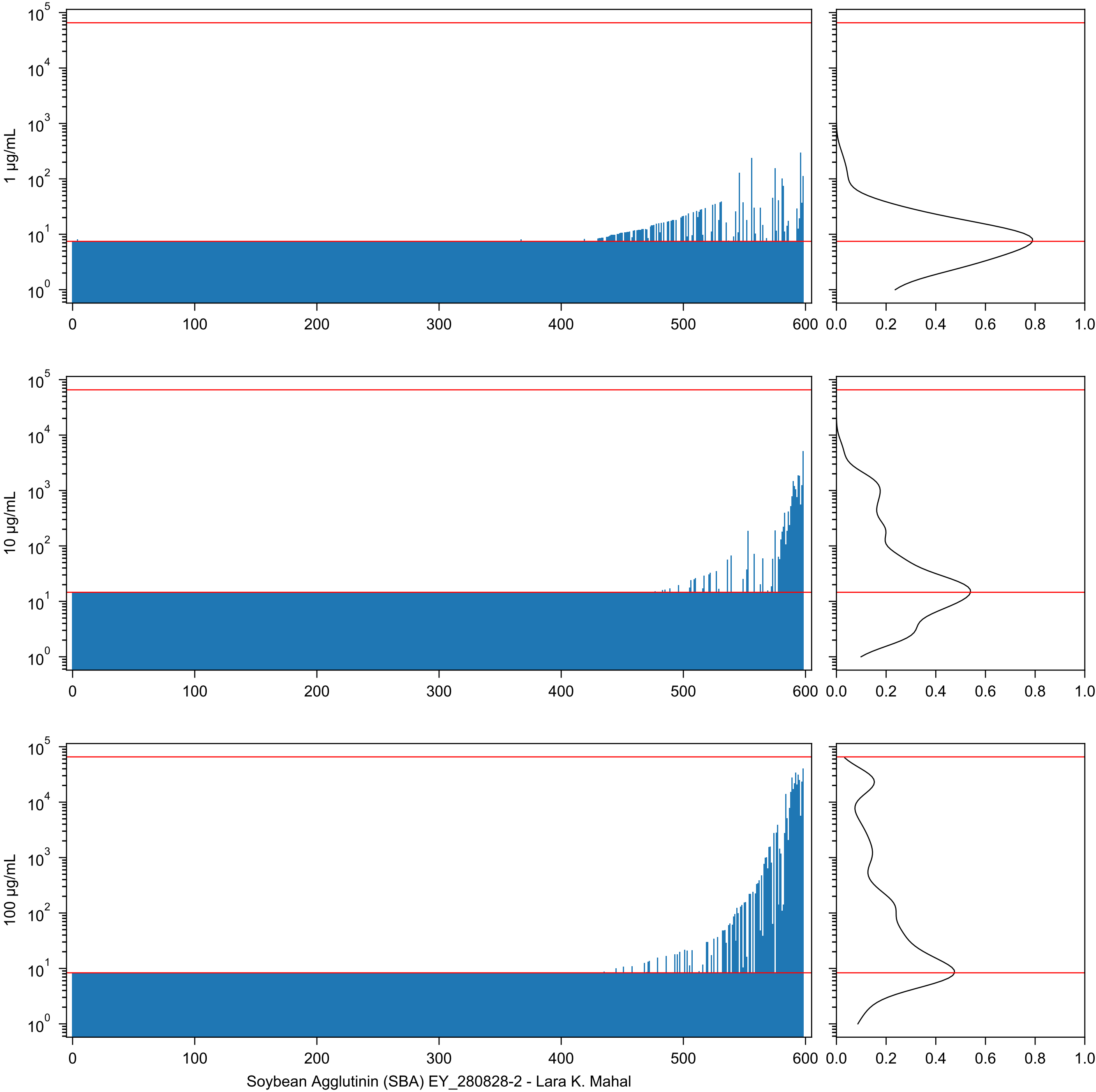

Soybean Agglutinin (SBA) EY_280828-2 - Lara K. Mahal

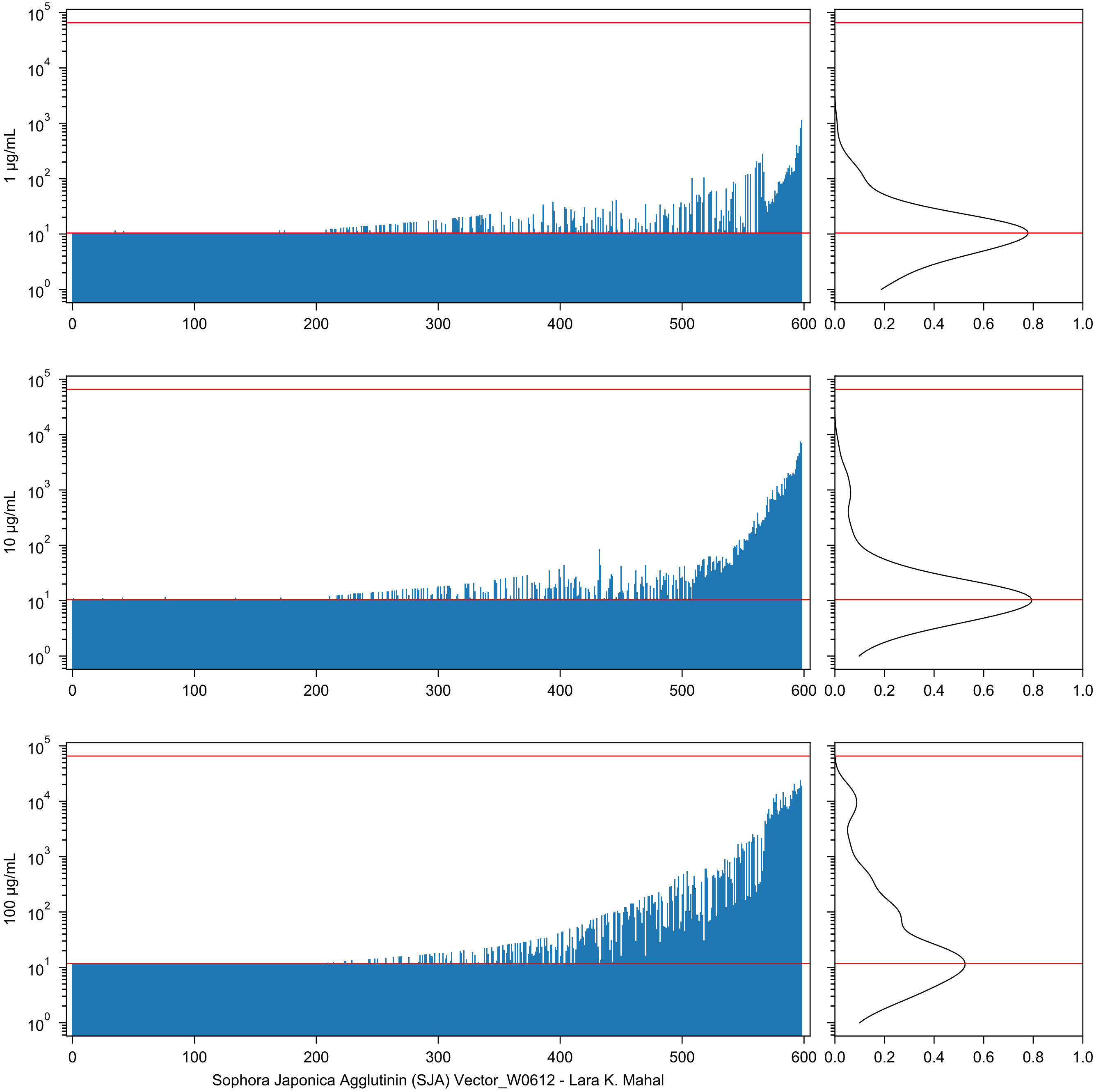
1 µg/mL
10 µg/mL
100 µg/mL
Sophora Japonica Agglutinin (SJA) Vector_W0612 - Lara K. Mahal

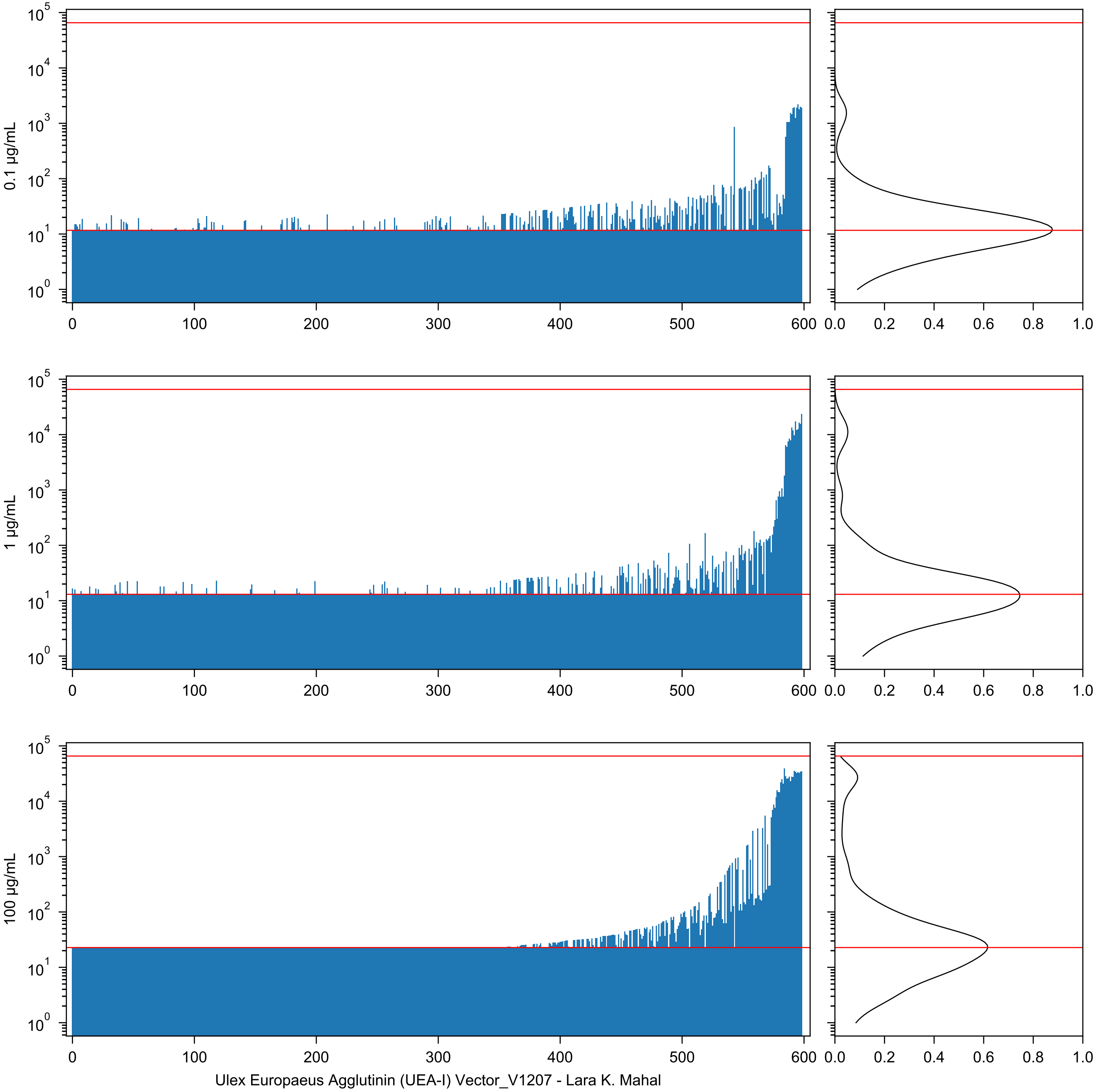
0.1 µg/mL
1 µg/mL
100 µg/mL
Ulex Europaeus Agglutinin (UEA-I) Vector_V1207 - Lara K. Mahal

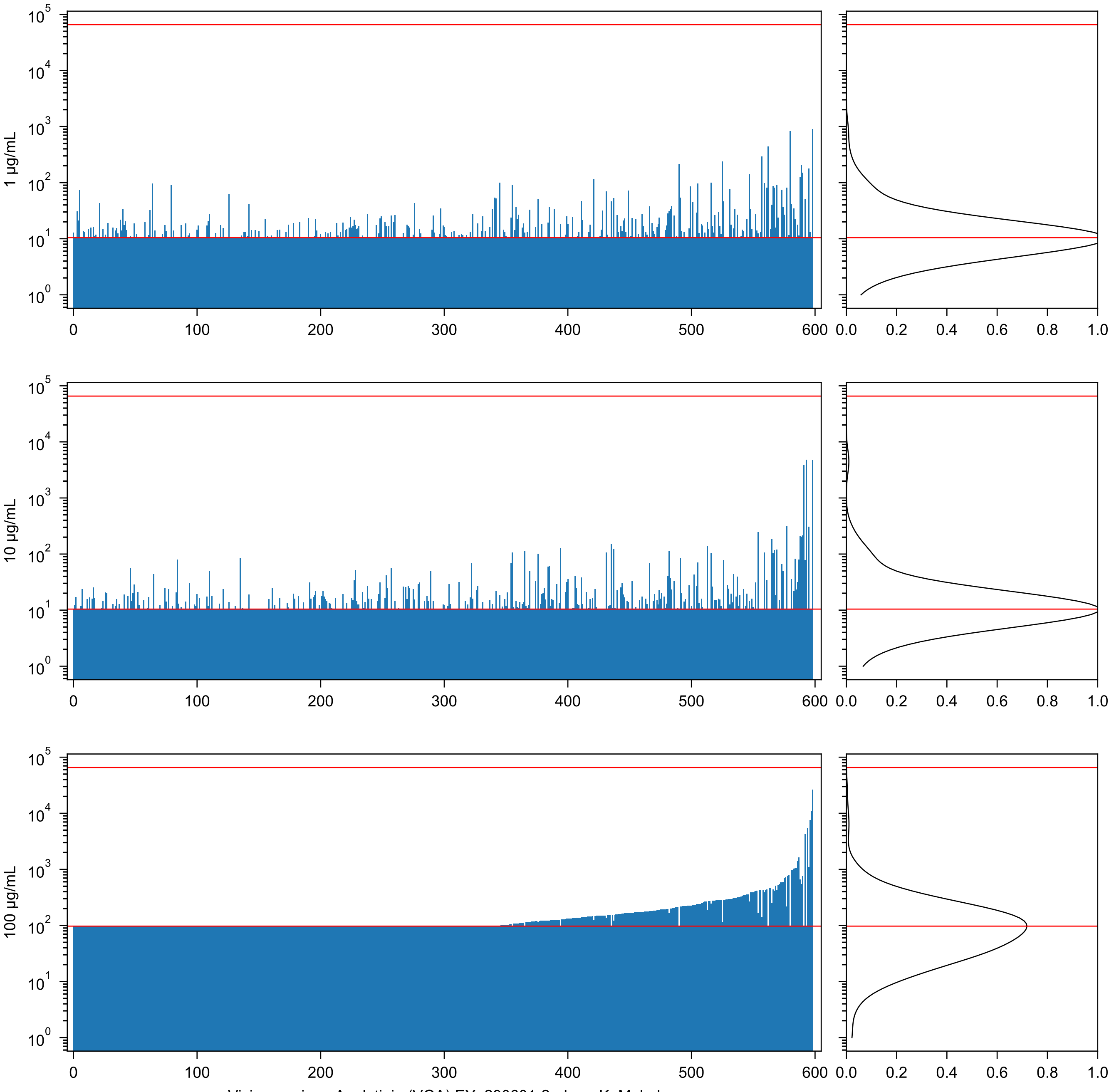
1 µg/mL
10 µg/mL
100 µg/mL
Vicia graminea Agglutinin (VGA) EY_290601-2 - Lara K. Mahal

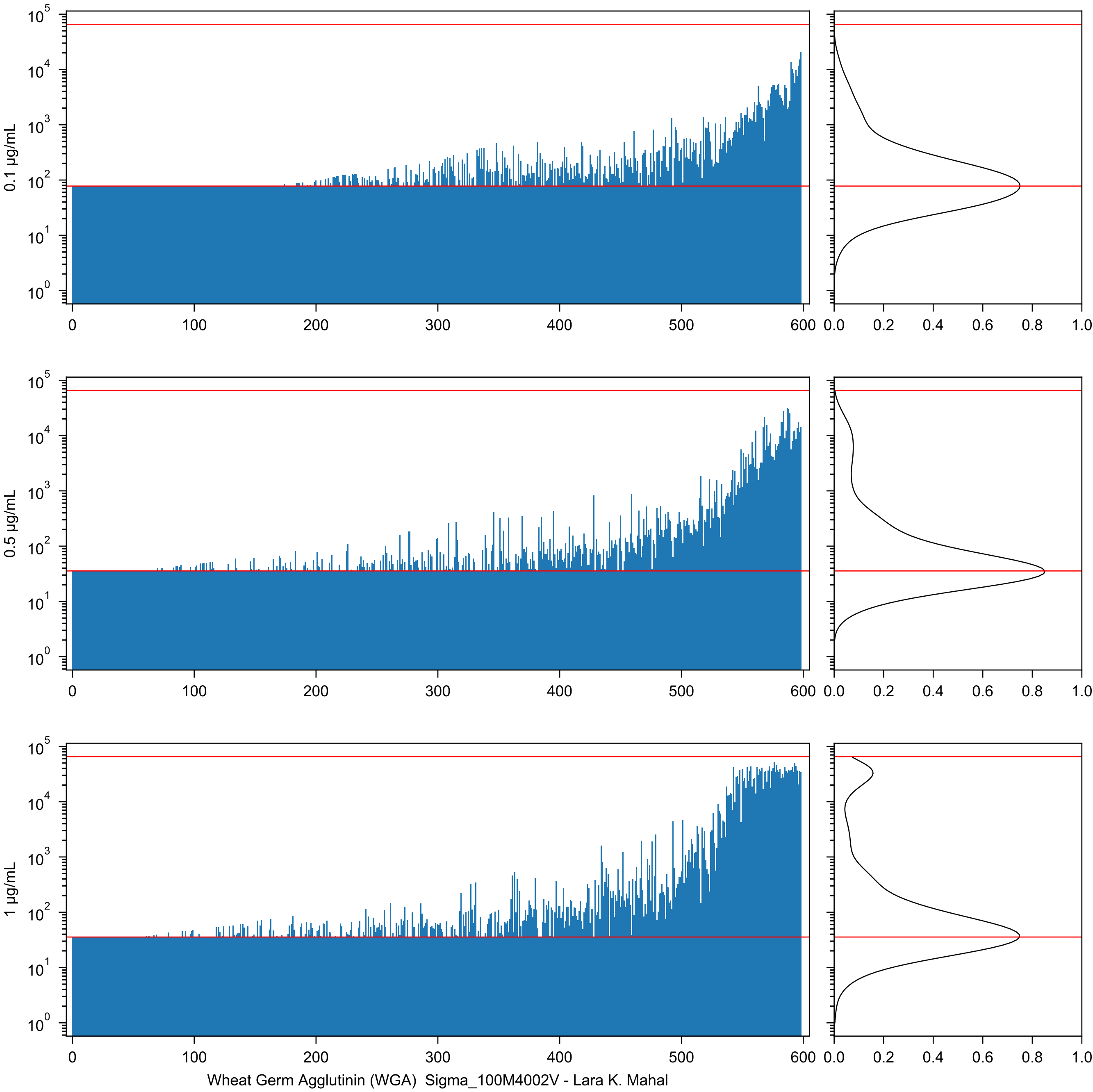

0.1 µg/mL
0.5 µg/mL
1 µg/mL
Wheat Germ Agglutinin (WGA) Sigma_100M4002V - Lara K. Mahal

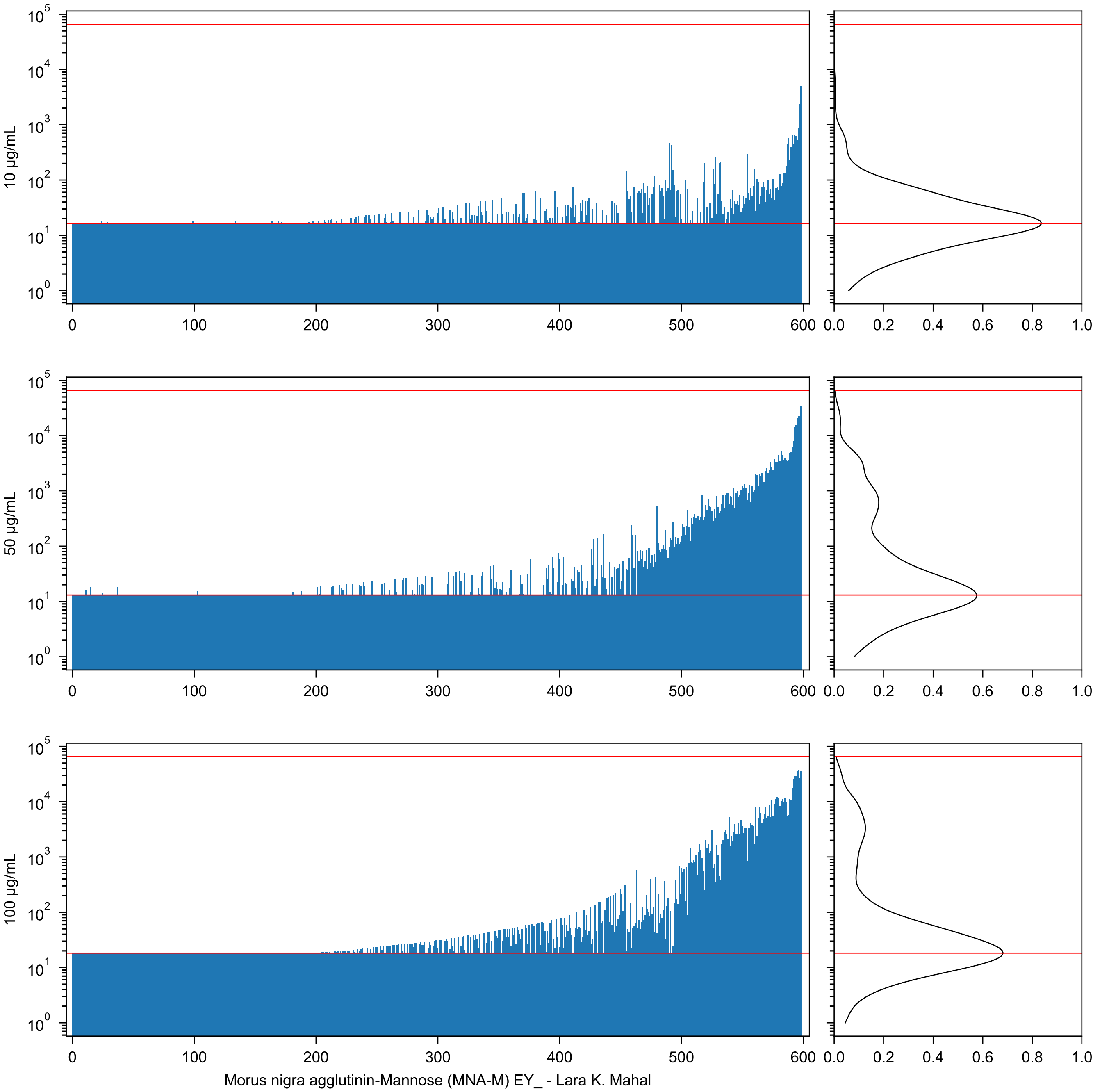

10 µg/mL
50 µg/mL
100 µg/mL
Morus nigra agglutinin-Mannose (MNA-M) EY_ - Lara K. Mahal

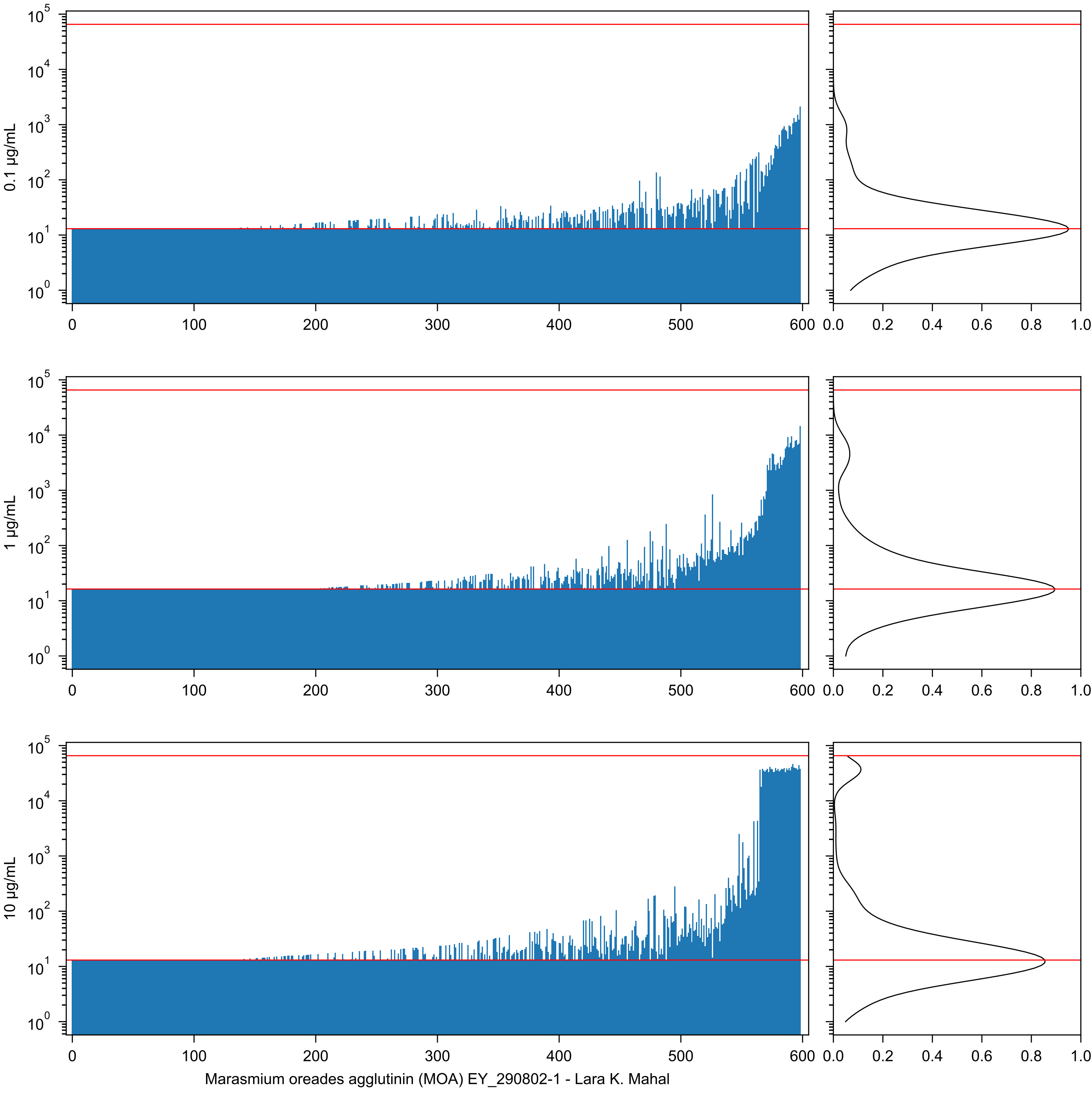

0.1 µg/mL
1 µg/mL
10 µg/mL
Marasmium oreades agglutinin (MOA) EY_290802-1 - Lara K. Mahal

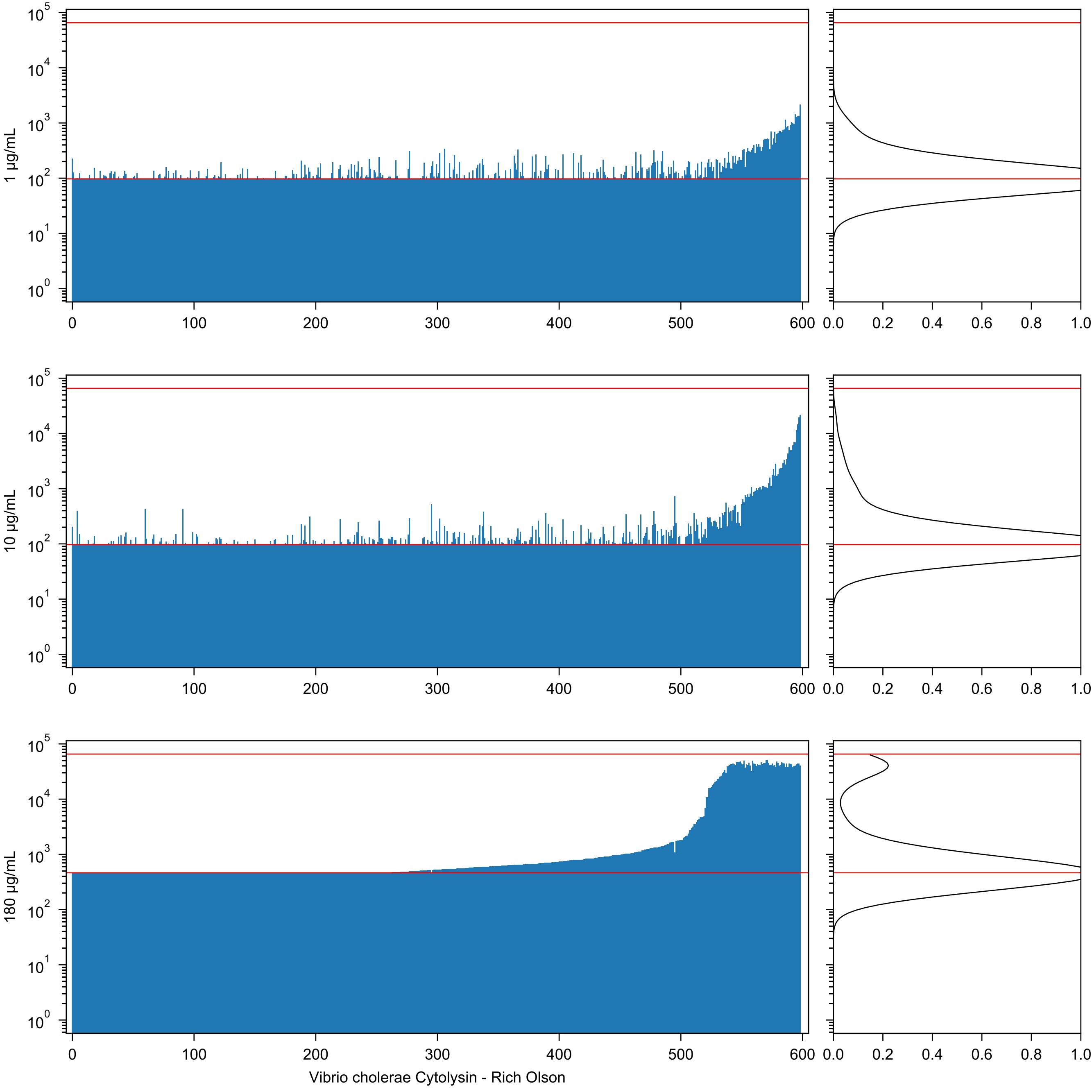

1 μg/mL
10 μg/mL
180 μg/mL
Vibrio cholerae Cytolysin - Rich Olson

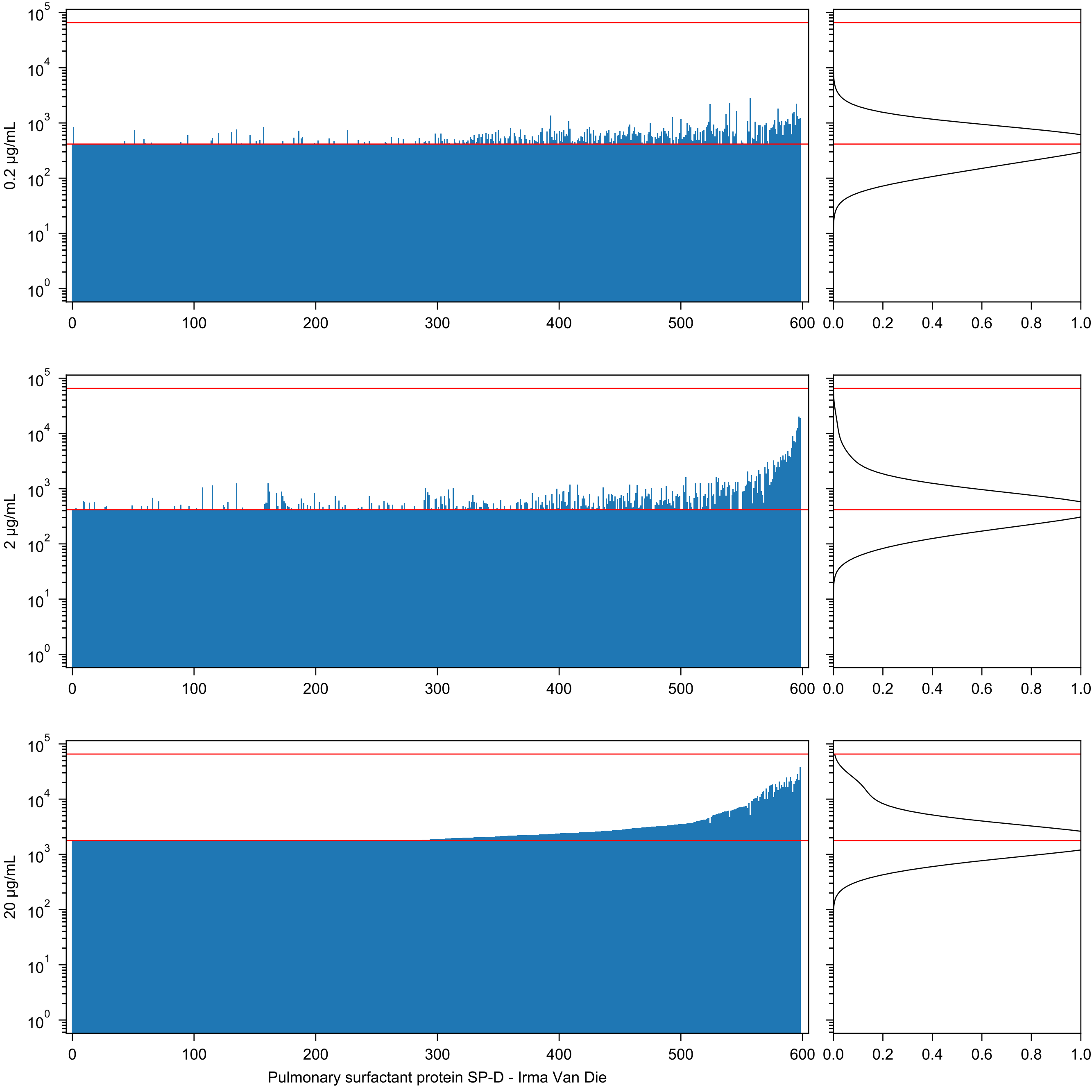

0.2 µg/mL
2 µg/mL
20 µg/mL
Pulmonary surfactant protein SP-D - Irma Van Die

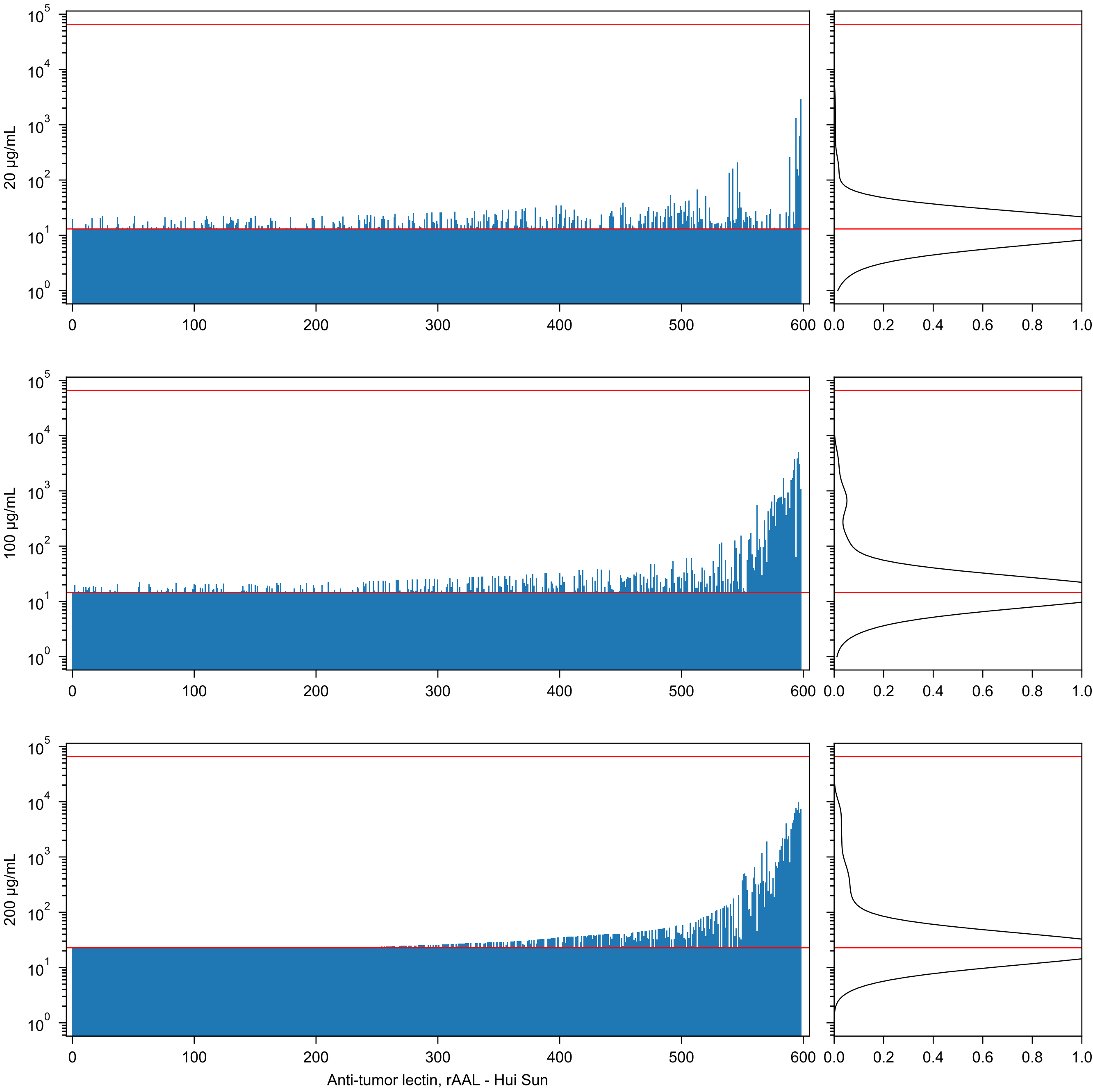

20 µg/mL
100 µg/mL
200 µg/mL
Anti-tumor lectin, rAAL - Hui Sun

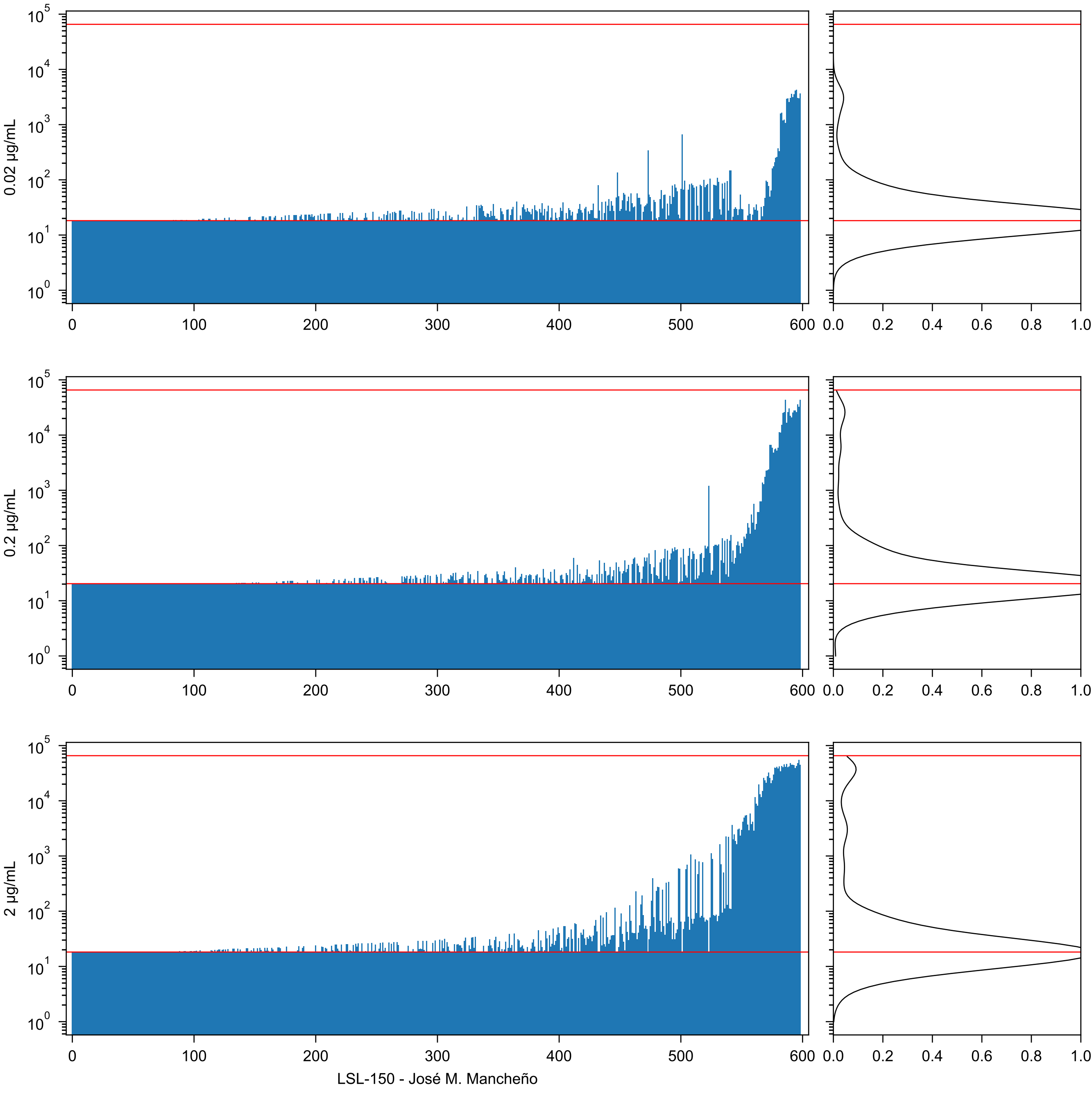
0.02 µg/mL
0.2 µg/mL
2 µg/mL
LSL-150 - José M. Mancheño

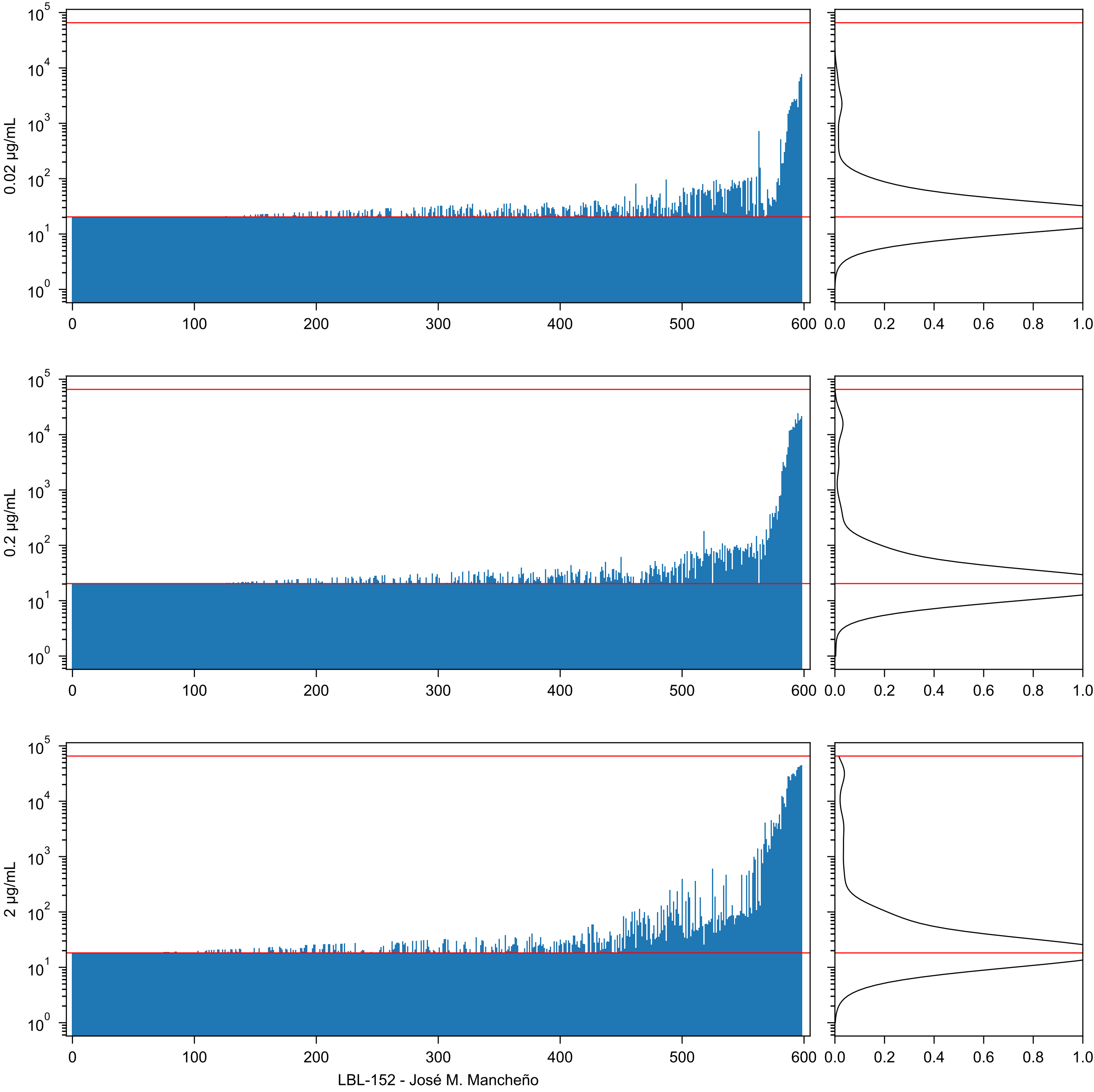
0.02 µg/mL
0.2 µg/mL
2 µg/mL
LBL-152 - José M. Mancheño

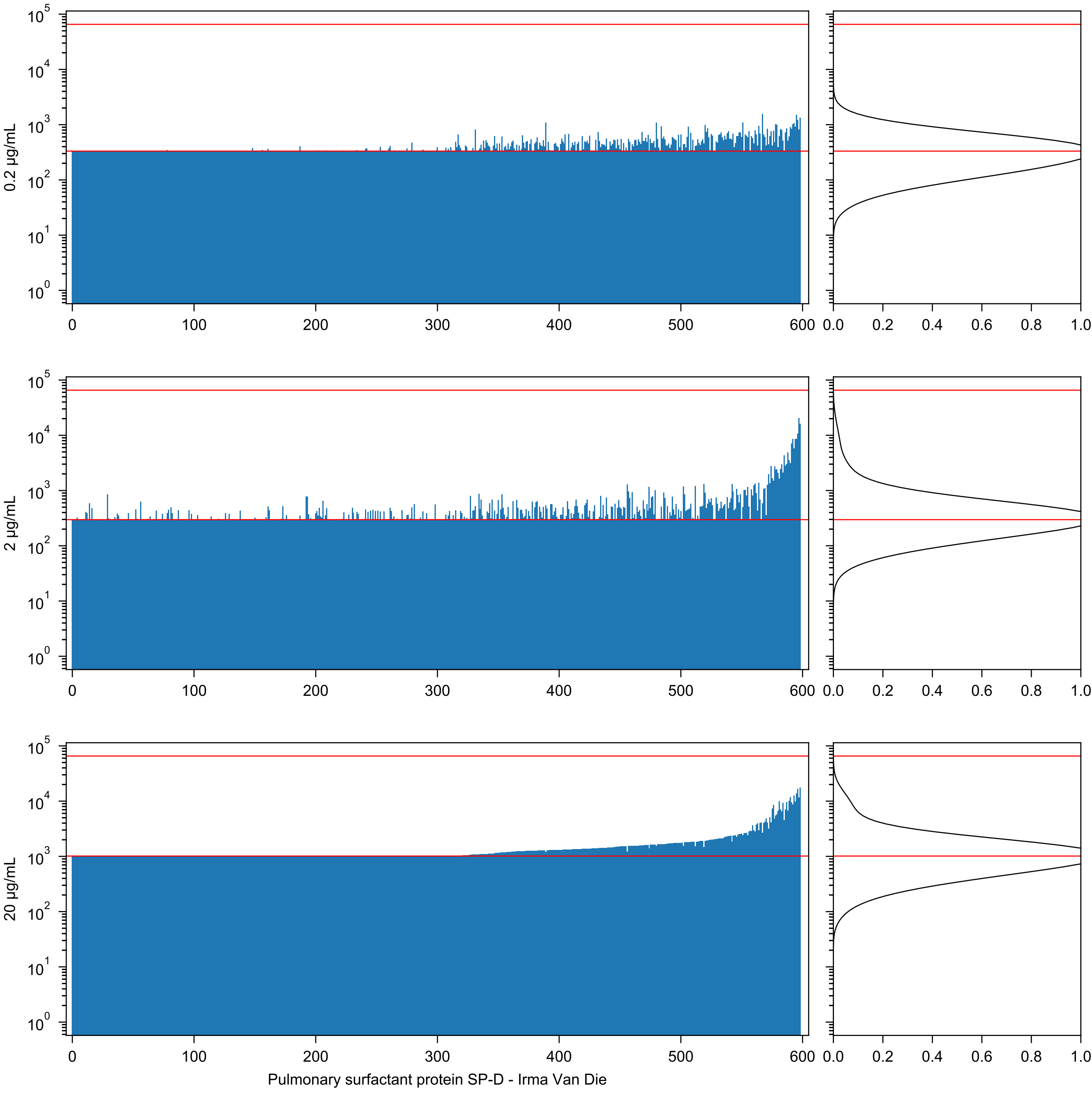
0.2 µg/mL
2 µg/mL
20 µg/mL
Pulmonary surfactant protein SP-D - Irma Van Die

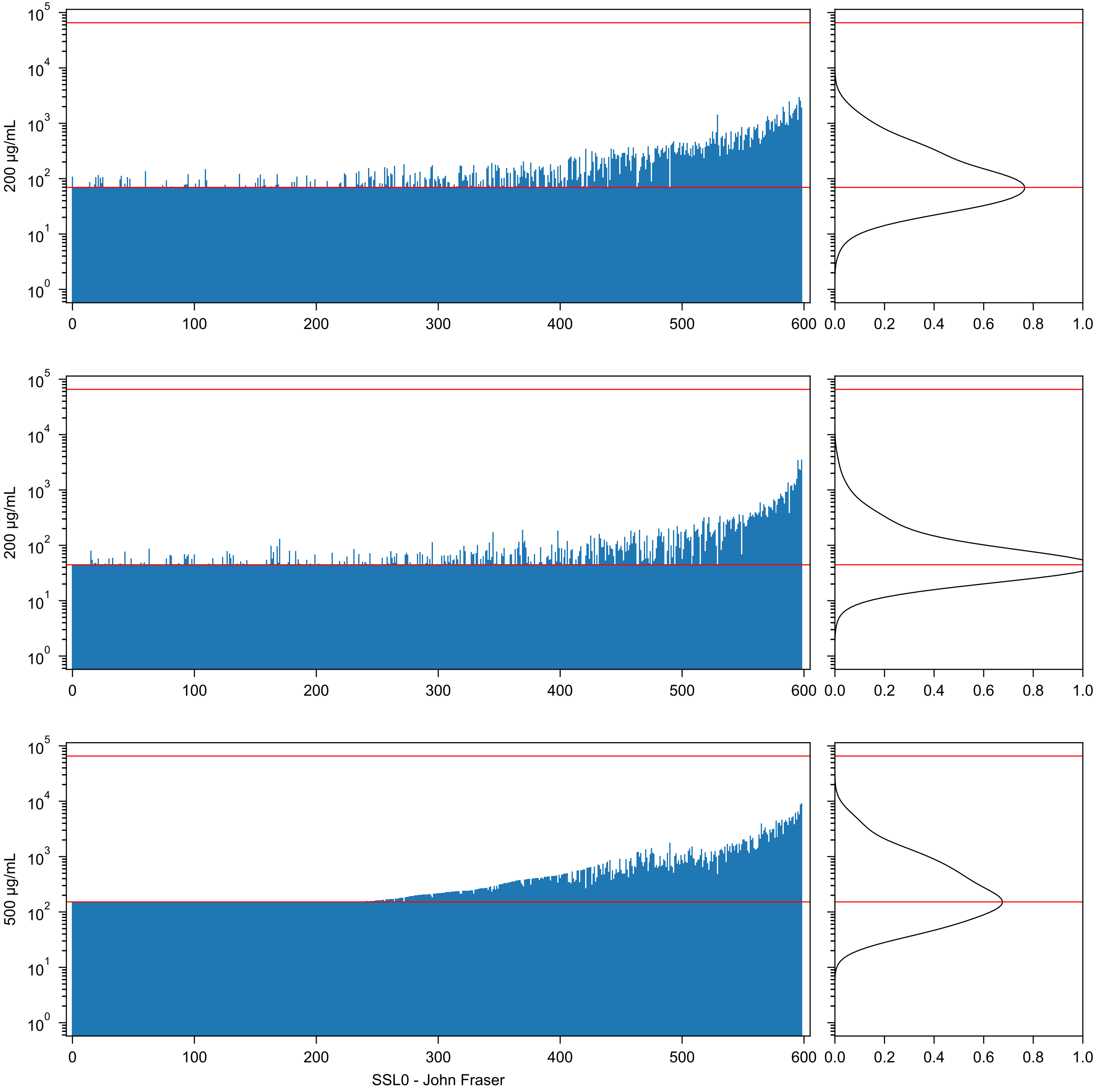
200 µg/mL
200 µg/mL
500 µg/mL
SSL0 - John Fraser

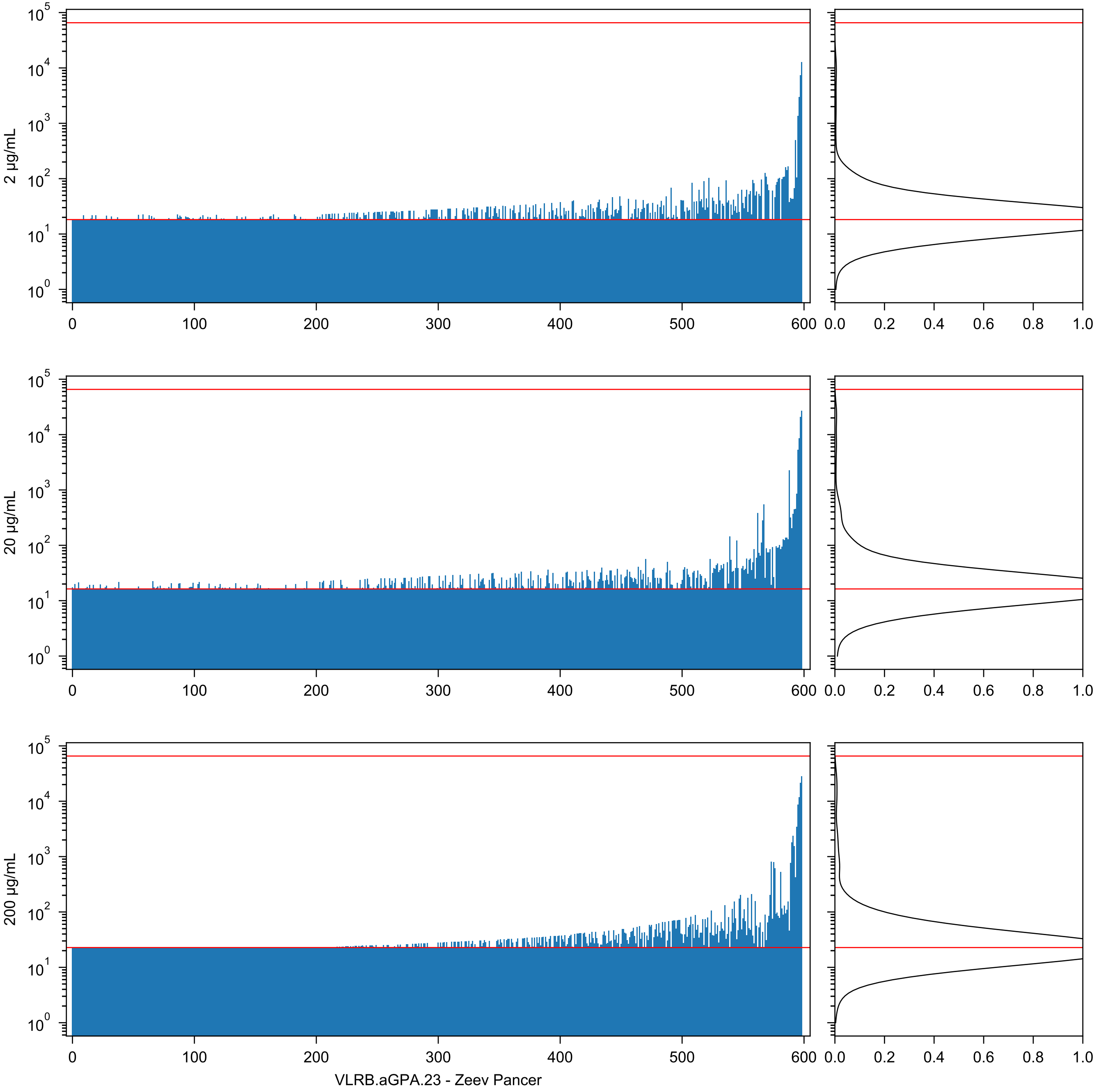
2 µg/mL
20 µg/mL
200 µg/mL
VLRB.aGPA.23 - Zeev Pancer

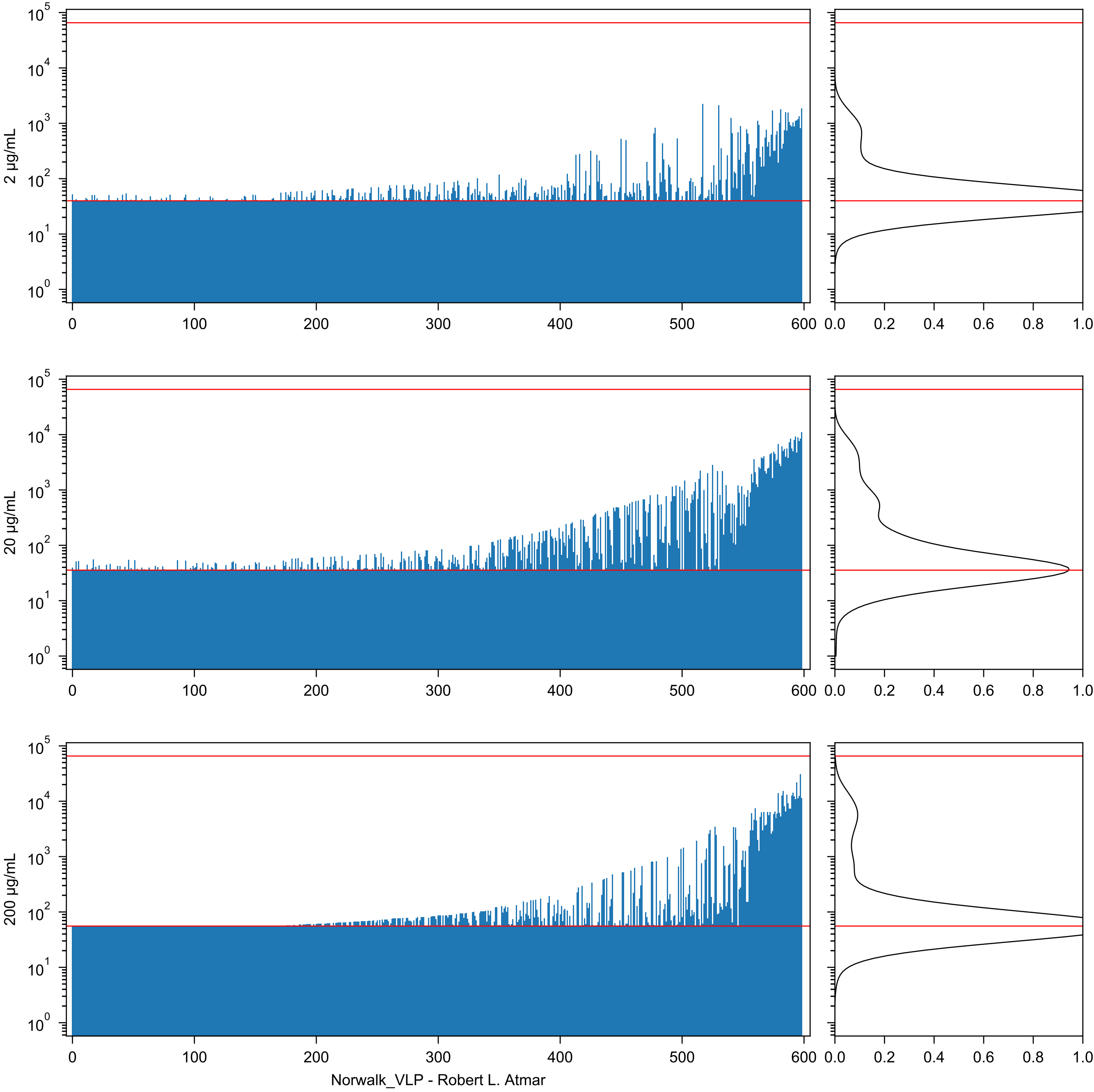

2 µg/mL
20 µg/mL
200 µg/mL
$10^0$
$10^1$
$10^2$
$10^3$
$10^4$
$10^5$
0
100
200
300
400
500
600
0.0
0.2
0.4
0.6
0.8
1.0
Norwalk_VLP - Robert L. Atmar

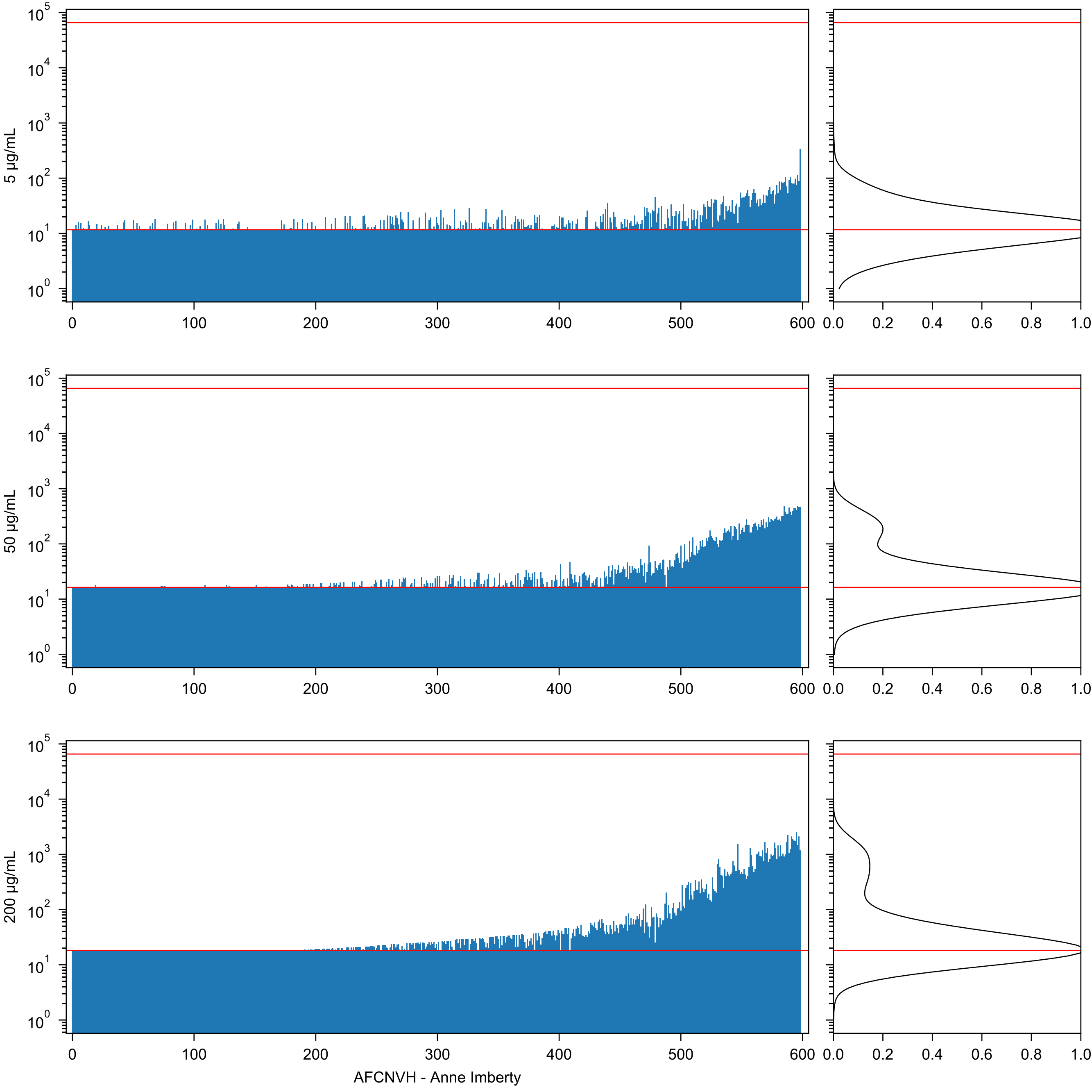
5 µg/mL
50 µg/mL
200 µg/mL
AFCNVH - Anne Imberty

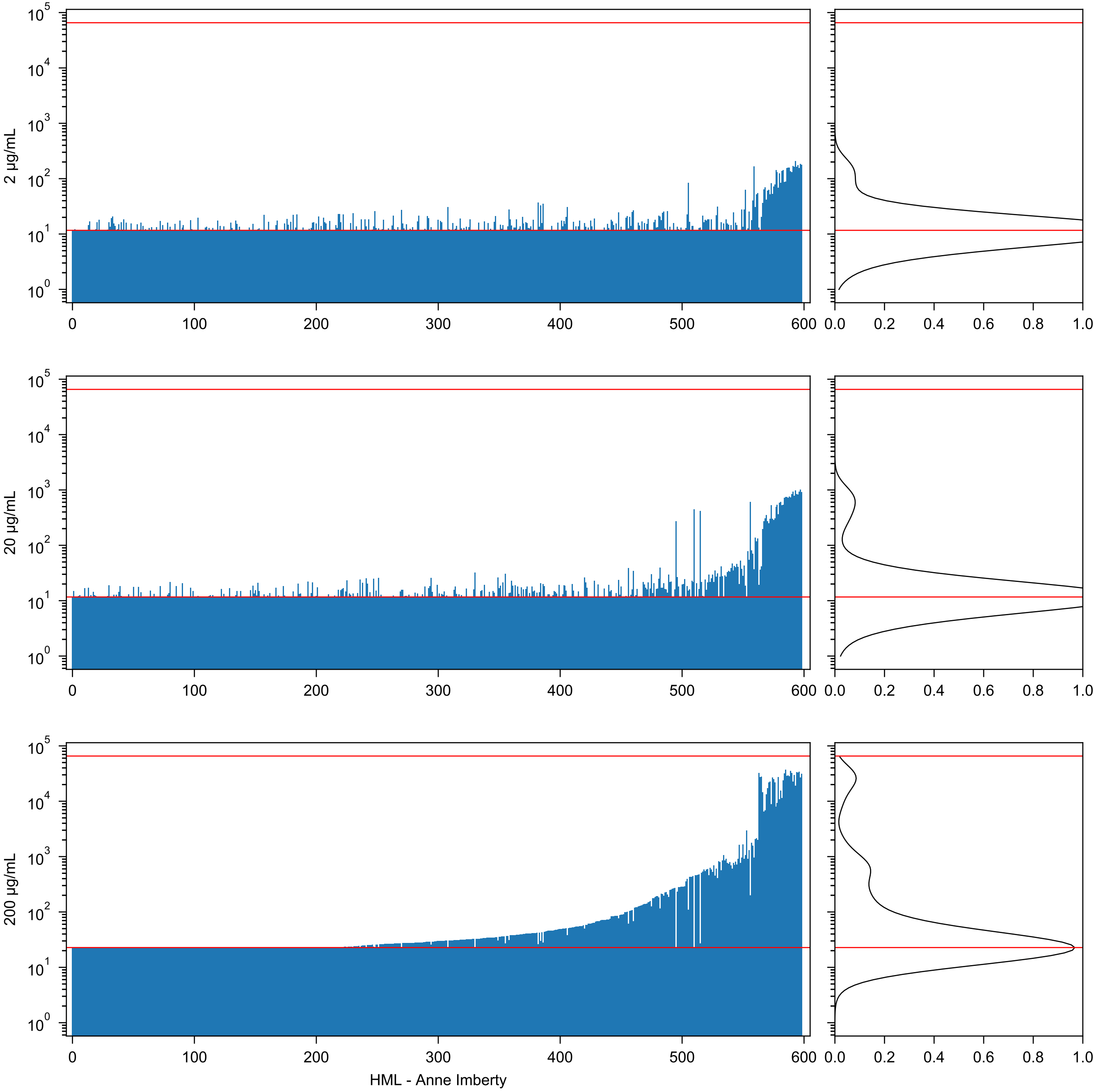

2 µg/mL
20 µg/mL
200 µg/mL
HML - Anne Imberty

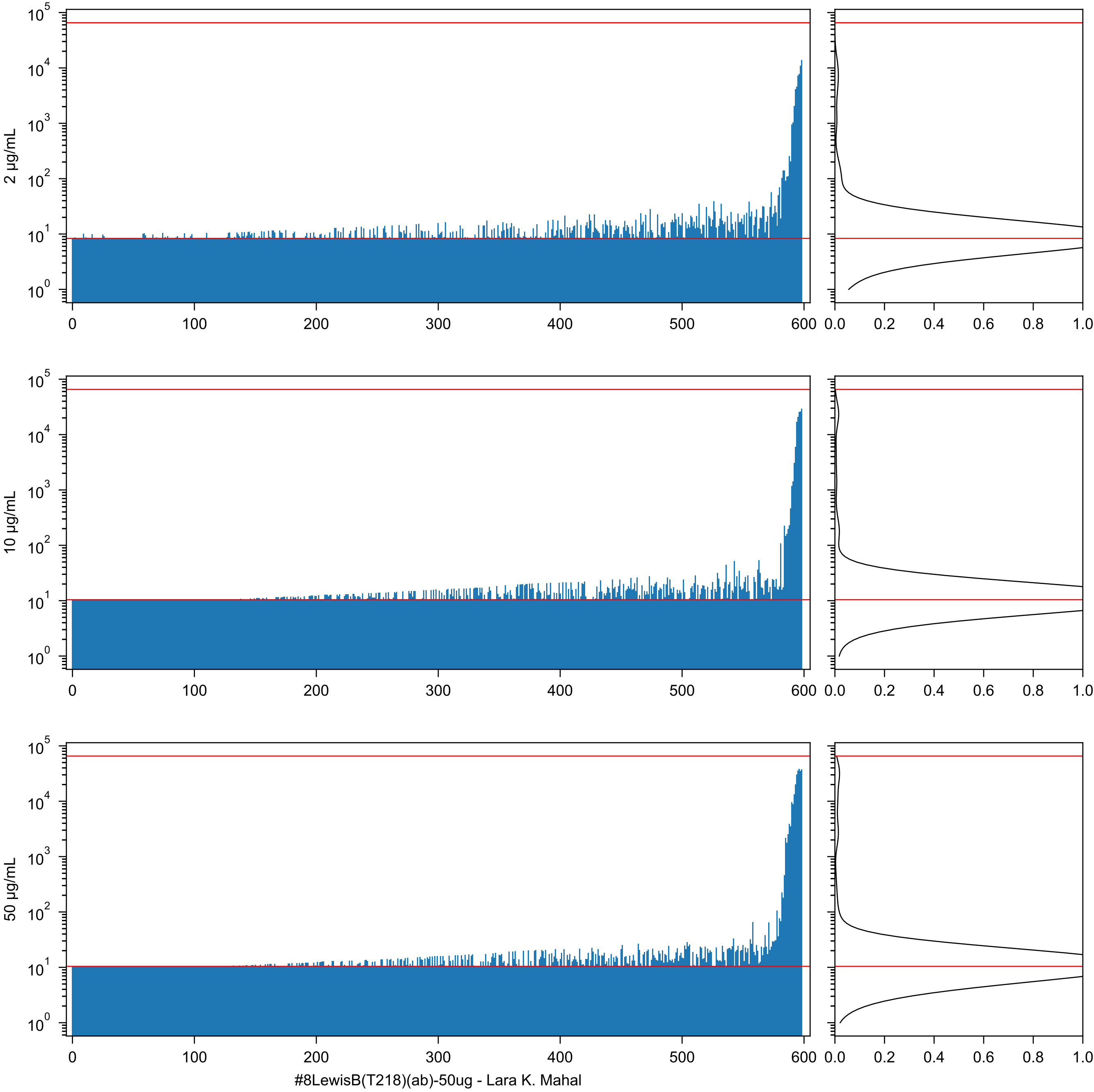
2 µg/mL
10 µg/mL
50 µg/mL
#8LewisB(T218)(ab)-50ug - Lara K. Mahal

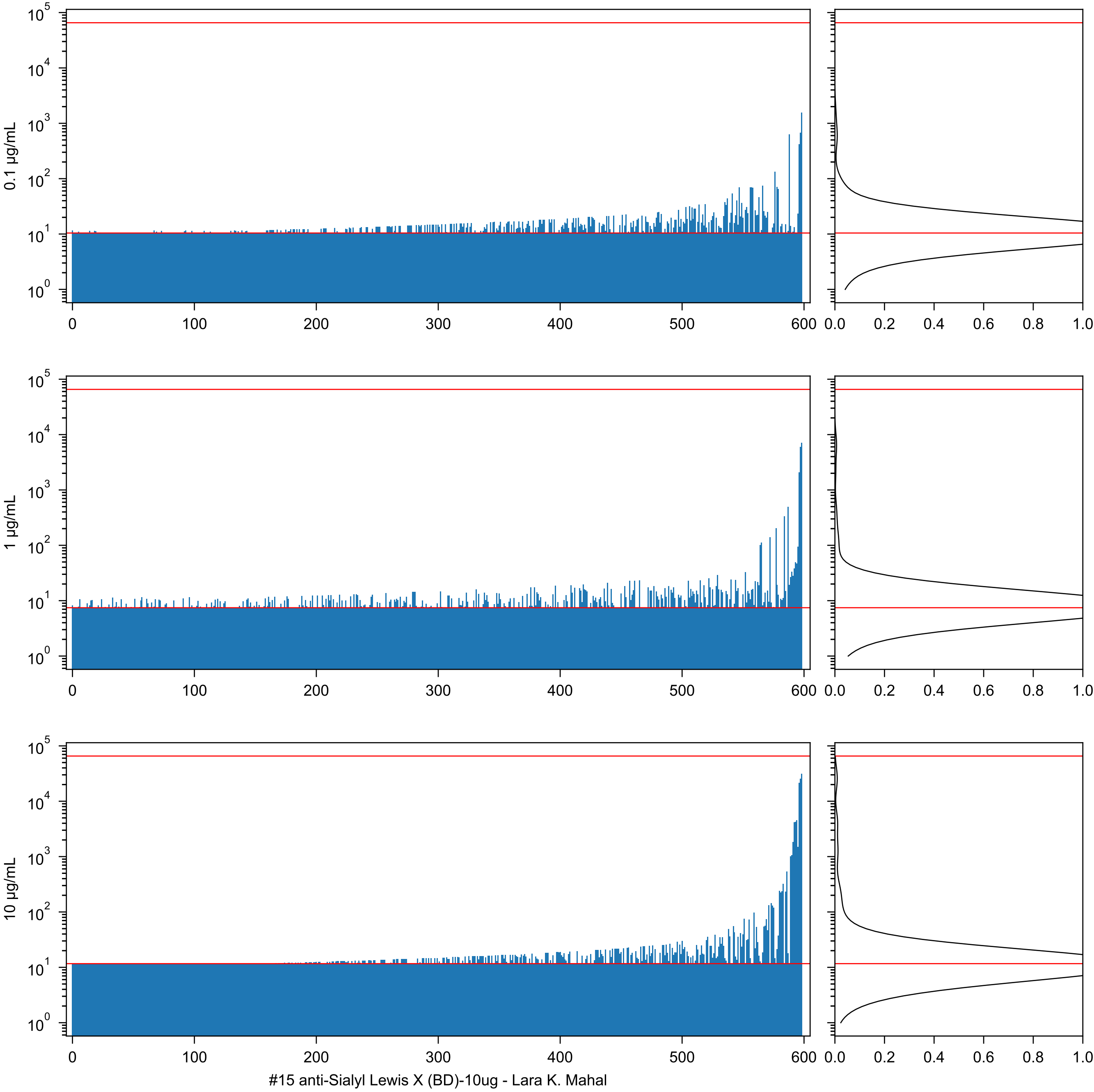
0.1 µg/mL
1 µg/mL
10 µg/mL
#15 anti-Sialyl Lewis X (BD)-10ug - Lara K. Mahal

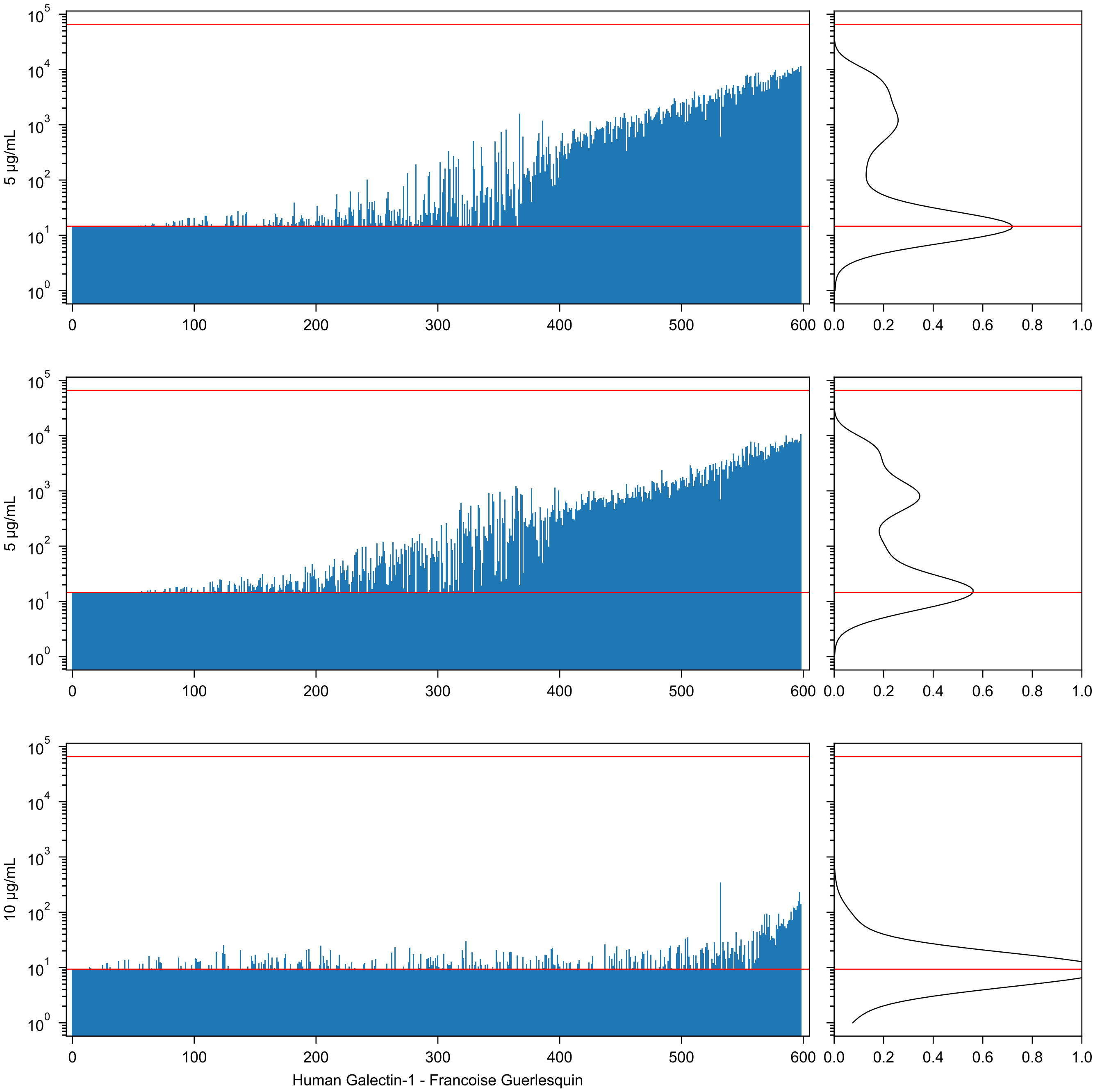

5 µg/mL
5 µg/mL
10 µg/mL
Human Galectin-1 - Francoise Guerlesquin

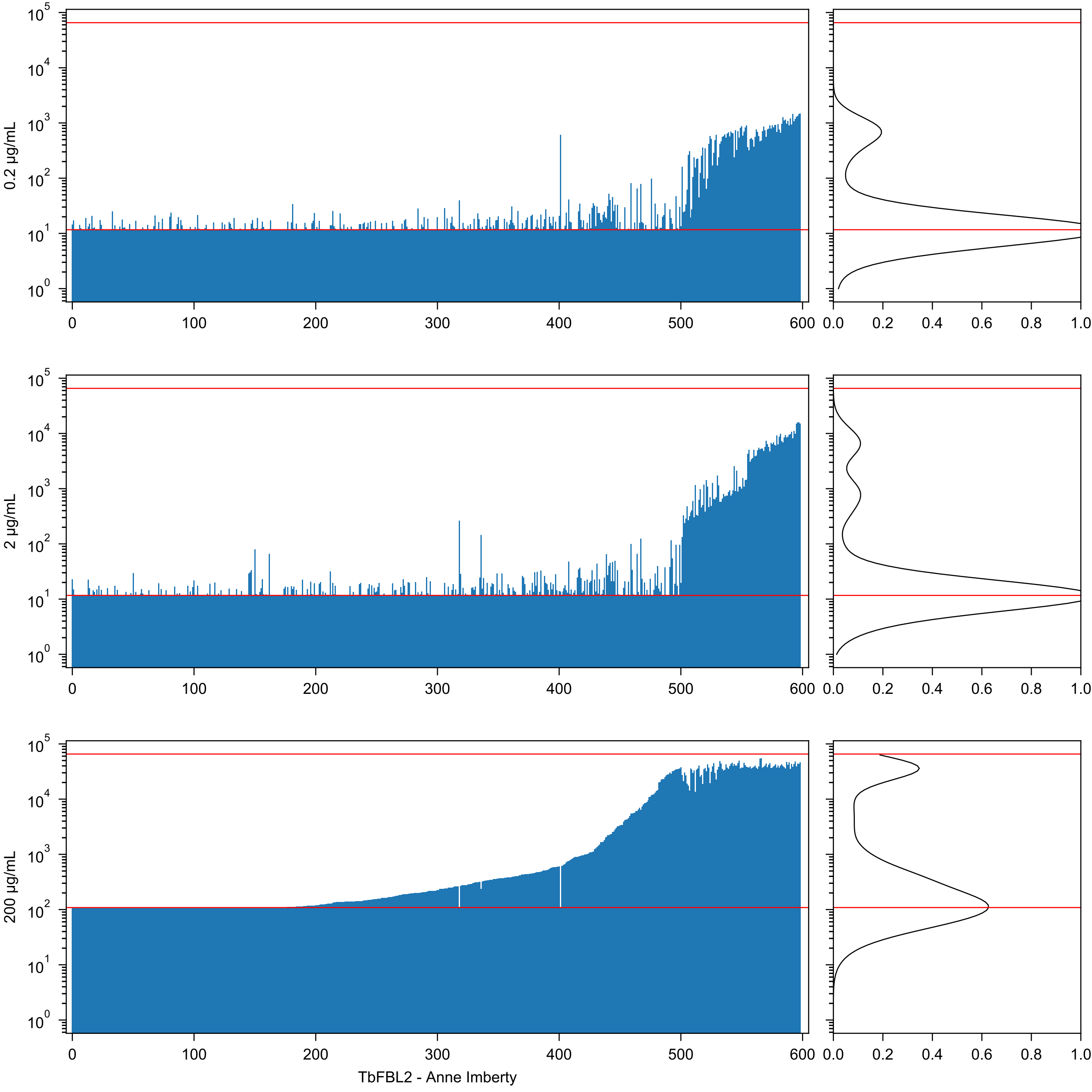
0.2 µg/mL
2 µg/mL
200 µg/mL
TbFBL2 - Anne Imberty

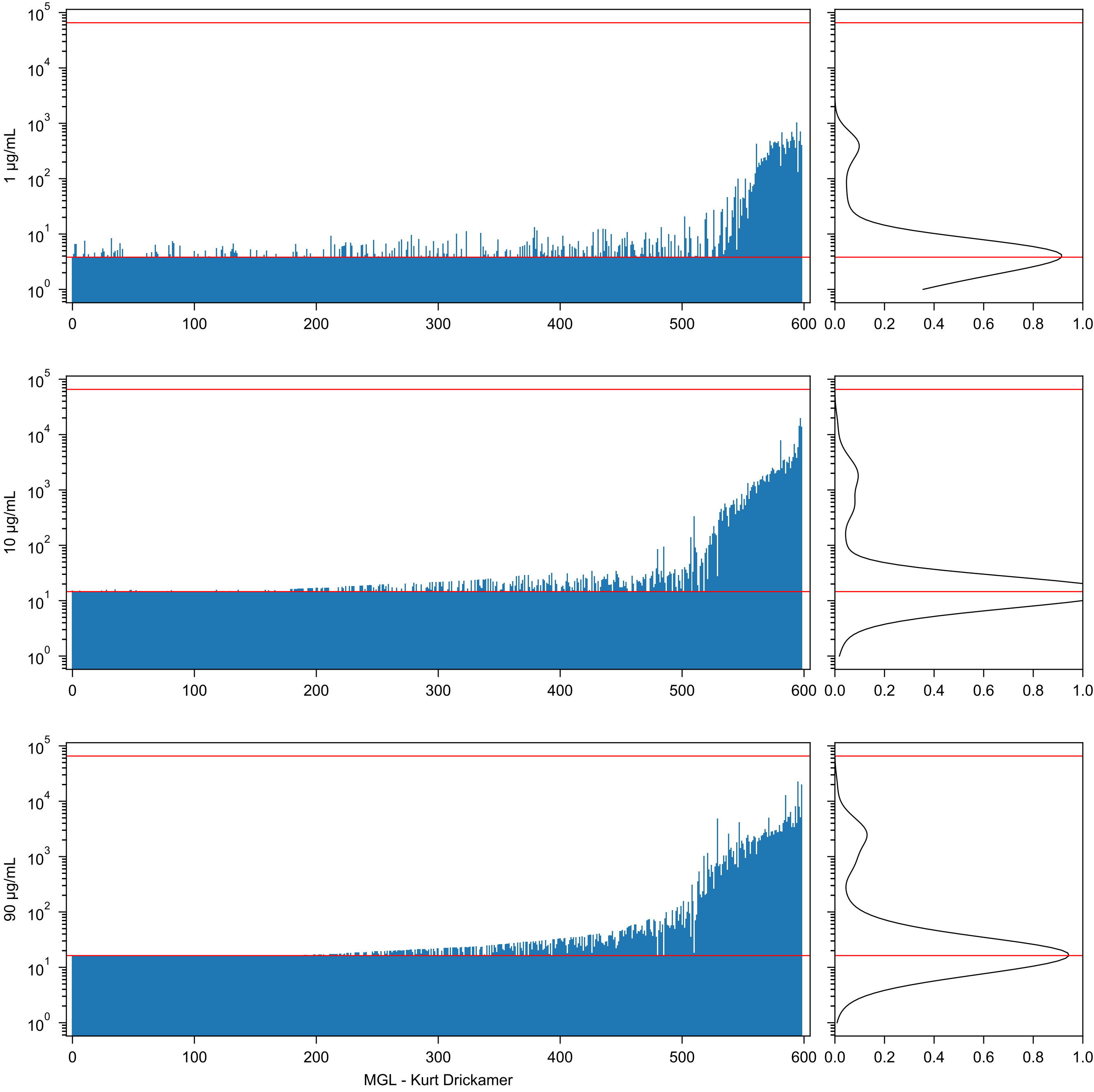

1 μg/mL
10 μg/mL
90 μg/mL
MGL - Kurt Drickamer

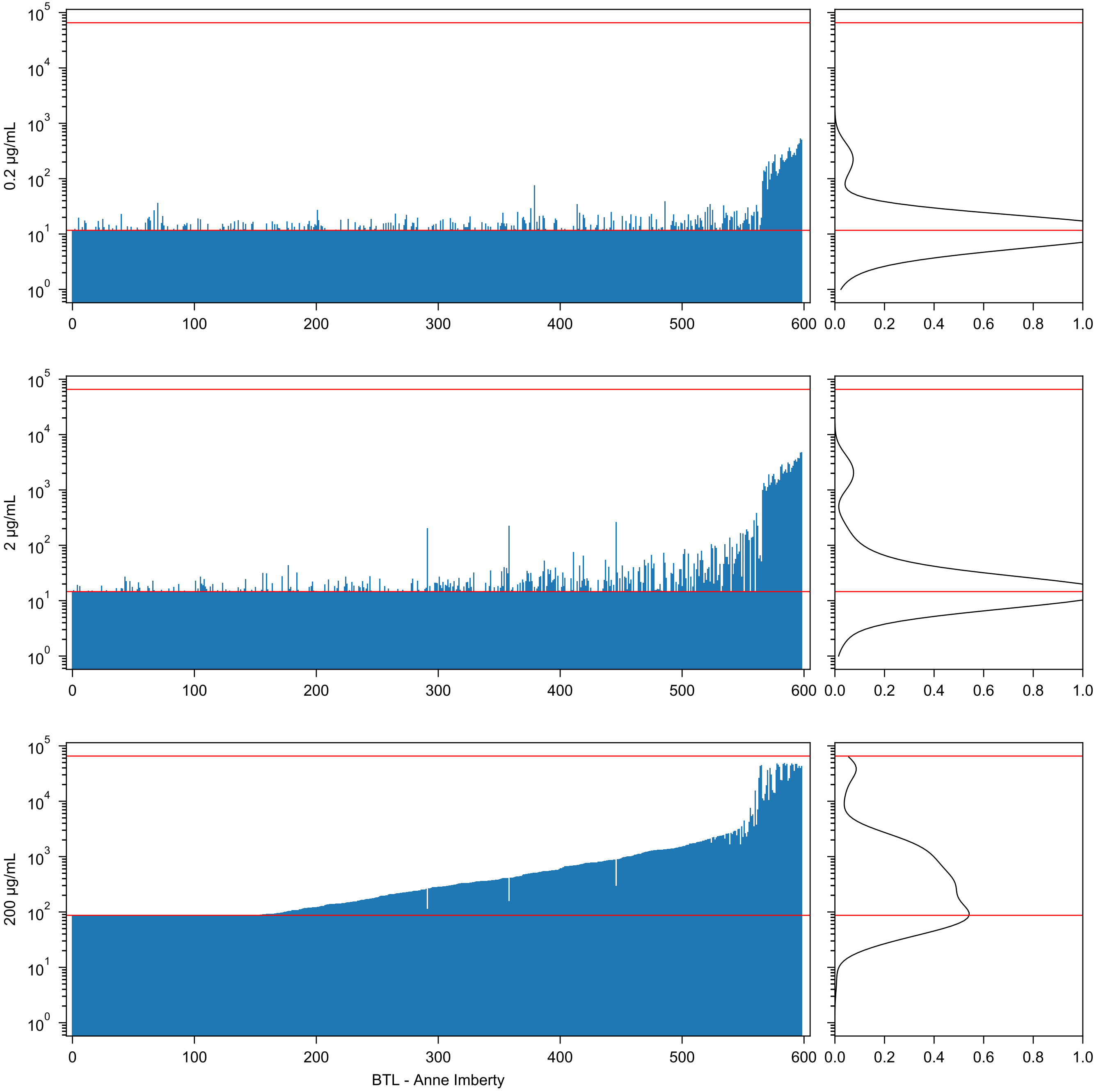

0.2 μg/mL
2 μg/mL
200 μg/mL
BTL - Anne Imberty

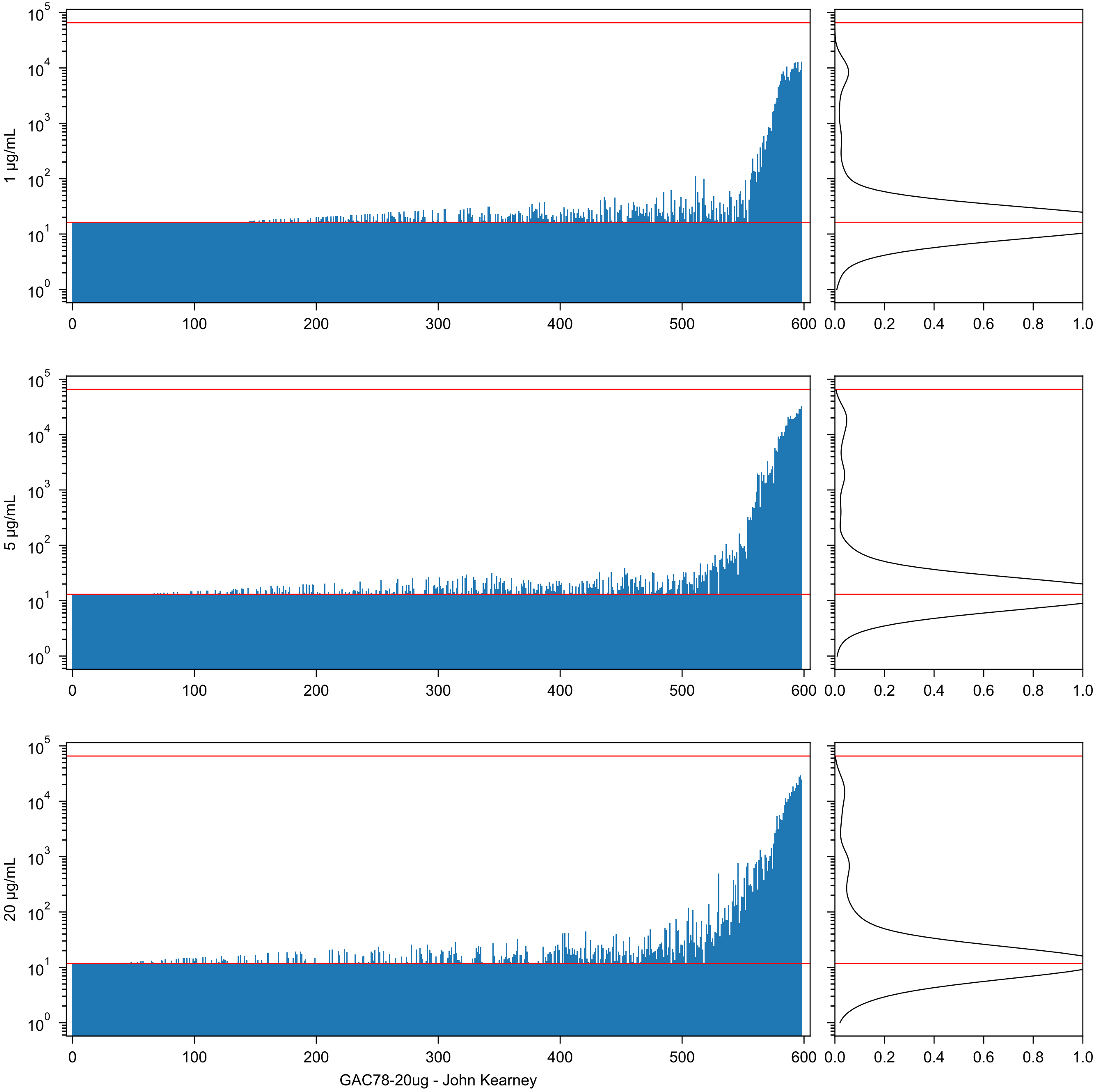
1 μg/mL
5 μg/mL
20 μg/mL
GAC78-20ug - John Kearney

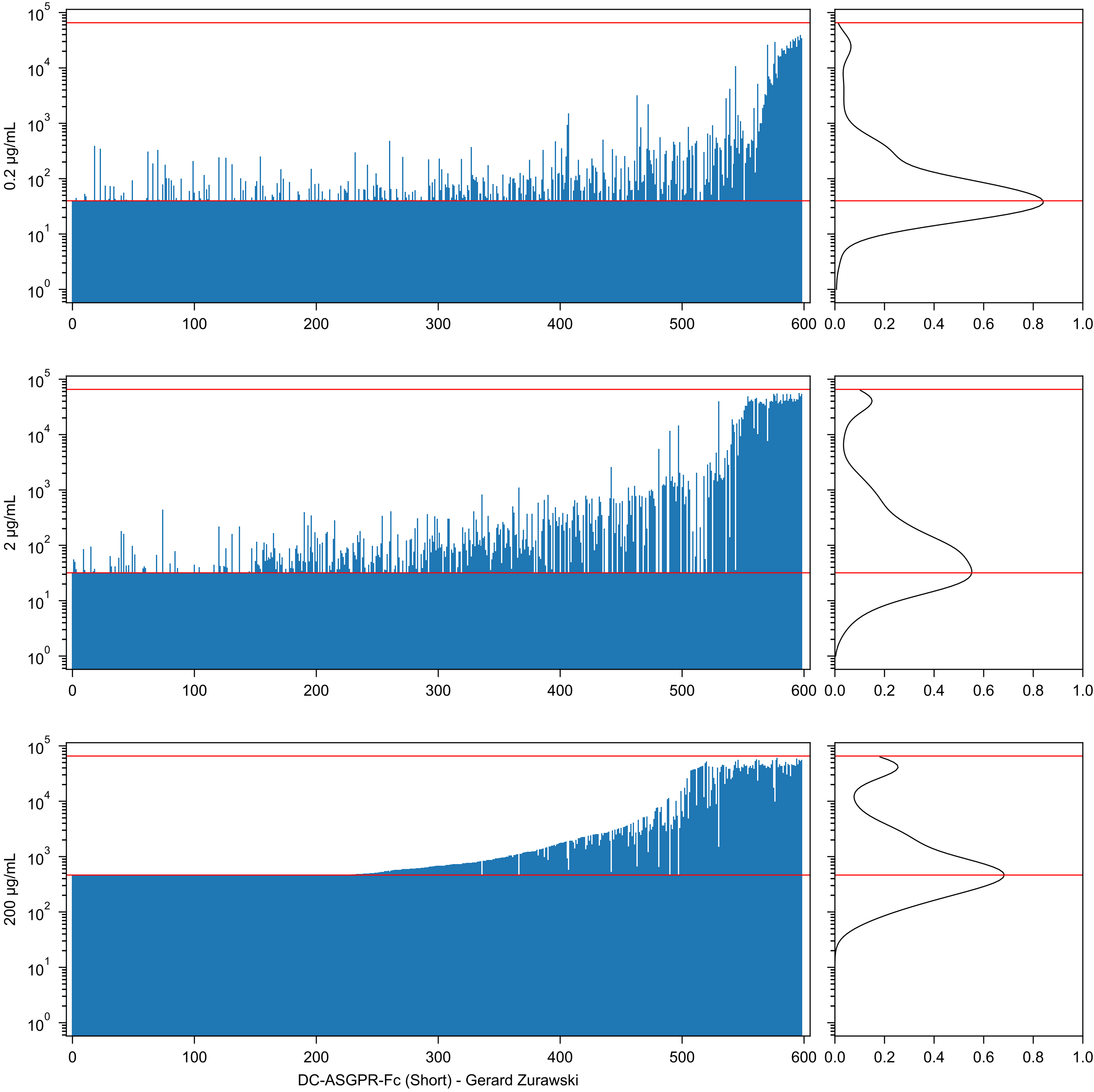
0.2 µg/mL
2 µg/mL
200 µg/mL
DC-ASGPR-Fc (Short) - Gerard Zurawski

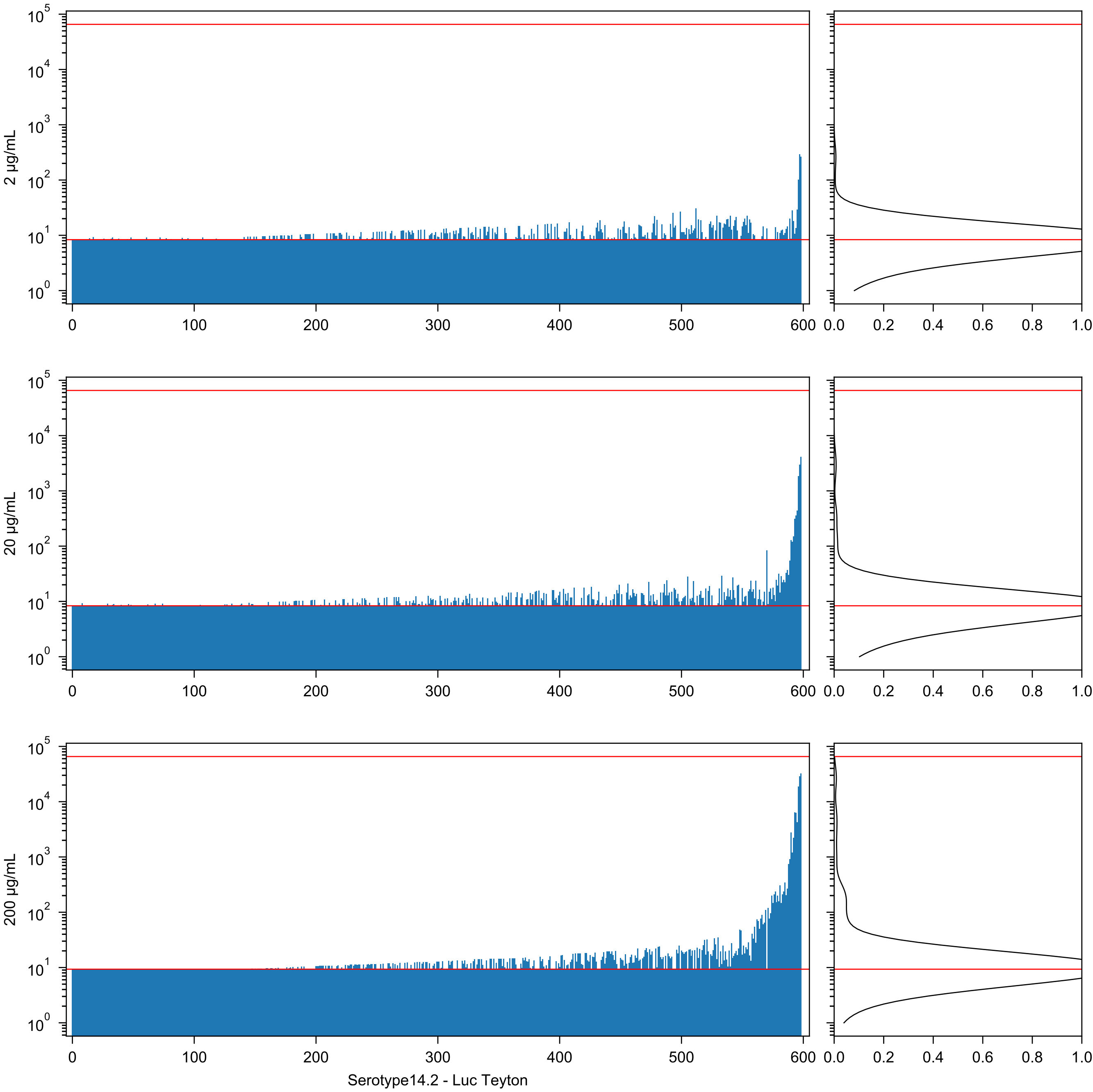
2 µg/mL
20 µg/mL
200 µg/mL
Serotype14.2 - Luc Teyton

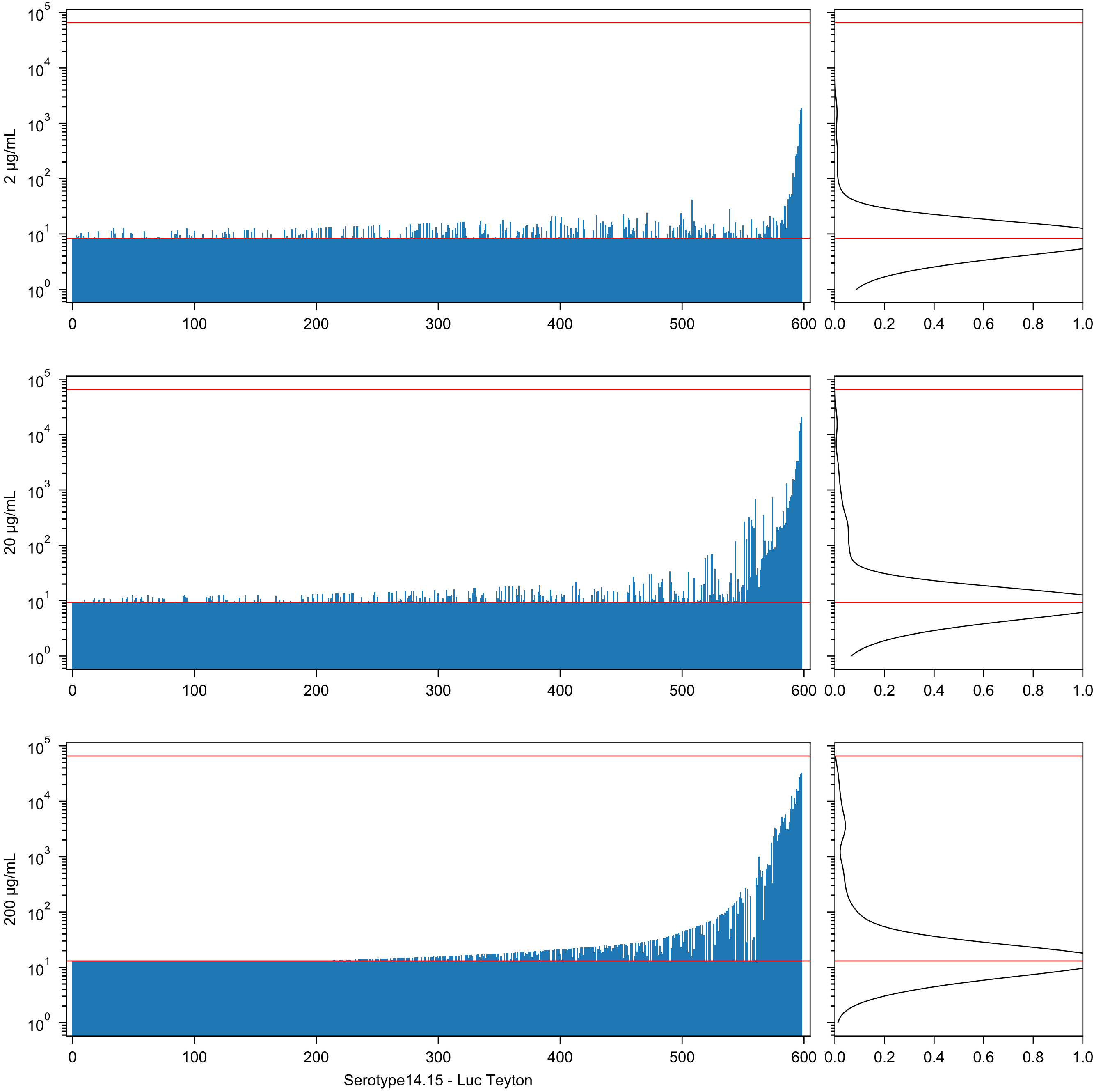
2 µg/mL
20 µg/mL
200 µg/mL
Serotype14.15 - Luc Teyton

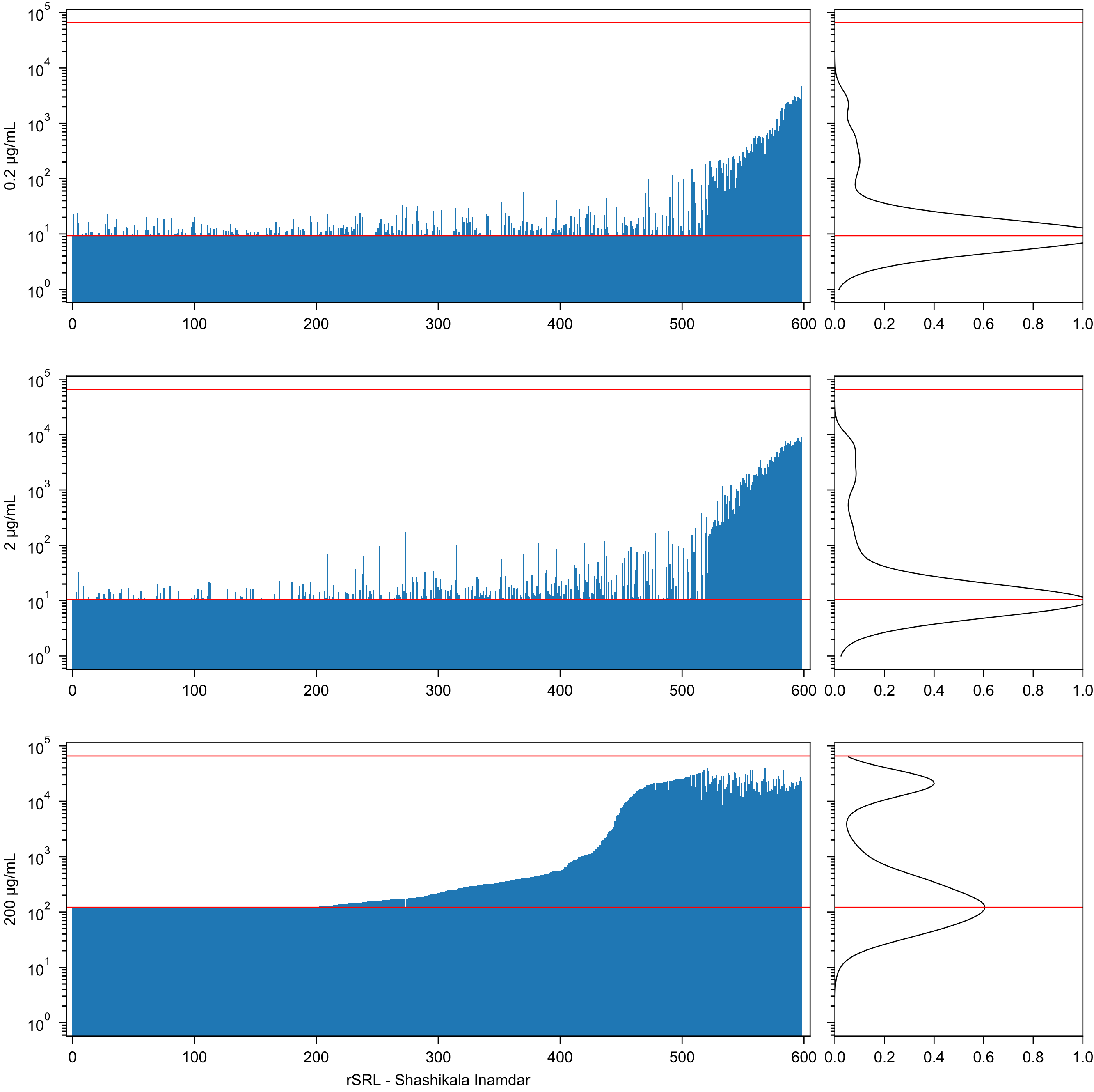
0.2 µg/mL
2 µg/mL
200 µg/mL
rSRL - Shashikala Inamdar

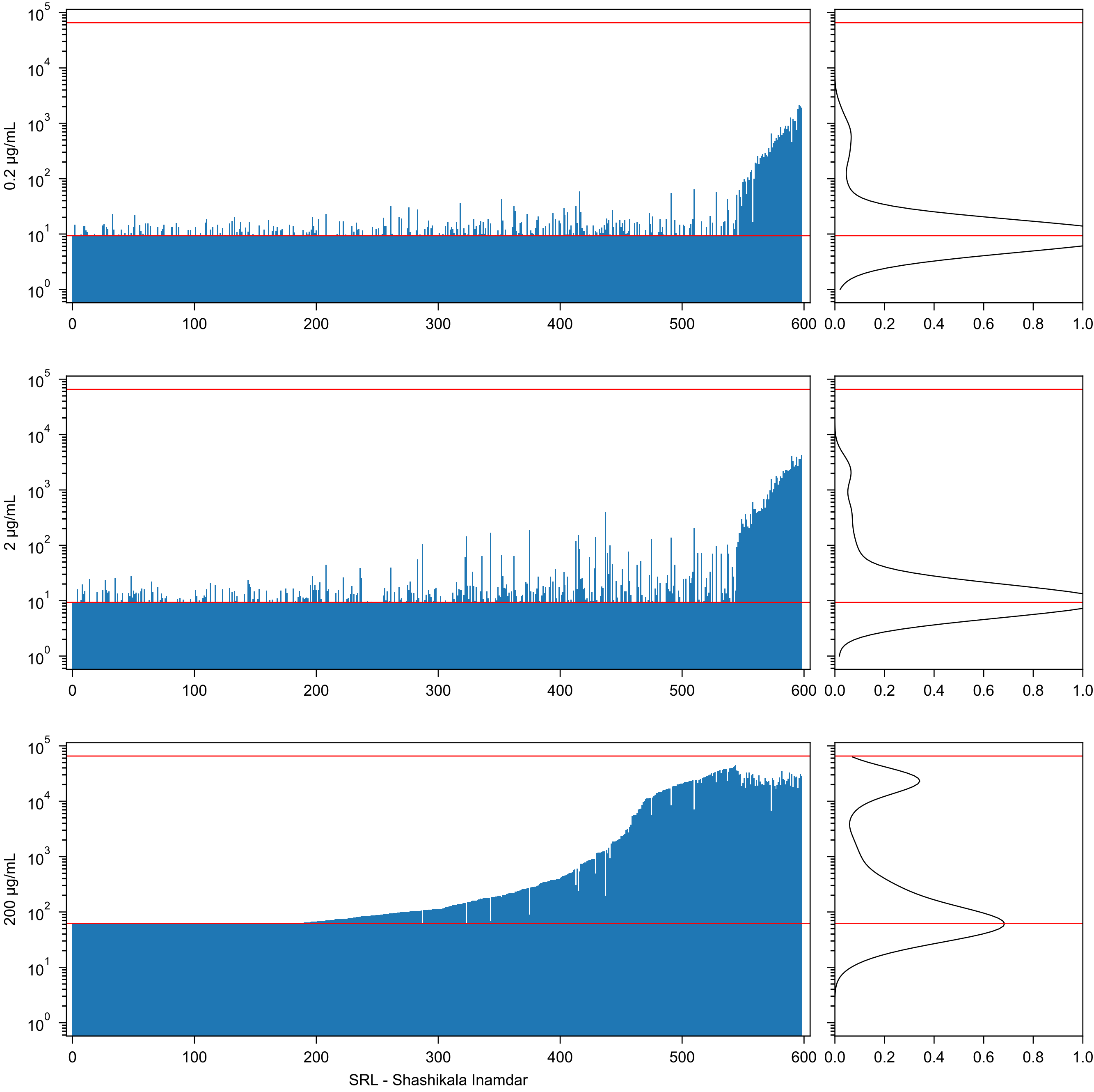

0.2 µg/mL
2 µg/mL
200 µg/mL
SRL - Shashikala Inamdar

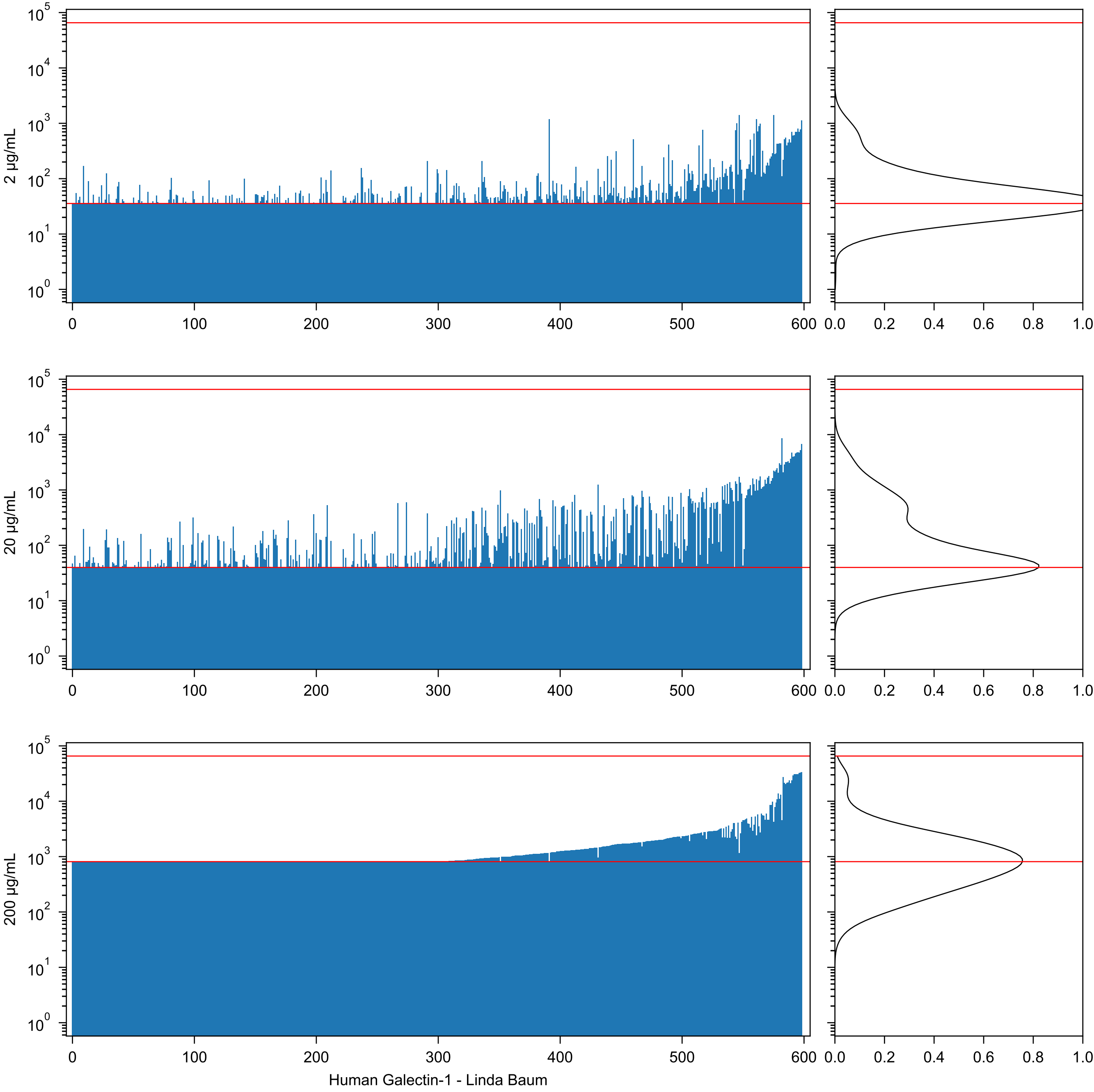
2 µg/mL
20 µg/mL
200 µg/mL
Human Galectin-1 - Linda Baum

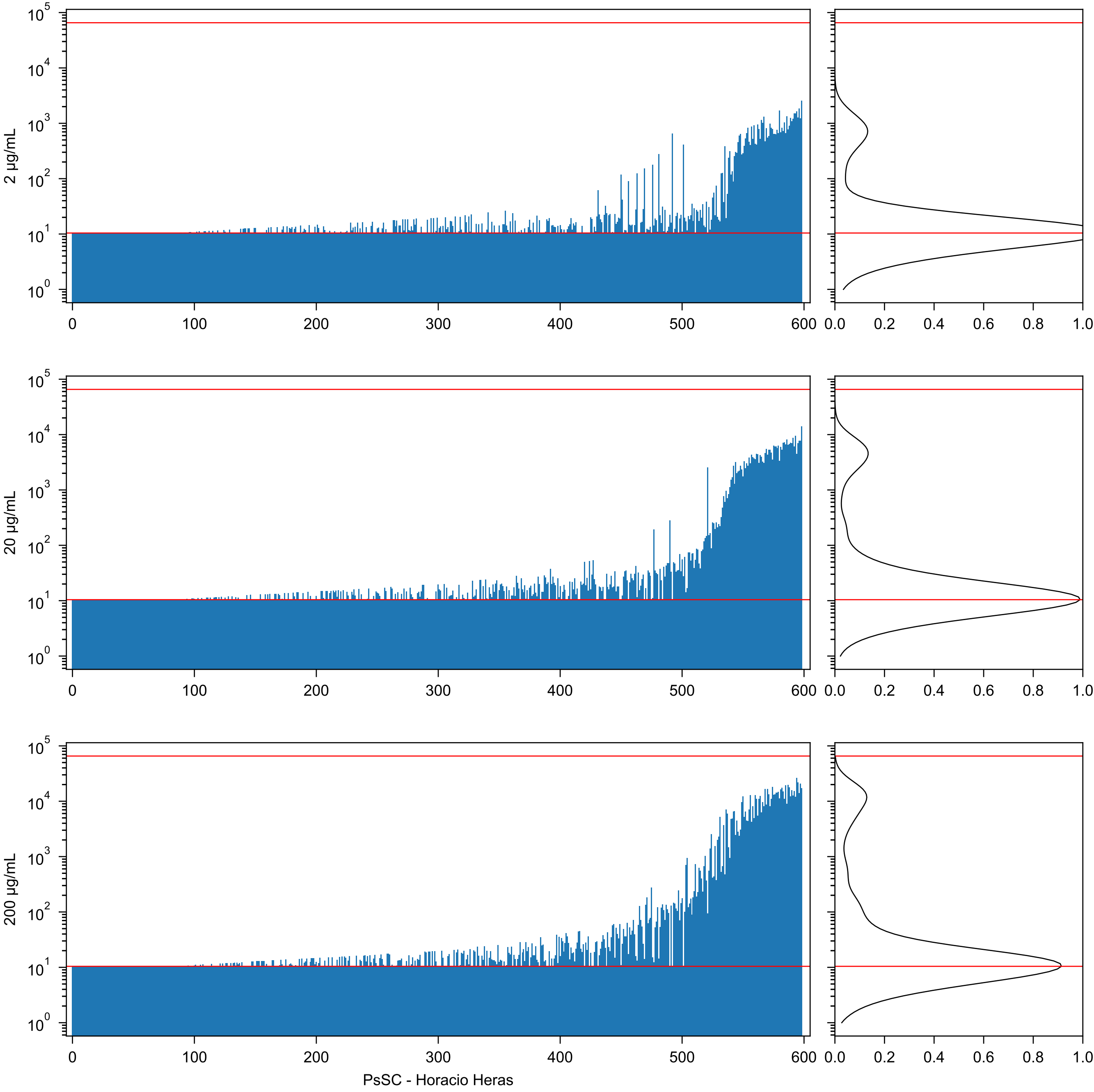
2 µg/mL
20 µg/mL
200 µg/mL
PsSC - Horacio Heras

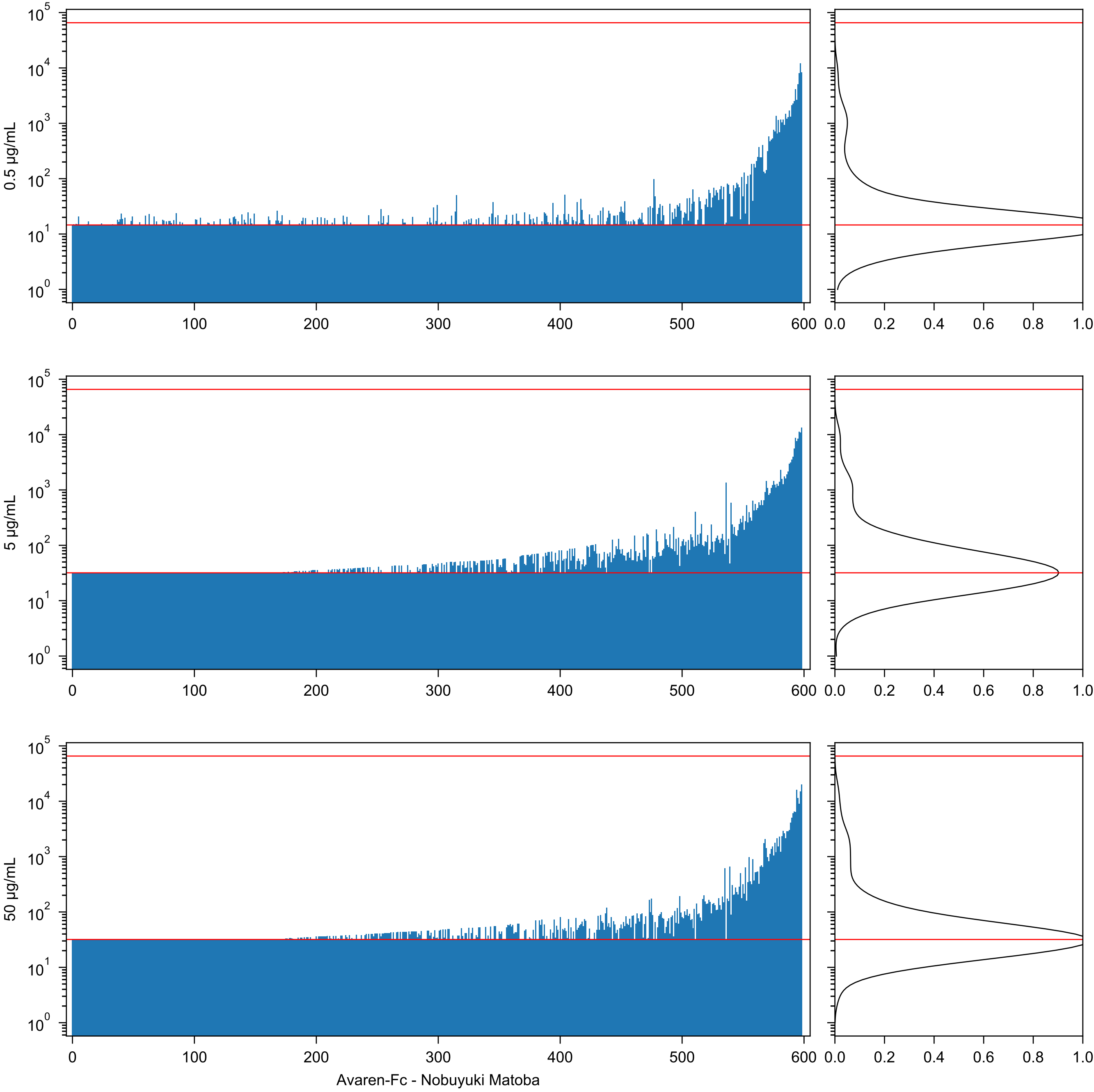

0.5 µg/mL
5 µg/mL
50 µg/mL
Avaren-Fc - Nobuyuki Matoba

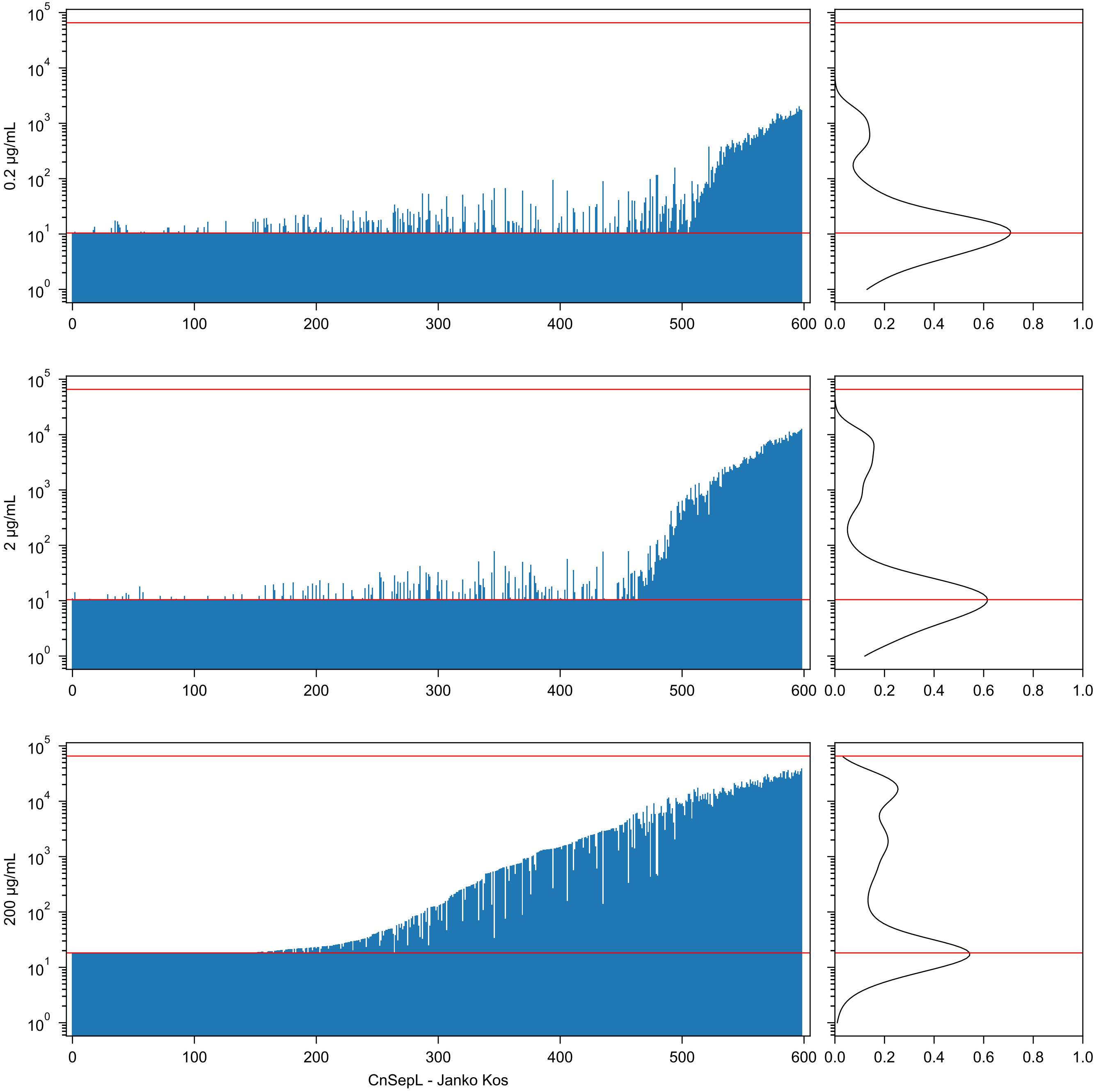

0.2 µg/mL
2 µg/mL
200 µg/mL
CnSepL - Janko Kos

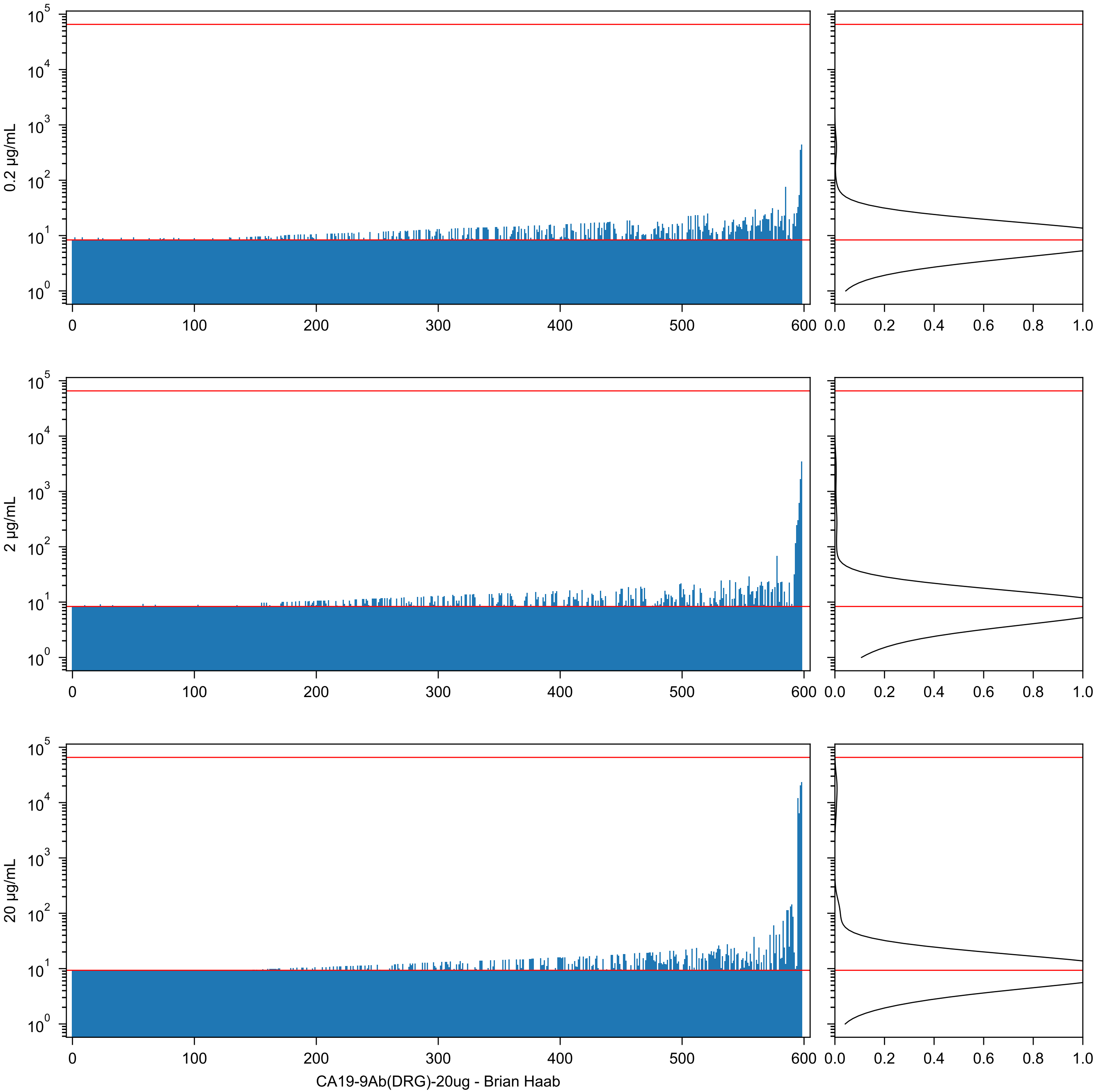

CA19-9Ab(DRG)-20ug - Brian Haab

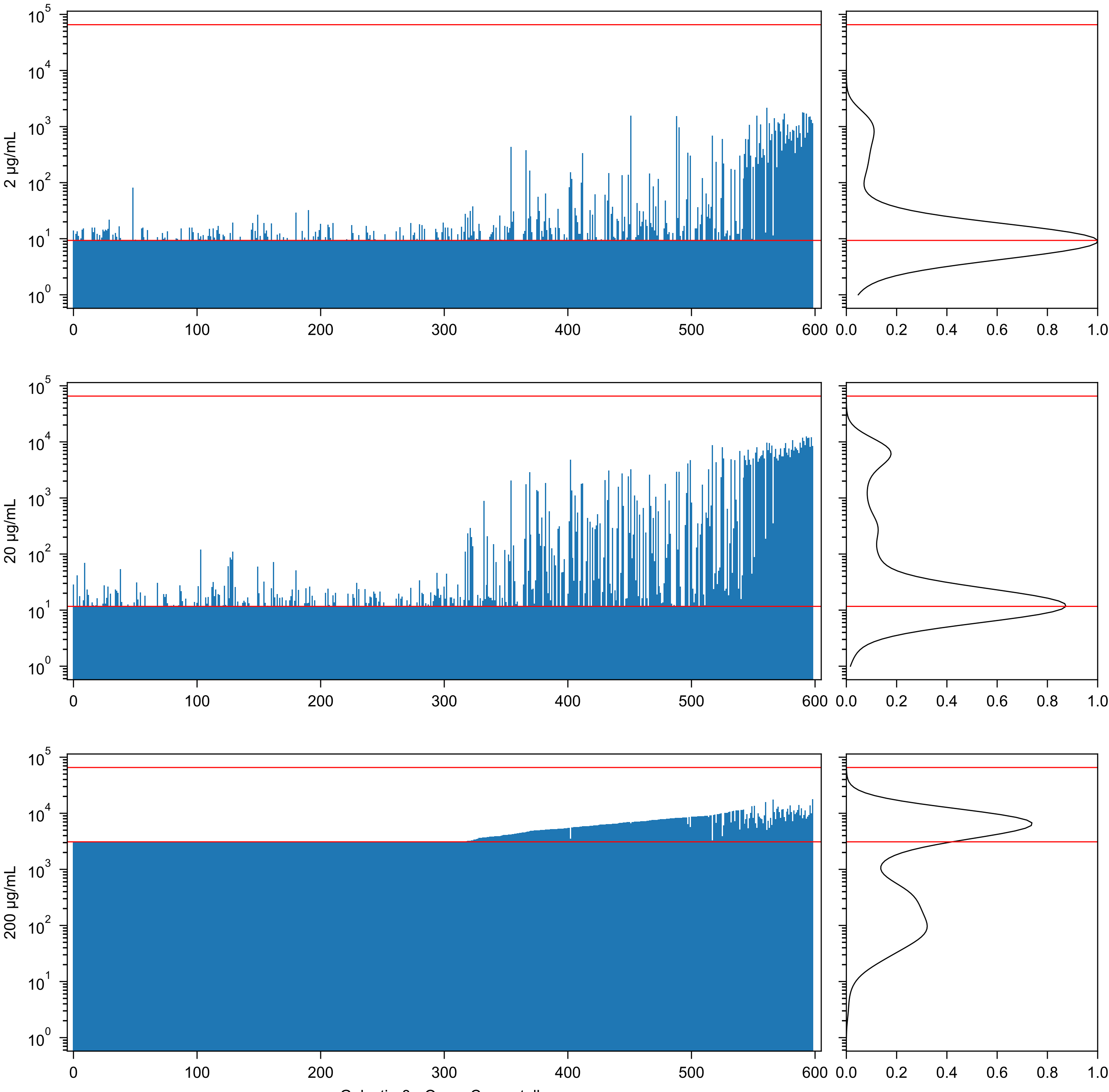

Galectin-8 - Oscar Campetella

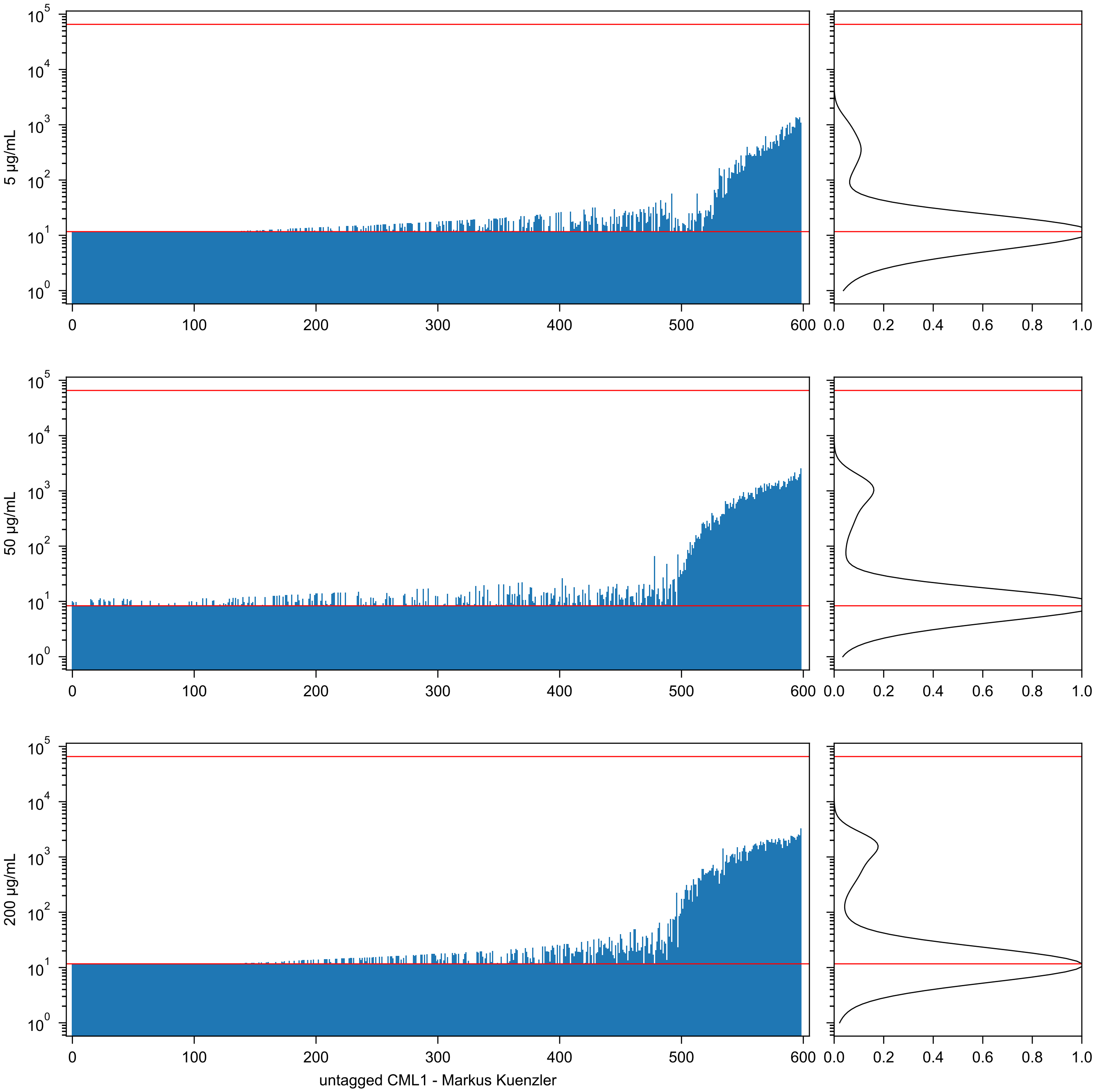

5 µg/mL
50 µg/mL
200 µg/mL
0
100
200
300
400
500
600
0.0
0.2
0.4
0.6
0.8
1.0
untagged CML1 - Markus Kuenzler

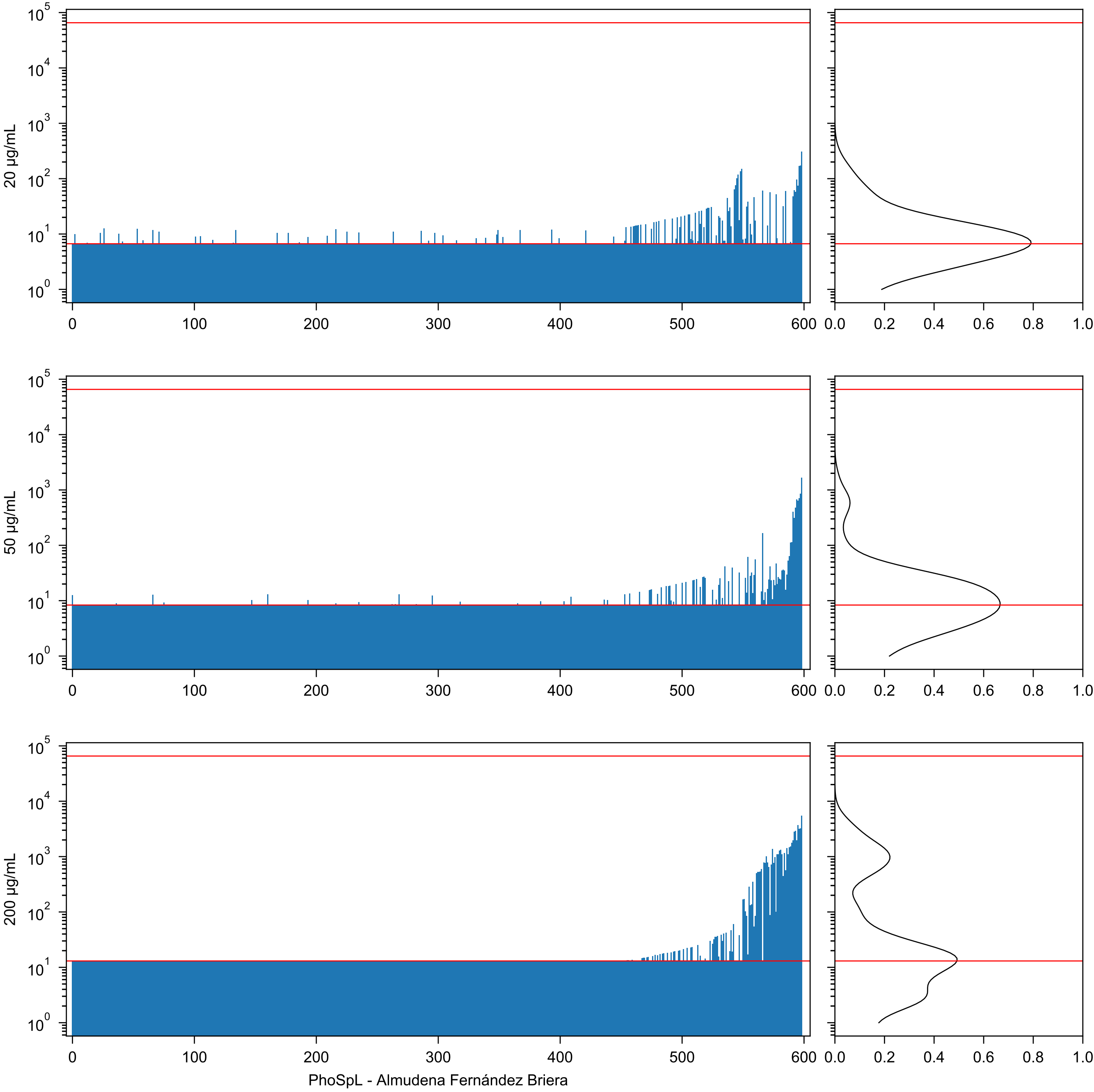
20 µg/mL
50 µg/mL
200 µg/mL
PhoSpL - Almudena Fernández Briera

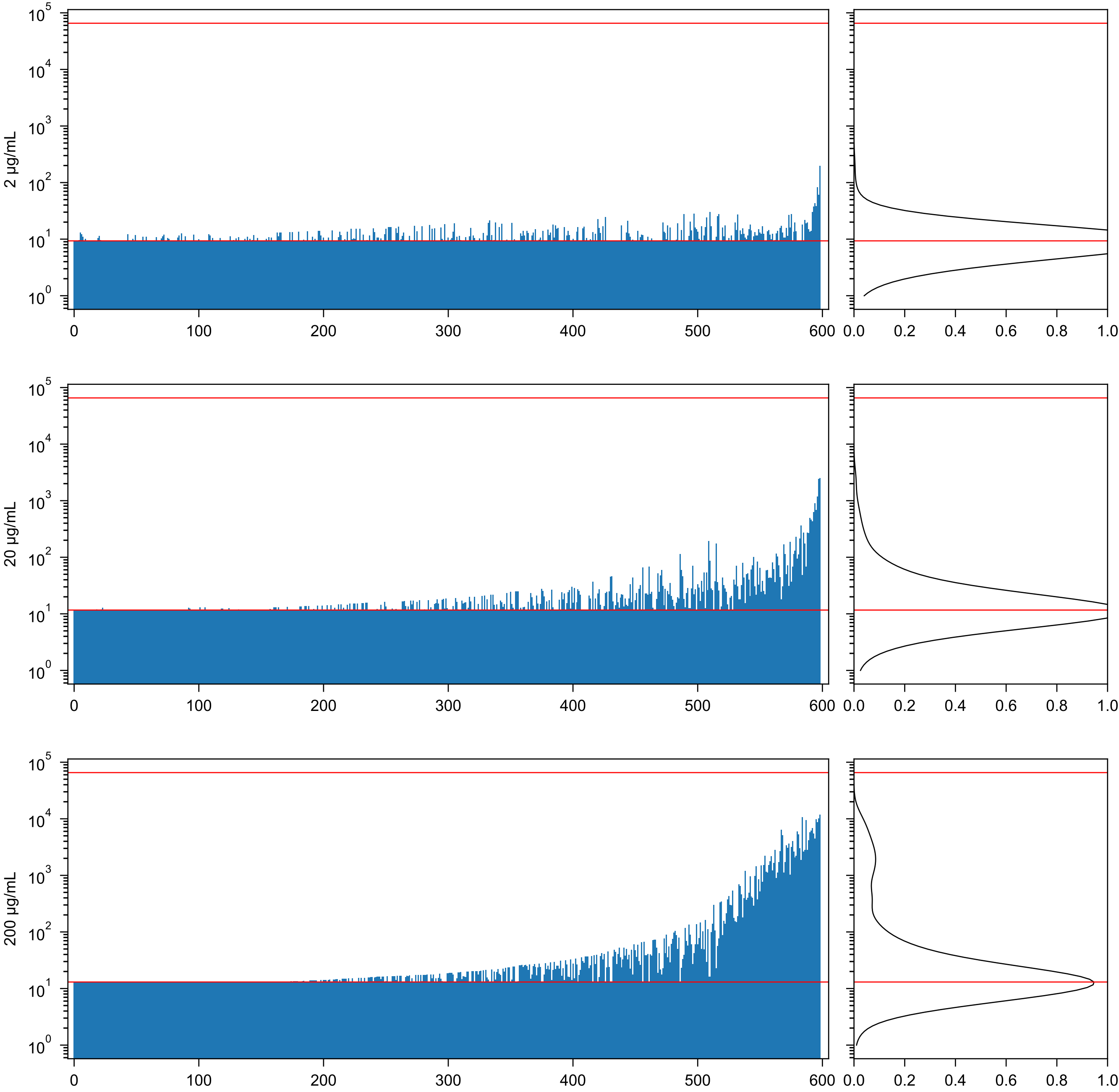

Epa-1 - Ronnie Willaert

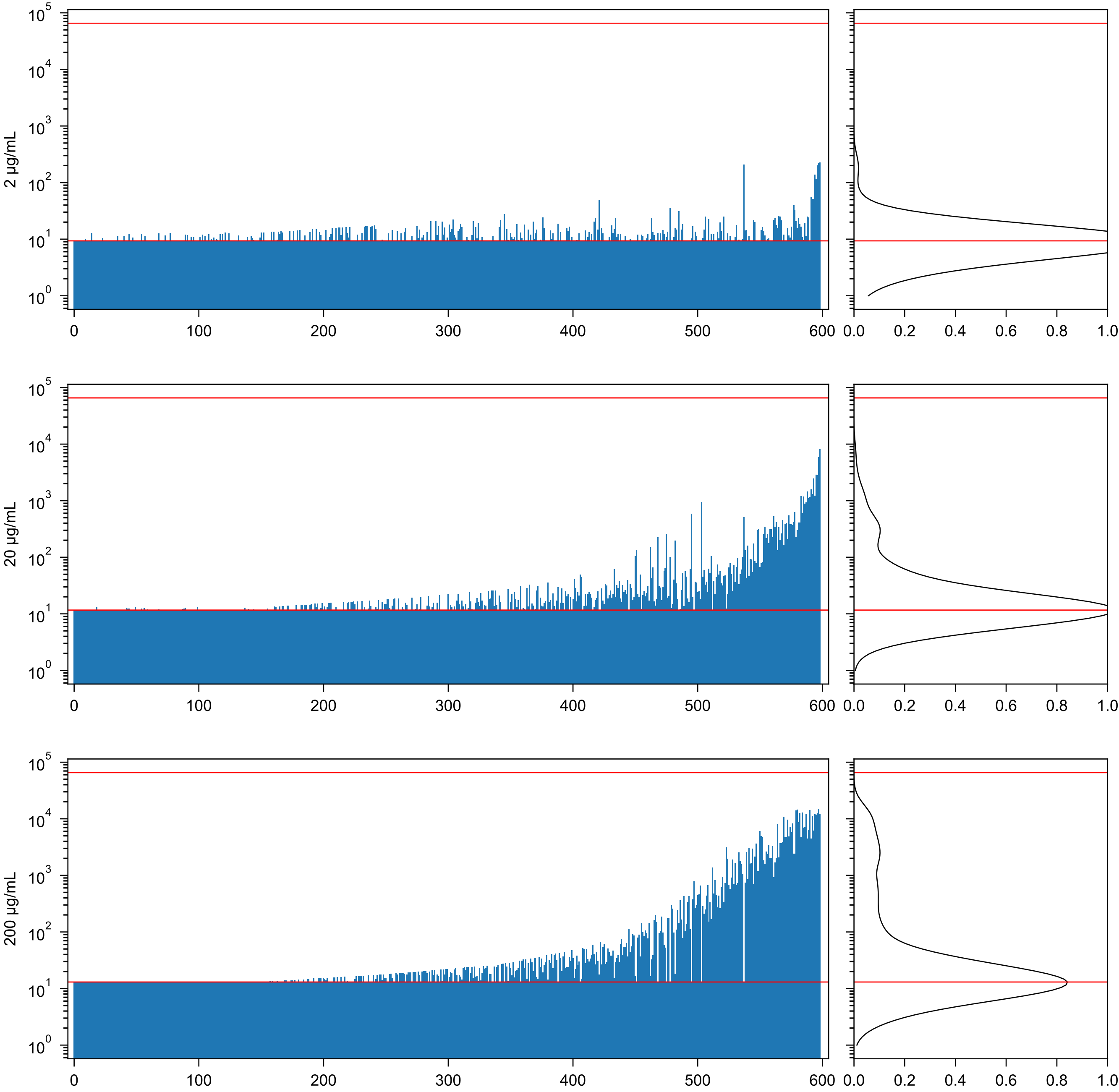


Epa-1 - Ronnie Willaert

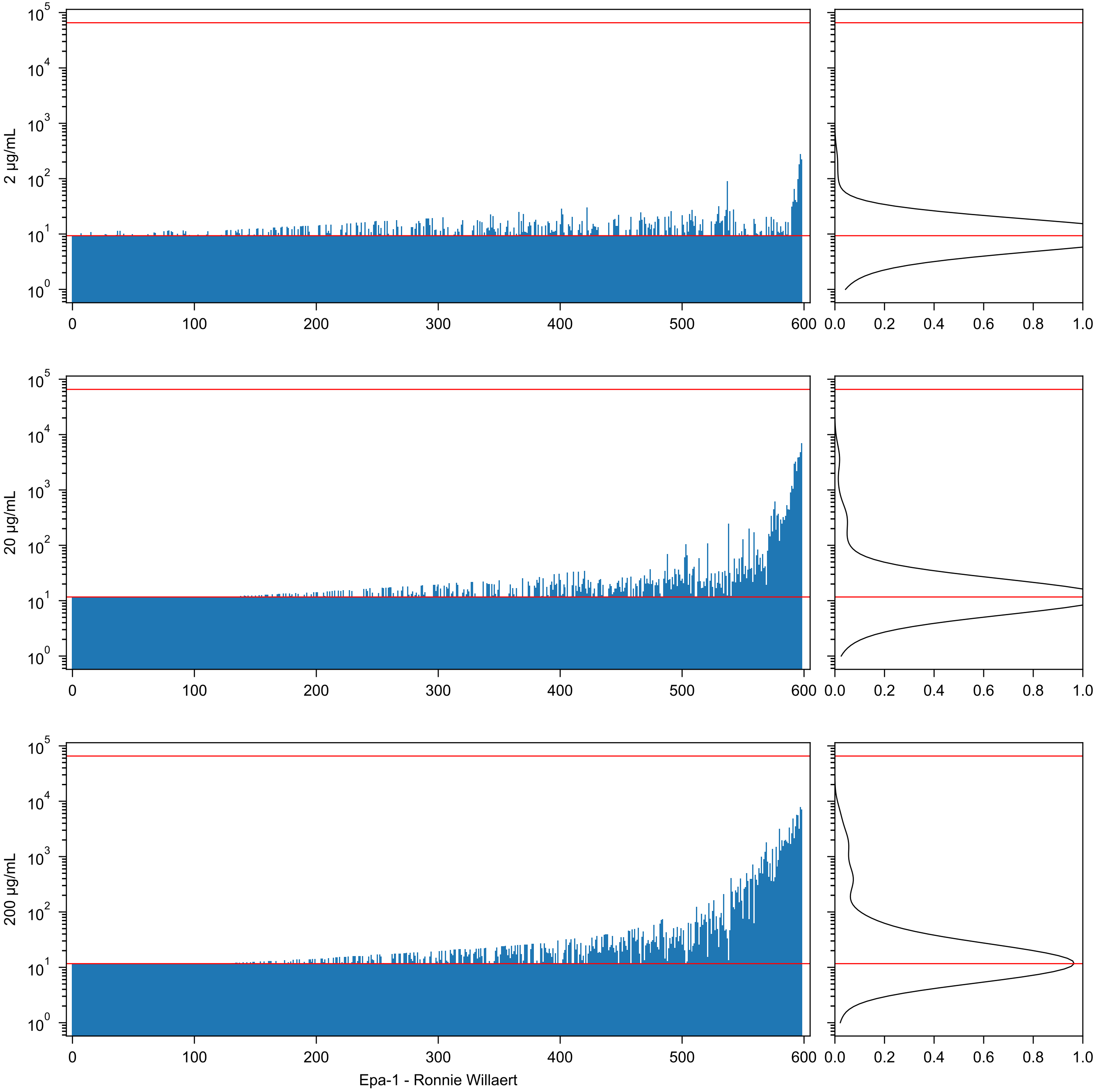

2 μg/mL
20 μg/mL
200 μg/mL
Epa-1 - Ronnie Willaert

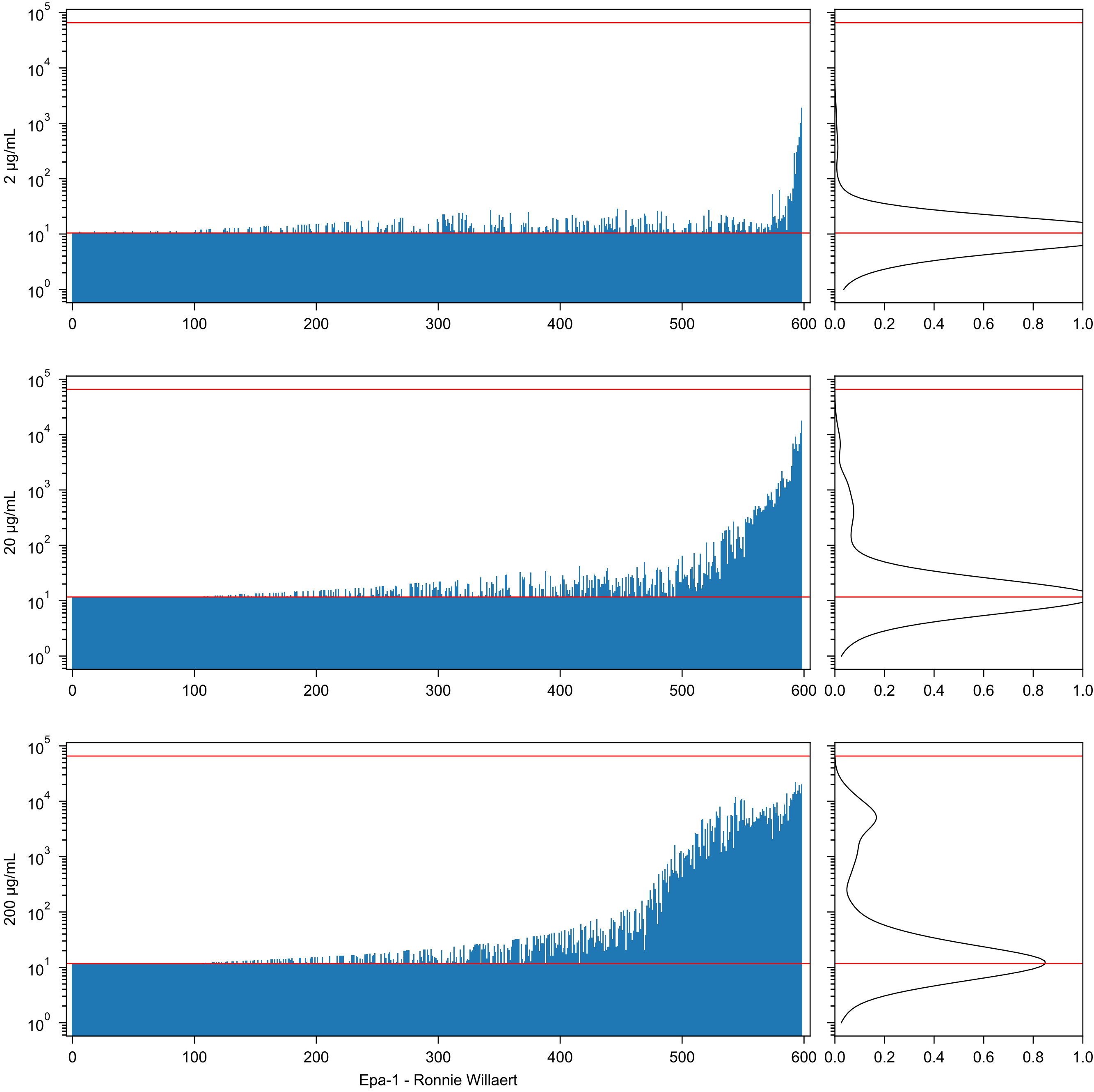

2 µg/mL
20 µg/mL
200 µg/mL
Epa-1 - Ronnie Willaert

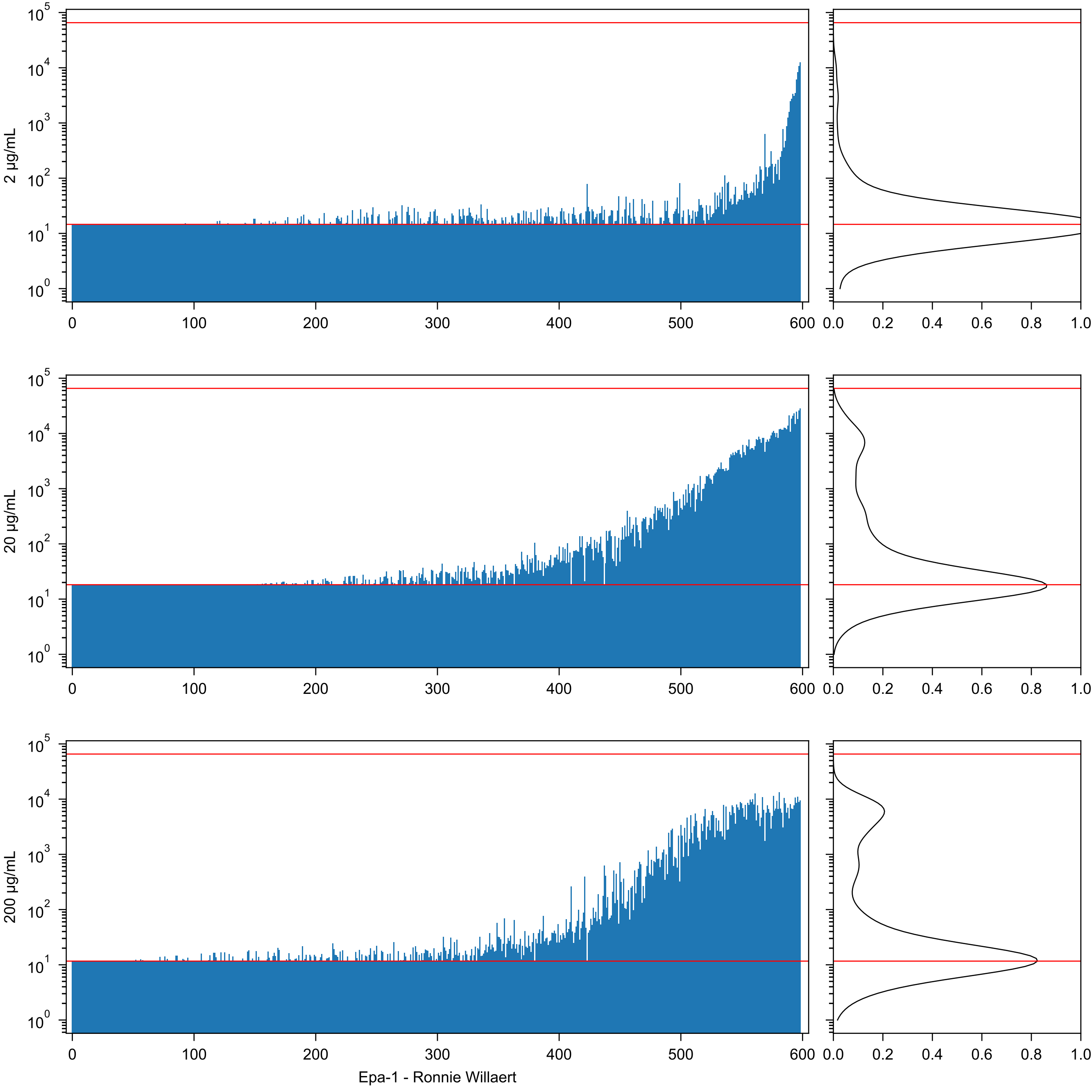
2 µg/mL
20 µg/mL
200 µg/mL
Epa-1 - Ronnie Willaert

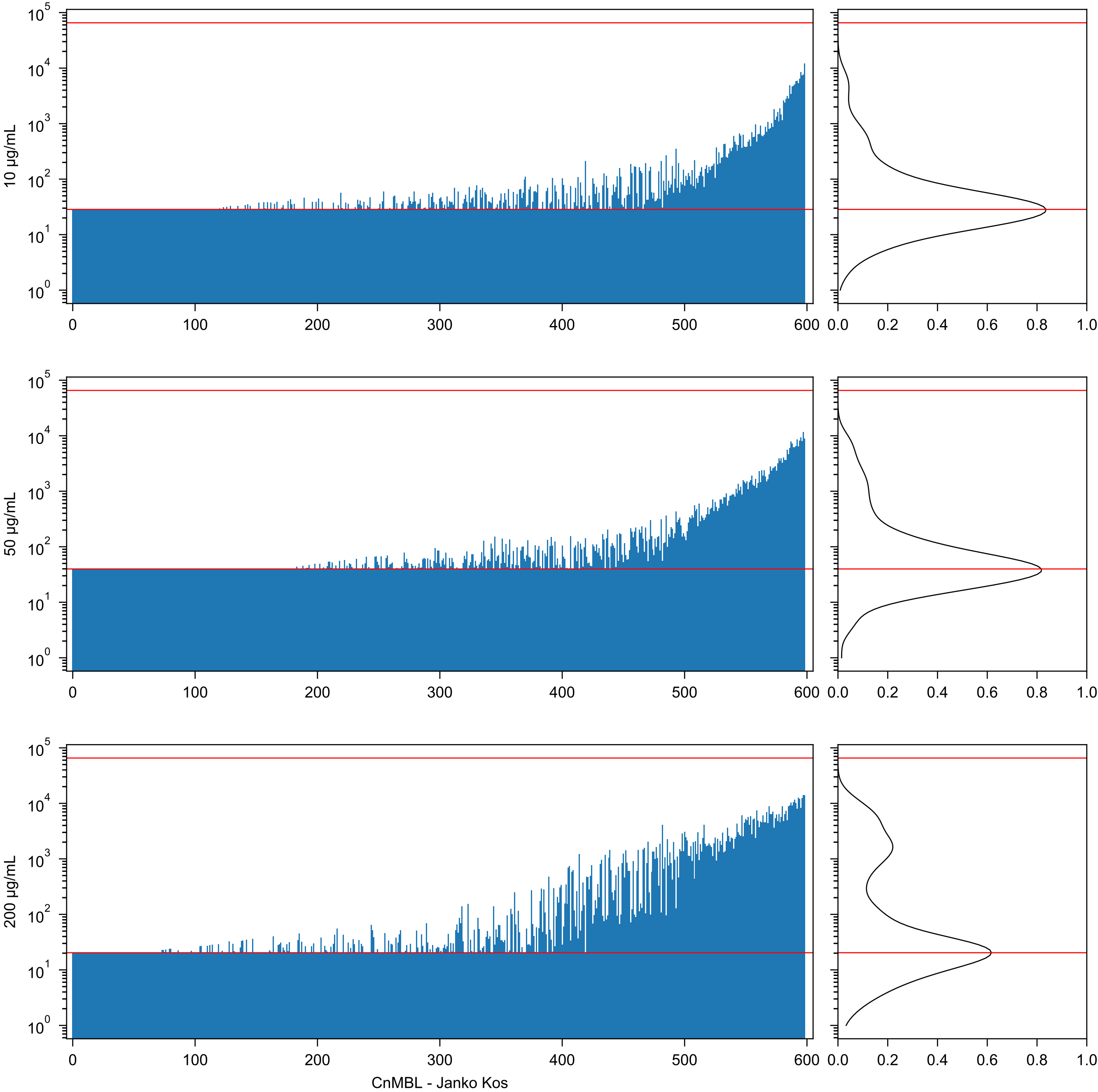

10 µg/mL
50 µg/mL
200 µg/mL
$10^5$
$10^4$
$10^3$
$10^2$
$10^1$
$10^0$
0
100
200
300
400
500
600
0.0
0.2
0.4
0.6
0.8
1.0
CnMBL - Janko Kos

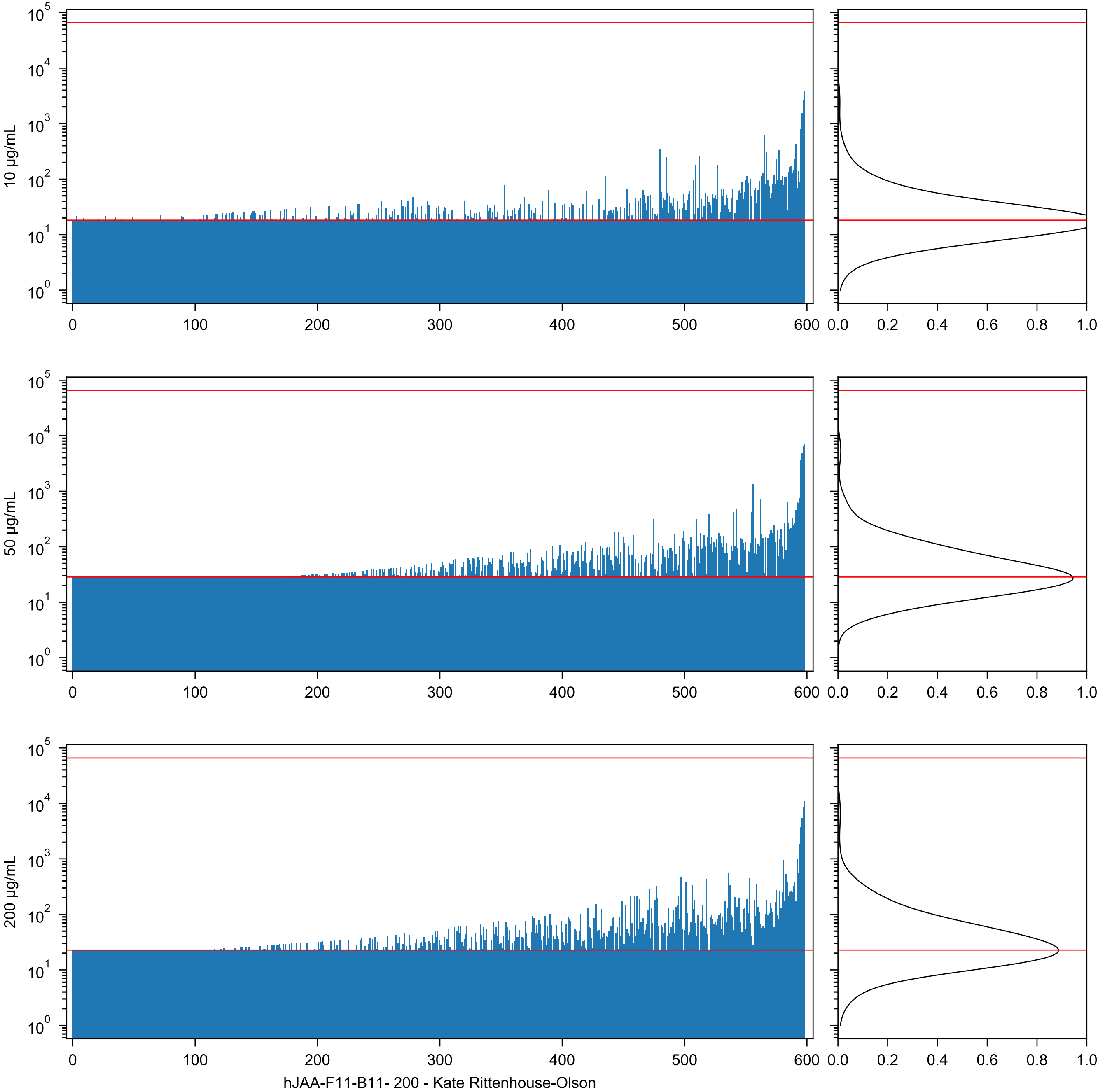

hJAA-F11-B11- 200 - Kate Rittenhouse-Olson
