## Supplementary material for "Atom-level Machine Learning of Protein-Glycan Interactions and Cross-chiral Recognition in Glycobiology": Per Glycan Prediction Plots

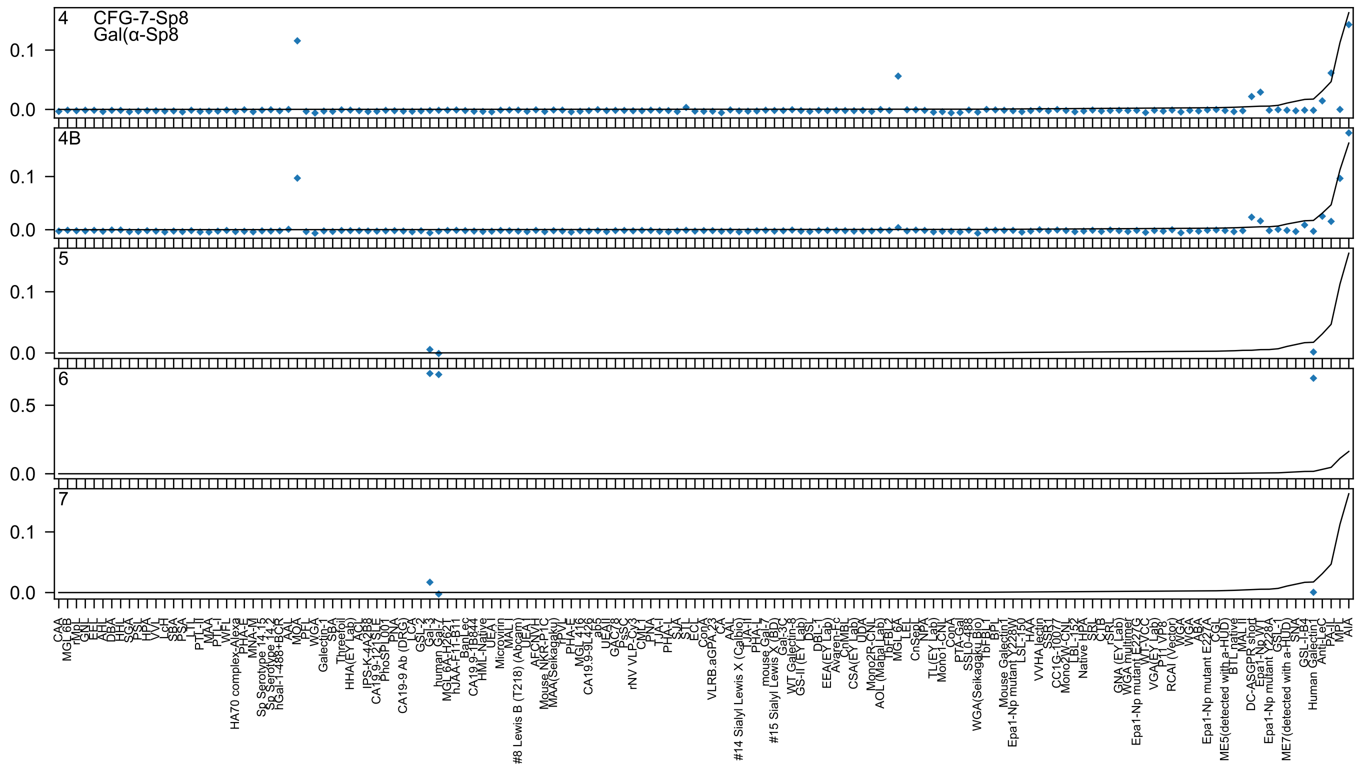

4
CFG-7-Sp8
Gal(α-Sp8
4B
5
6
7
0.0
0.1
0.5
CAA
MGL 6B
rMpL
GNL
EEL
AHL
DBA
HHL
SGA
PSL
HPA
VVL
LcH
SBA
PSA
LTL
PTL-II
MAA
PTL-I
WFL
HA70 complex-Alexa
PHA-E
MNA-M
Sp Serotype 14.15
Sp Serotype 14.2
hGal-1-488+BCR
AAL
MOA
PFL
WGA
Galectin-1
SBA
Threefoil
HHA(EY Lab)
ACL
IPS-K-4A2B8
CA19.9-121SLE
PhoSL001
PNA
CA19-9 Ab (DRG)
LCA
GSL-2
Gal-3
human Gal-7
MGL 6A-H262T
hJAA-F11-B11
BanLec
CA19.9-1B844
HML-Native
UEA I
Microvirin
MAL I
#8 Lewis B (T218) (Abcam)
UEA-I
AFCNVH
Mouse NKR-P1C
MAA(Seikagaku)
PVL
PHA-E
MGL 416
CA19.9-9L426
ab5
UEAII
GAC78
PsSC
rNV VLP-Cy3
CML1
PNA
TJA-I
PHA-L
SJA
STL
ECL
ConA
VLRB.aGPA.23
CA
AAL
#14 Sialyl Lewis X (Calbio)
TJA-II
PHA-L
mouse Gal-7
#15 Sialyl Lewis X (BD)
Gal-3C
WT Galectin-8
GS-II (EY Lab)
DSL
DBL-1
EEA(EY Lab)
Avaren-Fc
CnMBL
CSA(EY Lab)
UDA
Mono2R-CNL
AOL (Mana Lab)
TbFBL2
MGL 6A
LEL
CnSep1
NPA
TL(EY Lab)
Mono1-CNL
ConA
PTA-Gal
SSL0-488
WGA(Seikagaku Bio)
TbFBL1
BPL
Mouse Galectin1
Epa1-Np mutant Y228W
LSL-150
HAA
VVHA lectin
SSR1
CC1G-10077
Mono2W-CNL
LBL-152
Native HPA
SRL
CTB
rSRL
GNA (EY Lab)
WGA multimer
Epa1-Np mutant E227G
WT-YCC
VGA(EY Lab)
P11 VP8
RCAI (Vector)
WGA
ABA
Epa1-Np mutant E227A
CGL
ME5(detected with a-HUD)
BTL native
MAL II
DC-ASGPR short
Epa1-Np WT
Epa1-Np mutant Y228A
GSL-1
ME7(detected with a-HUD)
SNA
GSL-I-B4
Human Galectin1
Anti-LeC
PA-IL
MPL
AIA

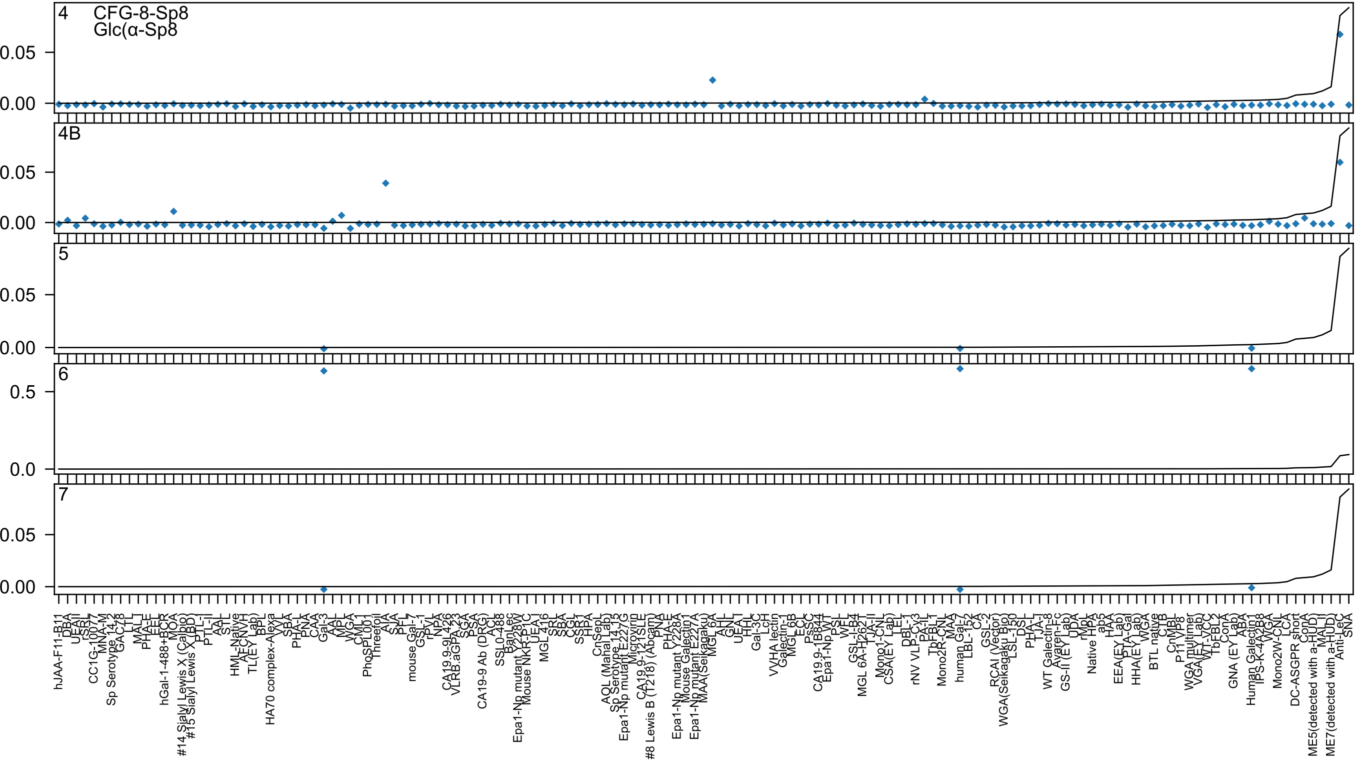
4
CFG-8-Sp8
Glc(α-Sp8
0.05
0.00
4B
0.05
0.00
5
0.05
0.00
6
0.5
0.0
7
0.05
0.00

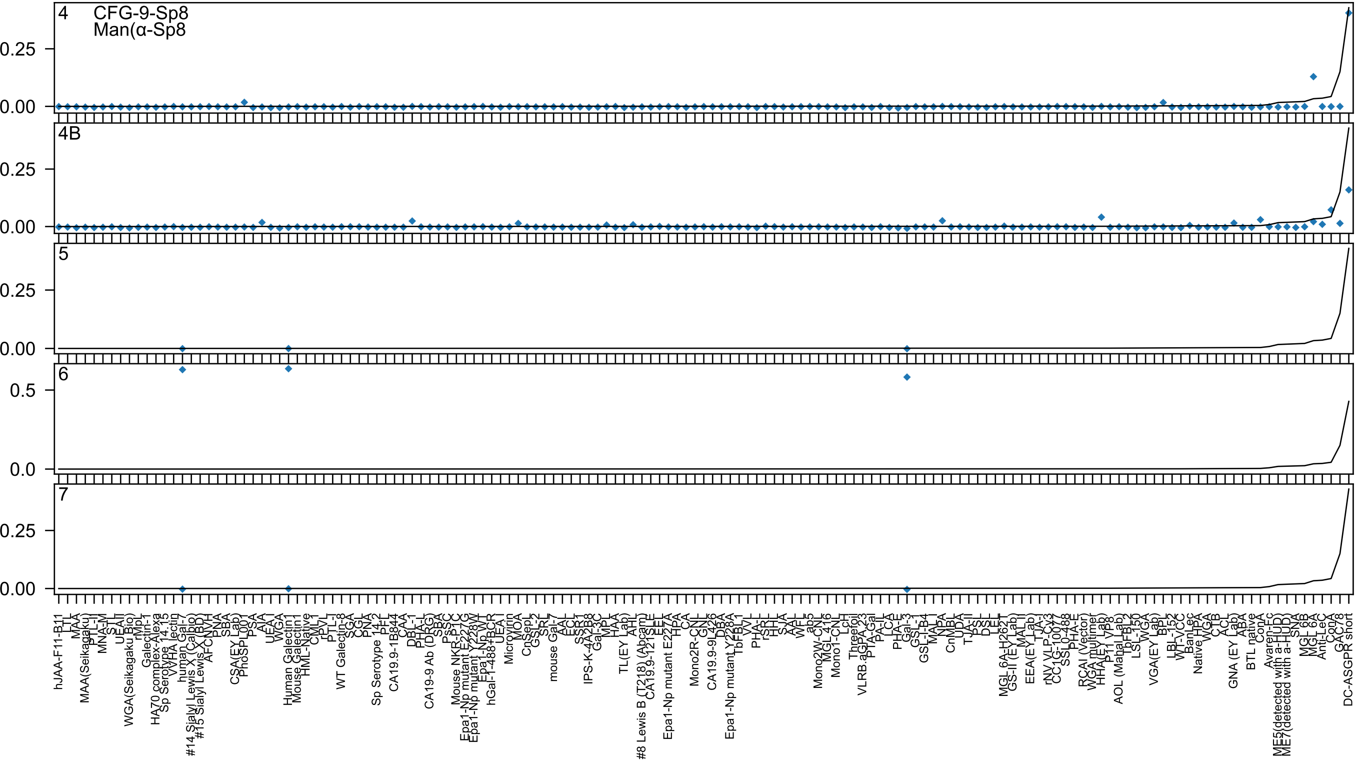
4
CFG-9-Sp8
Man(α-Sp8
4B
5
6
7
0.25
0.00
0.5
0.0

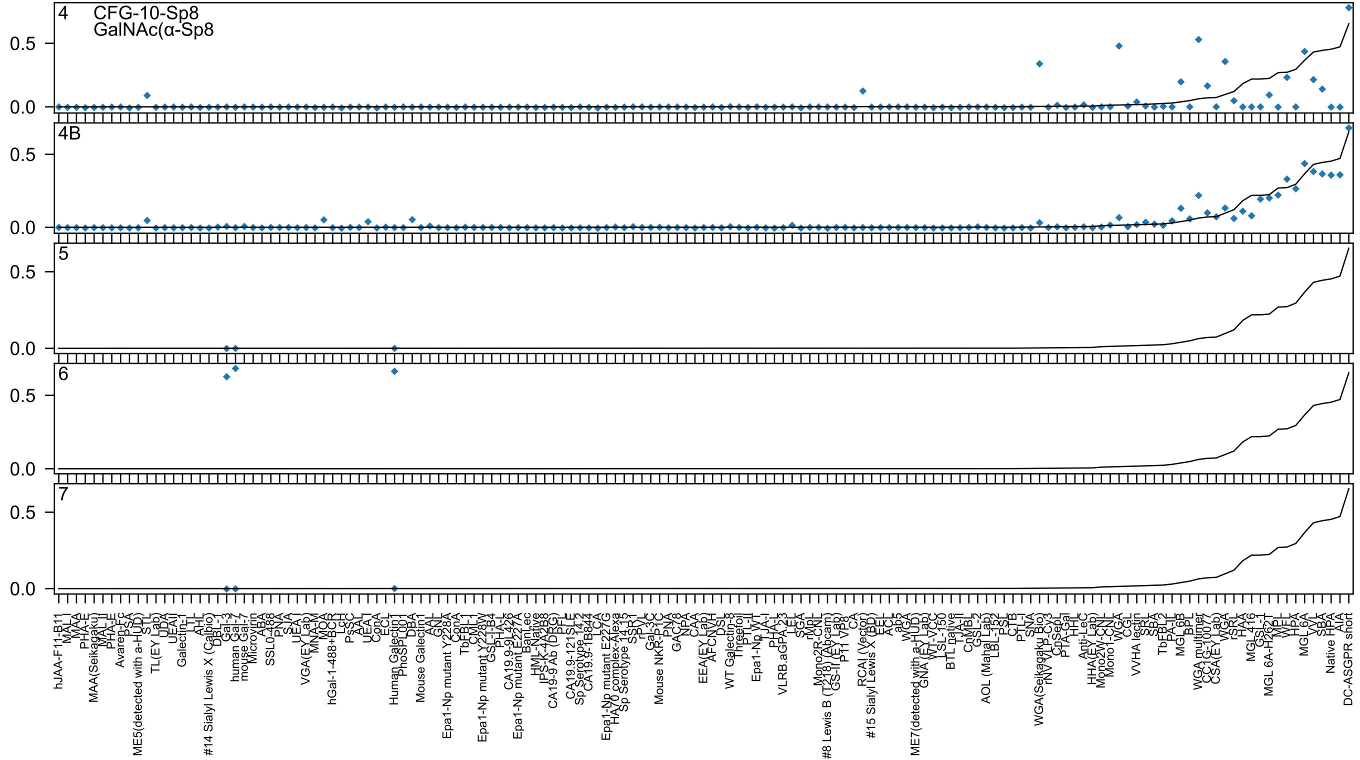
4
CFG-10-Sp8
GalNAc(α-Sp8
4B
5
6
7
0.5
0.0
hJAA-F11-B11
MAL_I
MAA
PHA-E
MAA(Seikagaku)
MAL_I
PHA-E
Avaren-Fc
PSA
ME5(detected with a-HUD)
STL
TL(EY Lab)
UDA
UEAII
Galectin-1
LTL
AHL
#14 Sialyl Lewis X (Calbio)
DBL-1
Gal-3
human Gal-7
mouse Gal-7
Microvirin
ABA
SSL0-488
PNA
SJA
UEA-I
VGA(EY Lab)
MNA-M
MOA
hGal-1-488+BCR
LcH
PsSC
AAL
UEA-I
ConA
ECL
Human Galectin1
PhoSL001
DBA
Mouse Galectin1
AAL
GNL
Epa1-Np mutant Y228A
ConA
TbFBL1
CML1
Epa1-Np mutant Y228W
GSL-I-B4
PHA-L
CA19.9-9L426
Epa1-Np mutant E227A
BanLec
HML-Native
IPS-K-4A2B8
CA19-9 Ab (DRG)
PFL
CA19.9-121SLE
Sp Serotype 14.2
CA19.9-1B844
LCA
Epa1-Np mutant E227G
HA70 complex-Alexa
Sp Serotype 14.15
SSR1
rPVL
Gal-3C
Mouse NKR-P1C
PNA
GAC78
NPA
CAA
EEA(EY Lab)
AFCNVH
DSL
WT Galectin-8
Threefoil
PTL-II
Epa1-Np WT
TJA-I
PHA-L
VLRB.aGPA.23
LEL
SGA
rMpL
Mono2R-CNL
#8 Lewis B (T218) (Abcam)
GS-II (EY Lab)
P11 VP8
CA
RCAI (Vector)
#15 Sialyl Lewis X (BD)
EEL
ACL
ab5
WGA
ME7(detected with a-HUD)
GNA (EY Lab)
WT-VCC
LSL-150
BTL native
TJA-II
CnMBL
GSL-2
AOL (Mahal Lab)
LBL-152
PSL
CTB
PTL-I
SNA
WGA(Seikagaku Bio)
rNV VLP-Cy3
CpSepL
PTA-Gal
HHL
Anti-LeC
HHA(EY Lab)
Mono2W-CNL
Mono1-CNL
WGA
CGL
VVHA lectin
SRA
SBA
TbFBL2
PA-IL
MGL 6B
BPL
WGA multimer
CC1G-10077
CSA(EY Lab)
WGA
rSRL
HAA
MGL 416
GSL-1
MGL 6A-H262T
MPL
WFL
HPA
MGL 6A
VVL
SBA
Native HPA
AIA
DC-ASGPR short

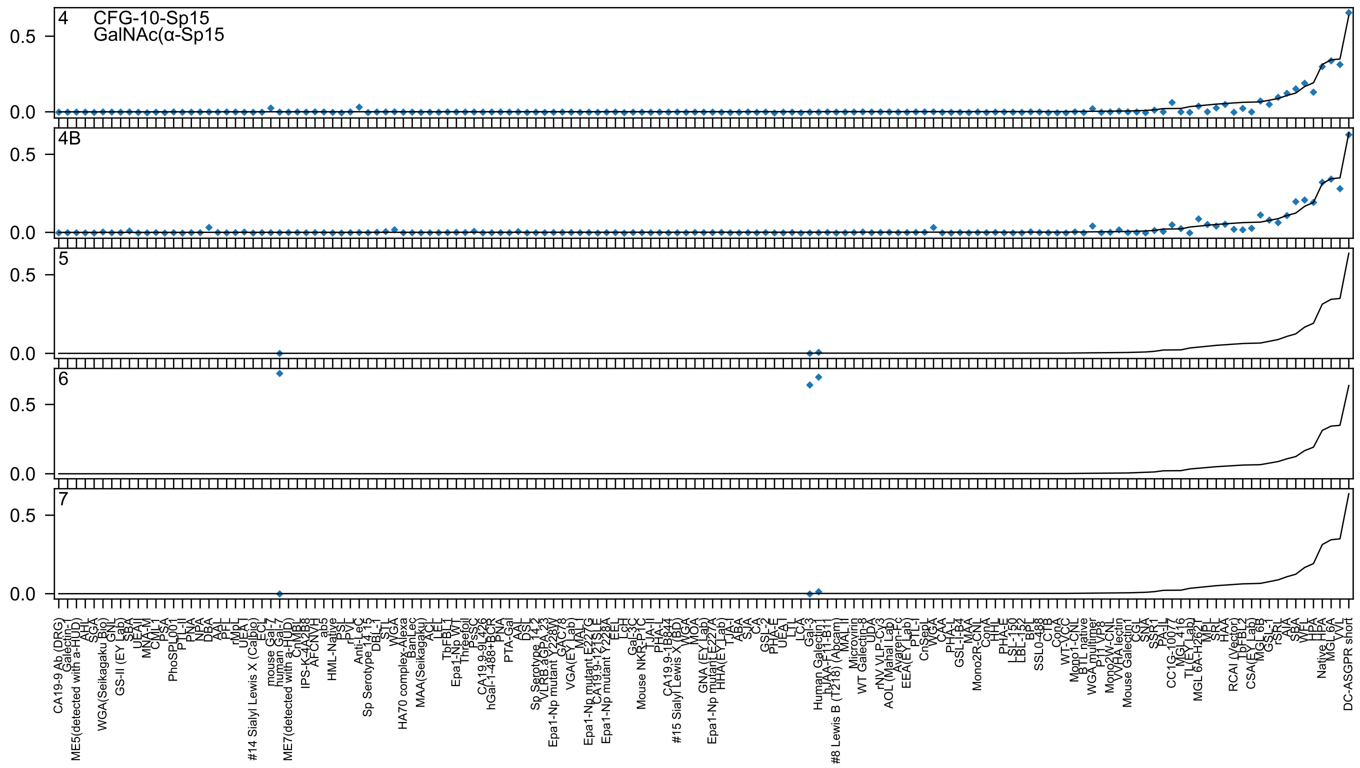
4
CFG-10-Sp15
GalNAc(α-Sp15
4B
5
6
7
0.5
0.0
CA19-9 Ab (DRG)
Galectin-1
ME5(detected with a-HUD)
AHL
SGA
WGA(Seikagaku Bio)
GNL
GS-II (EY Lab)
SBA
UEAII
MNA-M
CML-1
PSA
PhoSPL001
PTL-II
PNA
NPA
DBA
AAL
PFL
rMpL
UEA-I
ECL
#14 Sialyl Lewis X (Calbio)
mouse Gal-7
human Gal-7
ME7(detected with a-HUD)
CnMBL
IPS-K-4A2B8
AFCNVH
ab5
HML-Native
PSL
rPVL
Anti-LeC
Sp Serotype 14 15
DBL-I
STL
WGA
HA70 complex-Alexa
BanLec
MAA(Seikagaku)
ACL
LEL
TbFBL1
Epa1-Np WT
Threefoil
PSSC
CA19-9 9L426
hGal-1-488+BCR
PNA
PTA-Gal
AAL
DSL
Sp Serotype 14 2
VLRB aGPA 23
Epa1-Np mutant Y228W
GAC78
VGA(EY Lab)
MAL I
Epa1-Np mutant E227G
CA19-9 121SLE
Epa1-Np mutant Y228A
EEL
LcH
Gal-3C
Mouse NKR-P1C
TJA-II
PHA-L
CA19-9 1B844
#15 Sialyl Lewis X (BD)
WGA
MOA
GNA (EY Lab)
Epa1-Np mutant E227A
HHA(EY Lab)
TJA-I
ABA
SJA
CA
GSL-2
PHA-E
UEA
LTL
LCA
Gal-3
Human Galectin1
hJAA-F11-B11
#8 Lewis B (T218) (Abcam)
MAL II
Microvirin
WT Galectin-8
UDA
rNV VLP-Cy3
AOL (Manal Lab)
Avaren-Fc
EEA(EY Lab)
PTL-I
CnSepL
WGA
CAA
PHA-L
GSL-I-B4
MAA
Mono2R-CNL
ConA
HHL
PHA-E
LSL-150
LBL-152
BPL
SSL0-488
CTB
ConA
WT-VCC
Mono1-CNL
BTL native
WGA multimer
P11 VP8
Mono2W-CNL
VVHA lectin
Mouse Galectin1
CGL
SNA
SSR1
PA-IL
CC1G-10077
MGL 416
TL(EY Lab)
MGL 6A-H262T
MPL
SRL
HAA
RCAI (Vector)
TbFBL2
CSA(EY Lab)
MGL 6B
GSL-1
rSRL
AIA
SBA
WFL
Native HPA
MGL 6A
VVL
DC-ASGPR short

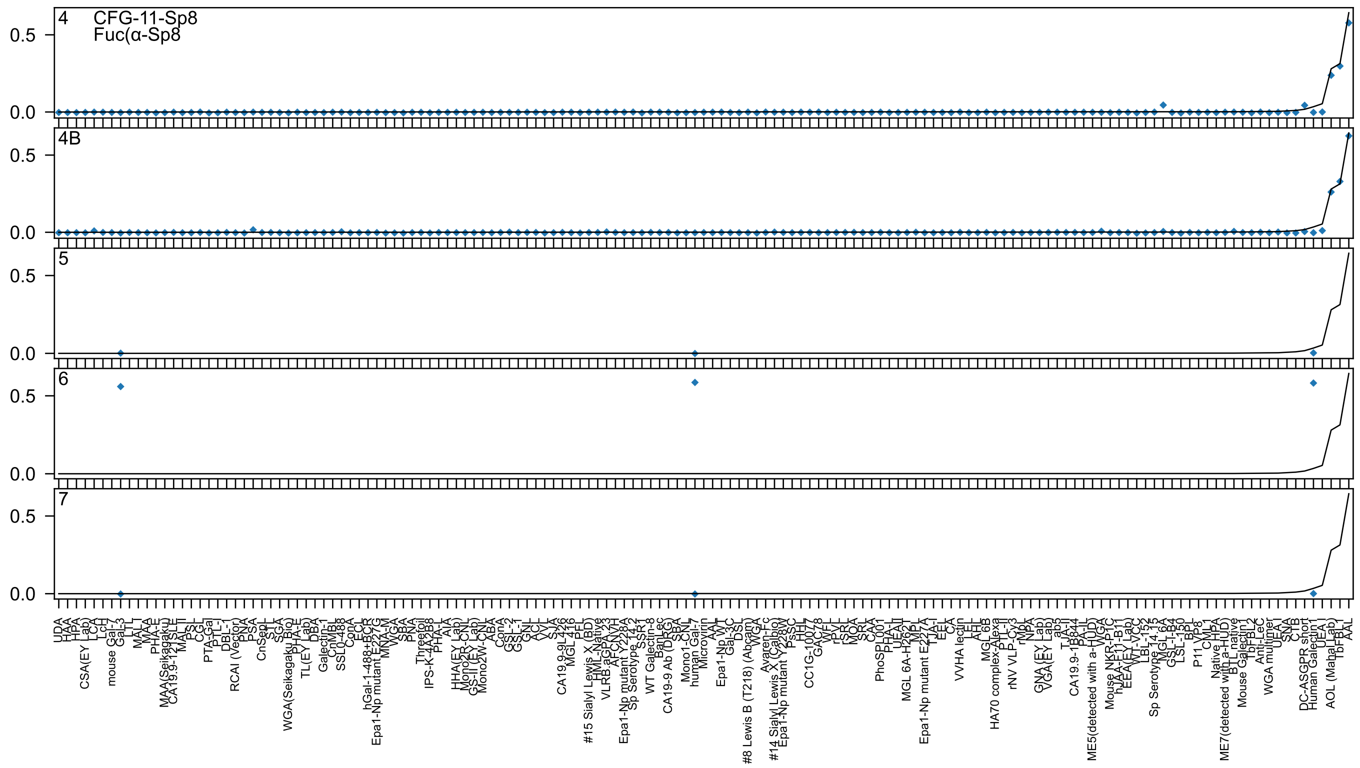
4 CFG-11-Sp8
Fuc(α-Sp8
4B
5
6
7
0.5
0.0

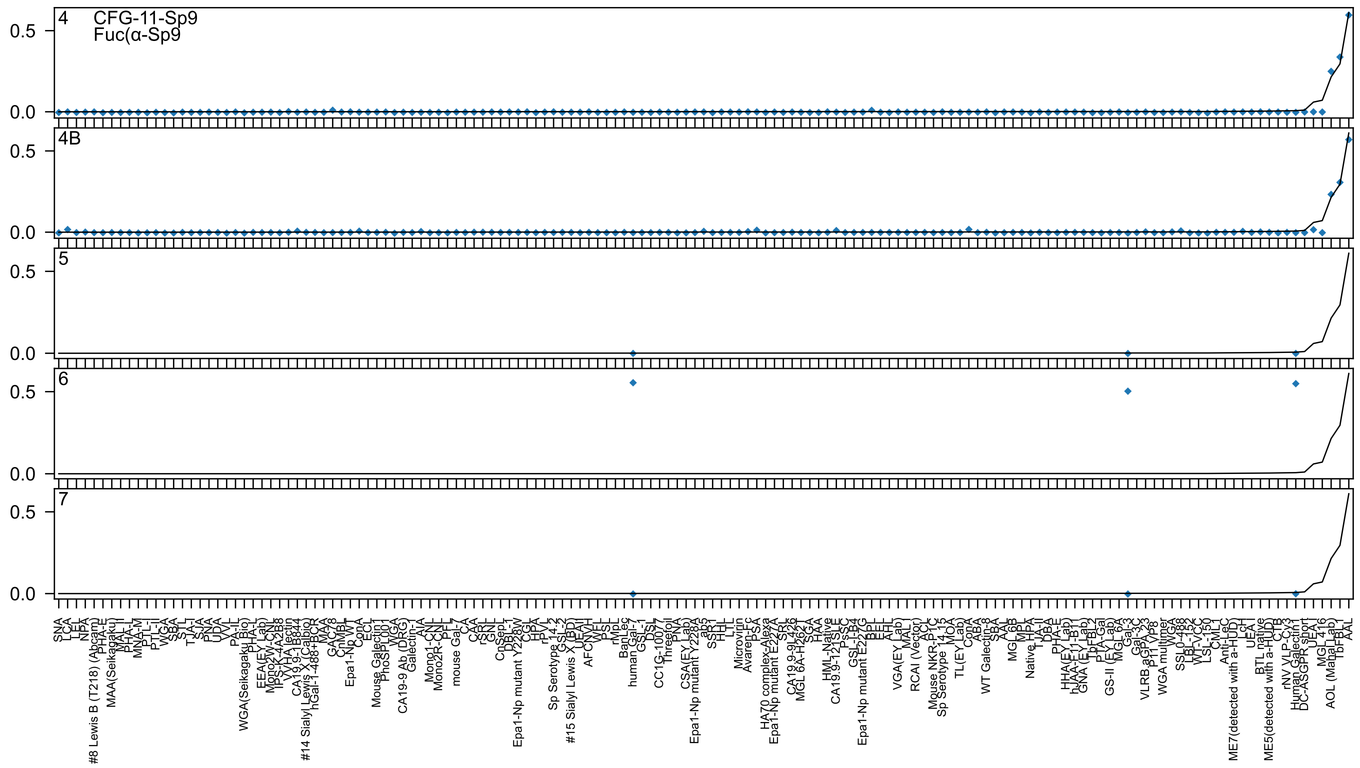

4
CFG-11-Sp9
Fuc(α-Sp9
4B
5
6
7

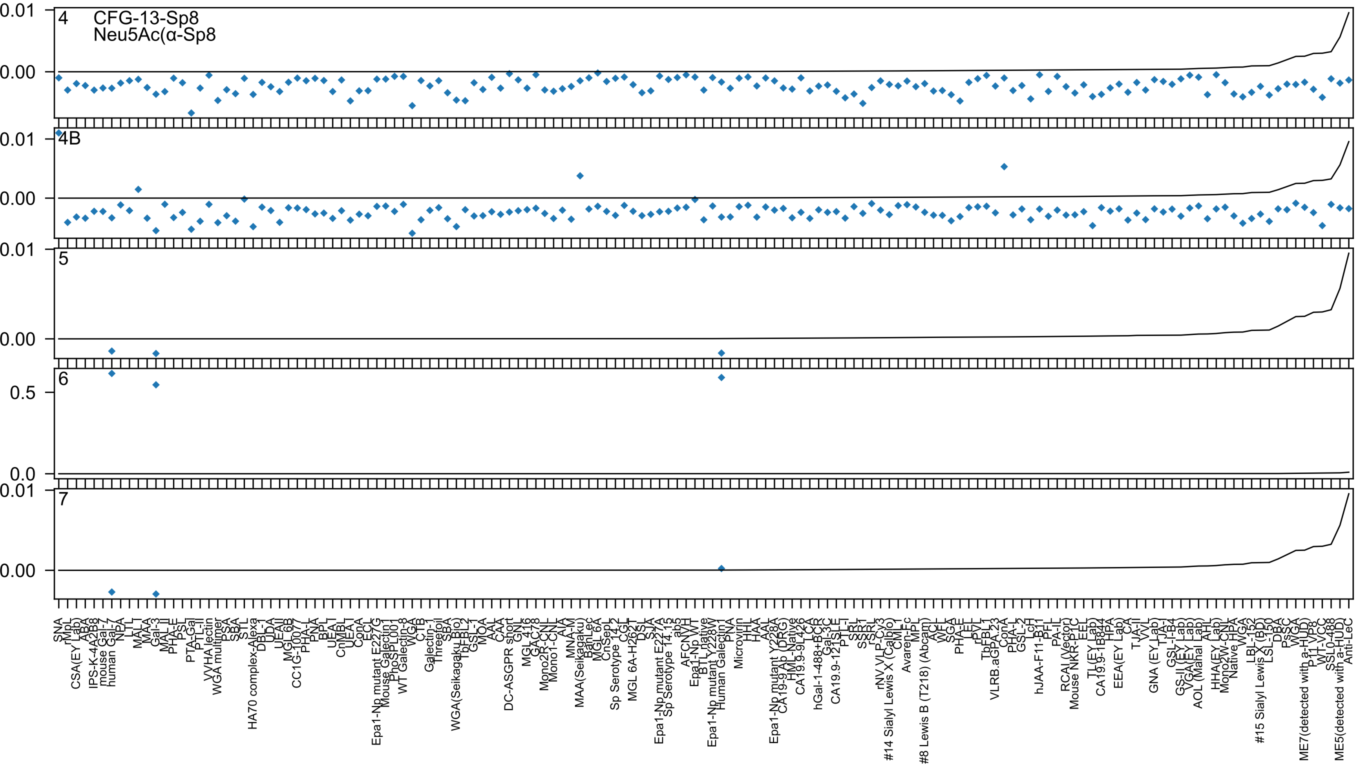

4 CFG-13-Sp8
Neu5Ac(α-Sp8
4B
5
6
7
0.01
0.00
0.5
0.0
SNA
rMpL
CSA(EY Lab)
ABA
IPS-K-4A2B8
mouse Gal-7
human Gal-7
NPA
LTL
MALT
MAA
Gal-3
MAL_II
PHA-E
PSL
PTA-Gal
PTL-II
VVHA lectin
WGA multimer
PSA
SBA
STL
HA70 complex-Alexa
DBL-1
UDA
UEAII
MGL 6B
CC1G-10077
PHA-L
PNA
BPL
UEA-I
CnMBL
UEA-I
ConA
ECL
Epa1-Np mutant E227G
Mouse Galectin1
PhoSL001
WT Galectin-8
WGA
CTB
Galectin-1
Threefoil
SBA
WGA(Seikagaku Bio)
TbFBL2
GSL-I
MOA
AAL
CAA
DC-ASGPR short
GNL
MGL 416
GAC78
Mono2R-CNL
Mono1-CNL
AIA
MNA-M
MAA(Seikagaku)
BanLec
MGL 6A
CnSep1
Sp Serotype 14.2
CGL
MGL 6A-H262T
DSL
SJA
Epa1-Np mutant E227A
Sp Serotype 14.15
ab5
AFCNVH
Epa1-Np WT
BTL native
Epa1-Np mutant Y228W
Human Galectin1
PNA
Microvirin
HHL
HAA
AAL
Epa1-Np mutant Y228A
CA19-9Ab (DRG)
HML-Native
CA19.9-9L426
LCA
hGal-1-488+BCR
Gal-3C
CA19.9-121SLE
PTL-I
SRL
SSR1
rSRL
rNV VLP-Cy3
#14 Sialyl Lewis X (Calbio)
CML
Avaren-Fc
MPL
#8 Lewis B (T218) (Abcam)
ACL
WFL
SGA
PHA-E
LEL
rPVL
TbFBL1
VLRB.aGPA.23
ConA
PHA-L
GSL-2
LcH
hJAA-F11-B11
PFL
PA-IL
RCAI (Vector)
Mouse NKR-P1C
EEL
TL(EY Lab)
CA19.9-1B844
HPA
EEA(EY Lab)
CA
TJA-II
VVL
GNA (EY Lab)
TJA-I
GSL-I-B4
GS-II (EY Lab)
VGA(EY Lab)
AOL (Mahal Lab)
AHL
HHA(EY Lab)
Mono2W-CNL
Native HPA
WGA
LBL-152
#15 Sialyl Lewis X (BD)
LSL-150
DBA
PSSC
WGA
ME7(detected with a-HUD)
P11 VP8
WT-VCC
SSL0-488
ME5(detected with a-HUD)
Anti-LeC

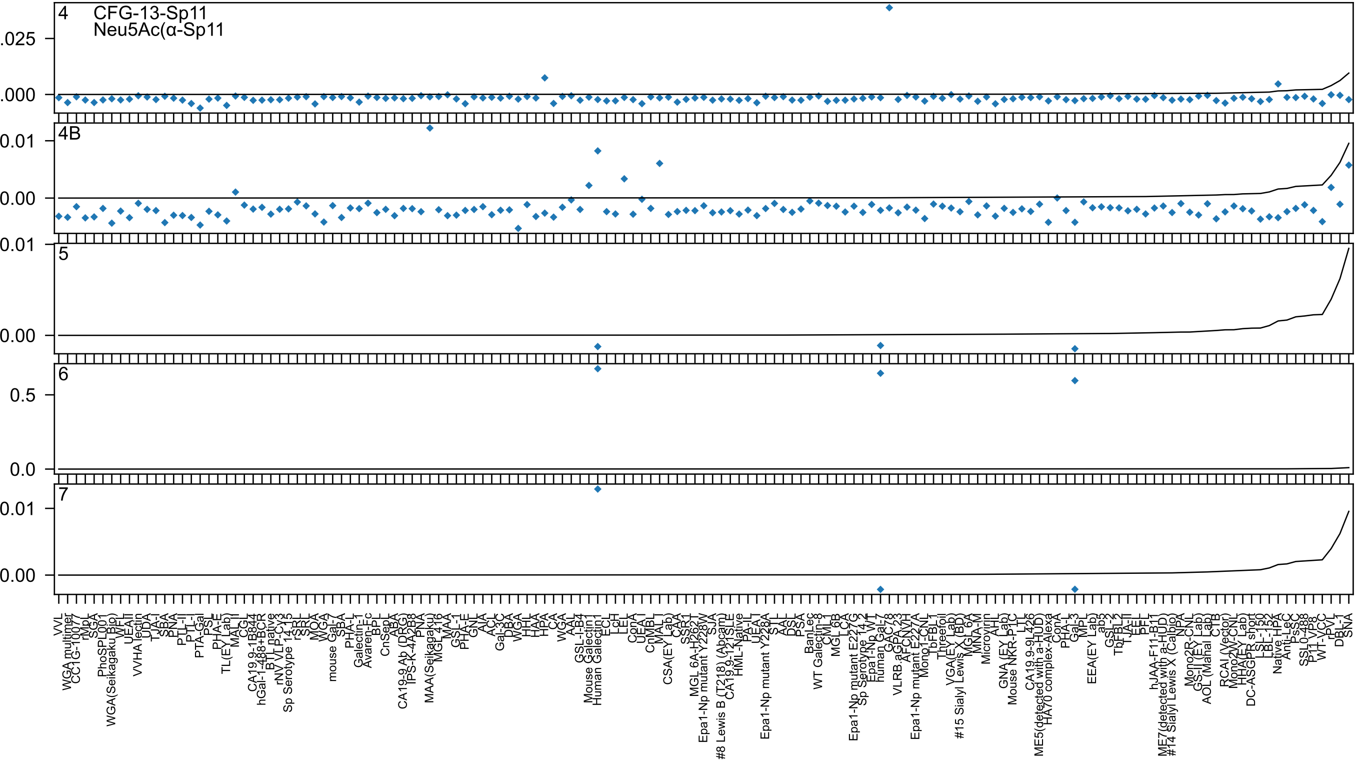
4
CFG-13-Sp11
Neu5Ac(α-Sp11
.025
.000
4B
0.01
0.00
5
0.01
0.00
6
0.5
0.0
7
0.01
0.00

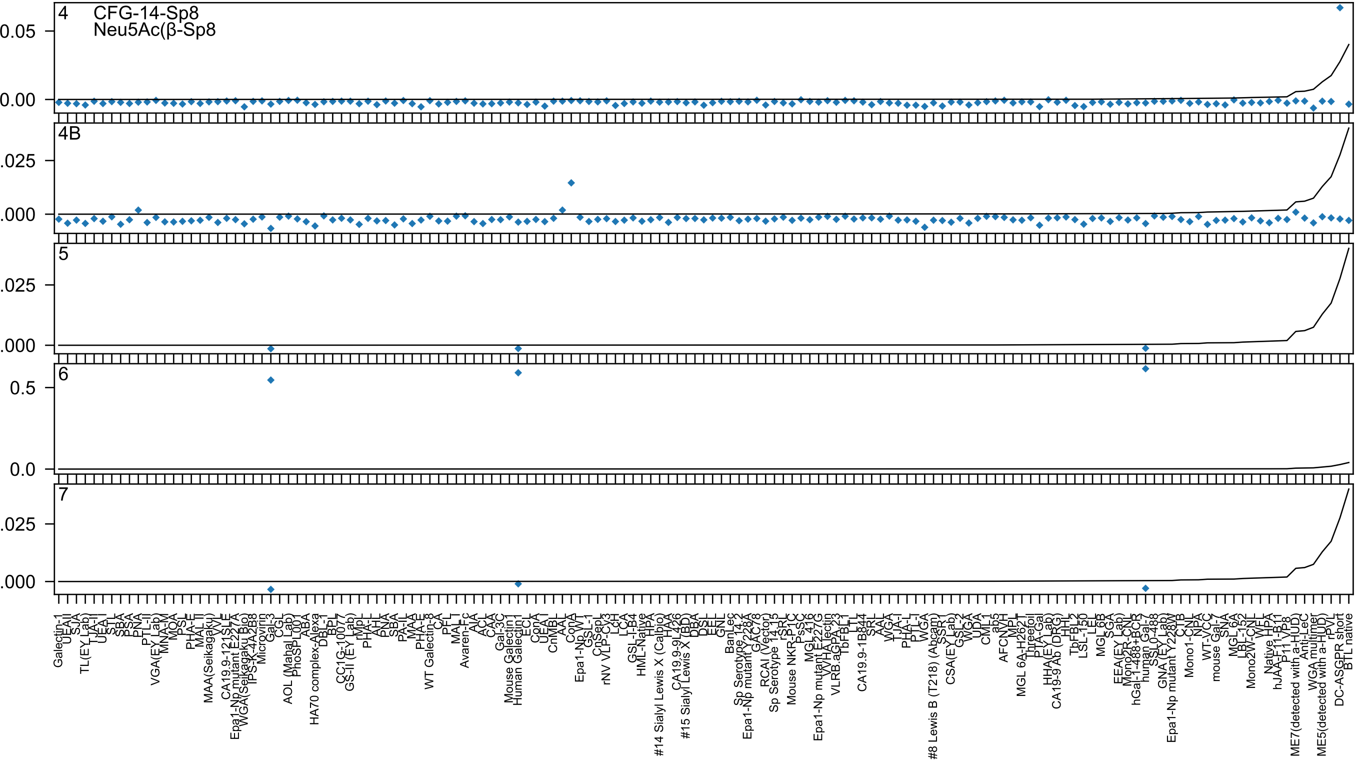
4
CFG-14-Sp8
Neu5Ac(β-Sp8
0.05
0.00
4B
.025
.000
5
.025
.000
6
0.5
0.0
7
.025
.000

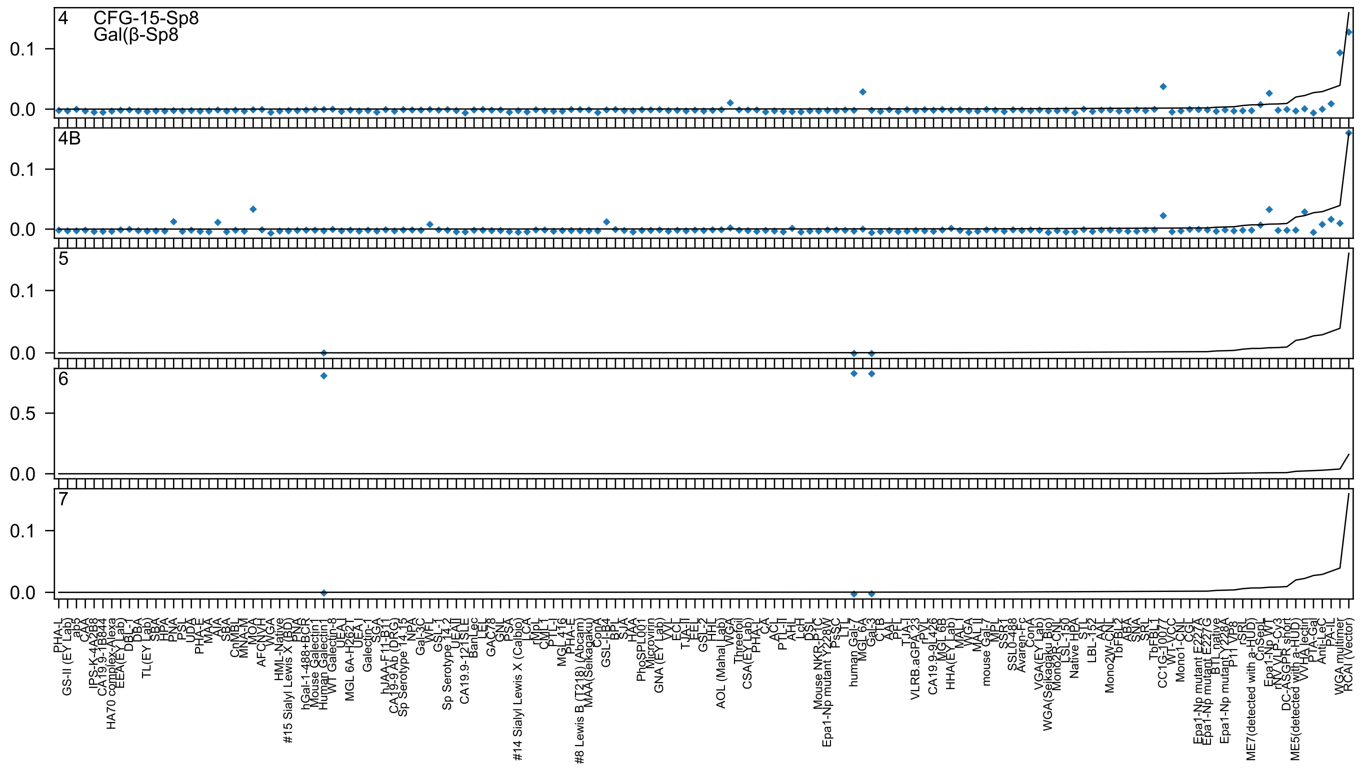
4
CFG-15-Sp8
Gal(β-Sp8
4B
5
6
7
0.0
0.1
0.5
PHA-L
GS-II (EY Lab)
ab5
CAA
IPS-K-4A2B8
CA19.9-1B844
HA70 complex-Alexa
EEA(EY Lab)
DBL-1
DBA
TL(EY Lab)
SBA
HPA
PNA
PSL
UDA
PHA-E
MAA
AIA
SBA
CnMBL
MNA-M
MOA
AFCNVH
WGA
HML-Native
#15 Sialyl Lewis X (BD)
PNA
hGal-1-488+BCR
Mouse Galectin1
Human Galectin1
WT Galectin-8
UEA-I
MGL 6A-H262T
UEA-I
Galectin-1
SGA
hJAA-F11-B11
CA19-9 Ab (DRG)
Sp Serotype 14.15
NPA
Gal-3C
WFL
GSL-1
Sp Serotype 14.2
UEAII
CA19.9-121SLE
BanLec
LEL
GAC78
GNL
PSA
#14 Sialyl Lewis X (Calbio)
LCA
rMpL
CML-1
PTL-I
MGL416
PHA-E
#8 Lewis B (T218) (Abcam)
MAA(Seikagaku)
ConA
GSL-I-B4
BPL
SJA
HAA
PhoSPL001
Microvirin
GNA (EY Lab)
VVL
ECL
TJA-II
LEL
GSL-2
HHL
AOL (Mahal Lab)
WGA
Threefoil
CSA(EY Lab)
PHA-L
CA
ACL
PTL-II
AHL
LcH
DSL
Mouse NKR-P1C
Epa1-Np mutant Y228W
PsSC
LTL
human Gal-7
MGL 6A
Gal-3
CTB
AAL
PFL
TJA-I
VLRB.aGPA.23
rPVL
CA19.9-9L426
MGL 6B
HHA(EY Lab)
MAL-I
WGA
MAL II
mouse Gal-7
MPL
SSR1
SSL0-488
Avaren-Fc
ConA
VGA(EY Lab)
WGA(Seikagaku Bio)
Mono2R-CNL
LSL-150
Native HPA
STL
LBL-152
AAL
Mono2W-CNL
TbFBL2
ABA
SNA
SRL
TbFBL1
CC1G-10077
WT-VCC
Mono1-CNL
CGL
Epa1-Np mutant E227A
Epa1-Np mutant E227G
BTL native
Epa1-Np mutant Y228A
P11 VP8
rSRL
ME7(detected with a-HUD)
CnSepL
Epa1-Np WT
rNV VLP-Cy3
DC-ASGPR short
ME5(detected with a-HUD)
WHA lectin
PTA-Gal
Anti-LeC
PA-IL
WGA multimer
RCAI (Vector)

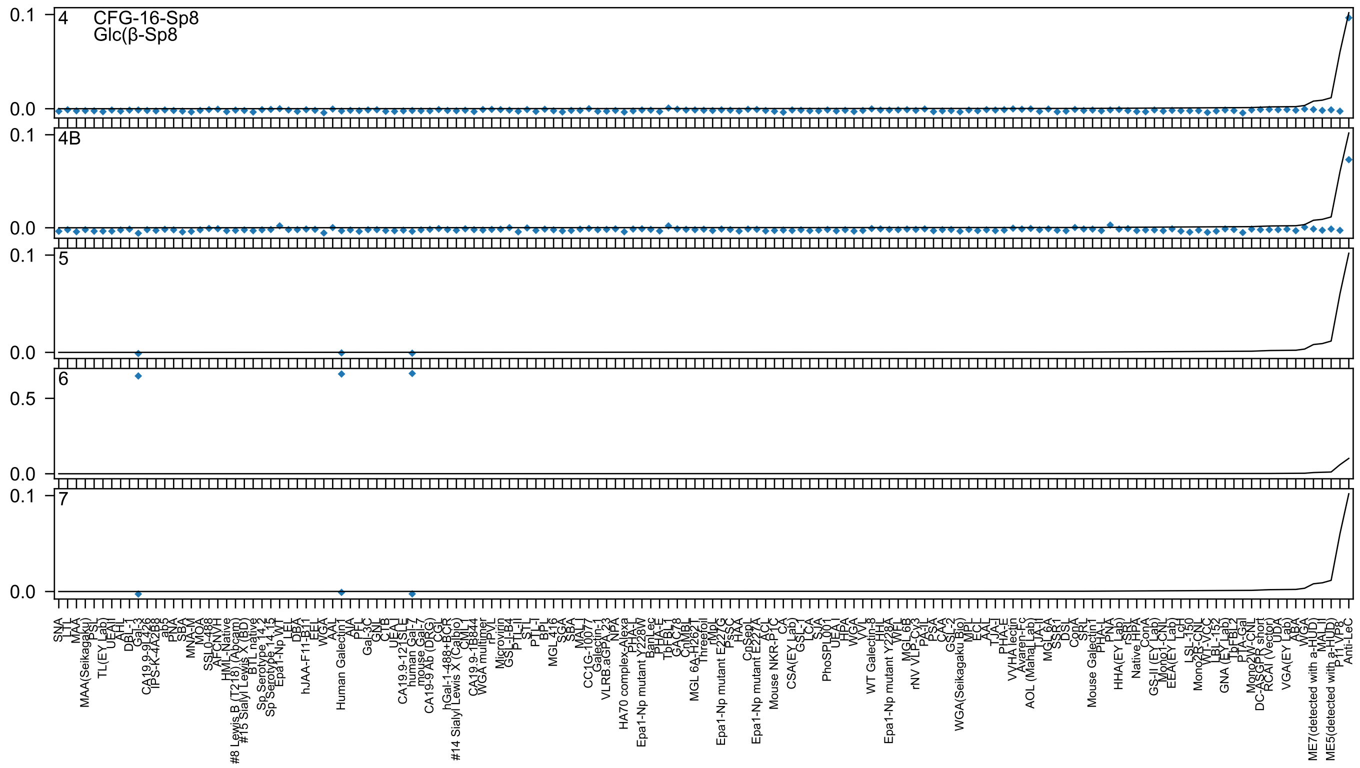
4
CFG-16-Sp8
Glc(β-Sp8
4B
5
6
7
0.1
0.0
0.5
SNA
Anti-LeC

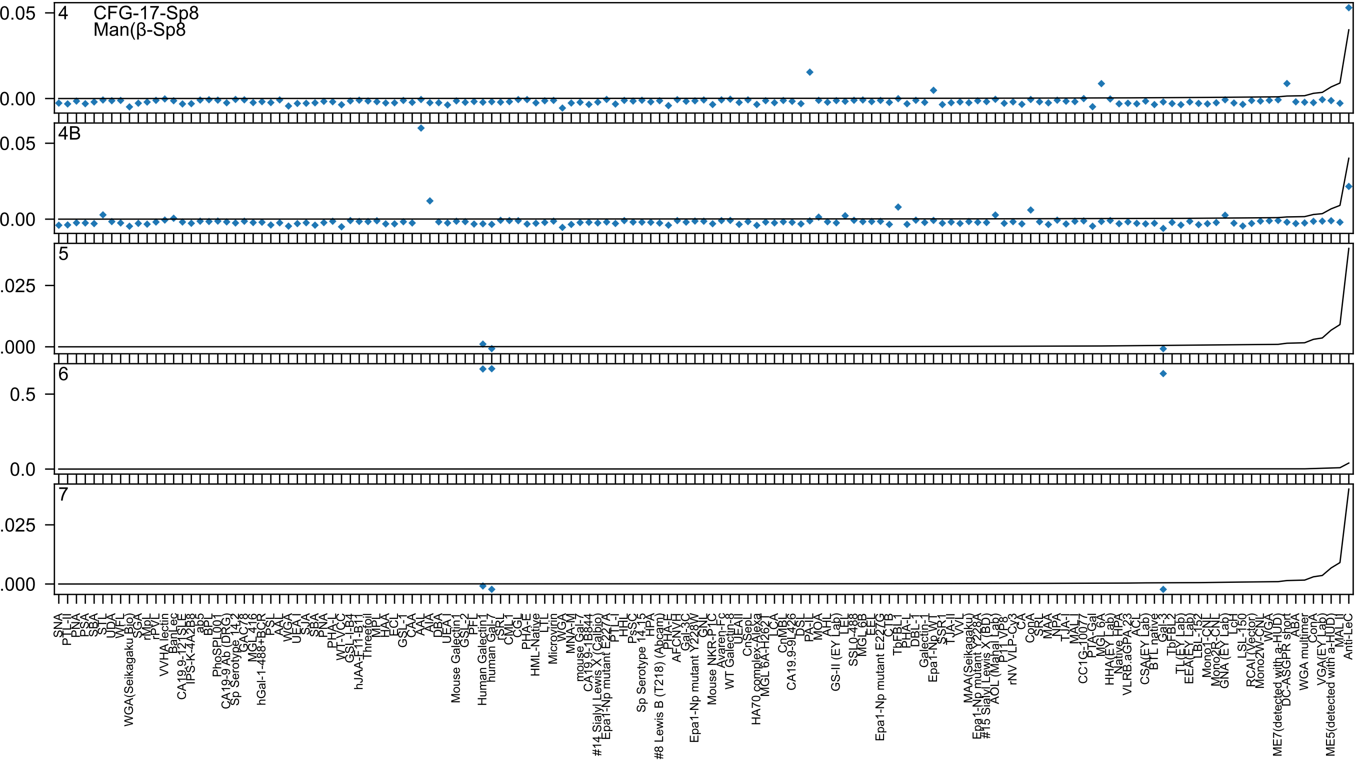

4
CFG-17-Sp8
Man(β-Sp8
4B
5
6
7

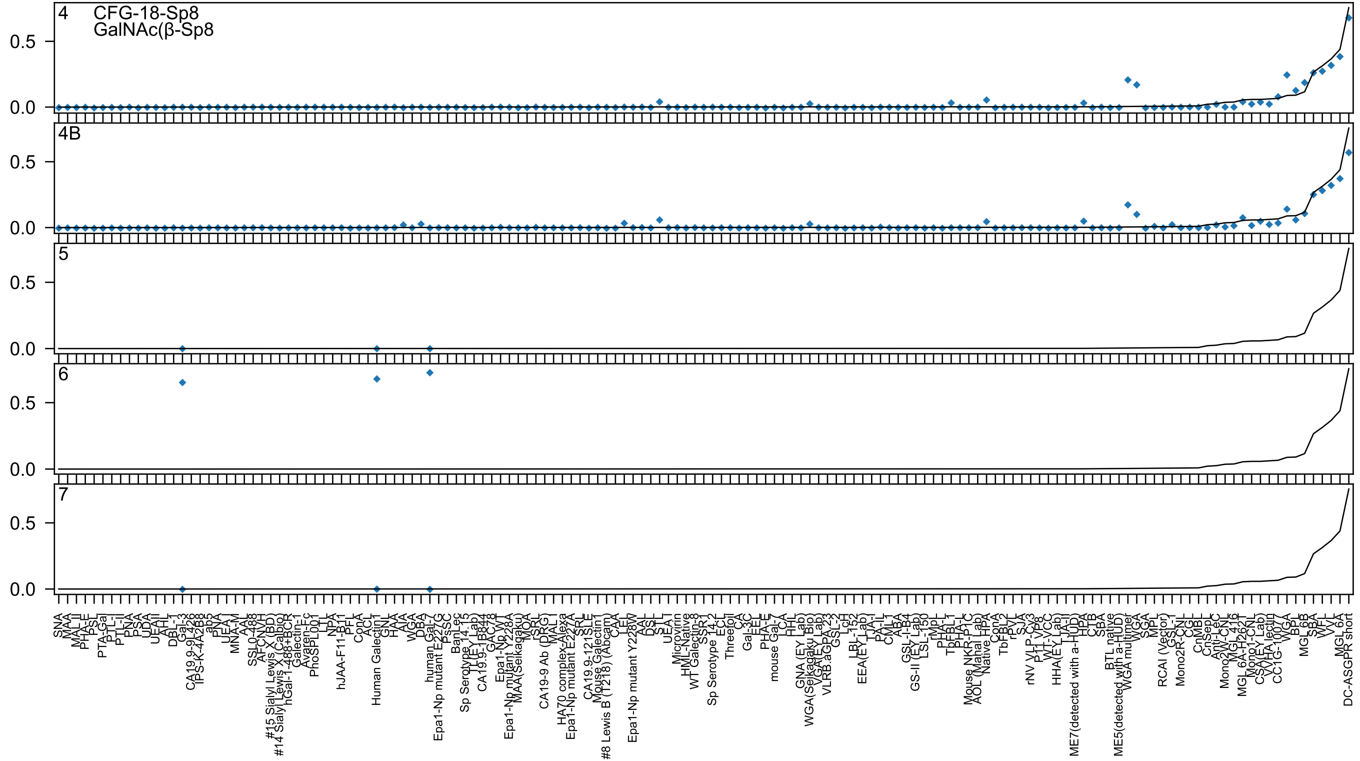
4
CFG-18-Sp8
GalNAc(β-Sp8
4B
5
6
7
0.0
0.5
SNA
MAA
MAL_II
PHA-E
PSL
PTA-Gal
PTL-I
PTL-II
PNA
PSA
UDA
UEAII
AHL
DBL-1
Gal-3
CA19.9-9L426
IPS-K-4A2B8
ab5
PNA
UEA-I
MNA-M
AAL
SSL0-488
AFCNVH
#15 Sialyl Lewis X (BD)
#14 Sialyl Lewis X (Calbio)
hGal-1-488+BCR
Galectin-1
Avaren-Fc
PhoSPL001
LTL
NPA
hJAA-F11-B1
PFL
ConA
ACL
Human Galectin1
GNL
HAA
AIA
WGA
DBA
human Gal-7
Epa1-Np mutant E227G
PsSC
BanLec
Sp Serotype 14.15
TJL(EY Lab)
CA19.9-1B844
GAC78
Epa1-Np WT
Epa1-Np mutant Y228A
MAA(Seikagaku)
MOA
rSRL
CA19-9 Ab (DRG)
MAL-I
HA70 complex-Alexa
Epa1-Np mutant E227A
SRL
CA19.9-121SLE
Mouse Galectin1
#8 Lewis B (T218) (Abcam)
CAA
LEL
Epa1-Np mutant Y228W
AAL
DSL
STL
UEA I
Microvirin
HML-Native
WT Galectin-8
SSR1
Sp Serotype 14.2
ECL
Threefoil
CA
Gal-3C
EEL
PHA-E
mouse Gal-7
LCA
HHL
GNA (EY Lab)
WGA(Seikagaku Bio)
VGA(EY Lab)
VLRB.aGPA.23
GSL-2
LcH
LBL-152
EEA(EY Lab)
TJA-I
PA-IL
CML-1
ABA
GSL-I-B4
GS-II (EY Lab)
LSL-150
rMpL
PHA-L
TbFBL1
PHA-L
Mouse NKR-P1C
AOL (Manal Lab)
Native HPA
ConA
TbFBL2
rPVL
SJA
rNV VLP-Cv3
P11 VP8
WT-VCC
HHA(EY Lab)
TJA-II
ME7(detected with a-HUD)
HPA
CTB
SBA
BTL native
ME5(detected with a-HUD)
WGA multimer
WGA
SGA
MPL
RCAI (Vector)
GSL-I
Mono2R-CNL
CGL
CnMBL
CnSepL
Anti-LeC
Mono2W-CNL
MGL416
MGL 6A-H262T
Mono1-CNL
CSA(EY Lab)
VVHA lectin
CC1G-10077
WGA
BPL
MGL 6B
SBA
WFL
VVL
MGL 6A
DC-ASGPR short

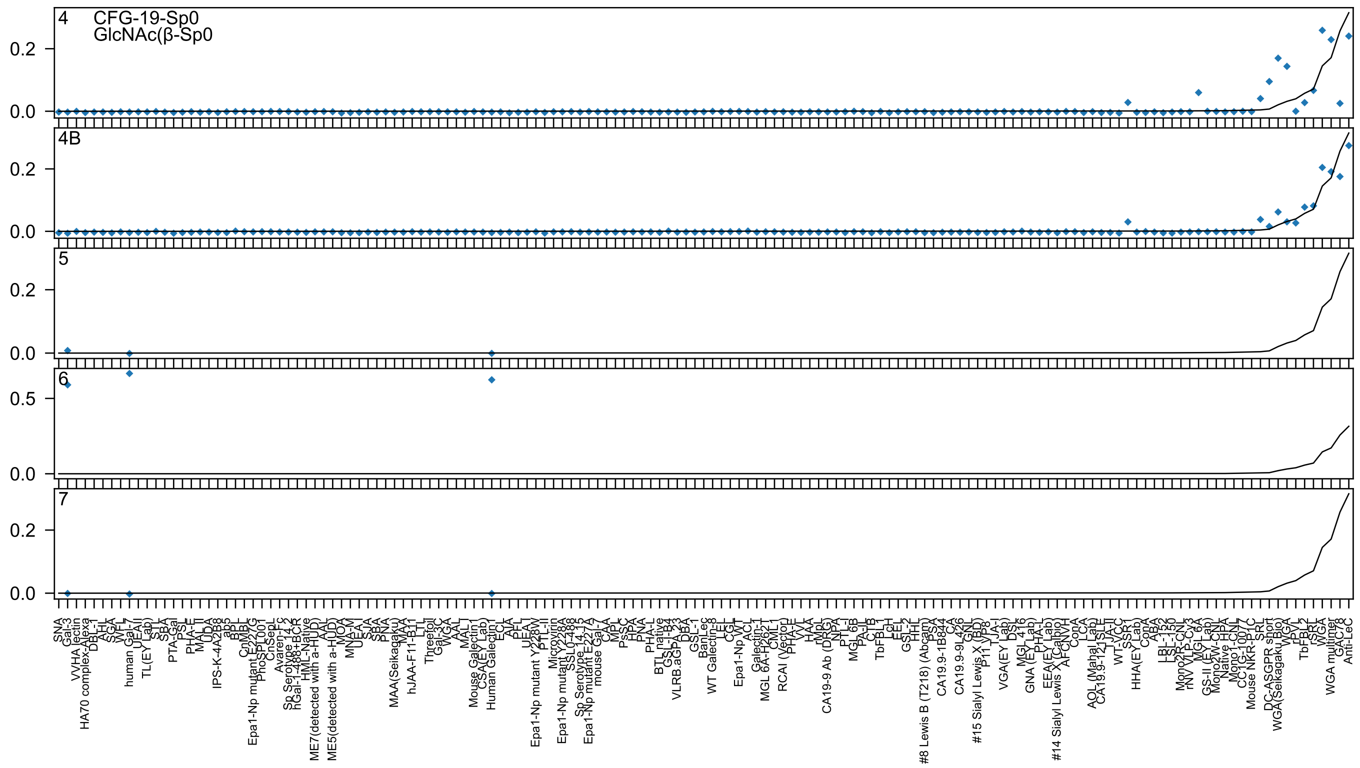
4
CFG-19-Sp0
GlcNAc(β-Sp0
4B
5
6
7

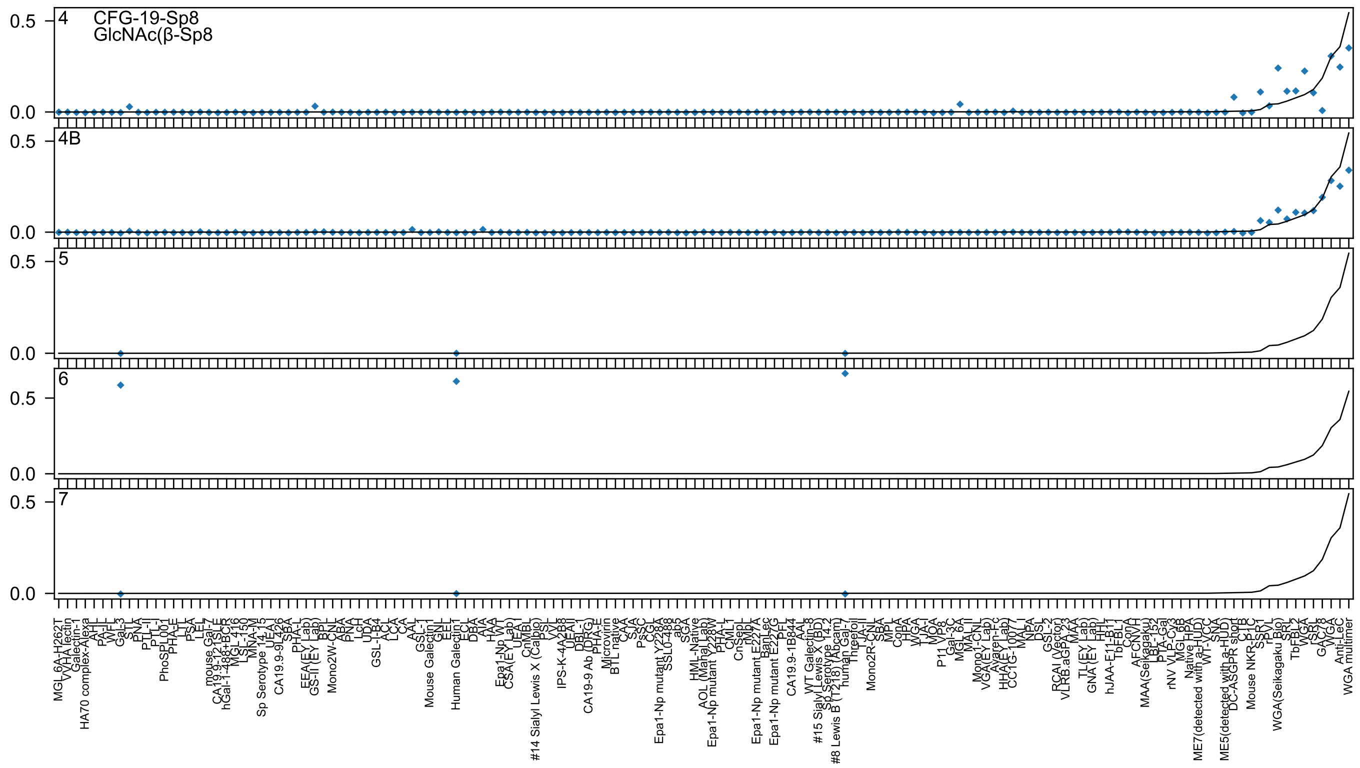
4
CFG-19-Sp8
GlcNAc(β-Sp8
4B
5
6
7

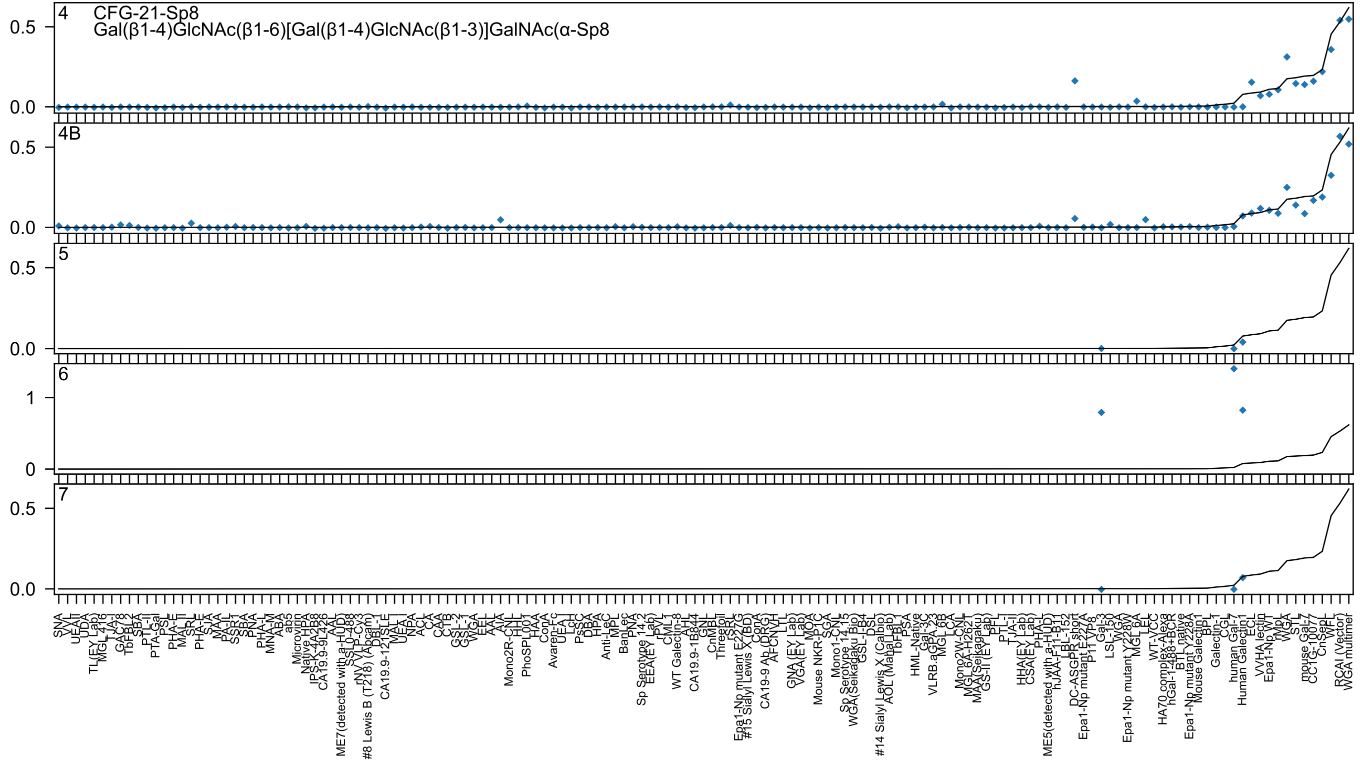
4
CFG-21-Sp8
Gal(β1-4)GlcNAc(β1-6)[Gal(β1-4)GlcNAc(β1-3)]GalNAc(α-Sp8
4B
5
6
7
0.5
0.0
1
0

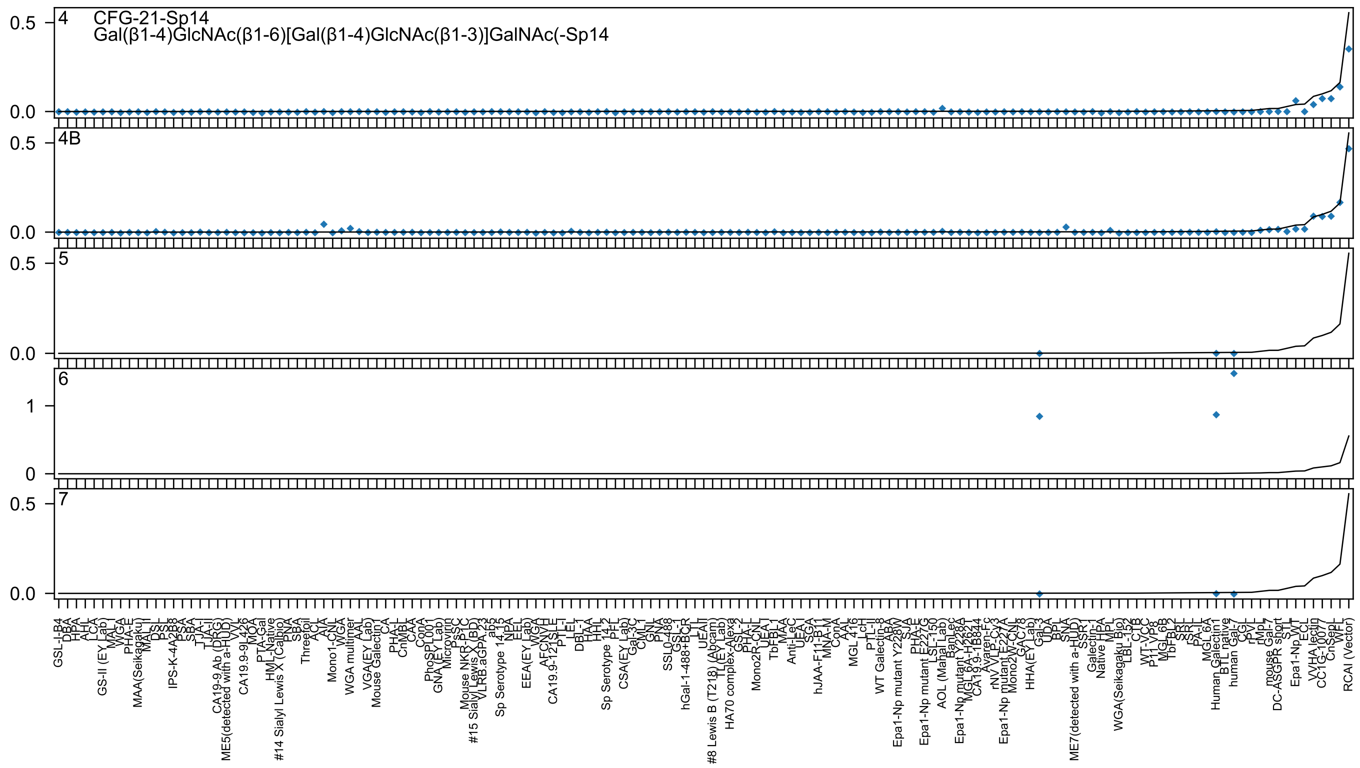

4 CFG-21-Sp14
Gal(β1-4)GlcNAc(β1-6)[Gal(β1-4)GlcNAc(β1-3)]GalNAc(-Sp14
4B
5
6
7

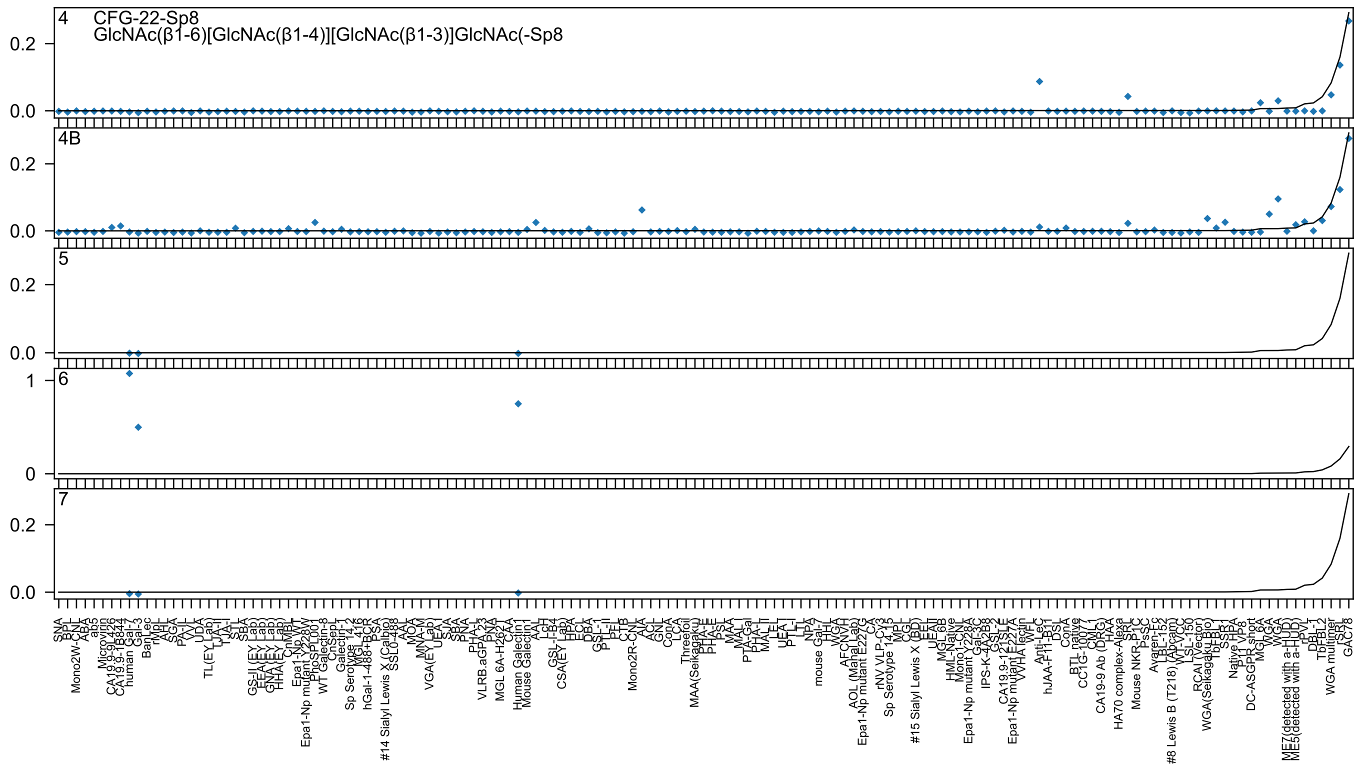

4 CFG-22-Sp8
GlcNAc(β1-6)[GlcNAc(β1-4)][GlcNAc(β1-3)]GlcNAc(-Sp8
4B
5
6
7
0.0
0.2
0
1
SNA
BPL
Mono2W-CNL
ABA
ab5
Microvirin
CA19.9-9L426
CA19.9-1B844
human Gal-7
Gal-3
BanLec
rMpL
AHL
SGA
PA-IL
VVL
UDA
TL(EY Lab)
TJA-II
TJA-I
STL
SBA
GS-II (EY Lab)
EEA(EY Lab)
GNA(EY Lab)
HHA(EY Lab)
CnMBL
Epa1-Np WT
Epa1-Np mutant Y228W
PhoSL001
WT Galectin-8
CnSepL
Galectin-1
Sp Serotype 14.2
MGL 416
hGal-1-488+BCR
PSA
#14 Sialyl Lewis X (Calbio)
SSL0-488
AAL
MOA
MNA-M
VGA(EY Lab)
UEA-I
SJA
SBA
PNA
PHA-L
VLRB.aGPA.23
PNA
MGL 6A-H262
CAA
Human Galectin1
Mouse Galectin1
AAL
LcH
GSL-I-B4
CSA(EY Lab)
HPA
ECL
DBA
GSL-I
PTL-II
PFL
CTB
Mono2R-CNL
AIA
ACL
GNL
ConA
LCA
Threefoil
MAA(Seikagaku)
PHA-E
PHA-E
PSL
MAA
MAL-I
PTA-Gal
PHA-L
MAL-II
LEL
UEA-I
PTL-I
LTL
NPA
mouse Gal-7
HHL
WGA
AFCNVH
AOL (Manal Lab)
Epa1-Np mutant E227G
CA
rNV VLP-Cy3
Sp Serotype 14.15
MPL
CGL
#15 Sialyl Lewis X (BD)
EEL
UEA-II
MGL 6B
HML-Native
Mono1-CNL
Epa1-Np mutant Y228A
Gal-3C
IPS-K-4A2B8
GSL-2
CA19.9-121SLE
Epa1-Np mutant E227A
VVHA lectin
WFL
Anti-LeC
hJAA-F11-B11
DSL
ConA
BTL native
CC1G-10077
CML-1
CA19-9 Ab (DRG)
HAA
HA70 complex-Alexa
SRL
Mouse NKR-P1C
PsSC
Ayaren-Fc
LBL-152
#8 Lewis B (T218) (Abcam)
WT-VCC
LSL-150
RCAI (Vector)
WGA(Seikagaku Bio)
TbFBL1
SSR1
Native HPA
P11 VP8
DC-ASGPR short
MGL 6A
WGA
WGA
ME7(detected with a-HUD)
ME5(detected with a-HUD)
rPVL
DBL-1
TbFBL2
WGA multimer
rSRL
GAC78

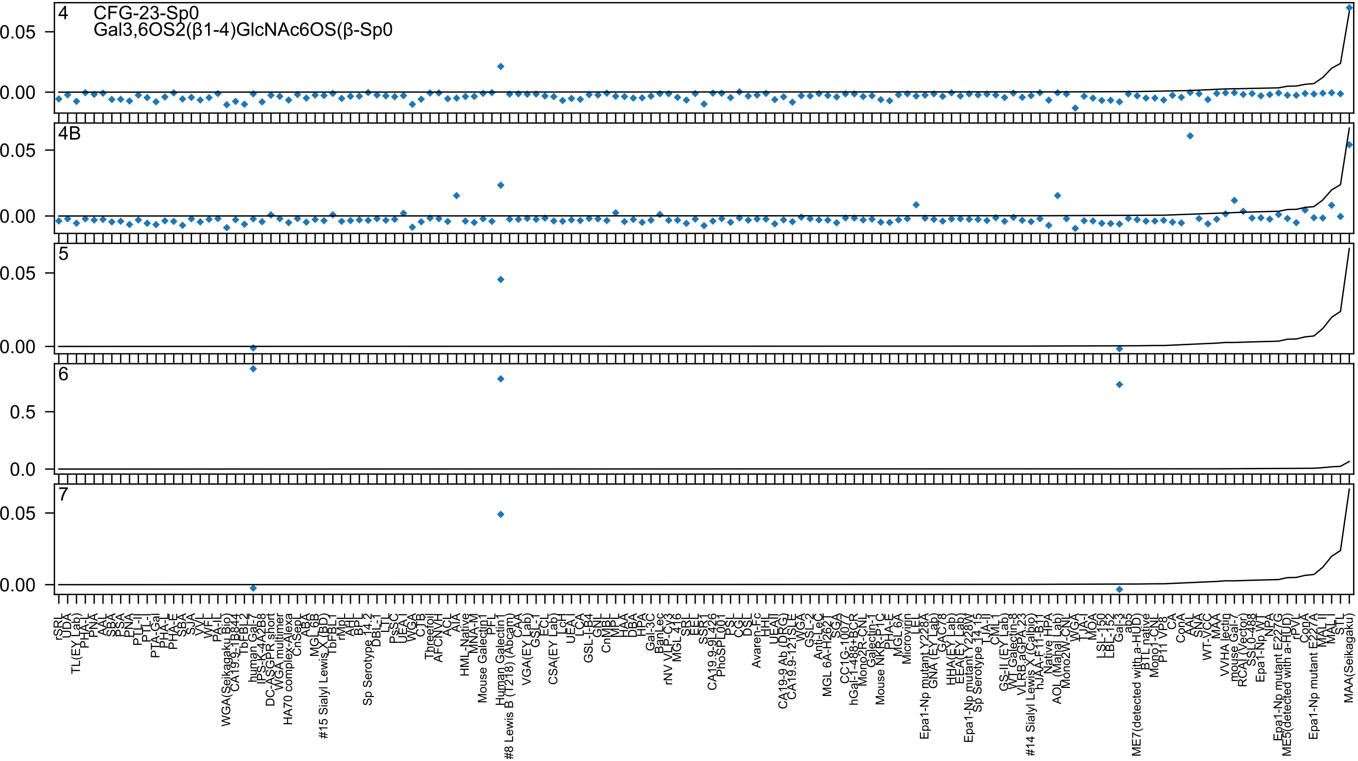

4
CFG-23-Sp0
Gal3,6OS2(β1-4)GlcNAc6OS(β-Sp0
4B
5
6
7
0.05
0.00
0.5
0.0

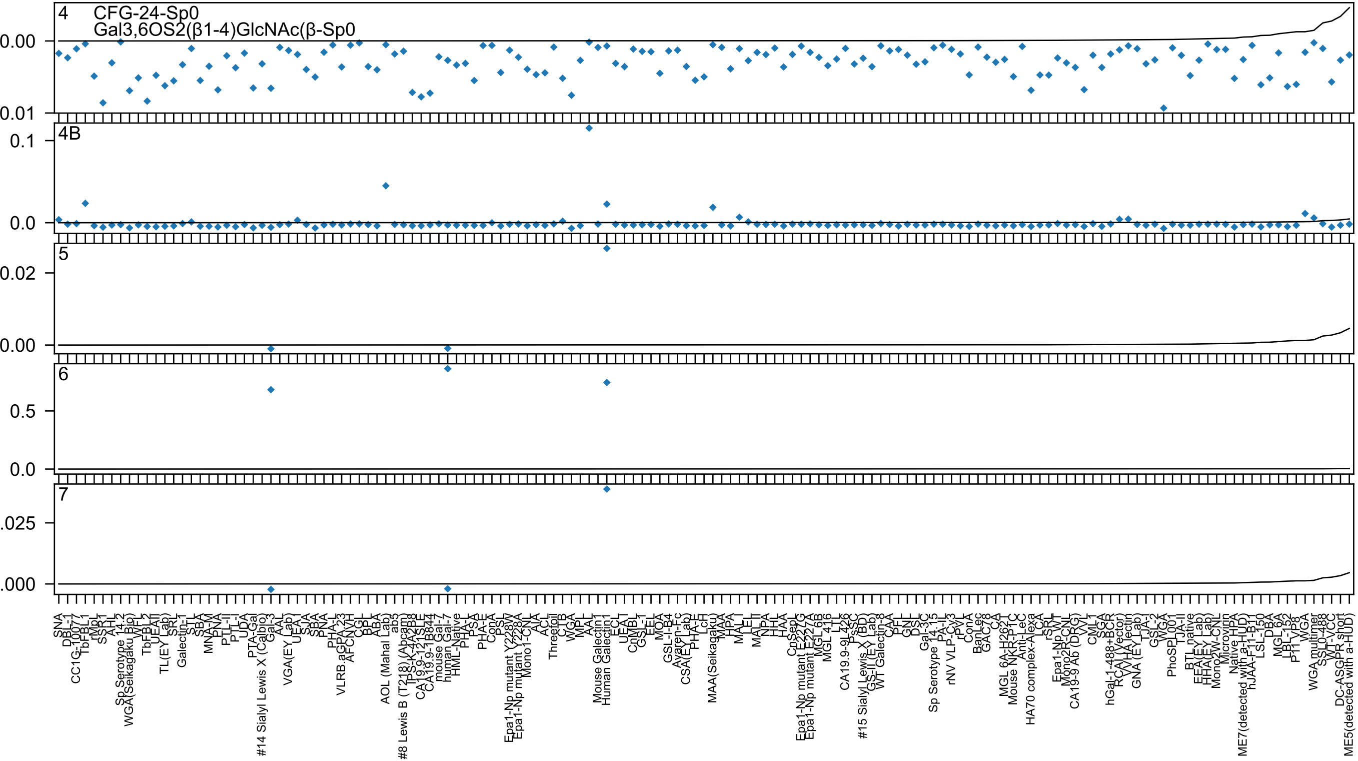
4 CFG-24-Sp0
Gal3,6OS2(β1-4)GlcNAc(β-Sp0
0.00
0.01
4B
0.1
0.0
5
0.02
0.00
6
0.5
0.0
7
.025
.000

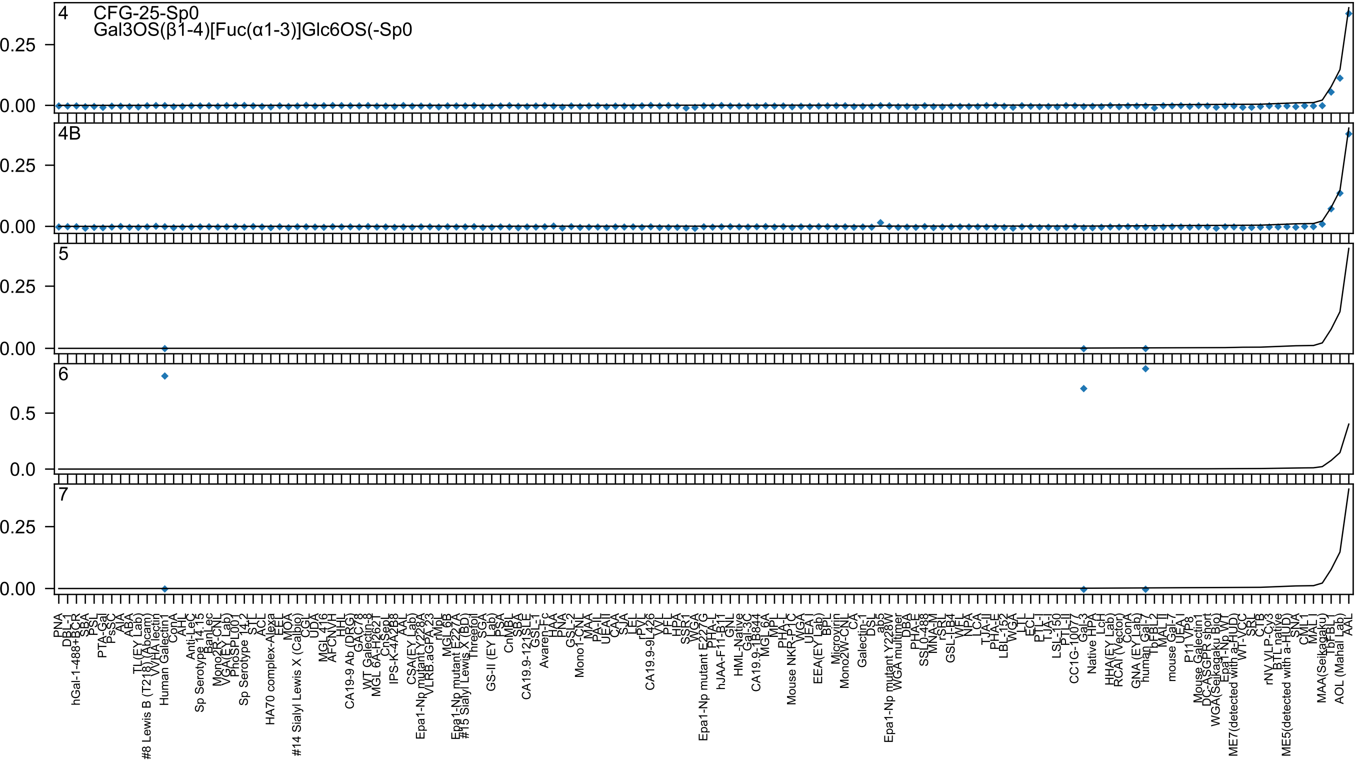

4 CFG-25-Sp0
Gal3OS(β1-4)[Fuc(α1-3)]Glc6OS(-Sp0
4B
5
6
7

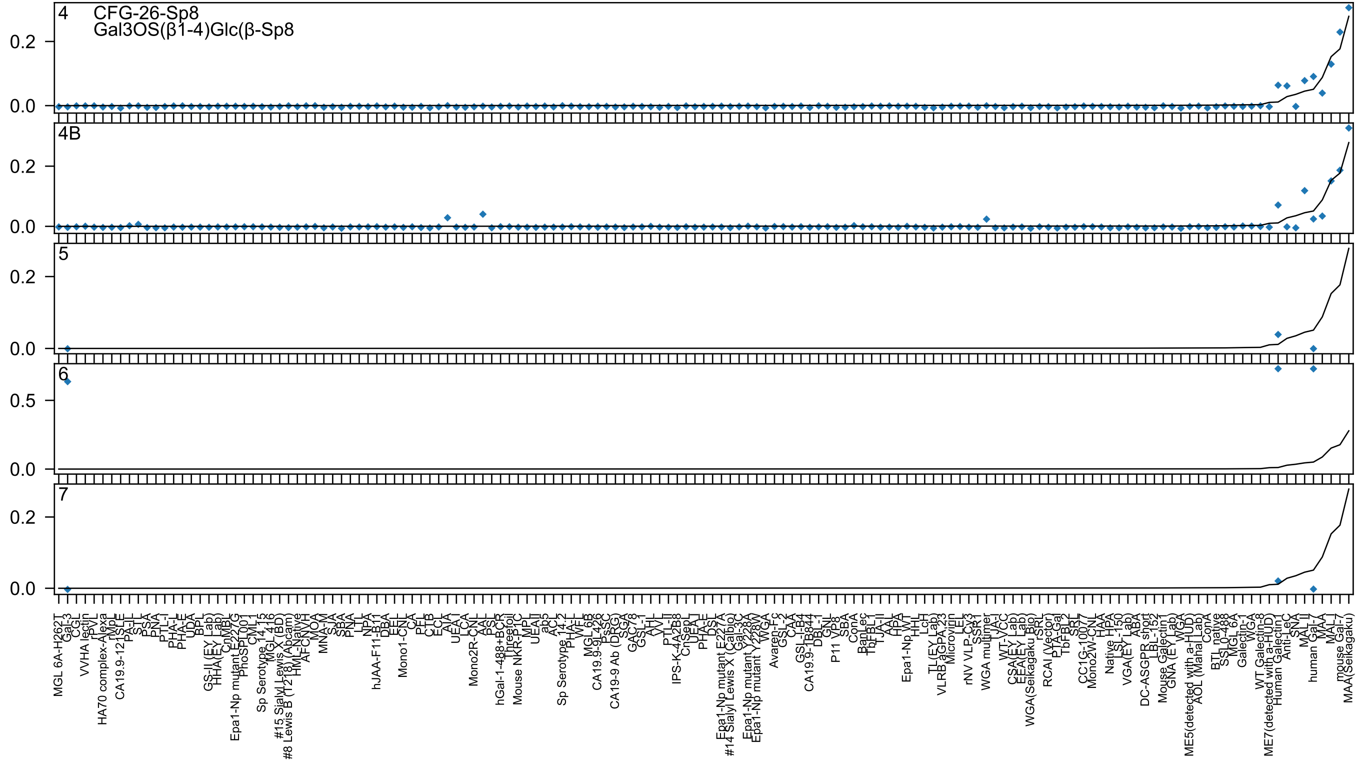

4
CFG-26-Sp8
Gal3OS(β1-4)Glc(β-Sp8
4B
5
6
7

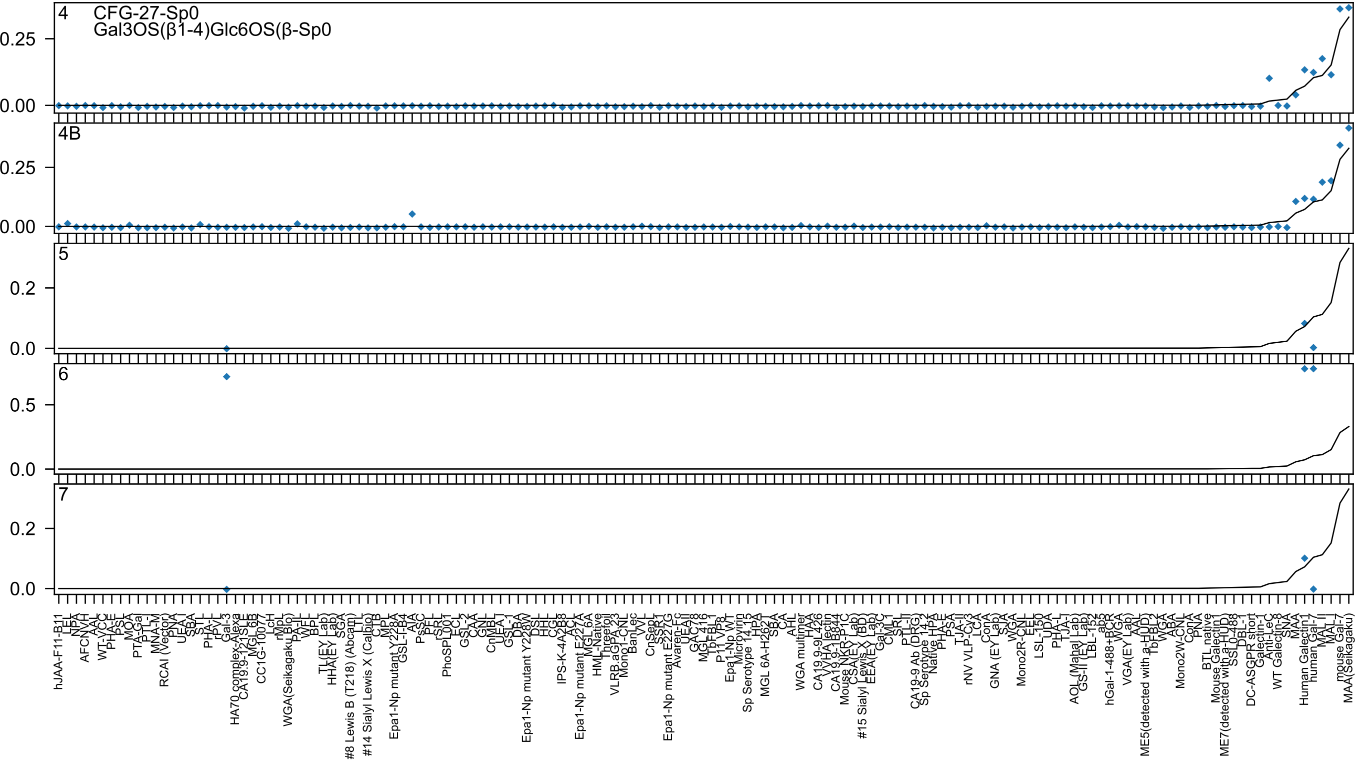

4
CFG-27-Sp0
Gal3OS(β1-4)Glc6OS(β-Sp0
0.25
0.00
4B
0.25
0.00
5
0.2
0.0
6
0.5
0.0
7
0.2
0.0
hJAA-F11-B11
LEL
NPA
AFCNVH
AAL
WT-VCC
PHA-E
PSL
MOA
PTA-Gal
PTL-I
MNA-M
RCAI (Vector)
PNA
UEA-I
SBA
STL
PHA-L
rPVL
Gal-3
HA70 complex-Alexa
CA19.9-121SLE
MGL 6B
CC1G-10077
LcH
rMpL
WGA(Seikagaku Bio)
PA-IL
WFL
BPL
TL(EY Lab)
HHA(EY Lab)
SGA
#8 Lewis B (T218) (Abcam)
LTL
#14 Sialyl Lewis X (Calbio)
CTB
MPL
Epa1-Np mutant Y228A
GSL-I-B4
AIA
PsSC
PFL
rSRL
PhoSL
PL001
ECL
GSL-2
CAA
GNL
CnMBL
UEA-I
GSL-1
DBA
Epa1-Np mutant Y228W
DSL
HHL
CGL
IPS-K-4A2B8
ACL
Epa1-Np mutant E227A
MGL 6A
HML-Native
Threefoil
VLRB.aGPA.23
Mono1-CNL
BanLec
VVL
CnSepL
SSR1
Epa1-Np mutant E227G
Avaren-Fc
UEAII
GAC78
MGL 416
TbFBL1
P11 VP8
Epa1-Np WT
Microvirin
Sp Serotype 14_15
HPA
MGL 6A-H262T
SBA
CA
AHL
WGA multimer
HAA
CA19.9-9L426
VVHA lectin
CA19.9-1B844
Mouse NKR-P1C
CSA(EY Lab)
#15 Sialyl Lewis X (BD)
EEA(EY Lab)
Gal-3C
CML-1
SRL
PTL-II
CA19-9 Ab (DRG)
Sp Serotype 14_2
Native HPA
PHA-E
PSA
TJA-II
rNV VLP-Cy3
LCA
ConA
GNA (EY Lab)
SJA
WGA
Mono2R-CNL
EEL
LSL-150
UDA
PHA-L
TJA-I
AOL (Mahal Lab)
GS-II (EY Lab)
LBL-152
ab5
hGal-1-488+BCR
WGA
VGA(EY Lab)
AAL
ME5(detected with a-HUD)
TbFBL2
WGA
ABA
Mono2W-CNL
ConA
PNA
BTL native
Mouse Galectin1
ME7(detected with a-HUD)
SSL0-488
DBL-1
DC-ASGPR short
Galectin-1
Anti-LeC
WT Galectin-8
SNA
MAA
Human Galectin1
human Gal-7
MAL-II
MAL-I
mouse Gal-7
MAA(Seikagaku)

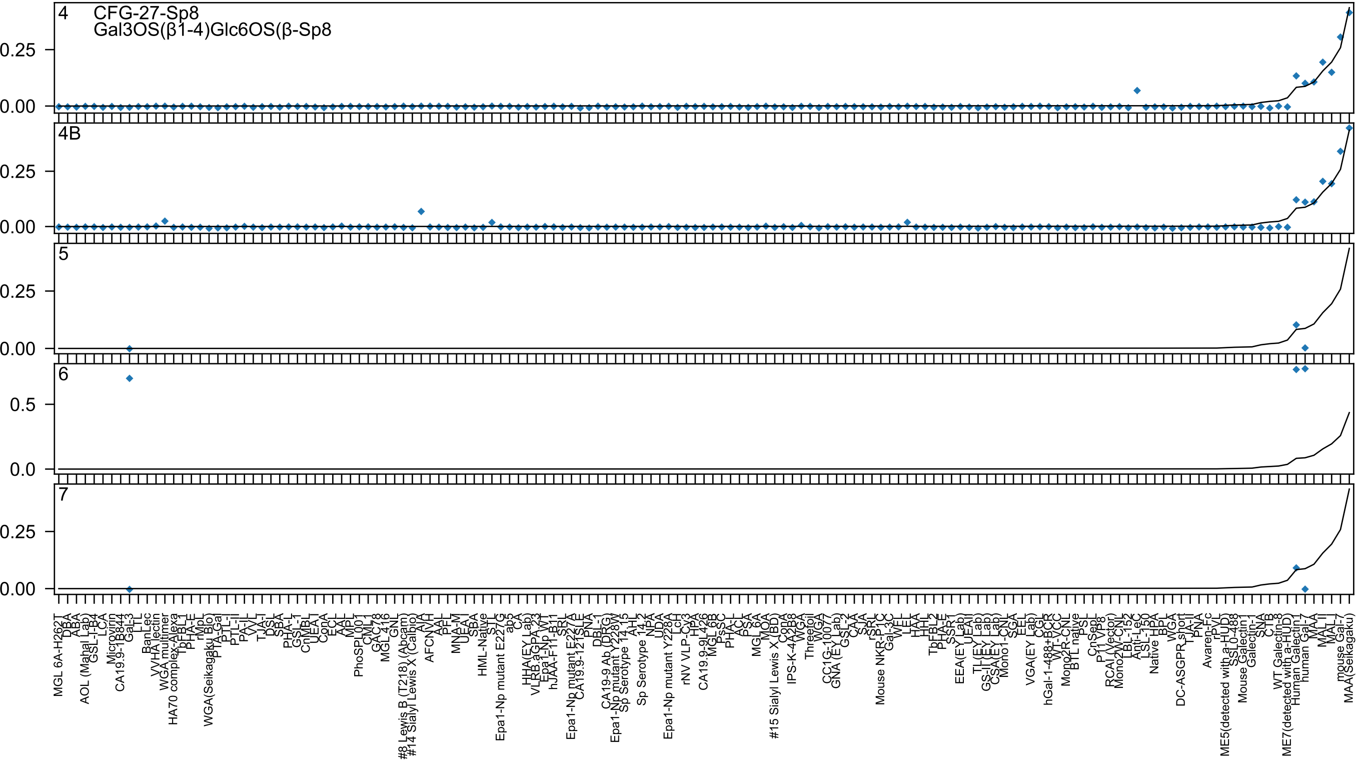

4
CFG-27-Sp8
Gal3OS(β1-4)Glc6OS(β-Sp8
4B
5
6
7
0.25
0.00
0.5
0.0

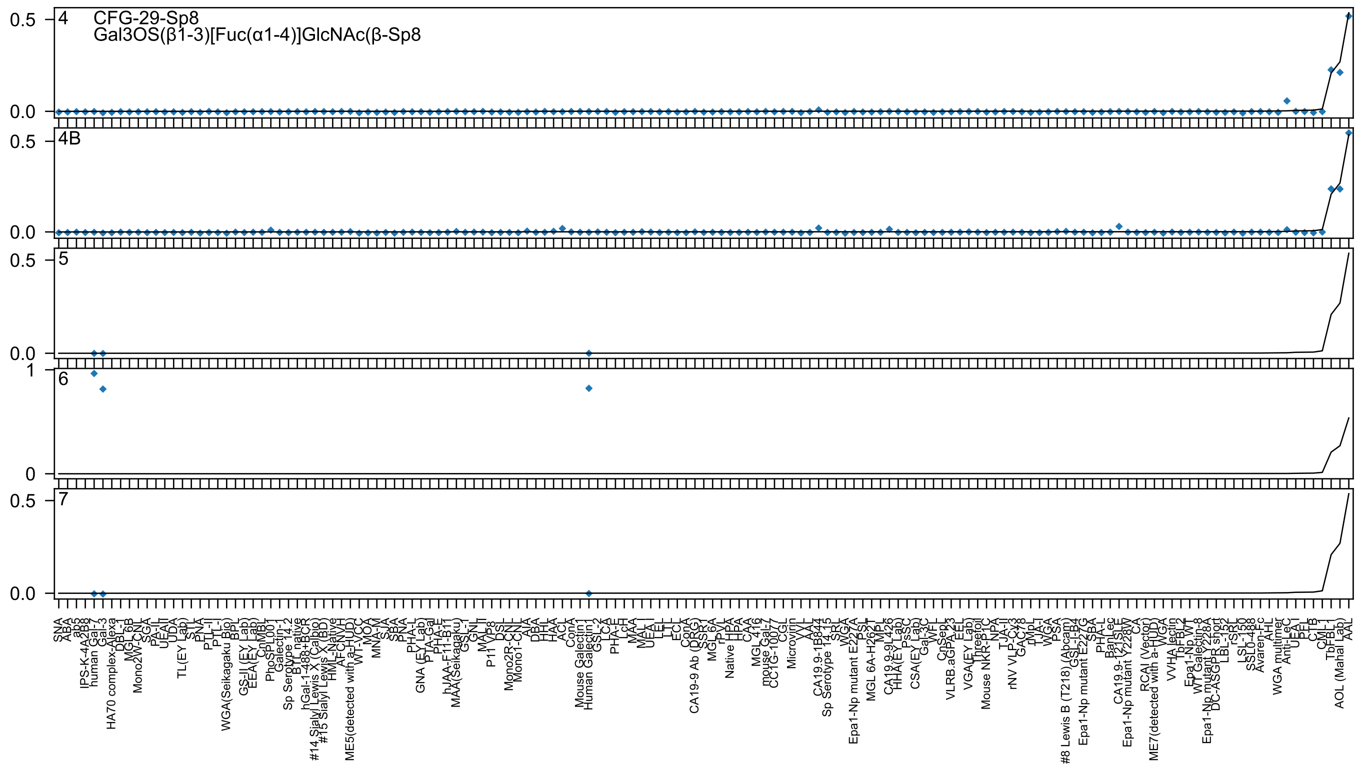
4
CFG-29-Sp8
Gal3OS(β1-3)[Fuc(α1-4)]GlcNAc(β-Sp8
4B
5
6
7
0.5
0.0
1
0

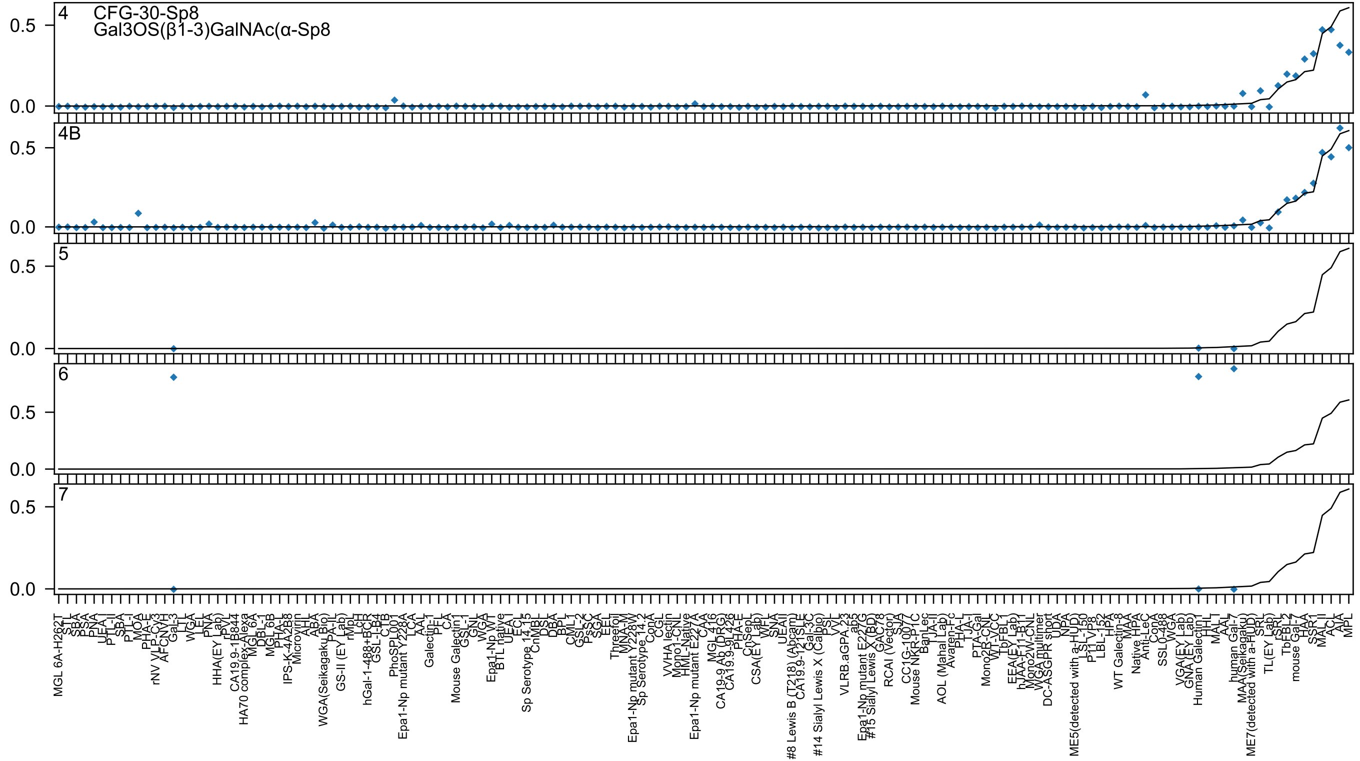
4
CFG-30-Sp8
Gal3OS(β1-3)GalNAc(α-Sp8
4B
5
6
7
0.5
0.0
MGL 6A-H262T
STL
SBA
PSA
PNA
UEA-I
PTL-II
SBA
PTL-I
MOA
PHA-E
rNV VLP-Cy3
AFCNYH
Gal-3
LTL
WGA
LEL
PNA
HHA(EY Lab)
rPVL
CA19.9-1B844
HA70 complex-Alexa
MGL 6A
DBL-1
MGL 6B
PHA-L
IPS-K-4A2B8
Microvirin
AHL
ABA
WGA(Seikagaku Bio)
PA-IL
GS-II (EY Lab)
rMpL
LcH
hGal-1-488+BCR
GSL-I-B4
CTB
PhoSPL001
Epa1-Np mutant Y228A
LCA
AAL
Galectin-1
PFL
CA
Mouse Galectin1
GSL-I
GNL
WGA
Epa1-Np WT
BTL native
UEA-I
ECL
Sp Serotype 14.15
CnMBL
DSL
DBA
BPL
CML-1
GSL-2
PsSC
SGA
EEL
Threefoil
MNA-M
Epa1-Np mutant Y228W
Sp Serotype 14.2
ConA
CGL
VVHA lectin
Mono1-CNL
HML-Native
Epa1-Np mutant E227A
CAA
MGL 416
CA19-9 Ab (DRG)
CA19.9-9L426
PHA-E
CnSepL
CSA(EY Lab)
WFL
SNA
UEAII
#8 Lewis B (T218) (Abcam)
CA19.9-121SLE
Gal-3C
#14 Sialyl Lewis X (Calbio)
PSL
VVL
VLRB.aGPA.23
ab5
Epa1-Np mutant E227G
#15 Sialyl Lewis X (BD)
GAC78
RCAI (Vector)
SJA
CC1G-10077
Mouse NKR-P1C
BanLec
TJA-II
AOL (Mahal Lab)
Avaren-Fc
PHA-L
TJA-I
PTA-Gal
Mono2R-CNL
WT-VCC
TbFBL1
EEA(EY Lab)
hJAA-F11-B1T
Mono2W-CNL
WGA multimer
DC-ASGPR short
UDA
NPA
ME5(detected with a-HUD)
LSL-150
P11 VP8
LBL-152
HPA
WT Galectin-8
MAA
Native HPA
Anti-LeC
ConA
SSL0-488
WGA
VGA(EY Lab)
GNA(EY Lab)
Human Galectin1
HHL
MAL-I
AAL
human Gal-7
MAA(Seikagaku)
ME7(detected with a-HUD)
SRL
TL(EY Lab)
rSRL
TbFBL-2
mouse Gal-7
HAA
SSR1
MAL-II
ACT
AIA
MPL

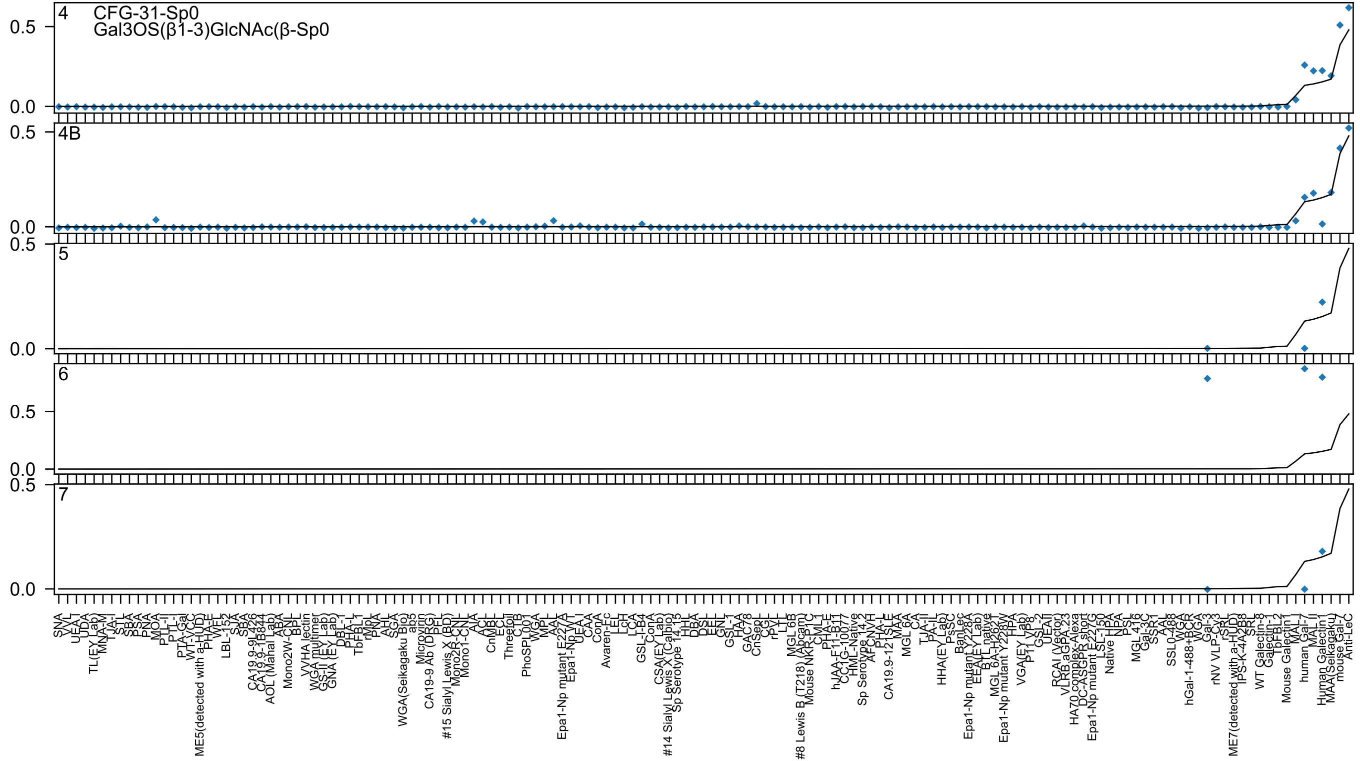

4
CFG-31-Sp0
Gal3OS(β1-3)GlcNAc(β-Sp0
4B
5
6
7
0.0
0.5
SNA
VVL
UEA-I
UDA
TL(EY Lab)
MNA-M
TJA-I
STL
SBA
PSA
PNA
MOA
PTL-II
PTL-I
PTA-Gal
WT-VCC
ME5(detected with a-HUD)
PHA-E
WFL
LBL-152
SJA
SBA
CA19.9-9L426
CA19.9-1B844
AOL (Mahal Lab)
ABA
Mono2W-CNL
BPL
VVHA lectin
WGA multimer
GS-II (EY Lab)
GNA (EY Lab)
DBL-1
PHA-L
TbFBL1
rMpL
PNA
AHL
SGA
WGA(Seikagaku Bio)
ab3
Microvirin
CA19-9 Ab (DRG)
PFL
#15 Sialyl Lewis X (BD)
Mono2R-CNL
Mono1-CNL
AIA
ACL
CnMBL
ECL
Threefoil
CLIB
PhoSPL001
WGA
MPL
AAL
Epa1-Np mutant E227A
Epa1-Np WT
UEA I
CAA
ConA
Avaren-Fc
LEL
LcH
LCA
GSL-I-B4
ConA
CSA(EY Lab)
#14 Sialyl Lewis X (Calbio)
Sp Serotype 14.13
HHL
DBA
DSL
EEL
GNL
GSL-I
HAA
GAC78
CnSepL
CGL
rPVL
LTL
MGL 6B
#8 Lewis B (T218) (Abcam)
Mouse NKR-P1C
CML1
PHA-E
hJAA-F11-B11
CC1G-10077
HML-Native
Sp Serotype 14.2
AFCNVH
PHA-L
CA19.9-121SLE
MAA
MGL 6A
CA
TJA-II
PA-IL
HHA(EY Lab)
PsSC
BanLec
Epa1-Np mutant Y228A
EEA(EY Lab)
BTL native
MGL 6A-H262T
Epa1-Np mutant Y228W
HPA
VGA(EY Lab)
P11 VP8
GSL-2
UEAII
RCAI (Vector)
VLRB.aGPA.23
HA70 complex-Alexa
DC-ASGPR short
Epa1-Np mutant E227G
LSL-150
Native HPA
NPA
PSL
MGL 416
Gal-3C
SSR1
AAL
SSL0-488
WGA
hGal-1-488+BCR
WGA
Gal-3
rNV VLP-Cy3
rSRL
ME7(detected with a-HUD)
IPS-K-4A2B8
SRL
WT Galectin-8
Galectin-1
TbFBL2
Mouse Galectin1
MAL-I
human Gal-7
MAL II
Human Galectin1
MAA(Seikagaku)
mouse Gal-7
Anti-LeC

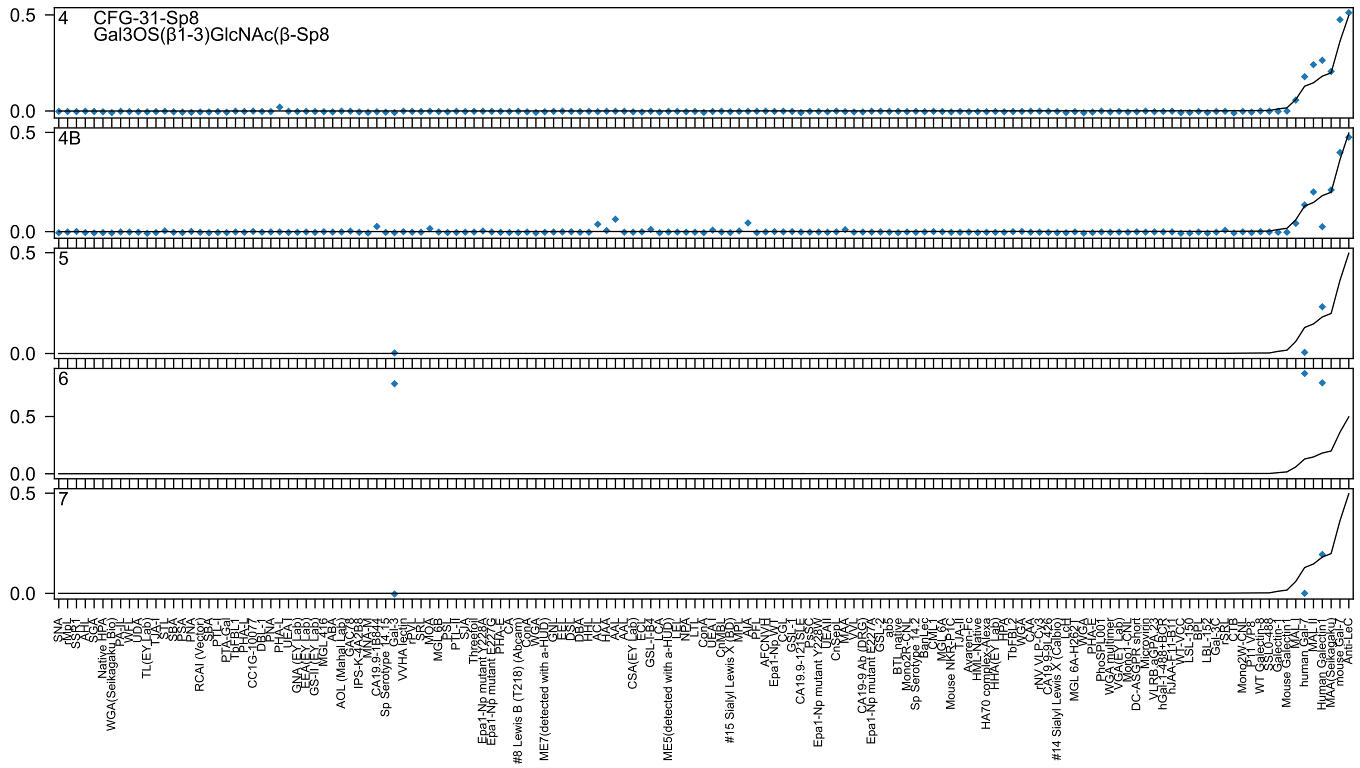
4
CFG-31-Sp8
Gal3OS(β1-3)GlcNAc(β-Sp8
4B
5
6
7
0.5
0.0

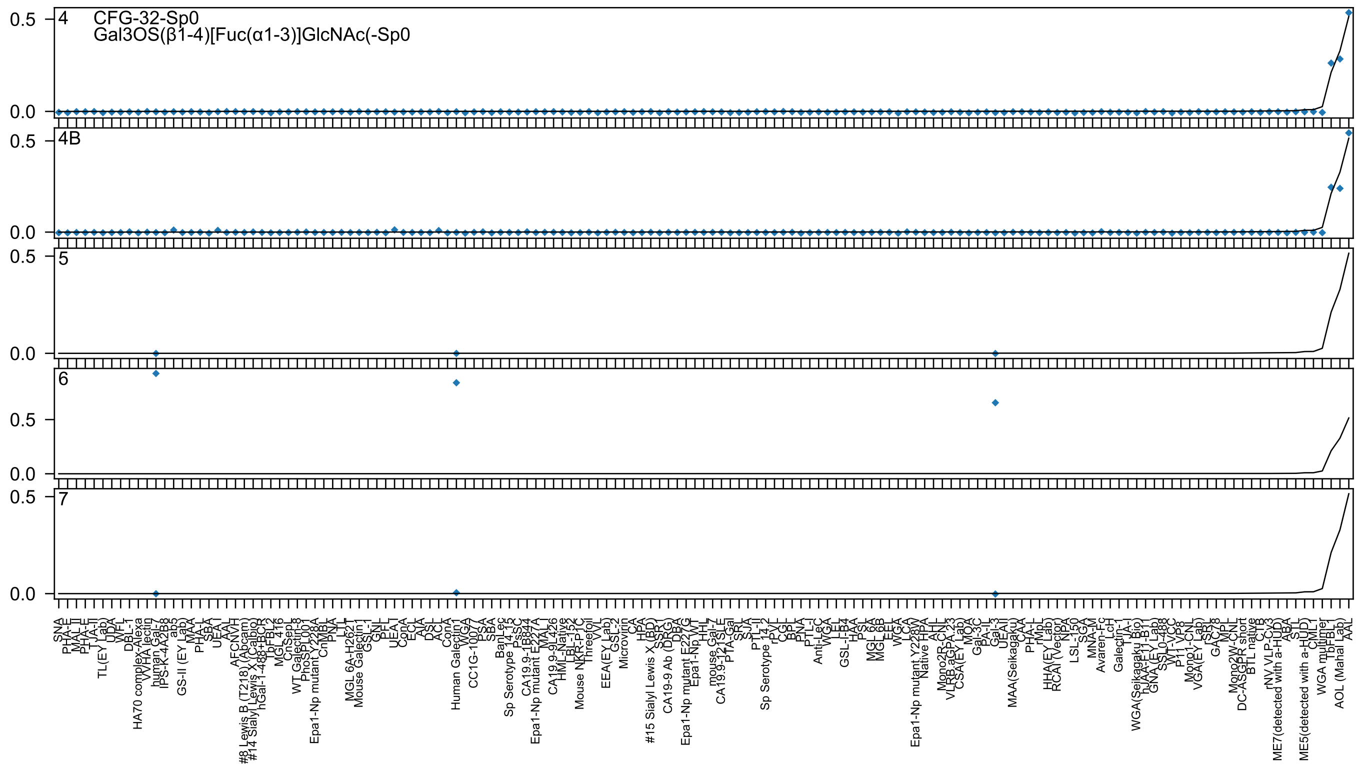

4
CFG-32-Sp0
Gal3OS(β1-4)[Fuc(α1-3)]GlcNAc(-Sp0
4B
5
6
7

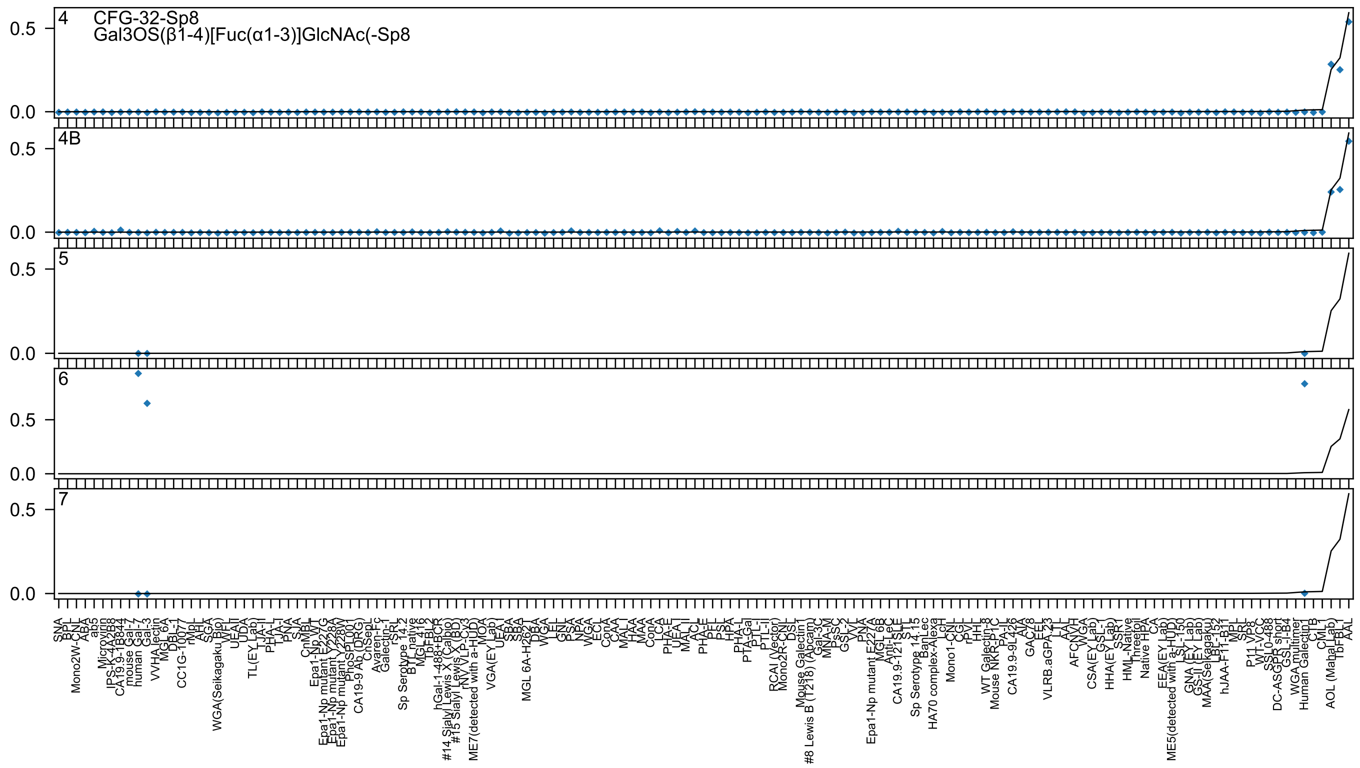
4
CFG-32-Sp8
Gal3OS(β1-4)[Fuc(α1-3)]GlcNAc(-Sp8
4B
5
6
7
0.5
0.0

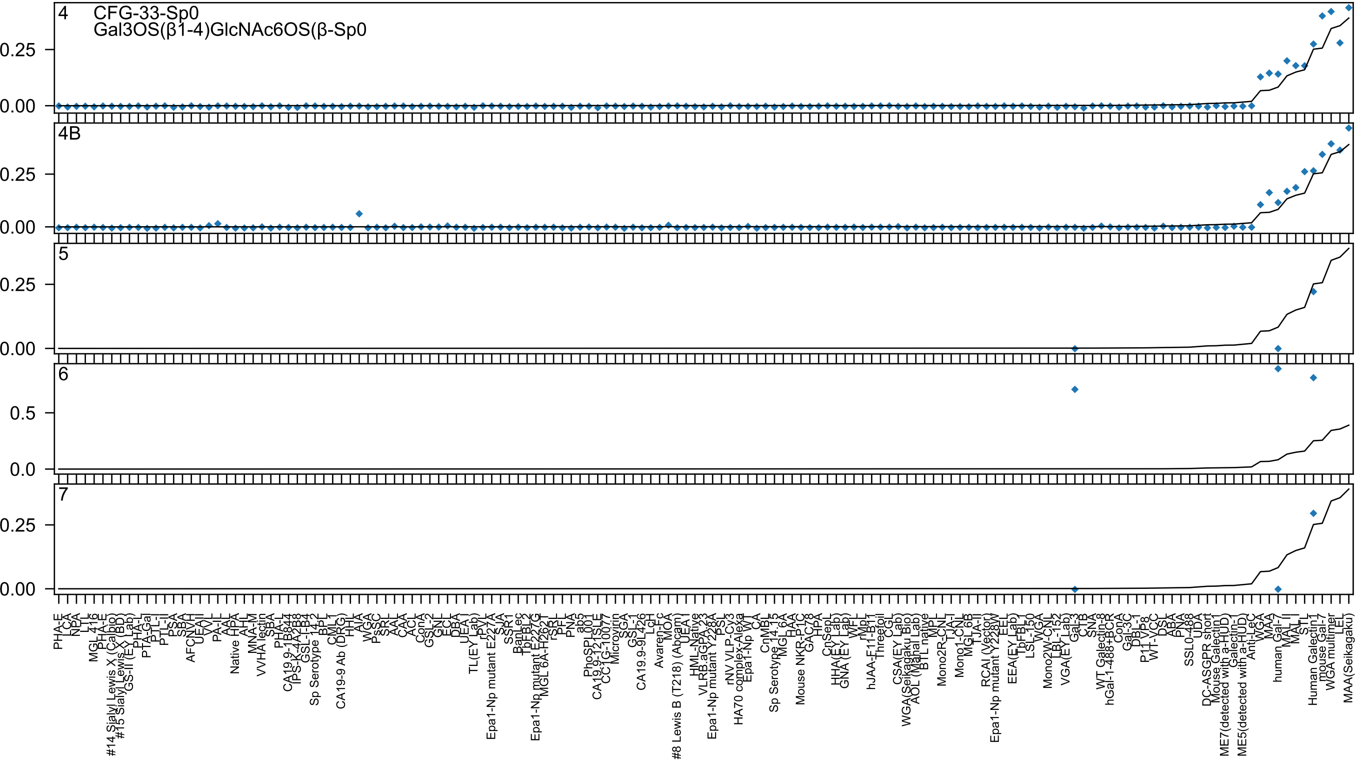
4
CFG-33-Sp0
Gal3OS(β1-4)GlcNAc6OS(β-Sp0
4B
5
6
7
0.25
0.00
0.5
0.0

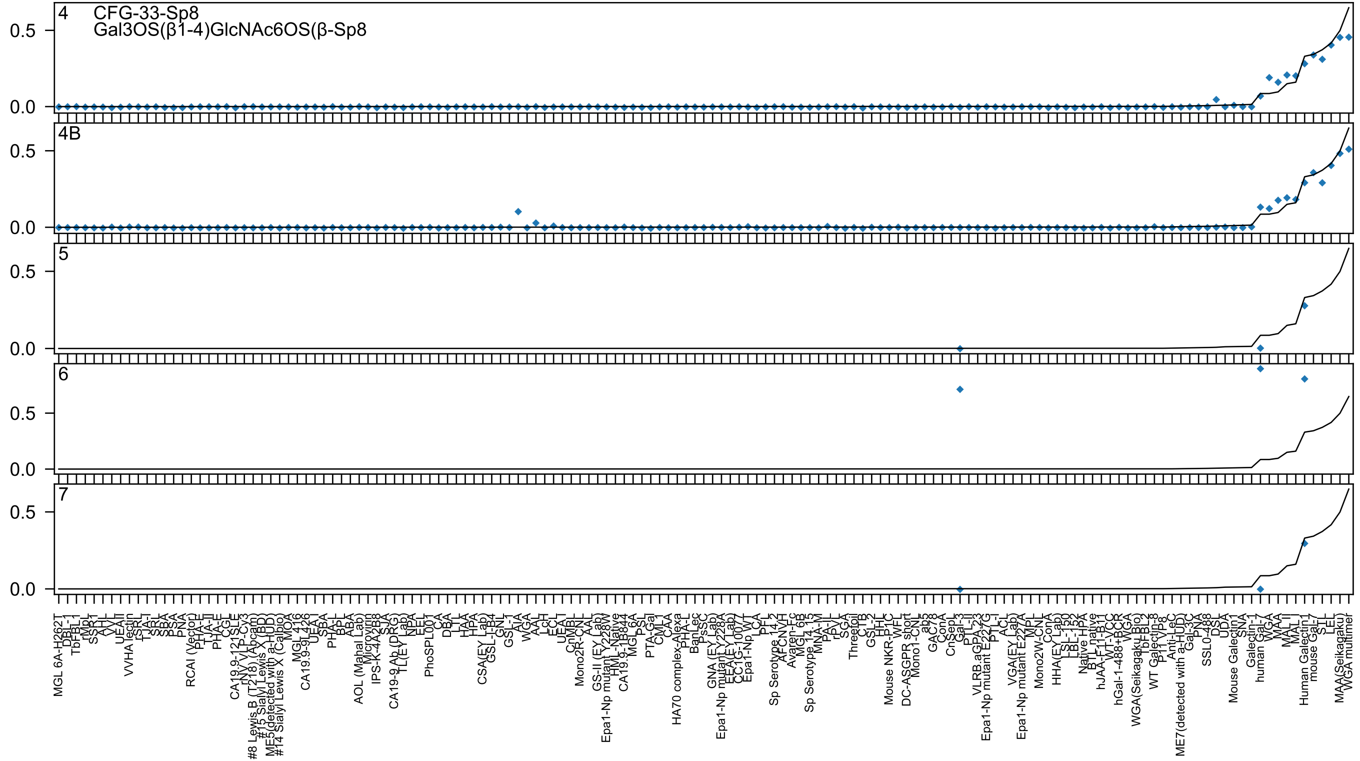

4 CFG-33-Sp8
Gal3OS(β1-4)GlcNAc6OS(β-Sp8
4B
5
6
7
0.5
0.0

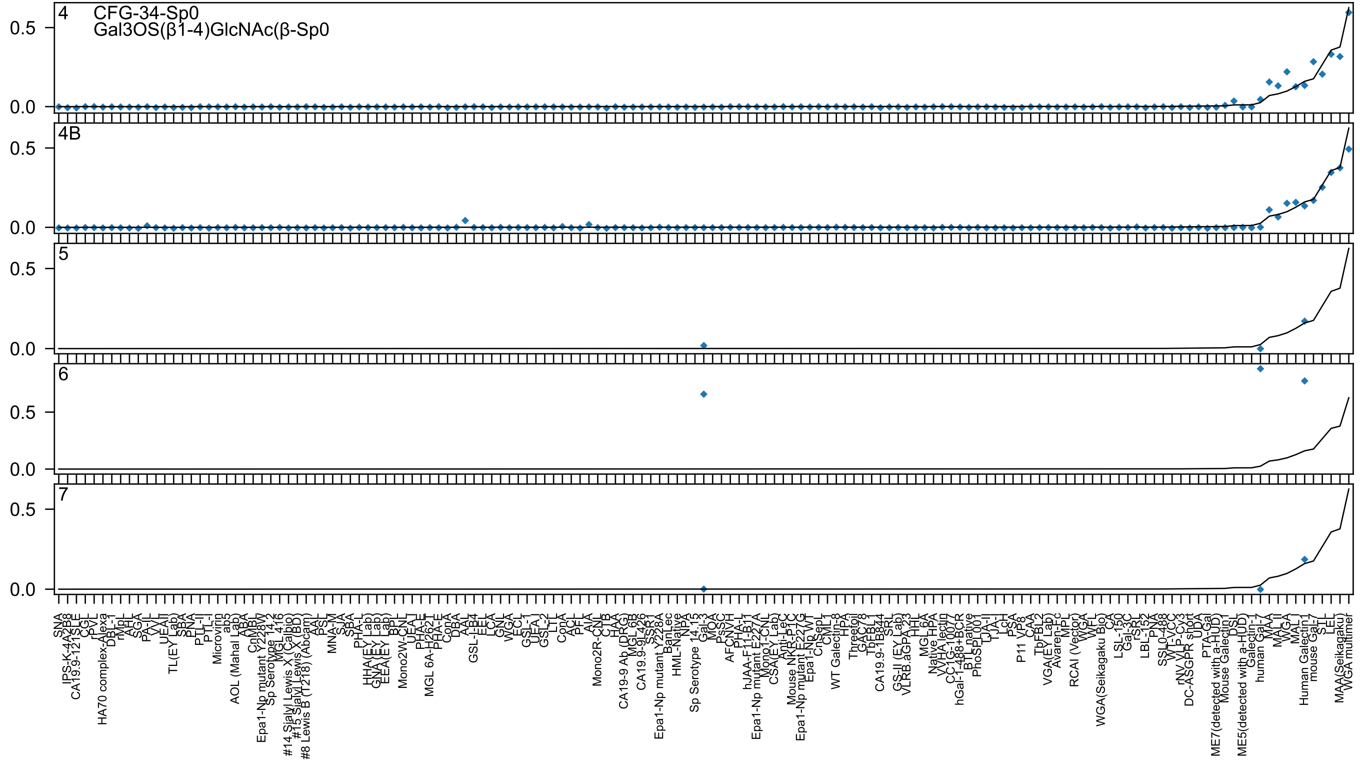

4
CFG-34-Sp0
Gal3OS(β1-4)GlcNAc(β-Sp0
4B
5
6
7
0.5
0.0
SNA
IPS-K-4A2B8
CA19.9-121SLE
CGL
rPVL
HA70 complex-Alexa
DBL-1
rMpL
AHL
SGA
PA-IL
VVL
UEA-II
TL(EY Lab)
SBA
PNA
PTL-II
PTL-I
Microvirin
ab5
AOL (Mahal Lab)
ABA
CnMBL
Epa1-Np mutant Y228W
Sp Serotype 14.2
MGL 416
#14 Sialyl Lewis X (Calbio)
#15 Sialyl Lewis X (BD)
#8 Lewis B (T218) (Abcam)
AAL
PSL
MNA-M
SJA
SBA
PHA-L
HHA(EY Lab)
GNA(EY Lab)
EEA(EY Lab)
BPL
Mono2W-CNL
UEA-I
PHA-E
MGL 6A-H262T
PHA-E
ConA
DBA
AAL
GSL-I-B4
EEL
LCA
GNL
WGA
ECL
GSL-1
UEA-I
GSL-2
LTL
ConA
ACL
PFL
AIA
Mono2R-CNL
CTB
HAA
CA19-9 Ab (DRG)
MGL 6B
CA19.9-9L426
SSR1
Epa1-Np mutant Y228A
BanLec
HML-Native
NPA
Sp Serotype 14.13
Gal-3
MOA
PsSC
AFCNVH
PHA-L
hJAA-F11-B11
Epa1-Np mutant E227A
Mono1-CNL
CSA(EY Lab)
Anti-LeC
Mouse NKR-P1C
Epa1-Np mutant E227G
Epa1-Np WT
CnSpL
CML
WT Galectin-8
HPA
Threefoil
GAC78
TbFBL1
CA19.9-1B844
SRL
GS-II (EY Lab)
VLRB.aGPA.23
HHL
MGL 6A
Native HPA
VVHA lectin
CC1G-10077
hGal-1-488+BCR
BTL native
PhoSL001
TJA-II
TJA-I
LcH
PSA
P11 VP8
CAA
TbFBL2
VGA(EY Lab)
Avaren-Fc
MPL
RCAI (Vector)
WGA
WGA(Seikagaku Bio)
WFL
CA
LSL-150
Gal-3C
rSRL
LBL-152
PNA
SSL0-488
WT-VCC
rNV VLP-Cy3
DC-ASGPR short
UDA
PTA-Gal
ME7(detected with a-HUD)
Mouse Galectin1
DSL
ME5(detected with a-HUD)
Galectin-1
human Gal-7
MAA
MAL-II
WGA
MAL-I
Human Galectin1
mouse Gal-7
STL
LEL
MAA(Seikagaku)
WGA multimer

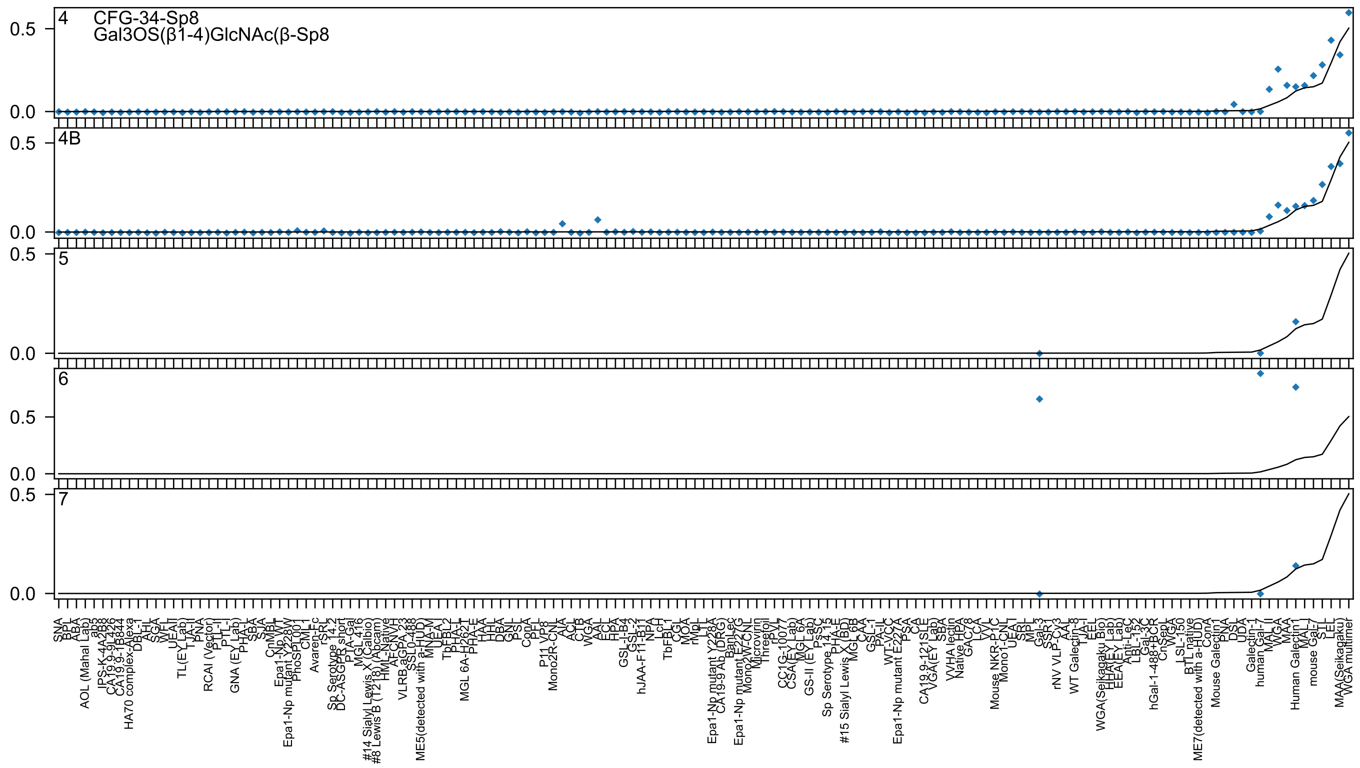
4
CFG-34-Sp8
Gal3OS(β1-4)GlcNAc(β-Sp8
4B
5
6
7

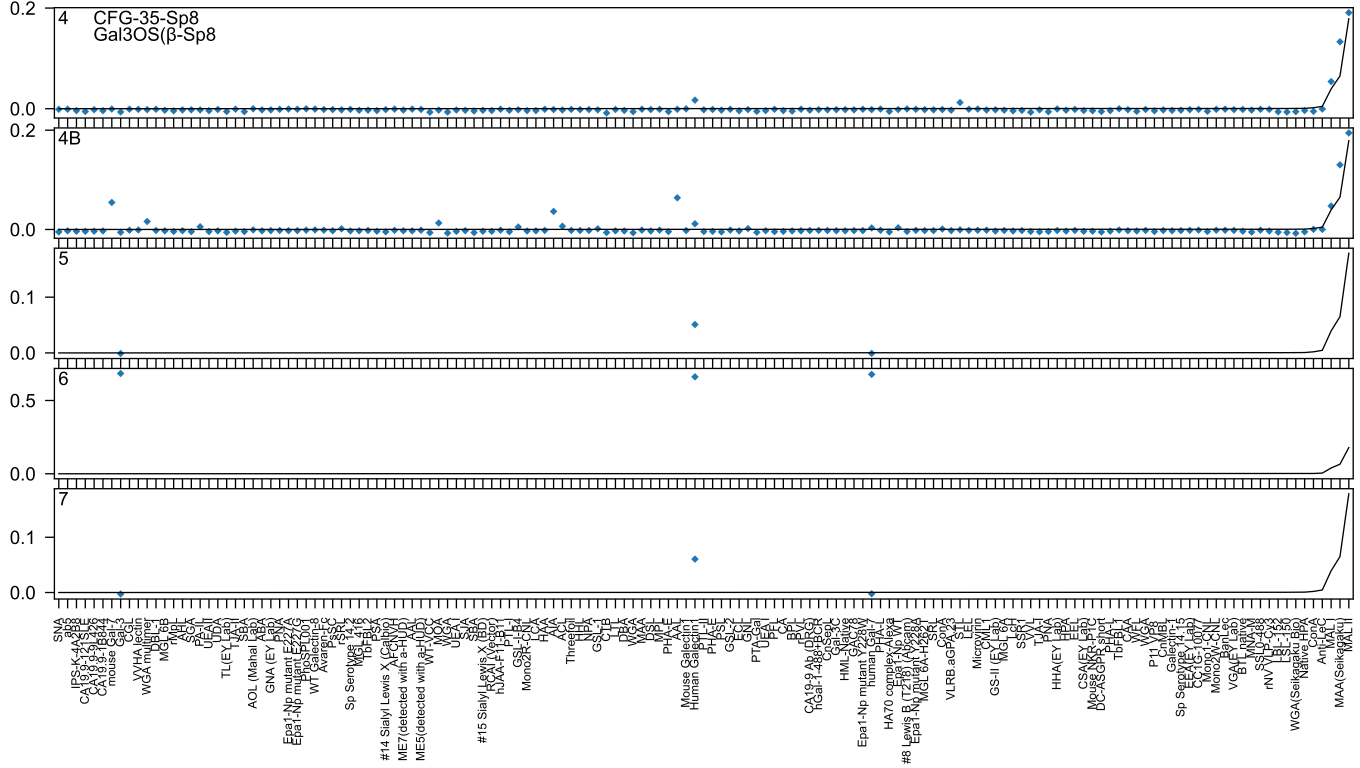

4
CFG-35-Sp8
Gal3OS(β-Sp8
4B
5
6
7
0.2
0.1
0.5
0.0

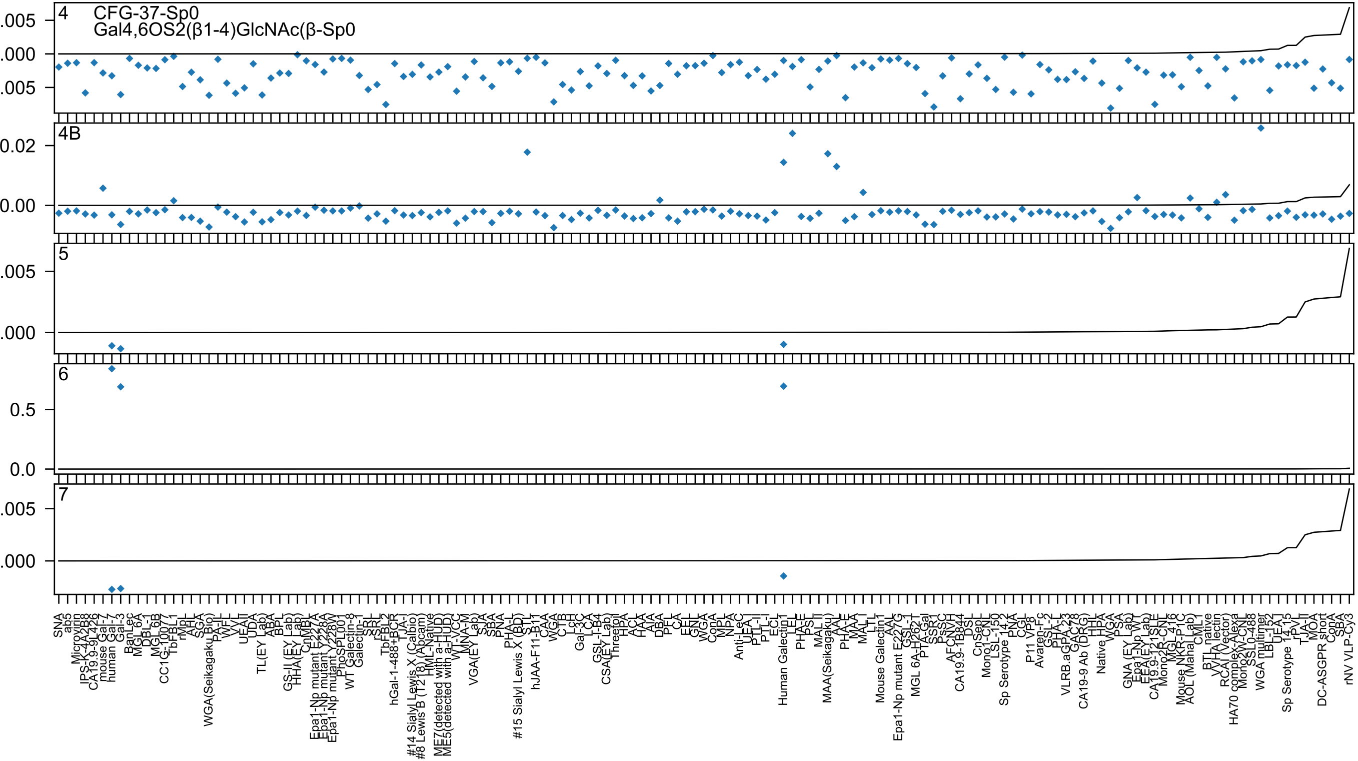

4
CFG-37-Sp0
Gal4,6OS2(β1-4)GlcNAc(β-Sp0
4B
5
6
7

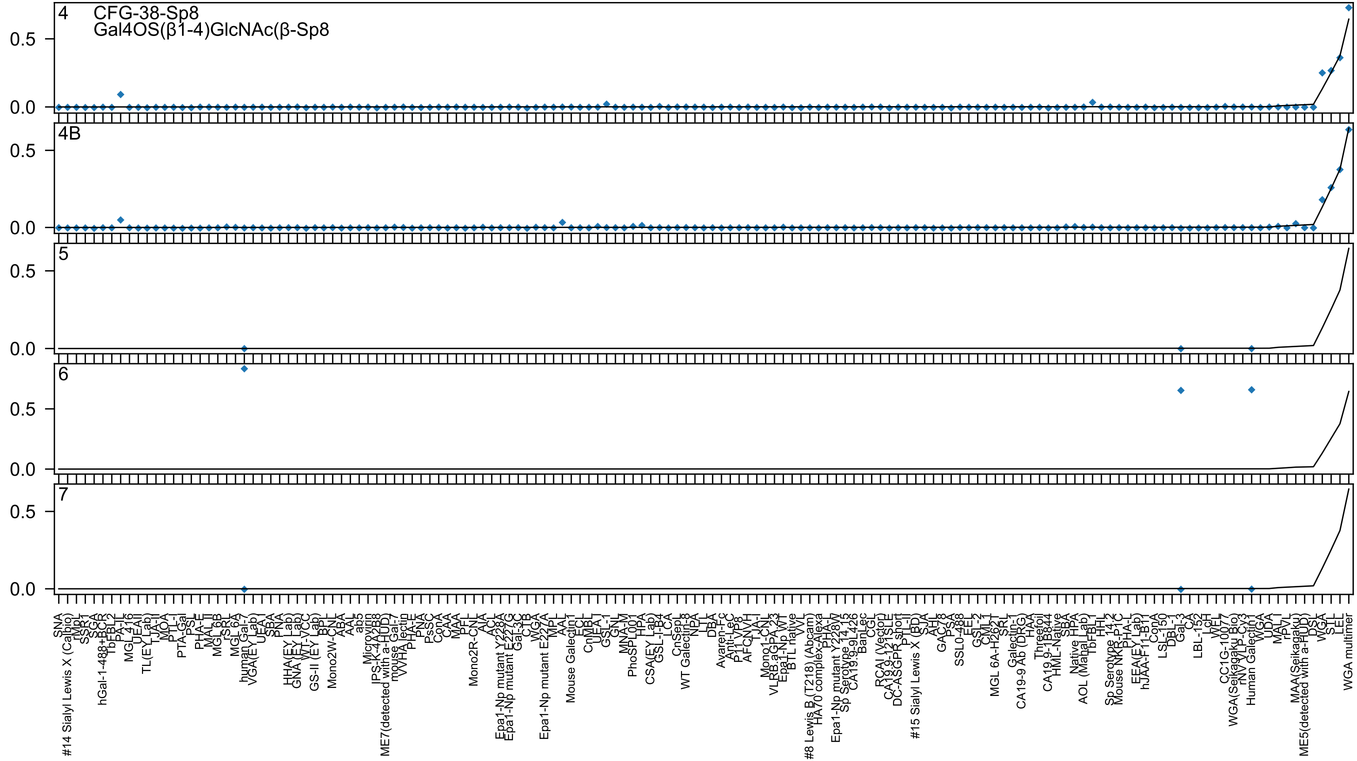

4
CFG-38-Sp8
Gal4OS(β1-4)GlcNAc(β-Sp8
4B
5
6
7
0.5
0.0

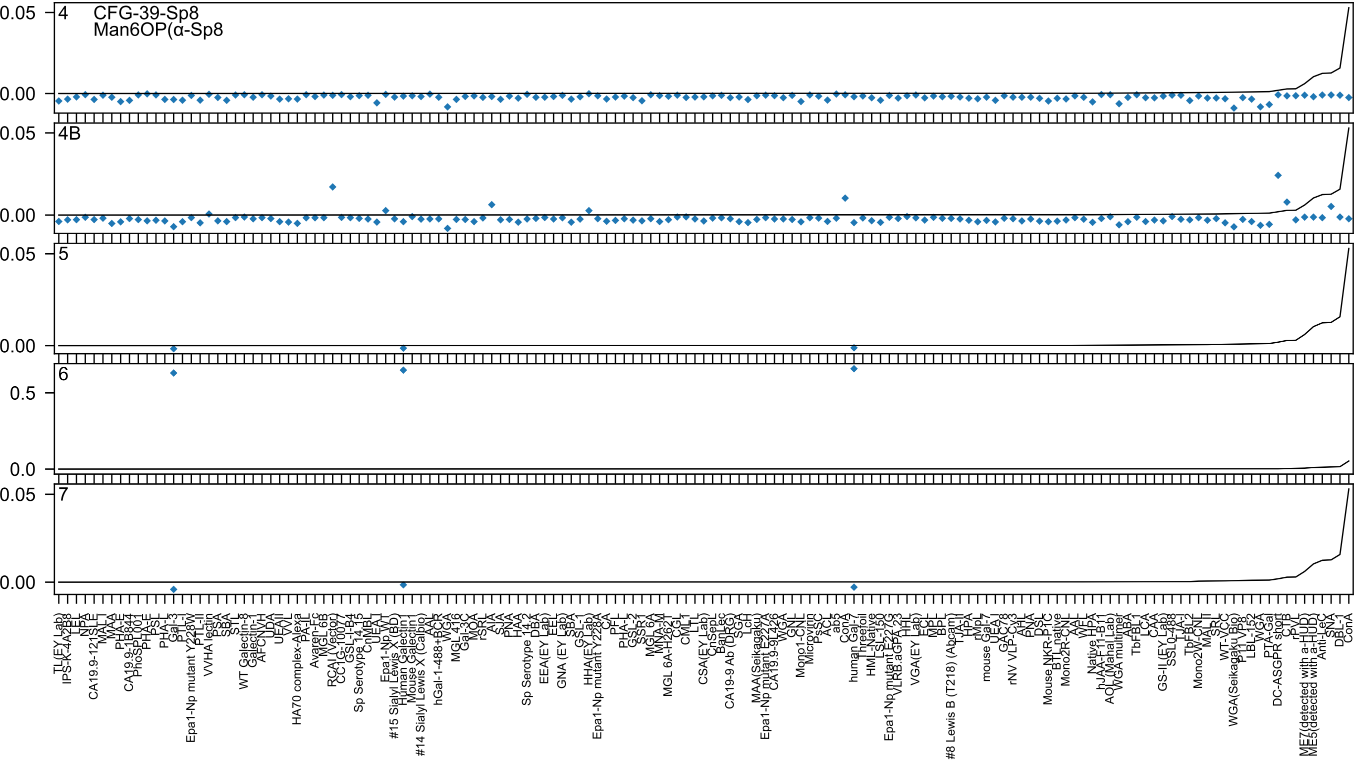

4
CFG-39-Sp8
Man6OP(α-Sp8
4B
5
6
7

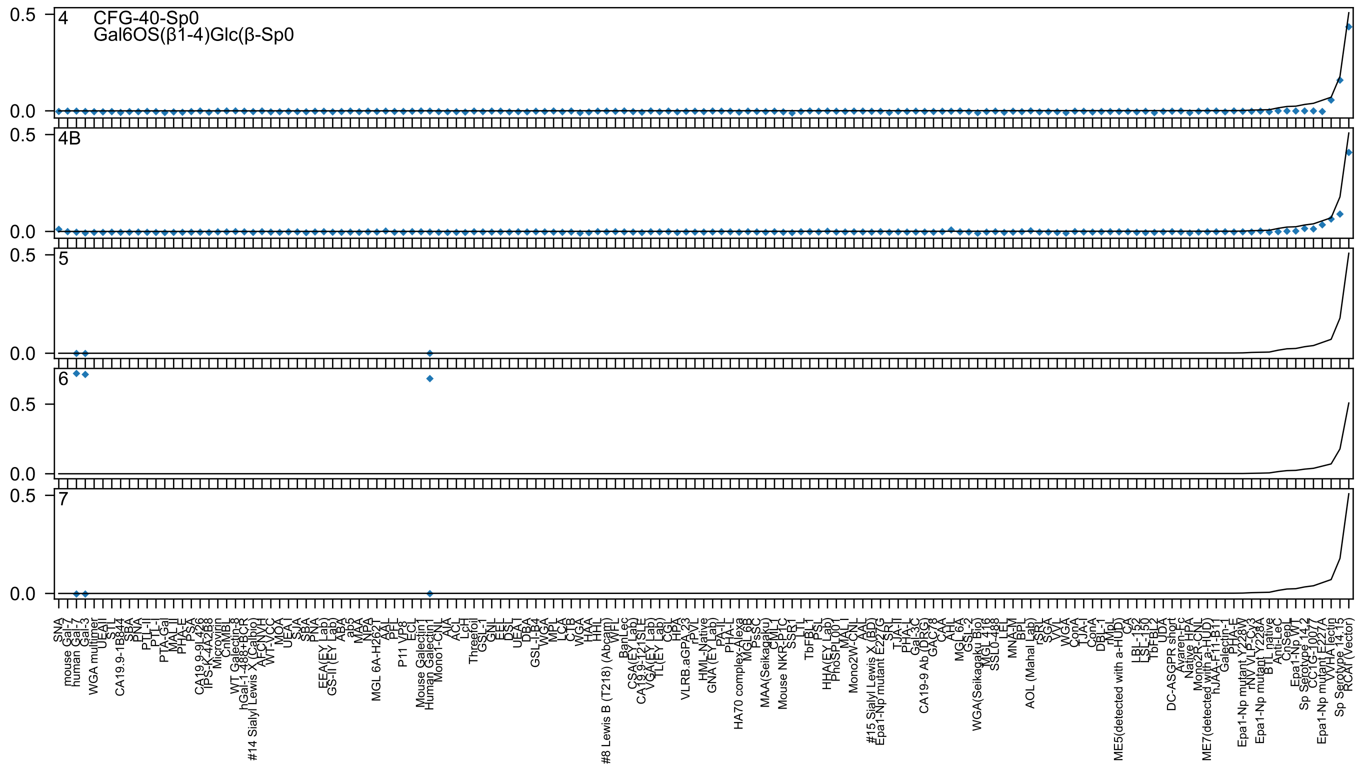

4
CFG-40-Sp0
Gal6OS(β1-4)Glc(β-Sp0
4B
5
6
7

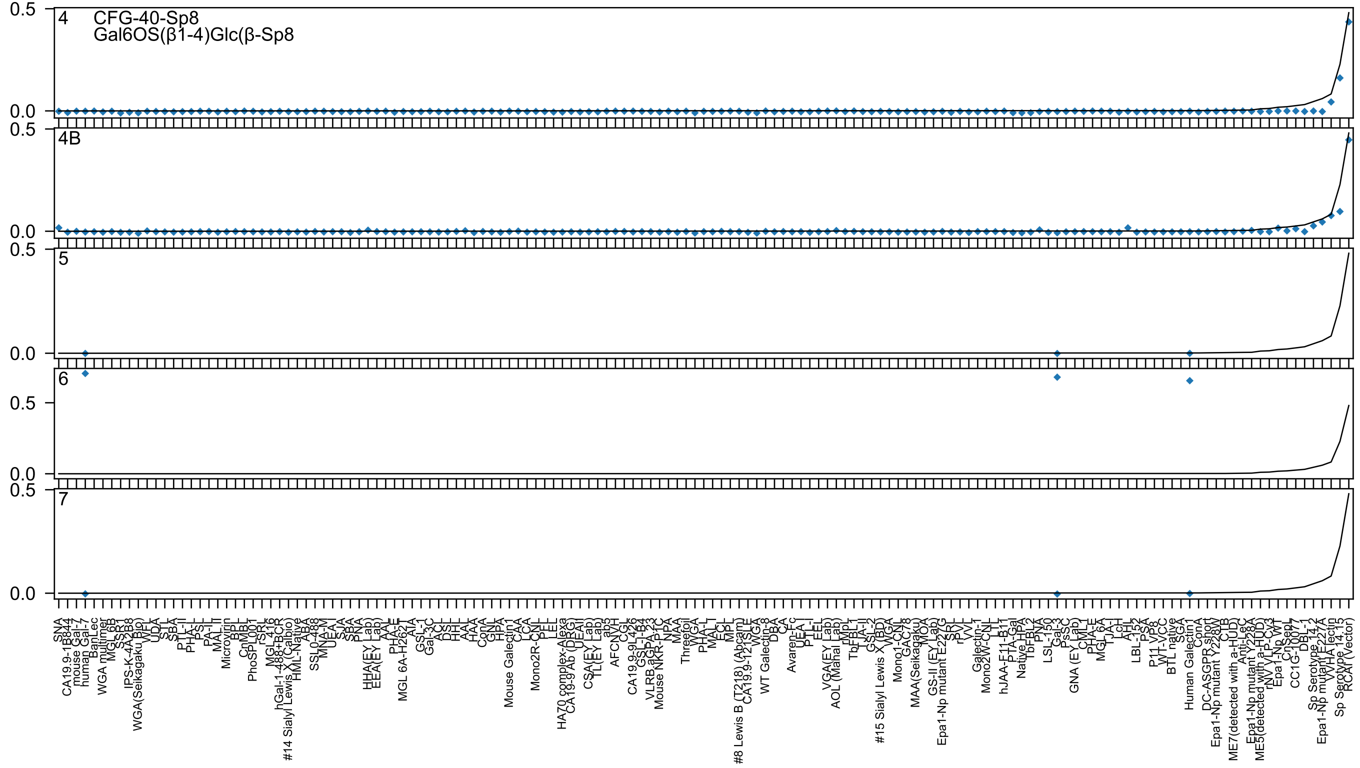
4
CFG-40-Sp8
Gal6OS(β1-4)Glc(β-Sp8
4B
5
6
7
0.5
0.0

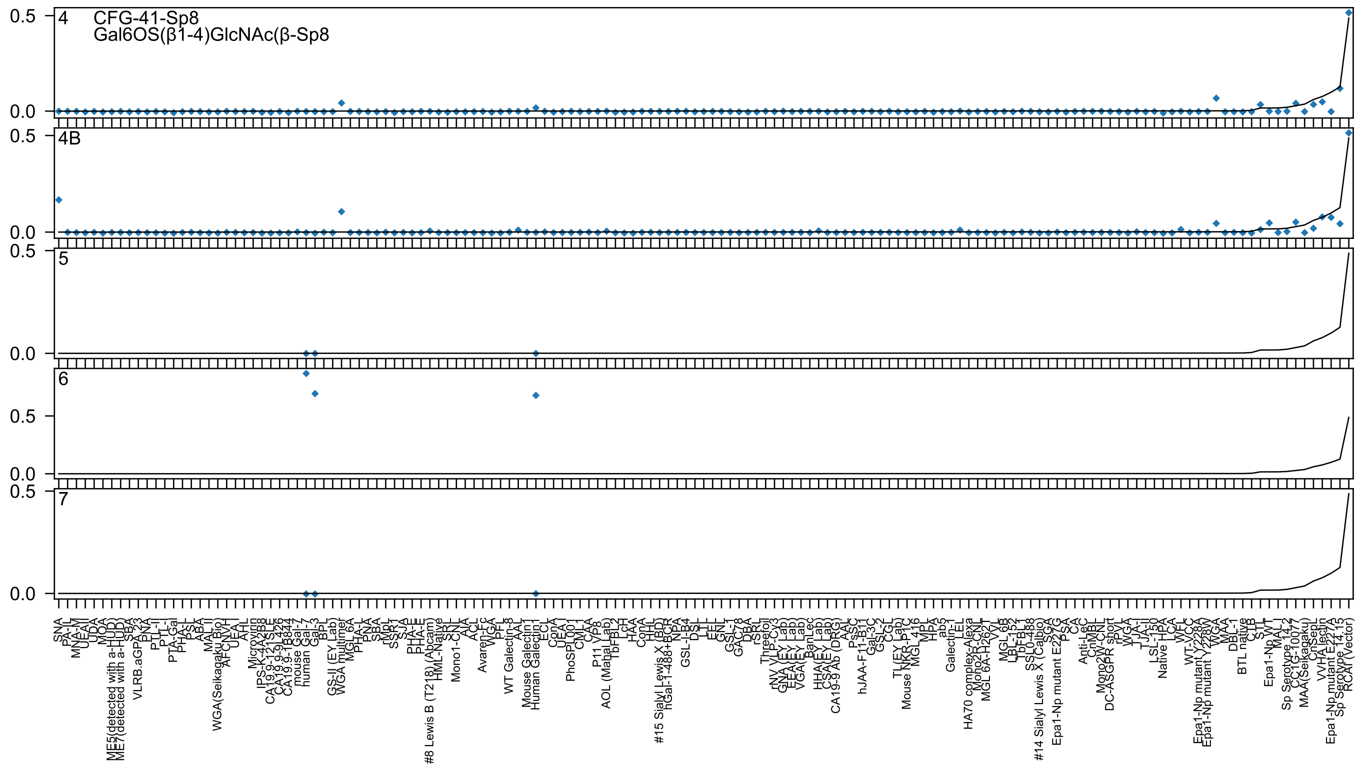

4
CFG-41-Sp8
Gal6OS(β1-4)GlcNAc(β-Sp8
4B
5
6
7
0.5
0.0
SNA
PA-IL
MNA-M
UEAII
UDA
MOA
ME5(detected with a-HUD)
ME7(detected with a-HUD)
SBA
VLRB.aGPA.23
PNA
PTL-II
PTL-I
PTA-Gal
PHA-L
PSL
ABA
MAL II
WGA(Seikagaku Bio)
AFCNVH
UEA I
AHL
Microvirin
IPS-K-4A2B8
CA19.9-121SLE
CA19.9-9L426
CA19.9-1B844
mouse Gal-7
human Gal-7
Gal-3
BPL
GS-II (EY Lab)
WGA multimer
MGL 6A
PHA-L
PNA
SBA
rMpL
SSR1
SJA
PHA-E
PHA-E
#8 Lewis B (T218) (Abcam)
HML-Native
SRL
Mono1-CNL
AIA
ACL
Avaren-Fc
WGA
PFL
WT Galectin-8
AAL
Mouse Galectin1
Human Galectin1
ECL
ConA
UEA I
PhoSL
PL001
CML1
CAA
P11 VP8
AOL (Mahal Lab)
TbFBL2
LcH
HAA
ConA
HHL
#15 Sialyl Lewis X (BD)
hGal-1-488+BCR
NPA
GSL-I-B4
DSL
LTL
EEL
GNL
GSL-I
GAC78
DBA
rSRL
Threefoil
rNV VLP-Cy3
GNA (EY Lab)
EEA(EY Lab)
VGA(EY Lab)
BanLec
HHA(EY Lab)
CSA(EY Lab)
CA19-9 Ab (DRG)
AAL
PsSC
hJAA-F11-B11
Gal-3C
GSL-2
CGL
TL(EY Lab)
Mouse NKR-P1C
MGL 416
MPL
HPA
ab5
Galectin-1
LEL
HA70 complex-Alexa
Mono2R-CNL
MGL 6A-H262T
VVL
MGL 6B
LBL-152
TbFBL1
SSL0-488
#14 Sialyl Lewis X (Calbio)
SGA
Epa1-Np mutant E227G
PSA
CA
Anti-LeC
CnMBL
Mono2W-CNL
DC-ASGPR short
rPVL
WGA
TJA-II
TJA-I
LSL-150
Native HPA
LCA
WFL
WT-VCC
Epa1-Np mutant Y228A
Epa1-Np mutant Y228W
WGA
MAA
DBL-1
BTL native
CTB
STL
Epa1-Np WT
MAL I
Sp Serotype 14.2
CC1G-10077
MAA(Seikagaku)
CnSeP
VVHA lectin
Epa1-Np mutant E227A
Sp Serotype 14.15
RCAI (Vector)

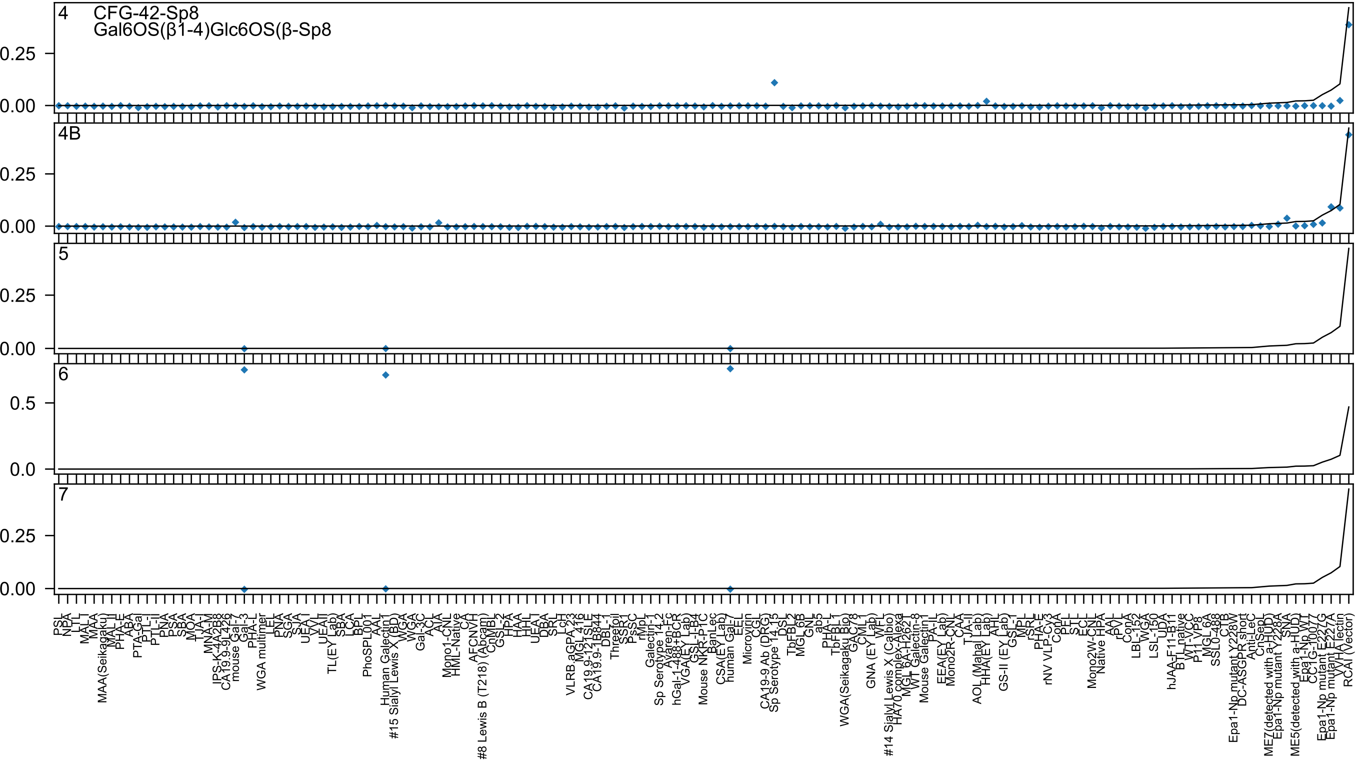

4
CFG-42-Sp8
Gal6OS(β1-4)Glc6OS(β-Sp8
4B
5
6
7

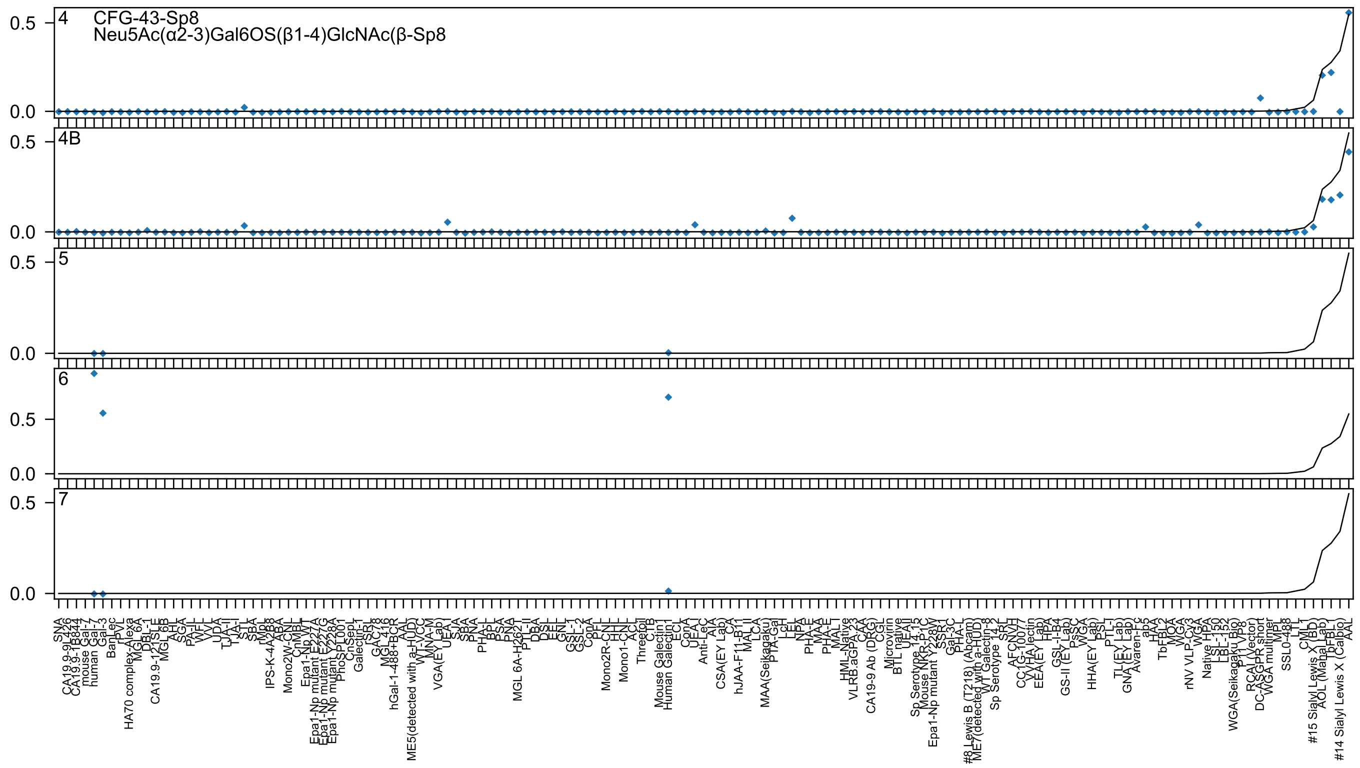

4
CFG-43-Sp8
Neu5Ac(α2-3)Gal6OS(β1-4)GlcNAc(β-Sp8
4B
5
6
7
0.5
0.0

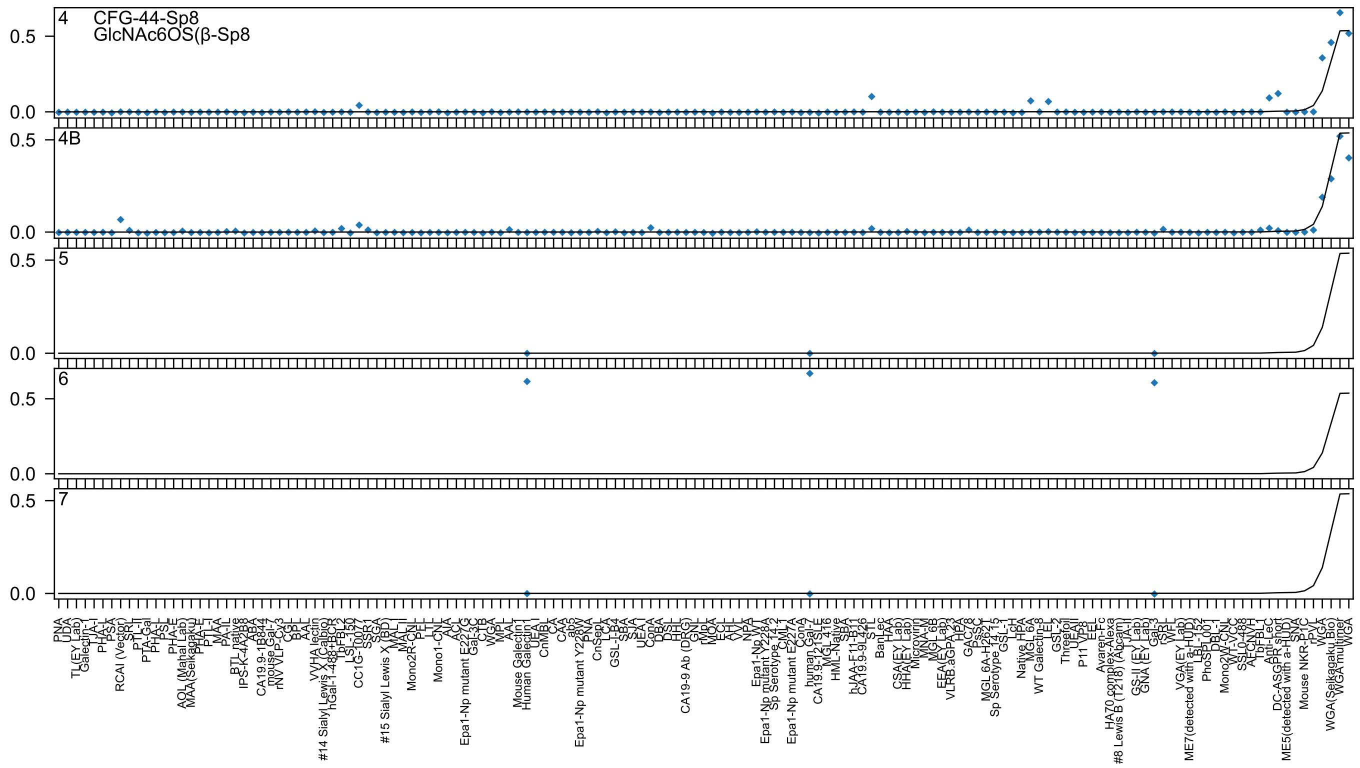
4
CFG-44-Sp8
GlcNAc6OS(β-Sp8
4B
5
6
7
0.5
0.0

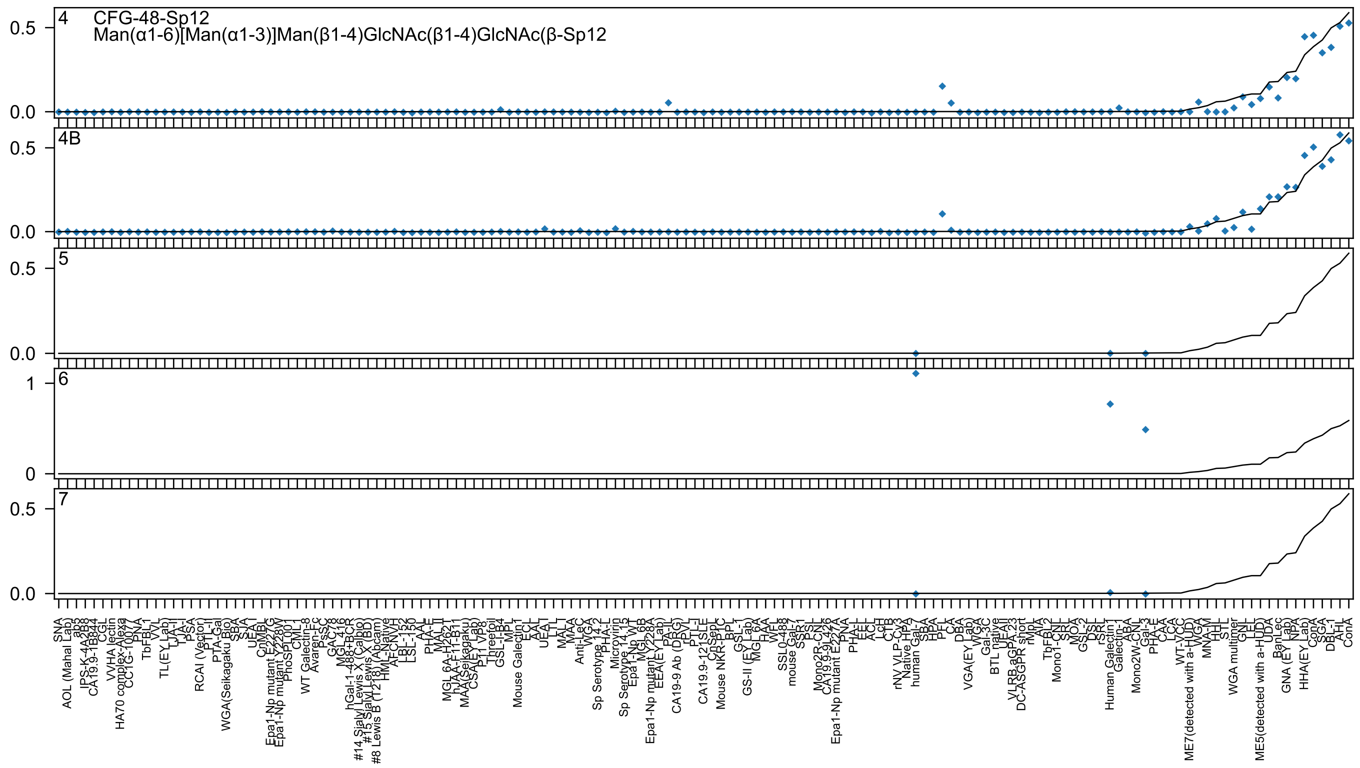

4
CFG-48-Sp12
Man(α1-6)[Man(α1-3)]Man(β1-4)GlcNAc(β1-4)GlcNAc(β-Sp12
4B
5
6
7

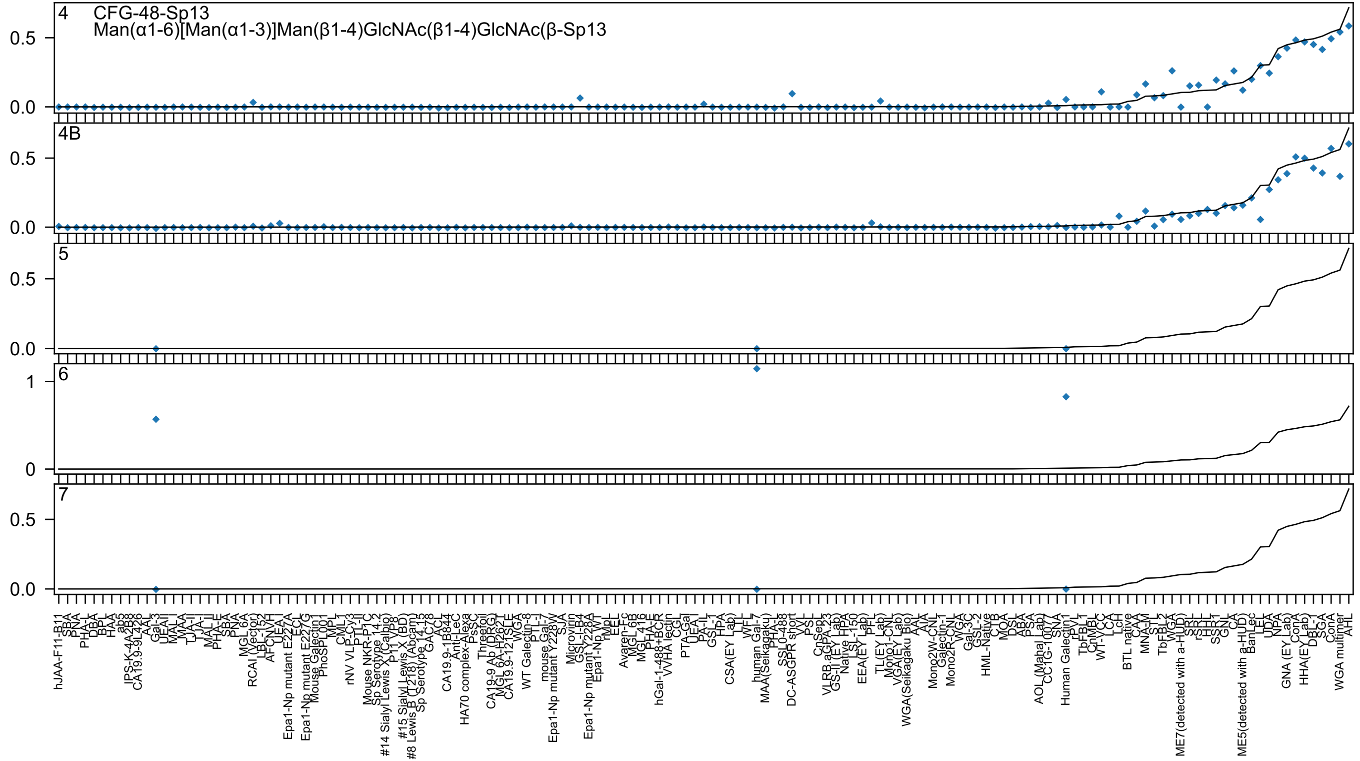
4
CFG-48-Sp13
Man(α1-6)[Man(α1-3)]Man(β1-4)GlcNAc(β1-4)GlcNAc(β-Sp13
4B
5
6
7
0.0
0.5
0
1
hJAA-F11-B11
SBA
PNA
PHA-L
DBA
BPL
HAA
ab5
IPS-K-4A2B8
CA19.9-9L426
AAL
Gal-3
UEAI
MAL-I
MAA
TJA-II
TJA-I
MAL-II
PHA-E
SBA
PNA
MGL 6A
RCAI (Vector)
LBL-152
AFCNVH
UEA-I
Epa1-Np mutant E227A
ECL
Epa1-Np mutant E227G
Mouse Galectin1
PhoSL001
MPL
CML-1
rNV VLP-Cy3
PTL-II
Mouse NKR-P1C
Sp Serotype 14.2
#14 Sialyl Lewis X (Calbio)
P11 VP8
#15 Sialyl Lewis X (BD)
#8 Lewis B (T218) (Abcam)
Sp Serotype 14.13
GAC-78
ACL
CA19.9-1B844
Anti-LeC
HA70 complex-Alexa
PsSC
Threefoil
CA19-9 Ab (DRG)
MGL 6A-H262T
CA19.9-121SLE
WGA
WT Galectin-8
PTL-I
mouse Gal-7
Epa1-Np mutant Y228W
SJA
Microvirin
GSL-I-B4
Epa1-Np mutant Y228A
Epa1-Np WT
rMpL
EEL
Avaren-Fc
MGL 6B
MGL 416
PHA-E
hGal-1-488+BCR
VVHA lectin
CGL
PTA-Gal
UEA-I
PA-IL
GSL-I
HPA
CSA(EY Lab)
LTL
WFL
human Gal-7
MAA(Seikagaku)
PHA-L
SSL0-488
DC-ASGPR short
VVL
PSL
CnSep
VLRB aGPA.23
GS-II (EY Lab)
Native HPA
LSL-150
EEA(EY Lab)
PFL
TL(EY Lab)
Mono1-CNL
VGA(EY Lab)
WGA(Seikagaku Bio)
AAL
AIA
Mono2W-CNL
Galectin-1
Mono2R-CNL
WGA
Gal-3C
GSL-2
HML-Native
CTB
MOA
DSL
ABA
PSA
AOL (Mahal Lab)
CC1G-10077
SNA
Human Galectin1
rPVL
TbFBL1
CnMBL
WT-YCC
LcH
BTL native
CAA
MNA-M
STL
TbFBL2
WGA
ME7(detected with a-HUD)
SRL
rSRL
HHL
SSR1
GNL
CA
ME5(detected with a-HUD)
BanLec
LEL
UDA
NPA
GNA (EY Lab)
ConA
HHA(EY Lab)
DBL-1
SGA
ConA
WGA multimer
AHL

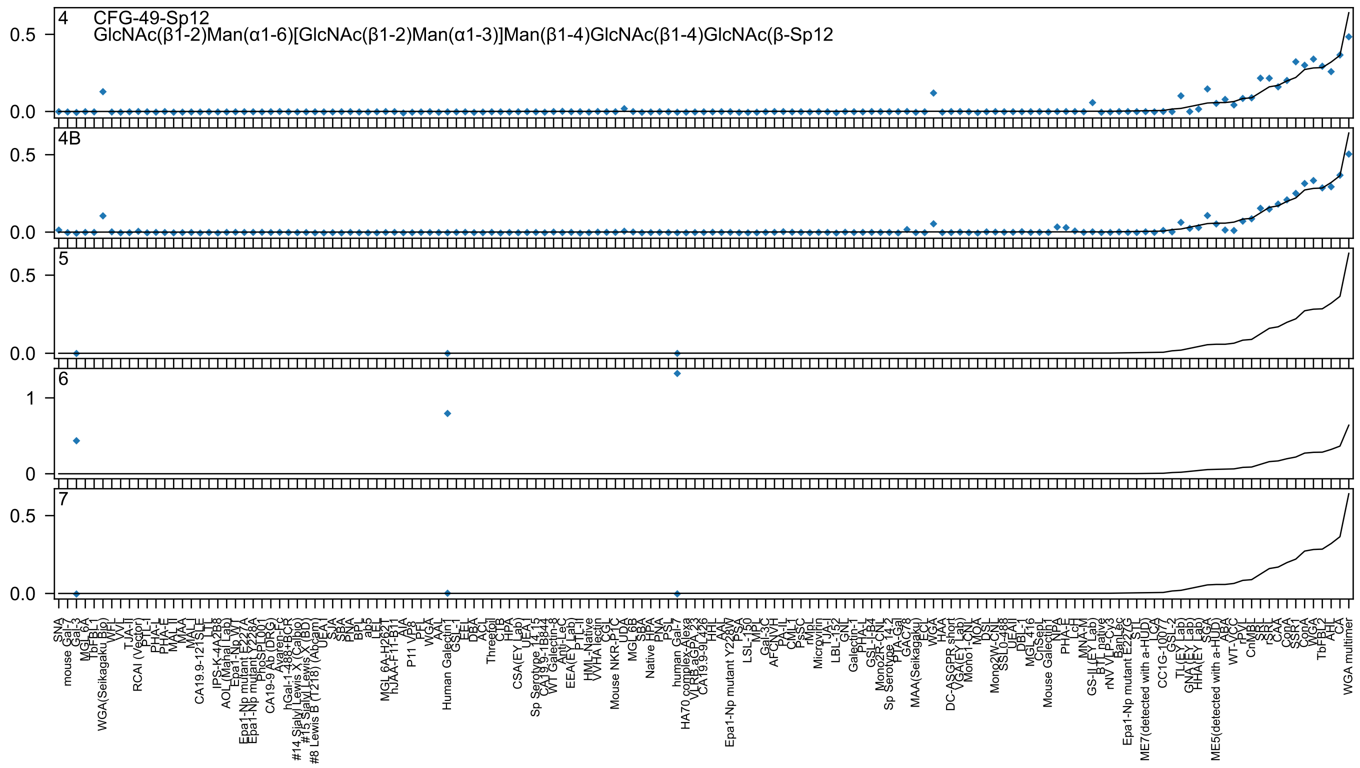

4 CFG-49-Sp12
GlcNAc(β1-2)Man(α1-6)[GlcNAc(β1-2)Man(α1-3)]Man(β1-4)GlcNAc(β1-4)GlcNAc(β-Sp12
4B
5
6
7

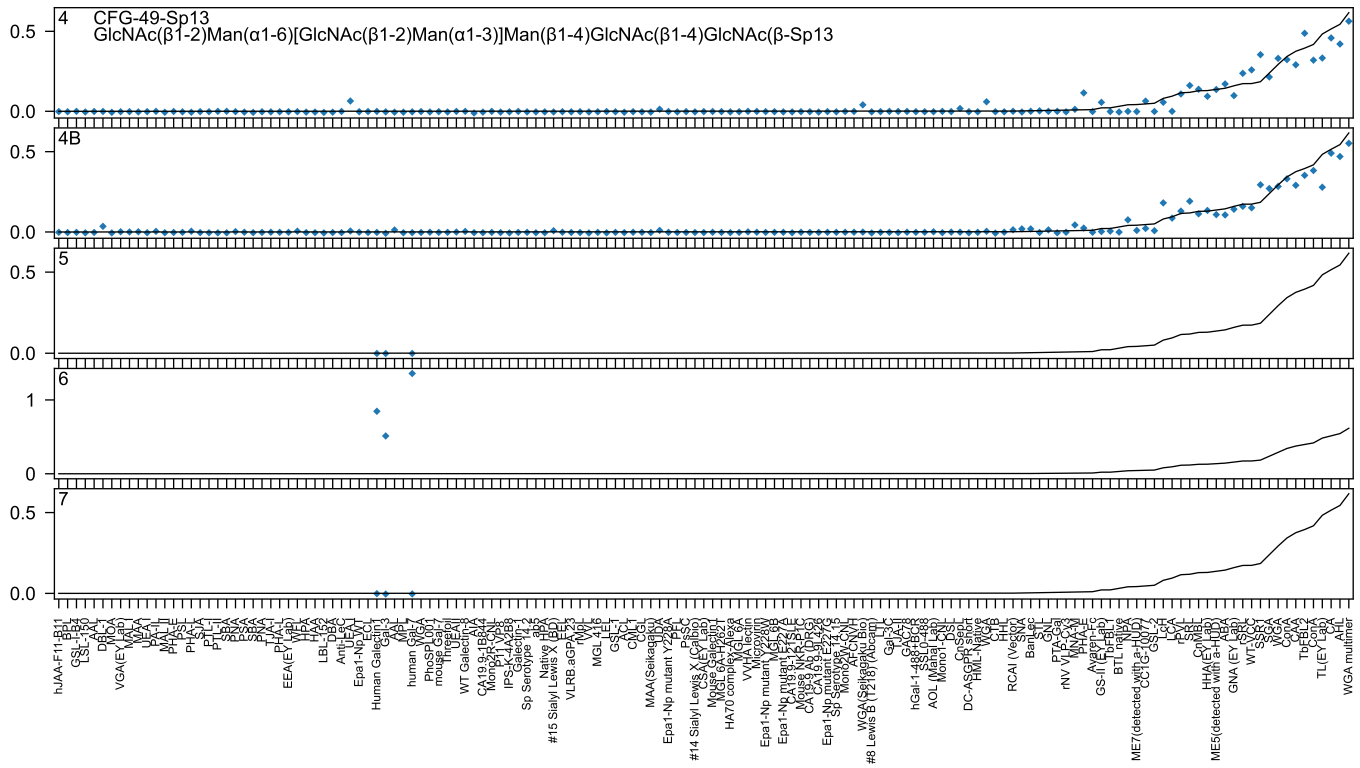
4
CFG-49-Sp13
GlcNAc(β1-2)Man(α1-6)[GlcNAc(β1-2)Man(α1-3)]Man(β1-4)GlcNAc(β1-4)GlcNAc(β-Sp13
4B
5
6
7

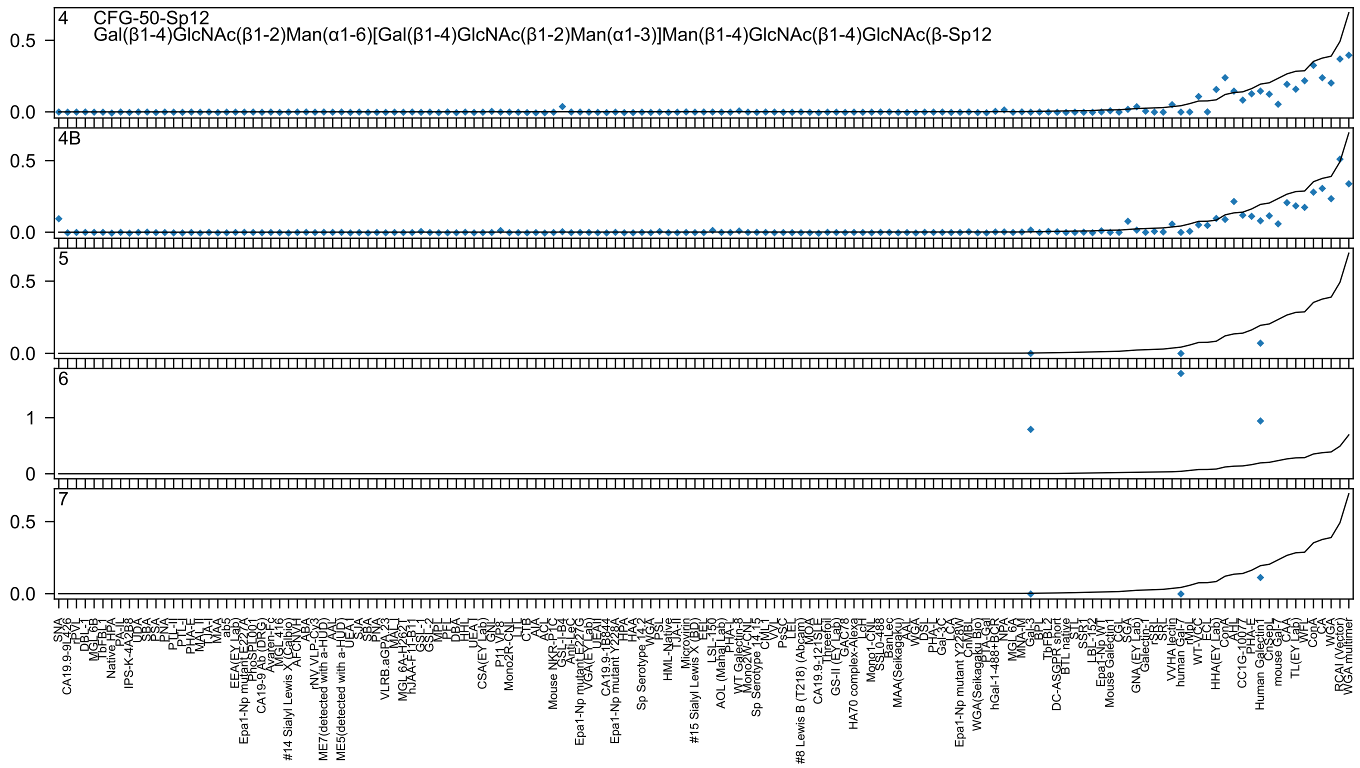
4
CFG-50-Sp12
Gal(β1-4)GlcNAc(β1-2)Man(α1-6)[Gal(β1-4)GlcNAc(β1-2)Man(α1-3)]Man(β1-4)GlcNAc(β1-4)GlcNAc(β-Sp12
4B
5
6
7

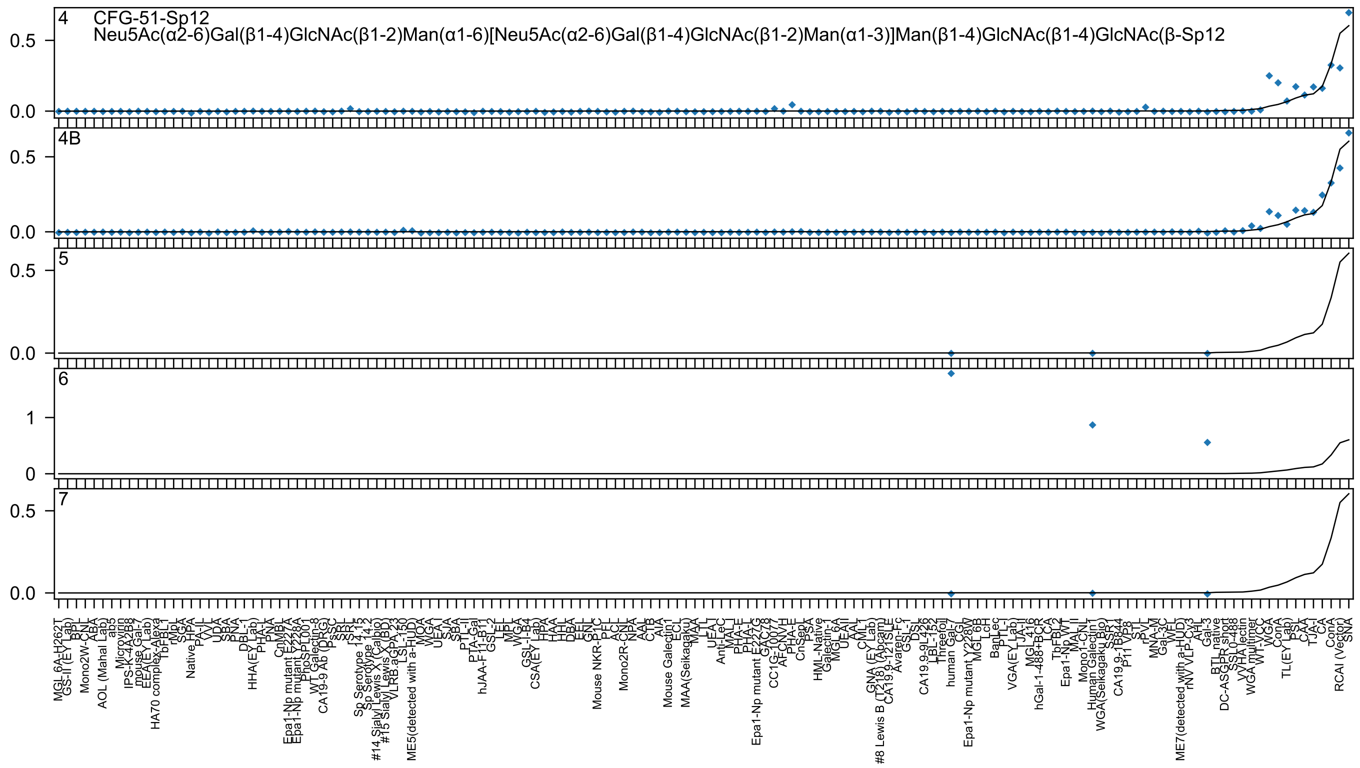

CFG-51-Sp12
Neu5Ac(α2-6)Gal(β1-4)GlcNAc(β1-2)Man(α1-6)[Neu5Ac(α2-6)Gal(β1-4)GlcNAc(β1-2)Man(α1-3)]Man(β1-4)GlcNAc(β1-4)GlcNAc(β-Sp12
4
4B
5
6
7

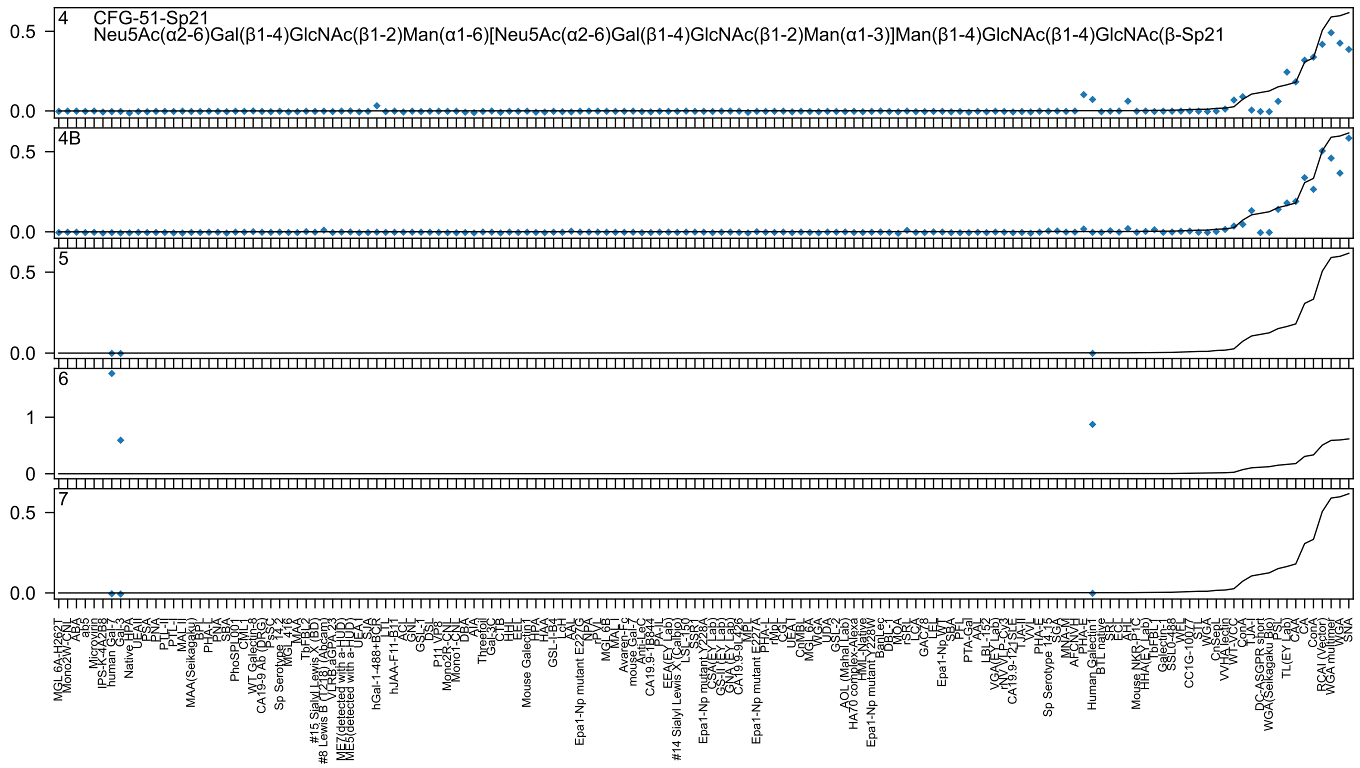

4 CFG-51-Sp21
Neu5Ac(α2-6)Gal(β1-4)GlcNAc(β1-2)Man(α1-6)[Neu5Ac(α2-6)Gal(β1-4)GlcNAc(β1-2)Man(α1-3)]Man(β1-4)GlcNAc(β1-4)GlcNAc(β-Sp21
4B
5
6
7

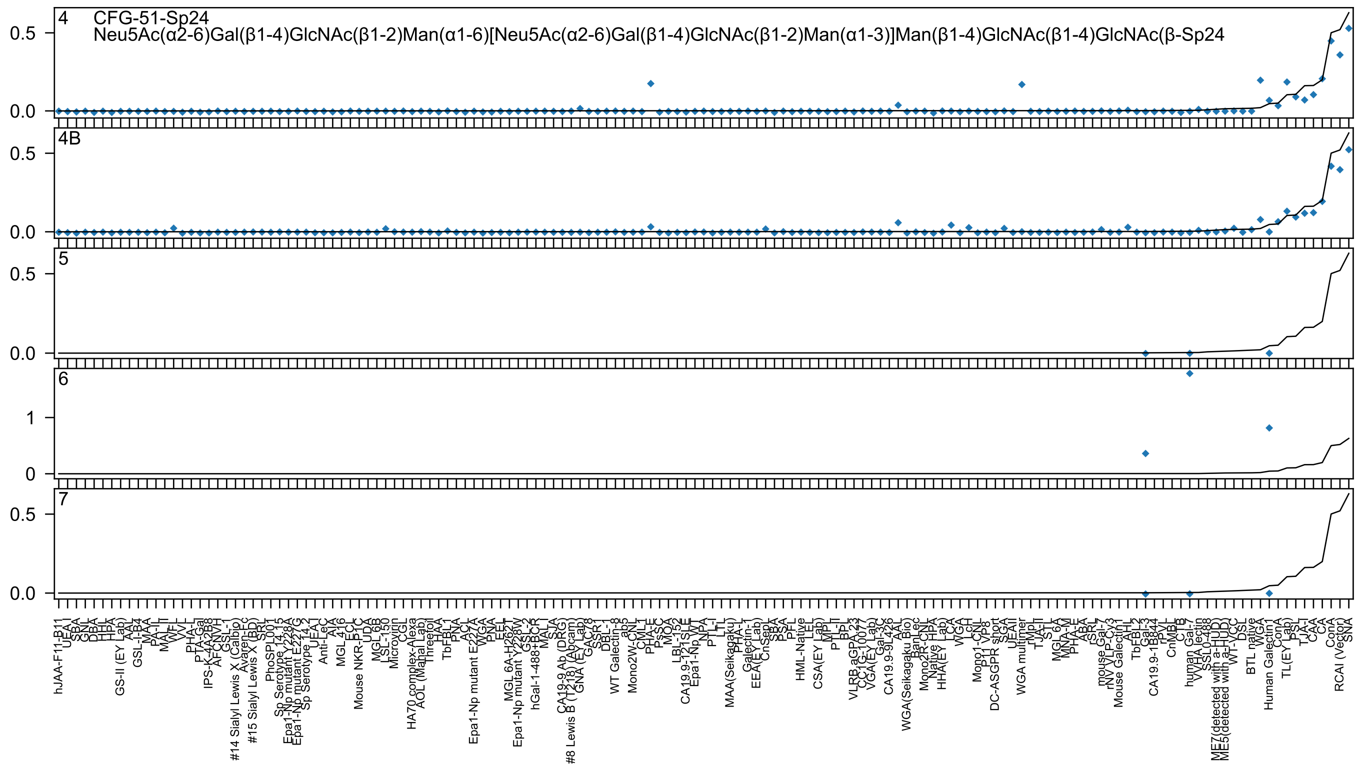
4
CFG-51-Sp24
Neu5Ac(α2-6)Gal(β1-4)GlcNAc(β1-2)Man(α1-6)[Neu5Ac(α2-6)Gal(β1-4)GlcNAc(β1-2)Man(α1-3)]Man(β1-4)GlcNAc(β1-4)GlcNAc(β-Sp24
4B
5
6
7
0.5
0.0
1
0
hJAA-F11-B11
UEA-I
SBA
GNL
DBA
HHL
HPA
GS-II (EY Lab)
AAL
GSL-I-B4
MAA
PA-IL
MAL-II
WFL
VVL
PHA-L
PTA-Gal
IPS-K-4A2B8
AFCNVH
GSL-1
#14 Sialyl Lewis X (Calbio)
Avaren-Fc
#15 Sialyl Lewis X (BD)
SRL
PhoSPL001
Sp Serotype 14.15
Epa1-Np mutant Y228A
Epa1-Np mutant E227G
Sp Serotype 14.2
UEA-I
Anti-LeC
AIA
MGL 4I6
ECL
Mouse NKR-P1C
UDA
MGL 6B
LSL-150
Microvirin
CGL
HA70 complex-Alexa
AOL (Mahal Lab)
Threefoil
HAA
TbFBL1
PNA
ACL
Epa1-Np mutant E227A
WGA
PNA
EEL
MGL 6A-H262T
Epa1-Np mutant Y228W
GSL-2
hGal-1-488+BCR
MAL-I
SJA
CA19-9 Ab (DRG)
#8 Lewis B (T218) (Abcam)
GNA (EY Lab)
GAC78
SSR-1
DBL-1
WT Galectin-8
ab5
Mono2W-CNL
CML1
PHA-E
PsSC
MOA
LBL-152
CA19.9-121SLE
Epa1-Np WT
NPA
PTL-I
LTL
MAA(Seikagaku)
PHA-L
Galectin-1
EEA(EY Lab)
CnSep
SBA
PSA
PFL
HML-Native
LEL
CSA(EY Lab)
MPL
PTL-II
BPL
VLRB.aGPA.23
CC1G-10077
VGA(EY Lab)
Gal-3C
CA19.9-9L426
AAL
WGA(Seikagaku Bio)
BanLec
Mono2R-CNL
Native HPA
HHA(EY Lab)
LCA
WGA
LcH
Mono1-CNL
P11 VP8
DC-ASGPR short
SGA
UEAII
WGA multimer
rMpL
TJA-II
STL
MGL 6A
MNA-M
PHA-E
ABA
rSRL
mouse Gal-7
rNV VLP-Cy3
Mouse Galectin1
AHL
TbFBL2
Gal-3
CA19.9-1B844
rPVL
CnMBL
CTB
human Gal-7
VVHA lectin
SSL0-488
ME7(detected with a-HUD)
ME5(detected with a-HUD)
WT-YCC
DSL
BTL native
WGA
Human Galectin1
ConA
TL(EY Lab)
PSL
TJA-I
CA
CAA
ConA
RCAI (Vector)
SNA

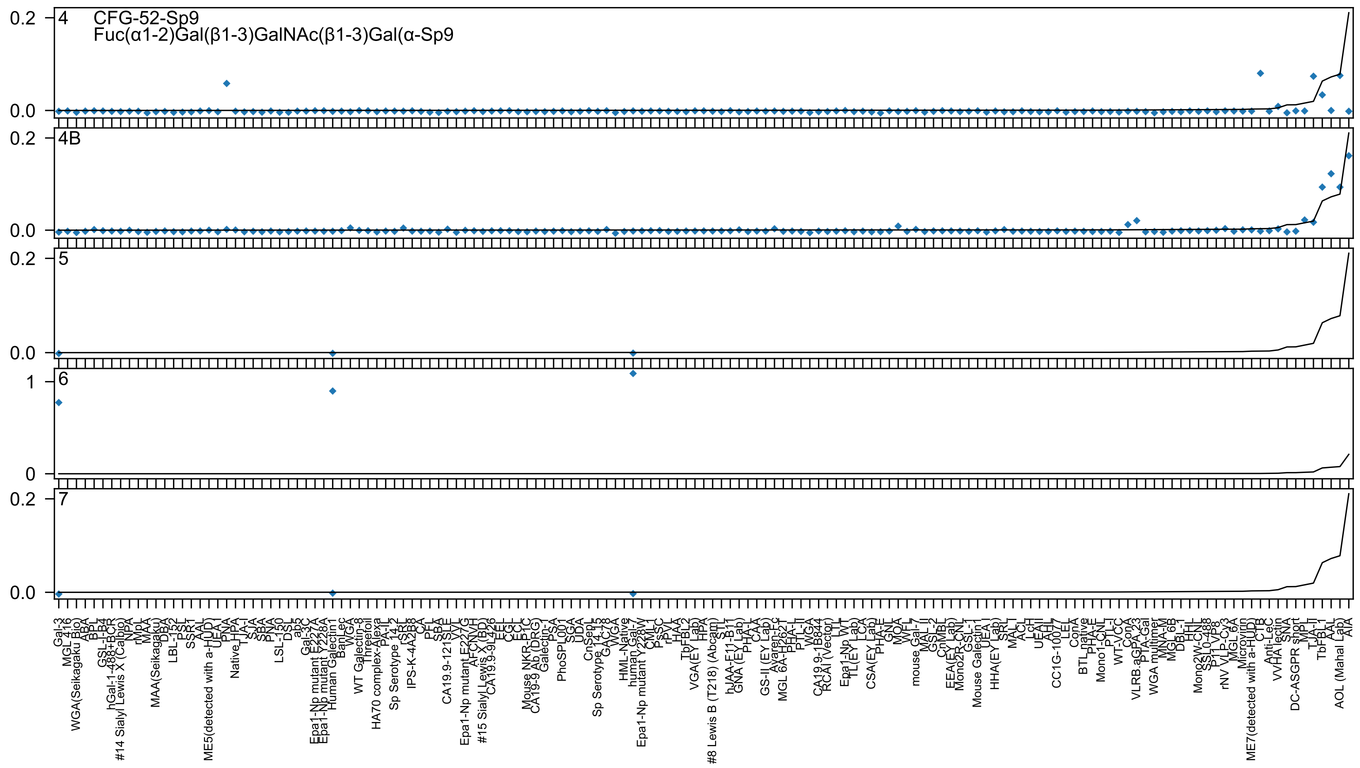
4
CFG-52-Sp9
Fuc(α1-2)Gal(β1-3)GalNAc(β1-3)Gal(α-Sp9
4B
5
6
7
0.2
0.0
1
0
Gal-3
MGL 4.16
WGA(Seikagaku Bio)
ABA
hGal-1-488+BCR
#14 Sialyl Lewis X (Calbio)
MAA(Seikagaku)
ME5(detected with a-HUD)
Native_HPA
Epa1-Np mutant E227A
Epa1-Np mutant Y228A
Human Galectin1
WT Galectin-8
HA70 complex-Alexa
Sp Serotype 14_2
IPS-K-4A2B8
CA19.9-121SLE
Epa1-Np mutant E227G
#15 Sialyl Lewis X (BD)
Mouse NKR-P1C
CA19-9 Ab (DRG)
PhoSPL001
Sp Serotype 14_3
HML-Native
human Gal-7
Epa1-Np mutant Y228W
VGA(EY Lab)
#8 Lewis B (T218) (Abcam)
hJAA-F11-B11
GNA (EY Lab)
GS-II (EY Lab)
MGL 6A-P262T
CA19.9-1B844
RCAI (Vector)
Epa1-Np WT
CSA(EY Lab)
mouse Gal-7
EEA(EY Lab)
Mouse Galectin1
HHA(EY Lab)
CC1G-10077
BTL native
VLRB.aGPA.23
WGA multimer
rNV VLP-Cy3
ME7(detected with a-HUD)
VVHA lectin
DC-ASGPR short
AOL (Mahal Lab)
AIA

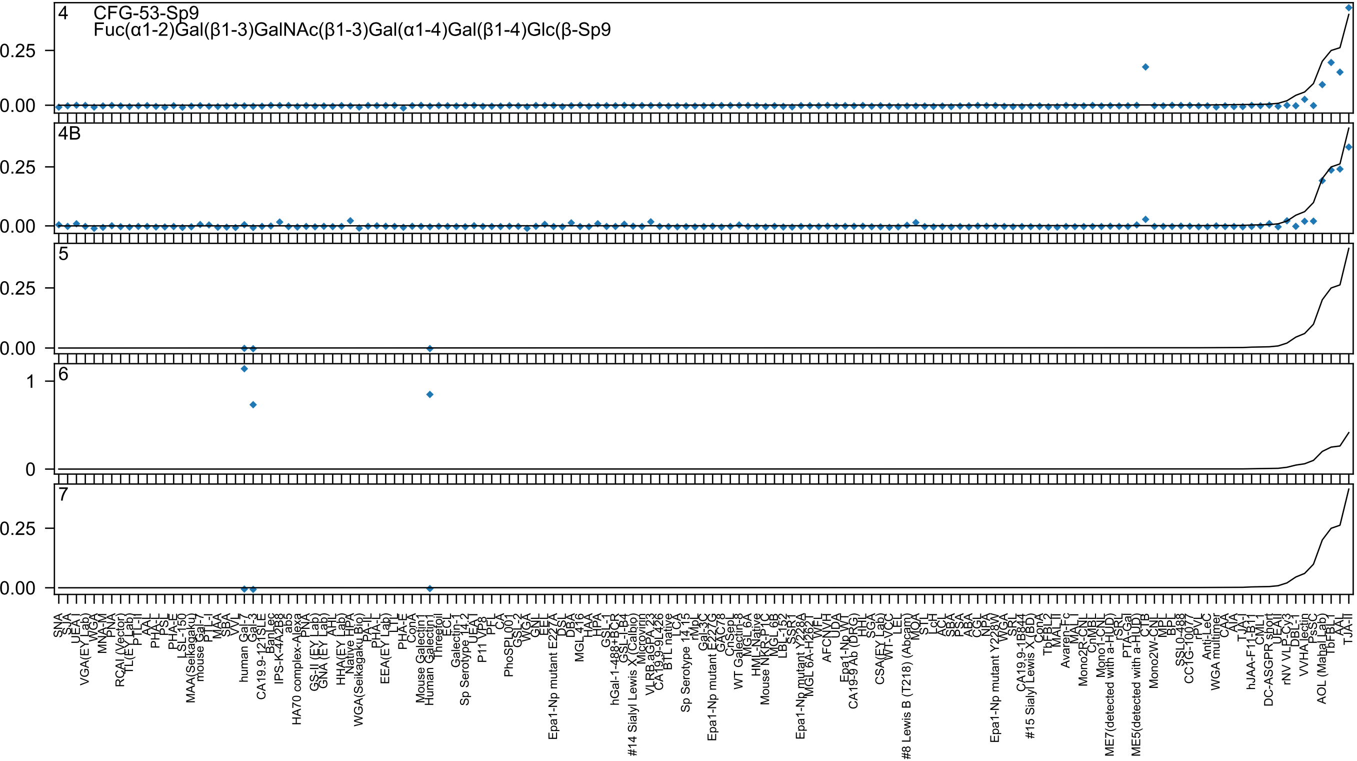

4
CFG-53-Sp9
Fuc(α1-2)Gal(β1-3)GalNAc(β1-3)Gal(α1-4)Gal(β1-4)Glc(β-Sp9
4B
5
6
7

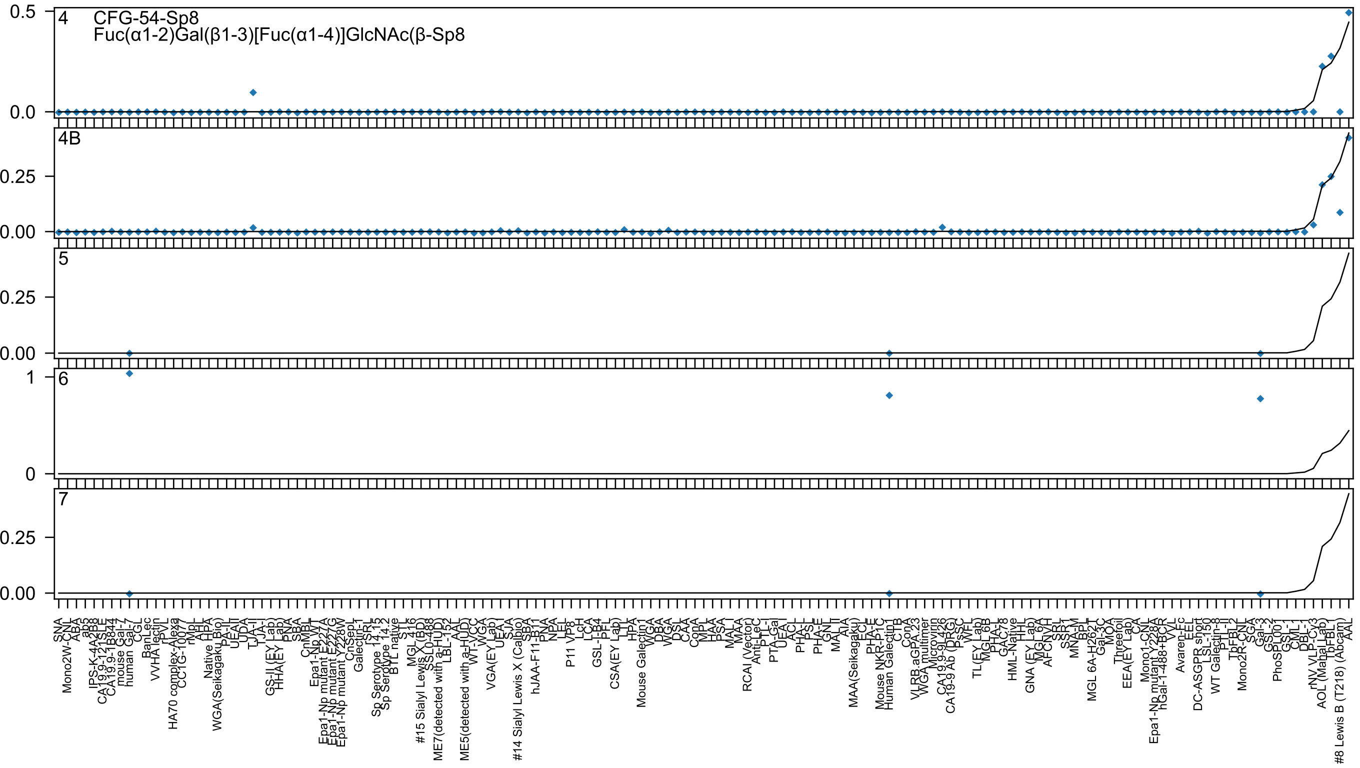
4
CFG-54-Sp8
Fuc(α1-2)Gal(β1-3)[Fuc(α1-4)]GlcNAc(β-Sp8
4B
5
6
7

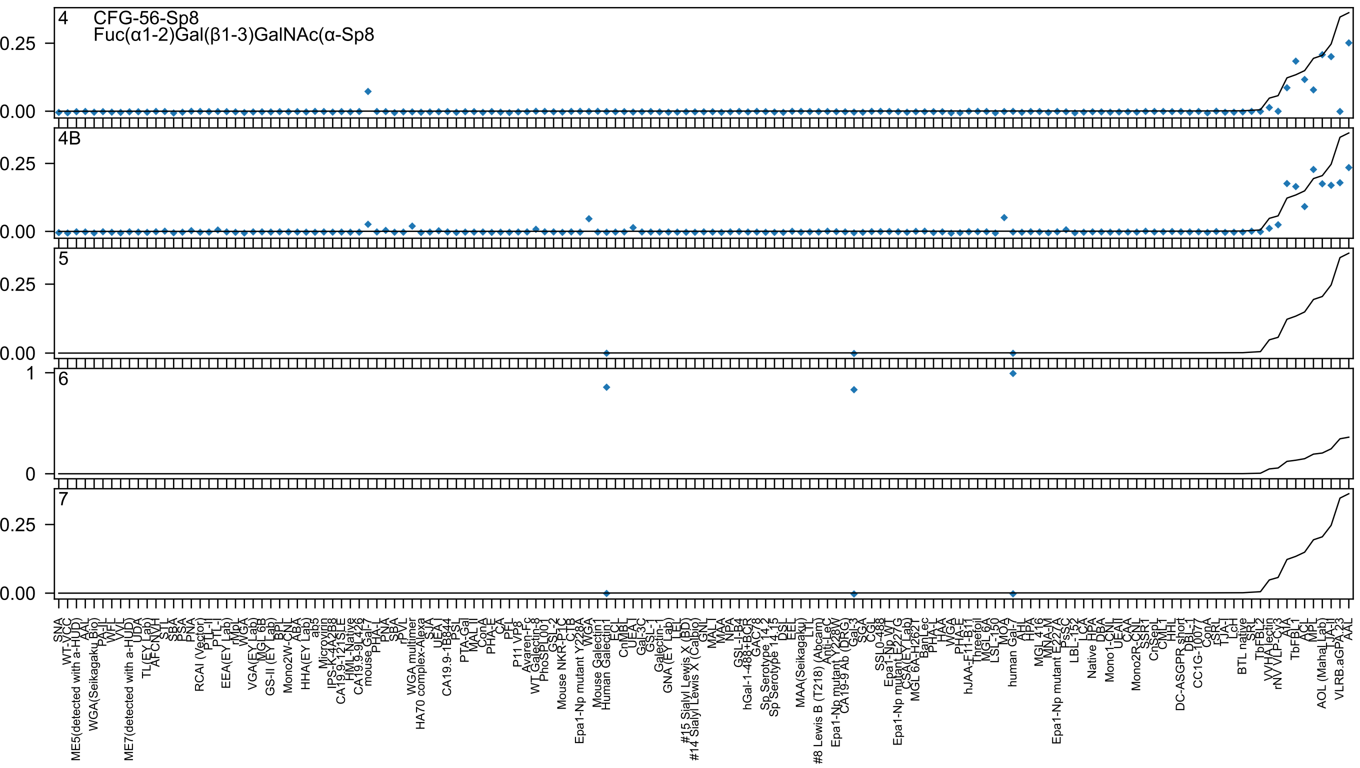
4
CFG-56-Sp8
Fuc(α1-2)Gal(β1-3)GalNAc(α-Sp8
4B
5
6
7
0.25
0.00
1
0
SNA
WT-VCC
ME5(detected with a-HUD)
AAL
WGA(Seikagaku Bio)
PA-II
WFL
VVL
ME7(detected with a-HUD)
UDA
TL(EY Lab)
AFCN-H
STL
SBA
PSA
PNA
RCAI (Vector)
PTL-II
PTL-I
EEA(EY Lab)
rMpL
WGA
VGA(EY Lab)
MGL 6B
GS-II (EY Lab)
BPL
Mono2W-CNL
ABA
HHA(EY Lab)
ab5
Microvirin
IPS-K-4A2B8
CA19.9-121SLE
HML-Native
CA19.9-9L426
mouse Gal-7
PHA-L
PNA
SBA
rPVL
WGA multimer
HA70 complex-Alexa
SJA
UEA-I
CA19.9-1B844
PSL
PTA-Gal
MAL II
ConA
PHA-E
CA
PFL
P11 VP8
Avaren-Fc
WT Galectin-8
PhoSL001
GSL-2
Mouse NKR-P1C
CTB
Epa1-Np mutant Y228A
WGA
Mouse Galectin1
Human Galectin1
ECL
CnMBL
UEA-I
Gal-3C
GSL-I
Galectin-1
GNA (EY Lab)
LEL
#15 Sialyl Lewis X (BD)
#14 Sialyl Lewis X (Calbio)
GNL
MAL-I
MAA
NPA
GSL-I-B4
hGal-1-488+BCR
GAC78
Sp Serotype 14.2
Sp Serotype 14.15
DSL
EEL
MAA(Seikagaku)
LTL
#8 Lewis B (T218) (Abcam)
Anti-LeC
Epa1-Np mutant Y228V
CA19-9 Ab (DRG)
Gal-3
SGA
CGL
SSL0-488
Epa1-Np WT
Epa1-Np mutant E227G
CSA(EY Lab)
MGL 6A-H262T
BanLec
PHA-L
HAA
WGA
PHA-E
hJAA-F11-B11
Threefoil
MGL 6A
LSL-150
MOA
human Gal-7
AHL
HPA
MGL 416
MNA-M
Epa1-Np mutant E227A
PsSC
LBL-152
LCA
Native HPA
DBA
Mono1-CNL
UEA-II
CAA
Mono2R-CNL
SSR1
CnSepL
CML1
HHL
DC-ASGPR short
DBL-1
CC1G-10077
ConA
rSRL
TJA-I
LcH
BTL native
SRL
TbFBL2
VVHA lectin
rNV VLP-Cy3
AIA
TbFBL1
ACL
MPL
AOL (Mahal Lab)
LTA-II
VLRB.aGPA.23
AAL

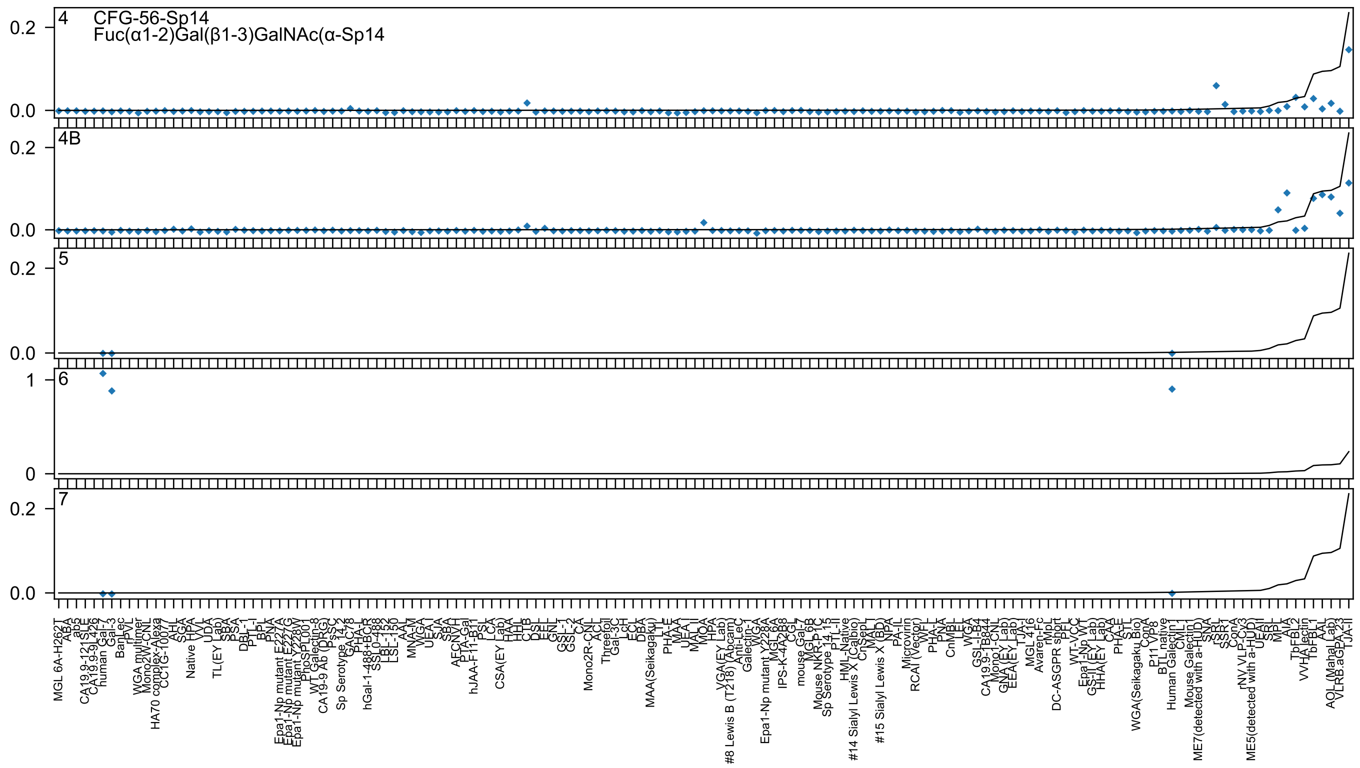
4 CFG-56-Sp14
Fuc(α1-2)Gal(β1-3)GalNAc(α-Sp14
4B
5
6
7
0.0
0.2
0
1

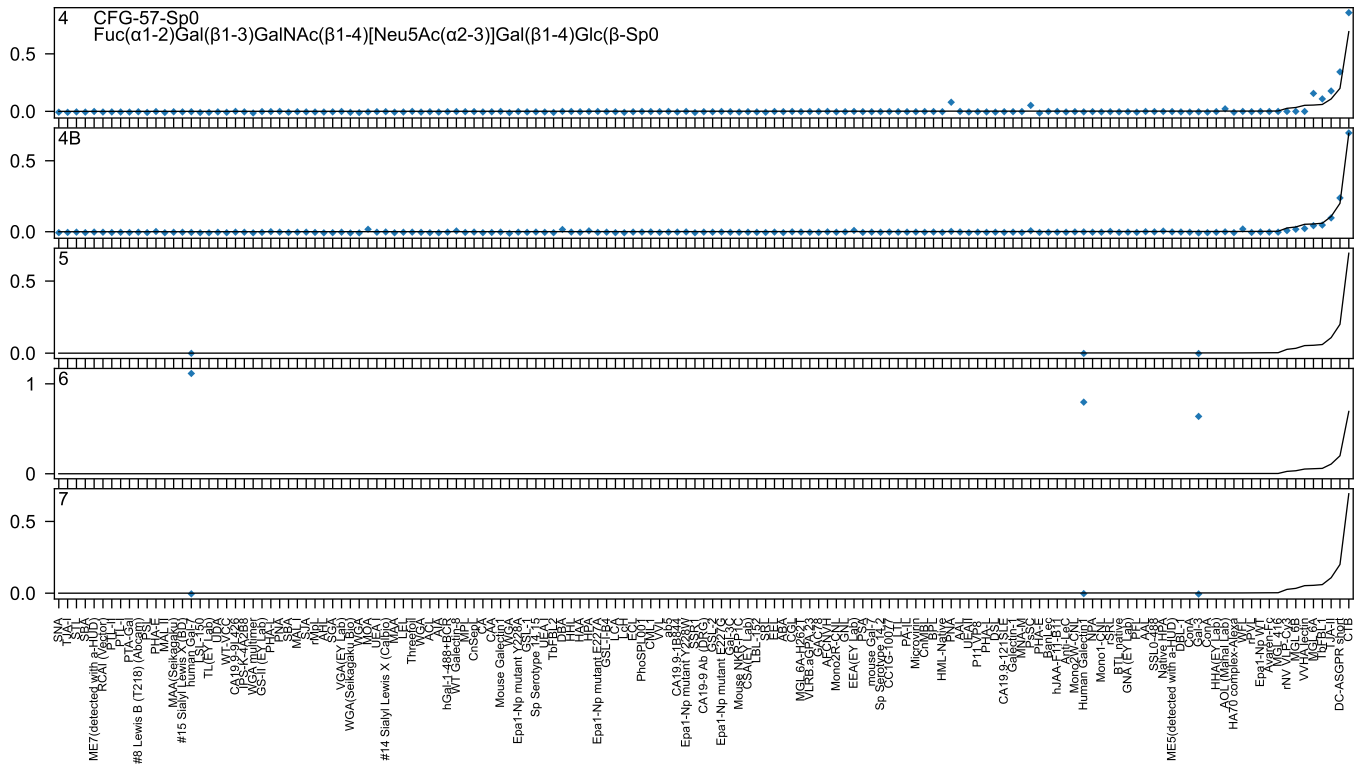

4
CFG-57-Sp0
Fuc(α1-2)Gal(β1-3)GalNAc(β1-4)[Neu5Ac(α2-3)]Gal(β1-4)Glc(β-Sp0
4B
5
6
7

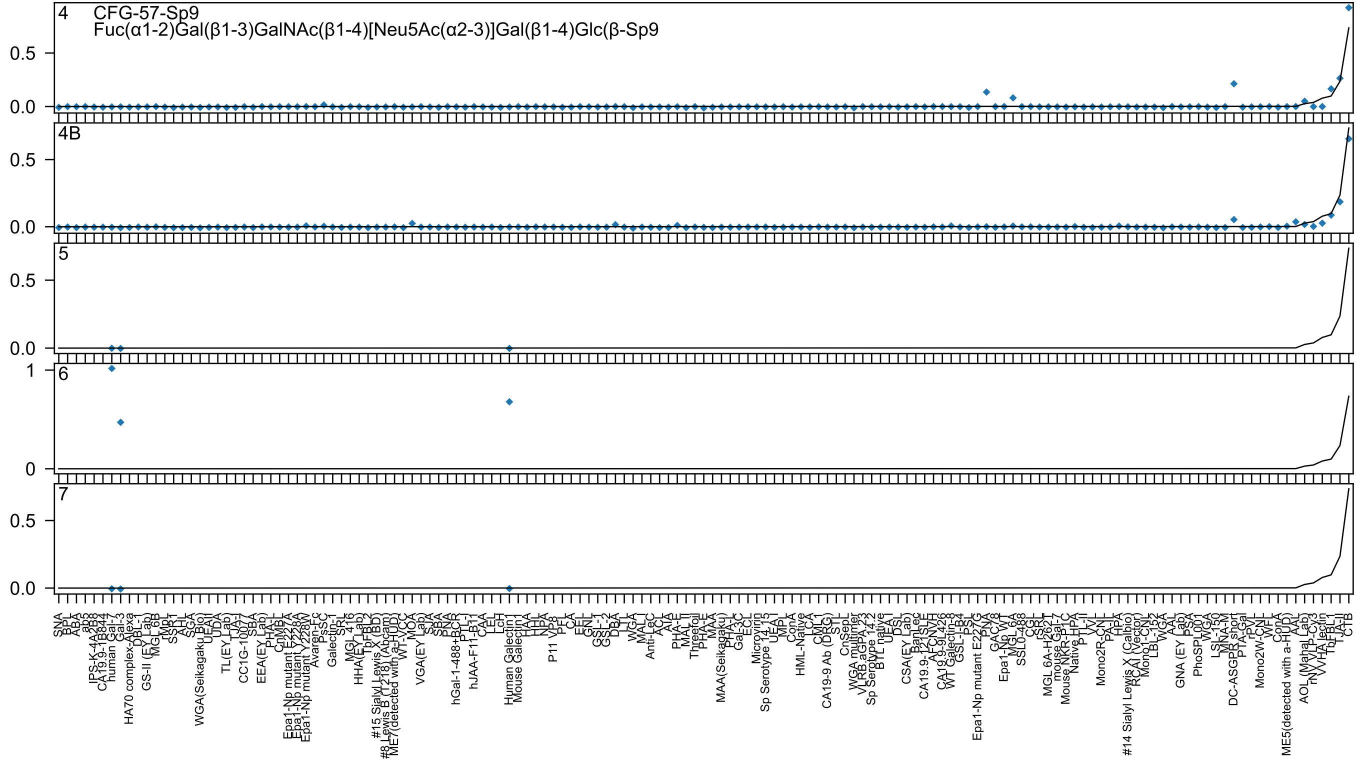

4
CFG-57-Sp9
Fuc(α1-2)Gal(β1-3)GalNAc(β1-4)[Neu5Ac(α2-3)]Gal(β1-4)Glc(β-Sp9
4B
5
6
7
0.0
0.5
1
0

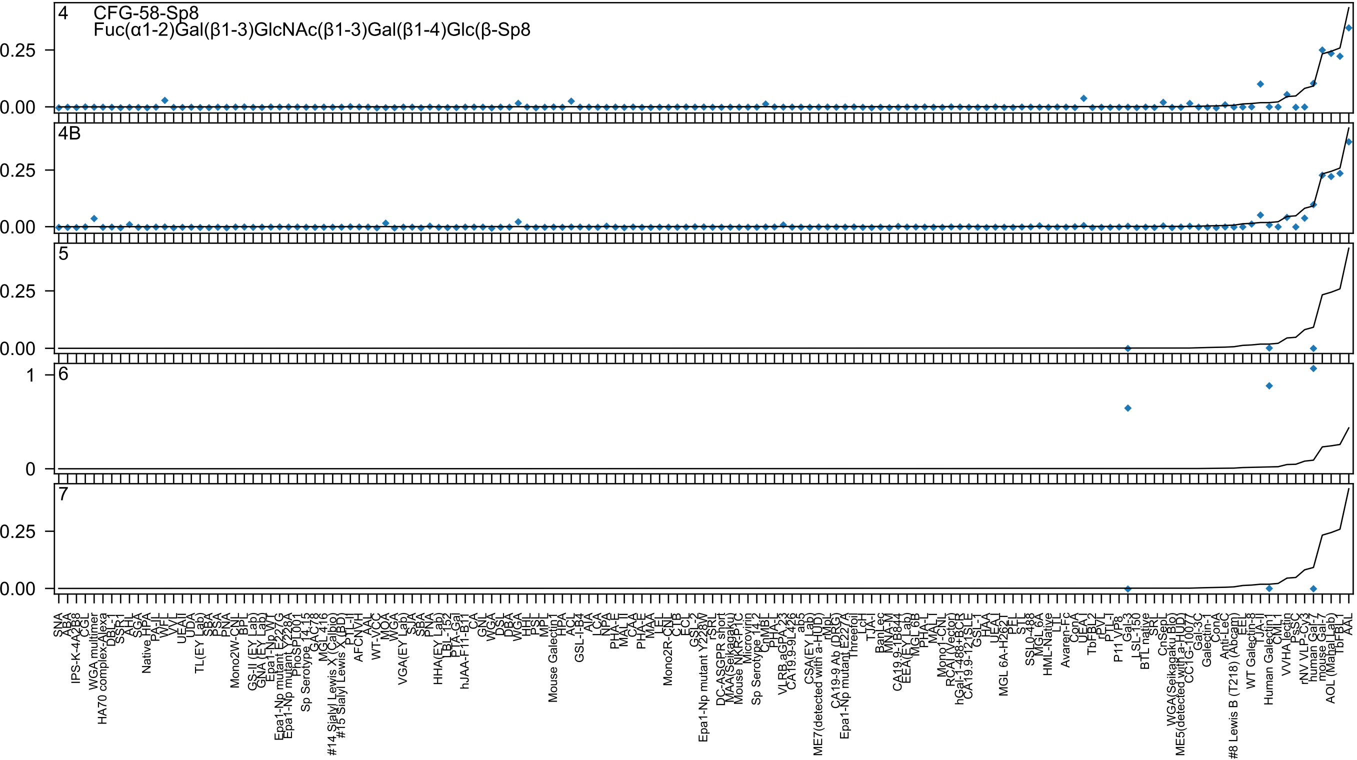
4
CFG-58-Sp8
Fuc(α1-2)Gal(β1-3)GlcNAc(β1-3)Gal(β1-4)Glc(β-Sp8
4B
5
6
7
0.25
0.00
1
0

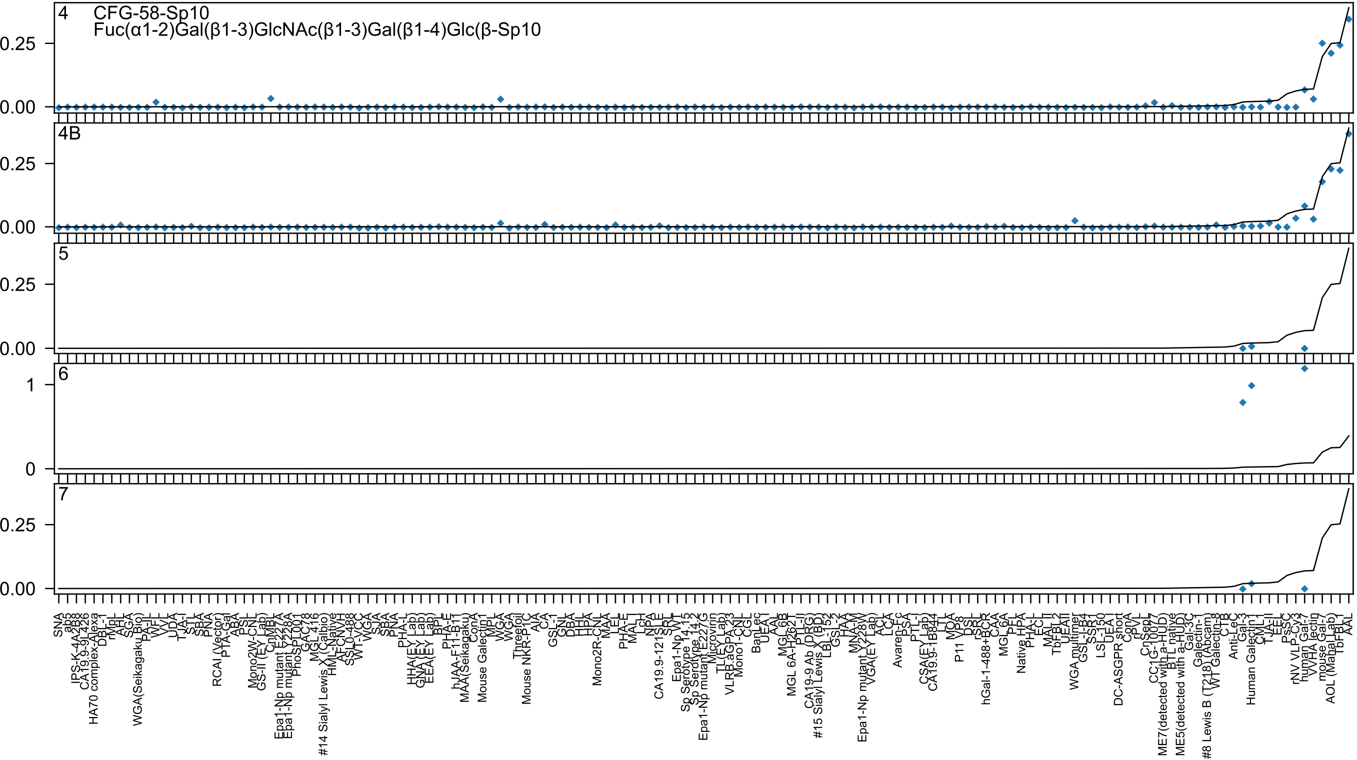

4 CFG-58-Sp10
Fuc(α1-2)Gal(β1-3)GlcNAc(β1-3)Gal(β1-4)Glc(β-Sp10
4B
5
6
7

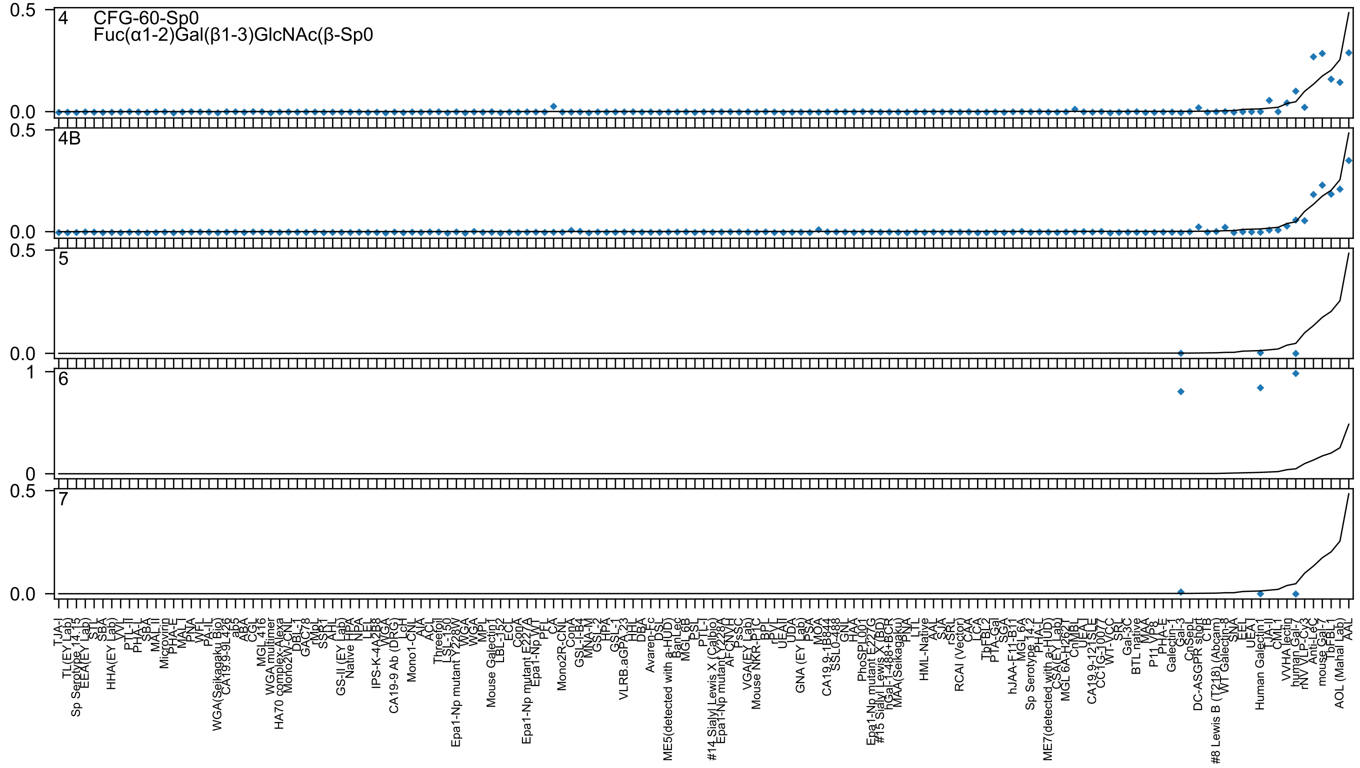
4 CFG-60-Sp0
Fuc(α1-2)Gal(β1-3)GlcNAc(β-Sp0
4B
5
6
7

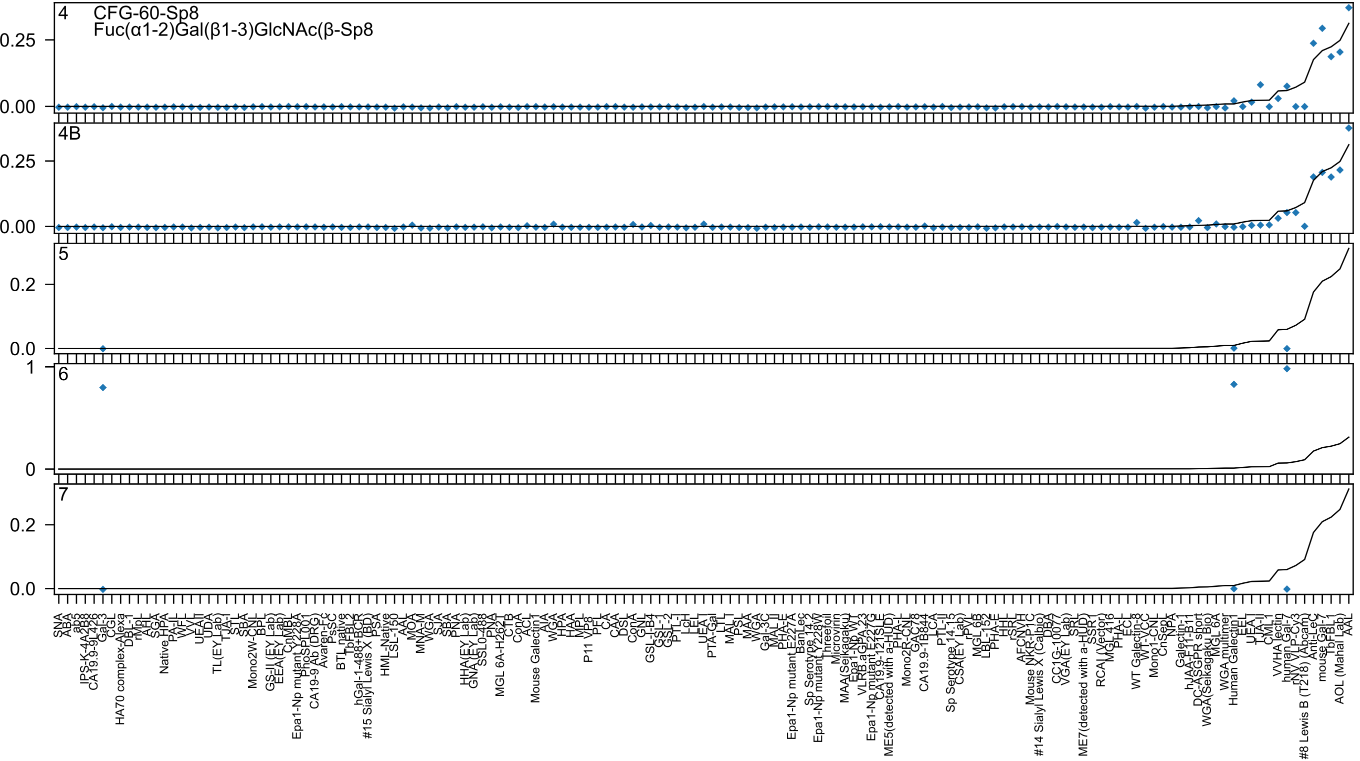

4 CFG-60-Sp8
Fuc(α1-2)Gal(β1-3)GlcNAc(β-Sp8
4B
5
6
7

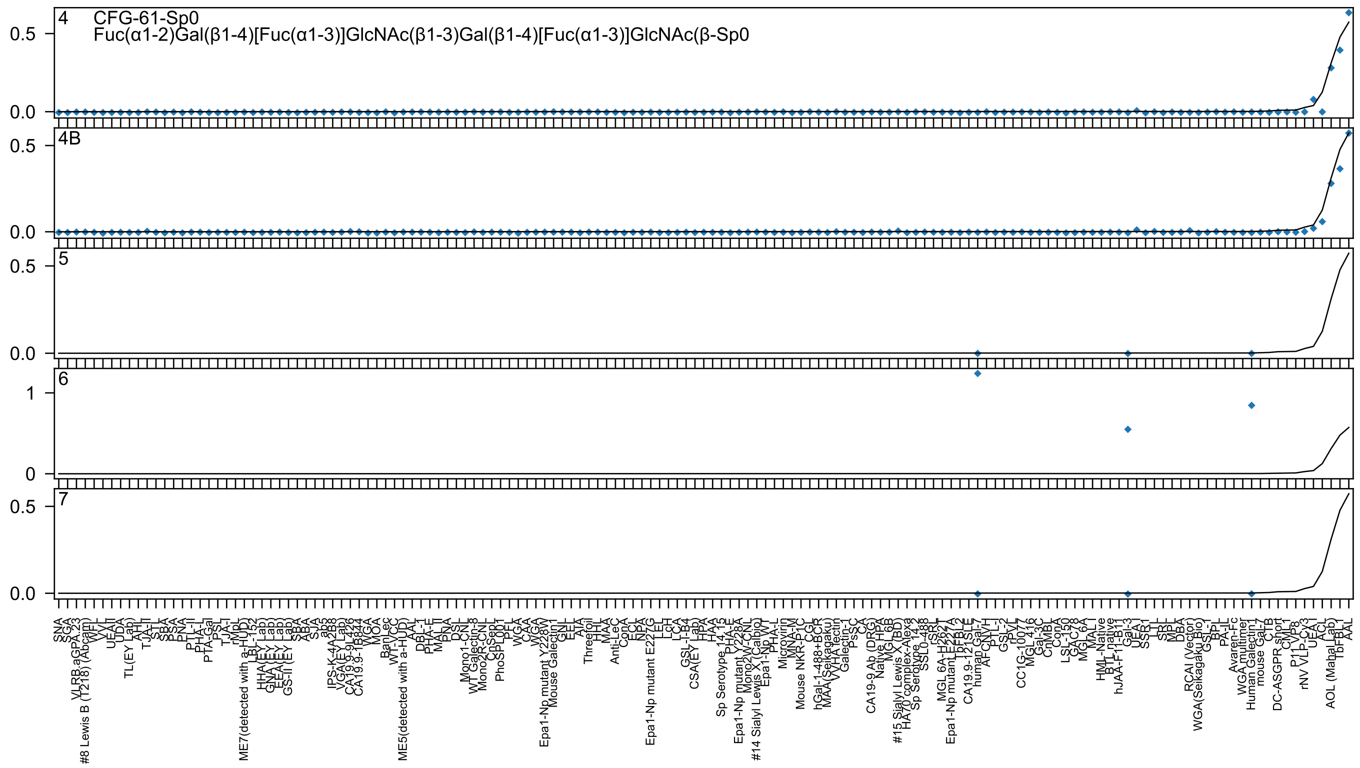
4
CFG-61-Sp0
Fuc(α1-2)Gal(β1-4)[Fuc(α1-3)]GlcNAc(β1-3)Gal(β1-4)[Fuc(α1-3)]GlcNAc(β-Sp0
0.5
0.0
4B
0.5
0.0
5
0.5
0.0
6
1
0
7
0.5
0.0

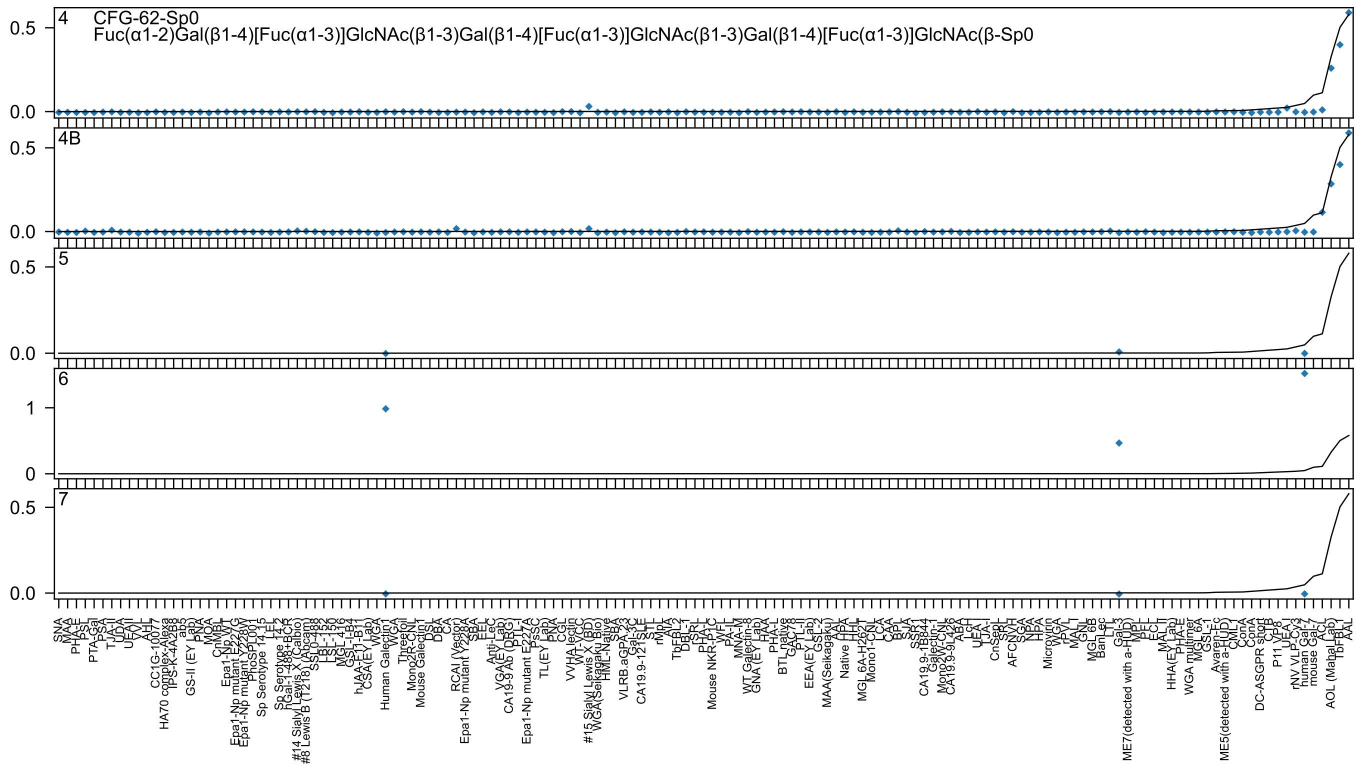

4
CFG-62-Sp0
Fuc(α1-2)Gal(β1-4)[Fuc(α1-3)]GlcNAc(β1-3)Gal(β1-4)[Fuc(α1-3)]GlcNAc(β1-3)Gal(β1-4)[Fuc(α1-3)]GlcNAc(β-Sp0
4B
5
6
7
0.0
0.5
0
1
SNA
MAA
PHA-E
PSL
PTA-Gal
PSA
TJA-II
UDA
UEAII
VV
AHL
CC1G-10077
HA70 complex-Alexa
IPS-K-4A2B8
ab5
GS-II (EY Lab)
PNA
MOA
CnMBL
Epa1-Np WT
Epa1-Np mutant E227G
Epa1-Np mutant Y228W
PhoSL001
Sp Serotype 14.15
LEL
Sp Serotype 14.2
hGal-1-488+BCR
#14 Sialyl Lewis X (Calbio)
#8 Lewis B (T218) (Abcam)
SSL0-488
LBL-152
LSL-150
MGL 416
GSL-I-B4
hJAA-F11-B11
CSA(EY Lab)
WGA
Human Galectin1
WGA
Threefoil
Mono2R-CNL
Mouse Galectin1
DSL
DBA
CA
RCAI (Vector)
Epa1-Np mutant Y228A
SBA
EEL
Anti-LeC
VGA(EY Lab)
CA19-9 Ab (DRG)
PTL-I
Epa1-Np mutant E227A
PsSC
TL(EY Lab)
PNA
CGL
VVHA lectin
WT-VCC
#15 Sialyl Lewis X (BD)
WGA(Seikagaku Bio)
HML-Native
SBA
VLRB.aGPA.23
Gal-3C
CA19.9-121SLE
STL
rMpL
AIA
TbFBL2
DBL-1
rSRL
PHA-L
Mouse NKR-P1C
WFL
PA-IL
MNA-M
WT Galectin-8
GNA (EY Lab)
HAA
PHA-L
BTL native
GAC78
PTL-II
EEA(EY Lab)
GSL-2
MAA(Seikagaku)
AAL
Native HPA
HHL
MGL 6A-H262T
Mono1-CNL
LCA
CAA
BPL
SJA
SSR1
CA19.9-1B844
Galectin-1
Mono2W-CNL
CA19.9-9L426
ABA
LcH
UEA-I
TJA-I
CnSepL
SRL
AFCNVH
SGA
NPA
HPA
Microvirin
WGA
rPVL
MAL-I
GNL
MGL 6B
BanLec
LTL
Gal-3
ME7(detected with a-HUD)
MPL
PFL
ECL
MAL-II
HHA(EY Lab)
PHA-E
WGA multimer
MGL 6A
GSL-1
Avaren-Fc
ME5(detected with a-HUD)
CML
ConA
ConA
DC-ASGPR short
CTB
P11 VP8
UEA-I
rNV VLP-Cy3
human Gal-7
mouse Gal-7
ACL
AOL (Mahal Lab)
TbFBL1
AAL

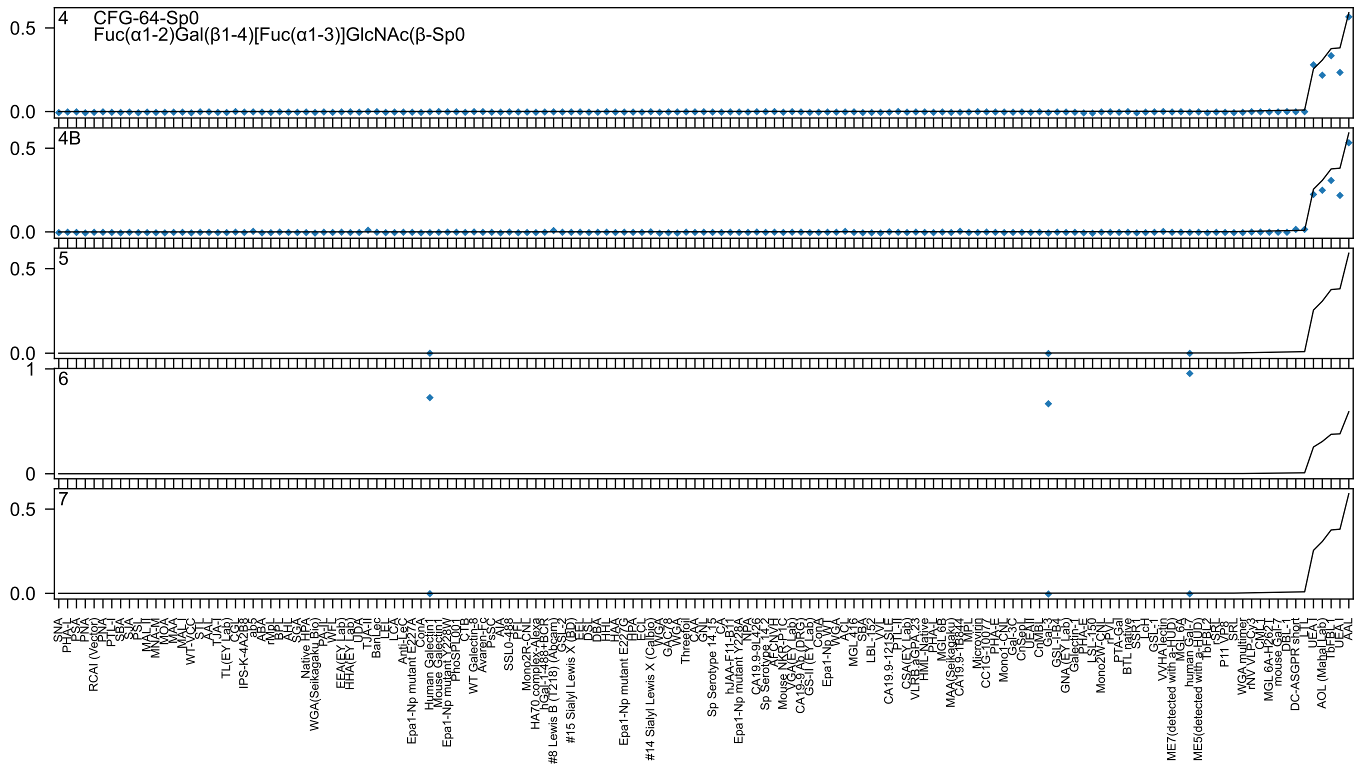

4 CFG-64-Sp0
Fuc(α1-2)Gal(β1-4)[Fuc(α1-3)]GlcNAc(β-Sp0
4B
5
6
7
0.5
0.0
1
0
SNA
PHA-L
PSA
PNA
RCAI (Vector)
PNA
PTL-I
SBA
SJA
PSL
MAL-I
MNA-M
MOA
MAA
MAL-I
WT-VCC
STL
AAL
TJA-I
TL(EY Lab)
CGL
IPS-K-4A2B8
ab5
ABA
rMpL
BPL
AHL
SGA
Native HPA
WGA(Seikagaku Bio)
PA-IL
WFL
EEA(EY Lab)
HHA(EY Lab)
UDA
TJA-II
BanLec
LEL
LCA
Anti-LeC
Epa1-Np mutant E227A
ConA
Human Galectin1
Mouse Galectin1
Epa1-Np mutant Y228W
PhoSL
PL001
CTB
WT Galectin-8
Avaren-Fc
PsSC
AIA
SSL0-488
PFL
Mono2R-CNL
HA70 complex-Alexa
hGal-1-488+BCR
#8 Lewis B (T218) (Abcam)
GSL-2
#15 Sialyl Lewis X (BD)
EEL
DSL
DBA
HHL
HAA
Epa1-Np mutant E227G
HPA
ECL
#14 Sialyl Lewis X (Calbio)
WGA
GAC78
WGA
Threefoil
CAA
CNL
Sp Serotype 14_15
CA
hJAA-F11-B11
Epa1-Np mutant Y228A
NPA
CA19.9-9L426
Sp Serotype 14_2
AFCNVH
Mouse NKR-P1C
VGA(EY Lab)
CA19-9 Ab (DRG)
GS-II (EY Lab)
ConA
Epa1-Np WT
WGA
ACL
MGL 416
SBA
LBL-152
VVL
CA19.9-121SLE
PTL-II
CSA(EY Lab)
VLRB aGPA 23
HML-Native
PHA-L
MGL 6B
MAA(Seikagaku)
CA19.9-1B844
MPL
Microvirin
CC1G-10077
PHA-E
Mono1-CNL
Gal-3C
CnSep
UEAII
CnMBL
Gal-3
GSL-I-B4
GNA (EY Lab)
Galectin-1
PHA-E
LSL-150
Mono2W-CNL
rPVL
PTA-Gal
BTL native
SSR1
LCH
GSL-1
VVHA lectin
ME7(detected with a-HUD)
MGL 6A
human Gal-7
ME5(detected with a-HUD)
TbFBL2
rSRL
P11 VP8
SRL
WGA multimer
rNV VLP-Cy3
CML
MGL 6A-H262T
mouse Gal-7
DBL-1
DC-ASGPR short
LTL
UEA I
AOL (Mahal Lab)
TbFBL1
UEA I
AAL

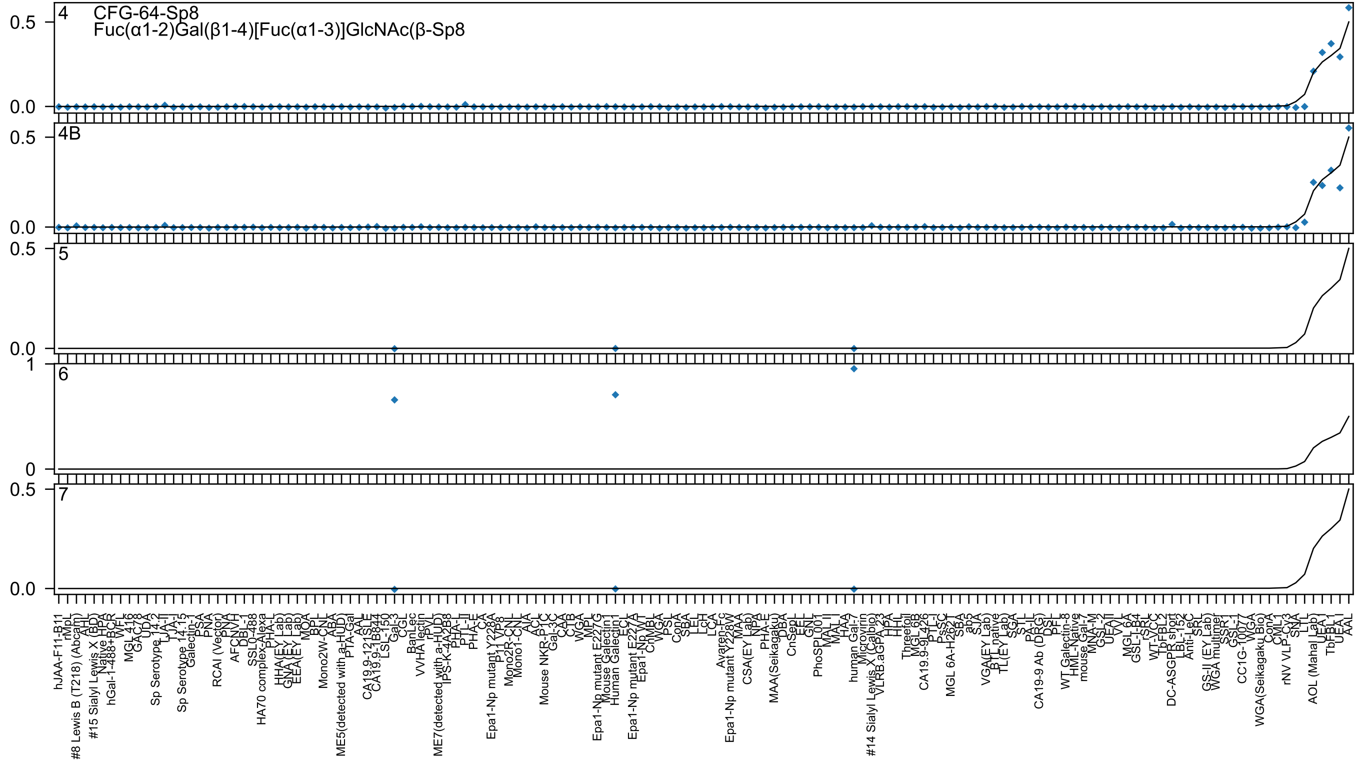

4
CFG-64-Sp8
Fuc(α1-2)Gal(β1-4)[Fuc(α1-3)]GlcNAc(β-Sp8
4B
5
6
7
0.5
0.0
1
0

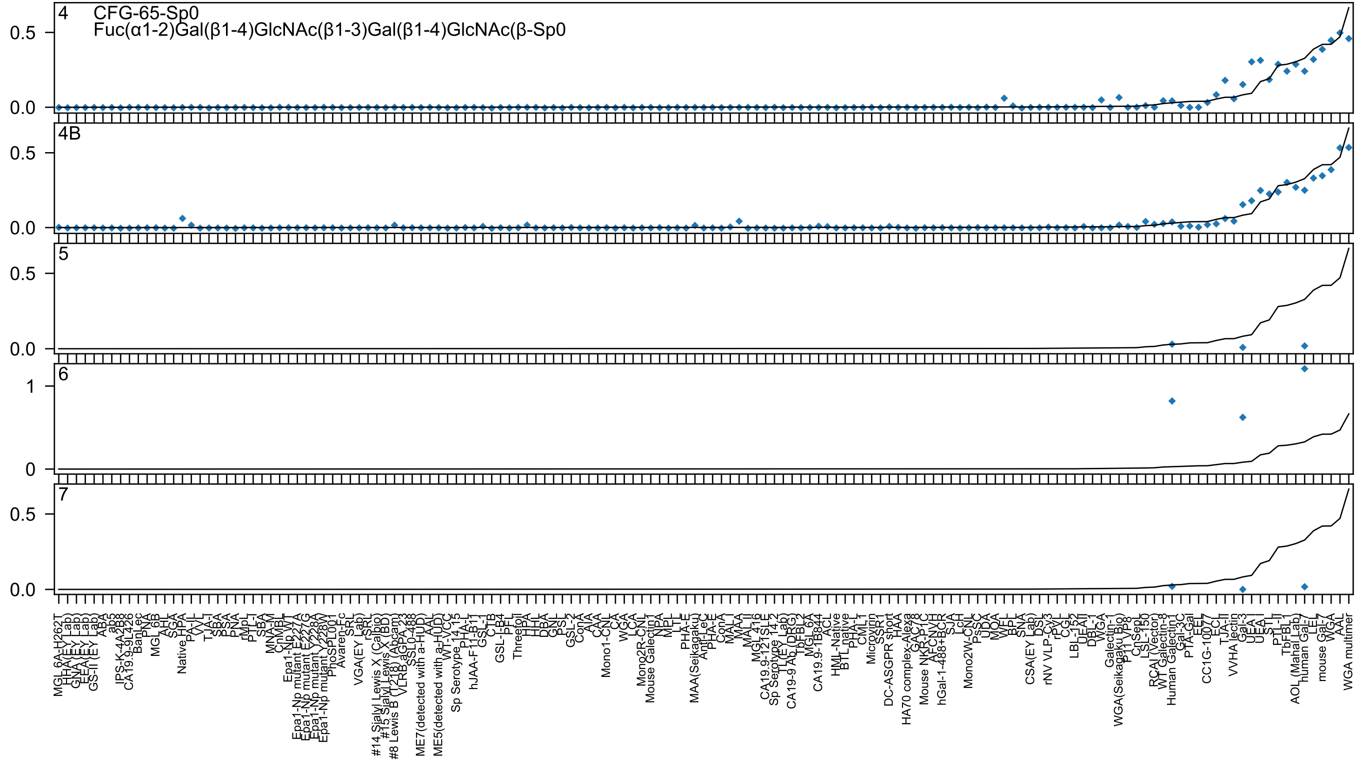
4 CFG-65-Sp0
Fuc(α1-2)Gal(β1-4)GlcNAc(β1-3)Gal(β1-4)GlcNAc(β-Sp0
4B
5
6
7

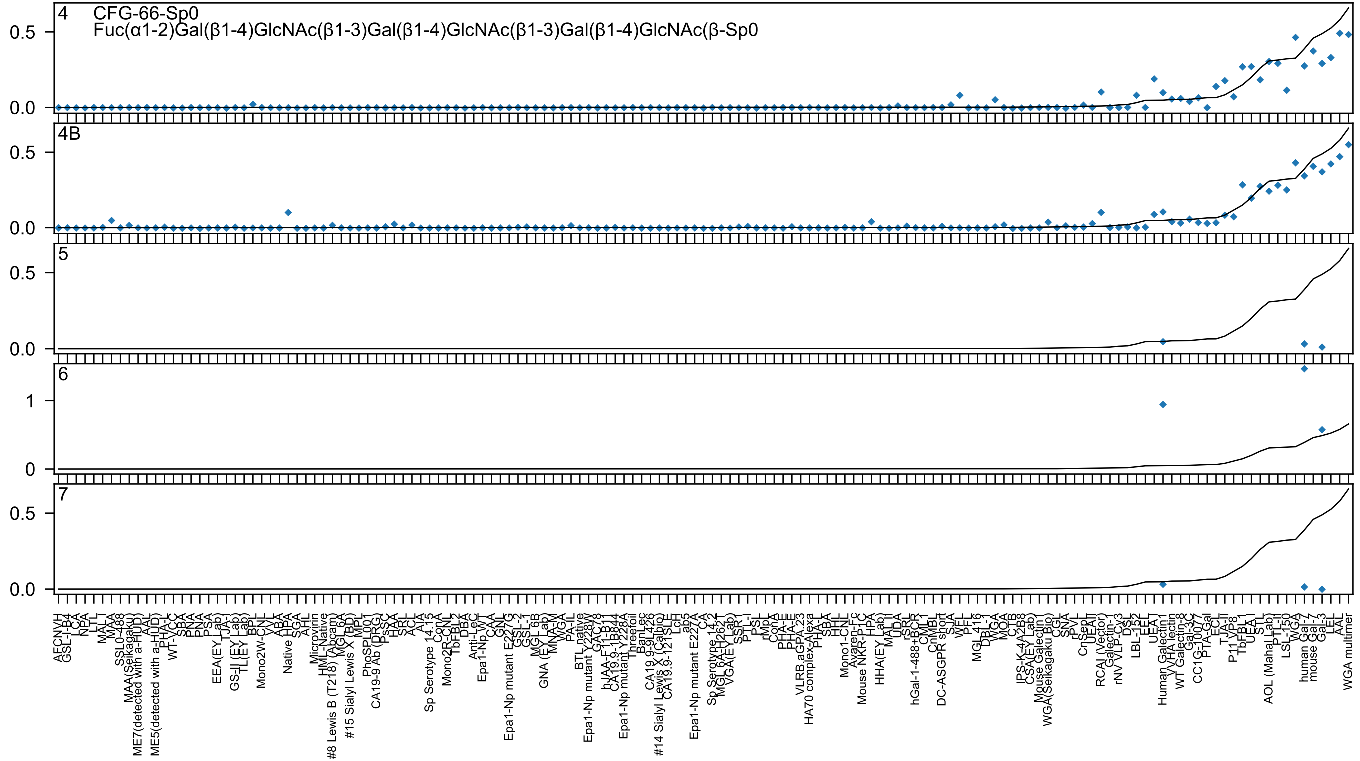
4 CFG-66-Sp0
Fuc(α1-2)Gal(β1-4)GlcNAc(β1-3)Gal(β1-4)GlcNAc(β1-3)Gal(β1-4)GlcNAc(β-Sp0
4B
5
6
7

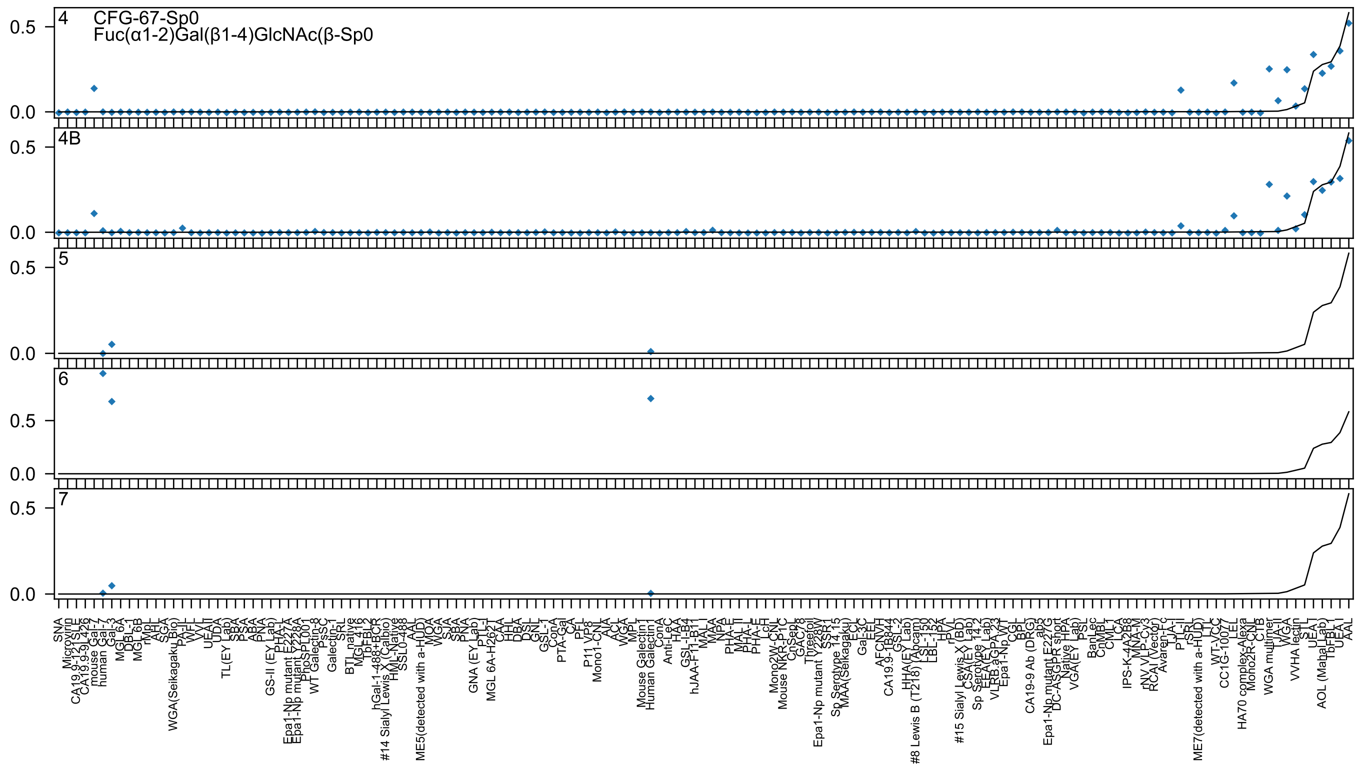
4
CFG-67-Sp0
Fuc(α1-2)Gal(β1-4)GlcNAc(β-Sp0
4B
5
6
7
0.5
0.0

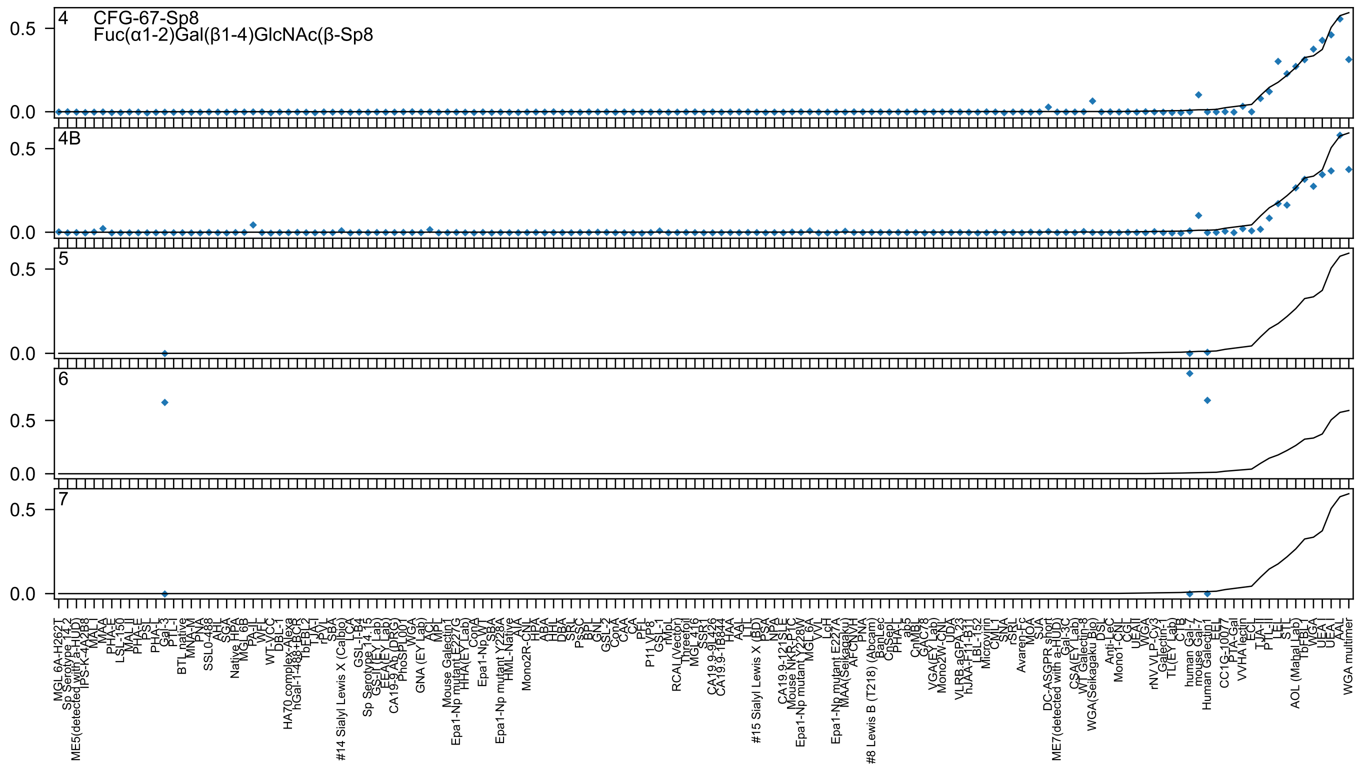

4
CFG-67-Sp8
Fuc(α1-2)Gal(β1-4)GlcNAc(β-Sp8
4B
5
6
7
0.5
0.0

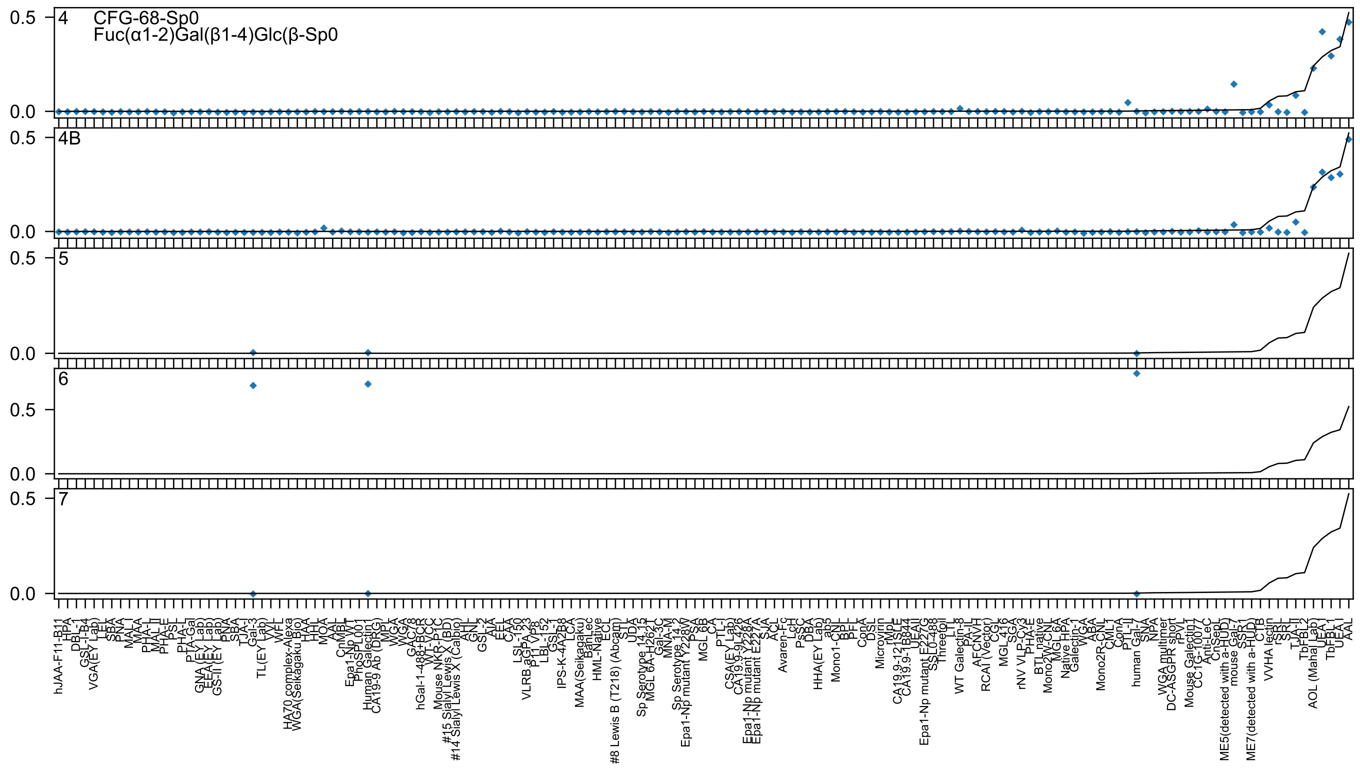

CFG-68-Sp0
Fuc(α1-2)Gal(β1-4)Glc(β-Sp0
4
4B
5
6
7
0.5
0.0

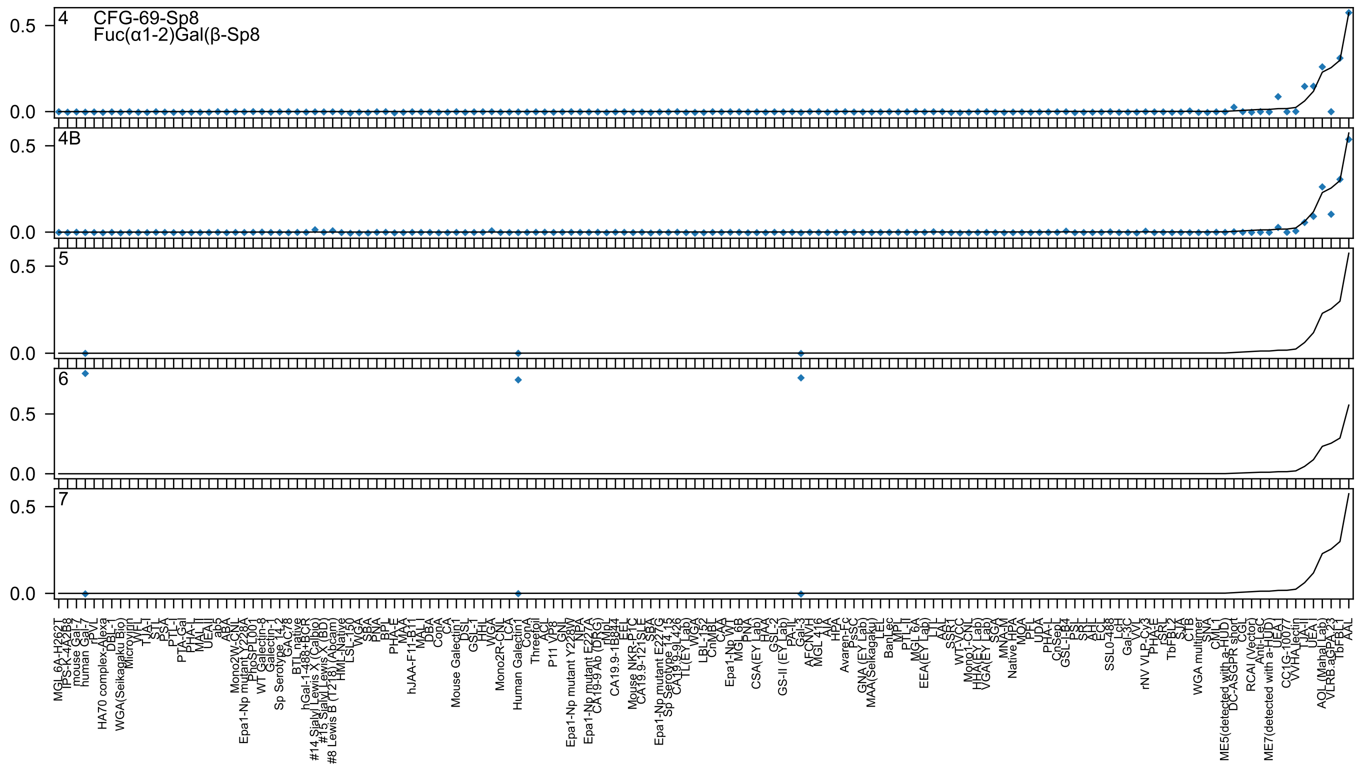
4
CFG-69-Sp8
Fuc(α1-2)Gal(β-Sp8
4B
5
6
7

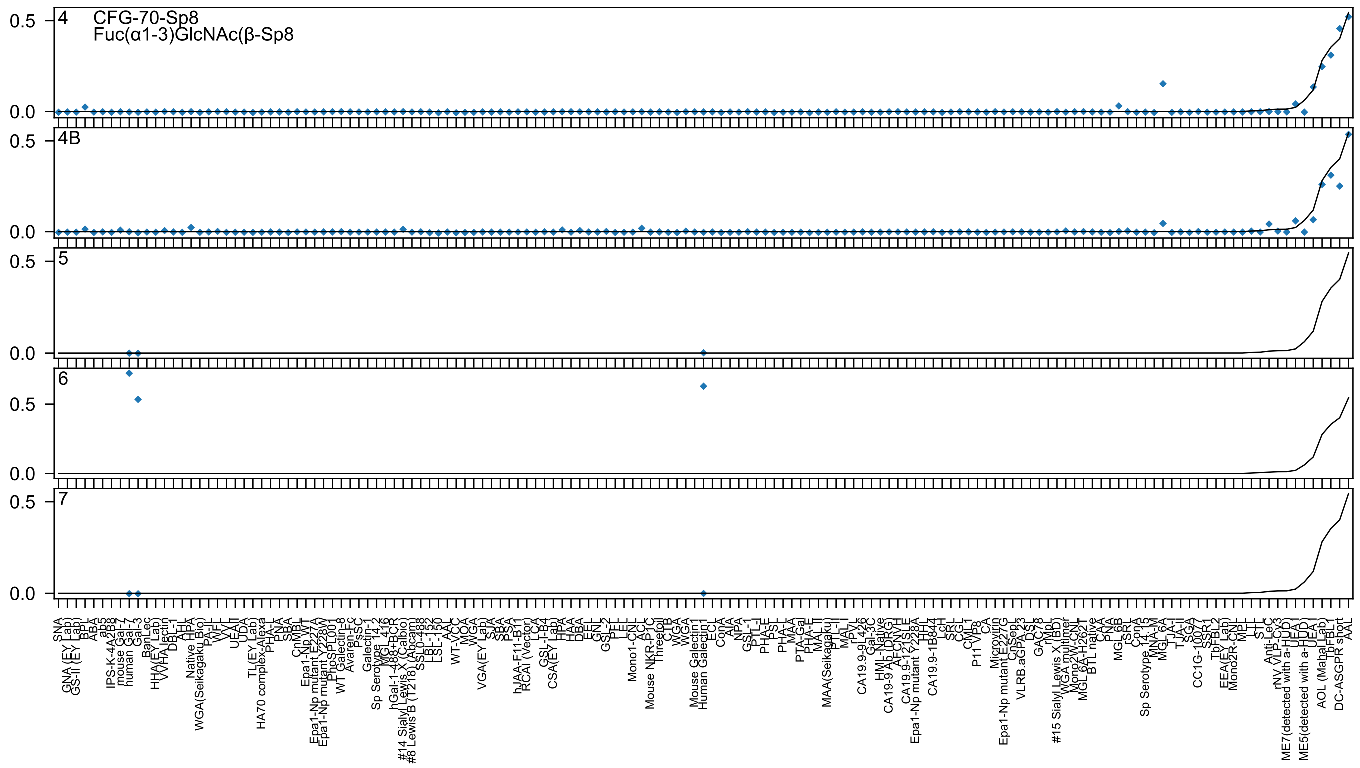

4
CFG-70-Sp8
Fuc(α1-3)GlcNAc(β-Sp8
4B
5
6
7

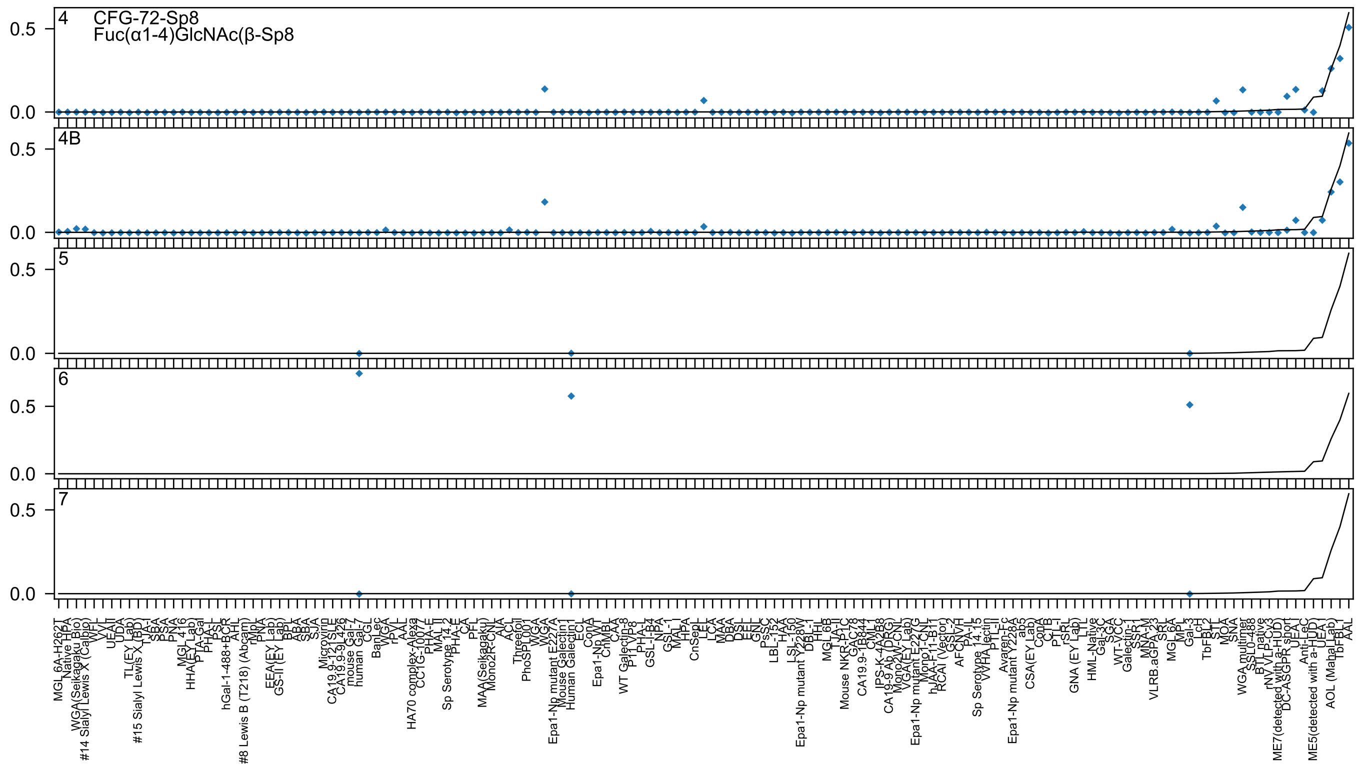
4
CFG-72-Sp8
Fuc(α1-4)GlcNAc(β-Sp8
4B
5
6
7

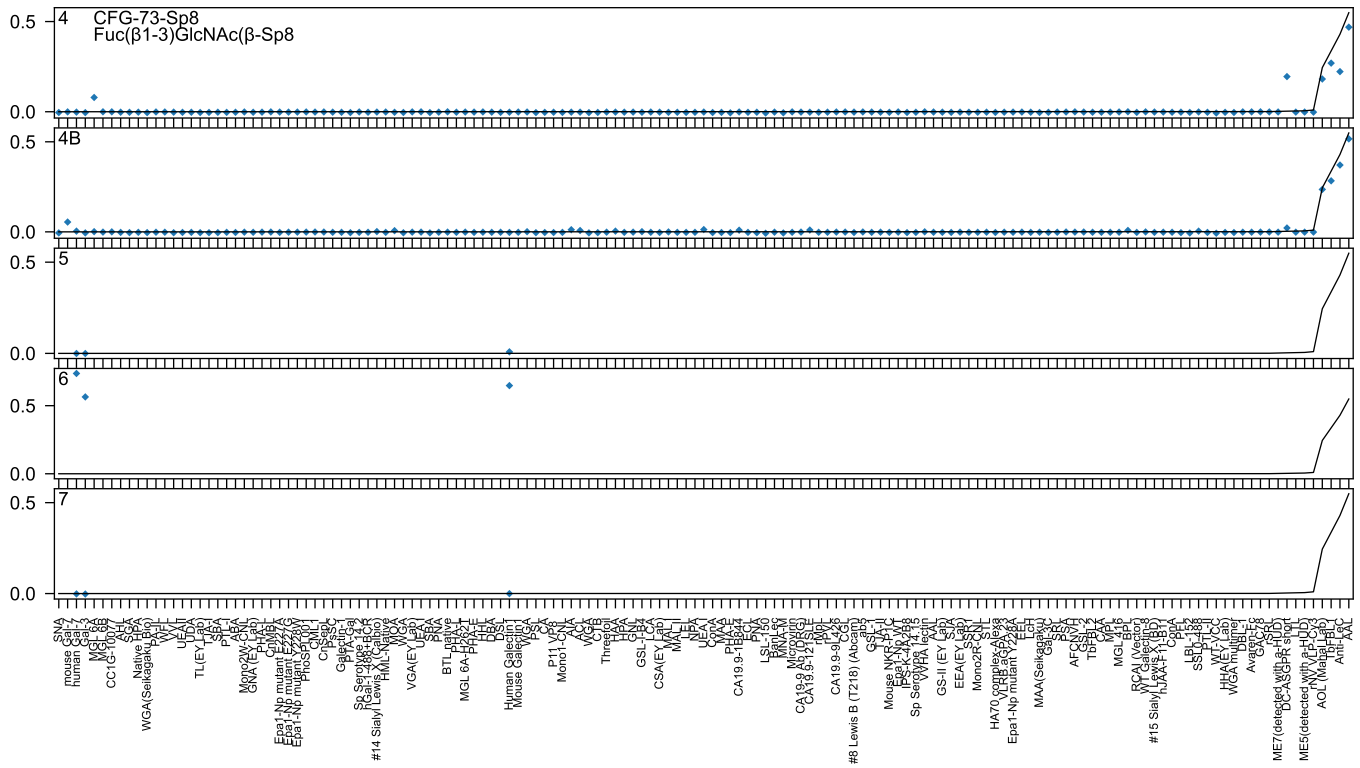
4
CFG-73-Sp8
Fuc(β1-3)GlcNAc(β-Sp8
4B
5
6
7
0.5
0.0
SNA
mouse Gal-7
human Gal-7
Gal-3
MGL 6A
MGL 6B
CC1G-10077
AHL
SGA
Native HPA
WGA(Seikagaku Bio)
PA-IL
WFL
VVL
UEAII
UDA
TL(EY Lab)
TJA-I
SBA
PTL-I
ABA
Mono2W-CNL
GNA (EY Lab)
PHA-L
CnMBL
Epa1-Np mutant E227A
Epa1-Np mutant E227G
Epa1-Np mutant Y228W
PhoSL001
CML1
CnSep
PsSC
Galectin-1
PTA-Gal
Sp Serotype 14.2
hGal-1-488+BCR
#14 Sialyl Lewis X (Calbio)
HML-Native
MOA
WGA
VGA(EY Lab)
UEA I
SBA
PNA
BTL native
PHA-L
MGL 6A-H262T
PHA-E
HHL
DBA
DSL
Human Galectin1
Mouse Galectin1
WGA
PSL
CA
P11 VP8
Mono1-CNL
AIA
ACL
WGA
CTB
Threefoil
HAA
HPA
GNL
GSL-I-B4
LCA
CSA(EY Lab)
MAL II
MAL I
LEL
NPA
UEA I
ConA
MAA
PHA-E
CA19.9-1B844
ECL
PNA
LSL-150
BanLec
MNA-M
Microvirin
CA19-9 Ab (DRG)
CA19.9-121SLE
rMpL
rPVL
CA19.9-9L426
CGL
#8 Lewis B (T218) (Abcam)
ab3
GSL-I
TJA-II
Mouse NKR-P1C
Epa1-Np WT
IPS-K-4A2B8
Sp Serotype 14.15
WHA lectin
AAL
GS-II (EY Lab)
SJA
EEA(EY Lab)
SSA
Mono2R-CNL
STL
HA70 complex-Alexa
VLRB.aGPA.23
Epa1-Np mutant Y228A
EEL
LcH
MAA(Seikagaku)
Gal-3C
SRL
PSA
AFCNVH
GSL-2
TbFBL2
CAA
MPL
MGL 416
BPL
RCAI (Vector)
WT Galectin-8
#15 Sialyl Lewis X (BD)
hJAA-F11-B11
ConA
PFL
LBL-152
SSL0-488
PTL-II
WT-VCC
HHA(EY Lab)
WGA multimer
DBL-I
Avaren-Fc
GAC78
rSRL
ME7(detected with a-HUD)
DC-ASGPR short
LTL
ME5(detected with a-HUD)
rNV VLP-Cy3
AOL (Mahal Lab)
TbFBL1
Anti-LeC
AAL

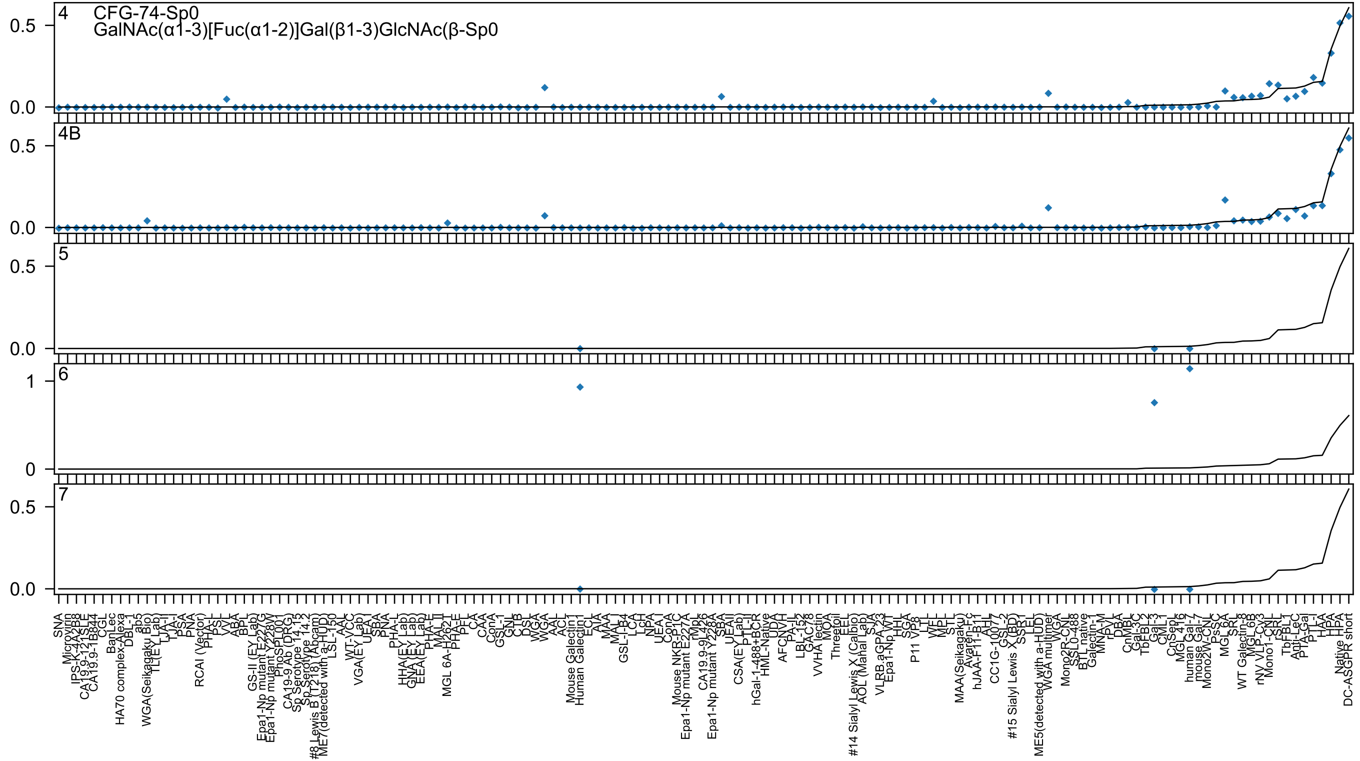
4 CFG-74-Sp0
GalNAc(α1-3)[Fuc(α1-2)]Gal(β1-3)GlcNAc(β-Sp0
4B
5
6
7

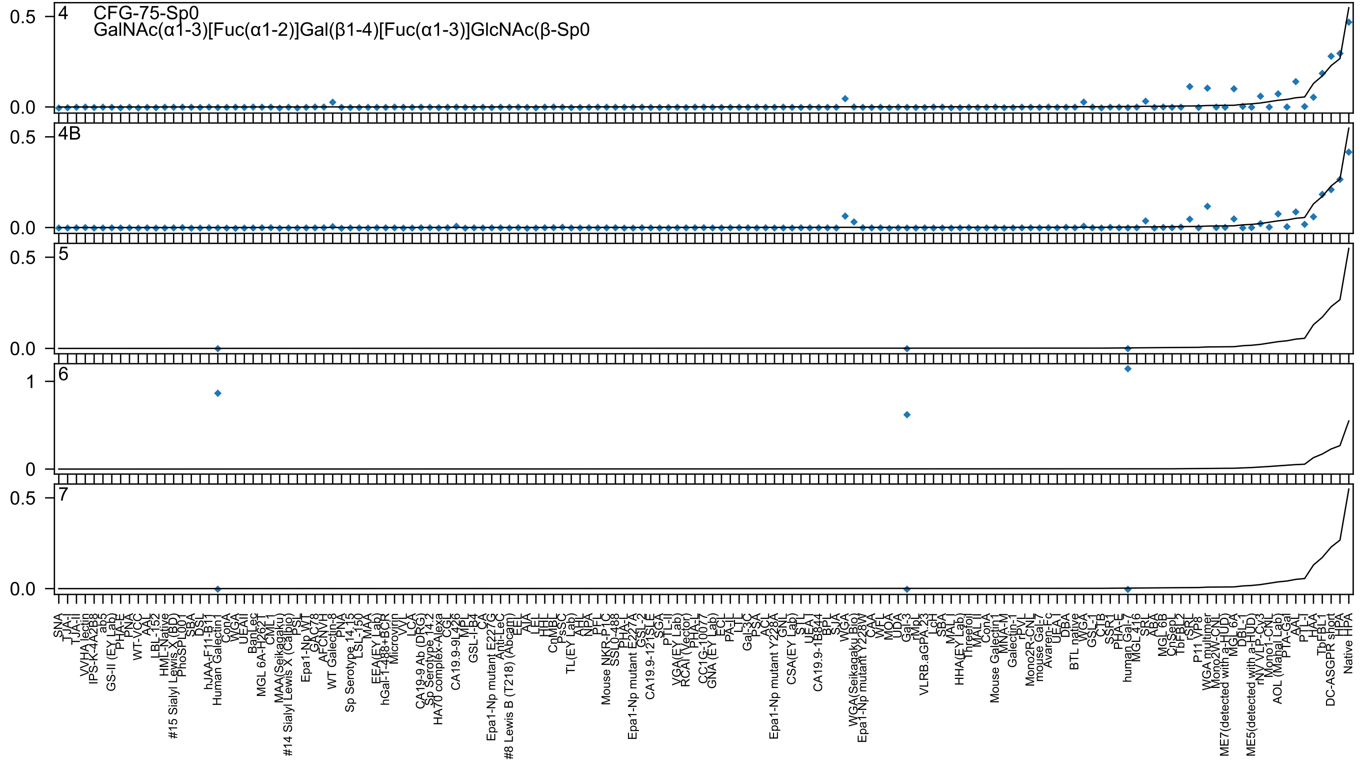
4
CFG-75-Sp0
GalNAc(α1-3)[Fuc(α1-2)]Gal(β1-4)[Fuc(α1-3)]GlcNAc(β-Sp0
4B
5
6
7

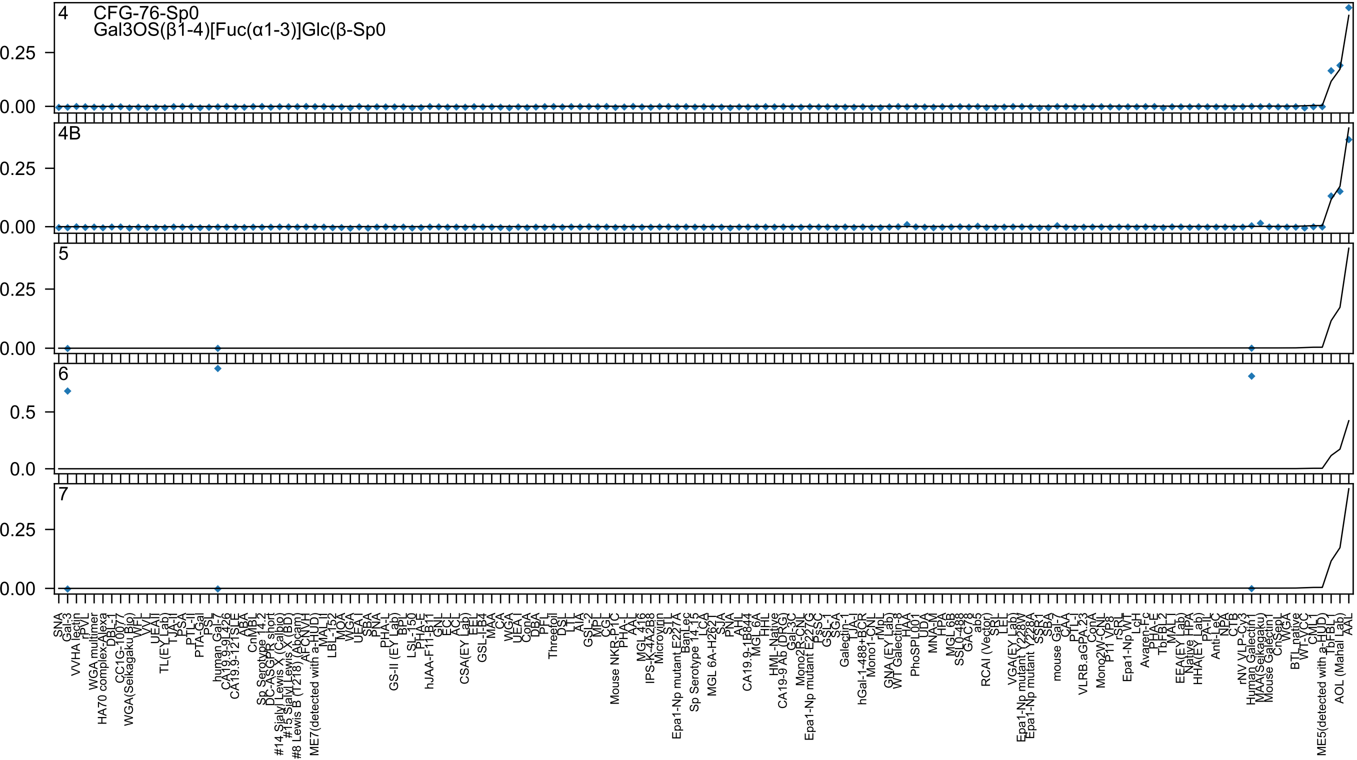

4
CFG-76-Sp0
Gal3OS(β1-4)[Fuc(α1-3)]Glc(β-Sp0
0.25
0.00
4B
0.25
0.00
5
0.25
0.00
6
0.5
0.0
7
0.25
0.00

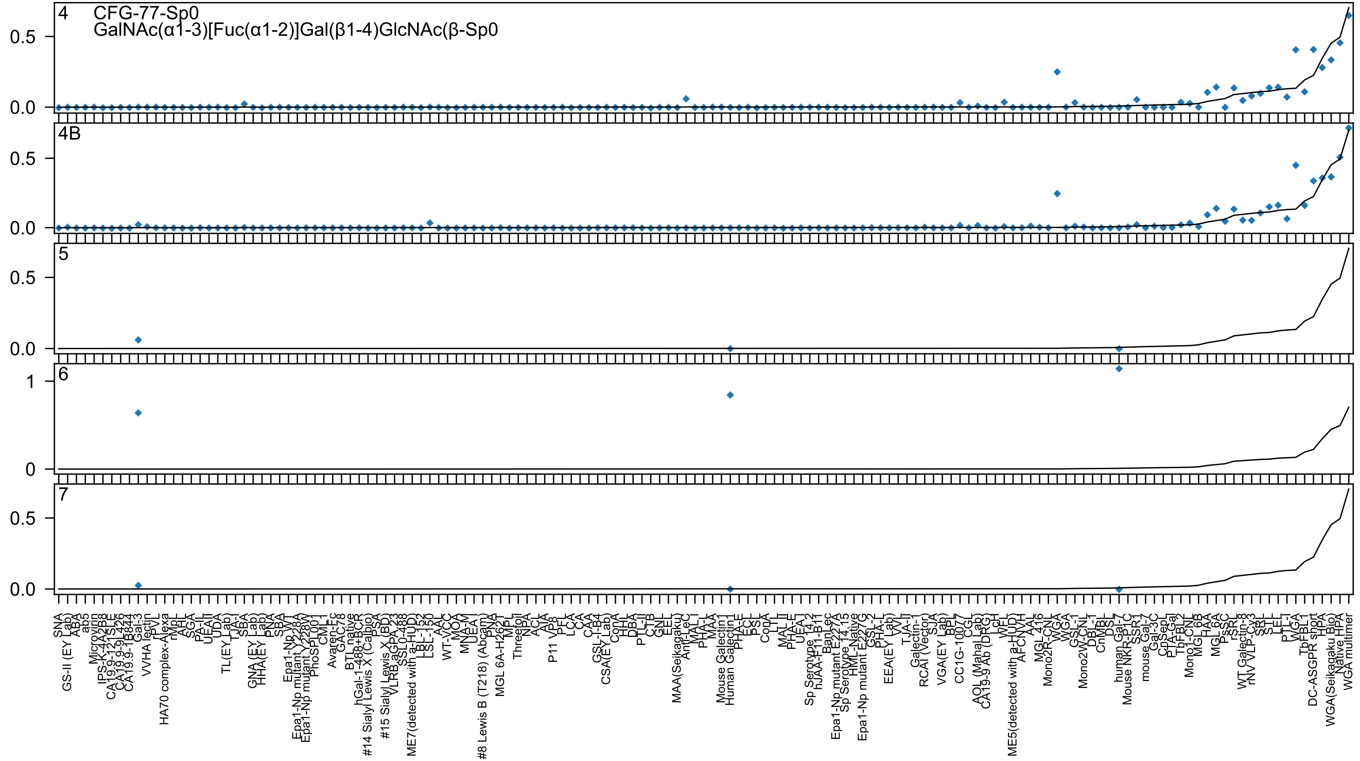
4 CFG-77-Sp0
GalNAc(α1-3)[Fuc(α1-2)]Gal(β1-4)GlcNAc(β-Sp0
4B
5
6
7

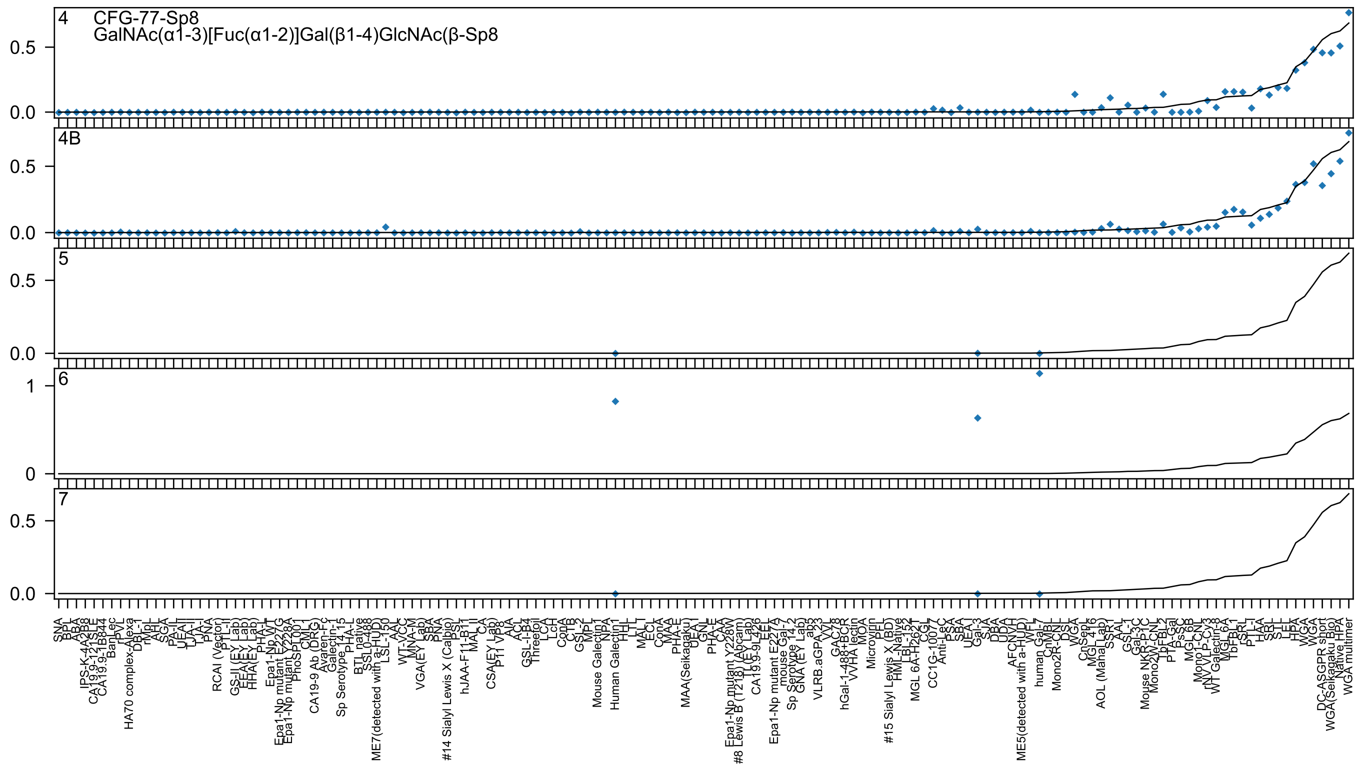
4
CFG-77-Sp8
GalNAc(α1-3)[Fuc(α1-2)]Gal(β1-4)GlcNAc(β-Sp8
4B
5
6
7

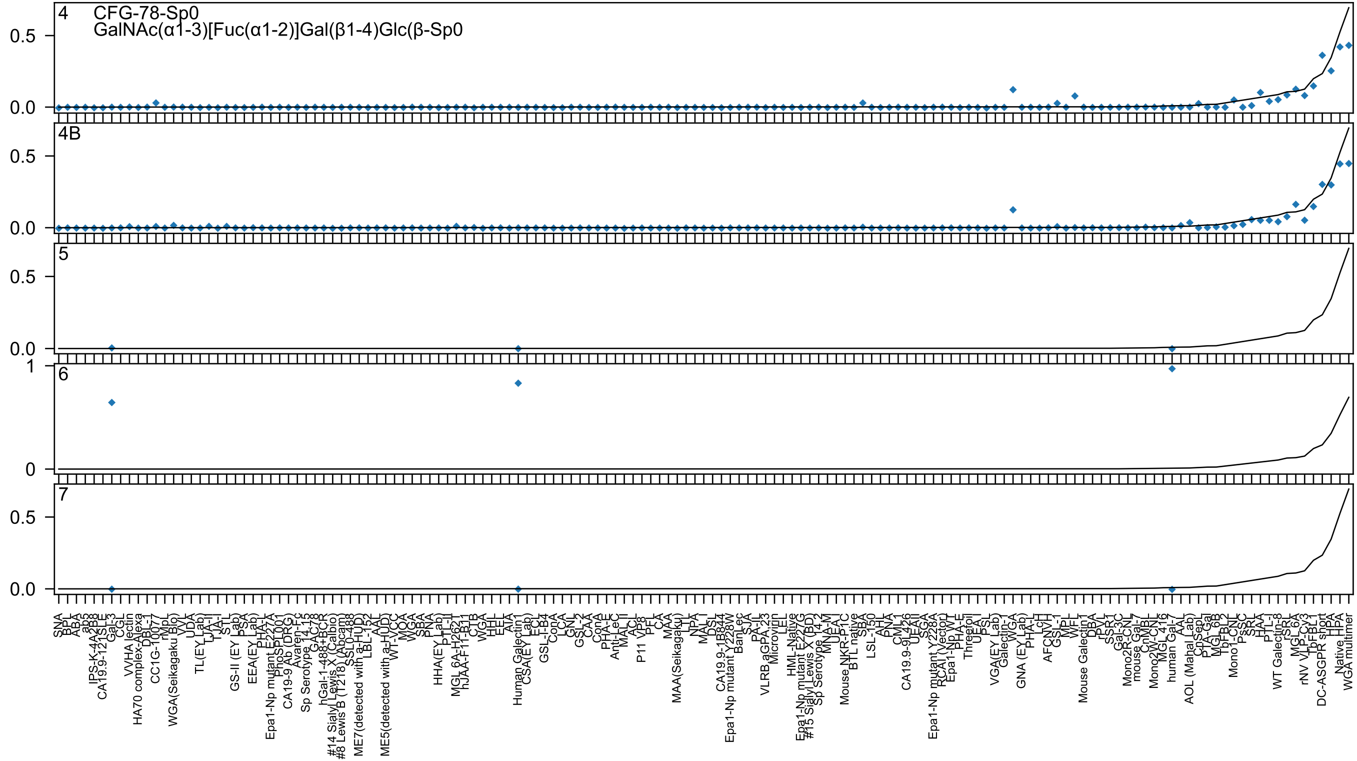
4
CFG-78-Sp0
GalNAc(α1-3)[Fuc(α1-2)]Gal(β1-4)Glc(β-Sp0
4B
5
6
7

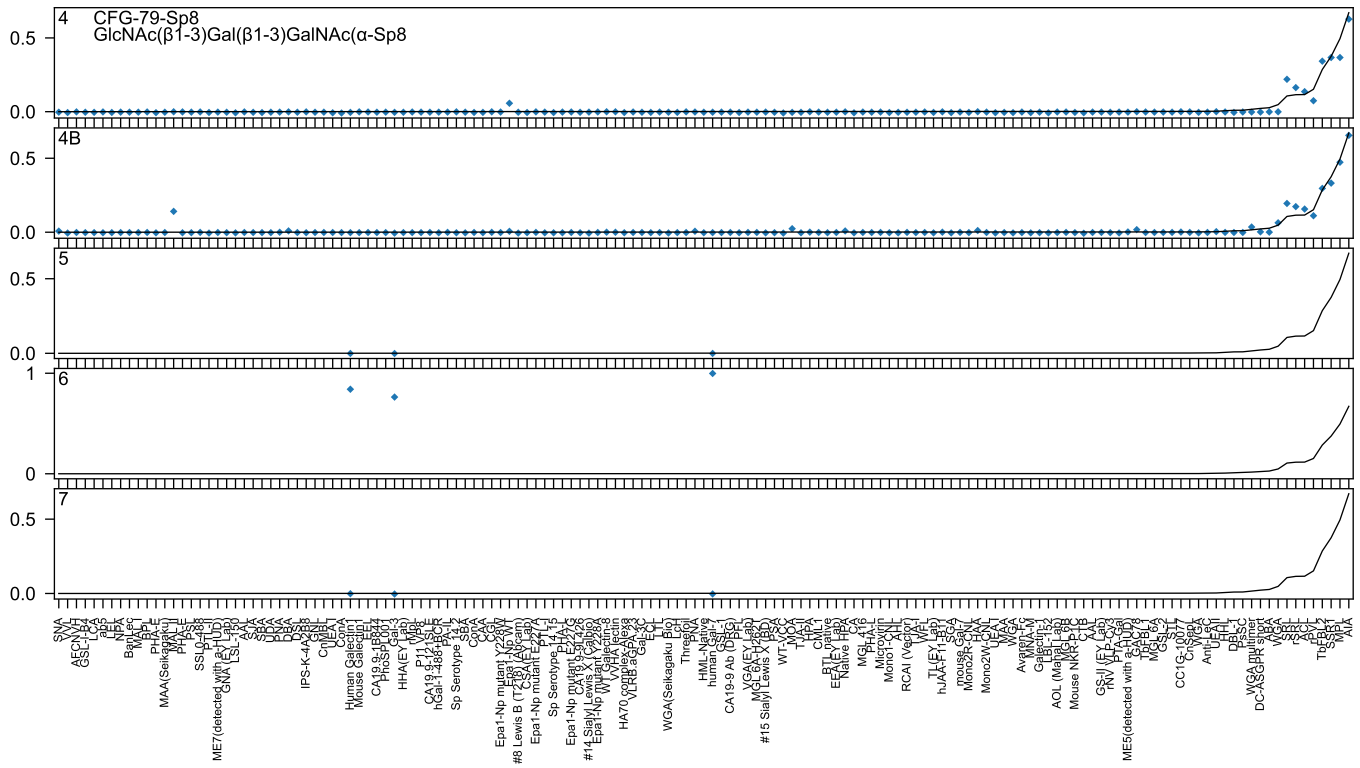
4
CFG-79-Sp8
GlcNAc(β1-3)Gal(β1-3)GalNAc(α-Sp8
4B
5
6
7
0.5
0.0
1
0

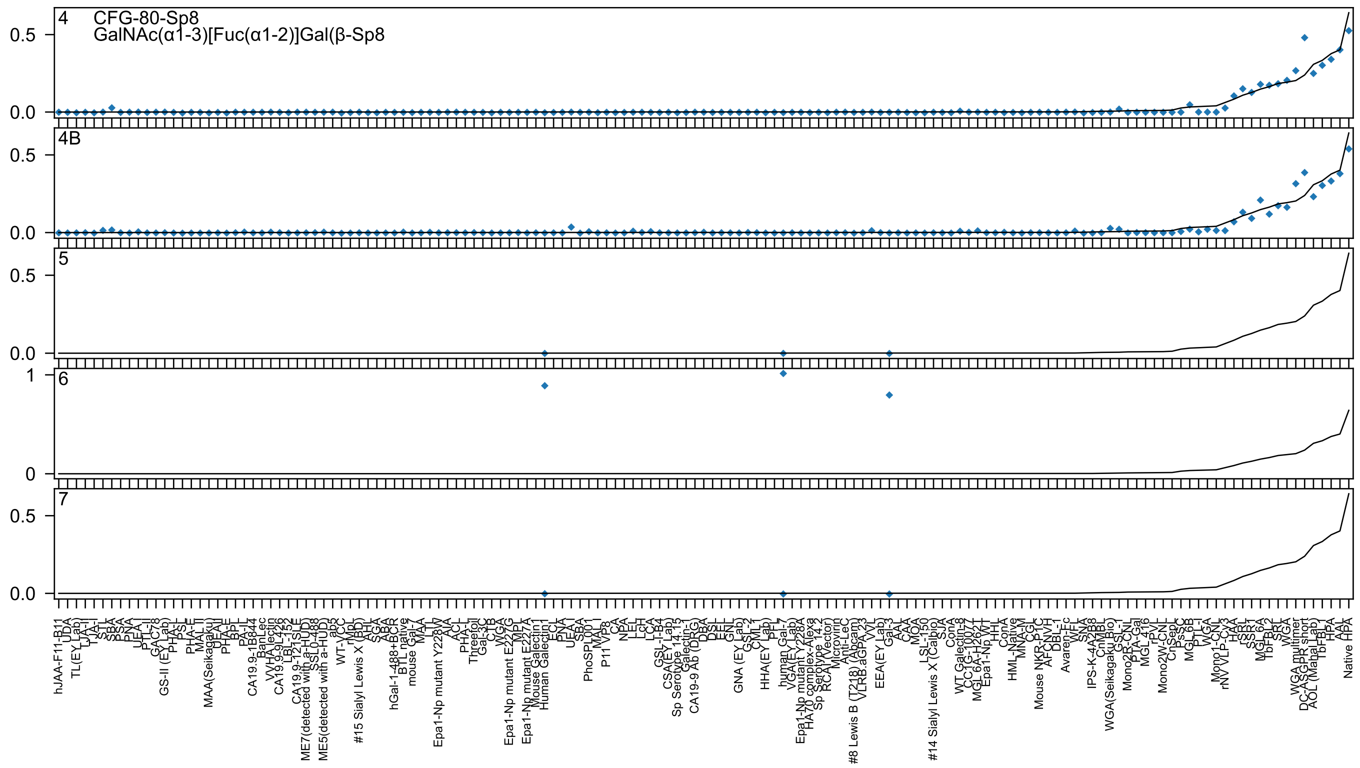
4
CFG-80-Sp8
GalNAc(α1-3)[Fuc(α1-2)]Gal(β-Sp8
4B
5
6
7

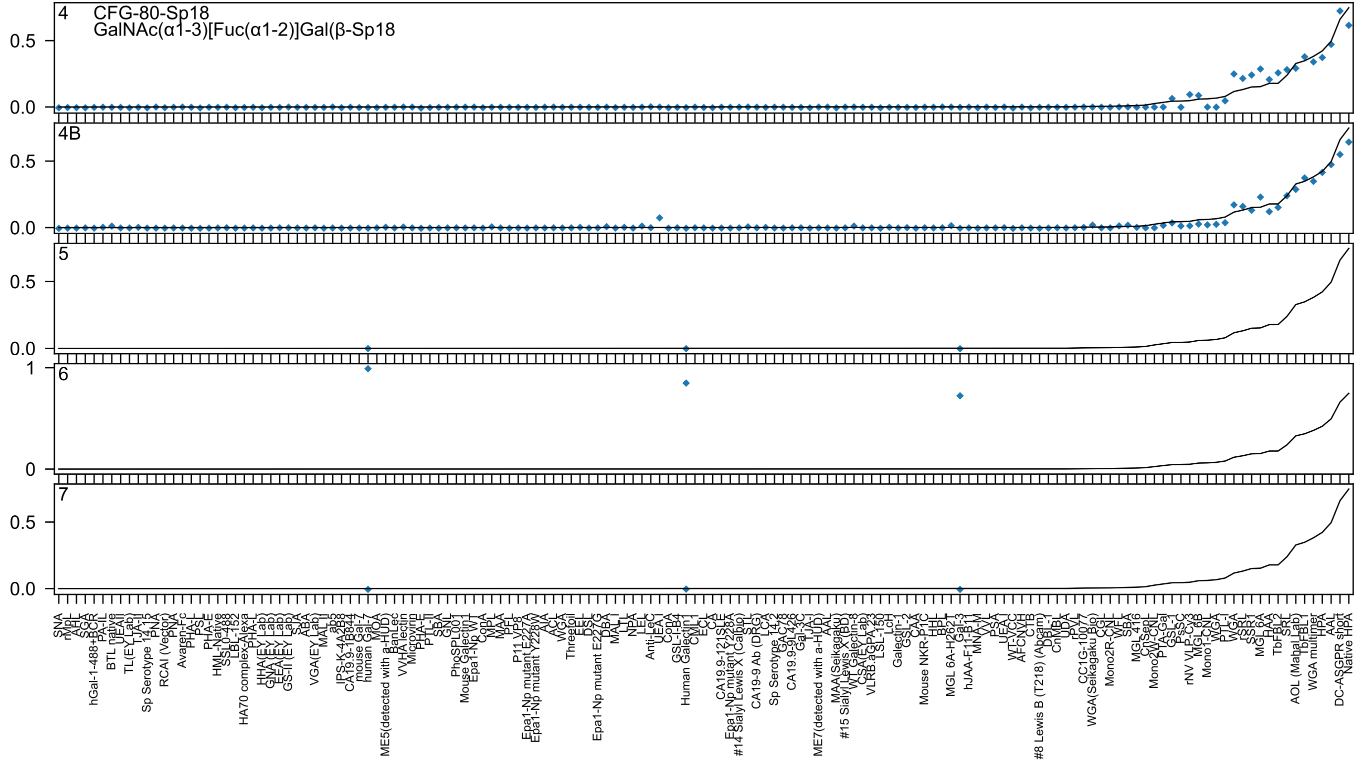
4
CFG-80-Sp18
GalNAc(α1-3)[Fuc(α1-2)]Gal(β-Sp18
4B
5
6
7

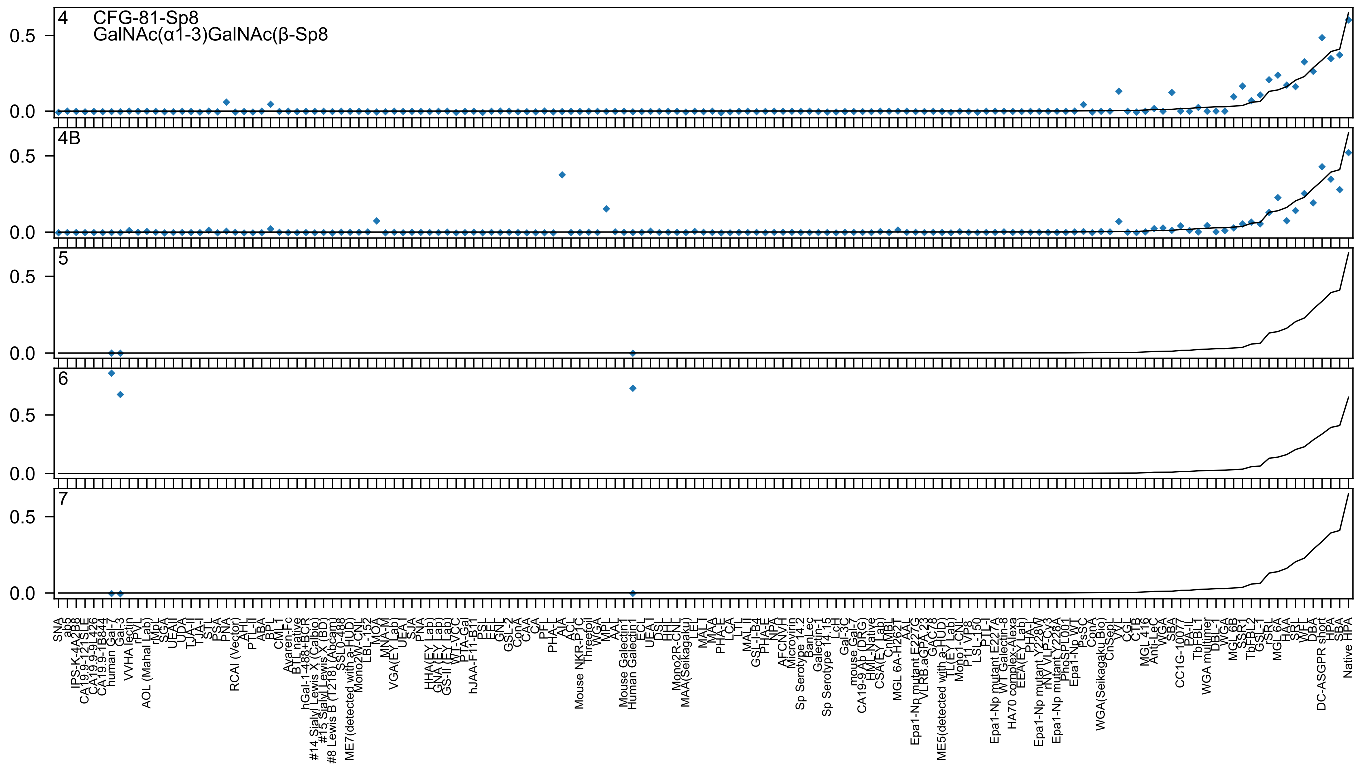

4 CFG-81-Sp8
GalNAc(α1-3)GalNAc(β-Sp8
4B
5
6
7
0.5
0.0

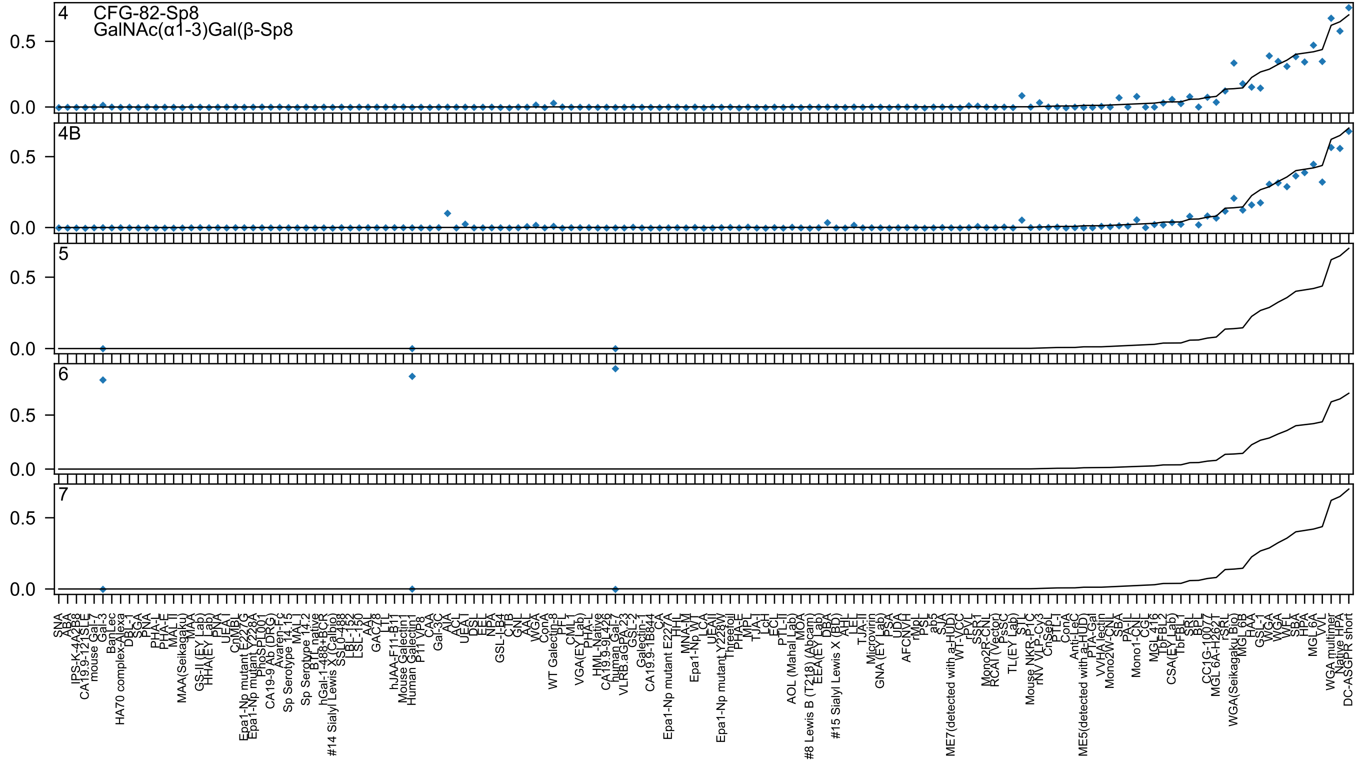

4
CFG-82-Sp8
GalNAc(α1-3)Gal(β-Sp8
4B
5
6
7
0.5
0.0
SNA
ABA
IPS-K-4A2B8
CA19.9-121SLE
mouse Gal-7
Gal-3
BanLec
HA70 complex-Alexa
DBL-1
SGA
PNA
PHA-L
PHA-E
MAL II
MAA(Seikagaku)
MAA
GS-II (EY Lab)
HHA(EY Lab)
PNA
UEA I
CnMBL
Epa1-Np mutant E227G
Epa1-Np mutant Y228A
PhoSL001
CA19-9 Ab (DRG)
Avaren-Fc
Sp Serotype 14.15
MAL I
Sp Serotype 14.2
BTL native
hGal-1-488+BCR
#14 Sialyl Lewis X (Calbio
SSL0-488
LBL-152
LBL-150
LSL-150
AAL
GAC78
LTL
hJAA-F11-B11
Mouse Galectin1
Human Galectin1
P11 VP8
CAA
Gal-3C
AIA
ACL
UEA I
DSL
EEL
NPA
GSL-I-B4
CTB
GNL
AAL
WGA
ConA
WT Galectin-8
PFL
CML1
VGA(EY Lab)
PHA-L
HML-Native
CA19.9-9L426
human Gal-7
VLRB.aGPA.23
GSL-2
Galectin-1
CA19.9-1B844
CA
Epa1-Np mutant E227A
HHL
MNA-M
Epa1-Np WT
LCA
UEA II
Epa1-Np mutant Y228W
Threefoil
PHA-E
MPL
TJA-I
LcH
ECL
PTL-II
AOL (Mahal Lab)
MOA
#8 Lewis B (T218) (Abcam)
EEA(EY Lab)
DBA
#15 Sialyl Lewis X (BD)
AHL
LEL
TJA-II
Microvirin
GNA (EY Lab)
PSA
UDA
AFCNVH
rMpL
PSL
ab5
SJA
ME7(detected with a-HUD)
WT-VCC
rPVL
SSR1
Mono2R-CNL
RCAI (Vector)
PsSC
TL(EY Lab)
STL
Mouse NKR-P1C
rNV VLP-Cy3
CnSepL
PTL-I
ConA
Anti-LeC
ME5(detected with a-HUD)
PTA-Gal
VVHA lectin
Mono2W-CNL
SBA
PA-IL
Mono1-CNL
CGL
MGL 416
TbFBL2
CSA(EY Lab)
TbFBL1
SRL
BPL
CC1G-10077
MGL 6A-H262T
rSRL
WGA(Seikagaku Bio)
MGL 6B
HAA
GSL-1
WGA
WFL
SBA
HPA
MGL 6A
VVL
WGA multimer
Native HPA
DC-ASGPR short

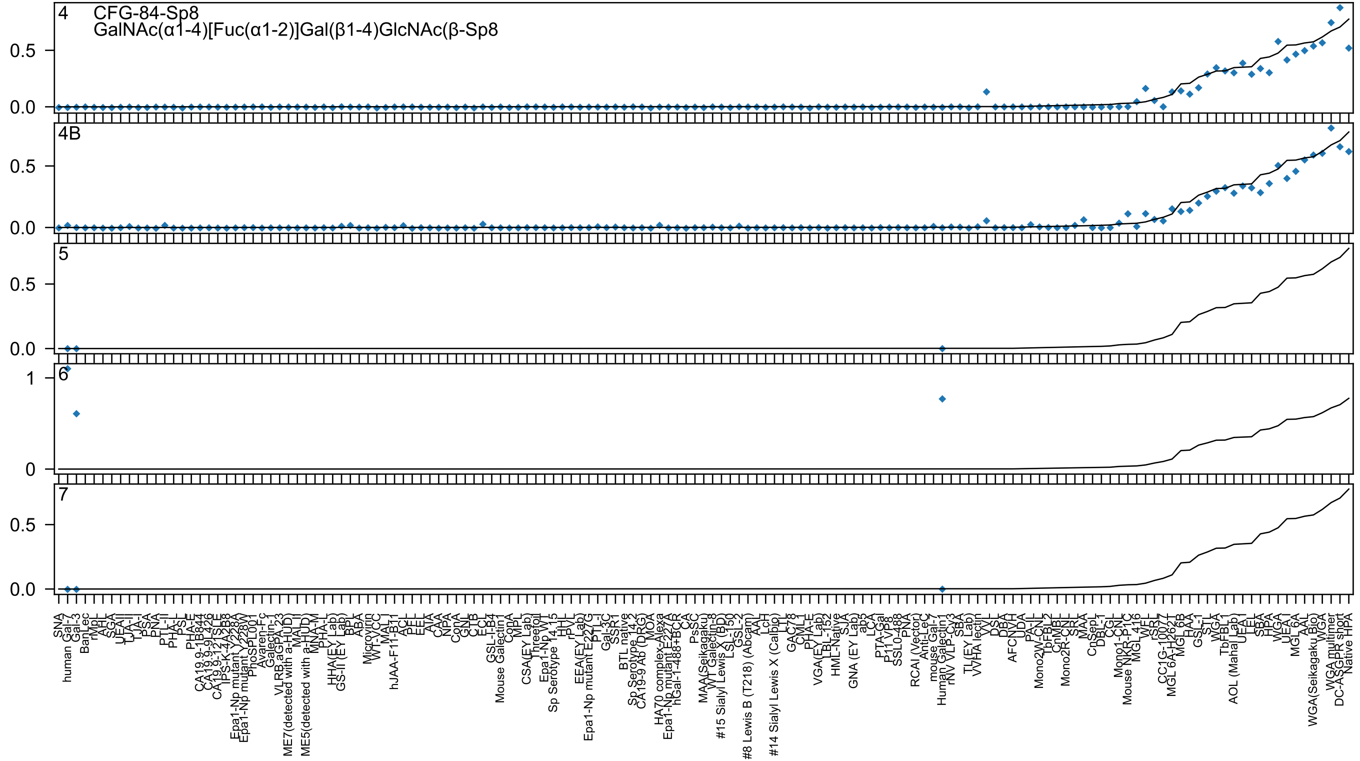

4
CFG-84-Sp8
GalNAc(α1-4)[Fuc(α1-2)]Gal(β1-4)GlcNAc(β-Sp8
4B
5
6
7

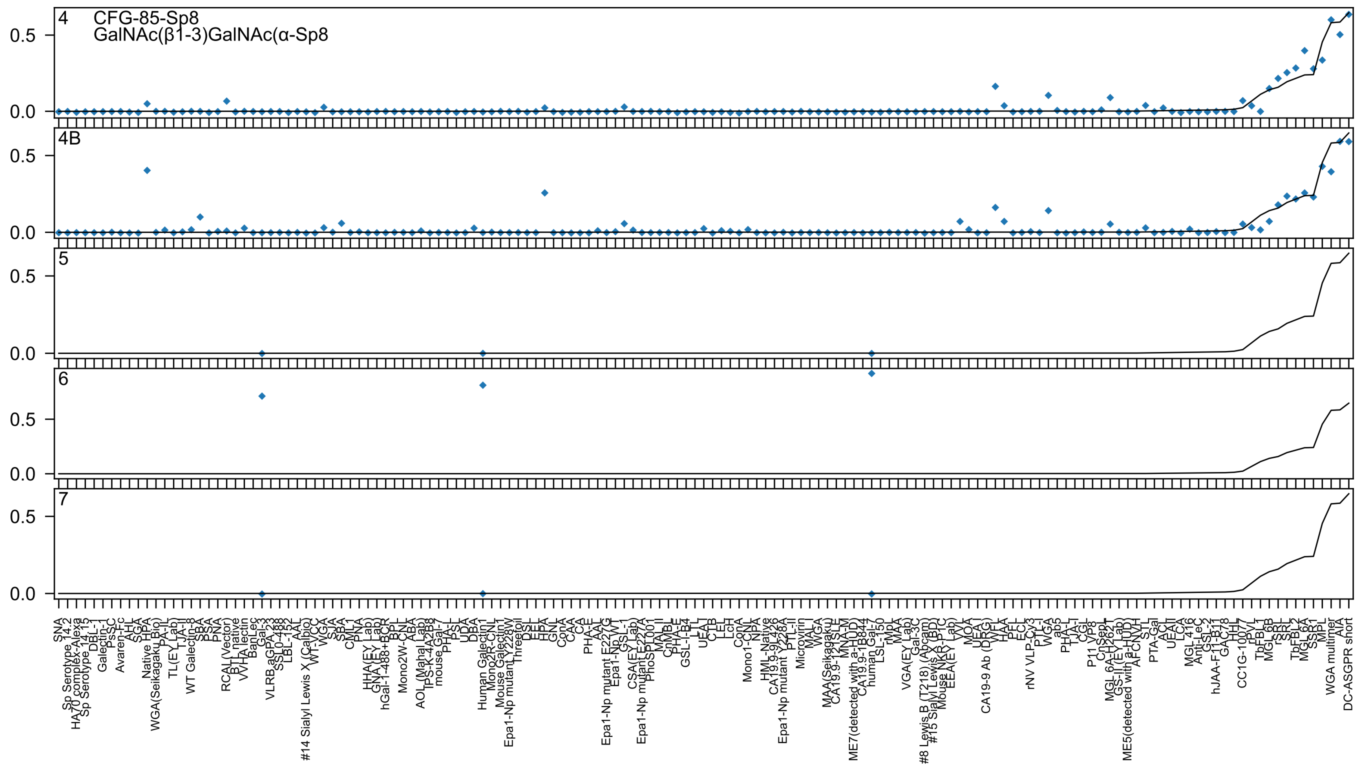
4
CFG-85-Sp8
GalNAc(β1-3)GalNAc(α-Sp8
4B
5
6
7
0.5
0.0

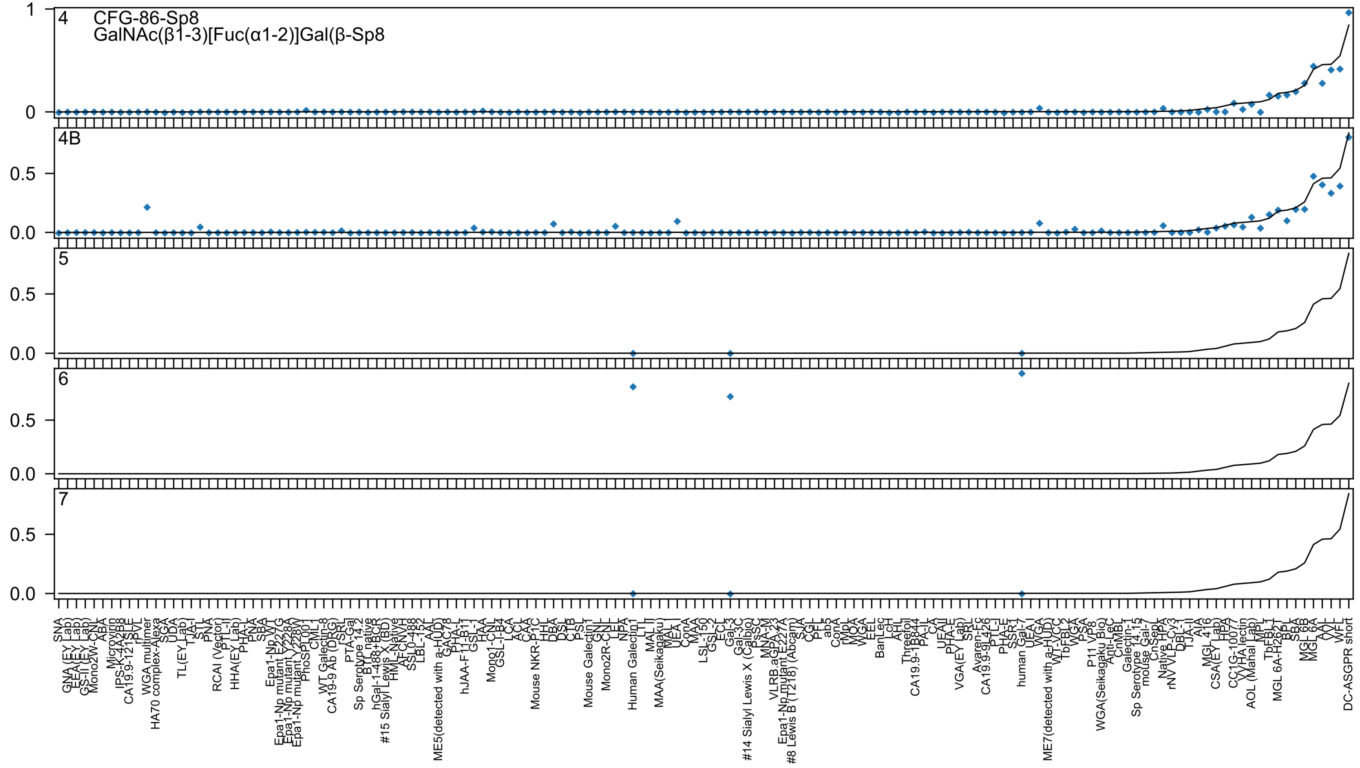
CFG-86-Sp8
GalNAc(β1-3)[Fuc(α1-2)]Gal(β-Sp8
4
4B
5
6
7
1
0
0.5
0.0

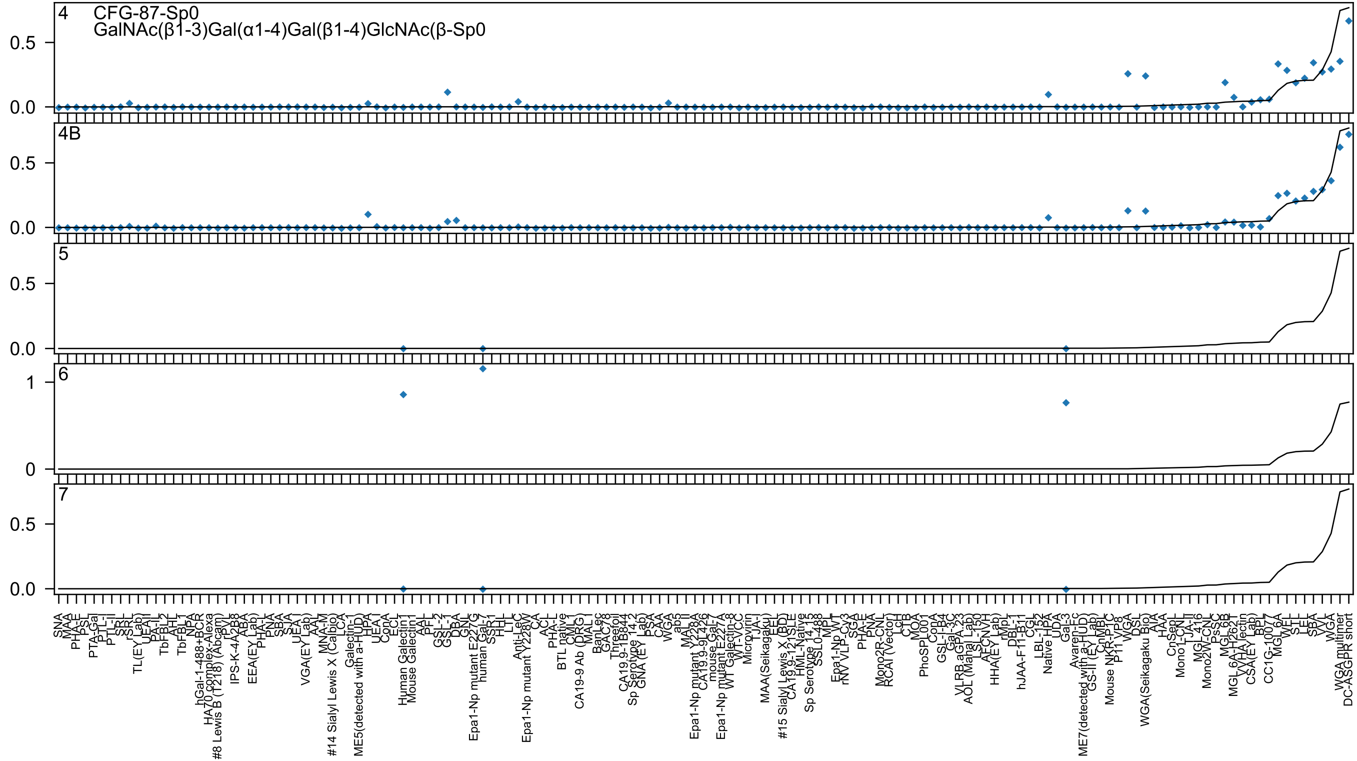

4 CFG-87-Sp0
GalNAc(β1-3)Gal(α1-4)Gal(β1-4)GlcNAc(β-Sp0
4B
5
6
7

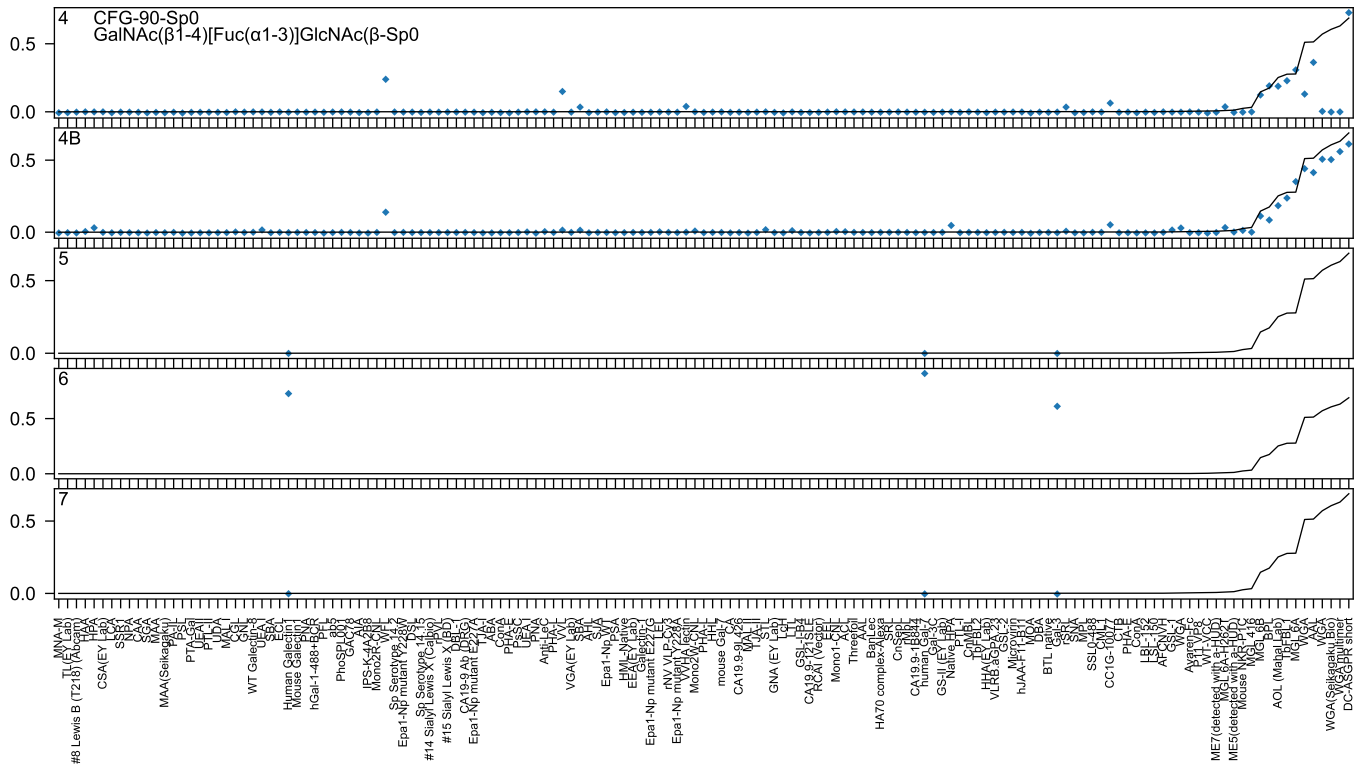
4
CFG-90-Sp0
GalNAc(β1-4)[Fuc(α1-3)]GlcNAc(β-Sp0
4B
5
6
7
0.5
0.0

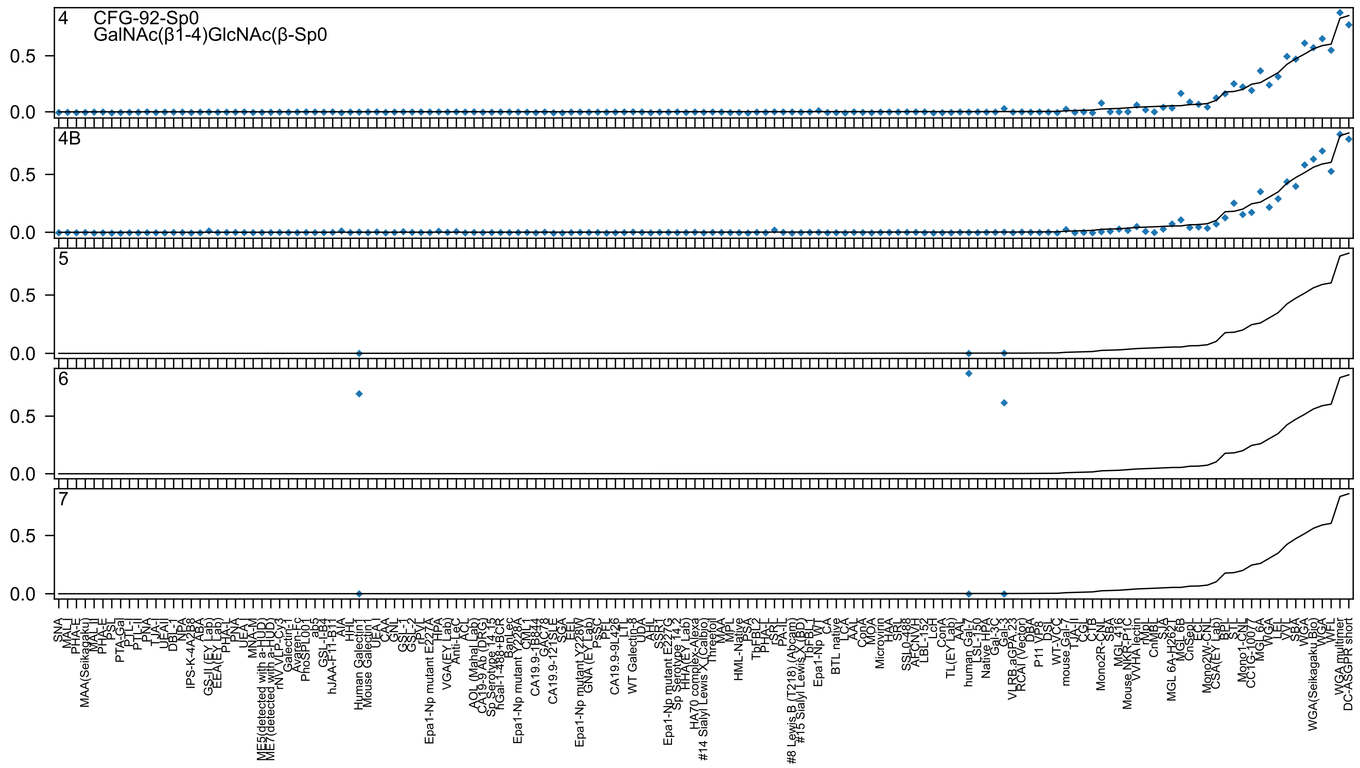
4
CFG-92-Sp0
GalNAc(β1-4)GlcNAc(β-Sp0
4B
5
6
7
0.0
0.5

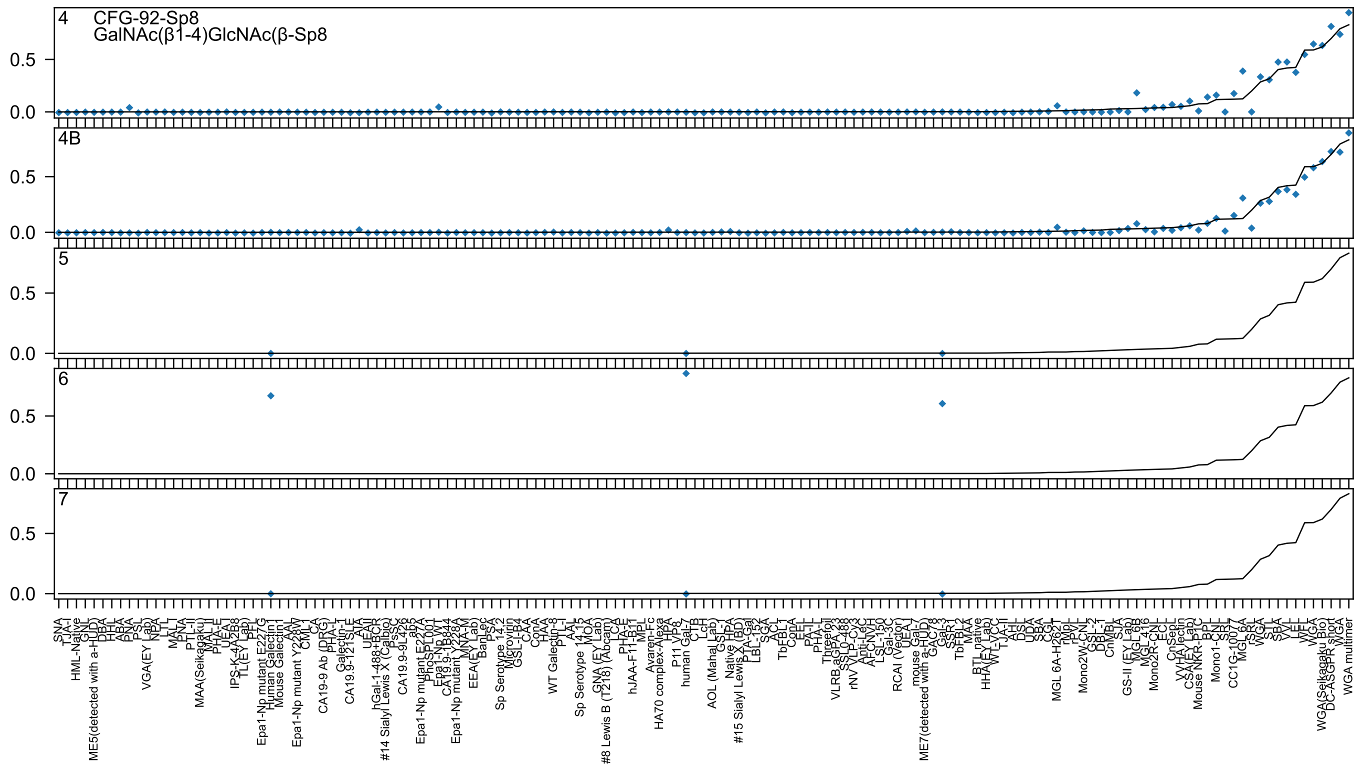
4
CFG-92-Sp8
GalNAc(β1-4)GlcNAc(β-Sp8
4B
5
6
7

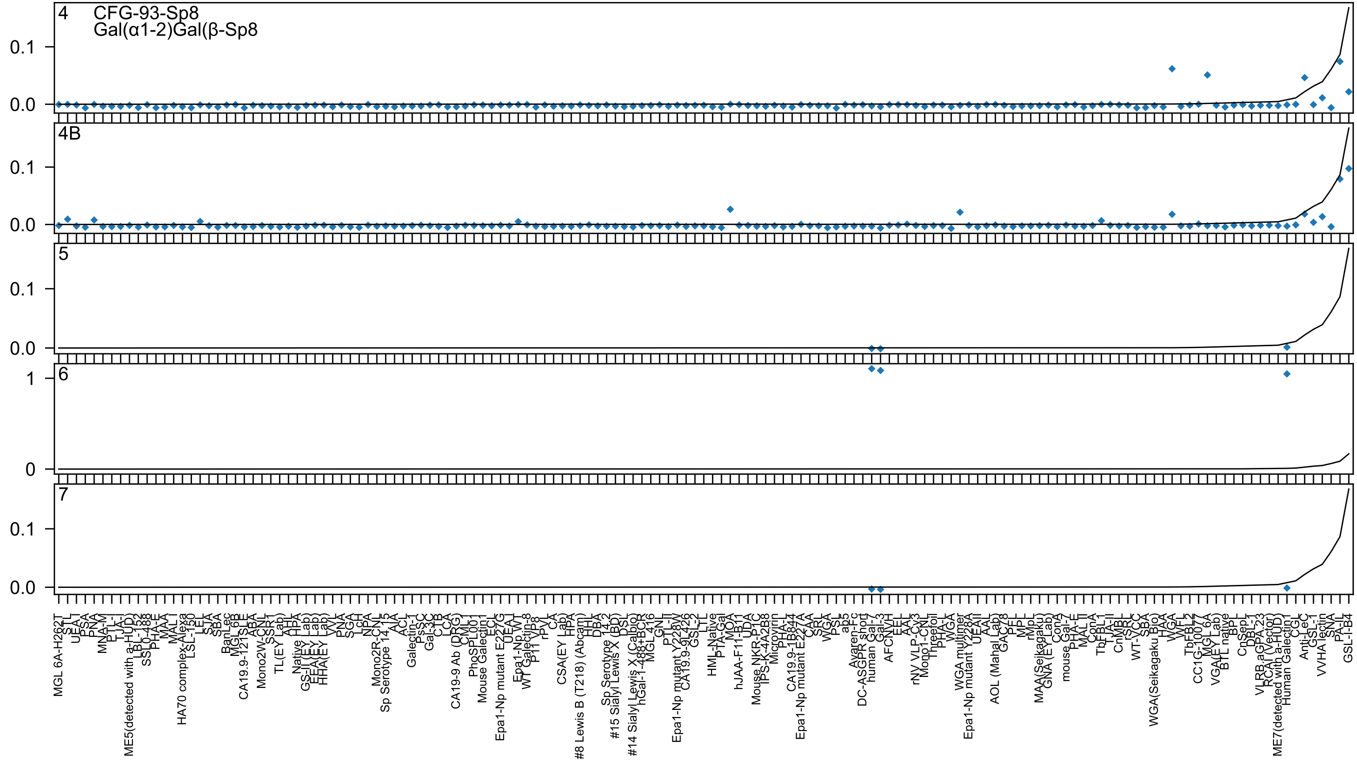

4
CFG-93-Sp8
Gal(α1-2)Gal(β-Sp8
4B
5
6
7
0.1
0.0
1
0
MGL 6A-H262T
STL
UEA-I
PSA
PNA
MNA-M
PTL-I
TJA-I
ME5(detected with a-HUD)
LBL-152
SSL0-488
PHA-E
MAA
MAL I
HA70 complex-Alexa
LSL-150
LEL
SJA
SBA
BanLec
MGL 6B
CA19.9-121SLE
ABA
Mono2W-CNL
SSR1
TL(EY Lab)
AHL
Native HPA
GS-II (EY Lab)
EEA(EY Lab)
HHA(EY Lab)
VVL
PNA
SGA
LcH
NPA
Mono2R-CNL
Sp Serotype 14_15
AIA
ACL
Galectin-1
PsSC
Gal-3C
CTB
LCA
CA19-9 Ab (DRG)
CML
PhoSL
PL001
Mouse Galectin1
ECL
Epa1-Np mutant E227G
UEA-I
Epa1-Np WT
WT Galectin-8
P11 VP8
rPVL
CA
CSA(EY Lab)
HPA
#8 Lewis B (T218) (Abcam)
HHL
DBA
Sp Serotype 14_2
#15 Sialyl Lewis X (BD)
DSL
#14 Sialyl Lewis X (Calbio)
hGal-1-488+BCR
MGL 416
GNL
PTL-II
Epa1-Np mutant Y228W
CA19.9-9L426
GSL-2
LTL
HML-Native
PTA-Gal
MOA
hJAA-F11-B11
UDA
Mouse NKR-P1C
IPS-K-4A2B8
Microvirin
PHA-L
CA19.9-1B844
Epa1-Np mutant E227A
CAA
SRL
WGA
PSL
ab5
Avaren-Fc
DC-ASGPR short
human Gal-7
Gal-3
AFCNVH
EEL
AAL
rNV VLP-Cy3
Mono1-CNL
Threefoil
PHA-L
WGA
WGA multimer
Epa1-Np mutant Y228A
UEAII
AAL
AOL (Mahal Lab)
GAC78
PFL
MPL
rMpL
MAA(Seikagaku)
GNA (EY Lab)
ConA
mouse Gal-7
PHA-E
MAL II
ConA
TbFBL1
TJA-II
CnMBL
rSRL
WT-VCC
SBA
WGA(Seikagaku Bio)
HAA
WGA
WFL
TbFBL2
CC1G-10077
MGL 6A
VGA(EY Lab)
BTL native
BPL
CnSep1
DBL-1
VLRB.aGPA.23
RCAI (Vector)
ME7(detected with a-HUD)
Human Galectin1
CGL
Anti-LeC
GSL-1
VVHA lectin
SNA
PA-IL
GSL-I-B4

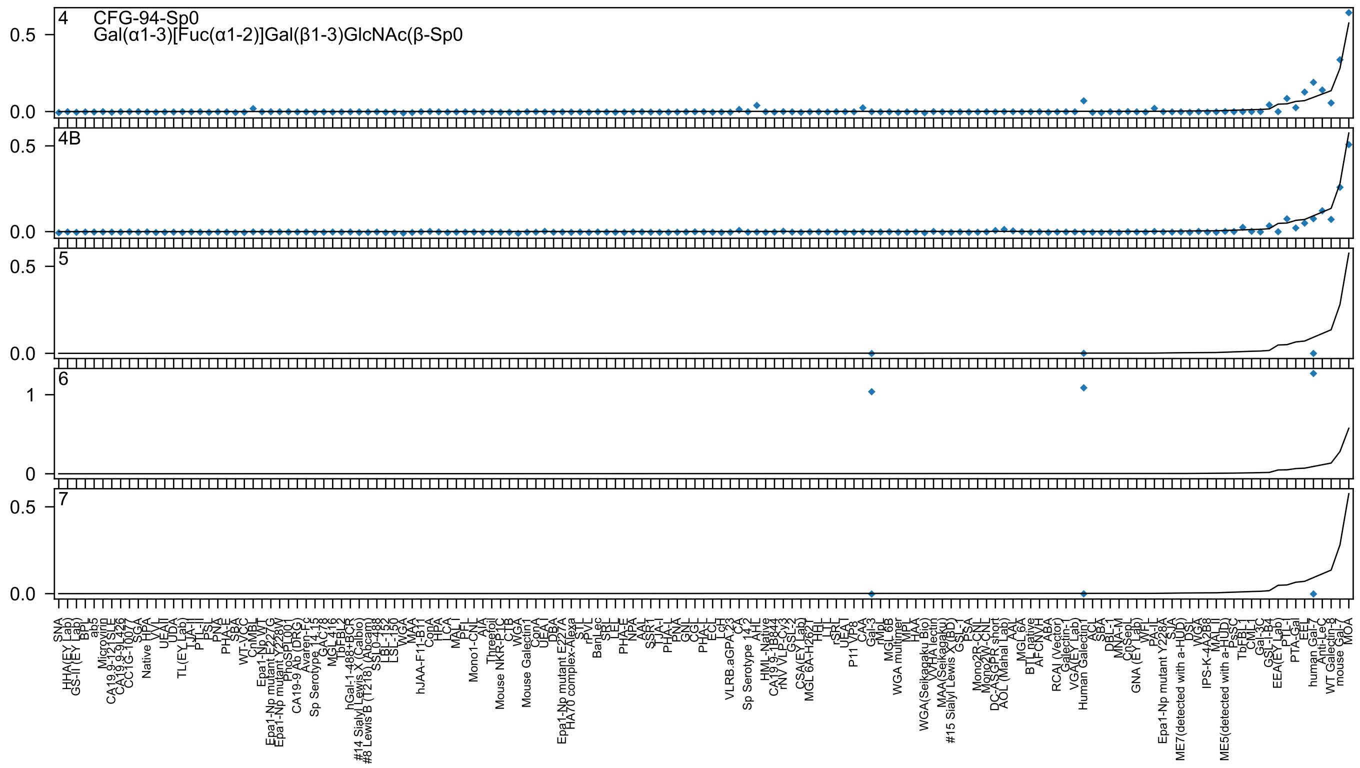
4 CFG-94-Sp0
Gal(α1-3)[Fuc(α1-2)]Gal(β1-3)GlcNAc(β-Sp0
4B
5
6
7

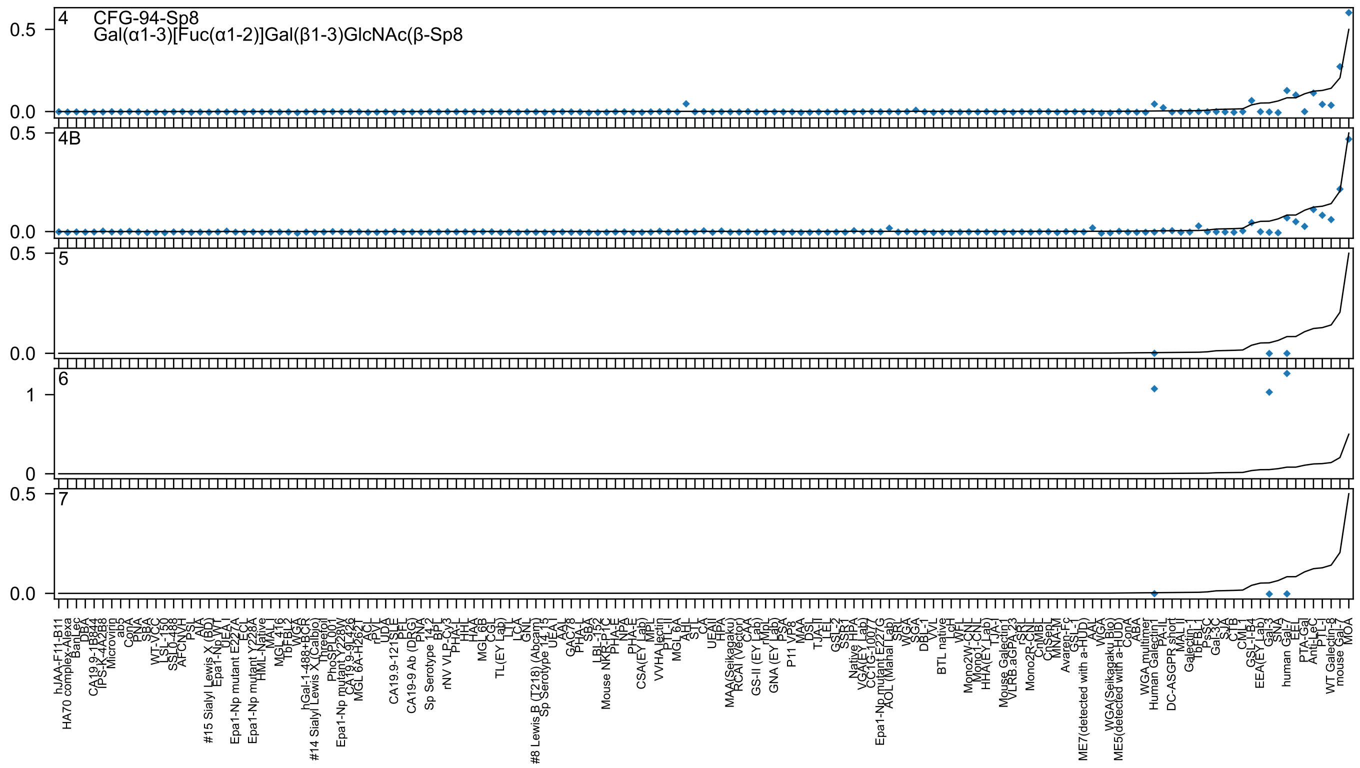

4 CFG-94-Sp8
Gal(α1-3)[Fuc(α1-2)]Gal(β1-3)GlcNAc(β-Sp8
4B
5
6
7

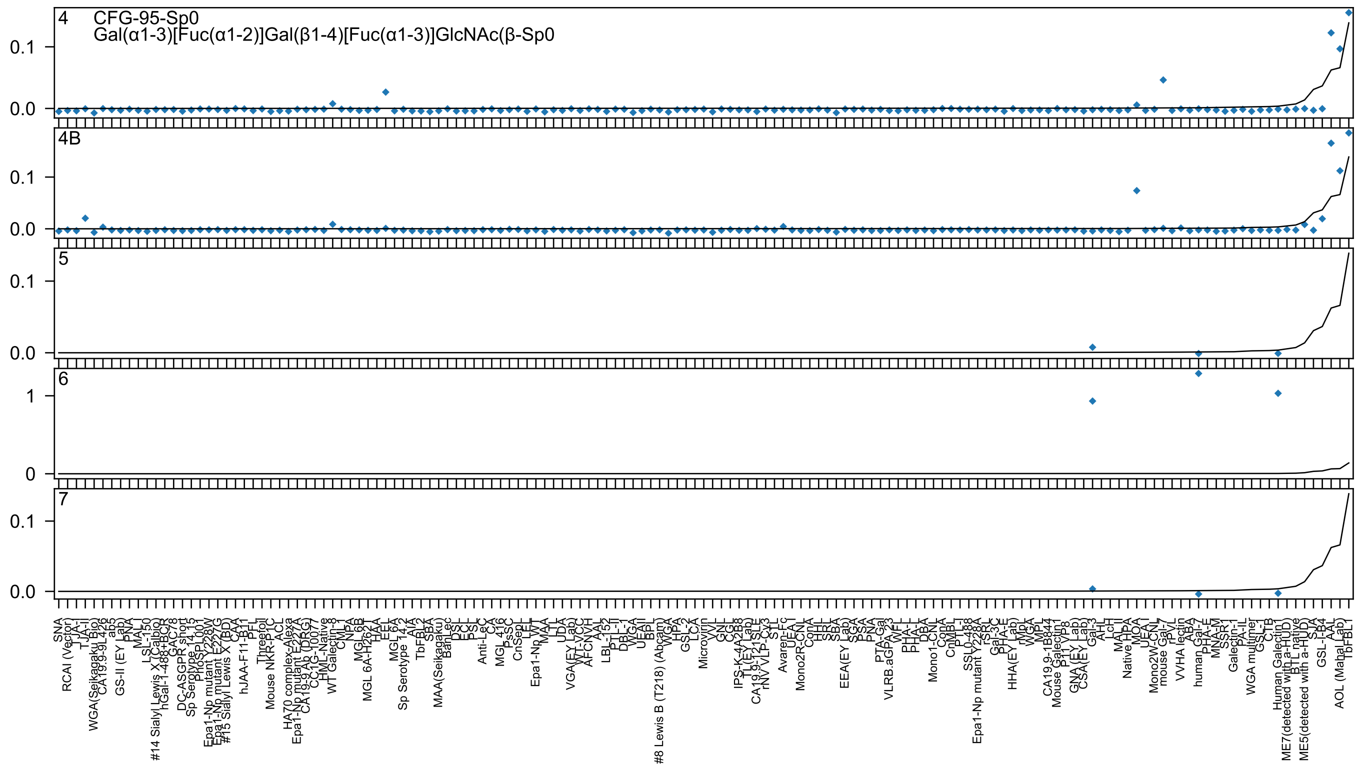

4
CFG-95-Sp0
Gal(α1-3)[Fuc(α1-2)]Gal(β1-4)[Fuc(α1-3)]GlcNAc(β-Sp0
0.1
0.0
4B
0.1
0.0
5
0.1
0.0
6
1
0
7
0.1
0.0
SNA
RCAI (Vector)
TJA-I
TJA-II
WGA(Seikagaku Bio)
CA19.9-9L426
ab5
GS-II (EY Lab)
PNA
MAL I
LSL-150
#14 Sialyl Lewis X (Calbio)
hGal-1-488+BCR
GAC78
DC-ASGPR short
Sp Serotype 14.15
PhoSL007
Epa1-Np mutant Y228W
Epa1-Np mutant E227G
#15 Sialyl Lewis X (BD)
CAA
hJAA-F11-B11
PFL
Threefoil
Mouse NKR-P1C
ACL
HA70 complex-Alexa
Epa1-Np mutant E227A
CA19-9 Ab (DRG)
CC1G-10077
HML-Native
WT Galectin-8
CML-1
NPA
MGL 6B
MGL 6A-H262T
HAA
EEL
MGL 6A
Sp Serotype 14.2
AIA
TbFBL2
SBA
MAA(Seikagaku)
BanLec
DSL
ECL
PSL
Anti-LeC
CA
MGL 416
PsSC
CnSepL
LEL
Epa1-Np WT
MAA
LTL
UDA
VGA(EY Lab)
WT-VCC
AFCNVH
AAL
LBL-152
PTL-II
DBL-1
WGA
UEAII
BPL
#8 Lewis B (T218) (Abcam)
WGA
HPA
GSL-2
LCA
Microvirin
VVL
GNL
CGL
IPS-K-4A2B8
TL(EY Lab)
CA19.9-121SLE
rNV VLP-Cy3
STL
Avaren-Fc
UEA I
Mono2R-CNL
ConA
HHL
SRL
SBA
EEA(EY Lab)
SGA
PSA
PNA
PTA-Gal
VLRB.aGPA.23
WFL
PHA-L
PHA-L
DBA
Mono1-CNL
ConA
CnMBL
PTL-I
SSL0-488
Epa1-Np mutant Y228A
rSRL
Gal-3C
PHA-E
HHA(EY Lab)
rMpL
WGA
MPL
CA19.9-1B844
Mouse Galectin1
P11 VP8
GNA (EY Lab)
CSA(EY Lab)
Gal-3
AH-I
LcH
MAL II
Native HPA
MOA
UEA-I
Mono2W-CNL
mouse Gal-7
rPVL
VVHA lectin
ABA
human Gal-7
PHA-E
MNA-M
SSR1
Galectin-1
PA-IL
WGA multimer
GSL-I
CTB
Human Galectin1
ME7(detected with a-HUD)
BTL native
ME5(detected with a-HUD)
SJA
GSL-I-B4
AAL
AOL (Mahal_ab)
TbFBL1

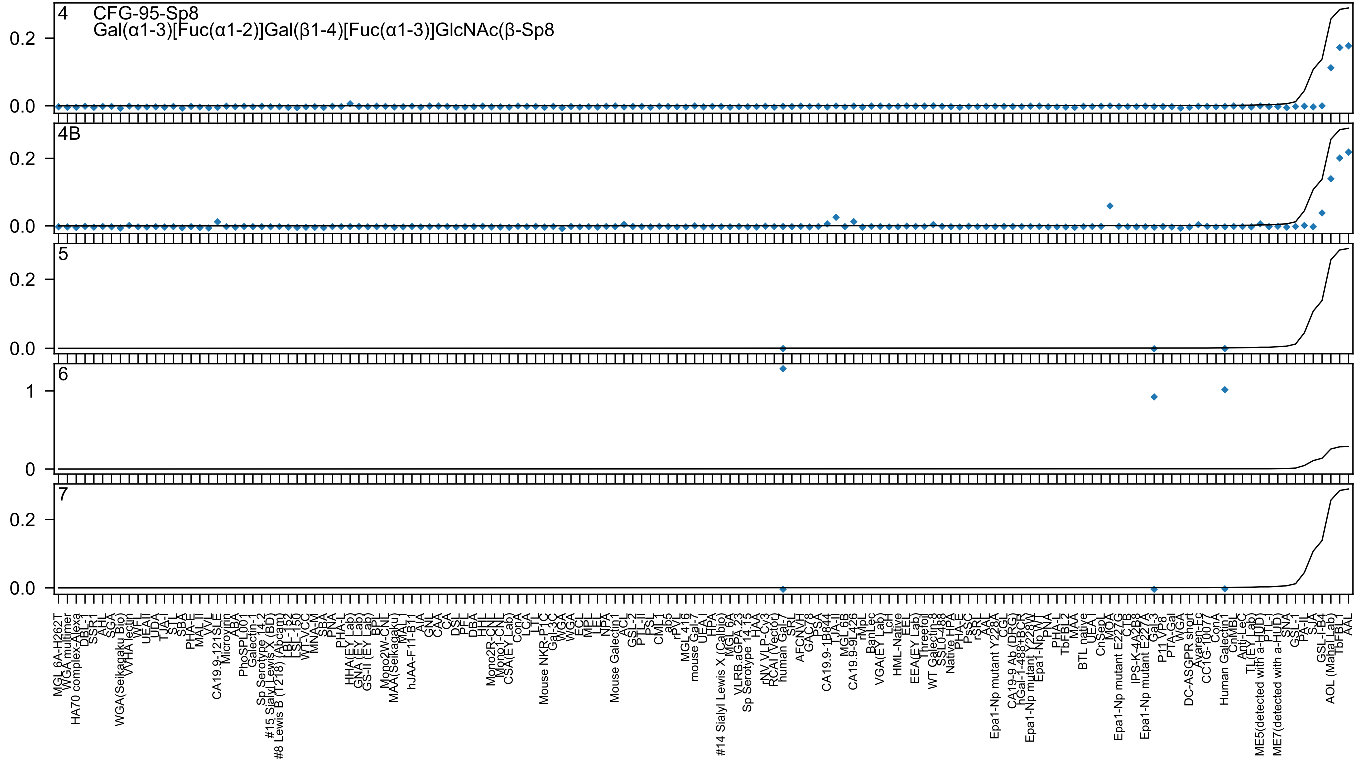

4 CFG-95-Sp8
Gal(α1-3)[Fuc(α1-2)]Gal(β1-4)[Fuc(α1-3)]GlcNAc(β-Sp8
4B
5
6
7

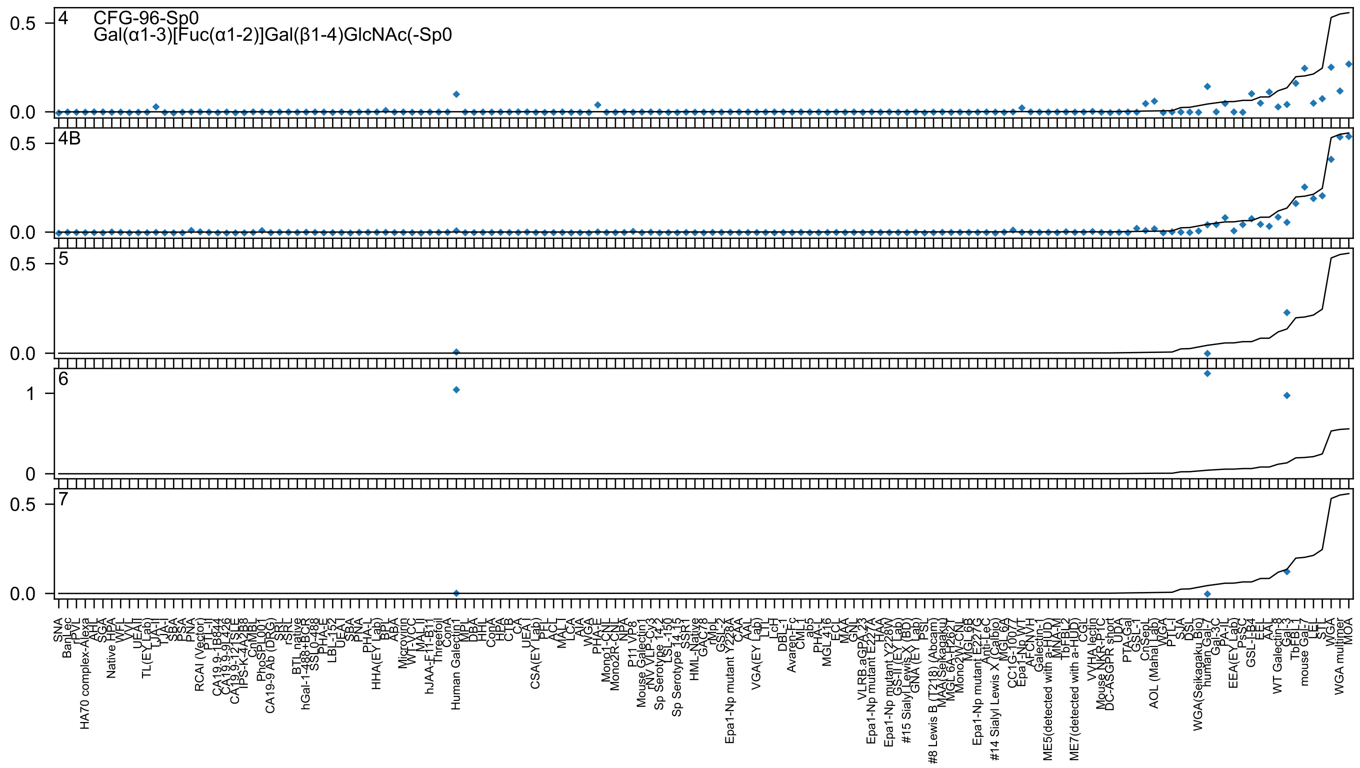

4
CFG-96-Sp0
Gal(α1-3)[Fuc(α1-2)]Gal(β1-4)GlcNAc(-Sp0
4B
5
6
7

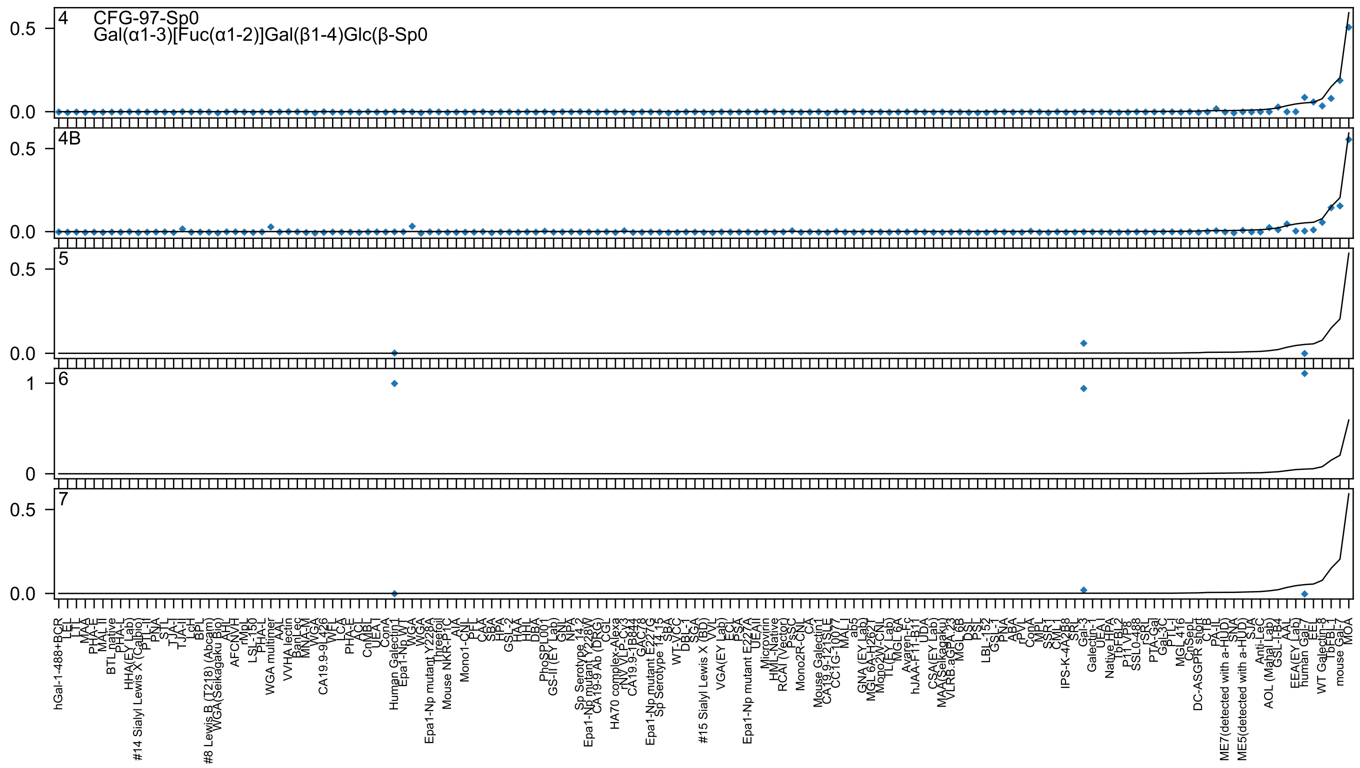

4 CFG-97-Sp0
Gal(α1-3)[Fuc(α1-2)]Gal(β1-4)Glc(β-Sp0
4B
5
6
7

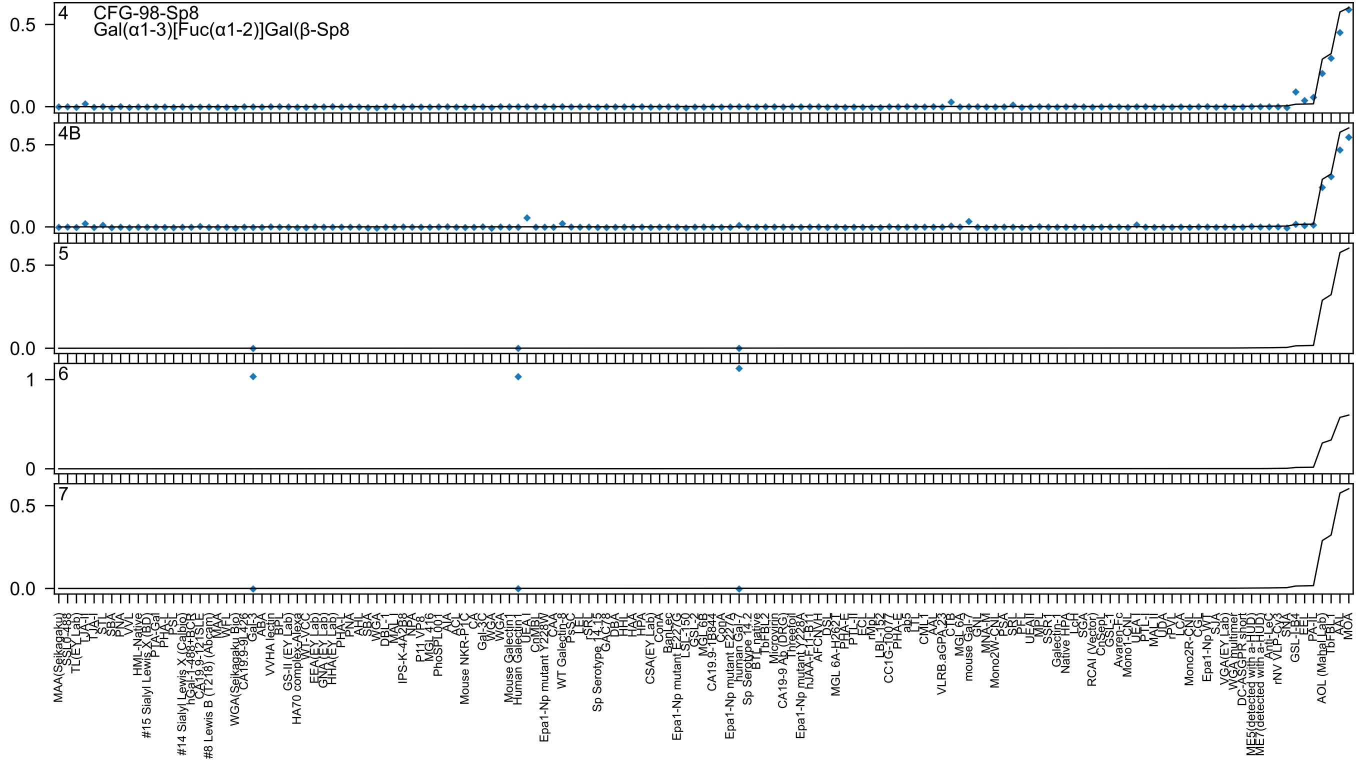
4
CFG-98-Sp8
Gal(α1-3)[Fuc(α1-2)]Gal(β-Sp8
4B
5
6
7

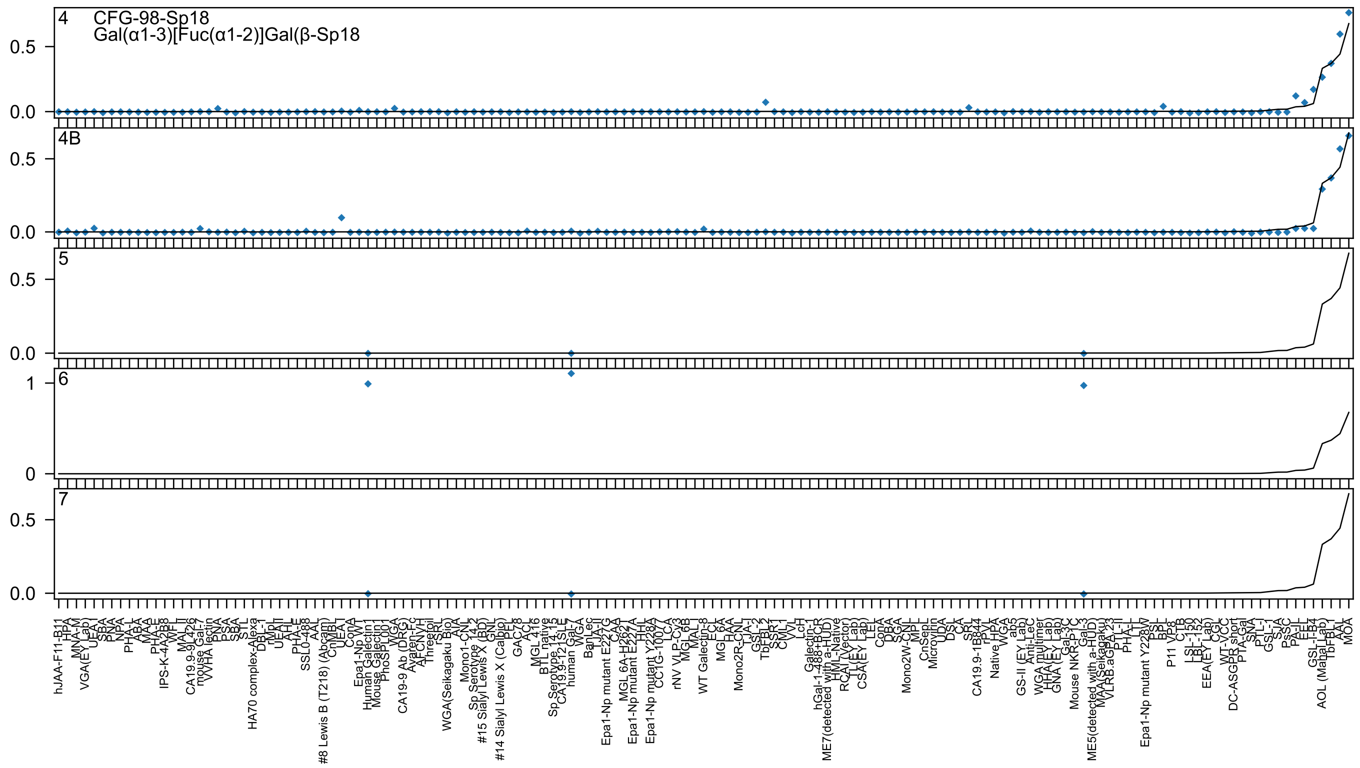

4
CFG-98-Sp18
Gal(α1-3)[Fuc(α1-2)]Gal(β-Sp18
4B
5
6
7
0.5
0.0
1
0

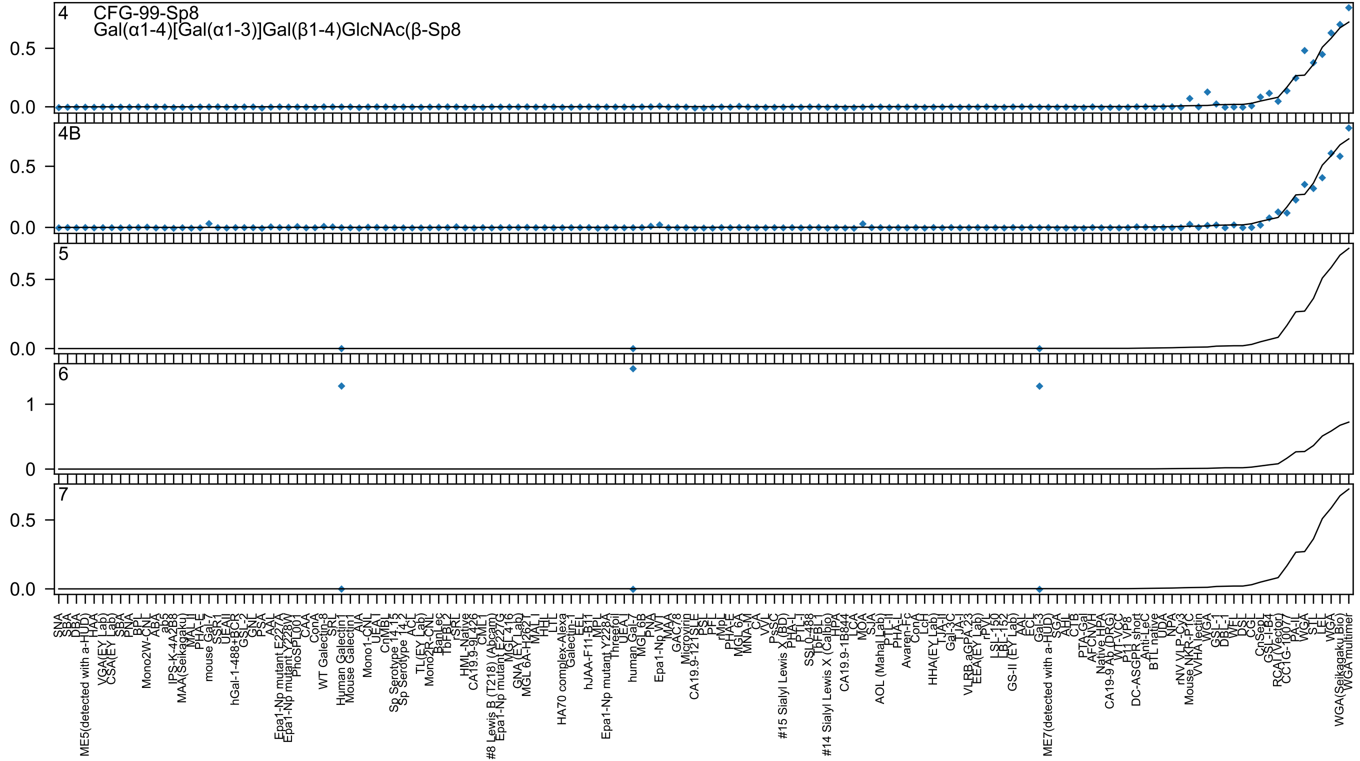
4
CFG-99-Sp8
Gal(α1-4)[Gal(α1-3)]Gal(β1-4)GlcNAc(β-Sp8
4B
5
6
7
0.0
0.5
1
0

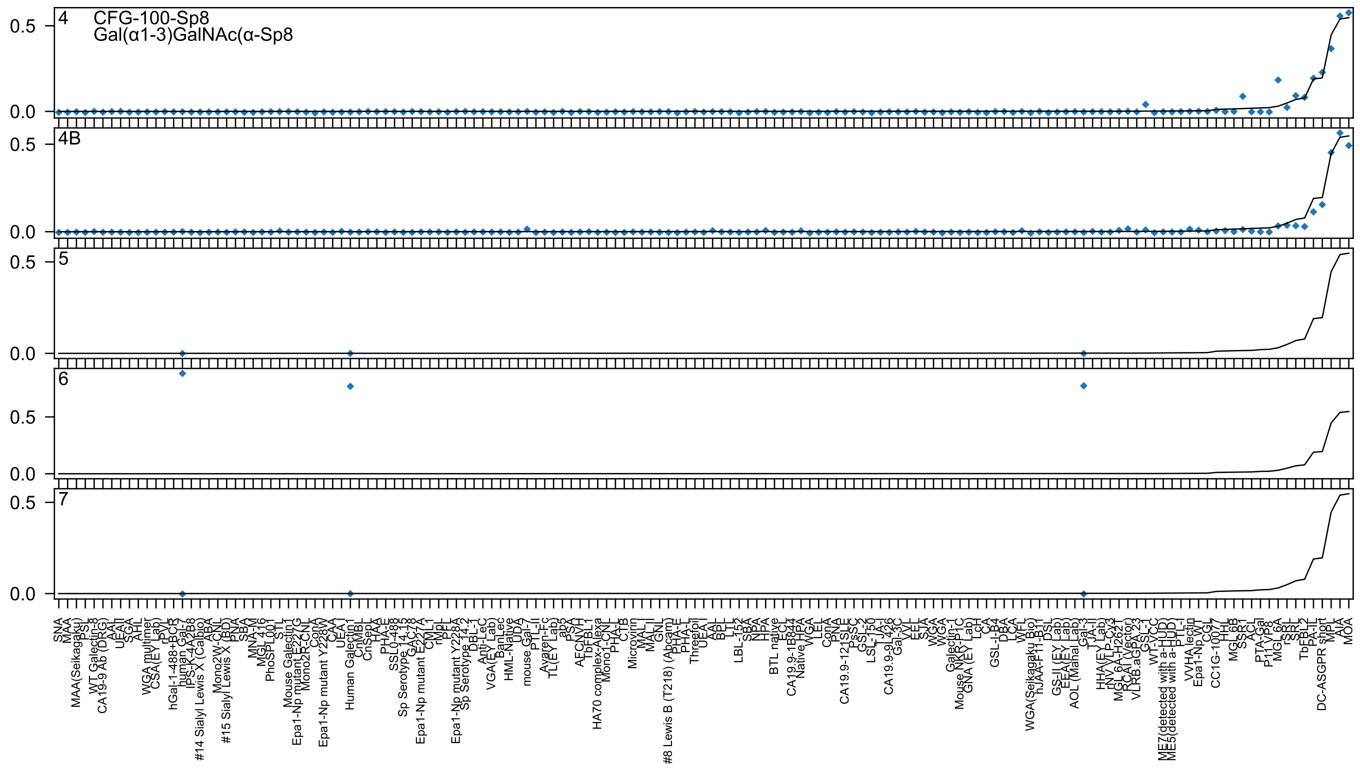
4
CFG-100-Sp8
Gal(α1-3)GalNAc(α-Sp8
4B
5
6
7
0.0
0.5

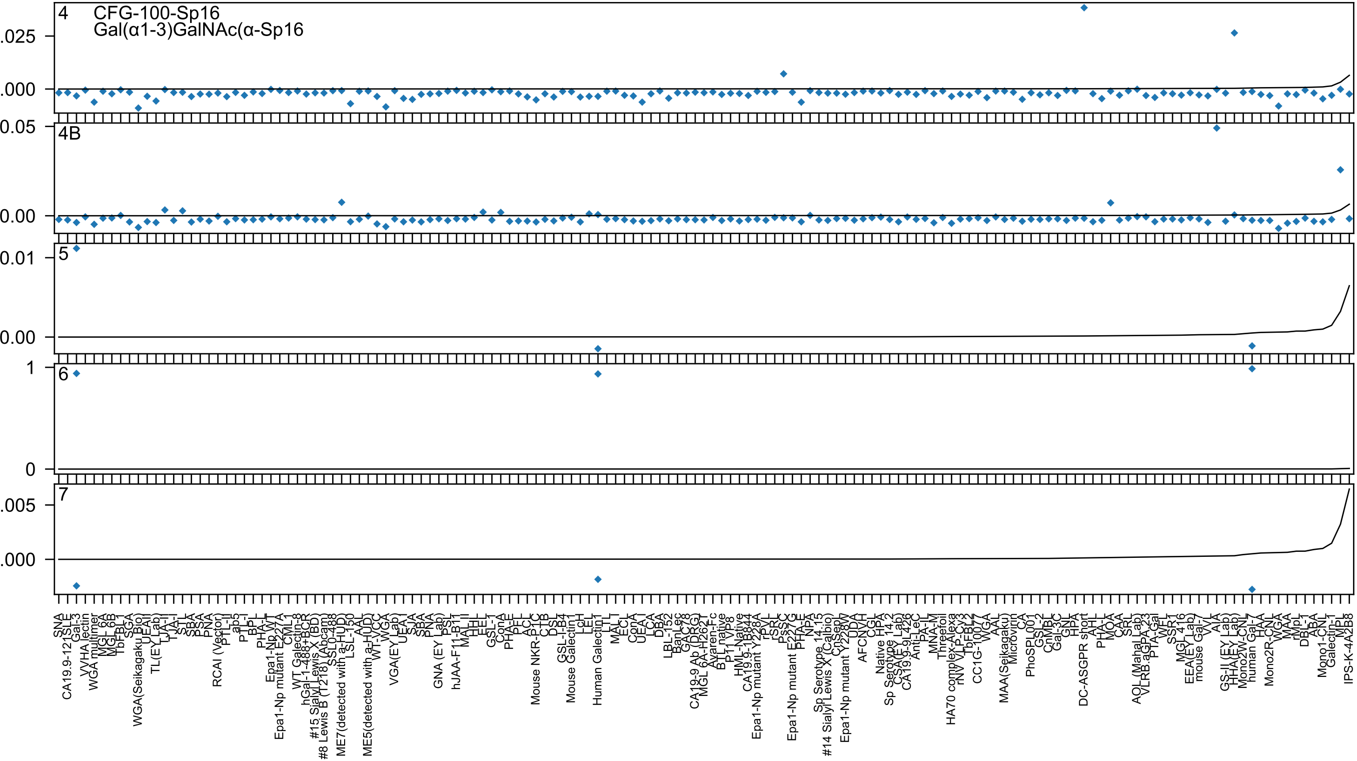
4
CFG-100-Sp16
Gal(α1-3)GalNAc(α-Sp16
.025
.000
4B
0.05
0.00
5
0.01
0.00
6
1
0
7
.005
.000
SNA
CA19.9-121
SLE
Gal-3
VVHA lectin
WGA multimer
MGL 6A
MGL 6B
TbFBL1
SGA
WGA(Seikagaku Bio)
UEAI
TL(EY Lab)
TJA-II
TJA-I
STL
SBA
PSA
PNA
RCAI (Vector)
PTL-II
ab5
PTL-I
BPL
PHA-L
Epa1-Np WT
Epa1-Np mutant E227A
CML1
WT Galectin-8
hGal-1-488+BCR
#15 Sialyl Lewis X (BD)
#8 Lewis B (T218) (Abcam)
SSL0-488
ME7(detected with a-HUD)
LSL-150
AAL
ME5(detected with a-HUD)
WT-VCC
WGA
VGA(EY Lab)
UEA-I
SJA
SBA
PNA
GNA (EY Lab)
PSL
hJAA-F11-B11
MAL_II
HHL
EEL
GSL-1
ConA
PHA-E
PEL
ACL
Mouse NKR-P1C
CTB
DSL
GSL-I-B4
Mouse Galectin1
LcH
LEL
Human Galectin1
LTL
MAL_I
ECL
ConA
UEA-I
LCA
DBA
LBL-152
BanLec
GAC78
CA19-9 Ab (DRG)
MGL 6A-H262T
Avaren-Fc
BTL native
P11 VP8
HML-Native
CA19.9-1B844
Epa1-Np mutant Y228A
rPVL
rSRL
PSSC
Epa1-Np mutant E227G
PHA-E
NPA
Sp Serotype 14.15
#14 Sialyl Lewis X (Calbio)
CnSep
Epa1-Np mutant Y228W
UDA
AFCNVH
CGL
Native HPA
Sp Serotype 14.2
CSA(EY Lab)
CA19.9-9L426
Anti-LeC
PA-IL
MNA-M
Threefoil
HA70 complex-Alexa
rNV VLP-Cy3
TbFBL2
CC1G-10077
WGA
AAL
MAA(Seikagaku)
Microvirin
CA
PhoSPL001
GSL-2
CnMBL
Gal-3C
GNL
HPA
DC-ASGPR short
AHL
PHA-L
MOA
CAA
SRL
AOL (Mahal Lab)
VLRB.aGPA.23
PTA-Gal
WFL
SSR1
MGL 416
EEA(EY Lab)
mouse Gal-7
VVL
AIA
GS-II (EY Lab)
HHA(EY Lab)
Mono2W-CNL
human Gal-7
HAA
Mono2R-CNL
WGA
MAA
rMpL
DBL-1
ABA
Mono1-CNL
Galectin-1
MPL
IPS-K-4A2B8

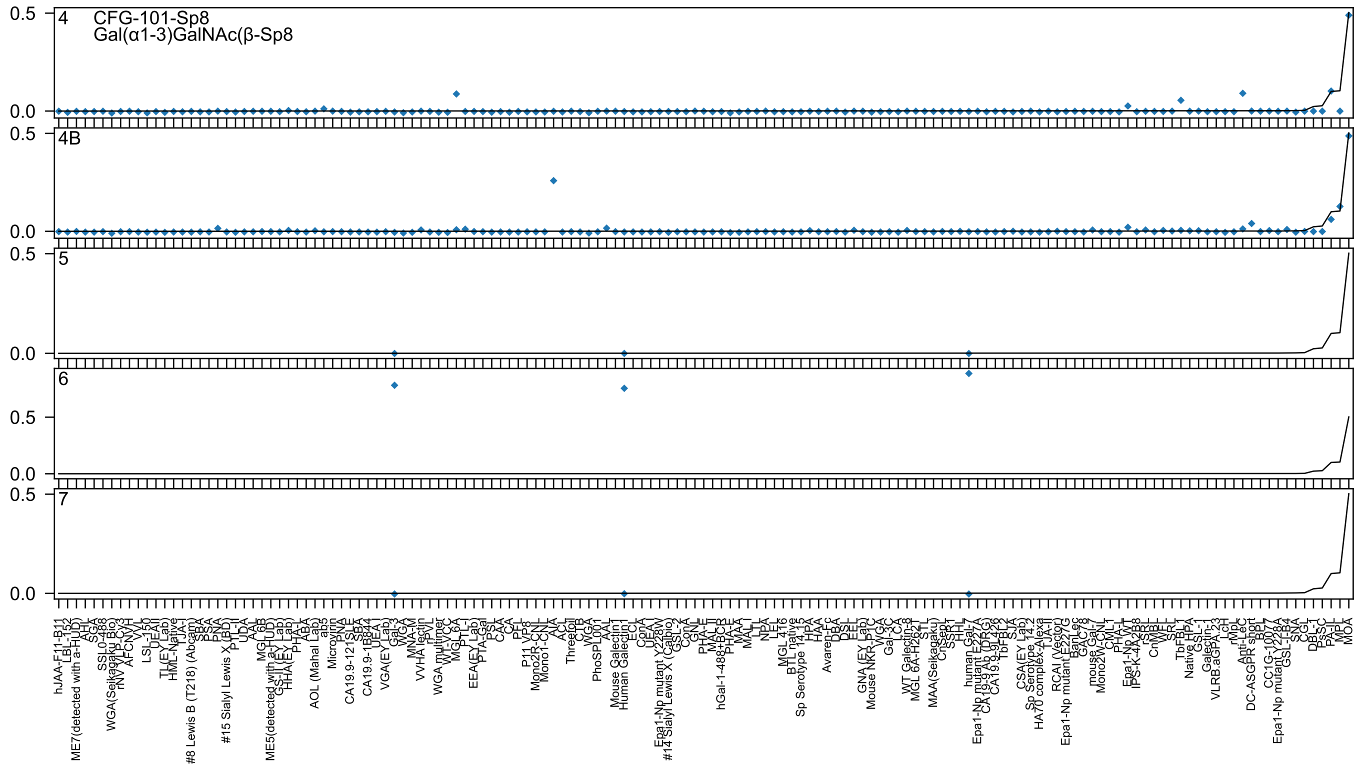

4
CFG-101-Sp8
Gal(α1-3)GalNAc(β-Sp8
4B
5
6
7
0.5
0.0

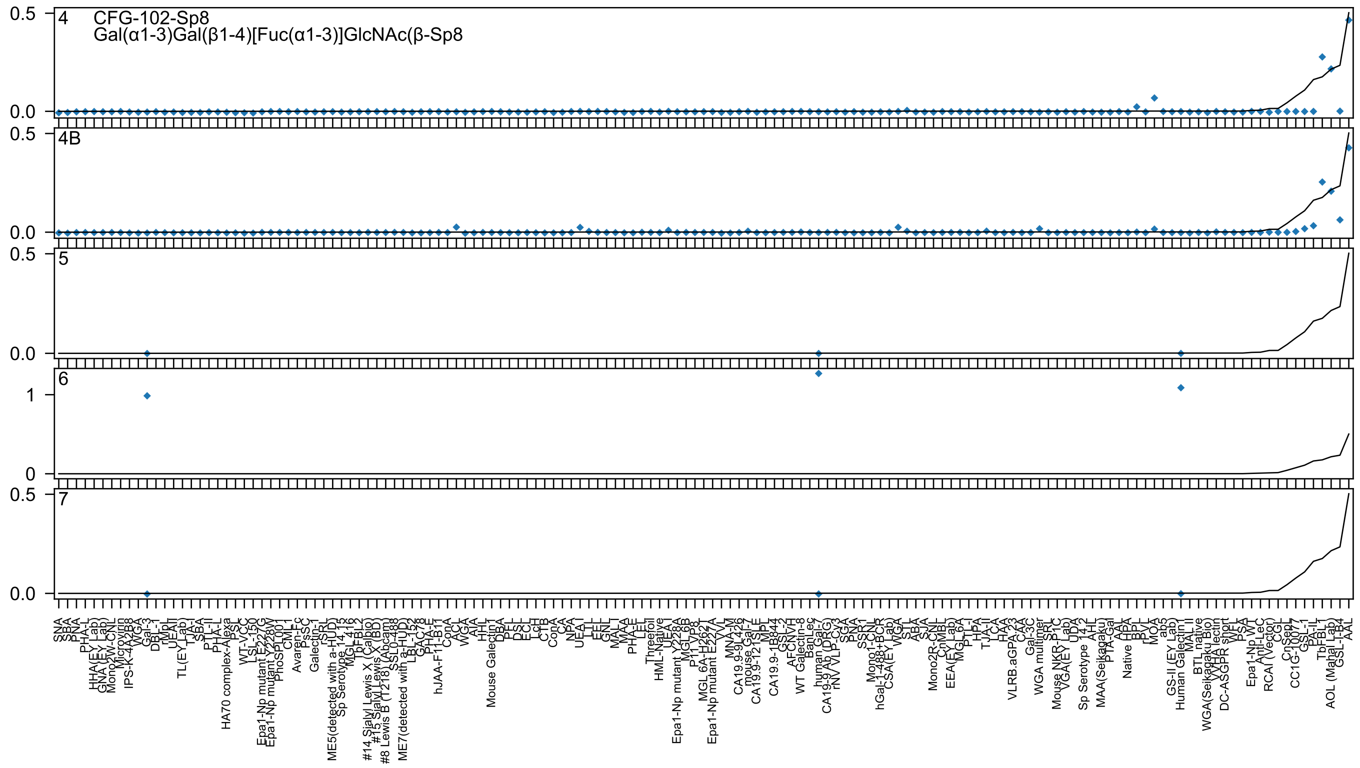

4
CFG-102-Sp8
Gal(α1-3)Gal(β1-4)[Fuc(α1-3)]GlcNAc(β-Sp8
4B
5
6
7
0.5
0.0
1
0

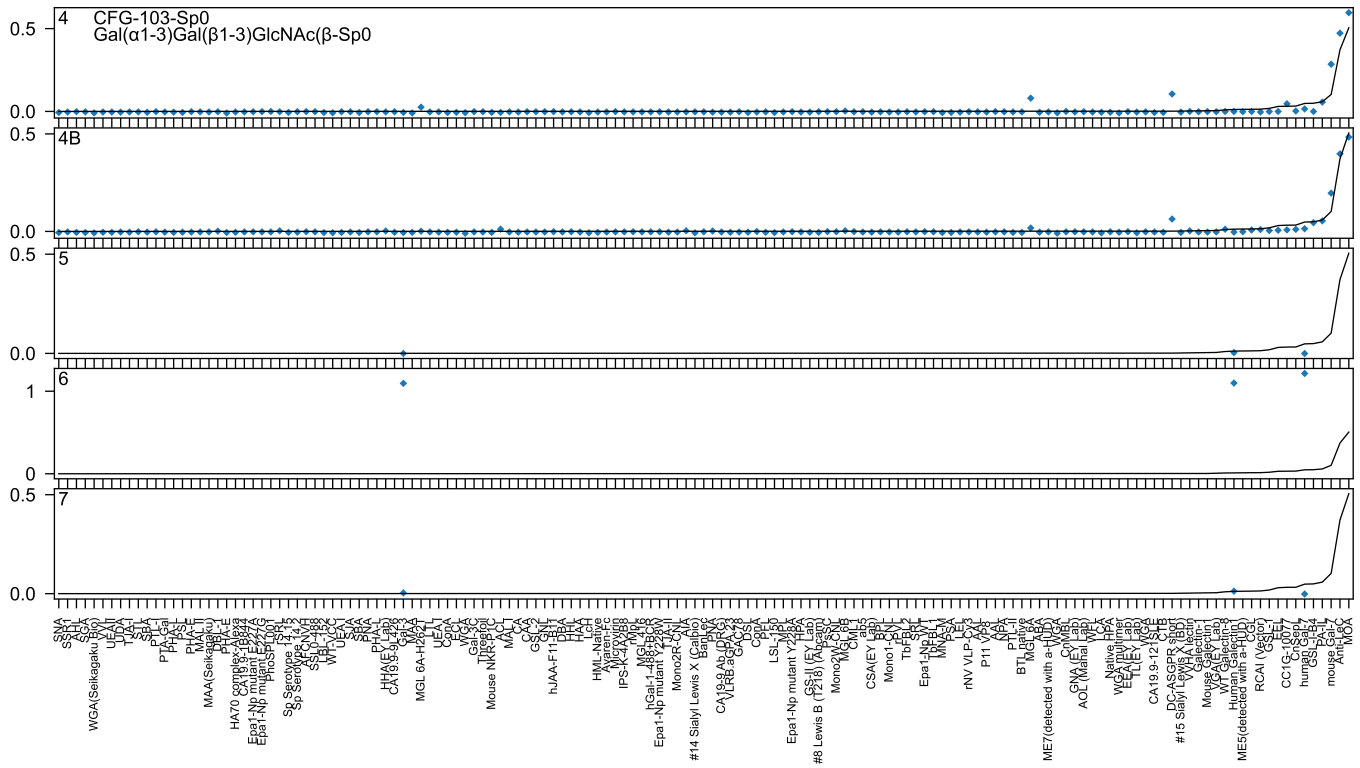
4 CFG-103-Sp0
Gal(α1-3)Gal(β1-3)GlcNAc(β-Sp0
4B
5
6
7
0.5
0.0
1
0

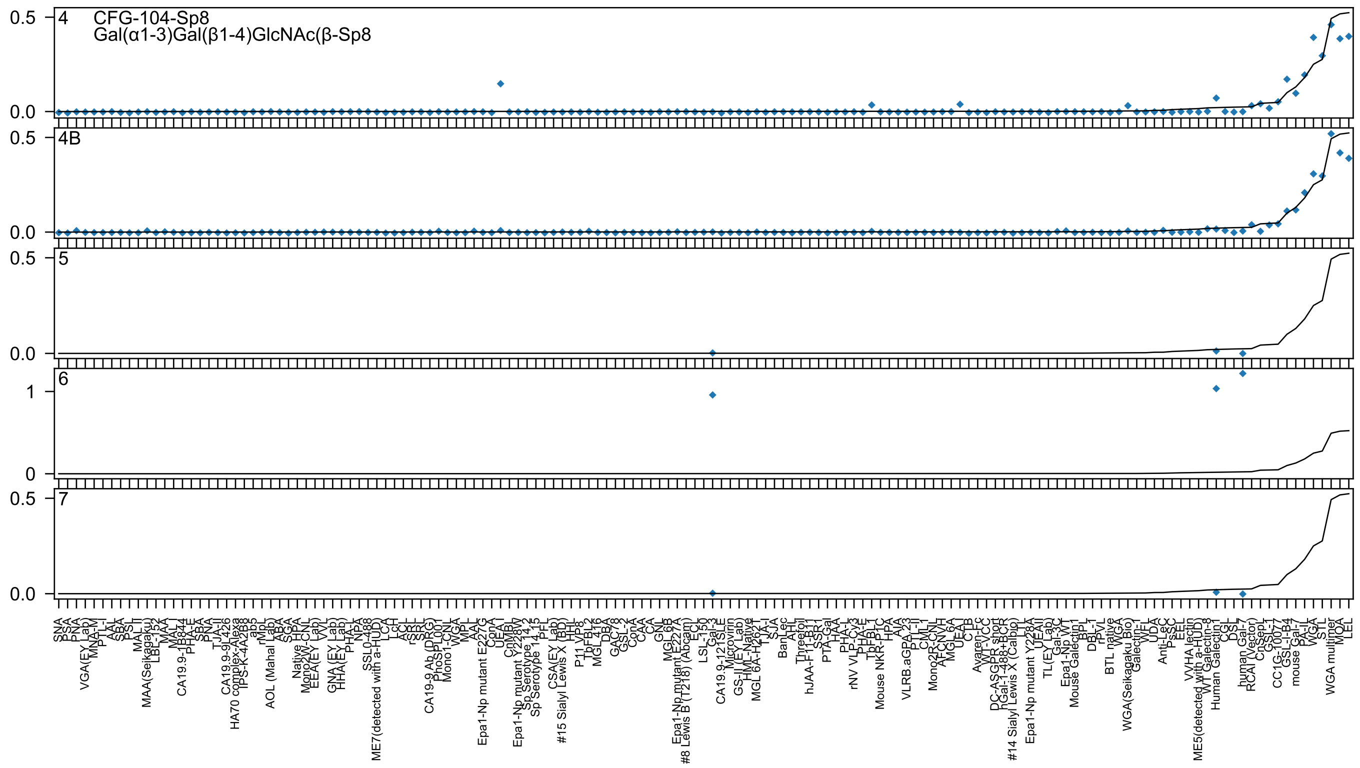
4
CFG-104-Sp8
Gal(α1-3)Gal(β1-4)GlcNAc(β-Sp8
4B
5
6
7
0.5
0.0
1
0

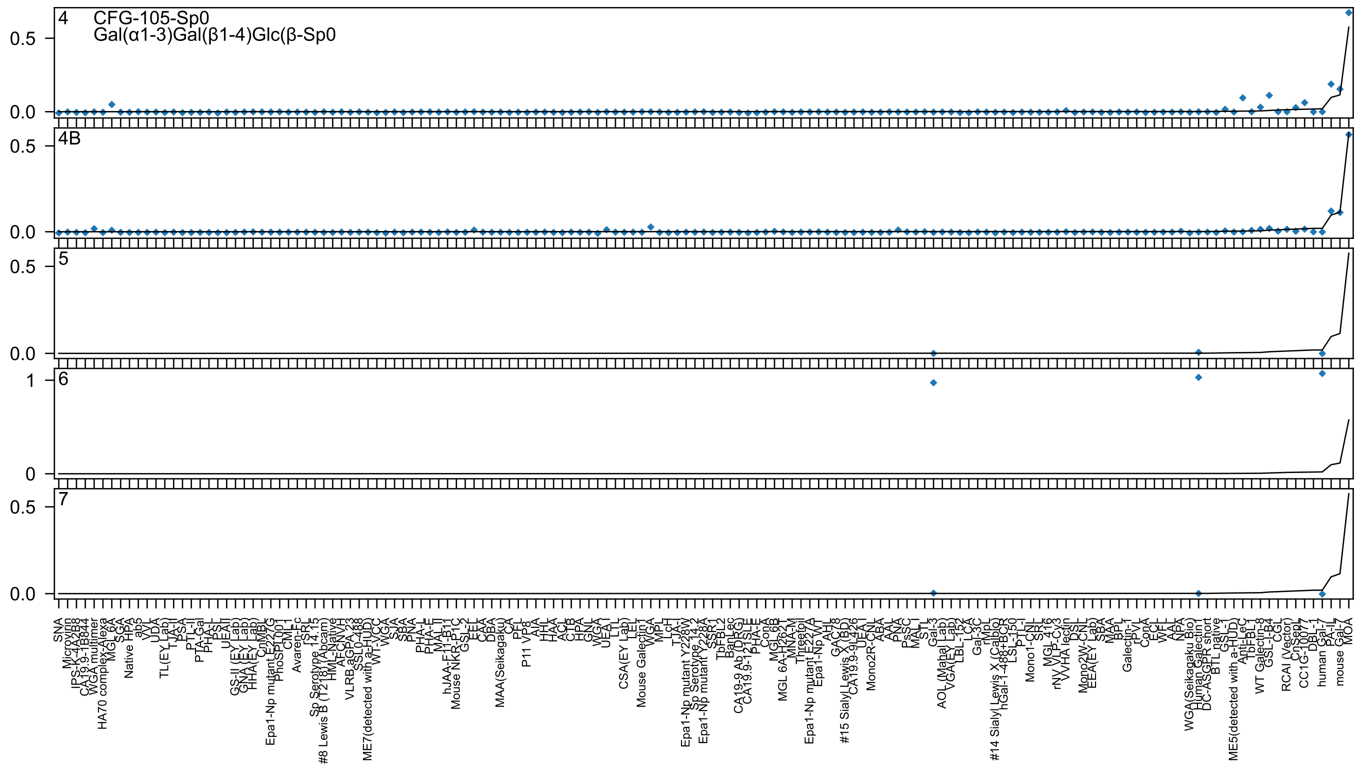
4
CFG-105-Sp0
Gal(α1-3)Gal(β1-4)Glc(β-Sp0
4B
5
6
7
0.5
0.0
1
0
SNA
Microvirin
IPS-K-4A2B8
CA19.9-1B844
WGA multimer
HA70 complex-Alexa
MGL_6A
SGA
Native HPA
ab5
VVL
UDA
TL(EY Lab)
TJA-II
PSA
PTL-II
PTA-Gal
PHA-L
PSL
UEAII
GS-II (EY Lab)
GNA (EY Lab)
HHA(EY Lab)
CnMBL
Epa1-Np mutant E227G
PhoSL001
CML-I
Avaren-Fc
rSRL
Sp Serotype 14.15
#8 Lewis B (T218) (Abcam)
HML-Native
AFCNVH
VLRB.aGPA.23
SSL0-488
ME7(detected with a-HUD)
WT-VCC
WGA
SJA
SBA
PNA
PHA-L
PHA-E
MAL II
hJAA-F11-B11
Mouse NKR-P1C
GSL-2
EEL
CAA
DBA
MAA(Seikagaku)
CA
PFL
P11 VP8
AIA
HHL
HAA
ACL
CTB
HPA
GNL
WGA
UEA I
LTL
CSA(EY Lab)
LEL
Mouse Galectin1
WGA
MPL
LcH
TJA-I
Epa1-Np mutant Y228W
Sp Serotype 14.2
Epa1-Np mutant Y228A
SSR1
TbFBL2
BanLec
CA19-9 Ab (DRG)
CA19.9-121SLE
PHA-E
ConA
MGL 6B
MGL 6A-H262T
MNA-M
Threetoil
Epa1-Np mutant E227A
Epa1-Np WT
AAL
GAC78
#15 Sialyl Lewis X (BD)
CA19.9-9L426
UEA I
Mono2R-CNL
ABA
AAL
PNA
PsSC
MAL I
STL
Gal-3
AOL (Mahal Lab)
VGA(EY Lab)
LBL-152
LCA
Gal-3C
rMpL
#14 Sialyl Lewis X (Calbio)
hGal-1-488+BCR
LSL-150
PTL-I
Mono1-CNL
SRL
MGL 416
rNV VLP-Cy3
VVHA lectin
DSL
Mono2W-CNL
EEA(EY Lab)
SBA
MAA
BPL
Galectin-1
rPVL
ConA
ECL
WFL
AAL
NPA
WGA(Seikagaku Bio)
Human Galectin1
DC-ASGPR short
BTL native
GSL-1
ME5(detected with a-HUD)
Anti-LeC
TbFBL1
WT Galectin-8
GSL-I-B4
CGL
RCAI (Vector)
CnSepL
CC1G-10077
DBL-1
human Gal-7
PA-IL
mouse Gal-7
MOA

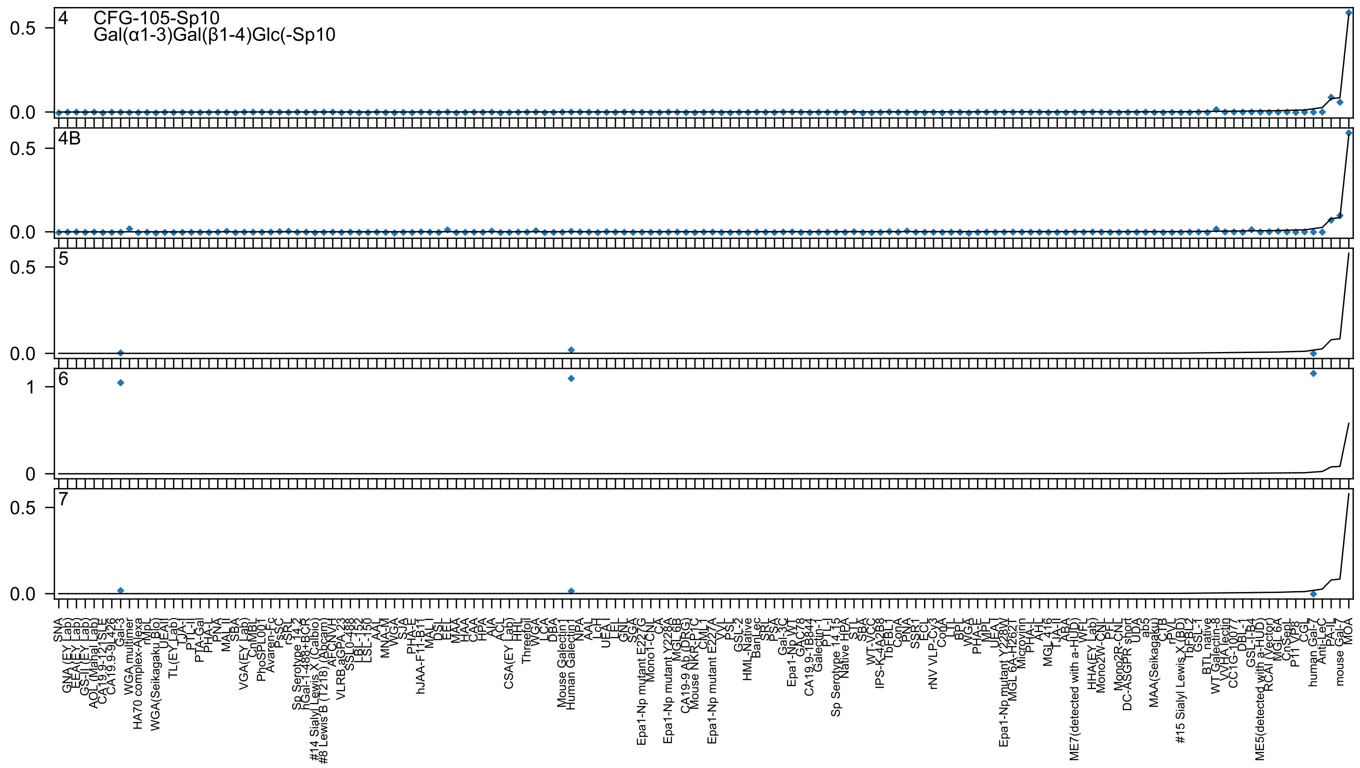

4 CFG-105-Sp10
Gal(α1-3)Gal(β1-4)Glc(-Sp10
4B
5
6
7

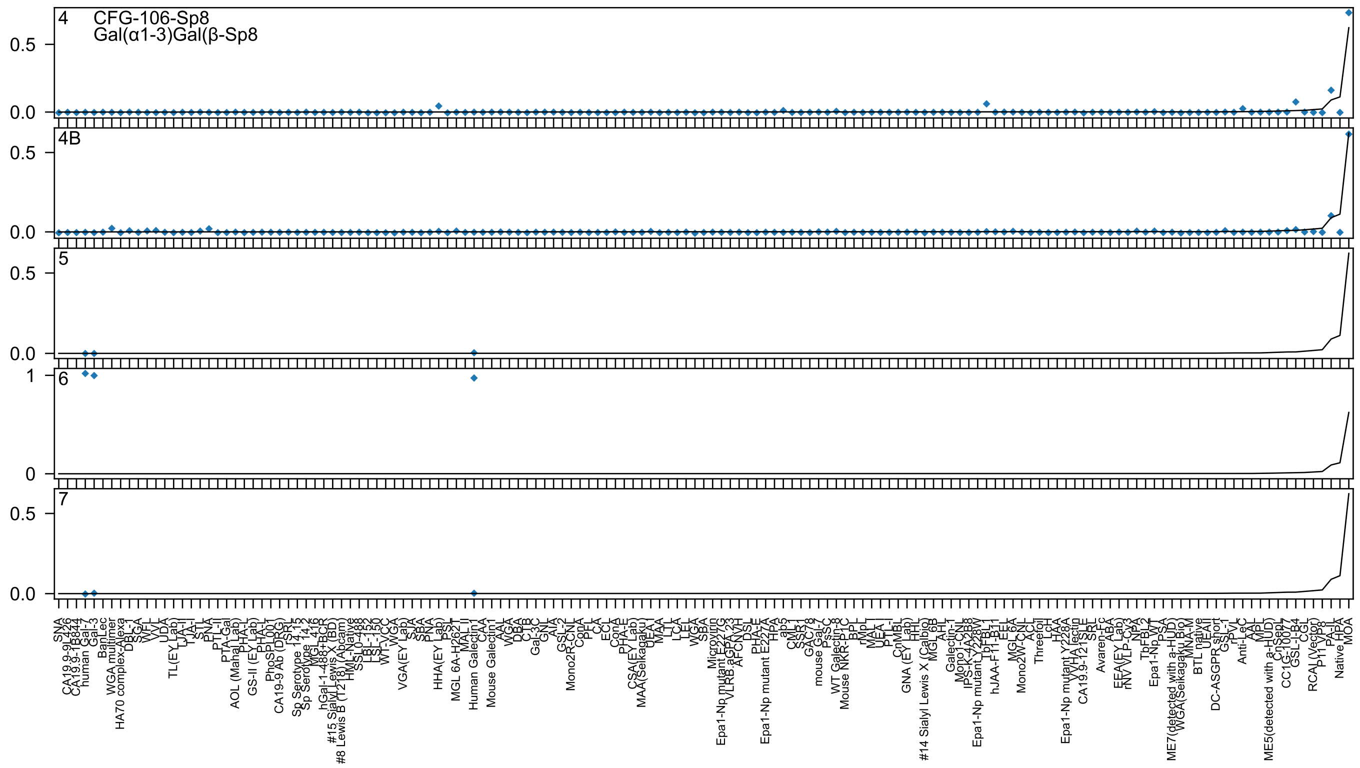

4
CFG-106-Sp8
Gal(α1-3)Gal(β-Sp8
4B
5
6
7
0.5
0.0
1
0

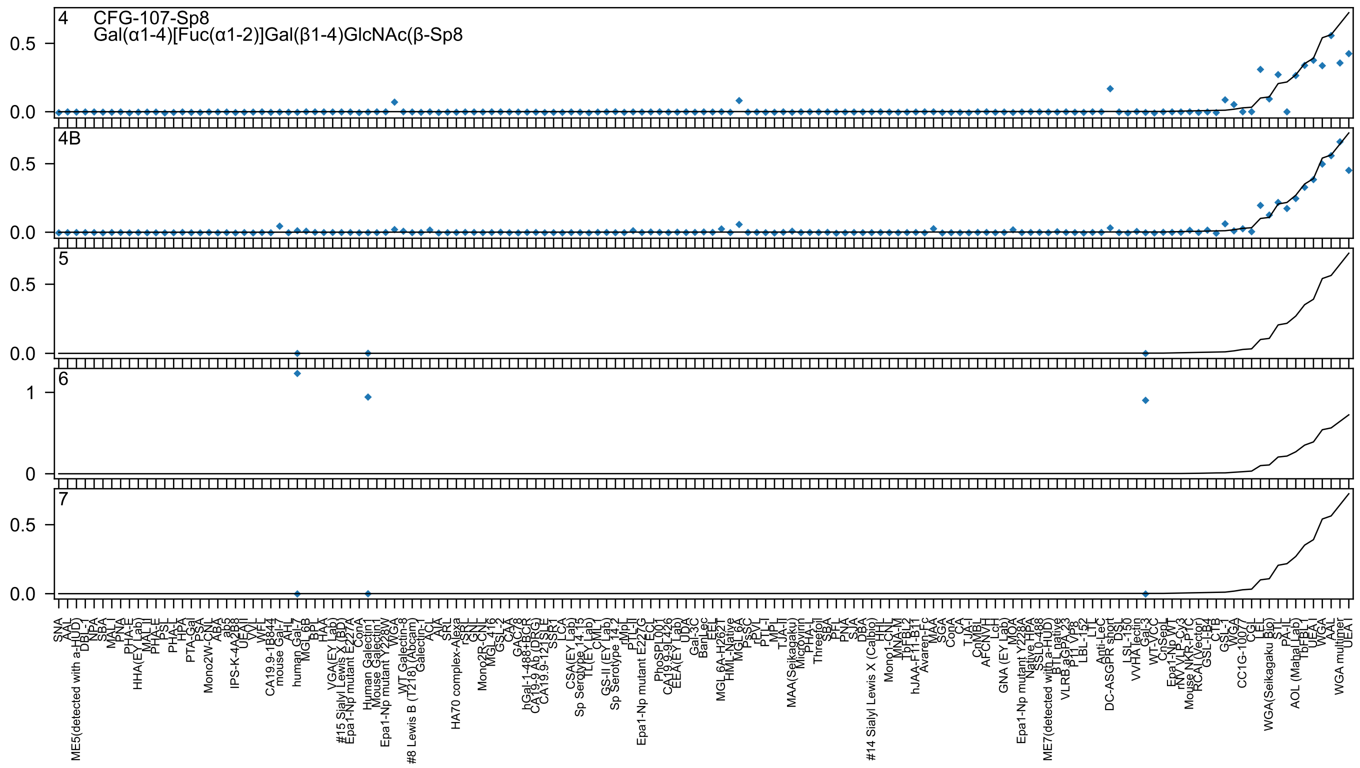
4
CFG-107-Sp8
Gal(α1-4)[Fuc(α1-2)]Gal(β1-4)GlcNAc(β-Sp8
4B
5
6
7

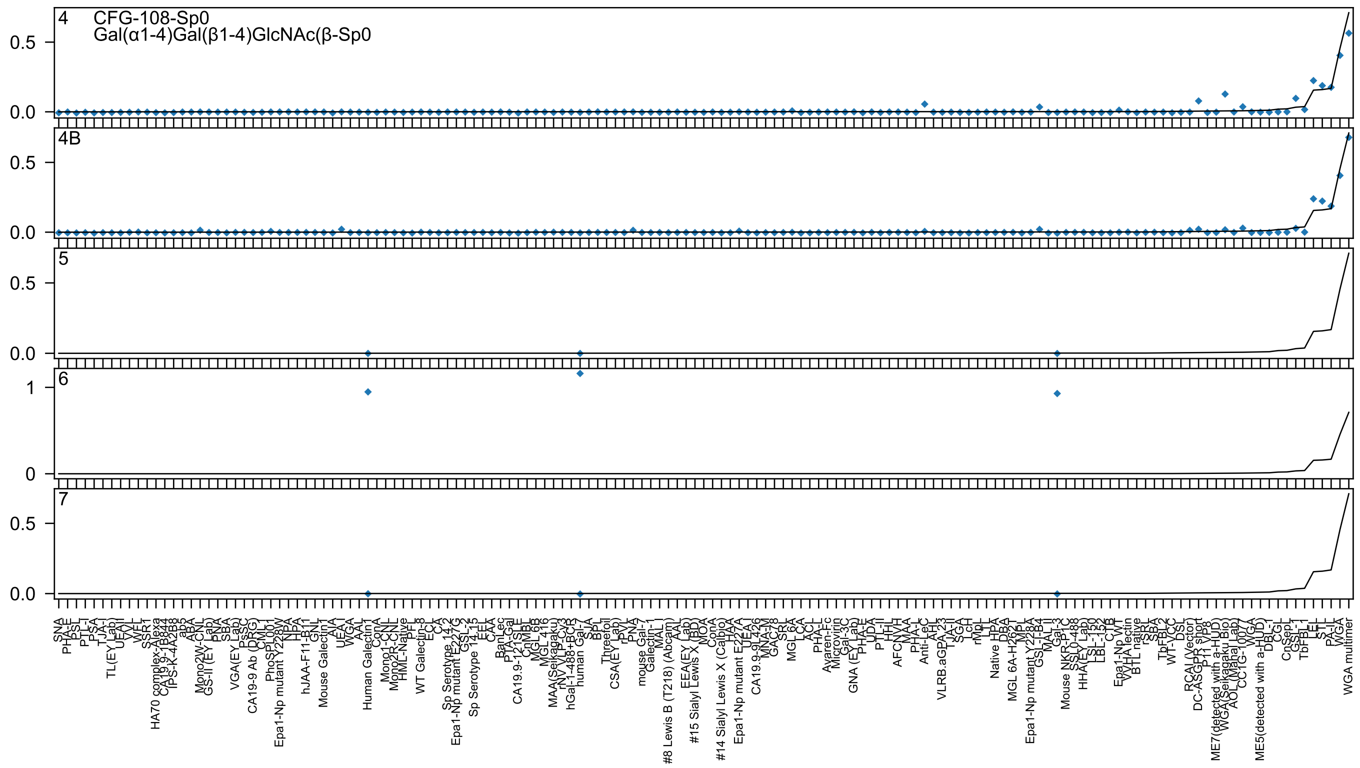

4
CFG-108-Sp0
Gal(α1-4)Gal(β1-4)GlcNAc(β-Sp0
4B
5
6
7
0.0
0.5
0
1

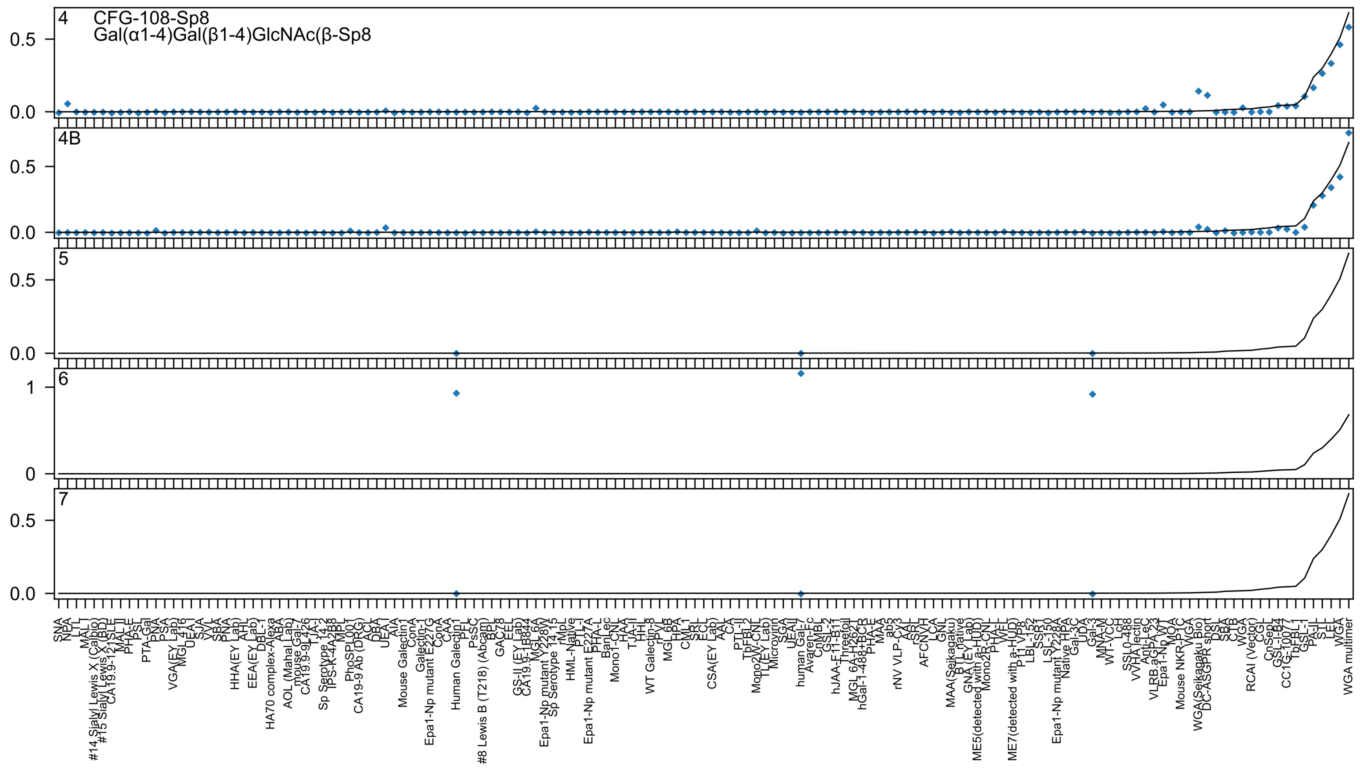

4
CFG-108-Sp8
Gal(α1-4)Gal(β1-4)GlcNAc(β-Sp8
4B
5
6
7

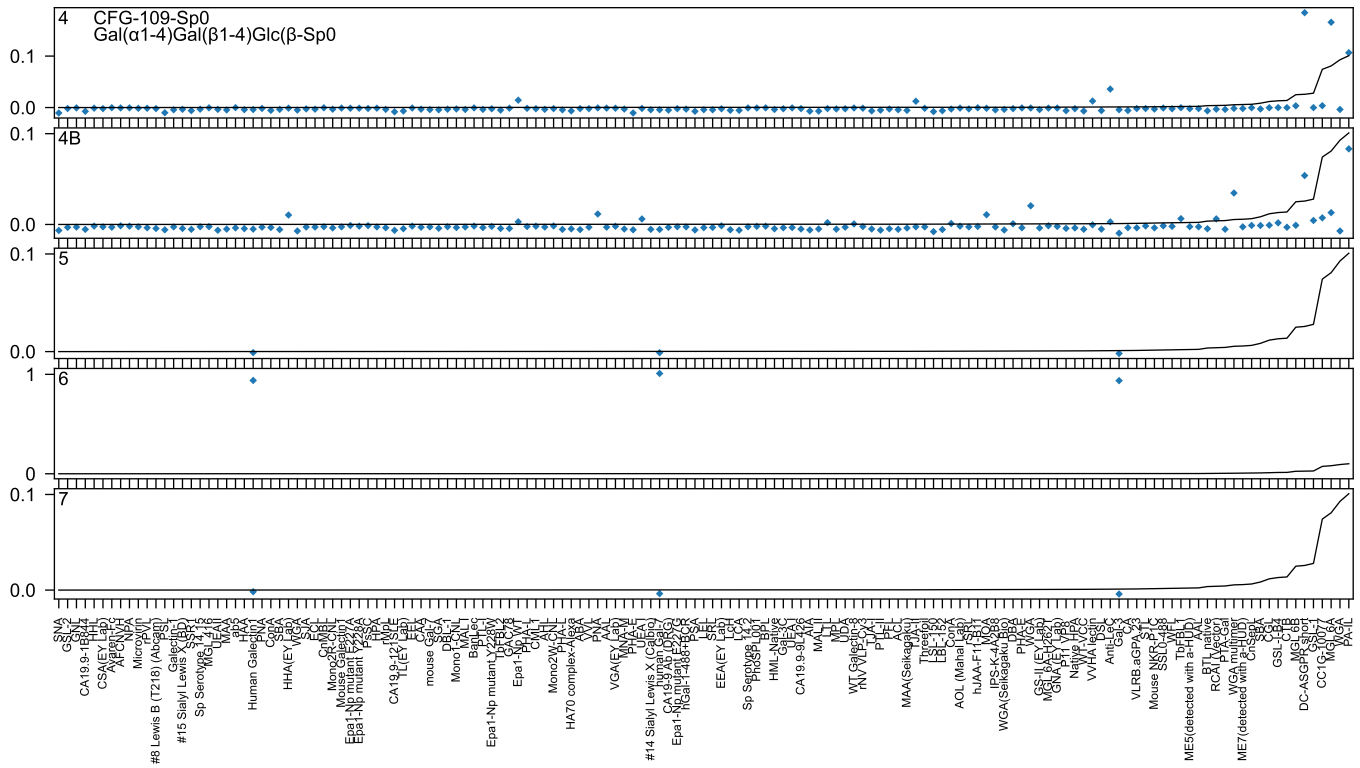
4
CFG-109-Sp0
Gal(α1-4)Gal(β1-4)Glc(β-Sp0
4B
5
6
7

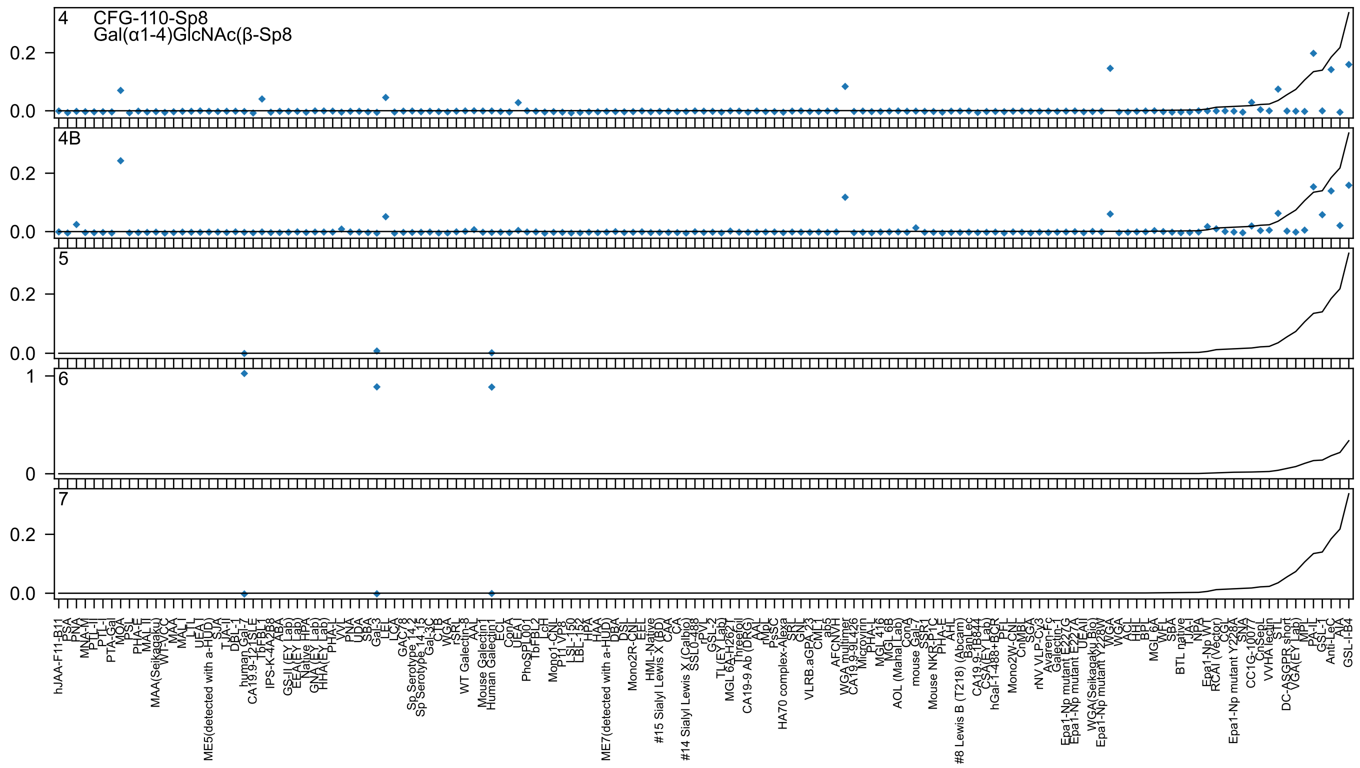

4
CFG-110-Sp8
Gal(α1-4)GlcNAc(β-Sp8
4B
5
6
7
0.2
0.0
1
0
hJAA-F11-B11
PSA
PNA
MNA-M
PTL-II
PTL-I
PTA-Gal
MOA
PSL
PHA-E
MAL II
MAA(Seikagaku)
WT-VCC
MAA
MAL I
LTL
UEA-I
ME5(detected with a-HUD)
SJA
TJA-II
DBL-1
human Gal-7
CA19.9-121SLE
TbFBL1
IPS-K-4A2B8
ABA
GS-II (EY Lab)
EEA(EY Lab)
Native HPA
GNA(EY Lab)
HHA(EY Lab)
PHA-L
VVL
PNA
UDA
SBA
Gal-3
LEL
LCA
GAC78
Sp Serotype 14.2
Sp Serotype 14.15
Gal-3C
CTB
WGA
rSRL
WT Galectin-8
AAL
Mouse Galectin1
Human Galectin1
ECL
ConA
UEA I
PhoSPL001
TbFBL2
LcH
Mono1-CNL
P11 VP8
LSL-150
LBL-152
HPA
HAA
ME7(detected with a-HUD)
DBA
DSL
Mono2R-CNL
EEL
HML-Native
#15 Sialyl Lewis X (BD)
CAA
CA
#14 Sialyl Lewis X (Calbio)
SSL0-488
rPVL
GSL-2
TL(EY Lab)
MGL 6A-H262T
Threefoil
CA19-9 Ab (DRG)
AAL
rMpL
PsSC
HA70 complex-Alexa
SRL
GNL
VLRB.aGPA.23
CML1
ab5
AFCNVH
WGA multimer
CA19.9-9L426
Microvirin
PHA-L
MGL 416
MGL 6B
AOL (Mahal Lab)
ConA
mouse Gal-7
SSR1
Mouse NKR-P1C
PHA-E
AHL
#8 Lewis B (T218) (Abcam)
BanLec
CA19.9-1B844
CSA(EY Lab)
hGal-1-488+BCR
PFL
Mono2W-CNL
CnMBL
SGA
rNV VLP-Cy3
Avaren-Fc
Galectin-1
Epa1-Np mutant E227G
Epa1-Np mutant E227A
UEAII
WGA(Seikagaku Bio)
Epa1-Np mutant Y228W
WGA
WGA
ACL
HHL
BPL
MGL 6A
WFL
SBA
BTL native
TJA-I
NPA
Epa1-Np WT
RCAI (Vector)
CGL
Epa1-Np mutant Y228A
SNA
CC1G-10077
CnSepL
VVHA lectin
STL
DC-ASGPR short
VGA(EY Lab)
MPL
PA-IL
GSL-1
Anti-LeC
AIA
GSL-I-B4

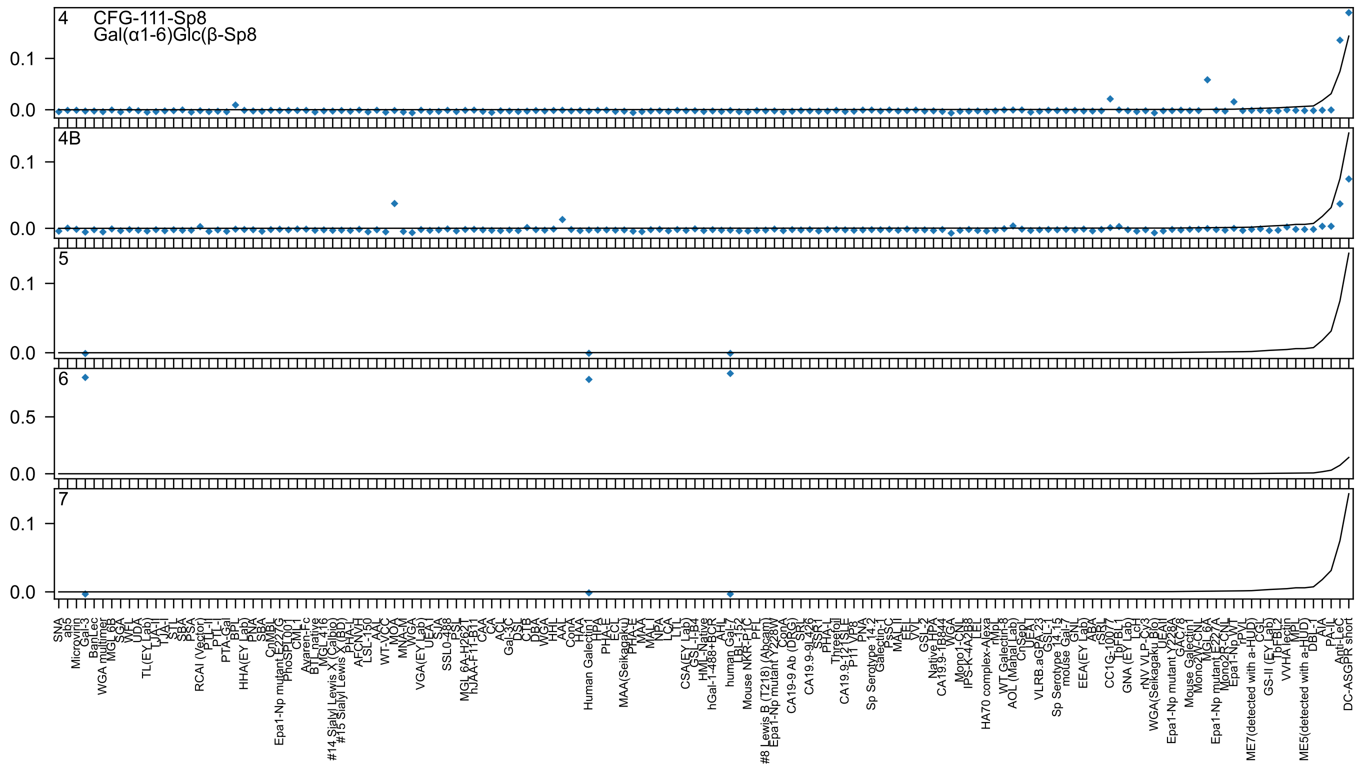
4
CFG-111-Sp8
Gal(α1-6)Glc(β-Sp8
4B
5
6
7

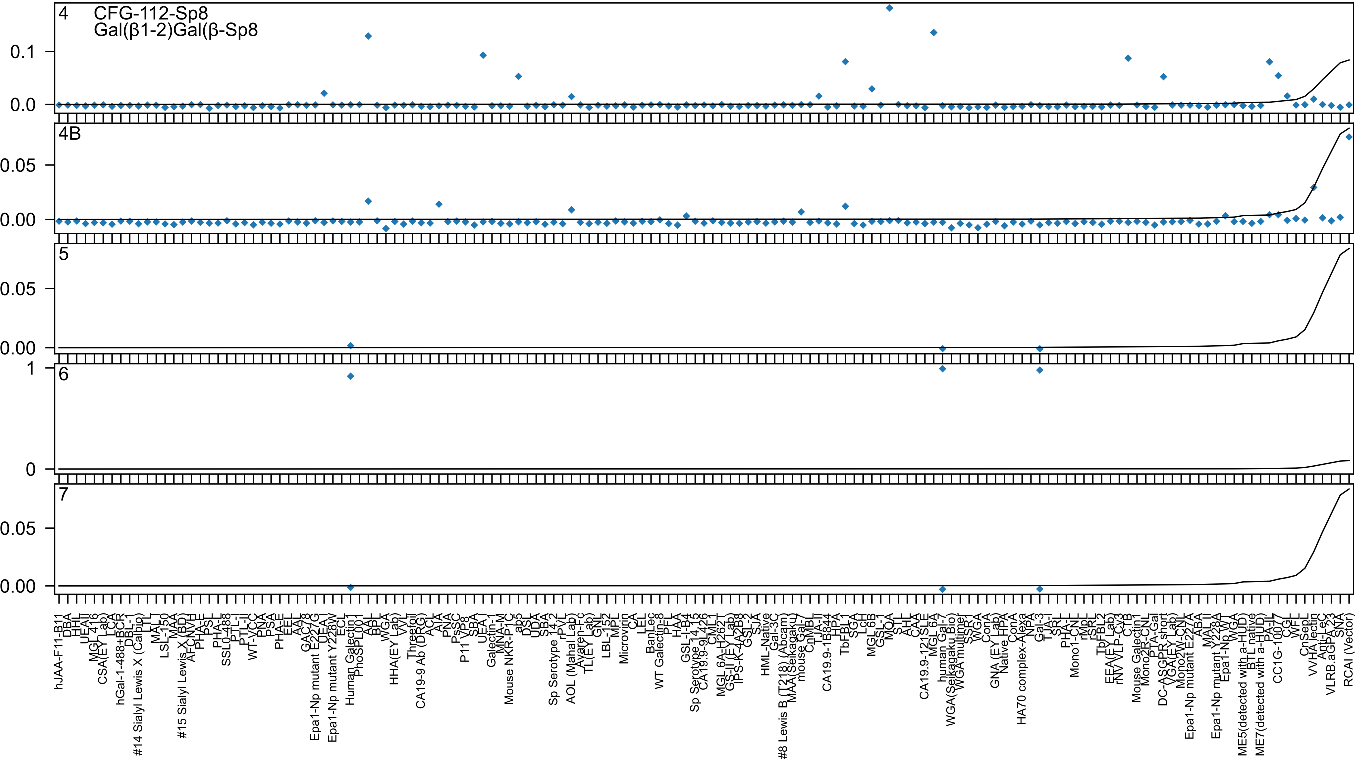
4
CFG-112-Sp8
Gal(β1-2)Gal(β-Sp8
4B
5
6
7

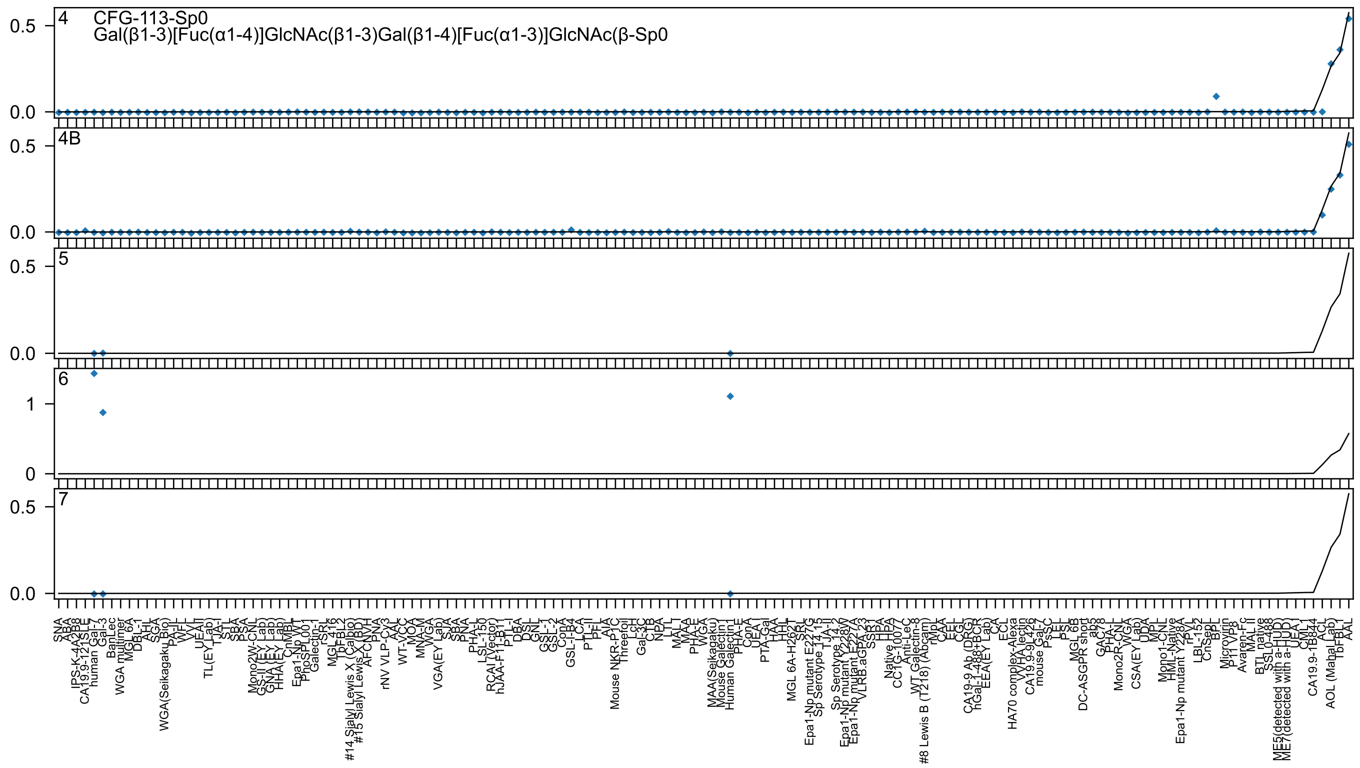

4
CFG-113-Sp0
Gal(β1-3)[Fuc(α1-4)]GlcNAc(β1-3)Gal(β1-4)[Fuc(α1-3)]GlcNAc(β-Sp0
4B
5
6
7
0.5
0.0
1
0
SNA
ABA
IPS-K-4A2B8
CA19.9-121SLE
human Gal-7
Gal-3
BanLec
WGA multimer
MGL 6A
DBL-1
AHL
SGA
WGA(Seikagaku Bio)
PA-IL
WFL
VVL
UEAII
TL(EY Lab)
TJA-I
STL
SBA
PSA
Mono2W-CNL
GS-II (EY Lab)
GNA (EY Lab)
HHA (EY Lab)
CnMBL
Epa1-Np WT
PhoSL001
Galectin-1
rSRL
MGL 416
TbFBL2
#14 Sialyl Lewis X (Calbio)
#15 Sialyl Lewis X (BD)
AFCNVH
PNA
rNV VLP-Cv3
AAL
WT-VCC
MOA
MNA-M
WGA
VGA(EY Lab)
SJA
SBA
PNA
PHA-L
LSL-150
RCAI (Vector)
hJAA-F11-B11
PTL-I
DBA
DSL
GNL
GSL-1
GSL-2
ConA
GSL-I-B4
LCA
PTL-II
PFL
AIA
Mouse NKR-P1C
Threefoil
LcH
Gal-3C
CTB
NPA
LTL
MAL-I
MAA
PHA-E
WGA
MAA(Seikagaku)
Mouse Galectin1
Human Galectin1
PHA-E
ConA
UEA-I
PTA-Gal
HAA
HHL
MGL 6A-H262T
SRL
Epa1-Np mutant E227G
Sp Serotype 14 15
TJA-II
Sp Serotype 14 2
Epa1-Np mutant Y228W
Epa1-Np mutant E227A
VLRB.aGPA.23
SSA
HPA
Native HPA
CC1G-10077
Anti-LeC
WT Galectin-8
#8 Lewis B (T218) (Abcam)
rMpL
CAA
EEL
CGL
CA19-9 Ab (DRG)
hGal-1-488+BCR
EEA(EY Lab)
CA
ECL
HA70 complex-Alexa
VVA lectin
CA19.9-9L426
mouse Gal-7
PsSC
PEL
PSL
MGL 6B
DC-ASGPR short
ab5
GAC78
PHA-L
Mono2R-CNL
WGA
CSA(EY Lab)
UDA
MPL
Mono1-CNL
HML-Native
Epa1-Np mutant Y228A
rPVL
LBL-152
CnSepL
BPL
Microvirin
P11 VP8
Avaren-Fc
MAL II
BTL native
SSL0-488
ME5(detected with a-HUD)
ME7(detected with a-HUD)
UEA I
CML1
CA19.9-1B844
ACL
AOL (Mahal Lab)
TbFBL1
AAL

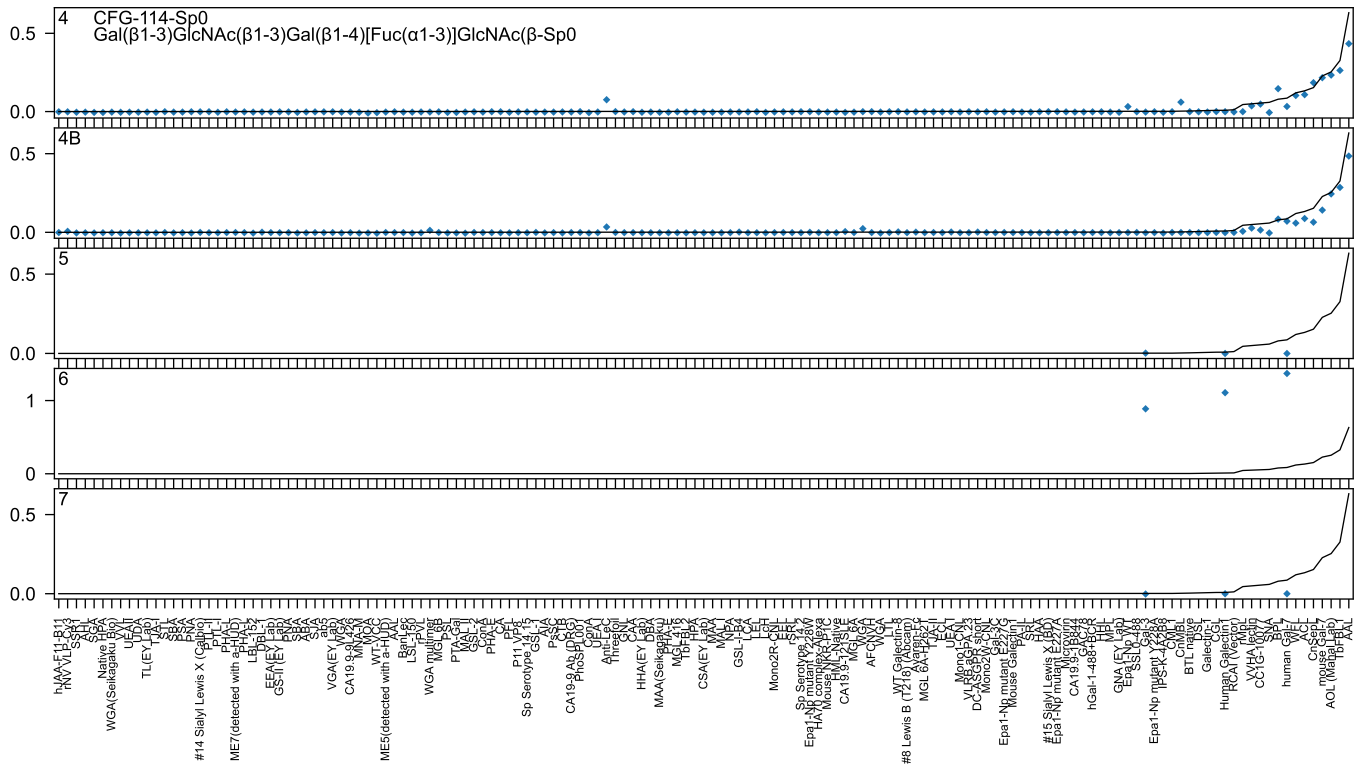

4 CFG-114-Sp0
Gal(β1-3)GlcNAc(β1-3)Gal(β1-4)[Fuc(α1-3)]GlcNAc(β-Sp0
4B
5
6
7

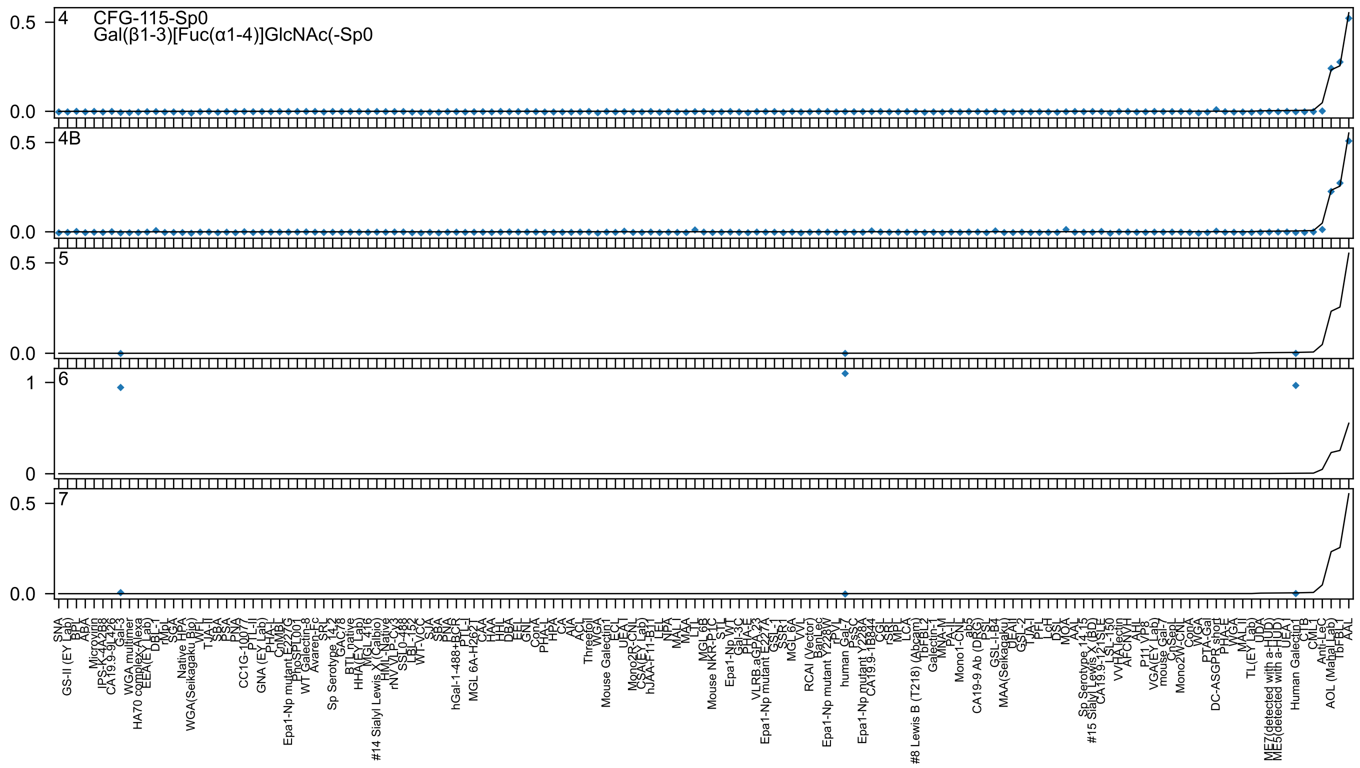
4
CFG-115-Sp0
Gal(β1-3)[Fuc(α1-4)]GlcNAc(-Sp0
4B
5
6
7

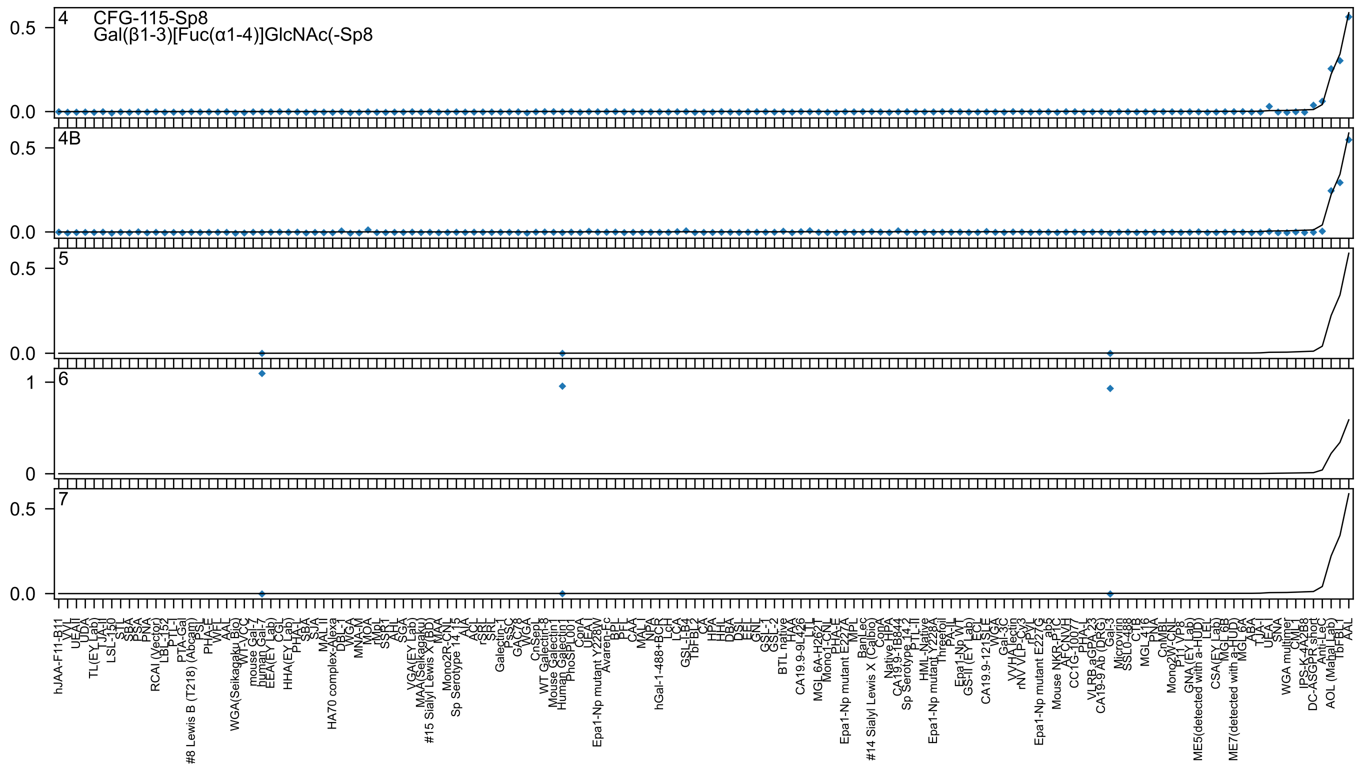

4
CFG-115-Sp8
Gal(β1-3)[Fuc(α1-4)]GlcNAc(-Sp8
4B
5
6
7

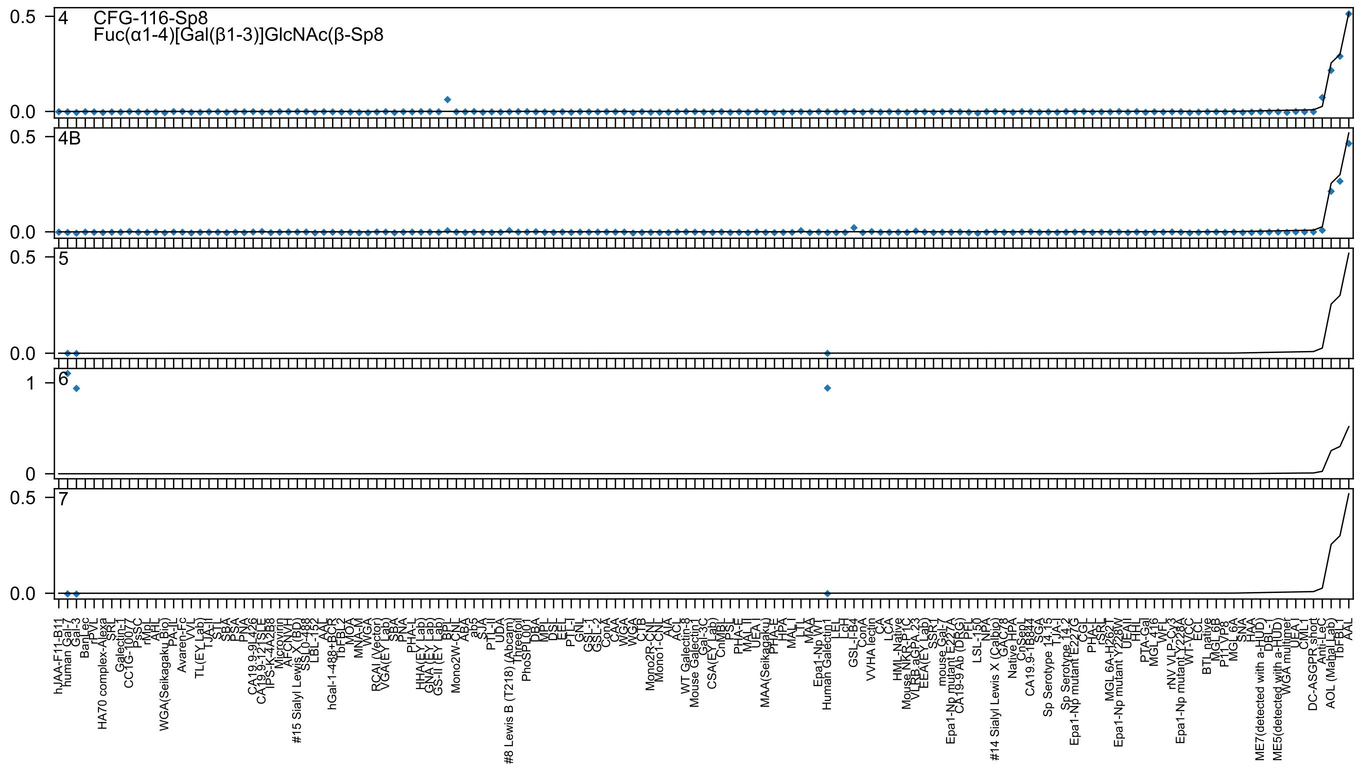
4
CFG-116-Sp8
Fuc(α1-4)[Gal(β1-3)]GlcNAc(β-Sp8
4B
5
6
7
0.5
0.0
1
0
hJAA-F11-B11
human Gal-7
Gal-3
BanLec
rPVL
HA70 complex-Alexa
SRT
Galectin-1
CC1G-10077
PsSC
rMpL
AHL
WGA(Seikagaku Bio)
PA-IL
Avaren-Fc
VVL
TL(EY Lab)
TJA-II
STL
SBA
PSA
PNA
CA19.9-9L426
CA19.9-121SLE
IPS-K-4A2B8
Microvirin
AFCNVH
#15 Sialyl Lewis X (BD)
SSL0-488
LBL-152
AAL
hGal-1-488+BCR
TbFBL2
MOA
MNA-M
WGA
RCAI (Vector)
VGA(EY Lab)
SBA
PNA
PHA-L
HHA(EY Lab)
GNA (EY Lab)
GS-II (EY Lab)
BPL
Mono2W-CNL
ABA
ab5
SJA
PTL-II
UDA
#8 Lewis B (T218) (Abcam)
Threefoil
PhoSL001
DBA
MPL
DSL
EEL
PTL-I
GNL
GSL-1
GSL-2
ConA
CAA
WGA
WGA
CTB
Mono2R-CNL
Mono1-CNL
AIA
ACL
WT Galectin-8
Mouse Galectin1
Gal-3C
CSA(EY Lab)
CnMBL
PSL
PHA-E
MAL-II
UEA-I
MAA(Seikagaku)
PHA-E
HPA
MAL-I
LTL
MAA
Epa1-Np WT
Human Galectin1
LEL
LcH
GSL-I-B4
ConA
VVHA lectin
CA
LCA
HML-Native
Mouse NKR-P1C
VLRB aGPA.23
EEA(EY Lab)
SSR1
mouse Gal-7
Epa1-Np mutant E227A
CA19-9 Ab (DRG)
PFL
LSL-150
NPA
#14 Sialyl Lewis X (Calbio)
GAC78
Native HPA
CnSepL
CA19.9-1B844
SGA
Sp Serotype 14.15
TJA-I
Sp Serotype 14.2
Epa1-Np mutant E227G
CGL
PHA-L
rSRL
MGL 6A-H262T
Epa1-Np mutant Y228W
UEAII
HHL
PTA-Gal
MGL 416
WFL
rNV VLP-Cy3
Epa1-Np mutant Y228A
WT-VCC
ECL
BTL native
MGL 6B
P11 VP8
MGL 6A
SNA
HAA
ME7(detected with a-HUD)
DBL
ME5(detected with a-HUD)
WGA multimer
UEA-I
CML1
DC-ASGPR short
Anti-LeC
AOL (Manal Lab)
TbFBL1
AAL

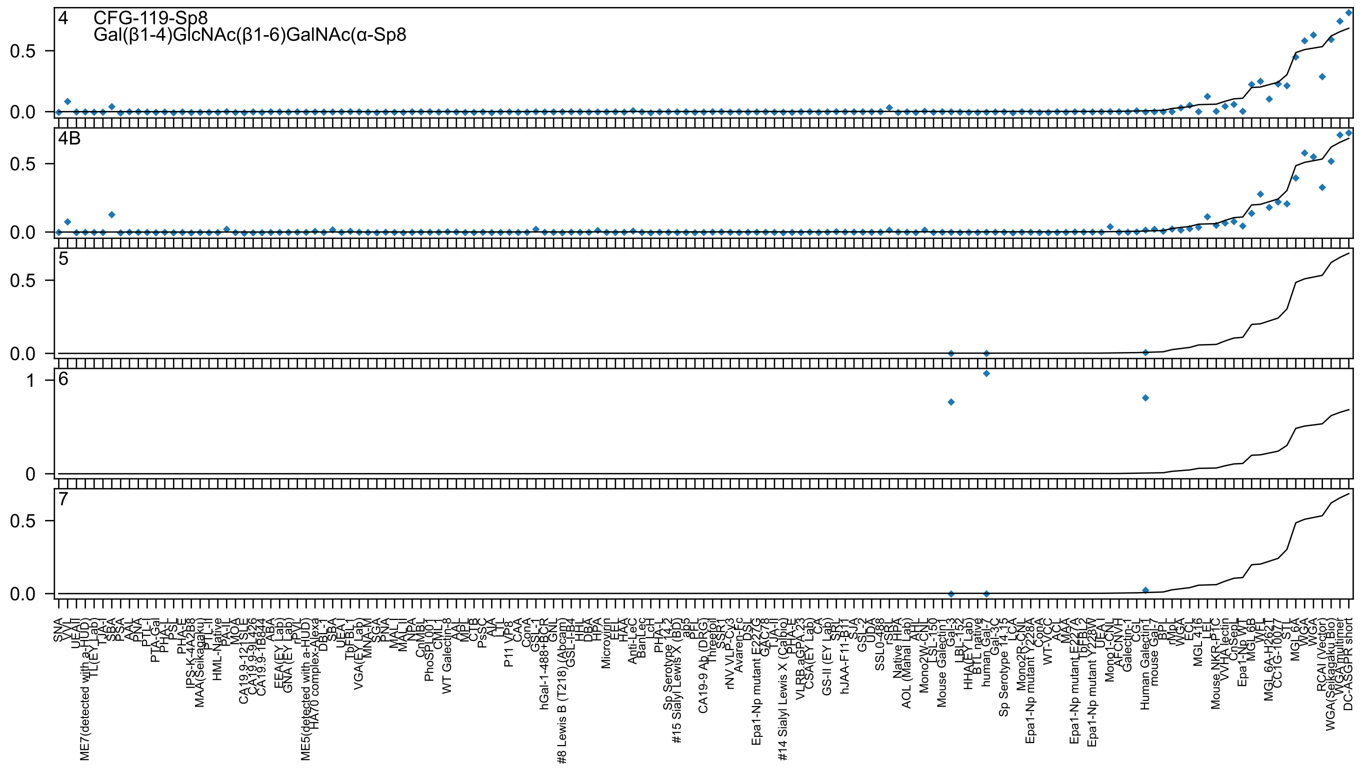
4
CFG-119-Sp8
Gal(β1-4)GlcNAc(β1-6)GalNAc(α-Sp8
4B
5
6
7
0.5
0.0
1
0

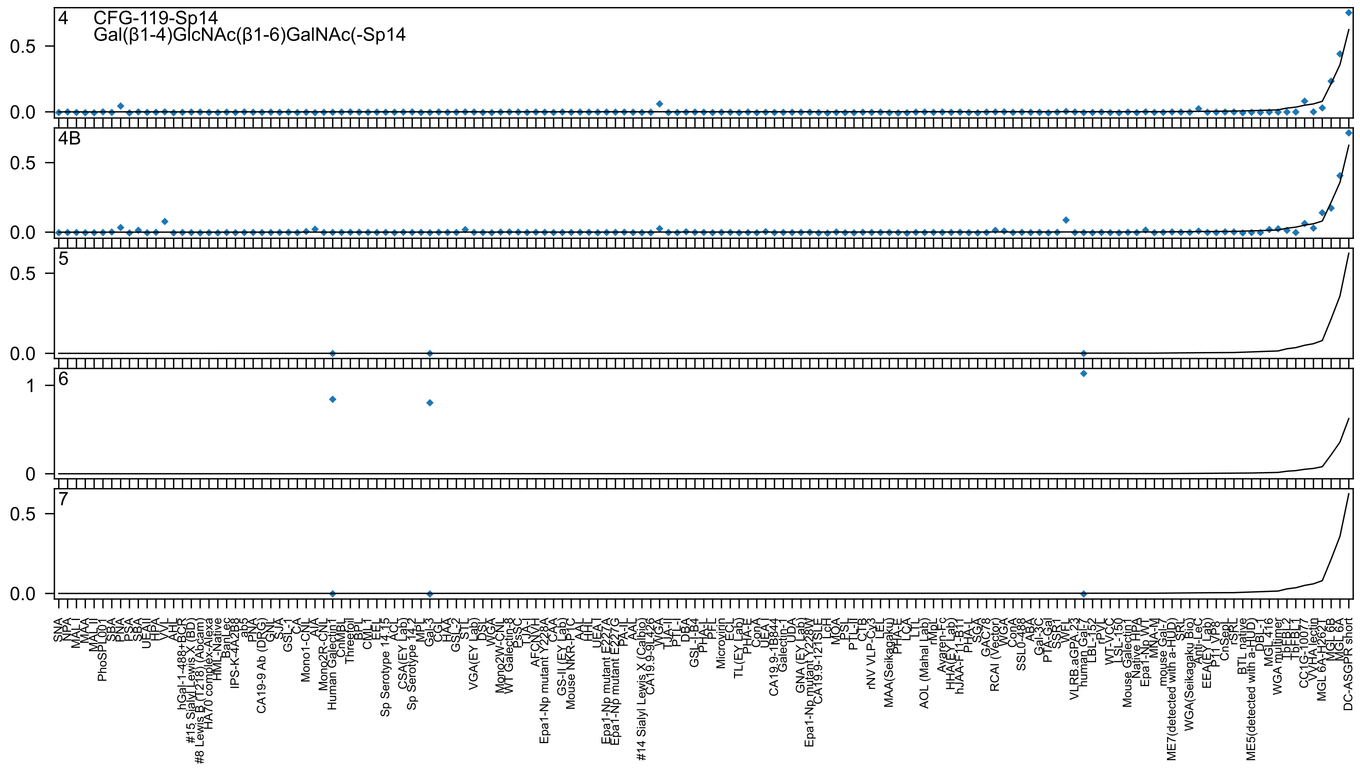
4 CFG-119-Sp14
Gal(β1-4)GlcNAc(β1-6)GalNAc(-Sp14
4B
5
6
7

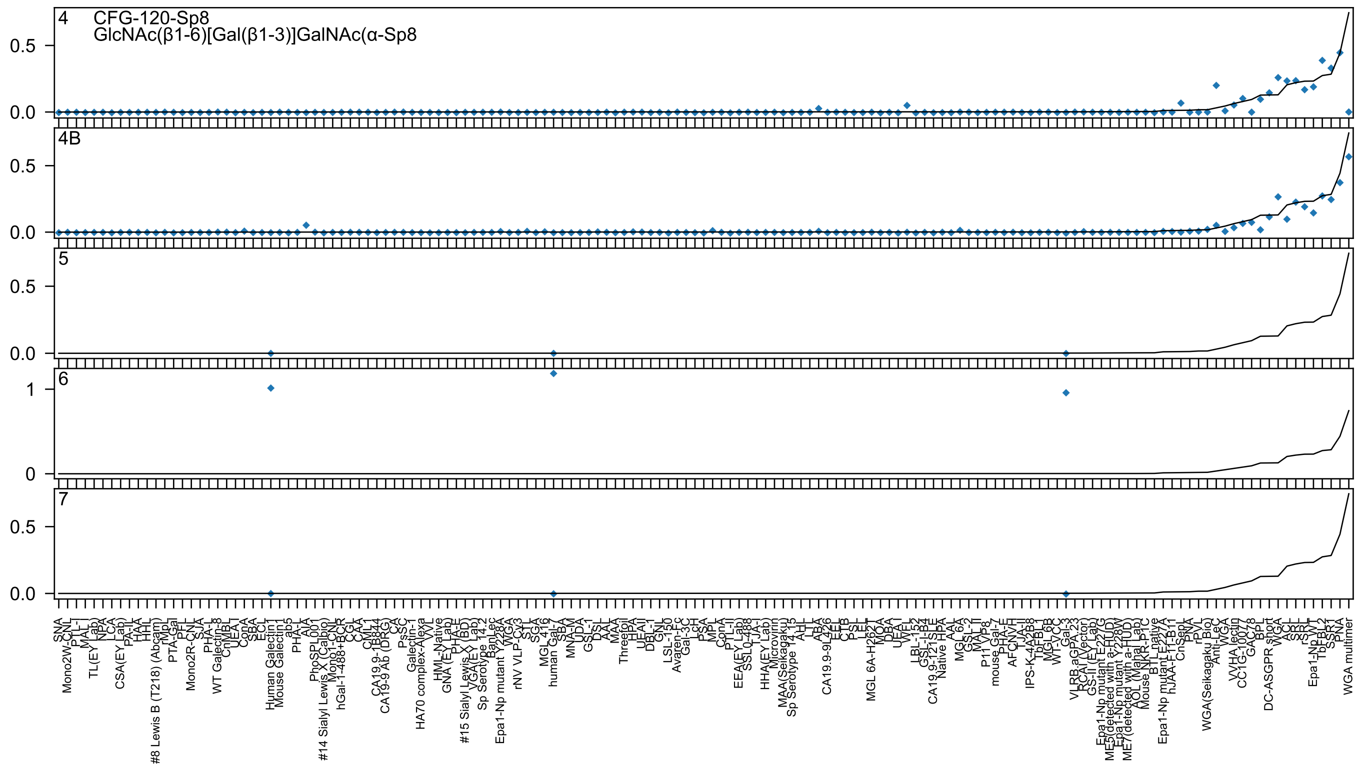

4
CFG-120-Sp8
GlcNAc(β1-6)[Gal(β1-3)]GalNAc(α-Sp8
4B
5
6
7

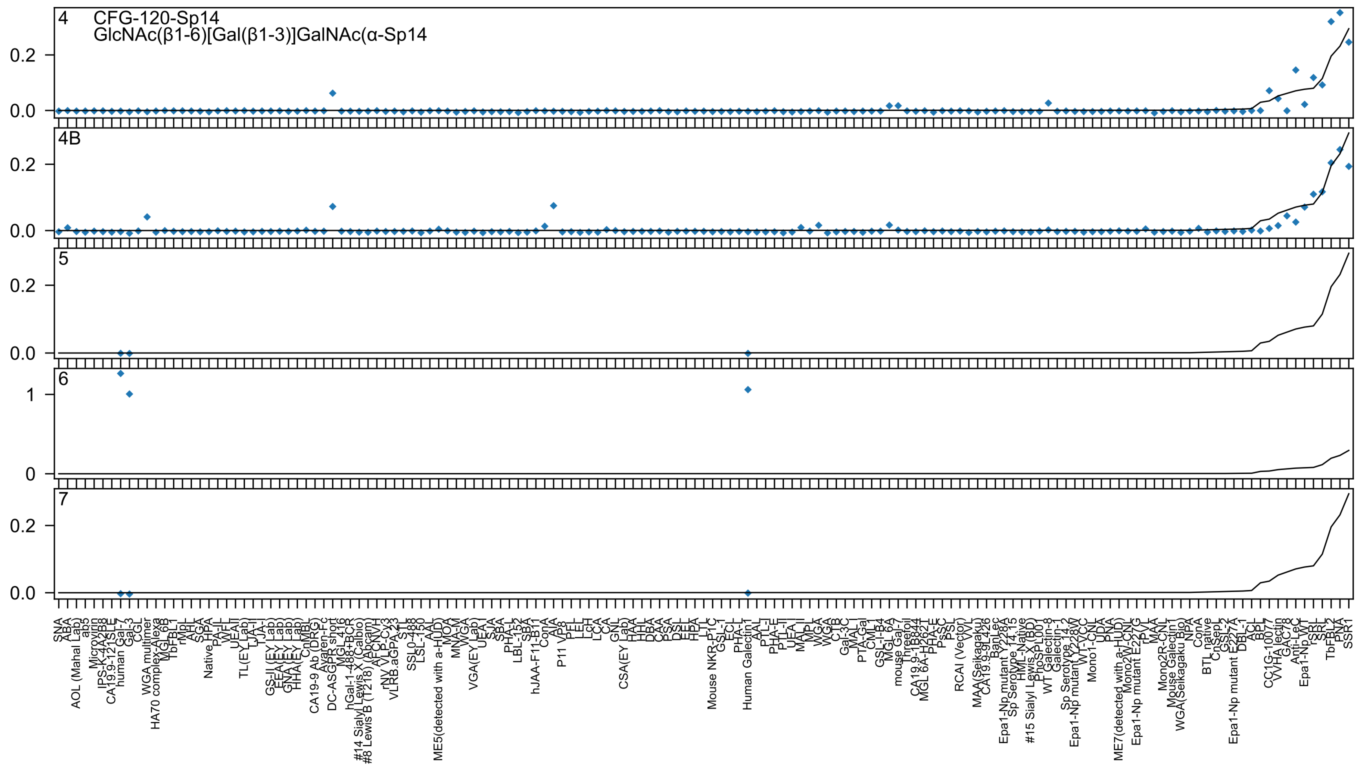
4 CFG-120-Sp14
GlcNAc(β1-6)[Gal(β1-3)]GalNAc(α-Sp14
4B
5
6
7

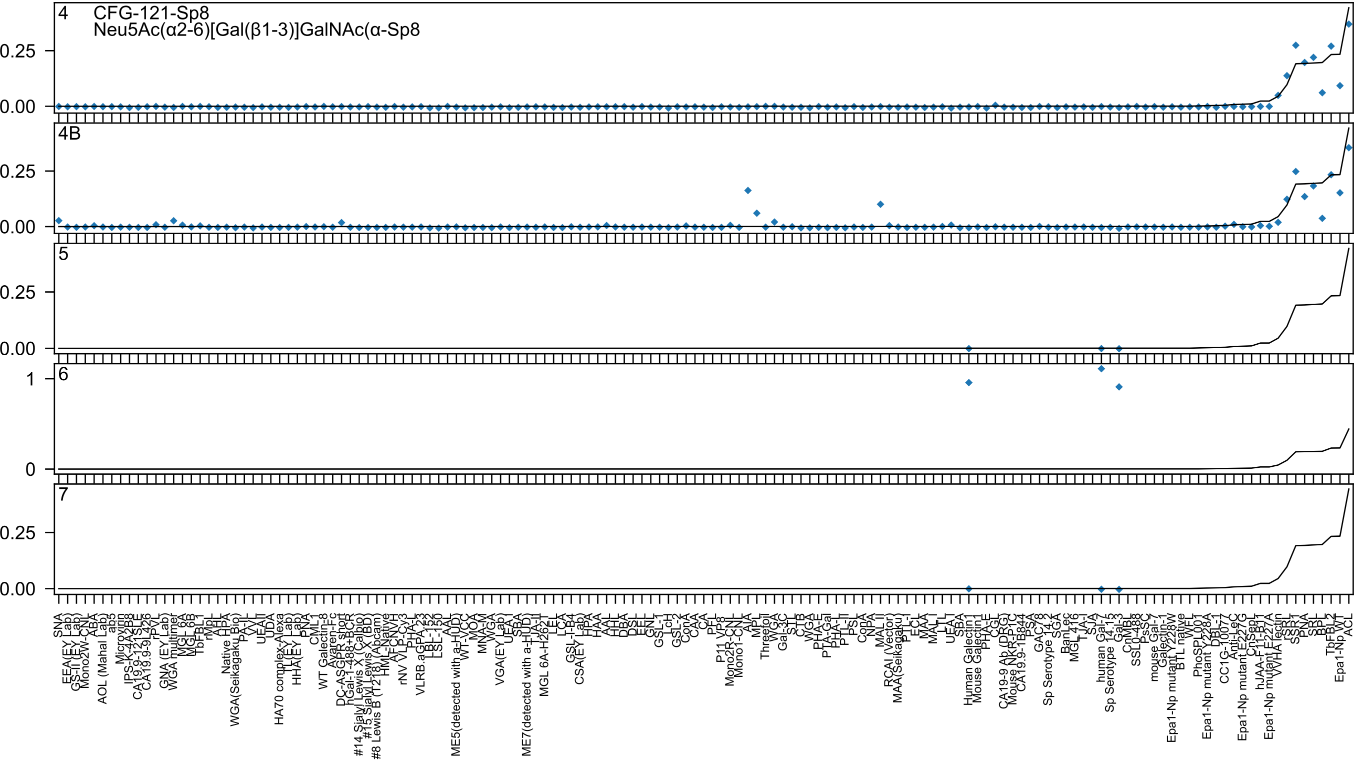
4
CFG-121-Sp8
Neu5Ac(α2-6)[Gal(β1-3)]GalNAc(α-Sp8
4B
5
6
7

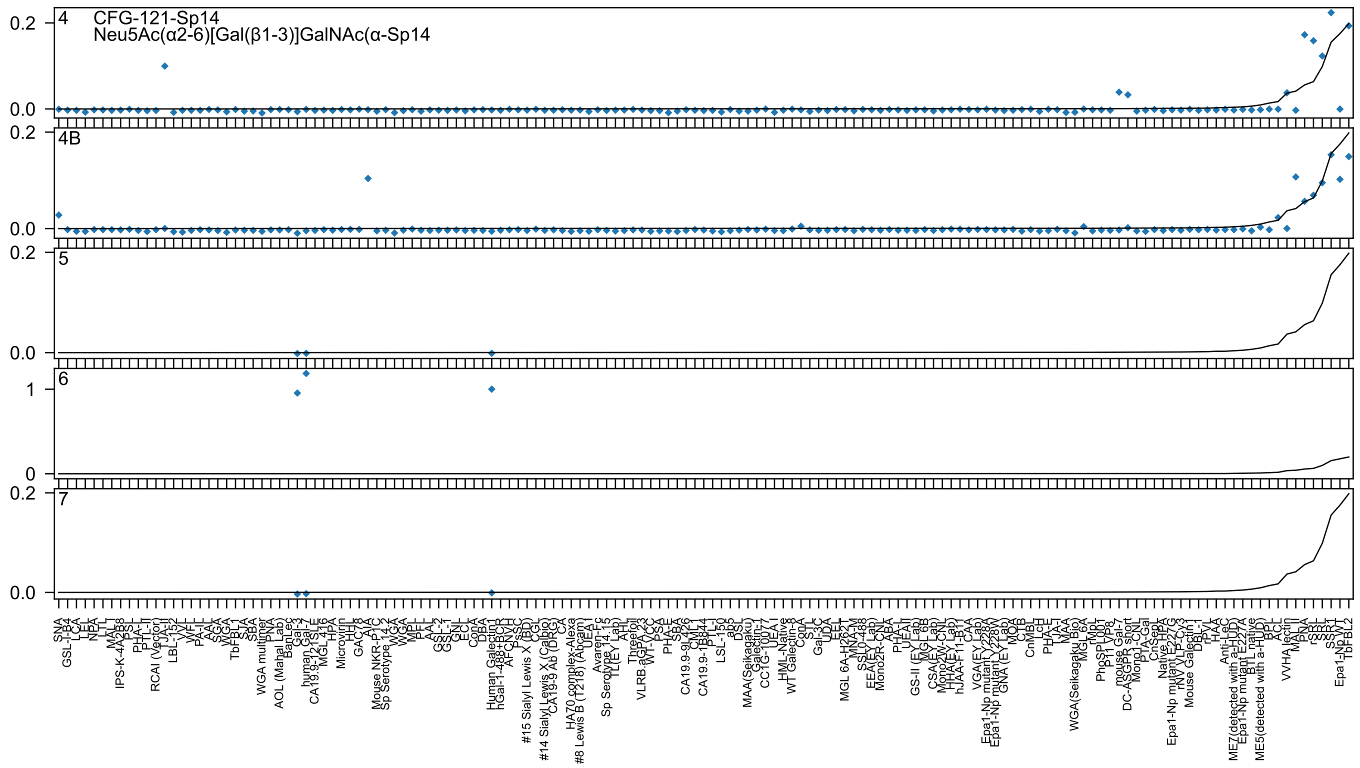
4
CFG-121-Sp14
Neu5Ac(α2-6)[Gal(β1-3)]GalNAc(α-Sp14
4B
5
6
7

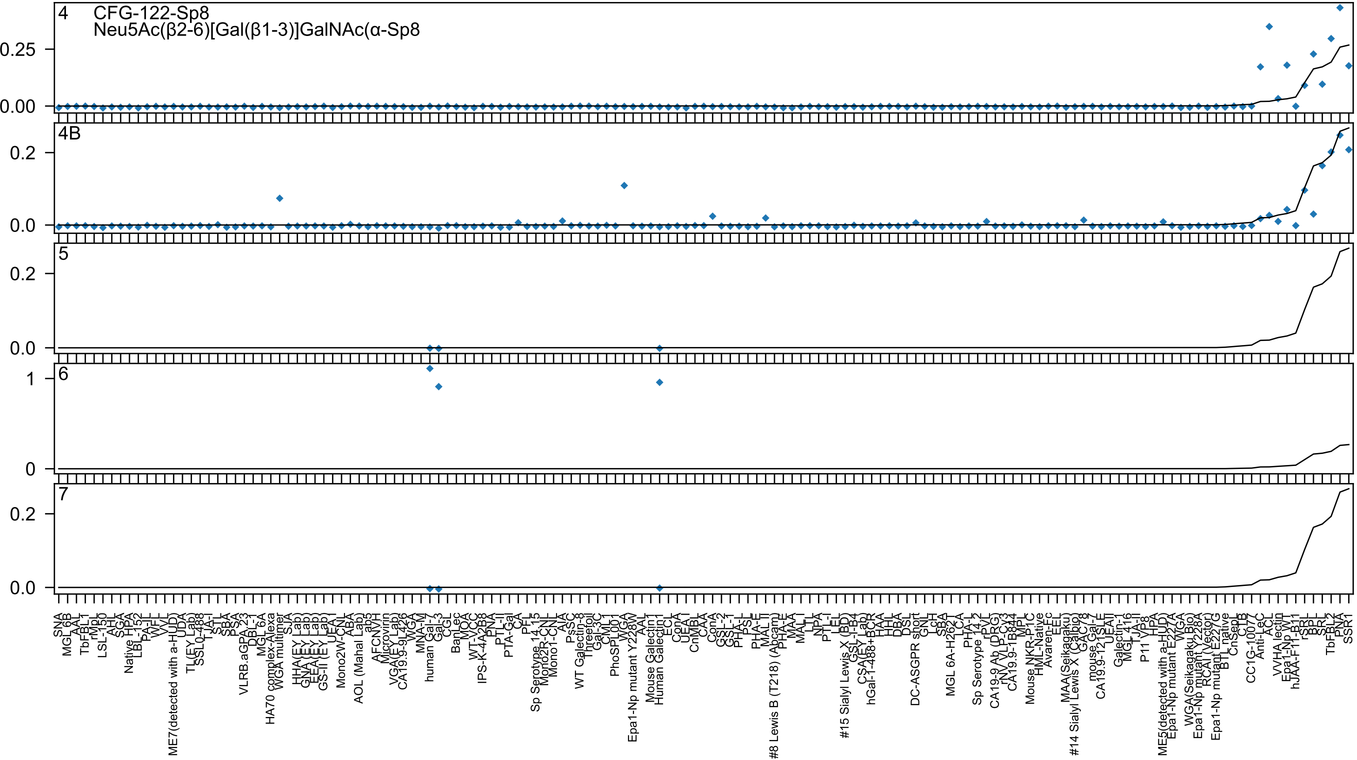

4
CFG-122-Sp8
Neu5Ac(β2-6)[Gal(β1-3)]GalNAc(α-Sp8
4B
5
6
7

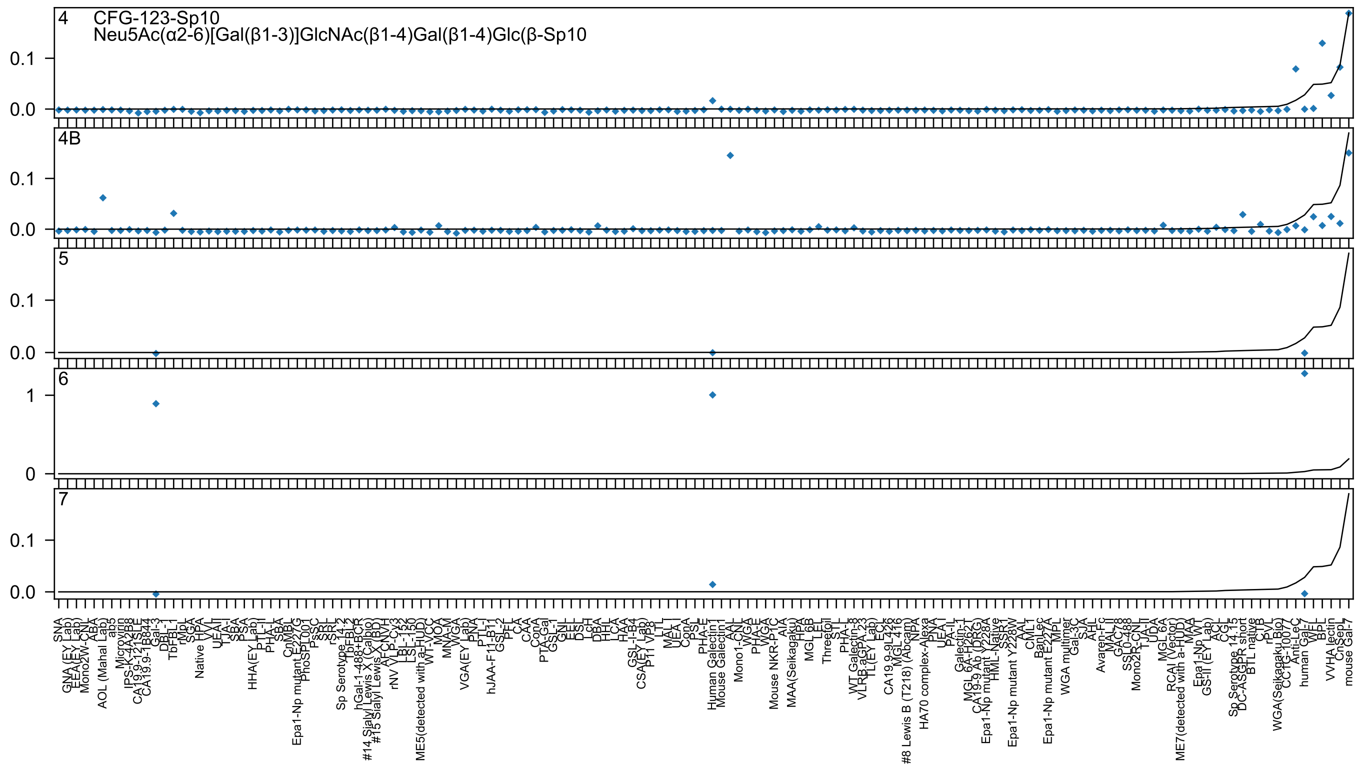

4 CFG-123-Sp10
Neu5Ac(α2-6)[Gal(β1-3)]GlcNAc(β1-4)Gal(β1-4)Glc(β-Sp10
4B
5
6
7

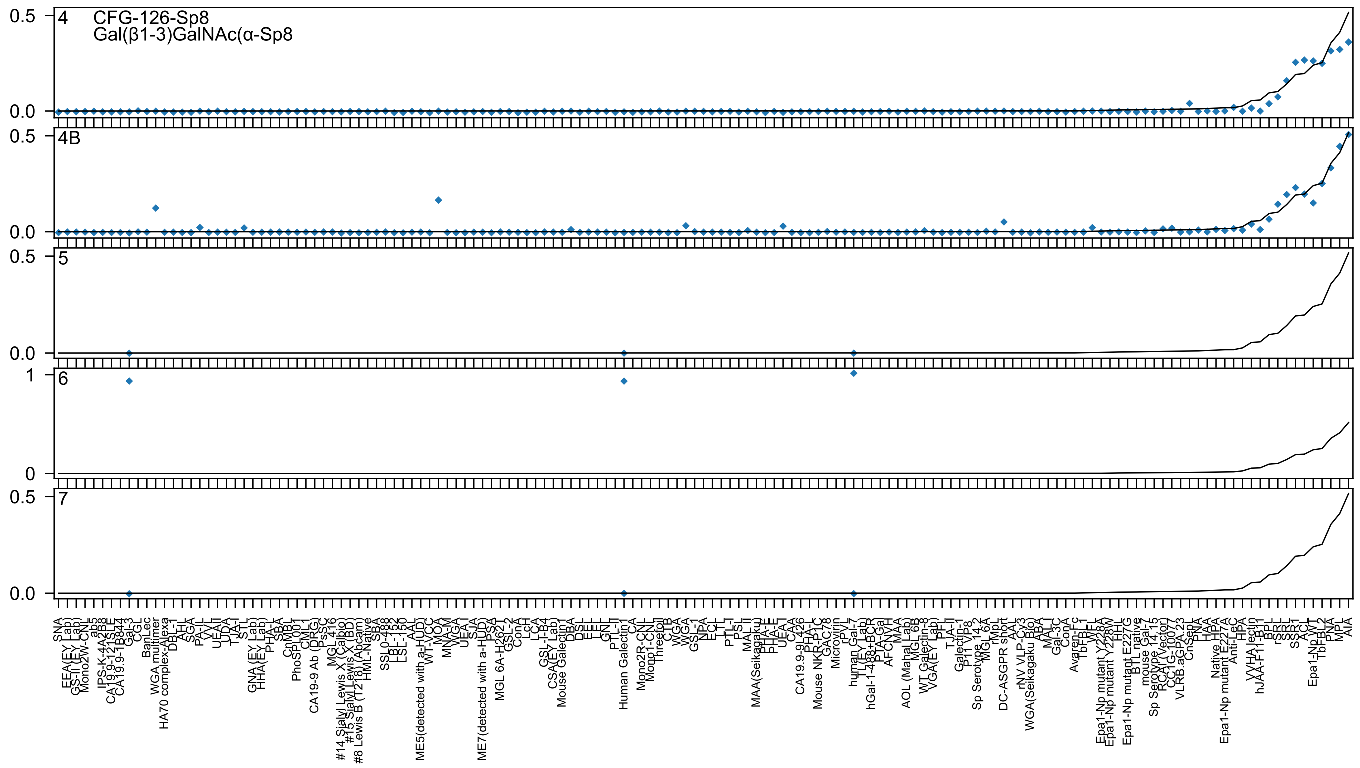

CFG-126-Sp8
Gal(β1-3)GalNAc(α-Sp8

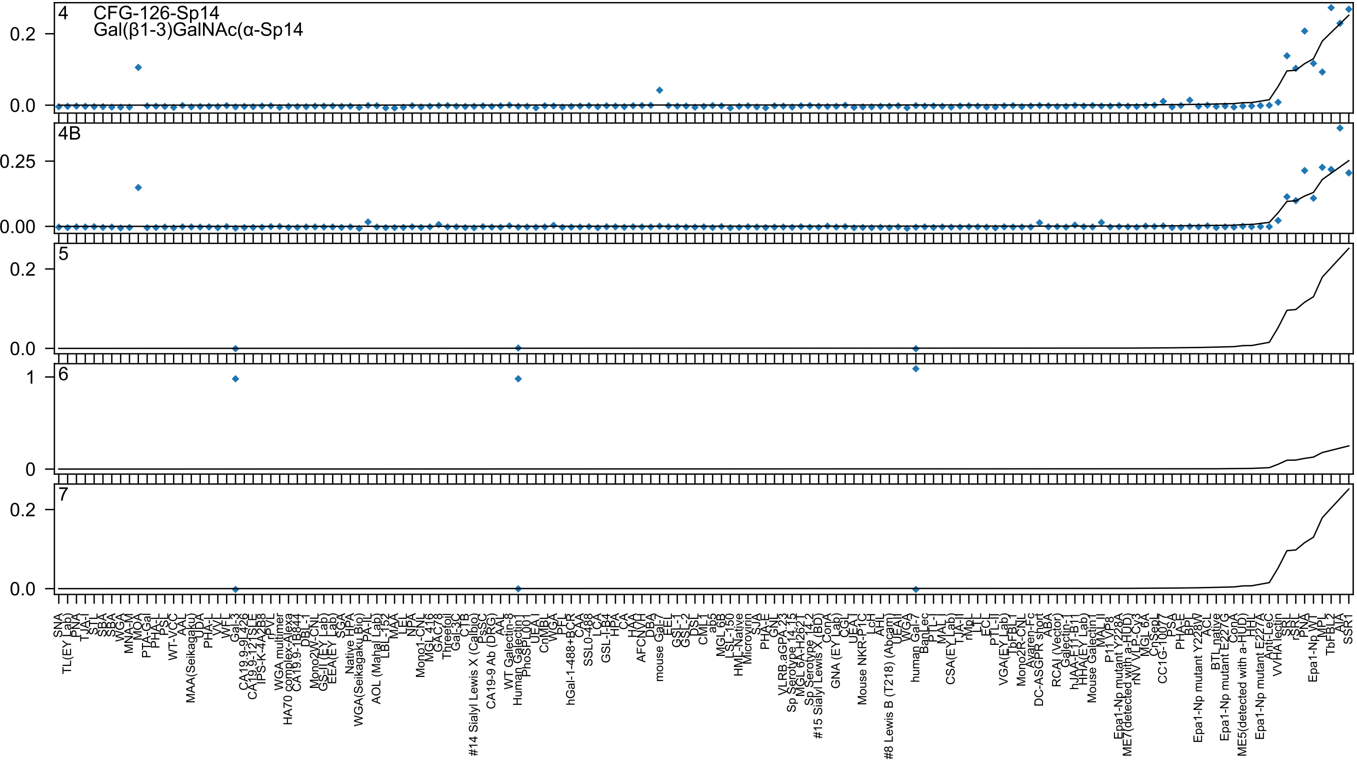

4
CFG-126-Sp14
Gal(β1-3)GalNAc(α-Sp14
0.2
0.0
4B
0.25
0.00
5
0.2
0.0
6
1
0
7
0.2
0.0

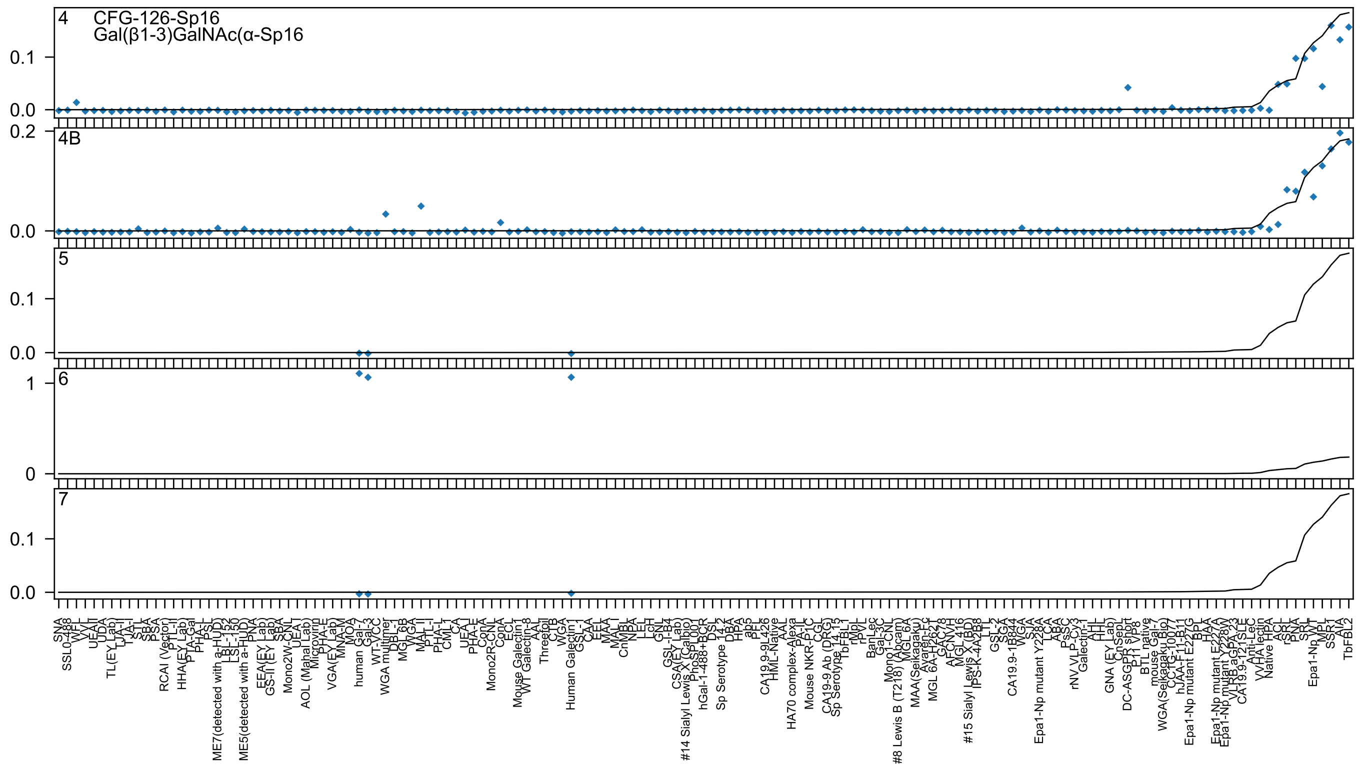
4
CFG-126-Sp16
Gal(β1-3)GalNAc(α-Sp16
0.1
0.0
0.2
4B
0.0
5
0.1
0.0
6
1
0
7
0.1
0.0
SNA
SSL0-488
WFL
VVL
UEA-I
UDA
TL(EY Lab)
TJA-II
TJA-I
STL
SBA
PSA
RCAI (Vector)
PTL-II
HHA(EY Lab)
PTA-Gal
PHA-L
PSL
ME7(detected with a-HUD)
LBL-152
LSL-150
ME5(detected with a-HUD)
PNA
EEA(EY Lab)
GS-II (EY Lab)
SBA
Mono2W-CNL
UEA-I
AOL (Manal Lab)
Microvirin
PHA-E
VGA(EY Lab)
MNA-M
MOA
human Gal-7
Gal-3
WT-YCC
WGA multimer
DBL-1
MGL 6B
WGA
MAL-II
PTL-I
PHA-L
CML-1
CA
UEA-I
PHA-E
ConA
Mono2R-CNL
ConA
ECL
Mouse Galectin1
WT Galectin-8
AAL
Threefoil
CTB
WGA
Human Galectin1
GSL-1
CAA
EEL
MAA
MAL-I
CnMBL
NPA
LEL
LcH
GNL
GSL-I-B4
CSA(EY Lab)
#14 Sialyl Lewis X (Calbio)
PhoSL
hGal-1-488+BCR
DSL
Sp Serotype 14.2
DBA
HPA
ab5
PFL
CA19.9-9L426
HML-Native
AAL
HA70 complex-Alexa
PA-IL
Mouse NKR-P1C
CGL
CA19-9 Ab (DRG)
Sp Serotype 14.15
TbFBL1
rMpL
rPVL
BanLec
Gal-3C
Mono1-CNL
#8 Lewis B (T218) (Abcam)
MGL 6A
MAA(Seikagaku)
Avaren-Fc
MGL 6A-H262T
GAC78
AFCNVH
MGL 416
#15 Sialyl Lewis X (BD)
IPS-K-4A2B8
LTL
GSL-2
SGA
CA19.9-1B844
WGA
SJA
Epa1-Np mutant Y228A
LCA
ABA
PsSC
rNV VLP-Cy3
Galectin-1
AHL
IHL
GNA (EY Lab)
CnSepL
DC-ASGPR short
P11 VP8
BTL native
mouse Gal-7
WGA(Seikagaku Bio)
CCTG-1007
hJAA-F11-B11
Epa1-Np mutant E227G
BPL
HAA
Epa1-Np mutant E227A
Epa1-Np mutant Y228W
VLRB aGPA 23
CA19.9-121SLE
Anti-LeC
VVHA lectin
Native HPA
ACL
rSRL
PNA
SRL
Epa1-Np WT
MPL
SSR1
AIA
TbFBL2

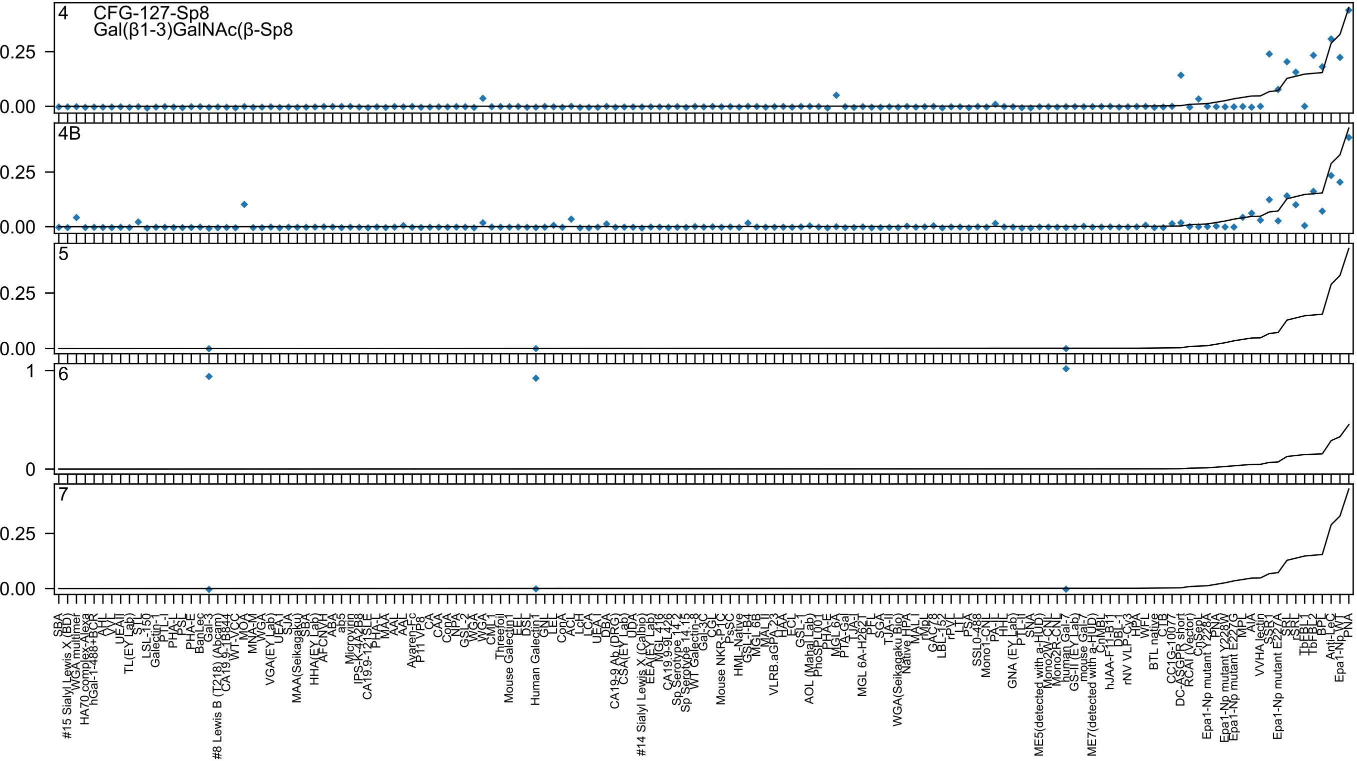
4
CFG-127-Sp8
Gal(β1-3)GalNAc(β-Sp8
4B
5
6
7

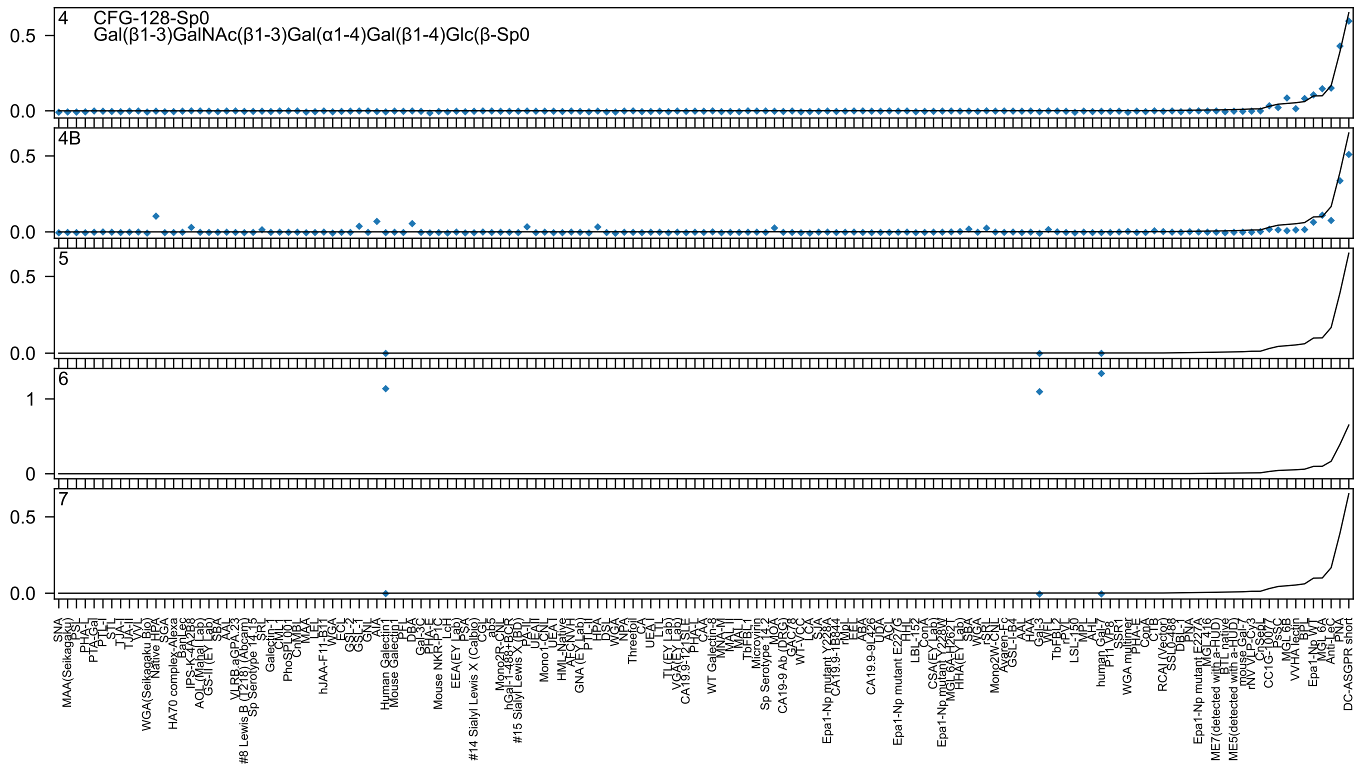
4
CFG-128-Sp0
Gal(β1-3)GalNAc(β1-3)Gal(α1-4)Gal(β1-4)Glc(β-Sp0
4B
5
6
7
0.0
0.5
0
1
SNA
MAA(Seikagaku)
PSL
PHA-L
PTA-Gal
PTL-I
STL
TJA-I
TJA-II
VVL
WGA(Seikagaku Bio)
Native HPA
SGA
HA70 complex-Alexa
BanLec
IPS-K-4A2B8
AOL (Mahal Lab)
GS-II (EY Lab)
SBA
AAL
VLRB aGPA.23
#8 Lewis B (T218) (Abcam)
Sp Serotype 14.15
SRL
Galectin-1
CML-I
PhoSL001
CnMBL
MAA
LEL
hJAA-F11-B11
WGA
ECL
GSL-2
GSL-1
GNL
AIA
Human Galectin1
Mouse Galectin1
PFL
DBA
Gal-3C
PHA-E
Mouse NKR-P1C
LcH
EEA(EY Lab)
PSA
#14 Sialyl Lewis X (Calbio)
CGL
ab5
Mono2R-CNL
hGal-1-488+BCR
#15 Sialyl Lewis X (BD)
PA-II
UEAII
Mono1-CNL
UEA-I
HML-Native
AFCNVH
GNA (EY Lab)
PTL-II
HPA
DSL
WGA
NPA
Threefoil
CA
UEA-I
LTL
TL(EY Lab)
VGA(EY Lab)
CA19.9-121SLE
PHA-L
CAA
WT Galectin-8
MNA-M
MAL-II
MAL-I
TbFBL1
Microvirin
Sp Serotype 14.2
MOA
CA19-9 Ab (DRG)
GAC78
WT-VCC
LCA
SJA
Epa1-Np mutant Y228A
CA19.9-1B844
rMpL
EEL
ABA
CA19.9-9L426
UDA
ACL
Epa1-Np mutant E227G
HHL
LBL-152
ConA
CSA(EY Lab)
Epa1-Np mutant Y228W
MGL 6A-H262T
HHA(EY Lab)
SBA
WGA
rSRL
Mono2W-CNL
Avaren-Fc
GSL-I-B4
AAL
HAA
Gal-3
WFL
TbFBL2
rPVL
LSL-150
MPL
AHL
human Gal-7
P11
VP8
SSR1
WGA multimer
PHA-E
ConA
CTB
RCAI (Vector)
SSL0-488
DBL-1
PNA
Epa1-Np mutant E227A
MGL 416
ME7(detected with a-HUD)
BTL native
ME5(detected with a-HUD)
mouse Gal-7
rNV VLP-Cy3
CnSepL
CC1G-10077
PsSC
MGL 6B
VVHA lectin
BPL
Epa1-Np WT
MGL 6A
Anti-LeC
PNA
DC-ASGPR short

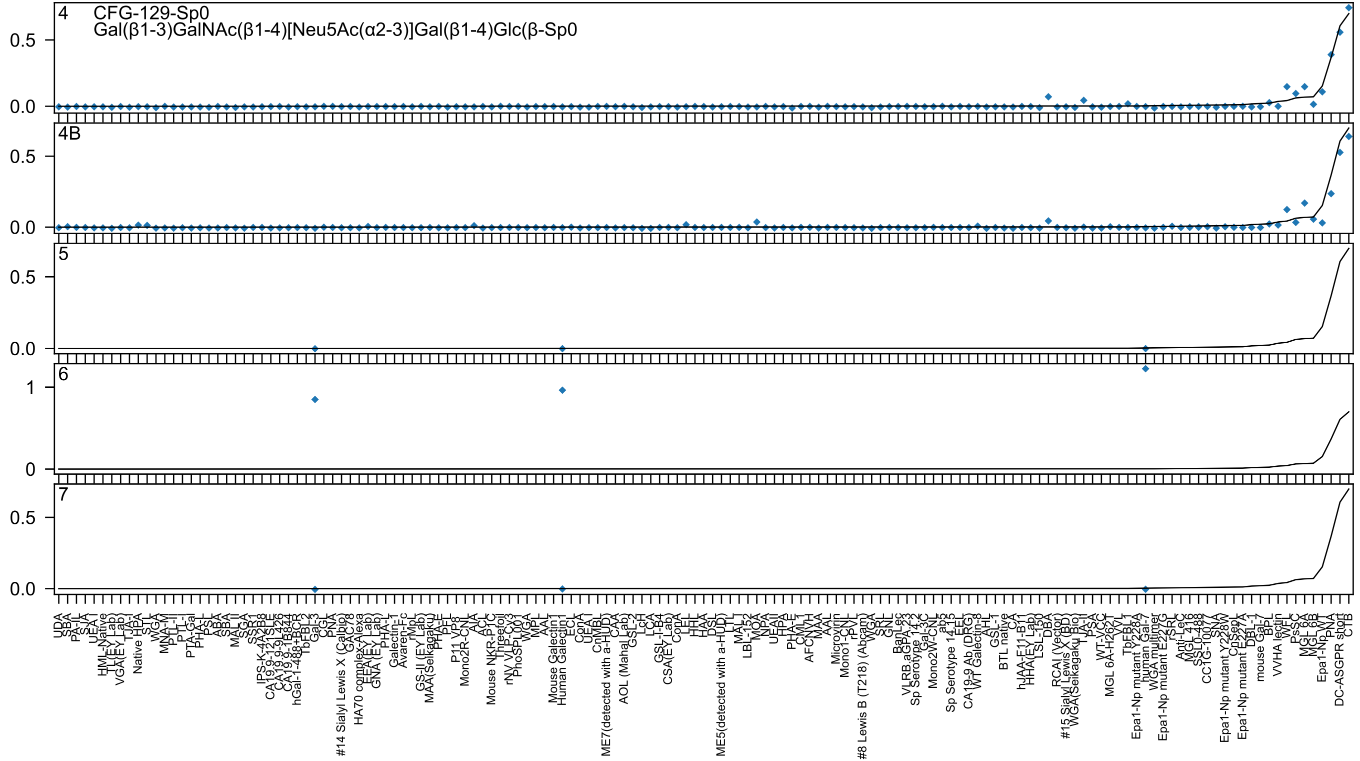

4
CFG-129-Sp0
Gal(β1-3)GalNAc(β1-4)[Neu5Ac(α2-3)]Gal(β1-4)Glc(β-Sp0
4B
5
6
7

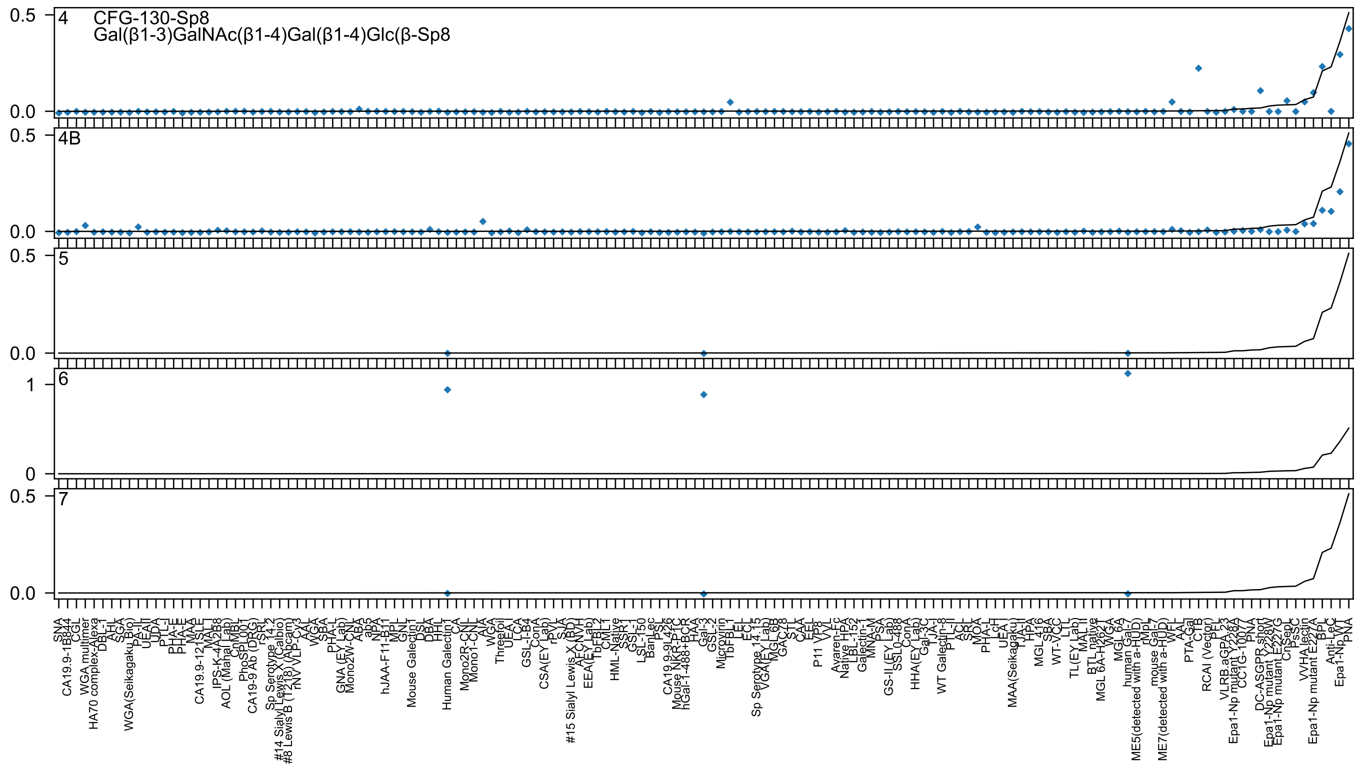

CFG-130-Sp8
Gal(β1-3)GalNAc(β1-4)Gal(β1-4)Glc(β-Sp8
4
4B
5
6
7
0.5
0.0
1
0
SNA
CA19.9-1B844
CGL
WGA multimer
HA70 complex-Alexa
DBL-I
AHL
SGA
WGA(Seikagaku Bio)
PA-II
UEAII
UDA
PTL-I
PHA-E
PHA-E
MAA
CA19.9-121SLE
MAL-I
IPS-K-4A2B8
AOL (Mahal Lab)
CnMBL
PhoSL001
CA19-9 Ab (DRG)
rSRL
Sp Serotype 14.2
#14 Sialyl Lewis X (Calbio)
#8 Lewis B (T218) (Abcam)
rNV VLP-Cy3
AAL
WGA
SBA
PHA-L
GNA (EY Lab)
Mono2W-CNL
ABA
ab5
NPA
hJAA-F11-B11
MPL
GNL
Mouse Galectin1
DSL
DBA
HHL
Human Galectin1
CA
Mono2R-CNL
Mono1-CNL
AIA
WGA
Threefoil
UEA-I
LCA
GSL-I-B4
ConA
CSA(EY Lab)
rPVL
SJA
#15 Sialyl Lewis X (BD)
AECNVH
EEA(EY Lab)
TbFBL2
CML1
HML-Native
SSR1
GSL-I
LSL-150
BanLec
PSL
CA19.9-9L426
Mouse NKR-P1C
hGal-1-488+BCR
HAA
Gal-3
GSL-2
Microvirin
TbFBL1
LEL
ECL
Sp Serotype 14.15
VGA(EY Lab)
MGL 6B
GAC78
STL
CAA
EEL
P11 VP8
VVL
Avaren-Fc
Native HPA
LBL-152
Galectin-1
MNA-M
PSA
GS-II (EY Lab)
SSL0-488
ConA
HHA(EY Lab)
Gal-3C
TJA-I
WT Galectin-8
PTL-II
ACL
SRL
MOA
PHA-L
LcH
UEA-I
MAA(Seikagaku)
TJA-II
HPA
MGL 416
SBA
WT-VCC
LTL
TL(EY Lab)
MAL-II
BTL native
MGL 6A-H262T
WGA
MGL 6A
human Gal-7
ME5(detected with a-HUD)
rMpL
mouse Gal-7
ME7(detected with a-HUD)
WFL
AAL
PTA-Gal
CTB
RCAI (Vector)
PFL
VLRB.aGPA.23
Epa1-Np mutant Y228A
CC1G-10077
PNA
DC-ASGPR short
Epa1-Np mutant Y228W
Epa1-Np mutant E227G
CnSep
PsSC
VVHA lectin
Epa1-Np mutant E227A
BPL
Anti-LeC
Epa1-Np WT
PNA

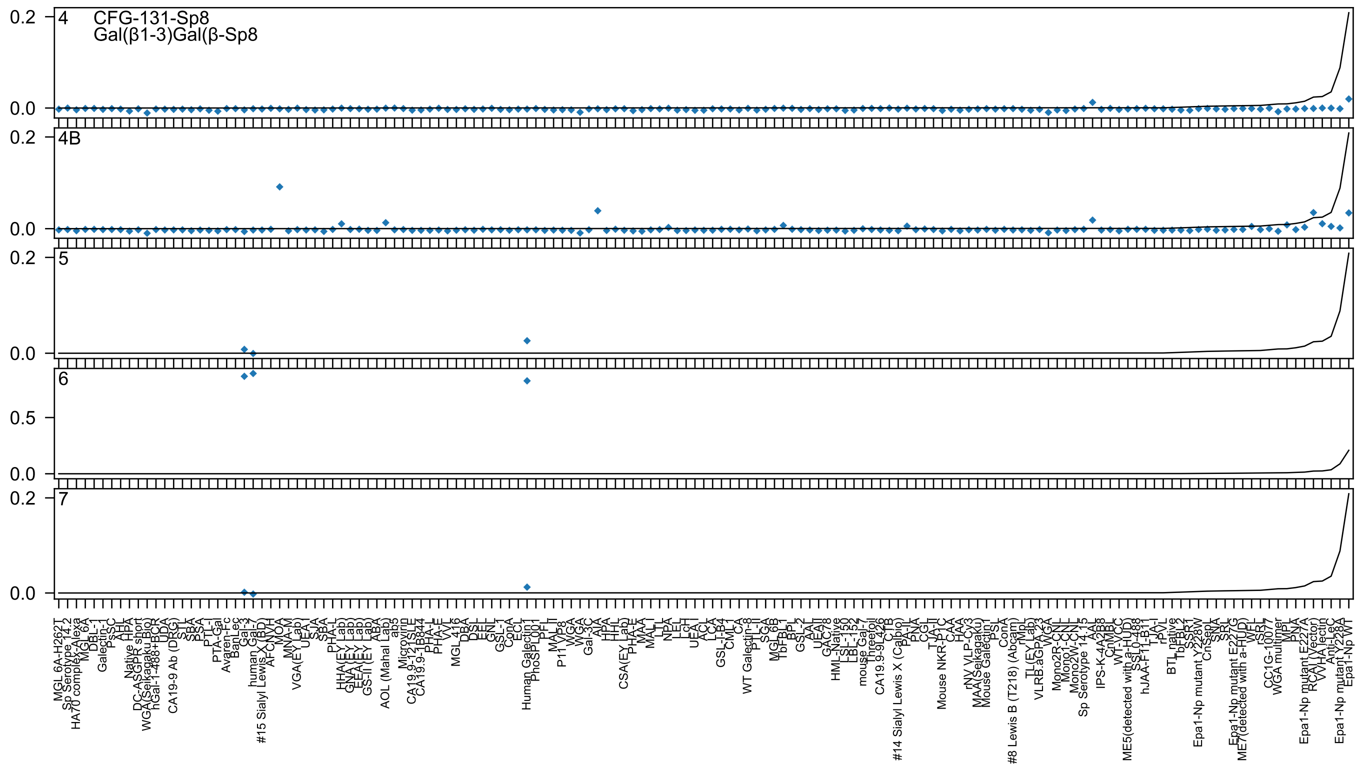

4
CFG-131-Sp8
Gal(β1-3)Gal(β-Sp8
4B
5
6
7
0.2
0.0
0.5
MGL 6A-H262T
Sp Serotype 14.2
HA70 complex-Alexa
MGL 6A
DBL-1
Galectin-1
PsSC
AHL
Native HPA
DC-ASGPR short
WGA(Seikagaku Bio)
hGal-1-488+BCR
UDA
CA19-9 Ab (DRG)
STL
SBA
PSA
PTL-I
PTA-Gal
Avaren-Fc
BanLec
Gal-3
human Gal-7
#15 Sialyl Lewis X (BD)
AFCNVH
MOA
MNA-M
VGA(EY Lab)
UEA-I
SJA
SBA
PHA-L
HHA(EY Lab)
GNA(EY Lab)
EEA(EY Lab)
GS-II (EY Lab)
ABA
AOL (Mahal Lab)
ab5
Microvirin
CA19.9-121SLE
CA19.9-1B844
PHA-L
PHA-E
VVL
MGL 416
DBA
DSL
EEL
GNL
GSL-1
ConA
ECL
Human Galectin1
PhoSL001
PFL
MAL-II
P11 VP8
WGA
WGA
Gal-3C
AIA
HPA
HHL
CSA(EY Lab)
PHA-E
MAA
MAL-I
LTL
NPA
LEL
LcH
UEA-I
ACL
LCA
GSL-I-B4
CML-1
CA
WT Galectin-8
PTL-II
SGA
MGL 6B
TbFBL1
BPL
GSL-2
AAL
UEAII
GAC78
HML-Native
LSL-150
LBL-152
mouse Gal-7
Threefoil
CA19.9-9L426
CTB
#14 Sialyl Lewis X (Calbio)
PA-IL
PNA
CGL
TJA-II
Mouse NKR-P1C
CAA
HAA
rNV VLP-Cy3
MAA(Seikagaku)
Mouse Galectin1
PSL
ConA
#8 Lewis B (T218) (Abcam)
rMpL
TL(EY Lab)
VLRB.aGPA.23
WGA
Mono2R-CNL
Mono1-CNL
Mono2W-CNL
Sp Serotype 14.15
AAL
IPS-K-4A2B8
CnMBL
WT-VCC
ME5(detected with a-HUD)
SSL0-488
hJAA-F11-B11
TJA-I
rPVL
BTL native
TbFBL2
SSR1
Epa1-Np mutant Y228W
CnSepL
SNA
SRL
Epa1-Np mutant E227G
ME7(detected with a-HUD)
WFL
rSRL
CC1G-10077
WGA multimer
MPL
PNA
Epa1-Np mutant E227A
RCAI (Vector)
VVHA lectin
Anti-LeC
Epa1-Np mutant Y228A
Epa1-Np WT

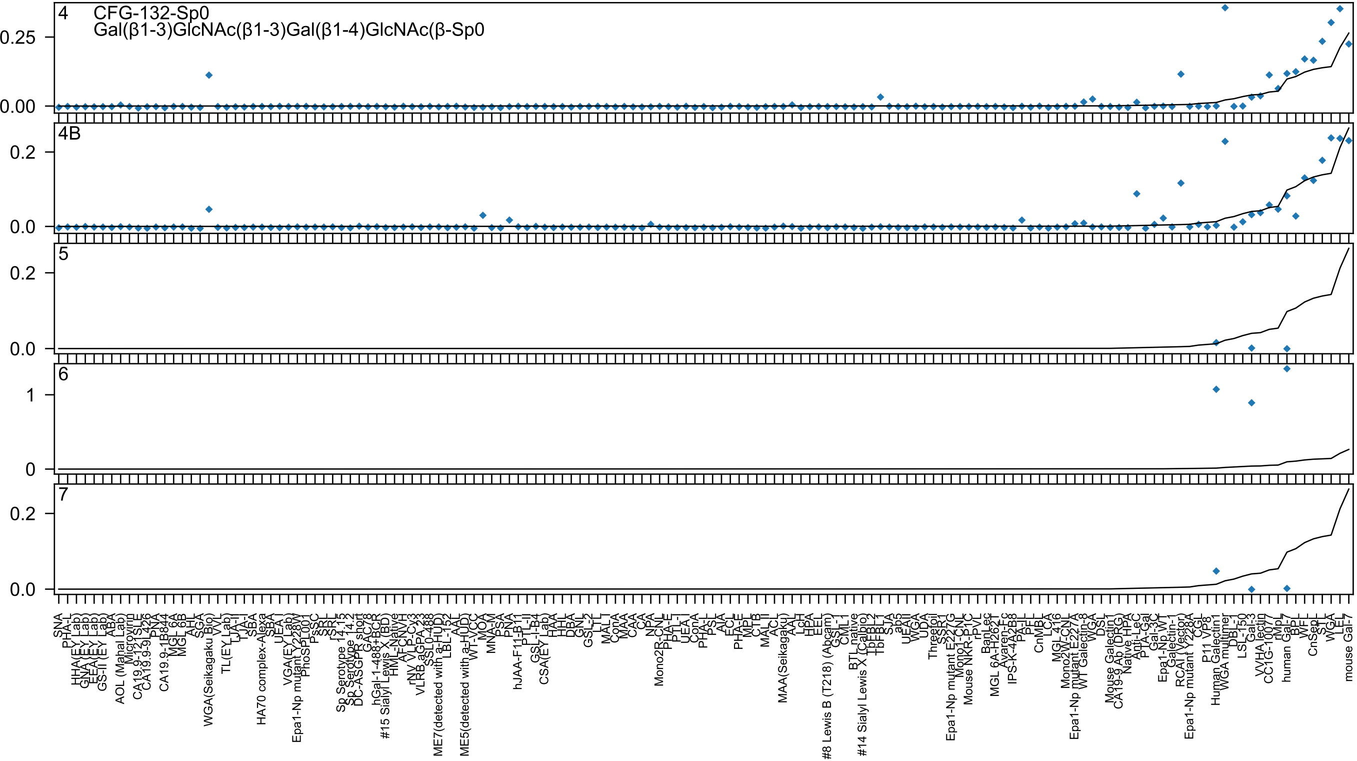

4
CFG-132-Sp0
Gal(β1-3)GlcNAc(β1-3)Gal(β1-4)GlcNAc(β-Sp0
4B
5
6
7

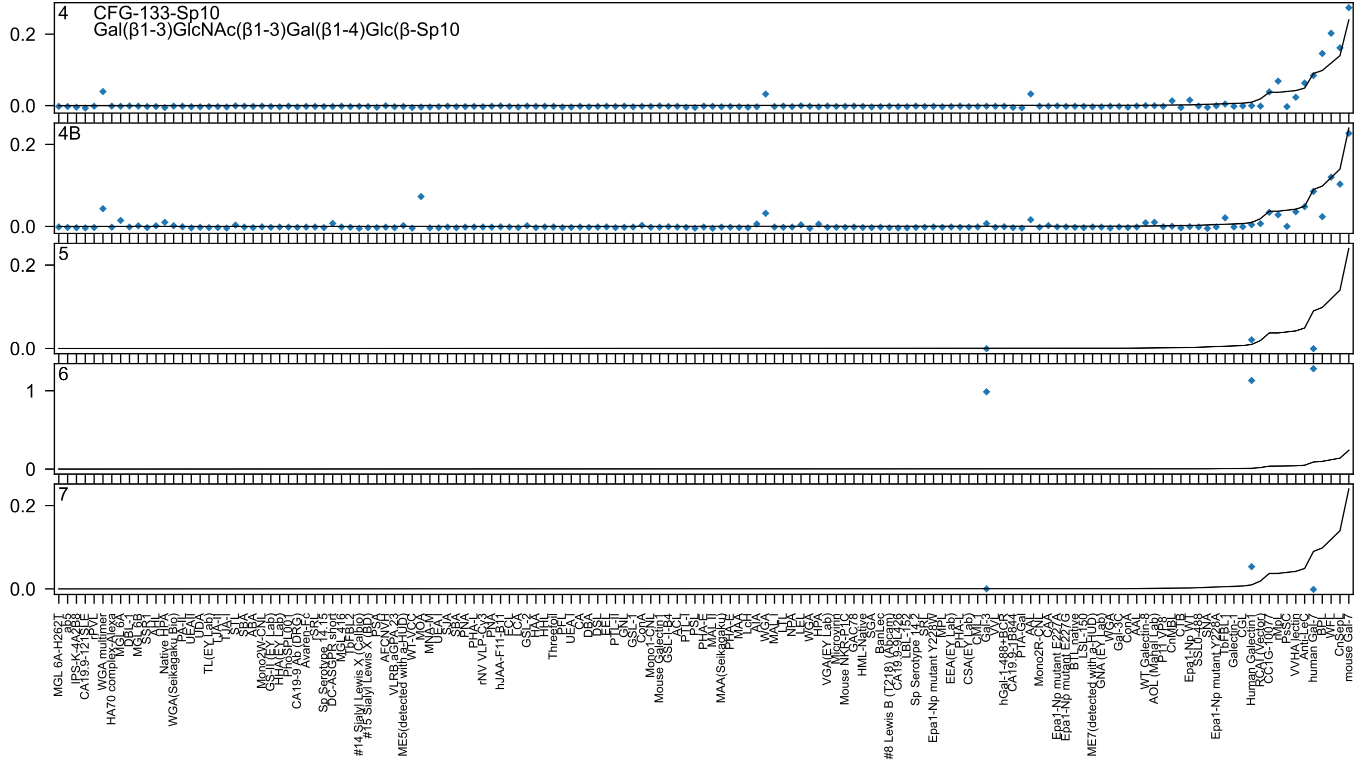
4 CFG-133-Sp10
Gal(β1-3)GlcNAc(β1-3)Gal(β1-4)Glc(β-Sp10
4B
5
6
7

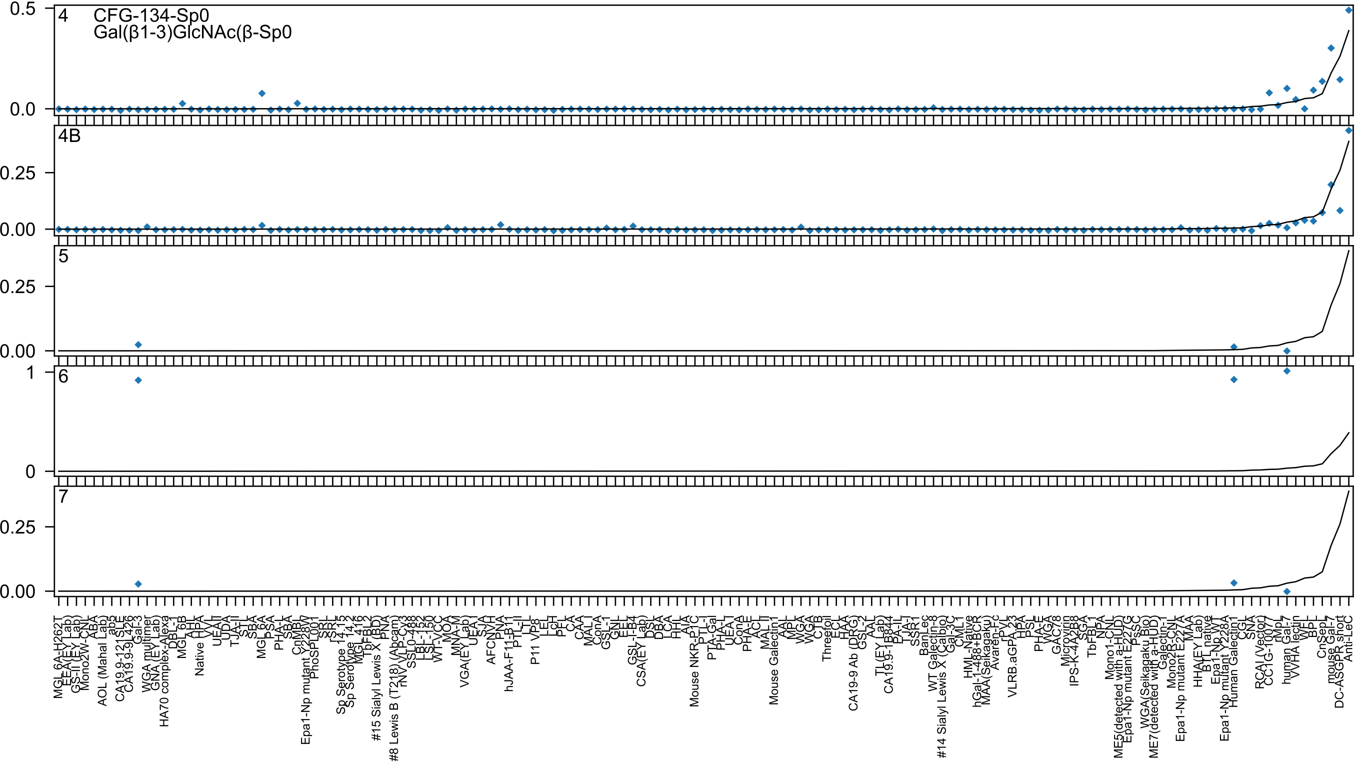
4
CFG-134-Sp0
Gal(β1-3)GlcNAc(β-Sp0
4B
5
6
7
0.5
0.0
0.25
0.00
1
0

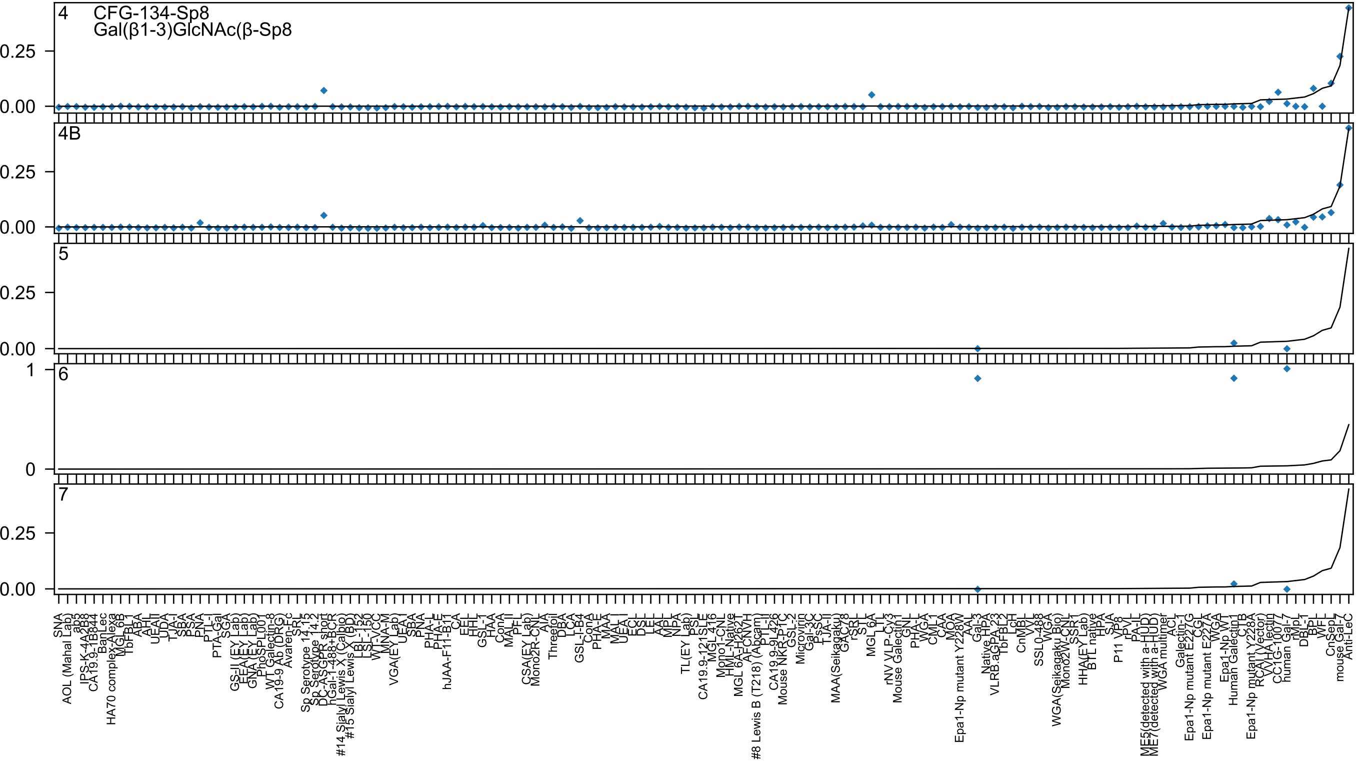
4
CFG-134-Sp8
Gal(β1-3)GlcNAc(β-Sp8
4B
5
6
7
0.25
0.00
1
0

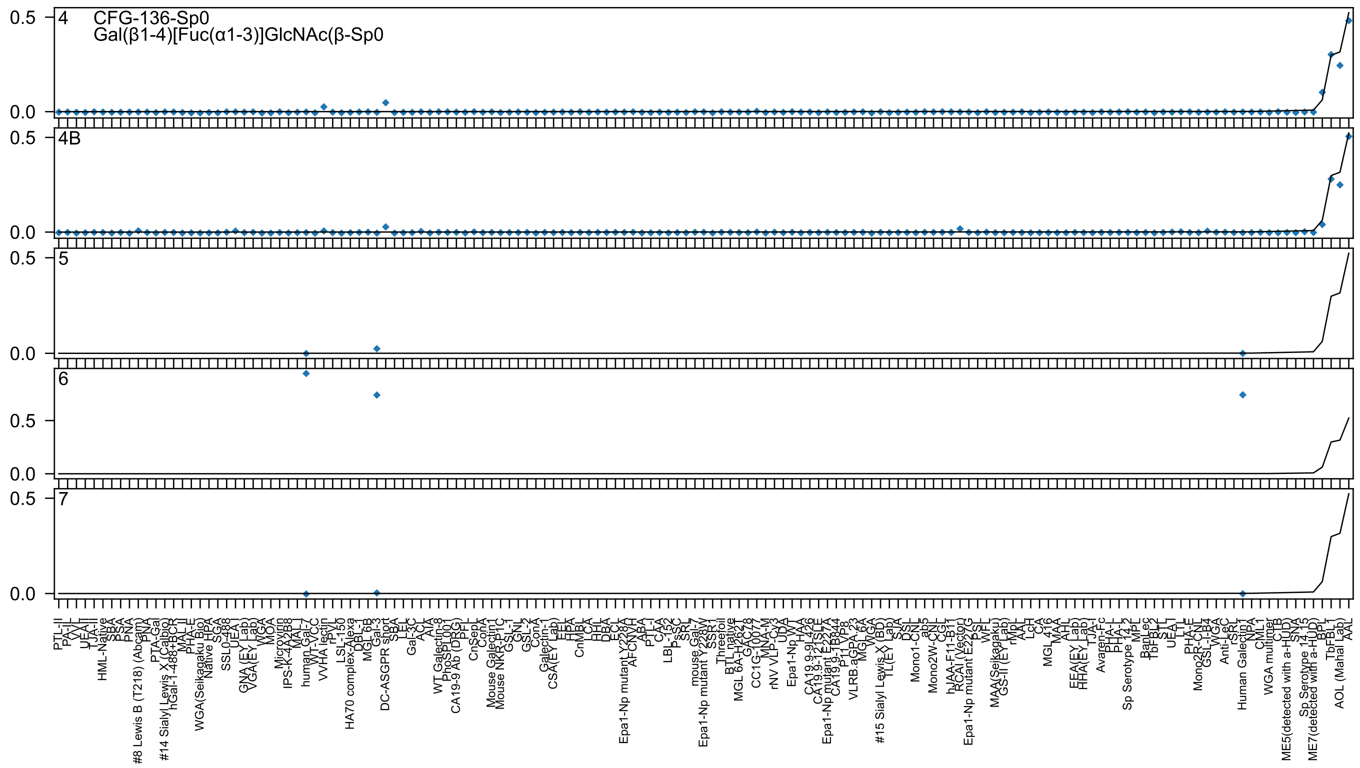

4
CFG-136-Sp0
Gal(β1-4)[Fuc(α1-3)]GlcNAc(β-Sp0
4B
5
6
7

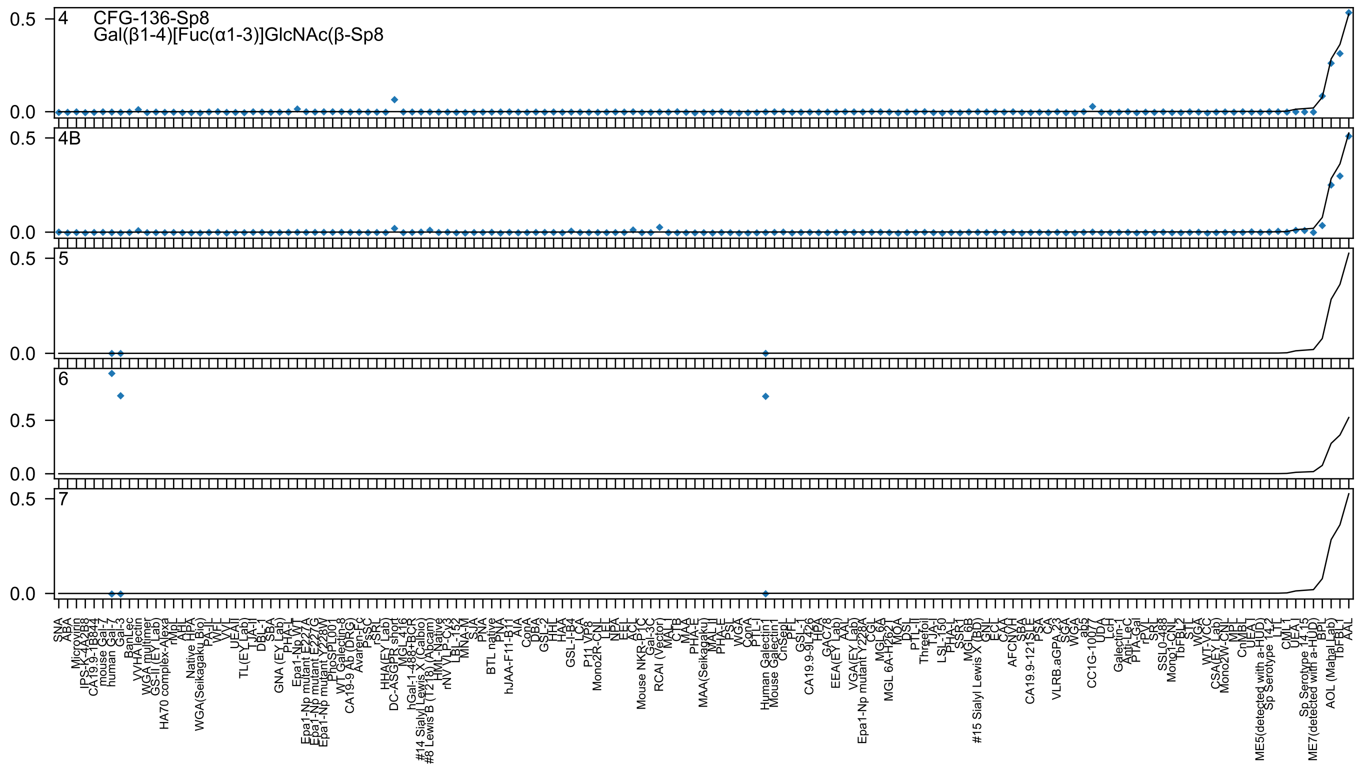
4 CFG-136-Sp8
Gal(β1-4)[Fuc(α1-3)]GlcNAc(β-Sp8
4B
5
6
7
0.5
0.0

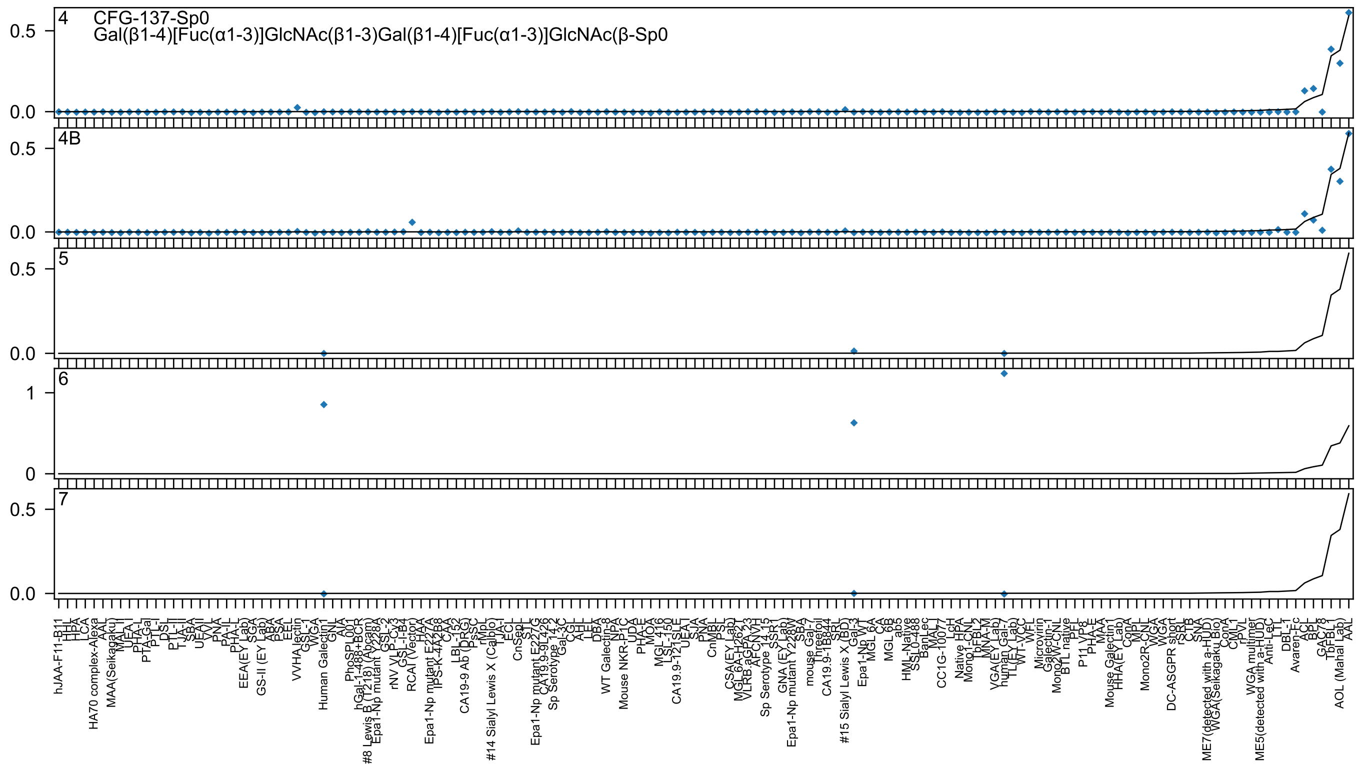

4
CFG-137-Sp0
Gal(β1-4)[Fuc(α1-3)]GlcNAc(β1-3)Gal(β1-4)[Fuc(α1-3)]GlcNAc(β-Sp0
4B
5
6
7
0.5
0.0
1
0
hJAA-F11-B11
HA70 complex-Alexa
MAA(Seikagaku)
EEA(EY Lab)
GS-II (EY Lab)
VVHA lectin
Human Galectin1
PhoSPL001
RCAI (Vector)
CA19-9 Ab (DRG)
#14 Sialyl Lewis X (Calbo)
Sp Serotype 14.2
WT Galectin-8
Mouse NKR-P1C
CA19.9-121SLE
Sp Serotype 14.15
mouse Gal-7
#15 Sialyl Lewis X (BD)
HML-Native
CC1G-10077
Native HPA
Microvirin
Mouse Galectin1
DC-ASGPR short
WGA multimer
Avaren-Fc
AOL (Mahal Lab)
AAL

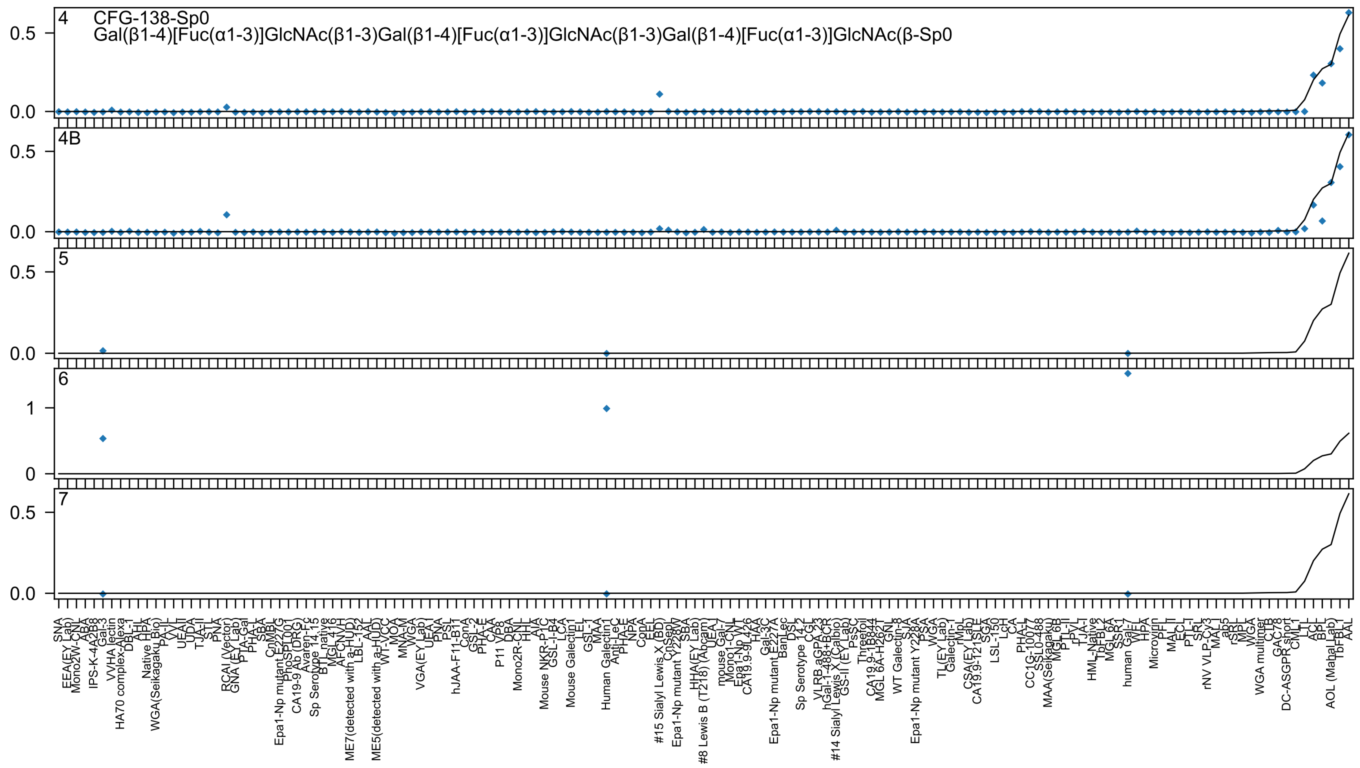

4
CFG-138-Sp0
Gal(β1-4)[Fuc(α1-3)]GlcNAc(β1-3)Gal(β1-4)[Fuc(α1-3)]GlcNAc(β1-3)Gal(β1-4)[Fuc(α1-3)]GlcNAc(β-Sp0
4B
5
6
7

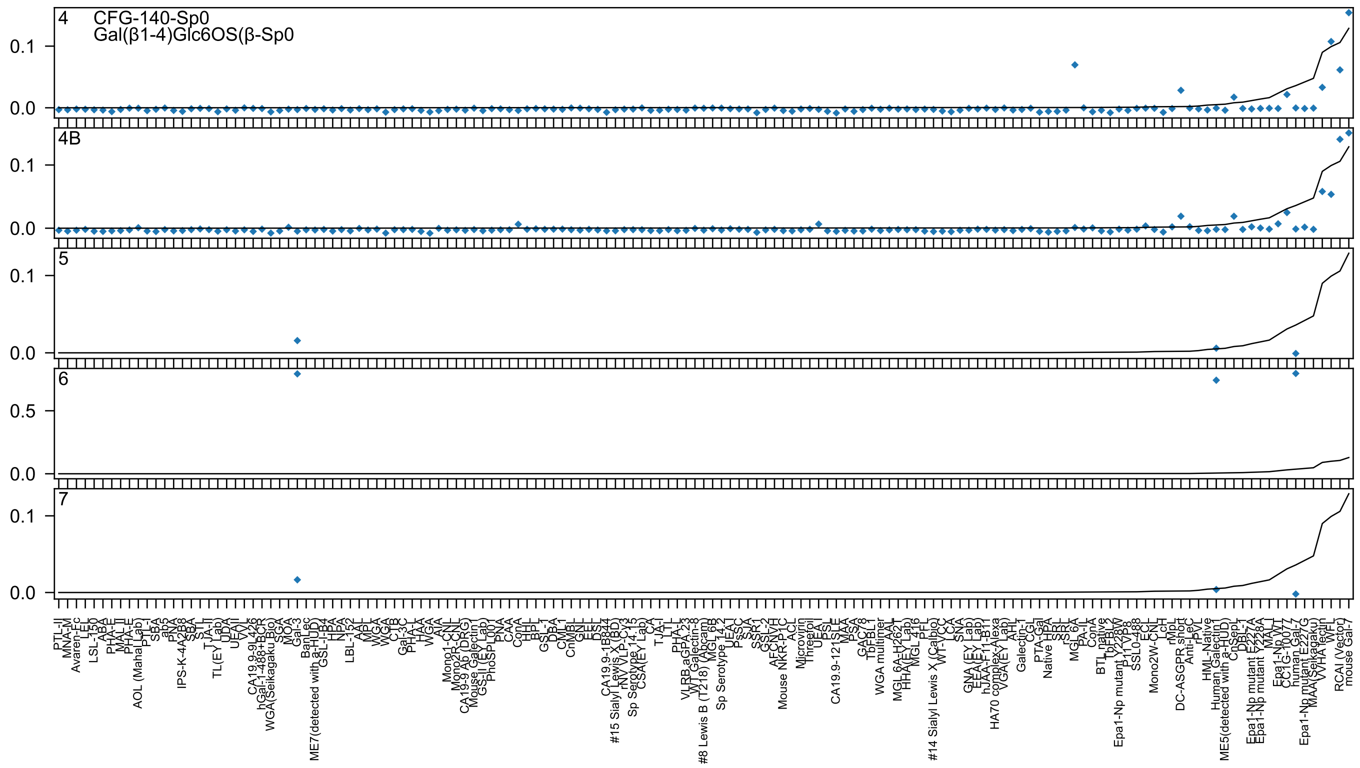

4
CFG-140-Sp0
Gal(β1-4)Glc6OS(β-Sp0
4B
5
6
7
0.1
0.0
0.5

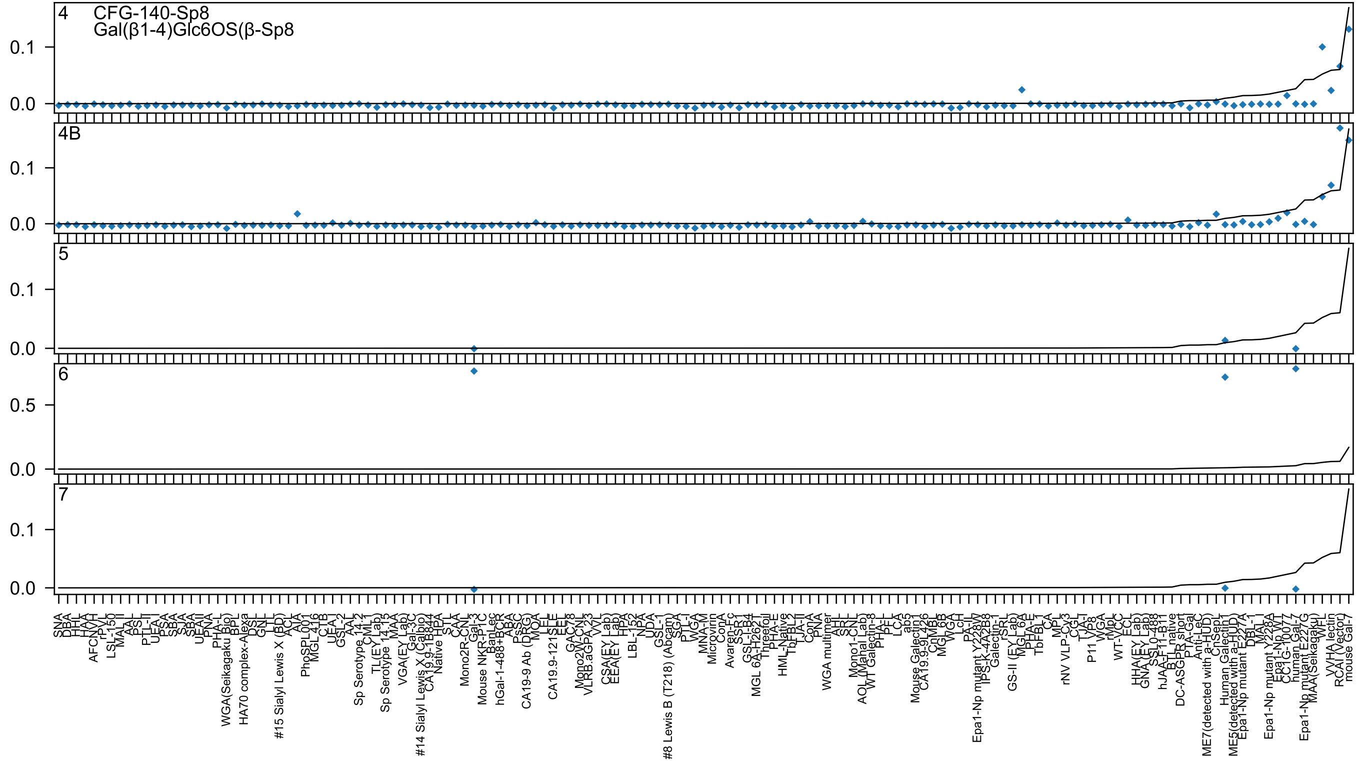

4
CFG-140-Sp8
Gal(β1-4)Glc6OS(β-Sp8
4B
5
6
7
0.1
0.0
0.5

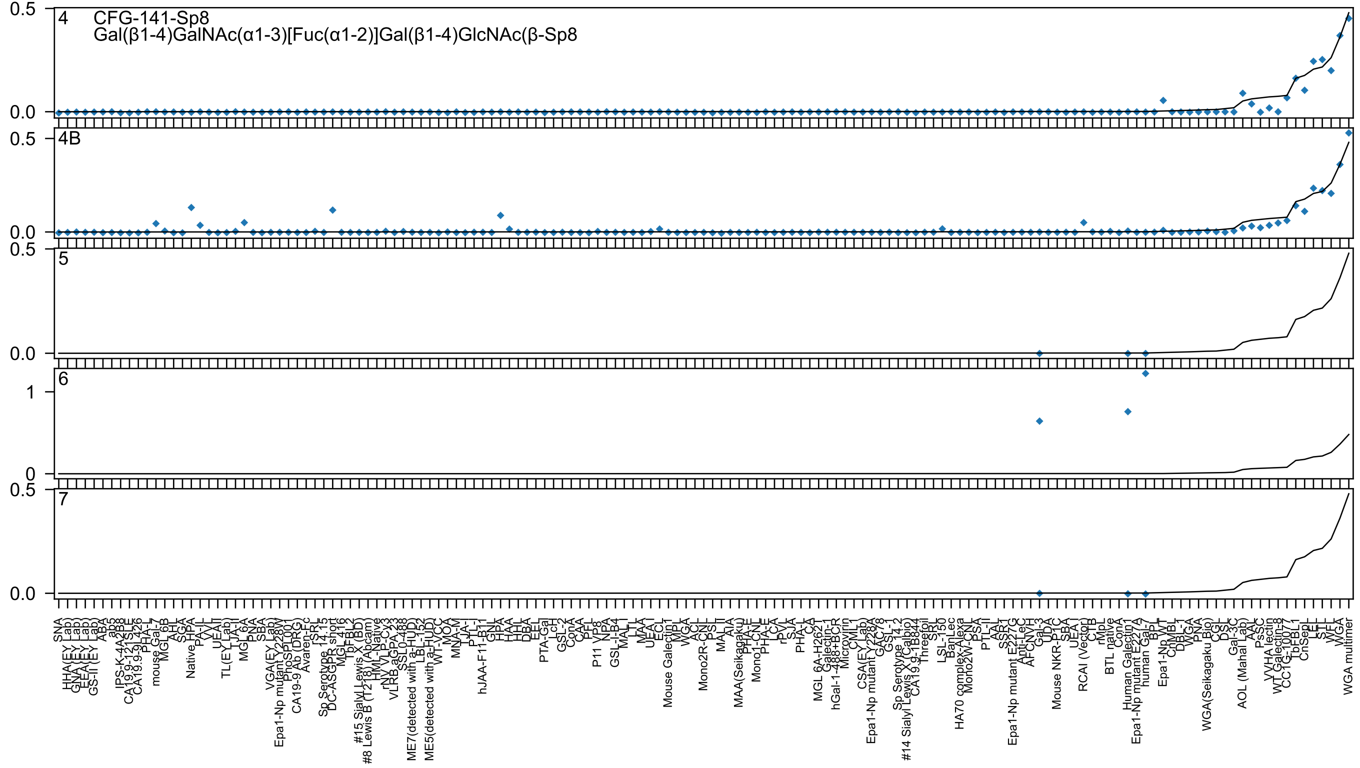
4
CFG-141-Sp8
Gal(β1-4)GalNAc(α1-3)[Fuc(α1-2)]Gal(β1-4)GlcNAc(β-Sp8
4B
5
6
7

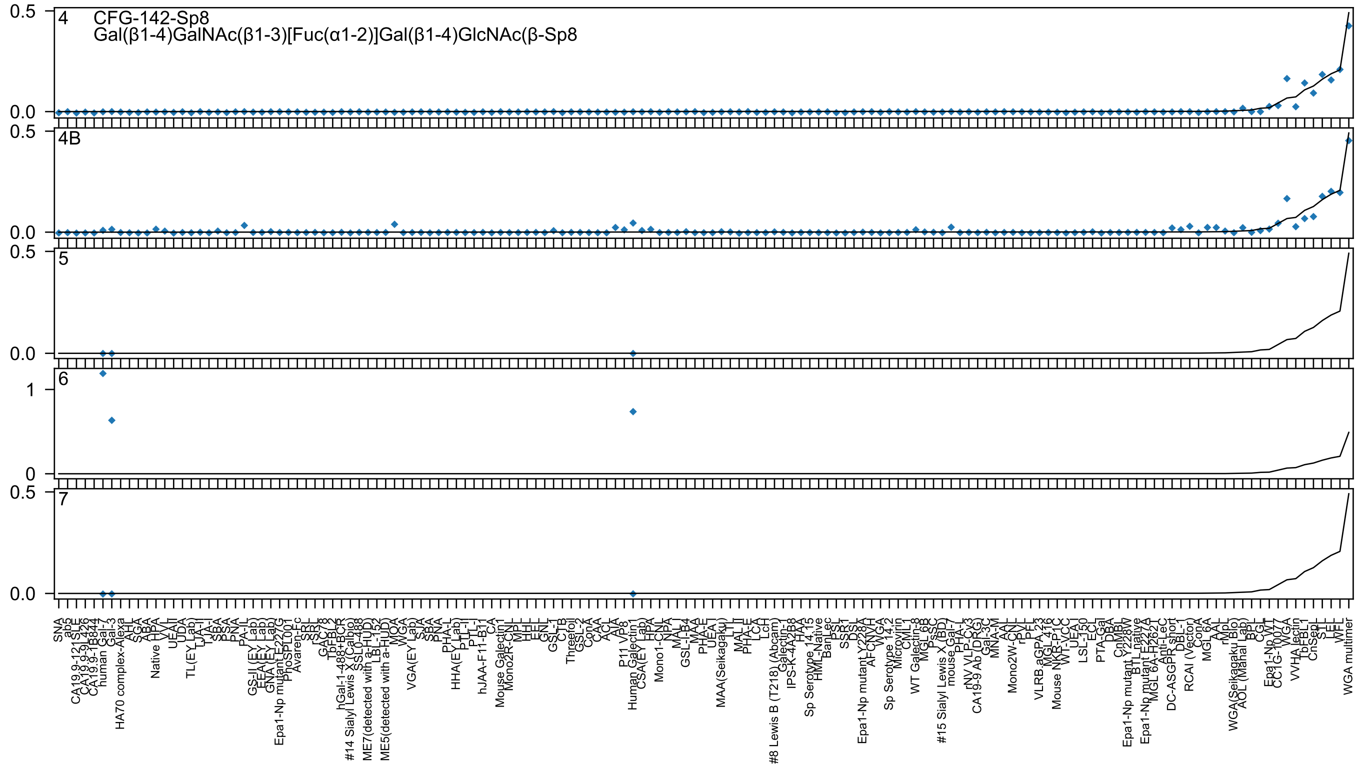

4 CFG-142-Sp8
Gal(β1-4)GalNAc(β1-3)[Fuc(α1-2)]Gal(β1-4)GlcNAc(β-Sp8
4B
5
6
7
0.5
0.0
1
0

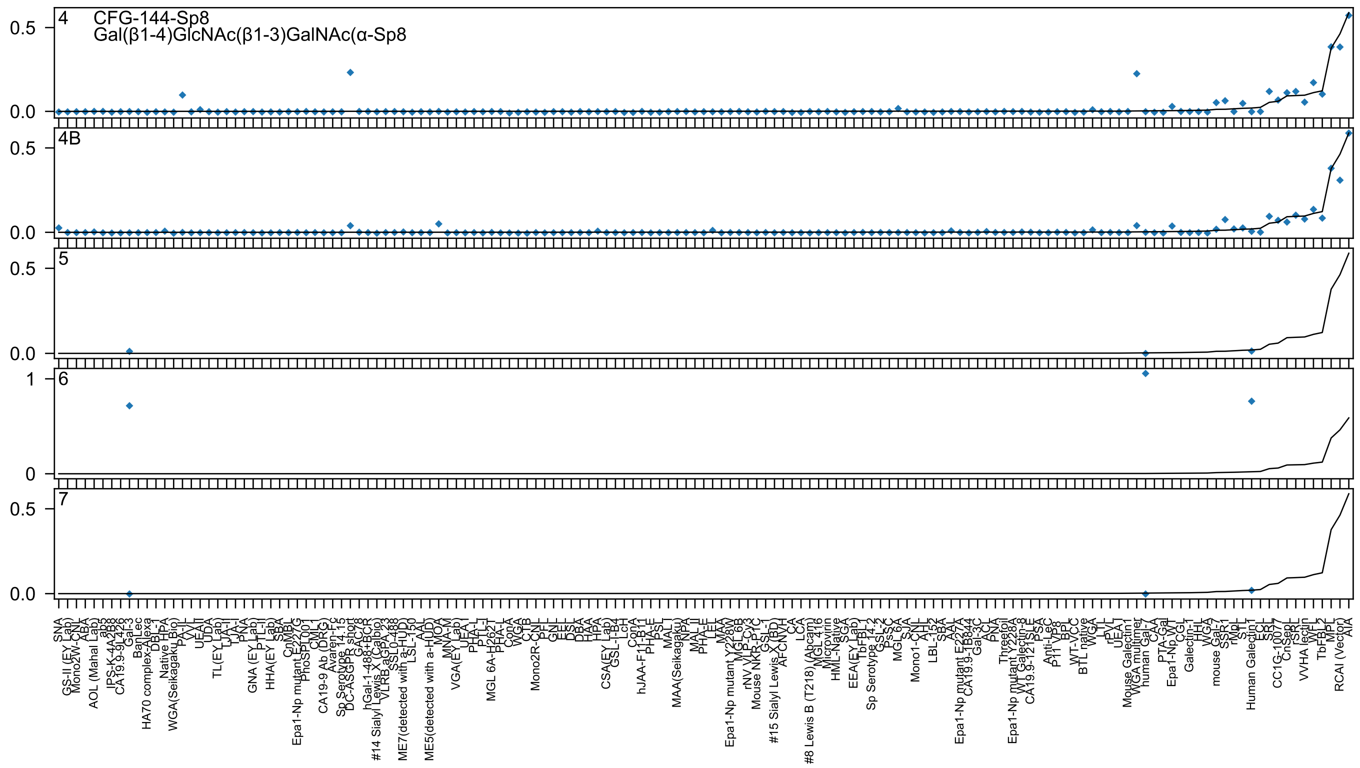

4
CFG-144-Sp8
Gal(β1-4)GlcNAc(β1-3)GalNAc(α-Sp8
4B
5
6
7

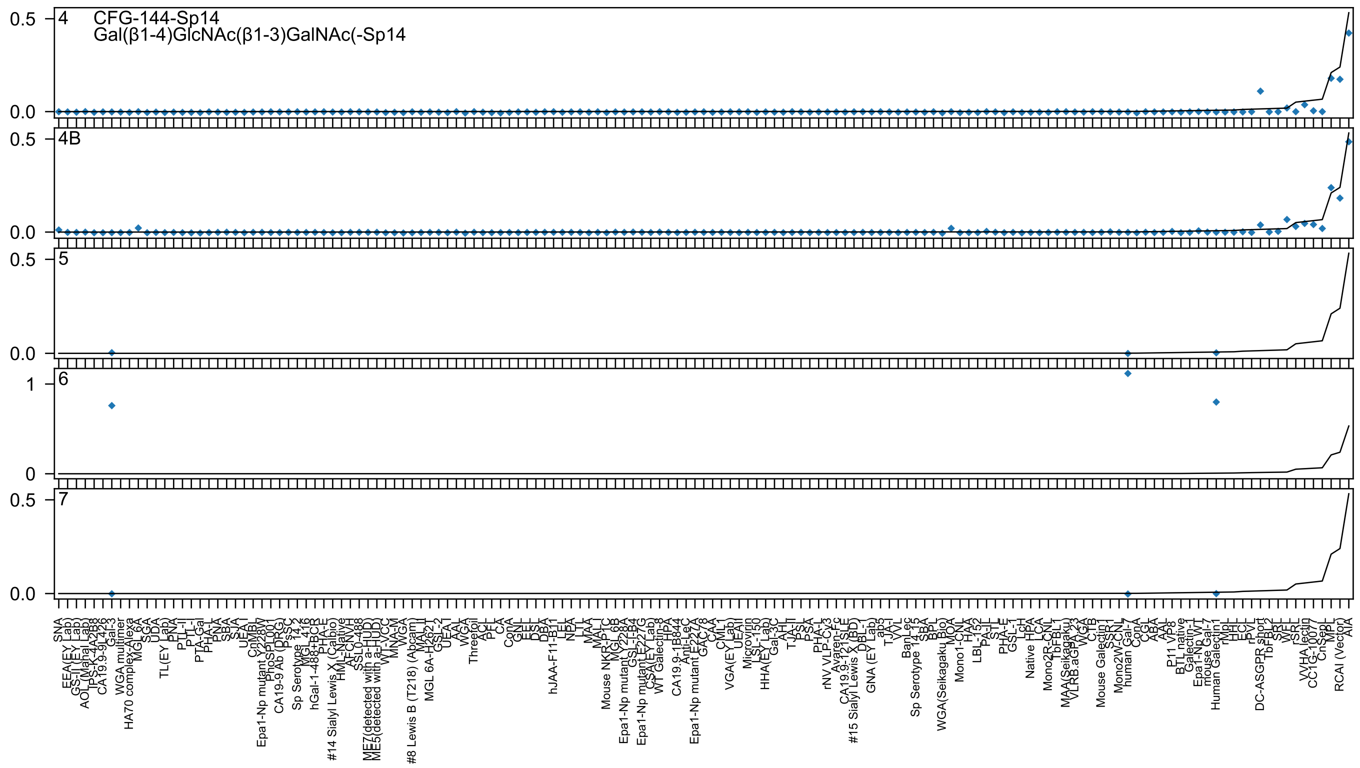
CFG-144-Sp14
Gal(β1-4)GlcNAc(β1-3)GalNAc(-Sp14
4
4B
5
6
7

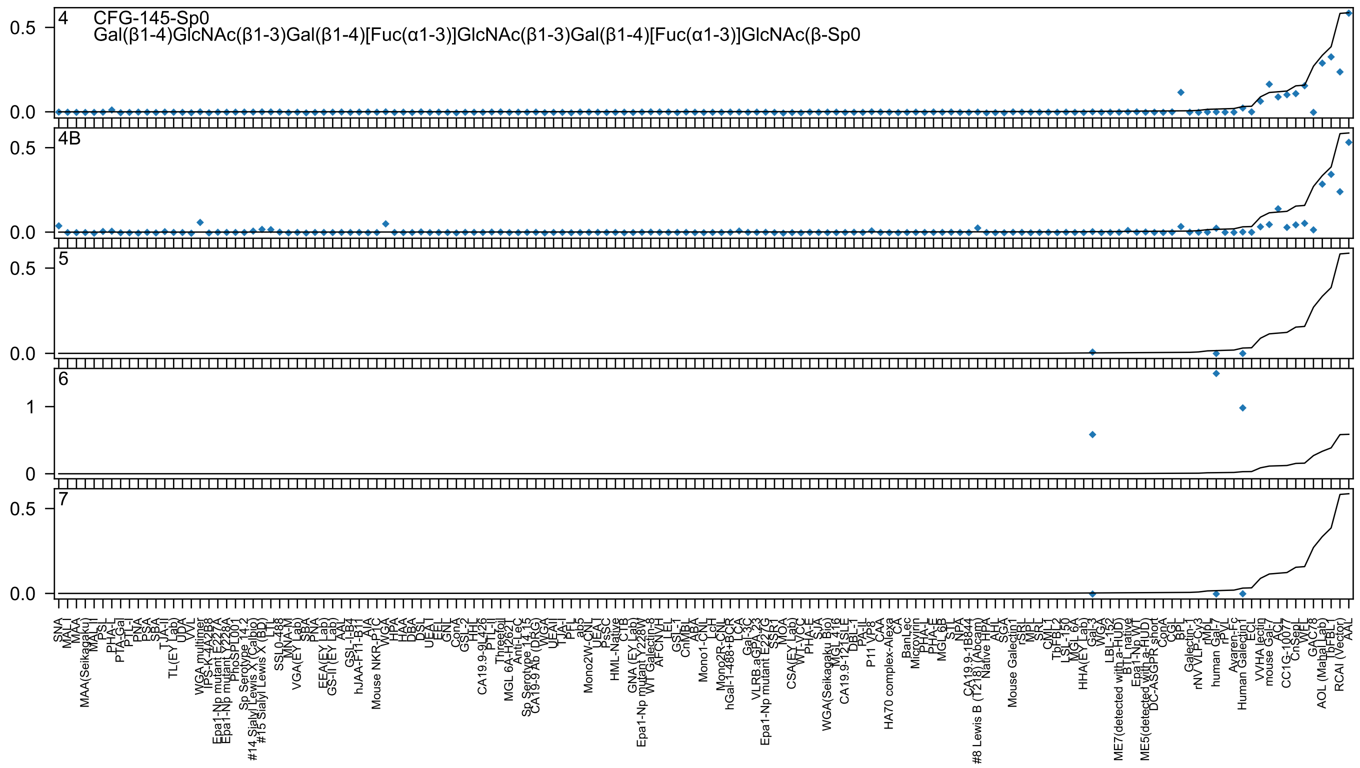

4 CFG-145-Sp0
Gal(β1-4)GlcNAc(β1-3)Gal(β1-4)[Fuc(α1-3)]GlcNAc(β1-3)Gal(β1-4)[Fuc(α1-3)]GlcNAc(β-Sp0
4B
5
6
7

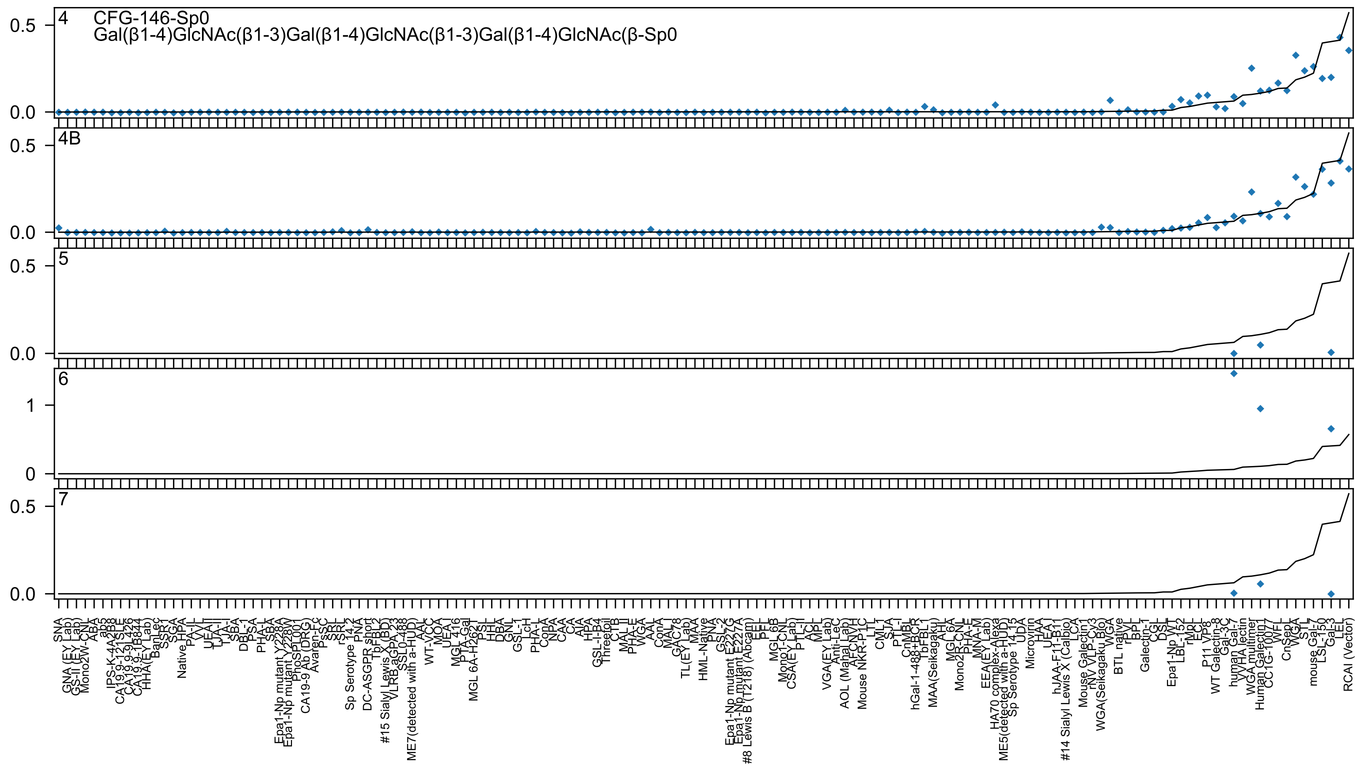

4
CFG-146-Sp0
Gal(β1-4)GlcNAc(β1-3)Gal(β1-4)GlcNAc(β1-3)Gal(β1-4)GlcNAc(β-Sp0
4B
5
6
7
0.5
0.0
1
0
SNA
GNA (EY Lab)
GS-II (EY Lab)
Mono2W-CNL
ABA
ab5
IPS-K-4A2B8
CA19.9-121SLE
CA19.9-9L426
CA19.9-1B844
HHA(EY Lab)
BanLec
SSR1
SGA
Native HPA
PA-IL
VVL
UEA-II
TJA-II
TJA-I
SBA
DBL-1
PSA
PHA-L
SBA
Epa1-Np mutant Y228A
Epa1-Np mutant Y228W
PhoSL001
CA19-9 Ab (DRG)
Avaren-Fc
PsSC
SRL
rSRL
Sp Serotype 14_2
PNA
DC-ASGPR short
TbFBL2
#15 Sialyl Lewis X (BD)
VLRB.aGPA.23
SSL0-488
ME7(detected with a-HUD)
AAL
WT-VCC
MOA
UEA-I
MGL 416
PTA-Gal
MGL 6A-H262L
PSL
HHL
DBA
GNL
GSL-1
LcH
PHA-L
ConA
NPA
CAA
CA
AIA
HPA
GSL-I-B4
Threetoil
CTB
MAL II
PHA-E
WGA
AAL
ConA
MAL-I
GAC78
TL(EY Lab)
MAA
HML-Native
PNA
GSL-2
Epa1-Np mutant E227G
Epa1-Np mutant E227A
#8 Lewis B (T218) (Abcam)
EEL
PTL
MGL 6B
Mono1-CNL
CSA(EY Lab)
PTL-II
ACL
MPL
VGA(EY Lab)
Anti-LeC
AOL (Mahal Lab)
AFCNVH
Mouse NKR-P1C
LTL
CML1
SJA
PTL-I
CnMBL
hGal-1-488+BCR
TbFBL1
MAA(Seikagaku)
AHL
MGL 6A
Mono2R-CNL
PHA-E
MNA-M
EEA(EY Lab)
HA70 complex-Alexa
ME5(detected with a-HUD)
Sp Serotype 14_3
UDA
Microvirin
HAA
UEA
hJAA-F11-B11
#14 Sialyl Lewis X (Calbio)
LCA
Mouse Galectin1
rNV VLP-Cy3
WGA(Seikagaku Bio)
WGA
BTL native
rPVL
BPL
Galectin-1
CGL
DSL
Epa1-Np WT
LBL-152
rMpL
ECL
P11 VP8
WT Galectin-8
Gal-3C
human Gal-7
VVHA lectin
WGA multimer
Human Galectin1
CC1G-10077
WFL
CnSep1
WGA
STL
mouse Gal-7
LSL-150
Gal-3
LEL
RCAI (Vector)

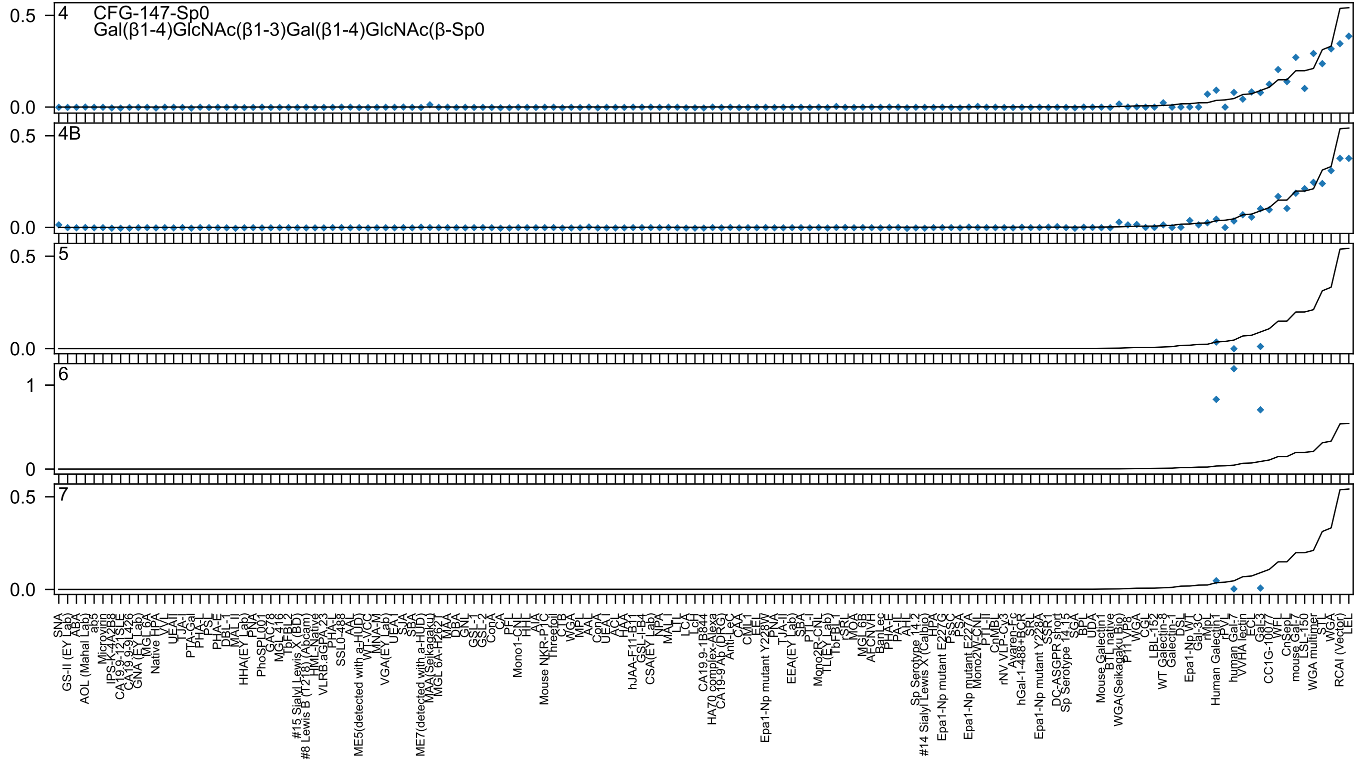
4
CFG-147-Sp0
Gal(β1-4)GlcNAc(β1-3)Gal(β1-4)GlcNAc(β-Sp0
4B
5
6
7

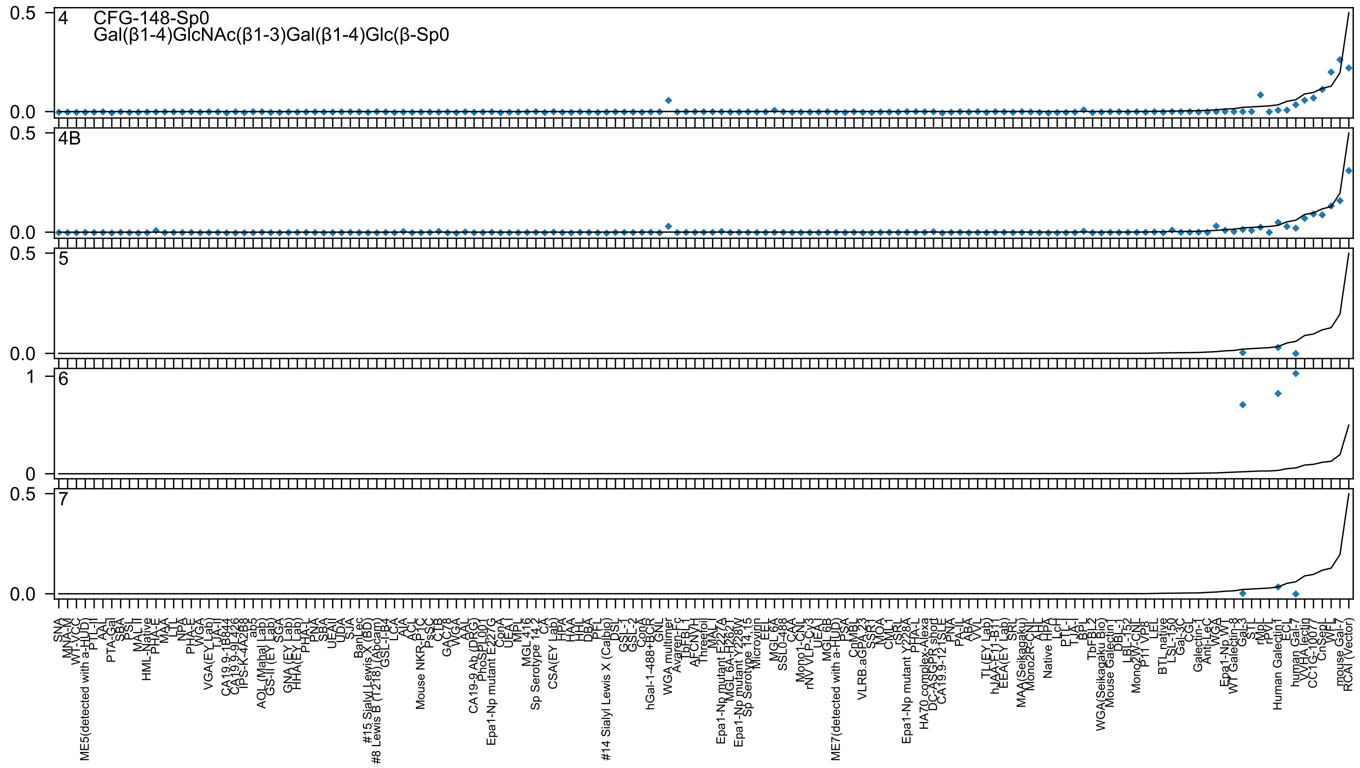

4
CFG-148-Sp0
Gal(β1-4)GlcNAc(β1-3)Gal(β1-4)Glc(β-Sp0
4B
5
6
7
0.5
0.0
1
0

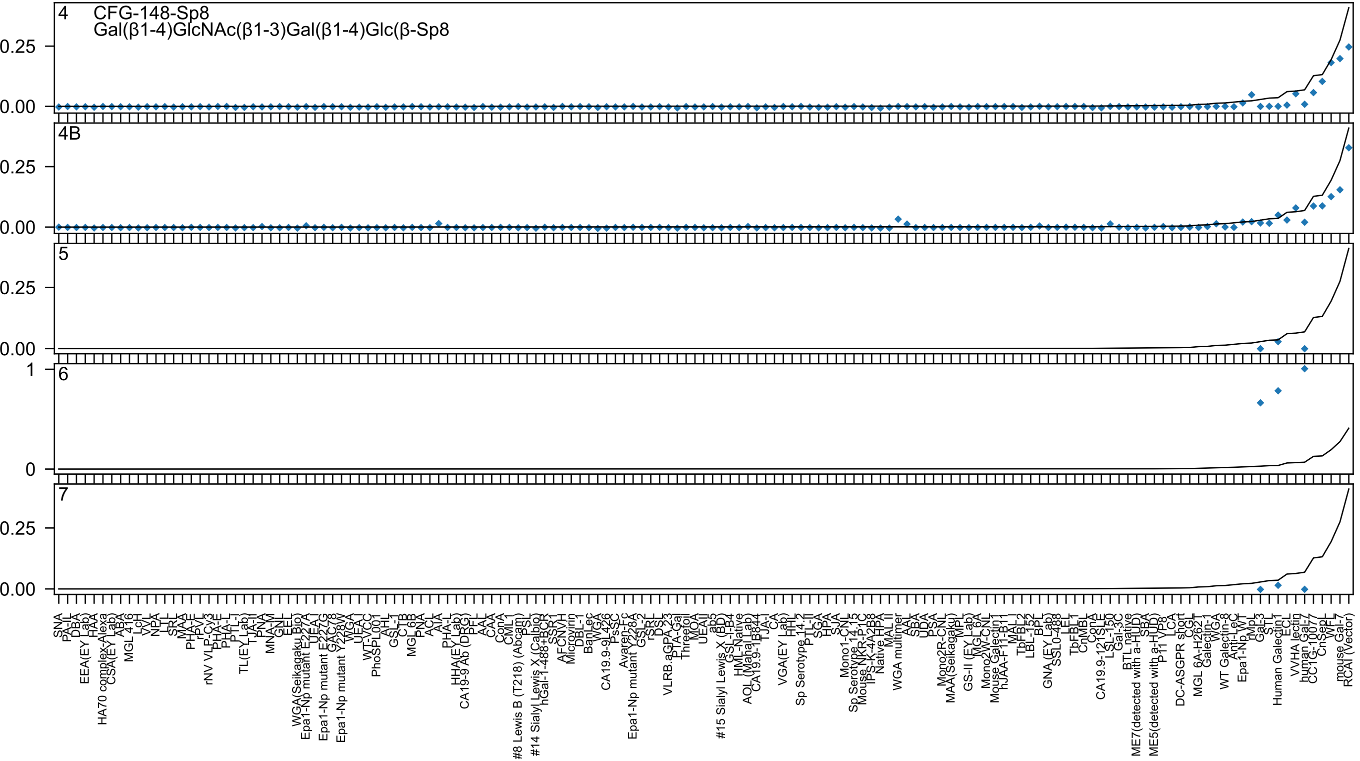
4 CFG-148-Sp8
Gal(β1-4)GlcNAc(β1-3)Gal(β1-4)Glc(β-Sp8
4B
5
6
7
0.25
0.00
1
0
SNA
PA-IL
DBA
EEA(EY Lab)
HAA
HA70 complex-Alexa
CSA(EY Lab)
ABA
MGL 416
LcH
VVL
NPA
LTL
SRL
MAA
PHA-E
rPVL
rNV VLP-Cy3
PHA-E
PHA-L
PTL-I
TL(EY Lab)
TJA-II
PNA
MNA-M
GNL
EEL
WGA(Seikagaku Bio)
Epa1-Np mutant E227A
UEA-I
Epa1-Np mutant E227G
GAC78
Epa1-Np mutant Y228W
WGA
UEA-I
WT-VCC
PhoSL001
AAL
GSL-I
CTB
MGL 6B
PNA
ACL
AIA
PHA-L
HHA(EY Lab)
CA19-9 Ab (DRG)
PFL
AAL
CAA
ConA
CML1
#8 Lewis B (T218) (Abcam)
PSL
#14 Sialyl Lewis X (Calbio)
hGal-1-488+BCR
SSR1
AFCNVH
Microvirin
DBL-1
BanLec
WGA
CA19.9-9L426
PsSC
Avaren-Fc
Epa1-Np mutant Y228A
GSL-2
rSRL
DSL
VLRB.aGPA.23
PTA-Gal
Threefoil
MOA
UEAII
ab5
#15 Sialyl Lewis X (BD)
GSL-I-B4
HML-Native
AOL (Mahal Lab)
CA19.9-1B844
TJA-I
CA
VGA(EY Lab)
HHL
Sp Serotype 14.2
PTL-II
SGA
HPA
SJA
Mono1-CNL
Sp Serotype 14.15
Mouse NKR-P1C
IPS-K-4A2B8
Native HPA
MAL II
WGA multimer
AAL
SBA
UDA
PSA
Mono2R-CNL
MAA(Seikagaku)
MPL
GS-II (EY Lab)
MGL 6A
Mono2W-CNL
Mouse Galectin1
hJAA-F11-B11
MAL I
TbFBL-2
LBL-152
BPL
GNA (EY Lab)
SSL0-488
LEL
TbFBL-1
CnMBL
ConA
CA19.9-121SLE
LSL-150
Gal-3C
BTL native
ME7(detected with a-HUD)
SBA
ME5(detected with a-HUD)
P11 VP8
LCA
DC-ASGPR short
CGL
MGL 6A-H262T
Galectin-1
WGA
WT Galectin-8
Anti-LeC
Epa1-Np WT
rMpL
Gal-3
STL
Human Galectin1
ECL
VVHA lectin
human Gal-7
CC1G-10077
CnSepL
WFL
mouse Gal-7
RCAI (Vector)

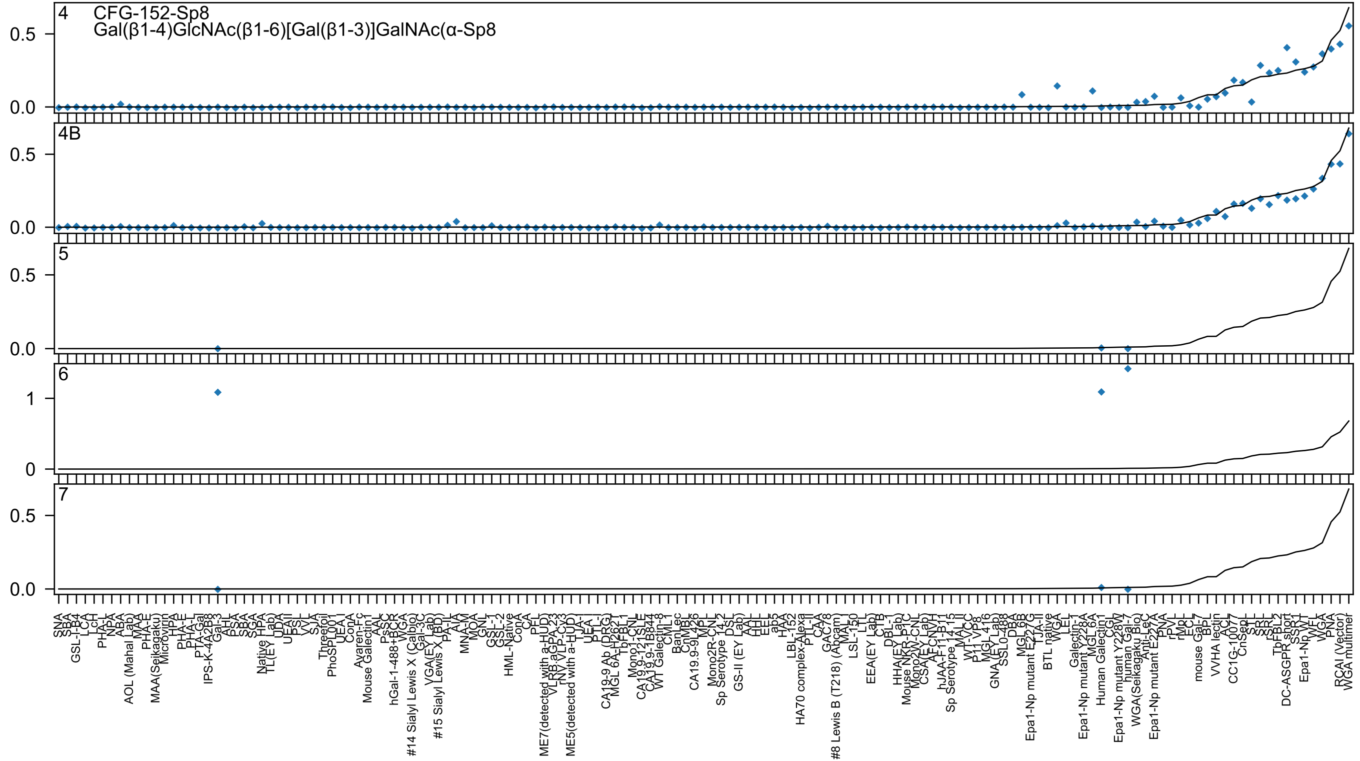
4
CFG-152-Sp8
Gal(β1-4)GlcNAc(β1-6)[Gal(β1-3)]GalNAc(α-Sp8
4B
5
6
7

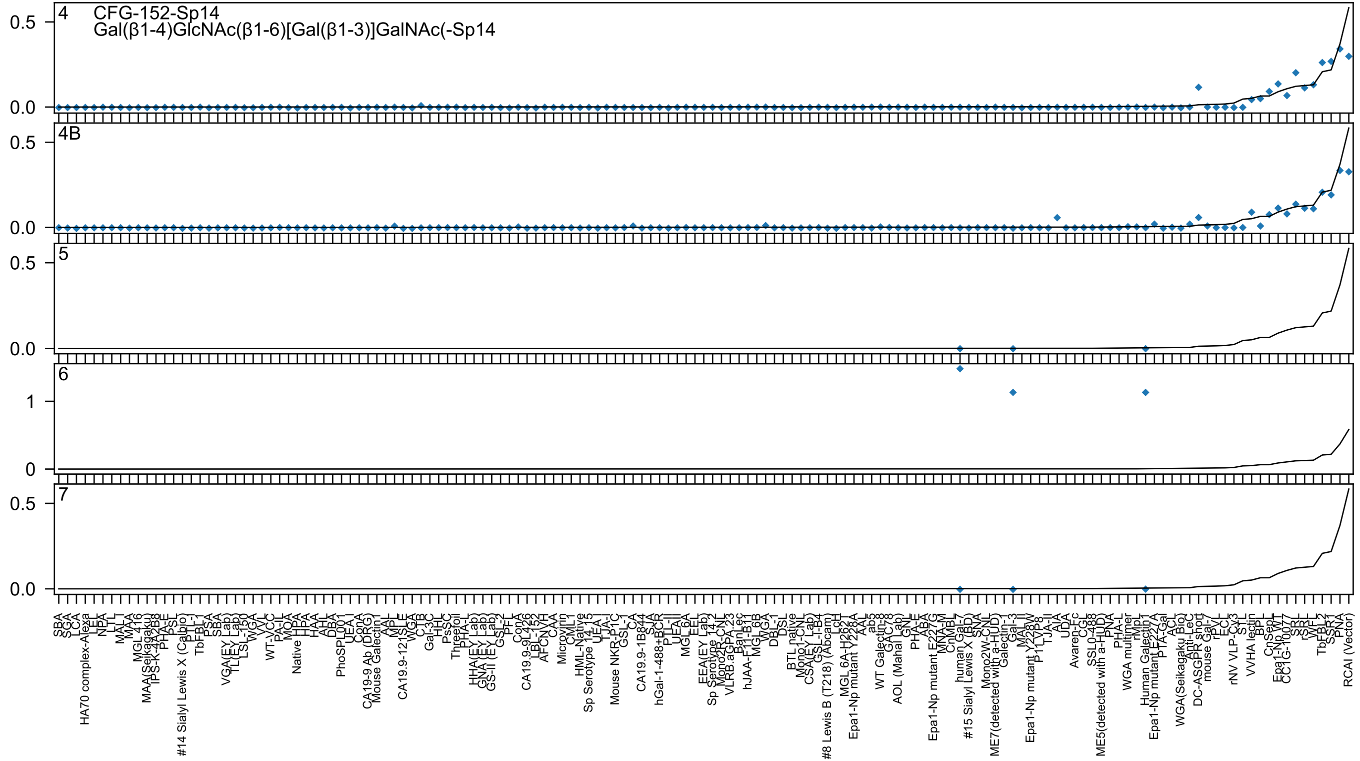
CFG-152-Sp14
Gal(β1-4)GlcNAc(β1-6)[Gal(β1-3)]GalNAc(-Sp14
4
4B
5
6
7

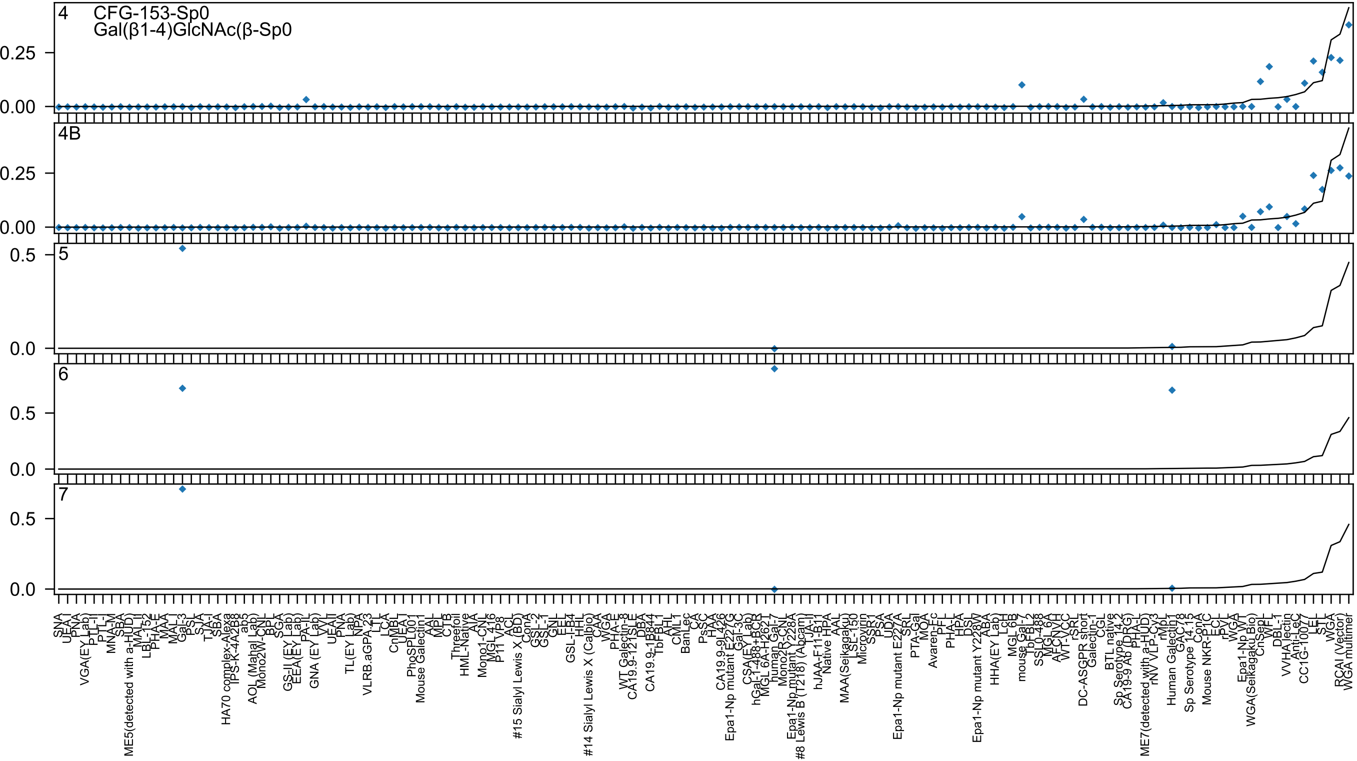
4
CFG-153-Sp0
Gal(β1-4)GlcNAc(β-Sp0
4B
5
6
7
0.25
0.00
0.5
0.0

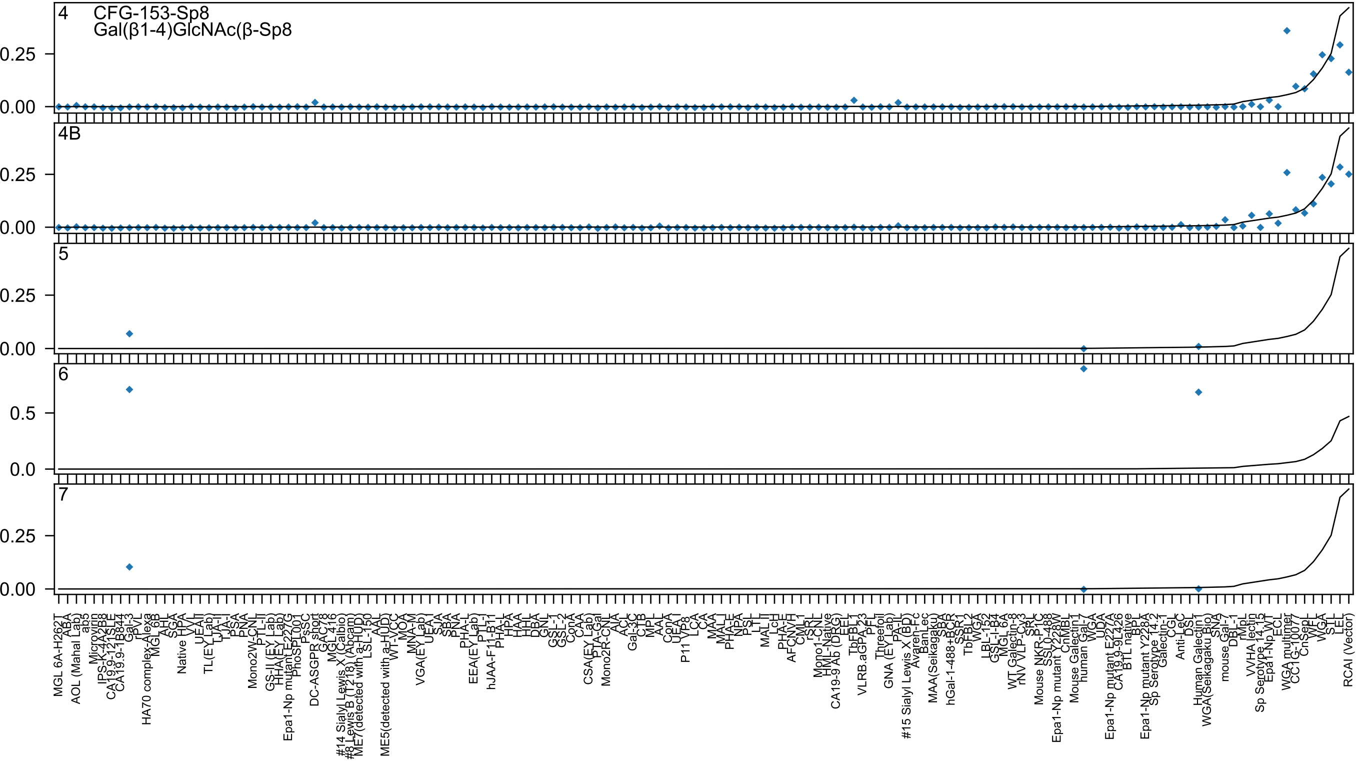

4
CFG-153-Sp8
Gal(β1-4)GlcNAc(β-Sp8
4B
5
6
7
0.25
0.00
0.5
0.0
MGL 6A-H262T
ABA
AOL (Mahal Lab)
ab5
Microvirin
IPS-K-4A2B8
CA19.9-121SLE
CA19.9-1B844
Gal-3
rPVL
HA70 complex-Alexa
MGL 6B
AHL
SGA
Native HPA
VVL
UEAII
TL(EY Lab)
TJA-II
TJA-I
PSA
PNA
Mono2W-CNL
PTL-II
GS-II (EY Lab)
HHA(EY Lab)
Epa1-Np mutant E227G
PhoSPL001
PsSC
DC-ASGPR short
GAC78
MGL 416
#14 Sialyl Lewis X (Calbio)
#8 Lewis B (T218) (Abcam)
ME7(detected with a-HUD)
LSL-150
AAL
ME5(detected with a-HUD)
WT-VCC
MOA
MNA-M
VGA(EY Lab)
UEA-I
SJA
SBA
PNA
PHA-L
EEA(EY Lab)
PTL-I
hJAA-F11-B11
PHA-L
HPA
HAA
HHL
DBA
GNL
GSL-1
GSL-2
ConA
CAA
CSA(EY Lab)
PTA-Gal
Mono2R-CNL
AIA
ACL
Gal-3C
CTB
MPL
AAL
ConA
UEA-I
P11 VP8
LCA
CA
MAA
MAL-I
PHA-E
NPA
PSL
LTL
MAL II
LcH
PHA-E
AFCNVH
CML-1
rSRL
Mono1-CNL
HML-Native
CA19-9 Ab (DRG)
EEL
TbFBL1
VLRB.aGPA.23
PFL
Threefoil
GNA (EY Lab)
PA-IL
#15 Sialyl Lewis X (BD)
Avaren-Fc
BanLec
MAA(Seikagaku)
SBA
hGal-1-488+BCR
SSR1
TbFBL2
WGA
LBL-152
GSL-I-B4
MGL 6A
WT Galectin-8
rNV VLP-Cy3
SRL
Mouse NKR-P1C
SSL0-488
Epa1-Np mutant Y228W
CnMBL
Mouse Galectin1
human Gal-7
WGA
UDA
Epa1-Np mutant E227A
CA19.9-9L426
BTL native
BPL
Epa1-Np mutant Y228A
Sp Serotype 14.2
Galectin-1
CGL
Anti-LeC
DSL
Human Galectin1
WGA(Seikagaku Bio)
SNA
mouse Gal-7
DBL-1
rMpL
VVHA lectin
Sp Serotype 14.15
Epa1-Np WT
ECL
WGA multimer
CC1G-10077
CnSepL
WFL
WGA
STL
LEL
RCAI (Vector)

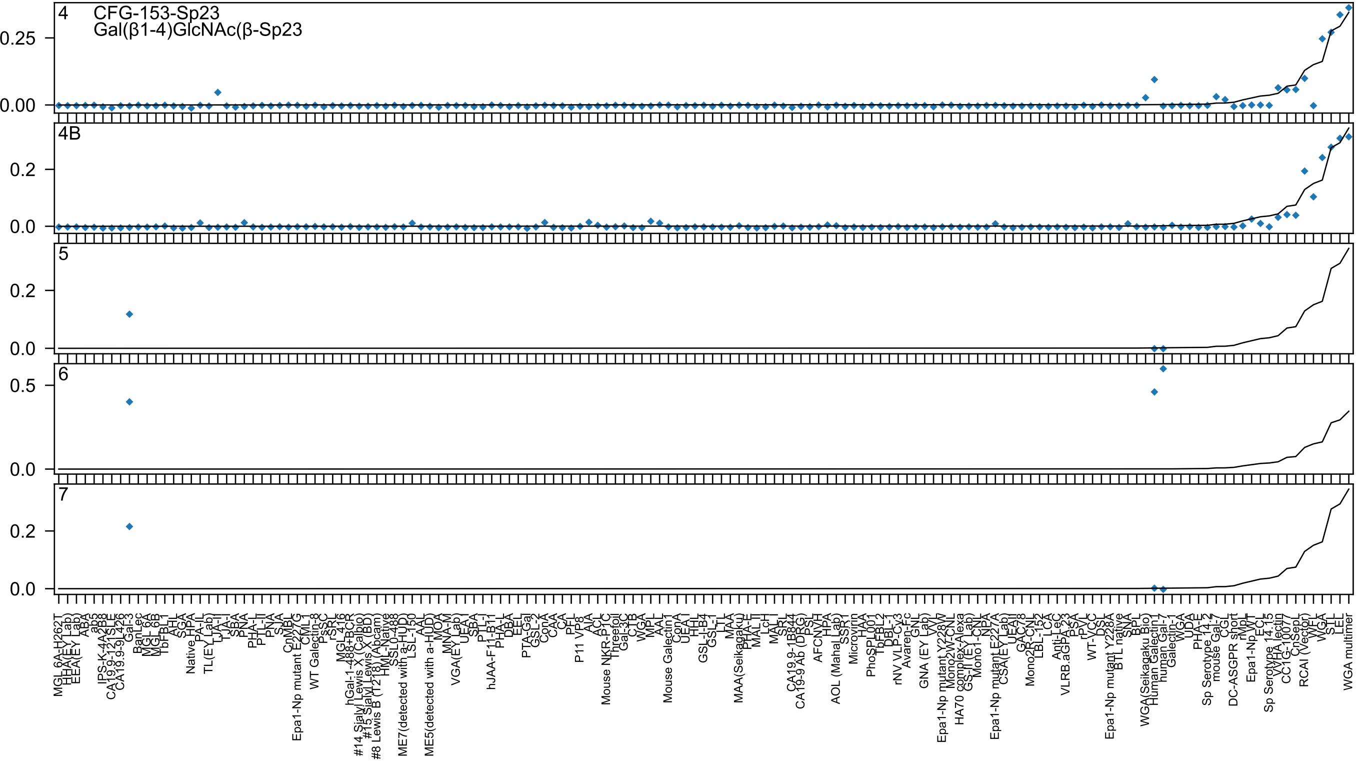

4
CFG-153-Sp23
Gal(β1-4)GlcNAc(β-Sp23
4B
5
6
7

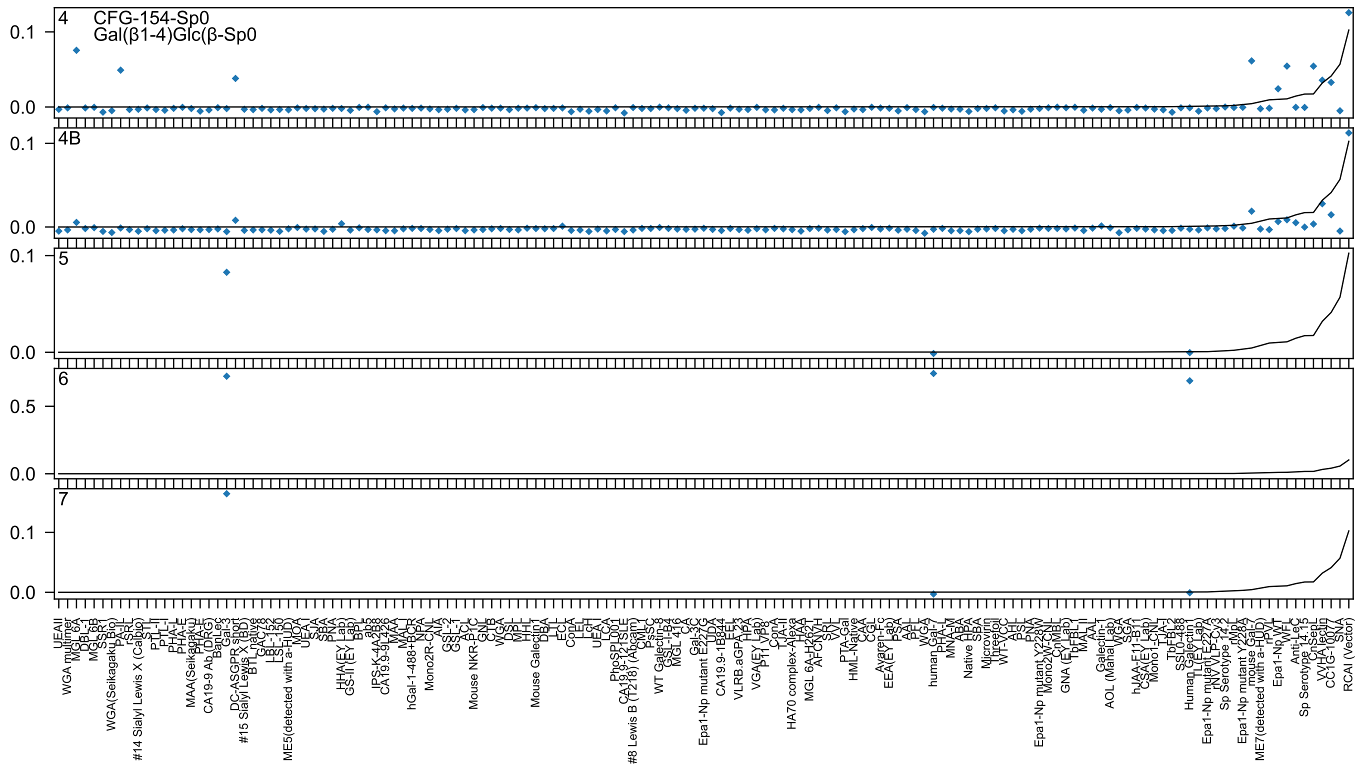

4
CFG-154-Sp0
Gal(β1-4)Glc(β-Sp0
4B
5
6
7

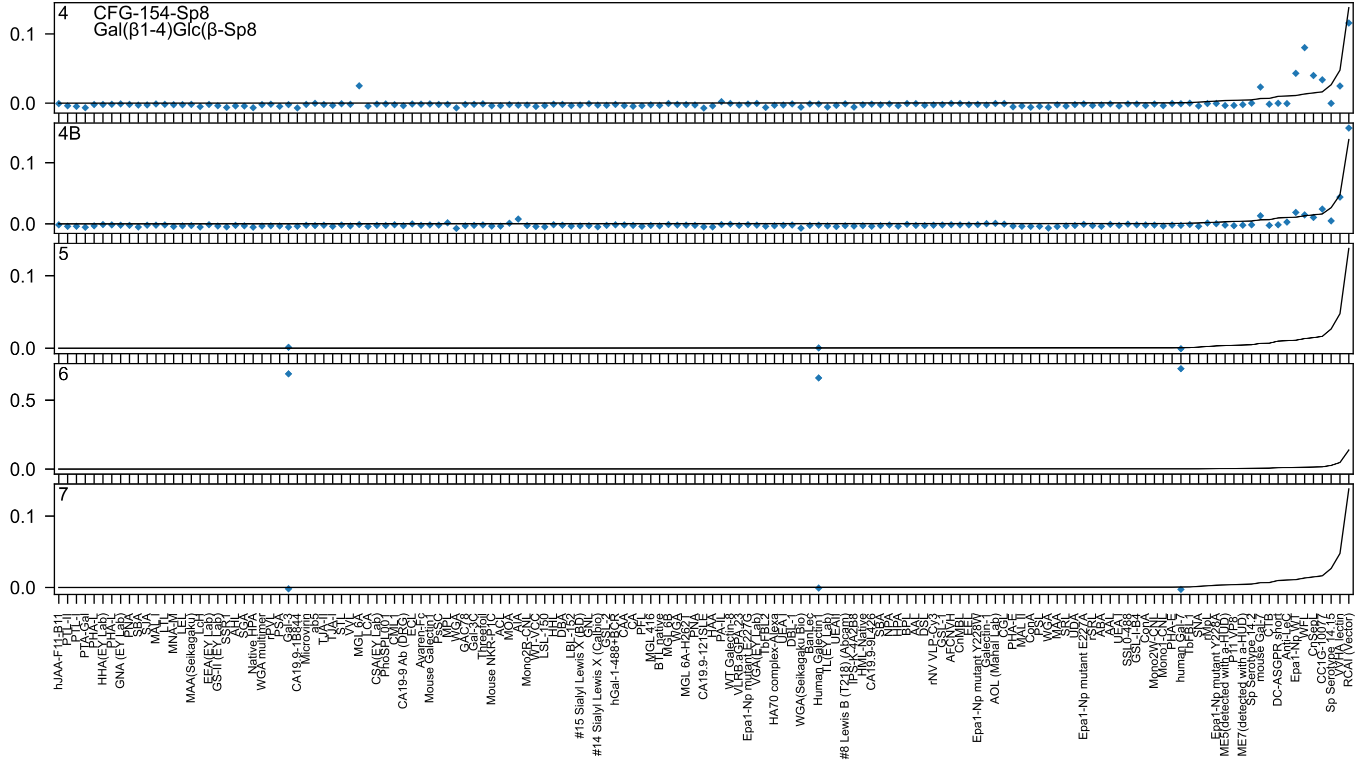

4 CFG-154-Sp8
Gal(β1-4)Glc(β-Sp8
4B
5
6
7
0.0
0.1
0.5
hJAA-F11-B11
PTL-II
PTL-I
PTA-Gal
PHA-L
HHA(EY Lab)
PHA-L
GNA (EY Lab)
PNA
SBA
SJA
MAL_I
LTL
MNA-M
LEL
MAA(Seikagaku)
LcH
EEA(EY Lab)
GS-II (EY Lab)
SSR1
AHL
SGA
Native HPA
WGA multimer
rPVL
PSA
Gal-3
CA19.9-1B844
Microvirin
ab5
TJA-II
TJA-I
STL
VVL
MGL 6A
LCA
CSA(EY Lab)
PhoSL001
CML1
CA19-9 Ab (DRG)
ECL
Avaren-Fc
Mouse Galectin1
PsSC
MPL
WGA
GAC78
Gal-3C
Threefoil
Mouse NKR-P1C
ACL
MOA
AIA
Mono2R-CNL
WT-VCC
LSL-150
HHL
DBA
LBL-152
#15 Sialyl Lewis X (BD)
GNL
#14 Sialyl Lewis X (Calbio)
GSL-2
hGal-1-488+BCR
CAA
CA
PFL
MGL 416
BTL native
MGL 6B
WGA
MGL 6A-H262T
PNA
CA19.9-121SLE
HAA
PA-IL
WT Galectin-8
VLRB.aGPA.23
Epa1-Np mutant E227G
VGA(EY Lab)
TbFBL2
HA70 complex-Alexa
UEA
DBL-1
WGA(Seikagaku Bio)
BanLec
Human Galectin1
TL(EY Lab)
UEAII
#8 Lewis B (T218) (Abcam)
TPS-K-4A2B8
HML-Native
CA19.9-9L426
SBA
NPA
HPA
BPL
AAL
DSL
rNV VLP-Cy3
GSL-1
AFCNVH
CnMBL
EEL
Epa1-Np mutant Y228W
Galectin-1
AOL (Mahal Lab)
CGL
PHA-E
MAL II
ConA
PSL
WGA
MAA
SRL
UDA
Epa1-Np mutant E227A
rSRL
ABA
AAL
UEA I
SSL0-488
GSL-I-B4
ConA
Mono2W-CNL
Mono1-CNL
PHA-E
human Gal-7
TbFBL1
SNA
rMpL
Epa1-Np mutant Y228A
ME5(detected with a-HUD)
P11 VP8
ME7(detected with a-HUD)
Sp Serotype 14.2
mouse Gal-7
CTB
DC-ASGPR short
Anti-LeC
Epa1-Np WT
WFL
CnSepL
CC1G-10077
Sp Serotype 14.15
VVHA lectin
RCAI (Vector)

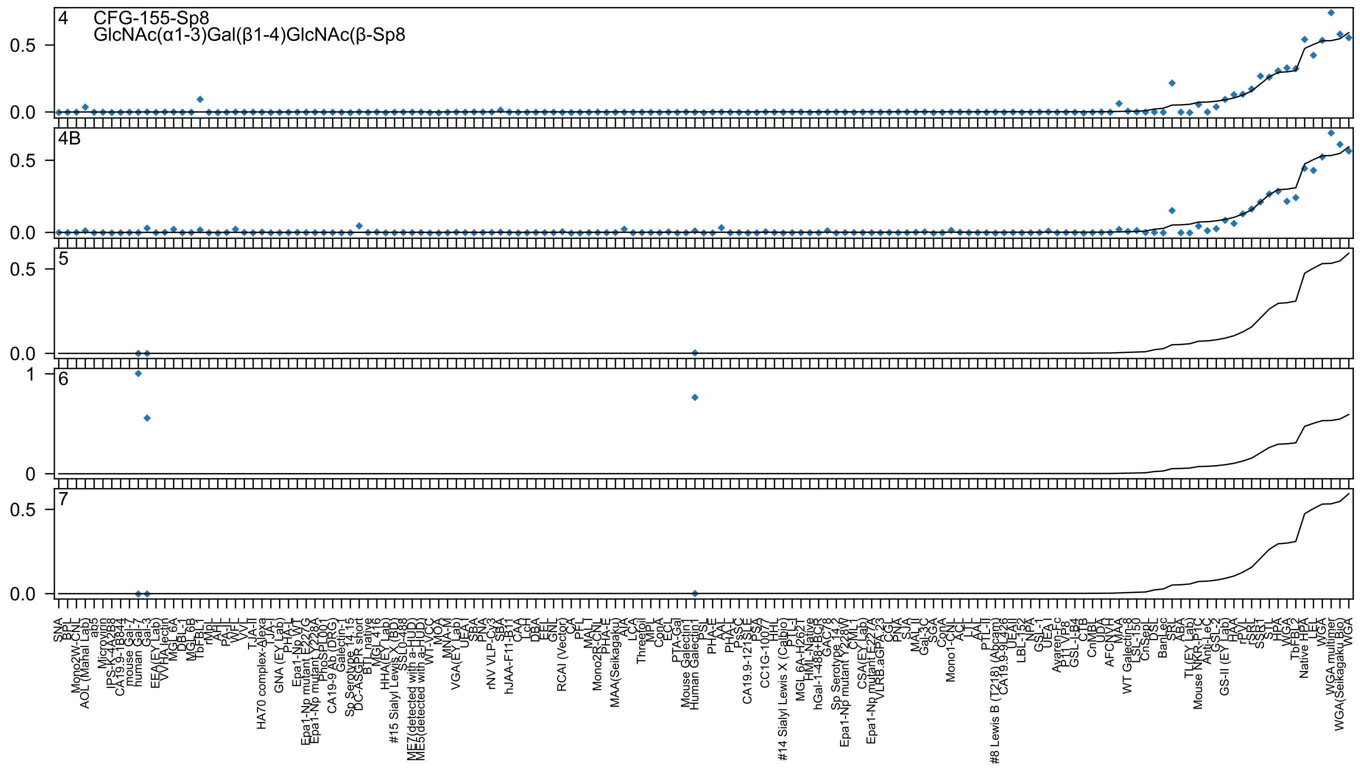
4
CFG-155-Sp8
GlcNAc(α1-3)Gal(β1-4)GlcNAc(β-Sp8
4B
5
6
7

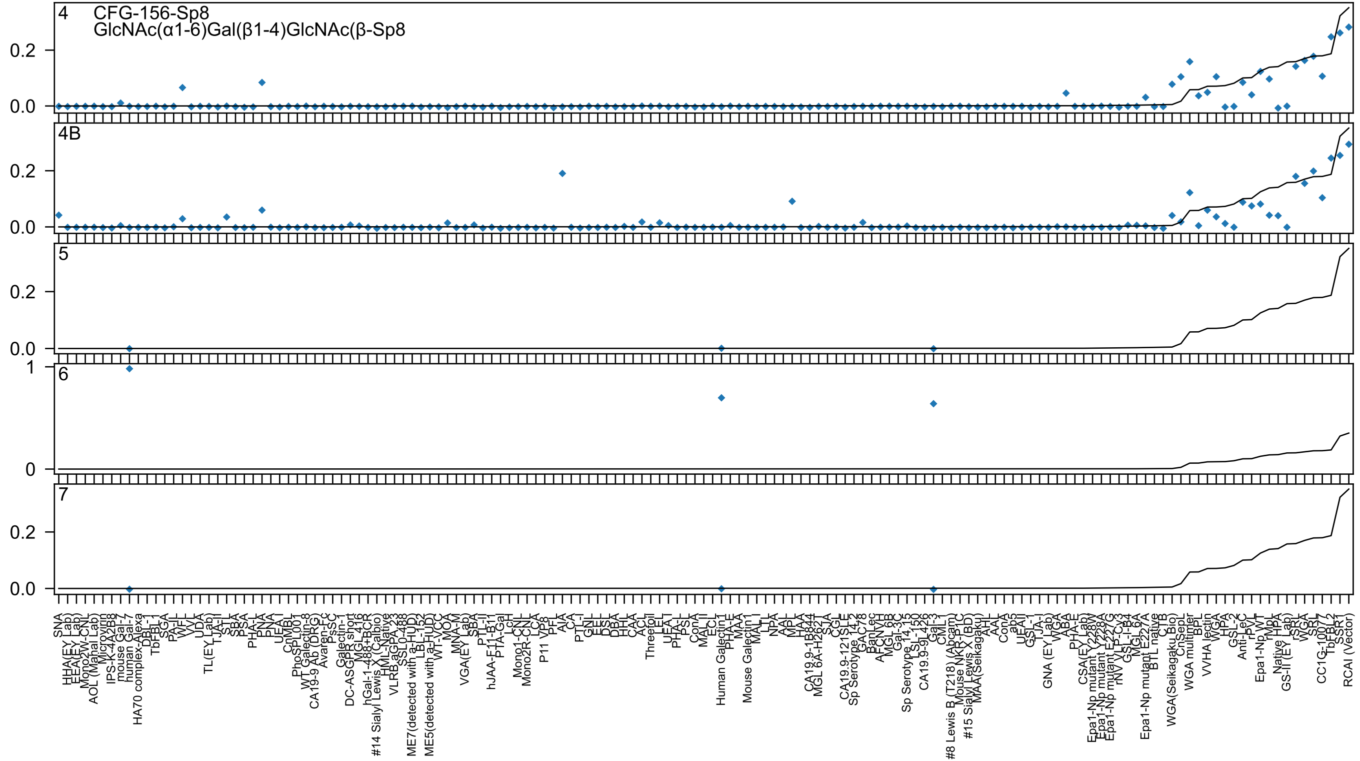
4
CFG-156-Sp8
GlcNAc(α1-6)Gal(β1-4)GlcNAc(β-Sp8
4B
5
6
7

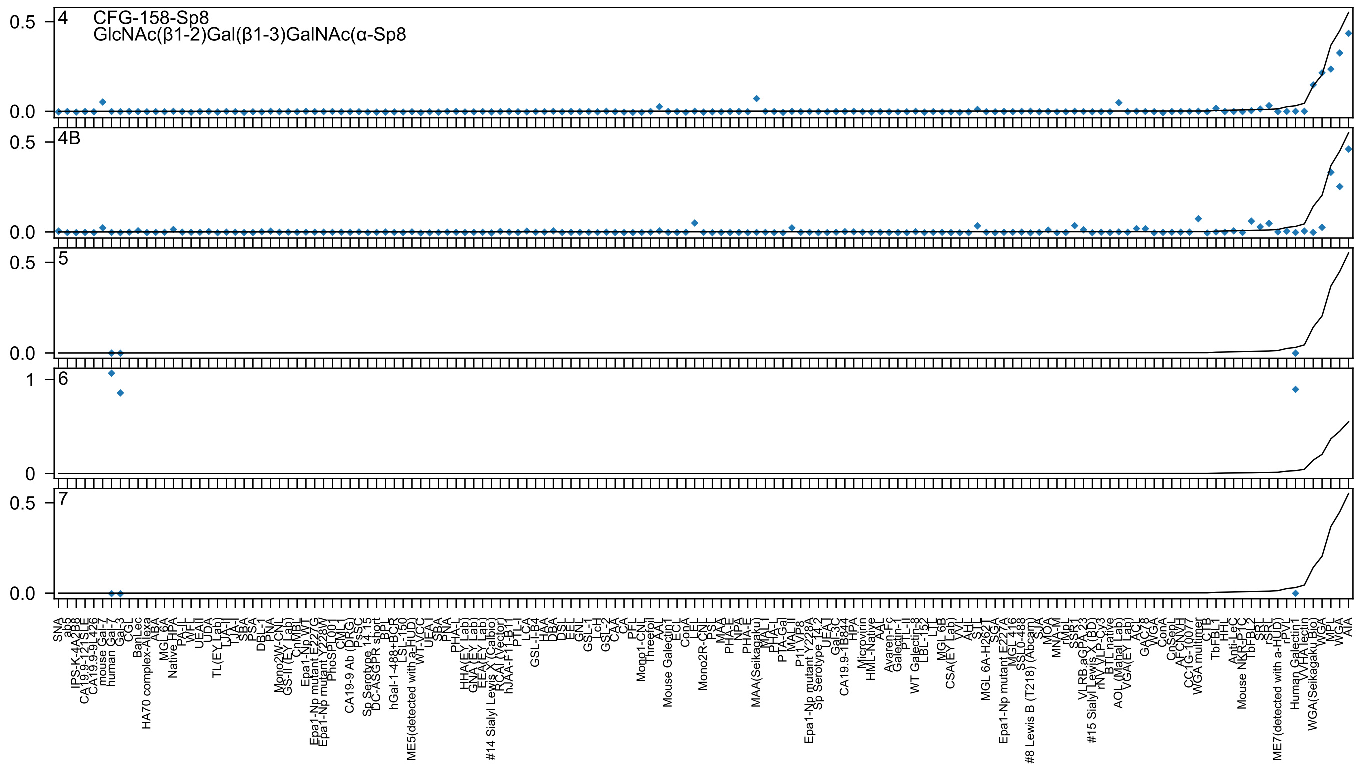

4
CFG-158-Sp8
GlcNAc(β1-2)Gal(β1-3)GalNAc(α-Sp8
4B
5
6
7
0.5
0.0
1
0

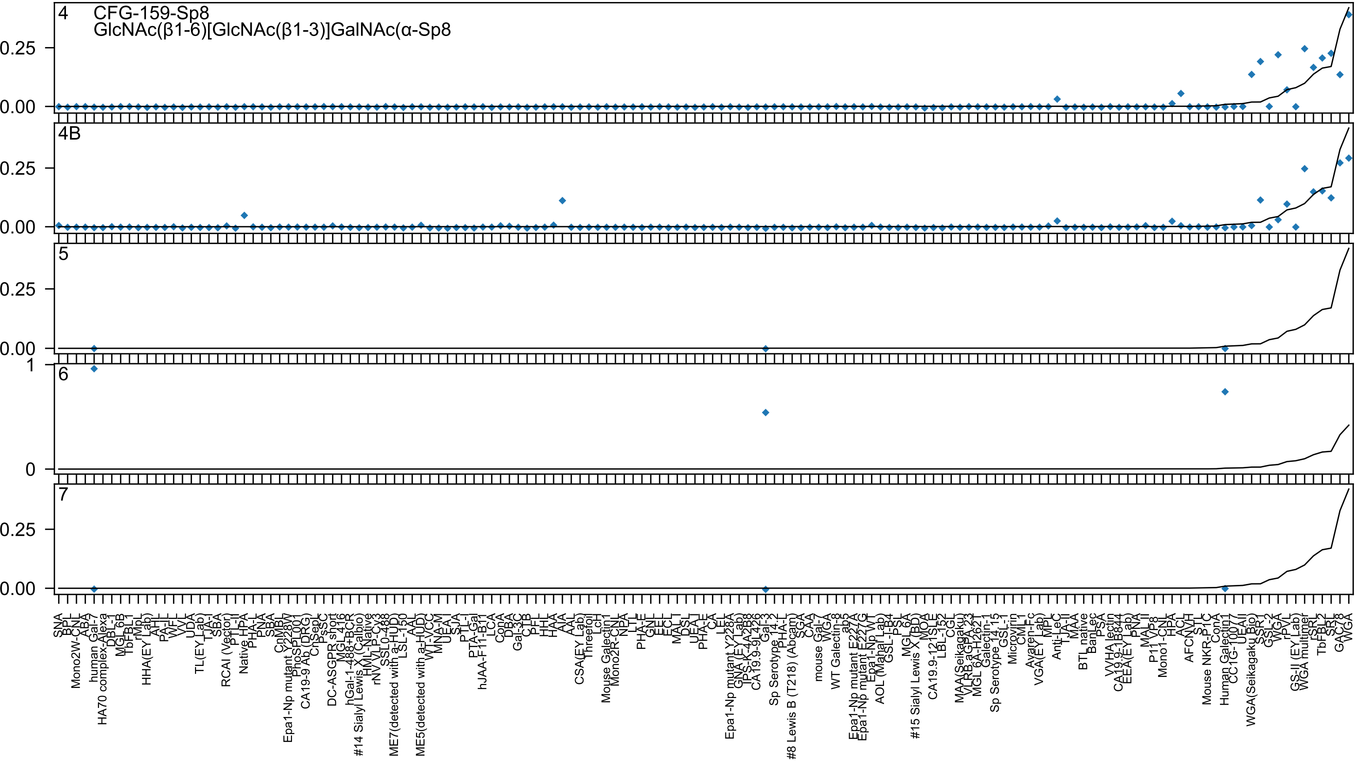
4
CFG-159-Sp8
GlcNAc(β1-6)[GlcNAc(β1-3)]GalNAc(α-Sp8
4B
5
6
7
0.25
0.00
1
0
SNA
BPL
Mono2W-CNL
ABA
human Gal-7
HA70 complex-Alexa
DBL-1
MGL 6B
TbFBL1
rMpL
HHA(EY Lab)
AHL
PA-IL
WFL
VVL
UDA
TL(EY Lab)
TJA-I
SBA
RCAI (Vector)
PTL-II
Native HPA
PHA-L
PNA
SBA
CnMBL
Epa1-Np mutant Y228V
PhoSL001
CA19-9 Ab (DRG)
ChSep
PsSC
DC-ASGPR short
MGL 416
hGal-1-488+BCR
#14 Sialyl Lewis X (Calbio)
HML-Native
rNV VLP-Cy3
SSL0-488
ME7(detected with a-HUD)
LSL-150
AAL
ME5(detected with a-HUD)
WT-VCC
MNA-M
UEA-I
SJA
PTL-I
PTA-Gal
hJAA-F11-B11
LCA
ConA
DBA
Gal-3C
CTB
PFL
HHL
HAA
AIA
AAL
CSA(EY Lab)
Threefoil
LcH
Mouse Galectin1
Mono2R-CNL
NPA
LTL
PHA-E
GNL
EEL
ECL
MAL I
DSL
UEA-I
PHA-E
CA
LEL
Epa1-Np mutant Y228A
GNA (EY Lab)
TPS-K-4A2B8
CA19.9-9L426
Gal-3
Sp Serotype 14.2
PHA-L
#8 Lewis B (T218) (Abcam)
SGA
CAA
mouse Gal-7
WGA
WT Galectin-8
ab5
Epa1-Np mutant E227A
Epa1-Np mutant E227G
Epa1-Np WT
AOL (Mahal Lab)
GSL-I-B4
PSL
MGL 6A
#15 Sialyl Lewis X (BD)
MOA
CA19.9-121SLE
LBL-152
CGL
MAA(Seikagaku)
VLRB.aGPA.23
MGL 6A-H262T
Galectin-1
Sp Serotype 14.15
GSL-I
Microvirin
CML1
Ayaren-Fc
VGA(EY Lab)
MPL
Anti-LeC
TJA-II
MAA
BTL native
BanLec
PSA
VVHA lectin
CA19.9-1B844
EEA(EY Lab)
PNA
MAL II
P11 VP8
Mono1-CNL
HPA
ACL
AFCNVH
STL
Mouse NKR-P1C
ConA
Human Galectin1
CC1G-10077
UEA II
WGA(Seikagaku Bio)
SSR1
GSL-2
WGA
rPVL
GS-II (EY Lab)
WGA multimer
rSRL
TbFBL2
SRL
GAC78
WGA

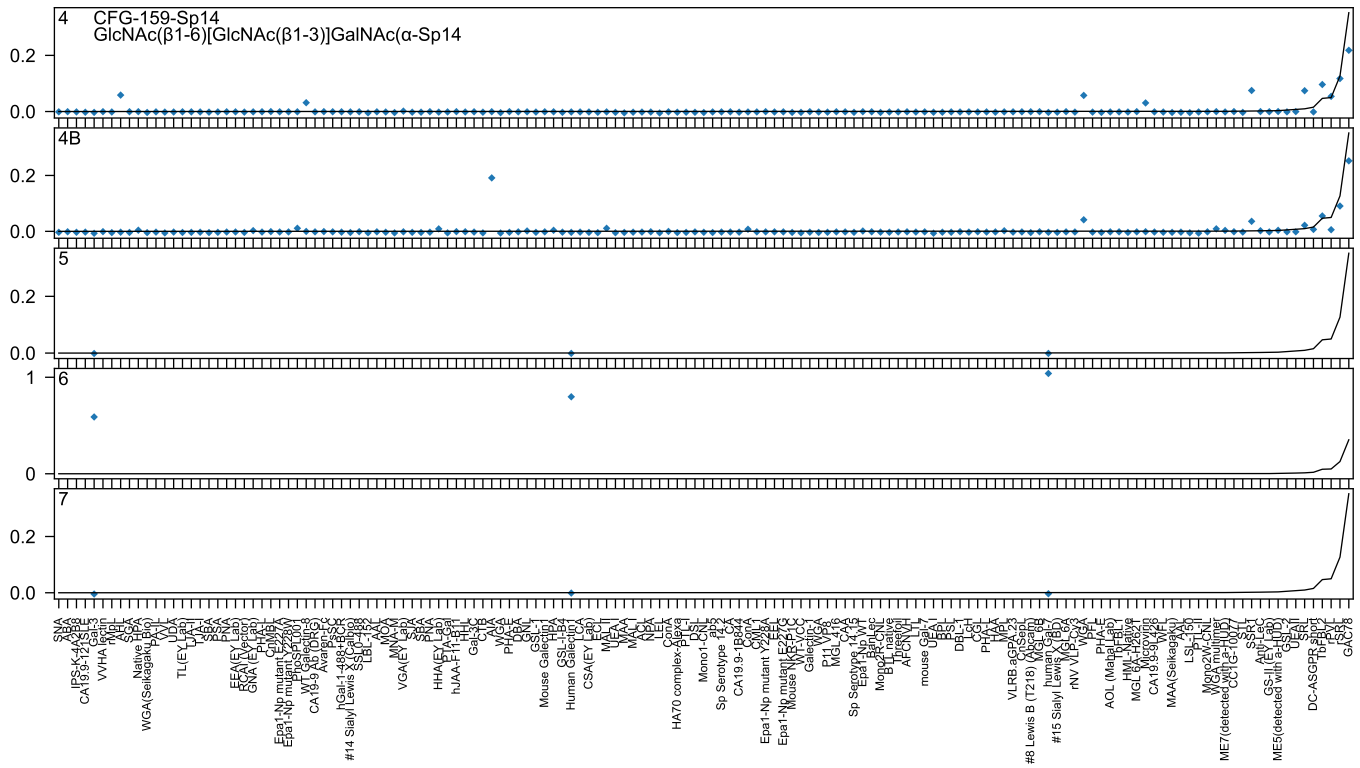
4
CFG-159-Sp14
GlcNAc(β1-6)[GlcNAc(β1-3)]GalNAc(α-Sp14
4B
5
6
7

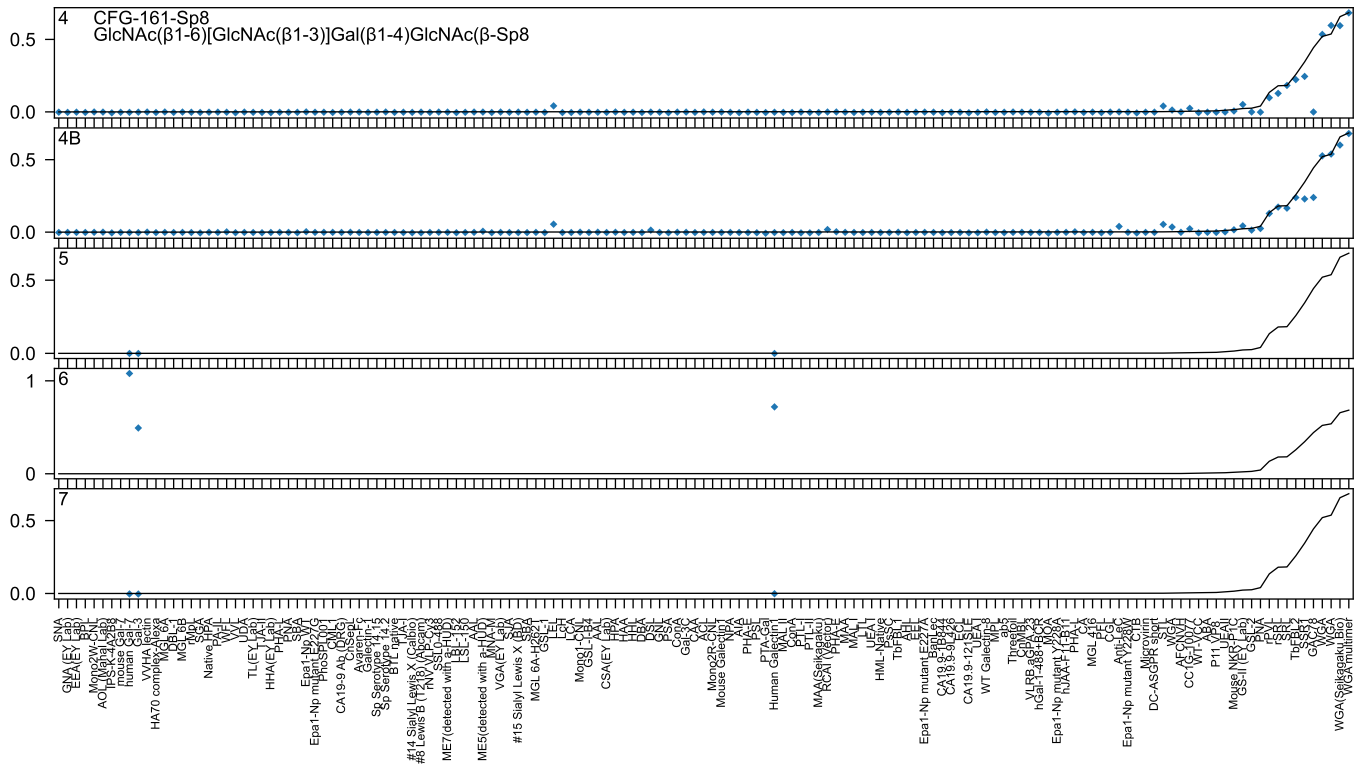

4
CFG-161-Sp8
GlcNAc(β1-6)[GlcNAc(β1-3)]Gal(β1-4)GlcNAc(β-Sp8
4B
5
6
7
0.0
0.5
0
1
SNA
GNA(EY Lab)
EEA(EY Lab)
BPL
Mono2W-CNL
AOL(Mahal Lab)
TPS-K-4A2B8
mouse Gal-7
human Gal-7
Gal-3
VVHA lectin
HA70 complex-Alexa
MGL 6A
DBL-1
MGL 6B
rMpL
SGA
Native HPA
PA-IL
WFL
VVL
UDA
TL(EY Lab)
TJA-II
HHA(EY Lab)
PHA-L
PNA
SBA
Epa1-Np WT
Epa1-Np mutant E227G
PhoSL001
CML1
CA19-9 Ab (DRG)
ChSepL
Avaren-Fc
Galectin-1
Sp Serotype 14.13
Sp Serotype 14.2
BTL native
TJA-I
#14 Sialyl Lewis X (Calbio)
#8 Lewis B (T218) (Abcam)
rNVL-CV3
SSL0-488
ME7(detected with a-HUD)
LBL-152
LSL-150
AAL
ME5(detected with a-HUD)
MNA-M
VGA(EY Lab)
SJA
#15 Sialyl Lewis X (BD)
SBA
MGL 6A-H262T
GSL-1
LEL
LcH
LCA
Mono1-CNL
GSL-I-B4
AAL
CSA(EY Lab)
HPA
HAA
HHL
DBA
DSL
GNL
PSA
ConA
Gal-3C
CAA
ACL
Mono2R-CNL
Mouse Galectin1
NPA
AIA
PHA-E
PSL
PTA-Gal
Human Galectin1
MAL II
ConA
PTL-I
PTL-II
MAA(Seikagaku)
RCAI (Vector)
PHA-E
MAA
MAL I
LTL
UEA I
HML-Native
PsSC
TbFBL1
AHL
EEL
Epa1-Np mutant E227A
BanLec
CA19.9-1B844
CA19.9-9L426
ECL
CA19.9-121SLE
UEA I
WT Galectin-8
MPL
ab5
Threefoil
CnMBL
VLRB.aGPA.23
hGal-1-488+BCR
MOA
Epa1-Np mutant Y228A
hJAA-F11-B11
PHA-L
CA
MGL 416
PFL
CGL
Anti-LeC
Epa1-Np mutant Y228W
CTB
Microvirin
DC-ASGPR short
STL
WGA
AFCNVH
CC1G-10077
WT-VCC
ABA
P11 VP8
UEAII
Mouse NKR-P1C
GS-II (EY Lab)
GSL-2
PNA
rPVL
rSRL
SRL
TbFBL2
SSR1
GAC78
WGA
VGA
WGA(Seikagaku Bio)
WGA multimer

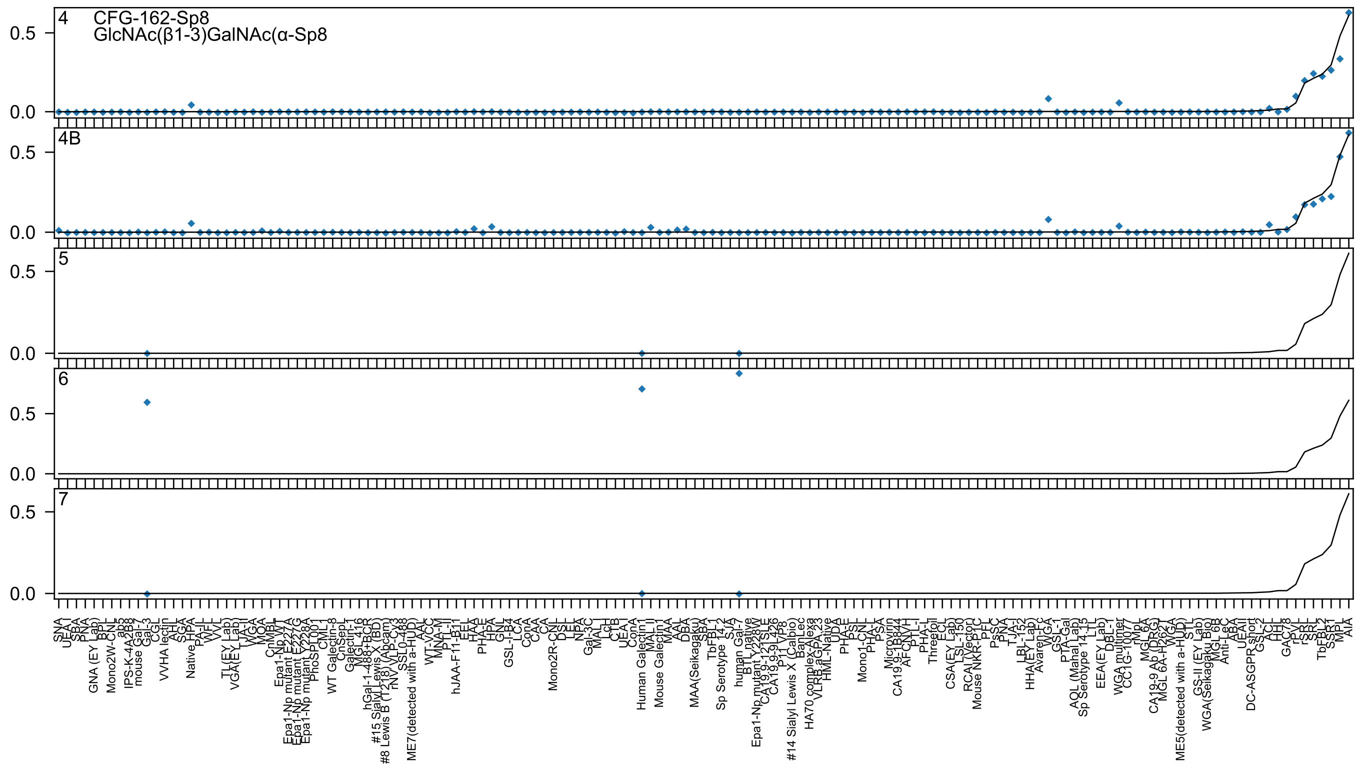

4
CFG-162-Sp8
GlcNAc(β1-3)GalNAc(α-Sp8
4B
5
6
7
0.5
0.0

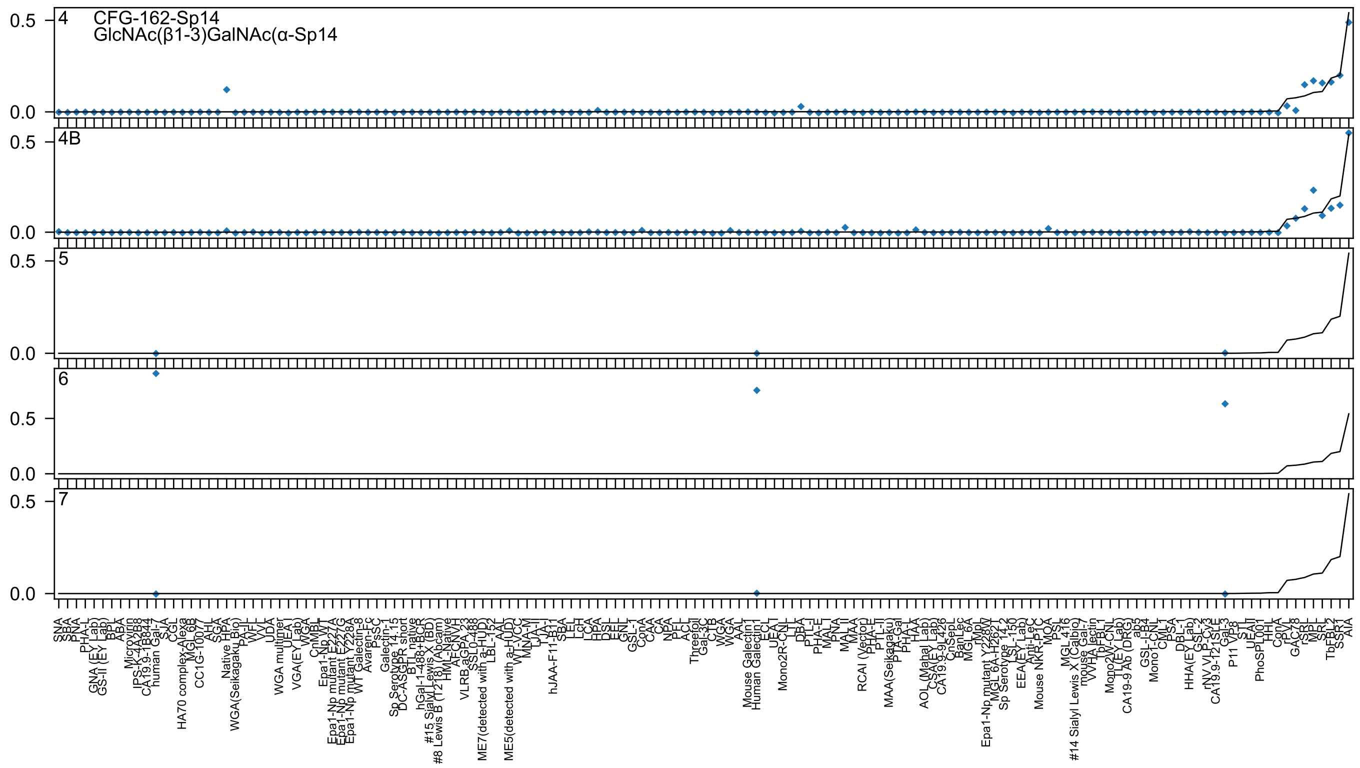

4
CFG-162-Sp14
GlcNAc(β1-3)GalNAc(α-Sp14
4B
5
6
7

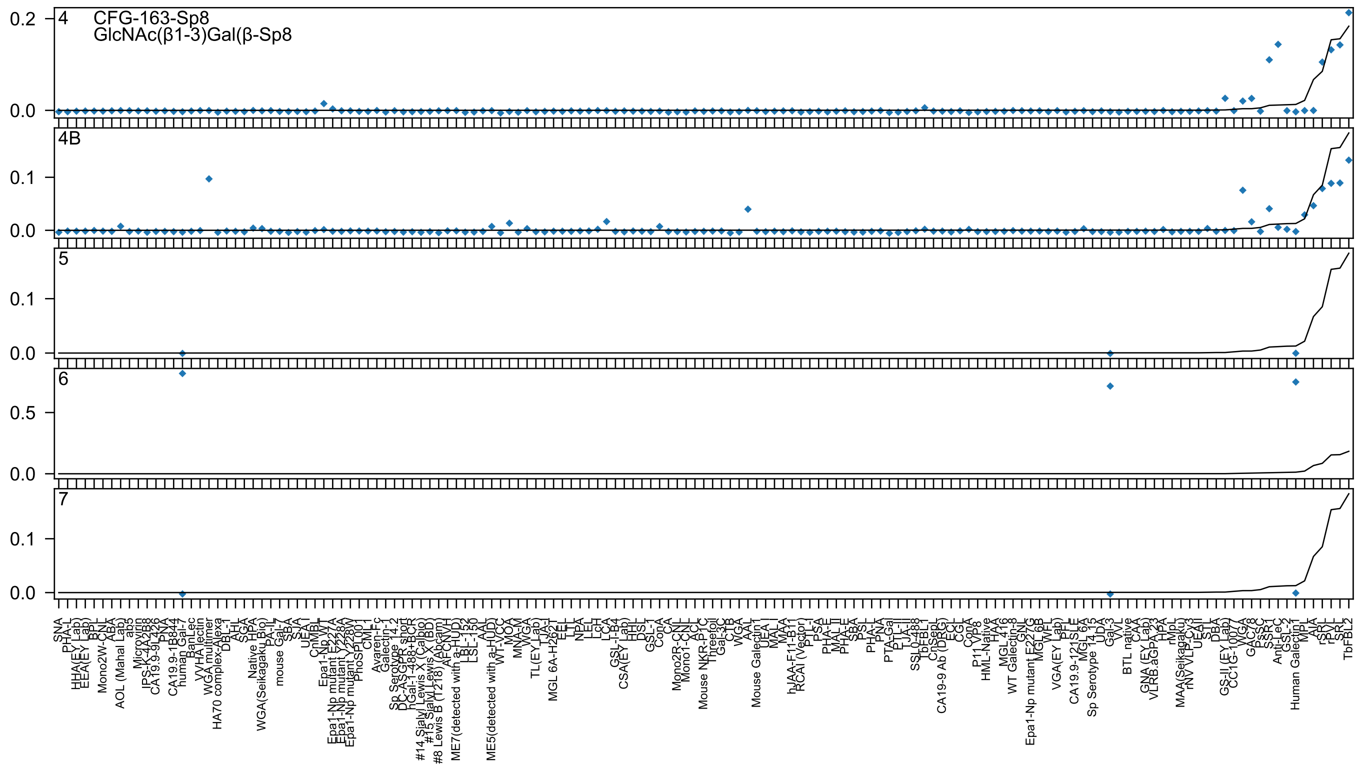

CFG-163-Sp8
GlcNAc(β1-3)Gal(β-Sp8
4
4B
5
6
7
0.0
0.1
0.2
0.5

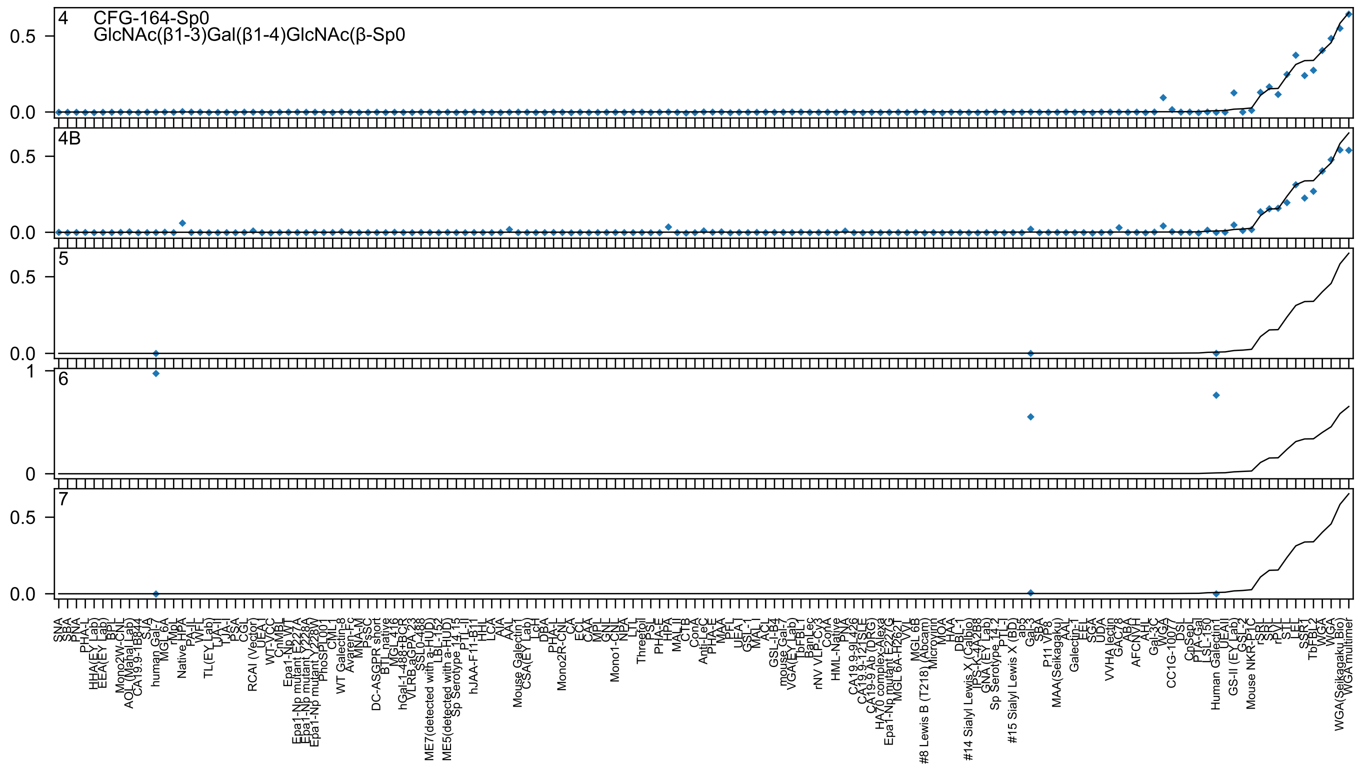

4
CFG-164-Sp0
GlcNAc(β1-3)Gal(β1-4)GlcNAc(β-Sp0
4B
5
6
7
0.5
0.0
1
0

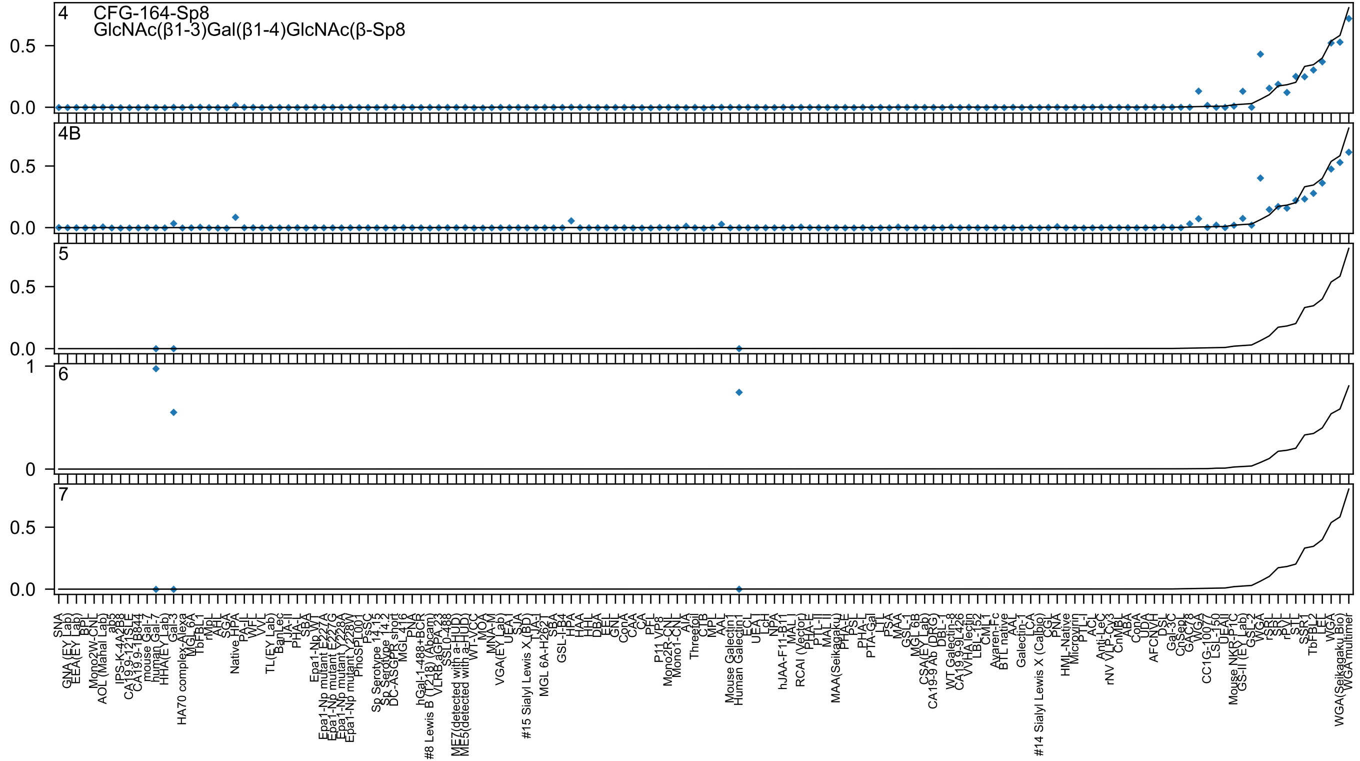
4
CFG-164-Sp8
GlcNAc(β1-3)Gal(β1-4)GlcNAc(β-Sp8
4B
5
6
7
0.5
0.0
1
0

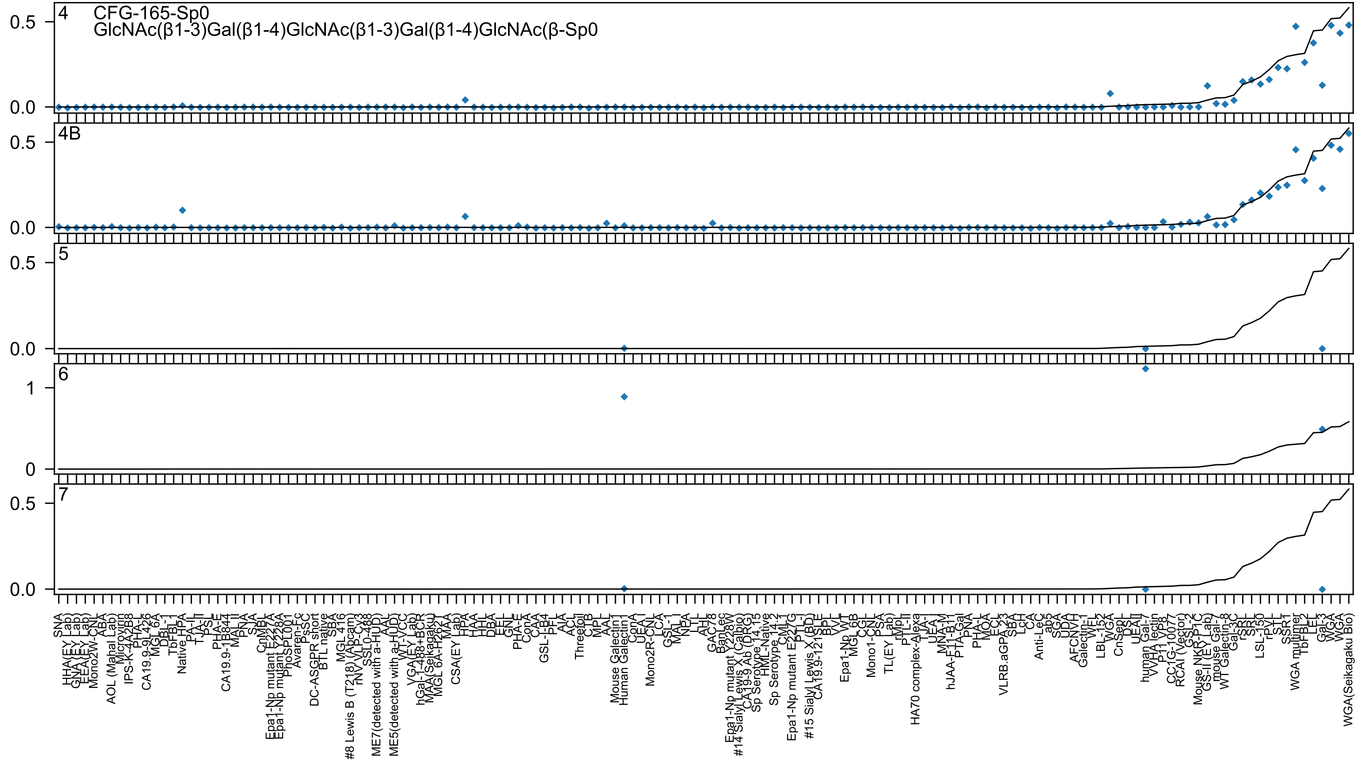
4
CFG-165-Sp0
GlcNAc(β1-3)Gal(β1-4)GlcNAc(β1-3)Gal(β1-4)GlcNAc(β-Sp0
4B
5
6
7
0.0
0.5
0
1
SNA
HHA(EY Lab)
GNA (EY Lab)
EEA(EY Lab)
Mono2W-CNL
ABA
AOL (Mahal Lab)
Microvirin
IPS-K-4A2B8
PHA-L
CA19.9-9L426
MGL 6A
DBL-1
TbFBL-1
Native HPA
PA-II
TJA-II
PSL
PHA-E
CA19.9-1B844
MAL II
PNA
SJA
CnMBL
Epa1-Np mutant E227A
Epa1-Np mutant Y228A
PhoSL001
Avaren-Fc
PsSC
DC-ASGPR short
BTL native
SBA
MGL 416
#8 Lewis B (T218) (Abcam)
rNV VLP-Cy3
SSL0-488
ME7(detected with a-HUD)
AAL
ME5(detected with a-HUD)
WT-VCC
VGA(EY Lab)
hGal-1-488+BCR
MAA(Seikagaku)
MGL 6A-H262T
MAA
CSA(EY Lab)
HPA
HAA
HHL
DBA
EEL
GNL
PHA-E
ConA
CAA
GSL-I-B4
PFL
AIA
ACL
Threefoil
CTB
MPL
AAL
Mouse Galectin1
Human Galectin1
ConA
UEA-I
Mono2R-CNL
LCA
GSL-1
MAL-I
NPA
LTL
AH
GAC78
BanLec
Epa1-Np mutant Y228W
#14 Sialyl Lewis X (Calbio)
CA19-9 Ab (DRG)
Sp Serotype 14.15
HML-Native
Sp Serotype 14.2
CML-1
Epa1-Np mutant E227G
PTL-I
#15 Sialyl Lewis X (BD)
CA19.9-121SLE
BPL
VVL
Epa1-Np WT
MGL 6B
CGL
Mono1-CNL
PSA
TL(EY Lab)
rMpL
PTL-II
HA70 complex-Alexa
TJA-I
UEA-I
MNA-M
hJAA-F11-B11
PTA-Gal
PNA
PHA-L
MOA
ECL
VLRB.aGPA.23
SBA
LcH
CA
Anti-LeC
ab5
SGA
UDA
AFCNVH
Galectin-1
WFL
LBL-152
WGA
CnSepL
DSL
UEA-II
human Gal-7
VVHA lectin
P11 VP8
CC1G-10077
RCAI (Vector)
GSL-2
Mouse NKR-P1C
GS-II (EY Lab)
mouse Gal-7
WT Galectin-8
Gal-3C
rSRL
SRL
LSL-150
rPVL
STL
SSR1
WGA multimer
TbFBL2
LEL
Gal-3
WGA
WGA
WGA(Seikagaku Bio)

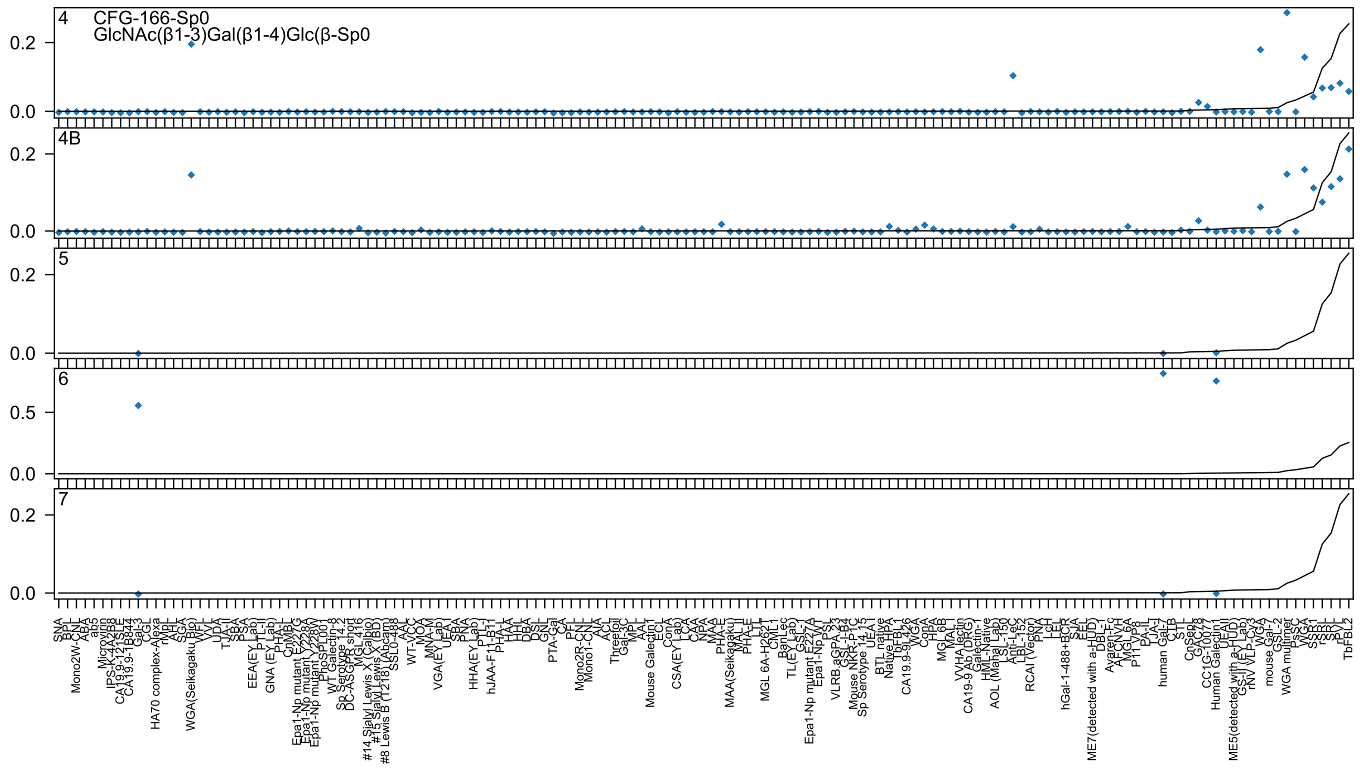

4
CFG-166-Sp0
GlcNAc(β1-3)Gal(β1-4)Glc(β-Sp0
4B
5
6
7
0.0
0.2
0.5

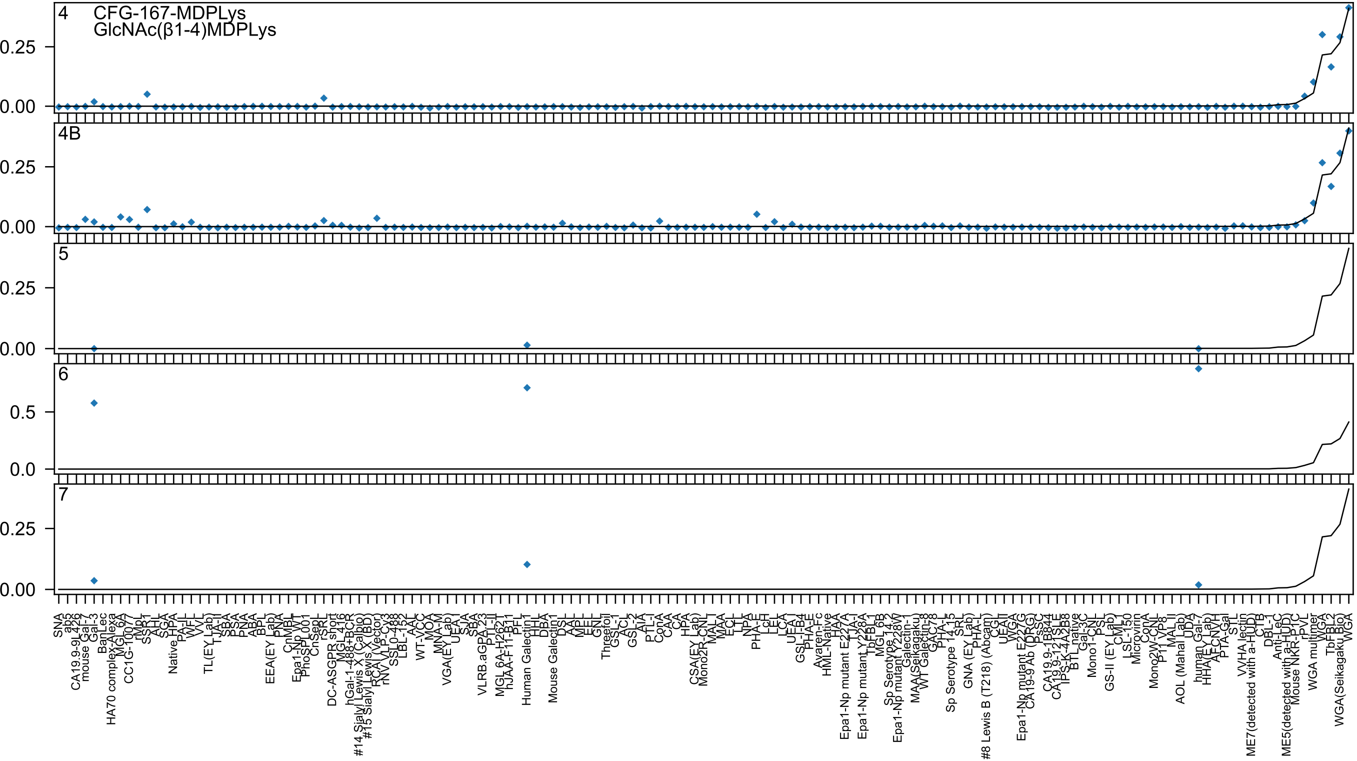

4
CFG-167-MDPLys
GlcNAc(β1-4)MDPLys
4B
5
6
7

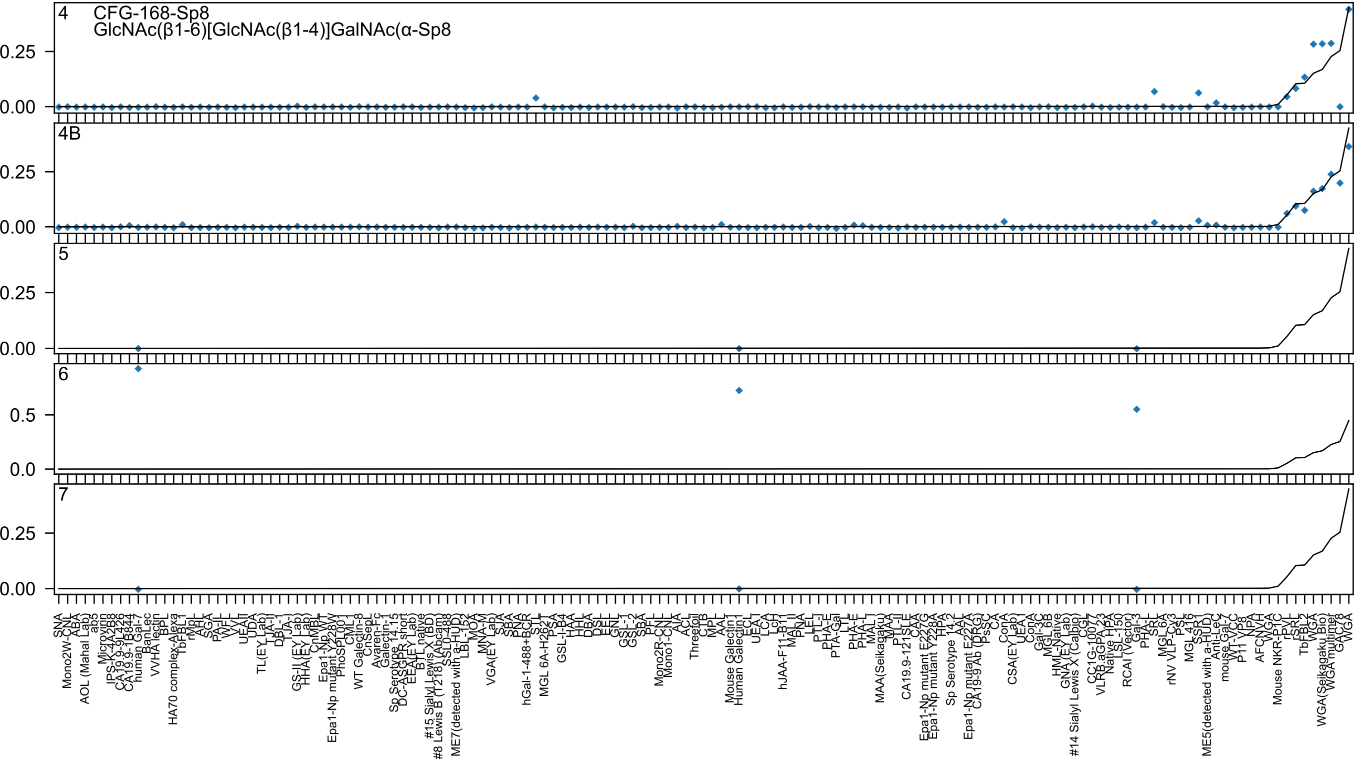
4
CFG-168-Sp8
GlcNAc(β1-6)[GlcNAc(β1-4)]GalNAc(α-Sp8
4B
5
6
7
0.25
0.00
0.5
0.0
SNA
Mono2W-CNL
ABA
AOL (Mahal Lab)
ab5
Microvirin
IPS-K-4A2B8
CA19.9-9L426
CA19.9-1B844
human Gal-7
BanLec
VVHA lectin
BPL
HA70 complex-Alexa
TbFBL1
rMpL
AHL
SGA
PA-IL
WFL
VVL
UEAII
UDA
TL(EY Lab)
TJA-II
DBL-I
TJA-I
GS-II (EY Lab)
HHA(EY Lab)
CnMBL
Epa1-Np WT
Epa1-Np mutant Y228W
PhoSL001
CML1
WT Galectin-8
CnSepL
Avaren-Fc
Galectin-1
Sp Serotype 14.15
DC-ASGPR short
EEA(EY Lab)
BTL native
#15 Sialyl Lewis X (BD)
#8 Lewis B (T218) (Abcam)
SSL0-488
ME7(detected with a-HUD)
LBL-152
MOA
MNA-M
VGA(EY Lab)
SJA
SBA
PNA
hGal-1-488+BCR
STL
MGL 6A-H262T
PSA
GSL-I-B4
HAA
HHL
DBA
DSL
EEL
GNL
GSL-1
GSL-2
SBA
PFL
Mono2R-CNL
Mono1-CNL
AIA
ACL
Threefoil
CTB
MPL
AAL
Mouse Galectin1
Human Galectin1
ECL
UEA I
LCA
LcH
hJAA-F11-B1
MAL II
PNA
LEL
PTL-I
PHA-E
PTA-Gal
LTL
PHA-E
PHA-L
MAL I
MAA(Seikagaku)
MAA
PTL-II
CA19.9-121SLE
CAA
Epa1-Np mutant E227G
Epa1-Np mutant Y228A
HPA
Sp Serotype 14.2
AAA
Epa1-Np mutant E227A
CA19-9 Ab (DRG)
PsSC
CA
ConA
CSA(EY Lab)
UEA I
ConA
Gal-3C
MGL 6B
HML-Native
GNA (EY Lab)
#14 Sialyl Lewis X (Calbo)
CGL
CC1G-1007
VLRB.aGPA.23
Native HPA
LSL-150
RCAI (Vector)
Gal-3
PHA-L
SRL
MGL 6A
rNV VLP-Cy3
PSL
MGL 416
SSP1
ME5(detected with a-HUD)
Anti-LeC
mouse Gal-7
WT-VCC
P11 VP8
NPA
AFCNVH
WGA
Mouse NKR-P1C
rPVL
rSRL
TbFBL2
WGA
WGA(Seikagaku Bio)
WGA multimer
GAC78
WGA

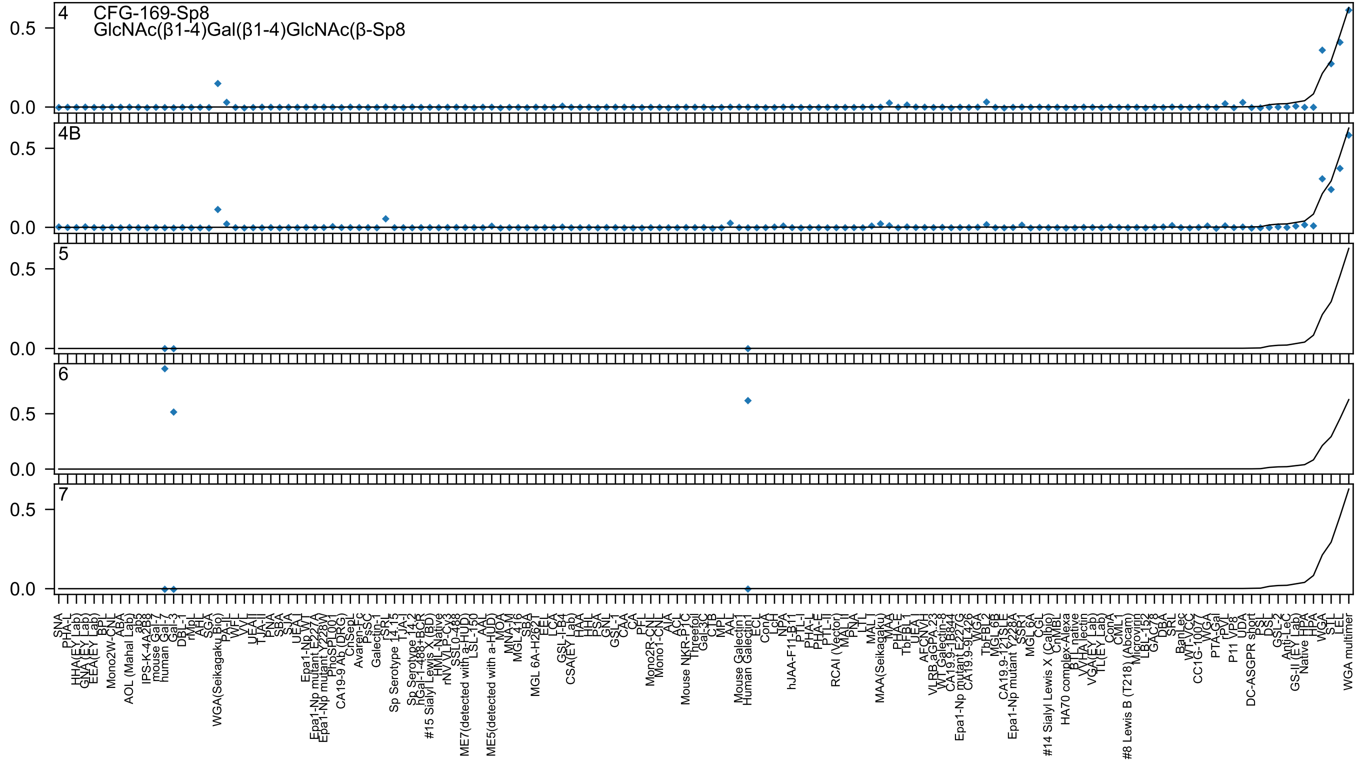
4
CFG-169-Sp8
GlcNAc(β1-4)Gal(β1-4)GlcNAc(β-Sp8
4B
5
6
7
0.5
0.0

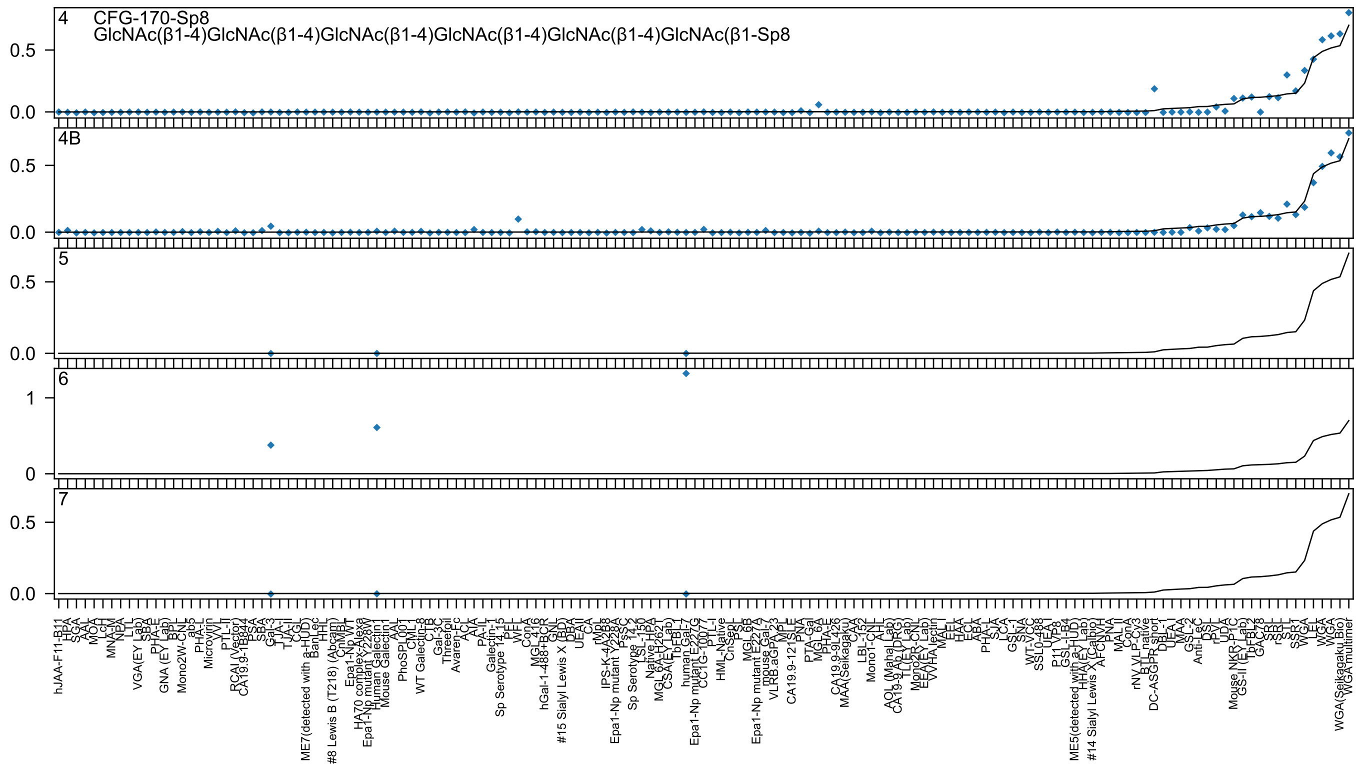
4 CFG-170-Sp8
GlcNAc(β1-4)GlcNAc(β1-4)GlcNAc(β1-4)GlcNAc(β1-4)GlcNAc(β1-4)GlcNAc(β1-Sp8
4B
5
6
7

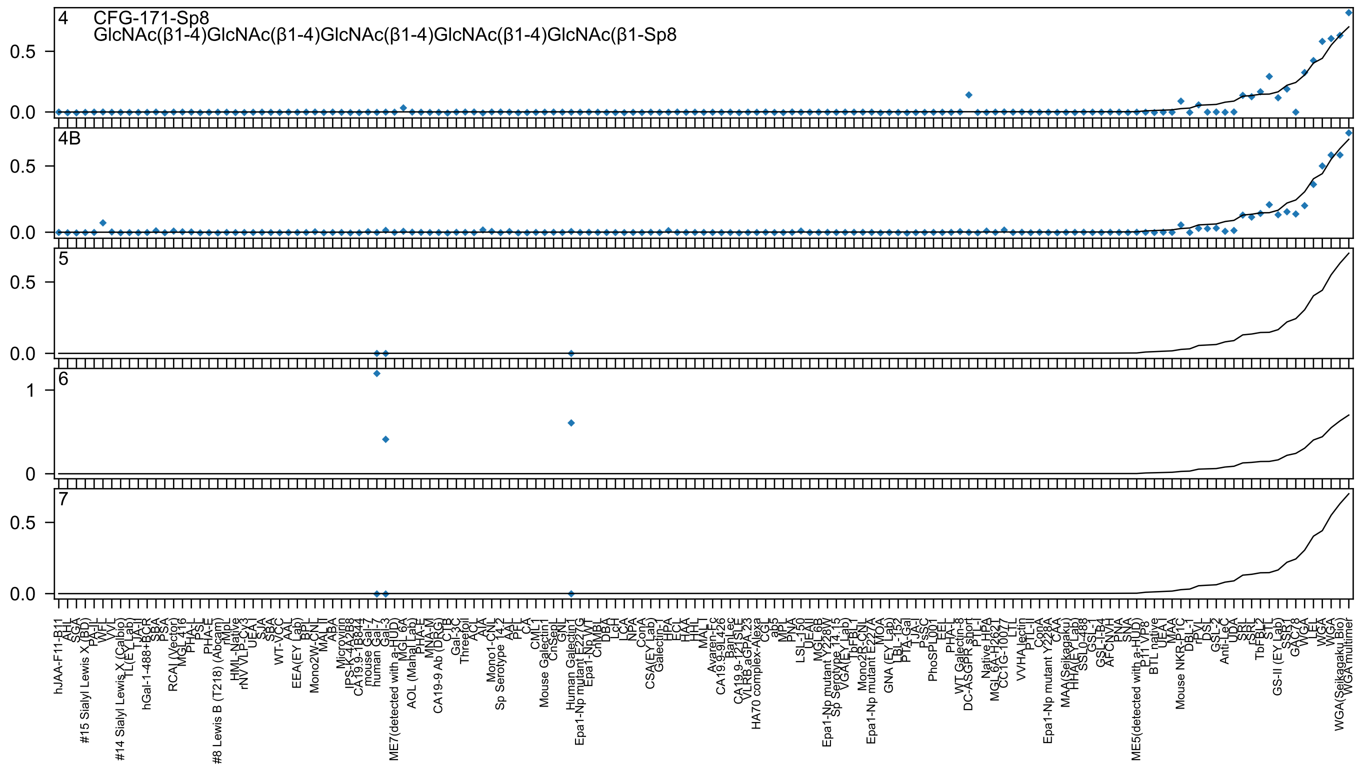

4 CFG-171-Sp8
GlcNAc(β1-4)GlcNAc(β1-4)GlcNAc(β1-4)GlcNAc(β1-4)GlcNAc(β1-Sp8
4B
5
6
7

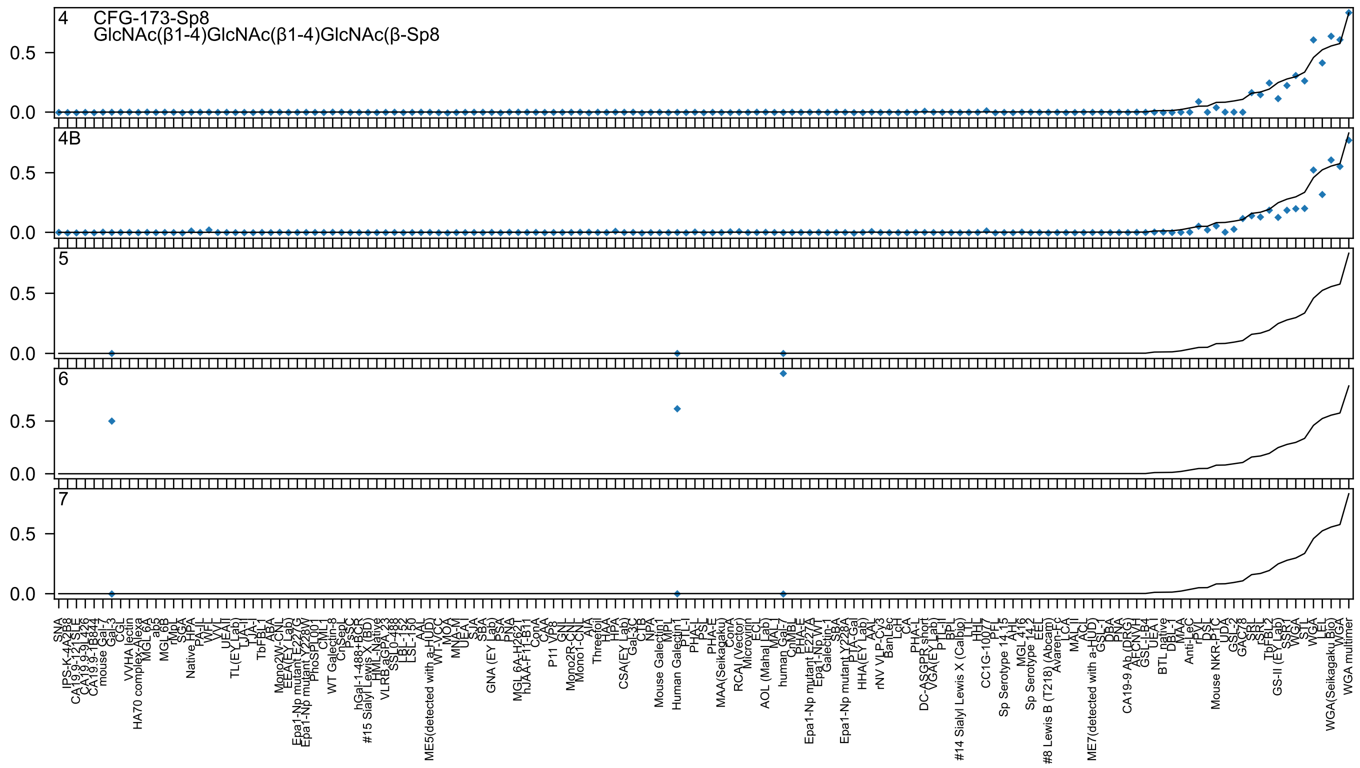

4
CFG-173-Sp8
GlcNAc(β1-4)GlcNAc(β1-4)GlcNAc(β-Sp8
4B
5
6
7
0.5
0.0

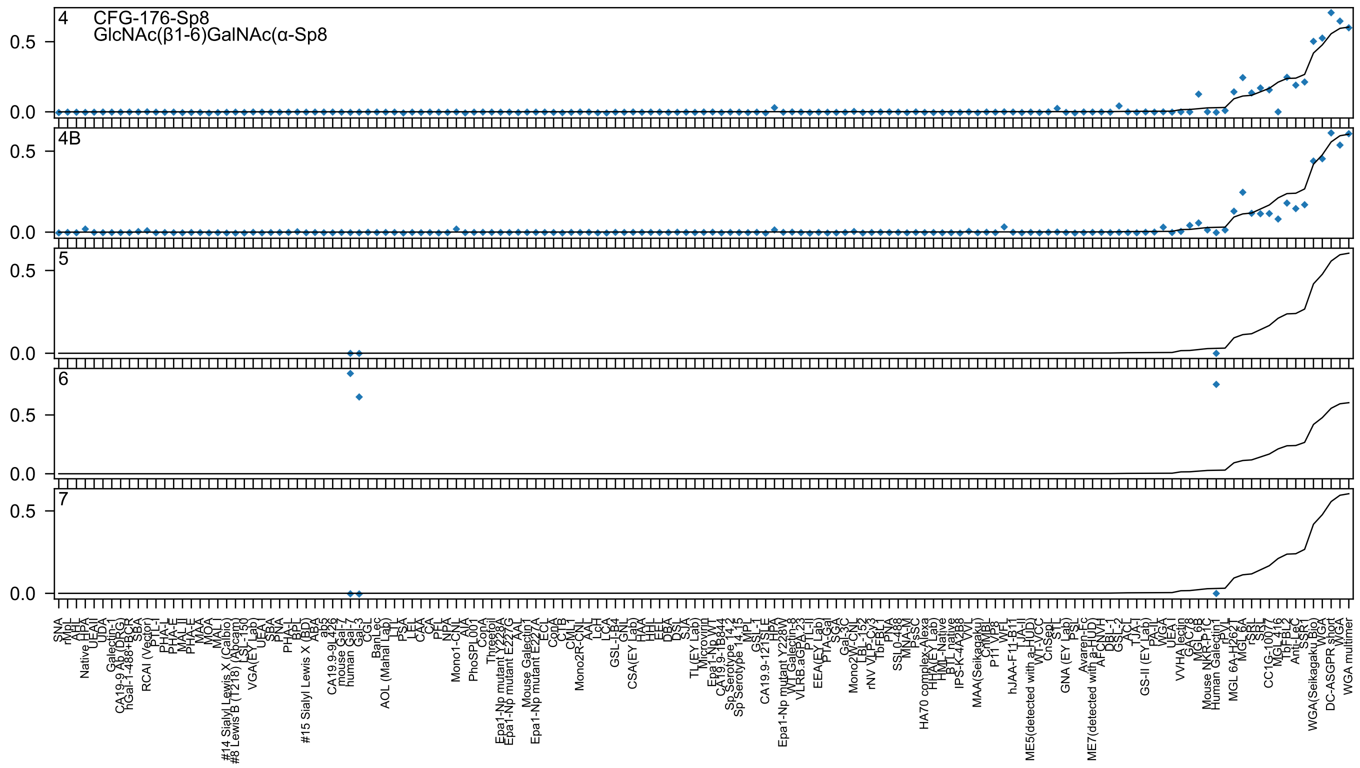
4 CFG-176-Sp8
GlcNAc(β1-6)GalNAc(α-Sp8
4B
5
6
7
0.5
0.0

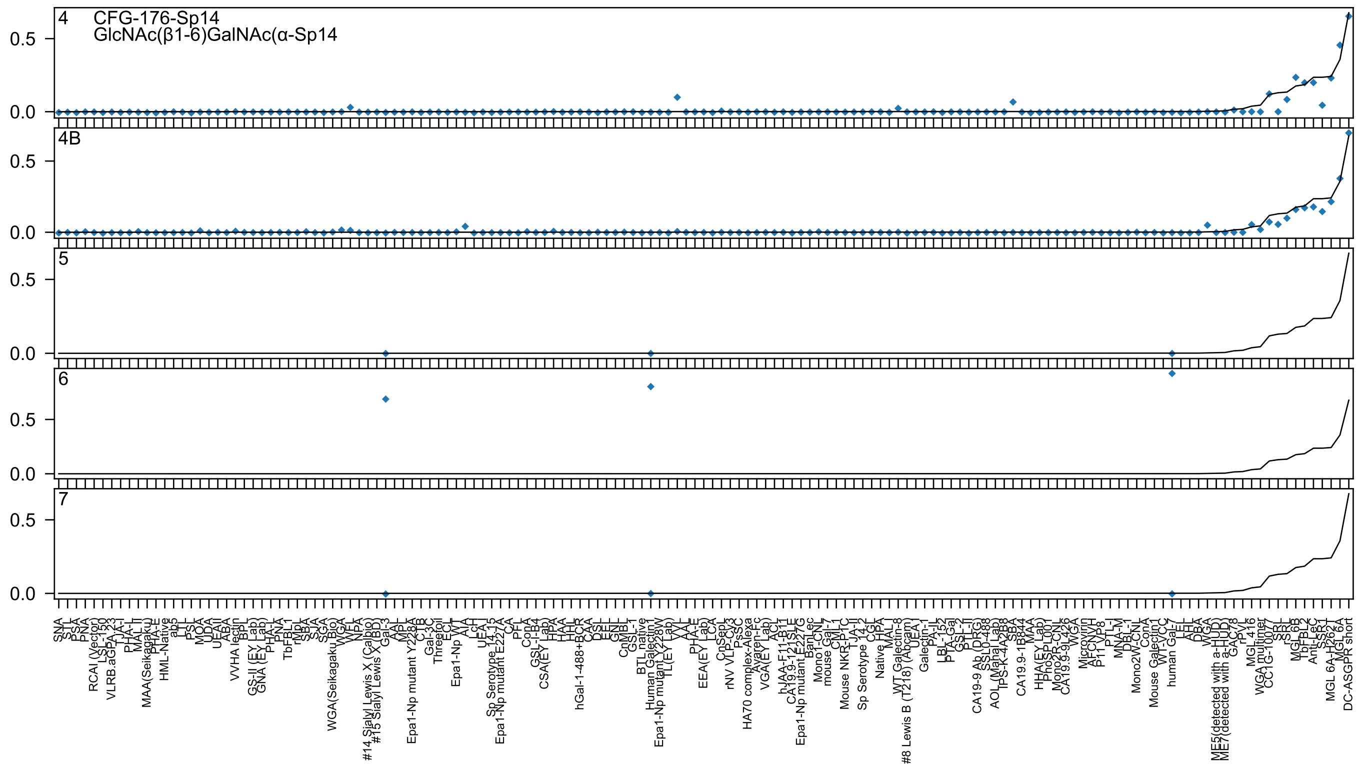
4 CFG-176-Sp14
GlcNAc(β1-6)GalNAc(α-Sp14
4B
5
6
7
0.5
0.0

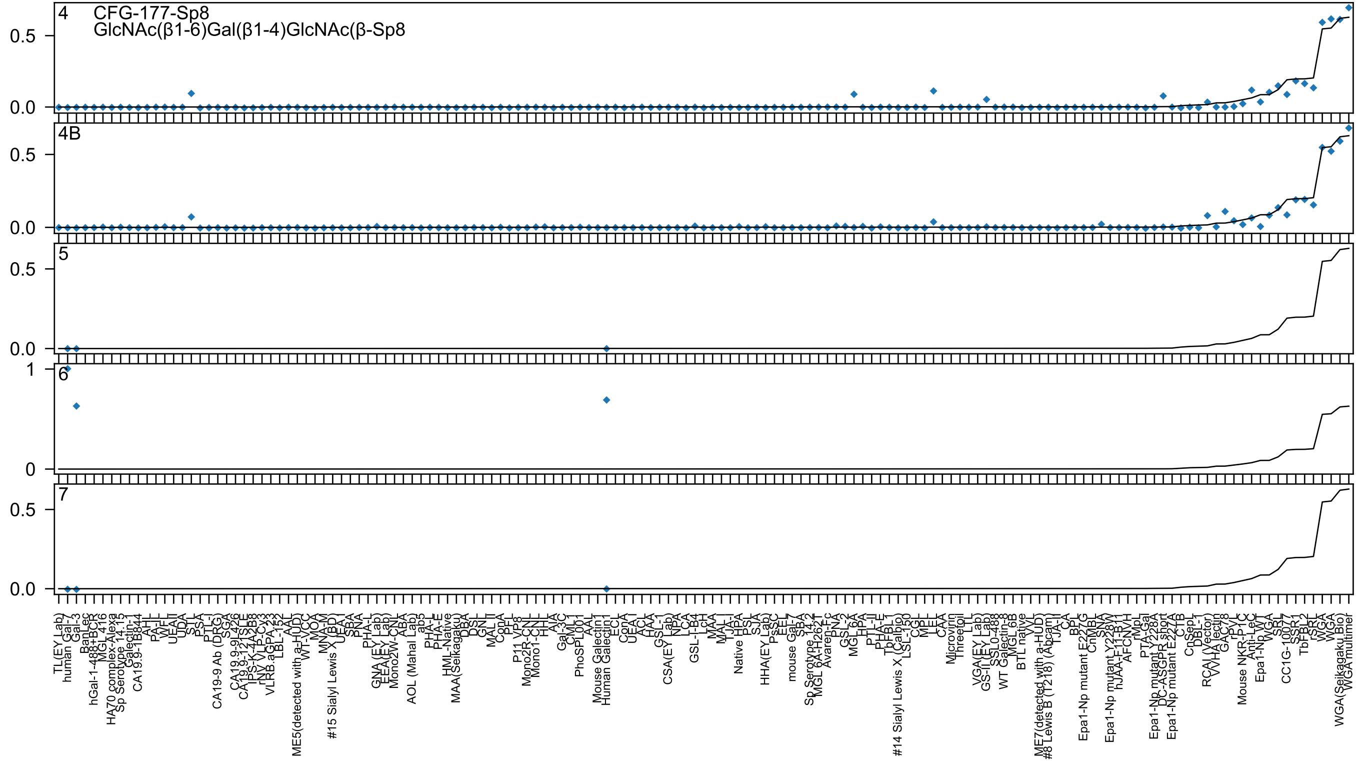
4 CFG-177-Sp8
GlcNAc(β1-6)Gal(β1-4)GlcNAc(β-Sp8
4B
5
6
7

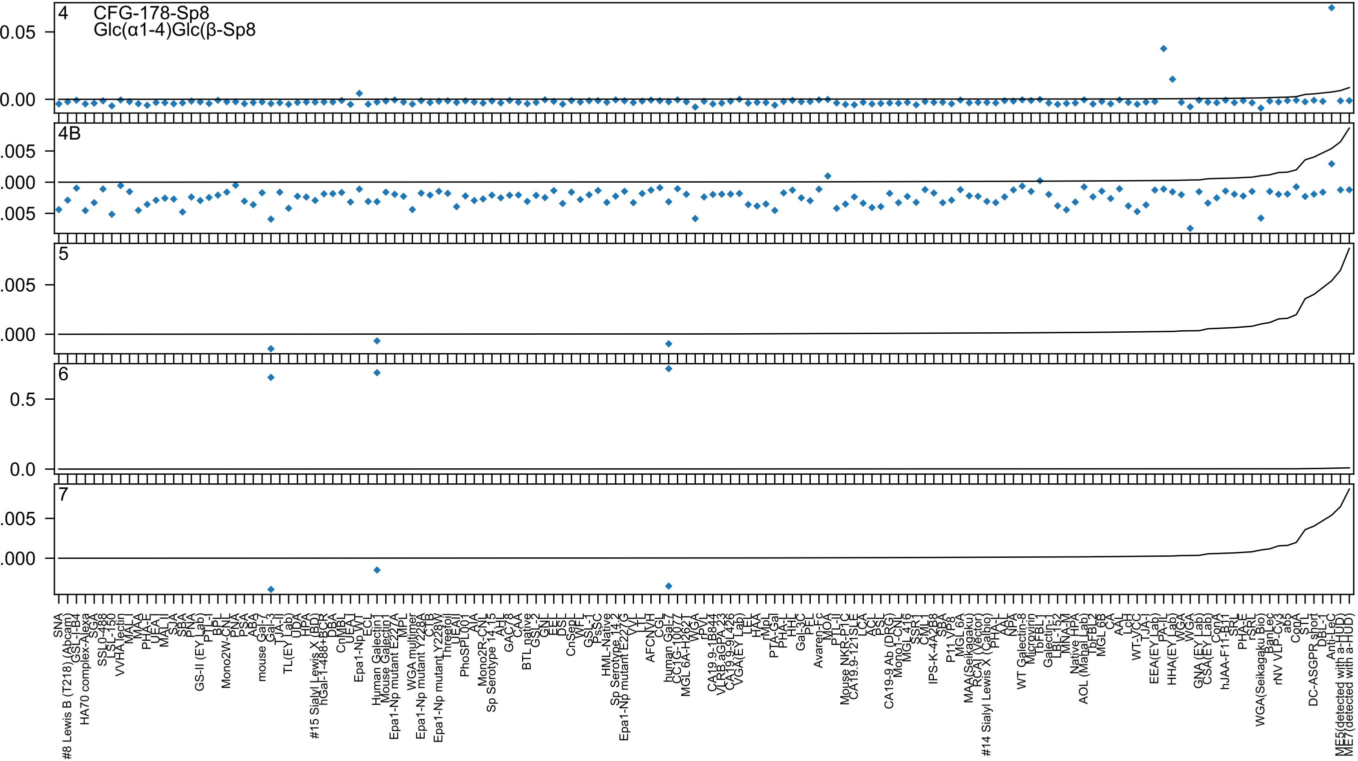

4
CFG-178-Sp8
Glc(α1-4)Glc(β-Sp8
4B
5
6
7

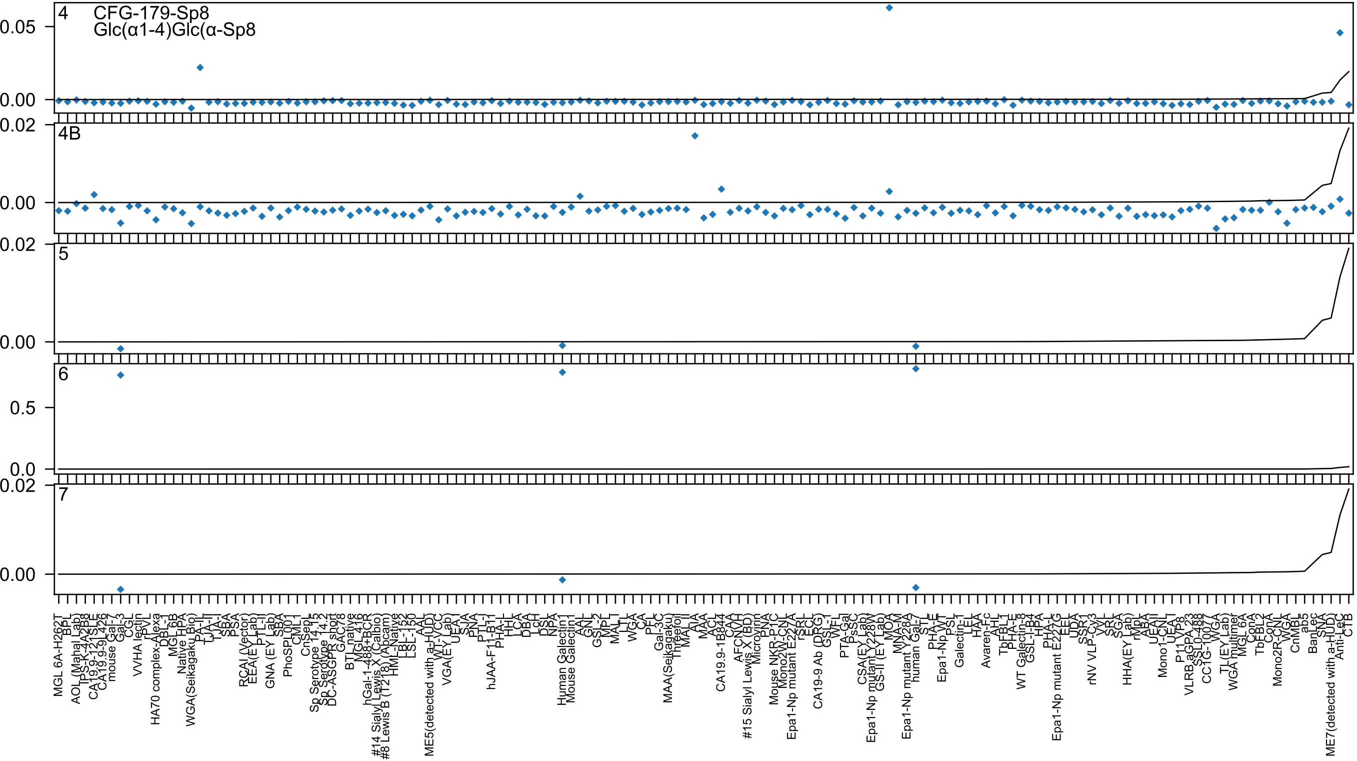
4 CFG-179-Sp8
Glc(α1-4)Glc(α-Sp8
4B
5
6
7

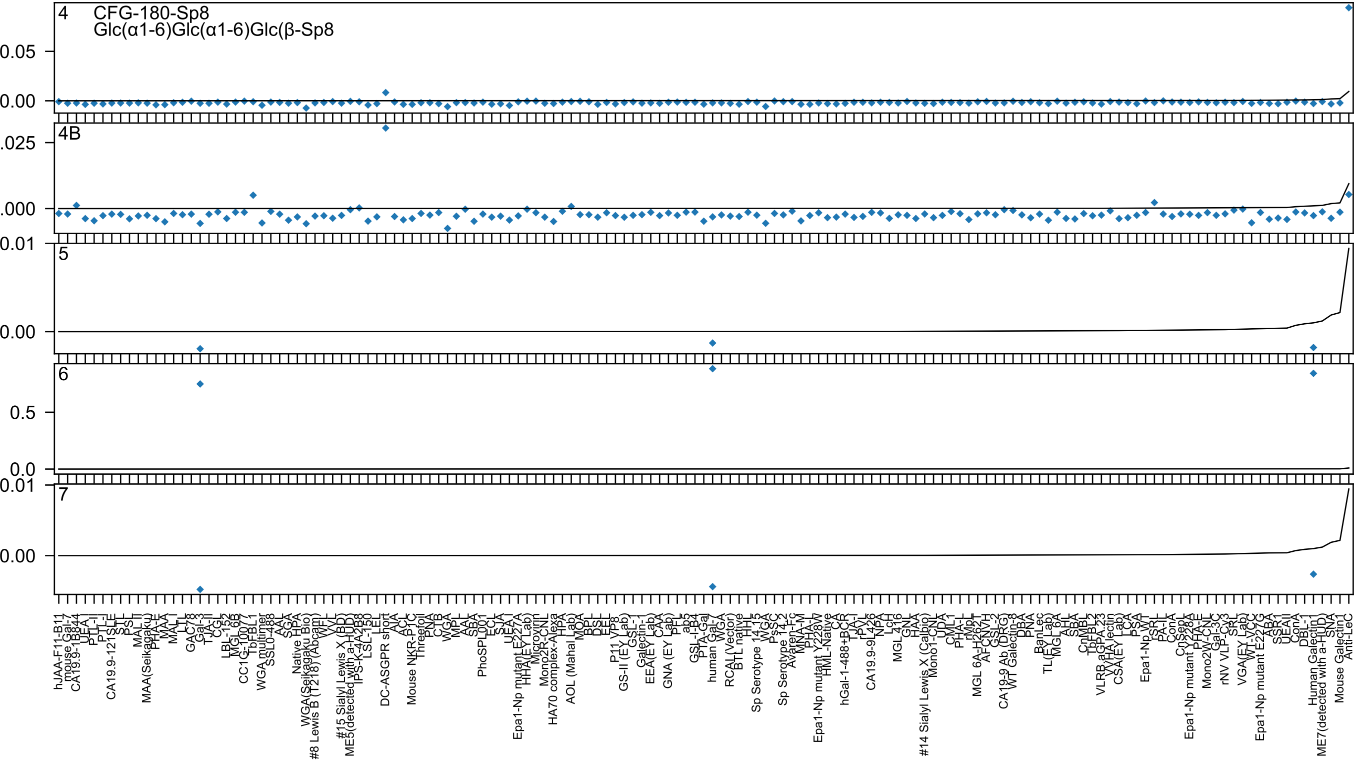
4
CFG-180-Sp8
Glc(α1-6)Glc(α1-6)Glc(β-Sp8
4B
5
6
7
0.05
0.00
.025
.000
0.01
0.5
0.0

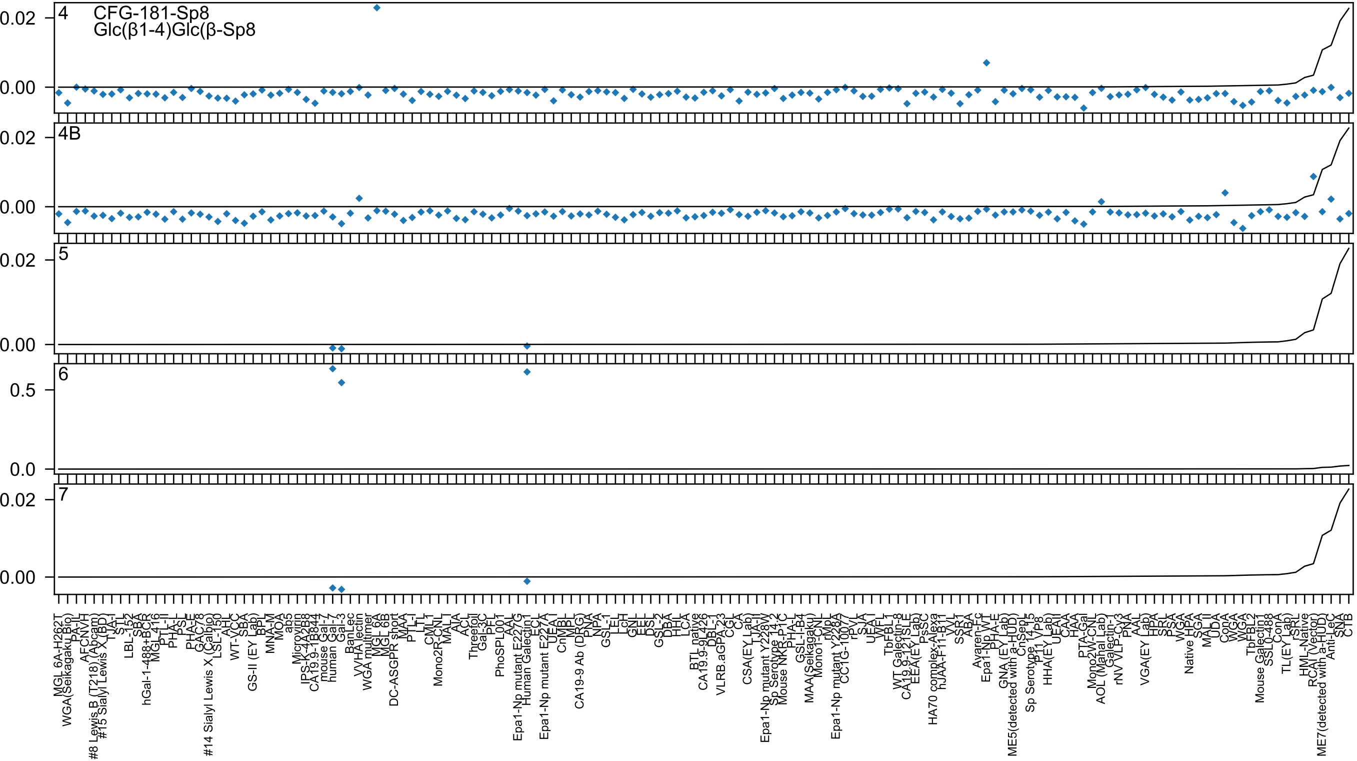

4
CFG-181-Sp8
Glc(β1-4)Glc(β-Sp8
4B
5
6
7

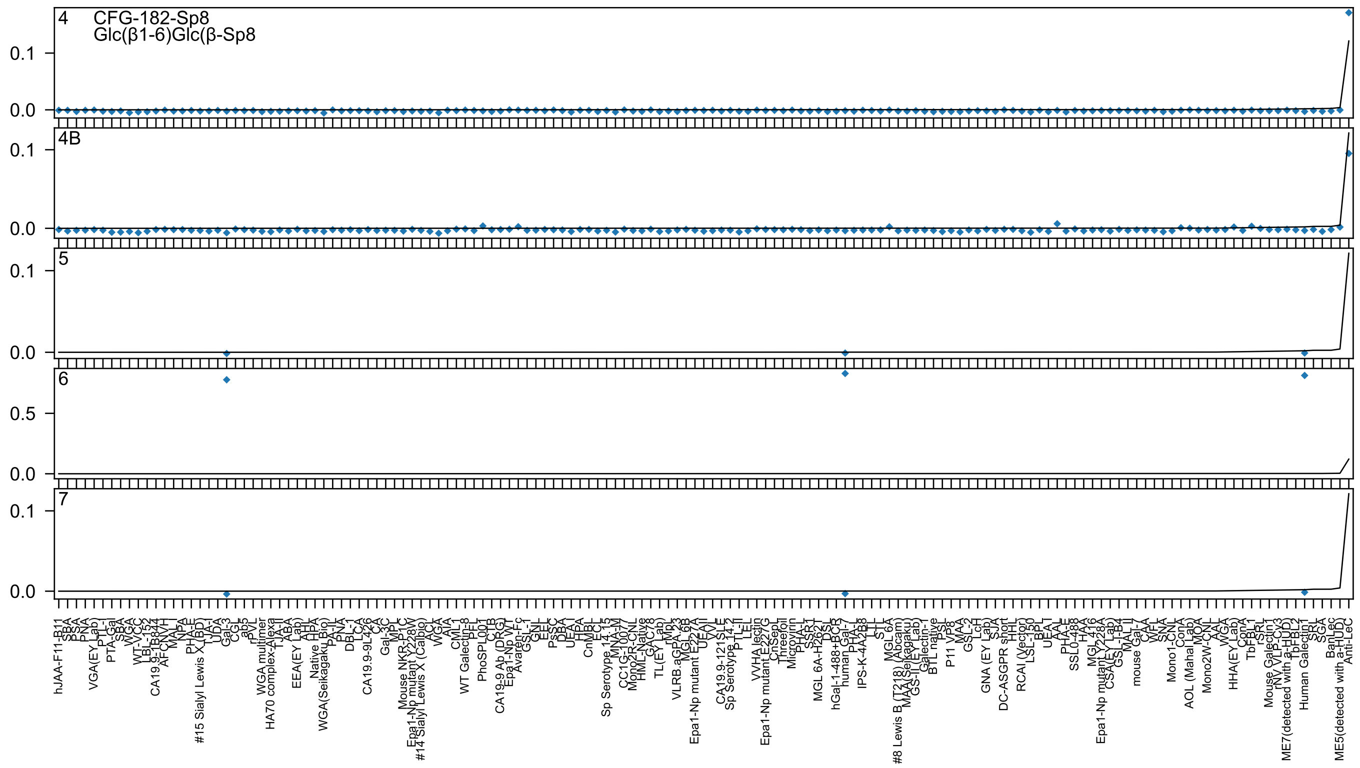

4 CFG-182-Sp8
Glc(β1-6)Glc(β-Sp8
4B
5
6
7

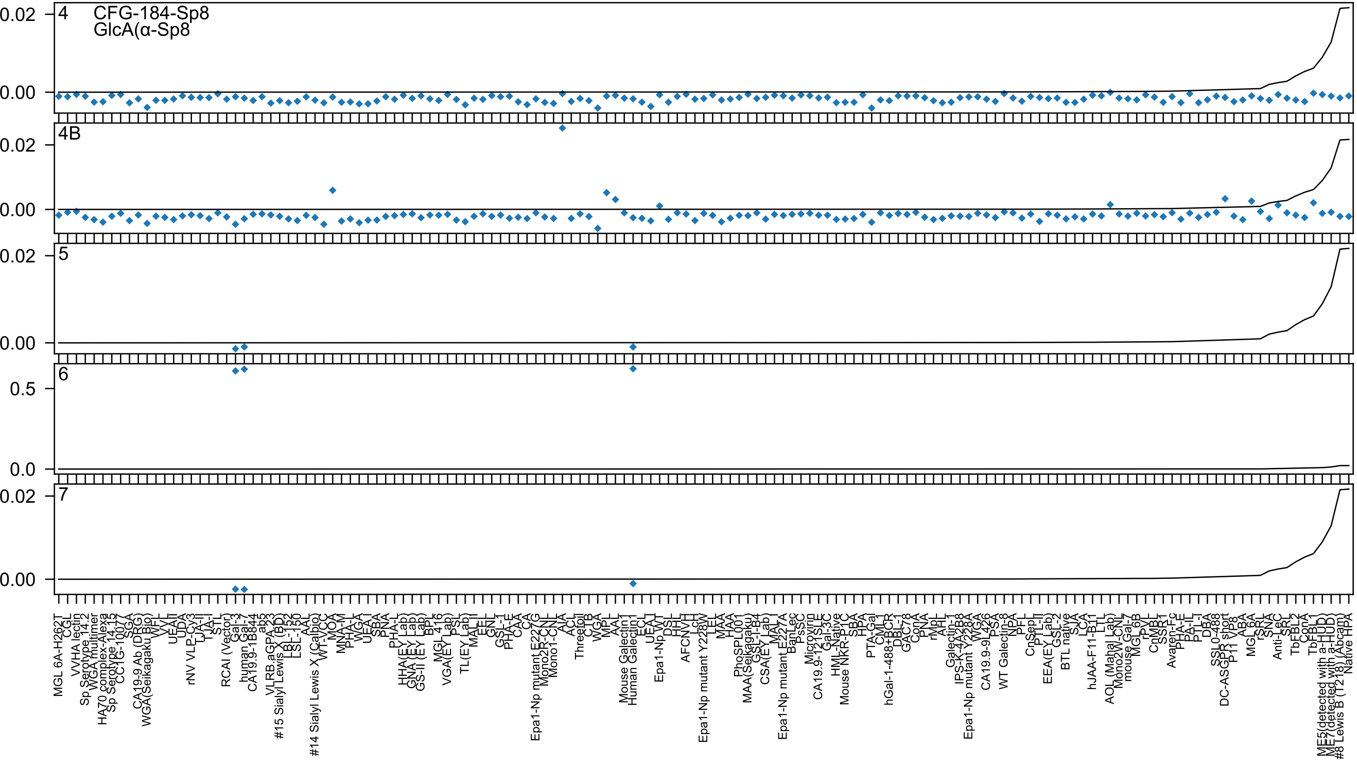
4
CFG-184-Sp8
GlcA(α-Sp8
4B
5
6
7

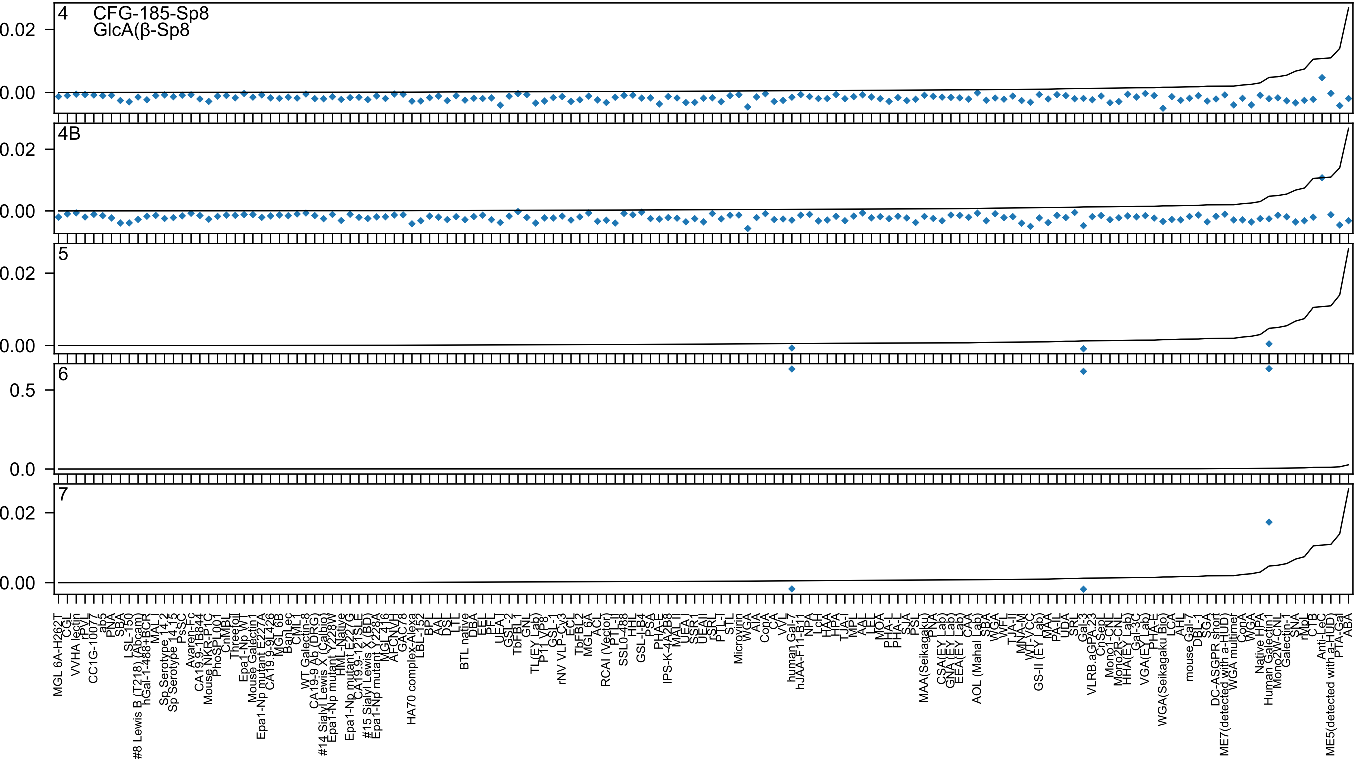
4
CFG-185-Sp8
GlcA(β-Sp8
4B
5
6
7
0.02
0.00
0.5
0.0
MGL 6A-H262T
CGL
VVHA lectin
rPY
CC1G-10077
ab5
PNA
SBA
LSL-150
#8 Lewis B (T218) (Abcam)
hGal-1-488+BCR
MAL-I
Sp Serotype 14.2
Sp Serotype 14.15
PsSC
Avaren-Fc
CA19.9-1B844
Mouse NKR-P1C
PhoSL001
CnMBL
Threefoil
Epa1-Np WT
Mouse Galectin1
Epa1-Np mutant E227A
CA19.9-9L426
MGL 6B
BanLec
CML-I
WT Galectin-8
CA19-9 Ab (DRG)
#14 Sialyl Lewis X (Calbio)
Epa1-Np mutant Y228W
HML-Native
Epa1-Np mutant E227G
CA19.9-121SLE
#15 Sialyl Lewis X (BD)
Epa1-Np mutant Y228A
MGL 4.16
AFCNVH
GAC78
HA70 complex-Alexa
LBL-152
BPL
AAL
DSL
LTL
BTL native
DBA
EEL
PFL
UEA-I
GSL-2
TbFBL1
GNL
TL(EY Lab)
P11 VP8
GSL-1
rNV VLP-Cy3
ECL
TbFBL2
MGL 6A
ACL
RCAI (Vector)
PTL-II
SSL0-488
HHL
GSL-I-B4
PSA
PHA-E
IPS-K-4A2B8
MAL-II
UEA-I
SSR1
UEAII
rSRL
PTL-I
STL
Microvirin
WGA
AIA
ConA
CA
VVL
human Gal-7
hJAA-F11-B11
NPA
LcH
HAA
HPA
TJA-I
MPL
AAL
LEL
MOA
PHA-L
PHA-L
SJA
PSL
MAA(Seikagaku)
PNA
CSA(EY Lab)
GNA(EY Lab)
EEA(EY Lab)
CAA
AOL (Mahal Lab)
SBA
WGA
WFL
TJA-II
MNA-M
WT-VCC
GS-II (EY Lab)
MAA
PA-IL
UDA
SRL
Gal-3
VLRB.aGPA.23
CnSepL
Mono1-CNL
Mono2R-CNL
HHA(EY Lab)
Gal-3C
VGA(EY Lab)
PHA-E
WGA(Seikagaku Bio)
LCA
AHL
mouse Gal-7
DBL-1
SGA
DC-ASGPR short
ME7(detected with a-HUD)
WGA multimer
ConA
WGA
Native HPA
Human Galectin1
Mono2W-CNL
Galectin-1
SNA
rMpL
CTB
Anti-LeC
ME5(detected with a-HUD)
PTA-Gal
ABA

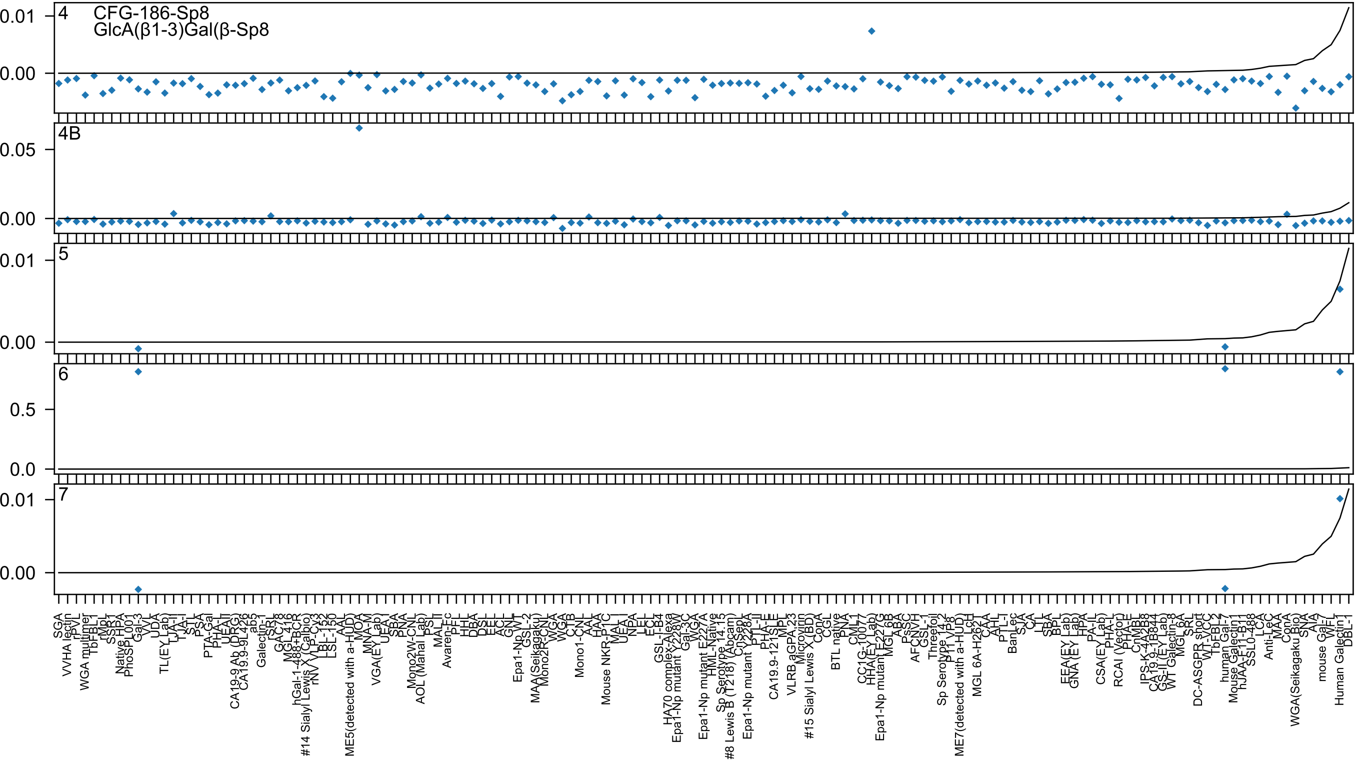
4
CFG-186-Sp8
GlcA(β1-3)Gal(β-Sp8
4B
5
6
7

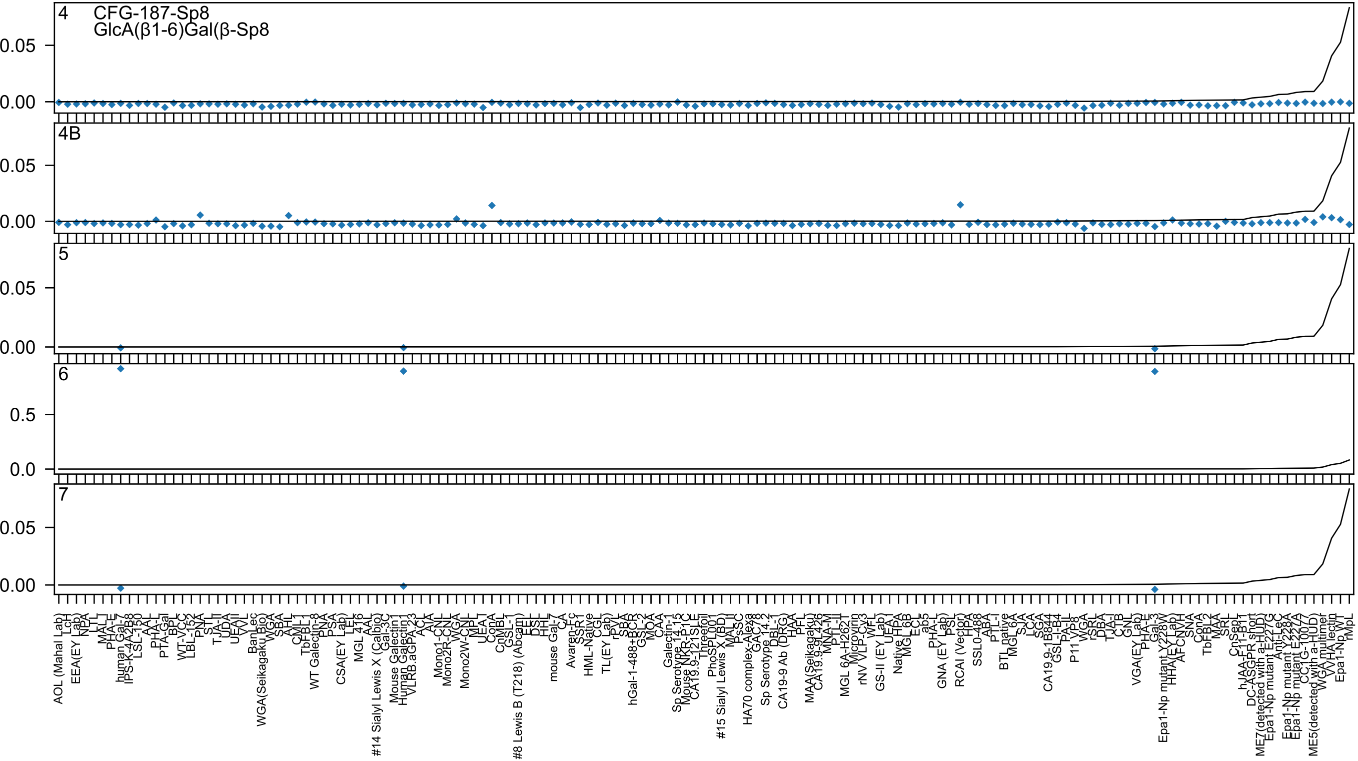
4
CFG-187-Sp8
GlcA(β1-6)Gal(β-Sp8
4B
5
6
7

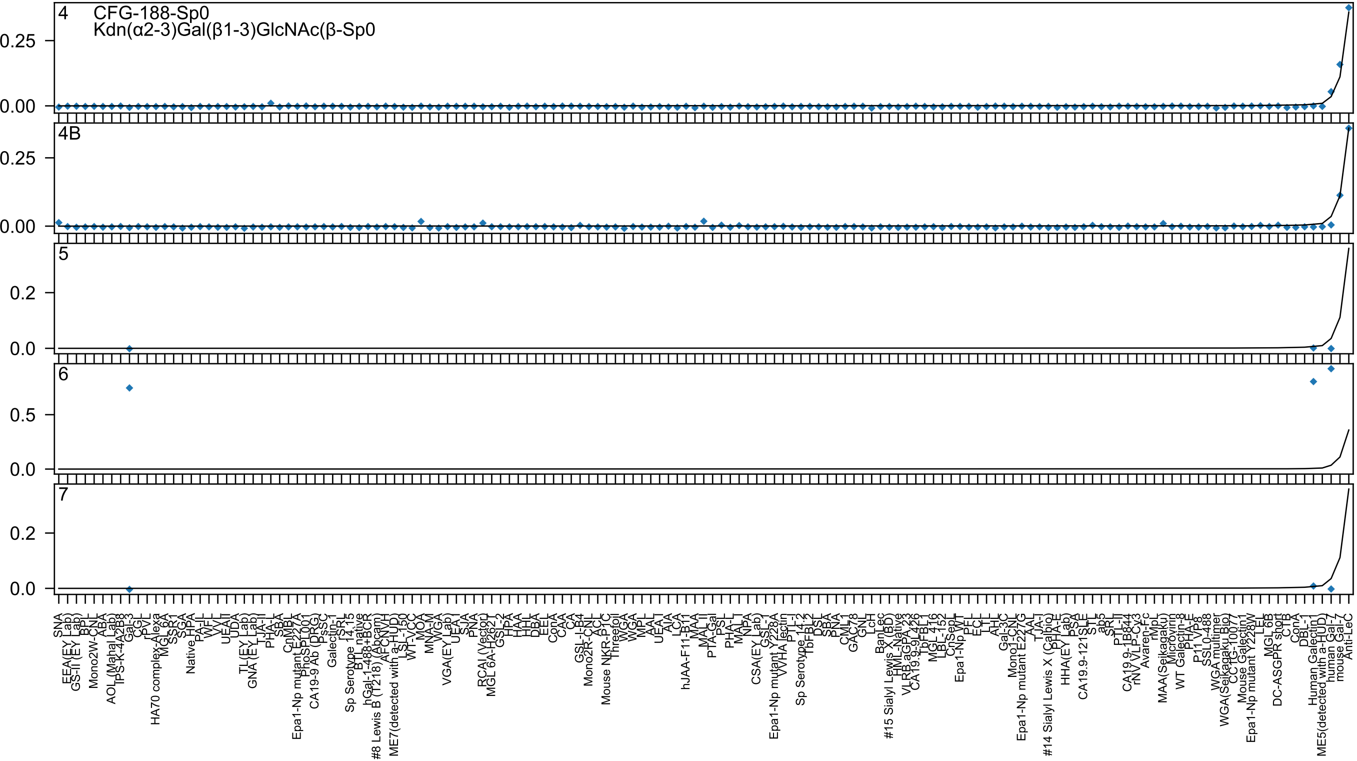

CFG-188-Sp0
Kdn(α2-3)Gal(β1-3)GlcNAc(β-Sp0
4
4B
5
6
7
SNA
Anti-LeC

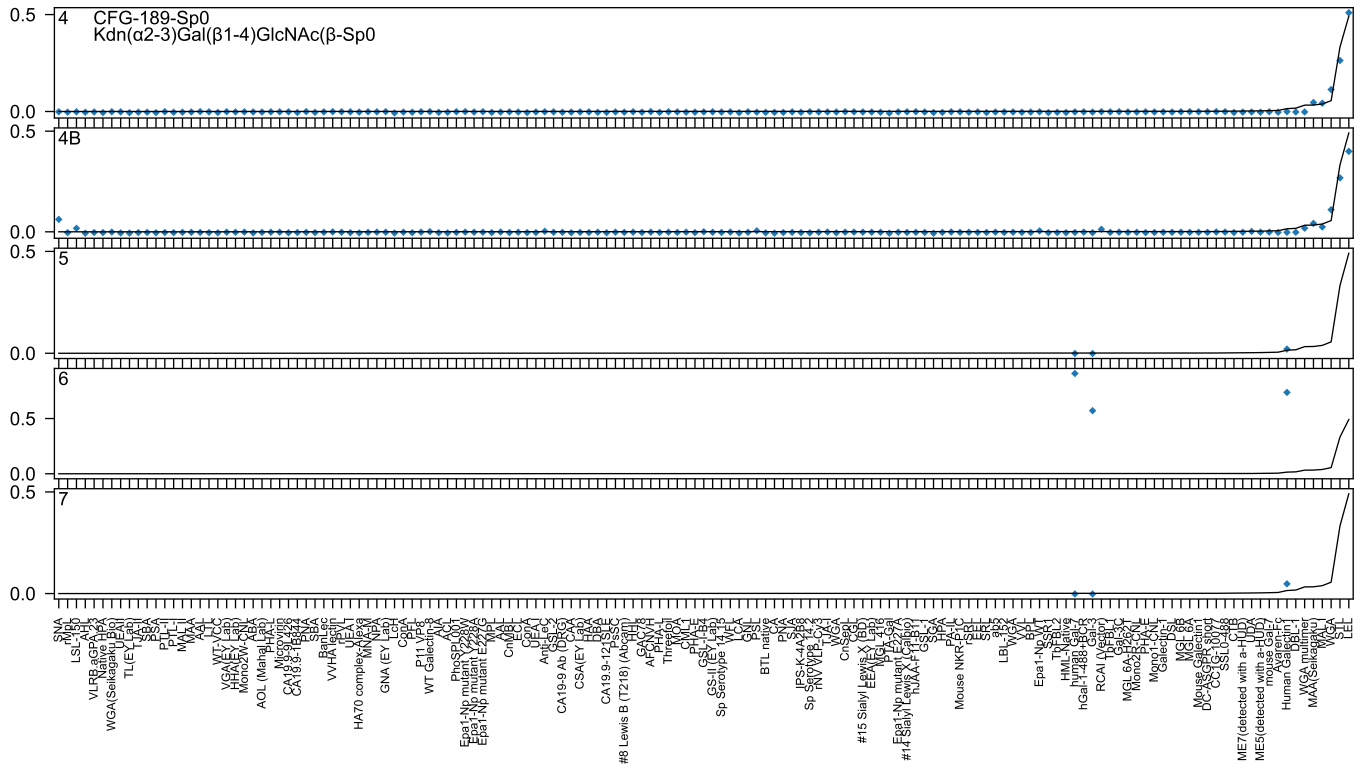
4
CFG-189-Sp0
Kdn(α2-3)Gal(β1-4)GlcNAc(β-Sp0
4B
5
6
7

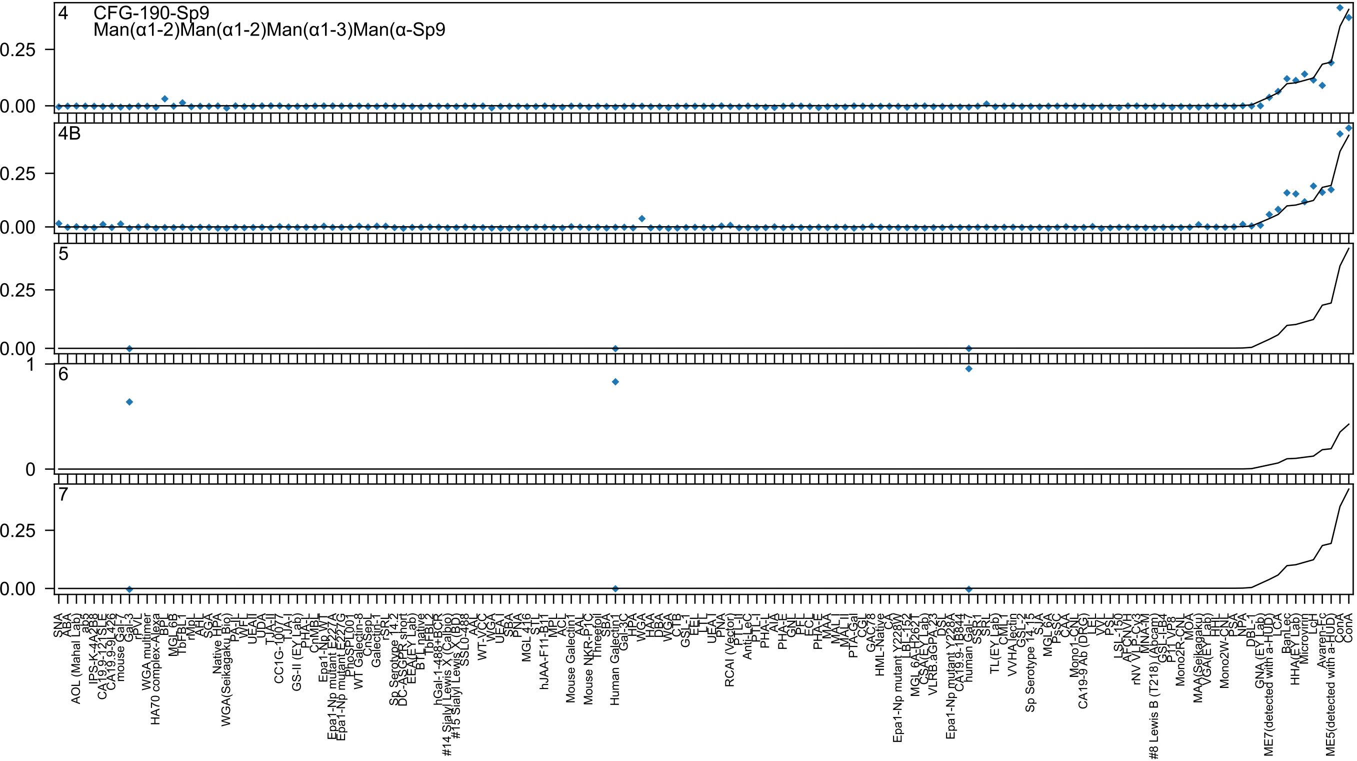
4
CFG-190-Sp9
Man(α1-2)Man(α1-2)Man(α1-3)Man(α-Sp9
4B
5
6
7
0.25
0.00
1
0
SNA
ABA
AOL (Mahal Lab)
ab5
IPS-K-4A2B8
CA19.9-121SLE
CA19.9-9L426
mouse Gal-7
Gal-3
rPVL
WGA multimer
HA70 complex-Alexa
BPL
MGL 6B
TbFBL1
rMpL
AHL
SGA
Native HPA
WGA(Seikagaku Bio)
PA-II
WFL
UEA II
UDA
TJA-II
CC1G-10077
TJA-I
GS-II (EY Lab)
PHA-L
CnMBL
Epa1-Np WT
Epa1-Np mutant E227A
Epa1-Np mutant E227G
PhoSL001
WT Galectin-8
CnSepL
Galectin-1
rSRL
Sp Serotype 14.2
DC-ASGPR short
EEA(EY Lab)
BTL native
TbFBL2
hGal-1-488+BCR
#14 Sialyl Lewis X (Calbio)
#15 Sialyl Lewis X (BD)
SSL0-488
AAL
WT-VCC
WGA
UEA I
SBA
PNA
MGL 416
STL
hJAA-F11-B11
MPL
ACL
Mouse Galectin1
AAL
Mouse NKR-P1C
Threefoil
SBA
Human Galectin1
Gal-3C
HPA
WGA
HAA
DBA
WGA
CTB
GSL-I
EEL
LTL
UEA I
PNA
RCAI (Vector)
PTL-II
Anti-LeC
PTL-I
PHA-L
AIA
PHA-E
GNL
PFL
ECL
PHA-E
MAA
MAL I
MAL II
PTA-Gal
CGL
GAC78
HML-Native
CA
Epa1-Np mutant Y228W
LBL-152
MGL 6A-H262L
CSA(EY Lab)
VLRB.aGPA.23
DSL
Epa1-Np mutant Y228A
CA19.9-1B844
human Gal-7
SSR1
SRL
TL(EY Lab)
CML
VVHA lectin
GSL-2
Sp Serotype 14.15
SJA
MGL 6A
PsSC
CAA
Mono1-CNL
CA19-9 Ab (DRG)
LEL
VVL
PSL
LSL-150
AFCNVH
rNV VLP-Cy3
MNA-M
#8 Lewis B (T218) (Abcam)
GSL-I-B4
P11 VP8
Mono2R-CNL
MOA
MAA(Seikagaku)
VGA(EY Lab)
HHL
Mono2W-CNL
PSA
NPA
DBL-1
GNA (EY Lab)
ME7(detected with a-HUD)
LCA
BanLec
HHA(EY Lab)
Microvirin
LcH
Avaren-Fc
ME5(detected with a-HUD)
ConA
ConA

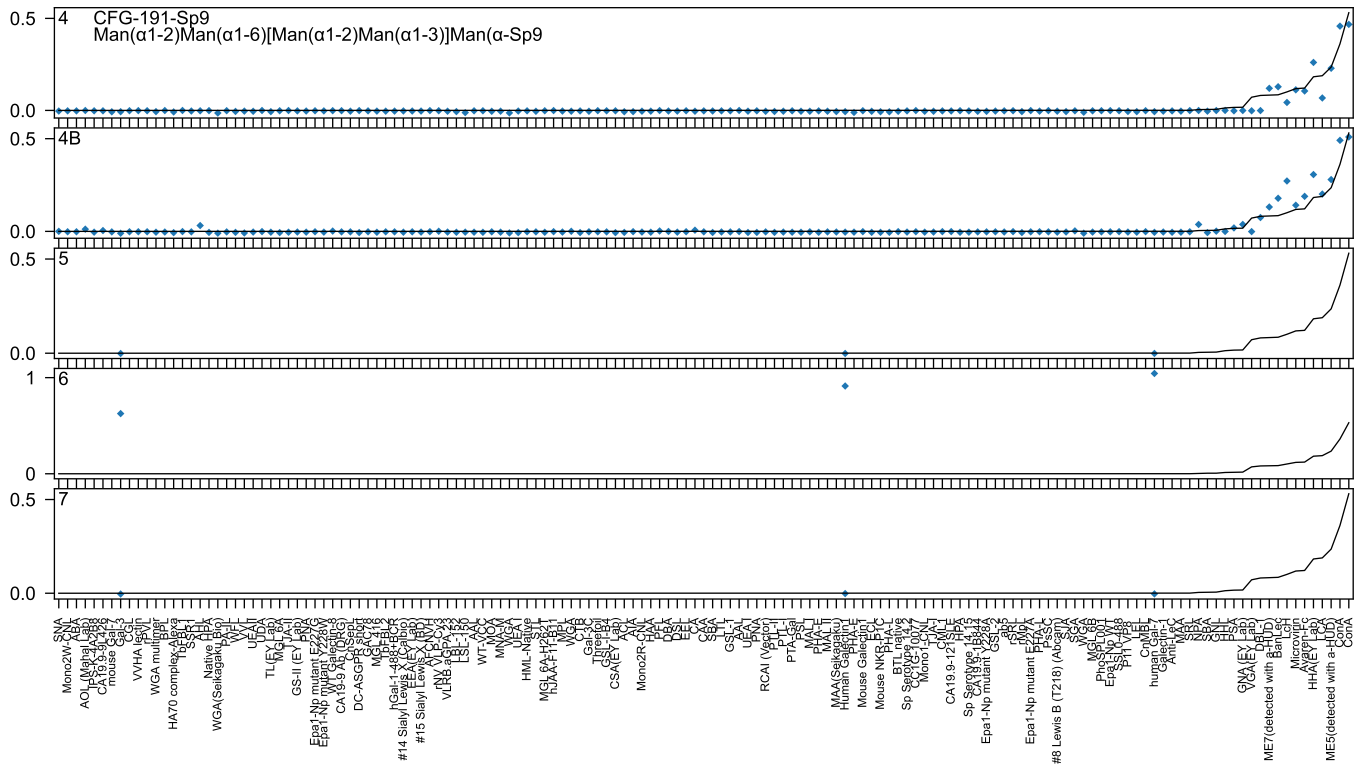

CFG-191-Sp9
Man(α1-2)Man(α1-6)[Man(α1-2)Man(α1-3)]Man(α-Sp9

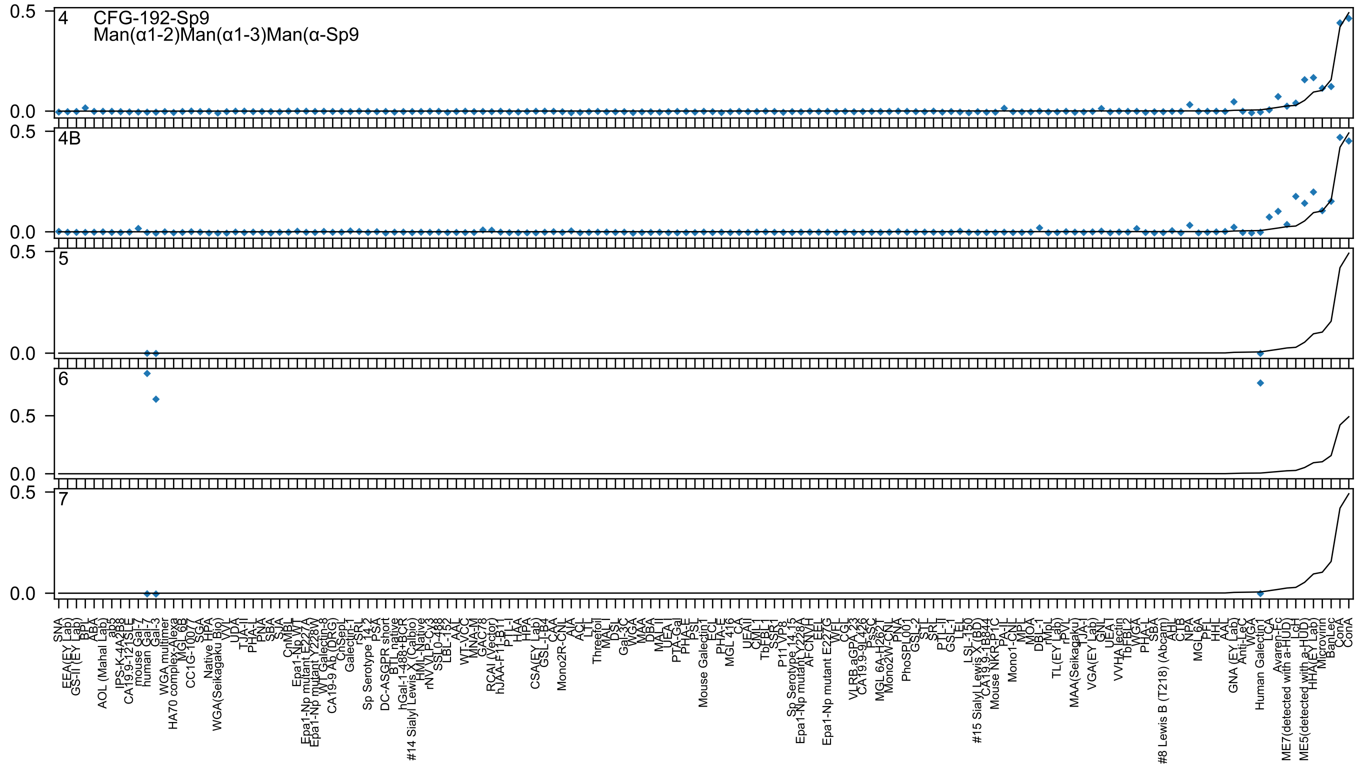
4
CFG-192-Sp9
Man(α1-2)Man(α1-3)Man(α-Sp9
4B
5
6
7
0.5
0.0

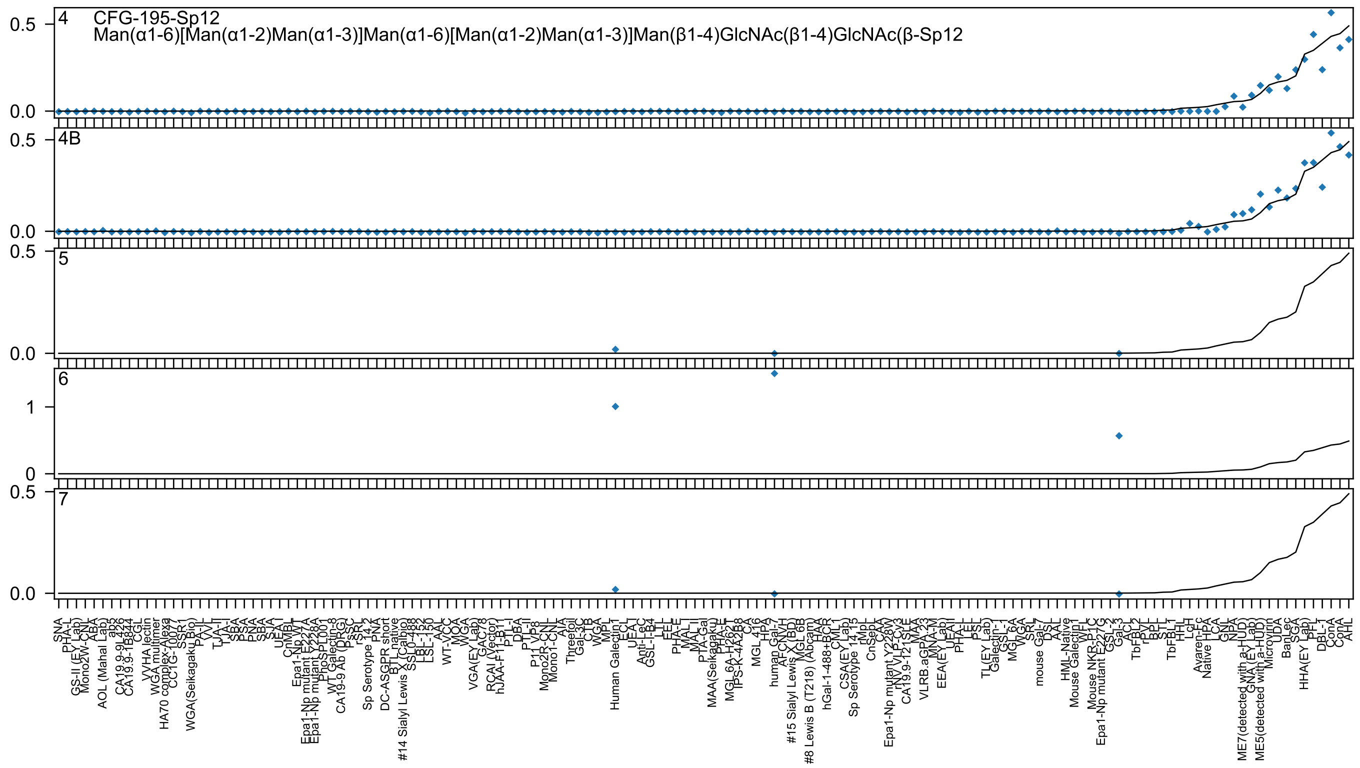
4
CFG-195-Sp12
Man(α1-6)[Man(α1-2)Man(α1-3)]Man(α1-6)[Man(α1-2)Man(α1-3)]Man(β1-4)GlcNAc(β1-4)GlcNAc(β-Sp12
4B
5
6
7

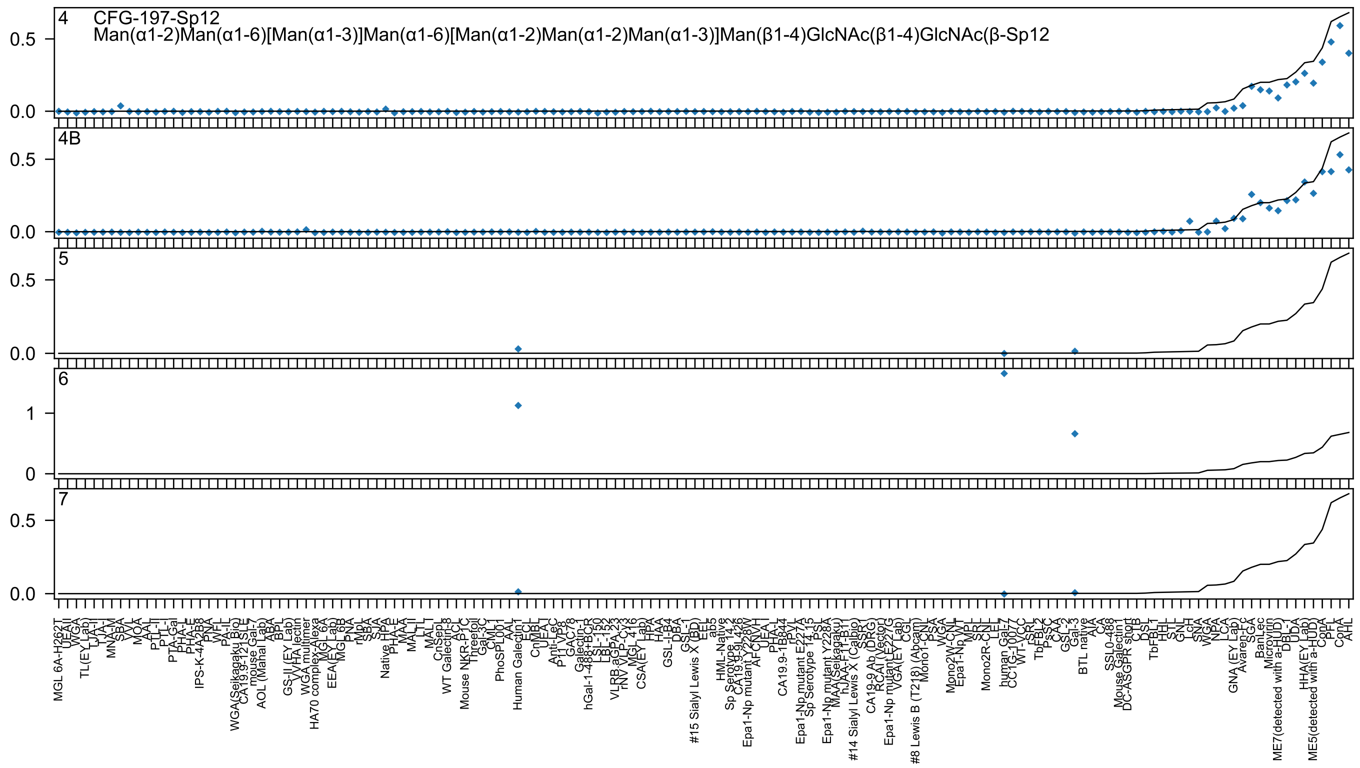

4 CFG-197-Sp12
Man(α1-2)Man(α1-6)[Man(α1-3)]Man(α1-6)[Man(α1-2)Man(α1-2)Man(α1-3)]Man(β1-4)GlcNAc(β1-4)GlcNAc(β-Sp12
4B
5
6
7

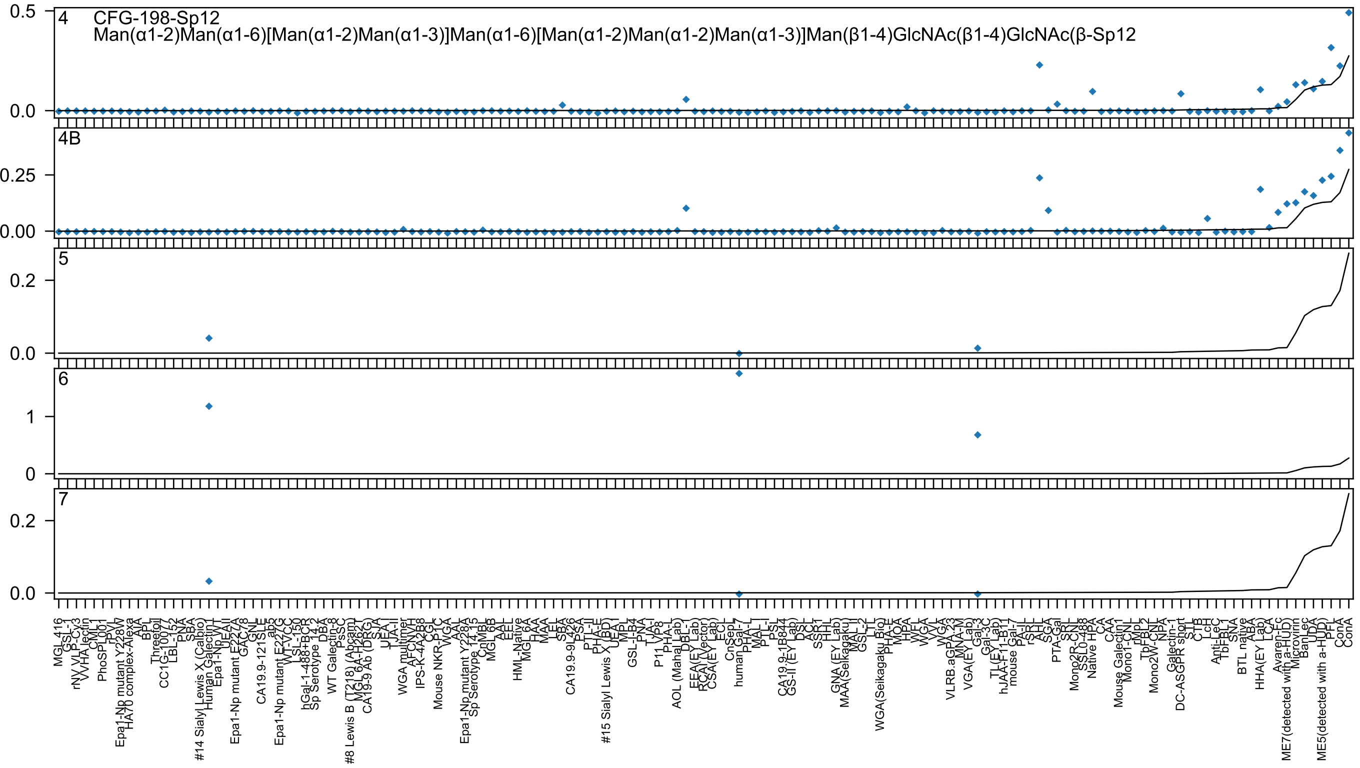
CFG-198-Sp12
Man(α1-2)Man(α1-6)[Man(α1-2)Man(α1-3)]Man(α1-6)[Man(α1-2)Man(α1-2)Man(α1-3)]Man(β1-4)GlcNAc(β1-4)GlcNAc(β-Sp12
4
4B
5
6
7

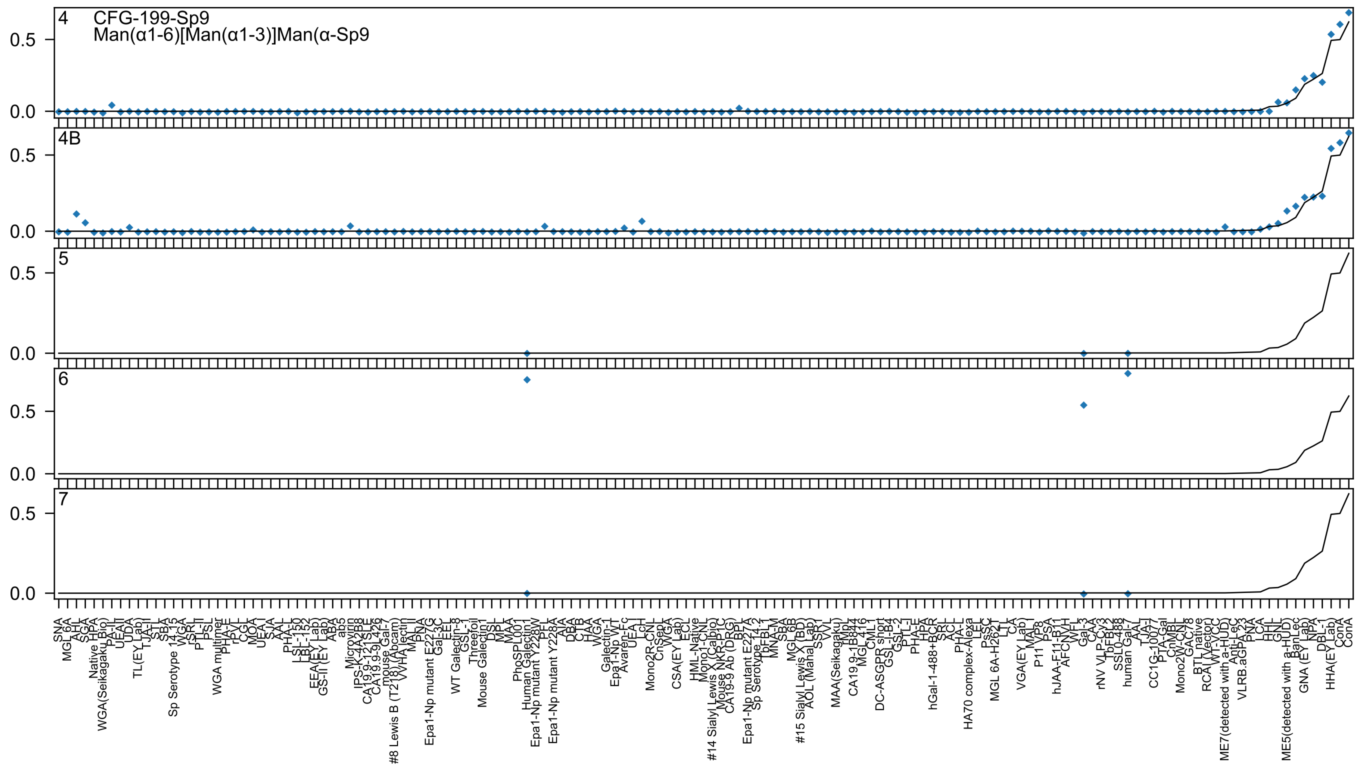
4
CFG-199-Sp9
Man(α1-6)[Man(α1-3)]Man(α-Sp9
4B
5
6
7
0.5
0.0

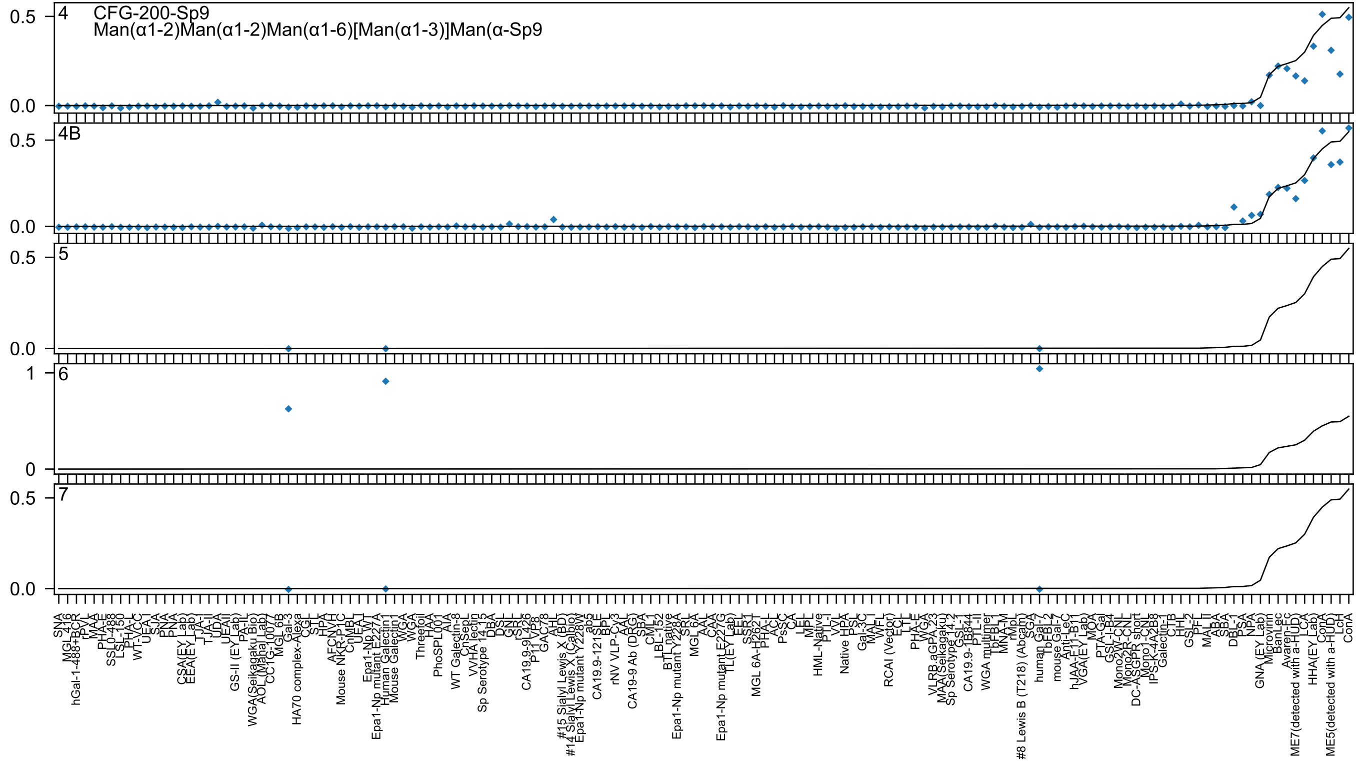

4
CFG-200-Sp9
Man(α1-2)Man(α1-2)Man(α1-6)[Man(α1-3)]Man(α-Sp9
4B
5
6
7

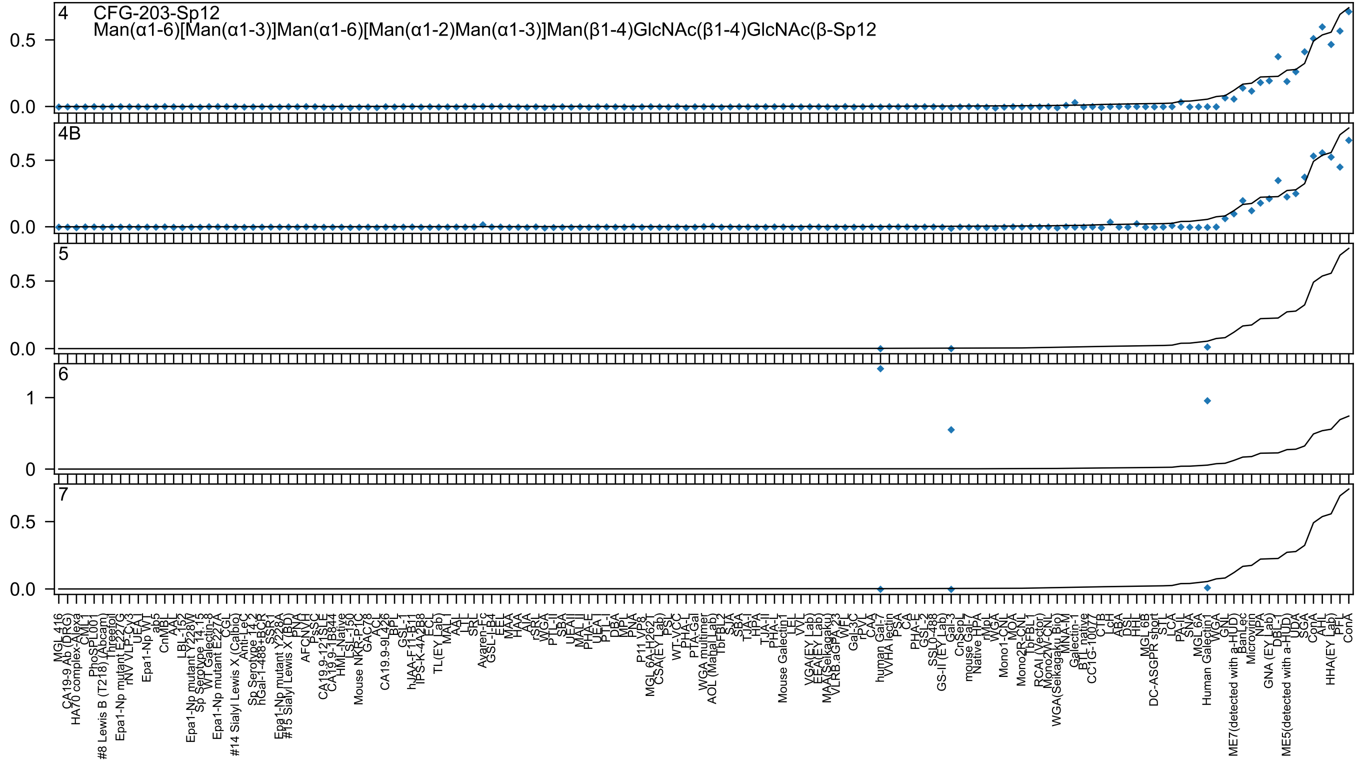

4 CFG-203-Sp12
Man(α1-6)[Man(α1-3)]Man(α1-6)[Man(α1-2)Man(α1-3)]Man(β1-4)GlcNAc(β1-4)GlcNAc(β-Sp12
4B
5
6
7

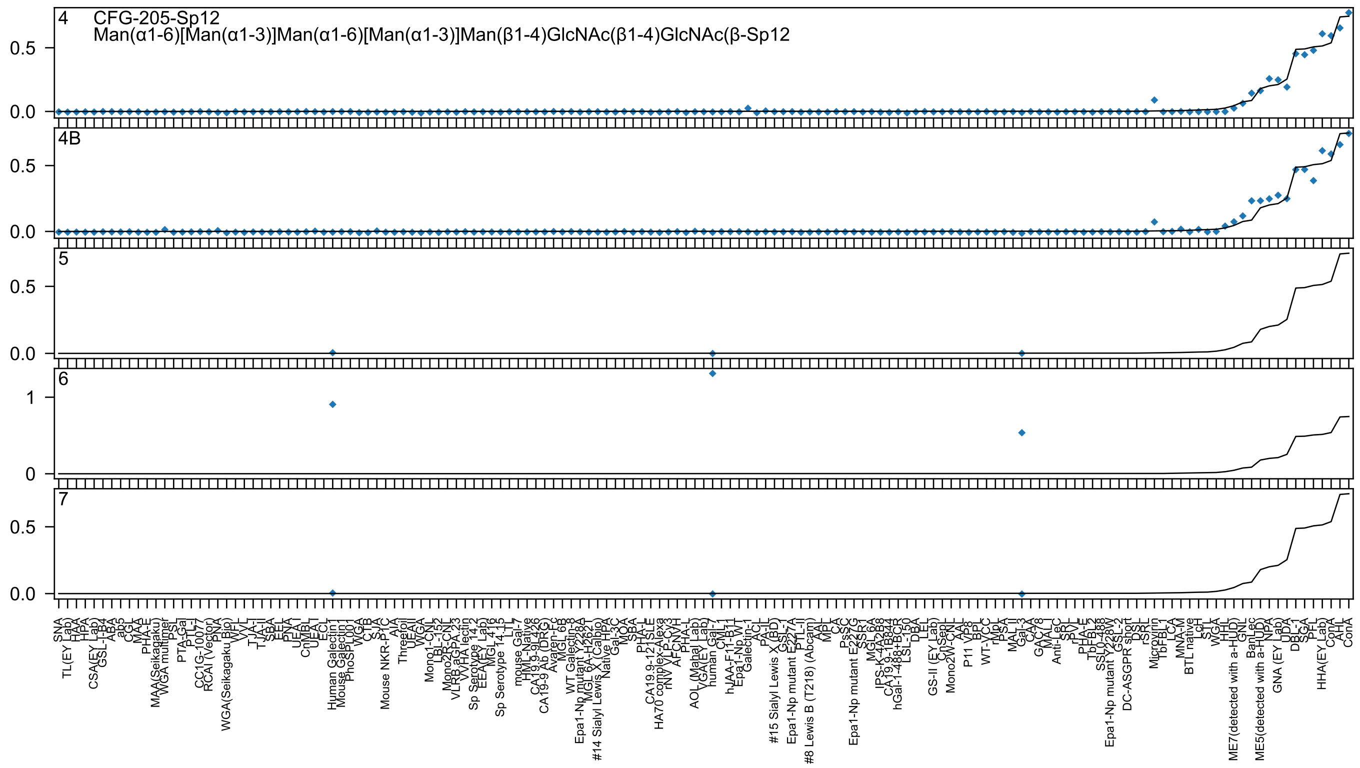

4
CFG-205-Sp12
Man(α1-6)[Man(α1-3)]Man(α1-6)[Man(α1-3)]Man(β1-4)GlcNAc(β1-4)GlcNAc(β-Sp12
4B
5
6
7

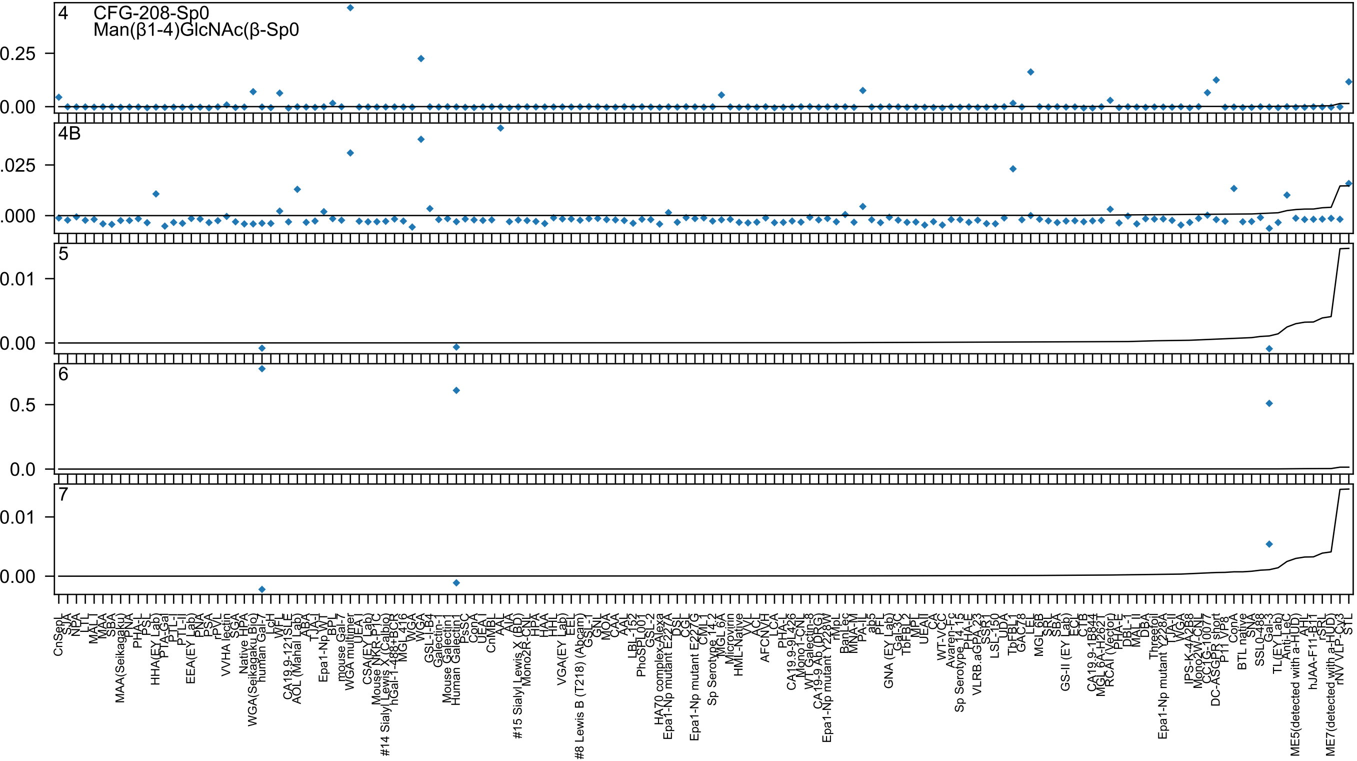

4
CFG-208-Sp0
Man(β1-4)GlcNAc(β-Sp0
4B
5
6
7

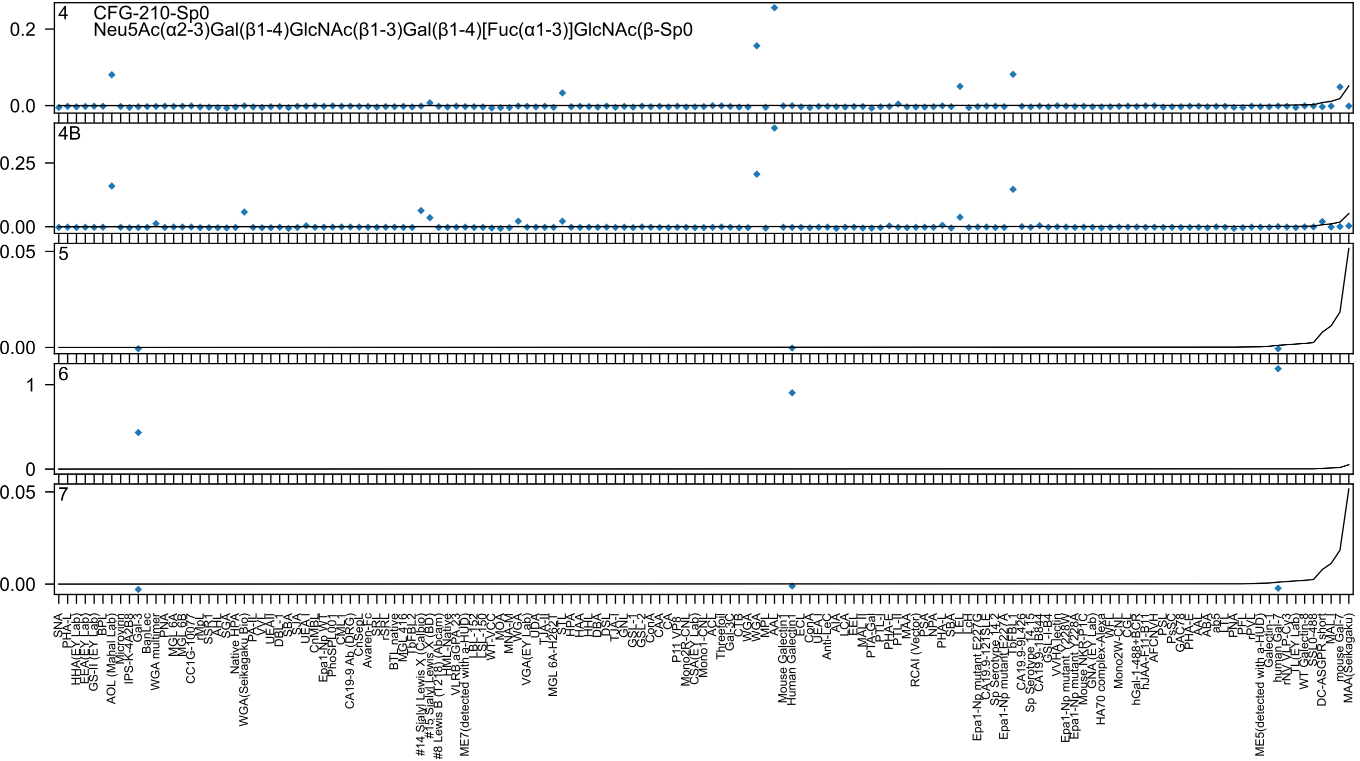

4
CFG-210-Sp0
Neu5Ac(α2-3)Gal(β1-4)GlcNAc(β1-3)Gal(β1-4)[Fuc(α1-3)]GlcNAc(β-Sp0
4B
5
6
7

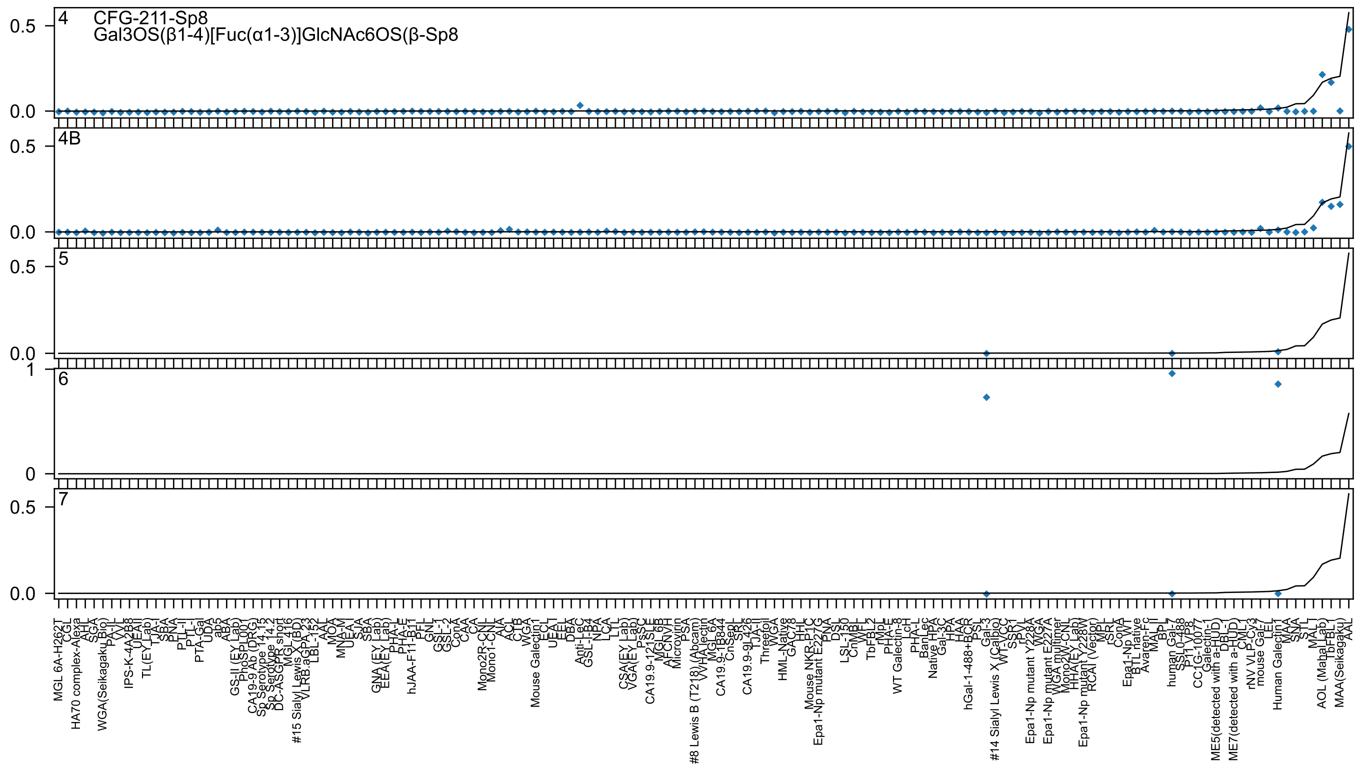

4
CFG-211-Sp8
Gal3OS(β1-4)[Fuc(α1-3)]GlcNAc6OS(β-Sp8
4B
5
6
7

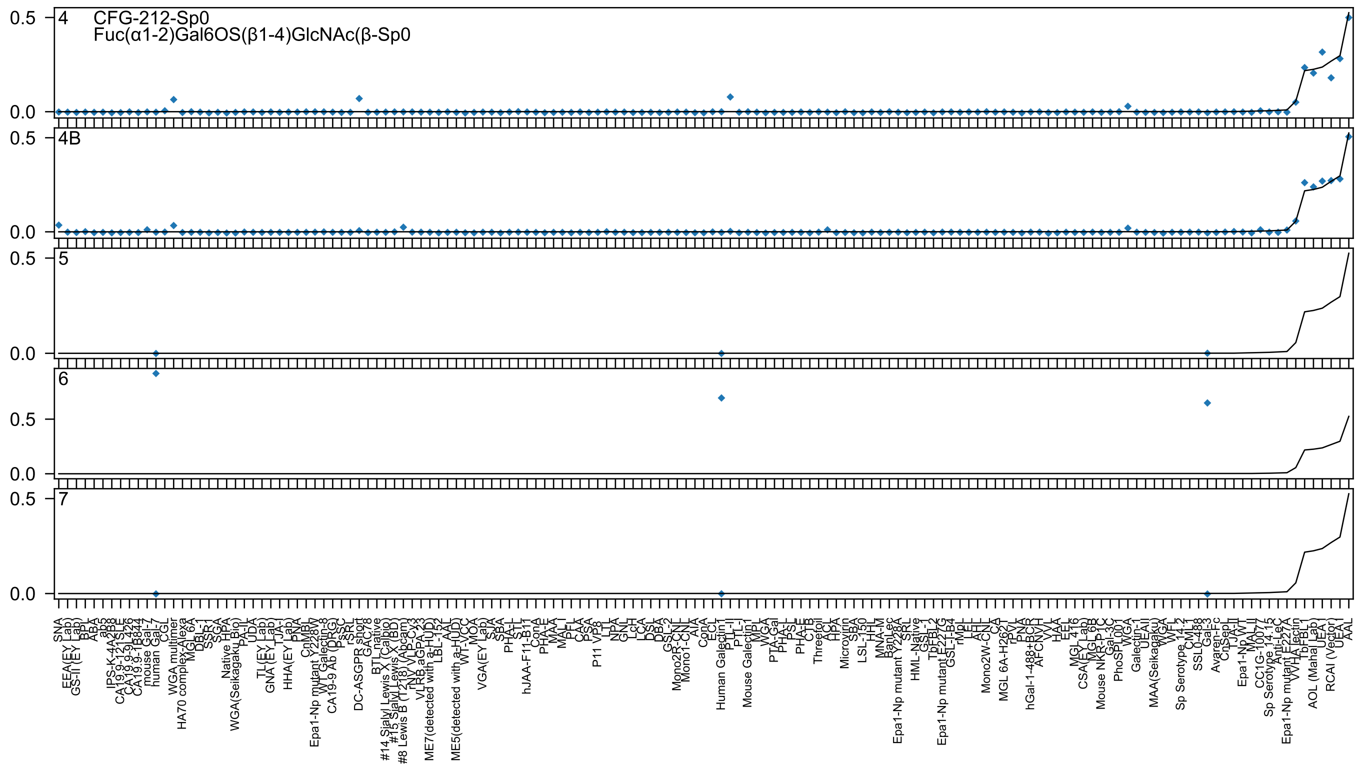

4
CFG-212-Sp0
Fuc(α1-2)Gal6OS(β1-4)GlcNAc(β-Sp0
4B
5
6
7
0.5
0.0

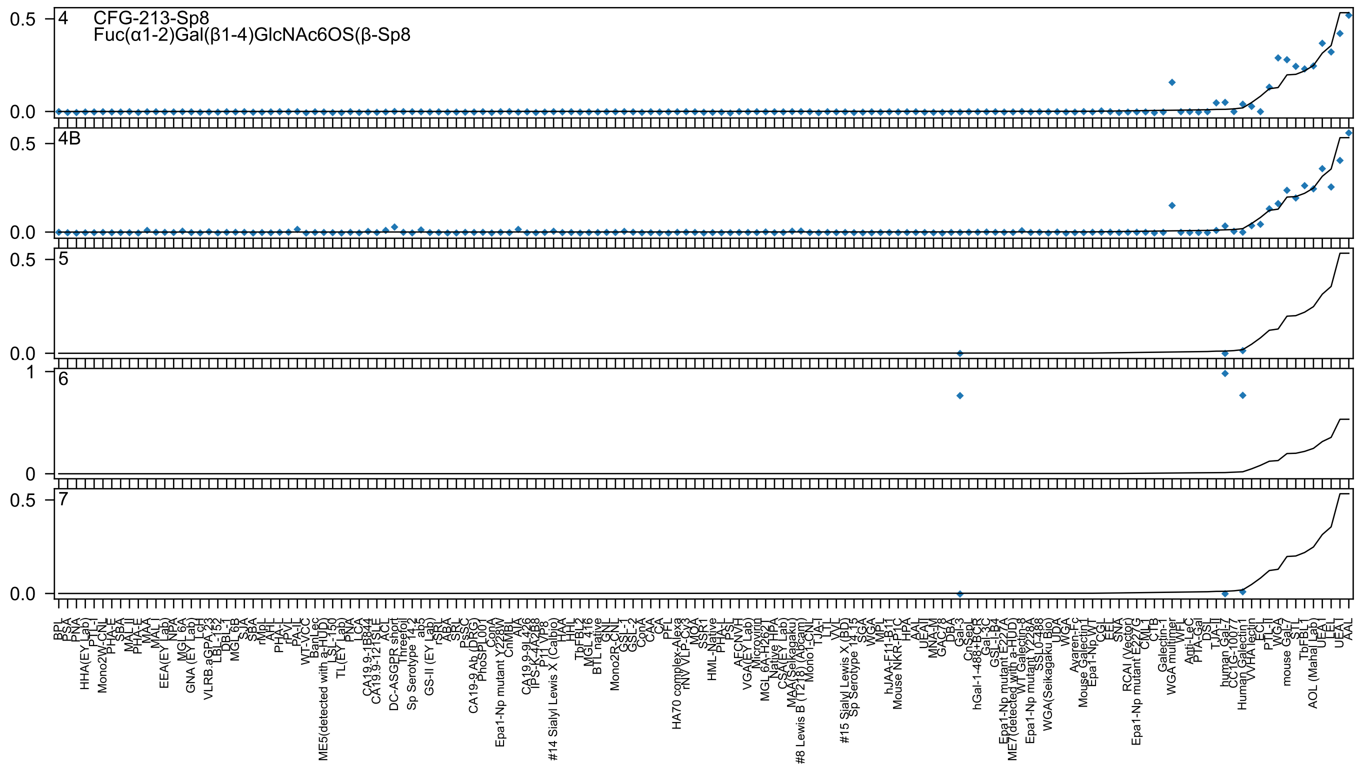

4 CFG-213-Sp8
Fuc(α1-2)Gal(β1-4)GlcNAc6OS(β-Sp8
4B
5
6
7
0.5
0.0
1
0
BPL
PSA
PNA
HHA(EY Lab)
PTL-I
Mono2W-CNL
PHA-E
SBA
MAL_II
PHA-E
MAA
MAL_I
EEA(EY Lab)
NPA
MGL 6A
GNA (EY Lab)
LcH
VLRB.aGPA.23
LBL-152
DBL-1
MGL 6B
SJA
SBA
rMpL
AAL
PHA-L
rPVL
PA-IL
WT-VCC
BanLec
ME5(detected with a-HUD)
LSL-150
TL(EY Lab)
PNA
LCA
CA19.9-1B844
CA19.9-121SLE
ACL
DC-ASGPR short
Threefoil
Sp Serotype 14.2
ab5
GS-II (EY Lab)
rSRL
ABA
SRL
PSSC
CA19-9 Ab (DRG)
PhoSL001
ConA
Epa1-Np mutant Y228W
CnMBL
AIA
CA19.9-9L426
IPS-K-4A2B8
P11 VP8
#14 Sialyl Lewis X (Calbio)
HAA
HHL
TbFBL2
MGL 416
BTL native
GNL
Mono2R-CNL
GSL-1
GSL-2
ConA
CAA
CA
PFL
HA70 complex-Alexa
rNV VLP-Cy3
MOA
SSR1
HML-Native
PHA-L
PSL
AFCNVH
VGA(EY Lab)
Microvirin
MGL 6A-H262
Native HPA
CSA(EY Lab)
MAA(Seikagaku)
#8 Lewis B (T218) (Abcam)
Mono1-CNL
TJA-I
LTL
VVL
#15 Sialyl Lewis X (BD)
Sp Serotype 14.15
SGA
WGA
MPL
hJAA-F11-B1
Mouse NKR-P1C
HPA
AAA
UEAII
MNA-M
GAC78
DBA
Gal-3
CnSep
hGal-1-488+BCR
Gal-3C
GSL-I-B4
Epa1-Np mutant E227A
ME7(detected with a-HUD)
WT Galectin-8
Epa1-Np mutant Y228A
SSL0-488
WGA(Seikagaku Bio)
UDA
WGA
Avaren-Fc
Mouse Galectin1
Epa1-Np WT
CGL
EEL
SNA
RCAI (Vector)
Epa1-Np mutant E227G
CVL1
CTB
Galectin-1
WGA multimer
WFL
Anti-LeC
PTA-Gal
DSL
TJA-II
human Gal-7
CC1G-10077
Human Galectin1
WHA lectin
ECL
PTL-II
WGA
mouse Gal-7
STL
TbFBL1
AOL (Mahal Lab)
UEA I
LEL
UEA I
AAL

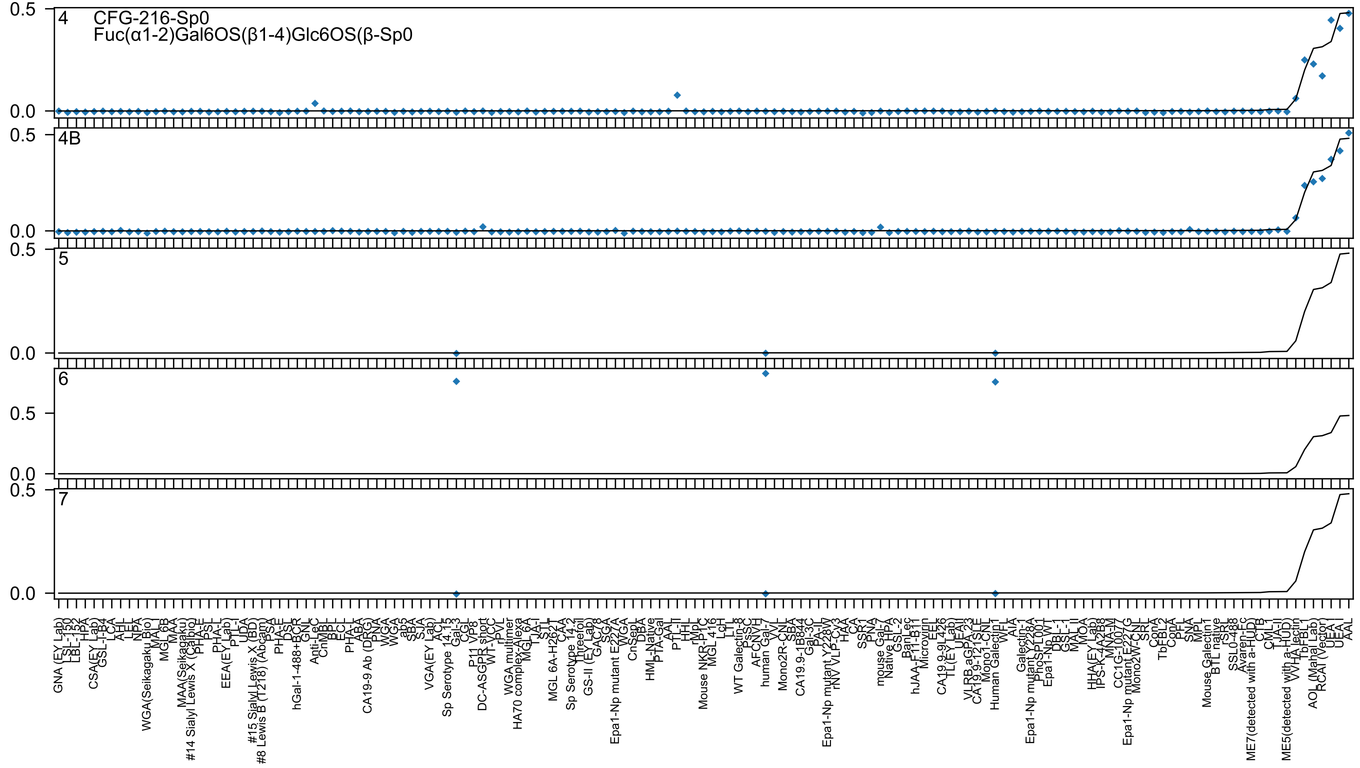

CFG-216-Sp0
Fuc(α1-2)Gal6OS(β1-4)Glc6OS(β-Sp0
4
4B
5
6
7
0.5
0.0
GNA (EY Lab)
LSL-150
LBL-152
HPA
CSA(EY Lab)
GSL-I-B4
LCA
AHL
LEL
NPA
WGA(Seikagaku Bio)
MAL
MGL 6B
MAA
MAA(Seikagaku)
#14 Sialyl Lewis X (Calbio)
PHA-E
PSL
PHA-L
EEA(EY Lab)
PTL-I
UDA
#15 Sialyl Lewis X (BD)
#8 Lewis B (T218) (Abcam)
PSA
PHA-E
DSL
hGal-1-488+BCR
GNL
Anti-LeC
CnMBL
BPL
ECL
PHA-L
ABA
CA19-9 Ab (DRG)
PNA
WGA
WGA
ab5
SBA
SJA
VGA(EY Lab)
ACL
Sp Serotype 14.15
Gal-3
CGL
VP8
P11 VP8
DC-ASGPR short
WT-VCC
rPVL
WGA multimer
HA70 complex-Alexa
MGL 6A
TJA-I
STL
MGL 6A-H262T
CAA
Sp Serotype 14.2
Threefoil
GS-II (EY Lab)
GAC78
SGA
Epa1-Np mutant E227A
WGA
CnSep
DBA
HML-Native
PTA-Gal
AAL
PTL-II
HHL
rMpL
Mouse NKR-P1C
MGL 416
LcH
LTL
WT Galectin-8
PsSC
AFCNVH
human Gal-7
VVL
Mono2R-CNL
SBA
CA19.9-1B844
Gal-3C
PA-IL
Epa1-Np mutant Y228W
rNV VLP-CV3
HAA
CA
SSR1
PNA
mouse Gal-7
Native HPA
GSL-2
BanLec
hJAA-F11-B11
Microvirin
EEL
CA19.9-9L426
TL(EY Lab)
UEAII
VLRB.aGPA.23
CA19.9-121SLE
Mono1-CNL
Human Galectin1
WFL
AIA
Galectin-1
Epa1-Np mutant Y228A
PhoSL001
Epa1-Np WT
DBL-1
GSL-I
MAL_II
MOA
HHA(EY Lab)
IPS-K-4A2B8
MNA-M
CC1G-10077
Epa1-Np mutant E227G
Mono2W-CNL
SRL
ConA
TbFBL2
ConA
PFL
SNA
MPL
Mouse Galectin1
BTL native
rSRL
SSL0-488
Avaren-Fc
ME7(detected with a-HUD)
CTB
CML-1
TJA-II
ME5(detected with a-HUD)
WHA lectin
TbFBL1
AOL (Mahal Lab)
RCA I (Vector)
UEA I
UEA I
AAL

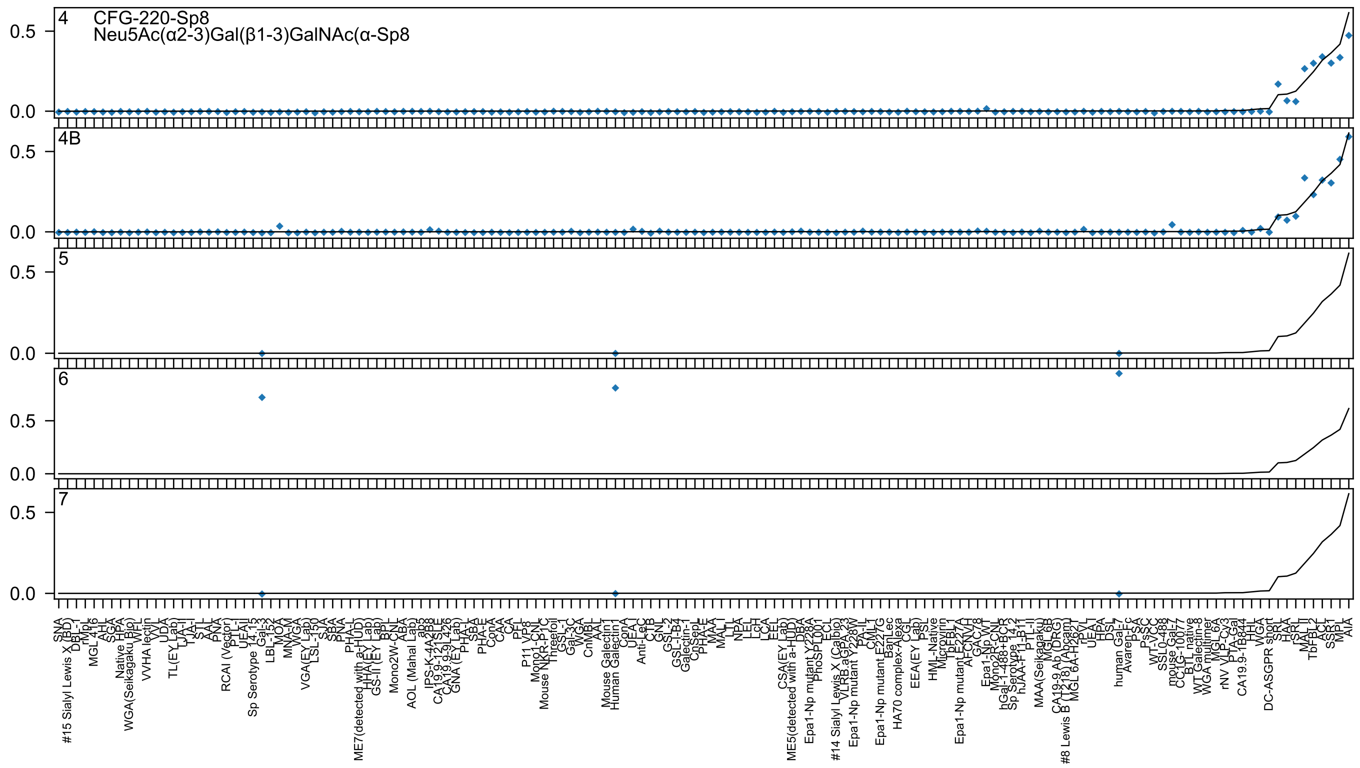

4 CFG-220-Sp8
Neu5Ac(α2-3)Gal(β1-3)GalNAc(α-Sp8
4B
5
6
7

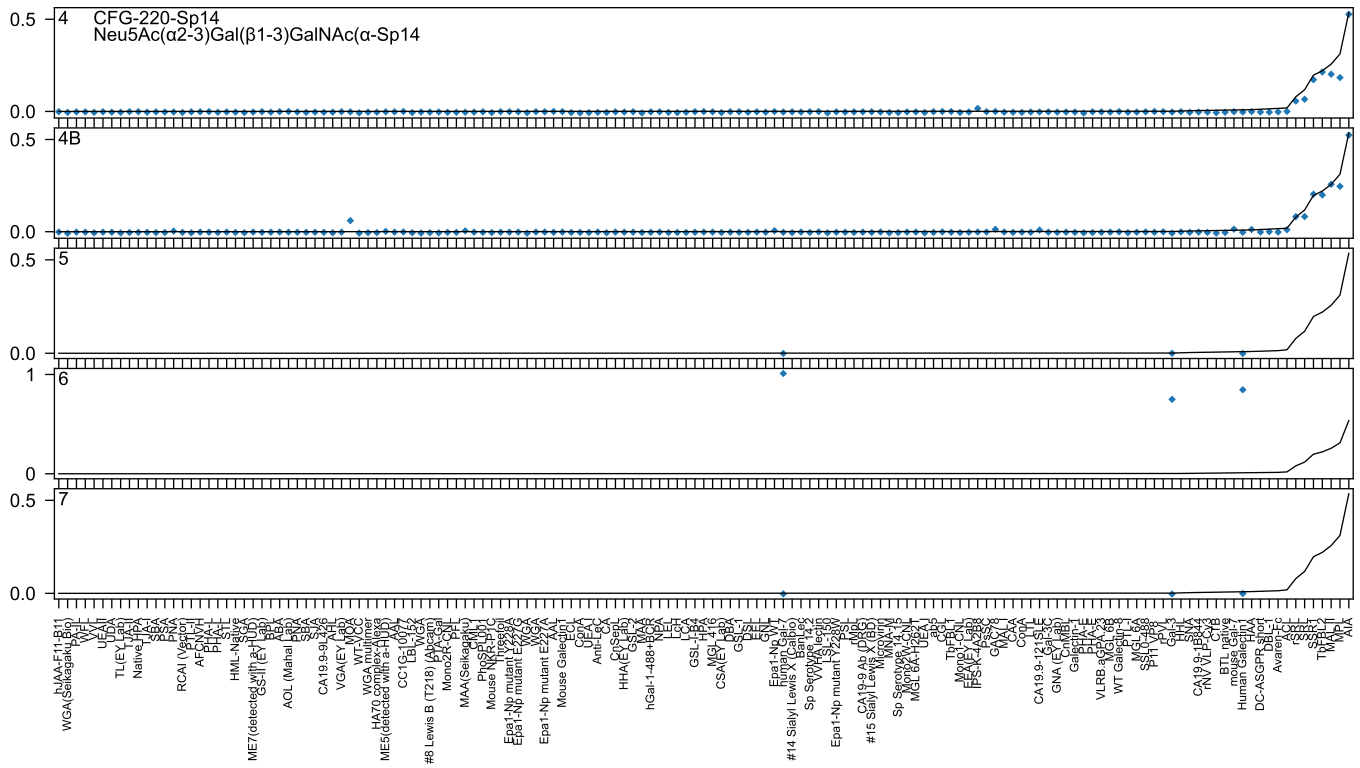
CFG-220-Sp14
Neu5Ac(α2-3)Gal(β1-3)GalNAc(α-Sp14
4
4B
5
6
7

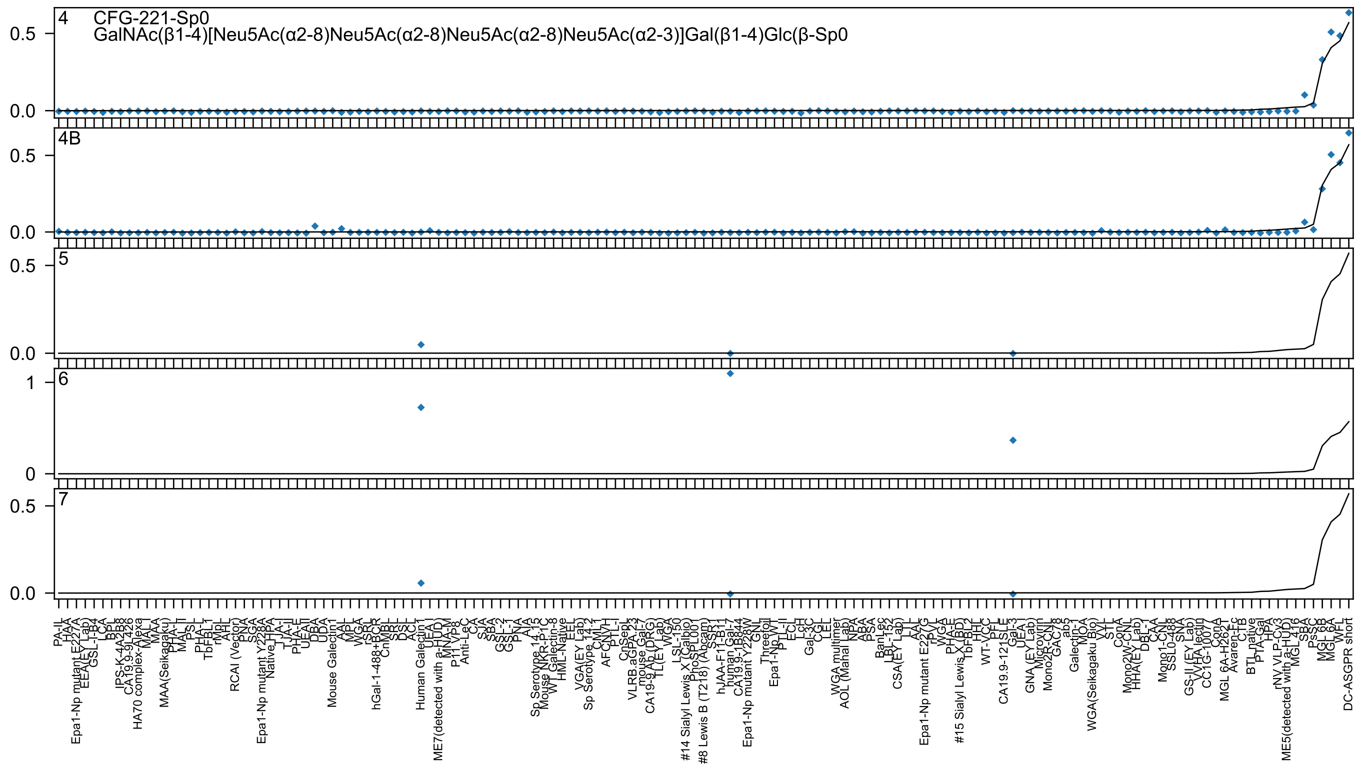

4 CFG-221-Sp0
GalNAc(β1-4)[Neu5Ac(α2-8)Neu5Ac(α2-8)Neu5Ac(α2-8)Neu5Ac(α2-3)]Gal(β1-4)Glc(β-Sp0
4B
5
6
7

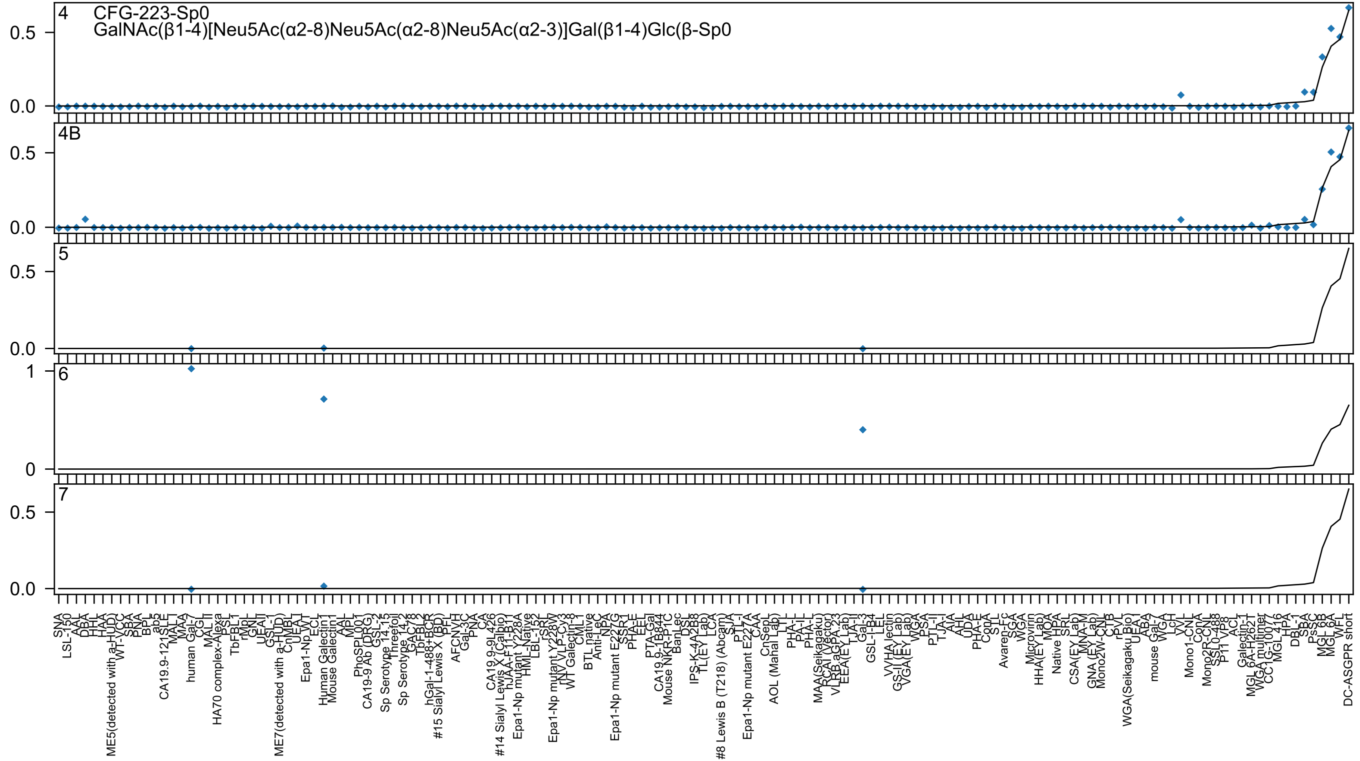

4 CFG-223-Sp0
GalNAc(β1-4)[Neu5Ac(α2-8)Neu5Ac(α2-8)Neu5Ac(α2-3)]Gal(β1-4)Glc(β-Sp0
4B
5
6
7
0.5
0.0
1
0
SNA
LSL-150
AAL
DBA
HHL
HAA
ME5(detected with a-HUD)
WT-VCC
SBA
PNA
BPL
ab5
CA19.9-121SLE
MAL1
MAA
human Gal-7
CGL
MAL II
HA70 complex-Alexa
PSL
TbFBL1
rMpL
GNL
UEAII
GSL-1
ME7(detected with a-HUD)
CnMBL
UEA1
Epa1-Np WT
ECL
Human Galectin1
Mouse Galectin1
AAL
MPL
PhoSL
CA19-9 Ab (DRG)
GSL-2
Sp Serotype 14.15
Threefoil
Sp Serotype 14.2
GAC78
TbFBL1
hGal-1-488+BCR
#15 Sialyl Lewis X (BD)
PFL
AFCNVH
Gal-3C
PNA
CA
CA19.9-9L426
#14 Sialyl Lewis X (Calbio)
hJAA-F11-B11
Epa1-Np mutant Y228A
HML-Native
LBL-152
rSRL
Epa1-Np mutant Y228W
rNV VLP-Cy3
WT Galectin-8
CML1
BTL native
Anti-LeC
NPA
Epa1-Np mutant E227G
SSR1
PHA-E
EEL
PTA-Gal
CA19.9-1B844
Mouse NKR-P1C
BanLec
DSL
IPS-K-4A2B8
TL(EY Lab)
LCA
#8 Lewis B (T218) (Abcam)
SJA
PTL-I
Epa1-Np mutant E227A
CAA
CnSepL
AOL (Mahal Lab)
LTL
PHA-L
PHA-L
PHA-L
MAA(Seikagaku)
RCAI (Vector)
VLRB.aGPA.23
EEA(EY Lab)
TJA-II
Gal-3
GSL-I-B4
LEL
VVHA lectin
GS-II (EY Lab)
VGA(EY Lab)
WGA
PSA
PTL-II
TJA-I
AIA
AHL
UDA
PHA-E
ConA
STL
Avaren-Fc
SGA
WGA
Microvirin
HHA(EY Lab)
MOA
Native HPA
SRL
CSA(EY Lab)
MNA-M
GNA (EY Lab)
Mono2W-CNL
CTB
rPVL
WGA(Seikagaku Bio)
UEA1
ABA
mouse Gal-7
WGA
LcH
VVL
Mono1-CNL
ConA
Mono2R-CNL
SSL0-488
P11 VP8
ACL
Galectin-1
MGL 6A-H262T
WGA multimer
CC1G-10077
MGL 416
HPA
DBL-1
SBA
PsSC
MGL 6B
MGL 6A
WFL
DC-ASGPR short

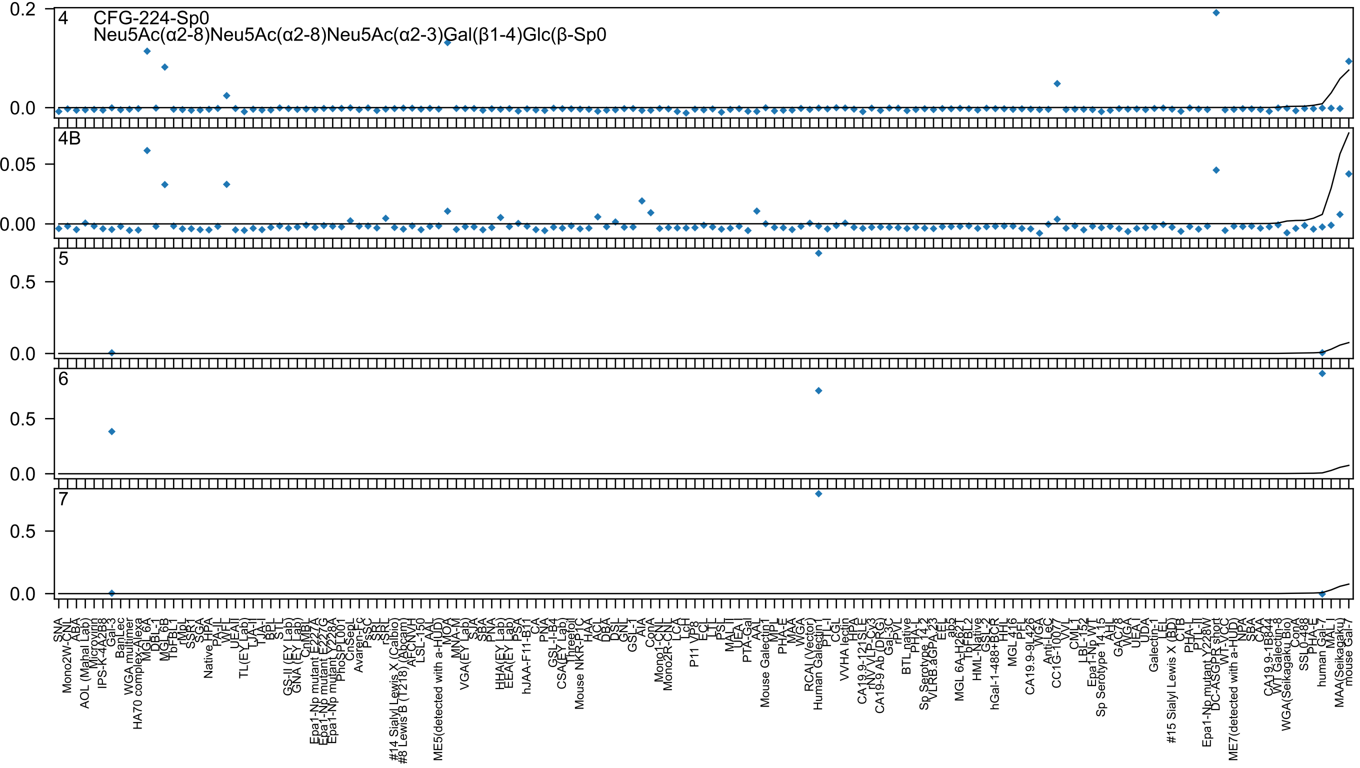
4
CFG-224-Sp0
Neu5Ac(α2-8)Neu5Ac(α2-8)Neu5Ac(α2-3)Gal(β1-4)Glc(β-Sp0
4B
5
6
7

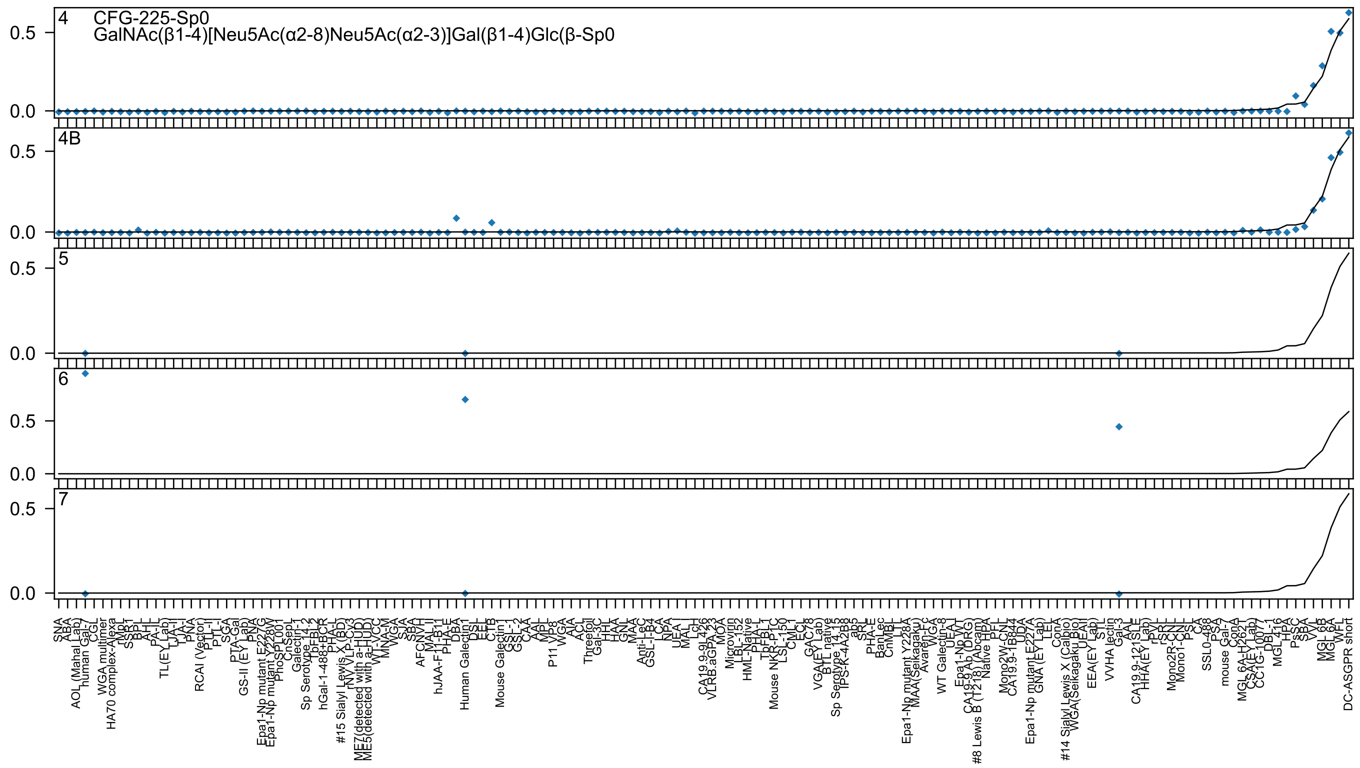
4 CFG-225-Sp0
GalNAc(β1-4)[Neu5Ac(α2-8)Neu5Ac(α2-3)]Gal(β1-4)Glc(β-Sp0
4B
5
6
7
0.5
0.0
SNA
ABA
AOL (Mahal Lab)
human Gal-7
CGL
WGA multimer
HA70 complex-Alexa
rMpL
SSR1
BPL
AAL
PA-IL
TL(EY Lab)
TJA-II
TJA-I
PNA
RCAI (Vector)
PTL-II
PTL-I
SGA
PTA-Gal
GS-II (EY Lab)
PNA
Epa1-Np mutant E227G
Epa1-Np mutant Y228W
PhoSL001
CnSep1
Galectin-1
Sp Serotype 14_2
TbFBL2
hGal-1-488+BCR
PHA-L
#15 Sialyl Lewis X (BD)
rNV VLP-Cy3
ME7(detected with a-HUD)
ME5(detected with a-HUD)
WT-VCC
MNA-M
WGA
SJA
SBA
AFCNVH
MAL-II
hJAA-F11-B11
PHA-E
DBA
Human Galectin1
DSL
EEL
CTB
Mouse Galectin1
GSL-I
GSL-2
CAA
AAL
MPL
P11 VP8
WGA
AIA
ACL
Threefoil
Gal-3C
HHL
HAA
GNL
MAA
Anti-LeC
GSL-I-B4
LCA
NPA
UEA-I
MAL-I
LcH
CA19.9-9L426
VLRB.aGPA.23
MOA
Microvirin
LBL-152
HML-Native
PHA-L
TbFBL1
Mouse NKR-P1C
LSL-150
CML-1
ECL
GAC78
VGA(EY Lab)
BTL native
Sp Serotype 14_3
IPS-K-4A2B8
ab5
SRL
PHA-E
BanLec
CnMBL
LTL
Epa1-Np mutant Y228A
MAA(Seikagaku)
Avaren-Fc
WGA
WT Galectin-8
UEA-I
Epa1-Np WT
CA19-9 Ab (DRG)
#8 Lewis B (T218) (Abcam)
Native HPA
PFL
Mono2W-CN
CA19.9-1B844
UDA
Epa1-Np mutant E227A
GNA (EY Lab)
LEL
ConA
#14 Sialyl Lewis X (Calbio)
WGA(Seikagaku Bio)
UEAII
EEA(EY Lab)
STL
VVHA lectin
Gal-3
AAL
CA19.9-121SLE
HHA(EY Lab)
rPVL
rSRL
Mono2R-CN
Mono1-CN
PSL
CA
SSL0-488
PSA
mouse Gal-7
ConA
MGL 6A-H262T
CSA(EY Lab)
CC1G-1007
DBL-1
MGL 416
HPA
HPL
PsSC
SBA
VVL
MGL 6B
MGL 6A
WFL
DC-ASGPR short

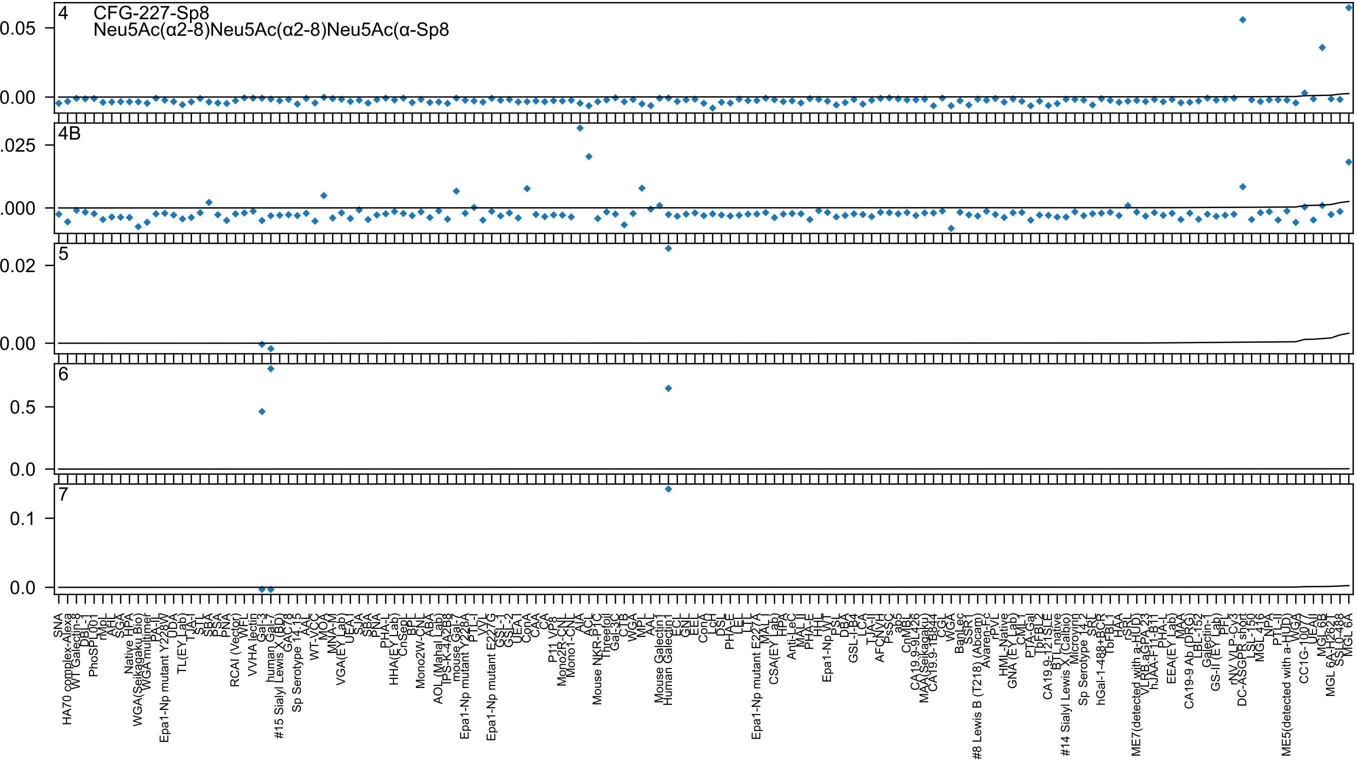

4
CFG-227-Sp8
Neu5Ac(α2-8)Neu5Ac(α2-8)Neu5Ac(α-Sp8
4B
5
6
7

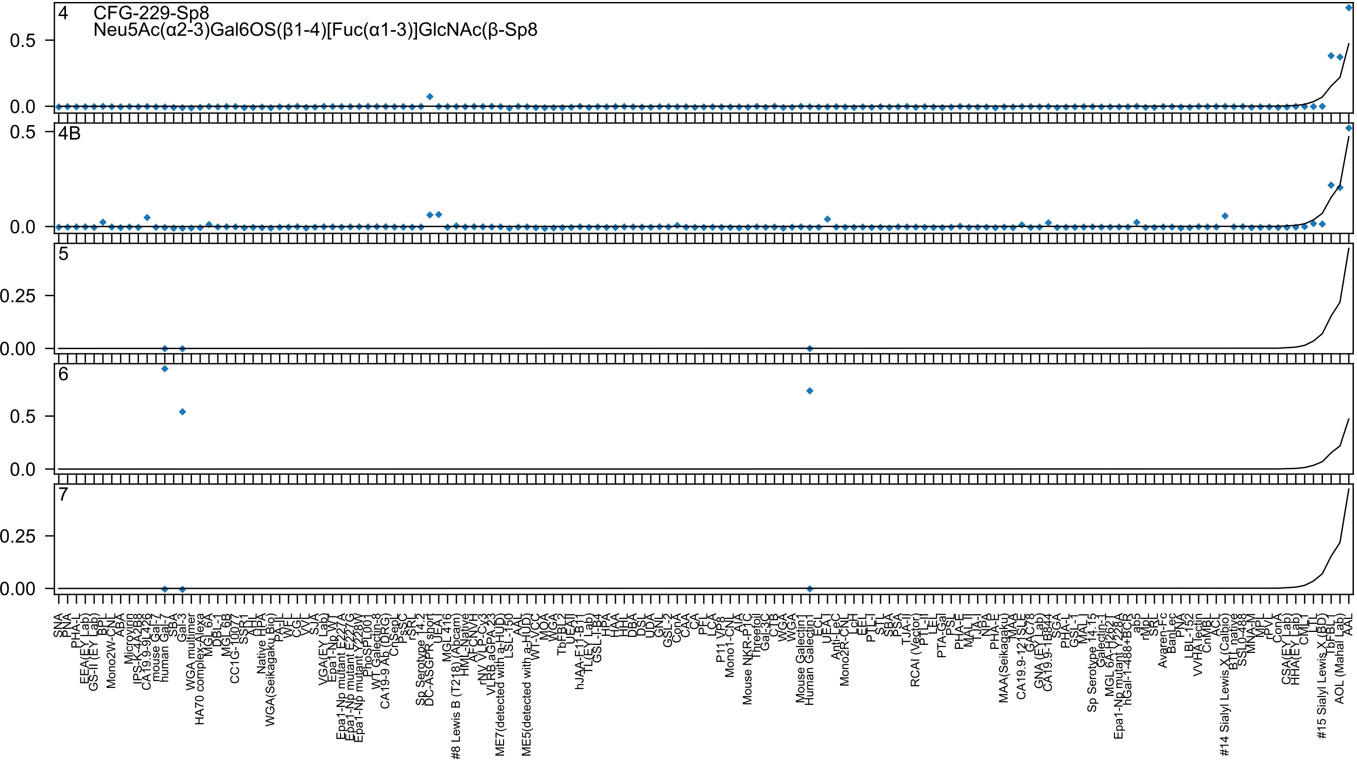
4
CFG-229-Sp8
Neu5Ac(α2-3)Gal6OS(β1-4)[Fuc(α1-3)]GlcNAc(β-Sp8
4B
5
6
7

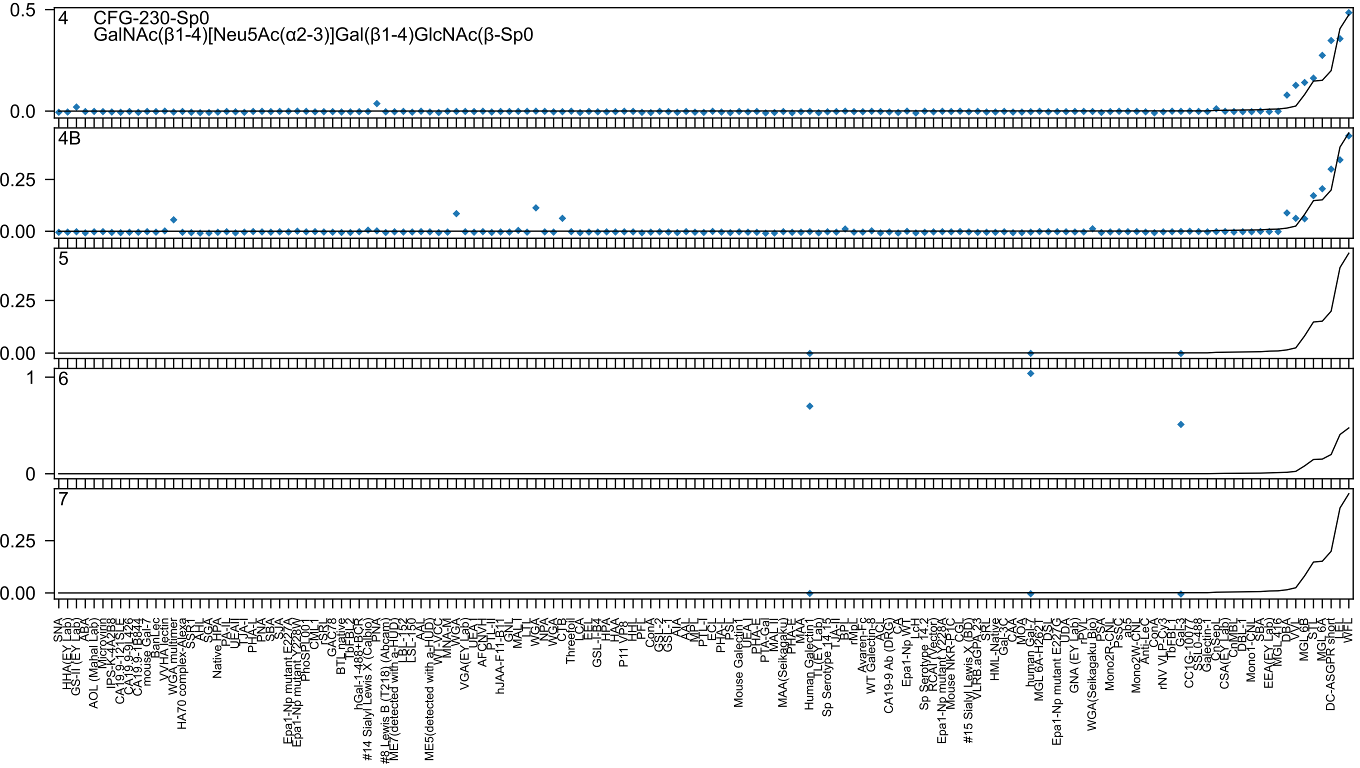

CFG-230-Sp0
GalNAc(β1-4)[Neu5Ac(α2-3)]Gal(β1-4)GlcNAc(β-Sp0
4
4B
5
6
7

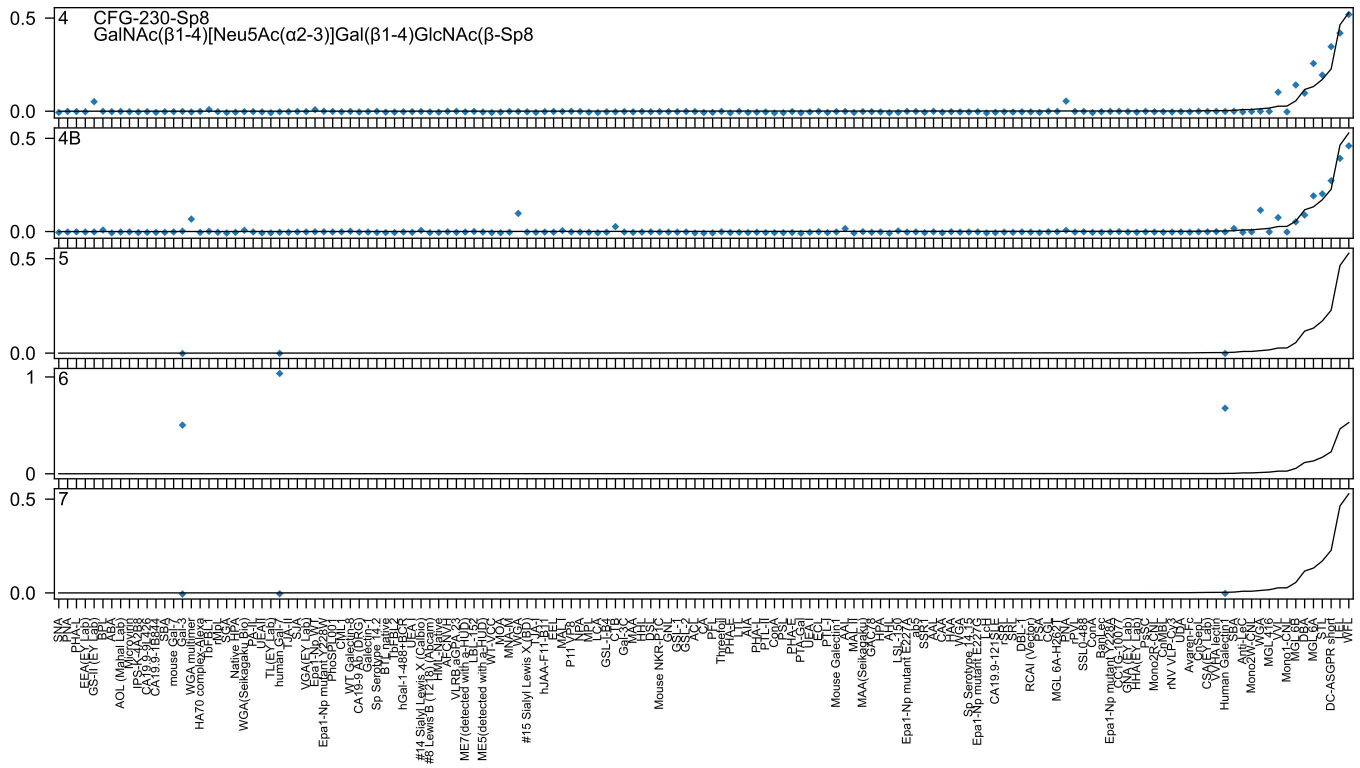
4
CFG-230-Sp8
GalNAc(β1-4)[Neu5Ac(α2-3)]Gal(β1-4)GlcNAc(β-Sp8
4B
5
6
7
0.5
0.0
1
0

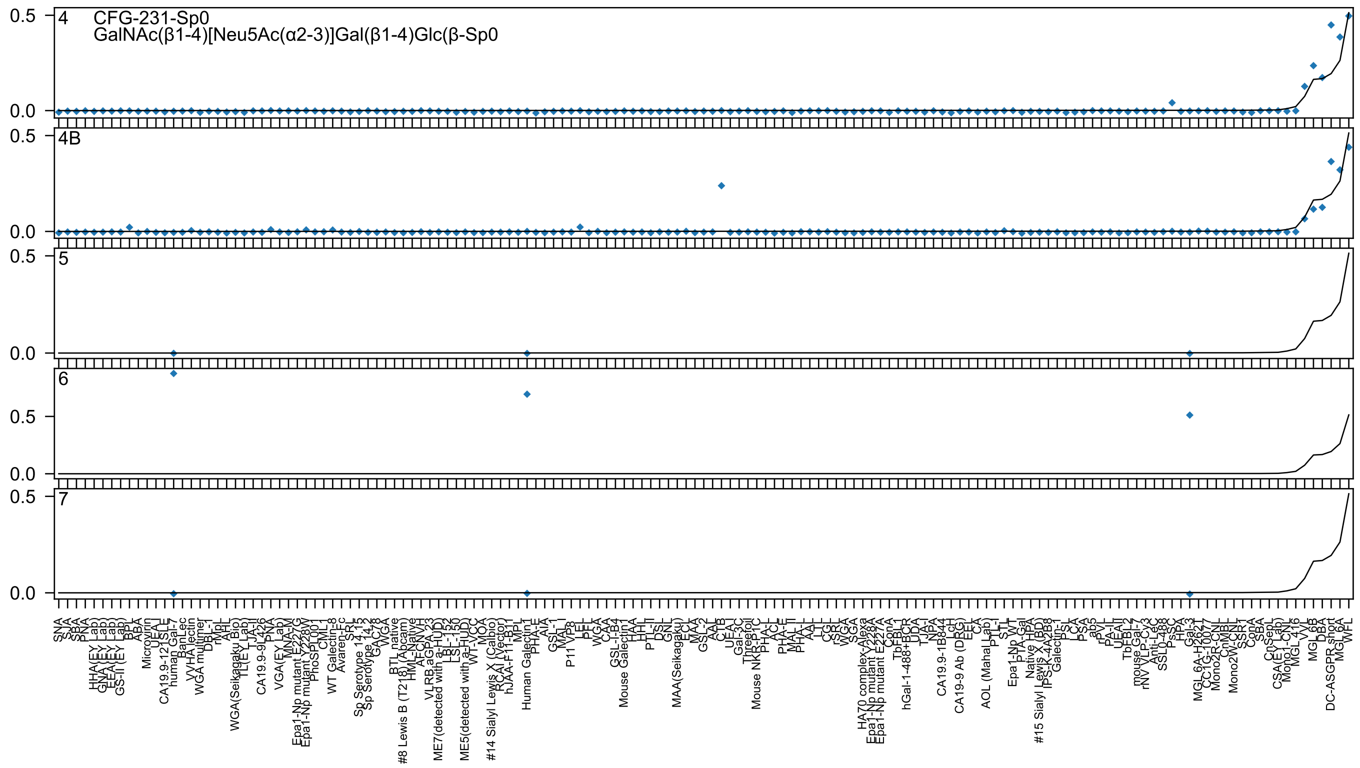

4
CFG-231-Sp0
GalNAc(β1-4)[Neu5Ac(α2-3)]Gal(β1-4)Glc(β-Sp0
4B
5
6
7
0.5
0.0

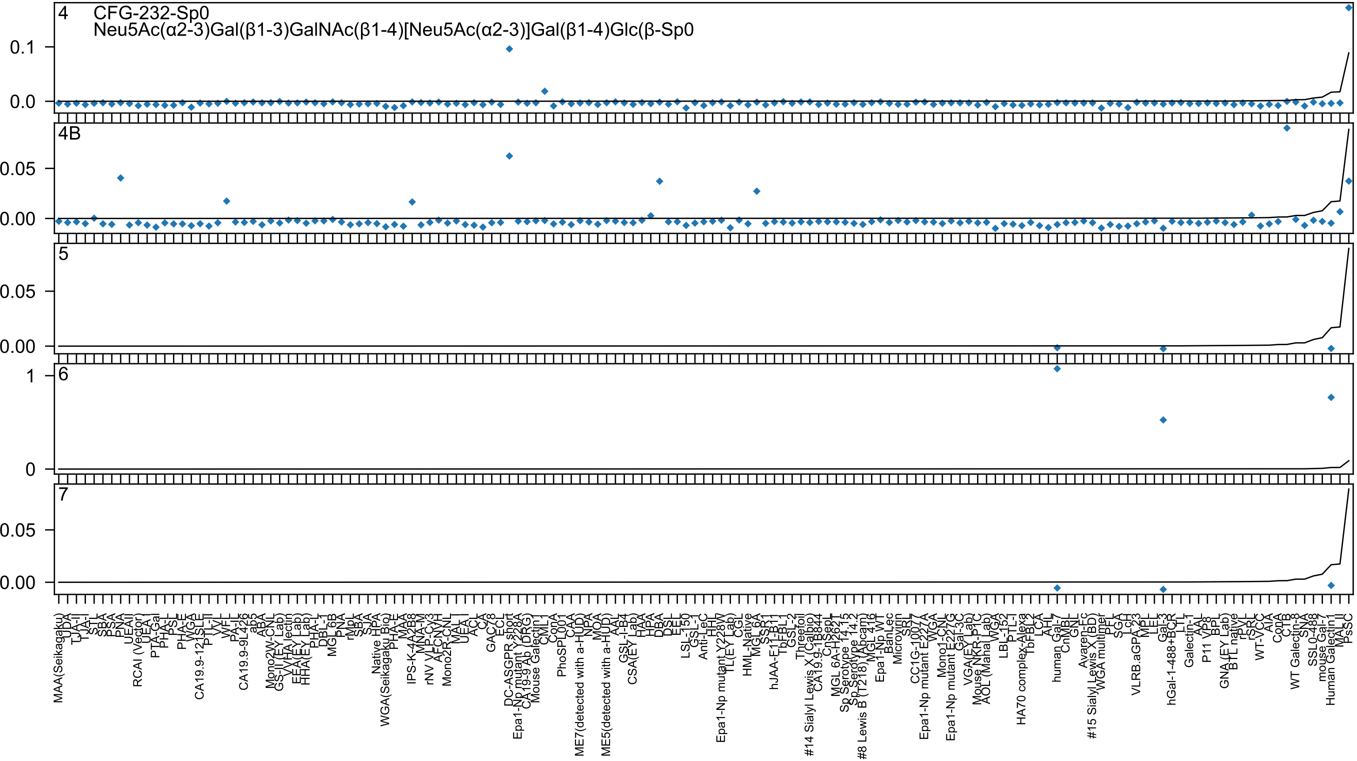
4
CFG-232-Sp0
Neu5Ac(α2-3)Gal(β1-3)GalNAc(β1-4)[Neu5Ac(α2-3)]Gal(β1-4)Glc(β-Sp0
0.1
0.0
4B
0.05
0.00
5
0.05
0.00
6
1
0
7
0.05
0.00

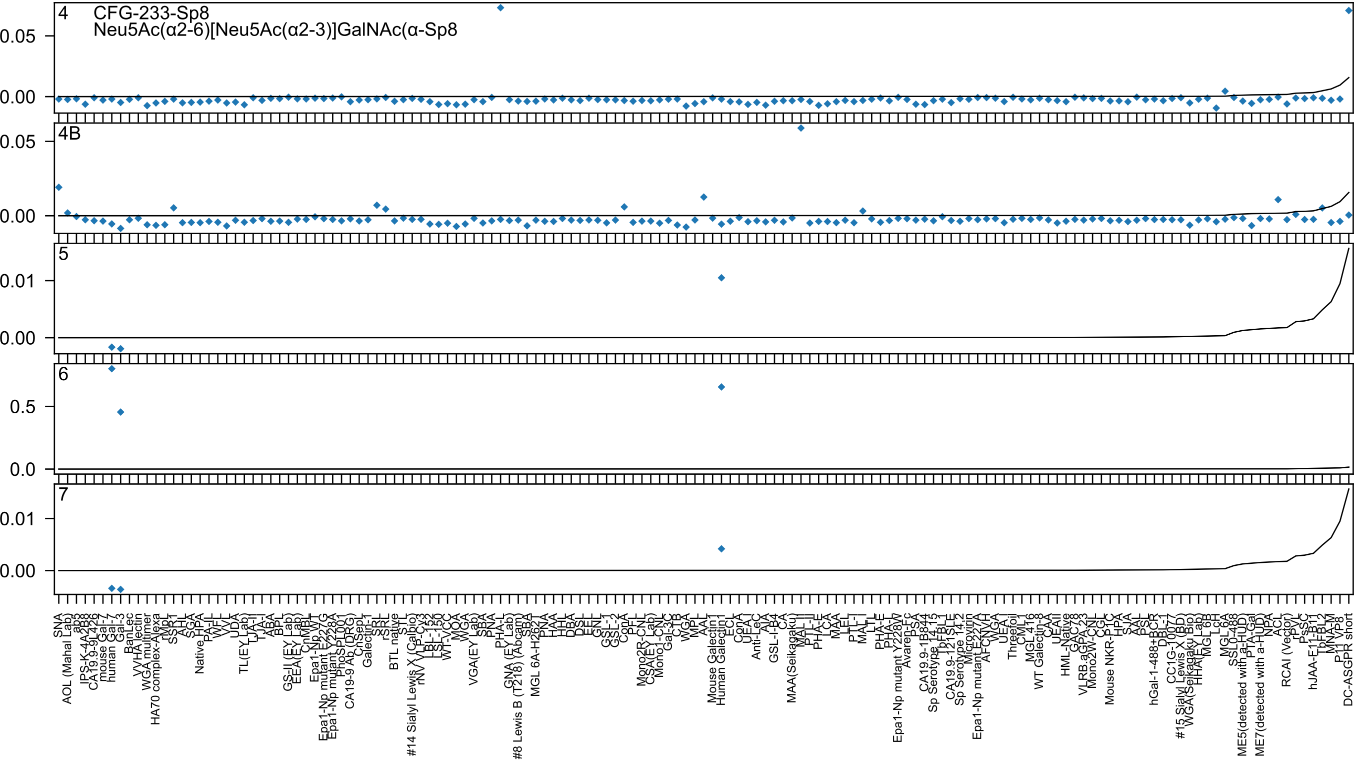
4
CFG-233-Sp8
Neu5Ac(α2-6)[Neu5Ac(α2-3)]GalNAc(α-Sp8
0.05
0.00
4B
0.05
0.00
5
0.01
0.00
6
0.5
0.0
7
0.01
0.00
SNA
AOL (Mahal Lab)
ab5
IPS-K-4A2B8
CA19.9-9L426
mouse Gal-7
human Gal-7
Gal-3
BanLec
VVHA lectin
WGA multimer
HA70 complex-Alexa
rMpL
SSR1
AHL
SGA
Native HPA
PA-IL
WFL
VVL
UDA
TL(EY Lab)
TJA-II
TJA-I
ABA
BPL
GS-II (EY Lab)
EEA(EY Lab)
CnMBL
Epa1-Np WT
Epa1-Np mutant E227G
Epa1-Np mutant Y228A
PhoSL001
CA19-9 Ab (DRG)
ChSep
Galectin-1
SRL
rSRL
BTL native
STL
#14 Sialyl Lewis X (Calbio)
rNV VLP-Cy3
LBL-152
LSL-150
WT-VCC
MOA
WGA
VGA(EY Lab)
SBA
PNA
PHA-L
GNA (EY Lab)
#8 Lewis B (T218) (Abcam)
SBA
MGL 6A-H262T
PNA
HAA
HHL
DBA
DSL
EEL
GNL
GSL-1
GSL-2
ConA
PFL
Mono2R-CNL
CSA(EY Lab)
Mono1-CNL
Gal-3C
CTB
WGA
MPL
AAL
Mouse Galectin1
Human Galectin1
ECL
ConA
UEA-I
Anti-LeC
AIA
GSL-I-B4
CA
MAA(Seikagaku)
MAL-II
PTL-II
PHA-E
LCA
MAA
LEL
PTL-I
MAL-I
LTL
PHA-E
PHA-L
Epa1-Np mutant Y228W
Avaren-Fc
PSA
CA19.9-1B844
Sp Serotype 14.15
TbFBL1
CA19.9-121SLE
Sp Serotype 14.2
Microvirin
Epa1-Np mutant E227A
AFCNVH
WGA
UEA
Threefoil
CML1
MGL 416
WT Galectin-8
CAA
UEAII
HML-Native
GAC78
VLRB.aGPA.23
Mono2W-CNL
CGL
Mouse NKR-P1C
HPA
SJA
AAL
PSL
hGal-1-488+BCR
DBL-1
CC1G-10077
#15 Sialyl Lewis X (BD)
WGA(Seikagaku Bio)
HHA(EY Lab)
MGL 6B
LcH
MGL 6A
SSL0-488
ME5(detected with a-HUD)
PTA-Gal
ME7(detected with a-HUD)
NPA
ACL
RCAI (Vector)
rPVL
PsSC
hJAA-E11-B11
TbFBL2
MNA-M
P11 VP8
DC-ASGPR short

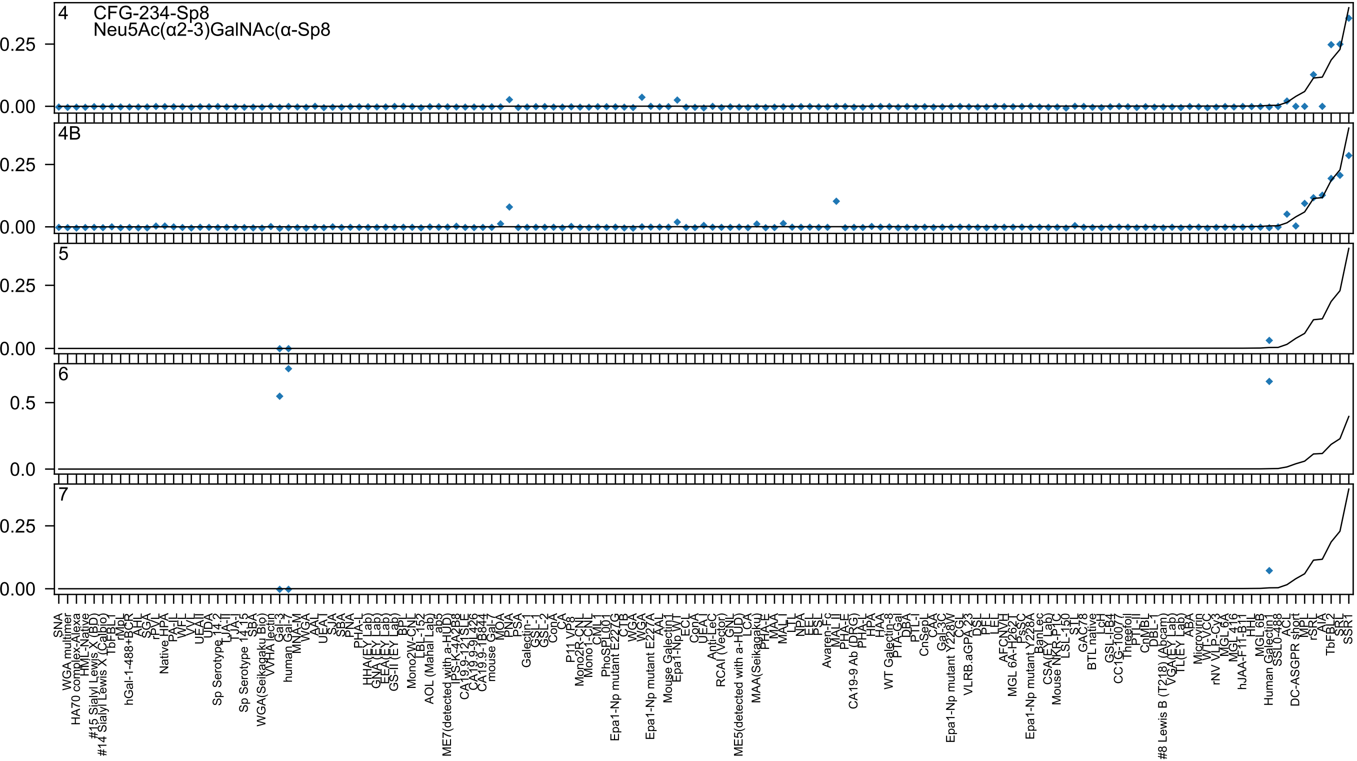
4
CFG-234-Sp8
Neu5Ac(α2-3)GalNAc(α-Sp8
4B
5
6
7
0.25
0.00
0.5
0.0

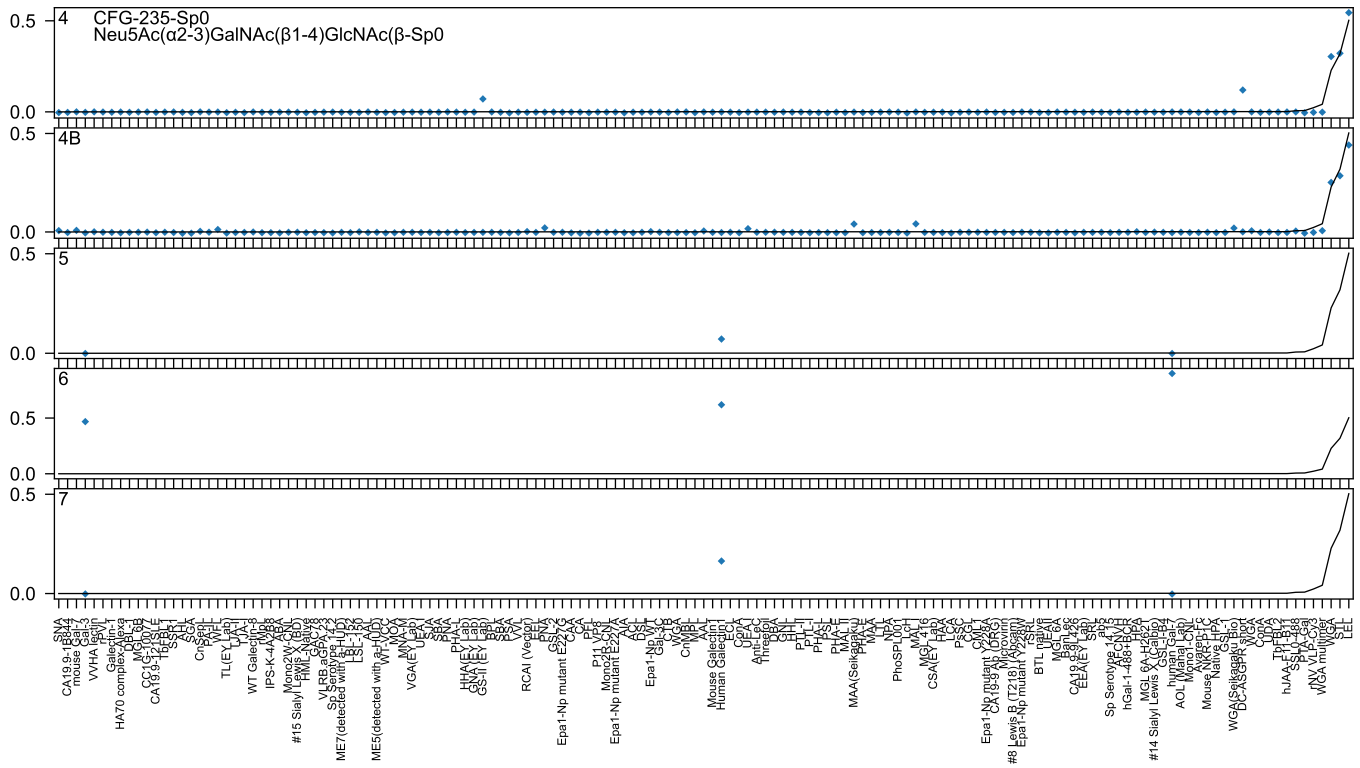

4
CFG-235-Sp0
Neu5Ac(α2-3)GalNAc(β1-4)GlcNAc(β-Sp0
4B
5
6
7
0.5
0.0

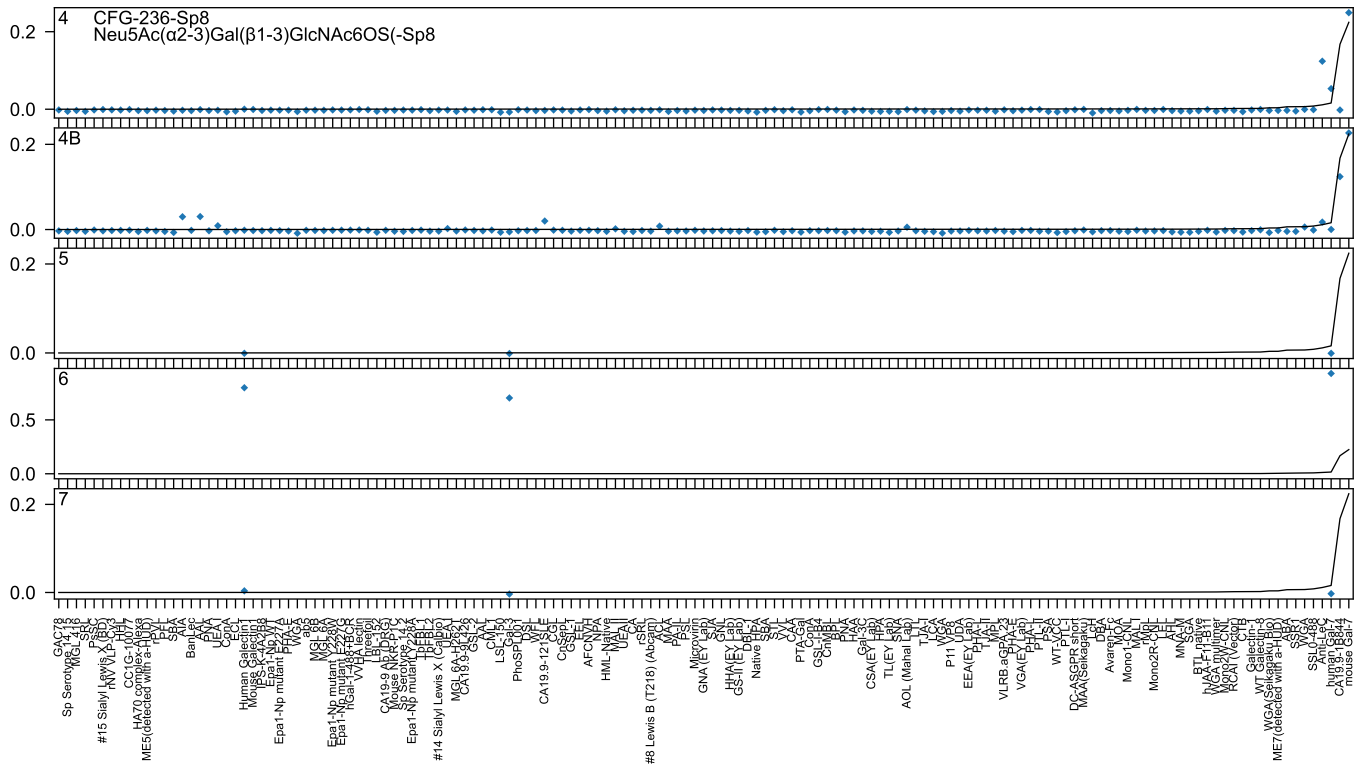

4
CFG-236-Sp8
Neu5Ac(α2-3)Gal(β1-3)GlcNAc6OS(-Sp8
4B
5
6
7

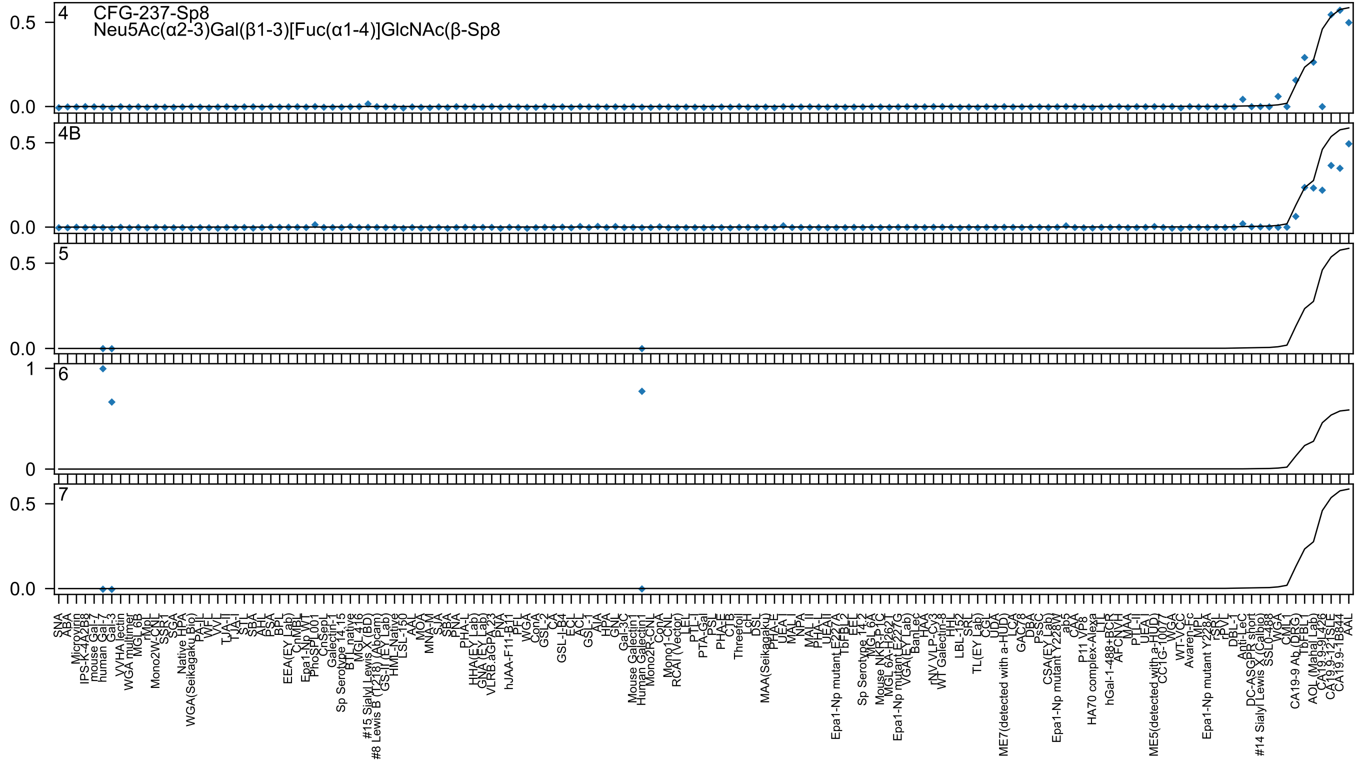

4
CFG-237-Sp8
Neu5Ac(α2-3)Gal(β1-3)[Fuc(α1-4)]GlcNAc(β-Sp8
4B
5
6
7

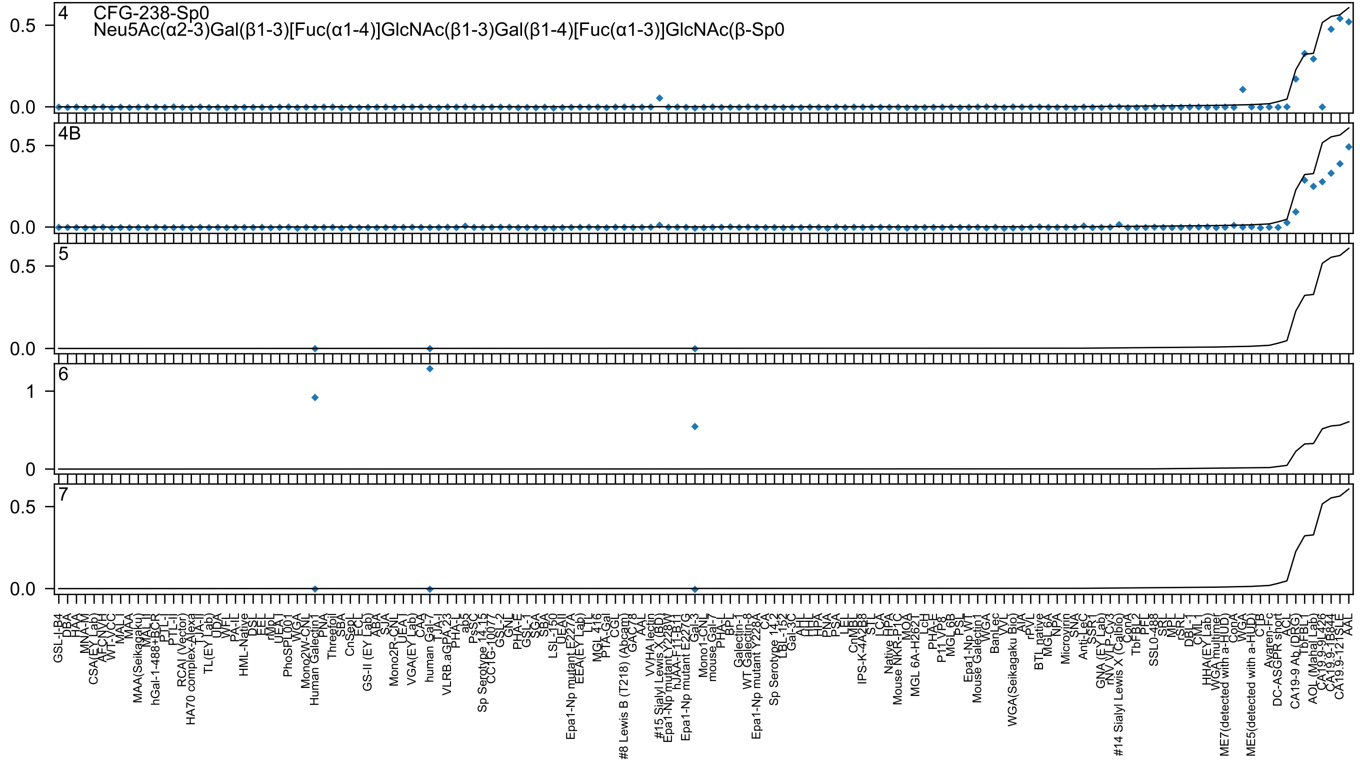
4
CFG-238-Sp0
Neu5Ac(α2-3)Gal(β1-3)[Fuc(α1-4)]GlcNAc(β1-3)Gal(β1-4)[Fuc(α1-3)]GlcNAc(β-Sp0
4B
5
6
7

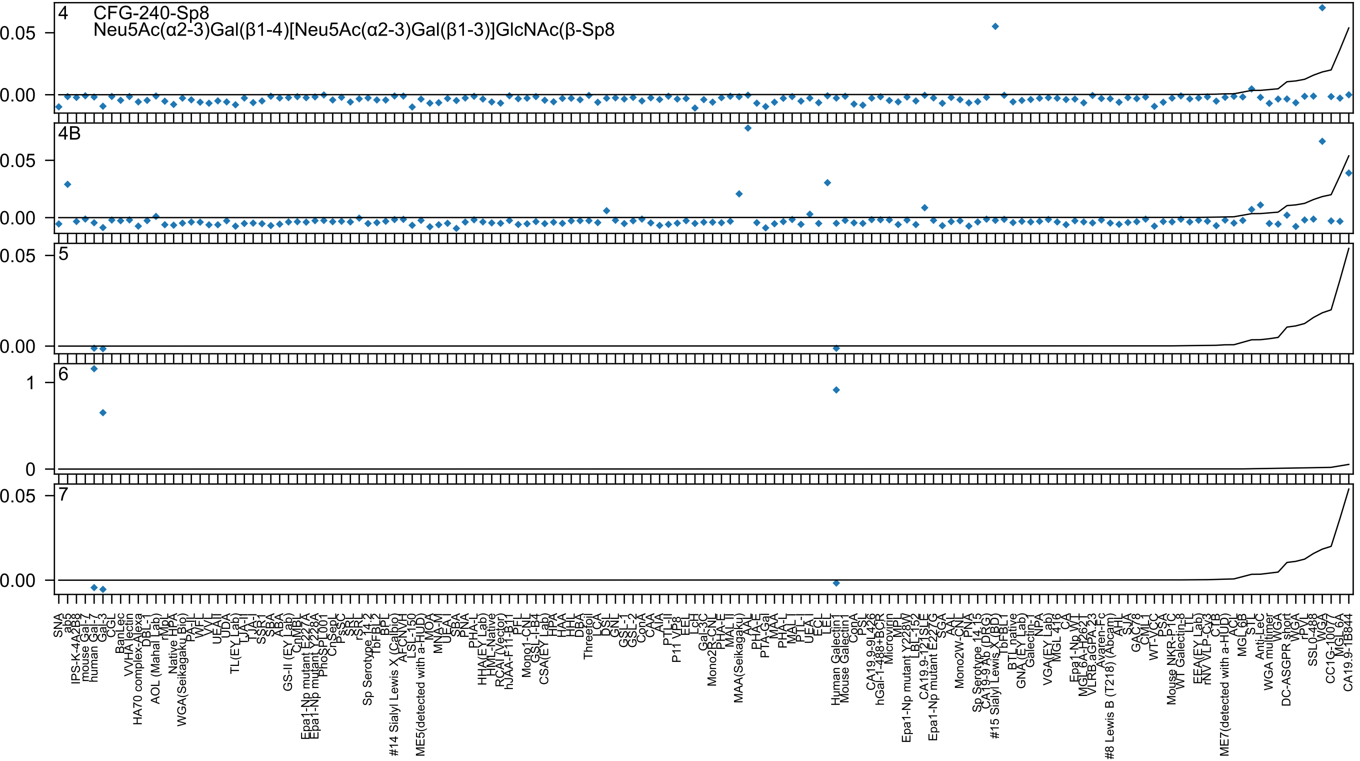
4
CFG-240-Sp8
Neu5Ac(α2-3)Gal(β1-4)[Neu5Ac(α2-3)Gal(β1-3)]GlcNAc(β-Sp8
4B
5
6
7
0.05
0.00
1
0
SNA
ab5
IPS-K-4A2B8
mouse Gal-7
human Gal-3
CGL
BanLec
VVHA lectin
HA70 complex-Alexa
DBL-I
AOL (Mahal Lab)
rMpL
Native HPA
WGA(Seikagaku Bio)
PA-II
WFL
VVL
UEA-II
UDA
TL(EY Lab)
LJA-I
TJA-I
SSR1
SBA
ABA
GS-II (EY Lab)
CnMBL
Epa1-Np mutant E227A
Epa1-Np mutant Y228A
PhoSL001
CnSeo
PsSC
SRL
rSRL
Sp Serotype 14.2
TbFBL2
BPL
#14 Sialyl Lewis X (Calbio)
AFCNVH
LSL-150
ME5(detected with a-HUD)
MOA
MNA-M
UEA-I
SBA
PNA
PHA-L
HHA(EY Lab)
HML-Native
RCAI (Vector)
hJAA-F11-B1
PFL
Mono1-CNL
GSL-I-B4
CSA(EY Lab)
HPA
HAA
HHL
DBA
Threefoil
LCA
DSL
GNL
GSL-1
GSL-2
ConA
CAA
AIA
PTL-II
P11 VP8
EEL
LcH
Gal-3C
Mono2R-CNL
PHA-E
MAL II
MAA(Seikagaku)
AAL
PHA-E
PTA-Gal
MAA
PHA-L
MAL I
PTL-I
UEA-I
ECL
LEL
Human Galectin1
Mouse Galectin1
ConA
PSL
CA19.9-9L426
hGal-1-488+BCR
Microvirin
MPL
Epa1-Np mutant Y228W
LBL-152
CA19.9-121SLE
Epa1-Np mutant E227G
SGA
AAL
Mono2W-CNL
PNA
Sp Serotype 14.15
CA19-9 Ab (DRG)
#15 Sialyl Lewis X (BD)
TbFBL1
BTL native
GNA (EY Lab)
Galectin-1
NPA
VGA(EY Lab)
MGL 4.16
CA
Epa1-Np WT
MGL6A-H262T
VLRB.aGPA.23
Avaren-Fc
#8 Lewis B (T218) (Abcam)
AHL
SJA
GAC78
CML1
WT-VCC
PSA
Mouse NKR-P1C
WT Galectin-8
LTL
EEA(EY Lab)
rNV VLP-Cy3
CTB
ME7(detected with a-HUD)
ACL
MGL 6B
STL
Anti-LeC
WGA multimer
WGA
DC-ASGPR short
WGA
rPVL
SSL0-488
WGA
CC1G-10077
MGL 6A
CA19.9-1B844

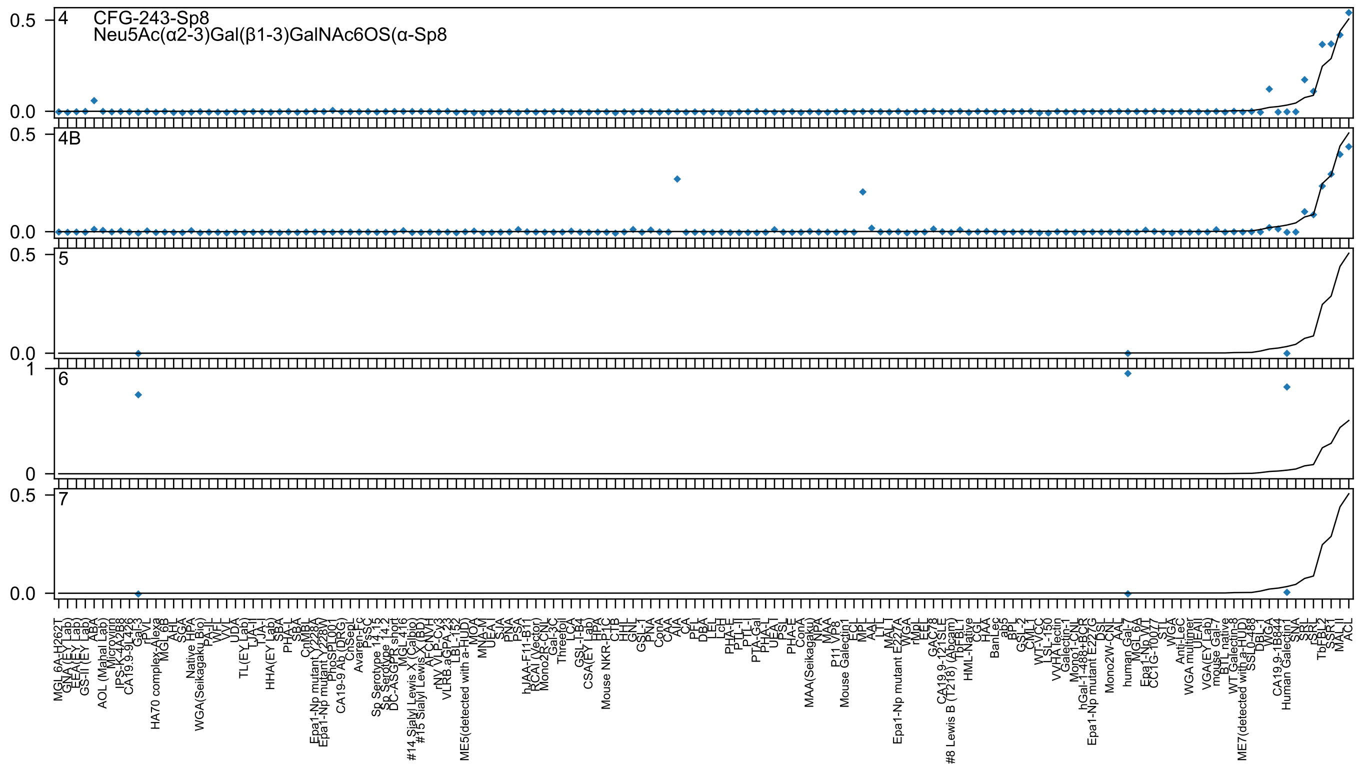
4
CFG-243-Sp8
Neu5Ac(α2-3)Gal(β1-3)GalNAc6OS(α-Sp8
4B
5
6
7

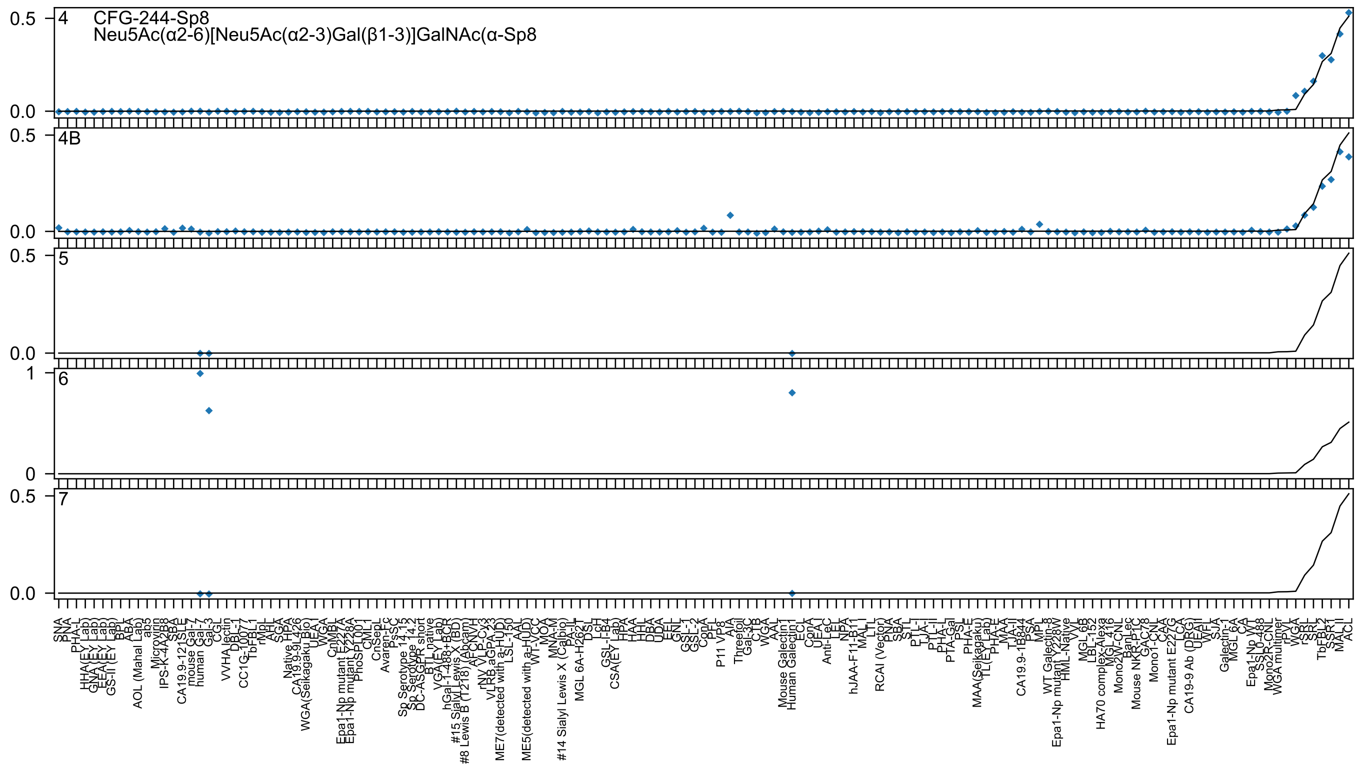

CFG-244-Sp8
Neu5Ac(α2-6)[Neu5Ac(α2-3)Gal(β1-3)]GalNAc(α-Sp8
4
4B
5
6
7
0.5
0.0
1
0

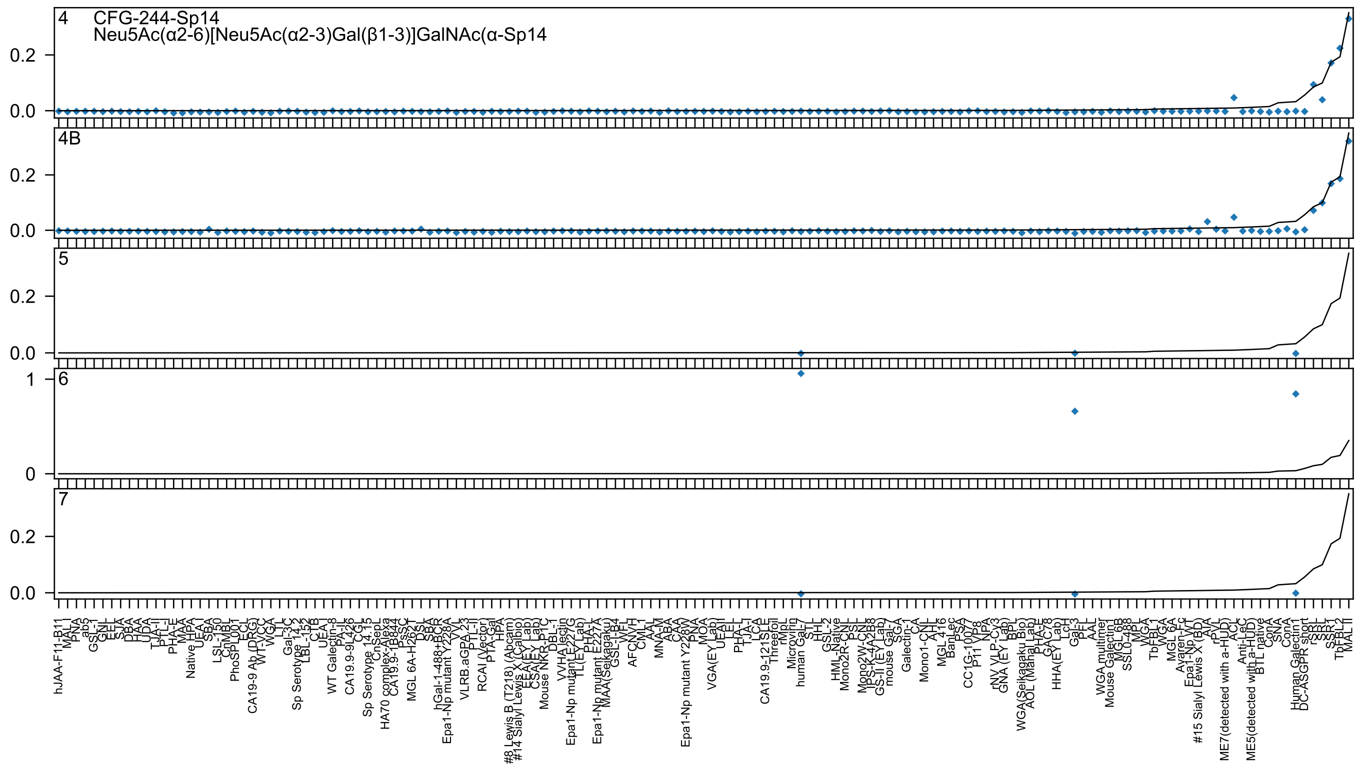
4 CFG-244-Sp14
Neu5Ac(α2-6)[Neu5Ac(α2-3)Gal(β1-3)]GalNAc(α-Sp14
4B
5
6
7

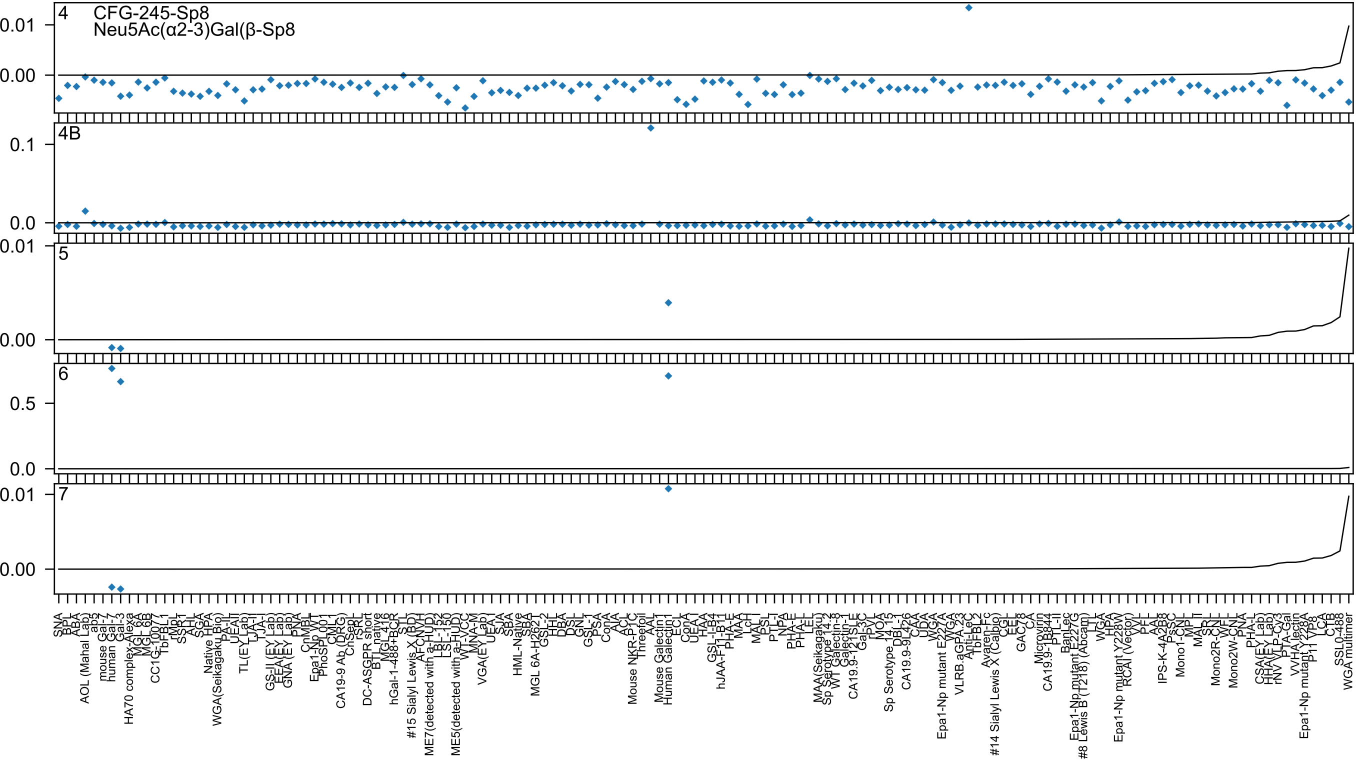

4
CFG-245-Sp8
Neu5Ac(α2-3)Gal(β-Sp8
4B
5
6
7

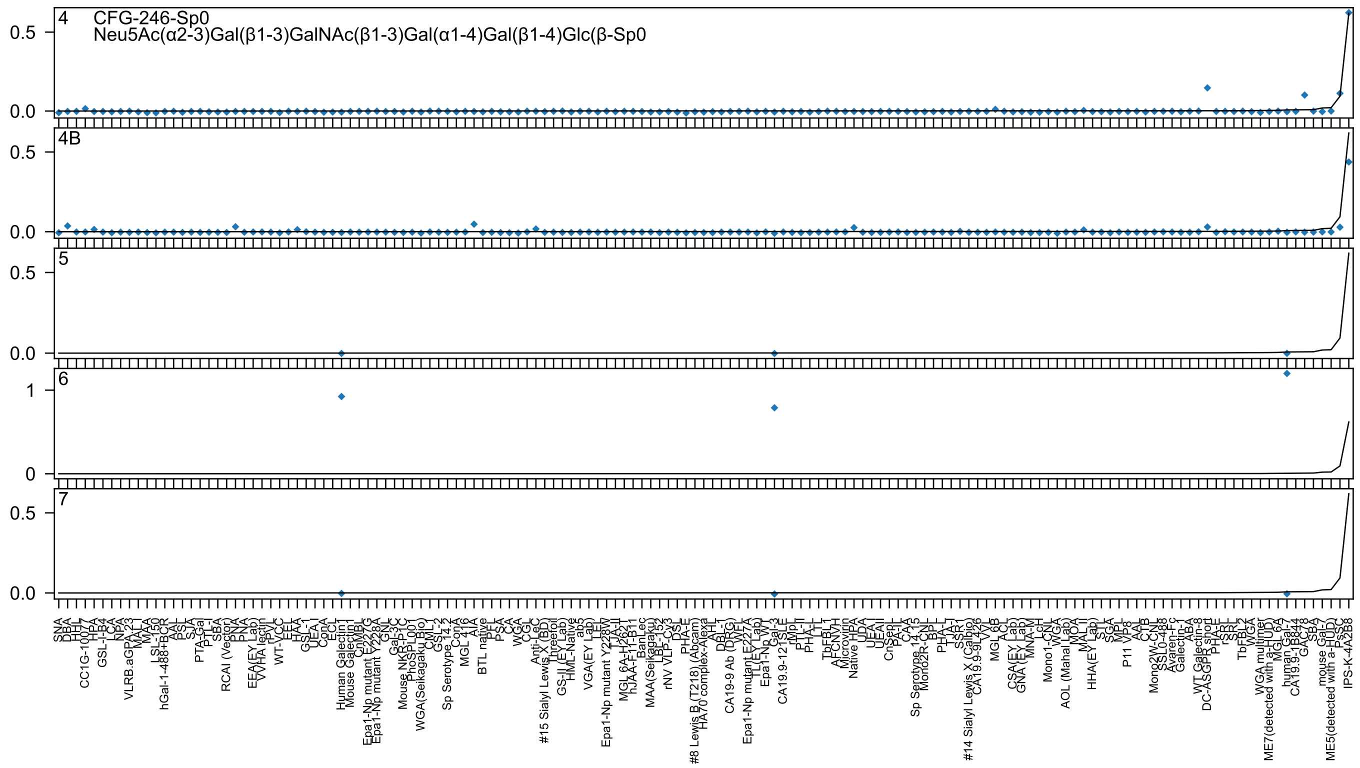

4 CFG-246-Sp0
Neu5Ac(α2-3)Gal(β1-3)GalNAc(β1-3)Gal(α1-4)Gal(β1-4)Glc(β-Sp0
4B
5
6
7
0.5
0.0
1
0
SNA
DBA
HHL
CC1G-10077
HPA
GSL-I-B4
LCA
NPA
VLRB.aGPA.23
MAL-I
MAA
LSL-150
hGal-1-488+BCR
AAL
PSL
SJA
PTA-Gal
PTL-I
SBA
RCAI (Vector)
PNA
PNA
EEA(EY Lab)
VVHA lectin
rPVL
WT-VCC
EEL
HAA
GSL-1
UEA-I
ConA
ECL
Human Galectin1
Mouse Galectin1
CnMBL
Epa1-Np mutant E227G
Epa1-Np mutant Y228A
GNL
Gal-3C
Mouse NKR-P1C
PhoSL001
WGA(Seikagaku Bio)
CML-1
GSL-2
Sp Serotype 14.2
ConA
MGL 416
AIA
BTL native
PFL
PSA
CA
WGA
CGL
Anti-LeC
#15 Sialyl Lewis X (BD)
Threefoil
GS-II (EY Lab)
HML-Native
ab5
VGA(EY Lab)
LEL
Epa1-Np mutant Y228W
TJA-II
MGL 6A-H262T
hJAA-F11-B11
BanLec
MAA(Seikagaku)
LBL-152
rNV VLP-Cy3
DSL
PHA-E
#8 Lewis B (T218) (Abcam)
HA70 complex-Alexa
AHL
DBL-1
CA19-9 Ab (DRG)
WFL
Epa1-Np mutant E227A
TL(EY Lab)
Epa1-Np WT
Gal-3
CA19.9-121SLE
rMpL
PTL-II
PHA-L
LTL
TbFBL1
AFCNVH
Microvirin
Native HPA
UDA
UEA-II
UEA-II
CnSep
PA-IL
CAA
Sp Serotype 14.15
Mono2R-CNL
BPL
PHA-L
TJA-I
SSR1
#14 Sialyl Lewis X (Calbio
CA19.9-9L426
VVL
MGL 6B
ACL
CSA(EY Lab)
GNA (EY Lab)
MNA-M
LcH
Mono1-CNL
WGA
AOL (Mahal Lab)
MOA
MAL-II
HHA(EY Lab)
STL
SGA
MPL
P11 VP8
AAL
CTB
Mono2W-CNL
SSL0-488
Avaren-Fc
Galectin-1
ABA
WT Galectin-8
DC-ASGPR short
PHA-E
rSRL
SRL
TbFBL2
WGA
WGA multimer
ME7(detected with a-HUD)
MGL 6A
human Gal-7
CA19.9-1B844
GAC78
SBA
mouse Gal-7
ME5(detected with a-HUD)
PsSC
IPS-K-4A2B8

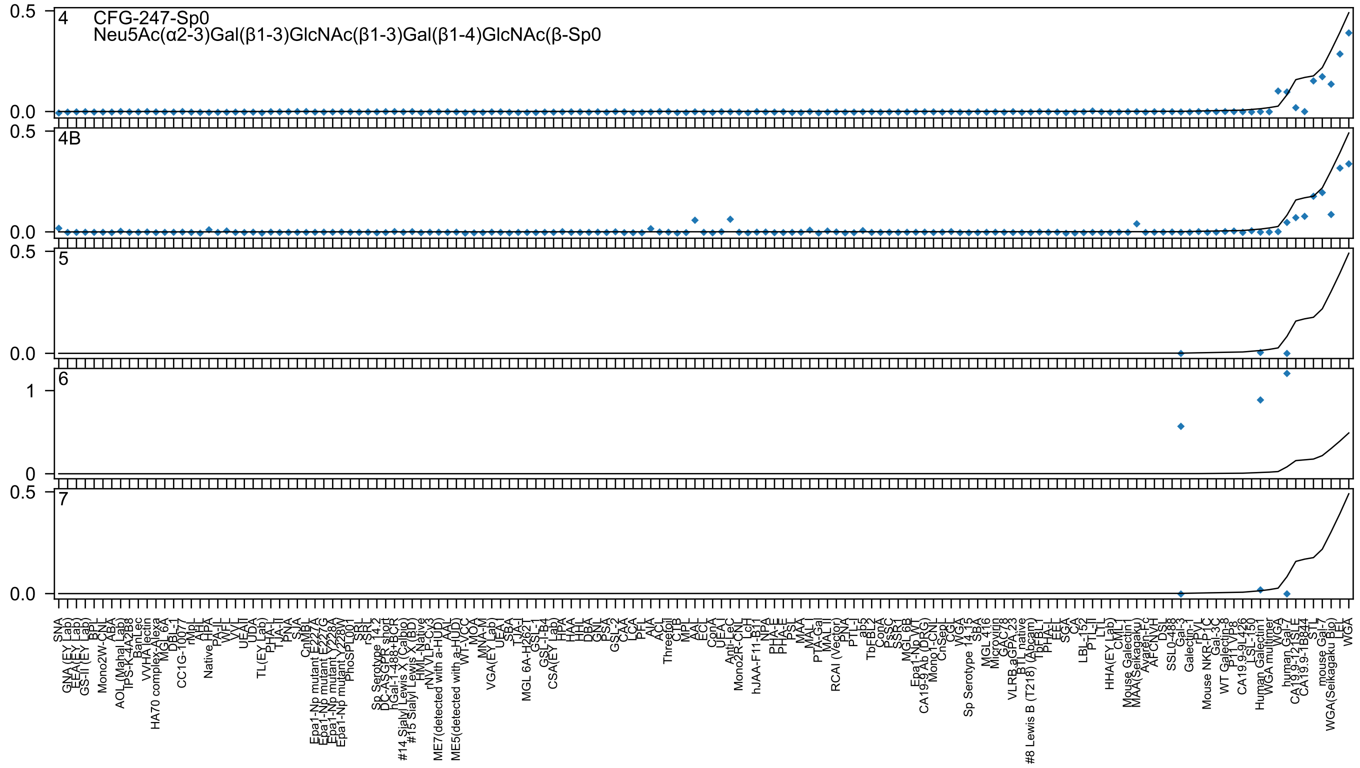
4
CFG-247-Sp0
Neu5Ac(α2-3)Gal(β1-3)GlcNAc(β1-3)Gal(β1-4)GlcNAc(β-Sp0
4B
5
6
7
0.5
0.0
1
0
SNA
GNA (EY Lab)
EEA(EY Lab)
GS-II (EY Lab)
BPL
Mono2W-CNL
ABA
AOL (Mahal Lab)
TPS-K-4A2B8
BanLec
VVHA lectin
HA70 complex-Alexa
MGL 6A
DBL-1
CC1G-10077
rMpL
AHL
Native HPA
PA-IL
WFL
VVL
UEA-II
UDA
TL(EY Lab)
PHA-L
TJA-II
PNA
SJA
CnMBL
Epa1-Np mutant E227A
Epa1-Np mutant E227G
Epa1-Np mutant Y228A
Epa1-Np mutant Y228W
PhoSL001
SRL
rSRL
Sp Serotype 14 2
DC-ASGPR short
hGal-1-488+BCR
#14 Sialyl Lewis X (Calbio)
#15 Sialyl Lewis X (BD)
HML-Native
rNV VLP-Cy3
ME7(detected with a-HUD)
AAL
ME5(detected with a-HUD)
WT-VCC
MOA
MNA-M
VGA(EY Lab)
UEA-I
SBA
TJA-I
MGL 6A-H262T
GSL-I
GSL-I-B4
CSA(EY Lab)
HPA
HAA
HHL
DBA
GNL
PSA
GSL-2
CAA
LCA
PEL
AIA
ACL
Threetoil
CTB
MPL
AAL
ECL
ConA
UEA-I
Anti-LeC
Mono2R-CNL
LcH
hJAA-F11-B11
NPA
PHA-E
PHA-E
PSL
MAA
MAL-I
PTA-Gal
MAL-II
RCAI (Vector)
PNA
PTL-I
ab5
TbFBL2
ConA
PsSC
SSR1
MGL 6B
Epa1-Np WT
CA19-9 Ab (DRG)
Mono1-CNL
CnSep
CGL
WGA
Sp Serotype 14 15
SBA
MGL 416
Microvirin
GAC78
VLRB aGPA 23
BTL native
#8 Lewis B (T218) (Abcam)
TbFBL1
PHA-L
EEL
SGA
CA
LBL-152
PTL-II
LTL
HHA(EY Lab)
CML1
Mouse Galectin1
MAA(Seikagaku)
Avaren-Fc
AFCN-H
DSL
SSL0-488
Gal-3
Galectin-1
TPL
Mouse NKR-P1C
Gal-3C
WT Galectin-8
P1 VP8
CA19.9-9L426
LSL-150
Human Galectin1
WGA multimer
WGA
human Gal-7
CA19.9-121SLE
CA19.9-1B844
STL
mouse Gal-7
WGA(Seikagaku Bio)
LEL
WGA

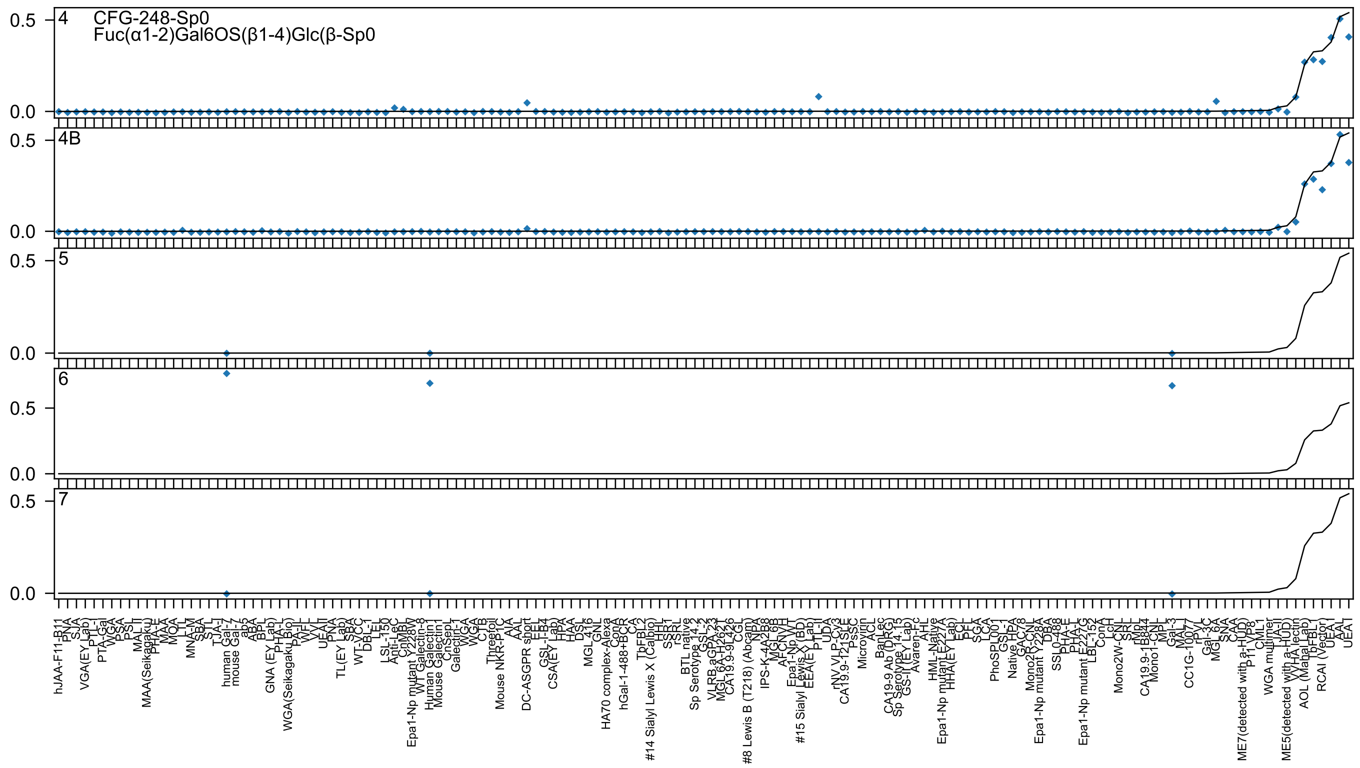
4
CFG-248-Sp0
Fuc(α1-2)Gal6OS(β1-4)Glc(β-Sp0
4B
5
6
7
0.5
0.0

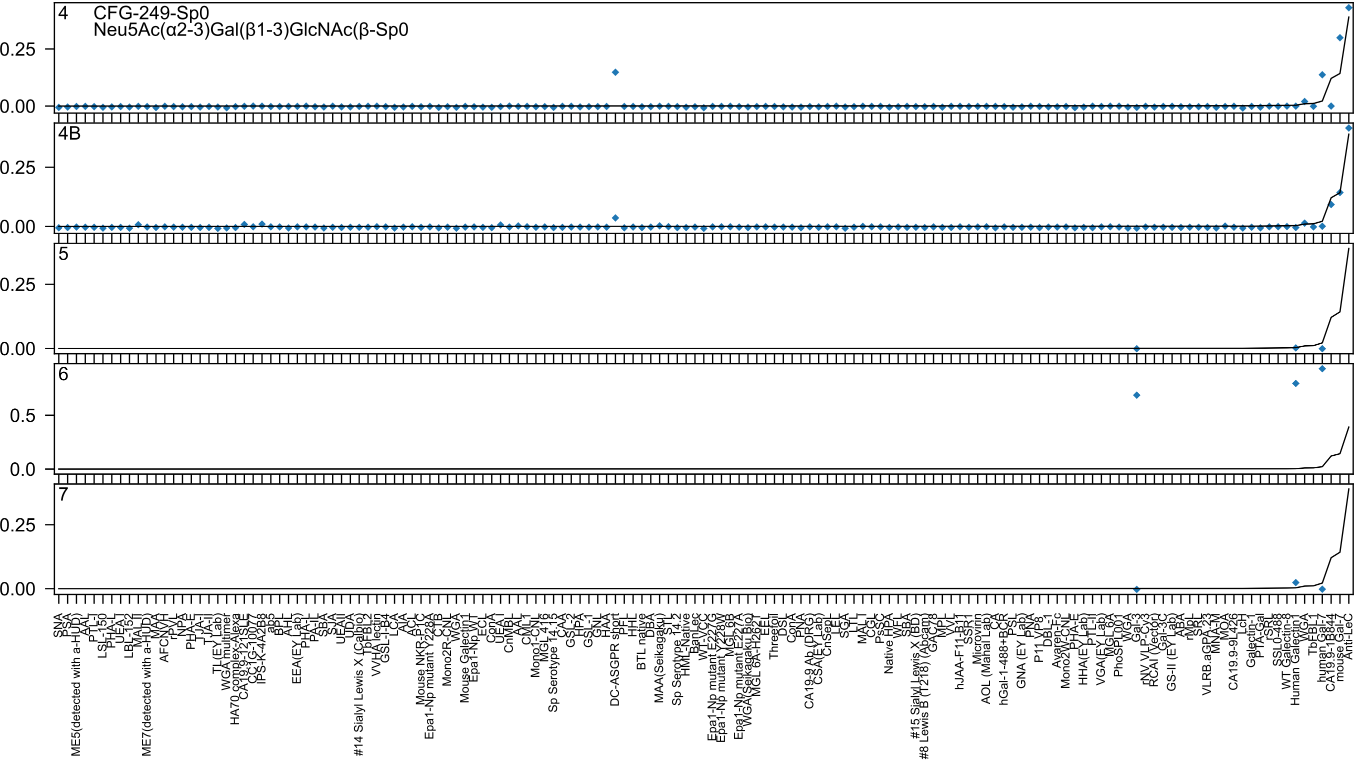
4 CFG-249-Sp0
Neu5Ac(α2-3)Gal(β1-3)GlcNAc(β-Sp0
4B
5
6
7
0.25
0.00
0.5
0.0

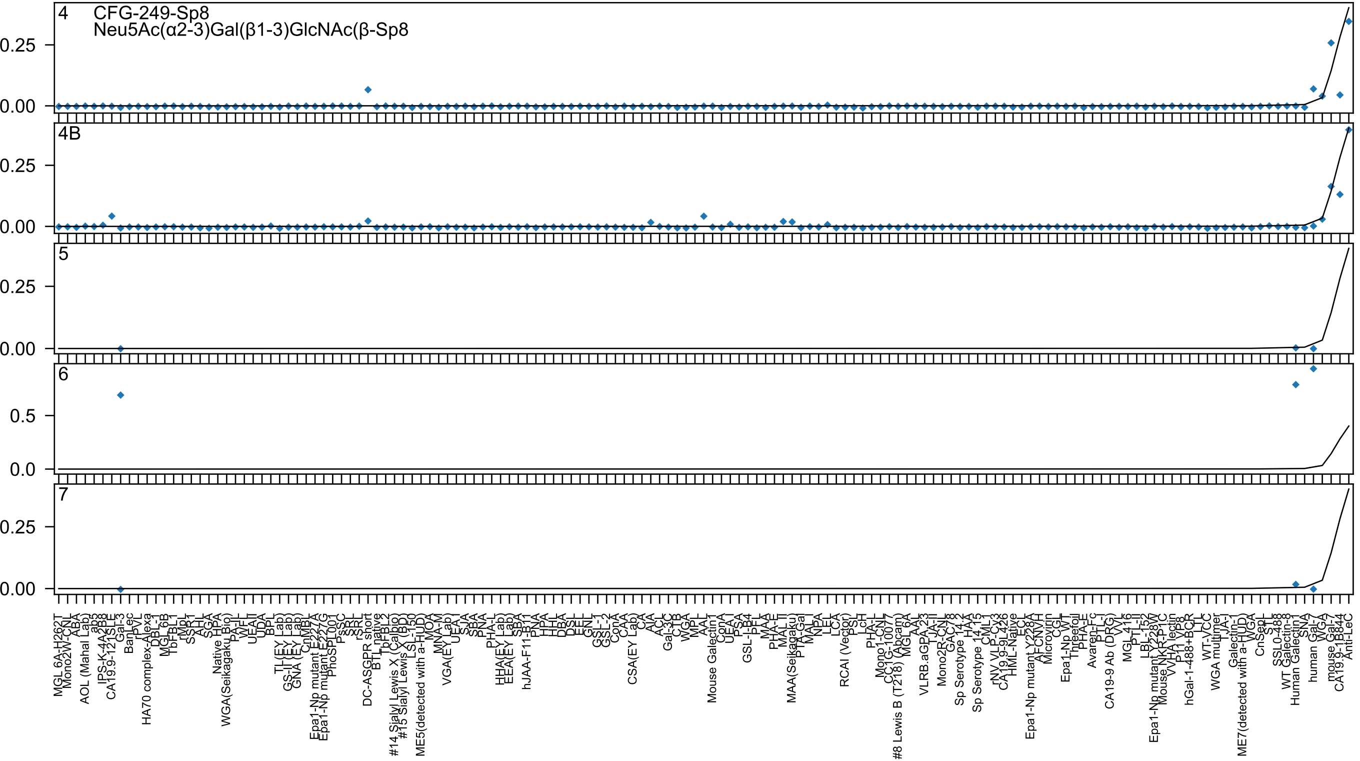
CFG-249-Sp8
Neu5Ac(α2-3)Gal(β1-3)GlcNAc(β-Sp8
4
4B
5
6
7

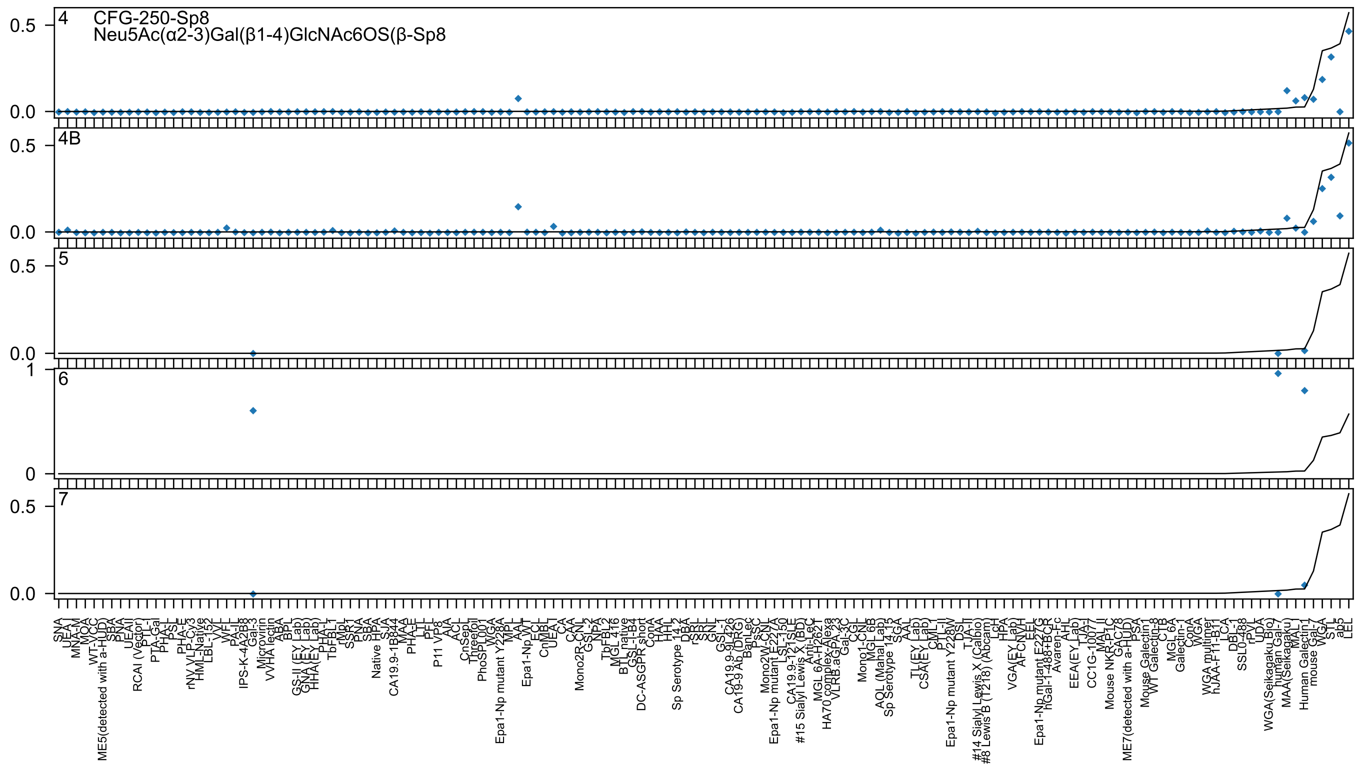

4
CFG-250-Sp8
Neu5Ac(α2-3)Gal(β1-4)GlcNAc6OS(β-Sp8
4B
5
6
7

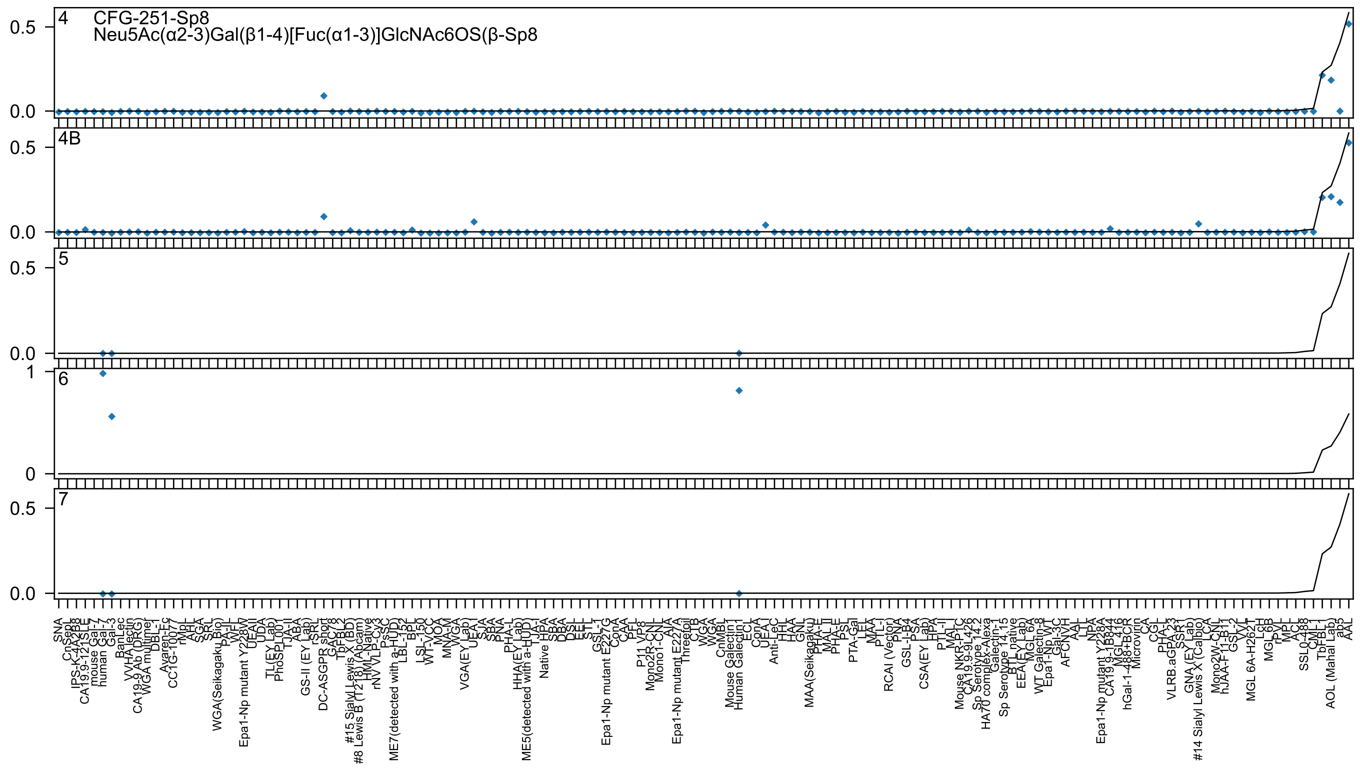
4
CFG-251-Sp8
Neu5Ac(α2-3)Gal(β1-4)[Fuc(α1-3)]GlcNAc6OS(β-Sp8
4B
5
6
7
0.5
0.0
1
0
SNA
ChSeql
IPS-K-4A2B8
CA19.9-121SLE
mouse Gal-7
human Gal-7
Gal-3
BanLec
VVHA lectin
CA19-9 Ab (DRG)
WGA multimer
DBL-1
Avaren-Fc
CC1G-1007
rMpL
AHL
SGA
SRL
WGA(Seikagaku Bio)
PA-IL
WFL
Epa1-Np mutant Y228W
UEAII
UDA
TL(EY Lab)
PhoSL001
TJA-II
ABA
GS-II (EY Lab)
rSRL
DC-ASGPR short
GAC78
TbFBL2
#15 Sialyl Lewis X (BD)
#8 Lewis B (T218) (Abcam)
HML-Native
rNV VLP-Cy3
PsSC
ME7(detected with a-HUD)
LBL-152
BPL
LSL-150
WT-VCC
MOA
MNA-M
WGA
VGA(EY Lab)
UEA-I
SJA
SBA
PNA
PHA-L
HHA(EY Lab)
ME5(detected with a-HUD)
TJA-I
Native HPA
SBA
DBA
DSL
EEL
STL
GSL-I
Epa1-Np mutant E227G
ConA
CAA
PFL
P11 VP8
Mono2R-CNL
Mono1-CNL
AIA
Epa1-Np mutant E227A
Threefoil
CTB
WGA
WGA
CnMBL
Mouse Galectin1
Human Galectin1
ECL
ConA
UEA-I
Anti-LeC
HHL
HAA
GNL
MAA(Seikagaku)
PHA-E
MAL_II
PHA-E
PSL
PTA-Gal
LEL
MAA
PTL-I
RCAI (Vector)
PNA
GSL-I-B4
PSA
CSA(EY Lab)
HPA
PTL-II
MAL-I
Mouse NKR-P1C
CA19.9-9L426
Sp Serotype 14.2
HA70 complex-Alexa
Galectin-1
Sp Serotype 14.15
BTL native
EEA(EY Lab)
MGL 6A
WT Galectin-8
Epa1-Np WT
Gal-3C
AFCNVH
AAL
LTL
NPA
Epa1-Np mutant Y228A
CA19.9-1B844
MGL 416
hGal-1-488+BCR
Microvirin
CA
CGL
PHA-L
VLRB.aGPA.23
SSR1
GNA (EY Lab)
#14 Sialyl Lewis X (Calbio)
LCA
Mono2W-CNL
hJAA-F11-B11
GSL-2
VVL
MGL 6A-H262T
LcH
MGL 6B
rPVL
MPL
ACL
SSL0-488
CML
TbFBL1
AOL (Mahal Lab)
ab5
AAL

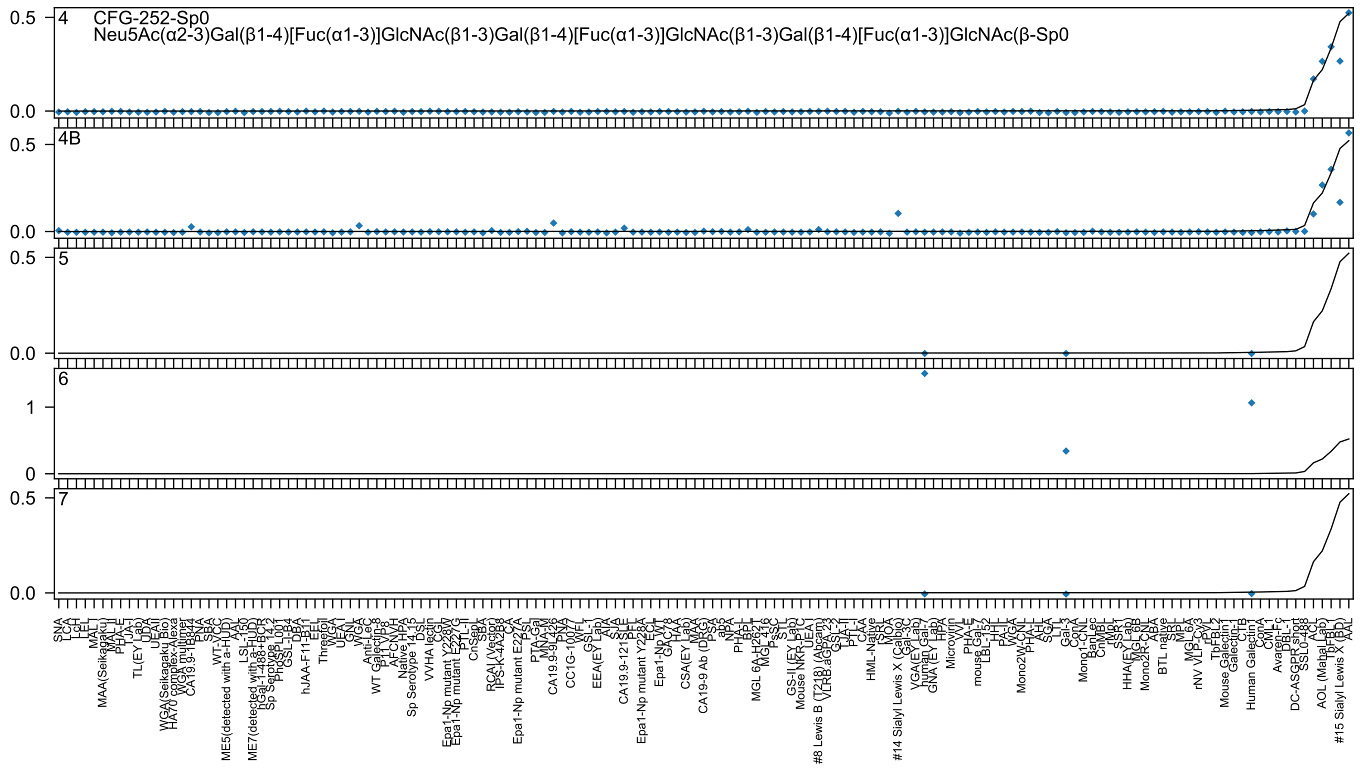

4
CFG-252-Sp0
Neu5Ac(α2-3)Gal(β1-4)[Fuc(α1-3)]GlcNAc(β1-3)Gal(β1-4)[Fuc(α1-3)]GlcNAc(β1-3)Gal(β1-4)[Fuc(α1-3)]GlcNAc(β-Sp0
4B
5
6
7

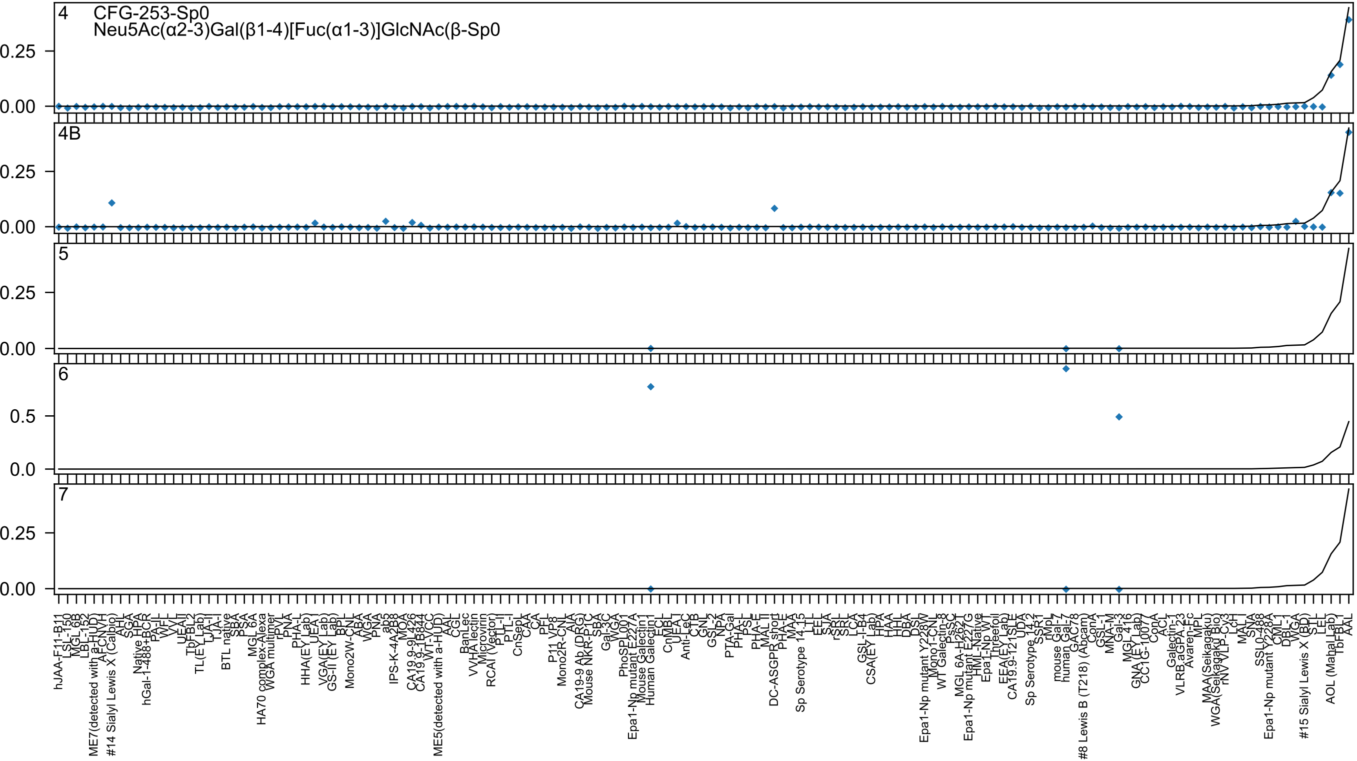
4 CFG-253-Sp0
Neu5Ac(α2-3)Gal(β1-4)[Fuc(α1-3)]GlcNAc(β-Sp0
4B
5
6
7

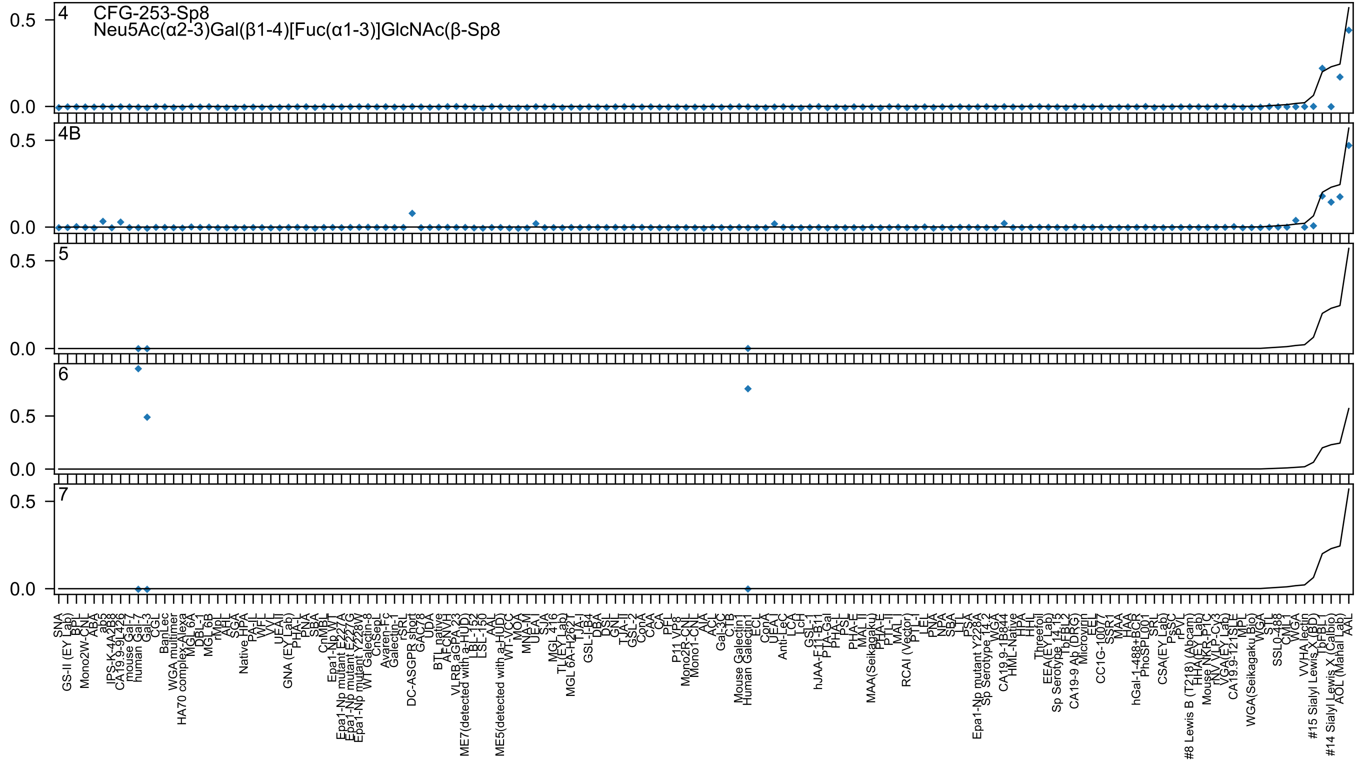
4
CFG-253-Sp8
Neu5Ac(α2-3)Gal(β1-4)[Fuc(α1-3)]GlcNAc(β-Sp8
4B
5
6
7
0.0
0.5
SNA
GS-II (EY Lab)
BPL
Mono2W-CNL
ABA
ab5
IPS-K-4A2B8
CA19.9-9L426
mouse Gal-7
human Gal-3
CGL
BanLec
WGA multimer
HA70 complex-Alexa
MGL 6A
DBL-1
MGL 6B
rMpL
AHL
SGA
Native HPA
PA-IL
WFL
VVL
UEAII
GNA (EY Lab)
PHA-L
PNA
SBA
CnMBL
Epa1-Np WT
Epa1-Np mutant E227G
Epa1-Np mutant E227A
Epa1-Np mutant Y228W
WT Galectin-8
CnSepL
Avaren-Fc
Galectin-1
rSRL
DC-ASGPR short
GAC78
UDA
BTL native
AFCNVH
VLRB aGPA.23
ME7(detected with a-HUD)
LBL-152
LSL-150
AAL
ME5(detected with a-HUD)
WT-VCC
MOA
MNA-M
UEA-I
SJA
MGL 416
TL(EY Lab)
MGL 6A-H262T
TJA-I
GSL-I-B4
DBA
DSL
GNL
TJA-II
GSL-2
ConA
CAA
CA
PFL
P11 VP8
Mono2R-CNL
Mono1-CNL
AIA
ACL
Gal-3C
CTB
Mouse Galectin1
Human Galectin1
ECL
ConA
UEAI
Anti-LeC
LCA
LcH
GSL-1
hJAA-F11-B11
PTA-Gal
PHA-L
PSL
PHA-E
MAL II
MAA(Seikagaku)
PHA-E
PTL-II
MAL I
RCAI (Vector)
PTL-I
LEL
PNA
NPA
SBA
LTL
PSA
Epa1-Np mutant Y228A
Sp Serotype 14.2
WGA
CA19.9-1B844
HML-Native
HPA
HHL
Threefoil
EEA(EY Lab)
Sp Serotype 14.13
TbFBL2
CA19-9 Ab (DRG)
Microvirin
EEL
CC1G-10077
SSR1
MAA
HAA
hGal-1-488+BCR
PhoSL001
SRL
CSA(EY Lab)
PsSC
rPVL
#8 Lewis B (T218) (Abcam)
HHA(EY Lab)
Mouse NKR-P1C
rNV VLP-Cy3
VGA(EY Lab)
CA19.9-121SLE
MPL
WGA(Seikagaku Bio)
WGA
STL
SSL0-488
CML1
WGA
VVHA lectin
#15 Sialyl Lewis X (BD)
TbFBL1
#14 Sialyl Lewis X (Calbio)
AOL (Manal Lab)
AAL

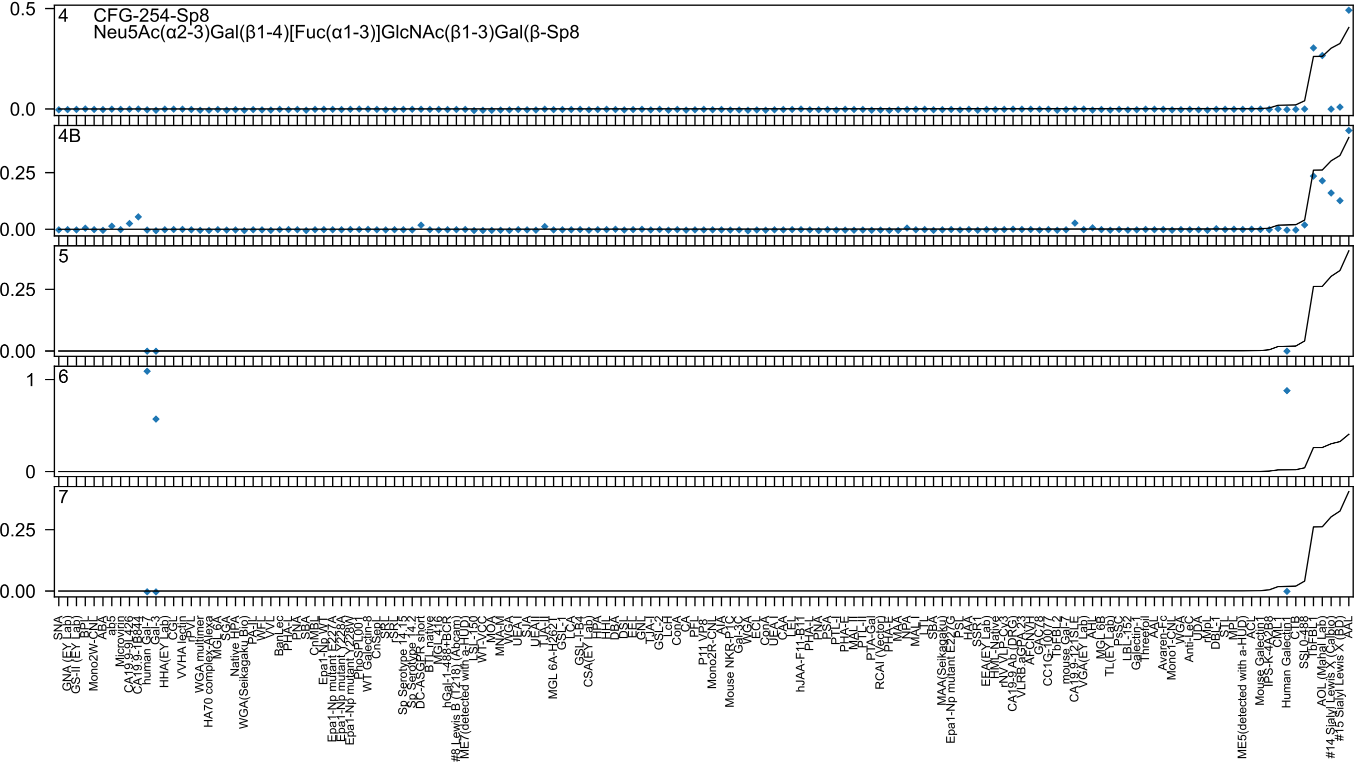
CFG-254-Sp8
Neu5Ac(α2-3)Gal(β1-4)[Fuc(α1-3)]GlcNAc(β1-3)Gal(β-Sp8
4
4B
5
6
7
0.5
0.0
0.25
0.00
1
0

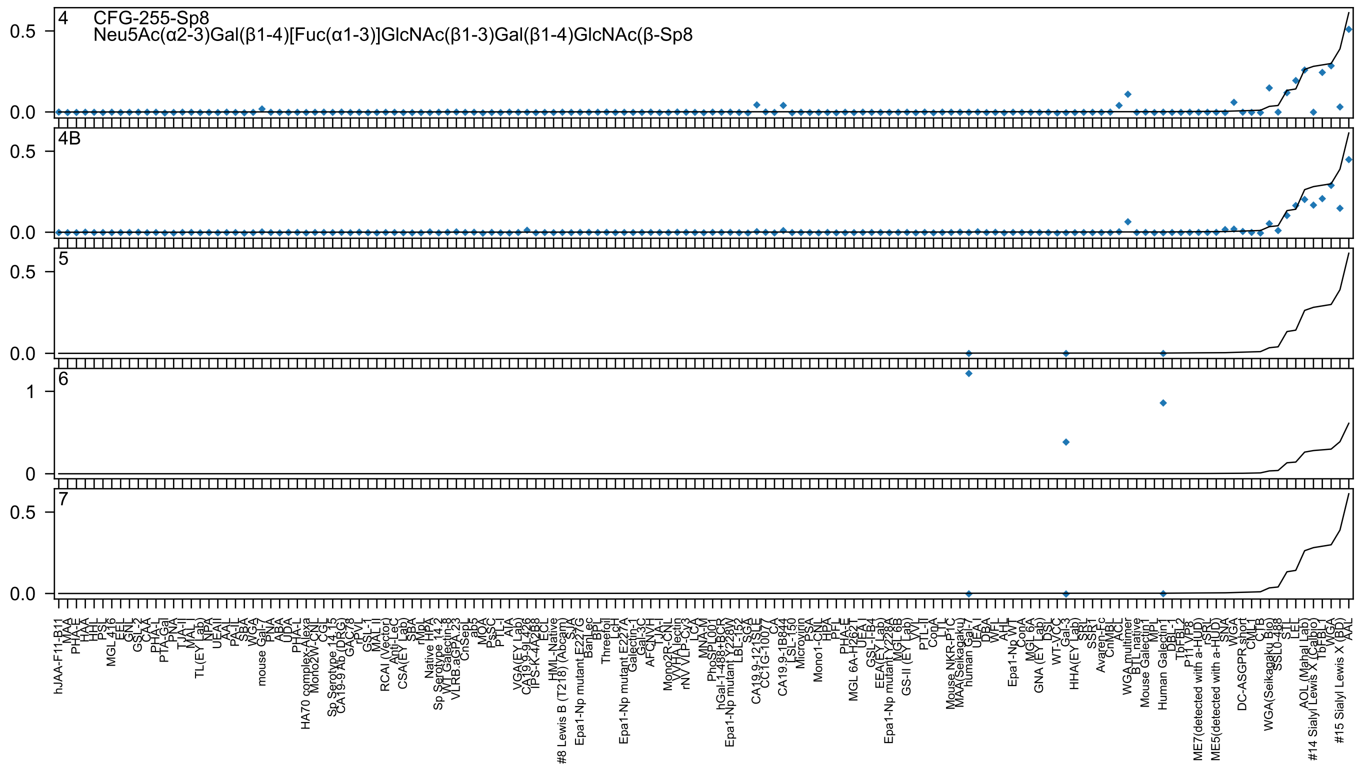

4
CFG-255-Sp8
Neu5Ac(α2-3)Gal(β1-4)[Fuc(α1-3)]GlcNAc(β1-3)Gal(β1-4)GlcNAc(β-Sp8
4B
5
6
7

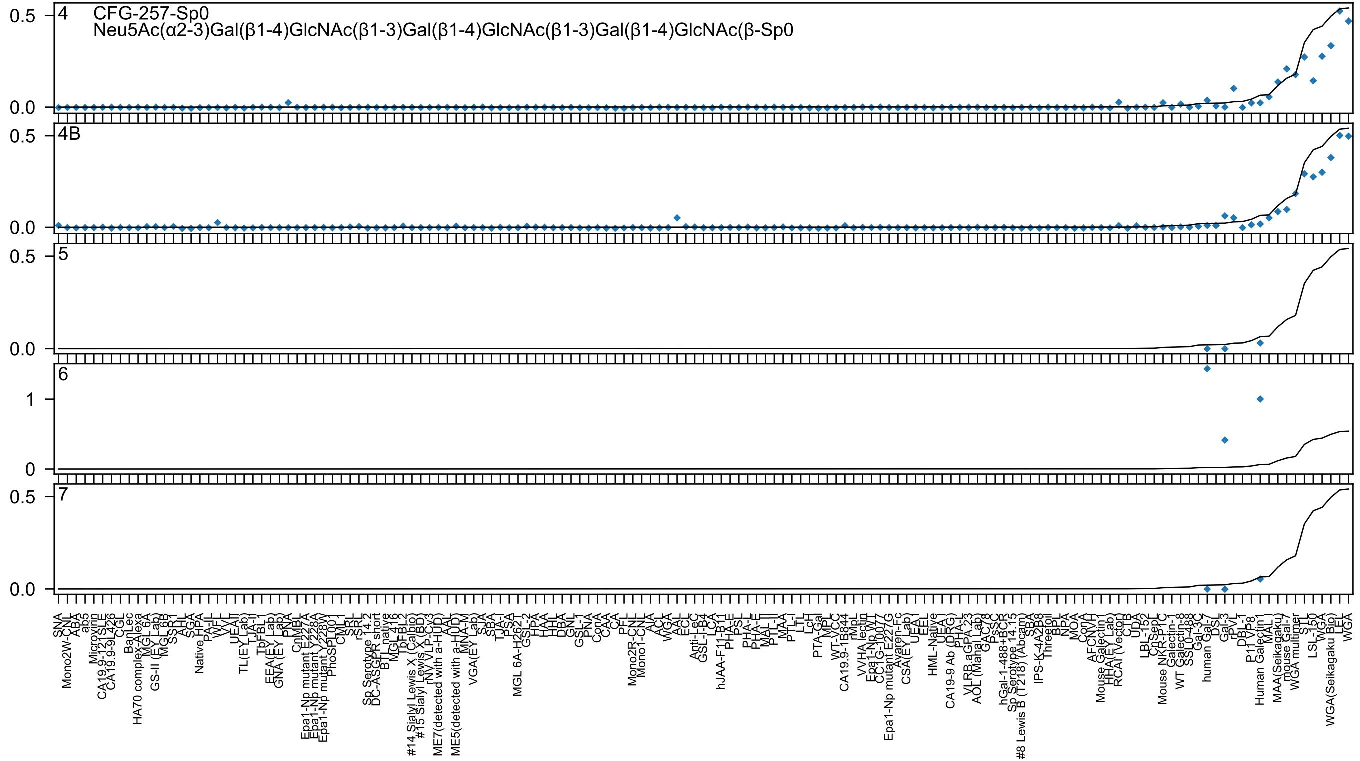

4
CFG-257-Sp0
Neu5Ac(α2-3)Gal(β1-4)GlcNAc(β1-3)Gal(β1-4)GlcNAc(β1-3)Gal(β1-4)GlcNAc(β-Sp0
4B
5
6
7

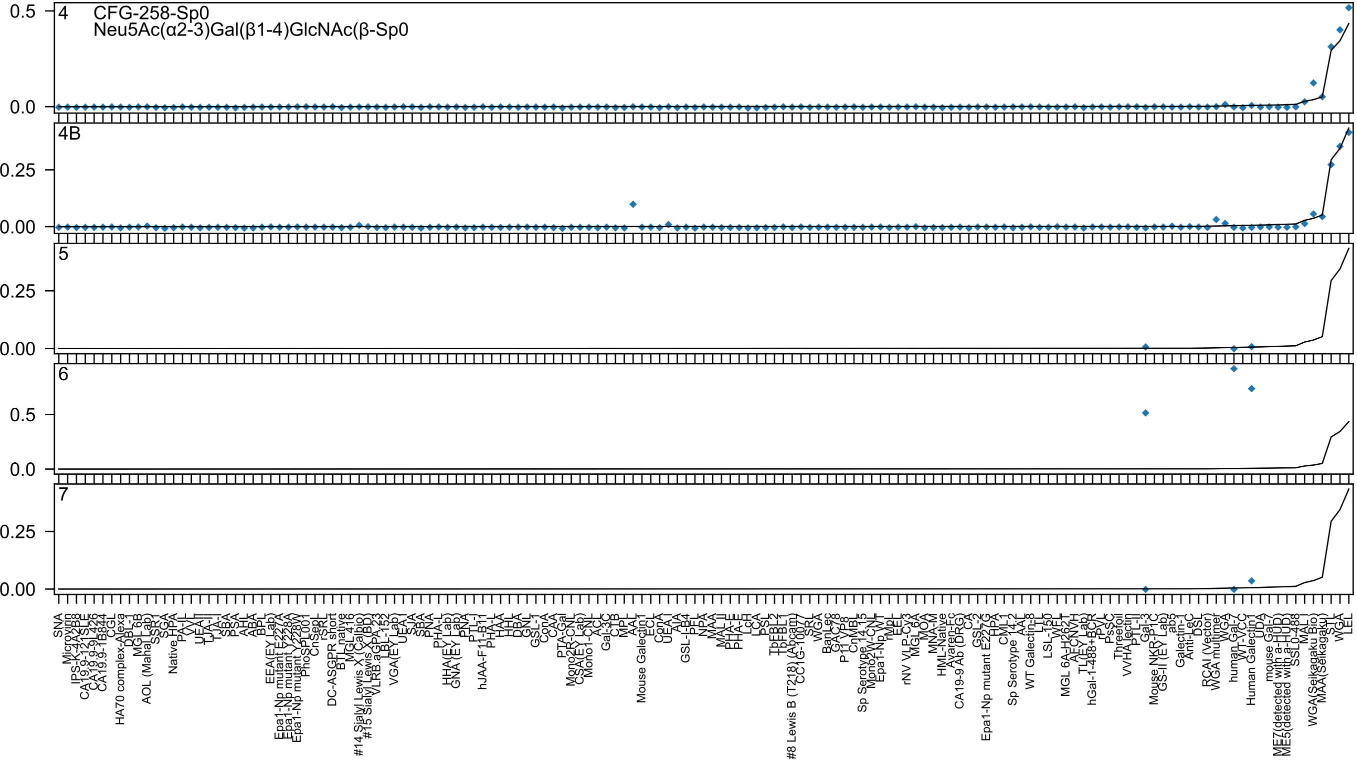
4
CFG-258-Sp0
Neu5Ac(α2-3)Gal(β1-4)GlcNAc(β-Sp0
0.5
0.0
4B
0.25
0.00
5
0.25
0.00
6
0.5
0.0
7
0.25
0.00

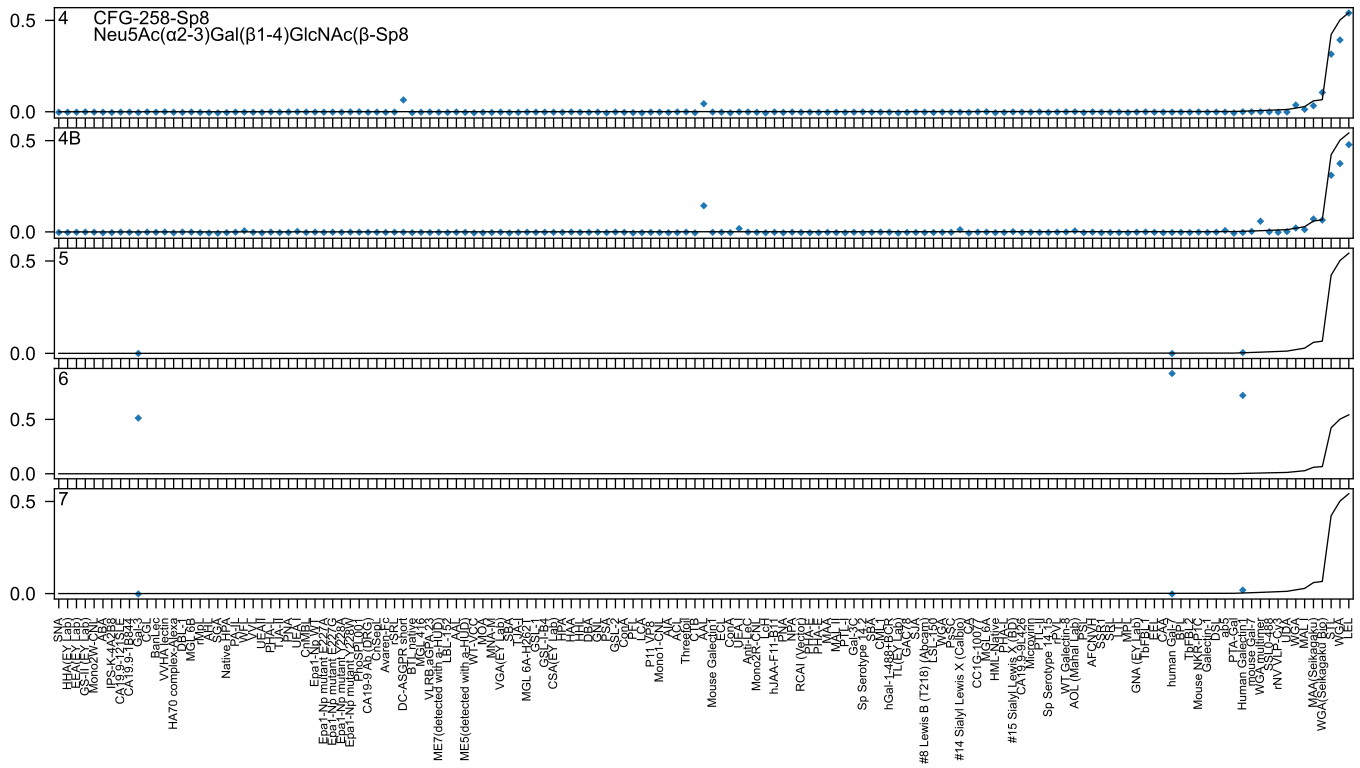
4
CFG-258-Sp8
Neu5Ac(α2-3)Gal(β1-4)GlcNAc(β-Sp8
4B
5
6
7
0.5
0.0

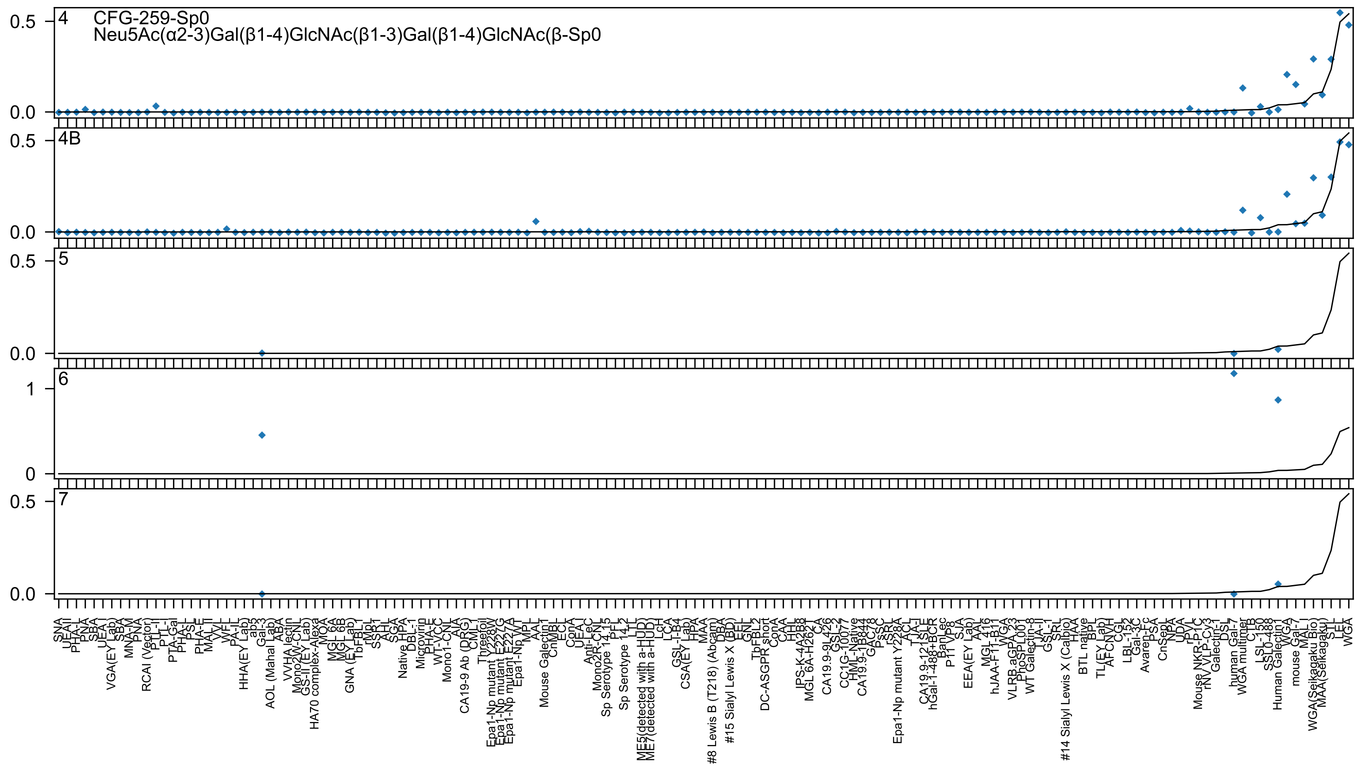

4 CFG-259-Sp0
Neu5Ac(α2-3)Gal(β1-4)GlcNAc(β1-3)Gal(β1-4)GlcNAc(β-Sp0
4B
5
6
7

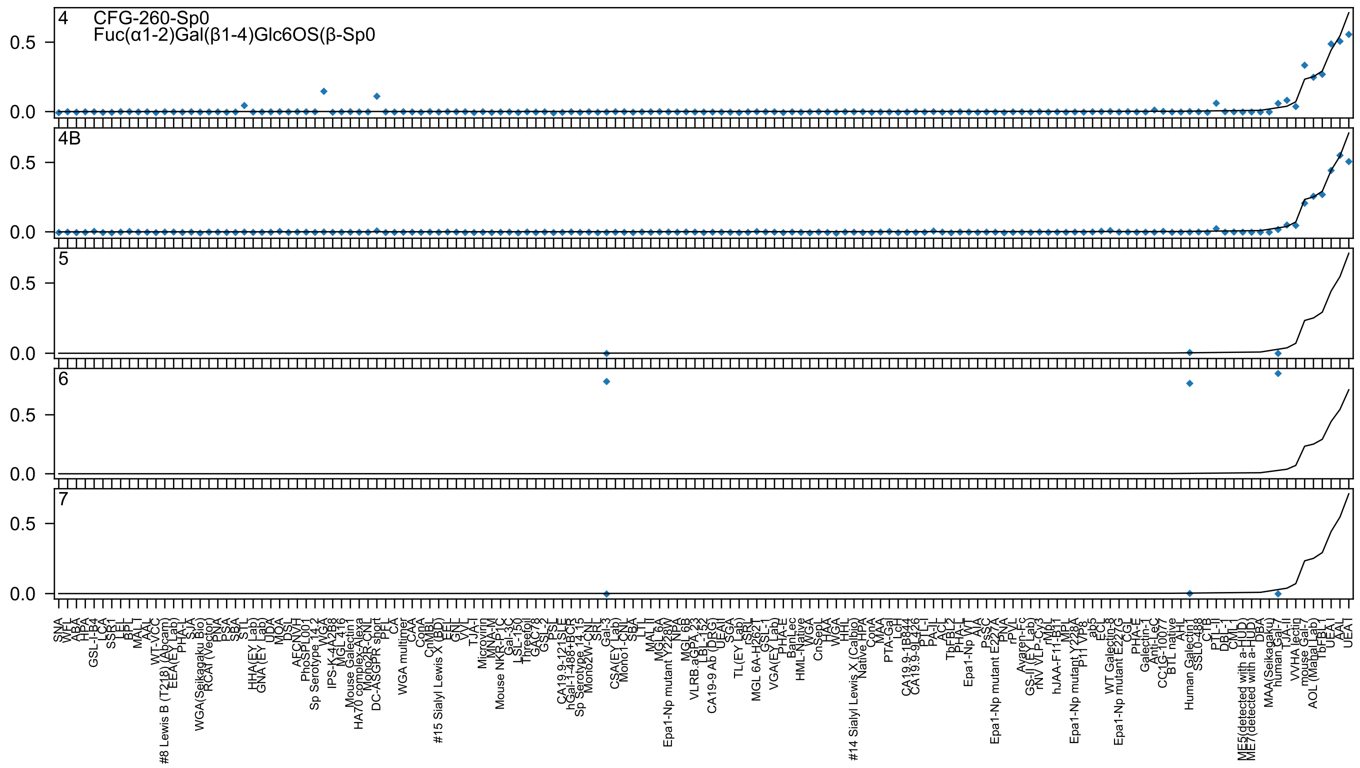
4 CFG-260-Sp0
Fuc(α1-2)Gal(β1-4)Glc6OS(β-Sp0
4B
5
6
7

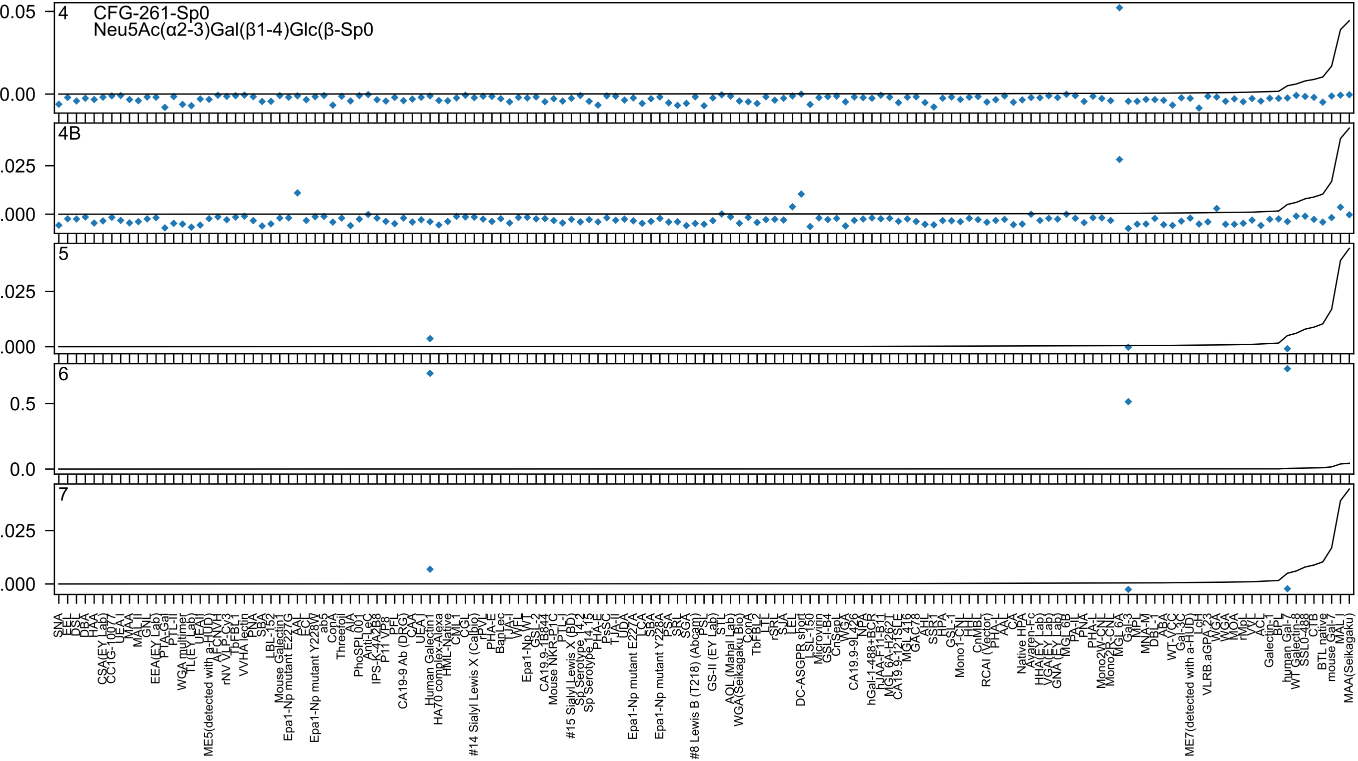
4
CFG-261-Sp0
Neu5Ac(α2-3)Gal(β1-4)Glc(β-Sp0
0.05
0.00
4B
.025
.000
5
.025
.000
6
0.5
0.0
7
.025
.000
SNA
EEL
DSL
DBA
HAA
CSA(EY Lab)
CC1G-10077
UEA-I
MAA
MAL-II
GNL
EEA(EY Lab)
PTA-Gal
PTL-II
WGA multimer
TL(EY Lab)
UEA-I
ME5(detected with a-HUD)
AFCNVH
rNV VLP-Cy3
TbFBL1
VVHA lectin
PNA
SBA
LBL-152
Mouse Galectin1
Epa1-Np mutant E227G
AAL
ECL
Epa1-Np mutant Y228W
ab5
ConA
Threefoil
AIA
PhoSPL001
Anti-LeC
IPS-K-4A2B8
P11 VP8
PFL
CA19-9 Ab (DRG)
CAA
UEA-I
Human Galectin1
HA70 complex-Alexa
HML-Native
CML1
CGL
#14 Sialyl Lewis X (Calbio)
rPVL
PHA-E
BanLec
LTA-I
WFL
Epa1-Np WT
GSL-2
CA19.9-1B844
Mouse NKR-P1C
PTL-I
#15 Sialyl Lewis X (BD)
Sp Serotype 14.2
Sp Serotype 14.15
PHA-E
PsSC
TJA-II
UDA
Epa1-Np mutant E227A
LCA
SBA
Epa1-Np mutant Y228A
PSA
SRL
SGA
#8 Lewis B (T218) (Abcam)
PSL
GS-II (EY Lab)
STL
AOL (Mahal Lab)
WGA(Seikagaku Bio)
ConA
TbFBL2
LTL
rSRL
SJA
LEL
DC-ASGPR short
LSL-150
Microvirin
GSL-I-B4
CnSept
WGA
CA19.9-9L426
NPA
hGal-1-488+BCR
hJAA-F11-B11
MGL 6A-H262T
CA19.9-121SLE
MGL 416
GAC78
AHL
SSA1
HPA
GSL-I
Mono1-CNL
HHL
CnMBL
RCAI (Vector)
PHA-L
AAL
CA
Native HPA
Avaren-Fc
HHA(EY Lab)
VGA(EY Lab)
GNA (EY Lab)
MGL 6B
PA-IL
PNA
PHA-L
Mono2W-CNL
Mono2R-CNL
MGL 6A
Gal-3
MPL
MNA-M
DBL-1
ABA
WT-VCC
Gal-3C
ME7(detected with a-HUD)
LcH
VLRB.aGPA.23
WGA
MOA
rMpL
VVL
ACL
Galectin-1
BPL
human Gal-7
WT Galectin-8
SSL0-488
CTB
BTL native
mouse Gal-7
MAL I
MAA(Seikagaku)

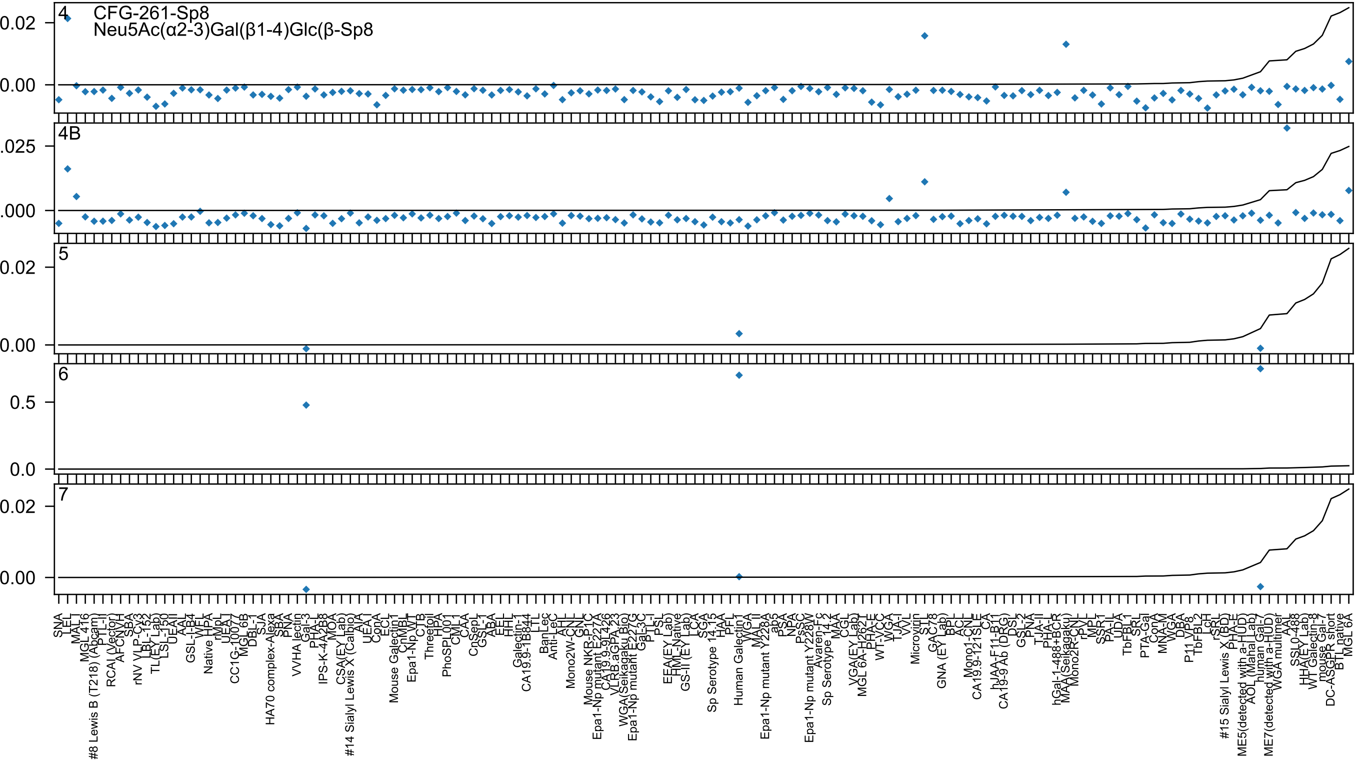
4
CFG-261-Sp8
Neu5Ac(α2-3)Gal(β1-4)Glc(β-Sp8
4B
5
6
7

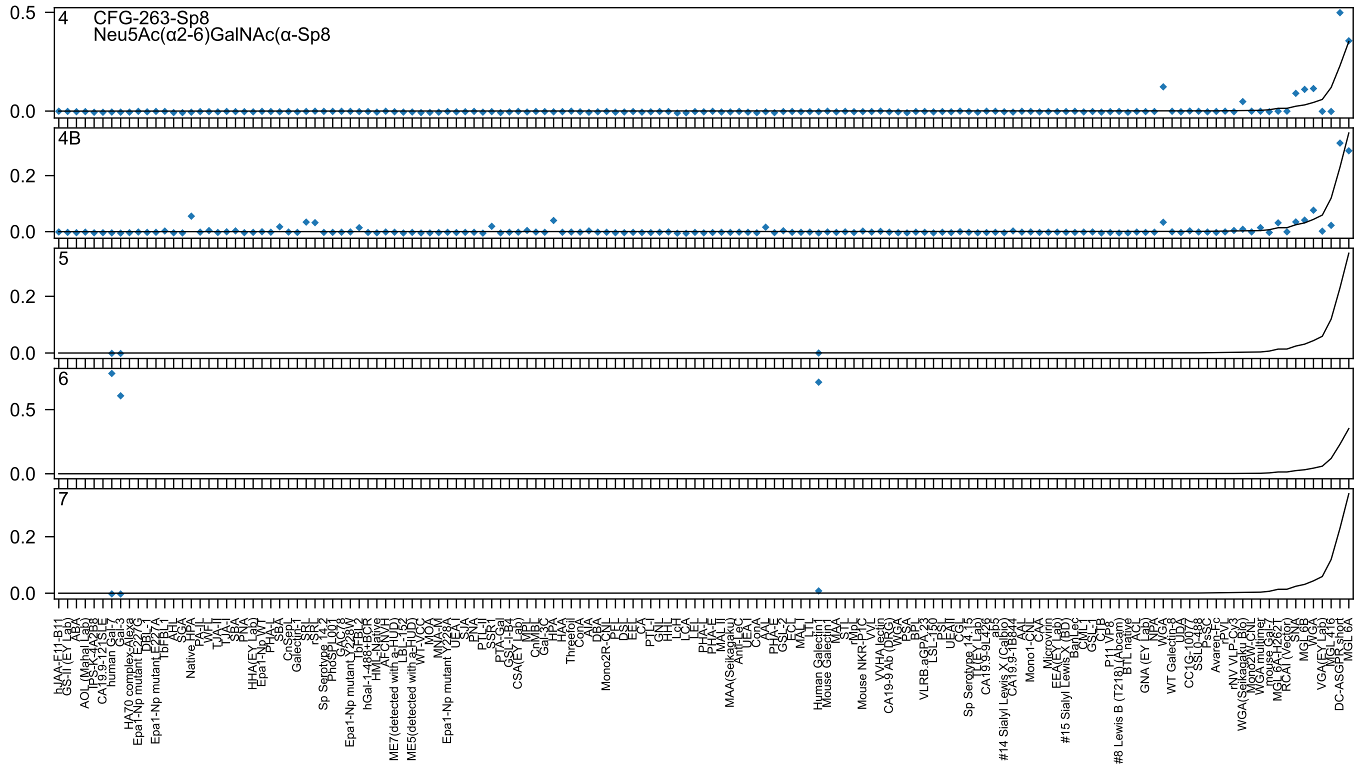

4
CFG-263-Sp8
Neu5Ac(α2-6)GalNAc(α-Sp8
4B
5
6
7

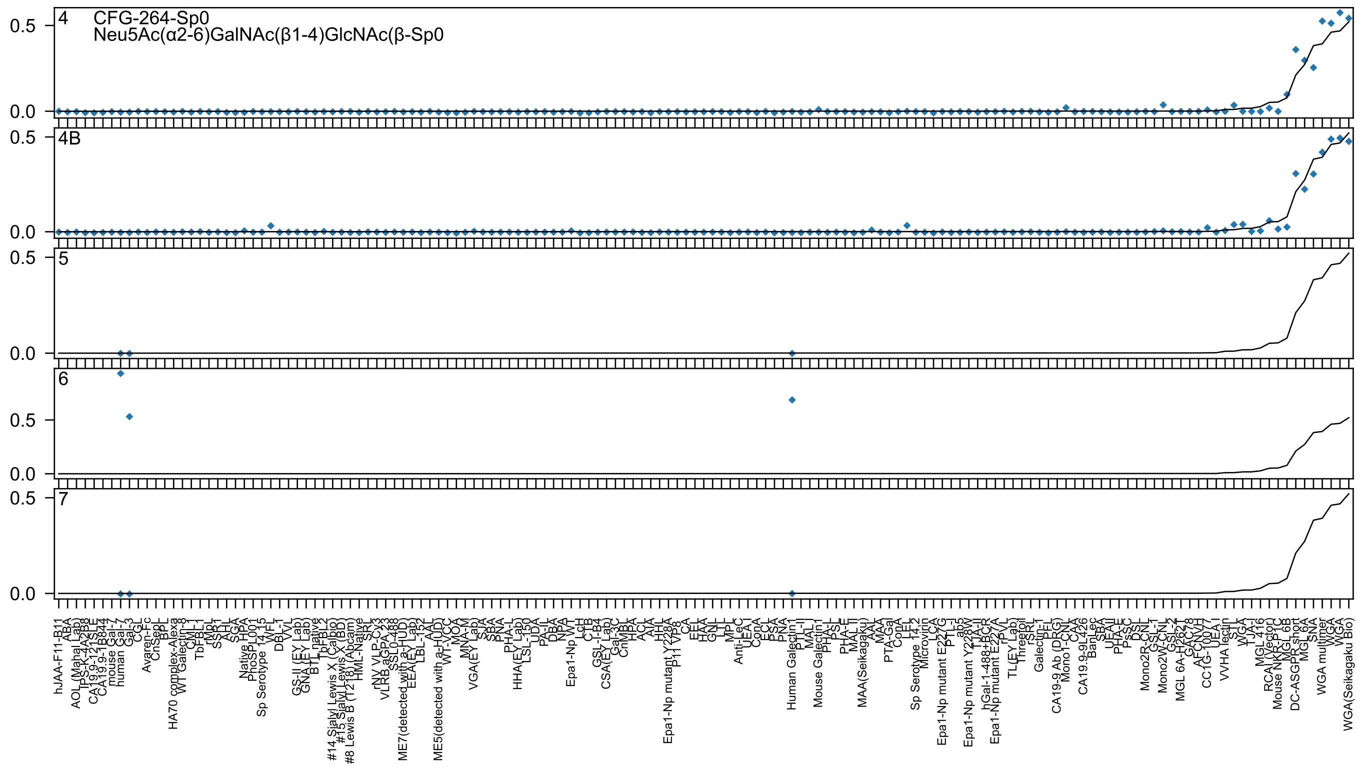

4
CFG-264-Sp0
Neu5Ac(α2-6)GalNAc(β1-4)GlcNAc(β-Sp0
4B
5
6
7
0.5
0.0

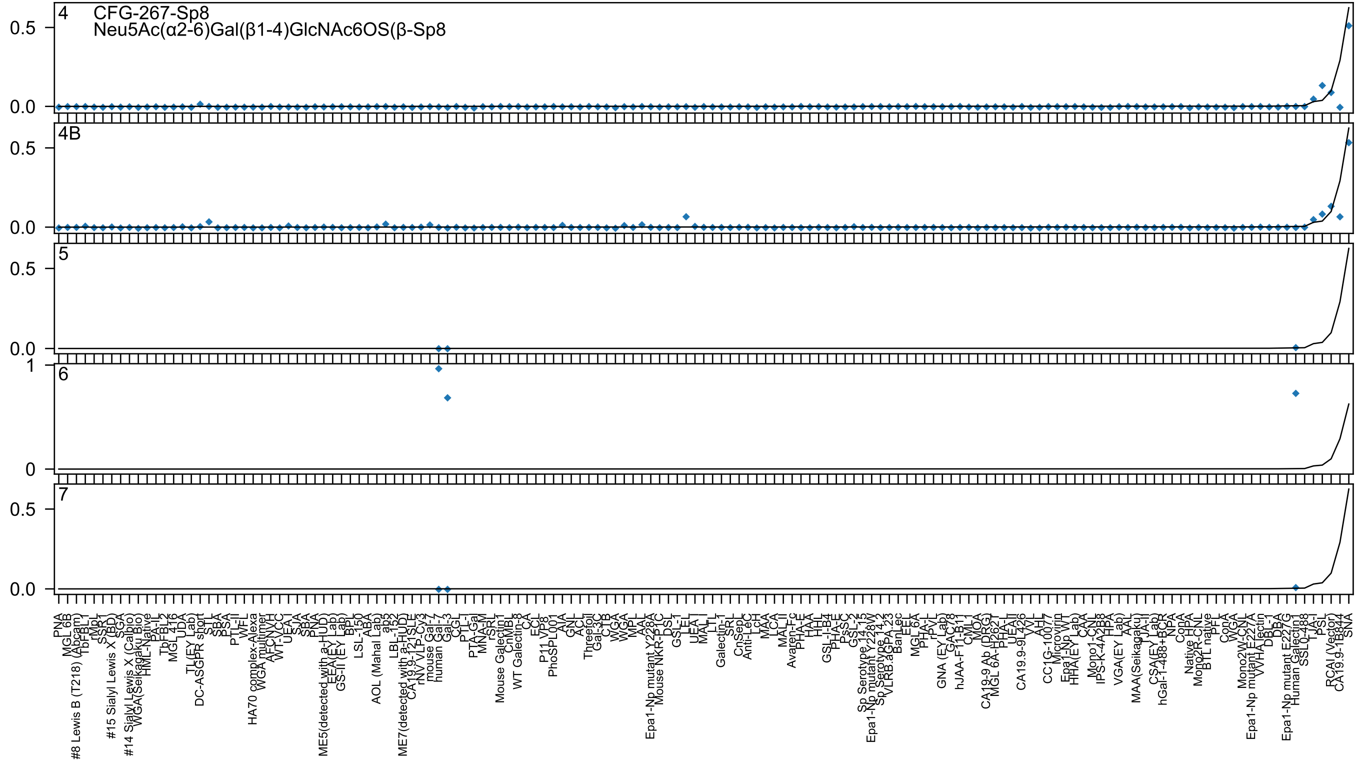
4
CFG-267-Sp8
Neu5Ac(α2-6)Gal(β1-4)GlcNAc6OS(β-Sp8
4B
5
6
7
0.5
0.0
1
0

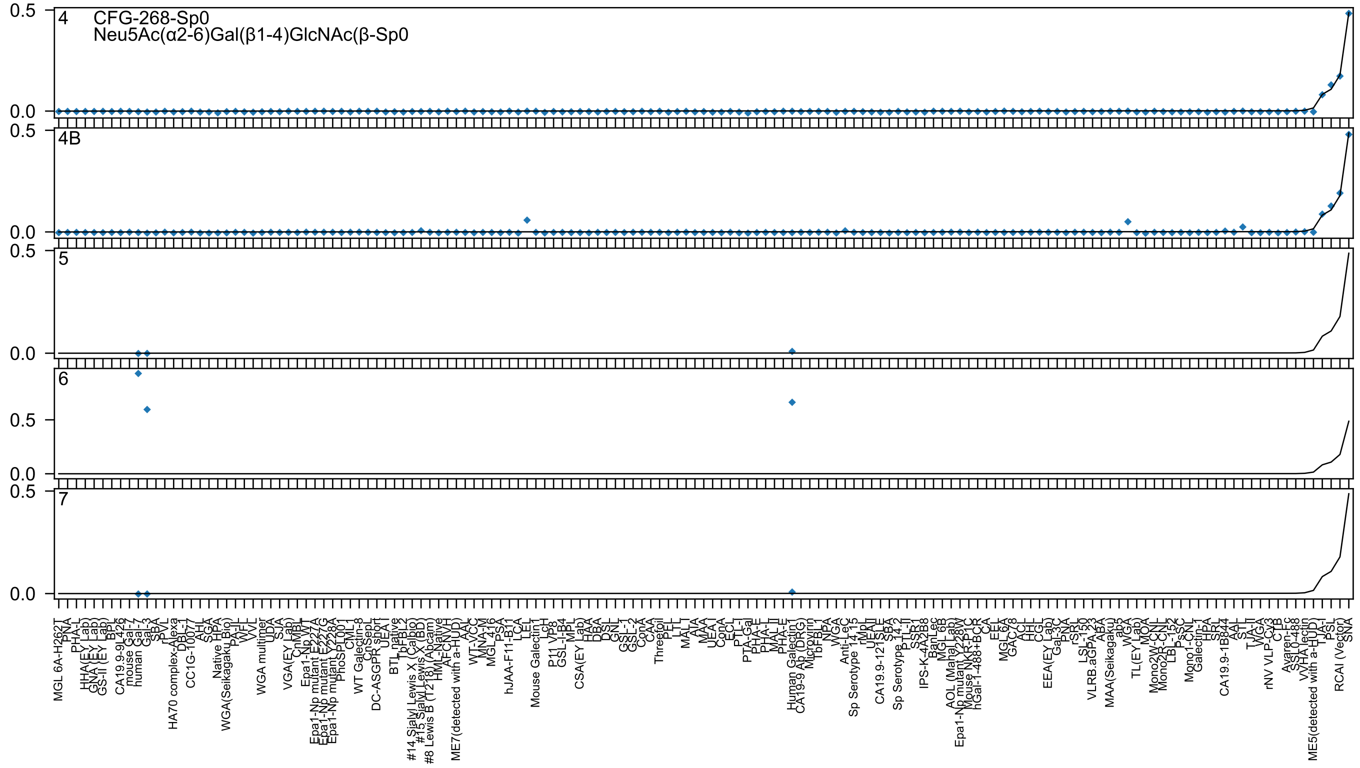

CFG-268-Sp0
Neu5Ac(α2-6)Gal(β1-4)GlcNAc(β-Sp0
4
4B
5
6
7
0.5
0.0

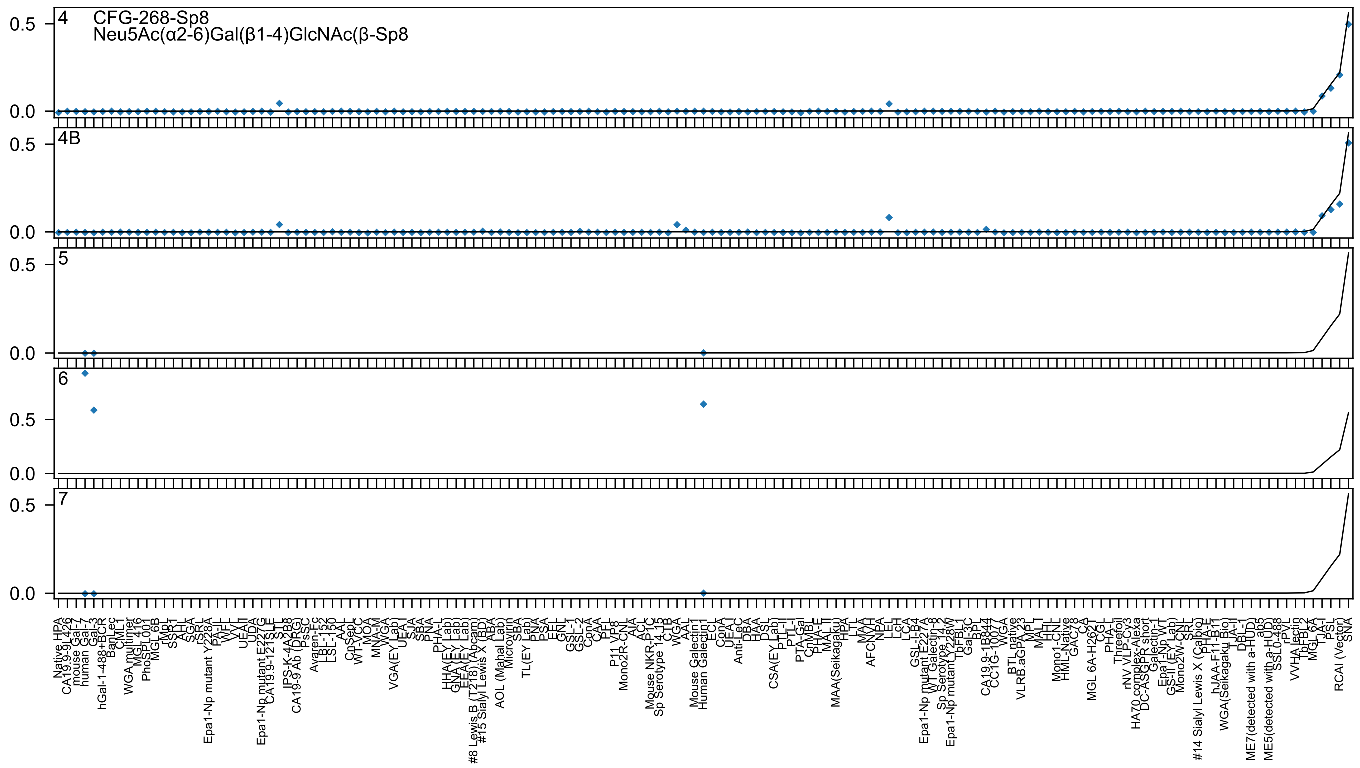
4
CFG-268-Sp8
Neu5Ac(α2-6)Gal(β1-4)GlcNAc(β-Sp8
4B
5
6
7
0.5
0.0

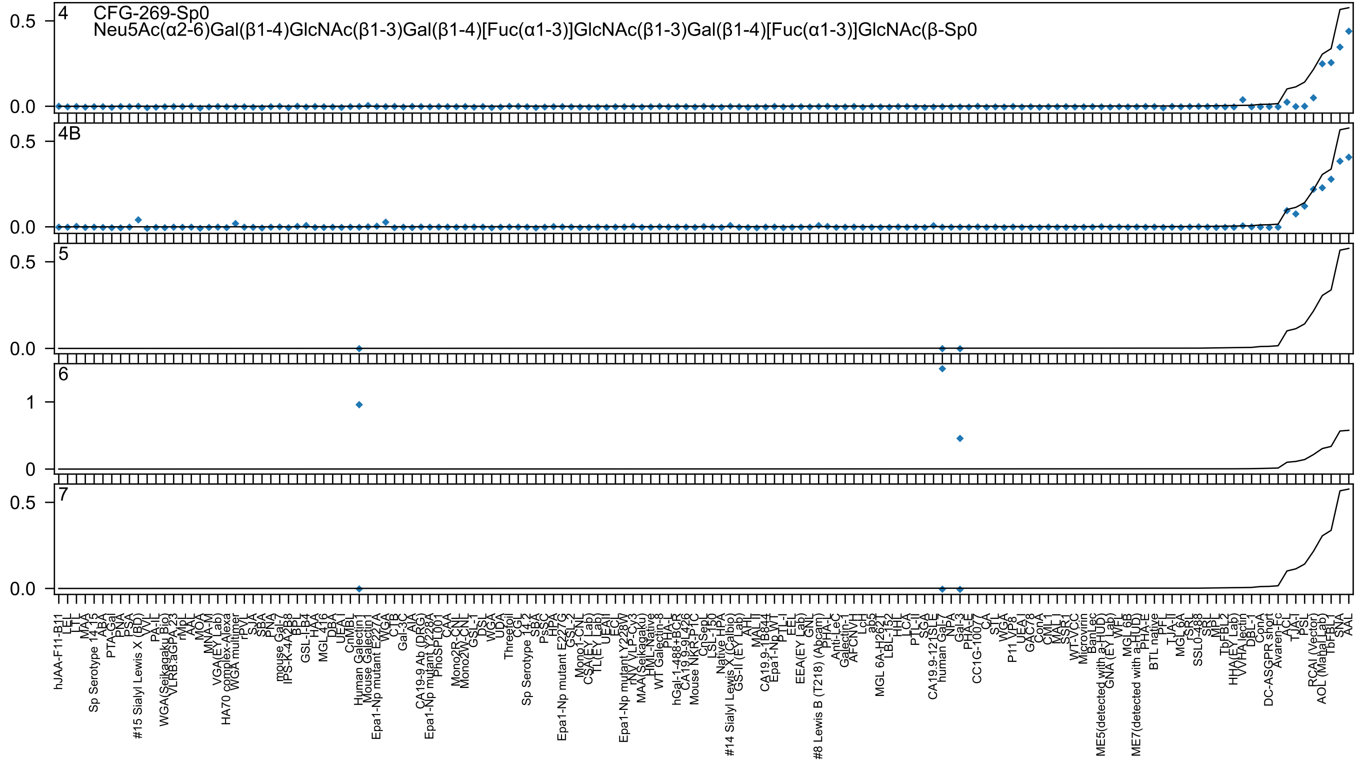
CFG-269-Sp0
Neu5Ac(α2-6)Gal(β1-4)GlcNAc(β1-3)Gal(β1-4)[Fuc(α1-3)]GlcNAc(β1-3)Gal(β1-4)[Fuc(α1-3)]GlcNAc(β-Sp0

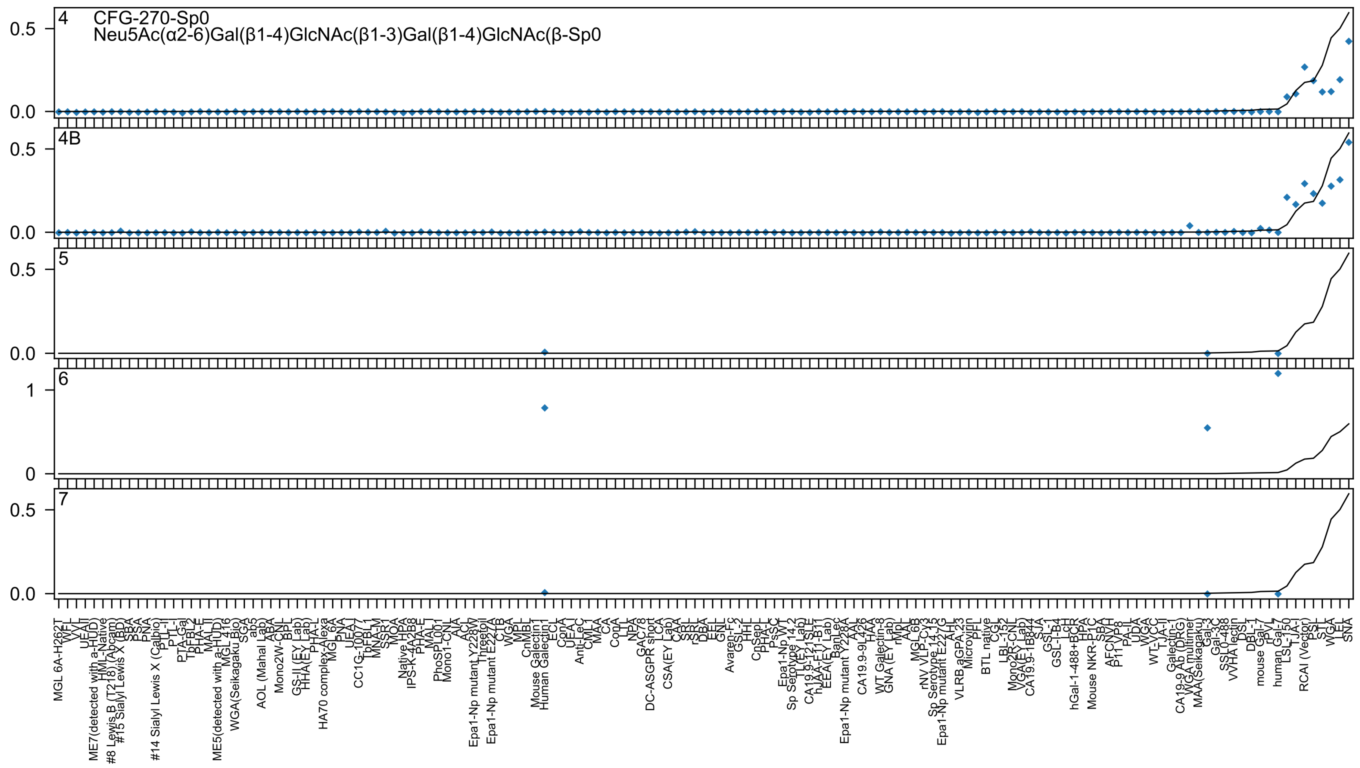
CFG-270-Sp0
Neu5Ac(α2-6)Gal(β1-4)GlcNAc(β1-3)Gal(β1-4)GlcNAc(β-Sp0
4
4B
5
6
7

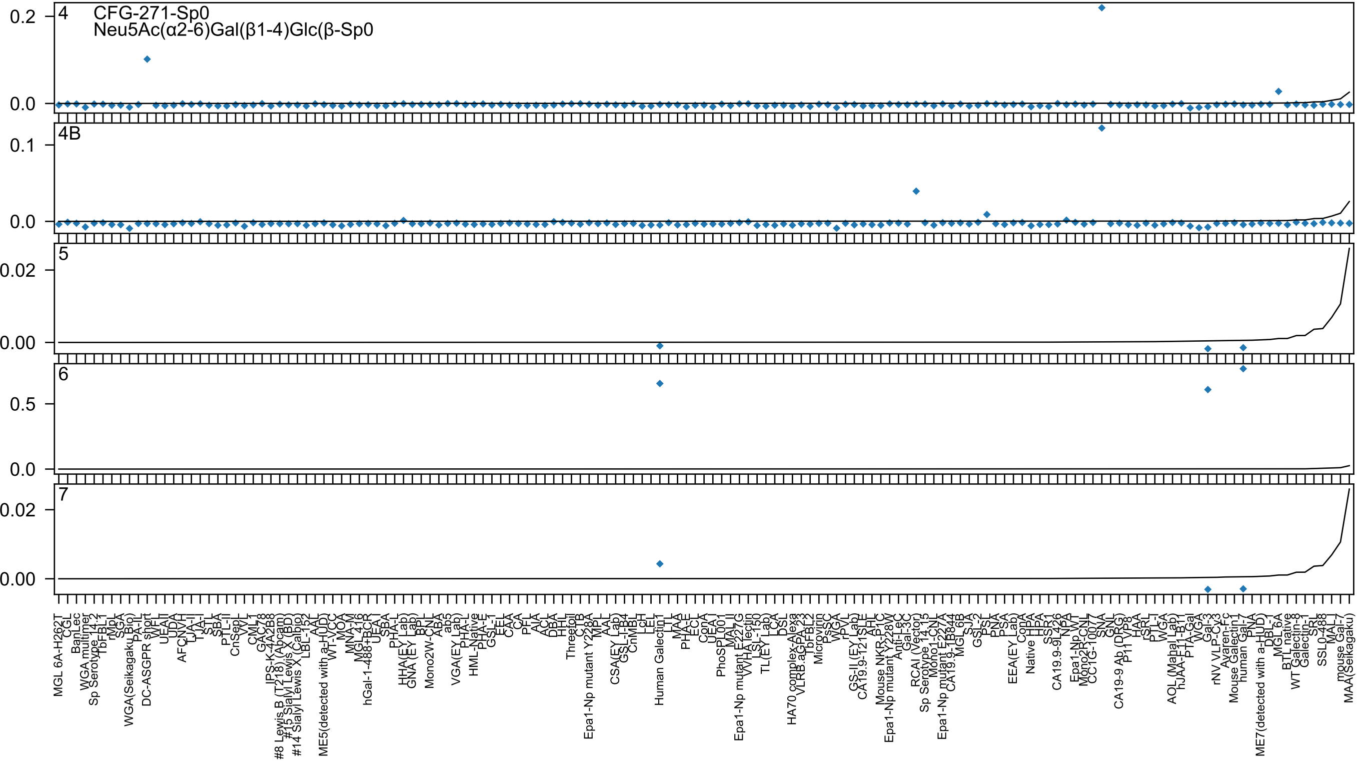

CFG-271-Sp0
Neu5Ac(α2-6)Gal(β1-4)Glc(β-Sp0
4
4B
5
6
7

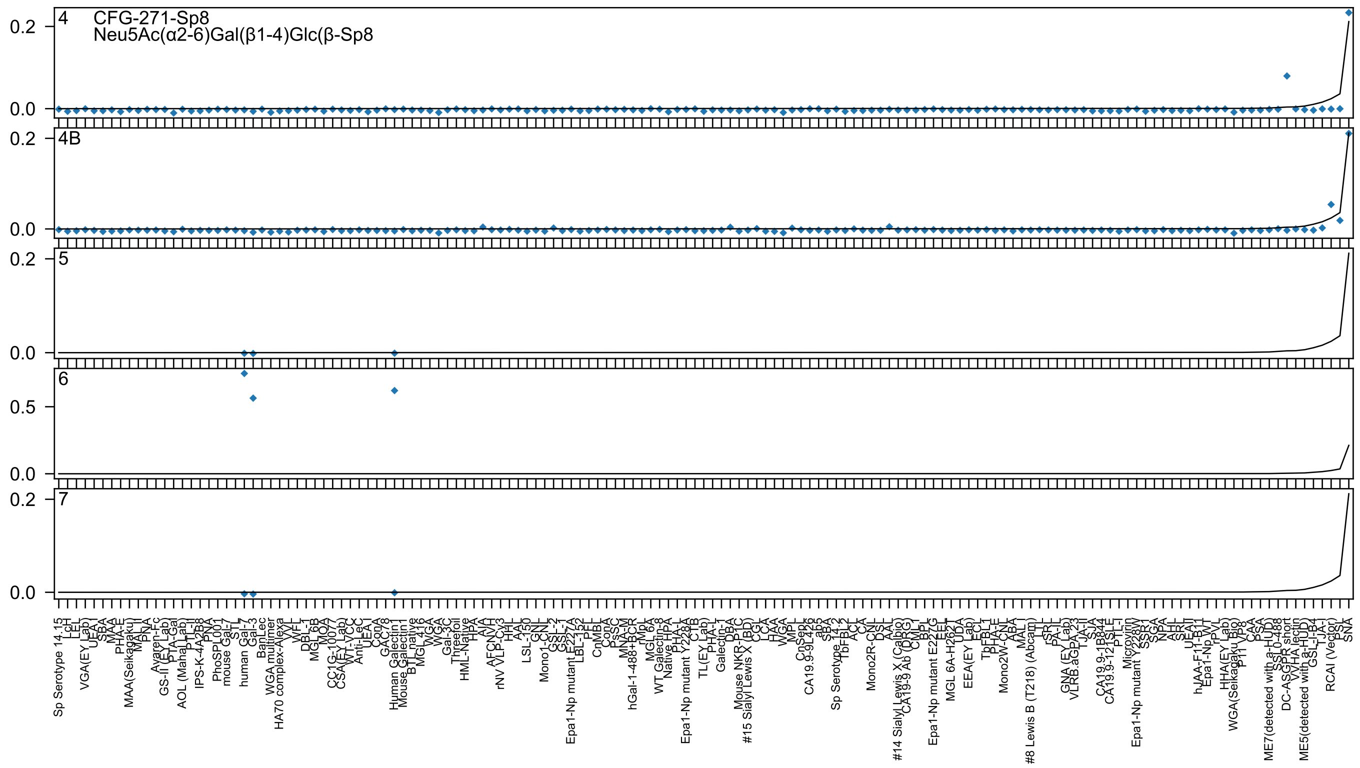

CFG-271-Sp8
Neu5Ac(α2-6)Gal(β1-4)Glc(β-Sp8
4
4B
5
6
7

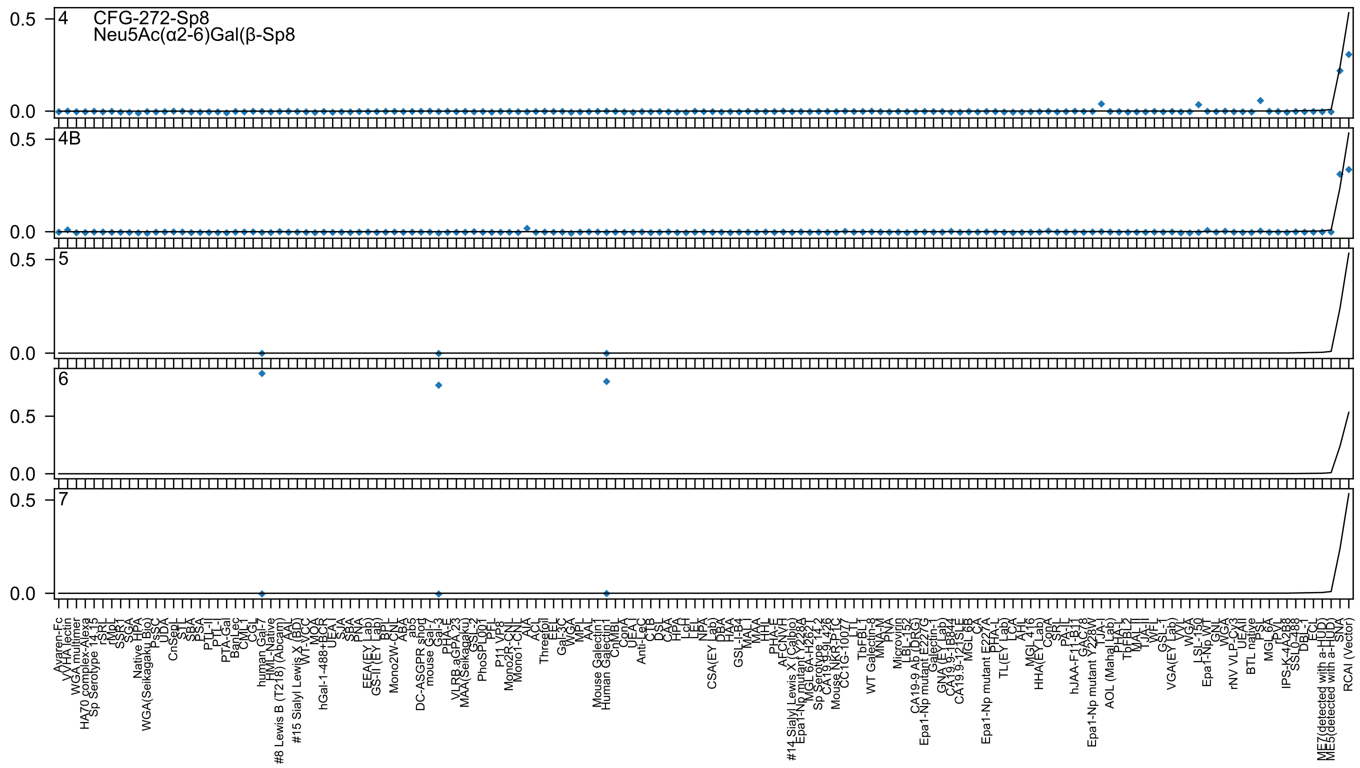

4 CFG-272-Sp8
Neu5Ac(α2-6)Gal(β-Sp8
4B
5
6
7
0.5
0.0

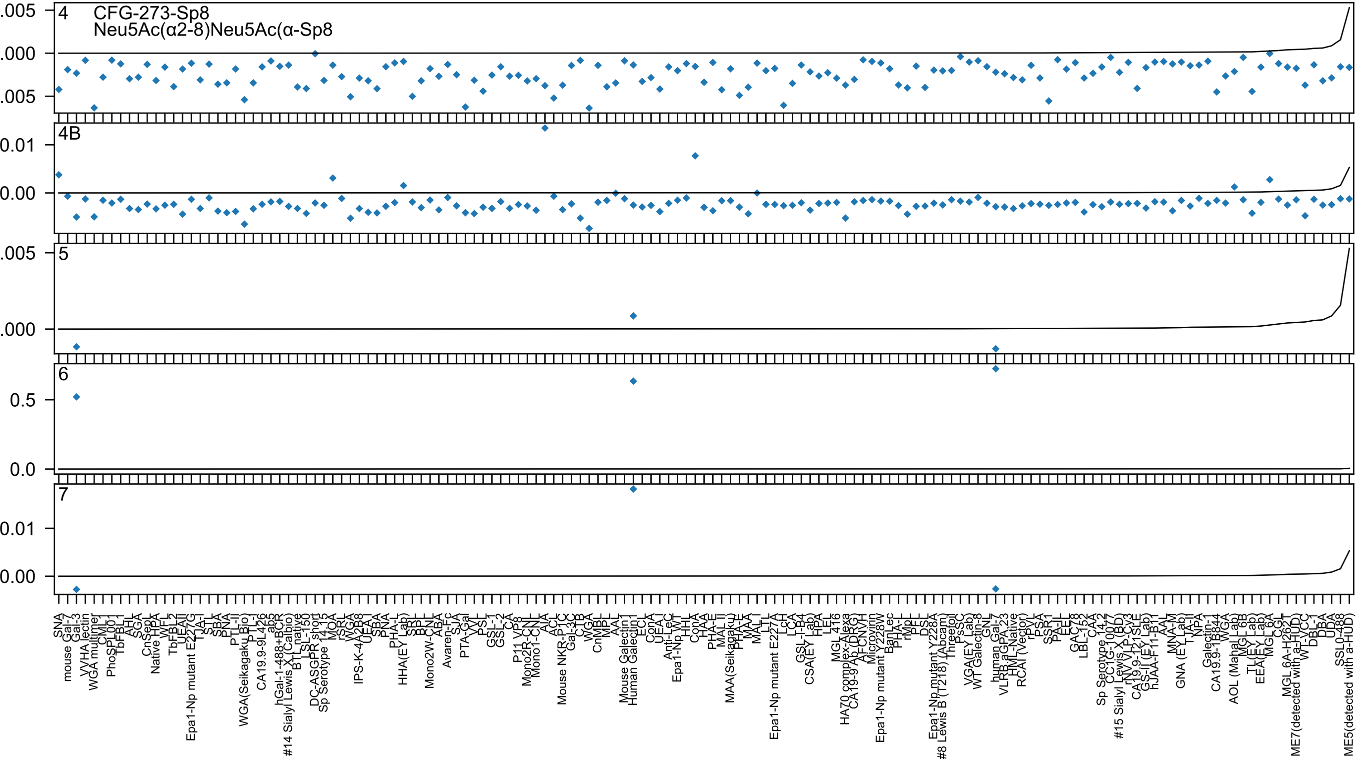

4
CFG-273-Sp8
Neu5Ac(α2-8)Neu5Ac(α-Sp8
.005
.000
.005
4B
0.01
0.00
5
.005
.000
6
0.5
0.0
7
0.01
0.00
SNA
mouse Gal-7
Gal-3
VVHA lectin
WGA multimer
CML1
PhoSL001
TbFBL1
AHL
SGA
CnSepL
Native HPA
WFL
TbFBL2
UEAII
Epa1-Np mutant E227G
TJA-I
STL
SBA
PNA
PTL-II
WGA(Seikagaku Bio)
PTL-I
CA19.9-9L426
ab5
hGal-1-488+BCR
#14 Sialyl Lewis X (Calbio)
BTL native
LSL-150
DC-ASGPR short
Sp Serotype 14.15
MOA
rSRL
WGA
IPS-K-4A2B8
UEA I
SBA
PNA
PHA-L
HHA(EY Lab)
SRL
BPL
Mono2W-CNL
ABA
Avaren-Fc
SJA
PTA-Gal
VVL
PSL
GSL-1
GSL-2
CA
P11 VP8
Mono2R-CNL
Mono1-CNL
AIA
ACL
Mouse NKR-P1C
Gal-3C
CTB
WGA
CnMBL
AAL
Mouse Galectin1
Human Galectin1
ECL
ConA
UEA-I
Anti-LeC
Epa1-Np WT
HHL
ConA
HAA
PHA-E
MAL II
MAA(Seikagaku)
PHA-E
MAA
MAL I
LTL
Epa1-Np mutant E227A
LcH
LCA
GSL-I-B4
CSA(EY Lab)
HPA
LEL
MGL 416
HA70 complex-Alexa
CA19-9 Ab (DRG)
AFCNVH
Microvirin
Epa1-Np mutant Y228W
BanLec
PHA-L
rMpL
PEL
DSL
Epa1-Np mutant Y228A
#8 Lewis B (T218) (Abcam)
Threefoil
PsSC
VGA(EY Lab)
WT Galectin-8
GNL
human Gal-7
VLRB aGPA.23
HML-Native
RCAI (Vector)
rPVL
PSA
SSR1
PA-IL
EEL
GAC78
LBL-152
CAA
Sp Serotype 14.2
CC1G-10077
#15 Sialyl Lewis X (BD)
rNV VLP-Cy3
CA19.9-121SLE
GS-II (EY Lab)
hJAA-F11-B11
AAL
MNA-M
GNA (EY Lab)
TJA-I
NPA
Galectin-1
CA19.9-1B844
WGA
AOL (Mahal Lab)
MGL 6B
TL(EY Lab)
EEA(EY Lab)
MGL 6A
CGL
MGL 6A-H262T
ME7(detected with a-HUD)
WT-VCC
DBL-1
DBA
UDA
SSL0-488
ME5(detected with a-HUD)

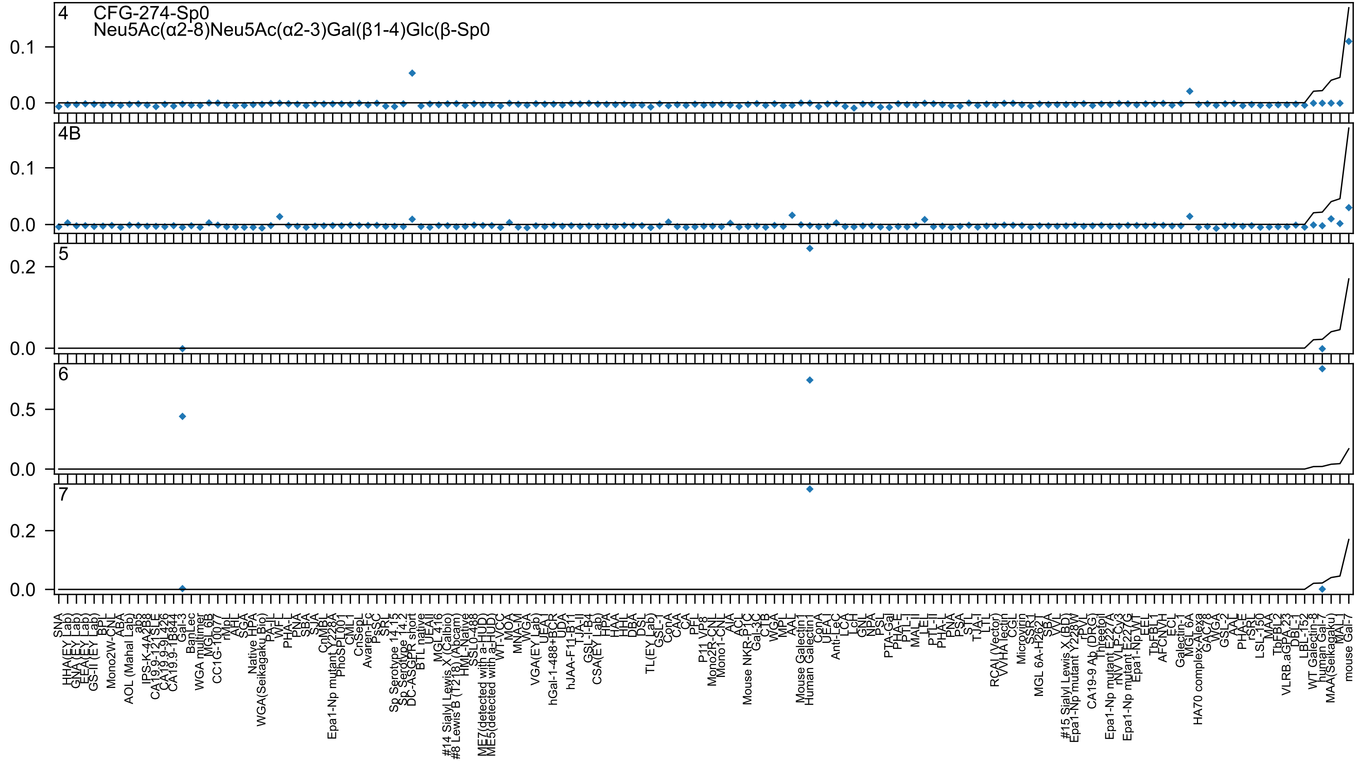

4
CFG-274-Sp0
Neu5Ac(α2-8)Neu5Ac(α2-3)Gal(β1-4)Glc(β-Sp0
0.1
0.0
4B
0.1
0.0
5
0.2
0.0
6
0.5
0.0
7
0.2
0.0

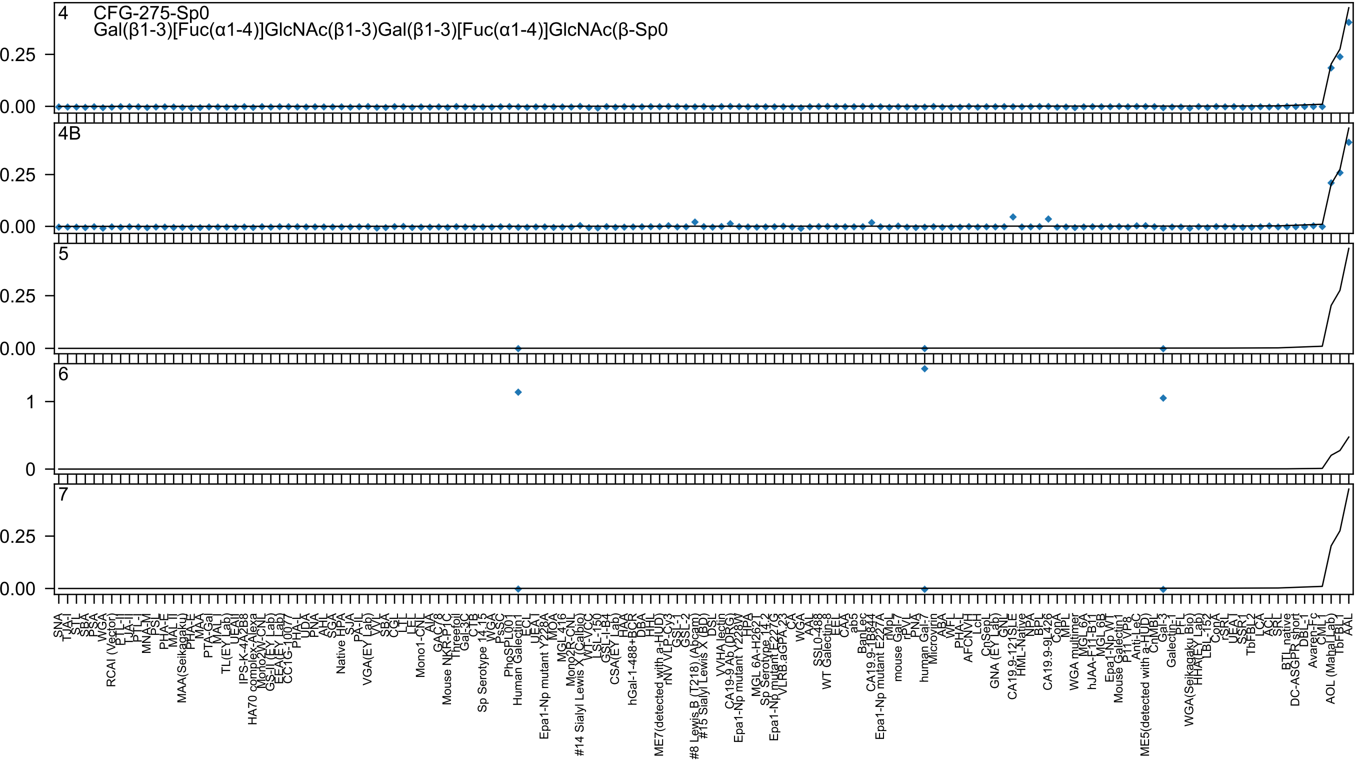

4 CFG-275-Sp0
Gal(β1-3)[Fuc(α1-4)]GlcNAc(β1-3)Gal(β1-3)[Fuc(α1-4)]GlcNAc(β-Sp0
4B
5
6
7
0.25
0.00
1
0

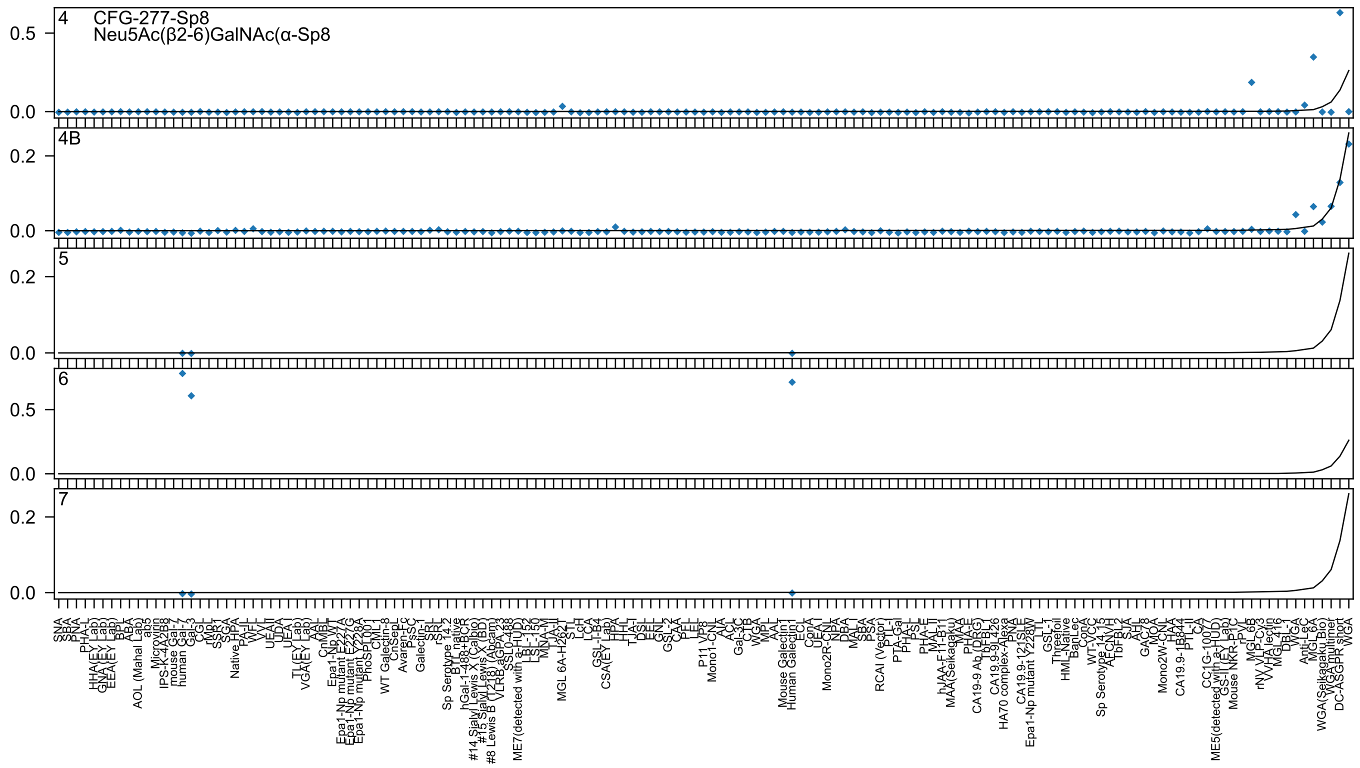
4 CFG-277-Sp8
Neu5Ac(β2-6)GalNAc(α-Sp8
4B
5
6
7

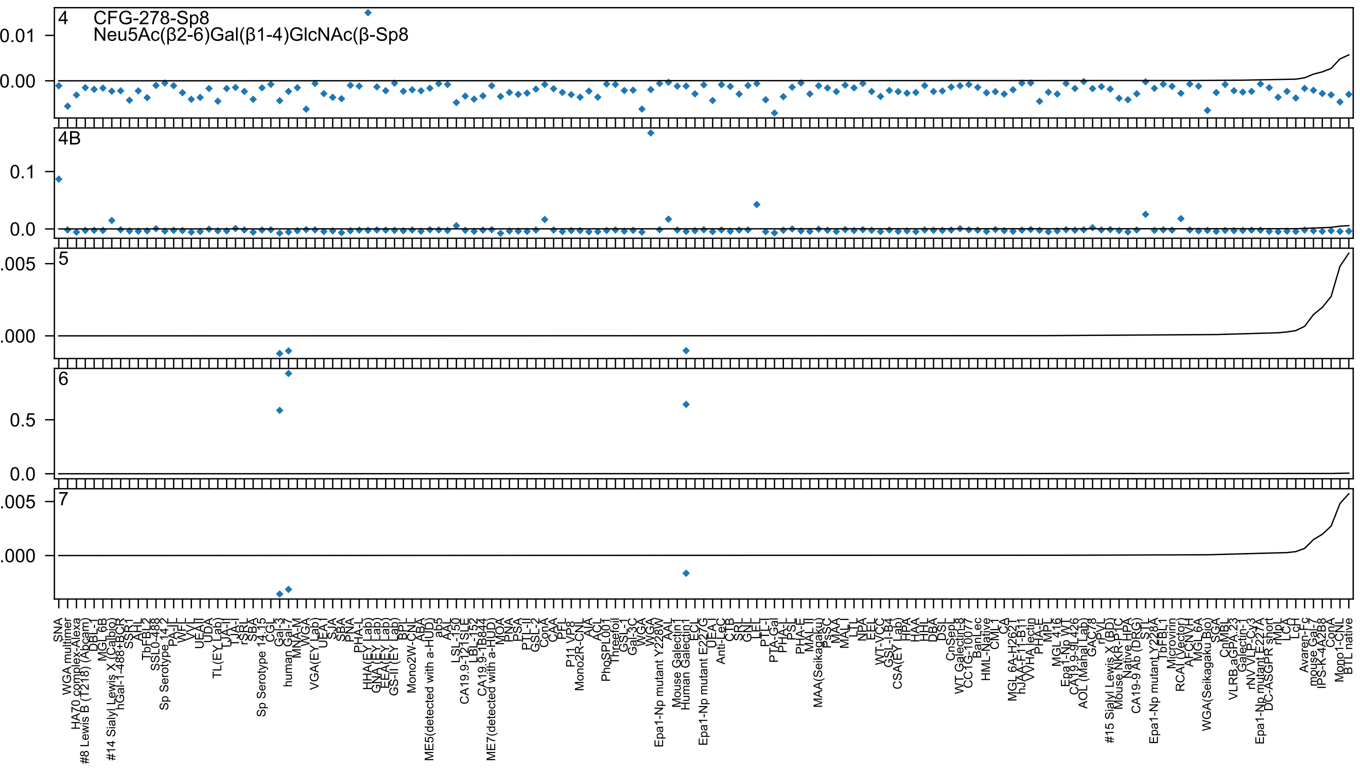
4
CFG-278-Sp8
Neu5Ac(β2-6)Gal(β1-4)GlcNAc(β-Sp8
4B
5
6
7

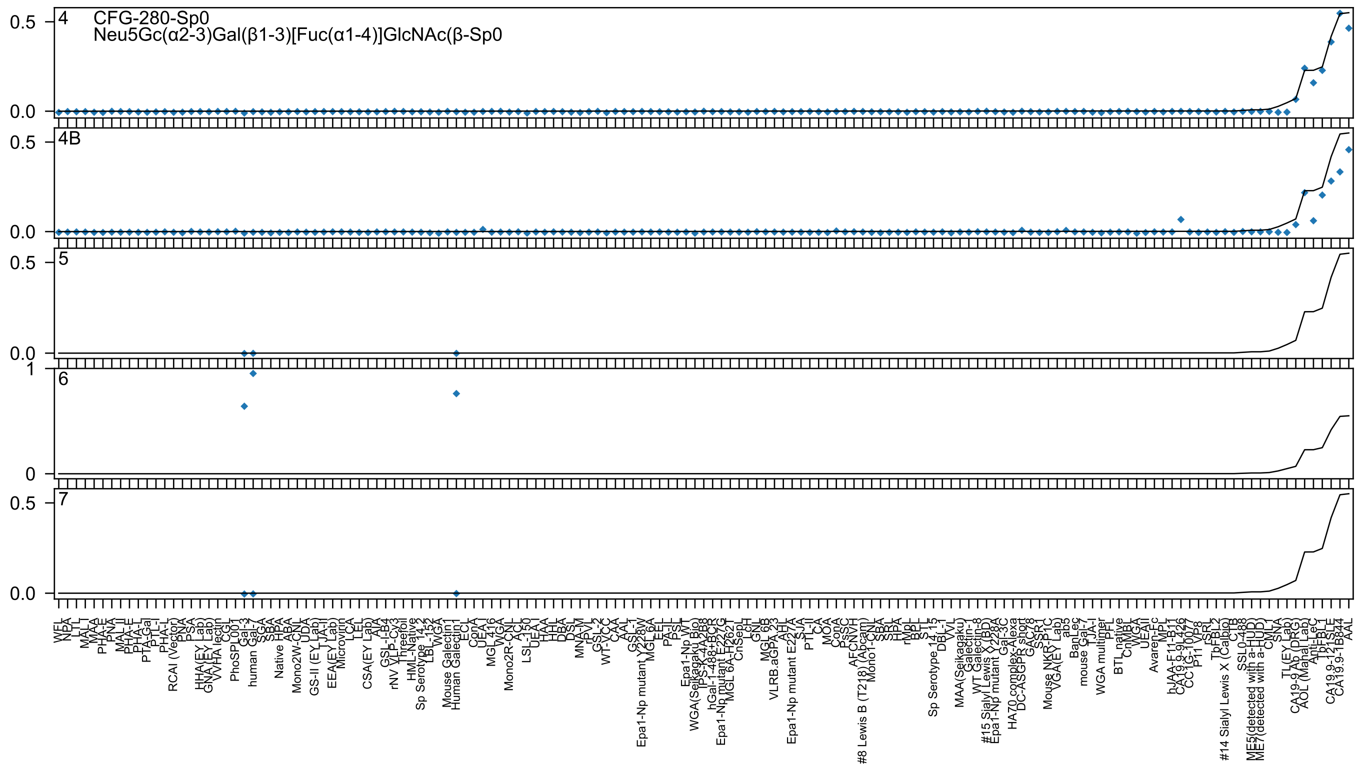
CFG-280-Sp0
Neu5Gc(α2-3)Gal(β1-3)[Fuc(α1-4)]GlcNAc(β-Sp0
4
4B
5
6
7

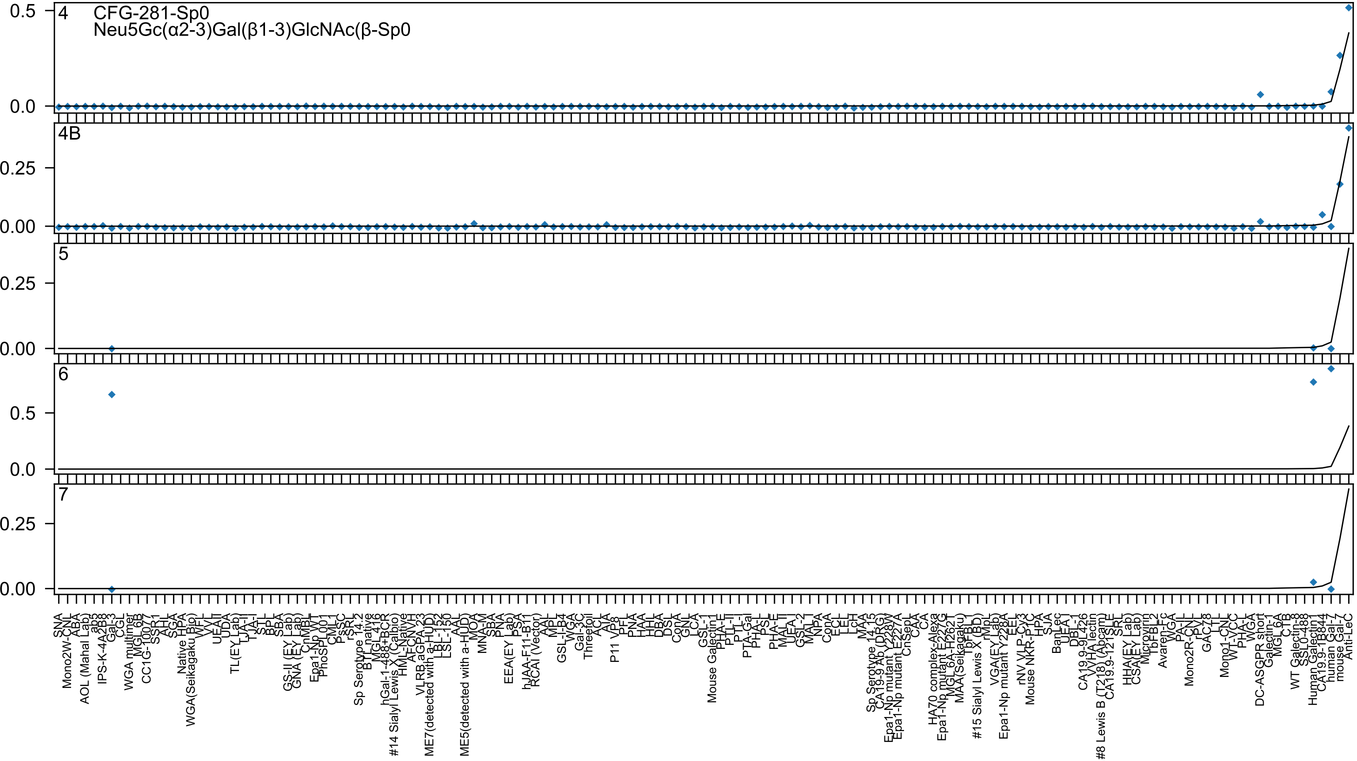
CFG-281-Sp0
Neu5Gc(α2-3)Gal(β1-3)GlcNAc(β-Sp0
4
4B
5
6
7

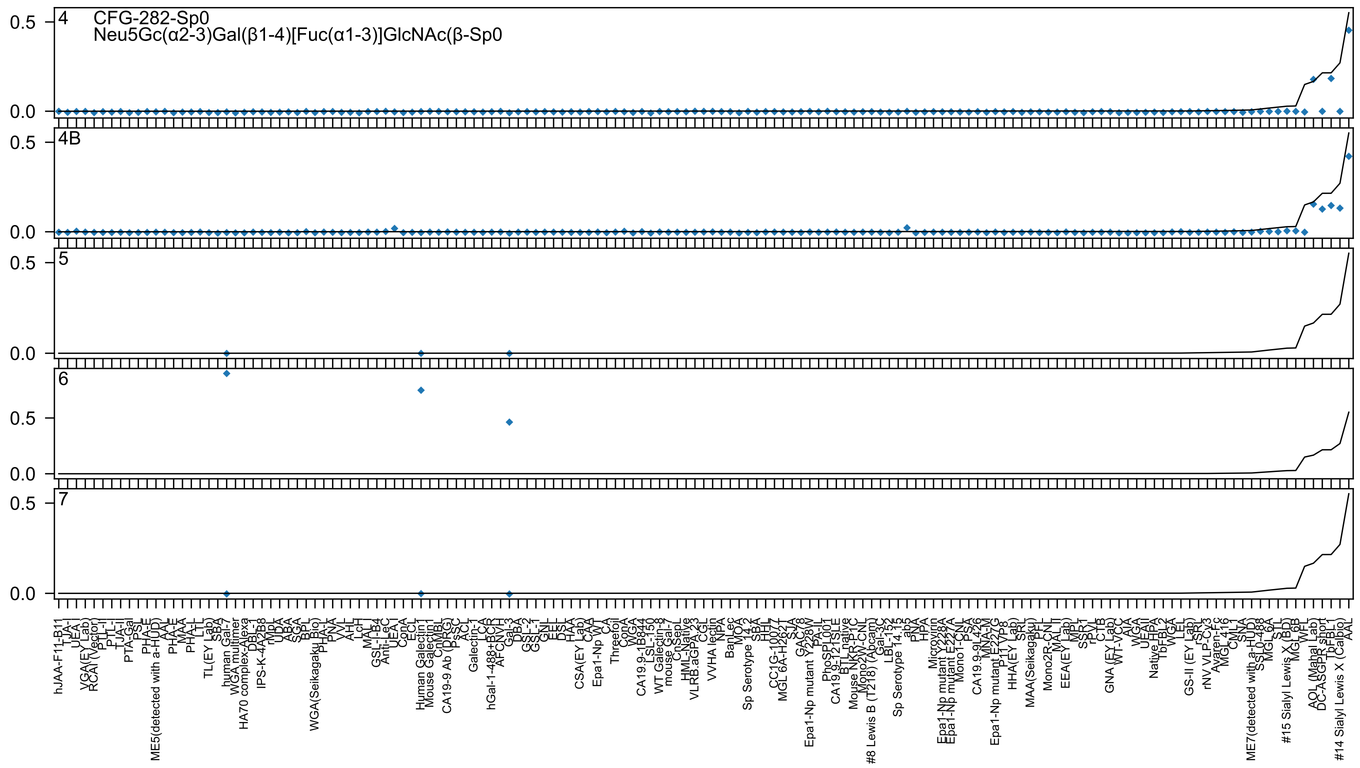

4
CFG-282-Sp0
Neu5Gc(α2-3)Gal(β1-4)[Fuc(α1-3)]GlcNAc(β-Sp0
4B
5
6
7
0.5
0.0

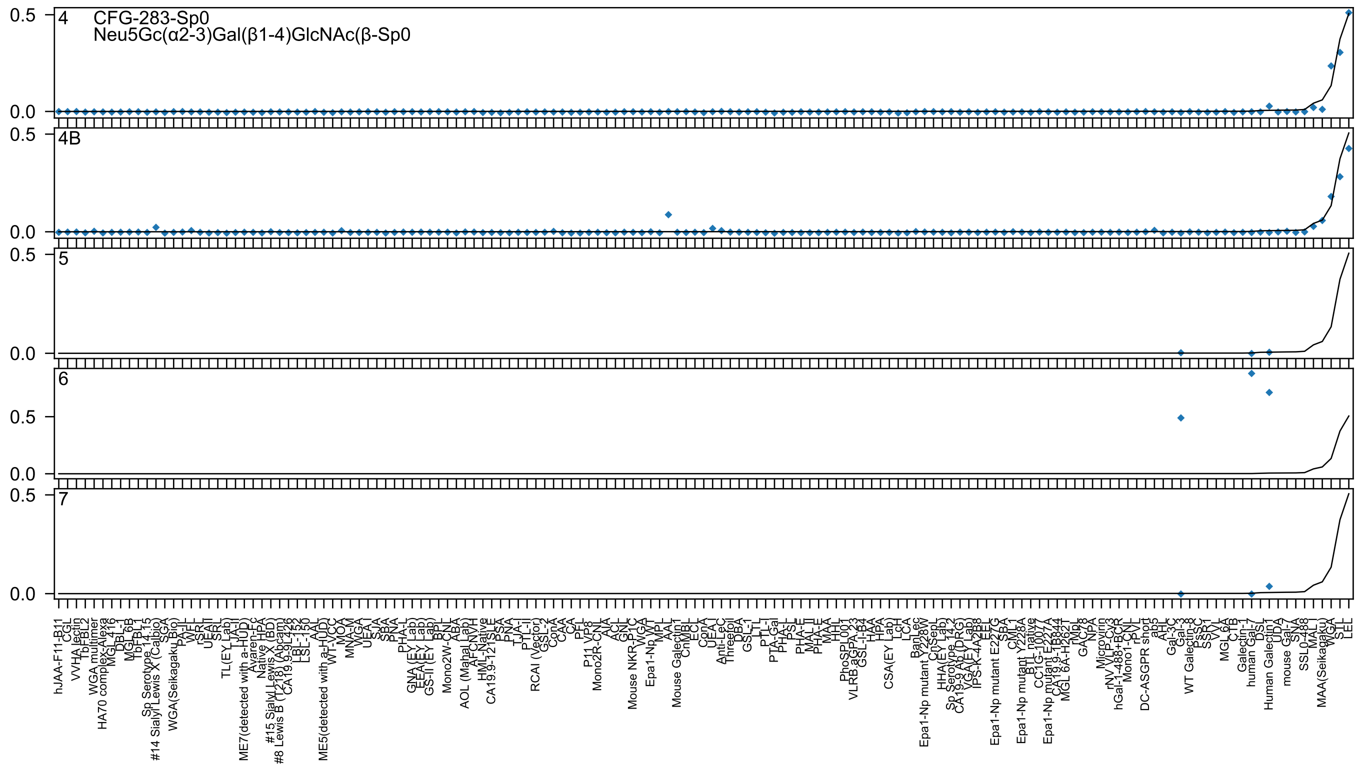
4
CFG-283-Sp0
Neu5Gc(α2-3)Gal(β1-4)GlcNAc(β-Sp0
4B
5
6
7
0.5
0.0

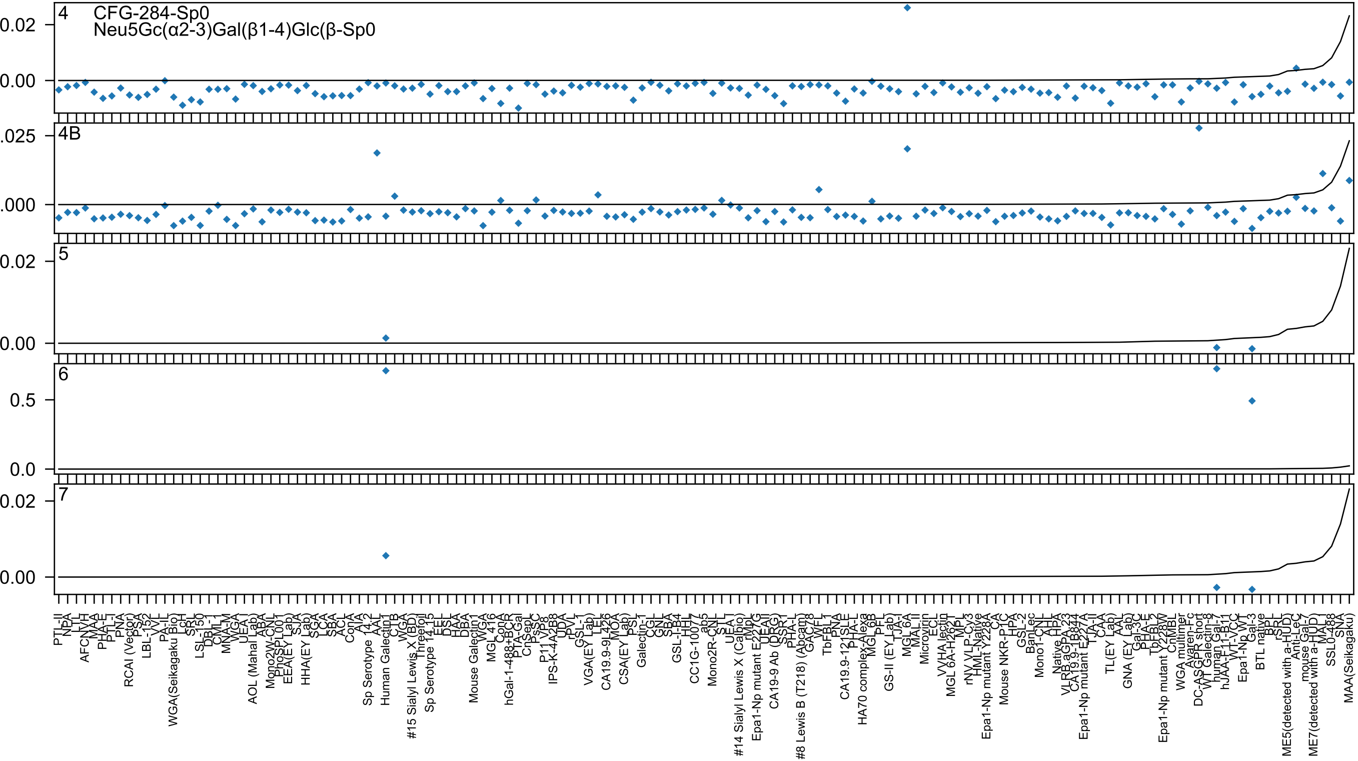
4
CFG-284-Sp0
Neu5Gc(α2-3)Gal(β1-4)Glc(β-Sp0
4B
5
6
7
0.02
0.00
.025
.000
0.5
0.0

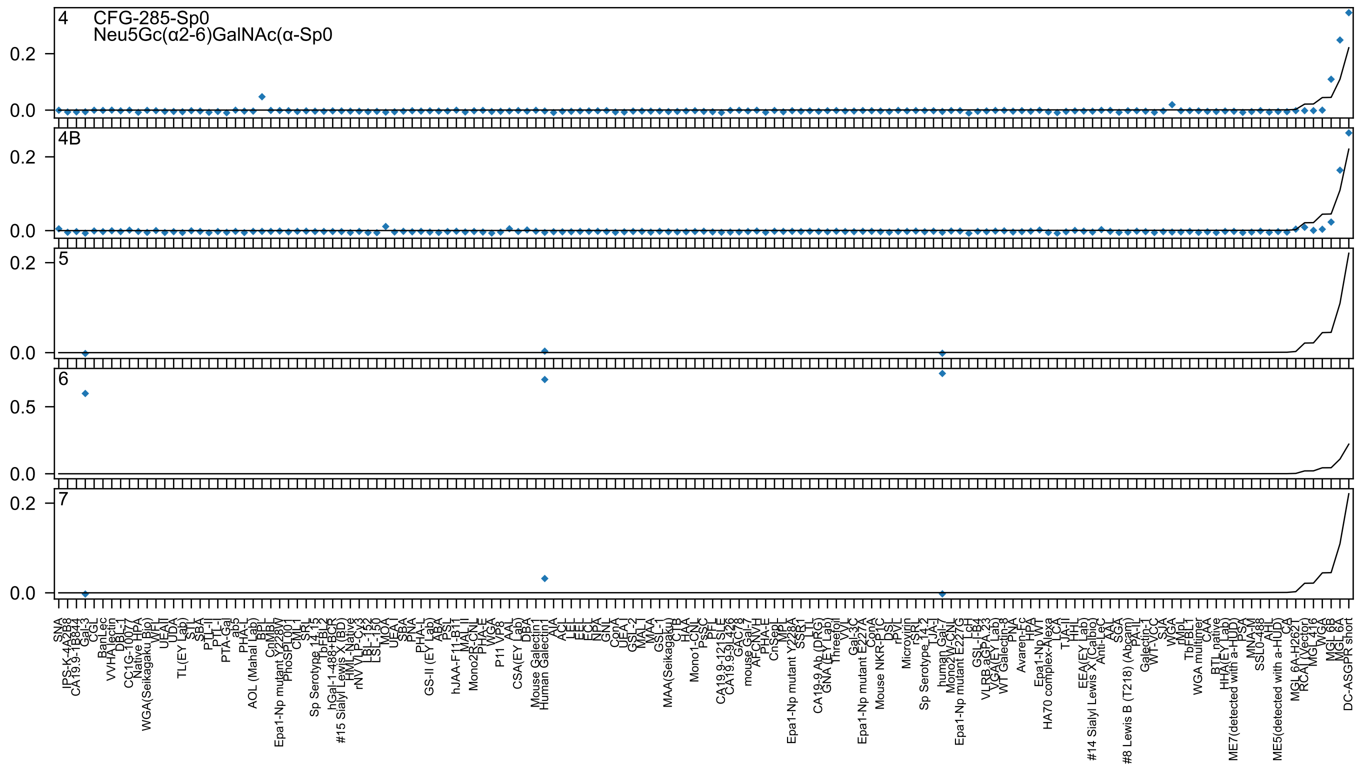
4
CFG-285-Sp0
Neu5Gc(α2-6)GalNAc(α-Sp0
4B
5
6
7
0.0
0.2
0.5

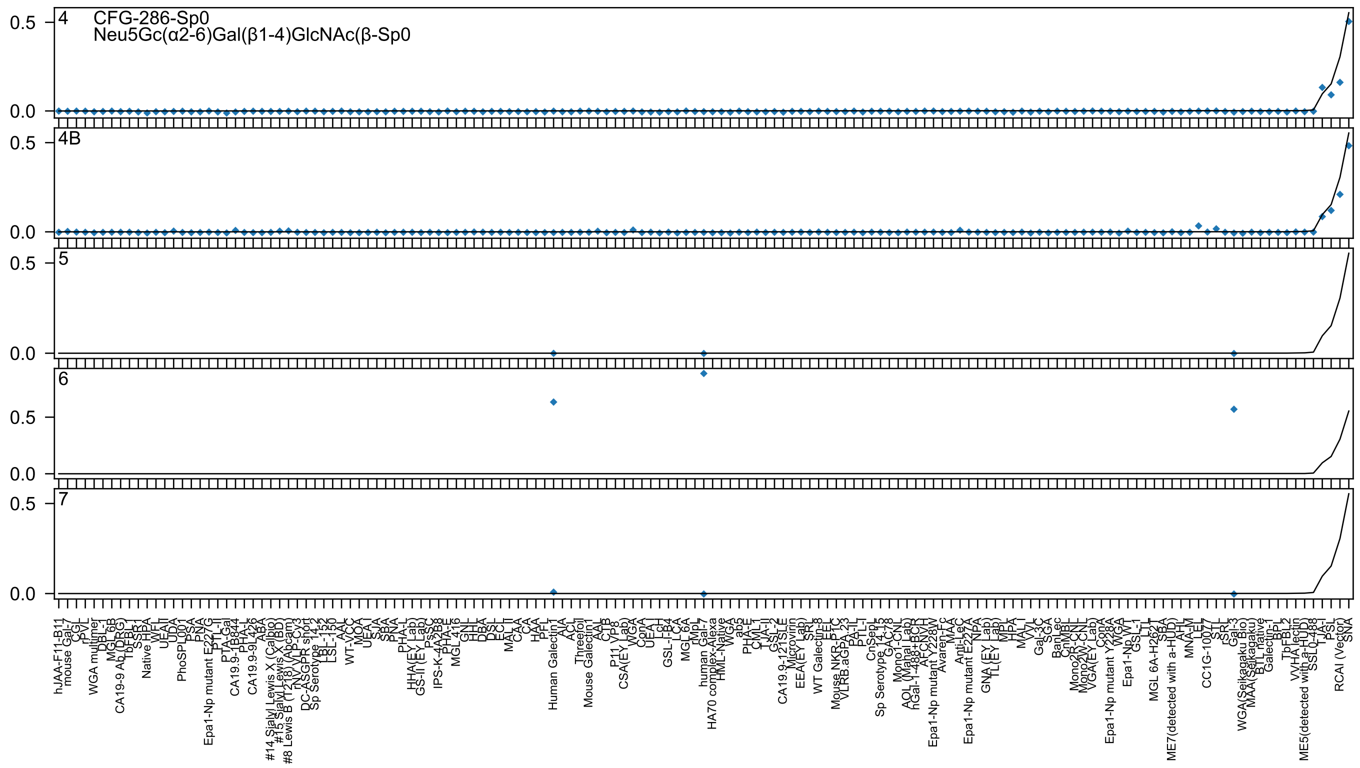
4
CFG-286-Sp0
Neu5Gc(α2-6)Gal(β1-4)GlcNAc(β-Sp0
4B
5
6
7
0.5
0.0
hJAA-F11-B11
mouse Gal-7
CGL
rPVL
WGA multimer
DBL-1
MGL 6B
CA19-9 Ab (DRG)
TbFBL1
SSR1
Native HPA
WFL
UEA-II
UDA
PhoSPL001
PSA
PNA
Epa1-Np mutant E227G
PTL-II
PTA-Gal
CA19.9-1B844
PHA-L
CA19.9-9L426
ABA
#14 Sialyl Lewis X (Calbio)
#15 Sialyl Lewis X (BD)
#8 Lewis B (T218) (Abcam)
rNV VLP-Cy3
DC-ASGPR short
Sp Serotype 14.2
LBL-152
LSL-150
AAL
WT-VCC
MOA
UEA-I
SJA
SBA
PNA
PHA-L
HHA(EY Lab)
GS-II (EY Lab)
PsSC
IPS-K-4A2B8
PHA-E
MGL 416
GNL
HHL
DBA
DSL
ECL
MAL II
CAA
CA
HAA
PFL
Human Galectin1
AIA
ACL
Threefoil
Mouse Galectin1
AAL
CTB
P11 VP8
CSA(EY Lab)
WGA
ConA
UEA-I
LcH
GSL-I-B4
LCA
MGL 6A
rMpL
human Gal-7
HA70 complex-Alexa
HML-Native
WGA
ab2
PHA-E
CML-I
TJA-II
GSL-2
CA19.9-121SLE
Microvirin
EEA(EY Lab)
SRL
EEL
WT Galectin-8
Mouse NKR-P1C
VLRB.aGPA.23
PA-IL
PTL-I
CnSepL
Sp Serotype 14-78
GAC-78
Mono1-CNL
AOL (Manal Lab)
hGal-1-488+BCR
AFCNVH
Epa1-Np mutant Y228W
Avaren-Fc
MAA
Anti-LeC
Epa1-Np mutant E227A
NPA
GNA (EY Lab)
TL(EY Lab)
MPL
HPA
MAL-I
VVL
Gal-3C
SGA
BanLec
CnMBL
Mono2R-CNL
Mono2W-CNL
VGA(EY Lab)
ConA
Epa1-Np mutant Y228A
WGA
Epa1-Np WT
GSL-1
LTL
MGL 6A-H262T
SBA
ME7(detected with a-HUD)
AHL
MNA-M
LEL
CC1G-10077
STL
rSRL
Gal-3
WGA(Seikagaku Bio)
MAA(Seikagaku)
BTL native
Galectin-1
BPL
TbFBL2
VVHA lectin
ME5(detected with a-HUD)
SSL0-488
TJA-I
PSL
RCAI (Vector)
SNA

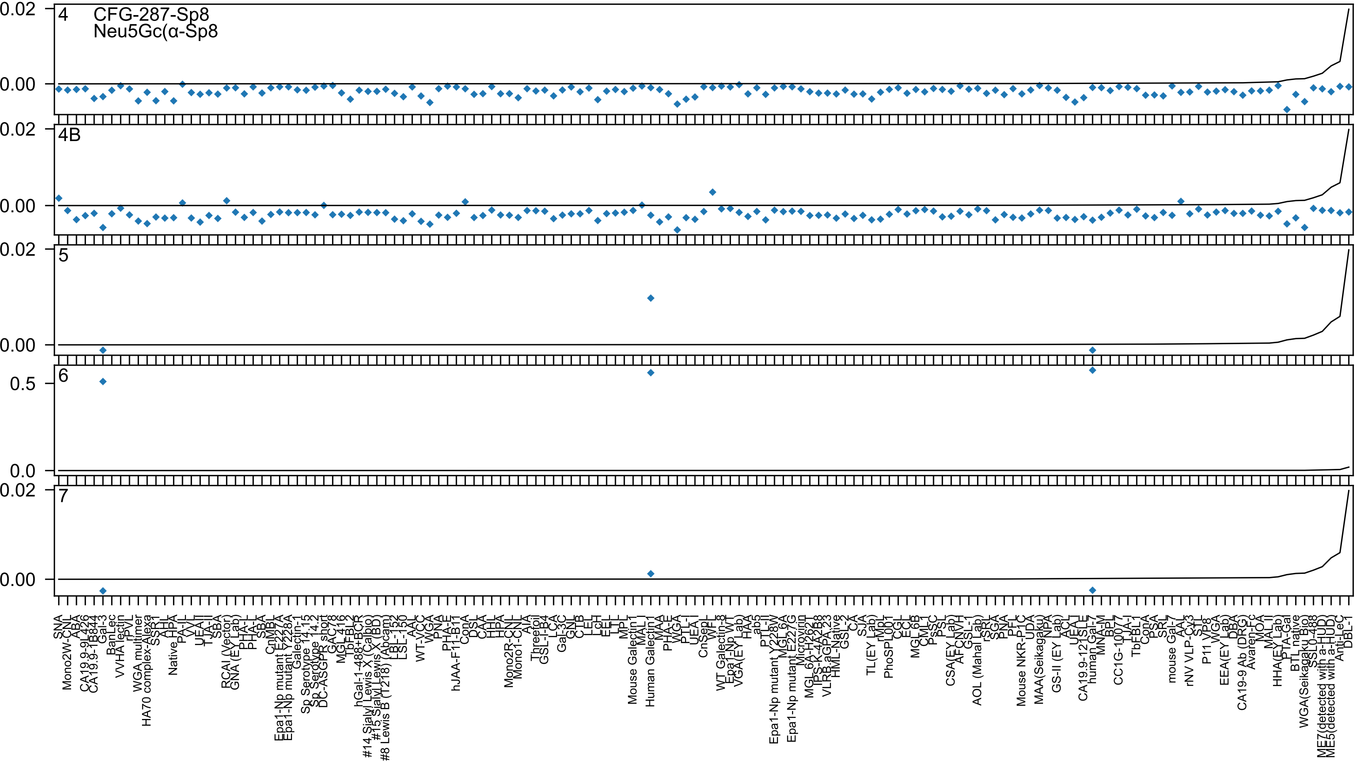
4
CFG-287-Sp8
Neu5Gc(α-Sp8
4B
5
6
7

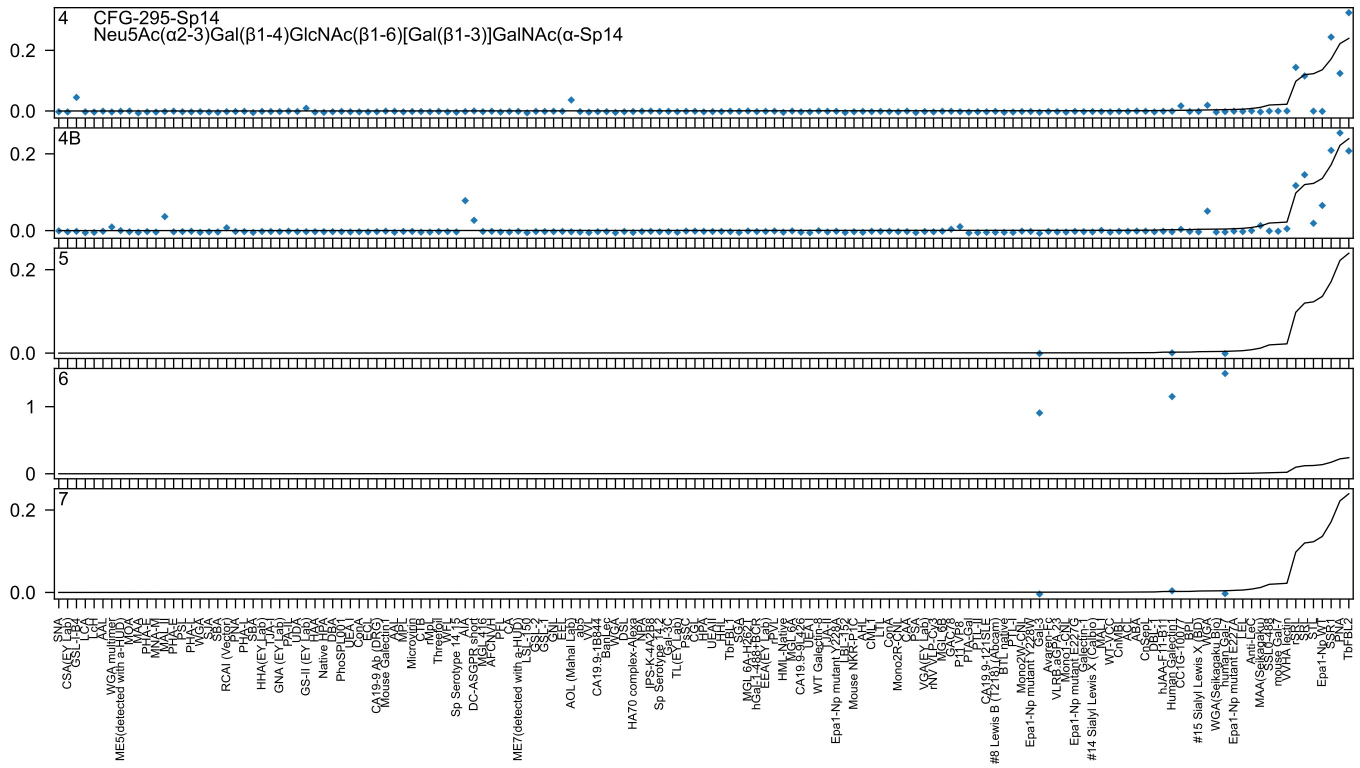

4
CFG-295-Sp14
Neu5Ac(α2-3)Gal(β1-4)GlcNAc(β1-6)[Gal(β1-3)]GalNAc(α-Sp14
4B
5
6
7
0.0
0.2
0
1
SNA
CSA(EY Lab)
GSL-I-B4
LCA
LcH
AAL
WGA multimer
ME5(detected with a-HUD)
MOA
MAA
PHA-E
MNA-M
MAL II
PHA-E
PSL
PHA-L
WGA
SJA
SBA
RCAI (Vector)
PNA
PHA-L
SBA
HHA(EY Lab)
TJA-I
GNA (EY Lab)
PA-IL
UDA
GS-II (EY Lab)
HAA
Native HPA
DBA
PhoSL001
UEA I
ConA
ECL
CA19-9 Ab (DRG)
Mouse Galectin1
AAL
MPL
Microvirin
CTB
rMpL
Threefoil
WFL
Sp Serotype 14.15
AIA
DC-ASGPR short
MGL 416
AFCNVH
PFL
CA
ME7(detected with a-HUD)
LSL-150
GSL-2
GSL-1
GNL
EEL
ab5
VVL
CA19.9-1B844
BanLec
WGA
DSL
HA70 complex-Alexa
NPA
IPS-K-4A2B8
Sp Serotype 14.2
Gal-3C
TL(EY Lab)
PsSC
CGL
HPA
UEAII
HHL
TbFBL1
SGA
MGL 6A-H262T
hGal-1-488+BCR
EEA(EY Lab)
rPVL
HML-Native
MGL 6A
CA19.9-9L426
UEA I
WT Galectin-8
TJA-II
Epa1-Np mutant Y228A
LBL-152
Mouse NKR-P1C
AHL
CML1
LTL
ConA
Mono2R-CNL
CAA
PSA
VGA(EY Lab)
rNV VLP-Cy3
MGL 6B
GAC78
P11 VP8
PTA-Gal
PTL-II
CA19.9-121SLE
#8 Lewis B (T218) (Abcam)
BTL native
PTL-I
Mono2W-CNL
Epa1-Np mutant Y228W
Gal-3
Avaren-Fc
VLRB.aGPA.23
Mono1-CNL
Epa1-Np mutant E227G
Galectin-1
#14 Sialyl Lewis X (Calbio)
MAL-I
WT-VCC
CnMBL
ACL
ABA
CnSepL
DBL-1
hJAA-F11-B11
Human Galectin1
CC1G-10077
BPL
#15 Sialyl Lewis X (BD)
WGA
WGA(Seikagaku Bio)
human Gal-7
Epa1-Np mutant E227A
LEL
Anti-LeC
MAA(Seikagaku)
SSL0-488
mouse Gal-7
WWHA lectin
rSRL
SRL
STL
Epa1-Np WT
SSR1
PNA
TbFBL2

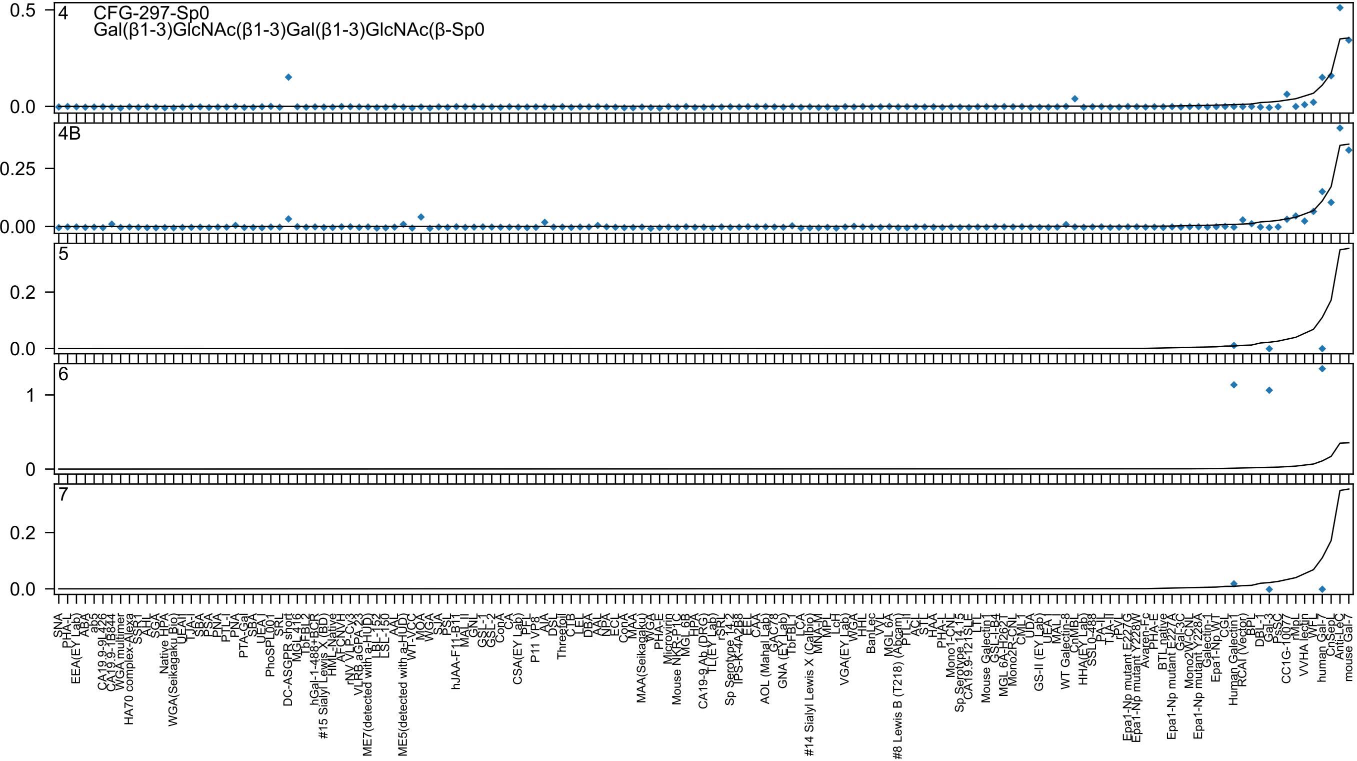
4
CFG-297-Sp0
Gal(β1-3)GlcNAc(β1-3)Gal(β1-3)GlcNAc(β-Sp0
4B
5
6
7
0.5
0.0
0.25
0.00
0.2
0.0
1
0
0.2
0.0

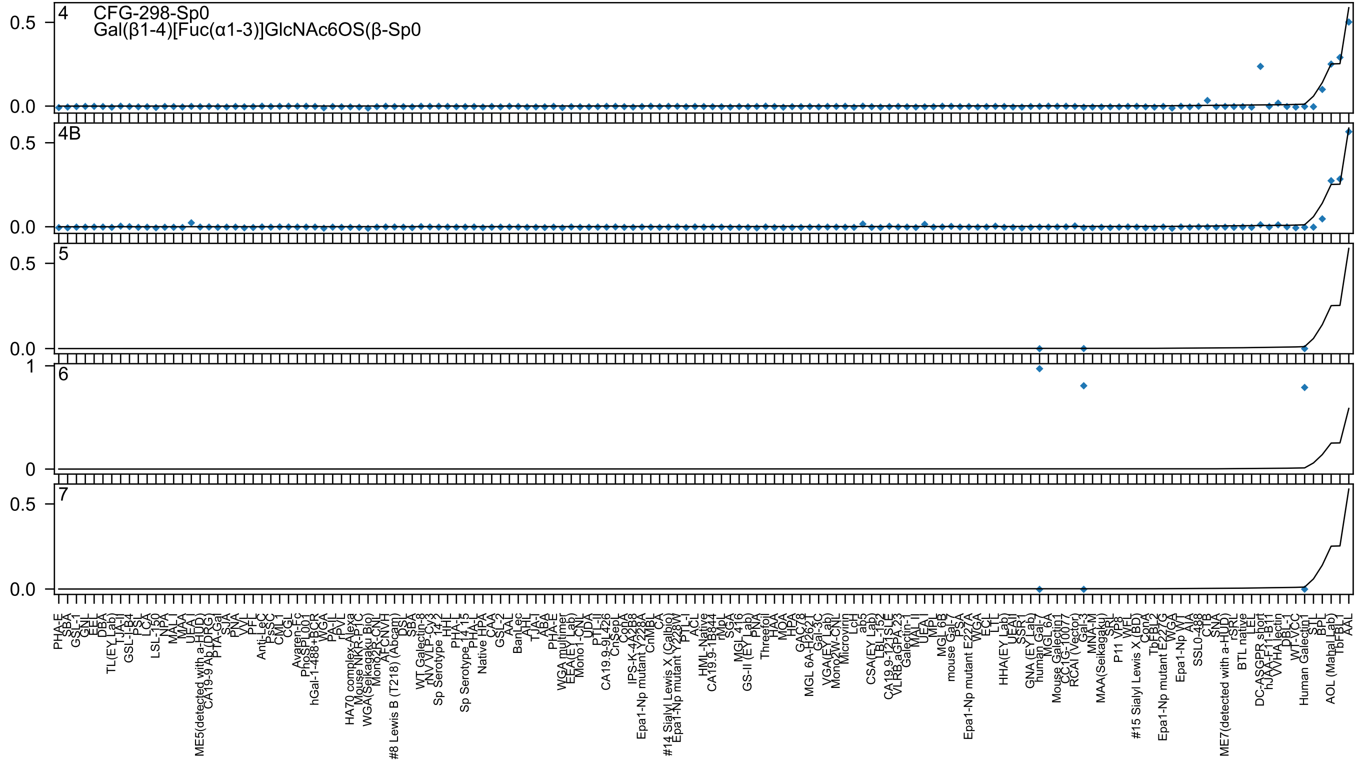
4
CFG-298-Sp0
Gal(β1-4)[Fuc(α1-3)]GlcNAc6OS(β-Sp0
4B
5
6
7

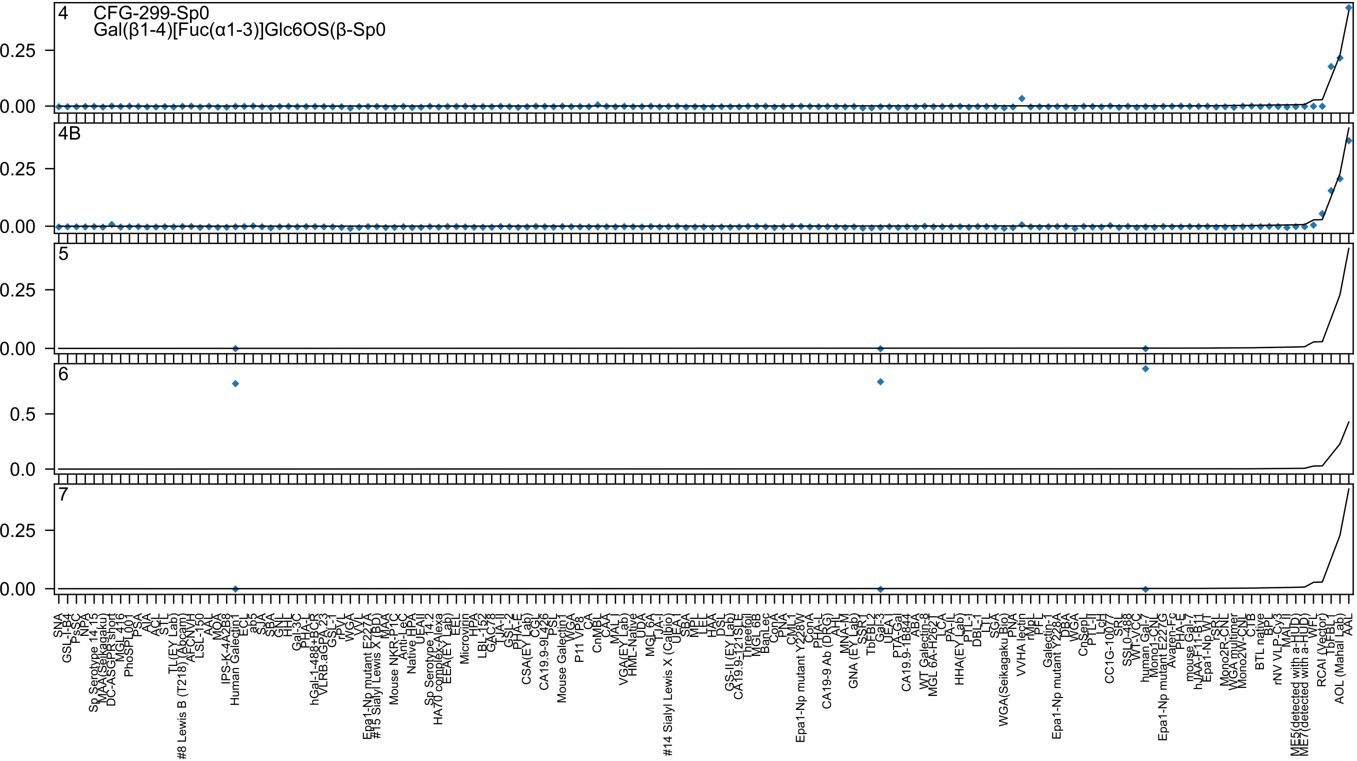

4 CFG-299-Sp0
Gal(β1-4)[Fuc(α1-3)]Glc6OS(β-Sp0
4B
5
6
7

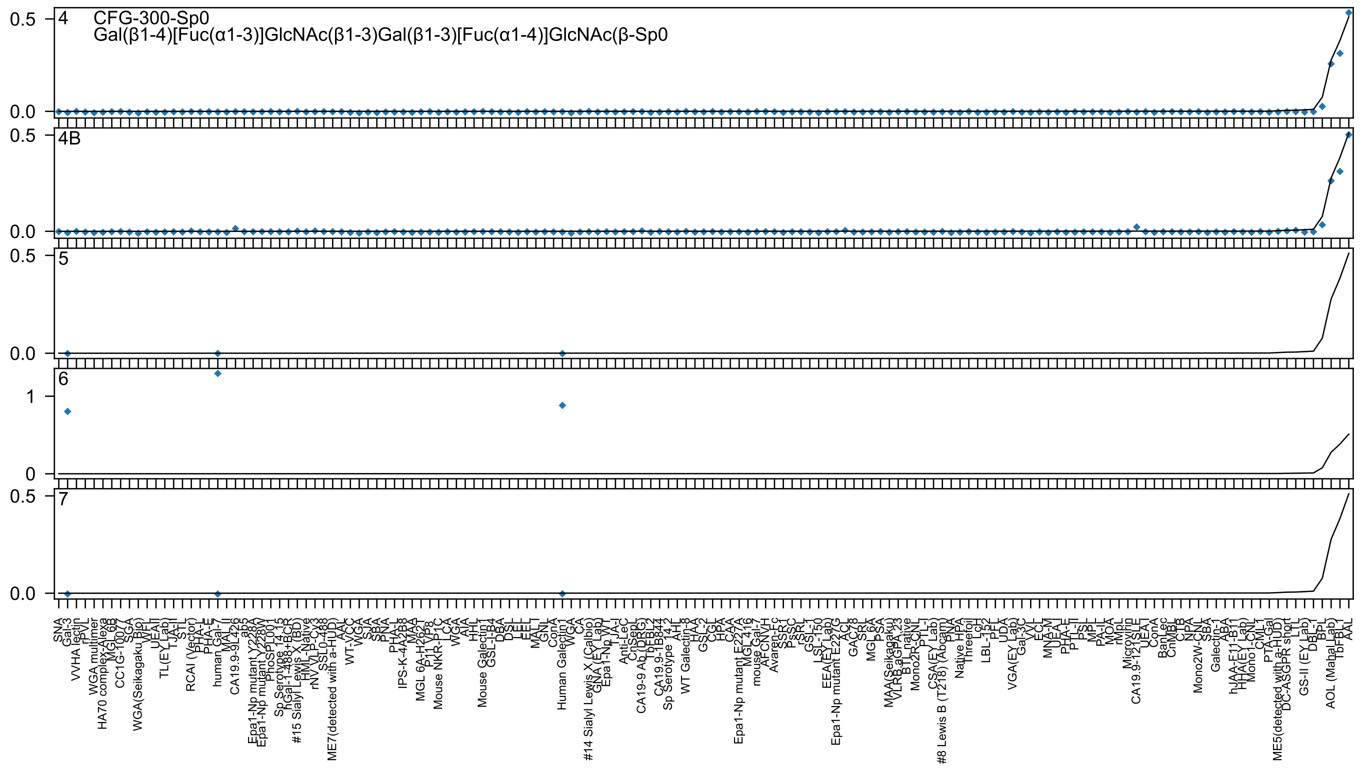

4
CFG-300-Sp0
Gal(β1-4)[Fuc(α1-3)]GlcNAc(β1-3)Gal(β1-3)[Fuc(α1-4)]GlcNAc(β-Sp0
4B
5
6
7

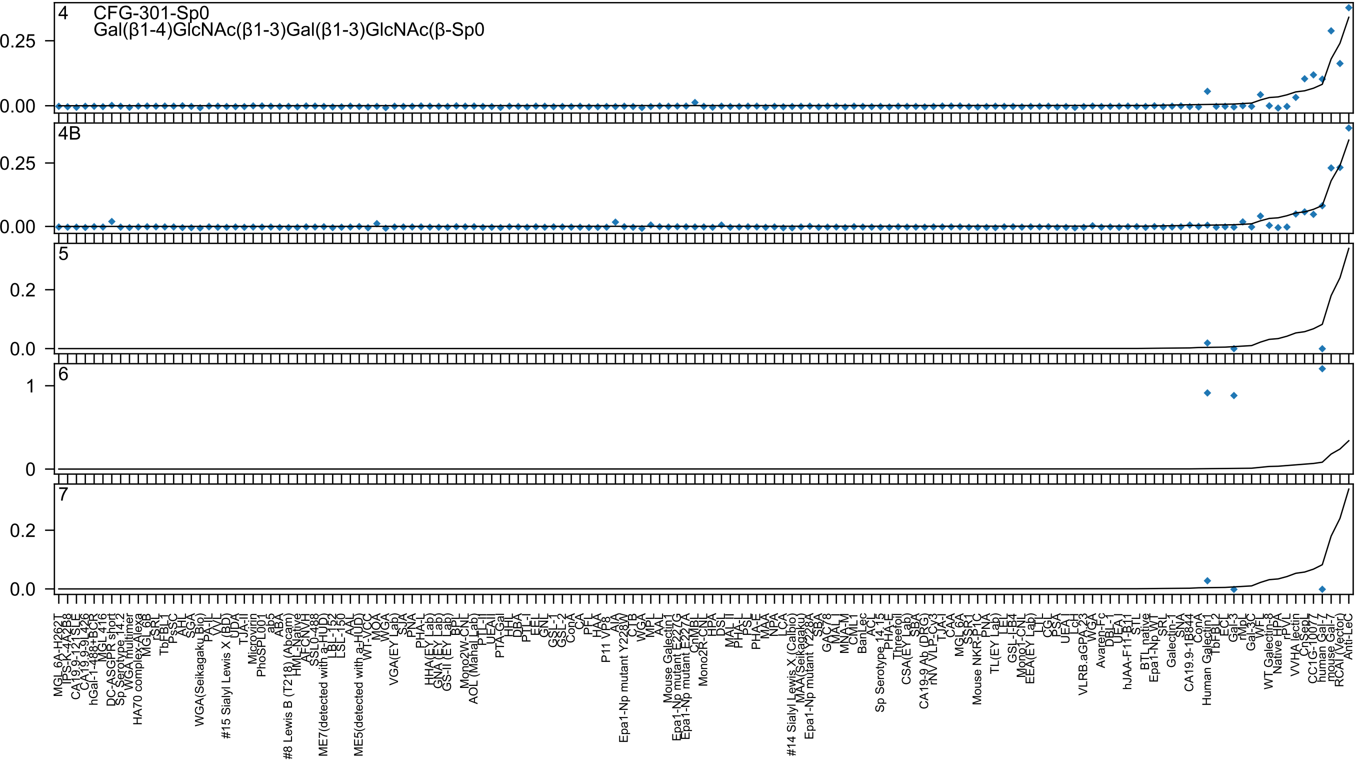
4
CFG-301-Sp0
Gal(β1-4)GlcNAc(β1-3)Gal(β1-3)GlcNAc(β-Sp0
0.25
0.00
4B
0.25
0.00
5
0.2
0.0
6
1
0
7
0.2
0.0
MGL_6A-H262T
IPs-K-4A2B8
CA19.9-121SLE
CA19.9-9L426
hGal-1-488+BCR
MGL_416
DC-ASGPR short
Sp Serotype_14.2
WGA multimer
HA70 complex-Alexa
MGL_6B
rSRL
TbFBL1
PsSC
AHL
SGA
WGA(Seikagaku_Bio)
PA-IL
VVL
#15 Sialyl Lewis X (BD)
UDA
TJA-II
Microvirin
PhoSPL001
ab5
ABA
#8 Lewis B (T218) (Abcam)
HML-Native
AFCNVH
SSL0-488
ME7(detected with a-HUD)
LBL-152
LSL-150
AAL
ME5(detected with a-HUD)
WT-VCC
MOA
WGA
VGA(EY Lab)
SJA
PNA
PHA-L
HHA(EY Lab)
GNA (EY Lab)
GS-II (EY Lab)
BPL
Mono2W-CNL
AOL (ManaLab)
PTL-II
UEAII
PTA-Gal
HHL
DBA
PTL-I
EEL
GNL
GSL-I
GSL-2
ConA
CA
PFL
HAA
P11 VP8
AIA
Epa1-Np mutant Y228W
CTB
WGA
MPL
AAL
Mouse Galectin1
Epa1-Np mutant E227G
Epa1-Np mutant E227A
CnMBL
Mono2R-CNL
HPA
DSL
MAL_II
PHA-L
PSL
PHA-E
MAA
NPA
LCA
#14 Sialyl Lewis X (Calbio)
MAA(Seikagaku)
Epa1-Np mutant Y228A
SBA
GAC78
MAL_I
MNA-M
CML1
BanLec
ACL
Sp Serotype_14.15
PHA-E
Threefoil
CSA(EY Lab)
SBA
CA19-9 Ab (DRG)
rNV VLP-CV3
TJA-I
CAA
MGL_6A
SSR1
Mouse NKR-P1C
PNA
TL(EY Lab)
LEL
GSL-I-B4
Mono1-CNL
EEA(EY Lab)
LTL
CGL
PSA
UEA I
LcH
VLRB.aGPA.23
WGA
Avaren-Fc
DBL-1
UEA I
hJAA-F11-B11
STL
BTL native
Epa1-Np WT
SRL
Galectin-1
SNA
CA19.9-1B844
ConA
Human Galectin1
TbFBL2
ECL
Gal-3
rMpL
Gal-3C
WFL
WT Galectin-8
Native HPA
rPVL
VVHA lectin
CnSep
CC1G-1007
human Gal-7
mouse Gal-7
RCAI (Vector)
Anti-LeC

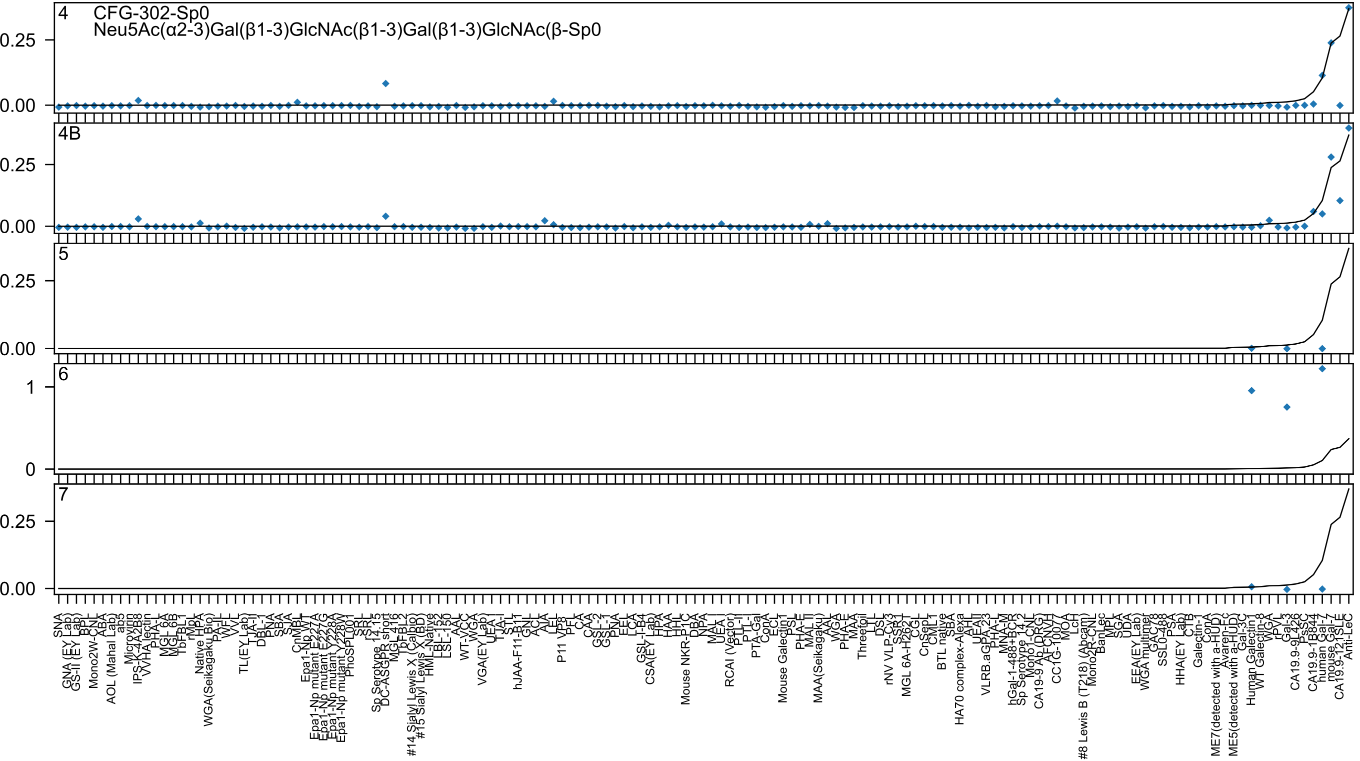
4
CFG-302-Sp0
Neu5Ac(α2-3)Gal(β1-3)GlcNAc(β1-3)Gal(β1-3)GlcNAc(β-Sp0
0.25
0.00
4B
0.25
0.00
5
0.25
0.00
6
1
0
7
0.25
0.00

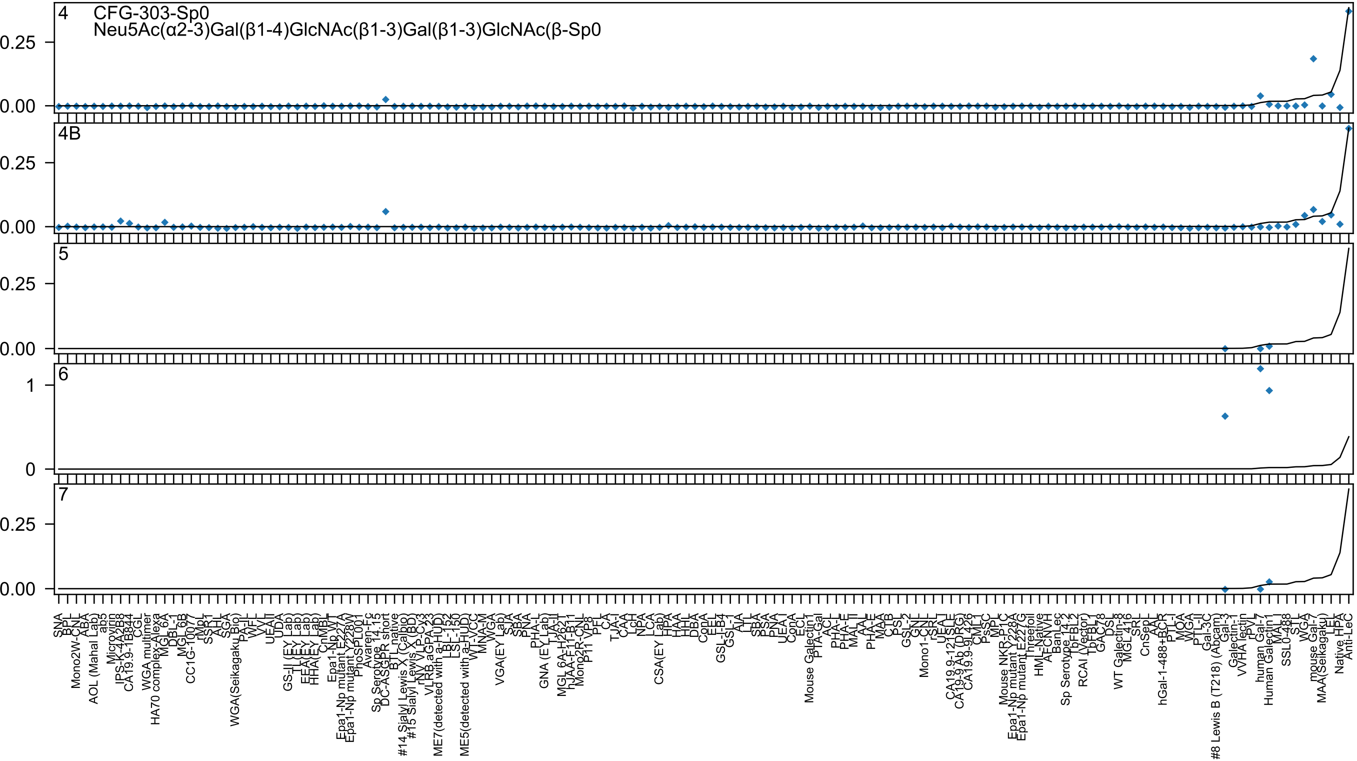
CFG-303-Sp0
Neu5Ac(α2-3)Gal(β1-4)GlcNAc(β1-3)Gal(β1-3)GlcNAc(β-Sp0

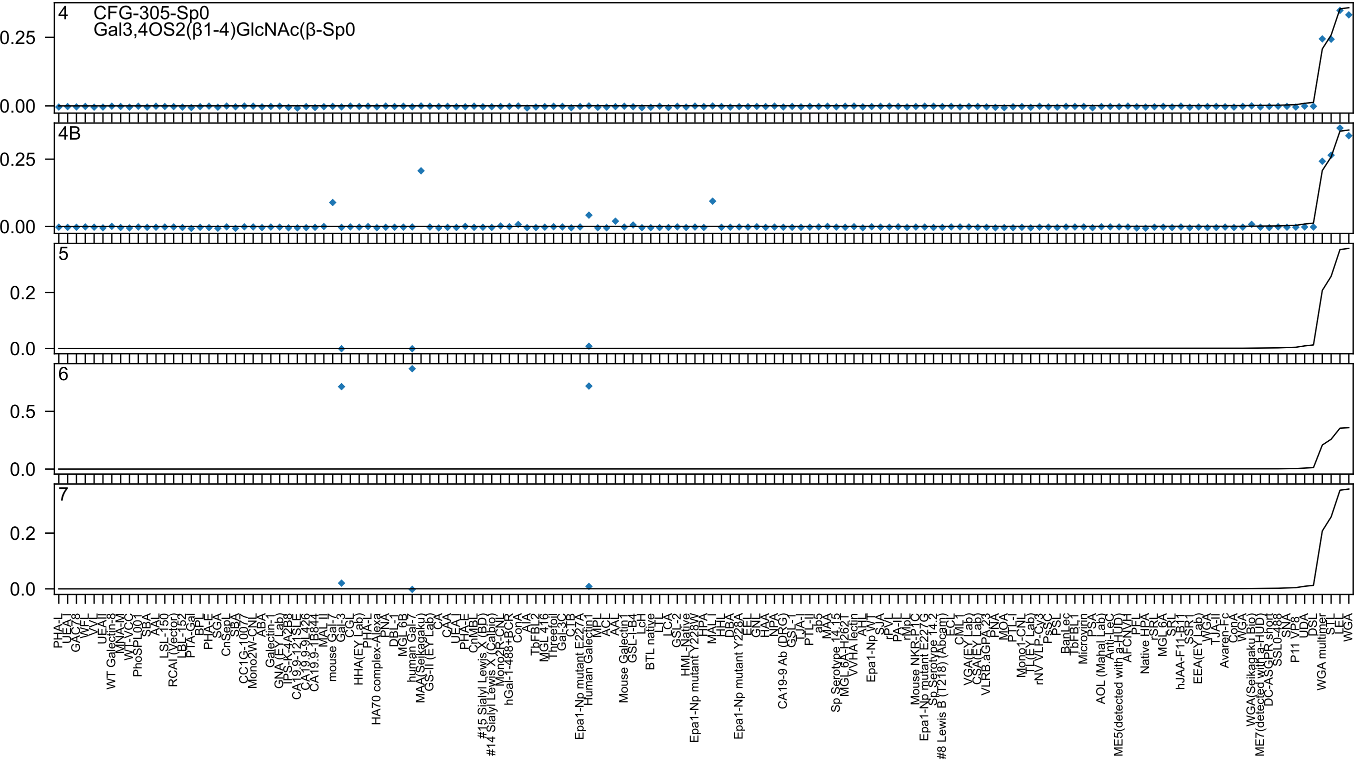
4
CFG-305-Sp0
Gal3,4OS2(β1-4)GlcNAc(β-Sp0
0.25
0.00
4B
0.25
0.00
5
0.2
0.0
6
0.5
0.0
7
0.2
0.0

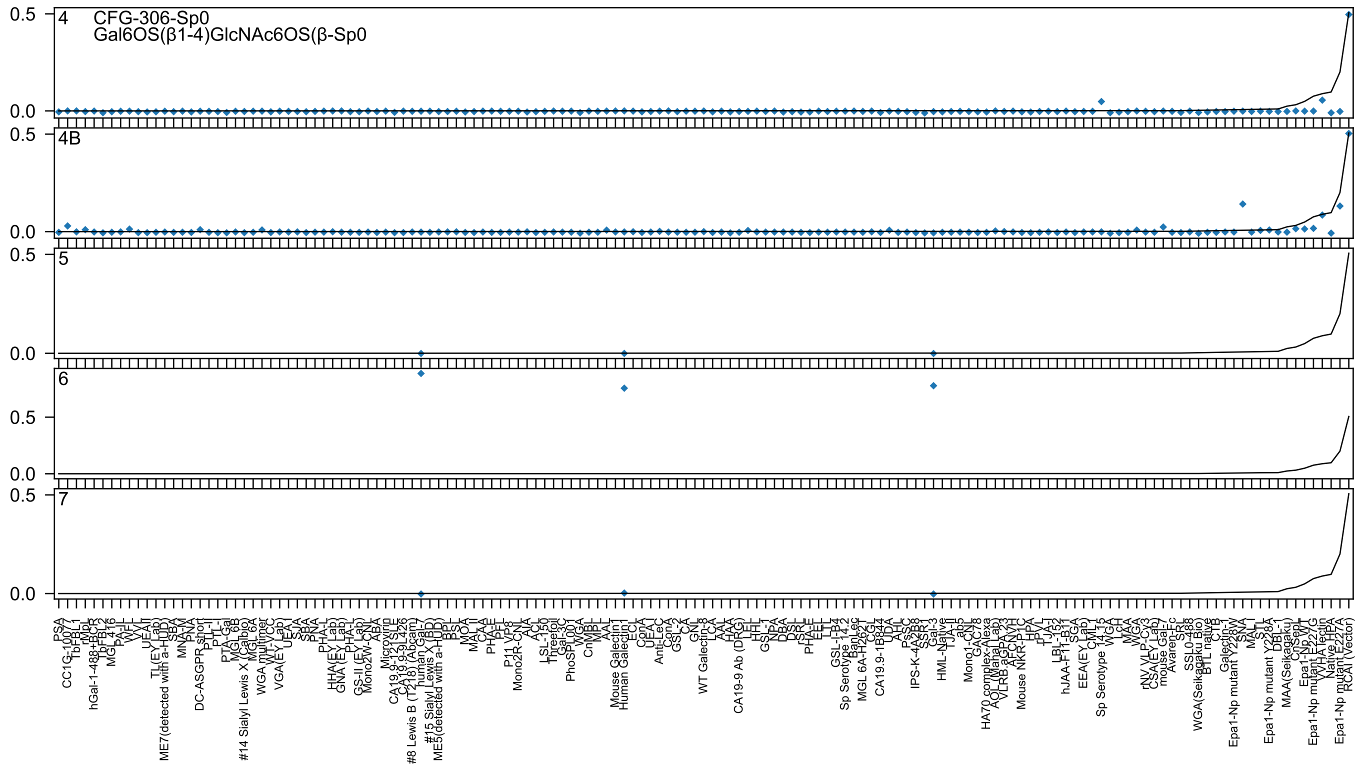

4
CFG-306-Sp0
Gal6OS(β1-4)GlcNAc6OS(β-Sp0
4B
5
6
7
0.5
0.0

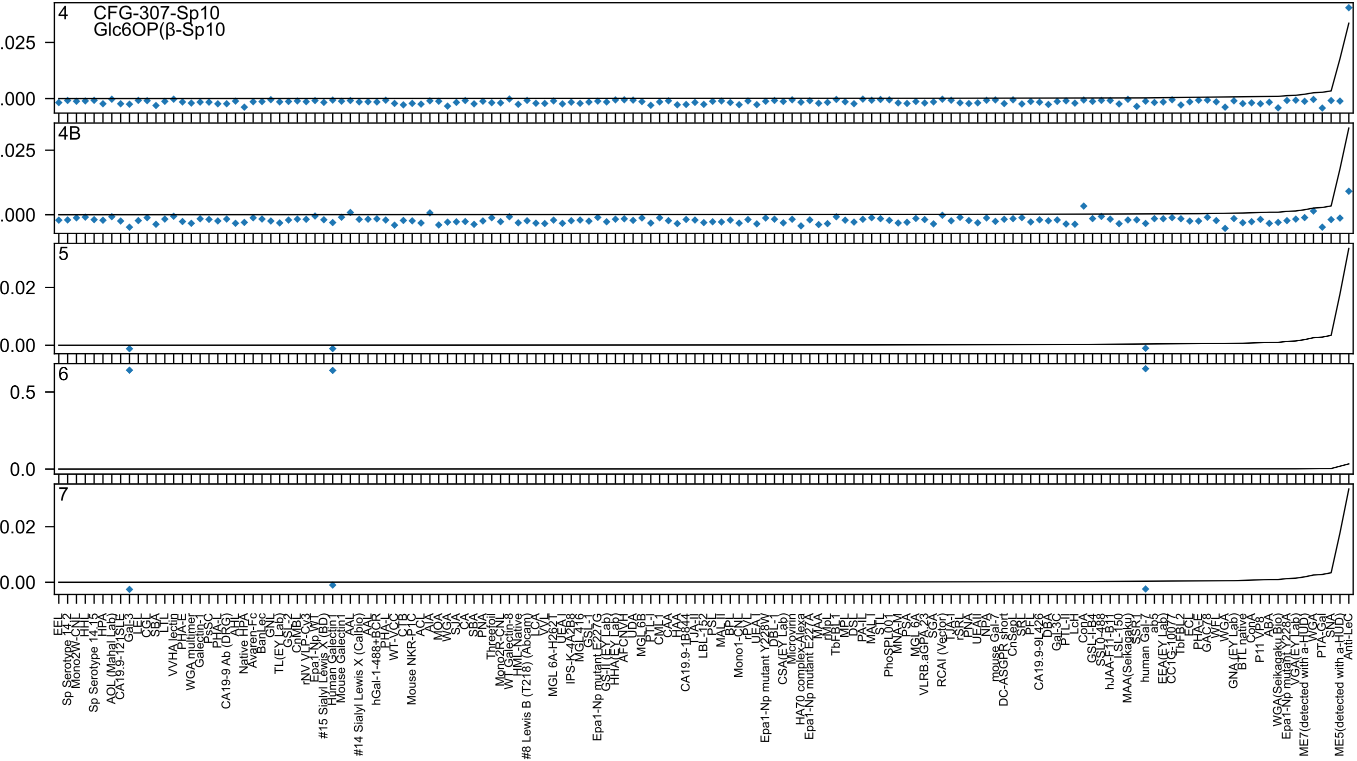
4
CFG-307-Sp10
Glc6OP(β-Sp10
4B
5
6
7

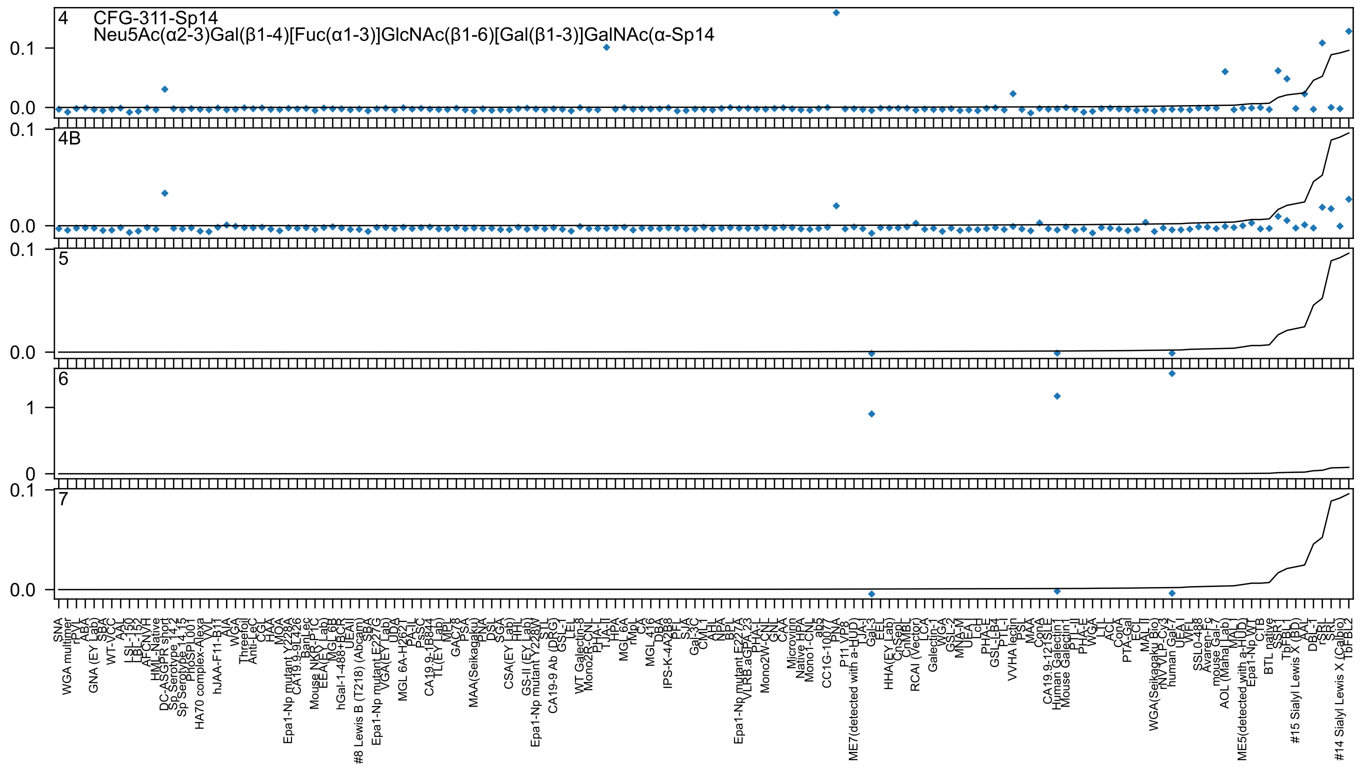

4
CFG-311-Sp14
Neu5Ac(α2-3)Gal(β1-4)[Fuc(α1-3)]GlcNAc(β1-6)[Gal(β1-3)]GalNAc(α-Sp14
4B
5
6
7

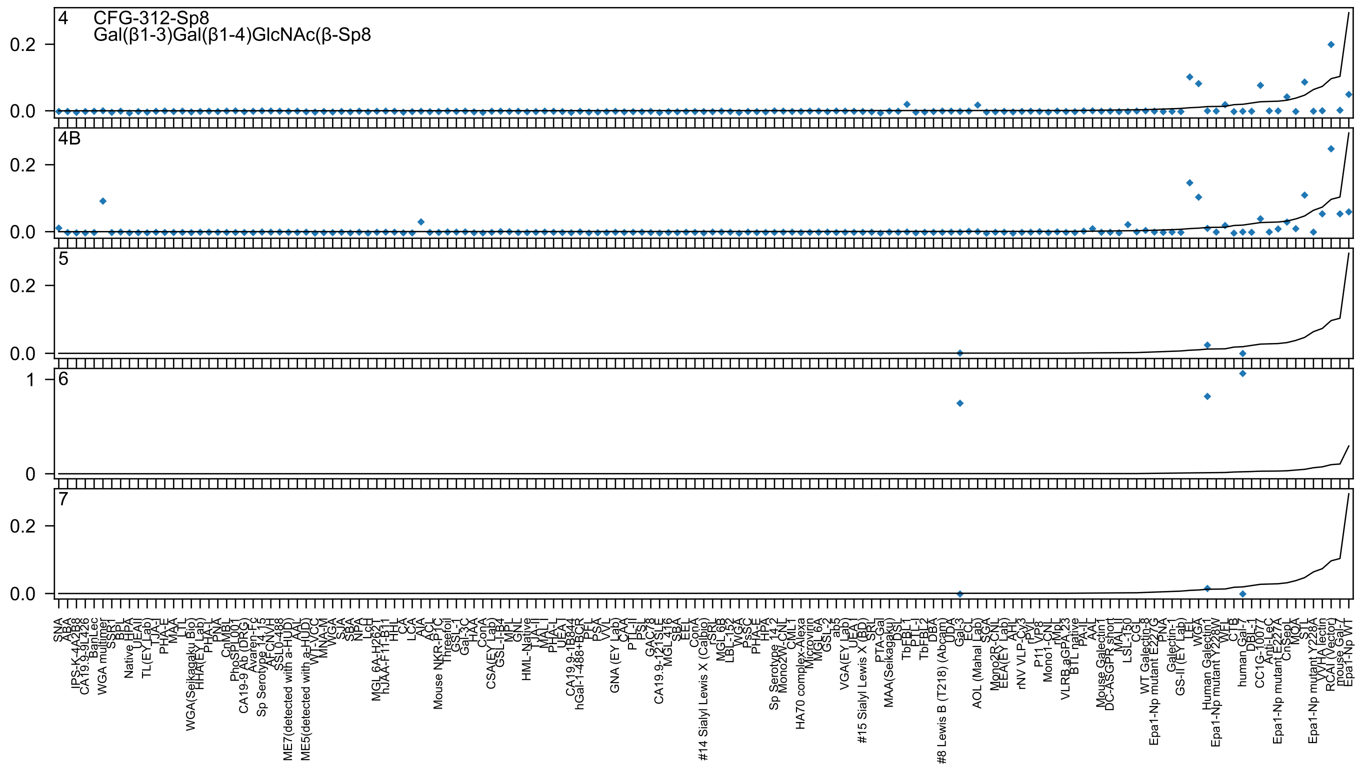

4 CFG-312-Sp8
Gal(β1-3)Gal(β1-4)GlcNAc(β-Sp8
4B
5
6
7

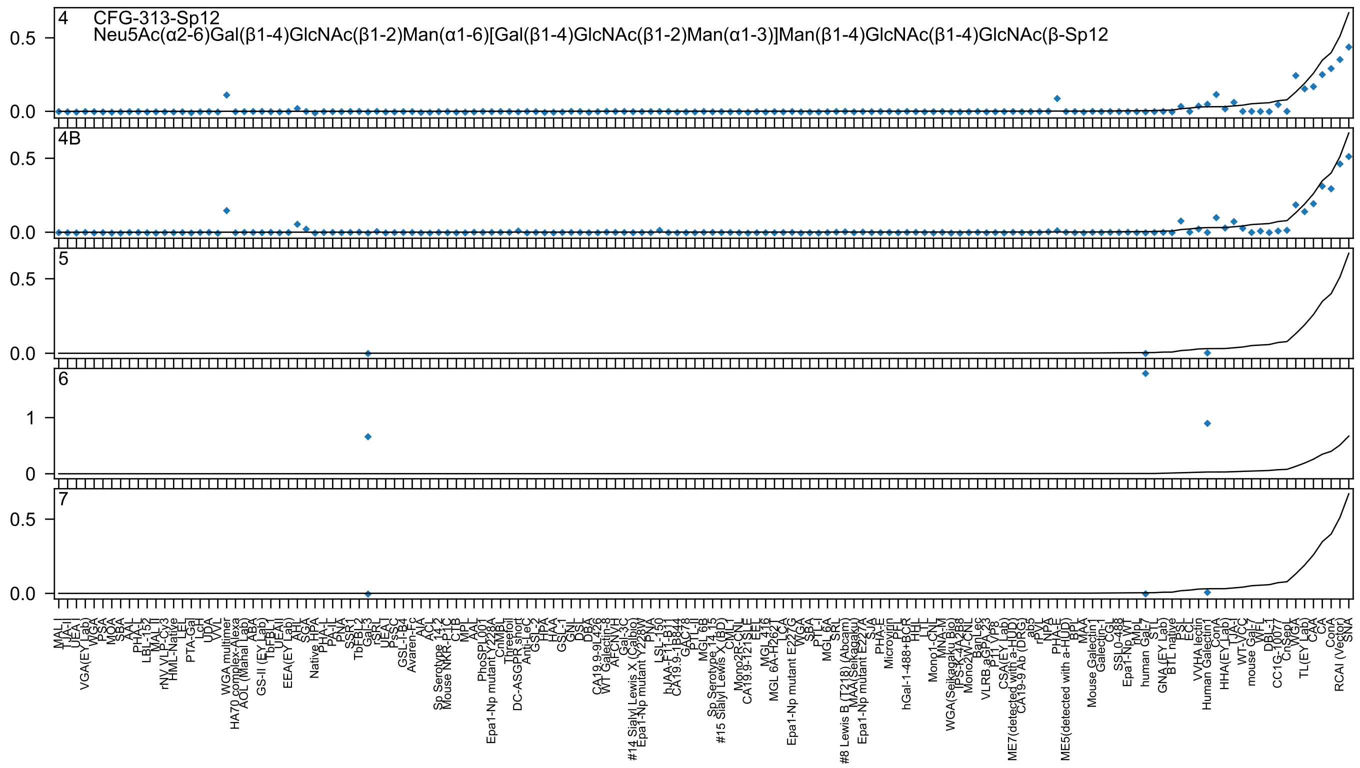
4
CFG-313-Sp12
Neu5Ac(α2-6)Gal(β1-4)GlcNAc(β1-2)Man(α1-6)[Gal(β1-4)GlcNAc(β1-2)Man(α1-3)]Man(β1-4)GlcNAc(β1-4)GlcNAc(β-Sp12
4B
5
6
7
0.5
0.0
1
0

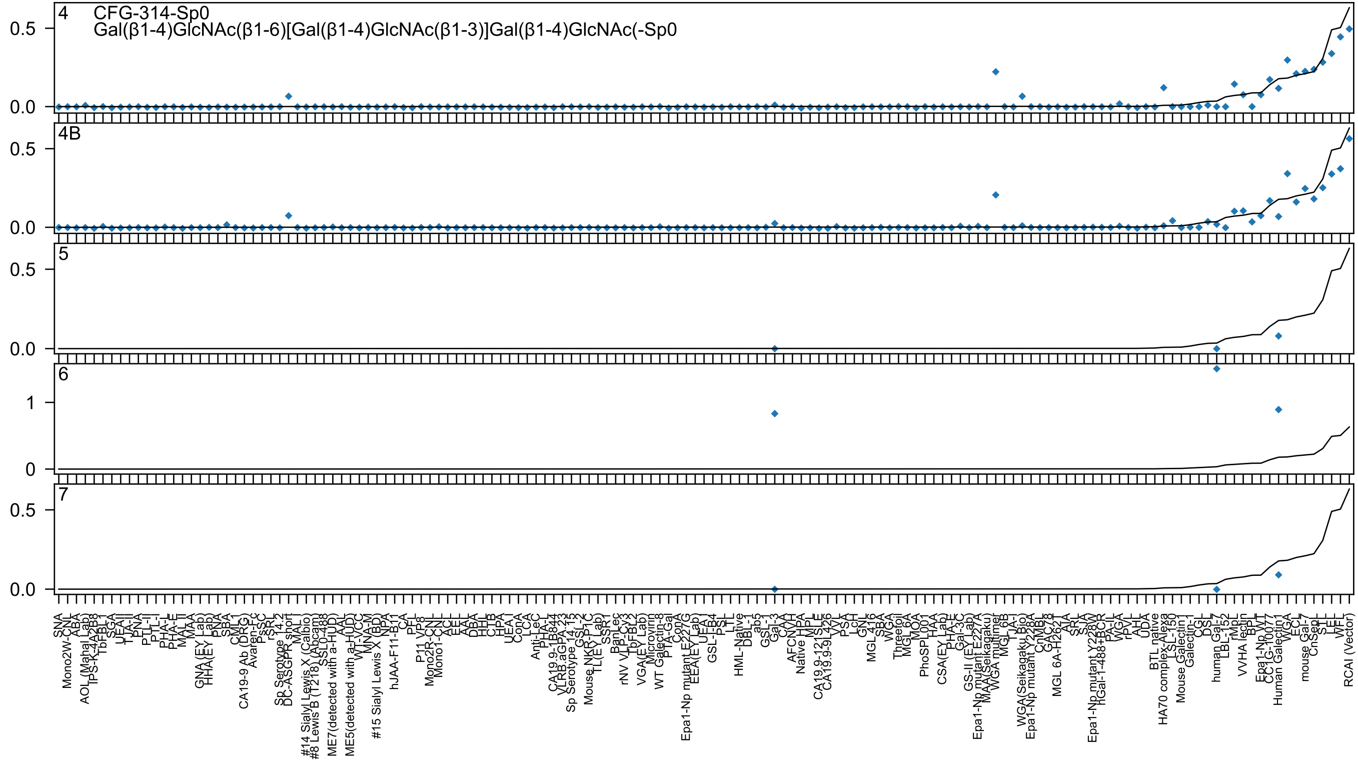
4 CFG-314-Sp0
Gal(β1-4)GlcNAc(β1-6)[Gal(β1-4)GlcNAc(β1-3)]Gal(β1-4)GlcNAc(-Sp0
4B
5
6
7

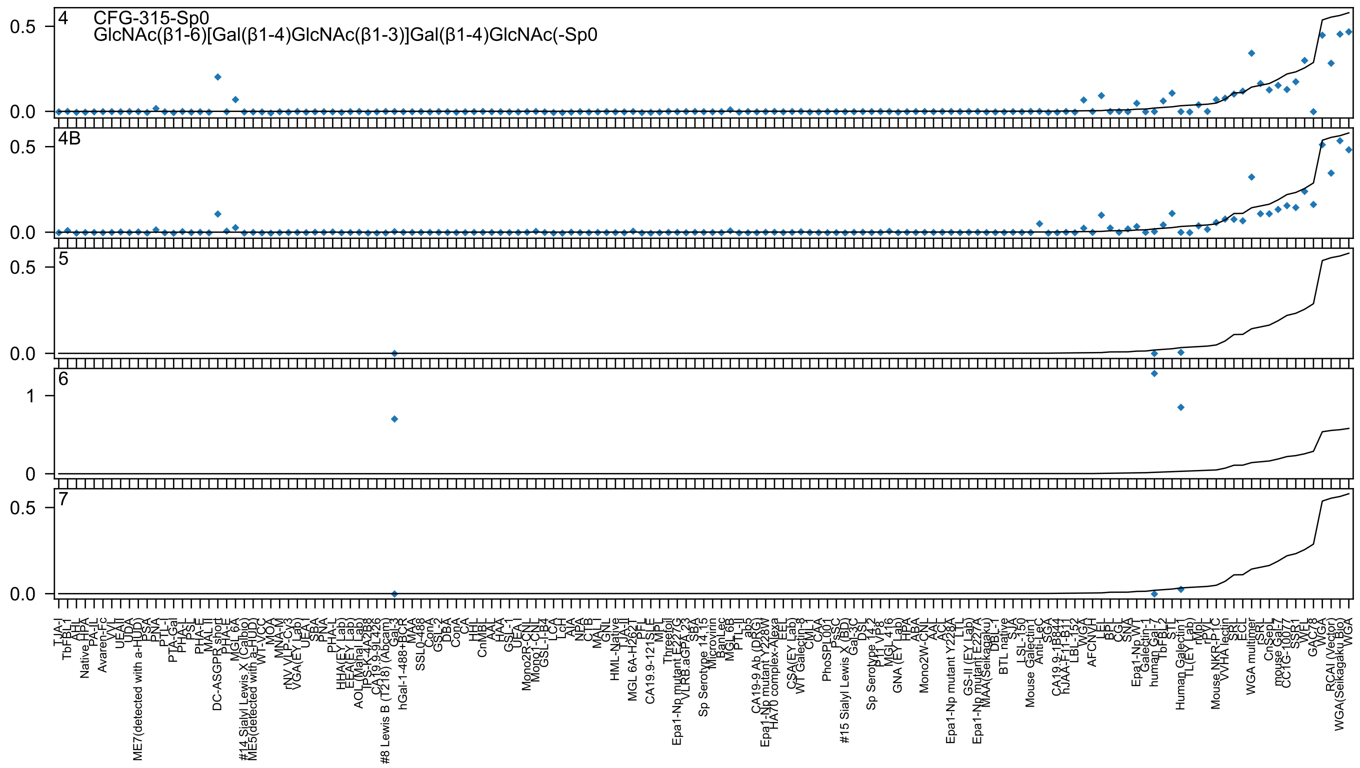
4
CFG-315-Sp0
GlcNAc(β1-6)[Gal(β1-4)GlcNAc(β1-3)]Gal(β1-4)GlcNAc(-Sp0
4B
5
6
7
0.0
0.5
0
1
TJA-I
TbFBL-1
AHL
HPA
Native HPA
PA-IL
Avaren-Fc
VVL
UEA-II
UDA
ME7(detected with a-HUD)
PSA
PNA
PTL-I
PTA-Gal
PHA-L
PSL
PHA-E
MAL II
DC-ASGPR_short
PHA-E
MGL 6A
#14 Sialyl Lewis X (Calbio)
ME5(detected with a-HUD)
WT-VCC
MOA
MNA-M
rNV VLP-Cy3
VGA(EY Lab)
UEA-I
SBA
PNA
PHA-L
HHA(EY Lab)
EEA(EY Lab)
AOL (Mahal Lab)
TPS-K-4A2B8
CA19.9-9L426
#8 Lewis B (T218) (Abcam)
Gal-3
hGal-1-488+BCR
MAA
SSL0-488
ConA
GSL-2
DBA
ConA
CA
HHL
CnMBL
AAL
HAA
GSL-1
UEA-I
Mono2R-CNL
Mono1-CNL
GSL-I-B4
LCA
LcH
AIA
NPA
CTB
MAL-I
GNL
HML-Native
TJA-II
MGL 6A-H262T
PFL
CA19.9-121SLE
MPL
Threefoil
Epa1-Np mutant E227G
VLRB.aGPA.23
SBA
Sp Serotype 14.15
Microvirin
BanLec
MGL 6B
PTL-II
ab5
CA19-9 Ab (DRG)
Epa1-Np mutant Y228W
HA70 complex-Alexa
EEL
CSA(EY Lab)
WT Galectin-8
CML-1
CAA
PhoSL001
PsSC
#15 Sialyl Lewis X (BD)
Gal-3C
DSL
Sp Serotype 14.2
P1 VP8
MGL 416
GNA (EY Lab)
HPA
ABA
Mono2W-CNL
AAL
ACL
Epa1-Np mutant Y228A
LTL
GS-II (EY Lab)
Epa1-Np mutant E227A
MAA(Seikagaku)
DBL-1
BTL native
SJA
LSL-150
Mouse Galectin1
Anti-LeC
SGA
CA19.9-1B844
hJAA-F11-B11
LBL-152
WGA
AFCNVH
LEL
BPL
CGL
SNA
Epa1-Np WT
Galectin-1
human Gal-7
TbFBL-2
STL
Human Galectin1
TL(EY Lab)
rMpL
rPVL
Mouse NKR-P1C
VVHA lectin
SRL
ECL
WGA multimer
rSRL
CnSepL
mouse Gal-7
CC1G-10077
SSR1
WFL
GAC78
WGA
RCAI (Vector)
WGA(Seikagaku Bio)
WGA

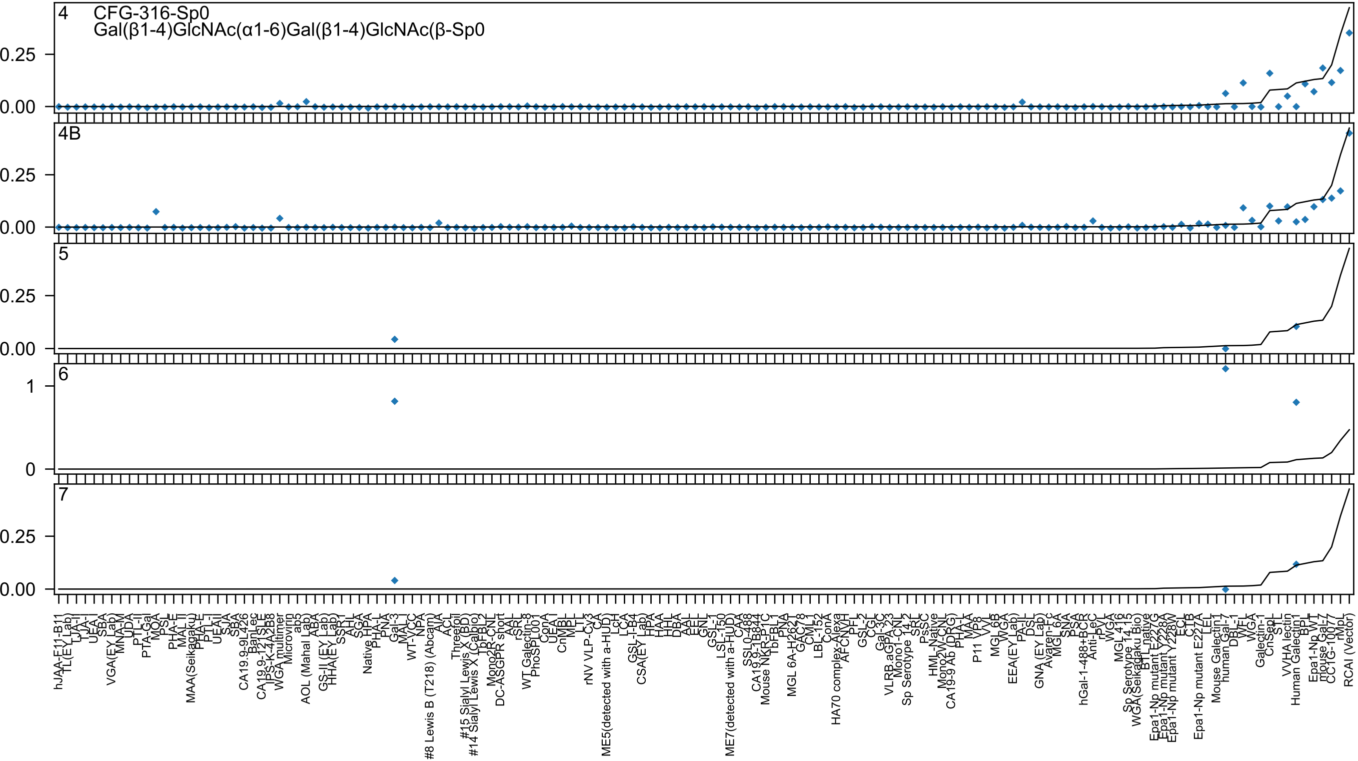
4
CFG-316-Sp0
Gal(β1-4)GlcNAc(α1-6)Gal(β1-4)GlcNAc(β-Sp0
4B
5
6
7
0.25
0.00
1
0

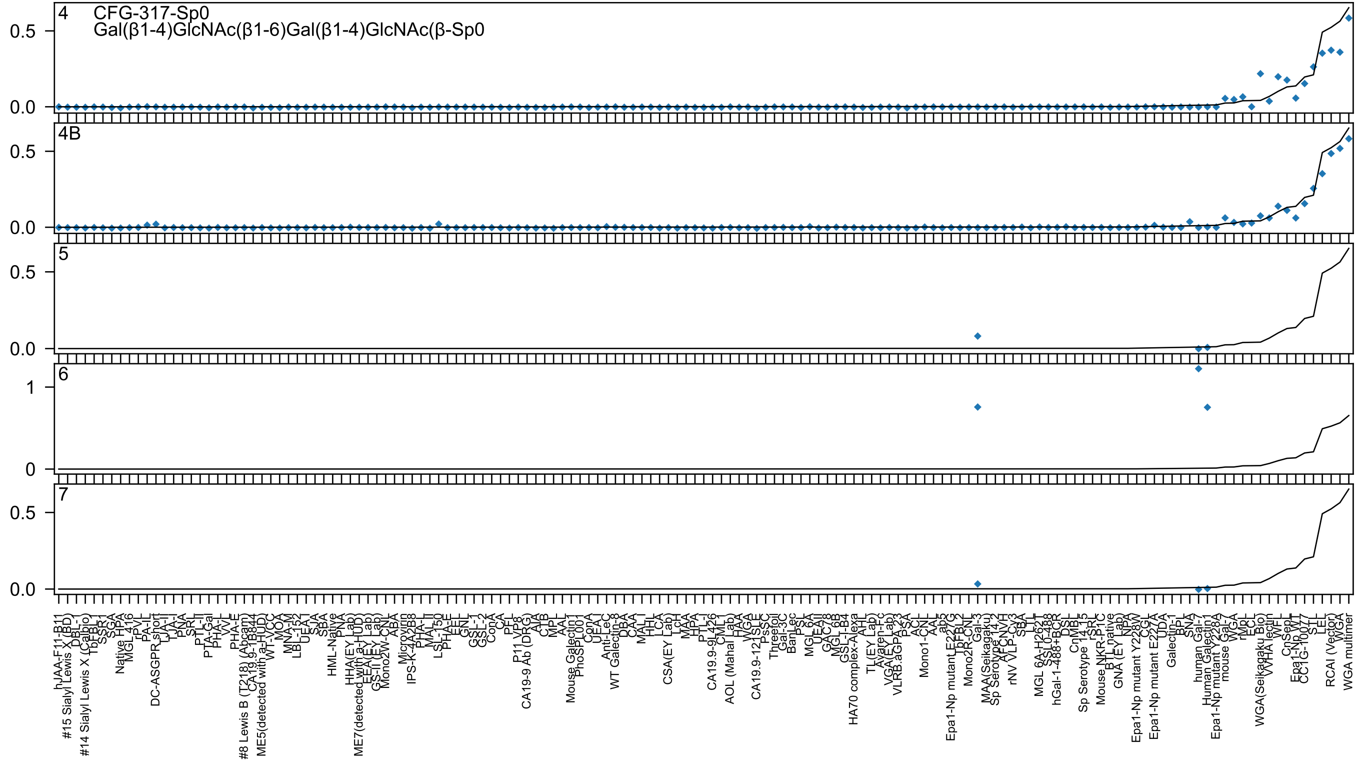
4 CFG-317-Sp0
Gal(β1-4)GlcNAc(β1-6)Gal(β1-4)GlcNAc(β-Sp0
4B
5
6
7

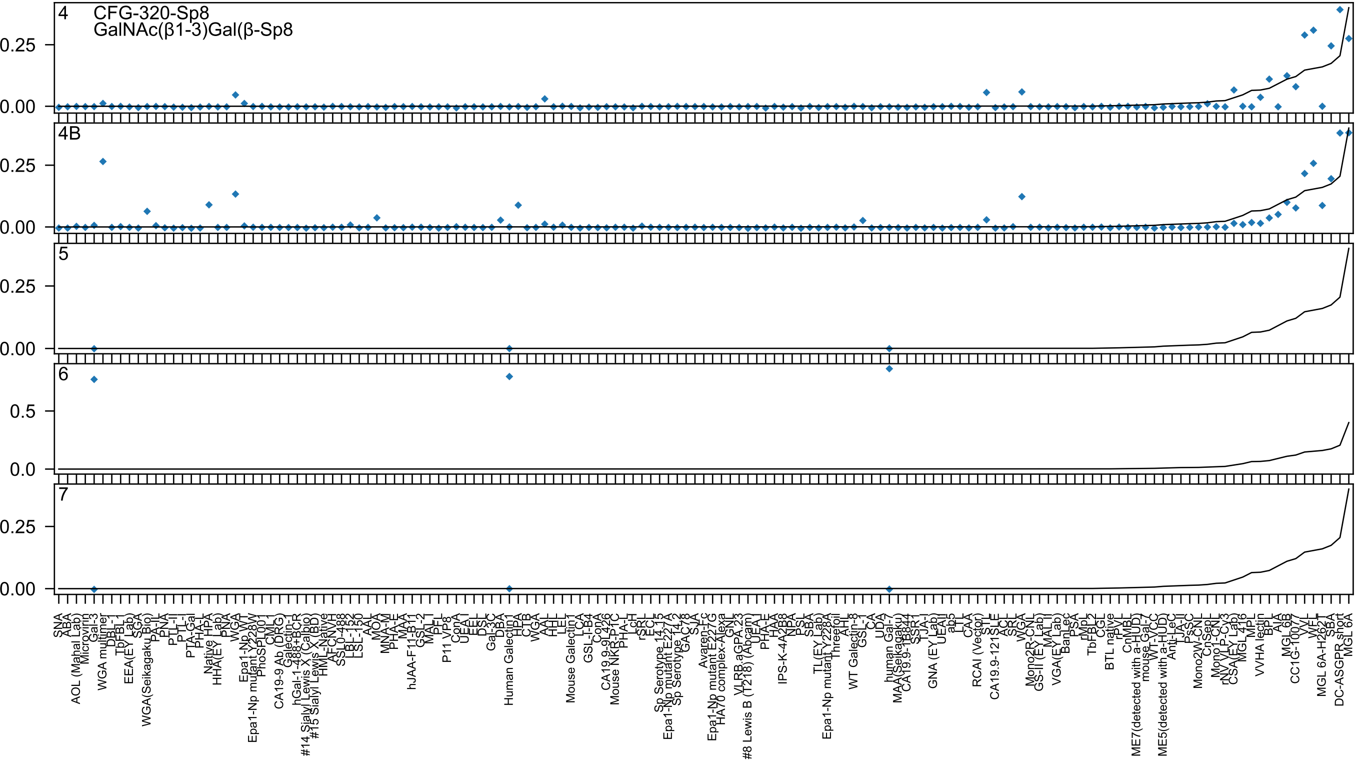

4
CFG-320-Sp8
GalNAc(β1-3)Gal(β-Sp8
4B
5
6
7

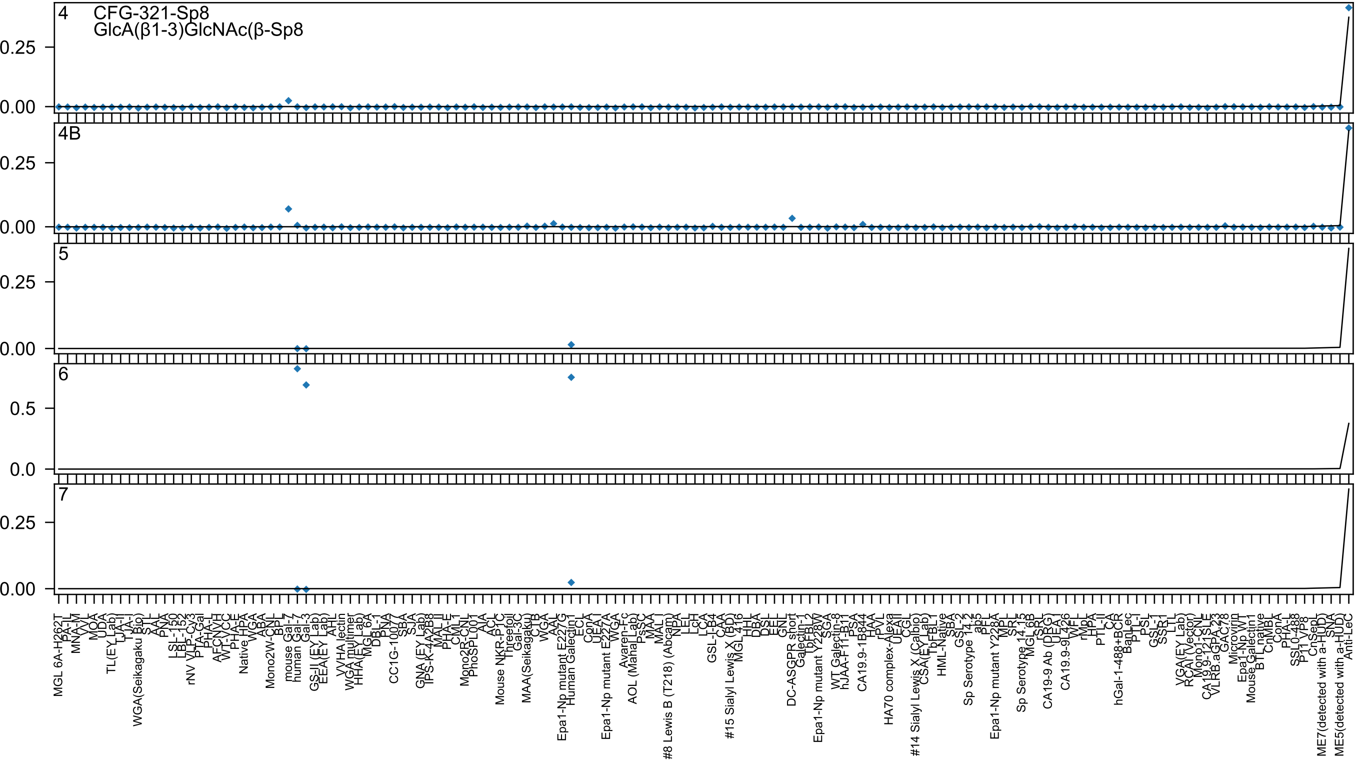
4
CFG-321-Sp8
GlcA(β1-3)GlcNAc(β-Sp8
4B
5
6
7

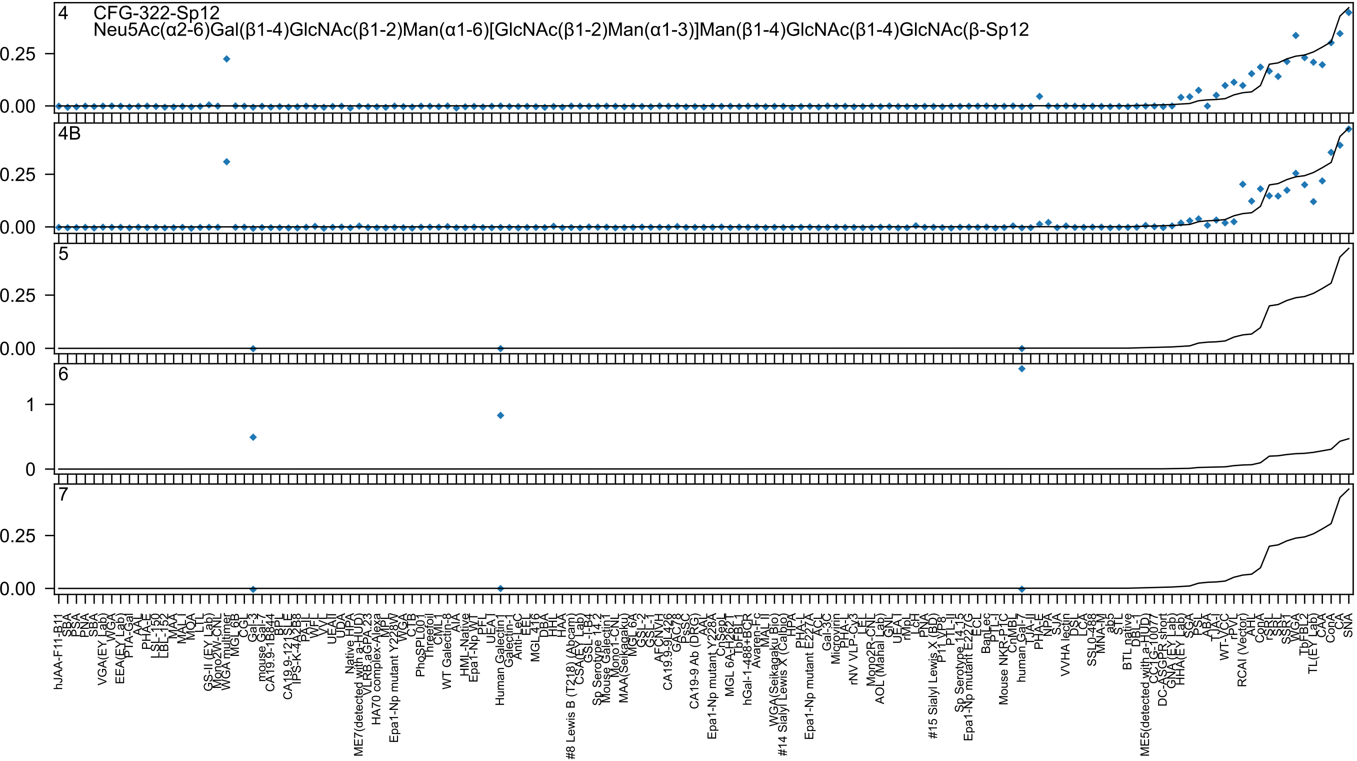
4
CFG-322-Sp12
Neu5Ac(α2-6)Gal(β1-4)GlcNAc(β1-2)Man(α1-6)[GlcNAc(β1-2)Man(α1-3)]Man(β1-4)GlcNAc(β1-4)GlcNAc(β-Sp12
4B
5
6
7
0.25
0.00
1
0
hJAA-F11-B11
SBA
PSA
PNA
SBA
VGA(EY Lab)
WGA
EEA(EY Lab)
PTA-Gal
AAL
PHA-E
LSL-150
LBL-152
MAA
MAL
MOA
LTL
GS-II (EY Lab)
Mono2W-CNL
WGA multimer
MGL 6B
CGL
Gal-3
mouse Gal-7
CA19.9-1B844
BPL
SLE
CA19.9-121SLE
IPS-K-4A2B8
PA-IL
WFL
VVL
UEA-II
UDA
HPA
Native
ME7(detected with a-HUD)
VLRB aGPA.23
HA70 complex-Alexa
MPL
Epa1-Np mutant Y228W
WGA
CTB
PhoSL
PPL001
Threefoil
CML1
WT Galectin-8
AIA
HML-Native
Epa1-Np WT
PFL
UEA-I
Human Galectin1
Galectin-1
Anti-LeC
EEL
MGL 416
DBA
HHL
HAA
#8 Lewis B (T218) (Abcam)
CSA(EY Lab)
GSL-I-B4
Sp Serotype 14.2
Mouse Galectin1
Mono1-CNL
MAA(Seikagaku)
MGL 6A
GSL-2
GSL-1
AFCN-H
CA19.9-9L426
GAC78
PsSC
CA19-9 Ab (DRG)
AAL
Epa1-Np mutant Y228A
CnSep
MGL 6A-H262T
TbFBL1
hGal-1-488+BCR
Avaren-Fc
MAL-II
WGA(Seikagaku Bio)
#14 Sialyl Lewis X (Calbio)
HPA
PHA-L
Epa1-Np mutant E227A
ACL
Gal-3C
Microvirin
PHA-L
rNV VLP-Cy3
LEL
Mono2R-CNL
AOL (Manal Lab)
GNL
UEA-I
rMpL
LcH
PNA
#15 Sialyl Lewis X (BD)
P11 VP8
PTL-II
Sp Serotype 14.15
Epa1-Np mutant E227G
ECL
BanLec
PTL-I
Mouse NKR-P1C
CnMBL
human Gal-7
TJA-II
PHA-E
NPA
SJA
VVHA lectin
DSL
LCA
SSL0-488
MNA-M
ab5
STL
BTL native
DBL-I
ME5(detected with a-HUD)
CCL2G-10077
DC-ASGPR short
GNA (EY Lab)
HHA(EY Lab)
SGA
PSL
ABA
TJA-I
WT-VCC
rPVL
RCAI (Vector)
AHL
ConA
rSRL
SRL
SSR1
WGA
TbFBL2
TL(EY Lab)
CAA
ConA
CA
SNA

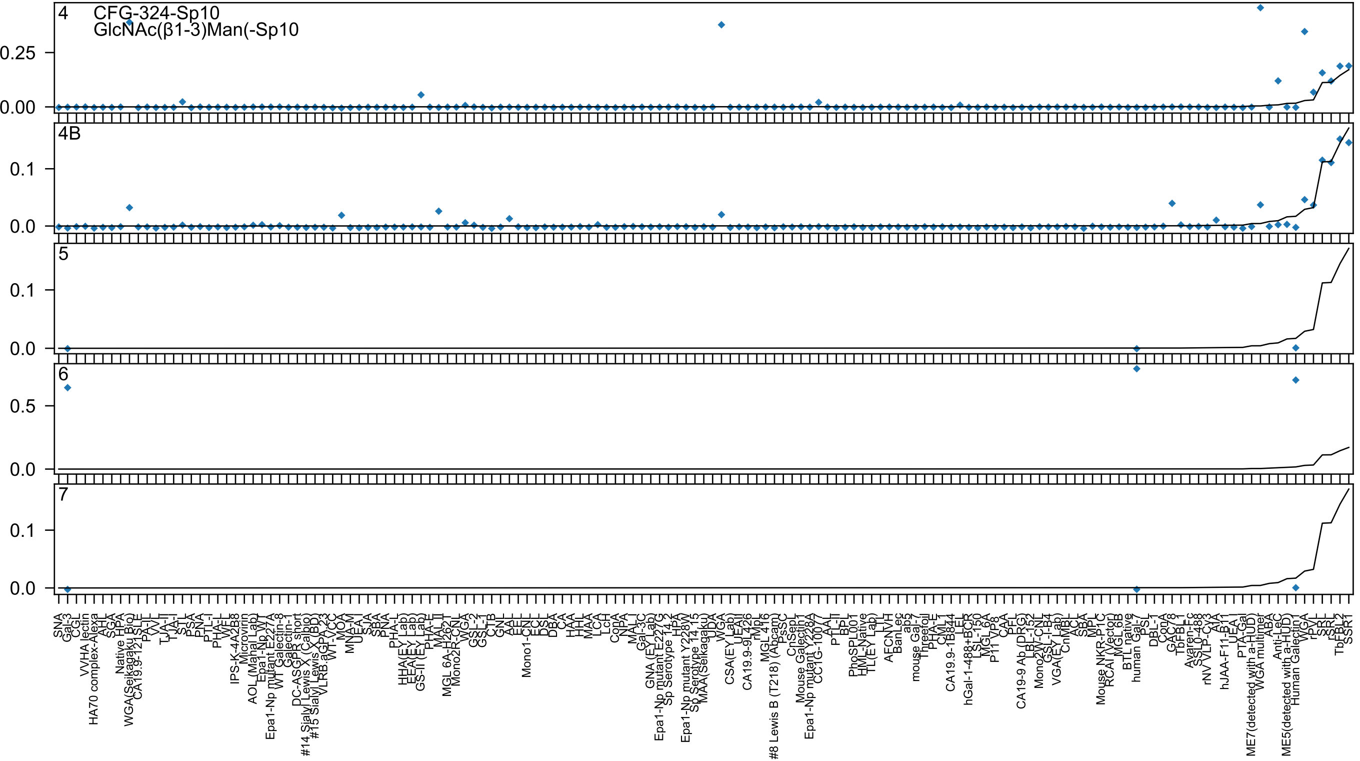
4
CFG-324-Sp10
GlcNAc(β1-3)Man(-Sp10
0.25
0.00
4B
0.1
0.0
5
0.1
0.0
6
0.5
0.0
7
0.1
0.0

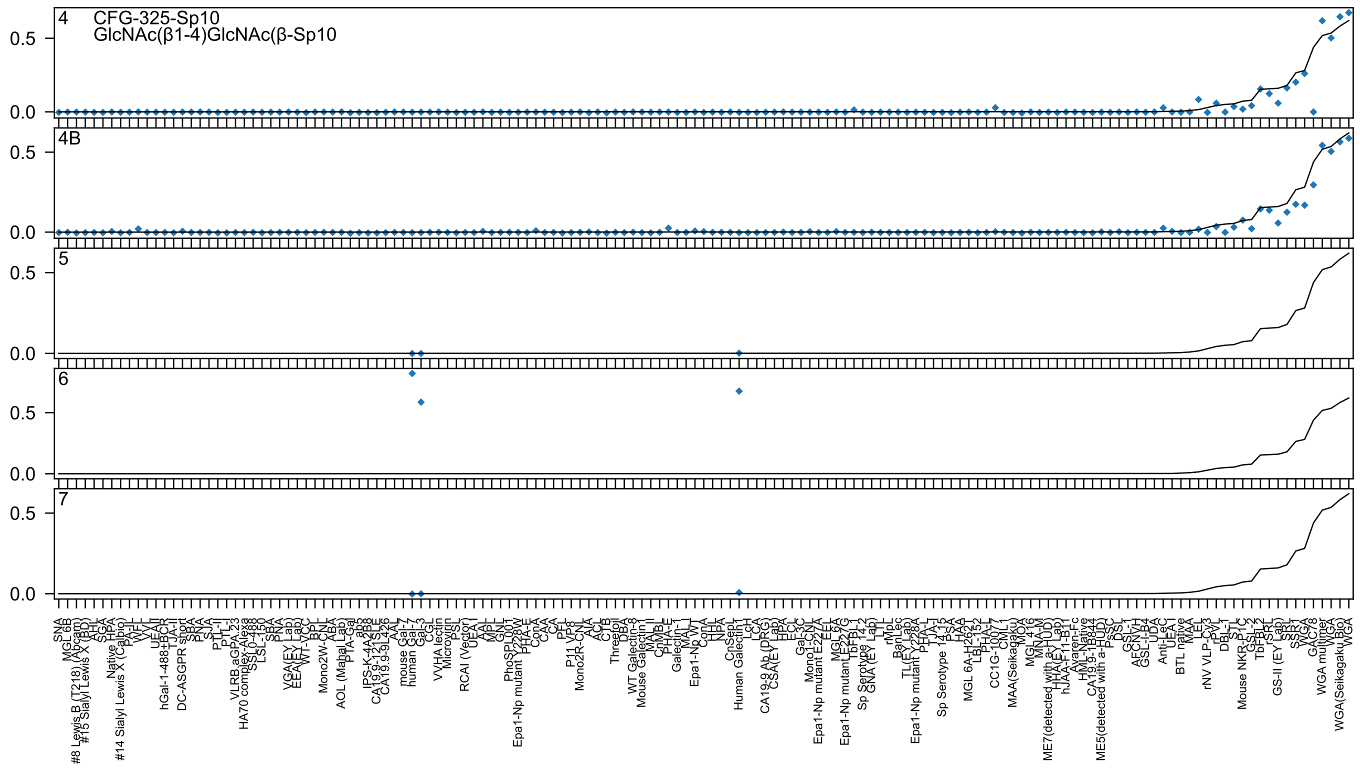
4 CFG-325-Sp10
GlcNAc(β1-4)GlcNAc(β-Sp10
4B
5
6
7
0.5
0.0

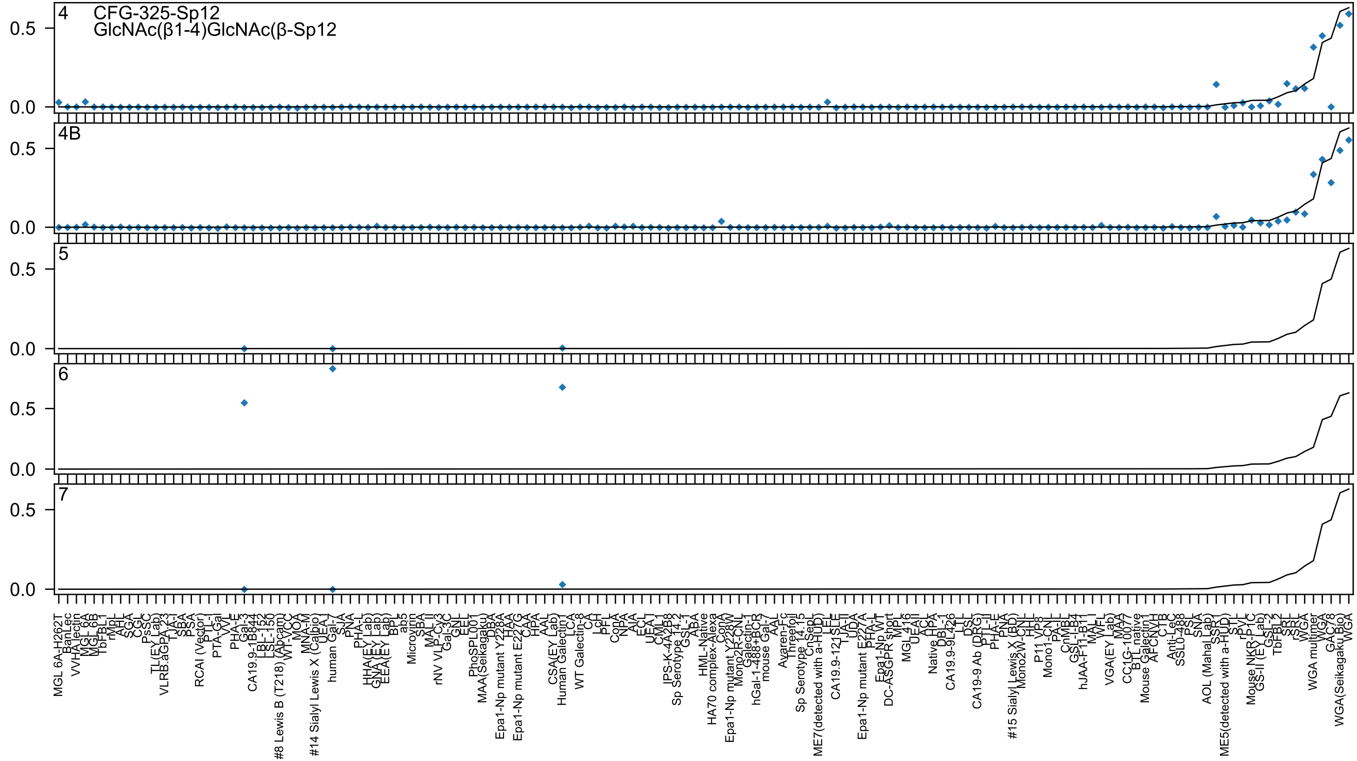

4
CFG-325-Sp12
GlcNAc(β1-4)GlcNAc(β-Sp12
4B
5
6
7

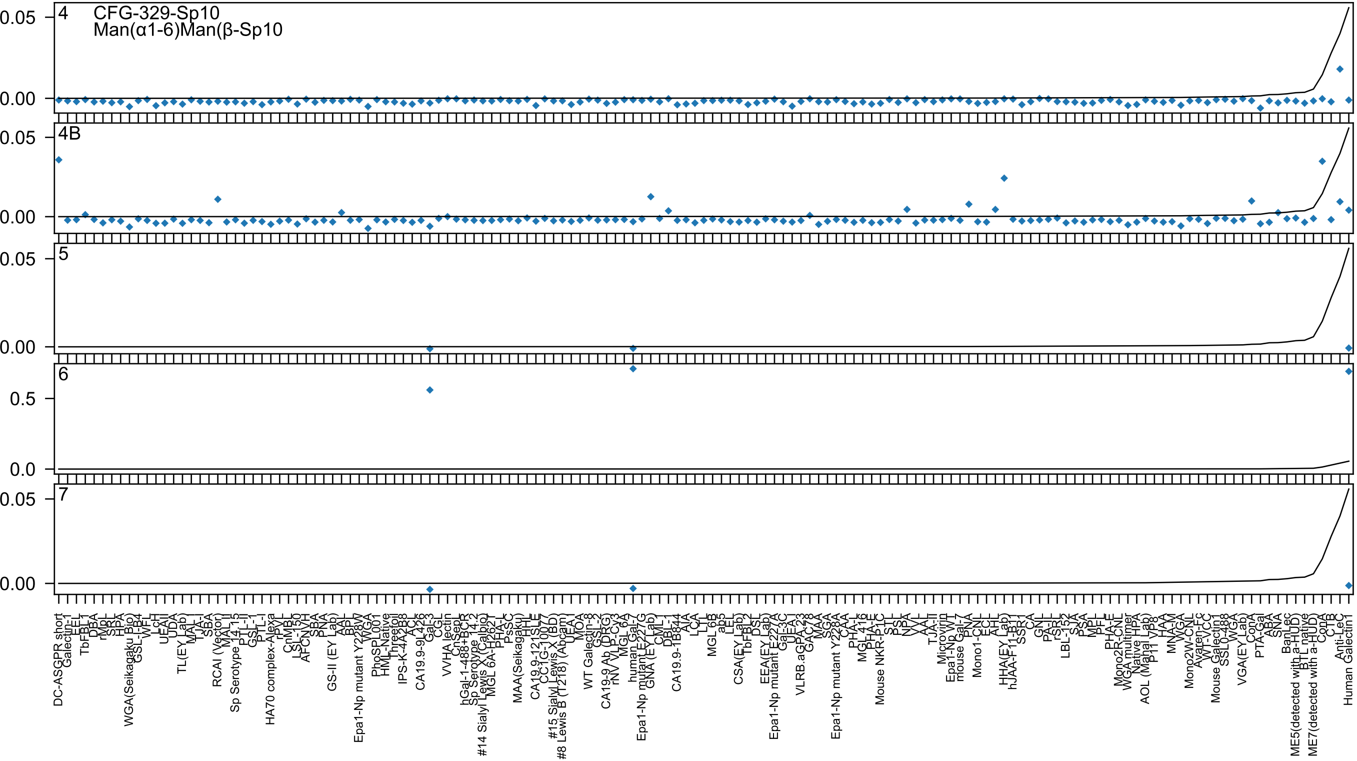

4
CFG-329-Sp10
Man(α1-6)Man(β-Sp10
4B
5
6
7
0.05
0.00
0.5
0.0

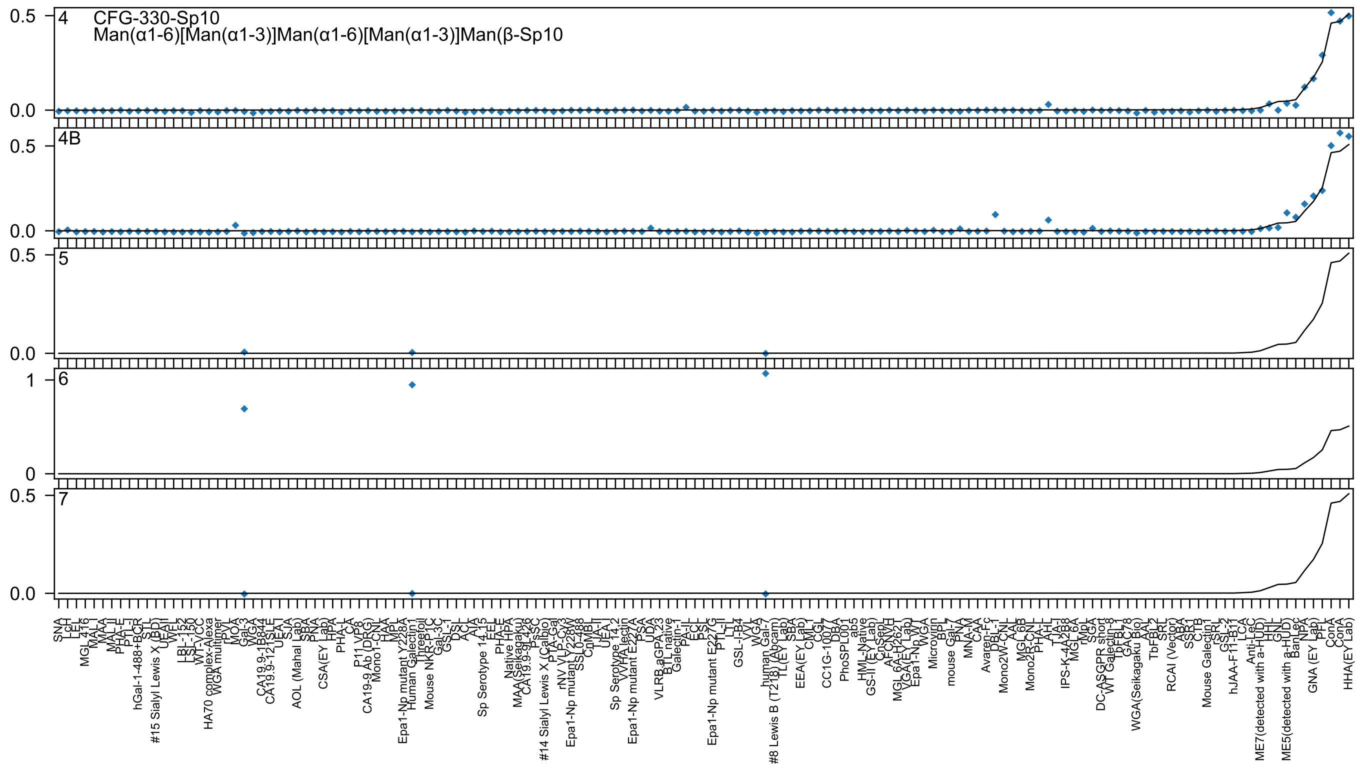

4
CFG-330-Sp10
Man(α1-6)[Man(α1-3)]Man(α1-6)[Man(α1-3)]Man(β-Sp10
4B
5
6
7

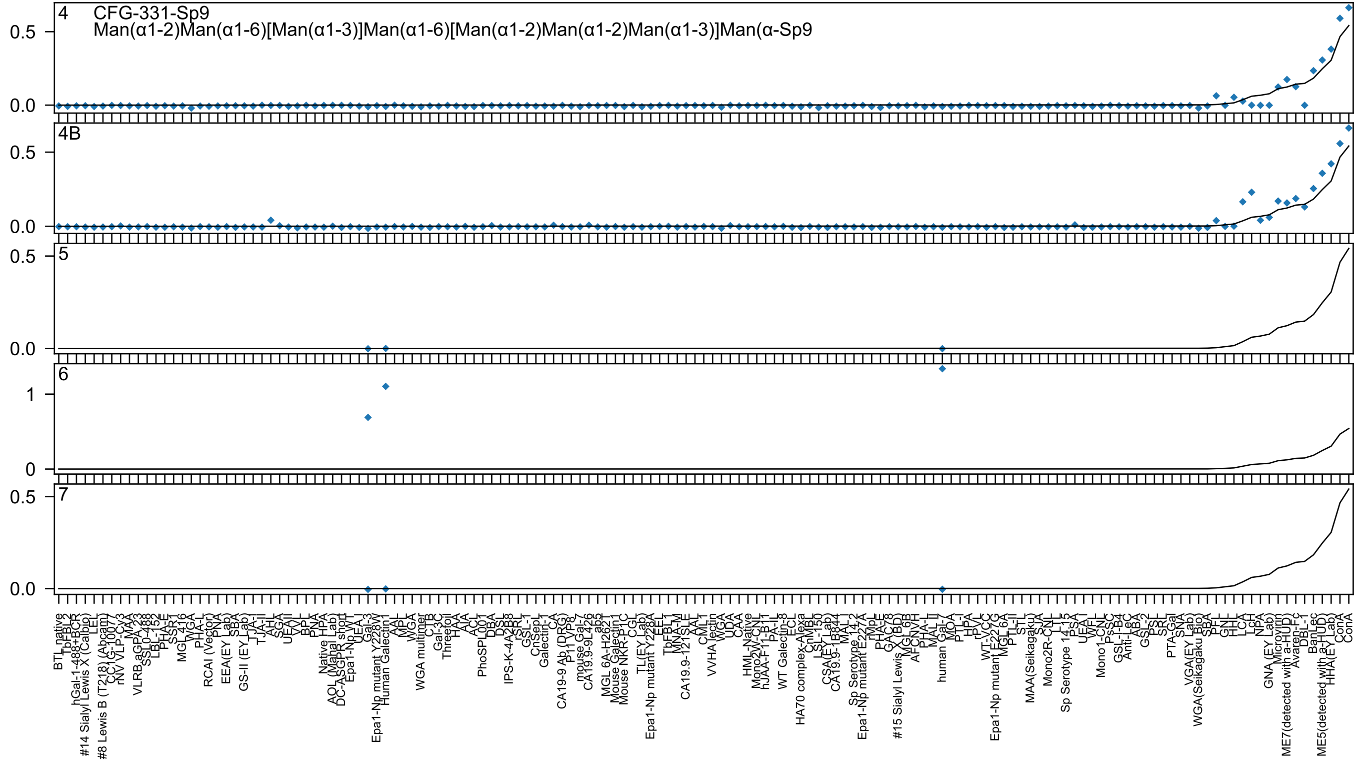

4 CFG-331-Sp9
Man(α1-2)Man(α1-6)[Man(α1-3)]Man(α1-6)[Man(α1-2)Man(α1-2)Man(α1-3)]Man(α-Sp9
4B
5
6
7

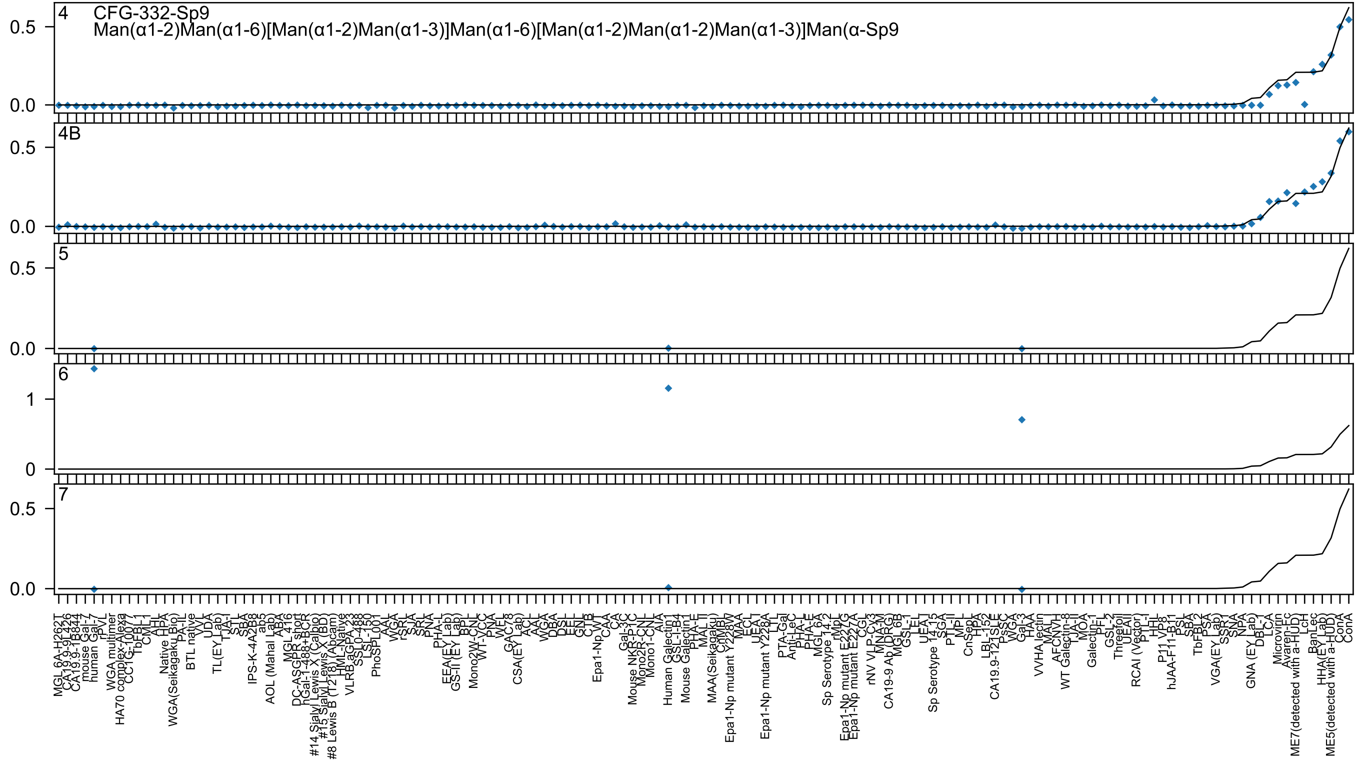

4
CFG-332-Sp9
Man(α1-2)Man(α1-6)[Man(α1-2)Man(α1-3)]Man(α1-6)[Man(α1-2)Man(α1-2)Man(α1-3)]Man(α-Sp9
4B
5
6
7

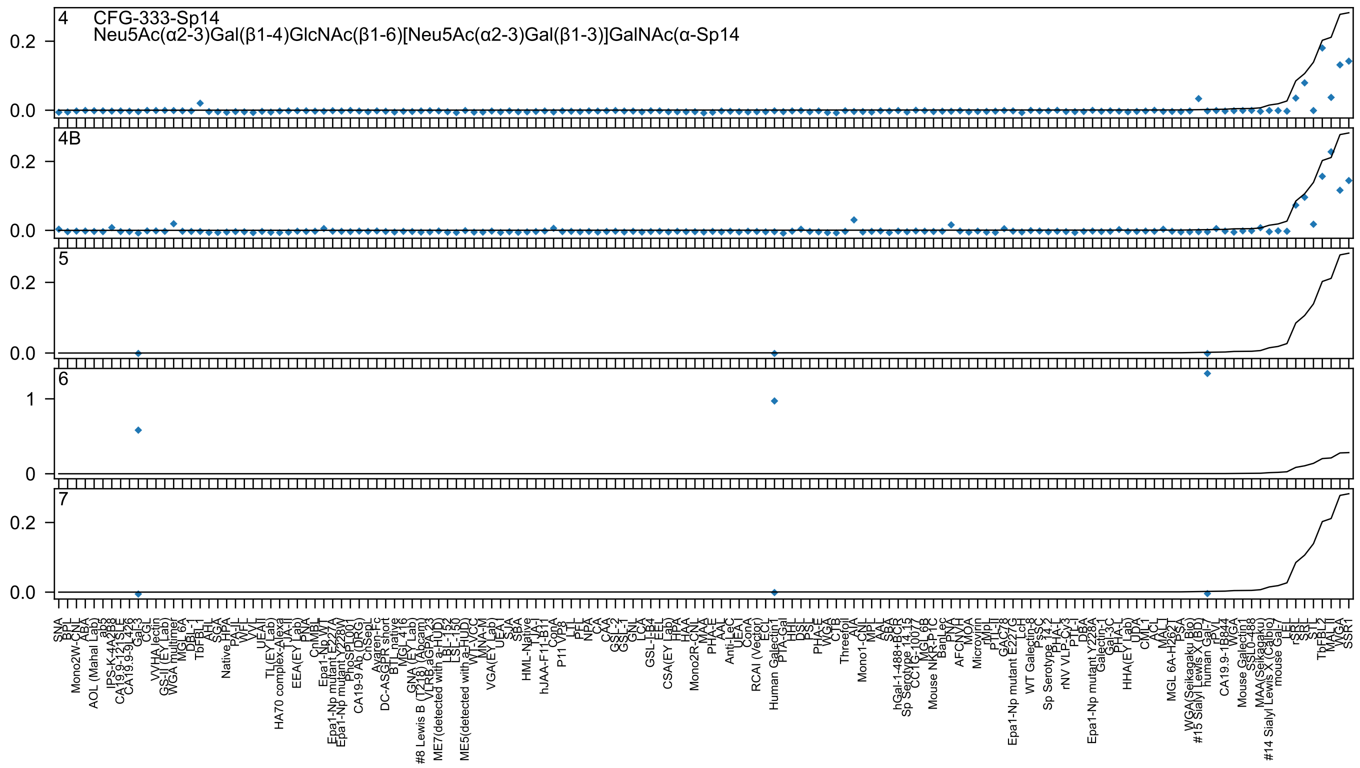

4 CFG-333-Sp14
Neu5Ac(α2-3)Gal(β1-4)GlcNAc(β1-6)[Neu5Ac(α2-3)Gal(β1-3)]GalNAc(α-Sp14
4B
5
6
7
0.2
0.0
1
0

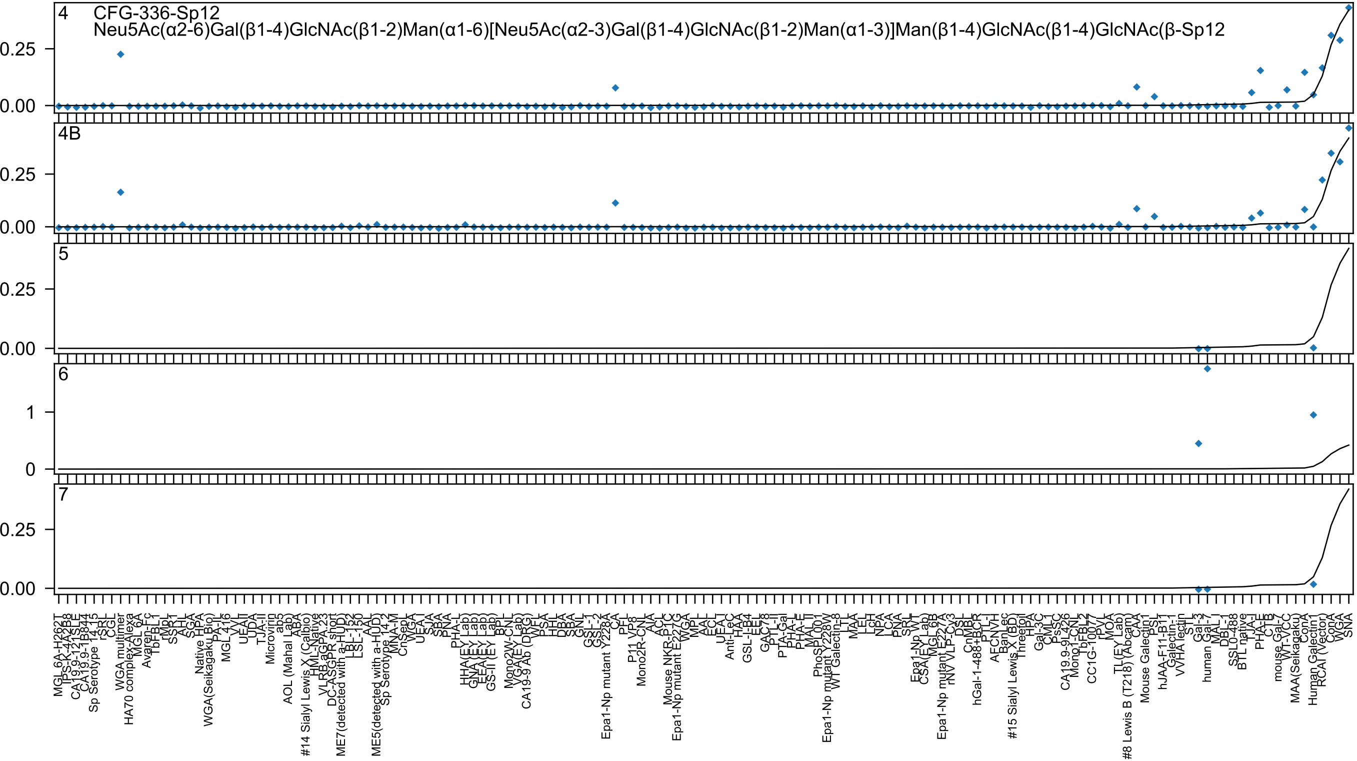

CFG-336-Sp12
Neu5Ac(α2-6)Gal(β1-4)GlcNAc(β1-2)Man(α1-6)[Neu5Ac(α2-3)Gal(β1-4)GlcNAc(β1-2)Man(α1-3)]Man(β1-4)GlcNAc(β1-4)GlcNAc(β-Sp12
4
4B
5
6
7
0.25
0.00
1
0

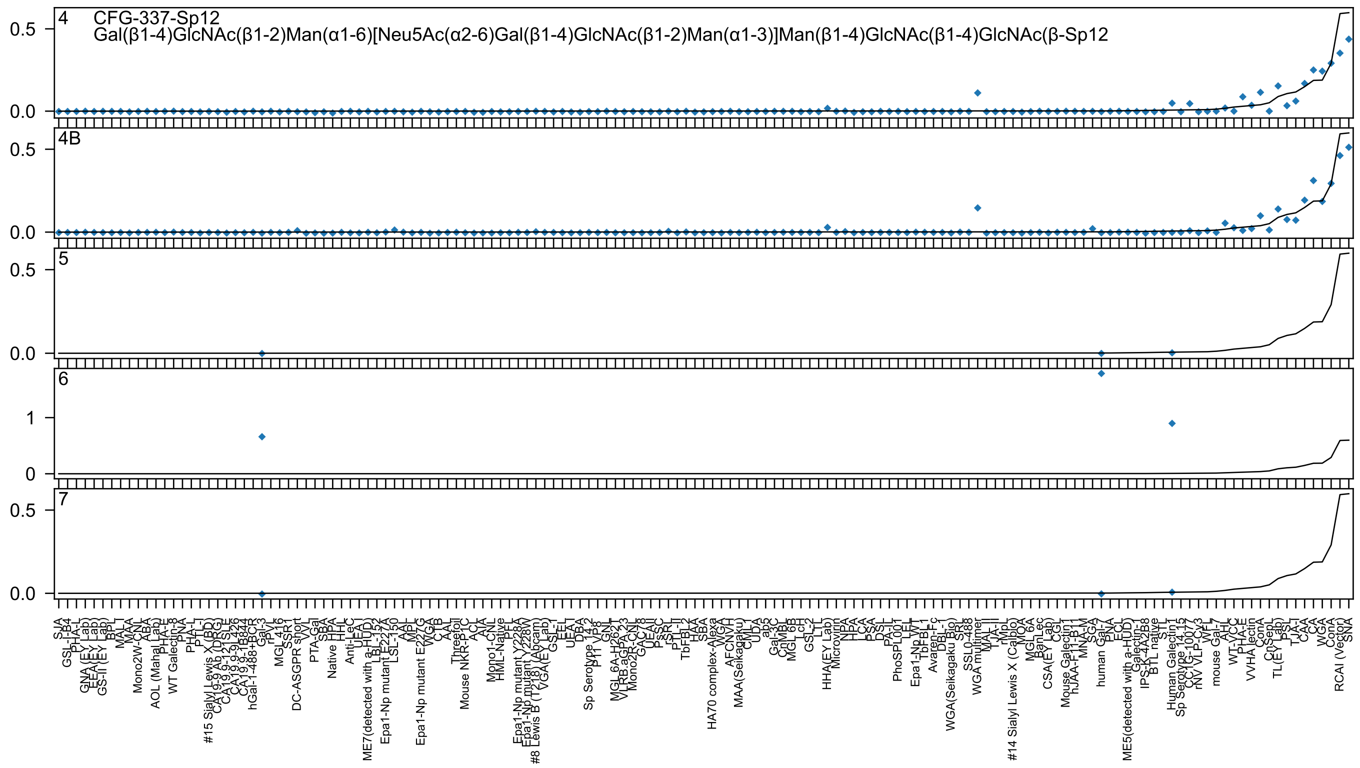
CFG-337-Sp12
Gal(β1-4)GlcNAc(β1-2)Man(α1-6)[Neu5Ac(α2-6)Gal(β1-4)GlcNAc(β1-2)Man(α1-3)]Man(β1-4)GlcNAc(β1-4)GlcNAc(β-Sp12
4
4B
5
6
7

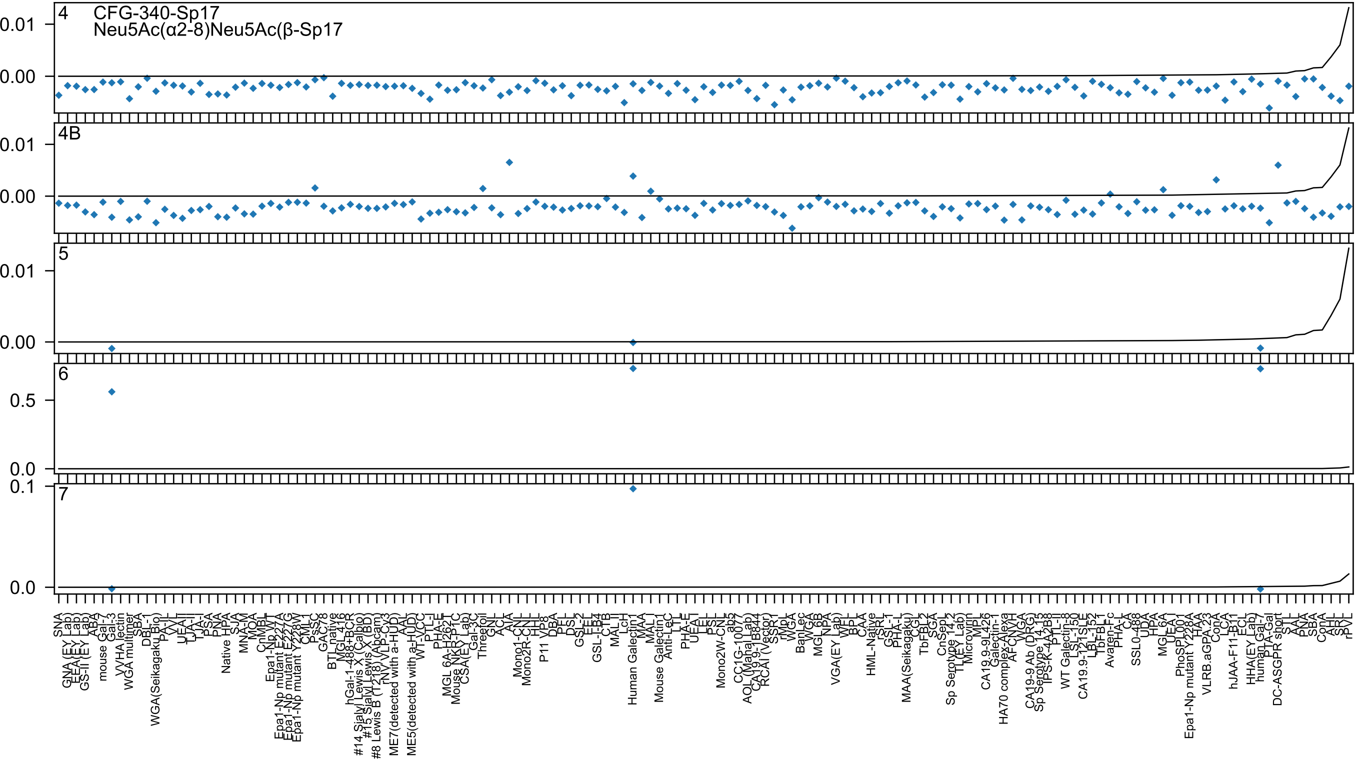
4
CFG-340-Sp17
Neu5Ac(α2-8)Neu5Ac(β-Sp17
0.01
0.00
4B
0.01
0.00
5
0.01
0.00
6
0.5
0.0
0.1
7
0.0

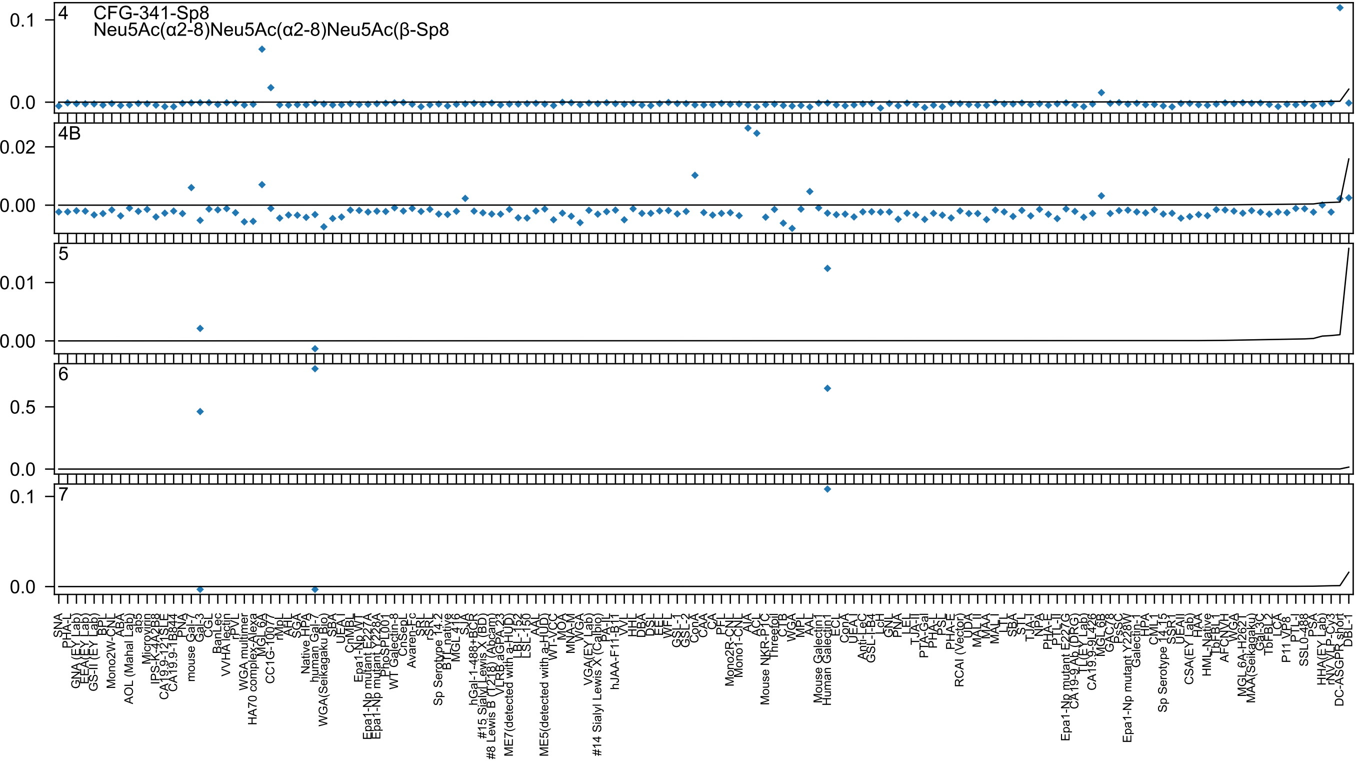

CFG-341-Sp8
Neu5Ac(α2-8)Neu5Ac(α2-8)Neu5Ac(β-Sp8
4
4B
5
6
7

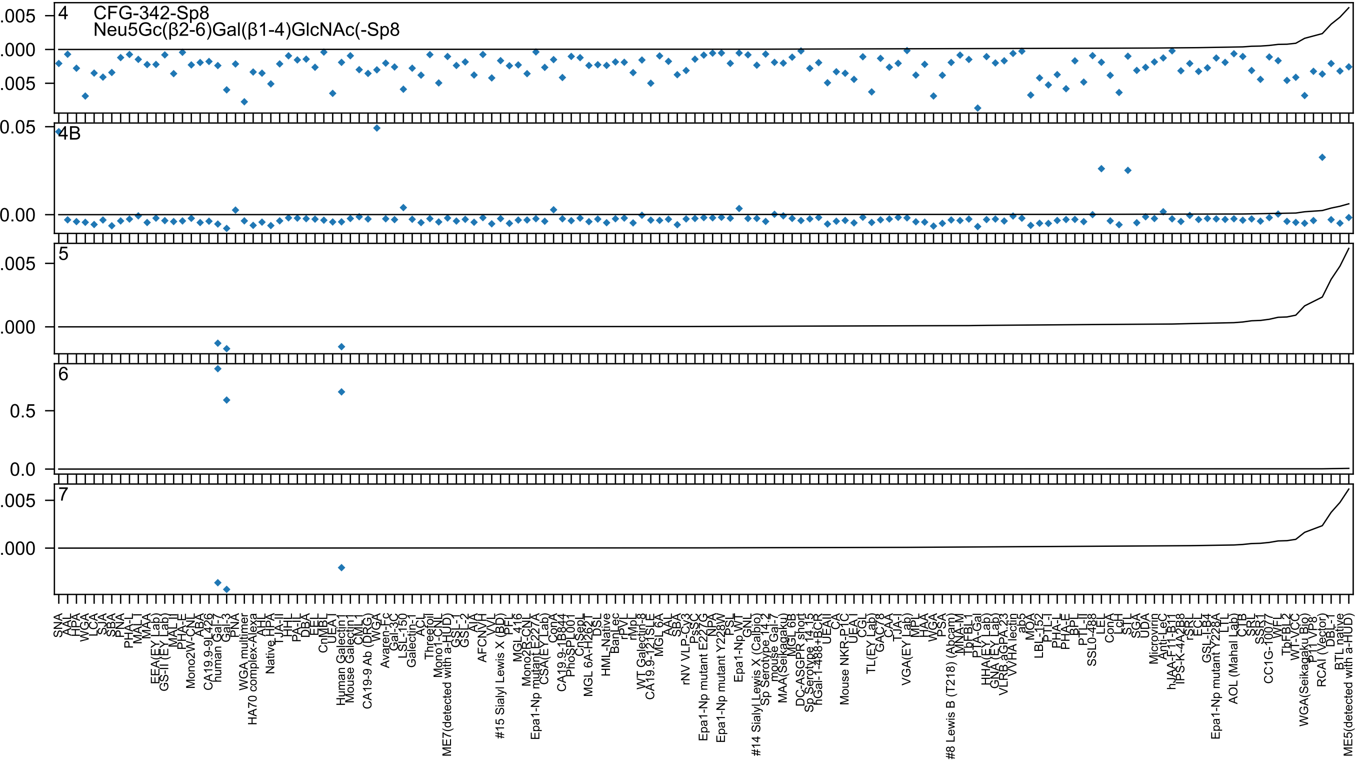

4
CFG-342-Sp8
Neu5Gc(β2-6)Gal(β1-4)GlcNAc(-Sp8
4B
5
6
7

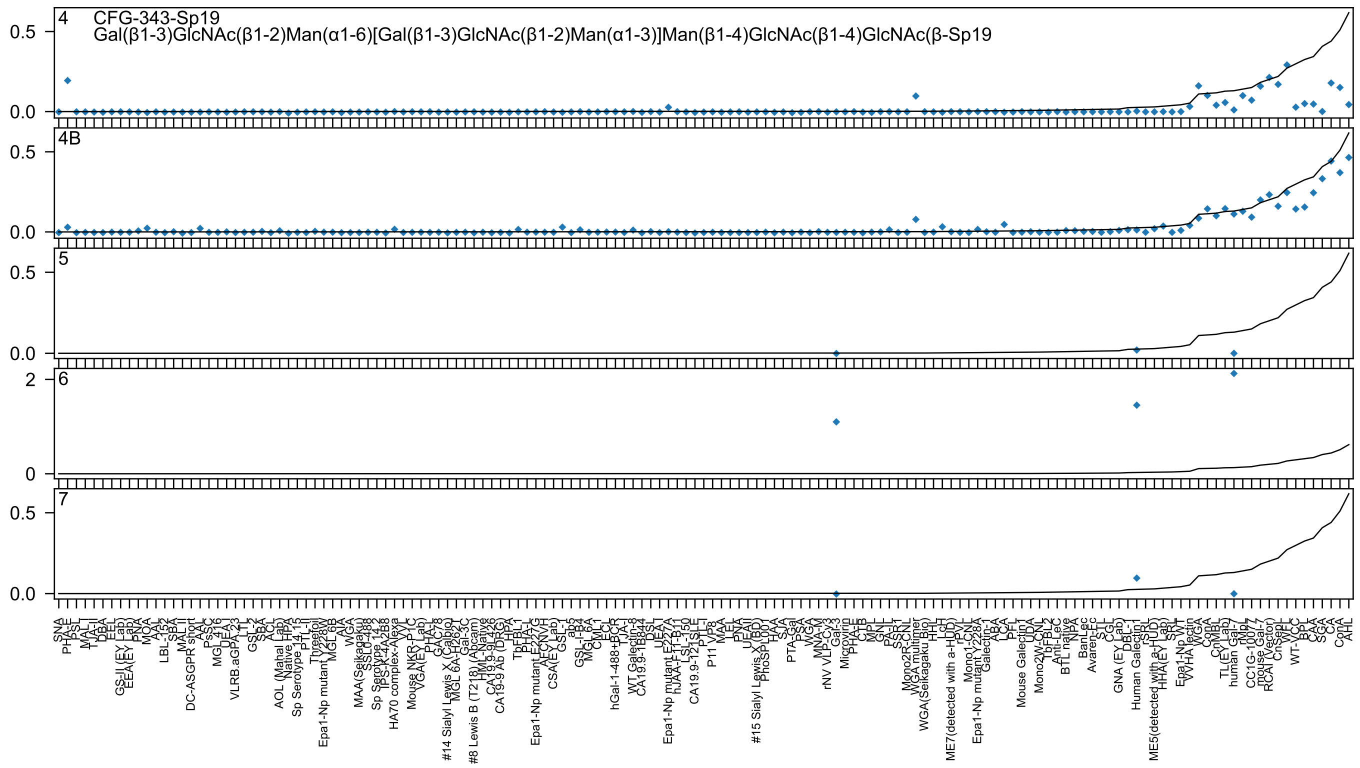
4
CFG-343-Sp19
Gal(β1-3)GlcNAc(β1-2)Man(α1-6)[Gal(β1-3)GlcNAc(β1-2)Man(α1-3)]Man(β1-4)GlcNAc(β1-4)GlcNAc(β-Sp19
4B
5
6
7

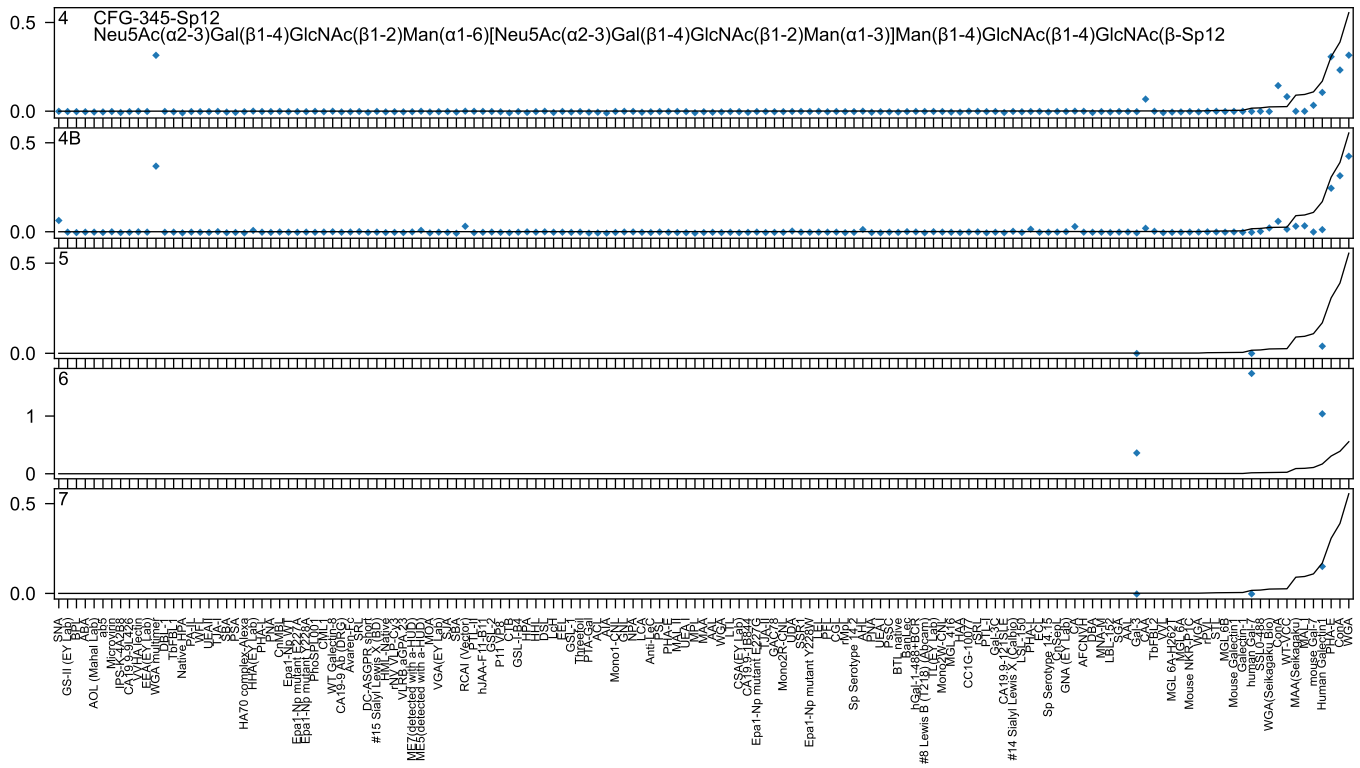

4 CFG-345-Sp12
Neu5Ac(α2-3)Gal(β1-4)GlcNAc(β1-2)Man(α1-6)[Neu5Ac(α2-3)Gal(β1-4)GlcNAc(β1-2)Man(α1-3)]Man(β1-4)GlcNAc(β1-4)GlcNAc(β-Sp12
4B
5
6
7

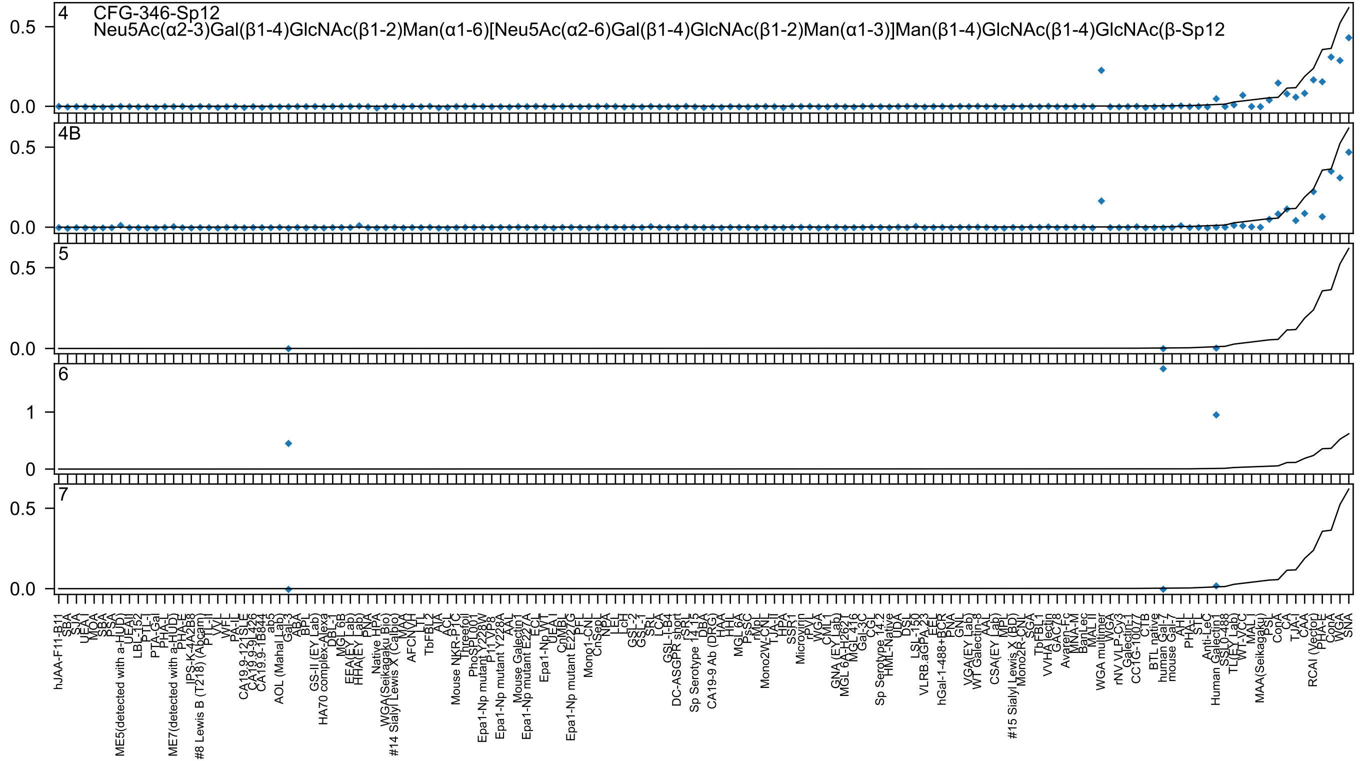

4 CFG-346-Sp12
Neu5Ac(α2-3)Gal(β1-4)GlcNAc(β1-2)Man(α1-6)[Neu5Ac(α2-6)Gal(β1-4)GlcNAc(β1-2)Man(α1-3)]Man(β1-4)GlcNAc(β1-4)GlcNAc(β-Sp12
4B
5
6
7

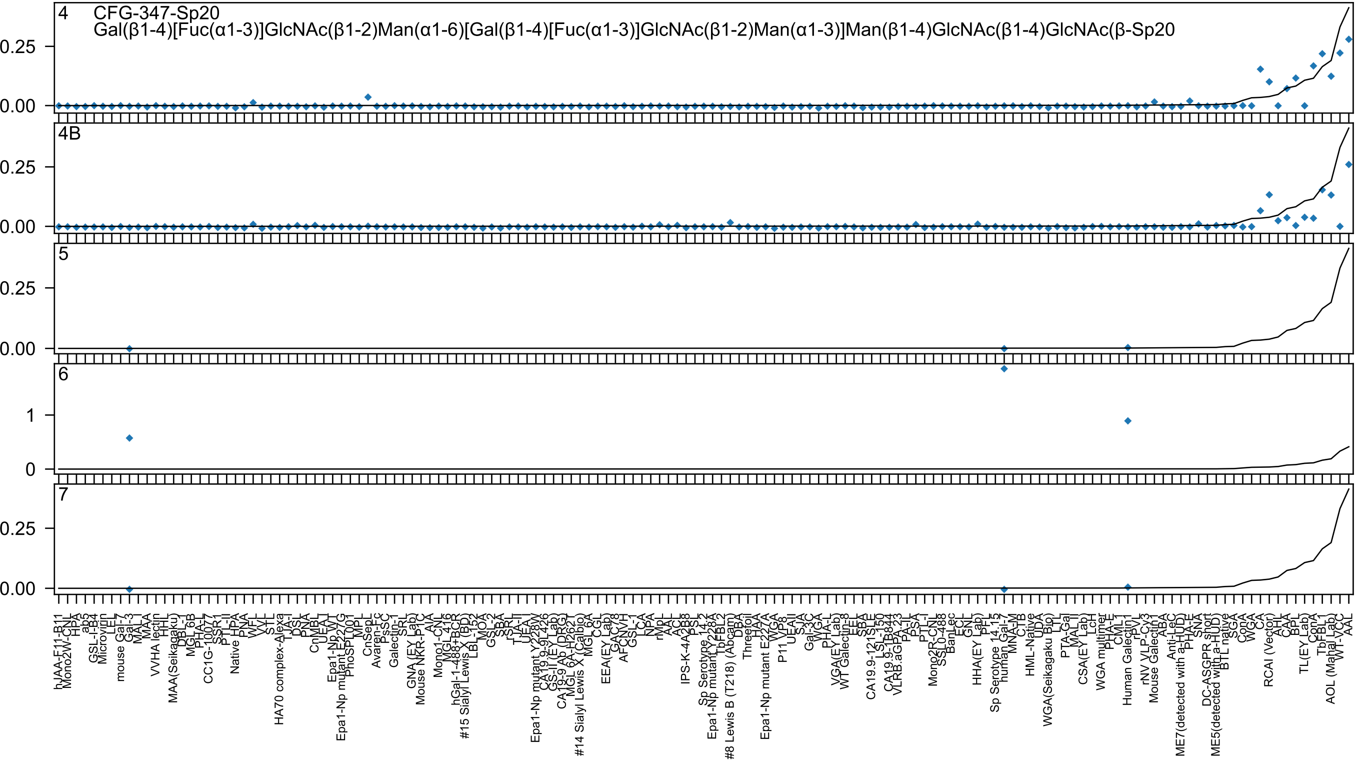

4
CFG-347-Sp20
Gal(β1-4)[Fuc(α1-3)]GlcNAc(β1-2)Man(α1-6)[Gal(β1-4)[Fuc(α1-3)]GlcNAc(β1-2)Man(α1-3)]Man(β1-4)GlcNAc(β1-4)GlcNAc(β-Sp20
4B
5
6
7
0.25
0.00
1
0

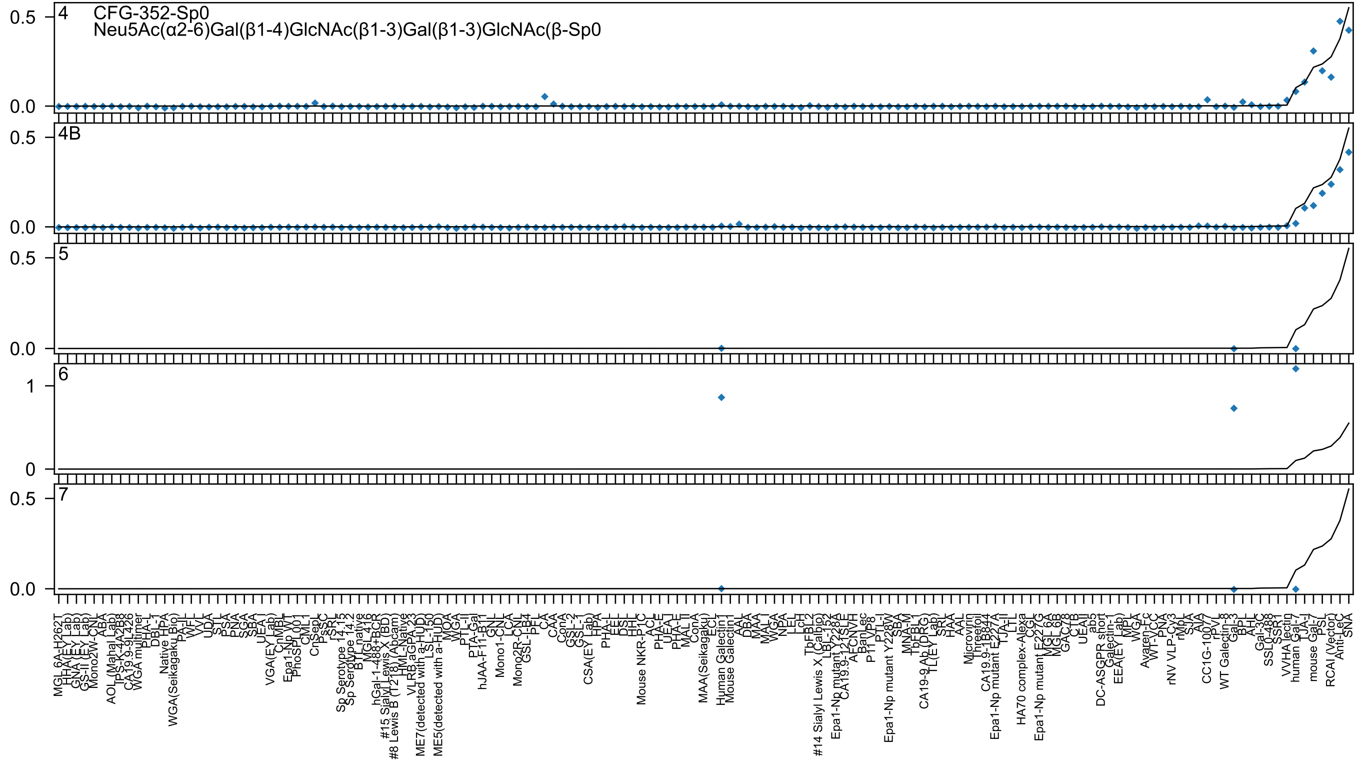

CFG-352-Sp0
Neu5Ac(α2-6)Gal(β1-4)GlcNAc(β1-3)Gal(β1-3)GlcNAc(β-Sp0
4
4B
5
6
7

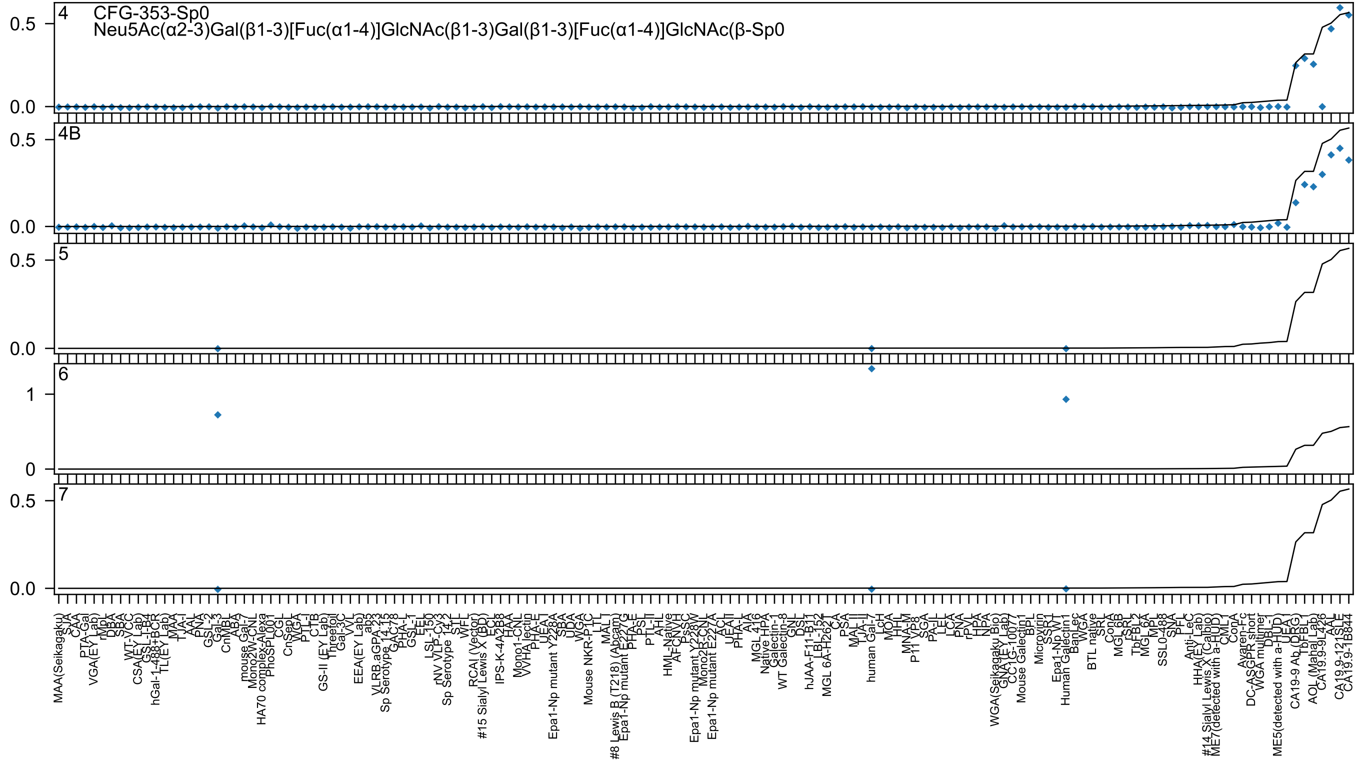

4
CFG-353-Sp0
Neu5Ac(α2-3)Gal(β1-3)[Fuc(α1-4)]GlcNAc(β1-3)Gal(β1-3)[Fuc(α1-4)]GlcNAc(β-Sp0
4B
5
6
7

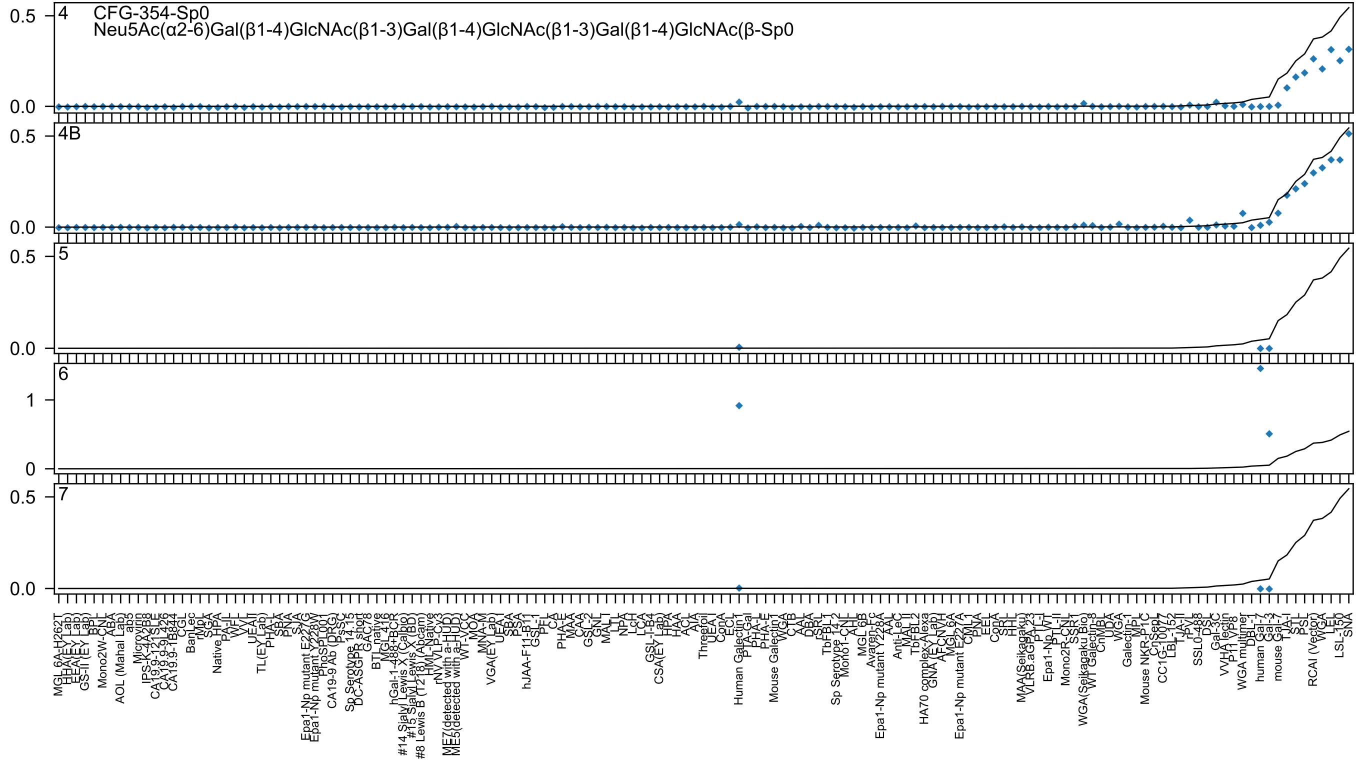
4 CFG-354-Sp0
Neu5Ac(α2-6)Gal(β1-4)GlcNAc(β1-3)Gal(β1-4)GlcNAc(β1-3)Gal(β1-4)GlcNAc(β-Sp0
4B
5
6
7
0.5
0.0
1
0
Human Galectin1
human Gal-7
Gal-3
mouse Gal-7
RCAI (Vector)
WGA
LEL
LSL-150
SNA

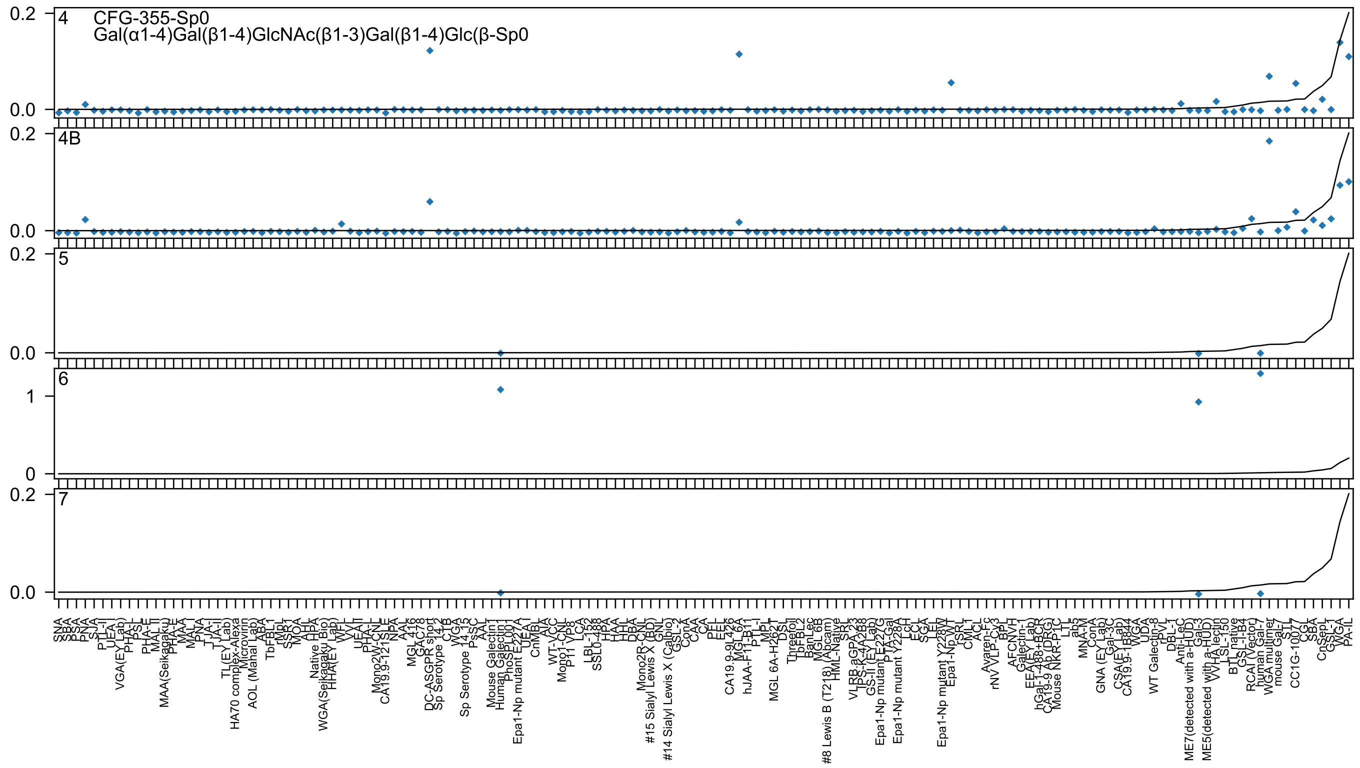

4
CFG-355-Sp0
Gal(α1-4)Gal(β1-4)GlcNAc(β1-3)Gal(β1-4)Glc(β-Sp0
4B
5
6
7
0.2
0.0
1
0

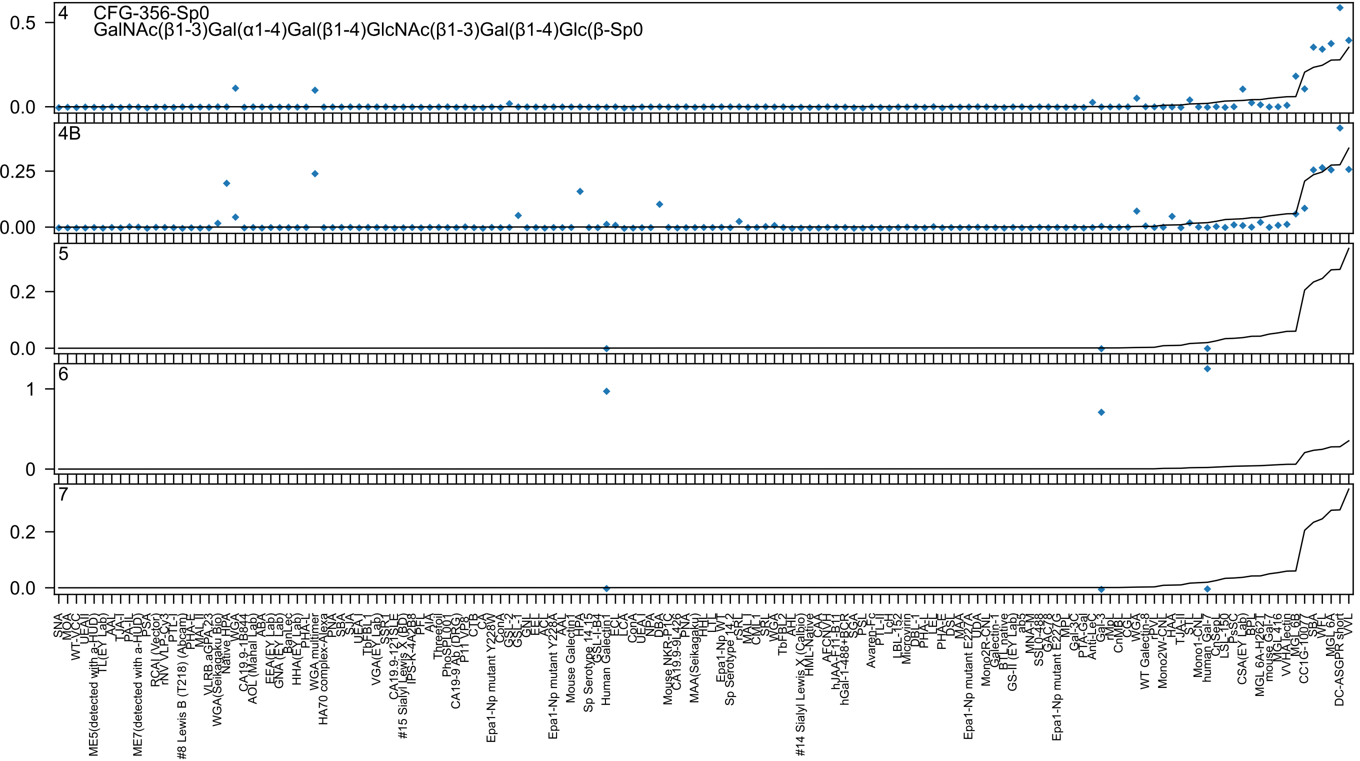

CFG-356-Sp0
GalNAc(β1-3)Gal(α1-4)Gal(β1-4)GlcNAc(β1-3)Gal(β1-4)Glc(β-Sp0
4
4B
5
6
7

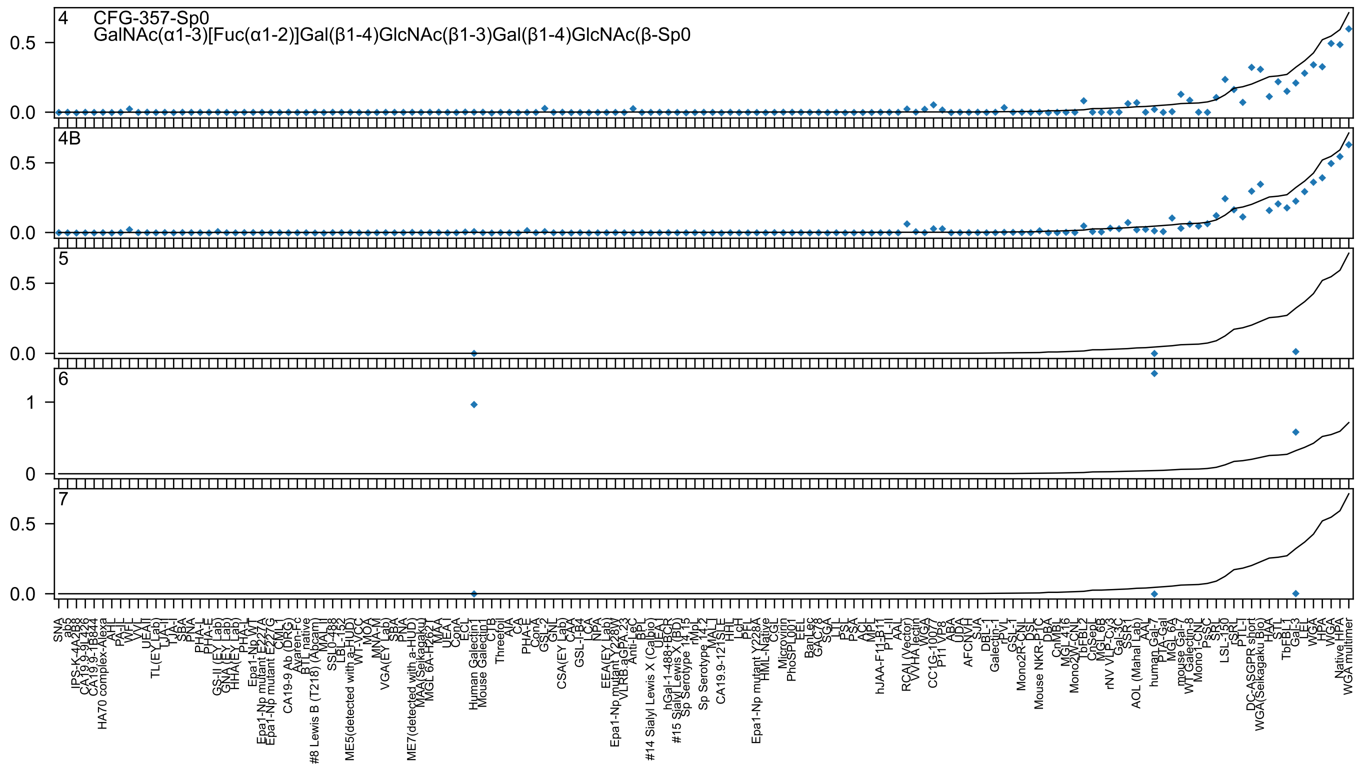
4 CFG-357-Sp0
GalNAc(α1-3)[Fuc(α1-2)]Gal(β1-4)GlcNAc(β1-3)Gal(β1-4)GlcNAc(β-Sp0
4B
5
6
7
0.0
0.5
0
1
SNA
ab5
IPS-K-4A2B8
CA19.9-9L426
CA19.9-1B844
HA70 complex-Alexa
AAL
PA-IL
WFL
VVL
UEA-II
TL(EY Lab)
TJA-II
TJA-I
SBA
PNA
PHA-L
PHA-E
GS-II (EY Lab)
GNA (EY Lab)
HHA(EY Lab)
PHA-L
Epa1-Np WT
Epa1-Np mutant E227A
Epa1-Np mutant E227G
CML-I
CA19-9 Ab (DRG)
Avaren-Fc
BTL native
#8 Lewis B (T218) (Abcam)
MAL-I
SSL0-488
LBL-152
ME5(detected with a-HUD)
WT-VCC
MOA
MNA-M
VGA(EY Lab)
SBA
PNA
ME7(detected with a-HUD)
MAA(Seikagaku)
MGL 6A-H262T
MAA
UEA-I
ConA
ECL
Human Galectin1
Mouse Galectin1
CTB
Threefoil
AIA
CA
PHA-E
ConA
GSL-2
GNL
CSA(EY Lab)
CAA
GSL-I-B4
LCA
NPA
EEA(EY Lab)
Epa1-Np mutant Y228W
VLRB.aGPA.23
Anti-LeC
BPL
UEA-I
#14 Sialyl Lewis X (Calbio)
hGal-1-488+BCR
#15 Sialyl Lewis X (BD)
Sp Serotype 14.15
rMpL
Sp Serotype 14.2
CA19.9-121SLE
MAL-I
HHL
LcH
PEL
Epa1-Np mutant Y228A
HML-Native
CGL
Microvirin
PhoSL001
EEL
BanLec
GAC78
SGA
LTL
PSL
PSA
ACL
MPL
hJAA-F11-B11
PTL-II
AAL
RCAI (Vector)
VVHA lectin
WGA
CC1G-10077
P11 VP8
ABA
UDA
AFCNVH
SJA
DBL-1
Galectin-1
rPVL
GSL-1
Mono2R-CNL
DSL
Mouse NKR-P1C
DBA
CnMBL
MGL 416
Mono2W-CNL
TbFBL2
CnSep1
MGL 6B
rNV VLP-Cy3
Gal-3C
SSR1
AOL (Mahal Lab)
AAL
human Gal-7
PTA-Gal
MGL 6A
mouse Gal-7
WT Galectin-8
Mono1-CNL
PsSC
SRL
LSL-150
rSRL
PTL-I
DC-ASGPR short
WGA(Seikagaku Bio)
HAA
STL
TbFBL1
Gal-3
LEL
WGA
HPA
WGA
Native HPA
WGA multimer

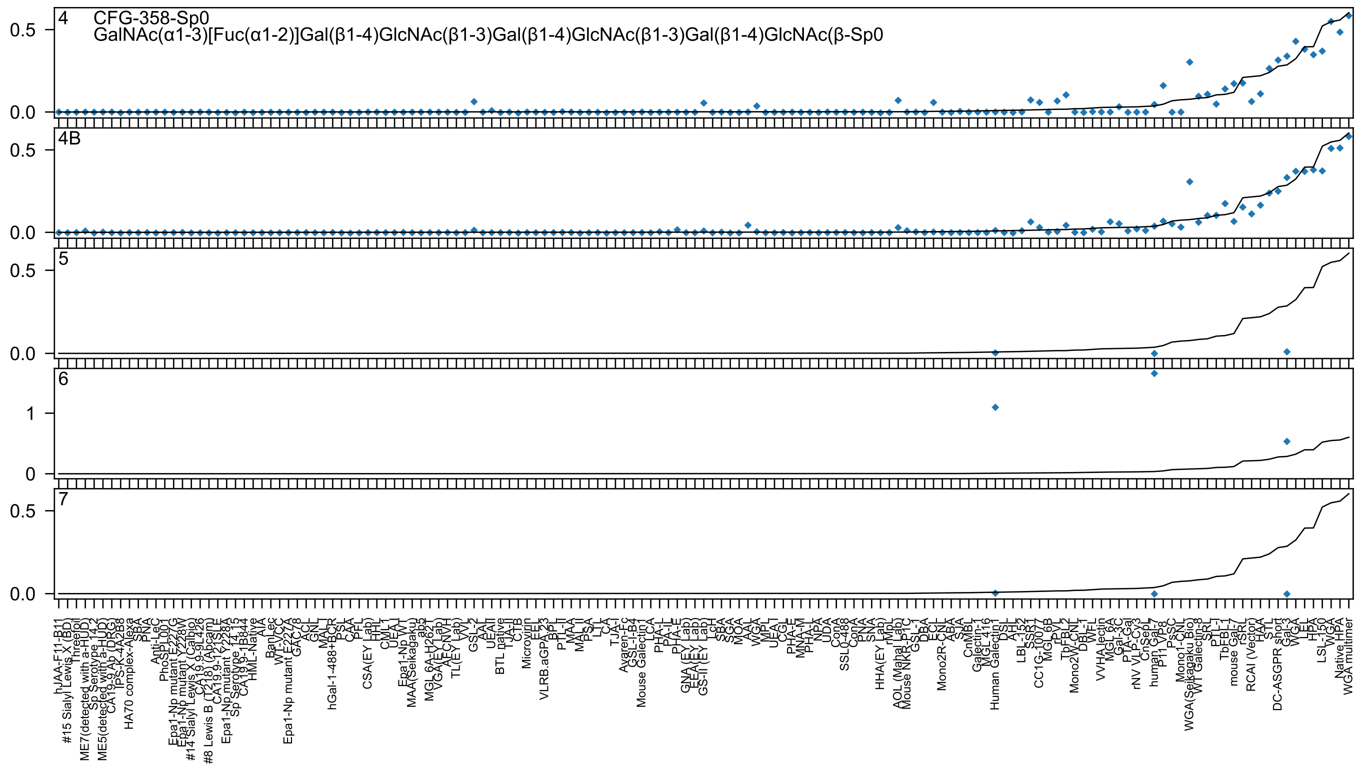

CFG-358-Sp0
GalNAc(α1-3)[Fuc(α1-2)]Gal(β1-4)GlcNAc(β1-3)Gal(β1-4)GlcNAc(β1-3)Gal(β1-4)GlcNAc(β-Sp0
4
4B
5
6
7

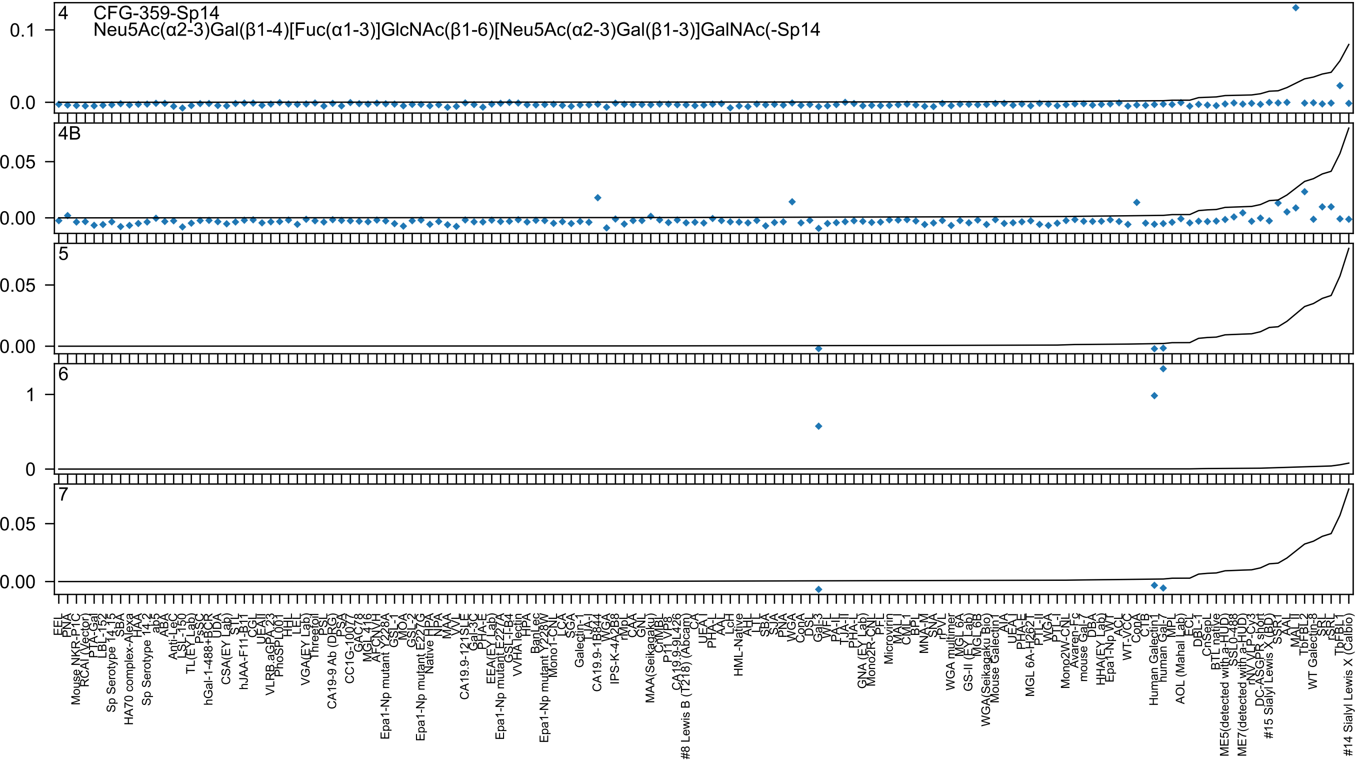
4
CFG-359-Sp14
Neu5Ac(α2-3)Gal(β1-4)[Fuc(α1-3)]GlcNAc(β1-6)[Neu5Ac(α2-3)Gal(β1-3)]GalNAc(-Sp14
0.1
0.0
4B
0.05
0.00
5
0.05
0.00
6
1
0
7
0.05
0.00

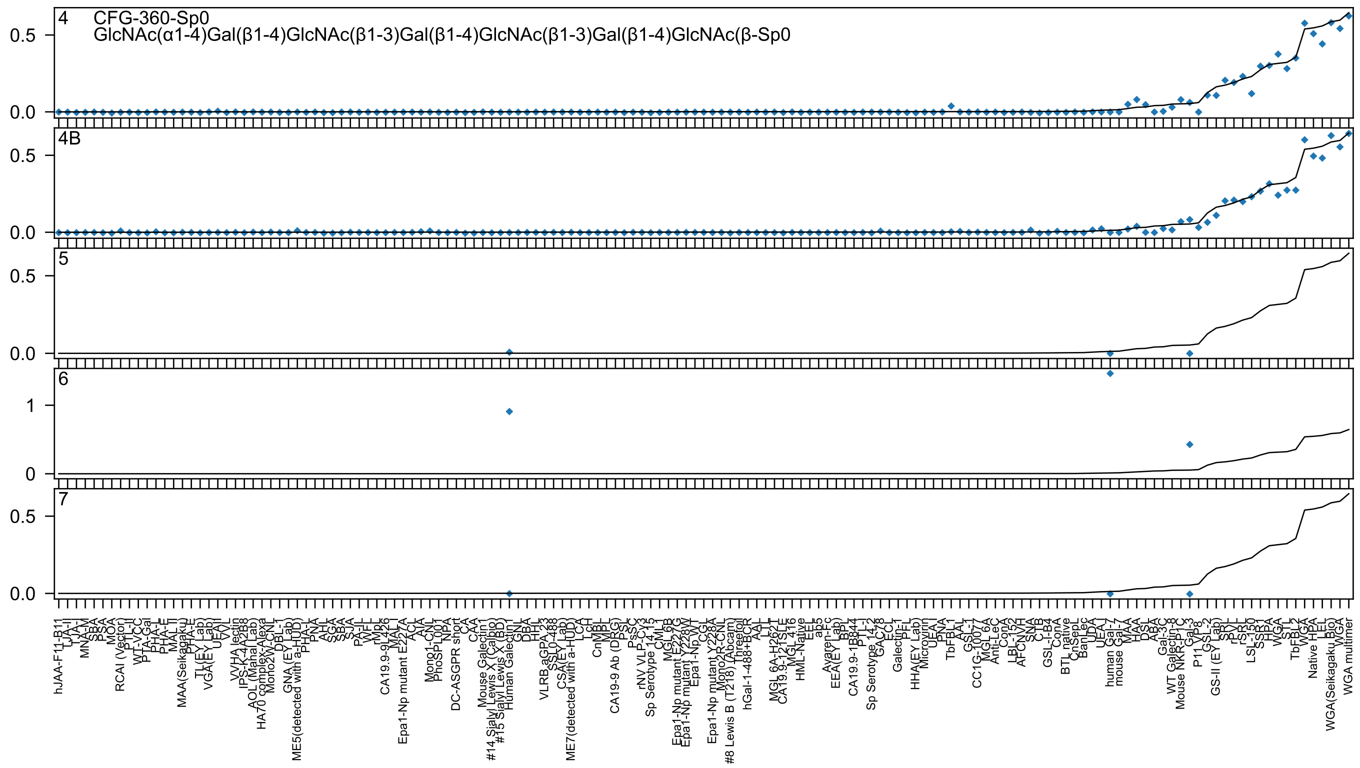

4
CFG-360-Sp0
GlcNAc(α1-4)Gal(β1-4)GlcNAc(β1-3)Gal(β1-4)GlcNAc(β1-3)Gal(β1-4)GlcNAc(β-Sp0
4B
5
6
7

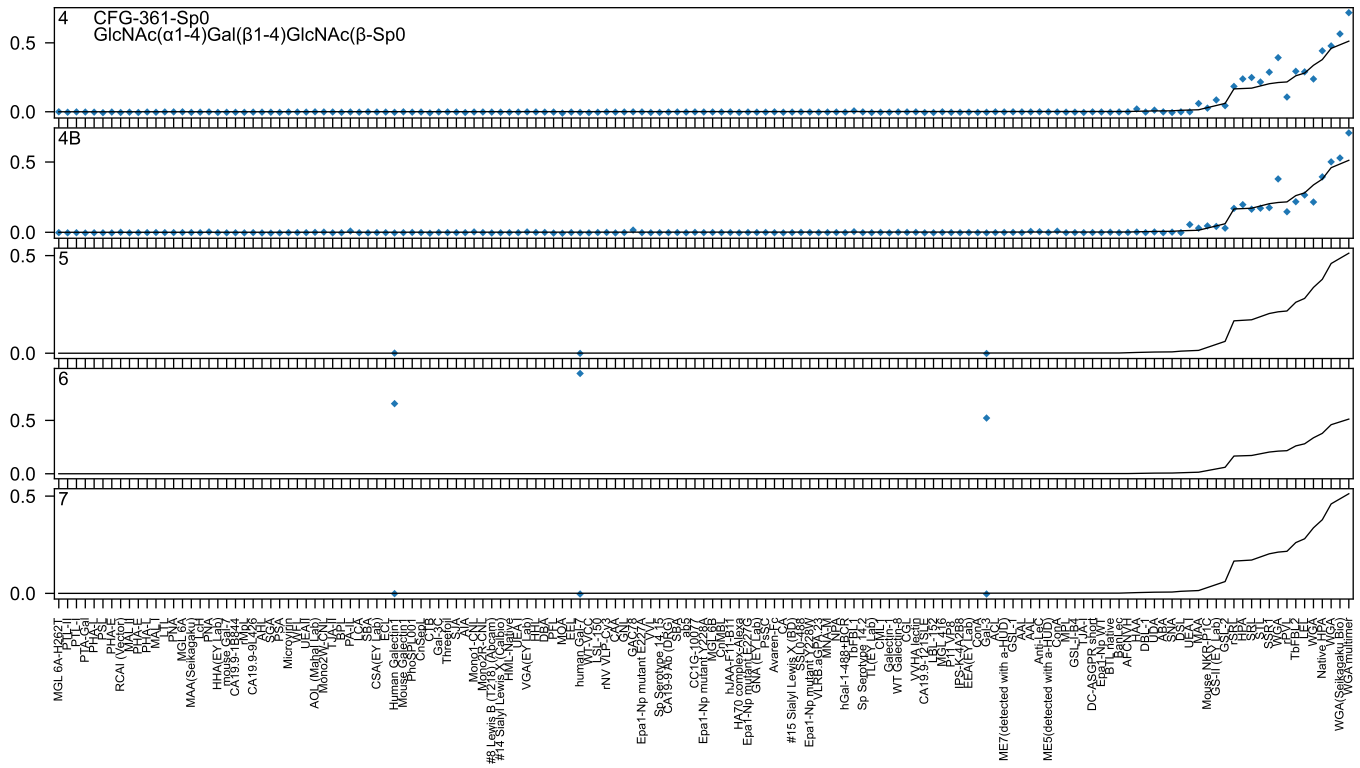
4
CFG-361-Sp0
GlcNAc(α1-4)Gal(β1-4)GlcNAc(β-Sp0
4B
5
6
7
0.5
0.0

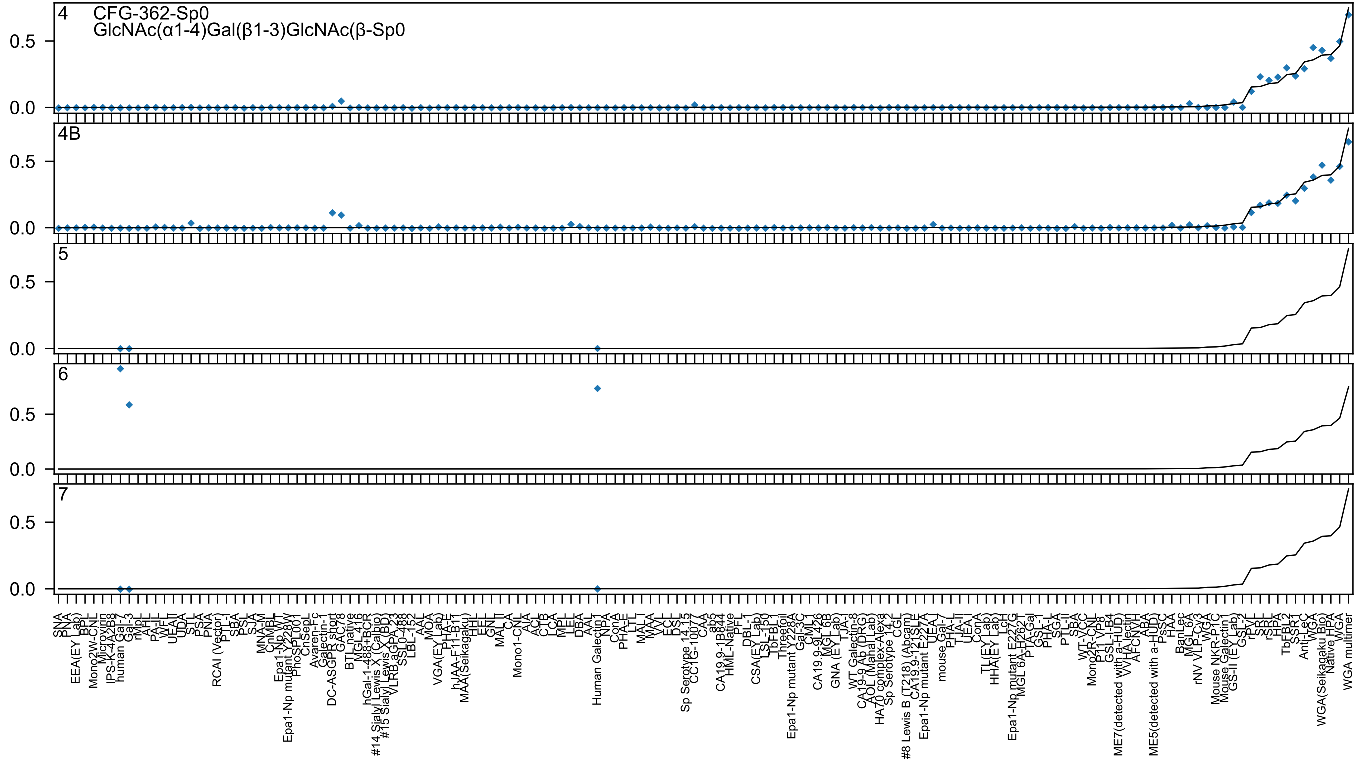
4
CFG-362-Sp0
GlcNAc(α1-4)Gal(β1-3)GlcNAc(β-Sp0
4B
5
6
7
0.5
0.0

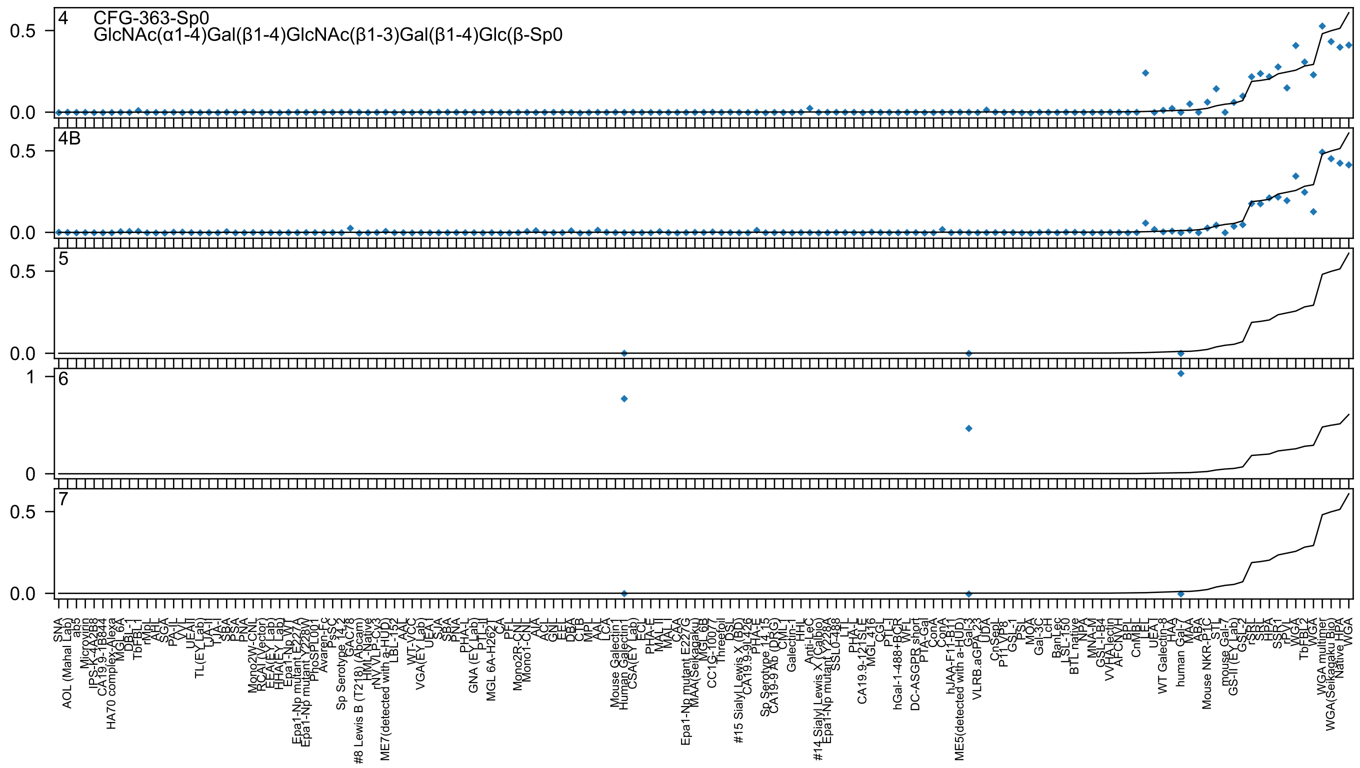

CFG-363-Sp0
GlcNAc(α1-4)Gal(β1-4)GlcNAc(β1-3)Gal(β1-4)Glc(β-Sp0
4
4B
5
6
7

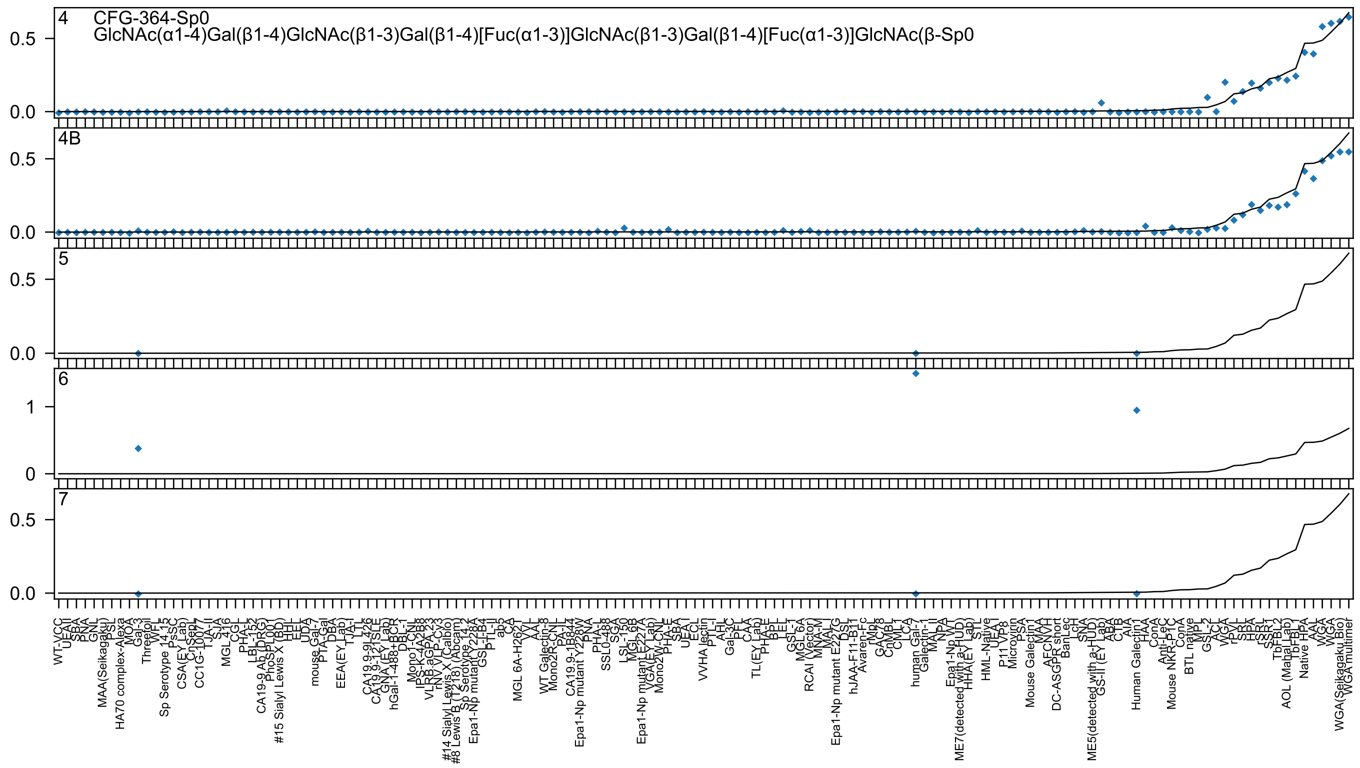
4 CFG-364-Sp0
GlcNAc(α1-4)Gal(β1-4)GlcNAc(β1-3)Gal(β1-4)[Fuc(α1-3)]GlcNAc(β1-3)Gal(β1-4)[Fuc(α1-3)]GlcNAc(β-Sp0
4B
5
6
7

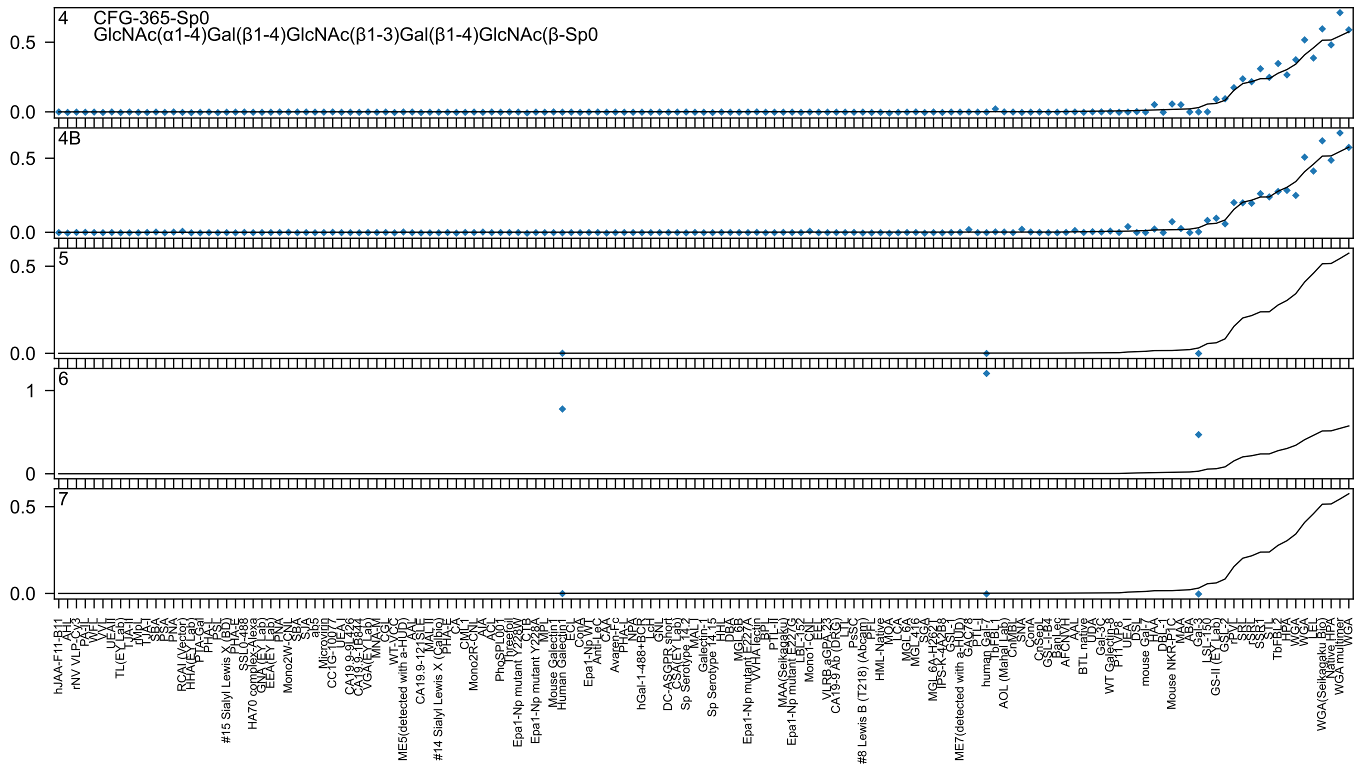

4 CFG-365-Sp0
GlcNAc(α1-4)Gal(β1-4)GlcNAc(β1-3)Gal(β1-4)GlcNAc(β-Sp0
4B
5
6
7

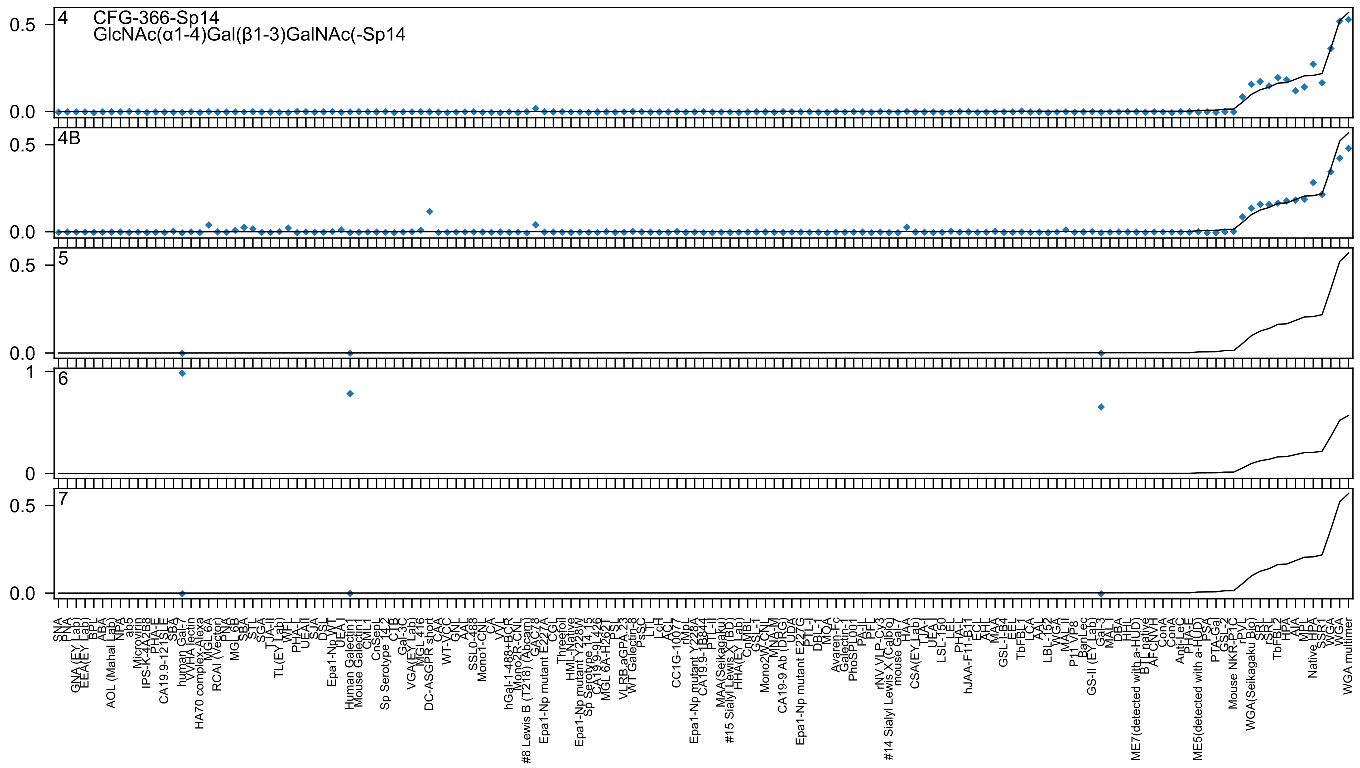

4
CFG-366-Sp14
GlcNAc(α1-4)Gal(β1-3)GalNAc(-Sp14
4B
5
6
7

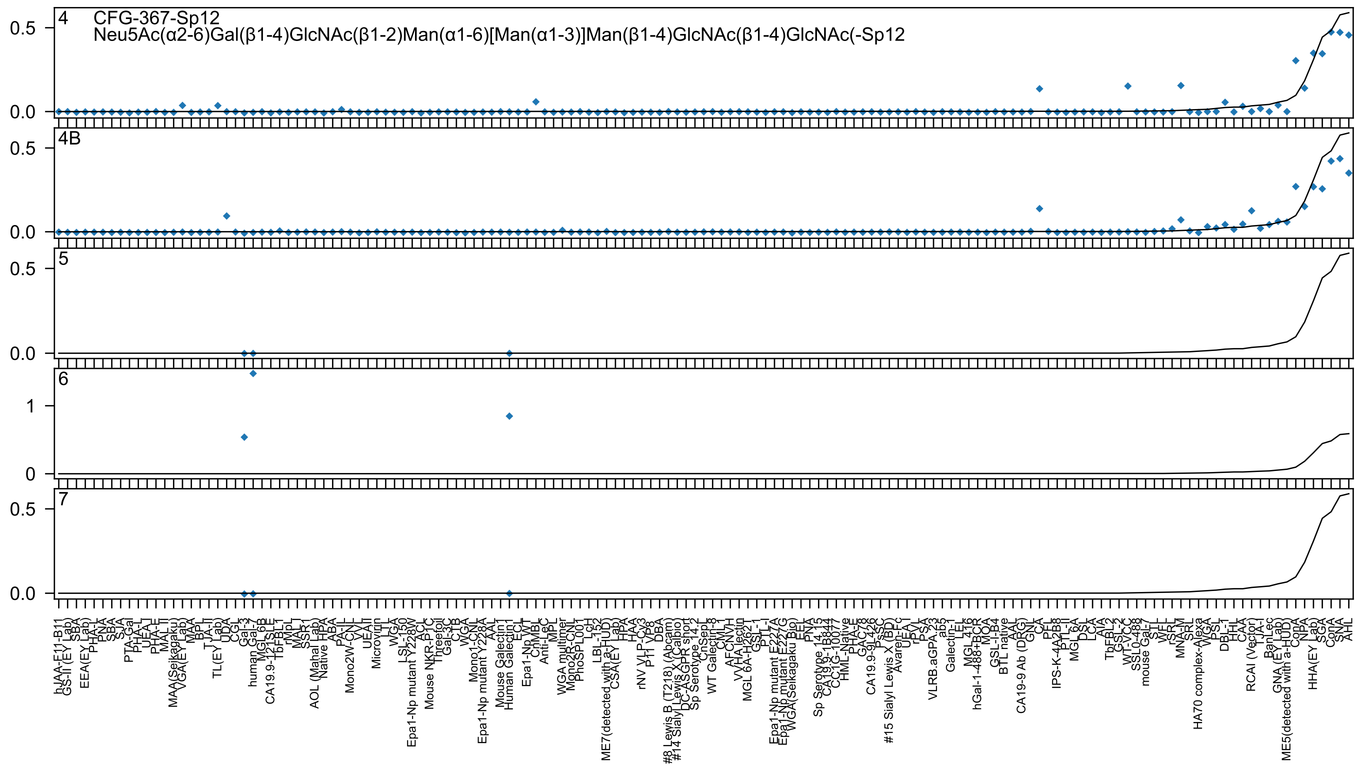
4
CFG-367-Sp12
Neu5Ac(α2-6)Gal(β1-4)GlcNAc(β1-2)Man(α1-6)[Man(α1-3)]Man(β1-4)GlcNAc(β1-4)GlcNAc(-Sp12
4B
5
6
7

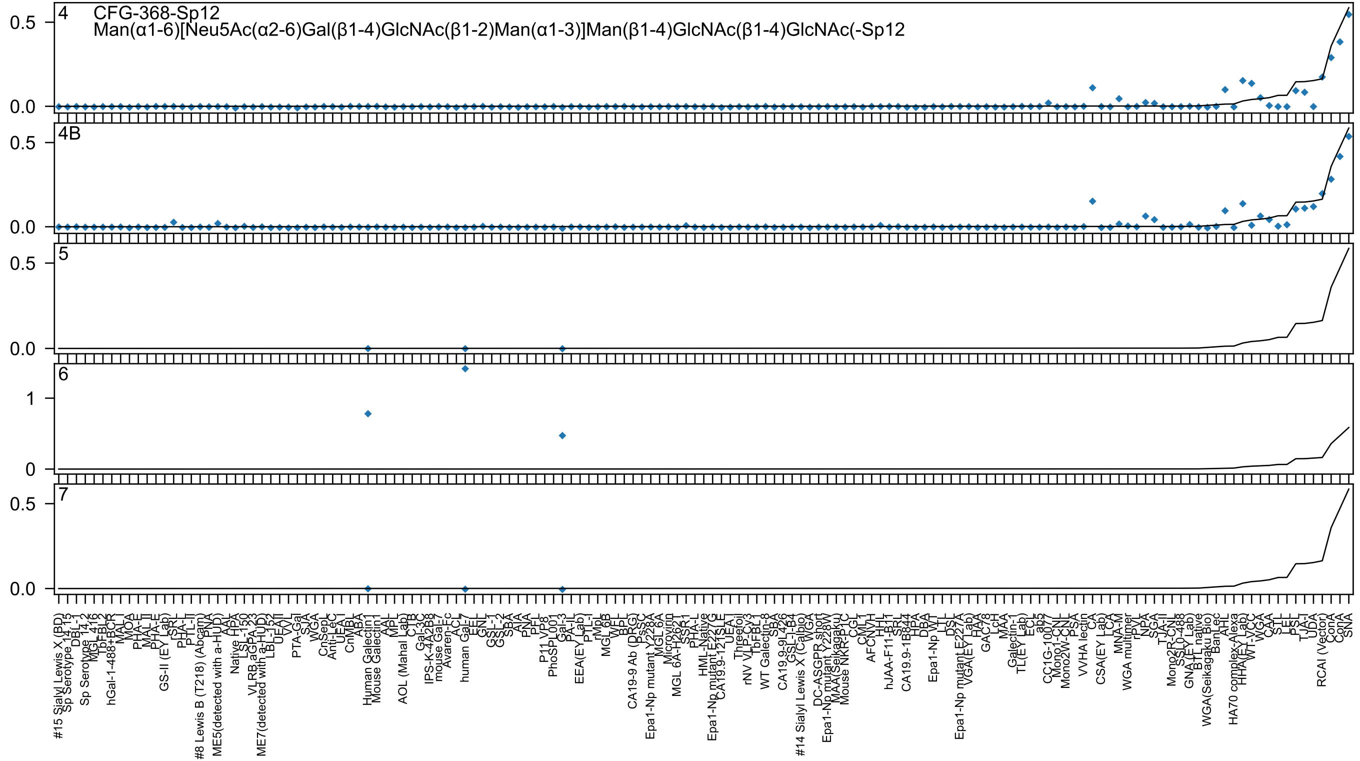
4
CFG-368-Sp12
Man(α1-6)[Neu5Ac(α2-6)Gal(β1-4)GlcNAc(β1-2)Man(α1-3)]Man(β1-4)GlcNAc(β1-4)GlcNAc(-Sp12
4B
5
6
7

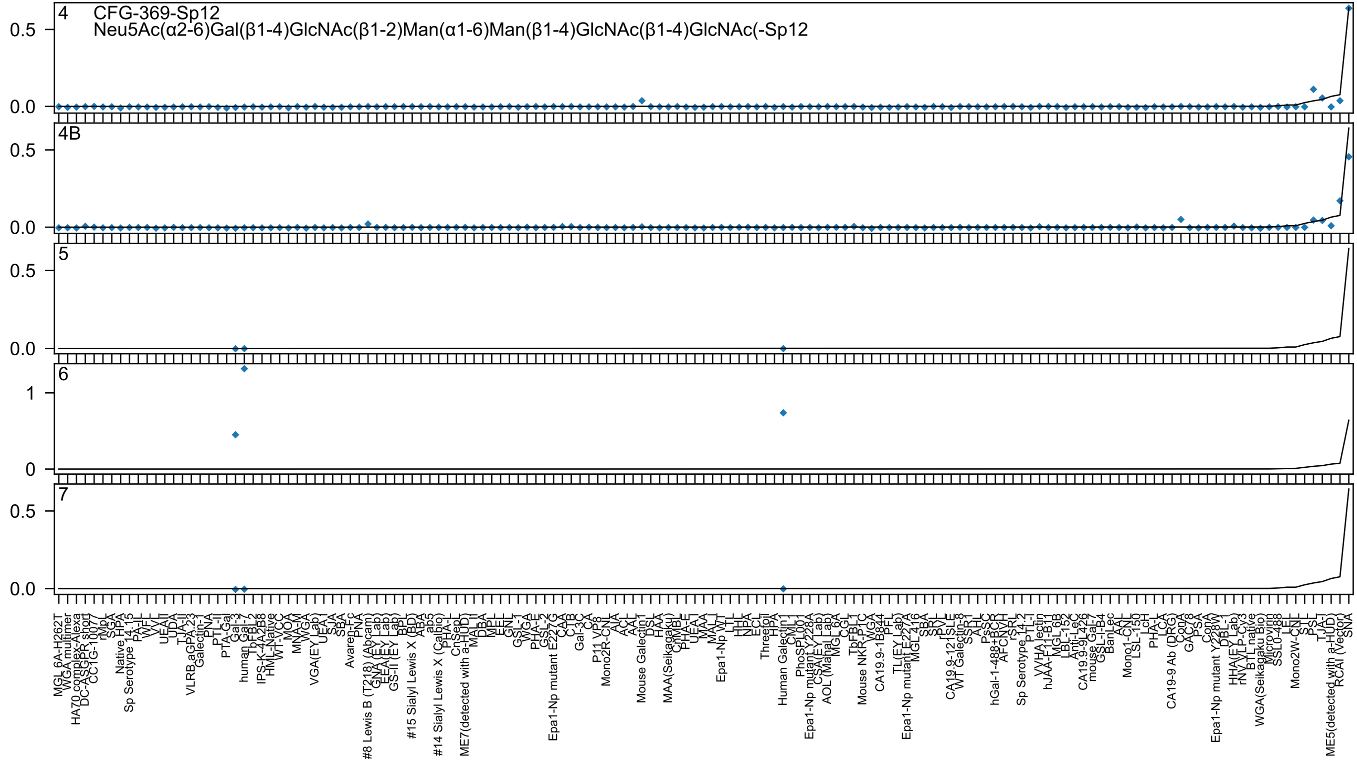
4
CFG-369-Sp12
Neu5Ac(α2-6)Gal(β1-4)GlcNAc(β1-2)Man(α1-6)Man(β1-4)GlcNAc(β1-4)GlcNAc(-Sp12
4B
5
6
7
0.5
0.0
1
0

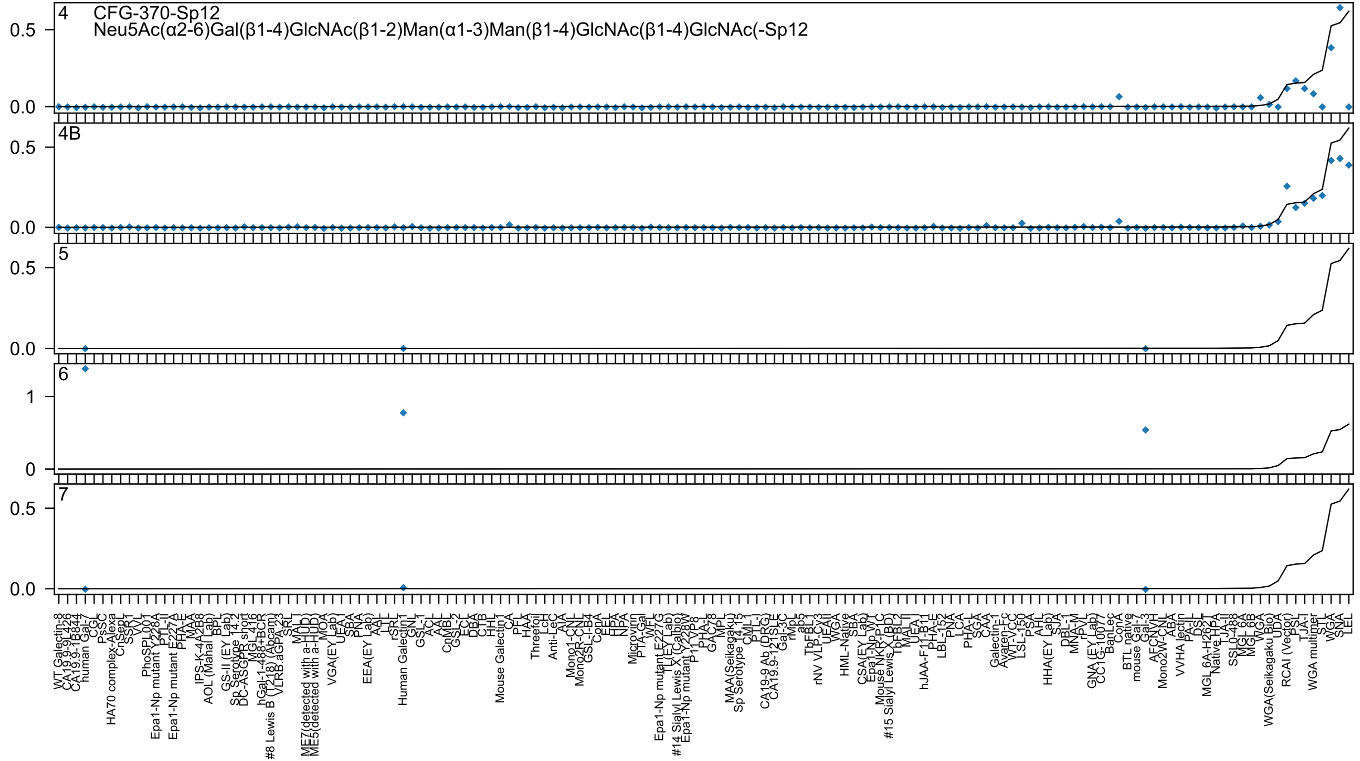
4
CFG-370-Sp12
Neu5Ac(α2-6)Gal(β1-4)GlcNAc(β1-2)Man(α1-3)Man(β1-4)GlcNAc(β1-4)GlcNAc(-Sp12
4B
5
6
7

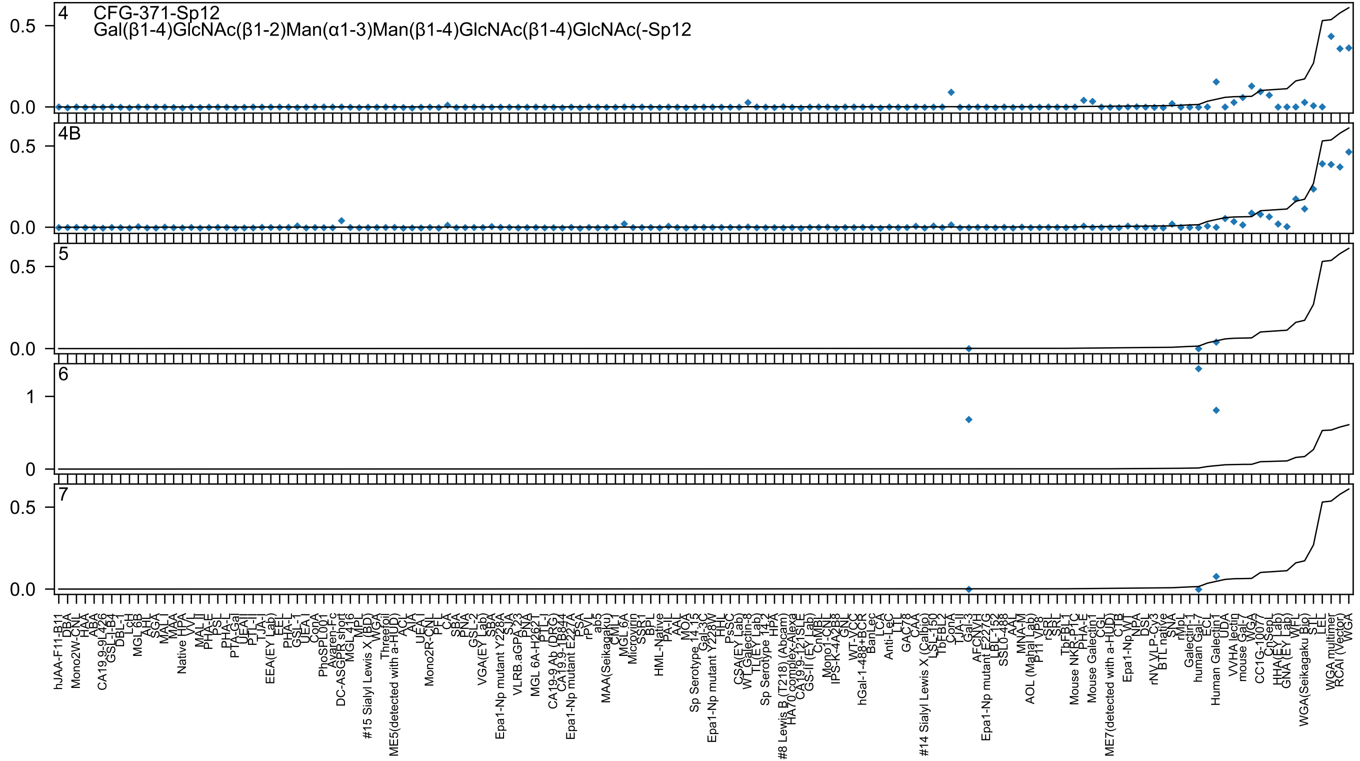

4 CFG-371-Sp12
Gal(β1-4)GlcNAc(β1-2)Man(α1-3)Man(β1-4)GlcNAc(β1-4)GlcNAc(-Sp12
4B
5
6
7

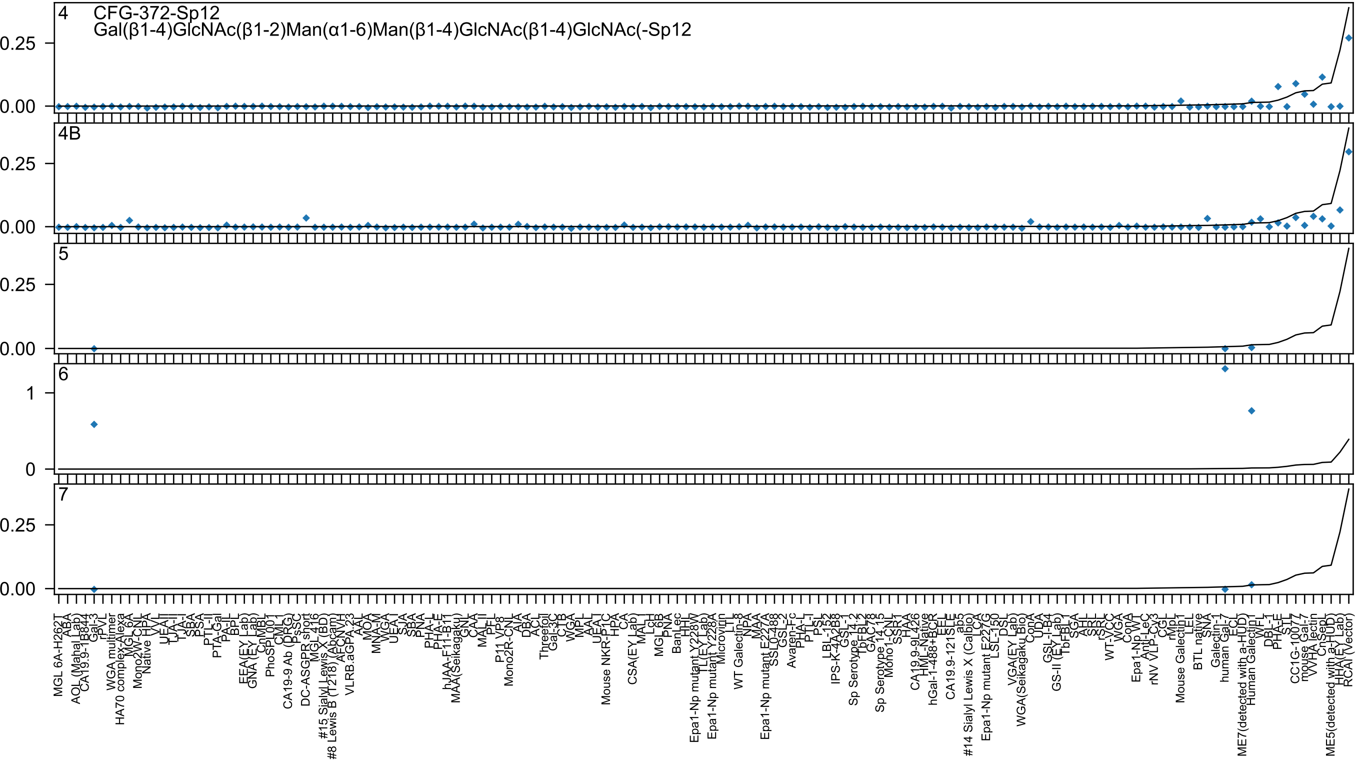

4
CFG-372-Sp12
Gal(β1-4)GlcNAc(β1-2)Man(α1-6)Man(β1-4)GlcNAc(β1-4)GlcNAc(-Sp12
4B
5
6
7
0.25
0.00
1
0

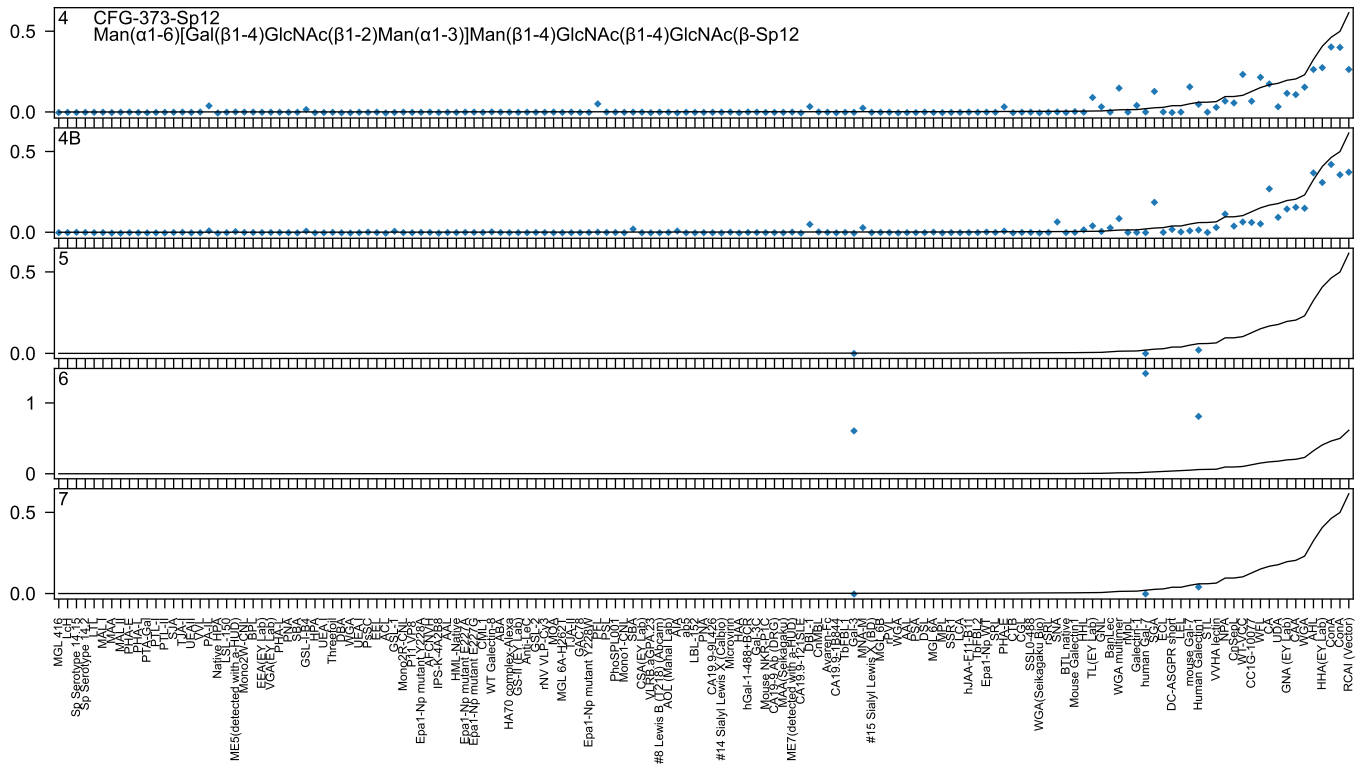
4
CFG-373-Sp12
Man(α1-6)[Gal(β1-4)GlcNAc(β1-2)Man(α1-3)]Man(β1-4)GlcNAc(β1-4)GlcNAc(β-Sp12
4B
5
6
7

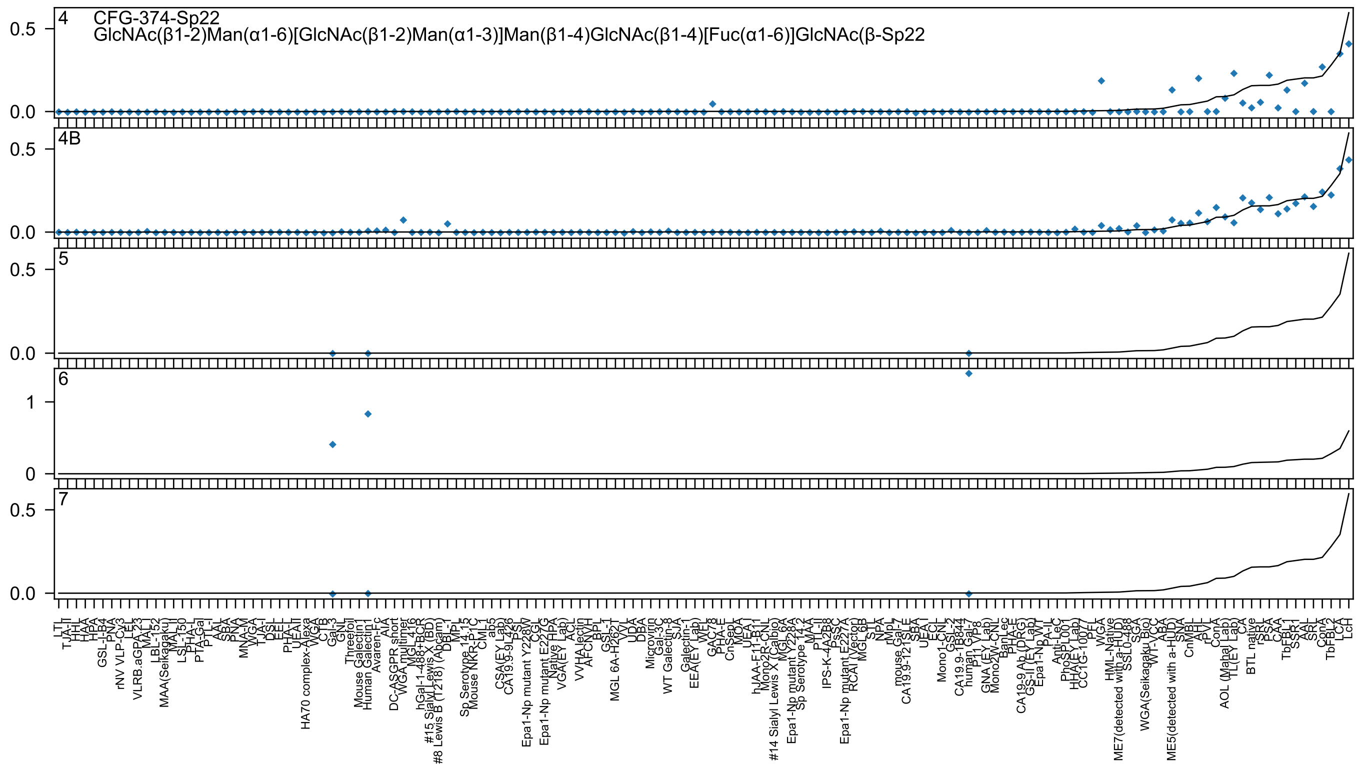
4 CFG-374-Sp22
GlcNAc(β1-2)Man(α1-6)[GlcNAc(β1-2)Man(α1-3)]Man(β1-4)GlcNAc(β1-4)[Fuc(α1-6)]GlcNAc(β-Sp22
4B
5
6
7

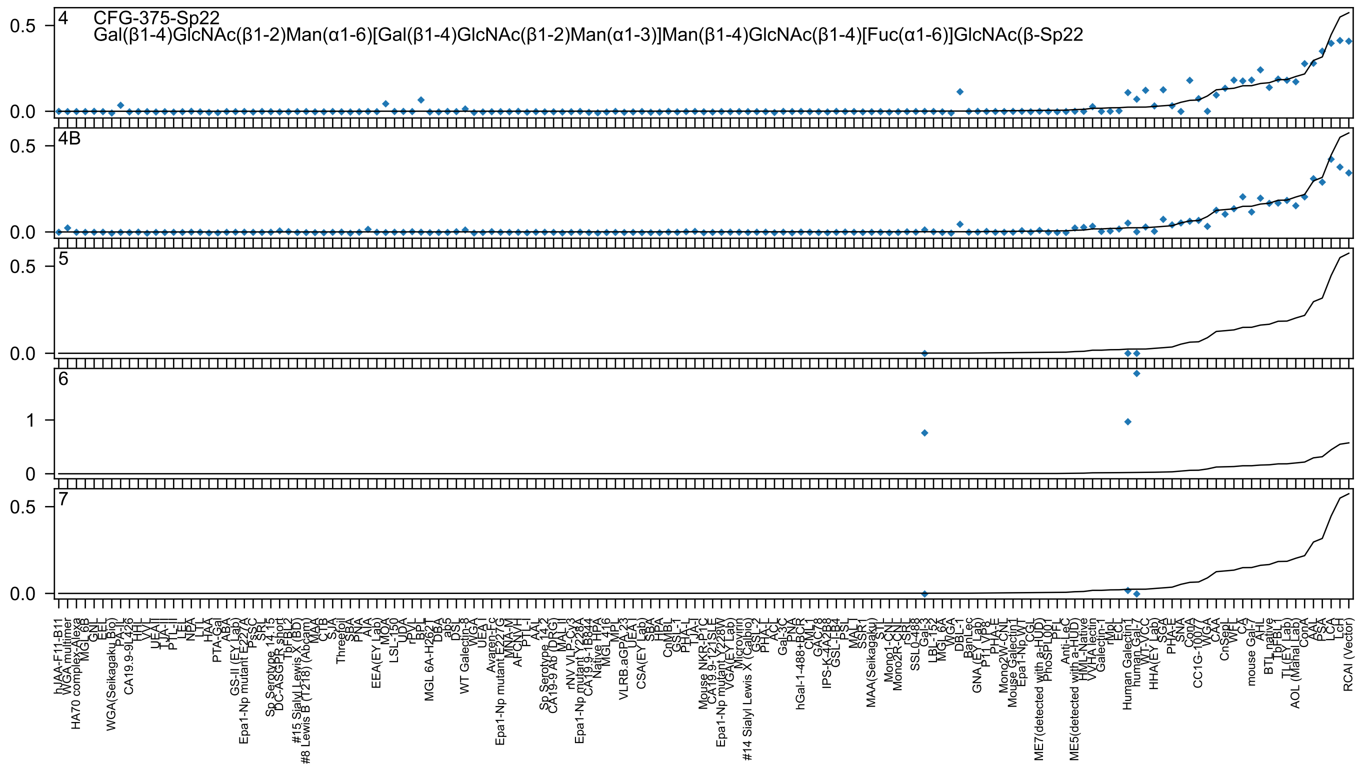

4
CFG-375-Sp22
Gal(β1-4)GlcNAc(β1-2)Man(α1-6)[Gal(β1-4)GlcNAc(β1-2)Man(α1-3)]Man(β1-4)GlcNAc(β1-4)[Fuc(α1-6)]GlcNAc(β-Sp22
0.5
0.0
4B
0.5
0.0
5
0.5
0.0
6
1
0
7
0.5
0.0
hJAA-F11-B11
WGA multimer
HA70 complex-Alexa
MGL 6B
GNL
EEL
WGA(Seikagaku_Bio)
PA-IL
CA19.9-9L426
HHL
VVL
UEAII
TJA-II
PTL-II
LEL
NPA
LTL
HAA
PTA-Gal
ABA
GS-II (EY Lab)
Epa1-Np mutant E227A
PsSC
SRL
Sp Serotype 14.15
DC-ASGPR short
TbFBL2
#15 Sialyl Lewis X (BD)
#8 Lewis B (T218) (Abcam)
MAA
CTB
SJA
Threefoil
SBA
PNA
AIA
EEA(EY Lab)
MOA
LSL-150
UDA
rPVL
BPL
MGL 6A-H262T
DBA
ab5
DSL
WT Galectin-8
WGA
UEA I
Avaren-Fc
Epa1-Np mutant E227G
MNA-M
AFCNVH
PTL-I
AAL
Sp Serotype 14.2
CA19-9 Ab (DRG)
MAL II
rNV VLP-Cy3
Epa1-Np mutant Y228A
CA19.9-1B844
Native HPA
MGL 416
MPL
VLRB.aGPA.23
UEA I
CSA(EY Lab)
SBA
HPA
CnMBL
GSL-I
PHA-L
TJA-I
Mouse NKR-P1C
CA19.9-121SLE
Epa1-Np mutant Y228W
VGA(EY Lab)
Microvirin
#14 Sialyl Lewis X (Calbio)
GSL-2
PHA-L
ACL
Gal-3C
PNA
hGal-1-488+BCR
CML1
GAC78
IPS-K-4A2B8
GSL-I-B4
PSL
MAL I
SSR1
MAA(Seikagaku)
STL
Mono1-CNL
Mono2R-CNL
rSRL
SSL0-488
Gal-3
LBL-152
MGL 6A
WGA
DBL-1
BanLec
GNA (EY Lab)
P11 VP8
PHA-E
Mono2W-CNL
Mouse Galectin1
Epa1-Np WT
CGL
ME7(detected with a-HUD)
PhoSL00
PFL
Anti-LeC
ME5(detected with a-HUD)
HML-Native
VVHA lectin
Galectin-1
rMpL
ECL
Human Galectin1
human Gal-7
WT-YCC
HHA(EY Lab)
SGA
PHA-E
SNA
ConA
CC1G-10077
WGA
CAA
CnSepL
WFL
CA
mouse Gal-7
AHL
BTL native
TbFBL1
TL(EY Lab)
AOL (Manal Lab)
ConA
AAL
PSA
LCA
LcH
RCAI (Vector)

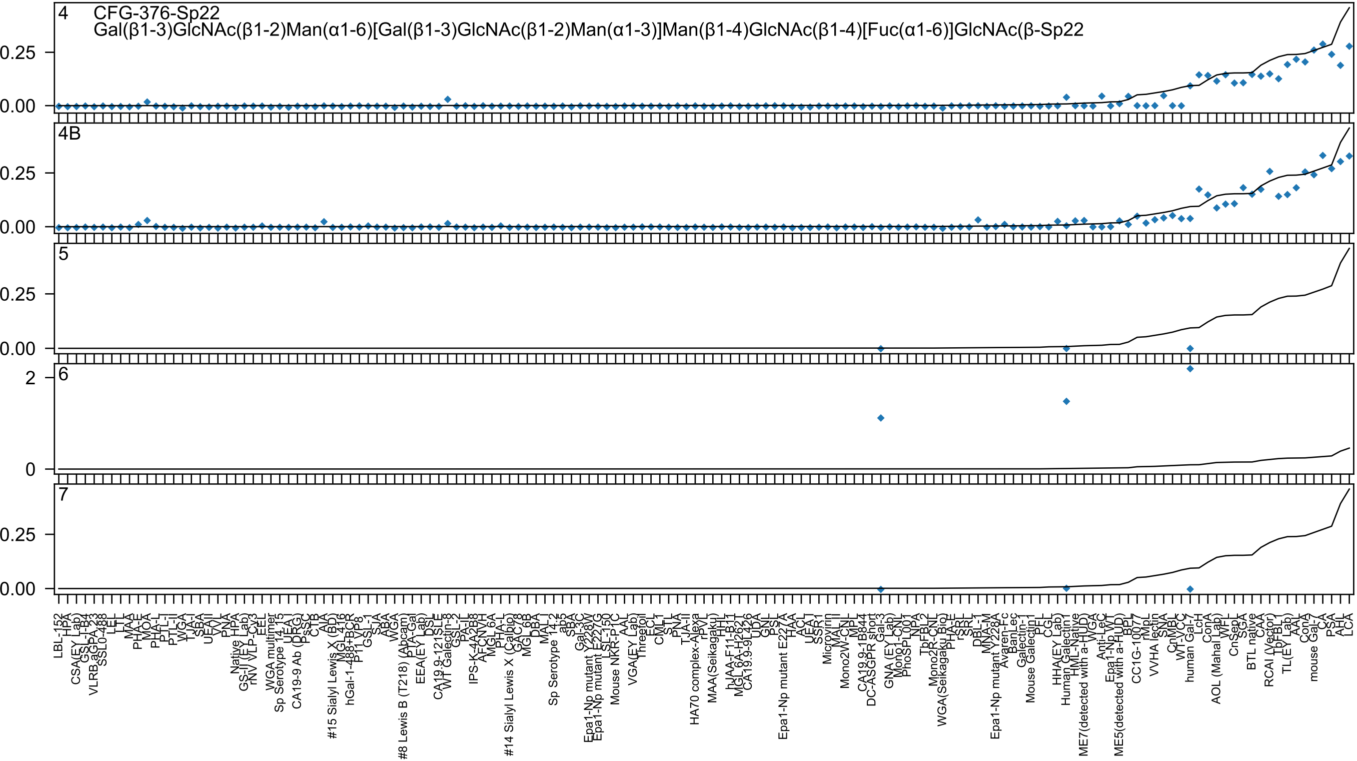
4 CFG-376-Sp22
Gal(β1-3)GlcNAc(β1-2)Man(α1-6)[Gal(β1-3)GlcNAc(β1-2)Man(α1-3)]Man(β1-4)GlcNAc(β1-4)[Fuc(α1-6)]GlcNAc(β-Sp22
4B
5
6
7
0.25
0.00
2
0

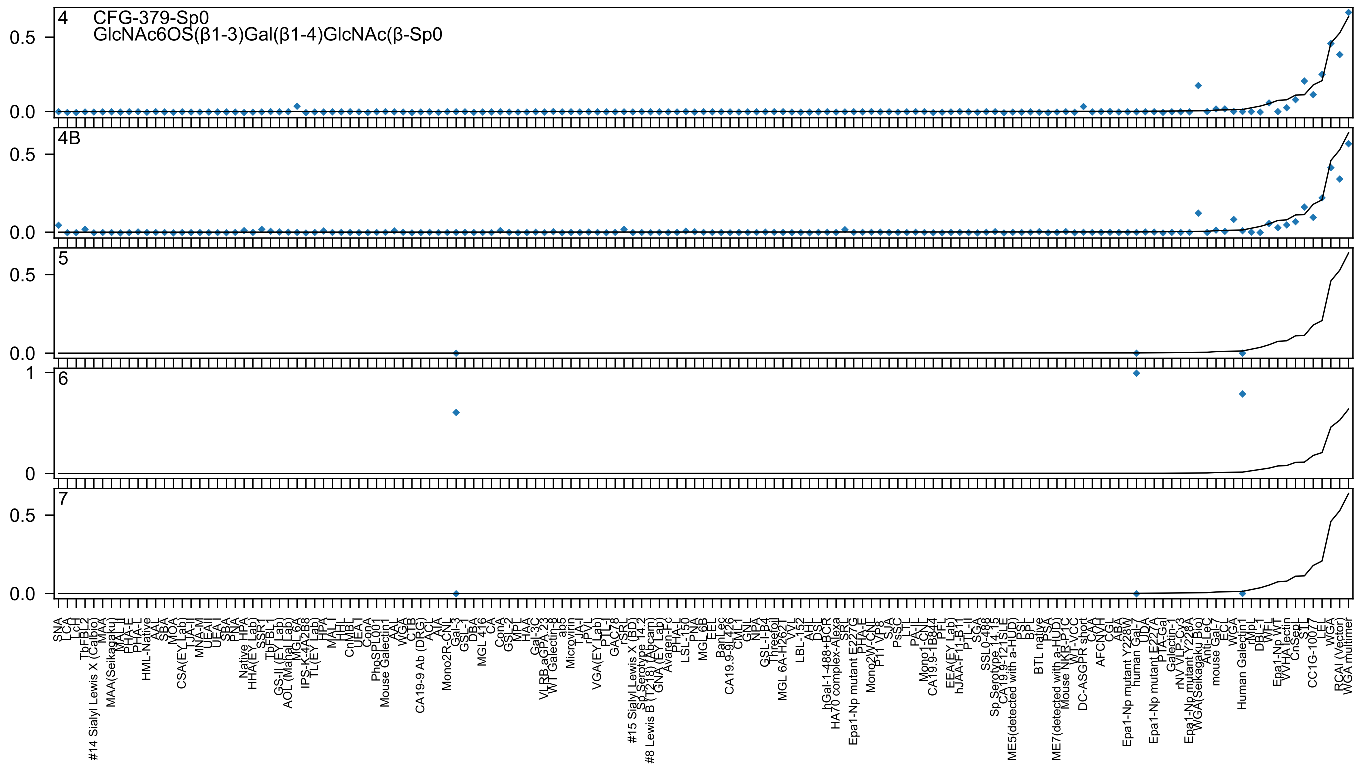
4
CFG-379-Sp0
GlcNAc6OS(β1-3)Gal(β1-4)GlcNAc(β-Sp0
4B
5
6
7
0.5
0.0
1
0
SNA
LCA
LcH
TbFBL2
#14 Sialyl Lewis X (Calbio)
MAA
MAA(Seikagaku)
MAL II
PHA-E
PHA-L
HML-Native
AAL
SBA
MOA
CSA(EY Lab)
TJA-II
MNA-M
UEAII
UEA I
SBA
PNA
Native HPA
HHA(EY Lab)
SSR1
TbFBL1
GS-II (EY Lab)
AOL (Mahal Lab)
MGL 6A
IPS-K-4A2B8
TL(EY Lab)
HPA
MAL I
HHL
CnMBL
UEA1
ConA
PhoSL001
Mouse Galectin1
AAL
WGA
CTB
CA19-9 Ab (DRG)
ACL
AIA
Mono2R-CNL
Gal-3
GSL-I
DBA
MGL 416
CA
ConA
GSL-2
MPL
HAA
Gal-3C
VLRB aGPA 23
WT Galectin-8
ab5
Microvirin
TJA-I
rPVL
VGA(EY Lab)
PTL-I
GAC78
rSRL
#15 Sialyl Lewis X (BD)
Sp Serotype 14.2
#8 Lewis B (T218) (Abcam)
GNA (EY Lab)
Avaren-Fc
PHA-L
LSL-150
PNA
MGL 6B
EEL
BanLec
CA19.9-9L426
CML1
GNL
NPA
GSL-I-B4
Threefoil
MGL 6A-H262T
VVL
LBL-152
AHL
DSL
hGal-1-488+BCR
HA70 complex-Alexa
SRL
Epa1-Np mutant E227G
PHA-E
Mono2W-CNL
P11 VP8
SJA
PsSC
LTL
PA-IL
Mono1-CNL
CA19.9-1B844
PFL
EEA(EY Lab)
hJAA-F11-B1
PTL-II
SGA
SSL0-488
Sp Serotype 14.15
CA19.9-121SLE
ME5(detected with a-HUD)
DSL
BPL
BTL native
PSA
ME7(detected with a-HUD)
Mouse NKR-P1C
WT-VCC
DC-ASGPR short
CAA
AFCNYH
CGL
ABA
Epa1-Np mutant Y228W
human Gal-7
UDA
Epa1-Np mutant E227A
PTA-Gal
Galectin-1
rNV VLP-Cy3
Epa1-Np mutant Y228A
WGA(Seikagaku Bio)
Anti-LeC
mouse Gal-7
ECL
WGA
Human Galectin1
rMpL
DBL-1
WFL
Epa1-Np WT
VVHA lectin
CnSepL
STL
CC1G-10077
LEL
WGA
RCAI (Vector)
WGA multimer

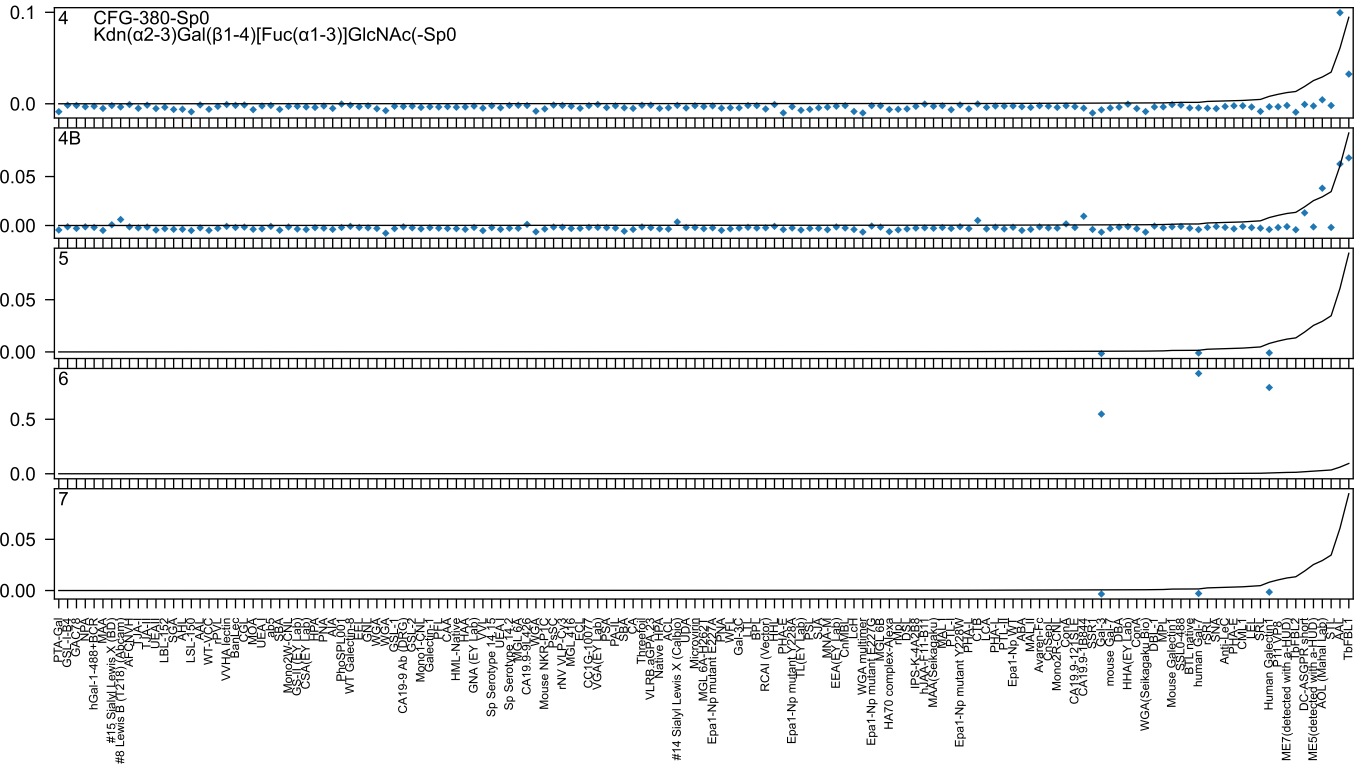

4
CFG-380-Sp0
Kdn(α2-3)Gal(β1-4)[Fuc(α1-3)]GlcNAc(-Sp0
4B
5
6
7
0.1
0.0
0.05
0.00
0.5
0.0
PTA-Gal
GSL-I-B4
GAC78
NPA
hGal-1-488+BCR
MAA
#15 Sialyl Lewis X (BD)
#8 Lewis B (T218) (Abcam)
AFCNVH
TJA-I
TJA-II
UEA-II
LBL-152
SGA
AHL
LSL-150
AAL
WT-VCC
rPVL
VVHA lectin
BanLec
CGL
MOA
UEA-I
ab5
SBA
Mono2W-CNL
GS-II (EY Lab)
CSA(EY Lab)
HPA
PNA
AIA
PhoSPL001
WT Galectin-8
EEL
GNL
WGA
WGA
GSL-1
CA19-9 Ab (DRG)
GSL-2
Mono1-CNL
Galectin-1
PFL
CAA
HML-Native
HAA
GNA (EY Lab)
VVL
Sp Serotype 14.15
UEA-I
Sp Serotype 14.2
MGL 6A
CA19.9-9L426
WGA
Mouse NKR-P1C
PsSC
rNV VLP-Cy3
MGL 416
ECL
CC1G-10077
VGA(EY Lab)
PSA
PA-IL
SBA
CA
Threefoil
VLRB.aGPA.23
Native HPA
ACL
#14 Sialyl Lewis X (Calbio)
UDA
Microvirin
MGL 6A-H262T
Epa1-Np mutant E227A
PNA
WFL
Gal-3C
LTL
BPL
RCAI (Vector)
HHL
PHA-E
Epa1-Np mutant Y228A
TL(EY Lab)
PSL
SJA
MNA-M
EEA(EY Lab)
CnMBL
LcH
WGA multimer
Epa1-Np mutant E227G
MGL 6B
HA70 complex-Alexa
rMpL
DSL
IPS-K-4A2B8
hJAA-F11-B11
MAA(Seikagaku)
MAL-I
PTL-I
Epa1-Np mutant Y228W
PHA-L
CTB
LCA
PHA-L
PTL-II
Epa1-Np WT
ABA
MAL-II
Avaren-Fc
CnSeL
Mono2R-CNL
ConA
CA19.9-121SLE
CA19.9-1B844
SSR1
Gal-3
mouse Gal-7
DBA
HHA(EY Lab)
ConA
WGA(Seikagaku Bio)
DBL-1
MPL
Mouse Galectin1
SSL0-488
BTL native
human Gal-7
rSRL
SNA
Anti-LeC
PHA-E
CML1
EEL
SRL
Human Galectin1
P11 VP8
ME7(detected with a-HUD)
TbFBL2
DC-ASGPR short
ME5(detected with a-HUD)
AOL (Mahal Lab)
STL
AAL
TbFBL1

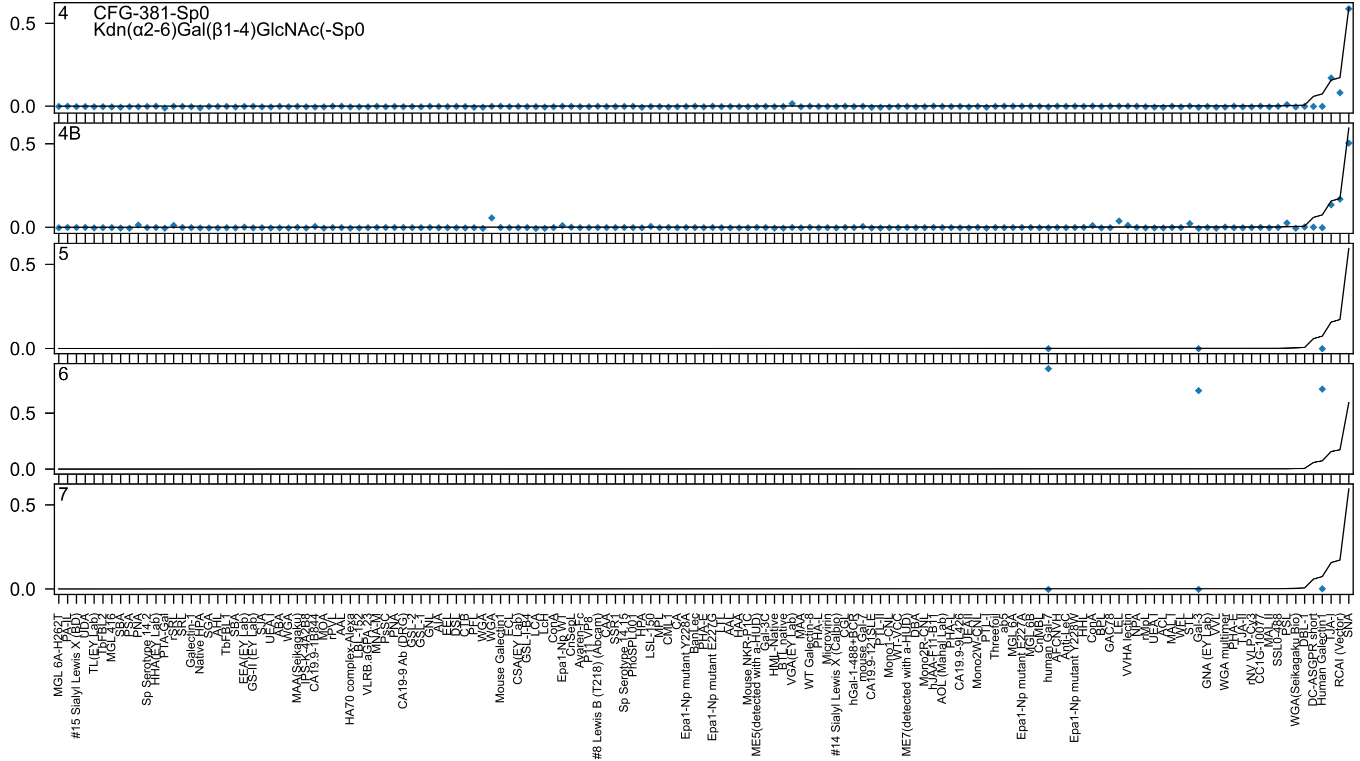

4
CFG-381-Sp0
Kdn(α2-6)Gal(β1-4)GlcNAc(-Sp0
4B
5
6
7

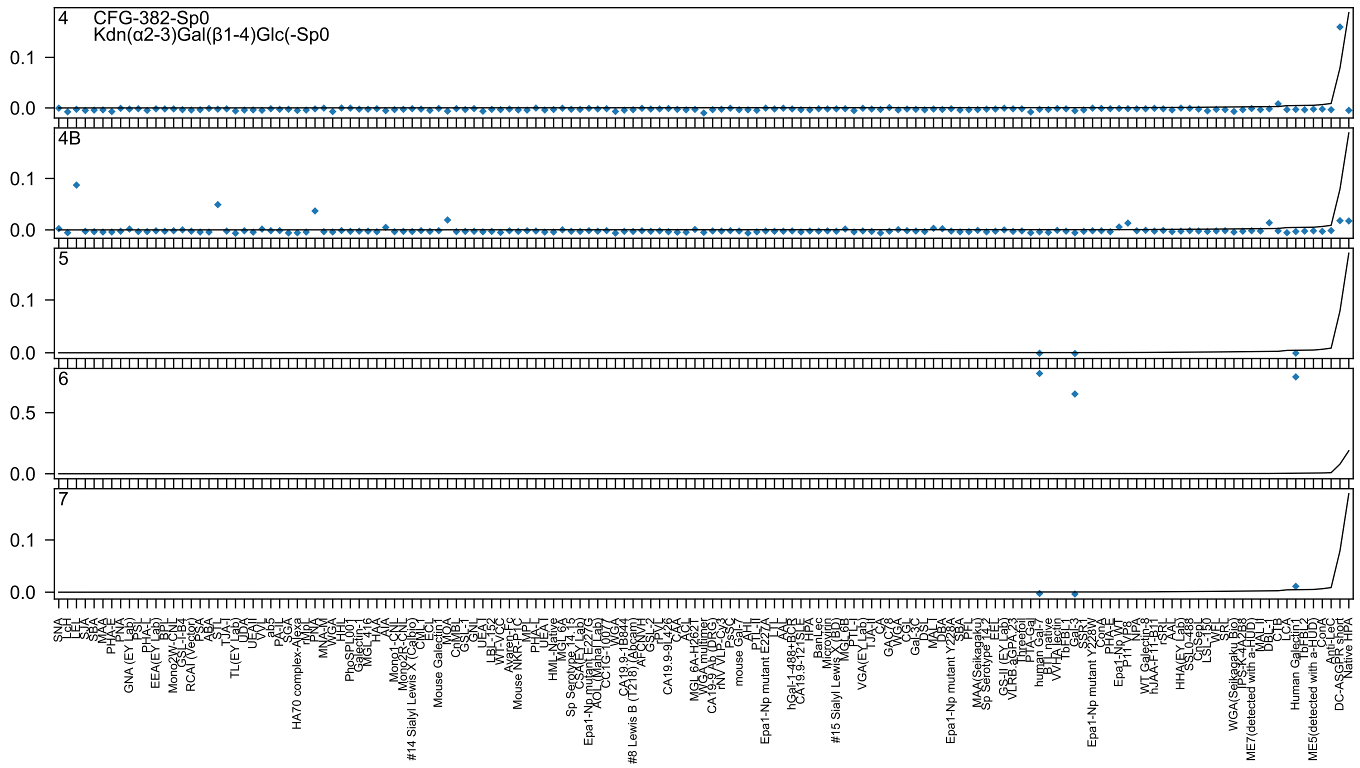

4 CFG-382-Sp0
Kdn(α2-3)Gal(β1-4)Glc(-Sp0
4B
5
6
7

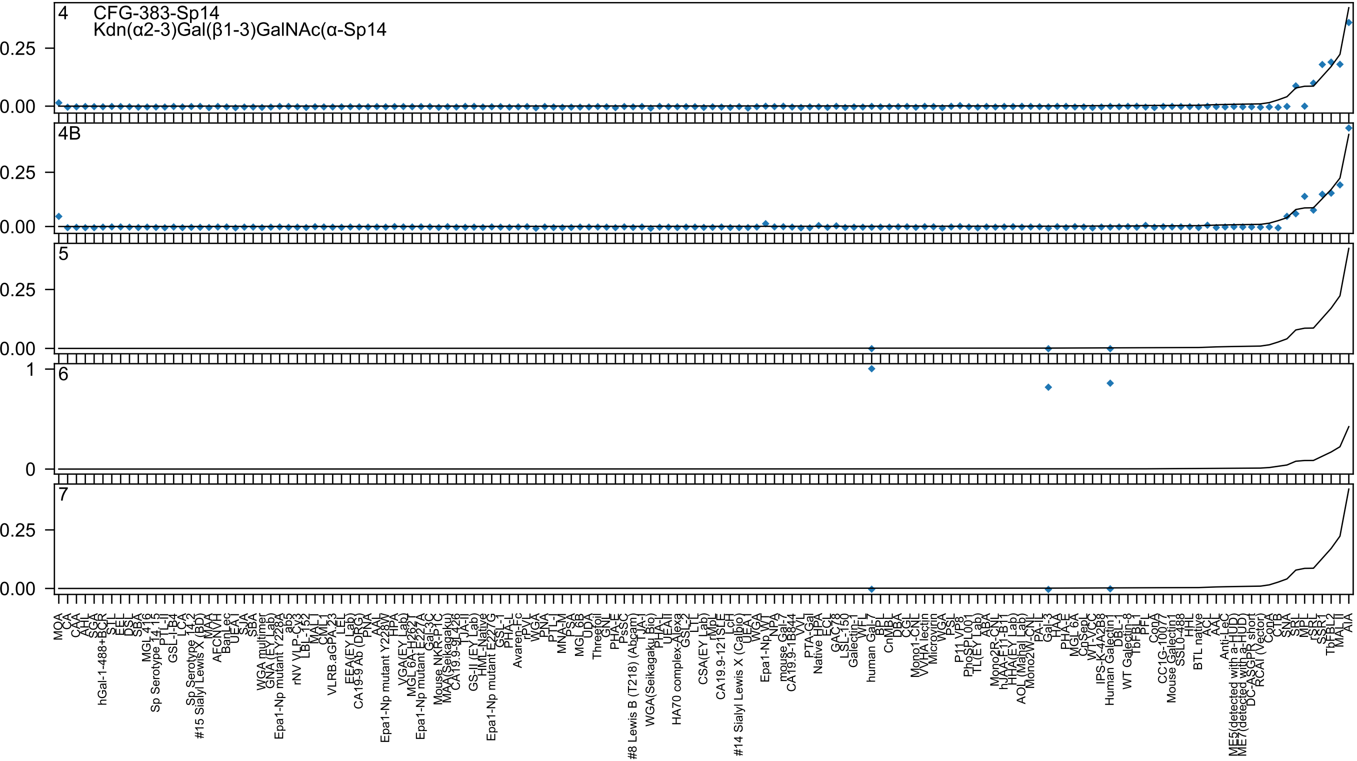
4
CFG-383-Sp14
Kdn(α2-3)Gal(β1-3)GalNAc(α-Sp14
4B
5
6
7
0.25
0.00
1
0

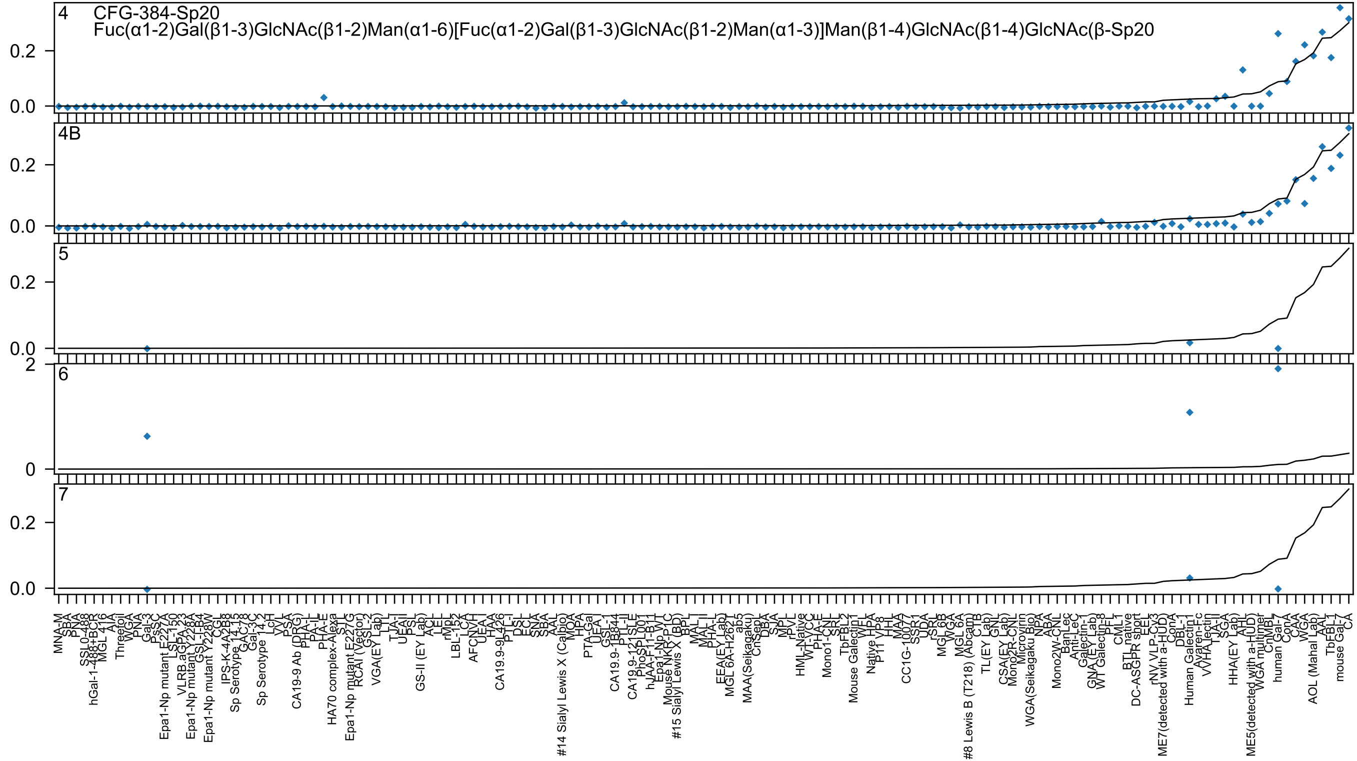

4 CFG-384-Sp20
Fuc(α1-2)Gal(β1-3)GlcNAc(β1-2)Man(α1-6)[Fuc(α1-2)Gal(β1-3)GlcNAc(β1-2)Man(α1-3)]Man(β1-4)GlcNAc(β1-4)GlcNAc(β-Sp20
4B
5
6
7

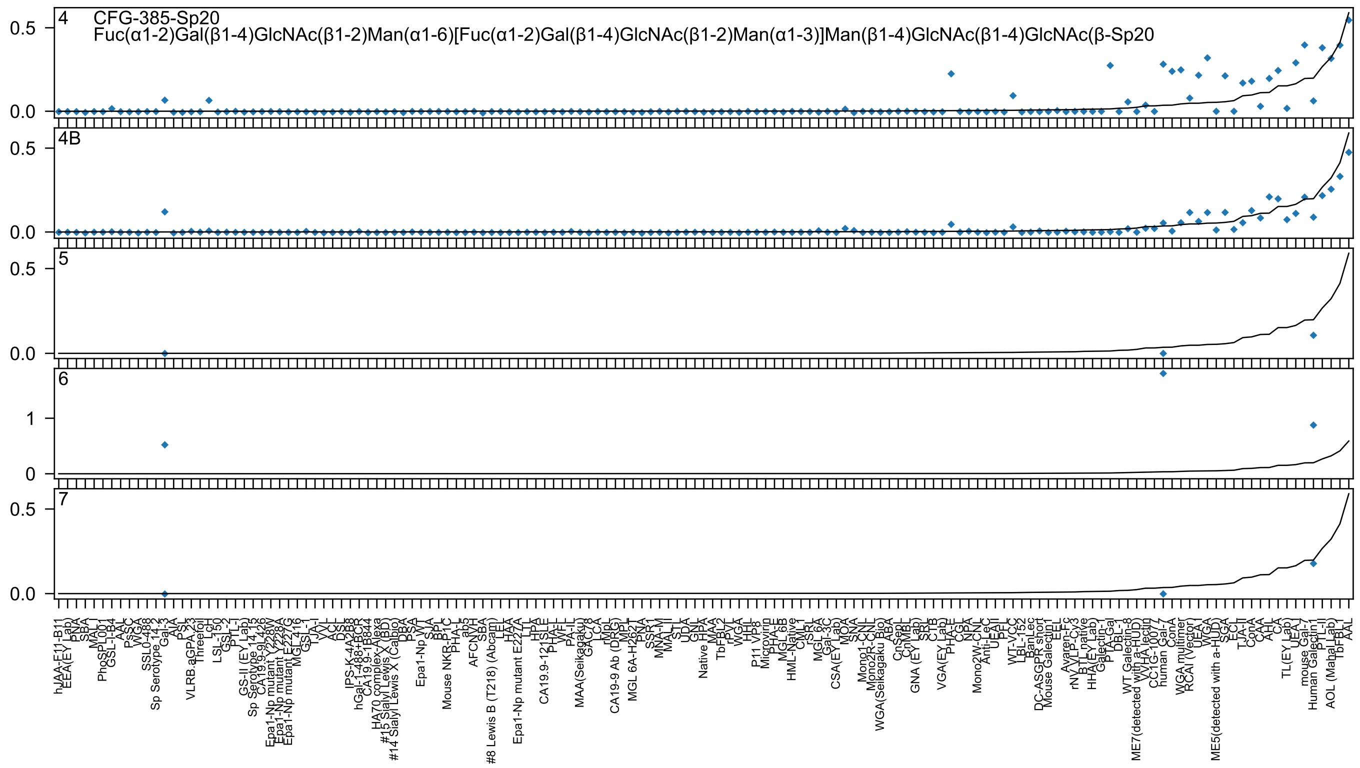
CFG-385-Sp20
Fuc(α1-2)Gal(β1-4)GlcNAc(β1-2)Man(α1-6)[Fuc(α1-2)Gal(β1-4)GlcNAc(β1-2)Man(α1-3)]Man(β1-4)GlcNAc(β1-4)GlcNAc(β-Sp20
4
4B
5
6
7

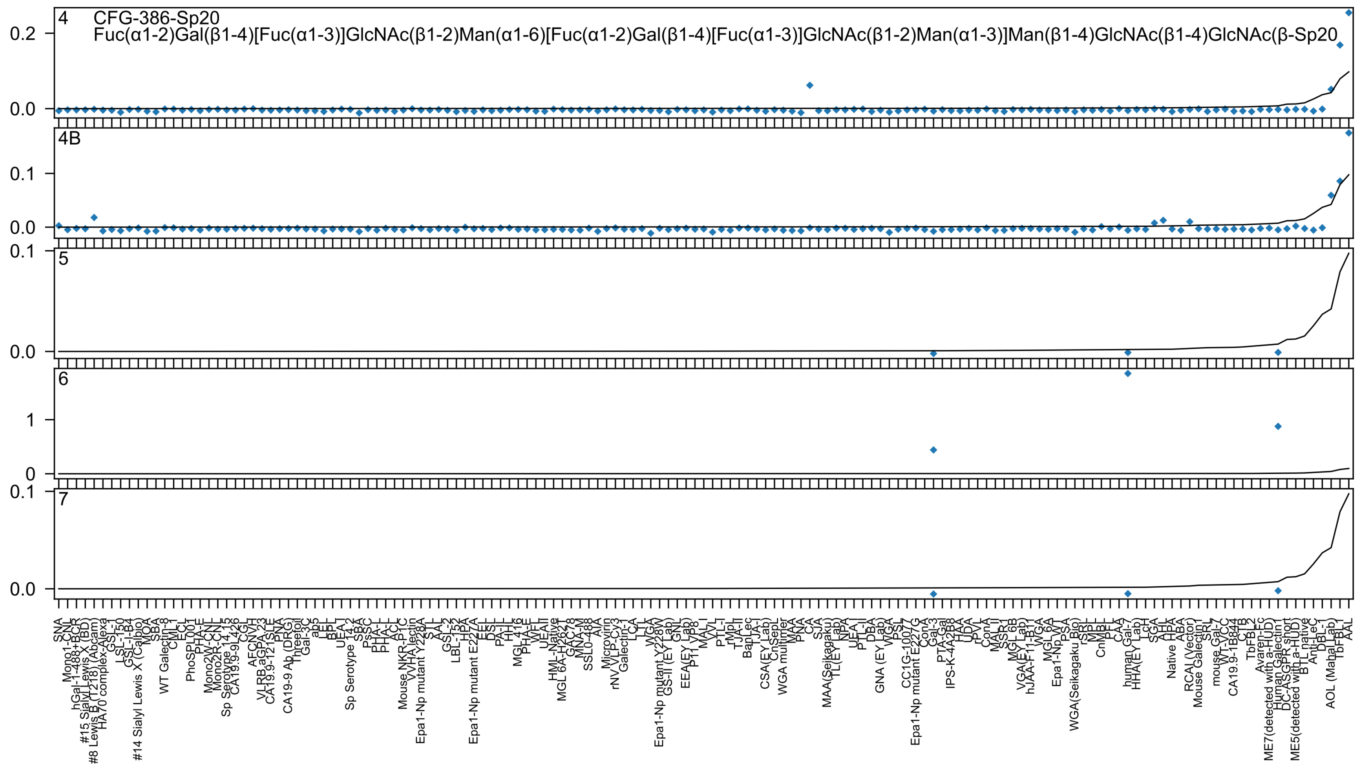

CFG-386-Sp20
Fuc(α1-2)Gal(β1-4)[Fuc(α1-3)]GlcNAc(β1-2)Man(α1-6)[Fuc(α1-2)Gal(β1-4)[Fuc(α1-3)]GlcNAc(β1-2)Man(α1-3)]Man(β1-4)GlcNAc(β1-4)GlcNAc(β-Sp20
4
4B
5
6
7

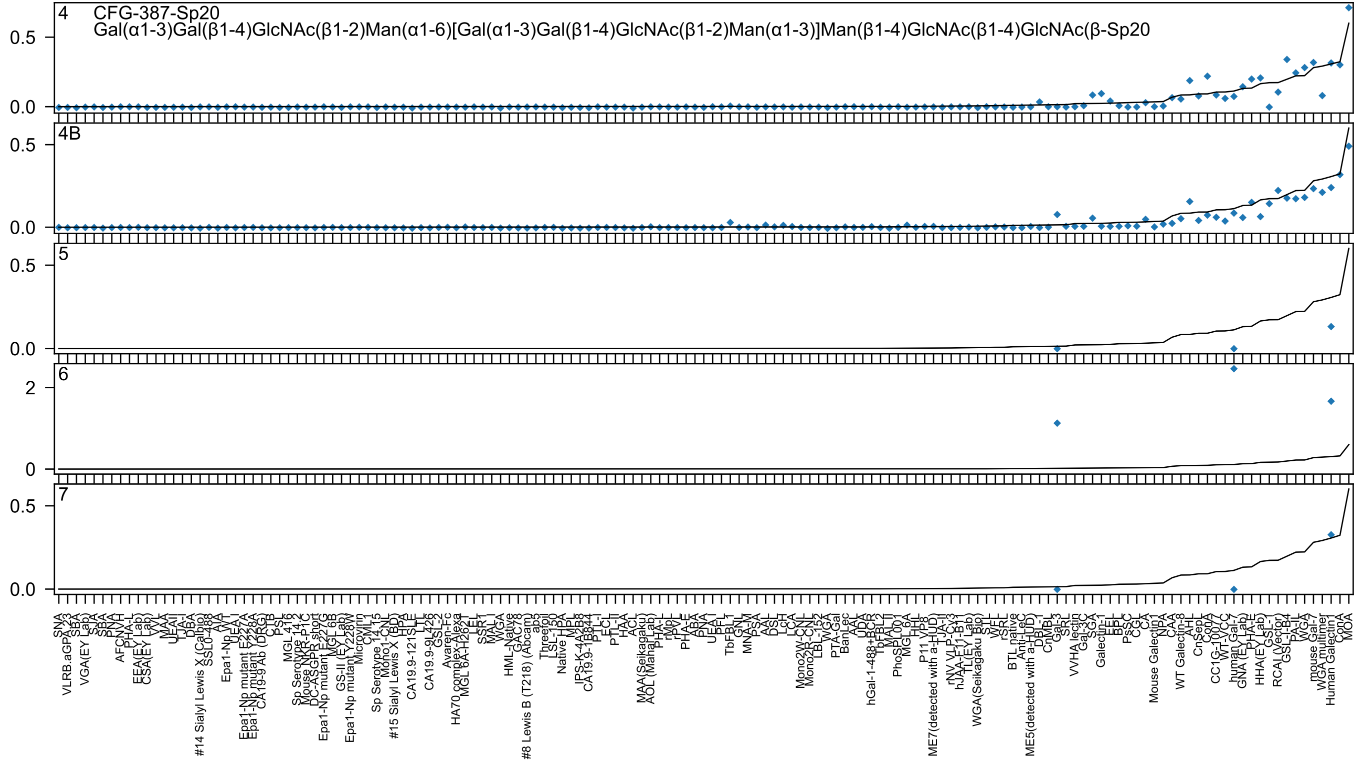

4 CFG-387-Sp20
Gal(α1-3)Gal(β1-4)GlcNAc(β1-2)Man(α1-6)[Gal(α1-3)Gal(β1-4)GlcNAc(β1-2)Man(α1-3)]Man(β1-4)GlcNAc(β1-4)GlcNAc(β-Sp20
4B
5
6
7

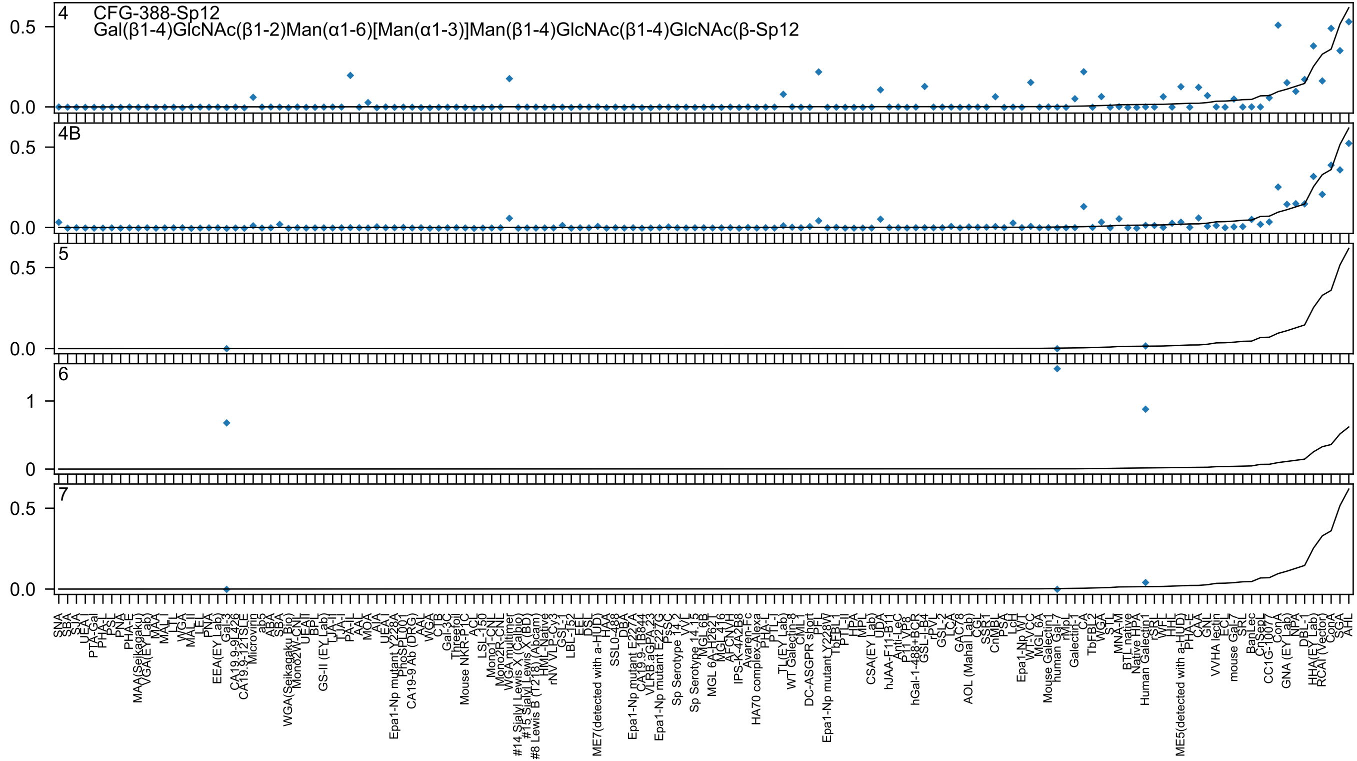

4
CFG-388-Sp12
Gal(β1-4)GlcNAc(β1-2)Man(α1-6)[Man(α1-3)]Man(β1-4)GlcNAc(β1-4)GlcNAc(β-Sp12
4B
5
6
7

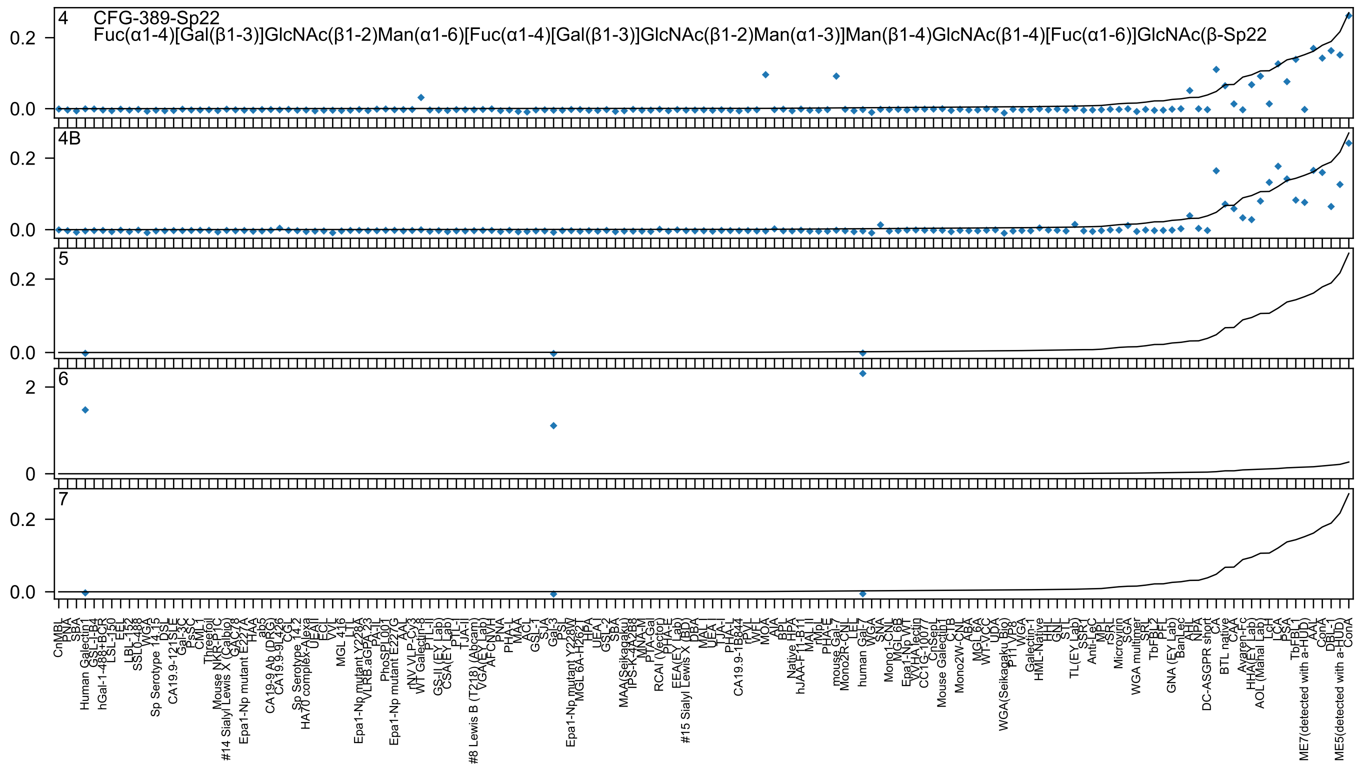

4 CFG-389-Sp22
Fuc(α1-4)[Gal(β1-3)]GlcNAc(β1-2)Man(α1-6)[Fuc(α1-4)[Gal(β1-3)]GlcNAc(β1-2)Man(α1-3)]Man(β1-4)GlcNAc(β1-4)[Fuc(α1-6)]GlcNAc(β-Sp22
4B
5
6
7

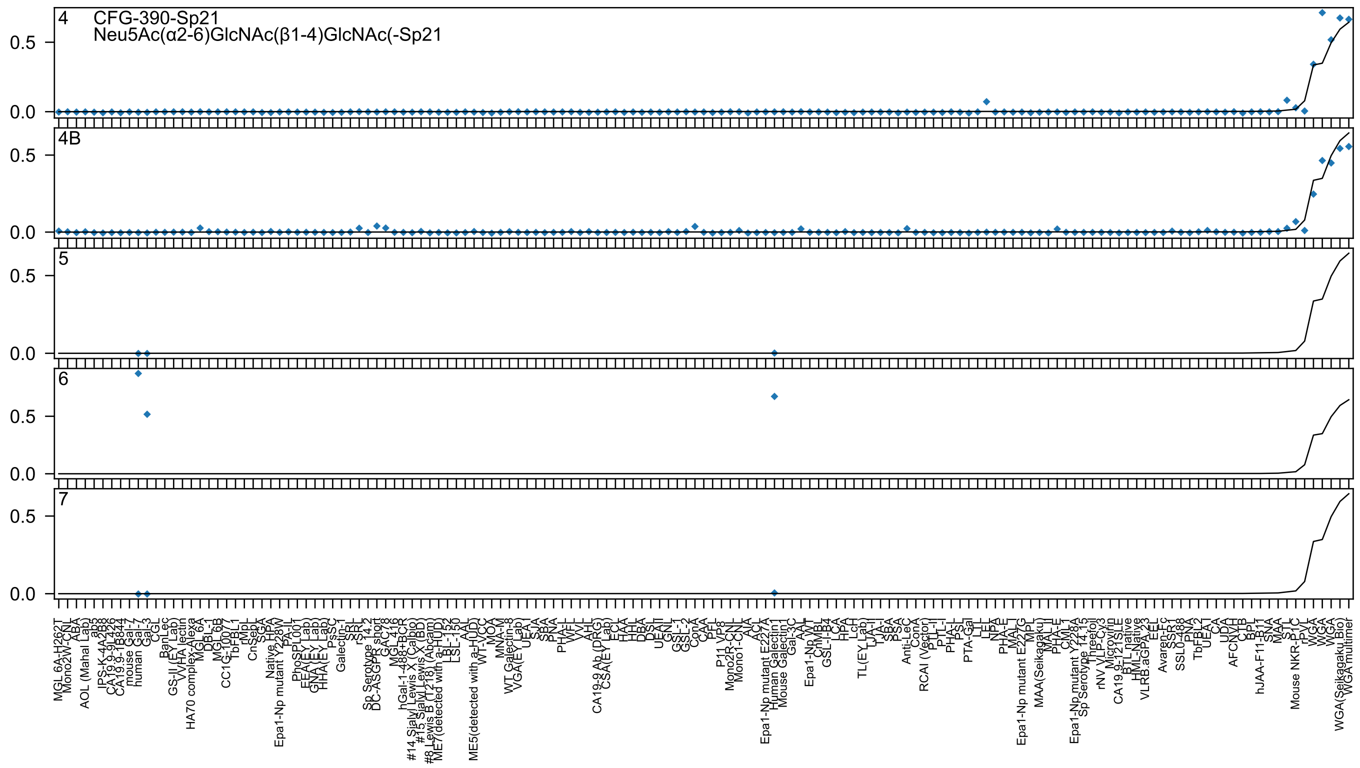

4 CFG-390-Sp21
Neu5Ac(α2-6)GlcNAc(β1-4)GlcNAc(-Sp21
4B
5
6
7

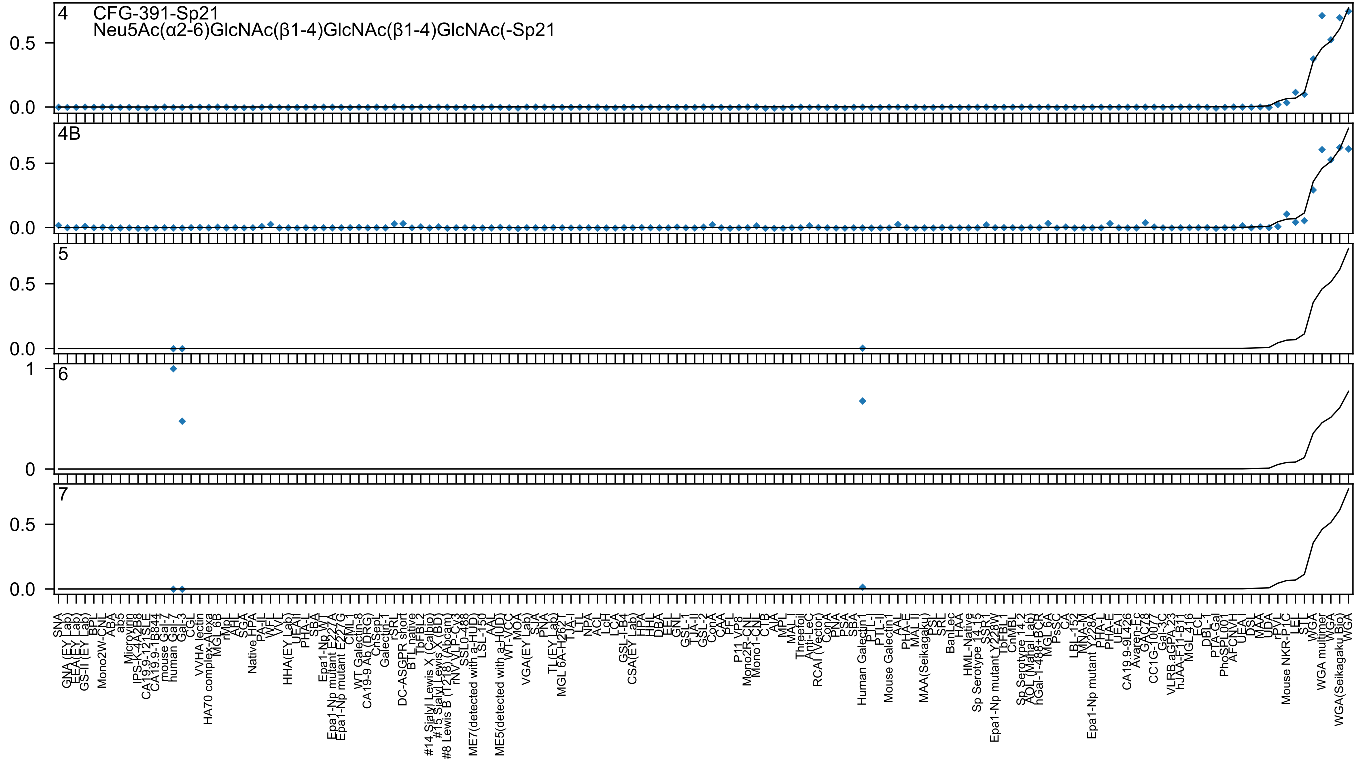

4 CFG-391-Sp21
Neu5Ac(α2-6)GlcNAc(β1-4)GlcNAc(β1-4)GlcNAc(-Sp21
4B
5
6
7

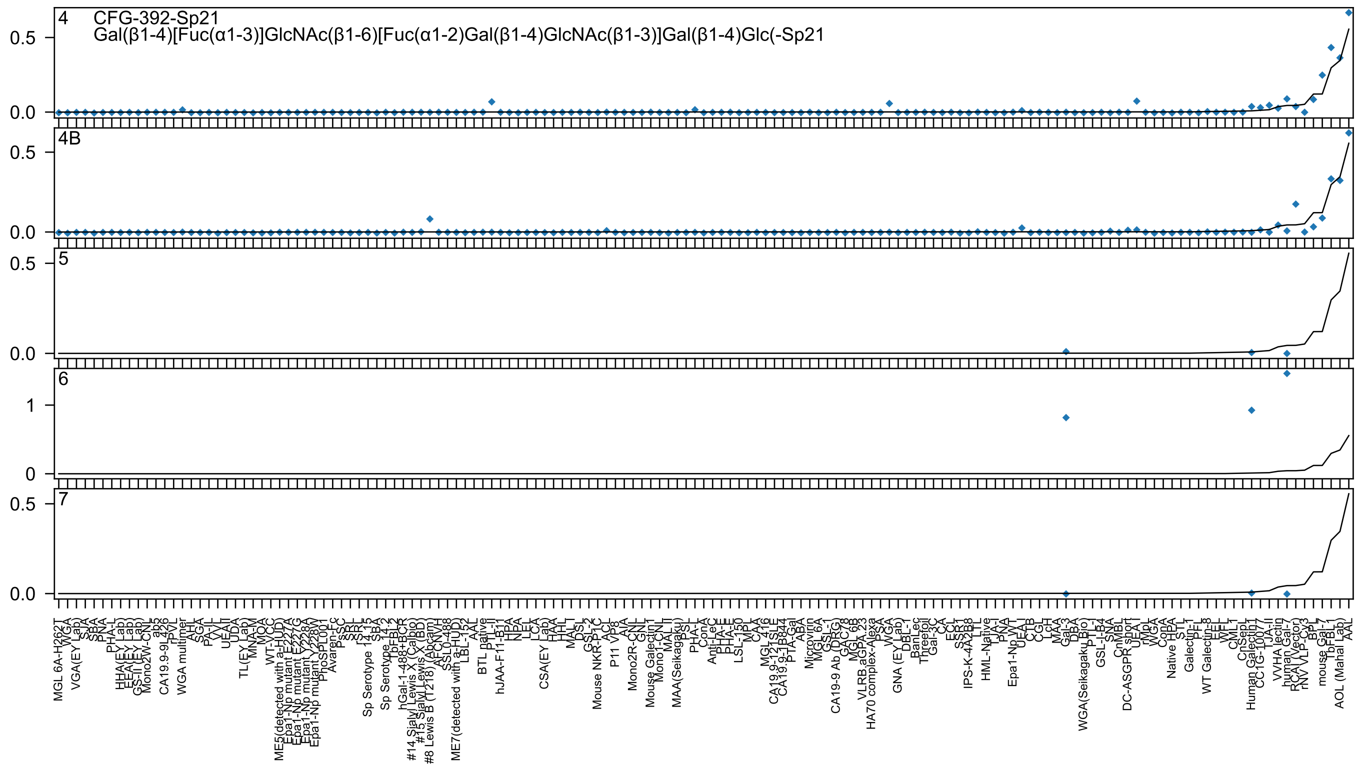

4
CFG-392-Sp21
Gal(β1-4)[Fuc(α1-3)]GlcNAc(β1-6)[Fuc(α1-2)Gal(β1-4)GlcNAc(β1-3)]Gal(β1-4)Glc(-Sp21
4B
5
6
7

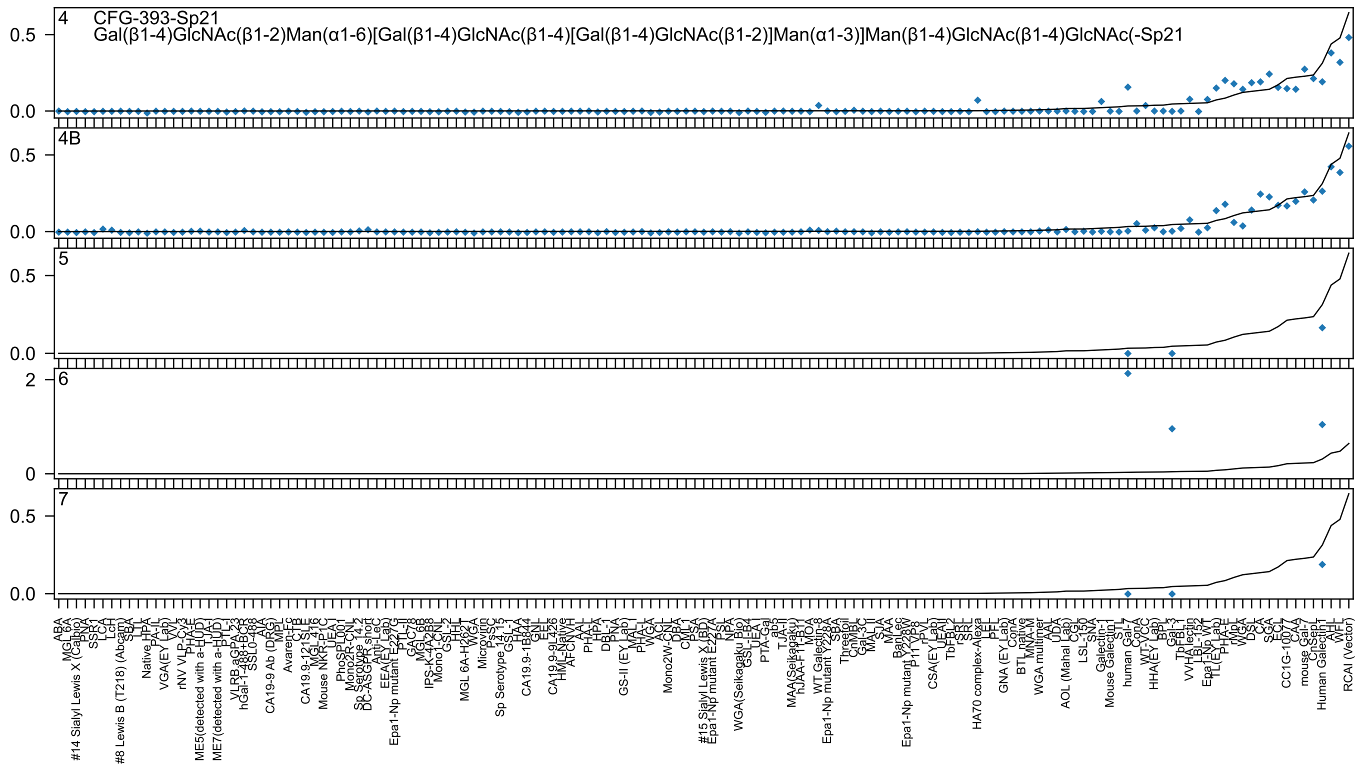

4 CFG-393-Sp21
Gal(β1-4)GlcNAc(β1-2)Man(α1-6)[Gal(β1-4)GlcNAc(β1-4)[Gal(β1-4)GlcNAc(β1-2)]Man(α1-3)]Man(β1-4)GlcNAc(β1-4)GlcNAc(-Sp21
0.5
0.0
4B
0.5
0.0
5
0.5
0.0
6
2
0
7
0.5
0.0
ABA
MGL 6A
#14 Sialyl Lewis X (Calbio)
PNA
SSR1
LCA
LcH
#8 Lewis B (T218) (Abcam)
SBA
LTL
Native HPA
PA-IL
VGA(EY Lab)
VVL
rNV VLP-Cy3
PHA-E
ME5(detected with a-HUD)
TJA-I
ME7(detected with a-HUD)
PTL-I
VLRB aGPA 23
hGal-1-488+BCR
SSL0-488
AIA
CA19-9 Ab (DRG)
MPL
Avaren-Fc
CTB
CA19.9-121SLE
MGL 416
Mouse NKR-P1C
UEA-I
PhoSL001
Mono2R-CNL
Sp Serotype 14.2
DC-ASGPR short
Anti-LeC
EEA(EY Lab)
Epa1-Np mutant E227G
PTL-II
GAC78
MGL 6B
IPS-K-4A2B8
Mono1-CNL
GSL-2
HHL
MGL 6A-H262T
WGA
Microvirin
PsSC
Sp Serotype 14.15
GSL-1
HAA
CA19.9-1B844
GNL
EEL
CA19.9-9L426
HML-Native
AFCNVH
AAL
PHA-L
HPA
DBL-1
PNA
GS-II (EY Lab)
MAL I
PHA-L
WGA
ACL
Mono2W-CNL
DBA
CML1
PSA
#15 Sialyl Lewis X (BD)
Epa1-Np mutant E227A
PSL
NPA
WGA(Seikagaku Bio)
GSL-I-B4
UEA I
PTA-Gal
ab5
TJA-II
MAA(Seikagaku)
hJAA-F11-B11
MOA
WT Galectin-8
Epa1-Np mutant Y228A
SBA
Threetoil
CnMBL
Gal-3C
MAL II
SJA
MAA
BanLec
Epa1-Np mutant Y228W
P11 VP8
rPVL
CSA(EY Lab)
UEAII
TbFBL2
rSRL
SRL
HA70 complex-Alexa
LEL
PFL
GNA (EY Lab)
ConA
BTL native
MNA-M
WGA multimer
AAL
UDA
AOL (Mahal Lab)
CGL
LSL-150
SNA
Galectin-1
Mouse Galectin-1
STL
human Gal-7
ConA
WT-YCC
HHA(EY Lab)
BPL
Gal-3
TbFBL1
VVHA lectin
LBL-152
Epa1-Np WT
TL(EY Lab)
PHA-E
rMpL
WGA
DSL
CA
SGA
ECL
CC1G-1007
CAA
mouse Gal-7
CnSepL
Human Galectin1
AHL
WFL
RCAI (Vector)

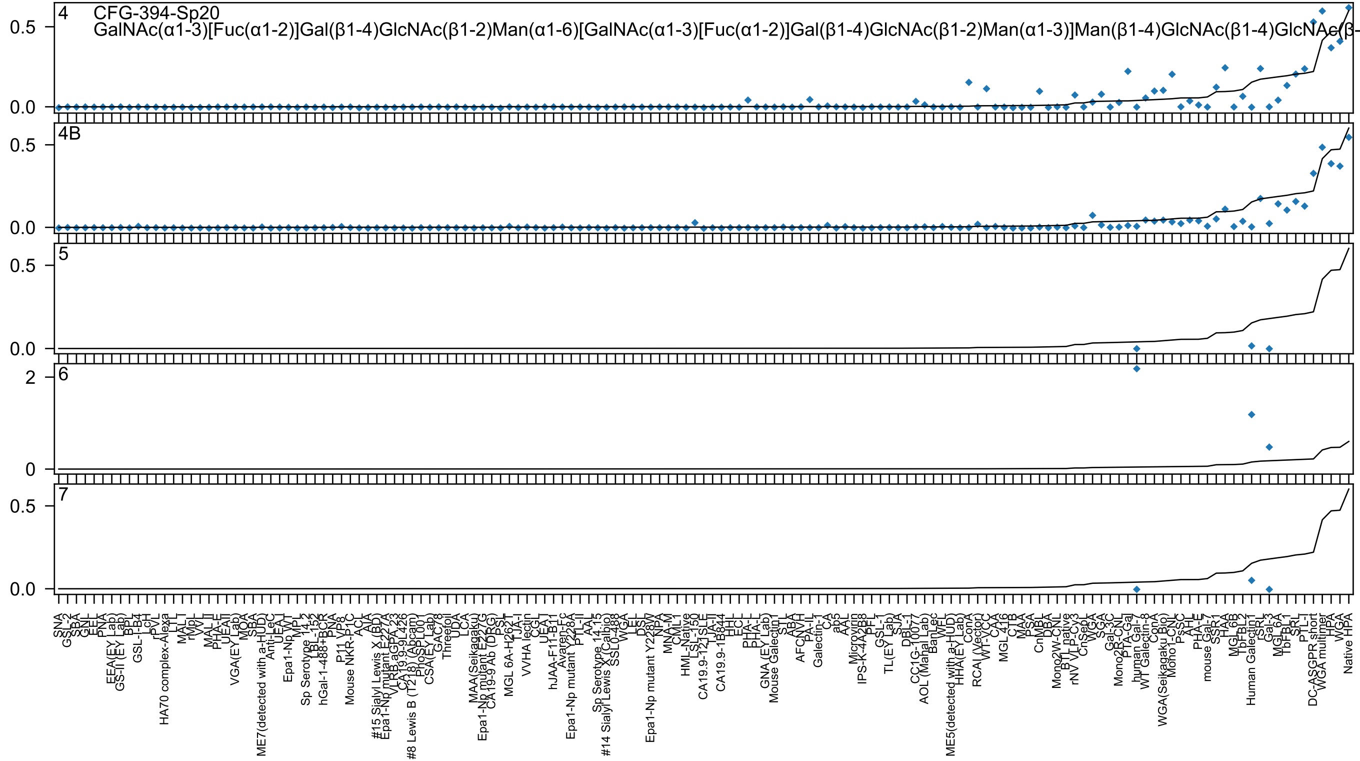
4
CFG-394-Sp20
GalNAc(α1-3)[Fuc(α1-2)]Gal(β1-4)GlcNAc(β1-2)Man(α1-6)[GalNAc(α1-3)[Fuc(α1-2)]Gal(β1-4)GlcNAc(β1-2)Man(α1-3)]Man(β1-4)GlcNAc(β1-4)GlcNAc(β-
4B
5
6
7

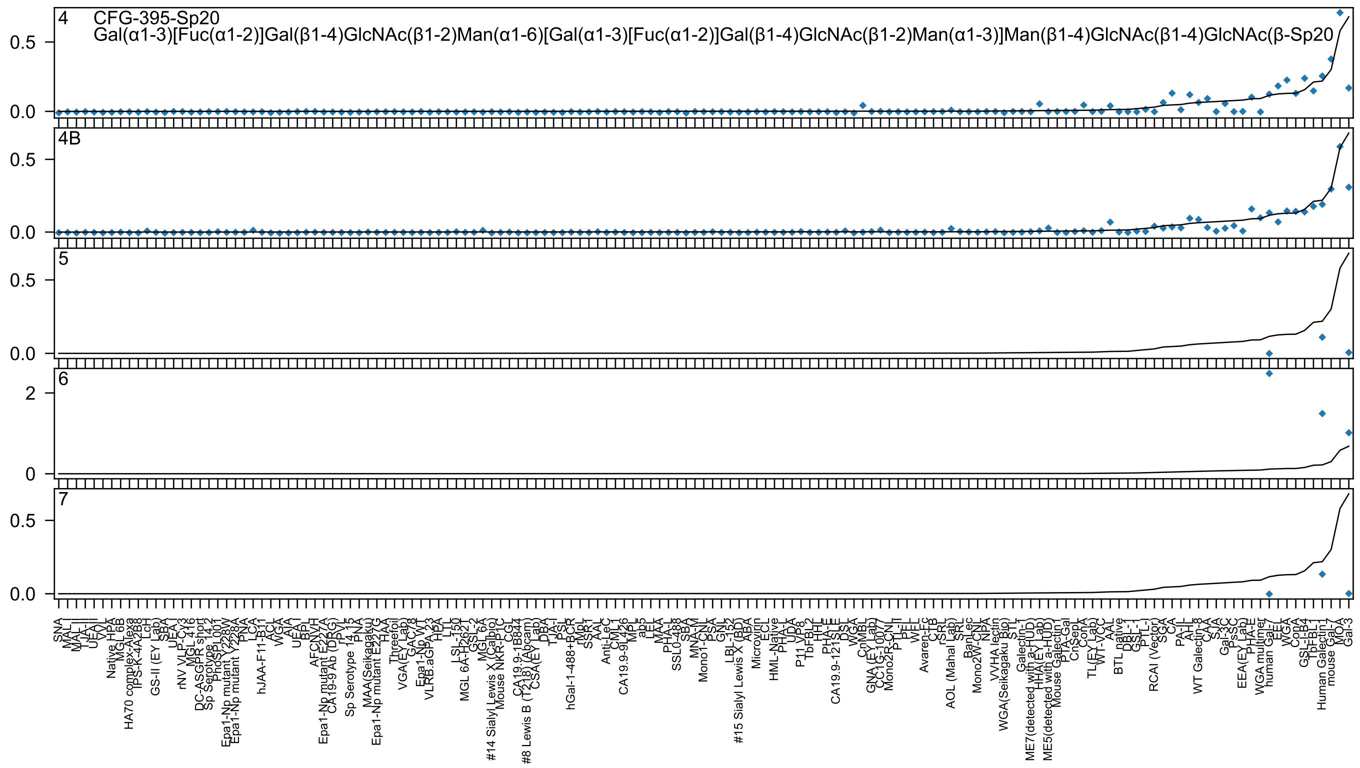
4 CFG-395-Sp20
Gal(α1-3)[Fuc(α1-2)]Gal(β1-4)GlcNAc(β1-2)Man(α1-6)[Gal(α1-3)[Fuc(α1-2)]Gal(β1-4)GlcNAc(β1-2)Man(α1-3)]Man(β1-4)GlcNAc(β1-4)GlcNAc(β-Sp20
4B
5
6
7

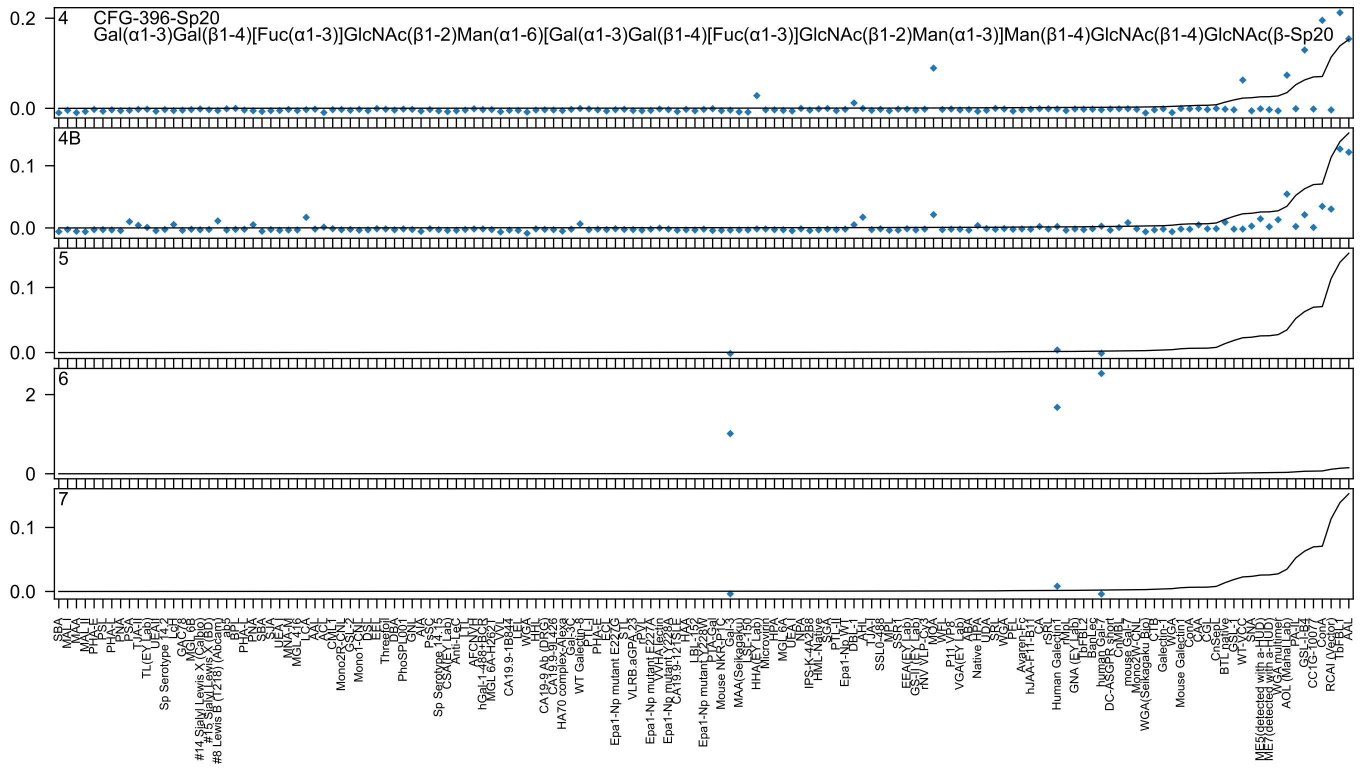
4
CFG-396-Sp20
Gal(α1-3)Gal(β1-4)[Fuc(α1-3)]GlcNAc(β1-2)Man(α1-6)[Gal(α1-3)Gal(β1-4)[Fuc(α1-3)]GlcNAc(β1-2)Man(α1-3)]Man(β1-4)GlcNAc(β1-4)GlcNAc(β-Sp20
4B
5
6
7

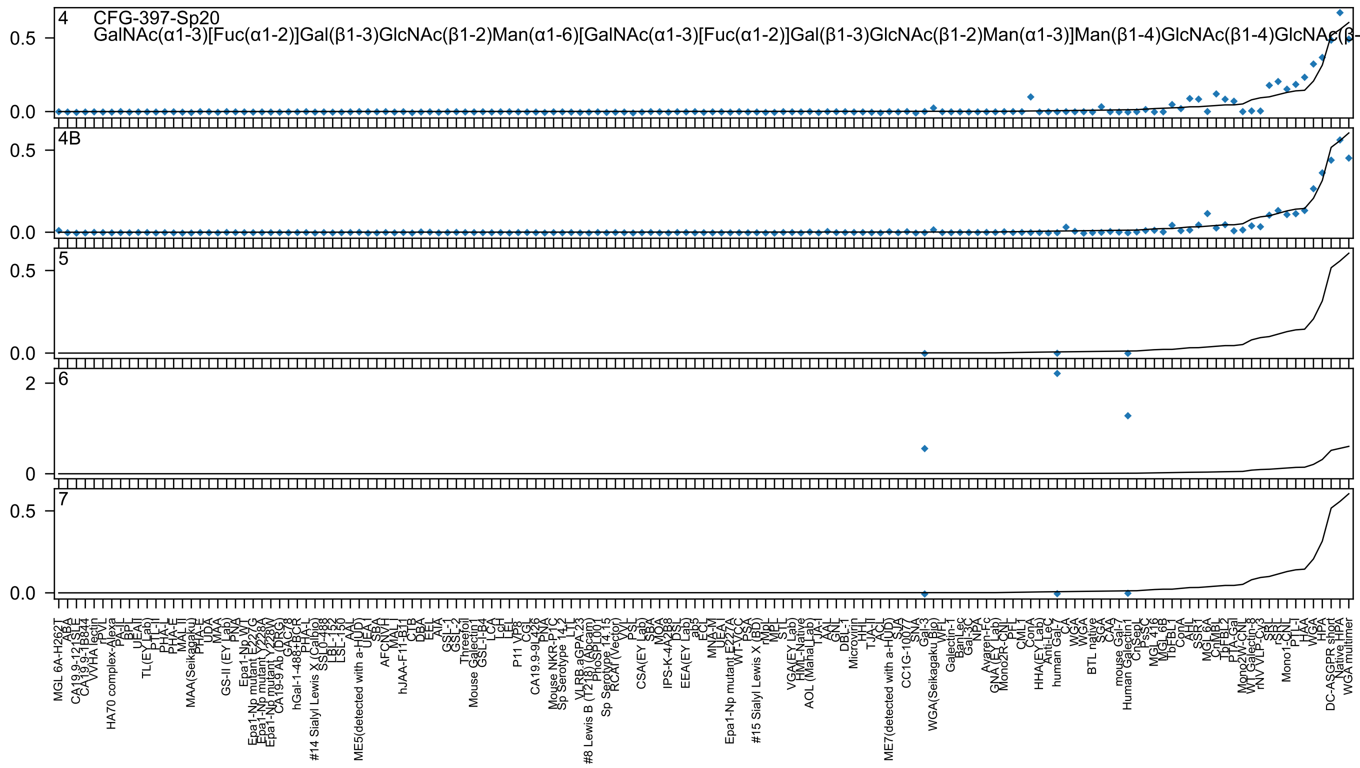

4 CFG-397-Sp20
GalNAc(α1-3)[Fuc(α1-2)]Gal(β1-3)GlcNAc(β1-2)Man(α1-6)[GalNAc(α1-3)[Fuc(α1-2)]Gal(β1-3)GlcNAc(β1-2)Man(α1-3)]Man(β1-4)GlcNAc(β1-4)GlcNAc(β-
4B
5
6
7
0.5
0.0
2
0

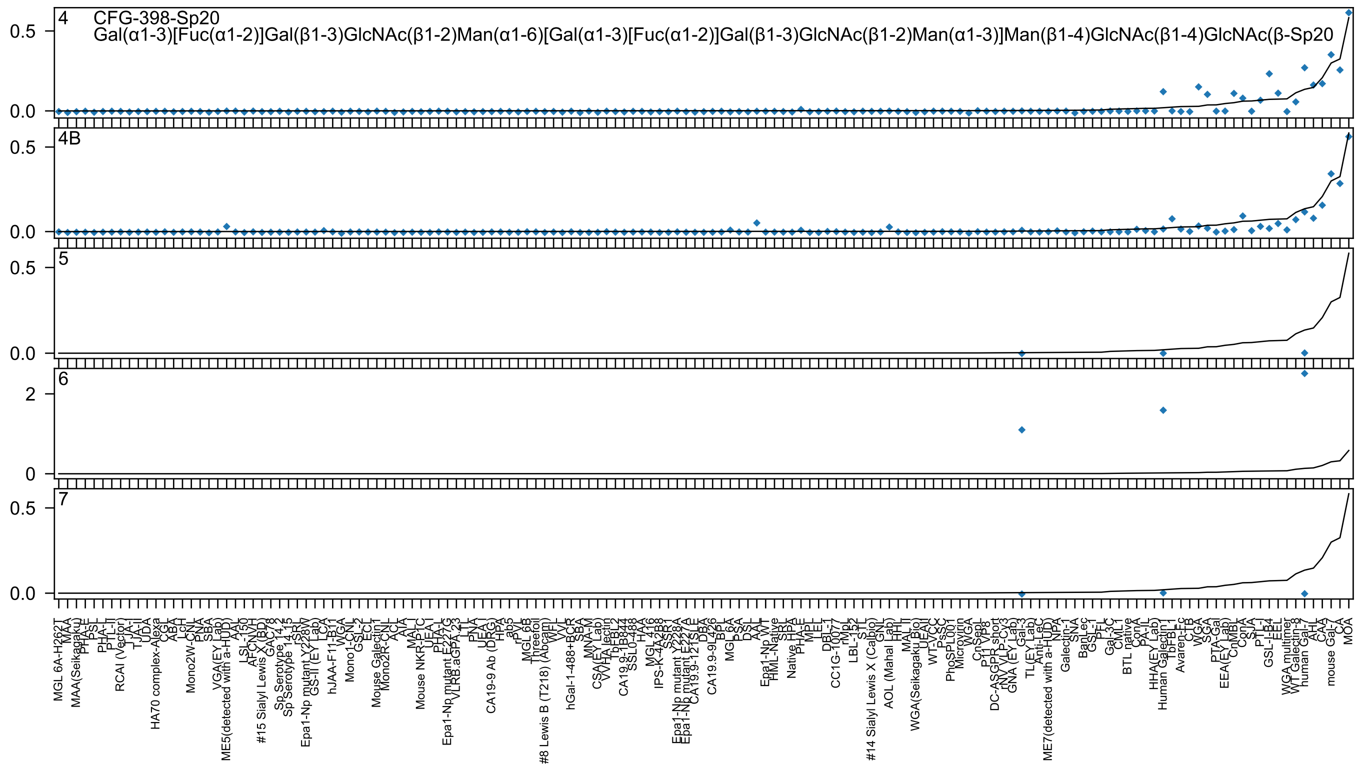

4
CFG-398-Sp20
Gal(α1-3)[Fuc(α1-2)]Gal(β1-3)GlcNAc(β1-2)Man(α1-6)[Gal(α1-3)[Fuc(α1-2)]Gal(β1-3)GlcNAc(β1-2)Man(α1-3)]Man(β1-4)GlcNAc(β1-4)GlcNAc(β-Sp20
4B
5
6
7

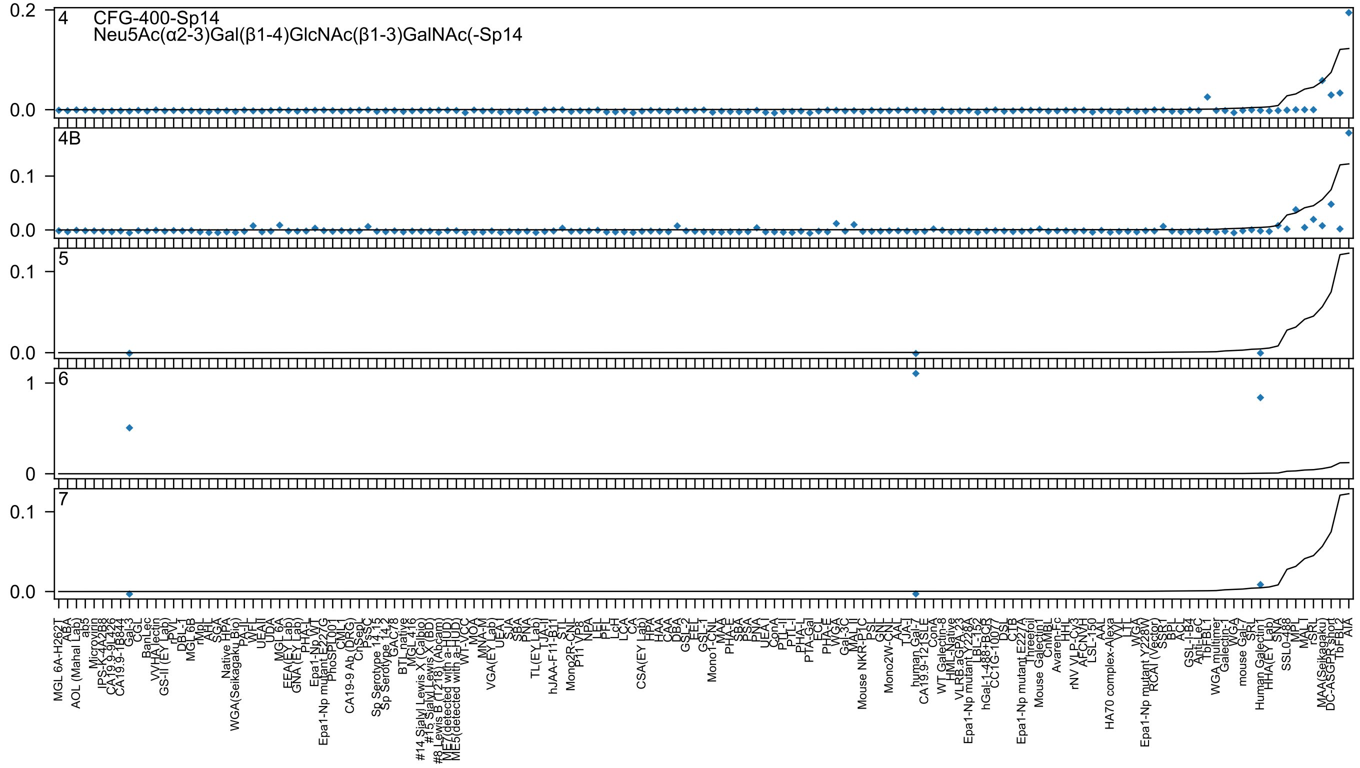
4
CFG-400-Sp14
Neu5Ac(α2-3)Gal(β1-4)GlcNAc(β1-3)GalNAc(-Sp14
4B
5
6
7

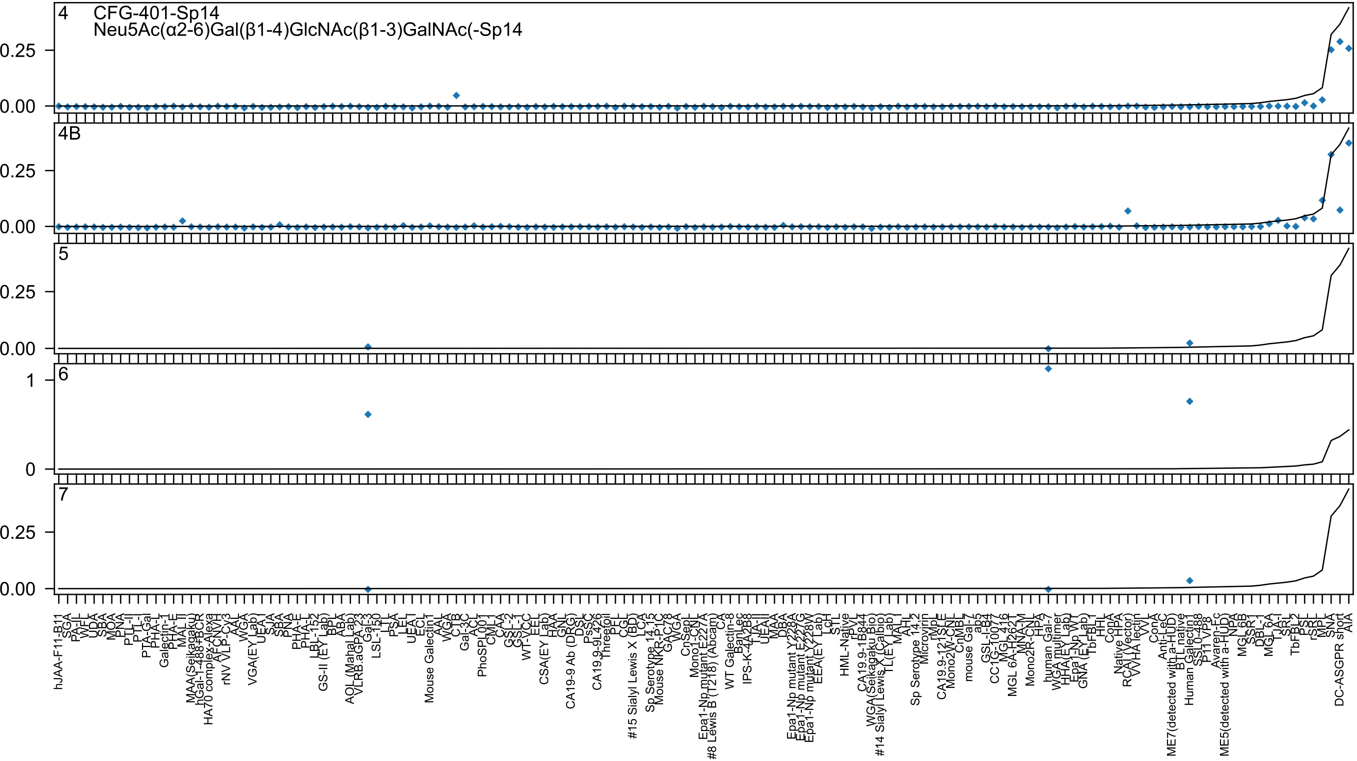
4
CFG-401-Sp14
Neu5Ac(α2-6)Gal(β1-4)GlcNAc(β1-3)GalNAc(-Sp14
4B
5
6
7
0.25
0.00
1
0

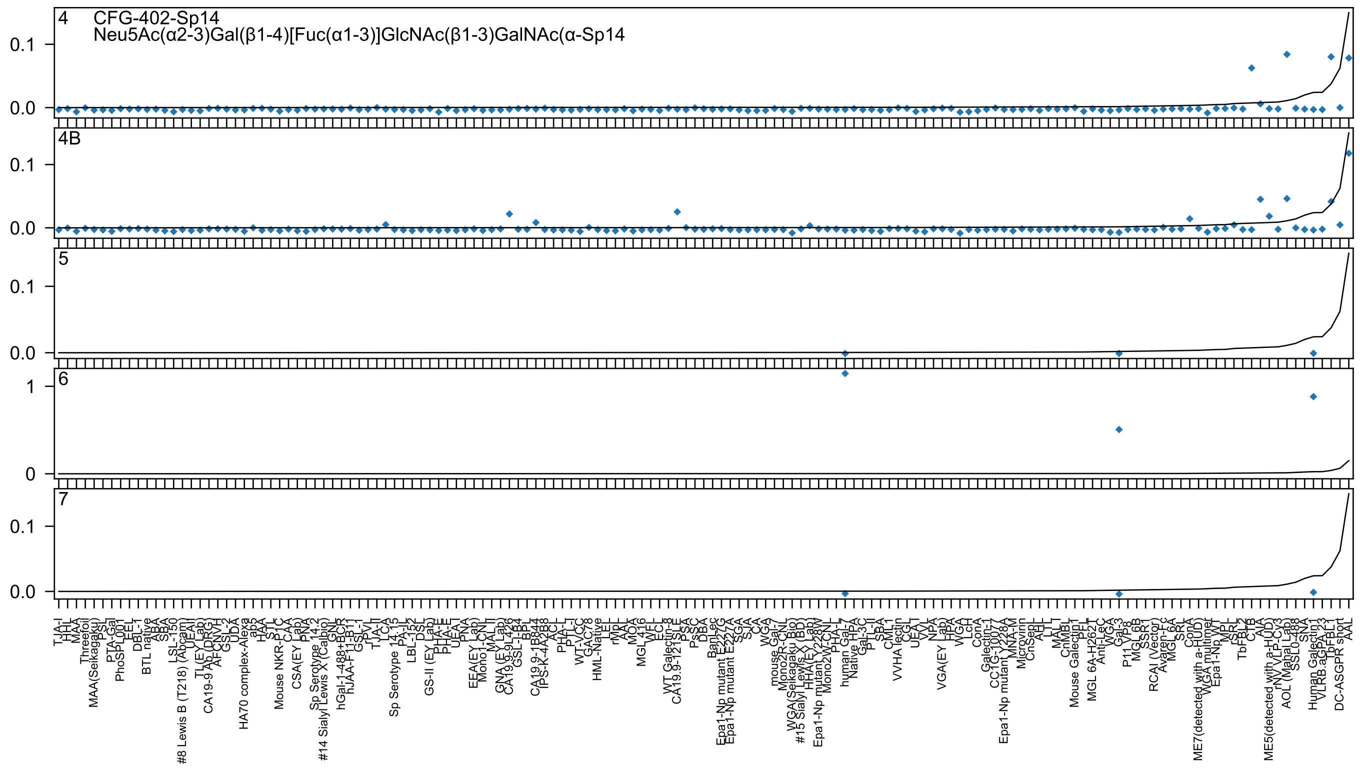

4
CFG-402-Sp14
Neu5Ac(α2-3)Gal(β1-4)[Fuc(α1-3)]GlcNAc(β1-3)GalNAc(α-Sp14
4B
5
6
7

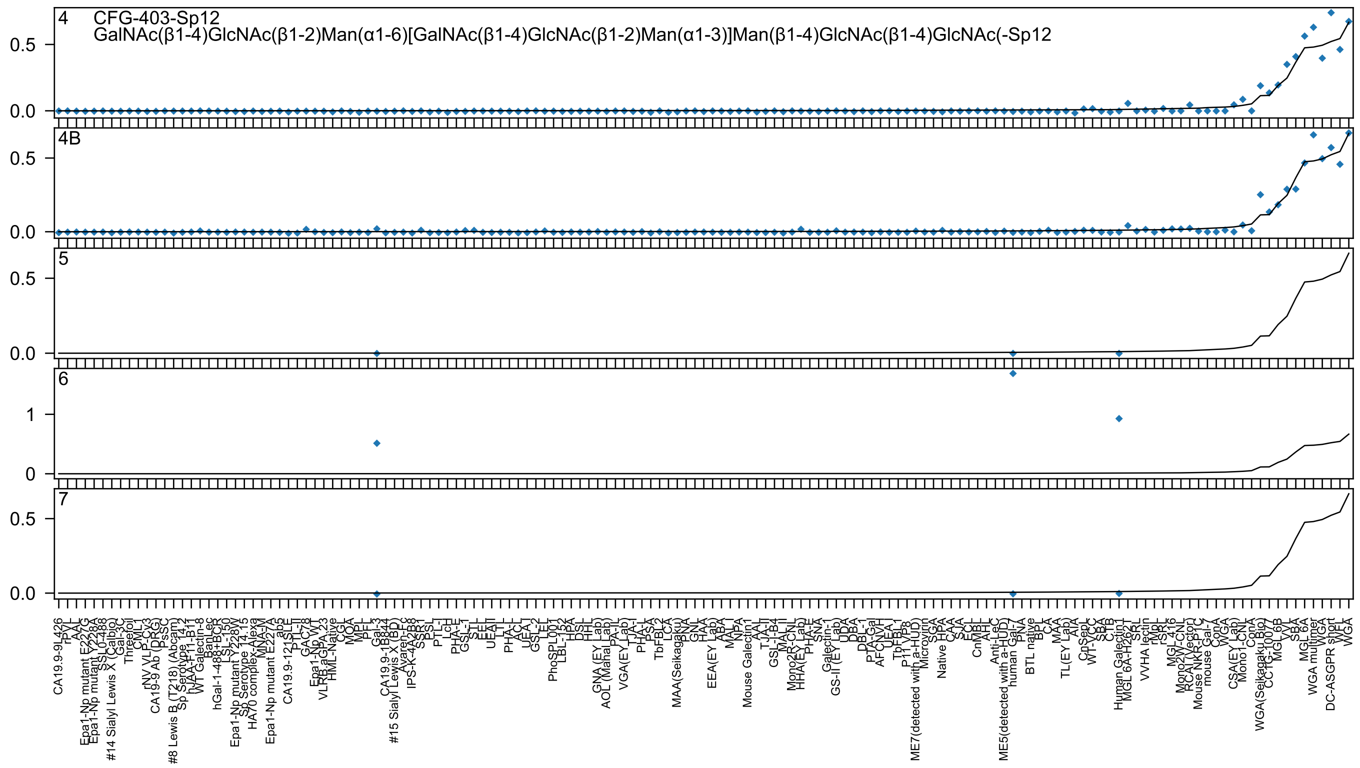

4 CFG-403-Sp12
GalNAc(β1-4)GlcNAc(β1-2)Man(α1-6)[GalNAc(β1-4)GlcNAc(β1-2)Man(α1-3)]Man(β1-4)GlcNAc(β1-4)GlcNAc(-Sp12
4B
5
6
7

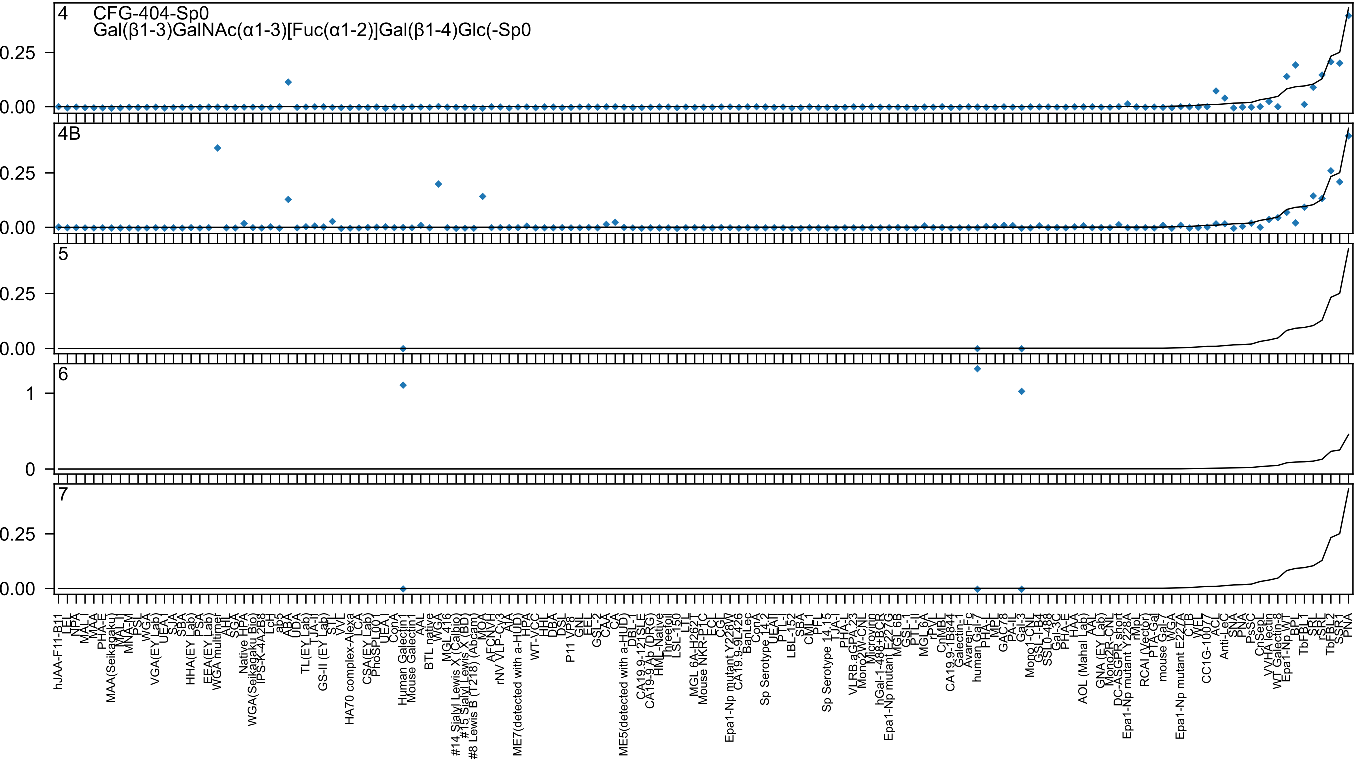
4
CFG-404-Sp0
Gal(β1-3)GalNAc(α1-3)[Fuc(α1-2)]Gal(β1-4)Glc(-Sp0
4B
5
6
7
0.25
0.00
1
0
hJAA-F11-B11
LEL
NPA
MAL-I
MAA
PHA-E
MAA(Seikagaku)
MAL-II
MNA-M
PSL
WGA
VGA(EY Lab)
UEA-I
SJA
SBA
HHA(EY Lab)
PSA
EEA(EY Lab)
WGA multimer
AHL
SGA
Native HPA
WGA(Seikagaku Bio)
IPS-K-4A2B8
LcH
ab5
ABA
UDA
TL(EY Lab)
TJA-II
GS-II (EY Lab)
STL
WVL
HA70 complex-Alexa
LCA
CSA(EY Lab)
PhoSL001
UEA-I
ConA
Human Galectin1
Mouse Galectin1
AAL
BTL native
WGA
MGL 416
#14 Sialyl Lewis X (Calbio)
#15 Sialyl Lewis X (BD)
#8 Lewis B (T218) (Abcam)
MOA
AFCNVH
rNV VLP-Cy3
AIA
ME7(detected with a-HUD)
HPA
WT-VCC
HHL
DBA
DSL
P11 VP8
GNL
EEL
GSL-2
CAA
CA
ME5(detected with a-HUD)
DBL-1
CA19.9-121SLE
CA19-9 Ab (DRG)
HML-Native
Threefoil
LSL-150
LTL
MGL 6A-H262T
Mouse NKR-P1C
ECL
CGL
Epa1-Np mutant Y228W
CA19.9-9L426
BanLec
ConA
Sp Serotype 14.2
UEAII
PTL-I
LBL-152
SBA
CML1
PFL
Sp Serotype 14.15
TJA-I
PHA-L
VLRB.aGPA.23
Mono2W-CNL
Microvirin
hGal-1-488+BCR
Epa1-Np mutant E227G
MGL 6B
GSL-I
PTL-II
MGL 6A
rPVL
CnMBL
CA19.9-1B844
Galectin-1
Avaren-Fc
human Gal-7
PHA-L
MPL
GAC78
PA-IL
Gal-3
Mono1-CNL
GSL-I-B4
SSL0-488
Gal-3C
PHA-E
HAA
AOL (Mahal Lab)
AAL
GNA (EY Lab)
Mono2R-CNL
DC-ASGPR short
Epa1-Np mutant Y228A
rMpL
RCAI (Vector)
PTA-Gal
mouse Gal-7
WGA
Epa1-Np mutant E227A
CTB
WFL
CC1G-10077
ACL
Anti-LeC
SNA
PsSC
CnSepL
VVHA lectin
WT Galectin-8
Epa1-Np WT
BPL
TbFBL1
SRL
TbFBL2
SSR1
PNA

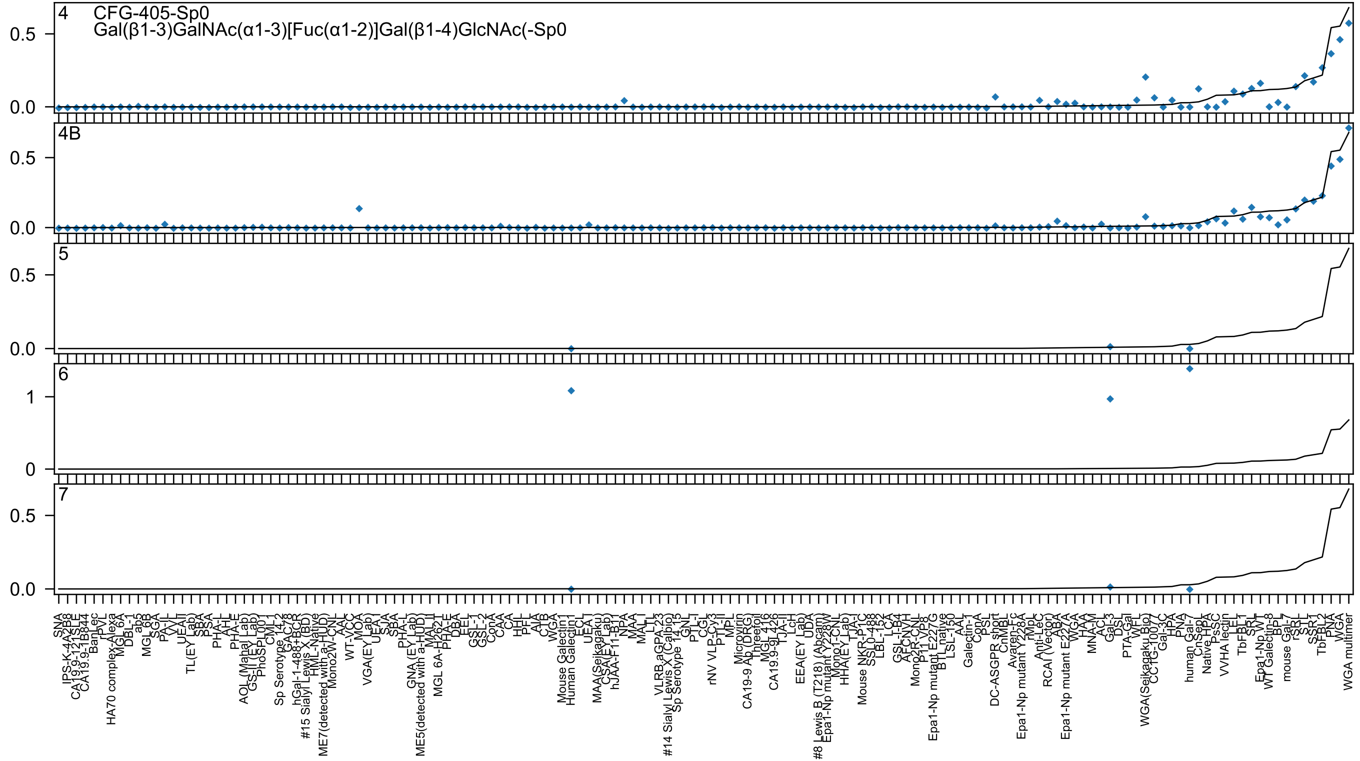

4 CFG-405-Sp0
Gal(β1-3)GalNAc(α1-3)[Fuc(α1-2)]Gal(β1-4)GlcNAc(-Sp0
4B
5
6
7

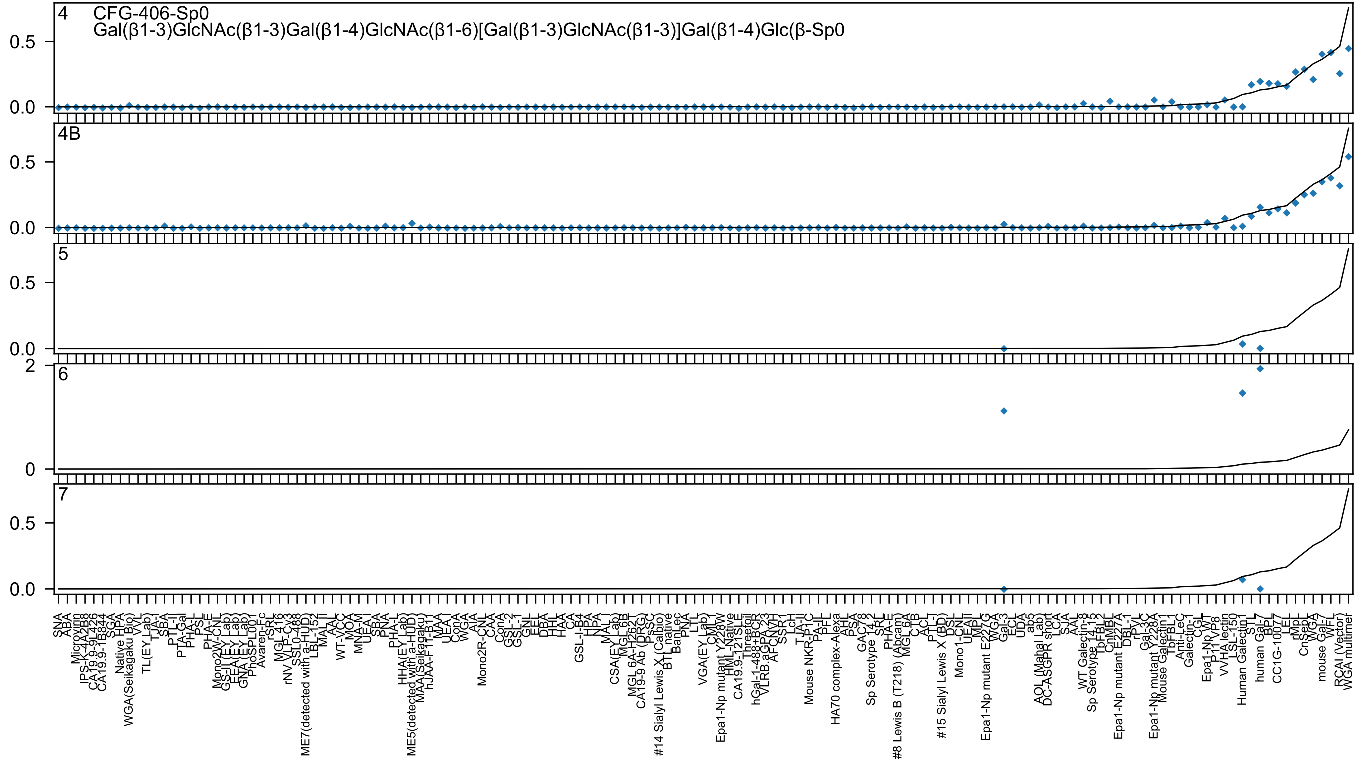

4 CFG-406-Sp0
Gal(β1-3)GlcNAc(β1-3)Gal(β1-4)GlcNAc(β1-6)[Gal(β1-3)GlcNAc(β1-3)]Gal(β1-4)Glc(β-Sp0
4B
5
6
7

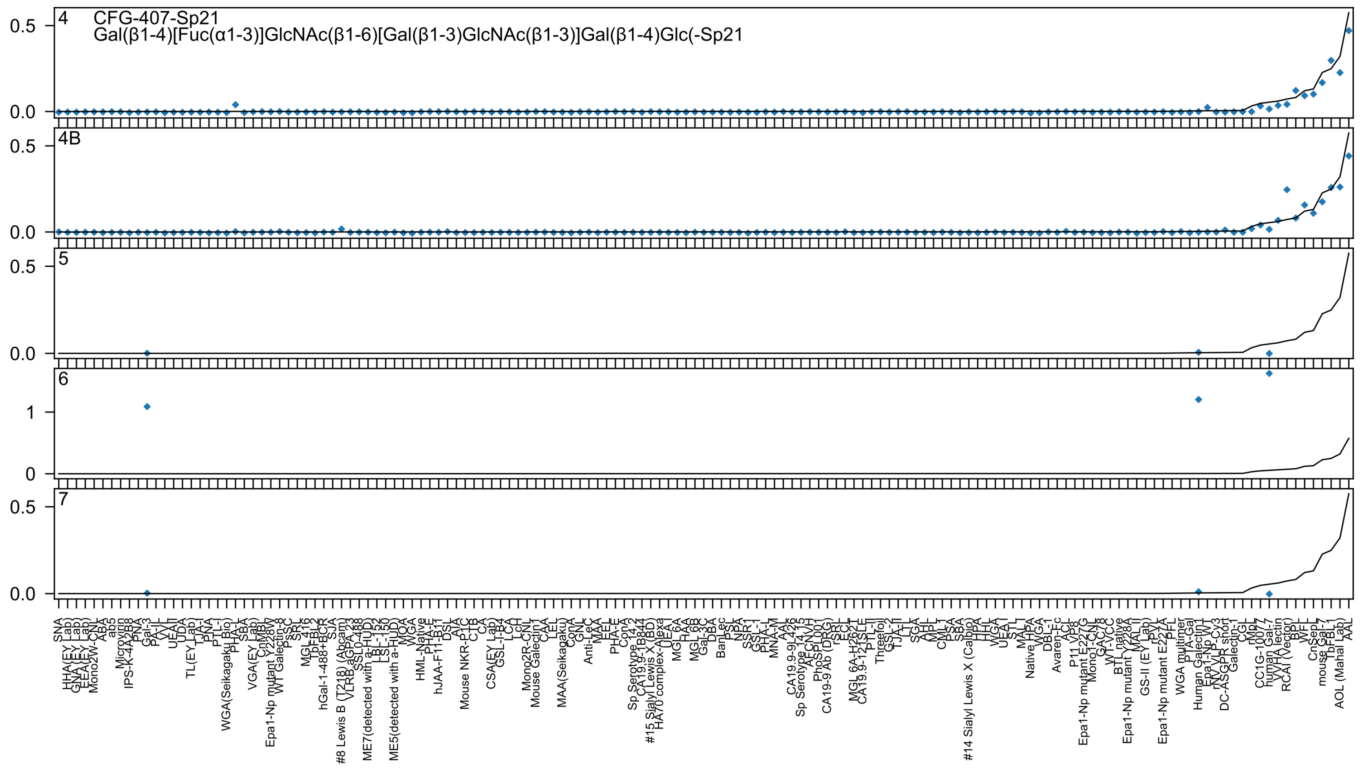
4
CFG-407-Sp21
Gal(β1-4)[Fuc(α1-3)]GlcNAc(β1-6)[Gal(β1-3)GlcNAc(β1-3)]Gal(β1-4)Glc(-Sp21
4B
5
6
7

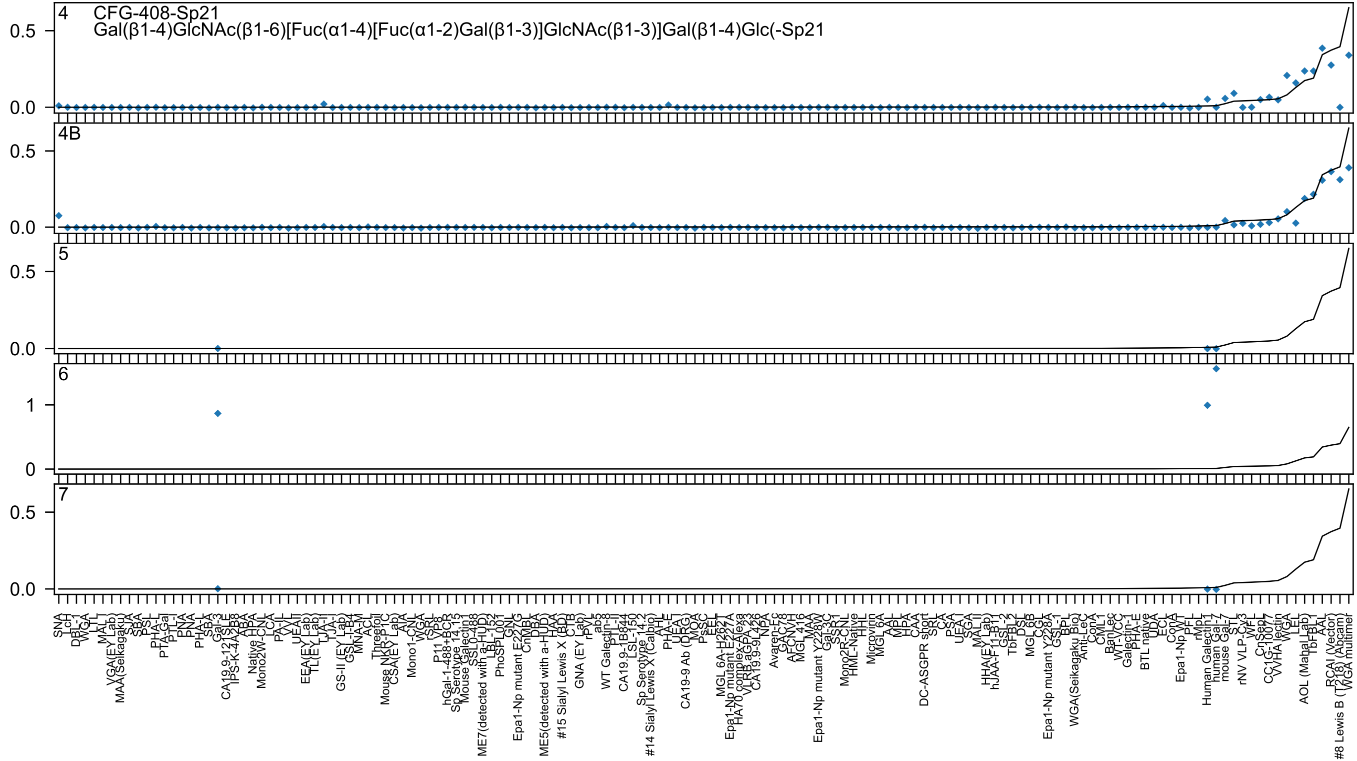
4 CFG-408-Sp21
Gal(β1-4)GlcNAc(β1-6)[Fuc(α1-4)[Fuc(α1-2)Gal(β1-3)]GlcNAc(β1-3)]Gal(β1-4)Glc(-Sp21
4B
5
6
7

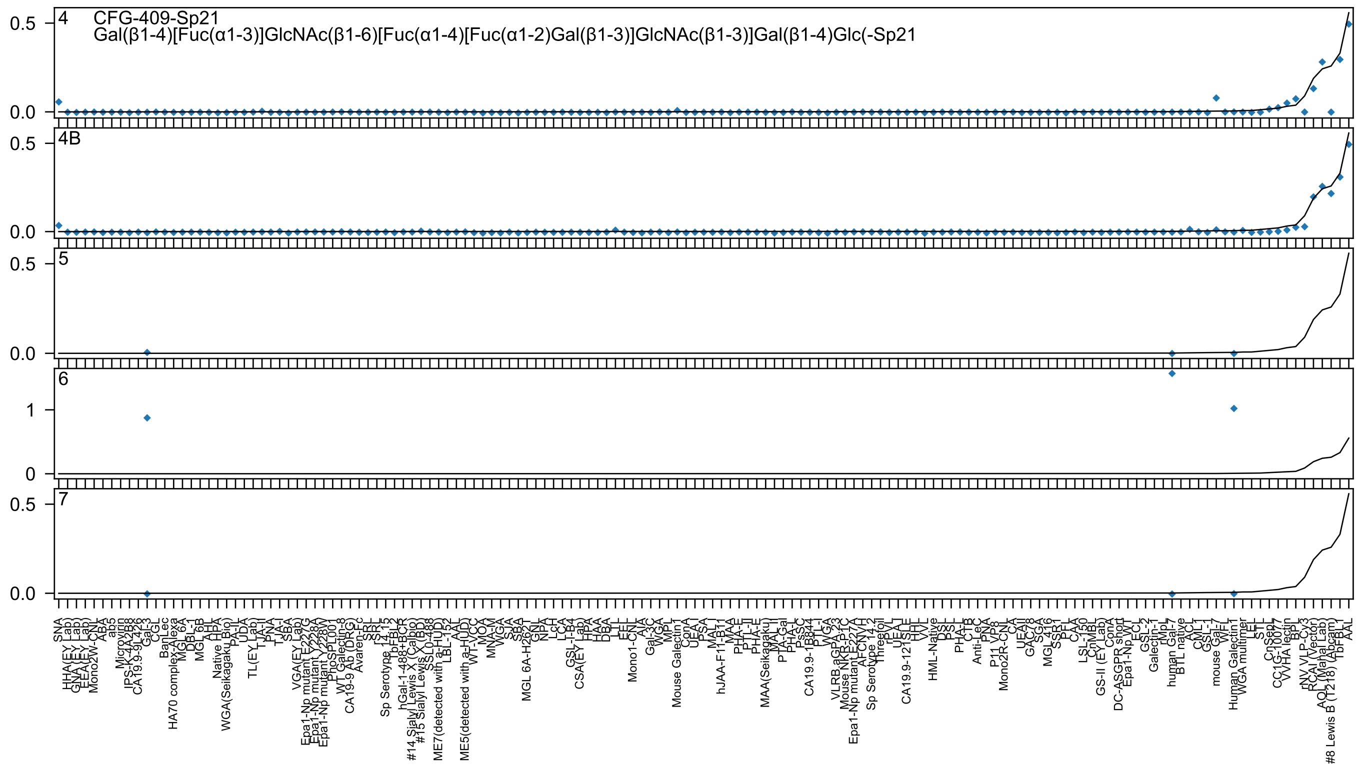
4
CFG-409-Sp21
Gal(β1-4)[Fuc(α1-3)]GlcNAc(β1-6)[Fuc(α1-4)[Fuc(α1-2)Gal(β1-3)]GlcNAc(β1-3)]Gal(β1-4)Glc(-Sp21
4B
5
6
7
0.5
0.0
1
0

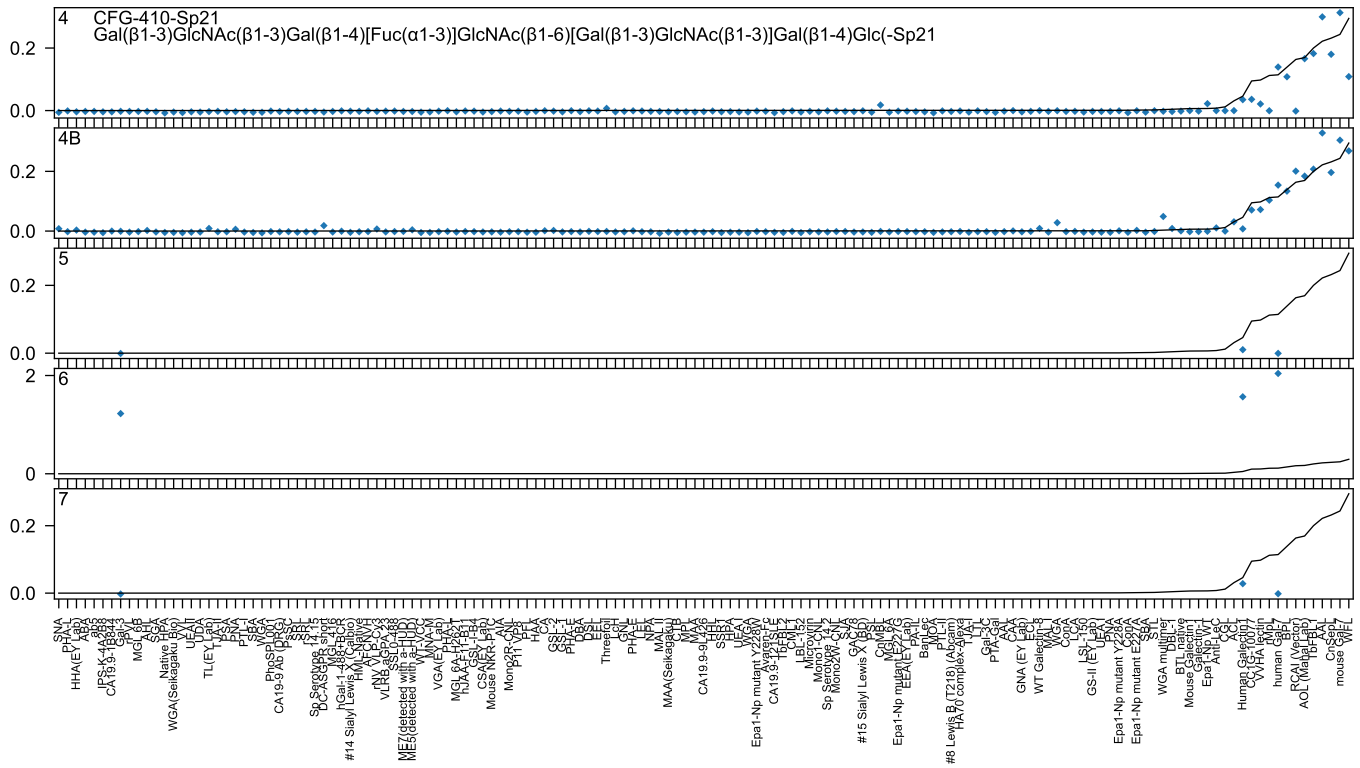
4
CFG-410-Sp21
Gal(β1-3)GlcNAc(β1-3)Gal(β1-4)[Fuc(α1-3)]GlcNAc(β1-6)[Gal(β1-3)GlcNAc(β1-3)]Gal(β1-4)Glc(-Sp21
4B
5
6
7
0.0
0.2
0
2

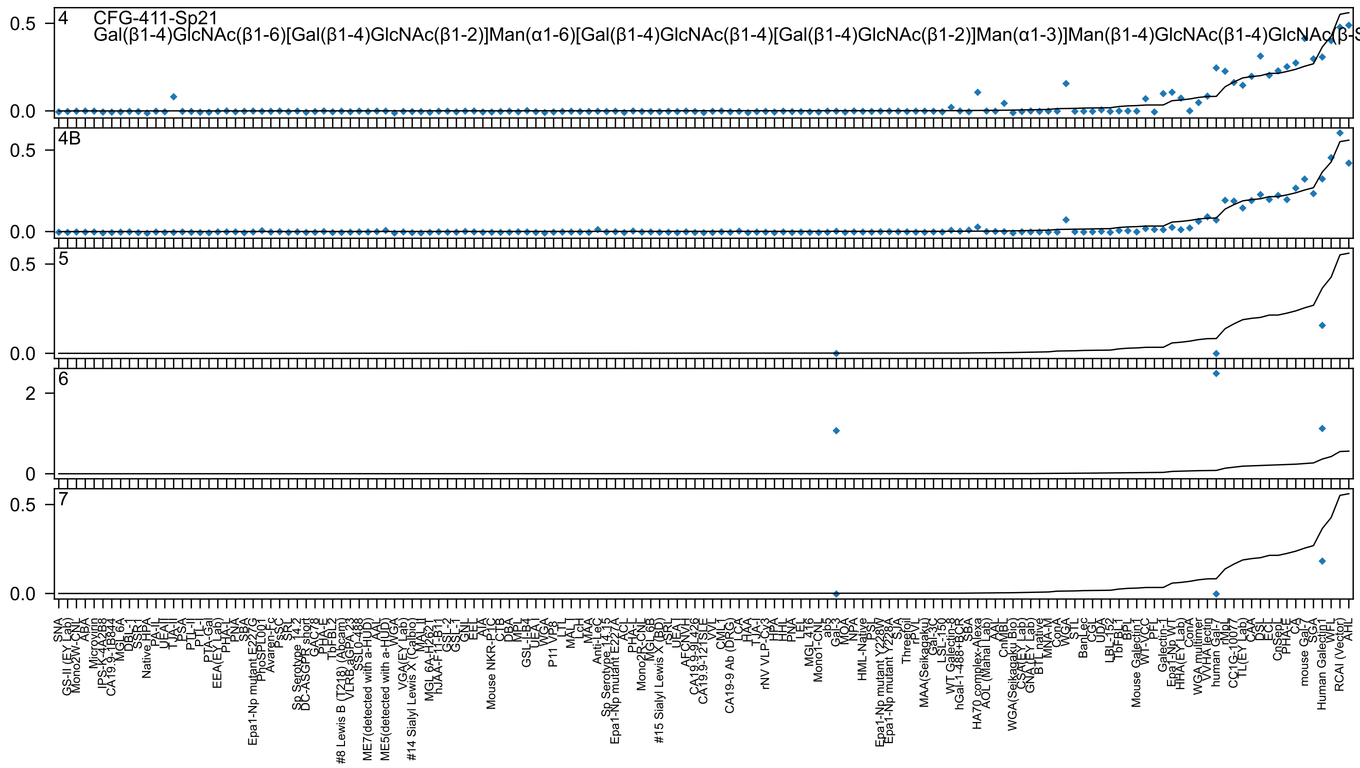

CFG-411-Sp21
Gal(β1-4)GlcNAc(β1-6)[Gal(β1-4)GlcNAc(β1-2)]Man(α1-6)[Gal(β1-4)GlcNAc(β1-4)[Gal(β1-4)GlcNAc(β1-2)]Man(α1-3)]Man(β1-4)GlcNAc(β1-4)GlcNAc(β-S
4
4B
5
6
7

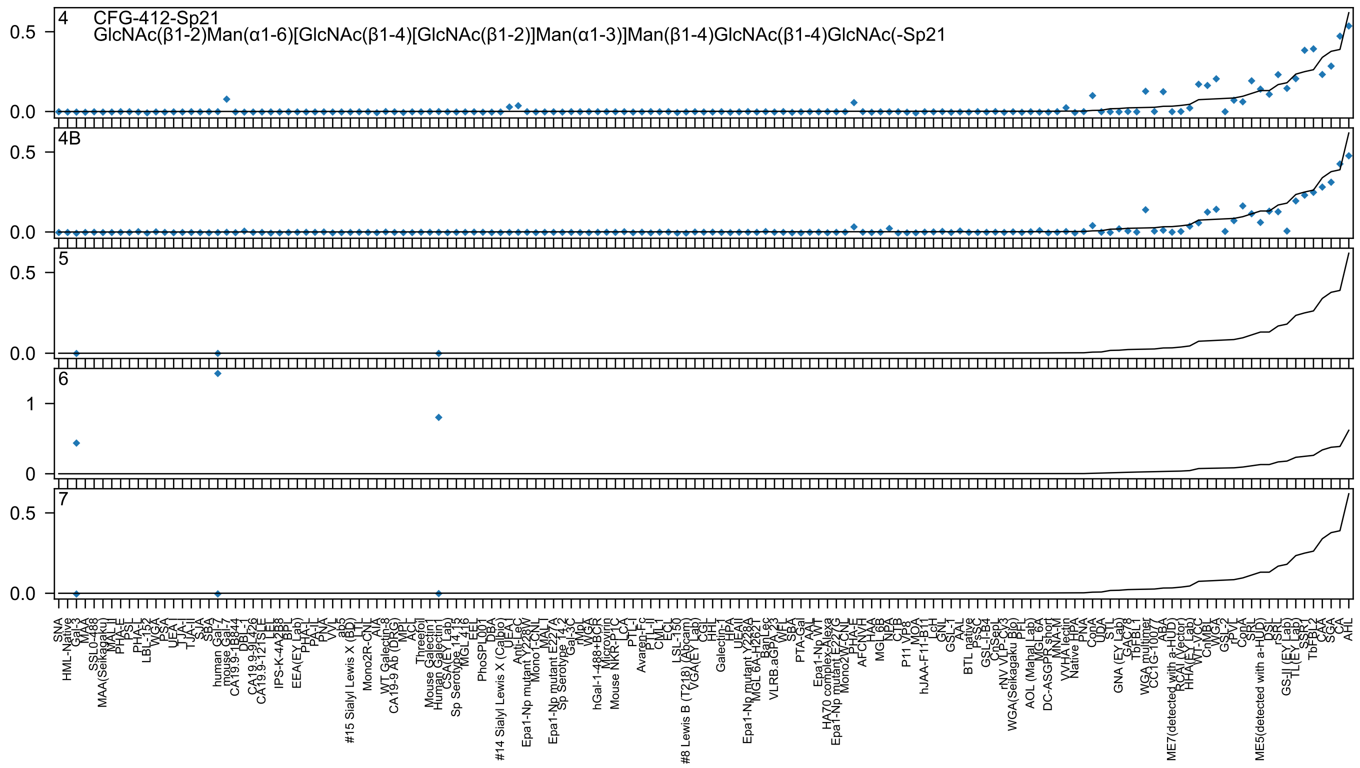
4
CFG-412-Sp21
GlcNAc(β1-2)Man(α1-6)[GlcNAc(β1-4)[GlcNAc(β1-2)]Man(α1-3)]Man(β1-4)GlcNAc(β1-4)GlcNAc(-Sp21
4B
5
6
7
0.0
0.5
0
1
SNA
HML-Native
Gal-3
MAA
SSL0-488
MAA(Seikagaku)
MAL_II
PHA-E
PSL
PHA-L
LBL-152
WGA
PSA
UEA-I
TJA-I
TJA-II
SJA
SBA
human Gal-7
mouse Gal-7
CA19.9-1B844
DBL-1
CA19.9-9L426
CA19.9-121SLE
LEL
IPS-K-4A2B8
BPL
EEA(EY Lab)
PHA-L
PA-IL
PNA
VVL
ab5
#15 Sialyl Lewis X (BD)
LTL
Mono2R-CNL
AIA
WT Galectin-8
CA19-9 Ab (DRG)
MPL
ACL
Threefoil
Mouse Galectin1
Human Galectin1
CSA(EY Lab)
Sp Serotype 14.15
MGL 4.16
EEL
PhoSPL001
DBA
#14 Sialyl Lewis X (Calbio)
UEA-I
Anti-LeC
Epa1-Np mutant Y228W
Mono1-CNL
MAL-I
Epa1-Np mutant E227A
Sp Serotype 14.2
Gal-3C
rMpL
WGA
hGal-1-488+BCR
Microvirin
Mouse NKR-P1C
LCA
PTL-I
Avaren-Fc
PTL-II
CML-1
ECL
LSL-150
#8 Lewis B (T218) (Abcam)
VGA(EY Lab)
CGL
HHL
Galectin-1
HPA
UEA-II
Epa1-Np mutant Y228A
MGL 6A-H262T
BanLec
VLRB.aGPA.23
WFL
SBA
PTA-Gal
AAL
Epa1-Np WT
HA70 complex-Alexa
Epa1-Np mutant E227G
Mono2-WT-CNL
PHA-E
AFCNVH
HAA
MGL 6B
NPA
CTB
P11 VP8
MOA
hJAA-F11-B11
LcH
GNL
GSL-1
AAL
BTL native
PsSC
GSL-I-B4
CnSepL
rNV VLP-Cv3
WGA(Seikagaku Bio)
PFL
AOL (Manal Lab)
MGL 6A
DC-ASGPR short
MNA-M
VVHA lectin
Native HPA
PNA
ConA
UDA
STL
GNA (EY Lab)
GAC78
TbFBL1
WGA multimer
CC1G-10077
ABA
ME7(detected with a-HUD)
RCAI (Vector)
HHA(EY Lab)
WT-VCC
CnMBL
WGA
GSL-2
rPVL
ConA
SRL
ME5(detected with a-HUD)
DSL
rSRL
GS-II (EY Lab)
TL(EY Lab)
SSR1
TbFBL2
CAA
SGA
CA
AHL

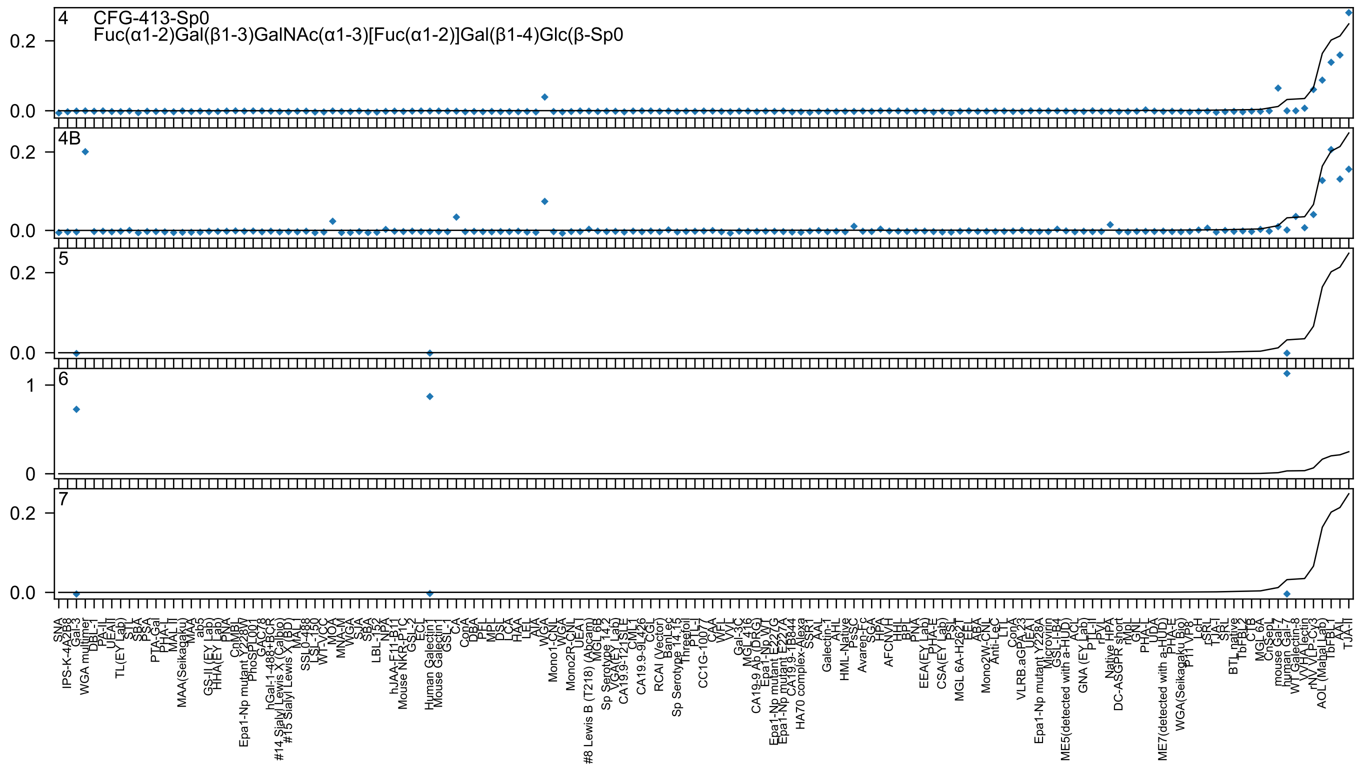
4
CFG-413-Sp0
Fuc(α1-2)Gal(β1-3)GalNAc(α1-3)[Fuc(α1-2)]Gal(β1-4)Glc(β-Sp0
4B
5
6
7
0.2
0.0
1
0

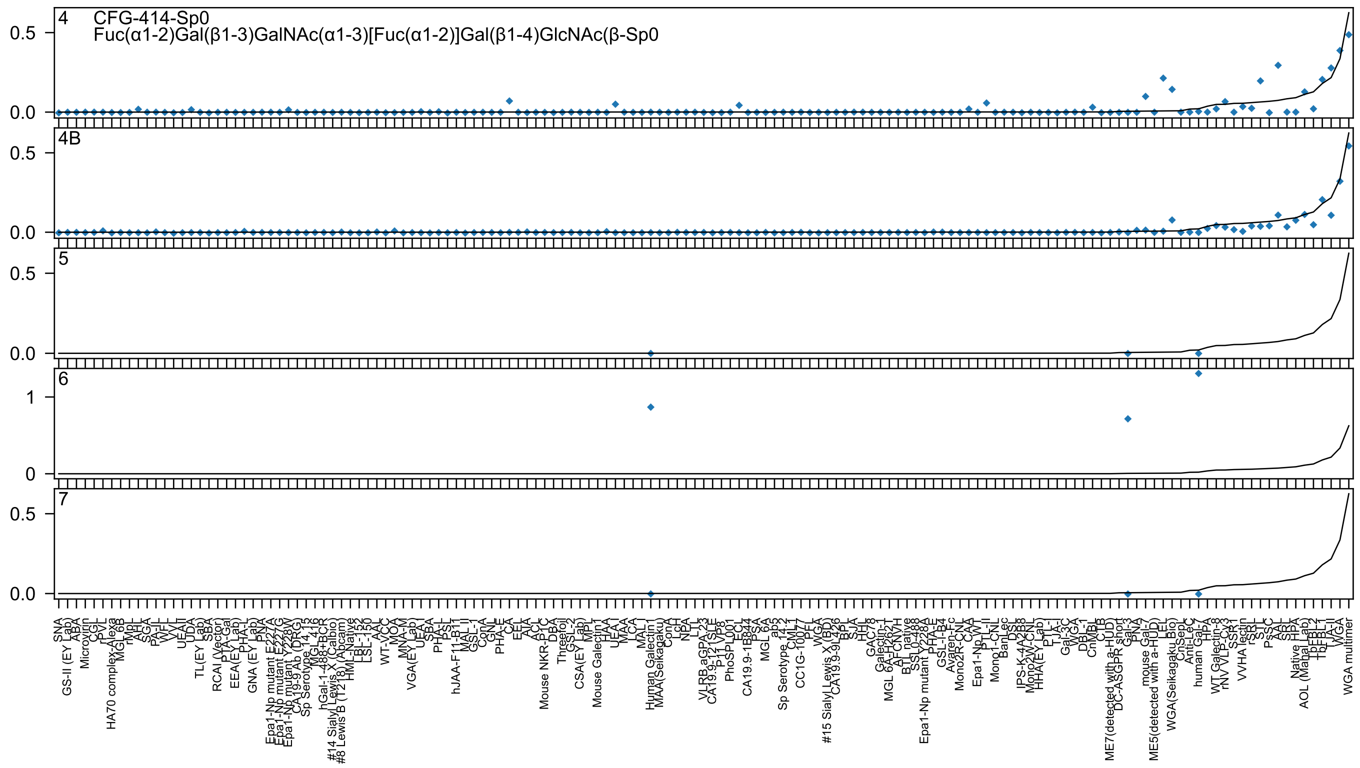

4 CFG-414-Sp0
Fuc(α1-2)Gal(β1-3)GalNAc(α1-3)[Fuc(α1-2)]Gal(β1-4)GlcNAc(β-Sp0
4B
5
6
7

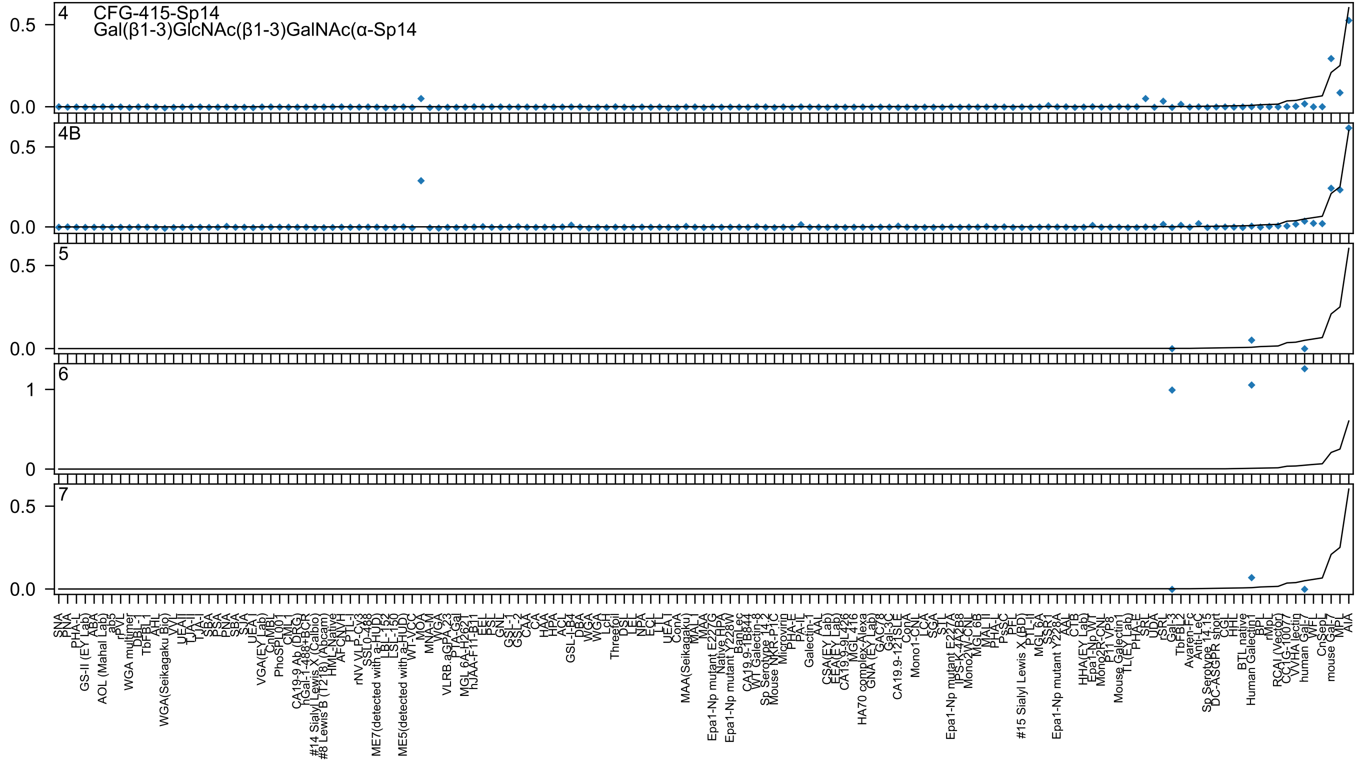
4
CFG-415-Sp14
Gal(β1-3)GlcNAc(β1-3)GalNAc(α-Sp14
0.5
0.0
4B
0.5
0.0
5
0.5
0.0
6
1
0
7
0.5
0.0

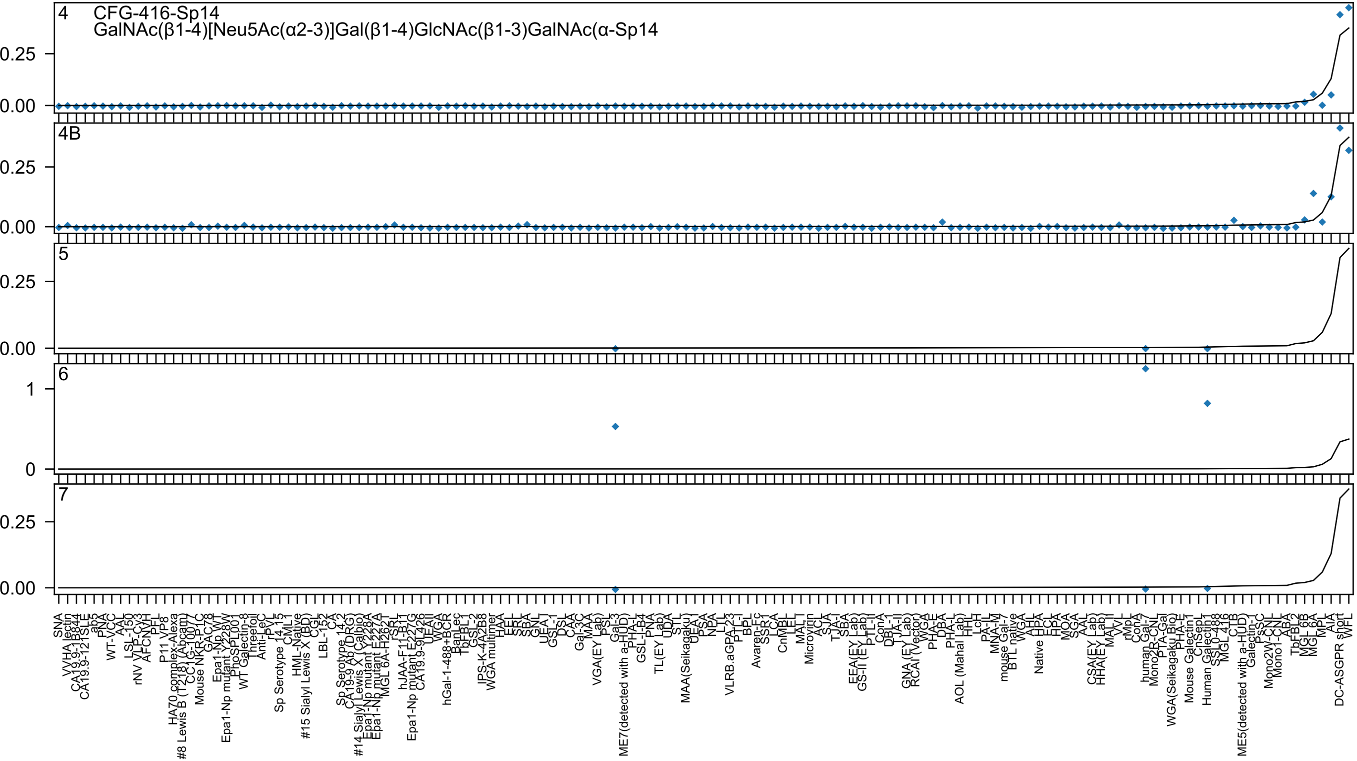
4 CFG-416-Sp14
GalNAc(β1-4)[Neu5Ac(α2-3)]Gal(β1-4)GlcNAc(β1-3)GalNAc(α-Sp14
4B
5
6
7
0.25
0.00
1
0

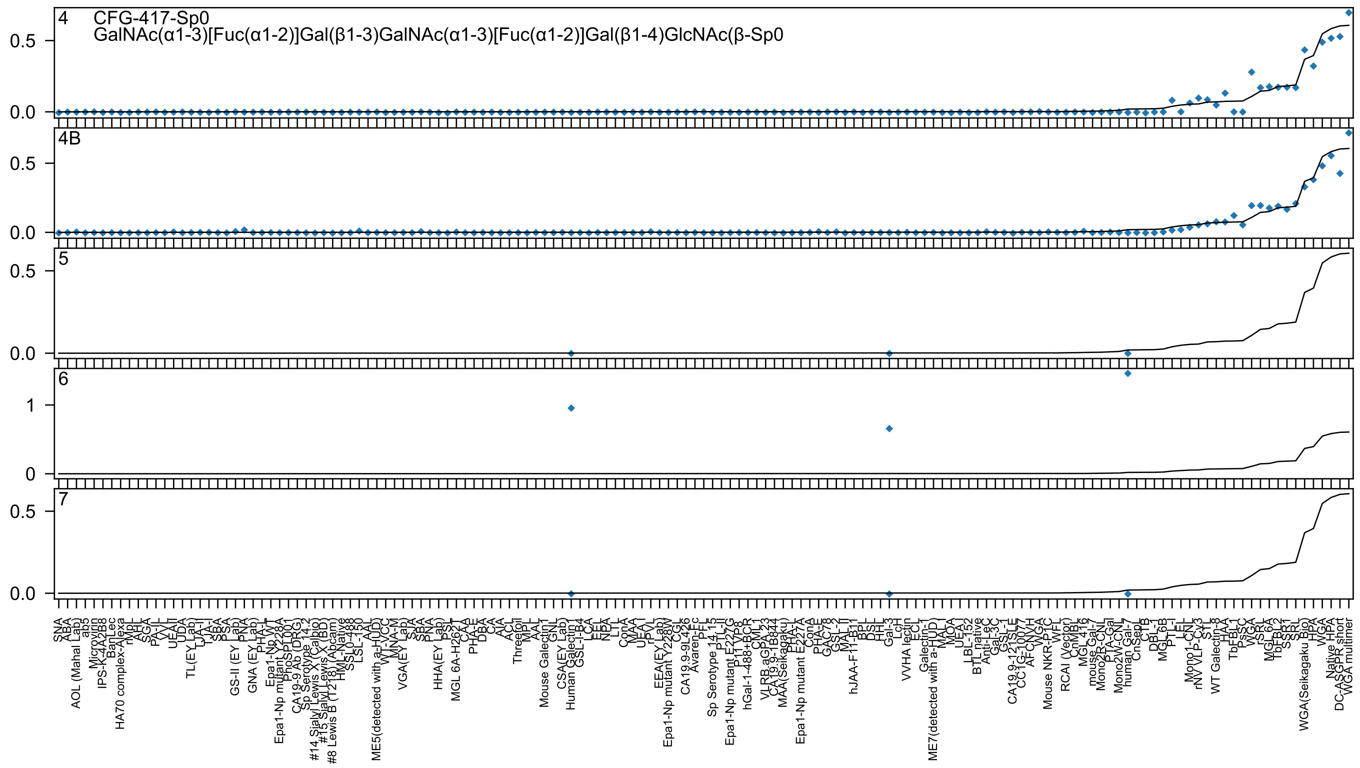

4
CFG-417-Sp0
GalNAc(α1-3)[Fuc(α1-2)]Gal(β1-3)GalNAc(α1-3)[Fuc(α1-2)]Gal(β1-4)GlcNAc(β-Sp0
4B
5
6
7
0.0
0.5
0
1
SNA
ABA
AOL (Mahal Lab)
ab5
Microvirin
IPS-K-4A2B8
BanLec
HA70 complex-Alexa
rMpL
AHL
SGA
PA-IL
VVL
UEAII
UDA
TL(EY Lab)
LTA-I
TJA-I
SBA
PSA
GS-II (EY Lab)
PNA
GNA (EY Lab)
PHA-L
Epa1-Np WT
Epa1-Np mutant Y228A
PhoSL001
CA19-9 Ab (DRG)
Sp Serotype 14.2
#14 Sialyl Lewis X (Calbio)
#15 Sialyl Lewis X (BD)
#8 Lewis B (T218) (Abcam)
HML-Native
SSL0-488
LSL-150
AAL
ME5(detected with a-HUD)
WT-VCC
MNA-M
VGA(EY Lab)
SJA
SBA
PNA
HHA(EY Lab)
PSL
MGL 6A-H262T
CAA
PHA-E
DBA
CA
AIA
ACL
Threefoil
MPL
AAL
Mouse Galectin1
GNL
CSA(EY Lab)
Human Galectin1
GSL-I-B4
LCA
EEL
NPA
LTL
ConA
MAA
UEA1
rPVL
EEA(EY Lab)
Epa1-Np mutant Y228W
CGL
CA19.9-9L426
Avaren-Fc
PFL
Sp Serotype 14.15
PTL-II
Epa1-Np mutant E227G
P11VP8
hGal-1-488+BCR
CML-1
VLRB.aGPA.23
CA19.9-1B844
MAA(Seikagaku)
PHA-L
Epa1-Np mutant E227A
ConA
PHA-E
GAC78
GSL-2
MAL II
hJAA-F11-B11
BPL
DSL
HHL
Gal-3
LcH
VVHA lectin
ECL
Galectin-1
ME7(detected with a-HUD)
MAL I
MOA
UEA-I
LBL-152
BTL native
Anti-LeC
Gal-3C
GSL-I
CA19.9-121SLE
CC1G-10077
AFC
NVH
WGA
Mouse NKR-P1C
WFL
RCAI (Vector)
CNMBL
MGL 416
mouse Gal-7
Mono2R-CNL
PTA-Gal
Mono2W-CNL
human Gal-7
CnSep
CTB
DBL-1
MGL 6B
PTL-I
LEL
Mono1-CNL
rNV VLP-Cy3
STL
WT Galectin-8
HAA
TbFBL1
PsSC
WGA
rSRL
MGL 6A
TbFBL2
SSR1
SRL
WGA(Seikagaku Bio)
HPA
WGA
Native HPA
DC-ASGPR short
WGA multimer

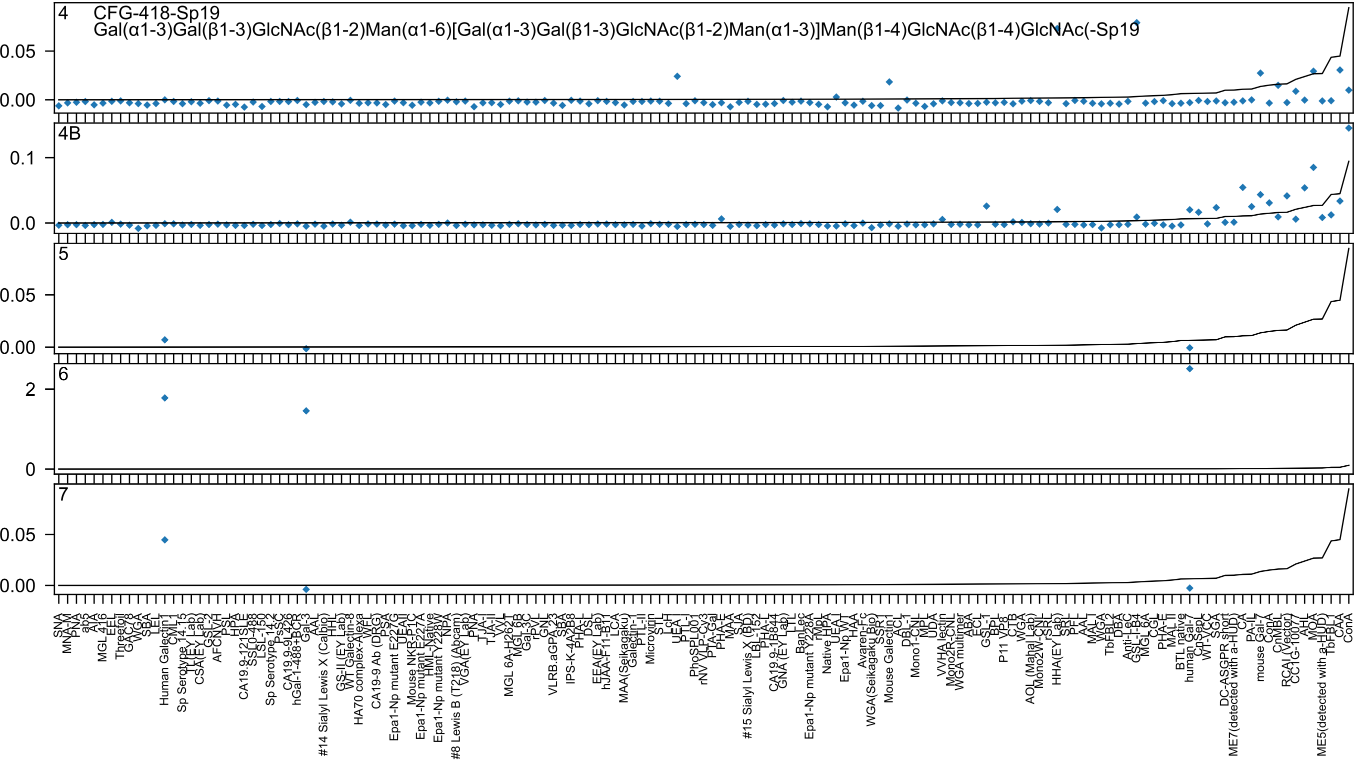
4
CFG-418-Sp19
Gal(α1-3)Gal(β1-3)GlcNAc(β1-2)Man(α1-6)[Gal(α1-3)Gal(β1-3)GlcNAc(β1-2)Man(α1-3)]Man(β1-4)GlcNAc(β1-4)GlcNAc(-Sp19
0.05
0.00
4B
0.1
0.0
5
0.05
0.00
6
2
0
7
0.05
0.00

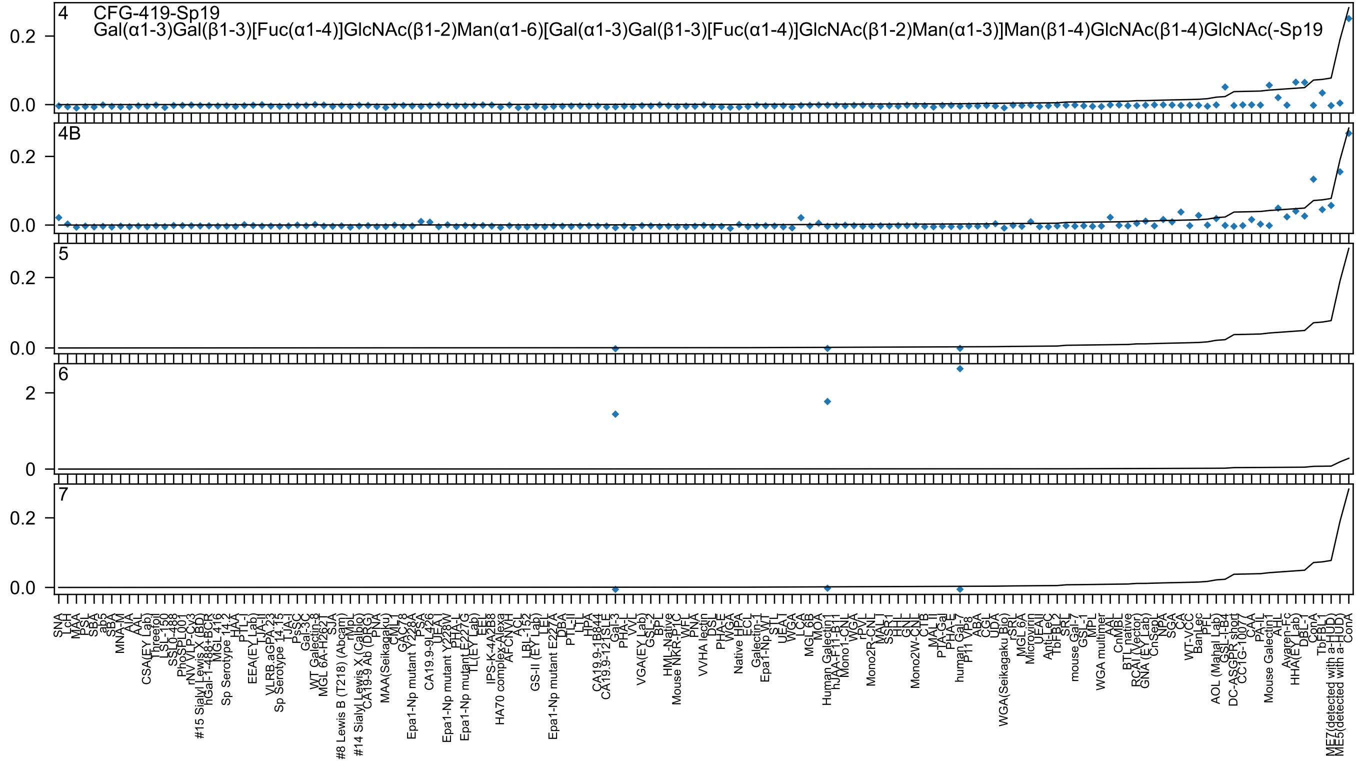

4
CFG-419-Sp19
Gal(α1-3)Gal(β1-3)[Fuc(α1-4)]GlcNAc(β1-2)Man(α1-6)[Gal(α1-3)Gal(β1-3)[Fuc(α1-4)]GlcNAc(β1-2)Man(α1-3)]Man(β1-4)GlcNAc(β1-4)GlcNAc(-Sp19
4B
5
6
7

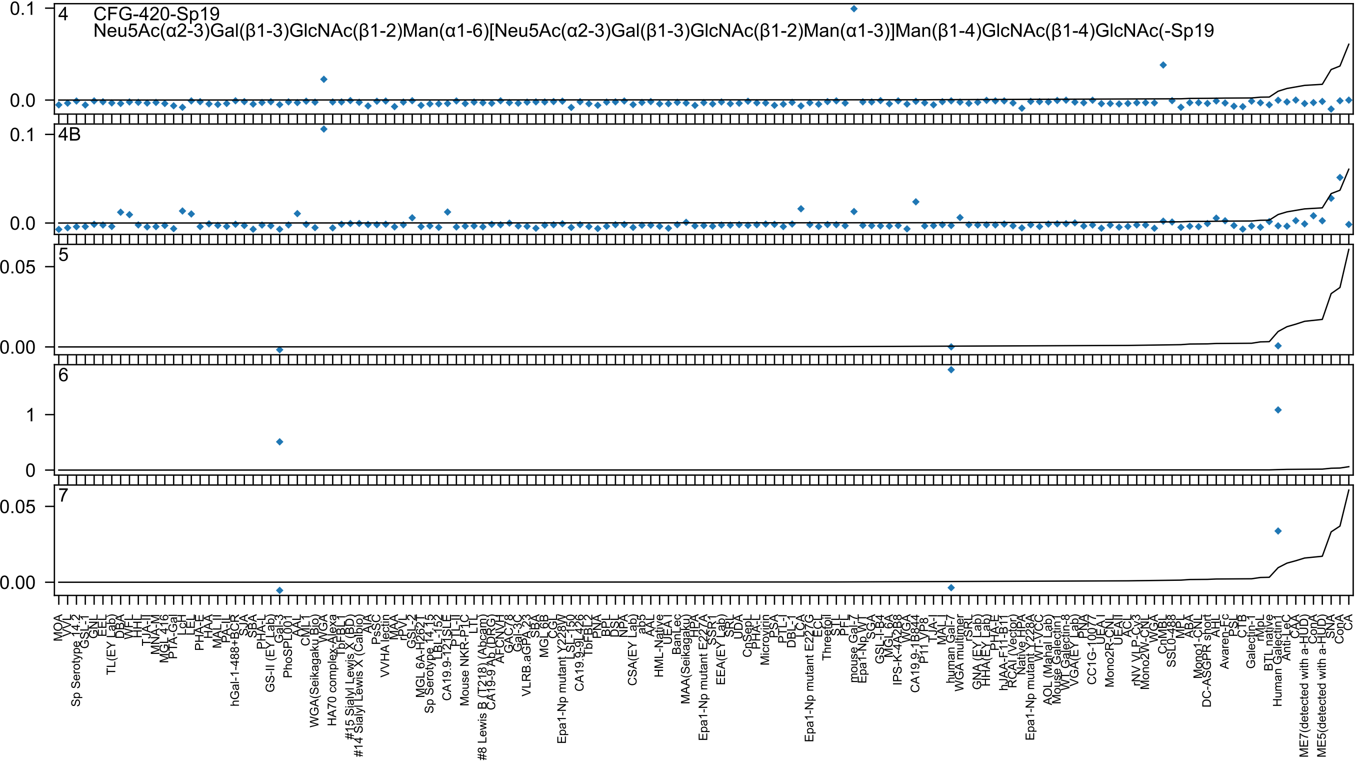

CFG-420-Sp19
Neu5Ac(α2-3)Gal(β1-3)GlcNAc(β1-2)Man(α1-6)[Neu5Ac(α2-3)Gal(β1-3)GlcNAc(β1-2)Man(α1-3)]Man(β1-4)GlcNAc(β1-4)GlcNAc(-Sp19
4
4B
5
6
7

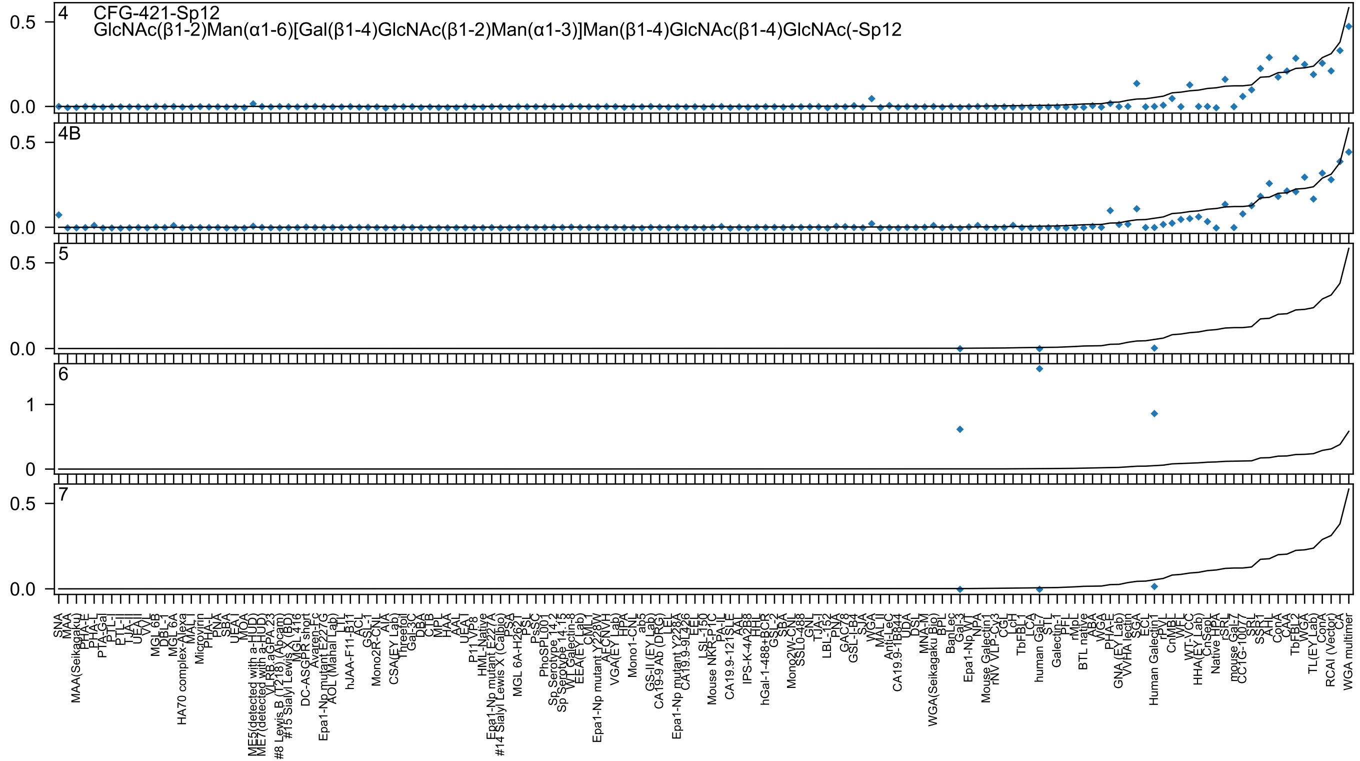
4
CFG-421-Sp12
GlcNAc(β1-2)Man(α1-6)[Gal(β1-4)GlcNAc(β1-2)Man(α1-3)]Man(β1-4)GlcNAc(β1-4)GlcNAc(-Sp12
4B
5
6
7
0.0
0.5
0
1
SNA
MAA
MAA(Seikagaku)
PHA-E
PHA-L
PTA-Gal
PTL-I
PTL-II
TJA-II
UEA-I
VVL
MGL 6B
DBL-1
MGL 6A
HA70 complex-Alexa
MAL-I
Microvirin
PHA-L
PNA
SBA
UEA-I
MOA
ME5(detected with a-HUD)
ME7(detected with a-HUD)
VLRB.aGPA.23
#8 Lewis B (T218) (Abcam)
#15 Sialyl Lewis X (BD)
MGL 416
DC-ASGPR short
Avaren-Fc
Epa1-Np mutant E227G
AOL (Mahal Lab)
LTL
hJAA-F11-B11
ACL
GSL-I
Mono2R-CNL
AIA
CSA(EY Lab)
Threefoil
Gal-3C
DBA
CTB
MPL
HAA
AAL
UEA-I
VP8
P11 VP8
HML-Native
Epa1-Np mutant E227A
#14 Sialyl Lewis X (Calbio)
PSA
MGL 6A-H262T
PSL
PsSC
PhoSL
PL001
Sp Serotype 14.2
Sp Serotype 14.15
WT Galectin-8
EEA(EY Lab)
CML
Epa1-Np mutant Y228W
AFCNVH
VGA(EY Lab)
HPA
Mono1-CNL
ab5
GS-II (EY Lab)
CA19-9 Ab (DRG)
LEL
Epa1-Np mutant Y228A
CA19.9-9L426
EEL
LSL-150
Mouse NKR-P1C
PA-IL
CA19.9-121SLE
AAL
IPS-K-4A2B8
HHL
hGal-1-488+BCR
GSL-2
SBA
Mono2W-CNL
SSL0-488
GNL
TJA-I
LBL-152
PNA
GAC78
GSL-I-B4
SJA
WGA
MAL II
Anti-LeC
CA19.9-1B844
UDA
DSL
MNA-M
WGA(Seikagaku Bio)
BPL
BanLec
Gal-3
Epa1-Np WT
NPA
Mouse Galectin1
rNV VLP-Cy3
CGL
LcH
TbF-BL1
LCA
human Gal-7
STL
Galectin-1
PFL
rMpL
BTL native
ABA
WGA
PHA-E
GNA (EY Lab)
WHA lectin
SGA
ECL
Human Galectin1
rPVL
CnMBL
WFL
WT-VCC
HHA(EY Lab)
CnSept
Native HPA
rSRL
mouse Gal-7
CC1G-10077
SRL
SSR1
AHL
ConA
CAA
TbF-BL2
WGA
TL(EY Lab)
ConA
RCAI (Vector)
CA
WGA multimer

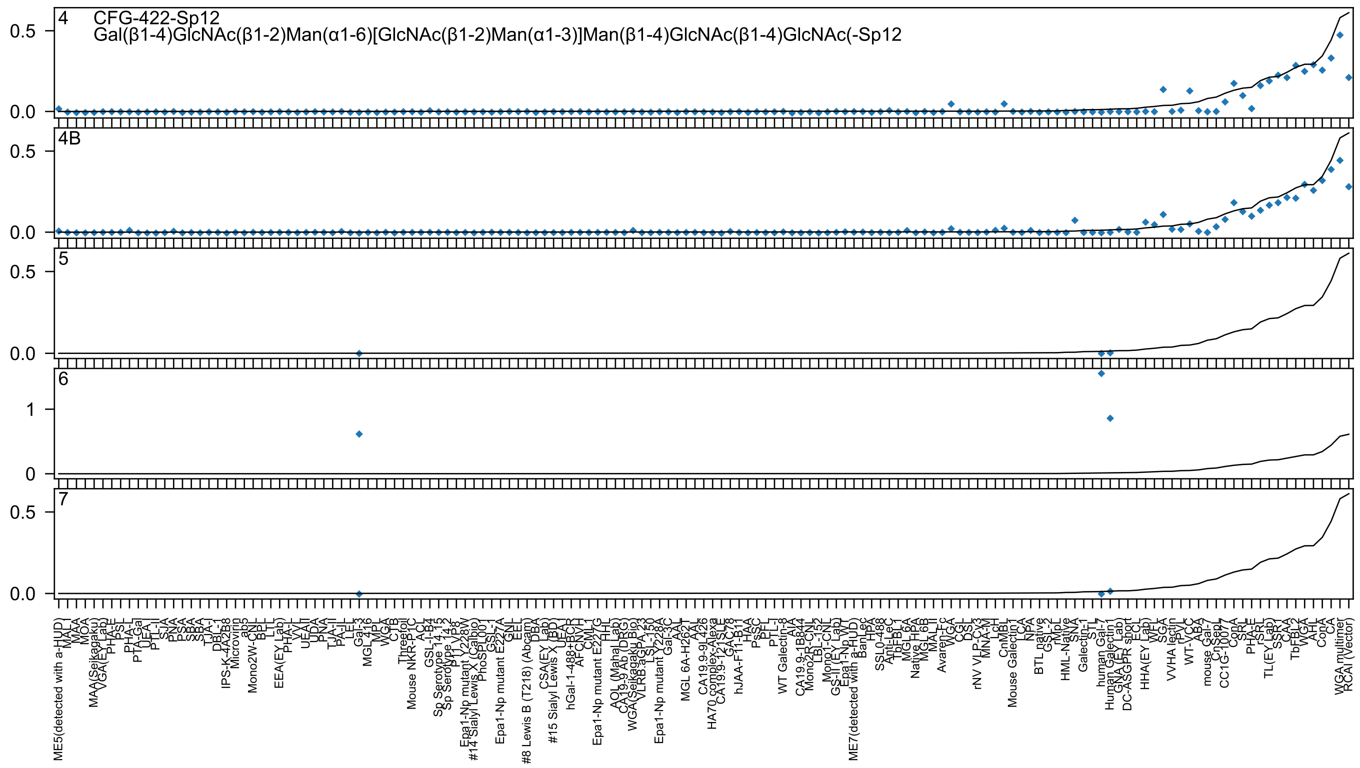

4 CFG-422-Sp12
Gal(β1-4)GlcNAc(β1-2)Man(α1-6)[GlcNAc(β1-2)Man(α1-3)]Man(β1-4)GlcNAc(β1-4)GlcNAc(-Sp12
4B
5
6
7

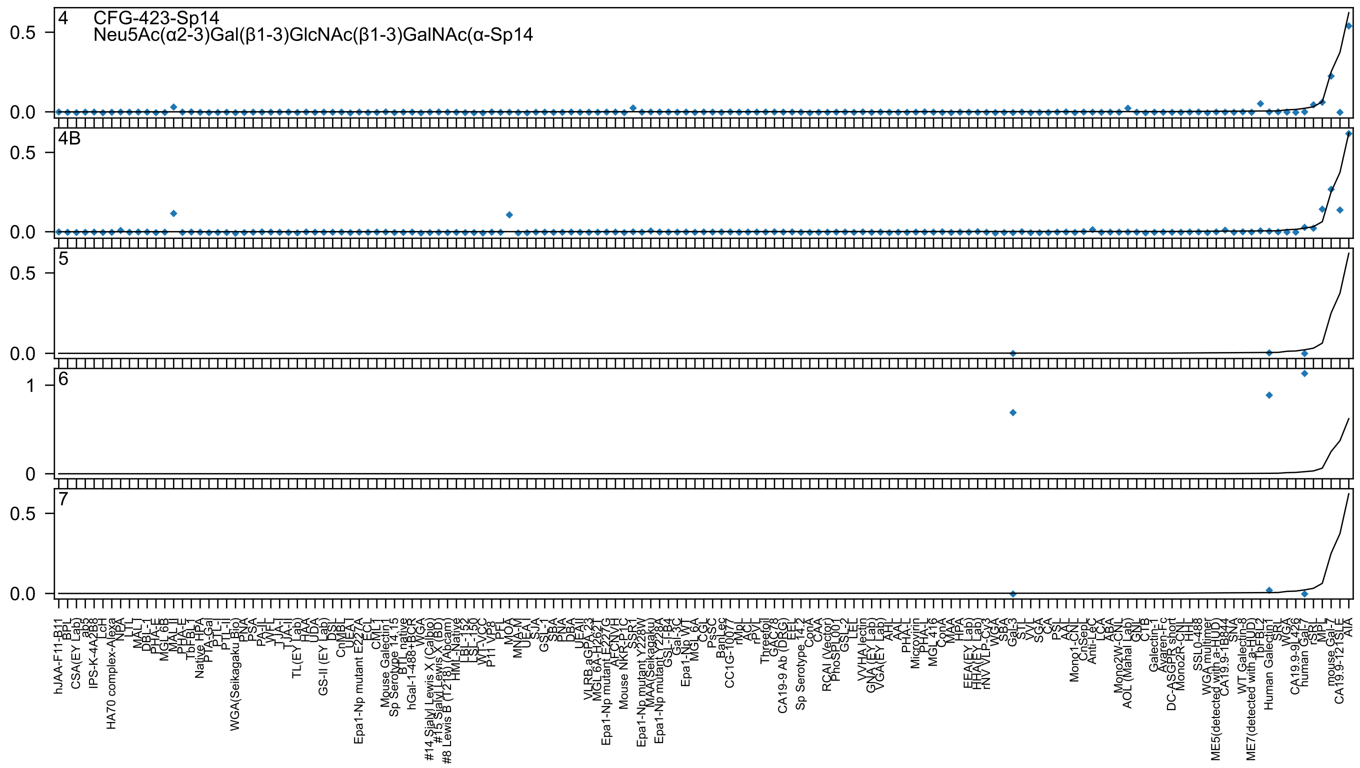
4 CFG-423-Sp14
Neu5Ac(α2-3)Gal(β1-3)GlcNAc(β1-3)GalNAc(α-Sp14
4B
5
6
7
0.0
0.5
0
1
hJAA-F11-B11
BPL
CSA(EY Lab)
ab5
IPS-K-4A2B8
LcH
HA70 complex-Alexa
NPA
LTL
MAL-I
DBL-I
PHA-E
MGL 6B
MAL II
PHA-E
TbFBL1
Native HPA
PTA-Gal
PTL-I
PTL-II
WGA(Seikagaku Bio)
PNA
PSA
PA-IL
WFL
TJA-I
TJA-II
TL(EY Lab)
HAA
UDA
GS-II (EY Lab)
DSL
CnMBL
UEA-I
Epa1-Np mutant E227A
ECL
CML-I
Mouse Galectin1
Sp Serotype 14.15
BTL native
hGal-1-488+BCR
WGA
#14 Sialyl Lewis X (Calbio)
#15 Sialyl Lewis X (BD)
#8 Lewis B (T218) (Abcam)
HML-Native
LBL-152
LSL-150
WT-VCC
P11 VP8
PFL
MOA
MNA-M
UEA-I
SJA
GSL-I
SBA
PNA
DBA
UEAII
VLRB.aGPA.23
MGL 6A-H262T
Epa1-Np mutant E227G
AFCNVH
Mouse NKR-P1C
SSR1
Epa1-Np mutant Y228W
MAA(Seikagaku)
Epa1-Np mutant Y228A
GSL-I-B4
Gal-3C
Epa1-Np WT
MGL 6A
CGL
PsSC
BanLec
CC1G-10077
rMpL
ACL
rPVL
Threefoil
GAC78
CA19-9 Ab (DRG)
EEL
Sp Serotype 14.2
ConA
CAA
RCAI (Vector)
PhoSL001
GSL-2
LEL
VVHA lectin
GNA (EY Lab)
VGA(EY Lab)
AHL
AAL
PHA-L
Microvirin
PHA-L
MGL 416
ConA
MAA
HPA
EEA(EY Lab)
HHA(EY Lab)
rNV VLP-Cy3
WGA
SBA
Gal-3
STL
VVL
SGA
CA
PSL
AAL
Mono1-CNL
CnSep
Anti-LeC
LCA
ABA
Mono2W-CNL
AOL (Mahal Lab)
GNL
CTB
Galectin-1
Avaren-Fc
DC-ASGPR short
Mono2R-CNL
HHL
SSL0-488
WGA multimer
ME5(detected with a-HUD)
CA19.9-1B844
SNA
WT Galectin-8
ME7(detected with a-HUD)
TbFBL2
Human Galectin1
SRL
WGA
CA19.9-9L426
human Gal-7
rSRL
MPL
mouse Gal-7
CA19.9-121SLE
AIA

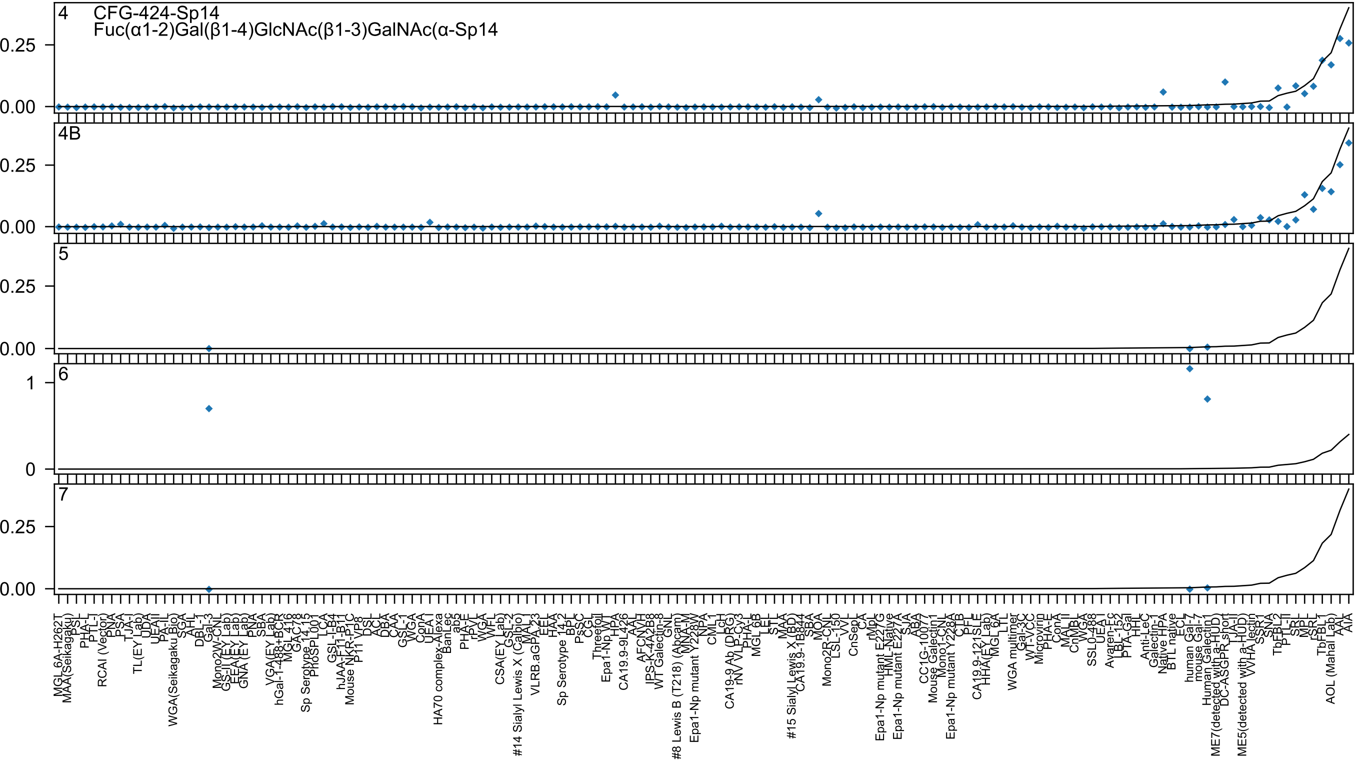

4 CFG-424-Sp14
Fuc(α1-2)Gal(β1-4)GlcNAc(β1-3)GalNAc(α-Sp14
4B
5
6
7

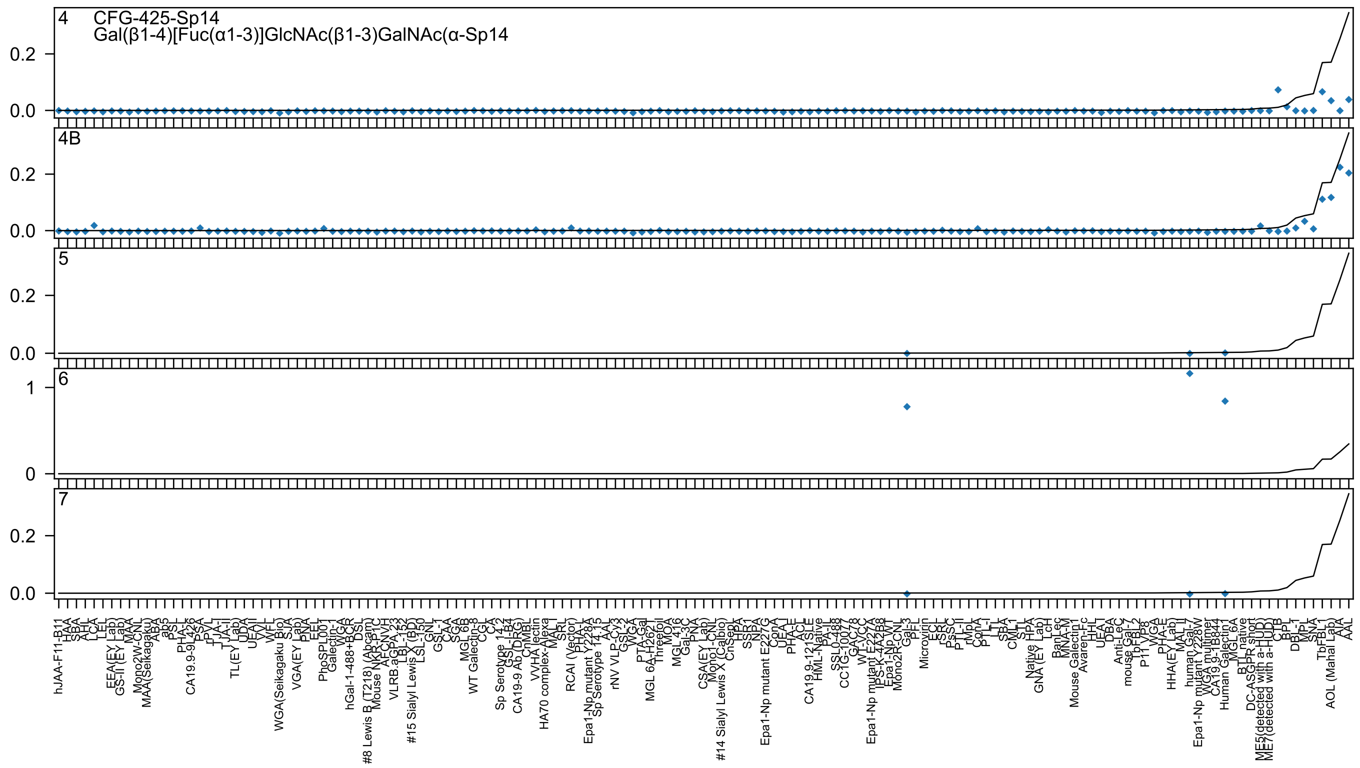

4
CFG-425-Sp14
Gal(β1-4)[Fuc(α1-3)]GlcNAc(β1-3)GalNAc(α-Sp14
4B
5
6
7
0.0
0.2
0
1
hJAA-F11-B11
HAA
SBA
AHL
LCA
LEL
EEA(EY Lab)
GS-II (EY Lab)
MAA
Mono2W-CNL
MAA(Seikagaku)
ABA
ab5
PSL
PHA-L
CA19.9-9L426
PSA
rPVL
TJA-II
TJA-I
TL(EY Lab)
UDA
UEAII
VVL
WFL
WGA(Seikagaku Bio)
SJA
VGA(EY Lab)
PNA
EEL
PhoSPL001
Galectin-1
WGA
hGal-1-488+BCR
DSL
#8 Lewis B (T218) (Abcam)
Mouse NKR-P1C
AFCNVH
VLRB.aGPA.23
LBL-152
#15 Sialyl Lewis X (BD)
LSL-150
GNL
GSL-1
CAA
SGA
MGL 6B
WT Galectin-8
CGL
CA
Sp Serotype 14.2
GSL-I-B4
CA19-9 Ab (DRG)
CnMBL
VVHA lectin
HA70 complex-Alexa
MAL
SRL
RCAI (Vector)
PHA-L
Epa1-Np mutant Y228A
Sp Serotype 14.15
AAL
rNV VLP-Cy3
GSL-2
WGA
PTA-Gal
MGL 6A-H262I
Threefoil
MOA
MGL 416
Gal-3C
PNA
CSA(EY Lab)
Mono1-CNL
#14 Sialyl Lewis X (Calbio)
CnSepL
HPA
SSA
NPA
Epa1-Np mutant E227G
ConA
UEA-I
PHA-E
ACL
CA19.9-121SLE
HML-Native
PA-II
SSL0-488
CC1G-10077
GAC78
WT-VCC
Epa1-Np mutant E227A
IPS-K-4A2B8
Epa1-Np WT
Mono2R-CNL
Gal-3
PFL
Microvirin
ECL
rSRL
PsSC
PTL-II
rMpL
ConA
PTL-I
LTL
SBA
CML1
STL
Native HPA
GNA (EY Lab)
LcH
BanLec
MNA-M
Mouse Galectin1
Avaren-Fc
HHL
UEA-I
DBA
Anti-LeC
mouse Gal-7
TbFBL2
P11 VP8
VGA
PHA-E
HHA(EY Lab)
MAL I
human Gal-7
Epa1-Np mutant Y228W
WGA multimer
CA19.9-1B844
Human Galectin1
MGL 6A
BTL native
DC-ASGPR short
ME5(detected with a-HUD)
ME7(detected with a-HUD)
CTB
BPL
DBL-1
MPL
SNA
TbFBL1
AOL (Mahal Lab)
AIA
AAL

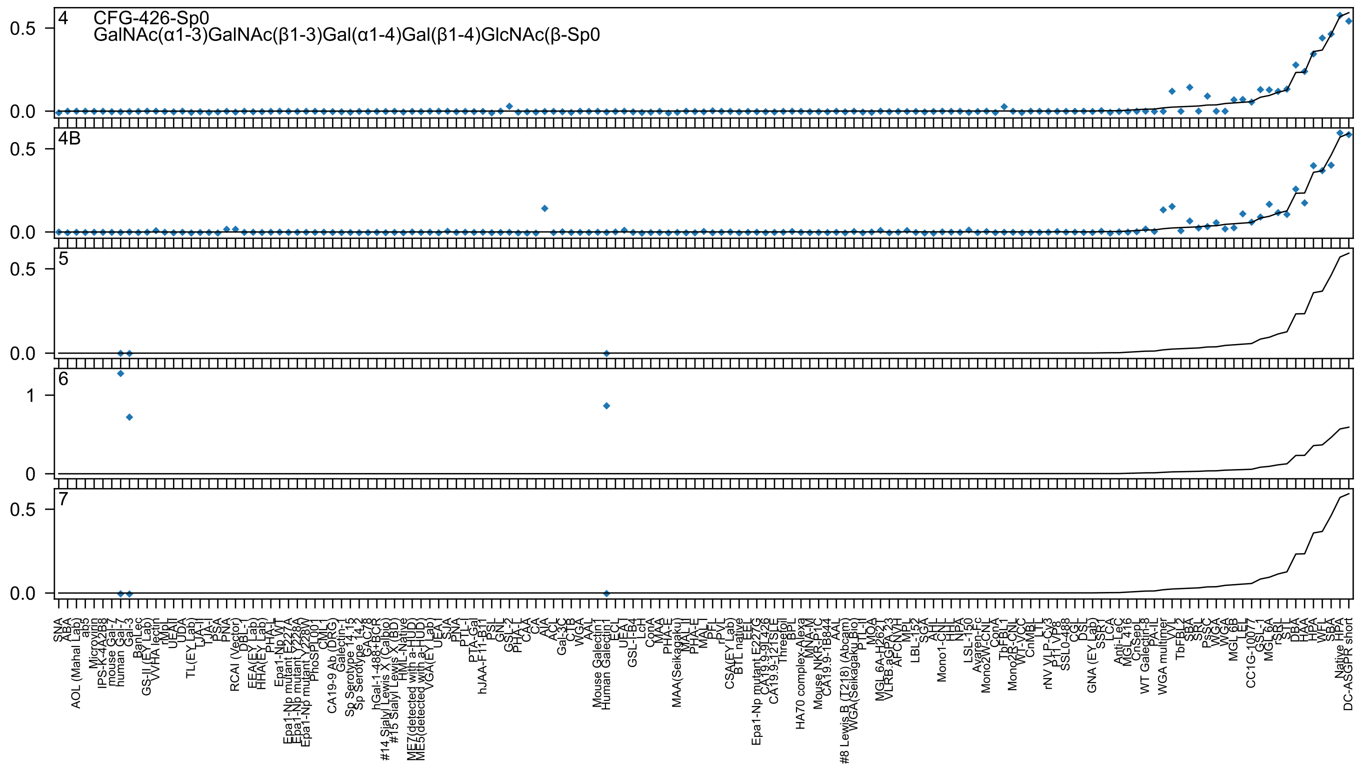

CFG-426-Sp0
GalNAc(α1-3)GalNAc(β1-3)Gal(α1-4)Gal(β1-4)GlcNAc(β-Sp0

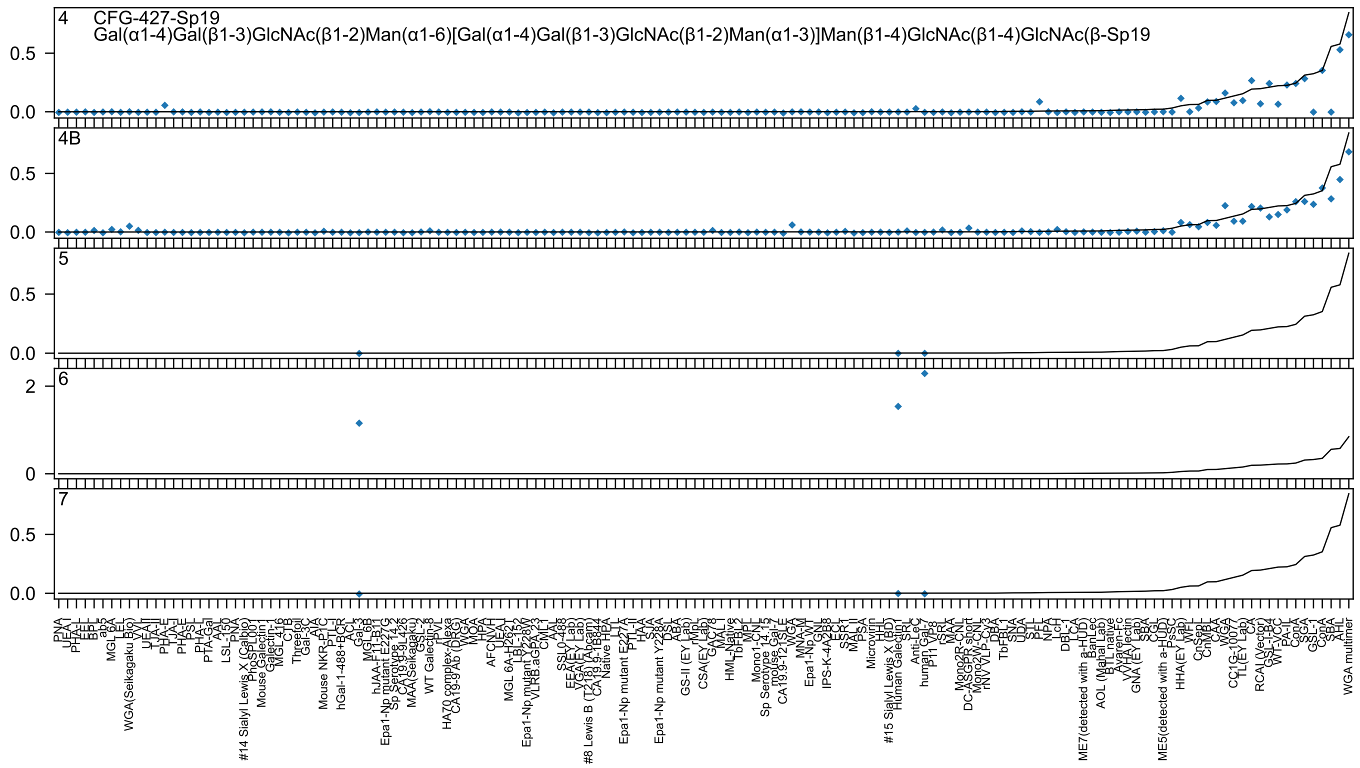

CFG-427-Sp19
Gal(α1-4)Gal(β1-3)GlcNAc(β1-2)Man(α1-6)[Gal(α1-4)Gal(β1-3)GlcNAc(β1-2)Man(α1-3)]Man(β1-4)GlcNAc(β1-4)GlcNAc(β-Sp19
4
4B
5
6
7

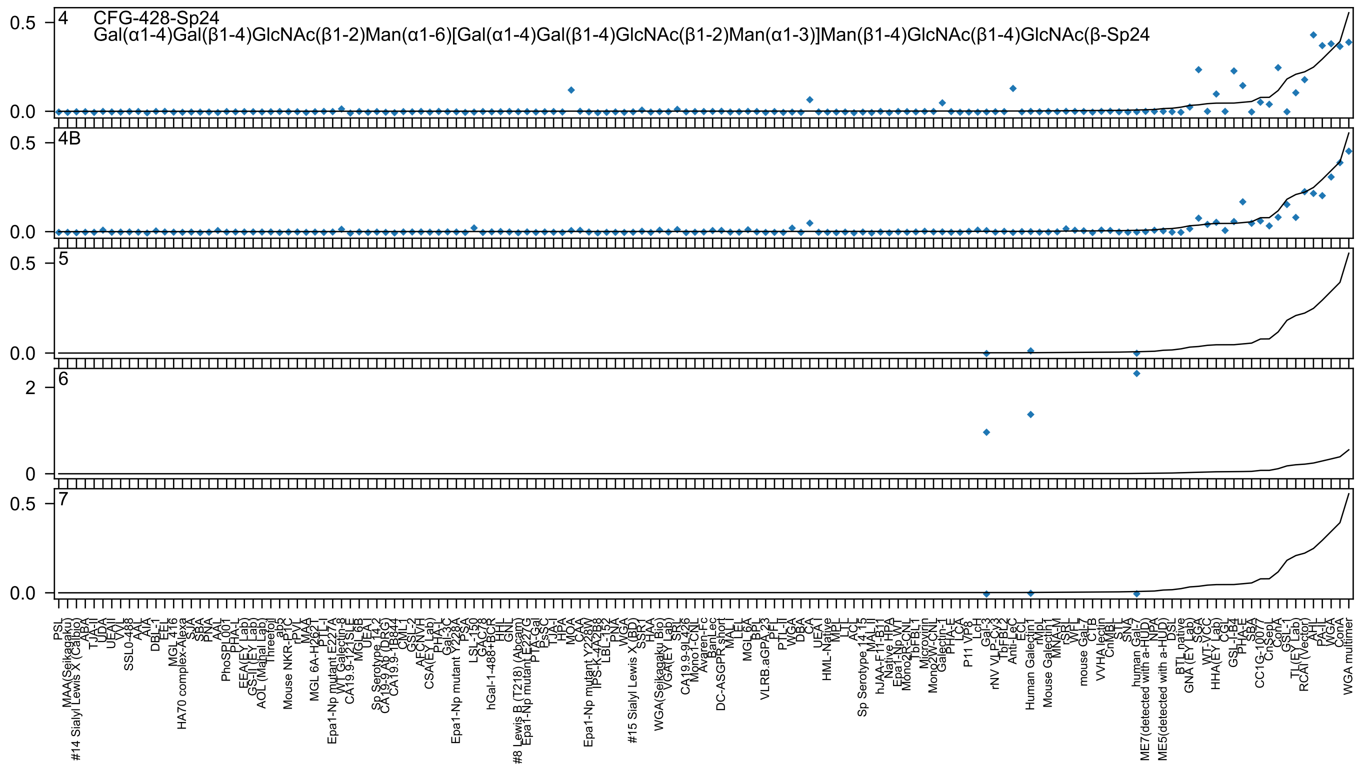
4 CFG-428-Sp24
Gal(α1-4)Gal(β1-4)GlcNAc(β1-2)Man(α1-6)[Gal(α1-4)Gal(β1-4)GlcNAc(β1-2)Man(α1-3)]Man(β1-4)GlcNAc(β1-4)GlcNAc(β-Sp24
4B
5
6
7
0.5
0.0
2
0

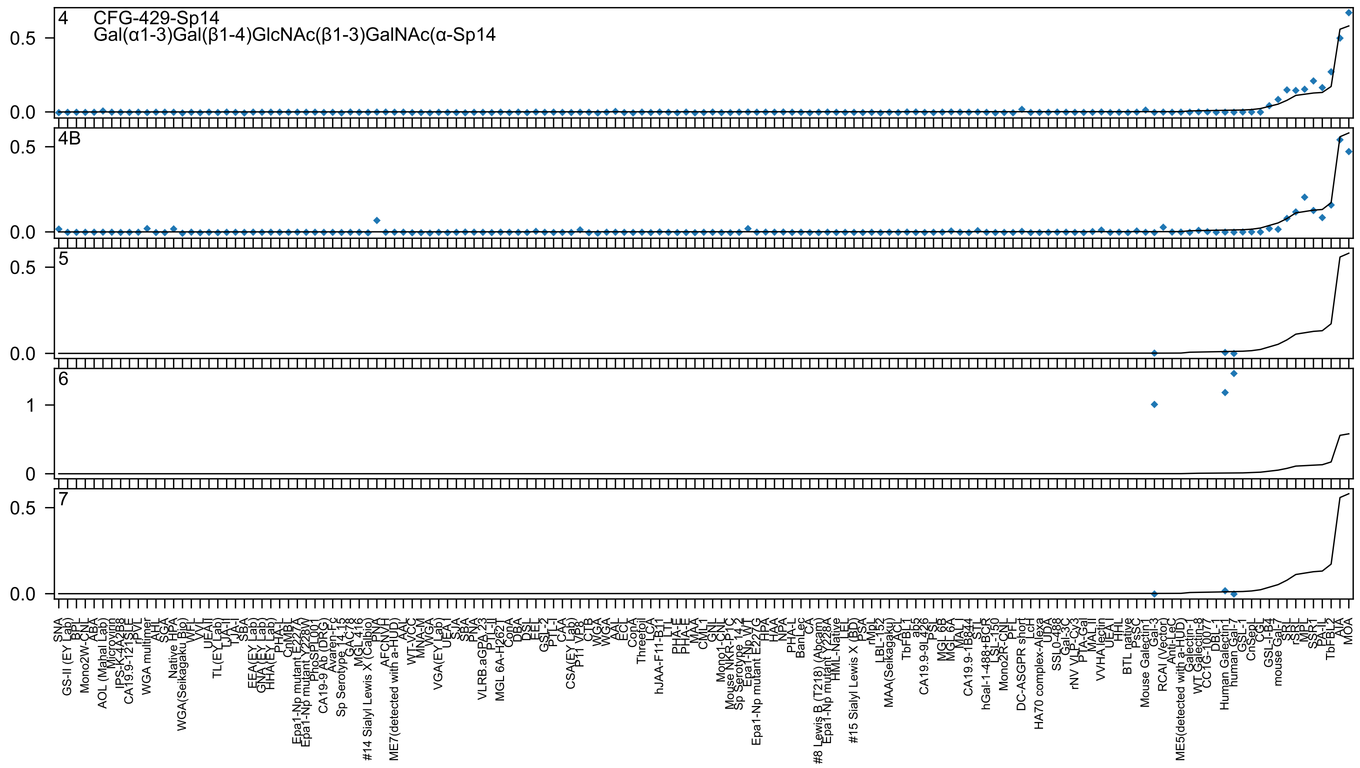
4 CFG-429-Sp14
Gal(α1-3)Gal(β1-4)GlcNAc(β1-3)GalNAc(α-Sp14
4B
5
6
7
0.0
0.5
0
1

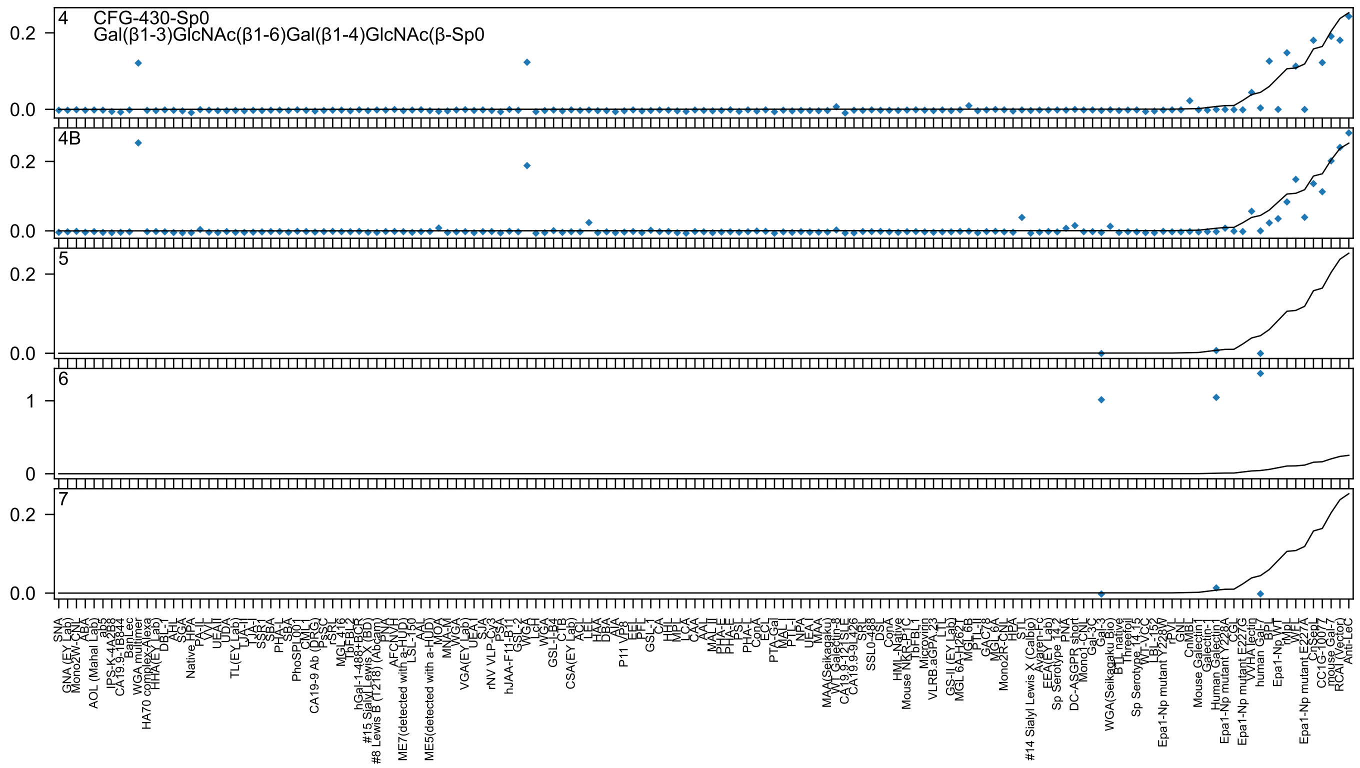
4
CFG-430-Sp0
Gal(β1-3)GlcNAc(β1-6)Gal(β1-4)GlcNAc(β-Sp0
4B
5
6
7

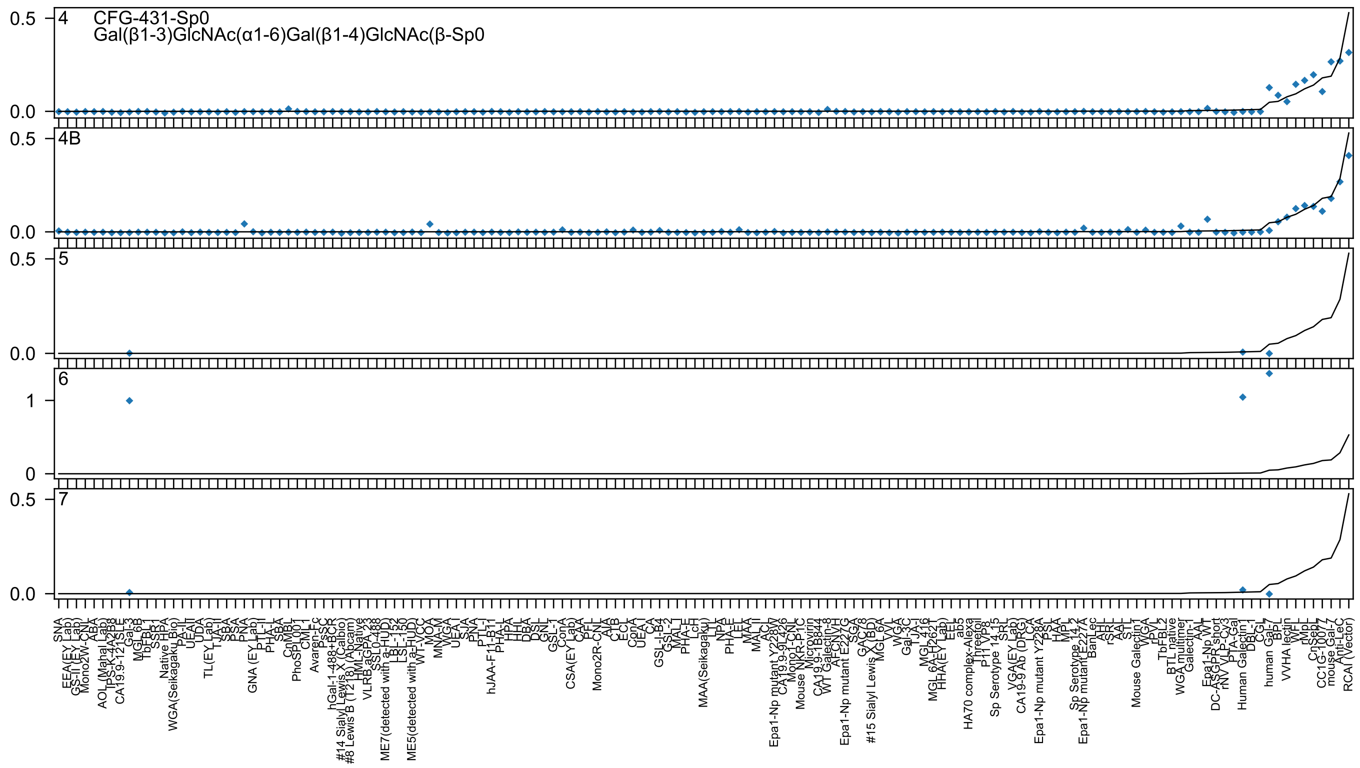

CFG-431-Sp0
Gal(β1-3)GlcNAc(α1-6)Gal(β1-4)GlcNAc(β-Sp0

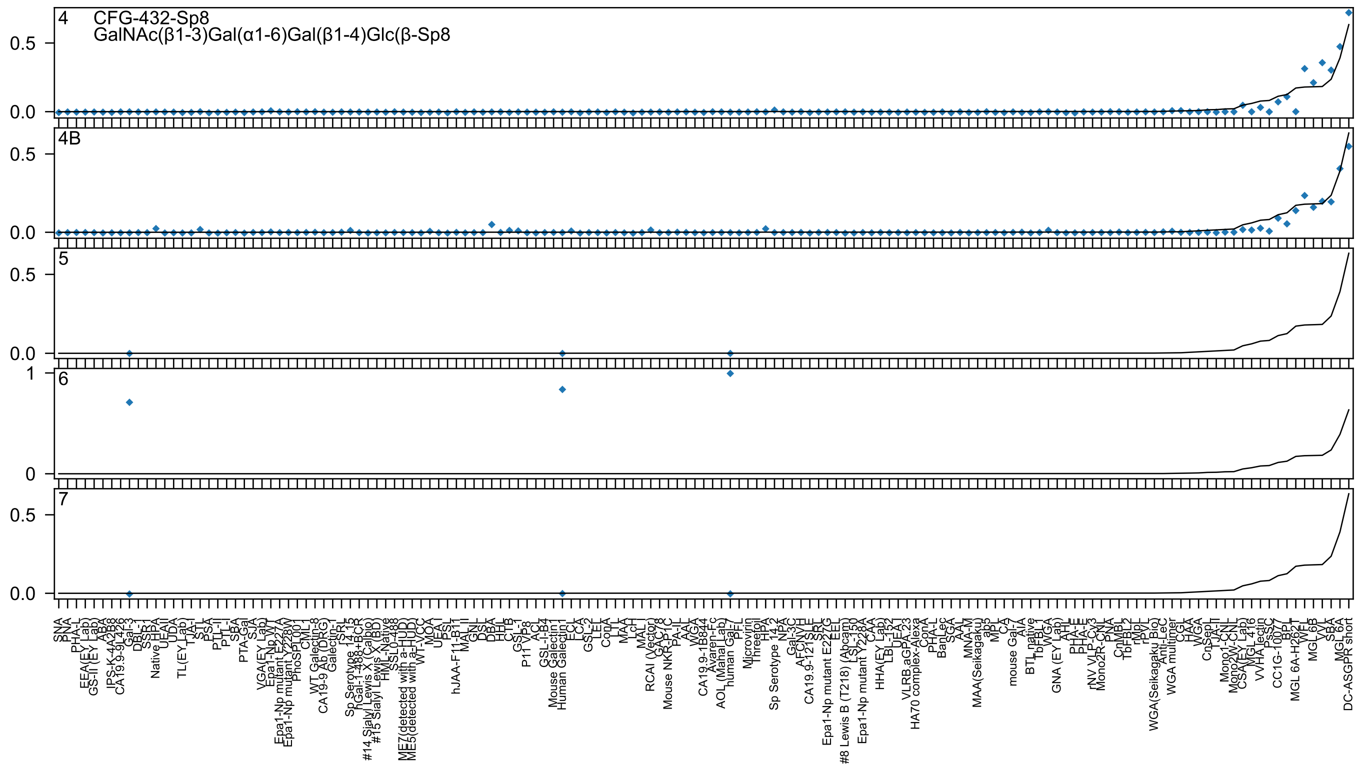
4
CFG-432-Sp8
GalNAc(β1-3)Gal(α1-6)Gal(β1-4)Glc(β-Sp8
4B
5
6
7
0.0
0.5
0
1
SNA
PNA
PHA-L
EEA(EY Lab)
GS-II (EY Lab)
ABA
IPS-K-4A2B8
CA19.9-9L426
Gal-3
DBL-1
SSR-1
Native HPA
UEAII
UDA
TL(EY Lab)
TJA-I
STL
PSA
PTL-II
PTL-I
SBA
PTA-Gal
SJA
VGA(EY Lab)
Epa1-Np WT
Epa1-Np mutant E227A
Epa1-Np mutant Y228W
PhoSL
CML1
WT Galectin-8
CA19-9 Ab (DRG)
Galectin-1
TSRL
Sp Serotype 14.15
hGal-1-488+BCR
#14 Sialyl Lewis X (Calbio)
#15 Sialyl Lewis X (BD)
HML-Native
SSL0-488
ME7(detected with a-HUD)
ME5(detected with a-HUD)
WT-VCC
MOA
UEA I
PSL
hJAA-F11-B11
MAL II
GNL
DSL
DBA
HHL
CTB
GSL-I
P11 VP8
ACL
GSL-I-B4
Mouse Galectin1
Human Galectin1
ECL
LCA
GSL-2
LEL
ConA
LTL
MAA
LcH
MAL I
RCAI (Vector)
GAC78
Mouse NKR-P1C
PA-IL
AAL
WGA
CA19.9-1B844
Avaren-Fc
AOL (Manal Lab)
human Gal-7
PFL
Microvirin
Threefoil
HPA
Sp Serotype 14.2
NPA
Gal-3C
AFCNYH
CA19.9-121SLE
SRL
Epa1-Np mutant E227G
EEL
#8 Lewis B (T218) (Abcam)
LSL-150
Epa1-Np mutant Y228A
CAA
HHA(EY Lab)
LBL-152
UEA1
VLRB.aGPA.23
HA70 complex-Alexa
ConA
PHA-L
BanLec
SGA
AAL
MNA-M
MAA(Seikagaku)
ab5
MPL
CA
mouse Gal-7
AIA
BTL native
TbFBL1
WGA
GNA (EY Lab)
AHL
PHA-E
PHA-E
rNV VLP-Cy3
Mono2R-CNL
PNA
CnMBL
TbFBL2
rMpL
rPyL
WGA(Seikagaku_Bio)
Anti-LeC
WGA multimer
CGL
HAA
WGA
CnSepL
TJA-II
Mono1-CNL
Mono2W-CNL
CSA(EY Lab)
MGL 416
VVHA lectin
PsSC
CC1G-10077
BPL
MGL 6A-H262T
WFL
MGL 6B
VVL
SBA
MGL 6A
DC-ASGPR short

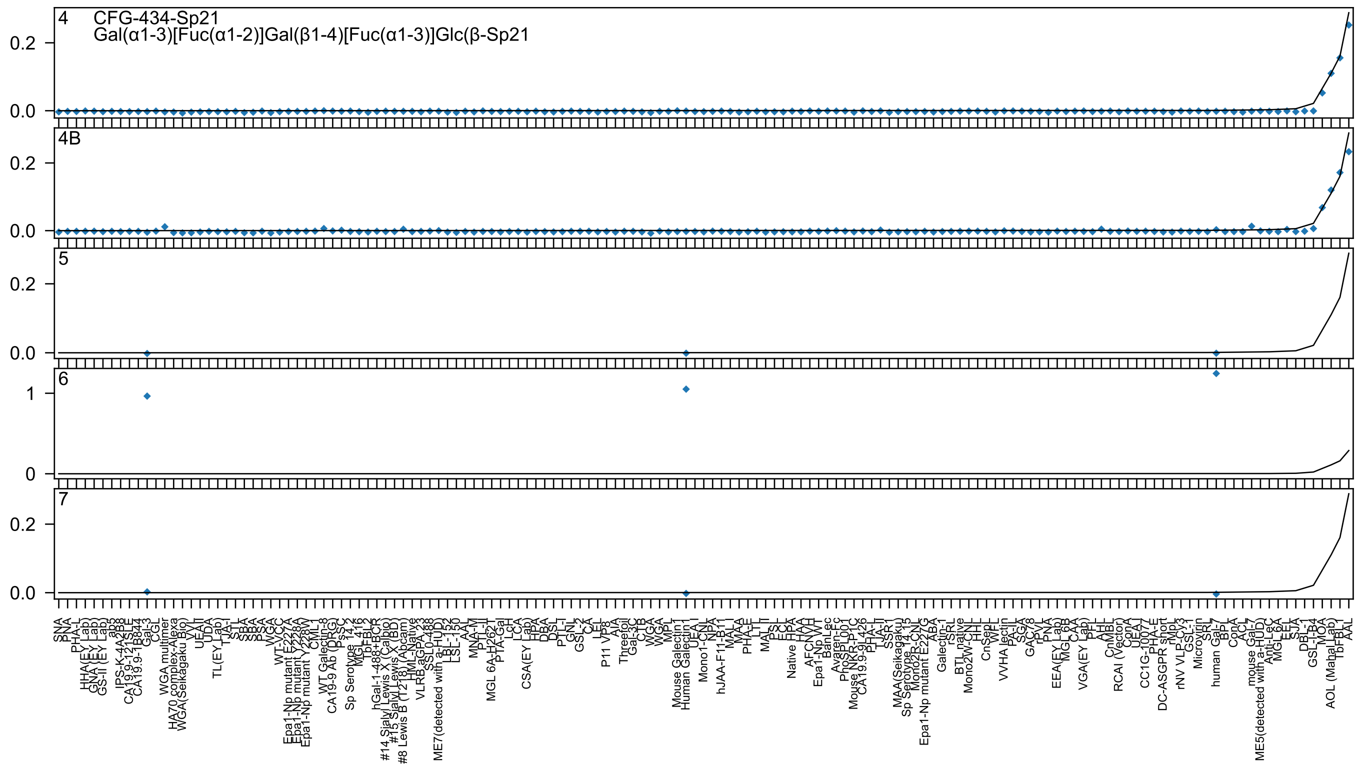
4
CFG-434-Sp21
Gal(α1-3)[Fuc(α1-2)]Gal(β1-4)[Fuc(α1-3)]Glc(β-Sp21
4B
5
6
7
0.2
0.0
1
0
SNA
PNA
PHA-L
HHA(EY Lab)
GNA (EY Lab)
GS-II (EY Lab)
ab3
IPS-K-4A2B8
CA19.9-121SLE
CA19.9-1B844
Gal-3
CGL
WGA multimer
HA70 complex-Alexa
WGA(Seikagaku Bio)
VVL
UEAII
UDA
TL(EY Lab)
TJA-I
STL
SBA
SBA
PSA
WGA
WT-VCC
Epa1-Np mutant E227A
Epa1-Np mutant Y228A
Epa1-Np mutant Y228W
CML
WT Galectin-8
CA19-9 Ab (DRG)
PsSC
Sp Serotype 14_2
MGL 416
TbFBL2
hGal-1-488+BCR
#14 Sialyl Lewis X (Calbio)
#15 Sialyl Lewis X (BD)
#8 Lewis B (T218) (Abcam)
HML-Native
VLRB aGPA.23
SSL0-488
ME7(detected with a-HUD)
LBL-152
LSL-150
AAL
MNA-M
PTL-II
MGL 6A-H262T
PTA-Gal
LcH
LCA
CSA(EY Lab)
HPA
DBA
DSL
PTL-I
GNL
GSL-2
CA
LEL
P11 VP8
AIA
Threefoil
Gal-3C
CTB
WGA
WGA
MPL
Mouse Galectin1
Human Galectin1
UEA-I
Mono1-CNL
NPA
hJAA-F11-B11
MAL-I
MAA
PHA-E
LTL
MAL-II
PSL
ECL
Native FPA
HAA
AFCN-VH
Epa1-Np WT
BanLec
Avaren-Fc
PhoSL001
Mouse NKR-P1C
CA19.9-9L426
PHA-L
TJA-II
SSR1
MAA(Seikagaku)
Sp Serotype 14_15
Mono2R-CNL
Epa1-Np mutant E227G
ABA
Galectin-1
rSRL
BTL native
Mono2W-CNL
HHL
CnSepL
WFL
VVHA lectin
PA-IL
SGA
GAC78
rPVL
PNA
EEA(EY Lab)
MGL 6B
CAA
VGA(EY Lab)
PFL
AHL
CnMBL
RCAI (Vector)
ConA
UEA-I
CC1G-10077
PHA-E
DC-ASGPR short
rMpL
rNV VLP-Cy3
GSL-I
Microvirin
SRL
human Gal-7
BPL
ConA
ACL
mouse Gal-7
ME5(detected with a-HUD)
Anti-LeC
MGL 6A
EEL
SJA
DBL-1
GSL-I-B4
MOA
AOL (Mahal Lab)
TbFBL1
AAL

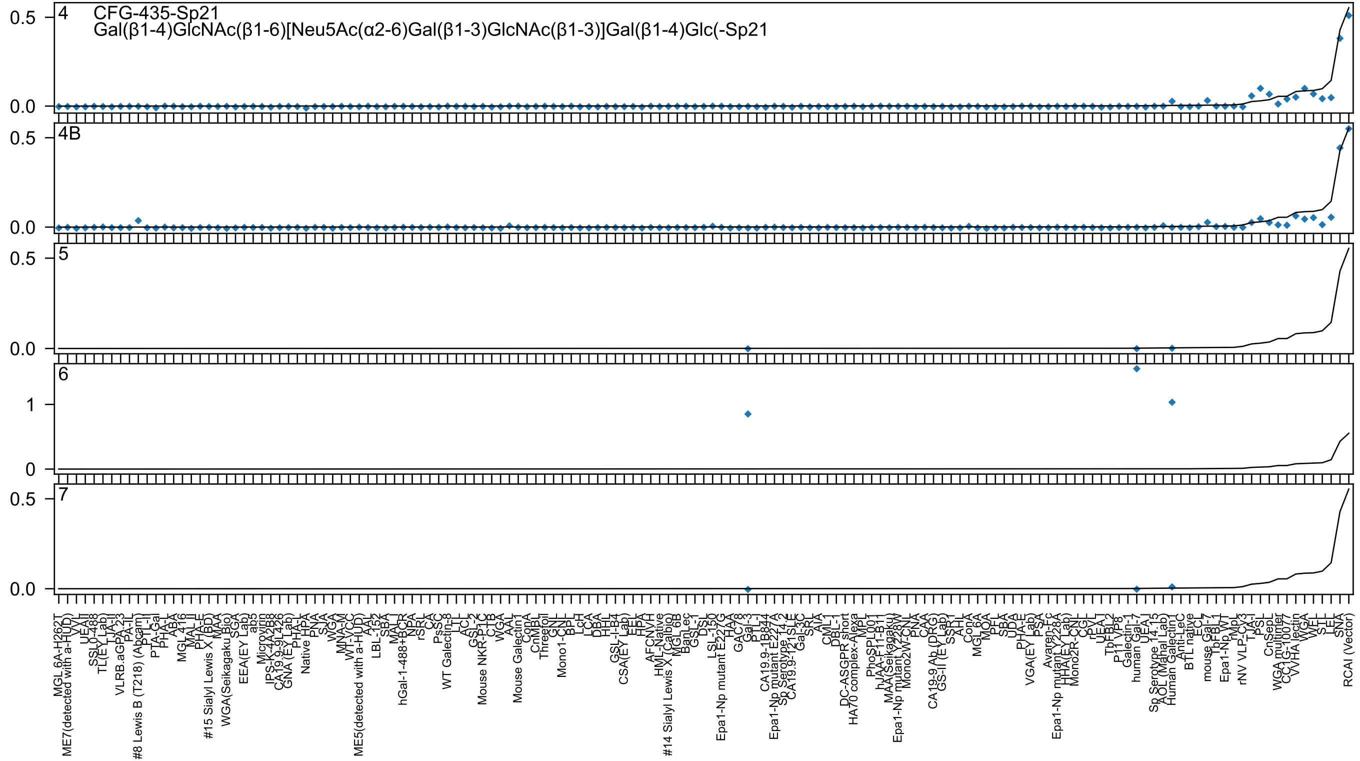

4 CFG-435-Sp21
Gal(β1-4)GlcNAc(β1-6)[Neu5Ac(α2-6)Gal(β1-3)GlcNAc(β1-3)]Gal(β1-4)Glc(-Sp21
4B
5
6
7

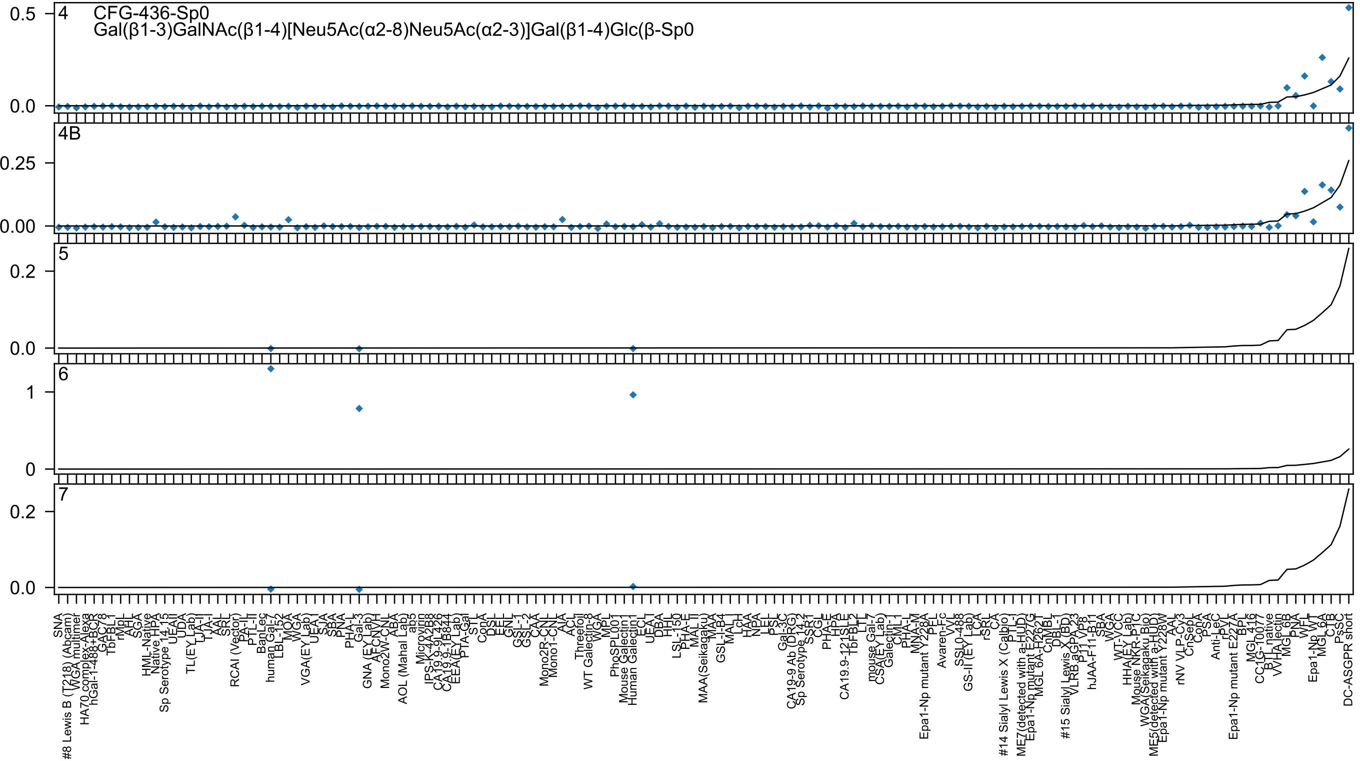

4 CFG-436-Sp0
Gal(β1-3)GalNAc(β1-4)[Neu5Ac(α2-8)Neu5Ac(α2-3)]Gal(β1-4)Glc(β-Sp0
4B
5
6
7

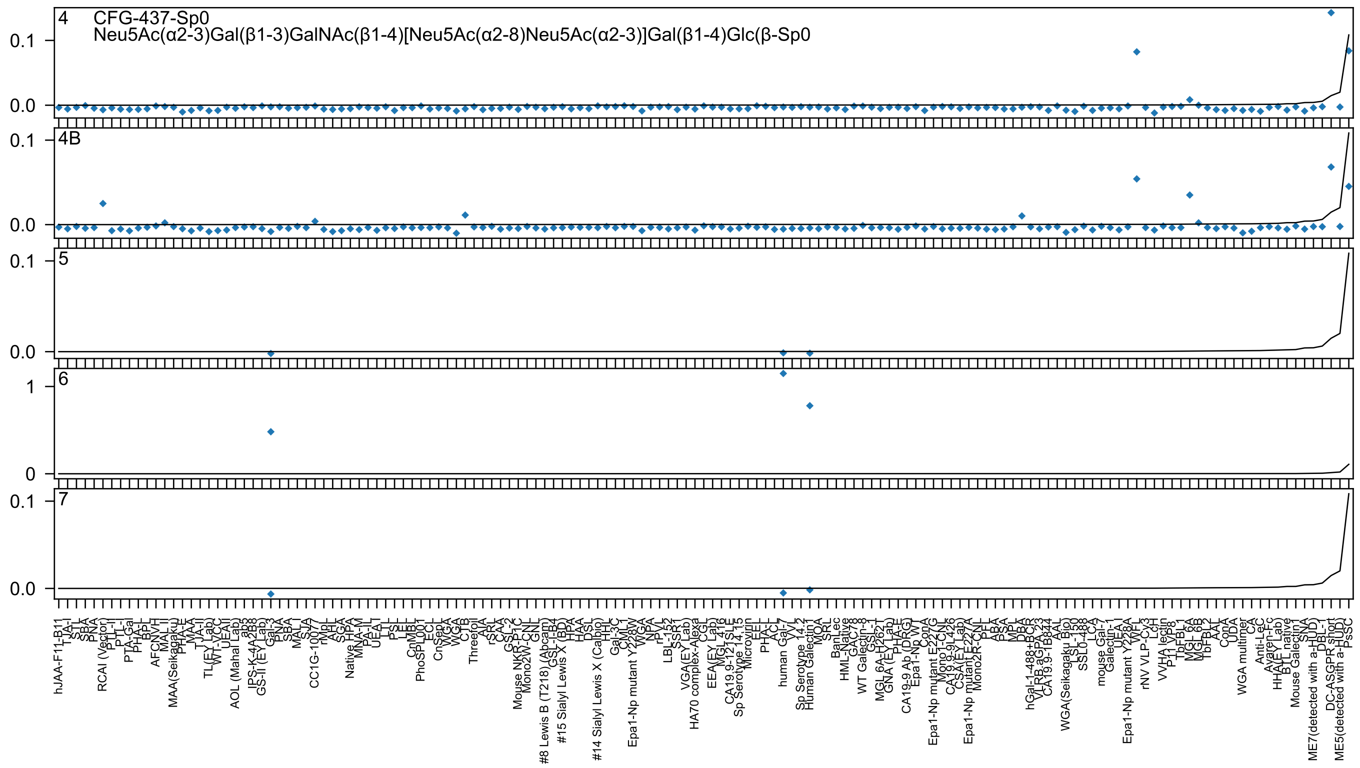

4
CFG-437-Sp0
Neu5Ac(α2-3)Gal(β1-3)GalNAc(β1-4)[Neu5Ac(α2-8)Neu5Ac(α2-3)]Gal(β1-4)Glc(β-Sp0
4B
5
6
7

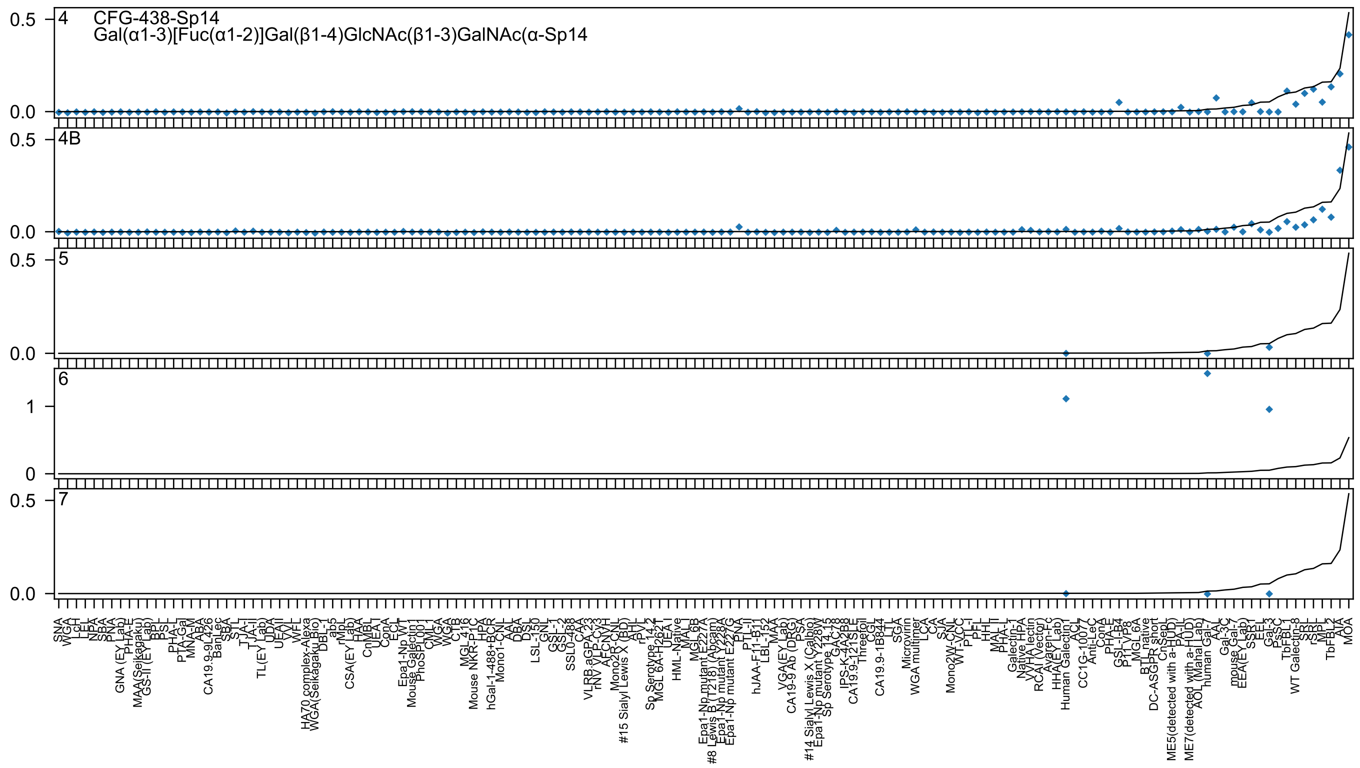
4 CFG-438-Sp14
Gal(α1-3)[Fuc(α1-2)]Gal(β1-4)GlcNAc(β1-3)GalNAc(α-Sp14
4B
5
6
7
0.5
0.0
1
0

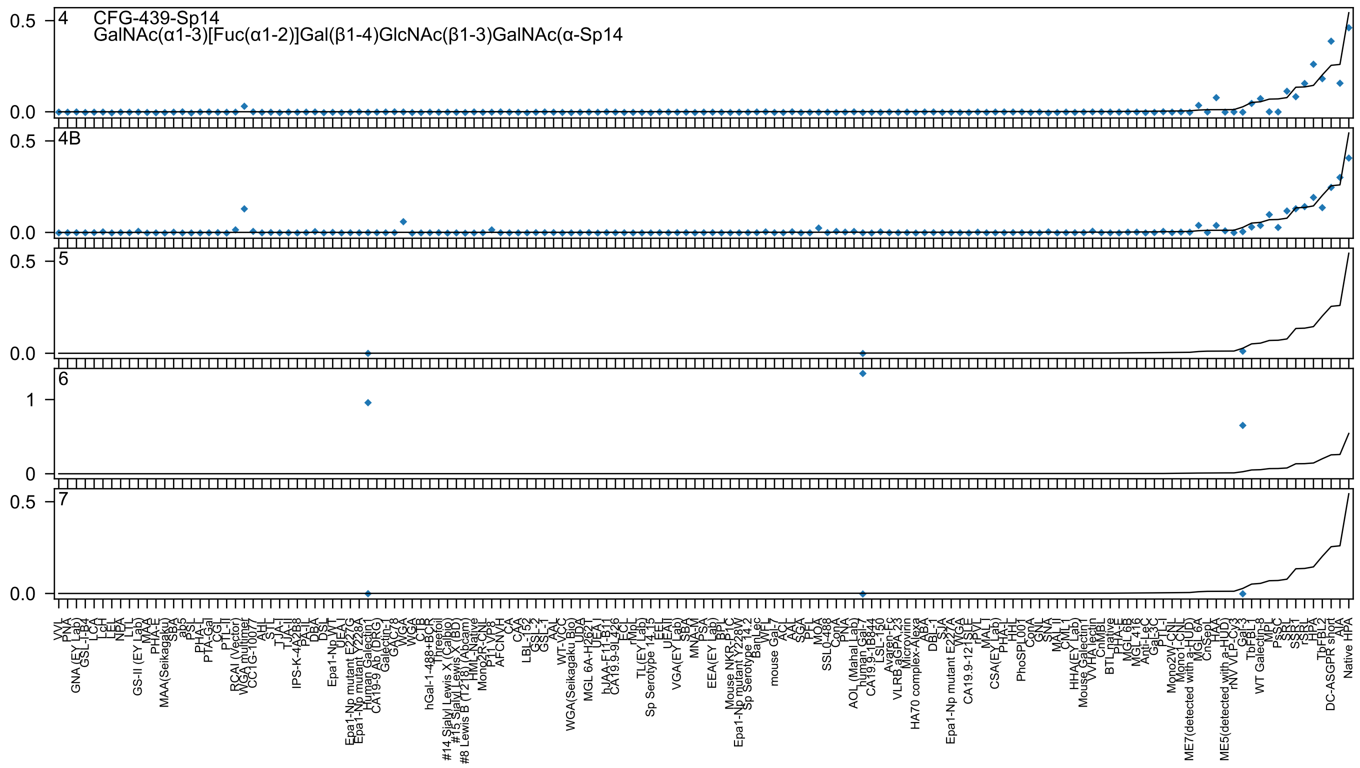

CFG-439-Sp14
GalNAc(α1-3)[Fuc(α1-2)]Gal(β1-4)GlcNAc(β1-3)GalNAc(α-Sp14
4
4B
5
6
7

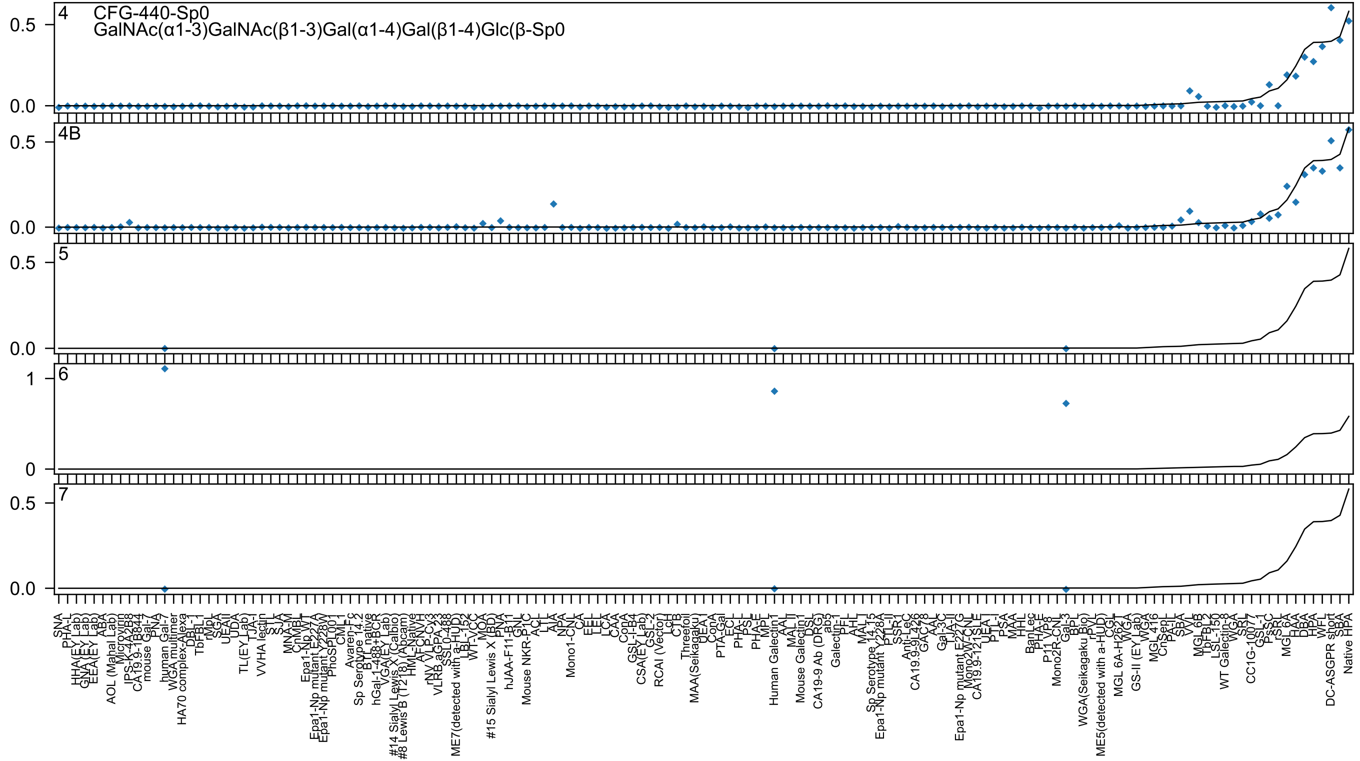

4
CFG-440-Sp0
GalNAc(α1-3)GalNAc(β1-3)Gal(α1-4)Gal(β1-4)Glc(β-Sp0
4B
5
6
7

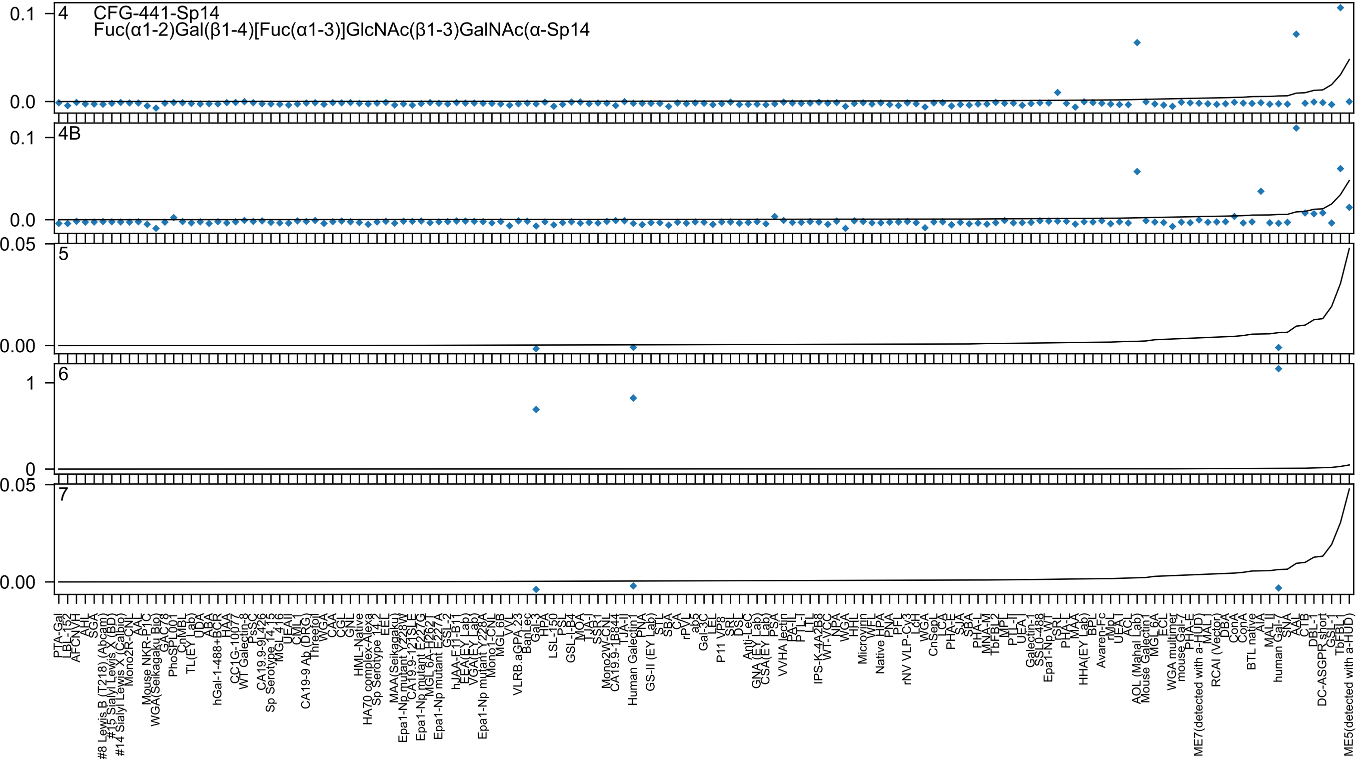
CFG-441-Sp14
Fuc(α1-2)Gal(β1-4)[Fuc(α1-3)]GlcNAc(β1-3)GalNAc(α-Sp14
4
4B
5
6
7

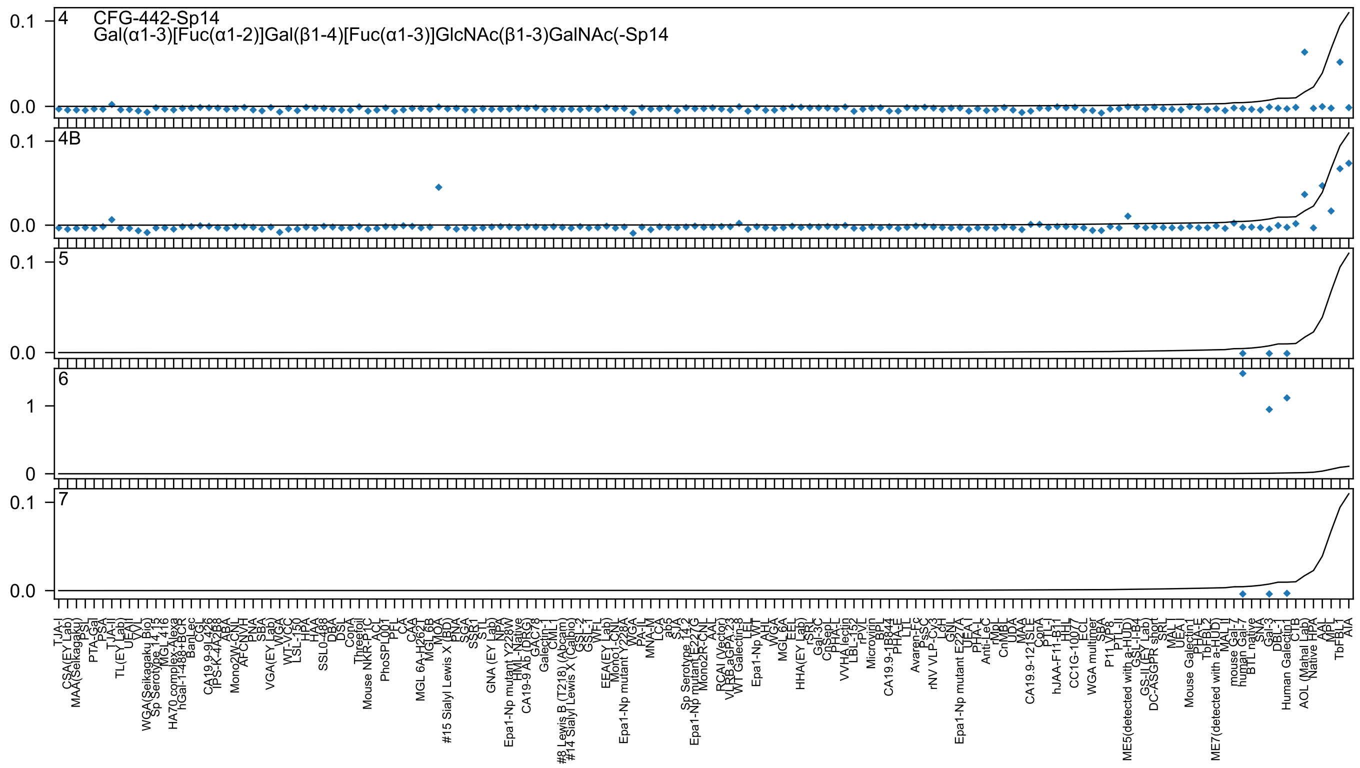

CFG-442-Sp14
Gal(α1-3)[Fuc(α1-2)]Gal(β1-4)[Fuc(α1-3)]GlcNAc(β1-3)GalNAc(-Sp14
4
4B
5
6
7

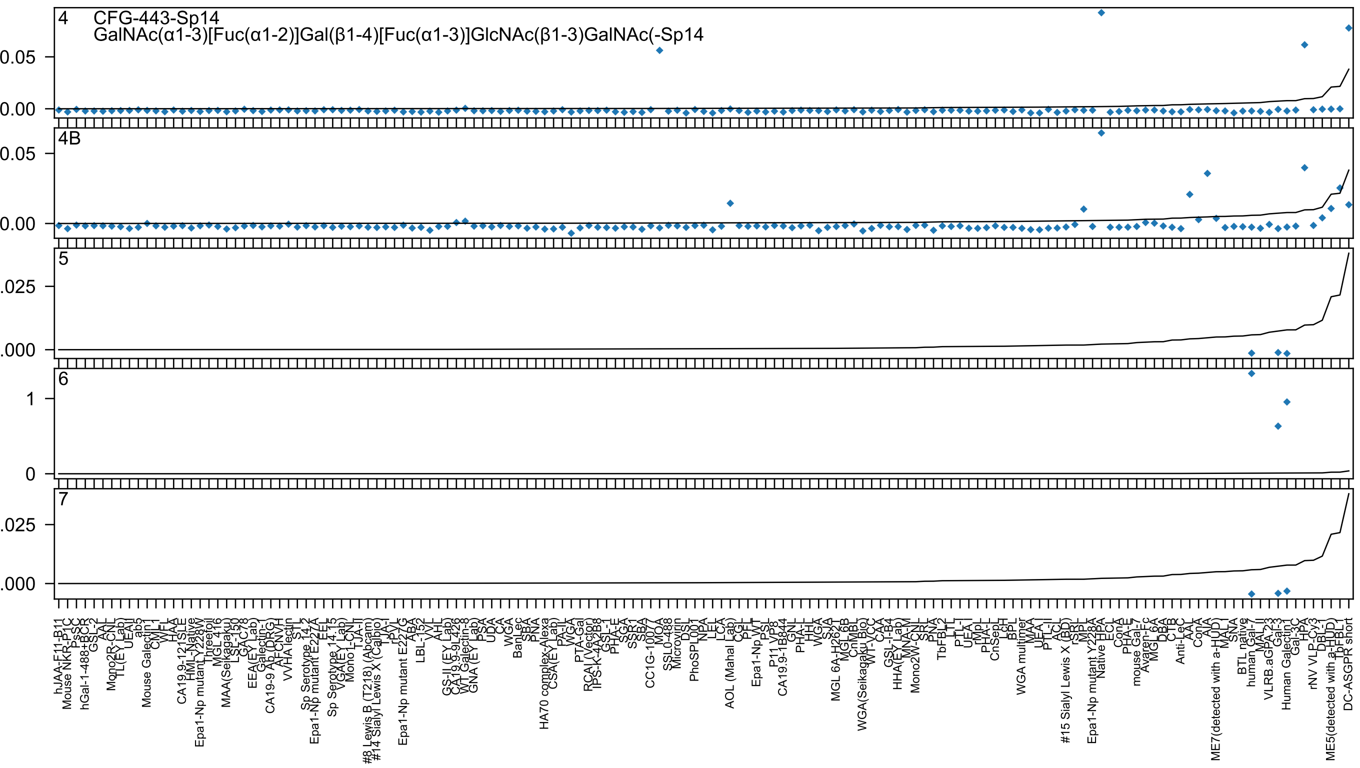
4 CFG-443-Sp14
GalNAc(α1-3)[Fuc(α1-2)]Gal(β1-4)[Fuc(α1-3)]GlcNAc(β1-3)GalNAc(-Sp14
4B
5
6
7

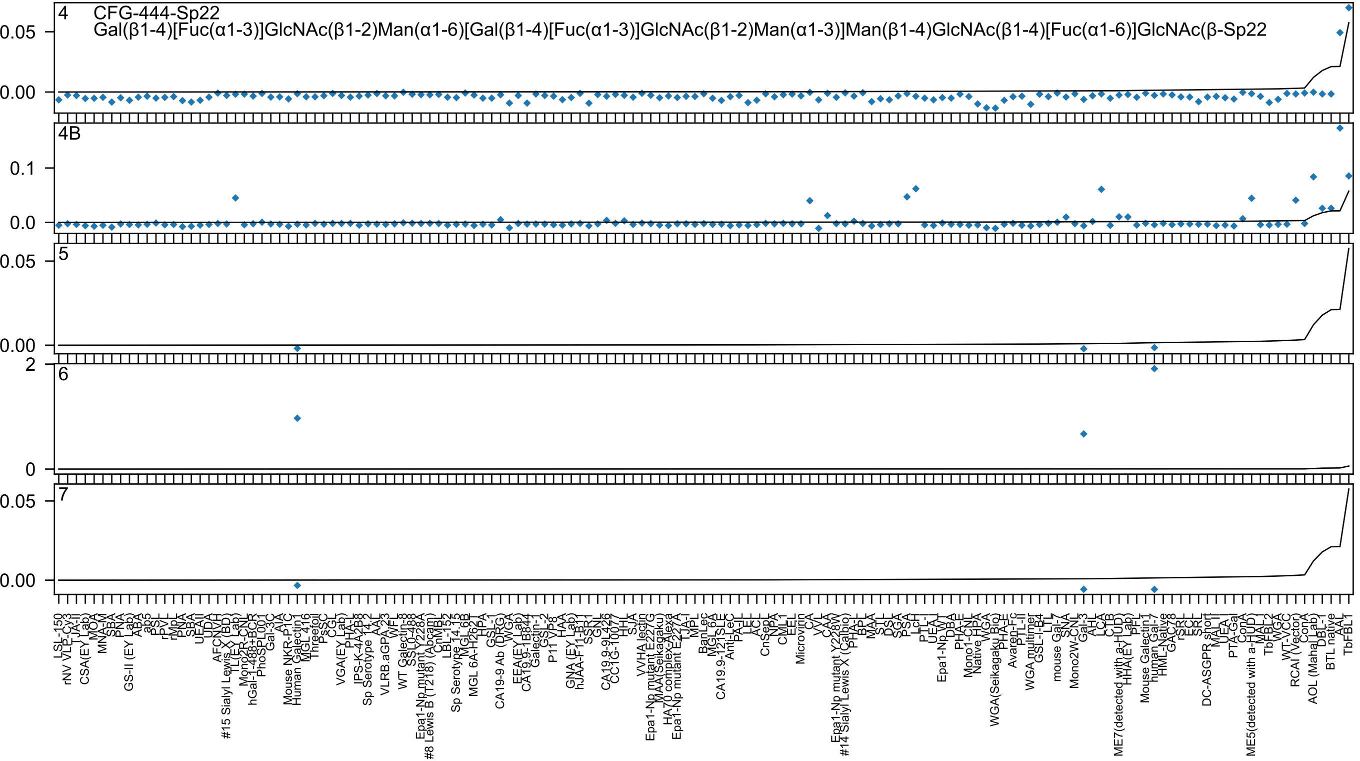
4 CFG-444-Sp22
Gal(β1-4)[Fuc(α1-3)]GlcNAc(β1-2)Man(α1-6)[Gal(β1-4)[Fuc(α1-3)]GlcNAc(β1-2)Man(α1-3)]Man(β1-4)GlcNAc(β1-4)[Fuc(α1-6)]GlcNAc(β-Sp22
0.05
0.00
4B
0.1
0.0
5
0.05
0.00
6
2
0
7
0.05
0.00
LSL-150
rNV VLP-Cy3
TJA-II
CSA(EY Lab)
MOA
MNA-M
SBA
PNA
GS-II (EY Lab)
ABA
ab5
PSL
r-PVL
rMpL
PNA
SBA
UEAII
UDA
AFCNVH
#15 Sialyl Lewis X (BD)
TL(EY Lab)
Mono2R-CNL
hGal-1-488+BCR
PhoSL
PPL001
Gal-3C
AIA
Mouse NKR-P1C
Human Galectin1
MGL 416
Threefoil
PsSC
CGL
VGA(EY Lab)
PHA-L
IPS-K-4A2B8
Sp Serotype 14.2
AAL
VLRB.aGPA.23
WFL
WT Galectin-8
SSL0-488
Epa1-Np mutant Y228A
#8 Lewis B (T218) (Abcam)
CnMBL
LBL-152
Sp Serotype 14.15
MGL 6B
MGL 6A-H262T
HPA
GSL-1
CA19-9 Ab (DRG)
WGA
EEA(EY Lab)
CA19.9-1B844
Galectin-1
GSL-2
P11 VP8
HAA
GNA (EY Lab)
hJAA-F11-B11
SSR1
GNL
CA19.9-9L426
CC1G-10077
HHL
SJA
VVHA lectin
Epa1-Np mutant E227G
MAA(Seikagaku)
HA70 complex-Alexa
Epa1-Np mutant E227A
LTA-I
MPL
BanLec
MGL 6A
CA19.9-121SLE
Anti-LeC
PA-IL
LEL
ACL
CnSep
NPA
CML1
EEL
Microvirin
CA
VVL
CAA
Epa1-Np mutant Y228W
#14 Sialyl Lewis X (Calbio)
PHA-L
BPL
MAA
STL
DSL
SGA
PSA
LcH
PTL-I
UEA-I
Epa1-Np WT
DBA
PHA-E
Mono1-CNL
Native HPA
WGA
WGA(Seikagaku Bio)
PHA-E
Avaren-Fc
PTL-II
WGA multimer
GSL-I-B4
LTL
mouse Gal-7
SNA
Mono2W-CNL
Gal-3
AAL
LCA
CTB
ME7(detected with a-HUD)
HHA(EY Lab)
PFL
Mouse Galectin1
human Gal-7
HML-Native
GAC78
rSRL
ECL
SRL
DC-ASGPR short
MAL-II
UEA-I
PTA-Gal
ConA
ME5(detected with a-HUD)
MAL-I
TbFBL2
WGA
WT-VCC
RCAI (Vector)
ConA
AOL (Mahal Lab)
DBL-I
BTL native
AAL
TbFBL1

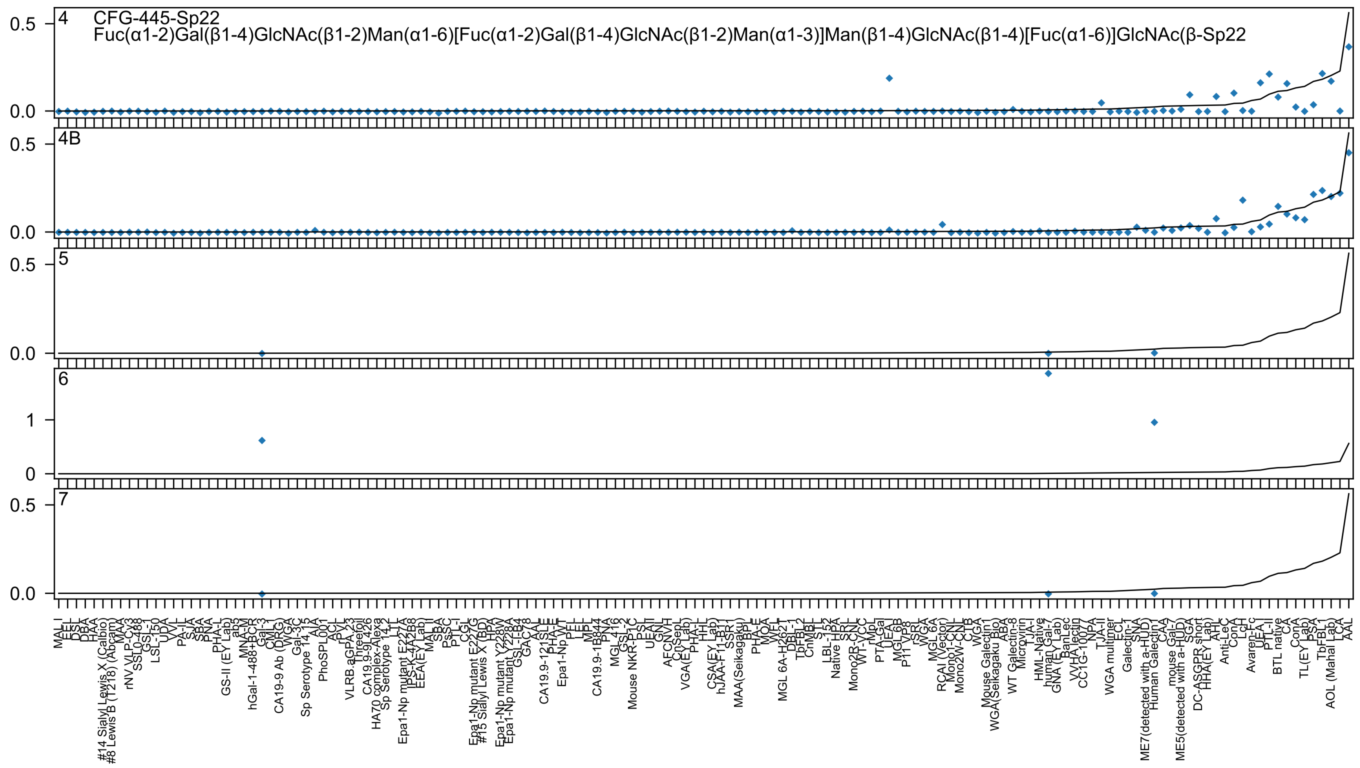
CFG-445-Sp22
Fuc(α1-2)Gal(β1-4)GlcNAc(β1-2)Man(α1-6)[Fuc(α1-2)Gal(β1-4)GlcNAc(β1-2)Man(α1-3)]Man(β1-4)GlcNAc(β1-4)[Fuc(α1-6)]GlcNAc(β-Sp22

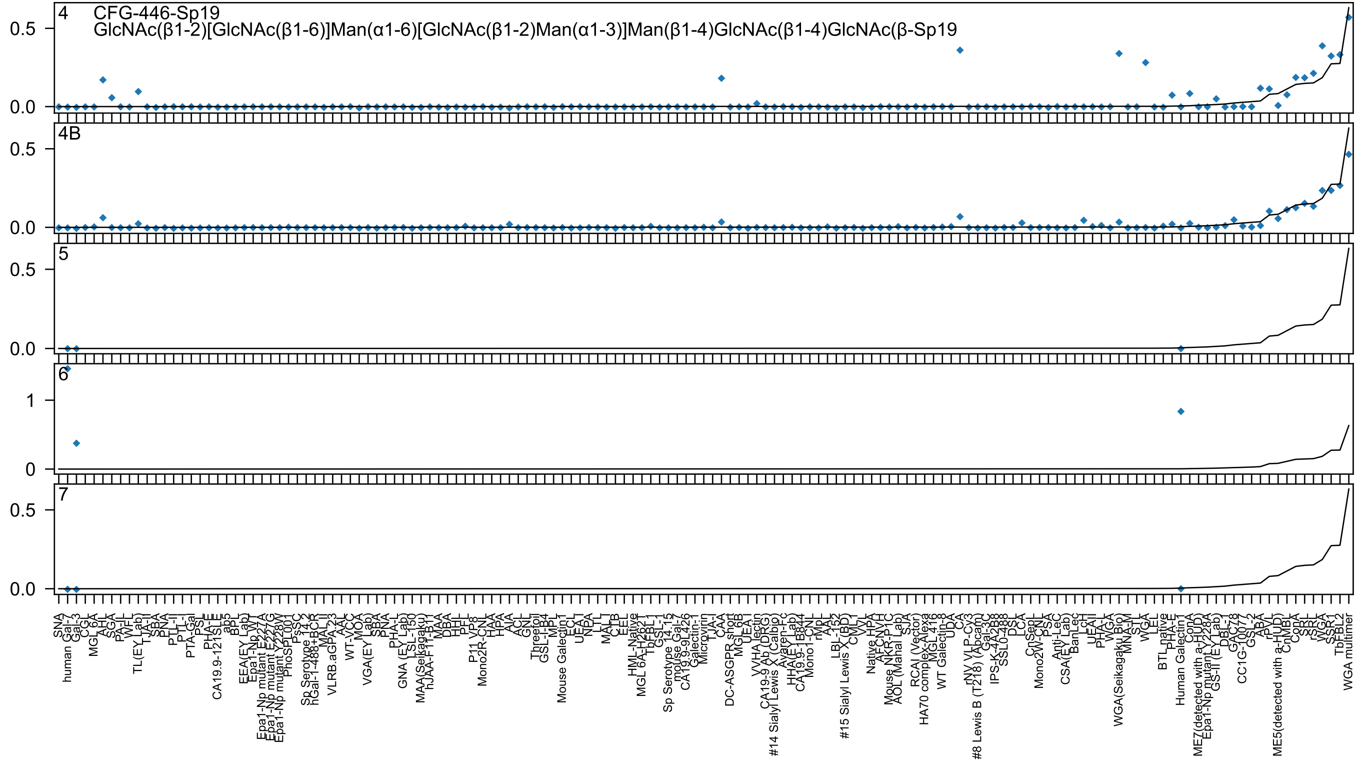

4
CFG-446-Sp19
GlcNAc(β1-2)[GlcNAc(β1-6)]Man(α1-6)[GlcNAc(β1-2)Man(α1-3)]Man(β1-4)GlcNAc(β1-4)GlcNAc(β-Sp19
4B
5
6
7

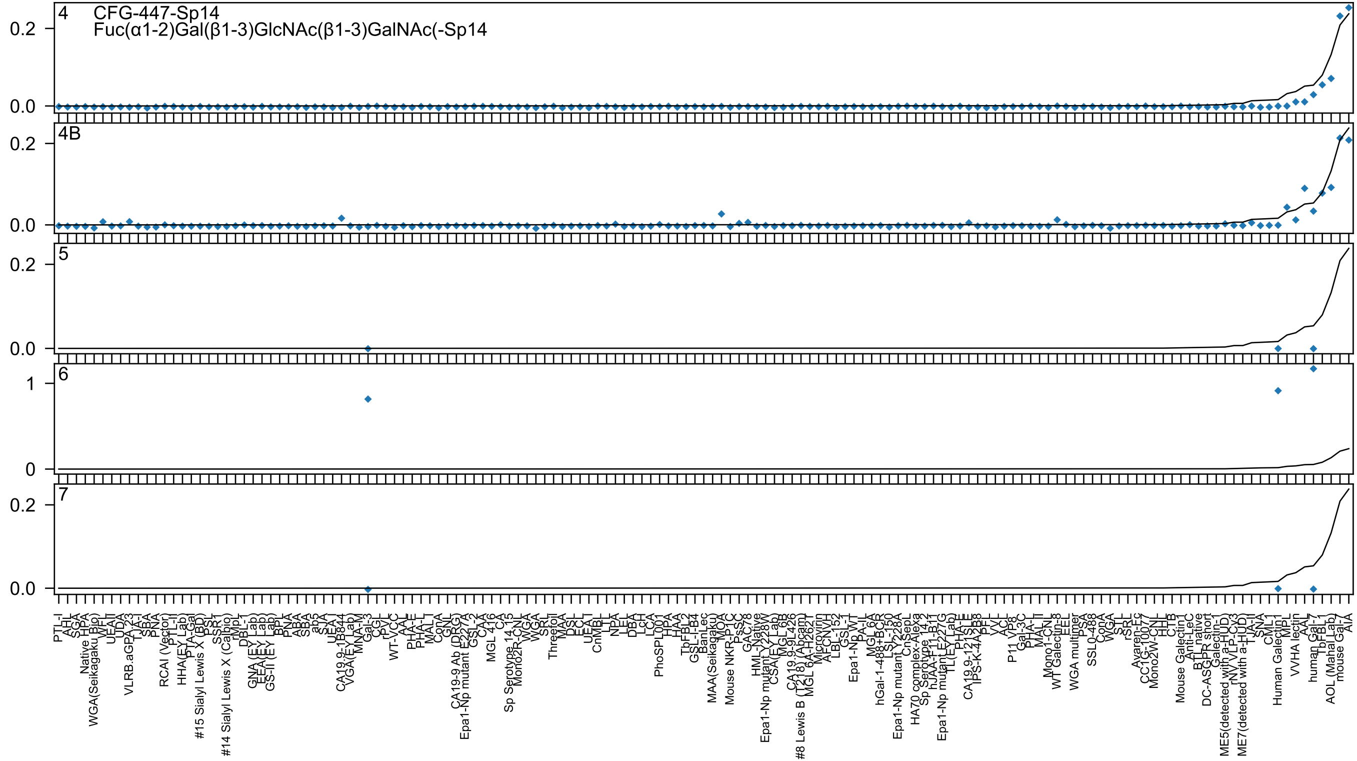

4 CFG-447-Sp14
Fuc(α1-2)Gal(β1-3)GlcNAc(β1-3)GalNAc(-Sp14
4B
5
6
7

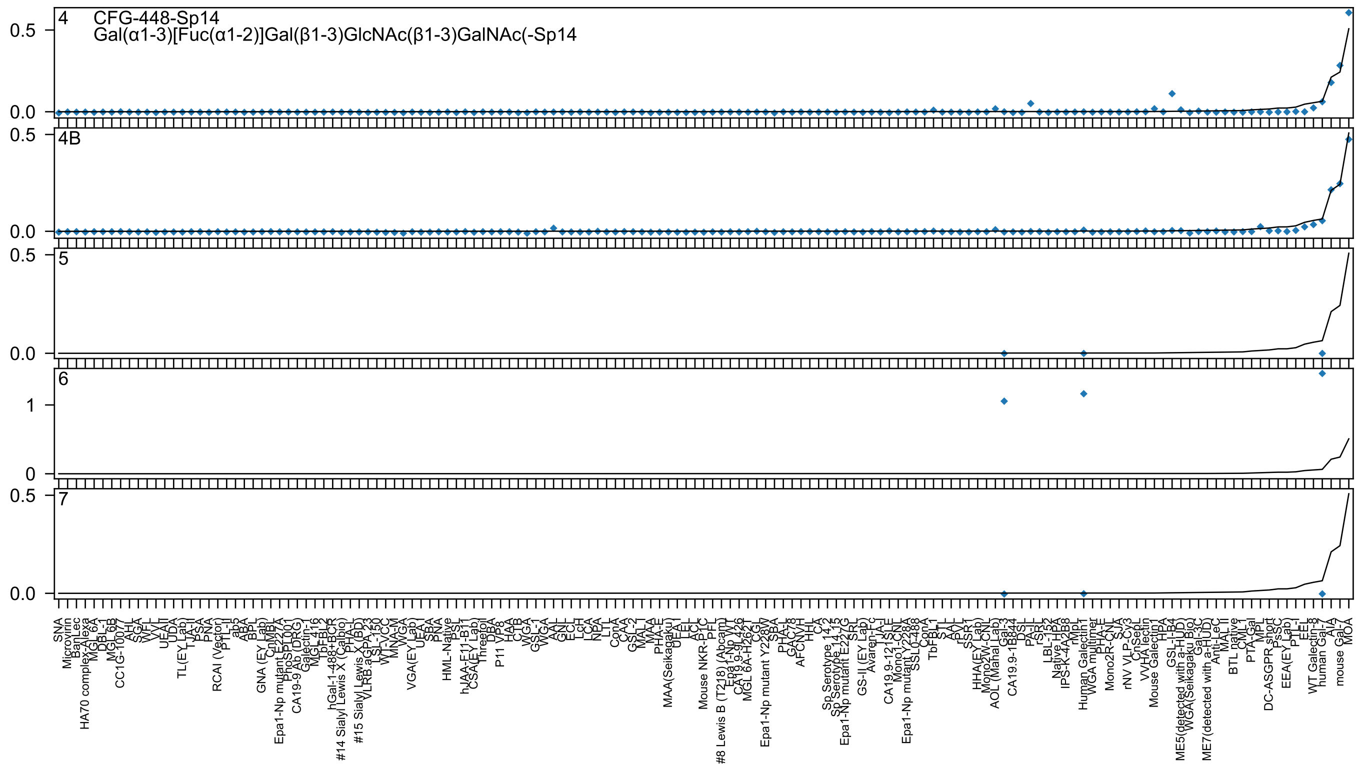

4 CFG-448-Sp14
Gal(α1-3)[Fuc(α1-2)]Gal(β1-3)GlcNAc(β1-3)GalNAc(-Sp14
4B
5
6
7

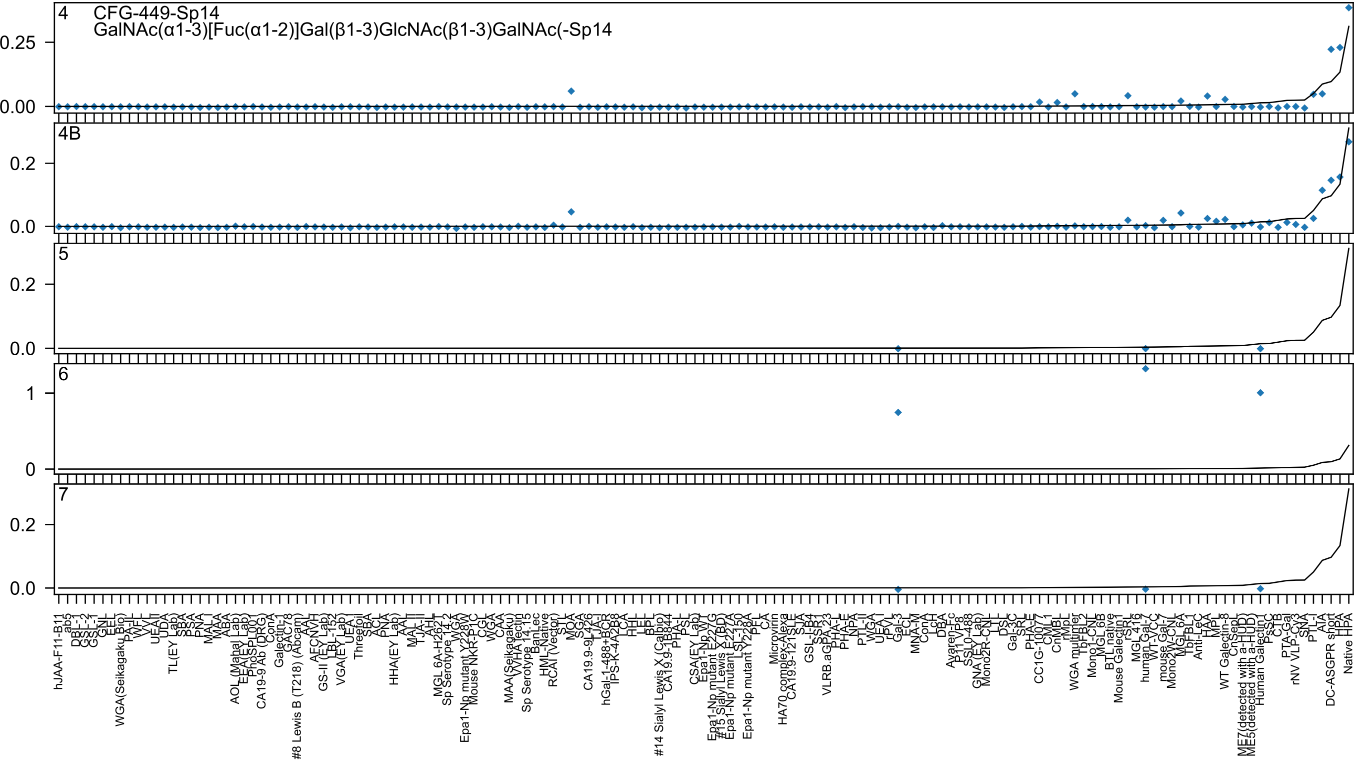

4
CFG-449-Sp14
GalNAc(α1-3)[Fuc(α1-2)]Gal(β1-3)GlcNAc(β1-3)GalNAc(-Sp14
0.25
0.00
4B
0.2
0.0
5
0.2
0.0
6
1
0
7
0.2
0.0

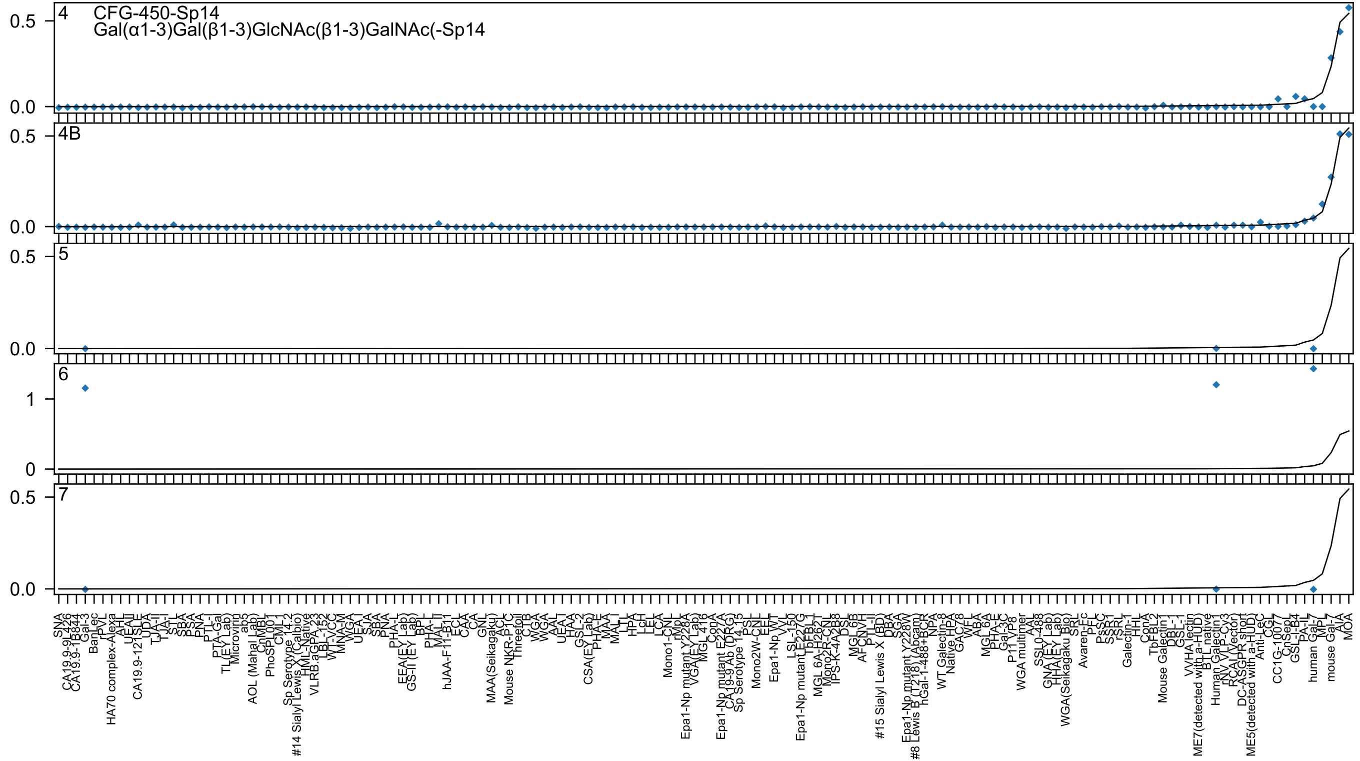
4
CFG-450-Sp14
Gal(α1-3)Gal(β1-3)GlcNAc(β1-3)GalNAc(-Sp14
4B
5
6
7

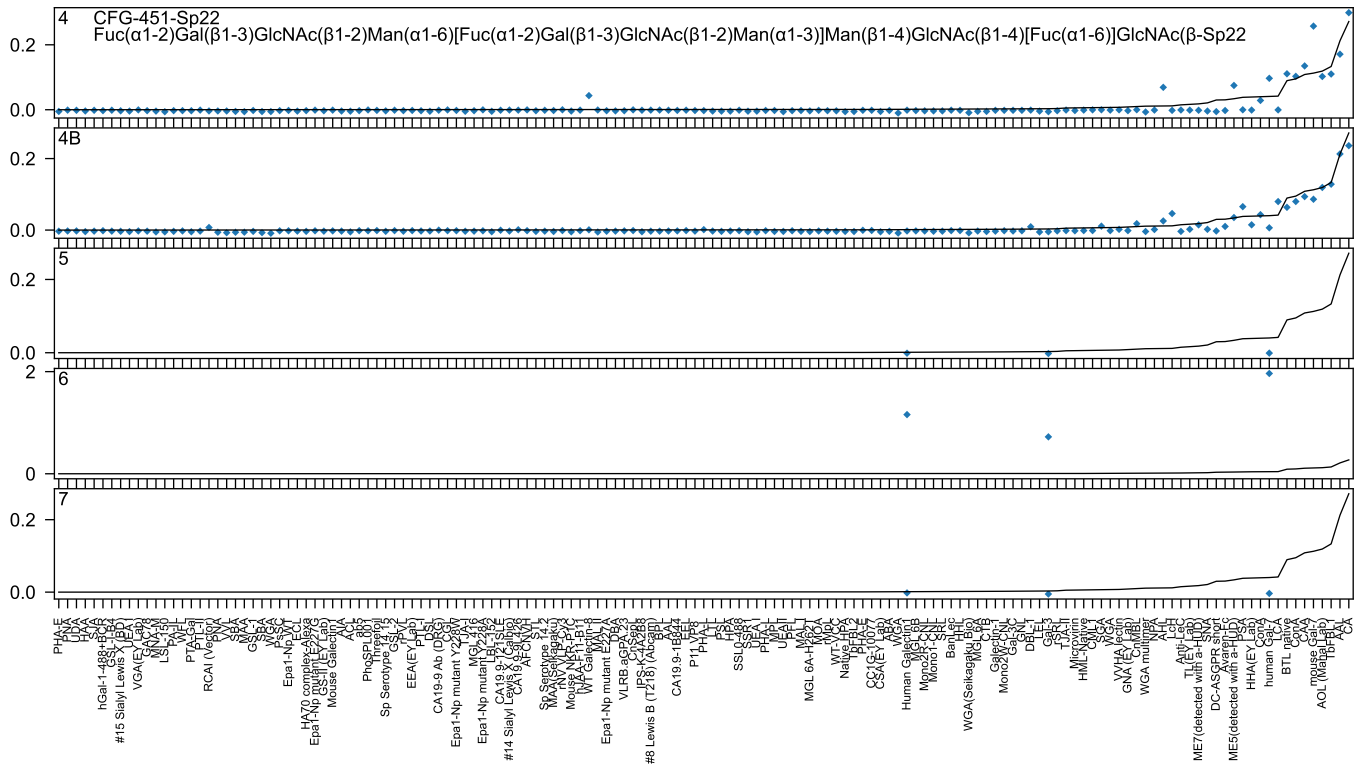
4 CFG-451-Sp22
Fuc(α1-2)Gal(β1-3)GlcNAc(β1-2)Man(α1-6)[Fuc(α1-2)Gal(β1-3)GlcNAc(β1-2)Man(α1-3)]Man(β1-4)GlcNAc(β1-4)[Fuc(α1-6)]GlcNAc(β-Sp22
4B
5
6
7
0.0
0.2
0
2

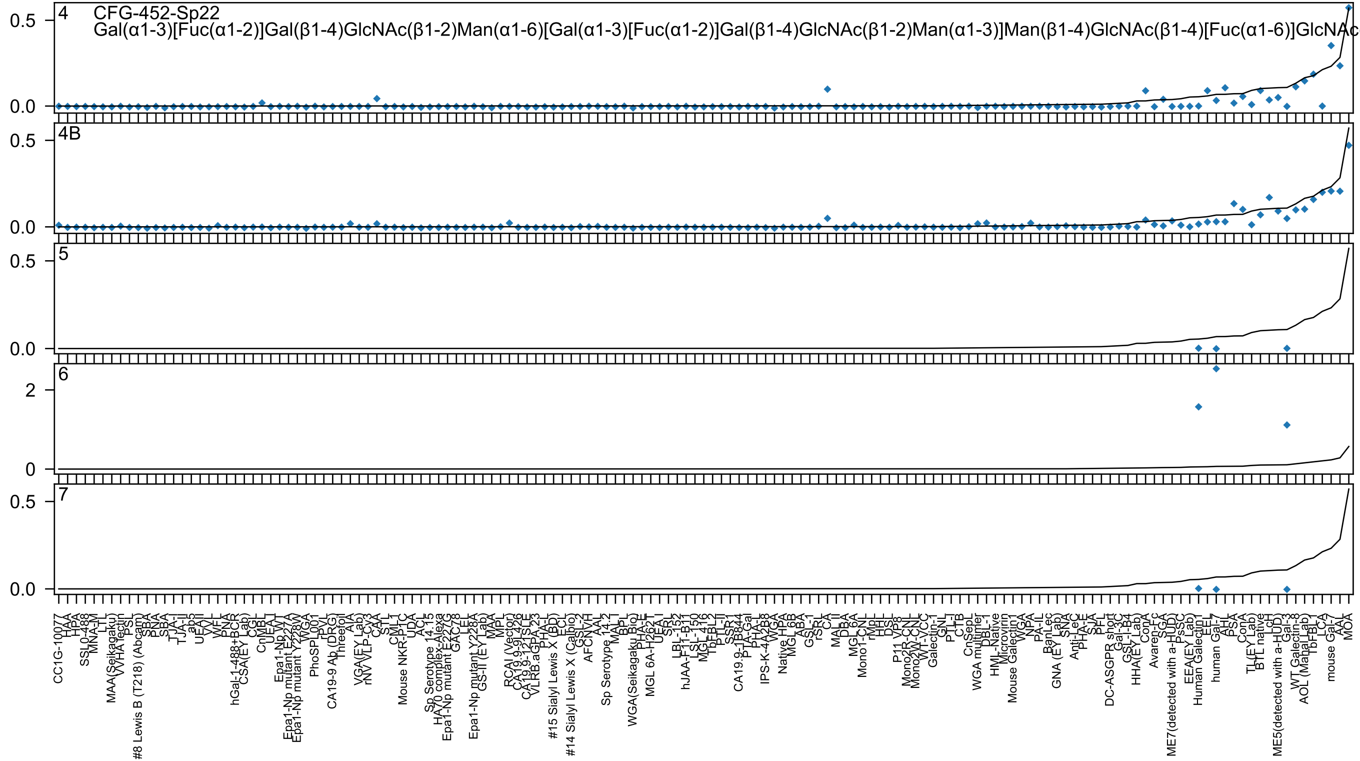
4
CFG-452-Sp22
Gal(α1-3)[Fuc(α1-2)]Gal(β1-4)GlcNAc(β1-2)Man(α1-6)[Gal(α1-3)[Fuc(α1-2)]Gal(β1-4)GlcNAc(β1-2)Man(α1-3)]Man(β1-4)GlcNAc(β1-4)[Fuc(α1-6)]GlcNAc
4B
5
6
7

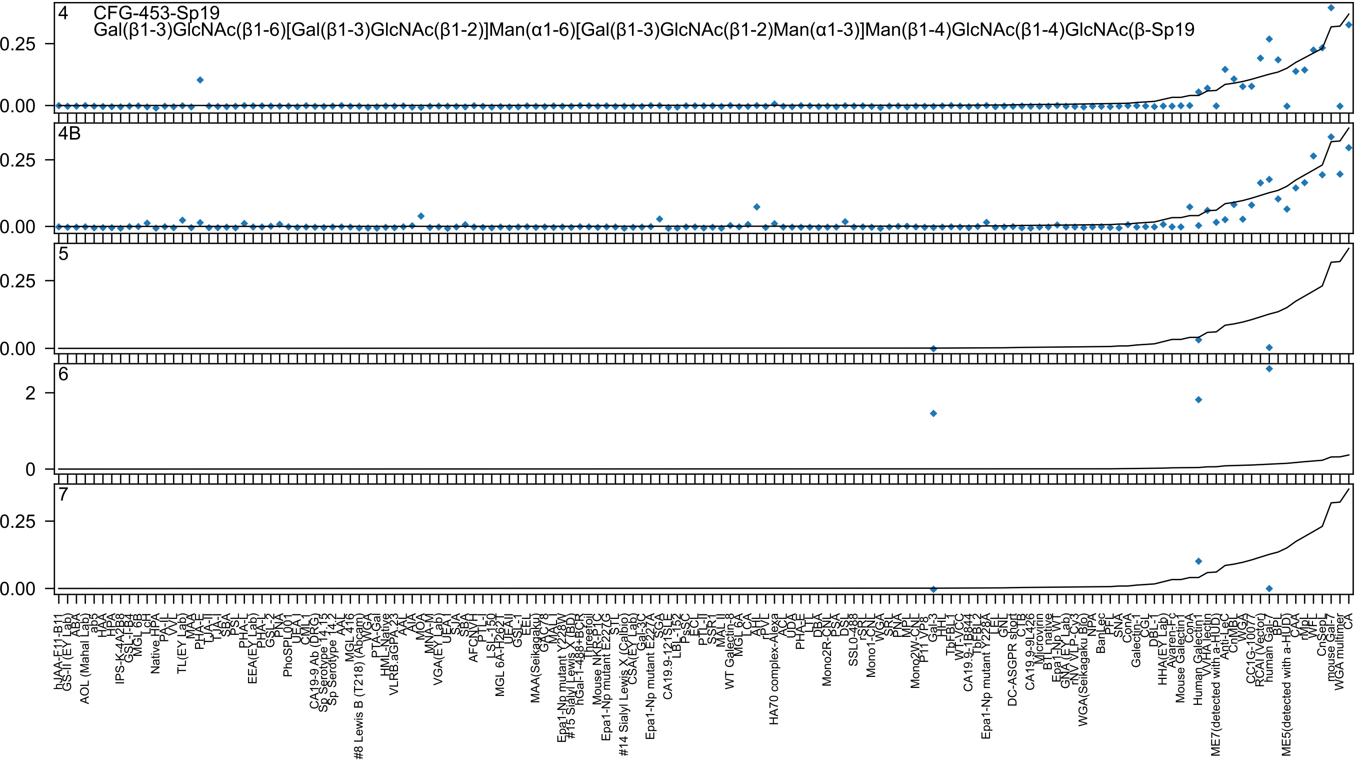

CFG-453-Sp19
Gal(β1-3)GlcNAc(β1-6)[Gal(β1-3)GlcNAc(β1-2)]Man(α1-6)[Gal(β1-3)GlcNAc(β1-2)Man(α1-3)]Man(β1-4)GlcNAc(β1-4)GlcNAc(β-Sp19
4
4B
5
6
7

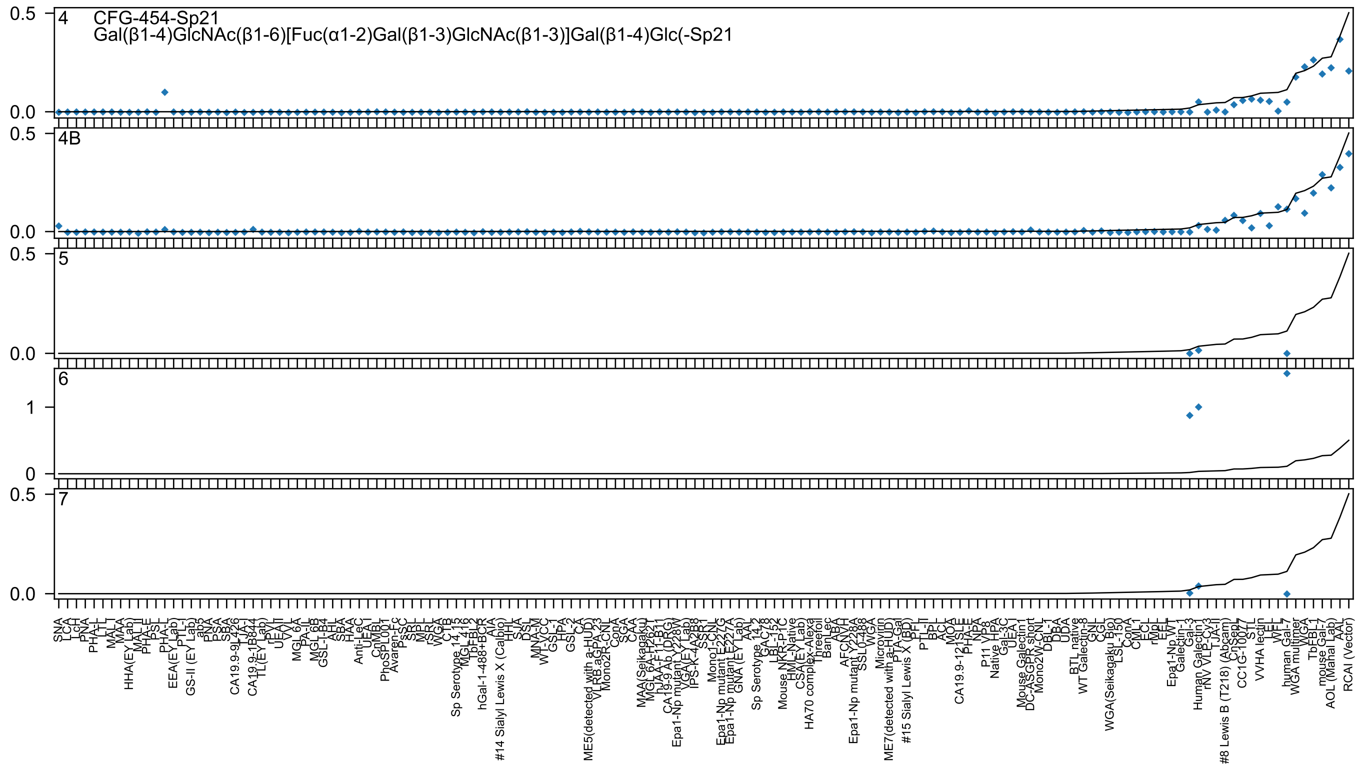
CFG-454-Sp21
Gal(β1-4)GlcNAc(β1-6)[Fuc(α1-2)Gal(β1-3)GlcNAc(β1-3)]Gal(β1-4)Glc(-Sp21

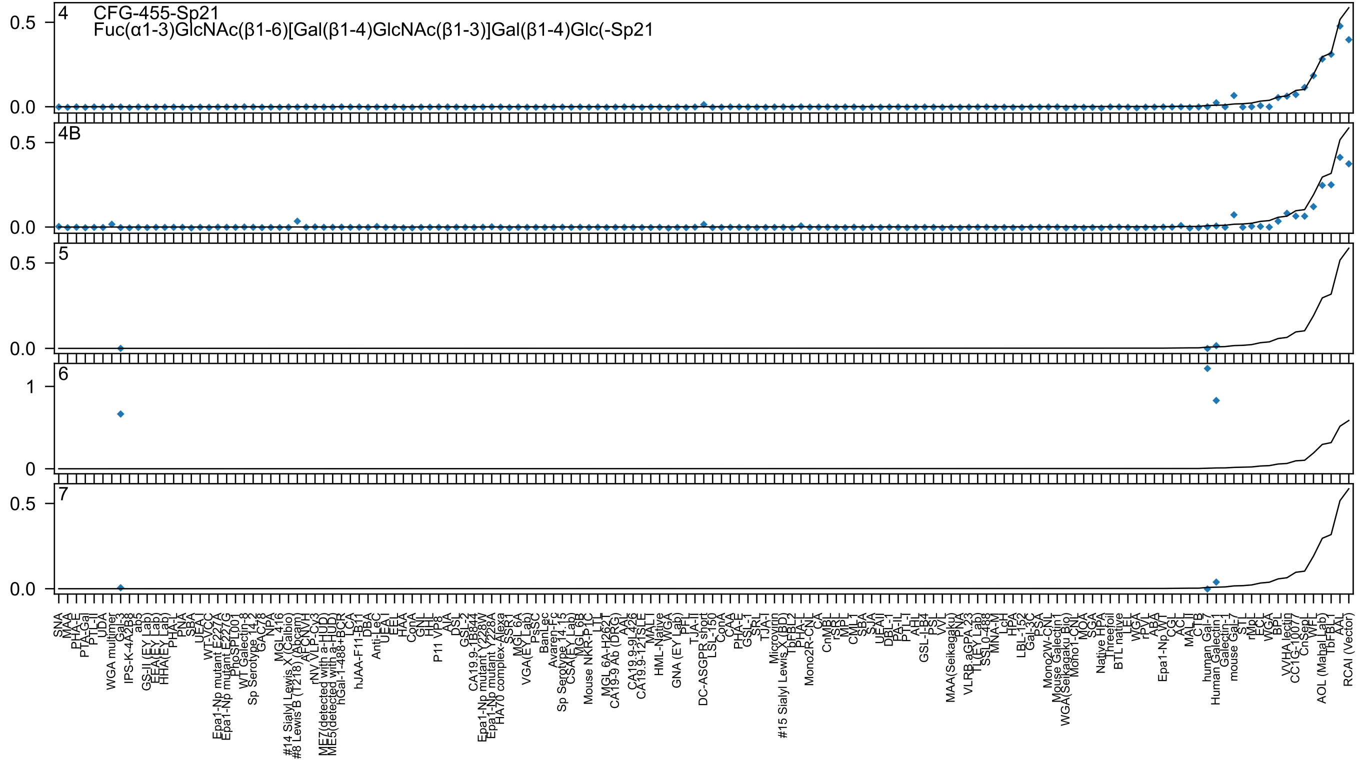
4 CFG-455-Sp21
Fuc(α1-3)GlcNAc(β1-6)[Gal(β1-4)GlcNAc(β1-3)]Gal(β1-4)Glc(-Sp21
4B
5
6
7
SNA
MAA
PHA-E
PTA-Gal
PTL-II
UDA
WGA multimer
Gal-3
IPS-K-4A2B8
ab5
GS-II (EY Lab)
EEA(EY Lab)
HHA(EY Lab)
PHA-L
PNA
SBA
UEA-I
WT-VCC
Epa1-Np mutant E227A
Epa1-Np mutant E227G
PhoSL
WT Galectin-8
Sp Serotype 14.2
GAC78
NPA
MGL 416
#14 Sialyl Lewis X (Calbio)
#8 Lewis B (T218) (Abcam)
AFCNVH
rNV VLP-Cy3
ME7(detected with a-HUD)
ME5(detected with a-HUD)
hGal-1-488+BCR
LCA
hJAA-F11-B11
DBA
Anti-LeC
UEA-I
EEL
HAA
ConA
GNL
HHL
P11 VP8
AIA
DSL
GSL-2
CA19.9-1B844
Epa1-Np mutant Y228W
Epa1-Np mutant Y228A
HA70 complex-Alexa
SSR1
MGL 6A
VGA(EY Lab)
PsSC
BanLec
Avaren-Fc
Sp Serotype 14.15
CSA(EY Lab)
MGL 6B
Mouse NKR-P1C
LTL
MGL 6A-H262T
CA19-9 Ab (DRG)
AAL
CA19.9-9L426
CA19.9-121SLE
MAL-I
HML-Native
WGA
GNA (EY Lab)
PFL
TJA-II
DC-ASGPR short
LSL-150
ConA
CAA
PHA-E
GSL-1
SRL
TJA-I
Microvirin
#15 Sialyl Lewis X (BD)
TbFBL2
PHA-L
Mono2R-CNL
CA
CnMBL
rSRL
MPL
CML1
SBA
SJA
UEAII
DBL-1
PA-IL
PTL-I
AHL
GSL-I-B4
PSL
VVL
MAA(Seikagaku)
RNA
VLRB.aGPA.23
TL(EY Lab)
SSL0-488
MNA-M
LcH
HPA
LBL-152
Gal-3C
PSA
Mono2W-CNL
Mouse Galectin1
WGA(Seikagaku Bio)
Mono1-CNL
MOA
SGA
Native HPA
Threefoil
BTL native
LEL
WGA
rPVL
ABA
Epa1-Np WT
CGL
ACL
MAL-II
CTB
human Gal-7
Human Galectin-1
Galectin-1
mouse Gal-7
STL
rMpL
ECL
WGA
BPL
VVHA lectin
CC1G-1007
CnSep1
WFL
AOL (Mahal Lab)
TbFBL1
AAL
RCAI (Vector)

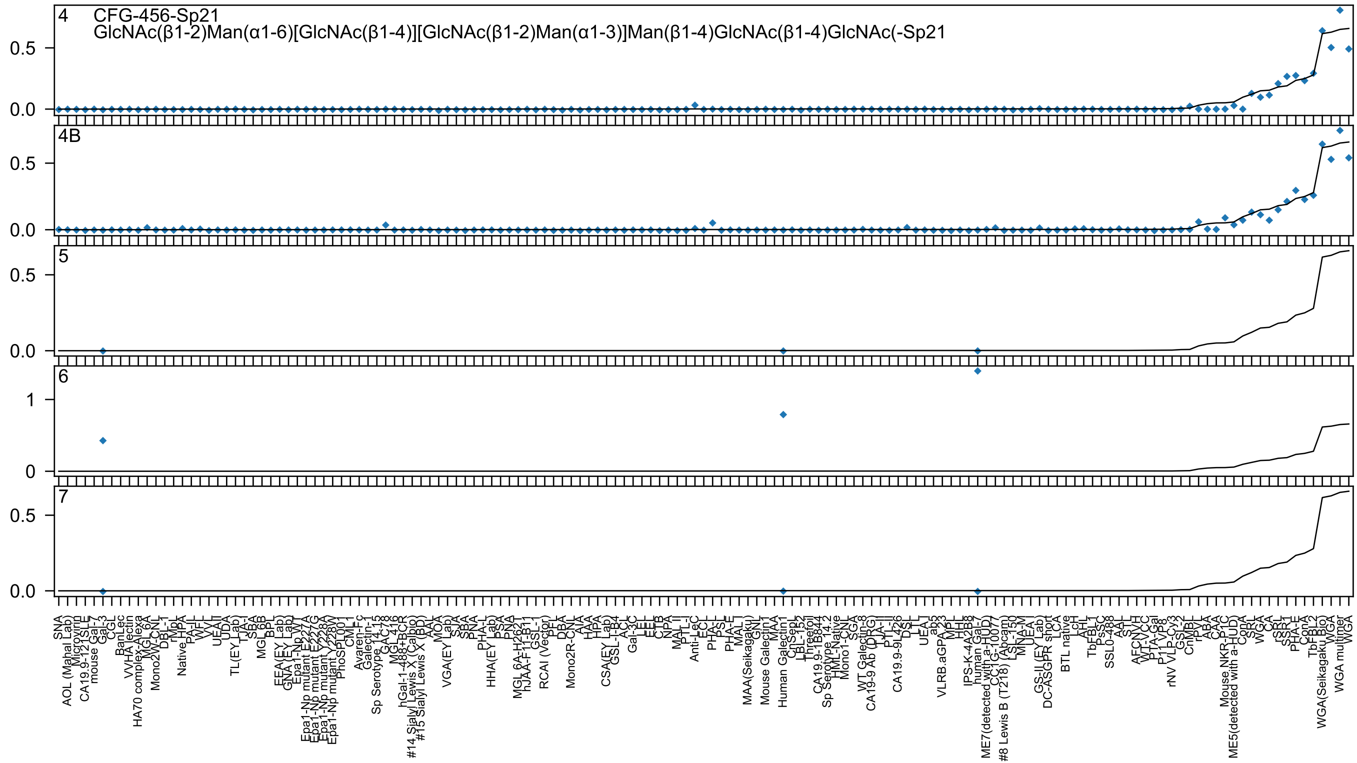
4
CFG-456-Sp21
GlcNAc(β1-2)Man(α1-6)[GlcNAc(β1-4)][GlcNAc(β1-2)Man(α1-3)]Man(β1-4)GlcNAc(β1-4)GlcNAc(-Sp21
4B
5
6
7
0.0
0.5
0
1

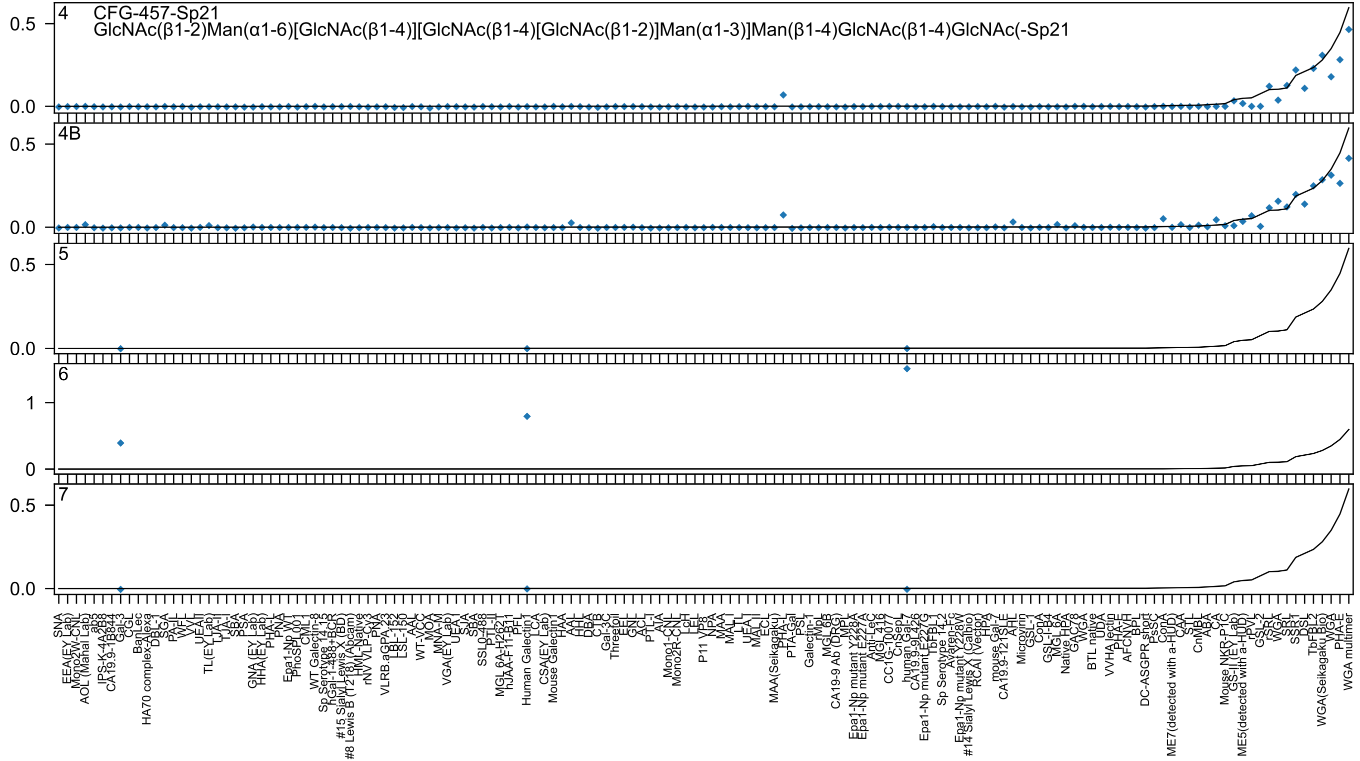

4 CFG-457-Sp21
GlcNAc(β1-2)Man(α1-6)[GlcNAc(β1-4)][GlcNAc(β1-4)[GlcNAc(β1-2)]Man(α1-3)]Man(β1-4)GlcNAc(β1-4)GlcNAc(-Sp21
4B
5
6
7

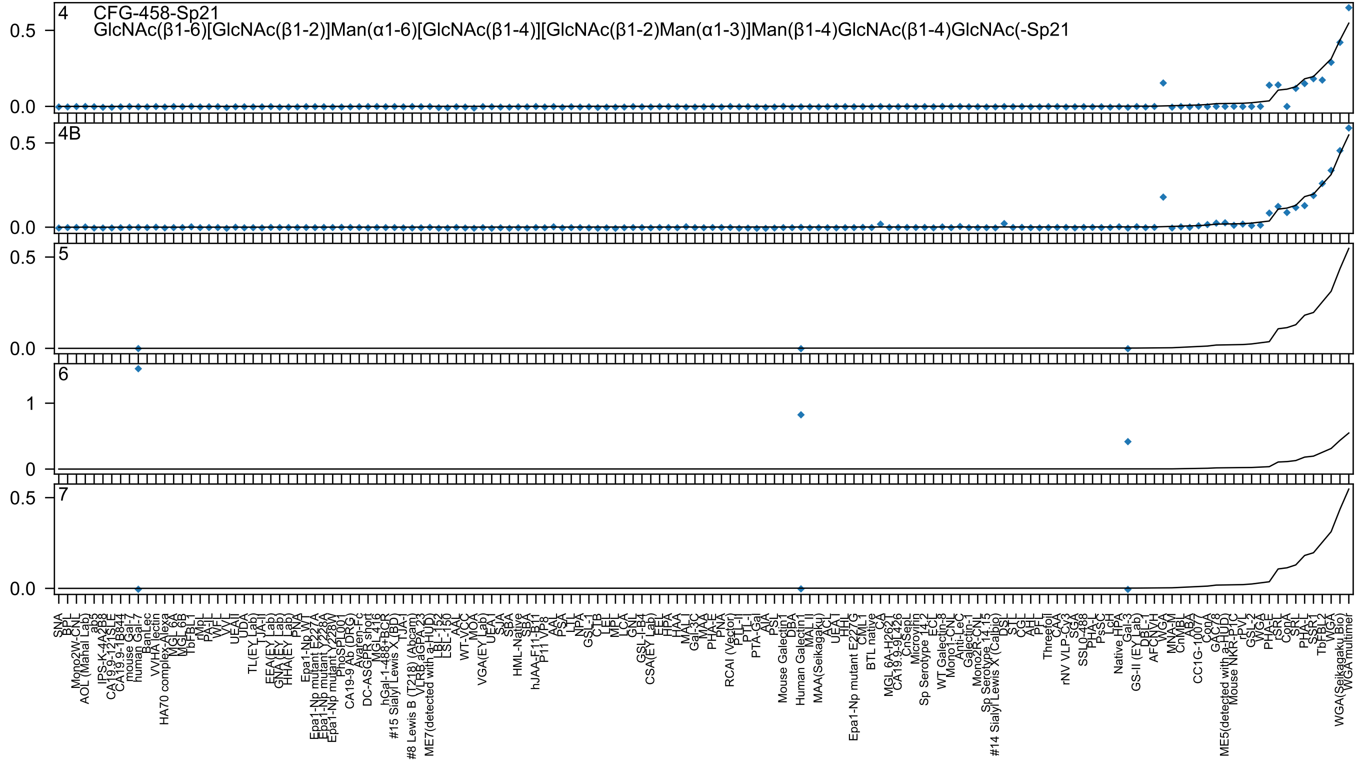

4 CFG-458-Sp21
GlcNAc(β1-6)[GlcNAc(β1-2)]Man(α1-6)[GlcNAc(β1-4)][GlcNAc(β1-2)Man(α1-3)]Man(β1-4)GlcNAc(β1-4)GlcNAc(-Sp21
4B
5
6
7

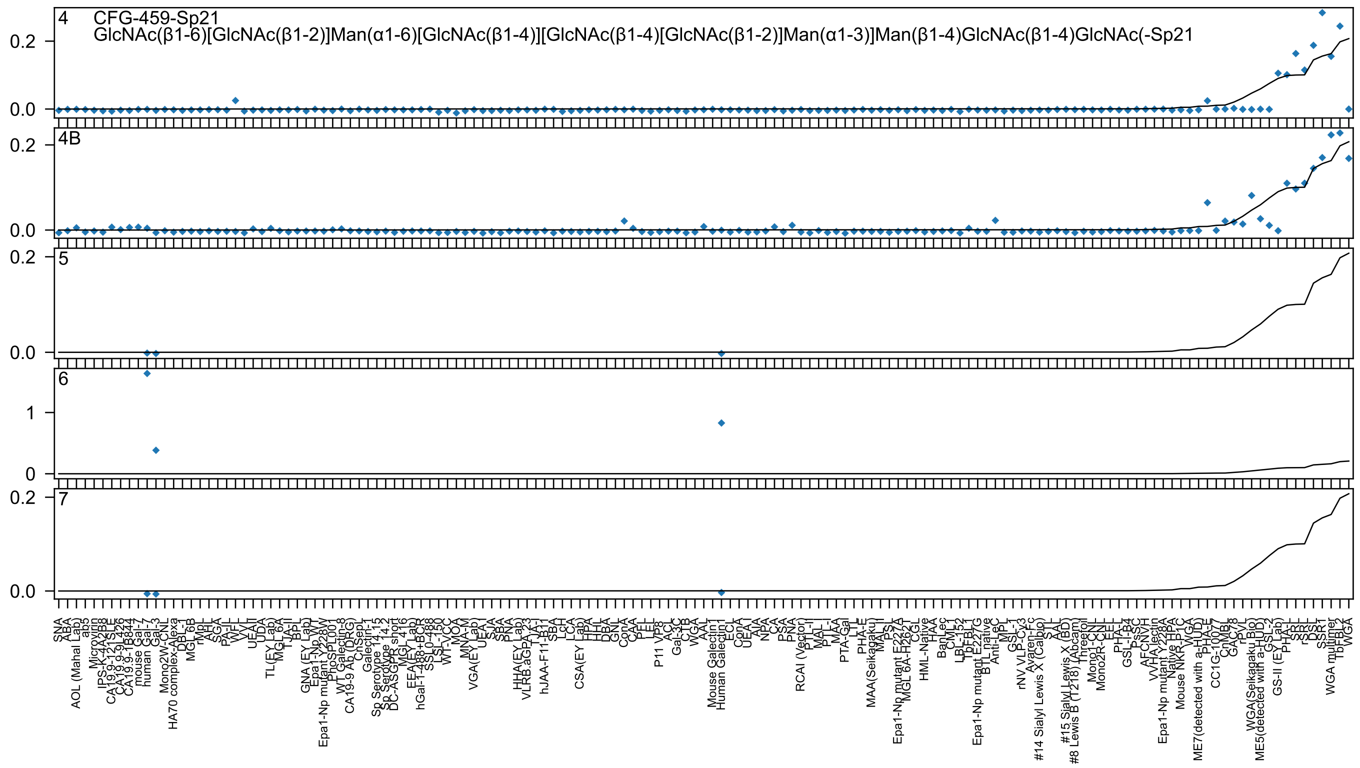

4
CFG-459-Sp21
GlcNAc(β1-6)[GlcNAc(β1-2)]Man(α1-6)[GlcNAc(β1-4)][GlcNAc(β1-4)[GlcNAc(β1-2)]Man(α1-3)]Man(β1-4)GlcNAc(β1-4)GlcNAc(-Sp21
4B
5
6
7

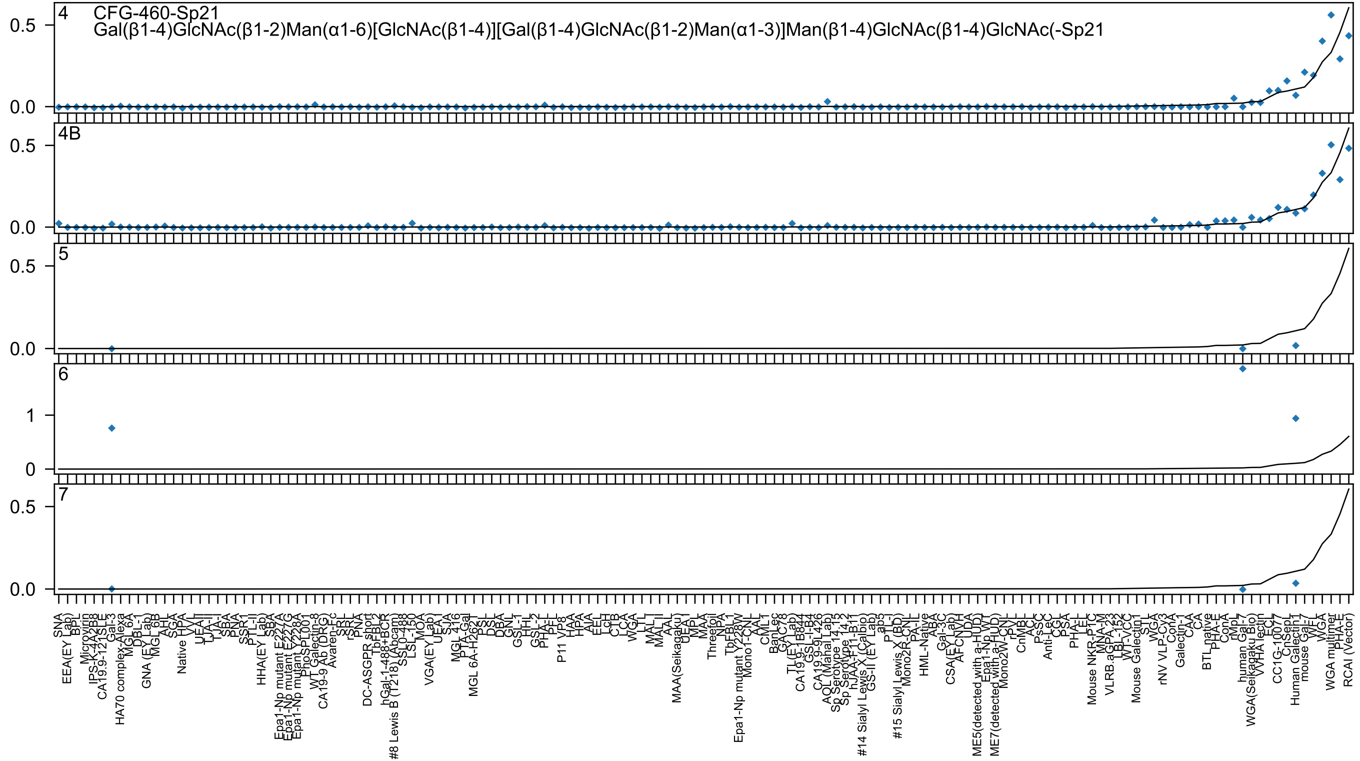
4
CFG-460-Sp21
Gal(β1-4)GlcNAc(β1-2)Man(α1-6)[GlcNAc(β1-4)][Gal(β1-4)GlcNAc(β1-2)Man(α1-3)]Man(β1-4)GlcNAc(β1-4)GlcNAc(-Sp21
4B
5
6
7
0.0
0.5
0
1

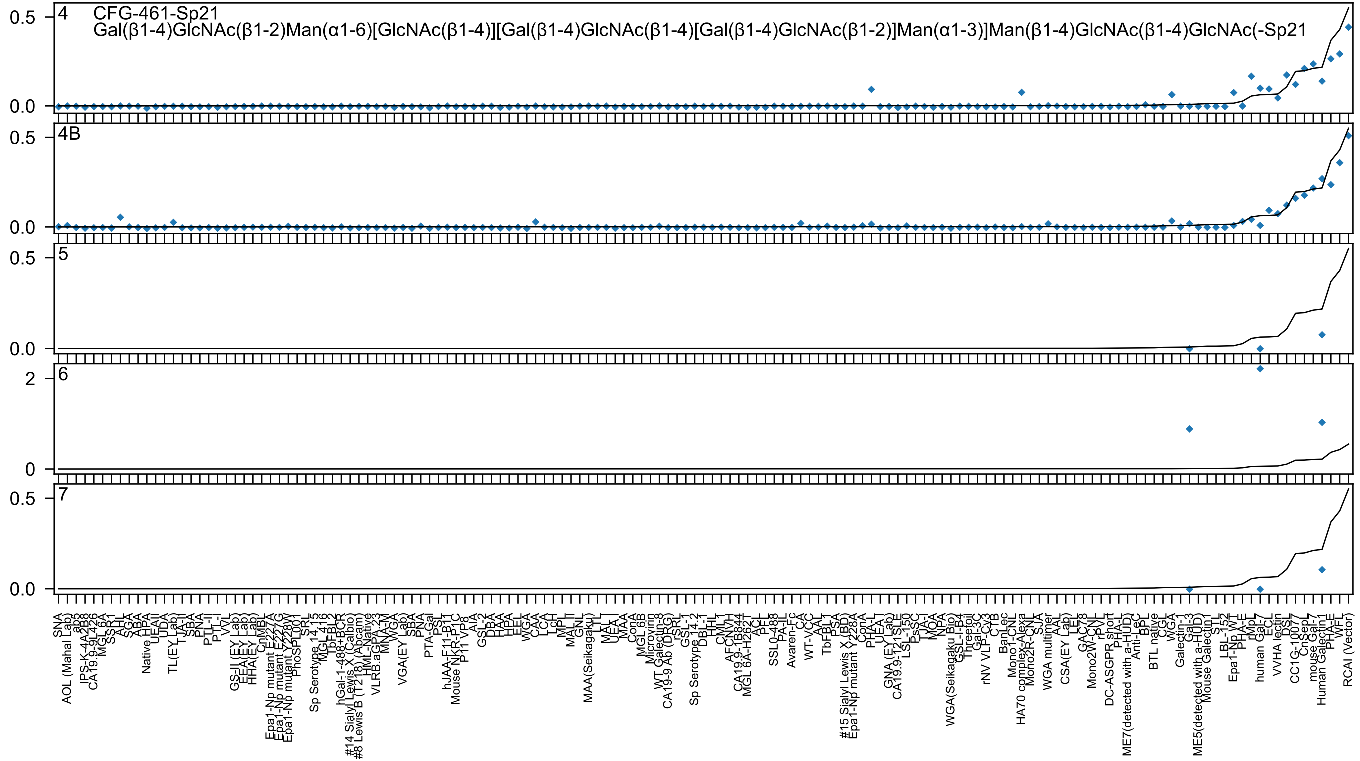

4
CFG-461-Sp21
Gal(β1-4)GlcNAc(β1-2)Man(α1-6)[GlcNAc(β1-4)][Gal(β1-4)GlcNAc(β1-4)[Gal(β1-4)GlcNAc(β1-2)]Man(α1-3)]Man(β1-4)GlcNAc(β1-4)GlcNAc(-Sp21
4B
5
6
7
0.5
0.0
2
0
SNA
AOL (Mahal Lab)
ab5
IPS-K-4A2B8
CA19.9-9L426
MGL 6A
SSR1
AHL
SGA
ABA
Native HPA
UEAII
UDA
TL(EY Lab)
TJA-II
SBA
PNA
PTL-II
PTL-I
VVL
GS-II (EY Lab)
EEA(EY Lab)
HHA(EY Lab)
CnMBL
Epa1-Np mutant E227A
Epa1-Np mutant E227G
Epa1-Np mutant Y228W
PhoSL001
SRL
Sp Serotype 14.15
MGL 416
TbFBL2
hGal-1-488+BCR
#14 Sialyl Lewis X (Calbio)
#8 Lewis B (T218) (Abcam)
HML-Native
VLRB.aGPA.23
MNA-M
WGA
VGA(EY Lab)
SBA
PNA
PTA-Gal
PSL
hJAA-F11-B11
Mouse NKR-P1C
P11 VP8
AIA
GSL-2
DBA
HAA
HPA
EEL
WGA
CAA
LCA
LcH
MPL
MAL II
GNL
MAA(Seikagaku)
LTL
MAL I
UEA I
MAA
ConA
MGL 6B
Microvirin
WT Galectin-8
CA19-9 Ab (DRG)
rSRL
GSL-I
Sp Serotype 14.2
DBL-1
HHL
CML1
AFCNVH
CA19.9-1B844
MGL 6A-H262T
ACL
PFL
SSL0-488
PA-IL
Avaren-Fc
CA
WT-VCC
AAL
TbFBL1
PSA
#15 Sialyl Lewis X (BD)
Epa1-Np mutant Y228A
ConA
PHA-L
UEA-I
GNA (EY Lab)
CA19.9-121SLE
LSL-150
PsSC
TJA-I
MOA
NPA
WGA(Seikagaku_Bio)
GSL-I-B4
Threefoil
Gal-3C
rNV VLP-Cy3
CTB
BanLec
Mono1-CNL
HA70 complex-Alexa
Mono2R-CNL
SJA
WGA multimer
AAL
CSA(EY Lab)
LEL
GAC78
Mono2W-CNL
rPVL
DC-ASGPR short
PHA-L
ME7(detected with a-HUD)
Anti-LeC
BPL
BTL native
CGL
WGA
Galectin-1
Gal-3
ME5(detected with a-HUD)
Mouse Galectin1
STL
LBL-152
Epa1-Np WT
PHA-E
rMpL
human Gal-7
ECL
VVHA lectin
DSL
CC1G-10077
CnSepL
mouse Gal-7
Human Galectin1
PHA-E
WFL
RCAI (Vector)

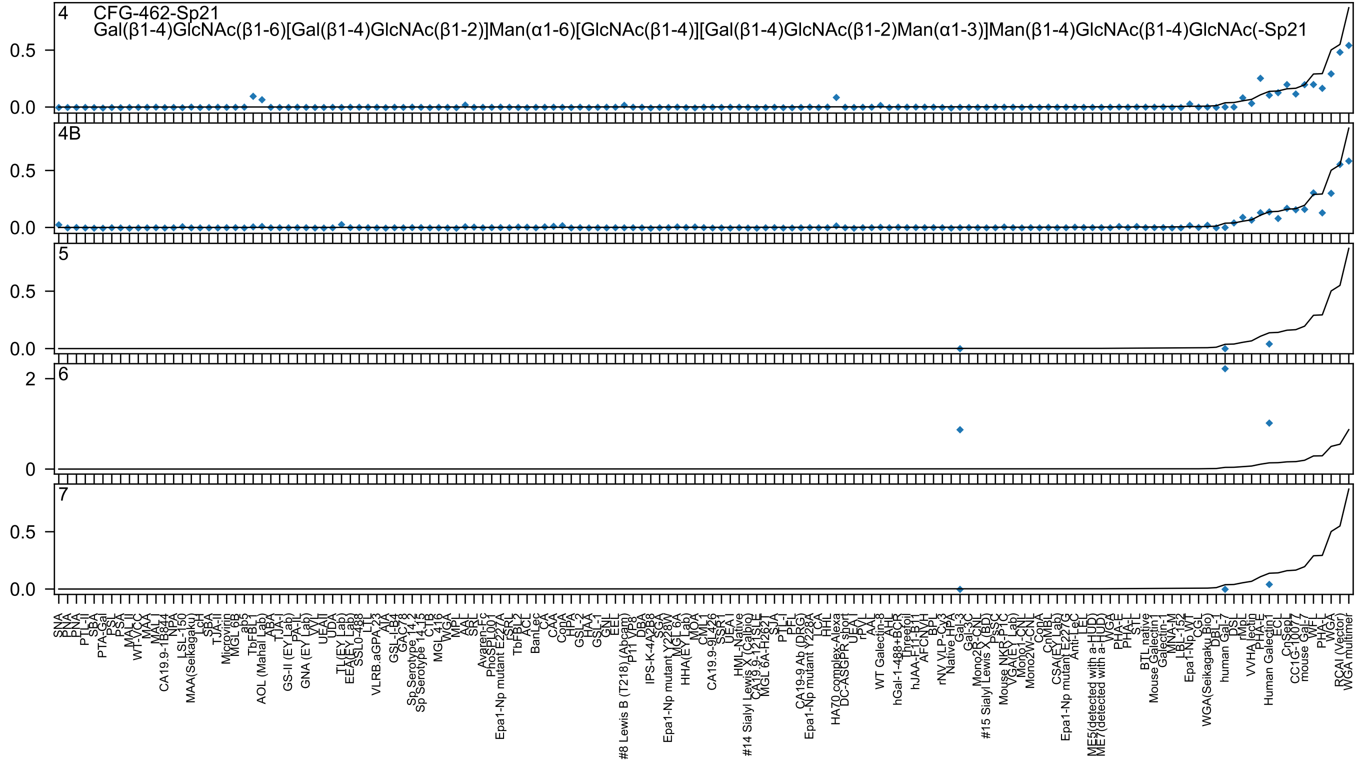
4
CFG-462-Sp21
Gal(β1-4)GlcNAc(β1-6)[Gal(β1-4)GlcNAc(β1-2)]Man(α1-6)[GlcNAc(β1-4)][Gal(β1-4)GlcNAc(β1-2)Man(α1-3)]Man(β1-4)GlcNAc(β1-4)GlcNAc(-Sp21
4B
5
6
7
0.0
0.5
0
2

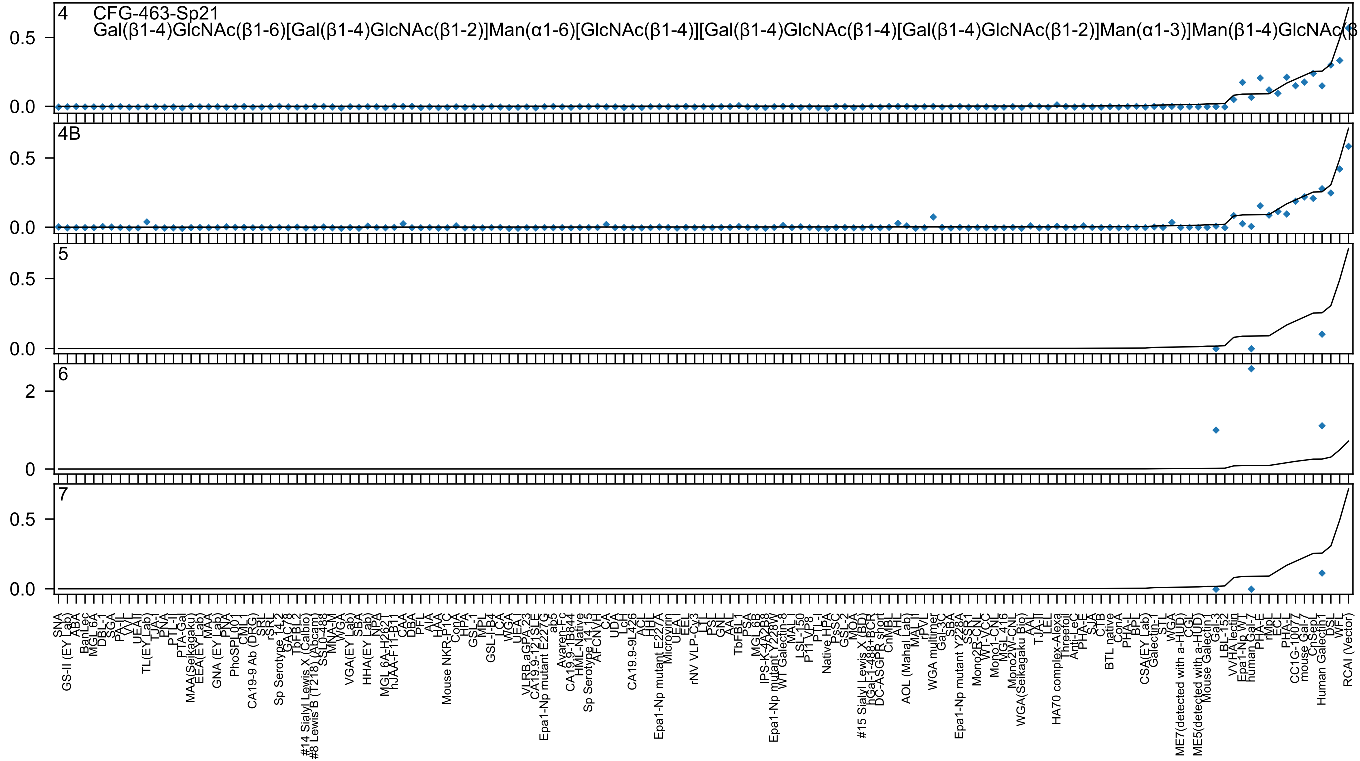

4 CFG-463-Sp21
Gal(β1-4)GlcNAc(β1-6)[Gal(β1-4)GlcNAc(β1-2)]Man(α1-6)[GlcNAc(β1-4)][Gal(β1-4)GlcNAc(β1-4)[Gal(β1-4)GlcNAc(β1-2)]Man(α1-3)]Man(β1-4)GlcNAc(β
4B
5
6
7

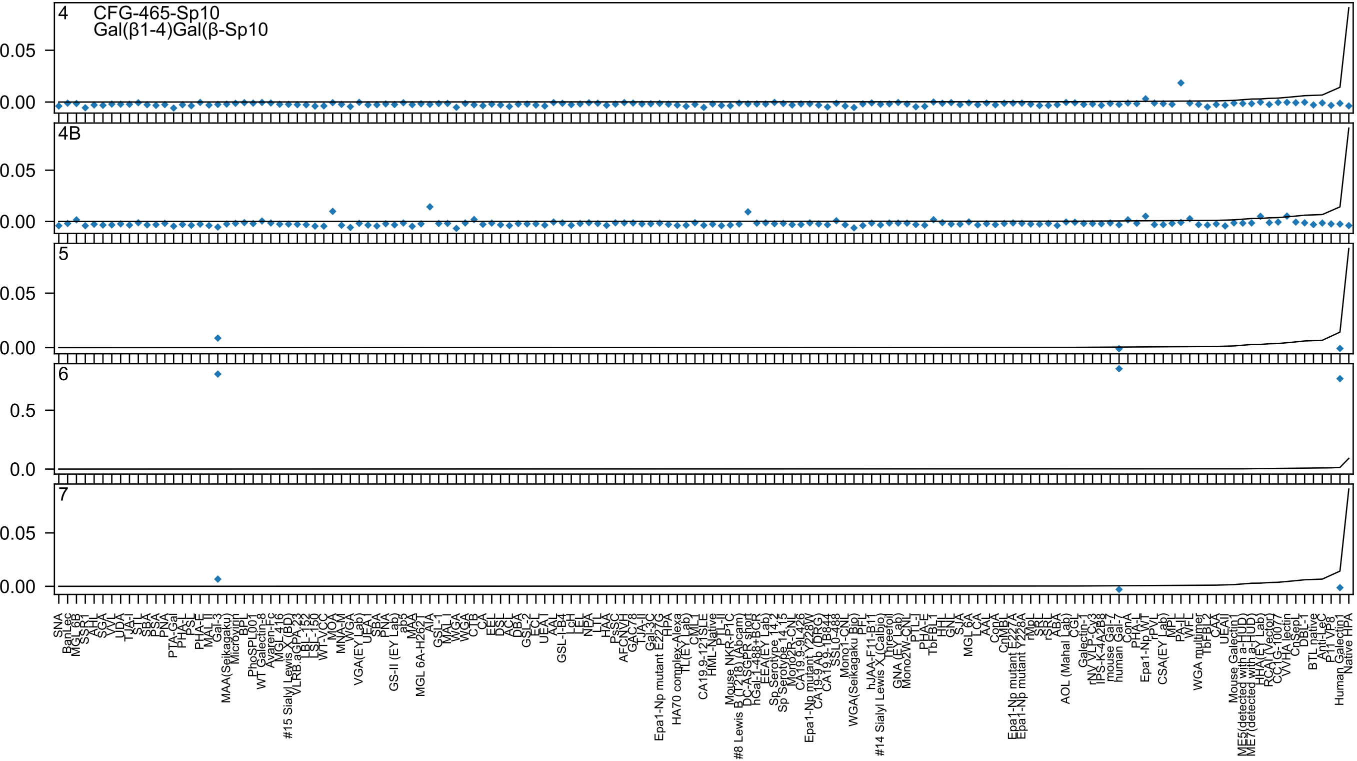
4
CFG-465-Sp10
Gal(β1-4)Gal(β-Sp10
4B
5
6
7
0.05
0.00
0.5
0.0

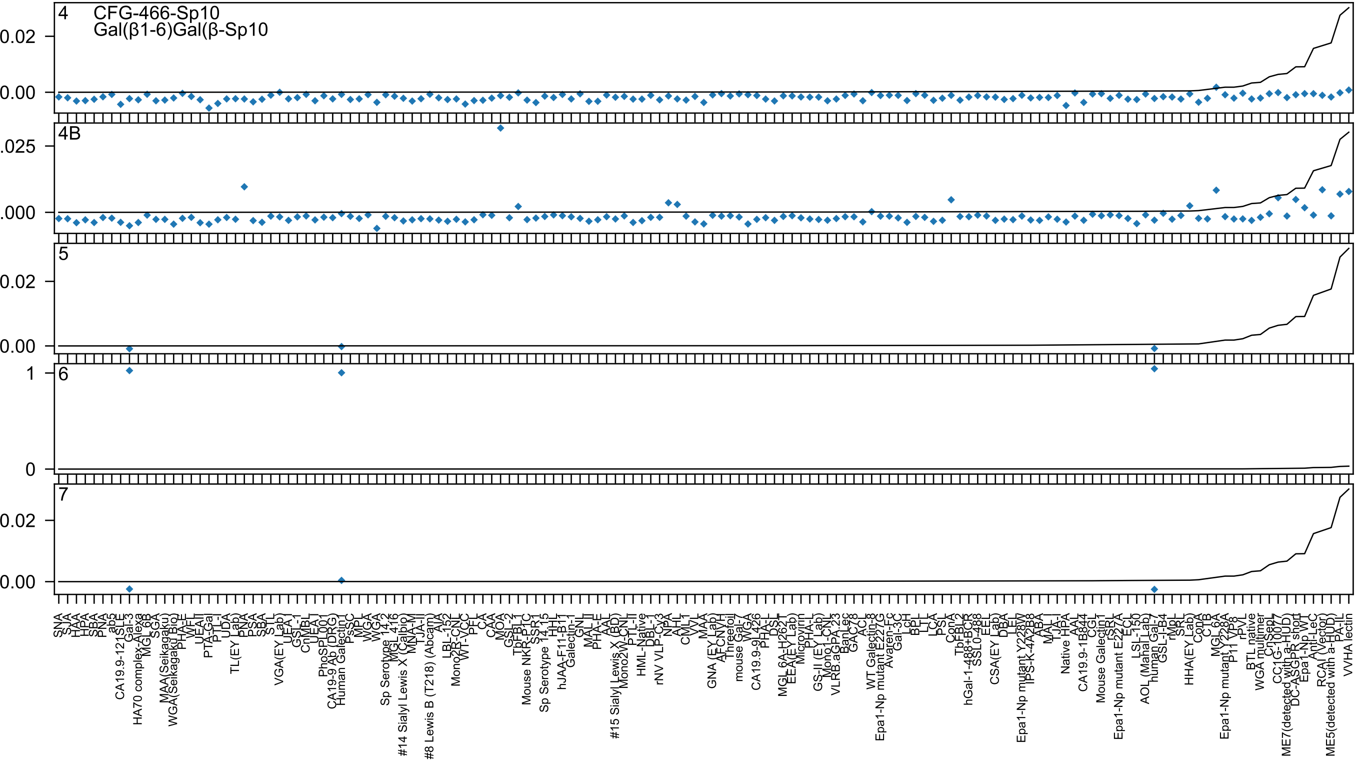
4
CFG-466-Sp10
Gal(β1-6)Gal(β-Sp10
4B
5
6
7

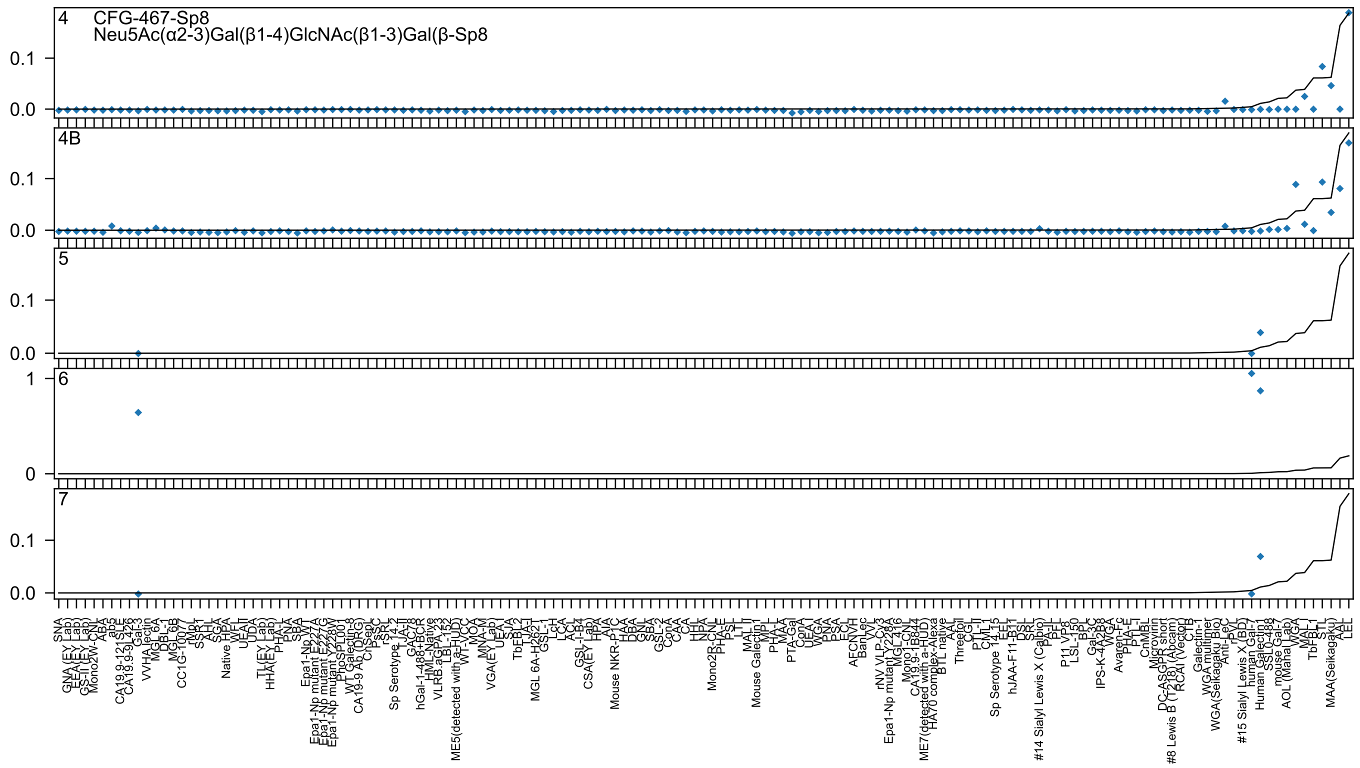

4 CFG-467-Sp8
Neu5Ac(α2-3)Gal(β1-4)GlcNAc(β1-3)Gal(β-Sp8
4B
5
6
7

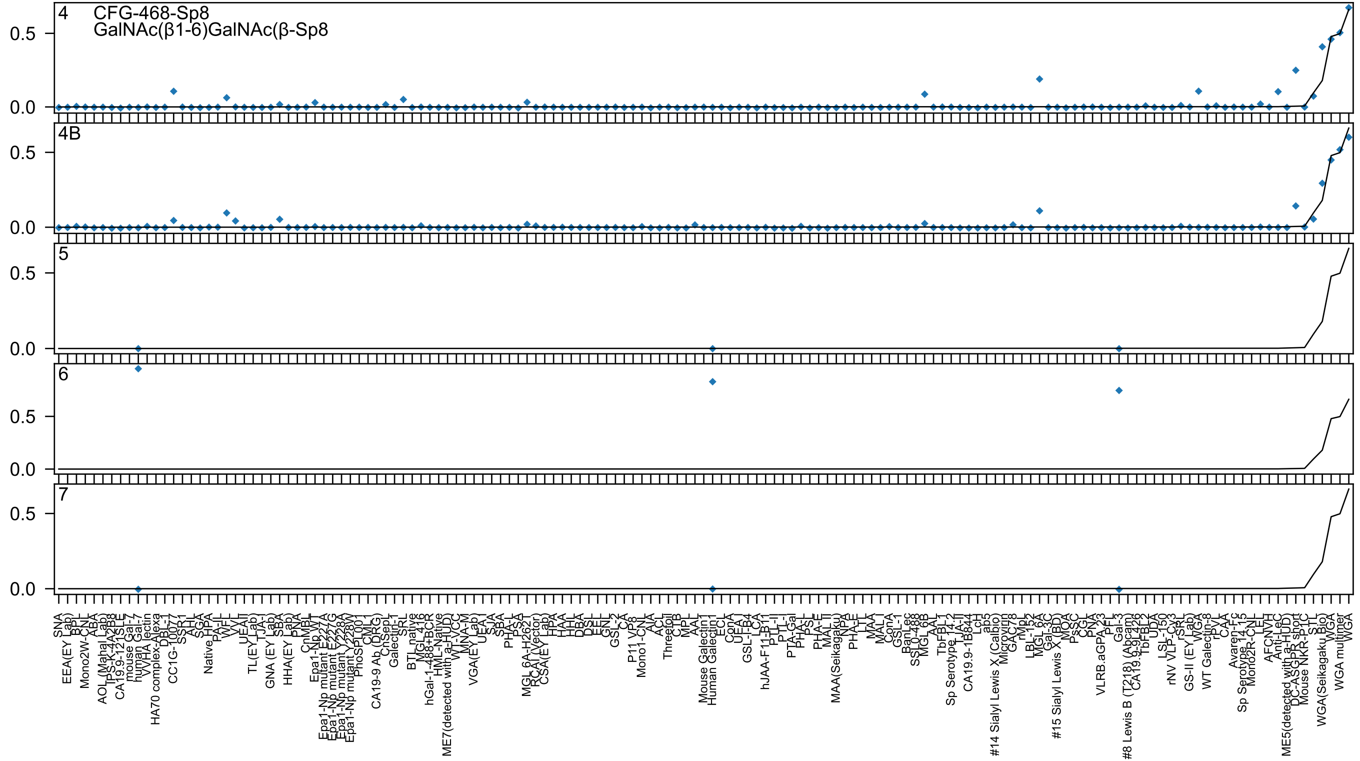

4
CFG-468-Sp8
GalNAc(β1-6)GalNAc(β-Sp8
4B
5
6
7
0.5
0.0

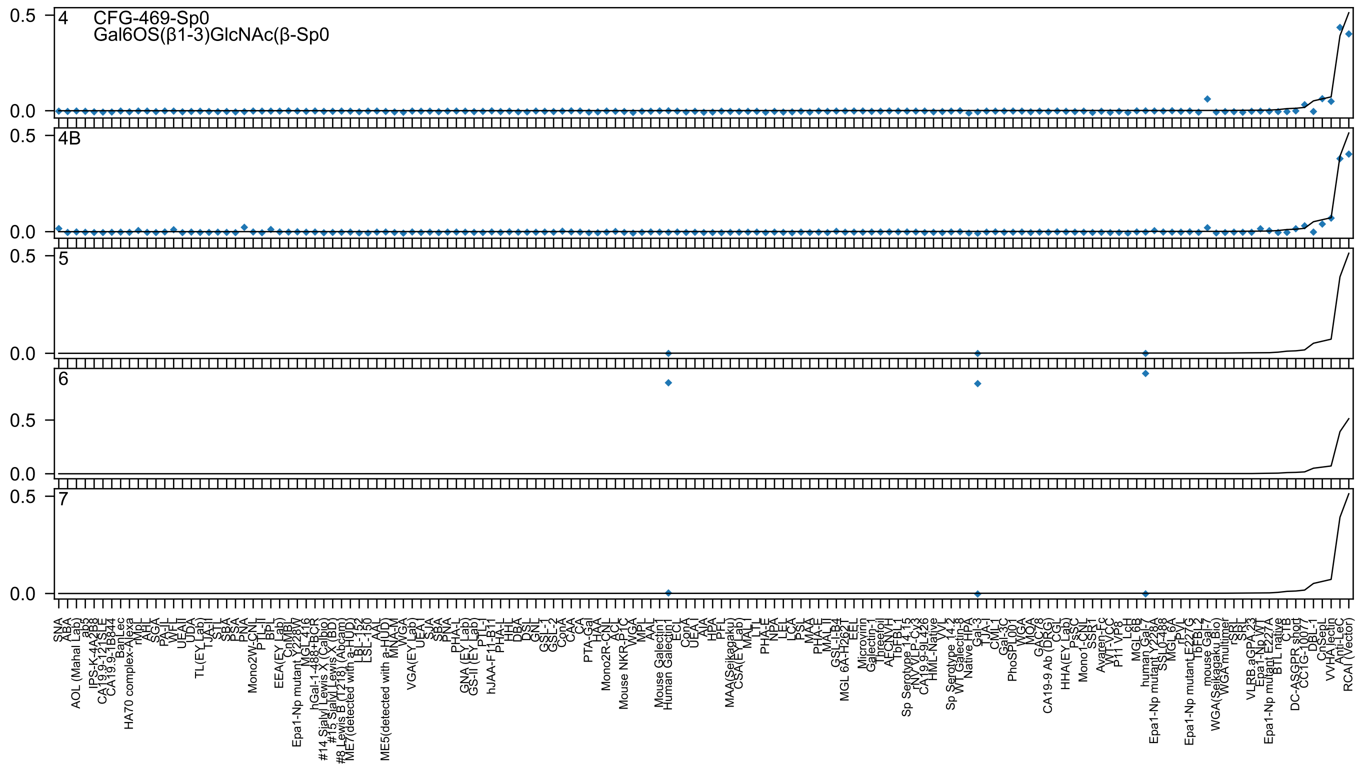
4
CFG-469-Sp0
Gal6OS(β1-3)GlcNAc(β-Sp0
4B
5
6
7
0.5
0.0

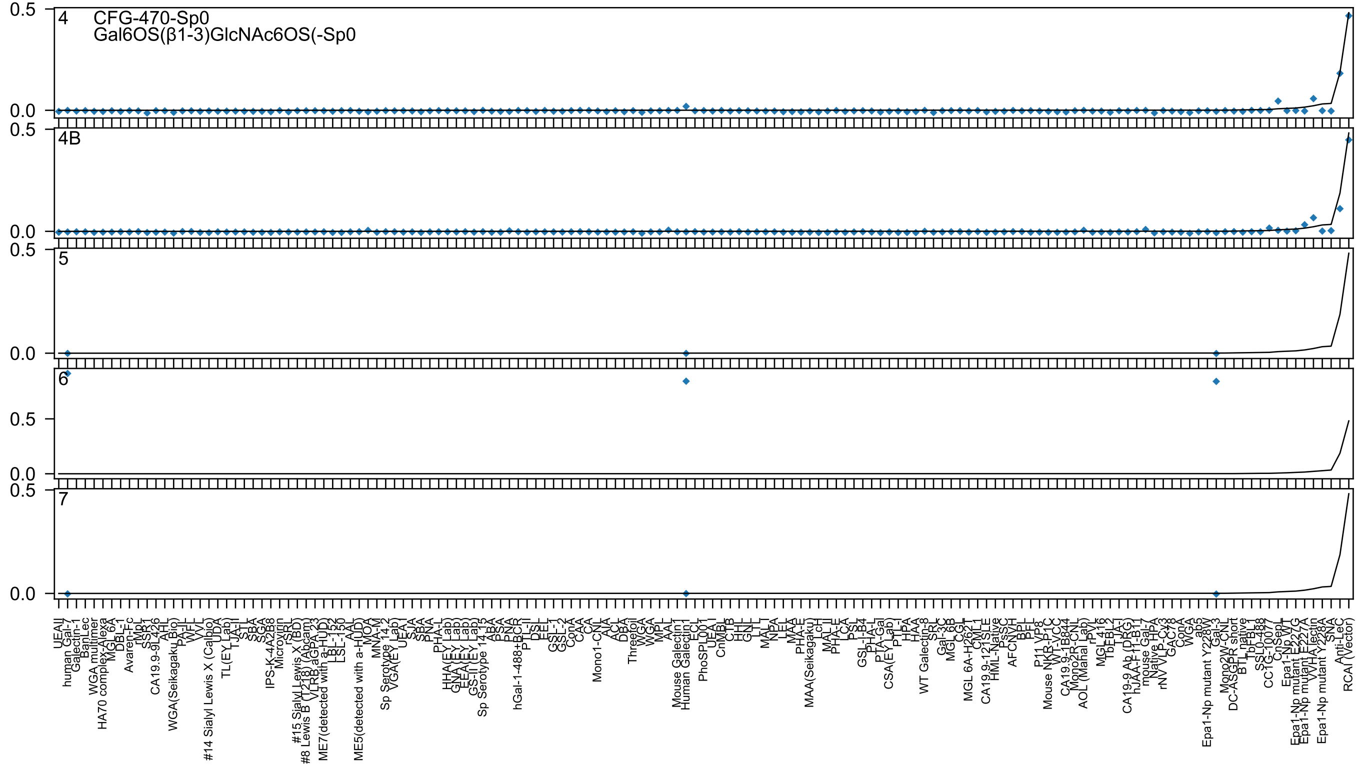

4
CFG-470-Sp0
Gal6OS(β1-3)GlcNAc6OS(-Sp0
4B
5
6
7
0.5
0.0
UEAII
human Gal-7
Galectin-1
BanLec
WGA multimer
HA70 complex-Alexa
MGL 6A
DBL-1
Avaren-Fc
rMpL
SSB1
CA19.9-9L426
AHL
WGA(Seikagaku Bio)
PA-IL
WFL
VVL
#14 Sialyl Lewis X (Calbio)
UDA
TL(EY Lab)
TJA-II
STL
SBA
SGA
IPS-K-4A2B8
Microvirin
rSRL
#15 Sialyl Lewis X (BD)
#8 Lewis B (T218) (Abcam)
VLRB aGPA.23
ME7(detected with a-HUD)
LBL-152
LSL-150
AAL
ME5(detected with a-HUD)
MOA
MNA-M
Sp Serotype 14.2
VGA(EY Lab)
UEA I
SJA
SBA
PNA
PHA-L
HHA(EY Lab)
GNA(EY Lab)
EEA(EY Lab)
GS-II (EY Lab)
Sp Serotype 14.15
ABA
PSA
PNA
hGal-1-488+BCR
PTL-II
DSL
ECL
GSL-I
GSL-2
ConA
CAA
CA
Mono1-CNL
AIA
ACL
DBA
Threefoil
WGA
WGA
MPL
AAL
Mouse Galectin1
Human Galectin1
ECL
PhoSL001
UEA I
CnMBL
CTB
HHL
GNL
LTL
MAL I
NPA
LEL
MAA
PHA-E
MAA(Seikagaku)
LcH
MAL II
PHA-E
LCA
PSL
GSL-I-B4
PHA-L
PTA-Gal
CSA(EY Lab)
PTL-I
HPA
HAA
WT Galectin-8
SRL
Gal-3C
MGL 6B
CGL
MGL 6A-H262T
CML-1
CA19.9-121SLE
HML-Native
PsSC
AFCNVH
BPL
PFL
P11 VP8
Mouse NKR-P1C
WT-VCC
CA19.9-1B844
Mono2R-CNL
AOL (Mahal Lab)
rPVL
MGL416
TbFBL2
TJA-I
CA19-9 Ab (DRG)
hJAA-F11-B11
mouse Gal-7
Native HPA
rNV VLP-Cy3
GAC78
ConA
WGA
ab5
Epa1-Np mutant Y228W
Gal-3
Mono2W-CNL
DC-ASGPR short
BTL native
TbFBL1
SSL0-488
CC1G-1007
CnSep
Epa1-Np WT
Epa1-Np mutant E227G
Epa1-Np mutant E227A
WVHA lectin
Epa1-Np mutant Y228A
SNA
Anti-LeC
RCAI (Vector)

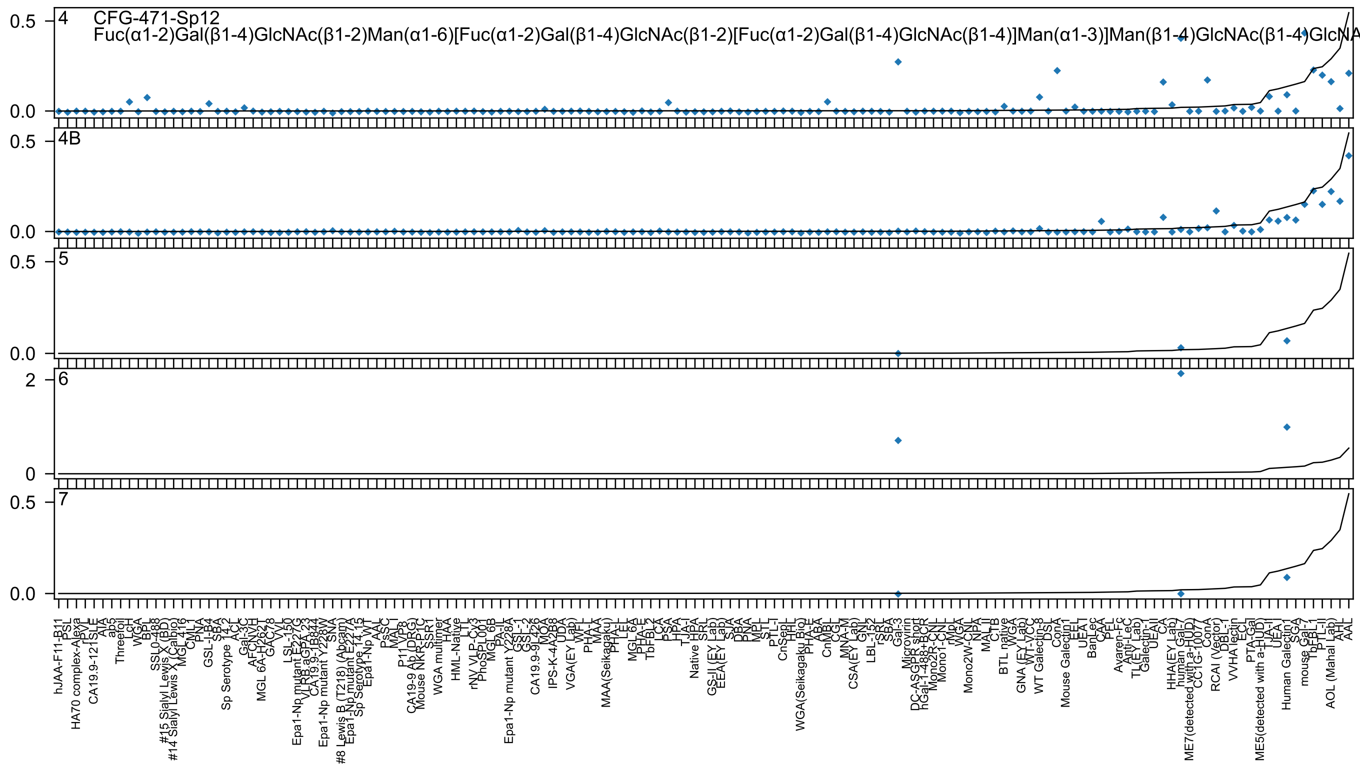
4 CFG-471-Sp12
Fuc(α1-2)Gal(β1-4)GlcNAc(β1-2)Man(α1-6)[Fuc(α1-2)Gal(β1-4)GlcNAc(β1-2)[Fuc(α1-2)Gal(β1-4)GlcNAc(β1-4)]Man(α1-3)]Man(β1-4)GlcNAc(β1-4)GlcNA
4B
5
6
7

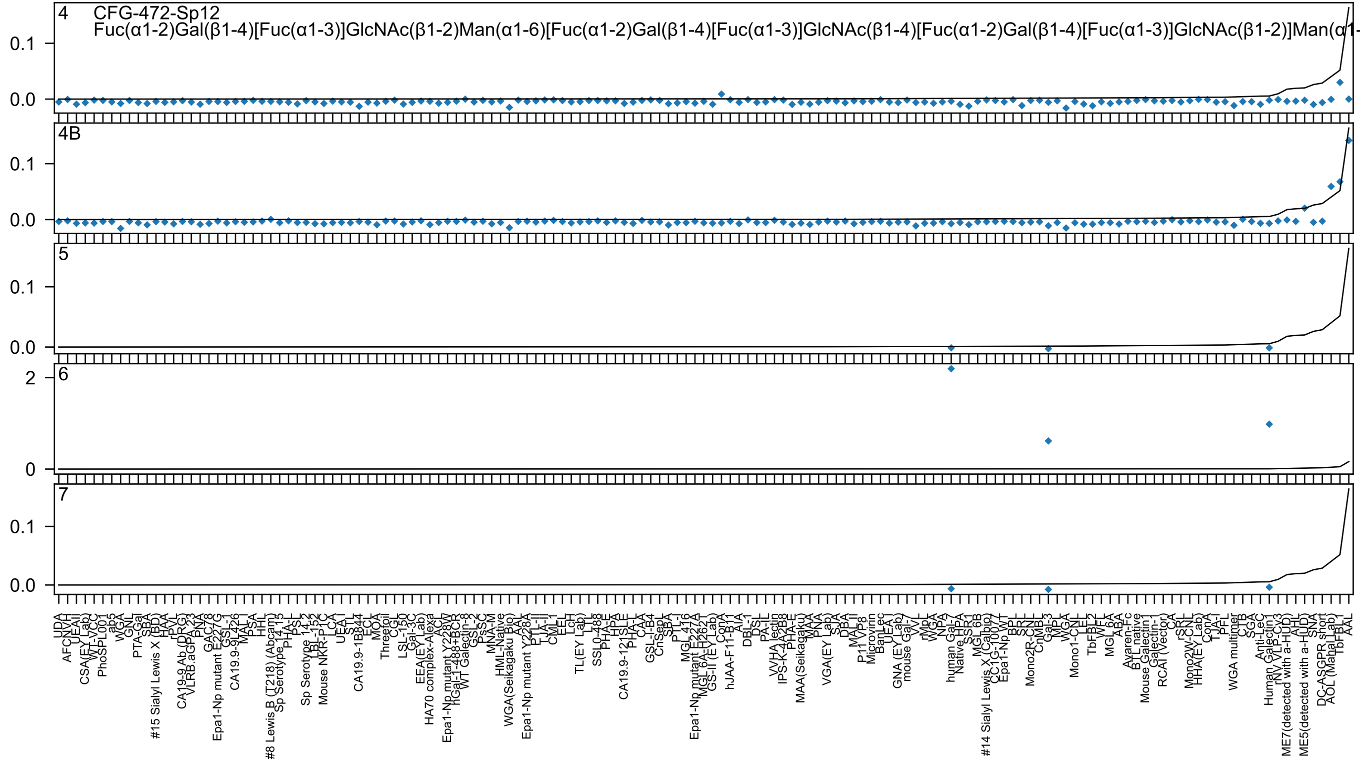

4 CFG-472-Sp12
Fuc(α1-2)Gal(β1-4)[Fuc(α1-3)]GlcNAc(β1-2)Man(α1-6)[Fuc(α1-2)Gal(β1-4)[Fuc(α1-3)]GlcNAc(β1-4)[Fuc(α1-2)Gal(β1-4)[Fuc(α1-3)]GlcNAc(β1-2)]Man(α1-
4B
5
6
7

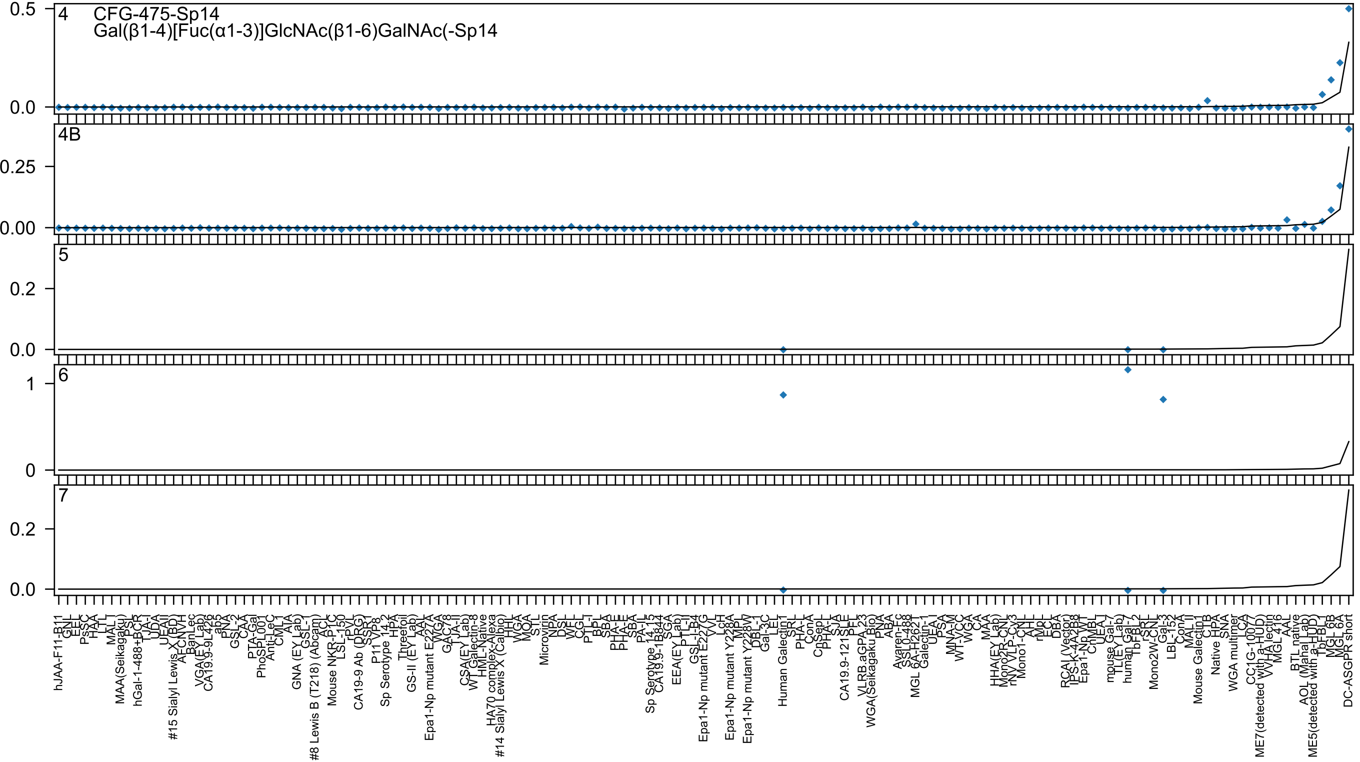

CFG-475-Sp14
Gal(β1-4)[Fuc(α1-3)]GlcNAc(β1-6)GalNAc(-Sp14
4
4B
5
6
7

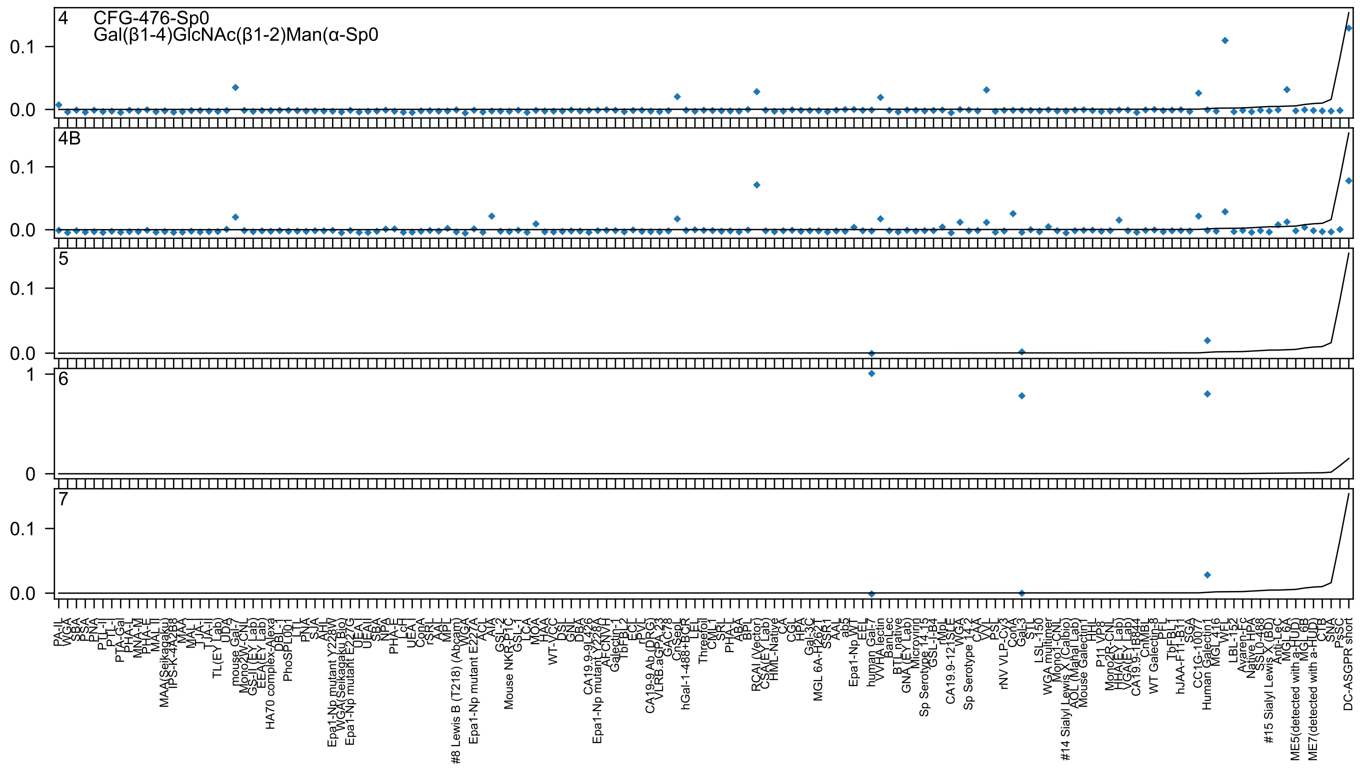
4
CFG-476-Sp0
Gal(β1-4)GlcNAc(β1-2)Man(α-Sp0
4B
5
6
7
0.0
0.1
0
1
PA-IL
WGA
SBA
PSA
PNA
PTL-II
PTL-I
PTA-Gal
PHA-L
MNA-M
PHA-E
MAL-II
MAA(Seikagaku)
IPS-K-4A2B8
MAA
MAL-I
TJA-I
TJA-II
TL(EY Lab)
UDA
mouse Gal-7
Mono2W-CNL
GS-II (EY Lab)
EEA(EY Lab)
HA70 complex-Alexa
DBL-1
PhoSPL001
LTL
PNA
SJA
AHL
Epa1-Np mutant Y228W
WGA(Seikagaku Bio)
Epa1-Np mutant E227G
UEA-I
UEA-II
SBA
NPA
PHA-E
LcH
UEA-I
ConA
rSRL
AAL
MPL
#8 Lewis B (T218) (Abcam)
WGA
Epa1-Np mutant E227A
ACL
AIA
GSL-2
Mouse NKR-P1C
GSL-1
LCA
MOA
HAA
WT-VCC
DSL
GNL
DBA
CA19.9-9L426
Epa1-Np mutant Y228A
AFCNVH
Galectin-1
TbFBL2
ECL
rPVL
CA19-9 Ab (DRG)
VLRB.aGPA.23
GAC78
CnSep
hGal-1-488+BCR
LEL
Threefoil
CML-1
SRL
PHA-L
ABA
BPL
RCAI (Vector)
CSA(EY Lab)
HML-Native
CA
CGL
HPA
Gal-3C
MGL 6A-H2621
SSR1
AAL
ab5
Epa1-Np WT
EEL
human Gal-7
VVHA lectin
BanLec
BTL native
GNA (EY Lab)
Microvirin
Sp Serotype 14.15
GSL-I-B4
rMpL
CA19.9-121SLE
WGA
Sp Serotype 14.2
CAA
VVL
PSL
rNV VLP-Cy3
ConA
Gal-3
STL
LSL-150
WGA multimer
Mono1-CNL
#14 Sialyl Lewis X (Calbio)
AOL (Mahal Lab)
Mouse Galectin1
HHL
P11 VP8
Mono2R-CNL
HHA(EY Lab)
VGA(EY Lab)
CA19.9-1B844
CnMBL
WT Galectin-8
PFL
TbFBL1
hJAA-F11-B11
SGA
CC1G-10077
Human Galectin1
MGL 416
WFL
LBL-152
Avaren-Fc
Native HPA
SSL0-488
#15 Sialyl Lewis X (BD)
Anti-LeC
MGL6A
ME5(detected with a-HUD)
MGL6B
ME7(detected with a-HUD)
CTB
SNA
PsSC
DC-ASGPR short

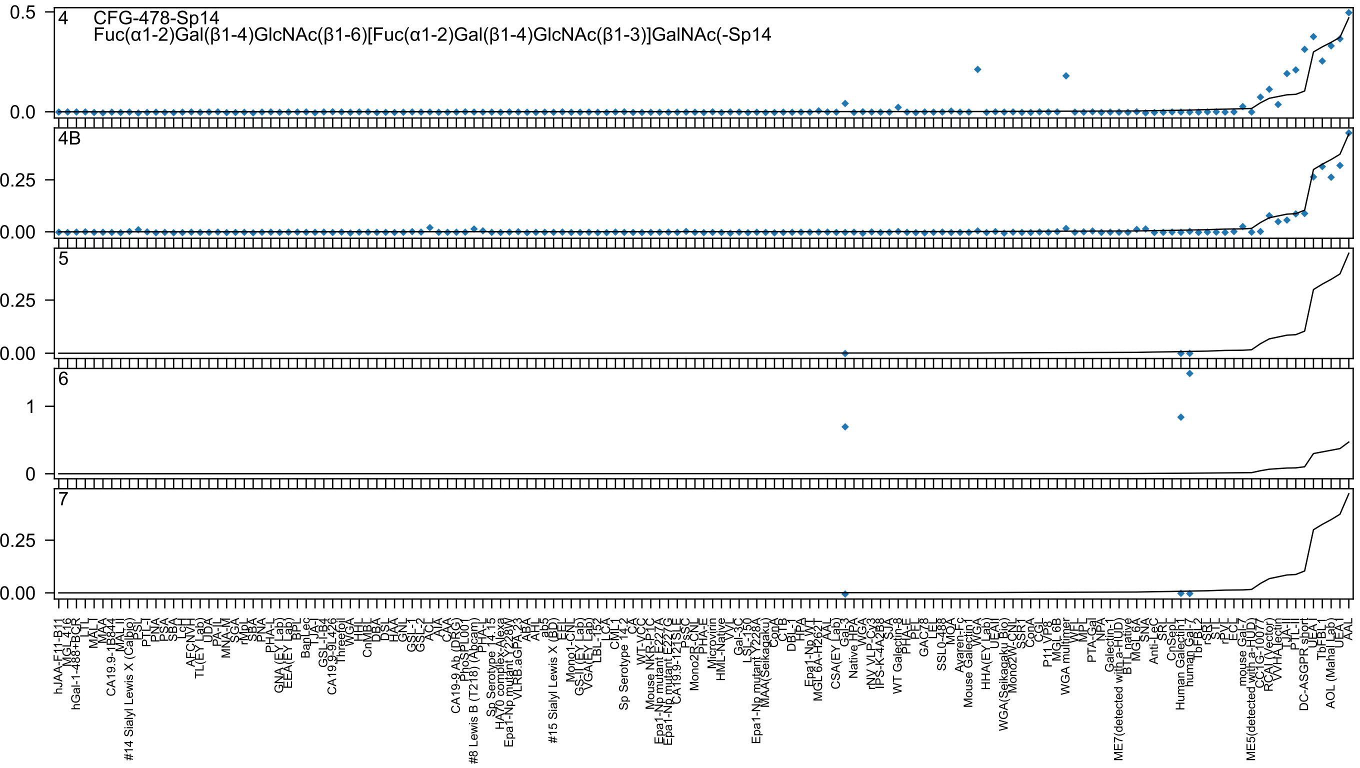
CFG-478-Sp14
Fuc(α1-2)Gal(β1-4)GlcNAc(β1-6)[Fuc(α1-2)Gal(β1-4)GlcNAc(β1-3)]GalNAc(-Sp14
4
4B
5
6
7

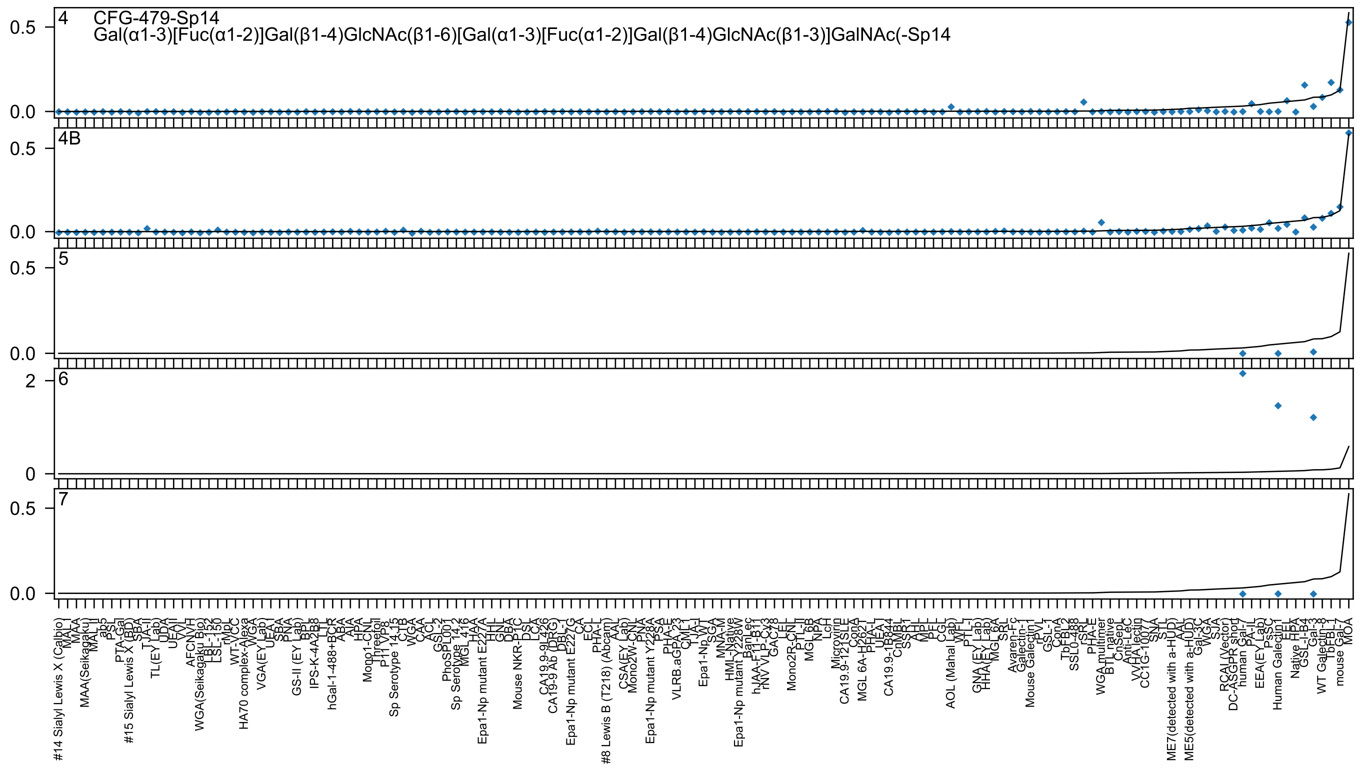
CFG-479-Sp14
Gal(α1-3)[Fuc(α1-2)]Gal(β1-4)GlcNAc(β1-6)[Gal(α1-3)[Fuc(α1-2)]Gal(β1-4)GlcNAc(β1-3)]GalNAc(-Sp14

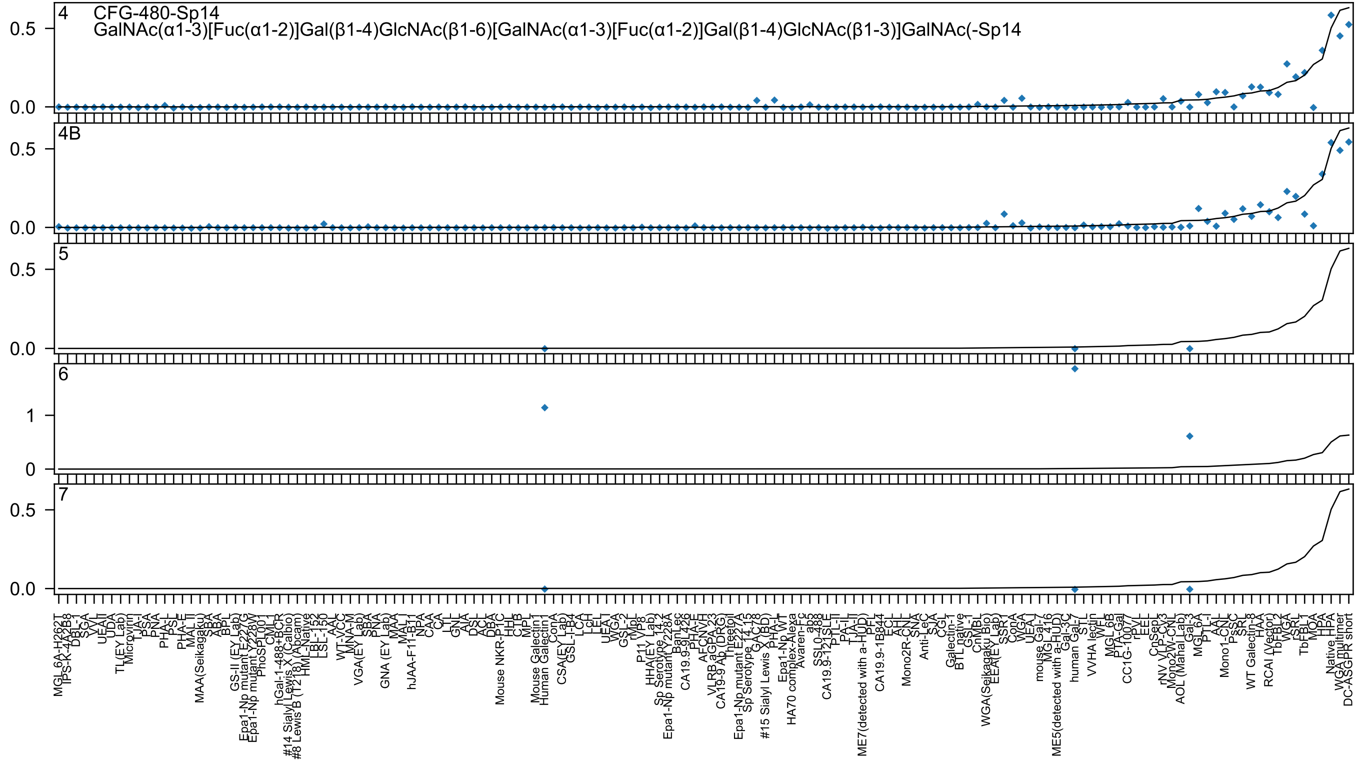
4 CFG-480-Sp14
GalNAc(α1-3)[Fuc(α1-2)]Gal(β1-4)GlcNAc(β1-6)[GalNAc(α1-3)[Fuc(α1-2)]Gal(β1-4)GlcNAc(β1-3)]GalNAc(-Sp14
4B
5
6
7
0.0
0.5
0
1
MGL_6A-H262T
Mouse Galectin1
Human Galectin1
human Gal-7
Gal-3
WGA multimer
DC-ASGPR short

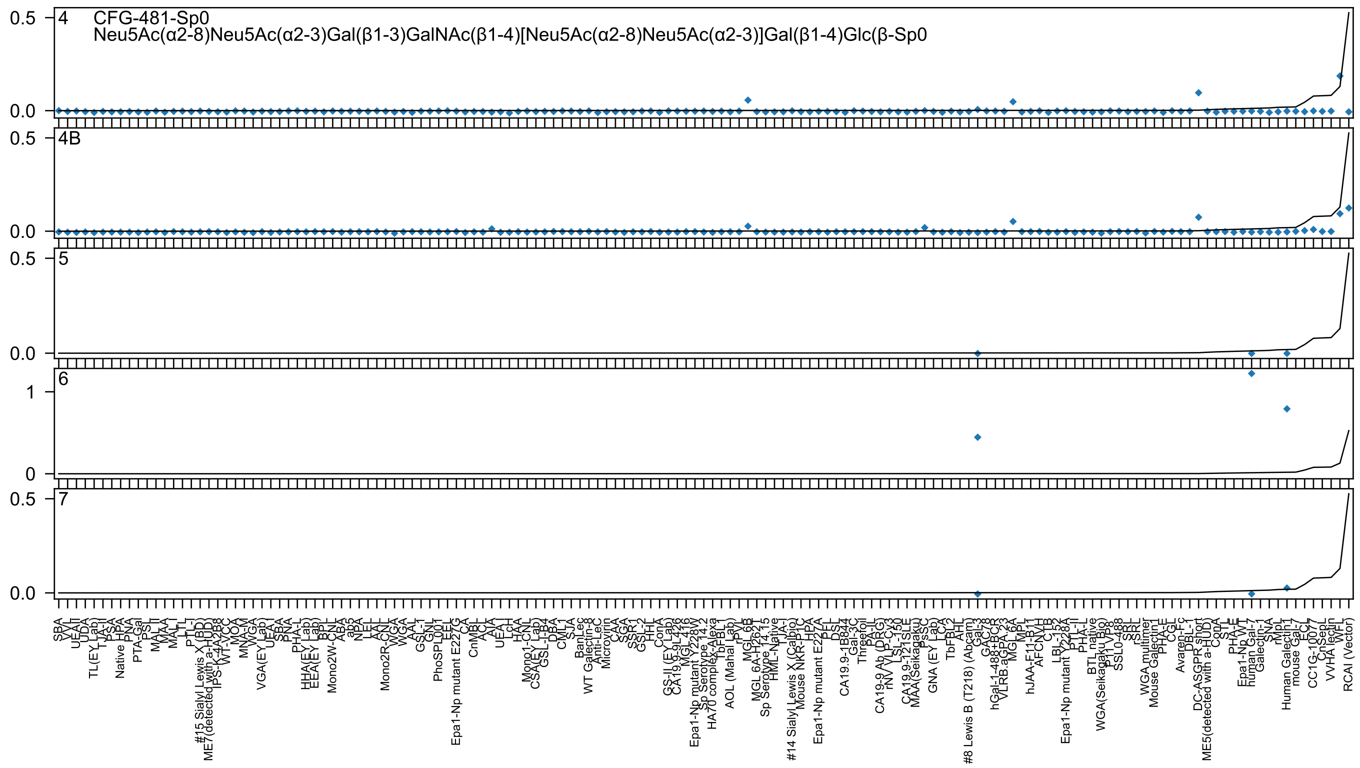

CFG-481-Sp0
Neu5Ac(α2-8)Neu5Ac(α2-3)Gal(β1-3)GalNAc(β1-4)[Neu5Ac(α2-8)Neu5Ac(α2-3)]Gal(β1-4)Glc(β-Sp0
4
4B
5
6
7

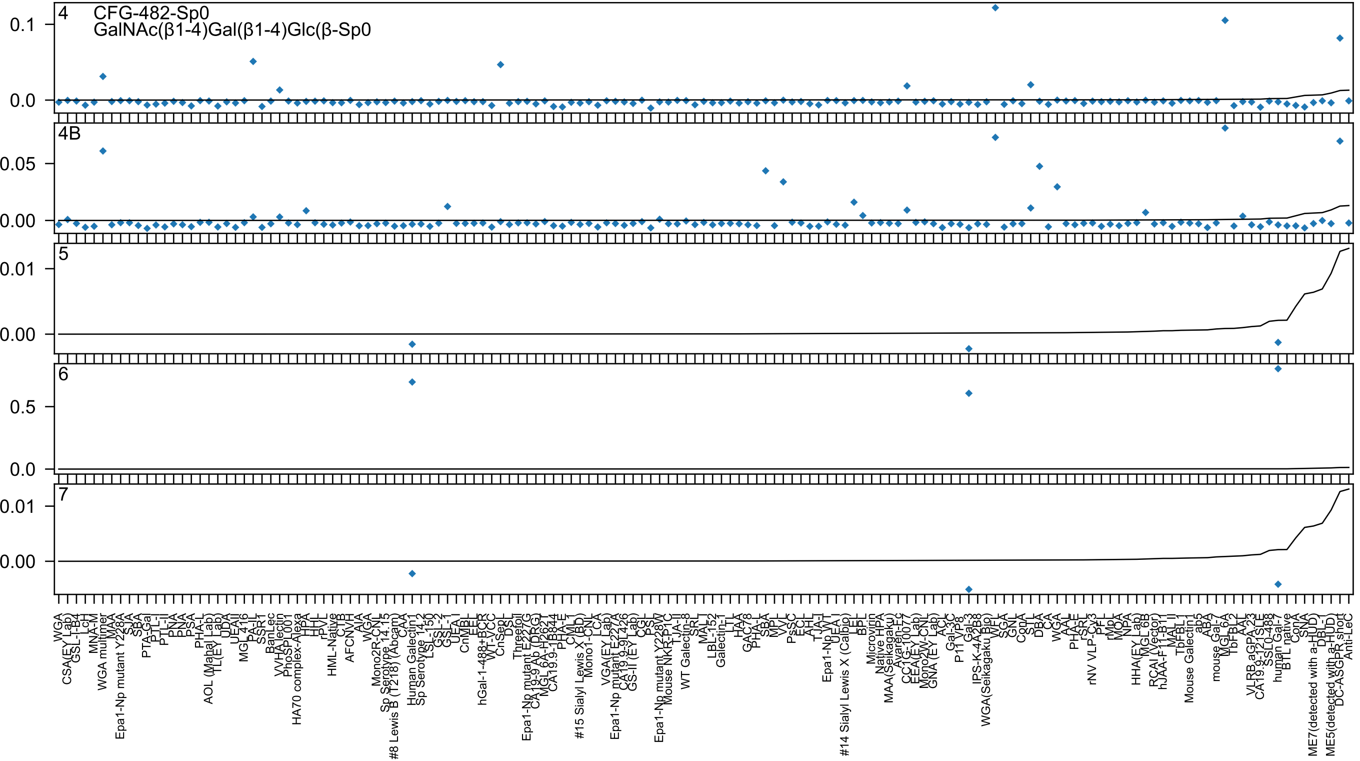
CFG-482-Sp0
GalNAc(β1-4)Gal(β1-4)Glc(β-Sp0
4
4B
5
6
7

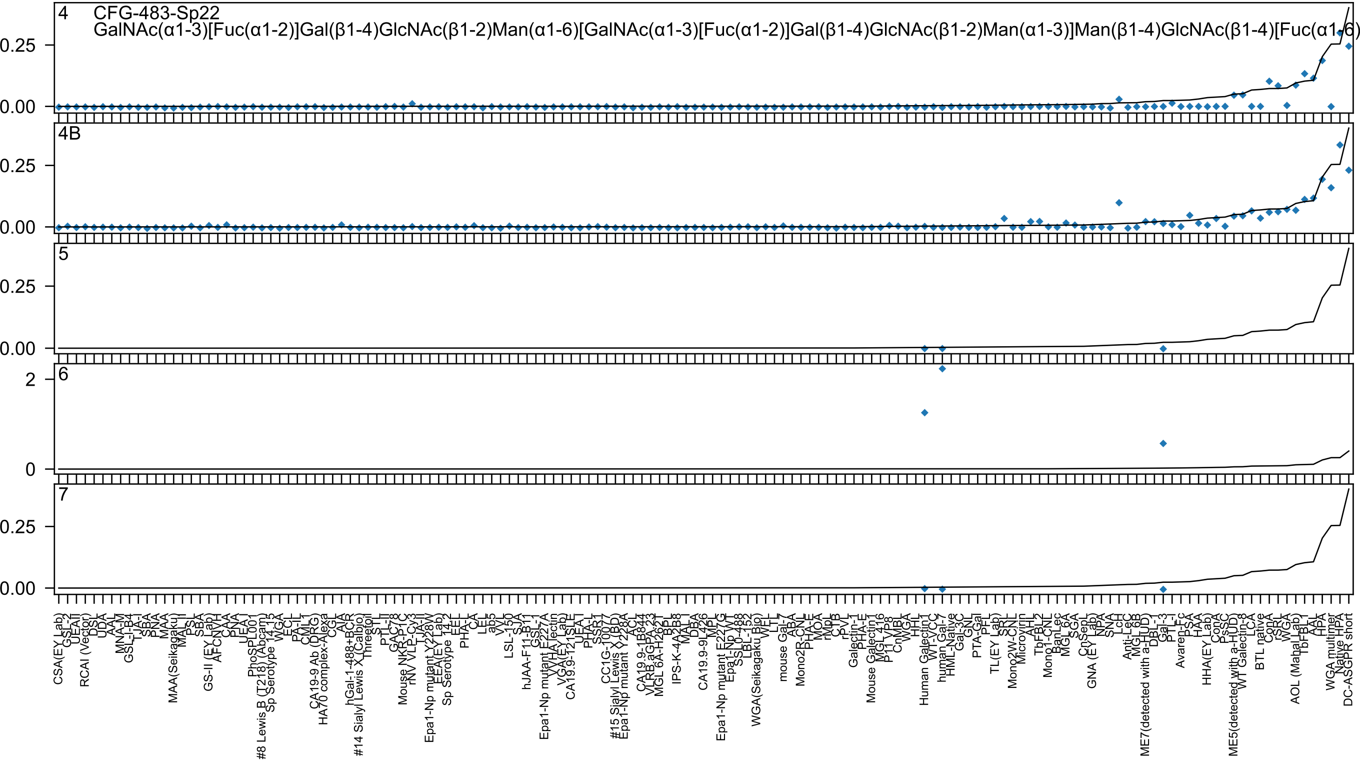
4 CFG-483-Sp22
GalNAc(α1-3)[Fuc(α1-2)]Gal(β1-4)GlcNAc(β1-2)Man(α1-6)[GalNAc(α1-3)[Fuc(α1-2)]Gal(β1-4)GlcNAc(β1-2)Man(α1-3)]Man(β1-4)GlcNAc(β1-4)[Fuc(α1-6)
4B
5
6
7
0.25
0.00
2
0

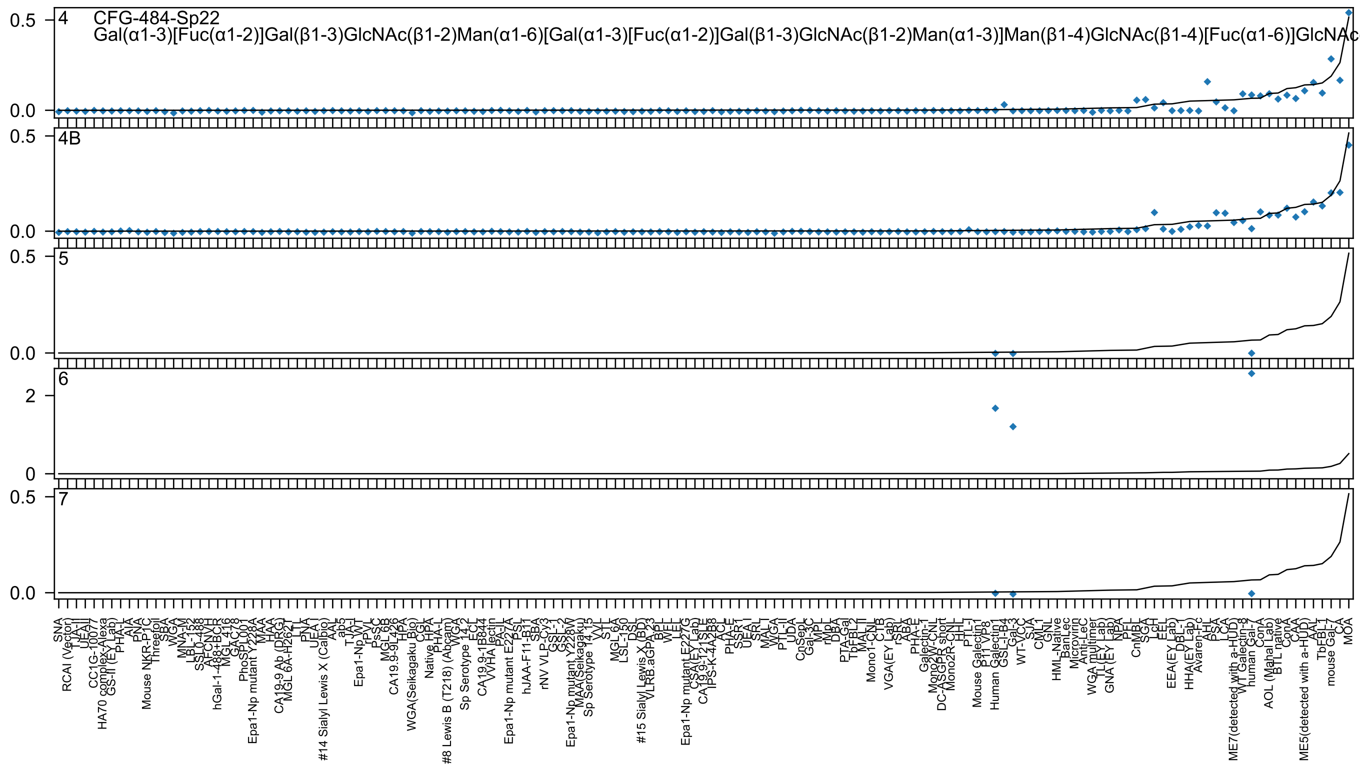

4
CFG-484-Sp22
Gal(α1-3)[Fuc(α1-2)]Gal(β1-3)GlcNAc(β1-2)Man(α1-6)[Gal(α1-3)[Fuc(α1-2)]Gal(β1-3)GlcNAc(β1-2)Man(α1-3)]Man(β1-4)GlcNAc(β1-4)[Fuc(α1-6)]GlcNAc
4B
5
6
7

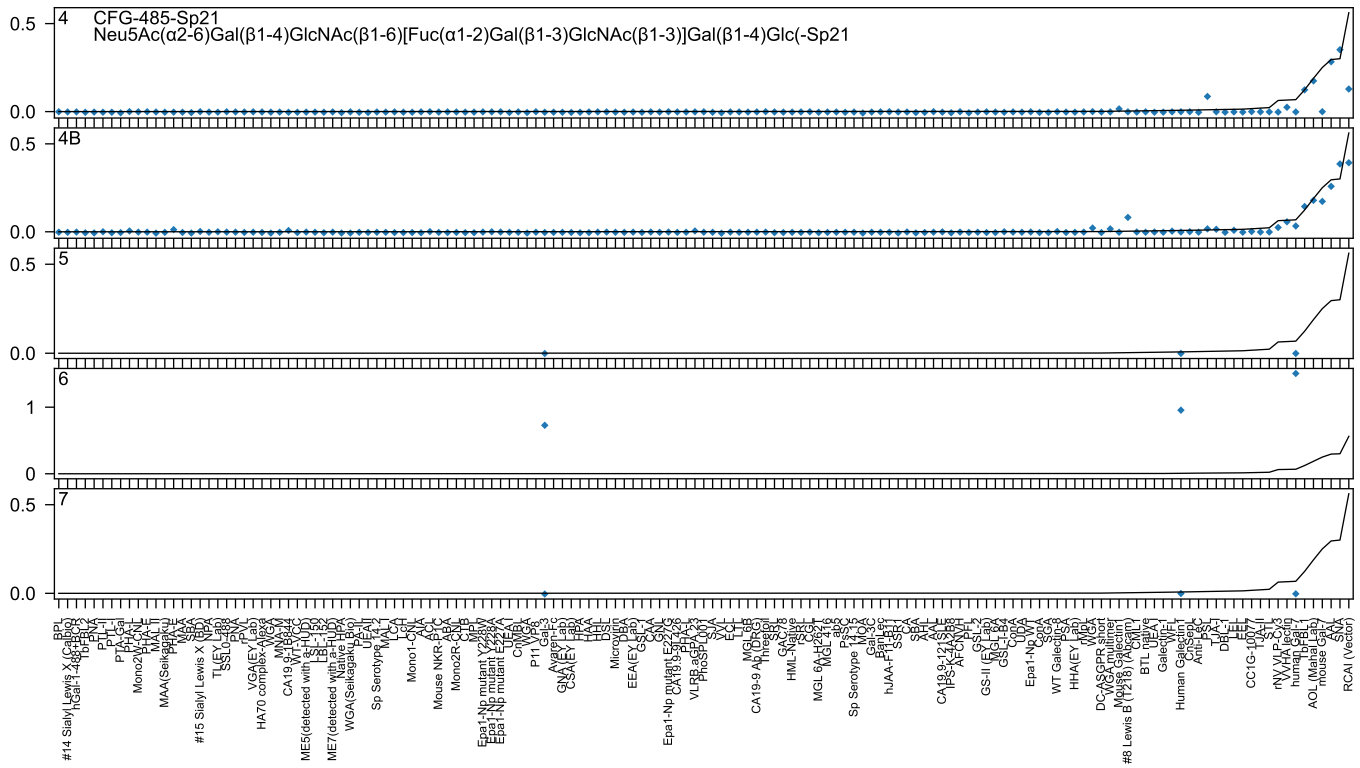

4 CFG-485-Sp21
Neu5Ac(α2-6)Gal(β1-4)GlcNAc(β1-6)[Fuc(α1-2)Gal(β1-3)GlcNAc(β1-3)]Gal(β1-4)Glc(-Sp21
4B
5
6
7

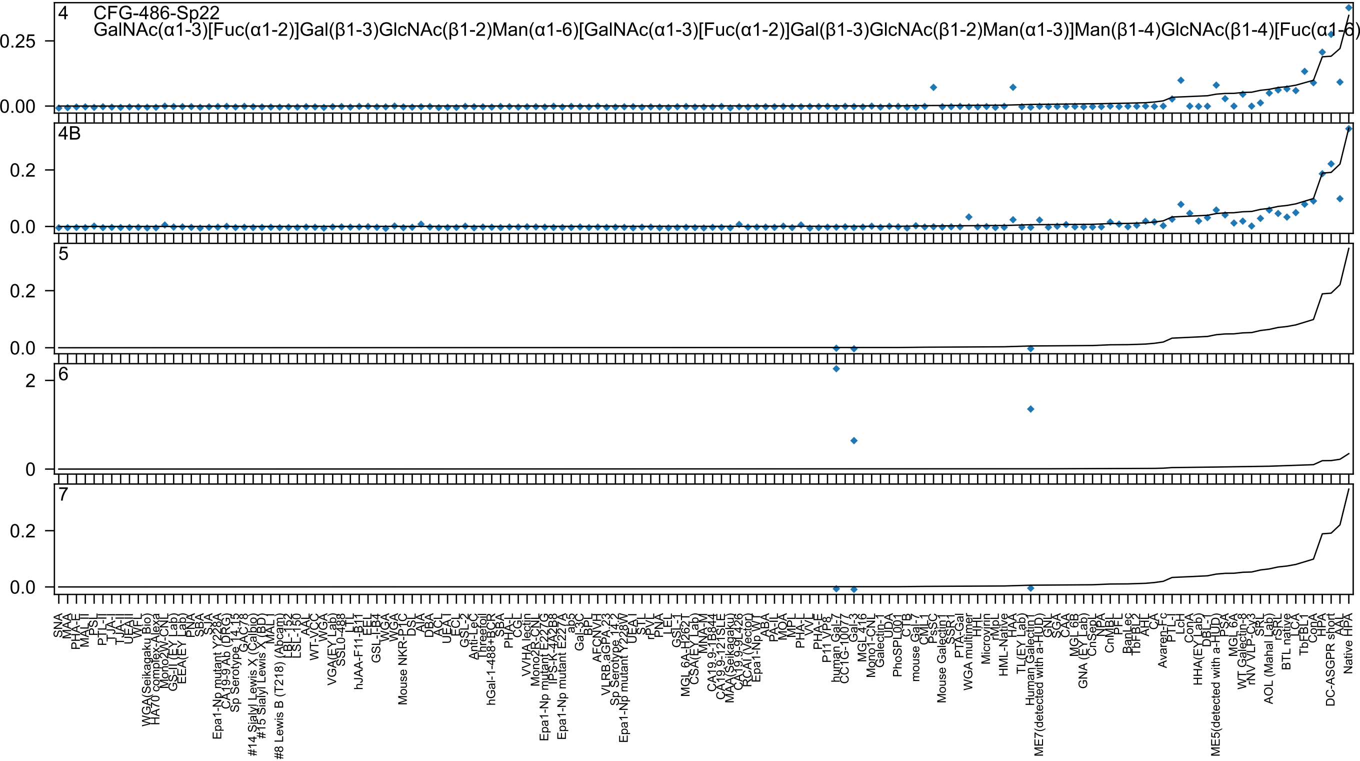
4
CFG-486-Sp22
GalNAc(α1-3)[Fuc(α1-2)]Gal(β1-3)GlcNAc(β1-2)Man(α1-6)[GalNAc(α1-3)[Fuc(α1-2)]Gal(β1-3)GlcNAc(β1-2)Man(α1-3)]Man(β1-4)GlcNAc(β1-4)[Fuc(α1-6)
4B
5
6
7

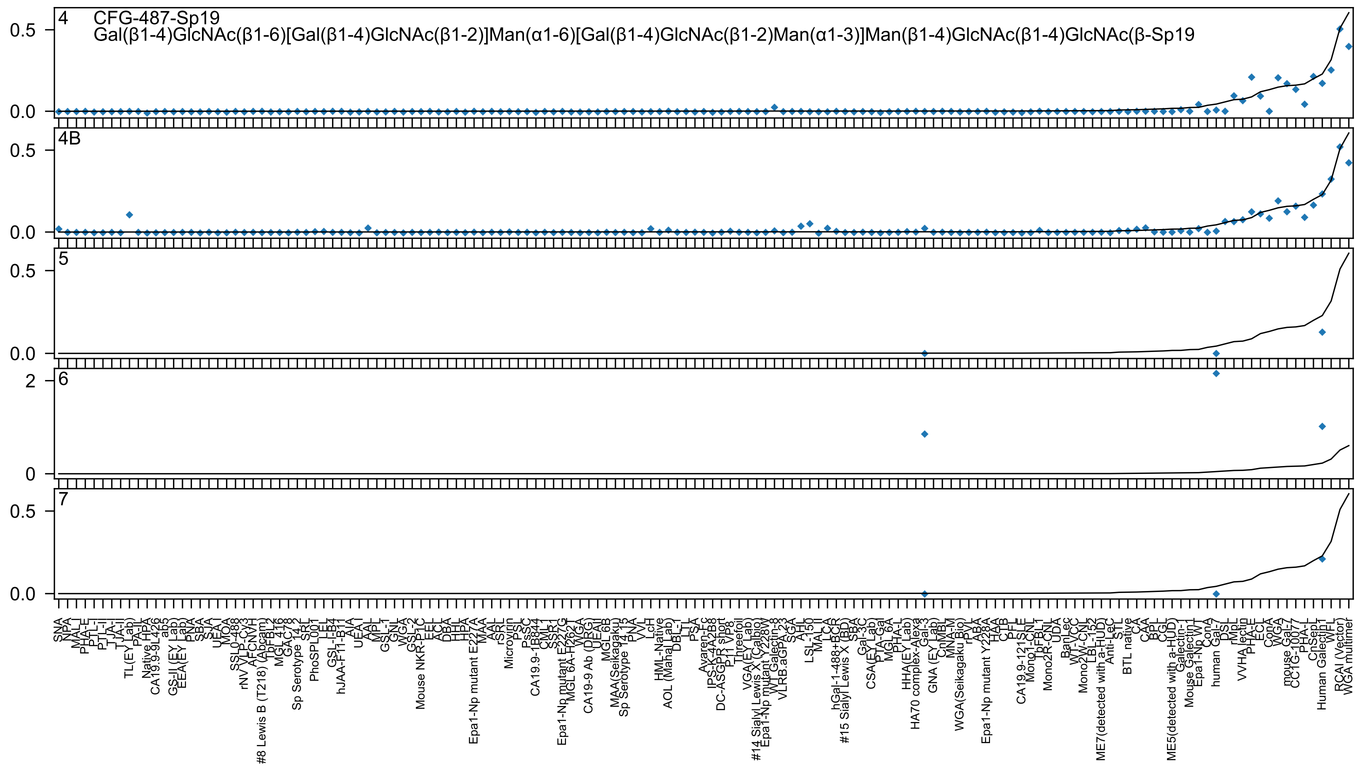
4 CFG-487-Sp19
Gal(β1-4)GlcNAc(β1-6)[Gal(β1-4)GlcNAc(β1-2)]Man(α1-6)[Gal(β1-4)GlcNAc(β1-2)Man(α1-3)]Man(β1-4)GlcNAc(β1-4)GlcNAc(β-Sp19
4B
5
6
7
0.5
0.0
2
0
SNA
NPA
MAL-I
PHA-E
PTL-I
PTL-II
TJA-I
TJA-II
TL(EY Lab)
PA-IL
Native HPA
CA19.9-9L426
ab5
GS-II (EY Lab)
EEA(EY Lab)
PNA
SBA
SJA
UEA-I
MOA
SSL0-488
rNV VLP-Cy3
AFCNVH
#8 Lewis B (T218) (Abcam)
TbFBL2
MGL416
GAC78
Sp Serotype 14.2
SRL
PhoSPL001
LEL
GSL-I-B4
hJAA-F11-B11
AIA
UEA-I
AAL
MPL
GSL-I
GNL
WGA
GSL-2
Mouse NKR-P1C
EEL
ACL
DBA
HHL
HPA
Epa1-Np mutant E227A
MAA
AAL
rSRL
Microvirin
PSL
PsSC
CA19.9-1B844
CML1
SSR1
Epa1-Np mutant E227G
MGL 6A-H262T
WGA
CA19-9 Ab (DRG)
UEAII
MGL 6B
MAA(Seikagaku)
Sp Serotype 14.15
PNA
VVL
LcH
HML-Native
AOL (Manal Lab)
DBL-I
LTL
PSA
Avaren-Fc
IPS-K-4A2B8
DC-ASGPR short
P11 VP8
Threefoil
VGA(EY Lab)
#14 Sialyl Lewis X (Calbio)
Epa1-Np mutant Y228W
WT Galectin-8
VLRB.aGPA.23
SGA
AHL
LSL-150
MAL-II
LCA
hGal-1-488+BCR
#15 Sialyl Lewis X (BD)
SBA
Gal-3C
CSA(EY Lab)
PTA-Gal
MGL 6A
PHA-L
HHA(EY Lab)
HA70 complex-Alexa
Gal-3
GNA (EY Lab)
CnMBL
MNA-M
WGA(Seikagaku Bio)
rPVL
ABA
Epa1-Np mutant Y228A
HAA
CTB
PFL
CA19.9-121SLE
Mono1-CNL
TbFBL1
Mono2R-CNL
UDA
BanLec
WT-CCL
Mono2W-CNL
LBL-152
ME7(detected with a-HUD)
Anti-LeC
STL
BTL native
CA
CAA
BPL
CGL
ME5(detected with a-HUD)
Galectin-1
Mouse Galectin-1
Epa1-Np WT
ConA
human Gal-7
DSL
rMpL
VVHA lectin
PHA-E
ECL
ConA
WGA
mouse Gal-7
CC1G-10077
PHA-L
CnSep1
Human Galectin-1
WFL
RCAI (Vector)
WGA multimer

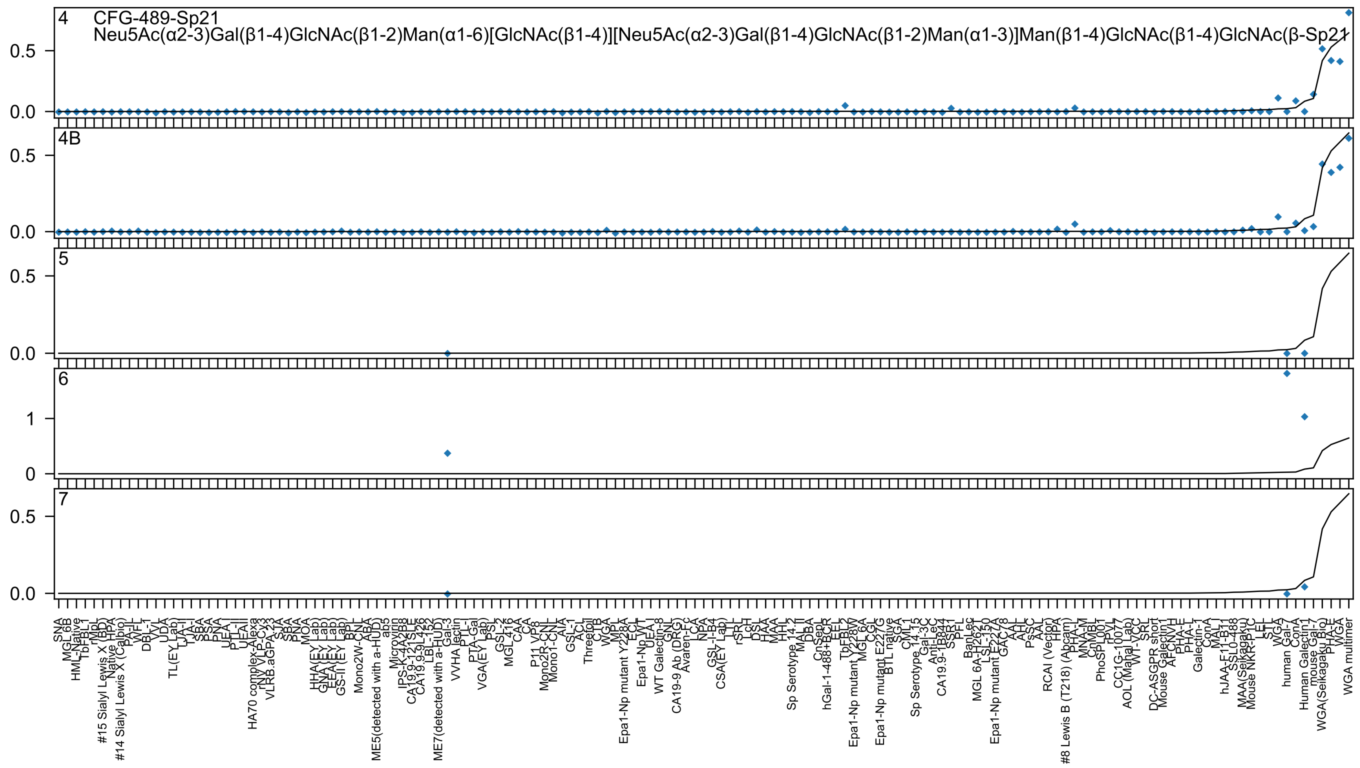
CFG-489-Sp21
Neu5Ac(α2-3)Gal(β1-4)GlcNAc(β1-2)Man(α1-6)[GlcNAc(β1-4)][Neu5Ac(α2-3)Gal(β1-4)GlcNAc(β1-2)Man(α1-3)]Man(β1-4)GlcNAc(β1-4)GlcNAc(β-Sp21
4
4B
5
6
7

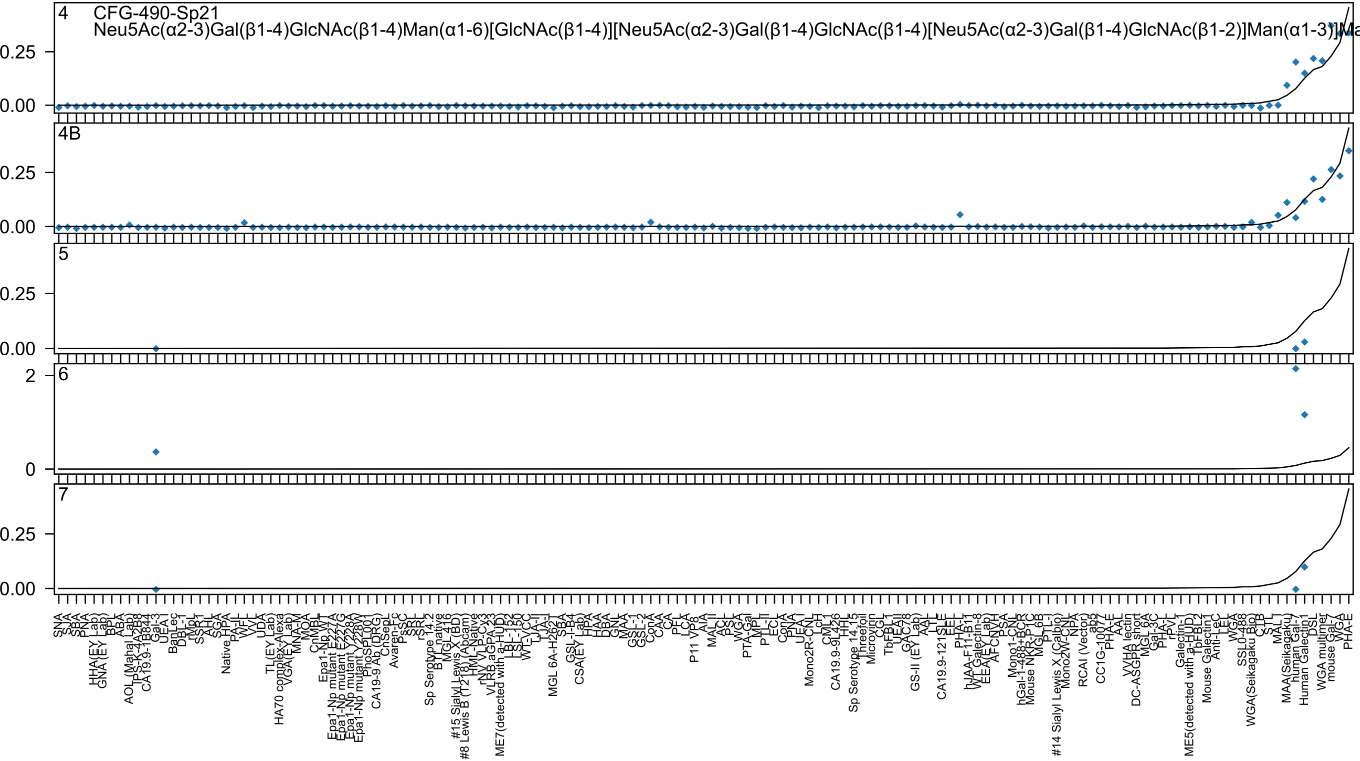
4 CFG-490-Sp21
Neu5Ac(α2-3)Gal(β1-4)GlcNAc(β1-4)Man(α1-6)[GlcNAc(β1-4)][Neu5Ac(α2-3)Gal(β1-4)GlcNAc(β1-4)[Neu5Ac(α2-3)Gal(β1-4)GlcNAc(β1-2)]Man(α1-3)]Ma
0.25
0.00
4B
0.25
0.00
5
0.25
0.00
6
2
0
7
0.25
0.00
SNA
SJA
SBA
PNA
HHA(EY Lab)
GNA (EY Lab)
BPL
ABA
AOL (Mahal Lab)
TPS-K-4A2B8
CA19.9-1B844
Gal-3
UEA-I
BanLec
DBL-1
rMpL
SSBT
AHL
SGA
Native HPA
PA-IL
WFL
VVL
UDA
TL(EY Lab)
HA70 complex-Alexa
VGA(EY Lab)
MNA-M
MOA
CnMBL
Epa1-Np WT
Epa1-Np mutant E227A
Epa1-Np mutant E227G
Epa1-Np mutant Y228A
Epa1-Np mutant Y228W
PhoSL001
CA19-9 Ab (DRG)
ChSepL
Avaren-Fc
PsSC
SRL
rSRL
Sp Serotype 14.2
BTL native
MGL 416
#15 Sialyl Lewis X (BD)
#8 Lewis B (T218) (Abcam)
HML-Native
rNV VLP-Cy3
VLRB.aGPA.23
ME7(detected with a-HUD)
LBL-152
LSL-150
WT-VCC
TJA-II
TJA-I
MGL 6A-H262T
SBA
GSL-I-B4
CSA(EY Lab)
HPA
HAA
DBA
GNL
MAA
GSL-1
GSL-2
ConA
CAA
CA
PFL
LCA
P11 VP8
AIA
MAL II
ACL
PSL
WGA
PTA-Gal
MPL
PTL-II
ECL
ConA
PNA
UEA-I
Mono2R-CNL
LcH
CML1
CA19.9-9L426
HHL
Sp Serotype 14.15
Threefoil
Microvirin
CGL
TbFBL1
UEAII
GAC78
GS-II (EY Lab)
AAL
LTL
CA19.9-121SLE
EEL
PHA-L
hJAA-F11-B11
WT Galectin-8
EEA(EY Lab)
AFCNVH
PSA
Mono1-CNL
hGal-1-488+BCR
Mouse NKR-P1C
MGL 6B
PTL-I
#14 Sialyl Lewis X (Calbio)
Mono2W-CNL
NPA
RCAI (Vector)
ab5
CC1G-1007
PHA-E
AAL
VVHA lectin
DC-ASGPR short
MGL 6A
Gal-3C
PHA-L
rPVL
Galectin-1
ME5(detected with a-HUD)
TbFBL2
Mouse Galectin1
Anti-LeC
LEL
WGA
SSL0-488
WGA(Seikagaku Bio)
CTB
STL
MAL I
MAA(Seikagaku)
human Gal-7
Human Galectin1
DSL
WGA multimer
mouse Gal-7
WGA
PHA-E

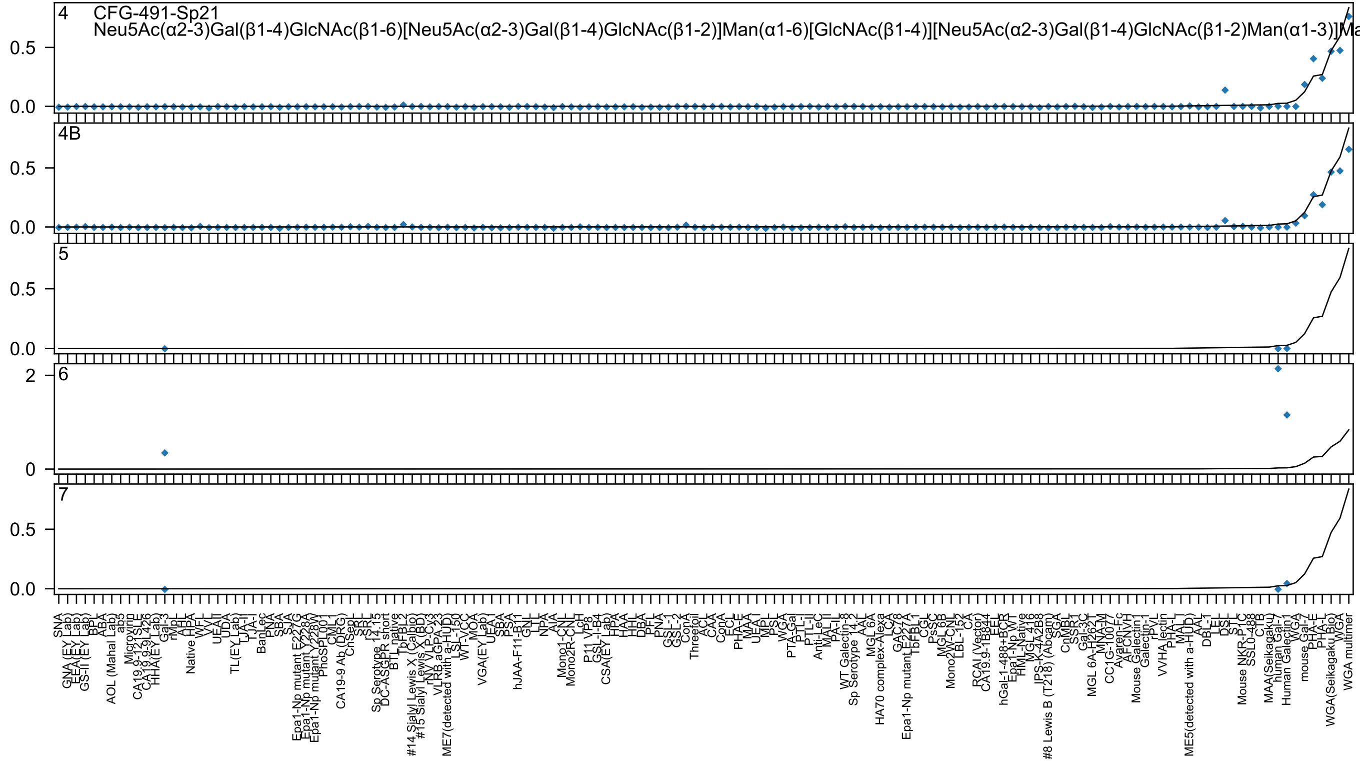
4
CFG-491-Sp21
Neu5Ac(α2-3)Gal(β1-4)GlcNAc(β1-6)[Neu5Ac(α2-3)Gal(β1-4)GlcNAc(β1-2)]Man(α1-6)[GlcNAc(β1-4)][Neu5Ac(α2-3)Gal(β1-4)GlcNAc(β1-2)Man(α1-3)]Ma
4B
5
6
7

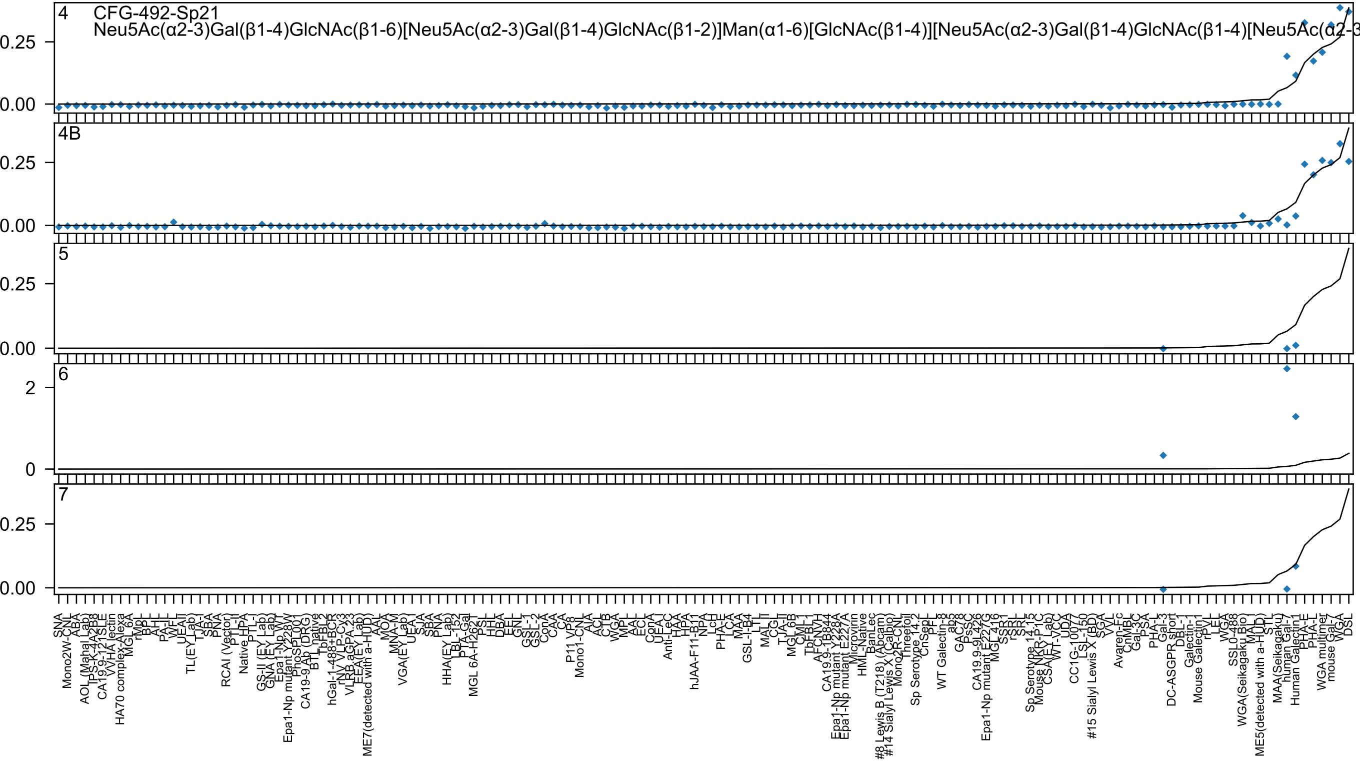
4 CFG-492-Sp21
Neu5Ac(α2-3)Gal(β1-4)GlcNAc(β1-6)[Neu5Ac(α2-3)Gal(β1-4)GlcNAc(β1-2)]Man(α1-6)[GlcNAc(β1-4)][Neu5Ac(α2-3)Gal(β1-4)GlcNAc(β1-4)[Neu5Ac(α2-3
4B
5
6
7

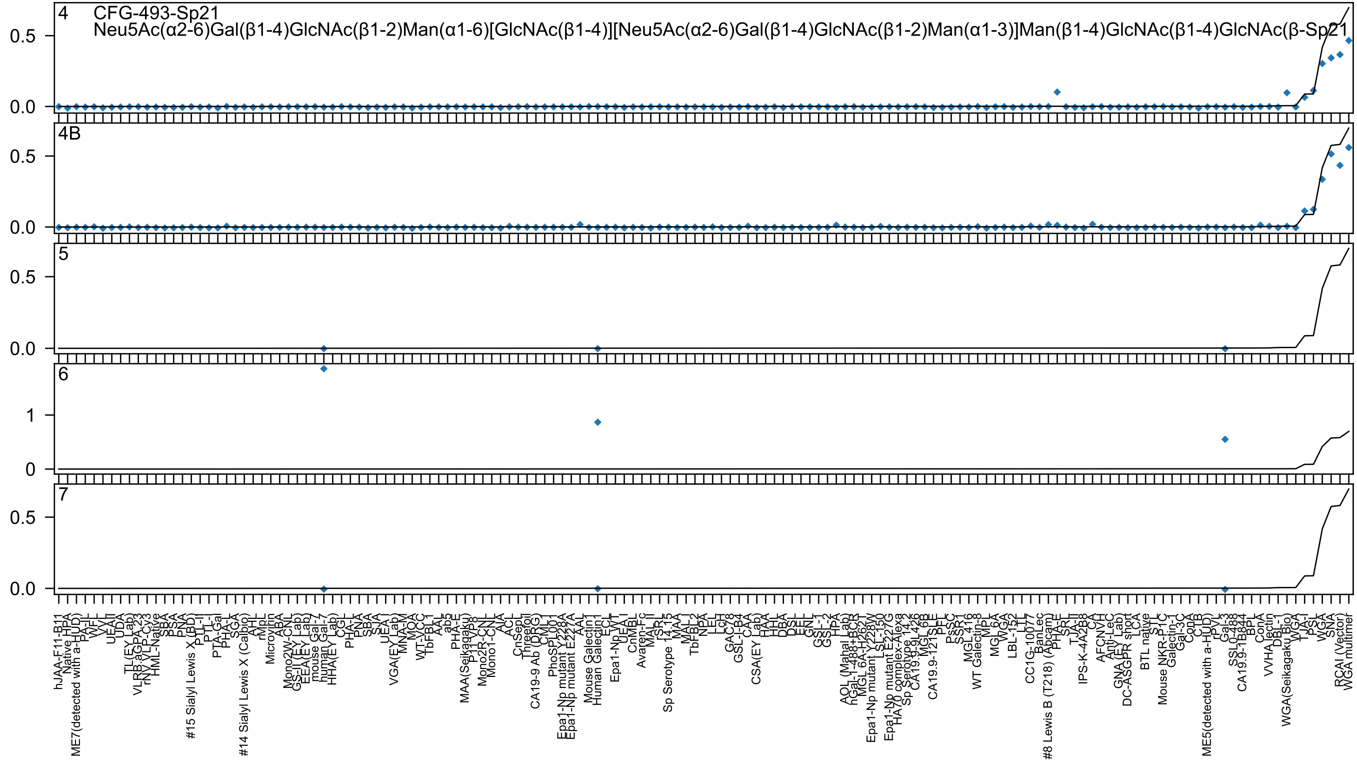

4 CFG-493-Sp21
Neu5Ac(α2-6)Gal(β1-4)GlcNAc(β1-2)Man(α1-6)[GlcNAc(β1-4)][Neu5Ac(α2-6)Gal(β1-4)GlcNAc(β1-2)Man(α1-3)]Man(β1-4)GlcNAc(β1-4)GlcNAc(β-Sp21
4B
5
6
7
0.0
0.5
0
1
hJAA-F11-B11
Native HPA
ME7(detected with a-HUD)
PA-IL
WFL
VVL
UEA-II
UDA
TL(EY Lab)
VLRB.aGPA.23
rNY VLP-Cy3
HML-Native
SBA
PSA
PNA
#15 Sialyl Lewis X (BD)
PTL-II
PTL-I
PTA-Gal
PHA-L
SGA
#14 Sialyl Lewis X (Calbio)
AHL
rMpL
Microvirin
ABA
Mono2W-CNL
GS-II (EY Lab)
EEA(EY Lab)
mouse Gal-7
human Gal-7
HHA(EY Lab)
CGL
PHA-L
PNA
SBA
SJA
UEA I
VGA(EY Lab)
MNA-M
MOA
WT-VCC
TbFBL1
AAL
ab5
PHA-E
MAA(Seikagaku)
P1P8
Mono2R-CNL
Mono1-CNL
AIA
ACL
CnSepL
Threefoil
CA19-9 Ab (DRG)
CML
PhoSL001
Epa1-Np mutant Y228A
Epa1-Np mutant E227A
AAL
Mouse Galectin1
Human Galectin1
ECL
Epa1-Np WT
UEA I
CnMBL
Avaren-Fc
MAL II
PSL
Sp Serotype 14.15
MAA
MAL I
TbFBL2
NPA
LEL
LcH
GAC78
GSL-I B4
CAA
CSA(EY Lab)
HAA
HHL
DBA
DSL
EEL
GNL
GSL-1
GSL-2
HPA
AOL (Mahal Lab)
hGal-1-488+BCR
MGL 6A-H262L
Epa1-Np mutant Y228W
LSL-150
Epa1-Np mutant E227G
HA70 complex-Alexa
Sp Serotype 14.2
CA19.9-9L426
MGL 6B
CA19.9-121SLE
PFL
PsSC
SSR1
MGL 416
WT Galectin-8
MPL
MGL 6A
WGA
LBL-152
LTL
CC1G-10077
BanLec
#8 Lewis B (T218) (Abcam)
PHA-E
SRT
TJA-II
IPS-K-4A2B8
CA
AFCNVH
Anti-LeC
GNA (EY Lab)
DC-ASGPR short
LCA
BTL native
STL
Mouse NKR-P1C
Galectin-1
Gal-3C
ConA
CTB
rPVL
ME5(detected with a-HUD)
Gal-3
SSL0-488
CA19.9-1B844
BPL
ConA
VVHA lectin
DBL-1
WGA(Seikagaku Bio)
WGA
TJA-I
PSL
WGA
SNA
RCAI (Vector)
WGA multimer

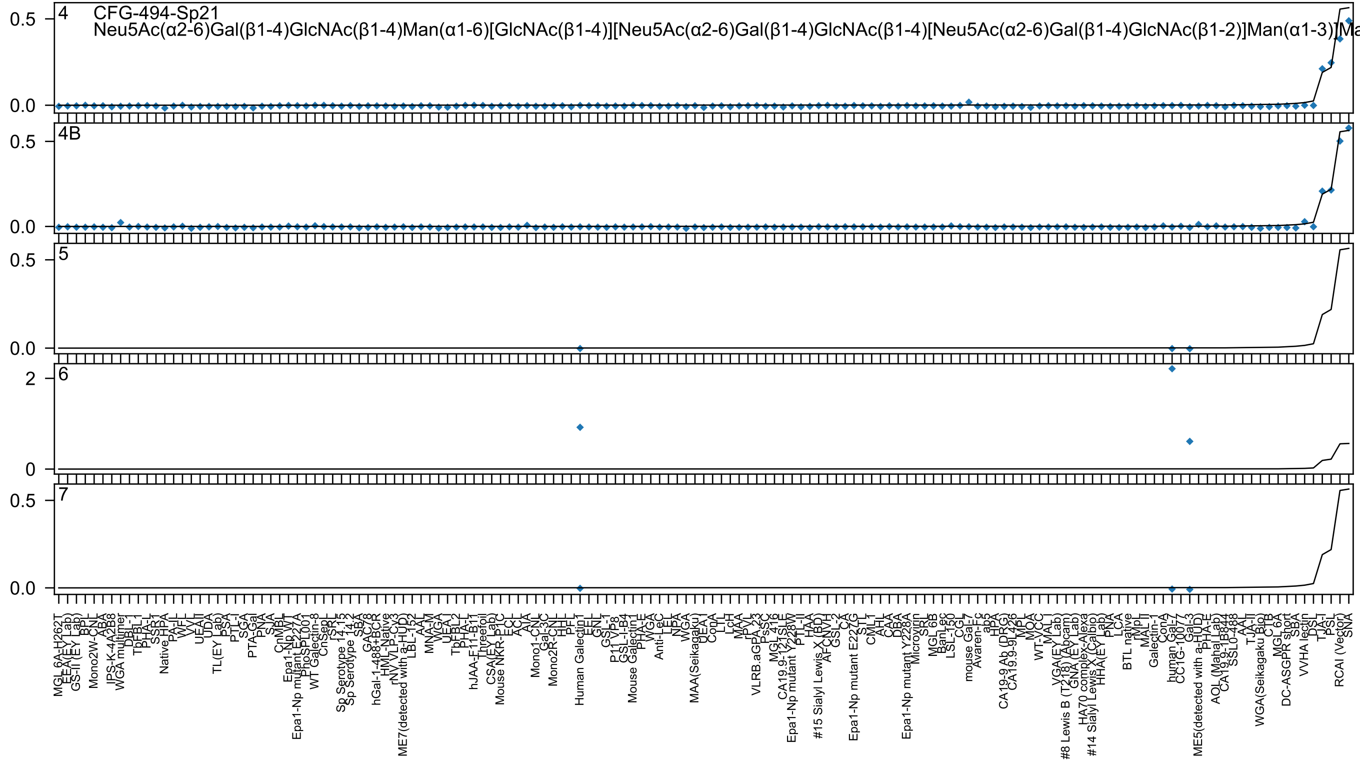

CFG-494-Sp21
Neu5Ac(α2-6)Gal(β1-4)GlcNAc(β1-4)Man(α1-6)[GlcNAc(β1-4)][Neu5Ac(α2-6)Gal(β1-4)GlcNAc(β1-4)[Neu5Ac(α2-6)Gal(β1-4)GlcNAc(β1-2)]Man(α1-3)]Ma

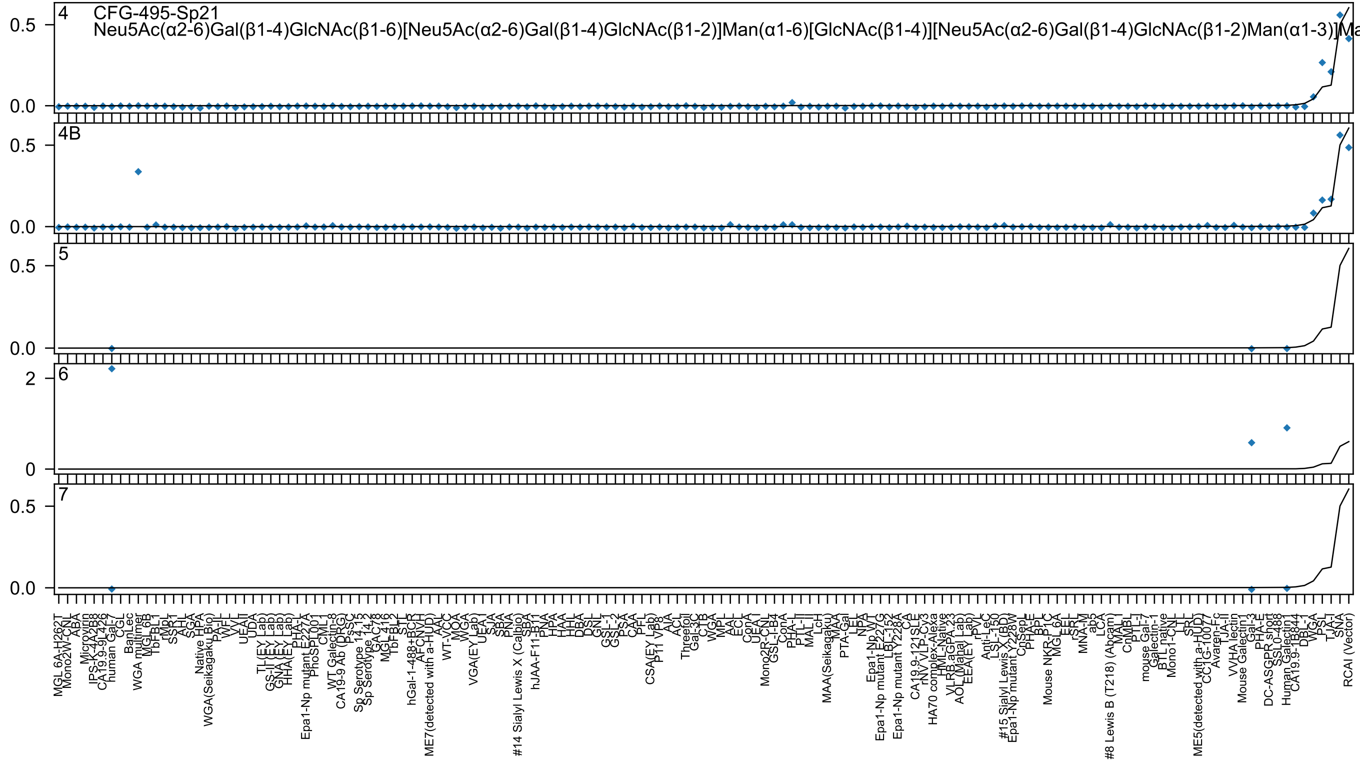
CFG-495-Sp21
Neu5Ac(α2-6)Gal(β1-4)GlcNAc(β1-6)[Neu5Ac(α2-6)Gal(β1-4)GlcNAc(β1-2)]Man(α1-6)[GlcNAc(β1-4)][Neu5Ac(α2-6)Gal(β1-4)GlcNAc(β1-2)Man(α1-3)]Ma
4
4B
5
6
7

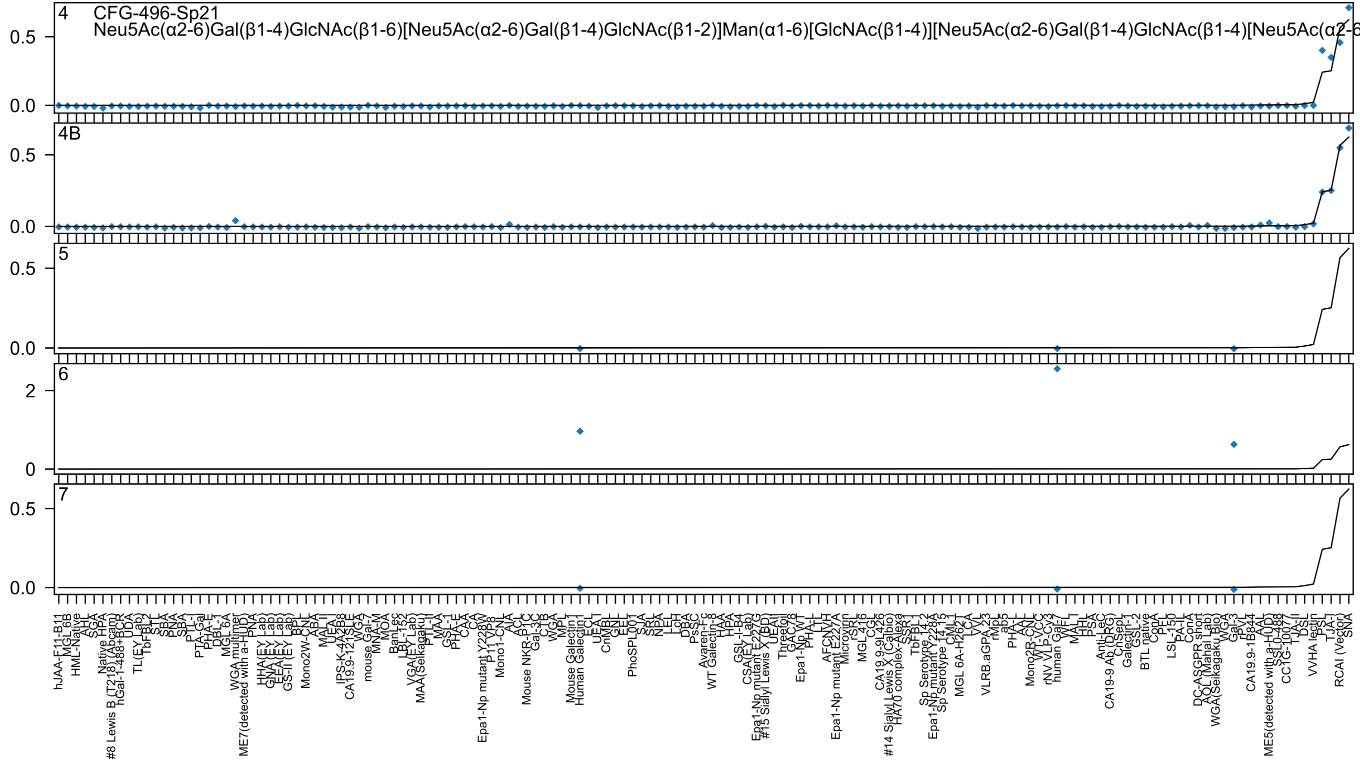

4
CFG-496-Sp21
Neu5Ac(α2-6)Gal(β1-4)GlcNAc(β1-6)[Neu5Ac(α2-6)Gal(β1-4)GlcNAc(β1-2)]Man(α1-6)[GlcNAc(β1-4)][Neu5Ac(α2-6)Gal(β1-4)GlcNAc(β1-4)[Neu5Ac(α2-6
4B
5
6
7
0.5
0.0
2
0

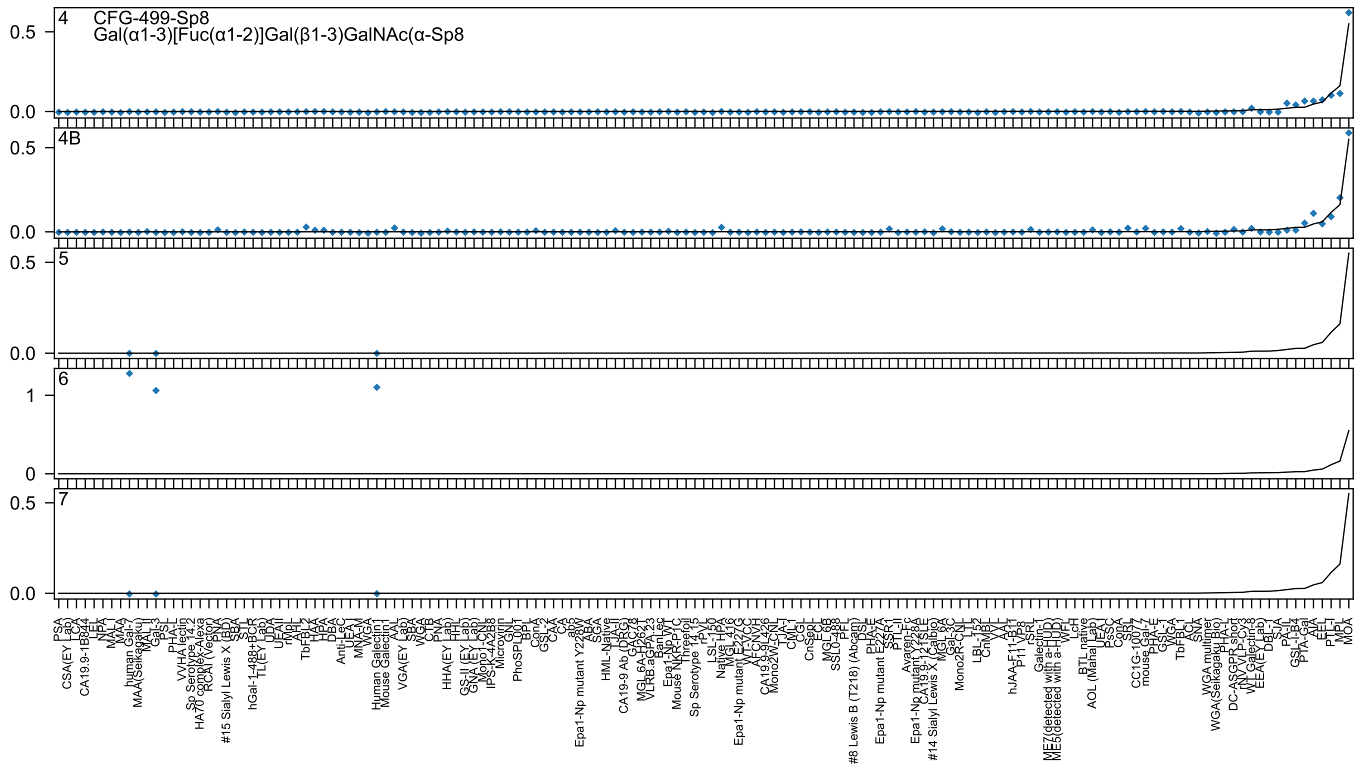

4
CFG-499-Sp8
Gal(α1-3)[Fuc(α1-2)]Gal(β1-3)GalNAc(α-Sp8
4B
5
6
7

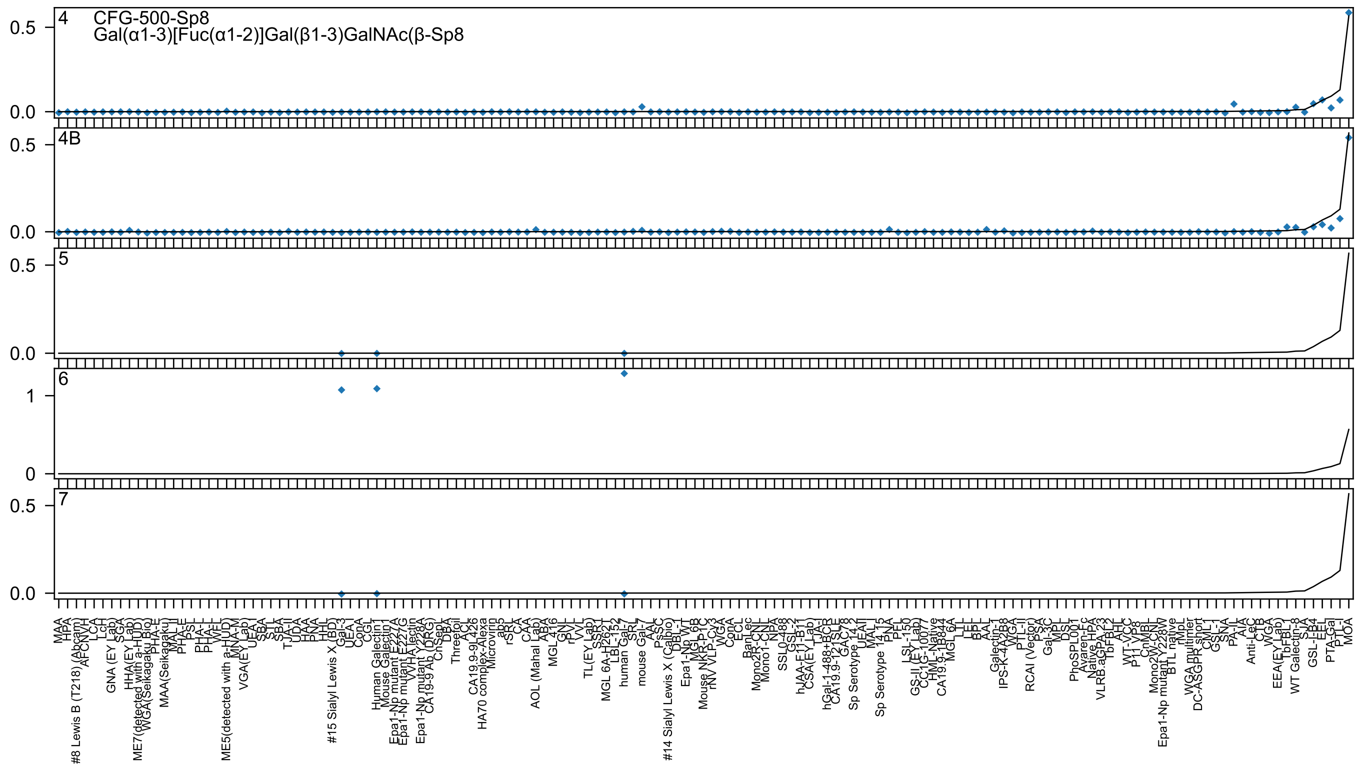

4 CFG-500-Sp8
Gal(α1-3)[Fuc(α1-2)]Gal(β1-3)GalNAc(β-Sp8
4B
5
6
7

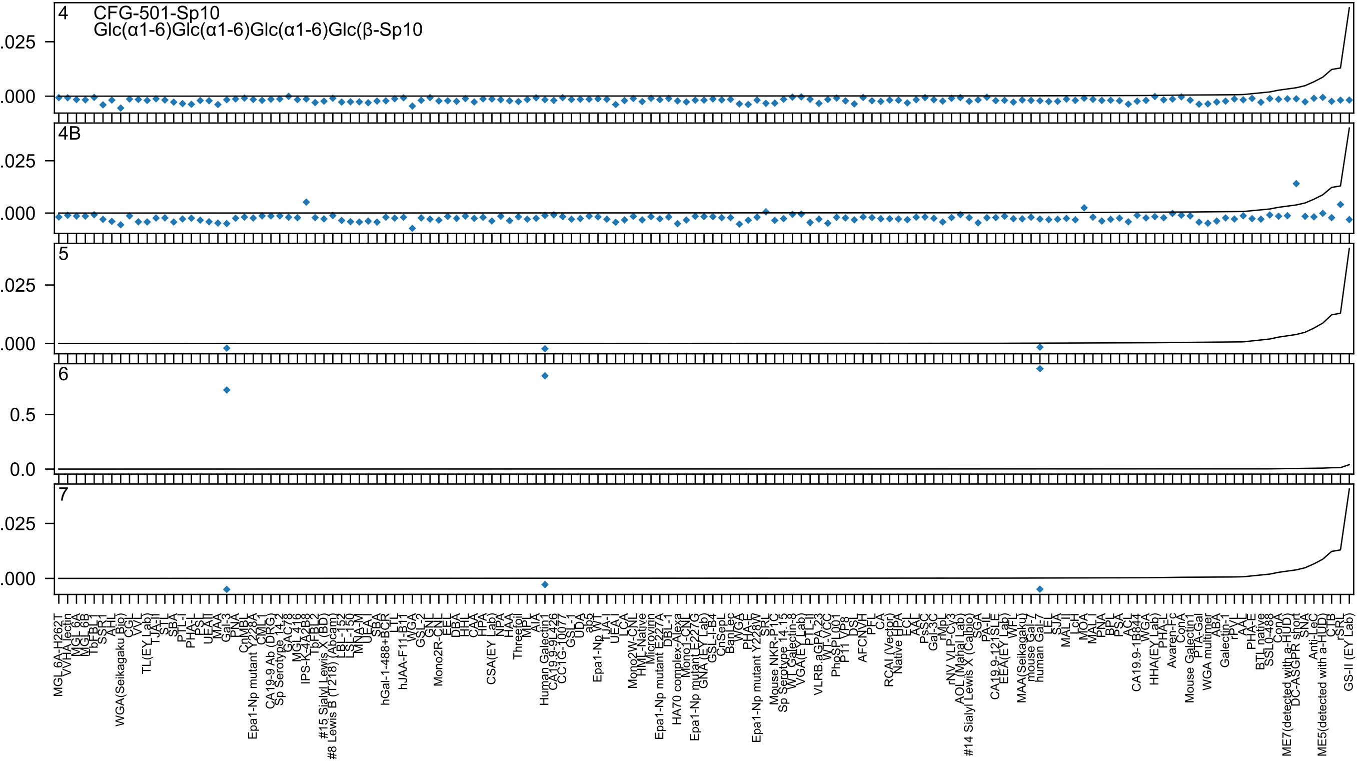
4
CFG-501-Sp10
Glc(α1-6)Glc(α1-6)Glc(α1-6)Glc(β-Sp10
4B
5
6
7
.025
.000
0.5
0.0

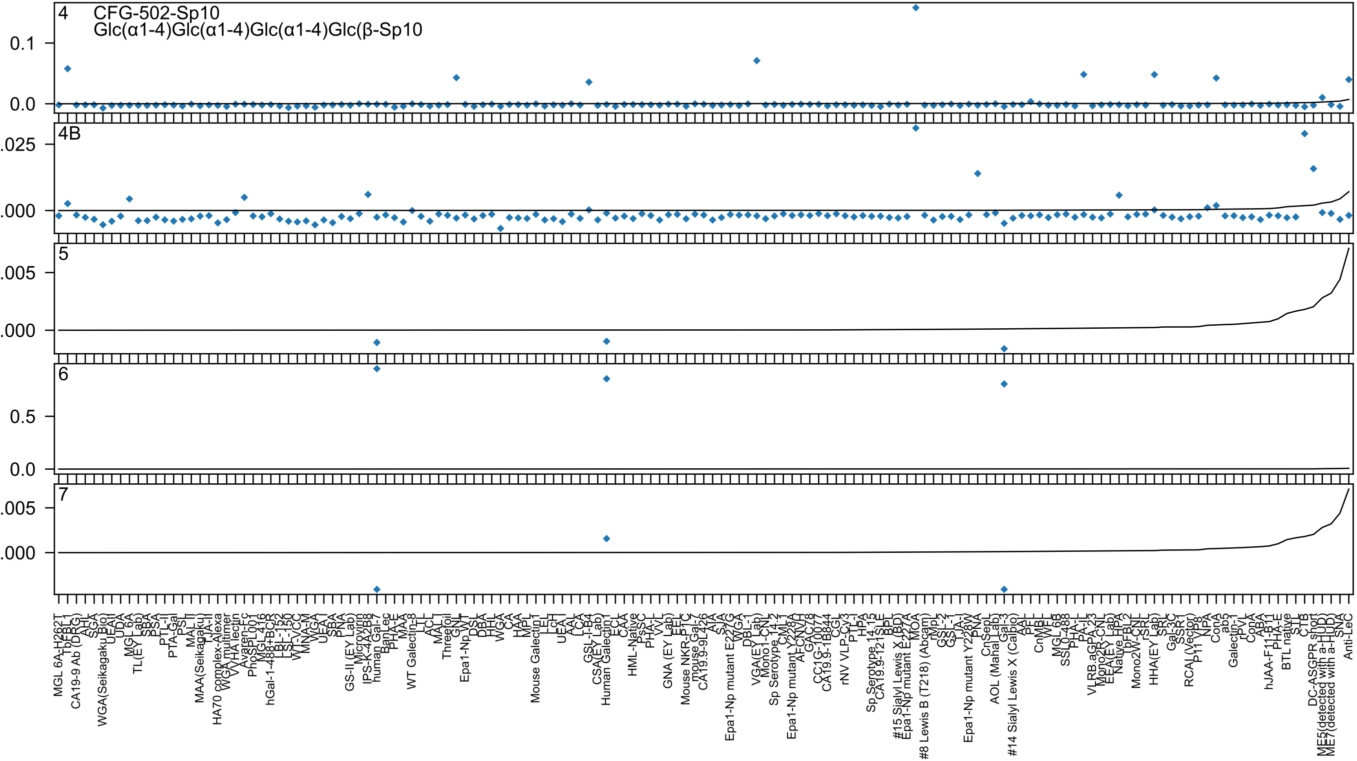

4
CFG-502-Sp10
Glc(α1-4)Glc(α1-4)Glc(α1-4)Glc(β-Sp10
4B
5
6
7

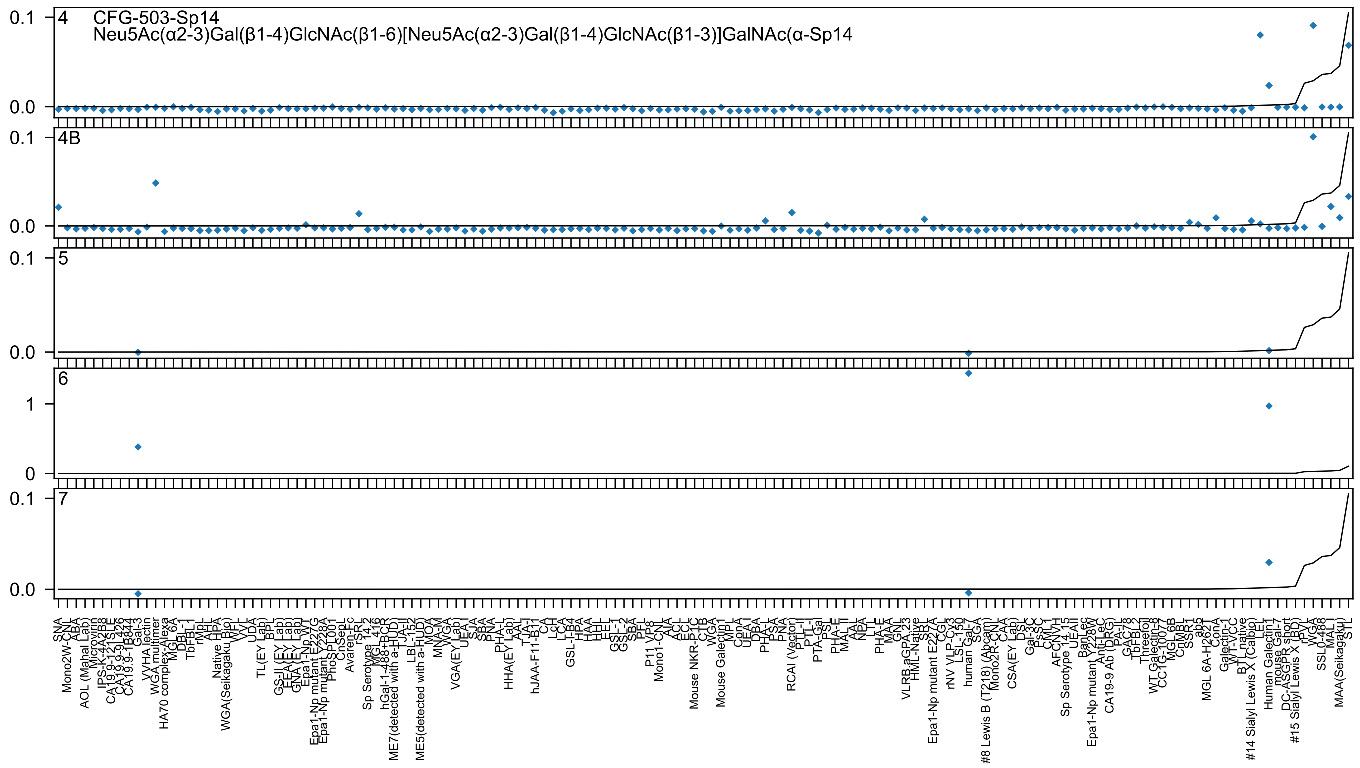
4
CFG-503-Sp14
Neu5Ac(α2-3)Gal(β1-4)GlcNAc(β1-6)[Neu5Ac(α2-3)Gal(β1-4)GlcNAc(β1-3)]GalNAc(α-Sp14
4B
5
6
7

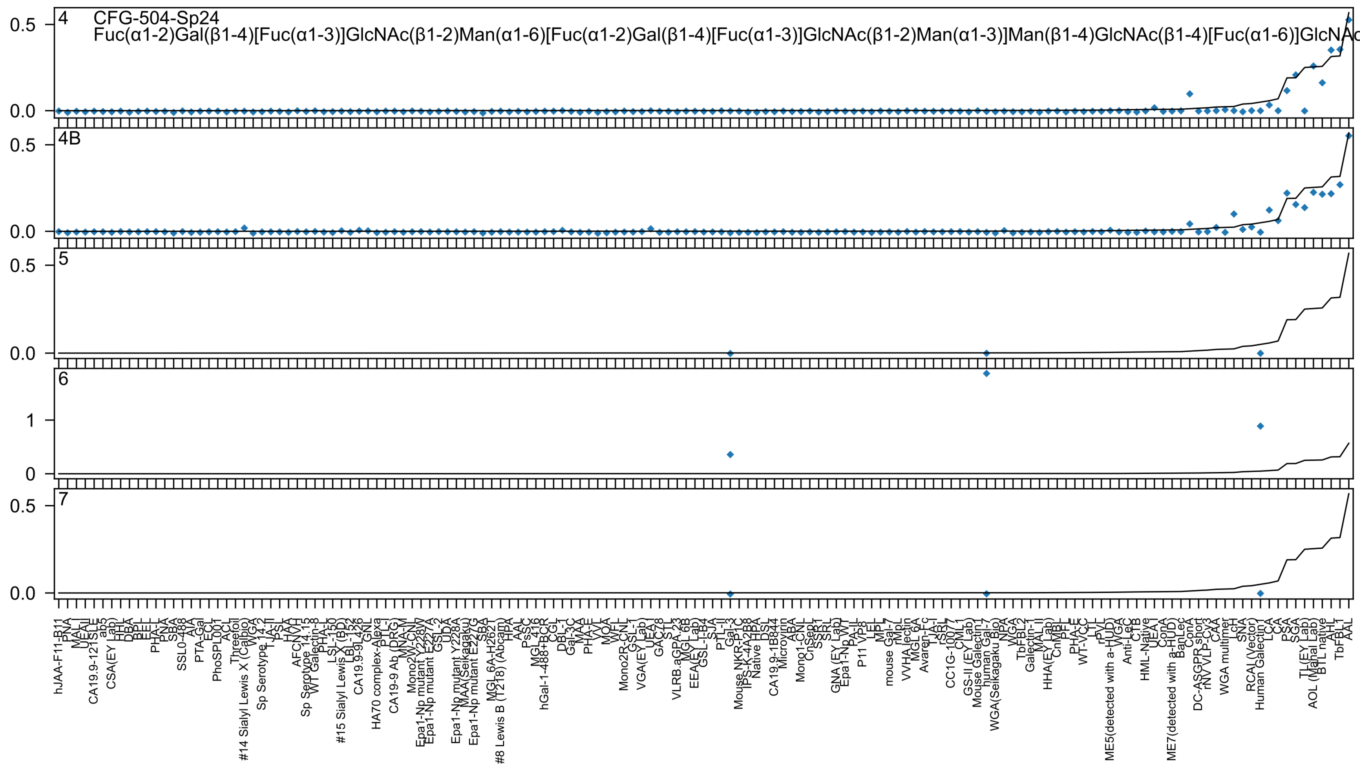
4 CFG-504-Sp24
Fuc(α1-2)Gal(β1-4)[Fuc(α1-3)]GlcNAc(β1-2)Man(α1-6)[Fuc(α1-2)Gal(β1-4)[Fuc(α1-3)]GlcNAc(β1-2)Man(α1-3)]Man(β1-4)GlcNAc(β1-4)[Fuc(α1-6)]GlcNAc
4B
5
6
7

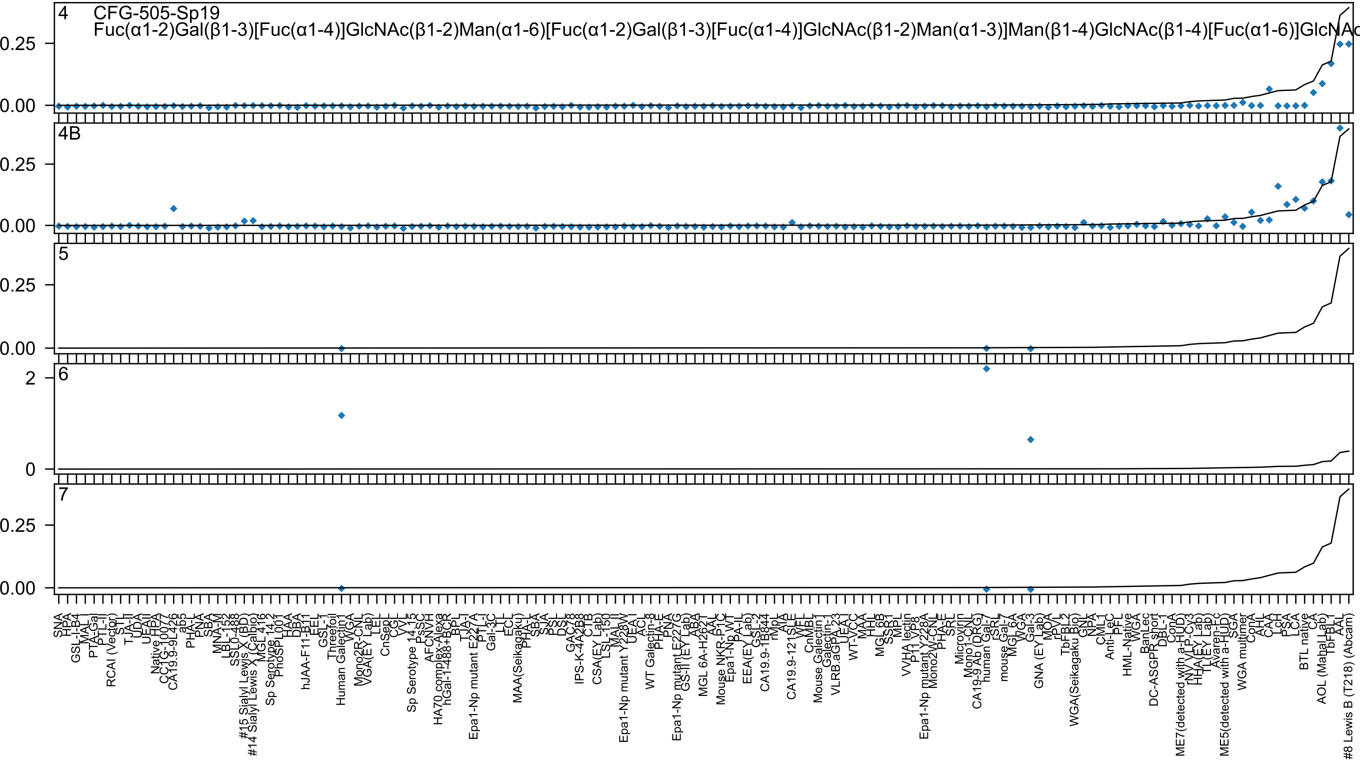

4
CFG-505-Sp19
Fuc(α1-2)Gal(β1-3)[Fuc(α1-4)]GlcNAc(β1-2)Man(α1-6)[Fuc(α1-2)Gal(β1-3)[Fuc(α1-4)]GlcNAc(β1-2)Man(α1-3)]Man(β1-4)GlcNAc(β1-4)[Fuc(α1-6)]GlcNAc
4B
5
6
7

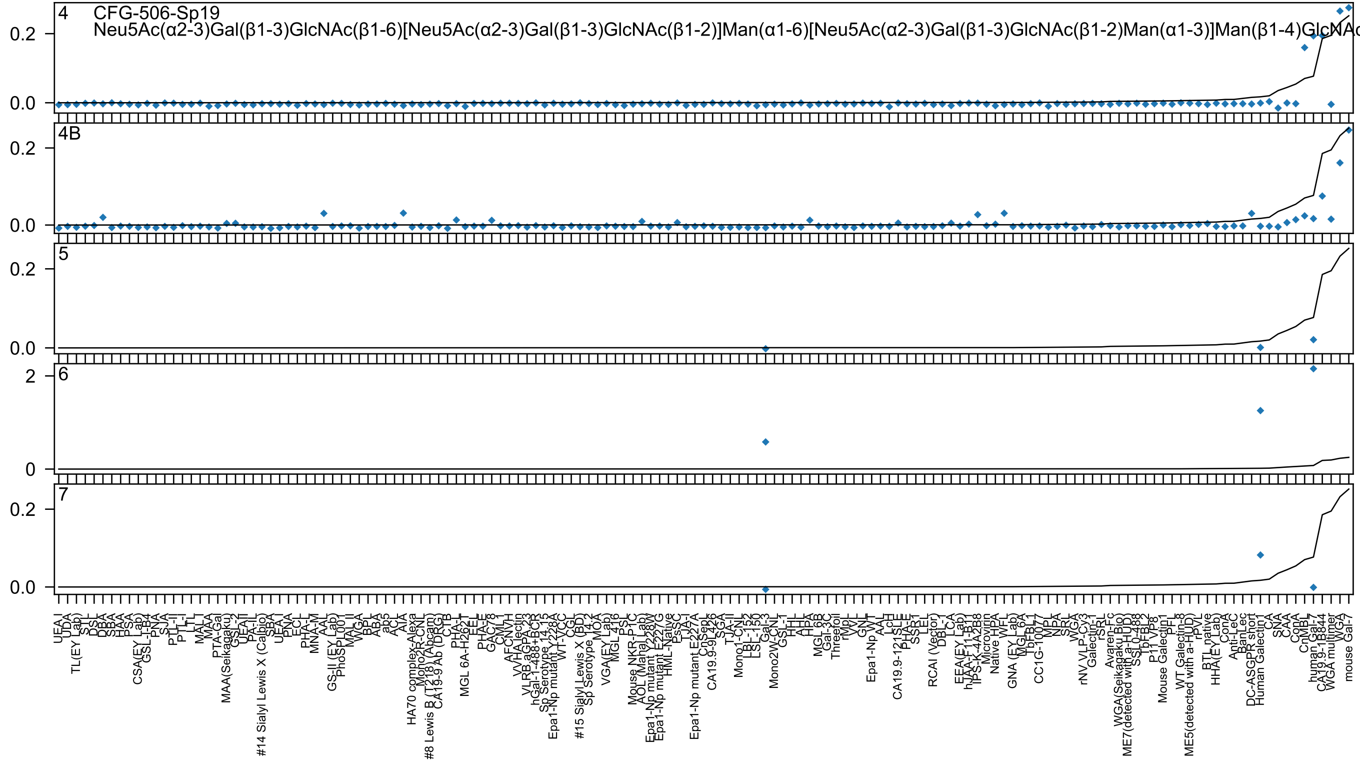
4
CFG-506-Sp19
Neu5Ac(α2-3)Gal(β1-3)GlcNAc(β1-6)[Neu5Ac(α2-3)Gal(β1-3)GlcNAc(β1-2)]Man(α1-6)[Neu5Ac(α2-3)Gal(β1-3)GlcNAc(β1-2)Man(α1-3)]Man(β1-4)GlcNAc
4B
5
6
7

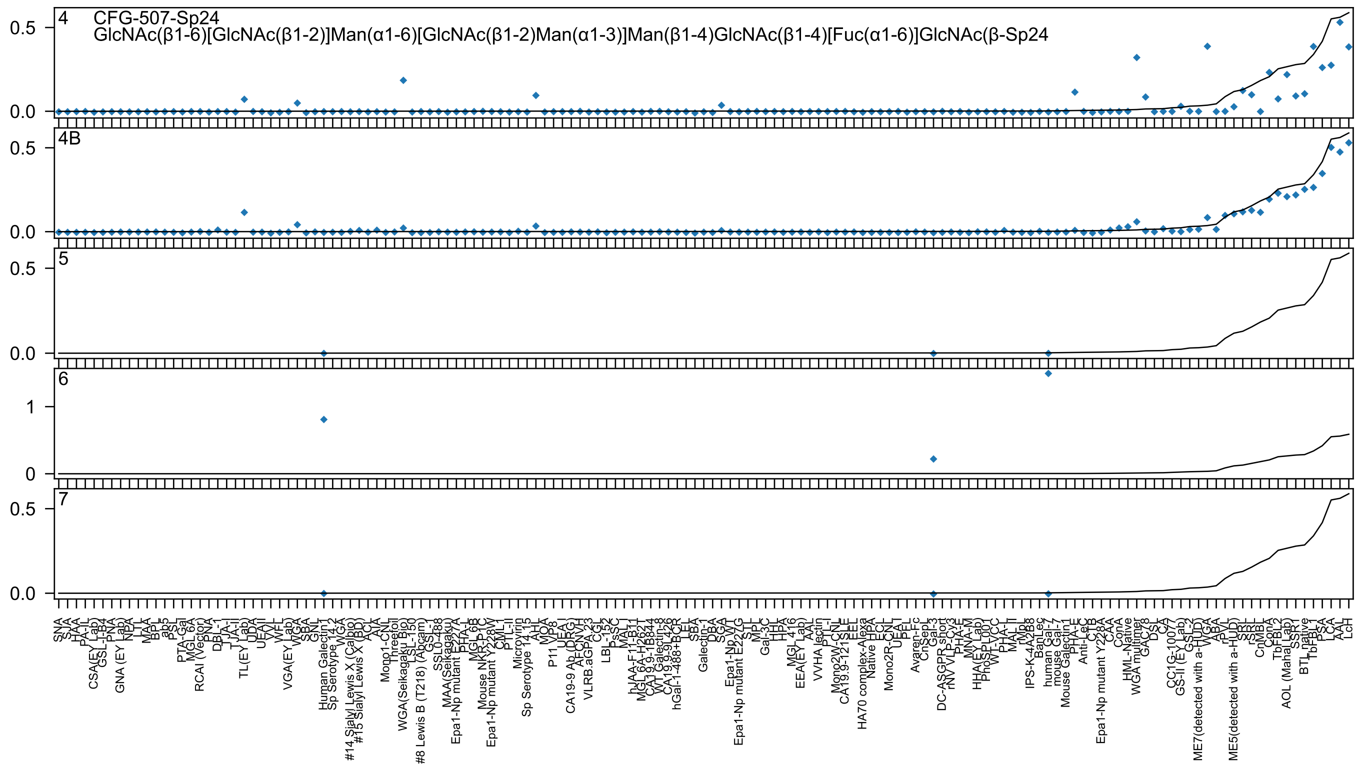
4
CFG-507-Sp24
GlcNAc(β1-6)[GlcNAc(β1-2)]Man(α1-6)[GlcNAc(β1-2)Man(α1-3)]Man(β1-4)GlcNAc(β1-4)[Fuc(α1-6)]GlcNAc(β-Sp24
4B
5
6
7
0.5
0.0
1
0
SNA
SJA
HAA
PA-IL
CSA(EY Lab)
GSL-I-B4
PNA
GNA (EY Lab)
NPA
LTL
MAA
BPL
ab5
PSL
PTA-Gal
MGL 6A
RCAI (Vector)
PNA
DBL-1
TJA-I
TJA-II
TL(EY Lab)
UDA
UEAII
VVL
WFL
VGA(EY Lab)
WGA
SBA
GNL
Human Galectin1
Sp Serotype 14.2
VVA
#14 Sialyl Lewis X (Calbio)
#15 Sialyl Lewis X (BD)
ACL
AIA
Mono1-CNL
Threefoil
WGA(Seikagaku Bio)
LSL-150
#8 Lewis B (T218) (Abcam)
GSL-I
SSL0-488
MAA(Seikagaku)
Epa1-Np mutant E227A
PHA-L
MGL 6B
Mouse NKR-P1C
Epa1-Np mutant Y228W
CML-I
PTL-II
Microvirin
Sp Serotype 14.15
AHL
MOA
P11 VP8
UEA-I
CA19-9 Ab (DRG)
AFCNVH
VLRB.aGPA.23
CGL
LBL-152
PsSC
MAL-I
hJAA-F11-B11
MGL 6A-H2621
CA19.9-1B844
WT Galectin-8
CA19.9-9L426
hGal-1-488+BCR
LEL
SBA
Galectin-1
DBA
SGA
Epa1-Np WT
Epa1-Np mutant E227G
STL
MPL
Gal-3C
HHL
HPA
MGL 416
EEA(EY Lab)
AAL
VVHA lectin
PTL-I
Mono2W-CNL
CA19.9-121SLE
EEL
HA70 complex-Alexa
Native HPA
ECL
Mono2R-CNL
UEA-I
PFL
Avaren-Fc
CnSep1
Gal-3
DC-ASGPR short
rNV VLP-Cy3
PHA-E
MNA-M
HHA(EY Lab)
PhoSL001
WT-LVCC
PHA-L
MAL-II
rMpL
IPS-K-4A2B8
BanLec
human Gal-7
mouse Gal-7
Mouse Galectin1
PHA-E
Anti-LeC
CTB
Epa1-Np mutant Y228A
CAA
ConA
HML-Native
WGA multimer
GAC78
DSL
CA
CC1G-10077
GS-II (EY Lab)
GSL-2
ME7(detected with a-HUD)
WGA
ABA
rPVL
ME5(detected with a-HUD)
SRL
rSRL
CnMBL
ConA
TbFBL2
AOL (Maha Lab)
SSR1
BTL native
TbFBL1
PSA
LCA
AAL
LcH

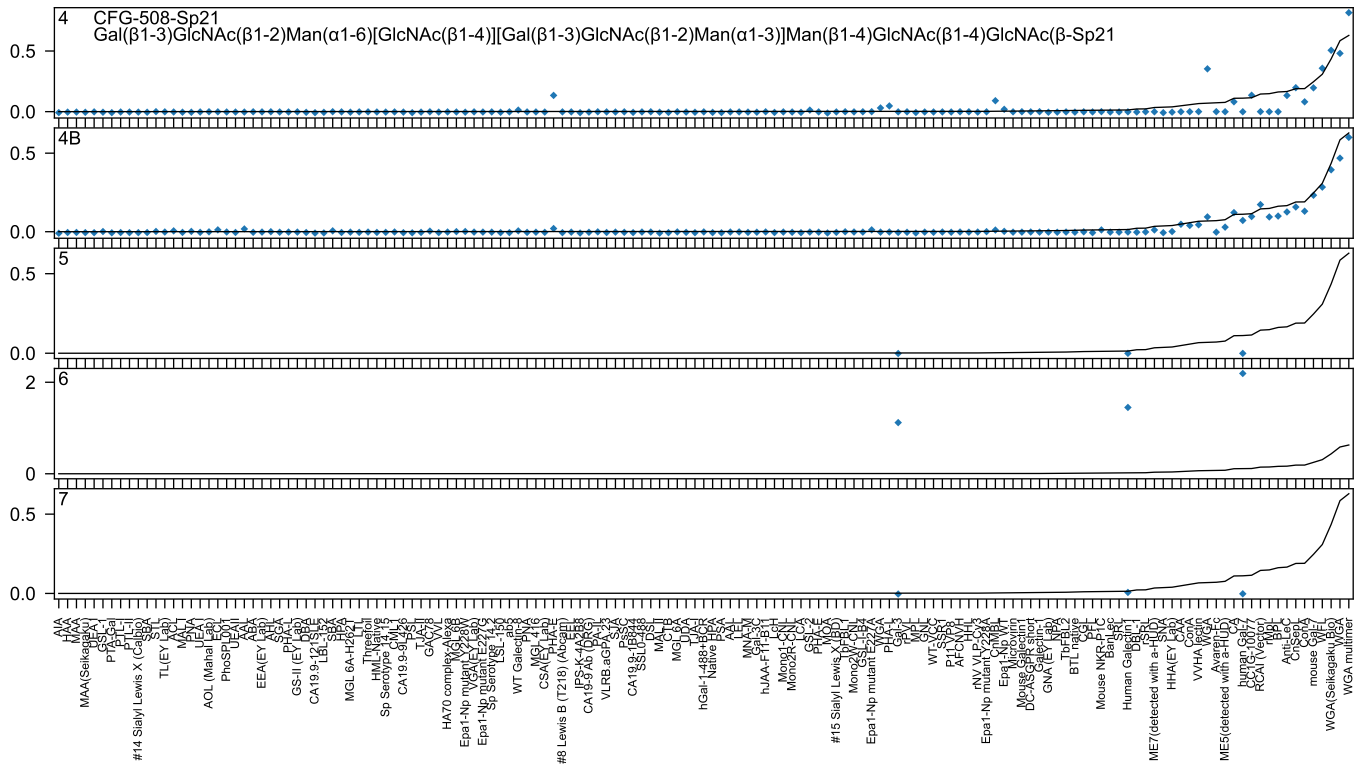

4 CFG-508-Sp21
Gal(β1-3)GlcNAc(β1-2)Man(α1-6)[GlcNAc(β1-4)][Gal(β1-3)GlcNAc(β1-2)Man(α1-3)]Man(β1-4)GlcNAc(β1-4)GlcNAc(β-Sp21
4B
5
6
7
0.0
0.5
0
2
AIA
HAA
MAA
MAA(Seikagaku)
UEA-I
GSL-I
PTA-Gal
PTL-I
PTL-II
#14 Sialyl Lewis X (Calbio)
SBA
STL
TL(EY Lab)
ACL
MAL-I
PNA
UEA-I
AOL (Mahal Lab)
ECL
PhoSL001
UEAII
AAL
ABA
EEA(EY Lab)
AHL
SGA
PHA-L
GS-II (EY Lab)
DBA
CA19.9-121SLE
LBL-152
SBA
HPA
MGL 6A-H262T
LTL
Threefoil
HML-Native
Sp Serotype 14_15
CML1
CA19.9-9L426
PSL
TJA-II
GAC78
VVL
HA70 complex-Alexa
MGL 6B
Epa1-Np mutant Y228W
VGA(EY Lab)
Epa1-Np mutant E227G
Sp Serotype 14_2
LSL-150
ab5
WT Galectin-8
PNA
MGL 416
CSA(EY Lab)
PHA-E
#8 Lewis B (T218) (Abcam)
EEL
IPS-K-4A2B8
CA19-9 Ab (DRG)
PA-IL
VLRB.aGPA.23
SJA
PsSC
CA19.9-1B844
SSL0-488
DSL
MAL-II
CTB
MGL 6A
UDA
TJA-I
hGal-1-488+BCR
Native HPA
PSA
AAL
LEL
MNA-M
Gal-3C
hJAA-F11-B11
LcH
Mono1-CNL
Mono2R-CNL
LCA
GSL-2
PHA-E
MOA
#15 Sialyl Lewis X (BD)
TbFBL1
Mono2W-CNL
GSL-I-B4
Epa1-Np mutant E227A
WGA
PHA-L
Gal-3
rPVL
MPL
GNL
WT-VCC
SSR1
P11 VP8
AFCNVH
HHL
rNV VLP-Cy3
Epa1-Np mutant Y228A
CnMBL
Epa1-Np WT
Microvirin
Mouse Galectin1
DC-ASGPR short
Galectin-1
GNA (EY Lab)
NPA
TbFBL2
BTL native
CGL
PFL
Mouse NKR-P1C
BanLec
SRL
Human Galectin1
DBL-1
rSRL
ME7(detected with a-HUD)
SNA
HHA(EY Lab)
CAA
ConA
VVHA lectin
WGA
Avaren-Fc
ME5(detected with a-HUD)
CA
human Gal-7
CC1G-10077
RCAI (Vector)
rMpL
BPL
Anti-LeC
CnSep1
ConA
mouse Gal-7
WFL
WGA(Seikagaku Bio)
WGA
WGA multimer

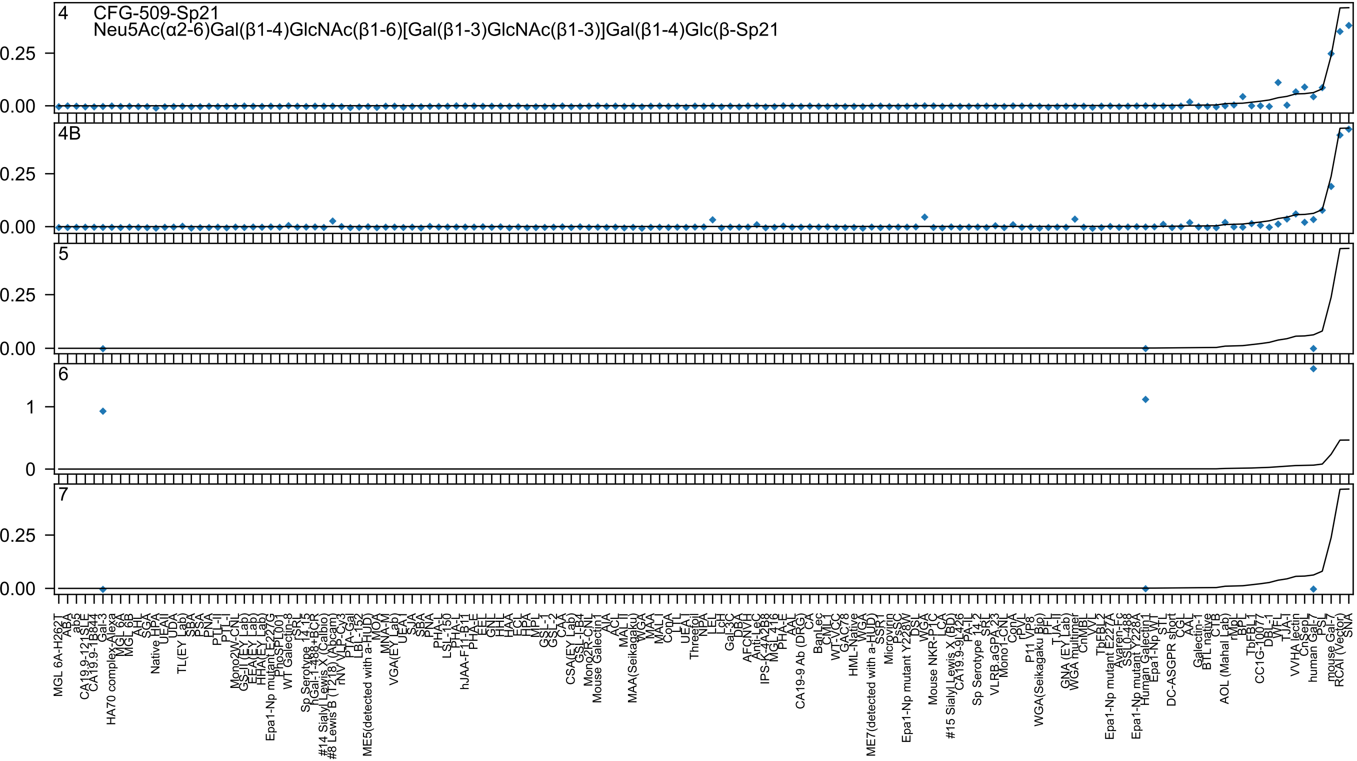
4
CFG-509-Sp21
Neu5Ac(α2-6)Gal(β1-4)GlcNAc(β1-6)[Gal(β1-3)GlcNAc(β1-3)]Gal(β1-4)Glc(β-Sp21
4B
5
6
7

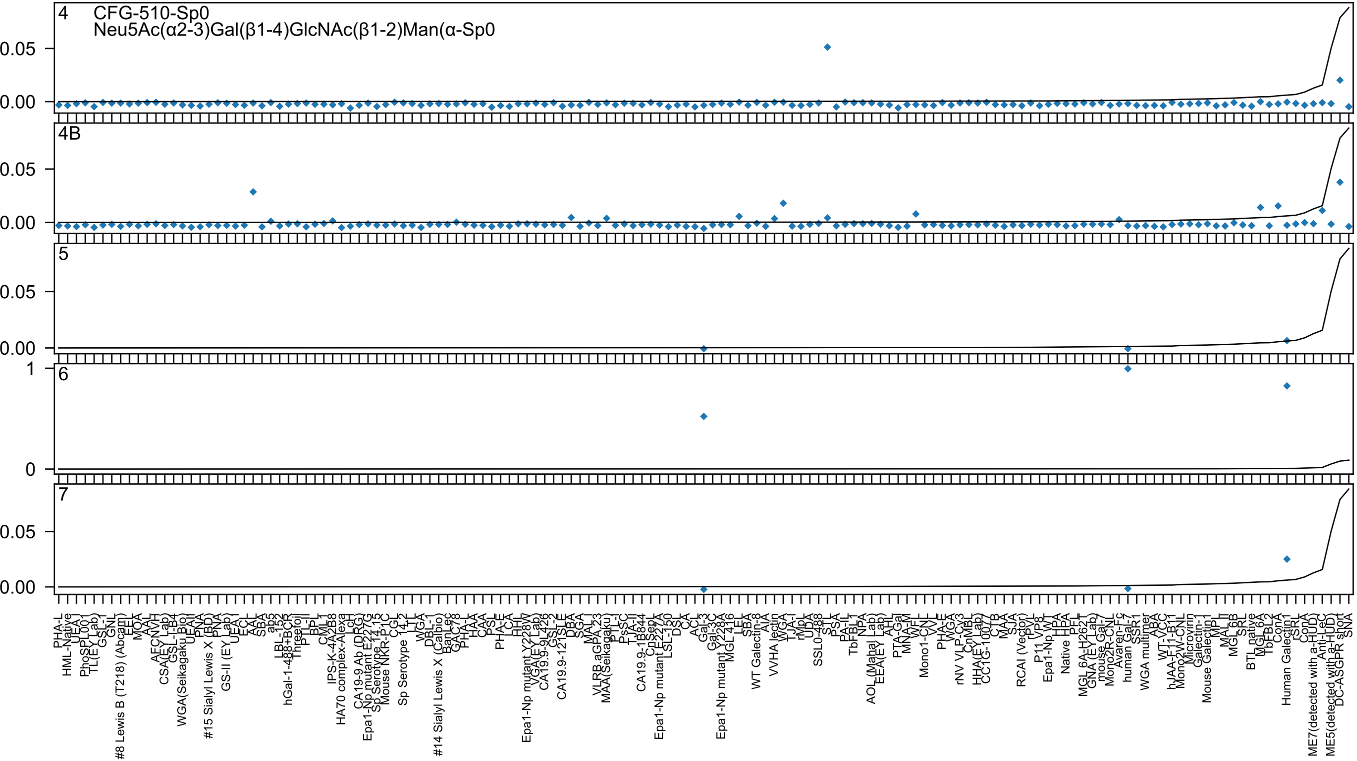

4 CFG-510-Sp0
Neu5Ac(α2-3)Gal(β1-4)GlcNAc(β1-2)Man(α-Sp0
4B
5
6
7
0.05
0.00
1
0

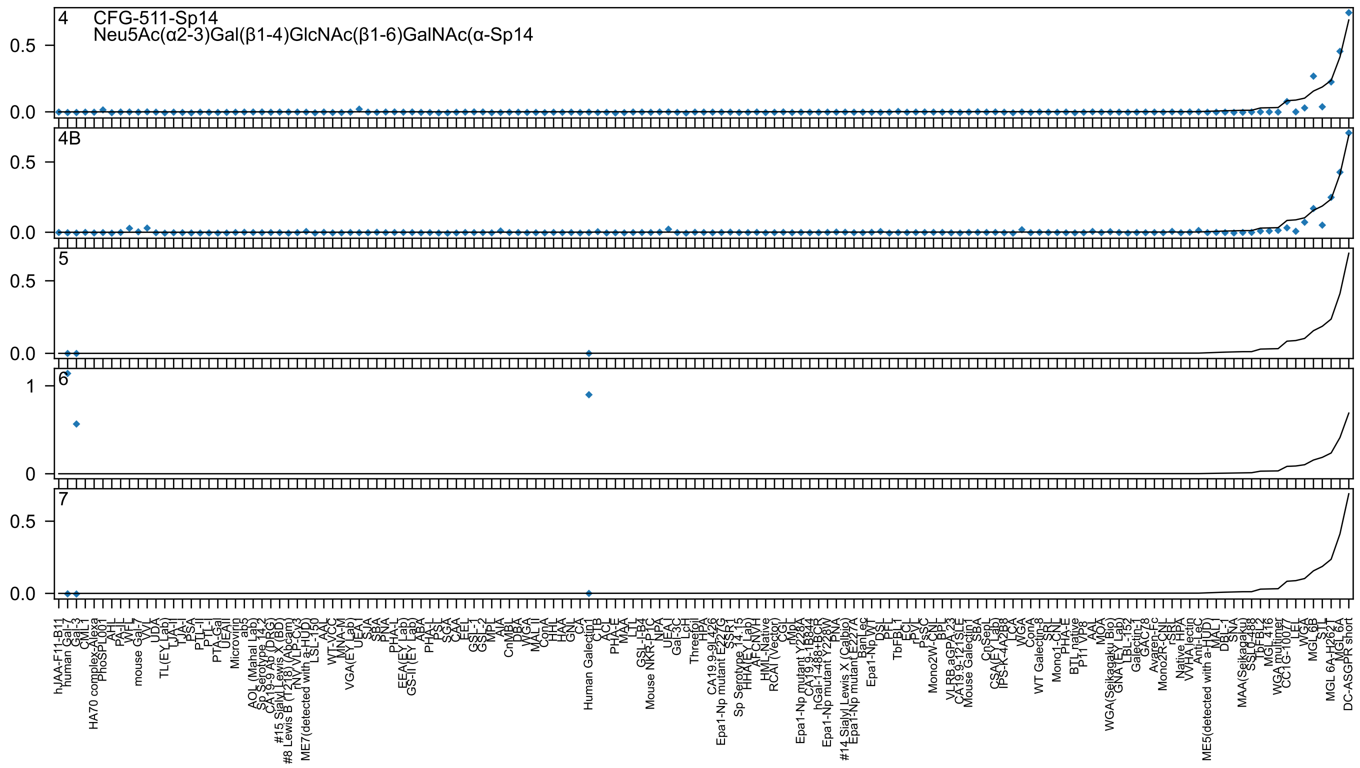

4
CFG-511-Sp14
Neu5Ac(α2-3)Gal(β1-4)GlcNAc(β1-6)GalNAc(α-Sp14
4B
5
6
7

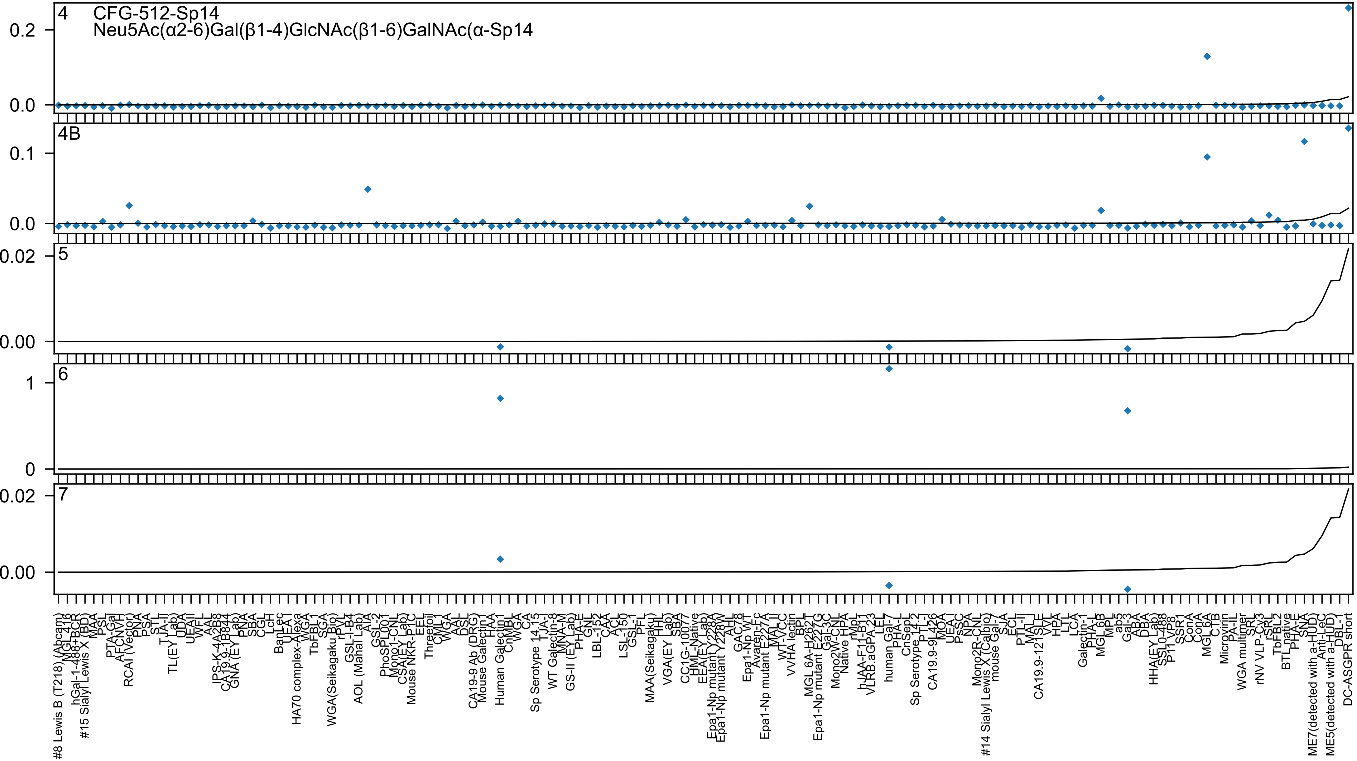

4 CFG-512-Sp14
Neu5Ac(α2-6)Gal(β1-4)GlcNAc(β1-6)GalNAc(α-Sp14
0.2
0.0
4B
0.1
0.0
5
0.02
0.00
6
1
0
7
0.02
0.00
#8 Lewis B (T218) (Abcam)
hGal-1-488+BCR
#15 Sialyl Lewis X (BD)
RCAI (Vector)
TL(EY Lab)
HA70 complex-Alexa
WGA(Seikagaku Bio)
AOL (Mahal Lab)
Mouse NKR-P1C
CA19-9 Ab (DRG)
Mouse Galectin1
Human Galectin1
Sp Serotype 14.15
WT Galectin-8
GS-II (EY Lab)
MAA(Seikagaku)
VGA(EY Lab)
CC1G-10077
Epa1-Np mutant Y228A
Epa1-Np mutant Y228W
Epa1-Np WT
Epa1-Np mutant E227A
VVHA lectin
MGL 6A-H262T
Epa1-Np mutant E227G
Mono2W-CNL
hJAA-F11-B11
VLRB.aGPA.23
human Gal-7
Sp Serotype 14.2
CA19.9-9L426
#14 Sialyl Lewis X (Calbio)
mouse Gal-7
CA19.9-121SLE
Galectin-1
MGL 6B
Gal-3
HHA(EY Lab)
SSL0-488
P11 VP8
SSR1
MGL 6A
Microvirin
WGA multimer
rNV VLP-Cy3
TbFBL2
BTL native
PHA-E
SNA
ME7(detected with a-HUD)
Anti-LeC
ME5(detected with a-HUD)
DBL-1
DC-ASGPR short

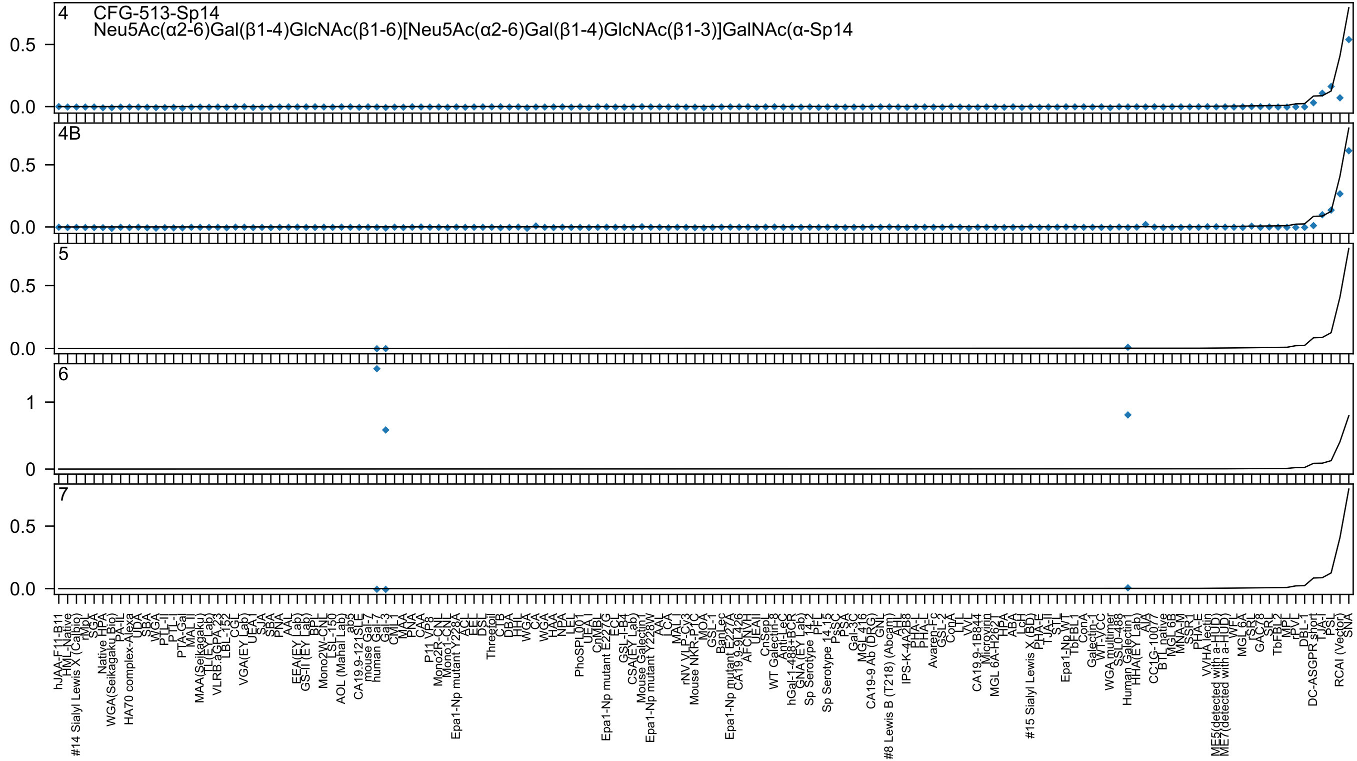
4 CFG-513-Sp14
Neu5Ac(α2-6)Gal(β1-4)GlcNAc(β1-6)[Neu5Ac(α2-6)Gal(β1-4)GlcNAc(β1-3)]GalNAc(α-Sp14
4B
5
6
7

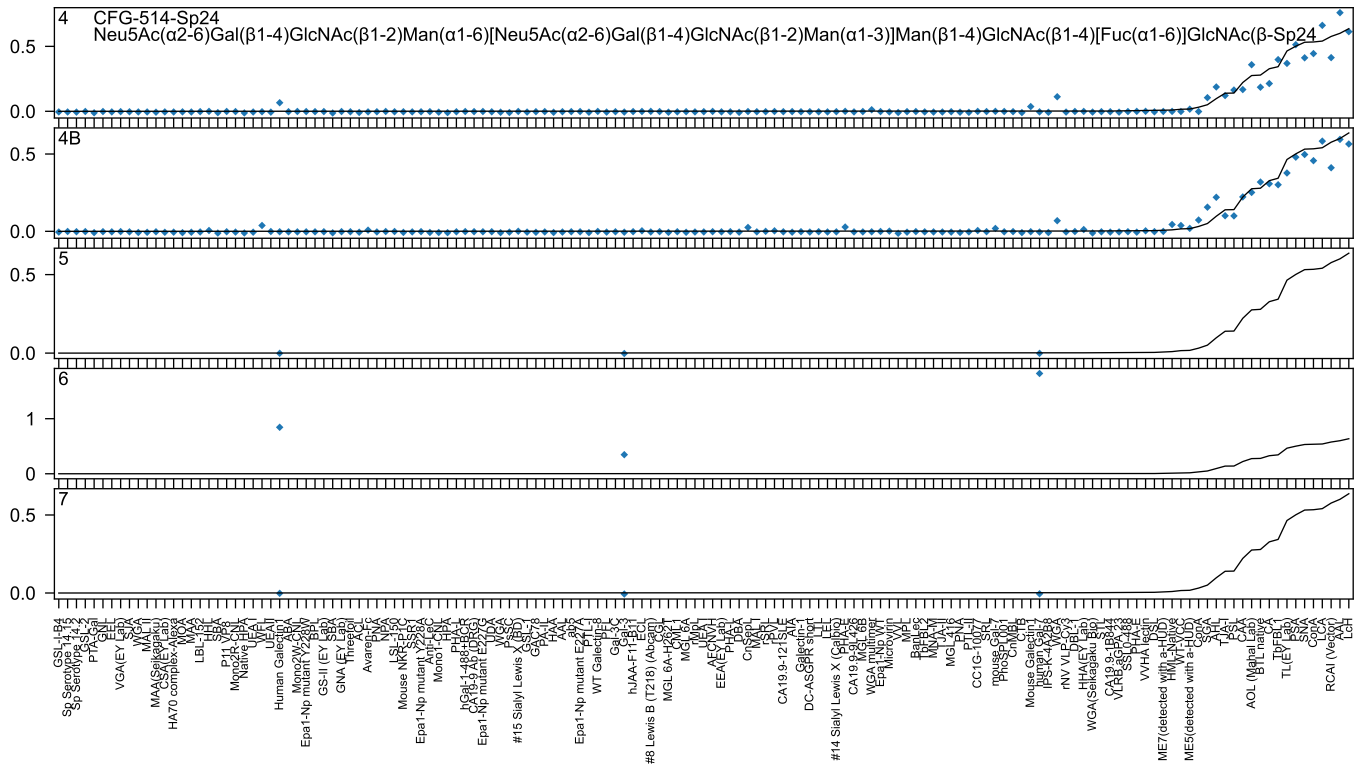
4
CFG-514-Sp24
Neu5Ac(α2-6)Gal(β1-4)GlcNAc(β1-2)Man(α1-6)[Neu5Ac(α2-6)Gal(β1-4)GlcNAc(β1-2)Man(α1-3)]Man(β1-4)GlcNAc(β1-4)[Fuc(α1-6)]GlcNAc(β-Sp24
0.5
0.0
4B
0.5
0.0
5
0.5
0.0
6
1
0
7
0.5
0.0
GSL-I-B4
Sp Serotype 14.15
Sp Serotype 14.2
GSL-2
PTA-Gal
GNL
EEL
VGA(EY Lab)
SJA
WGA
MAL II
MAA(Seikagaku)
CSA(EY Lab)
HA70 complex-Alexa
MOA
MAA
LBL-152
HHL
SBA
P11 VP8
Mono2R-CNL
Native HPA
UEA-I
WFL
UEAII
Human Galectin1
ABA
Mono2W-CNL
Epa1-Np mutant Y228W
BPL
GS-II (EY Lab)
SBA
GNA (EY Lab)
Threefoil
ACL
Avaren-Fc
PNA
NPA
LSL-150
Mouse NKR-P1C
SSR1
Epa1-Np mutant Y228A
Anti-LeC
Mono1-CNL
HPA
PHA-L
hGal-1-488+BCR
CA19-9 Ab (DRG)
Epa1-Np mutant E227G
UDA
WGA
PsSC
#15 Sialyl Lewis X (BD)
GSL-1
GAC78
PA-IL
HAA
AAL
ab5
Epa1-Np mutant E227A
PTL-I
WT Galectin-8
PFL
Gal-3C
Gal-3
hJAA-F11-B11
ECL
#8 Lewis B (T218) (Abcam)
CGL
MGL 6A-H262T
CML1
MGL 6A
rMpL
UEA I
AECNVH
EEA(EY Lab)
PHA-L
DBA
CnSepL
MAL I
rSRL
rPVL
CA19.9-121SLE
AIA
Galectin-1
DC-ASGPR short
LTL
LEL
#14 Sialyl Lewis X (Calbio)
PHA-E
CA19.9-9L426
MGL 6B
WGA multimer
Epa1-Np WT
Microvirin
VVL
MPL
BanLec
TbFBL2
MNA-M
TJA-II
MGL416
PNA
PTL-II
CC1G-10077
SRL
mouse Gal-7
PhoSL001
CnMBL
CTB
Mouse Galectin1
human Gal-7
IPS-K-4A2B8
WGA
rNV VLP-Cy3
DBL-1
HHA(EY Lab)
WGA(Seikagaku Bio)
STL
CA19.9-1B844
VLRB.aGPA.23
SSL0-488
PHA-E
VVHA lectin
DSL
ME7(detected with a-HUD)
HML-Native
WT-VCC
ME5(detected with a-HUD)
ConA
SGA
AHL
TJA-I
PSL
CAA
AOL (Mahal Lab)
BTL native
CA
TbFBL1
TL(EY Lab)
PSA
SNA
ConA
LCA
RCAI (Vector)
AAL
LcH

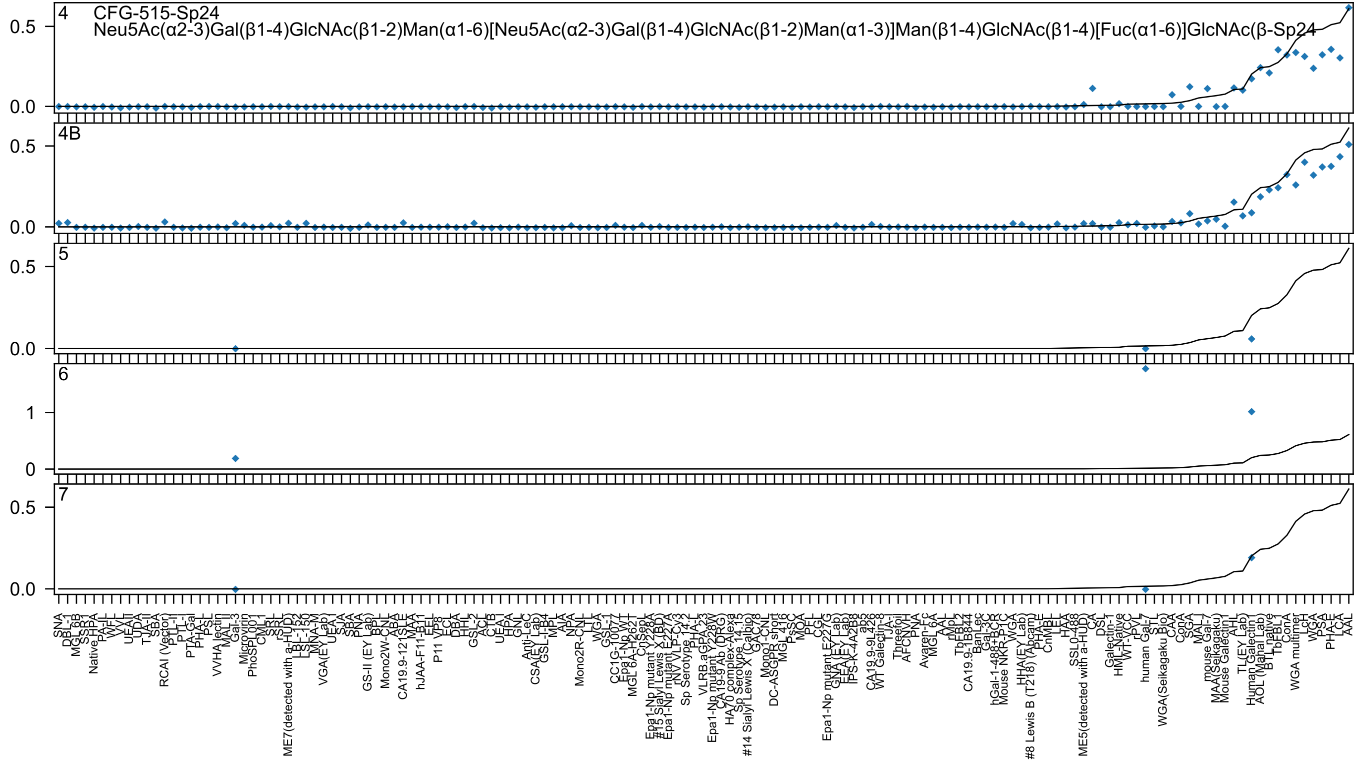

4
CFG-515-Sp24
Neu5Ac(α2-3)Gal(β1-4)GlcNAc(β1-2)Man(α1-6)[Neu5Ac(α2-3)Gal(β1-4)GlcNAc(β1-2)Man(α1-3)]Man(β1-4)GlcNAc(β1-4)[Fuc(α1-6)]GlcNAc(β-Sp24
4B
5
6
7

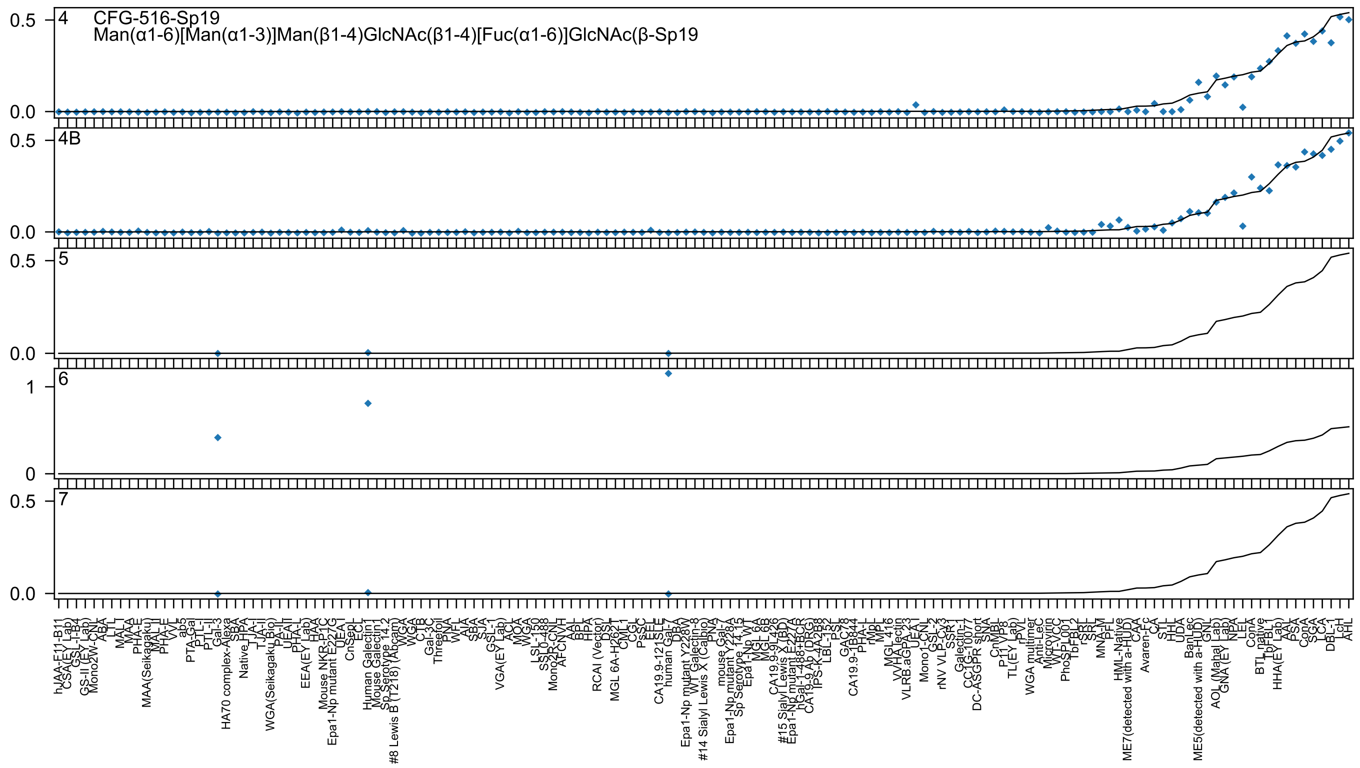

4
CFG-516-Sp19
Man(α1-6)[Man(α1-3)]Man(β1-4)GlcNAc(β1-4)[Fuc(α1-6)]GlcNAc(β-Sp19
4B
5
6
7
0.5
0.0
1
0
hJAA-F11-B11
CSA(EY Lab)
GSL-I-B4
GS-II (EY Lab)
Mono2W-CNL
ABA
LTL
MAL-I
MAA
PHA-E
MAA(Seikagaku)
MAL-II
PHA-E
VVL
ab5
PTA-Gal
PTL-I
PTL-II
Gal-3
HA70 complex-Alexa
SBA
Native HPA
TJA-I
TJA-II
WGA(Seikagaku Bio)
PA-IL
UEA-II
PHA-L
EEA(EY Lab)
HAA
Mouse NKR-P1C
Epa1-Np mutant E227G
UEA-I
CnSepL
ECL
Human Galectin1
Mouse Galectin1
Sp Serotype 14.2
#8 Lewis B (T218) (Abcam)
WGA
WGA
CTB
Gal-3C
Threefoil
PNA
WFL
AIA
SBA
SJA
GSL-1
VGA(EY Lab)
ACL
MOA
WGA
LSL-150
SSL0-488
Mono2R-CNL
AFCNVH
AAL
BPL
HPA
RCAI (Vector)
DSL
MGL 6A-H262T
CML-1
CGL
PsSC
EEL
CA19.9-121SLE
human Gal-7
DBA
Epa1-Np mutant Y228W
WT Galectin-8
#14 Sialyl Lewis X (Calbio)
PNA
mouse Gal-7
Epa1-Np mutant Y228A
Sp Serotype 14.15
Epa1-Np WT
MGL 6A
MGL 6B
CA19.9-9L426
#15 Sialyl Lewis X (BD)
Epa1-Np mutant E227A
hGal-1-488+BCR
CA19-9 Ab (DRG)
IPS-K-4A2B8
LBL-152
PSL
GAC78
CA19.9-1B844
PHA-L
rMpL
MPL
MGL 416
VVHA lectin
VLRB.aGPA.23
UEA-I
Mono1-CNL
GSL-2
rNV VLP-Cy3
SSR1
Galectin-1
CC1G-10077
DC-ASGPR short
SNA
CnMBL
P11 VP8
TL(EY Lab)
rPVL
WGA multimer
Anti-LeC
Microvirin
WT-YCC
PhoSL001
TbFBL2
rSRL
SRL
MNA-M
PFL
HML-Native
ME7(detected with a-HUD)
CAA
Avaren-Fc
CA
STL
HHL
UDA
BanLec
ME5(detected with a-HUD)
GNL
AOL (Manal Lab)
GNA (EY Lab)
NPA
LEL
ConA
BTL native
TbFBL1
HHA(EY Lab)
AAL
PSA
ConA
SGA
LCA
DBL-1
LcH
AHL

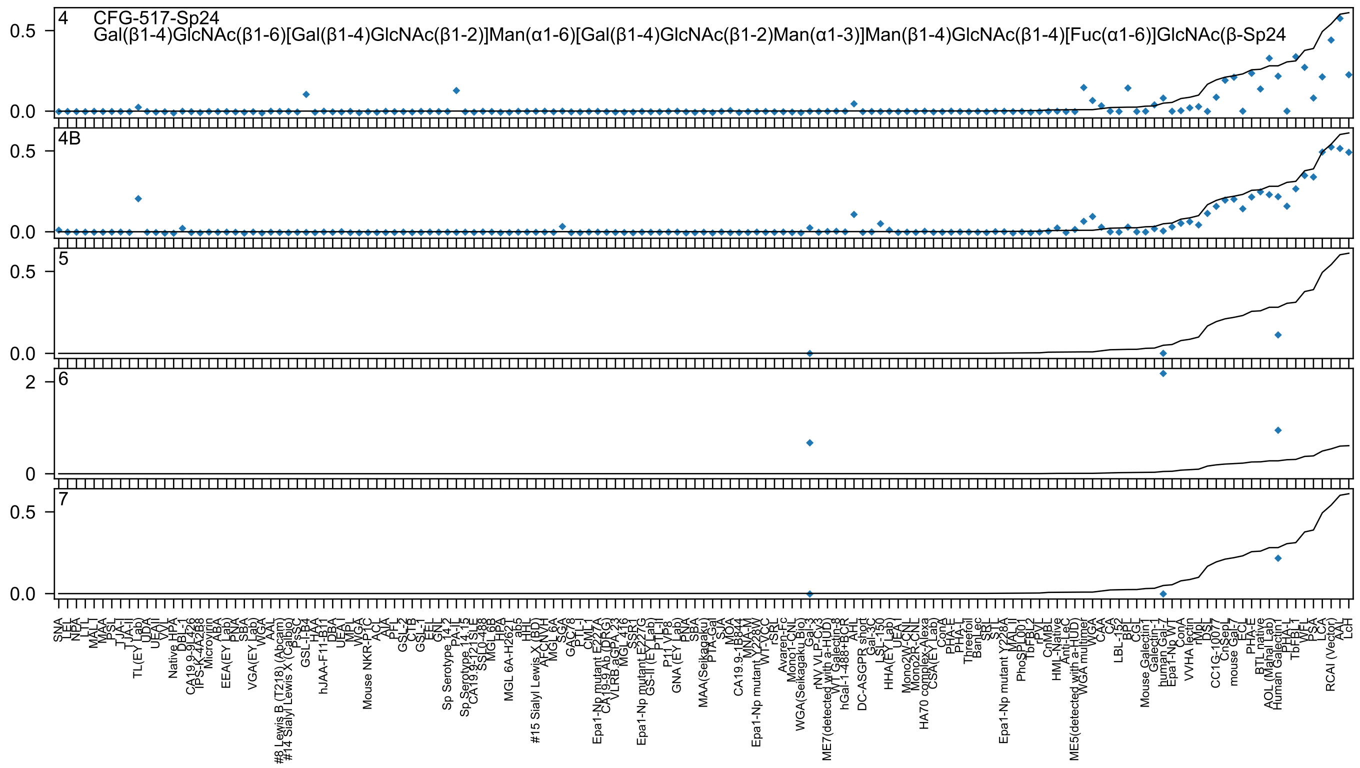

4 CFG-517-Sp24
Gal(β1-4)GlcNAc(β1-6)[Gal(β1-4)GlcNAc(β1-2)]Man(α1-6)[Gal(β1-4)GlcNAc(β1-2)Man(α1-3)]Man(β1-4)GlcNAc(β1-4)[Fuc(α1-6)]GlcNAc(β-Sp24
4B
5
6
7

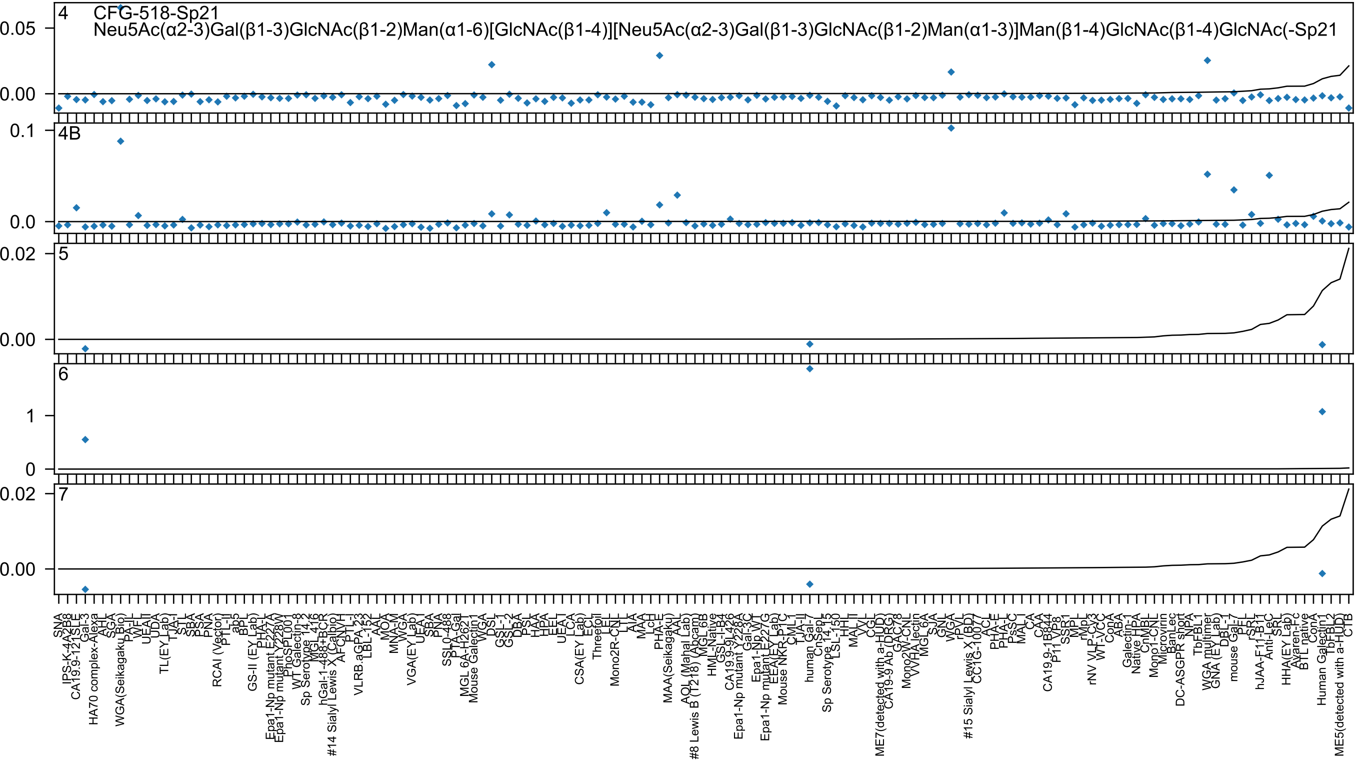

4 CFG-518-Sp21
Neu5Ac(α2-3)Gal(β1-3)GlcNAc(β1-2)Man(α1-6)[GlcNAc(β1-4)][Neu5Ac(α2-3)Gal(β1-3)GlcNAc(β1-2)Man(α1-3)]Man(β1-4)GlcNAc(β1-4)GlcNAc(-Sp21
4B
5
6
7

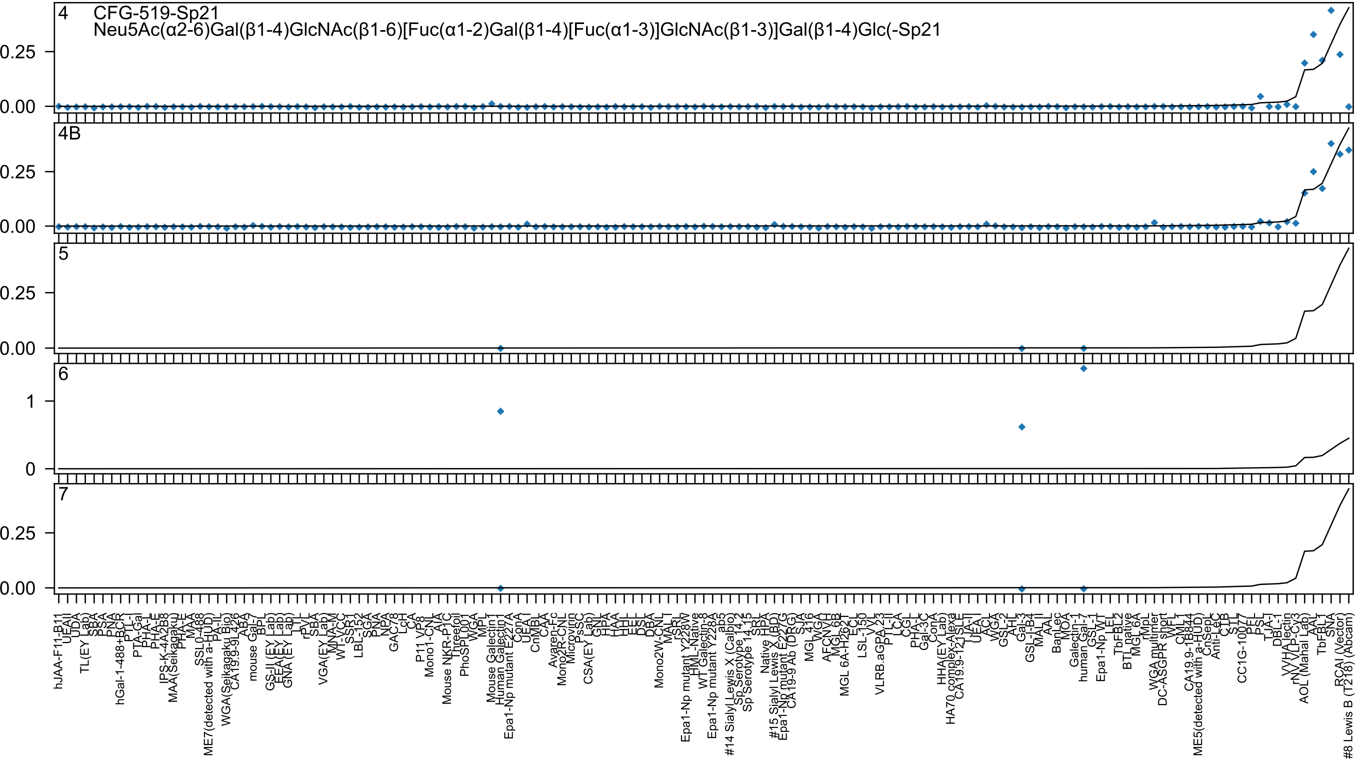

4
CFG-519-Sp21
Neu5Ac(α2-6)Gal(β1-4)GlcNAc(β1-6)[Fuc(α1-2)Gal(β1-4)[Fuc(α1-3)]GlcNAc(β1-3)]Gal(β1-4)Glc(-Sp21
4B
5
6
7

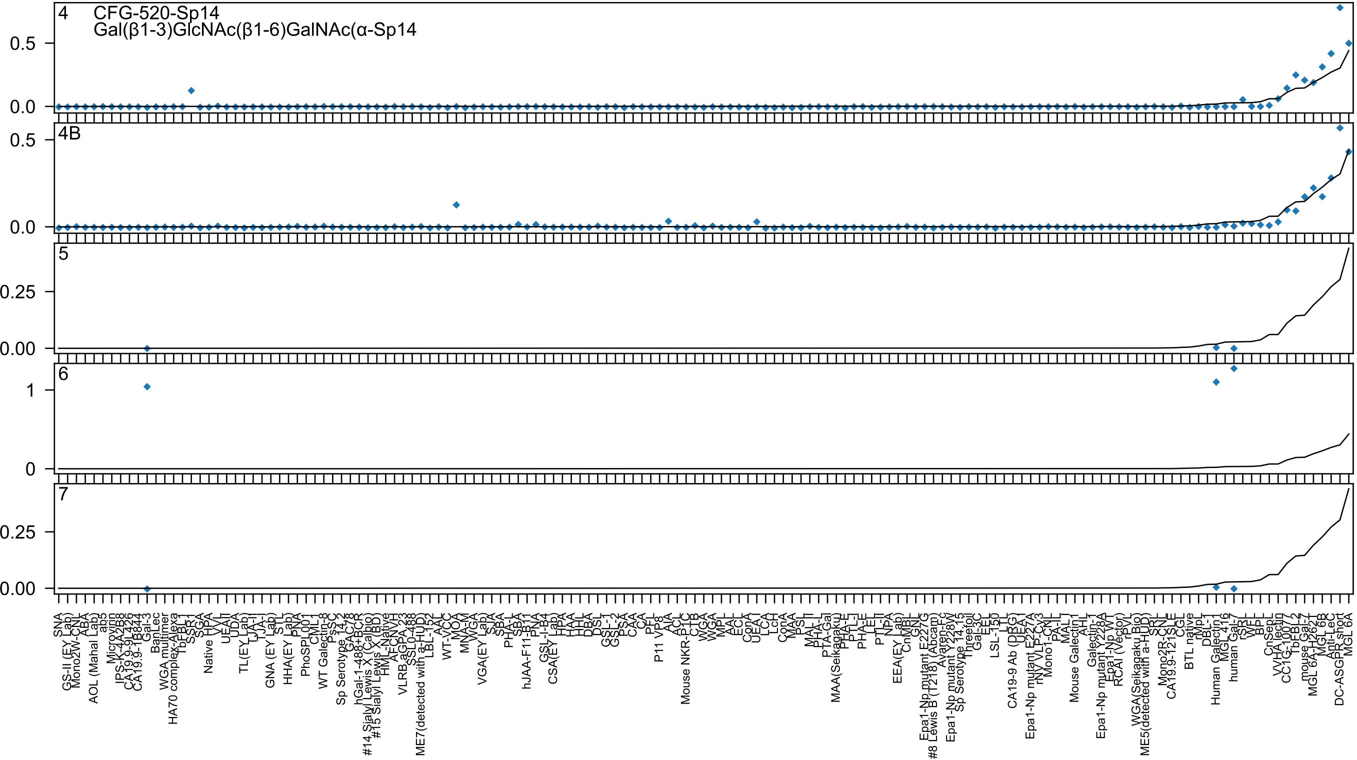

4
CFG-520-Sp14
Gal(β1-3)GlcNAc(β1-6)GalNAc(α-Sp14
0.5
0.0
4B
0.5
0.0
5
0.25
0.00
6
1
0
7
0.25
0.00
SNA
GS-II (EY Lab)
Mono2W-CNL
ABA
AOL (Mahal Lab)
ab5
Microvirin
IPS-K-4A2B8
CA19.9-9L426
CA19.9-1B844
Gal-3
BanLec
WGA multimer
HA70 complex-Alexa
TbFBL1
SSR1
SGA
Native HPA
VVL
UEAII
UDA
TL(EY Lab)
TJA-II
TJA-I
GNA (EY Lab)
STL
HHA(EY Lab)
PNA
PhoSL001
CML1
WT Galectin-8
PSSC
Sp Serotype 14.2
GAC78
hGal-1-488+BCR
#14 Sialyl Lewis X (Calbio)
#15 Sialyl Lewis X (BD)
HML-Native
AFCNVH
VLRB.aGPA.23
SSL0-488
ME7(detected with a-HUD)
LBL-152
AAL
WT-VCC
MOA
MNA-M
WGA
VGA(EY Lab)
SJA
SBA
PHA-L
SBA
hJAA-F11-B11
PNA
GSL-I-B4
CSA(EY Lab)
HPA
HAA
HHL
DBA
DSL
GSL-I
GSL-2
PSA
CAA
CA
PFL
P11 VP8
AIA
ACL
Mouse NKR-P1C
CTB
WGA
WGA
MPL
AAL
ECL
ConA
UEA-I
LCA
LcH
ConA
MAA
PSL
MAL-II
PHA-L
PTA-Gal
MAA(Seikagaku)
PHA-E
PTL-I
PHA-E
LEL
PTL-II
NPA
EEA(EY Lab)
CnMBL
GNL
Epa1-Np mutant E227G
#8 Lewis B (T218) (Abcam)
Avaren-Fc
Epa1-Np mutant Y228W
Sp Serotype 14.15
Threefoil
Gal-3C
EEL
LSL-150
LTL
CA19-9 Ab (DRG)
UEA-I
Epa1-Np mutant E227A
rNY VLP-Cy3
Mono1-CNL
PA-IL
MAL-I
Mouse Galectin1
AHL
Galectin-1
Epa1-Np mutant Y228A
Epa1-Np WT
RCAI (Vector)
rPVL
WGA(Seikagaku Bio)
ME5(detected with a-HUD)
SRL
Mono2R-CNL
CA19.9-121SLE
CGL
BTL native
rMpL
DBL-1
Human Galectin1
MGL 416
human Gal-7
rSRL
WFL
BPL
CnSep
VVHA lectin
CC1G-10077
TbFBL2
mouse Gal-7
MGL 6A-H262T
MGL 6B
Anti-LeC
DC-ASGPR short
MGL 6A

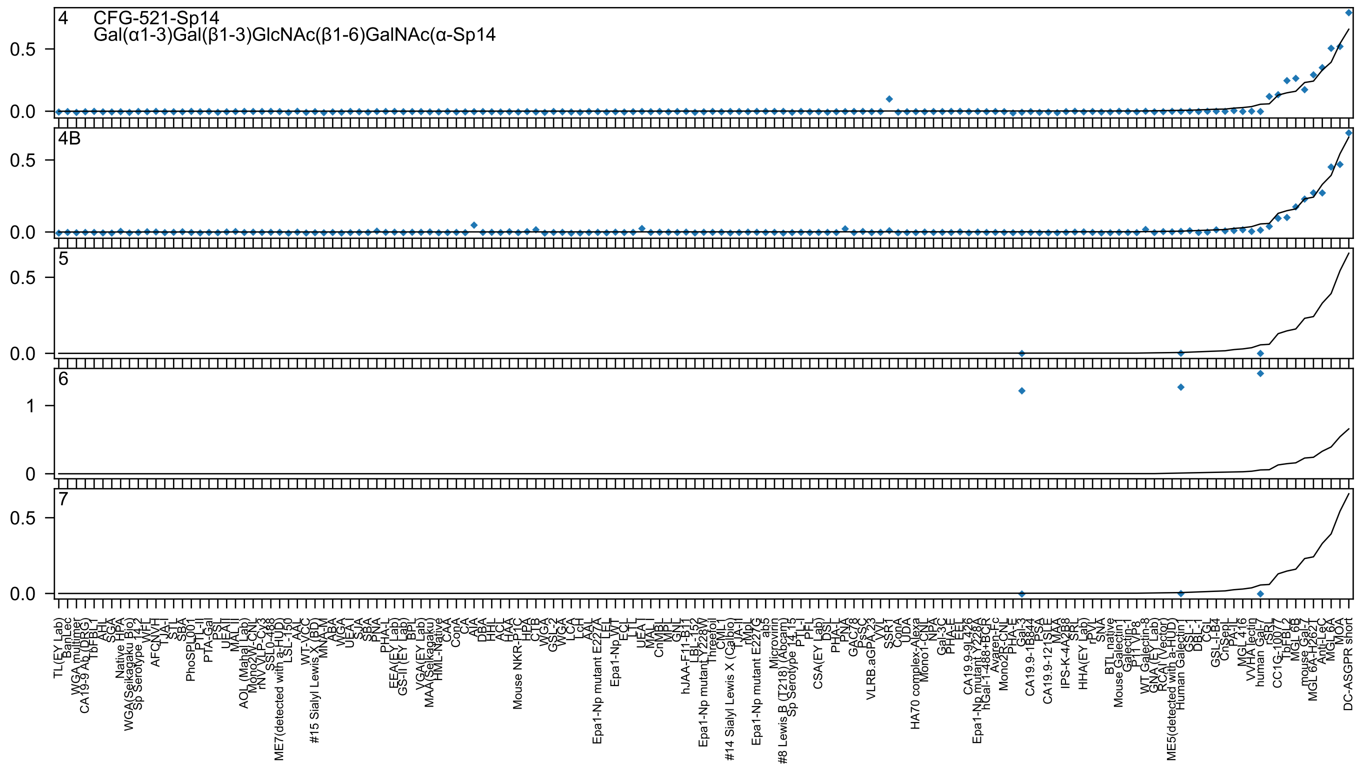
4 CFG-521-Sp14
Gal(α1-3)Gal(β1-3)GlcNAc(β1-6)GalNAc(α-Sp14
4B
5
6
7

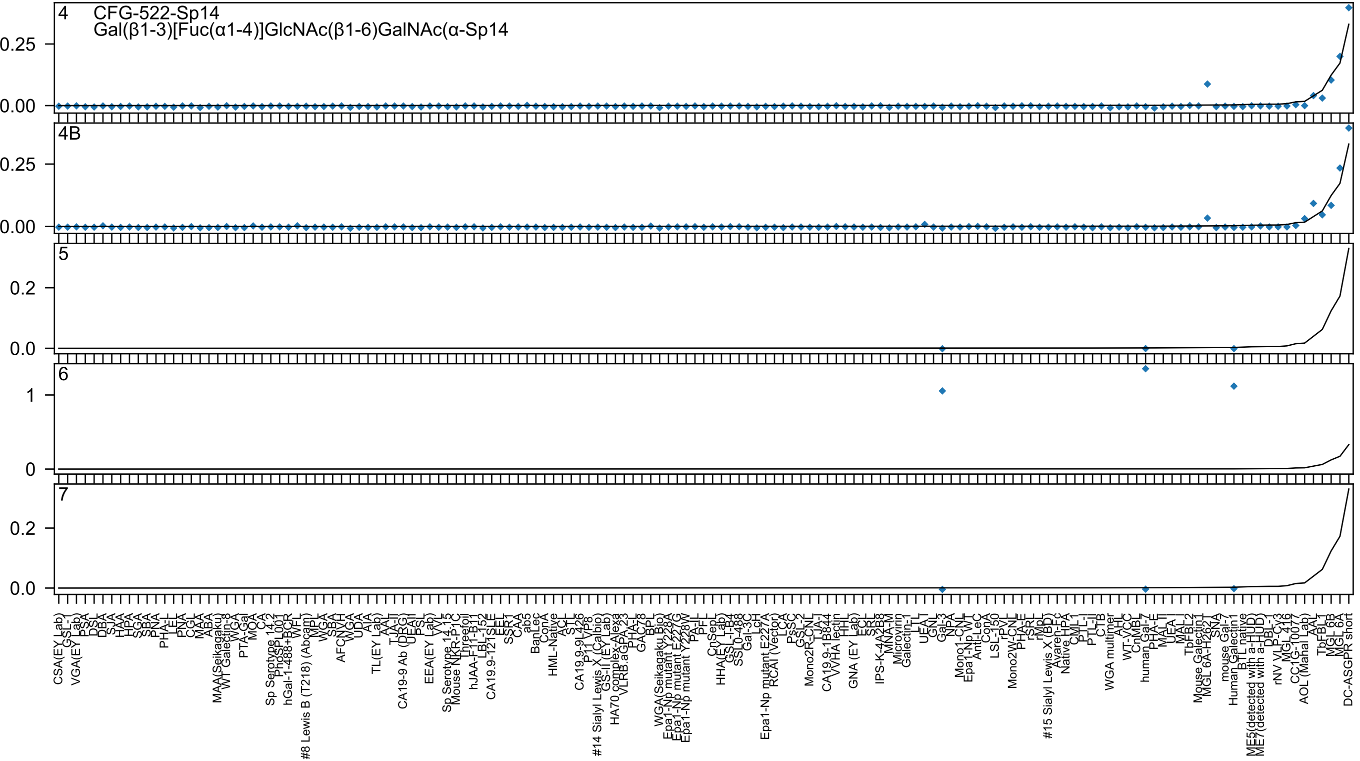
4
CFG-522-Sp14
Gal(β1-3)[Fuc(α1-4)]GlcNAc(β1-6)GalNAc(α-Sp14
0.25
0.00
4B
0.25
0.00
5
0.2
0.0
6
1
0
7
0.2
0.0

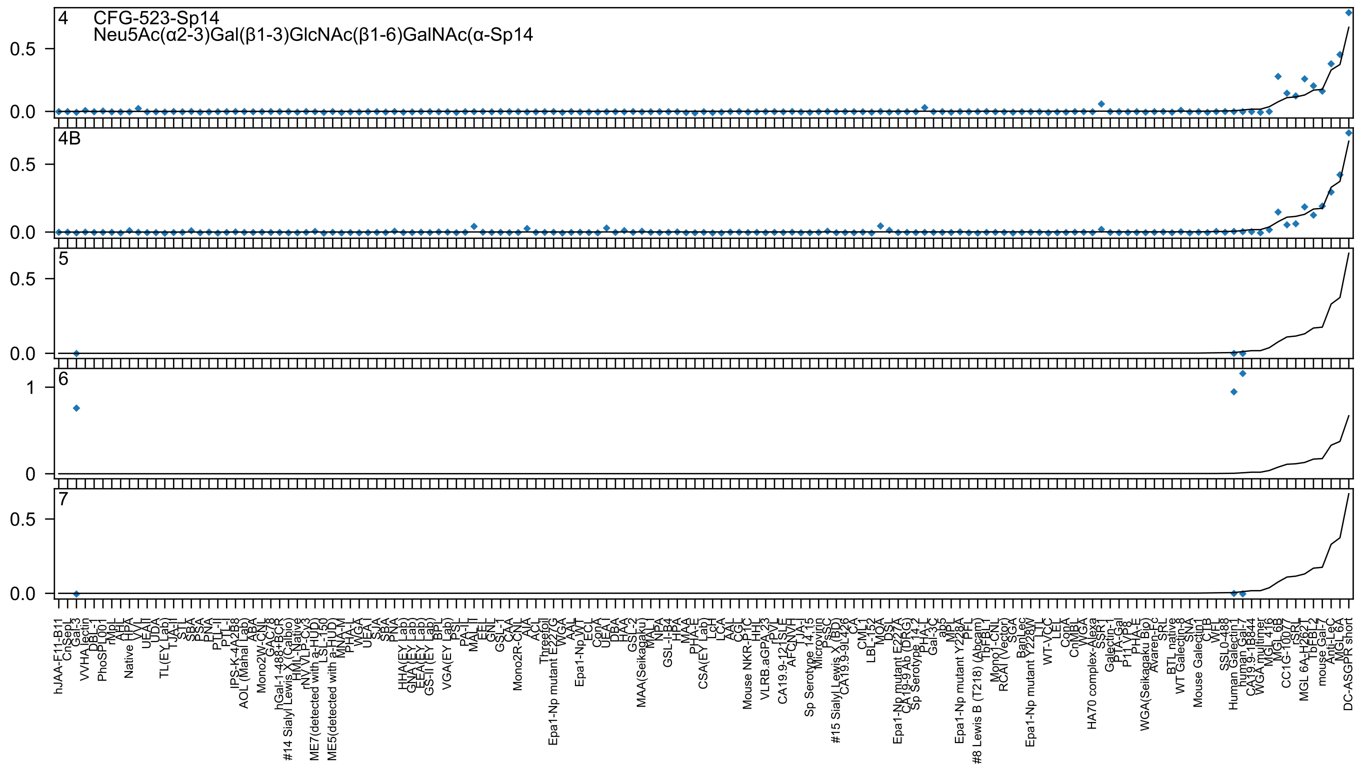
CFG-523-Sp14
Neu5Ac(α2-3)Gal(β1-3)GlcNAc(β1-6)GalNAc(α-Sp14
4
4B
5
6
7

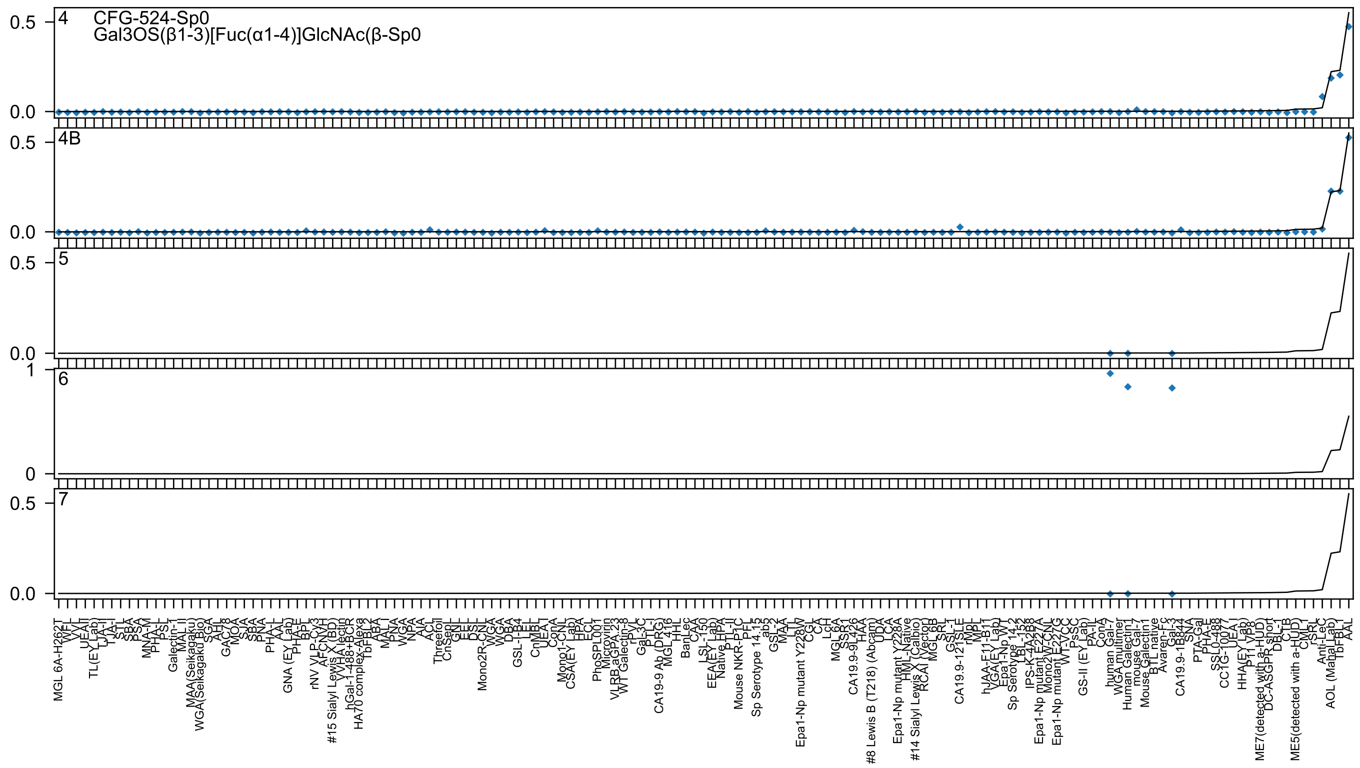

CFG-524-Sp0
Gal3OS(β1-3)[Fuc(α1-4)]GlcNAc(β-Sp0

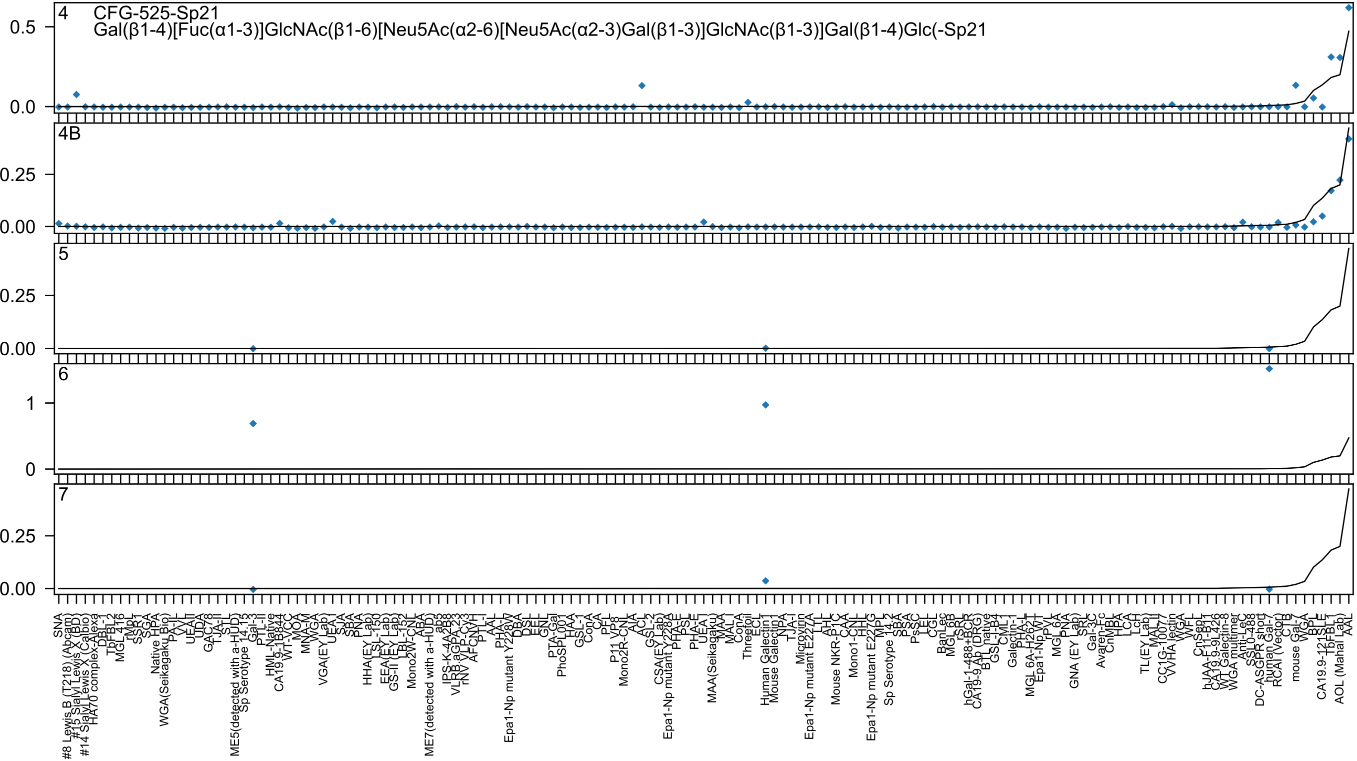
4
CFG-525-Sp21
Gal(β1-4)[Fuc(α1-3)]GlcNAc(β1-6)[Neu5Ac(α2-6)[Neu5Ac(α2-3)Gal(β1-3)]GlcNAc(β1-3)]Gal(β1-4)Glc(-Sp21
4B
5
6
7

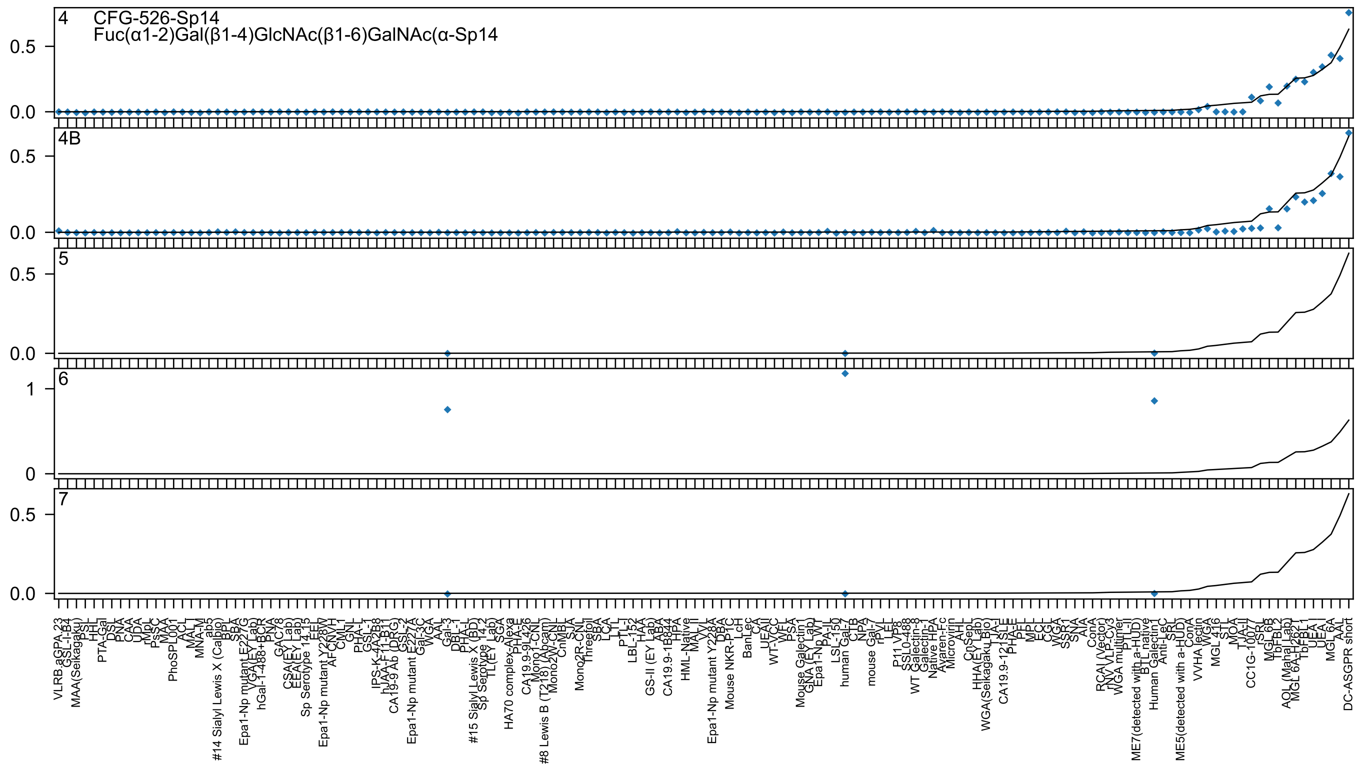

4 CFG-526-Sp14
Fuc(α1-2)Gal(β1-4)GlcNAc(β1-6)GalNAc(α-Sp14
4B
5
6
7

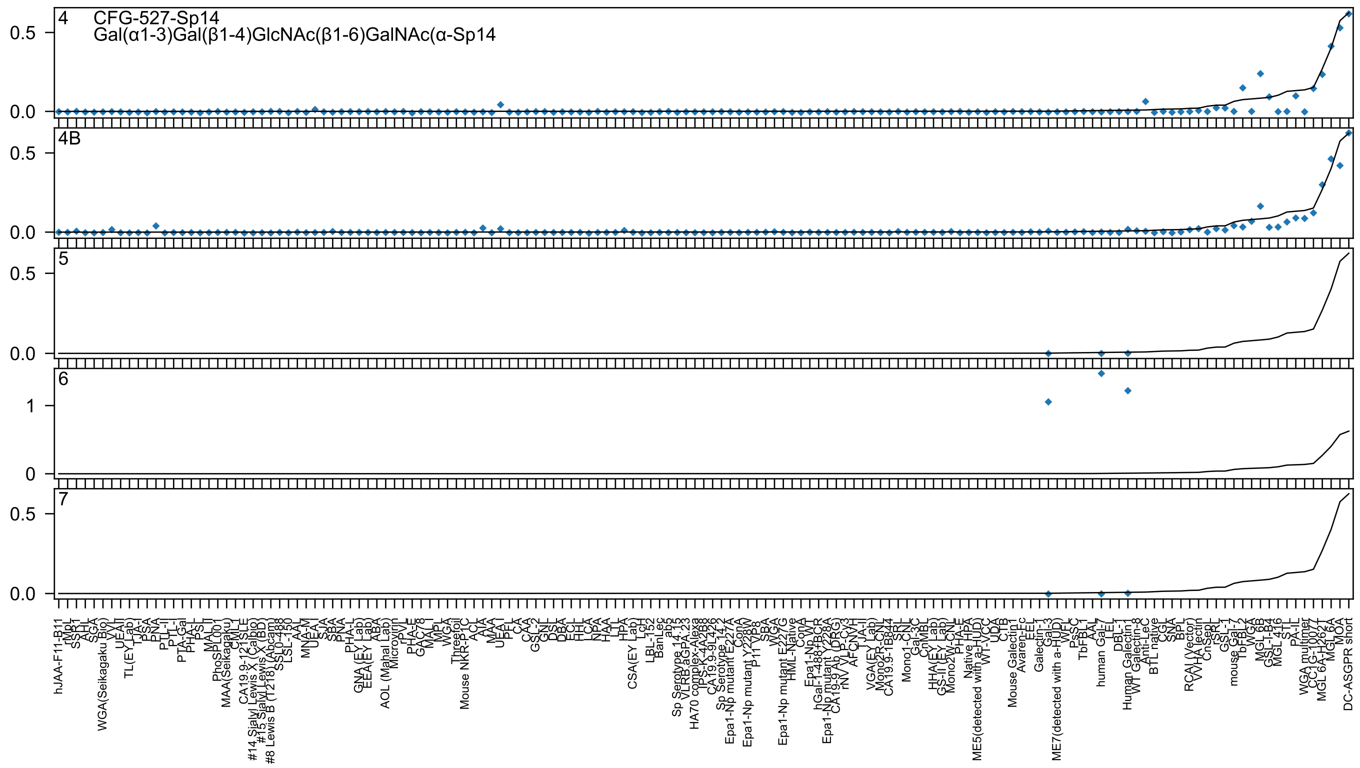
4
CFG-527-Sp14
Gal(α1-3)Gal(β1-4)GlcNAc(β1-6)GalNAc(α-Sp14
4B
5
6
7
0.5
0.0
1
0
hJAA-F11-B11
rMpL
SSR1
AHL
SGA
WGA(Seikagaku Bio)
VVL
UEA-II
TL(EY Lab)
TJA-I
PSA
PNA
PTL-II
PTL-I
PTA-Gal
PHA-L
PSL
MAL-II
PhoSL
MAA(Seikagaku)
CML
CA19.9-121SLE
#14 Sialyl Lewis X (Calbio)
#15 Sialyl Lewis X (BD)
#8 Lewis B (T218) (Abcam)
LSL-150
AAL
MNA-M
UEA-I
SJA
SBA
PNA
PHA-L
GNA(EY Lab)
EEA(EY Lab)
ABA
AOL (Mahal Lab)
Microvirin
rPVL
PHA-E
MAL-I
MPL
WGA
Threefoil
Mouse NKR-P1C
ACL
AIA
MAA
UEA-I
CA
CAA
GSL-2
GNL
DSL
DBA
ECL
HHL
LCA
NPA
HAA
LTL
HPA
CSA(EY Lab)
LcH
LBL-152
BanLec
ab5
Sp Serotype 14.15
VLRB aGPA.23
HA70 complex-Alexa
CA19.9-9L426
Sp Serotype 14.2
Epa1-Np mutant E227A
ConA
Epa1-Np mutant Y228W
P11 VP8
SBA
WGA
Epa1-Np mutant E227G
HML-Native
ConA
Epa1-Np WT
hGal-1-488+BCR
Epa1-Np mutant Y228A
CA19-9 Ab (DRG)
rNV VLP-CV3
AFCNVH
TJA-II
VGA(EY Lab)
Mono2R-CNL
CA19.9-1B844
SRL
Mono1-CNL
Gal-3C
CnMBL
HHA(EY Lab)
GS-II (EY Lab)
Mono2W-CNL
PHA-E
Native HPA
ME5(detected with a-HUD)
WT-VCC
UDA
CTB
Mouse Galectin1
Avaren-Fc
EEL
Galectin-1
Gal-3
ME7(detected with a-HUD)
WFL
TbFBL1
AAL
human Gal-7
LEL
DBL-1
Human Galectin-1
WT Galectin-8
Anti-LeC
BTL native
CGL
SNA
BPL
RCAI (Vector)
VVHA lectin
CnSepL
rSRL
GSL-1
mouse Gal-7
TbFBL2
WGA
MGL 6B
GSL-I-B4
MGL 416
STL
PA-IL
WGA multimer
CC1G-10077
MGL 6A-H262T
MGL 6A
MOA
DC-ASGPR short

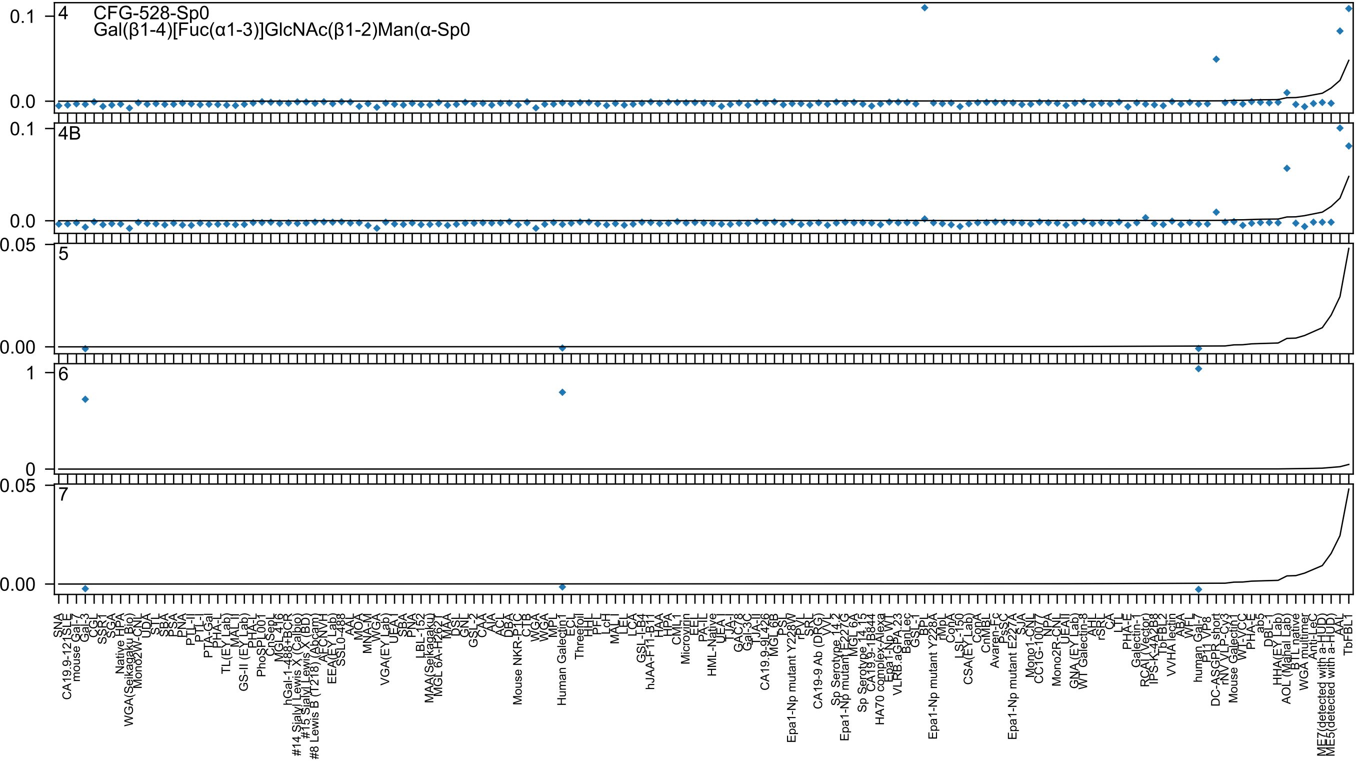

4
CFG-528-Sp0
Gal(β1-4)[Fuc(α1-3)]GlcNAc(β1-2)Man(α-Sp0
4B
5
6
7

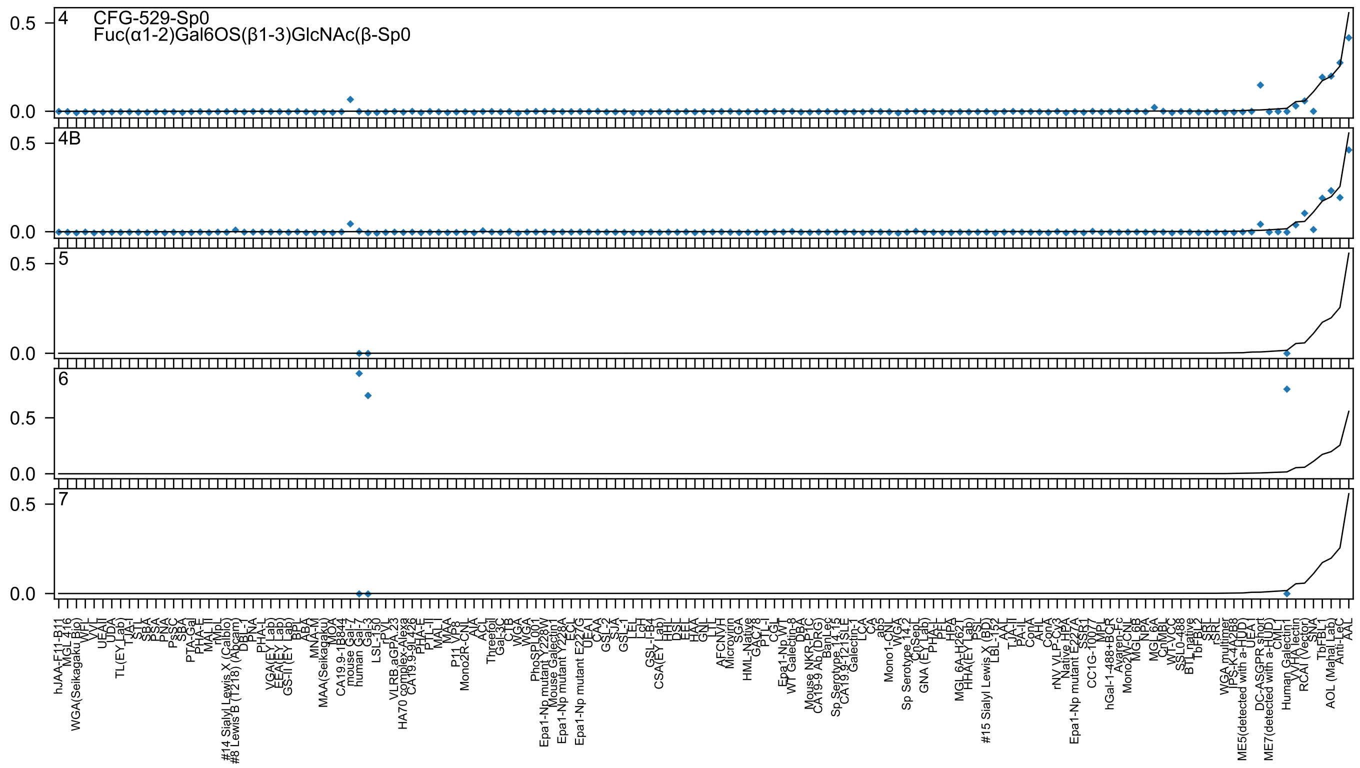

4 CFG-529-Sp0
Fuc(α1-2)Gal6OS(β1-3)GlcNAc(β-Sp0
4B
5
6
7

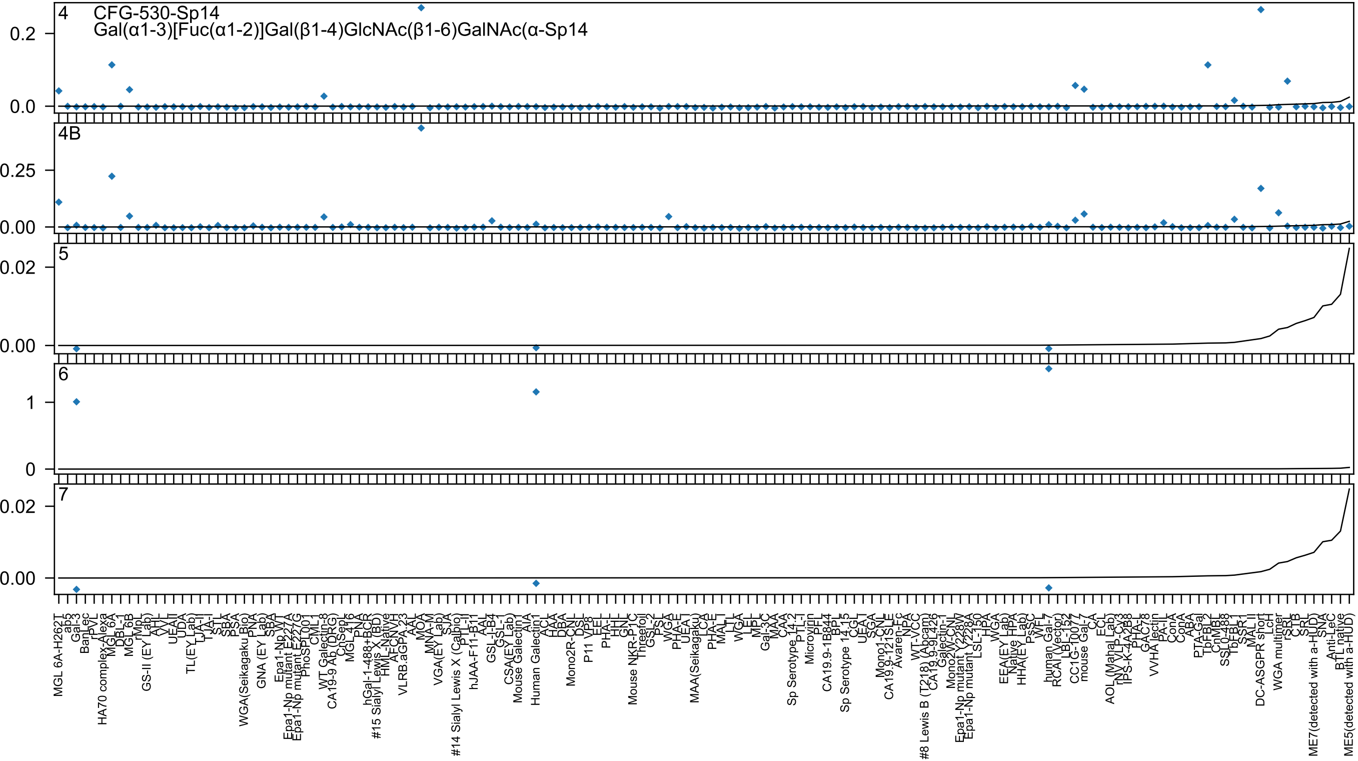
4 CFG-530-Sp14
Gal(α1-3)[Fuc(α1-2)]Gal(β1-4)GlcNAc(β1-6)GalNAc(α-Sp14
0.2
0.0
4B
0.25
0.00
5
0.02
0.00
6
1
0
7
0.02
0.00
MGL 6A-H262T
ab5
Gal-3
BanLec
rPVL
HA70 complex-Alexa
MGL 6A
DBL-1
MGL 6B
rMpL
GS-II (EY Lab)
AHL
UEA-II
UDA
TL(EY Lab)
LTA-I
TJA-I
STL
SBA
PSA
WGA(Seikagaku Bio)
PNA
GNA (EY Lab)
SBA
Epa1-Np WT
Epa1-Np mutant E227A
Epa1-Np mutant E227G
PhoSL001
CML1
WT Galectin-8
CA19-9 Ab (DRG)
ChSep
MGL 416
PNA
hGal-1-488+BCR
#15 Sialyl Lewis X (BD)
HML-Native
AFCNVH
VLRB.aGPA.23
AAL
MOA
MNA-M
VGA(EY Lab)
SJA
#14 Sialyl Lewis X (Calbio)
PTL-II
hJAA-F11-B11
AAL
GSL-I-B4
GSL-I
CSA(EY Lab)
Mouse Galectin1
AIA
Human Galectin1
ACL
HAA
DBA
Mono2R-CNL
DSL
P11 VP8
EEL
PHA-L
HHL
GNL
Mouse NKR-P1C
Threefoil
GSL-2
PSL
WGA
PHA-E
UEA-I
MAA(Seikagaku)
LCA
PHA-E
MAL-I
LTL
WGA
LEL
MPL
Gal-3C
MAA
CAA
Sp Serotype 14.2
PTL-I
Microvirin
PFL
CA19.9-1B844
BPL
Sp Serotype 14.15
CGL
UEA-I
SGA
Mono1-CNL
CA19.9-121SLE
Avaren-Fc
NPA
WT-VCC
#8 Lewis B (T218) (Abcam)
CA19.9-9L426
Galectin-1
Mono2W-CNL
Epa1-Np mutant Y228W
Epa1-Np mutant Y228A
LSL-150
HPA
WGA
EEA(EY Lab)
Native HPA
HHA(EY Lab)
PsSC
WFL
human Gal-7
RCAI (Vector)
LBL-152
CC1G-1007
mouse Gal-7
CA
ECL
AOL (Mahal Lab)
rNV VLP-CV3
IPS-K-4A2B8
PHA-L
GAC78
VVHA lectin
PA-IL
ConA
ConA
ABA
PTA-Gal
TbFBL2
CnMBL
SSL0-488
TbFBL1
SSR1
MAL II
DC-ASGPR short
LcH
WGA multimer
rSRL
CTB
SRL
SNA
Anti-LeC
BTL native
ME7(detected with a-HUD)
ME5(detected with a-HUD)

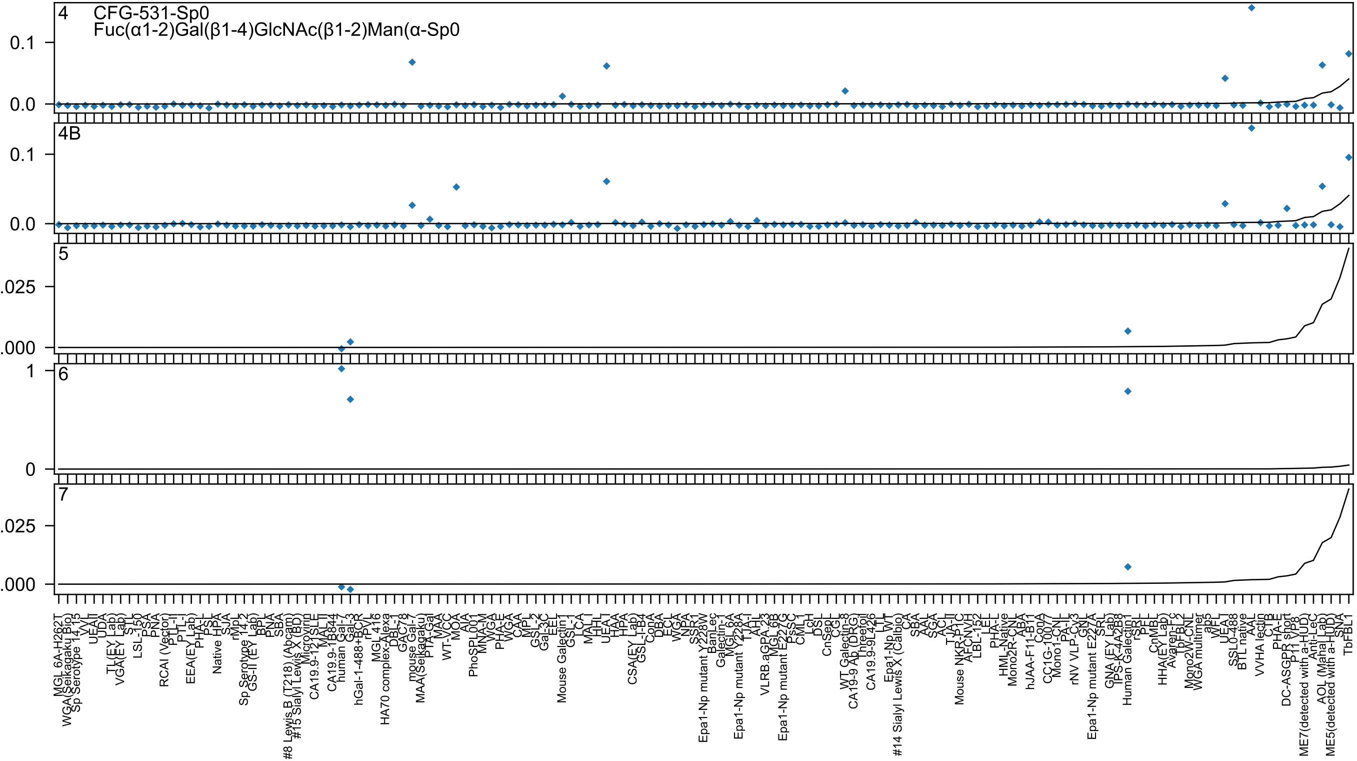

4 CFG-531-Sp0
Fuc(α1-2)Gal(β1-4)GlcNAc(β1-2)Man(α-Sp0
4B
5
6
7

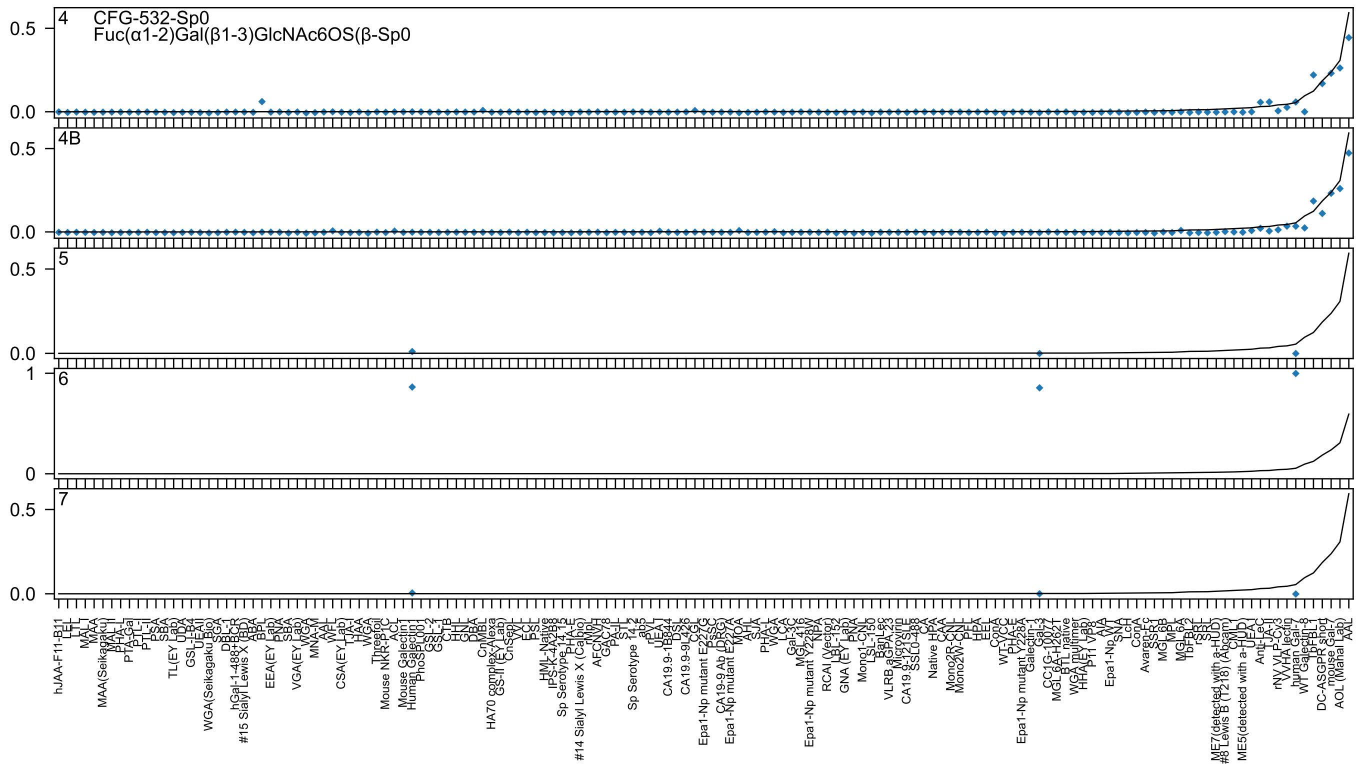

4 CFG-532-Sp0
Fuc(α1-2)Gal(β1-3)GlcNAc6OS(β-Sp0
4B
5
6
7
0.5
0.0
1
0
hJAA-F11-B11
AAL
AOL (Manal Lab)
mouse Gal-7
DC-ASGPR short
WT Galectin-8
human Gal-7

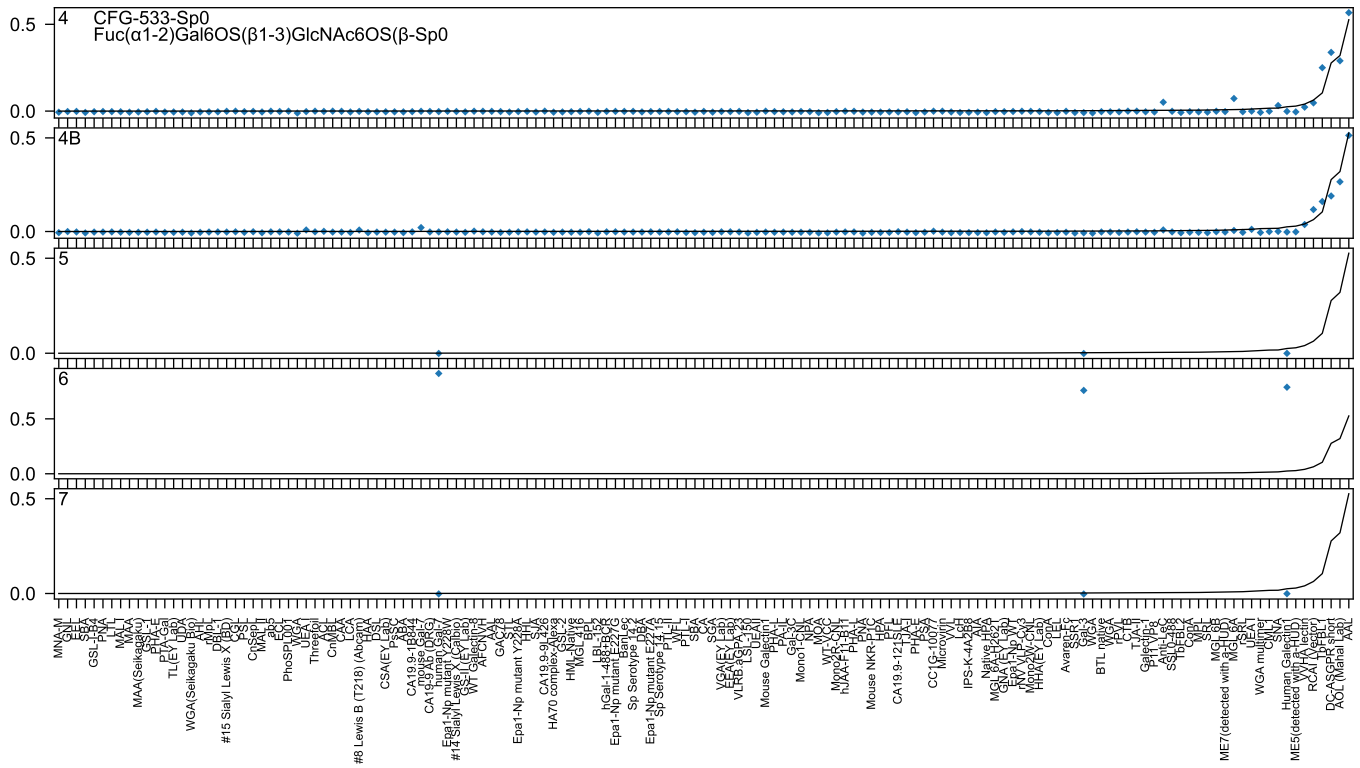
4
CFG-533-Sp0
Fuc(α1-2)Gal6OS(β1-3)GlcNAc6OS(β-Sp0
4B
5
6
7
0.5
0.0

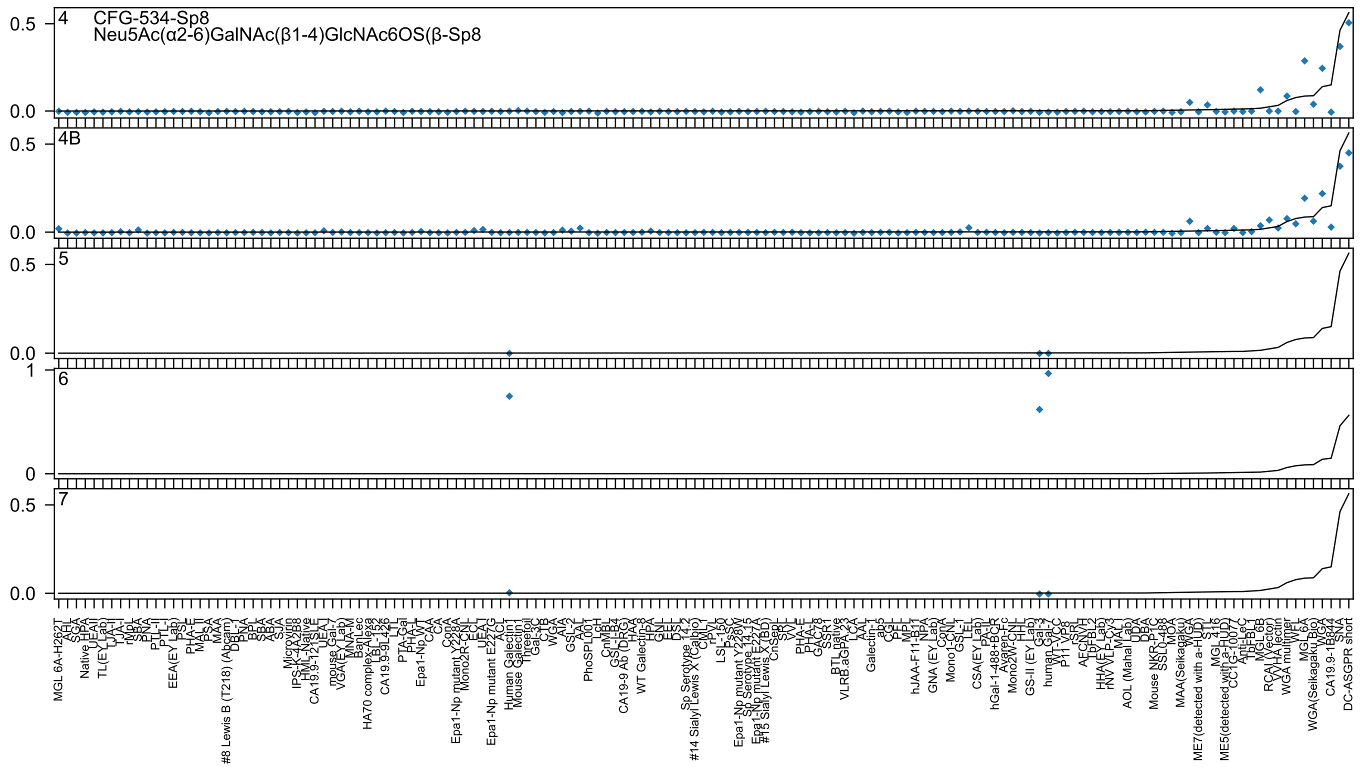
4
CFG-534-Sp8
Neu5Ac(α2-6)GalNAc(β1-4)GlcNAc6OS(β-Sp8
4B
5
6
7
0.5
0.0
1
0
MGL 6A-H262T
AHL
SGA
Native HPA
UEAII
TL(EY Lab)
LJA-II
TJA-I
rMpL
SBA
PNA
PTL-II
PTL-I
EEA(EY Lab)
PSL
PHA-E
MAL-II
PSA
MAA
#8 Lewis B (T218) (Abcam)
DBL-1
PNA
BPL
SBA
ABA
SJA
Microvirin
IPS-K-4A2B8
HML-Native
CA19.9-121SLE
UEA-I
mouse Gal-7
VGA(EY Lab)
MNA-M
BanLec
HA70 complex-Alexa
LBL-136
CA19.9-9L426
LTL
PTA-Gal
PHA-L
Epa1-Np WT
CAA
CA
ConA
Epa1-Np mutant Y228A
Mono2R-CNL
ECL
UEA-I
Epa1-Np mutant E227G
ACL
Human Galectin1
Mouse Galectin1
Threefoil
Gal-3C
CTB
WGA
AIA
GSL-2
AAL
PhoSPL001
LcH
CnMBL
GSL-I-B4
CA19-9 Ab (DRG)
HAA
WT Galectin-8
HPA
GNL
EEL
DSL
Sp Serotype 14_2
#14 Sialyl Lewis X (Calbio)
CML-I
rPVL
LSL-150
PsSC
Epa1-Np mutant Y228W
Sp Serotype 14_15
Epa1-Np mutant E227A
#15 Sialyl Lewis X (BD)
CnSep1
SRL
VVL
PHA-E
PHA-L
GAC78
SSR1
BTL native
VLRB.aGPA.23
LCA
AAL
Galectin-1
ab5
CGL
PFL
MPL
hJAA-F11-B1
NPA
GNA (EY Lab)
ConA
Mono1-CNL
GSL-I
LEL
CSA(EY Lab)
PA-IL
hGal-1-488+BCR
Avaren-Fc
Mono2W-CNL
HHL
GS-II (EY Lab)
Gal-3
human Gal-7
WT-VCC
P11 VP8
rSRL
AFCNVH
TbFBL2
HHA(EY Lab)
rNV VLP-Cy3
MAL-I
AOL (Mahal Lab)
UDA
DBA
Mouse NKR-P1C
SSL0-488
MOA
MAA(Seikagaku)
WGA
ME7(detected with a-HUD)
STL
MGL 416
ME5(detected with a-HUD)
CC1G-10077
Anti-LeC
TbFBL1
MGL 6B
RCAI (Vector)
VVHA lectin
WGA multimer
WFL
MGL 6A
WGA(Seikagaku Bio)
WGA
CA19.9-1B844
SNA
DC-ASGPR short

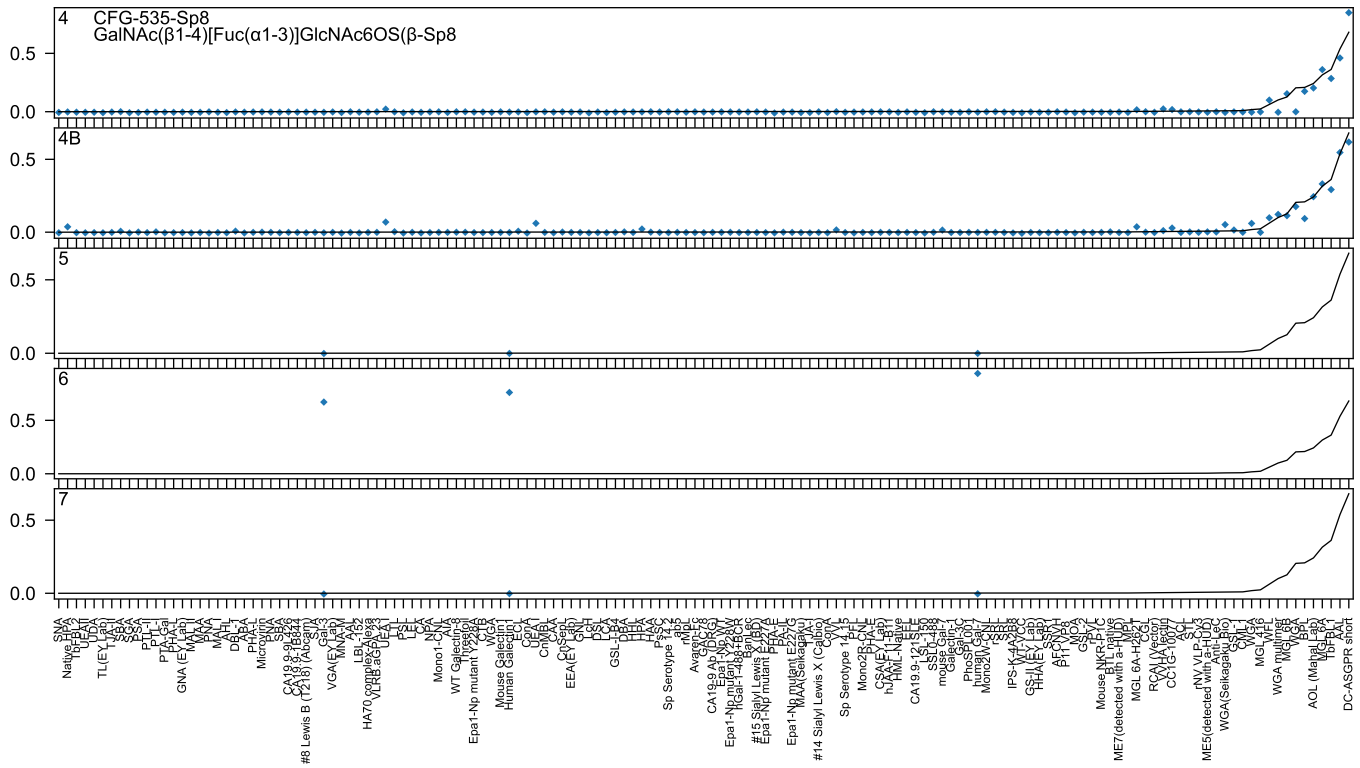

4 CFG-535-Sp8
GalNAc(β1-4)[Fuc(α1-3)]GlcNAc6OS(β-Sp8
4B
5
6
7
0.5
0.0

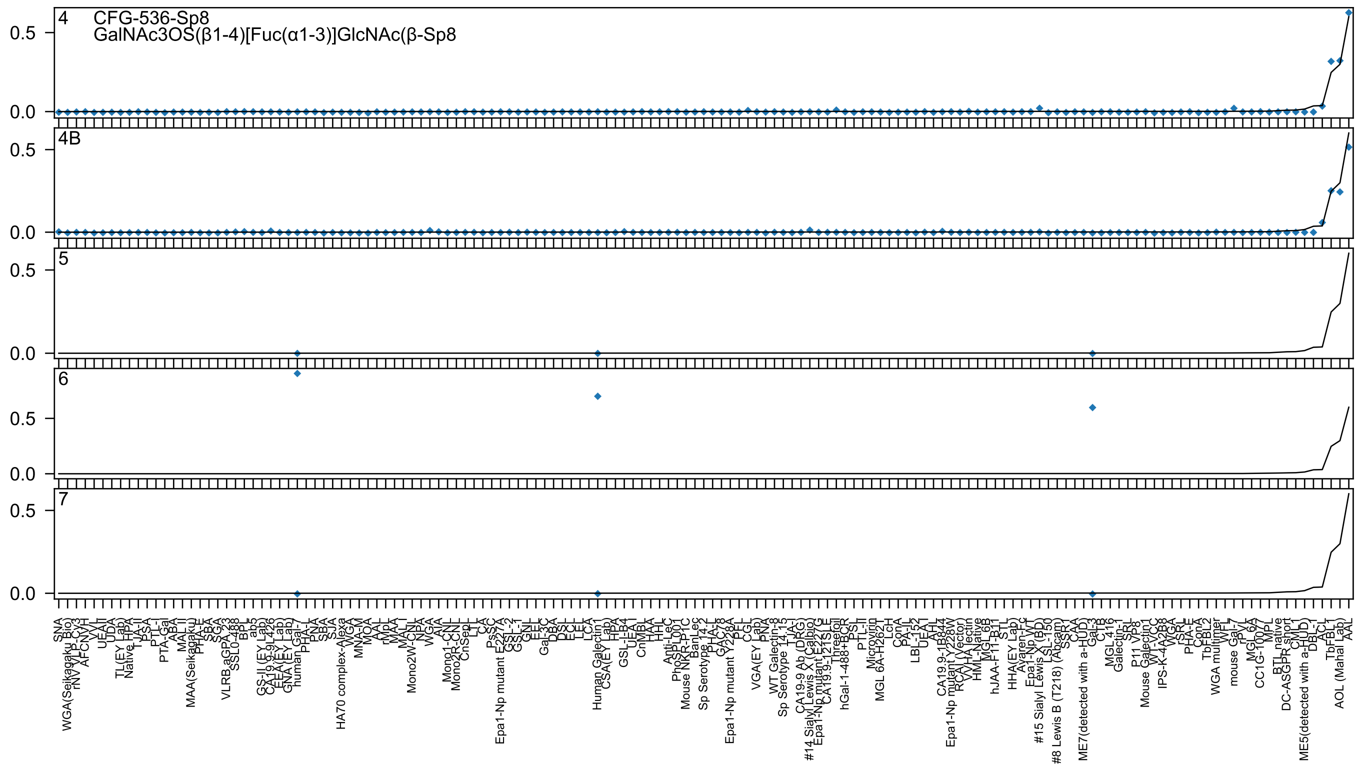
4
CFG-536-Sp8
GalNAc3OS(β1-4)[Fuc(α1-3)]GlcNAc(β-Sp8
4B
5
6
7

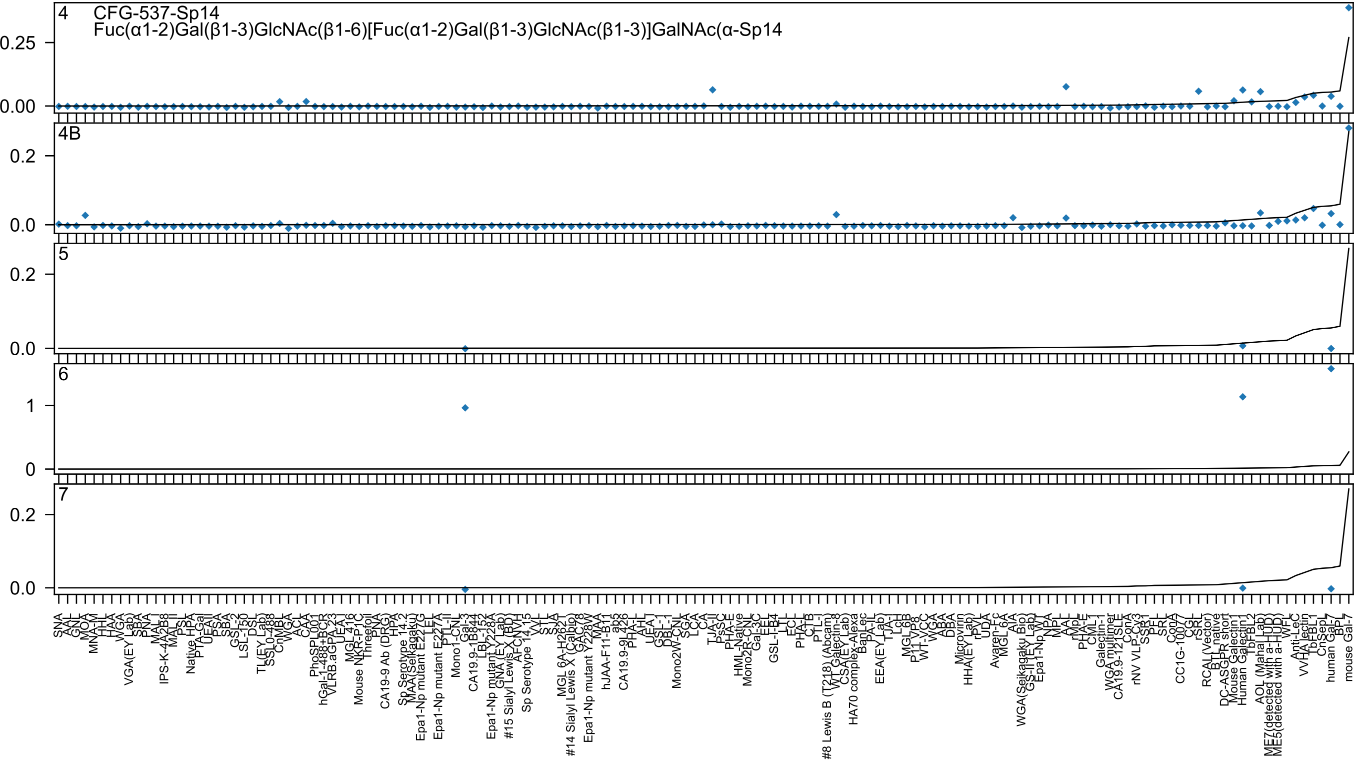

4 CFG-537-Sp14
Fuc(α1-2)Gal(β1-3)GlcNAc(β1-6)[Fuc(α1-2)Gal(β1-3)GlcNAc(β1-3)]GalNAc(α-Sp14
4B
5
6
7

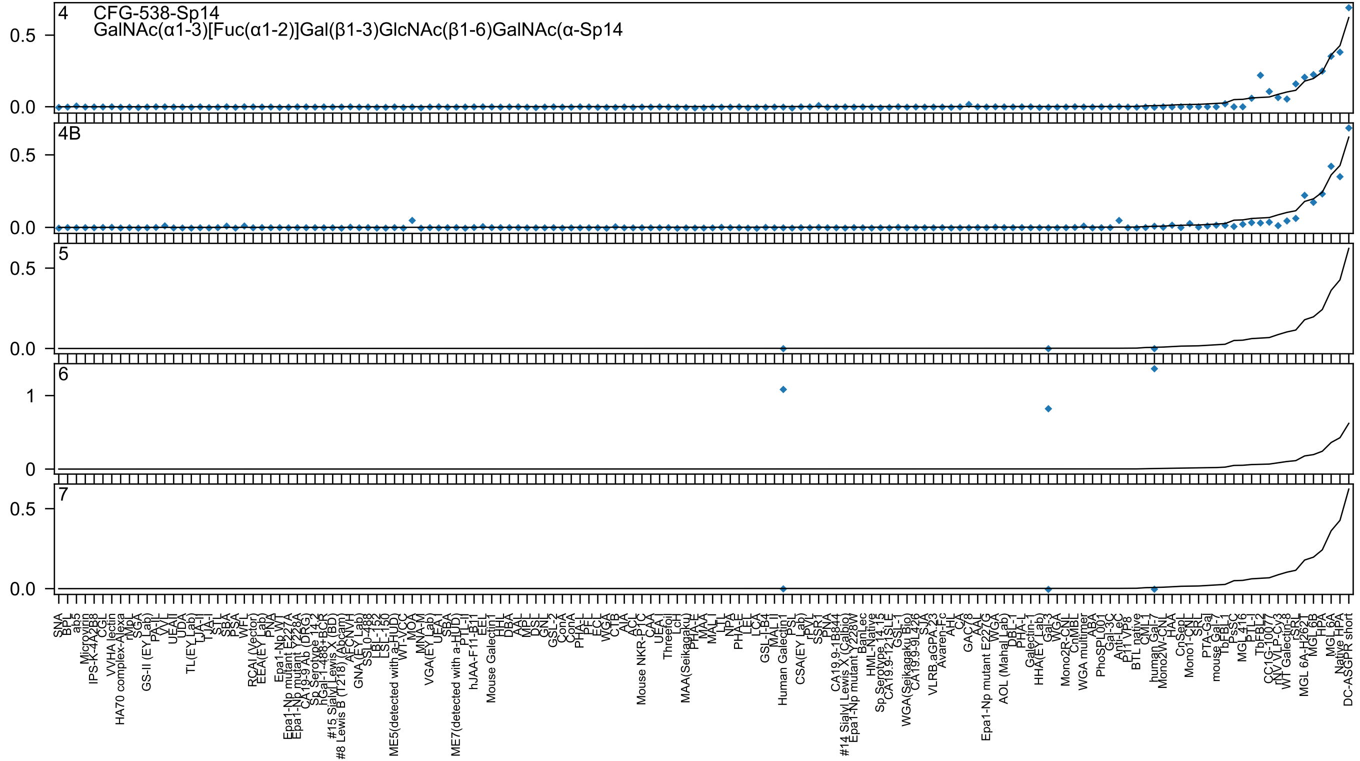
4
CFG-538-Sp14
GalNAc(α1-3)[Fuc(α1-2)]Gal(β1-3)GlcNAc(β1-6)GalNAc(α-Sp14
4B
5
6
7

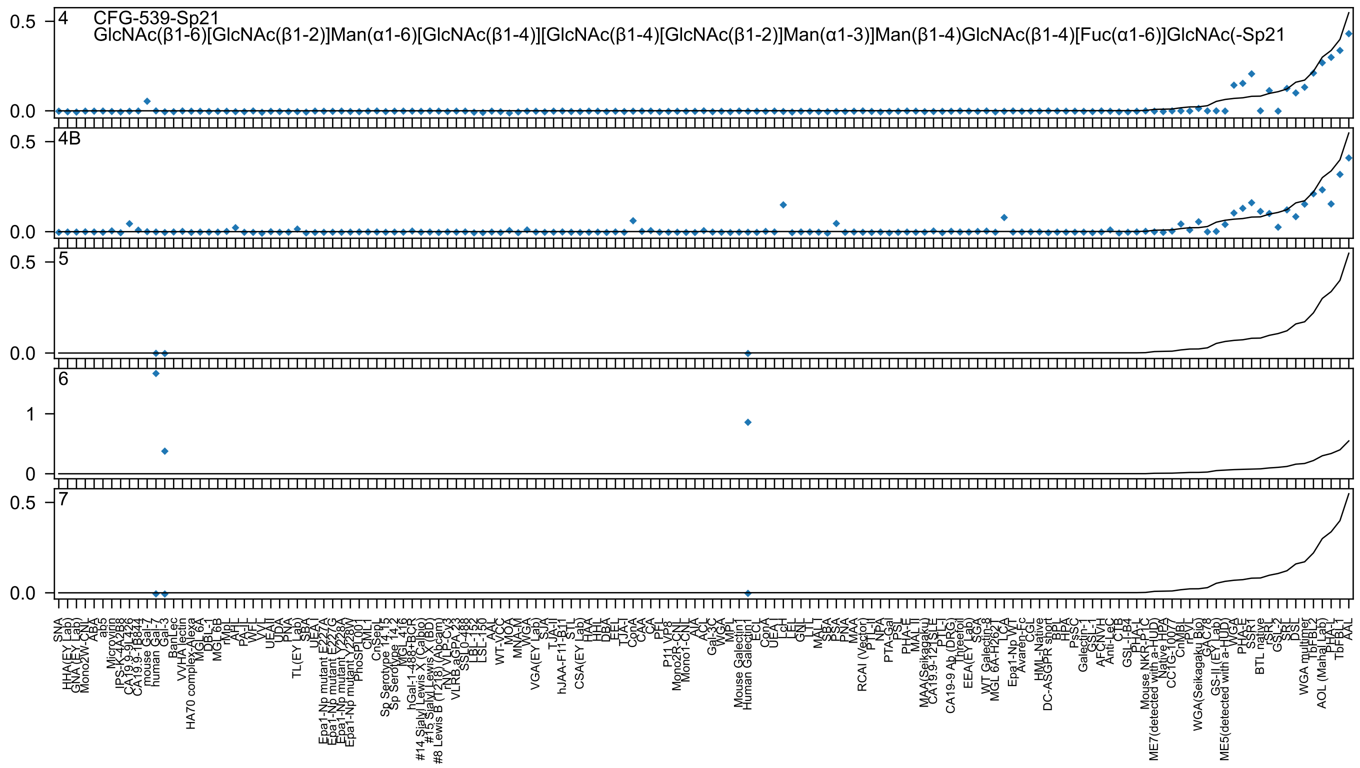

CFG-539-Sp21
GlcNAc(β1-6)[GlcNAc(β1-2)]Man(α1-6)[GlcNAc(β1-4)][GlcNAc(β1-4)[GlcNAc(β1-2)]Man(α1-3)]Man(β1-4)GlcNAc(β1-4)[Fuc(α1-6)]GlcNAc(-Sp21
4
4B
5
6
7
0.0
0.5
0
1
SNA
HHA(EY Lab)
GNA(EY Lab)
Mono2W-CNL
ABA
ab5
Microvirin
IPS-K-4A2B8
CA19.9-9L426
CA19.9-1B844
mouse Gal-7
human Gal-7
Gal-3
BanLec
VVHA lectin
HA70 complex-Alexa
MGL 6A
DBL-1
MGL 6B
rMpL
AHL
PA-IL
WFL
VVL
UEAII
UDA
PNA
TL(EY Lab)
SBA
UEAI
Epa1-Np mutant E227A
Epa1-Np mutant E227G
Epa1-Np mutant Y228A
Epa1-Np mutant Y228W
PhoSL001
CML1
CnSepL
Sp Serotype 14.15
Sp Serotype 14.2
MGL 416
hGal-1-488+BCR
#14 Sialyl Lewis X (Calbio)
#15 Sialyl Lewis X (BD)
#8 Lewis B (T218) (Abcam)
rNY VLP-Cy3
VLRB aGPA.23
SSL0-488
LBL-152
LSL-150
AAL
WT-VCC
MOA
MNA-M
WGA
VGA(EY Lab)
SJA
TJA-II
hJAA-F11-B11
STL
CSA(EY Lab)
HAA
HHL
DBA
EEL
TJA-I
ConA
CAA
CA
PFL
P11 VP8
Mono2R-CNL
Mono1-CNL
AIA
ACL
Gal-3C
WGA
MPL
Mouse Galectin1
Human Galectin1
ECL
ConA
UEA1
LcH
LEL
GNL
LTL
MAL I
SBA
PSA
PNA
MAA
RCAI (Vector)
PTL-II
NPA
PTA-Gal
PSL
PHA-E
MAL II
MAA(Seikagaku)
CA19.9-121SLE
PTL-I
CA19-9 Ab (DRG)
Threefoil
EEA(EY Lab)
SGA
WT Galectin-8
MGL 6A-H262
LCA
Epa1-Np WT
Avaren-Fc
CGL
HML-Native
DC-ASGPR short
BPL
HPA
PsSC
Galectin-1
GSL-1
AFCNVH
Anti-LeC
CTB
GSL-I-B4
PHA-L
Mouse NKR-P1C
ME7(detected with a-HUD)
Native HPA
CC1G-1007
CnMBL
rPVL
WGA(Seikagaku Bio)
GAC78
GS-II (EY Lab)
ME5(detected with a-HUD)
WGA
PHA-E
SSR1
BTL native
rSRL
GSL-2
SRL
DSL
WGA multimer
TbFBL2
AOL (Mahal Lab)
PHA-L
TbFBL1
AAL

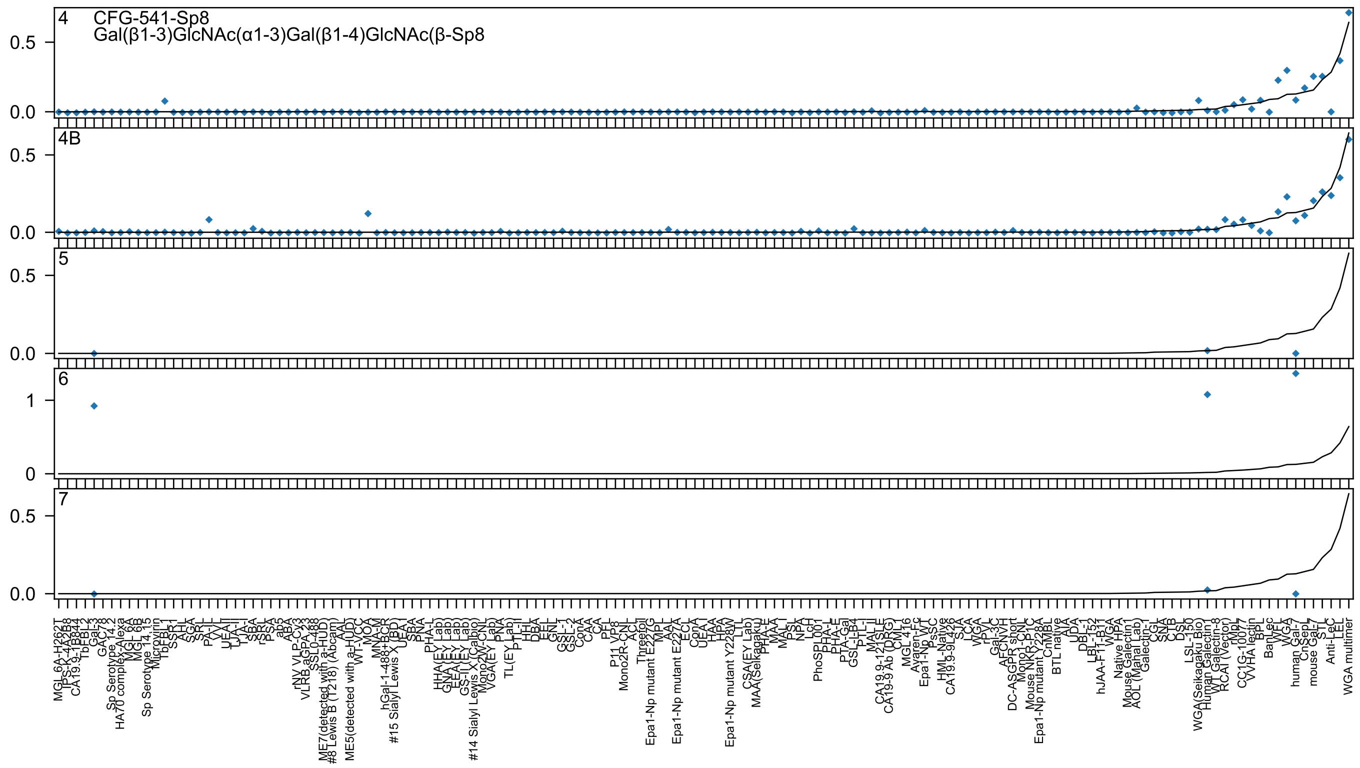
4 CFG-541-Sp8
Gal(β1-3)GlcNAc(α1-3)Gal(β1-4)GlcNAc(β-Sp8
4B
5
6
7
0.5
0.0
1
0
WGA multimer
LEL
Anti-LeC
STL
mouse Gal-7
CnSepL
human Gal-7
WGA
WFL
BanLec
BPL
VVHA lectin
CC1G-10077
rMpL
RCAI (Vector)
WT Galectin-8
Human Galectin1
WGA(Seikagaku Bio)
LSL-150
MGL_6A-H262T
Gal-3

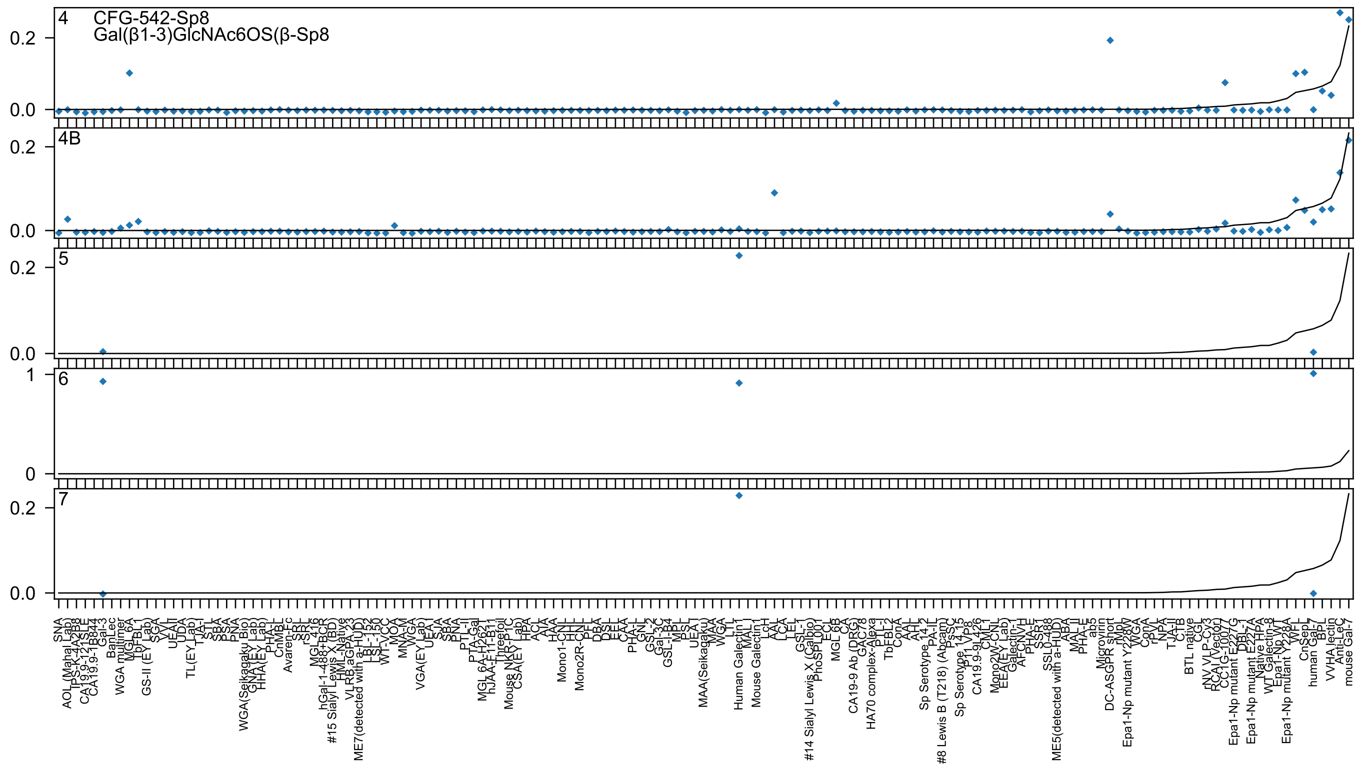
4
CFG-542-Sp8
Gal(β1-3)GlcNAc6OS(β-Sp8
4B
5
6
7
0.2
0.0
1
0

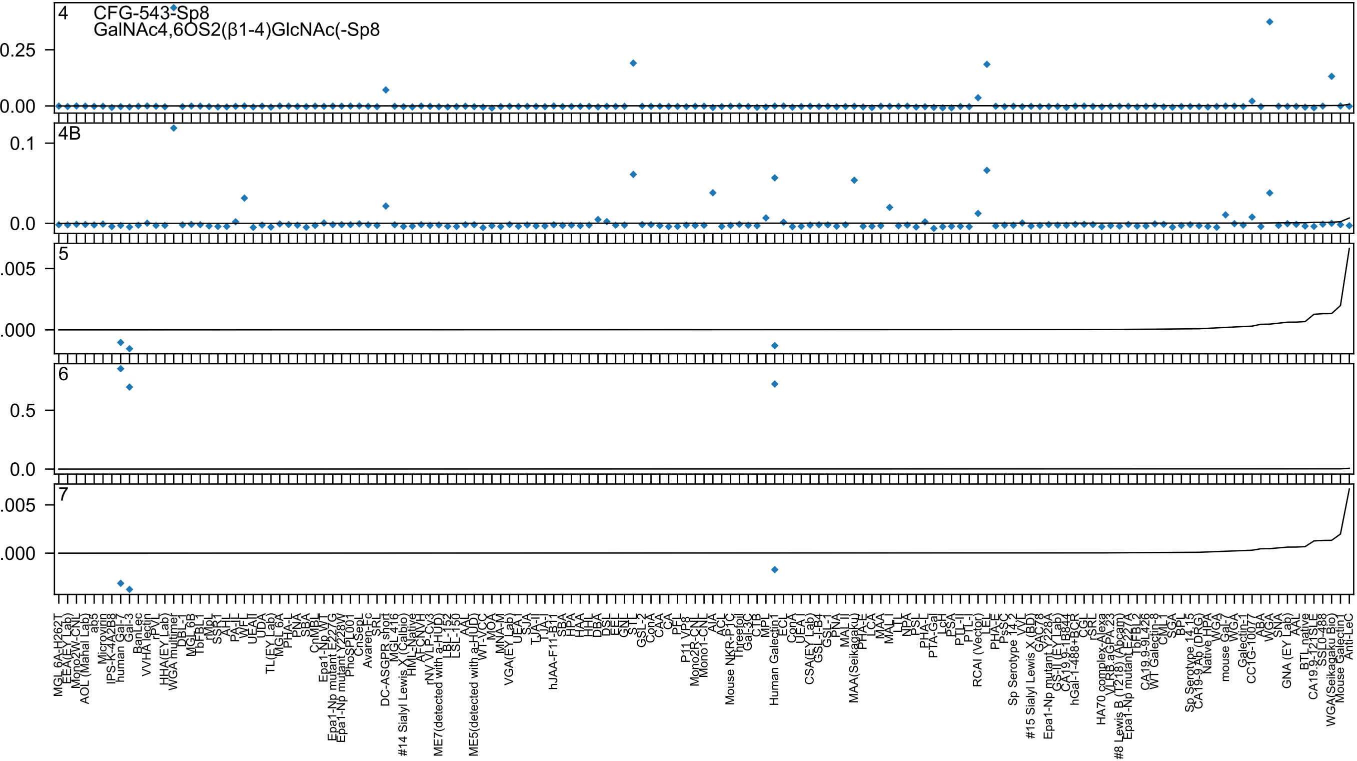

CFG-543-Sp8
GalNAc4,6OS2(β1-4)GlcNAc(-Sp8
4
4B
5
6
7

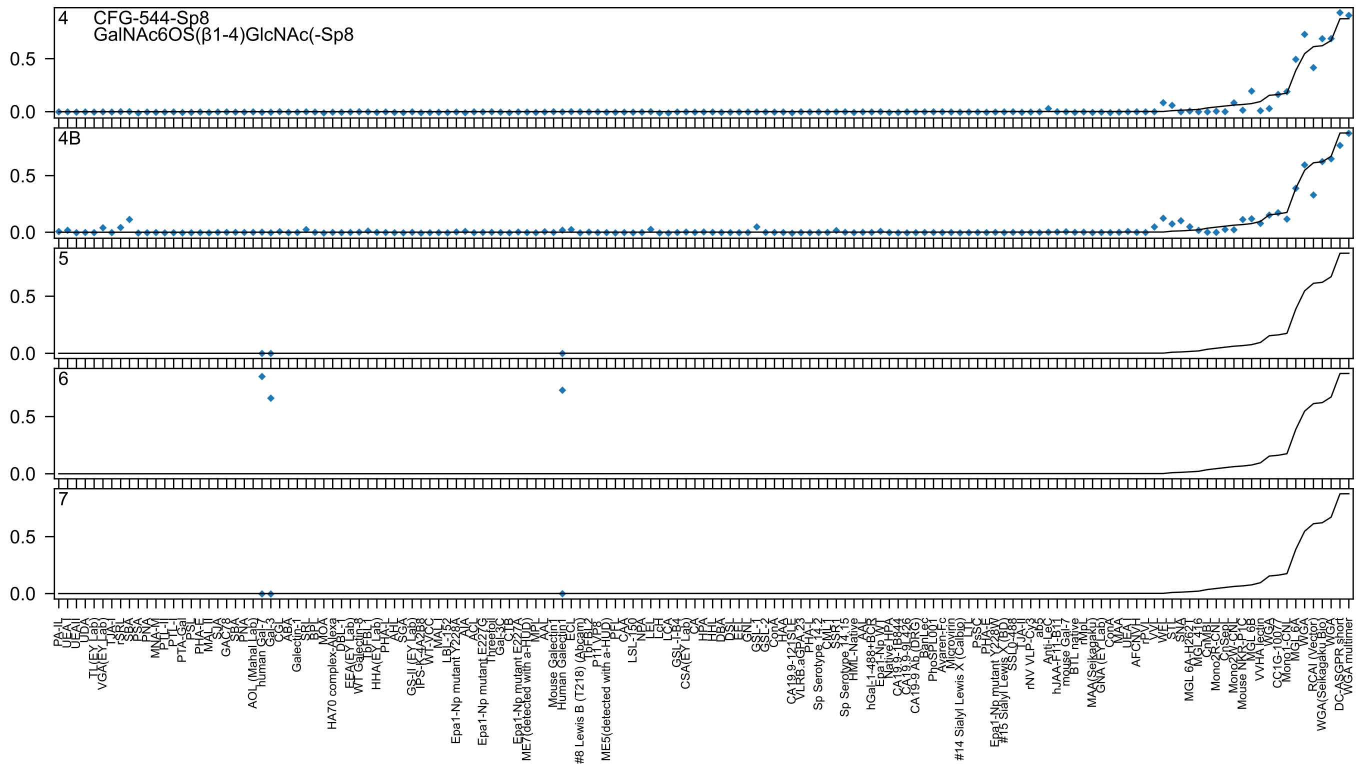

4
CFG-544-Sp8
GalNAc6OS(β1-4)GlcNAc(-Sp8
4B
5
6
7
0.0
0.5

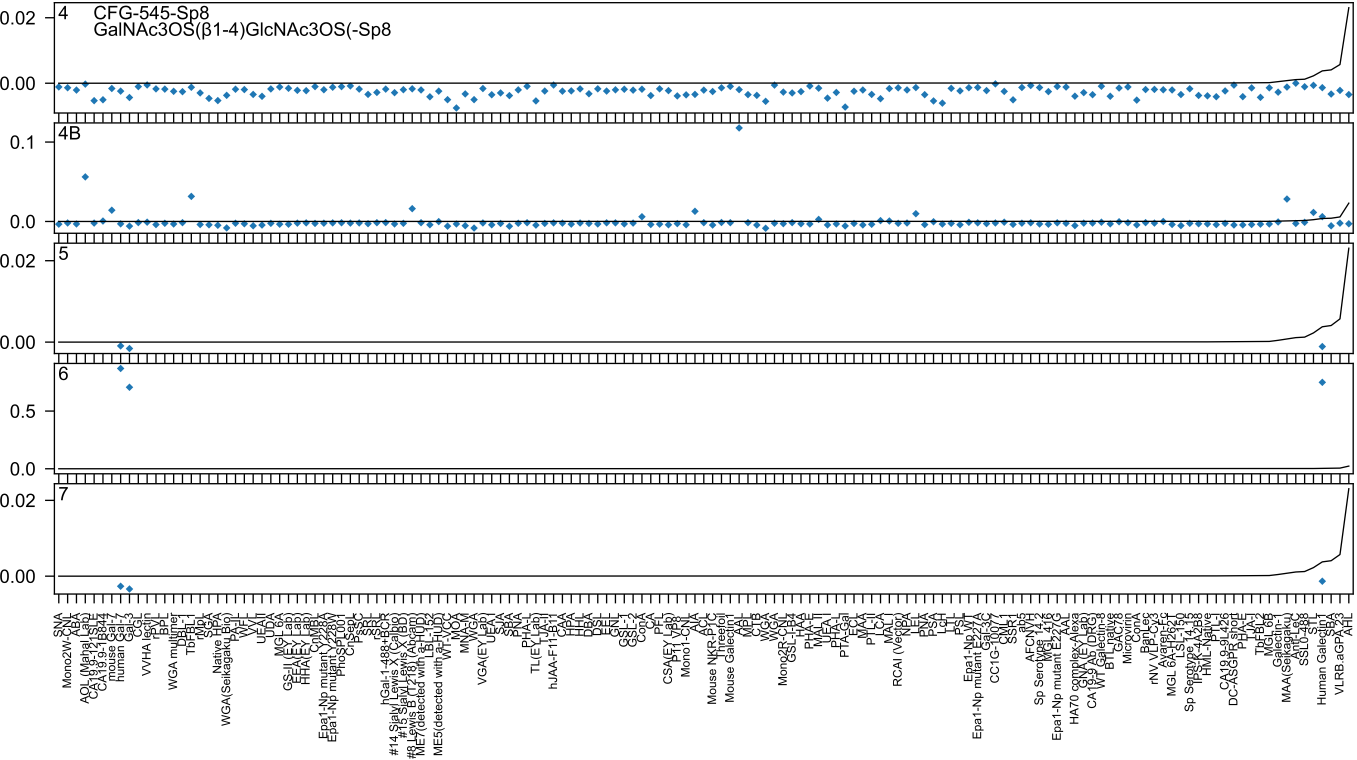

4
CFG-545-Sp8
GalNAc3OS(β1-4)GlcNAc3OS(-Sp8
0.02
0.00
4B
0.1
0.0
5
0.02
0.00
6
0.5
0.0
7
0.02
0.00

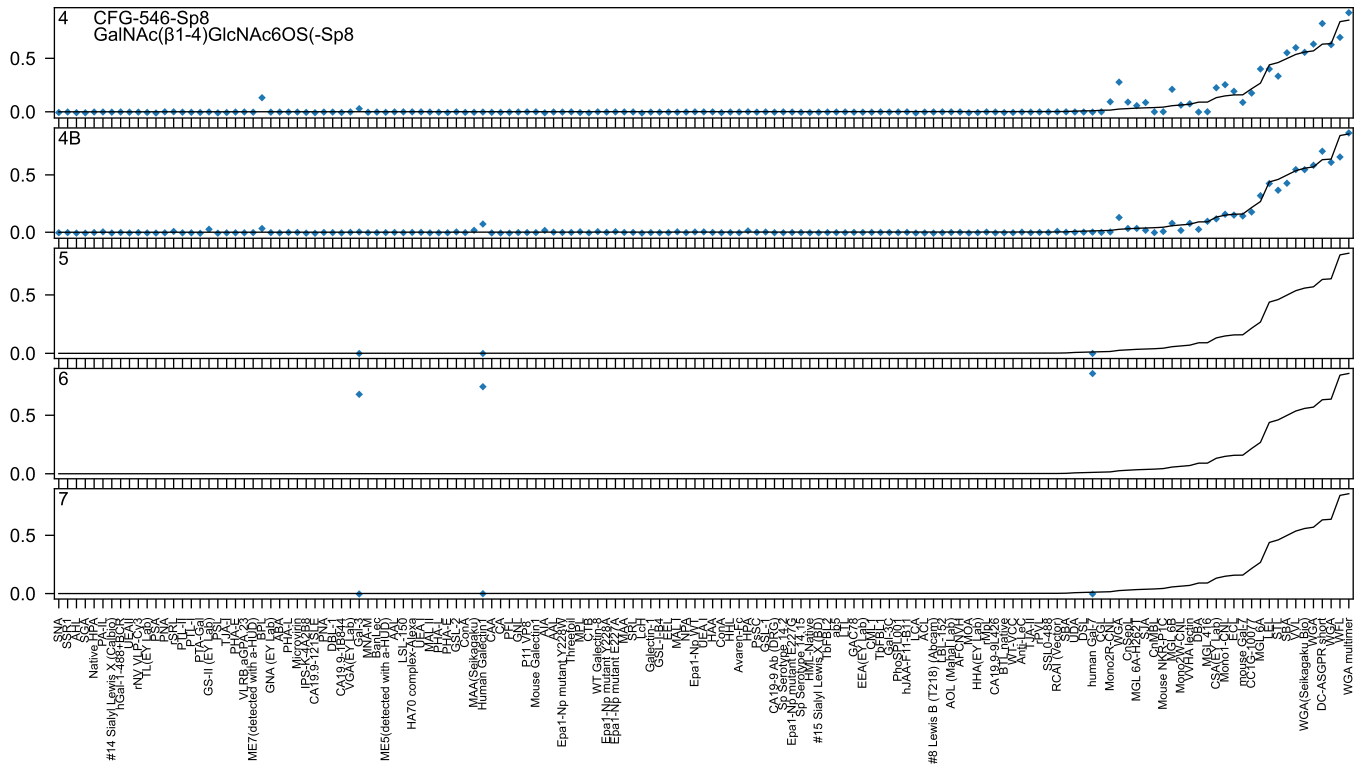

4
CFG-546-Sp8
GalNAc(β1-4)GlcNAc6OS(-Sp8
4B
5
6
7
0.5
0.0

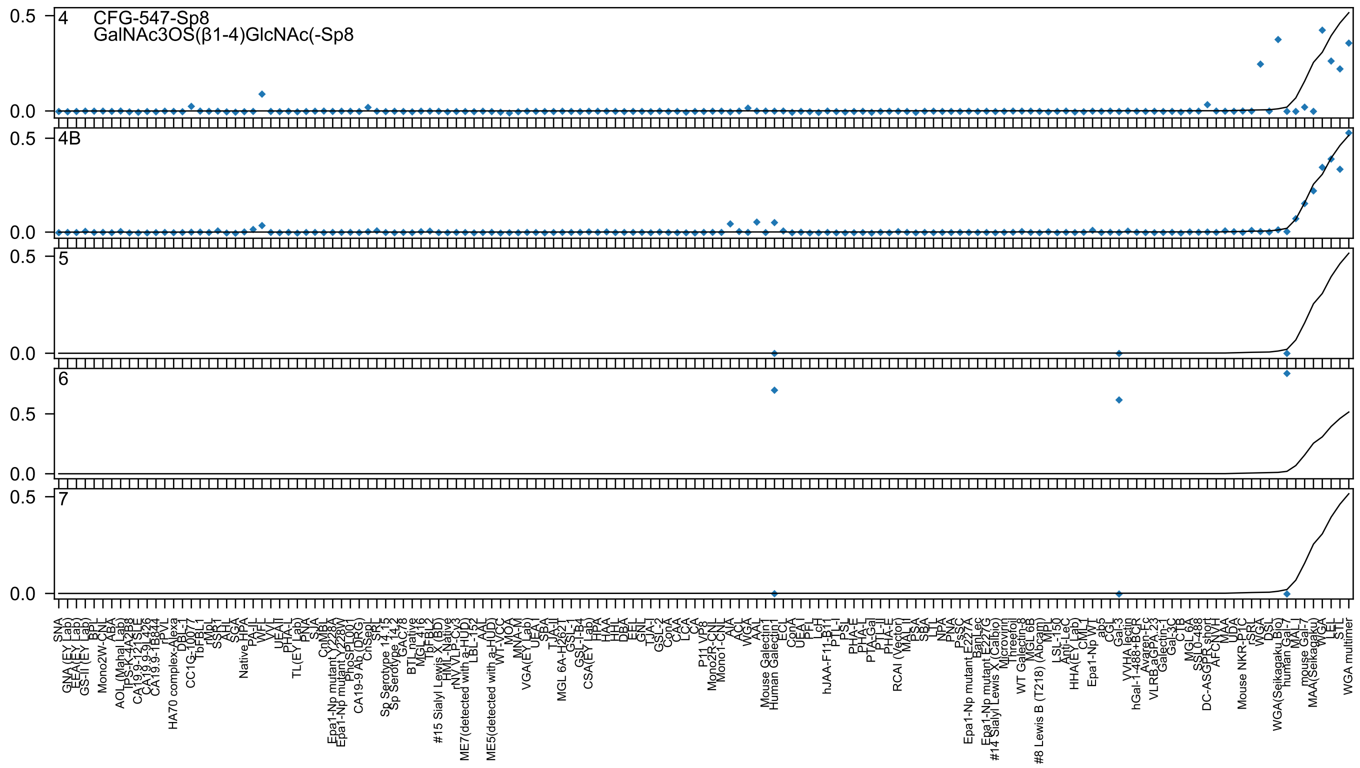
4
CFG-547-Sp8
GalNAc3OS(β1-4)GlcNAc(-Sp8
4B
5
6
7

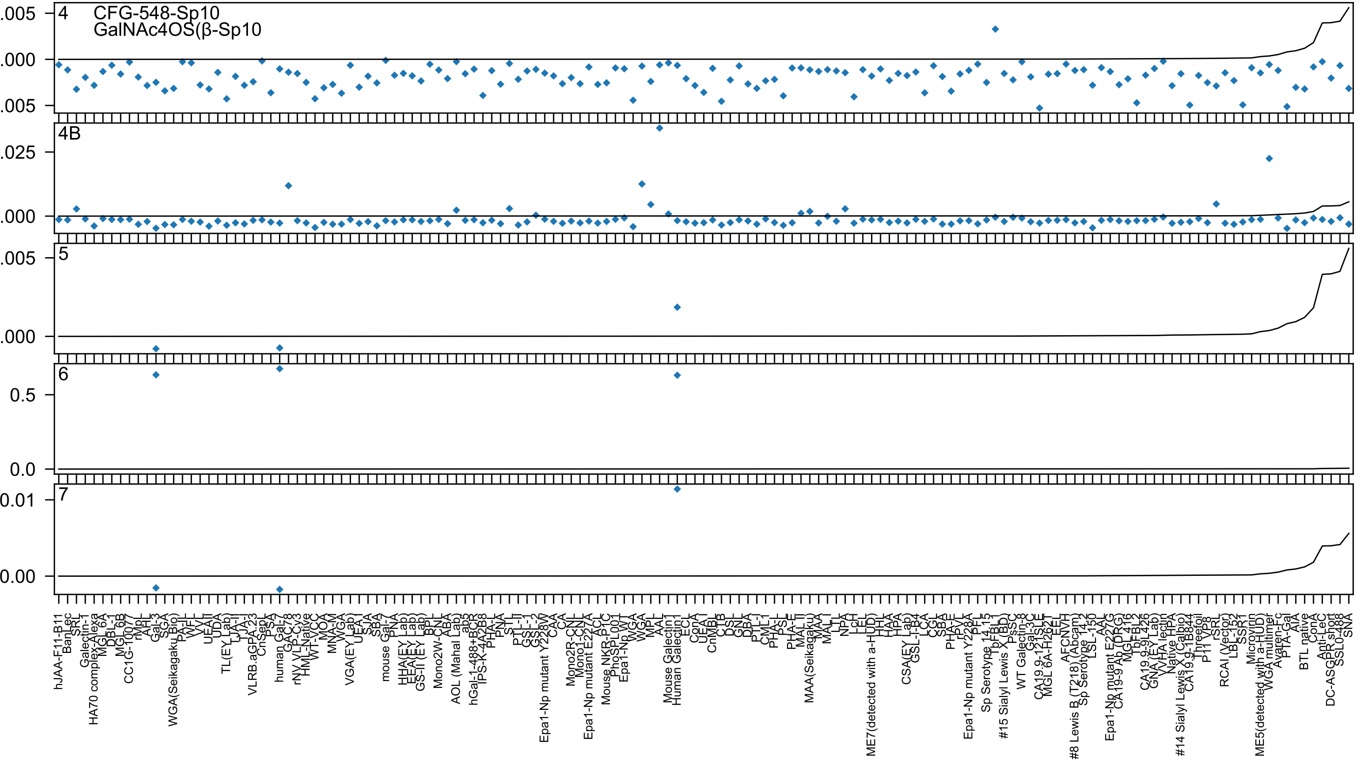
CFG-548-Sp10
GalNAc4OS(β-Sp10
4
4B
5
6
7

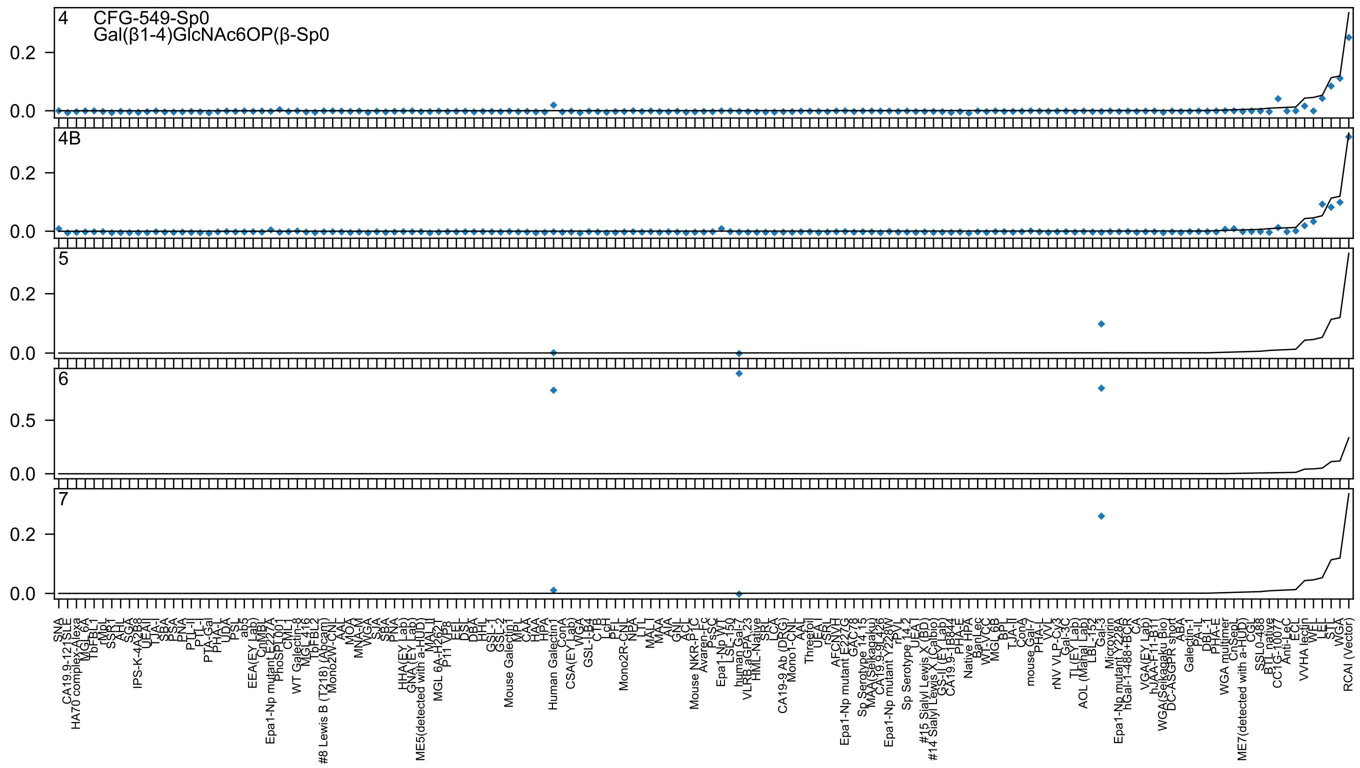
4 CFG-549-Sp0
Gal(β1-4)GlcNAc6OP(β-Sp0
4B
5
6
7
0.2
0.0
0.5
SNA
RCAI (Vector)
Human Galectin1
human Gal-7
Gal-3

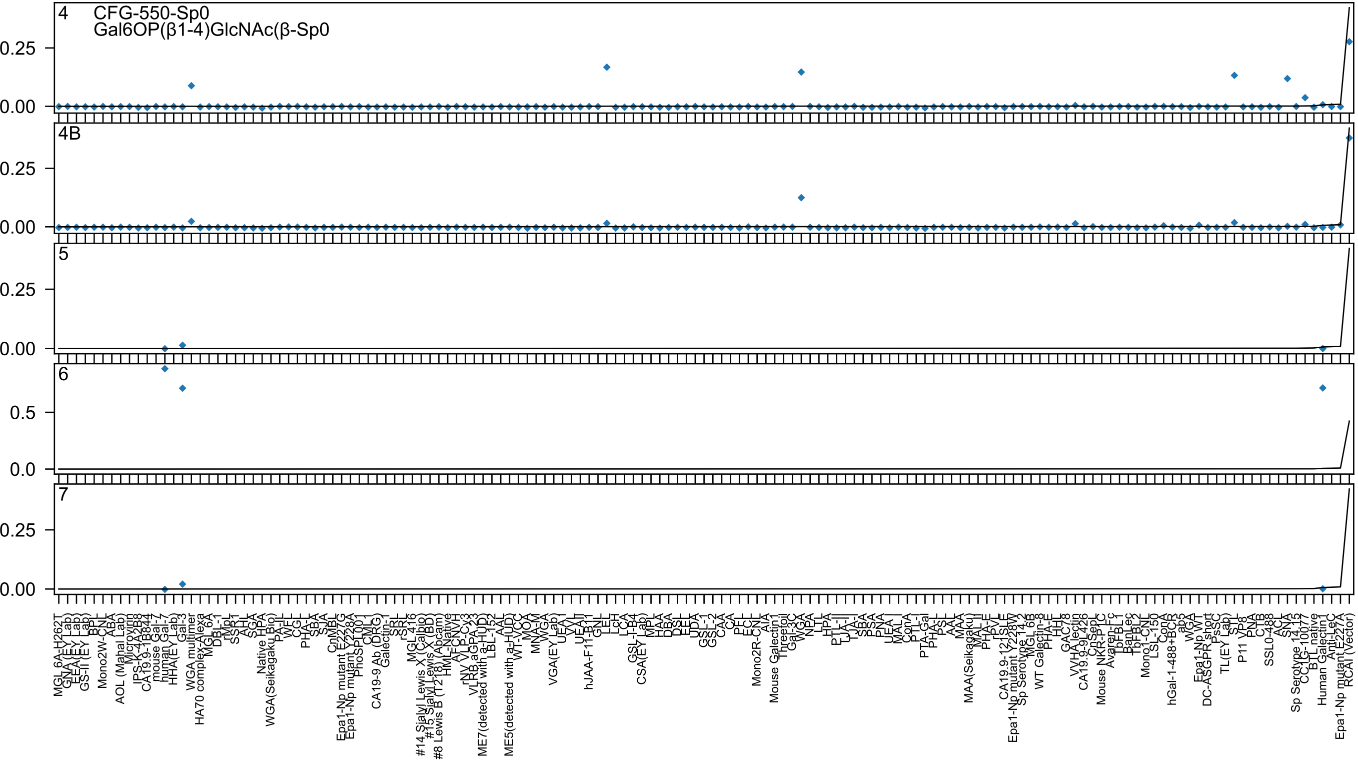
4
CFG-550-Sp0
Gal6OP(β1-4)GlcNAc(β-Sp0
4B
5
6
7

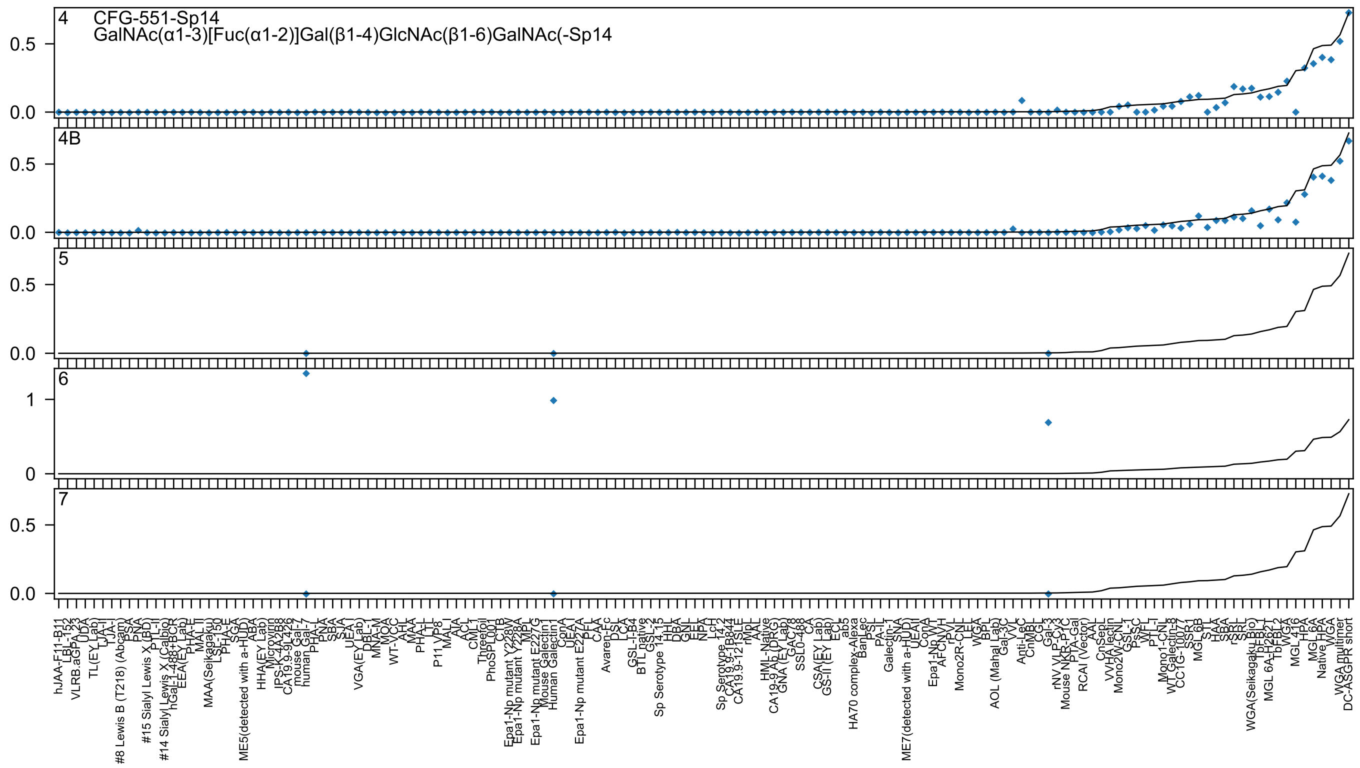
4
CFG-551-Sp14
GalNAc(α1-3)[Fuc(α1-2)]Gal(β1-4)GlcNAc(β1-6)GalNAc(-Sp14
4B
5
6
7
0.5
0.0
1
0

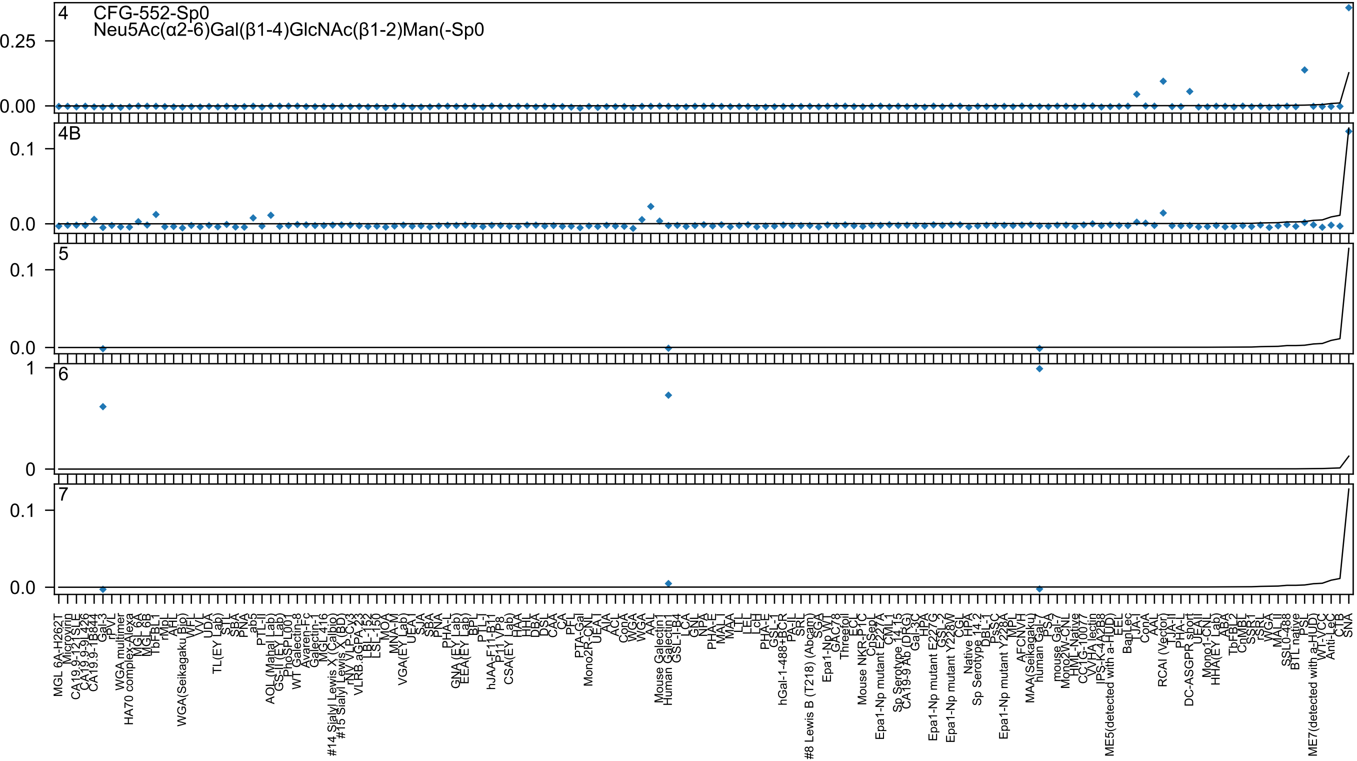
4 CFG-552-Sp0
Neu5Ac(α2-6)Gal(β1-4)GlcNAc(β1-2)Man(-Sp0
4B
5
6
7

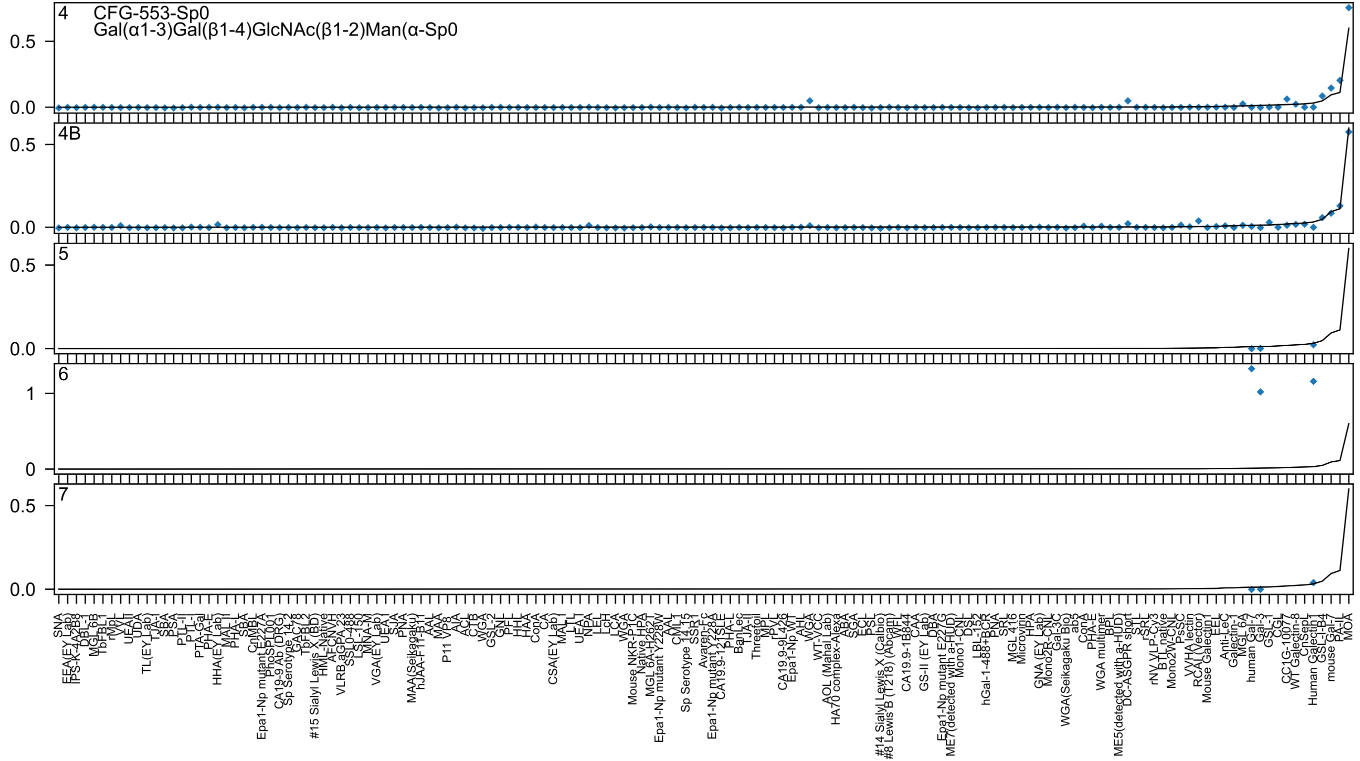

4
CFG-553-Sp0
Gal(α1-3)Gal(β1-4)GlcNAc(β1-2)Man(α-Sp0
4B
5
6
7
0.5
0.0
1
0

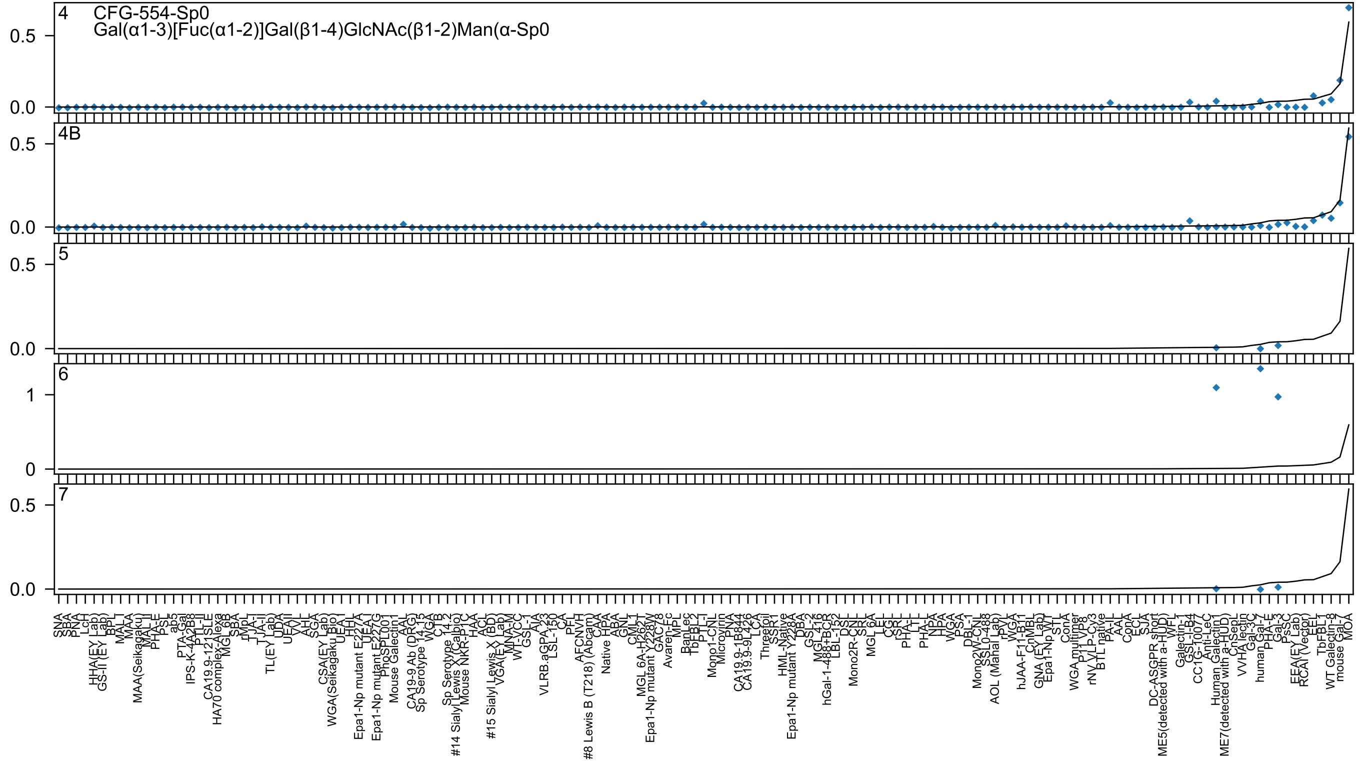

4
CFG-554-Sp0
Gal(α1-3)[Fuc(α1-2)]Gal(β1-4)GlcNAc(β1-2)Man(α-Sp0
4B
5
6
7

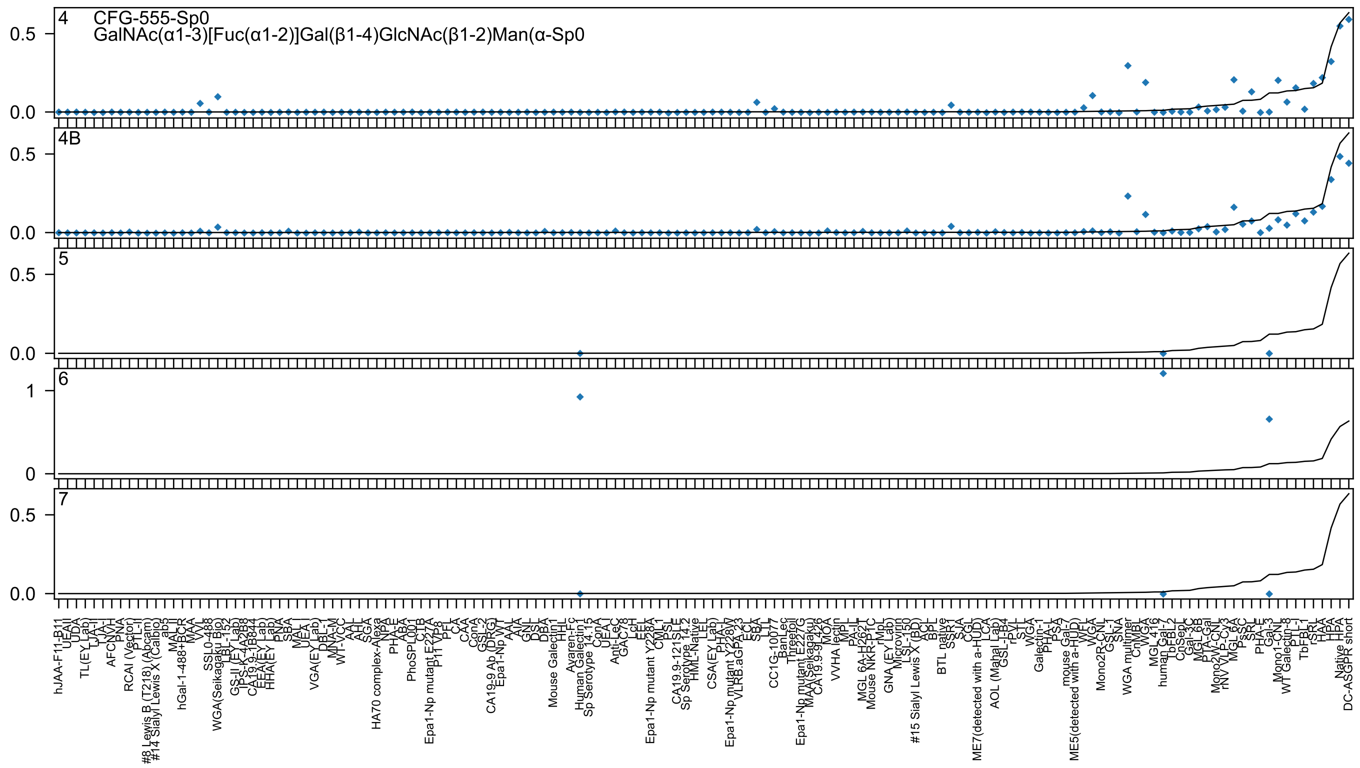

CFG-555-Sp0
GalNAc(α1-3)[Fuc(α1-2)]Gal(β1-4)GlcNAc(β1-2)Man(α-Sp0

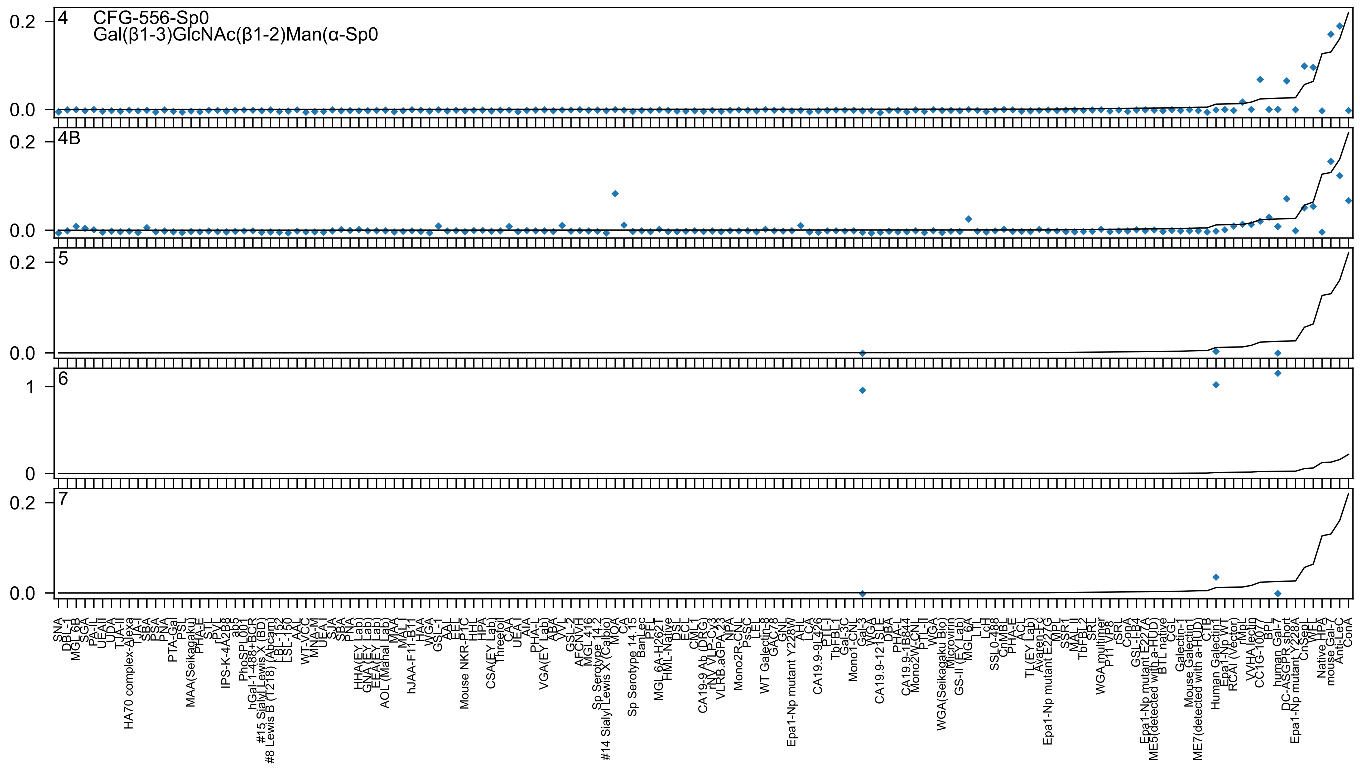
4
CFG-556-Sp0
Gal(β1-3)GlcNAc(β1-2)Man(α-Sp0
4B
5
6
7
0.2
0.0
1
0

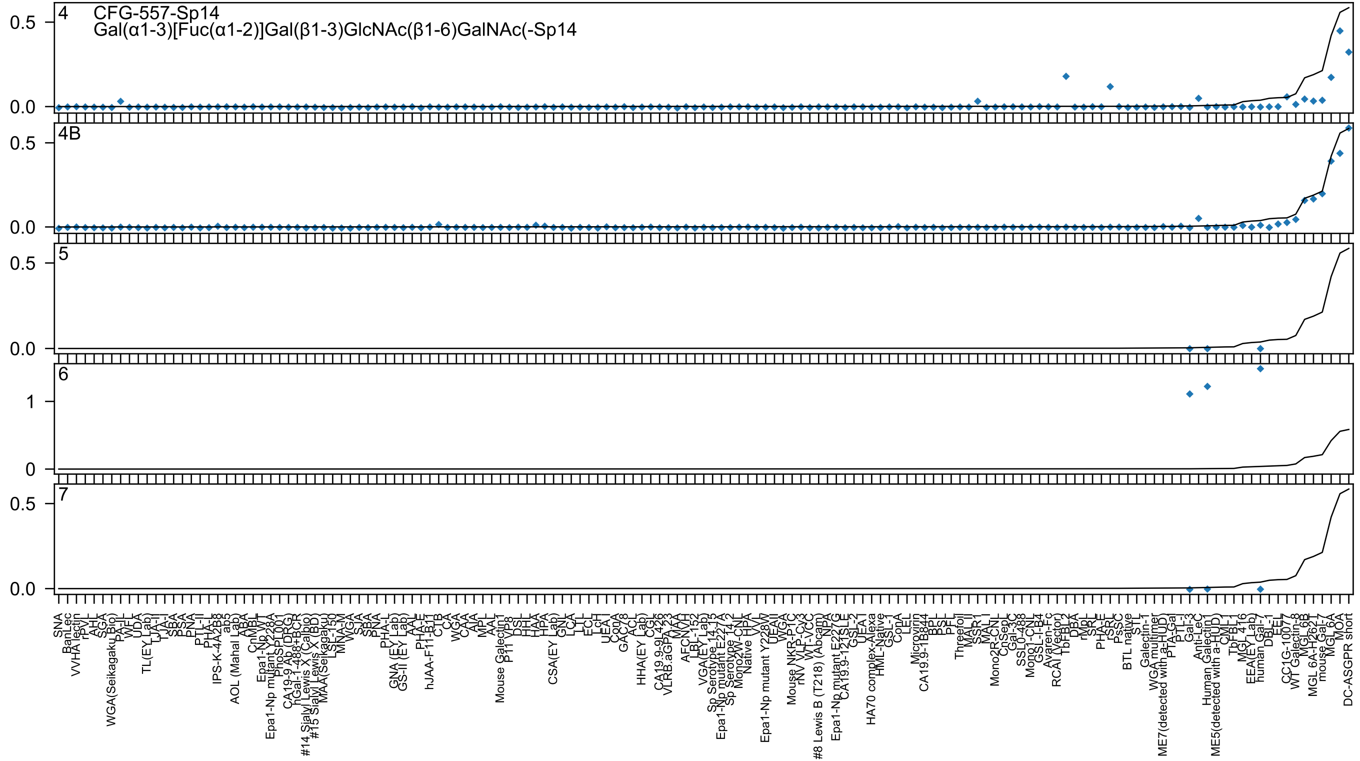
4
CFG-557-Sp14
Gal(α1-3)[Fuc(α1-2)]Gal(β1-3)GlcNAc(β1-6)GalNAc(-Sp14
4B
5
6
7

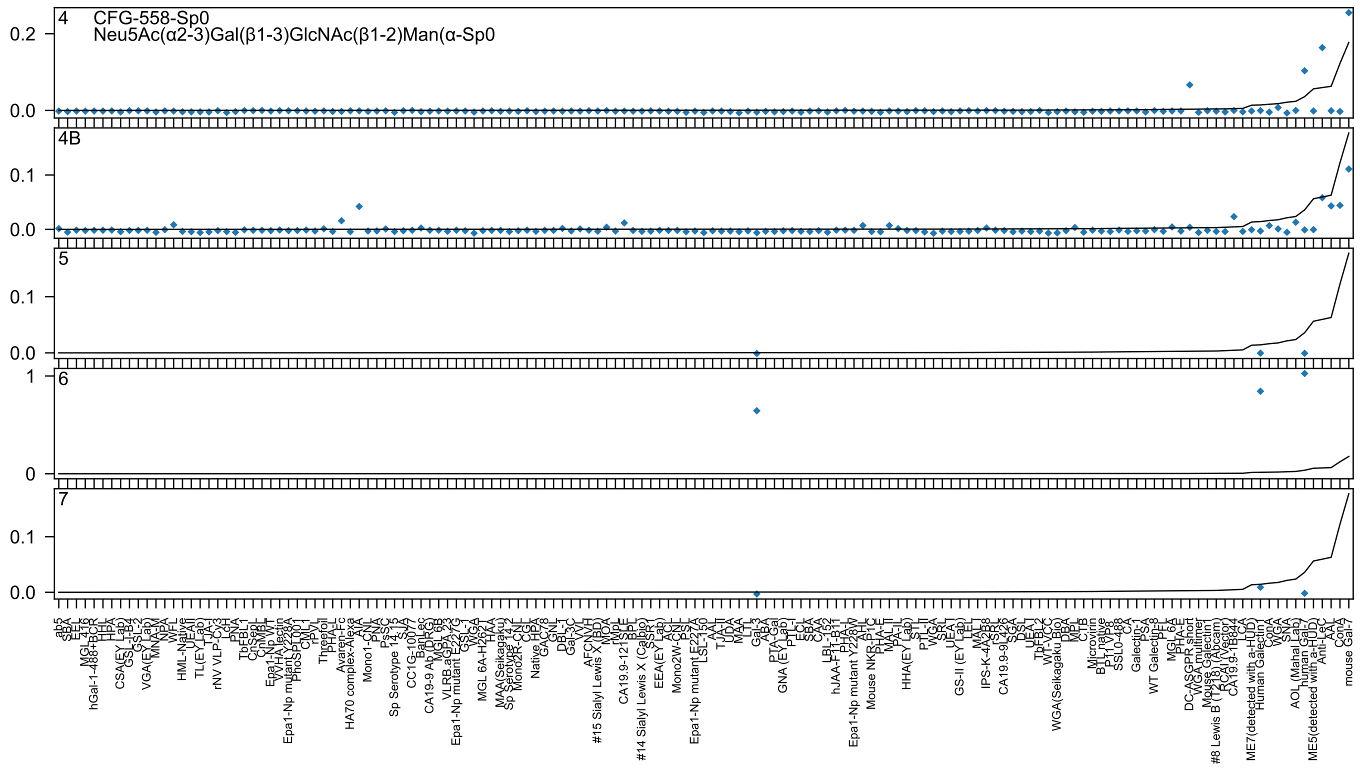
4
CFG-558-Sp0
Neu5Ac(α2-3)Gal(β1-3)GlcNAc(β1-2)Man(α-Sp0
4B
5
6
7
0.2
0.1
0.0
1
0
ab5
SBA
EEL
MGL 416
hGal-1-488+BCR
HHL
HPA
CSA(EY Lab)
GSL-I-B4
GSL-2
VGA(EY Lab)
MNA-M
NPA
WFL
HML-Native
UEAII
TL(EY Lab)
TJA-I
rNV VLP-Cy3
LcH
PNA
TbFBL1
CnSepL
CnMBL
Epa1-Np WT
VVHA lectin
Epa1-Np mutant Y228A
PhoSL001
CML1
rPVL
Threefoil
PHA-L
Avaren-Fc
HA70 complex-Alexa
AIA
Mono1-CNL
PNA
PsSC
Sp Serotype 14.15
SJA
CC1G-10077
BanLec
CA19-9 Ab (DRG)
MGL 6B
VLRB.aGPA.23
Epa1-Np mutant E227G
GSL-I
WGA
MGL 6A-H262T
HAA
MAA(Seikagaku)
Sp Serotype 14.2
Mono2R-CNL
CGL
Native HPA
GAC78
GNL
DBL-1
Gal-3C
VVL
AFCNVH
#15 Sialyl Lewis X (BD)
MOA
rMPL
CA19.9-121SLE
BPL
#14 Sialyl Lewis X (Calbio)
SSR1
EEA(EY Lab)
ACL
Mono2W-CNL
PSL
Epa1-Np mutant E227A
LSL-150
AAL
TJA-II
UDA
MAA
LTL
Gal-3
ABA
PTA-Gal
GNA (EY Lab)
PTL-I
ECL
SBA
CAA
LBL-152
hJAA-F11-B11
PHA-L
Epa1-Np mutant Y228W
AHL
Mouse NKR-P1C
PHA-E
MAL II
PA-IL
HHA(EY Lab)
STL
PTL-II
WGA
SRL
UEA I
GS-II (EY Lab)
LEL
MAL I
IPS-K-4A2B8
rSRL
CA19.9-9L426
SGA
DSL
UEA I
TbFBL2
WT-VCC
WGA(Seikagaku Bio)
DBA
MPL
CTB
Microvirin
BTL native
P11 VP8
SSL0-488
CA
Galectin-1
PSA
WT Galectin-8
PFL
MGL 6A
PHA-E
DC-ASGPR short
WGA multimer
Mouse Galectin1
#8 Lewis B (T218) (Abcam)
RCAI (Vector)
CA19.9-1B844
LCA
ME7(detected with a-HUD)
Human Galectin1
ConA
WGA
SNA
AOL (Mahal Lab)
human Gal-7
ME5(detected with a-HUD)
Anti-LeC
AAL
ConA
mouse Gal-7

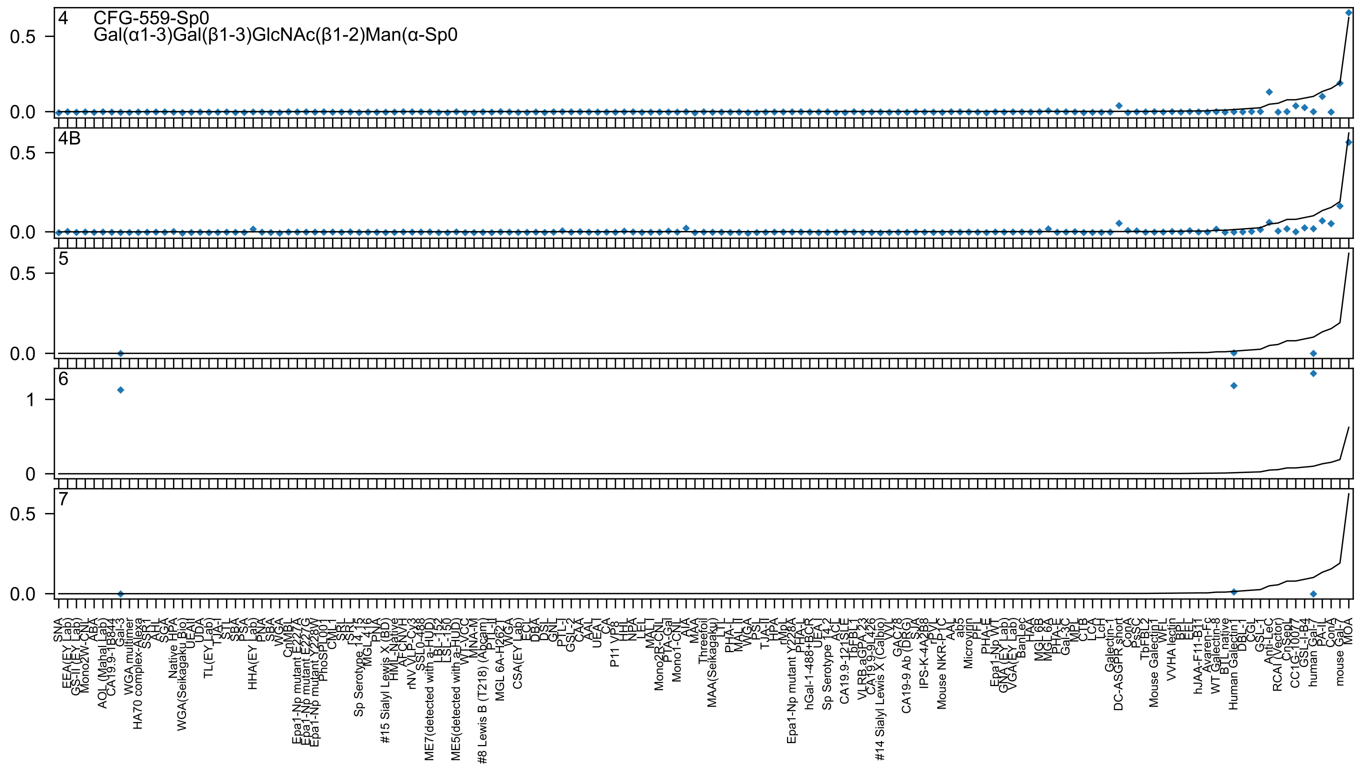
4 CFG-559-Sp0
Gal(α1-3)Gal(β1-3)GlcNAc(β1-2)Man(α-Sp0
4B
5
6
7

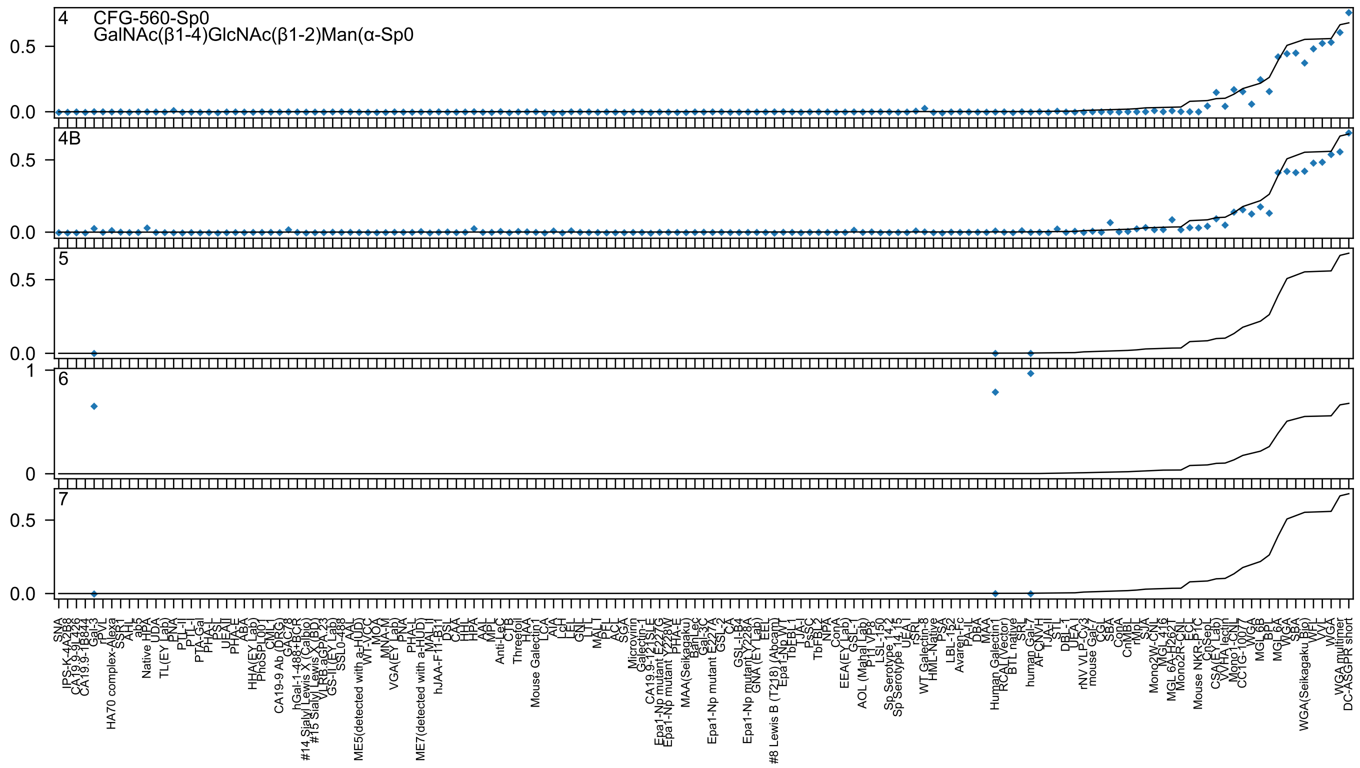

4
CFG-560-Sp0
GalNAc(β1-4)GlcNAc(β1-2)Man(α-Sp0
4B
5
6
7

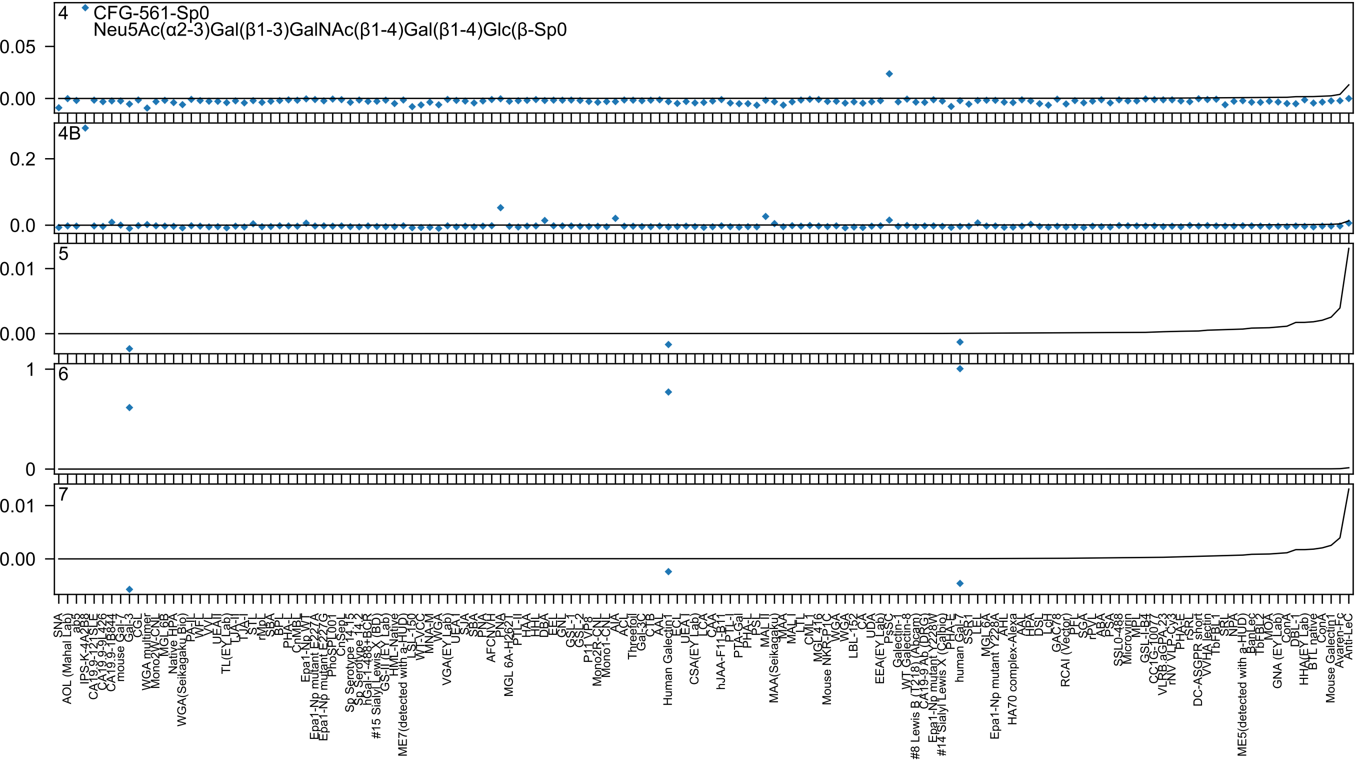
CFG-561-Sp0
Neu5Ac(α2-3)Gal(β1-3)GalNAc(β1-4)Gal(β1-4)Glc(β-Sp0
4
4B
5
6
7

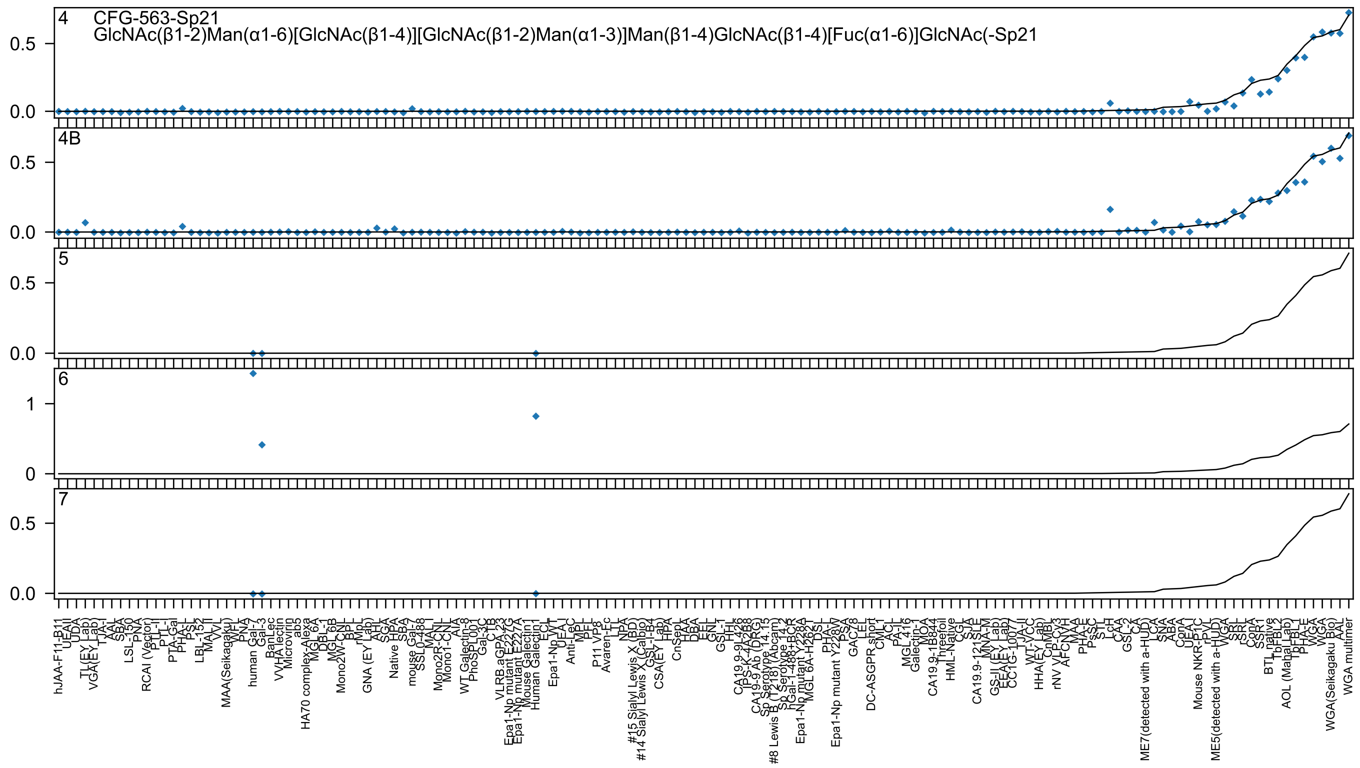
4 CFG-563-Sp21
GlcNAc(β1-2)Man(α1-6)[GlcNAc(β1-4)][GlcNAc(β1-2)Man(α1-3)]Man(β1-4)GlcNAc(β1-4)[Fuc(α1-6)]GlcNAc(-Sp21
4B
5
6
7
0.0
0.5
1
0

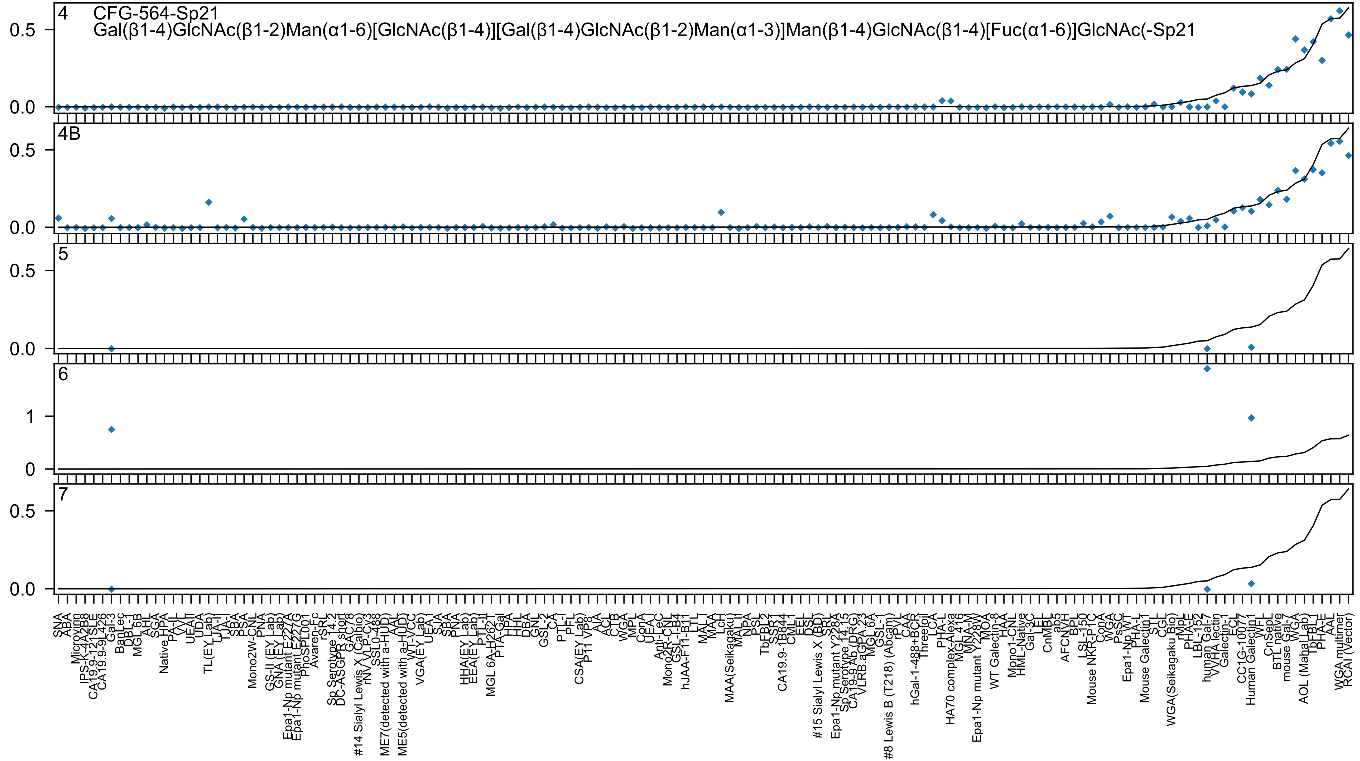
4 CFG-564-Sp21
Gal(β1-4)GlcNAc(β1-2)Man(α1-6)[GlcNAc(β1-4)][Gal(β1-4)GlcNAc(β1-2)Man(α1-3)]Man(β1-4)GlcNAc(β1-4)[Fuc(α1-6)]GlcNAc(-Sp21
4B
5
6
7

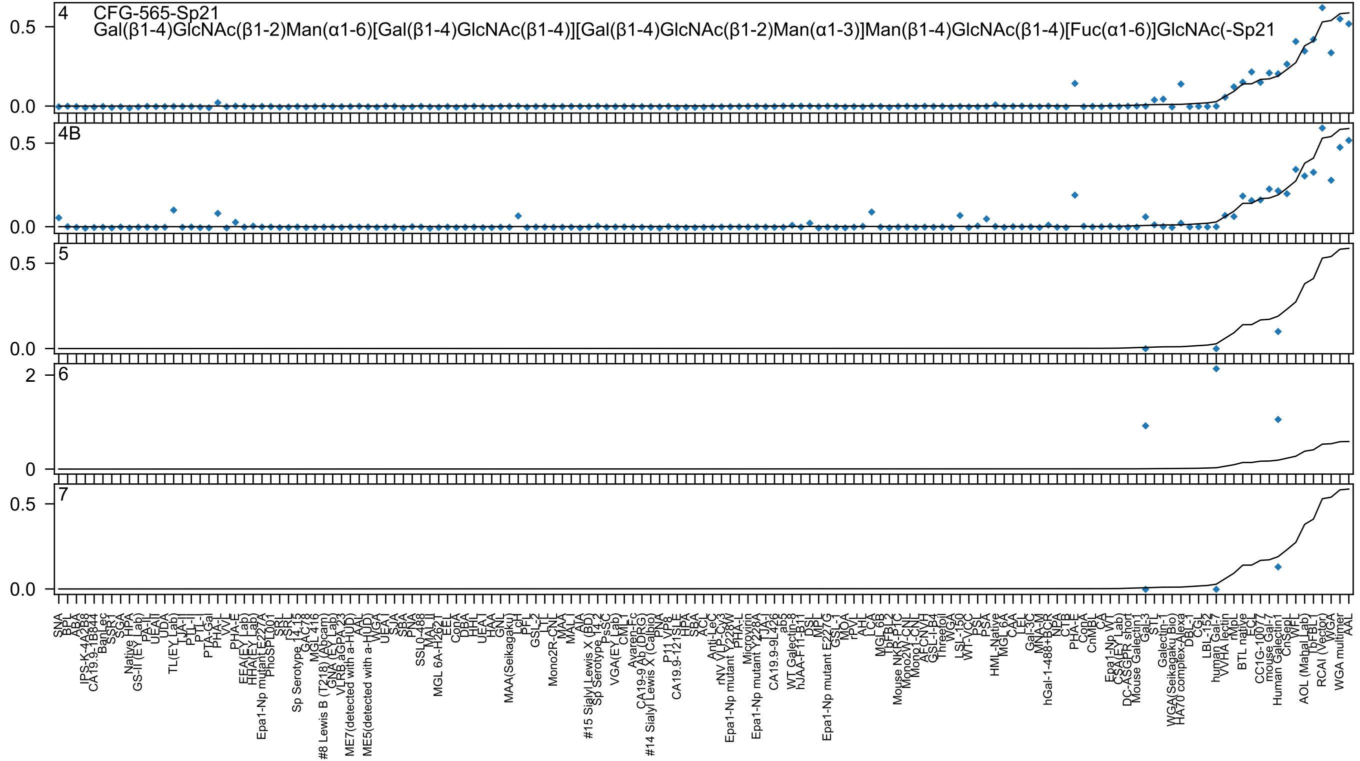

4
CFG-565-Sp21
Gal(β1-4)GlcNAc(β1-2)Man(α1-6)[Gal(β1-4)GlcNAc(β1-4)][Gal(β1-4)GlcNAc(β1-2)Man(α1-3)]Man(β1-4)GlcNAc(β1-4)[Fuc(α1-6)]GlcNAc(-Sp21
4B
5
6
7
0.0
0.5
0
2
SNA
BPL
ABA
IPS-K-4A2B8
CA19.9-1B844
BanLec
SSR1
SGA
Native HPA
GS-II(EY Lab)
PA-II
UEAII
UDA
TL(EY Lab)
TJA-II
PTL-II
PTL-I
PTA-Gal
PHA-L
VVL
PHA-E
EEA(EY Lab)
HHA(EY Lab)
Epa1-Np mutant E227A
PhoSL001
SRL
rSRL
Sp Serotype 14-15
GAC78
MGL 416
#8 Lewis B (T218) (Abcam)
GNA (EY Lab)
VLRB aGPA 23
ME7(detected with a-HUD)
AAL
ME5(detected with a-HUD)
WGA
UEA-I
SJA
SBA
PNA
SSL0-488
MAL-II
MGL 6A-H262T
EEL
ConA
DBA
HHL
UEA-I
HAA
GNL
MAA(Seikagaku)
PcL
PFL
GSL-2
LTL
Mono2R-CNL
MAA
MAL-I
AIA
#15 Sialyl Lewis X (BD)
Sp Serotype 14-2
PsSC
VGA(EY Lab)
CML
Avaren-Fc
CA19-9 Ab (DRG)
#14 Sialyl Lewis X (Calbio)
PNA
P11 VP8
CA19.9-121SLE
HPA
SBA
ACL
Anti-LeC
rNV VLP-Cy3
Epa1-Np mutant Y228W
PHA-L
Microvirin
Epa1-Np mutant Y228A
TJA-I
CA19.9-9L426
ab5
WT Galectin-8
hJAA-F11-B11
DSL
MPL
Epa1-Np mutant E227G
GSL-I
MOA
rPVL
AHL
LCA
MGL 6B
TbFBL2
Mouse NKR-P1C
Mono2W-CNL
Mono1-CNL
AFCNVH
GSL-I-B4
Threefoil
WGA
LSL-150
WT-VCC
PSL
PSA
HML-Native
MGL 6A
CAA
LEL
Gal-3C
MNA-M
hGal-1-488+BCR
NPA
CTB
PHA-E
ConA
CnMBL
CA
Epa1-Np WT
CSA(EY Lab)
DC-ASGPR short
Mouse Galectin1
Gal-3
STL
Galectin-1
WGA(Seikagaku Bio)
HA70 complex-Alexa
DBL-I
CGL
LBL-152
human Gal-7
VVHA lectin
rMpL
BTL native
ECL
CC1G-10077
mouse Gal-7
Human Galectin1
CnSepL
WFL
AOL (Mahal Lab)
TbFBL1
RCAI (Vector)
WGA
WGA multimer
AAL

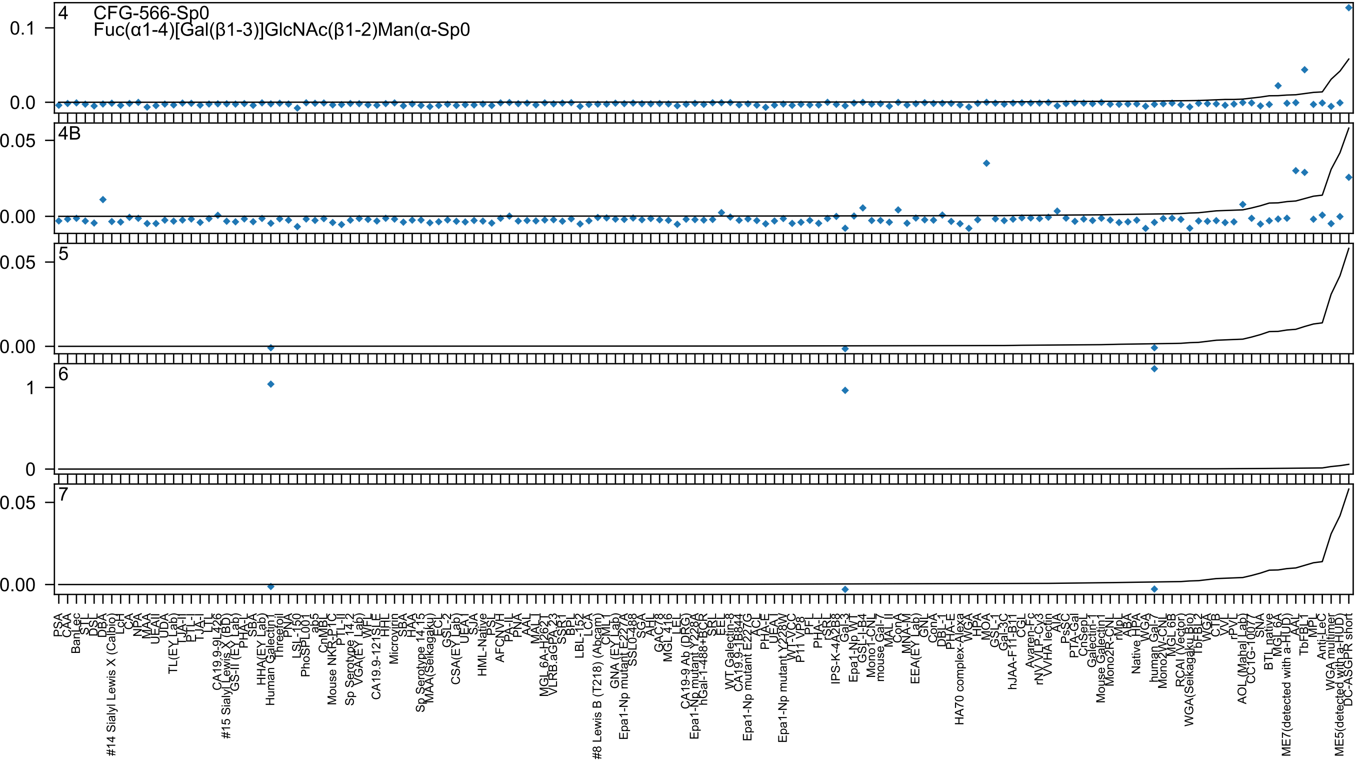

4
CFG-566-Sp0
Fuc(α1-4)[Gal(β1-3)]GlcNAc(β1-2)Man(α-Sp0
4B
5
6
7

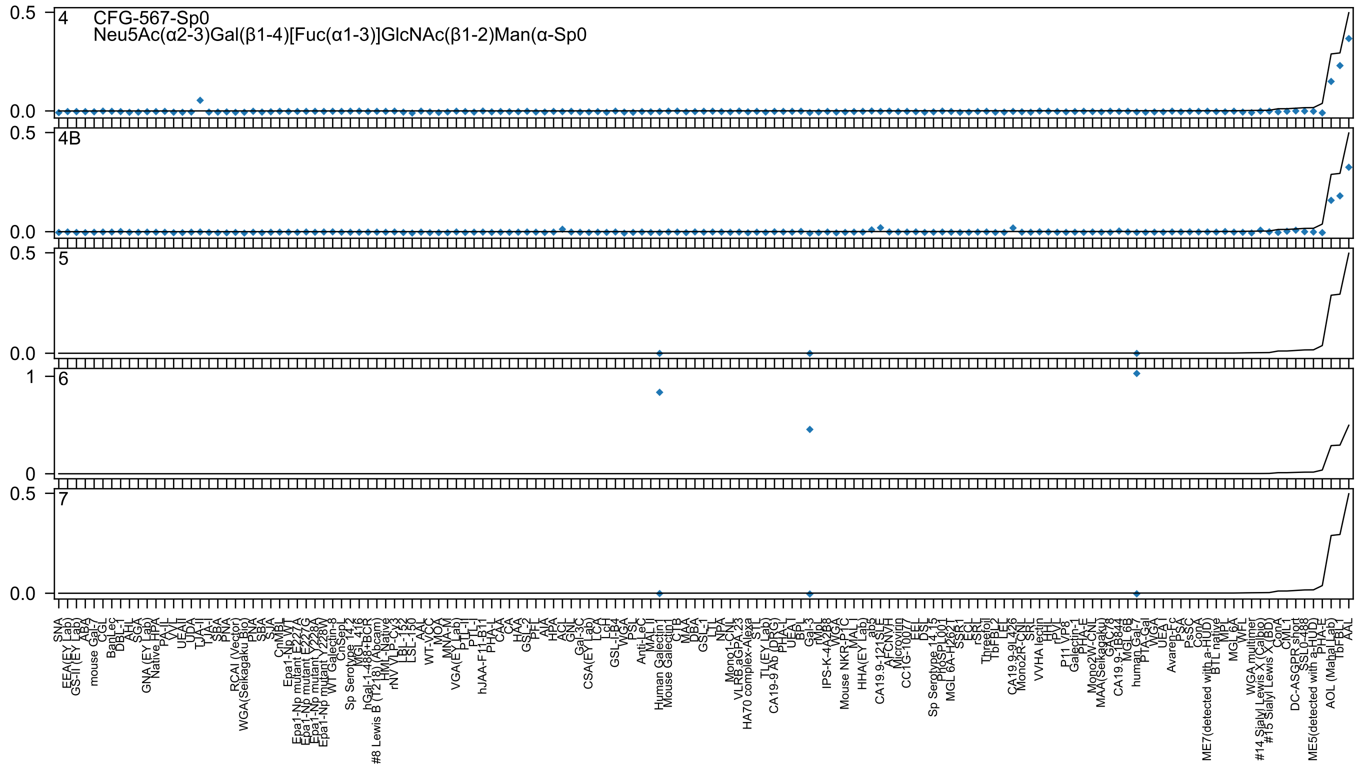
4
CFG-567-Sp0
Neu5Ac(α2-3)Gal(β1-4)[Fuc(α1-3)]GlcNAc(β1-2)Man(α-Sp0
4B
5
6
7
0.5
0.0
1
0
SNA
EEA(EY Lab)
GS-II (EY Lab)
ABA
mouse Gal-7
CGL
BanLec
DBL-1
AHL
SGA
GNA (EY Lab)
Native HPA
PA-IL
VVL
UEAII
UDA
TJA-II
TJA-I
SBA
PNA
RCAI (Vector)
WGA(Seikagaku Bio)
PNA
SBA
SJA
CnMBL
Epa1-Np WT
Epa1-Np mutant E227G
Epa1-Np mutant E227A
Epa1-Np mutant Y228A
Epa1-Np mutant Y228W
WT Galectin-8
CnSeP
Sp Serotype 14.2
MGL 416
hGal-1-488+BCR
#8 Lewis B (T218) (Abcam)
HML-Native
rNV VLP-Cy3
LBL-152
LSL-150
AAL
WT-VCC
MOA
MNA-M
VGA(EY Lab)
PTL-II
PTL-I
hJAA-F11-B11
PHA-L
CAA
CA
HAA
GSL-2
PFL
AIA
HPA
ACL
GNL
Gal-3C
CSA(EY Lab)
LCA
LcH
GSL-I-B4
WGA
PSL
Anti-LeC
MAL II
Human Galectin1
Mouse Galectin1
CTB
MAA
DBA
GSL-1
LTL
NPA
Mono1-CNL
VLRB aGPA.23
HA70 complex-Alexa
STL
TL(EY Lab)
CA19-9 Ab (DRG)
PHA-L
UEA I
BPL
Gal-3
rMpL
IPS-K-4A2B8
WGA
Mouse NKR-P1C
MAL I
HHA(EY Lab)
ab5
CA19.9-121SLE
AFCNVH
Microvirin
CC1G-10077
EEL
DSL
Sp Serotype 14.15
ProSPL001
MGL 6A-H262T
SSR1
ECL
rSRL
Threefoil
TbFBL2
LEL
CA19.9-9L426
Mono2R-CNL
SRL
VVHA lectin
HHL
rPVL
P11 VP8
Galectin-1
PHA-E
Mono2W-CNL
MAA(Seikagaku)
GAC78
CA19.9-1B844
MGL 6B
human Gal-7
PTA-Gal
WGA
UEA I
Avaren-Fc
PSA
PsSC
ConA
ME7(detected with a-HUD)
BTL native
MPL
MGL 6A
WFL
WGA multimer
#14 Sialyl Lewis X (Calbio)
#15 Sialyl Lewis X (BD)
ConA
CML1
DC-ASGPR short
SSL0-488
ME5(detected with a-HUD)
PHA-E
AOL (Mahal Lab)
TbFBL1
AAL

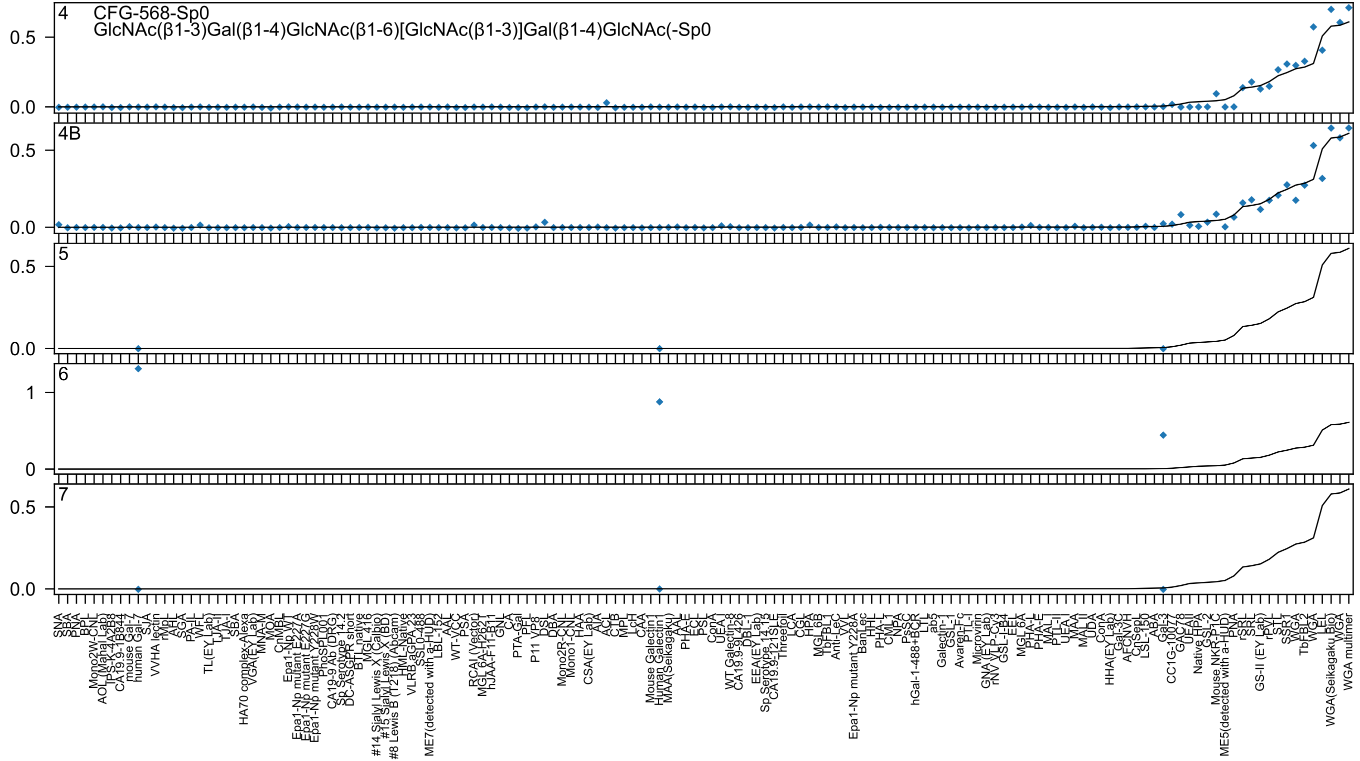
4 CFG-568-Sp0
GlcNAc(β1-3)Gal(β1-4)GlcNAc(β1-6)[GlcNAc(β1-3)]Gal(β1-4)GlcNAc(-Sp0
4B
5
6
7

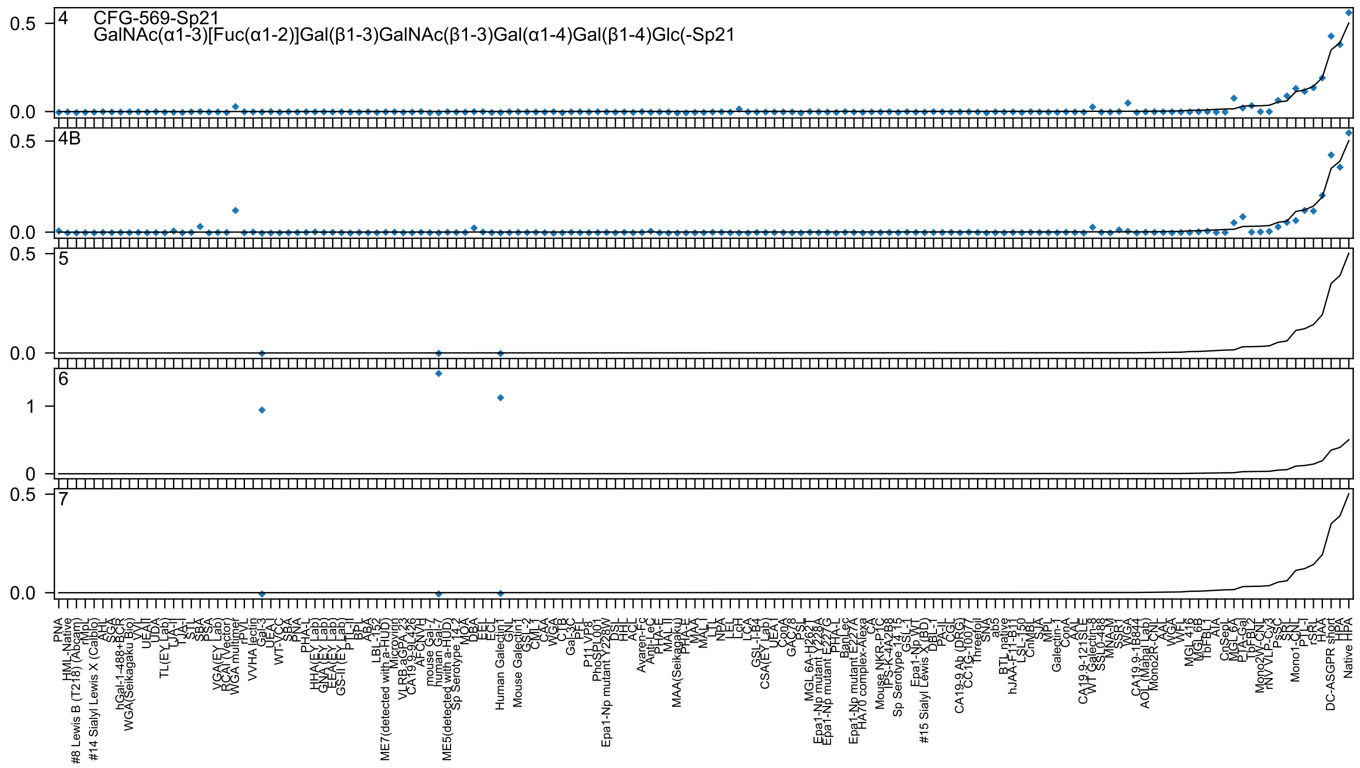
4 CFG-569-Sp21
GalNAc(α1-3)[Fuc(α1-2)]Gal(β1-3)GalNAc(β1-3)Gal(α1-4)Gal(β1-4)Glc(-Sp21
4B
5
6
7

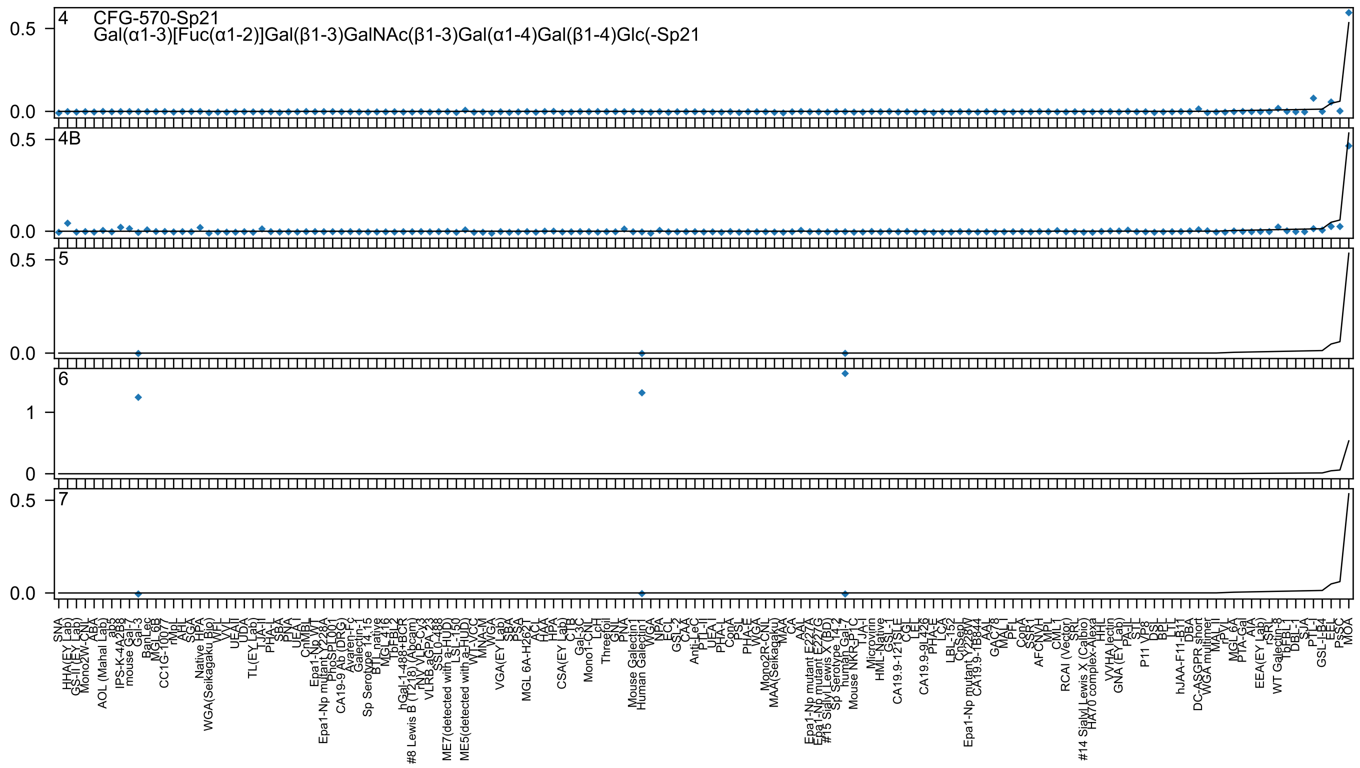

4 CFG-570-Sp21
Gal(α1-3)[Fuc(α1-2)]Gal(β1-3)GalNAc(β1-3)Gal(α1-4)Gal(β1-4)Glc(-Sp21
4B
5
6
7

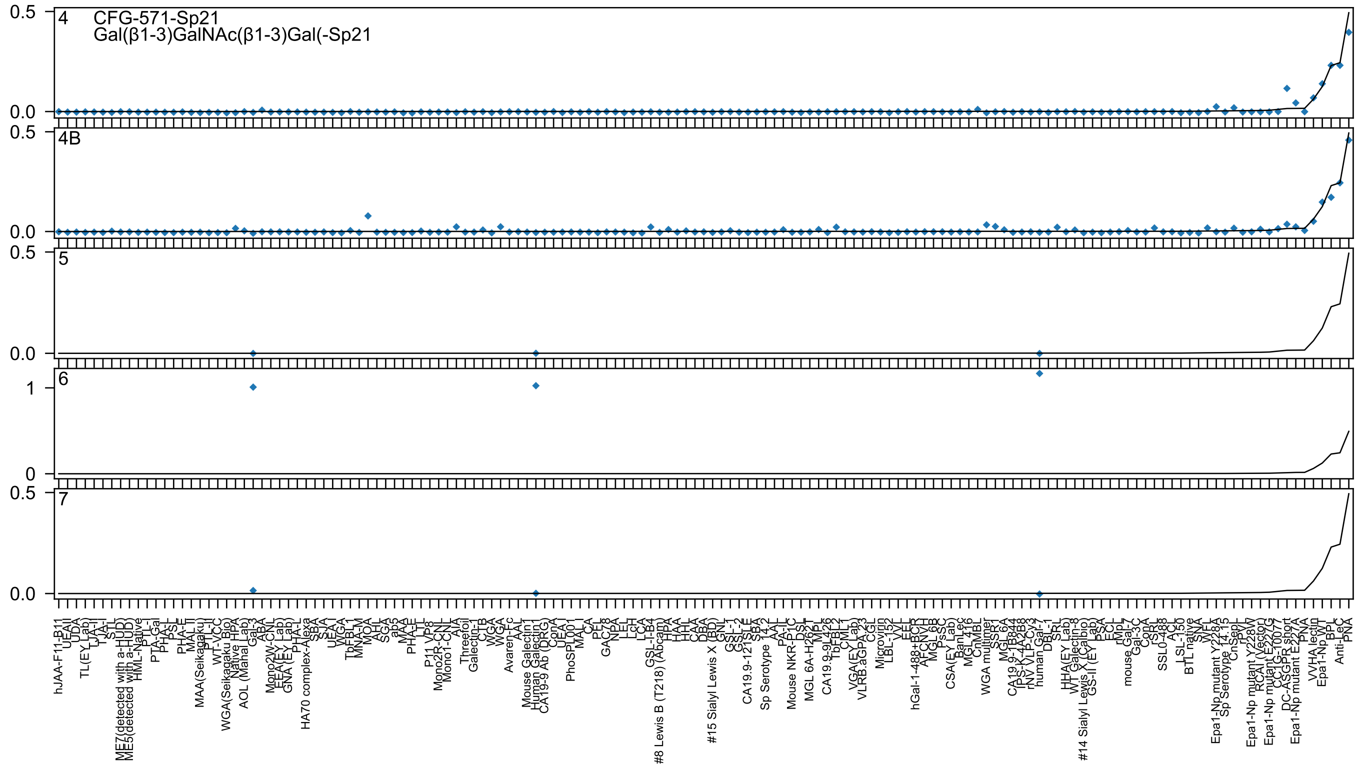
4
CFG-571-Sp21
Gal(β1-3)GalNAc(β1-3)Gal(-Sp21
4B
5
6
7

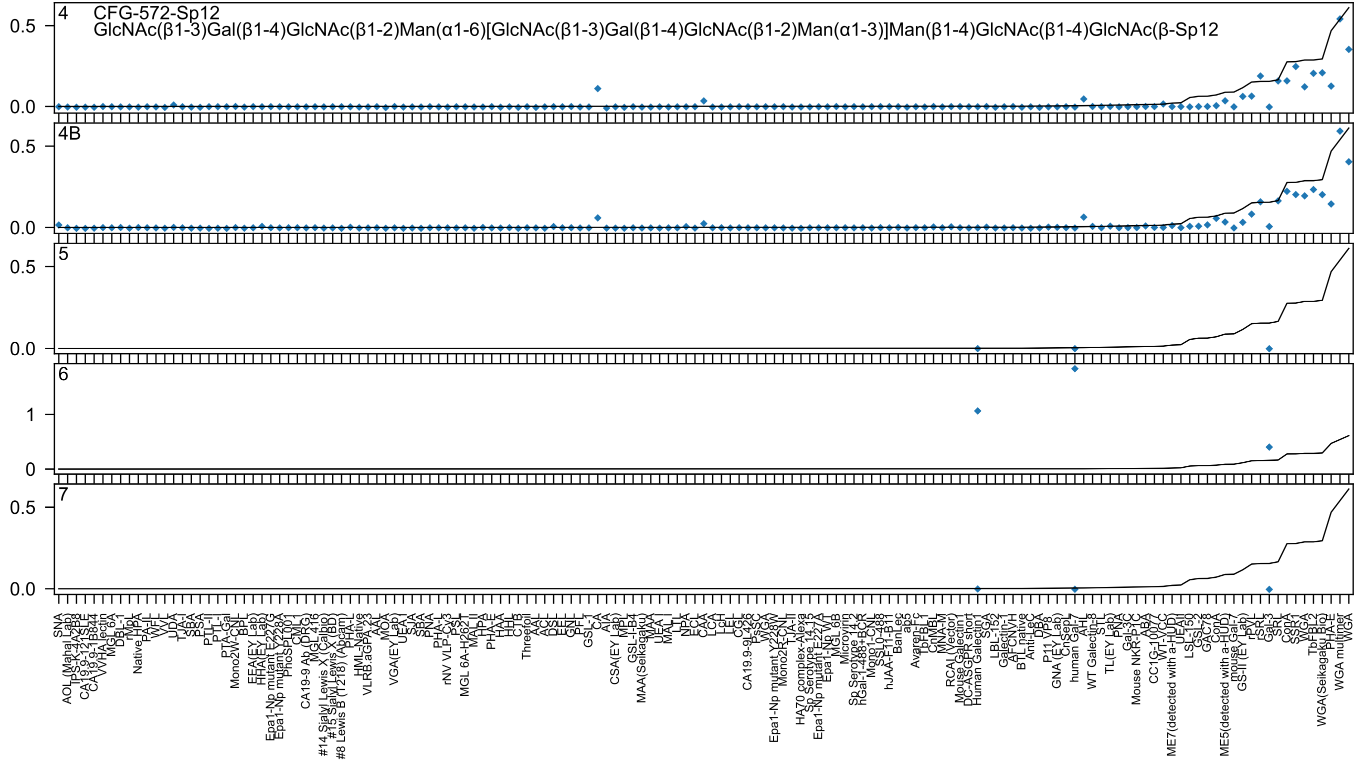

CFG-572-Sp12
GlcNAc(β1-3)Gal(β1-4)GlcNAc(β1-2)Man(α1-6)[GlcNAc(β1-3)Gal(β1-4)GlcNAc(β1-2)Man(α1-3)]Man(β1-4)GlcNAc(β1-4)GlcNAc(β-Sp12
4
4B
5
6
7

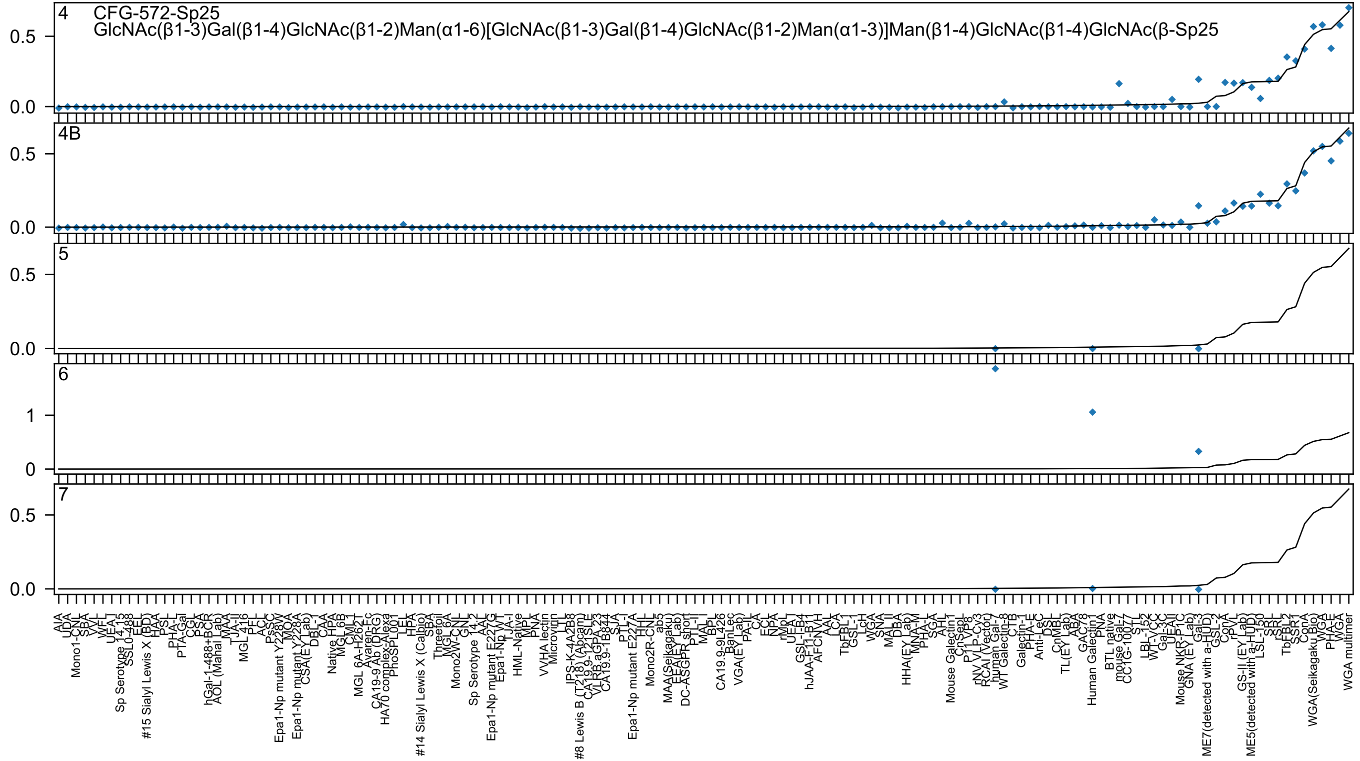

4
CFG-572-Sp25
GlcNAc(β1-3)Gal(β1-4)GlcNAc(β1-2)Man(α1-6)[GlcNAc(β1-3)Gal(β1-4)GlcNAc(β1-2)Man(α1-3)]Man(β1-4)GlcNAc(β1-4)GlcNAc(β-Sp25
4B
5
6
7

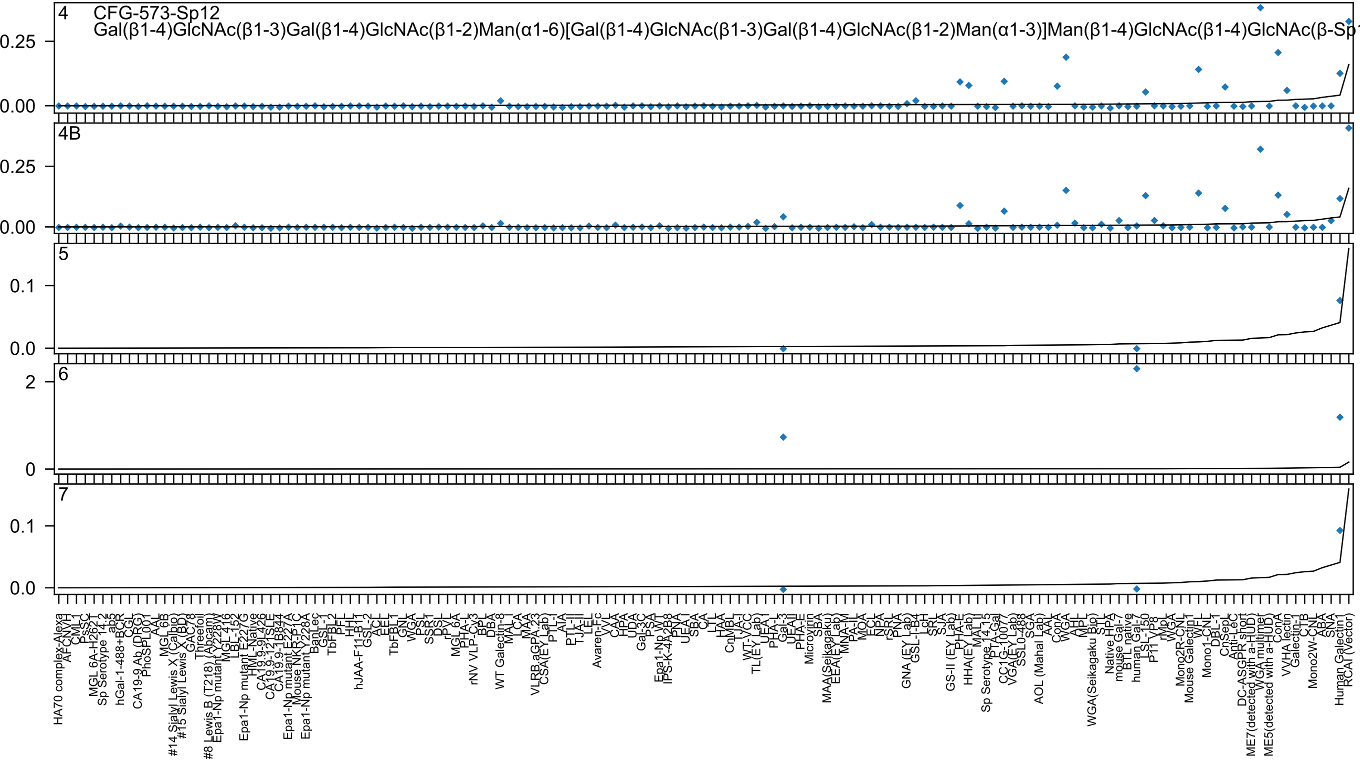
4 CFG-573-Sp12
Gal(β1-4)GlcNAc(β1-3)Gal(β1-4)GlcNAc(β1-2)Man(α1-6)[Gal(β1-4)GlcNAc(β1-3)Gal(β1-4)GlcNAc(β1-2)Man(α1-3)]Man(β1-4)GlcNAc(β1-4)GlcNAc(β-Sp1
4B
5
6
7

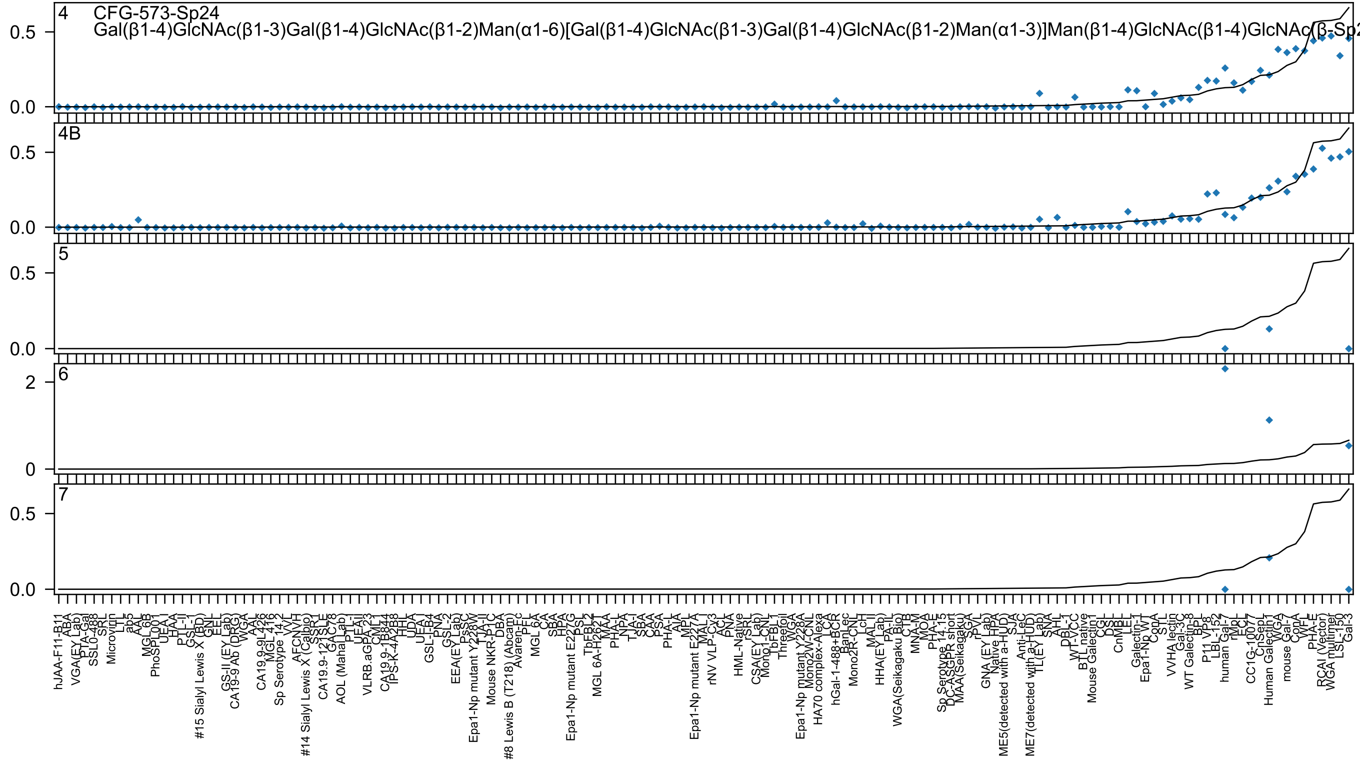
4 CFG-573-Sp24
Gal(β1-4)GlcNAc(β1-3)Gal(β1-4)GlcNAc(β1-2)Man(α1-6)[Gal(β1-4)GlcNAc(β1-3)Gal(β1-4)GlcNAc(β1-2)Man(α1-3)]Man(β1-4)GlcNAc(β1-4)GlcNAc(β-Sp2
4B
5
6
7
0.0
0.5
0
2

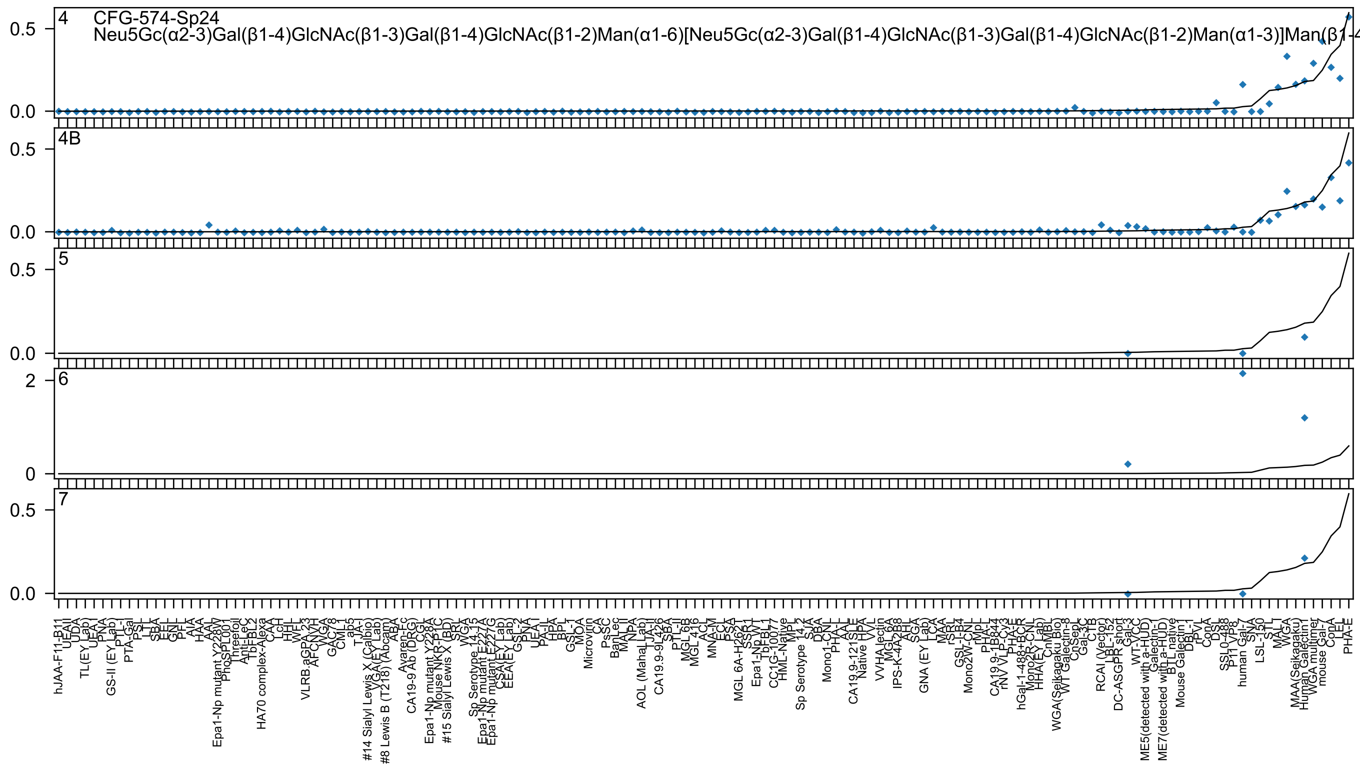

4
CFG-574-Sp24
Neu5Gc(α2-3)Gal(β1-4)GlcNAc(β1-3)Gal(β1-4)GlcNAc(β1-2)Man(α1-6)[Neu5Gc(α2-3)Gal(β1-4)GlcNAc(β1-3)Gal(β1-4)GlcNAc(β1-2)Man(α1-3)]Man(β1-4
4B
5
6
7
0.5
0.0
2
0
hJAA-F11-B11
UEA-II
UDA
TL(EY Lab)
UEA-I
PNA
GS-II (EY Lab)
PTL-I
PTA-Gal
PSL
LTL
SBA
EEL
GNL
PFL
AIA
HAA
AAL
Epa1-Np mutant Y228W
PhoSL
Threefoil
Anti-Lec
TbFBL2
HA70 complex-Alexa
CAA
LcH
HHL
WFL
VLRB.aGPA.23
AFCNVH
WGA
GAC78
CML-1
ab5
TJA-I
#14 Sialyl Lewis X (Calbio)
VGA(EY Lab)
#8 Lewis B (T218) (Abcam)
ABA
Avaren-Fc
CA19-9 Ab (DRG)
CGL
Epa1-Np mutant Y228A
Mouse NKR-P1C
#15 Sialyl Lewis X (BD)
SRL
WGA
Sp Serotype 14_15
Epa1-Np mutant E227A
Epa1-Np mutant E227G
CSA(EY Lab)
EEA(EY Lab)
GSL-2
PNA
UEA-I
PA-IL
HPA
BPL
GSL-1
MOA
Microvirin
CA
PsSC
BanLec
MAL-II
NPA
AOL (Mahal Lab)
LJA-II
CA19.9-9L426
SBA
PTL-II
MGL 6B
MGL 416
ACL
MNA-M
ECL
PSA
MGL 6A-H262
SSP-1
Epa1-Np WT
TbFBL1
CC1G-10077
HML-Native
MPL
Sp Serotype 14_2
SJA
DBA
Mono1-CNL
PHA-L
AAL
CA19.9-121SLE
Native HPA
VVL
VVHA lectin
MGL 6A
IPS-K-4A2B8
AAL
SGA
GNA (EY Lab)
LCA
MAA
rSRL
GSL-I-B4
Mono2W-CNL
rMpL
PHA-L
CA19.9-1B844
rNV VLP-Cy3
PHA-E
hGal-1-488+BCR
Mono2R-CNL
HHA(EY Lab)
CnMBL
WGA(Seikagaku Bio)
WT Galectin-8
CnSepL
Gal-3C
CTB
RCAI (Vector)
LBL-152
DC-ASGPR short
Gal-3
WT-YCC
ME5(detected with a-HUD)
Galectin-1
ME7(detected with a-HUD)
BTL native
Mouse Galectin1
DBL-1
rPVL
ConA
DSL
SSL0-488
P11 VP8
human Gal-7
SNA
LSL-150
STL
MAL-I
WGA
MAA(Seikagaku)
Human Galectin1
WGA multimer
mouse Gal-7
ConA
LEL
PHA-E

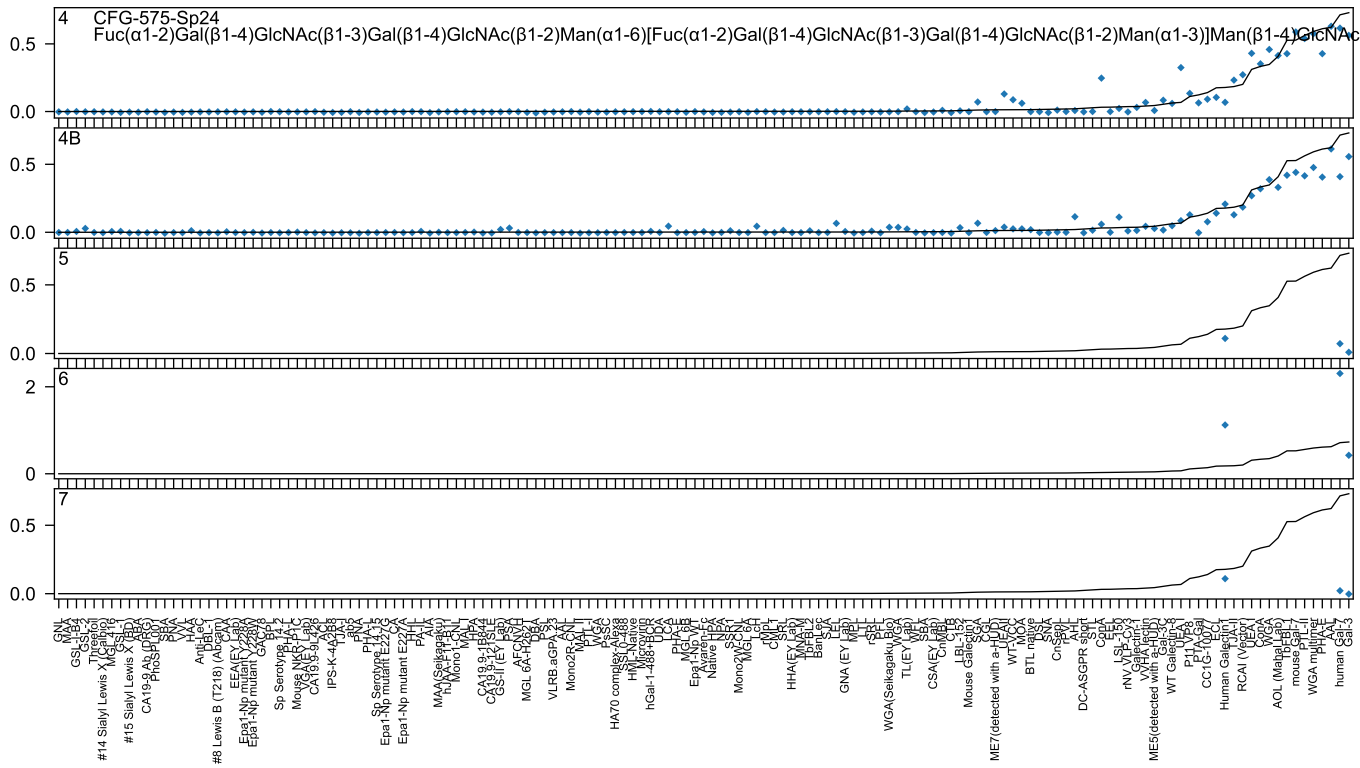

4 CFG-575-Sp24
Fuc(α1-2)Gal(β1-4)GlcNAc(β1-3)Gal(β1-4)GlcNAc(β1-2)Man(α1-6)[Fuc(α1-2)Gal(β1-4)GlcNAc(β1-3)Gal(β1-4)GlcNAc(β1-2)Man(α1-3)]Man(β1-4)GlcNAc
4B
5
6
7

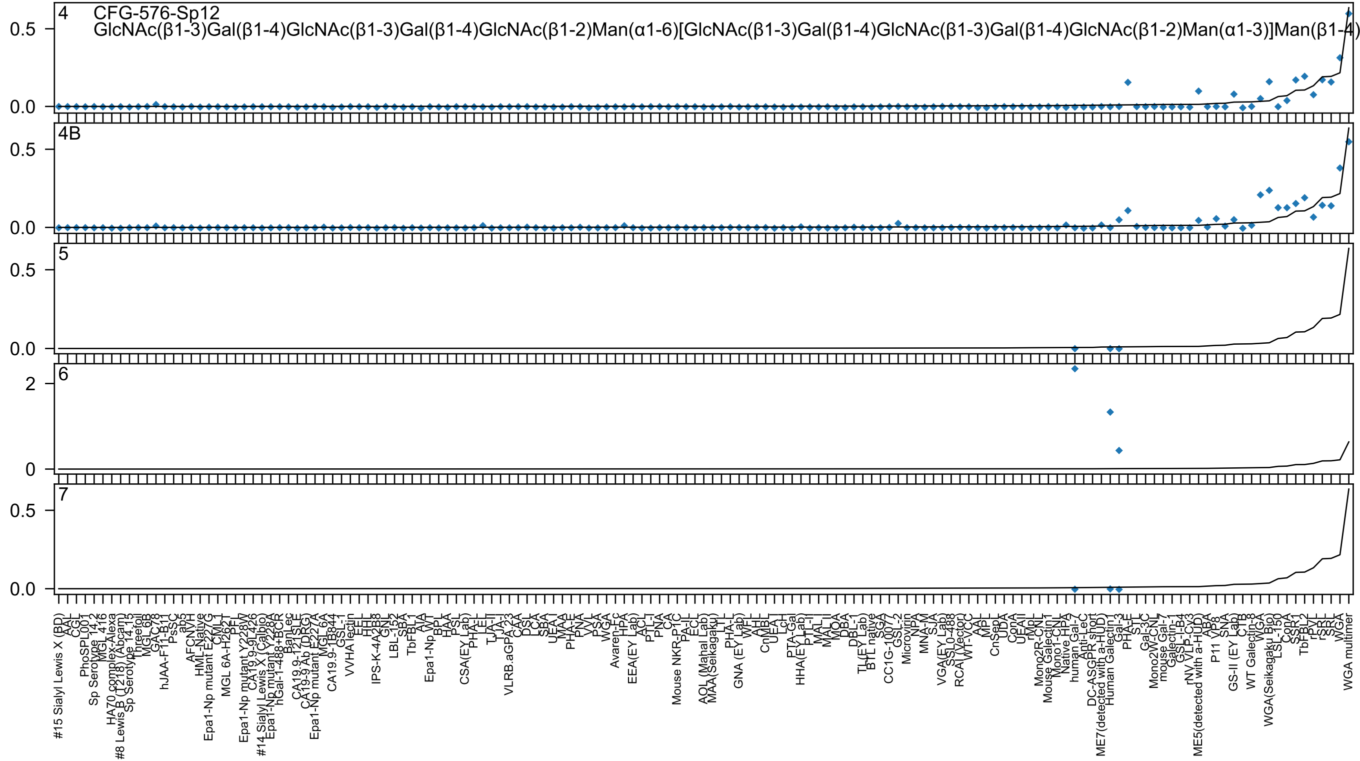
4
CFG-576-Sp12
GlcNAc(β1-3)Gal(β1-4)GlcNAc(β1-3)Gal(β1-4)GlcNAc(β1-2)Man(α1-6)[GlcNAc(β1-3)Gal(β1-4)GlcNAc(β1-3)Gal(β1-4)GlcNAc(β1-2)Man(α1-3)]Man(β1-4)
4B
5
6
7
0.5
0.0
2
0
#15 Sialyl Lewis X (BD)
WGA multimer

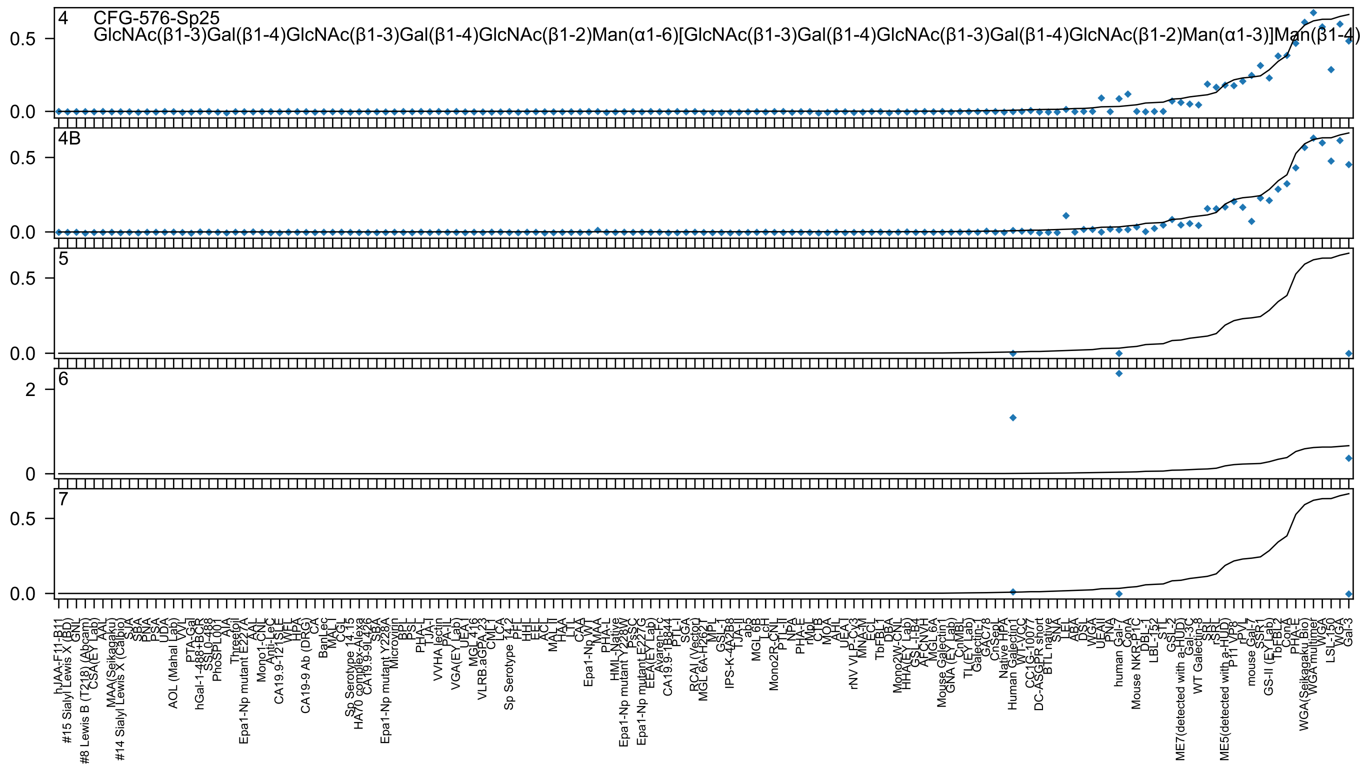
4 CFG-576-Sp25
GlcNAc(β1-3)Gal(β1-4)GlcNAc(β1-3)Gal(β1-4)GlcNAc(β1-2)Man(α1-6)[GlcNAc(β1-3)Gal(β1-4)GlcNAc(β1-3)Gal(β1-4)GlcNAc(β1-2)Man(α1-3)]Man(β1-4)
4B
5
6
7
0.0
0.5
0
2
hJAA-F11-B11
#15 Sialyl Lewis X (BD)
GNL
#8 Lewis B (T218) (Abcam)
CSA(EY Lab)
AAL
MAA(Seikagaku)
#14 Sialyl Lewis X (Calbio)
SJA
SBA
PNA
PSA
UDA
AOL (Mahal Lab)
WVL
PTA-Gal
hGal-1-488+BCR
SSL0-488
PhoSPL001
AIA
Threefoil
Epa1-Np mutant E227A
AAL
Mono1-CNL
Anti-LeC
CA19.9-121SLE
WFL
HPA
CA19-9 Ab (DRG)
CA
BanLec
MAL I
CGL
Sp Serotype 14.15
HA70 complex-Alexa
CA19.9-9L426
SBA
Epa1-Np mutant Y228A
Microvirin
BPL
PSL
PHA-L
TJA-I
VVHA lectin
PA-IL
VGA(EY Lab)
UEA-I
MGL416
VLRB.aGPA.23
CML-I
LCA
Sp Serotype 14.2
PFL
HHL
EEL
ACL
MAL II
HAA
LTL
CAA
Epa1-Np WT
MAA
PHA-L
HML-Native
Epa1-Np mutant Y228W
PsSC
Epa1-Np mutant E227G
EEA(EY Lab)
Avaren-Fc
CA19.9-1B844
PTL-I
SGA
RCAI (Vector)
MGL 6A-H262T
MPL
GSL-I
IPS-K-4A2B8
TJA-II
ab5
MGL 6B
LcH
Mono2R-CNL
PTL-II
NPA
PHA-E
rMpL
CTB
MOA
AAHL
UEA-I
rNV VLP-Cy3
MNA-M
ECL
TbFBL1
DBA
Mono2W-CNL
HHA(EY Lab)
GSL-I-B4
AFC-NVH
MGL 6A
Mouse Galectin1
GNA (EY Lab)
CnMBL
TL(EY Lab)
Galectin-1
GAC78
CnSep
Native HPA
Human Galectin1
WT-VCC
CC1G-10077
DC-ASGPR short
BTL native
SNA
LEL
ABA
DSL
WGA
UEAII
PNA
human Gal-7
ConA
Mouse NKR-P1C
DBL-1
LBL-152
STL
GSL-2
ME7(detected with a-HUD)
Gal-3C
WT Galectin-8
SRL
rSRL
ME5(detected with a-HUD)
P11 VP8
rPVL
mouse Gal-7
SSR1
GS-II (EY Lab)
TbFBL2
ConA
PHA-E
WGA(Seikagaku Bio)
WGA multimer
WGA
LSL-150
WGA
Gal-3

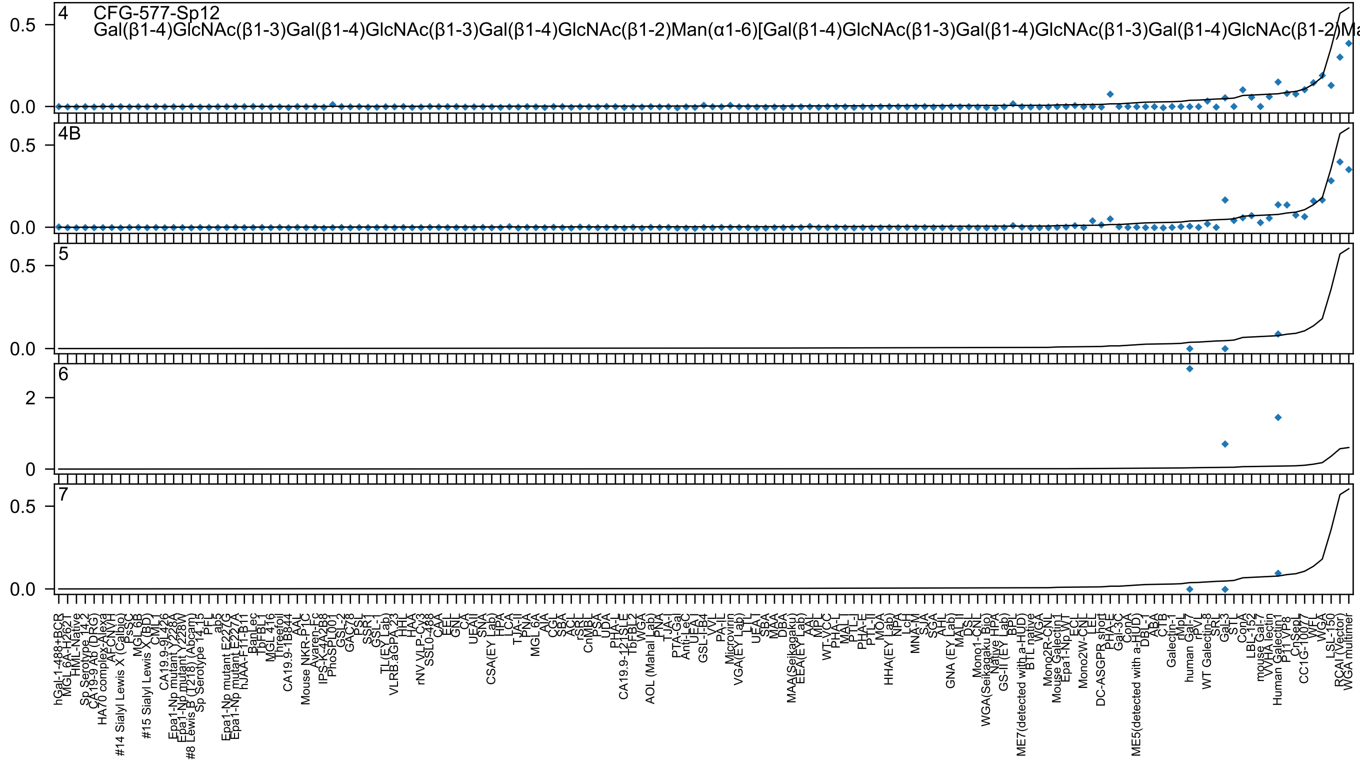
4
CFG-577-Sp12
Gal(β1-4)GlcNAc(β1-3)Gal(β1-4)GlcNAc(β1-3)Gal(β1-4)GlcNAc(β1-2)Man(α1-6)[Gal(β1-4)GlcNAc(β1-3)Gal(β1-4)GlcNAc(β1-3)Gal(β1-4)GlcNAc(β1-2)Ma
4B
5
6
7

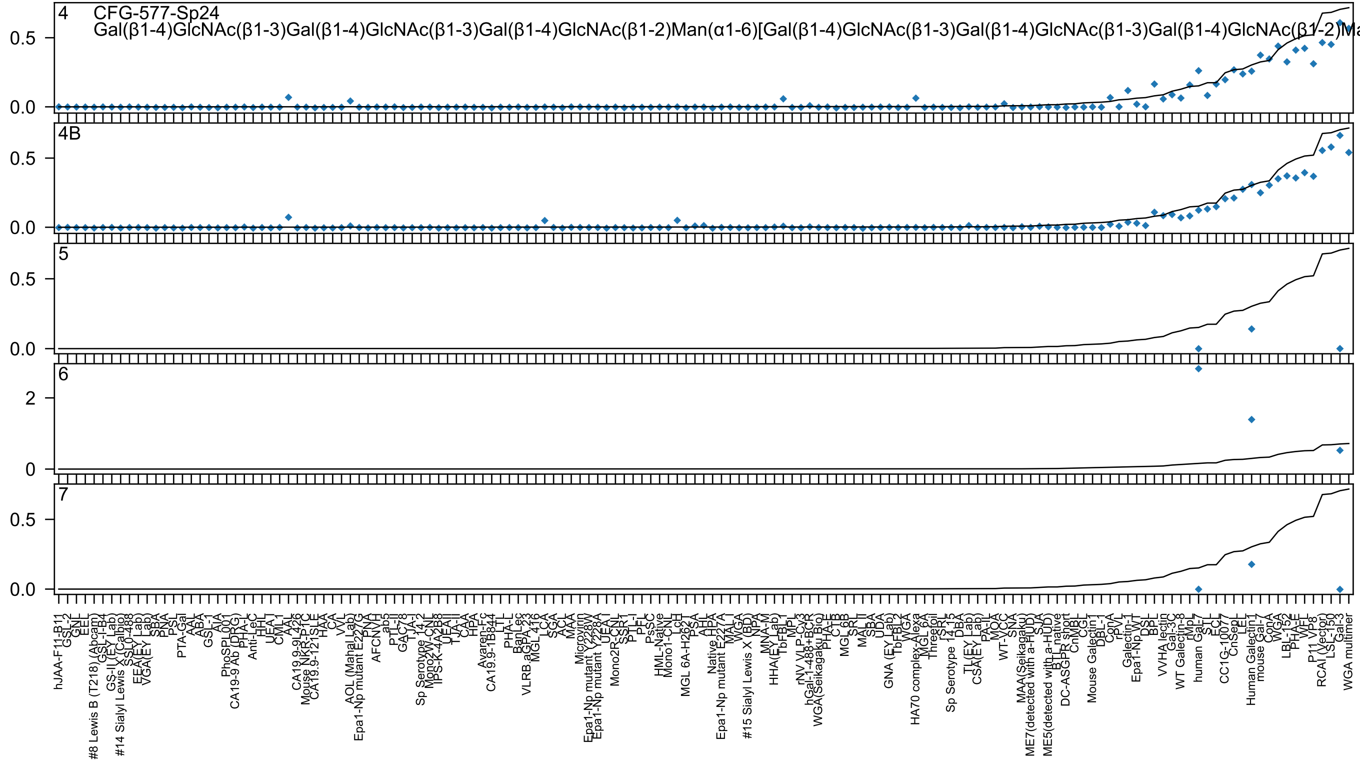
CFG-577-Sp24
Gal(β1-4)GlcNAc(β1-3)Gal(β1-4)GlcNAc(β1-3)Gal(β1-4)GlcNAc(β1-2)Man(α1-6)[Gal(β1-4)GlcNAc(β1-3)Gal(β1-4)GlcNAc(β1-3)Gal(β1-4)GlcNAc(β1-2)Ma

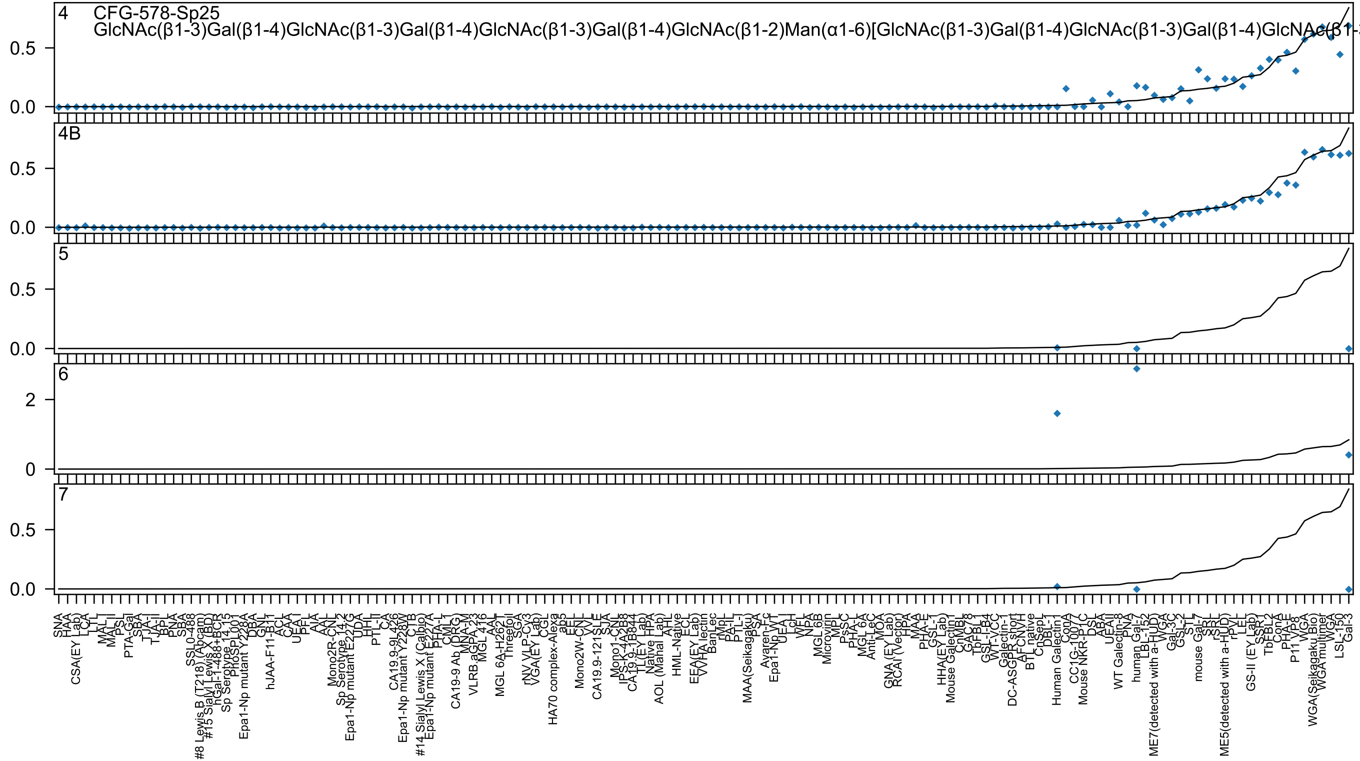

4
CFG-578-Sp25
GlcNAc(β1-3)Gal(β1-4)GlcNAc(β1-3)Gal(β1-4)GlcNAc(β1-3)Gal(β1-4)GlcNAc(β1-2)Man(α1-6)[GlcNAc(β1-3)Gal(β1-4)GlcNAc(β1-3)Gal(β1-4)GlcNAc(β1-
4B
5
6
7

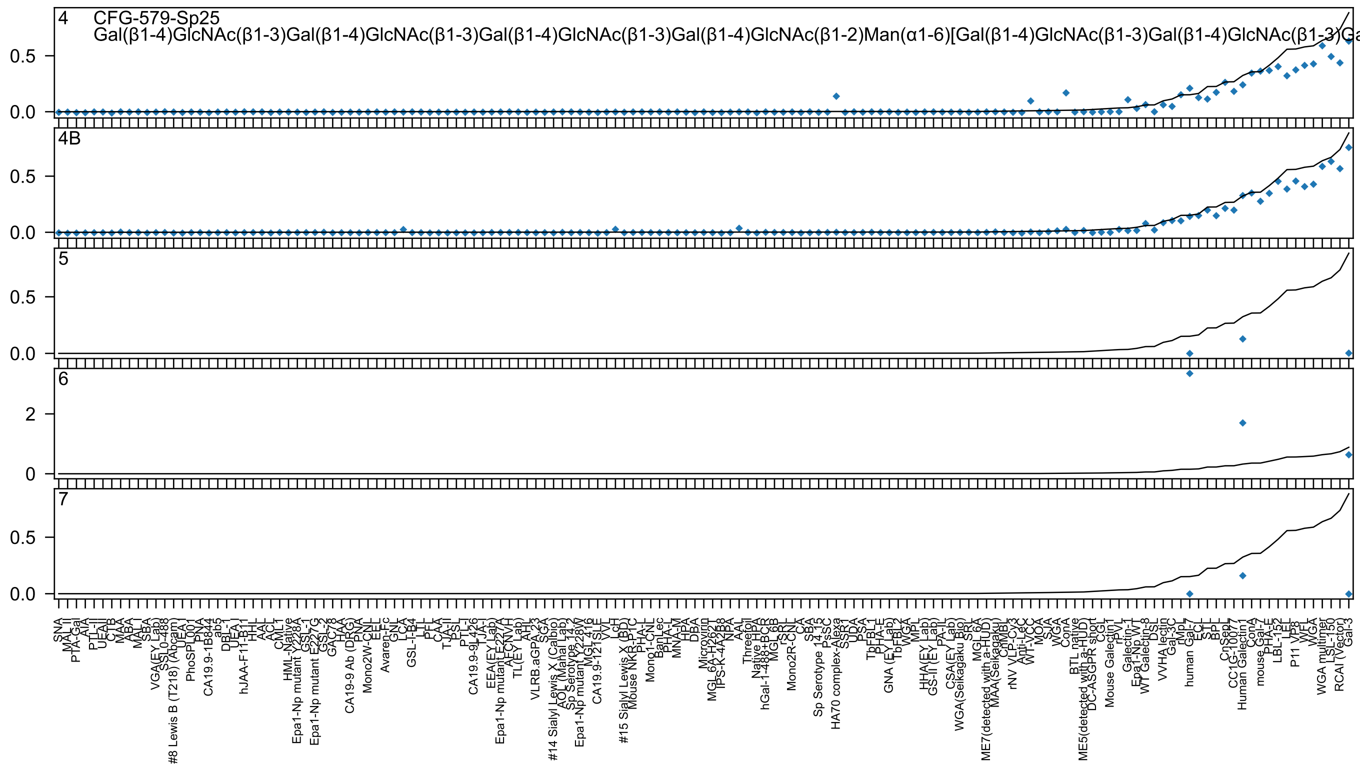

4
CFG-579-Sp25
Gal(β1-4)GlcNAc(β1-3)Gal(β1-4)GlcNAc(β1-3)Gal(β1-4)GlcNAc(β1-3)Gal(β1-4)GlcNAc(β1-2)Man(α1-6)[Gal(β1-4)GlcNAc(β1-3)Gal(β1-4)GlcNAc(β1-3)Ga
4B
5
6
7

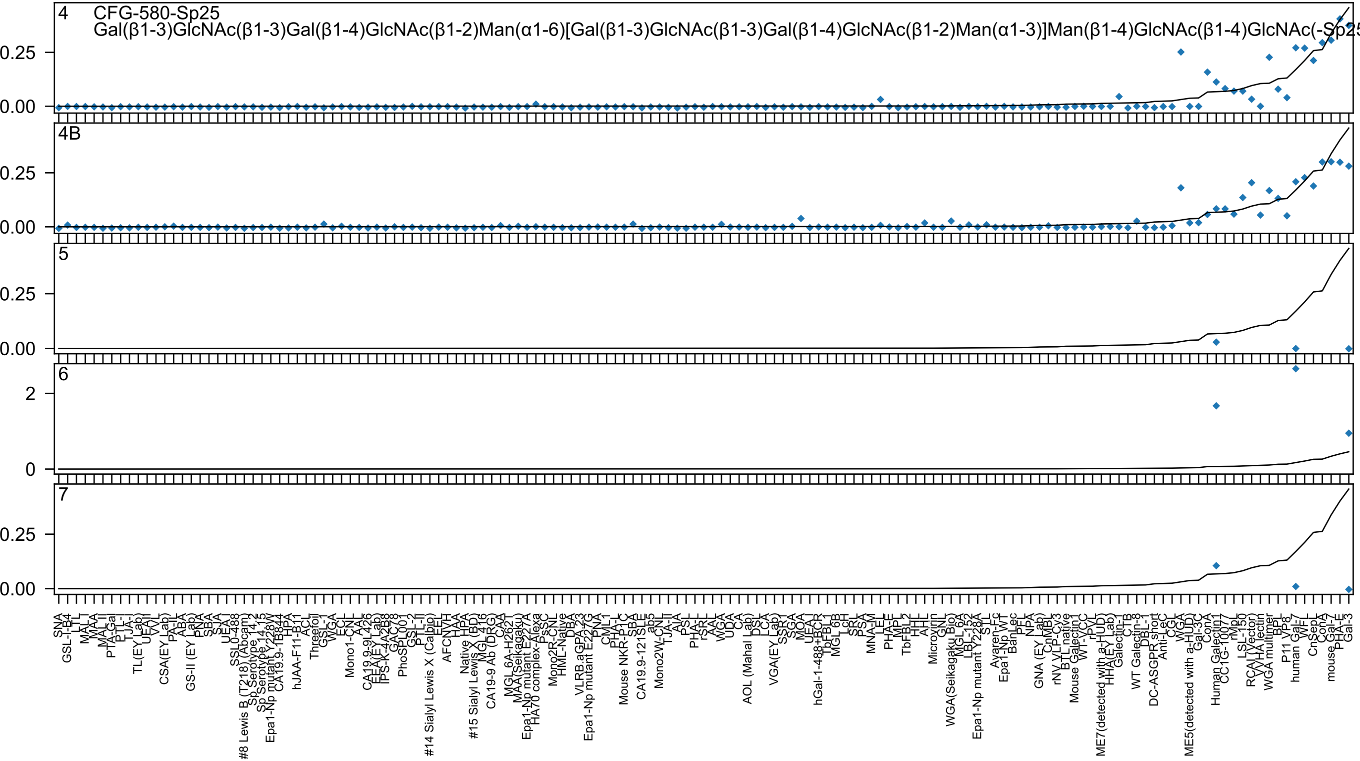
4
CFG-580-Sp25
Gal(β1-3)GlcNAc(β1-3)Gal(β1-4)GlcNAc(β1-2)Man(α1-6)[Gal(β1-3)GlcNAc(β1-3)Gal(β1-4)GlcNAc(β1-2)Man(α1-3)]Man(β1-4)GlcNAc(β1-4)GlcNAc(-Sp25
0.25
0.00
4B
0.25
0.00
5
0.25
0.00
6
2
0
7
0.25
0.00
SNA
GSL-I-B4
LTL
MAL-I
MAA
MAL_II
PTA-Gal
PTL-I
TJA-I
TL(EY Lab)
UEA-II
VVL
CSA(EY Lab)
PA-IL
ABA
GS-II (EY Lab)
PNA
SBA
SJA
UEA-I
SSL0-488
#8 Lewis B (T218) (Abcam)
Sp Serotype 14.2
Sp Serotype 14.15
Epa1-Np mutant Y228W
CA19.9-1B844
HPA
hJAA-F11-B11
ACL
Threefoil
GSL-I
WGA
ECL
Mono1-CNL
AAL
CA19.9-9L426
EEA(EY Lab)
IPS-K-4A2B8
GAC78
PhoSL001
GSL-2
PTL-II
#14 Sialyl Lewis X (Calbio)
EEL
AFCNVH
HAA
Native HPA
#15 Sialyl Lewis X (BD)
MGL 416
CA19-9 Ab (DRG)
CAA
MGL 6A-H2621
MAA(Seikagaku)
Epa1-Np mutant E227A
HA70 complex-Alexa
PsSC
Mono2R-CNL
HML-Native
DBA
VLRB.aGPA.23
Epa1-Np mutant E227G
PNA
CML1
PHA-L
Mouse NKR-P1C
SBA
CA19.9-121SLE
ab5
Mono2W-CNL
TJA-II
AIA
PSL
PHA-L
rSRL
AAL
WGA
UDA
CA
AOL (Mahal Lab)
DSL
LCA
VGA(EY Lab)
SSA
SGA
MOA
UEA-I
hGal-1-488+BCR
TbFBL1
MGL 6B
LcH
SRL
PSA
MNA-M
LEL
PHA-E
MPL
TbFBL2
HHL
AHL
Microvirin
GNL
WGA(Seikagaku Bio)
MGL 6A
LBL-152
Epa1-Np mutant Y228A
STL
Ayaren-Fc
Epa1-Np WT
BanLec
PEL
NPA
GNA (EY Lab)
CnMBL
rNV VLP-Cy3
BTL native
Mouse Galectin1
WT-VCC
rPVL
ME7(detected with a-HUD)
HHA(EY Lab)
Galectin-1
CTB
WT Galectin-8
DBL-1
DC-ASGPR short
Anti-LeC
CGL
WGA
ME5(detected with a-HUD)
Gal-3C
ConA
Human Galectin1
CC1G-10077
rMp
LSL-150
RCA I (Vector)
VVHA lectin
WGA multimer
BPL
P11 VP8
human Gal-7
WFL
CnSepL
ConA
mouse Gal-7
PHA-E
Gal-3

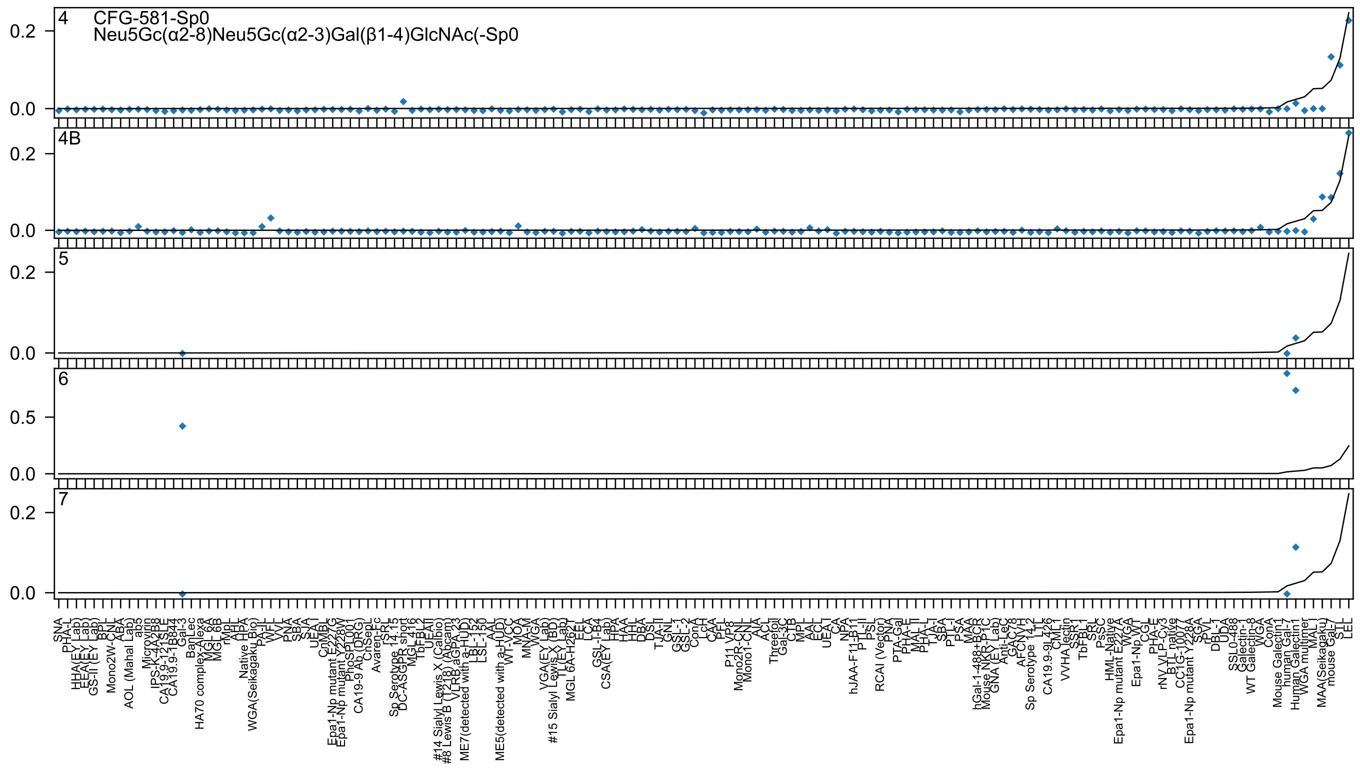
4
CFG-581-Sp0
Neu5Gc(α2-8)Neu5Gc(α2-3)Gal(β1-4)GlcNAc(-Sp0
4B
5
6
7
0.2
0.0
0.5
SNA
PHA-L
HHA(EY Lab)
EEA(EY Lab)
GS-II (EY Lab)
BPL
Mono2W-CNL
ABA
AOL (Mahal Lab)
ab5
Microvirin
IPS-K-4A2B8
CA19.9-121SLE
CA19.9-1B844
Gal-3
BanLec
HA70 complex-Alexa
MGL 6A
MGL 6B
rMpL
AHL
Native HPA
WGA(Seikagaku Bio)
PA-IL
WFL
VVL
PNA
SBA
SJA
UEA I
CnMBL
Epa1-Np mutant E227G
Epa1-Np mutant Y228W
PhoSL001
CA19-9 Ab (DRG)
ChSepL
Avaren-Fc
TSRL
Sp Serotype 14.15
DC-ASGPR short
MGL 416
TbFBL2
UEAII
#14 Sialyl Lewis X (Calbio)
#8 Lewis B (T218) (Abcam)
VLRB aGPA.23
ME7(detected with a-HUD)
LBL-152
LSL-150
AAL
ME5(detected with a-HUD)
WT-VCC
MOA
MNA-M
WGA
VGA(EY Lab)
#15 Sialyl Lewis X (BD)
TL(EY Lab)
MGL 6A-H262T
EEL
LCA
GSL-I-B4
CSA(EY Lab)
HPA
HAA
HHL
DBA
DSL
TJA-II
GNL
GSL-1
GSL-2
ConA
LcH
CAA
PFL
P11 VP8
Mono2R-CNL
Mono1-CNL
AIA
ACL
Threefoil
Gal-3C
CTB
MPL
AAL
ECL
UEA I
CA
NPA
hJAA-F11-B11
PTL-II
PSL
RCAI (Vector)
PNA
PTA-Gal
PHA-E
MAL II
PHA-L
TJA-I
SBA
PTL-I
PSA
MAA
hGal-1-488+BCR
Mouse NKR-P1C
GNA (EY Lab)
Anti-LeC
GAC78
AFCNYH
Sp Serotype 14.2
LTL
CA19.9-9L426
CMFL
VVHA lectin
SSR1
TbFBL1
SRL
PsSC
HML-Native
Epa1-Np mutant E227A
WGA
Epa1-Np WT
CGL
PHA-E
rNV VLP-Cy3
BTL native
CC1G-10077
Epa1-Np mutant Y228A
SGA
rPVL
DBL-1
UDA
SSL0-488
Galectin-1
WT Galectin-8
WGA
ConA
Mouse Galectin1
human Gal-7
Human Galectin1
WGA multimer
MAL I
MAA(Seikagaku)
mouse Gal-7
STL
LEL

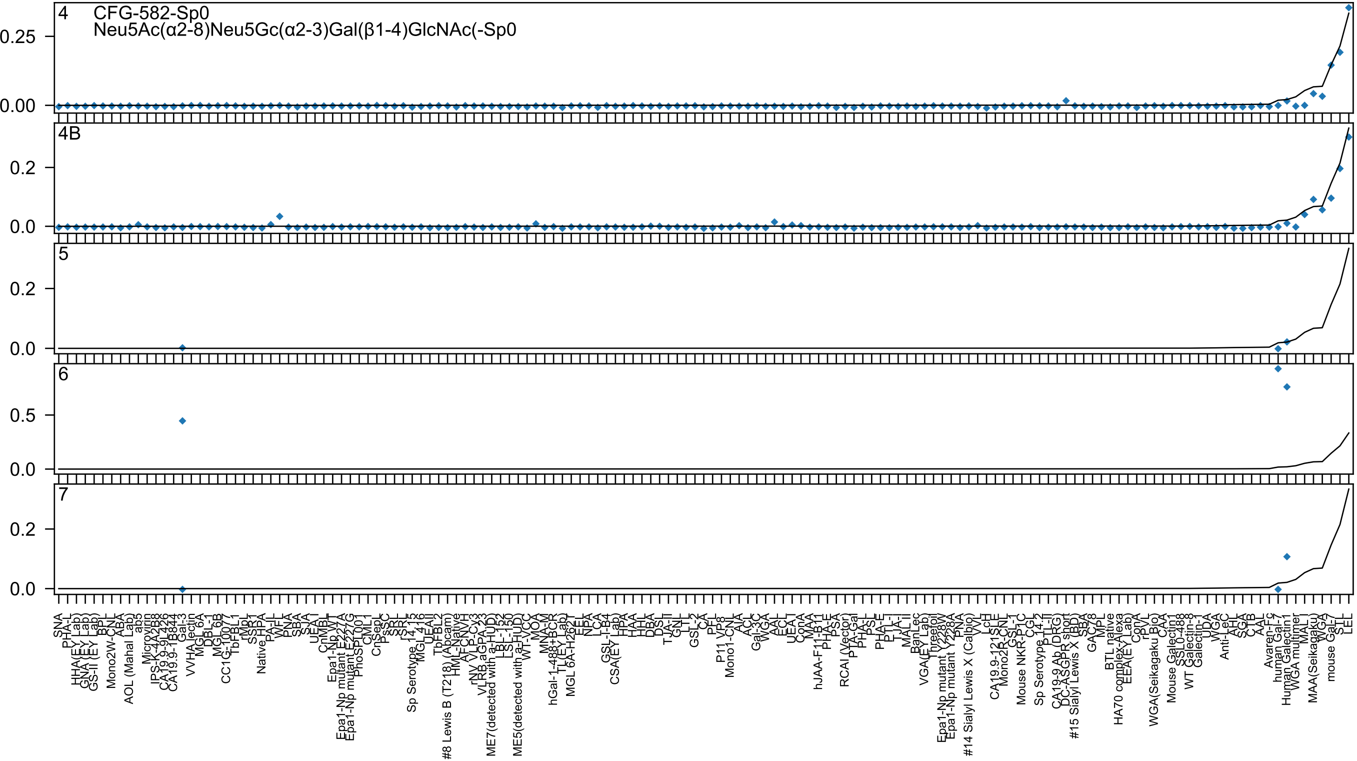

4
CFG-582-Sp0
Neu5Ac(α2-8)Neu5Gc(α2-3)Gal(β1-4)GlcNAc(-Sp0
0.25
0.00
4B
0.2
0.0
5
0.2
0.0
6
0.5
0.0
7
0.2
0.0

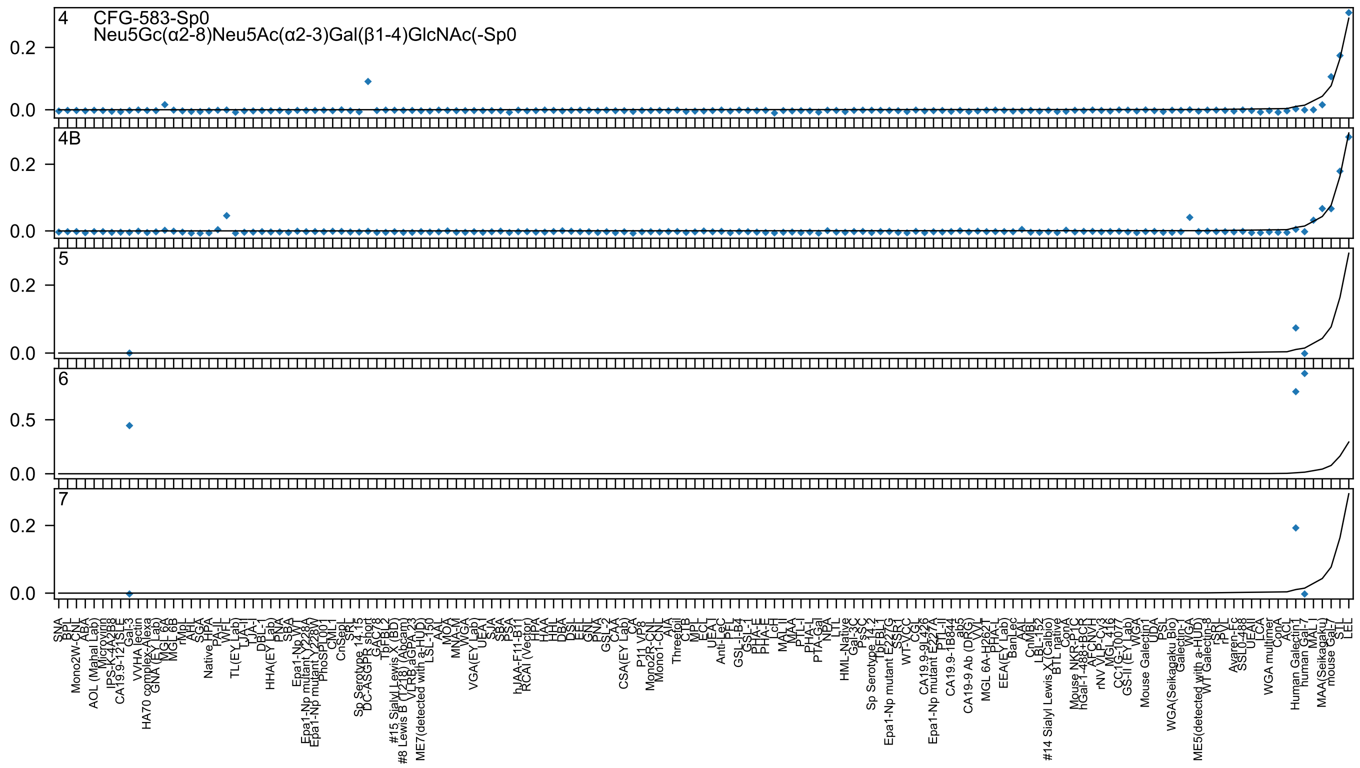

4
CFG-583-Sp0
Neu5Gc(α2-8)Neu5Ac(α2-3)Gal(β1-4)GlcNAc(-Sp0
0.2
0.0
4B
0.2
0.0
5
0.2
0.0
6
0.5
0.0
7
0.2
0.0

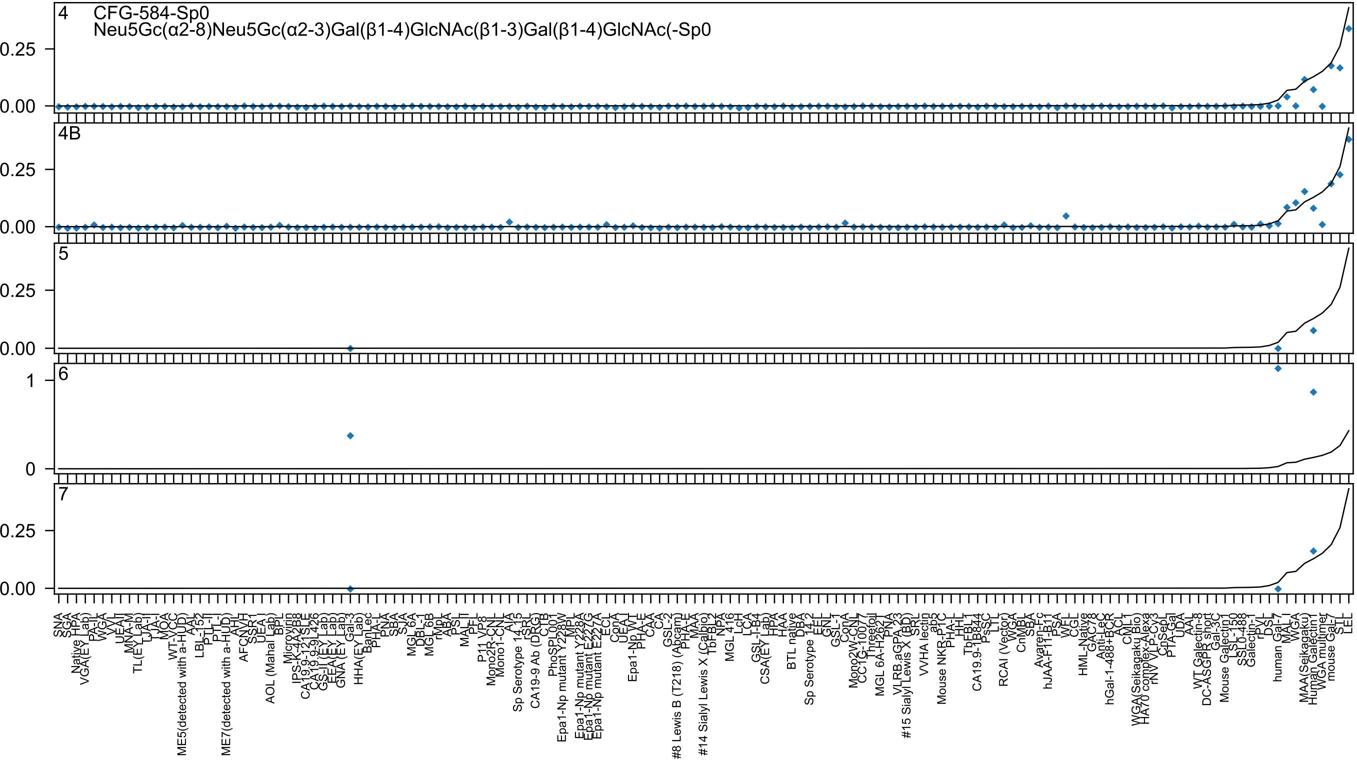
4
CFG-584-Sp0
Neu5Gc(α2-8)Neu5Gc(α2-3)Gal(β1-4)GlcNAc(β1-3)Gal(β1-4)GlcNAc(-Sp0
0.25
0.00
4B
0.25
0.00
5
0.25
0.00
6
1
0
7
0.25
0.00
SNA
SGA
Native HPA
VGA(EY Lab)
PA-IL
WGA
VVL
UEA-II
MNA-M
TL(EY Lab)
LTA-I
LJA-I
MOA
WT-VCC
ME5(detected with a-HUD)
AAL
LBL-152
PTL-II
PTL-I
ME7(detected with a-HUD)
AHL
AFCNVH
SSR1
UEA I
AOL (Mahal Lab)
BPL
Microvirin
IPS-K-4A2B8
CA19.9-121SLE
CA19.9-9L426
GS-II (EY Lab)
EEA(EY Lab)
GNA (EY Lab)
Gal-3
HHA(EY Lab)
BanLec
PHA-L
PNA
SBA
SJA
MGL 6A
DBL-1
MGL 6B
rMpL
ABA
PSL
MAL II
PFL
P11 VP8
Mono2R-CNL
Mono1-CNL
AIA
Sp Serotype 14_15
rSRL
CA19-9 Ab (DRG)
CTB
PhoSPL001
Epa1-Np mutant Y228W
MPL
Epa1-Np mutant Y228A
Epa1-Np mutant E227G
Epa1-Np mutant E227A
ECL
ConA
UEA-I
Epa1-Np WT
PHA-E
CAA
CA
GSL-2
#8 Lewis B (T218) (Abcam)
PHA-E
MAA
#14 Sialyl Lewis X (Calbio)
TbFBL2
NPA
MGL 416
LcH
LCA
GSL-I-B4
CSA(EY Lab)
HPA
HAA
BTL native
DBA
Sp Serotype 14_2
EEL
GNL
GSL-1
ConA
Mono2W-CNL
CC1G-10077
Threefoil
MGL 6A-H262T
PNA
VLRB.aGPA.23
#15 Sialyl Lewis X (BD)
SRL
VVHA lectin
ab5
Mouse NKR-P1C
PHA-L
HHL
TbFBL1
CA19.9-1B844
PsSC
LTL
RCAI (Vector)
WGA
CnMBL
SBA
Avaren-Fc
hJAA-F11-B11
PSA
WFL
CGL
HML-Native
GAC78
Anti-LeC
hGal-1-488+BCR
ACL
CML1
WGA(Seikagaku Bio)
HA70 complex-Alexa
rNV VLP-Cy3
CnSepL
PTA-Gal
UDA
AAL
WT Galectin-8
DC-ASGPR short
Gal-3C
Mouse Galectin1
LSL-150
SSL0.488
Galectin-1
rPyL
DSL
human Gal-7
MAL-I
WGA
MAA(Seikagaku)
Human Galectin1
WGA multimer
mouse Gal-7
STL
LEL

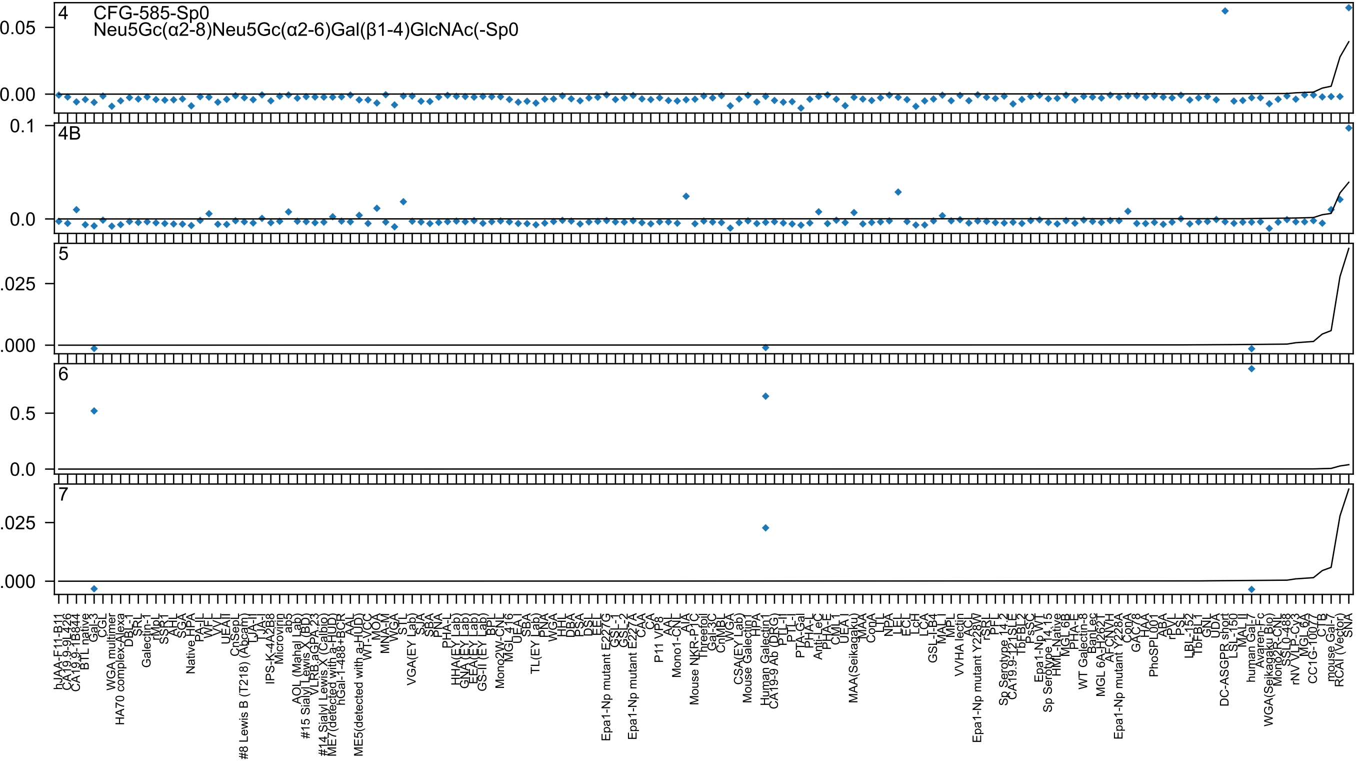

4
CFG-585-Sp0
Neu5Gc(α2-8)Neu5Gc(α2-6)Gal(β1-4)GlcNAc(-Sp0
4B
5
6
7

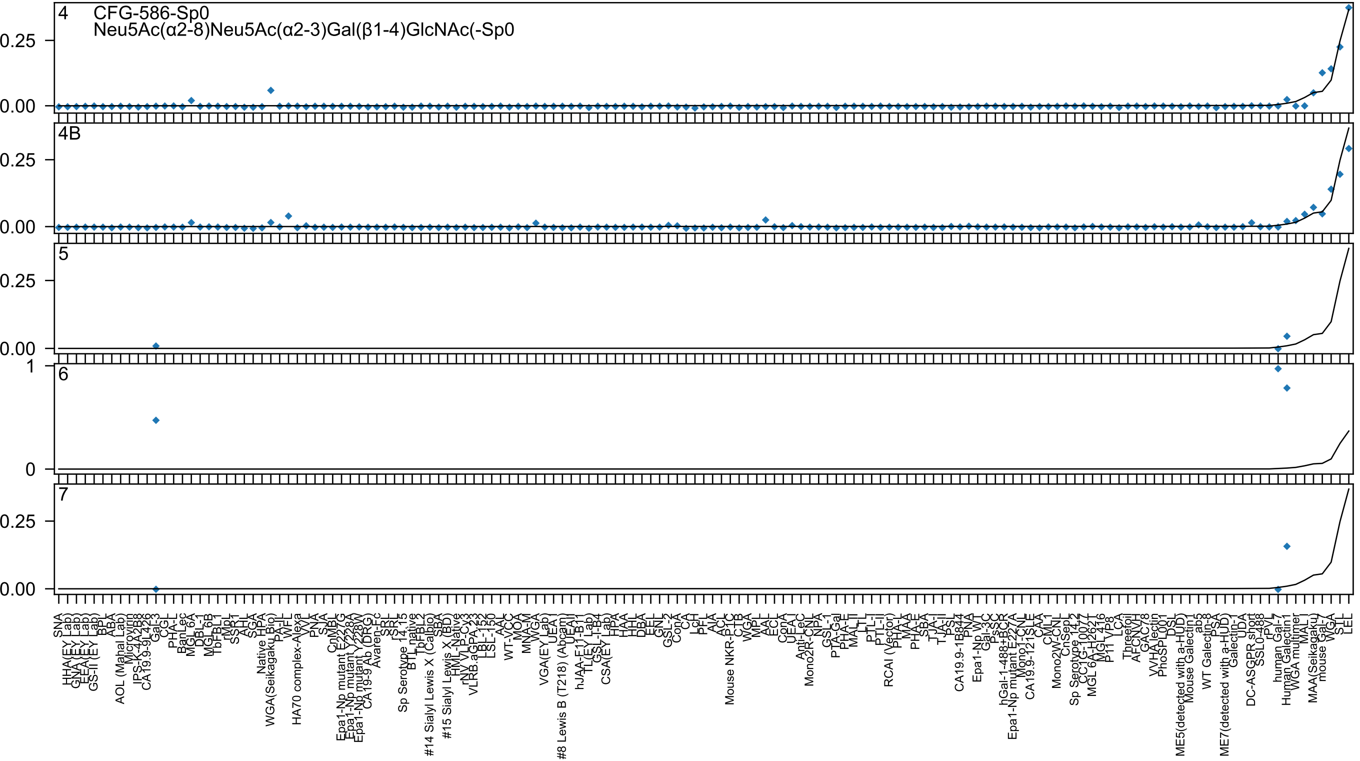
4
CFG-586-Sp0
Neu5Ac(α2-8)Neu5Ac(α2-3)Gal(β1-4)GlcNAc(-Sp0
4B
5
6
7
0.25
0.00
1
0

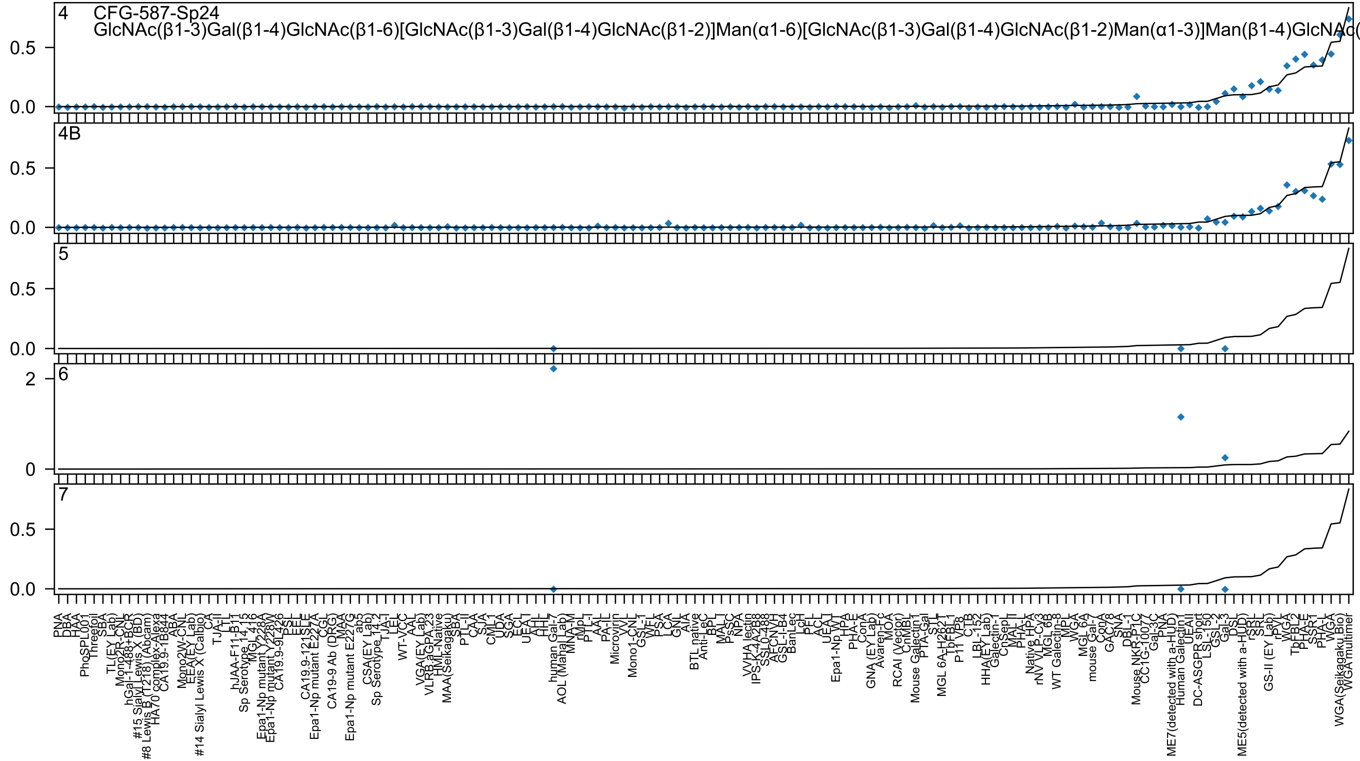
4 CFG-587-Sp24
GlcNAc(β1-3)Gal(β1-4)GlcNAc(β1-6)[GlcNAc(β1-3)Gal(β1-4)GlcNAc(β1-2)]Man(α1-6)[GlcNAc(β1-3)Gal(β1-4)GlcNAc(β1-2)Man(α1-3)]Man(β1-4)GlcNAc(
4B
5
6
7

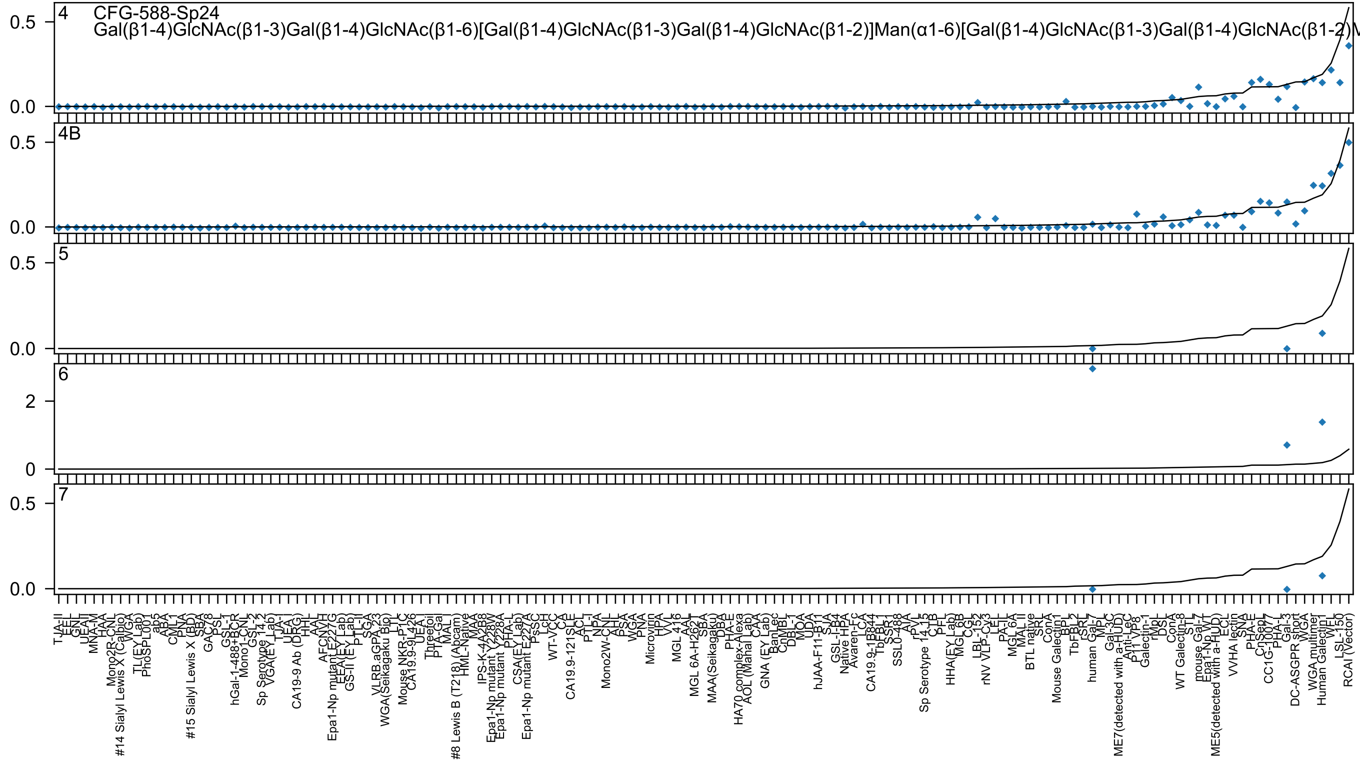
4
CFG-588-Sp24
Gal(β1-4)GlcNAc(β1-3)Gal(β1-4)GlcNAc(β1-6)[Gal(β1-4)GlcNAc(β1-3)Gal(β1-4)GlcNAc(β1-2)]Man(α1-6)[Gal(β1-4)GlcNAc(β1-3)Gal(β1-4)GlcNAc(β1-2)M
4B
5
6
7
0.0
0.5
0
2

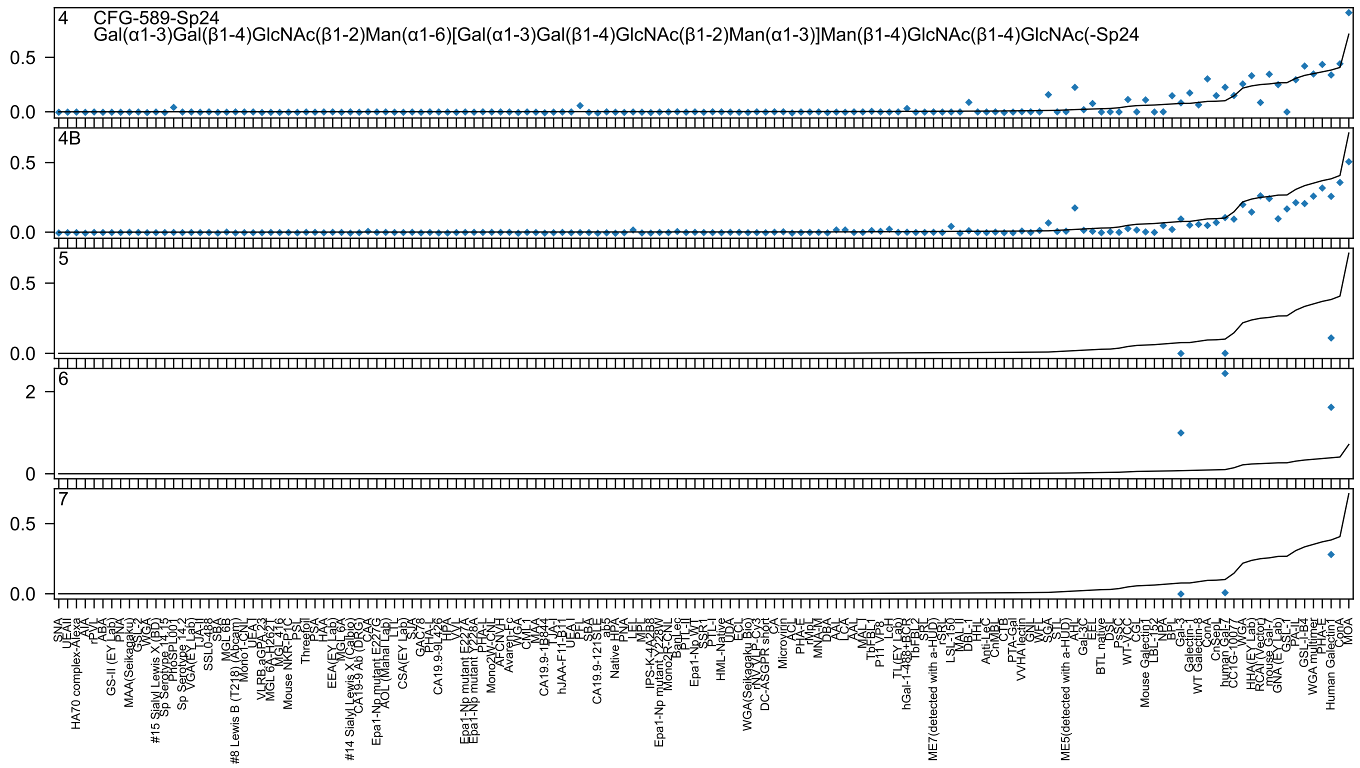
4
CFG-589-Sp24
Gal(α1-3)Gal(β1-4)GlcNAc(β1-2)Man(α1-6)[Gal(α1-3)Gal(β1-4)GlcNAc(β1-2)Man(α1-3)]Man(β1-4)GlcNAc(β1-4)GlcNAc(-Sp24
4B
5
6
7
0.0
0.5
0
2
SNA
UEAII
HA70 complex-Alexa
AIA
rPVL
ABA
GS-II (EY Lab)
PNA
MAA(Seikagaku)
GSL-2
WGA
#15 Sialyl Lewis X (BD)
Sp Serotype 14.15
PhoSPL001
Sp Serotype 14.2
VGA(EY Lab)
TJA-I
SSL0-488
SBA
MGL 6B
#8 Lewis B (T218) (Abcam)
Mono1-CNL
UEA-I
VLRB.aGPA.23
MGL 6A-H262T
MGL 416
Mouse NKR-P1C
PSL
Threefoil
PSA
HAA
EEA(EY Lab)
MGL 6A
#14 Sialyl Lewis X (Calbio)
CA19-9 Ab (DRG)
CAA
Epa1-Np mutant E227G
AOL (Manal Lab)
LTL
CSA(EY Lab)
SJA
GAC78
PHA-L
CA19.9-9L426
HPA
VVL
Epa1-Np mutant E227A
Epa1-Np mutant Y228A
PHA-L
Mono2W-CNL
AFCNVH
Avaren-Fc
WGA
CML-1
MAA
CA19.9-1B844
TJA-I
hJAA-F11-B1
UEA-I
PFL
SBA
CA19.9-121SLE
ab5
Native HPA
PNA
LEL
MPL
IPS-K-4A2B8
Epa1-Np mutant Y228W
Mono2R-CNL
BanLec
PTL-II
Epa1-Np WT
SSR1
PTL-I
HML-Native
UDA
ECL
WGA(Seikagaku Bio)
rNV VLP-Cy3
DC-ASGPR short
CA
Microvirin
ACL
PHA-E
rMpL
MNA-M
DBA
AAL
LCA
AAL
MAL-I
TbFBL-I
P11 VP8
LcH
TL(EY Lab)
hGal-1-488+BCR
TbFBL2
SRL
ME7(detected with a-HUD)
rSRL
LSL-150
MAL-II
DBL-I
HHL
Anti-LeC
CnMBL
CTB
PTA-Gal
VVHA lectin
GNL
WFL
SGA
STL
ME5(detected with a-HUD)
AHL
Gal-3C
EEL
BTL native
DSL
PsSC
WT-VCC
CGL
Mouse Galectin1
LBL-152
NPA
BPL
Gal-3
Galectin-8
WT Galectin-8
ConA
CnSepL
human Gal-7
CC1G-10077
WGA
HHA(EY Lab)
RCAI (Vector)
mouse Gal-7
GNA (EY Lab)
GSL-I
PA-IL
GSL-I-B4
WGA multimer
PHA-E
Human Galectin1
ConA
MOA

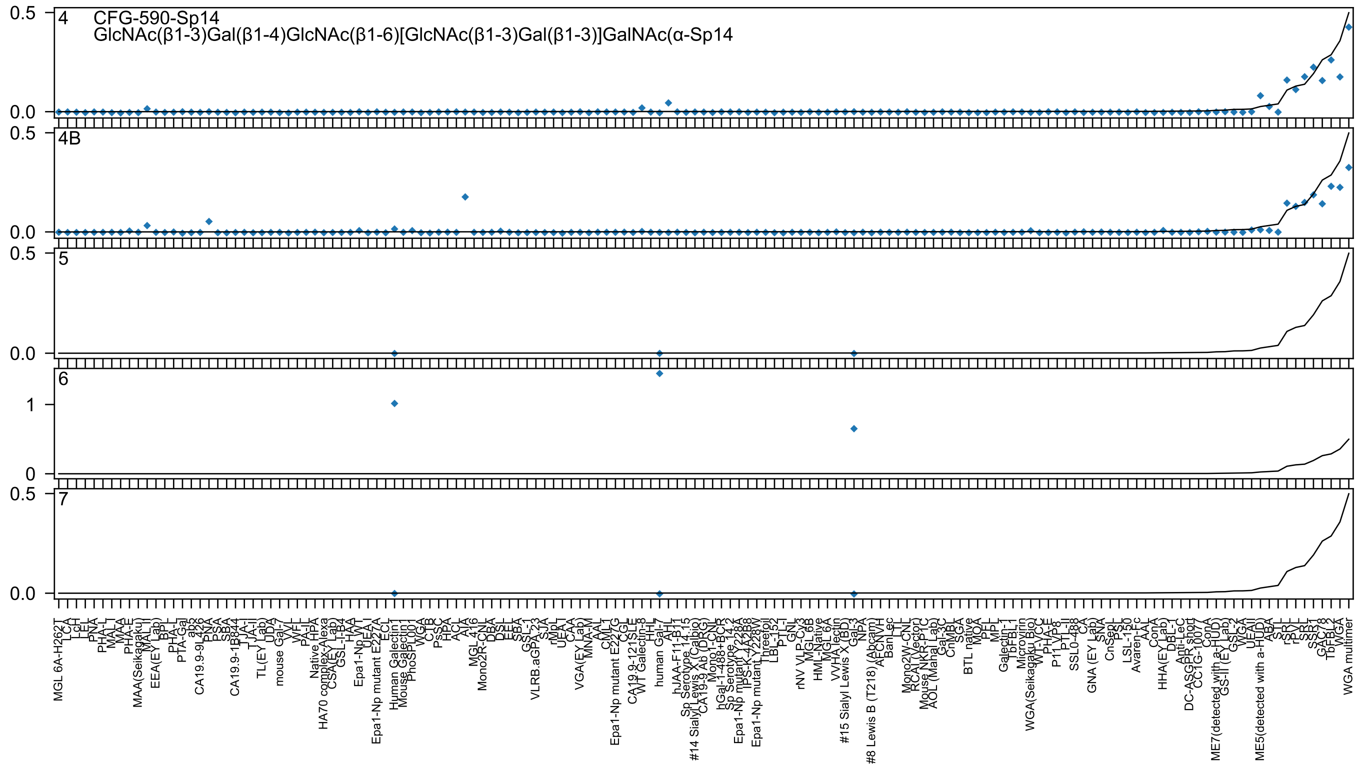
CFG-590-Sp14
GlcNAc(β1-3)Gal(β1-4)GlcNAc(β1-6)[GlcNAc(β1-3)Gal(β1-3)]GalNAc(α-Sp14
4
4B
5
6
7

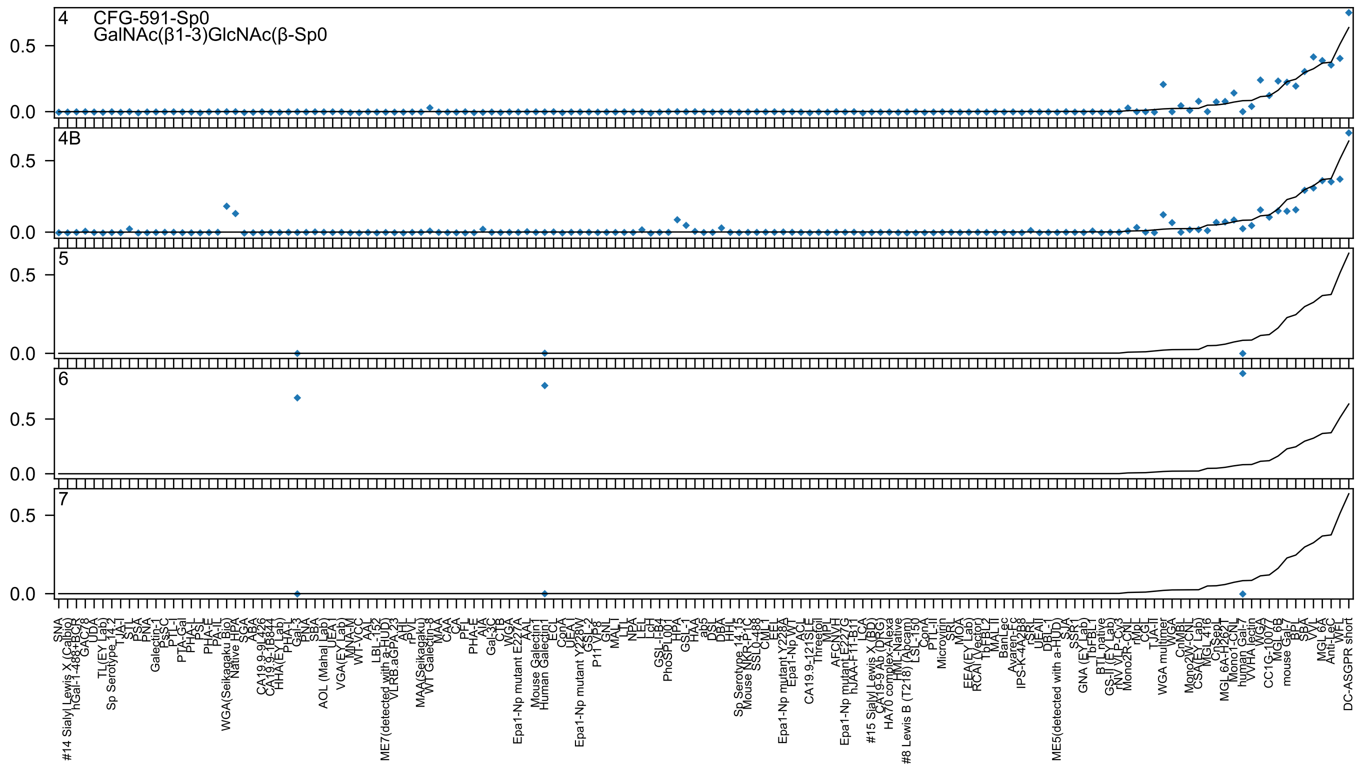
4 CFG-591-Sp0
GalNAc(β1-3)GlcNAc(β-Sp0
4B
5
6
7

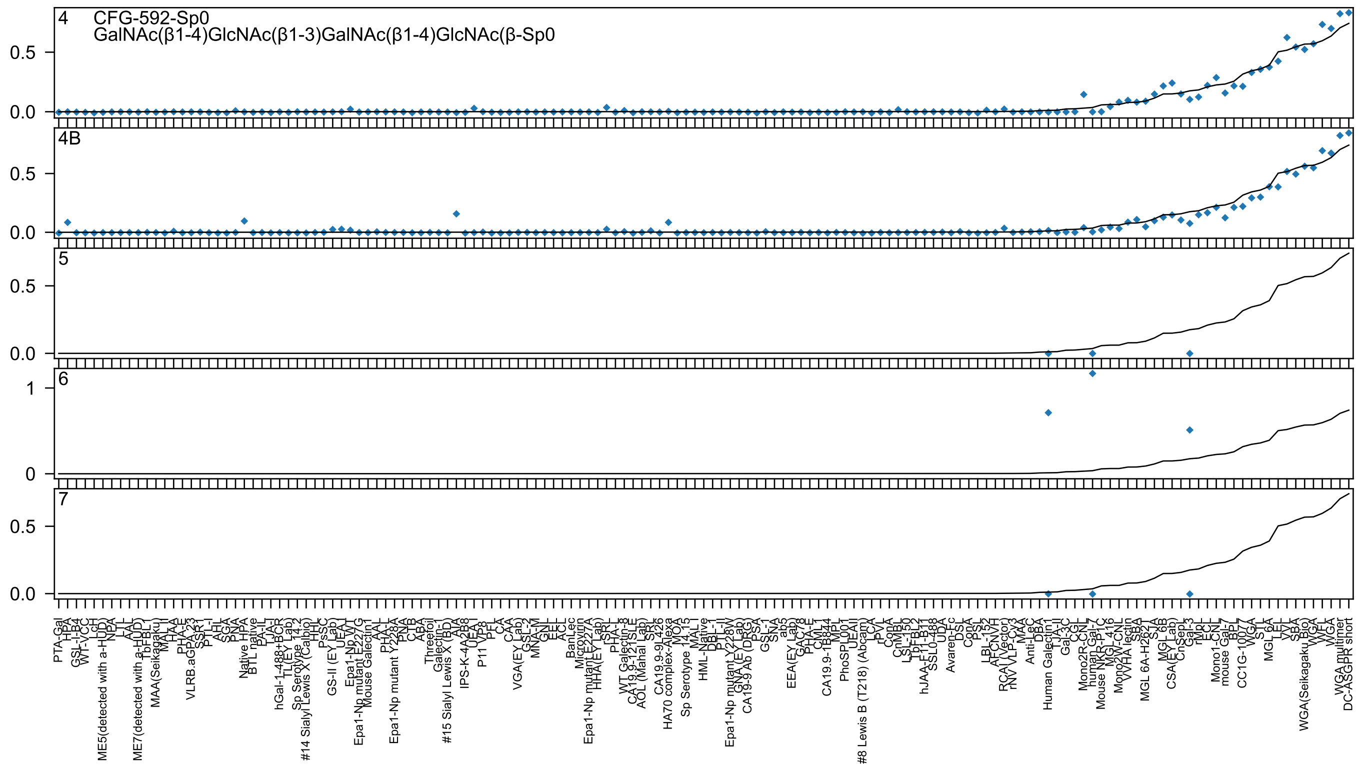
4
CFG-592-Sp0
GalNAc(β1-4)GlcNAc(β1-3)GalNAc(β1-4)GlcNAc(β-Sp0
0.5
0.0
4B
0.5
0.0
5
0.5
0.0
6
1
0
7
0.5
0.0

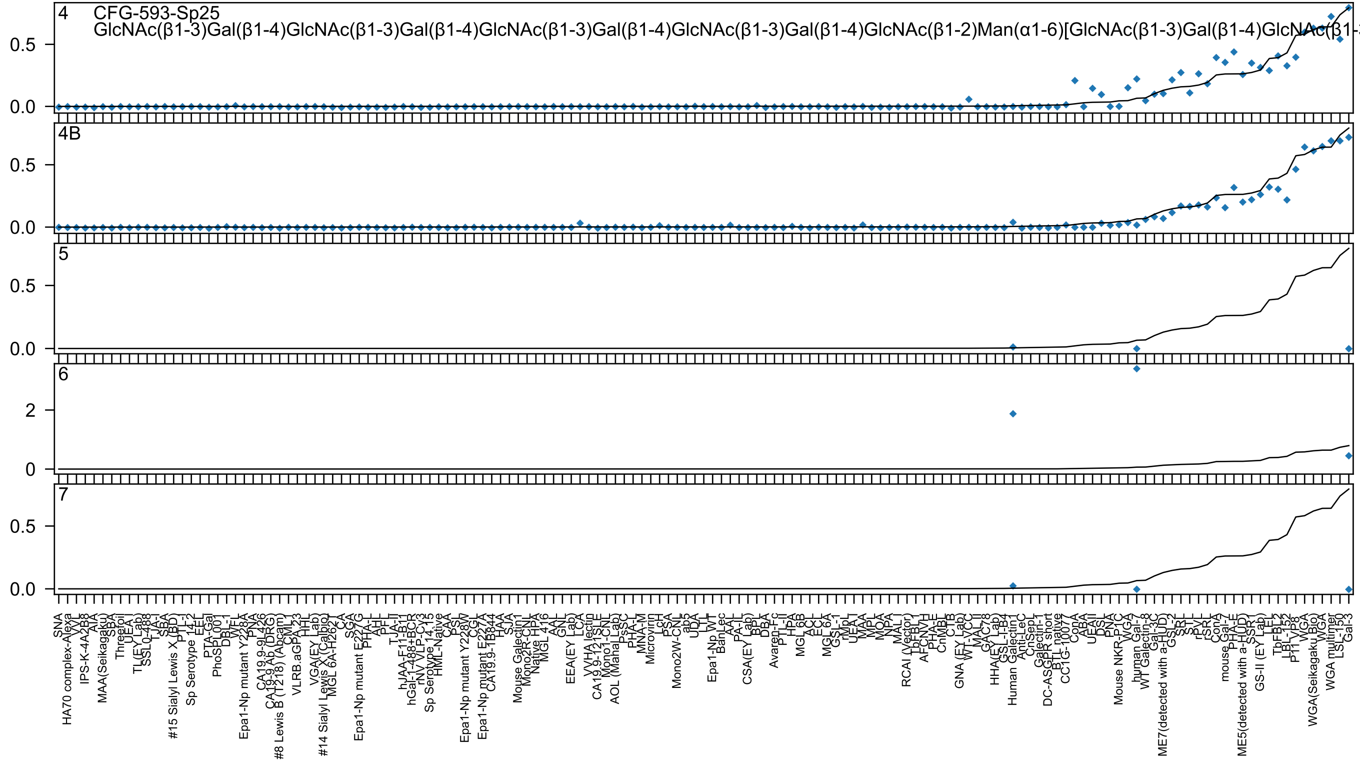

4 CFG-593-Sp25
GlcNAc(β1-3)Gal(β1-4)GlcNAc(β1-3)Gal(β1-4)GlcNAc(β1-3)Gal(β1-4)GlcNAc(β1-3)Gal(β1-4)GlcNAc(β1-2)Man(α1-6)[GlcNAc(β1-3)Gal(β1-4)GlcNAc(β1-
4B
5
6
7

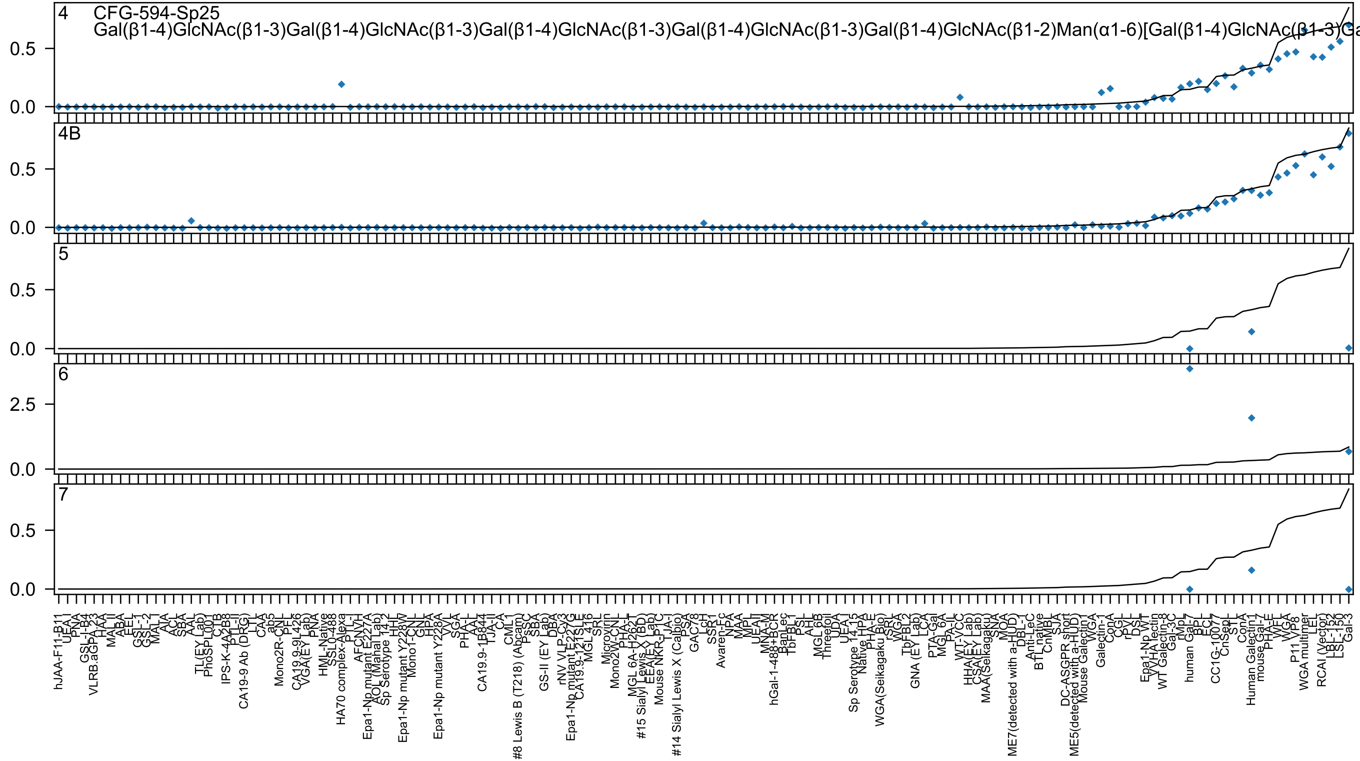
4 CFG-594-Sp25
Gal(β1-4)GlcNAc(β1-3)Gal(β1-4)GlcNAc(β1-3)Gal(β1-4)GlcNAc(β1-3)Gal(β1-4)GlcNAc(β1-3)Gal(β1-4)GlcNAc(β1-2)Man(α1-6)[Gal(β1-4)GlcNAc(β1-3)Ga
4B
5
6
7

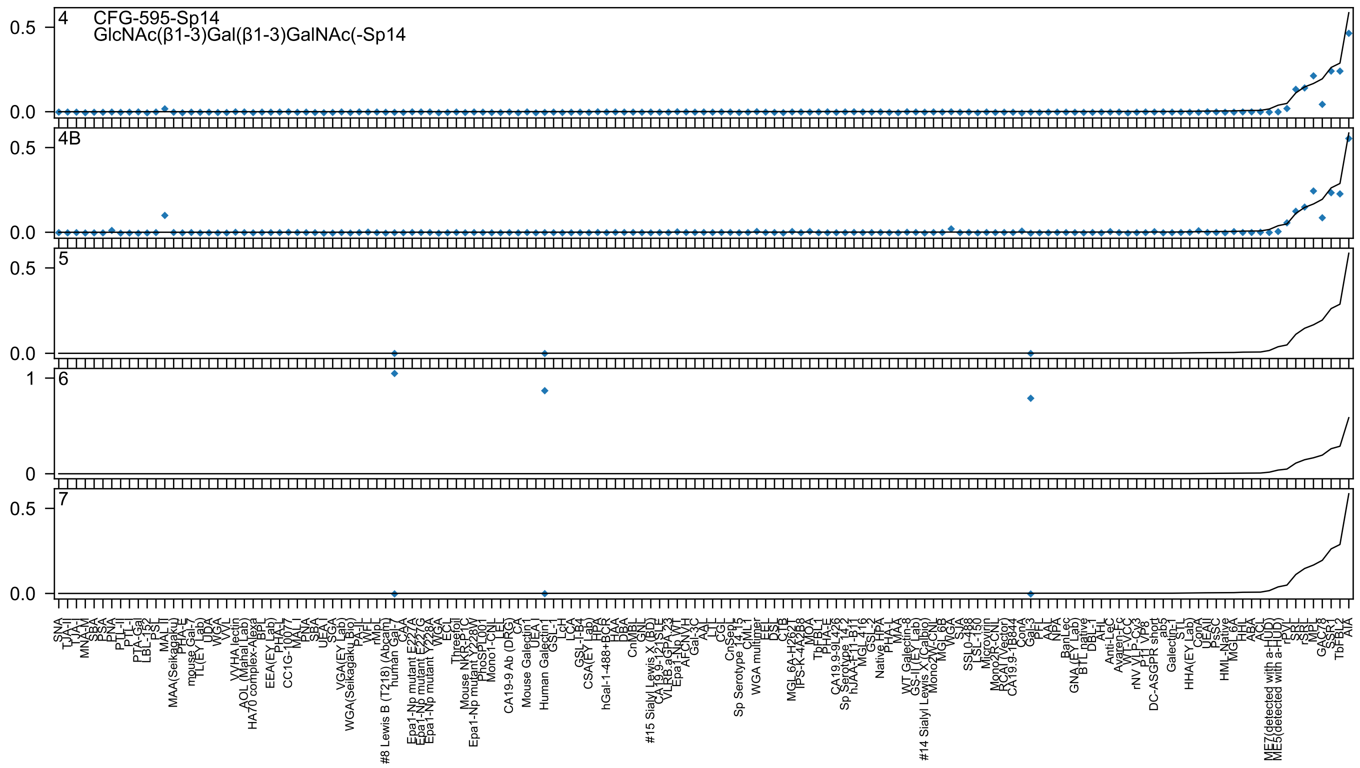

4
CFG-595-Sp14
GlcNAc(β1-3)Gal(β1-3)GalNAc(-Sp14
4B
5
6
7
0.5
0.0
1
0

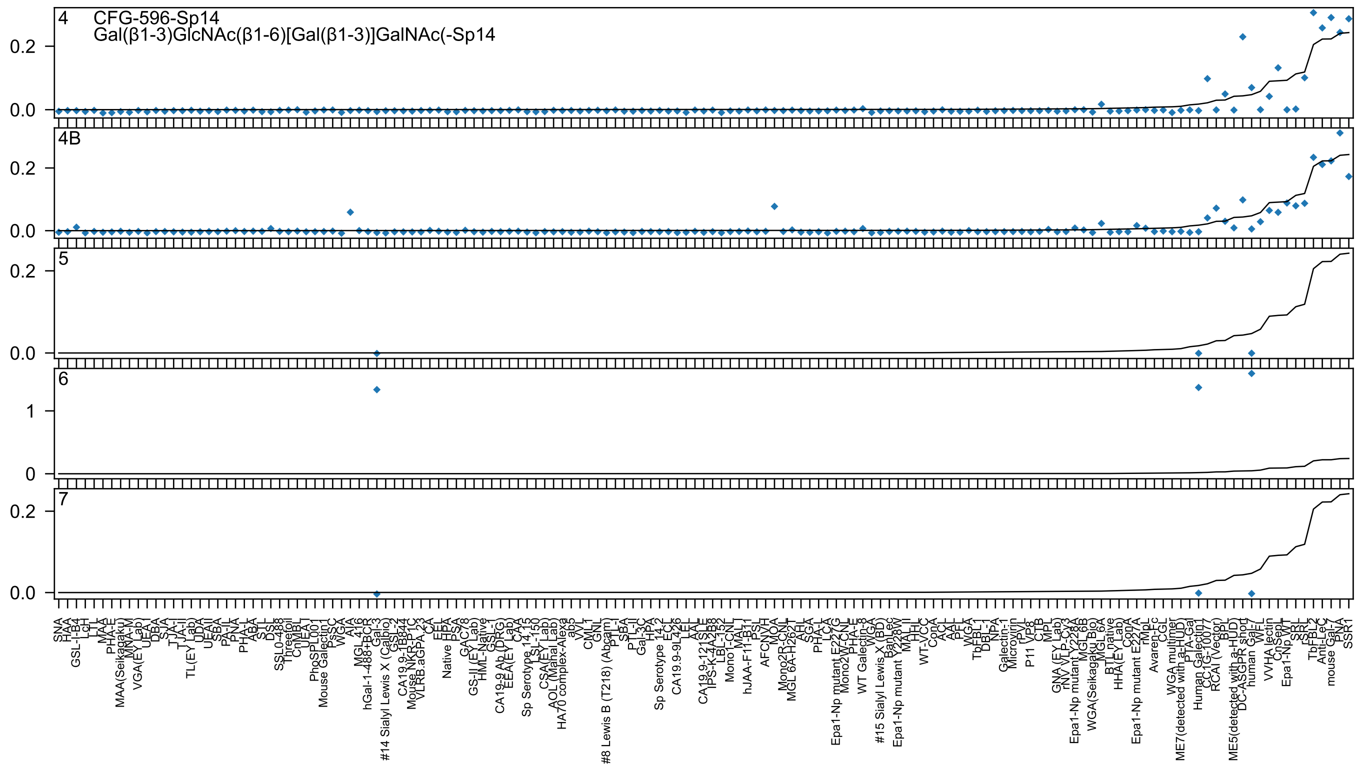

4 CFG-596-Sp14
Gal(β1-3)GlcNAc(β1-6)[Gal(β1-3)]GalNAc(-Sp14
4B
5
6
7

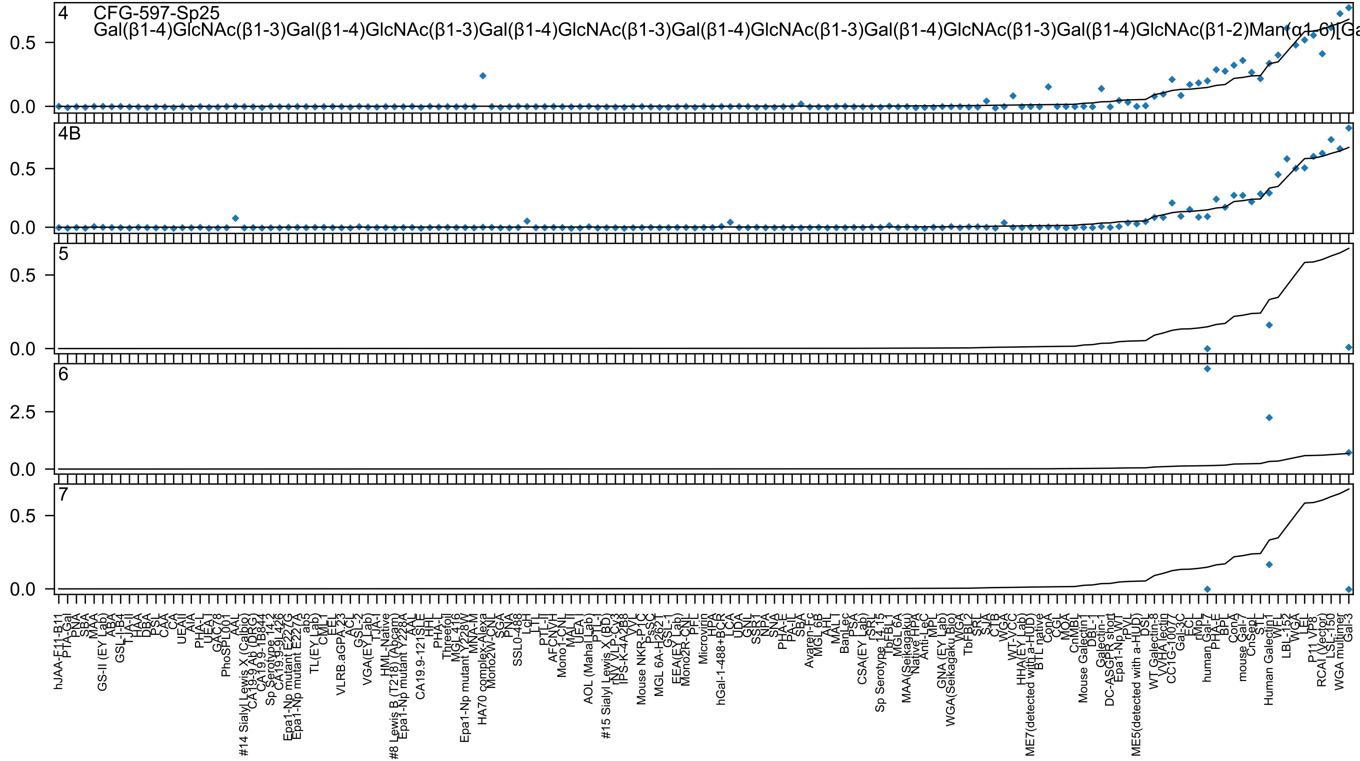

4 CFG-597-Sp25
Gal(β1-4)GlcNAc(β1-3)Gal(β1-4)GlcNAc(β1-3)Gal(β1-4)GlcNAc(β1-3)Gal(β1-4)GlcNAc(β1-3)Gal(β1-4)GlcNAc(β1-3)Gal(β1-4)GlcNAc(β1-2)Man(α1-6)[Ga
4B
5
6
7

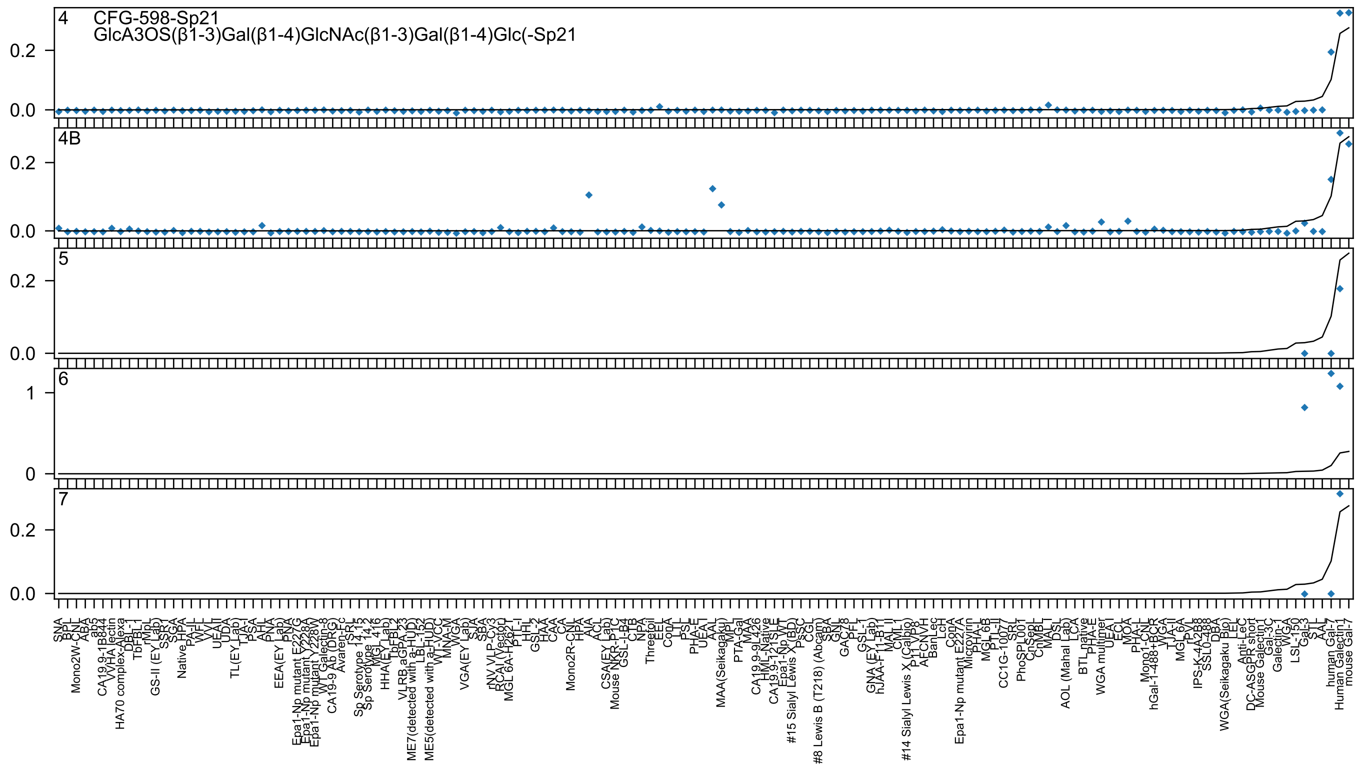
4
CFG-598-Sp21
GlcA3OS(β1-3)Gal(β1-4)GlcNAc(β1-3)Gal(β1-4)Glc(-Sp21
4B
5
6
7

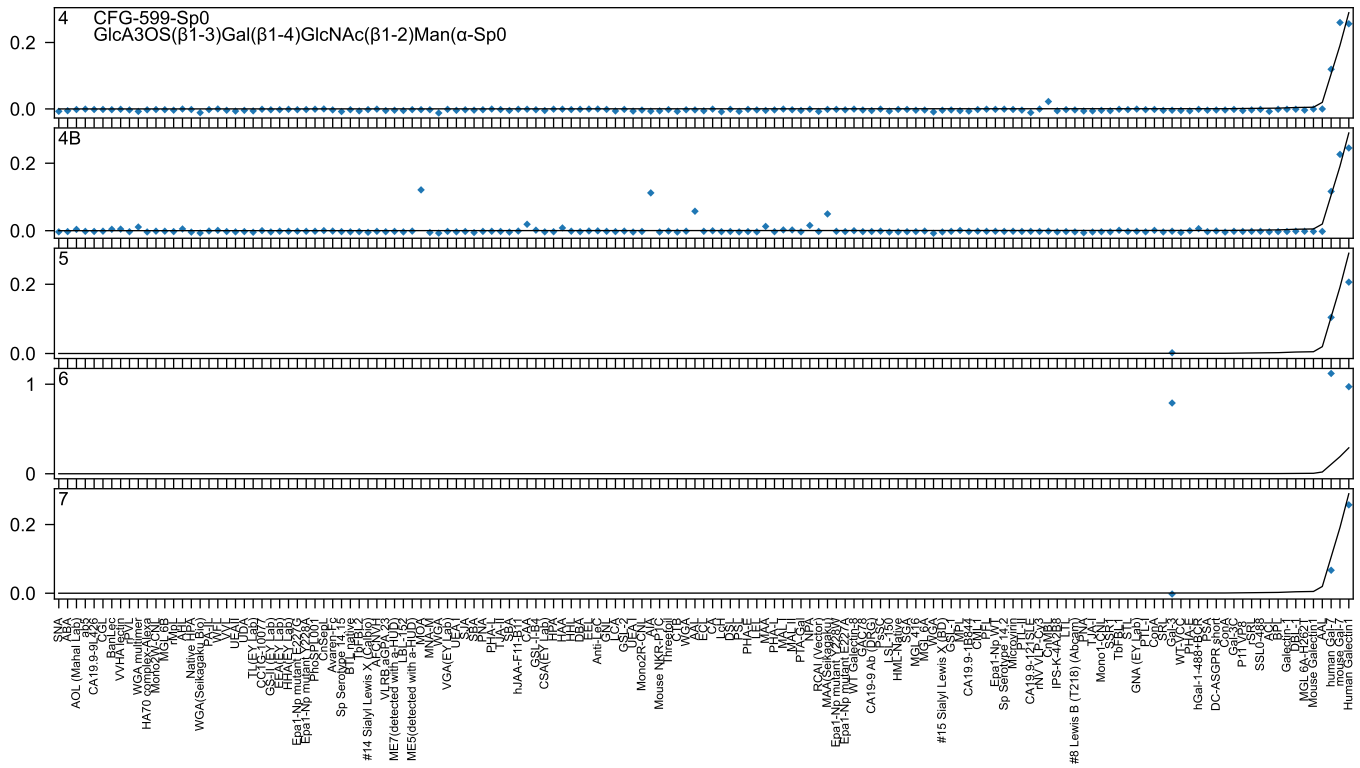
4
CFG-599-Sp0
GlcA3OS(β1-3)Gal(β1-4)GlcNAc(β1-2)Man(α-Sp0
0.2
0.0
4B
0.2
0.0
5
0.2
0.0
6
1
0
7
0.2
0.0

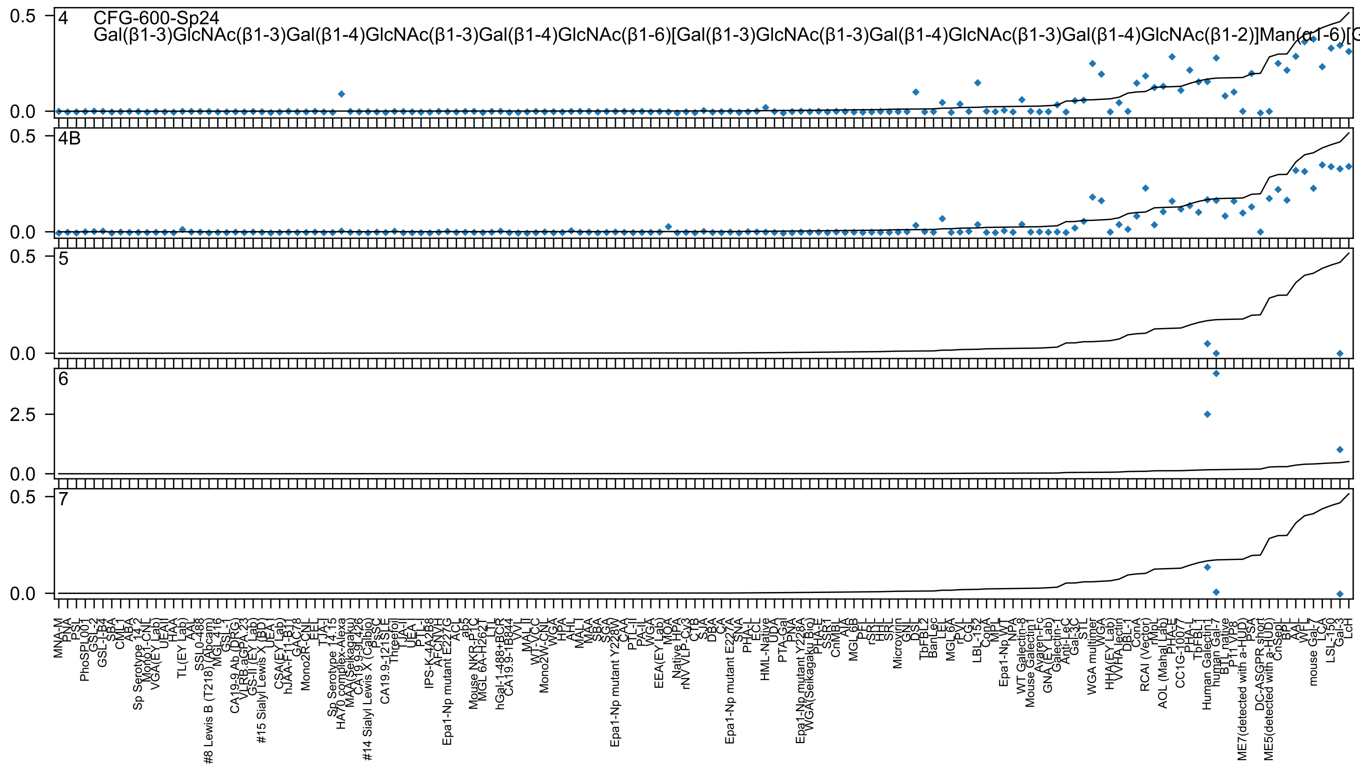

4 CFG-600-Sp24
Gal(β1-3)GlcNAc(β1-3)Gal(β1-4)GlcNAc(β1-3)Gal(β1-4)GlcNAc(β1-6)[Gal(β1-3)GlcNAc(β1-3)Gal(β1-4)GlcNAc(β1-3)Gal(β1-4)GlcNAc(β1-2)]Man(α1-6)[G
4B
5
6
7

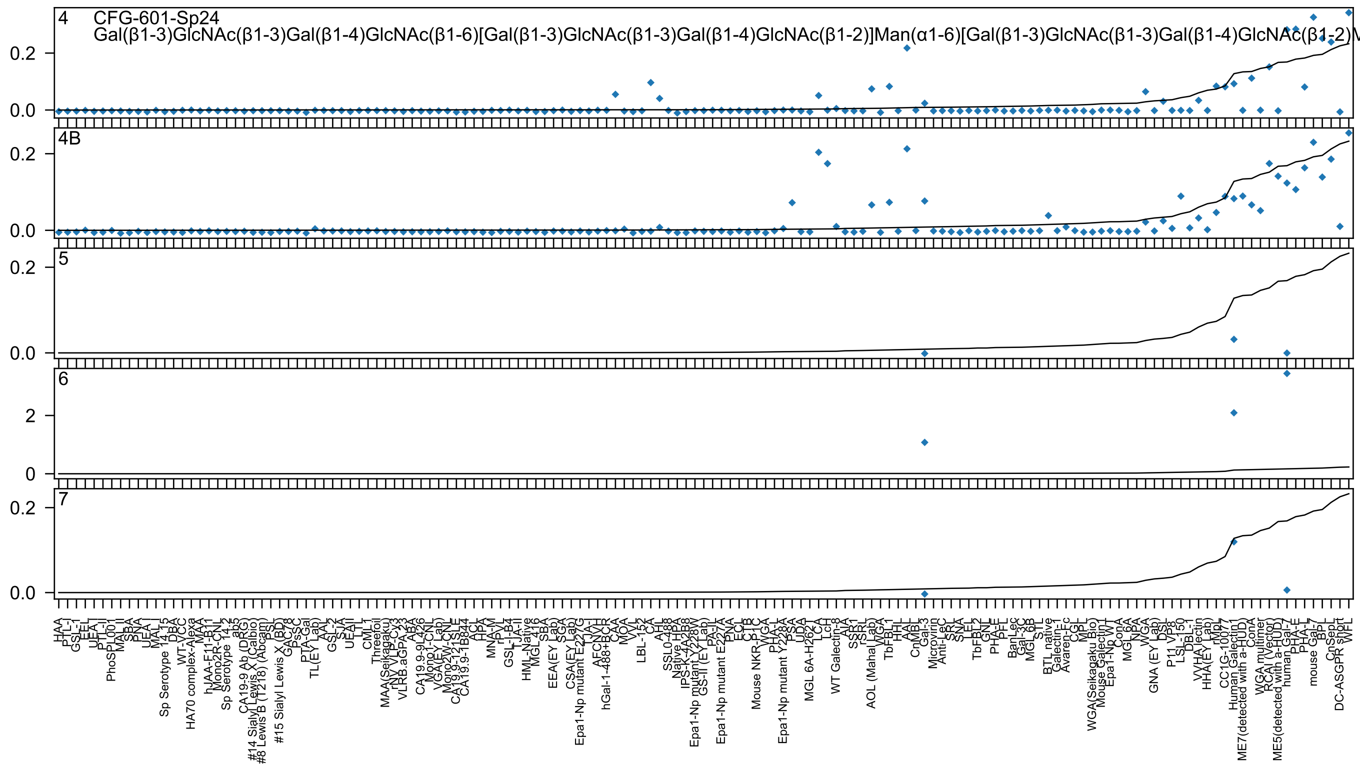

4 CFG-601-Sp24
Gal(β1-3)GlcNAc(β1-3)Gal(β1-4)GlcNAc(β1-6)[Gal(β1-3)GlcNAc(β1-3)Gal(β1-4)GlcNAc(β1-2)]Man(α1-6)[Gal(β1-3)GlcNAc(β1-3)Gal(β1-4)GlcNAc(β1-2)M
4B
5
6
7

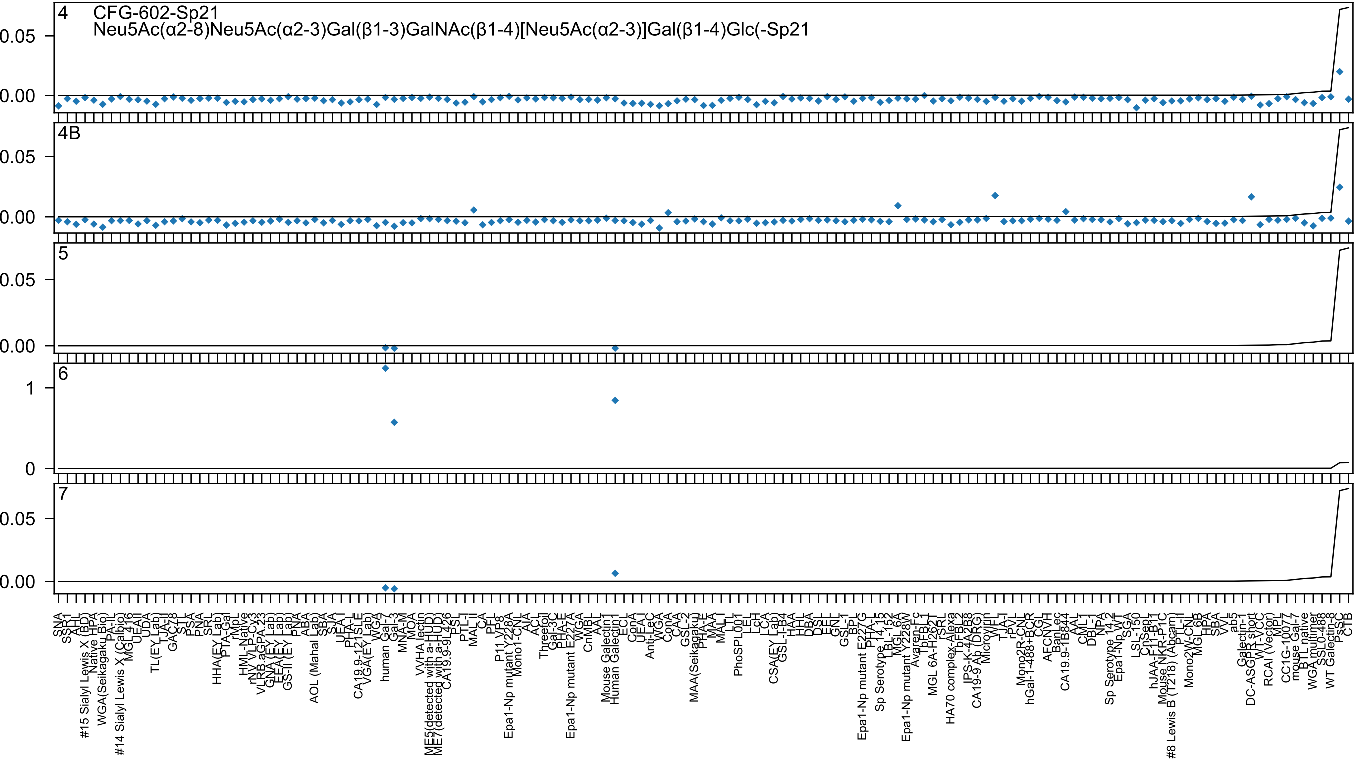
4 CFG-602-Sp21
Neu5Ac(α2-8)Neu5Ac(α2-3)Gal(β1-3)GalNAc(β1-4)[Neu5Ac(α2-3)]Gal(β1-4)Glc(-Sp21
4B
5
6
7
0.05
0.00
1
0

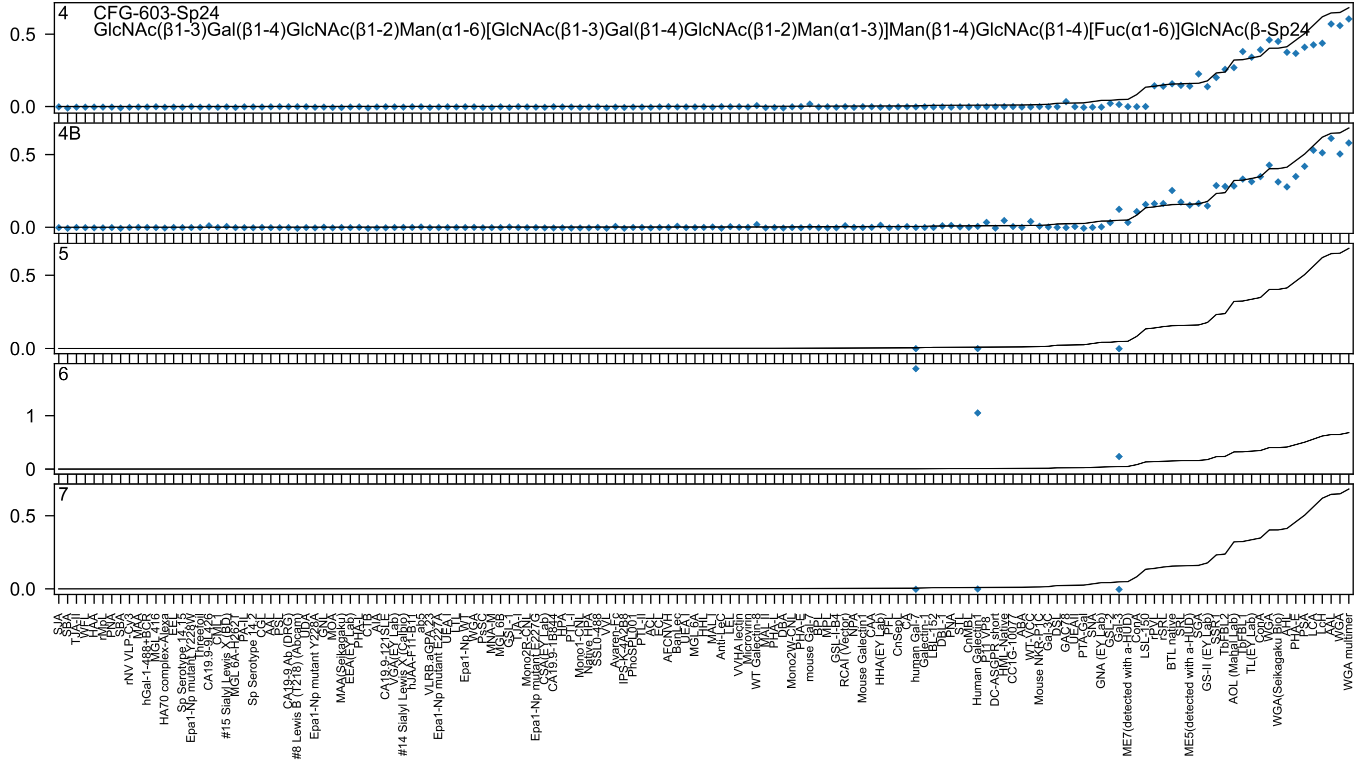
4 CFG-603-Sp24
GlcNAc(β1-3)Gal(β1-4)GlcNAc(β1-2)Man(α1-6)[GlcNAc(β1-3)Gal(β1-4)GlcNAc(β1-2)Man(α1-3)]Man(β1-4)GlcNAc(β1-4)[Fuc(α1-6)]GlcNAc(β-Sp24
4B
5
6
7

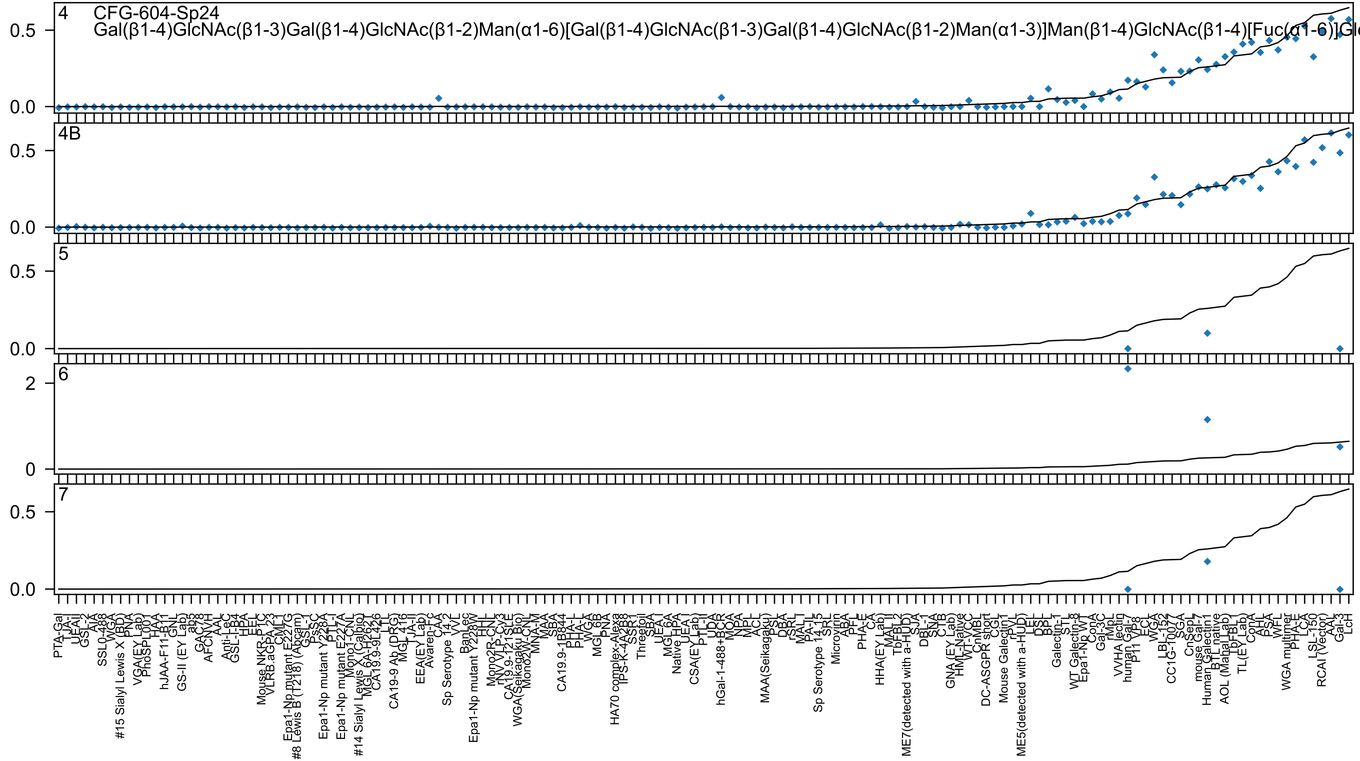
4 CFG-604-Sp24
Gal(β1-4)GlcNAc(β1-3)Gal(β1-4)GlcNAc(β1-2)Man(α1-6)[Gal(β1-4)GlcNAc(β1-3)Gal(β1-4)GlcNAc(β1-2)Man(α1-3)]Man(β1-4)GlcNAc(β1-4)[Fuc(α1-6)]Gl
4B
5
6
7
0.0
0.5
0
2
PTA-Gal
TJA-I
UEA-I
GSL-2
AIA
SSL0-488
WGA
#15 Sialyl Lewis X (BD)
PNA
VGA(EY Lab)
PhoSL001
HAA
hJAA-F11-B11
GNL
GS-II (EY Lab)
ab3
GAC78
AFCNVH
AAL
Anti-LeC
GSL-I-B4
HPA
EEL
Mouse NKR-P1C
VLRB.aGPA.23
CML1
Epa1-Np mutant E227G
#8 Lewis B (T218) (Abcam)
GSL-1
PsSC
Epa1-Np mutant Y228A
PTL-I
Epa1-Np mutant E227A
Mono1-CNL
#14 Sialyl Lewis X (Calbio)
MGL 6A-H262T
CA19.9-9L426
LTL
CA19-9 Ab (DRG)
MGL 416
TJA-II
EEA(EY Lab)
Avaren-Fc
CAA
Sp Serotype 14-2
VVL
BanLec
Epa1-Np mutant Y228W
HHL
Mono2R-CNL
rNV VLP-Cy3
CA19.9-121SLE
WGA(Seikagaku Bio)
Mono2W-CNL
MNA-M
MAA
SBA
CA19.9-1B844
PHA-L
PHA-L
WGA
MGL 6B
PNA
HA70 complex-Alexa
IPS-K-4A2B8
SSR1
Threefoil
SBA
UEA-I
MGL 6A
Native HPA
UEA-I
CSA(EY Lab)
PTL-II
UDA
hGal-1-488+BCR
MOA
NPA
MPL
ACL
MAA(Seikagaku)
PSL
DBA
rSRL
MAL-I
PA-IL
Sp Serotype 14-15
SRL
Microvirin
ABA
PFL
PHA-E
CA
HHA(EY Lab)
MAL-II
TbFBL2
ME7(detected with a-HUD)
SJA
DBL-1
SNA
CTB
GNA (EY Lab)
HML-Native
WT-VCC
CnMBL
DC-ASGPR short
CGL
Mouse Galectin1
rPVL
ME5(detected with a-HUD)
LEL
DSL
BPL
Galectin-1
STL
WT Galectin-8
Epa1-Np WT
ConA
Gal-3C
rMpL
VVHA lectin
human Gal-7
P11 VP8
ECL
WGA
LBL-152
CC1G-10077
SGA
CnSepL
mouse Gal-7
Human Galectin1
BTL native
AOL (Mahal Lab)
TbFBL1
TL(EY Lab)
ConA
AHL
PSA
WFL
WGA multimer
PHA-E
LCA
LSL-150
RCAI (Vector)
AAL
Gal-3
LcH

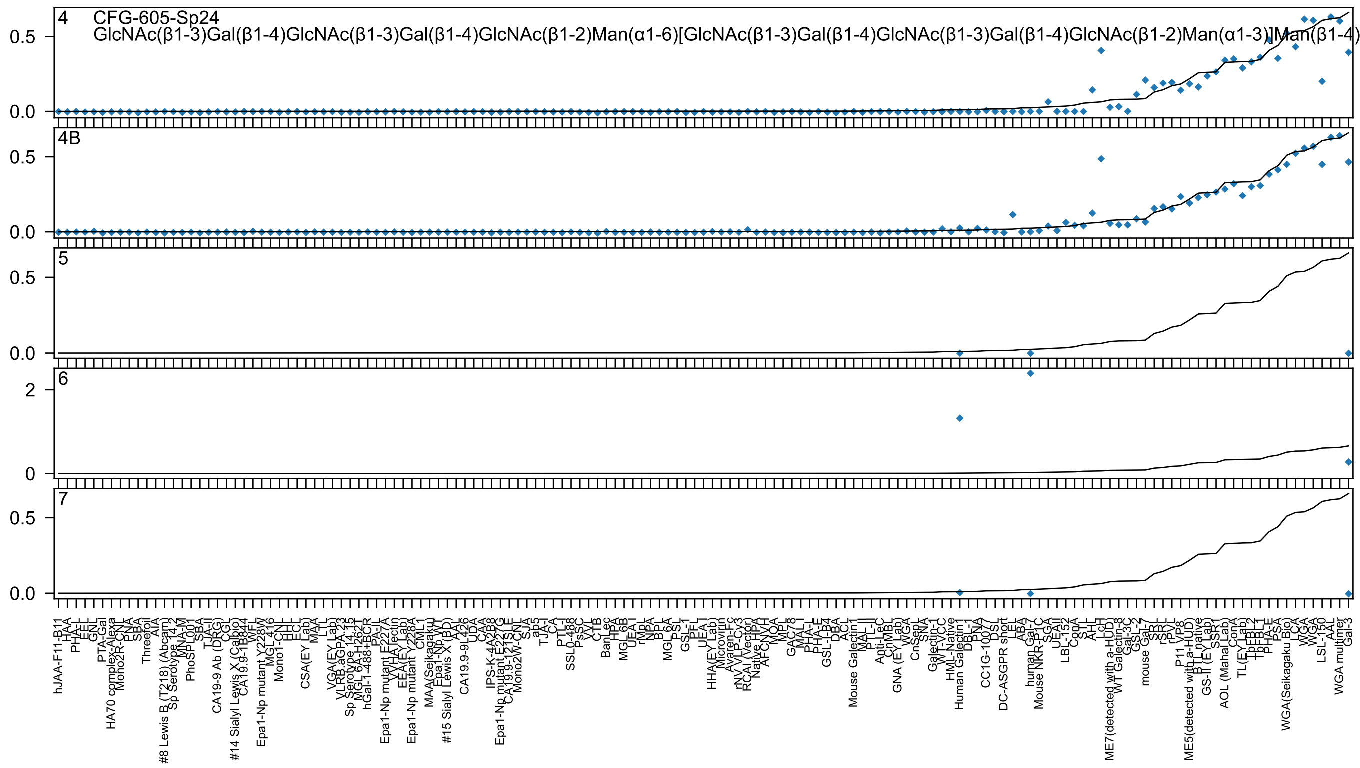

4
CFG-605-Sp24
GlcNAc(β1-3)Gal(β1-4)GlcNAc(β1-3)Gal(β1-4)GlcNAc(β1-2)Man(α1-6)[GlcNAc(β1-3)Gal(β1-4)GlcNAc(β1-3)Gal(β1-4)GlcNAc(β1-2)Man(α1-3)]Man(β1-4)
4B
5
6
7

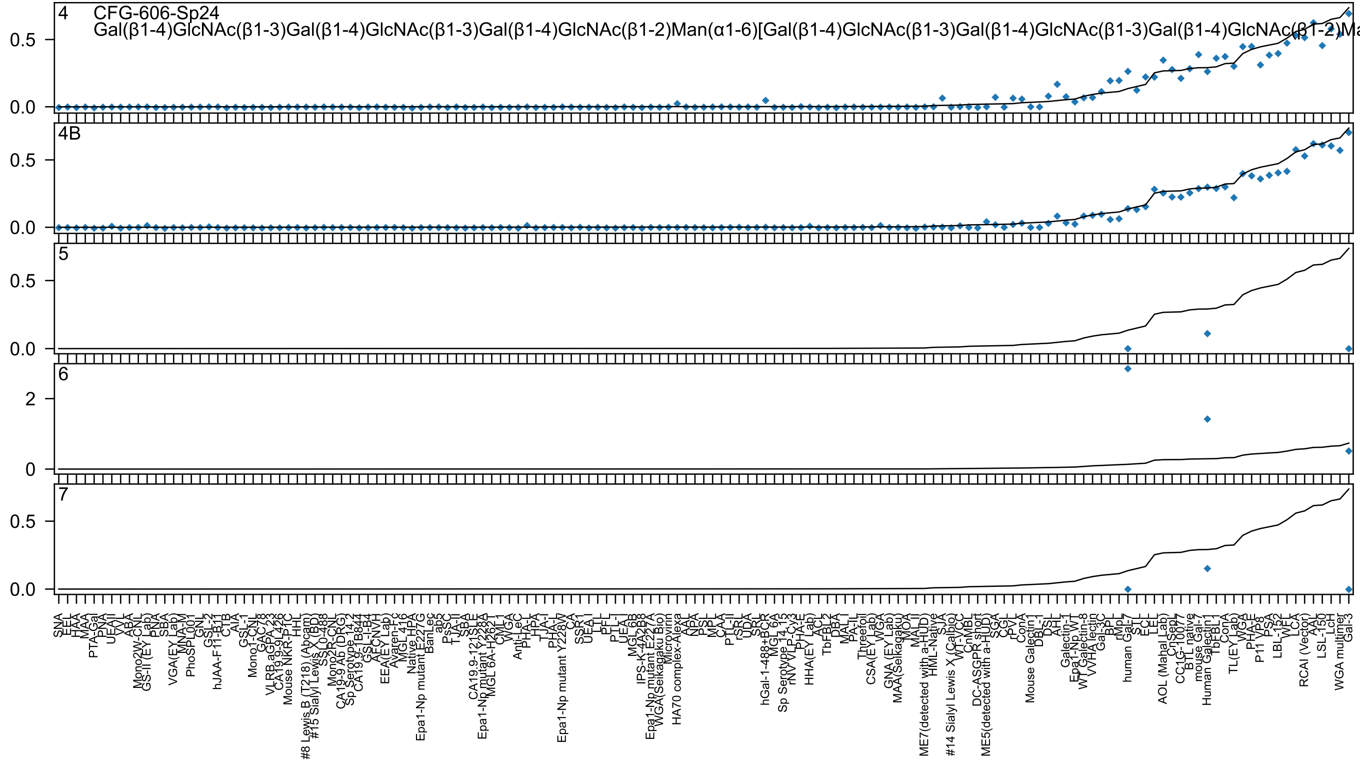
4 CFG-606-Sp24
Gal(β1-4)GlcNAc(β1-3)Gal(β1-4)GlcNAc(β1-3)Gal(β1-4)GlcNAc(β1-2)Man(α1-6)[Gal(β1-4)GlcNAc(β1-3)Gal(β1-4)GlcNAc(β1-3)Gal(β1-4)GlcNAc(β1-2)Ma
4B
5
6
7
0.0
0.5
0
2
SNA
EEL
HAA
MAA
PTA-Gal
PNA
UEA-II
VVL
ABA
Mono2W-CNL
GS-II (EY Lab)
PNA
SBA
VGA(EY Lab)
MNA-M
PhoSL001
GNL
GSL-2
hJAA-F11-B11
CTB
AIA
GSL-1
Mono1-CNL
GAC78
VLRB.aGPA.23
CA19.9-9L426
Mouse NKR-P1C
HHL
#8 Lewis B (T218) (Abcam)
#15 Sialyl Lewis X (BD)
SSL0-488
Mono2R-CNL
CA19-9 Ab (DRG)
Sp Serotype 14.2
CA19.9-1B844
GSL-I-B4
AFCNVH
EEA(EY Lab)
Avaren-Fc
MGL 416
Native HPA
Epa1-Np mutant E227G
BanLec
ab5
PsSC
TJA-II
SBA
CA19.9-121SLE
Epa1-Np mutant Y228A
MGL 6A-H262T
CML-1
WGA
Anti-LeC
PHA-L
HPA
TJA-I
PHA-L
Epa1-Np mutant Y228W
CA
SSR1
UEA-I
LTL
PFL
PTL-I
UEA-I
MGL 6B
IPS-K-4A2B8
Epa1-Np mutant E227A
WGA(Seikagaku Bio)
Microvirin
HA70 complex-Alexa
AAL
NPA
PSL
MPL
CAA
PTL-II
rSRL
UDA
SRL
hGal-1-488+BCR
MGL 6A
Sp Serotype 14.15
rNV VLP-Cy3
PHA-E
HHA(EY Lab)
ACL
TbFBL2
DBA
MAL-I
PA-IL
Threefoil
CSA(EY Lab)
WGA
GNA (EY Lab)
MAA(Selkagaku)
MOA
MAL-II
ME7(detected with a-HUD)
HML-Native
SJA
#14 Sialyl Lewis X (Calbio)
WT-VCC
CnMBL
DC-ASGPR short
ME5(detected with a-HUD)
SGA
CGL
rPVL
ConA
Mouse Galectin1
DBL-1
DSL
AHL
Galectin-1
Epa1-Np WT
WT Galectin-8
VVHA lectin
Gal-3C
BPL
rMpL
human Gal-7
STL
ECL
LEL
AOL (Mahal Lab)
CnSeP
CC1G-1007
BTL native
mouse Gal-7
Human Galectin1
TbFBL1
ConA
TL(EY Lab)
WGA
PHA-E
P11 VP8
PSA
LBL-152
WFL
LCA
RCAI (Vector)
AAL
LSL-150
LcH
WGA multimer
Gal-3

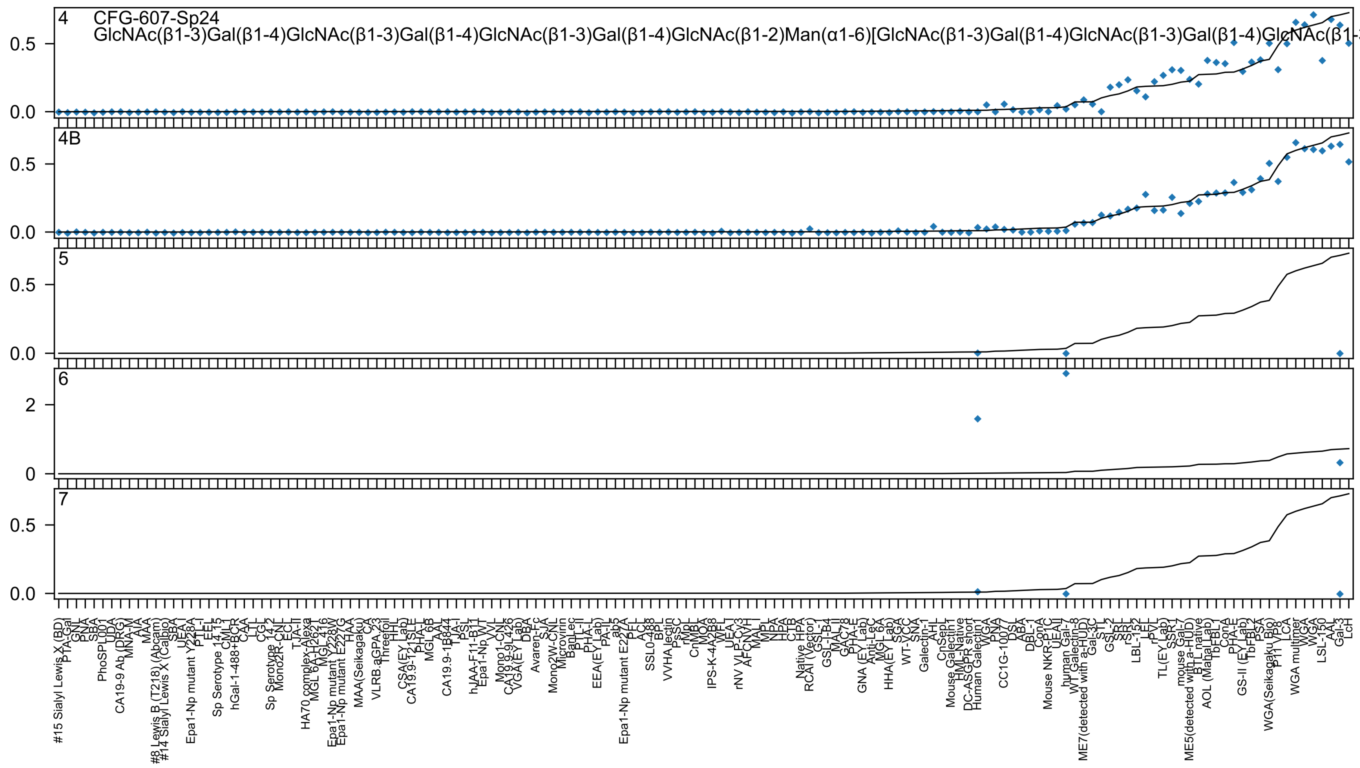
4
CFG-607-Sp24
GlcNAc(β1-3)Gal(β1-4)GlcNAc(β1-3)Gal(β1-4)GlcNAc(β1-3)Gal(β1-4)GlcNAc(β1-2)Man(α1-6)[GlcNAc(β1-3)Gal(β1-4)GlcNAc(β1-3)Gal(β1-4)GlcNAc(β1-
4B
5
6
7

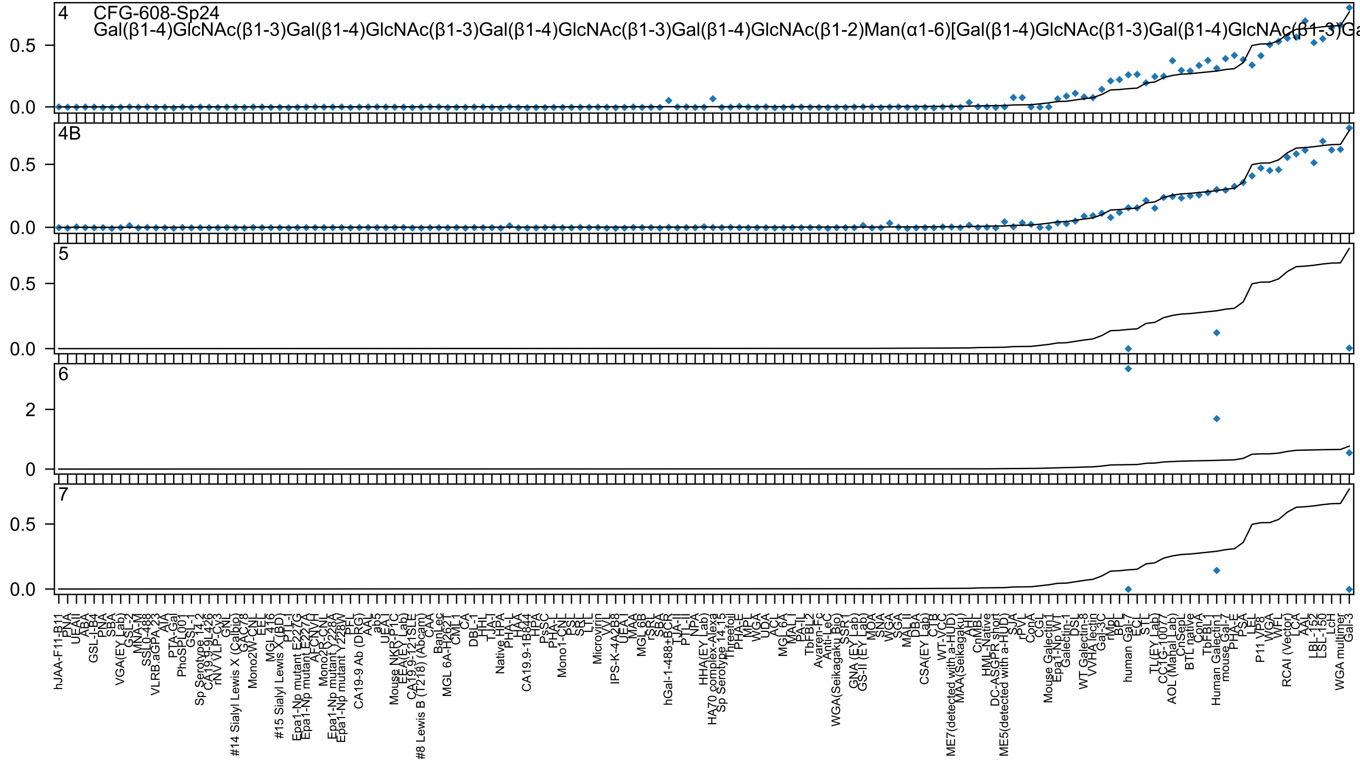

4 CFG-608-Sp24
Gal(β1-4)GlcNAc(β1-3)Gal(β1-4)GlcNAc(β1-3)Gal(β1-4)GlcNAc(β1-3)Gal(β1-4)GlcNAc(β1-2)Man(α1-6)[Gal(β1-4)GlcNAc(β1-3)Gal(β1-4)GlcNAc(β1-3)Ga
4B
5
6
7

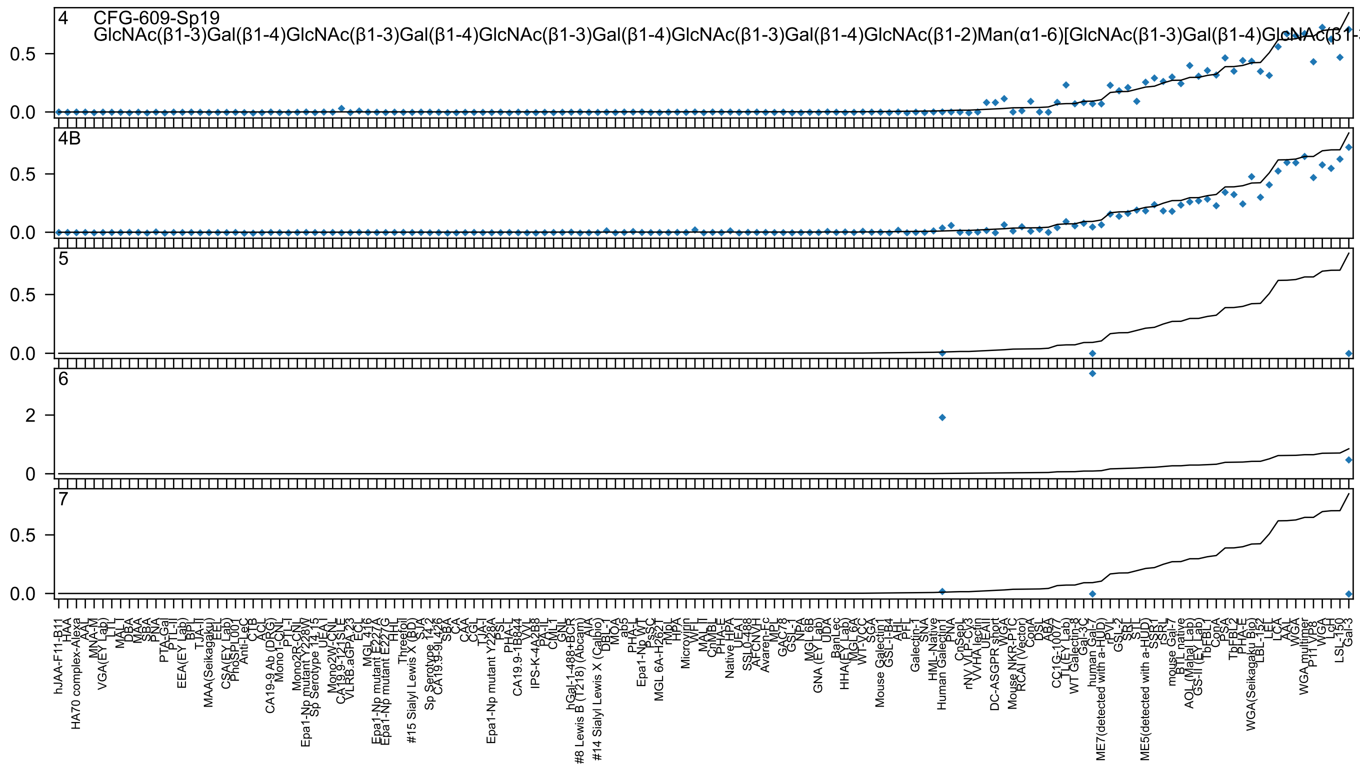
4
CFG-609-Sp19
GlcNAc(β1-3)Gal(β1-4)GlcNAc(β1-3)Gal(β1-4)GlcNAc(β1-3)Gal(β1-4)GlcNAc(β1-3)Gal(β1-4)GlcNAc(β1-2)Man(α1-6)[GlcNAc(β1-3)Gal(β1-4)GlcNAc(β1-
4B
5
6
7

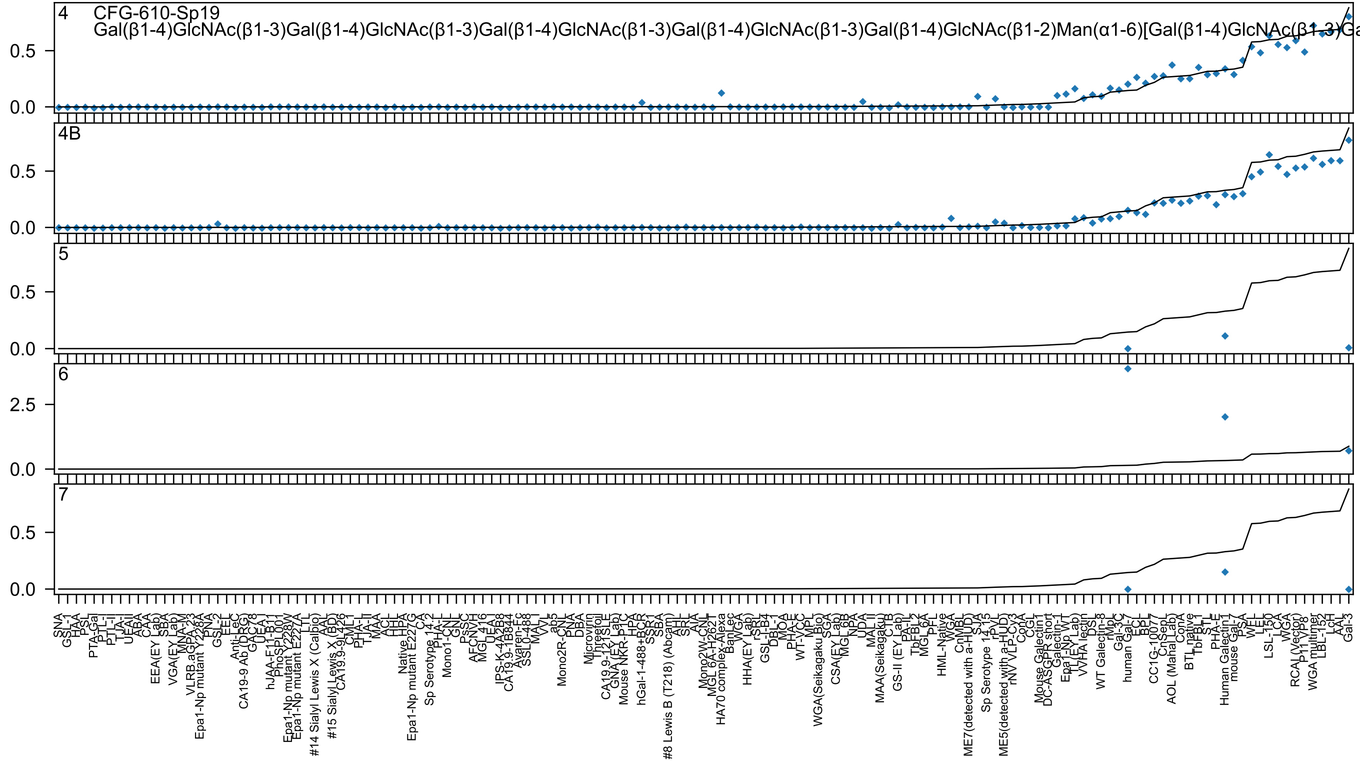
4 CFG-610-Sp19
Gal(β1-4)GlcNAc(β1-3)Gal(β1-4)GlcNAc(β1-3)Gal(β1-4)GlcNAc(β1-3)Gal(β1-4)GlcNAc(β1-3)Gal(β1-4)GlcNAc(β1-2)Man(α1-6)[Gal(β1-4)GlcNAc(β1-3)Ga
4B
5
6
7

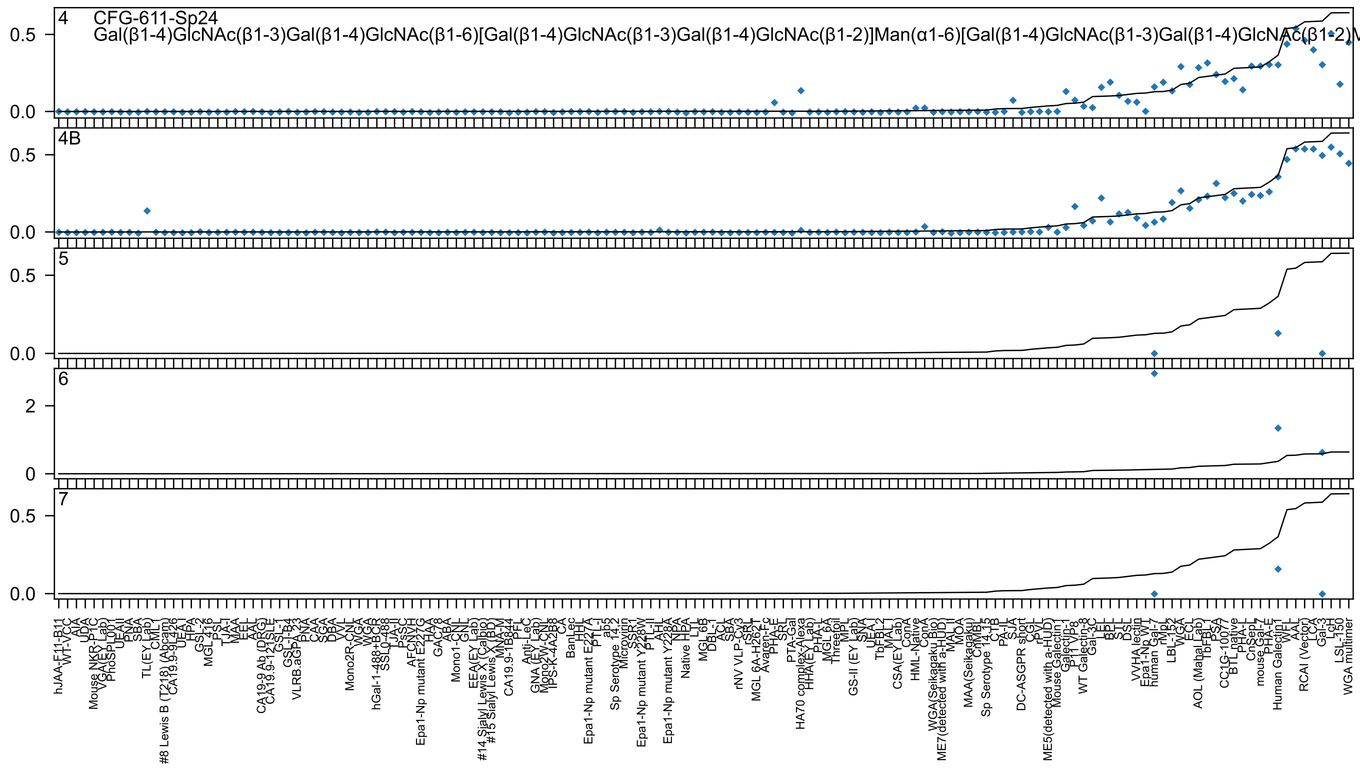

4 CFG-611-Sp24
Gal(β1-4)GlcNAc(β1-3)Gal(β1-4)GlcNAc(β1-6)[Gal(β1-4)GlcNAc(β1-3)Gal(β1-4)GlcNAc(β1-2)]Man(α1-6)[Gal(β1-4)GlcNAc(β1-3)Gal(β1-4)GlcNAc(β1-2)M
4B
5
6
7

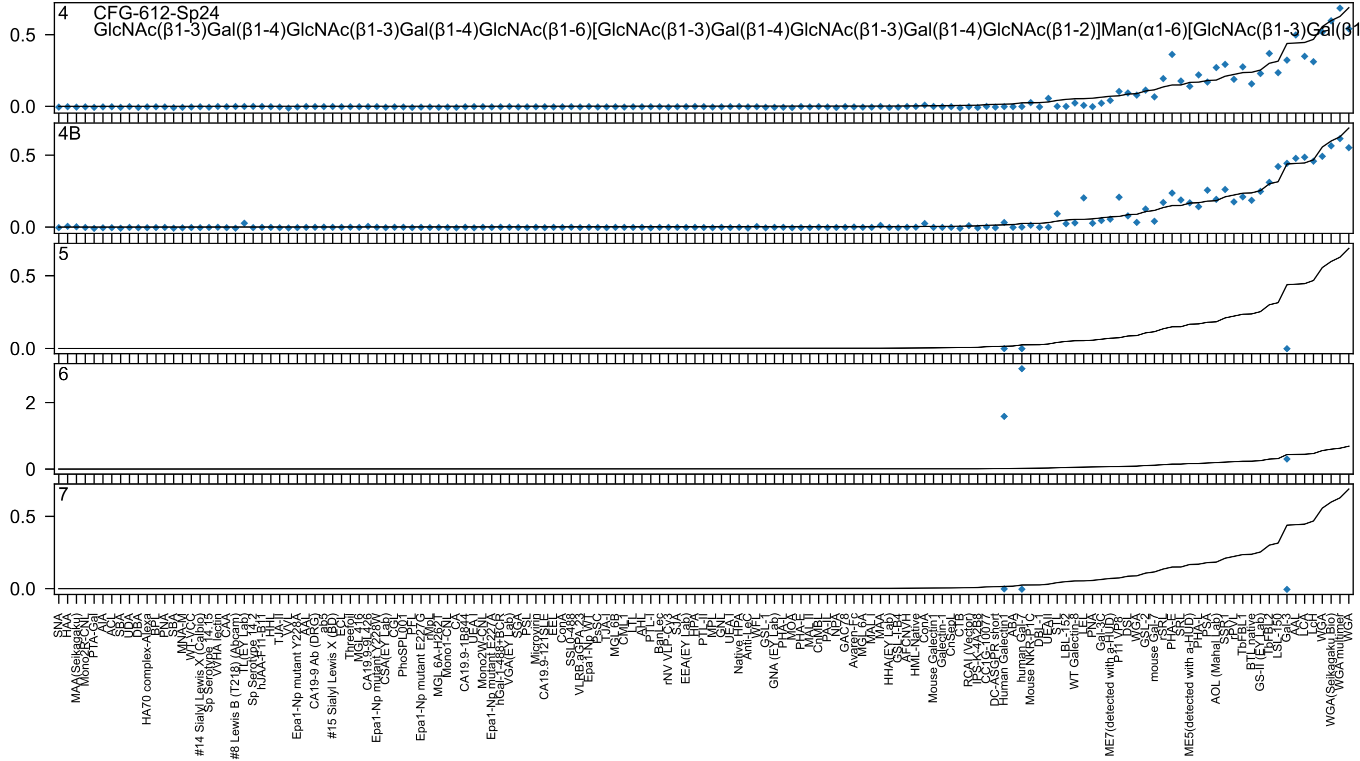

4
CFG-612-Sp24
GlcNAc(β1-3)Gal(β1-4)GlcNAc(β1-3)Gal(β1-4)GlcNAc(β1-6)[GlcNAc(β1-3)Gal(β1-4)GlcNAc(β1-3)Gal(β1-4)GlcNAc(β1-2)]Man(α1-6)[GlcNAc(β1-3)Gal(β1
4B
5
6
7
0.5
0.0
2
0

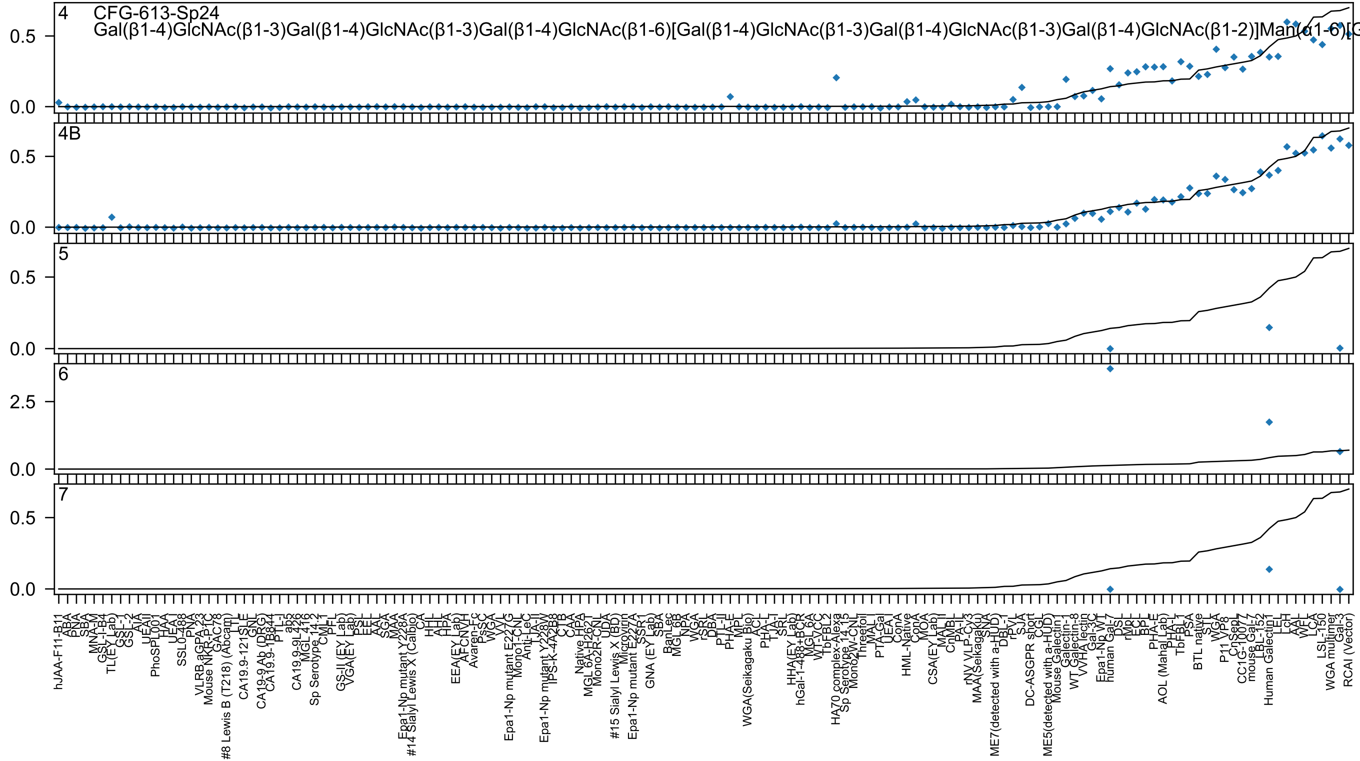
4 CFG-613-Sp24
Gal(β1-4)GlcNAc(β1-3)Gal(β1-4)GlcNAc(β1-3)Gal(β1-4)GlcNAc(β1-6)[Gal(β1-4)GlcNAc(β1-3)Gal(β1-4)GlcNAc(β1-3)Gal(β1-4)GlcNAc(β1-2)]Man(α1-6)[G
4B
5
6
7
0.0
0.5
2.5

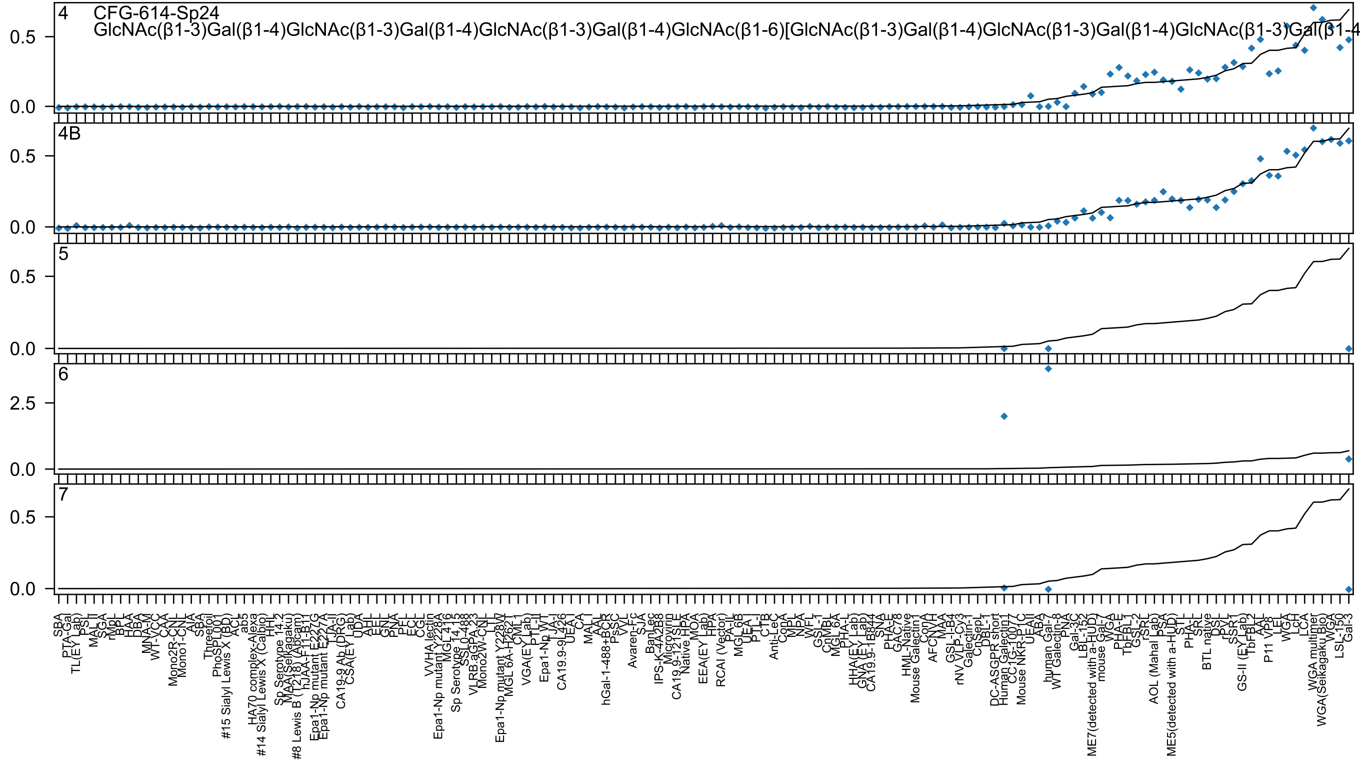

4 CFG-614-Sp24
GlcNAc(β1-3)Gal(β1-4)GlcNAc(β1-3)Gal(β1-4)GlcNAc(β1-3)Gal(β1-4)GlcNAc(β1-6)[GlcNAc(β1-3)Gal(β1-4)GlcNAc(β1-3)Gal(β1-4)GlcNAc(β1-3)Gal(β1-4
4B
5
6
7

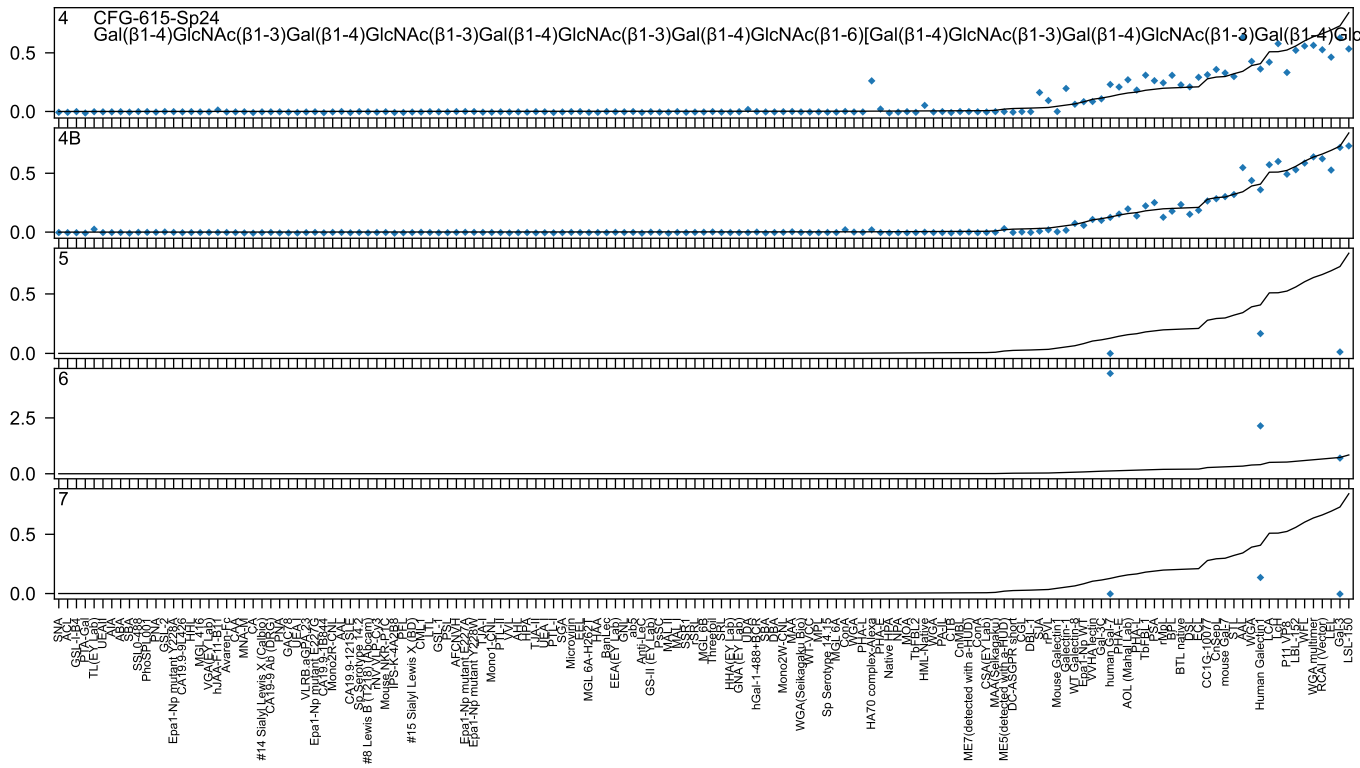
4
CFG-615-Sp24
Gal(β1-4)GlcNAc(β1-3)Gal(β1-4)GlcNAc(β1-3)Gal(β1-4)GlcNAc(β1-3)Gal(β1-4)GlcNAc(β1-6)[Gal(β1-4)GlcNAc(β1-3)Gal(β1-4)GlcNAc(β1-3)Gal(β1-4)Glc
4B
5
6
7

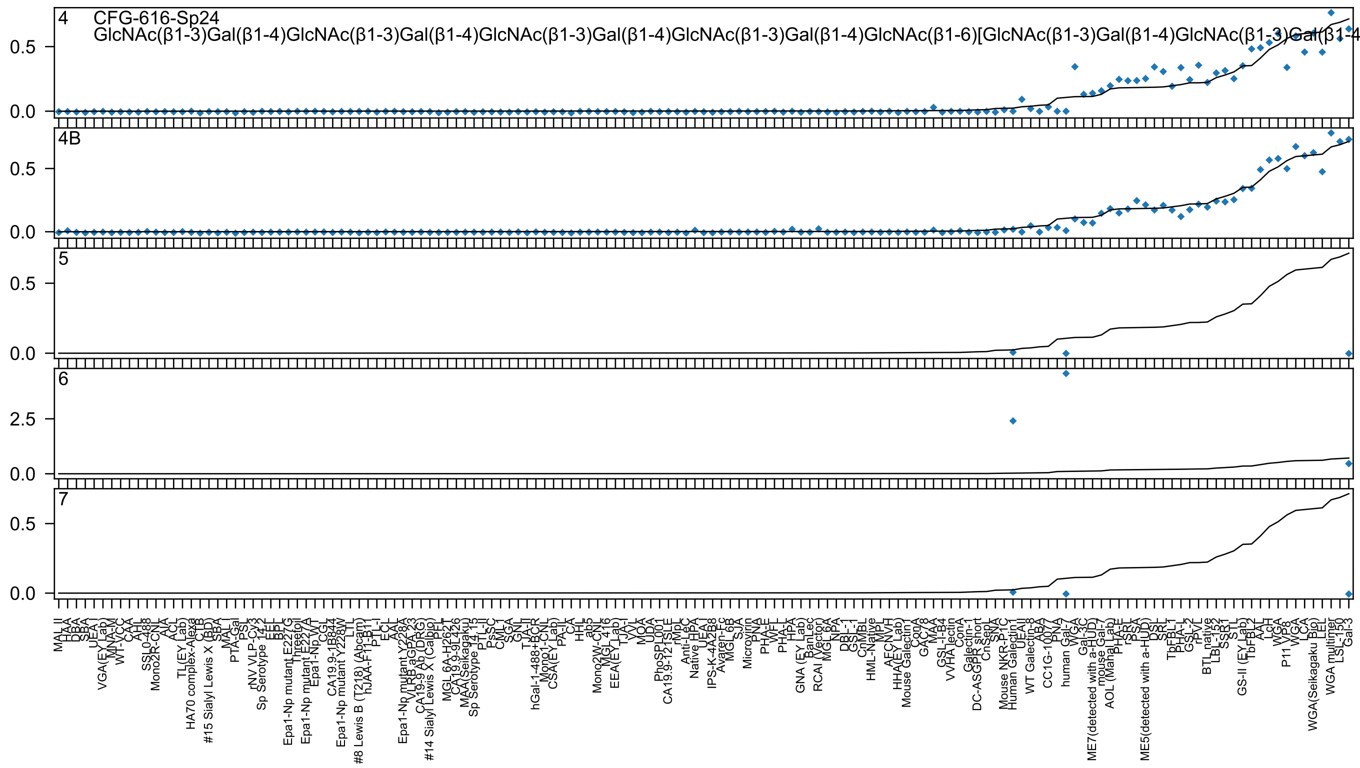
4
CFG-616-Sp24
GlcNAc(β1-3)Gal(β1-4)GlcNAc(β1-3)Gal(β1-4)GlcNAc(β1-3)Gal(β1-4)GlcNAc(β1-3)Gal(β1-4)GlcNAc(β1-6)[GlcNAc(β1-3)Gal(β1-4)GlcNAc(β1-3)Gal(β1-4
4B
5
6
7
0.0
0.5
2.5

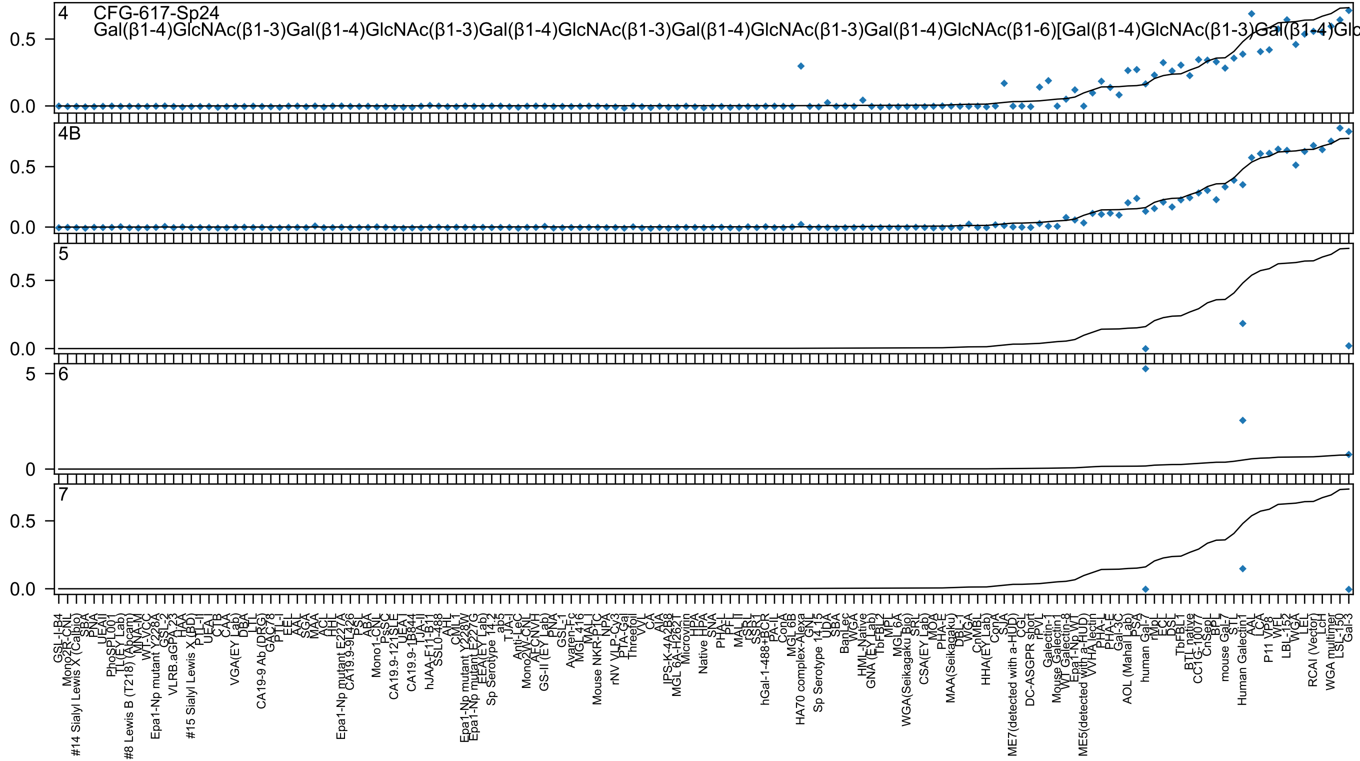
4 CFG-617-Sp24
Gal(β1-4)GlcNAc(β1-3)Gal(β1-4)GlcNAc(β1-3)Gal(β1-4)GlcNAc(β1-3)Gal(β1-4)GlcNAc(β1-3)Gal(β1-4)GlcNAc(β1-6)[Gal(β1-4)GlcNAc(β1-3)Gal(β1-4)Glc
4B
5
6
7

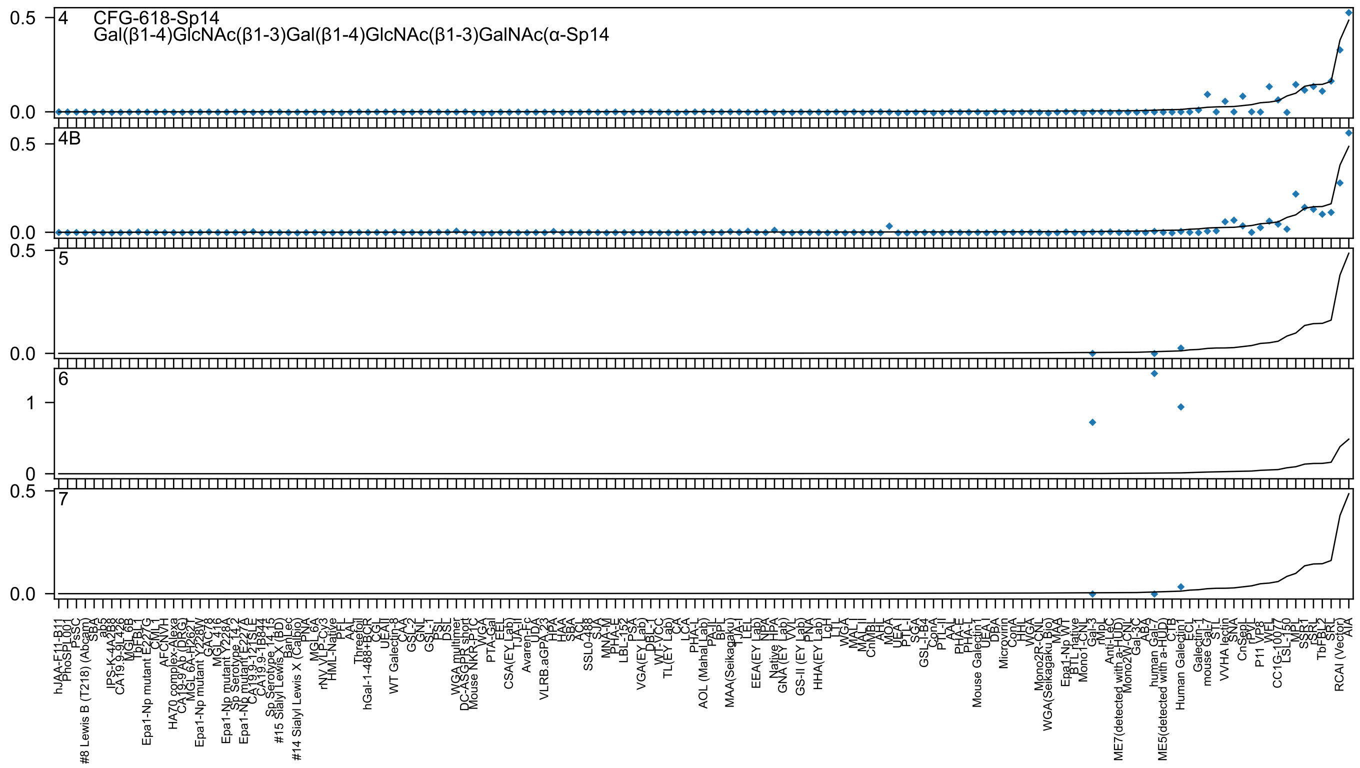

4 CFG-618-Sp14
Gal(β1-4)GlcNAc(β1-3)Gal(β1-4)GlcNAc(β1-3)GalNAc(α-Sp14
4B
5
6
7

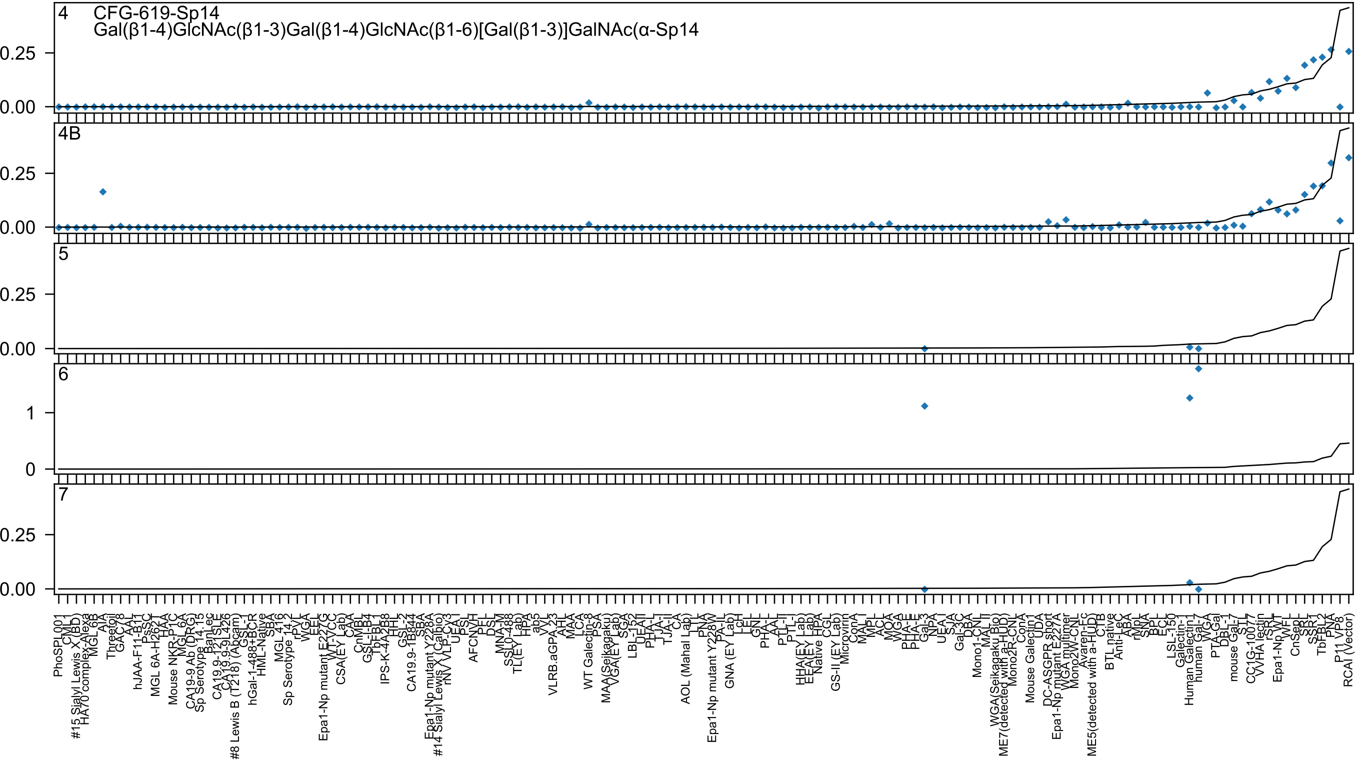

4 CFG-619-Sp14
Gal(β1-4)GlcNAc(β1-3)Gal(β1-4)GlcNAc(β1-6)[Gal(β1-3)]GalNAc(α-Sp14
4B
5
6
7

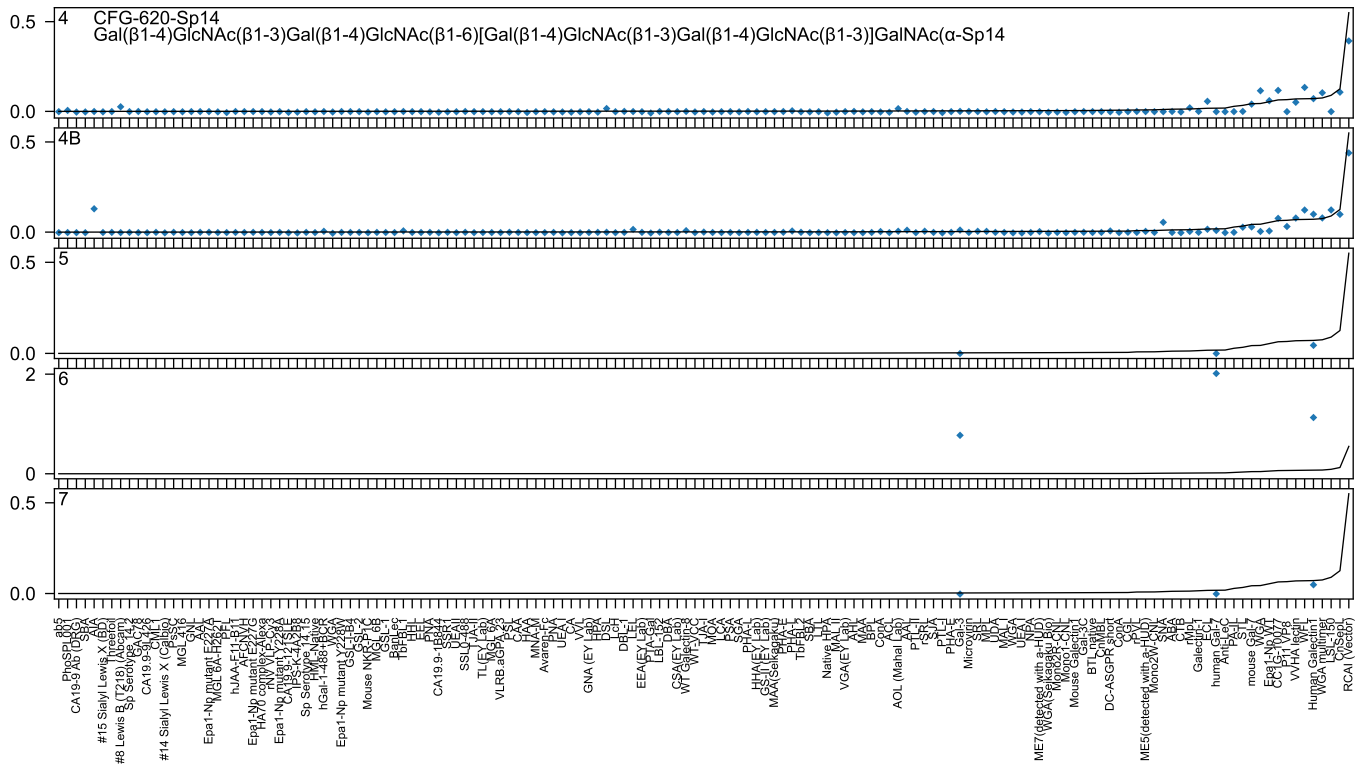

4
CFG-620-Sp14
Gal(β1-4)GlcNAc(β1-3)Gal(β1-4)GlcNAc(β1-6)[Gal(β1-4)GlcNAc(β1-3)Gal(β1-4)GlcNAc(β1-3)]GalNAc(α-Sp14
4B
5
6
7
0.5
0.0
2
0
ab5
PhoSL001
CA19-9 Ab (DRG)
SBA
AIA
#15 Sialyl Lewis X (BD)
Threefoil
#8 Lewis B (T218) (Abcam)
Sp Serotype 14.2
GAC78
CA19.9-9L426
CML1
#14 Sialyl Lewis X (Calbio)
PsSC
MGL 416
GNL
AAL
Epa1-Np mutant E227A
MGL 6A-H262T
PFL
hJAA-F11-B11
AFCNYH
Epa1-Np mutant E227G
HA70 complex-Alexa
rNV VLP-Cy3
Epa1-Np mutant Y228A
CA19.9-121SLE
IPS-K-4A2B8
Sp Serotype 14.15
HML-Native
hGal-1-488+BCR
WGA
Epa1-Np mutant Y228W
GSL-I-B4
GSL-2
Mouse NKR-P1C
MGL 6B
GSL-1
BanLec
TbFBL1
HHL
EEL
PNA
CA19.9-1B844
SSR1
UEAII
SSL0-488
TJA-II
TL(EY Lab)
MGL 6A
VLRB.aGPA.23
PSL
CAA
HAA
MNA-M
Avaren-Fc
PNA
UEA-I
CA
VVL
GNA (EY Lab)
HPA
DSL
LcH
DBL-1
LEL
EEA(EY Lab)
PTA-Gal
LBL-152
DBA
CSA(EY Lab)
WT Galectin-8
WT-VCC
TJA-I
MOA
LCA
PSA
SGA
PHA-L
HHA(EY Lab)
GS-II (EY Lab)
MAA(Seikagaku)
PHA-E
PHA-L
TbFBL2
SBA
LTL
Native HPA
MAL II
VGA(EY Lab)
AHL
MAA
BPL
ConA
ACL
AOL (Mahal Lab)
AAL
PTL-II
rSRL
SJA
PTL-I
PHA-E
Gal-3
Microvirin
SRL
MPL
UDA
MAL I
WGA
UEA I
NPA
ME7(detected with a-HUD)
WGA(Seikagaku Bio)
Mono2R-CNL
Mono1-CNL
Mouse Galectin1
Gal-3C
BTL native
CnMBL
DC-ASGPR short
ConA
CGL
rPVL
ME5(detected with a-HUD)
Mono2W-CNL
SNA
ABA
CTB
rMpL
Galectin-1
ECL
human Gal-7
Anti-LeC
PA-IL
STL
mouse Gal-7
WGA
Epa1-Np WT
CC1G-10077
P11 VP8
VVHA lectin
WFL
Human Galectin1
WGA multimer
LSL-150
CnSepL
RCAI (Vector)

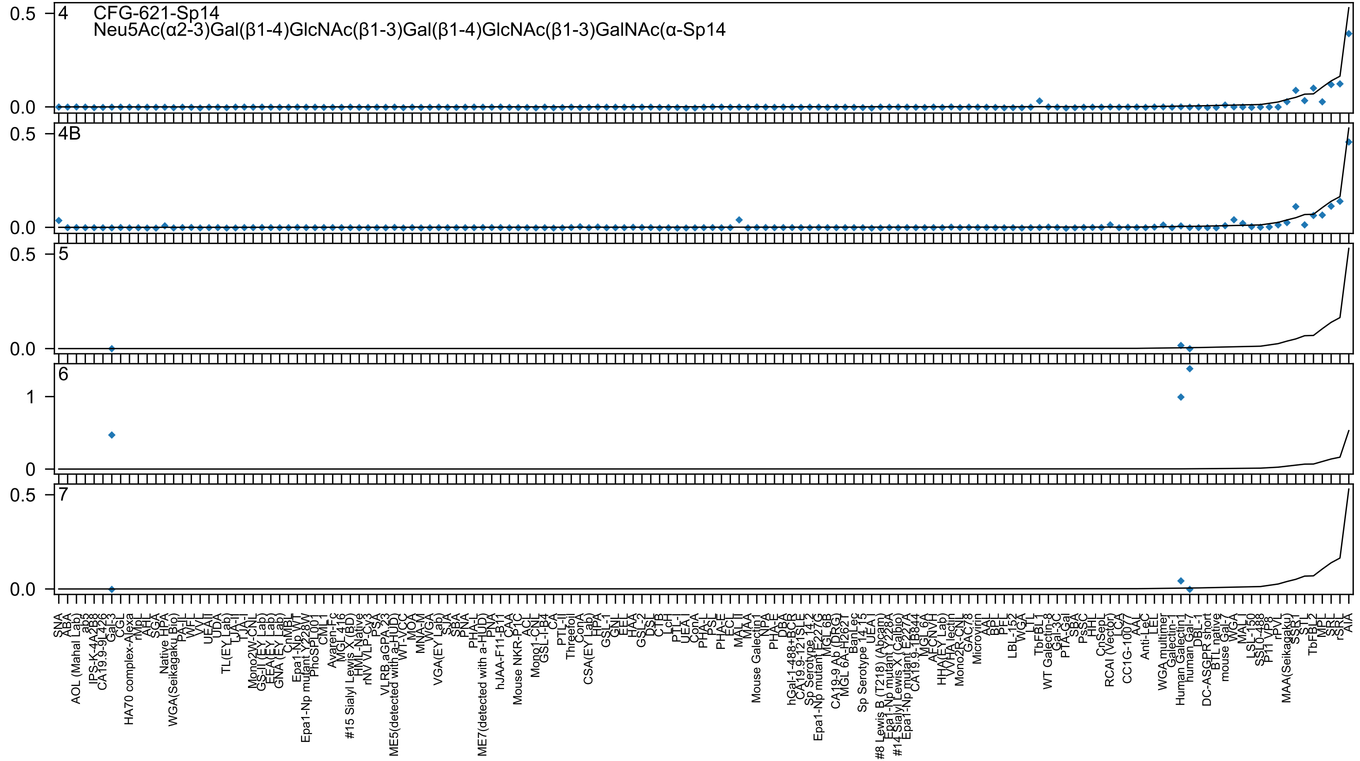
4
CFG-621-Sp14
Neu5Ac(α2-3)Gal(β1-4)GlcNAc(β1-3)Gal(β1-4)GlcNAc(β1-3)GalNAc(α-Sp14
4B
5
6
7

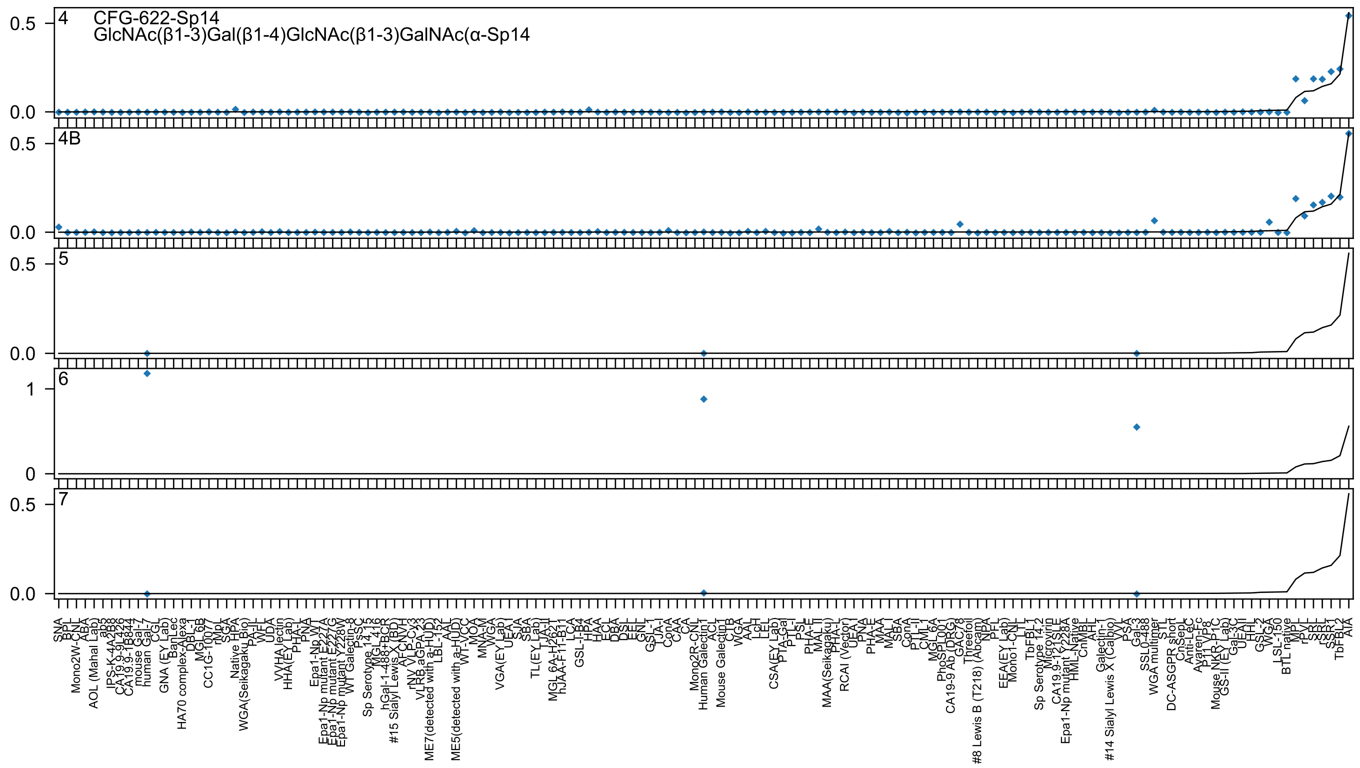
CFG-622-Sp14
GlcNAc(β1-3)Gal(β1-4)GlcNAc(β1-3)GalNAc(α-Sp14
4
4B
5
6
7

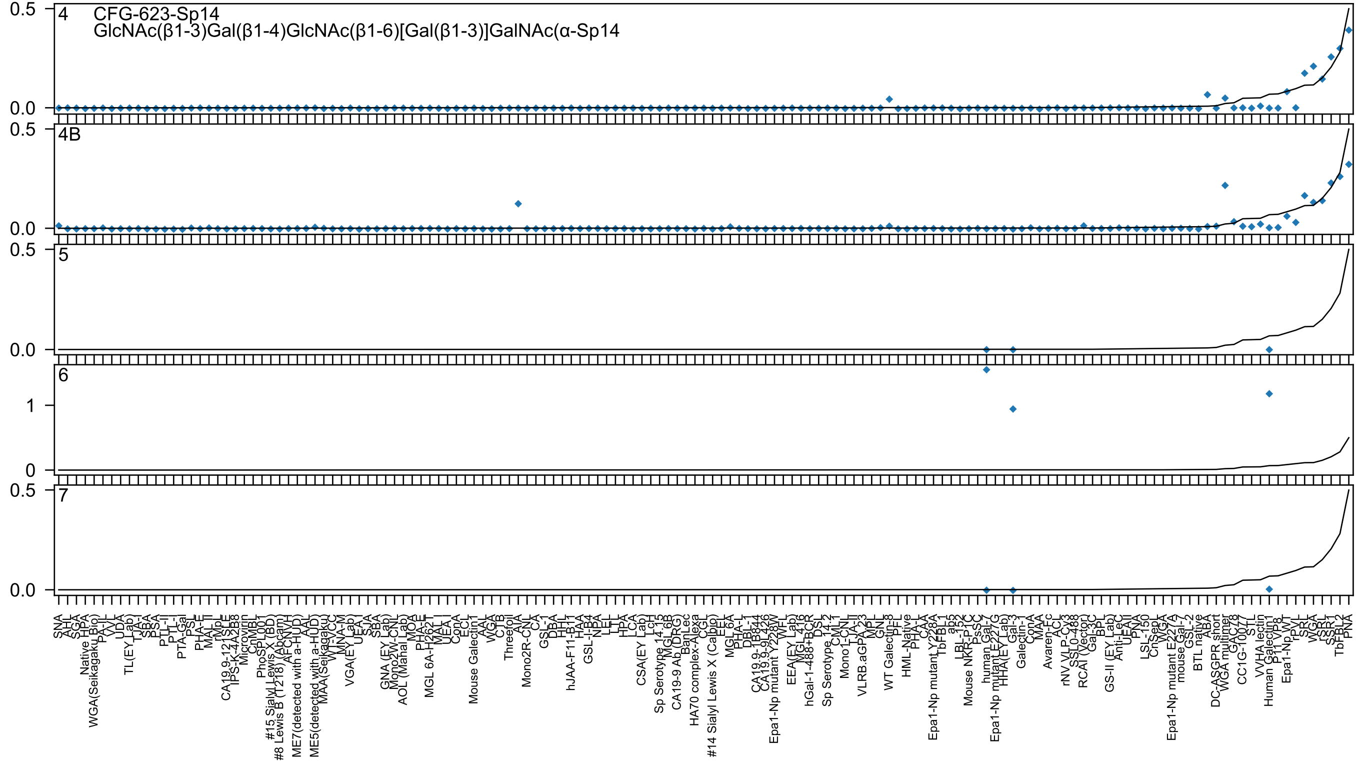

CFG-623-Sp14
GlcNAc(β1-3)Gal(β1-4)GlcNAc(β1-6)[Gal(β1-3)]GalNAc(α-Sp14
4
4B
5
6
7
0.5
0.0
1
0

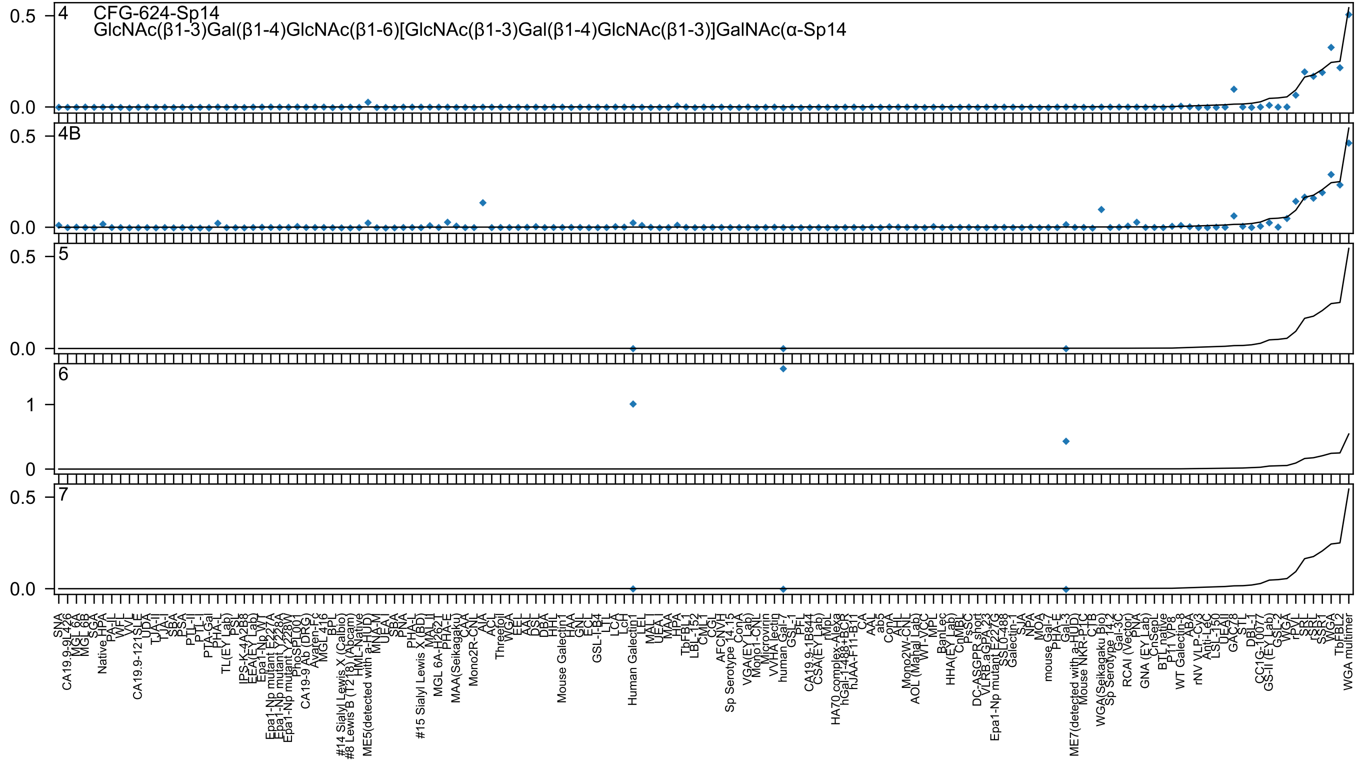

4 CFG-624-Sp14
GlcNAc(β1-3)Gal(β1-4)GlcNAc(β1-6)[GlcNAc(β1-3)Gal(β1-4)GlcNAc(β1-3)]GalNAc(α-Sp14
4B
5
6
7

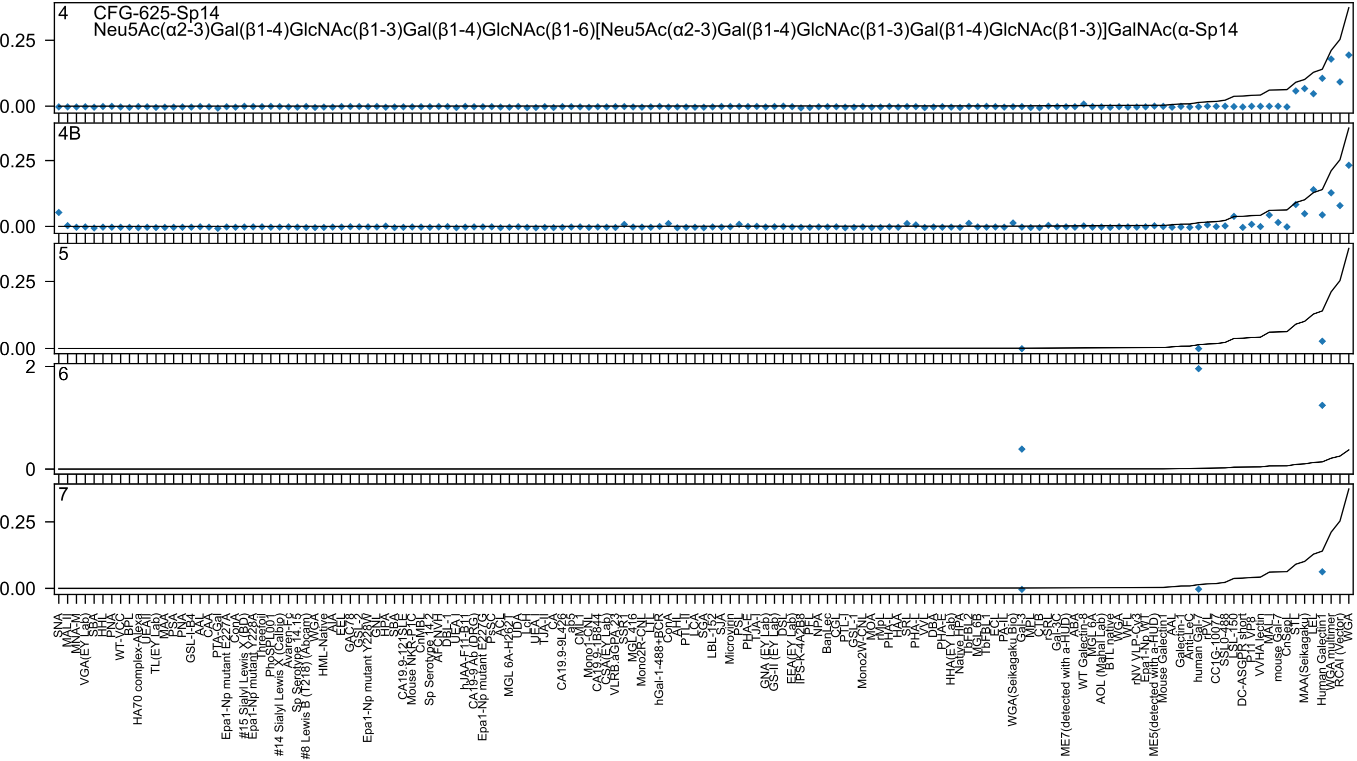

4
CFG-625-Sp14
Neu5Ac(α2-3)Gal(β1-4)GlcNAc(β1-3)Gal(β1-4)GlcNAc(β1-6)[Neu5Ac(α2-3)Gal(β1-4)GlcNAc(β1-3)Gal(β1-4)GlcNAc(β1-3)]GalNAc(α-Sp14
4B
5
6
7
0.25
0.00
2
0

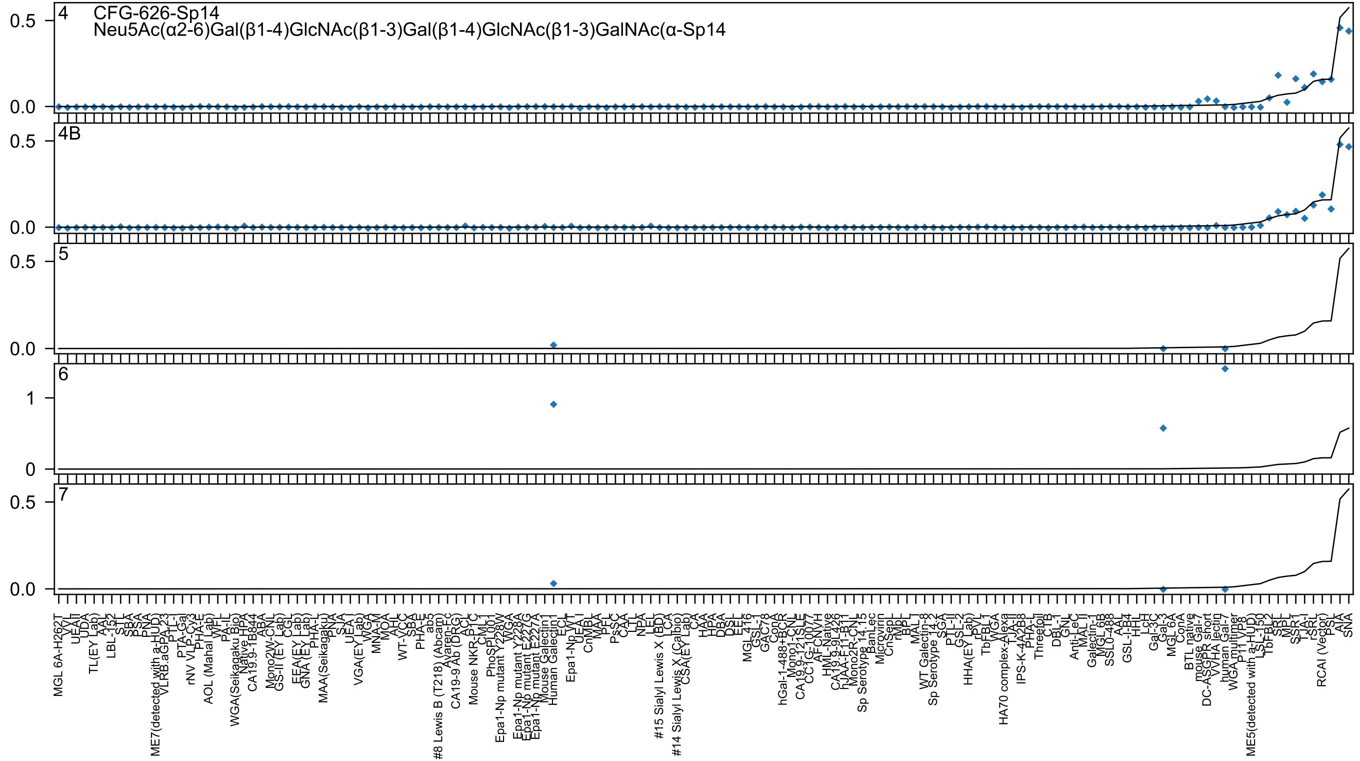

4
CFG-626-Sp14
Neu5Ac(α2-6)Gal(β1-4)GlcNAc(β1-3)Gal(β1-4)GlcNAc(β1-3)GalNAc(α-Sp14
4B
5
6
7
0.0
0.5
0
1
MGL 6A-H262T
VVL
UEA-II
UDA
TL(EY Lab)
AAL
LBL-152
STL
SBA
PSA
PNA
ME7(detected with a-HUD)
VLRB.aGPA.23
PTL-I
PTA-Gal
rNV VLP-Cy3
PHA-E
AOL (Mahal Lab)
WFL
PA-IL
WGA(Seikagaku Bio)
Native HPA
CA19.9-1B844
ABA
Mono2W-CNL
GS-II (EY Lab)
CGL
EEA(EY Lab)
GNA (EY Lab)
PHA-L
MAA(Seikagaku)
BPL
SJA
UEA-I
VGA(EY Lab)
WGA
MNA-M
MOA
AHL
WT-VCC
SBA
PHA-E
ab5
#8 Lewis B (T218) (Abcam)
Avaren-Fc
CA19-9 Ab (DRG)
ACL
Mouse NKR-P1C
CML-1
PhoSL001
Epa1-Np mutant Y228W
WGA
Epa1-Np mutant Y228A
Epa1-Np mutant E227G
Epa1-Np mutant E227A
Mouse Galectin1
Human Galectin1
ECL
Epa1-Np WT
UEA-I
CnMBL
MAA
PFL
PsSC
CAA
LTL
NPA
LEL
#15 Sialyl Lewis X (BD)
LCA
#14 Sialyl Lewis X (Calbio)
CSA(EY Lab)
CA
HAA
HPA
DBA
DSL
EEL
MGL 416
GSL-1
GAC78
ConA
hGal-1-488+BCR
Mono1-CNL
CA19.9-121SLE
CC1G-1007
AFCNVH
HML-Native
CA19.9-9L426
hJAA-F11-B11
Mono2R-CNL
Sp Serotype 14.15
BanLec
Microvirin
CnSepL
rMpL
BPL
MAL
WT Galectin-8
Sp Serotype 14.2
SGA
PTL-II
GSL-2
HHA(EY Lab)
rPVL
TbFBL1
WGA
HA70 complex-Alexa
TJA-II
IPS-K-4A2B8
PHA-L
Threefoil
CTB
DBL-1
GNL
Anti-LeC
MAL II
Galectin-1
MGL 6B
SSL0-488
AAL
GSL-I-B4
HHL
LcH
Gal-3C
Gal-3
MGL 6A
ConA
BTL native
mouse Gal-7
DC-ASGPR short
VVHA lectin
human Gal-7
WGA multimer
P11 VP8
ME5(detected with a-HUD)
LSL-150
TbFBL2
SRL
MPL
SSR1
TJA-I
rSRL
RCAI (Vector)
PSL
AIA
SNA

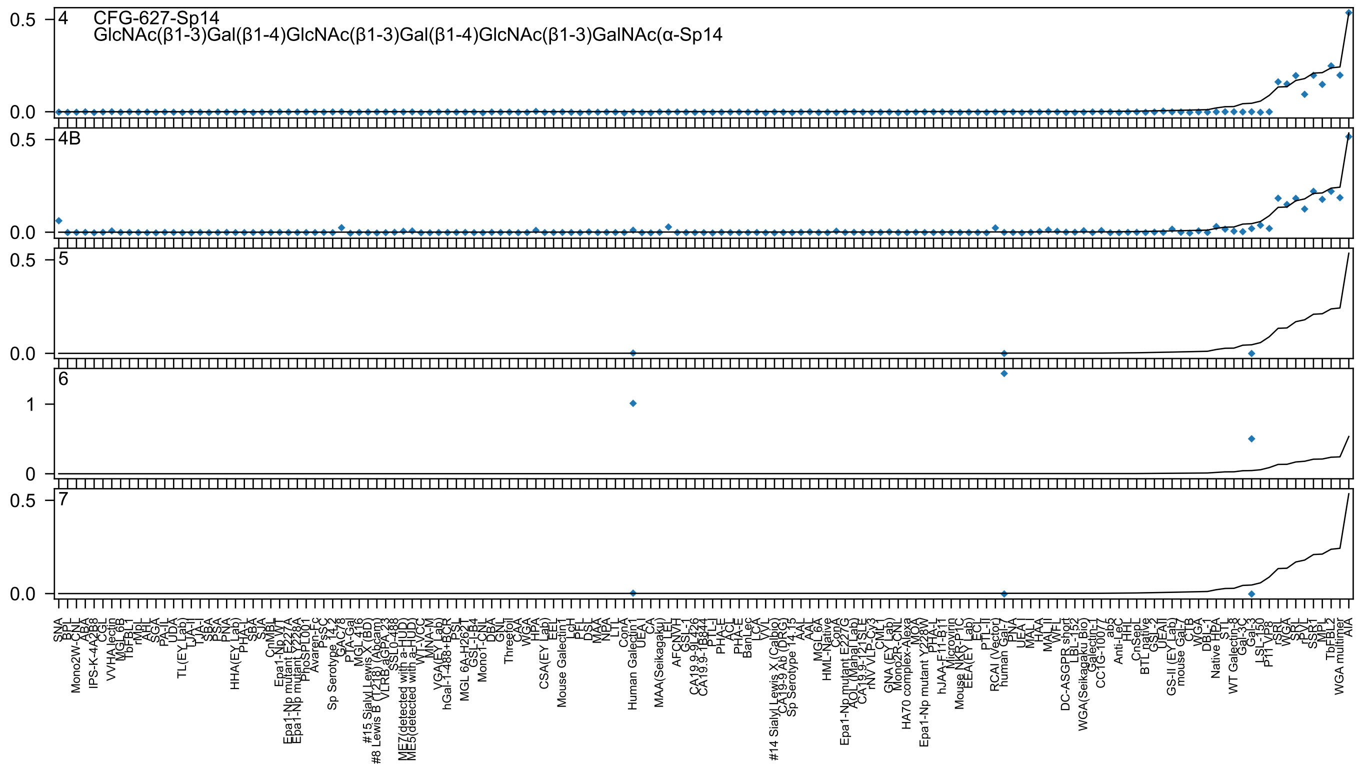

CFG-627-Sp14
GlcNAc(β1-3)Gal(β1-4)GlcNAc(β1-3)Gal(β1-4)GlcNAc(β1-3)GalNAc(α-Sp14
4
4B
5
6
7

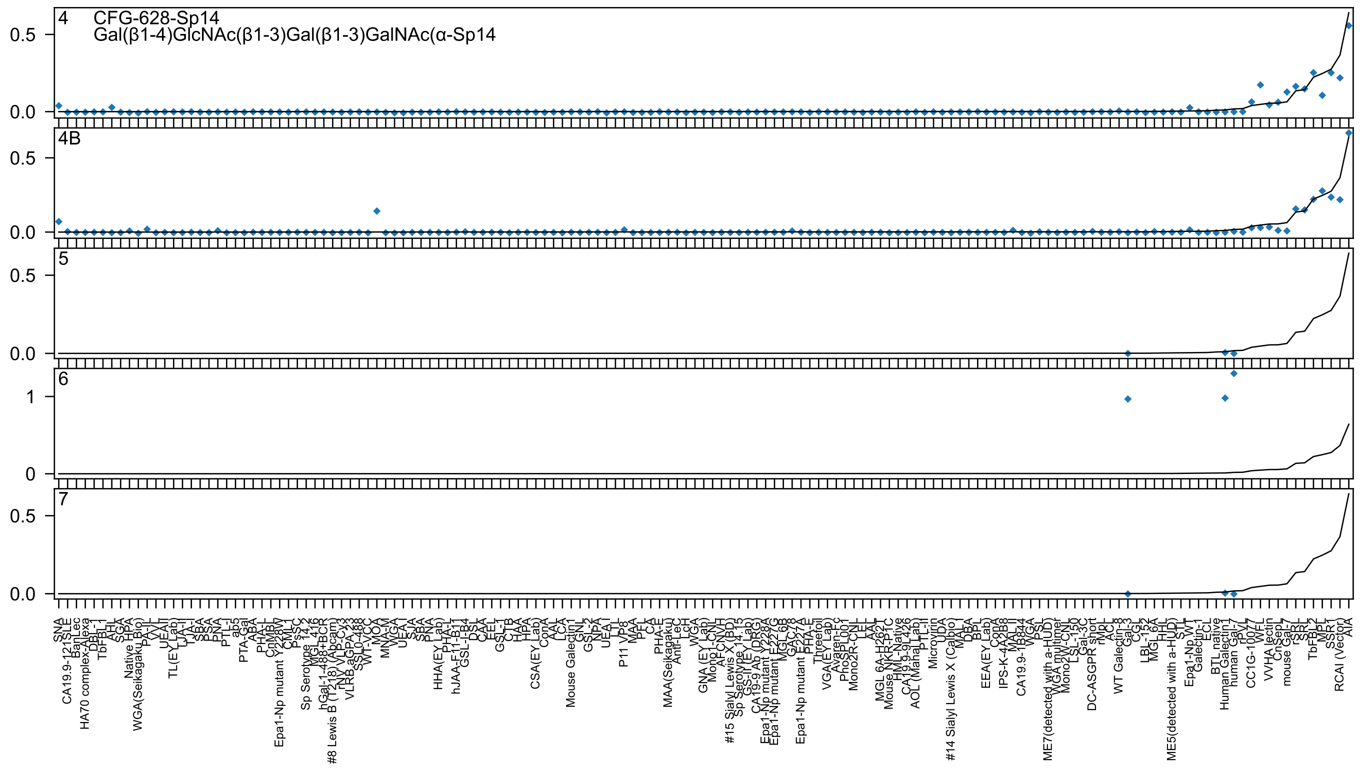

4 CFG-628-Sp14
Gal(β1-4)GlcNAc(β1-3)Gal(β1-3)GalNAc(α-Sp14
4B
5
6
7

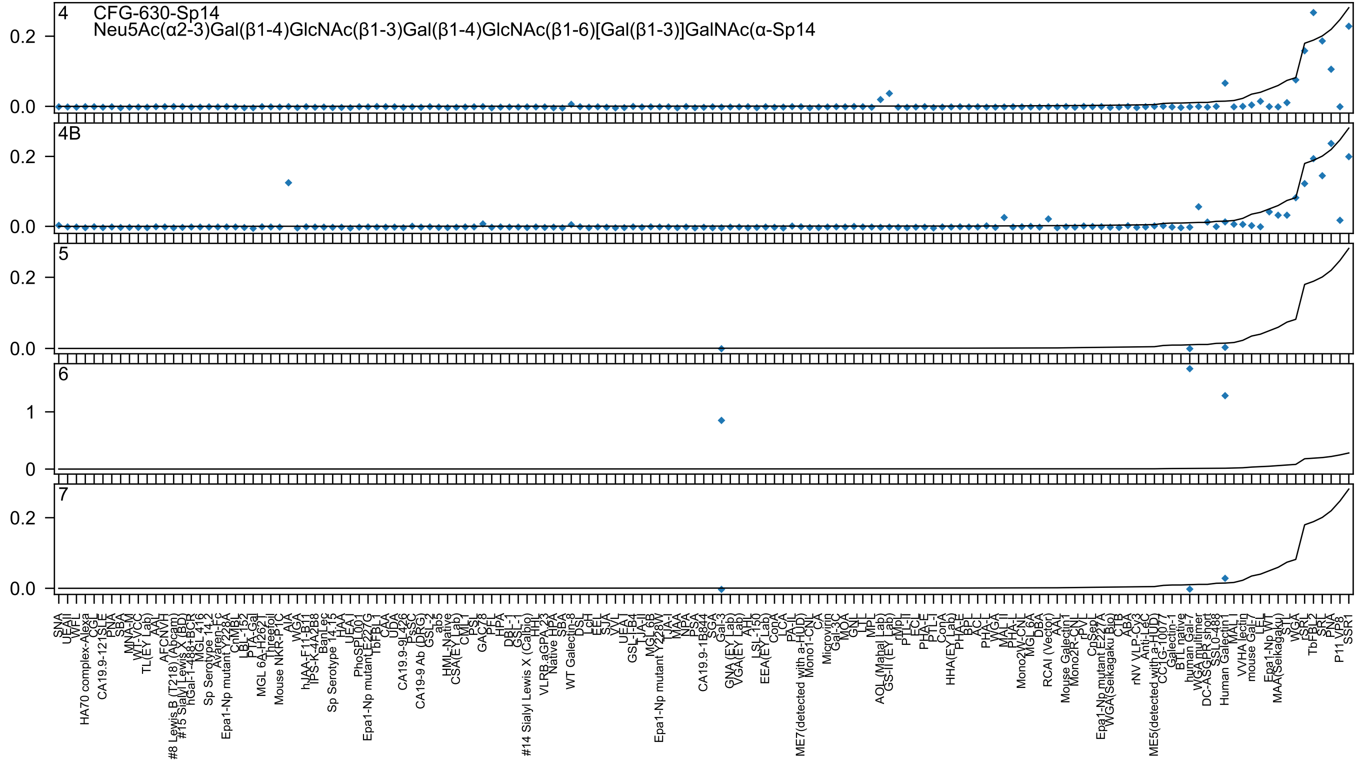

4 CFG-630-Sp14
Neu5Ac(α2-3)Gal(β1-4)GlcNAc(β1-3)Gal(β1-4)GlcNAc(β1-6)[Gal(β1-3)]GalNAc(α-Sp14
4B
5
6
7
0.0
0.2
0
1
SNA
UEAII
WFL
HA70 complex-Alexa
CGL
CA19.9-121SLE
PNA
SBA
MNA-M
WT-VCC
TL(EY Lab)
AAL
AFCNVH
#8 Lewis B (T218) (Abcam)
#15 Sialyl Lewis X (BD)
hGal-1-488+BCR
MGL416
Sp Serotype 14-2
Avaren-Fc
Epa1-Np mutant Y228A
CnMBL
LBL-152
PTA-Gal
MGL 6A-H262T
Threefoil
Mouse NKR-P1C
AIA
WGA
hJAA-F11-B11
IPS-K-4A2B8
BanLec
Sp Serotype 14-15
HAA
UEA-I
PhoSL001
Epa1-Np mutant E227G
TbFBL1
CAA
UDA
CA19.9-9L426
PsSC
CA19-9 Ab (DRG)
GSL-2
ab5
HML-Native
CSA(EY Lab)
CML-I
PSL
GAC78
PFL
HPA
DBL-1
GSL-1
#14 Sialyl Lewis X (Calbio)
HHL
VLRB.aGPA.23
Native HPA
SBA
WT Galectin-8
DSL
LcH
EEL
SJA
VVL
UEA-I
GSL-I-B4
TJA-II
MGL 6B
Epa1-Np mutant Y228W
TJA-I
MAA
NPA
PSA
CA19.9-1B844
SGA
Gal-3
GNA (EY Lab)
VGA(EY Lab)
AHL
LSL-150
EEA(EY Lab)
ConA
LCA
PA-IL
ME7(detected with a-HUD)
Mono1-CNL
CA
Microvirin
Gal-3C
MOA
GNL
LTL
MPL
AOL (Mahal Lab)
GS-II (EY Lab)
rMpL
PTL-II
ECL
PHA-E
PTL-I
ConA
HHA(EY Lab)
PHA-E
BPL
ACL
PHA-L
WGA
MAL_II
PHA-L
Mono2W-CNL
MGL 6A
DBA
RCAI (Vector)
AAL
Mouse Galectin1
Mono2R-CNL
rPVL
CnSep
Epa1-Np mutant E227A
WGA(Seikagaku Bio)
CTB
ABA
rNV VLP-Cy3
Anti-LeC
ME5(detected with a-HUD)
CC1G-1007
Galectin-1
BTL native
human Gal-7
WGA multimer
DC-ASGPR short
SSL0-488
Human Galectin1
MAL_I
VVHA lectin
mouse Gal-7
LEL
Epa1-Np WT
MAA(Seikagaku)
STL
WGA
rSRL
TbFBL2
SRL
PNA
P11 VP8
SSR1

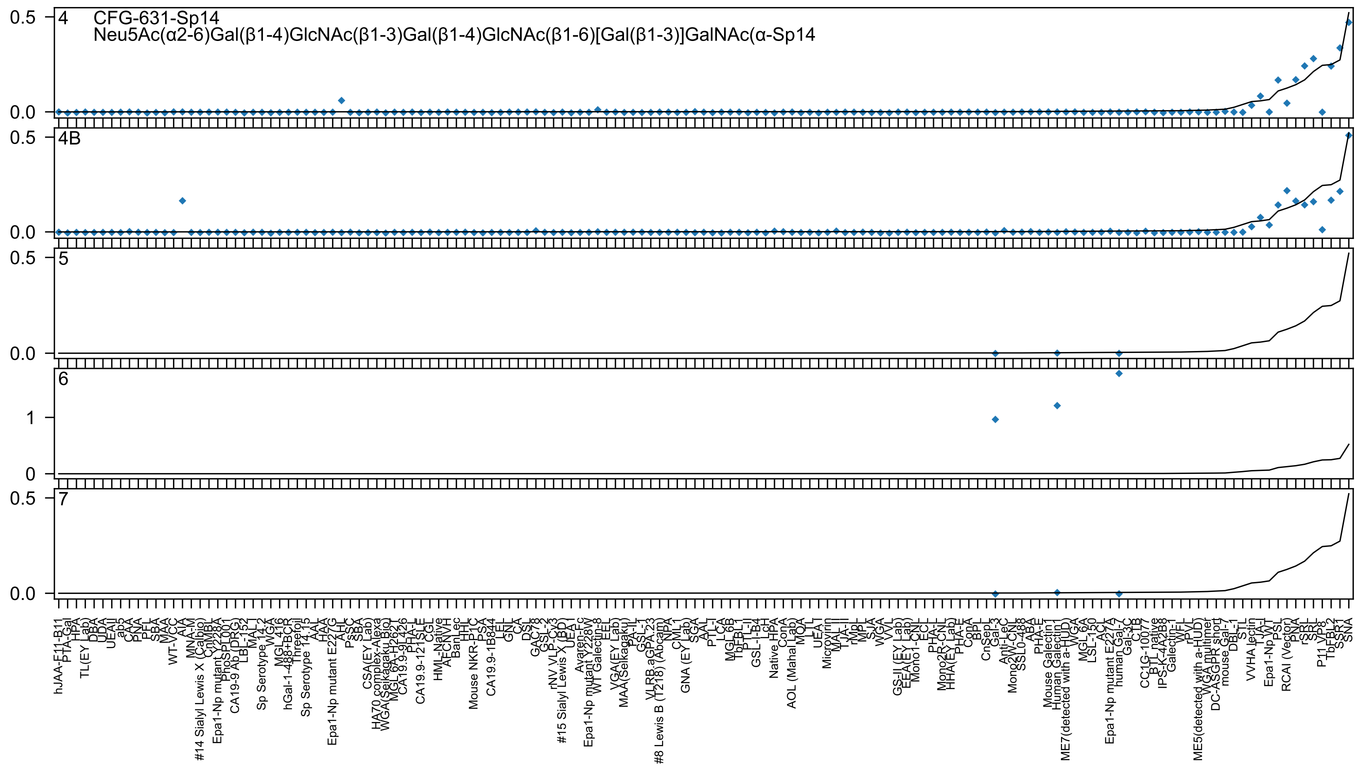

CFG-631-Sp14
Neu5Ac(α2-6)Gal(β1-4)GlcNAc(β1-3)Gal(β1-4)GlcNAc(β1-6)[Gal(β1-3)]GalNAc(α-Sp14
4
4B
5
6
7
0.5
0.0
1
0
hJAA-F11-B11
PTA-Gal
HPA
TL(EY Lab)
DBA
UDA
UEA-II
ab5
CAA
PNA
PFL
SBA
MAA
WT-VCC
AIA
MNA-M
#14 Sialyl Lewis X (Calbio)
CnMBL
Epa1-Np mutant Y228A
PhoSL001
CA19-9 Ab (DRG)
LBL-152
MAL-I
Sp Serotype 14.2
WGA
MGL416
hGal-1-488+BCR
Threefoil
Sp Serotype 14.15
AAL
HAA
Epa1-Np mutant E227G
AHL
PsSC
SBA
CSA(EY Lab)
HA70 complex-Alexa
WGA(Seikagaku Bio)
MGL 6A-H262T
CA19.9-9L426
PHA-L
CA19.9-121SLE
CGL
HML-Native
AFCNVH
BanLec
HHL
Mouse NKR-P1C
PSA
CA19.9-1B844
LEL
GNL
CA
DSL
GAC78
GSL-2
rNV VLP-Cy3
#15 Sialyl Lewis X (BD)
UEA-I
Avaren-Fc
Epa1-Np mutant Y228W
WT Galectin-8
EEL
VGA(EY Lab)
MAA(Seikagaku)
PA-IL
GSL-I
VLRB aGPA.23
#8 Lewis B (T218) (Abcam)
NPA
CML-I
GNA (EY Lab)
SGA
AAL
PTL-I
LCA
MGL 6B
TbFBL1
PTL-II
GSL-I-B4
LcH
Native HPA
ConA
AOL (Mahal Lab)
MOA
LTL
UEA-I
Microvirin
MAL-II
TJA-II
rMpL
MPL
SJA
WGA
VVL
GS-II (EY Lab)
EEA(EY Lab)
Mono1-CNL
ECL
PHA-L
Mono2R-CNL
HHA(EY Lab)
PHA-E
CobA
BPL
CnSepL
Gal-3
Anti-LeC
Mono2W-CNL
SSL0-488
ABA
PHA-E
Mouse Galectin1
Human Galectin1
ME7(detected with a-HUD)
WGA
MGL 6A
LSL-150
ACL
Epa1-Np mutant E227A
human Gal-7
Gal-3C
CTB
CC1G-10077
BTL native
IPS-K-4A2B8
Galectin-1
WFL
rPVL
ME5(detected with a-HUD)
WGA multimer
DC-ASGPR short
mouse Gal-7
DBL-1
STL
VVHA lectin
TJA-I
Epa1-Np WT
PSL
RCAI (Vector)
PNA
rSRL
SRL
P11 VP8
TbFBL2
SSR1
SNA

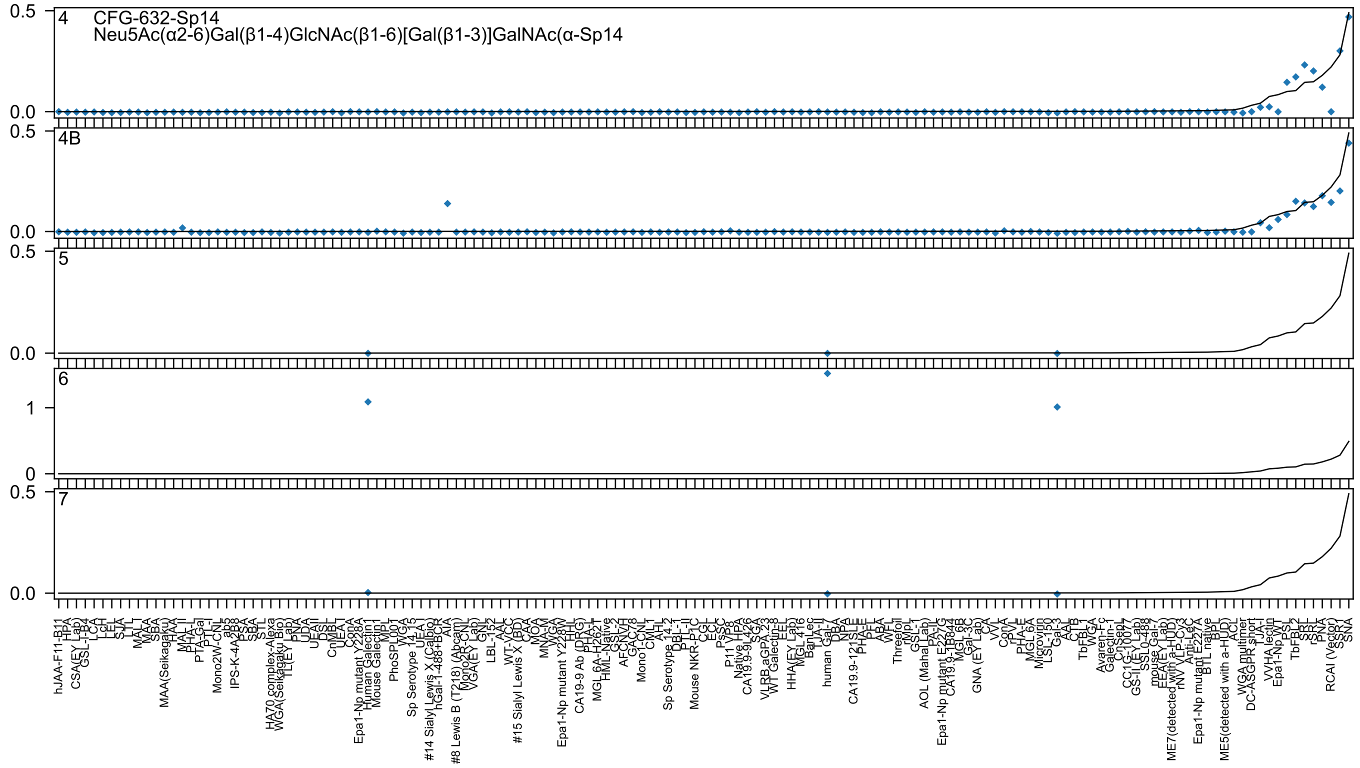
4 CFG-632-Sp14
Neu5Ac(α2-6)Gal(β1-4)GlcNAc(β1-6)[Gal(β1-3)]GalNAc(α-Sp14
4B
5
6
7

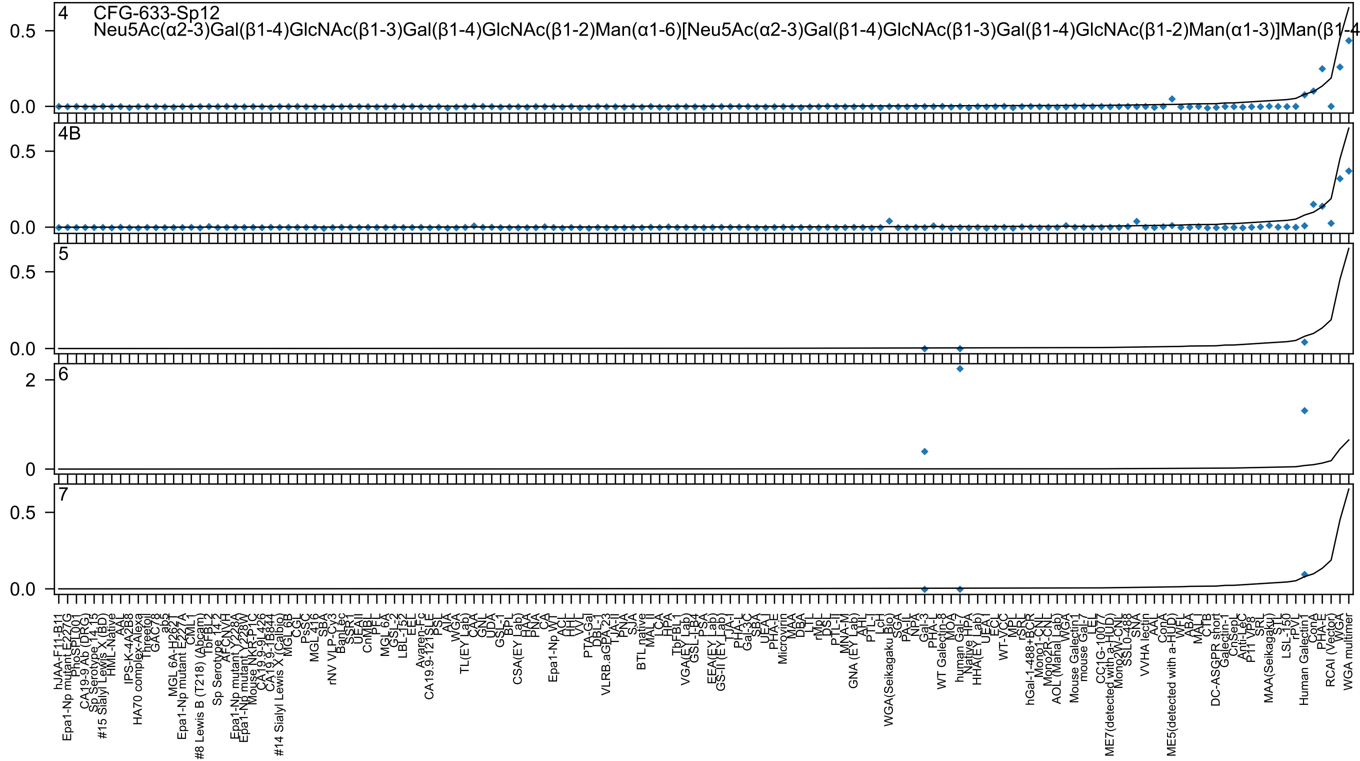
4 CFG-633-Sp12
Neu5Ac(α2-3)Gal(β1-4)GlcNAc(β1-3)Gal(β1-4)GlcNAc(β1-2)Man(α1-6)[Neu5Ac(α2-3)Gal(β1-4)GlcNAc(β1-3)Gal(β1-4)GlcNAc(β1-2)Man(α1-3)]Man(β1-4
4B
5
6
7
0.0
0.5
2
0

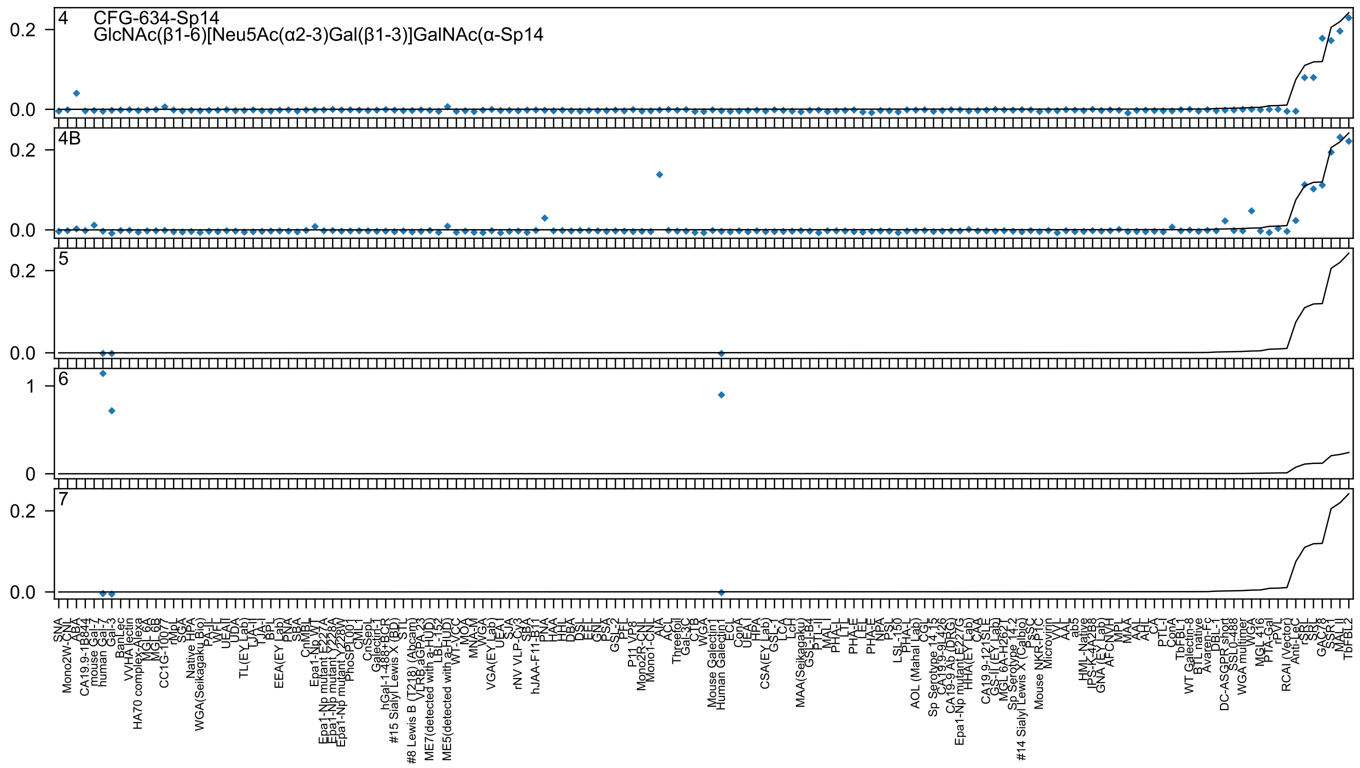
4
CFG-634-Sp14
GlcNAc(β1-6)[Neu5Ac(α2-3)Gal(β1-3)]GalNAc(α-Sp14
4B
5
6
7

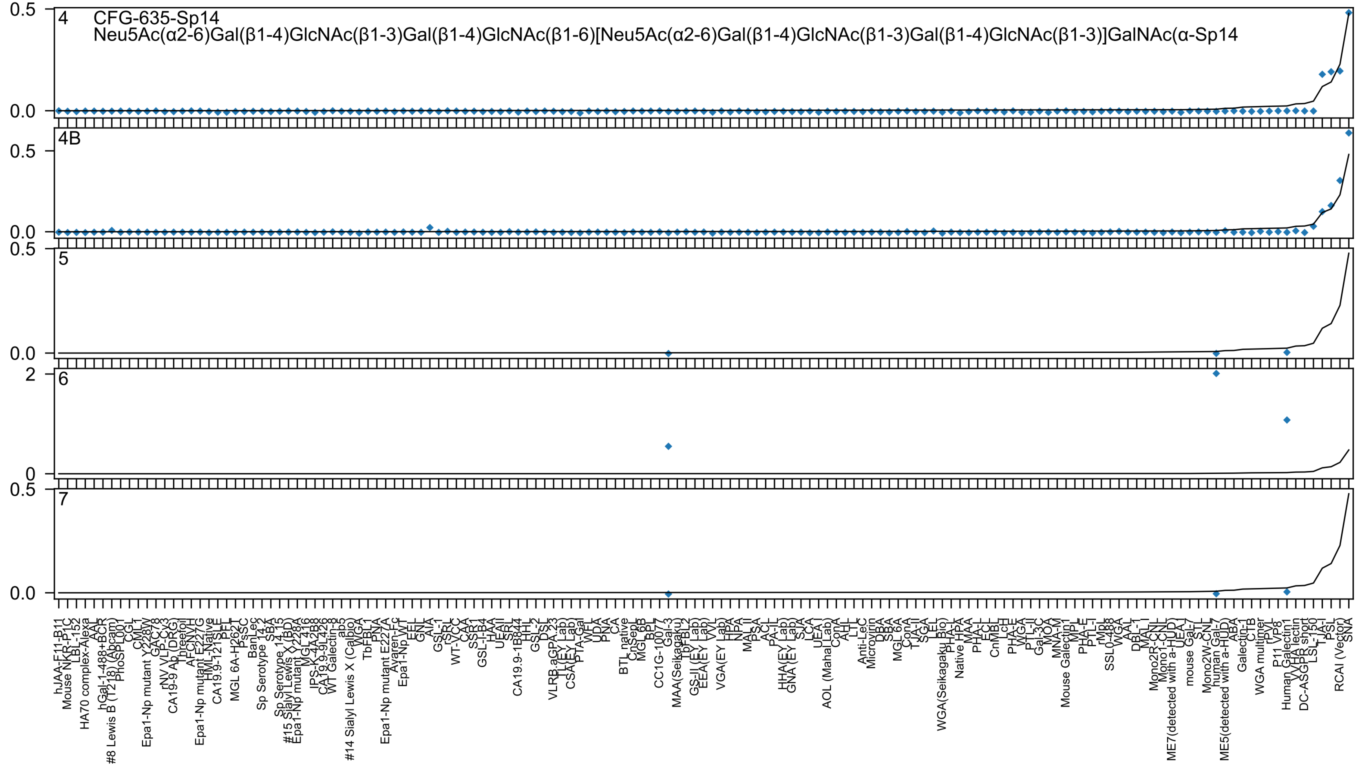

CFG-635-Sp14
Neu5Ac(α2-6)Gal(β1-4)GlcNAc(β1-3)Gal(β1-4)GlcNAc(β1-6)[Neu5Ac(α2-6)Gal(β1-4)GlcNAc(β1-3)Gal(β1-4)GlcNAc(β1-3)]GalNAc(α-Sp14
4
4B
5
6
7
0.5
0.0
2
0
hJAA-F11-B11
Mouse NKR-P1C
LBL-152
HA70 complex-Alexa
AAL
hGal-1-488+BCR
#8 Lewis B (T218) (Abcam)
PhoSL001
CGL
CML1
Epa1-Np mutant Y228W
GAC78
rNV VLP-Cy3
CA19-9 Ab (DRG)
Threefoil
AFCNVH
Epa1-Np mutant E227G
HML-Native
CA19.9-121SLE
PFL
MGL 6A-H262T
PsSC
BanLec
Sp Serotype 14.2
SBA
Sp Serotype 14.15
#15 Sialyl Lewis X (BD)
Epa1-Np mutant Y228A
MGL 416
IPS-K-4A2B8
CA19.9-9L426
WT Galectin-8
ab5
#14 Sialyl Lewis X (Calbio)
WGA
TbFBL1
PNA
Epa1-Np mutant E227A
Ayaren-Fc
Epa1-Np WT
EEL
GNL
AIA
GSL-I
rSRL
WT-VCC
CAA
SSR1
GSL-I-B4
HAA
UEAII
SRL
CA19.9-1B844
HHL
GSL-2
DSL
VLRB aGPA.23
TL(EY Lab)
CSA(EY Lab)
PTA-Gal
WFL
UDA
PNA
CA
BTL native
CnSep
MGL 6B
BPL
CC1G-10077
Gal-3
MAA(Seikagaku)
TbFBL2
GS-II (EY Lab)
EEA(EY Lab)
VVL
VGA(EY Lab)
HPA
NPA
MAL II
PSA
ACL
PA-IL
HHA(EY Lab)
GNA (EY Lab)
SJA
LCA
UEA I
AOL (Maha Lab)
ConA
AHL
LTL
Anti-LeC
Microvirin
DBA
SBA
MGL 6A
ConA
TJA-II
SGA
LEL
WGA(Seikagaku Bio)
PHA-L
Native HPA
MAA
PHA-L
ECL
CnMBL
LcH
PHA-E
WGA
PTL-II
Gal-3C
MOA
MNA-M
Mouse Galectin1
MPL
PHA-E
PTL-I
rMpL
SSL0-488
WGA
AAL
DBL-1
MAL I
Mono2R-CNL
Mono1-CNL
ME7(detected with a-HUD)
UEA I
mouse Gal-7
STL
Mono2W-CNL
human Gal-7
ME5(detected with a-HUD)
ABA
Galectin-1
CTB
WGA multimer
rPVL
P11 VP8
Human Galectin1
VVHA lectin
DC-ASGPR short
LSL-150
TJA-I
PSL
RCAI (Vector)
SNA

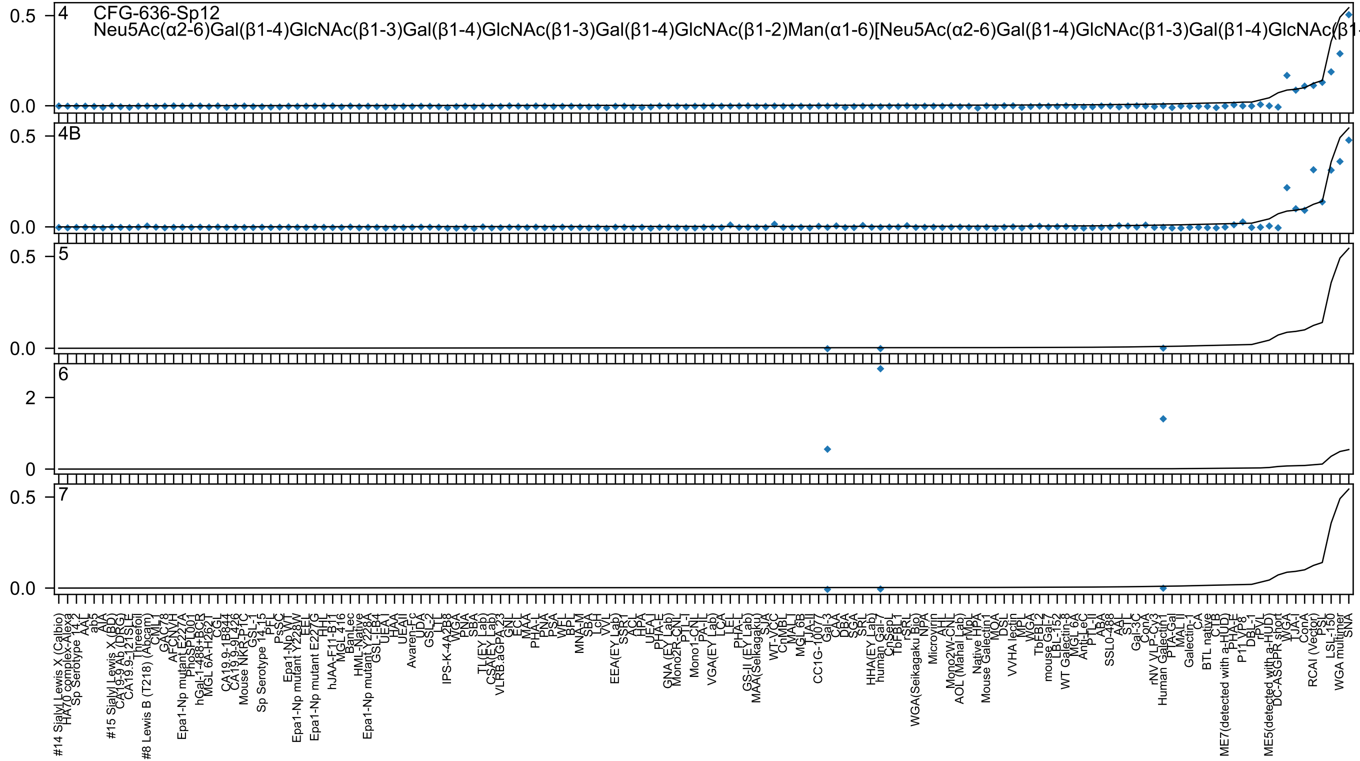
4
CFG-636-Sp12
Neu5Ac(α2-6)Gal(β1-4)GlcNAc(β1-3)Gal(β1-4)GlcNAc(β1-3)Gal(β1-4)GlcNAc(β1-2)Man(α1-6)[Neu5Ac(α2-6)Gal(β1-4)GlcNAc(β1-3)Gal(β1-4)GlcNAc(β1
4B
5
6
7

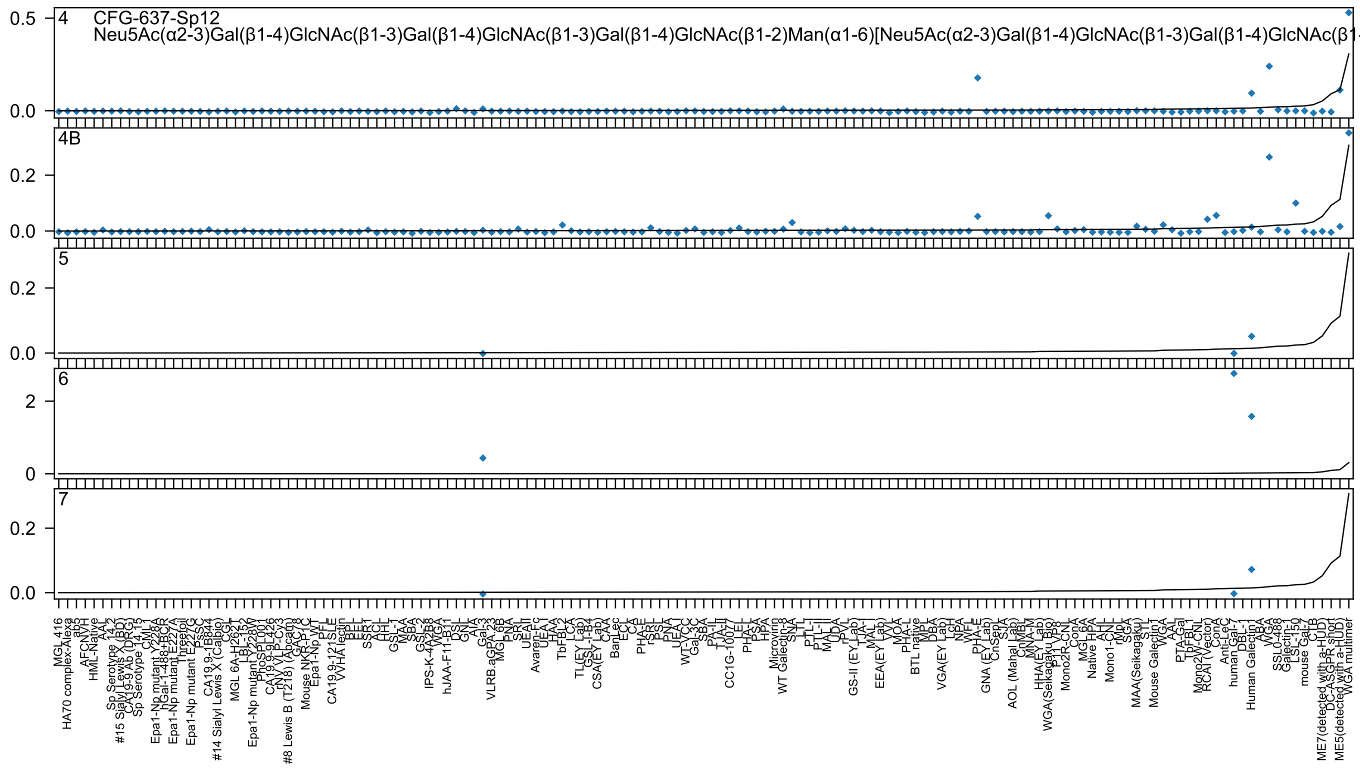

CFG-637-Sp12
Neu5Ac(α2-3)Gal(β1-4)GlcNAc(β1-3)Gal(β1-4)GlcNAc(β1-3)Gal(β1-4)GlcNAc(β1-2)Man(α1-6)[Neu5Ac(α2-3)Gal(β1-4)GlcNAc(β1-3)Gal(β1-4)GlcNAc(β1
4
4B
5
6
7

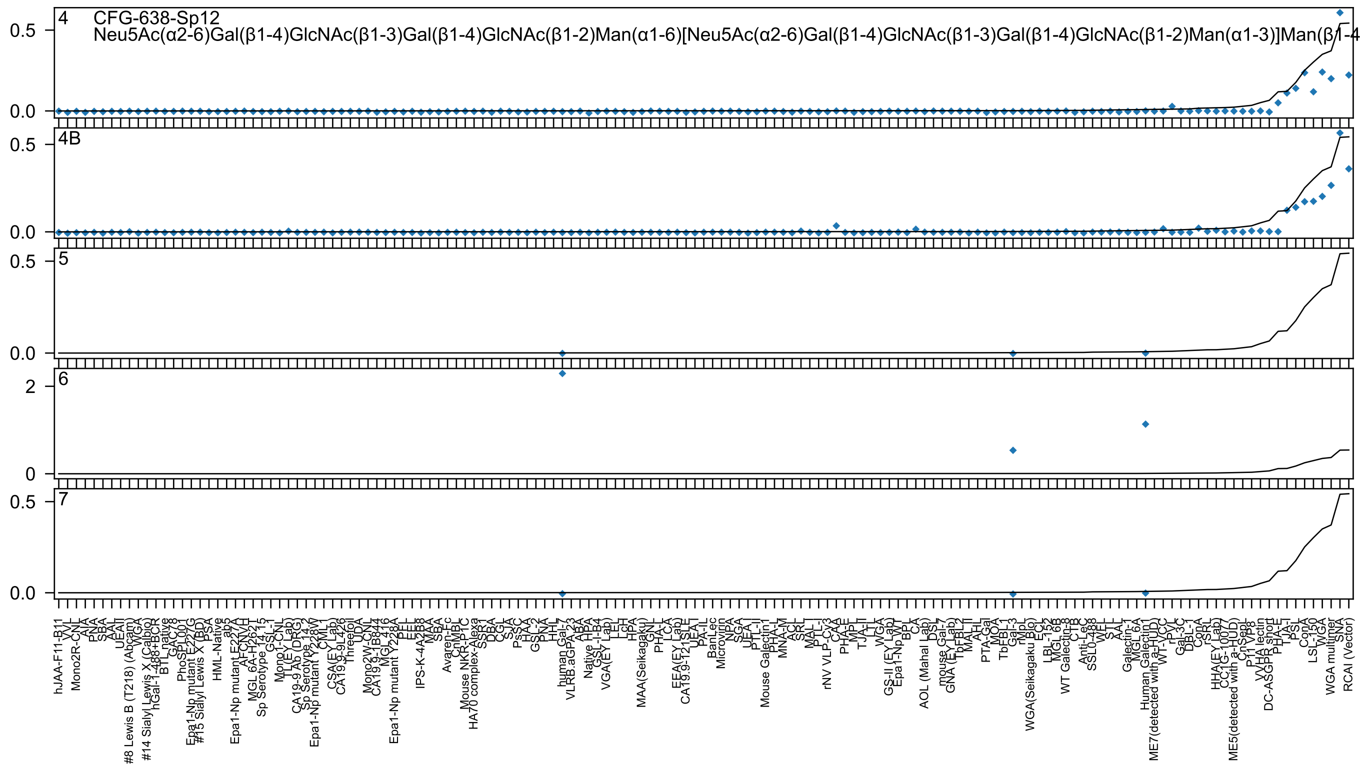

4
CFG-638-Sp12
Neu5Ac(α2-6)Gal(β1-4)GlcNAc(β1-3)Gal(β1-4)GlcNAc(β1-2)Man(α1-6)[Neu5Ac(α2-6)Gal(β1-4)GlcNAc(β1-3)Gal(β1-4)GlcNAc(β1-2)Man(α1-3)]Man(β1-4
4B
5
6
7
0.5
0.0
2
0

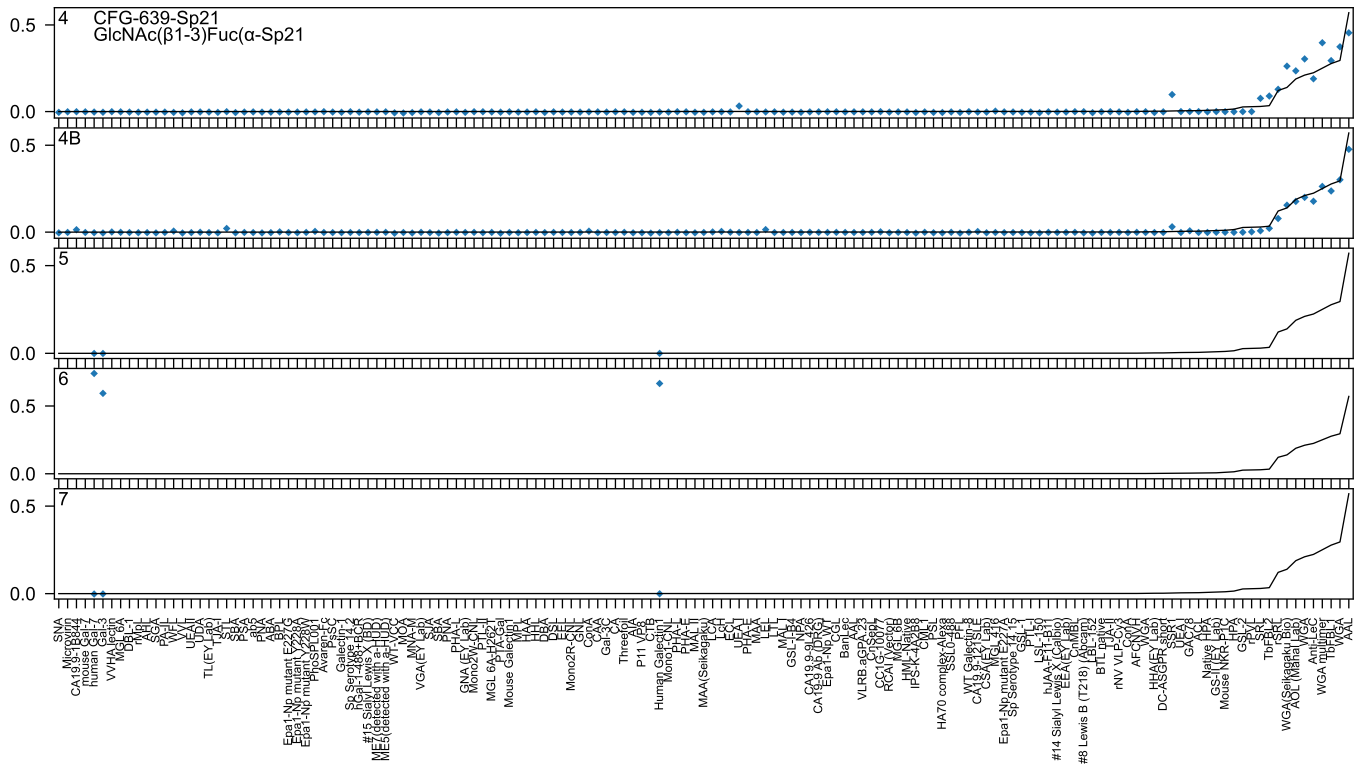
4
CFG-639-Sp21
GlcNAc(β1-3)Fuc(α-Sp21
4B
5
6
7
0.0
0.5

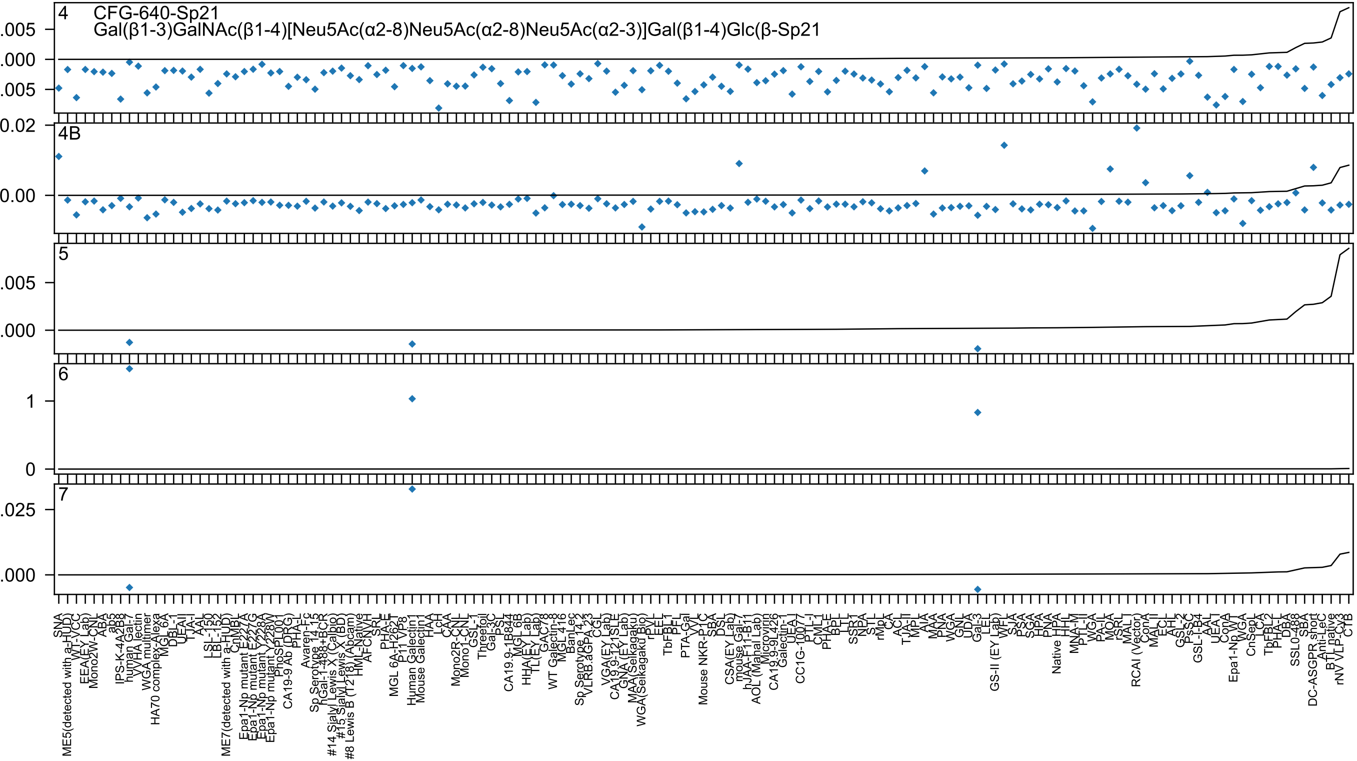

4
CFG-640-Sp21
Gal(β1-3)GalNAc(β1-4)[Neu5Ac(α2-8)Neu5Ac(α2-8)Neu5Ac(α2-3)]Gal(β1-4)Glc(β-Sp21
.005
.000
.005
0.02
4B
0.00
5
.005
.000
6
1
0
7
.025
.000
