## Supplementary material for "Atom-level Machine Learning of Protein-Glycan Interactions and Cross-chiral Recognition in Glycobiology": Predictions Across Concentrations

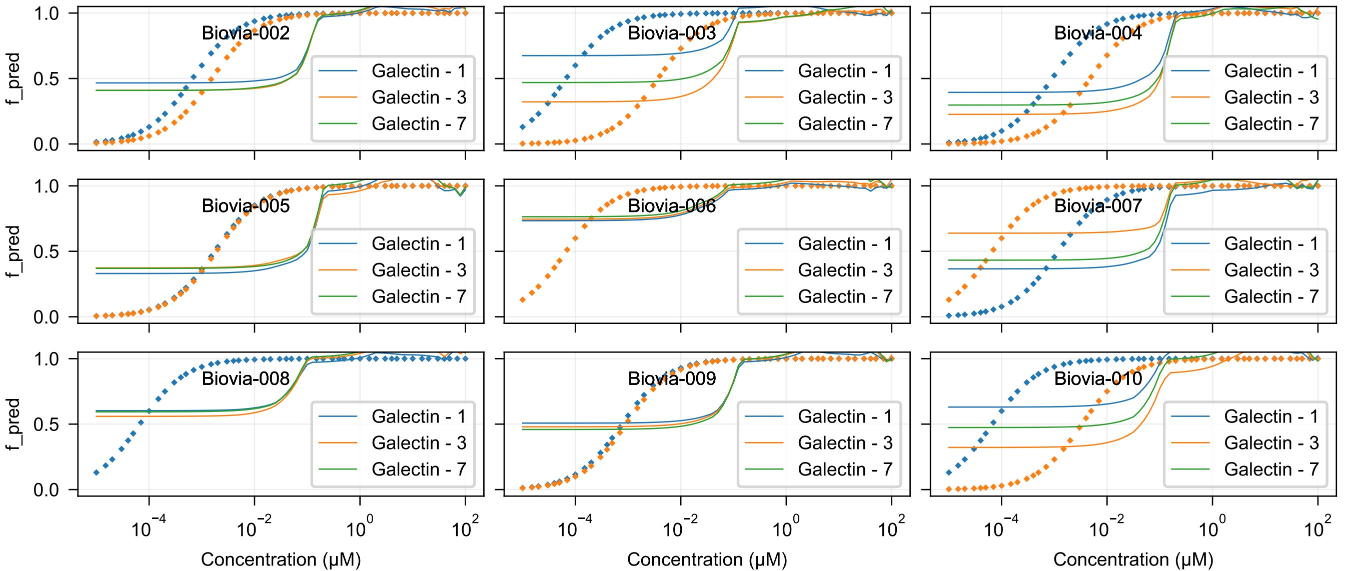
Biovia-002
Biovia-003
Biovia-004
Biovia-005
Biovia-006
Biovia-007
Biovia-008
Biovia-009
Biovia-010
Galectin - 1
Galectin - 3
Galectin - 7
f_pred
Concentration (μM)

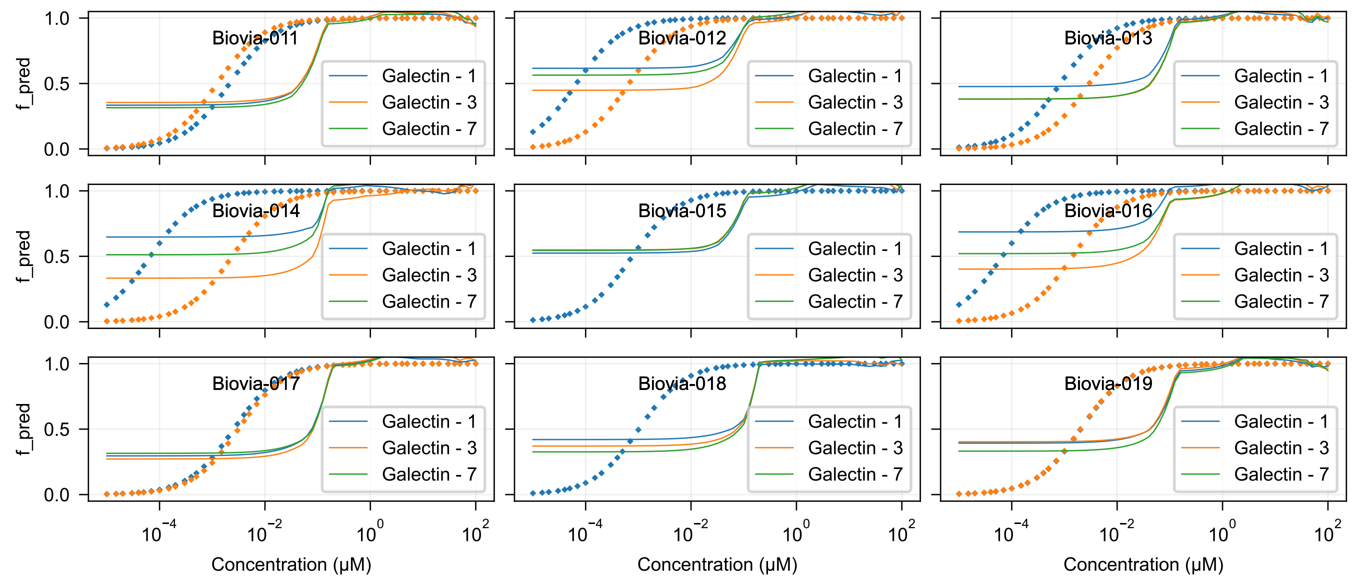

Biovia-011
Biovia-012
Biovia-013
Biovia-014
Biovia-015
Biovia-016
Biovia-017
Biovia-018
Biovia-019
Galectin - 1
Galectin - 3
Galectin - 7
f_pred
Concentration (μM)

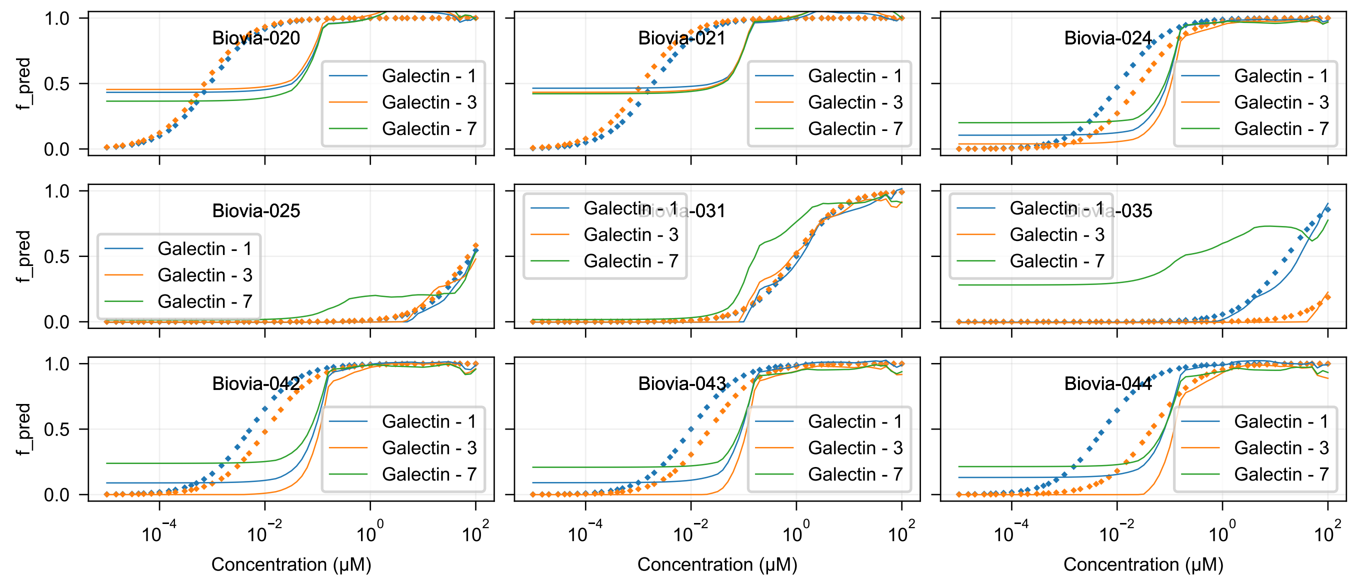
Biovia-020
Biovia-021
Biovia-024
Biovia-025
Biovia-031
Biovia-035
Biovia-042
Biovia-043
Biovia-044
Galectin - 1
Galectin - 3
Galectin - 7
f_pred
Concentration (μM)

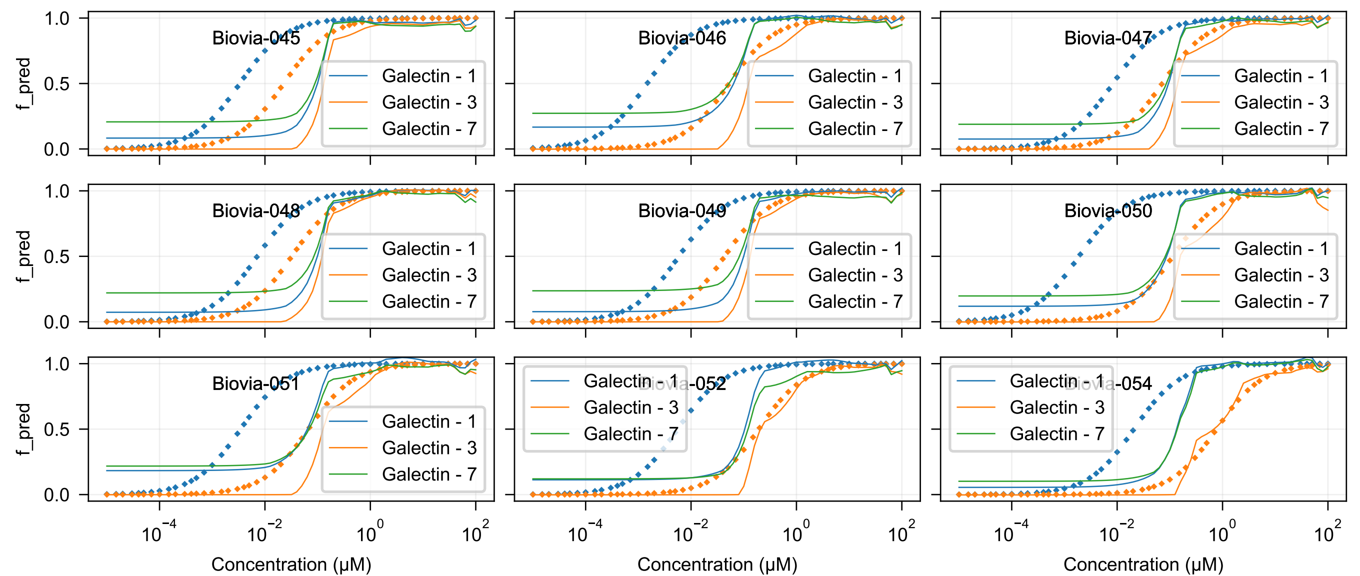

Biovia-045
Biovia-046
Biovia-047
Biovia-048
Biovia-049
Biovia-050
Biovia-051
Biovia-052
Biovia-054
Galectin - 1
Galectin - 3
Galectin - 7
f_pred
Concentration (μM)

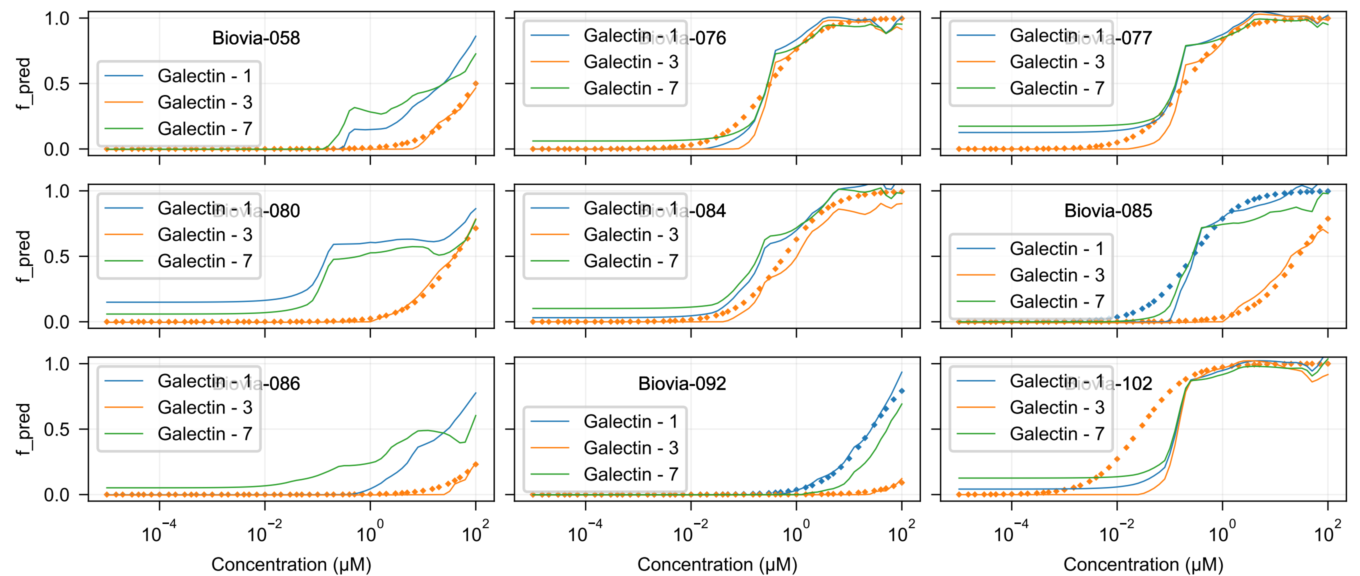

Biovia-058
Biovia-076
Biovia-077
Biovia-080
Biovia-084
Biovia-085
Biovia-086
Biovia-092
Biovia-102
Galectin - 1
Galectin - 3
Galectin - 7
f_pred
1.0
0.5
0.0
$10^{-4}$
$10^{-2}$
$10^{0}$
$10^{2}$
Concentration (μM)

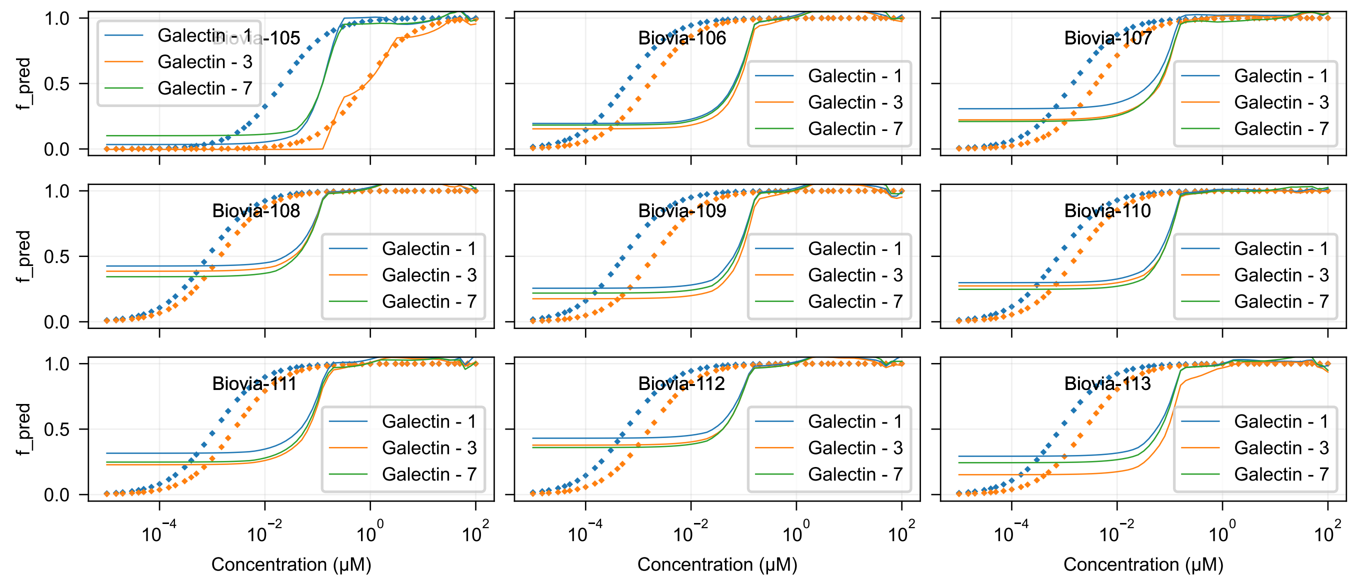
Biovia-105
Biovia-106
Biovia-107
Biovia-108
Biovia-109
Biovia-110
Biovia-111
Biovia-112
Biovia-113
Galectin - 1
Galectin - 3
Galectin - 7
f_pred
Concentration (μM)

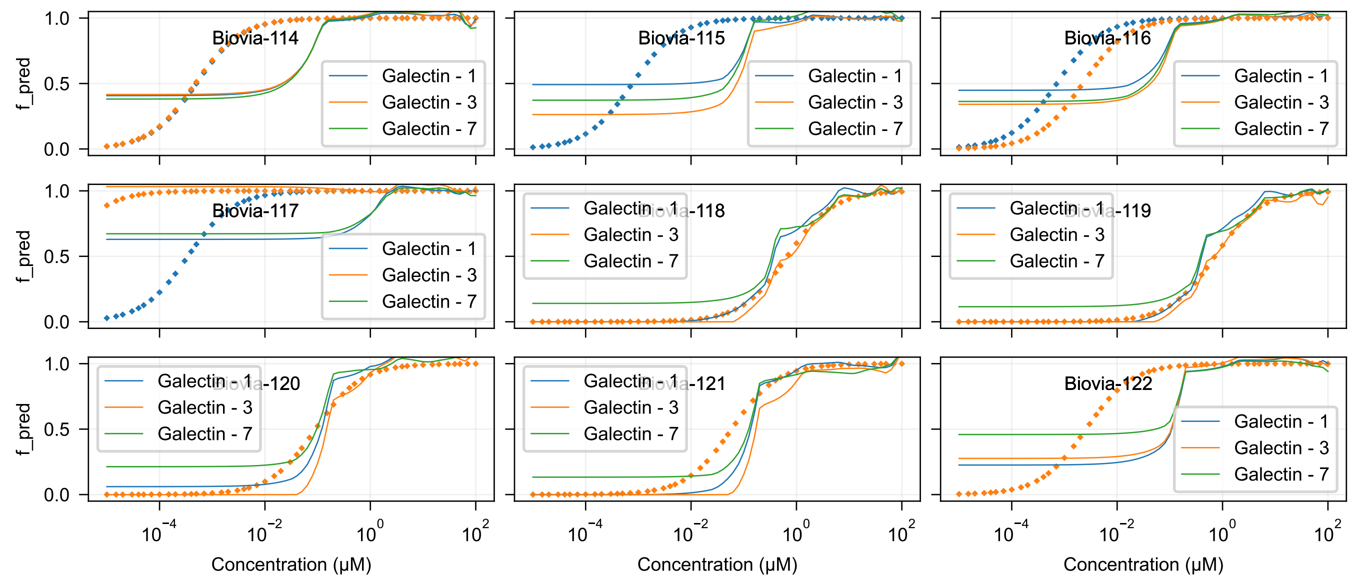

Biovia-114
Biovia-115
Biovia-116
Biovia-117
Biovia-118
Biovia-119
Biovia-120
Biovia-121
Biovia-122
Galectin - 1
Galectin - 3
Galectin - 7
f_pred
Concentration (μM)

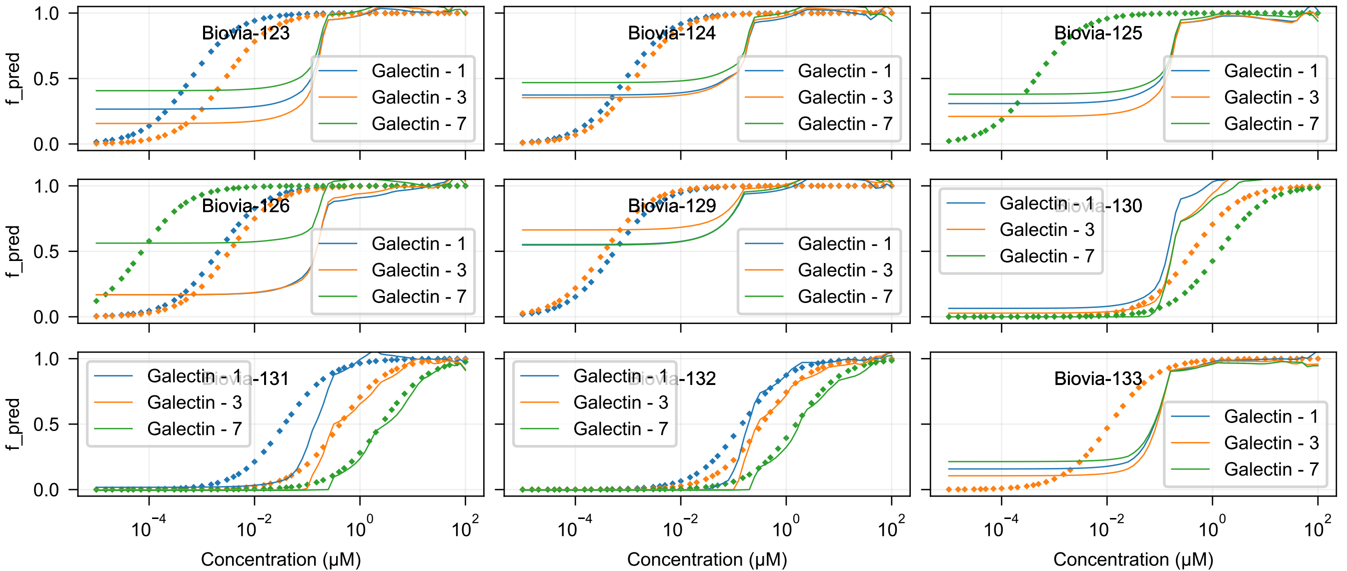
Biovia-123
Biovia-124
Biovia-125
Biovia-126
Biovia-129
Biovia-130
Biovia-131
Biovia-132
Biovia-133
Galectin - 1
Galectin - 3
Galectin - 7
f_pred
Concentration (μM)

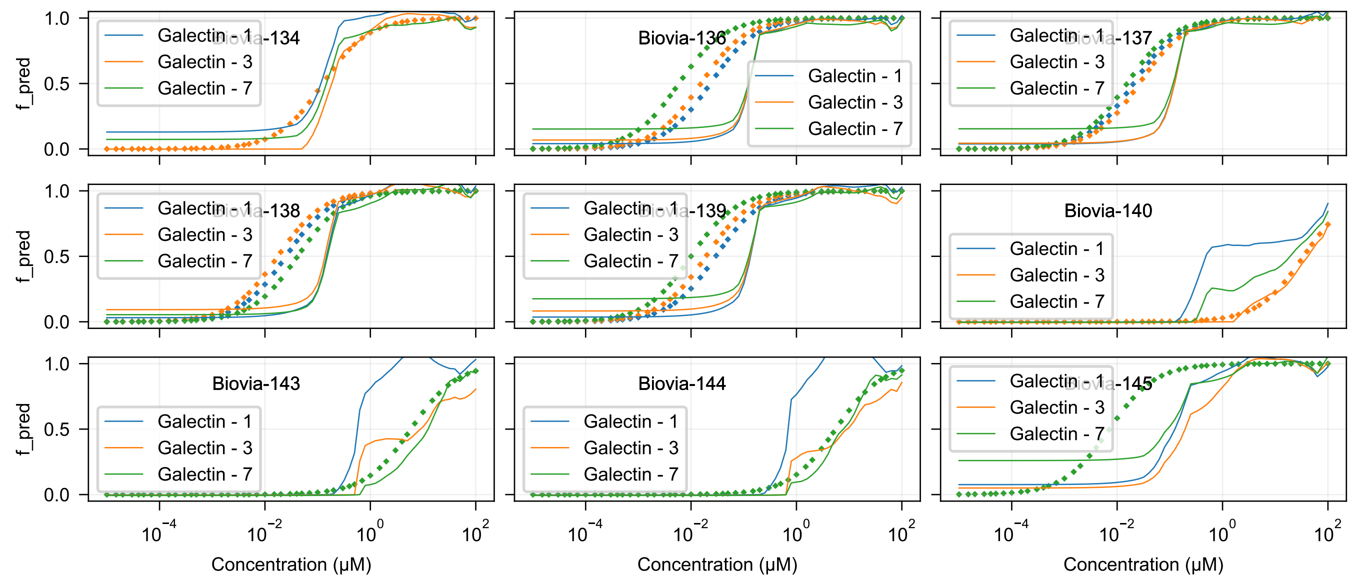

Biovia-134
Biovia-136
Biovia-137
Biovia-138
Biovia-139
Biovia-140
Biovia-143
Biovia-144
Biovia-145
Galectin - 1
Galectin - 3
Galectin - 7
f_pred
Concentration (μM)

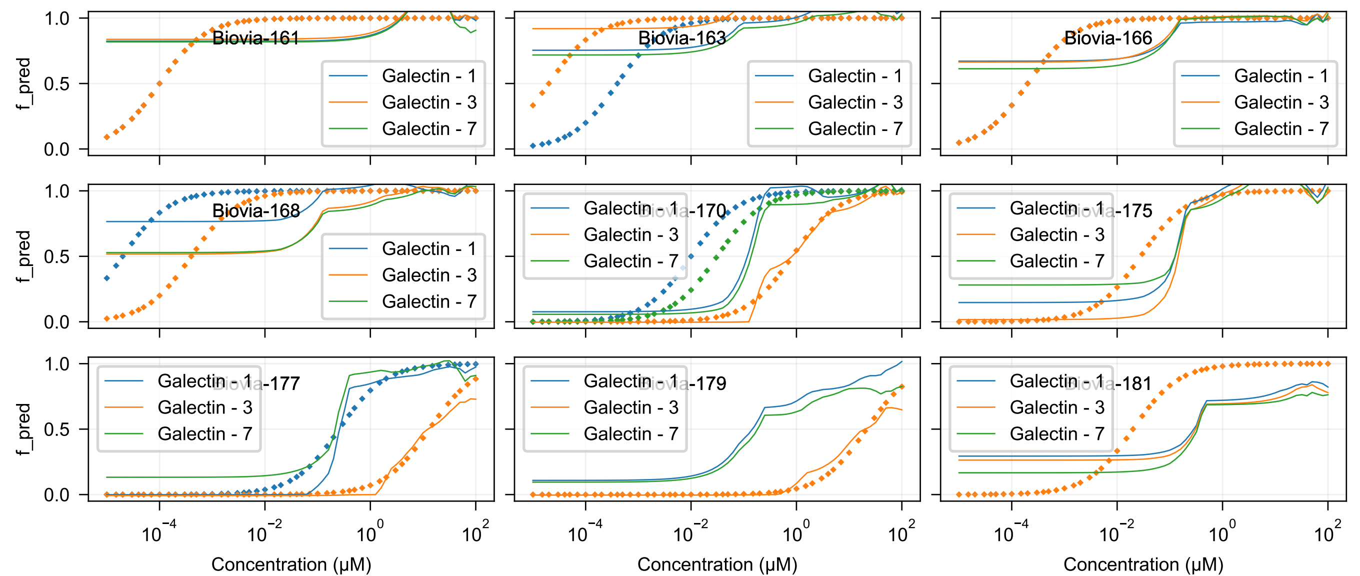
Biovia-161
Biovia-163
Biovia-166
Biovia-168
Biovia-170
Biovia-175
Biovia-177
Biovia-179
Biovia-181
Galectin - 1
Galectin - 3
Galectin - 7
f_pred
Concentration (μM)
0.0
0.5
1.0
$10^{-4}$
$10^{-2}$
$10^{0}$
$10^{2}$

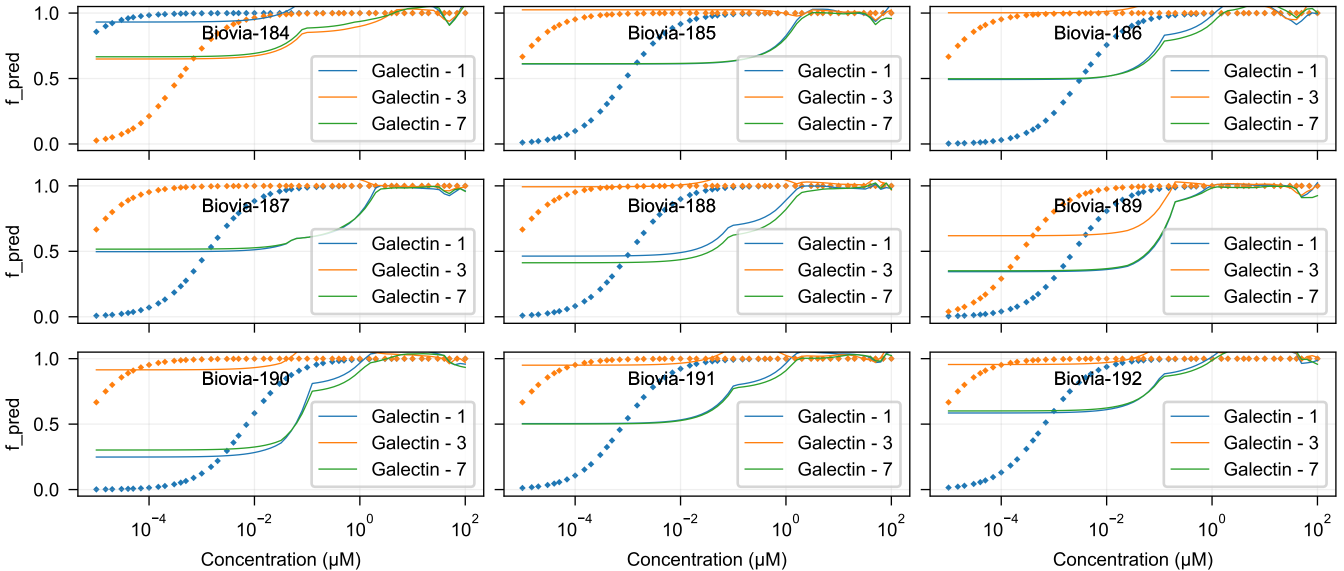
Biovia-184
Biovia-185
Biovia-186
Biovia-187
Biovia-188
Biovia-189
Biovia-190
Biovia-191
Biovia-192
Galectin - 1
Galectin - 3
Galectin - 7
f_pred
Concentration (μM)

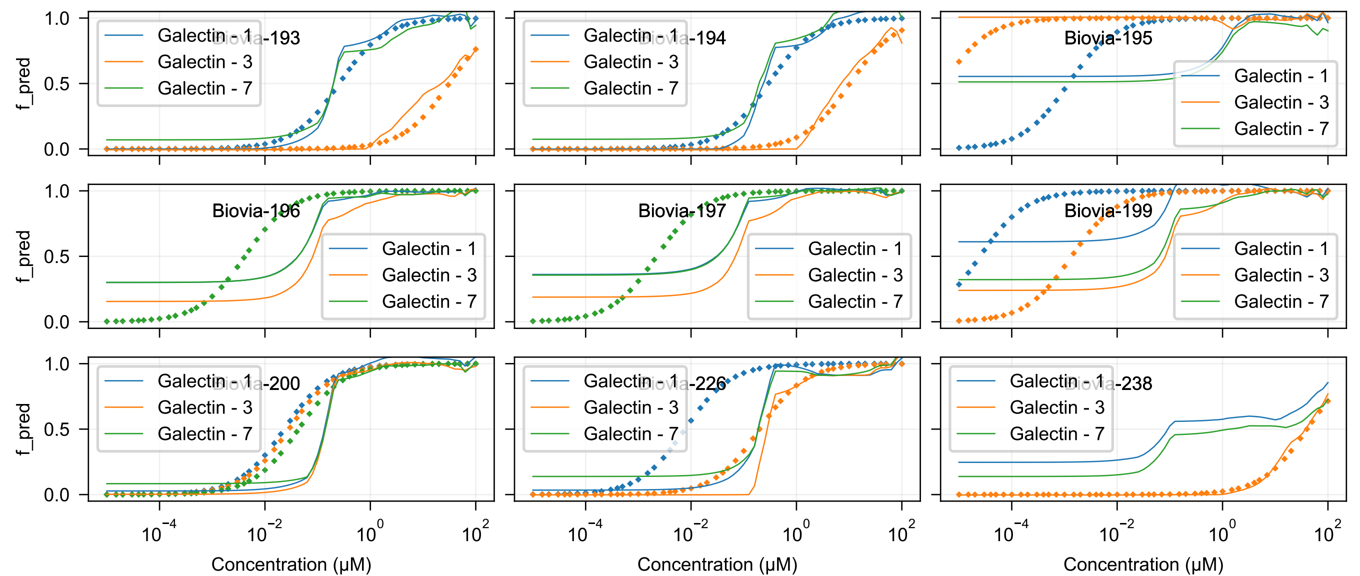
Biovia-193
Biovia-194
Biovia-195
Biovia-196
Biovia-197
Biovia-199
Biovia-200
Biovia-226
Biovia-238
Galectin - 1
Galectin - 3
Galectin - 7
f_pred
Concentration (μM)

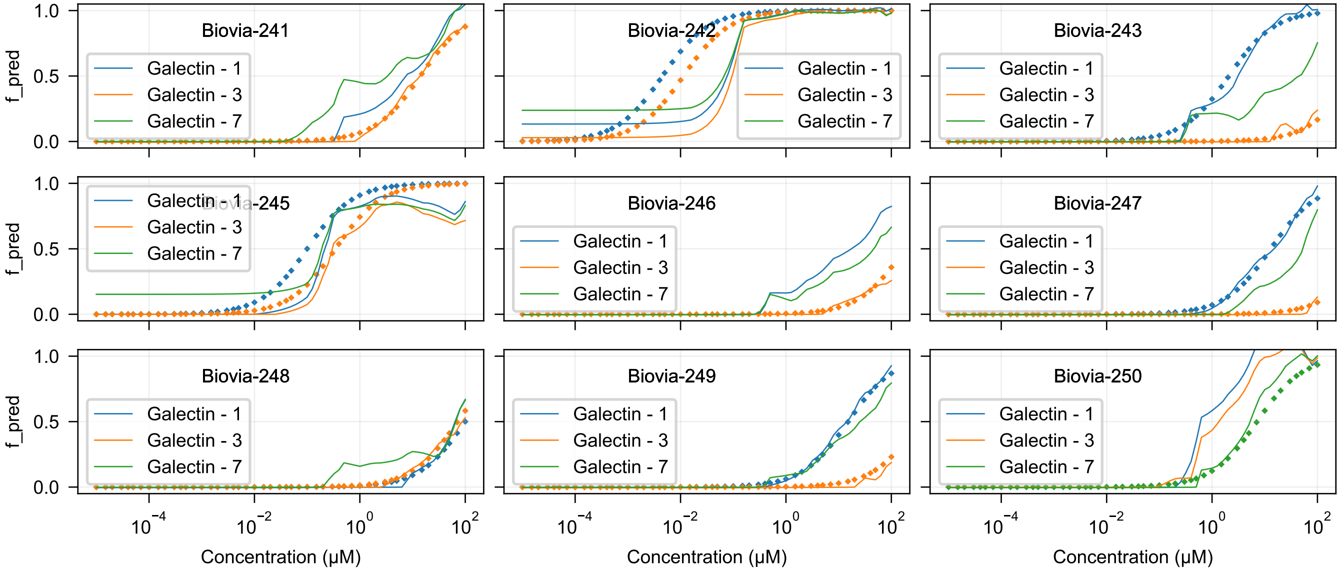

Biovia-241
Biovia-242
Biovia-243
Biovia-245
Biovia-246
Biovia-247
Biovia-248
Biovia-249
Biovia-250
Galectin - 1
Galectin - 3
Galectin - 7
f_pred
1.0
0.5
0.0
$10^{-4}$
$10^{-2}$
$10^{0}$
$10^{2}$
Concentration (μM)

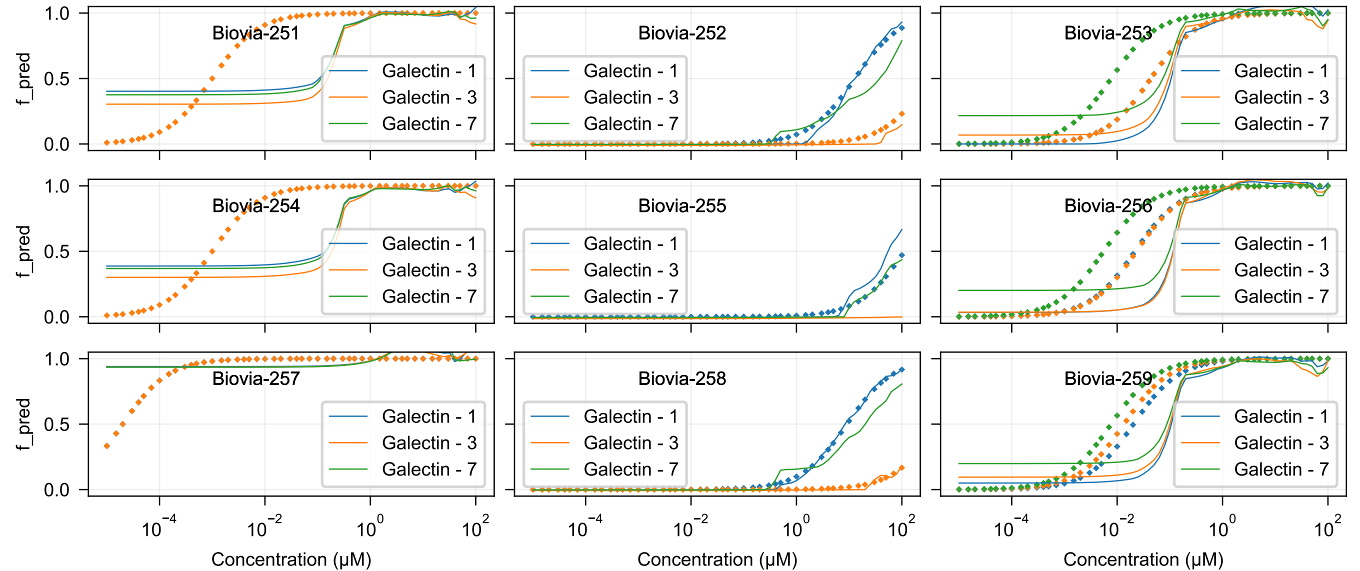

Biovia-251
Biovia-252
Biovia-253
Biovia-254
Biovia-255
Biovia-256
Biovia-257
Biovia-258
Biovia-259
Galectin - 1
Galectin - 3
Galectin - 7
f_pred
1.0
0.5
0.0
$10^{-4}$
$10^{-2}$
$10^{0}$
$10^{2}$
Concentration (μM)

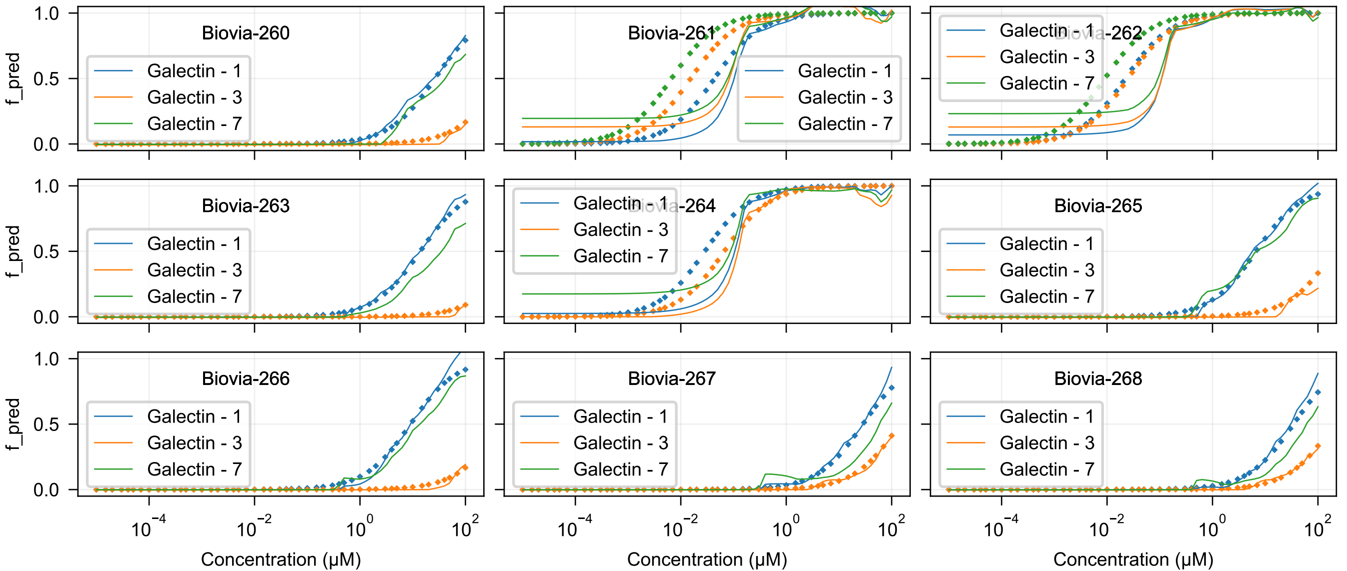
Biovia-260
Biovia-261
Biovia-262
Biovia-263
Biovia-264
Biovia-265
Biovia-266
Biovia-267
Biovia-268
Galectin - 1
Galectin - 3
Galectin - 7
f_pred
0.0
0.5
1.0
$10^{-4}$
$10^{-2}$
$10^{0}$
$10^{2}$
Concentration (μM)

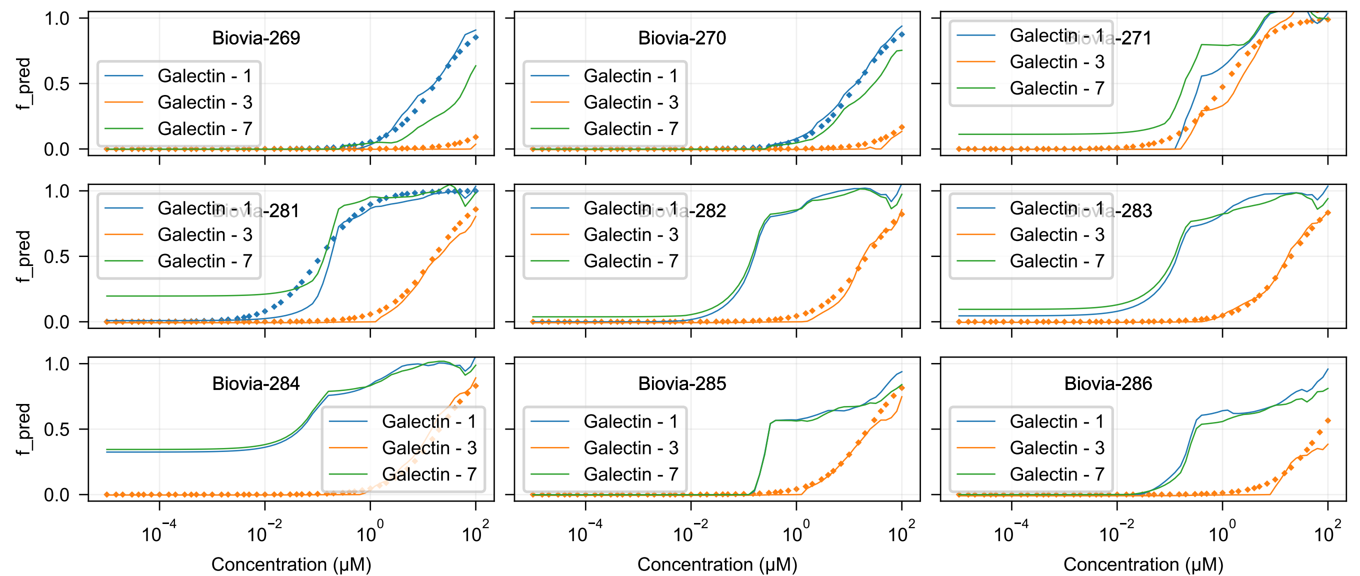
Biovia-269
Biovia-270
Biovia-271
Biovia-281
Biovia-282
Biovia-283
Biovia-284
Biovia-285
Biovia-286
Galectin - 1
Galectin - 3
Galectin - 7
f_pred
Concentration (μM)

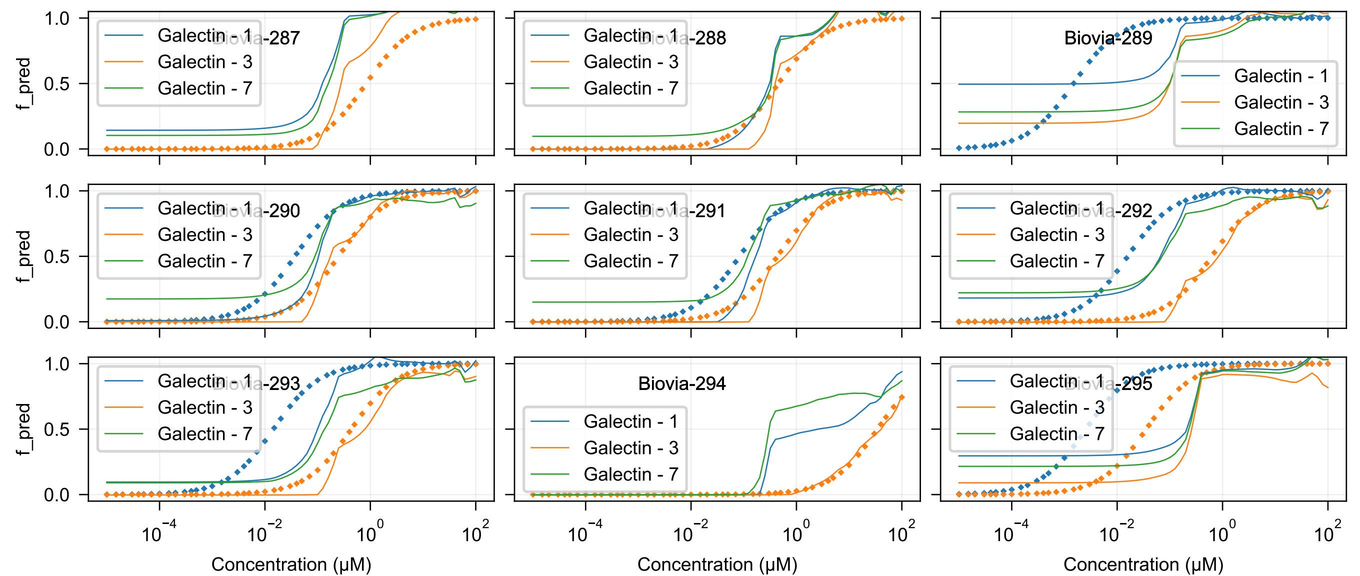
Biovia-287
Biovia-288
Biovia-289
Biovia-290
Biovia-291
Biovia-292
Biovia-293
Biovia-294
Biovia-295
Galectin - 1
Galectin - 3
Galectin - 7
f_pred
Concentration (μM)

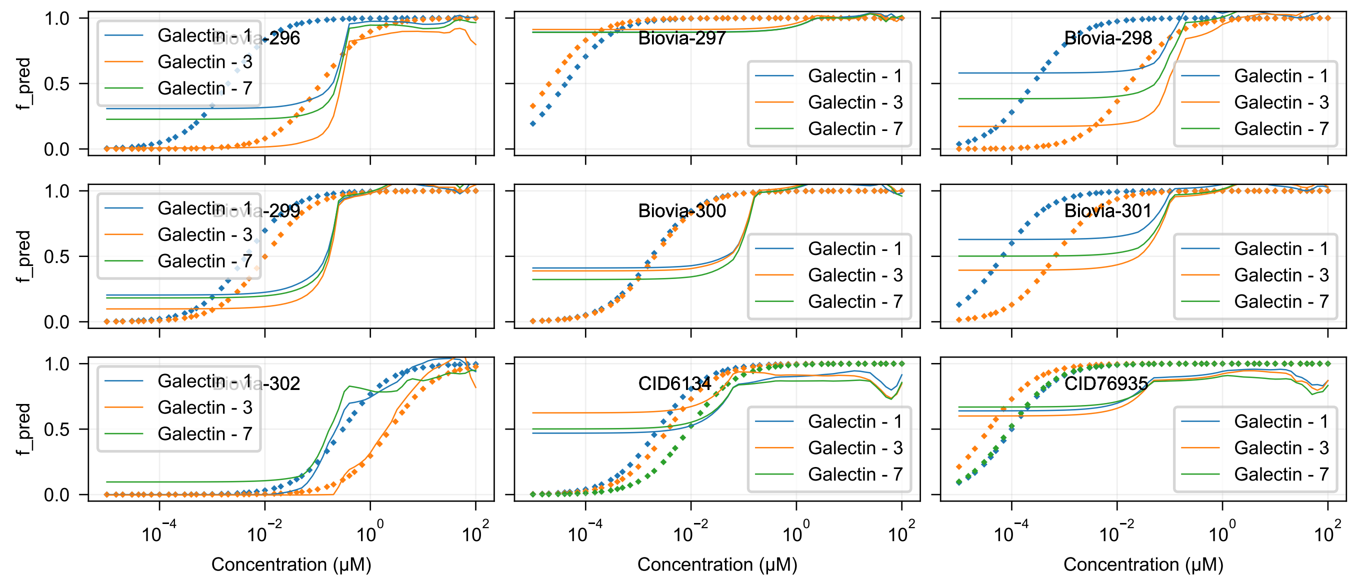

Biovia-296
Biovia-297
Biovia-298
Biovia-299
Biovia-300
Biovia-301
Biovia-302
CID6134
CID76935
Galectin - 1
Galectin - 3
Galectin - 7
f_pred
1.0
0.5
0.0
$10^{-4}$
$10^{-2}$
$10^{0}$
$10^{2}$
Concentration (μM)

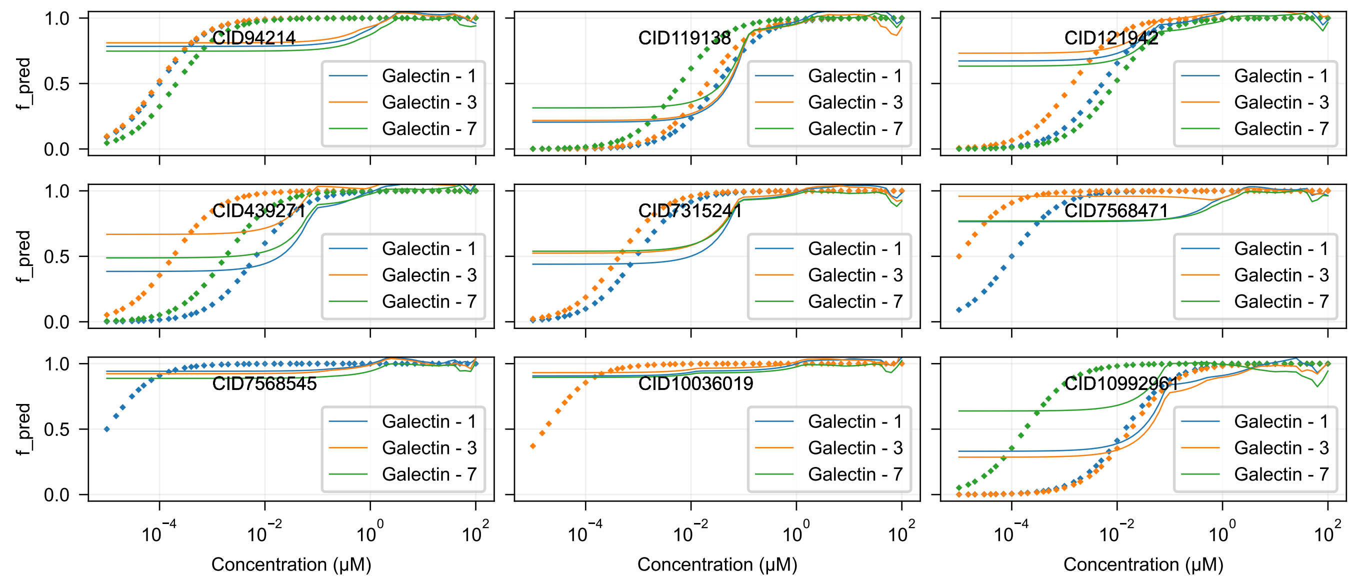
CID94214
CID119138
CID121942
CID439271
CID7315241
CID7568471
CID7568545
CID10036019
CID10992961
Galectin - 1
Galectin - 3
Galectin - 7
f_pred
Concentration (μM)

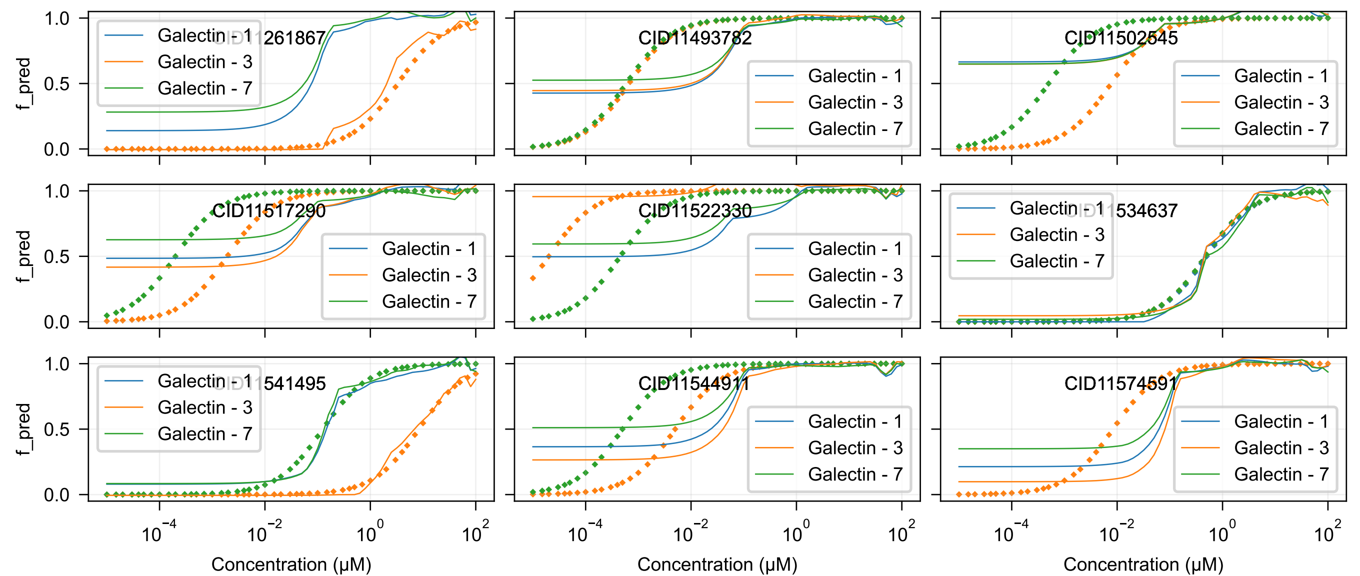

CID11261867
CID11493782
CID11502545
CID11517290
CID11522330
CID11534637
CID11541495
CID11544911
CID11574591
Galectin - 1
Galectin - 3
Galectin - 7
f_pred
Concentration (μM)

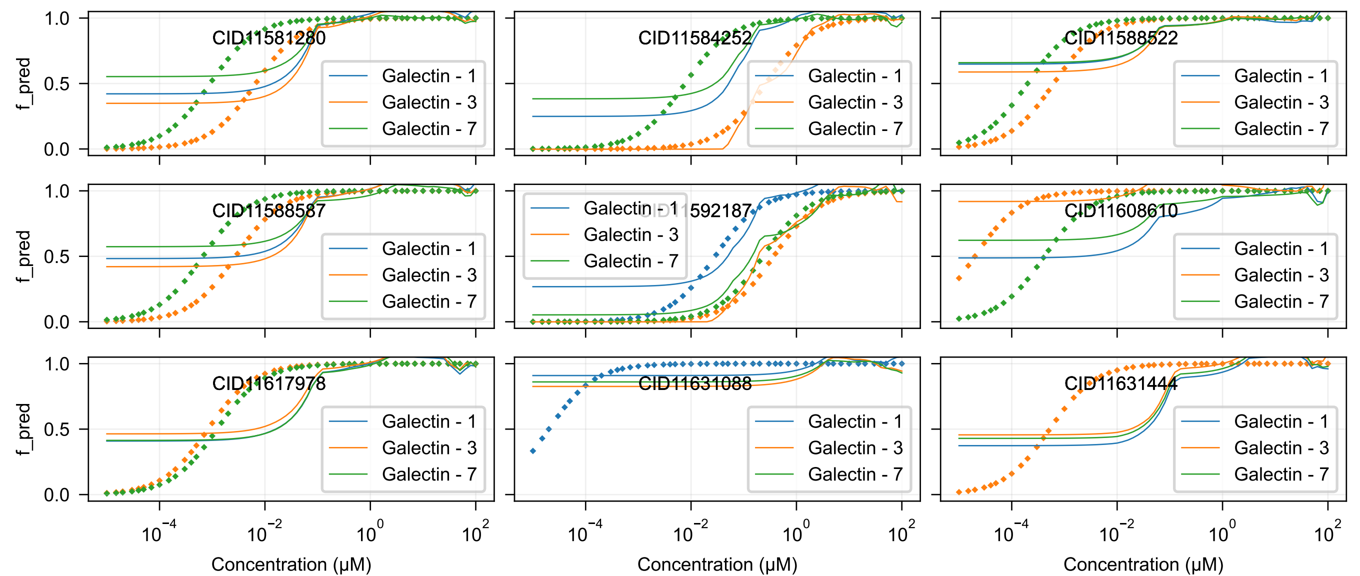
CID11581280
CID11584252
CID11588522
CID11588587
CID11592187
CID11608610
CID11617978
CID11631088
CID11631444
Galectin - 1
Galectin - 3
Galectin - 7
f_pred
Concentration (μM)

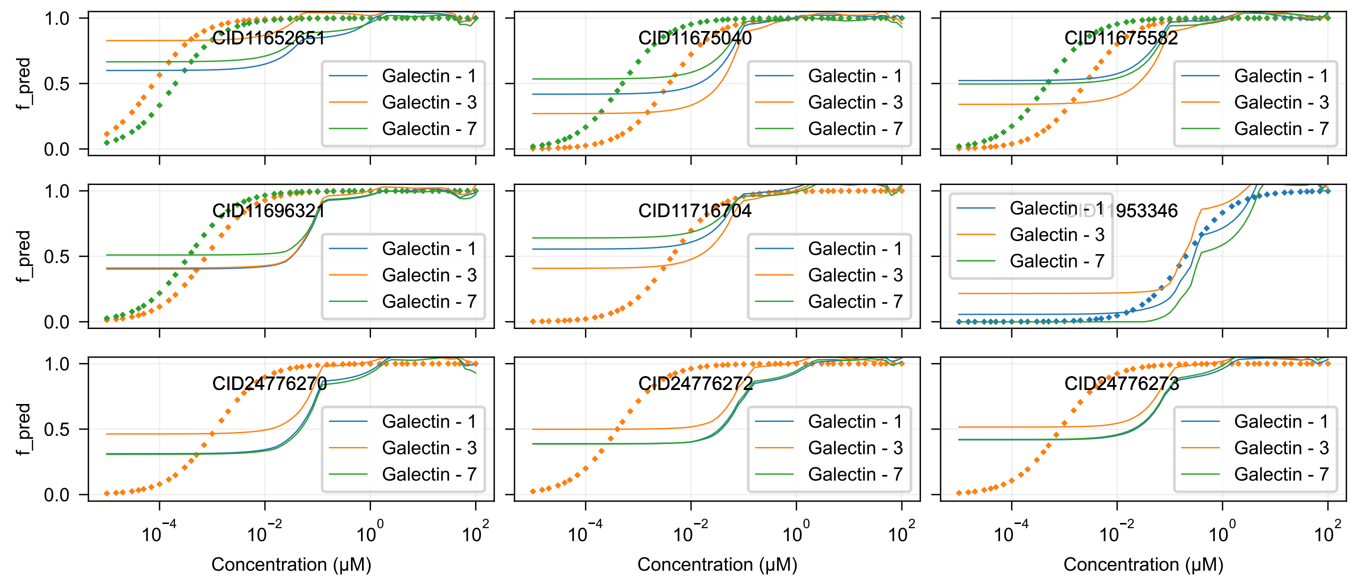

CID11652651
CID11675040
CID11675582
CID11696321
CID11716704
CID11953346
CID24776270
CID24776272
CID24776273
Galectin - 1
Galectin - 3
Galectin - 7
f_pred
Concentration (μM)

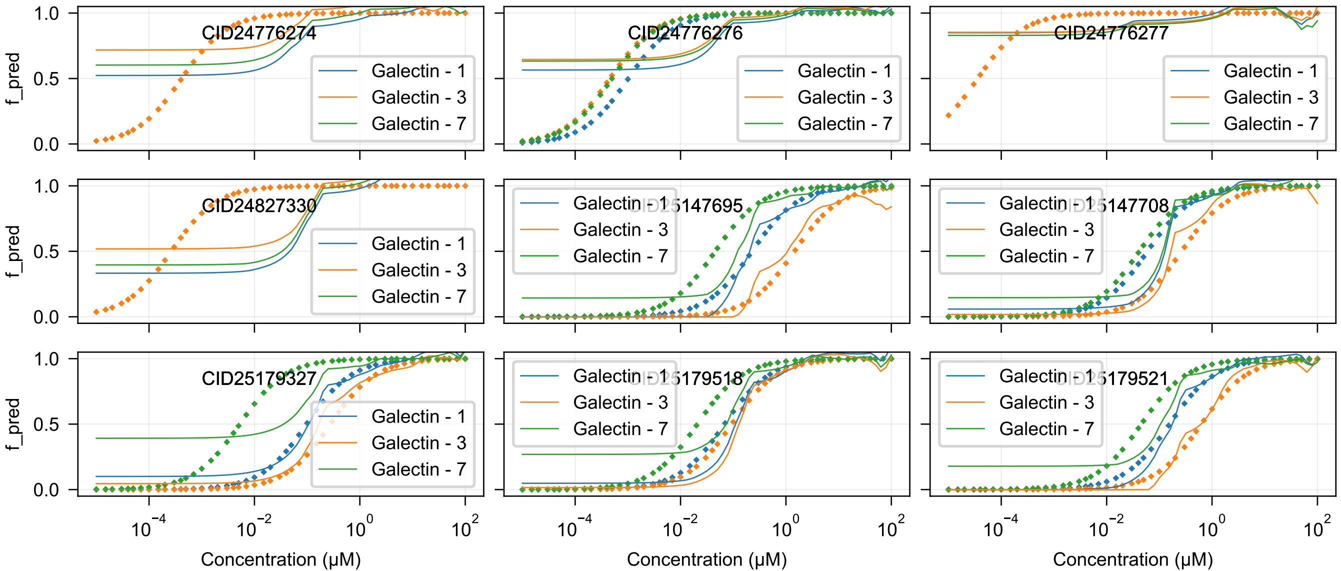

CID24776274
CID24776276
CID24776277
CID24827330
CID25147695
CID25147708
CID25179327
CID25179518
CID25179521
Galectin - 1
Galectin - 3
Galectin - 7
f_pred
Concentration (μM)

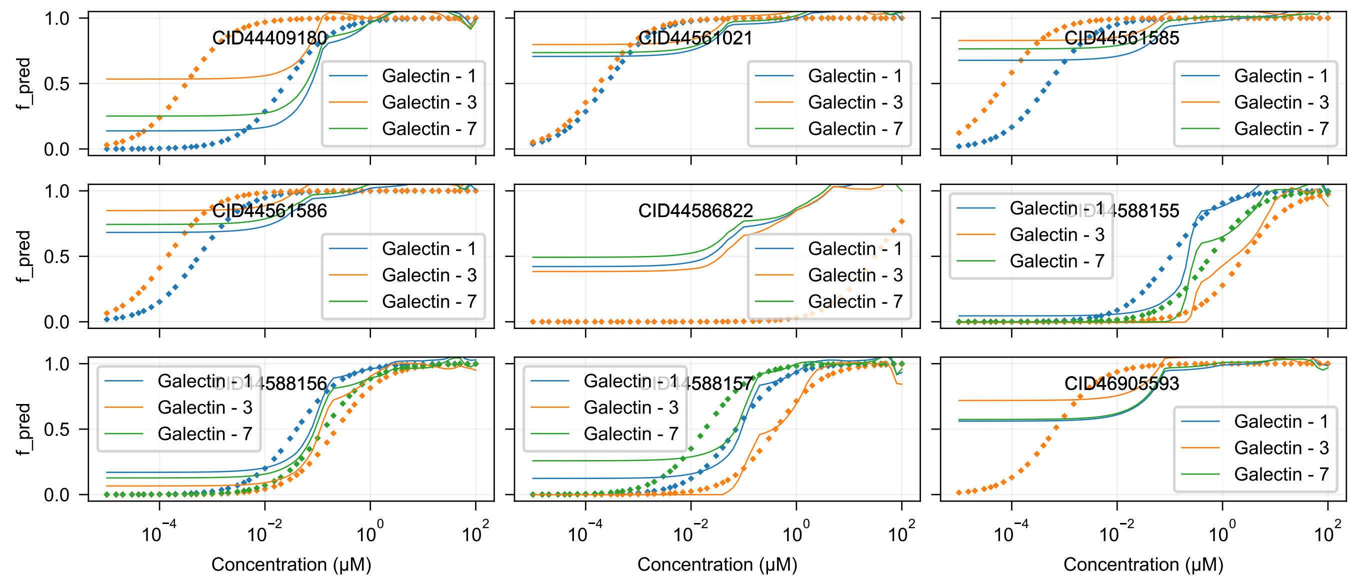
CID44409180
CID44561021
CID44561585
CID44561586
CID44586822
CID44588155
CID44588156
CID44588157
CID46905593
Galectin - 1
Galectin - 3
Galectin - 7
f_pred
Concentration (μM)

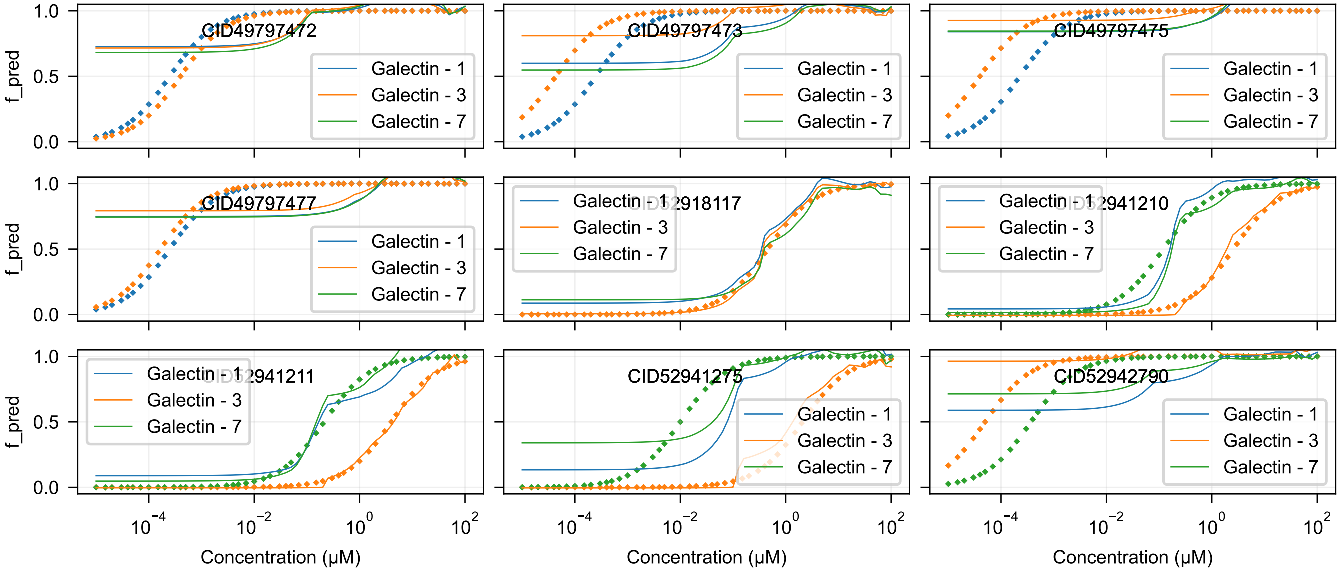

CID49797472
CID49797473
CID49797475
CID49797477
CID52918117
CID52941210
CID52941211
CID52941275
CID52942790
Galectin - 1
Galectin - 3
Galectin - 7
f_pred
Concentration (μM)

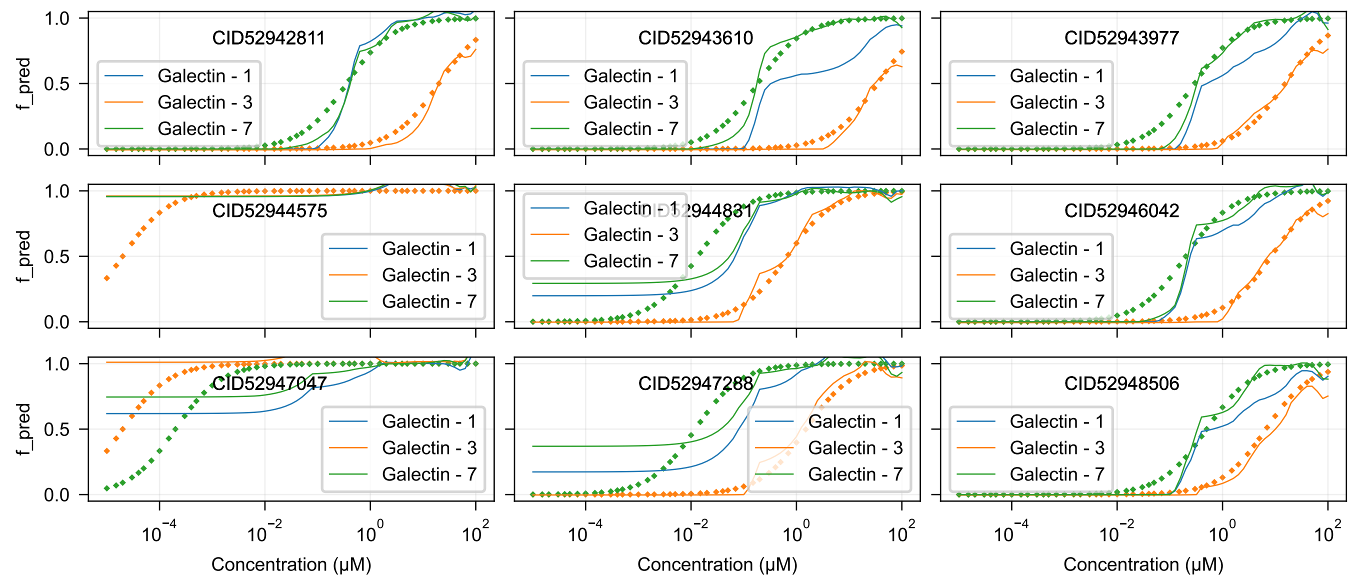

CID52942811
CID52943610
CID52943977
CID52944575
CID52944831
CID52946042
CID52947047
CID52947288
CID52948506
Galectin - 1
Galectin - 3
Galectin - 7
f_pred
1.0
0.5
0.0
$10^{-4}$
$10^{-2}$
$10^{0}$
$10^{2}$
Concentration (μM)

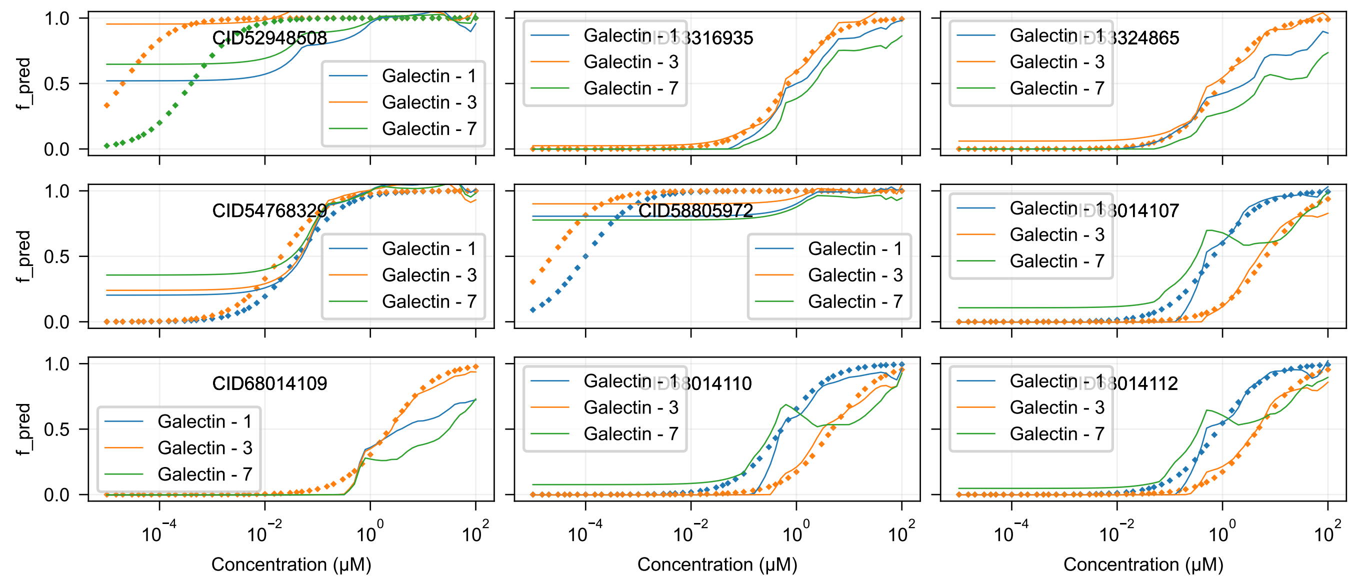

CID52948508
CID63316935
CID63324865
CID54768329
CID58805972
CID68014107
CID68014109
CID68014110
CID68014112
Galectin - 1
Galectin - 3
Galectin - 7
f_pred
Concentration (μM)

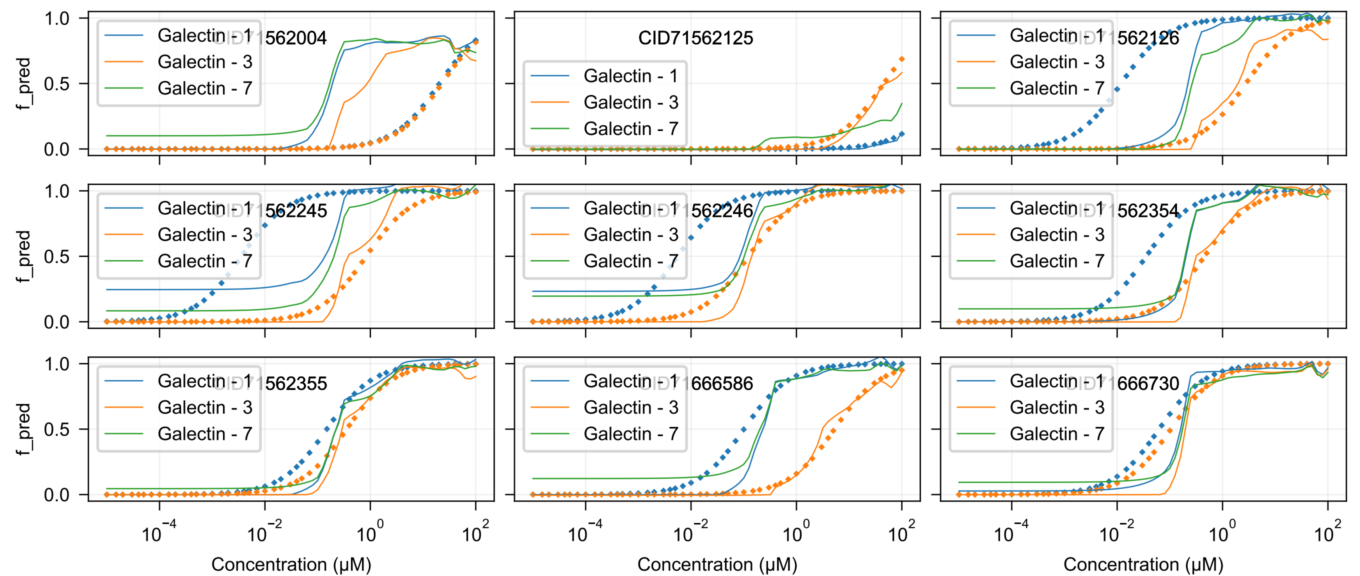

CID71562004
CID71562125
CID71562126
CID71562245
CID71562246
CID71562354
CID71562355
CID71666586
CID71666730
Galectin - 1
Galectin - 3
Galectin - 7
f_pred
0.0
0.5
1.0
$10^{-4}$
$10^{-2}$
$10^{0}$
$10^{2}$
Concentration (μM)

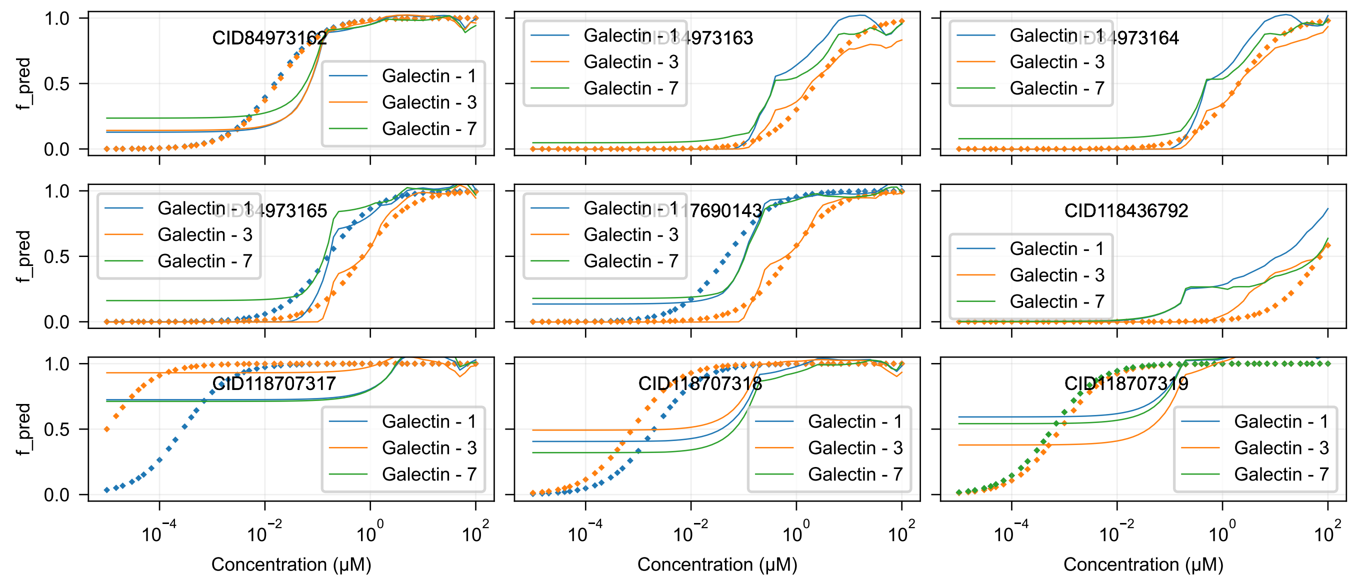

CID84973162
CID84973163
CID84973164
CID84973165
CID117690143
CID118436792
CID118707317
CID118707318
CID118707319
Galectin - 1
Galectin - 3
Galectin - 7
f_pred
1.0
0.5
0.0
$10^{-4}$
$10^{-2}$
$10^{0}$
$10^{2}$
Concentration (μM)

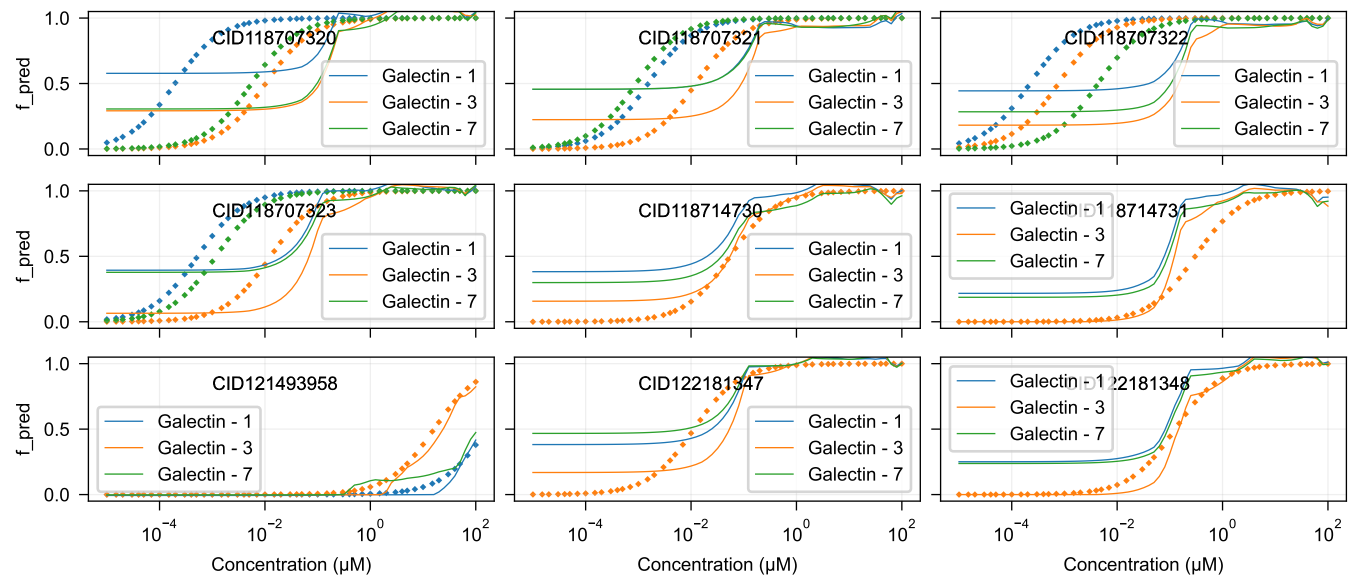
CID118707320
CID118707321
CID118707322
CID118707323
CID118714730
CID118714731
CID121493958
CID122181347
CID122181348
Galectin - 1
Galectin - 3
Galectin - 7
f_pred
Concentration (μM)

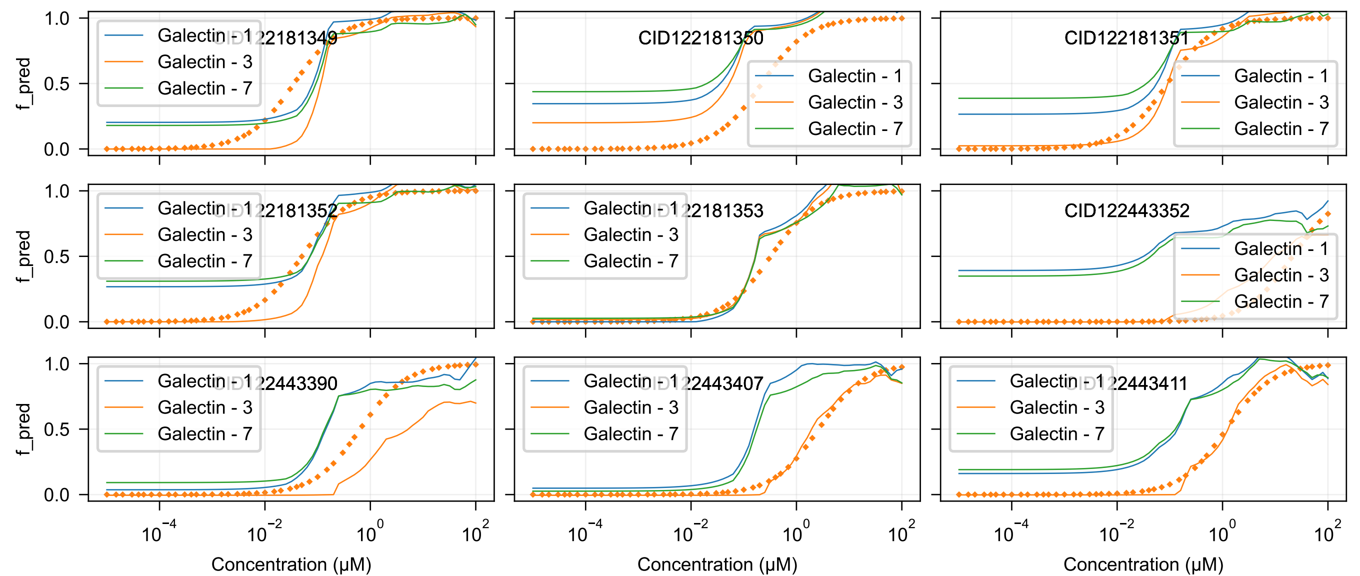

CID122181349
CID122181350
CID122181351
CID122181352
CID122181353
CID122443352
CID122443390
CID122443407
CID122443411
Galectin - 1
Galectin - 3
Galectin - 7
f_pred
Concentration (μM)

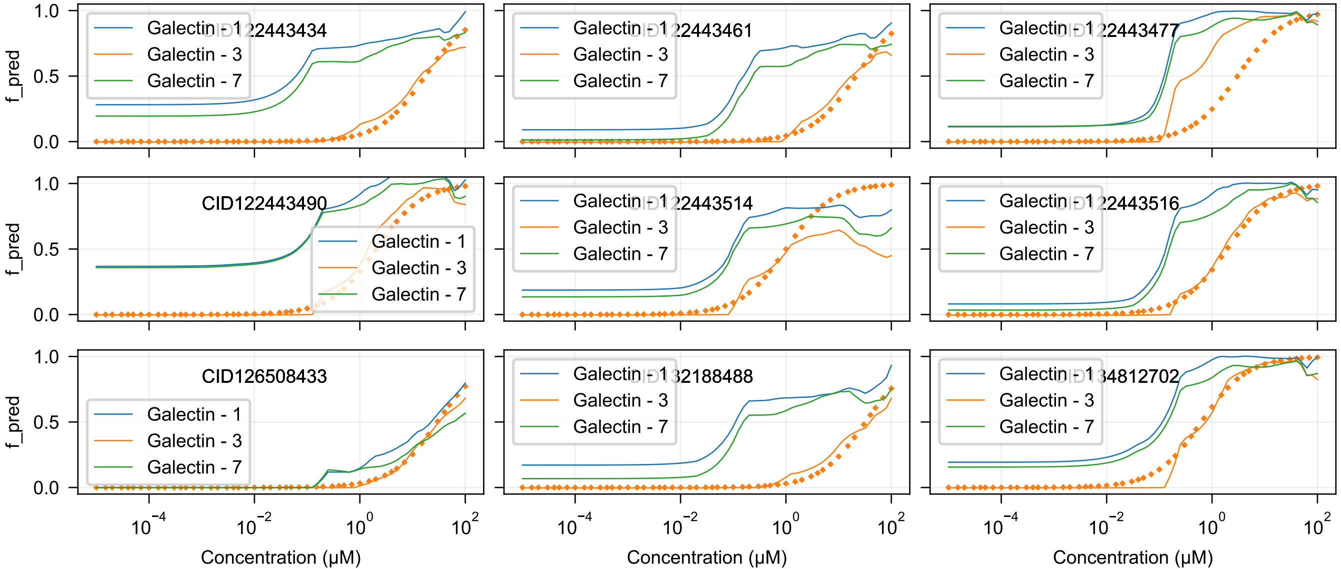
CID122443434
CID122443461
CID122443477
CID122443490
CID122443514
CID122443516
CID126508433
CID132188488
CID134812702
Galectin - 1
Galectin - 3
Galectin - 7
f_pred
Concentration (μM)
1.0
0.5
0.0
$10^{-4}$
$10^{-2}$
$10^{0}$
$10^{2}$

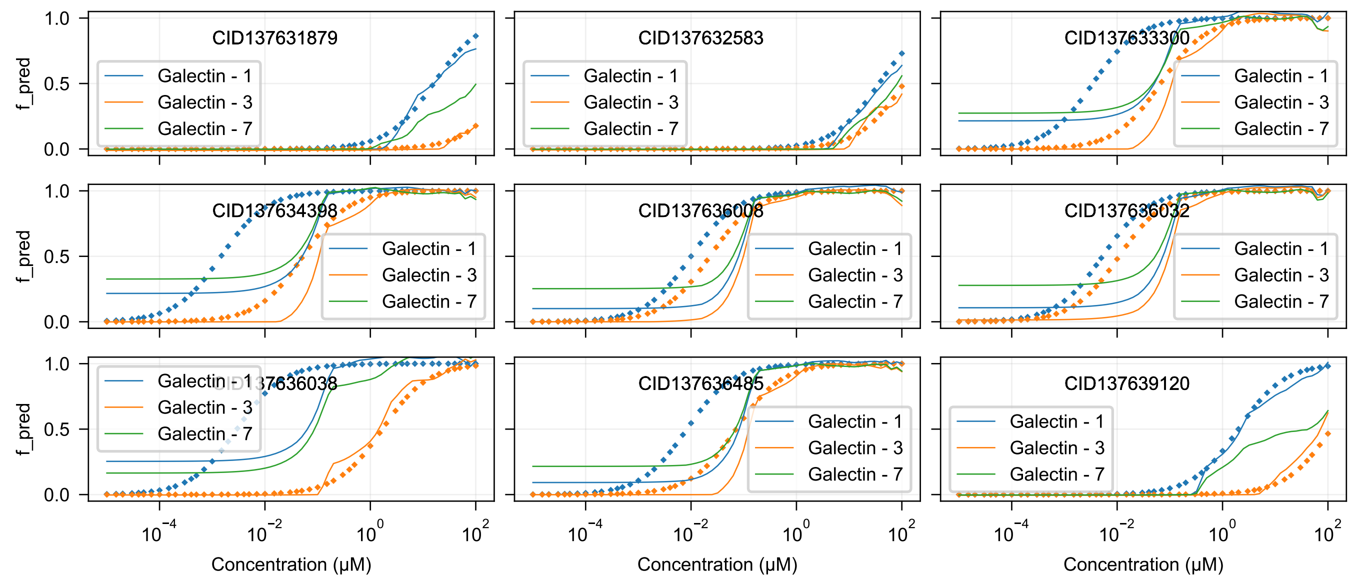
CID137631879
CID137632583
CID137633300
CID137634398
CID137636008
CID137636032
CID137636038
CID137636485
CID137639120
Galectin - 1
Galectin - 3
Galectin - 7
f_pred
1.0
0.5
0.0
$10^{-4}$
$10^{-2}$
$10^{0}$
$10^{2}$
Concentration (μM)

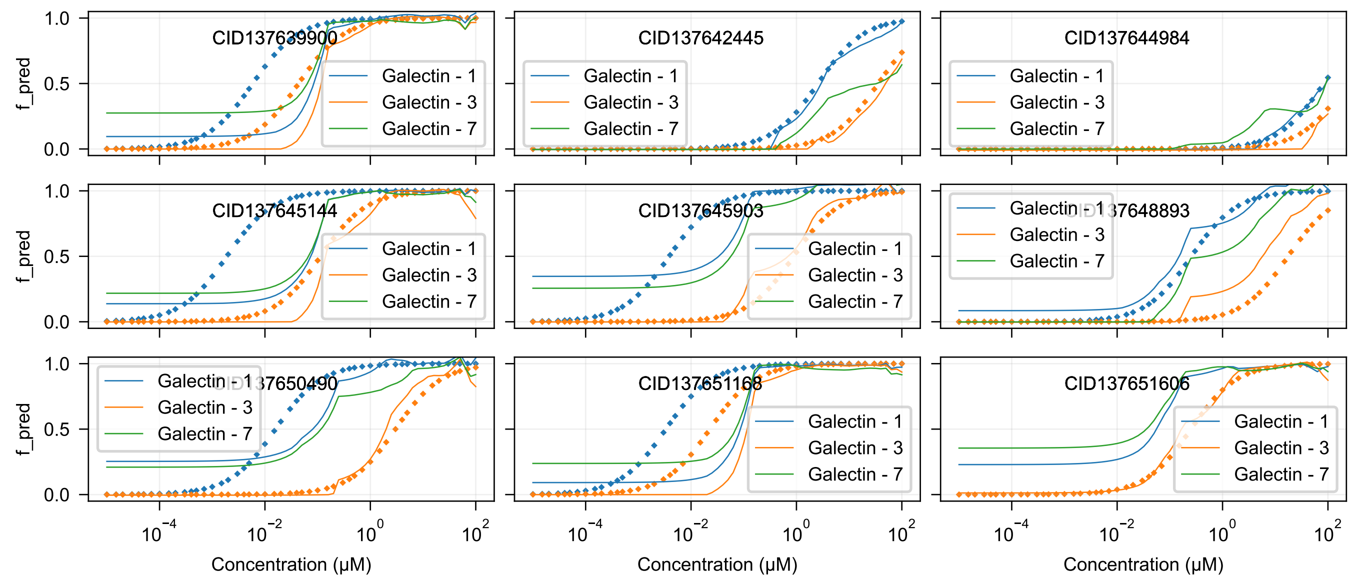

CID137639900
CID137642445
CID137644984
CID137645144
CID137645903
CID137648893
CID137650490
CID137651168
CID137651606
Galectin - 1
Galectin - 3
Galectin - 7
f_pred
Concentration (μM)

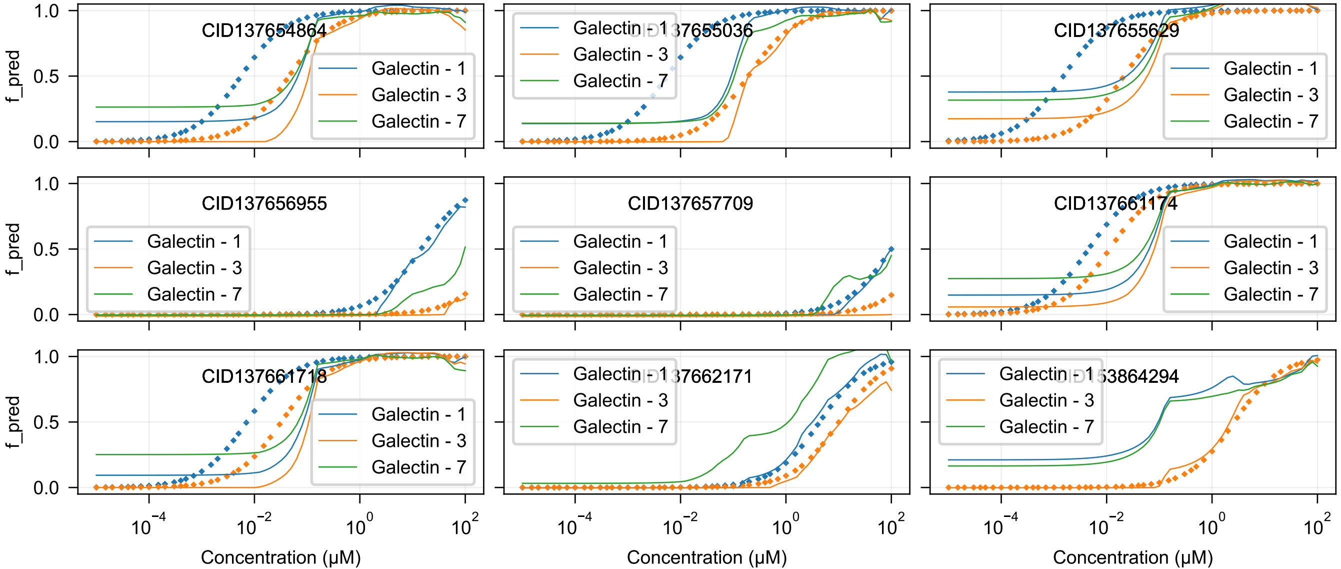

CID137654864
CID137655036
CID137655629
CID137656955
CID137657709
CID137661174
CID137661718
CID137662171
CID153864294
Galectin - 1
Galectin - 3
Galectin - 7
f_pred
1.0
0.5
0.0
$10^{-4}$
$10^{-2}$
$10^{0}$
$10^{2}$
Concentration (μM)

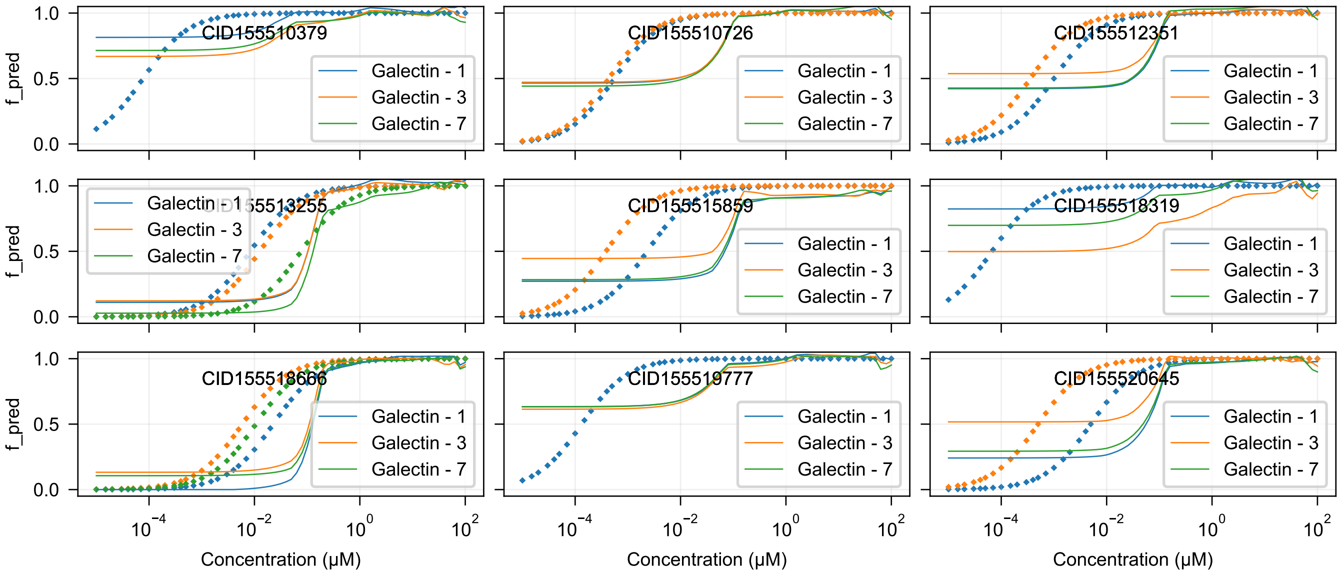
CID155510379
CID155510726
CID155512351
CID155513255
CID155515859
CID155518319
CID155518666
CID155519777
CID155520645
Galectin - 1
Galectin - 3
Galectin - 7
f_pred
Concentration (μM)

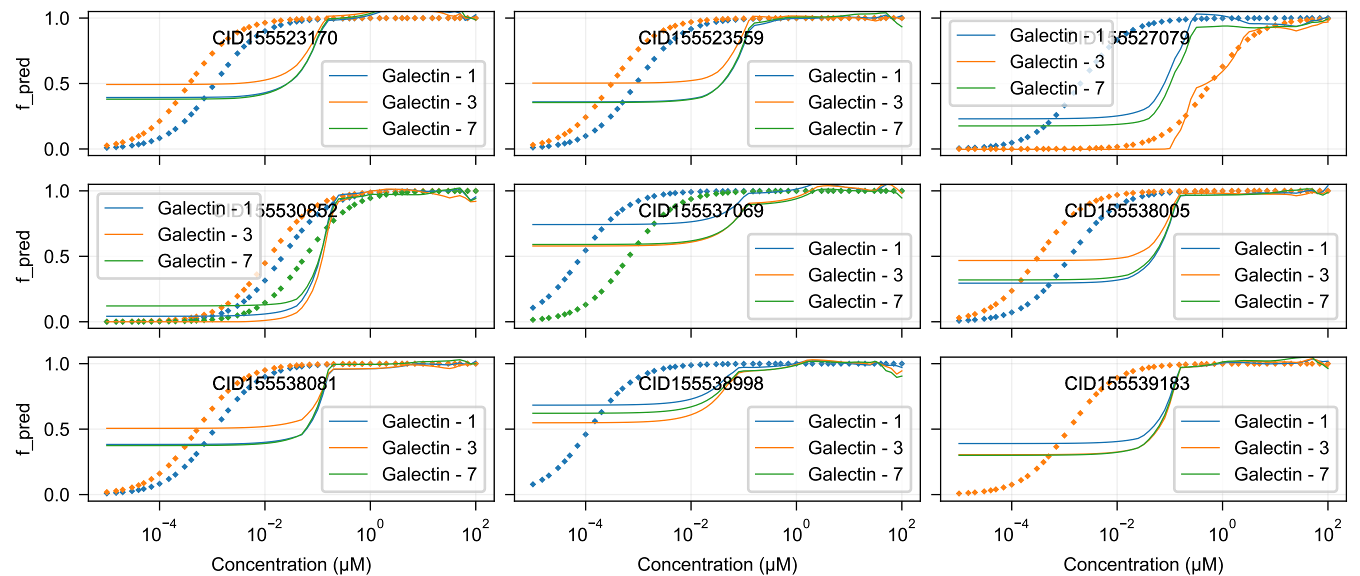
CID155523170
CID155523559
CID155527079
CID155530852
CID155537069
CID155538005
CID155538081
CID155538998
CID155539183
Galectin - 1
Galectin - 3
Galectin - 7
f_pred
Concentration (μM)

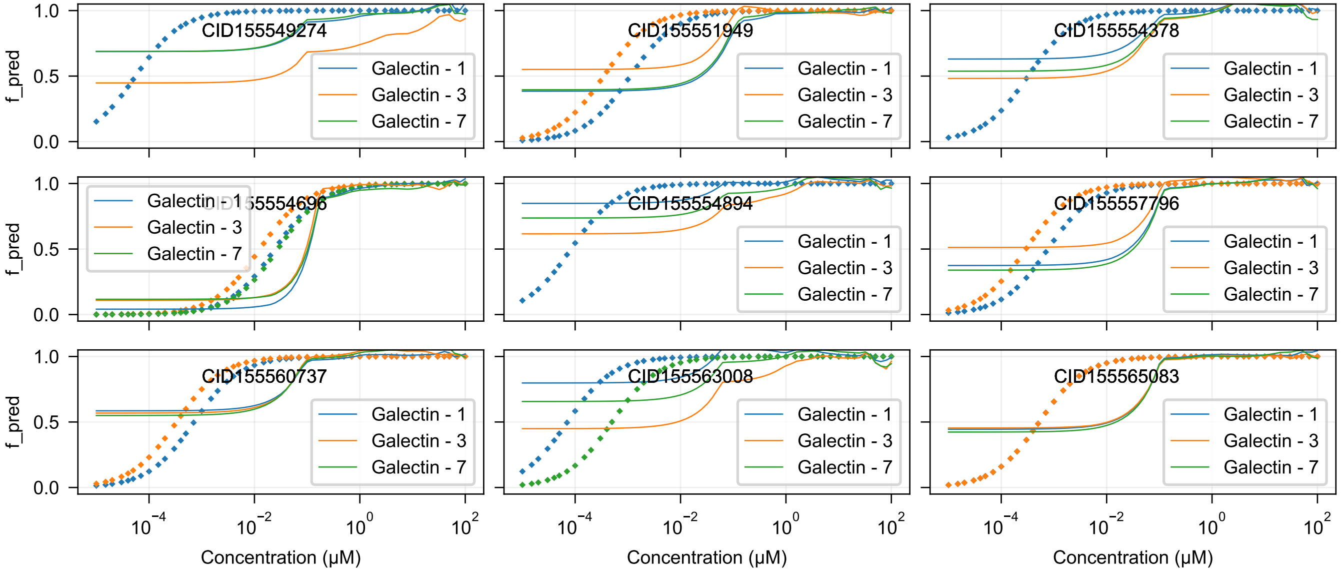
CID155549274
CID155551949
CID155554378
CID155554696
CID155554894
CID155557796
CID155560737
CID155563008
CID155565083
Galectin - 1
Galectin - 3
Galectin - 7
f_pred
1.0
0.5
0.0
$10^{-4}$
$10^{-2}$
$10^{0}$
$10^{2}$
Concentration (μM)

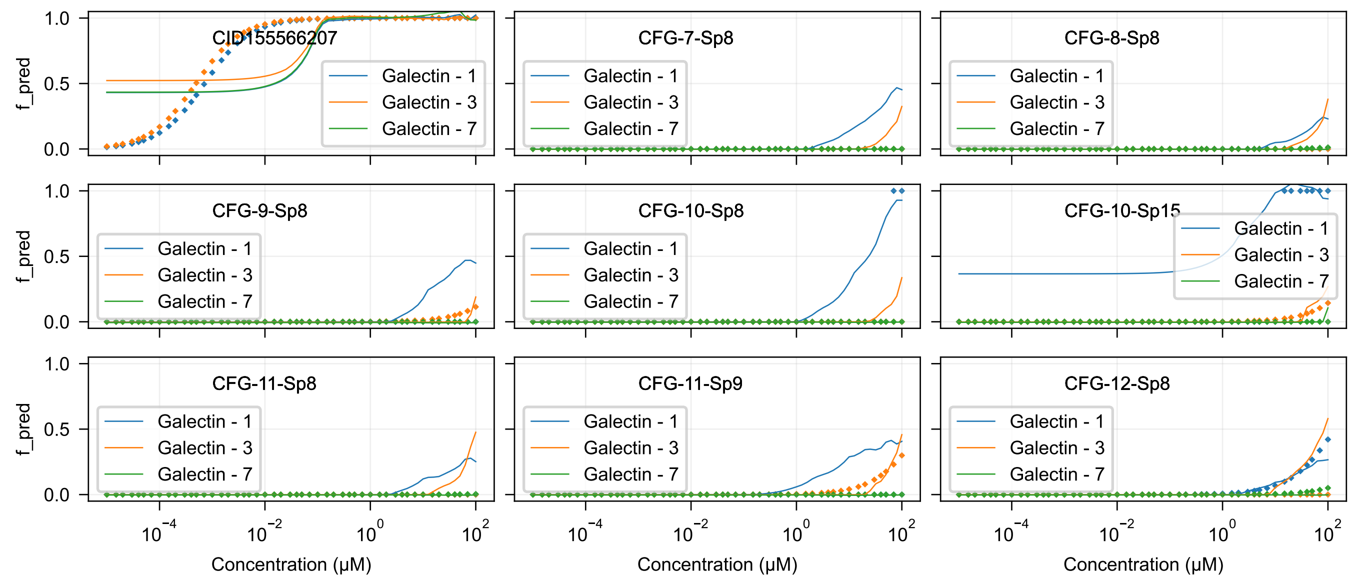
CID155566207
CFG-7-Sp8
CFG-8-Sp8
CFG-9-Sp8
CFG-10-Sp8
CFG-10-Sp15
CFG-11-Sp8
CFG-11-Sp9
CFG-12-Sp8
Galectin - 1
Galectin - 3
Galectin - 7
f_pred
Concentration (μM)

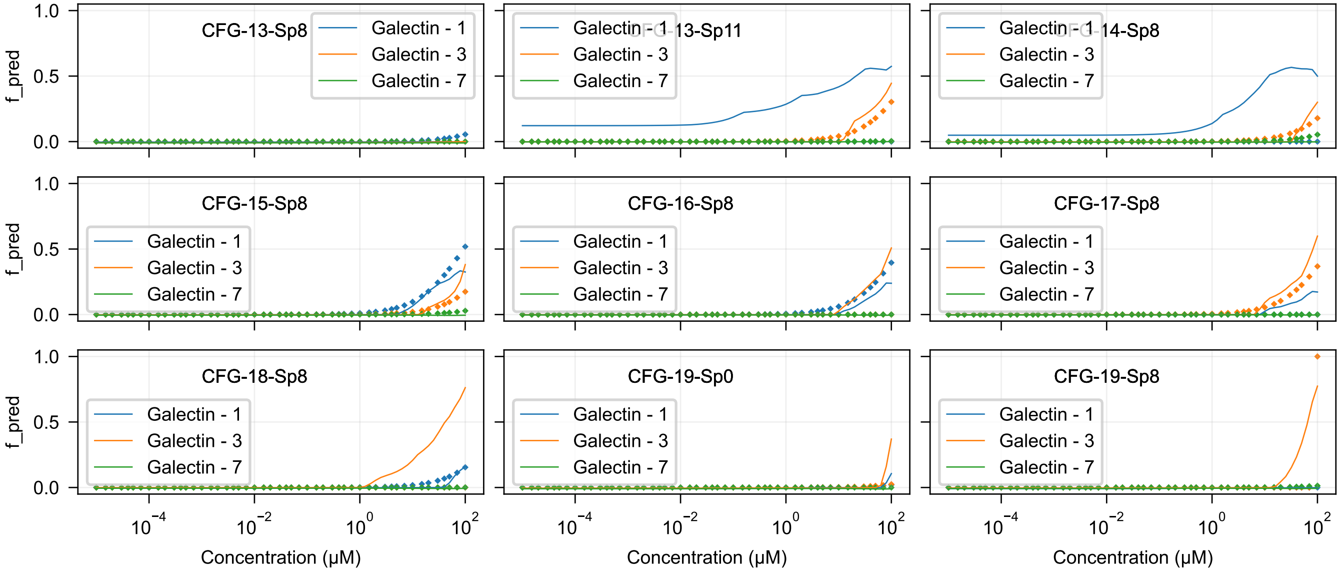

CFG-13-Sp8
CFG-13-Sp11
CFG-14-Sp8
CFG-15-Sp8
CFG-16-Sp8
CFG-17-Sp8
CFG-18-Sp8
CFG-19-Sp0
CFG-19-Sp8
Galectin - 1
Galectin - 3
Galectin - 7
f_pred
0.0
0.5
1.0
$10^{-4}$
$10^{-2}$
$10^{0}$
$10^{2}$
Concentration (μM)

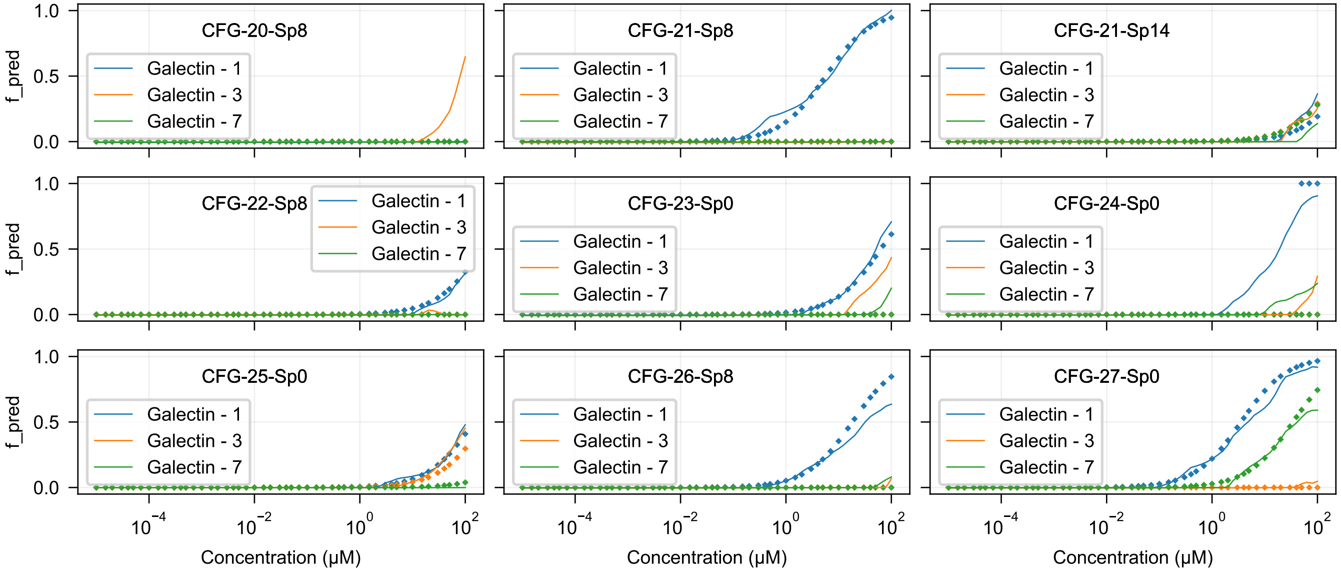

CFG-20-Sp8
CFG-21-Sp8
CFG-21-Sp14
CFG-22-Sp8
CFG-23-Sp0
CFG-24-Sp0
CFG-25-Sp0
CFG-26-Sp8
CFG-27-Sp0
Galectin - 1
Galectin - 3
Galectin - 7
f_pred
1.0
0.5
0.0
$10^{-4}$
$10^{-2}$
$10^{0}$
$10^{2}$
Concentration (μM)

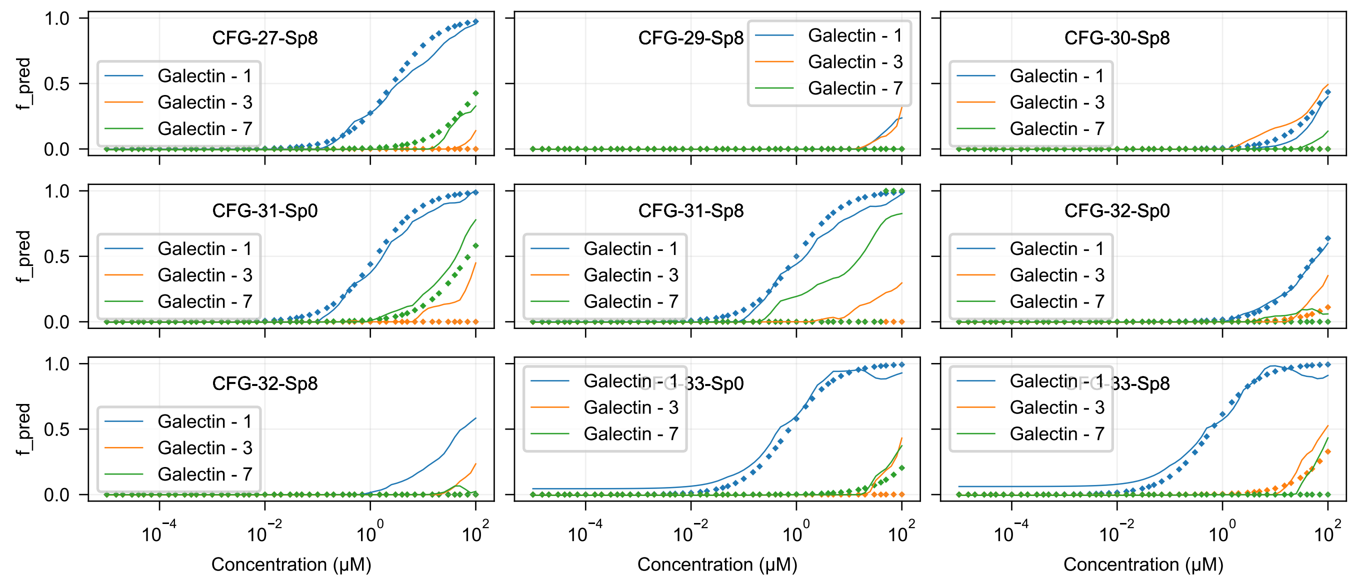
CFG-27-Sp8
CFG-29-Sp8
CFG-30-Sp8
CFG-31-Sp0
CFG-31-Sp8
CFG-32-Sp0
CFG-32-Sp8
CFG-33-Sp0
CFG-33-Sp8
Galectin - 1
Galectin - 3
Galectin - 7
f_pred
1.0
0.5
0.0
$10^{-4}$
$10^{-2}$
$10^{0}$
$10^{2}$
Concentration (μM)

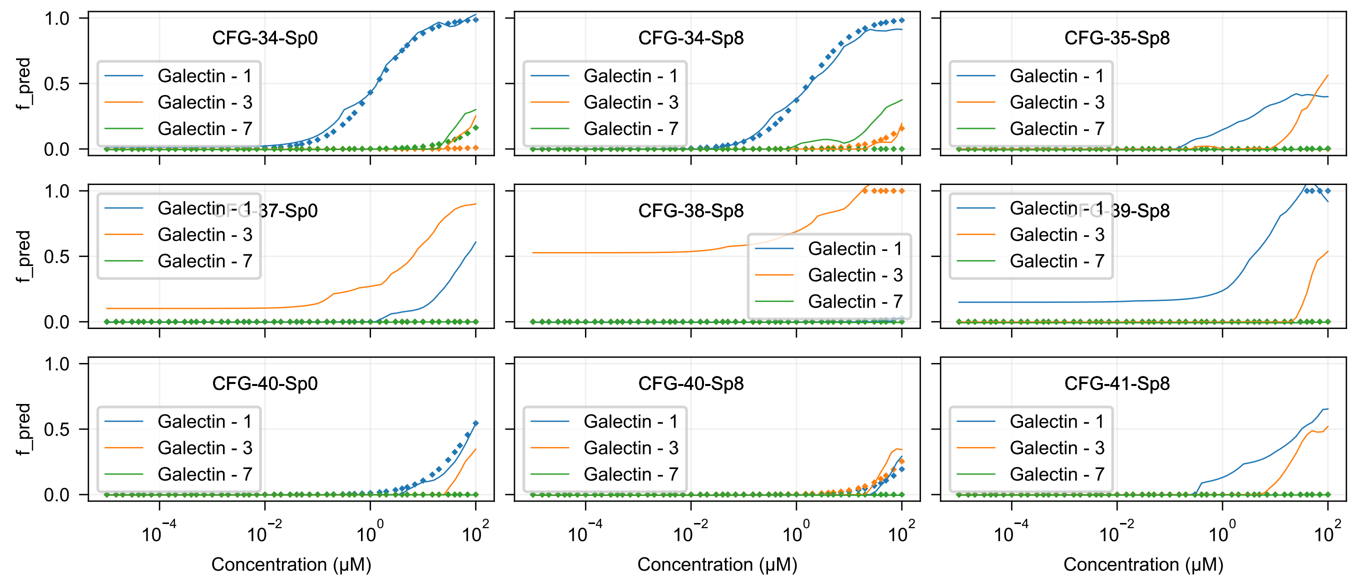

CFG-34-Sp0
CFG-34-Sp8
CFG-35-Sp8
CFG-37-Sp0
CFG-38-Sp8
CFG-39-Sp8
CFG-40-Sp0
CFG-40-Sp8
CFG-41-Sp8
Galectin - 1
Galectin - 3
Galectin - 7
f_pred
Concentration (μM)

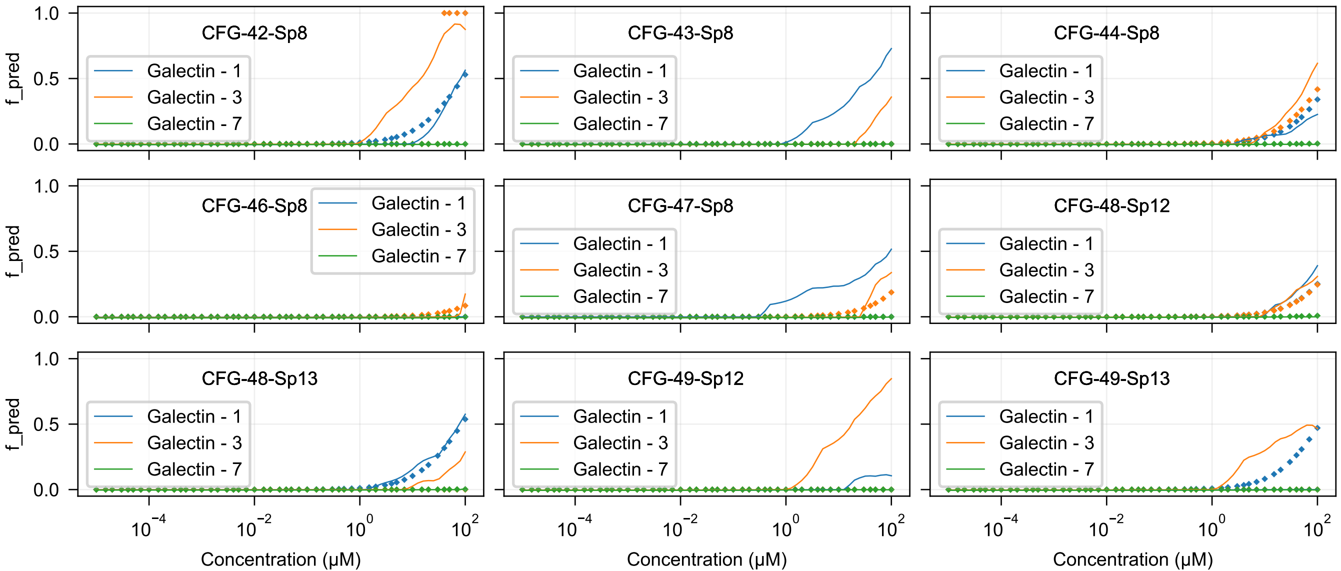

CFG-42-Sp8
CFG-43-Sp8
CFG-44-Sp8
CFG-46-Sp8
CFG-47-Sp8
CFG-48-Sp12
CFG-48-Sp13
CFG-49-Sp12
CFG-49-Sp13
Galectin - 1
Galectin - 3
Galectin - 7
f_pred
0.0
0.5
1.0
$10^{-4}$
$10^{-2}$
$10^{0}$
$10^{2}$
Concentration (μM)

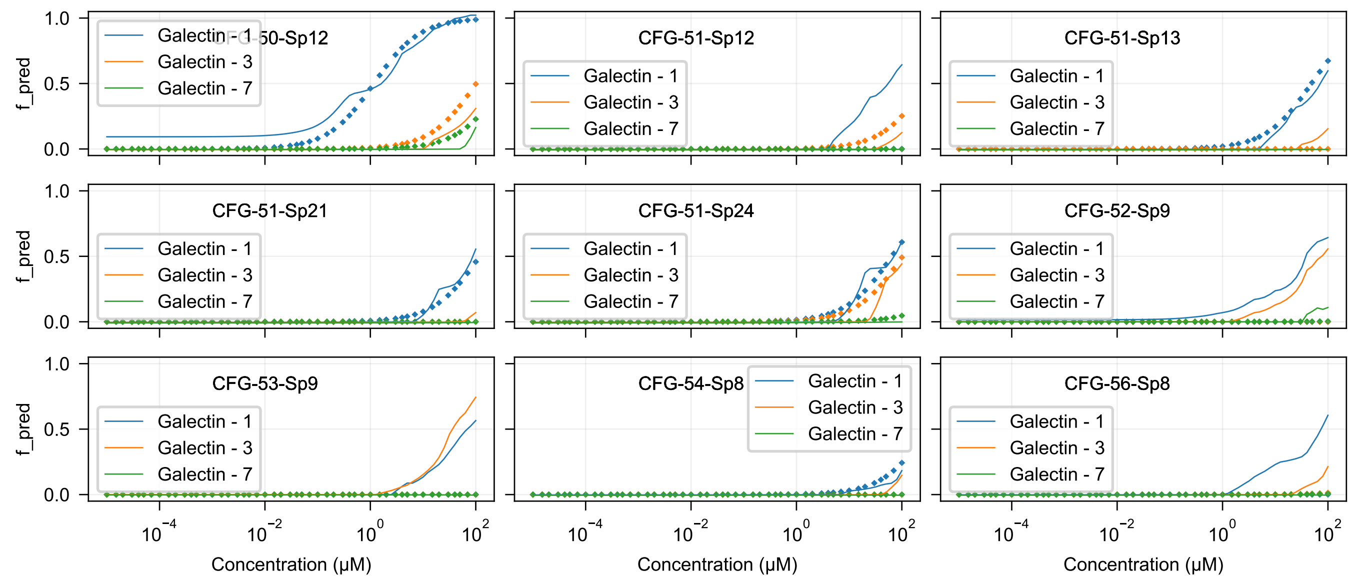

CFG-50-Sp12
CFG-51-Sp12
CFG-51-Sp13
CFG-51-Sp21
CFG-51-Sp24
CFG-52-Sp9
CFG-53-Sp9
CFG-54-Sp8
CFG-56-Sp8
Galectin - 1
Galectin - 3
Galectin - 7
f_pred
Concentration (μM)

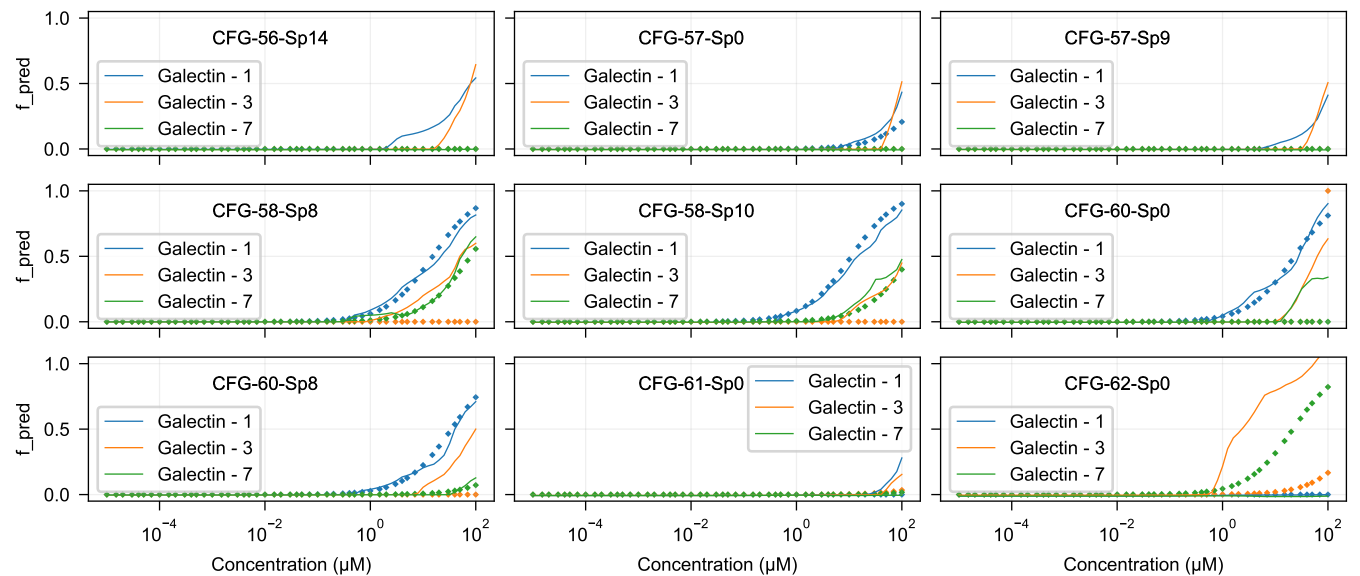

CFG-56-Sp14
CFG-57-Sp0
CFG-57-Sp9
CFG-58-Sp8
CFG-58-Sp10
CFG-60-Sp0
CFG-60-Sp8
CFG-61-Sp0
CFG-62-Sp0
Galectin - 1
Galectin - 3
Galectin - 7
f_pred
1.0
0.5
0.0
$10^{-4}$
$10^{-2}$
$10^{0}$
$10^{2}$
Concentration (μM)

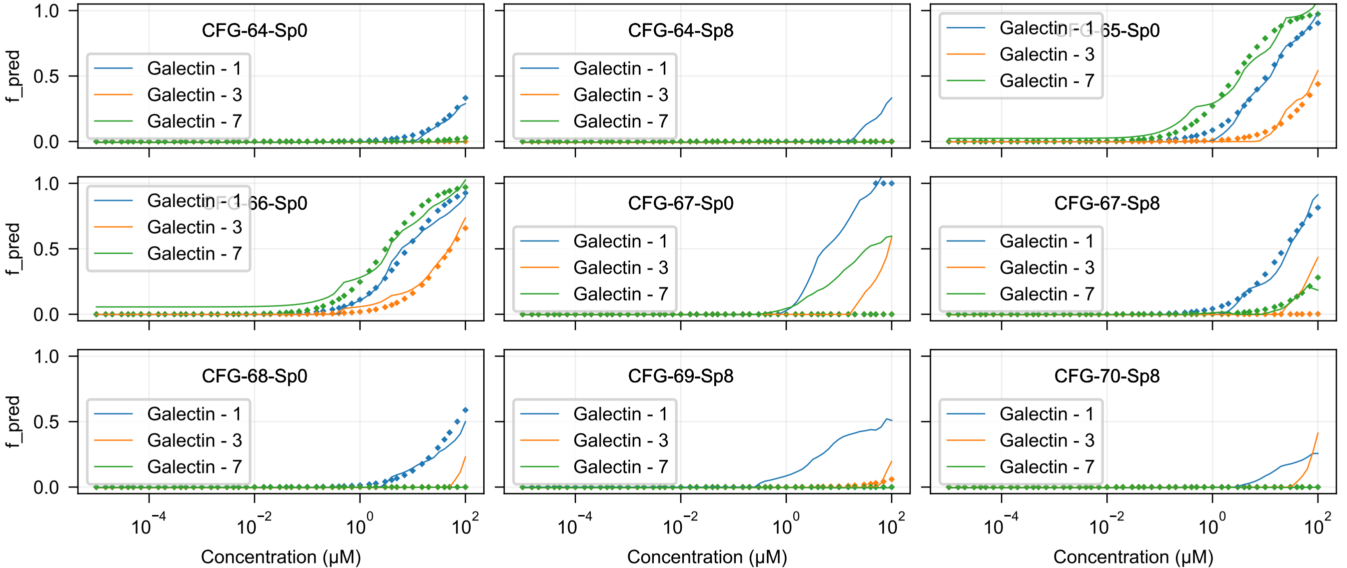
CFG-64-Sp0
CFG-64-Sp8
CFG-65-Sp0
CFG-66-Sp0
CFG-67-Sp0
CFG-67-Sp8
CFG-68-Sp0
CFG-69-Sp8
CFG-70-Sp8
Galectin - 1
Galectin - 3
Galectin - 7
f_pred
1.0
0.5
0.0
$10^{-4}$
$10^{-2}$
$10^{0}$
$10^{2}$
Concentration (μM)

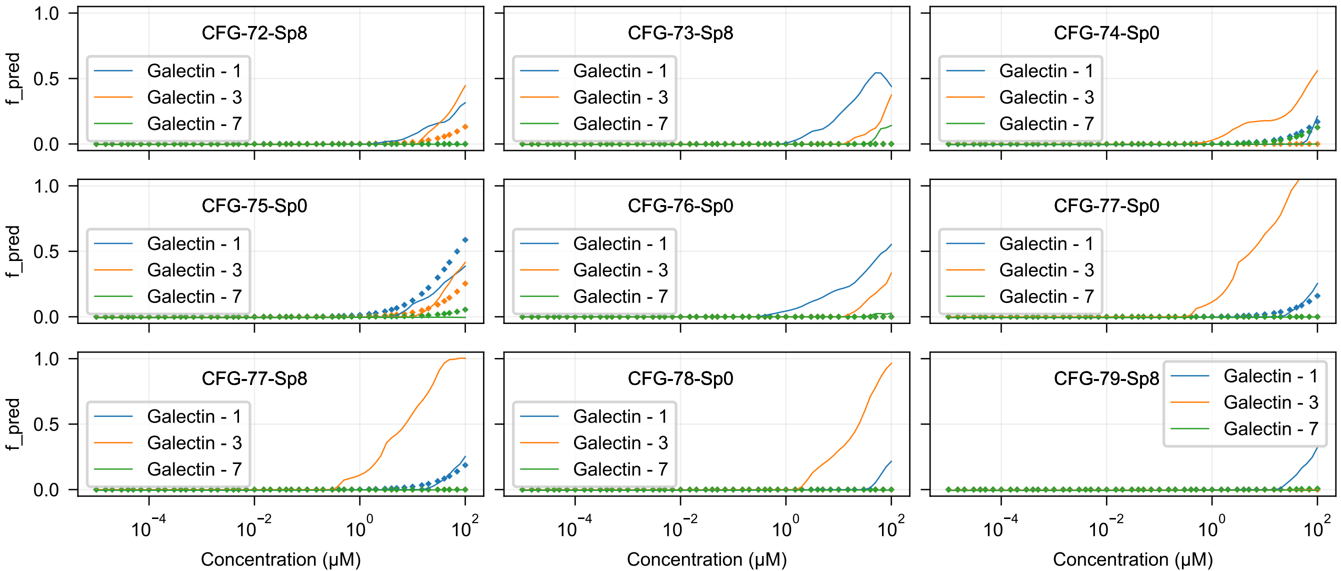
CFG-72-Sp8
CFG-73-Sp8
CFG-74-Sp0
CFG-75-Sp0
CFG-76-Sp0
CFG-77-Sp0
CFG-77-Sp8
CFG-78-Sp0
CFG-79-Sp8
Galectin - 1
Galectin - 3
Galectin - 7
f_pred
1.0
0.5
0.0
$10^{-4}$
$10^{-2}$
$10^{0}$
$10^{2}$
Concentration (μM)

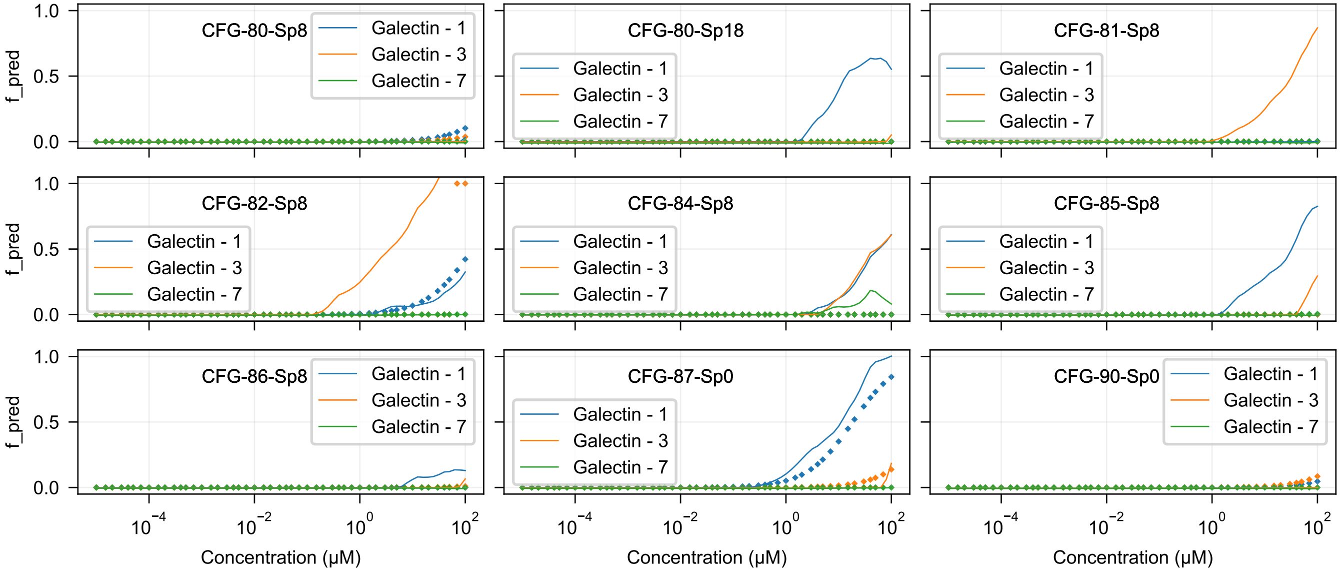

CFG-80-Sp8
CFG-80-Sp18
CFG-81-Sp8
CFG-82-Sp8
CFG-84-Sp8
CFG-85-Sp8
CFG-86-Sp8
CFG-87-Sp0
CFG-90-Sp0
Galectin - 1
Galectin - 3
Galectin - 7
f_pred
1.0
0.5
0.0
$10^{-4}$
$10^{-2}$
$10^{0}$
$10^{2}$
Concentration (μM)

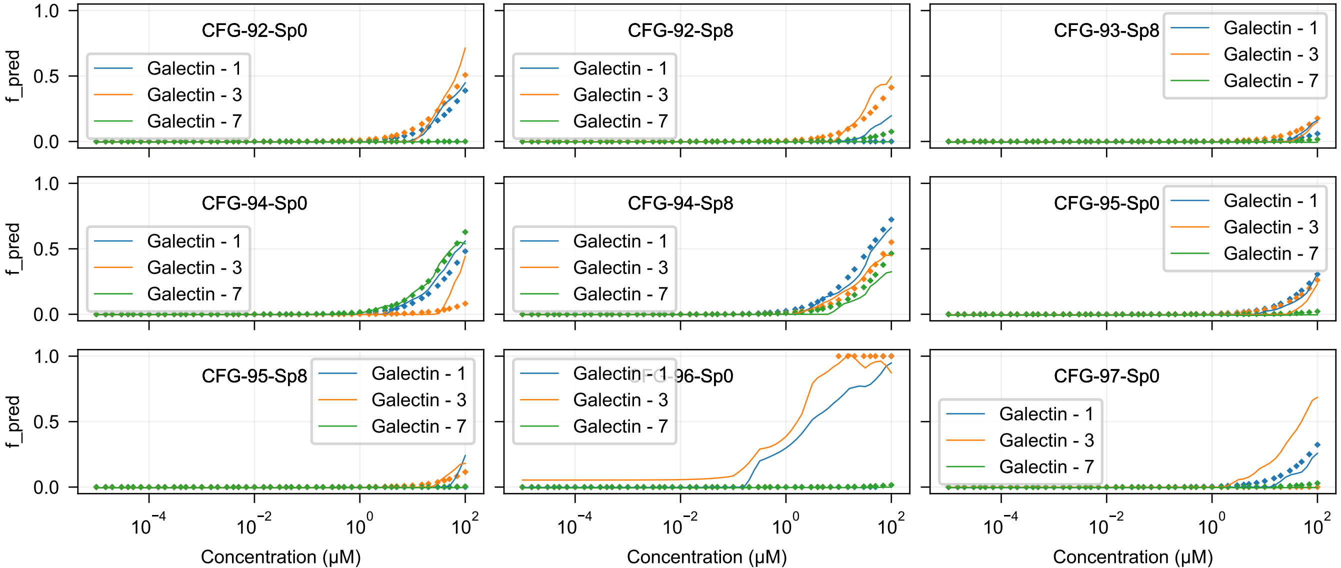
CFG-92-Sp0
CFG-92-Sp8
CFG-93-Sp8
CFG-94-Sp0
CFG-94-Sp8
CFG-95-Sp0
CFG-95-Sp8
CFG-96-Sp0
CFG-97-Sp0
Galectin - 1
Galectin - 3
Galectin - 7
f_pred
Concentration (μM)

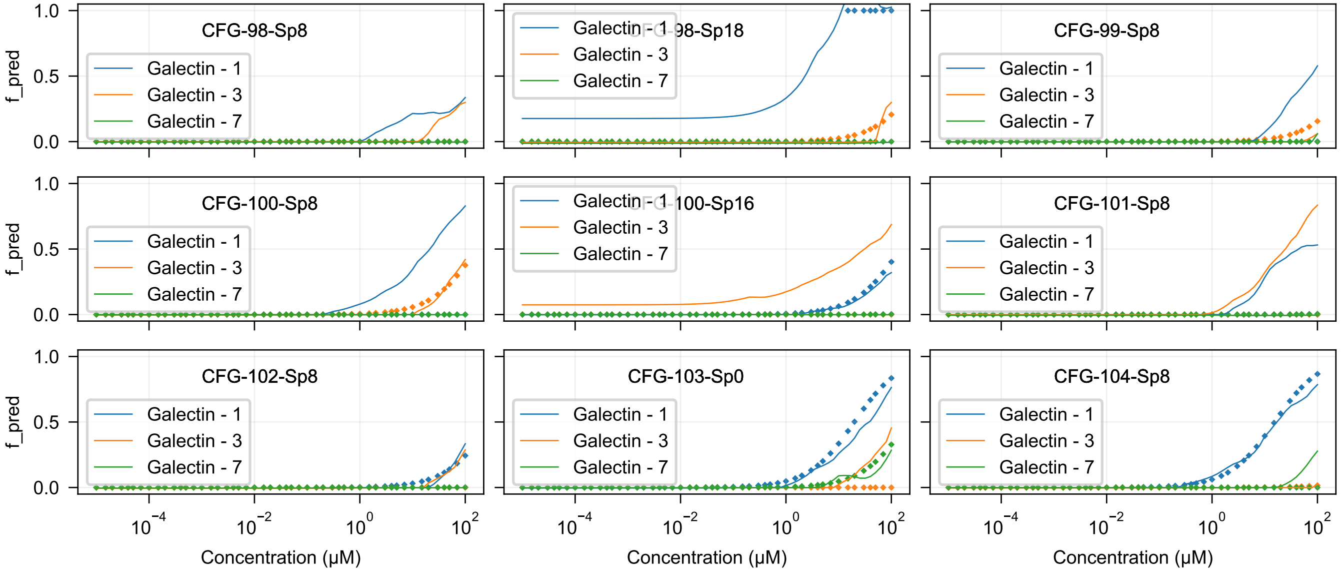
CFG-98-Sp8
CFG-98-Sp18
CFG-99-Sp8
CFG-100-Sp8
CFG-100-Sp16
CFG-101-Sp8
CFG-102-Sp8
CFG-103-Sp0
CFG-104-Sp8
Galectin - 1
Galectin - 3
Galectin - 7
f_pred
Concentration (μM)
0.0
0.5
1.0
$10^{-4}$
$10^{-2}$
$10^{0}$
$10^{2}$

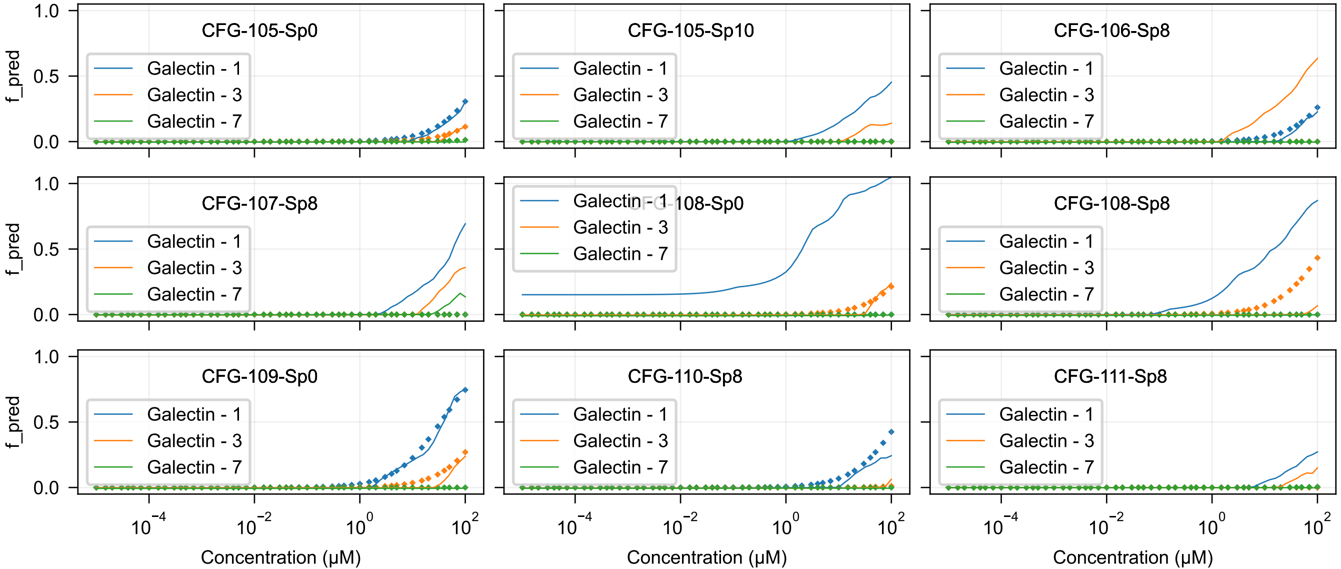

CFG-105-Sp0
CFG-105-Sp10
CFG-106-Sp8
CFG-107-Sp8
CFG-108-Sp0
CFG-108-Sp8
CFG-109-Sp0
CFG-110-Sp8
CFG-111-Sp8
Galectin - 1
Galectin - 3
Galectin - 7
f_pred
Concentration (μM)

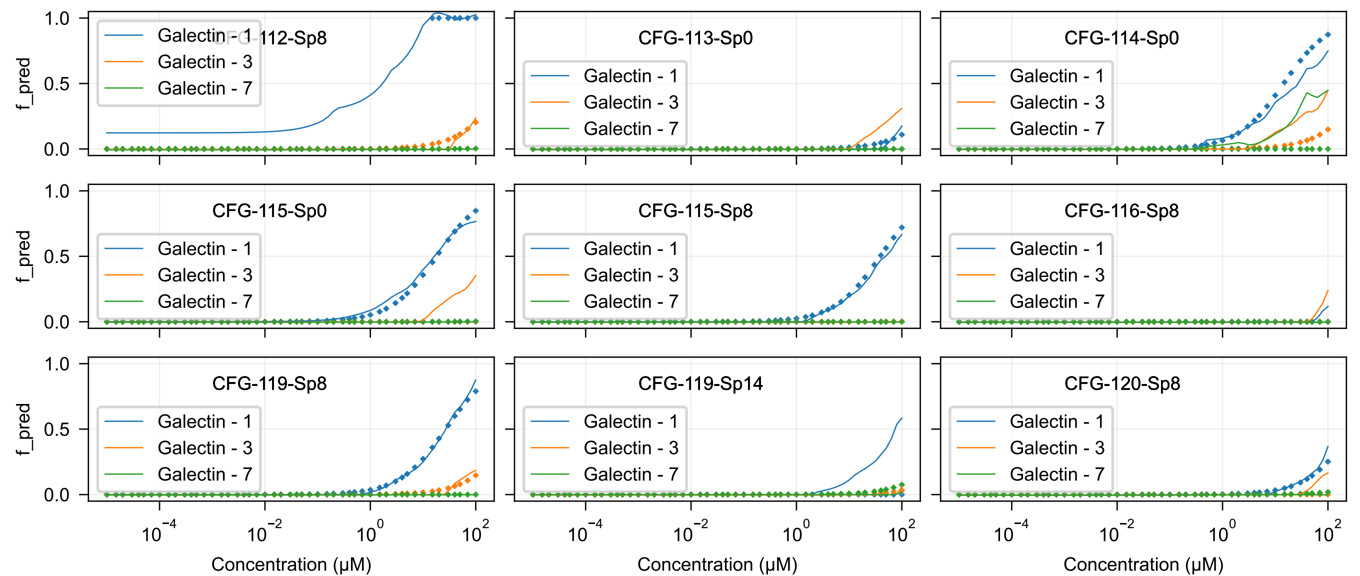

CFG-112-Sp8
CFG-113-Sp0
CFG-114-Sp0
CFG-115-Sp0
CFG-115-Sp8
CFG-116-Sp8
CFG-119-Sp8
CFG-119-Sp14
CFG-120-Sp8
Galectin - 1
Galectin - 3
Galectin - 7
f_pred
Concentration (μM)

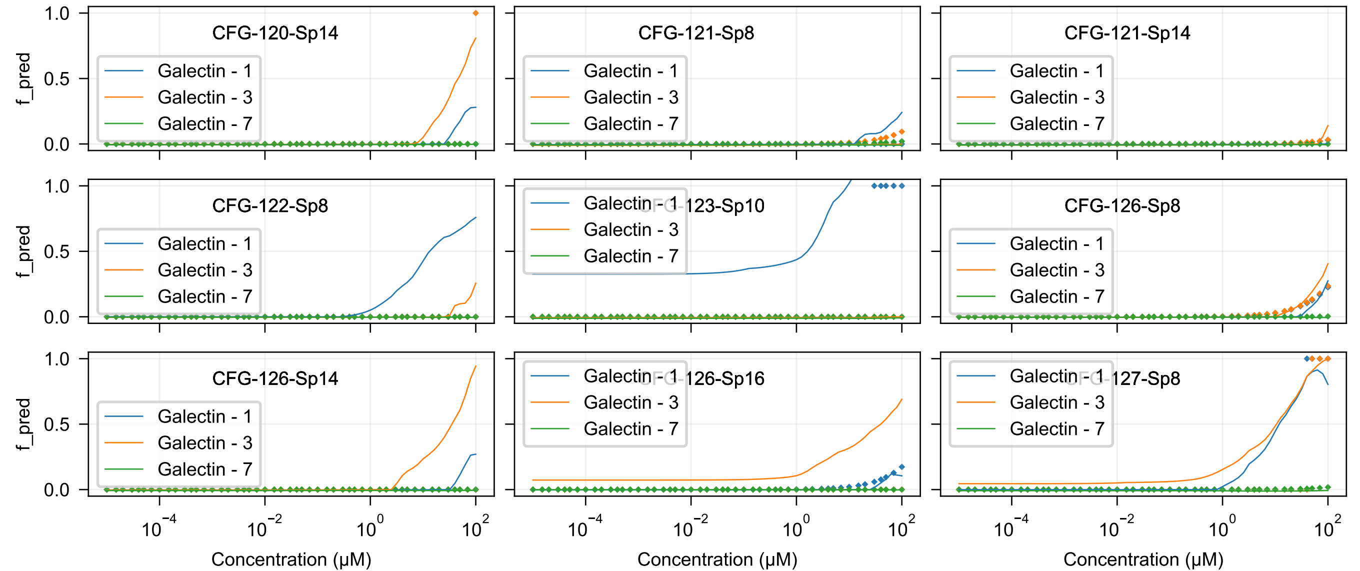

CFG-120-Sp14
CFG-121-Sp8
CFG-121-Sp14
CFG-122-Sp8
CFG-123-Sp10
CFG-126-Sp8
CFG-126-Sp14
CFG-126-Sp16
CFG-127-Sp8
Galectin - 1
Galectin - 3
Galectin - 7
f_pred
1.0
0.5
0.0
$10^{-4}$
$10^{-2}$
$10^{0}$
$10^{2}$
Concentration (μM)

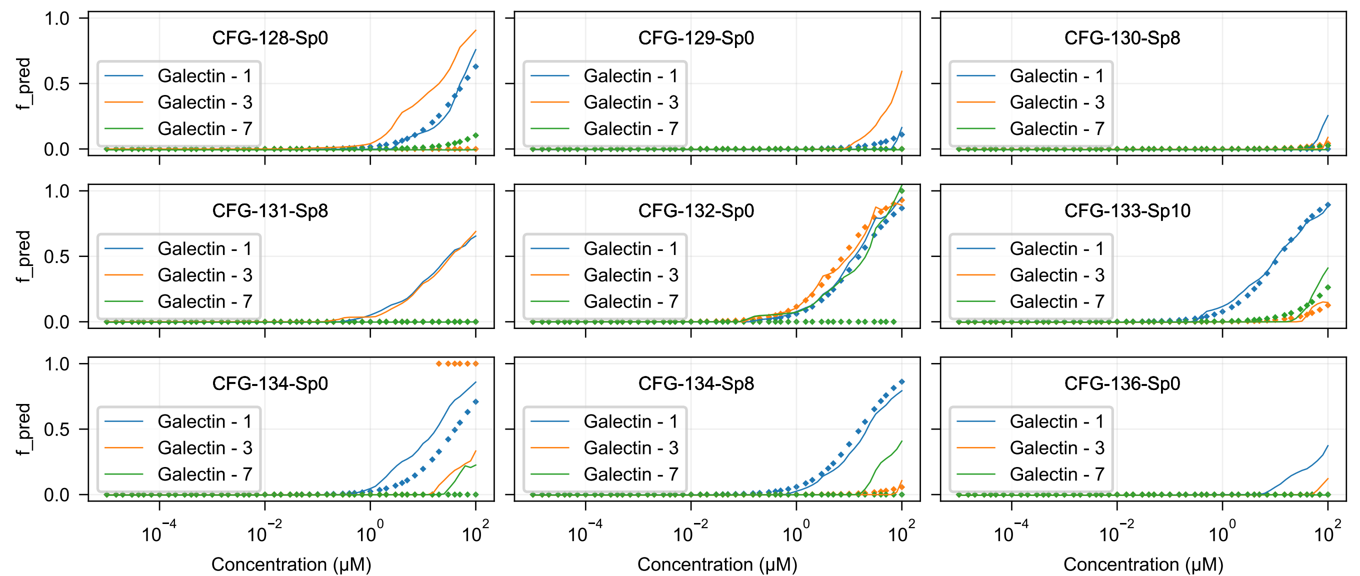
CFG-128-Sp0
CFG-129-Sp0
CFG-130-Sp8
CFG-131-Sp8
CFG-132-Sp0
CFG-133-Sp10
CFG-134-Sp0
CFG-134-Sp8
CFG-136-Sp0
Galectin - 1
Galectin - 3
Galectin - 7
f_pred
1.0
0.5
0.0
$10^{-4}$
$10^{-2}$
$10^{0}$
$10^{2}$
Concentration (μM)

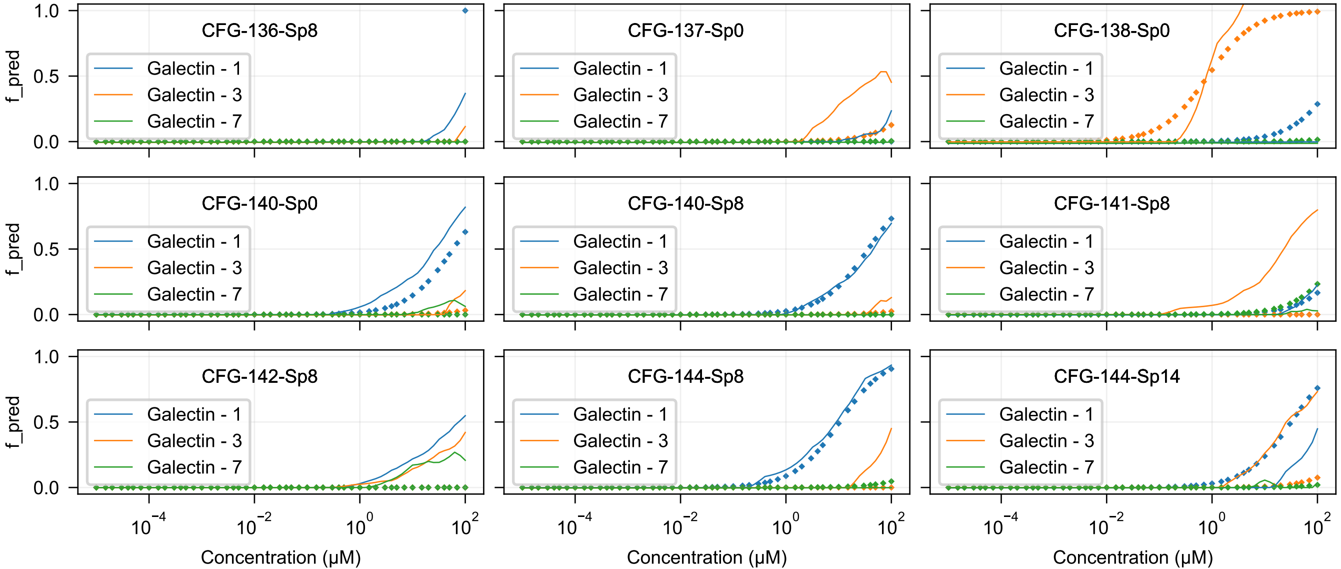
CFG-136-Sp8
CFG-137-Sp0
CFG-138-Sp0
CFG-140-Sp0
CFG-140-Sp8
CFG-141-Sp8
CFG-142-Sp8
CFG-144-Sp8
CFG-144-Sp14
Galectin - 1
Galectin - 3
Galectin - 7
f_pred
Concentration (μM)

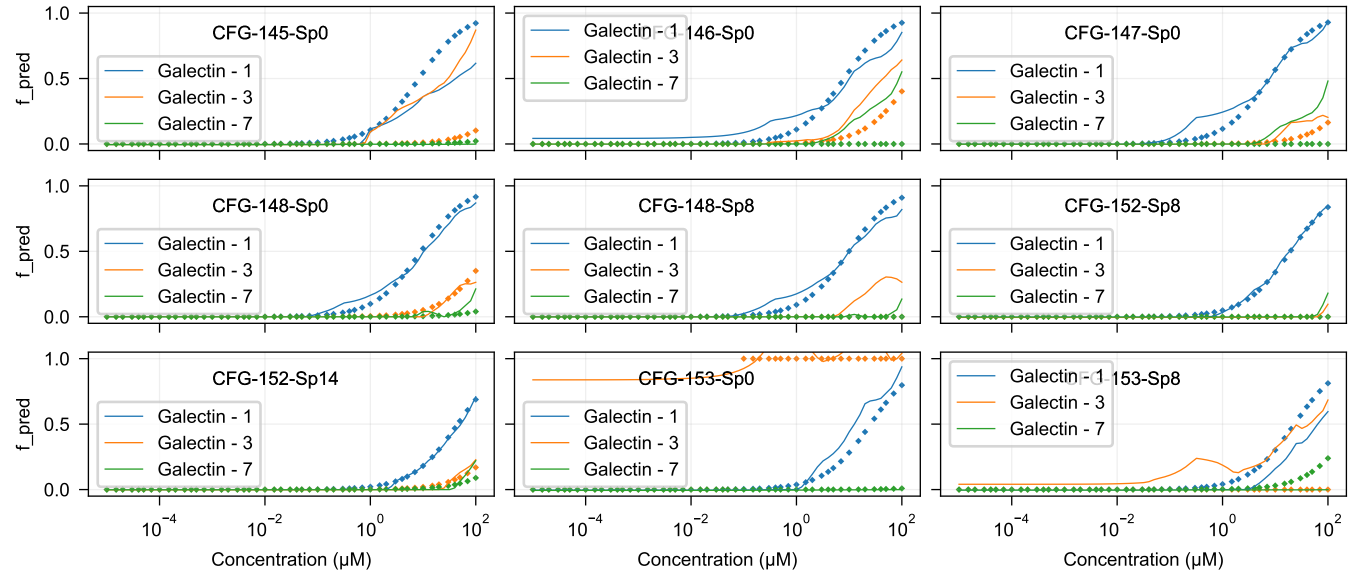
CFG-145-Sp0
CFG-146-Sp0
CFG-147-Sp0
CFG-148-Sp0
CFG-148-Sp8
CFG-152-Sp8
CFG-152-Sp14
CFG-153-Sp0
CFG-153-Sp8
Galectin - 1
Galectin - 3
Galectin - 7
f_pred
1.0
0.5
0.0
$10^{-4}$
$10^{-2}$
$10^{0}$
$10^{2}$
Concentration (μM)

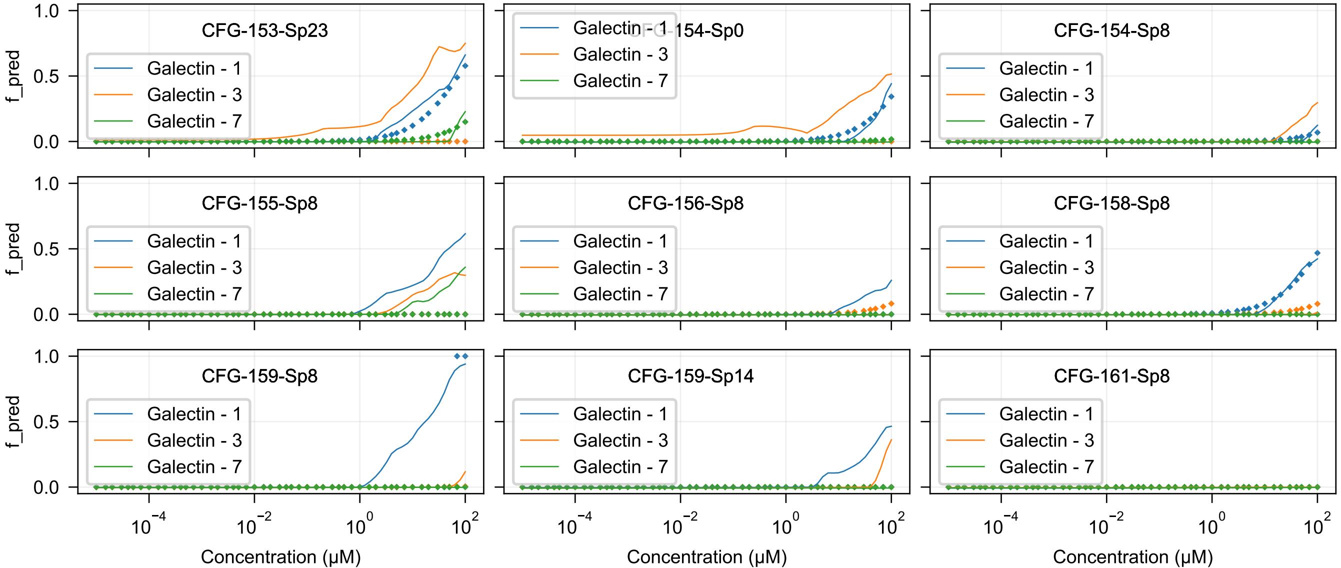

CFG-153-Sp23
CFG-154-Sp0
CFG-154-Sp8
CFG-155-Sp8
CFG-156-Sp8
CFG-158-Sp8
CFG-159-Sp8
CFG-159-Sp14
CFG-161-Sp8
Galectin - 1
Galectin - 3
Galectin - 7
f_pred
1.0
0.5
0.0
$10^{-4}$
$10^{-2}$
$10^{0}$
$10^{2}$
Concentration (μM)

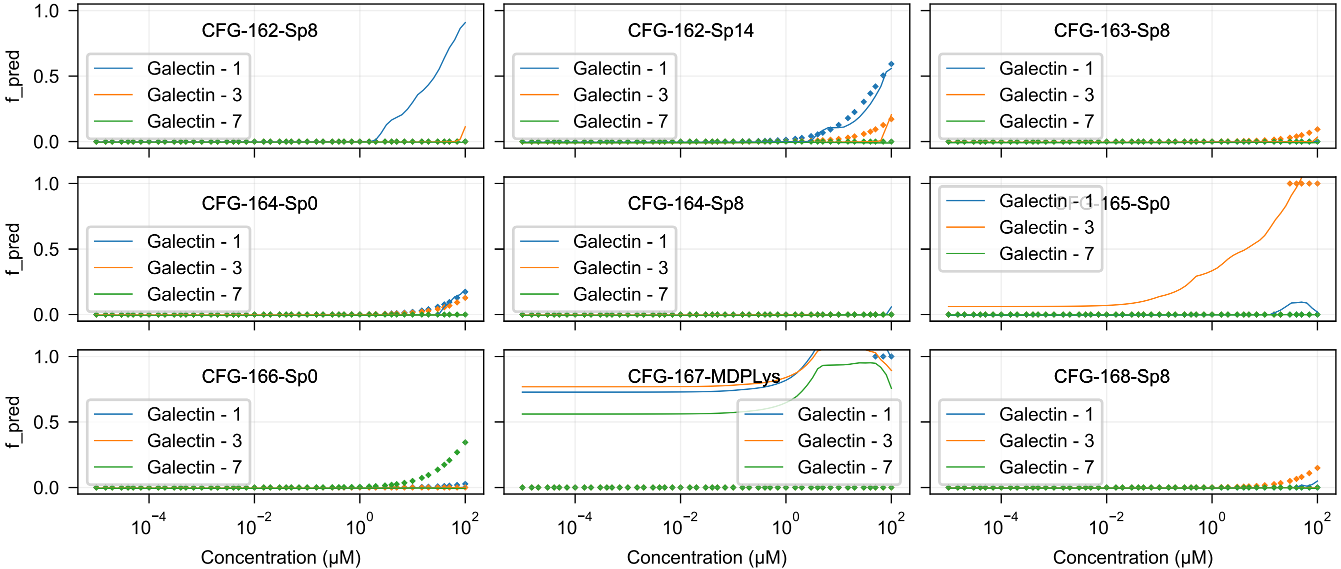
CFG-162-Sp8
CFG-162-Sp14
CFG-163-Sp8
CFG-164-Sp0
CFG-164-Sp8
CFG-165-Sp0
CFG-166-Sp0
CFG-167-MDPLys
CFG-168-Sp8
Galectin - 1
Galectin - 3
Galectin - 7
f_pred
1.0
0.5
0.0
$10^{-4}$
$10^{-2}$
$10^{0}$
$10^{2}$
Concentration (μM)

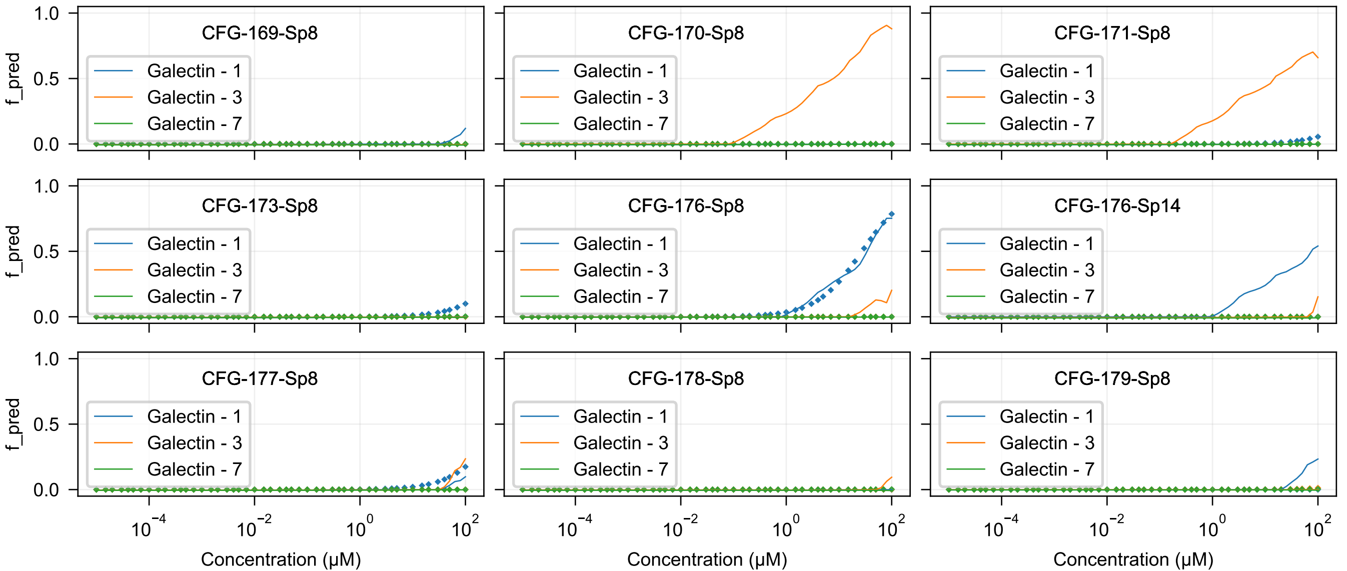

CFG-169-Sp8
CFG-170-Sp8
CFG-171-Sp8
CFG-173-Sp8
CFG-176-Sp8
CFG-176-Sp14
CFG-177-Sp8
CFG-178-Sp8
CFG-179-Sp8
Galectin - 1
Galectin - 3
Galectin - 7
f_pred
Concentration (μM)

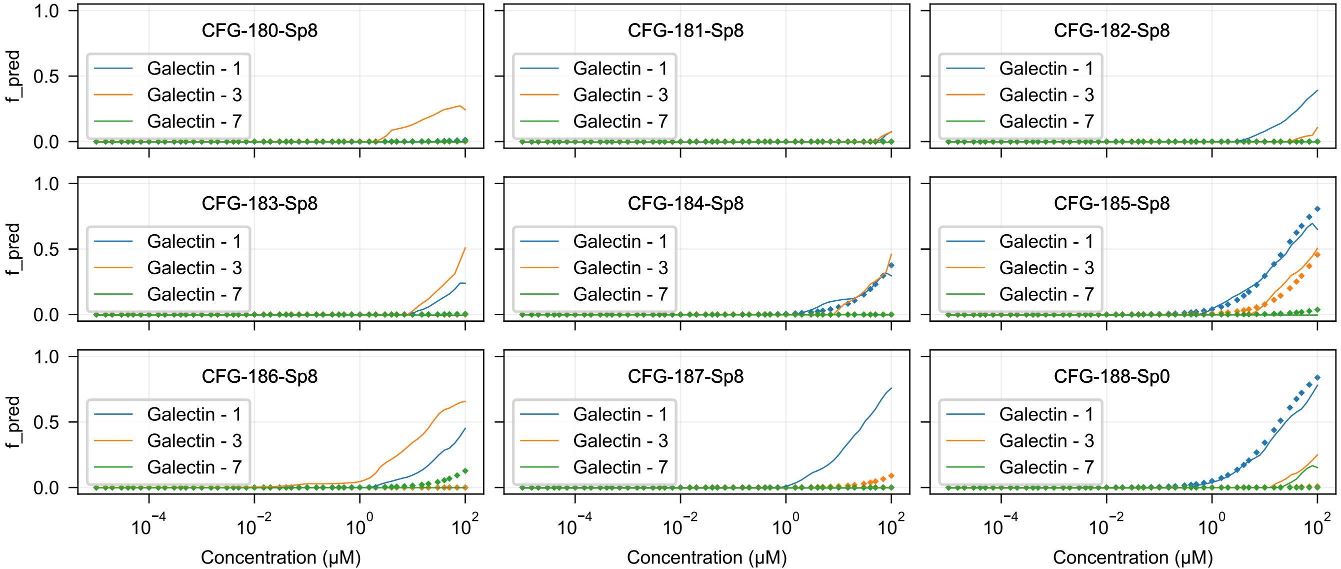
CFG-180-Sp8
CFG-181-Sp8
CFG-182-Sp8
CFG-183-Sp8
CFG-184-Sp8
CFG-185-Sp8
CFG-186-Sp8
CFG-187-Sp8
CFG-188-Sp0
Galectin - 1
Galectin - 3
Galectin - 7
f_pred
Concentration (μM)

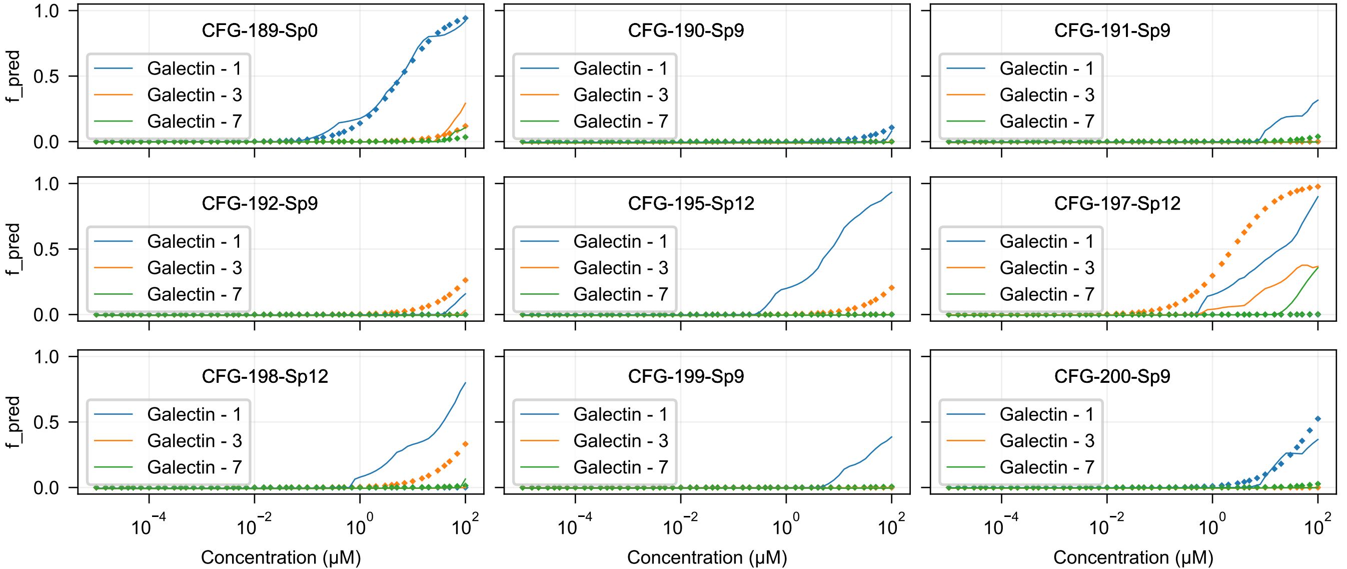
CFG-189-Sp0
CFG-190-Sp9
CFG-191-Sp9
CFG-192-Sp9
CFG-195-Sp12
CFG-197-Sp12
CFG-198-Sp12
CFG-199-Sp9
CFG-200-Sp9
Galectin - 1
Galectin - 3
Galectin - 7
f_pred
Concentration (μM)

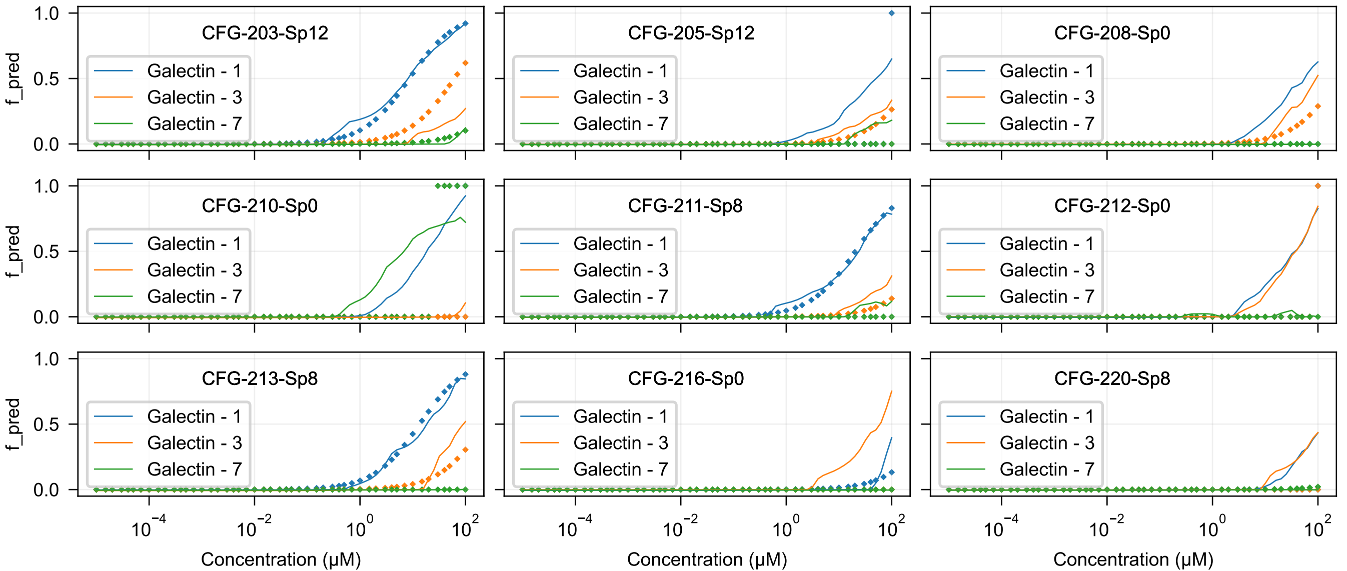

CFG-203-Sp12
CFG-205-Sp12
CFG-208-Sp0
CFG-210-Sp0
CFG-211-Sp8
CFG-212-Sp0
CFG-213-Sp8
CFG-216-Sp0
CFG-220-Sp8
Galectin - 1
Galectin - 3
Galectin - 7
f_pred
1.0
0.5
0.0
$10^{-4}$
$10^{-2}$
$10^{0}$
$10^{2}$
Concentration (μM)

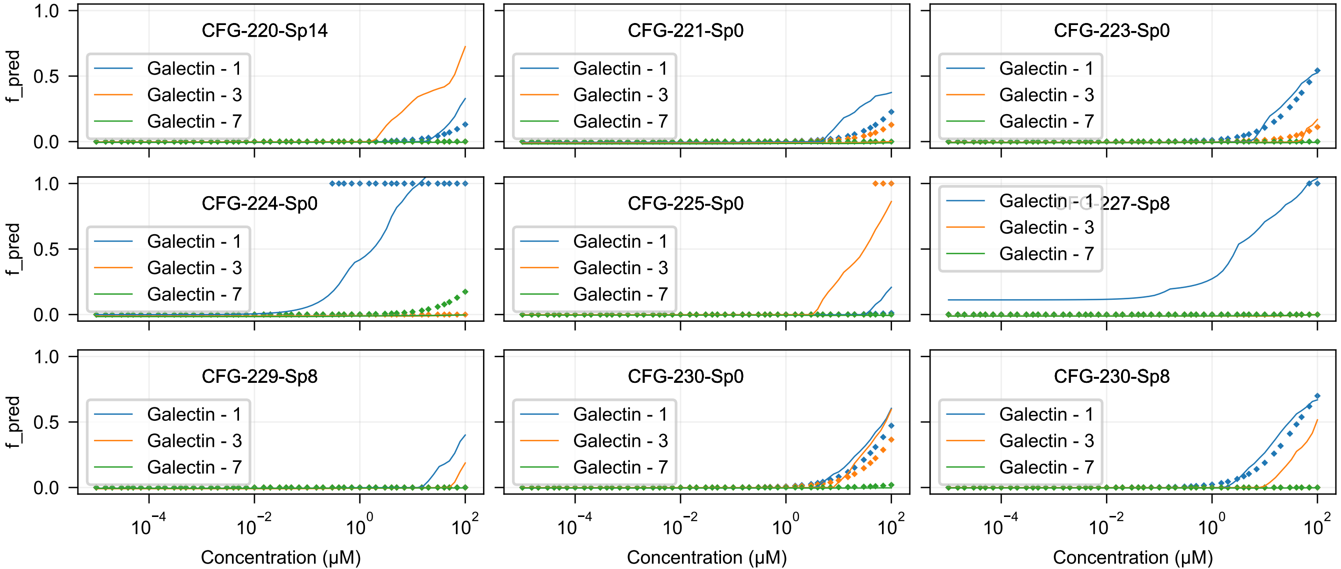
CFG-220-Sp14
CFG-221-Sp0
CFG-223-Sp0
CFG-224-Sp0
CFG-225-Sp0
CFG-227-Sp8
CFG-229-Sp8
CFG-230-Sp0
CFG-230-Sp8
Galectin - 1
Galectin - 3
Galectin - 7
f_pred
1.0
0.5
0.0
$10^{-4}$
$10^{-2}$
$10^{0}$
$10^{2}$
Concentration (μM)

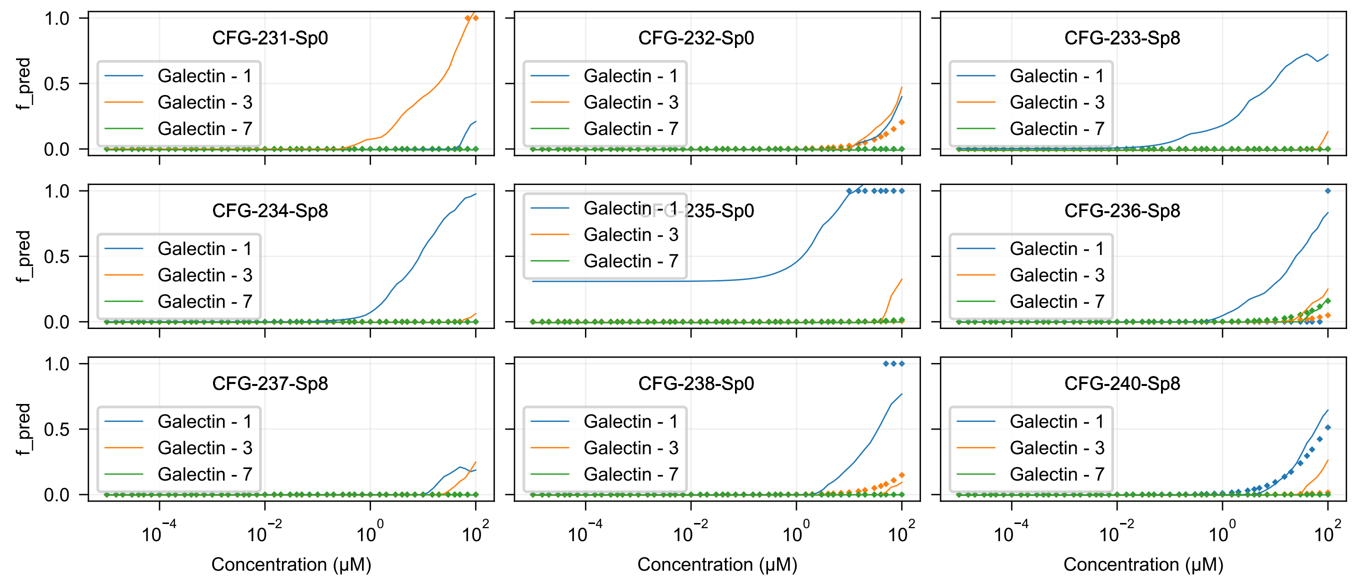
CFG-231-Sp0
CFG-232-Sp0
CFG-233-Sp8
CFG-234-Sp8
CFG-235-Sp0
CFG-236-Sp8
CFG-237-Sp8
CFG-238-Sp0
CFG-240-Sp8
Galectin - 1
Galectin - 3
Galectin - 7
f_pred
Concentration (μM)

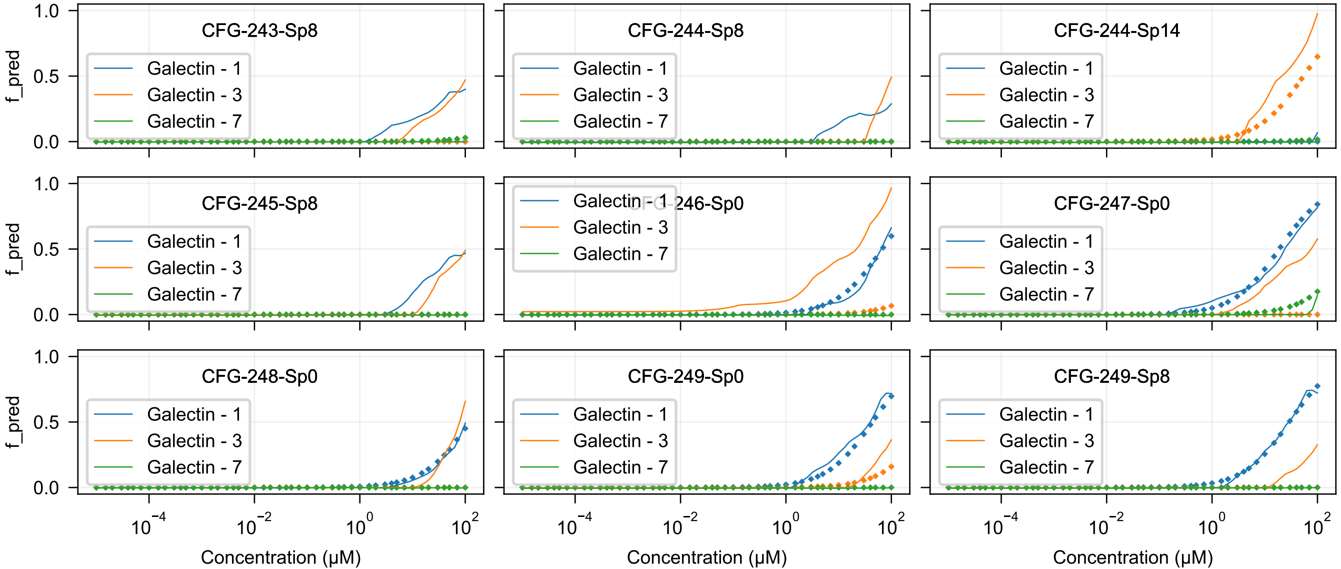
CFG-243-Sp8
CFG-244-Sp8
CFG-244-Sp14
CFG-245-Sp8
CFG-246-Sp0
CFG-247-Sp0
CFG-248-Sp0
CFG-249-Sp0
CFG-249-Sp8
Galectin - 1
Galectin - 3
Galectin - 7
f_pred
0.0
0.5
1.0
$10^{-4}$
$10^{-2}$
$10^{0}$
$10^{2}$
Concentration (μM)

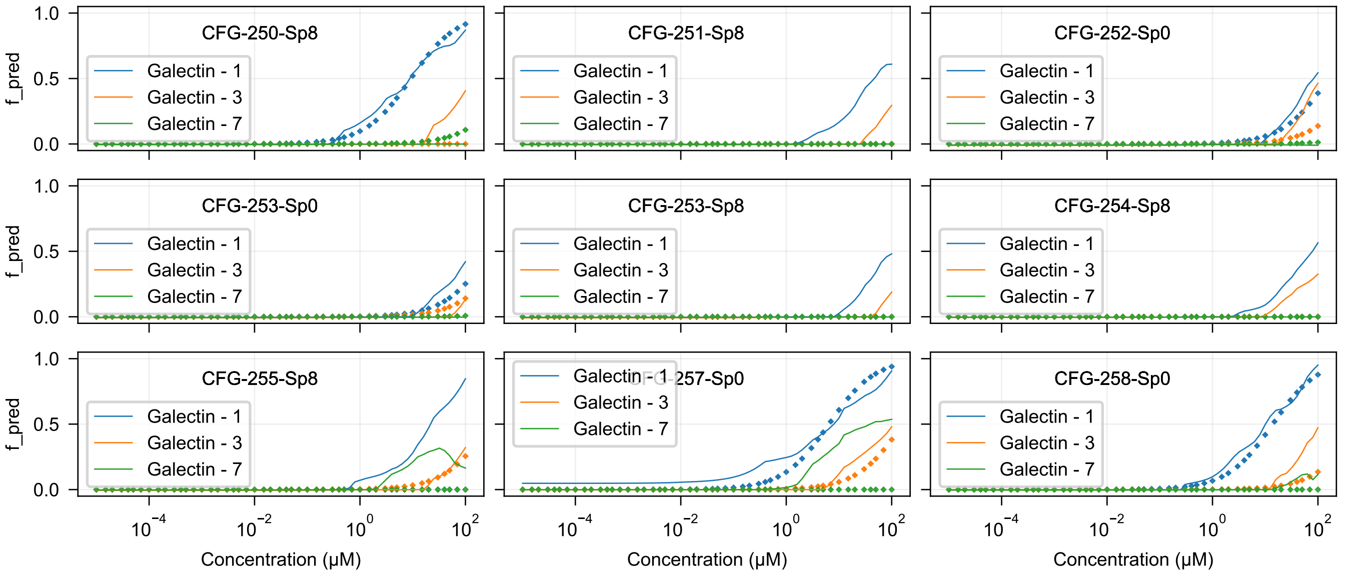

CFG-250-Sp8
CFG-251-Sp8
CFG-252-Sp0
CFG-253-Sp0
CFG-253-Sp8
CFG-254-Sp8
CFG-255-Sp8
CFG-257-Sp0
CFG-258-Sp0
Galectin - 1
Galectin - 3
Galectin - 7
f_pred
1.0
0.5
0.0
$10^{-4}$
$10^{-2}$
$10^{0}$
$10^{2}$
Concentration (μM)

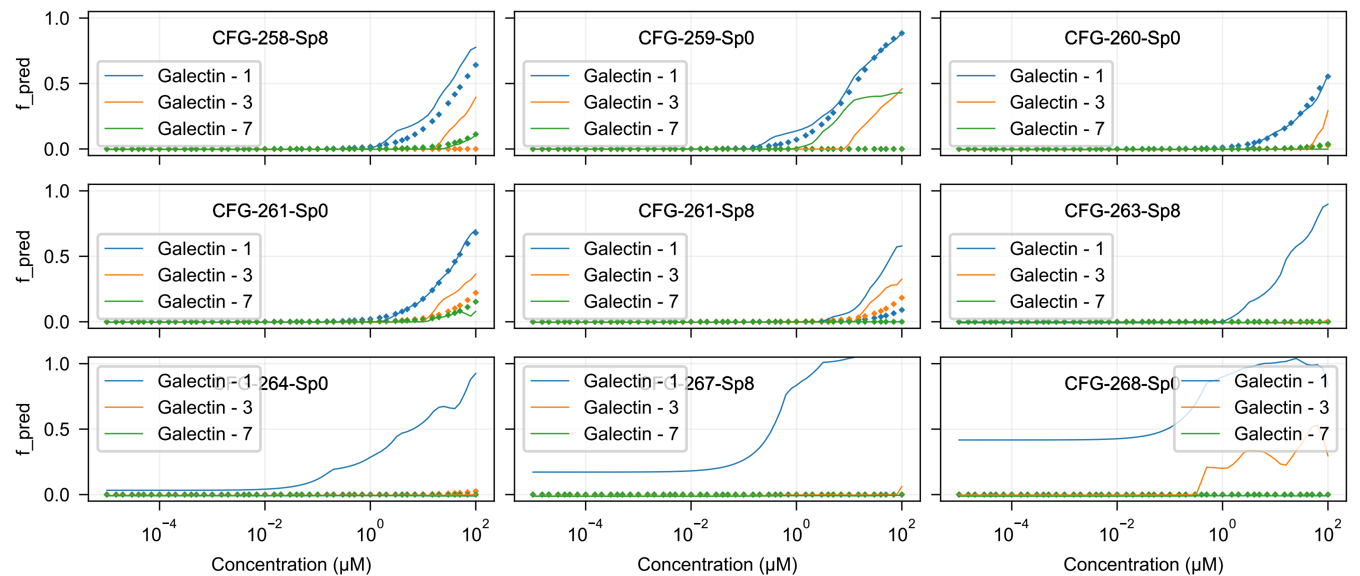

CFG-258-Sp8
CFG-259-Sp0
CFG-260-Sp0
CFG-261-Sp0
CFG-261-Sp8
CFG-263-Sp8
CFG-264-Sp0
CFG-267-Sp8
CFG-268-Sp0
Galectin - 1
Galectin - 3
Galectin - 7
f_pred
Concentration (μM)

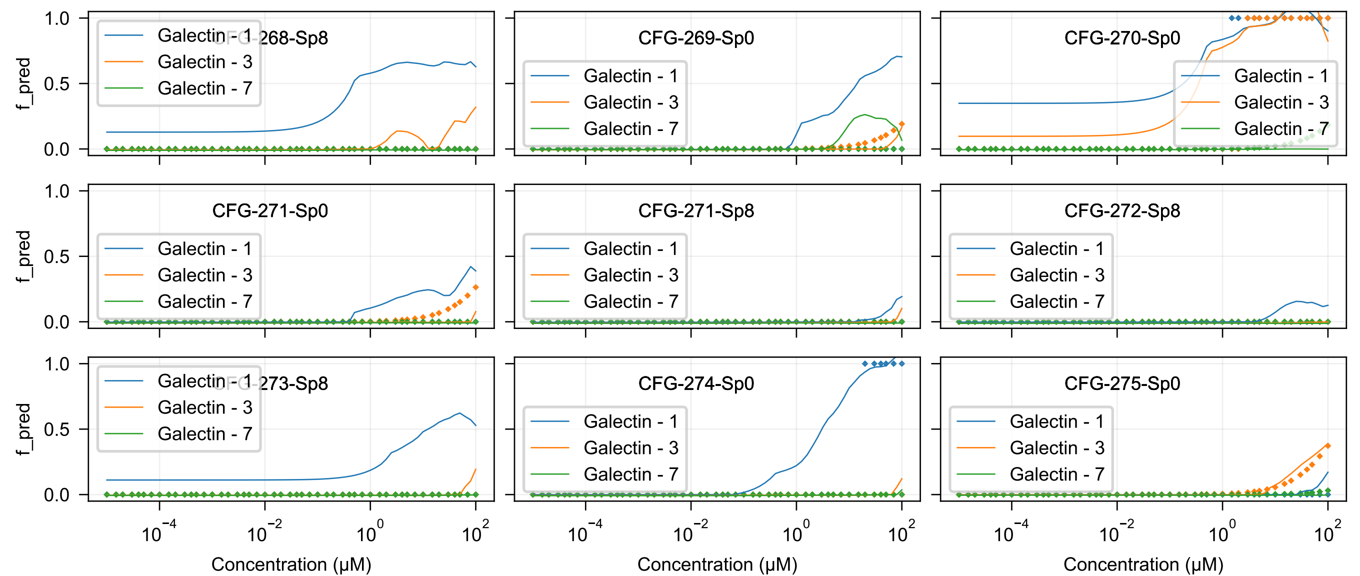

CFG-268-Sp8
CFG-269-Sp0
CFG-270-Sp0
CFG-271-Sp0
CFG-271-Sp8
CFG-272-Sp8
CFG-273-Sp8
CFG-274-Sp0
CFG-275-Sp0
Galectin - 1
Galectin - 3
Galectin - 7
f_pred
Concentration (μM)

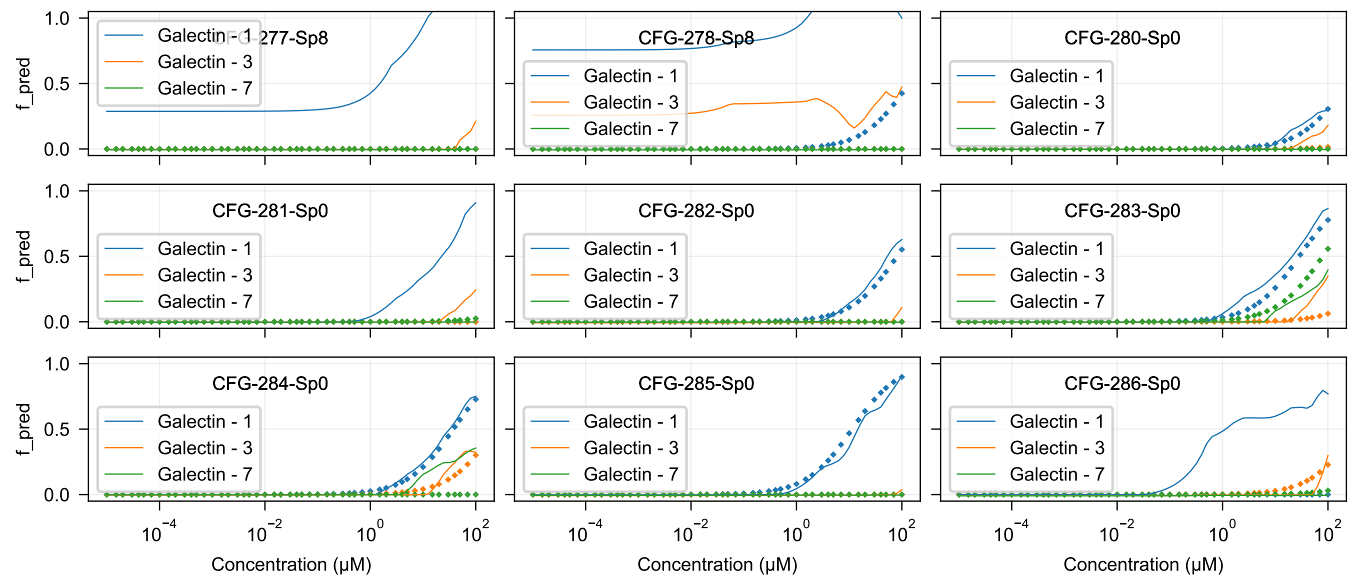
CFG-277-Sp8
CFG-278-Sp8
CFG-280-Sp0
CFG-281-Sp0
CFG-282-Sp0
CFG-283-Sp0
CFG-284-Sp0
CFG-285-Sp0
CFG-286-Sp0
Galectin - 1
Galectin - 3
Galectin - 7
f_pred
Concentration (μM)

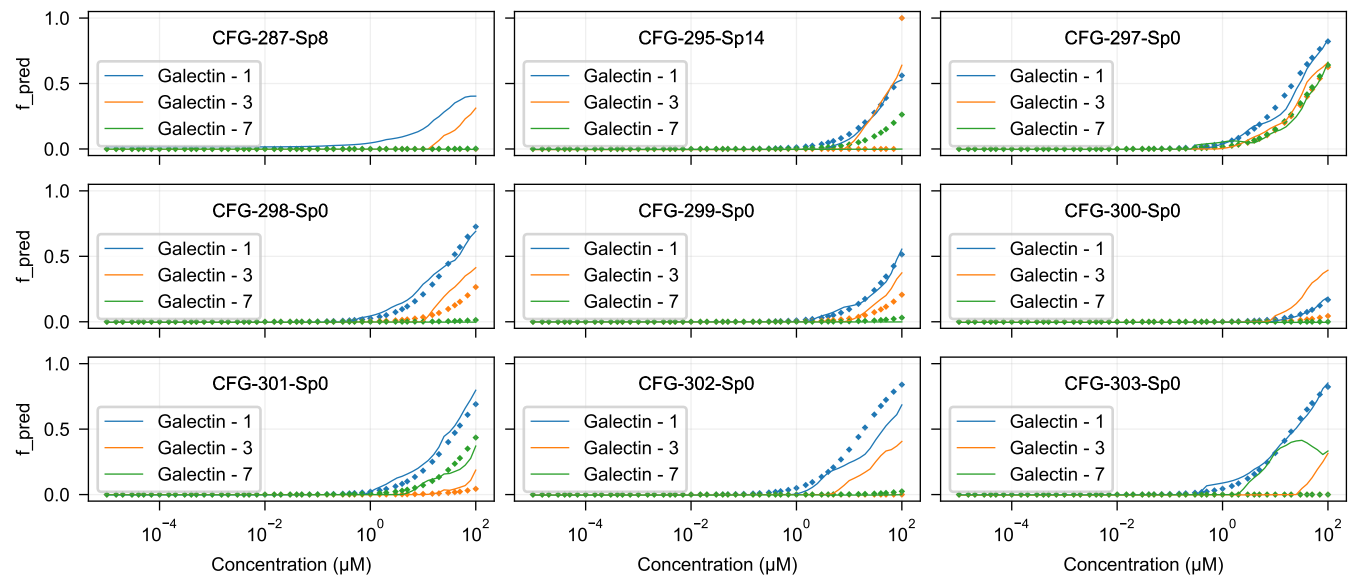
CFG-287-Sp8
CFG-295-Sp14
CFG-297-Sp0
CFG-298-Sp0
CFG-299-Sp0
CFG-300-Sp0
CFG-301-Sp0
CFG-302-Sp0
CFG-303-Sp0
Galectin - 1
Galectin - 3
Galectin - 7
f_pred
Concentration (μM)

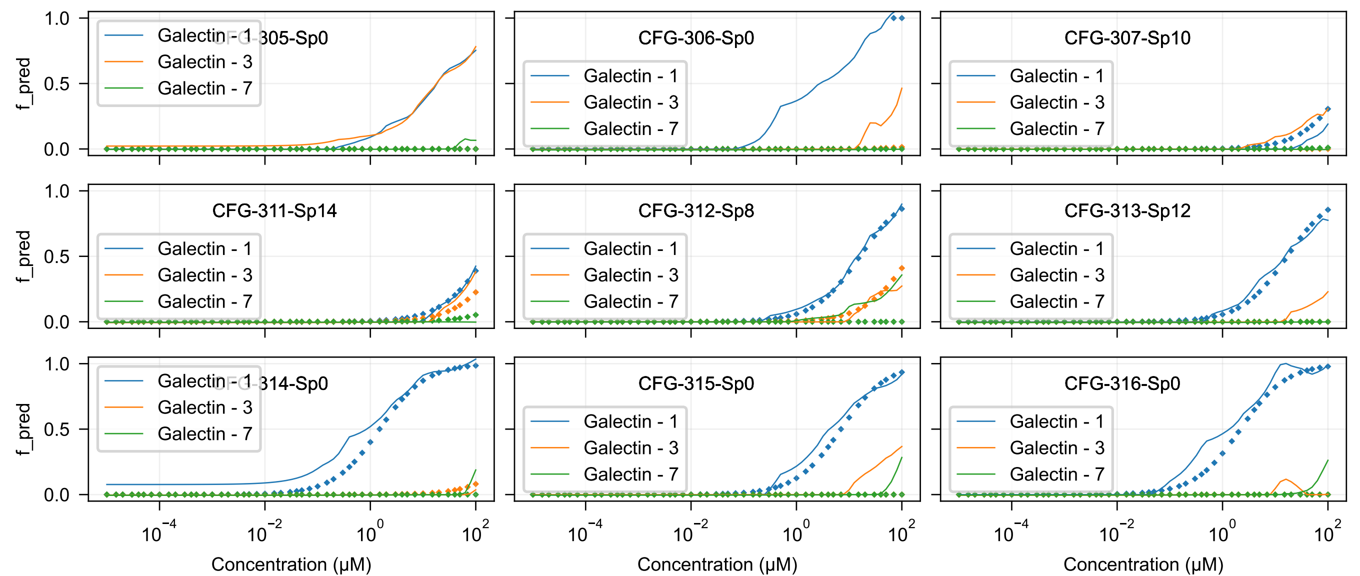
CFG-305-Sp0
CFG-306-Sp0
CFG-307-Sp10
CFG-311-Sp14
CFG-312-Sp8
CFG-313-Sp12
CFG-314-Sp0
CFG-315-Sp0
CFG-316-Sp0
Galectin - 1
Galectin - 3
Galectin - 7
f_pred
1.0
0.5
0.0
$10^{-4}$
$10^{-2}$
$10^{0}$
$10^{2}$
Concentration (μM)

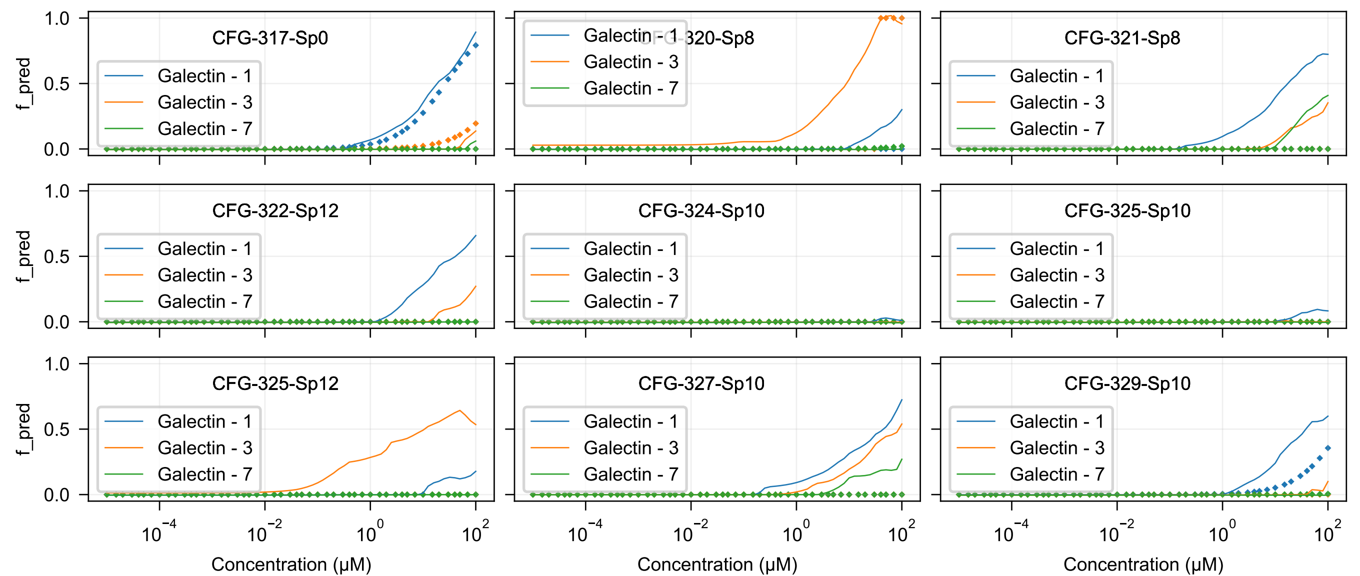

CFG-317-Sp0
CFG-320-Sp8
CFG-321-Sp8
CFG-322-Sp12
CFG-324-Sp10
CFG-325-Sp10
CFG-325-Sp12
CFG-327-Sp10
CFG-329-Sp10
Galectin - 1
Galectin - 3
Galectin - 7
f_pred
0.0
0.5
1.0
$10^{-4}$
$10^{-2}$
$10^{0}$
$10^{2}$
Concentration (μM)

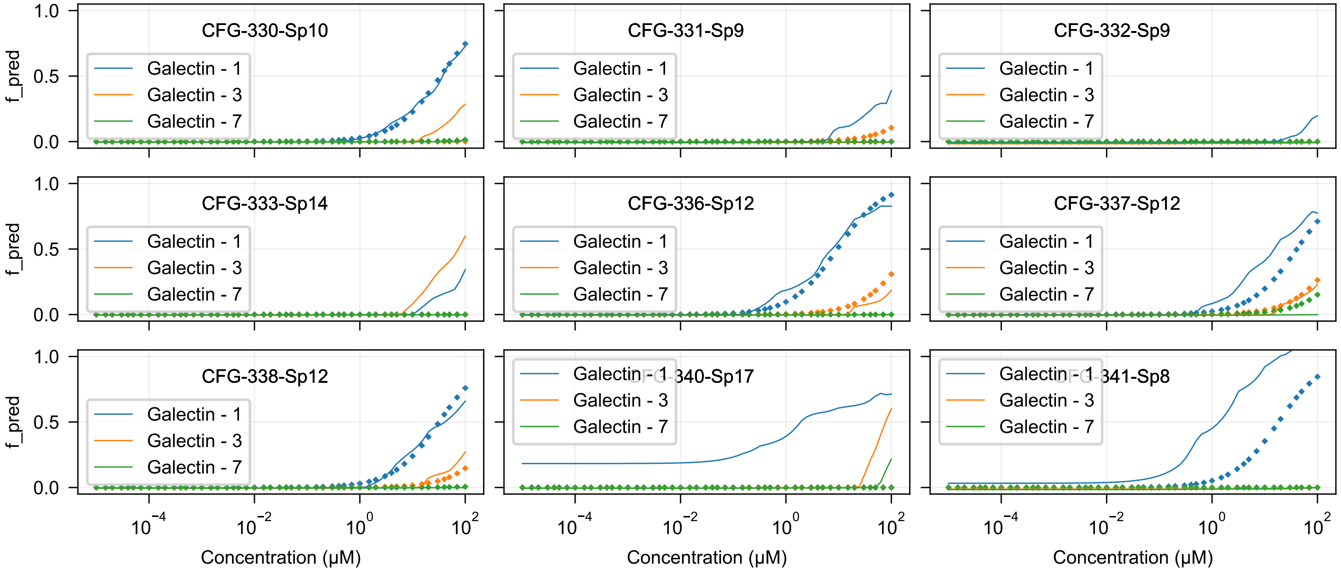
CFG-330-Sp10
CFG-331-Sp9
CFG-332-Sp9
CFG-333-Sp14
CFG-336-Sp12
CFG-337-Sp12
CFG-338-Sp12
CFG-340-Sp17
CFG-341-Sp8
Galectin - 1
Galectin - 3
Galectin - 7
f_pred
Concentration (μM)
1.0
0.5
0.0
$10^{-4}$
$10^{-2}$
$10^{0}$
$10^{2}$

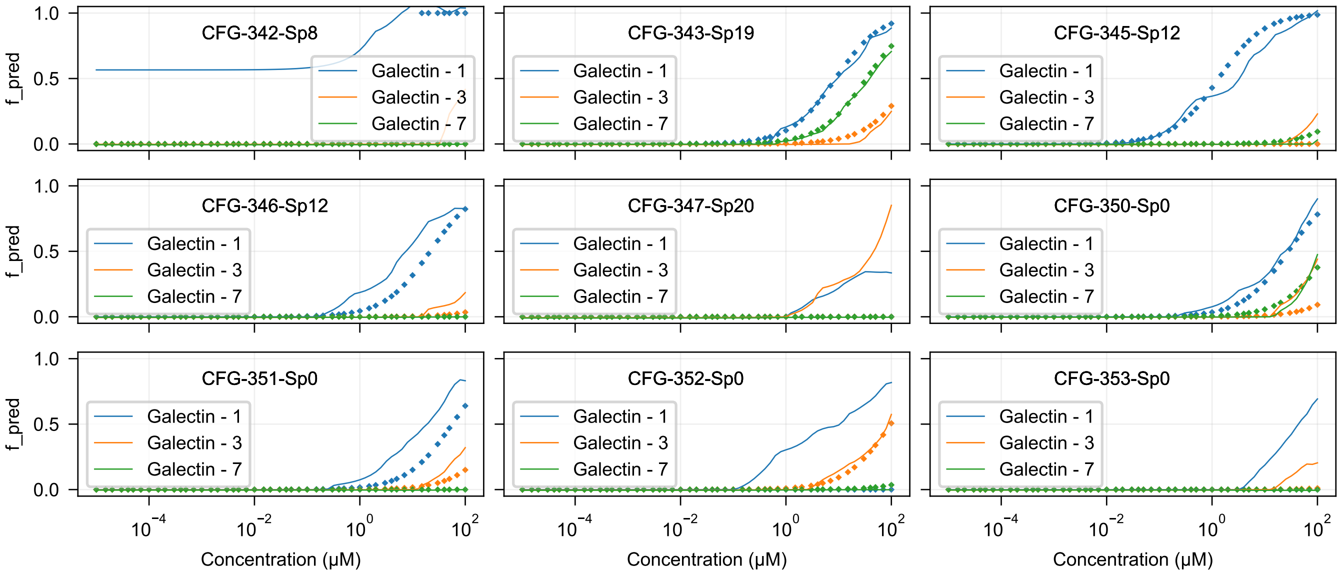
CFG-342-Sp8
CFG-343-Sp19
CFG-345-Sp12
CFG-346-Sp12
CFG-347-Sp20
CFG-350-Sp0
CFG-351-Sp0
CFG-352-Sp0
CFG-353-Sp0
Galectin - 1
Galectin - 3
Galectin - 7
f_pred
Concentration (μM)

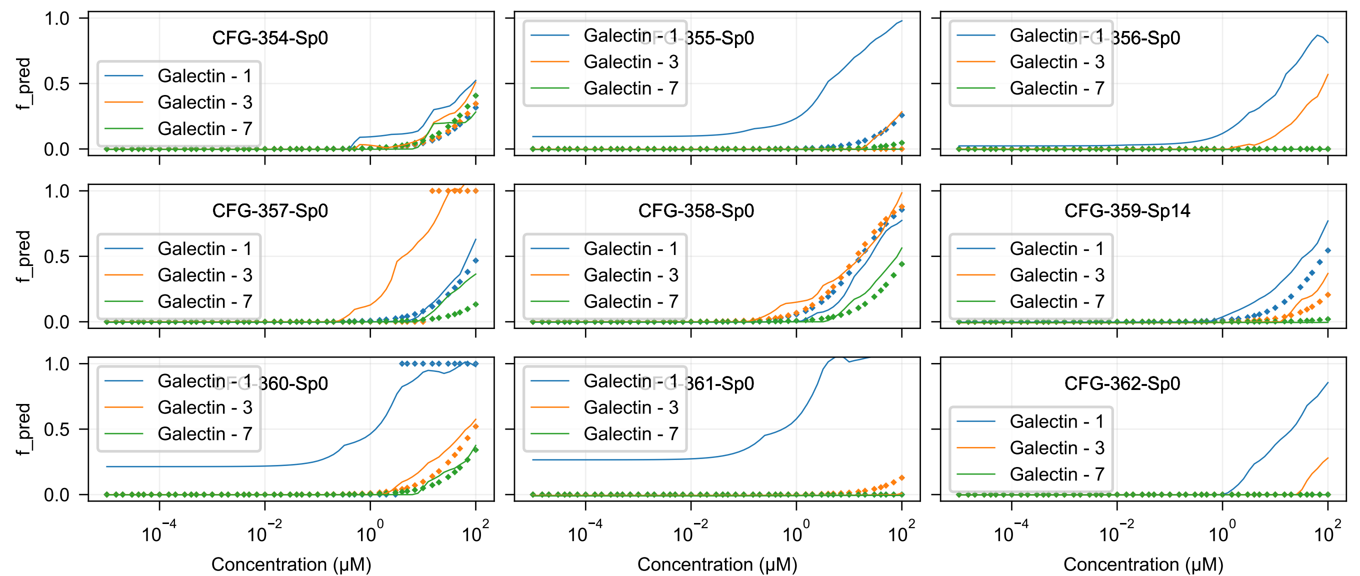

CFG-354-Sp0
CFG-355-Sp0
CFG-356-Sp0
CFG-357-Sp0
CFG-358-Sp0
CFG-359-Sp14
CFG-360-Sp0
CFG-361-Sp0
CFG-362-Sp0
Galectin - 1
Galectin - 3
Galectin - 7
f_pred
1.0
0.5
0.0
$10^{-4}$
$10^{-2}$
$10^{0}$
$10^{2}$
Concentration (μM)

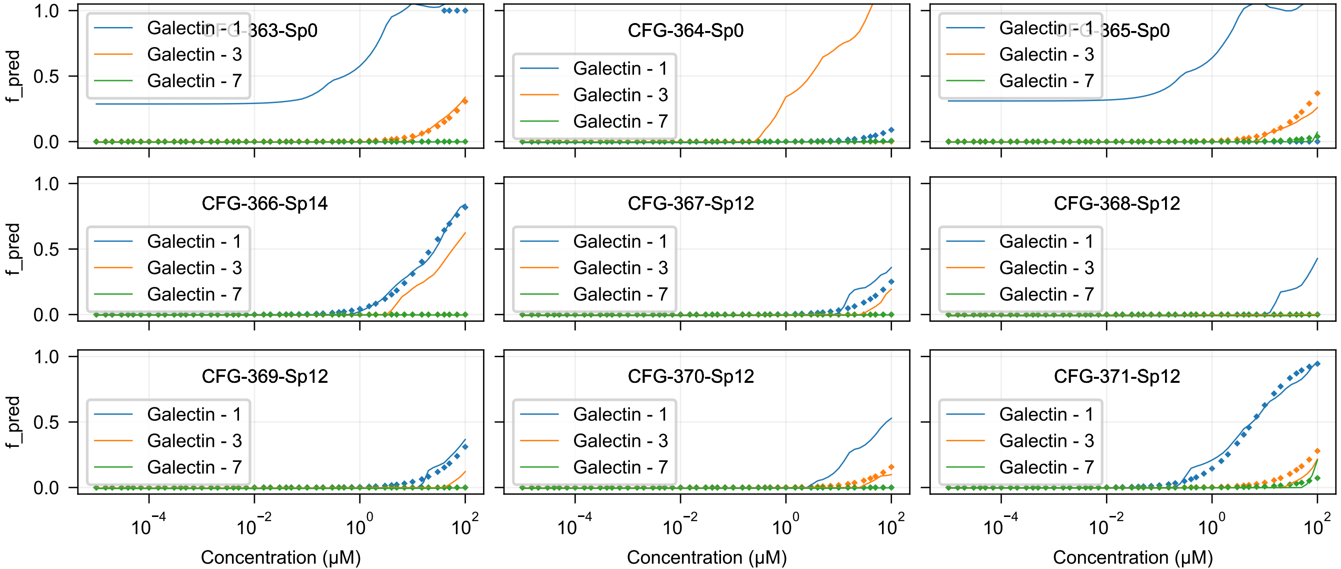
CFG-363-Sp0
CFG-364-Sp0
CFG-365-Sp0
CFG-366-Sp14
CFG-367-Sp12
CFG-368-Sp12
CFG-369-Sp12
CFG-370-Sp12
CFG-371-Sp12
Galectin - 1
Galectin - 3
Galectin - 7
f_pred
Concentration (μM)

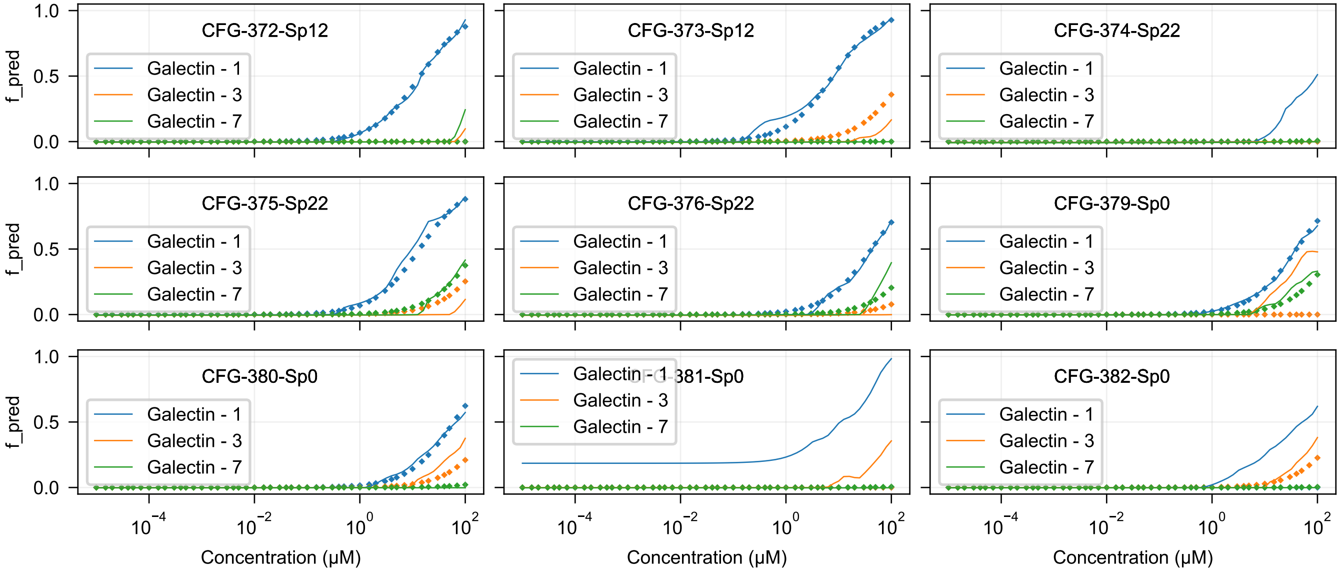
CFG-372-Sp12
CFG-373-Sp12
CFG-374-Sp22
CFG-375-Sp22
CFG-376-Sp22
CFG-379-Sp0
CFG-380-Sp0
CFG-381-Sp0
CFG-382-Sp0
Galectin - 1
Galectin - 3
Galectin - 7
f_pred
Concentration (μM)

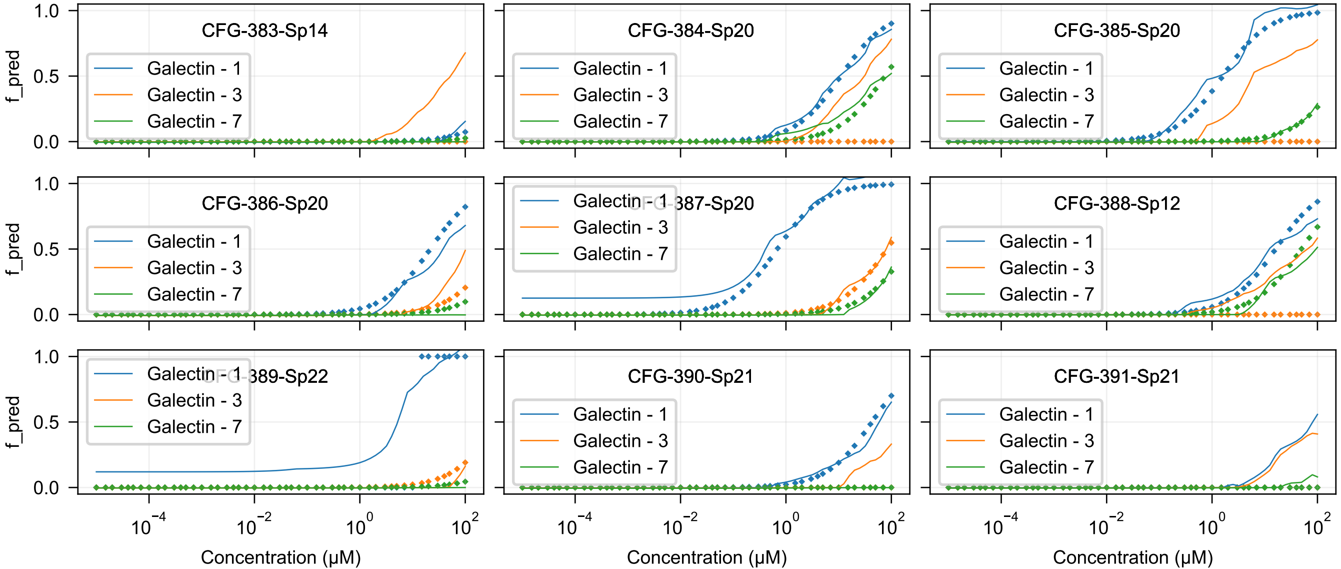
CFG-383-Sp14
CFG-384-Sp20
CFG-385-Sp20
CFG-386-Sp20
CFG-387-Sp20
CFG-388-Sp12
CFG-389-Sp22
CFG-390-Sp21
CFG-391-Sp21
Galectin - 1
Galectin - 3
Galectin - 7
f_pred
Concentration (μM)
1.0
0.5
0.0
$10^{-4}$
$10^{-2}$
$10^{0}$
$10^{2}$

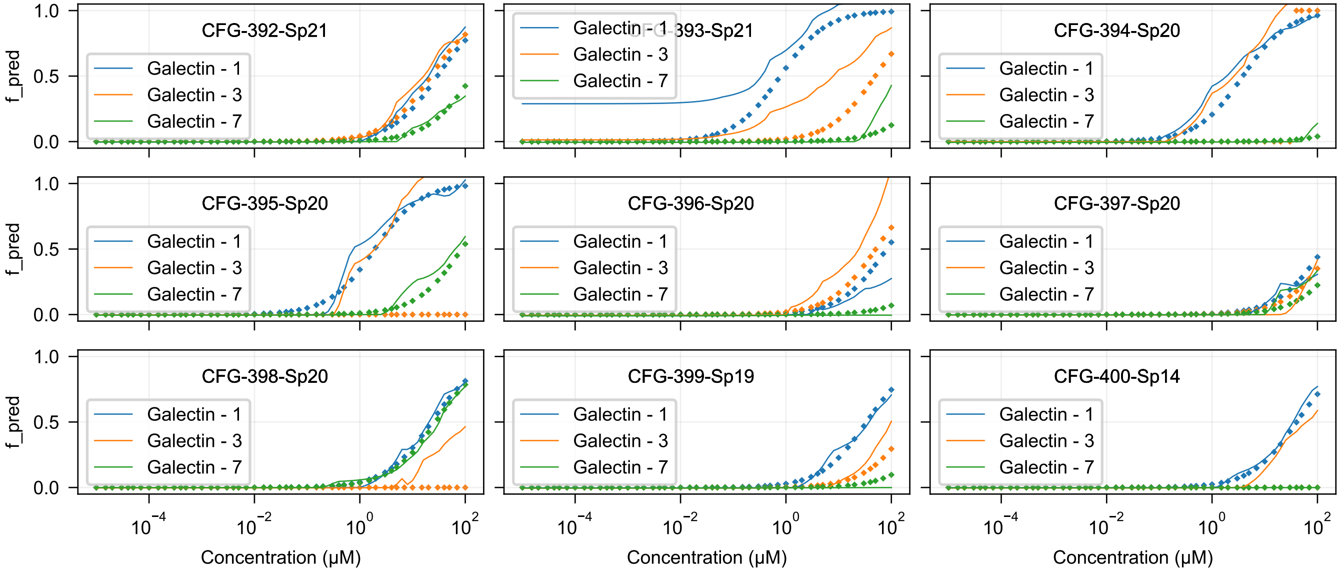

CFG-392-Sp21
CFG-393-Sp21
CFG-394-Sp20
CFG-395-Sp20
CFG-396-Sp20
CFG-397-Sp20
CFG-398-Sp20
CFG-399-Sp19
CFG-400-Sp14
Galectin - 1
Galectin - 3
Galectin - 7
f_pred
1.0
0.5
0.0
$10^{-4}$
$10^{-2}$
$10^{0}$
$10^{2}$
Concentration (μM)

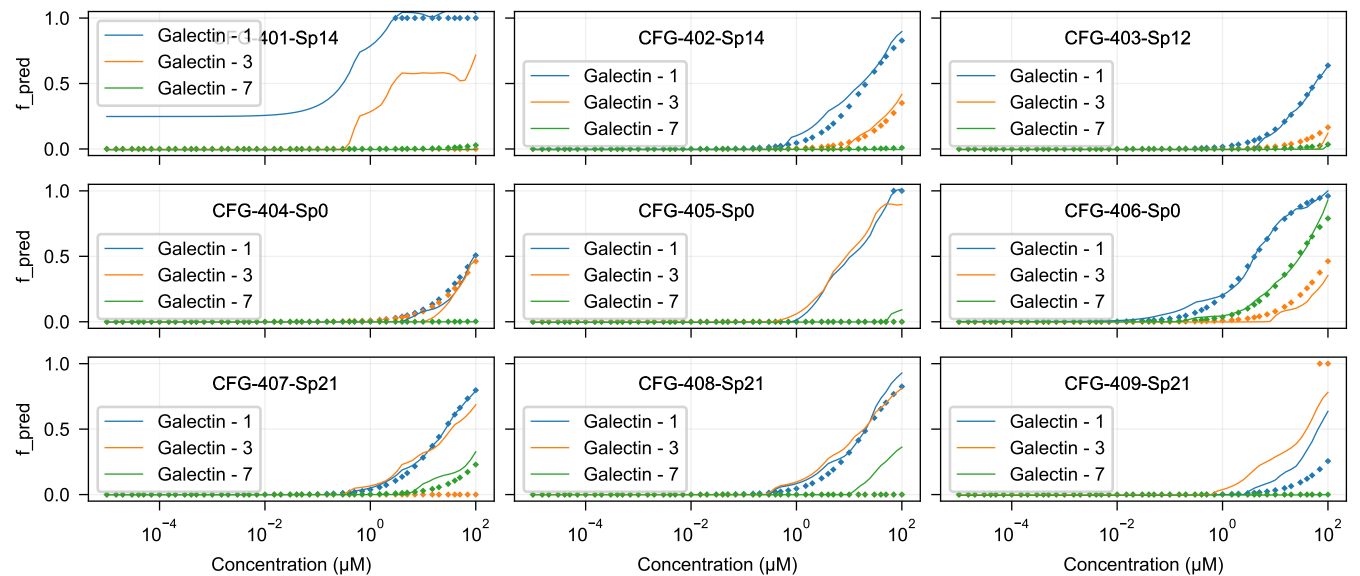

CFG-401-Sp14
CFG-402-Sp14
CFG-403-Sp12
CFG-404-Sp0
CFG-405-Sp0
CFG-406-Sp0
CFG-407-Sp21
CFG-408-Sp21
CFG-409-Sp21
Galectin - 1
Galectin - 3
Galectin - 7
f_pred
Concentration (μM)

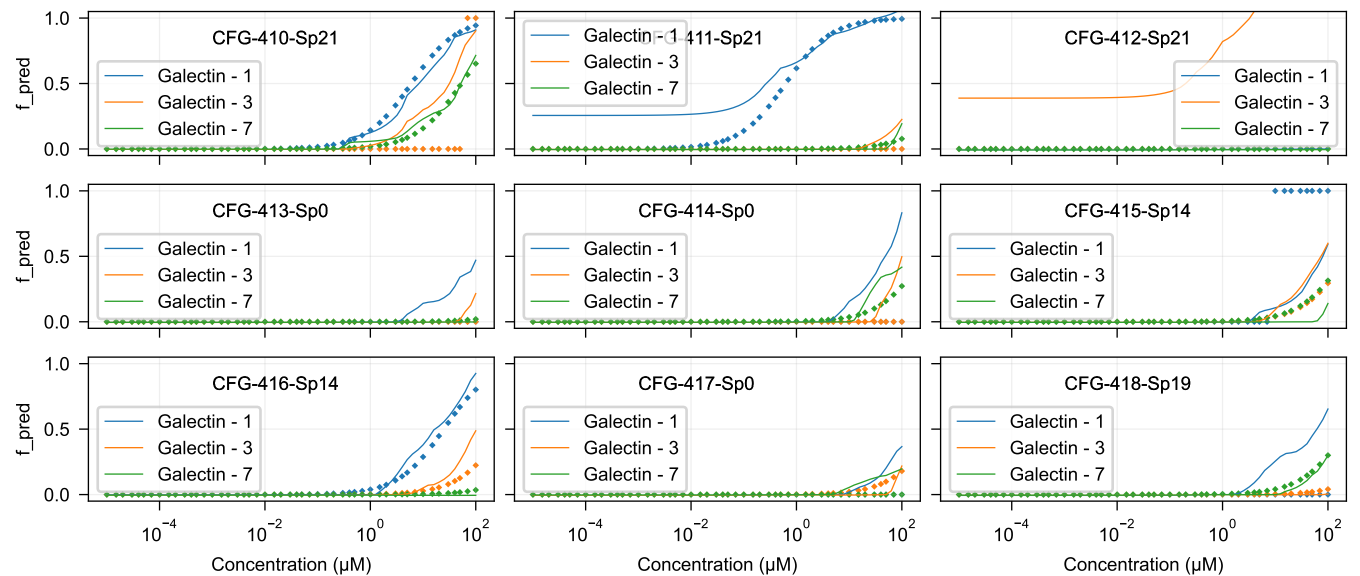

CFG-410-Sp21
CFG-411-Sp21
CFG-412-Sp21
CFG-413-Sp0
CFG-414-Sp0
CFG-415-Sp14
CFG-416-Sp14
CFG-417-Sp0
CFG-418-Sp19
Galectin - 1
Galectin - 3
Galectin - 7
f_pred
Concentration (μM)

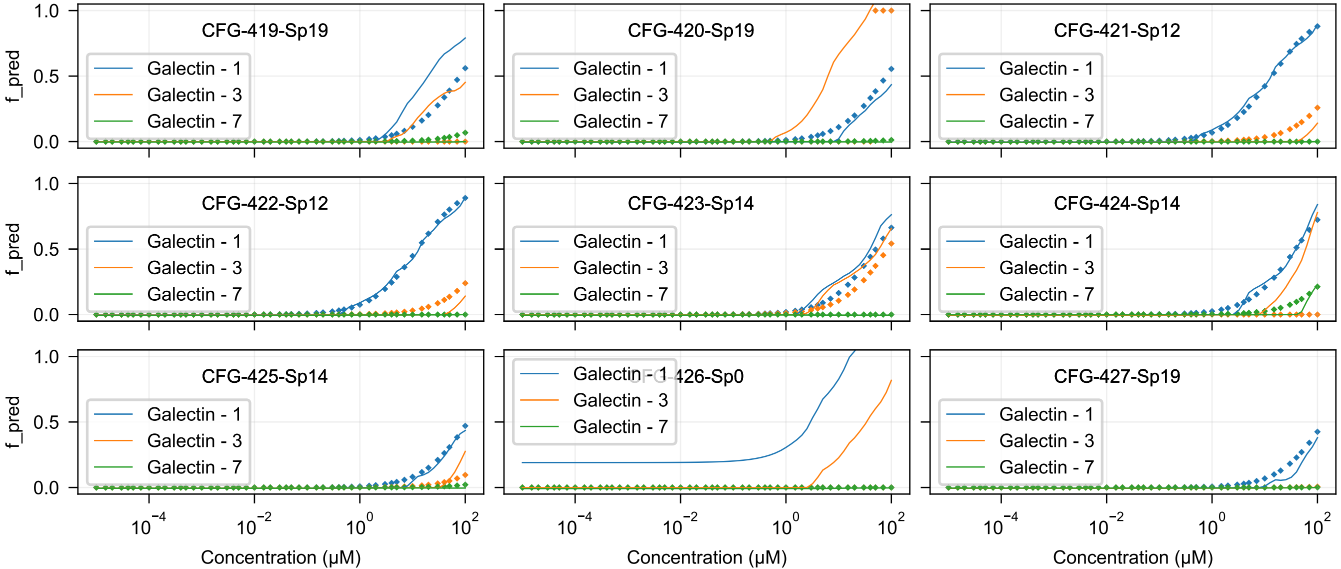
CFG-419-Sp19
CFG-420-Sp19
CFG-421-Sp12
CFG-422-Sp12
CFG-423-Sp14
CFG-424-Sp14
CFG-425-Sp14
CFG-426-Sp0
CFG-427-Sp19
Galectin - 1
Galectin - 3
Galectin - 7
f_pred
1.0
0.5
0.0
$10^{-4}$
$10^{-2}$
$10^{0}$
$10^{2}$
Concentration (μM)

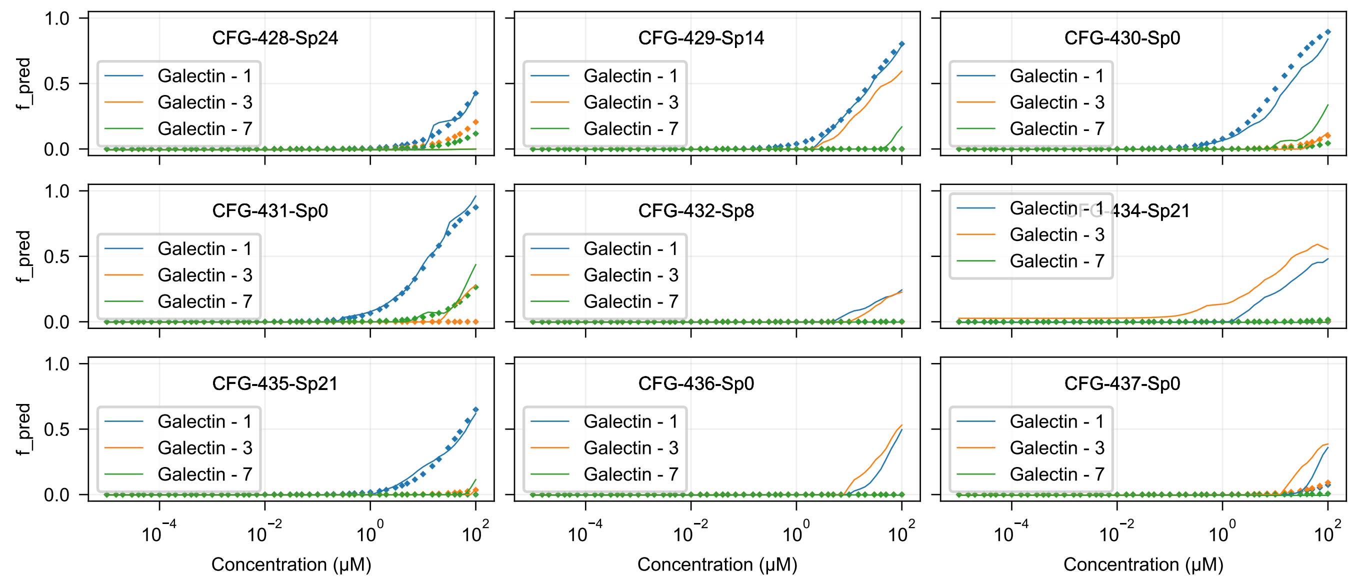
CFG-428-Sp24
CFG-429-Sp14
CFG-430-Sp0
CFG-431-Sp0
CFG-432-Sp8
CFG-434-Sp21
CFG-435-Sp21
CFG-436-Sp0
CFG-437-Sp0
Galectin - 1
Galectin - 3
Galectin - 7
f_pred
Concentration (μM)
0.0
0.5
1.0
$10^{-4}$
$10^{-2}$
$10^{0}$
$10^{2}$

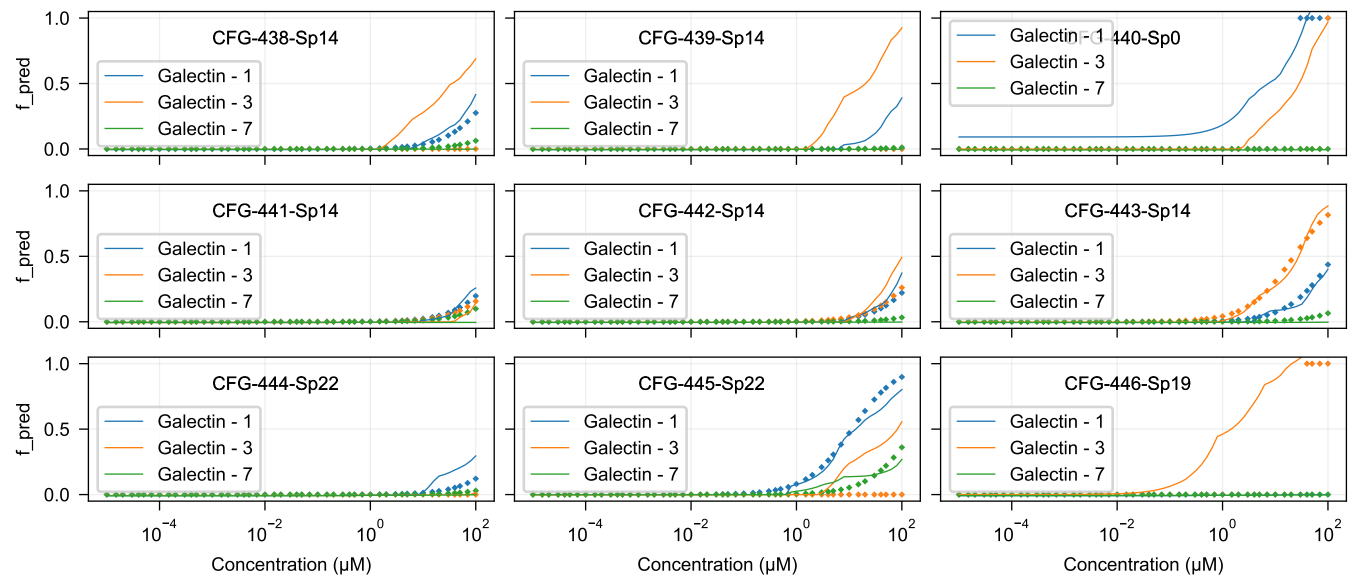

CFG-438-Sp14
CFG-439-Sp14
CFG-440-Sp0
CFG-441-Sp14
CFG-442-Sp14
CFG-443-Sp14
CFG-444-Sp22
CFG-445-Sp22
CFG-446-Sp19
Galectin - 1
Galectin - 3
Galectin - 7
f_pred
0.0
0.5
1.0
$10^{-4}$
$10^{-2}$
$10^{0}$
$10^{2}$
Concentration (μM)

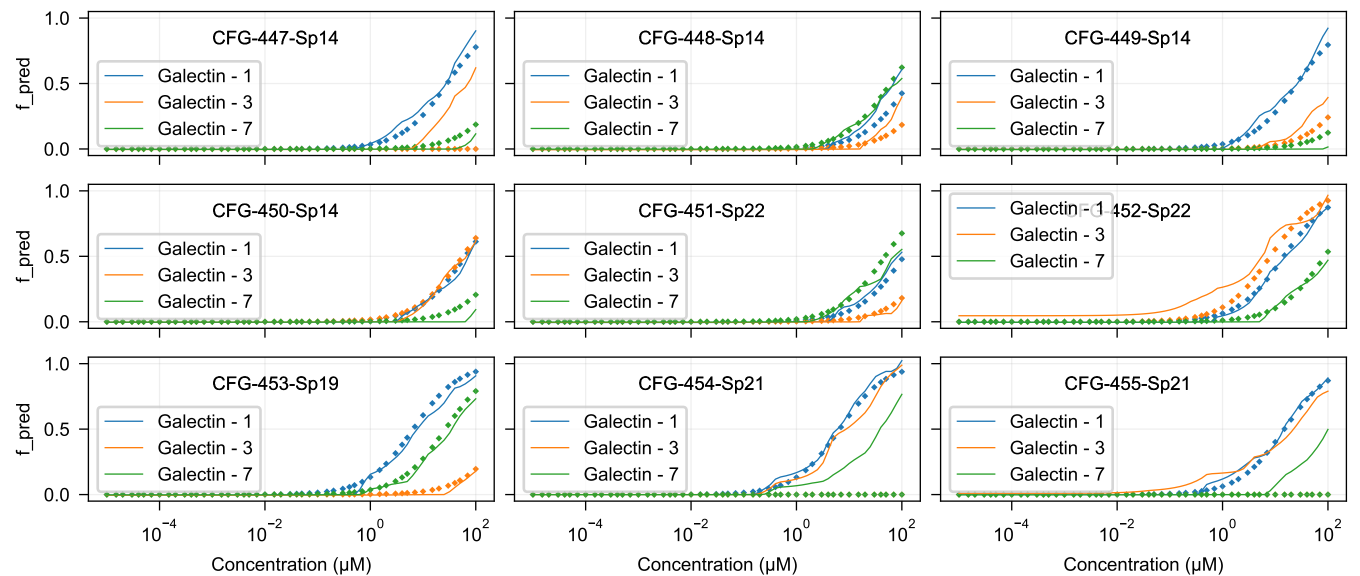
CFG-447-Sp14
CFG-448-Sp14
CFG-449-Sp14
CFG-450-Sp14
CFG-451-Sp22
CFG-452-Sp22
CFG-453-Sp19
CFG-454-Sp21
CFG-455-Sp21
Galectin - 1
Galectin - 3
Galectin - 7
f_pred
Concentration (μM)
1.0
0.5
0.0
$10^{-4}$
$10^{-2}$
$10^{0}$
$10^{2}$

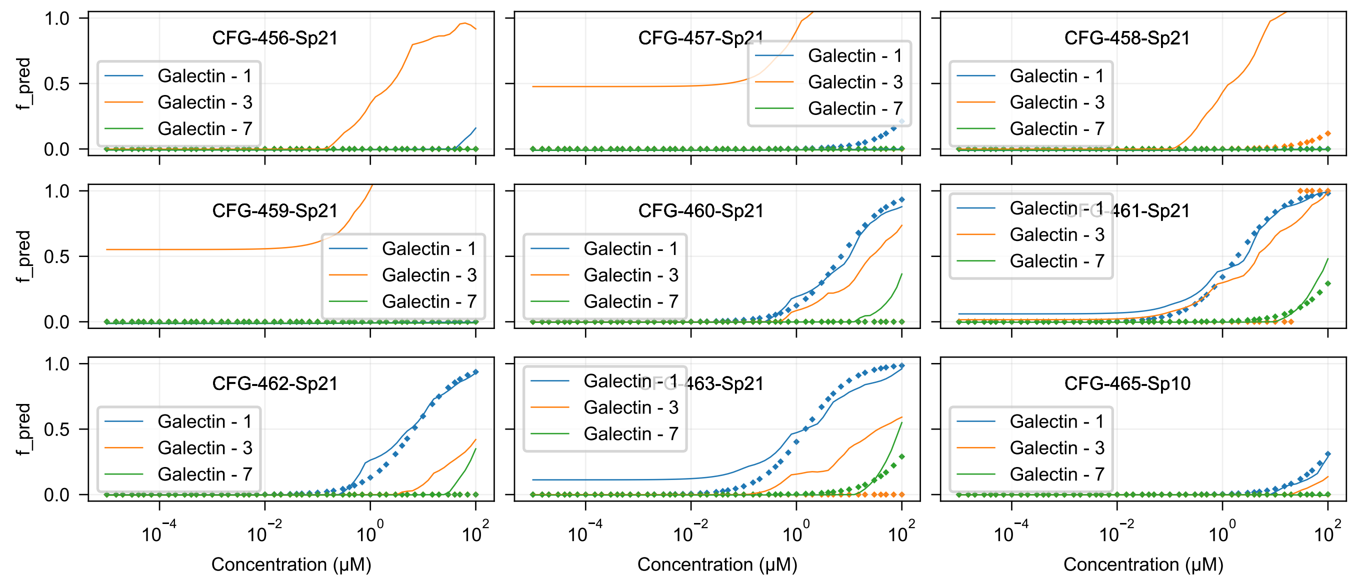
CFG-456-Sp21
CFG-457-Sp21
CFG-458-Sp21
CFG-459-Sp21
CFG-460-Sp21
CFG-461-Sp21
CFG-462-Sp21
CFG-463-Sp21
CFG-465-Sp10
Galectin - 1
Galectin - 3
Galectin - 7
f_pred
Concentration (μM)

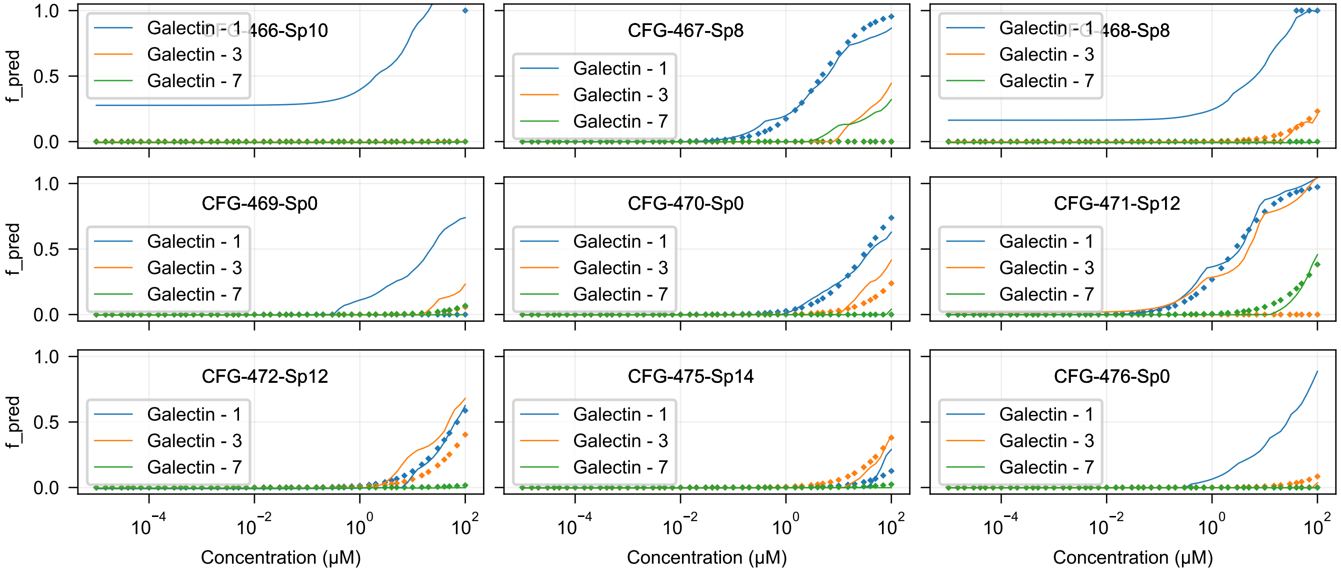
CFG-466-Sp10
CFG-467-Sp8
CFG-468-Sp8
CFG-469-Sp0
CFG-470-Sp0
CFG-471-Sp12
CFG-472-Sp12
CFG-475-Sp14
CFG-476-Sp0
Galectin - 1
Galectin - 3
Galectin - 7
f_pred
Concentration (μM)

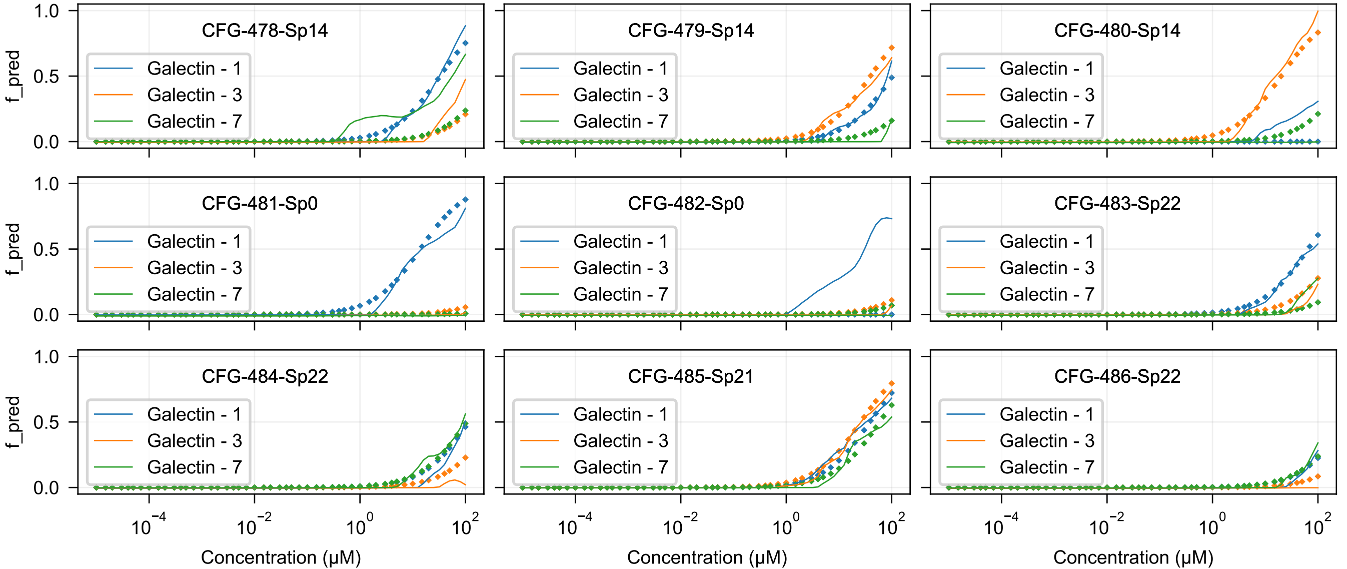
CFG-478-Sp14
CFG-479-Sp14
CFG-480-Sp14
CFG-481-Sp0
CFG-482-Sp0
CFG-483-Sp22
CFG-484-Sp22
CFG-485-Sp21
CFG-486-Sp22
Galectin - 1
Galectin - 3
Galectin - 7
f_pred
0.0
0.5
1.0
$10^{-4}$
$10^{-2}$
$10^{0}$
$10^{2}$
Concentration (μM)

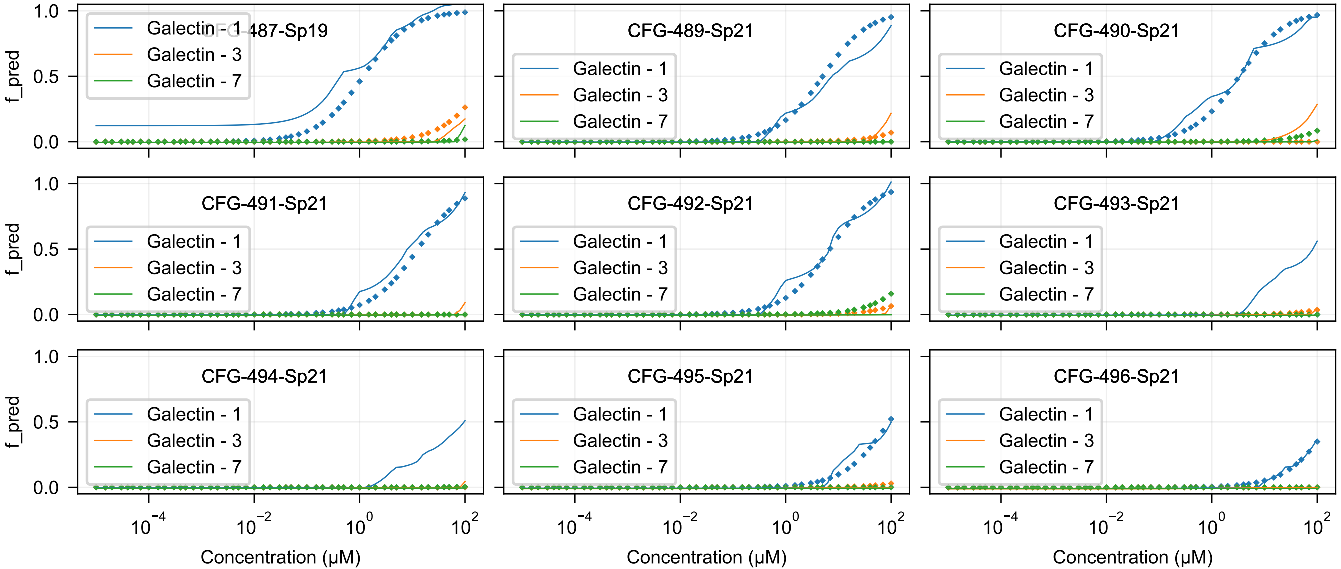
CFG-487-Sp19
CFG-489-Sp21
CFG-490-Sp21
CFG-491-Sp21
CFG-492-Sp21
CFG-493-Sp21
CFG-494-Sp21
CFG-495-Sp21
CFG-496-Sp21
Galectin - 1
Galectin - 3
Galectin - 7
f_pred
1.0
0.5
0.0
$10^{-4}$
$10^{-2}$
$10^{0}$
$10^{2}$
Concentration (μM)

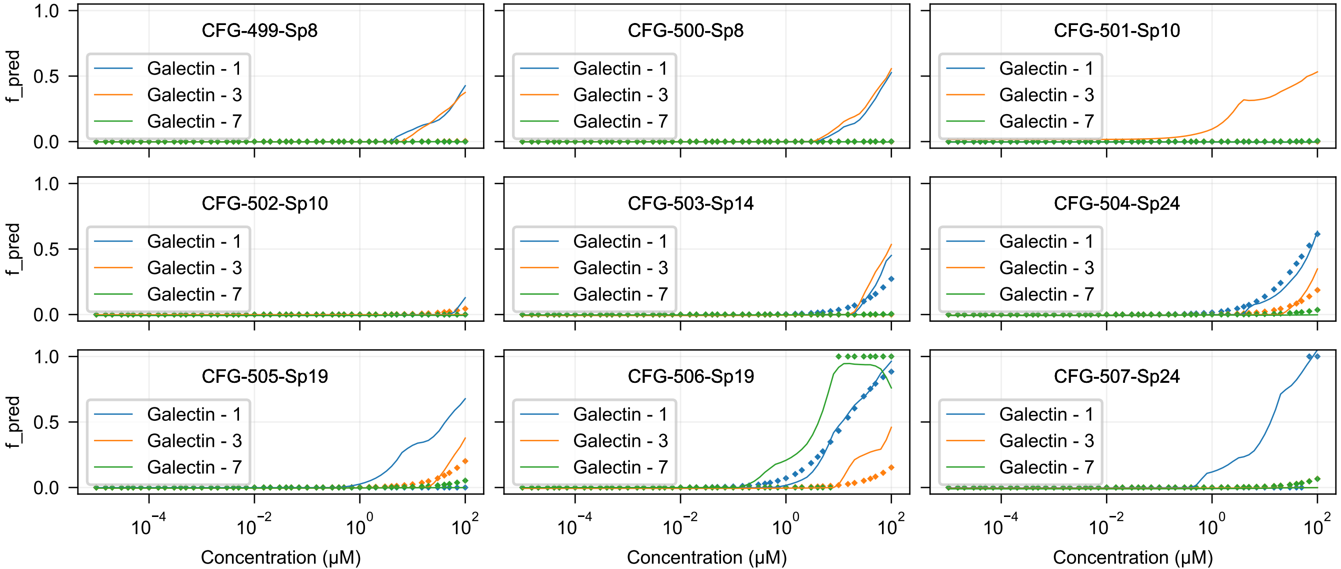

CFG-499-Sp8
CFG-500-Sp8
CFG-501-Sp10
CFG-502-Sp10
CFG-503-Sp14
CFG-504-Sp24
CFG-505-Sp19
CFG-506-Sp19
CFG-507-Sp24
Galectin - 1
Galectin - 3
Galectin - 7
f_pred
Concentration (μM)
1.0
0.5
0.0
$10^{-4}$
$10^{-2}$
$10^{0}$
$10^{2}$

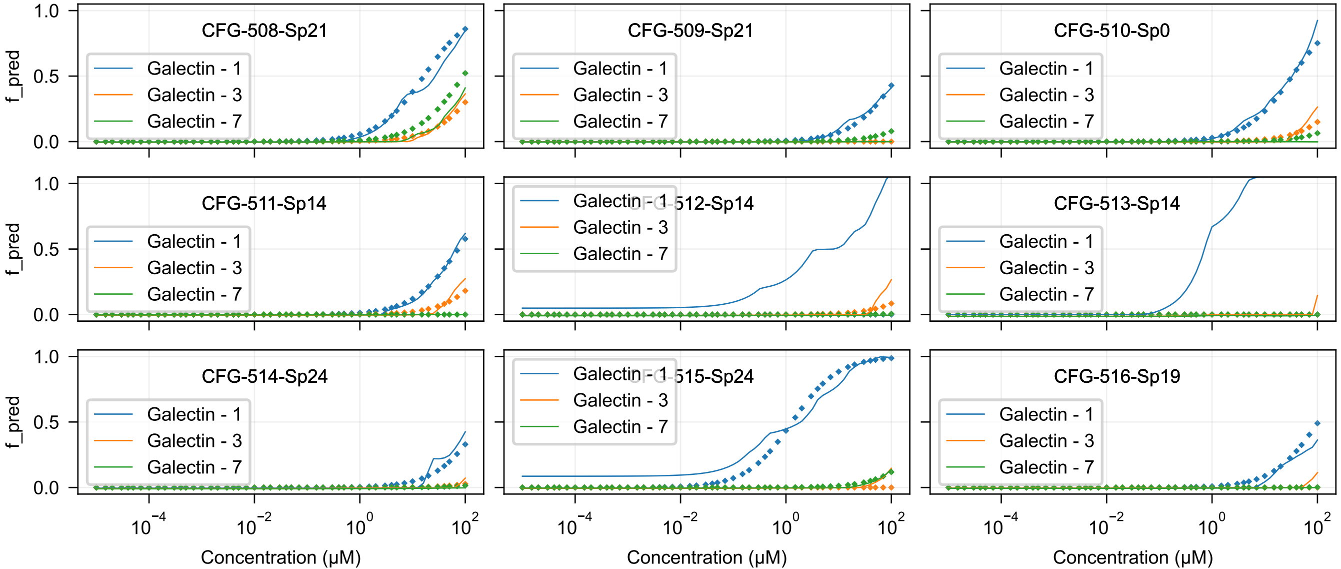
CFG-508-Sp21
CFG-509-Sp21
CFG-510-Sp0
CFG-511-Sp14
CFG-512-Sp14
CFG-513-Sp14
CFG-514-Sp24
CFG-515-Sp24
CFG-516-Sp19
Galectin - 1
Galectin - 3
Galectin - 7
f_pred
Concentration (μM)
1.0
0.5
0.0
$10^{-4}$
$10^{-2}$
$10^{0}$
$10^{2}$

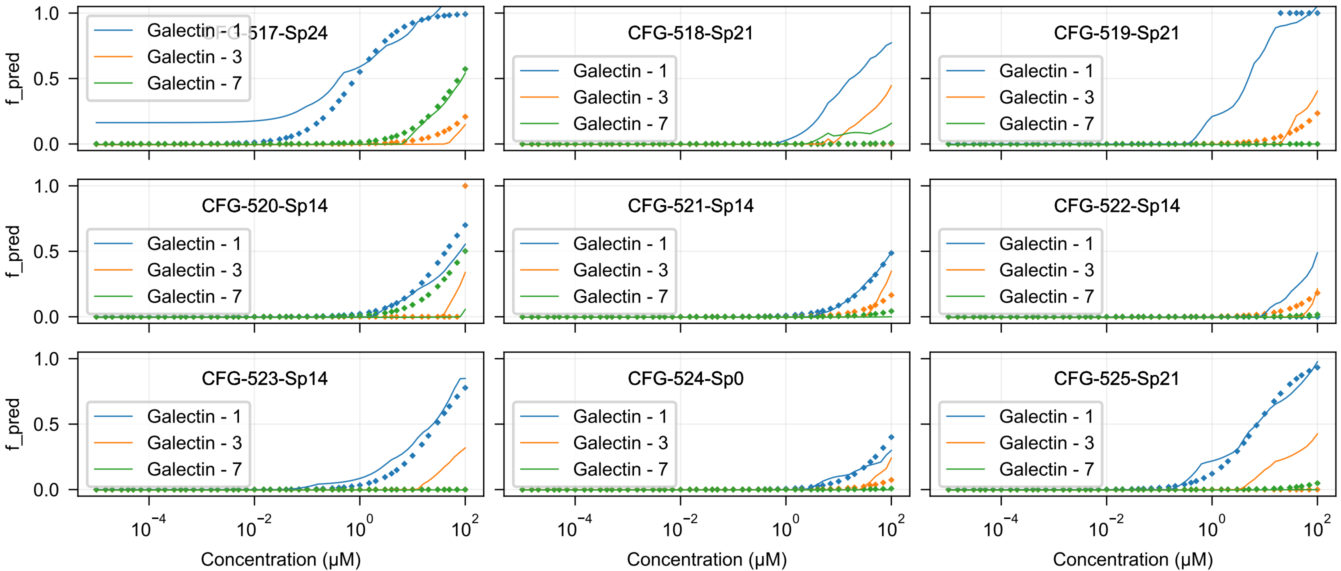
CFG-517-Sp24
CFG-518-Sp21
CFG-519-Sp21
CFG-520-Sp14
CFG-521-Sp14
CFG-522-Sp14
CFG-523-Sp14
CFG-524-Sp0
CFG-525-Sp21
Galectin - 1
Galectin - 3
Galectin - 7
f_pred
0.0
0.5
1.0
$10^{-4}$
$10^{-2}$
$10^{0}$
$10^{2}$
Concentration (μM)

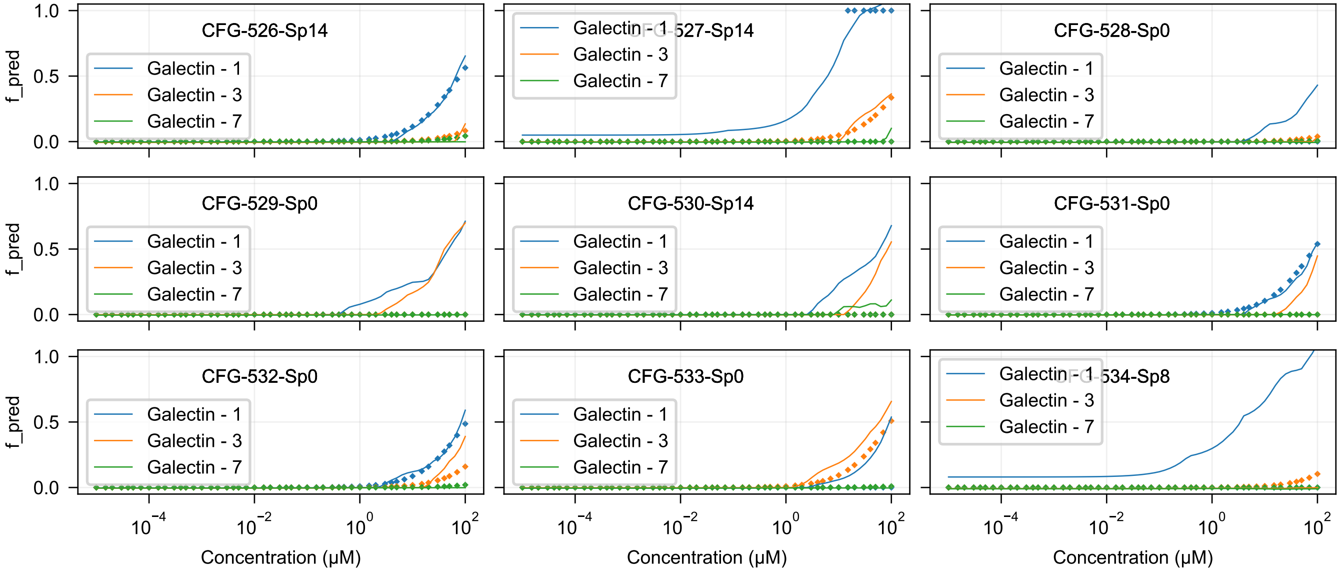
CFG-526-Sp14
CFG-527-Sp14
CFG-528-Sp0
CFG-529-Sp0
CFG-530-Sp14
CFG-531-Sp0
CFG-532-Sp0
CFG-533-Sp0
CFG-534-Sp8
Galectin - 1
Galectin - 3
Galectin - 7
f_pred
1.0
0.5
0.0
$10^{-4}$
$10^{-2}$
$10^{0}$
$10^{2}$
Concentration (μM)

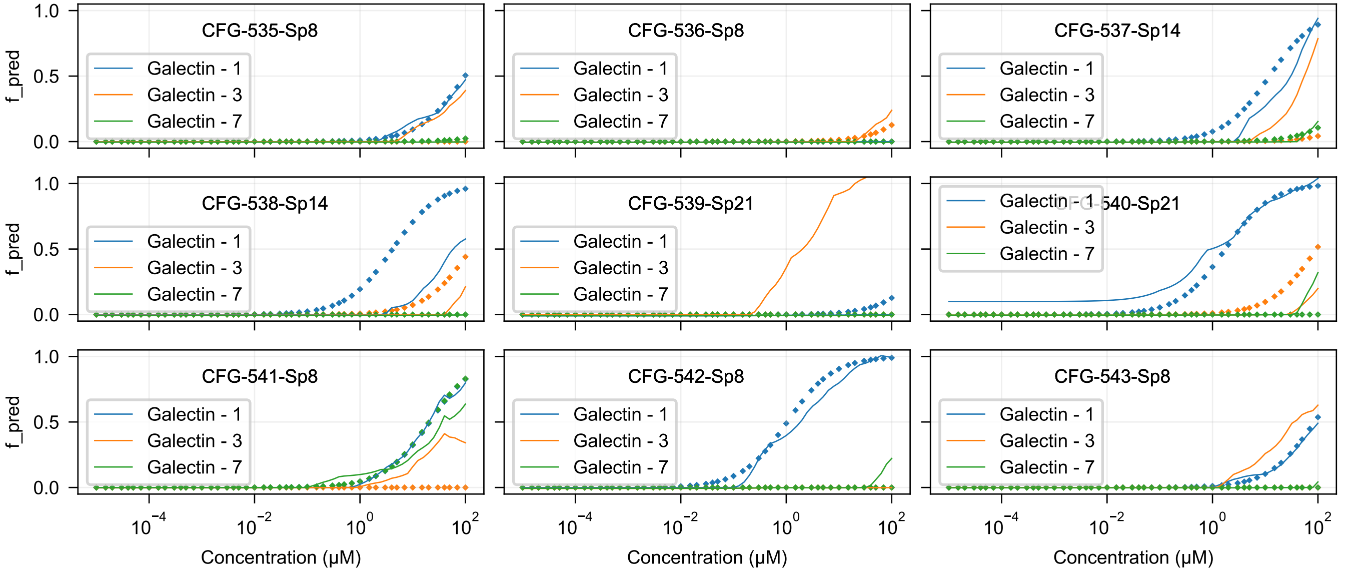

CFG-535-Sp8
CFG-536-Sp8
CFG-537-Sp14
CFG-538-Sp14
CFG-539-Sp21
CFG-540-Sp21
CFG-541-Sp8
CFG-542-Sp8
CFG-543-Sp8
Galectin - 1
Galectin - 3
Galectin - 7
f_pred
Concentration (μM)

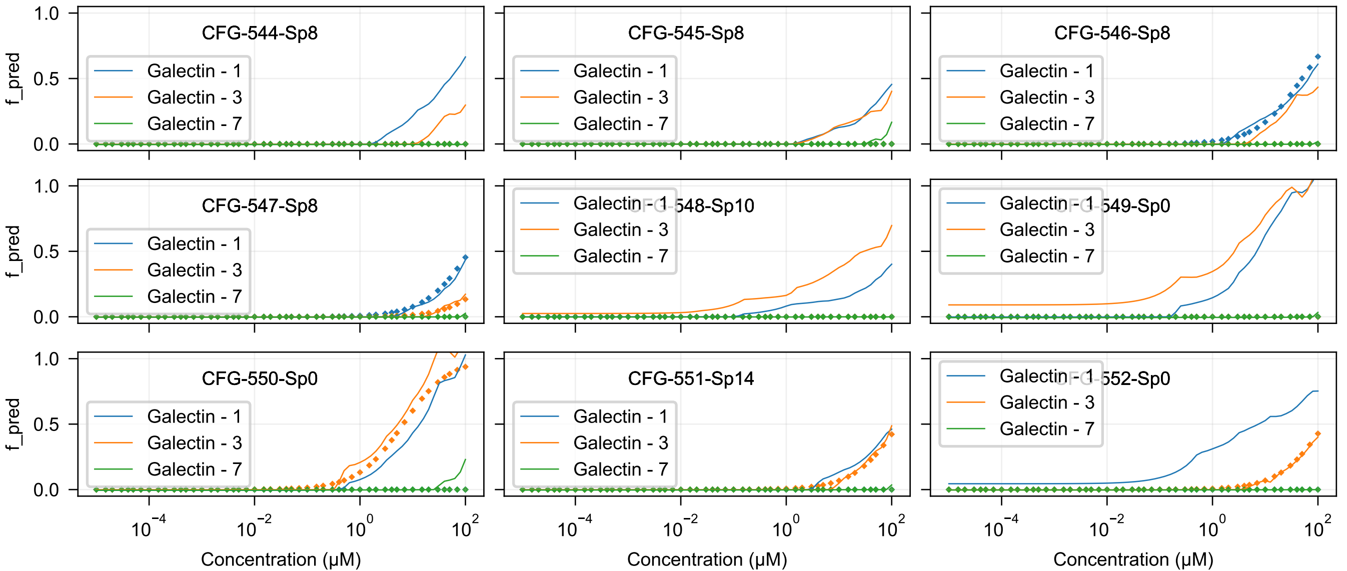
CFG-544-Sp8
CFG-545-Sp8
CFG-546-Sp8
CFG-547-Sp8
CFG-548-Sp10
CFG-549-Sp0
CFG-550-Sp0
CFG-551-Sp14
CFG-552-Sp0
Galectin - 1
Galectin - 3
Galectin - 7
f_pred
1.0
0.5
0.0
$10^{-4}$
$10^{-2}$
$10^{0}$
$10^{2}$
Concentration (μM)

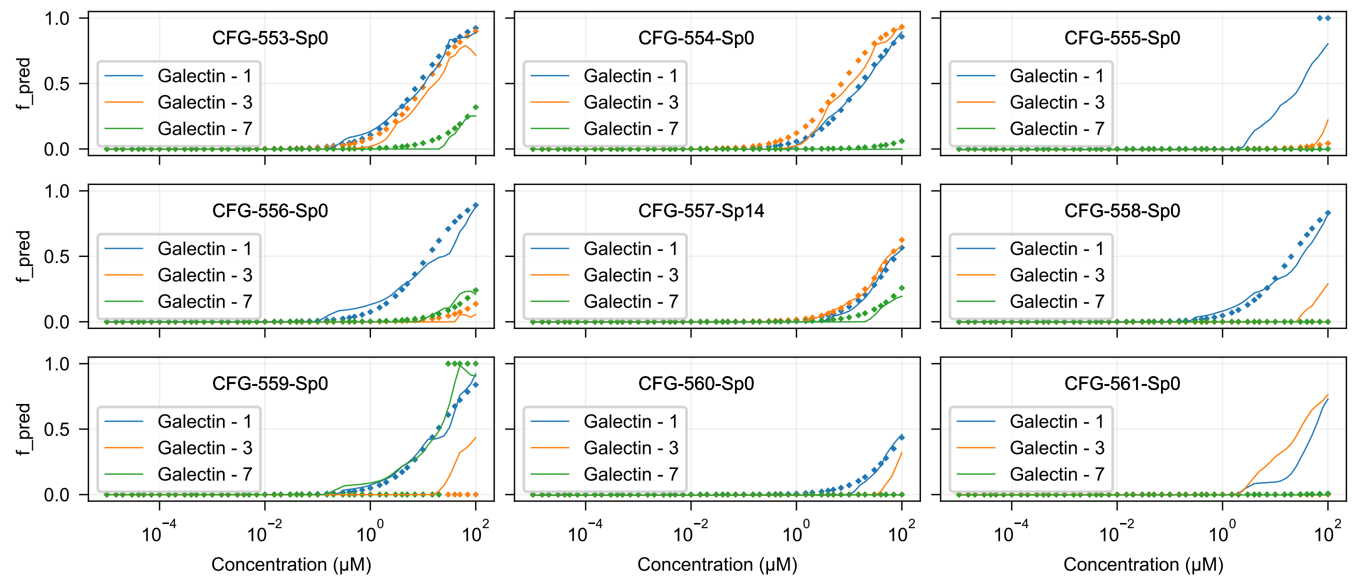
CFG-553-Sp0
CFG-554-Sp0
CFG-555-Sp0
CFG-556-Sp0
CFG-557-Sp14
CFG-558-Sp0
CFG-559-Sp0
CFG-560-Sp0
CFG-561-Sp0
Galectin - 1
Galectin - 3
Galectin - 7
f_pred
Concentration (μM)
1.0
0.5
0.0
$10^{-4}$
$10^{-2}$
$10^{0}$
$10^{2}$

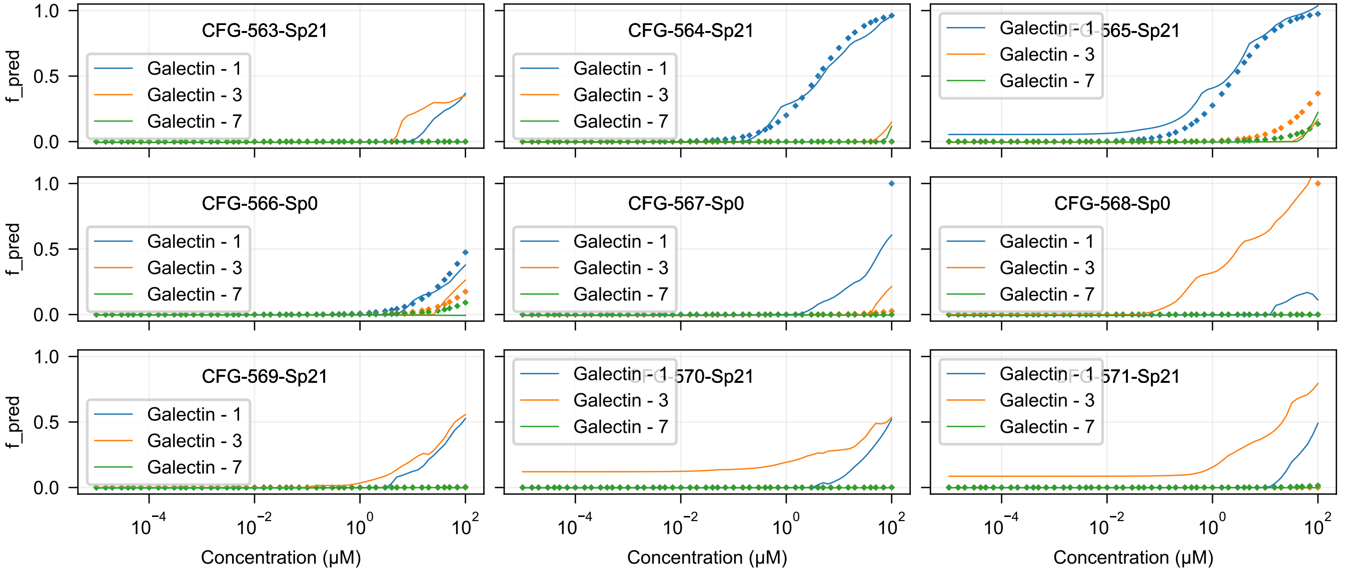
CFG-563-Sp21
CFG-564-Sp21
CFG-565-Sp21
CFG-566-Sp0
CFG-567-Sp0
CFG-568-Sp0
CFG-569-Sp21
CFG-570-Sp21
CFG-571-Sp21
Galectin - 1
Galectin - 3
Galectin - 7
f_pred
Concentration (μM)
1.0
0.5
0.0
$10^{-4}$
$10^{-2}$
$10^{0}$
$10^{2}$

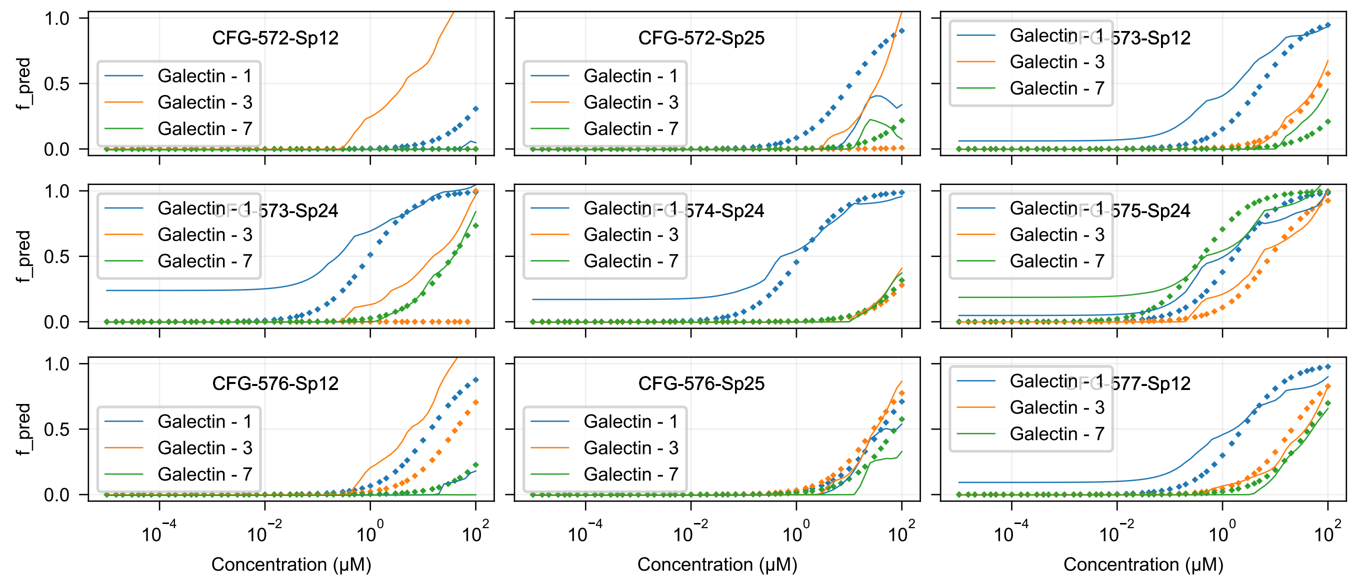
CFG-572-Sp12
CFG-572-Sp25
CFG-573-Sp12
CFG-573-Sp24
CFG-574-Sp24
CFG-575-Sp24
CFG-576-Sp12
CFG-576-Sp25
CFG-577-Sp12
Galectin - 1
Galectin - 3
Galectin - 7
f_pred
Concentration (μM)

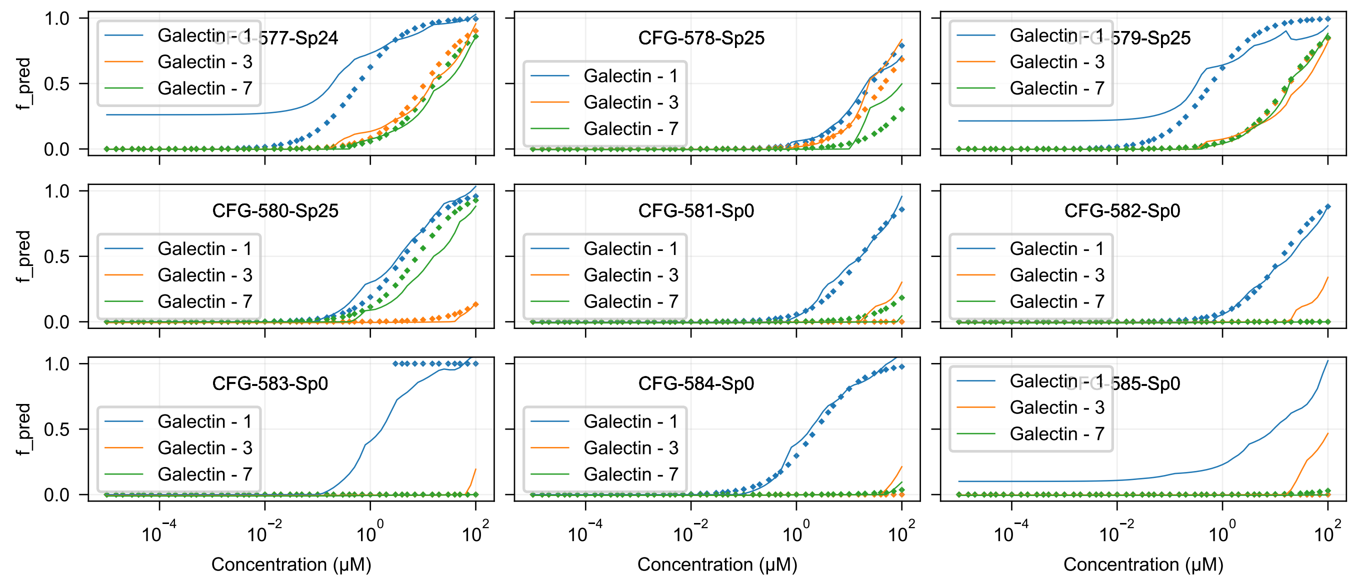
CFG-577-Sp24
CFG-578-Sp25
CFG-579-Sp25
CFG-580-Sp25
CFG-581-Sp0
CFG-582-Sp0
CFG-583-Sp0
CFG-584-Sp0
CFG-585-Sp0
Galectin - 1
Galectin - 3
Galectin - 7
f_pred
0.0
0.5
1.0
$10^{-4}$
$10^{-2}$
$10^{0}$
$10^{2}$
Concentration (μM)

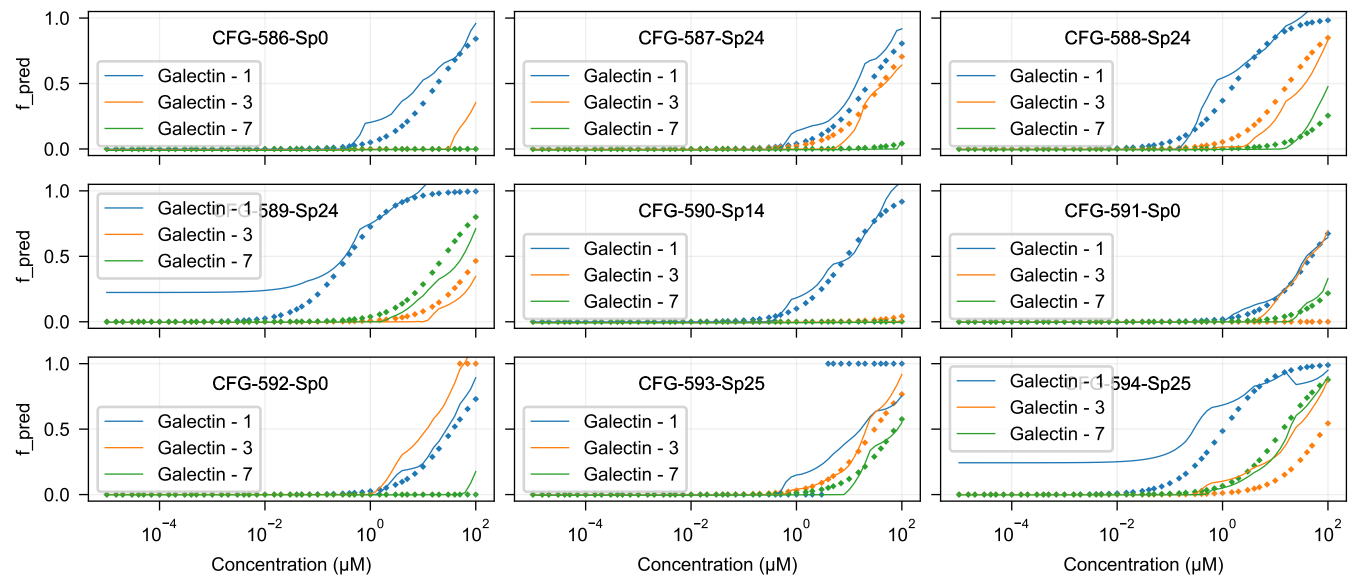

CFG-586-Sp0
CFG-587-Sp24
CFG-588-Sp24
CFG-589-Sp24
CFG-590-Sp14
CFG-591-Sp0
CFG-592-Sp0
CFG-593-Sp25
CFG-594-Sp25
Galectin - 1
Galectin - 3
Galectin - 7
f_pred
Concentration (μM)

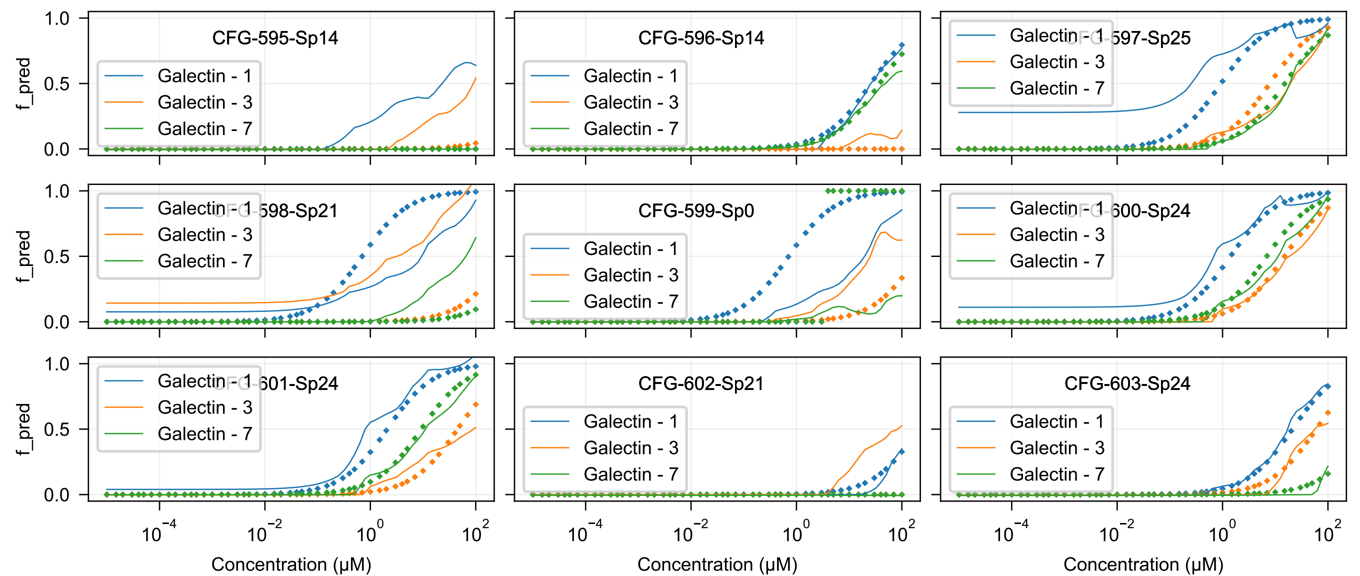
CFG-595-Sp14
CFG-596-Sp14
CFG-597-Sp25
CFG-598-Sp21
CFG-599-Sp0
CFG-600-Sp24
CFG-601-Sp24
CFG-602-Sp21
CFG-603-Sp24
Galectin - 1
Galectin - 3
Galectin - 7
f_pred
Concentration (μM)

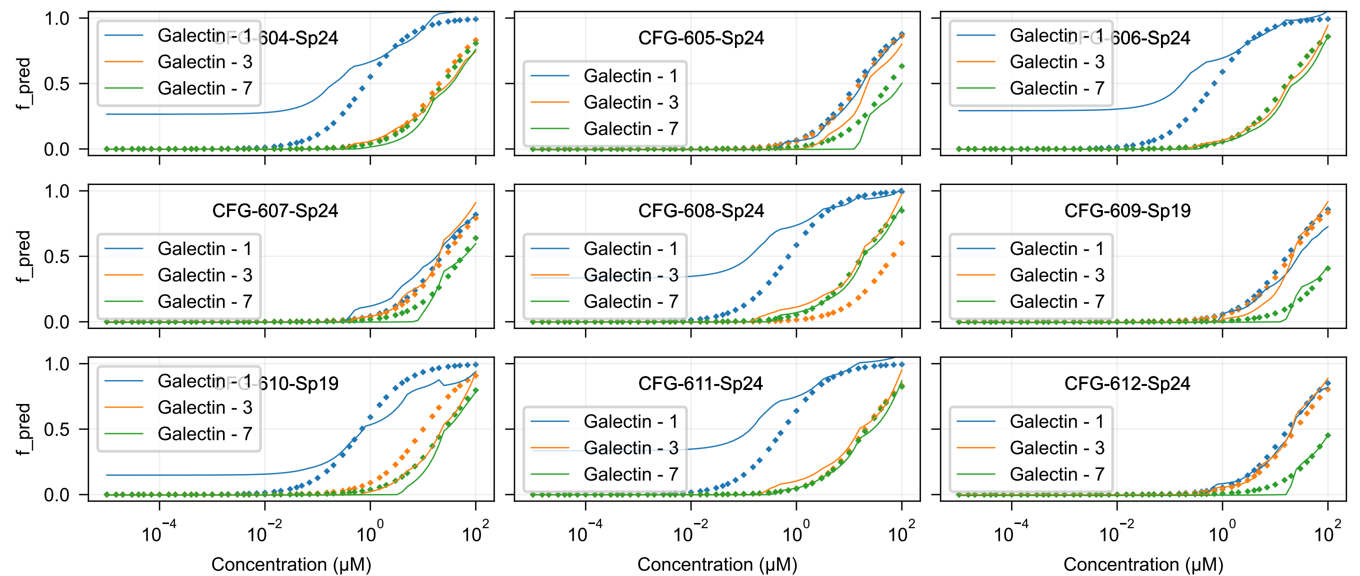

CFG-604-Sp24
CFG-605-Sp24
CFG-606-Sp24
CFG-607-Sp24
CFG-608-Sp24
CFG-609-Sp19
CFG-610-Sp19
CFG-611-Sp24
CFG-612-Sp24
Galectin - 1
Galectin - 3
Galectin - 7
f_pred
Concentration (μM)

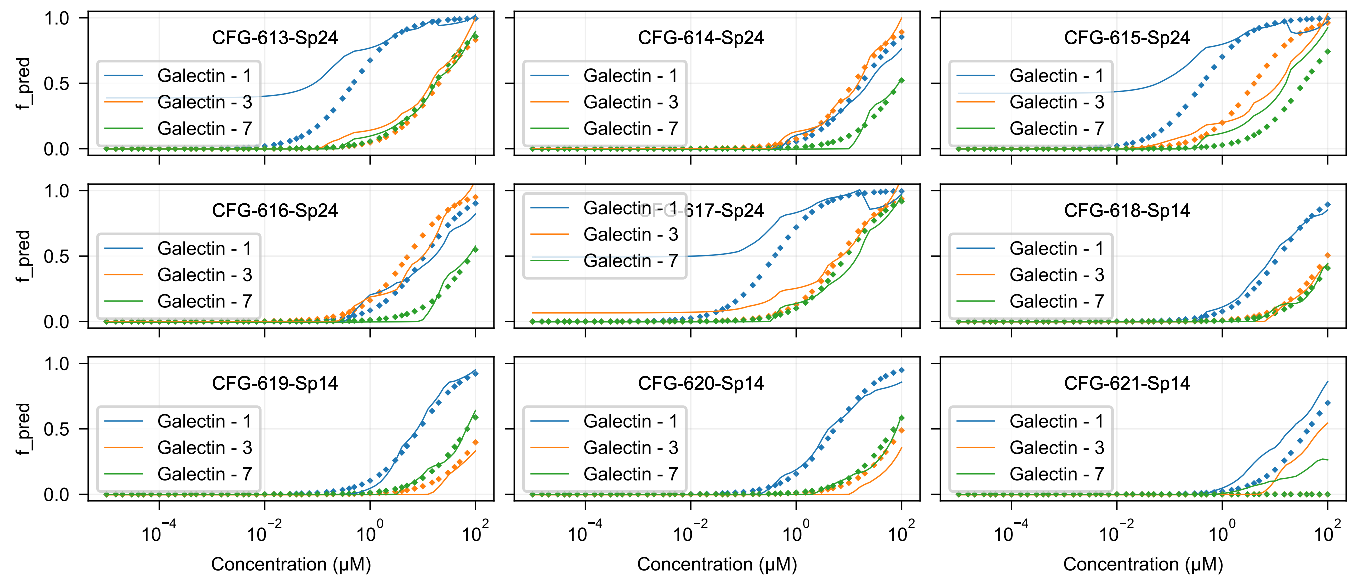
CFG-613-Sp24
CFG-614-Sp24
CFG-615-Sp24
CFG-616-Sp24
CFG-617-Sp24
CFG-618-Sp14
CFG-619-Sp14
CFG-620-Sp14
CFG-621-Sp14
Galectin - 1
Galectin - 3
Galectin - 7
f_pred
0.0
0.5
1.0
$10^{-4}$
$10^{-2}$
$10^{0}$
$10^{2}$
Concentration (μM)

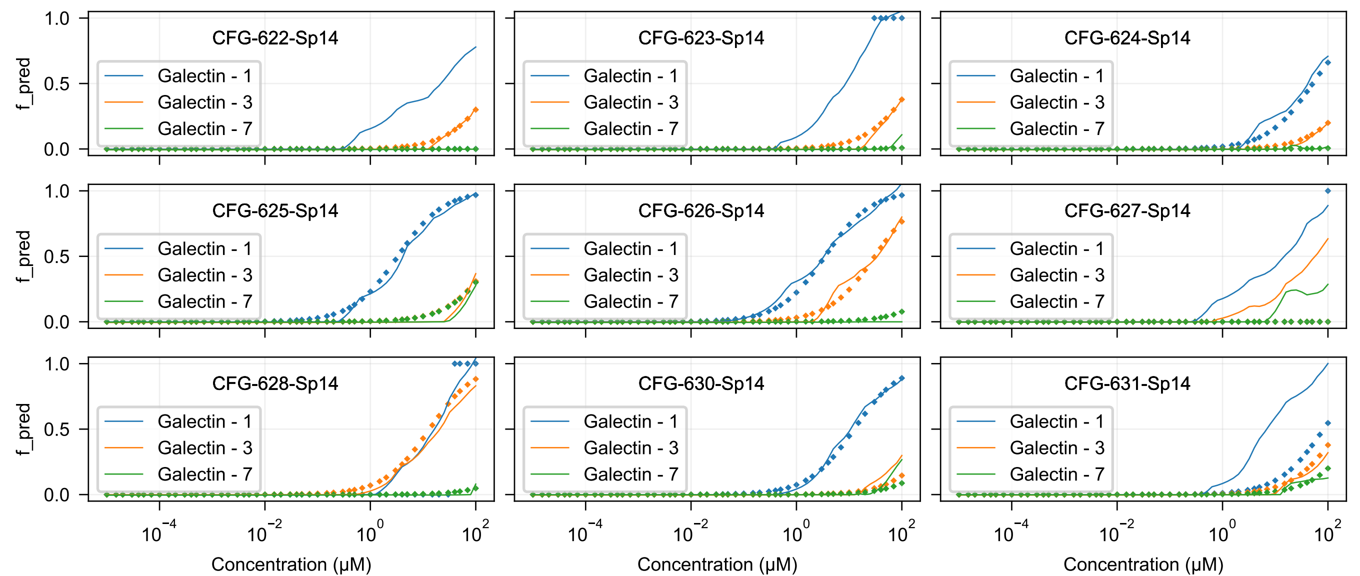

CFG-622-Sp14
CFG-623-Sp14
CFG-624-Sp14
CFG-625-Sp14
CFG-626-Sp14
CFG-627-Sp14
CFG-628-Sp14
CFG-630-Sp14
CFG-631-Sp14
Galectin - 1
Galectin - 3
Galectin - 7
f_pred
1.0
0.5
0.0
$10^{-4}$
$10^{-2}$
$10^{0}$
$10^{2}$
Concentration (μM)

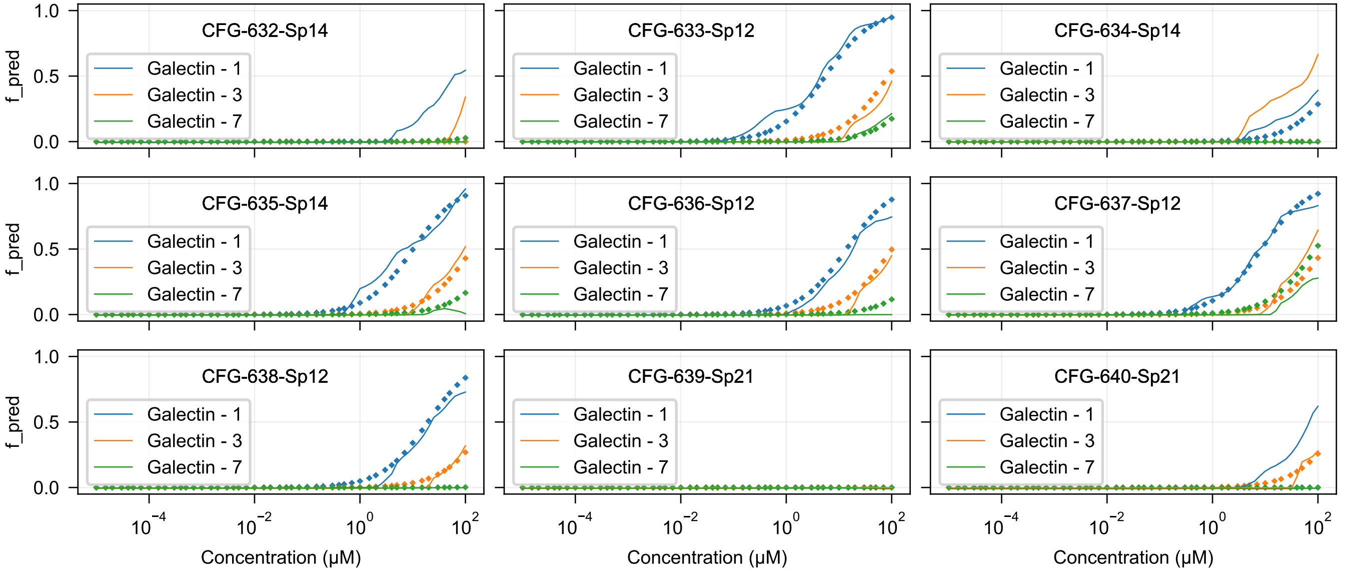

CFG-632-Sp14
CFG-633-Sp12
CFG-634-Sp14
CFG-635-Sp14
CFG-636-Sp12
CFG-637-Sp12
CFG-638-Sp12
CFG-639-Sp21
CFG-640-Sp21
Galectin - 1
Galectin - 3
Galectin - 7
f_pred
0.0
0.5
1.0
$10^{-4}$
$10^{-2}$
$10^{0}$
$10^{2}$
Concentration (μM)
