## Supplementary material for "Atom-level Machine Learning of Protein-Glycan Interactions and Cross-chiral Recognition in Glycobiology": Mannose Enantiomer Prediction Plots

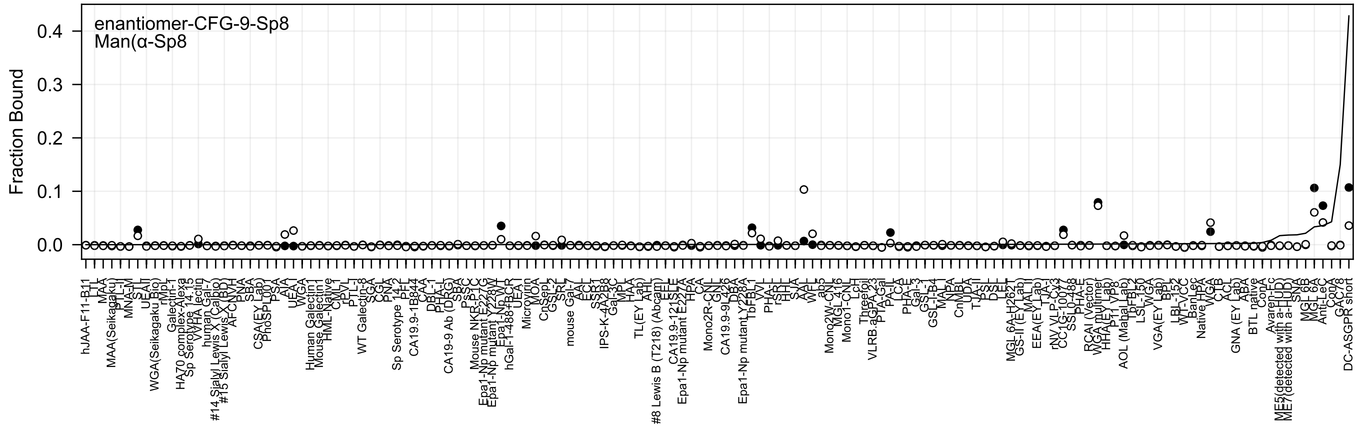
enantiomer-CFG-9-Sp8
Man(α-Sp8
Fraction Bound
0.4
0.3
0.2
0.1
0.0

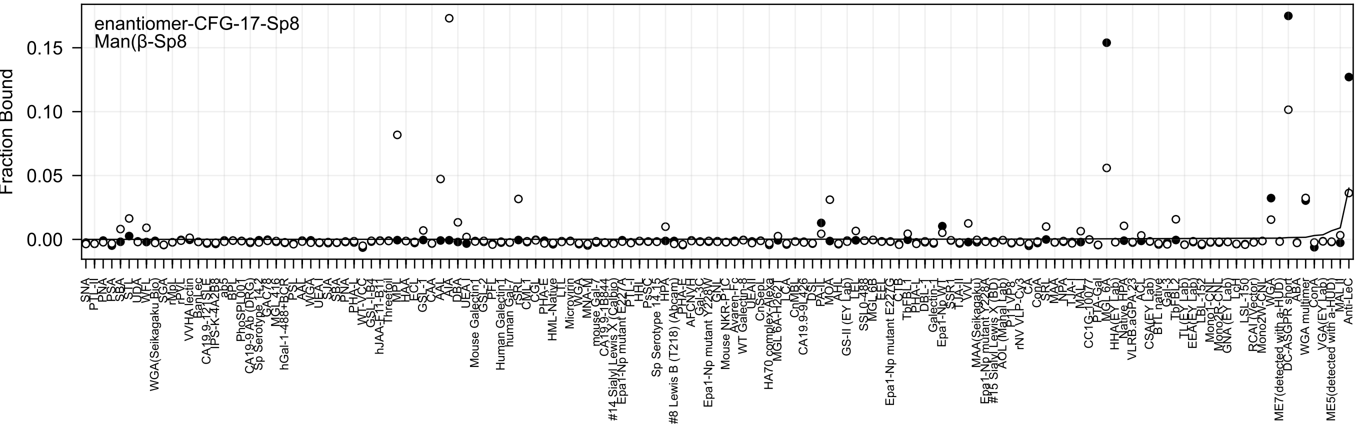
enantiomer-CFG-17-Sp8
Man(β-Sp8
Fraction Bound
0.15
0.10
0.05
0.00

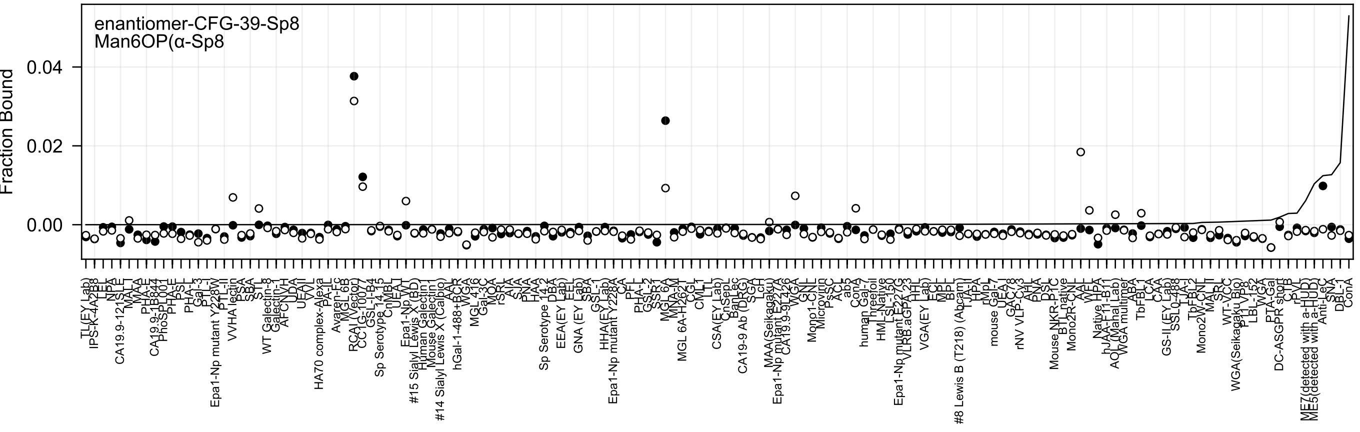

enantiomer-CFG-39-Sp8
Man6OP(α-Sp8
Fraction Bound
0.04
0.02
0.00

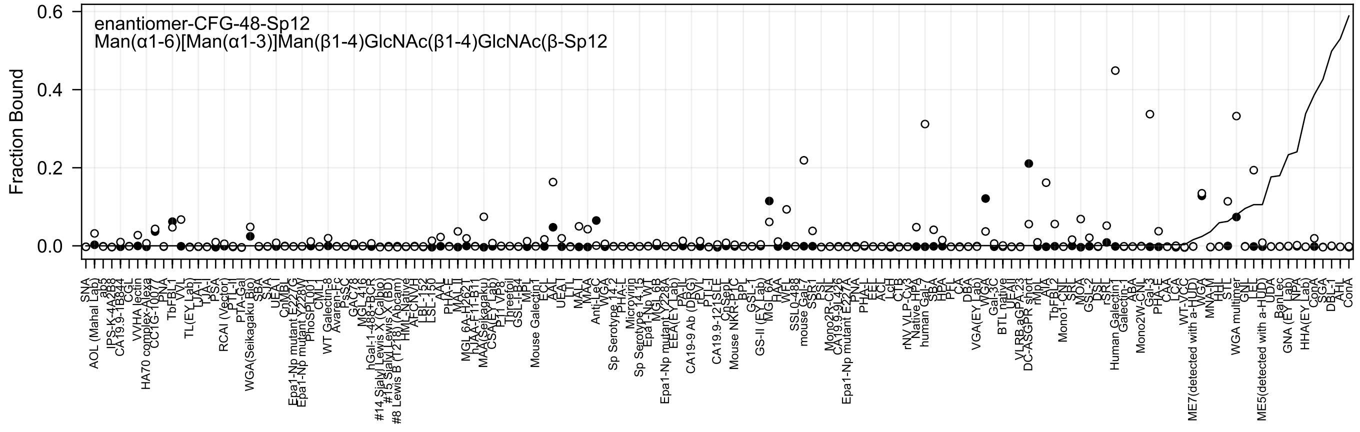
enantiomer-CFG-48-Sp12
Man(α1-6)[Man(α1-3)]Man(β1-4)GlcNAc(β1-4)GlcNAc(β-Sp12
Fraction Bound
0.6
0.4
0.2
0.0

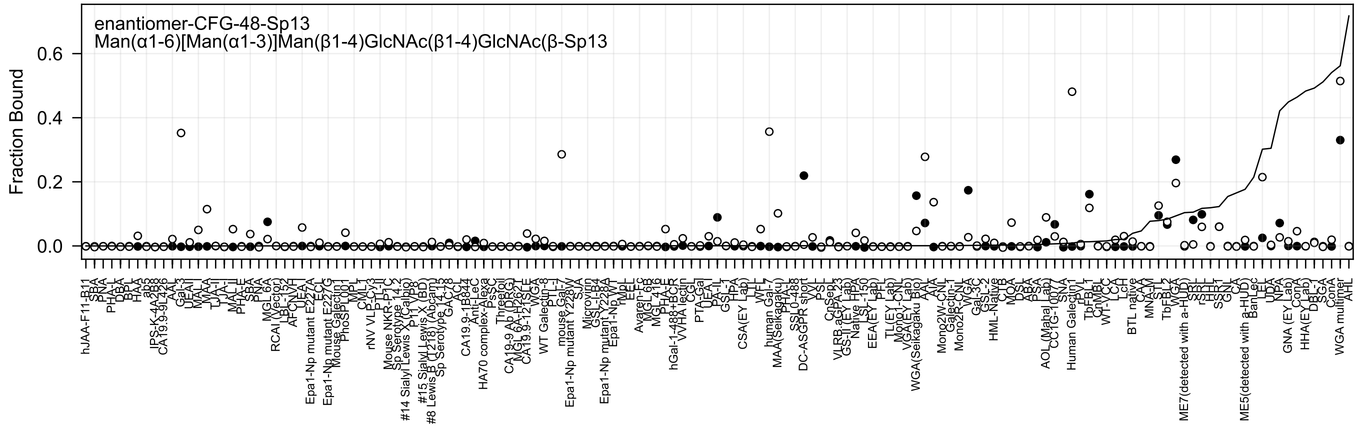
enantiomer-CFG-48-Sp13
Man(α1-6)[Man(α1-3)]Man(β1-4)GlcNAc(β1-4)GlcNAc(β-Sp13
Fraction Bound
0.6
0.4
0.2
0.0

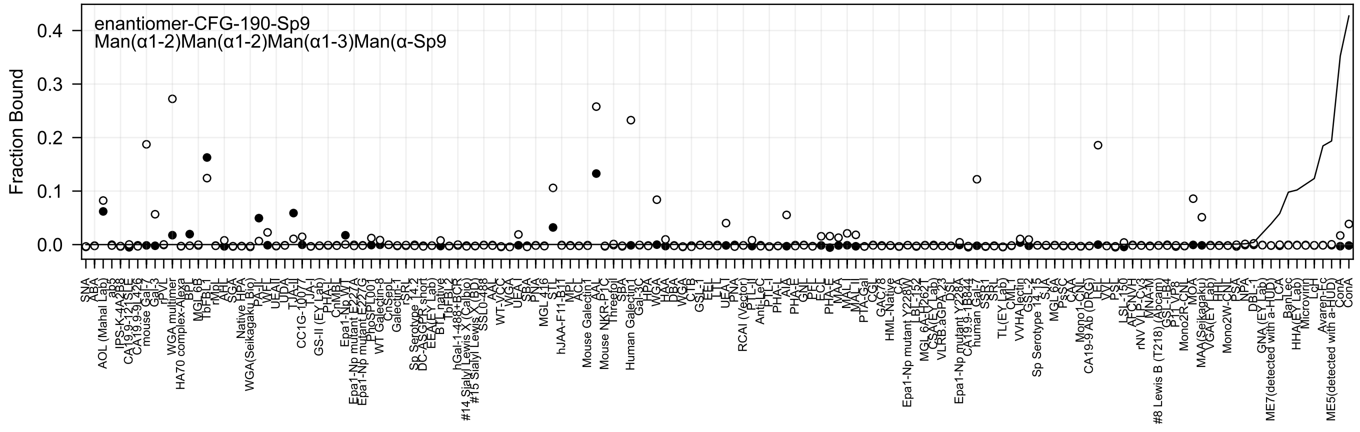
enantiomer-CFG-190-Sp9
Man(α1-2)Man(α1-2)Man(α1-3)Man(α-Sp9
Fraction Bound
0.4
0.3
0.2
0.1
0.0

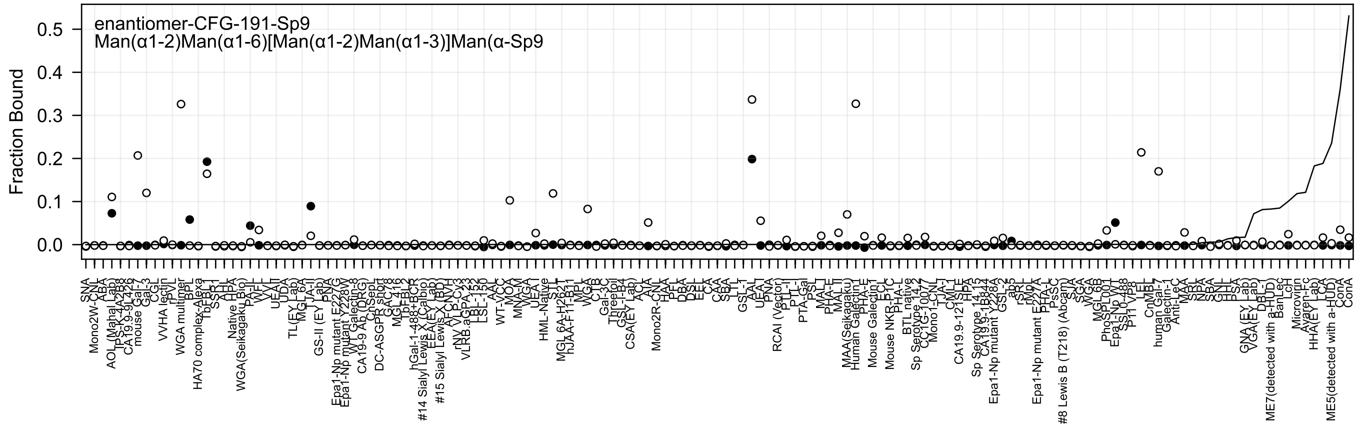
enantiomer-CFG-191-Sp9
Man(α1-2)Man(α1-6)[Man(α1-2)Man(α1-3)]Man(α-Sp9
Fraction Bound
0.5
0.4
0.3
0.2
0.1
0.0

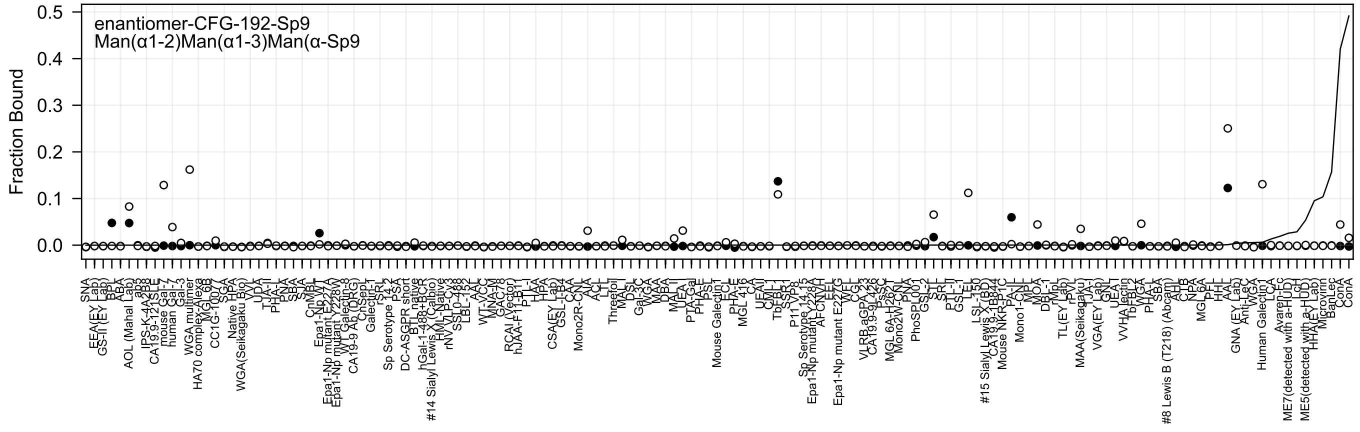
enantiomer-CFG-192-Sp9
Man(α1-2)Man(α1-3)Man(α-Sp9
Fraction Bound
0.5
0.4
0.3
0.2
0.1
0.0

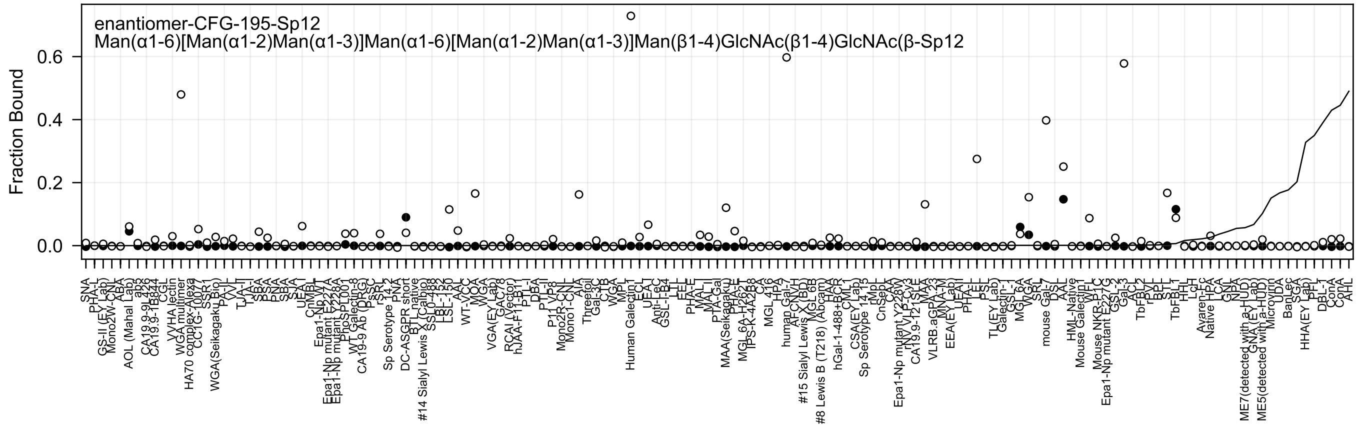
enantiomer-CFG-195-Sp12
Man(α1-6)[Man(α1-2)Man(α1-3)]Man(α1-6)[Man(α1-2)Man(α1-3)]Man(β1-4)GlcNAc(β1-4)GlcNAc(β-Sp12
Fraction Bound
0.6
0.4
0.2
0.0

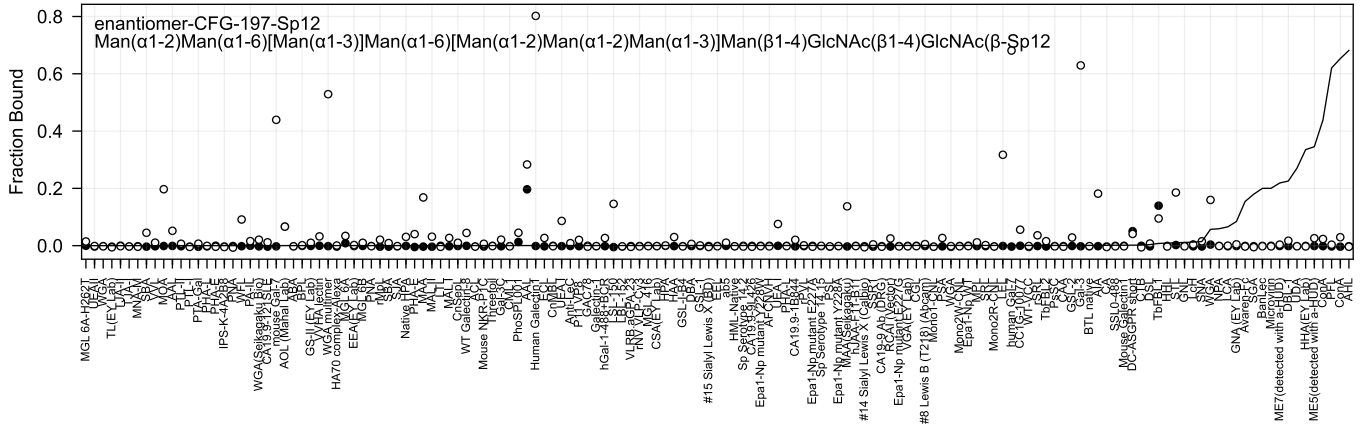

enantiomer-CFG-197-Sp12
Man(α1-2)Man(α1-6)[Man(α1-3)]Man(α1-6)[Man(α1-2)Man(α1-2)Man(α1-3)]Man(β1-4)GlcNAc(β1-4)GlcNAc(β-Sp12
Fraction Bound
0.0
0.2
0.4
0.6
0.8
MGL 6A-H262T
UEA-I
WGA
TL(EY Lab)
LTA-I
TJA-I
MNA-M
SBA
VVL
MOA
AAL
PTL-I
PTL-II
PTA-Gal
PHA-L
PHA-E
IPS-K-4A2B8
PNA
WFL
PA-IL
WGA(Seikagaku Bio)
CA19.9-121SLE
mouse Gal-7
AOL (Mahal Lab)
ABA
BPL
GS-II (EY Lab)
VVHA lectin
WGA multimer
HA70 complex-Alexa
MGL 6A
EEA(EY Lab)
MGL 6B
PNA
rVpL
SBA
SJA
Native HPA
PHA-E
MAA
MAL-II
LTL
MAL-I
CnSep-1
WT Galectin-3
ACL
Mouse NKR-P1C
Threefoil
Gal-3C
CML-1
PhoSL001
AAL
Human Galectin1
ECL
CnMBL
UEA-I
Anti-Lec
P11 VP8
GAC78
Galectin-1
hGal-1-488+BCR
LSL-150
LBL-152
VLRB.aGPA.23
rNV VLP-Cy3
MGL 416
CSA(EY Lab)
HPA
HAA
GSL-I-B4
DBA
GSL-I
#15 Sialyl Lewis X (BD)
EEL
ab5
HML-Native
Sp Serotype 14.2
CA19.9-9L426
Epa1-Np mutant Y228W
AFCNVH
UEA-I
PHA-L
CA19.9-1B844
rPVL
Epa1-Np mutant E227A
Sp Serotype 14.15
PSL
Epa1-Np mutant Y228A
MAA(Seikagaku)
#14 Sialyl Lewis X (Calbio)
CA19-9 Ab (DRG)
RCAI (Vector)
Epa1-Np mutant E227G
VGA(EY Lab)
CGL
#8 Lewis B (T218) (Abcam)
Mono1-CNL
PSA
WGA
Mono2W-CNL
Epa1-Np WT
MPL
SRL
Mono2R-CNL
LEL
human Gal-7
WT-VCC
rSRL
TbFBL2
PsSC
CAA
GSL-2
Gal-3
BTL native
AIA
CA
SSL0-488
Mouse Galectin1
DC-ASGPR short
CTB
DSL
TbFBL1
HHL
STL
GNL
LcH
SNA
WGA
NPA
LCA
GNA (EY Lab)
Avaren-Fc
SGA
BanLec
Microvirin
ME7(detected with a-HUD)
DBL-1
UDA
HHA(EY Lab)
ME5(detected with a-HUD)
CopA
PFL
ConA
AHL

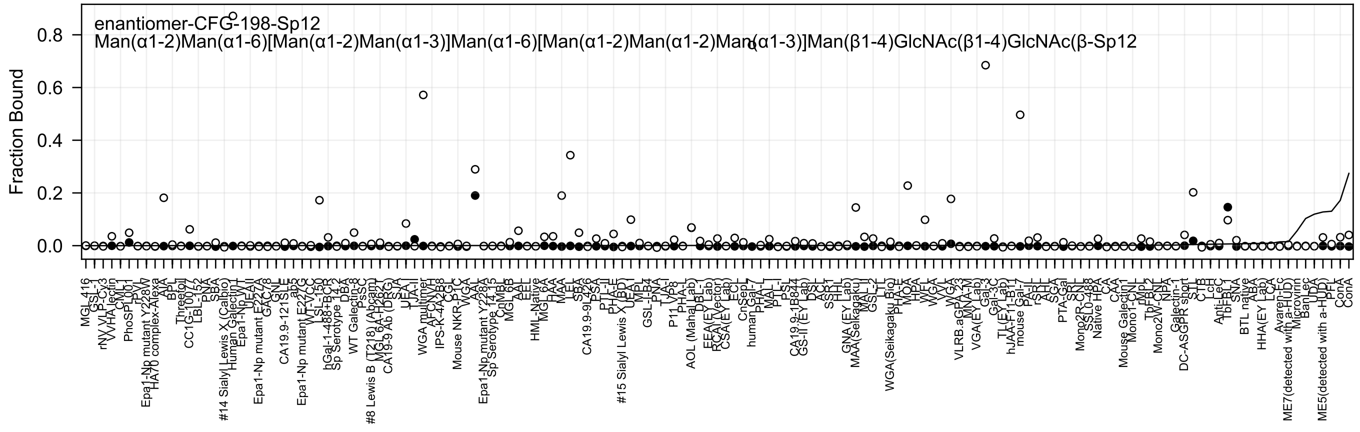
enantiomer-CFG-198-Sp12
Man(α1-2)Man(α1-6)[Man(α1-2)Man(α1-3)]Man(α1-6)[Man(α1-2)Man(α1-2)Man(α1-3)]Man(β1-4)GlcNAc(β1-4)GlcNAc(β-Sp12
Fraction Bound
0.0
0.2
0.4
0.6
0.8

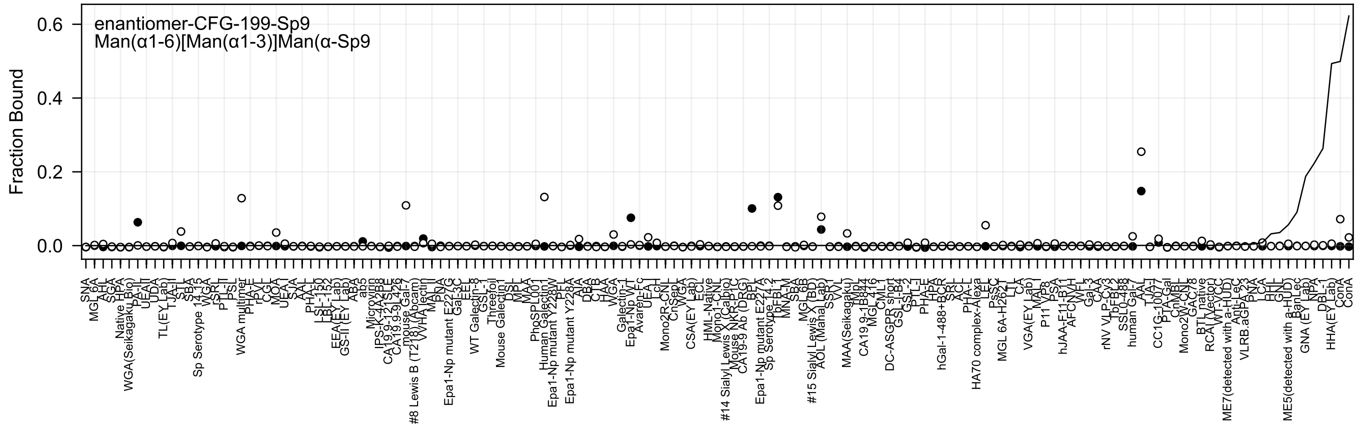

enantiomer-CFG-199-Sp9
Man(α1-6)[Man(α1-3)]Man(α-Sp9
Fraction Bound
0.0
0.2
0.4
0.6

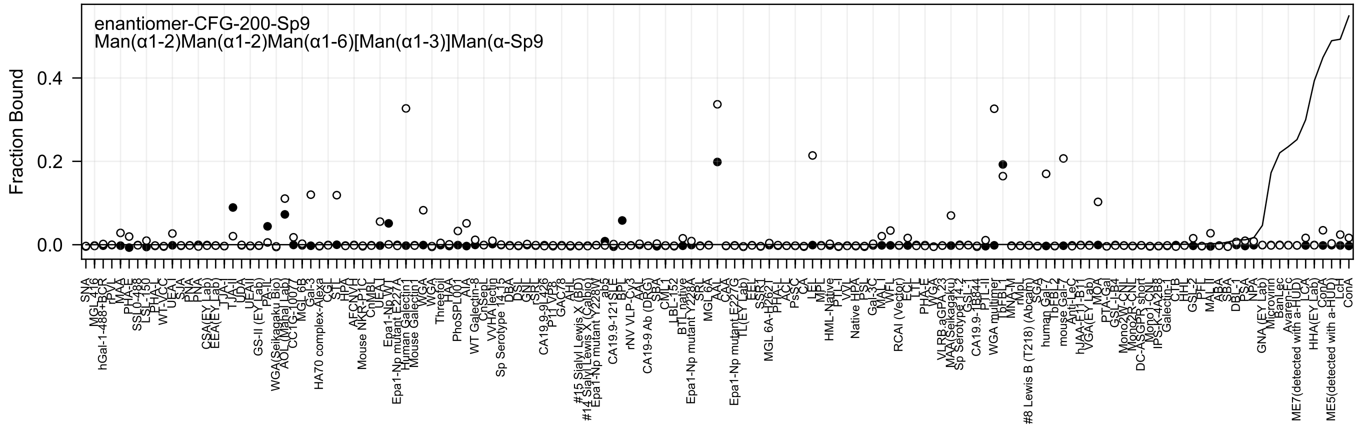

enantiomer-CFG-200-Sp9
Man(α1-2)Man(α1-2)Man(α1-6)[Man(α1-3)]Man(α-Sp9
Fraction Bound
0.0
0.2
0.4
SNA
MGL 416
hGal-1-488+BCR
rPVL
MAA
PHA-E
SSL0-488
LSL-150
PHA-L
WT-VCC
UEA-I
SJA
PNA
PNA
CSA(EY Lab)
EEA(EY Lab)
TJA-I
TJA-II
UDA
UEA-II
GS-II (EY Lab)
PA-IL
WGA(Seikagaku Bio)
AOL (Mahal Lab)
CC1G-1007
MGL 6B
Gal-3
HA70 complex-Alexa
CGL
STL
HPA
AFCNVH
Mouse NKR-P1C
CnMBL
UEA-I
Epa1-Np WT
Epa1-Np mutant E227A
Human Galectin1
Mouse Galectin1
WGA
Threefoil
HAA
PhoSPL001
AIA
WT Galectin-8
CnSepL
VVHA lectin
Sp Serotype 14.15
DBA
DSL
GNL
rSRL
CA19.9-9L426
P11 VP8
GAC78
AAL
#15 Sialyl Lewis X (BD)
#14 Sialyl Lewis X (Calbio)
Epa1-Np mutant Y228W
ab5
CA19.9-121 SLE
BPL
rNV VLP-Cy3
AAL
CA19-9 Ab (DRG)
SBA
CML1
LBL-152
BTL native
Epa1-Np mutant Y228A
SRL
MGL 6A
AAA
CAA
Epa1-Np mutant E227G
TL(EY Lab)
EEL
SSR1
MGL 6A-H262T
PHA-L
ACL
PsSC
CA
LEL
MPL
HML-Native
PTL-I
VVL
Native HPA
PSL
Gal-3C
MAL-I
WFL
RCAI (Vector)
ECL
LTL
PHA-E
WGA
VLRB.aGPA.23
MAA(Seikagaku)
Sp Serotype 14.2
GSL-I
CA19.9-1B844
PTL-II
WGA multimer
TbFBL1
MNA-M
#8 Lewis B (T218) (Abcam)
rMpL
SGA
human Gal-7
TbFBL2
mouse Gal-7
Anti-LeC
hJAA-E1-B11
VGA(EY Lab)
MOA
PTA-Gal
GSL-I-B4
Mono2W-CNL
Mono2R-CNL
DC-ASGPR short
Mono1-CNL
IPS-K-4A2B8
Galectin-3
CTB
HHL
GSL-2
PFL
MAL-II
ABA
SBA
DBL-1
PSA
NPA
GNA (EY Lab)
Microvirin
BanLec
Avaren-Fc
ME7(detected with a-HUD)
LCA
HHA(EY Lab)
ConA
ME5(detected with a-HUD)
LcH
ConA

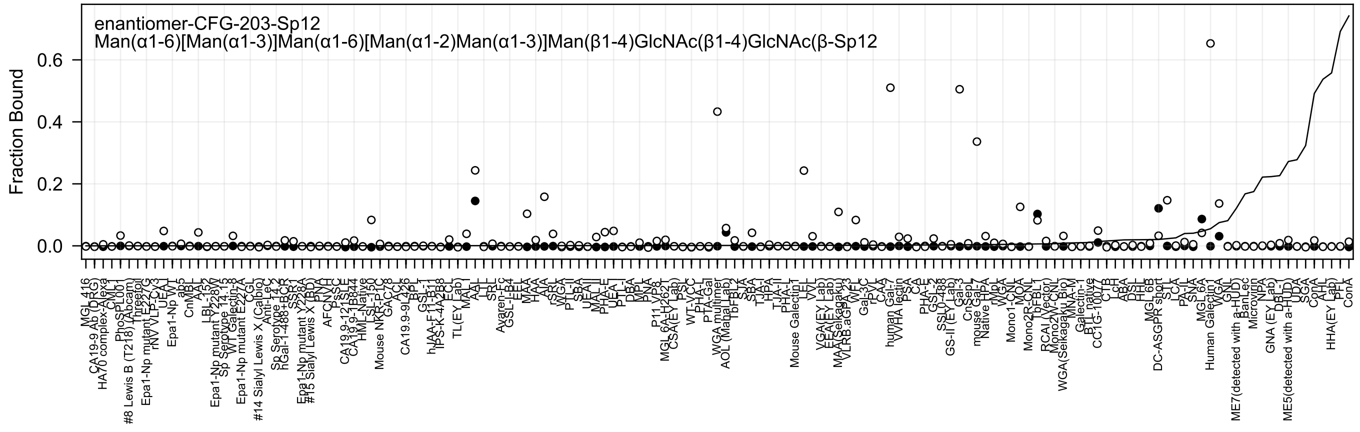
enantiomer-CFG-203-Sp12
Man(α1-6)[Man(α1-3)]Man(α1-6)[Man(α1-2)Man(α1-3)]Man(β1-4)GlcNAc(β1-4)GlcNAc(β-Sp12
Fraction Bound
0.0
0.2
0.4
0.6

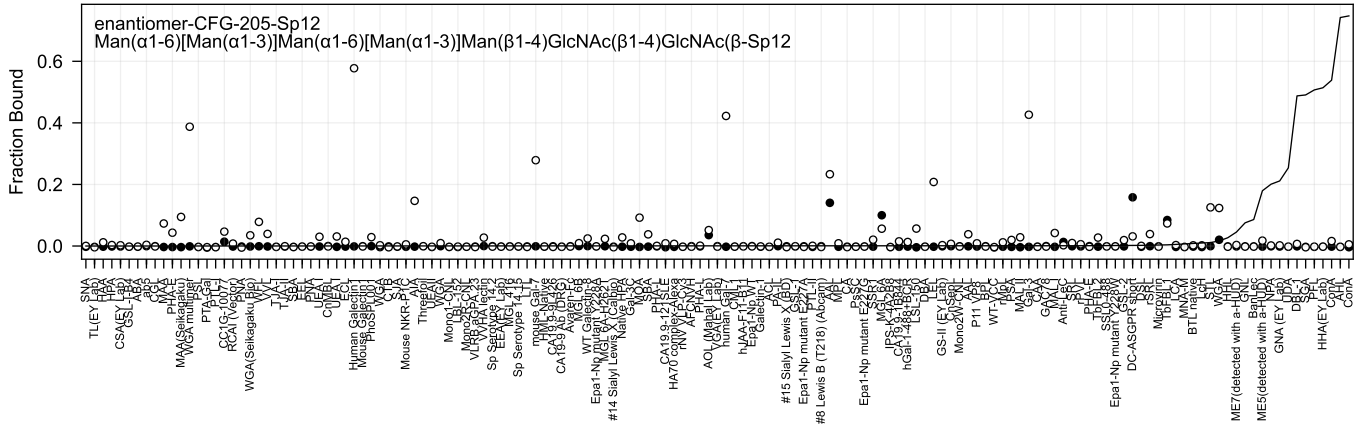
enantiomer-CFG-205-Sp12
Man(α1-6)[Man(α1-3)]Man(α1-6)[Man(α1-3)]Man(β1-4)GlcNAc(β1-4)GlcNAc(β-Sp12
Fraction Bound
0.0
0.2
0.4
0.6
SNA
TL(EY Lab)
HAA
HPA
CSA(EY Lab)
GSL-I-B4
ABA
ab5
CGL
MAA
PHA-E
MAA(Seikagaku)
WGA multimer
PSL
PTA-Gal
PTL-I
CC1G-10077
RCAI (Vector)
PNA
WGA(Seikagaku Bio)
WFL
VVL
TJA-II
TJA-I
SBA
EEL
PNA
UEA-I
CnMBL
UEA-I
ECL
Human Galectin-1
Mouse Galectin-1
PhoSL001
WGA
CTB
SJA
Mouse NKR-P1C
AIA
Threefoil
UEA-II
WGA
Mono1-CNL
LBL-152
Mono2H-CNL
VLRB aGPA.23
WHA lectin
Sp Serotype 14.2
EEA(EY Lab)
MGL 4.12
Sp Serotype 14.15
LTL
mouse Gal-7
HML-Native
CA19.9-9L426
CA19-9 Ab (DRG)
Avaren-Fc
MGL 6B
WT Galectin-8
Epa1-Np mutant Y228A
MGL 6A-H262
#14 Sialyl Lewis X (Calbio)
Native HPA
Gal-3C
MOA
SBA
PHA-L
CA19.9-121SLE
HA70 complex-Alexa
rNV VLP-Cy3
AFCNVH
PHA-L
AOL (Mahal Lab)
VGA(EY Lab)
human Gal-7
CML-1
hJAA-F11-B11
Epa1-Np WT
Galectin-1
ACL
PA-IL
#15 Sialyl Lewis X (BD)
GSL-I
Epa1-Np mutant E227A
PTL-II
#8 Lewis B (T218) (Abcam)
AAL
MPL
CA
PsSC
Epa1-Np mutant E227G
SSP-1
MGL 6A
IPS-K-4A2B8
CA19.9-1B844
hGal-1-488+BCR
LSL-150
DBA
LEL
GS-II (EY Lab)
CnSep
Mono2W-CNL
AAL
P11 VP8
BPL
WT-VCC
rMpL
PSA
MAL II
Gal-3
CAA
GAC78
MAL I
Anti-LeC
SRL
SVL
rPVL
PHA-E
TbFBL2
SSL0-488
Epa1-Np mutant Y228W
GSL-2
DC-ASGPR short
DSL
rSRL
Microvirin
TbFBL1
LCA
MNA-M
BTL native
LcH
STL
WGA
HHL
ME7(detected with a-HUD)
GNL
BanLec
ME5(detected with a-HUD)
NPA
GNA (EY Lab)
UDA
DBL-1
SGA
PFL
HHA(EY Lab)
ConA
AHL
ConA

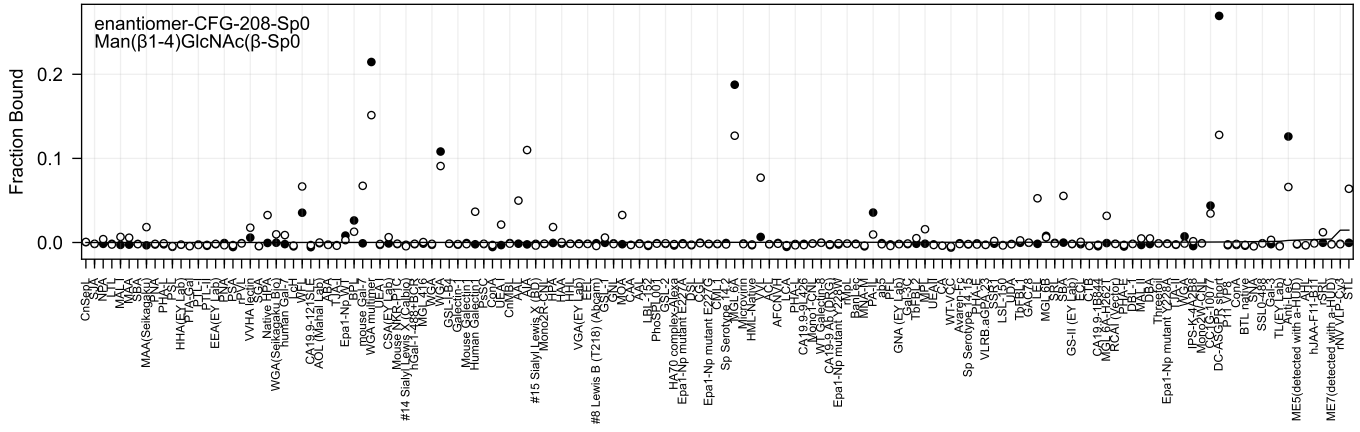
enantiomer-CFG-208-Sp0
Man(β1-4)GlcNAc(β-Sp0
Fraction Bound
0.2
0.1
0.0

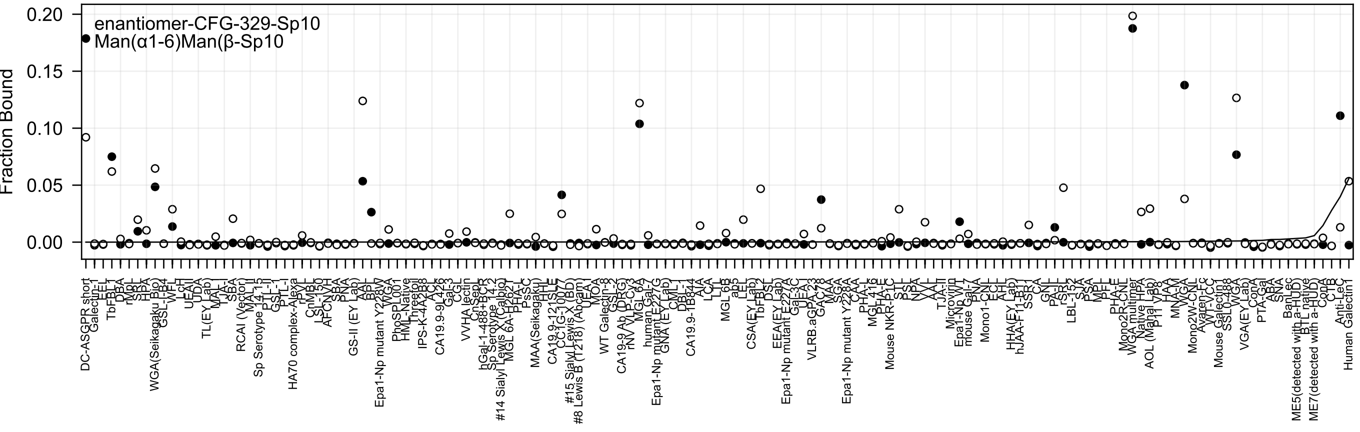

enantiomer-CFG-329-Sp10
Man(α1-6)Man(β-Sp10
Fraction Bound
0.20
0.15
0.10
0.05
0.00

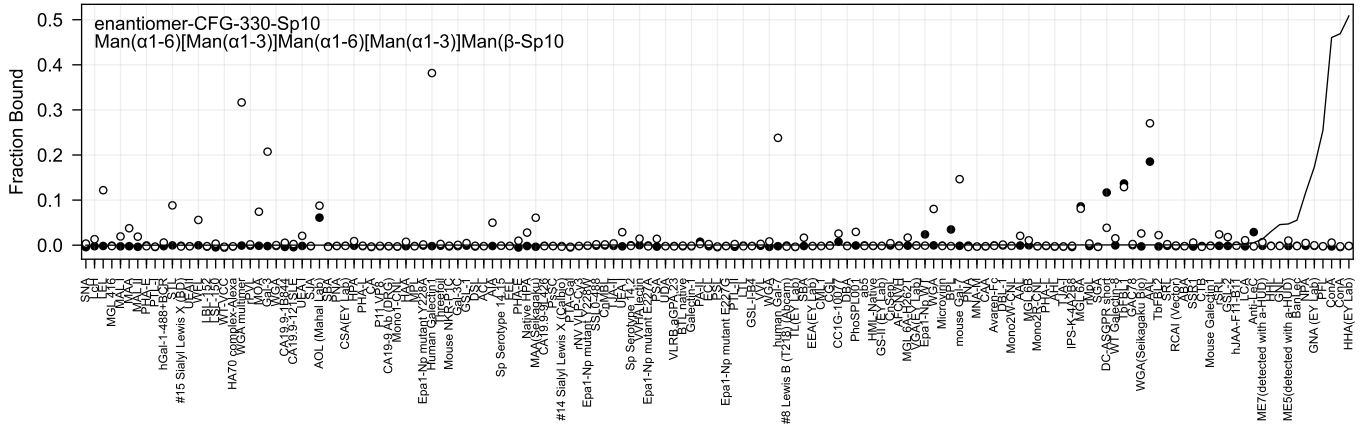
enantiomer-CFG-330-Sp10
Man(α1-6)[Man(α1-3)]Man(α1-6)[Man(α1-3)]Man(β-Sp10
Fraction Bound
0.0
0.1
0.2
0.3
0.4
0.5

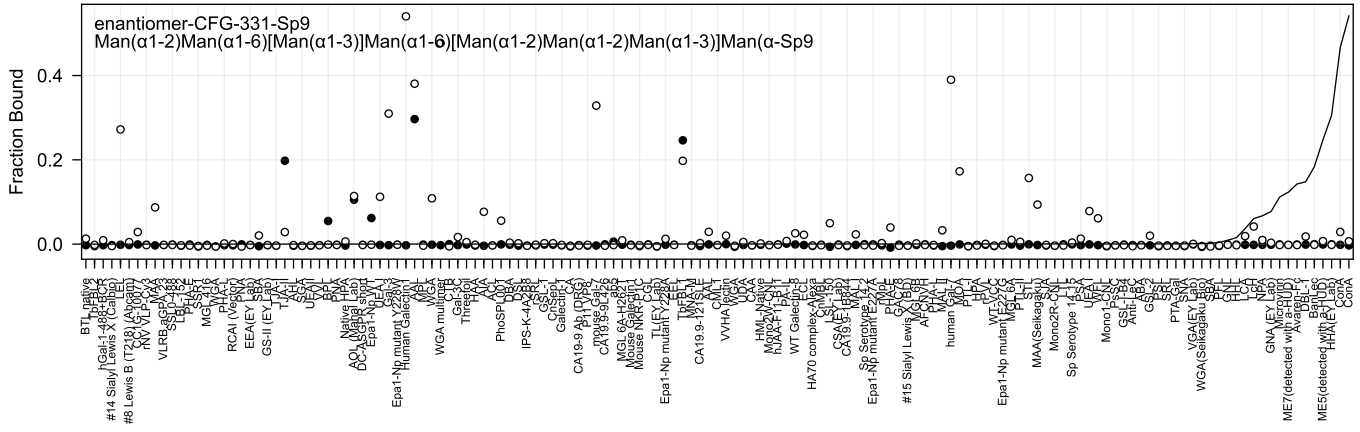

enantiomer-CFG-331-Sp9
Man(α1-2)Man(α1-6)[Man(α1-3)]Man(α1-6)[Man(α1-2)Man(α1-2)Man(α1-3)]Man(α-Sp9
Fraction Bound
0.0
0.2
0.4

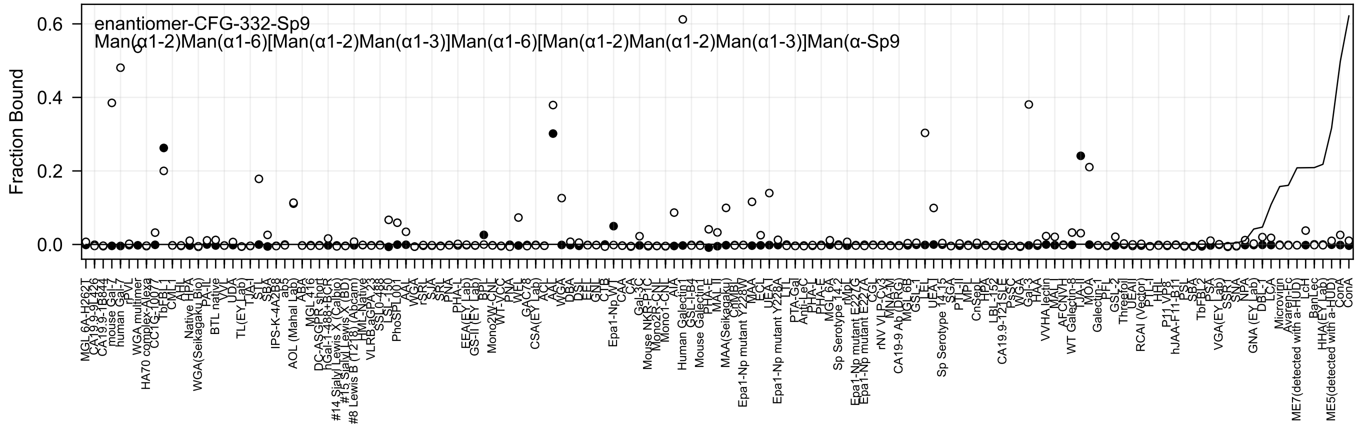

enantiomer-CFG-332-Sp9
Man(α1-2)Man(α1-6)[Man(α1-2)Man(α1-3)]Man(α1-6)[Man(α1-2)Man(α1-2)Man(α1-3)]Man(α-Sp9
Fraction Bound
0.6
0.4
0.2
0.0
MGL 6A-H262T
CA19.9-9L426
CA19.9-1B844
mouse Gal-7
human Gal-7
rPVL
WGA multimer
HA70 complex-Alexa
CC1G-10077
TbFBL-1
CML-I
AHL
Native HPA
WGA(Seikagaku Bio)
PA-IL
BTL native
VVL
UDA
TL(EY Lab)
TJA-I
STL
SBA
IPS-K-4A2B8
ab5
AOL (Mahal Lab)
ABA
MGL 416
DC-ASGPR short
hGal-1-488+BCR
#14 Sialyl Lewis X (Calbio)
#15 Sialyl Lewis X (BD)
#8 Lewis B (T218) (Abcam)
HML-Native
VLRB.aGPA.23
SSL0-488
LSL-150
PhoSL001
AAL
WGA
rSRL
SJA
SRL
PNA
PHA-L
EEA(EY Lab)
GS-II (EY Lab)
BPL
Mono2W-CNL
WT-VCC
PNA
WFL
GAC78
CSA(EY Lab)
ACL
AAL
WGA
DBA
DSL
EEL
GNL
CTB
Epa1-Np WT
CAA
CA
Gal-3C
Mouse NKR-P1C
Mono2R-CNL
Mono1-CNL
AIA
Human Galectin1
GSL-I-B4
Mouse Galectin1
PHA-E
MAL-II
MAA(Seikagaku)
ChMBL
Epa1-Np mutant Y228W
MAA
ECL
UEA-I
Epa1-Np mutant Y228A
LTL
PTA-Gal
Anti-LeC
PHA-E
MGL 6A
Sp Serotype 14.2
rMpL
Epa1-Np mutant E227G
Epa1-Np mutant E227A
CGL
rNV VLP-Cy3
MNA-M
CA19-9 Ab (DRG)
MGL 6B
GSL-I
LEL
UEA-I
Sp Serotype 14.15
SGA
PTL-II
MPL
CnSept
HPA
LBL-152
CA19.9-121SLE
PsSC
WGA
Gal-3
HAA
VVHA lectin
MAL-I
AFCNVH
WT Galectin-8
TJA-II
MOA
Galectin-1
PFL
GSL-2
Threefoil
UEAII
RCAI (Vector)
PTL-I
HHL
P11 VP8
hJAA-F11-B11
PSL
SBA
TbFBL2
PSA
VGA(EY Lab)
SSR1
SNA
NPA
GNA (EY Lab)
DBL-I
LCA
Microvirin
Avaren-Fc
ME7(detected with a-HUD)
LcH
BanLec
HHA(EY Lab)
ME5(detected with a-HUD)
ConA
ConA

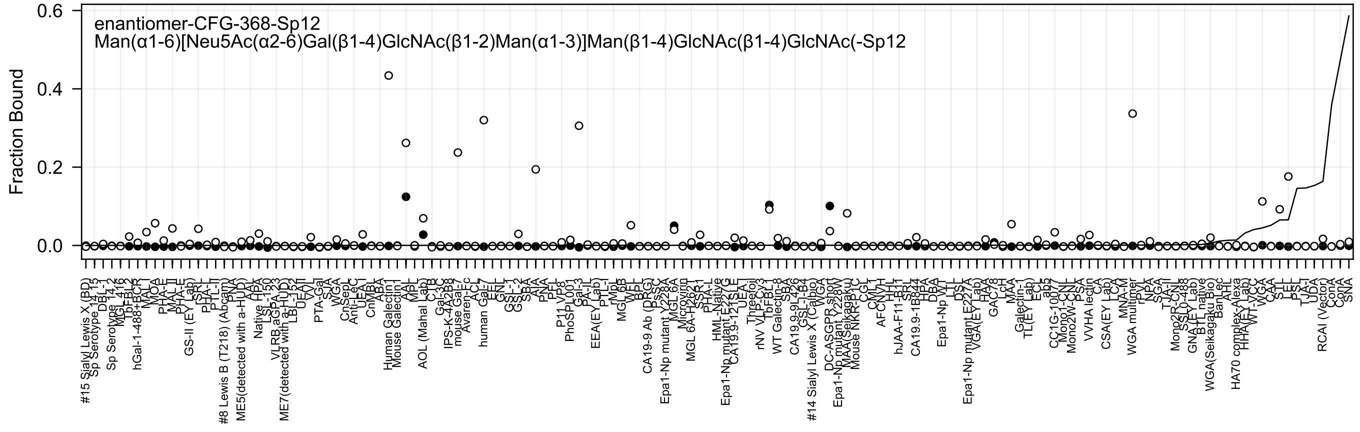
enantiomer-CFG-368-Sp12
Man(α1-6)[Neu5Ac(α2-6)Gal(β1-4)GlcNAc(β1-2)Man(α1-3)]Man(β1-4)GlcNAc(β1-4)GlcNAc(-Sp12
Fraction Bound
0.6
0.4
0.2
0.0

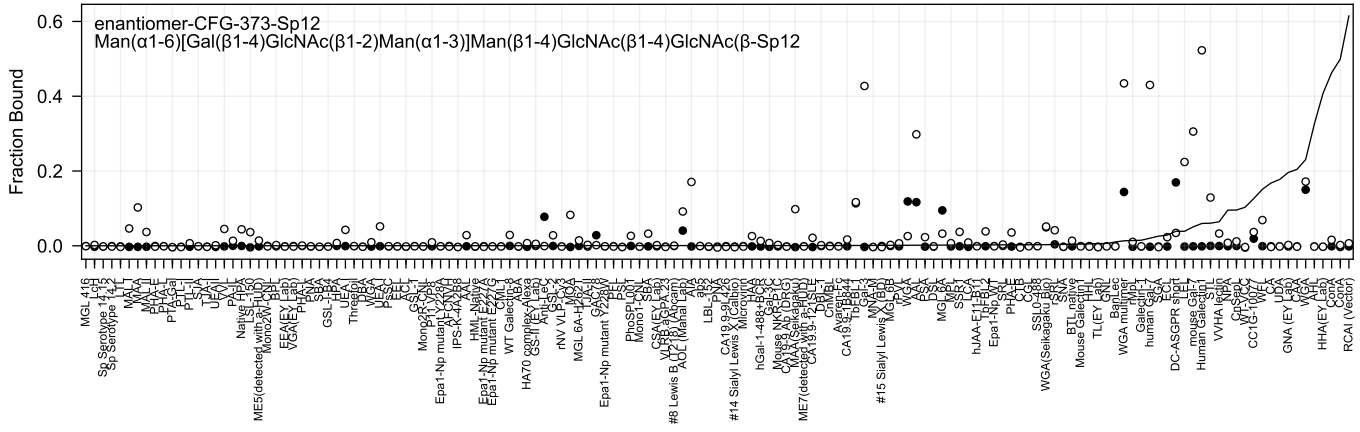
enantiomer-CFG-373-Sp12
Man(α1-6)[Gal(β1-4)GlcNAc(β1-2)Man(α1-3)]Man(β1-4)GlcNAc(β1-4)GlcNAc(β-Sp12
Fraction Bound
0.6
0.4
0.2
0.0

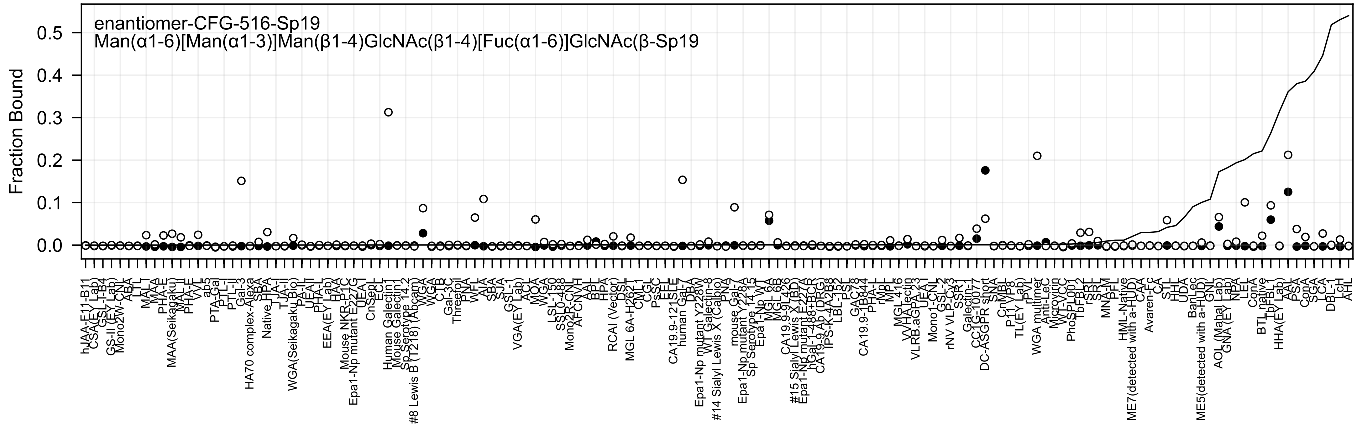
enantiomer-CFG-516-Sp19
Man(α1-6)[Man(α1-3)]Man(β1-4)GlcNAc(β1-4)[Fuc(α1-6)]GlcNAc(β-Sp19
Fraction Bound
0.5
0.4
0.3
0.2
0.1
0.0
