## Supplementary material for "Atom-level Machine Learning of Protein-Glycan Interactions and Cross-chiral Recognition in Glycobiology": All Enantiomer Prediction Plots

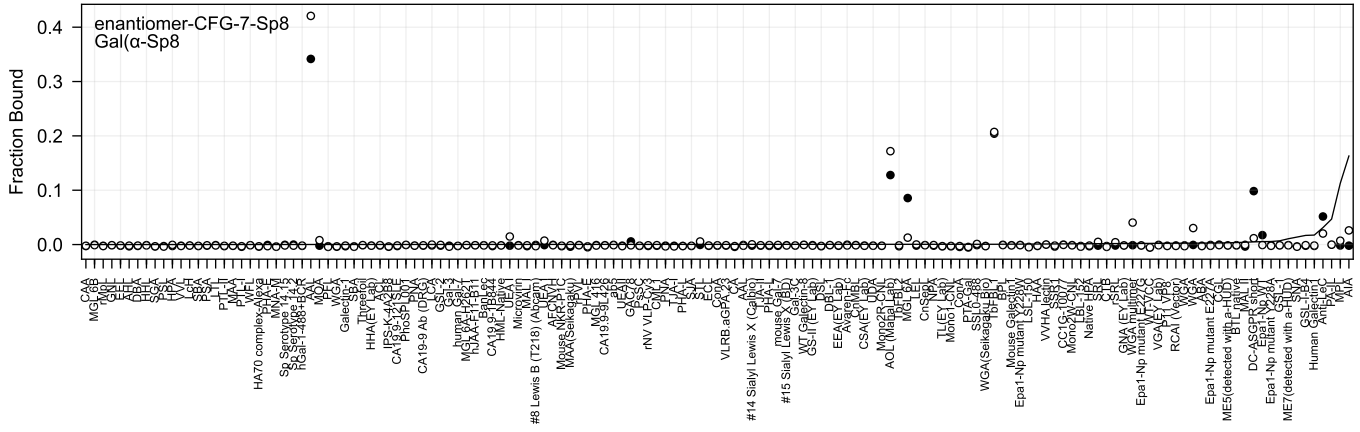
enantiomer-CFG-7-Sp8
Gal(α-Sp8
Fraction Bound
0.4
0.3
0.2
0.1
0.0

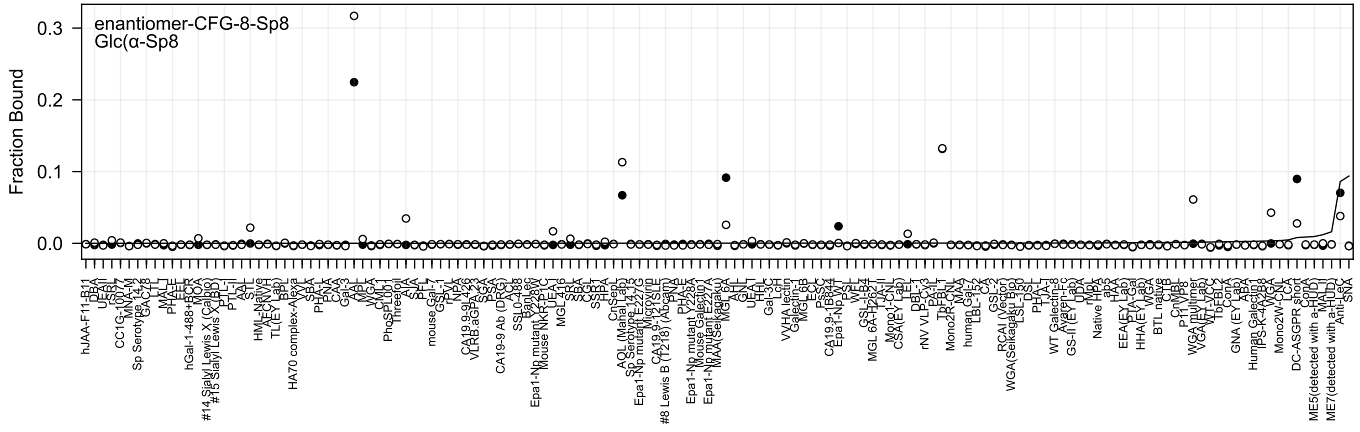
enantiomer-CFG-8-Sp8
Glc(α-Sp8
Fraction Bound
0.3
0.2
0.1
0.0

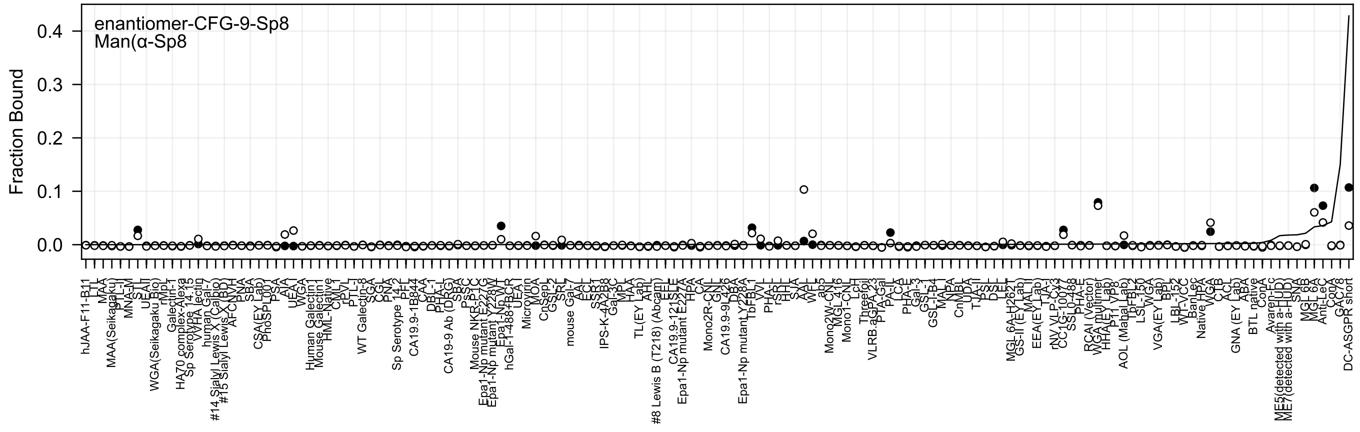
enantiomer-CFG-9-Sp8
Man(α-Sp8
Fraction Bound
0.0
0.1
0.2
0.3
0.4

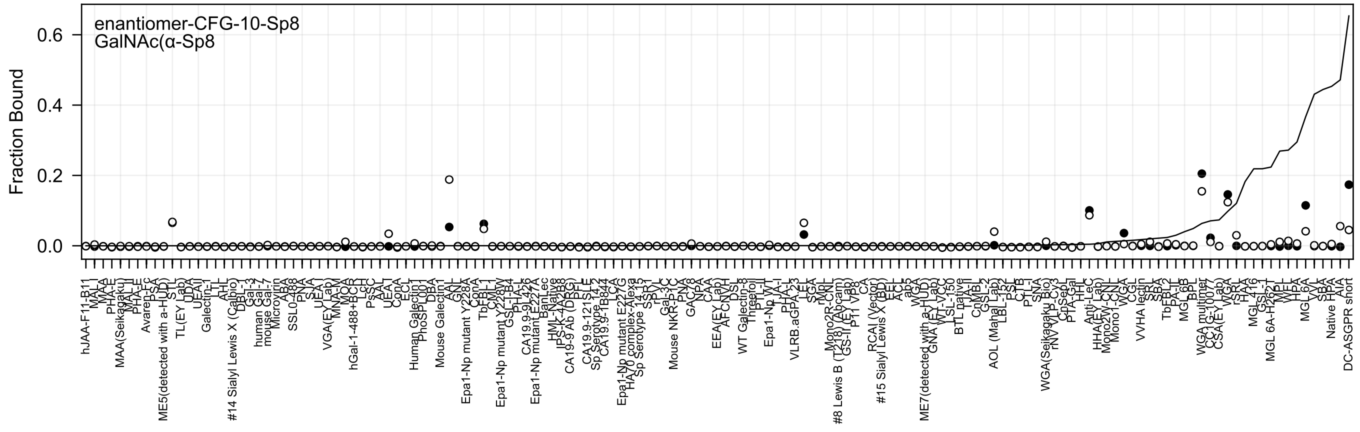

enantiomer-CFG-10-Sp8
GalNAc(α-Sp8
Fraction Bound
0.0
0.2
0.4
0.6

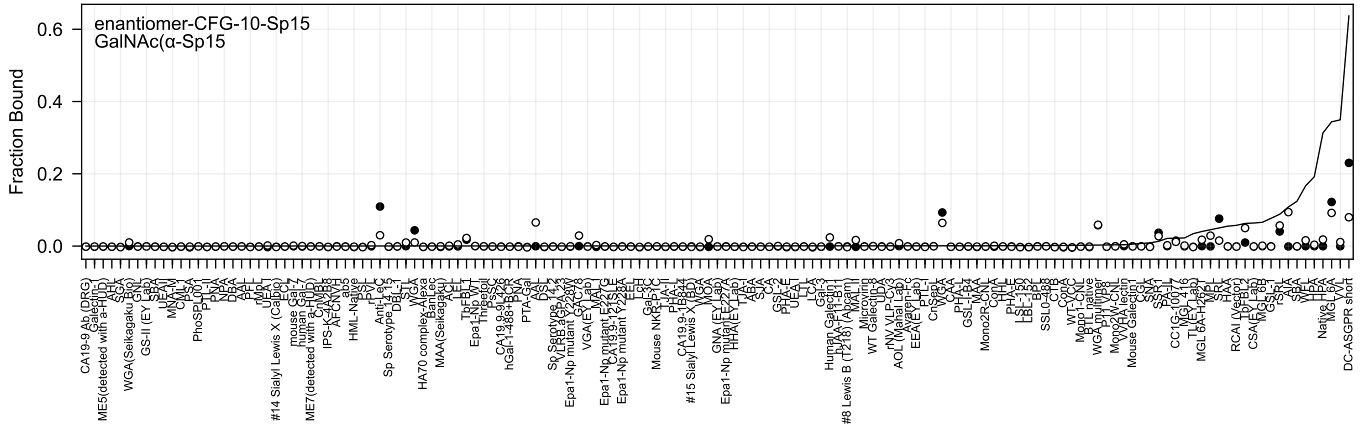
enantiomer-CFG-10-Sp15
GalNAc(α-Sp15
Fraction Bound
0.6
0.4
0.2
0.0

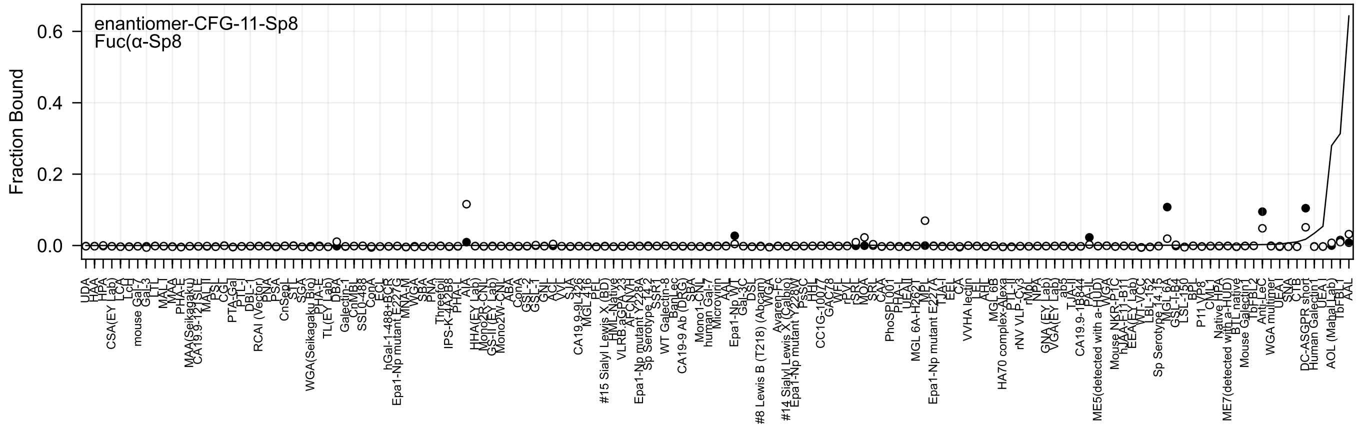
enantiomer-CFG-11-Sp8
Fuc(α-Sp8
Fraction Bound
0.0
0.2
0.4
0.6

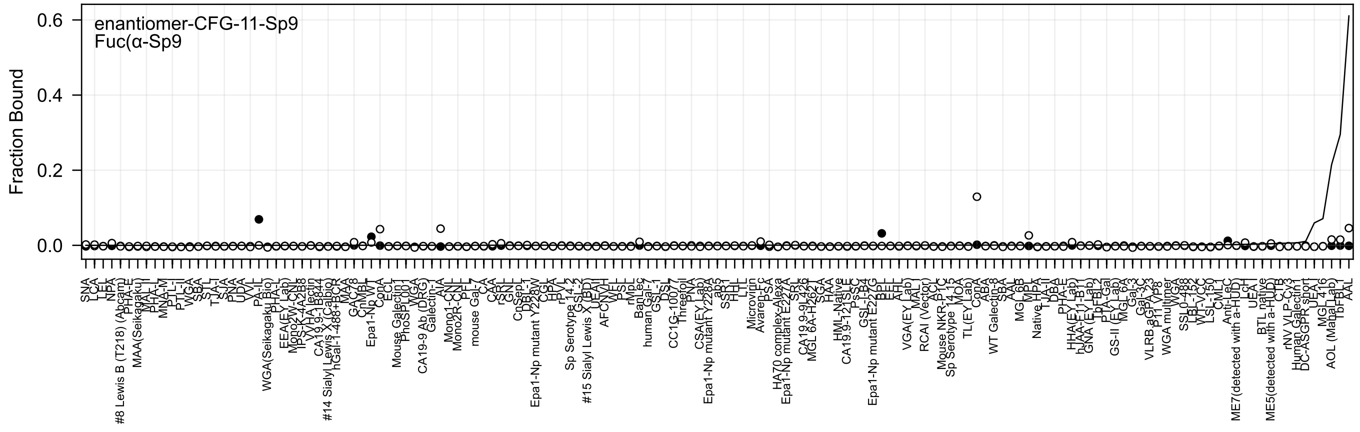
enantiomer-CFG-11-Sp9
Fuc(α-Sp9
Fraction Bound
0.6
0.4
0.2
0.0

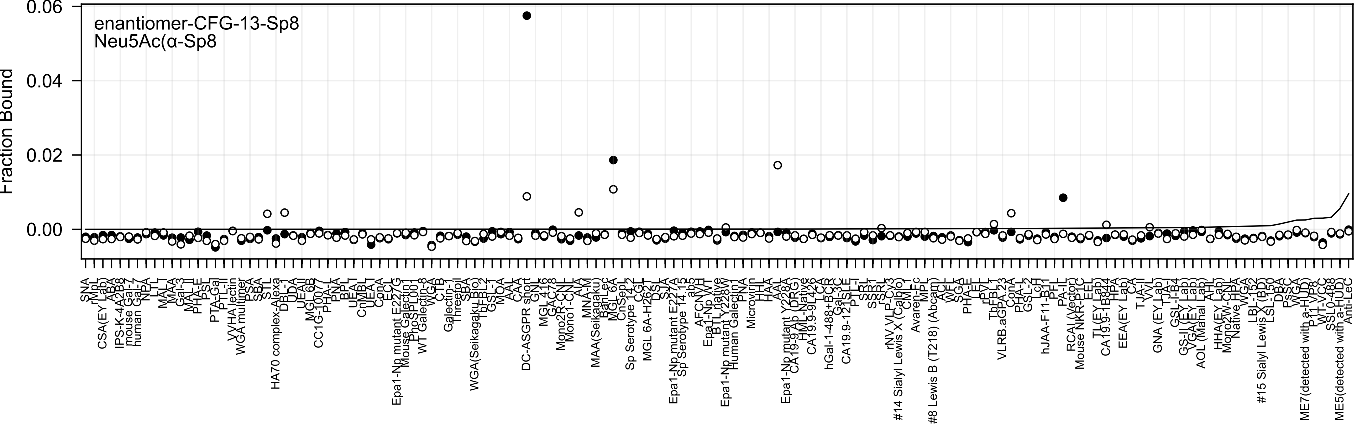
enantiomer-CFG-13-Sp8
Neu5Ac(α-Sp8
Fraction Bound
0.06
0.04
0.02
0.00

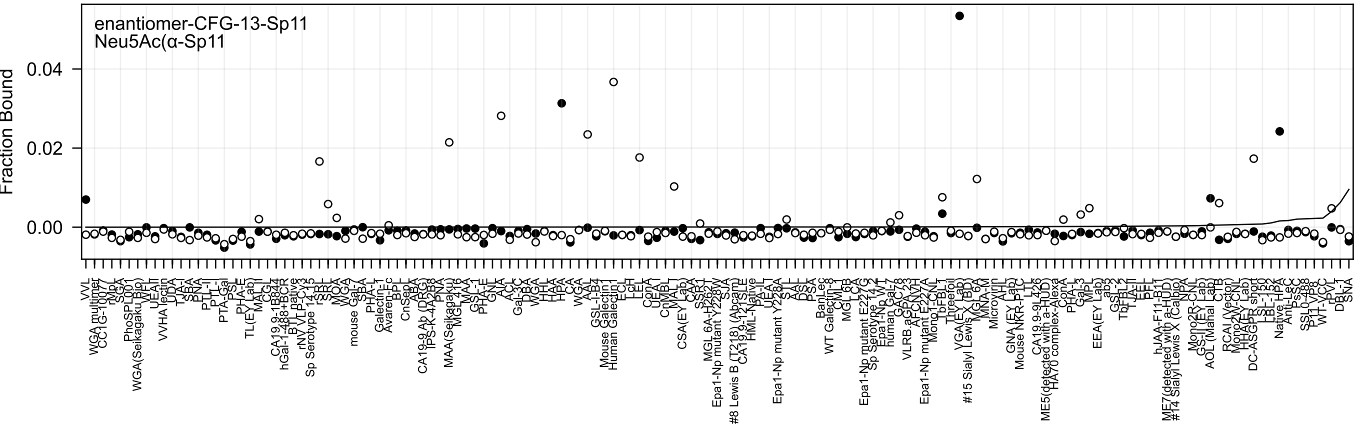
enantiomer-CFG-13-Sp11
Neu5Ac(α-Sp11
Fraction Bound
0.04
0.02
0.00

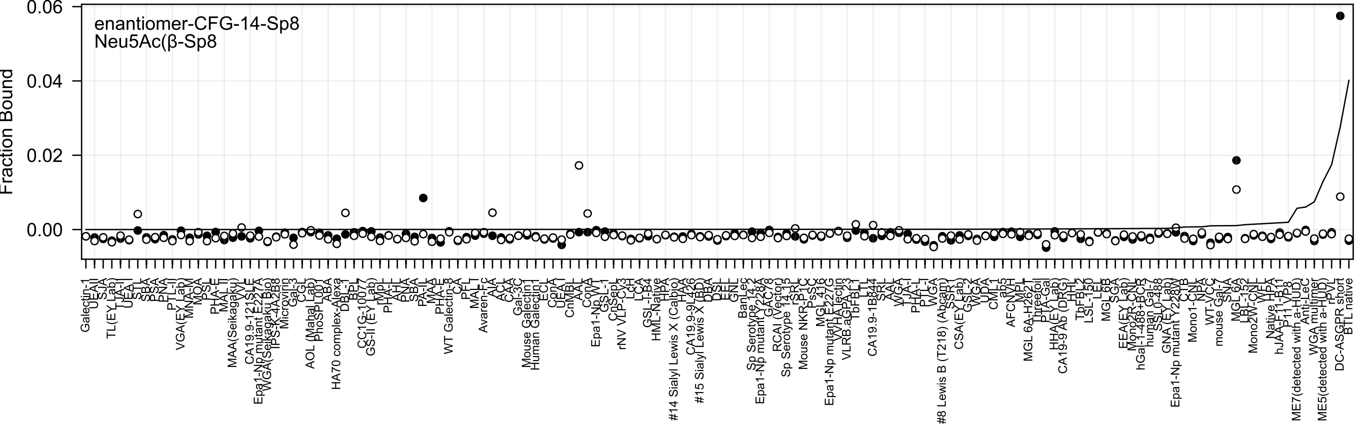
enantiomer-CFG-14-Sp8
Neu5Ac(β-Sp8
Fraction Bound
0.06
0.04
0.02
0.00

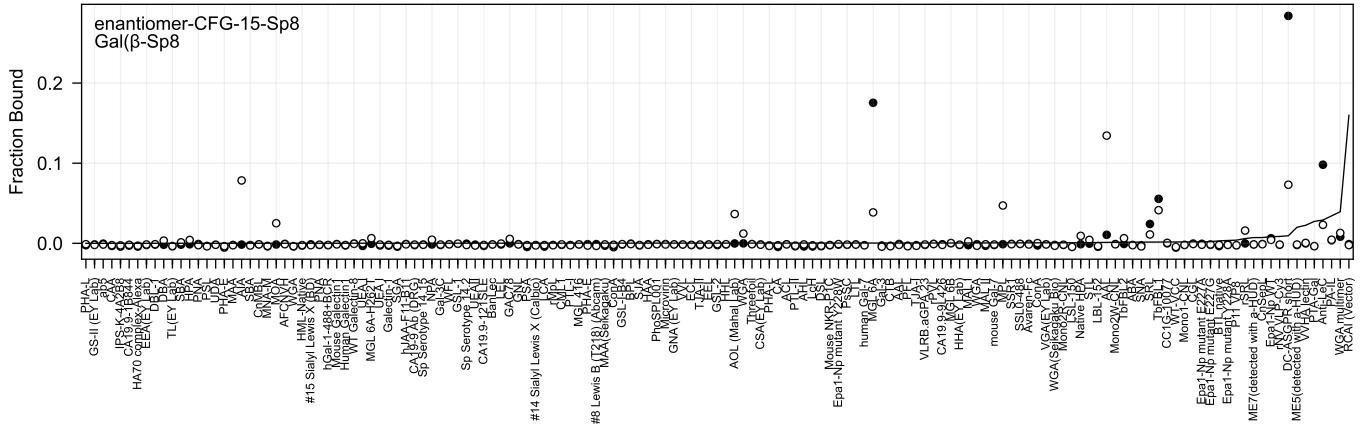
enantiomer-CFG-15-Sp8
Gal(β-Sp8
Fraction Bound
0.2
0.1
0.0

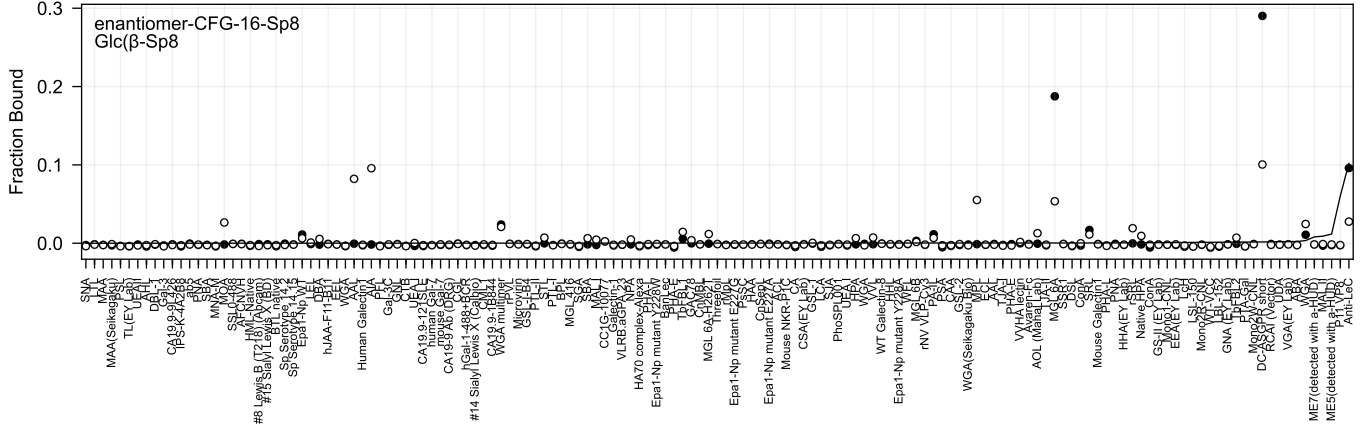
enantiomer-CFG-16-Sp8
Glc(β-Sp8
Fraction Bound
0.3
0.2
0.1
0.0

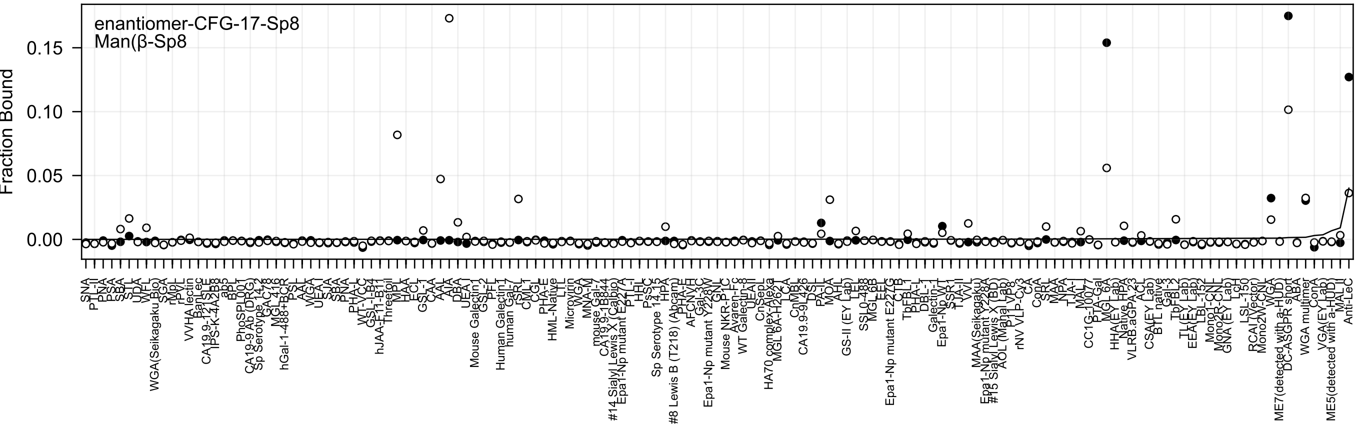
enantiomer-CFG-17-Sp8
Man(β-Sp8
Fraction Bound
0.15
0.10
0.05
0.00
SNA
PTL-II
PNA
PSA
SBA
STL
UDA
WFL
WGA(Seikagaku Bio)
SGA
rMpL
rPVL
VVHA lectin
BanLec
CA19.9-121SLE
IPS-K-4A2B8
ab5
BPL
PhoSL001
CA19-9 Ab (DRG)
Sp Serotype 14-2
GAC78
MGL416
hGal-1-488+BCR
PSL
AAL
WGA
UEA-I
SJA
SBA
PNA
PHA-L
WT-VCC
GSL-I-B4
hJAA-F11-B11
Threefoil
MPL
HAA
ECL
GSL-I
CAA
AAL
AIA
DBA
UEA-I
Mouse Galectin1
GSL-2
PFL
Human Galectin1
human Gal-7
rSRL
CML1
CGL
PHA-E
HML-Native
LTL
Microvirin
WGA
MNA-M
mouse Gal-7
CA19.9-1B844
#14 Sialyl Lewis X (Calbio)
Epa1-Np mutant E227A
PTL-I
HHL
PsSC
Sp Serotype 14-15
HPA
#8 Lewis B (T218) (Abcam)
PHA-E
AFCNYH
Gal-3C
Epa1-Np mutant Y228W
GNL
Mouse NKR-P1C
Avaren-Fc
WT Galectin-8
UEAII
CnSepL
HA70 complex-Alexa
MGL 6A-H262T
LCA
CnMBL
CA19.9-9L426
DSL
PA-IL
MOA
AHL
GS-II (EY Lab)
LEL
SSL0-488
MGL 6B
EEL
Epa1-Np mutant E227G
CTB
TbFBL1
PHA-L
DBL-1
Galectin-1
Epa1-Np WT
SSR1
TJA-II
VVL
MAA(Seikagaku)
Epa1-Np mutant Y228A
#15 Sialyl Lewis X (BD)
AOL (Mahal Lab)
P1 VP8
rNV VLP-Cy3
CA
ConA
SRL
MAA
NPA
TJA-I
MAL-I
CC1G-10077
PTA-Gal
MGL 6A
HHA(EY Lab)
Native HPA
VLRB.aGPA.23
ACL
CSA(EY Lab)
BTL native
Gal-3
TbFBL2
TL(EY Lab)
EEA(EY Lab)
LBL-152
Mono1-CNL
Mono2R-CNL
GNA (EY Lab)
LcH
LSL-150
RCAI (Vector)
Mono2W-CNL
WGA
ME7(detected with a-HUD)
DC-ASGPR short
ABA
WGA multimer
ConA
VGA(EY Lab)
ME5(detected with a-HUD)
MAL-II
Anti-LeC

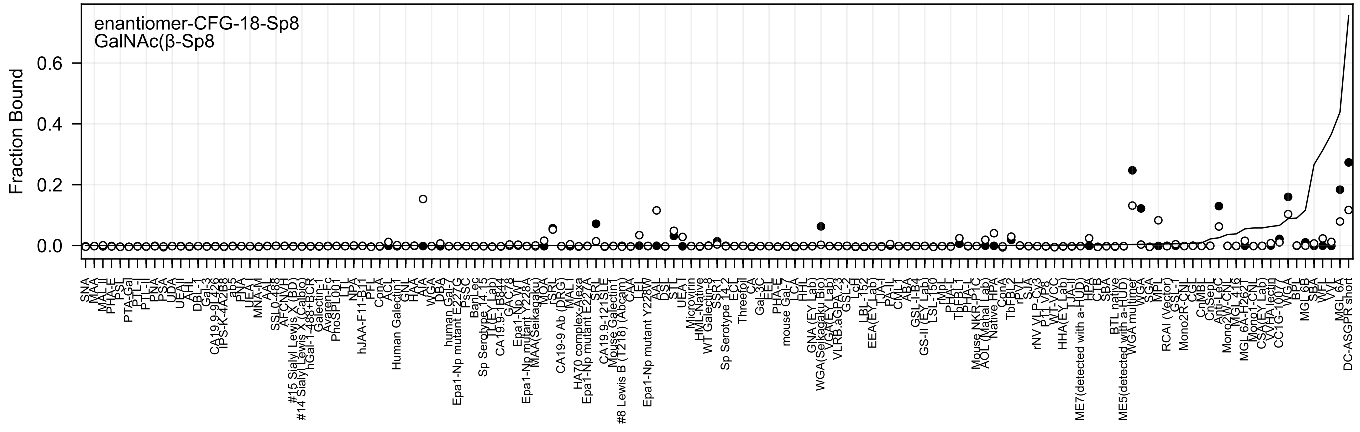
enantiomer-CFG-18-Sp8
GalNAc(β-Sp8
Fraction Bound
0.0
0.2
0.4
0.6
SNA
MAA
MAL-II
PHA-E
PSL
PTA-Gal
PTL-I
PTL-II
PNA
PSA
UDA
UEAII
AAL
DBL-1
Gal-3
CA19.9-9L426
TPS-K-4A2B8
ab5
PNA
UEA-I
MNA-M
AIA
SSL0-488
AFCNVH
#15 Sialyl Lewis X (BD)
#14 Sialyl Lewis X (Calbio)
hGal-1-488+BCR
Galectin-1
Avaren-Fc
PhoSPL001
LTL
NPA
hJAA-F11-B11
PFL
ConA
ACL
Human Galectin1
GNL
HAA
AIA
WGA
DBA
human Gal-7
Epa1-Np mutant E227G
PsSC
BanLec
Sp Serotype 14.15
TL(EY Lab)
CA19.9-1B844
GAC78
Epa1-Np WT
Epa1-Np mutant Y228A
MAA(Seikagaku)
MOA
rSRL
CA19-9 Ab (DRG)
MAL-I
HA70 complex-Alexa
Epa1-Np mutant E227A
SRL
CA19.9-121SLE
Mouse Galectin1
#8 Lewis B (T218) (Abcam)
CAA
LEL
Epa1-Np mutant Y228W
AAL
DSL
STL
UEA-I
Microvirin
HML-Native
WT Galectin-8
SSR1
Sp Serotype 14.2
ECL
Threefoil
CA
Gal-3C
EEL
PHA-E
mouse Gal-7
LCA
HHL
GNA (EY Lab)
WGA(Seikagaku Bio)
VGA(EY Lab)
VLRB.aGPA.23
GSL-2
LcH
LBL-152
EEA(EY Lab)
TJA-I
PA-IL
CML-1
ABA
GSL-I-B4
GS-II (EY Lab)
LSL-150
rMpL
PHA-L
TbFBL1
PHA-L
Mouse NKR-P1C
AOL (Nahal Lab)
Native HPA
ConA
TbFBL2
rPVL
SJA
rNV VLP-Cv3
P1 VP8
WT-VCC
HHA(EY Lab)
LTA-II
ME7(detected with a-HUD)
HPA
CTB
SBA
BTL native
ME5(detected with a-HUD)
WGA multimer
WGA
SGA
MPL
RCAI (Vector)
GSL-I
Mono2R-CNL
CGL
CnMBL
CnSept
Anti-LeC
Mono2WT-CNL
MGL 4.16
MGL 6A-H262T
Mono1-CNL
CSA(EY Lab)
VVHA lectin
CC1G-10077
WGA
BPL
MGL 6B
SBA
WFL
VVL
MGL 6A
DC-ASGPR short

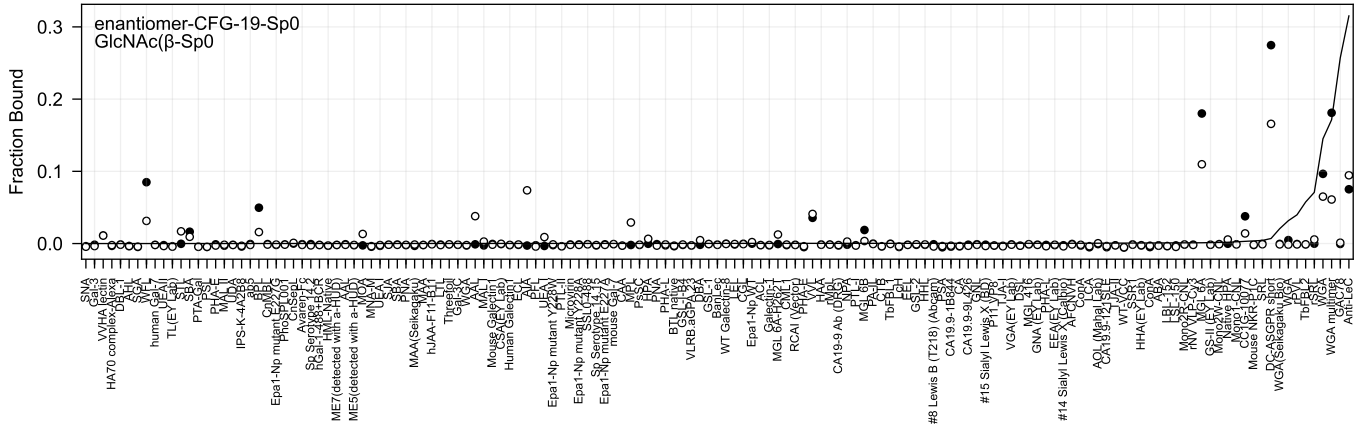
enantiomer-CFG-19-Sp0
GlcNAc(β-Sp0
Fraction Bound

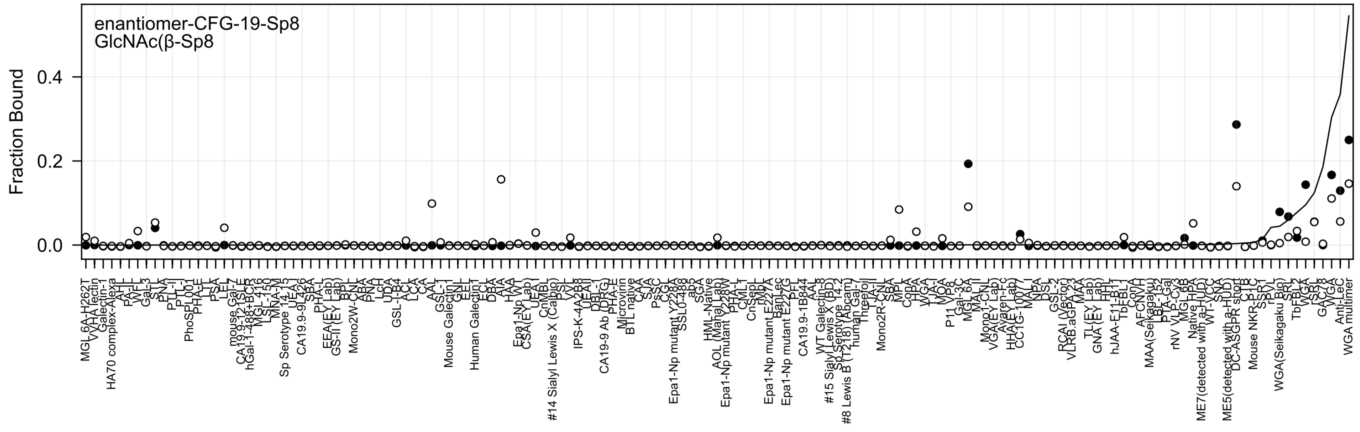
enantiomer-CFG-19-Sp8
GlcNAc(β-Sp8
Fraction Bound
0.0
0.2
0.4

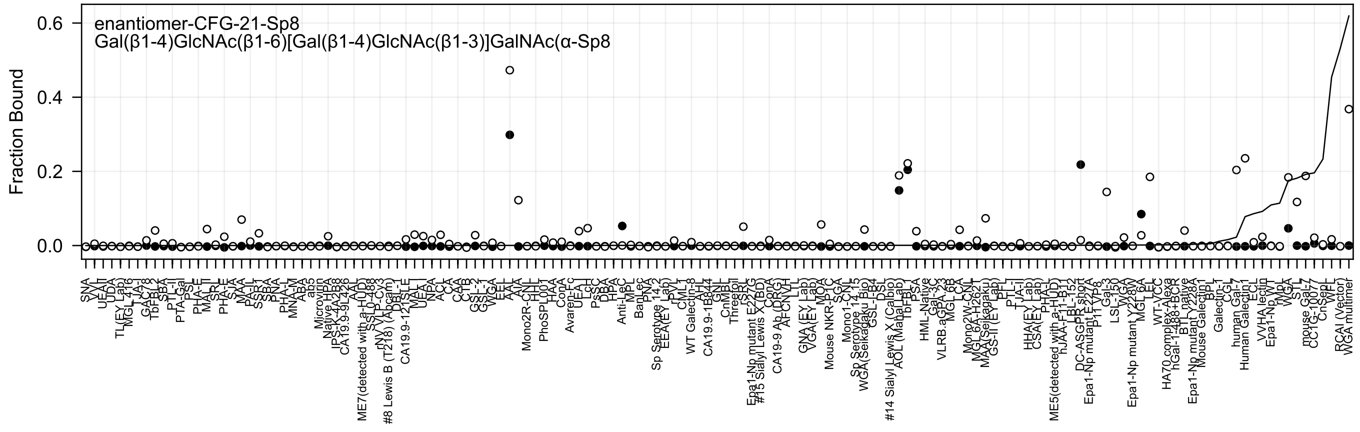

enantiomer-CFG-21-Sp8
Gal(β1-4)GlcNAc(β1-6)[Gal(β1-4)GlcNAc(β1-3)]GalNAc(α-Sp8
Fraction Bound
0.0
0.2
0.4
0.6

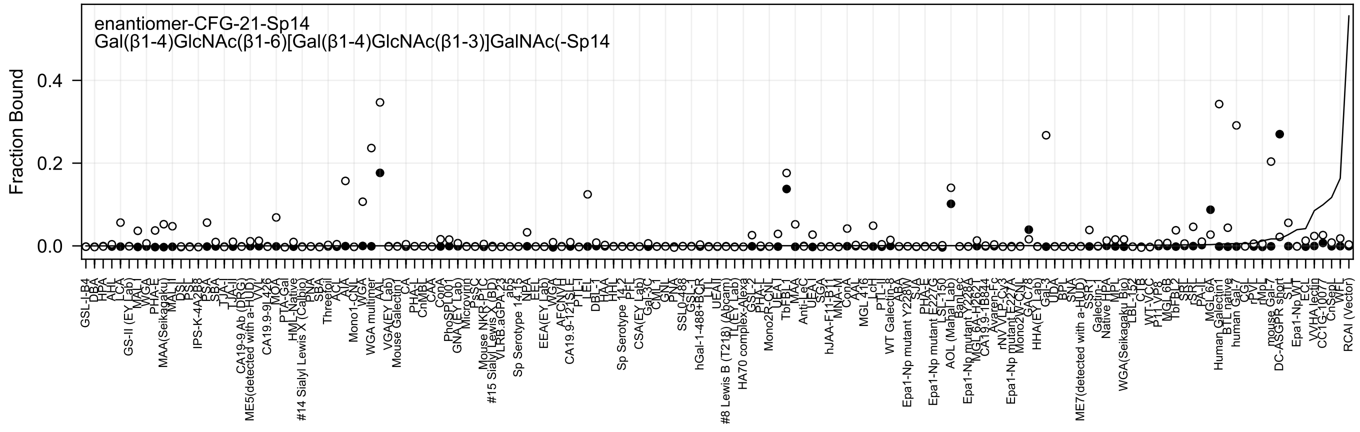
enantiomer-CFG-21-Sp14
Gal(β1-4)GlcNAc(β1-6)[Gal(β1-4)GlcNAc(β1-3)]GalNAc(-Sp14
Fraction Bound
0.4
0.2
0.0

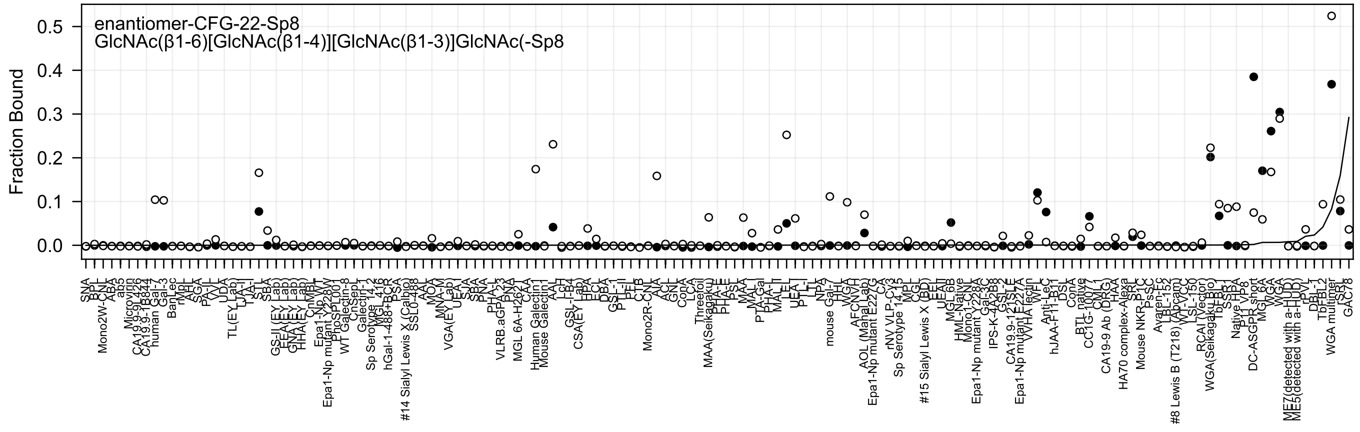

enantiomer-CFG-22-Sp8
GlcNAc(β1-6)[GlcNAc(β1-4)][GlcNAc(β1-3)]GlcNAc(-Sp8
Fraction Bound

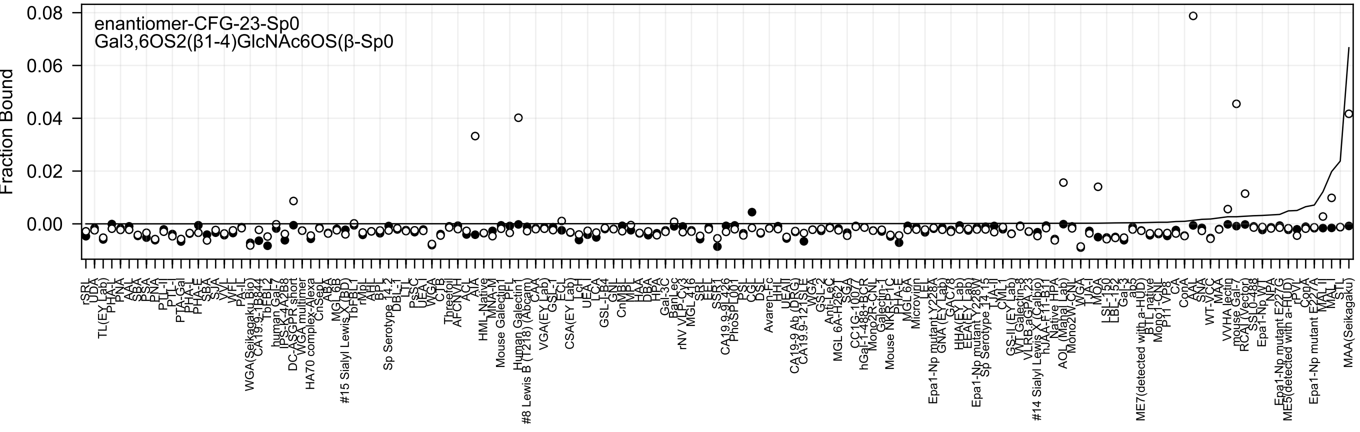
enantiomer-CFG-23-Sp0
Gal3,6OS2(β1-4)GlcNAc6OS(β-Sp0
Fraction Bound
0.08
0.06
0.04
0.02
0.00
rSRL
UDA
TL(EY Lab)
PHA-L
PNA
AAL
SBA
PSA
PNA
PTL-II
PTL-I
PTA-Gal
PHA-L
PHA-E
SBA
SJA
VVL
WFL
PA-IL
WGA(Seikagaku Bio)
CA19.9-1B844
TbBLL-2
human Gal-7
IPS-K-4A2B8
DC-ASGPR short
WGA multimer
HA70 complex-Alexa
CnSept
ABA
MGL6B
#15 Sialyl Lewis X (BD)
TbPBLL
rMpL
AAL
BPL
Sp Serotype 14.2
DBL-1
LTL
PsSC
UEA-I
WGA
CTB
Threefoil
AFCNVH
ACL
AIA
HML-Native
MNA-M
Mouse Galectin1
PFL
Human Galectin1
#8 Lewis B (T218) (Abcam)
CAA
VGA(EY Lab)
GSL-I
ECL
CSA(EY Lab)
LcH
UEA-I
LCA
GSL-I-B4
GNL
CnMBL
MPL
HAA
DBA
HPA
Gal-3C
BanLec
rNV VLP-Cy3
MGL 416
SBL
EEL
SSP1
CA19.9-9L426
PhoSL001
PSL
CGL
DSL
Avaren-Fc
HHL
UEA-II
CA19-9 Ab (DRG)
CA19.9-121SLE
WGA
GSL-2
Anti-LeC
MGL 6A-H262T
SGA
CC1G-1007
hGal-1-488+BCR
Mono2R-CNT
Galectin-1
Mouse NKR-P1C
PHA-E
MGL 6A
Microvirin
LEL
Epa1-Np mutant Y228A
GNA (EY Lab)
GAC78
HHA(EY Lab)
EEA(EY Lab)
Epa1-Np mutant Y228W
Sp Serotype 14.15
TJA-II
CML-I
GS-II (EY Lab)
WT-Galectin-8
VLRB.aGPA.23
#14 Sialyl Lewis X (Calbio)
hJAA-F11-B11
Native HPA
AOL (Mahal Lab)
Mono2W-CNT
WGA-I
TJA-I
MOA
LSL-150
LBL-152
Gal-3
ab5
ME7(detected with a-HUD)
BTL native
Mono1-CNT
P11 VP8
CA
ConA
AAL
SNA
WT-VCC
MAA
VVHA lectin
mouse Gal-7
RCAI (Vector)
SSL0-488
Epa1-Np WT
NPA
Epa1-Np mutant E227G
ME5(detected with a-HUD)
rPVL
ConA
Epa1-Np mutant E227A
MAL-II
MAL-I
STL
MAA(Seikagaku)

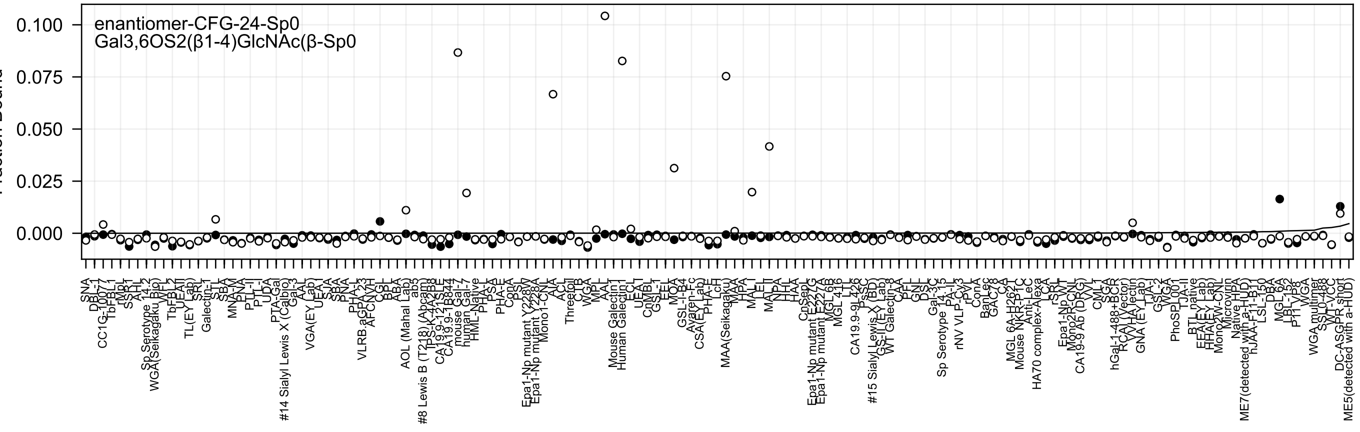
enantiomer-CFG-24-Sp0
Gal3,6OS2(β1-4)GlcNAc(β-Sp0

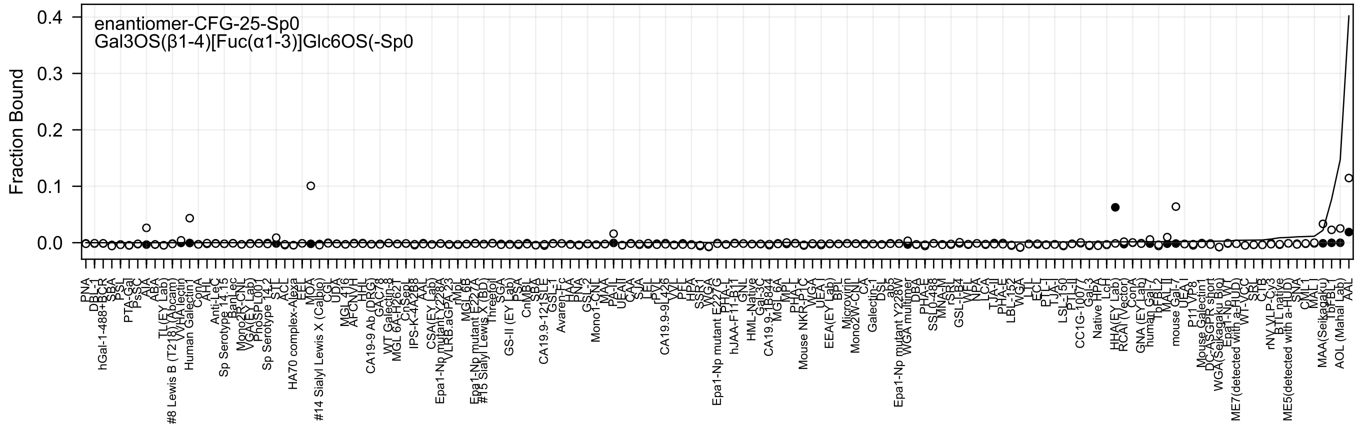
enantiomer-CFG-25-Sp0
Gal3OS(β1-4)[Fuc(α1-3)]Glc6OS(-Sp0
Fraction Bound
0.4
0.3
0.2
0.1
0.0

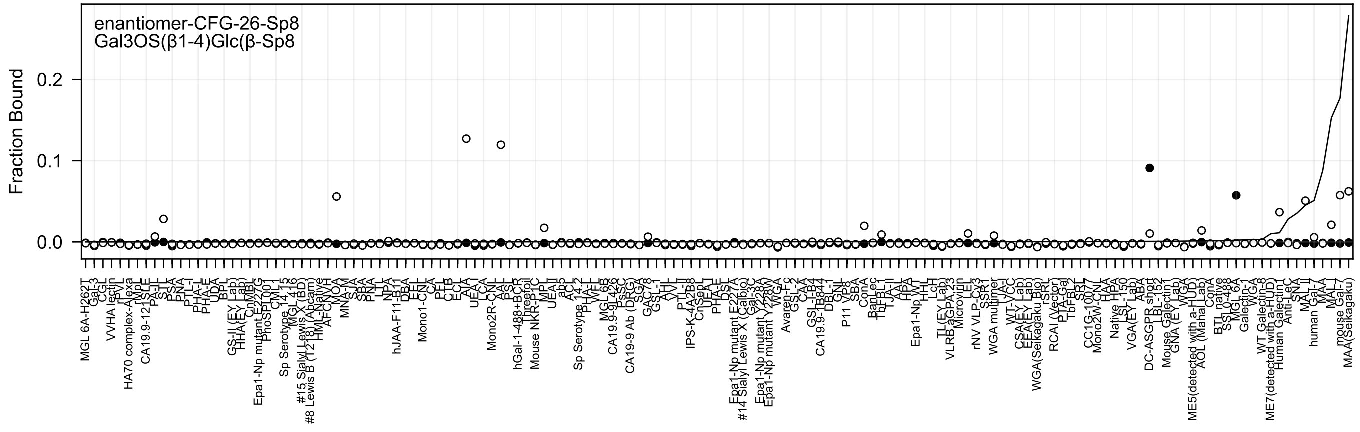
enantiomer-CFG-26-Sp8
Gal3OS(β1-4)Glc(β-Sp8
Fraction Bound
0.2
0.1
0.0

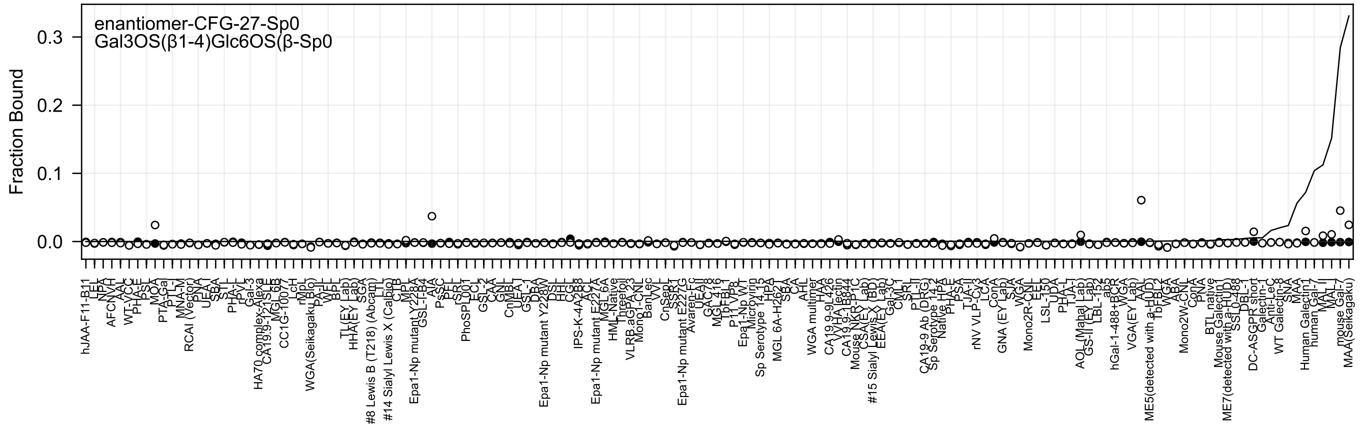
enantiomer-CFG-27-Sp0
Gal3OS(β1-4)Glc6OS(β-Sp0
Fraction Bound
0.3
0.2
0.1
0.0

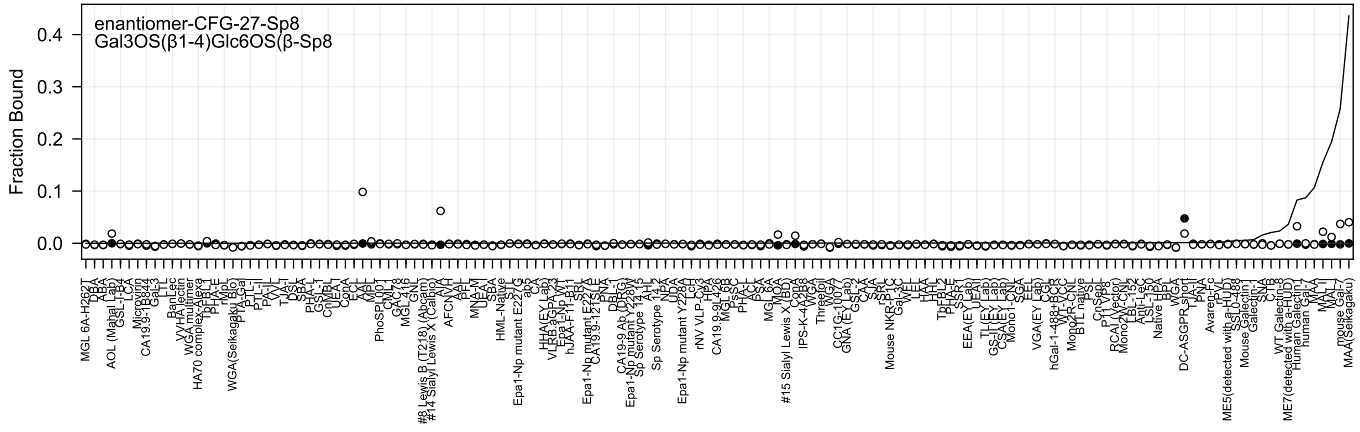

enantiomer-CFG-27-Sp8
Gal3OS(β1-4)Glc6OS(β-Sp8
Fraction Bound
0.4
0.3
0.2
0.1
0.0

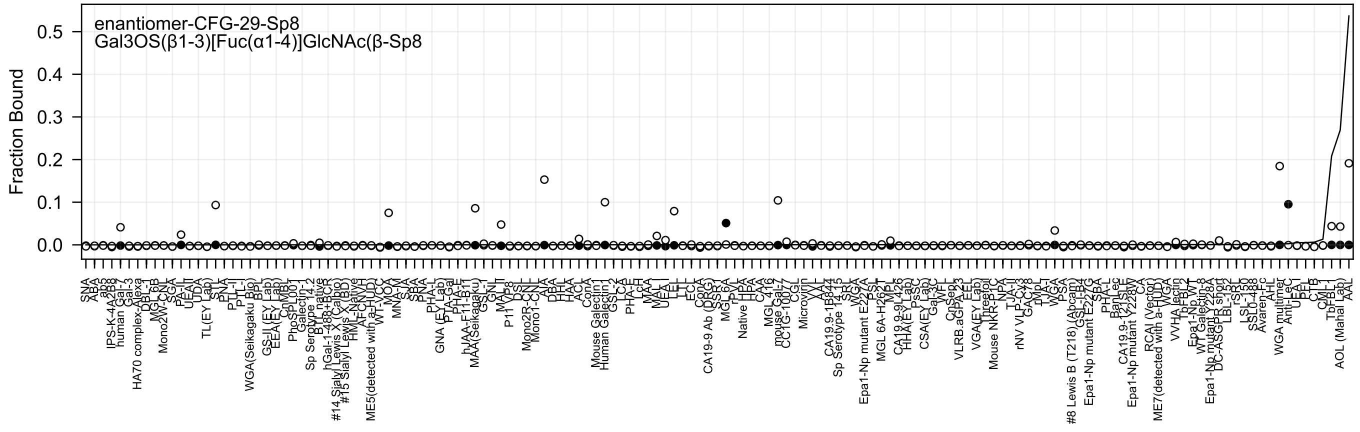
enantiomer-CFG-29-Sp8
Gal3OS(β1-3)[Fuc(α1-4)]GlcNAc(β-Sp8
Fraction Bound
0.5
0.4
0.3
0.2
0.1
0.0

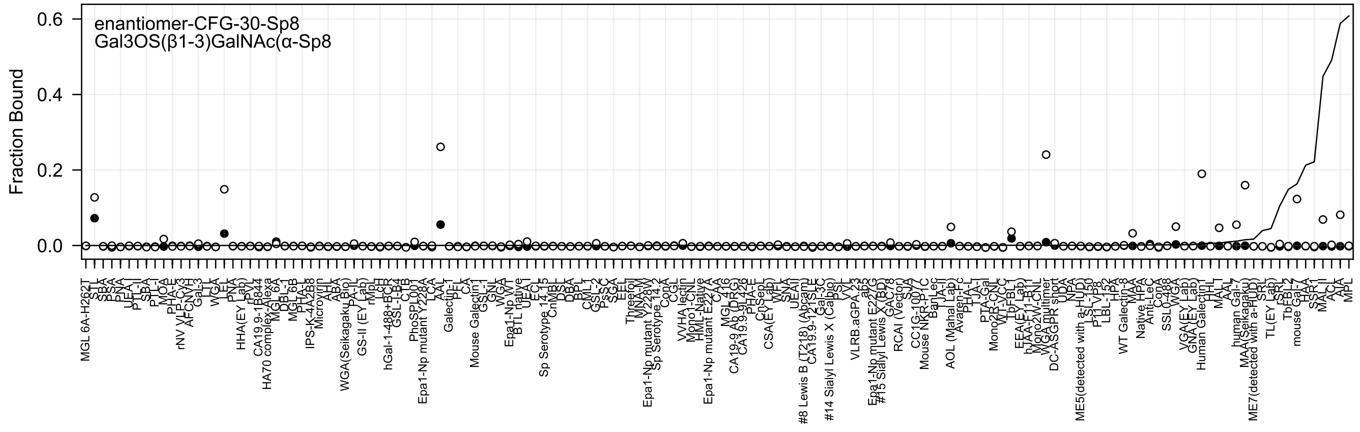

enantiomer-CFG-30-Sp8
Gal3OS(β1-3)GalNAc(α-Sp8
Fraction Bound
0.6
0.4
0.2
0.0

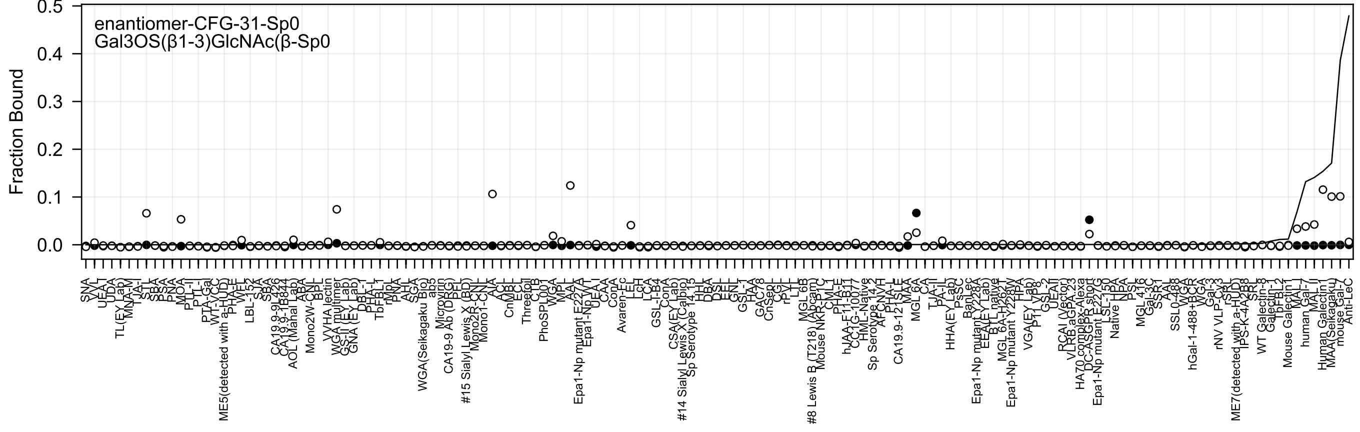

enantiomer-CFG-31-Sp0
Gal3OS(β1-3)GlcNAc(β-Sp0
Fraction Bound
0.0
0.1
0.2
0.3
0.4
0.5

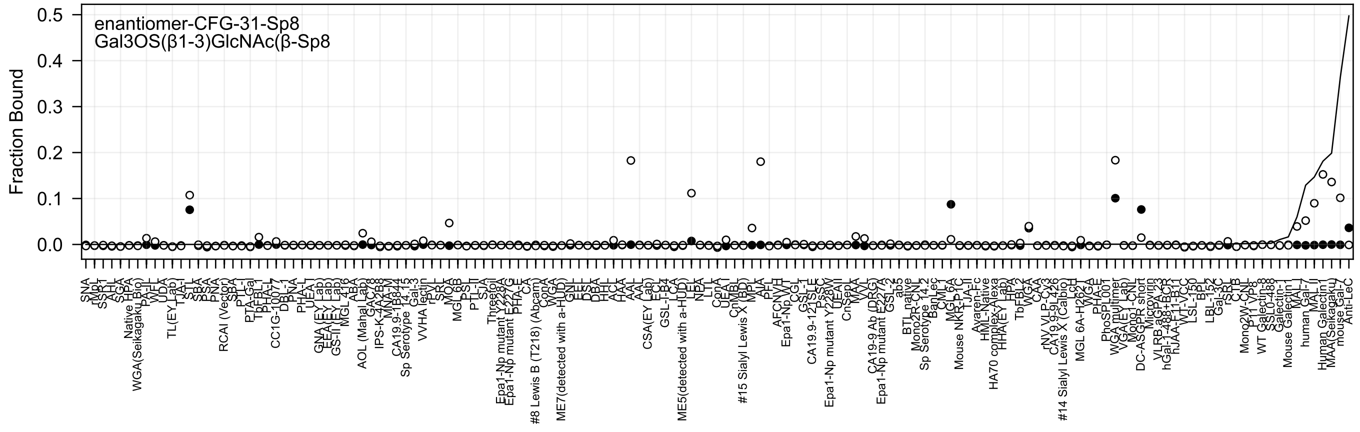

enantiomer-CFG-31-Sp8
Gal3OS(β1-3)GlcNAc(β-Sp8
Fraction Bound
0.5
0.4
0.3
0.2
0.1
0.0

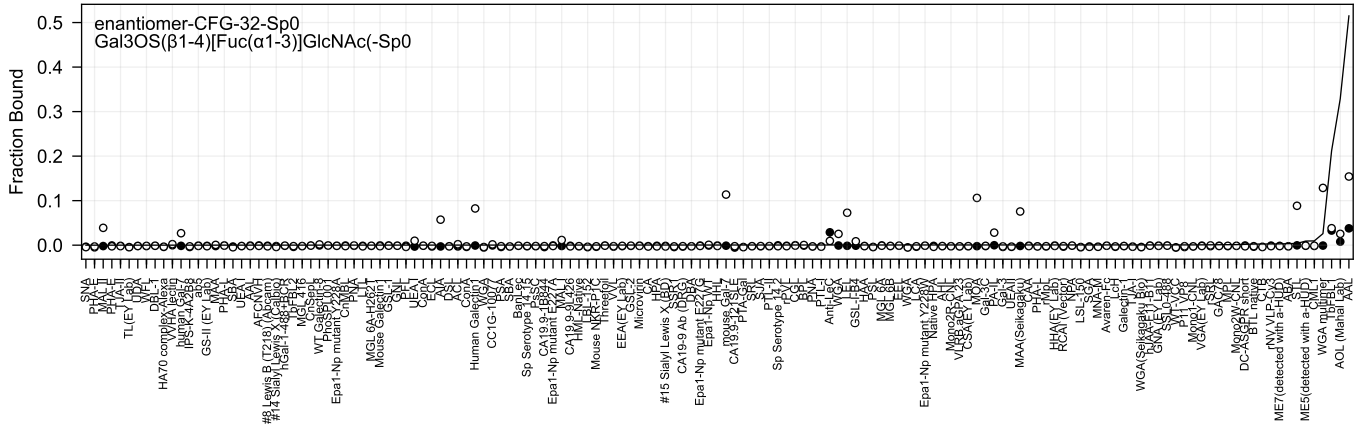
enantiomer-CFG-32-Sp0
Gal3OS(β1-4)[Fuc(α1-3)]GlcNAc(-Sp0
Fraction Bound
0.5
0.4
0.3
0.2
0.1
0.0

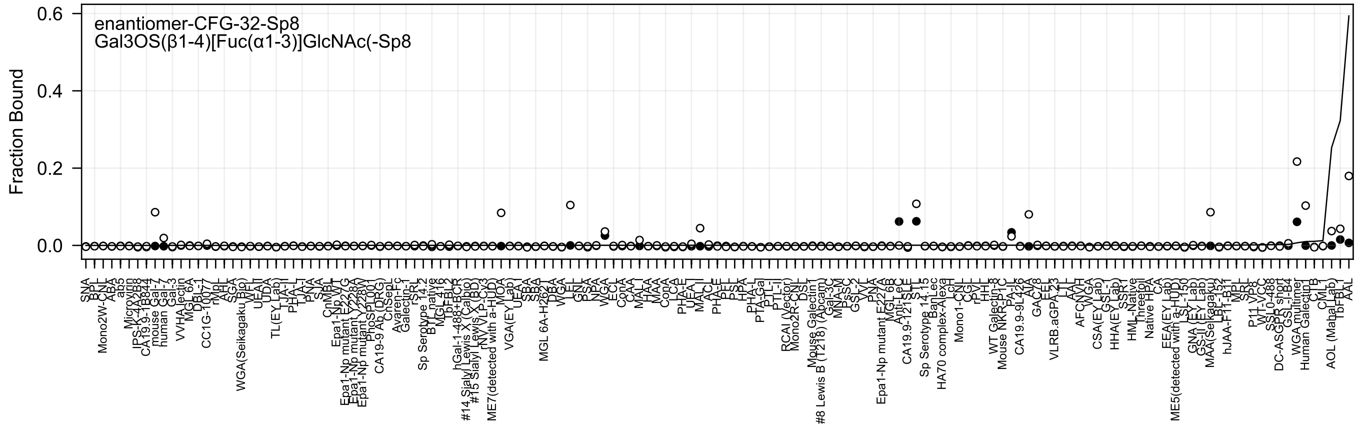

enantiomer-CFG-32-Sp8
Gal3OS(β1-4)[Fuc(α1-3)]GlcNAc(-Sp8
Fraction Bound
0.6
0.4
0.2
0.0

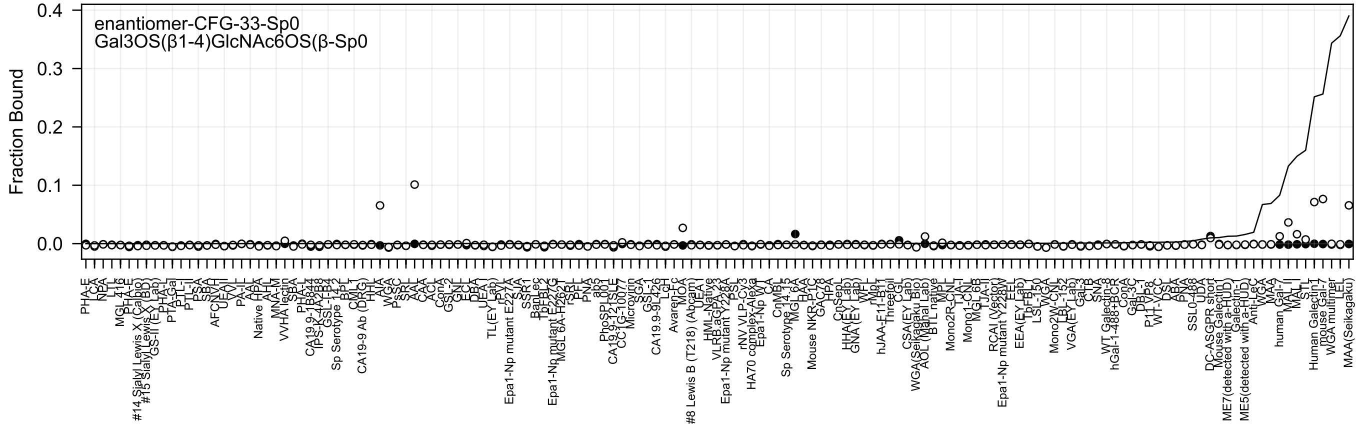
enantiomer-CFG-33-Sp0
Gal3OS(β1-4)GlcNAc6OS(β-Sp0
Fraction Bound
0.4
0.3
0.2
0.1
0.0

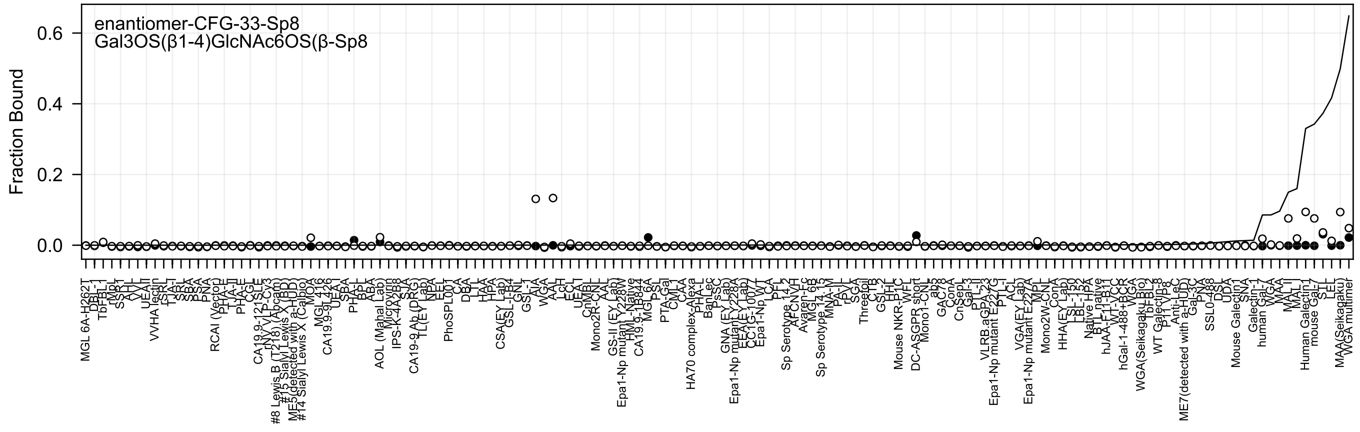
enantiomer-CFG-33-Sp8
Gal3OS(β1-4)GlcNAc6OS(β-Sp8
Fraction Bound

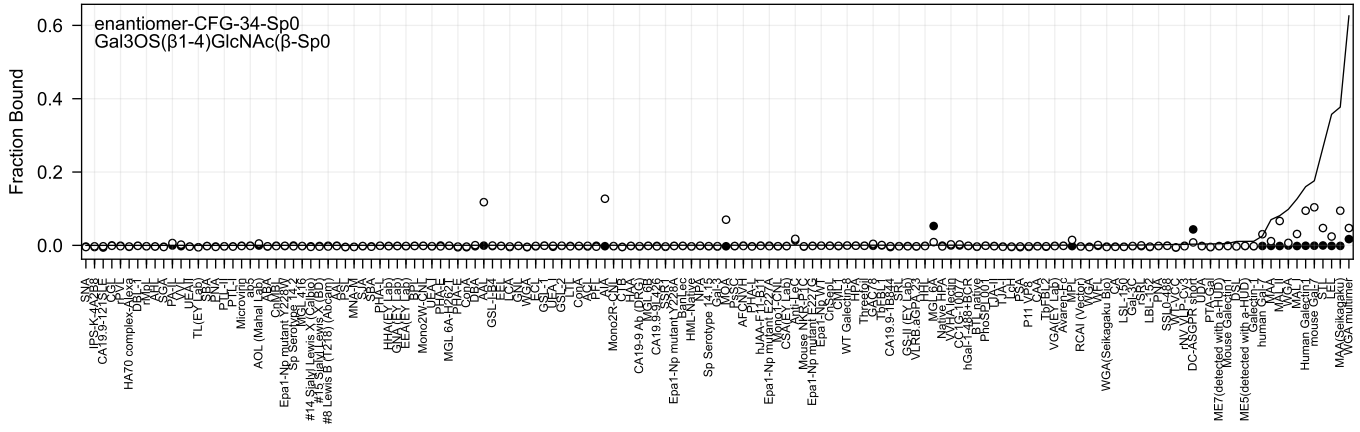

enantiomer-CFG-34-Sp0
Gal3OS(β1-4)GlcNAc(β-Sp0
Fraction Bound
0.6
0.4
0.2
0.0

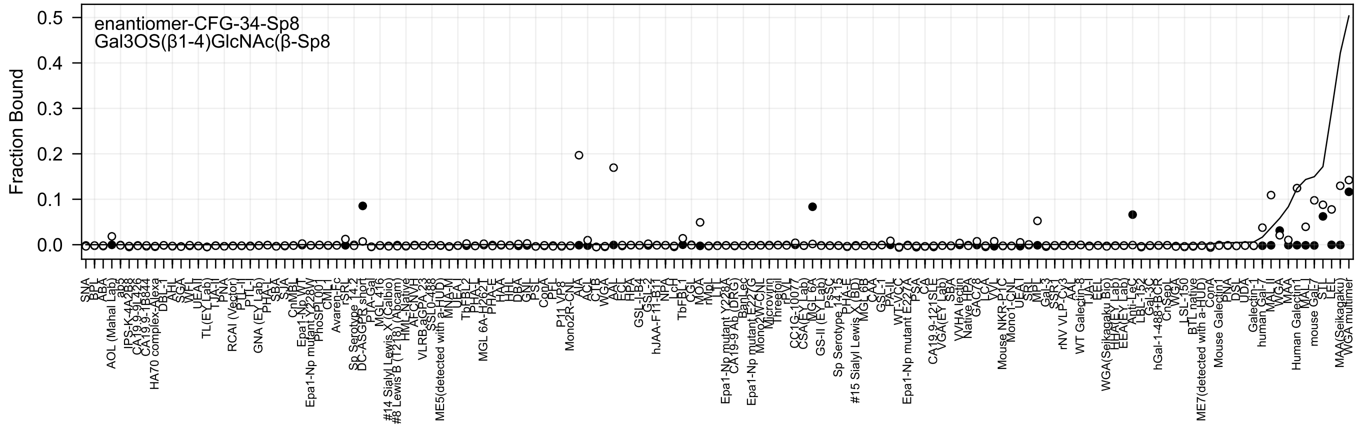
enantiomer-CFG-34-Sp8
Gal3OS(β1-4)GlcNAc(β-Sp8
Fraction Bound
0.0
0.1
0.2
0.3
0.4
0.5
SNA
BPL
ABA
AOL (Mahal Lab)
ab3
IPS-K-4A2B8
CA19.9-9L426
CA19.9-1B844
HA70 complex-Alexa
DBL-1
AHL
SGA
WFL
UEAII
TL(EY Lab)
TJA-I
PNA
RCAI (Vector)
PTL-I
PTL-II
GNA (EY Lab)
PHA-L
SBA
SJA
CnMBL
Epa1-Np WT
Epa1-Np mutant Y228W
PhoSL100
CML-1
Avaren-Fc
rSRL
Sp Serotype 14.2
DC-ASGPR short
PTA-Gal
MGL 416
#14 Sialyl Lewis X (Calbio)
#8 Lewis B (T218) (Abcam)
HML-Native
AFC-NVT
VLRB.aGPA.23
SSL0-488
ME5(detected with a-HUD)
MNA-M
UEA-I
TbFBL2
PHA-L
MGL 6A-H262T
PHA-E
HAA
HHL
DBA
GNL
PSL
ConA
PFL
P11 VP8
Mono2R-CNL
AIA
ACL
CTB
WGA
AAL
ECL
HPA
GSL-I-B4
GSL-2
hJAA-F11-B11
NPA
LcH
TbFBL1
CGL
MOA
rMpL
TJA-II
Epa1-Np mutant Y228A
CA19-9 Ab (DRG)
BanLec
Epa1-Np mutant E227G
Mono2W-CNL
Microvirin
Threefoil
rPVL
CC1G-10077
CSA(EY Lab)
MGL 6A
GS-II (EY Lab)
PsSC
Sp Serotype 14.15
PHA-E
#15 Sialyl Lewis X (BD)
MGL 6B
CAA
GSL-1
PA-IL
WT-VCC
Epa1-Np mutant E227A
PSA
CA
CA19.9-121SLE
VGA(EY Lab)
SBA
VVHA lectin
Native HPA
GAC78
LCA
VVL
Mouse NKR-P1C
Mono1-CNL
UEA-I
SRL
MPL
Gal-3
SSR1
rNV VLP-Cy3
AAL
WT Galectin-8
TJA-I
EEL
WGA(Seikagaku Bio)
HHA(EY Lab)
EEA(EY Lab)
Anti-LeC
LBL-152
Gal-3C
hGal-1-488+BCR
CnSepL
VGA
LSL-150
BTL native
ME7(detected with a-HUD)
ConA
Mouse Galectin1
PNA
DSL
UDA
Galectin-1
human Gal-7
MAL II
WGA
MAA
Human Galectin1
MAL I
mouse Gal-7
STL
LEL
MAA(Seikagaku)
WGA multimer

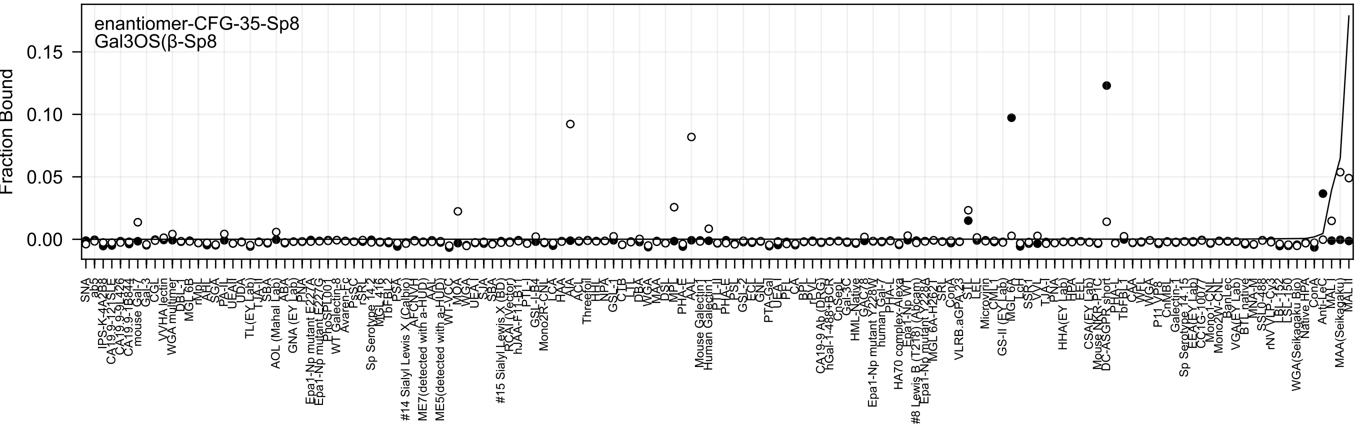
enantiomer-CFG-35-Sp8
Gal3OS(β-Sp8
Fraction Bound
0.15
0.10
0.05
0.00

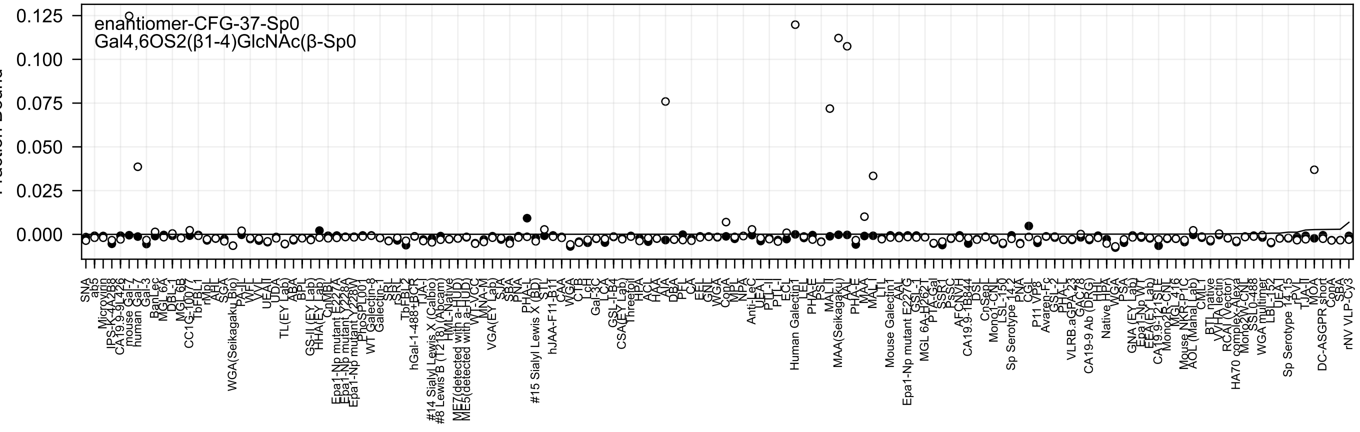
enantiomer-CFG-37-Sp0
Gal4,6OS2(β1-4)GlcNAc(β-Sp0
0.125
0.100
0.075
0.050
0.025
0.000
SNA
ab5
Microvirin
IPS-K-4A2B8
CA19.9-9L426
mouse Gal-7
human Gal-7
Gal-3
BanLec
MGL 6A
DBL-1
MGL 6B
CC1G-1007
TbFBL1
rMpL
AHT
SGA
WGA(Seikagaku Bio)
PA-IL
WFL
VVL
UEA-I
UDA
TL(EY Lab)
ABA
BPL
GS-II (EY Lab)
HHA(EY Lab)
CNMBL
Epa1-Np mutant E227A
Epa1-Np mutant Y228A
Epa1-Np mutant Y228W
PhoSL001
WT Galectin-8
Galectin-1
SRL
rSRL
TbFBL2
hGal-1-488+BCR
TJA-I
#14 Sialyl Lewis X (Calbio
#8 Lewis B (T218) (Abcam
HML-Native
ME7(detected with a-HUD)
ME5(detected with a-HUD)
WT-VCC
MNA-M
VGA(EY Lab)
SJA
SBA
PNA
PHA-L
#15 Sialyl Lewis X (BD)
STL
hJAA-F11-B11
CAA
WGA
CTB
LcH
Gal-3C
LCA
GSL-I-B4
CSA(EY Lab)
Threefoil
HPA
ACL
HAA
AIA
DBA
PFL
CA
EEL
GNL
WGA
ConA
MPL
NPA
Anti-LeC
UEA-I
PTL-I
PTL-II
ECL
Human Galectin1
LEL
PHA-E
PSL
MAL-II
MAA(Seikagaku)
AAL
PHA-E
MAA
MAL-I
LTL
Mouse Galectin1
AAL
Epa1-Np mutant E227G
GSL-II
MGL 6A-H262T
PTA-Gal
SSR1
PsSC
AFCNVH
CA19.9-1B844
DSL
CnSep
Mono1-CNL
LSL-150
Sp Serotype 14-2
PNA
CGL
P11 VP8
Avaren-Fc
GSL-2
PHA-L
VLRB.aGPA.23
GAC78
CA19-9 Ab (DRG)
HHL
Native HPA
WGA
PSA
GNA (EY Lab)
Epa1-Np WT
EEA (EY Lab)
CA19.9-121SLE
Mono2R-CNL
MGL 416
Mouse NKR-P1C
AOL (Manal Lab)
CML
BTL native
VHA lectin
RCA I (Vector)
HA70 complex-Alexa
Mono2W-CNL
SSL0-488
WGA multimer
LBL-152
UEA-I
Sp Serotype 14-15
rPVL
TJA-II
MOA
DC-ASGPR short
ConA
SBA
rNV VLP-Cy3

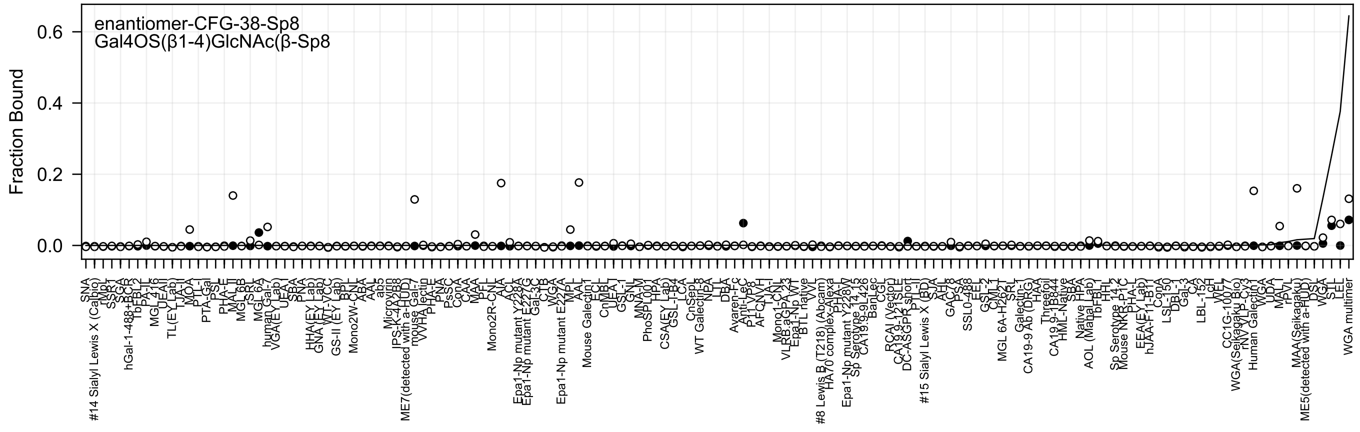
enantiomer-CFG-38-Sp8
Gal4OS(β1-4)GlcNAc(β-Sp8
Fraction Bound

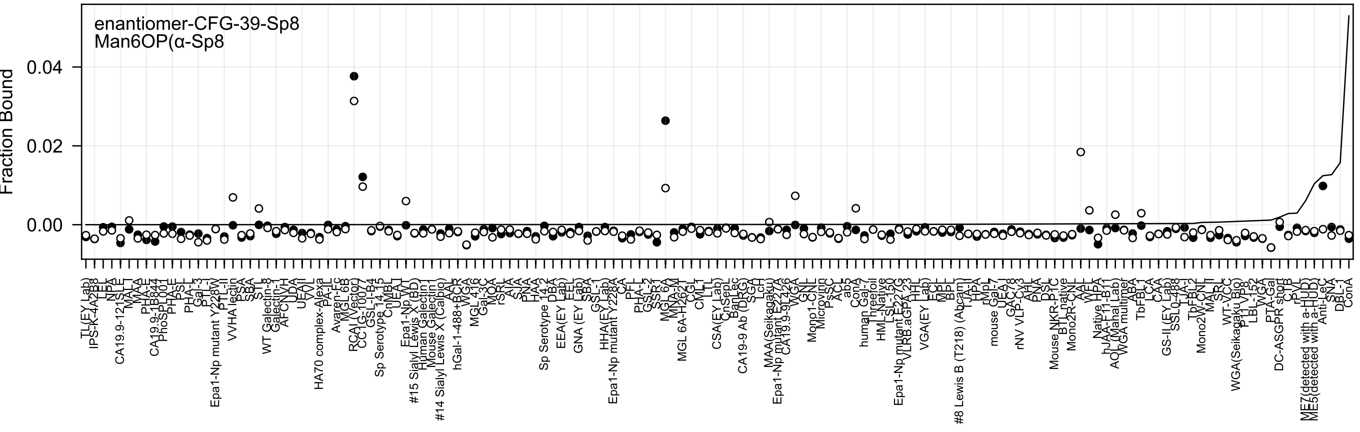

enantiomer-CFG-39-Sp8
Man6OP(α-Sp8
Fraction Bound
0.04
0.02
0.00

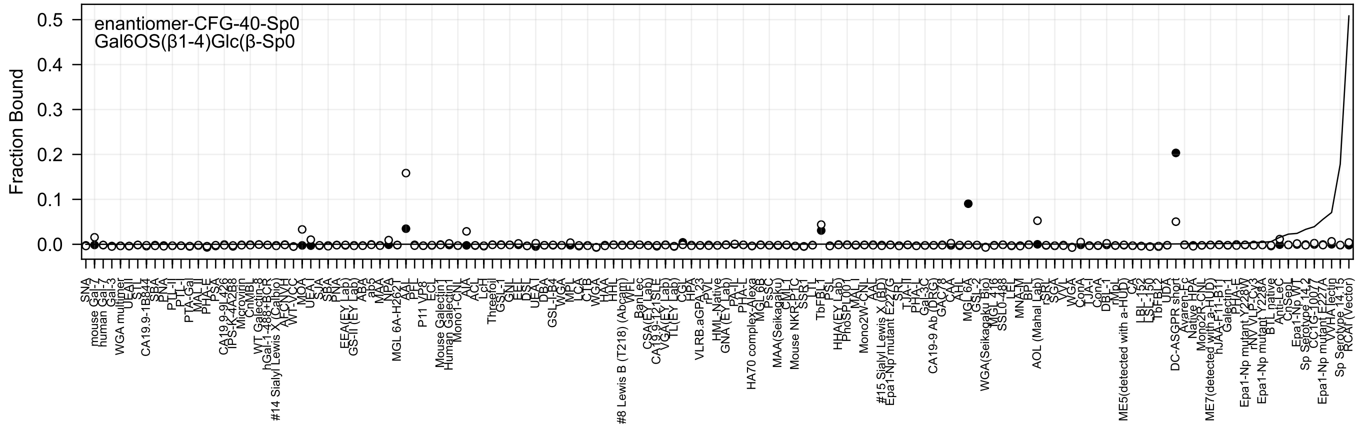
enantiomer-CFG-40-Sp0
Gal6OS(β1-4)Glc(β-Sp0
Fraction Bound

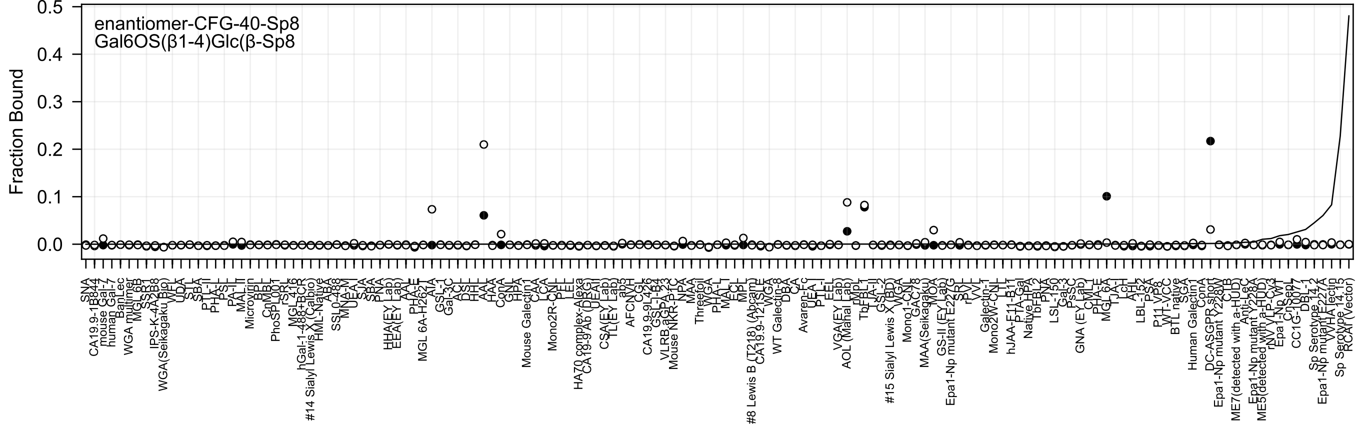

enantiomer-CFG-40-Sp8
Gal6OS(β1-4)Glc(β-Sp8
Fraction Bound
0.5
0.4
0.3
0.2
0.1
0.0

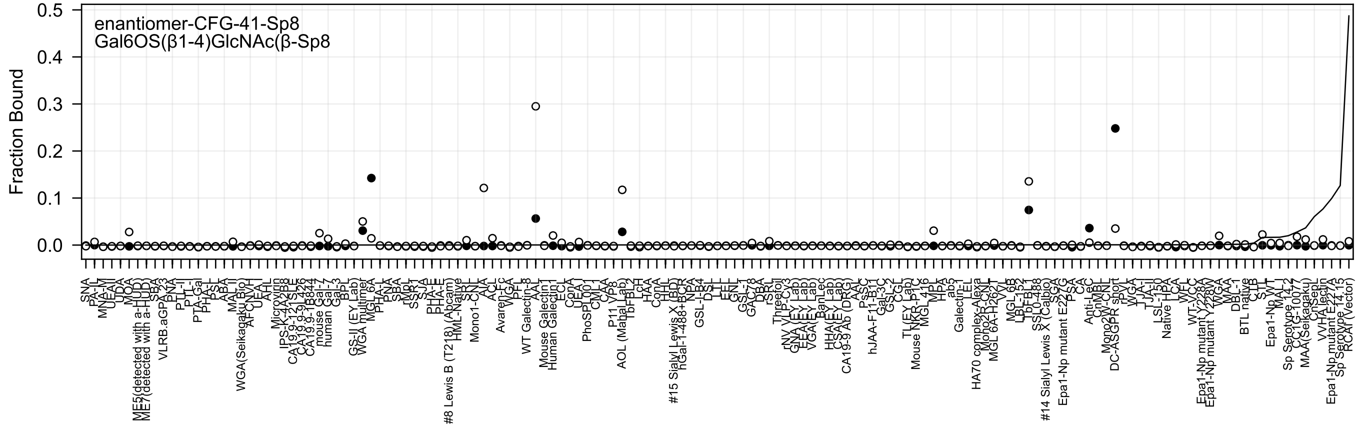
enantiomer-CFG-41-Sp8
Gal6OS(β1-4)GlcNAc(β-Sp8
Fraction Bound
0.5
0.4
0.3
0.2
0.1
0.0
SNA
PA-IL
MNA-M
UEA-I
UDA
MOA
ME5(detected with a-HUD)
ME7(detected with a-HUD)
SBA
VLRB.aGPA23
PNA
PTL-II
PTL-I
PTA-Gal
PHA-L
PSL
ABA
MAL II
WGA(Seikagaku Bio)
AFCNVH
UEA-I
AAL
Microvirin
IPS-K-4A2B8
CA19.9-121SLE
CA19.9-9L426
CA19.9-1B844
mouse Gal-7
human Gal-7
Gal-3
BPL
GS-II (EY Lab)
WGA multimer
MGL 6A
PHA-L
PNA
SBA
rMpL
SSR1
SJA
PHA-E
PHA-E
#8 Lewis B (T218) (Abcam)
HML-Native
SNL
Mono1-CNL
AIA
ACL
Avaren-Fc
WGA
PFL
WT Galectin-8
AAL
Mouse Galectin1
Human Galectin1
ECL
ConA
UEA-I
PhoSL001
CML-1
CAA
P11 VP8
AOL (Mahal Lab)
TbFBL2
LcH
HAA
ConA
HHL
#15 Sialyl Lewis X (BD)
hGal-1-488+BCR
NPA
GSL-I-B4
DSL
LEL
EEL
GNL
GSL-1
GAC78
DBA
rSRL
Threefoil
rNY VP-Cy3
GNA (EY Lab)
EEA(EY Lab)
VGA(EY Lab)
BanLec
HHA(EY Lab)
CSA(EY Lab)
CA19-9 Ab (DRG)
AAL
PsSC
hJAA-F11-B11
Gal-3C
GSL-2
CGL
TL(EY Lab)
Mouse NKR-P1C
MGL 416
MPL
HPA
ab5
Galectin-1
LEL
HA70 complex-Alexa
Mono2R-CNL
MGL 6A-H262T
VVL
MGL 6B
LBL-152
TbFBL1
SSL0-488
#14 Sialyl Lewis X (Calbio)
SGA
Epa1-Np mutant E227G
PSA
CA
Anti-LeC
CnMBL
Mono2W-CNL
DC-ASGPR short
rPVL
WGA
TJA-I
TJA-I
LSL-150
Native HPA
LCA
WFL
WT-VCC
Epa1-Np mutant Y228A
Epa1-Np mutant Y228W
WGA
MAA
DBL-1
BTL native
CTB
STL
Epa1-Np WT
MAL-I
Sp Serotype 14.2
CC1G-1007
MAA(Seikagaku)
CnSepL
VVHA lectin
Epa1-Np mutant E227A
Sp Serotype 14.15
RCA1 (Vector)

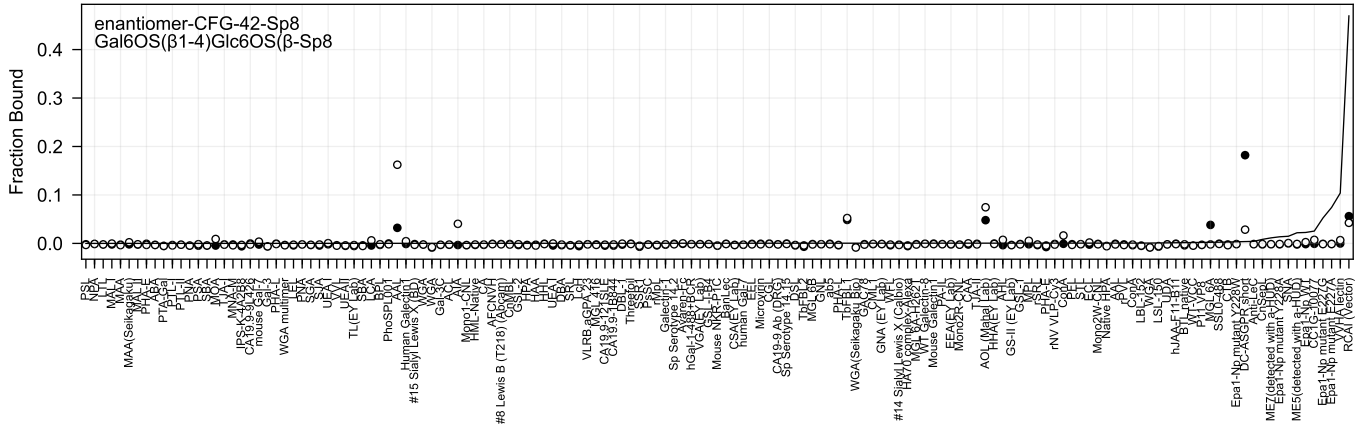
enantiomer-CFG-42-Sp8
Gal6OS(β1-4)Glc6OS(β-Sp8
Fraction Bound

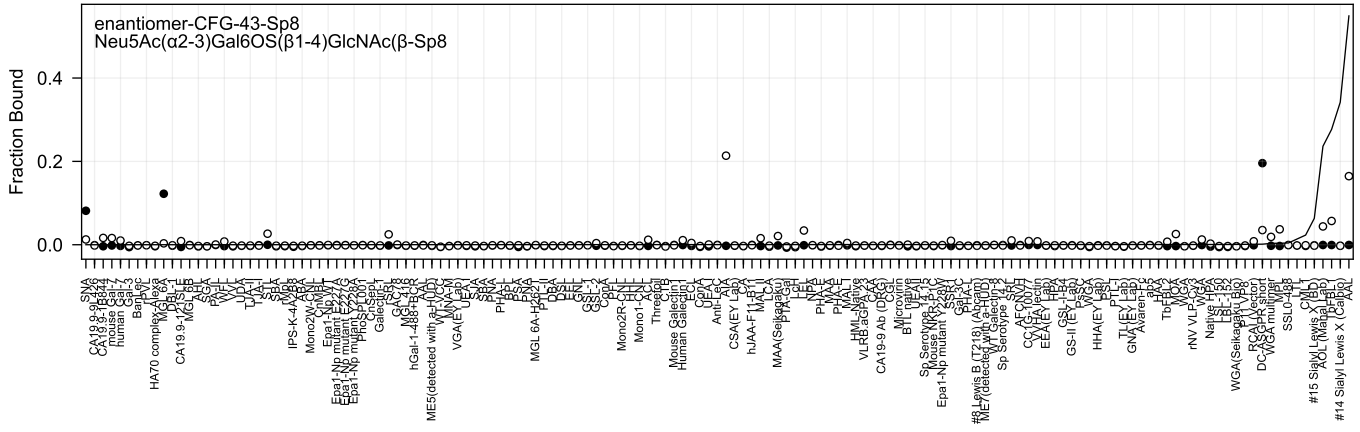
enantiomer-CFG-43-Sp8
Neu5Ac(α2-3)Gal6OS(β1-4)GlcNAc(β-Sp8
Fraction Bound
0.0
0.2
0.4

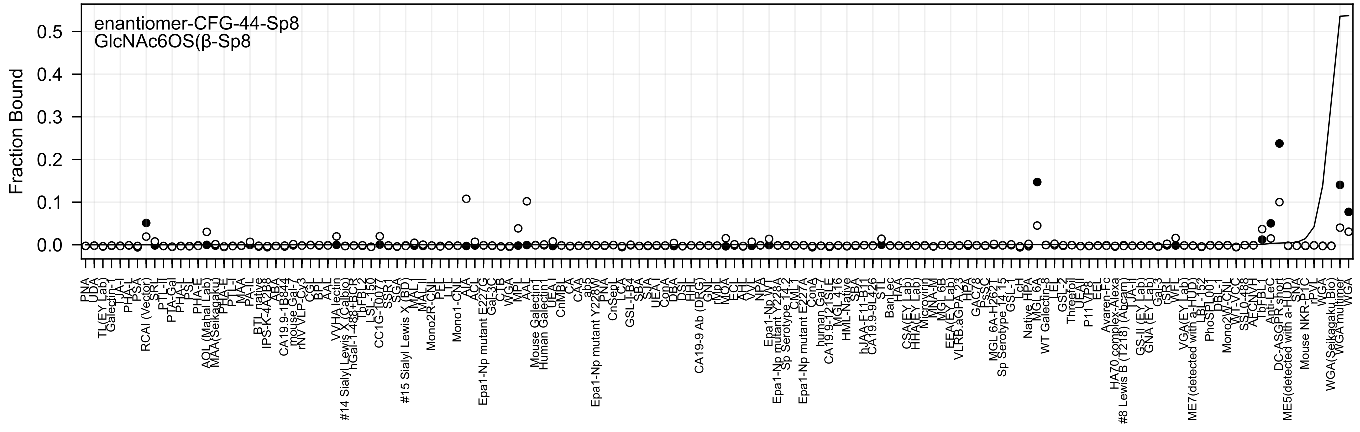
enantiomer-CFG-44-Sp8
GlcNAc6OS(β-Sp8
Fraction Bound
0.0
0.1
0.2
0.3
0.4
0.5
PNA
UDA
TL(EY Lab)
Galectin-1
TJA-I
PHA-L
PSA
RCAI (Vector)
SRL
PTL-II
PTA-Gal
PHA-L
PSL
PHA-E
AOL (Mahal Lab)
MAA(Seikagaku)
PHA-E
PTL-I
MAA
PA-IL
BTL native
IPS-K-4A2B8
ABA
CA19.9-1B844
mouse Gal-7
rNV VLP-Cy3
CGL
BPL
AAL
VVHA lectin
#14 Sialyl Lewis X (Calbio)
hGal-1-488+BCR
TbFBL2
LSL-150
CC1G-10077
SSR1
SGA
#15 Sialyl Lewis X (BD)
MAL-I
MAL-I
Mono2R-CNL
PFL
LTL
Mono1-CNL
AIA
ACL
Epa1-Np mutant E227G
Gal-3C
CTB
WGA
MPL
AAL
Mouse Galectin1
Human Galectin1
UEA-I
CnMBL
CAA
ab5
Epa1-Np mutant Y228W
PNA
CnSepL
LCA
GSL-I-B4
SBA
SJA
UEA
ConA
DBA
DSL
HHL
CA19-9 Ab (DRG)
GNL
rMpL
MOA
ECL
AHL
VVL
NPA
Epa1-Np WT
Epa1-Np mutant Y228A
Sp Serotype 14.2
CML1
Epa1-Np mutant E227A
ConA
human Gal-7
CA19.9-121SLE
MGL 416
HML-Native
SBA
hJAA-F11-B11
CA19.9-9L426
STL
BanLec
HAA
CSA(EY Lab)
HHA(EY Lab)
Microvirin
MNA-M
MGL 6B
EEA(EY Lab)
VLRB.aGPA.23
HPA
GAC78
PsSC
MGL 6A-H262T
Sp Serotype 14.15
GSL-I
LcH
Native HPA
MGL 6A
WT Galectin-8
LEL
GSL-2
Threefoil
UEAII
P11 VP8
EEL
Avaren-Fc
HA70 complex-Alexa
#8 Lewis B (T218) (Abcam)
TJA-II
GS-II (EY Lab)
GNA (EY Lab)
Gal-3
rSRL
WFL
VGA(EY Lab)
ME7(detected with a-HUD)
LBL-152
PhoSL001
DBL-I
Mono2W-CNL
WT-VCC
SSL0-488
AFCNVH
TbFBL1
Anti-LeC
DC-ASGPR short
ME5(detected with a-HUD)
SNA
Mouse NKR-P1C
rPVL
WGA
WGA(Seikagaku.Bio)
WGA multimer
WGA

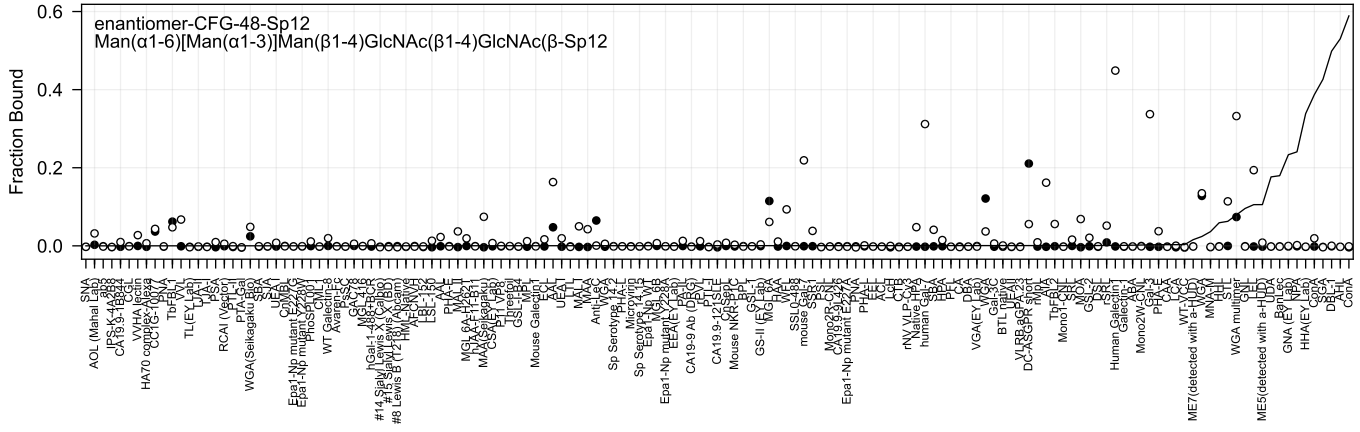
enantiomer-CFG-48-Sp12
Man(α1-6)[Man(α1-3)]Man(β1-4)GlcNAc(β1-4)GlcNAc(β-Sp12
Fraction Bound
0.6
0.4
0.2
0.0

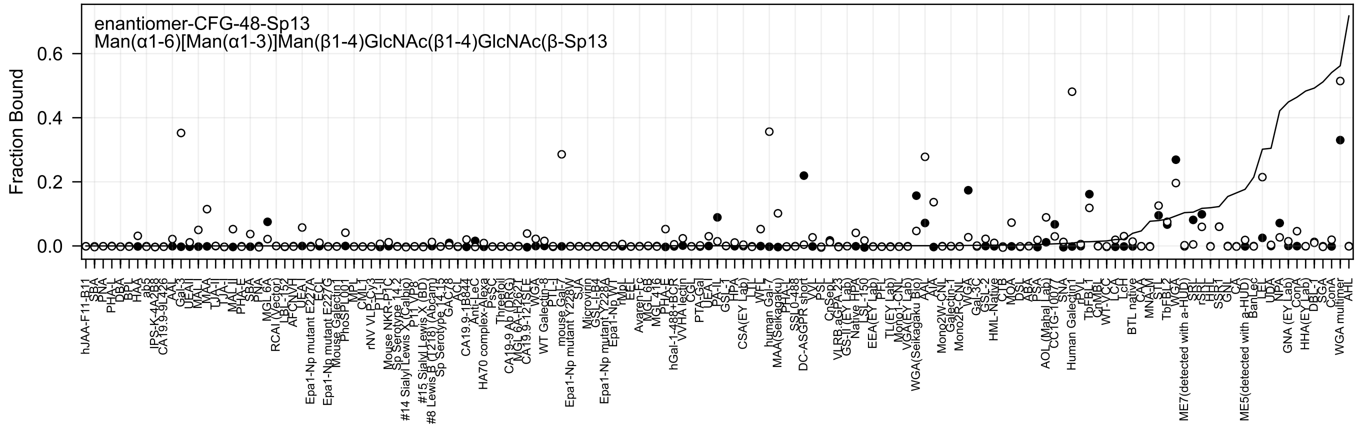
enantiomer-CFG-48-Sp13
Man(α1-6)[Man(α1-3)]Man(β1-4)GlcNAc(β1-4)GlcNAc(β-Sp13
Fraction Bound
0.0
0.2
0.4
0.6

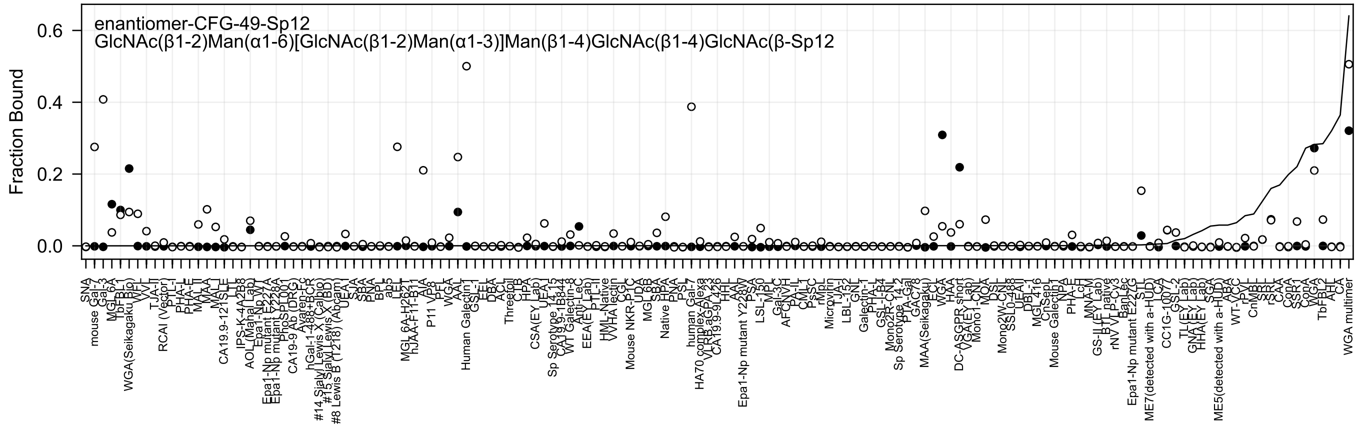
enantiomer-CFG-49-Sp12
GlcNAc(β1-2)Man(α1-6)[GlcNAc(β1-2)Man(α1-3)]Man(β1-4)GlcNAc(β1-4)GlcNAc(β-Sp12
Fraction Bound
0.6
0.4
0.2
0.0

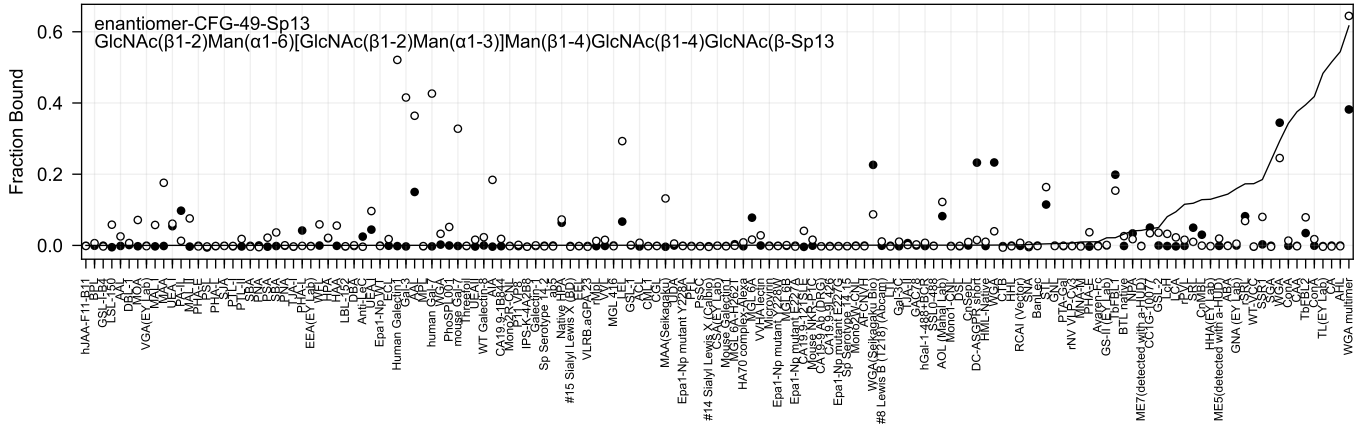
enantiomer-CFG-49-Sp13
GlcNAc(β1-2)Man(α1-6)[GlcNAc(β1-2)Man(α1-3)]Man(β1-4)GlcNAc(β1-4)GlcNAc(β-Sp13
Fraction Bound
0.0
0.2
0.4
0.6

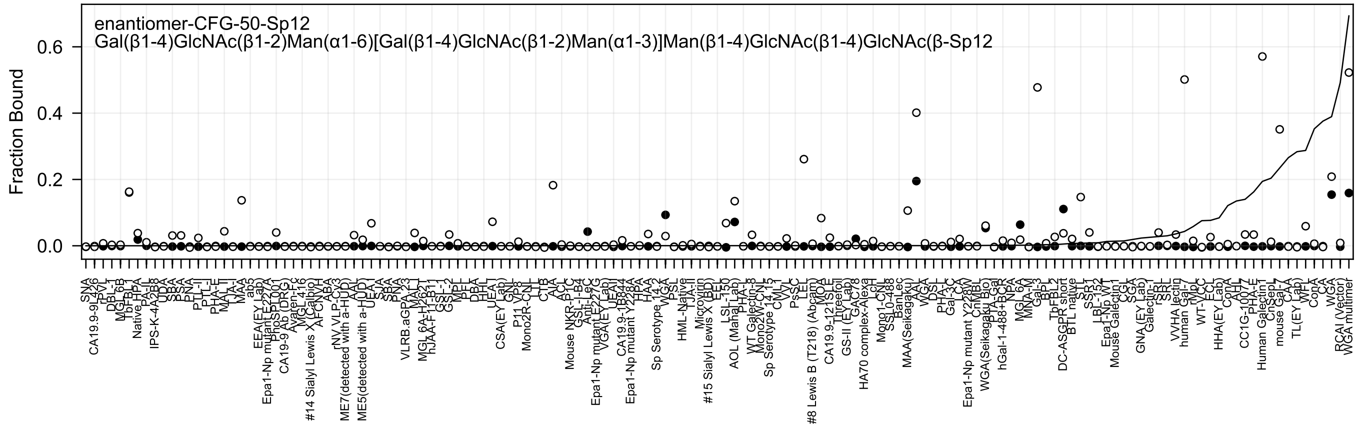
enantiomer-CFG-50-Sp12
Gal(β1-4)GlcNAc(β1-2)Man(α1-6)[Gal(β1-4)GlcNAc(β1-2)Man(α1-3)]Man(β1-4)GlcNAc(β1-4)GlcNAc(β-Sp12
Fraction Bound
0.6
0.4
0.2
0.0

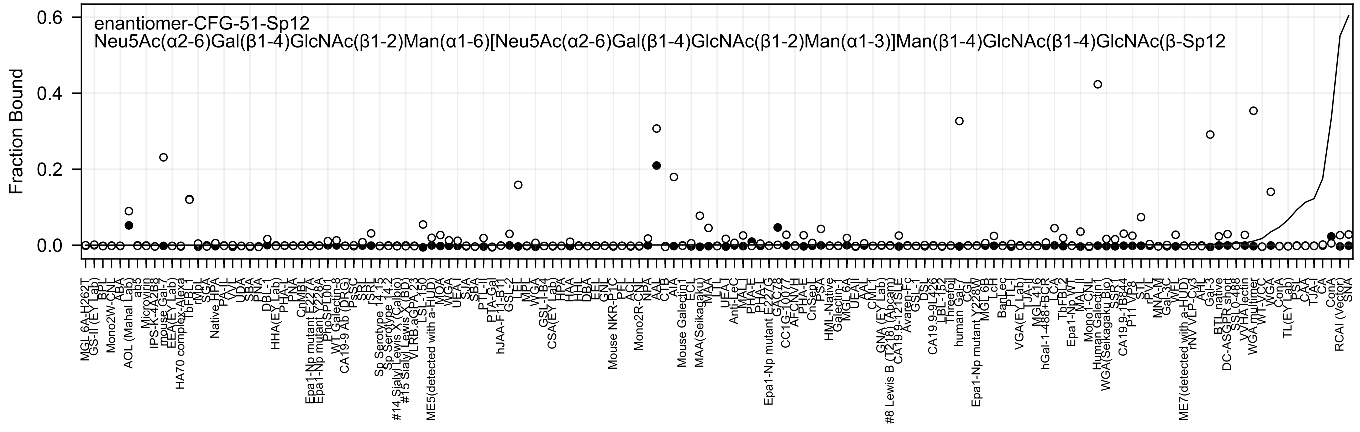
enantiomer-CFG-51-Sp12
Neu5Ac(α2-6)Gal(β1-4)GlcNAc(β1-2)Man(α1-6)[Neu5Ac(α2-6)Gal(β1-4)GlcNAc(β1-2)Man(α1-3)]Man(β1-4)GlcNAc(β1-4)GlcNAc(β-Sp12
Fraction Bound
0.6
0.4
0.2
0.0
MGL 6A-H262T
GS-II (EY Lab)
BPL
Mono2W-CNL
ABA
AOL (Mahal Lab)
ab5
Microvirin
IPS-K-4A2B8
mouse Gal-7
EEA(EY Lab)
HA70 complex-Alexa
TbFBL1
rMpL
SGA
Native HPA
PA-IL
VVL
UDA
SBA
PNA
DBL-1
HHA(EY Lab)
PHA-L
PNA
CnMBL
Epa1-Np mutant E227A
Epa1-Np mutant Y228A
PhoSL001
WT Galectin-8
CA19-9 Ab (DRG)
PsSC
SRL
Sp Serotype 14.15
Sp Serotype 14.2
#14 Sialyl Lewis X (Calbio)
#15 Sialyl Lewis X (BD)
VLRB.aGPA.23
LSL-150
ME5(detected with a-HUD)
MOA
WGA
UEA-I
SJA
SBA
PTL-II
PTA-Gal
hJAA-F11-B1.2
GSL-II
LEL
MPL
GSL-I-B4
CSA(EY Lab)
HPA
HAA
HHL
DBA
EEL
GNL
Mouse NKR-P1C
PEL
ACL
Mono2R-CNL
NPA
AAL
CTB
AIA
Mouse Galectin1
ECL
MAA(Seikagaku)
MAL
Anti-LeC
MAL-I
PHA-E
PHA-L
Epa1-Np mutant E227G
AFCNVH
PHA-E
CnSepL
PSA
HML-Native
Galectin-1
MGL 6A
UEA-II
AAL
GNA (EY Lab)
#8 Lewis B (T218) (Abcam)
CA19.9-121SLE
Avaren-Fc
GSL-I
DSL
CA19.9-9L426
LBL-152
Threefoil
human Gal-7
CGL
Epa1-Np mutant Y228W
MGL 6B
LcH
BanLec
PTL-I
VGA(EY Lab)
TJA-II
MGL
hGal-1-488+BCR
LCA
TbFBL2
Epa1-Np WT
MAL-II
Mono1-CNL
Human Galectin1
WGA(Seikagaku Bio)
SSA
CA19.9-1B8.4
P11 VP8
STL
rPVL
MNA-M
Gal-3C
WFL
ME7(detected with a-HUD)
rNV VLP-Cy3
AHL
Gal-3
BTL native
DC-ASGPR short
SSL0-488
VVHA lectin
WGA multimer
WT-VCC
WGA
ConA
TL(EY Lab)
PSL
CAA
TJA-I
CA
ConA
RCAI (Vector)
SNA

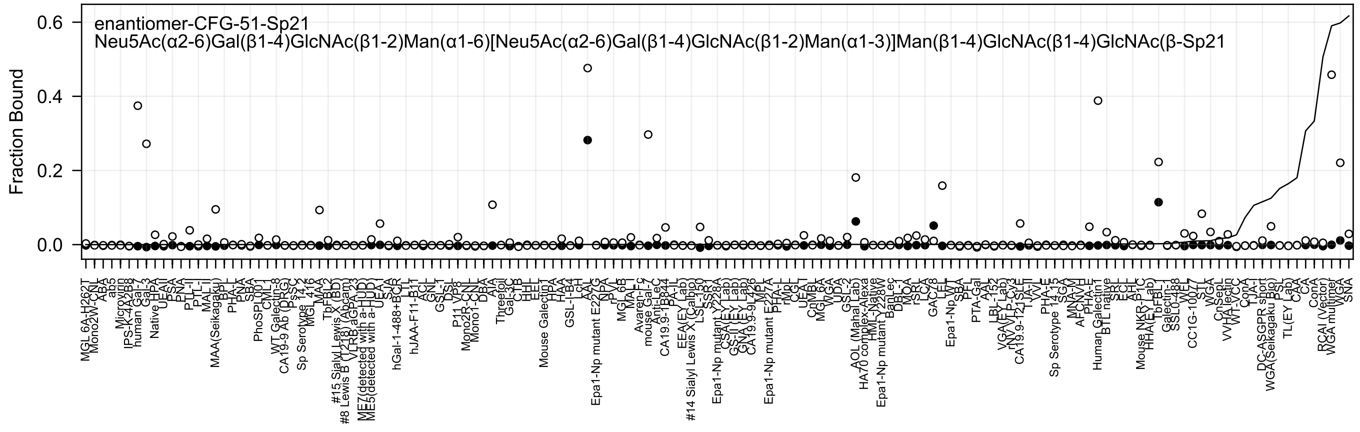
enantiomer-CFG-51-Sp21
Neu5Ac(α2-6)Gal(β1-4)GlcNAc(β1-2)Man(α1-6)[Neu5Ac(α2-6)Gal(β1-4)GlcNAc(β1-2)Man(α1-3)]Man(β1-4)GlcNAc(β1-4)GlcNAc(β-Sp21
Fraction Bound
0.6
0.4
0.2
0.0

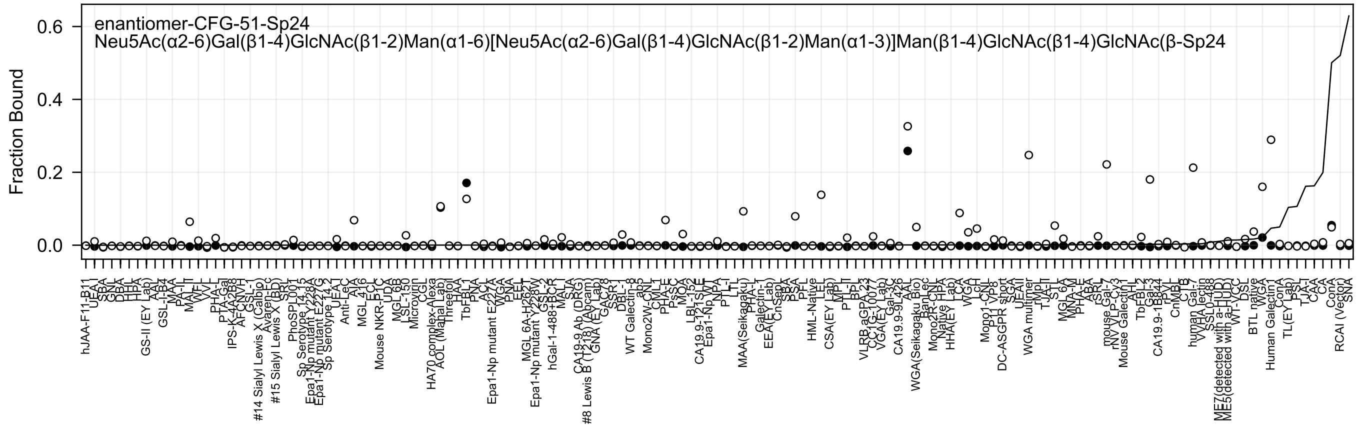
enantiomer-CFG-51-Sp24
Neu5Ac(α2-6)Gal(β1-4)GlcNAc(β1-2)Man(α1-6)[Neu5Ac(α2-6)Gal(β1-4)GlcNAc(β1-2)Man(α1-3)]Man(β1-4)GlcNAc(β1-4)GlcNAc(β-Sp24
Fraction Bound
0.0
0.2
0.4
0.6
hJAA-F11-B11
UEA-I
SBA
GNL
DBA
HHL
HPA
GS-II (EY Lab)
AAL
GSL-I-B4
MAA
PA-IL
MAL-II
WFL
VVL
PHA-L
PTA-Gal
IPS-K-4A2B8
AFCNVH
GSL-I
#14 Sialyl Lewis X (Calbio)
Avaren-Fc
#15 Sialyl Lewis X (BD)
SRL
PhoSPL001
Sp Serotype 14.15
Epa1-Np mutant Y228A
Epa1-Np mutant E227G
Sp Serotype 14.2
UEA-I
Anti-LeC
AIA
MGL 4-16
Mouse NKR-P1C
ECL
UDA
MGL 6B
LSL-150
Microvirin
CGL
HA70 complex-Alexa
AOL (Mahal Lab)
Threefoil
HAA
TbFBL1
PNA
ACL
Epa1-Np mutant E227A
WGA
PNA
EEL
MGL 6A-H262T
Epa1-Np mutant Y228W
GSL-2
hGal-1-488+BCR
MAL-I
SJA
CA19-9 Ab (DRG)
#8 Lewis B (T218) (Abcam)
GNA (EY Lab)
GAC78
SSR-1
DBL-1
WT Galectin-8
ab5
Mono2W-CNL
CML-1
PHA-E
PsSC
MOA
LBL-152
CA19.9-121SLE
Epa1-Np WT
NPA
PTL-I
LTL
MAA(Seikagaku)
PHA-L
Galectin-1
EEA(EY Lab)
CnSept
SBA
PSA
PFL
HML-Native
LEL
CSA(EY Lab)
MPL
PTL-II
BPL
VLRB aGPA.23
CC1G-10077
VGA(EY Lab)
Gal-3C
CA19.9-9L426
AAL
WGA(Seikagaku Bio)
BanLec
Mono2R-CNL
Native HPA
HHA(EY Lab)
LCA
WGA
LcH
Mono1-CNL
P11 VP8
DC-ASGPR short
SGA
UEAII
WGA multimer
rMpL
TJA-II
STL
MGL 6A
MNA-M
PHA-E
ABA
rSRL
mouse Gal-7
rNV VLP-Cy3
Mouse Galectin1
AHL
TbFBL2
Gal-3
CA19.9-1B844
rPVL
CnMBL
CTB
human Gal-7
VVHA lectin
SSL0-488
ME7(detected with a-HUD)
ME5(detected with a-HUD)
WT-VCC
DSL
BTL native
WGA
Human Galectin1
ConA
TL(EY Lab)
PSL
TJA-I
CAA
CA
ConA
RCAI (Vector)
SNA

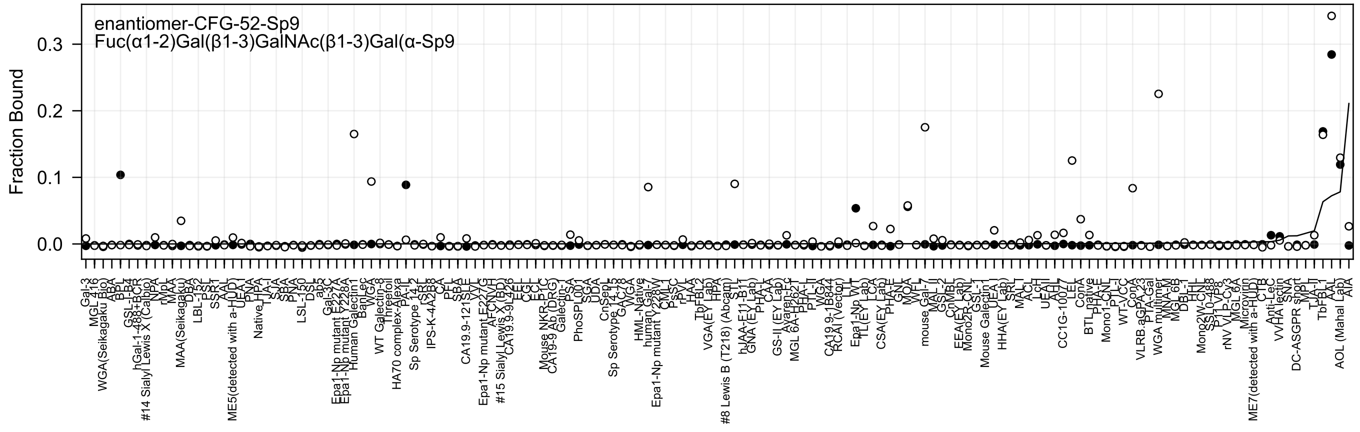
enantiomer-CFG-52-Sp9
Fuc(α1-2)Gal(β1-3)GalNAc(β1-3)Gal(α-Sp9
Fraction Bound
0.0
0.1
0.2
0.3

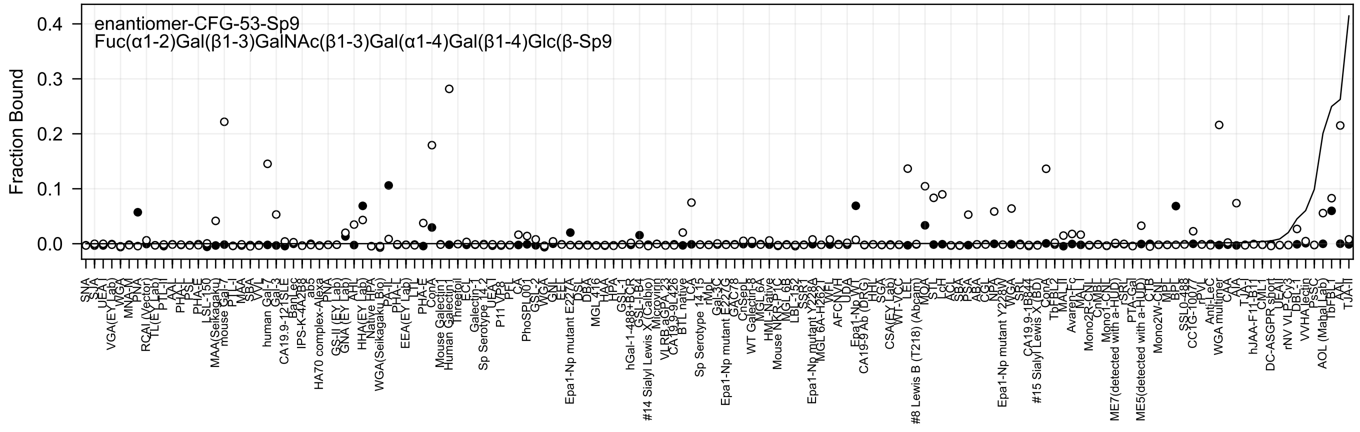
enantiomer-CFG-53-Sp9
Fuc(α1-2)Gal(β1-3)GalNAc(β1-3)Gal(α1-4)Gal(β1-4)Glc(β-Sp9
Fraction Bound

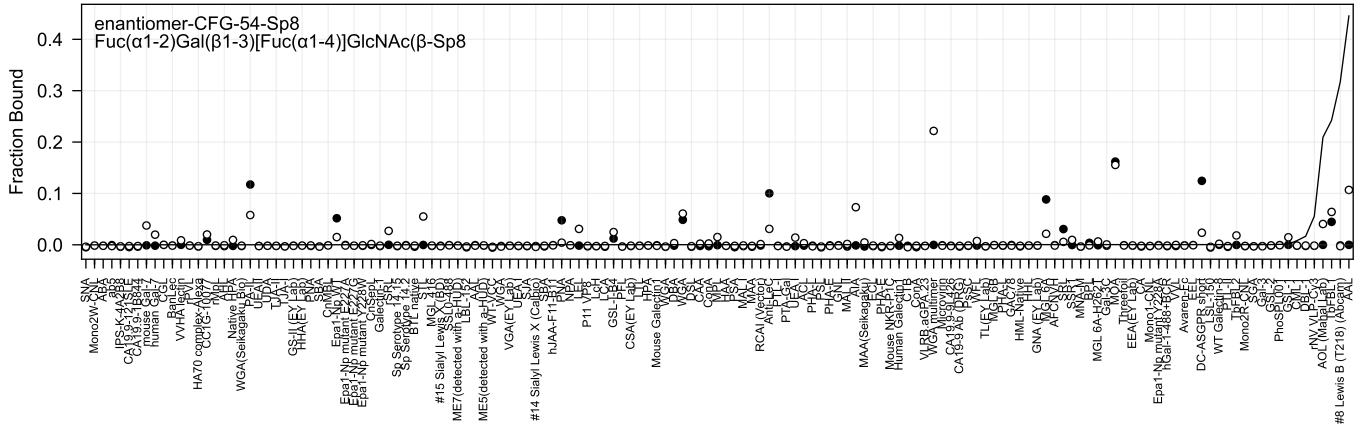

enantiomer-CFG-54-Sp8
Fuc(α1-2)Gal(β1-3)[Fuc(α1-4)]GlcNAc(β-Sp8
Fraction Bound

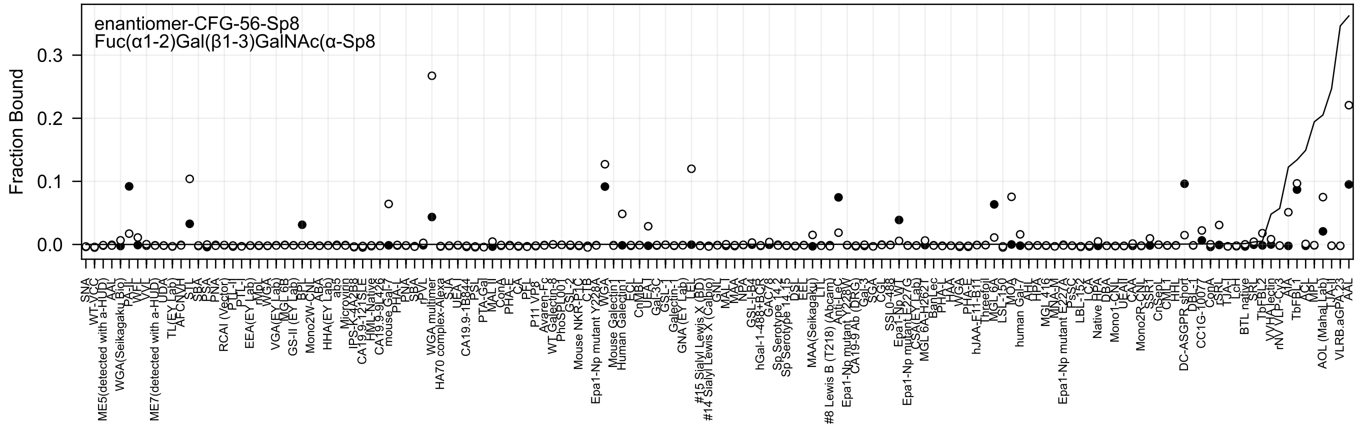
enantiomer-CFG-56-Sp8
Fuc(α1-2)Gal(β1-3)GalNAc(α-Sp8
Fraction Bound
0.3
0.2
0.1
0.0

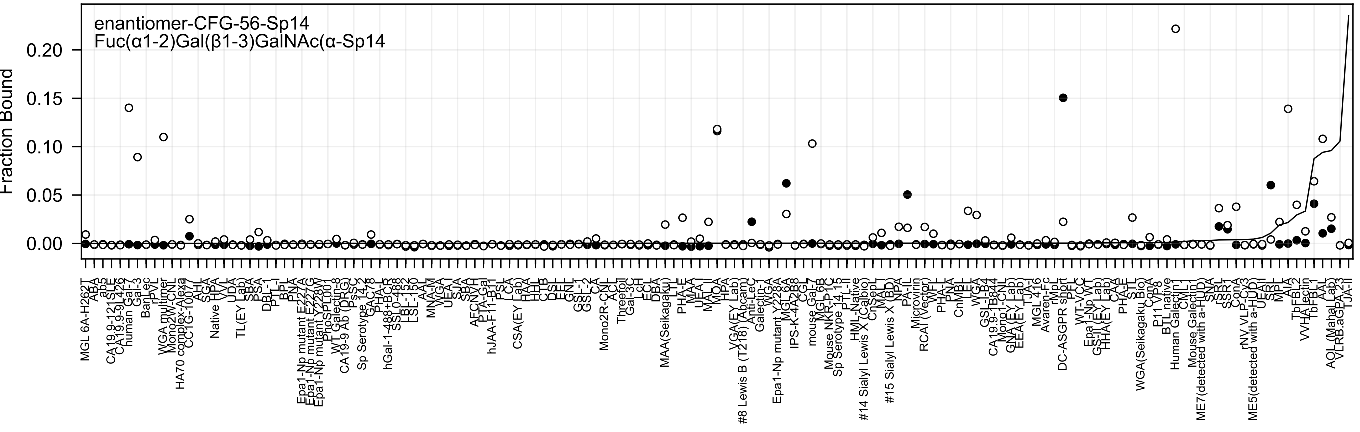
enantiomer-CFG-56-Sp14
Fuc(α1-2)Gal(β1-3)GalNAc(α-Sp14
Fraction Bound
0.00
0.05
0.10
0.15
0.20

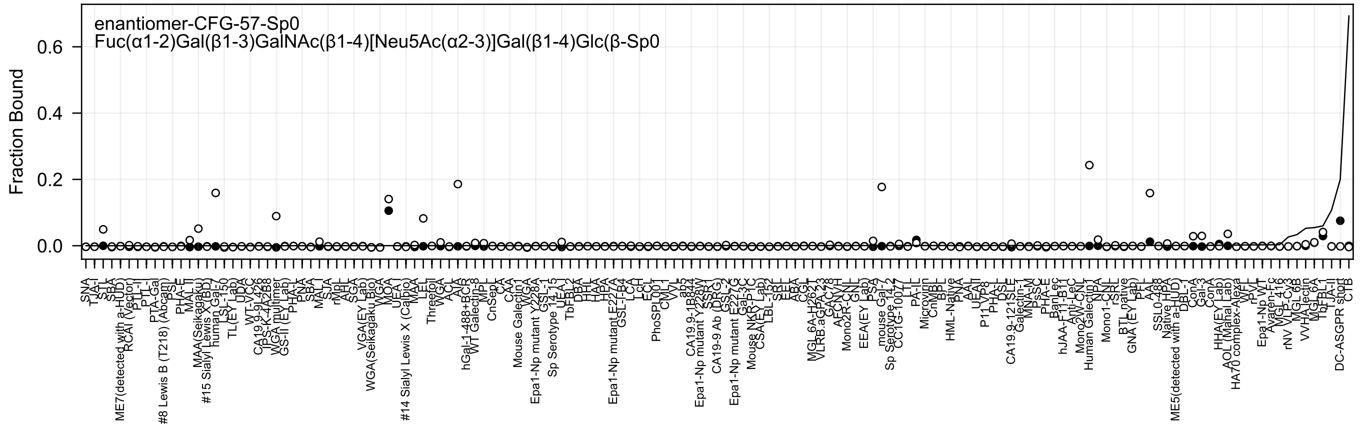

enantiomer-CFG-57-Sp0
Fuc(α1-2)Gal(β1-3)GalNAc(β1-4)[Neu5Ac(α2-3)]Gal(β1-4)Glc(β-Sp0
Fraction Bound
0.0
0.2
0.4
0.6

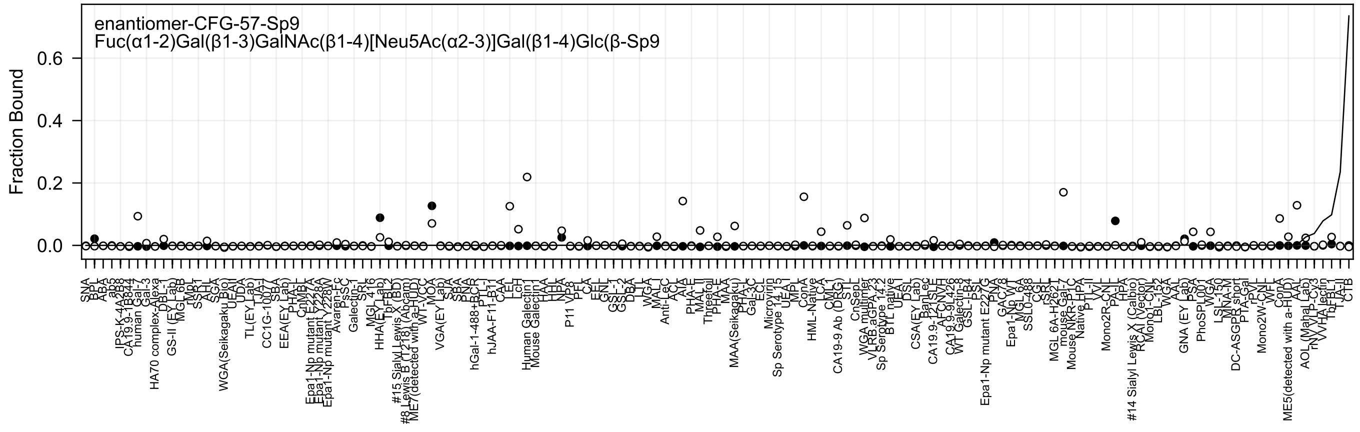

enantiomer-CFG-57-Sp9
Fuc(α1-2)Gal(β1-3)GalNAc(β1-4)[Neu5Ac(α2-3)]Gal(β1-4)Glc(β-Sp9
Fraction Bound
0.6
0.4
0.2
0.0
SNA
BPL
ABA
ab3
IPS-K-4A2B8
CA19.9-1B844
human Gal-7
Gal-3
HA70 complex-Alexa
DBL-1
GS-II (EY Lab)
MGL 6B
rMb
SSB1
AAL
SGA
WGA(Seikagaku Bio)
UEAII
UDA
TL(EY Lab)
TJA-I
CC1G-1007
SBA
EEA(EY Lab)
PHA-L
CnMBL
Epa1-Np mutant E227A
Epa1-Np mutant Y228A
Epa1-Np mutant Y228W
Avaren-Fc
PsSC
Galectin-1
SRL
MGL 416
HHA(EY Lab)
TbFBL2
#15 Sialyl Lewis X (BD)
#8 Lewis B (T218) (Abcam)
ME7(detected with a-HUD)
WT-VCC
MOA
VGA(EY Lab)
SJA
SBA
PNA
hGal-1-488+BCR
PTL-I
hJAA-F11-B11
CAA
LEL
LcH
Human Galectin1
Mouse Galectin1
HAA
HHL
NPA
P11 VP8
PFL
CA
EEL
GNL
GSL-I
GSL-2
DBA
LTL
WGA
MAL-I
Anti-LeC
ACL
AIA
PHA-E
MAL-II
Threefoil
PHA-E
MAA
MAA(Seikagaku)
PHA-L
Gal-3C
ECL
Microvirin
Sp Serotype 14.15
UEA-I
MPL
ConA
HML-Native
LCA
CML1
CA19-9 Ab (DRG)
STL
CnSepL
WGA multimer
VLRB.aGPA.23
Sp Serotype 14.2
BTL native
UEA-I
DSL
CSA(EY Lab)
BanLec
CA19.9-121SLE
AFCNVH
CA19.9-9L426
WT Galectin-8
GSL-IB4
PSL
Epa1-Np mutant E227G
PNA
GAC78
Epa1-Np WT
MGL 6A
SSL0-488
CGL
rSRL
MGL 6A-H262T
mouse Gal-7
Mouse NKR-P1C
Native HPA
PTL-II
VVL
Mono2R-CNL
PA-IL
HPA
#14 Sialyl Lewis X (Calbio)
RCAI (Vector)
Mono1-CNL
LBL-152
WGA
AAL
GNA (EY Lab)
PSA
PhoSL001
WGA
LSL-150
MNA-M
DC-ASGPR short
PTA-Gal
rPVL
Mono2W-CNL
WFL
ConA
ME5(detected with a-HUD)
AAL
AOL (Mahal Lab)
rNYVLP-Cy3
VVHA lectin
TbFBL1
TJA-II
CTB

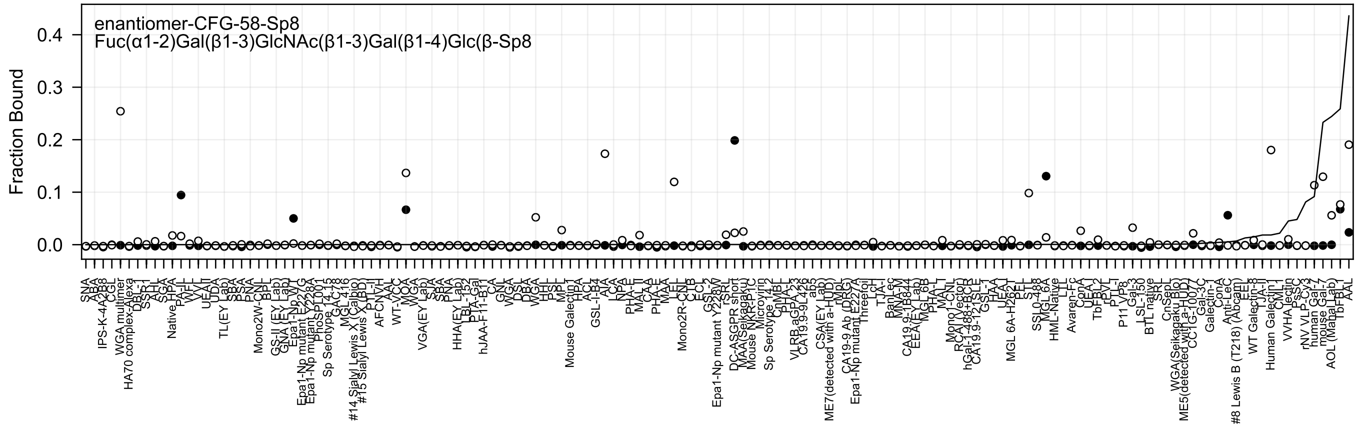

enantiomer-CFG-58-Sp8
Fuc(α1-2)Gal(β1-3)GlcNAc(β1-3)Gal(β1-4)Glc(β-Sp8
Fraction Bound
0.4
0.3
0.2
0.1
0.0

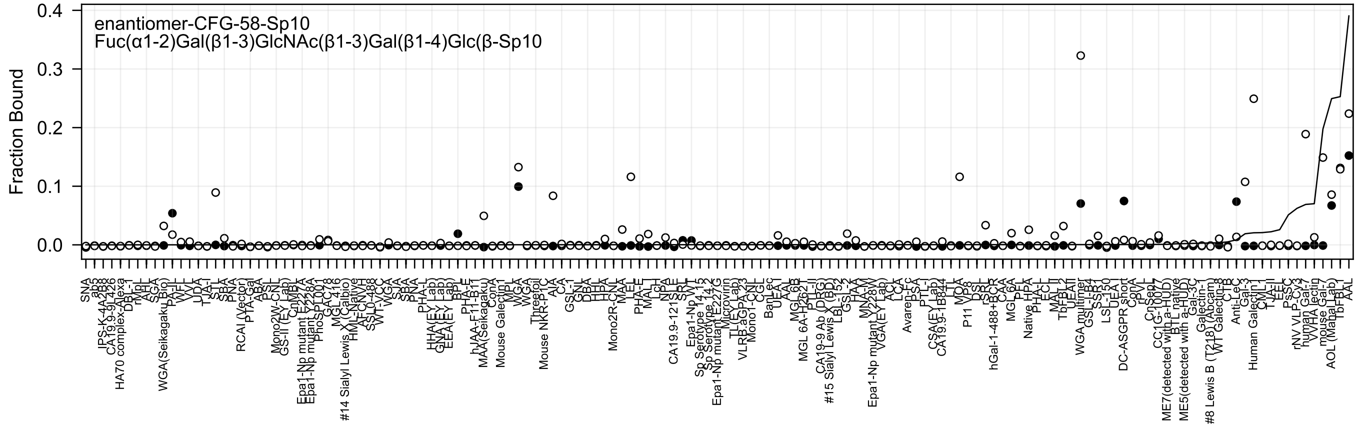

enantiomer-CFG-58-Sp10
Fuc(α1-2)Gal(β1-3)GlcNAc(β1-3)Gal(β1-4)Glc(β-Sp10
Fraction Bound
0.4
0.3
0.2
0.1
0.0

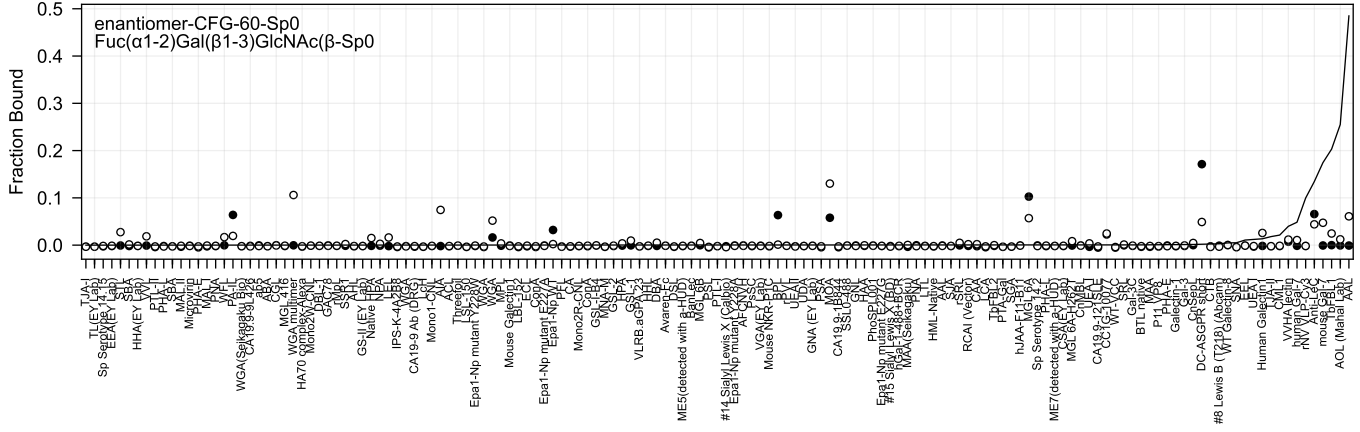
enantiomer-CFG-60-Sp0
Fuc(α1-2)Gal(β1-3)GlcNAc(β-Sp0
Fraction Bound
0.5
0.4
0.3
0.2
0.1
0.0

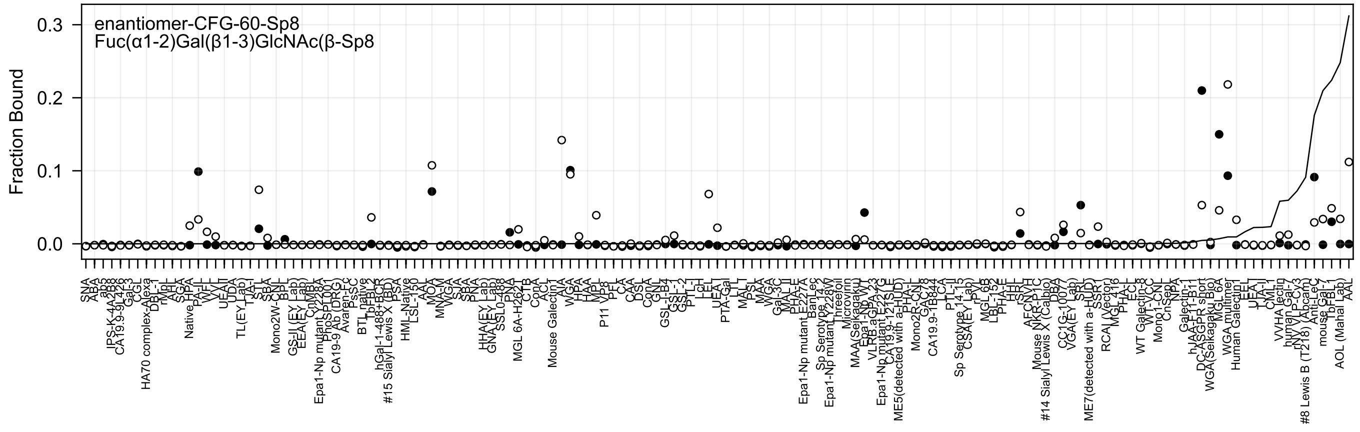
enantiomer-CFG-60-Sp8
Fuc(α1-2)Gal(β1-3)GlcNAc(β-Sp8
Fraction Bound
0.0
0.1
0.2
0.3

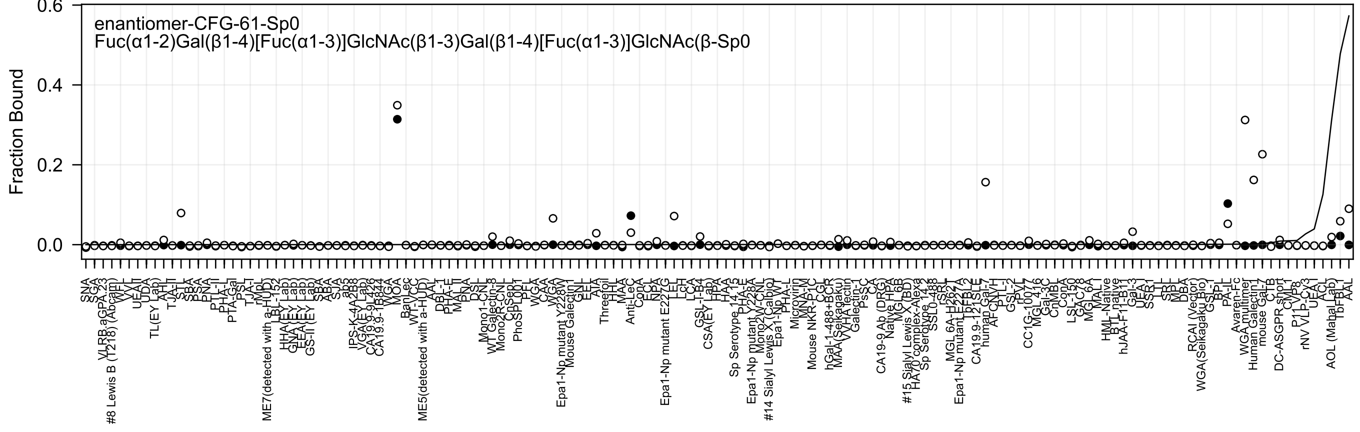
enantiomer-CFG-61-Sp0
Fuc(α1-2)Gal(β1-4)[Fuc(α1-3)]GlcNAc(β1-3)Gal(β1-4)[Fuc(α1-3)]GlcNAc(β-Sp0
Fraction Bound
0.6
0.4
0.2
0.0

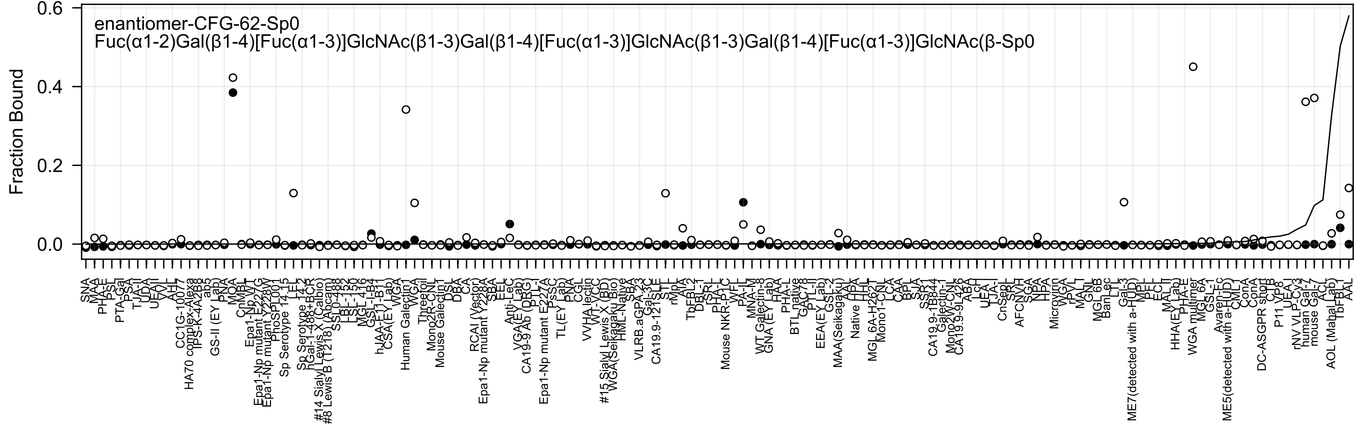
enantiomer-CFG-62-Sp0
Fuc(α1-2)Gal(β1-4)[Fuc(α1-3)]GlcNAc(β1-3)Gal(β1-4)[Fuc(α1-3)]GlcNAc(β1-3)Gal(β1-4)[Fuc(α1-3)]GlcNAc(β-Sp0
Fraction Bound
0.0
0.2
0.4
0.6

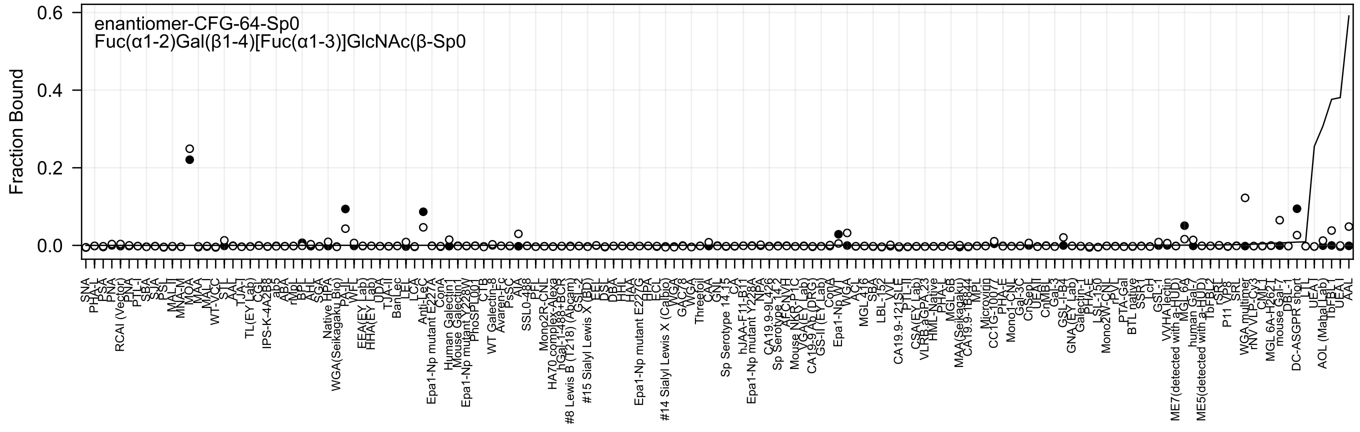
enantiomer-CFG-64-Sp0
Fuc(α1-2)Gal(β1-4)[Fuc(α1-3)]GlcNAc(β-Sp0
Fraction Bound
0.6
0.4
0.2
0.0

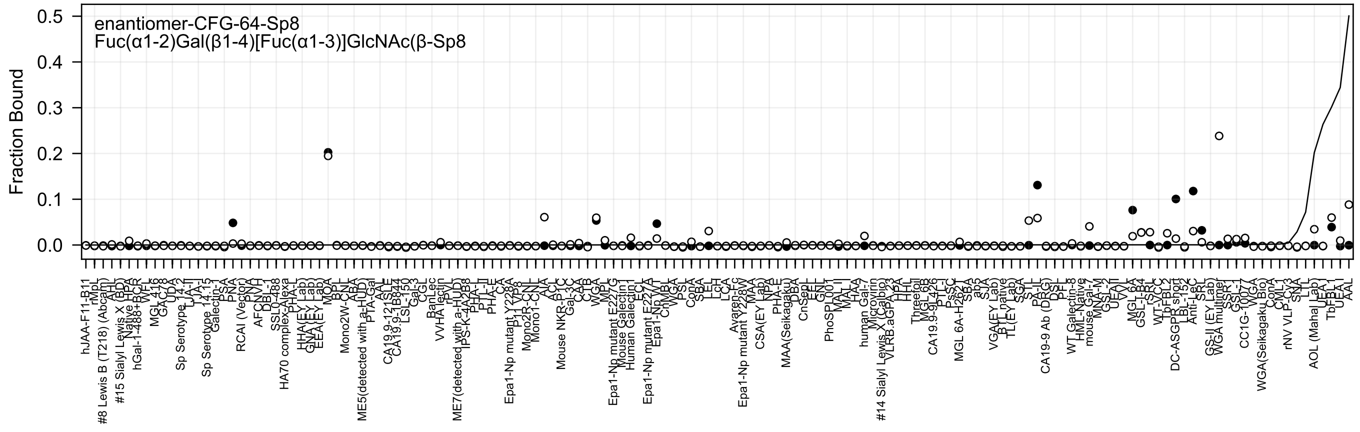

enantiomer-CFG-64-Sp8
Fuc(α1-2)Gal(β1-4)[Fuc(α1-3)]GlcNAc(β-Sp8
Fraction Bound
0.5
0.4
0.3
0.2
0.1
0.0

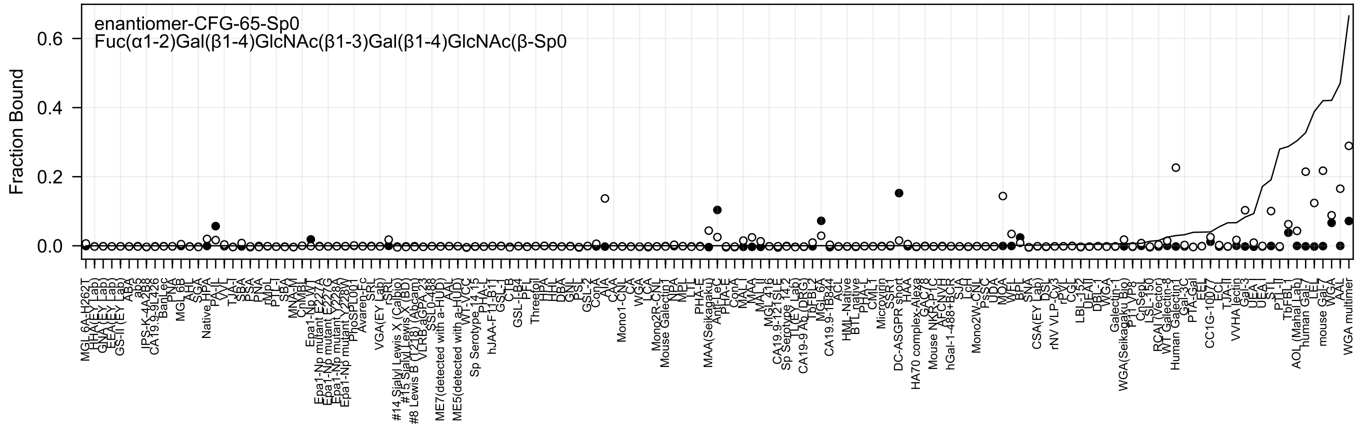
enantiomer-CFG-65-Sp0
Fuc(α1-2)Gal(β1-4)GlcNAc(β1-3)Gal(β1-4)GlcNAc(β-Sp0
Fraction Bound
0.6
0.4
0.2
0.0

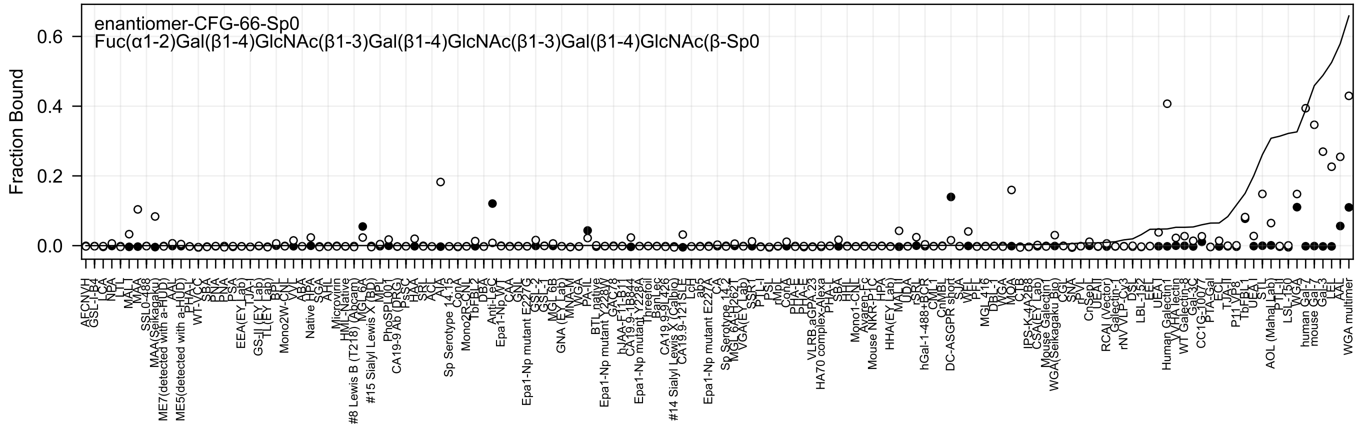

enantiomer-CFG-66-Sp0
Fuc(α1-2)Gal(β1-4)GlcNAc(β1-3)Gal(β1-4)GlcNAc(β1-3)Gal(β1-4)GlcNAc(β-Sp0
Fraction Bound

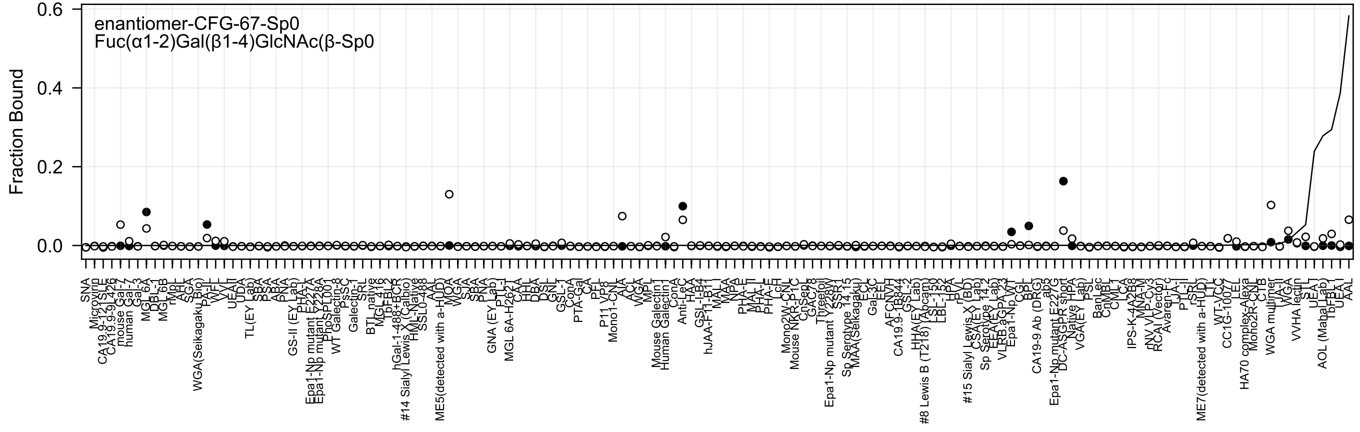
enantiomer-CFG-67-Sp0
Fuc(α1-2)Gal(β1-4)GlcNAc(β-Sp0
Fraction Bound
0.6
0.4
0.2
0.0

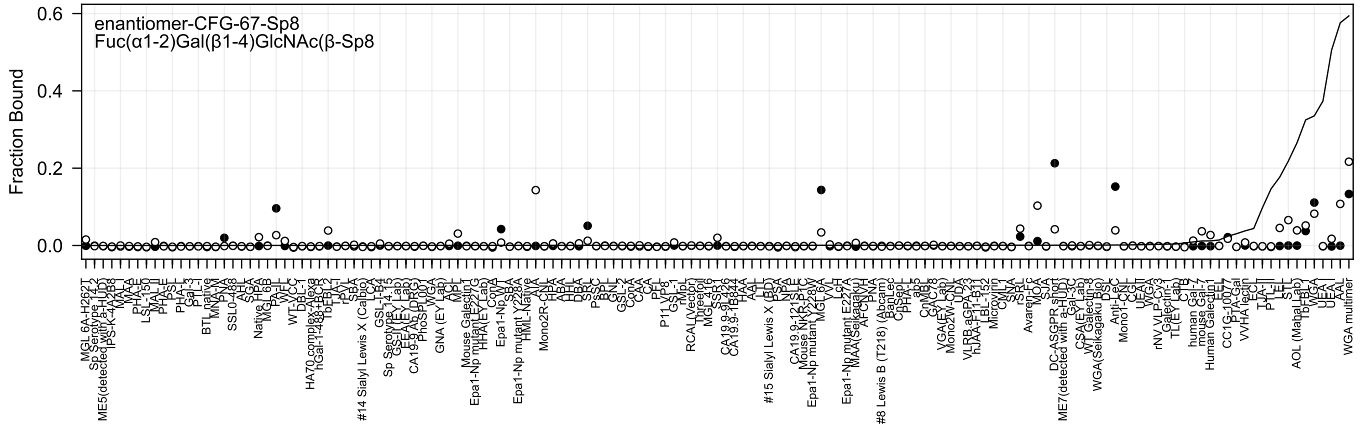
enantiomer-CFG-67-Sp8
Fuc(α1-2)Gal(β1-4)GlcNAc(β-Sp8
Fraction Bound
0.6
0.4
0.2
0.0

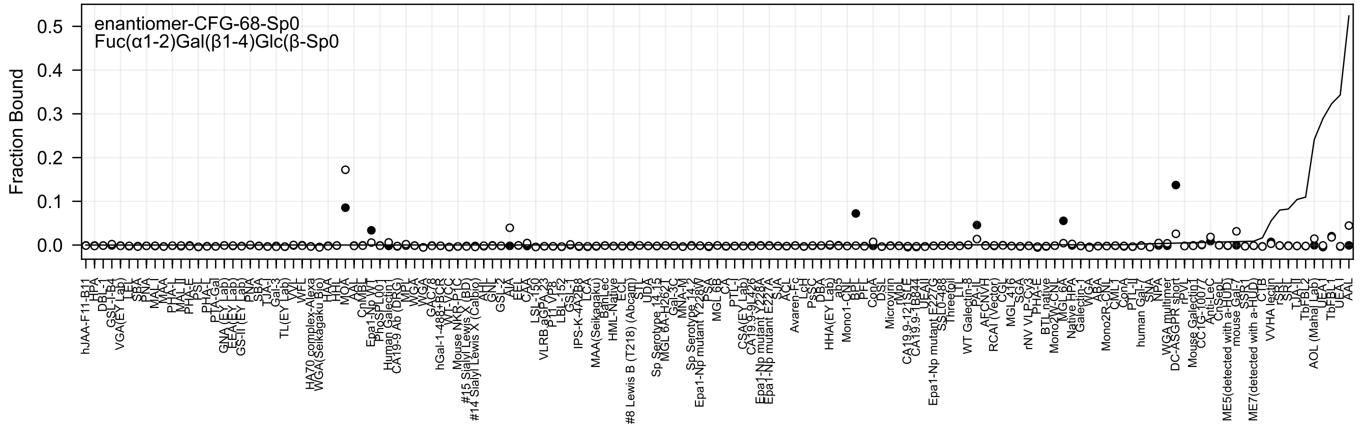
enantiomer-CFG-68-Sp0
Fuc(α1-2)Gal(β1-4)Glc(β-Sp0
Fraction Bound

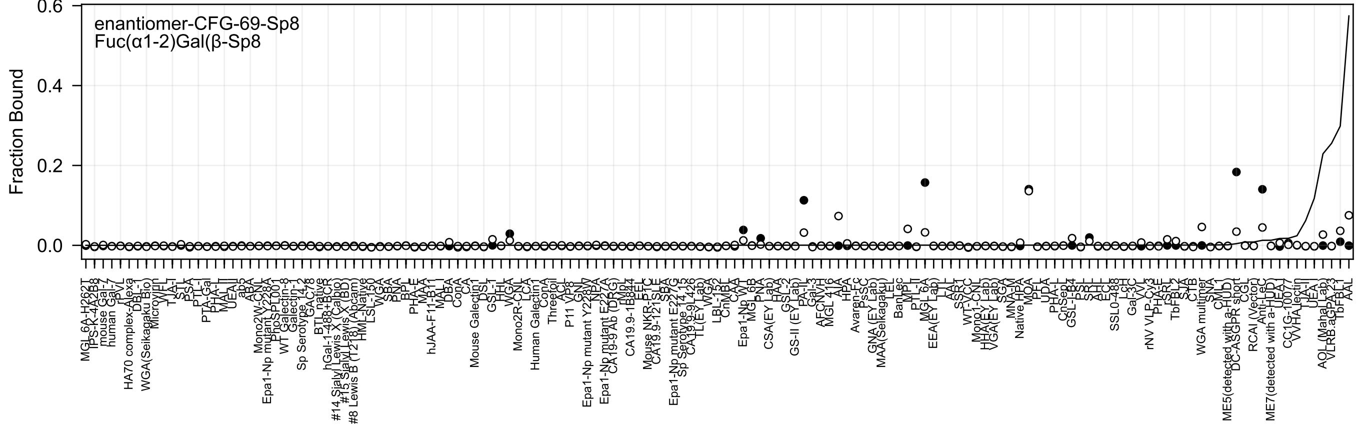

enantiomer-CFG-69-Sp8
Fuc(α1-2)Gal(β-Sp8
Fraction Bound
0.6
0.4
0.2
0.0

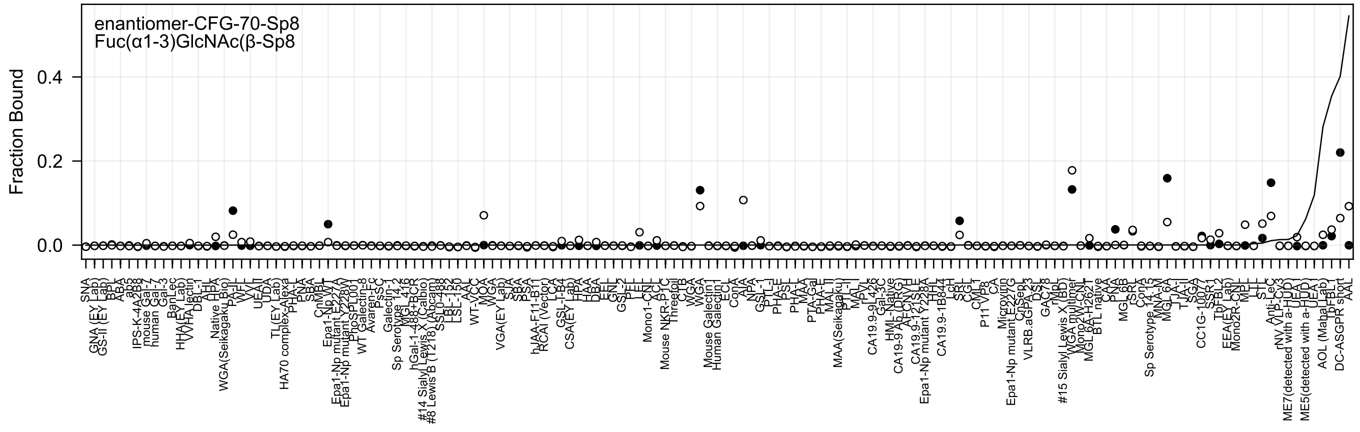
enantiomer-CFG-70-Sp8
Fuc(α1-3)GlcNAc(β-Sp8
Fraction Bound
0.4
0.2
0.0

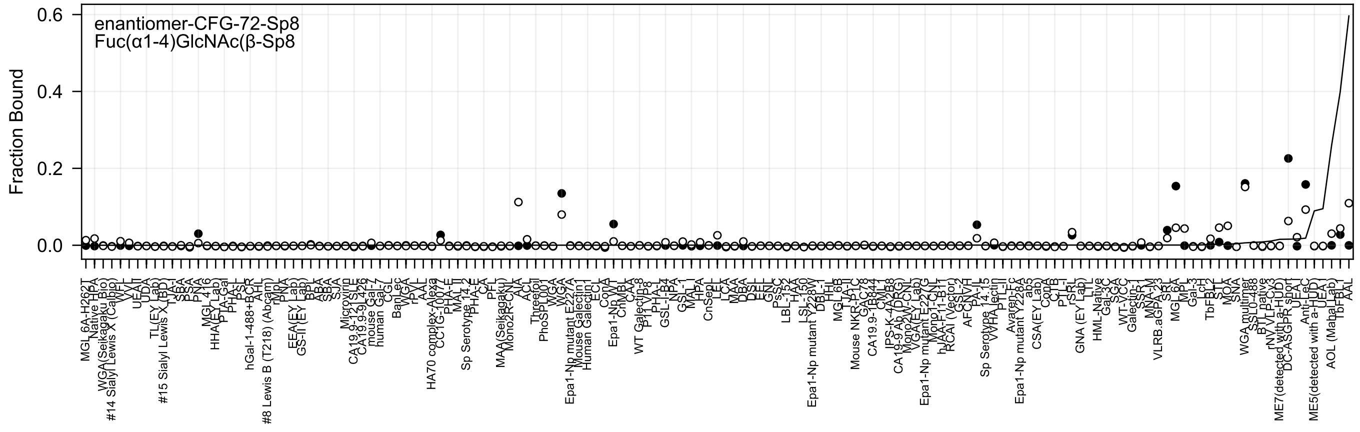
enantiomer-CFG-72-Sp8
Fuc(α1-4)GlcNAc(β-Sp8
Fraction Bound
0.6
0.4
0.2
0.0

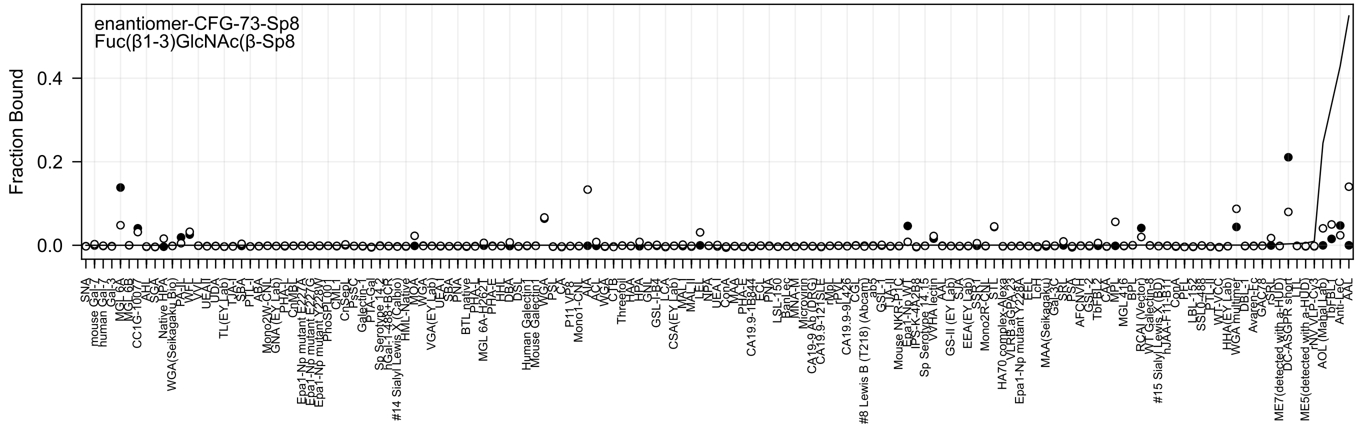
enantiomer-CFG-73-Sp8
Fuc(β1-3)GlcNAc(β-Sp8
Fraction Bound

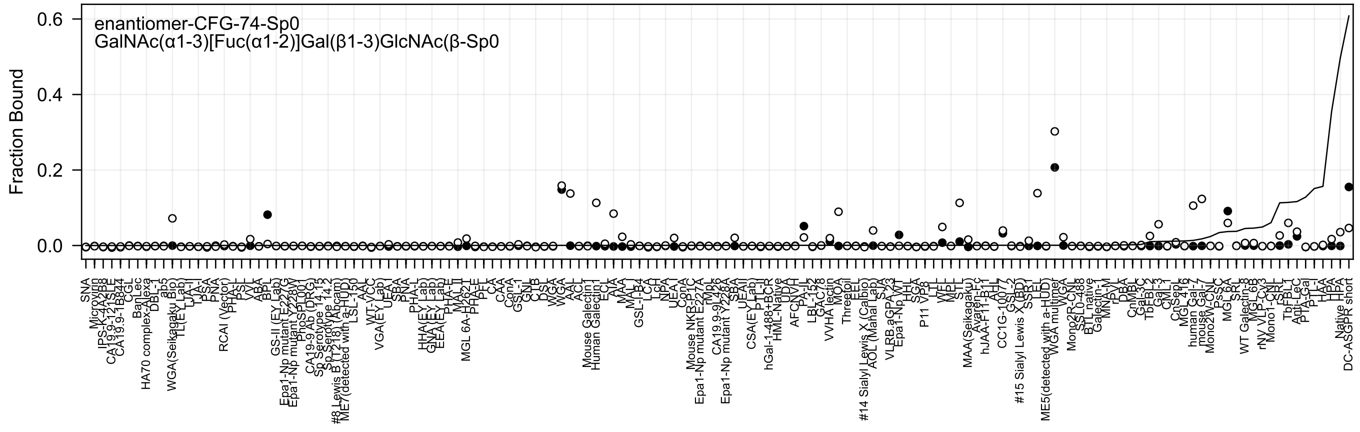
enantiomer-CFG-74-Sp0
GalNAc(α1-3)[Fuc(α1-2)]Gal(β1-3)GlcNAc(β-Sp0
Fraction Bound
0.6
0.4
0.2
0.0

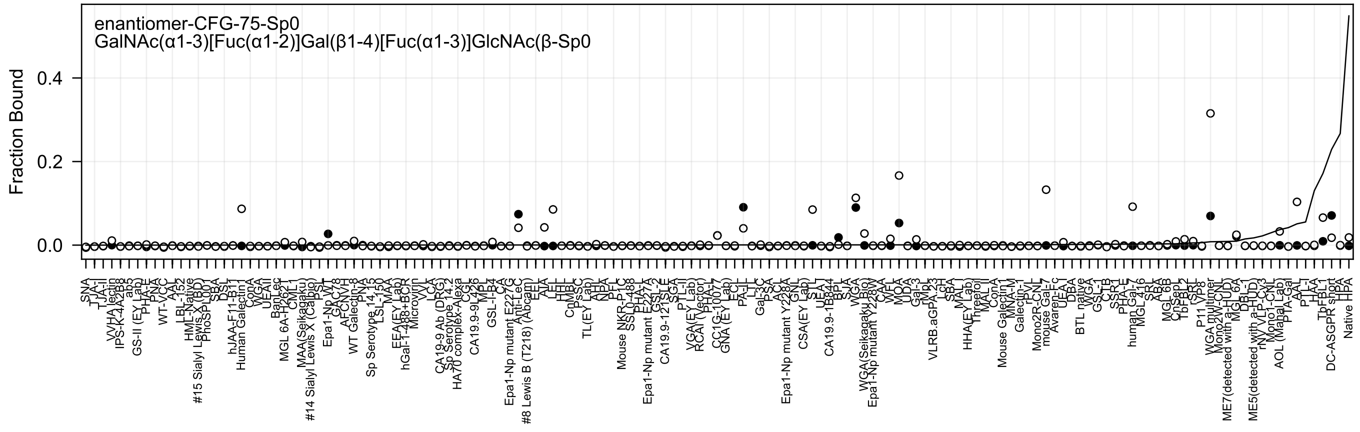
enantiomer-CFG-75-Sp0
GalNAc(α1-3)[Fuc(α1-2)]Gal(β1-4)[Fuc(α1-3)]GlcNAc(β-Sp0
Fraction Bound

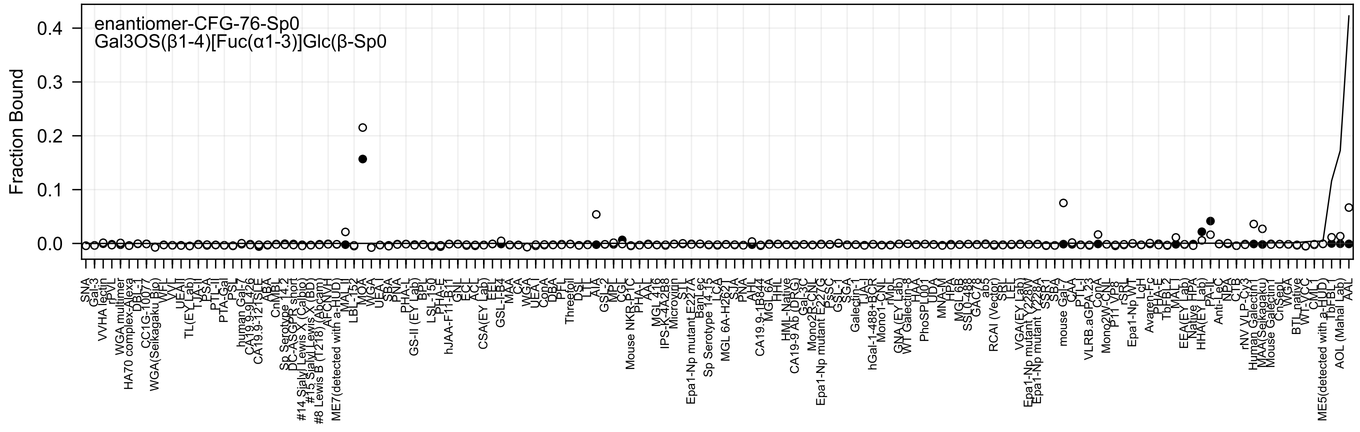
enantiomer-CFG-76-Sp0
Gal3OS(β1-4)[Fuc(α1-3)]Glc(β-Sp0
Fraction Bound
0.0
0.1
0.2
0.3
0.4

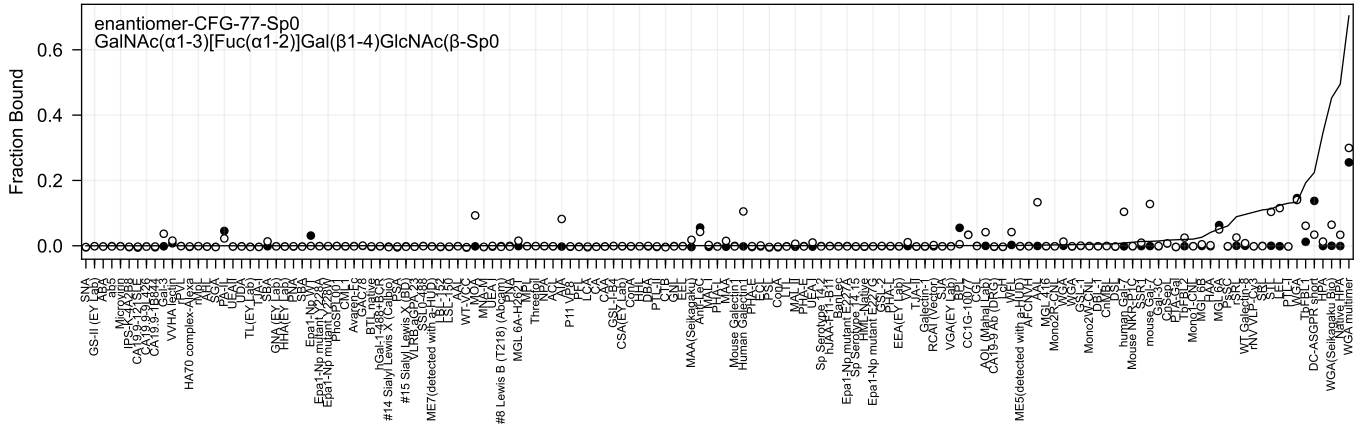
enantiomer-CFG-77-Sp0
GalNAc(α1-3)[Fuc(α1-2)]Gal(β1-4)GlcNAc(β-Sp0
Fraction Bound
0.6
0.4
0.2
0.0

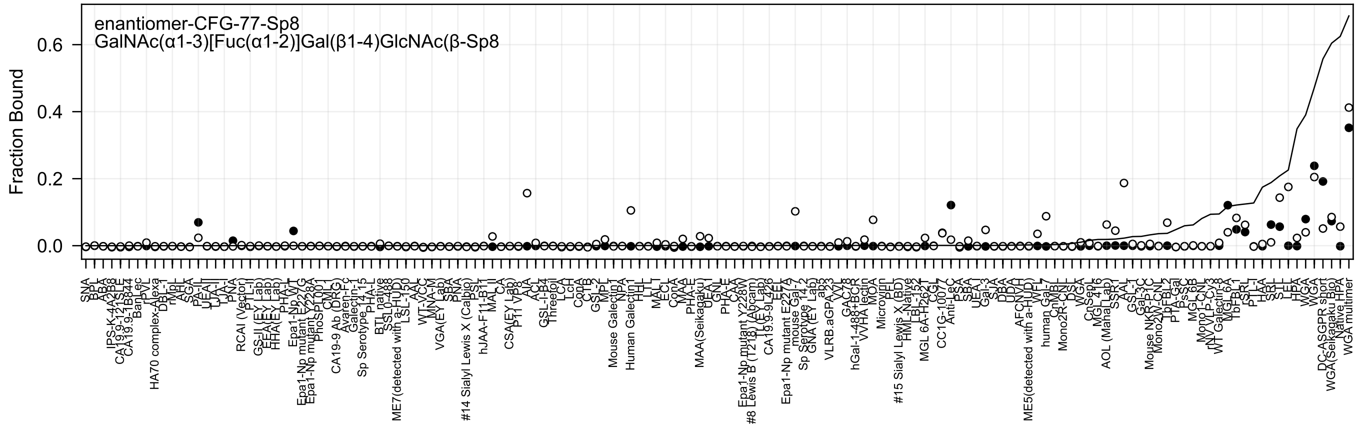
enantiomer-CFG-77-Sp8
GalNAc(α1-3)[Fuc(α1-2)]Gal(β1-4)GlcNAc(β-Sp8
Fraction Bound
0.6
0.4
0.2
0.0

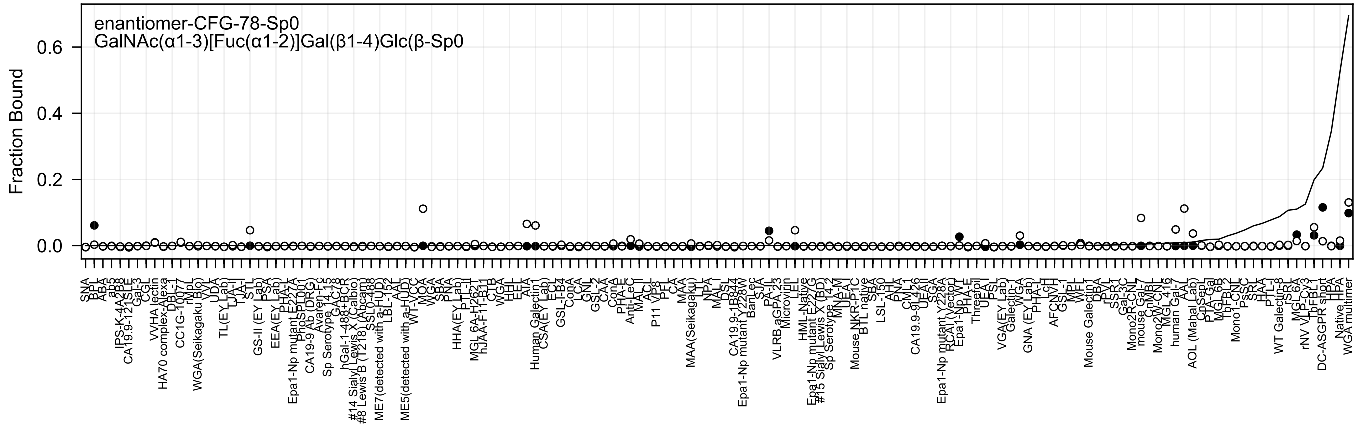
enantiomer-CFG-78-Sp0
GalNAc(α1-3)[Fuc(α1-2)]Gal(β1-4)Glc(β-Sp0
Fraction Bound
0.6
0.4
0.2
0.0
SNA
BPL
ABA
ab3
IPS-K-4A2B8
CA19.9-121SLE
Gal-3
CGL
VVHA lectin
HA70 complex-Alexa
DBL-1
CC1G-10077
rMpL
WGA(Seikagaku Bio)
VVL
UDA
TL(EY Lab)
LJA-II
TJA-I
STL
GS-II (EY Lab)
PSA
EEA(EY Lab)
PHA-L
Epa1-Np mutant E227A
PhoSL001
CA19-9 Ab (DRG)
Avaren-Fc
Sp Serotype 14-15
GAC78
hGal-1-488+BCR
#14 Sialyl Lewis X (Calbio)
#8 Lewis B (T218) (Abcam)
SSL0-488
ME7(detected with a-HUD)
LBL-152
AAL
ME5(detected with a-HUD)
WT-VCC
MOA
WGA
SBA
PNA
HHA(EY Lab)
PTL-II
MGL 6A-H262T
hJAA-F11-B11
CTB
WGA
HHL
EEL
AIA
Human Galectin1
CSA(EY Lab)
ECL
GSL-I-B4
ConA
LCA
GNL
GSL-2
CAA
ConA
PHA-E
Anti-LeC
MAL II
ACL
P11 VP8
PFL
CA
MAA
MAA(Seikagaku)
LTL
NPA
MAL I
DSL
CA19.9-1B844
Epa1-Np mutant Y228W
BanLec
SJA
PA-I
VLRB aGPA.23
Microvirin
LEL
HML-Native
Epa1-Np mutant E227S
#15 Sialyl Lewis X (BD)
Sp Serotype 14-2
MNA-M
UEA-I
Mouse NKR-P1C
BTL native
SBA
LSL-150
AHL
PNA
CML1
CA19.9-9L426
UEAII
SGA
Epa1-Np mutant Y228A
RCAI (Vector)
Epa1-Np WT
PHA-E
Threeoil
UEA I
PSL
VGA(EY Lab)
Galectin-1
WGA
GNA (EY Lab)
PHA-L
LcH
AFCNVH
GSL-1
MPL
WFL
Mouse Galectin1
DBA
rPVL
SSR1
Gal-3C
Mono2R-CNL
mouse Gal-7
CnMBL
Mono2W-CNL
MGL 416
human Gal-7
AAL
AOL (Mahal Lab)
CnSep
PTA-Gal
MGL 6B
TbFBL2
Mono1-CNL
PsSC
SRL
HAA
PTL-II
WT Galectin-8
rSRL
MGL 6A
rNV VLP-Cy3
TbFBL1
DC-ASGPR short
HPA
Native HPA
WGA multimer

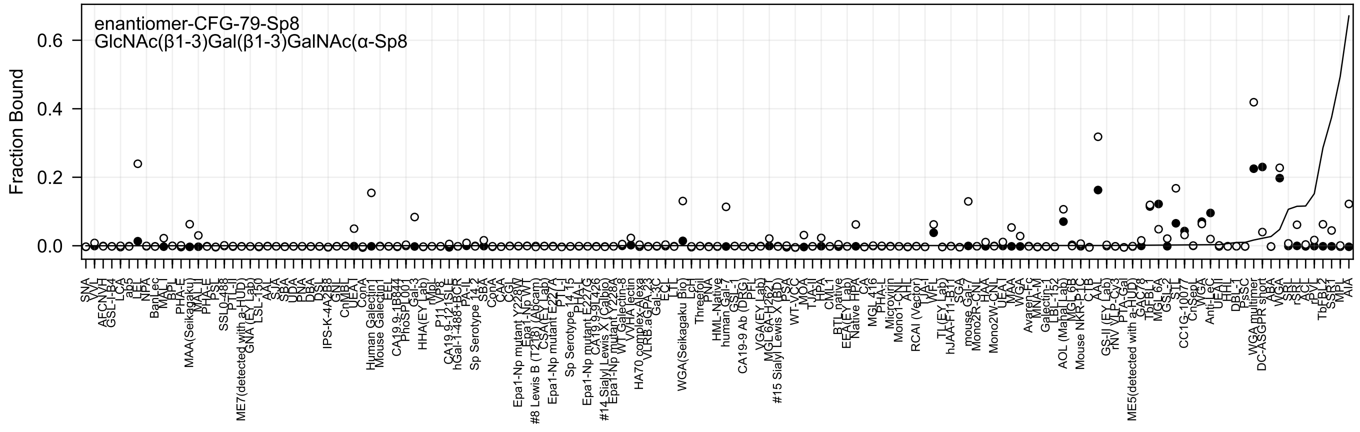

enantiomer-CFG-79-Sp8
GlcNAc(β1-3)Gal(β1-3)GalNAc(α-Sp8
Fraction Bound
0.0
0.2
0.4
0.6

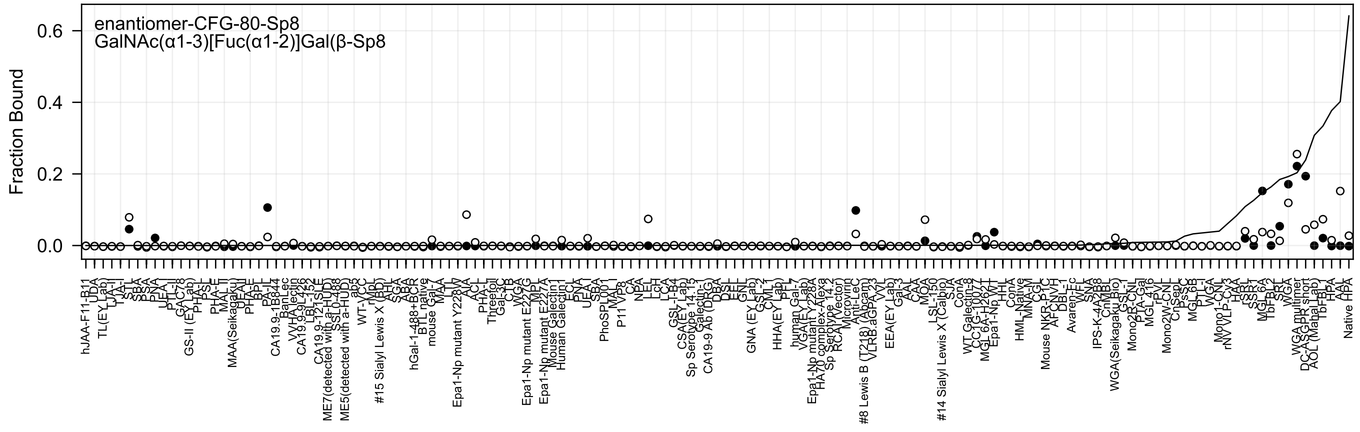

enantiomer-CFG-80-Sp8
GalNAc(α1-3)[Fuc(α1-2)]Gal(β-Sp8
Fraction Bound

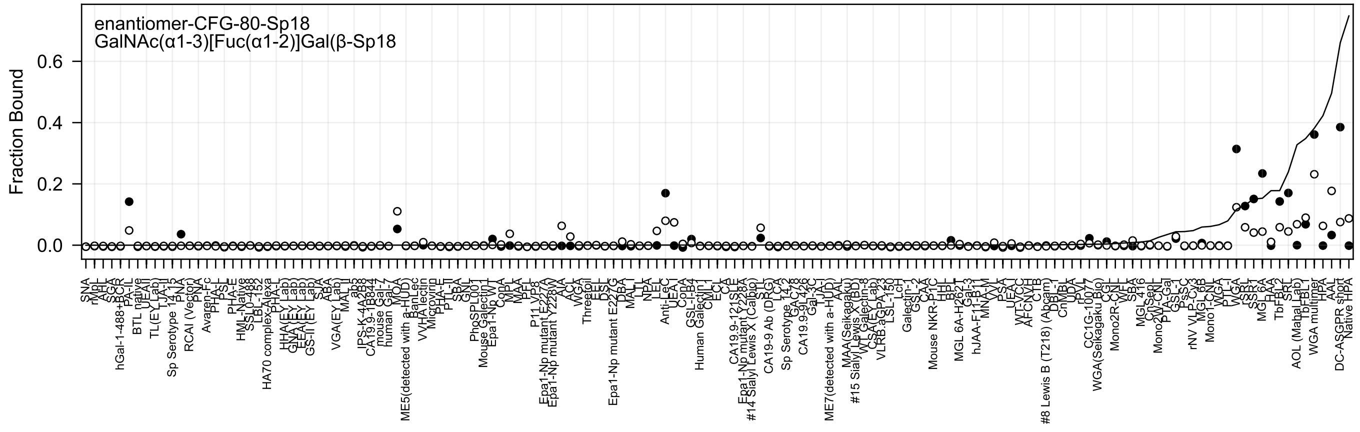

enantiomer-CFG-80-Sp18
GalNAc(α1-3)[Fuc(α1-2)]Gal(β-Sp18
Fraction Bound
0.0
0.2
0.4
0.6

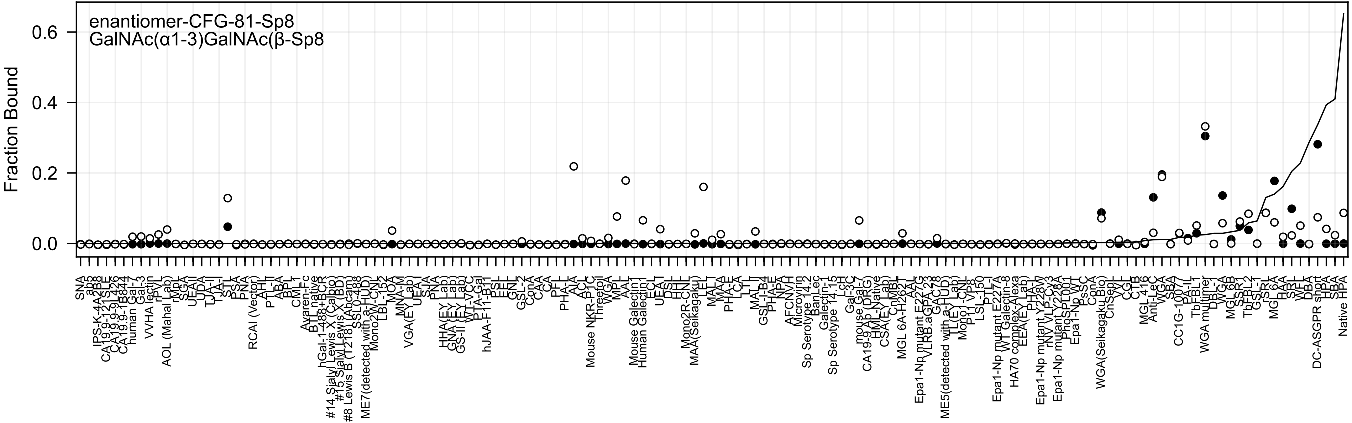
enantiomer-CFG-81-Sp8
GalNAc(α1-3)GalNAc(β-Sp8
Fraction Bound
0.6
0.4
0.2
0.0

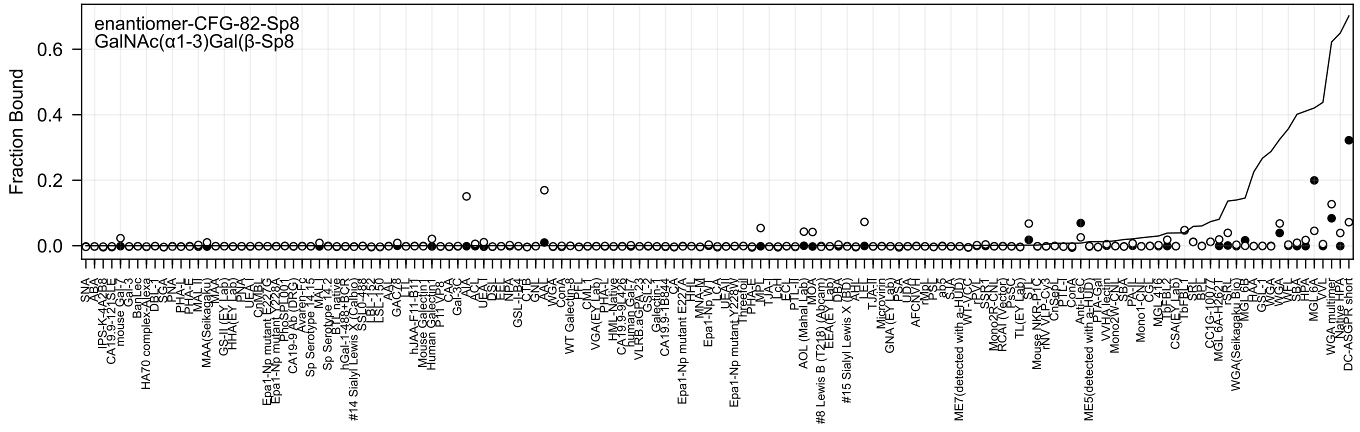
enantiomer-CFG-82-Sp8
GalNAc(α1-3)Gal(β-Sp8
Fraction Bound
0.6
0.4
0.2
0.0

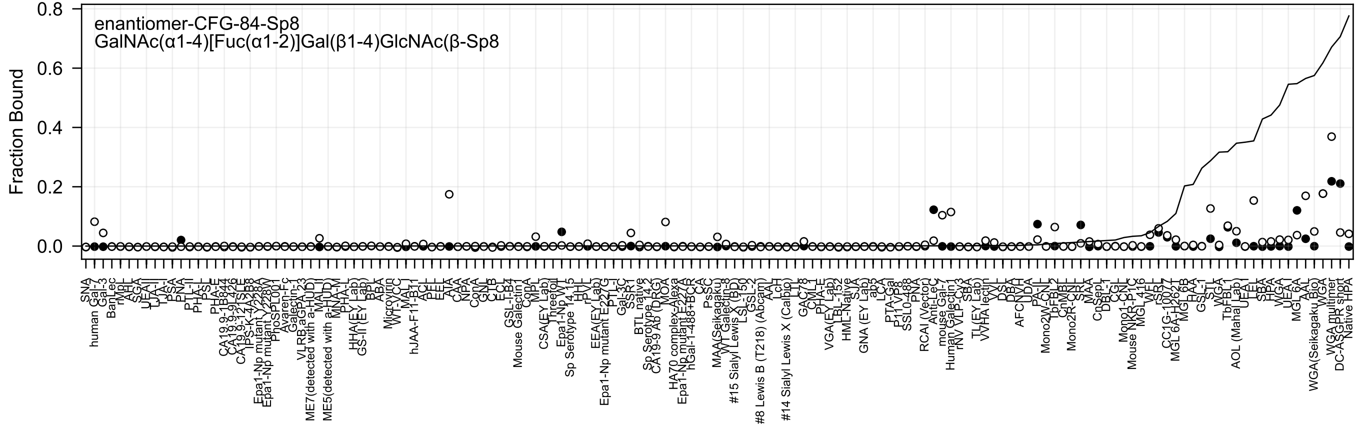
enantiomer-CFG-84-Sp8
GalNAc(α1-4)[Fuc(α1-2)]Gal(β1-4)GlcNAc(β-Sp8
Fraction Bound
0.8
0.6
0.4
0.2
0.0

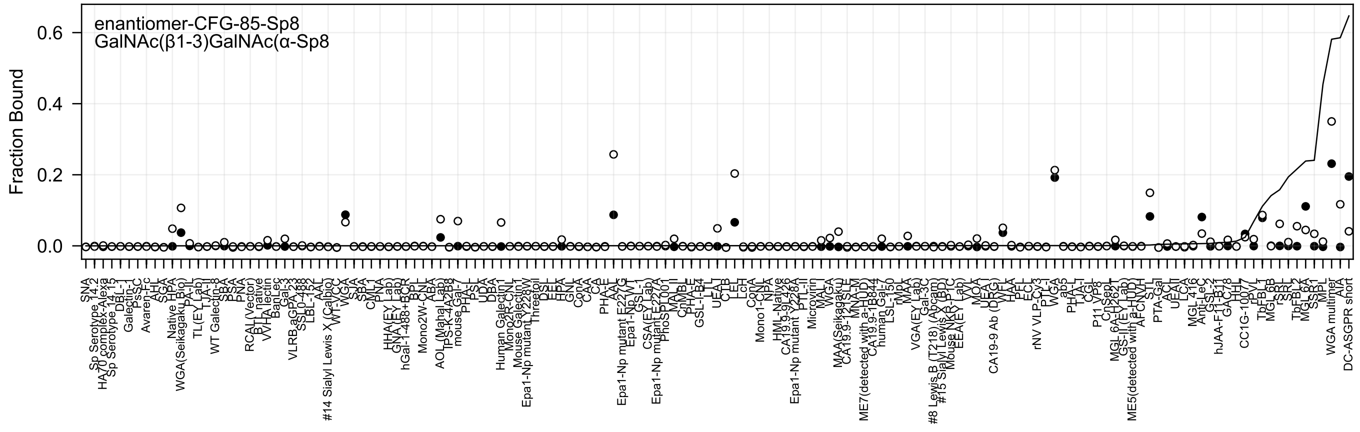
enantiomer-CFG-85-Sp8
GalNAc(β1-3)GalNAc(α-Sp8
Fraction Bound
0.6
0.4
0.2
0.0

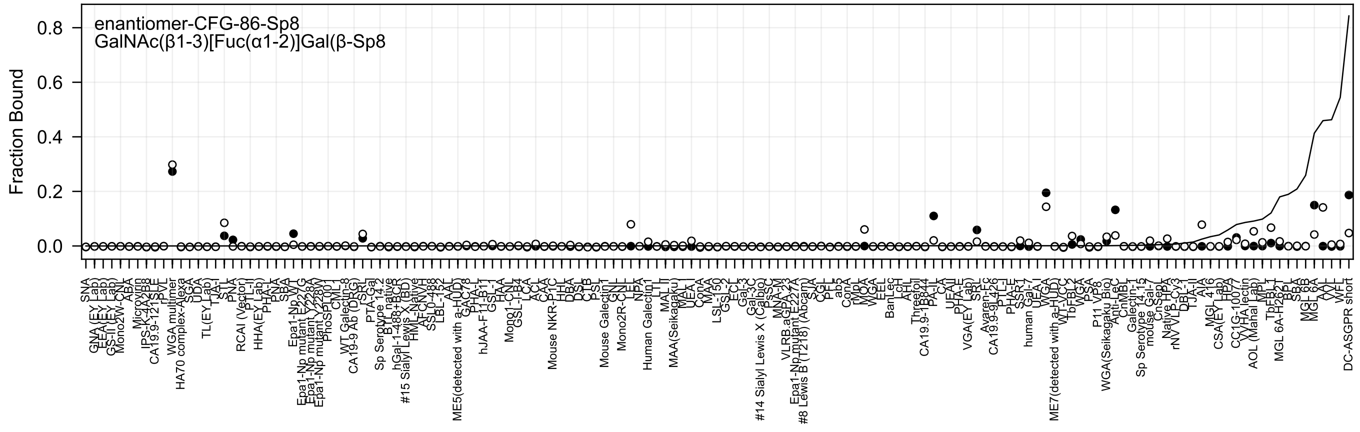
enantiomer-CFG-86-Sp8
GalNAc(β1-3)[Fuc(α1-2)]Gal(β-Sp8
Fraction Bound
0.8
0.6
0.4
0.2
0.0

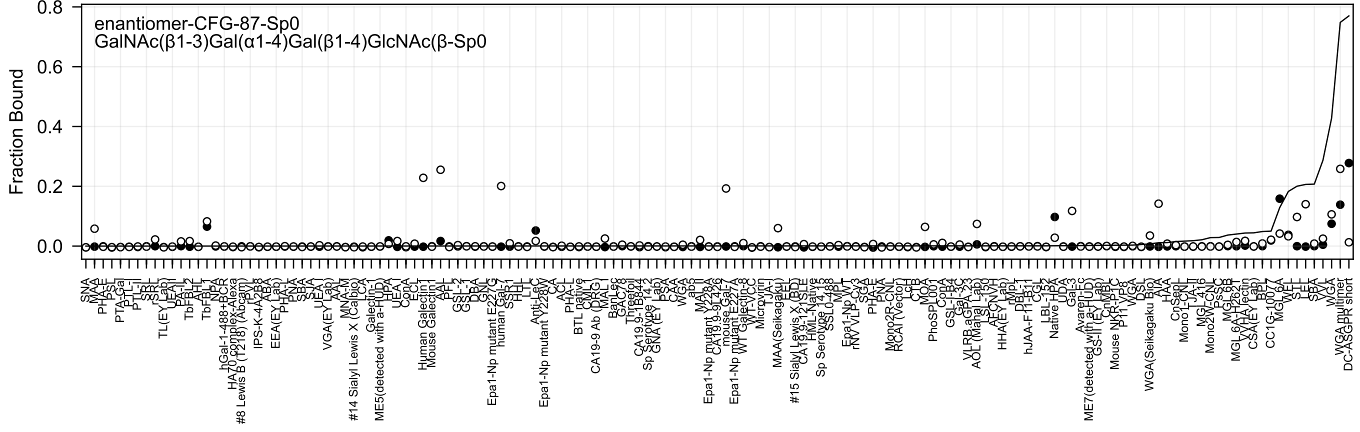

enantiomer-CFG-87-Sp0
GalNAc(β1-3)Gal(α1-4)Gal(β1-4)GlcNAc(β-Sp0
Fraction Bound
0.0
0.2
0.4
0.6
0.8

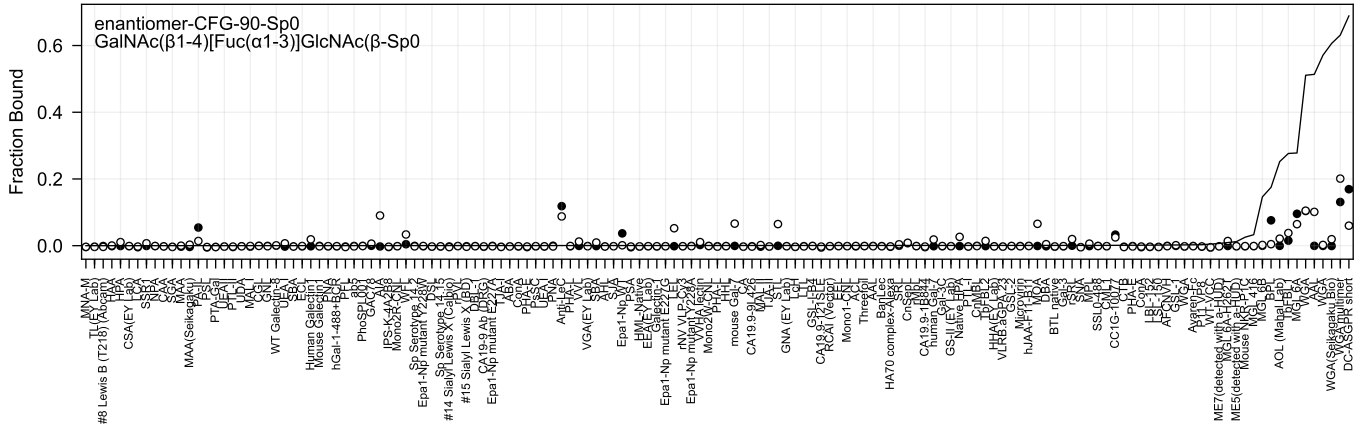
enantiomer-CFG-90-Sp0
GalNAc(β1-4)[Fuc(α1-3)]GlcNAc(β-Sp0
Fraction Bound
0.6
0.4
0.2
0.0

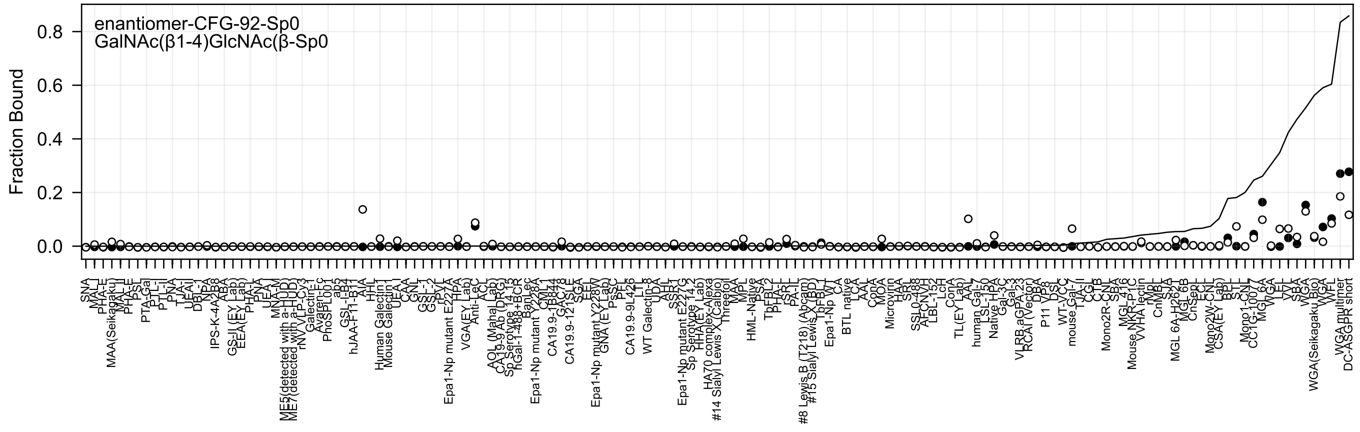

enantiomer-CFG-92-Sp0
GalNAc(β1-4)GlcNAc(β-Sp0
Fraction Bound
0.0
0.2
0.4
0.6
0.8

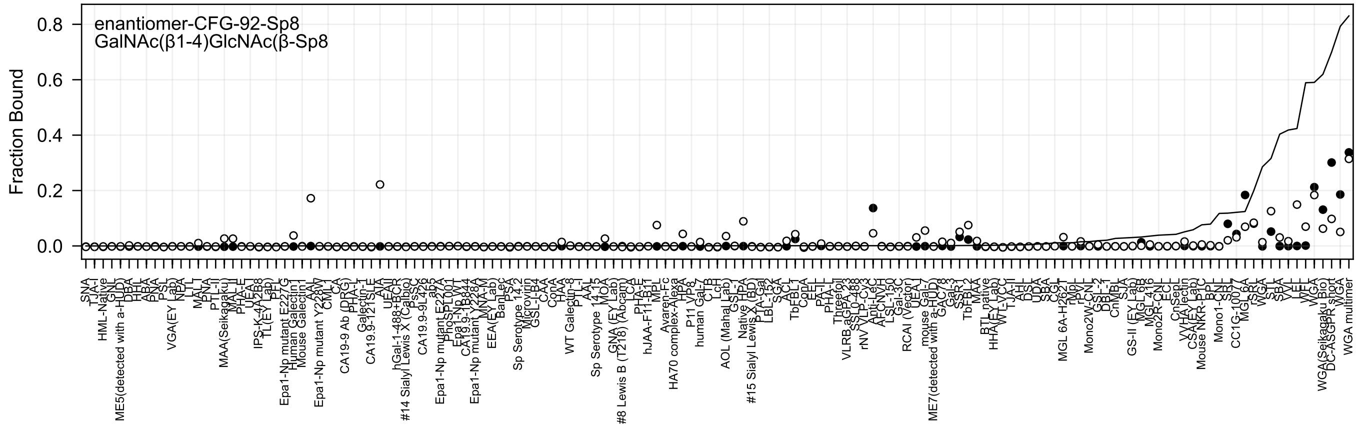
enantiomer-CFG-92-Sp8
GalNAc(β1-4)GlcNAc(β-Sp8
Fraction Bound
0.8
0.6
0.4
0.2
0.0

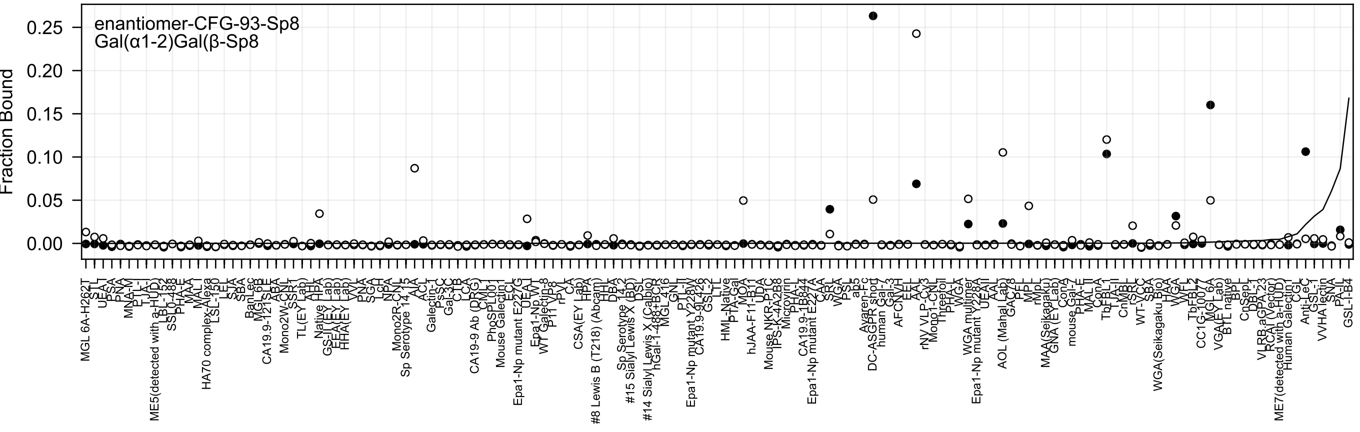

enantiomer-CFG-93-Sp8
Gal(α1-2)Gal(β-Sp8
Fraction Bound
0.25
0.20
0.15
0.10
0.05
0.00
MGL 6A-H262T
STL
UEA-I
PSA
PNA
MNA-M
PTL-I
TJA-I
ME5(detected with a-HUD)
LBL-152
SSL0-488
PHA-E
MAA
MAL-I
HA70 complex-Alexa
LSL-150
LEL
SJA
SBA
BanLec
MGL 6B
CA19.9-121SLE
ABA
Mono2W-CNL
SSR1
TL(EY Lab)
AHA
Native HPA
GS-II(EY Lab)
EEA(EY Lab)
HHA(EY Lab)
VVL
PNA
SGA
LcH
NPA
Mono2R-CNL
Sp Serotype 14.15
AIA
ACL
Galectin-1
PsSC
Gal-3C
CTB
LCA
CA19-9 Ab (DRG)
CML
PhoSPL001
Mouse Galectin1
ECL
Epa1-Np mutant E227G
UEA-I
Epa1-Np WT
WT Galectin-8
P11 VP8
rPVL
CA
CSA(EY Lab)
HPA
HHL
#8 Lewis B (T218) (Abcam)
DBA
Sp Serotype 14.2
#15 Sialyl Lewis X (BD)
DSL
#14 Sialyl Lewis X (Calbio)
hGal-1-488+BCR
MGL416
GNL
PTL-II
Epa1-Np mutant Y228W
CA19.9-9L426
GSL-2
LTL
HML-Native
PTA-Gal
MOA
hJAA-F11-B11
UDA
Mouse NKR-P1C
IPS-K-4A2B8
Microvirin
PHA-L
CA19.9-1B844
Epa1-Np mutant E227A
CAA
SRL
WGA
PSL
ab5
Avaren-Fc
DC-ASGPR short
human Gal-7
Gal-3
AFCNVH
EEL
AAL
rNV VLP-Cy3
Mono1-CNL
Threefoil
PHA-L
WGA
WGA multimer
Epa1-Np mutant Y228A
UEAII
AAL
AOL (Mahal Lab)
GAC78
PFL
MPL
rMpL
MAA(Seikagaku)
GNA (EY Lab)
ConA
mouse Gal-7
PHA-E
MAL II
ConA
TbFBL1
TJA-II
CnMBL
rSRL
WT-VCC
SBA
WGA(Seikagaku Bio)
HAA
WGA
WFL
TbFBL2
CC1G-1007
MGL 6A
VGA(EY Lab)
BTL native
BPL
CnSepL
DBL-1
VLRB.aGPA.23
RCAI (Vector)
ME7(detected with a-HUD)
Human Galectin1
CGL
Anti-LeC
GSL-I
VVHA lectin
SNA
PA-IL
GSL-I-B4

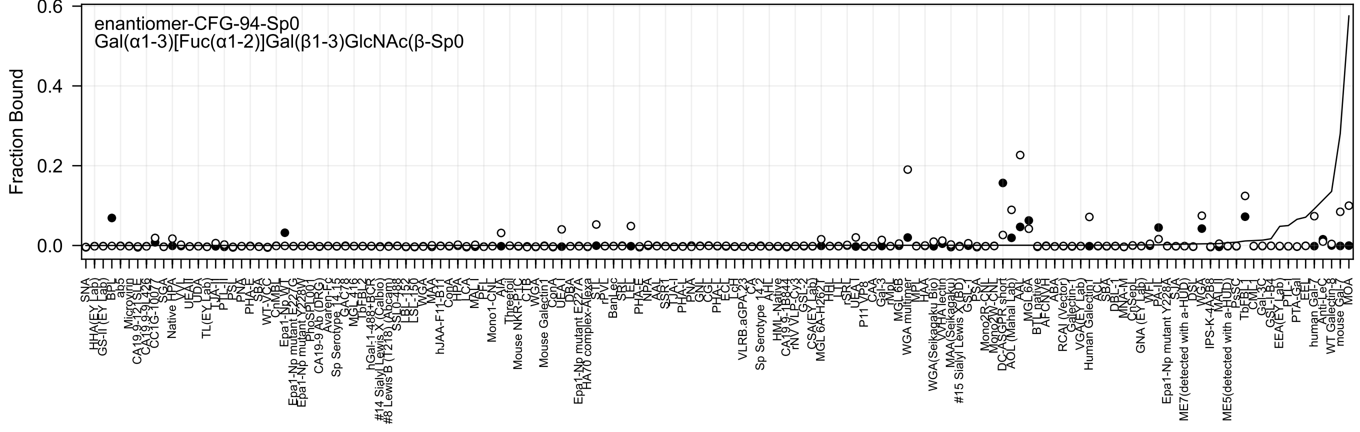
enantiomer-CFG-94-Sp0
Gal(α1-3)[Fuc(α1-2)]Gal(β1-3)GlcNAc(β-Sp0
Fraction Bound
0.0
0.2
0.4
0.6

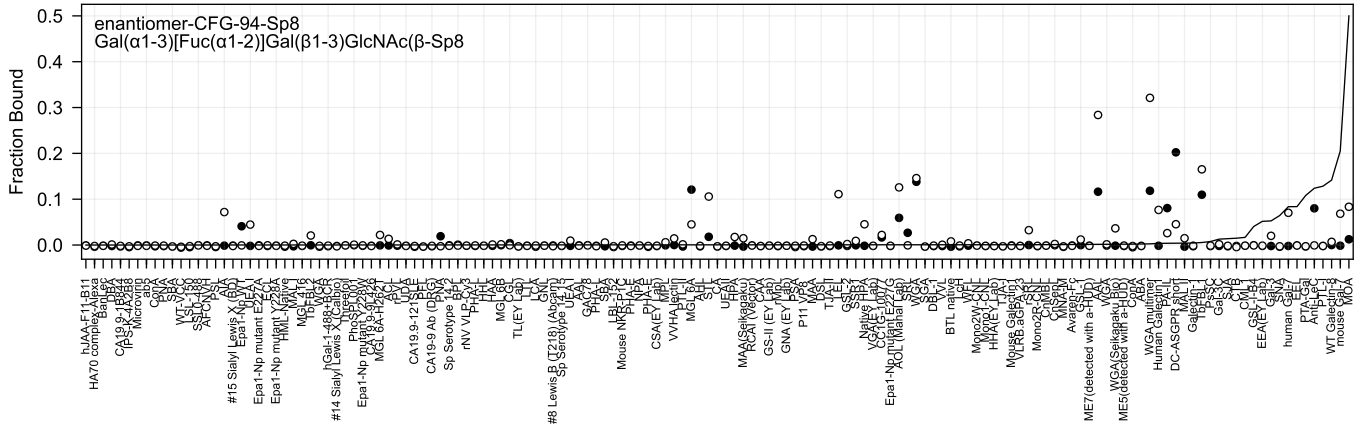

enantiomer-CFG-94-Sp8
Gal(α1-3)[Fuc(α1-2)]Gal(β1-3)GlcNAc(β-Sp8
Fraction Bound

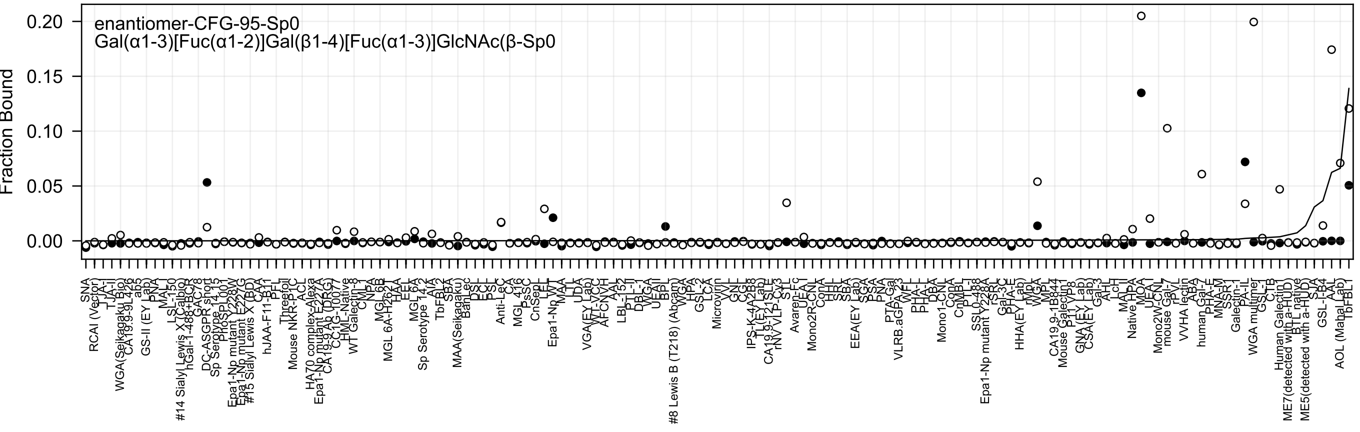
enantiomer-CFG-95-Sp0
Gal(α1-3)[Fuc(α1-2)]Gal(β1-4)[Fuc(α1-3)]GlcNAc(β-Sp0
Fraction Bound
0.20
0.15
0.10
0.05
0.00

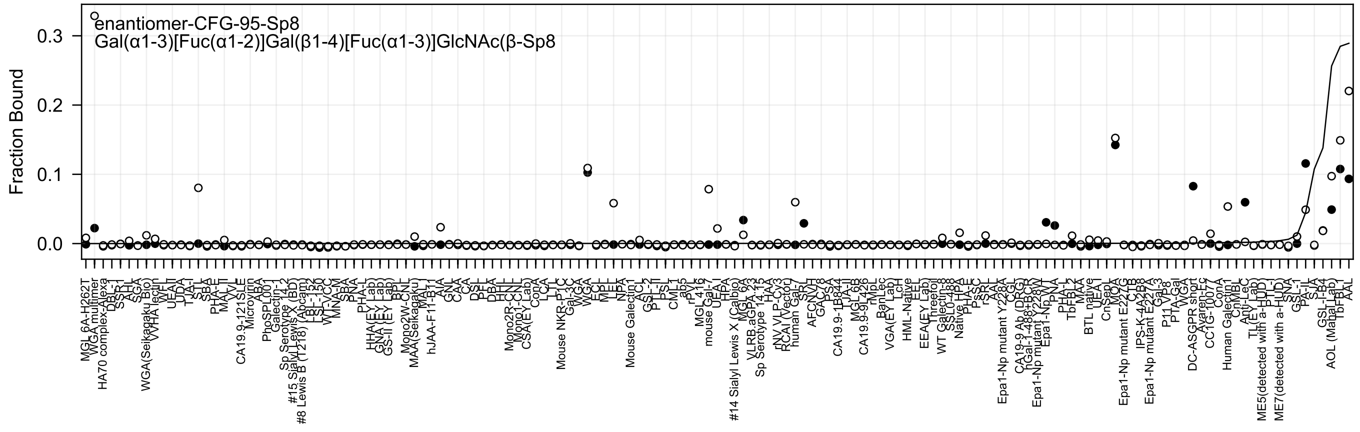
enantiomer-CFG-95-Sp8
Gal(α1-3)[Fuc(α1-2)]Gal(β1-4)[Fuc(α1-3)]GlcNAc(β-Sp8
Fraction Bound

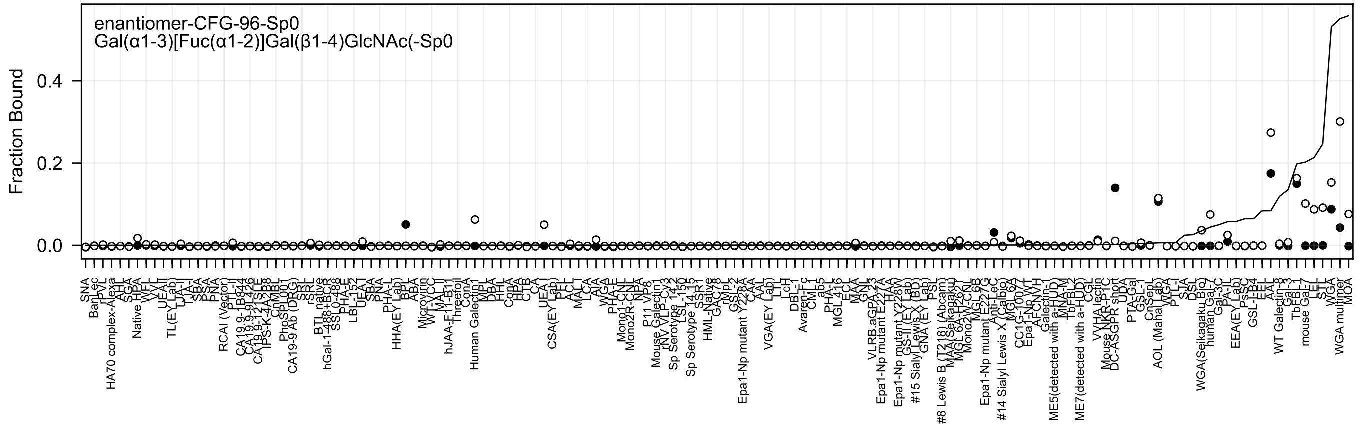
enantiomer-CFG-96-Sp0
Gal(α1-3)[Fuc(α1-2)]Gal(β1-4)GlcNAc(-Sp0
Fraction Bound
0.0
0.2
0.4

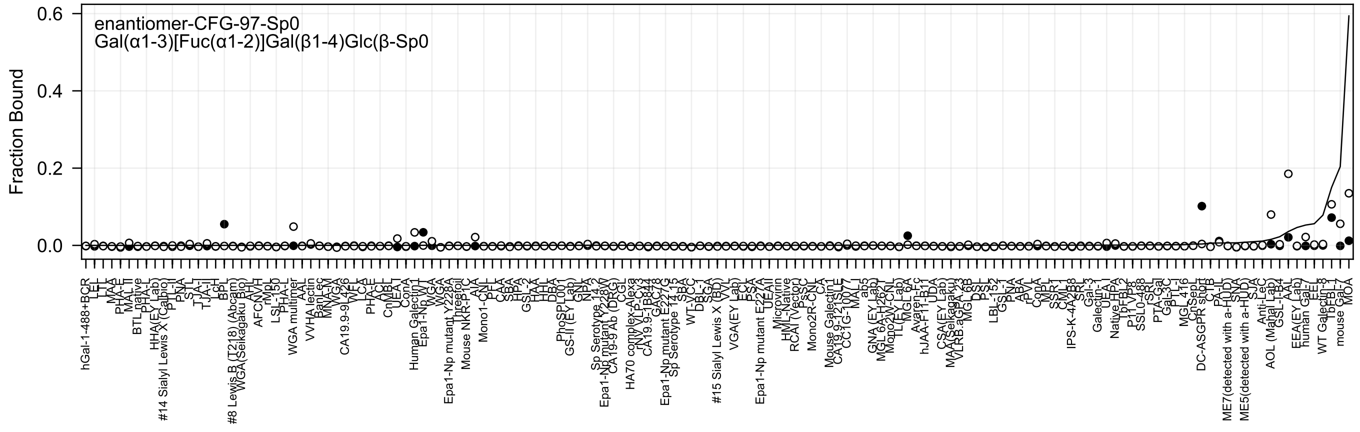
enantiomer-CFG-97-Sp0
Gal(α1-3)[Fuc(α1-2)]Gal(β1-4)Glc(β-Sp0
Fraction Bound

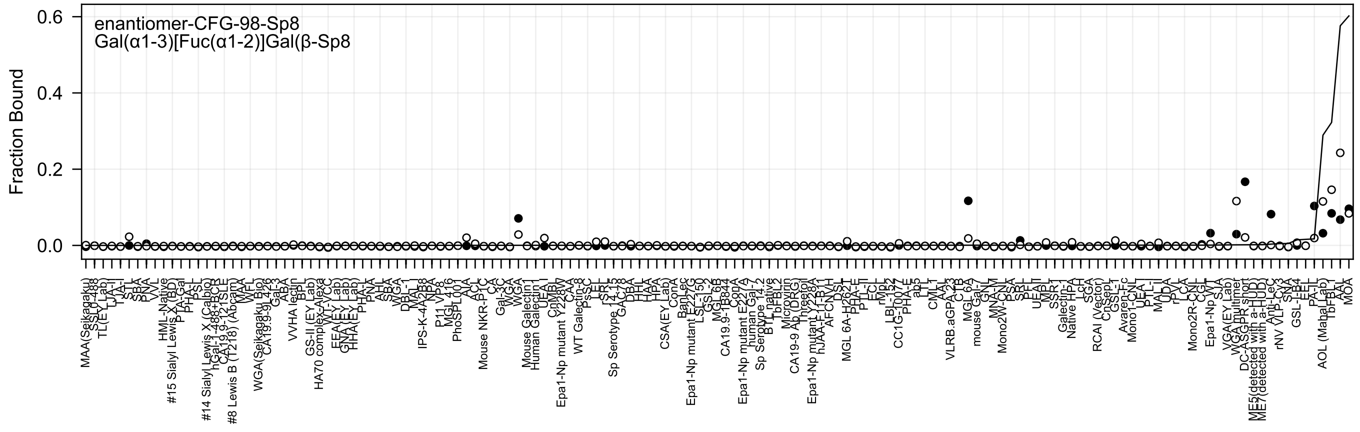
enantiomer-CFG-98-Sp8
Gal(α1-3)[Fuc(α1-2)]Gal(β-Sp8
Fraction Bound

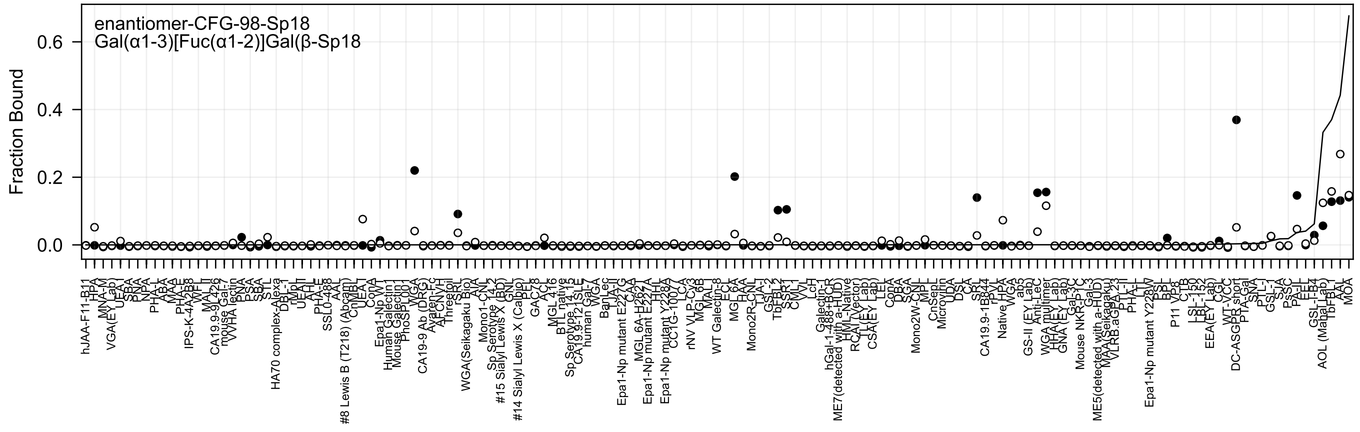

enantiomer-CFG-98-Sp18
Gal(α1-3)[Fuc(α1-2)]Gal(β-Sp18
Fraction Bound
0.0
0.2
0.4
0.6

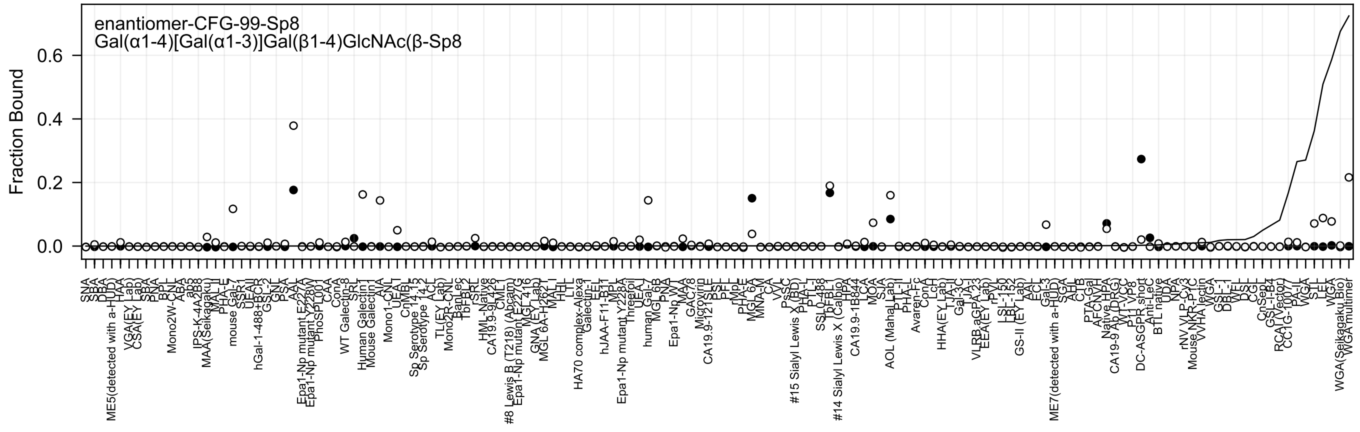
enantiomer-CFG-99-Sp8
Gal(α1-4)[Gal(α1-3)]Gal(β1-4)GlcNAc(β-Sp8
Fraction Bound
0.0
0.2
0.4
0.6

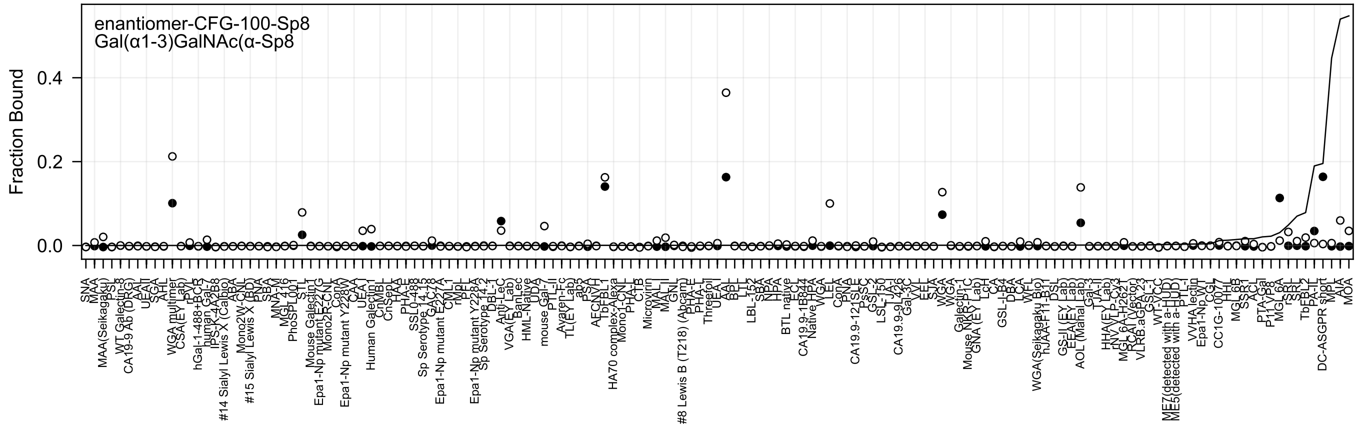
enantiomer-CFG-100-Sp8
Gal(α1-3)GalNAc(α-Sp8
Fraction Bound
0.0
0.2
0.4

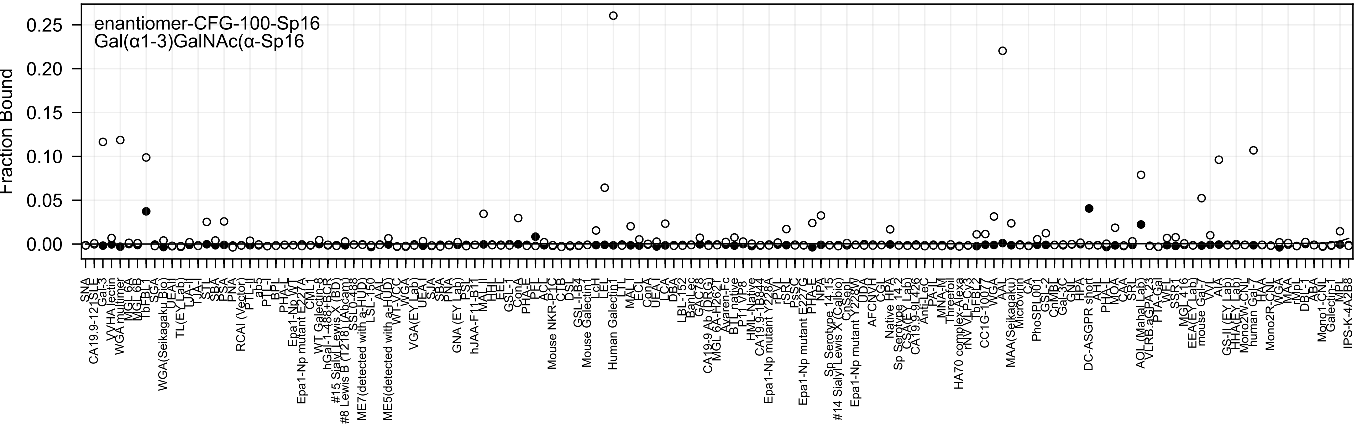
enantiomer-CFG-100-Sp16
Gal(α1-3)GalNAc(α-Sp16
Fraction Bound
0.25
0.20
0.15
0.10
0.05
0.00

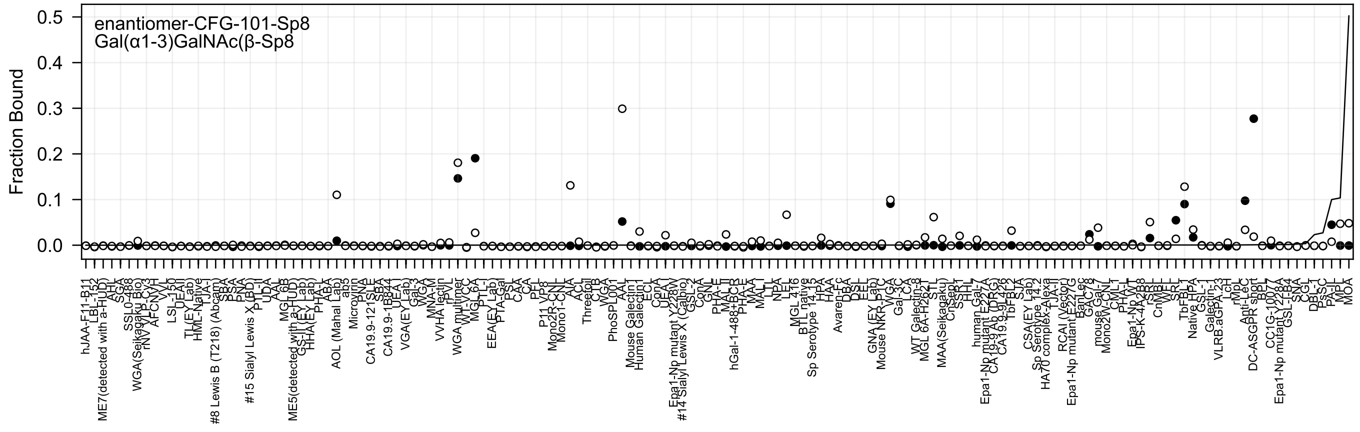

enantiomer-CFG-101-Sp8
Gal(α1-3)GalNAc(β-Sp8
Fraction Bound

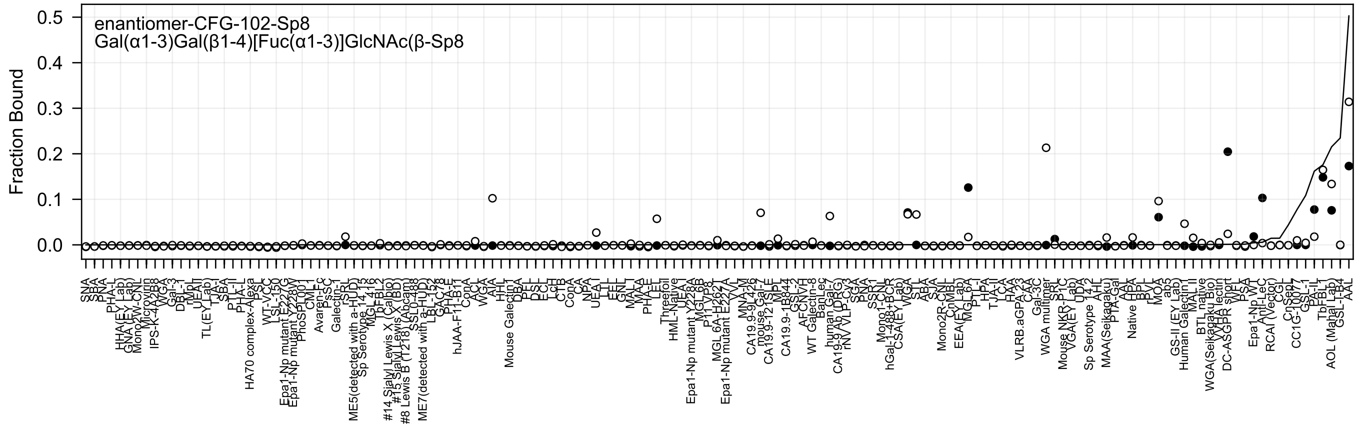
enantiomer-CFG-102-Sp8
Gal(α1-3)Gal(β1-4)[Fuc(α1-3)]GlcNAc(β-Sp8
Fraction Bound
0.0
0.1
0.2
0.3
0.4
0.5

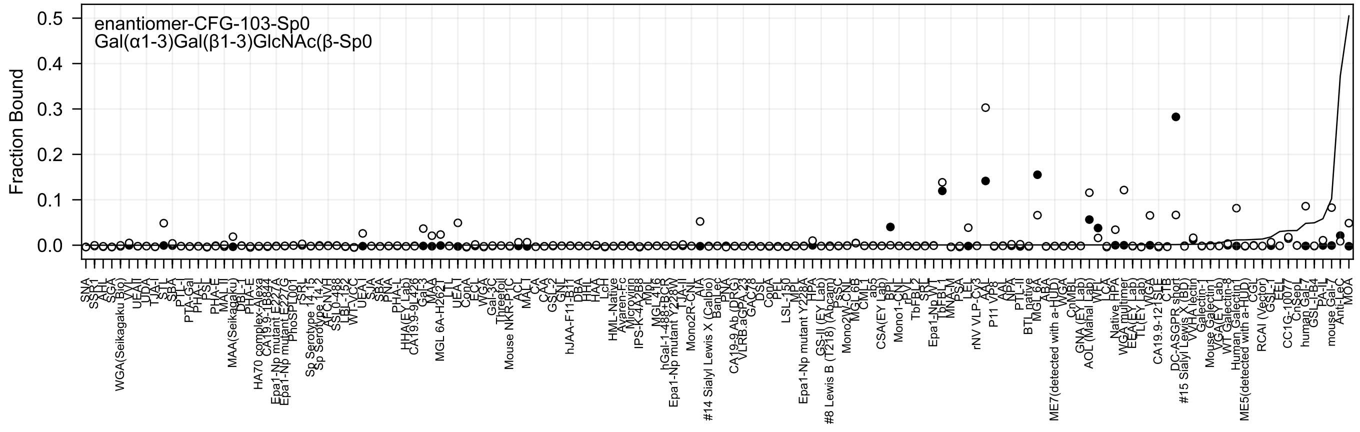
enantiomer-CFG-103-Sp0
Gal(α1-3)Gal(β1-3)GlcNAc(β-Sp0
Fraction Bound
0.5
0.4
0.3
0.2
0.1
0.0

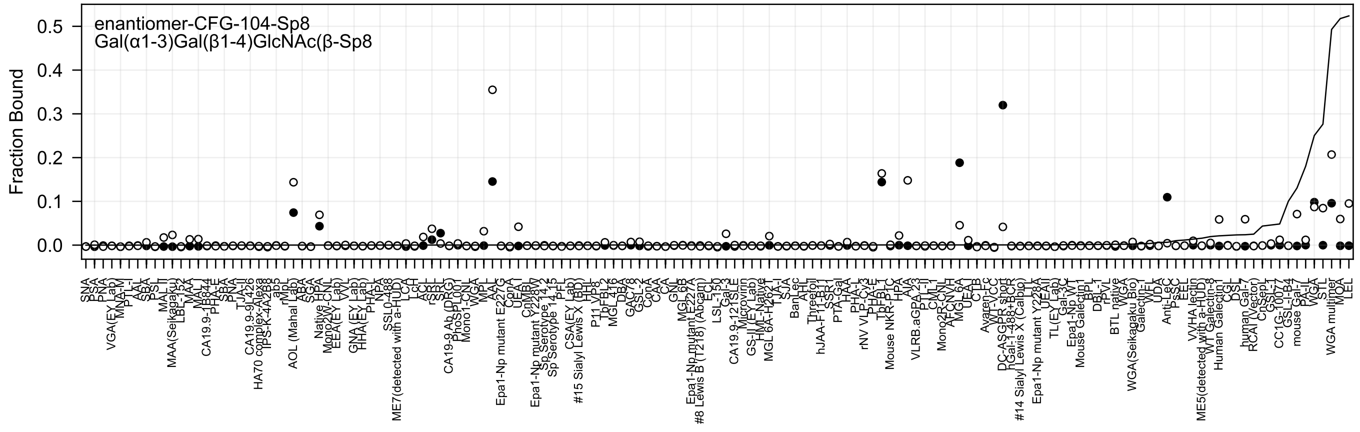

enantiomer-CFG-104-Sp8
Gal(α1-3)Gal(β1-4)GlcNAc(β-Sp8
Fraction Bound
0.0
0.1
0.2
0.3
0.4
0.5
SNA
PSA
PNA
VGA(EY Lab)
MNA-M
PTL-I
AAL
SBA
PSL
MAL II
MAA(Seikagaku)
LBL-152
MAA
MAL I
CA19.9-1B844
PHA-E
SBA
PNA
TJA-II
CA19.9-9L426
HA70 complex-Alexa
IPS-K-4A2B8
ab5
rMpL
AOL (Mahal Lab)
ABA
SGA
Native HPA
Mono2W-CNL
EEA(EY Lab)
VVL
GNA (EY Lab)
HHA(EY Lab)
PHA-L
NPA
SSL0-488
ME7(detected with a-HUD)
LCH
LCA
ACL
rSRL
SRL
CA19-9 Ab (DRG)
PhoSL001
Mono1-CNL
WGA
MPL
AAL
Epa1-Np mutant E227G
ConA
UEA-I
CNMBL
Epa1-Np mutant Y228W
Sp Serotype 14-2
Sp Serotype 14-15
PTL
CSA(EY Lab)
#15 Sialyl Lewis X (BD)
HHL
P11 VP8
TbFBL2
MGL 4-16
DBA
GAC78
GSL-2
ConA
CAA
CA
GNL
MGL 6B
Epa1-Np mutant E227A
#8 Lewis B (T218) (Abcam)
ECL
LSL-150
Gal-3
CA19.9-121SLE
Microvirin
GS-II (EY Lab)
HML-Native
MGL 6A-H262T
TJA-I
SJA
BanLec
AHL
Threefoil
hJAA-F113-BT
SSR1
PTA-Gal
HAA
PHA-L
rNV VLP-Cy3
PHA-E
TbFBL1
Mouse NKR-P1C
HPA
AIA
VLRB.aGPA.23
PTL-II
CML-1
Mono2R-CNL
AFCNVH
MGL 6A
UEA-I
CTB
Avaren-Fc
WT-VCC
DC-ASGPR short
hGal-1-488+BCR
#14 Sialyl Lewis X (Calbio)
LTL
Epa1-Np mutant Y228A
UEAII
TL(EY Lab)
Gal-3C
Epa1-Np WT
Mouse Galectin-1
BPL
DBL-1
rPVL
BTL native
VVA
WGA(Seikagaku Bio)
Galectin-1
WFL
UDA
Anti-LeC
PSSC
EEL
VVHA lectin
ME5(detected with a-HUD)
WT Galectin-8
Human Galectin-1
CSL
DSL
human Gal-7
RCAI (Vector)
CnSepL
GSL-I
CC1G-10077
GSL-I-B4
mouse Gal-7
PA-IL
WGA
STL
WGA multimer
MOA
LEL

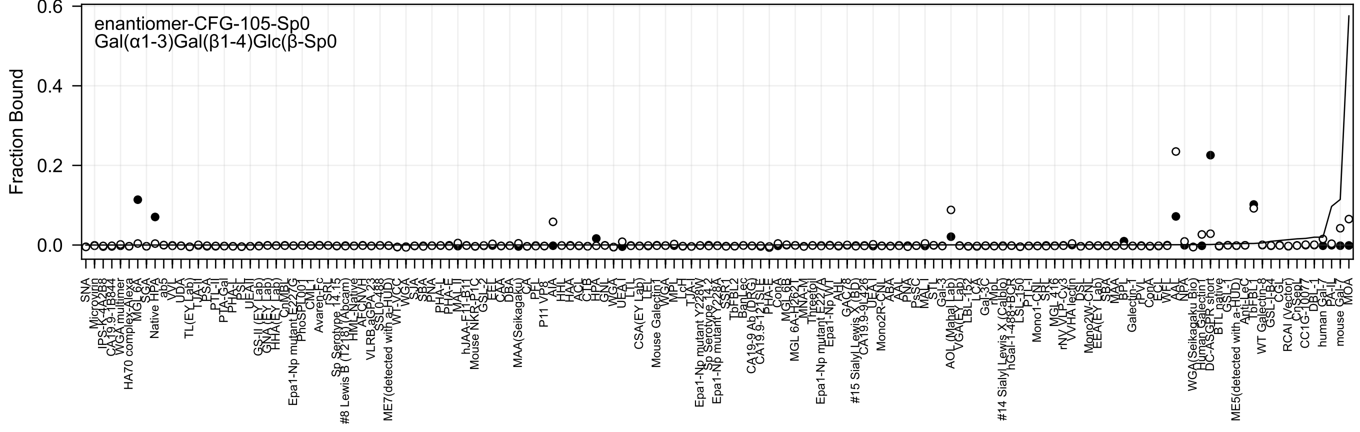

enantiomer-CFG-105-Sp0
Gal(α1-3)Gal(β1-4)Glc(β-Sp0
Fraction Bound
0.6
0.4
0.2
0.0

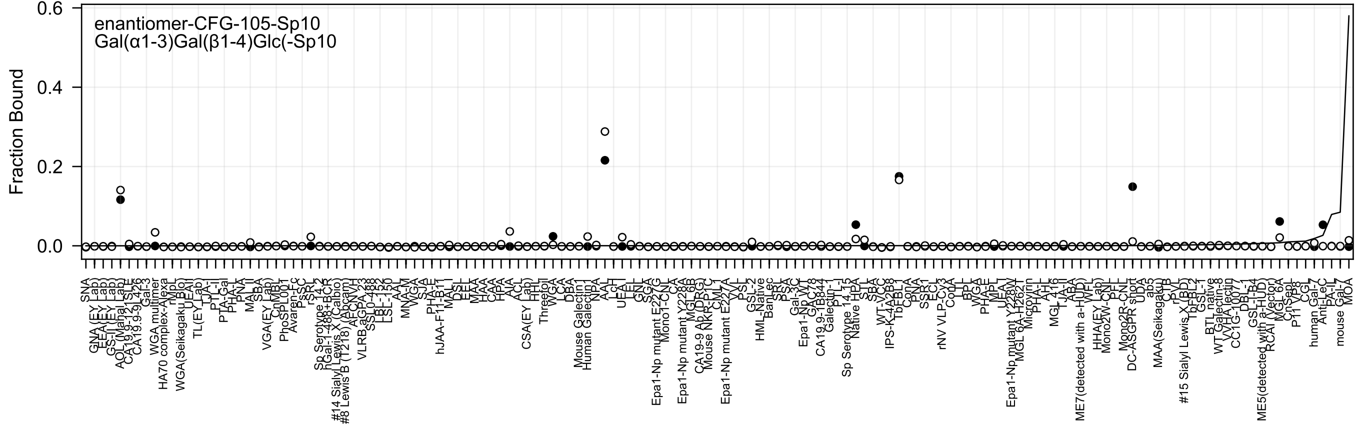
enantiomer-CFG-105-Sp10
Gal(α1-3)Gal(β1-4)Glc(-Sp10
Fraction Bound
0.6
0.4
0.2
0.0
SNA
GNA(EY Lab)
EEA(EY Lab)
GS-II(EY Lab)
AOL(Manal Lab)
CA19.9-121SLE
CA19.9-9L426
Gal-3
WGA multimer
HA70 complex-Alexa
rMpL
WGA(Seikagaku Bio)
UEAI
TL(EY Lab)
TJA-I
PTL-I
PTA-Gal
PHA-L
PNA
MAL-II
SBA
VGA(EY Lab)
CnMBL
PhoSL001
Avaren-Fc
PsSC
rSRL
Sp Serotype 14.2
hGal-1-488+BCR
#14 Sialyl Lewis X (Calbio)
#8 Lewis B (T218) (Abcam)
AFCNVH
VLRB.aGPA.23
SSL0-488
LBL-152
LSL-150
AAL
MNA-M
VVGA
SJA
PHA-E
hJAA-F11-B11
MAL-I
DSL
EEL
MAA
HAA
CAA
HPA
AIA
ACL
CSA(EY Lab)
HHL
Threefoil
WGA
LCA
DBA
Mouse Galectin1
Human Galectin1
NPA
AAL
LcH
UEA-I
LEL
GNL
SGA
Epa1-Np mutant E227G
Mono1-CNL
CA
Epa1-Np mutant Y228A
MGL 6B
CA19-9 Ab (DRG)
Mouse NKR-P1C
CML1
Epa1-Np mutant E227A
VVL
PSL
GSL-2
HML-Native
BanLec
SRL
PSA
Gal-3C
Epa1-Np WT
GAC78
CA19.9-1B844
Galectin-1
PTL-I
Sp Serotype 14.15
Native HPA
STL
SBA
WT-VCC
IPS-K-4A2B8
TbFBL1
ConA
PNA
SSR1
ECL
rNV VLP-Cy3
ConA
LTL
BPL
WGA
PHA-E
MPL
UEA-I
Epa1-Np mutant Y228W
MGL 6A-H262T
Microvirin
PHA-L
AH
MGL 416
TJA-II
ABA
ME7(detected with a-HUD)
WFL
HHA(EY Lab)
Mono2W-CNL
PFL
Mono2R-CNL
DC-ASGPR short
UDA
ab5
MAA(Seikagaku)
CTB
rPVL
#15 Sialyl Lewis X (BD)
TbFBL2
GSL-1
BTL native
WT Galectin-8
VVHA lectin
CC1G-10077
DBL-1
GSL-I-B4
ME5(detected with a-HUD)
RCAI (Vector)
MGL 6A
CnSepL
P11 VP8
CGL
human Gal-7
Anti-LeC
PA-IL
mouse Gal-7
MOA

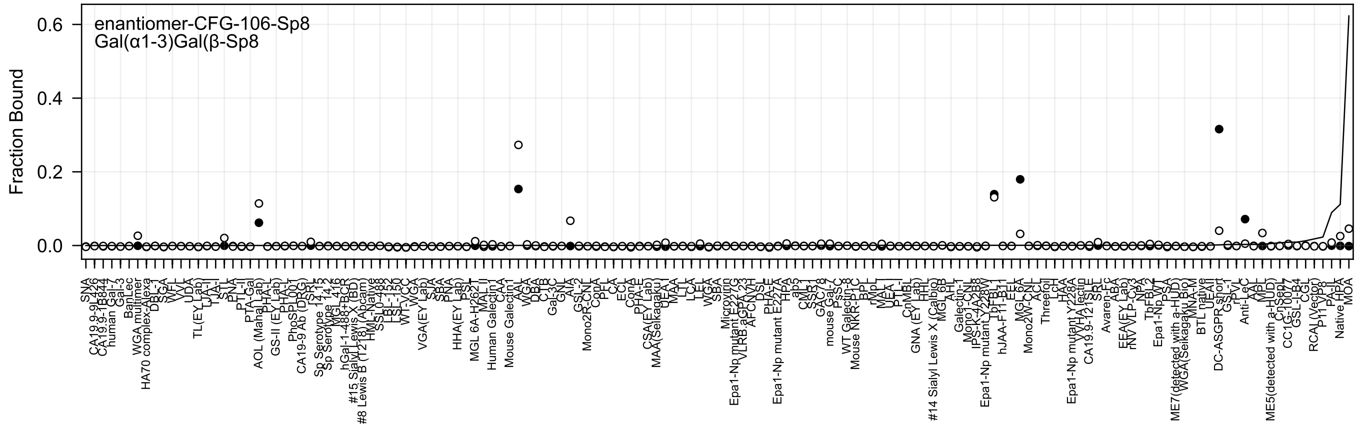
enantiomer-CFG-106-Sp8
Gal(α1-3)Gal(β-Sp8
Fraction Bound

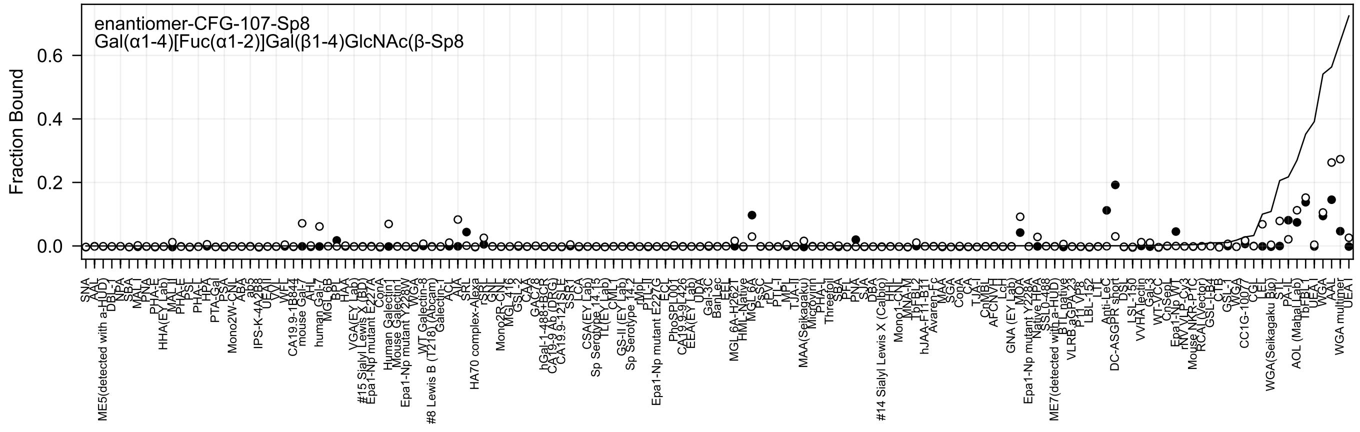
enantiomer-CFG-107-Sp8
Gal(α1-4)[Fuc(α1-2)]Gal(β1-4)GlcNAc(β-Sp8
Fraction Bound
0.0
0.2
0.4
0.6

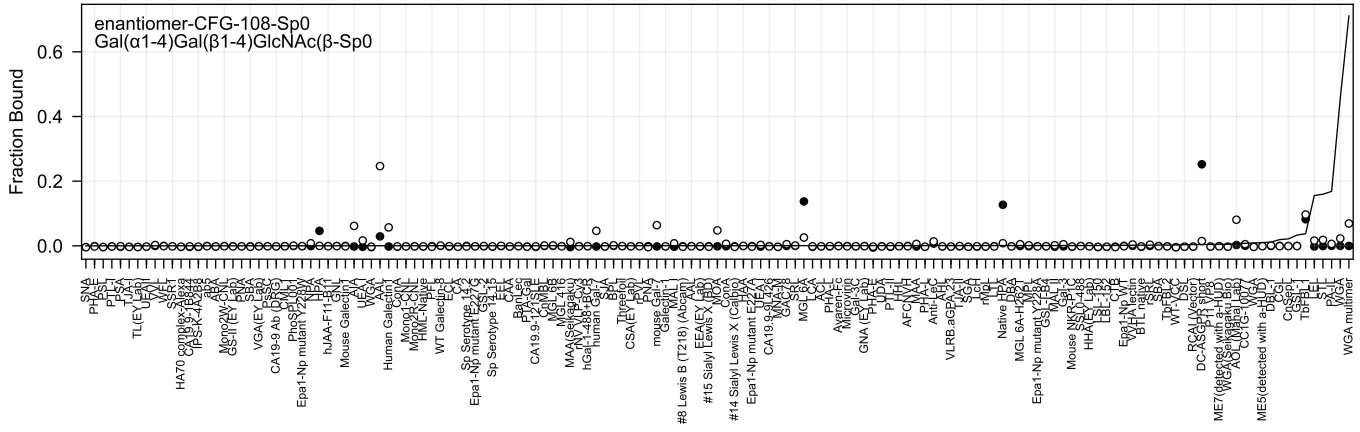
enantiomer-CFG-108-Sp0
Gal(α1-4)Gal(β1-4)GlcNAc(β-Sp0
Fraction Bound

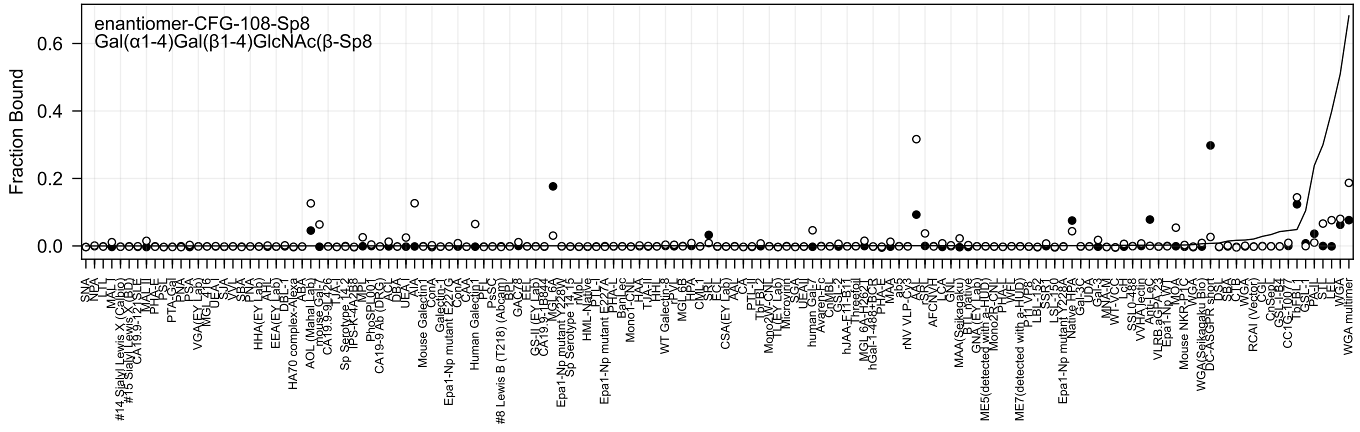
enantiomer-CFG-108-Sp8
Gal(α1-4)Gal(β1-4)GlcNAc(β-Sp8
Fraction Bound
0.6
0.4
0.2
0.0

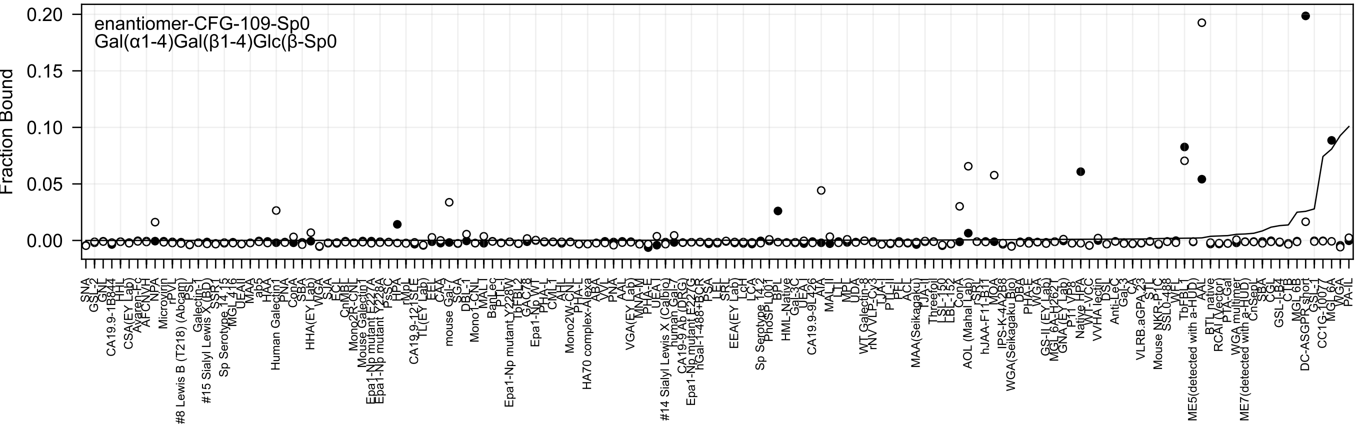
enantiomer-CFG-109-Sp0
Gal(α1-4)Gal(β1-4)Glc(β-Sp0
Fraction Bound
0.20
0.15
0.10
0.05
0.00
SNA
GSL-2
GNL
CA19.9-1B844
HHL
CSA(EY Lab)
Ayaren-Fc
AFCNVH
NPA
Microvirin
rPVL
#8 Lewis B (T218) (Abcam)
PSL
Galectin-1
#15 Sialyl Lewis X (BD)
SSR1
Sp Serotype 14.15
MGL 416
UEA-II
MAA
ab5
HAA
Human Galectin1
PNA
ConA
SBA
HHA(EY Lab)
WGA
SJA
ECL
CnMBL
Mono2R-CNL
Mouse Galectin1
Epa1-Np mutant E227A
Epa1-Np mutant Y228A
PsSC
HPA
rMpL
CA19.9-121SLE
TL(EY Lab)
EEL
CAA
mouse Gal-7
SGA
DBL-1
Mono1-CNL
MAL-I
BanLec
PTL-I
Epa1-Np mutant Y228W
TbFBL2
GAC78
Epa1-Np WT
PHA-L
CML1
AHL
Mono2W-CNL
PHA-L
HA70 complex-Alexa
ABA
VVL
PNA
AAL
VGA(EY Lab)
MNA-M
PHA-E
UEA-I
#14 Sialyl Lewis X (Calbio)
human Gal-7
CA19-9 Ab (DRG)
Epa1-Np mutant E227G
hGal-1-488+BCR
PSA
LEL
SRL
EEA(EY Lab)
LcH
LCA
Sp Serotype 14.2
PhoSL001
BPL
HML-Native
Gal-3C
UEA-I
CA19.9-9L426
AIA
MAL-II
LTL
MPL
UDA
WT Galectin-8
rNV VLP-CY3
TJA-II
PTL-II
PFL
ACL
MAA(Seikagaku)
TJA-I
Threefoil
LSL-150
LBL-152
ConA
AOL (Mahal Lab)
rSRT
hJAA-F11-B11
MOA
IPS-K-4A2B8
WGA(Seikagaku Bio)
DBA
PHA-E
WGA
GS-II (EY Lab)
MGL 6A-H262
GNA (EY Lab)
P11 VP8
Native HPA
WT-VCC
VVHA lectin
DSL
Anti-LeC
Gal-3
CA
VLRB.aGPA.23
STL
Mouse NKR-P1C
SSL0-488
WFL
TbFBL1
ME5(detected with a-HUD)
AAL
BTL native
RCAI (Vector)
PTA-Gal
WGA multimer
ME7(detected with a-HUD)
CnSep
SBA
CGL
GSL-I-B4
CTB
MGL 6B
DC-ASGPR short
GSL-I
CC1G-1007
MGL6A
WGA
PA-IL

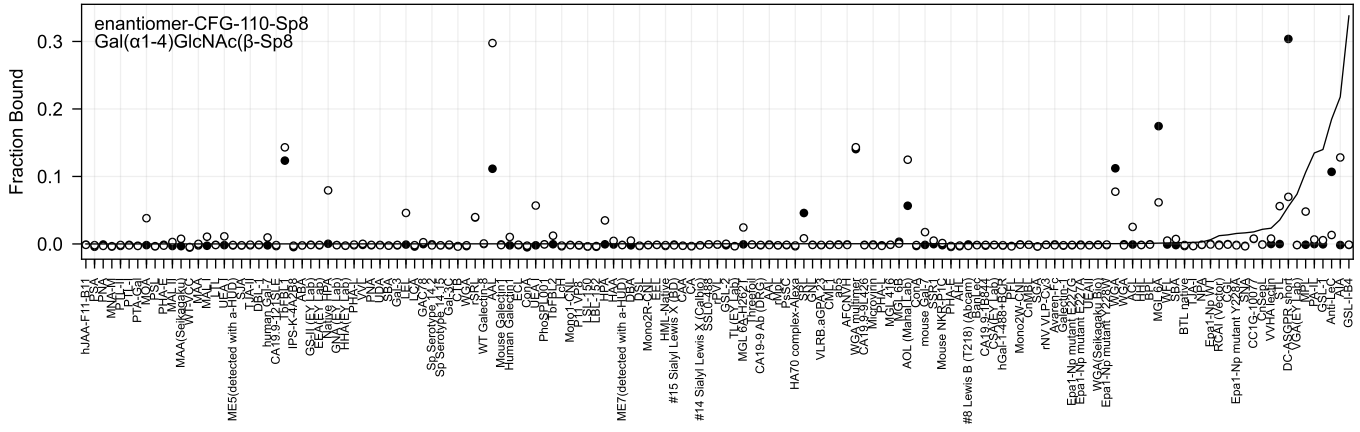
enantiomer-CFG-110-Sp8
Gal(α1-4)GlcNAc(β-Sp8
Fraction Bound
0.3
0.2
0.1
0.0

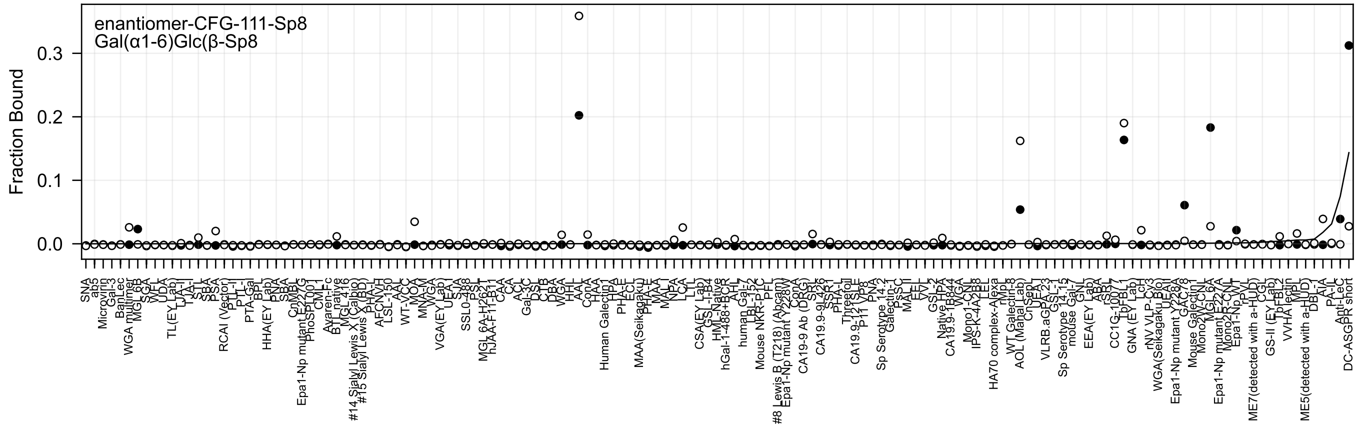
enantiomer-CFG-111-Sp8
Gal(α1-6)Glc(β-Sp8
Fraction Bound
0.0
0.1
0.2
0.3
SNA
ab5
Microvirin
Gal-3
BanLec
WGA multimer
MGL 6B
SGA
WFL
UDA
TL(EY Lab)
TJA-I
TJA-II
STL
SBA
PSA
RCAI (Vector)
PTL-II
PTL-I
PTA-Gal
BPL
HHA(EY Lab)
PNA
SBA
CnMBL
Epa1-Np mutant E227G
PhoSL
CML
Avaren-Fc
BTL native
MGL 416
#14 Sialyl Lewis X (Calbio)
#15 Sialyl Lewis X (BD)
PHA-L
AFCNVH
LSL-150
AAL
WT-VCC
MOA
MNA-M
WGA
VGA(EY Lab)
UEA I
SJA
SSL0-488
PSL
MGL 6A-H262T
hJAA-F11-B11
CAA
CA
ACL
Gal-3C
DSL
CTB
DBA
WGA
HHL
AAL
ConA
HAA
Human Galectin1
HPA
PHA-E
ECL
MAA(Seikagaku)
PHA-E
MAA
MAL
NPA
LCA
LTL
CSA(EY Lab)
GSL-I-B4
HML-Native
hGal-1-488+BCR
AAL
human Gal-7
LBL-152
Mouse NKR-P1C
PFL
#8 Lewis B (T218) (Abcam)
Epa1-Np mutant Y228W
ConA
CA19-9 Ab (DRG)
SRL
CA19.9-9L426
SSA
PHA-L
Threefoil
CA19.9-121SLE
P11
VP8
PNA
Sp Serotype 14.2
Galectin-1
PsSC
MAL II
EEL
VVL
GSL-2
Native HPA
CA19.9-1B844
WGA
Mono1-CNL
IPS-K-4A2B8
LEL
HA70 complex-Alexa
rVP
WT Galectin-8
AOL (Manal Lab)
CnSepL
UEA I
VLRB.aGPA.23
GSL-1
Sp Serotype 14.15
mouse Gal-7
GNL
EEA(EY Lab)
ABA
rSRL
CC1G-10077
TbFBL 1
GNA (EY Lab)
LcH
rNV VLP-Cy3
WGA(Seikagaku Bio)
UEA I
Epa1-Np mutant Y228A
GAC78
Mouse Galectin1
Mono2W-CNL
MGL 6A
Epa1-Np mutant E227A
Mono2R-CNL
Epa1-Np WT
rPVL
CGL
GS-II (EY Lab)
TbFBL 2
VVHA lectin
MPL
ME7(detected with a-HUD)
ME5(detected with a-HUD)
DBL
AIA
PA-IL
Anti-LeC
DC-ASGPR short

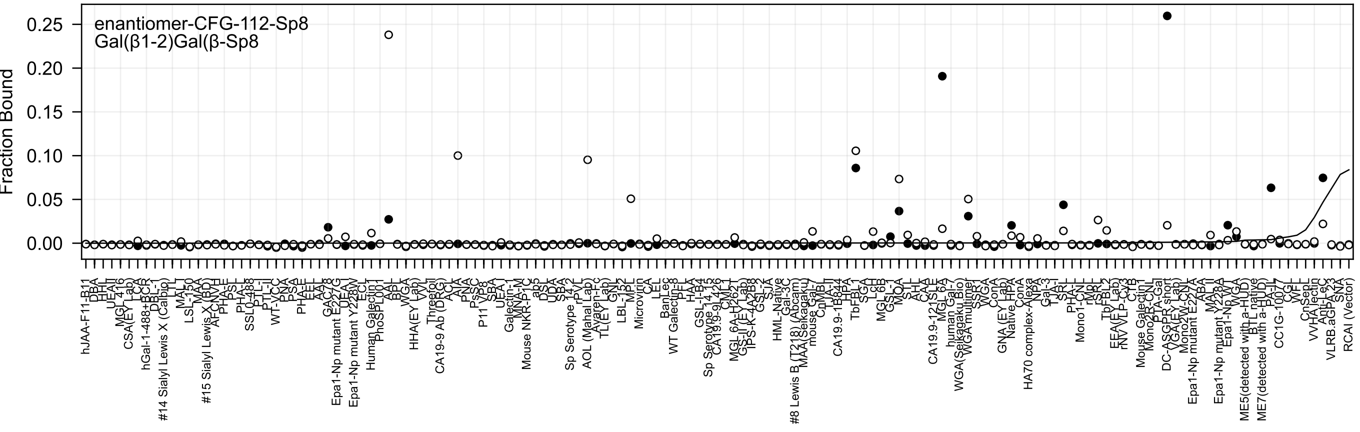

enantiomer-CFG-112-Sp8
Gal(β1-2)Gal(β-Sp8
Fraction Bound
0.25
0.20
0.15
0.10
0.05
0.00

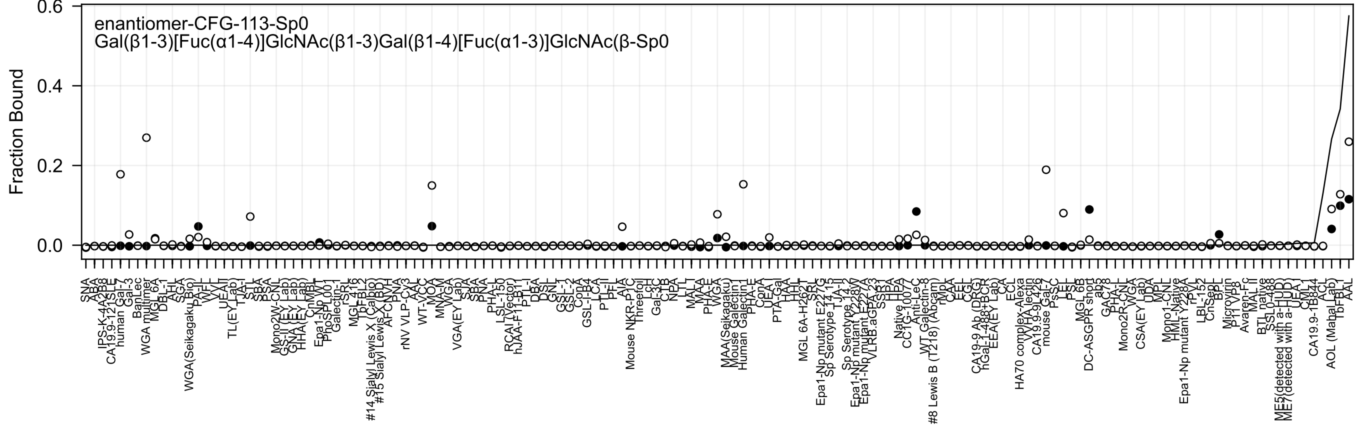

enantiomer-CFG-113-Sp0
Gal(β1-3)[Fuc(α1-4)]GlcNAc(β1-3)Gal(β1-4)[Fuc(α1-3)]GlcNAc(β-Sp0
Fraction Bound

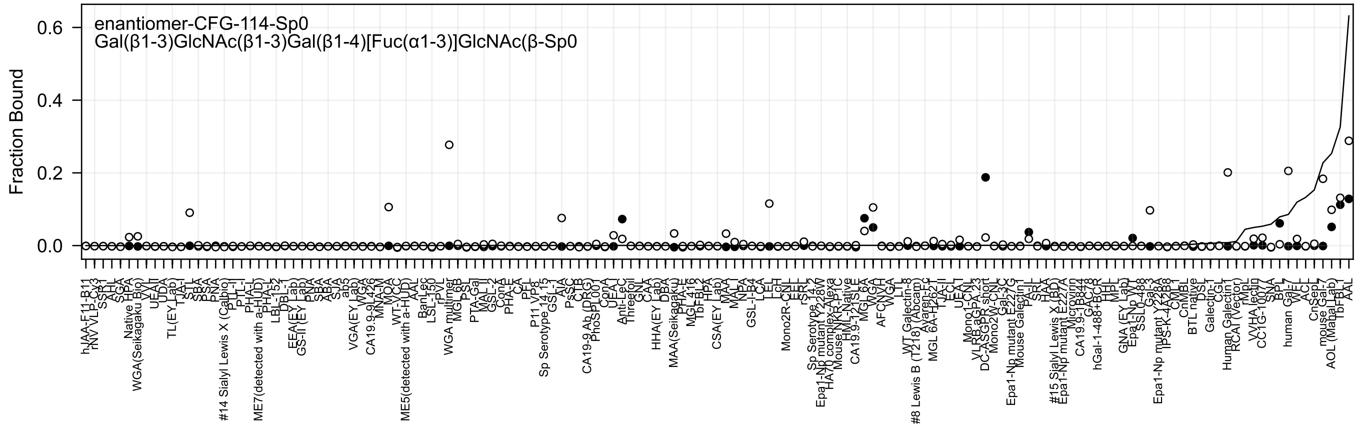
enantiomer-CFG-114-Sp0
Gal(β1-3)GlcNAc(β1-3)Gal(β1-4)[Fuc(α1-3)]GlcNAc(β-Sp0
Fraction Bound
0.6
0.4
0.2
0.0

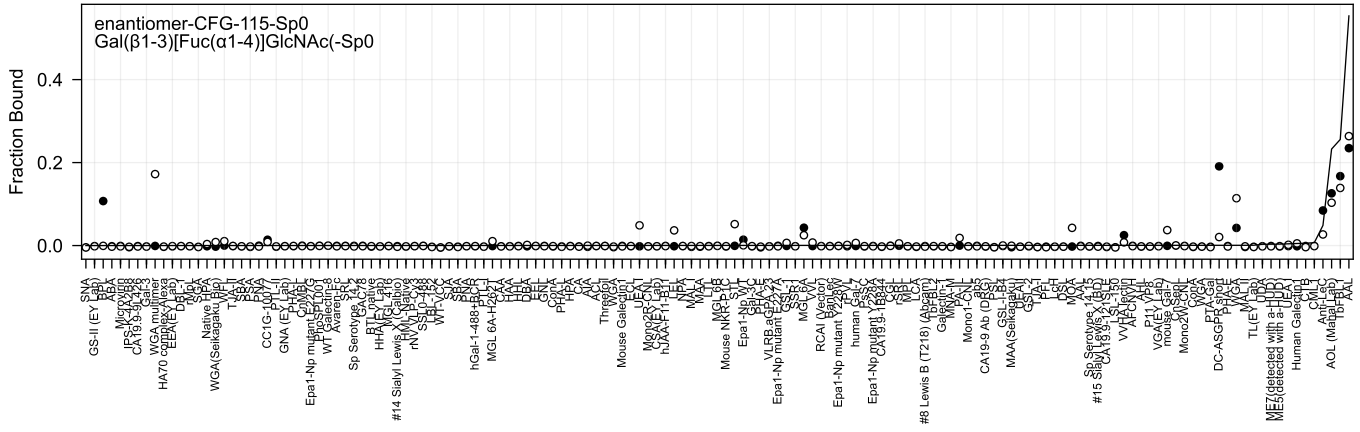

enantiomer-CFG-115-Sp0
Gal(β1-3)[Fuc(α1-4)]GlcNAc(-Sp0
Fraction Bound
0.0
0.2
0.4

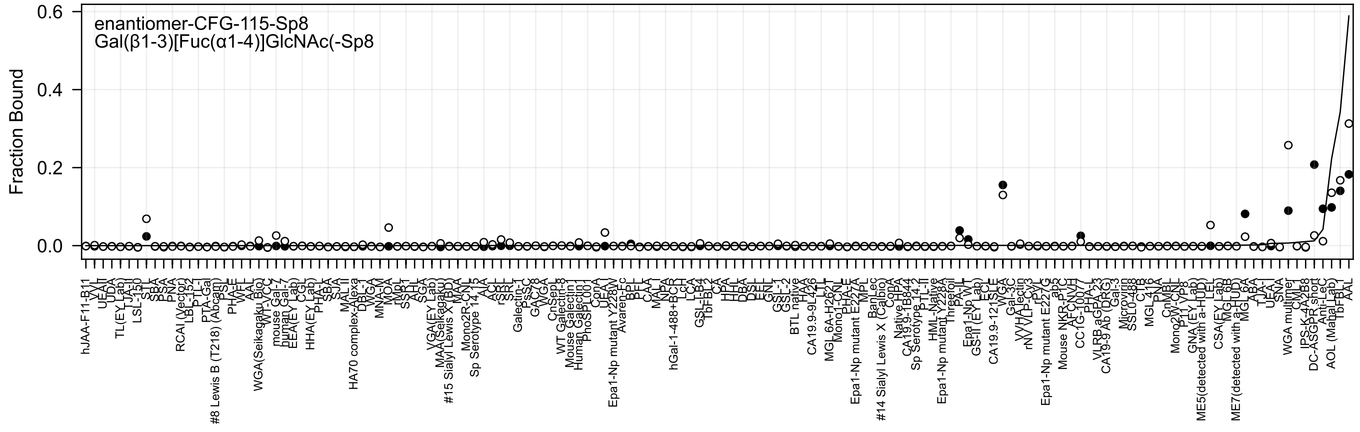
enantiomer-CFG-115-Sp8
Gal(β1-3)[Fuc(α1-4)]GlcNAc(-Sp8
Fraction Bound
0.6
0.4
0.2
0.0

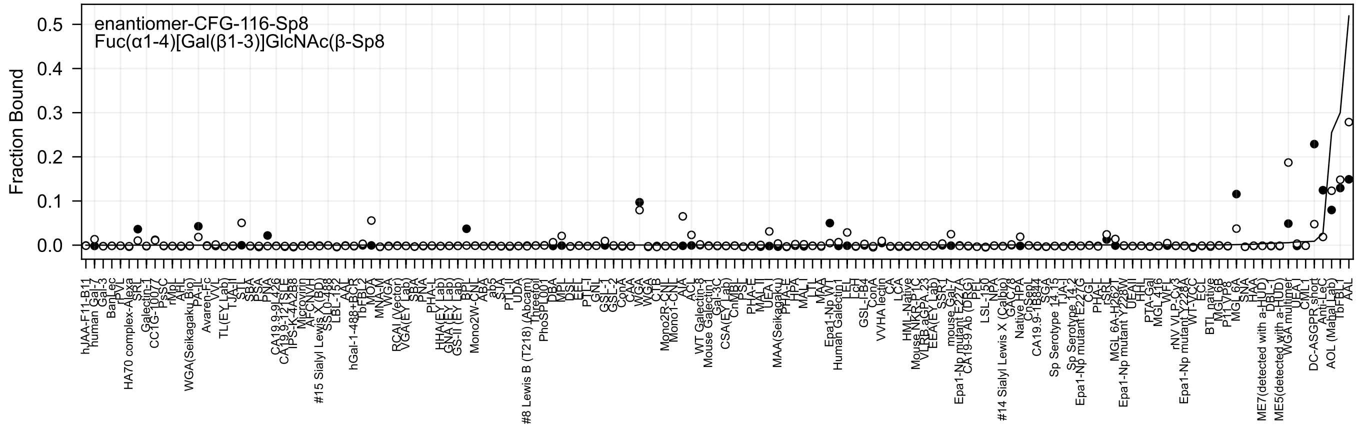
enantiomer-CFG-116-Sp8
Fuc(α1-4)[Gal(β1-3)]GlcNAc(β-Sp8
Fraction Bound
0.5
0.4
0.3
0.2
0.1
0.0

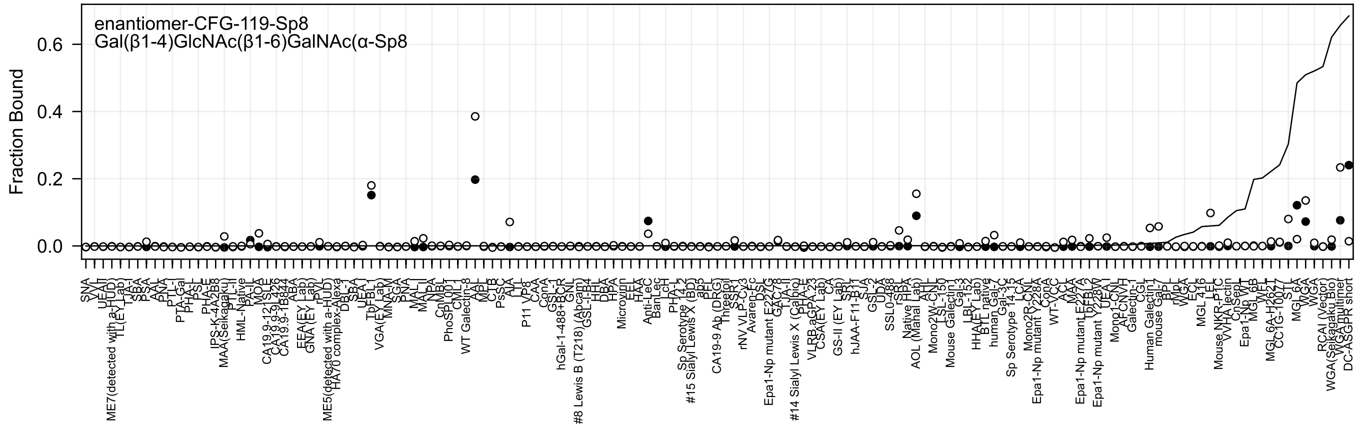

enantiomer-CFG-119-Sp8
Gal(β1-4)GlcNAc(β1-6)GalNAc(α-Sp8
Fraction Bound
0.6
0.4
0.2
0.0

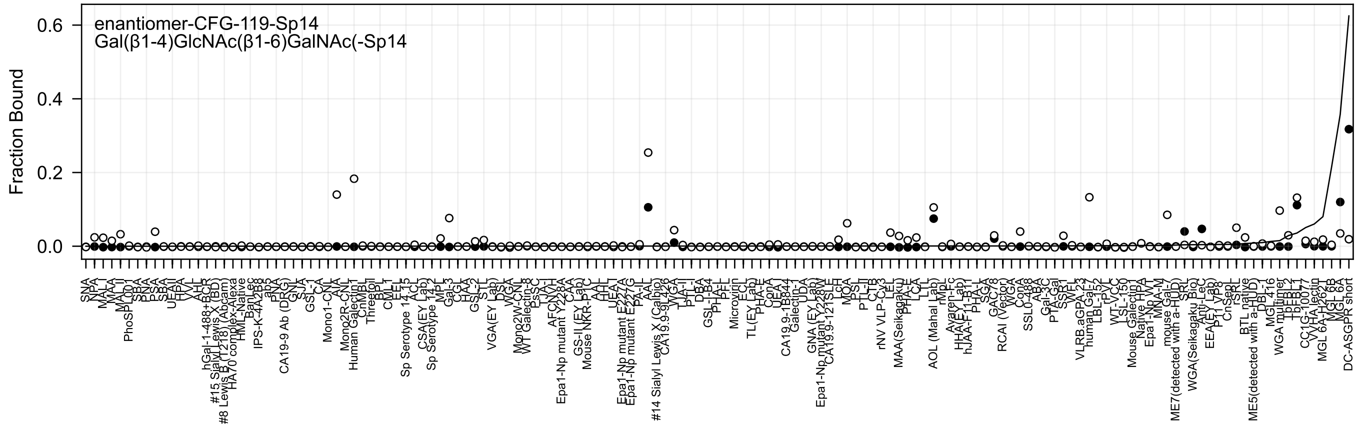
enantiomer-CFG-119-Sp14
Gal(β1-4)GlcNAc(β1-6)GalNAc(-Sp14
Fraction Bound

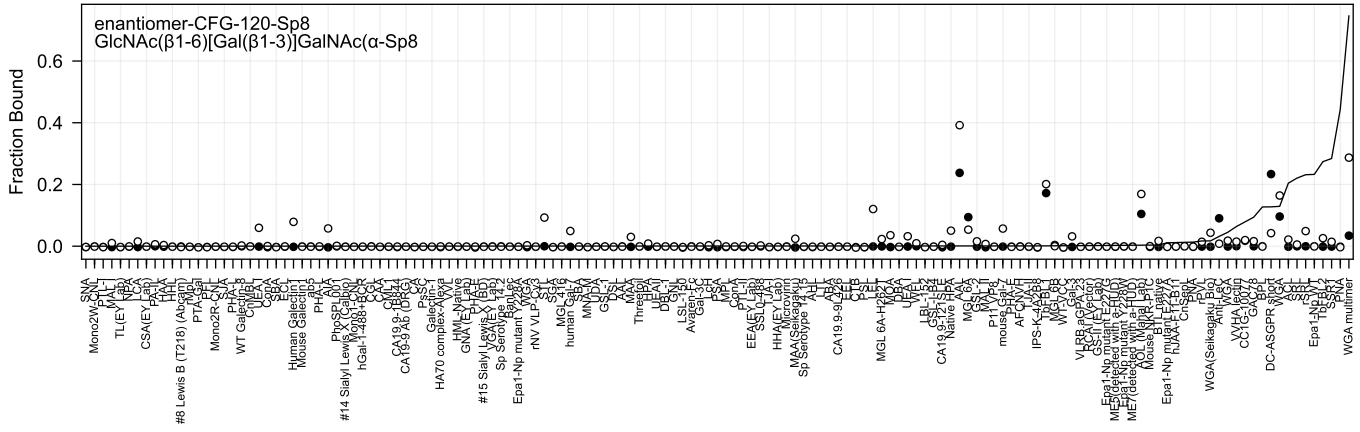
enantiomer-CFG-120-Sp8
GlcNAc(β1-6)[Gal(β1-3)]GalNAc(α-Sp8
Fraction Bound

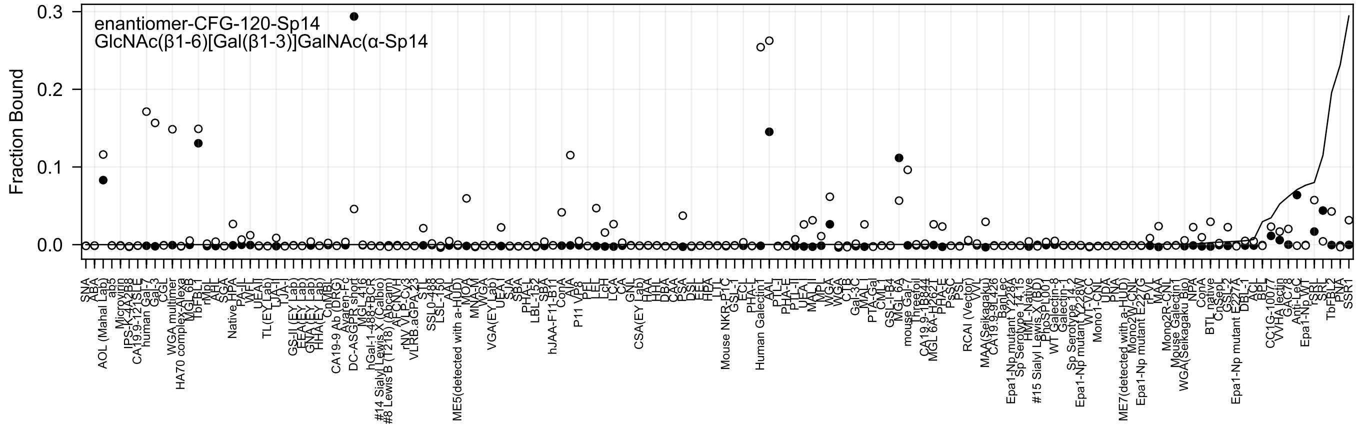
enantiomer-CFG-120-Sp14
GlcNAc(β1-6)[Gal(β1-3)]GalNAc(α-Sp14
Fraction Bound
0.3
0.2
0.1
0.0

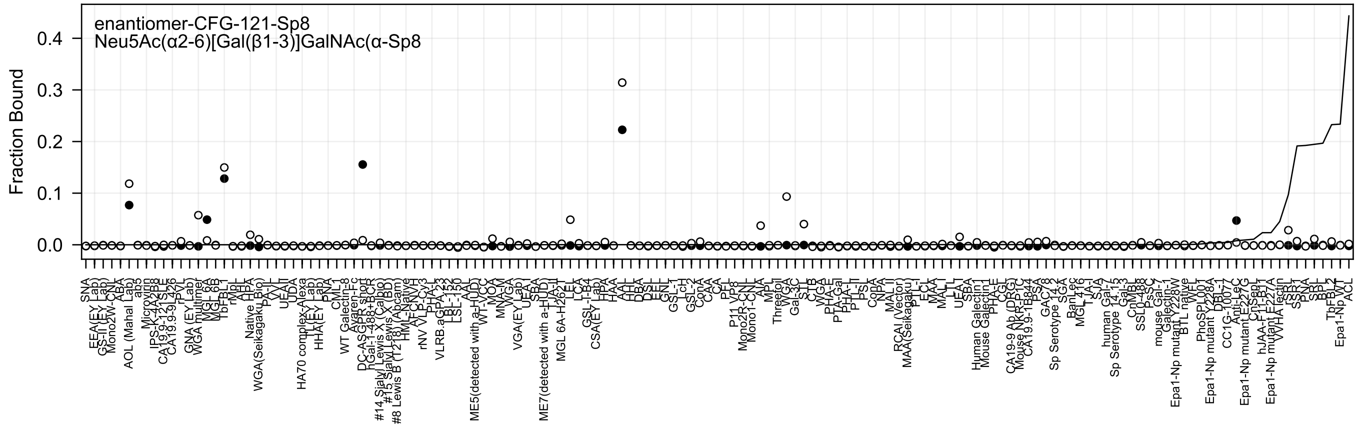
enantiomer-CFG-121-Sp8
Neu5Ac(α2-6)[Gal(β1-3)]GalNAc(α-Sp8
Fraction Bound
0.0
0.1
0.2
0.3
0.4

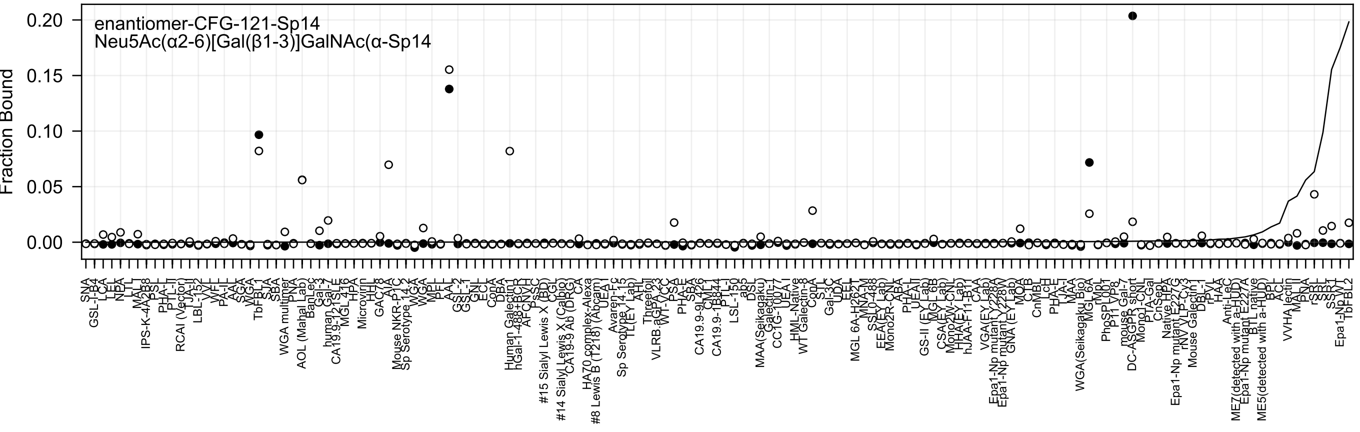
enantiomer-CFG-121-Sp14
Neu5Ac(α2-6)[Gal(β1-3)]GalNAc(α-Sp14
Fraction Bound
0.20
0.15
0.10
0.05
0.00

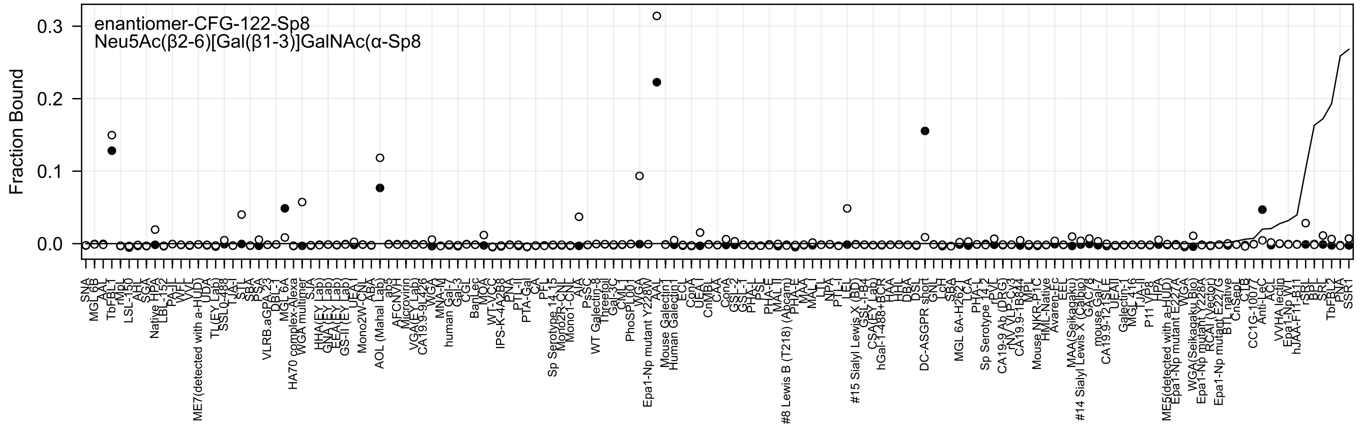
enantiomer-CFG-122-Sp8
Neu5Ac(β2-6)[Gal(β1-3)]GalNAc(α-Sp8
Fraction Bound
0.3
0.2
0.1
0.0

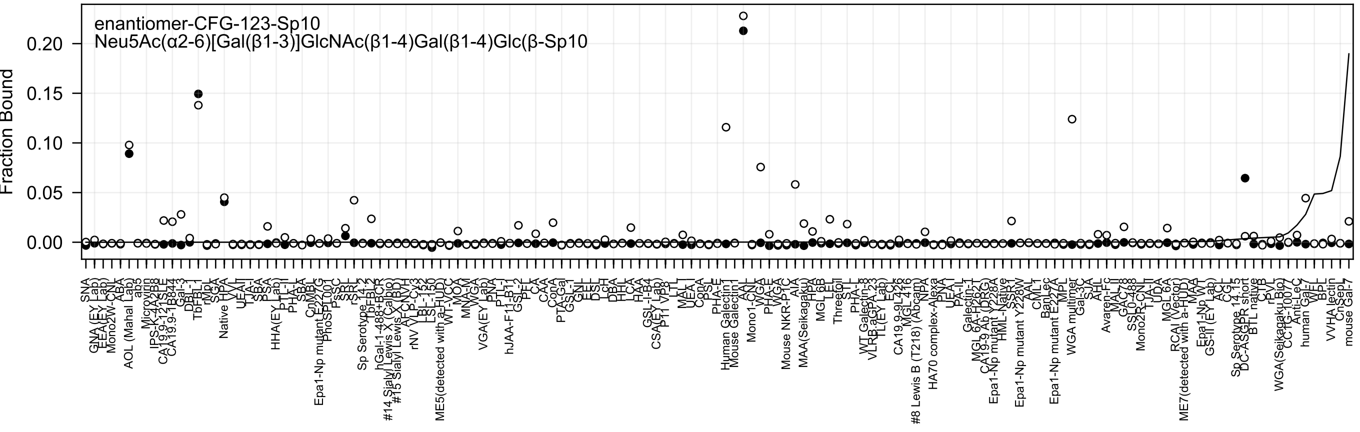

enantiomer-CFG-123-Sp10
Neu5Ac(α2-6)[Gal(β1-3)]GlcNAc(β1-4)Gal(β1-4)Glc(β-Sp10
Fraction Bound
0.00
0.05
0.10
0.15
0.20
SNA
GNA(EY Lab)
EEA(EY Lab)
Mono2W-CNL
ABA
AOL (Mahal Lab)
ab5
Microvirin
IPS-K-4A2B8
CA19.9-121SPE
CA19.9-1B844
Gal-3
DBL-1
TbFBL-1
rMpL
SGA
Native HPA
VVL
UEA-I
LTA-I
SBA
SSA
HHA(EY Lab)
PTL-II
PHA-L
SBA
CnMBL
Epa1-Np mutant E227G
PhoSL001
PsSC
SRL
rSRL
Sp Serotype 14.2
TbFBL-2
hGal-1-488+BCR
#14 Sialyl Lewis X (Calbio)
#15 Sialyl Lewis X (BD)
AFCNVH
rNV VLP-Cy3
LBL-152
LSL-150
WT-VCC
ME5(detected with a-HUD)
MOA
MNA-M
WGA
VGA(EY Lab)
PNA
PTL-I
hJAA-F11-B11
GSL-2
PFL
CA
CAA
ConA
PTA-Gal
GSL-1
GNL
EEL
DSL
LcH
DBA
HHL
LCA
HAA
GSL-I-B4
CSA(EY Lab)
P11 VP8
LTL
MAL-I
UEA-I
ConA
PSL
PHA-E
Human Galectin1
Mouse Galectin1
AAL
Mono1-CNL
WGA
PHA-E
VVA
Mouse NKR-P1C
AIA
MAA(Seikagaku)
HPA
MGL 6B
LEL
Threonil
STL
PHA-L
WT Galectin-3
VLRB.aGPA.23
TL(EY Lab)
ECL
CA19.9-9L426
MGL 4I6
#8 Lewis B (T218) (Abcam)
NPA
HA70 complex-Alexa
PNA
UEA-I
PA-IL
Galectin-1
MGL 6A-H26L
CA19.9 Ab (DRG)
Epa1-Np mutant Y228A
HML-Native
Epa1-Np mutant Y228W
SSA
AAL
CML1
BanLec
Epa1-Np mutant E227A
MPL
WGA multimer
Gal-3C
SJA
AHL
Avaren-Fc
MAL-II
GAC78
SSL0-488
Mono2R-CNL
TJA-II
UDA
MGL 6A
RCA I (Vector)
ME7(detected with a-HUD)
MAA
Epa1-Np WT
GS-II (EY Lab)
ACL
CGL
Sp Serotype 14.15
DC-ASGPR short
BTL native
CTB
rPVL
PVL
WGA(Seikagaku Bio)
CCTG-1007
Anti-LeC
human Gal-7
WFL
BPL
VVHA lectin
CnSep
mouse Gal-7

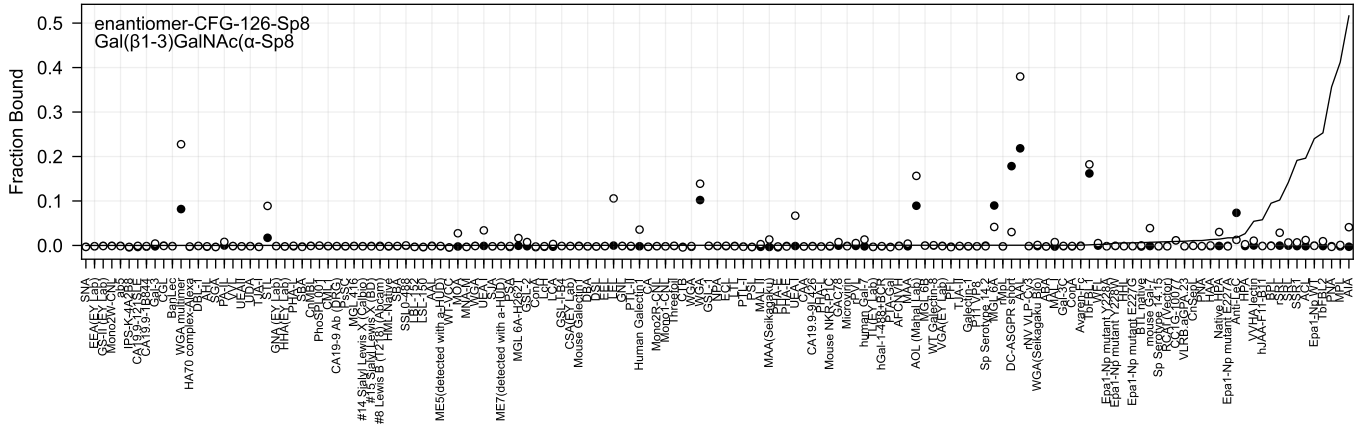
enantiomer-CFG-126-Sp8
Gal(β1-3)GalNAc(α-Sp8
Fraction Bound
0.0
0.1
0.2
0.3
0.4
0.5

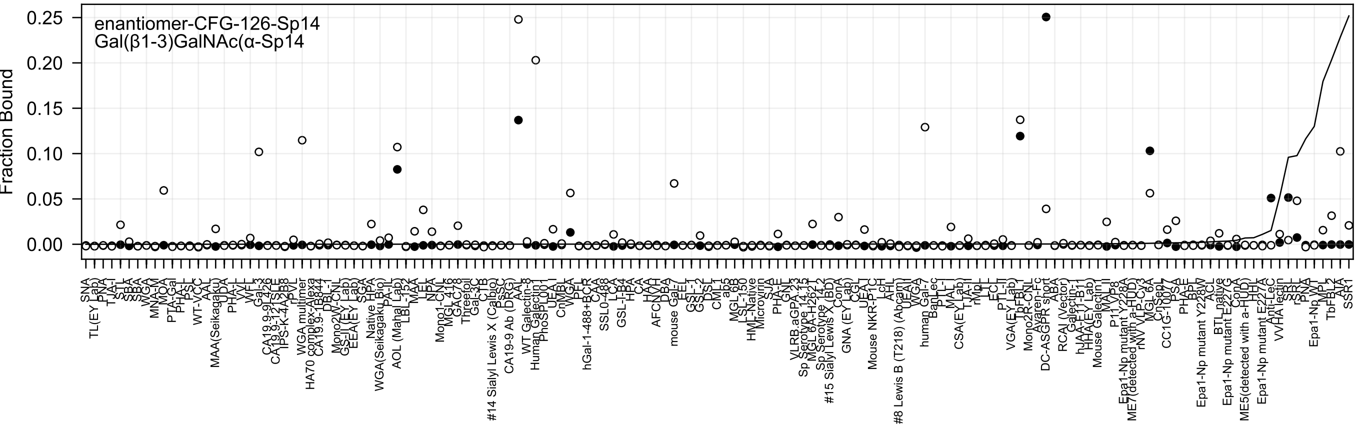

enantiomer-CFG-126-Sp14
Gal(β1-3)GalNAc(α-Sp14
Fraction Bound
0.25
0.20
0.15
0.10
0.05
0.00
SNA
TL(EY Lab)
PNA
TJA-I
STL
SBA
SBA
WGA
MNA-M
MOA
PTA-Gal
PHA-L
PSL
WT-VCC
AAL
MAA(Seikagaku)
UDA
PHA-L
VVL
WFL
Gal-3
CA19.9-9L426
CA19.9-12 SLE
IPS-K-4A2B8
rPVL
WGA multimer
HA70 complex-Alexa
CA19.9-1B844
DBL-1
Mono2W-CNL
GS-II(EY Lab)
EEA(EY Lab)
SGA
Native HPA
WGA(Seikagaku Bio)
PA-IL
AOL (Mahal Lab)
LBL-152
MAA
LEL
NPA
Mono1-CNL
MGL 416
GAC78
Threefoil
Gal-3C
CTB
#14 Sialyl Lewis X (Calbio)
PsSC
CA19-9 Ab (DRG)
AAL
WT Galectin-8
Human Galectin1
PhoSL001
UEA-I
CnMBL
WGA
PFL
hGal-1-488+BCR
CAA
SSL0-488
LCA
GSL-I-B4
HPA
CA
HAA
AFCN-H
DBA
mouse Gal-7
EEL
GSL-1
GSL-2
DSL
CML-1
ab5
MGL 6B
LSL-150
HML-Native
Microvirin
SJA
PHA-E
GNL
VLRB.aGPA.23
Sp Serotype 14.15
MGL 6A-H262T
Sp Serotype 14.2
#15 Sialyl Lewis X (BD)
ConA
GNA (EY Lab)
CGL
UEA-I
Mouse NKR-P1C
LcH
AHL
#8 Lewis B (T218) (Abcam)
UEAII
WGA
human Gal-7
BanLec
PTL-I
MAL-I
CSA(EY Lab)
TJA-II
rMpL
LTL
ECL
PTL-II
VGA(EY Lab)
TbFBL1
Mono2R-CNL
Ayaren-Fc
DC-ASGPR short
ABA
RCAI (Vector)
Galectin-1
hJAA-EY-B1
HHA(EY Lab)
Mouse Galectin1
MAL-II
P11 Y P8
Epa1-Np mutant Y228A
ME7(detected with a-HUD)
rNV VLP-Cy3
MGL 6A
CnSep7
CC1G-1007
PSA
PHA-E
BPL
Epa1-Np mutant Y228W
ACL
BTL native
Epa1-Np mutant E227G
ConA
ME5(detected with a-HUD)
HHL
Epa1-Np mutant E227A
Anti-LeC
VVHA lectin
SRL
rSRL
PNA
Epa1-Np WT
MPL
TbFBL2
AIA
SSR1

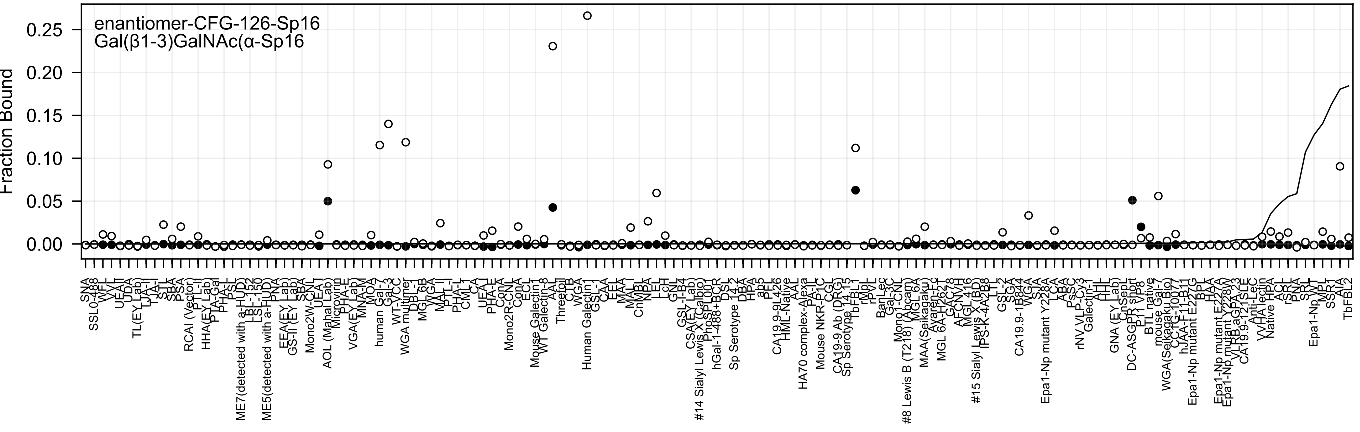
enantiomer-CFG-126-Sp16
Gal(β1-3)GalNAc(α-Sp16
Fraction Bound
0.25
0.20
0.15
0.10
0.05
0.00
SNA
SSL0-488
WFL
VVL
UEA-II
UDA
TL(EY Lab)
TJA-II
TJA-I
STL
SJA
SBA
PSA
RCAI (Vector)
PTL-II
HHA(EY Lab)
PTA-Gal
PHA-L
PSL
ME7(detected with a-HUD)
LBL-152
LSL-150
ME5(detected with a-HUD)
PNA
EEA(EY Lab)
GS-II (EY Lab)
SBA
Mono2W-CNL
UEA-I
AOL (Mahal Lab)
Microvirin
PHA-E
VGA(EY Lab)
MNA-M
MOA
human Gal-7
Gal-3
WT-VCC
WGA multimer
DBL-I
MGL 6B
VVGA
MAL-II
PTL-I
PHA-L
CML-I
CA
UEA-I
PHA-E
ConA
Mono2R-CNL
ConA
ECL
Mouse Galectin-1
WT Galectin-8
AAL
Threefoil
CLB
WGA
Human Galectin1
GSL-1
CAA
EEL
MAA
MAL-I
CnMBL
NPA
LcH
GNL
GSL-I-B4
CSA(EY Lab)
#14 Sialyl Lewis X (Calbio)
PhoSL007
hGal-1-488+BCR
DSL
Sp Serotype 14.2
DBA
HPA
ab5
PF
CA19.9-9L426
HML-Native
AAL
HA70 complex-Alexa
PA-IL
Mouse NKR-P1C
CGL
CA19-9 Ab (DRG)
Sp Serotype 14.15
TbFBL1
rMpL
PVL
BanLec
Gal-3C
Mono1-CNL
#8 Lewis B (T218) (Abcam)
MGL 6A
MAA(Seikagaku)
Ayaren-Fc
MGL 6A-H262T
GAC78
AFCNVH
MGL 416
#15 Sialyl Lewis X (BD)
IPS-K-4A2B8
LTL
GSL-2
SGA
CA19.9-1B844
WGA
SJA
Epa1-Np mutant Y228A
LCA
ABA
PsSC
rNV VLP-Cy3
Galectin-1
AHL
GNA (EY Lab)
CnSepL
DC-ASGPR short
P11 VP8
BTL native
mouse Gal-7
WGA(Seikagaku Bio)
hJAA-F11-B11
Epa1-Np mutant E227G
BPL
HAA
Epa1-Np mutant E227A
Epa1-Np mutant Y228W
VLRB.aGPA.23
CA19.9-121SLE
Anti-LeC
VVHA lectin
Native HPA
ACL
rSRL
PNA
SRL
Epa1-Np WT
MPL
AIA
TbFBL2

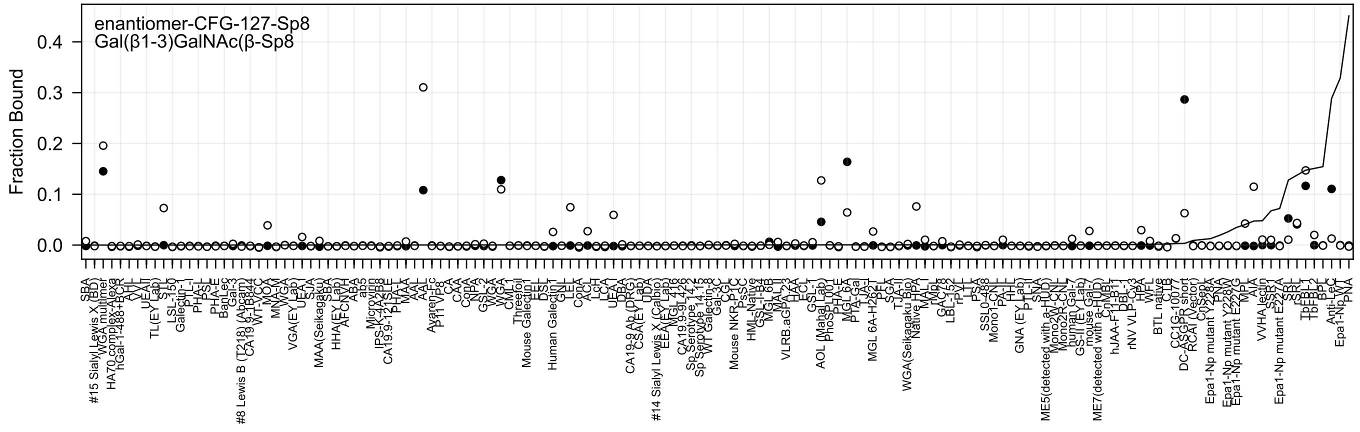
enantiomer-CFG-127-Sp8
Gal(β1-3)GalNAc(β-Sp8
Fraction Bound
0.4
0.3
0.2
0.1
0.0

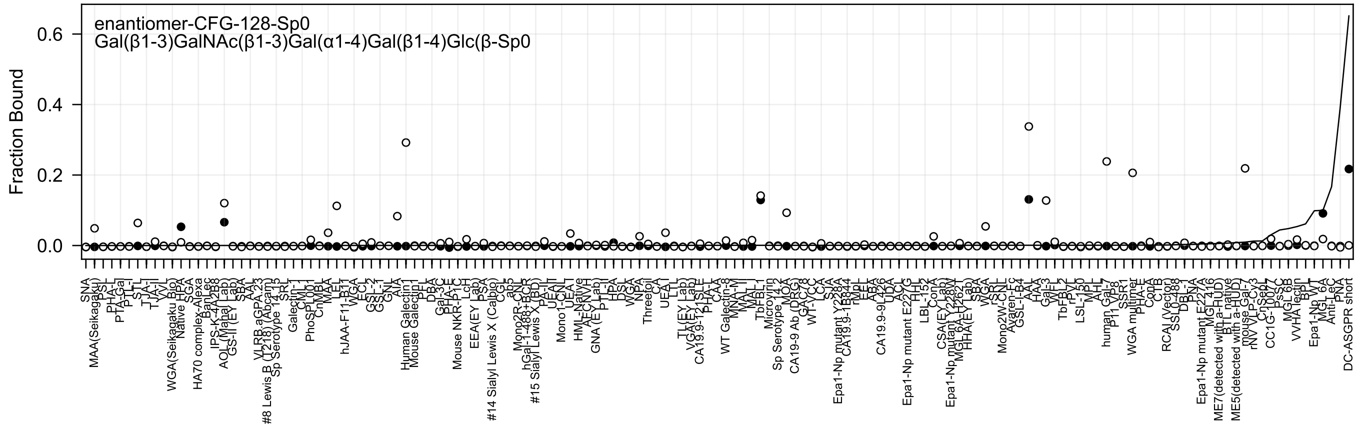
enantiomer-CFG-128-Sp0
Gal(β1-3)GalNAc(β1-3)Gal(α1-4)Gal(β1-4)Glc(β-Sp0
Fraction Bound
0.6
0.4
0.2
0.0

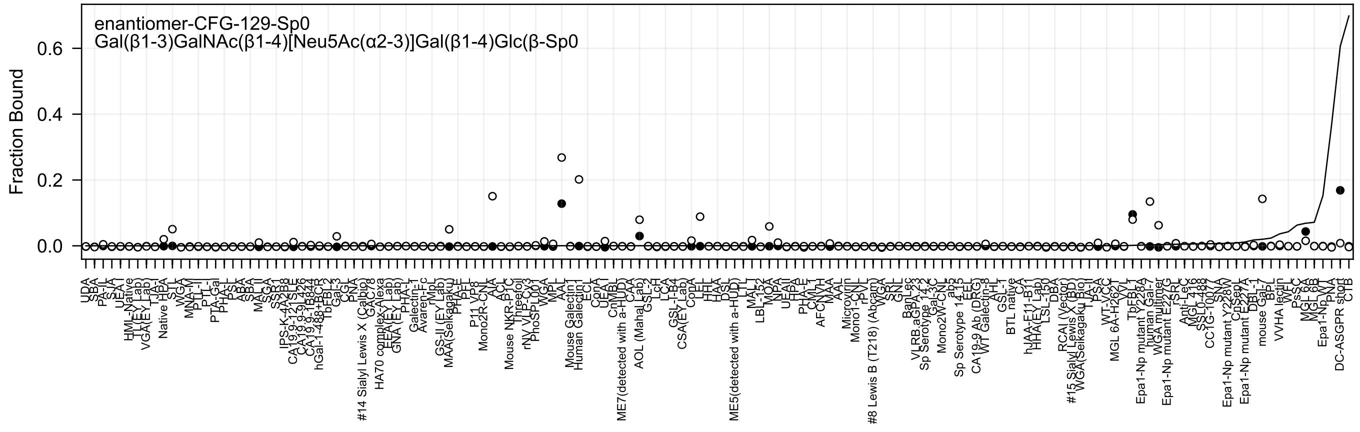

enantiomer-CFG-129-Sp0
Gal(β1-3)GalNAc(β1-4)[Neu5Ac(α2-3)]Gal(β1-4)Glc(β-Sp0
Fraction Bound
0.0
0.2
0.4
0.6

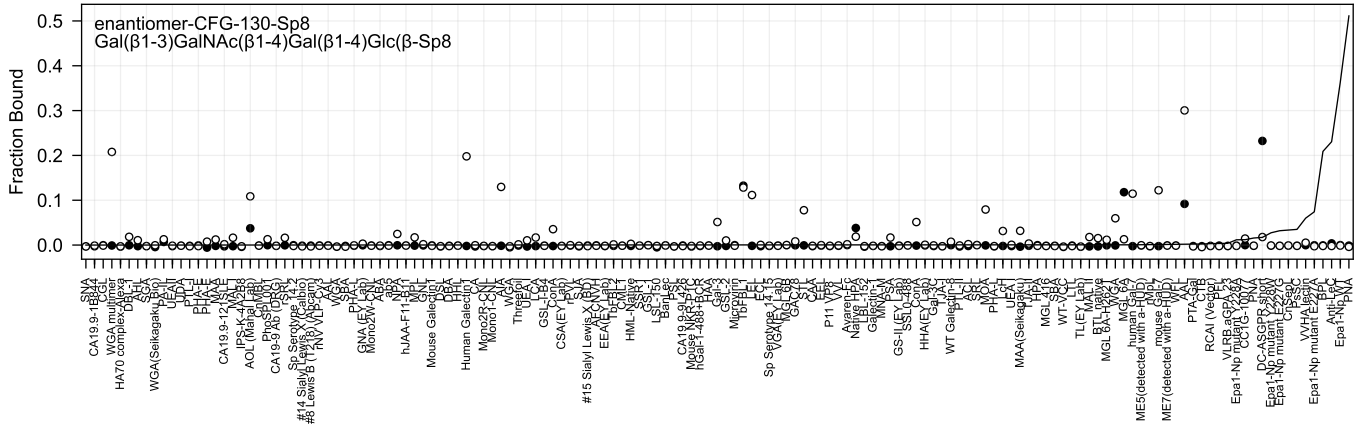
enantiomer-CFG-130-Sp8
Gal(β1-3)GalNAc(β1-4)Gal(β1-4)Glc(β-Sp8
Fraction Bound
0.5
0.4
0.3
0.2
0.1
0.0

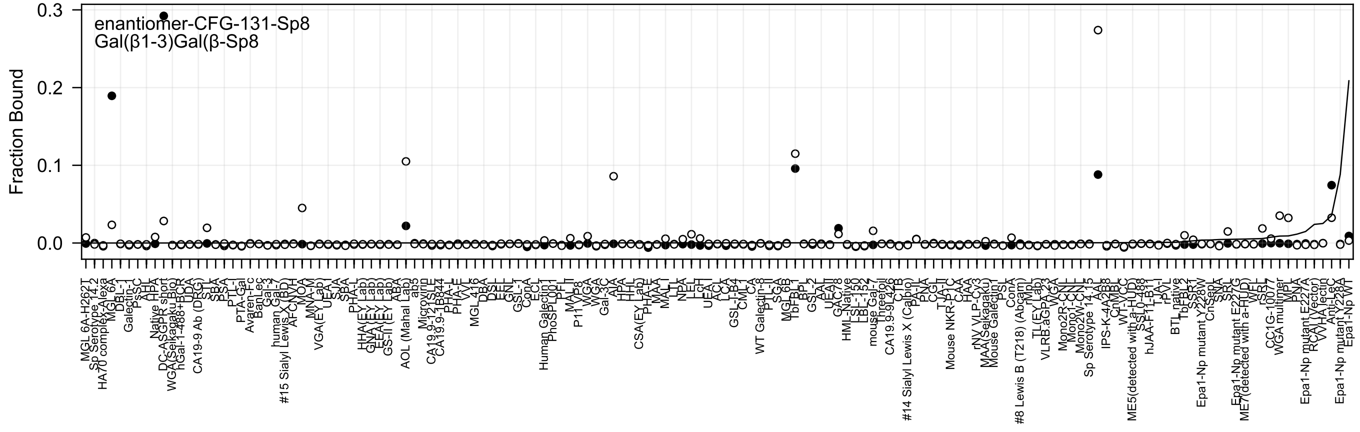

enantiomer-CFG-131-Sp8
Gal(β1-3)Gal(β-Sp8
Fraction Bound
0.3
0.2
0.1
0.0

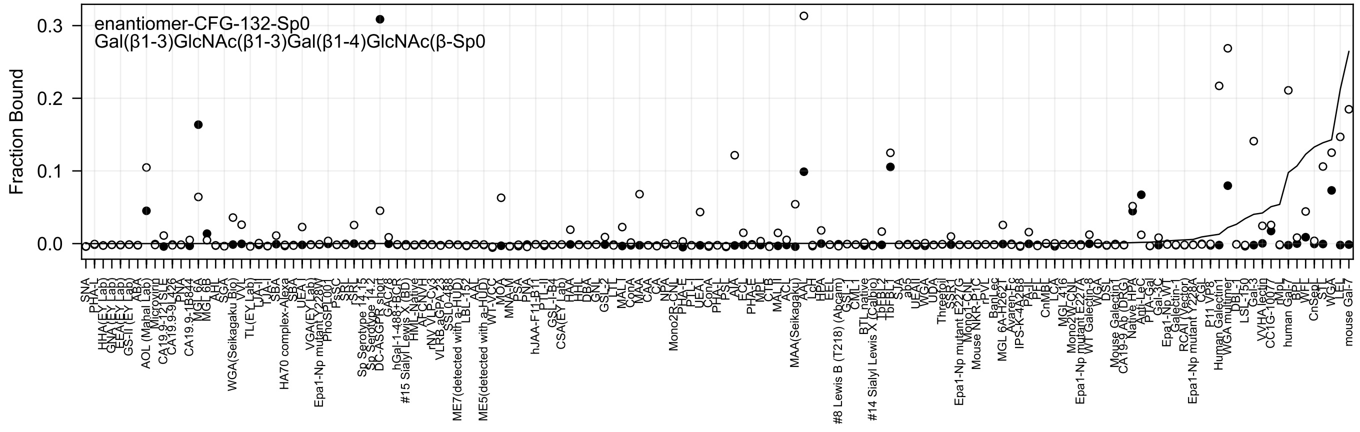
enantiomer-CFG-132-Sp0
Gal(β1-3)GlcNAc(β1-3)Gal(β1-4)GlcNAc(β-Sp0
Fraction Bound
0.3
0.2
0.1
0.0

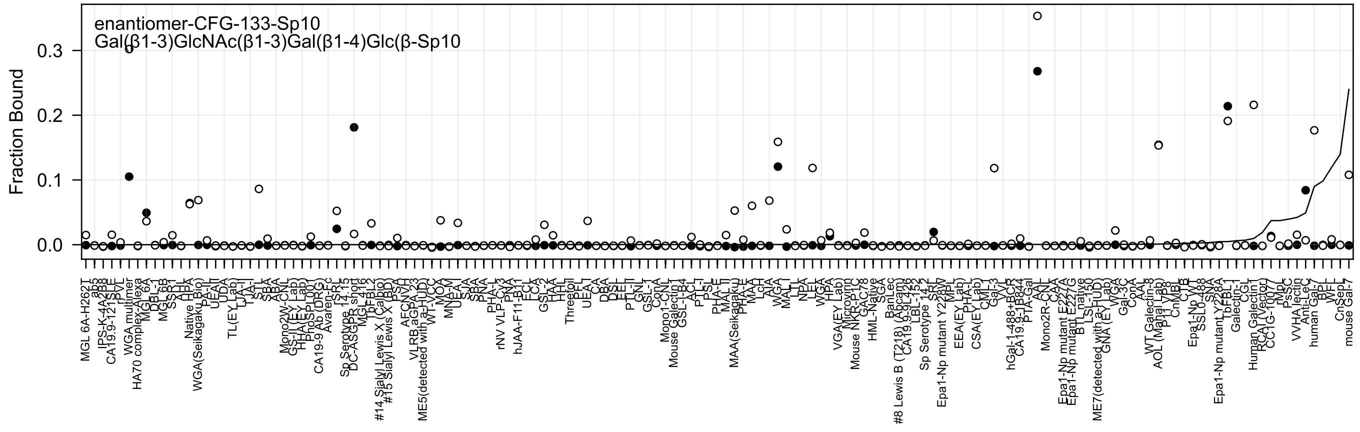
enantiomer-CFG-133-Sp10
Gal(β1-3)GlcNAc(β1-3)Gal(β1-4)Glc(β-Sp10
Fraction Bound
0.3
0.2
0.1
0.0

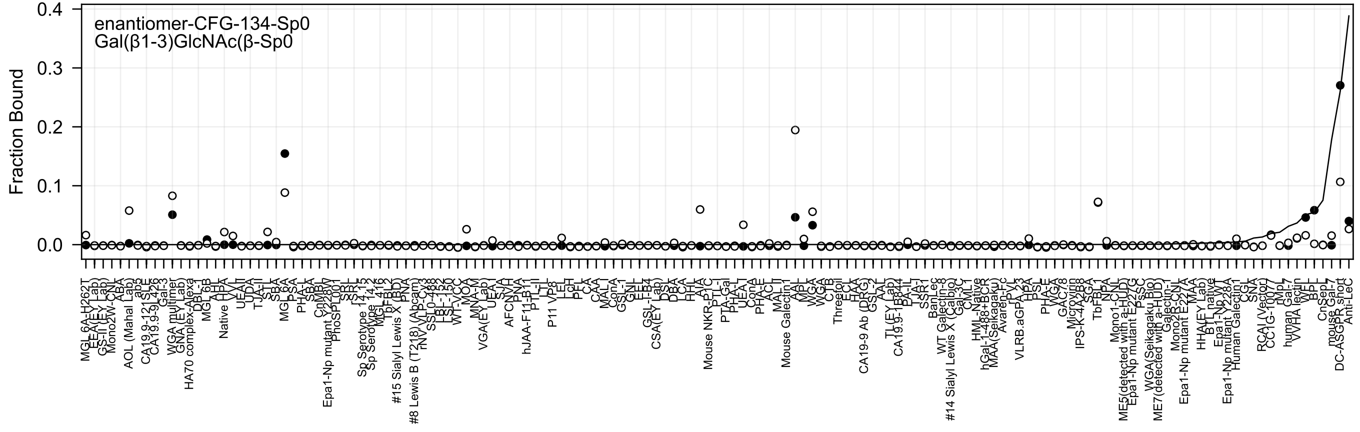
enantiomer-CFG-134-Sp0
Gal(β1-3)GlcNAc(β-Sp0
Fraction Bound
0.4
0.3
0.2
0.1
0.0
MGL 6A-H262T
AOL (Mahal Lab)
WGA multimer
HA70 complex-Alexa
Native
MGL 6A
Epa1-Np mutant Y228W
Sp Serotype 14,15
#15 Sialyl Lewis X (BD)
#8 Lewis B (T218) (Abcam)
VGA(EY Lab)
hJAA-F11-B11
P11 VP8
CSA(EY Lab)
Mouse NKR-P1C
Mouse Galectin1
Threefoil
CA19-9 Ab (DRG)
#14 Sialyl Lewis X (Calbio)
VLRB.aGPA.23
TbFBL1
ME5(detected with a-HUD)
ME7(detected with a-HUD)
Epa1-Np mutant E227A
Epa1-Np mutant Y228A
Human Galectin1
RCAI (Vector)
WVA lectin
DC-ASGPR short
Anti-LeC

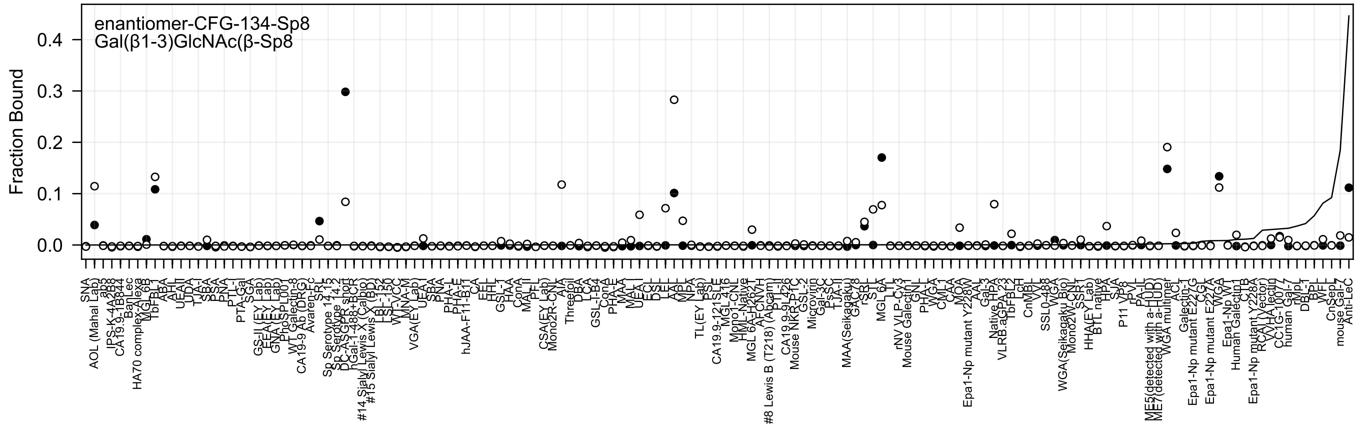
enantiomer-CFG-134-Sp8
Gal(β1-3)GlcNAc(β-Sp8
Fraction Bound

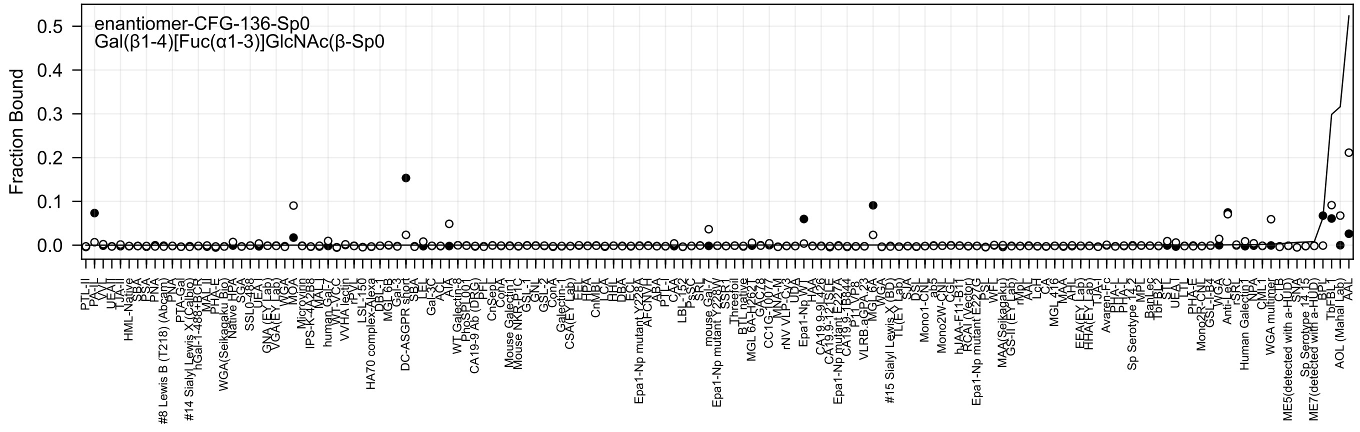

enantiomer-CFG-136-Sp0
Gal(β1-4)[Fuc(α1-3)]GlcNAc(β-Sp0
Fraction Bound
0.5
0.4
0.3
0.2
0.1
0.0

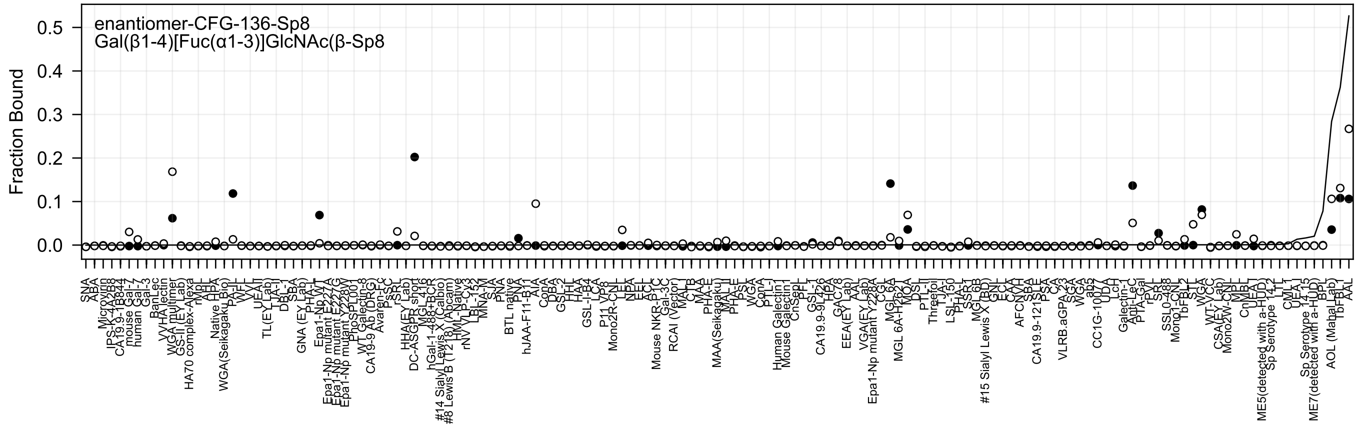

enantiomer-CFG-136-Sp8
Gal(β1-4)[Fuc(α1-3)]GlcNAc(β-Sp8
Fraction Bound
0.0
0.1
0.2
0.3
0.4
0.5

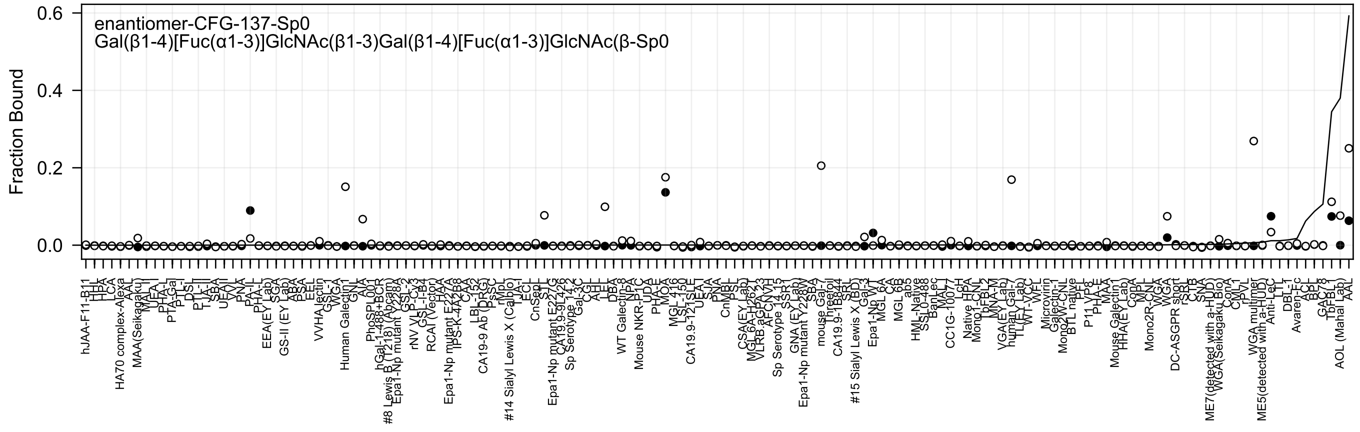

enantiomer-CFG-137-Sp0
Gal(β1-4)[Fuc(α1-3)]GlcNAc(β1-3)Gal(β1-4)[Fuc(α1-3)]GlcNAc(β-Sp0
Fraction Bound
0.6
0.4
0.2
0.0

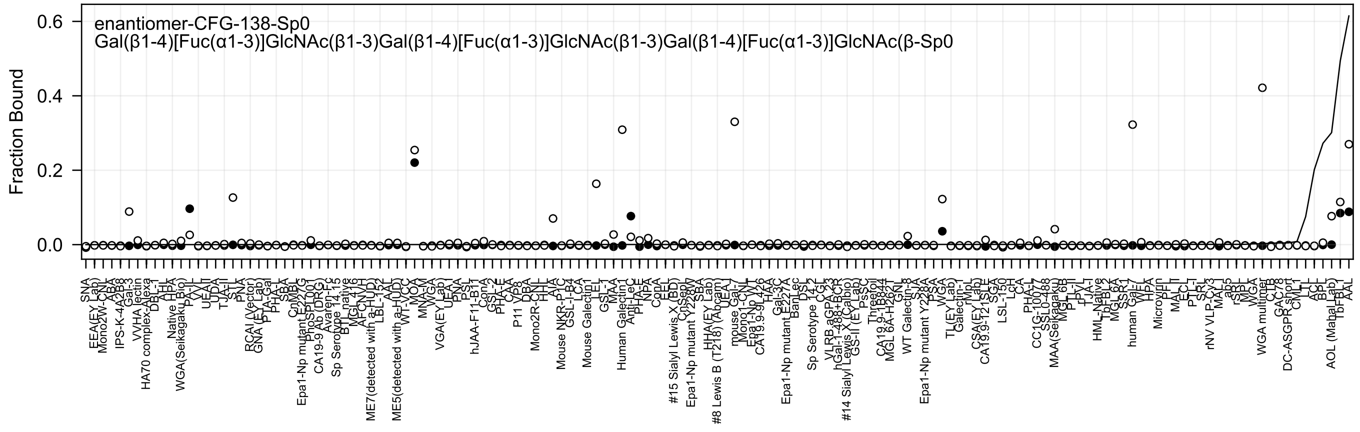
enantiomer-CFG-138-Sp0
Gal(β1-4)[Fuc(α1-3)]GlcNAc(β1-3)Gal(β1-4)[Fuc(α1-3)]GlcNAc(β1-3)Gal(β1-4)[Fuc(α1-3)]GlcNAc(β-Sp0
Fraction Bound
0.6
0.4
0.2
0.0
SNA
EEA(EY Lab)
Mono2W-CNL
ABA
IPS-K-4A2B8
Gal-3
VVHA lectin
HA70 complex-Alexa
DBL-I
HHL
Native HPA
WGA(Seikagaku Bio)
PA-II
VVL
UEA-II
UDA
TJA-II
STL
PNA
RCAI (Vector)
GNA (EY Lab)
PTA-Gal
PHA-L
SBA
CnMBL
Epa1-Np mutant E227G
PhoSL001
CA19-9 Ab (DRG)
Avaren-Fc
Sp Serotype 14.15
BTL native
MGL419
AFCN-H
ME7(detected with a-HUD)
LBL-152
AAL
ME5(detected with a-HUD)
WT-VCC
MOA
MNA-M
WGA
VGA(EY Lab)
UEA-I
PNA
PSL
hJAA-F11-B1
ConA
GSL-2
PHA-E
CAA
P11 VP8
DBA
Mono2R-CNL
HHL
AIA
Mouse NKR-P1C
GSL-I-B4
LCA
Mouse Galectin1
LEL
GSL-I
MAA
Human Galectin1
Anti-LeC
PHA-E
NPA
ConA
EEL
#15 Sialyl Lewis X (BD)
CnSep
Epa1-Np mutant Y228W
SBA
HHA(EY Lab)
#8 Lewis B (T218) (Abcam)
UEA-I
mouse Gal-1
Mono1-CNL
Epa1-Np WT
CA19.9-9L426
HAA
Gal-3C
Epa1-Np mutant E227A
BanLec
DSL
Sp Serotype 14.2
CGL
VLRB.aGPA.23
hGal-1-488+BCR
#14 Sialyl Lewis X (Calbio)
GS-II (EY Lab)
PsSC
Threefoil
CA19.9-1B844
MGL 6A-H262T
GNL
WT Galectin-8
STA
Epa1-Np mutant Y228A
PSA
WGA
TL(EY Lab)
Galectin-1
rMpL
CSA(EY Lab)
CA19.9-T21SLE
SGA
LSL-150
LcH
CA
PHA-L
CC1G-10077
SSL0-488
MAA(Seikagaku)
MGL6B
PTL-II
rPVL
TJA-I
HML-Native
TbFBL2
MGL6A
SSR1
human Gal-7
WFL
HPA
Microvirin
PFL
MAL-II
ECL
PTL-I
SRL
rNV VLP-Cy3
MAL-I
ab5
rSRL
MPL
WGA
WGA multimer
CTB
GAC78
DC-ASGPR short
CML1
LTL
ACL
BPL
AOL (Mahal Lab)
TbFBL1
AAL

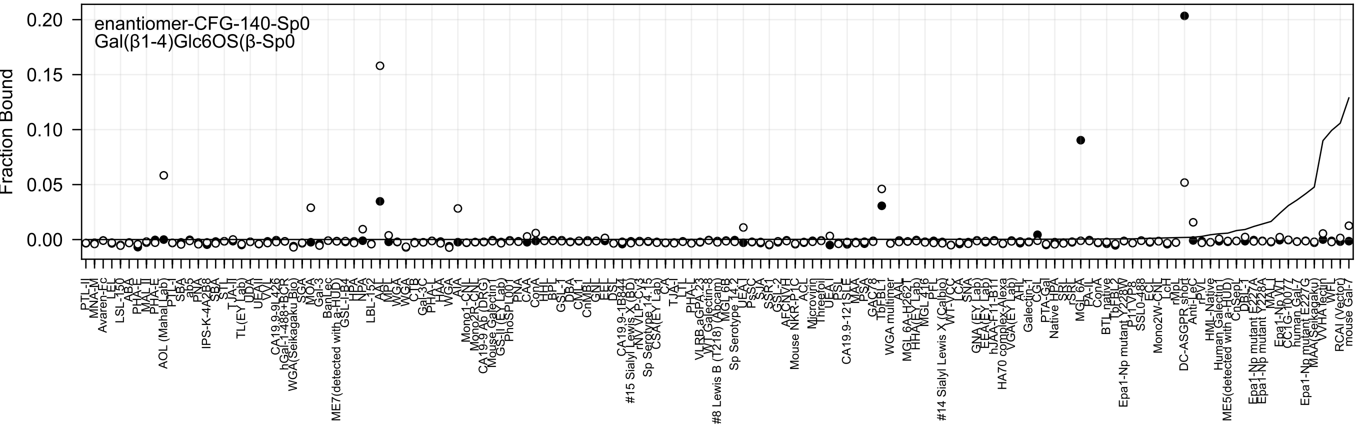
enantiomer-CFG-140-Sp0
Gal(β1-4)Glc6OS(β-Sp0
Fraction Bound
0.20
0.15
0.10
0.05
0.00

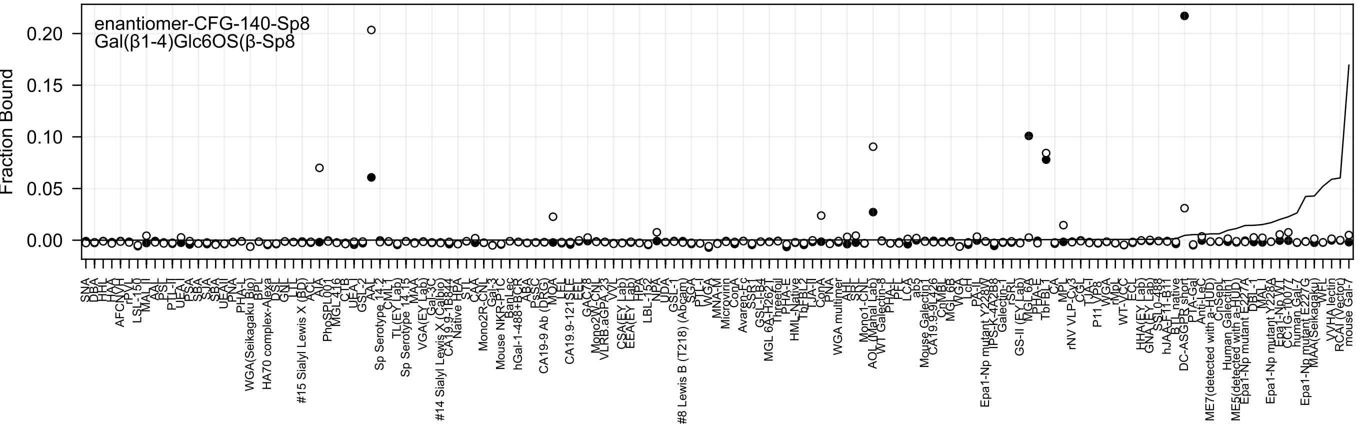
enantiomer-CFG-140-Sp8
Gal(β1-4)Glc6OS(β-Sp8
Fraction Bound
0.00
0.05
0.10
0.15
0.20

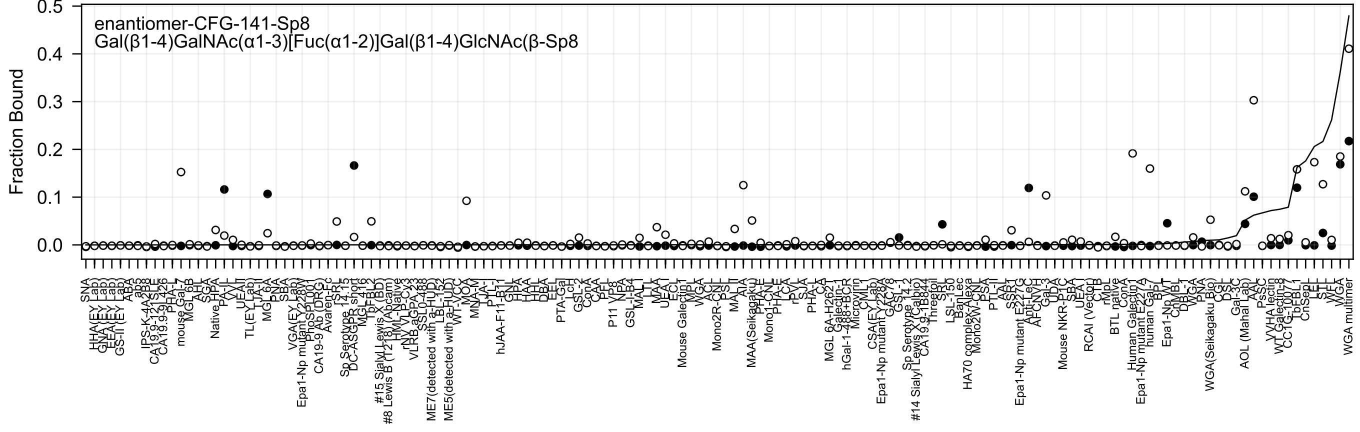
enantiomer-CFG-141-Sp8
Gal(β1-4)GalNAc(α1-3)[Fuc(α1-2)]Gal(β1-4)GlcNAc(β-Sp8
Fraction Bound

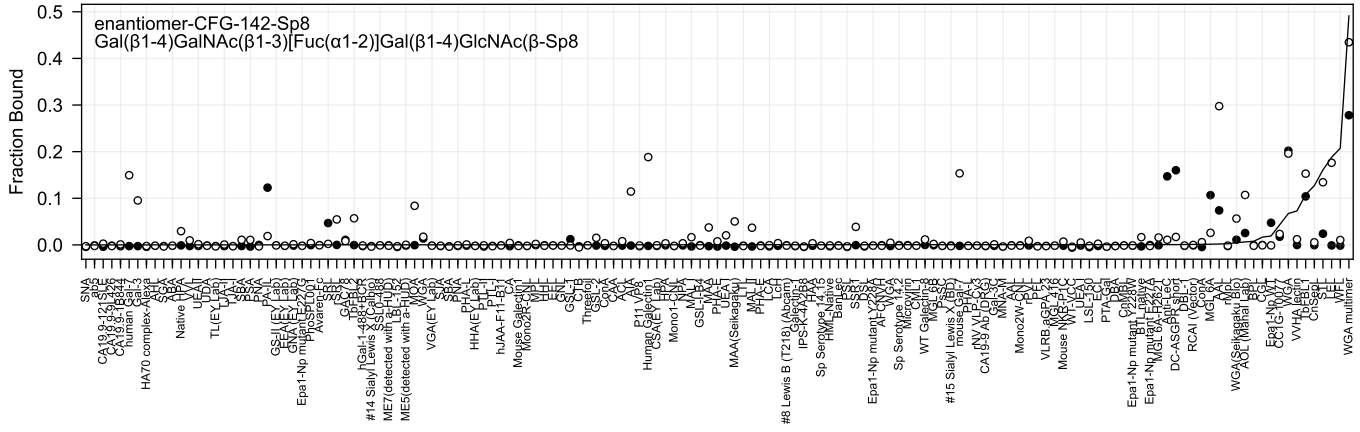
enantiomer-CFG-142-Sp8
Gal(β1-4)GalNAc(β1-3)[Fuc(α1-2)]Gal(β1-4)GlcNAc(β-Sp8
Fraction Bound
0.5
0.4
0.3
0.2
0.1
0.0

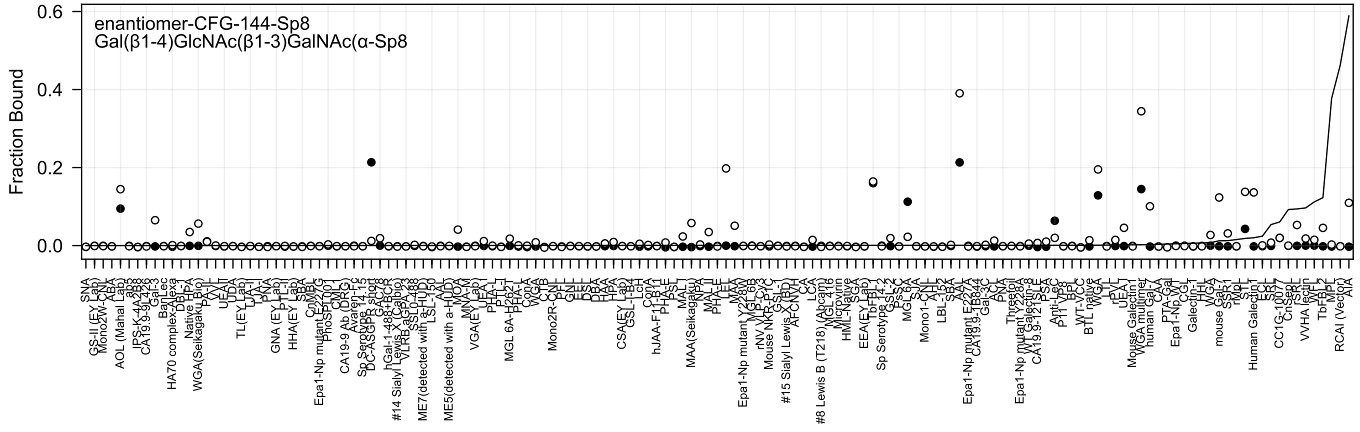
enantiomer-CFG-144-Sp8
Gal(β1-4)GlcNAc(β1-3)GalNAc(α-Sp8
Fraction Bound
0.6
0.4
0.2
0.0

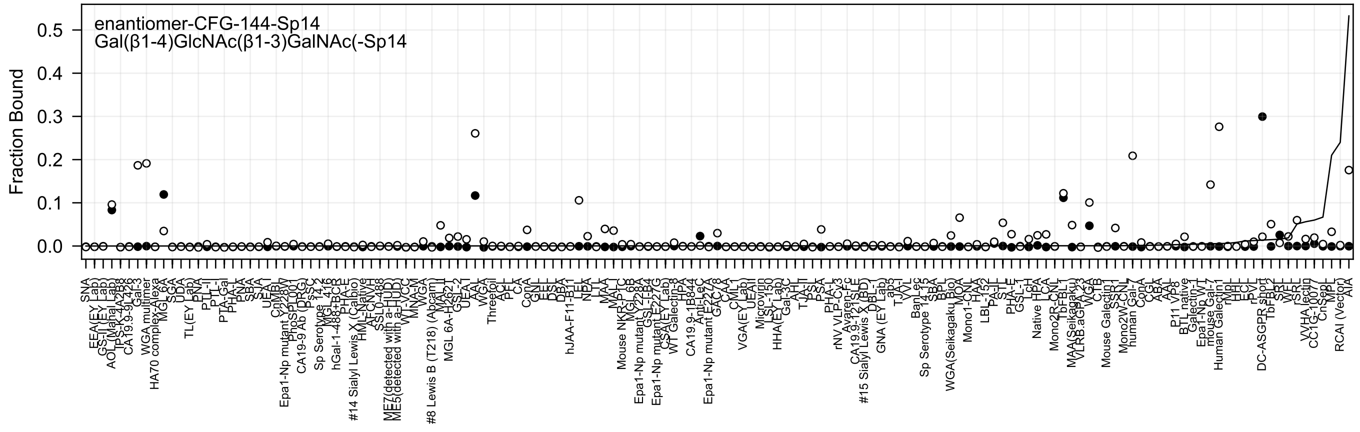
enantiomer-CFG-144-Sp14
Gal(β1-4)GlcNAc(β1-3)GalNAc(-Sp14
Fraction Bound
0.0
0.1
0.2
0.3
0.4
0.5

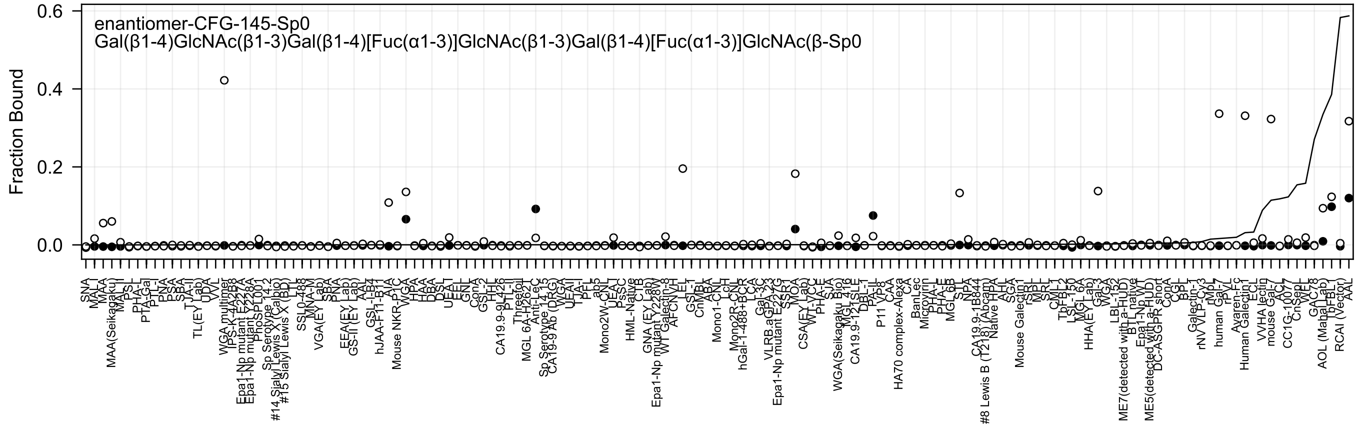
enantiomer-CFG-145-Sp0
Gal(β1-4)GlcNAc(β1-3)Gal(β1-4)[Fuc(α1-3)]GlcNAc(β1-3)Gal(β1-4)[Fuc(α1-3)]GlcNAc(β-Sp0
Fraction Bound
0.6
0.4
0.2
0.0

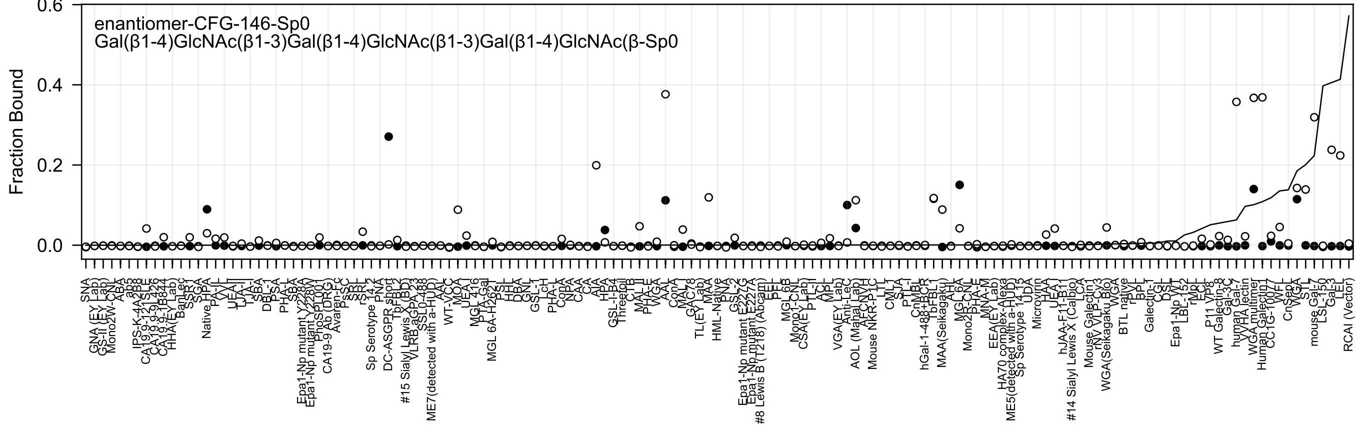
enantiomer-CFG-146-Sp0
Gal(β1-4)GlcNAc(β1-3)Gal(β1-4)GlcNAc(β1-3)Gal(β1-4)GlcNAc(β-Sp0
Fraction Bound
0.6
0.4
0.2
0.0

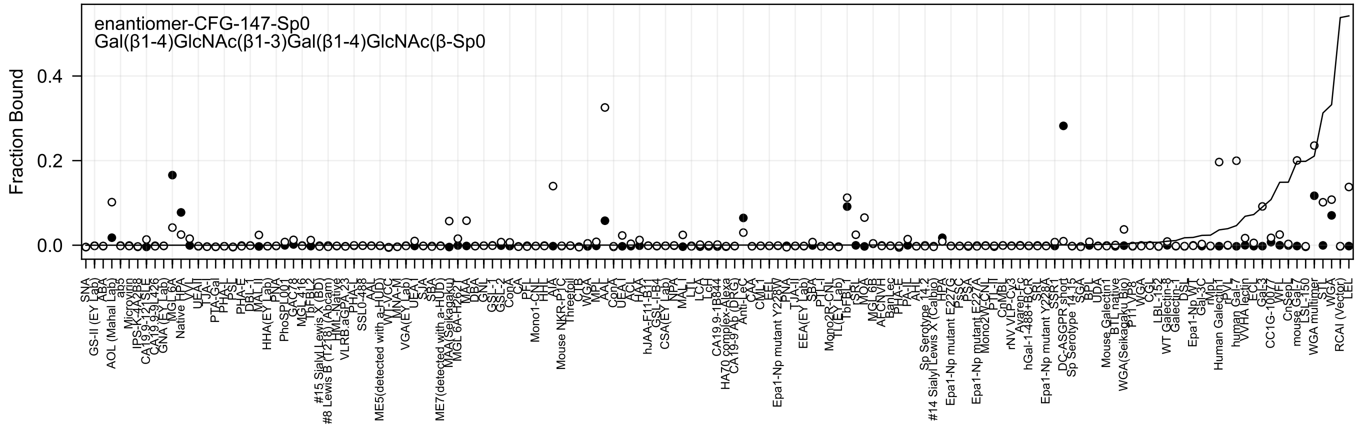
enantiomer-CFG-147-Sp0
Gal(β1-4)GlcNAc(β1-3)Gal(β1-4)GlcNAc(β-Sp0
Fraction Bound
0.0
0.2
0.4
SNA
GS-II (EY Lab)
ABA
AOL (Mahal Lab)
ab5
Microvirin
IPS-K-4A2B8
CA19-9-121SLE
CA19-9-9L426
GNA (EY Lab)
MGL 6A
Native HPA
VVL
UEA-I
TJA-I
PTA-Gal
PHA-L
PSL
PHA-E
DBL-1
MAL-II
HHA(EY Lab)
PNA
PhoSL001
GAC78
MGL 416
TbFBL2
#15 Sialyl Lewis X (BD)
#8 Lewis B (T218) (Abcam)
HML-Native
VLRB.aGPA.23
PHA-L
SSL0-488
AAL
ME5(detected with a-HUD)
WT-VCC
MNA-M
VGA(EY Lab)
UEA-I
SJA
SBA
ME7(detected with a-HUD)
MAA (Seikagaku)
MGL 6A-H262A
KAA
DBA
GNL
GSL-2
GSL-1
ConA
CA
PFL
Mono1-CNL
HHL
AIA
Mouse NKR-P1C
Threefoil
CTB
WGA
MPL
AAL
ConA
UEA-I
ACL
HAA
hJAA-F11-B11
GSL-I-B4
CSA(EY Lab)
NPA
MAL-I
LTL
LCA
LcH
CA19.9-1B844
HA70 complex-Alexa
CA 19-9 Ab (DRG)
Anti-LeC
CAA
CML-1
EEL
Epa1-Np mutant Y228W
PNA
TJA-II
EEA(EY Lab)
SBA
PTL-I
Mono2R-CNL
TL(EY Lab)
TbFBL1
rSRL
MOA
MGL 6B
AECNVH
BanLec
PHA-E
PA-IL
Sp Serotype 14.2
#14 Sialyl Lewis X (Calbio)
HPA
Epa1-Np mutant E227G
PSA
Epa1-Np mutant E227A
Mono2W-CNL
PTL-II
CnMBL
rNV VLP-Cy3
hGal-1-488+BCR
SRL
Epa1-Np mutant Y228A
SSR1
DC-ASGPR short
Sp Serotype 14.3
SGA
BPL
UDA
Mouse Galectin1
BTL native
WGA(Seikagaku Bio)
P11 VP8
WGA
CGL
LBL-152
WT Galectin-8
Galectin-1
DSL
Epa1-Np WT
Gal-3C
rMp
Human Galectin1
rPVL
human Gal-7
WHA lectin
ECL
Gal-3
CC1G-10077
WFL
CnSep
mouse Gal-7
LSL-150
WGA multimer
STL
WGA
RCAI (Vector)
LEL

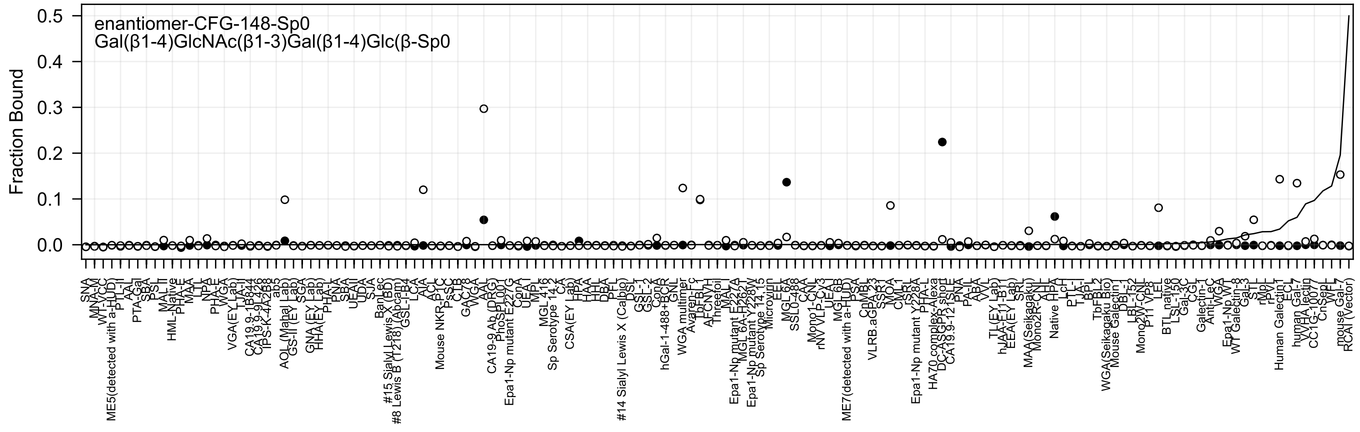
enantiomer-CFG-148-Sp0
Gal(β1-4)GlcNAc(β1-3)Gal(β1-4)Glc(β-Sp0
Fraction Bound
0.5
0.4
0.3
0.2
0.1
0.0

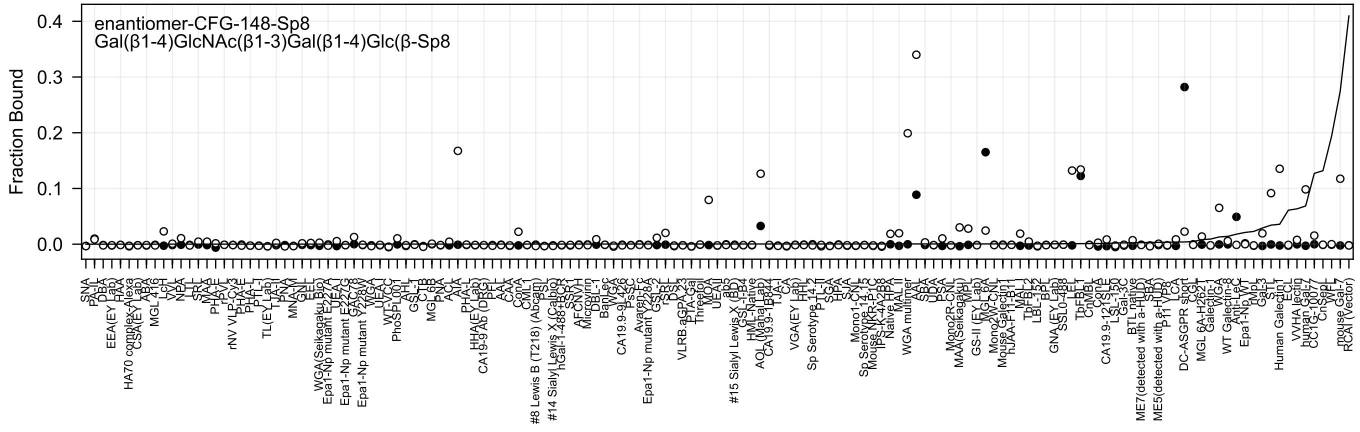
enantiomer-CFG-148-Sp8
Gal(β1-4)GlcNAc(β1-3)Gal(β1-4)Glc(β-Sp8
Fraction Bound
0.4
0.3
0.2
0.1
0.0

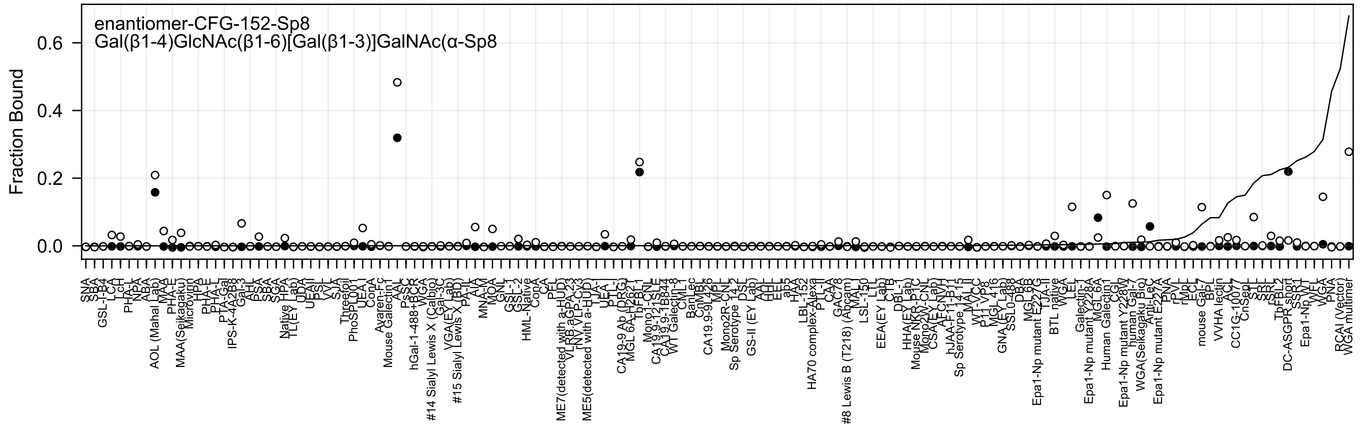
enantiomer-CFG-152-Sp8
Gal(β1-4)GlcNAc(β1-6)[Gal(β1-3)]GalNAc(α-Sp8
Fraction Bound
0.0
0.2
0.4
0.6

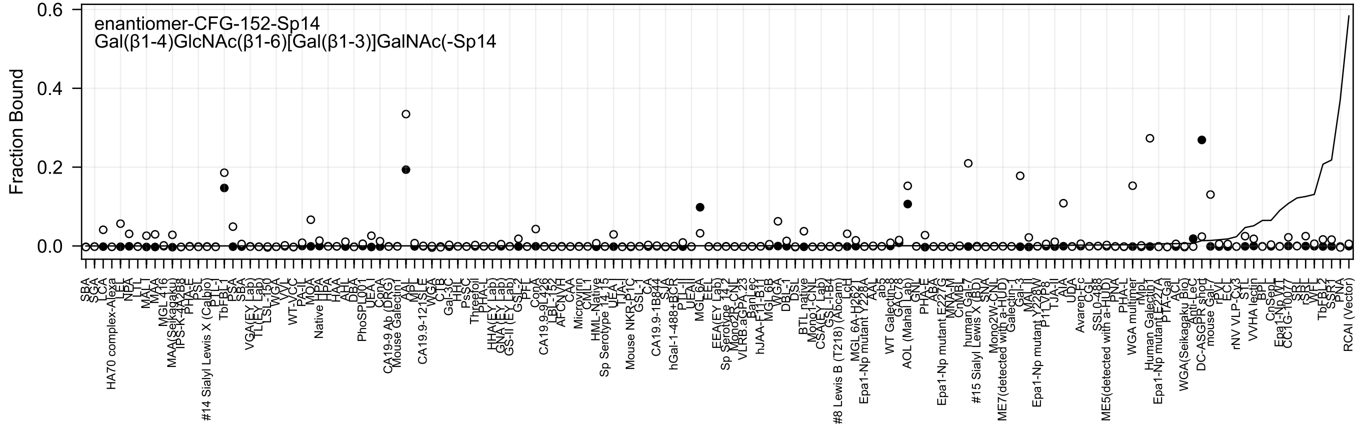

enantiomer-CFG-152-Sp14
Gal(β1-4)GlcNAc(β1-6)[Gal(β1-3)]GalNAc(-Sp14
Fraction Bound
0.6
0.4
0.2
0.0

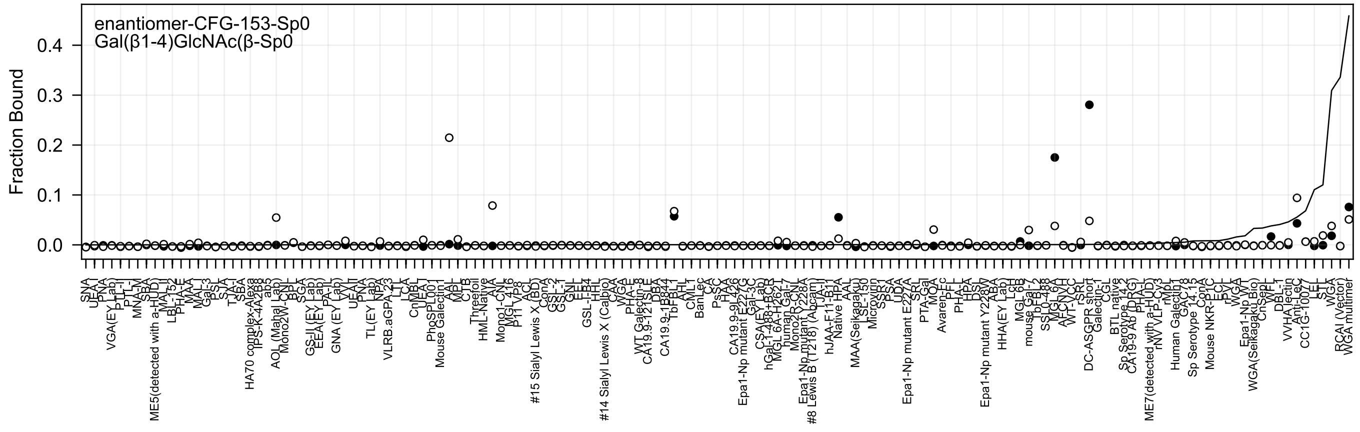

enantiomer-CFG-153-Sp0
Gal(β1-4)GlcNAc(β-Sp0
Fraction Bound
0.0
0.1
0.2
0.3
0.4

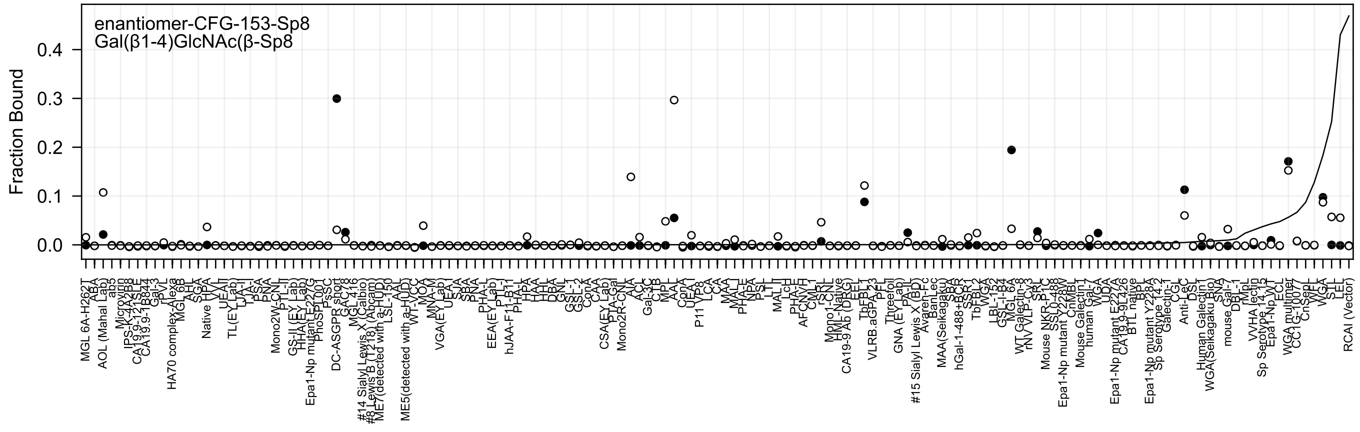
enantiomer-CFG-153-Sp8
Gal(β1-4)GlcNAc(β-Sp8
Fraction Bound

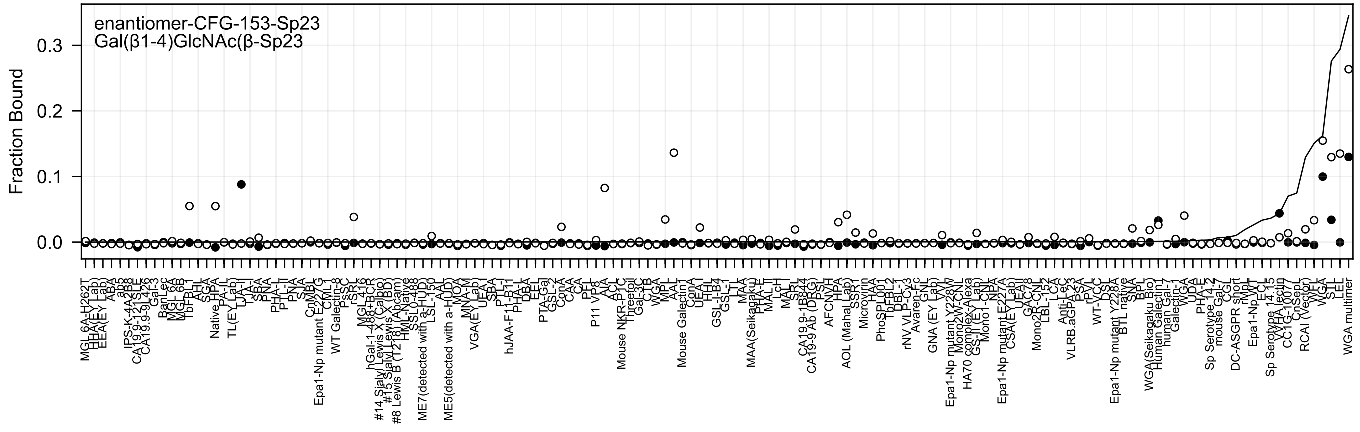

enantiomer-CFG-153-Sp23
Gal(β1-4)GlcNAc(β-Sp23
Fraction Bound
0.0
0.1
0.2
0.3

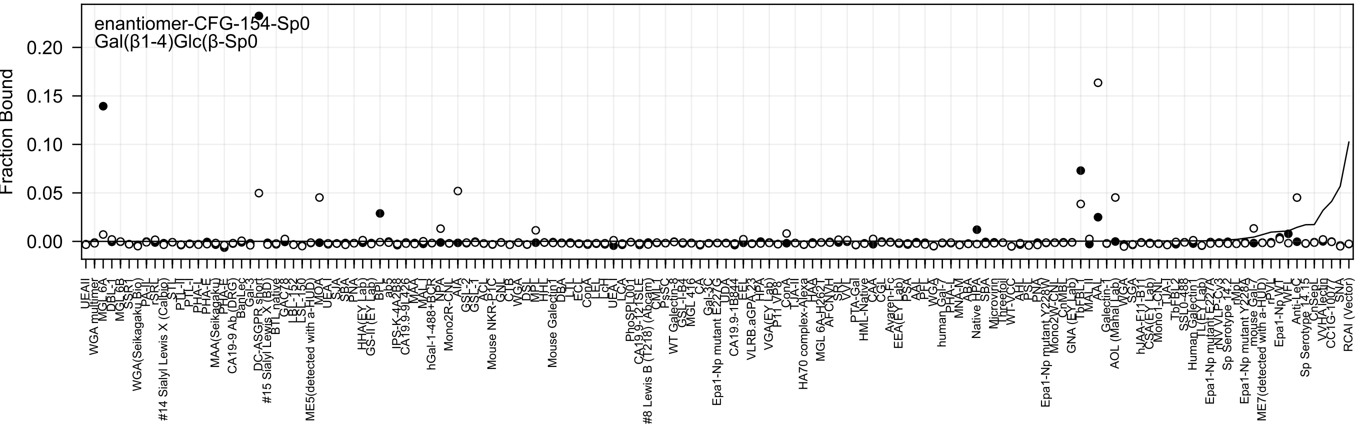
enantiomer-CFG-154-Sp0
Gal(β1-4)Glc(β-Sp0
Fraction Bound
0.00
0.05
0.10
0.15
0.20

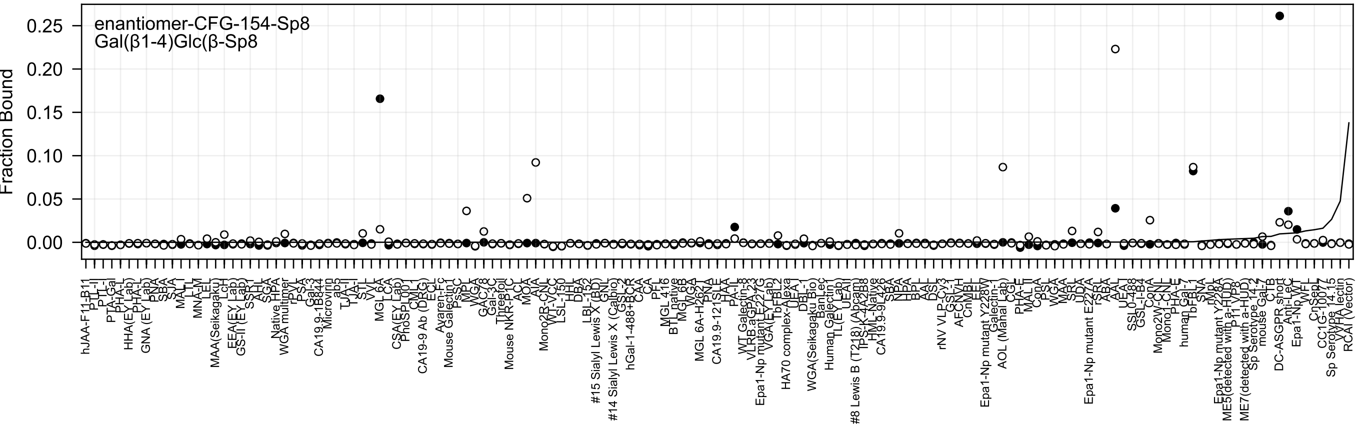
enantiomer-CFG-154-Sp8
Gal(β1-4)Glc(β-Sp8
Fraction Bound
0.25
0.20
0.15
0.10
0.05
0.00

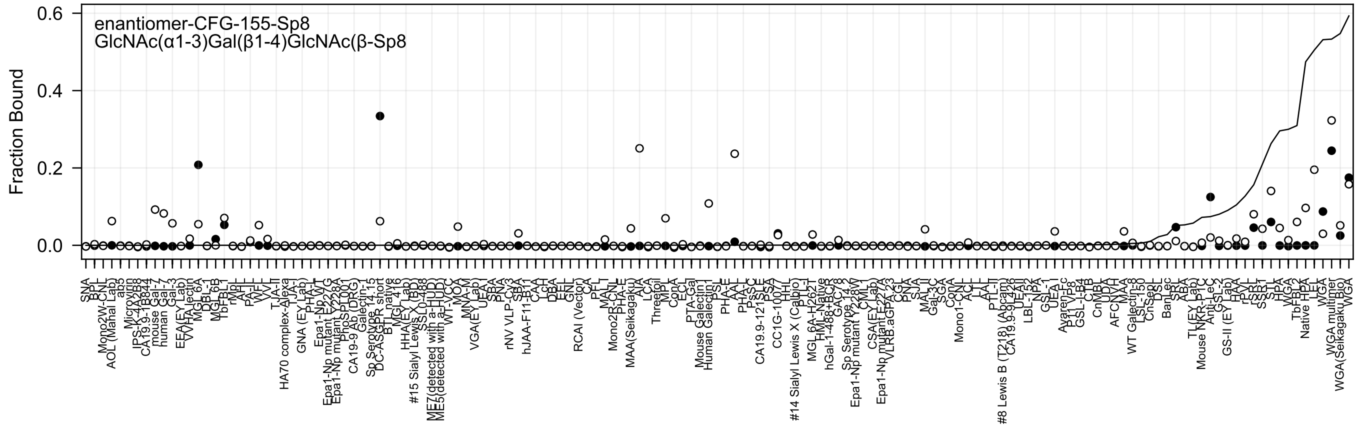
enantiomer-CFG-155-Sp8
GlcNAc(α1-3)Gal(β1-4)GlcNAc(β-Sp8
Fraction Bound
0.6
0.4
0.2
0.0

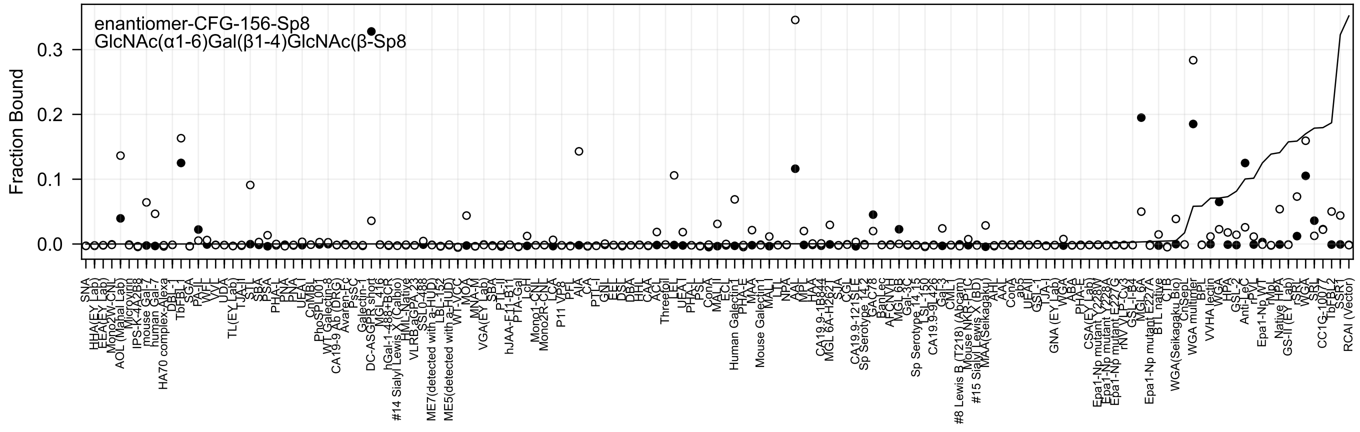
enantiomer-CFG-156-Sp8
GlcNAc(α1-6)Gal(β1-4)GlcNAc(β-Sp8
Fraction Bound
0.0
0.1
0.2
0.3

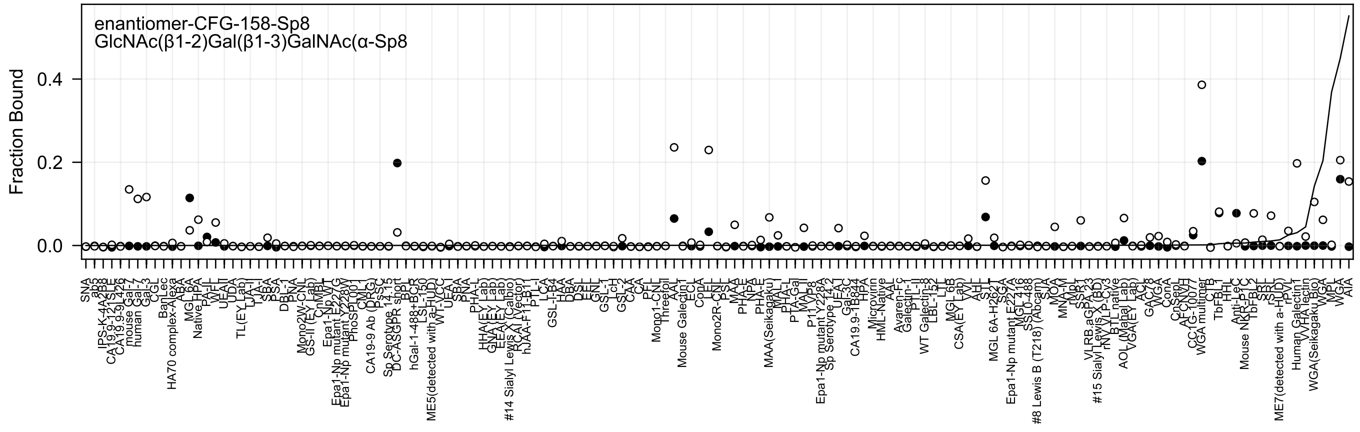
enantiomer-CFG-158-Sp8
GlcNAc(β1-2)Gal(β1-3)GalNAc(α-Sp8
Fraction Bound
0.4
0.2
0.0

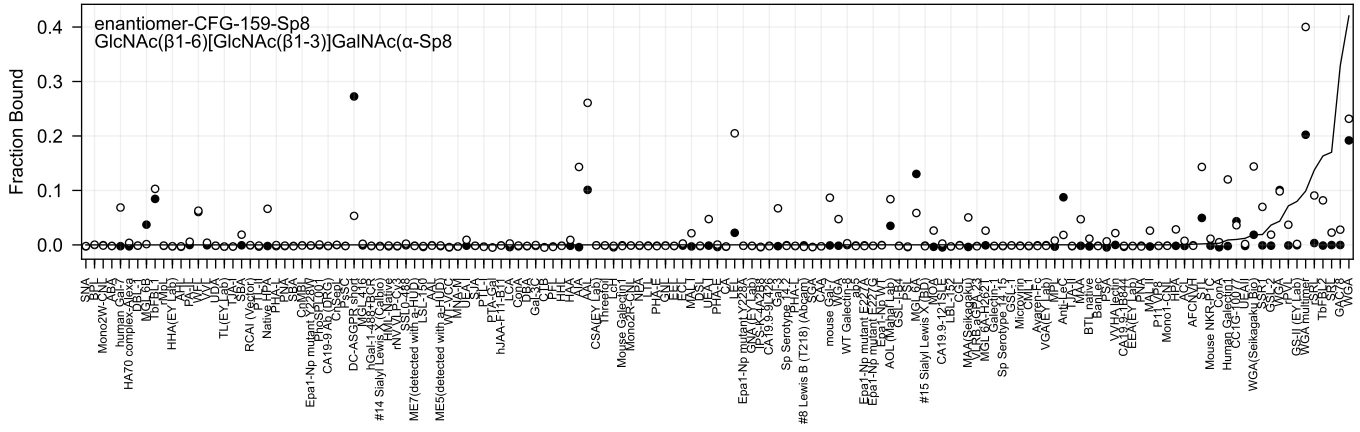
enantiomer-CFG-159-Sp8
GlcNAc(β1-6)[GlcNAc(β1-3)]GalNAc(α-Sp8
Fraction Bound
0.4
0.3
0.2
0.1
0.0

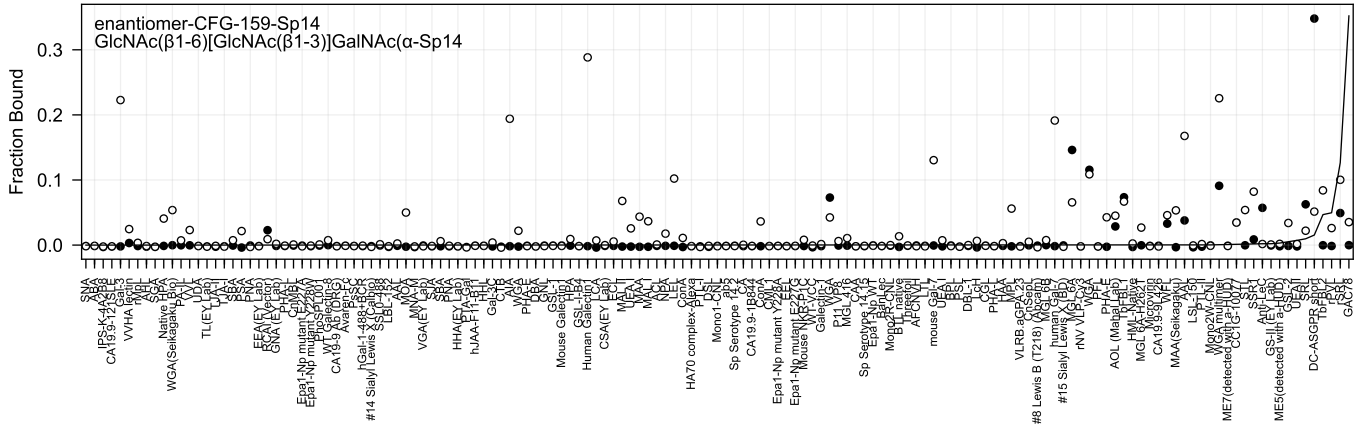
enantiomer-CFG-159-Sp14
GlcNAc(β1-6)[GlcNAc(β1-3)]GalNAc(α-Sp14
Fraction Bound
0.0
0.1
0.2
0.3

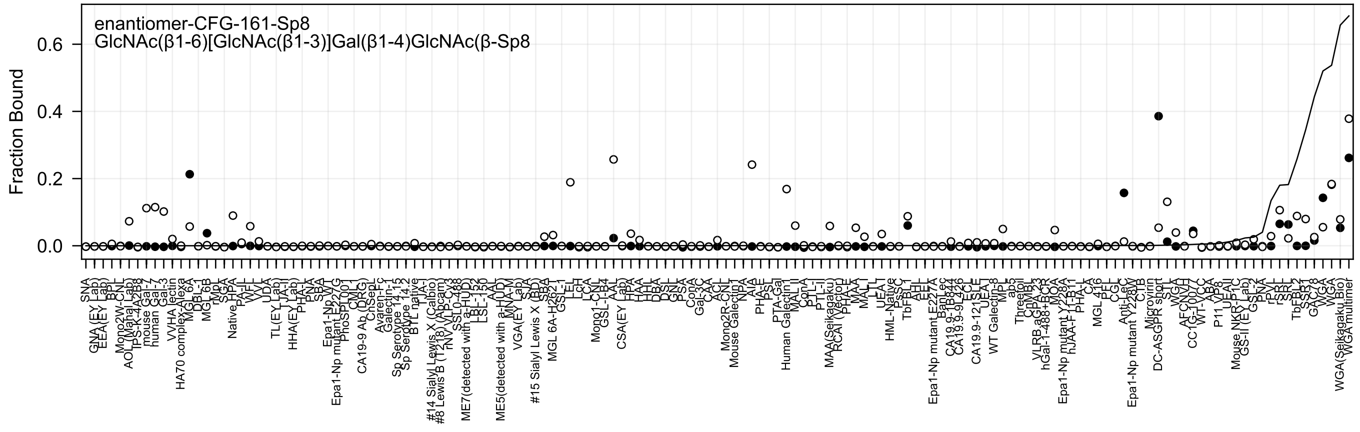
enantiomer-CFG-161-Sp8
GlcNAc(β1-6)[GlcNAc(β1-3)]Gal(β1-4)GlcNAc(β-Sp8
Fraction Bound
0.6
0.4
0.2
0.0

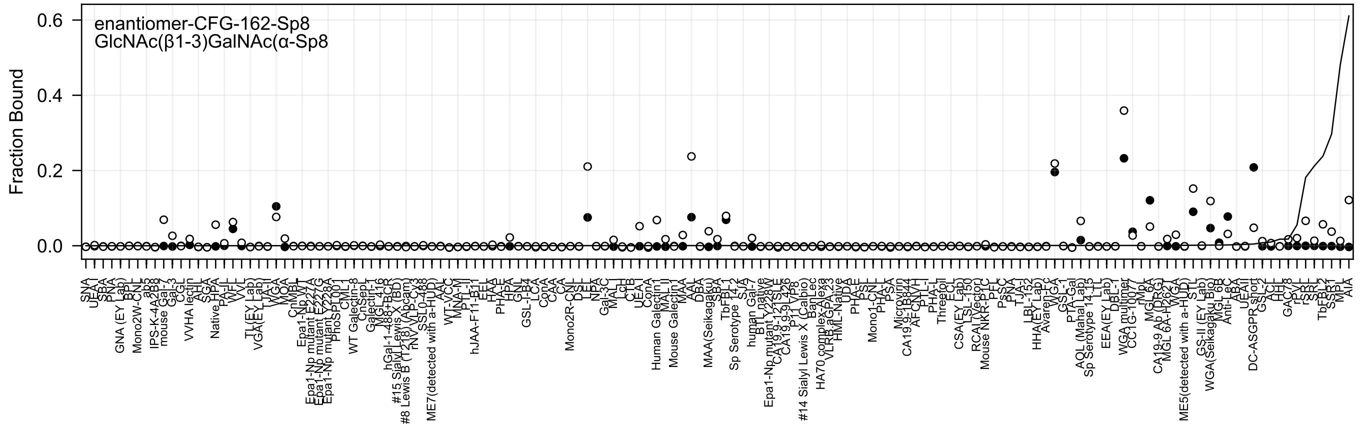
enantiomer-CFG-162-Sp8
GlcNAc(β1-3)GalNAc(α-Sp8
Fraction Bound
0.6
0.4
0.2
0.0

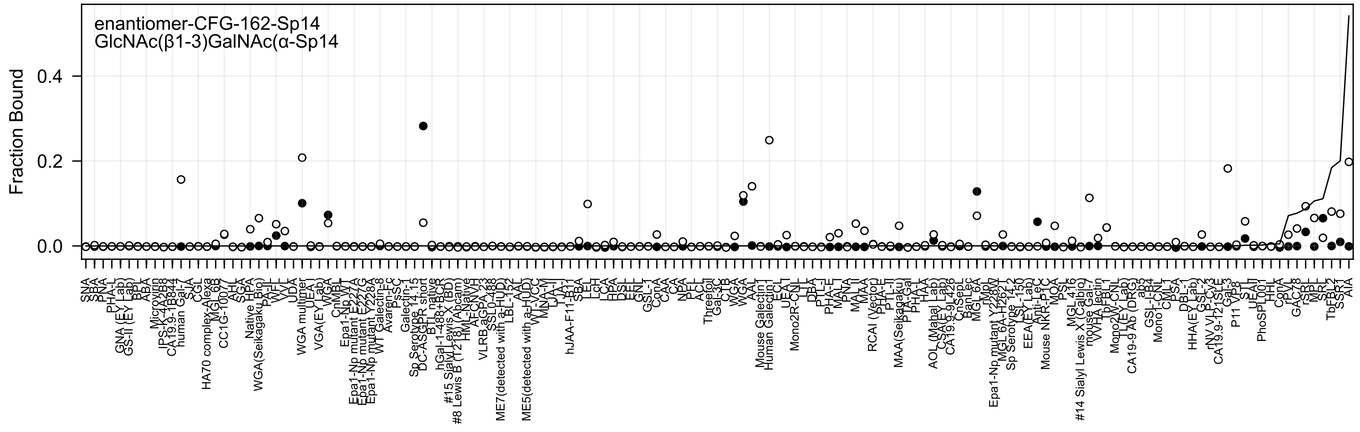
enantiomer-CFG-162-Sp14
GlcNAc(β1-3)GalNAc(α-Sp14
Fraction Bound
0.4
0.2
0.0

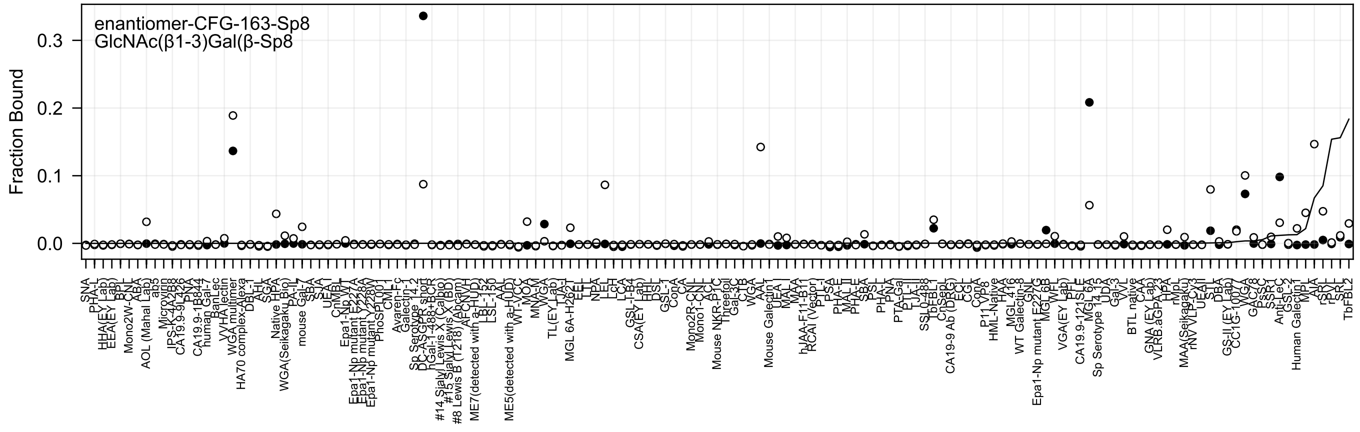
enantiomer-CFG-163-Sp8
GlcNAc(β1-3)Gal(β-Sp8
Fraction Bound
0.3
0.2
0.1
0.0

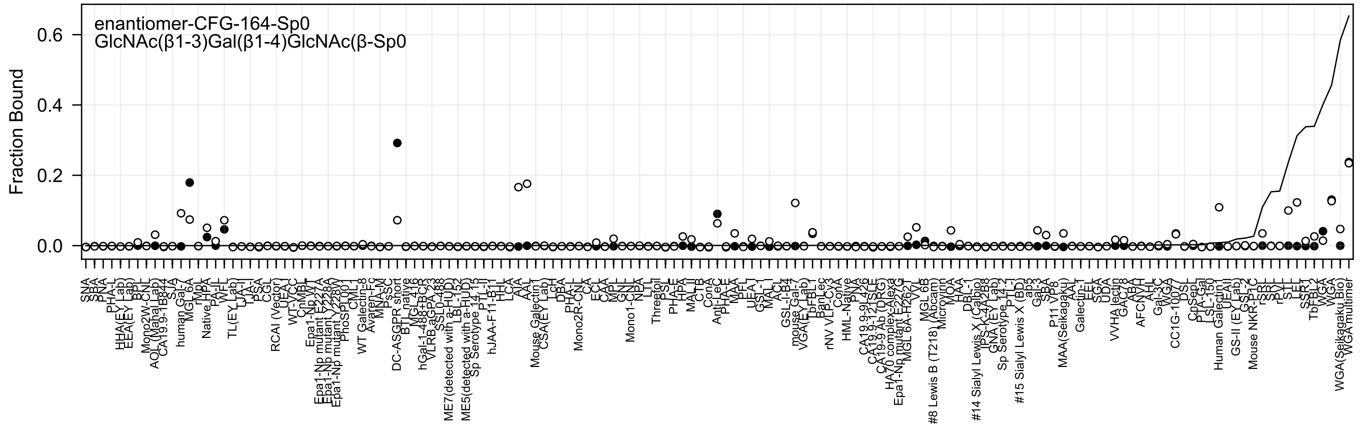
enantiomer-CFG-164-Sp0
GlcNAc(β1-3)Gal(β1-4)GlcNAc(β-Sp0
Fraction Bound
0.0
0.2
0.4
0.6

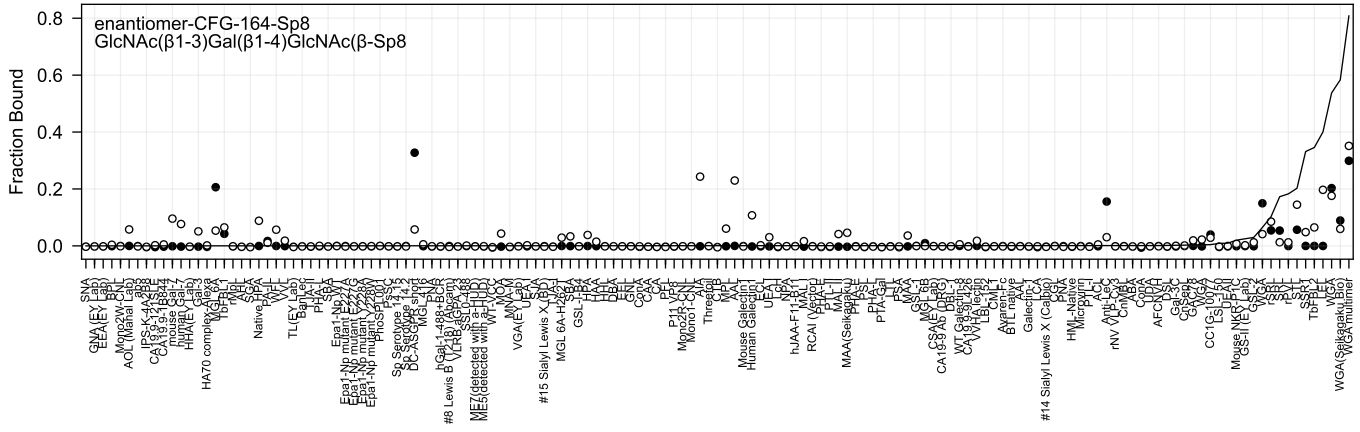
enantiomer-CFG-164-Sp8
GlcNAc(β1-3)Gal(β1-4)GlcNAc(β-Sp8
Fraction Bound

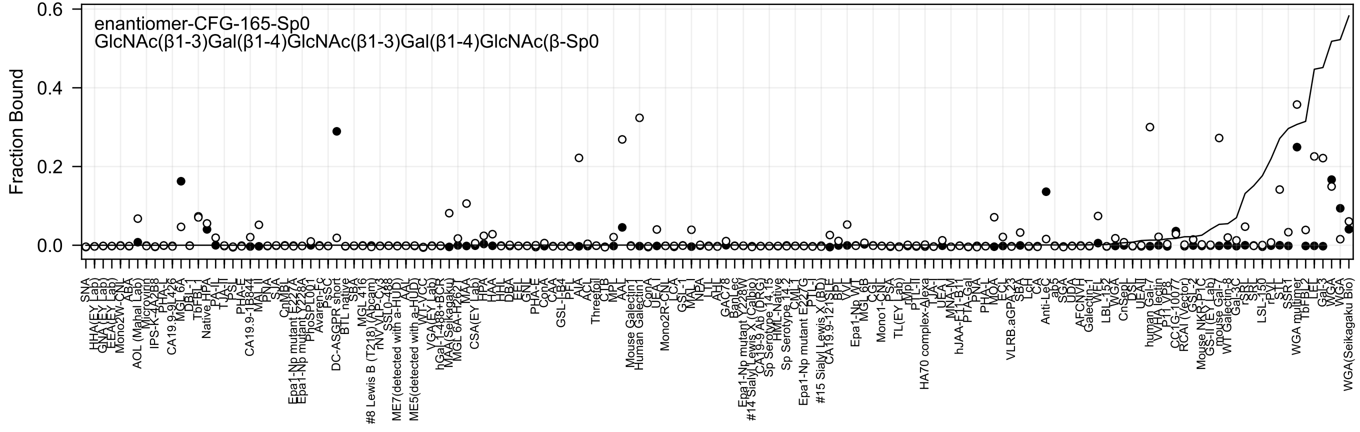
enantiomer-CFG-165-Sp0
GlcNAc(β1-3)Gal(β1-4)GlcNAc(β1-3)Gal(β1-4)GlcNAc(β-Sp0
Fraction Bound
0.6
0.4
0.2
0.0

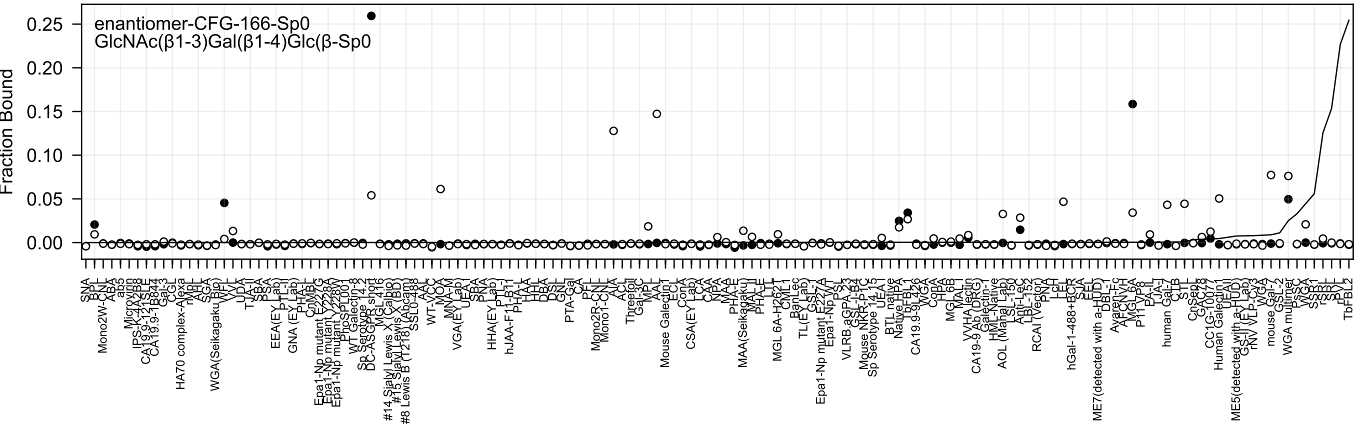

enantiomer-CFG-166-Sp0
GlcNAc(β1-3)Gal(β1-4)Glc(β-Sp0
Fraction Bound

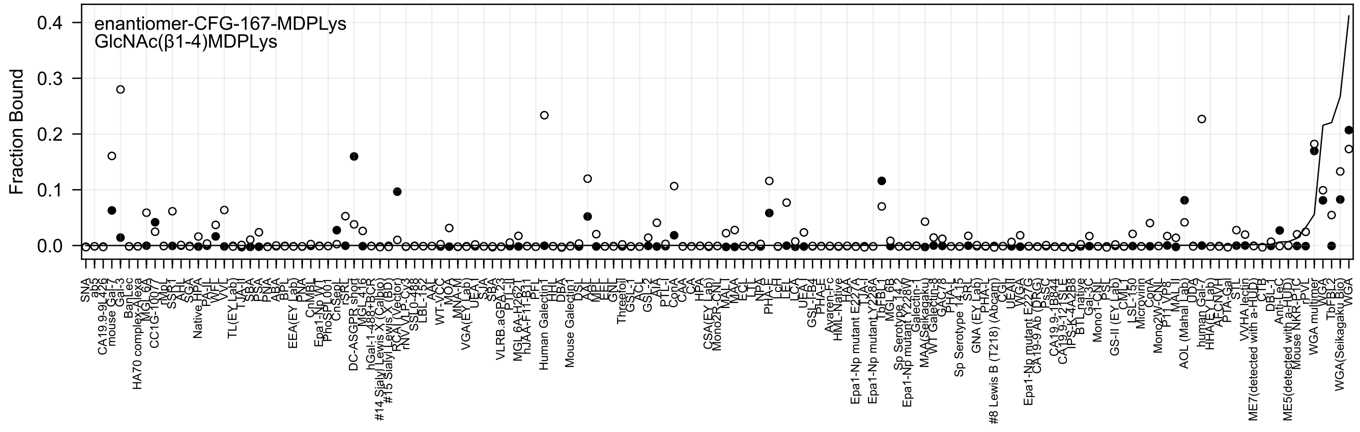
enantiomer-CFG-167-MDPLys
GlcNAc(β1-4)MDPLys
Fraction Bound
0.4
0.3
0.2
0.1
0.0

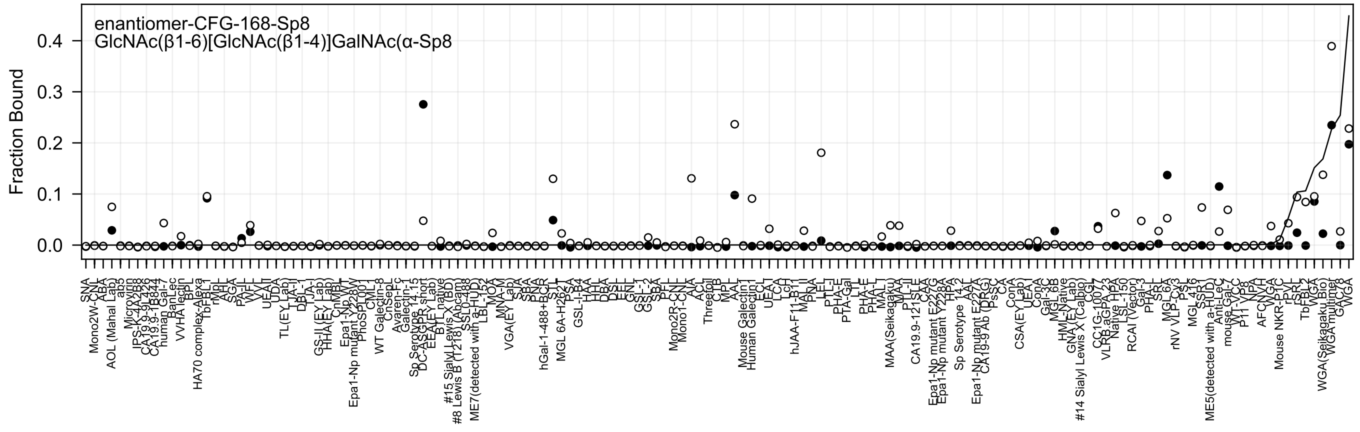
enantiomer-CFG-168-Sp8
GlcNAc(β1-6)[GlcNAc(β1-4)]GalNAc(α-Sp8
Fraction Bound
0.0
0.1
0.2
0.3
0.4

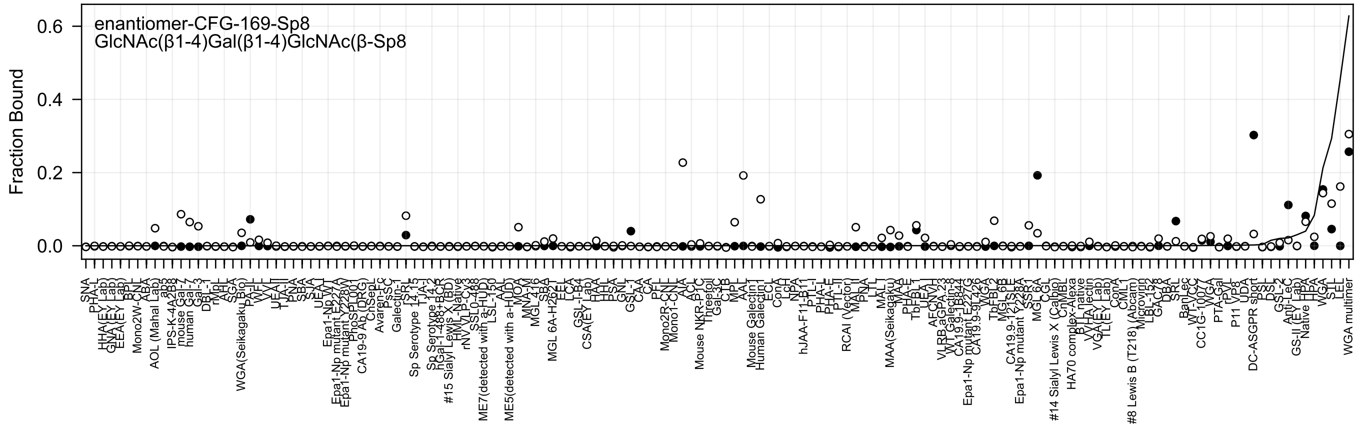
enantiomer-CFG-169-Sp8
GlcNAc(β1-4)Gal(β1-4)GlcNAc(β-Sp8
Fraction Bound
0.0
0.2
0.4
0.6
SNA
PHA-L
HHA(EY Lab)
GNA(EY Lab)
EEA(EY Lab)
BPL
Mono2W-CNL
ABA
AOL (Mahal Lab)
ab3
IPS-K-4A2B8
mouse Gal-7
human Gal-3
DBL-1
rMpL
AHL
SGA
WGA(Seikagaku Bio)
PA-IL
WFL
VVL
UEA-I
TJA-II
PNA
SBA
SJA
UEA-I
Epa1-Np WT
Epa1-Np mutant E227A
Epa1-Np mutant Y228W
PhoSL001
CA19-9 Ab (DRG)
ChSepL
Avaren-Fc
PsSC
Galectin-1
SRL
Sp Serotype 14.15
TJA-I
Sp Serotype 14.2
hGal-1-488+BCR
#15 Sialyl Lewis X (BD)
HML-Native
rNV VLP-GX3
SSL0-488
ME7(detected with a-HUD)
LSL-150
AAL
ME5(detected with a-HUD)
MOA
MNA-M
MGL 416
SBA
MGL 6A-H262T
LEL
ECA
GSL-I-B4
CSA(EY Lab)
HAA
HHL
PSA
GNL
GSL-1
CAA
CA
PFL
Mono2R-CNL
Mono1-CNL
AIA
ACL
Mouse NKR-P1C
Threefoil
Gal-3C
CTB
MPL
AAL
Mouse Galectin1
Human Galectin1
ECL
ConA
LcH
NPA
hJAA-F11-B11
PTL-I
PHA-L
PHA-E
PTL-II
RCAI (Vector)
MAL-II
PNA
LTL
MAL-I
MAA(Seikagaku)
MAA
PHA-E
TbFBL1
UEA-I
AFCNVL
VLRB.aGPA.23
WT Galectin-8
CA19.9-1B844
Epa1-Np mutant E227G
CA19.9-9L426
WGA
TbFBL2
MGL 6B
CA19.9-121SLE
Epa1-Np mutant Y228A
SSR1
MGL 6A
CGL
#14 Sialyl Lewis X (Calbio)
CnMBL
HA70 complex-Alexa
BTL native
VVHA lectin
VGA(EY Lab)
TL(EY Lab)
ConA
CML1
#8 Lewis B (T218) (Abcam)
Microvirin
LBL-152
GAC78
DBA
SRL
BanLec
WT-VCC
CC1G-1007
WGA
PTA-Gal
PVL
P11 VP8
UDA
DC-ASGPR short
PSL
DSL
GSL-2
Anti-LeC
GS-II (EY Lab)
Native HPA
HPA
WGA
STL
LEL
WGA multimer

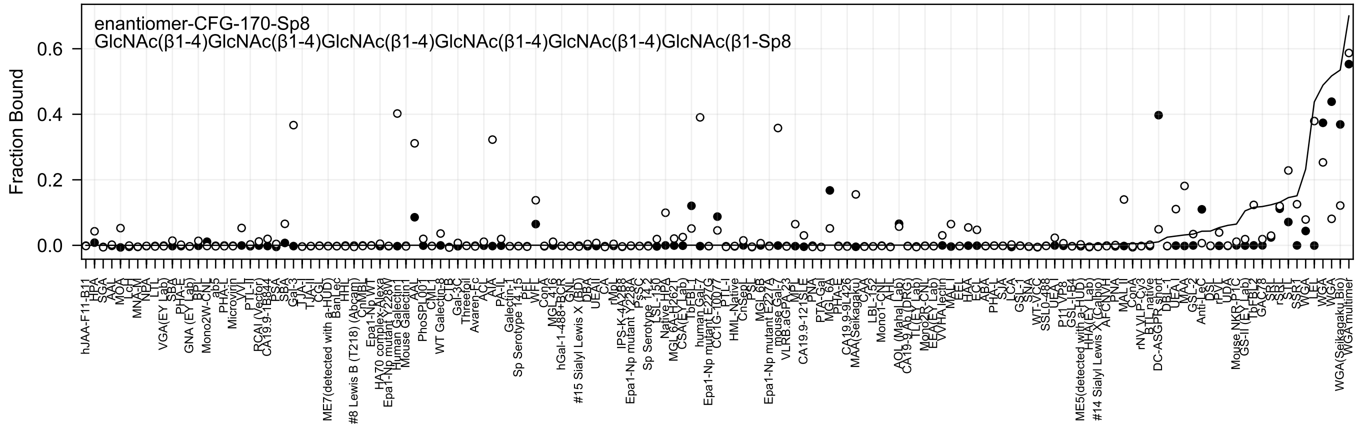
enantiomer-CFG-170-Sp8
GlcNAc(β1-4)GlcNAc(β1-4)GlcNAc(β1-4)GlcNAc(β1-4)GlcNAc(β1-4)GlcNAc(β1-Sp8
Fraction Bound
0.6
0.4
0.2
0.0

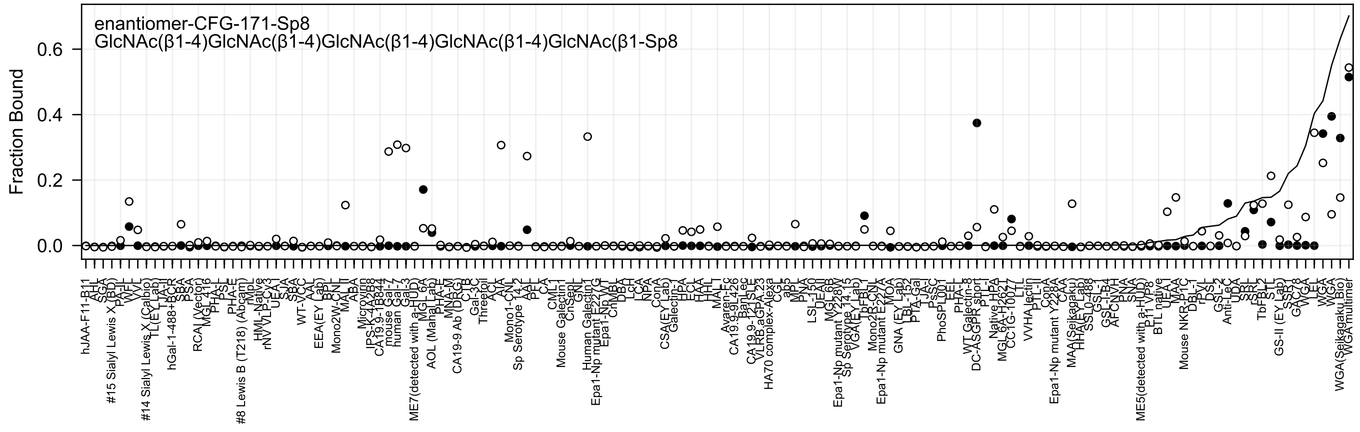
enantiomer-CFG-171-Sp8
GlcNAc(β1-4)GlcNAc(β1-4)GlcNAc(β1-4)GlcNAc(β1-4)GlcNAc(β1-Sp8
Fraction Bound

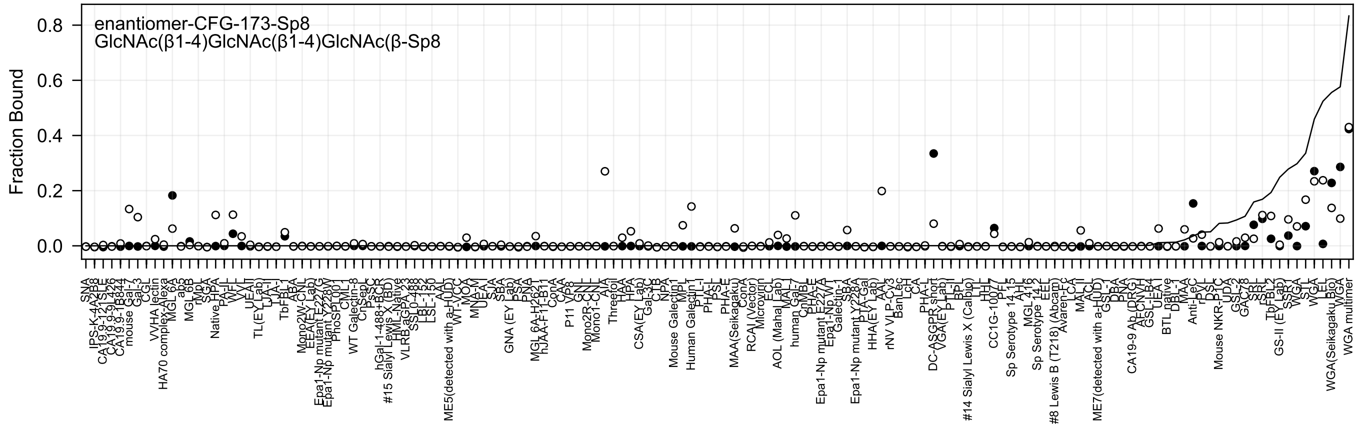

enantiomer-CFG-173-Sp8
GlcNAc(β1-4)GlcNAc(β1-4)GlcNAc(β-Sp8
Fraction Bound
0.8
0.6
0.4
0.2
0.0
SNA
IPS-K-4A2B8
mouse Gal-7
Gal-3
CGL
VVHA lectin
HA70 complex-Alexa
MGL 6A
ab5
MGL 6B
rMpL
SGA
Native HPA
PA-IL
WFL
VVL
UEAII
TL(EY Lab)
LTA-I
TJA-I
TbFBL-1
ABA
Mono2W-CNL
EEA(EY Lab)
Epa1-Np mutant E227G
Epa1-Np mutant Y228W
PhoSL001
CML-1
WT Galectin-8
CnSepL
PsSC
hGal-1-488+BCR
#15 Sialyl Lewis X (BD)
HML-Native
VLRB.aGPA.23
SSL0-488
LBL-152
LSL-150
AAL
ME5(detected with a-HUD)
WT-VCC
MOA
MNA-M
UEA-I
SJA
SBA
GNA (EY Lab)
PSA
PNA
MGL 6A-H262T
hJAA-F11-B11
ConA
CAA
P11 VP8
GNL
Mono2R-CNL
Mono1-CNL
AIA
Threefoil
HAA
HPA
CSA(EY Lab)
Gal-3C
CTB
NPA
Mouse Galectin1
MPL
Human Galectin1
PTL-I
PHA-L
PSL
PHA-E
MAA(Seikagaku)
ConA
RCAI (Vector)
Microvirin
ECL
AOL (Mahal Lab)
MAL-I
human Gal-7
CnMBL
PHA-E
Epa1-Np mutant E227A
Epa1-Np WT
Galectin-1
SBA
Epa1-Np mutant Y228A
PTA-Gal
HHA(EY Lab)
AAL
rNV VLP-Cy3
BanLec
LcH
CA
PHA-L
DC-ASGPR short
VGA(EY Lab)
PTL-II
BPL
#14 Sialyl Lewis X (Calbio)
LTL
HHL
CC1G-10077
PFL
Sp Serotype 14 15
AHL
MGL 416
Sp Serotype 14 2
EEL
#8 Lewis B (T218) (Abcam)
Avaren-Fc
LCA
MAL II
ACL
ME7(detected with a-HUD)
GSL-I
DBA
BPA
CA19-9 Ab (DRG)
AECNYH
GSL-I-B4
UEA-I
BTL native
DBL-1
MAA
Anti-LeC
rPVL
DSL
Mouse NKR-P1C
UDA
GSL-II
GAC78
SRL
rSRL
TbFBL2
GS-II (EY Lab)
SSR1
WGA
STL
WGA
LEL
WGA(Seikagaku Bio)
WGA
WGA multimer

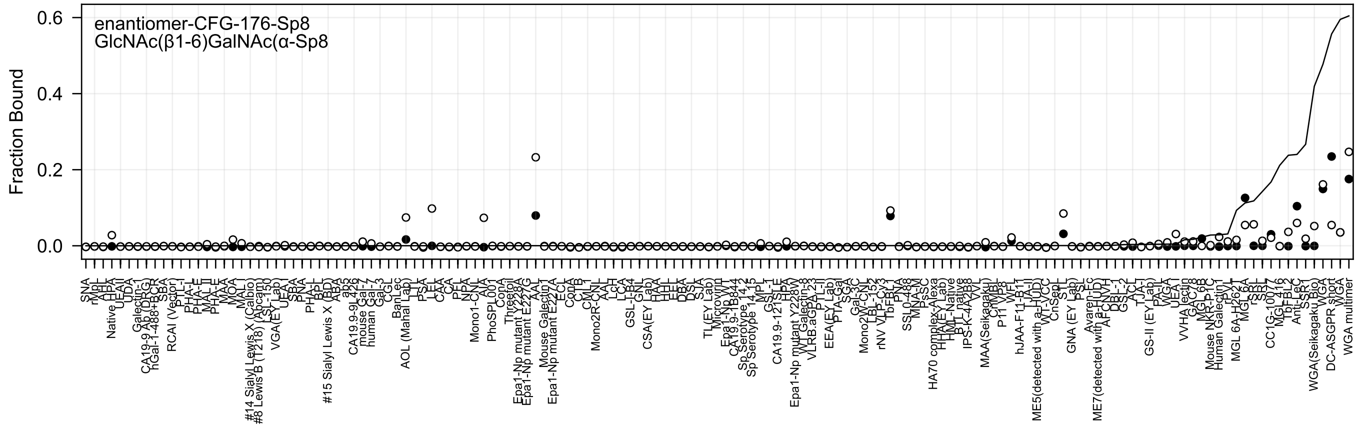
enantiomer-CFG-176-Sp8
GlcNAc(β1-6)GalNAc(α-Sp8
Fraction Bound
0.6
0.4
0.2
0.0

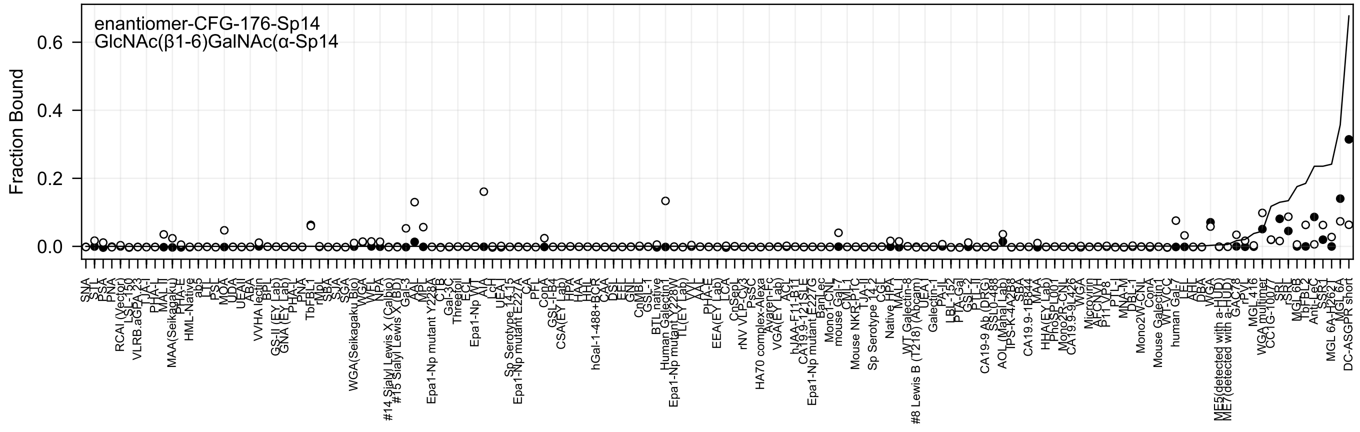

enantiomer-CFG-176-Sp14
GlcNAc(β1-6)GalNAc(α-Sp14
Fraction Bound
0.0
0.2
0.4
0.6

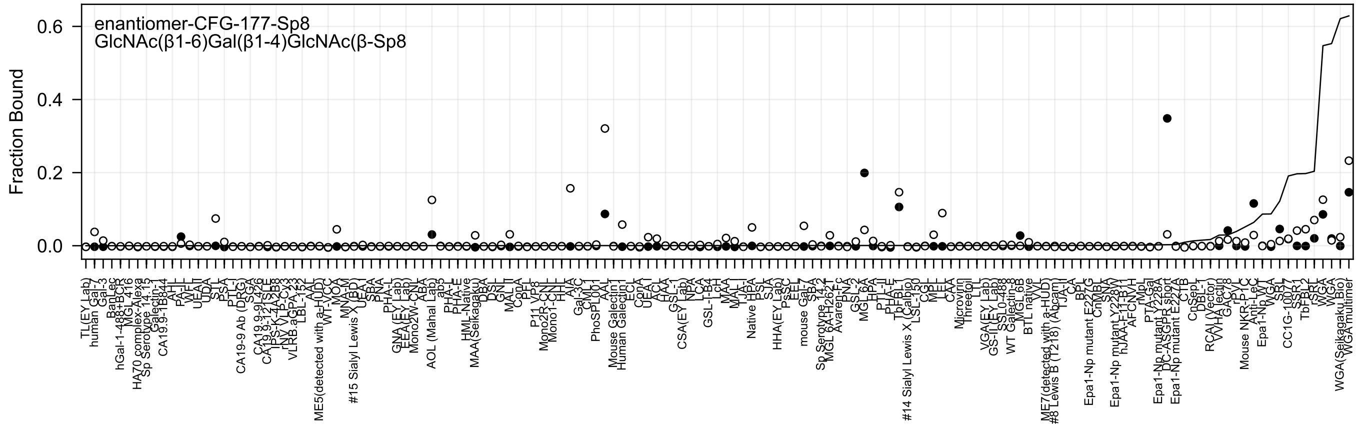
enantiomer-CFG-177-Sp8
GlcNAc(β1-6)Gal(β1-4)GlcNAc(β-Sp8
Fraction Bound
0.6
0.4
0.2
0.0

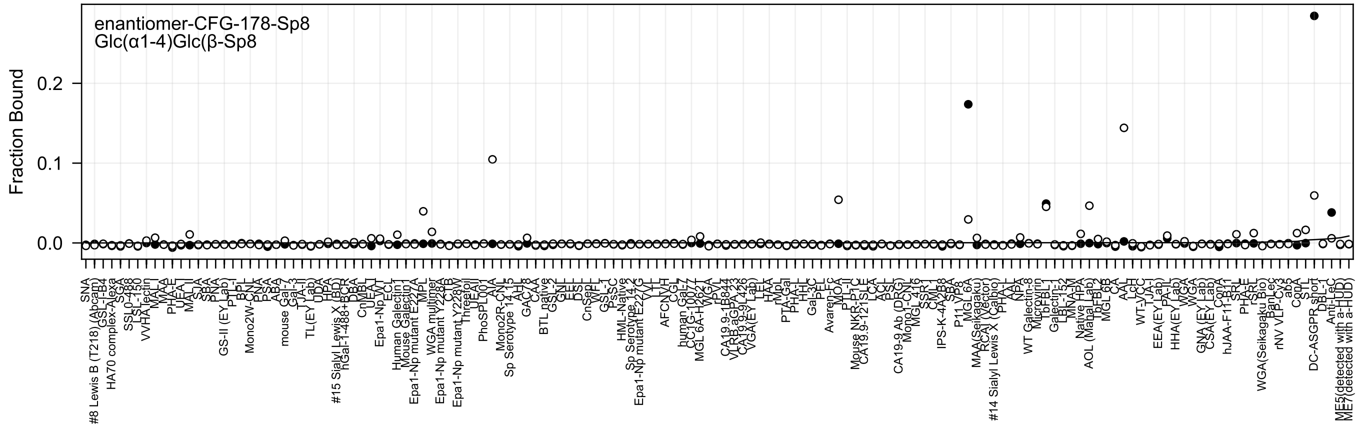
enantiomer-CFG-178-Sp8
Glc(α1-4)Glc(β-Sp8
Fraction Bound
0.2
0.1
0.0

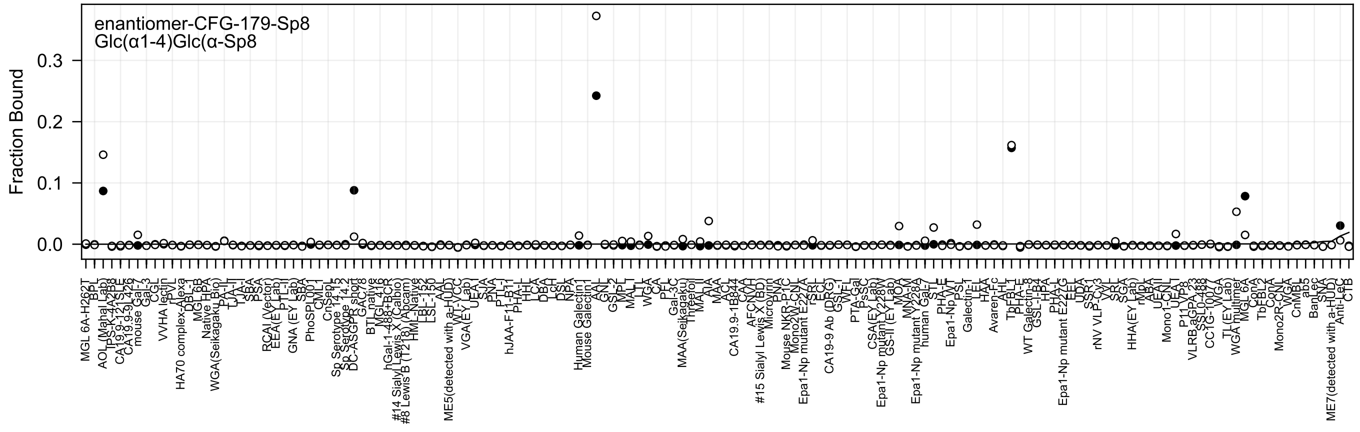
enantiomer-CFG-179-Sp8
Glc(α1-4)Glc(α-Sp8
Fraction Bound
0.3
0.2
0.1
0.0

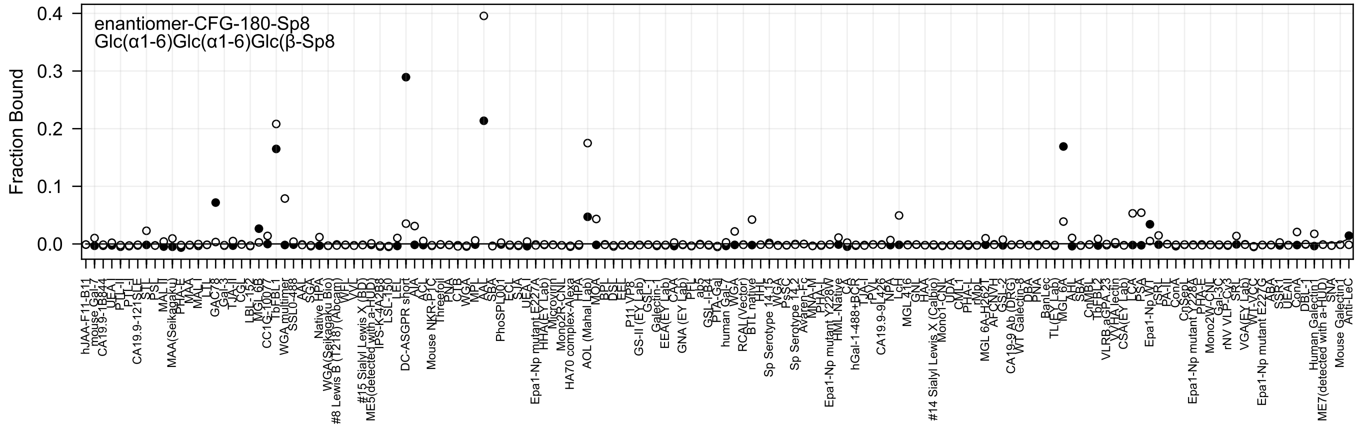

enantiomer-CFG-180-Sp8
Glc(α1-6)Glc(α1-6)Glc(β-Sp8
Fraction Bound
0.0
0.1
0.2
0.3
0.4

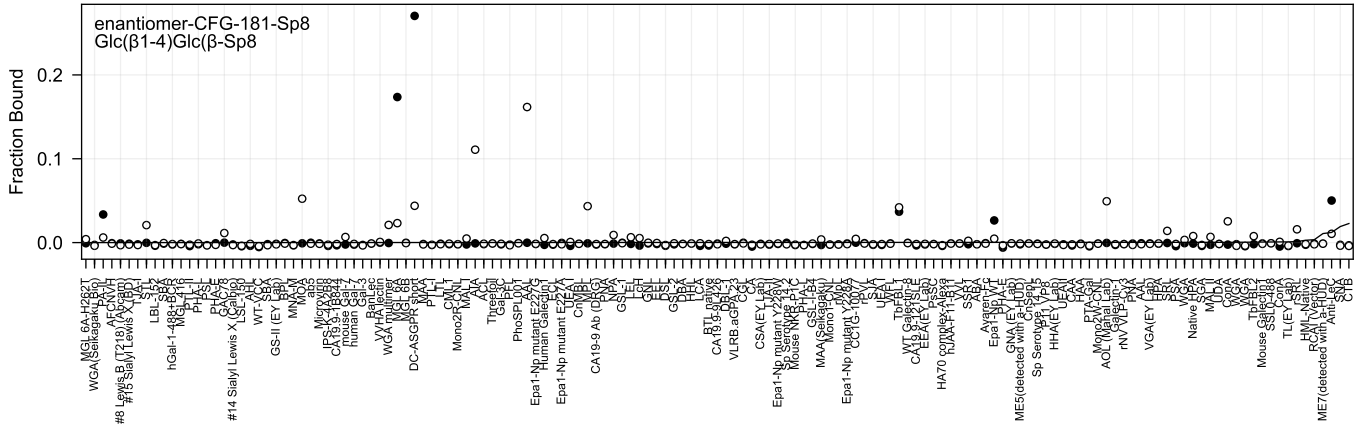

enantiomer-CFG-181-Sp8
Glc(β1-4)Glc(β-Sp8
Fraction Bound
0.2
0.1
0.0

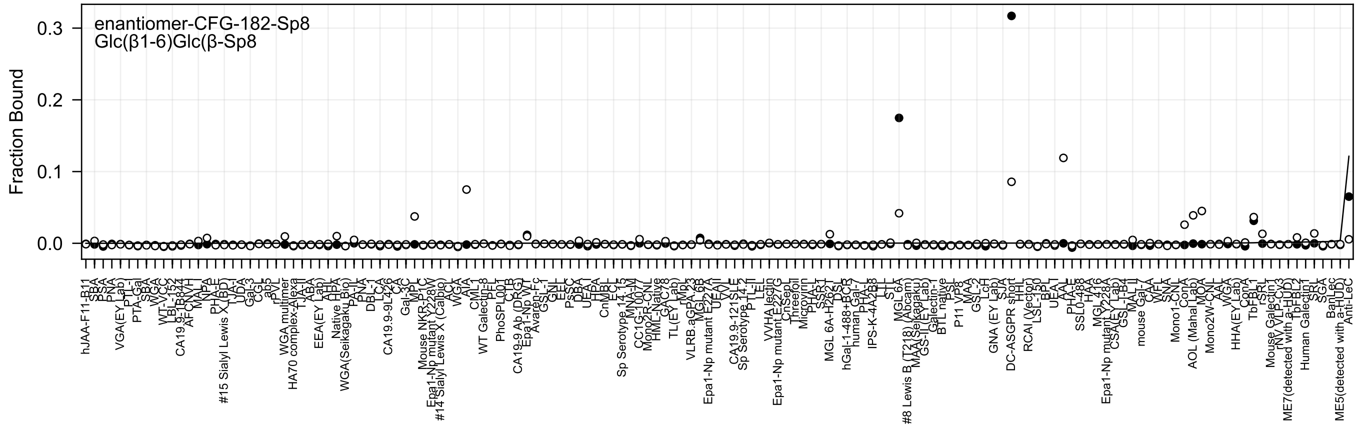
enantiomer-CFG-182-Sp8
Glc(β1-6)Glc(β-Sp8
Fraction Bound
0.3
0.2
0.1
0.0

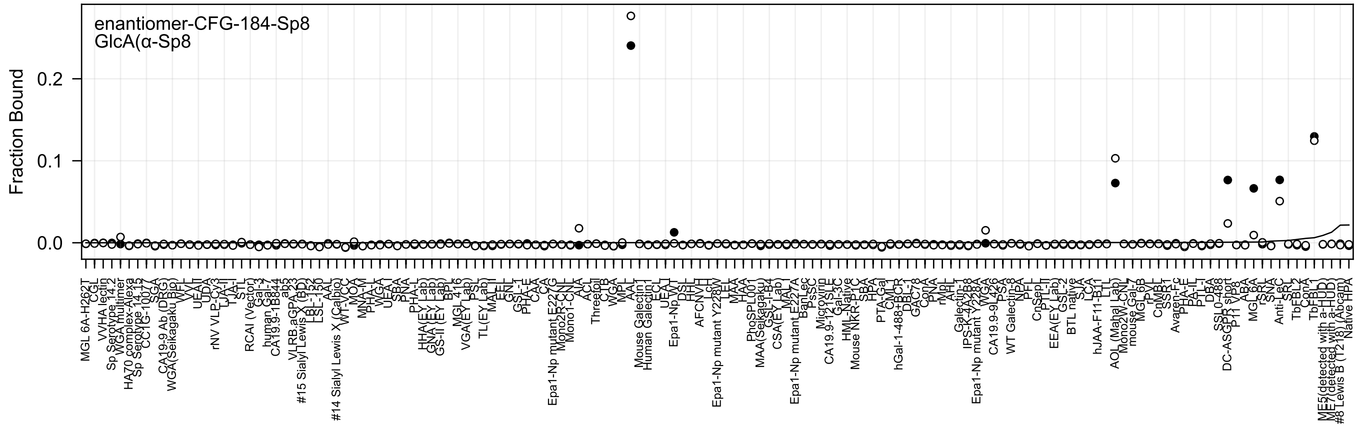
enantiomer-CFG-184-Sp8
GlcA(α-Sp8
Fraction Bound
0.0
0.1
0.2

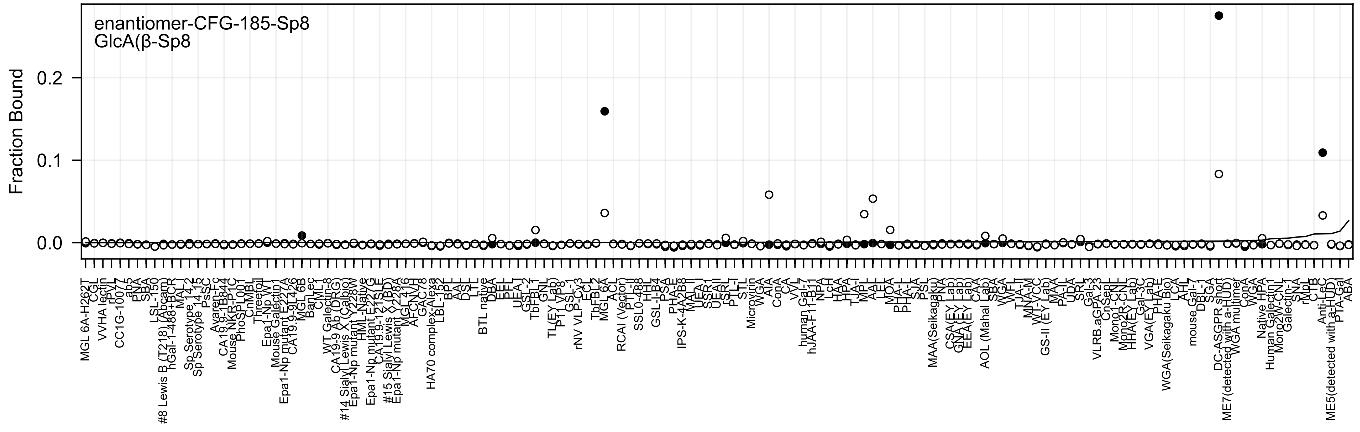
enantiomer-CFG-185-Sp8
GlcA(β-Sp8
Fraction Bound
0.2
0.1
0.0

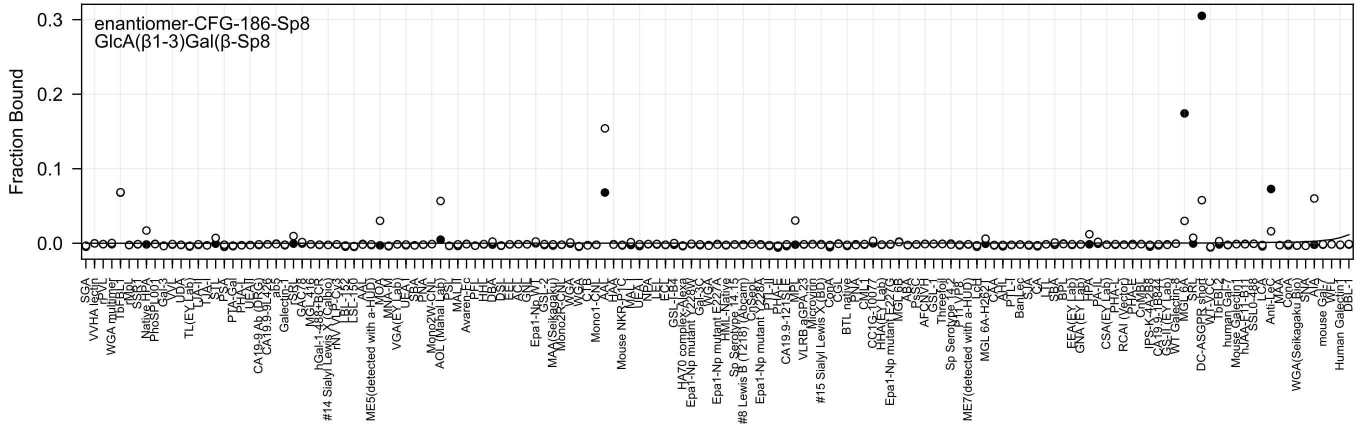
enantiomer-CFG-186-Sp8
GlcA(β1-3)Gal(β-Sp8
Fraction Bound
0.3
0.2
0.1
0.0
SGA
VVHA lectin
rPVL
WGA multimer
TbF-BL1
rMpL
SSR1
Native HPA
PhoSL001
Gal-3
VVL
UDA
TL(EY Lab)
LJA-I
TJA-I
STL
PSA
PTA-Gal
PHA-L
UEA-II
CA19-9 Ab (DRG)
CA19.9-9L426
ab5
Galectin-1
rSRL
GAL3R
MGL-416
hGal-1-488+BCR
#14 Sialyl Lewis X (Calbio)
rNV VLP-Cy3
LSL-152
LSL-150
AAL
ME5(detected with a-HUD)
MOA
MNA-M
VGA(EY Lab)
UEA-I
SBA
PNA
Mono2W-CNL
AOL (Mahal Lab)
PSL
MAL-I
Avaren-Fc
PFL
HHL
DBA
DSL
EEL
ACL
GNL
Epa1-Np WT
GSL-2
MAA(Seikagaku)
Mono2R-CNL
WGA
WGA
CTB
Mono1-CNL
AAL
HAA
Mouse NKR-P1C
MAL-I
UEA-I
NPA
LEL
ECL
GSL-I-B4
HA70 complex-Alexa
Epa1-Np mutant Y228W
Gal-3C
WGA
Epa1-Np mutant E227A
HML-Native
Sp Serotype 14.15
#8 Lewis B (T218) (Abcam)
CnSep
Epa1-Np mutant Y228A
PTL-II
PHA-E
CA19.9-121SLE
MPL
VLRB aGPA.23
Microvirin
#15 Sialyl Lewis X (BD)
ConA
CGL
BTL native
PNA
CML1
CC1G-10077
HHA(EY Lab)
Epa1-Np mutant E227G
MGL 6B
ABA
PsSC
AFCNVH
GSL-1
Threefoil
Sp Serotype 14.2
P11 VP8
ME7(detected with a-HUD)
LcH
MGL 6A-H262T
CAA
AHL
PTL-I
BanLec
SJA
CA
LTL
SBA
BPL
EEA(EY Lab)
GNA (EY Lab)
HPA
PA-IL
CSA(EY Lab)
PHA-L
RCAI (Vector)
PHA-E
CnMBL
IPS-K-4A2B8
CA19.9-1B844
GS-II (EY Lab)
WT Galectin-8
MGL 6A
SRL
DC-ASGPR short
WT-VCC
TbFBL2
human Gal-7
Mouse Galectin1
hJAA-F11-B11
SSL0-488
LCA
Anti-LeC
MAA
ConA
WGA(Seikagaku Bio)
SNA
AIA
mouse Gal-7
WFL
Human Galectin1
DBL-1

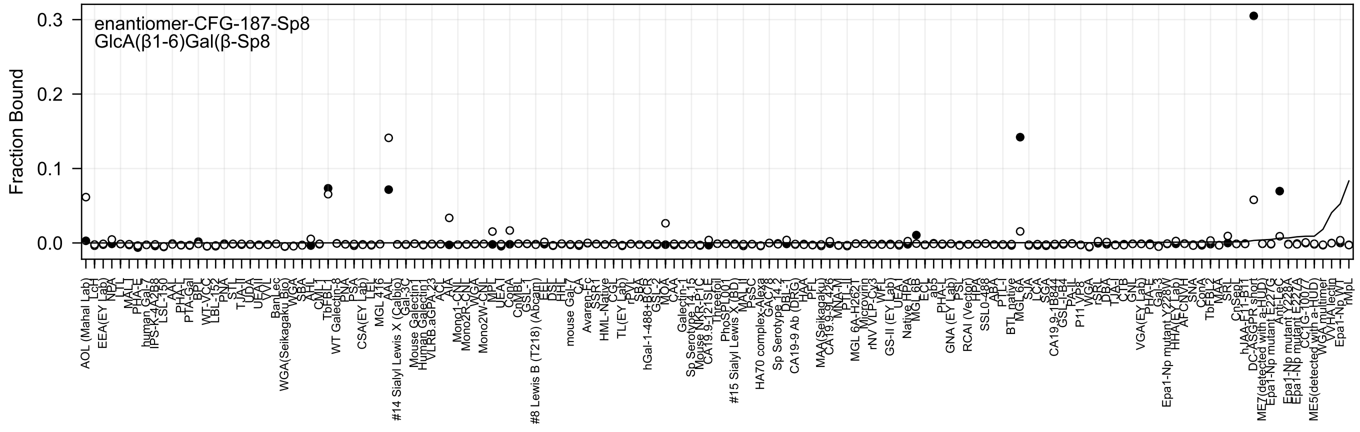
enantiomer-CFG-187-Sp8
GlcA(β1-6)Gal(β-Sp8
Fraction Bound

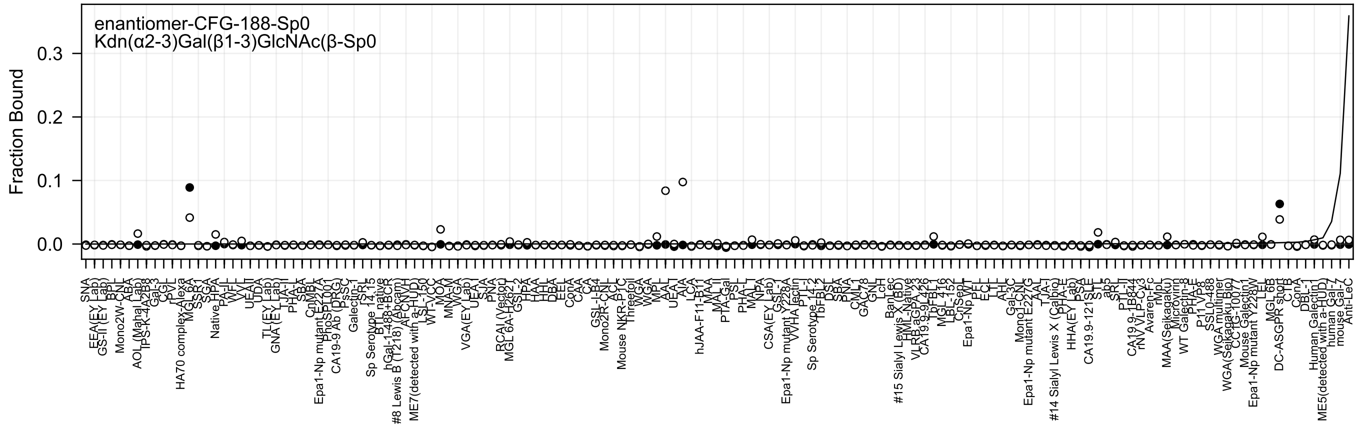

enantiomer-CFG-188-Sp0
Kdn(α2-3)Gal(β1-3)GlcNAc(β-Sp0
Fraction Bound
0.0
0.1
0.2
0.3

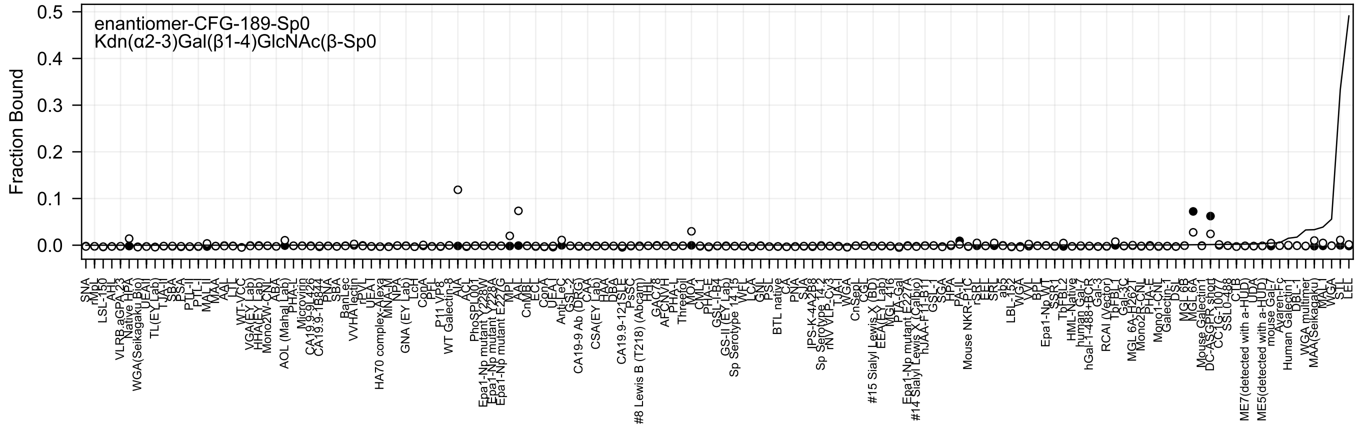
enantiomer-CFG-189-Sp0
Kdn(α2-3)Gal(β1-4)GlcNAc(β-Sp0
Fraction Bound
0.5
0.4
0.3
0.2
0.1
0.0

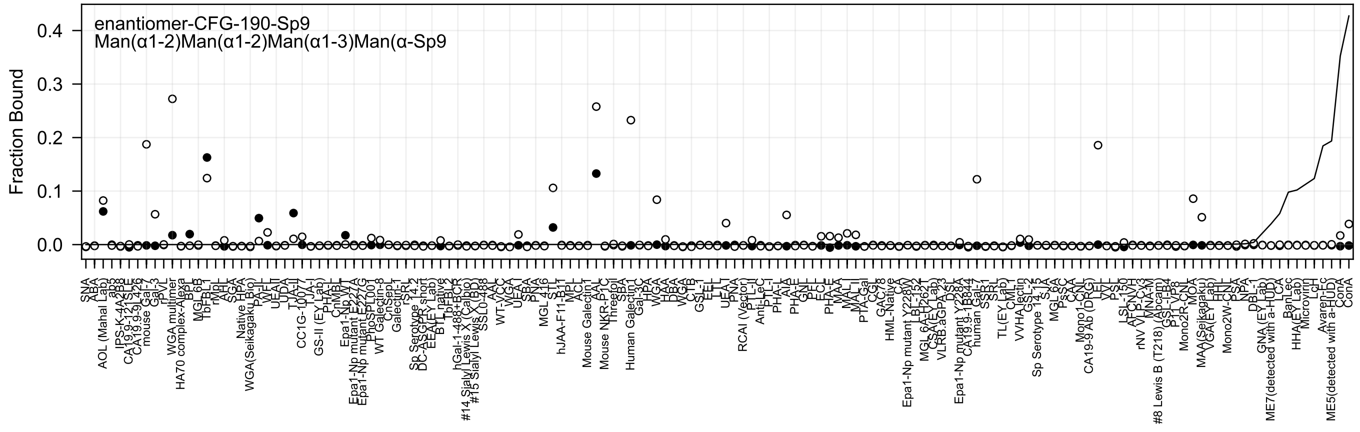
enantiomer-CFG-190-Sp9
Man(α1-2)Man(α1-2)Man(α1-3)Man(α-Sp9
Fraction Bound
0.4
0.3
0.2
0.1
0.0

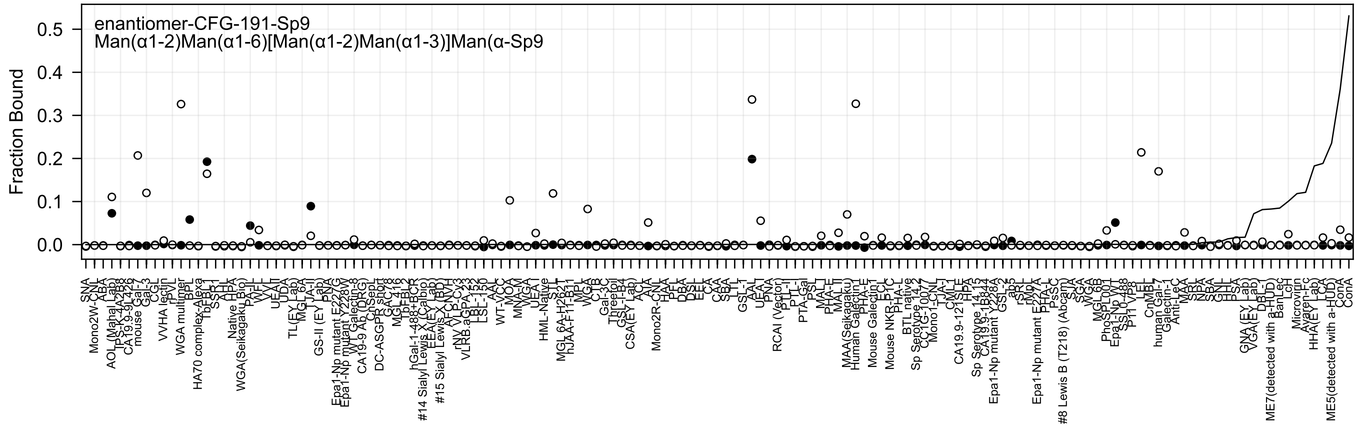
enantiomer-CFG-191-Sp9
Man(α1-2)Man(α1-6)[Man(α1-2)Man(α1-3)]Man(α-Sp9
Fraction Bound
0.5
0.4
0.3
0.2
0.1
0.0

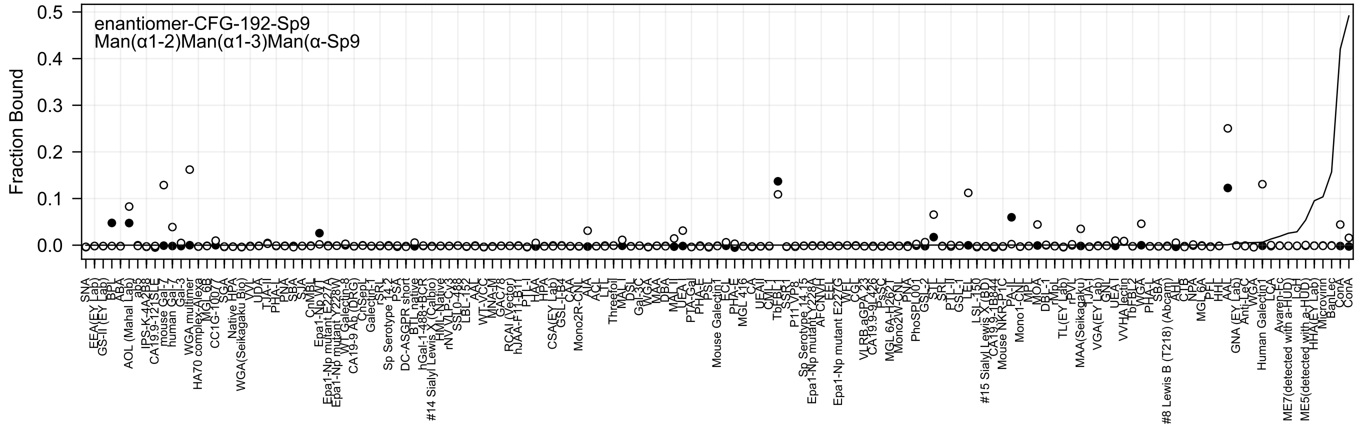
enantiomer-CFG-192-Sp9
Man(α1-2)Man(α1-3)Man(α-Sp9
Fraction Bound
0.5
0.4
0.3
0.2
0.1
0.0

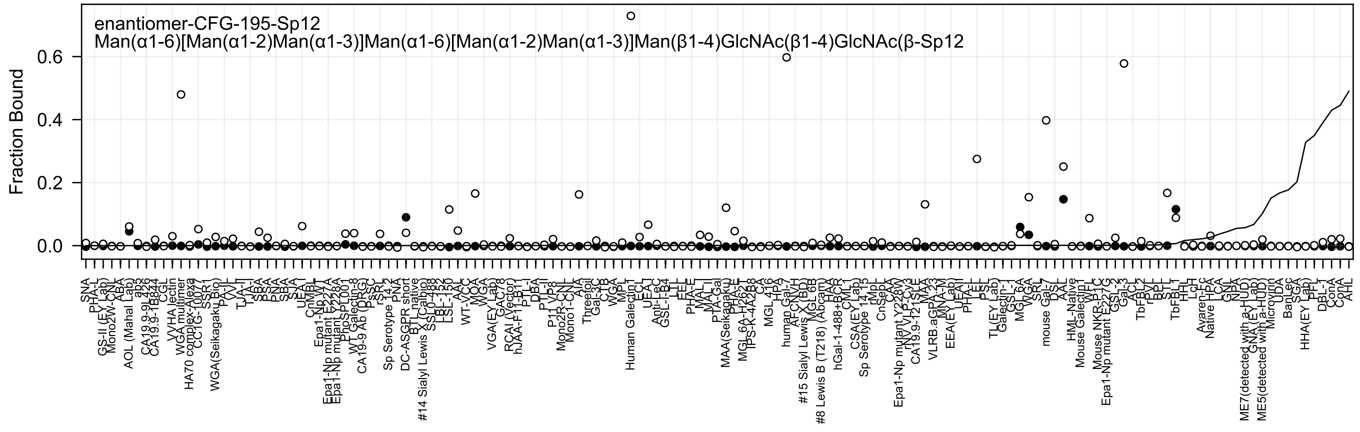

enantiomer-CFG-195-Sp12
Man(α1-6)[Man(α1-2)Man(α1-3)]Man(α1-6)[Man(α1-2)Man(α1-3)]Man(β1-4)GlcNAc(β1-4)GlcNAc(β-Sp12
Fraction Bound
0.0
0.2
0.4
0.6

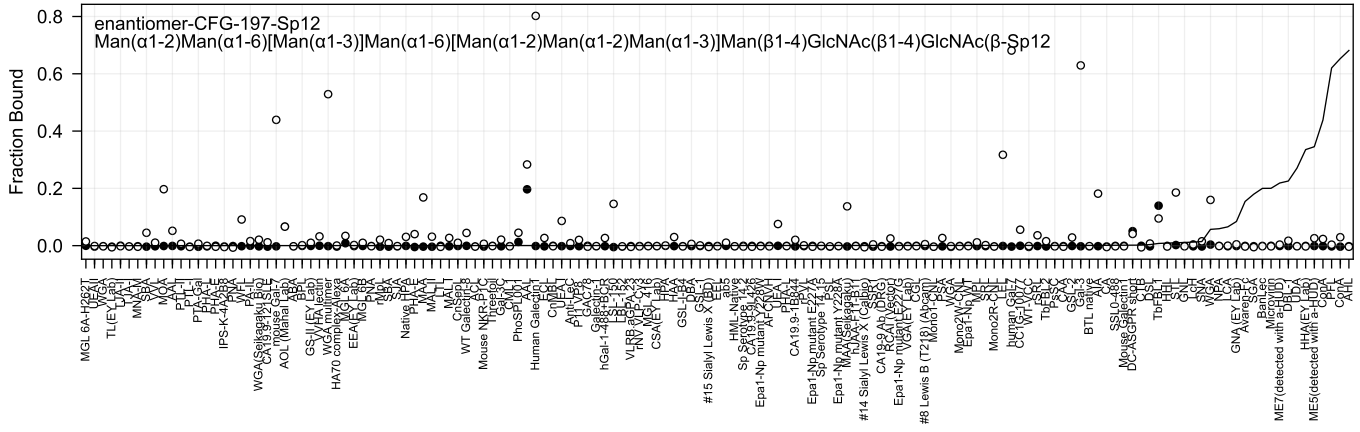
enantiomer-CFG-197-Sp12
Man(α1-2)Man(α1-6)[Man(α1-3)]Man(α1-6)[Man(α1-2)Man(α1-2)Man(α1-3)]Man(β1-4)GlcNAc(β1-4)GlcNAc(β-Sp12
Fraction Bound
0.8
0.6
0.4
0.2
0.0

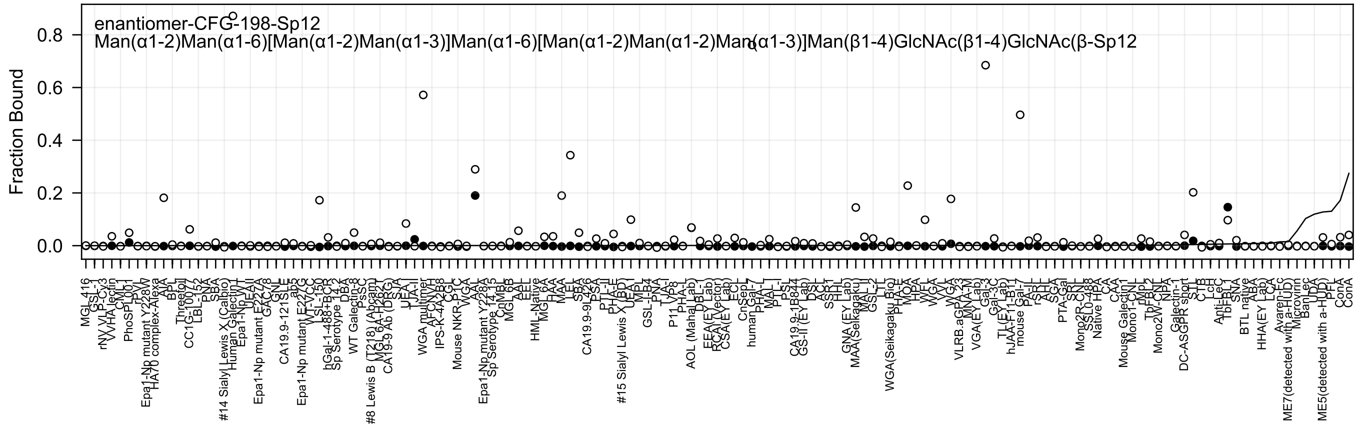
enantiomer-CFG-198-Sp12
Man(α1-2)Man(α1-6)[Man(α1-2)Man(α1-3)]Man(α1-6)[Man(α1-2)Man(α1-2)Man(α1-3)]Man(β1-4)GlcNAc(β1-4)GlcNAc(β-Sp12
Fraction Bound
0.0
0.2
0.4
0.6
0.8
MGL_416
GSL-1
rNV VLP-Cy3
VVHA lectin
CML1
PhoSPL001
rPV
Epa1-Np mutant Y228W
HA70 complex-Alexa
AIA
BPL
Threefoil
CC1G-10077
LBL-152
PNA
SBA
#14 Sialyl Lewis X (Calbio)
Human Galectin1
Epa1-Np WT
UEAII
Epa1-Np mutant E227A
GAC-78
GNL
CA19.9-121SLE
ab3
Epa1-Np mutant E227G
WT-VCC
LSL-150
hGal-1-488+BCR
Sp Serotype 14.2
DBA
WT Galectin-8
PsSC
#8 Lewis B (T218) (Abcam)
MGL6A-H262T
CA19-9 Ab (DRG)
SJA
UEA-I
LTA-II
WGA multimer
AFCNYH
IPS-K-4A2B8
CGL
Mouse NKR-P1C
WGA
AAL
Epa1-Np mutant Y228A
Sp Serotype 14.15
CnMBL
MGL 6B
AAL
EEL
HML-Native
MGL 6A
HAA
MAA
LEL
SBA
CA19.9-9L426
PSA
PTL-II
PHA-E
#15 Sialyl Lewis X (BD)
UEA-I
MPL
GSL-I-B4
PNA
TJA-I
P11_VP8
PHA-L
AOL (Mahal Lab)
DBL
EEA(EY Lab)
RCA(Vector)
CSA(EY Lab)
ECL
CnSepL
human Gal-7
PHA-L
MAL-I
PTL-I
PSL
CA19.9-1B844
GS-II (EY Lab)
DSL
ACL
SSP1
HHL
GNA (EY Lab)
MAA(Selkagaku)
MAL-I
GSL-2
LTL
WGA(Seikagaku Bio)
PHA-E
MOA
HPA
WFL
WGA
VVL
WGA
VLRB.aGPA.23
MNA-M
VGA(EY Lab)
Gal-3
Gal-3C
TL(EY Lab)
hJAA-F11-B11
mouse Gal-7
PA-IL
rSRL
AAH
SGA
PTA-Gal
SRL
Mono2R-CNL
SSL0-488
Native HPA
CA
CAA
Mouse Galectin1
Mono1-CNL
rMpL
TbFBL2
Mono2W-CNL
NPA
Galectin-1
DC-ASGPR short
STL
CTB
LcH
Anti-LeC
TbFBL1
SNA
BTL native
ABA
HHA(EY Lab)
LCA
Avaren-Fc
ME7(detected with a-HUD)
Microvirin
BanLec
UDA
ME5(detected with a-HUD)
PFL
ConA
ConA

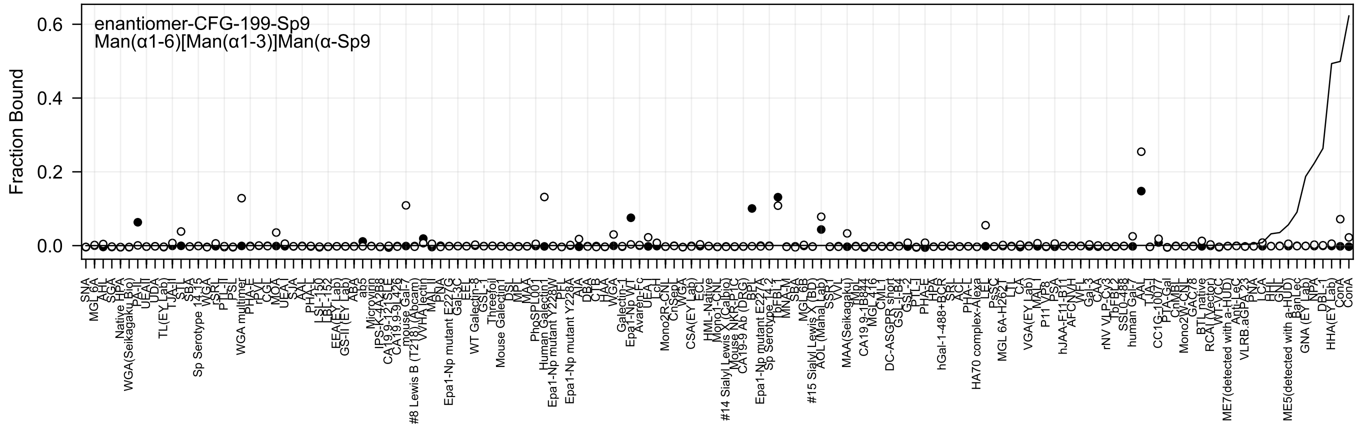
enantiomer-CFG-199-Sp9
Man(α1-6)[Man(α1-3)]Man(α-Sp9
Fraction Bound
0.6
0.4
0.2
0.0
SNA
MGL 6A
AHL
SGA
Native HPA
WGA(Seikagaku Bio)
PA-II
UEAI
UDA
TL(EY Lab)
TJA-II
STL
SBA
Sp Serotype 14.15
WGA
rSRL
PTL-II
PSL
WGA multimer
PHA-E
rPVL
CGL
MOA
UEA I
SJA
AAL
PHA-L
LSL-150
LBL-152
EEA(EY Lab)
GS-II (EY Lab)
ABA
ab5
Microvirin
IPS-K-4A2B8
CA19.9-121SLE
CA19.9-9L426
mouse Gal-7
#8 Lewis B (T218) (Abcam)
WHA lectin
MAL-II
PNA
Epa1-Np mutant E227G
Gal-3C
EEL
WT Galectin-8
GSL-1
Threefoil
Mouse Galectin1
DSL
MPL
MAA
PhoSPL001
Human Galectin1
Epa1-Np mutant Y228W
PPL
Epa1-Np mutant Y228A
AIA
DBA
CTB
HAA
WGA
Galectin-1
Epa1-Np WT
Avaren-Fc
UEA-I
LcH
Mono2R-CNL
CnSept
WGA
CSA(EY Lab)
ECL
HML-Native
Mono1-CNL
#14 Sialyl Lewis X (Calbio)
Mouse NKR-P1C
CA19-9 Ab (DRG)
BPL
Epa1-Np mutant E227A
Sp Serotype 14.2
TbF-BL2
MNA-M
SBA
MGL 6B
#15 Sialyl Lewis X (BD)
AOL (Mana Lab)
SSL1
VVL
MAA(Seikagaku)
TMpL
CA19.9-1B844
MGL 416
CML1
DC-ASGPR short
GSL-I-B4
GSL-2
PTL-I
PHA-E
HPA
hGal-1-488+BCR
SRL
ACL
PHA-L
HA70 complex-Alexa
LEL
PsSC
MGL 6A-H262
LTL
CA
VGA(EY Lab)
MAL-I
P11 VP8
PSA
hJAA-F11-B11
AFCNVH
WFL
Gal-3
CAA
rNV VLP-Cy3
TbFBL2
SSL0-488
human Gal-7
AAL
TJA-I
CC1G-10077
PTA-Gal
ChMBL
Mono2W-CNL
GAC78
BTL native
RCA I (Vector)
WT-YCC
ME7(detected with a-HUD)
Anti-Lec
VLRB.aGPA.23
PNA
LCA
HHL
GNL
ME5(detected with a-HUD)
BanLec
GNA (EY Lab)
NPA
DBL-1
HHA(EY Lab)
ConA
ConA

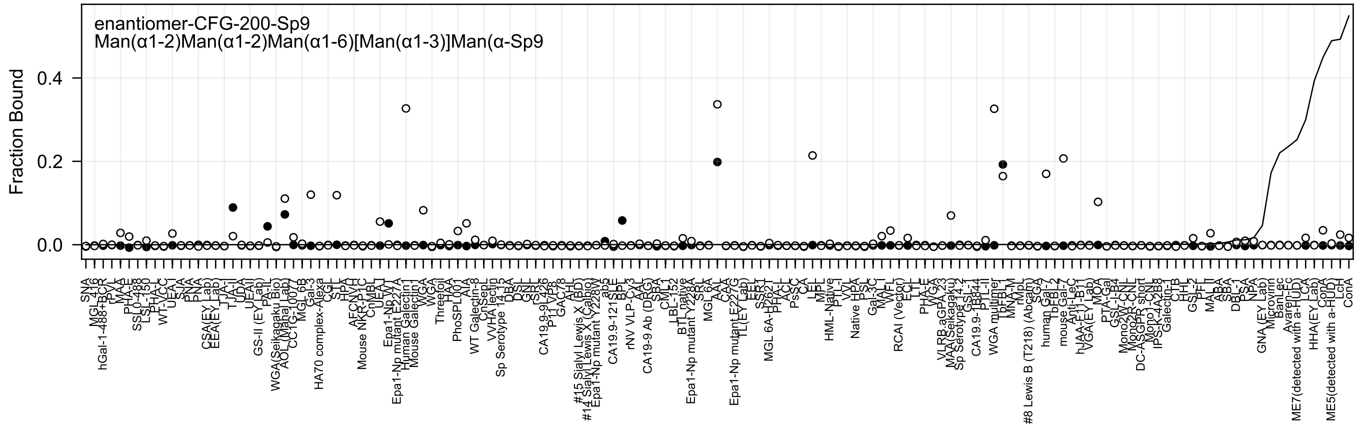

enantiomer-CFG-200-Sp9
Man(α1-2)Man(α1-2)Man(α1-6)[Man(α1-3)]Man(α-Sp9
Fraction Bound
0.0
0.2
0.4

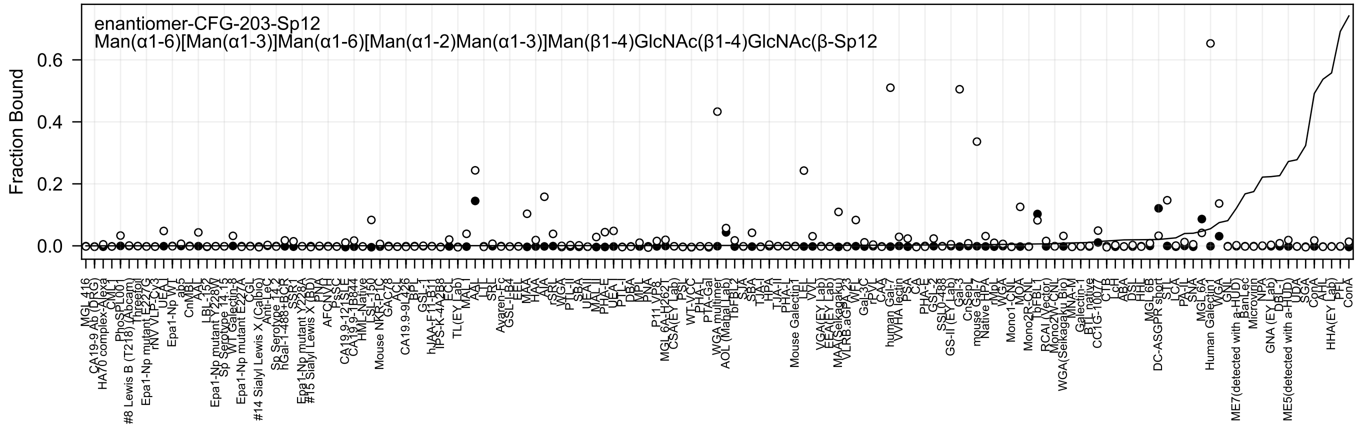
enantiomer-CFG-203-Sp12
Man(α1-6)[Man(α1-3)]Man(α1-6)[Man(α1-2)Man(α1-3)]Man(β1-4)GlcNAc(β1-4)GlcNAc(β-Sp12
Fraction Bound

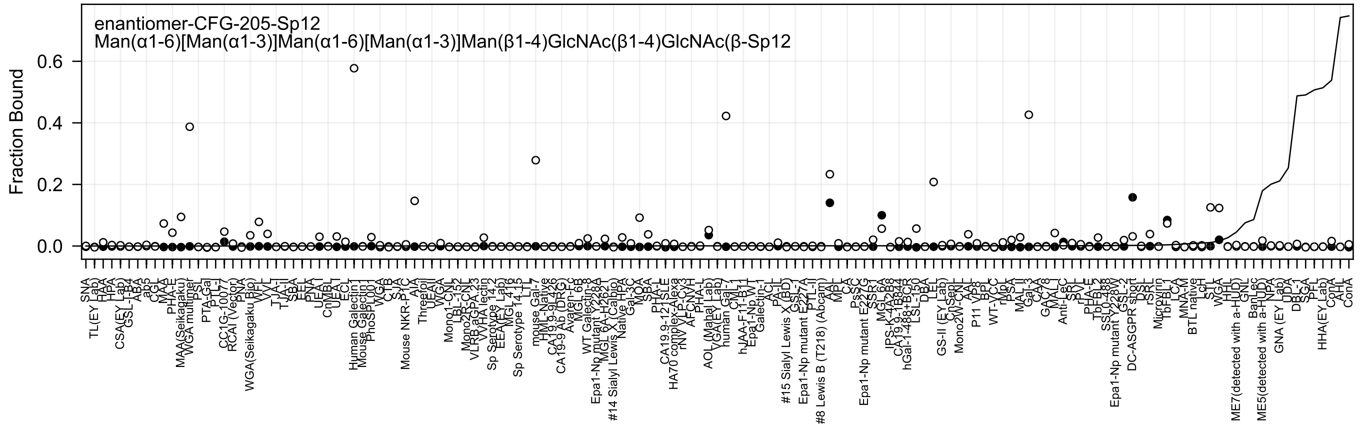
enantiomer-CFG-205-Sp12
Man(α1-6)[Man(α1-3)]Man(α1-6)[Man(α1-3)]Man(β1-4)GlcNAc(β1-4)GlcNAc(β-Sp12
Fraction Bound
0.0
0.2
0.4
0.6

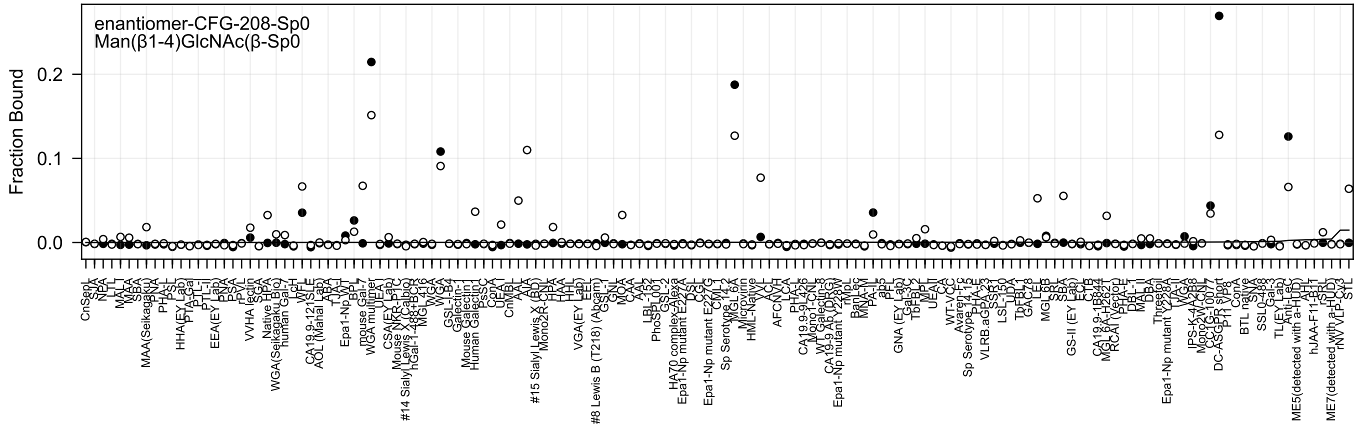
enantiomer-CFG-208-Sp0
Man(β1-4)GlcNAc(β-Sp0
Fraction Bound
0.0
0.1
0.2

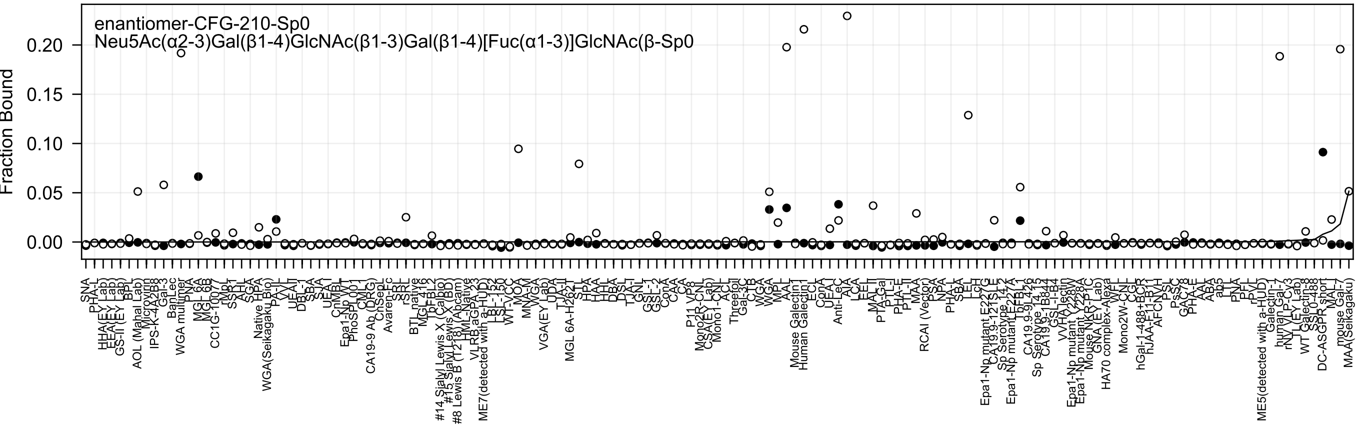

enantiomer-CFG-210-Sp0
Neu5Ac(α2-3)Gal(β1-4)GlcNAc(β1-3)Gal(β1-4)[Fuc(α1-3)]GlcNAc(β-Sp0
Fraction Bound

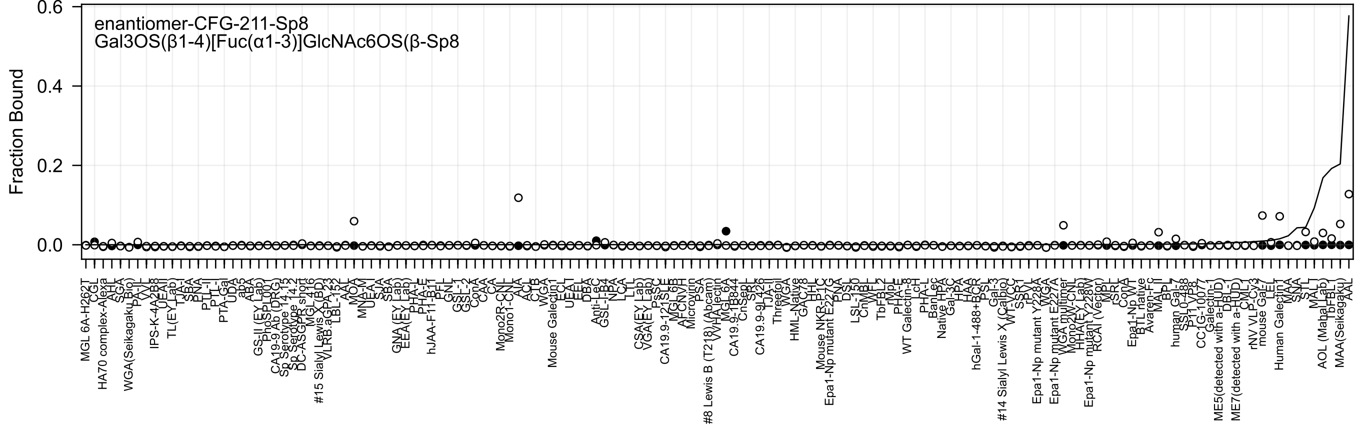
enantiomer-CFG-211-Sp8
Gal3OS(β1-4)[Fuc(α1-3)]GlcNAc6OS(β-Sp8
Fraction Bound

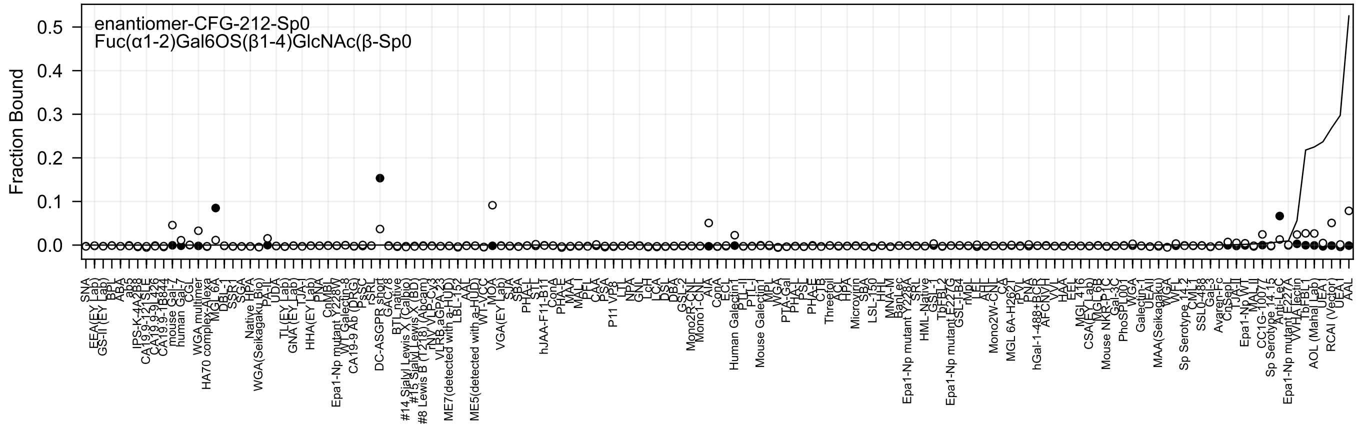
enantiomer-CFG-212-Sp0
Fuc(α1-2)Gal6OS(β1-4)GlcNAc(β-Sp0
Fraction Bound
0.0
0.1
0.2
0.3
0.4
0.5
SNA
EEA(EY Lab)
GS-II (EY Lab)
BPL
ABA
ab5
IPS-K-4A2B8
CA19.9-121SLE
CA19.9-9L426
CA19.9-1B844
mouse Gal-7
human Gal-7
CGL
WGA multimer
HA70 complex-Alexa
MGL 6A
DBL-1
SSR-1
SGA
Native HPA
WGA(Seikagaku Bio)
PA-II
UDA
TL(EY Lab)
GNA (EY Lab)
TJA-I
HHA(EY Lab)
PNA
CnMBL
Epa1-Np mutant Y228W
WT Galectin-8
CA19-9 Ab (DRG)
PsSC
rSRL
DC-ASGPR short
GAC78
BTL native
#14 Sialyl Lewis X (Calbio)
#15 Sialyl Lewis X (BD)
#8 Lewis B (T218) (Abcam)
rNY VLP-Cy3
VLRB.aGPA.23
ME7(detected with a-HUD)
LBL-152
AAL
ME5(detected with a-HUD)
WT-VCC
MOA
VGA(EY Lab)
SJA
SBA
PHA-L
STL
hJAA-F11-B11
ConA
PHA-E
MAA
MAL-I
PFL
CAA
PSA
P11 VP8
LTL
NPA
GNL
LcH
LCA
DSA
DBA
GSL-2
Mono2R-CNL
Mono1-CNL
AIA
ConA
ECL
Human Galectin1
PTL-II
PTL-I
Mouse Galectin1
MPL
WGA
PTA-Gal
PHA-L
PSL
PHA-E
CTB
Threefoil
ACL
HPA
Microvirin
SBA
LSL-150
HHL
MNA-M
BanLec
Epa1-Np mutant Y228A
SRL
HML-Native
GSL-1
TbFBL2
Epa1-Np mutant E227G
GSL-I-B4
rMpL
LEL
AHL
Mono2W-CNL
CA
MGL 6A-H262T
rPVL
PNA
hGal-1-488+BCR
AFCNVH
VVL
HAA
EEL
MGL 416
CSA(EY Lab)
MGL 6B
Mouse NKR-P1C
Gal-3C
PhoSL001
WGA
Galectin-1
UEAII
MAA(Seikagaku)
WGA
WFL
Sp Serotype 14.2
CML1
SSL0-488
Gal-3
Avaren-Fc
CnSep1
TJA-II
Epa1-Np WT
MAL-II
CC1G-10077
Sp Serotype 14.15
Anti-LeC
Epa1-Np mutant E227A
WHA lectin
TbFBL1
AOL (Mahal Lab)
UEA I
RCAI (Vector)
UEA-I
AAL

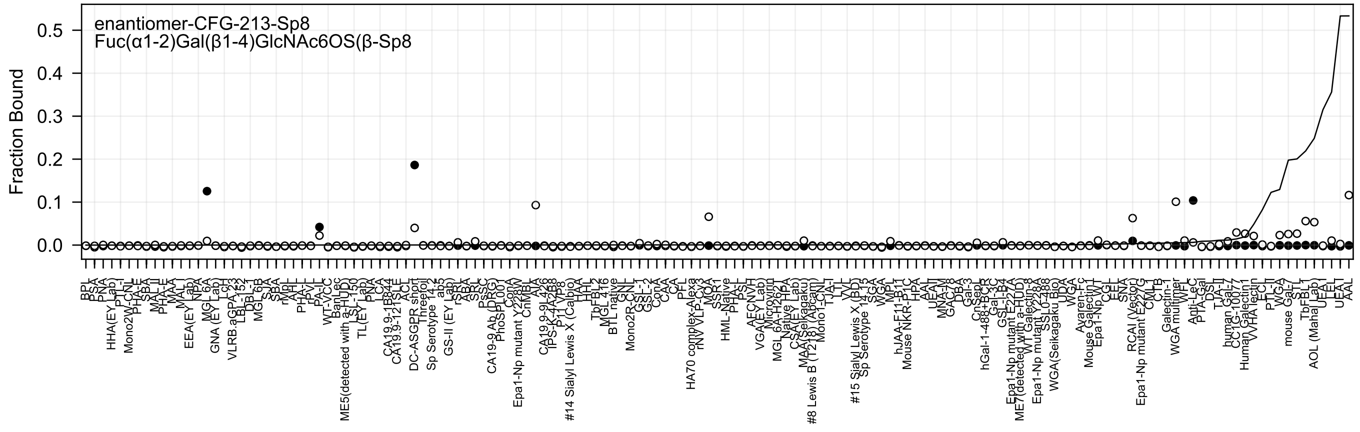
enantiomer-CFG-213-Sp8
Fuc(α1-2)Gal(β1-4)GlcNAc6OS(β-Sp8
Fraction Bound
0.5
0.4
0.3
0.2
0.1
0.0

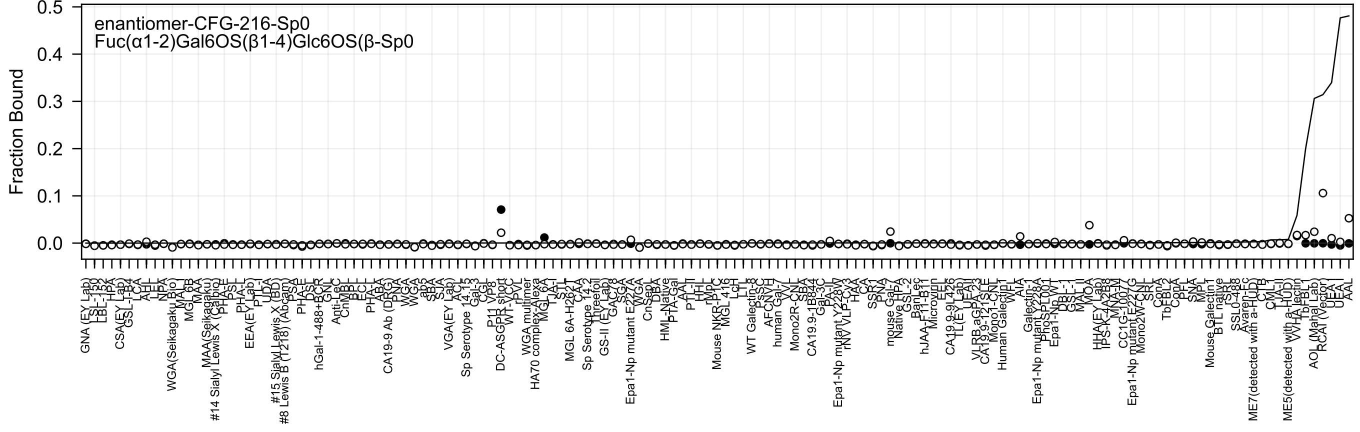

enantiomer-CFG-216-Sp0
Fuc(α1-2)Gal6OS(β1-4)Glc6OS(β-Sp0
Fraction Bound
0.5
0.4
0.3
0.2
0.1
0.0

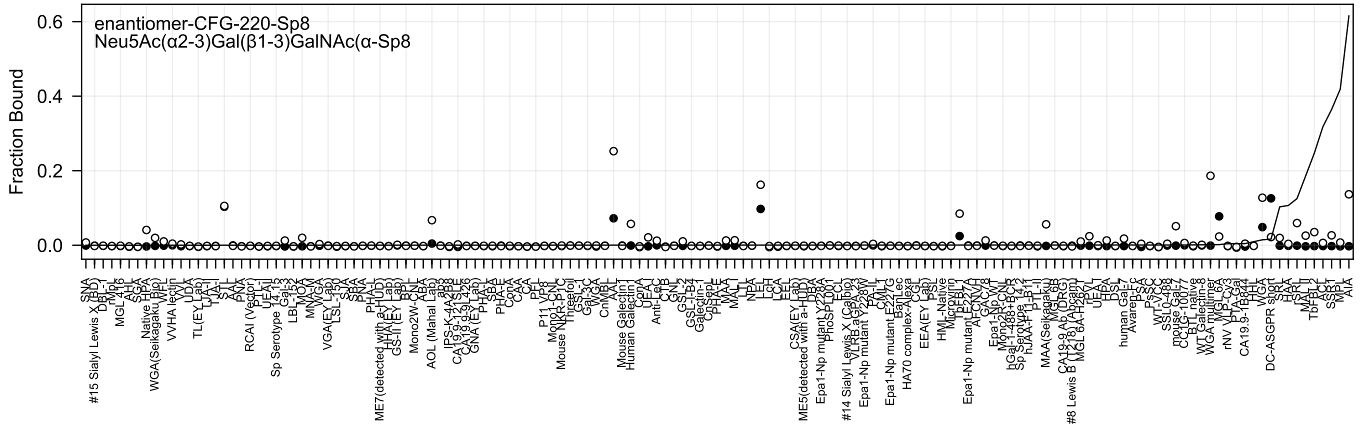
enantiomer-CFG-220-Sp8
Neu5Ac(α2-3)Gal(β1-3)GalNAc(α-Sp8
Fraction Bound
0.0
0.2
0.4
0.6

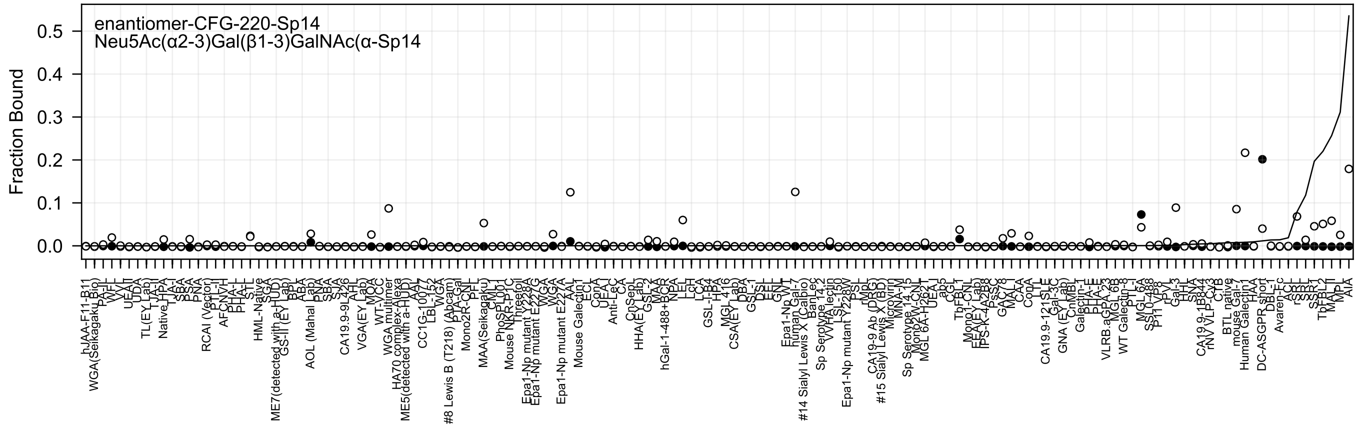

enantiomer-CFG-220-Sp14
Neu5Ac(α2-3)Gal(β1-3)GalNAc(α-Sp14
Fraction Bound
0.5
0.4
0.3
0.2
0.1
0.0

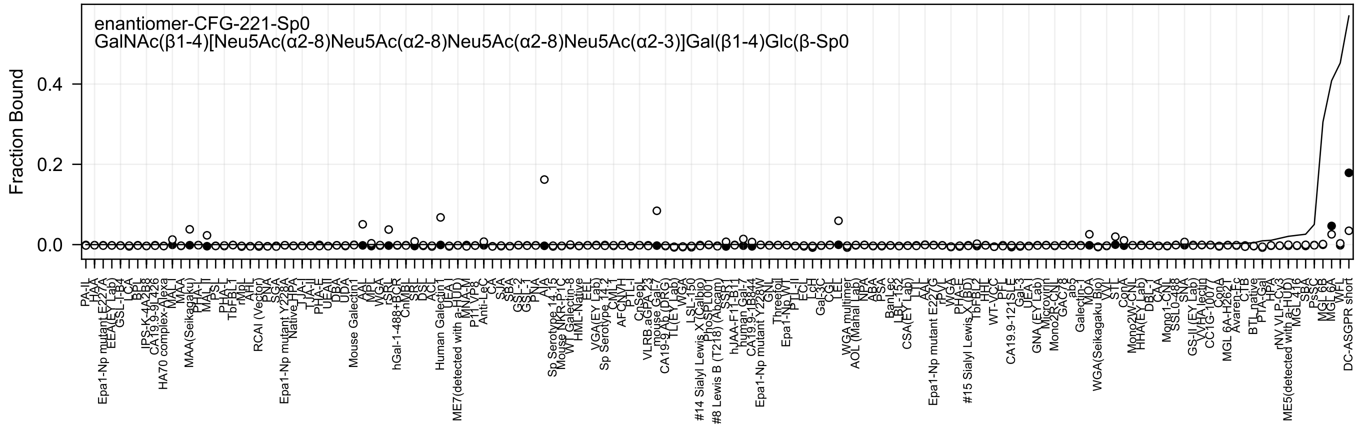

enantiomer-CFG-221-Sp0
GalNAc(β1-4)[Neu5Ac(α2-8)Neu5Ac(α2-8)Neu5Ac(α2-8)Neu5Ac(α2-3)]Gal(β1-4)Glc(β-Sp0
Fraction Bound
0.4
0.2
0.0

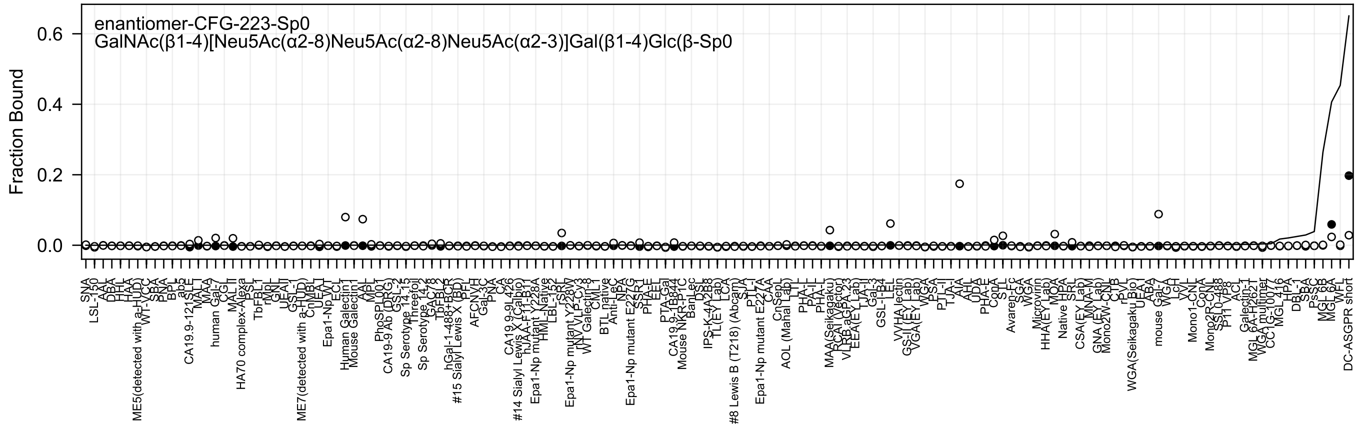

enantiomer-CFG-223-Sp0
GalNAc(β1-4)[Neu5Ac(α2-8)Neu5Ac(α2-8)Neu5Ac(α2-3)]Gal(β1-4)Glc(β-Sp0
Fraction Bound

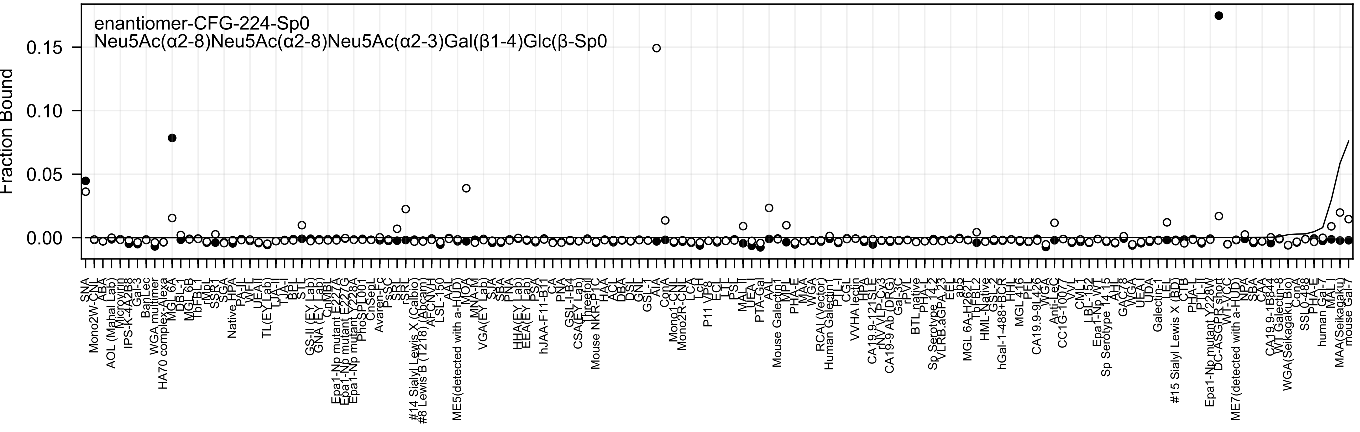
enantiomer-CFG-224-Sp0
Neu5Ac(α2-8)Neu5Ac(α2-8)Neu5Ac(α2-3)Gal(β1-4)Glc(β-Sp0
Fraction Bound
0.15
0.10
0.05
0.00

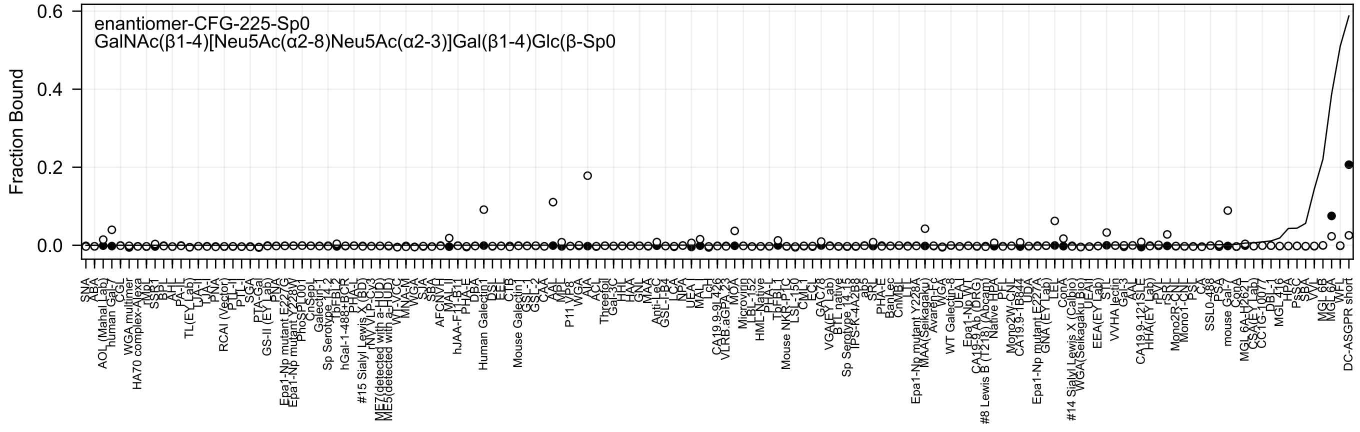
enantiomer-CFG-225-Sp0
GalNAc(β1-4)[Neu5Ac(α2-8)Neu5Ac(α2-3)]Gal(β1-4)Glc(β-Sp0
Fraction Bound
0.0
0.2
0.4
0.6

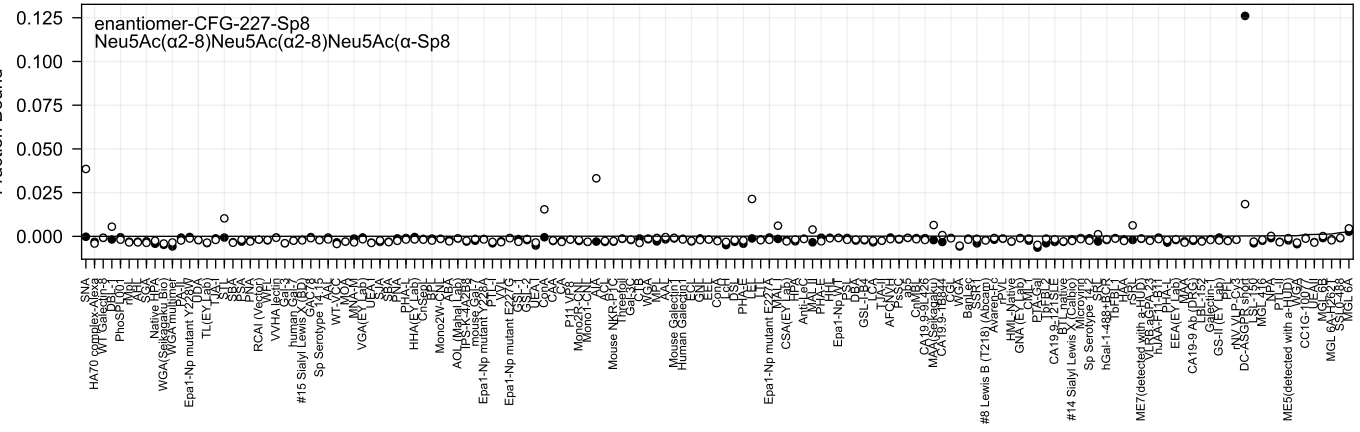

enantiomer-CFG-227-Sp8
Neu5Ac(α2-8)Neu5Ac(α2-8)Neu5Ac(α-Sp8

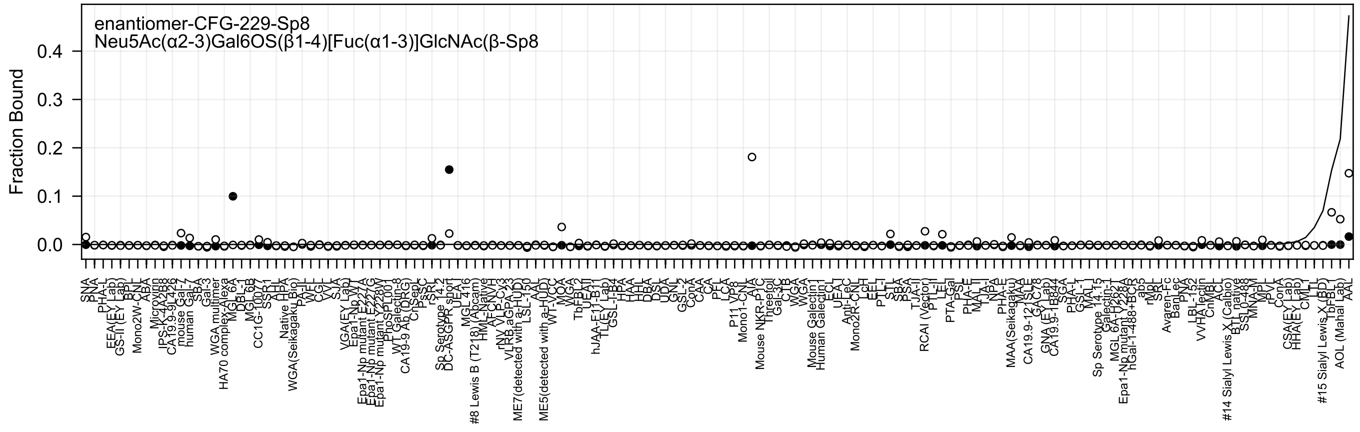

enantiomer-CFG-229-Sp8
Neu5Ac(α2-3)Gal6OS(β1-4)[Fuc(α1-3)]GlcNAc(β-Sp8
Fraction Bound

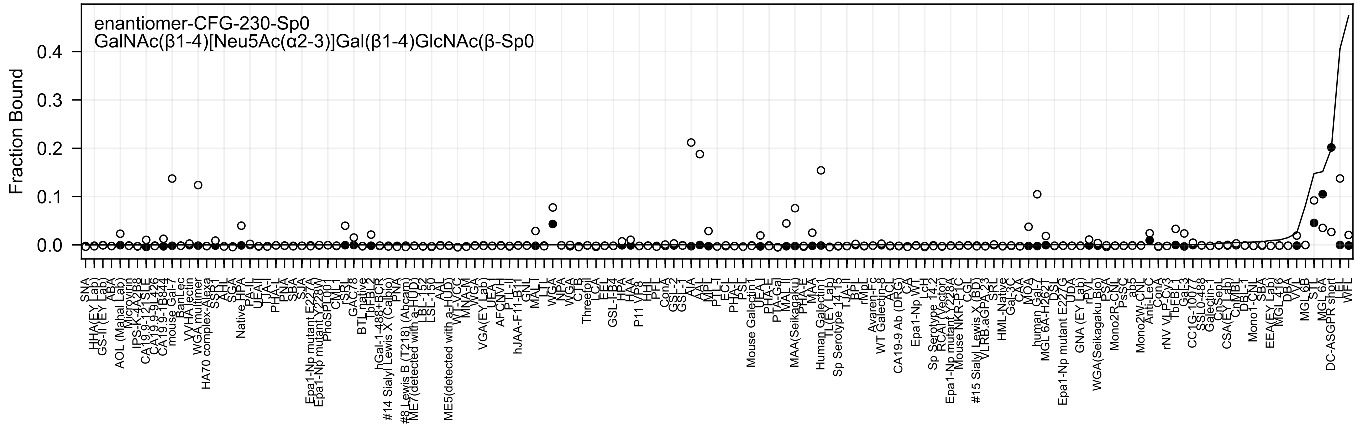
enantiomer-CFG-230-Sp0
GalNAc(β1-4)[Neu5Ac(α2-3)]Gal(β1-4)GlcNAc(β-Sp0
Fraction Bound
0.0
0.1
0.2
0.3
0.4

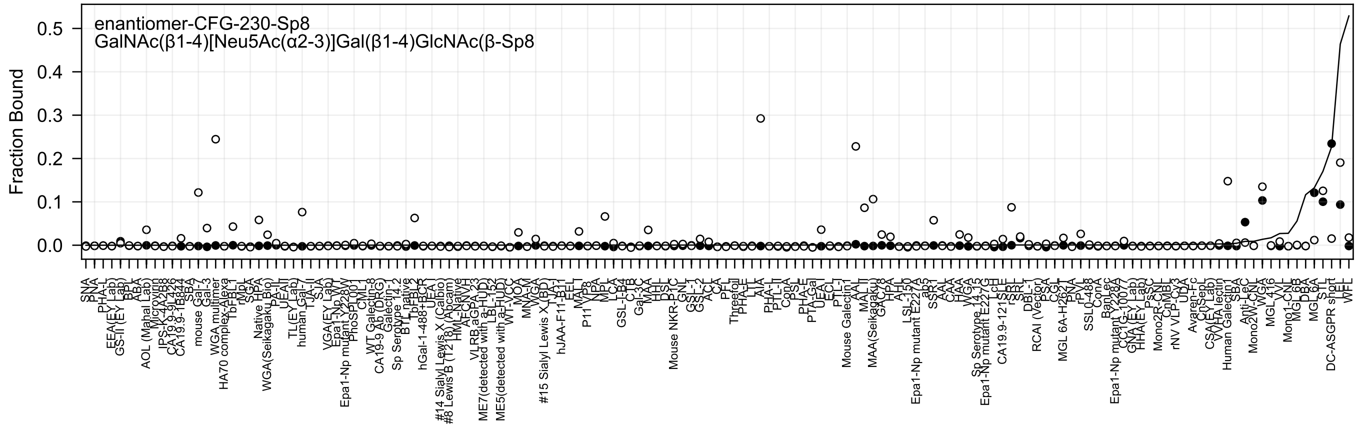
enantiomer-CFG-230-Sp8
GalNAc(β1-4)[Neu5Ac(α2-3)]Gal(β1-4)GlcNAc(β-Sp8
Fraction Bound
0.5
0.4
0.3
0.2
0.1
0.0

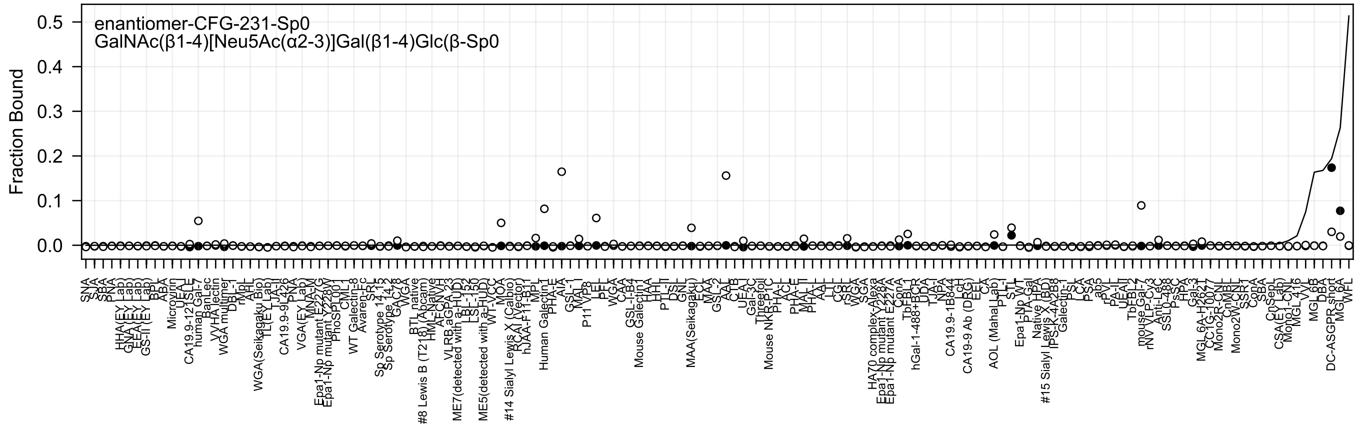
enantiomer-CFG-231-Sp0
GalNAc(β1-4)[Neu5Ac(α2-3)]Gal(β1-4)Glc(β-Sp0
Fraction Bound

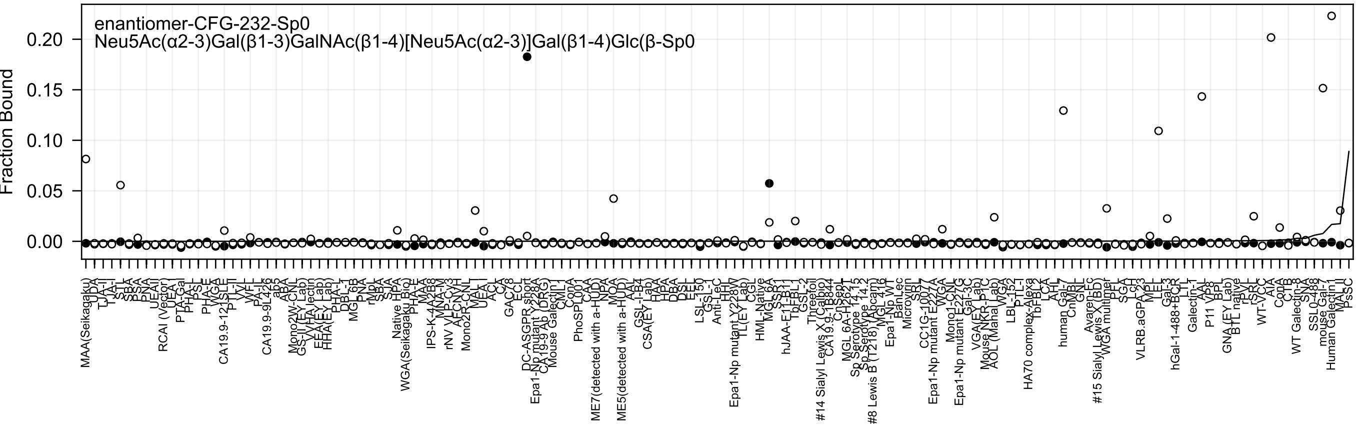
enantiomer-CFG-232-Sp0
Neu5Ac(α2-3)Gal(β1-3)GalNAc(β1-4)[Neu5Ac(α2-3)]Gal(β1-4)Glc(β-Sp0
Fraction Bound
0.00
0.05
0.10
0.15
0.20

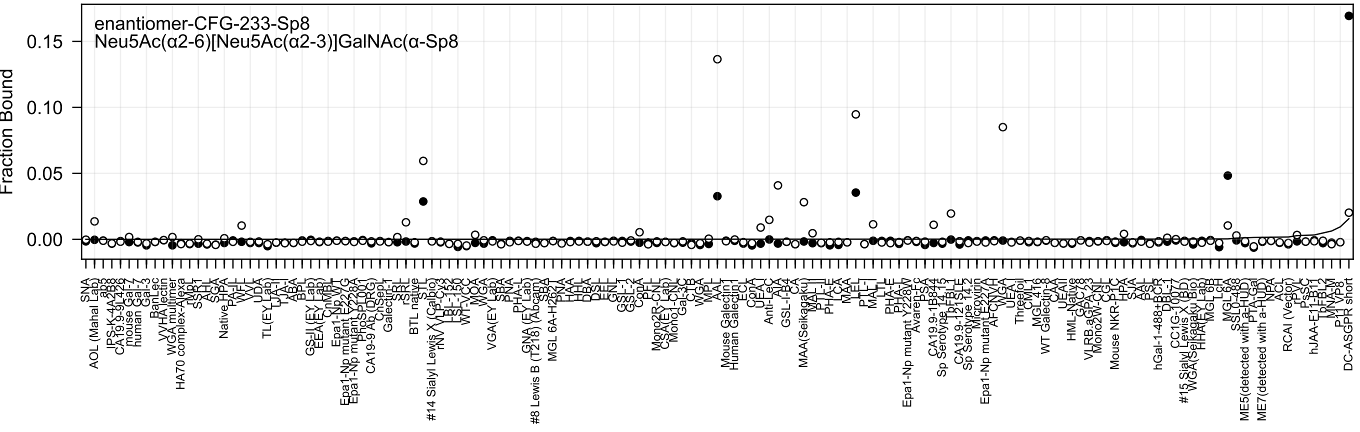
enantiomer-CFG-233-Sp8
Neu5Ac(α2-6)[Neu5Ac(α2-3)]GalNAc(α-Sp8
Fraction Bound
0.15
0.10
0.05
0.00

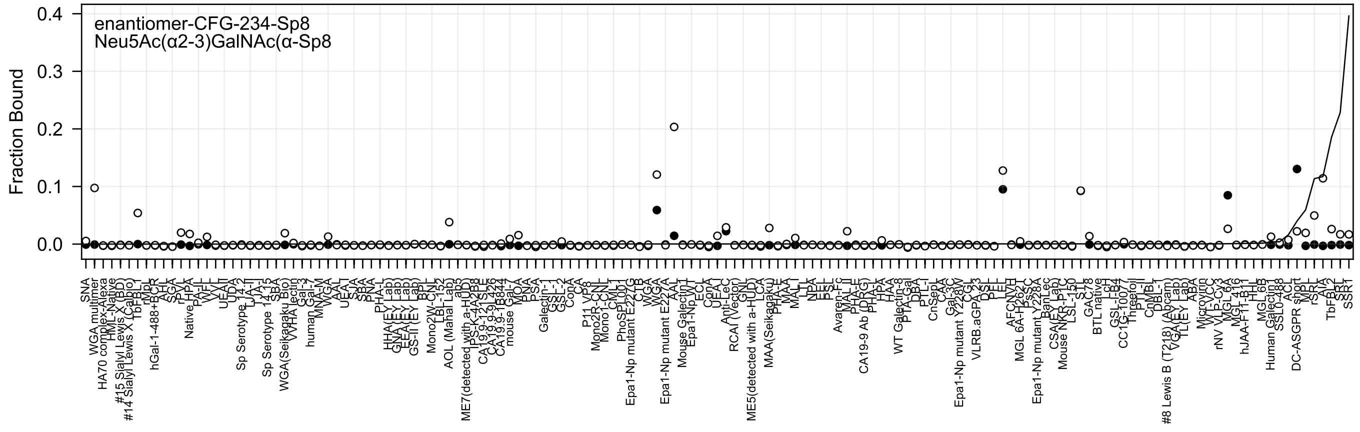
enantiomer-CFG-234-Sp8
Neu5Ac(α2-3)GalNAc(α-Sp8
Fraction Bound
0.4
0.3
0.2
0.1
0.0

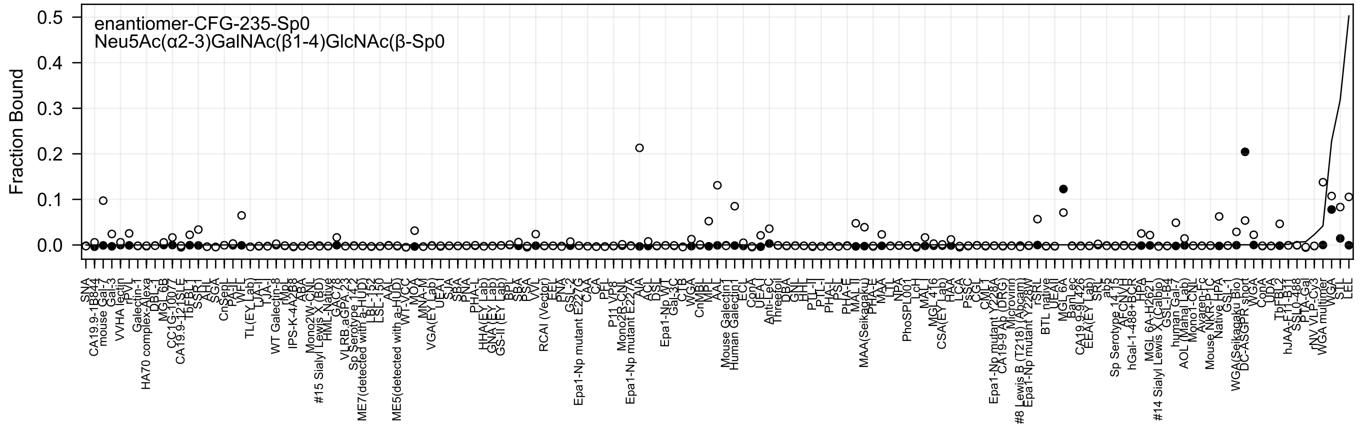
enantiomer-CFG-235-Sp0
Neu5Ac(α2-3)GalNAc(β1-4)GlcNAc(β-Sp0
Fraction Bound
0.0
0.1
0.2
0.3
0.4
0.5

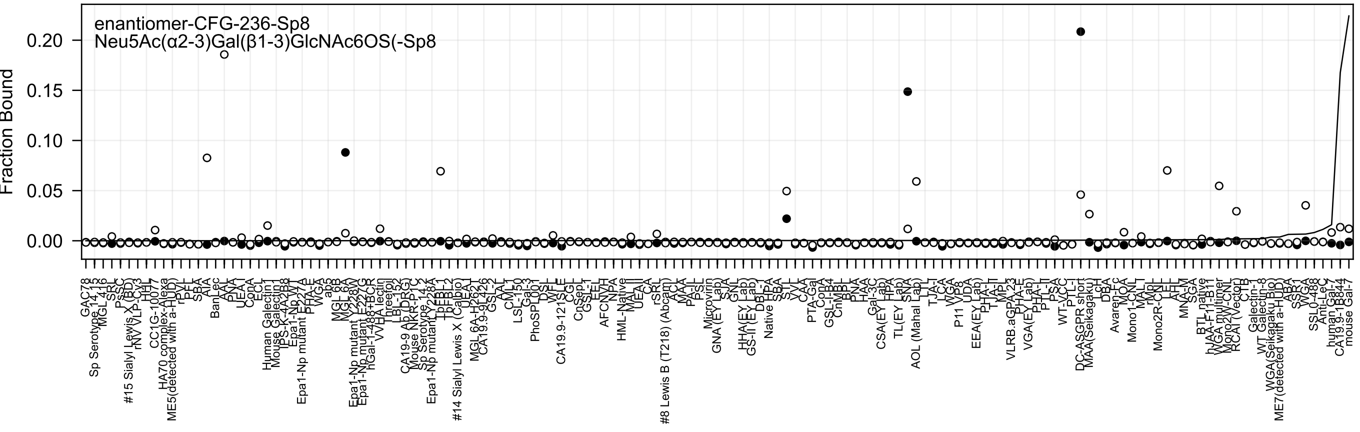
enantiomer-CFG-236-Sp8
Neu5Ac(α2-3)Gal(β1-3)GlcNAc6OS(-Sp8
Fraction Bound
0.00
0.05
0.10
0.15
0.20
GAC78
Sp Serotype 14.13
MGL 416
SRL
PsSC
#15 Sialyl Lewis X (BD)
rNV VLP-Cy3
HHL
CC1G-10077
HA70 complex-Alexa
ME5(detected with a-HuD)
rPVL
PFL
SBA
AIA
BanLec
AAL
PNA
UEA-I
ConA
ECL
Human Galectin1
Mouse Galectin1
IPS-K-4A2B8
Epa1-Np WT
Epa1-Np mutant E227A
PHA-E
WGA
ab5
MGL 6B
MGL 6A
Epa1-Np mutant Y228W
Epa1-Np mutant E227G
hGal-1-488+BCR
VVHA lectin
Threefoil
LBL-152
CA19-9 Ab (DRG)
Mouse NKR-P1C
Sp Serotype 14.2
Epa1-Np mutant Y228A
TbFBL-1
TbFBL-2
#14 Sialyl Lewis X (Calbio)
UEA-I
MGL 6A-H262T
CA19.9-9L426
GSL-2
AAL
CML-I
LSL-150
Gal-3
PhoSL001
DSL
WFL
CA19.9-121SLE
CGL
CnSepL
GSL-1
EEL
AFCNVH
NPA
HML-Native
MAL-II
UEA-II
CA
rSRL
#8 Lewis B (T218) (Abcam)
ACL
MAA
PA-IL
PSL
Microvirin
GNA (EY Lab)
SJA
GNL
HHA(EY Lab)
GS-II (EY Lab)
DBL
Native HPA
SBA
STL
VVL
CAA
PTA-Gal
ConA
GSL-I-B4
CnMBL
BPL
PNA
HAA
Gal-3C
CSA(EY Lab)
HPA
TL(EY Lab)
SNA
AOL (Mahal Lab)
LTL
TJA-I
LCA
WGA
P11 VP8
UDA
EEA(EY Lab)
PHA-L
TJA-II
MPL
VLRB.aGPA.23
PHA-E
VGA(EY Lab)
PHA-L
PTL-II
PSA
WT-VCC
PTL-I
DC-ASGPR short
MAA(Seikagaku)
LcH
DBA
Avaren-Fc
MOA
Mono1-CNL
MAL-I
rMpL
Mono2R-CNL
LEL
AAL
MNA-M
SGA
BTL native
hJAA-F11-B11
WGA multimer
Mono2W-CNL
RCAI (Vector)
CTB
Galectin-1
WT Galectin-8
WGA(Seikagaku Bio)
ME7(detected with a-HuD)
ABA
SSR1
WGA
SSL0-488
Anti-LeC
human Gal-7
CA19.9-1B844
mouse Gal-7

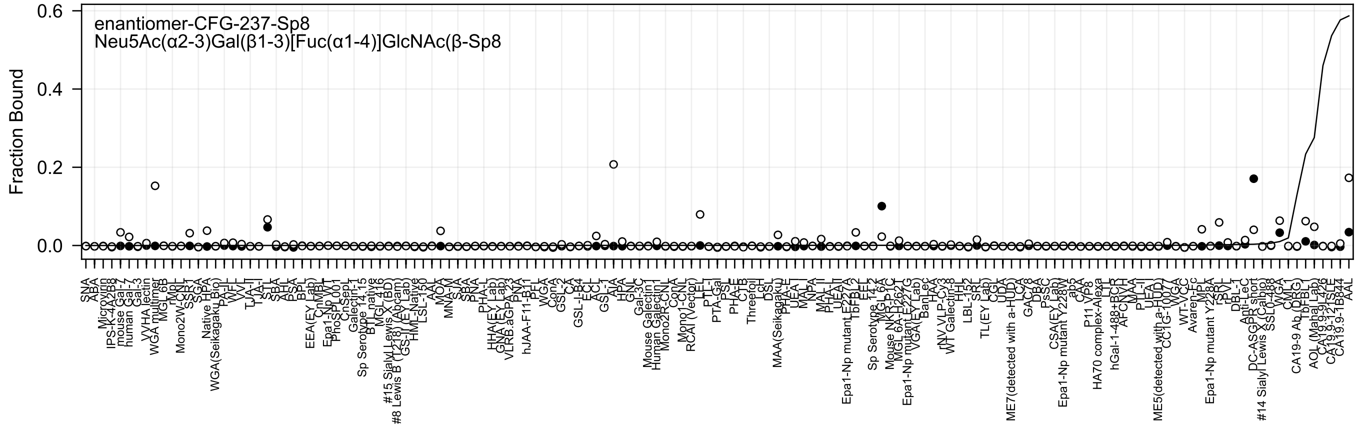

enantiomer-CFG-237-Sp8
Neu5Ac(α2-3)Gal(β1-3)[Fuc(α1-4)]GlcNAc(β-Sp8
Fraction Bound
0.6
0.4
0.2
0.0

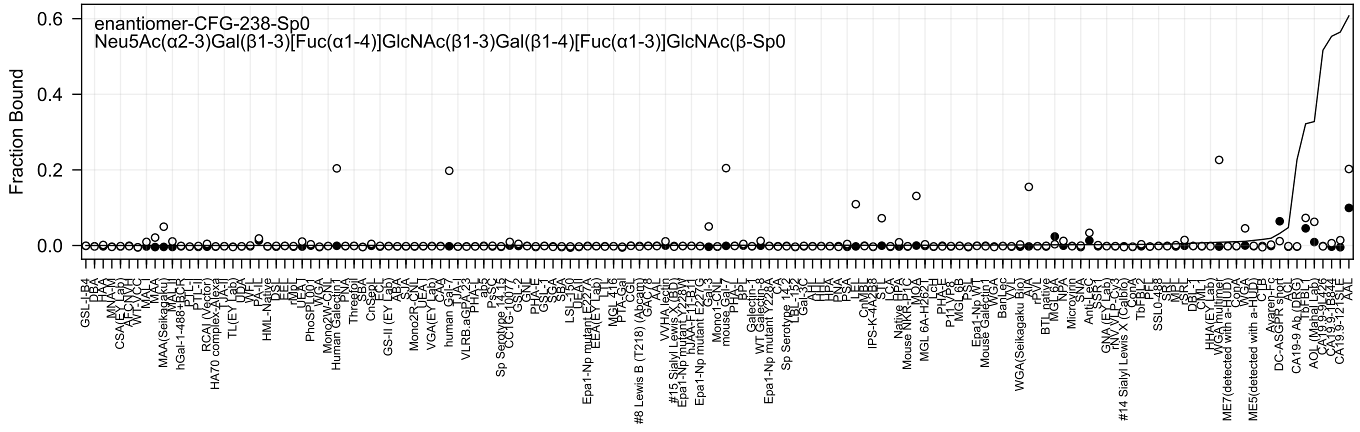
enantiomer-CFG-238-Sp0
Neu5Ac(α2-3)Gal(β1-3)[Fuc(α1-4)]GlcNAc(β1-3)Gal(β1-4)[Fuc(α1-3)]GlcNAc(β-Sp0
Fraction Bound
0.6
0.4
0.2
0.0

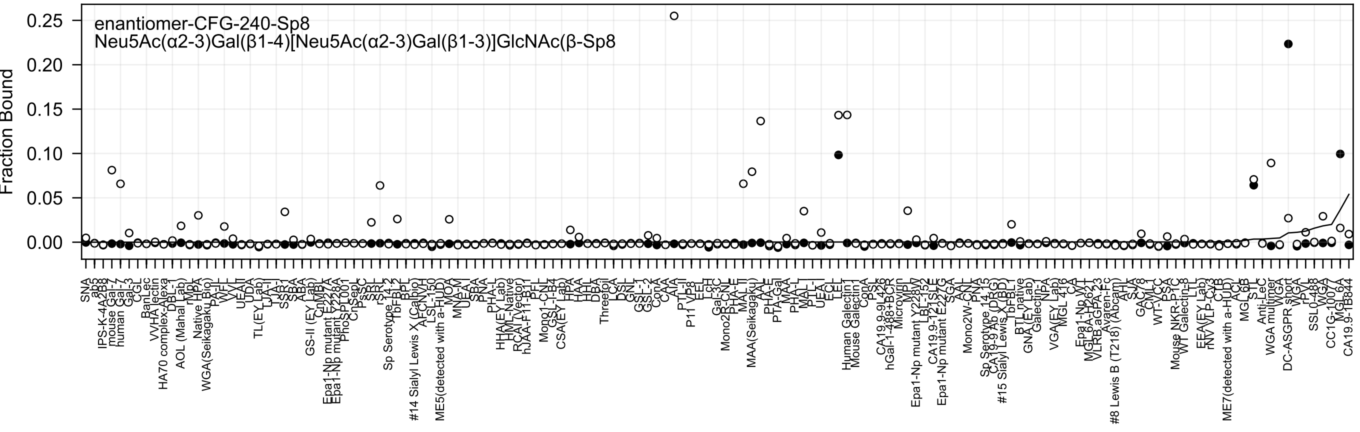
enantiomer-CFG-240-Sp8
Neu5Ac(α2-3)Gal(β1-4)[Neu5Ac(α2-3)Gal(β1-3)]GlcNAc(β-Sp8
Fraction Bound
0.25
0.20
0.15
0.10
0.05
0.00
SNA
ab5
IPS-K-4A2B8
mouse Gal-7
human Gal-3
Gal-3
CGL
BanLec
VVHA lectin
HA70 complex-Alexa
DBL-1
AOL (Mahal Lab)
rMpL
Native HPA
WGA(Seikagaku Bio)
PA-IL
WFL
VVL
UEA-II
UDA
TL(EY Lab)
LTA-I
TJA-I
SSR1
SBA
ABA
GS-II (EY Lab)
CnMBL
Epa1-Np mutant E227A
Epa1-Np mutant Y228A
PhoSL001
CnSeoL
PsSC
SRL
rSRL
Sp Serotype 14-2
TbFBL2
BPL
#14 Sialyl Lewis X (Calbio)
AFCNVH
LSL-150
ME5(detected with a-HUD)
MOA
MNA-M
UEA
SBA
PNA
PHA-L
HHA(EY Lab)
HM-Native
RCAI (Vector)
hJAA-F11-B11
PFL
Mono1-CNL
GSL-I-B4
CSA(EY Lab)
HPA
HAA
HHL
DBA
Threefoil
LCA
DSL
GNL
GSL-1
GSL-2
ConA
CAA
AIA
PTL-II
P11 VP8
EEL
LcH
Gal-3C
Mono2R-CNL
PHA-E
MAL II
MAA(Seikagaku)
AAL
PHA-E
PTA-Gal
MAA
PHA-L
MAL-I
PTL-I
UEA-I
ECL
LEL
Human Galectin1
Mouse Galectin1
ConA
PSL
CA19.9-9L426
hGal-1-488+BCR
Microvirin
MPL
Epa1-Np mutant Y228W
LBL-152
CA19.9-121SLE
Epa1-Np mutant E227G
SGA
AAL
Mono2W-CNL
PNA
Sp Serotype 14-15
CA19-9 Ab (DRG)
#15 Sialyl Lewis X (BD)
TbFBL1
BTL native
GNA (EY Lab)
Galectin-1
NPA
VGA(EY Lab)
MGL 41G
CA
Epa1-Np WT
MGL 6A-H262T
VLRB.aGPA.23
Avaren-Fc
#8 Lewis B (T218) (Abcam)
AIL
SJA
GAC78
CML1
WT-VCC
PSA
Mouse NKR-P1C
WT Galectin-8
LTL
EEA(EY Lab)
rNV VLP-Cv3
CTB
ME7(detected with a-HUD)
ACL
MGL 6B
STL
Anti-LeC
WGA multimer
WGA
DC-ASGPR short
WGA
rPVL
SSL0-488
WGA
CC1G-1007A
MGL 6A
CA19.9-1B844

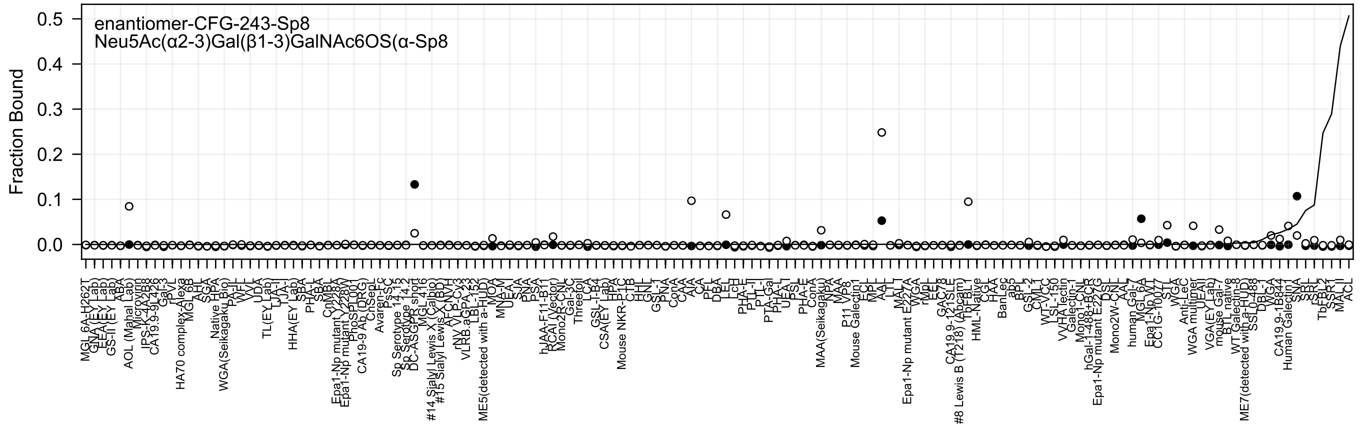

enantiomer-CFG-243-Sp8
Neu5Ac(α2-3)Gal(β1-3)GalNAc6OS(α-Sp8
Fraction Bound

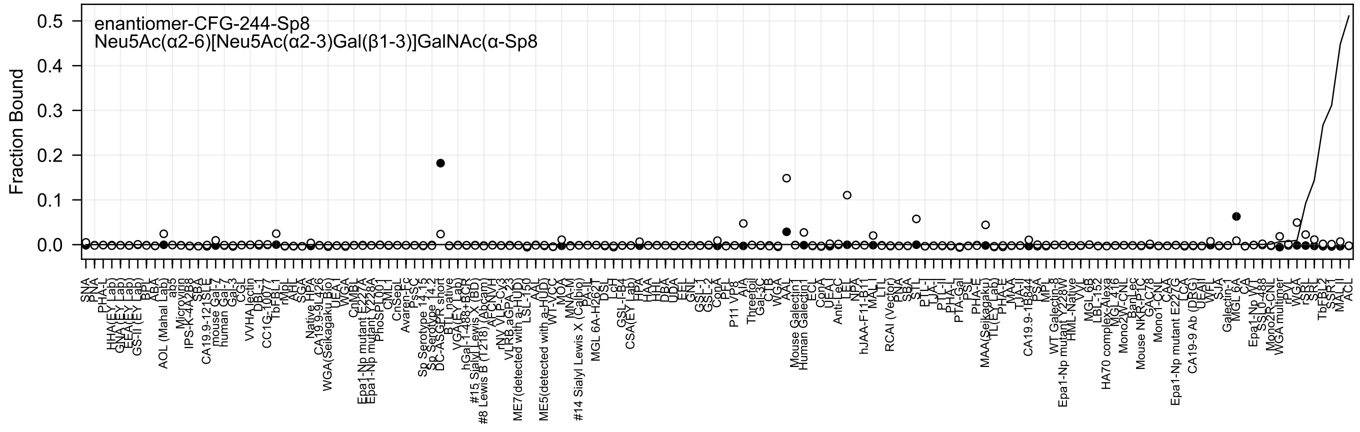
enantiomer-CFG-244-Sp8
Neu5Ac(α2-6)[Neu5Ac(α2-3)Gal(β1-3)]GalNAc(α-Sp8
Fraction Bound
0.5
0.4
0.3
0.2
0.1
0.0

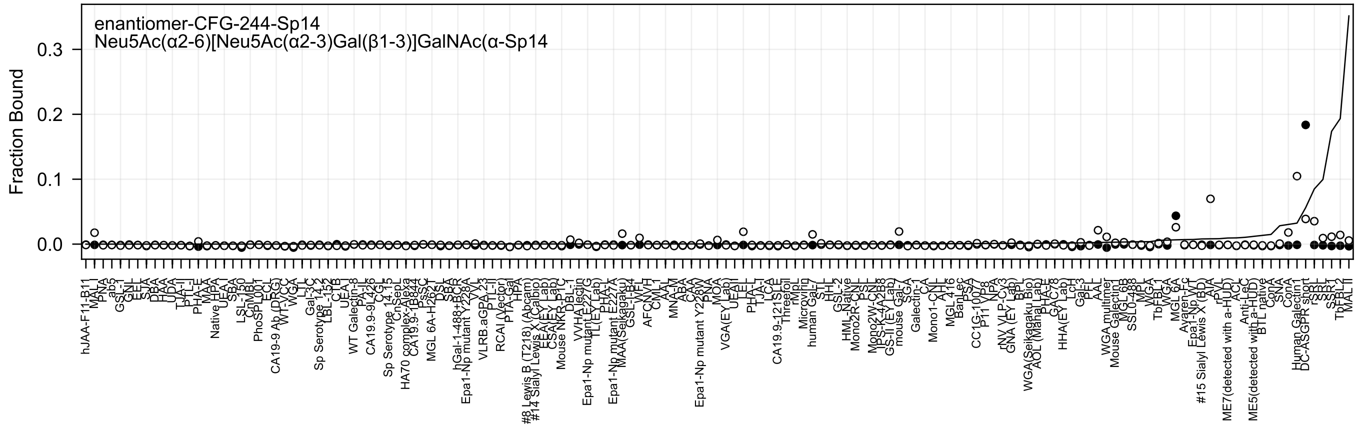
enantiomer-CFG-244-Sp14
Neu5Ac(α2-6)[Neu5Ac(α2-3)Gal(β1-3)]GalNAc(α-Sp14
Fraction Bound
0.3
0.2
0.1
0.0

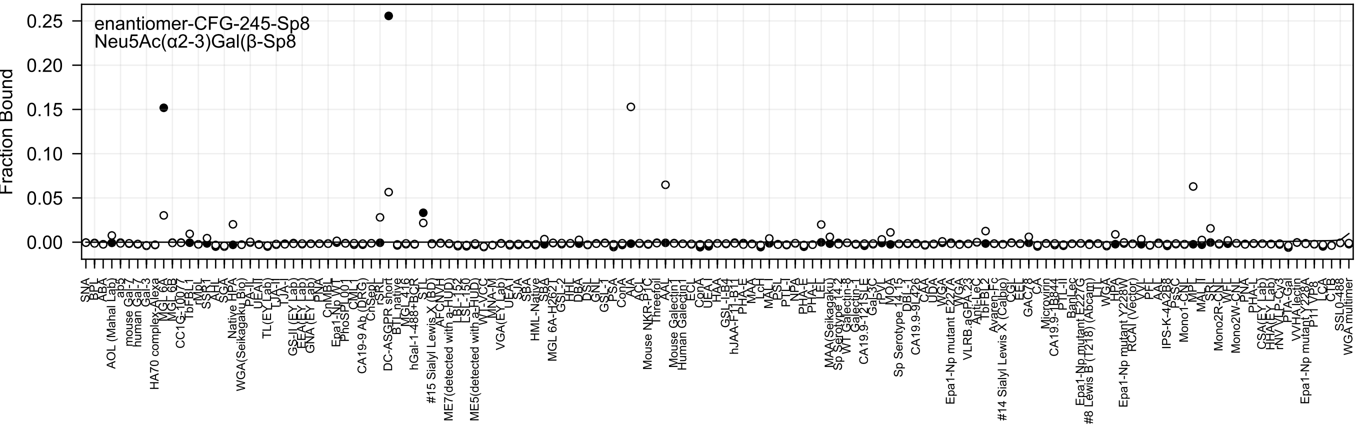
enantiomer-CFG-245-Sp8
Neu5Ac(α2-3)Gal(β-Sp8
Fraction Bound
0.25
0.20
0.15
0.10
0.05
0.00

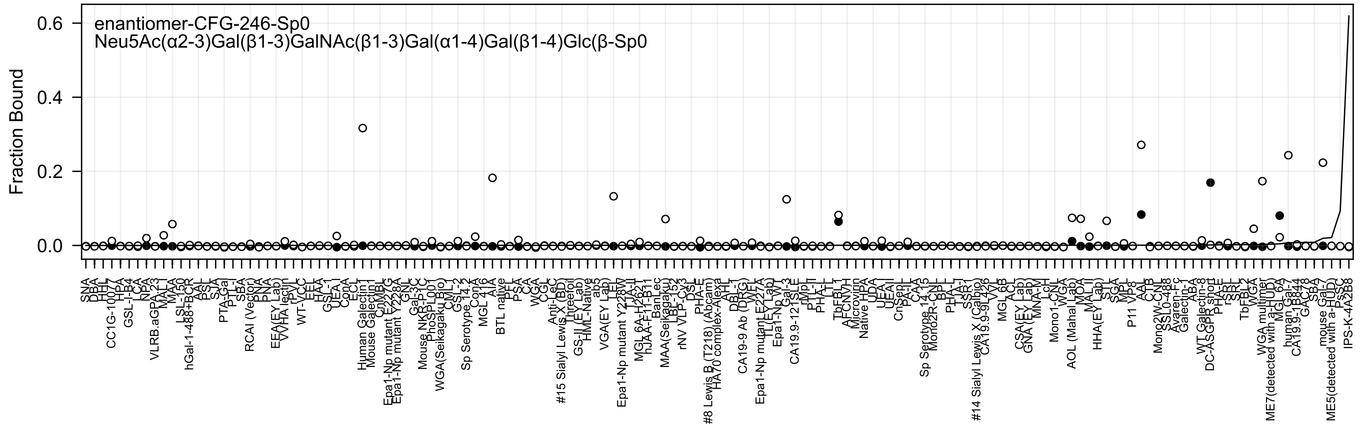
enantiomer-CFG-246-Sp0
Neu5Ac(α2-3)Gal(β1-3)GalNAc(β1-3)Gal(α1-4)Gal(β1-4)Glc(β-Sp0
Fraction Bound
0.6
0.4
0.2
0.0
SNA
DBA
HHL
CC1G-10077
HPA
GSL-I-B4
LCA
NPA
VLRB.aGPA.23
MAL-I
MAA
LSL-150
hGal-1-488+BCR
AAL
PSL
SJA
PTA-Gal
PTL-I
SBA
RCAI (Vector)
PNA
PNA
EEA(EY Lab)
WVHA lectin
rPVL
WT-VCC
EEL
HAA
GSL-I
UEA-I
ConA
ECL
Human Galectin1
Mouse Galectin1
CnMBL
Epa1-Np mutant E227G
Epa1-Np mutant Y228A
GNL
Gal-3C
Mouse NKR-P1C
PhoSL001
WGA(Seikagaku Bio)
CML-I
GSL-2
Sp Serotype 14.2
ConA
MGL 416
AIA
BTL native
PFL
PSA
CA
WGA
CGL
Anti-LeC
#15 Sialyl Lewis X (BD)
Threerol
GS-II (EY Lab)
HML-Native
ab5
VGA(EY Lab)
LEL
Epa1-Np mutant Y228W
TJA-II
MGL 6A-H262T
hJAA-F11-B11
BanLec
MAA(Seikagaku)
LBL-152
rNV VLP-Cy3
DSL
PHA-E
#8 Lewis B (T218) (Abcam)
HA70 complex-Alexa
AHL
DBL-1
CA19-9 Ab (DRG)
WFL
Epa1-Np mutant E227A
TL(EY Lab)
Epa1-Np WT
Gal-3
CA19.9-121SLE
rMpL
PTL-II
PHA-L
LTL
TbFBL1
AFCNVH
Microvirin
Native HPA
UDA
UEA-I
UEAII
CnSep
PA-IL
CAA
Sp Serotype 14.15
Mono2R-CN
BPL
PHA-L
TJA-I
SSR1
#14 Sialyl Lewis X (Calbio)
CA19.9-9L426
VVL
MGL 6B
ACL
CSA(EY Lab)
GNA (EY Lab)
MNA-M
LcH
Mono1-CN
WGA
AOL (Mahal Lab)
MOA
MAL-II
HHA(EY Lab)
STL
SGA
MPL
P11 VP8
AAL
CTB
Mono2W-CN
SSL0-488
Avaren-Fc
Galectin-1
ABA
WT Galectin-8
DC-ASGPR short
PHA-E
rSRL
SRL
TbFBL2
WGA
WGA multimer
ME7(detected with a-HUD)
MGL 6A
human Gal-7
CA19.9-1B844
GAC78
SBA
mouse Gal-7
ME5(detected with a-HUD)
PSSC
IPS-K-4A2B8

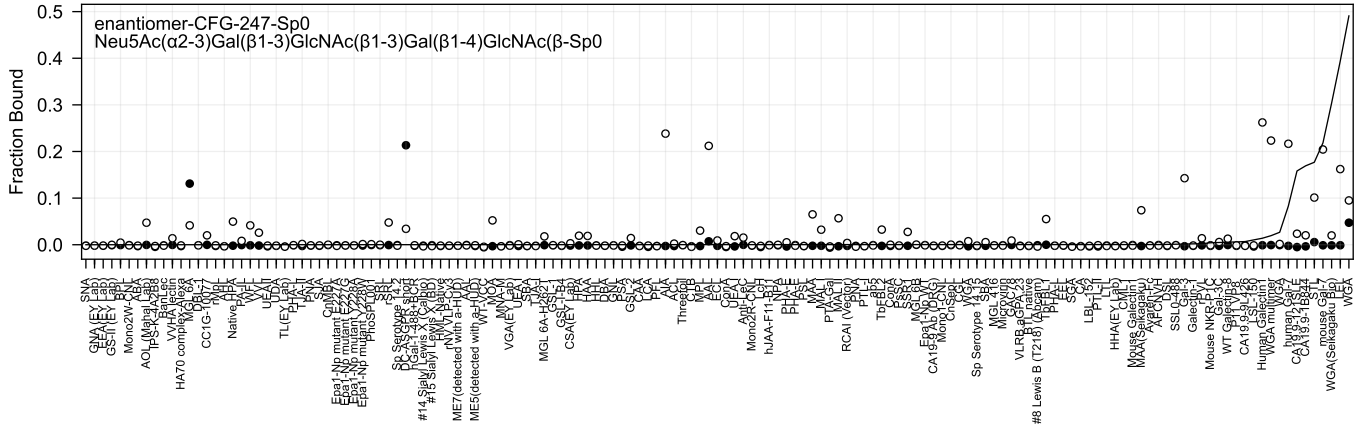

enantiomer-CFG-247-Sp0
Neu5Ac(α2-3)Gal(β1-3)GlcNAc(β1-3)Gal(β1-4)GlcNAc(β-Sp0
Fraction Bound

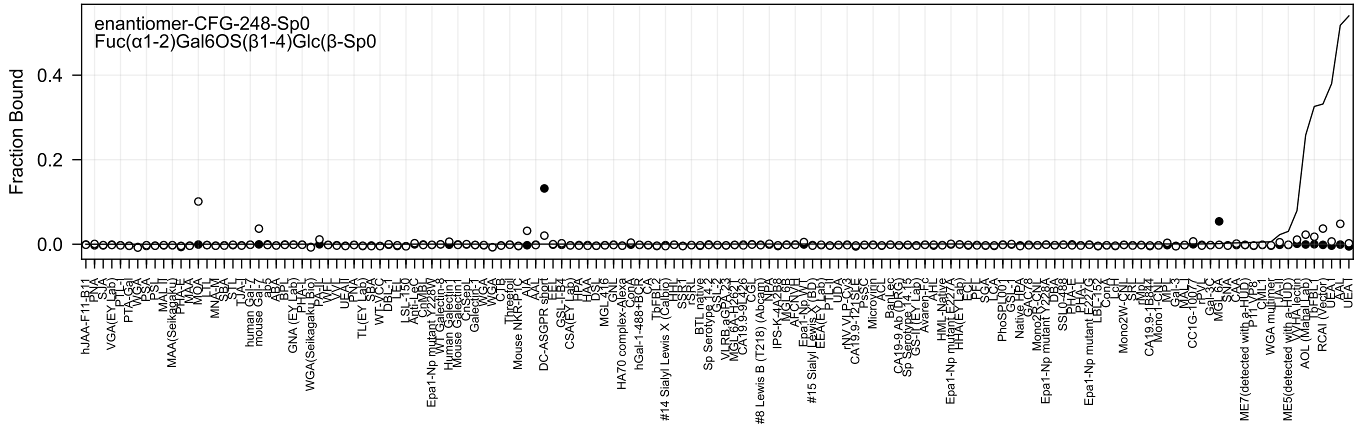
enantiomer-CFG-248-Sp0
Fuc(α1-2)Gal6OS(β1-4)Glc(β-Sp0
Fraction Bound
0.4
0.2
0.0

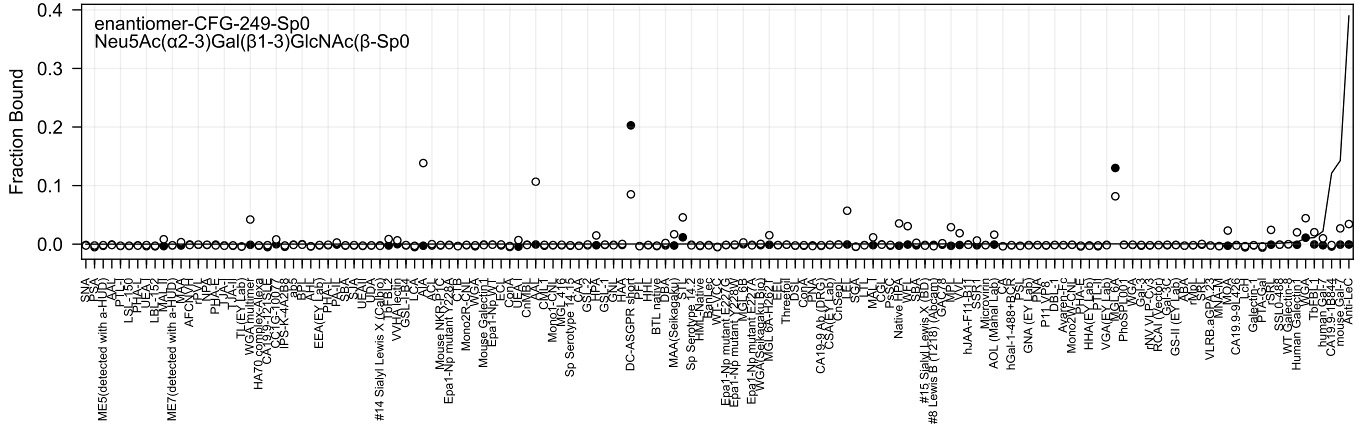

enantiomer-CFG-249-Sp0
Neu5Ac(α2-3)Gal(β1-3)GlcNAc(β-Sp0
Fraction Bound

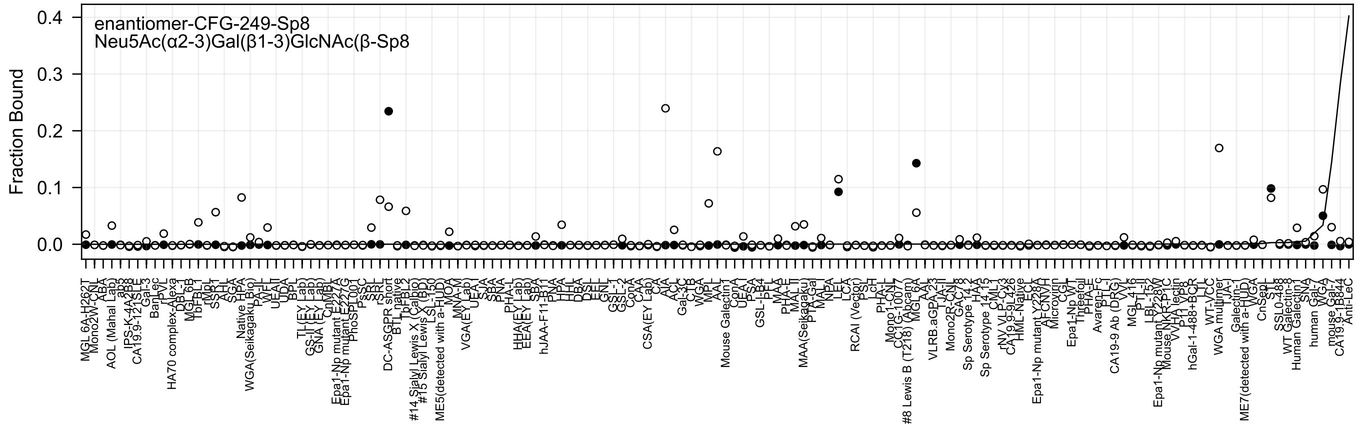
enantiomer-CFG-249-Sp8
Neu5Ac(α2-3)Gal(β1-3)GlcNAc(β-Sp8
Fraction Bound
0.4
0.3
0.2
0.1
0.0

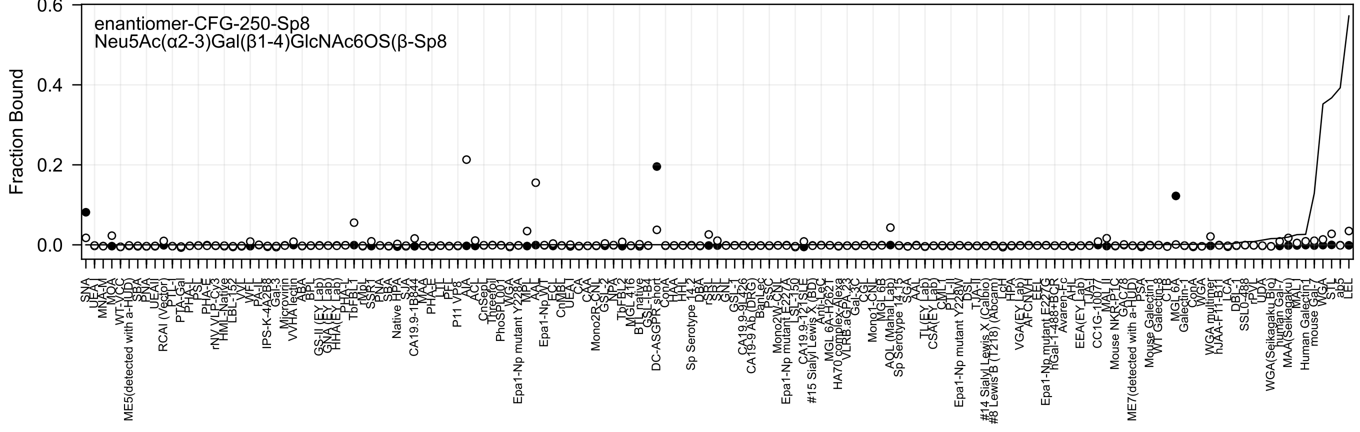
enantiomer-CFG-250-Sp8
Neu5Ac(α2-3)Gal(β1-4)GlcNAc6OS(β-Sp8
Fraction Bound
0.6
0.4
0.2
0.0

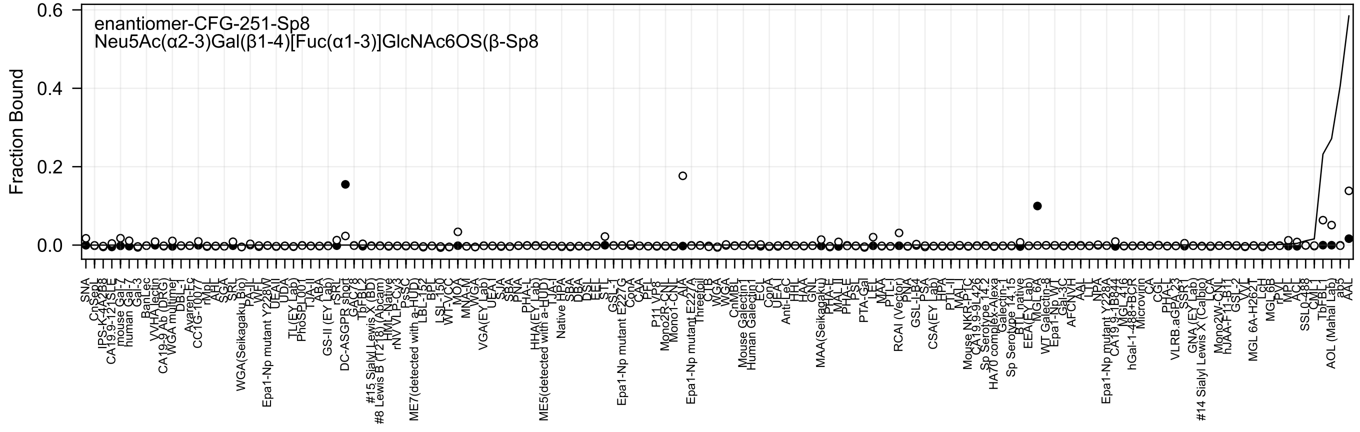

enantiomer-CFG-251-Sp8
Neu5Ac(α2-3)Gal(β1-4)[Fuc(α1-3)]GlcNAc6OS(β-Sp8
Fraction Bound
0.6
0.4
0.2
0.0

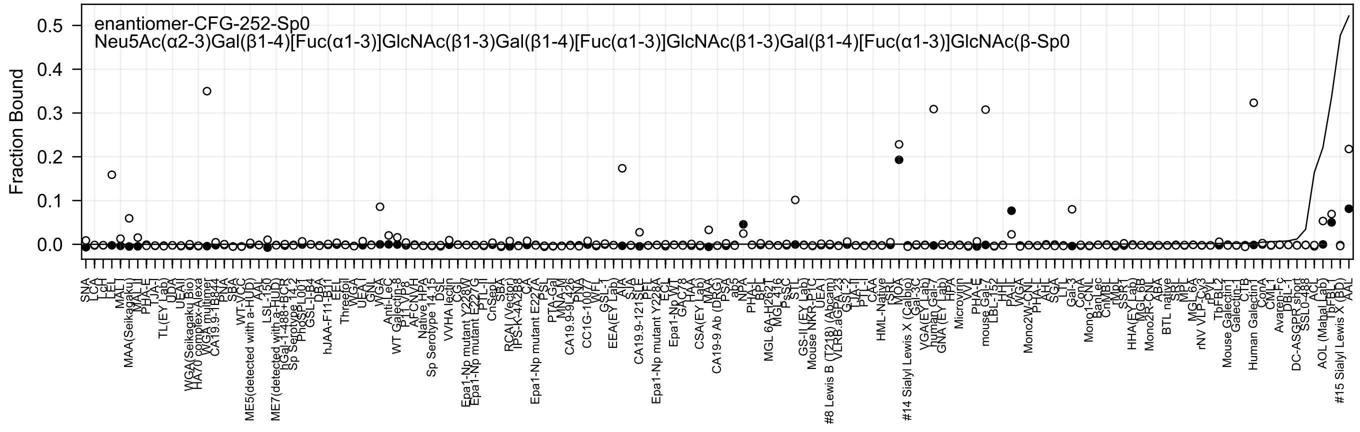
enantiomer-CFG-252-Sp0
Neu5Ac(α2-3)Gal(β1-4)[Fuc(α1-3)]GlcNAc(β1-3)Gal(β1-4)[Fuc(α1-3)]GlcNAc(β1-3)Gal(β1-4)[Fuc(α1-3)]GlcNAc(β-Sp0
Fraction Bound
0.5
0.4
0.3
0.2
0.1
0.0

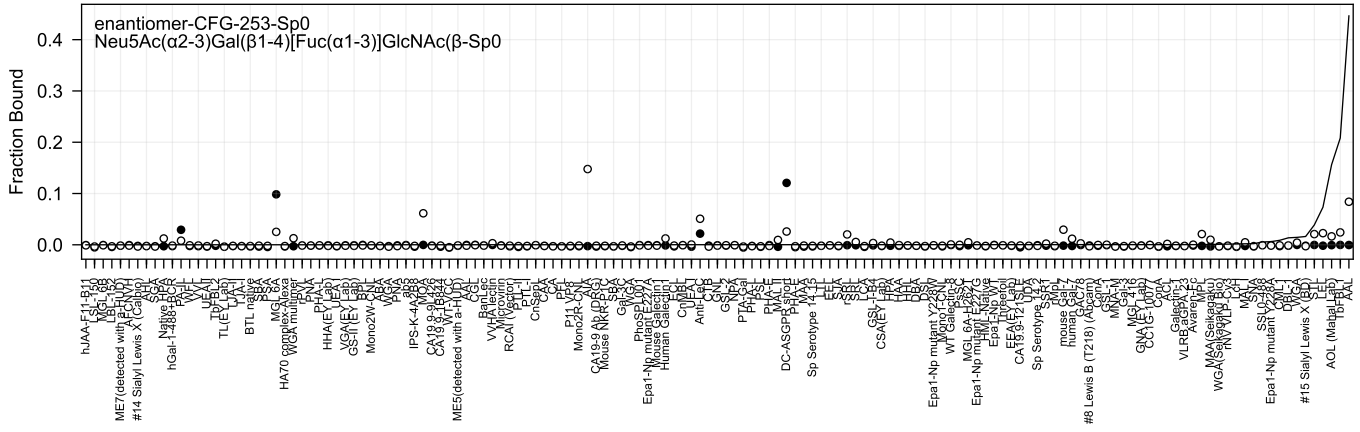
enantiomer-CFG-253-Sp0
Neu5Ac(α2-3)Gal(β1-4)[Fuc(α1-3)]GlcNAc(β-Sp0
Fraction Bound
0.4
0.3
0.2
0.1
0.0

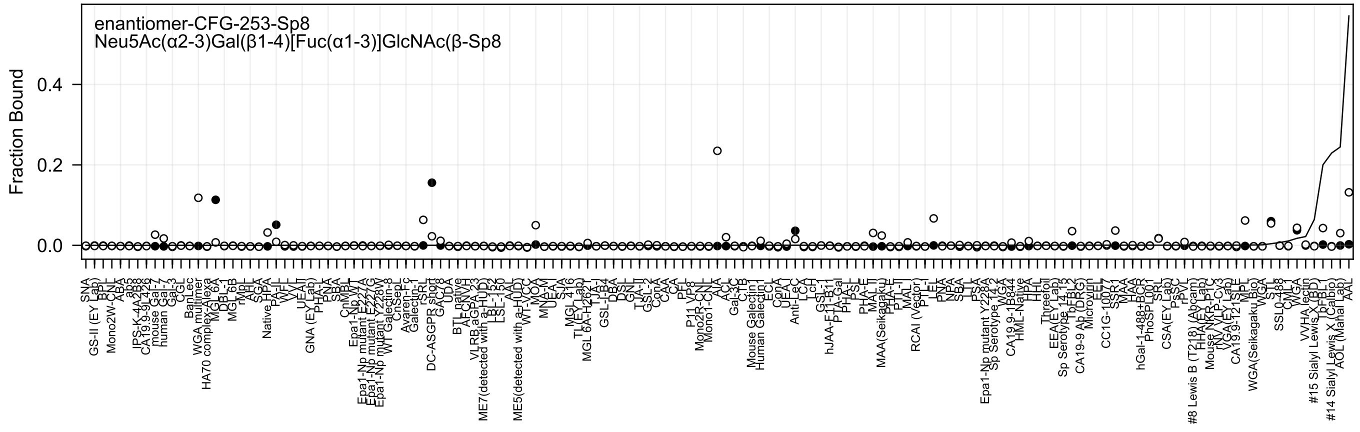
enantiomer-CFG-253-Sp8
Neu5Ac(α2-3)Gal(β1-4)[Fuc(α1-3)]GlcNAc(β-Sp8
Fraction Bound
0.0
0.2
0.4

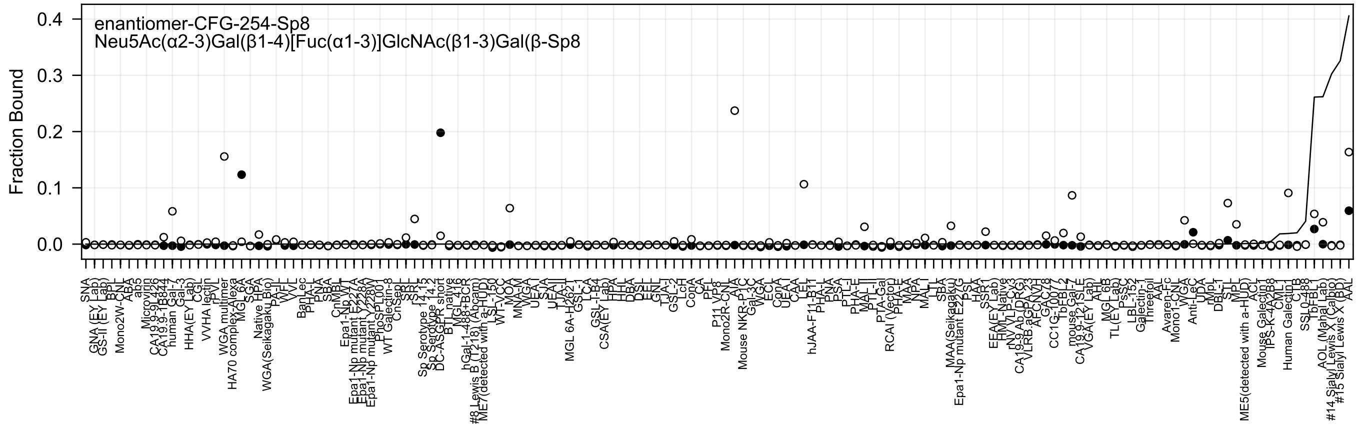

enantiomer-CFG-254-Sp8
Neu5Ac(α2-3)Gal(β1-4)[Fuc(α1-3)]GlcNAc(β1-3)Gal(β-Sp8
Fraction Bound
0.4
0.3
0.2
0.1
0.0

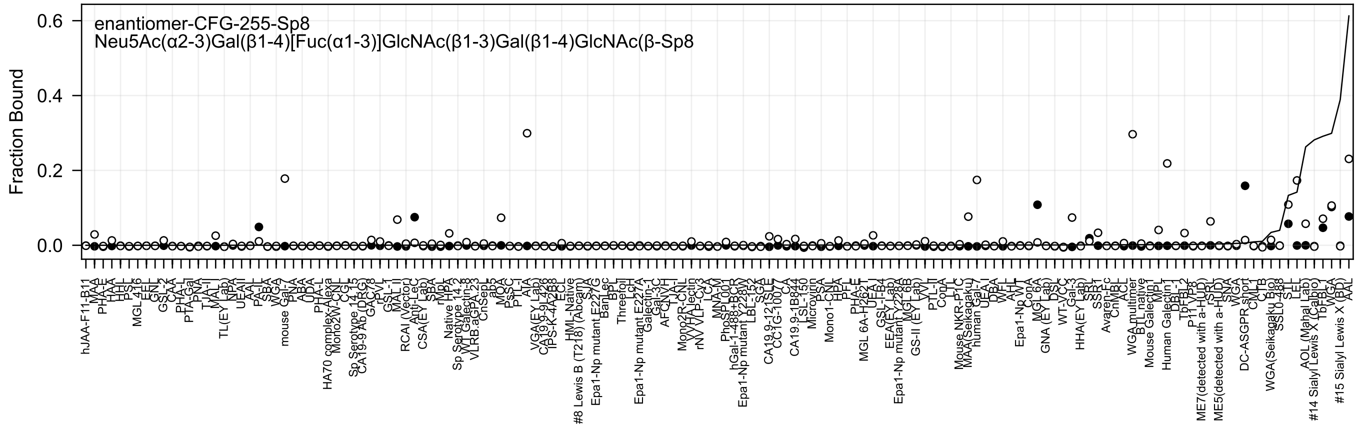
enantiomer-CFG-255-Sp8
Neu5Ac(α2-3)Gal(β1-4)[Fuc(α1-3)]GlcNAc(β1-3)Gal(β1-4)GlcNAc(β-Sp8
Fraction Bound
0.6
0.4
0.2
0.0

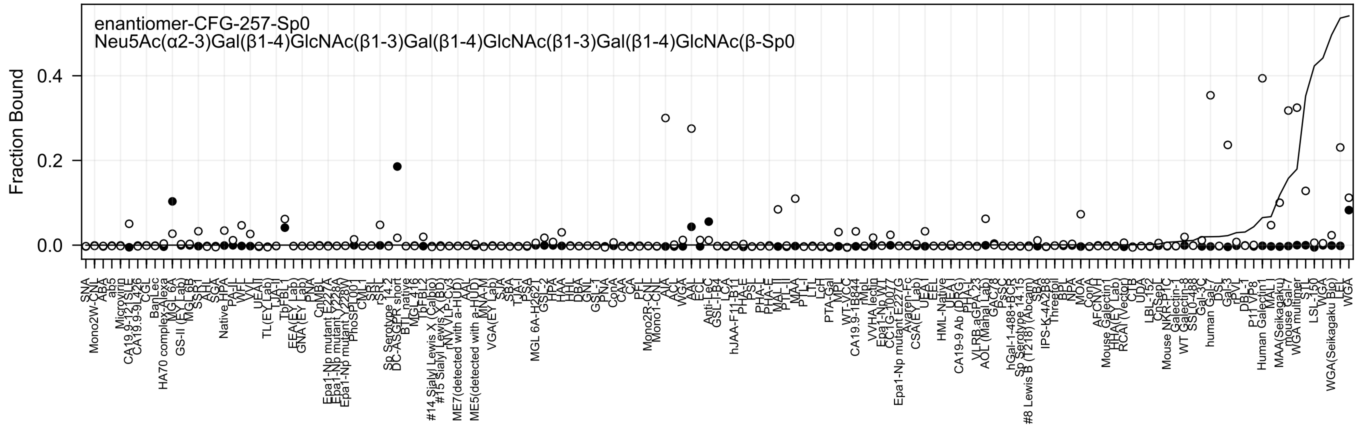

enantiomer-CFG-257-Sp0
Neu5Ac(α2-3)Gal(β1-4)GlcNAc(β1-3)Gal(β1-4)GlcNAc(β1-3)Gal(β1-4)GlcNAc(β-Sp0
Fraction Bound
0.0
0.2
0.4

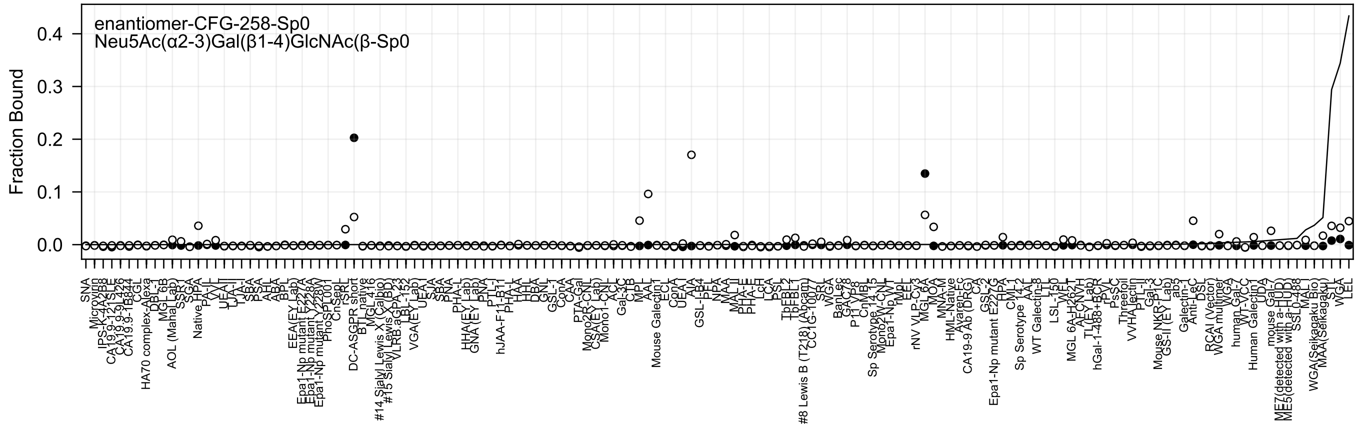
enantiomer-CFG-258-Sp0
Neu5Ac(α2-3)Gal(β1-4)GlcNAc(β-Sp0
Fraction Bound
0.4
0.3
0.2
0.1
0.0

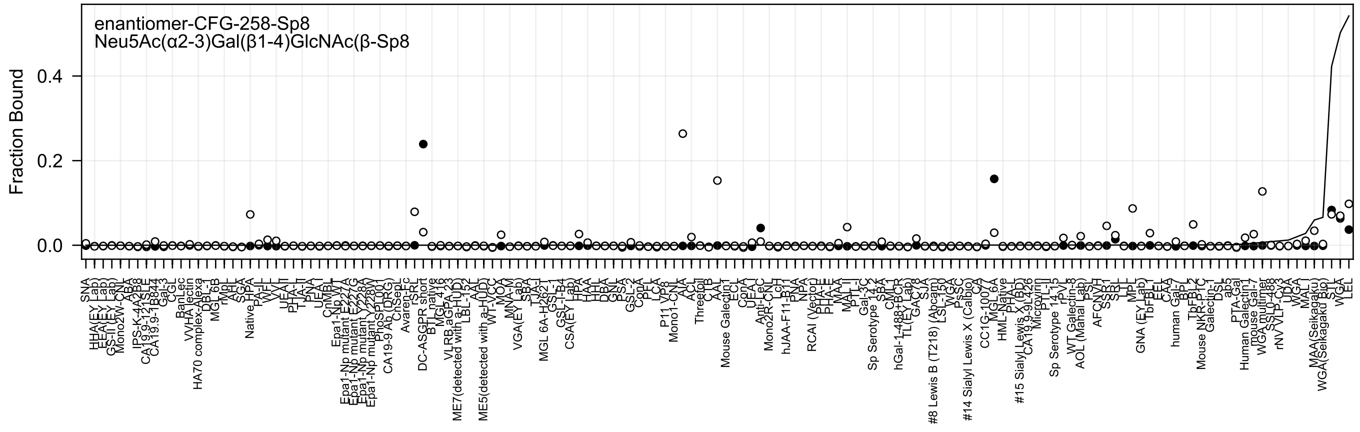
enantiomer-CFG-258-Sp8
Neu5Ac(α2-3)Gal(β1-4)GlcNAc(β-Sp8
Fraction Bound
0.0
0.2
0.4

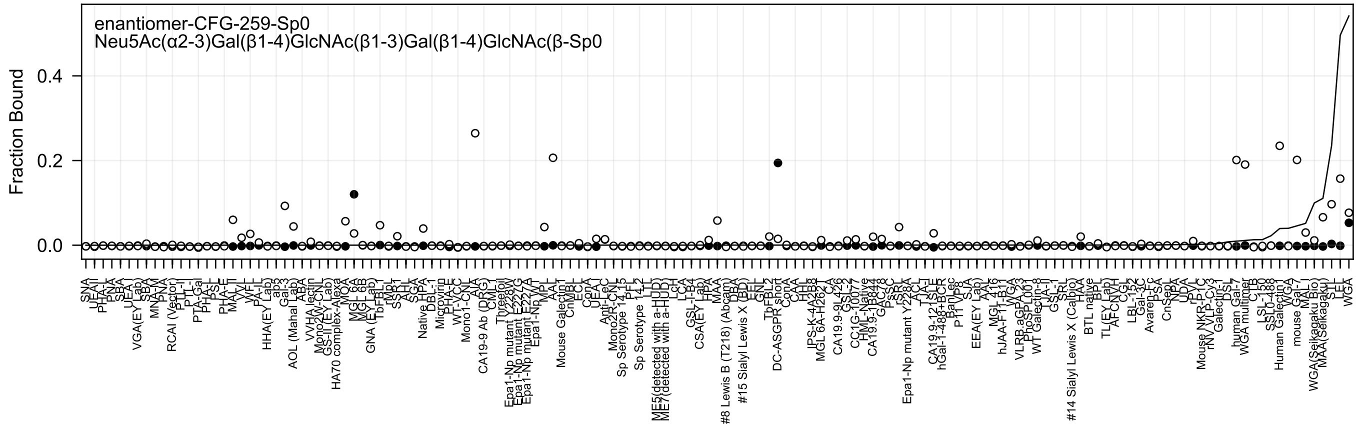
enantiomer-CFG-259-Sp0
Neu5Ac(α2-3)Gal(β1-4)GlcNAc(β1-3)Gal(β1-4)GlcNAc(β-Sp0
Fraction Bound
0.0
0.2
0.4

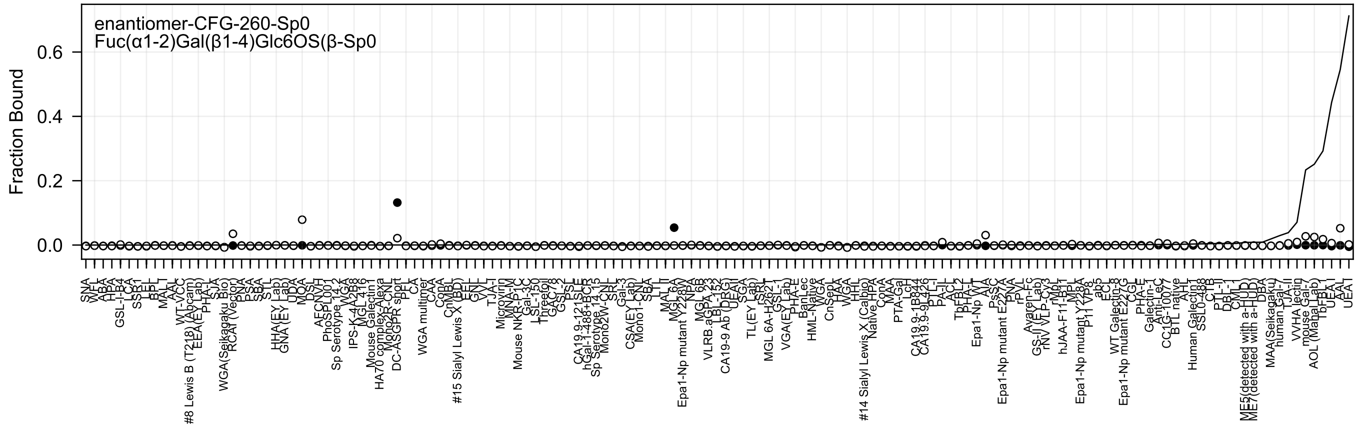
enantiomer-CFG-260-Sp0
Fuc(α1-2)Gal(β1-4)Glc6OS(β-Sp0
Fraction Bound
0.6
0.4
0.2
0.0

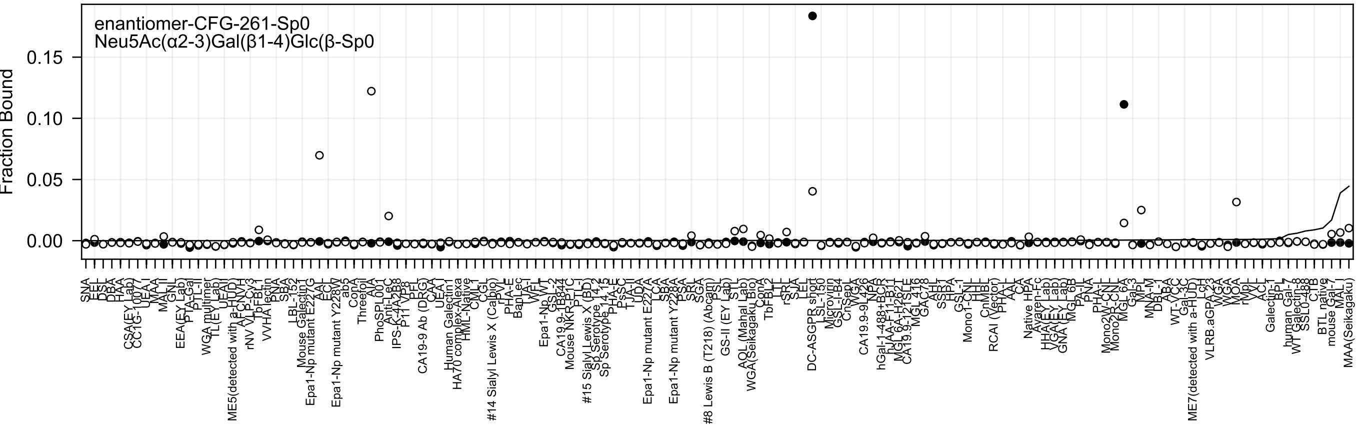
enantiomer-CFG-261-Sp0
Neu5Ac(α2-3)Gal(β1-4)Glc(β-Sp0
Fraction Bound
0.15
0.10
0.05
0.00

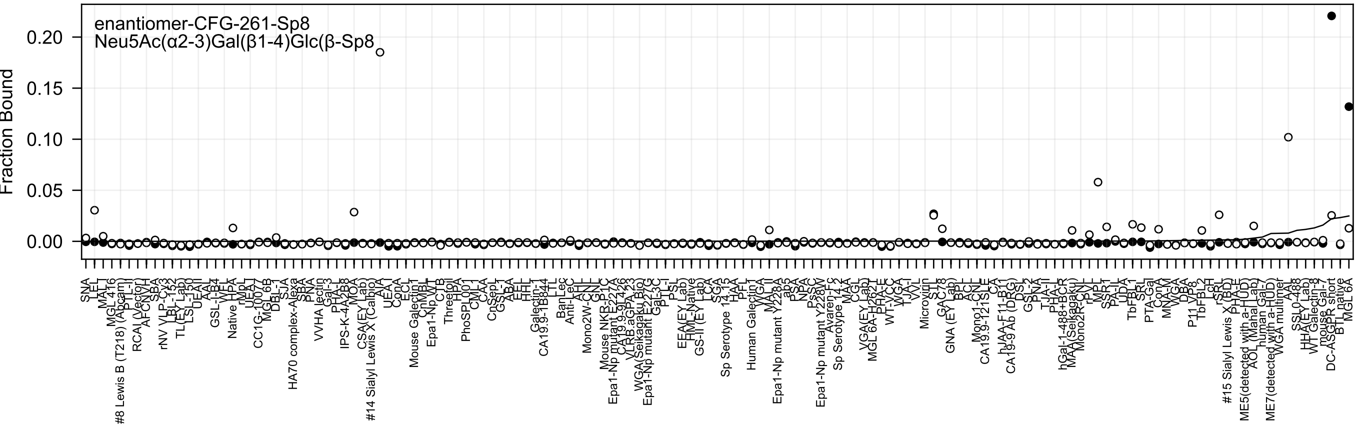

enantiomer-CFG-261-Sp8
Neu5Ac(α2-3)Gal(β1-4)Glc(β-Sp8
Fraction Bound
0.00
0.05
0.10
0.15
0.20

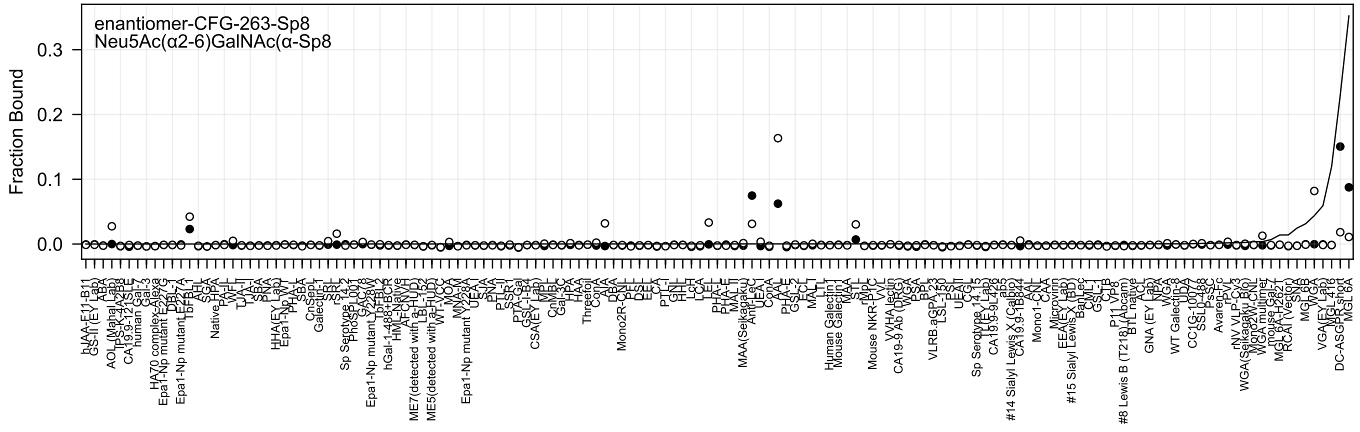
enantiomer-CFG-263-Sp8
Neu5Ac(α2-6)GalNAc(α-Sp8
Fraction Bound
0.0
0.1
0.2
0.3

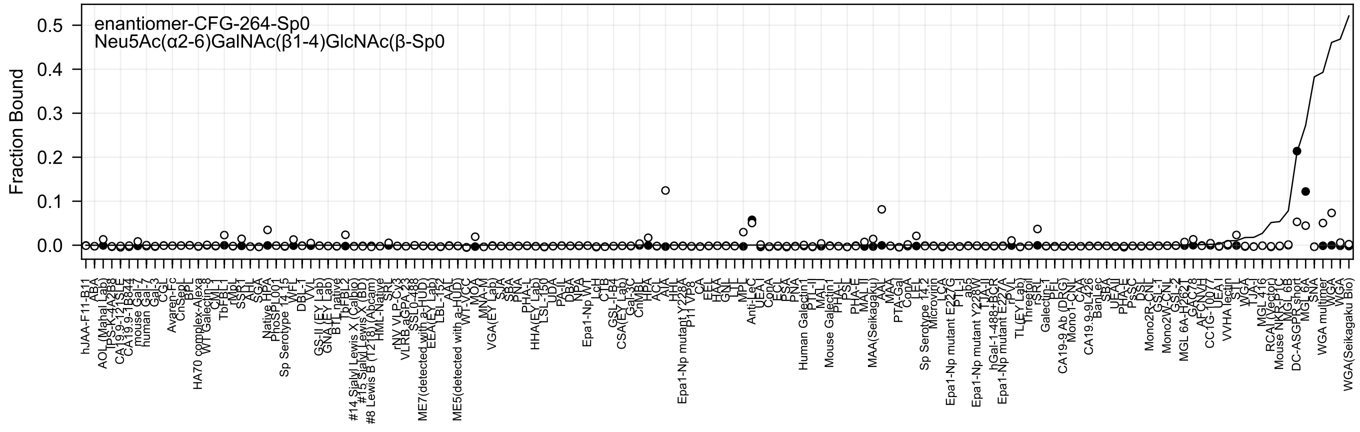
enantiomer-CFG-264-Sp0
Neu5Ac(α2-6)GalNAc(β1-4)GlcNAc(β-Sp0
Fraction Bound
0.5
0.4
0.3
0.2
0.1
0.0

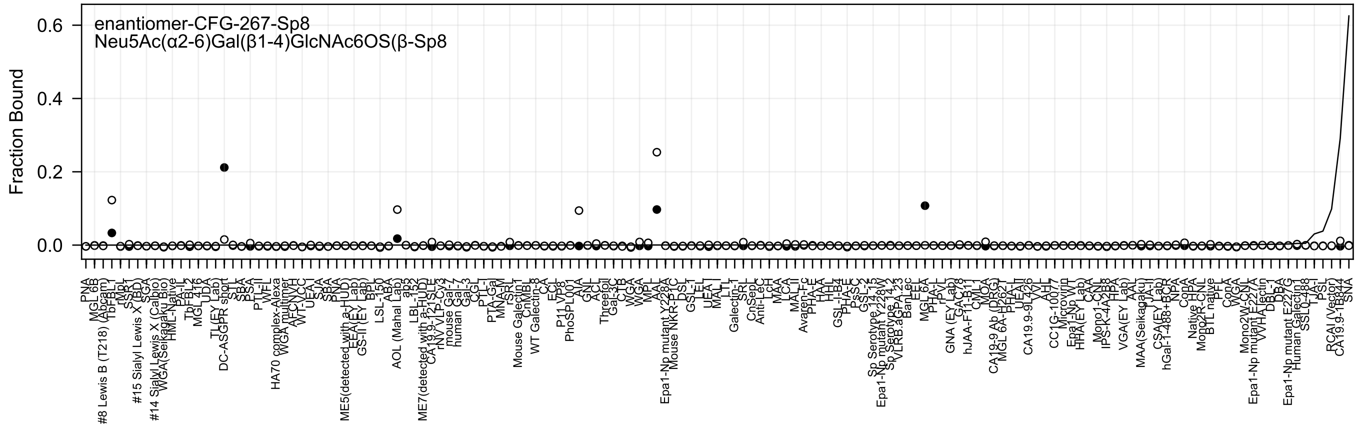

enantiomer-CFG-267-Sp8
Neu5Ac(α2-6)Gal(β1-4)GlcNAc6OS(β-Sp8
Fraction Bound
0.6
0.4
0.2
0.0

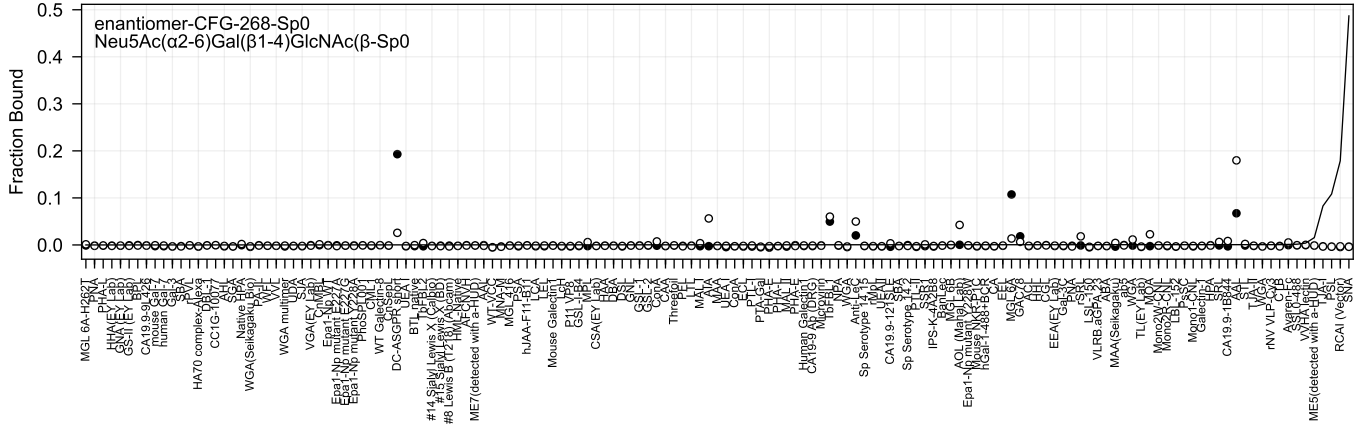

enantiomer-CFG-268-Sp0
Neu5Ac(α2-6)Gal(β1-4)GlcNAc(β-Sp0
Fraction Bound
0.5
0.4
0.3
0.2
0.1
0.0

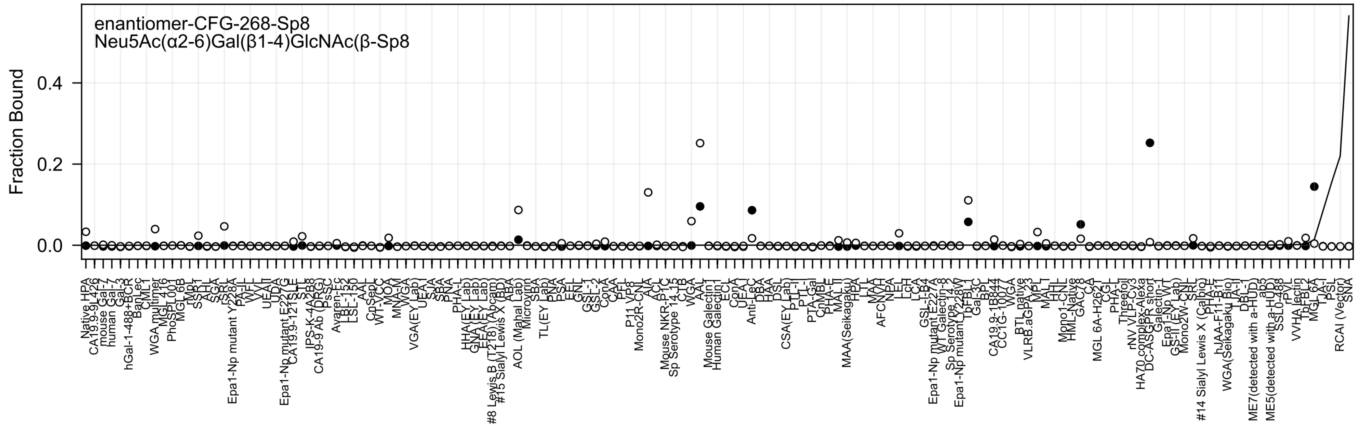
enantiomer-CFG-268-Sp8
Neu5Ac(α2-6)Gal(β1-4)GlcNAc(β-Sp8
Fraction Bound
0.4
0.2
0.0

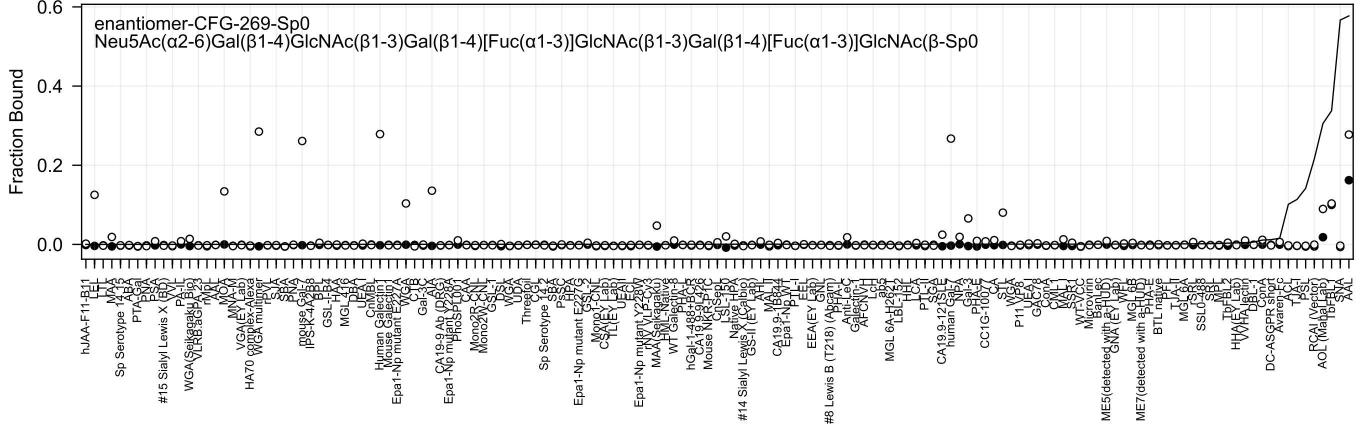
enantiomer-CFG-269-Sp0
Neu5Ac(α2-6)Gal(β1-4)GlcNAc(β1-3)Gal(β1-4)[Fuc(α1-3)]GlcNAc(β1-3)Gal(β1-4)[Fuc(α1-3)]GlcNAc(β-Sp0
Fraction Bound

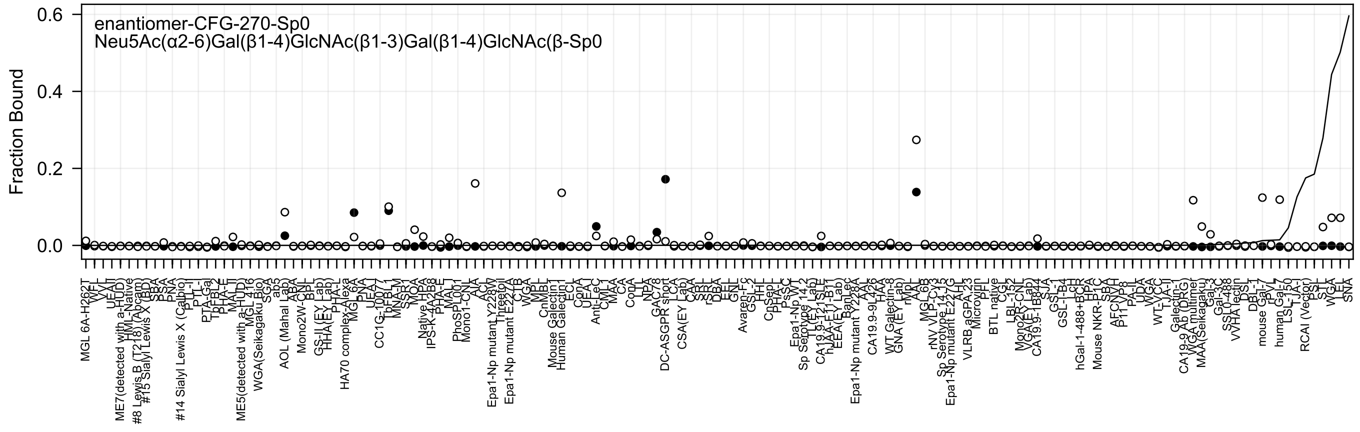
enantiomer-CFG-270-Sp0
Neu5Ac(α2-6)Gal(β1-4)GlcNAc(β1-3)Gal(β1-4)GlcNAc(β-Sp0
Fraction Bound
0.0
0.2
0.4
0.6

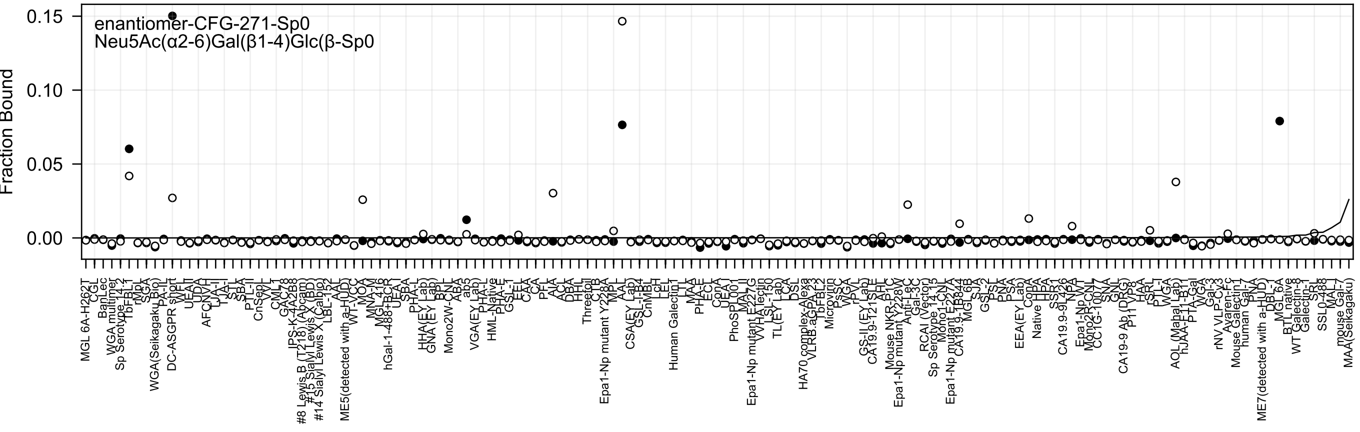
enantiomer-CFG-271-Sp0
Neu5Ac(α2-6)Gal(β1-4)Glc(β-Sp0
Fraction Bound
0.15
0.10
0.05
0.00

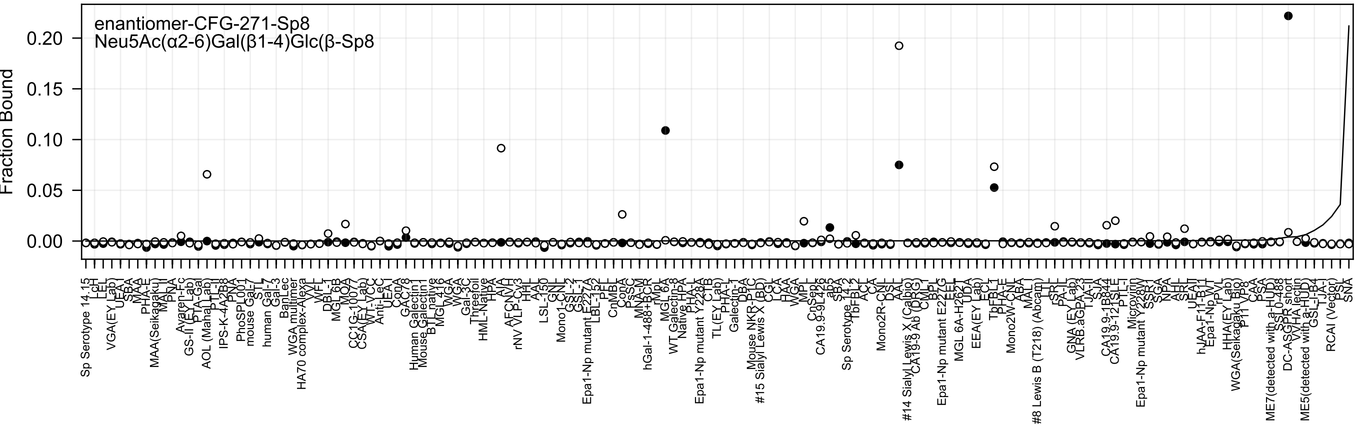
enantiomer-CFG-271-Sp8
Neu5Ac(α2-6)Gal(β1-4)Glc(β-Sp8
Fraction Bound
0.00
0.05
0.10
0.15
0.20

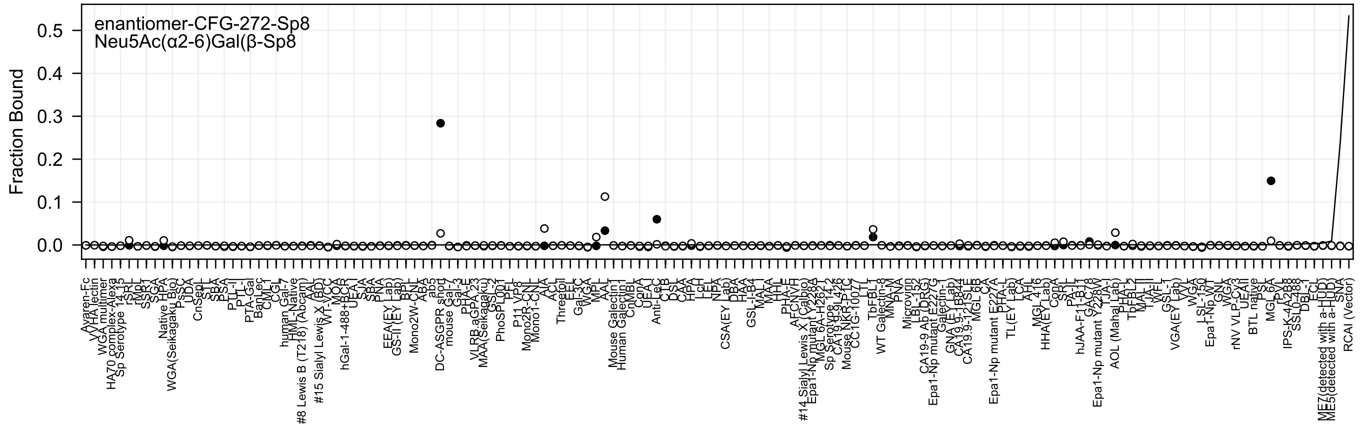

enantiomer-CFG-272-Sp8
Neu5Ac(α2-6)Gal(β-Sp8
Fraction Bound
0.5
0.4
0.3
0.2
0.1
0.0

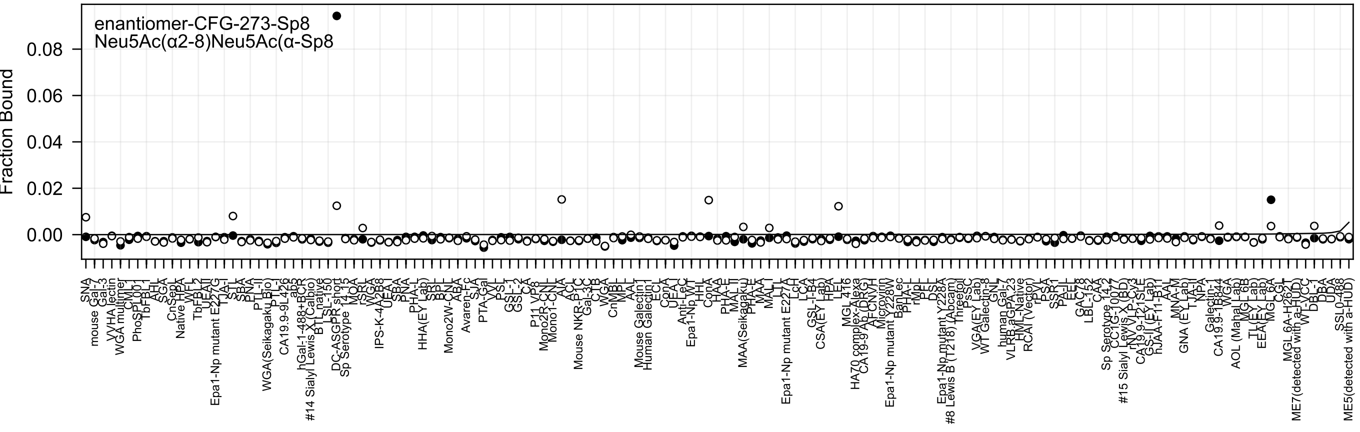

enantiomer-CFG-273-Sp8
Neu5Ac(α2-8)Neu5Ac(α-Sp8
Fraction Bound
0.08
0.06
0.04
0.02
0.00

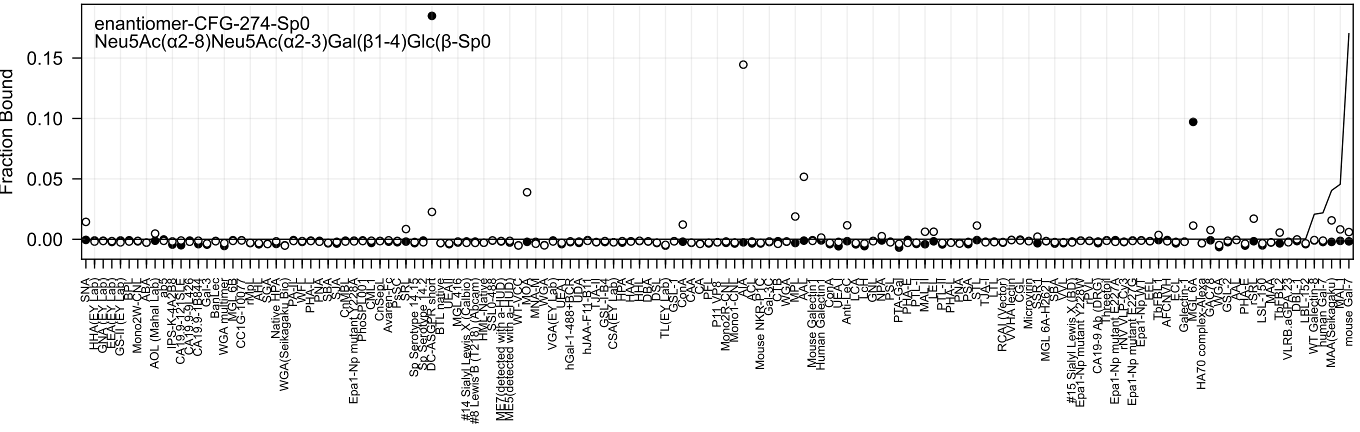
enantiomer-CFG-274-Sp0
Neu5Ac(α2-8)Neu5Ac(α2-3)Gal(β1-4)Glc(β-Sp0
Fraction Bound
0.15
0.10
0.05
0.00

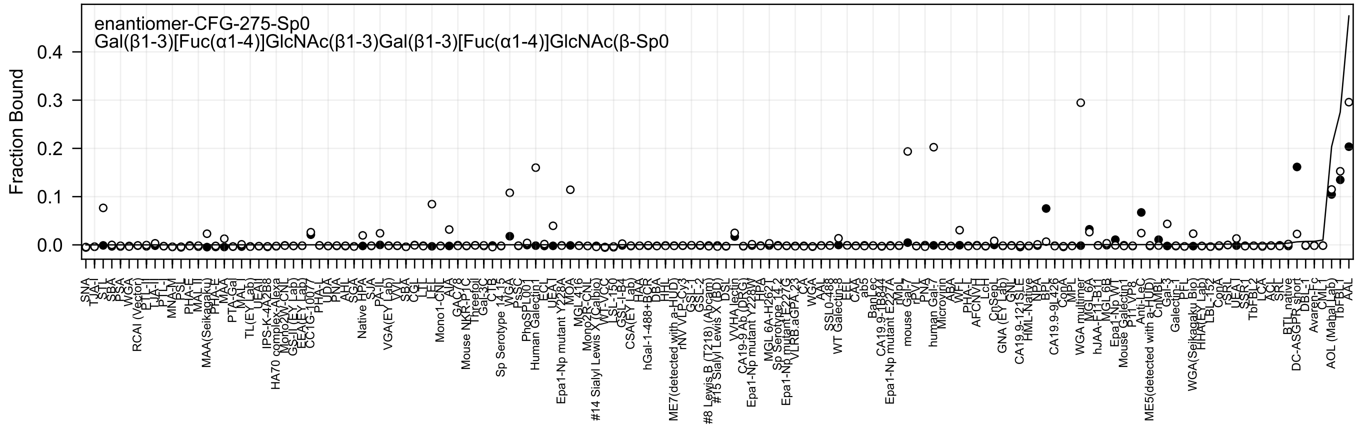
enantiomer-CFG-275-Sp0
Gal(β1-3)[Fuc(α1-4)]GlcNAc(β1-3)Gal(β1-3)[Fuc(α1-4)]GlcNAc(β-Sp0
Fraction Bound

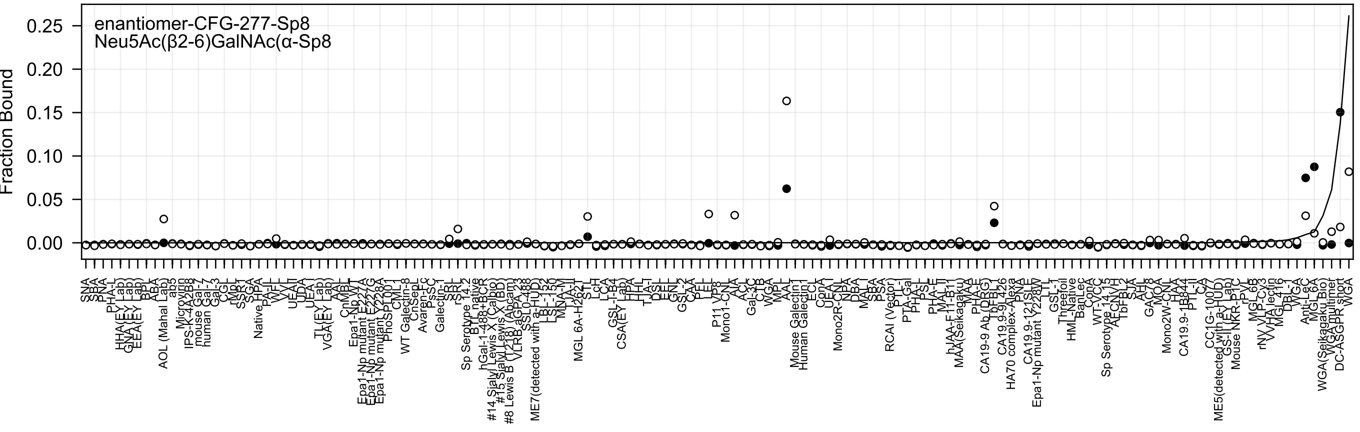
enantiomer-CFG-277-Sp8
Neu5Ac(β2-6)GalNAc(α-Sp8
Fraction Bound
0.25
0.20
0.15
0.10
0.05
0.00

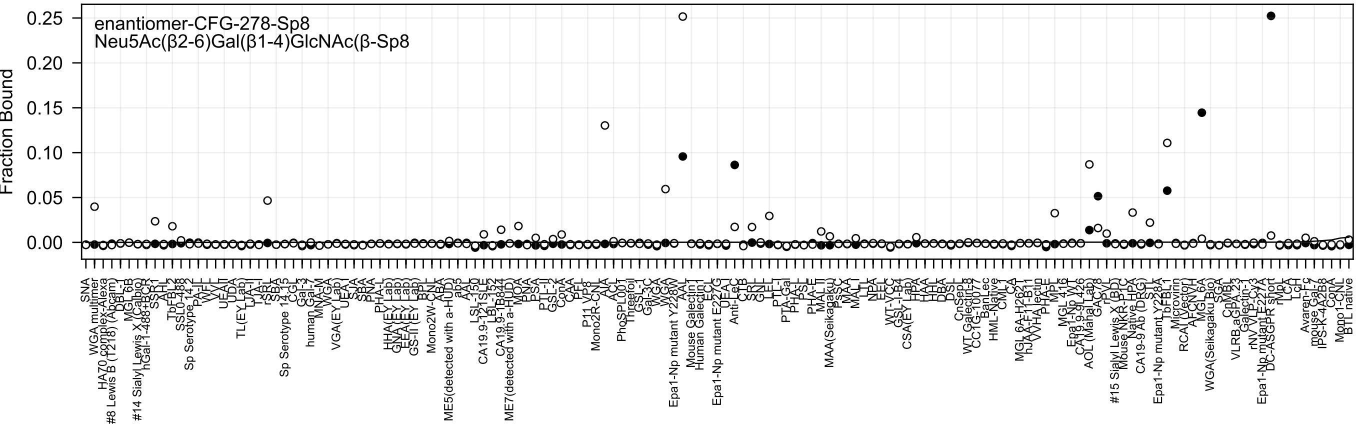
enantiomer-CFG-278-Sp8
Neu5Ac(β2-6)Gal(β1-4)GlcNAc(β-Sp8
Fraction Bound
0.25
0.20
0.15
0.10
0.05
0.00

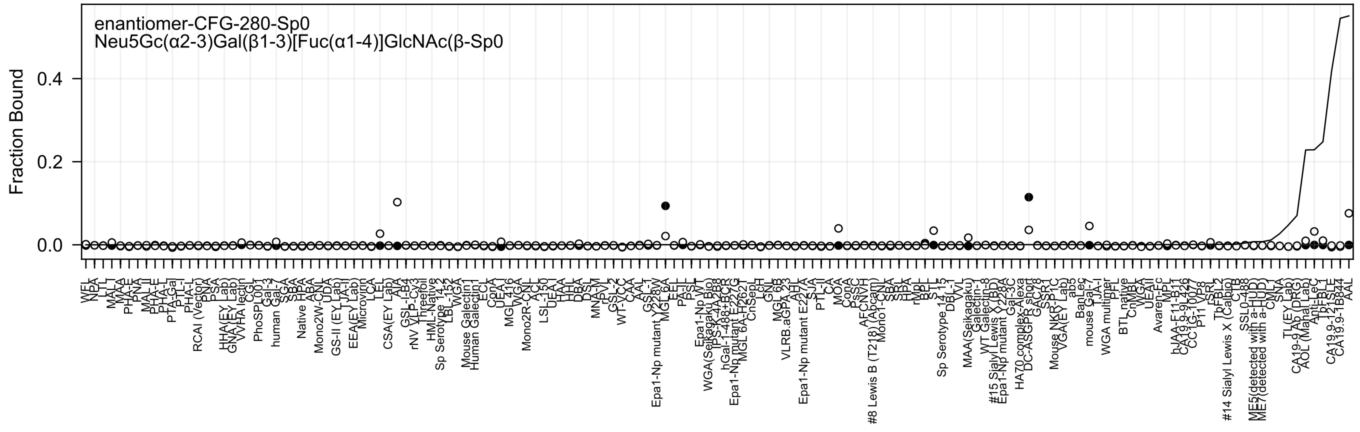
enantiomer-CFG-280-Sp0
Neu5Gc(α2-3)Gal(β1-3)[Fuc(α1-4)]GlcNAc(β-Sp0
Fraction Bound

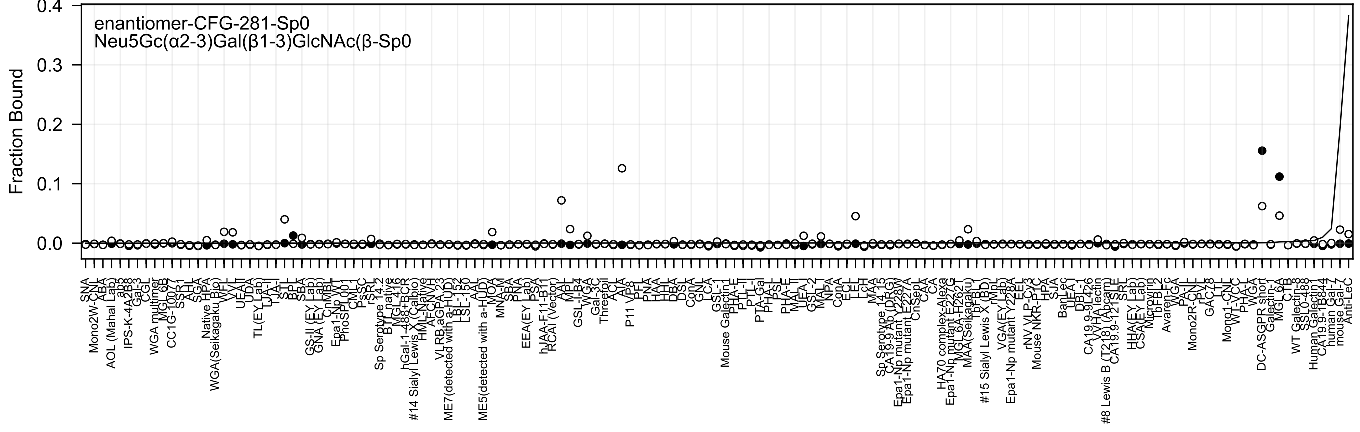
enantiomer-CFG-281-Sp0
Neu5Gc(α2-3)Gal(β1-3)GlcNAc(β-Sp0
Fraction Bound
0.4
0.3
0.2
0.1
0.0

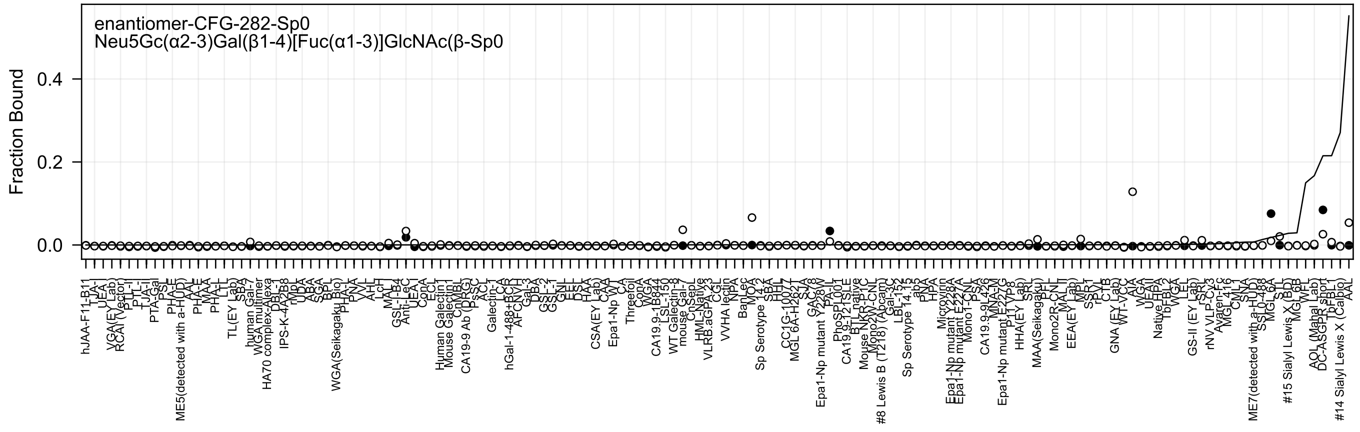
enantiomer-CFG-282-Sp0
Neu5Gc(α2-3)Gal(β1-4)[Fuc(α1-3)]GlcNAc(β-Sp0
Fraction Bound
0.4
0.2
0.0
hJAA-F11-B11
TJA-I
UEA-I
VGA(EY Lab)
RCAI (Vector)
PTL-II
PTL-I
TJA-II
PTA-Gal
PSL
PHA-E
ME5(detected with a-HUD)
AAL
PHA-E
MAA
PHA-L
LTL
TL(EY Lab)
SBA
human Gal-7
WGA multimer
HA70 complex-Alexa
DBL-1
IPS-K-4A2B8
rIMpL
UDA
ABA
SGA
BPL
WGA(Seikagaku Bio)
PHA-L
PNA
VVL
AHL
LcH
MAL-I
GSL-I-B4
Anti-LeC
UEA-I
ConA
ECL
Human Galectin1
Mouse Galectin1
CnMBL
CA19-9 Ab (DRG)
PsSC
ACL
Galectin-1
LCA
hGal-1-488+BCR
AFCNVH
Gal-3
DBA
GSL-2
GSL-1
GNL
EEL
DSL
HAA
CSA(EY Lab)
CAA
Epa1-Np WT
CA
Threefoil
ConA
WGA
CA19.9-1B844
LSL-150
WT Galectin-8
mouse Gal-7
CnSepL
HML-Native
VLRB.aGPA.23
CGL
VVHA lectin
NPA
BanLec
MOA
Sp Serotype 14.2
SBA
HHL
CC1G-10077
MGL 6A-H262T
SJA
GAC48
Epa1-Np mutant Y228W
PA-IL
PhoSL001
CA19.9-121SLE
BTL native
Mouse NKR-P1C
Mono2WT-CNL
#8 Lewis B (T218) (Abcam)
Gal-3C
LBL-152
Sp Serotype 14.15
ab5
PNA
HPA
Microvirin
Epa1-Np mutant Y228A
Epa1-Np mutant E227A
Mono1-CNL
PSA
CA19.9-9L426
MNA-M
Epa1-Np mutant E227G
P11VP8
HHA(EY Lab)
SRL
MAA(Seikagaku)
PFL
Mono2R-CNL
MAL II
EEA(EY Lab)
MPL
SSR1
rPVL
CTB
GNA (EY Lab)
WT-VacC
AIA
WGA
UEAII
Native HPA
TbFBL2
WGA
LEL
GS-II (EY Lab)
rSRL
rNV VLP-Cy3
Avaren-Fc
MGL416
CML1
SNA
ME7(detected with a-HUD)
SSL0-488
MGL 6A
STL
#15 Sialyl Lewis X (BD)
MGL6B
WFL
AOL (Mahal Lab)
DC-ASGPR short
TbFBL1
#14 Sialyl Lewis X (Calbio)
AAL

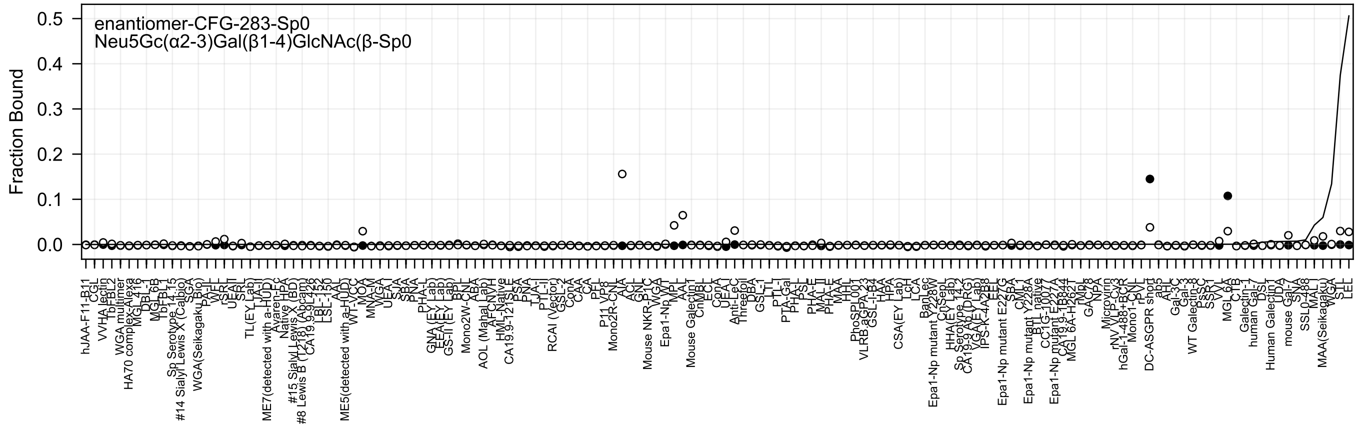
enantiomer-CFG-283-Sp0
Neu5Gc(α2-3)Gal(β1-4)GlcNAc(β-Sp0
Fraction Bound
0.5
0.4
0.3
0.2
0.1
0.0
hJAA-F11-B11
CGL
VVHA lectin
TbFBL2
WGA multimer
HA70 complex-Alexa
MGL 416
DBL-1
MGL 6B
TbFBL1
Sp Serotype 14.15
#14 Sialyl Lewis X (Calbio)
SGA
WGA(Seikagaku Bio)
PA-IL
WFL
rSRL
UEAII
SRL
TL(EY Lab)
TJA-I
ME7(detected with a-HuD)
Avaren-Fc
Native HPA
#15 Sialyl Lewis X (BD)
#8 Lewis B (T218) (Abcam)
CA19.9_9L426
LBL-152
LSL-150
AAL
ME5(detected with a-HuD)
WT-VCC
MOA
MNA-M
WGA
UEA-I
SJA
SBA
PNA
PHA-L
GNA (EY Lab)
EEA (EY Lab)
GS-II (EY Lab)
BPL
Mono2W-CNL
ABA
AOL (Mahal Lab)
AFCNVH
HML-Native
CA19.9-121SLE
PSA
PNA
TJA-II
PTL-II
RCAI (Vector)
GSL-2
ConA
CAA
CA
PFL
P11 VP8
Mono2R-CNL
AIA
ACL
GNL
Mouse NKR-P1C
WGA
Epa1-Np WT
MPL
AAL
Mouse Galectin1
CnMBL
ECL
ConA
UEA-I
Anti-LeC
Threefoil
DBA
GSL-I
LTL
PTL-I
PTA-Gal
PHA-L
PSL
PHA-E
MAL-II
PHA-E
MAA
HHL
PhoSL001
VLRB.aGPA.23
GSL-I-B4
HAA
HPA
CSA(EY Lab)
LCH
LCA
BanLec
Epa1-Np mutant Y228W
CnSepL
HHA(EY Lab)
Sp Serotype 14.2
CA19.9 Ab (DRG)
VGA (EY Lab)
IPS-K-4A2B8
EEL
Epa1-Np mutant E227G
SBA
CML1
Epa1-Np mutant Y228A
BTL native
CC1G-1007
Epa1-Np mutant E227A
CA19.9-1B844
MGL 6A-H362I
rNp
GAC78
NPA
Microvirin
rNV VLP-Cy3
hGal-1-488+BCR
Mono1-CNL
rPVL
DC-ASGPR short
ab5
AHL
Gal-3C
Gal-3
WT Galectin-8
PsSC
SSR1
VVL
MGL 6A
CTB
Galectin-1
human Gal-7
DSL
Human Galectin1
UDA
mouse Gal-7
SNA
SSL0-488
MAL-I
MAA(Seikagaku)
WGA
STL
LEL

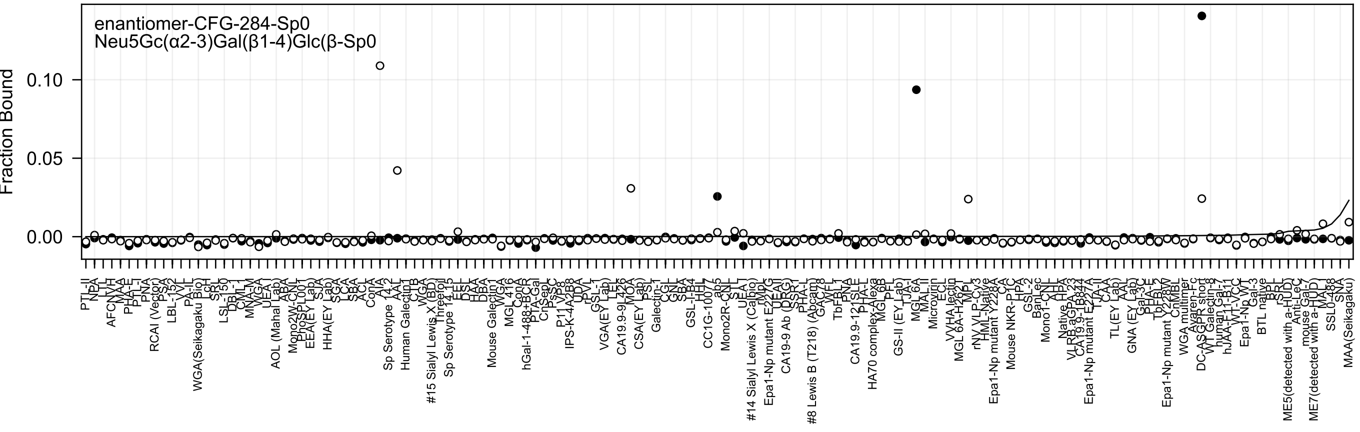
enantiomer-CFG-284-Sp0
Neu5Gc(α2-3)Gal(β1-4)Glc(β-Sp0
Fraction Bound
0.10
0.05
0.00

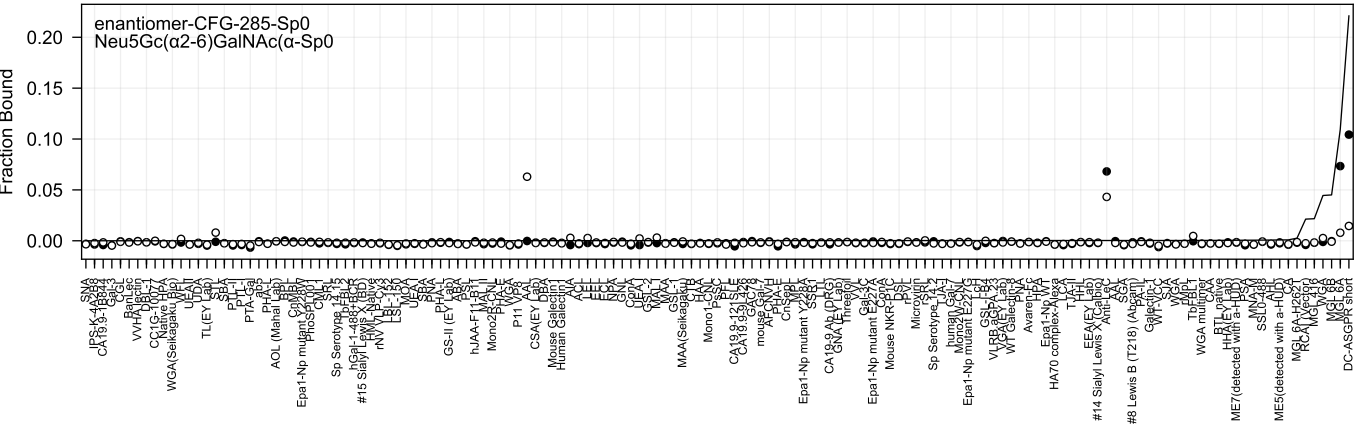

enantiomer-CFG-285-Sp0
Neu5Gc(α2-6)GalNAc(α-Sp0
Fraction Bound
0.20
0.15
0.10
0.05
0.00

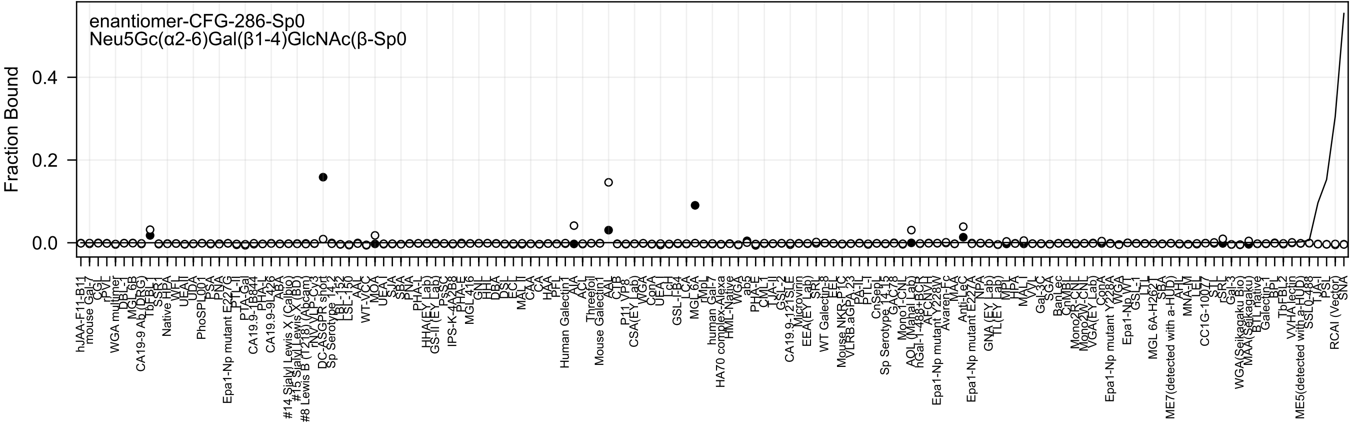

enantiomer-CFG-286-Sp0
Neu5Gc(α2-6)Gal(β1-4)GlcNAc(β-Sp0
Fraction Bound
0.4
0.2
0.0

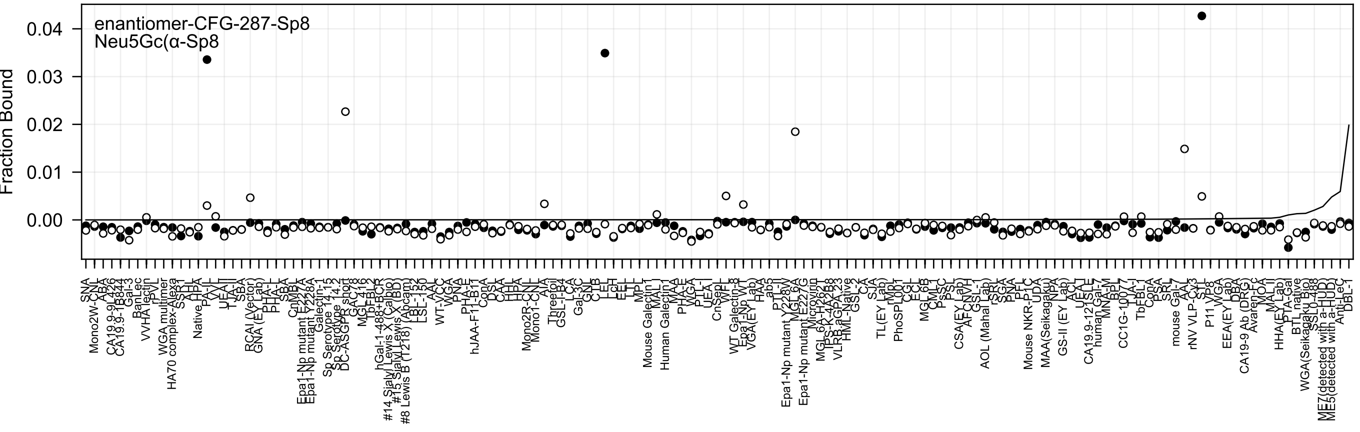

enantiomer-CFG-287-Sp8
Neu5Gc(α-Sp8
Fraction Bound
0.04
0.03
0.02
0.01
0.00

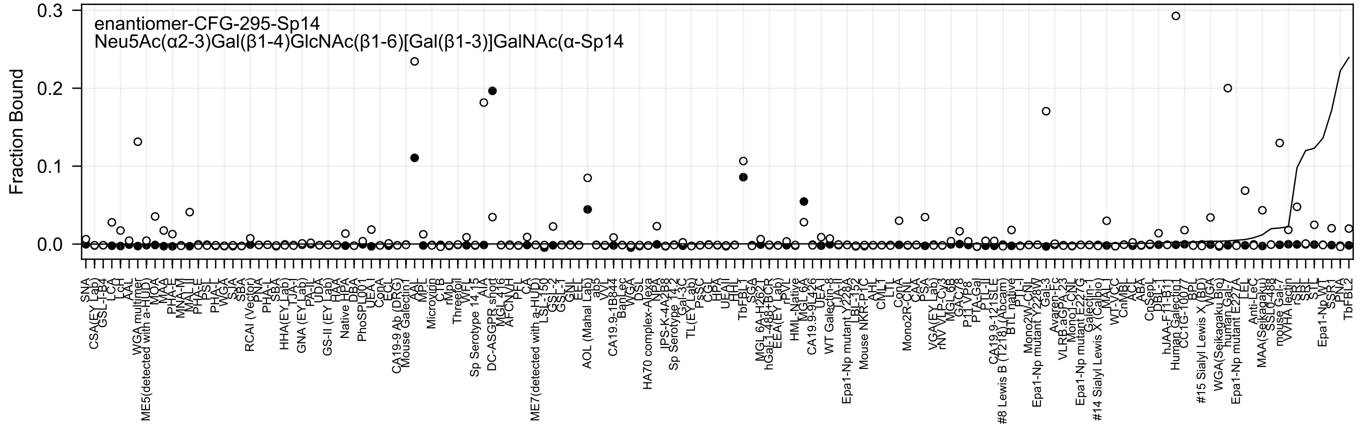

enantiomer-CFG-295-Sp14
Neu5Ac(α2-3)Gal(β1-4)GlcNAc(β1-6)[Gal(β1-3)]GalNAc(α-Sp14
Fraction Bound
0.3
0.2
0.1
0.0

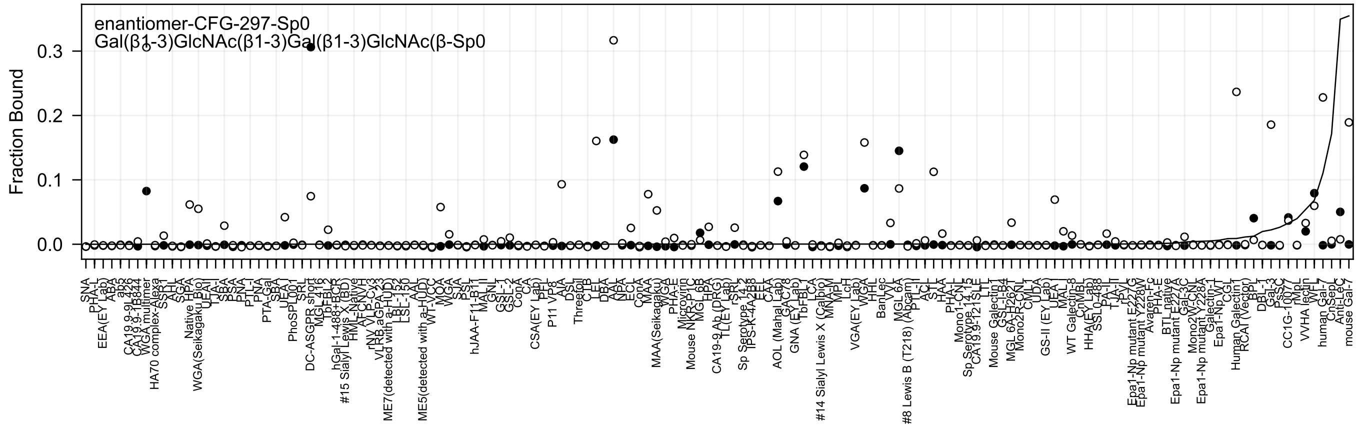

enantiomer-CFG-297-Sp0
Gal(β1-3)GlcNAc(β1-3)Gal(β1-3)GlcNAc(β-Sp0
Fraction Bound
0.3
0.2
0.1
0.0

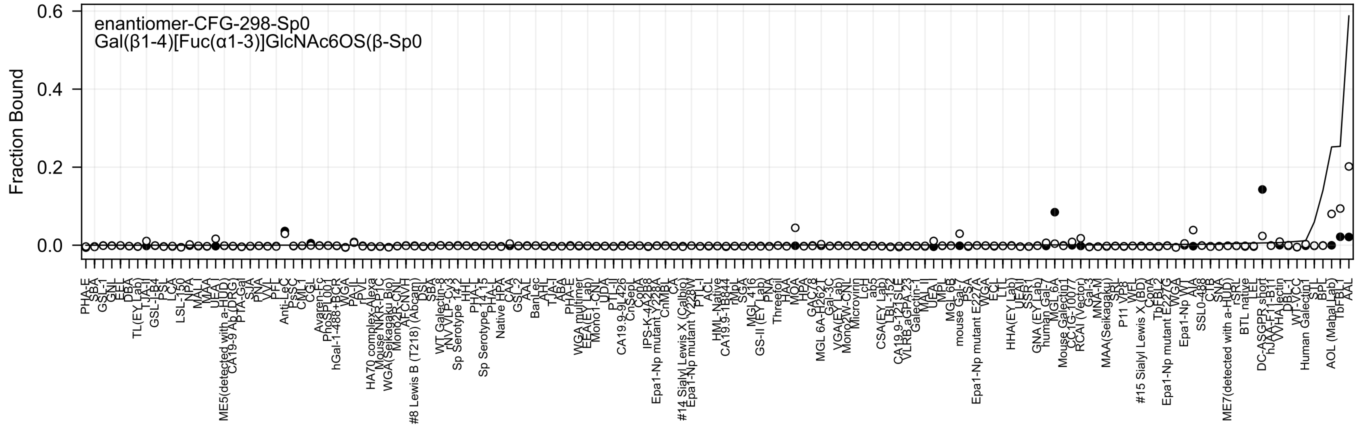
enantiomer-CFG-298-Sp0
Gal(β1-4)[Fuc(α1-3)]GlcNAc6OS(β-Sp0
Fraction Bound
0.6
0.4
0.2
0.0

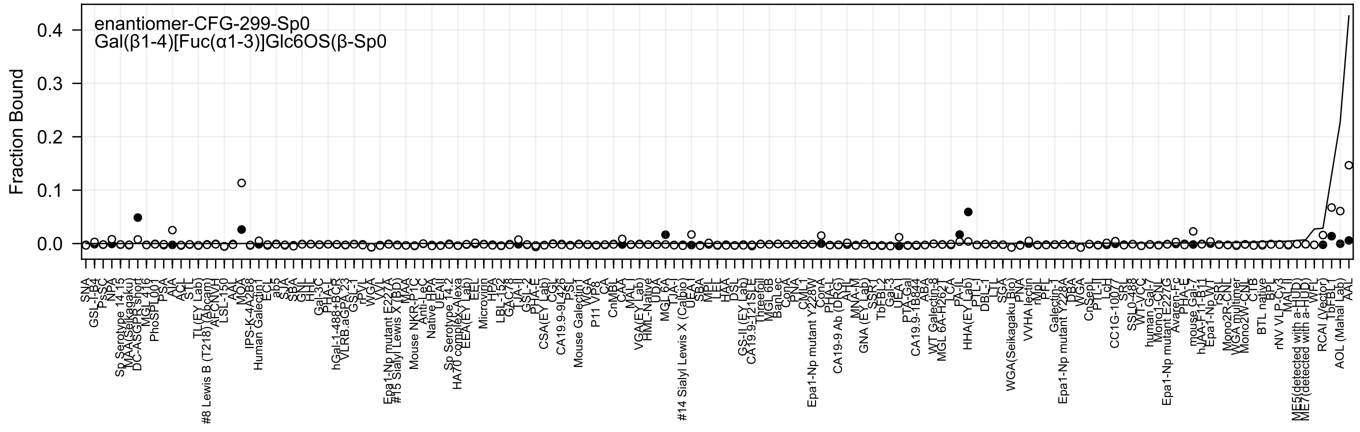
enantiomer-CFG-299-Sp0
Gal(β1-4)[Fuc(α1-3)]Glc6OS(β-Sp0
Fraction Bound
0.4
0.3
0.2
0.1
0.0

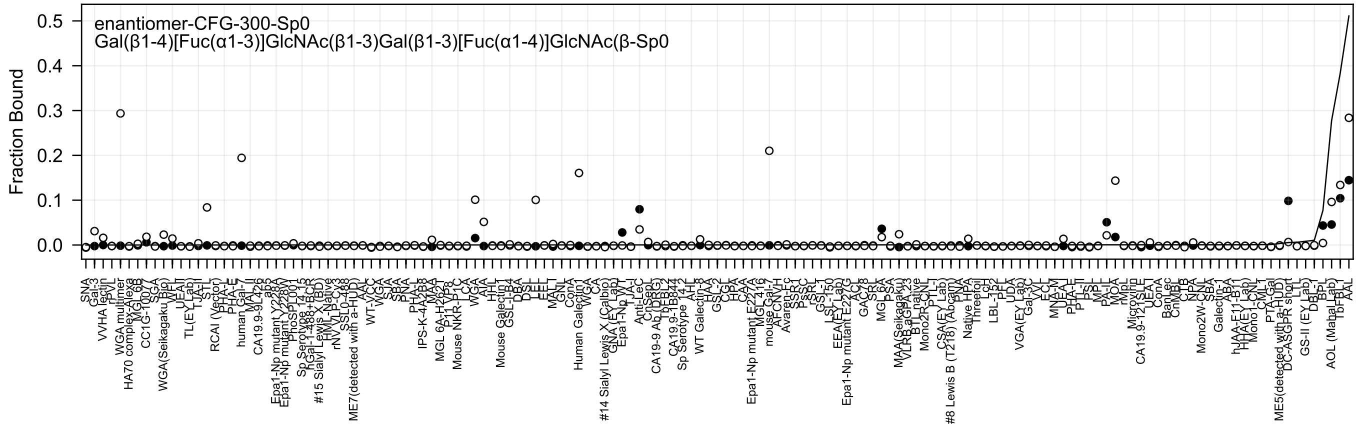

enantiomer-CFG-300-Sp0
Gal(β1-4)[Fuc(α1-3)]GlcNAc(β1-3)Gal(β1-3)[Fuc(α1-4)]GlcNAc(β-Sp0
Fraction Bound
0.0
0.1
0.2
0.3
0.4
0.5
SNA
Gal-3
VVHA lectin
rPVL
WGA multimer
HA70 complex-Alexa
MGL 6B
CC1G-1007
SGA
WGA(Seikagaku Bio)
WFL
UEAI
TL(EY Lab)
TJA-II
STL
RCAI (Vector)
PHA-L
PHA-E
human Gal-7
MAL II
CA19.9-9L426
ab5
Epa1-Np mutant Y228A
Epa1-Np mutant Y228W
PhoSL001
Sp Serotype 14-15
hGal-1-488+BCR
#15 Sialyl Lewis X (BD)
HML-Native
rNV VLP-Cy3
SSL0-488
ME7(detected with a-HUD)
AAL
WT-VCC
WGA
SJA
SBA
PNA
PHA-L
IPS-K-4A2B8
MAA
MGL 6A-H262T
P11 VP8
Mouse NKR-P1C
LCA
WGA
AIA
HHL
Mouse Galectin1
GSL-I-B4
DBA
DSL
LEL
EEL
MAL I
GNL
GNA
ConA
Human Galectin1
WGA
CA
#14 Sialyl Lewis X (Calbio)
GNA (EY Lab)
Epa1-Np WT
TJA-I
Anti-LeC
ChSep
CA19-9 Ab (DRG)
TbFBL2
CA19.9-1B844
Sp Serotype 14-2
AH
WT Galectin-8
HAA
GSL-2
CGL
HPA
CAA
Epa1-Np mutant E227A
MGL416
mouse Gal-7
AFCNVH
Avaren-Fc
SSR1
PsSC
rSRL
GSL-1
LSL-150
EEA(EY Lab)
Epa1-Np mutant E227G
ACL
GAC78
SRL
MGL 6A
PSA
MAA(Seikagaku)
VLRB aGPA.23
BTL native
Mono2R-CNL
PTL-I
CSA(EY Lab)
#8 Lewis B (T218) (Abcam)
PNA
Native HPA
Threefoil
LcH
LBL-152
PFL
UDA
VGA(EY Lab)
Gal-3C
VVL
ECL
MNA-M
UEA-I
PHA-E
PTL-II
PSL
MPL
PA-IL
MOA
rMpL
Microvirin
CA19.9-121SLE
UEA1
ConA
BanLec
CnMBL
CTB
NPA
Mono2W-CNL
SBA
Galectin-1
ABA
hJAA-E11-B11
HHA(EY Lab)
Mono1-CNL
CML1
PTA-Gal
ME5(detected with a-HUD)
DC-ASGPR short
LTL
GS-II (EY Lab)
DBL-I
BPL
AOL (Mahal Lab)
TbFBL1
AAL

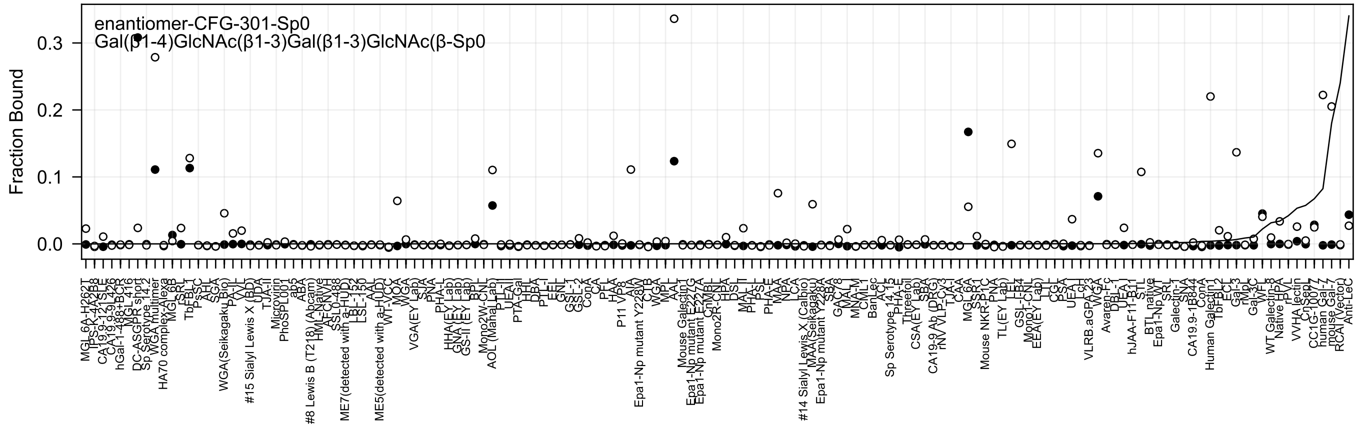
enantiomer-CFG-301-Sp0
Gal(β1-4)GlcNAc(β1-3)Gal(β1-3)GlcNAc(β-Sp0
Fraction Bound
0.0
0.1
0.2
0.3
MGL 6A-H262T
TPS-K-4A2B8
CA19.9-121SLE
CA19.9-9L426
hGal-1-488+BCR
MGL 416
DC-ASGPR short
Sp Serotype 14.2
WGA multimer
HA70 complex-Alexa
MGL 6B
rSRL
TbFBL1
PsSC
AHL
SGA
WGA(Seikagaku Bio)
PA-IL
#15 Sialyl Lewis X (BD)
UDA
TJA-II
Microvirin
PhoSPL001
ab5
ABA
#8 Lewis B (T218) (Abcam)
HML-Native
AFCNVH
SSL0-488
ME7(detected with a-HUD)
LBL-152
LSL-150
AAL
ME5(detected with a-HUD)
WT-VCC
MOA
WGA
VGA(EY Lab)
SJA
PNA
PHA-L
HHA(EY Lab)
GNA(EY Lab)
GS-II (EY Lab)
BPL
Mono2W-CNL
AOL (Mahal Lab)
PTL-I
UEA-II
PTA-Gal
HHL
DBA
PTL-I
EEL
GNL
GSL-1
GSL-2
ConA
CA
PFL
HAA
P11 VP8
AIA
Epa1-Np mutant Y228W
CTB
WGA
MPL
AAL
Mouse Galectin1
Epa1-Np mutant E227G
Epa1-Np mutant E227A
CnMBL
Mono2R-CNL
HPA
DSL
MAL-II
PHA-L
PSL
PHA-E
MAA
NPA
LCA
#14 Sialyl Lewis X (Calbio)
MAA (Seikagaku)
Epa1-Np mutant Y228A
SBA
GAG78
MAL-I
MNA-M
CML1
BanLec
ACL
Sp Serotype 14.15
PHA-E
Threefoil
CSA(EY Lab)
SBA
CA19-9 Ab (DRG)
rNV VLP-Cy3
TJA-I
CAA
MGL 6A
SSR1
Mouse NKR-P1C
PNA
TL(EY Lab)
LEL
GSL-I-B4
Mono1-CNL
EEA(EY Lab)
LTL
CGL
PSA
UEA-I
Lcl
VLRB.aGPA.23
WGA
Avaren-Fc
DBL-1
UEA-I
hJAA-F11-B11
STL
BTL native
Epa1-Np WT
SRL
Galectin-1
SNA
CA19.9-1B844
ConA
Human Galectin1
TbFBL2
ECL
Gal-3
rMpL
Gal-3C
WFL
WT Galectin-8
Native HPA
rPVL
VVHA lectin
CnSep
CC1G-10077
human Gal-7
mouse Gal-7
RCA I (Vector)
Anti-LeC

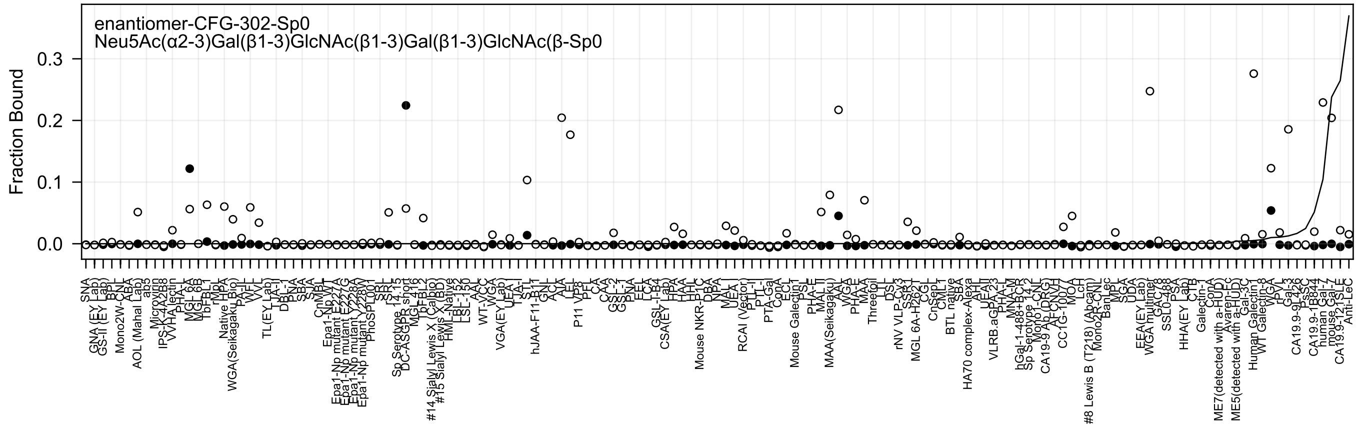

enantiomer-CFG-302-Sp0
Neu5Ac(α2-3)Gal(β1-3)GlcNAc(β1-3)Gal(β1-3)GlcNAc(β-Sp0
Fraction Bound
0.0
0.1
0.2
0.3

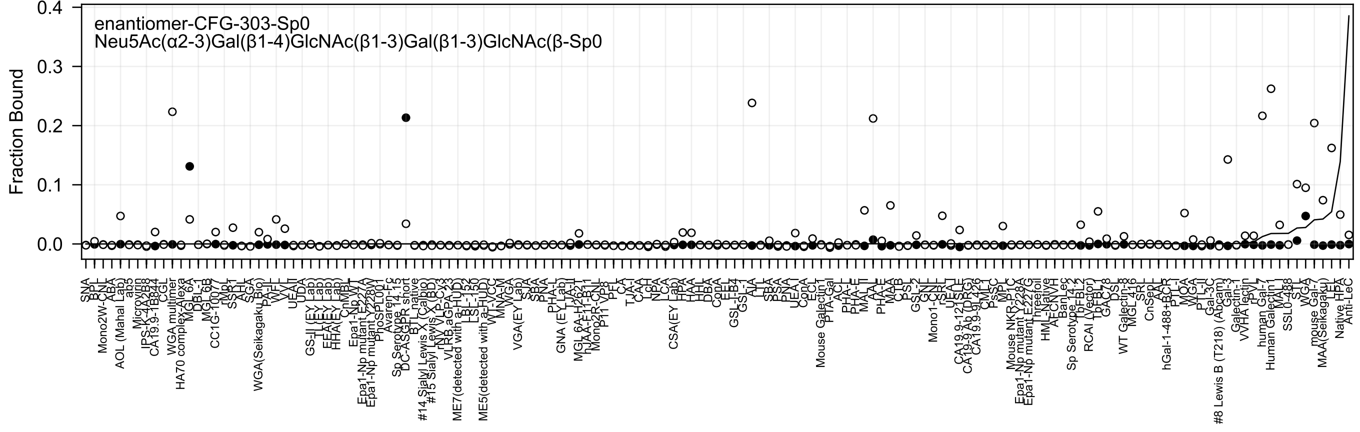
enantiomer-CFG-303-Sp0
Neu5Ac(α2-3)Gal(β1-4)GlcNAc(β1-3)Gal(β1-3)GlcNAc(β-Sp0
Fraction Bound

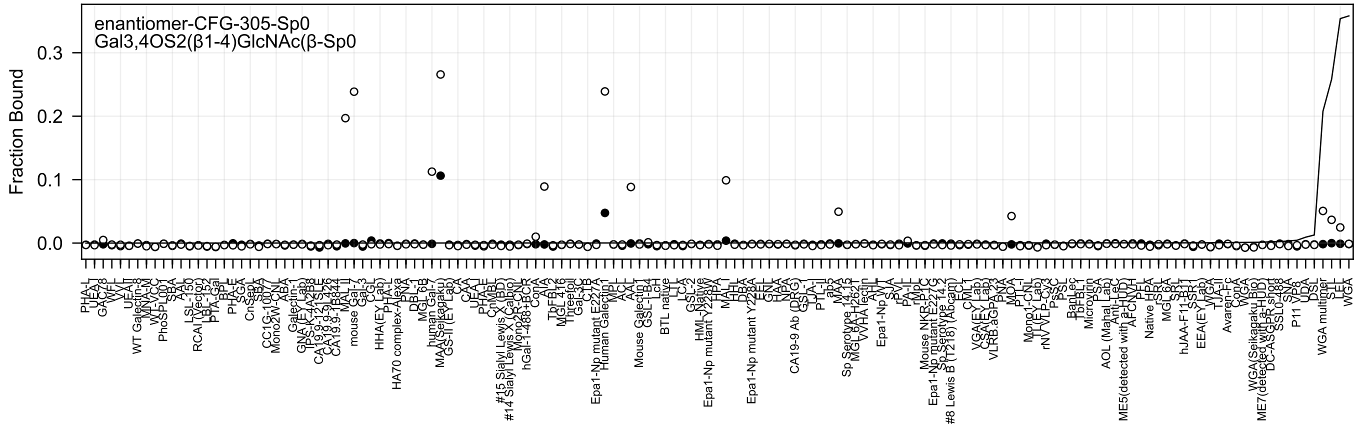

enantiomer-CFG-305-Sp0
Gal3,4OS2(β1-4)GlcNAc(β-Sp0
Fraction Bound
0.0
0.1
0.2
0.3

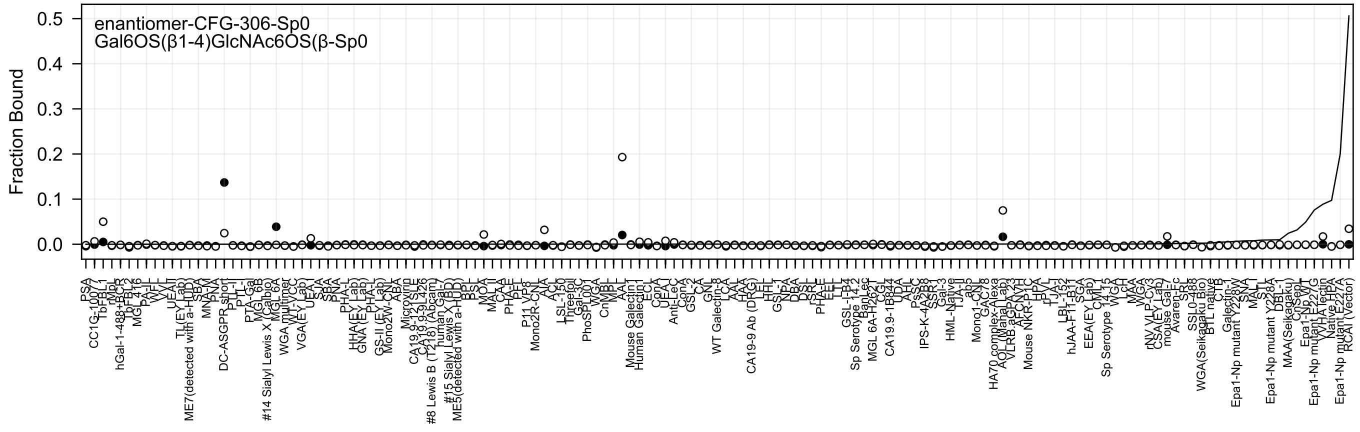
enantiomer-CFG-306-Sp0
Gal6OS(β1-4)GlcNAc6OS(β-Sp0
Fraction Bound
0.5
0.4
0.3
0.2
0.1
0.0

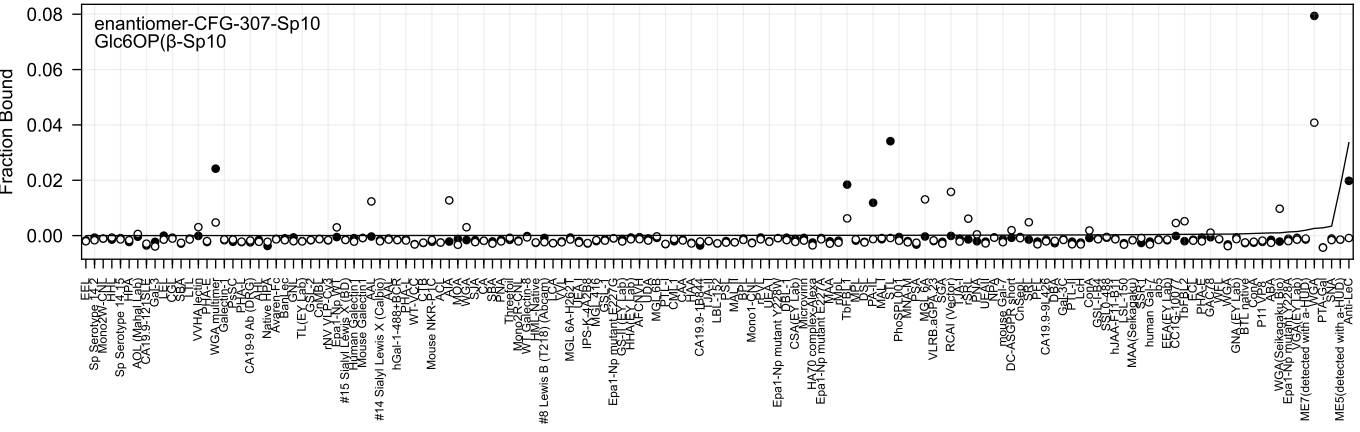
enantiomer-CFG-307-Sp10
Glc6OP(β-Sp10
Fraction Bound
0.08
0.06
0.04
0.02
0.00

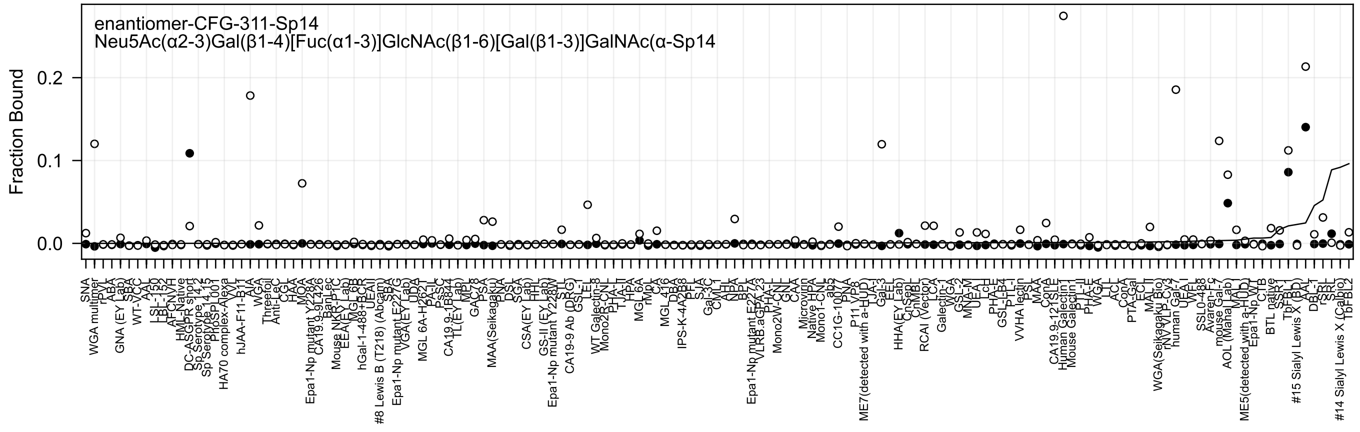
enantiomer-CFG-311-Sp14
Neu5Ac(α2-3)Gal(β1-4)[Fuc(α1-3)]GlcNAc(β1-6)[Gal(β1-3)]GalNAc(α-Sp14
Fraction Bound
0.0
0.1
0.2

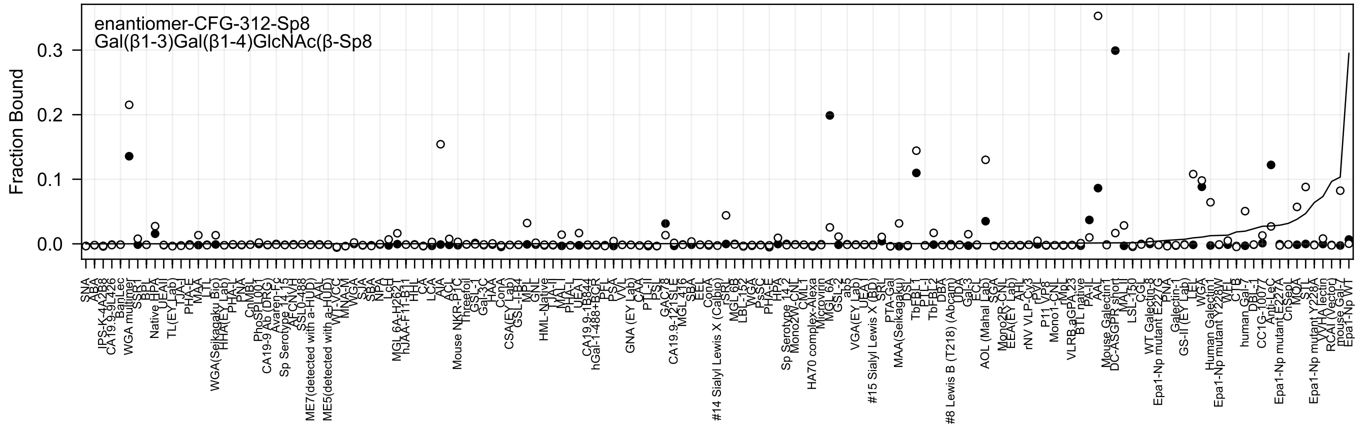
enantiomer-CFG-312-Sp8
Gal(β1-3)Gal(β1-4)GlcNAc(β-Sp8
Fraction Bound
0.0
0.1
0.2
0.3

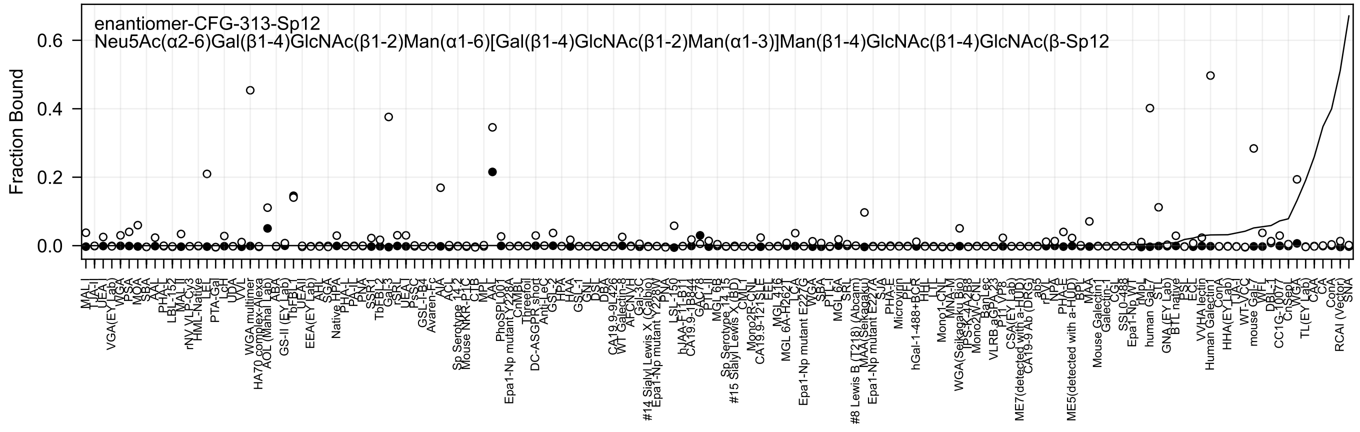
enantiomer-CFG-313-Sp12
Neu5Ac(α2-6)Gal(β1-4)GlcNAc(β1-2)Man(α1-6)[Gal(β1-4)GlcNAc(β1-2)Man(α1-3)]Man(β1-4)GlcNAc(β1-4)GlcNAc(β-Sp12
Fraction Bound
0.0
0.2
0.4
0.6

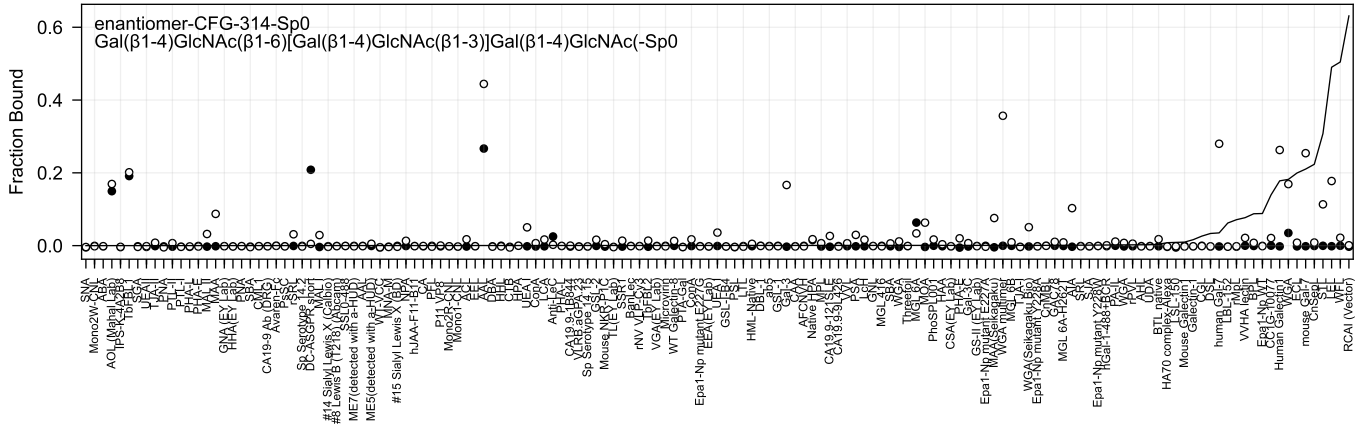

enantiomer-CFG-314-Sp0
Gal(β1-4)GlcNAc(β1-6)[Gal(β1-4)GlcNAc(β1-3)]Gal(β1-4)GlcNAc(-Sp0
Fraction Bound

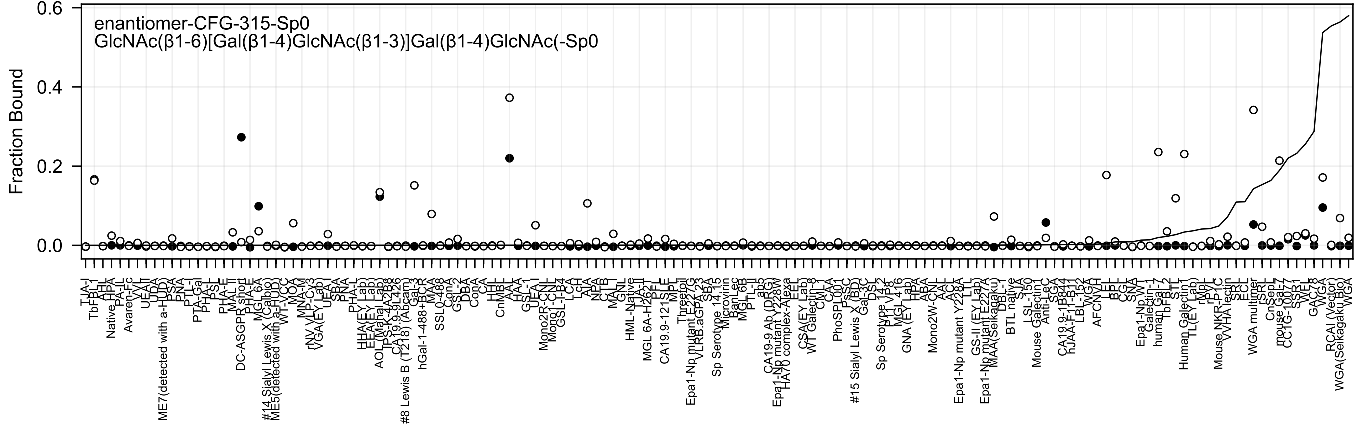
enantiomer-CFG-315-Sp0
GlcNAc(β1-6)[Gal(β1-4)GlcNAc(β1-3)]Gal(β1-4)GlcNAc(-Sp0
Fraction Bound
0.6
0.4
0.2
0.0

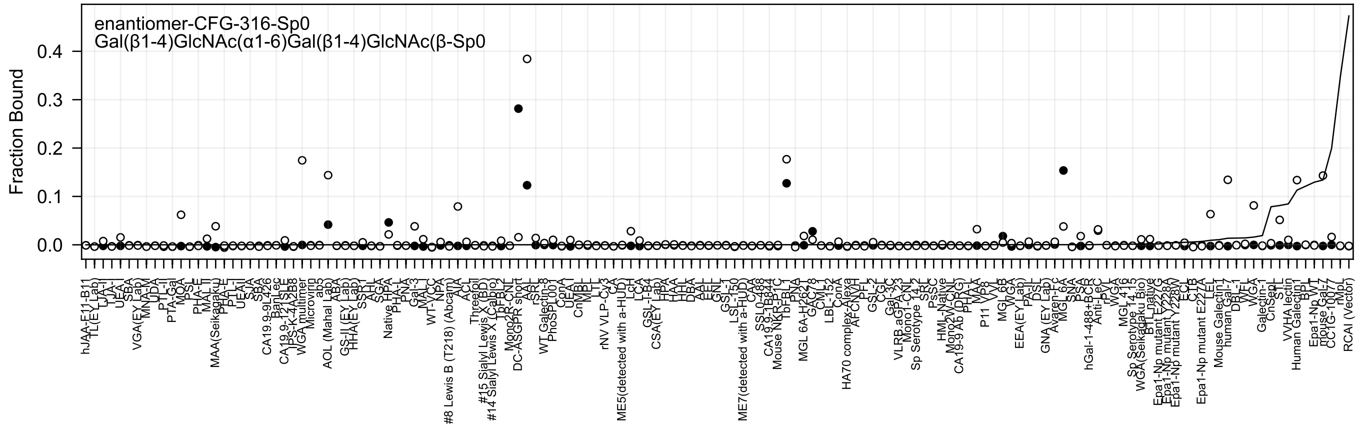
enantiomer-CFG-316-Sp0
Gal(β1-4)GlcNAc(α1-6)Gal(β1-4)GlcNAc(β-Sp0
Fraction Bound

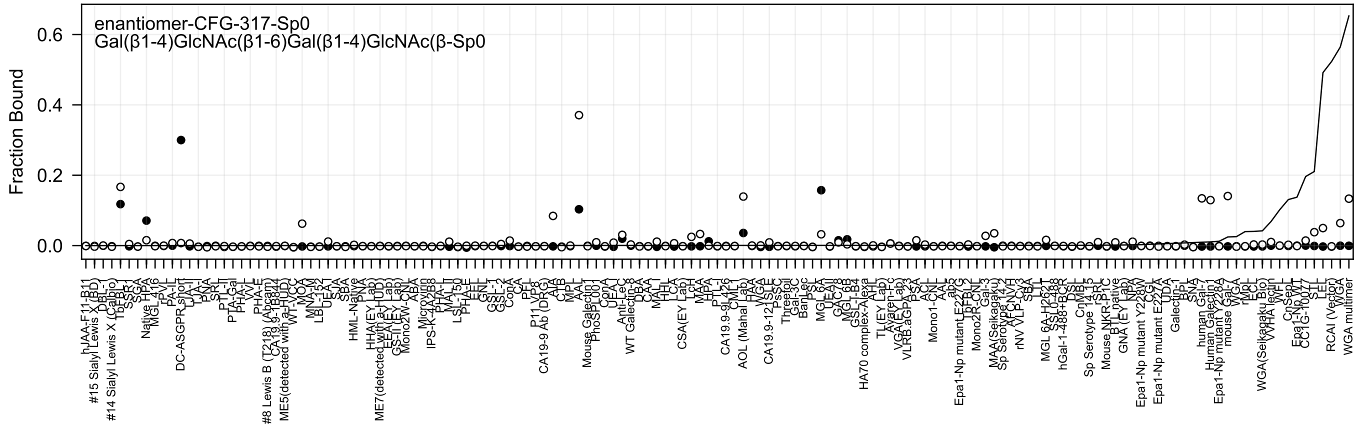

enantiomer-CFG-317-Sp0
Gal(β1-4)GlcNAc(β1-6)Gal(β1-4)GlcNAc(β-Sp0
Fraction Bound

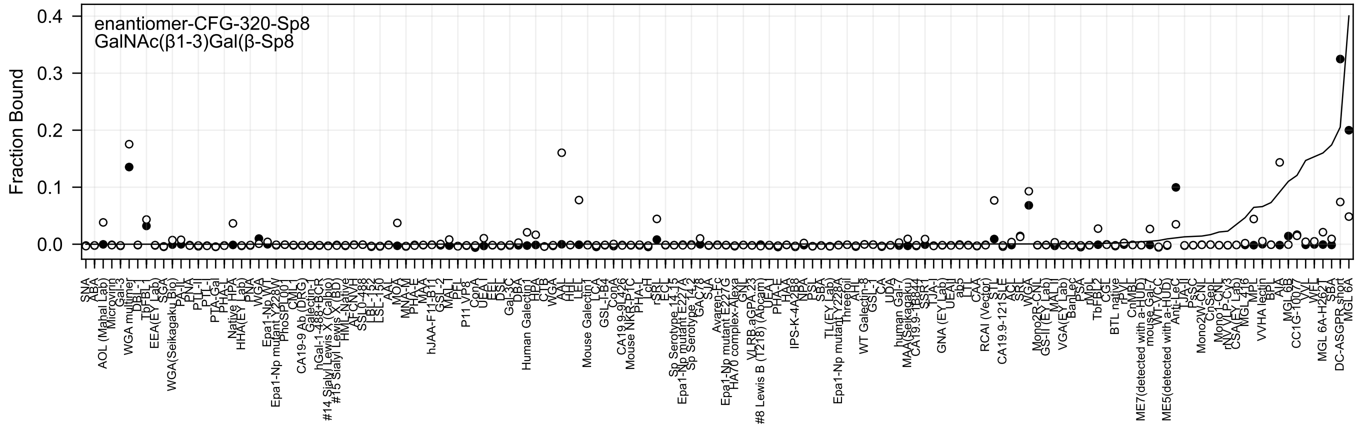

enantiomer-CFG-320-Sp8
GalNAc(β1-3)Gal(β-Sp8
Fraction Bound
0.4
0.3
0.2
0.1
0.0

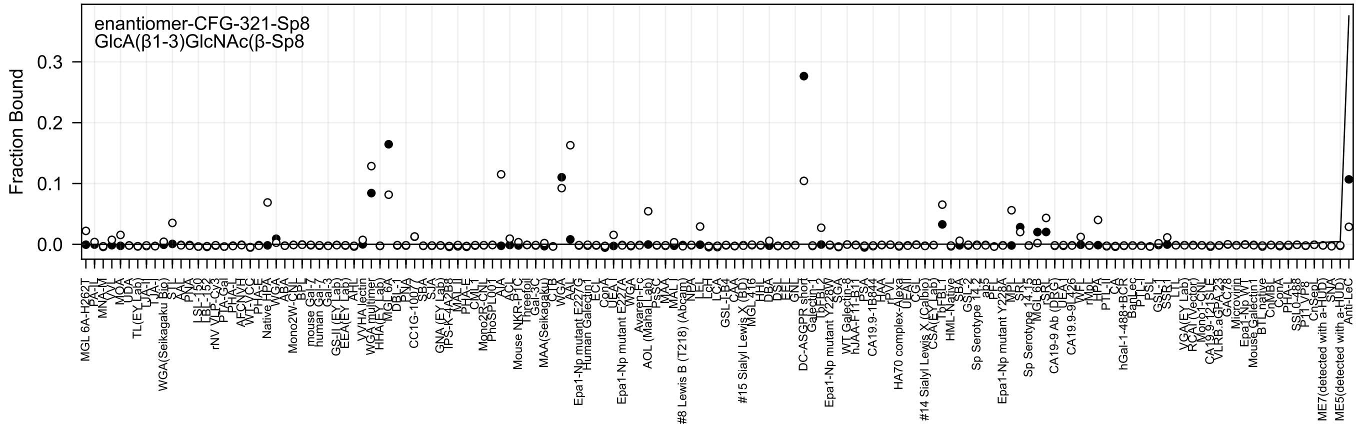
enantiomer-CFG-321-Sp8
GlcA(β1-3)GlcNAc(β-Sp8
Fraction Bound
0.3
0.2
0.1
0.0

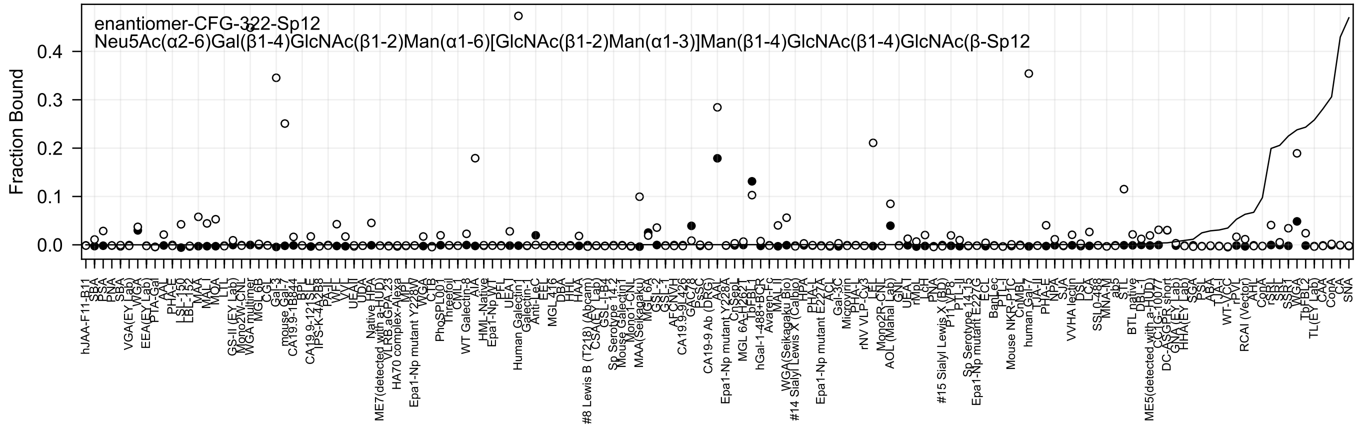
enantiomer-CFG-322-Sp12
Neu5Ac(α2-6)Gal(β1-4)GlcNAc(β1-2)Man(α1-6)[GlcNAc(β1-2)Man(α1-3)]Man(β1-4)GlcNAc(β1-4)GlcNAc(β-Sp12
Fraction Bound
0.0
0.1
0.2
0.3
0.4

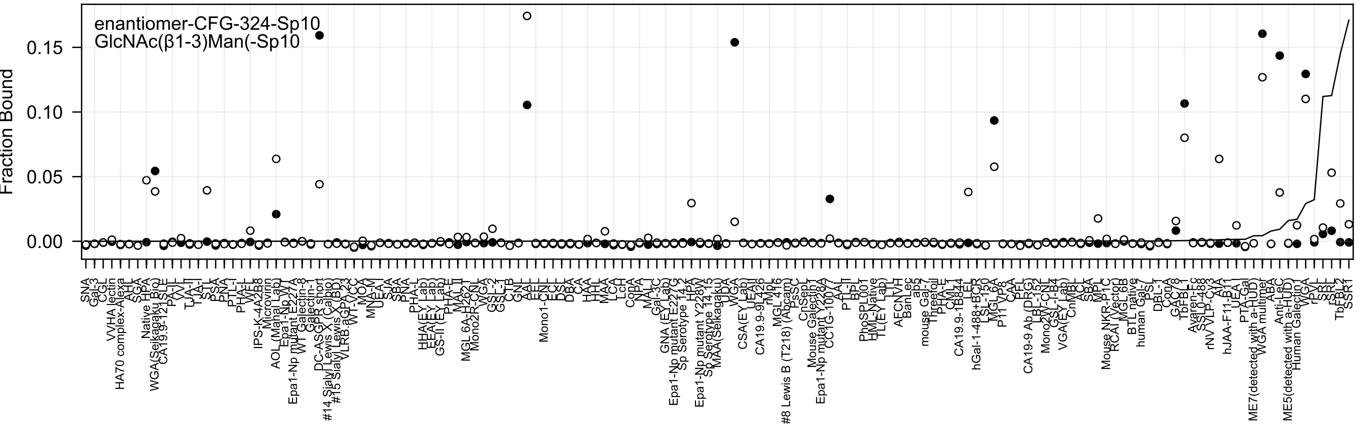
enantiomer-CFG-324-Sp10
GlcNAc(β1-3)Man(-Sp10
Fraction Bound
0.00
0.05
0.10
0.15

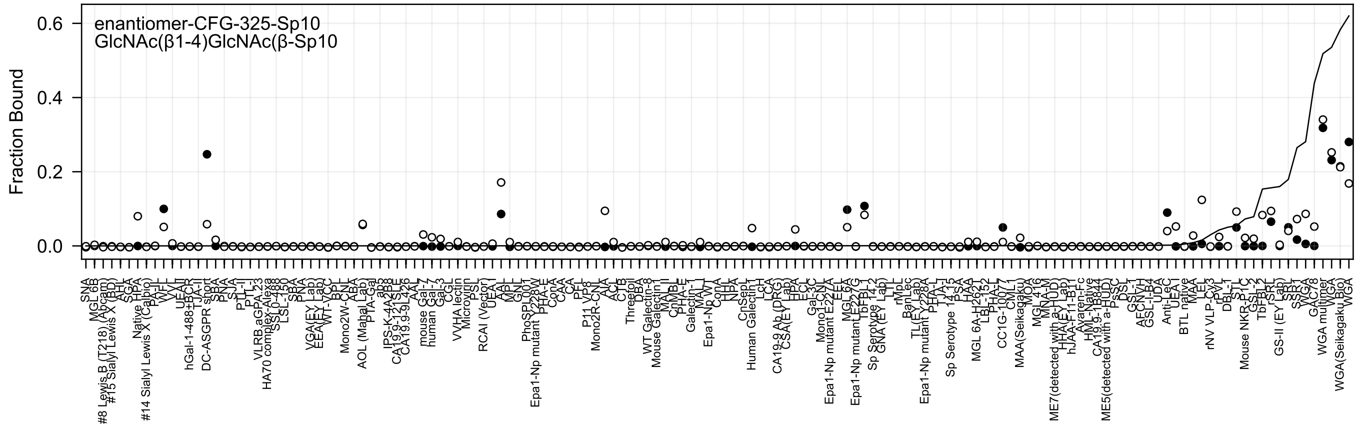
enantiomer-CFG-325-Sp10
GlcNAc(β1-4)GlcNAc(β-Sp10
Fraction Bound
0.6
0.4
0.2
0.0
SNA
MGL 6B
#8 Lewis B (T218) (Abcam)
#15 Sialyl Lewis X (BD)
SGA
Native HPA
#14 Sialyl Lewis X (Calbio)
WFL
VVL
UEA-II
hGal-1-488+BCR
TJA-II
DC-ASGPR short
SBA
PNA
SJA
PTL-II
PTL-I
VLRB.aGPA.23
HA70 complex-Alexa
SSL0-488
LSL-150
SBA
RNA
VGA(EY Lab)
EEA(EY Lab)
WT-YCC
BPL
Mono2W-CNL
ABA
AOL (Mahal Lab)
PTA-Gal
ab5
IPS-K-4A2B8
CA19.9-121SLE
CA19.9-9L426
AAL
mouse Gal-7
human Gal-7
Gal-3
CGL
VVHA lectin
Microvirin
PSL
RCAI (Vector)
UEA-I
AAL
MPL
GNL
PhoSL001
Epa1-Np mutant Y228W
PHA-E
ConA
CAA
CA
PFL
P11 VP8
Mono2R-CNL
AIA
ACL
CTB
Threefoil
DBA
WT Galectin-8
Mouse Galectin-1
MAL-II
CnMBL
PHA-E
Galectin-1
MAL-I
Epa1-Np WT
ConA
NHL
NPA
CnSept
Human Galectin1
LcH
LCA
CA19-9 Ab (DRG)
CSA(EY Lab)
HPA
ECL
Gal-3C
Mono1-CNL
Epa1-Np mutant E227A
EEL
MGL 6A
Epa1-Np mutant E227G
TbFBL1
Sp Serotype 14-2
GNA (EY Lab)
LTL
rMpL
BanLec
TL(EY Lab)
Epa1-Np mutant Y228A
PHA-L
TJA-I
Sp Serotype 14-15
PSA
HAA
MGL 6A-H262T
LBL-152
PHA-L
CC1G-1007
CML1
MAA(Seikagaku)
MOA
MGL 416
MNA-M
ME7(detected with a-HUD)
HHA(EY Lab)
hJAA-F11-B1
Avaren-Fc
HML-Native
CA19.9-1B844
ME5(detected with a-HUD)
PsSC
DSL
GSL-1
AECNYH
GSL-I-B4
UDA
Anti-LeC
UEA-I
BTL native
MAA
LEL
rNV VLP-Cy3
rPVL
DBL-1
STL
Mouse NKR-P1C
GSL-2
TbFBL2
rSRT
GS-II (EY Lab)
SSR1
SSR1
WGA
GAC78
WGA multimer
WGA
WGA(Seikagaku Bio)
WGA

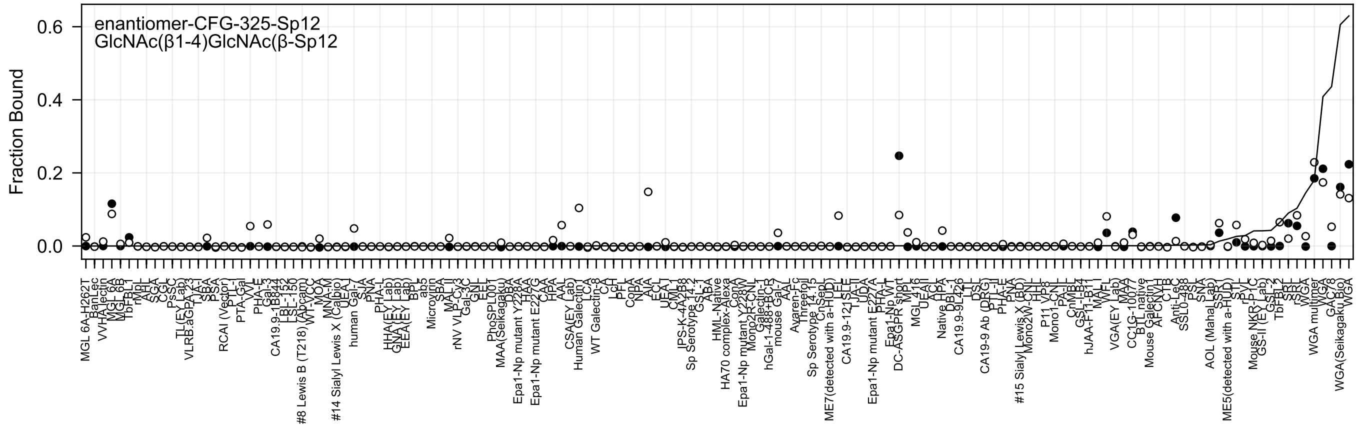
enantiomer-CFG-325-Sp12
GlcNAc(β1-4)GlcNAc(β-Sp12
Fraction Bound

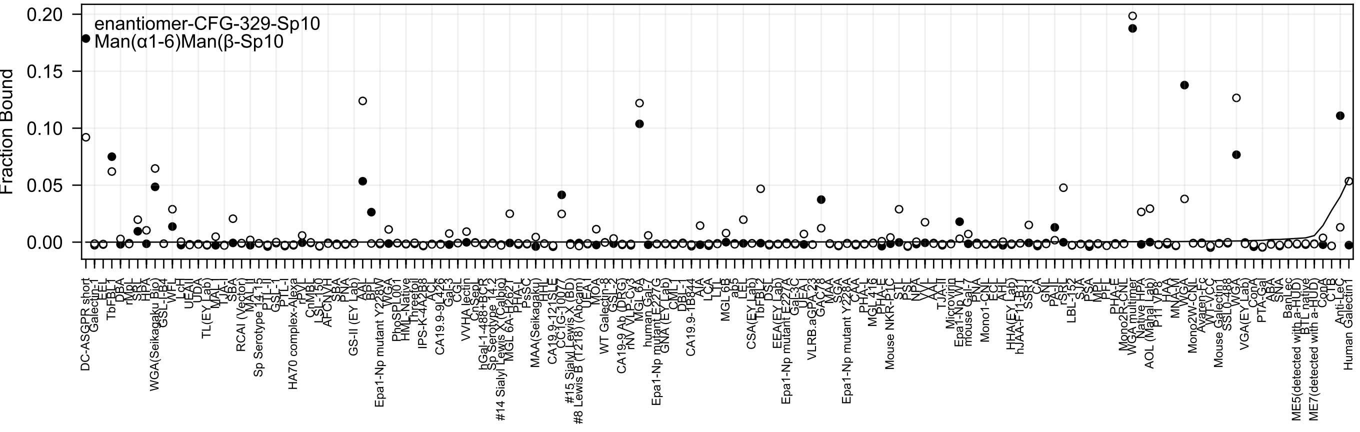
Fraction Bound
enantiomer-CFG-329-Sp10
Man(α1-6)Man(β-Sp10

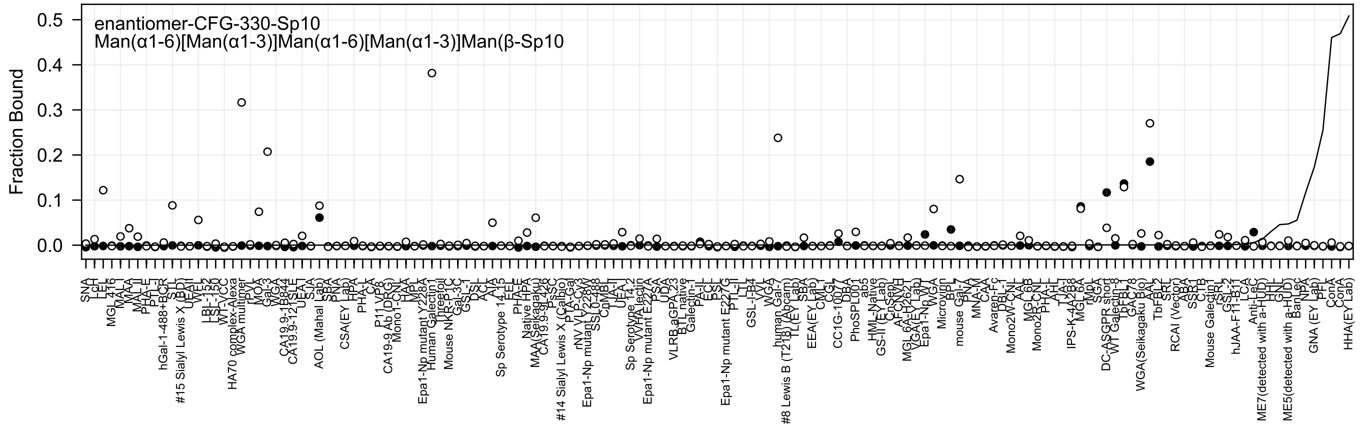
enantiomer-CFG-330-Sp10
Man(α1-6)[Man(α1-3)]Man(α1-6)[Man(α1-3)]Man(β-Sp10
Fraction Bound
0.5
0.4
0.3
0.2
0.1
0.0

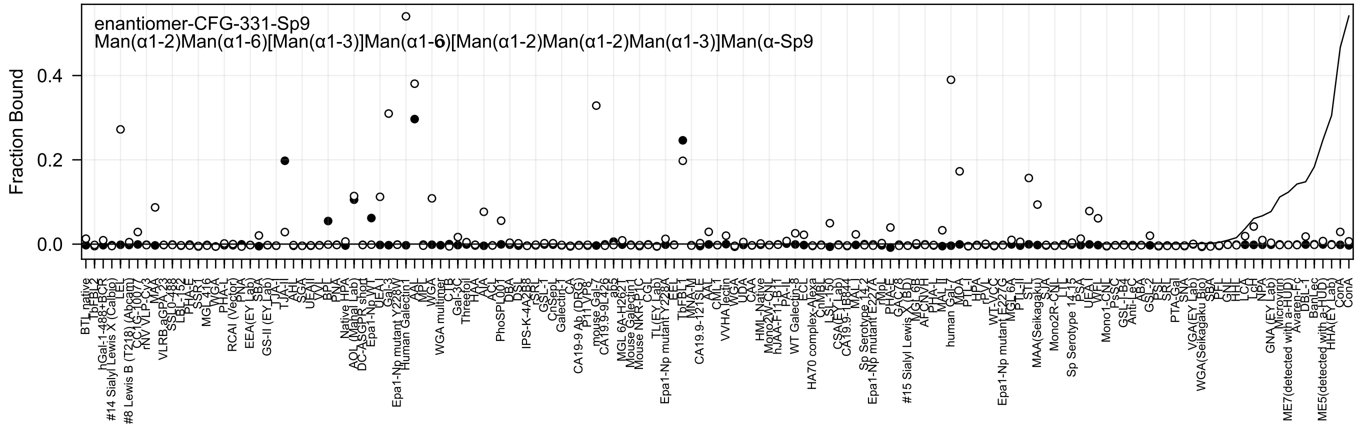
enantiomer-CFG-331-Sp9
Man(α1-2)Man(α1-6)[Man(α1-3)]Man(α1-6)[Man(α1-2)Man(α1-2)Man(α1-3)]Man(α-Sp9
Fraction Bound
0.0
0.2
0.4

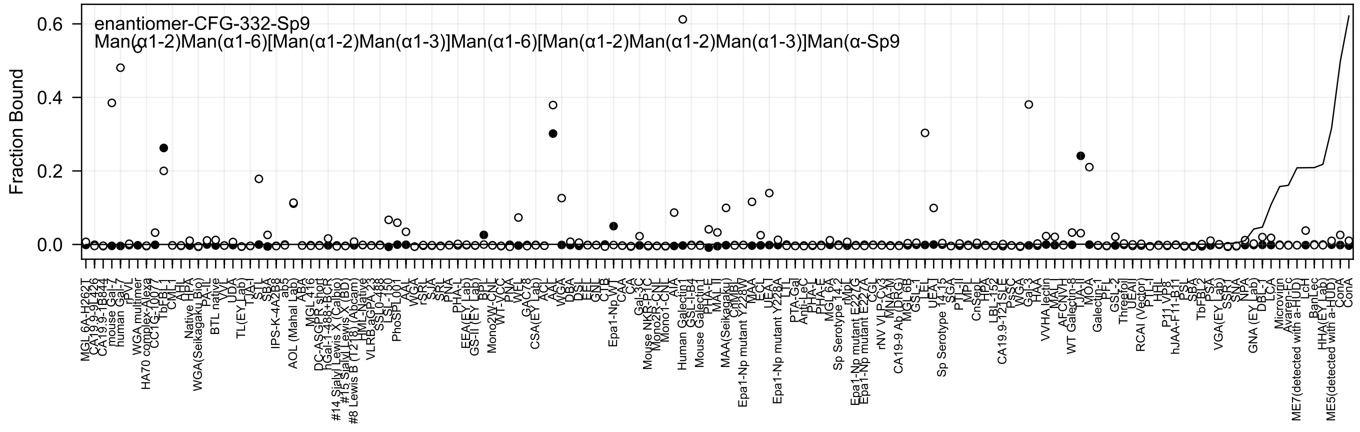
enantiomer-CFG-332-Sp9
Man(α1-2)Man(α1-6)[Man(α1-2)Man(α1-3)]Man(α1-6)[Man(α1-2)Man(α1-2)Man(α1-3)]Man(α-Sp9
Fraction Bound
0.6
0.4
0.2
0.0

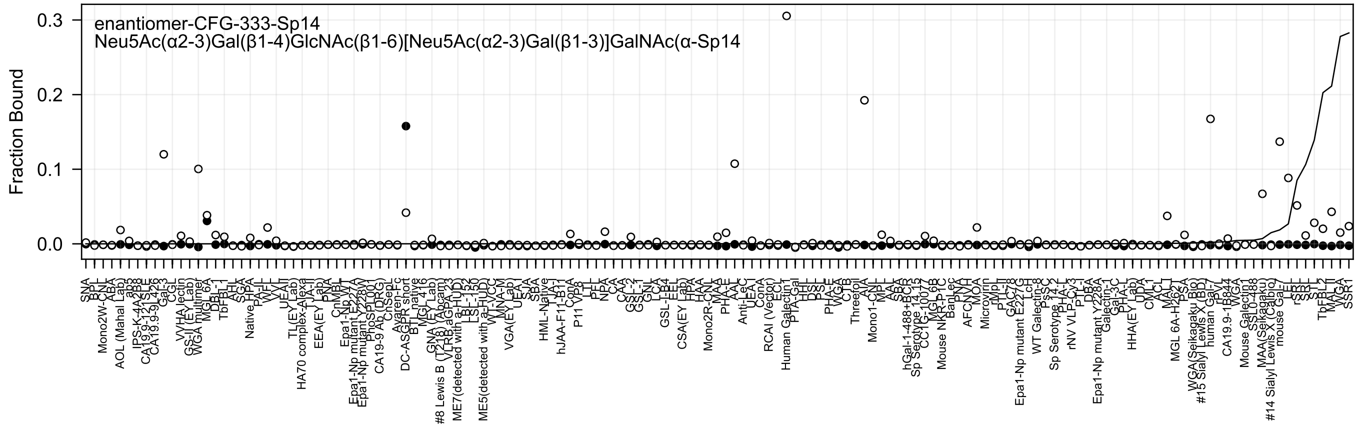
enantiomer-CFG-333-Sp14
Neu5Ac(α2-3)Gal(β1-4)GlcNAc(β1-6)[Neu5Ac(α2-3)Gal(β1-3)]GalNAc(α-Sp14
Fraction Bound
0.3
0.2
0.1
0.0

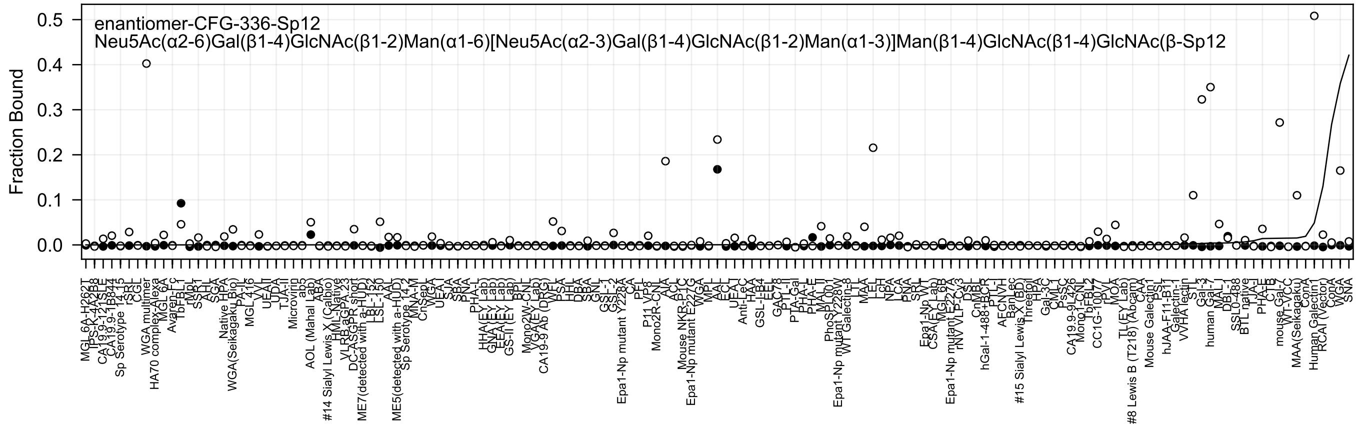
enantiomer-CFG-336-Sp12
Neu5Ac(α2-6)Gal(β1-4)GlcNAc(β1-2)Man(α1-6)[Neu5Ac(α2-3)Gal(β1-4)GlcNAc(β1-2)Man(α1-3)]Man(β1-4)GlcNAc(β1-4)GlcNAc(β-Sp12
Fraction Bound
0.0
0.1
0.2
0.3
0.4
0.5

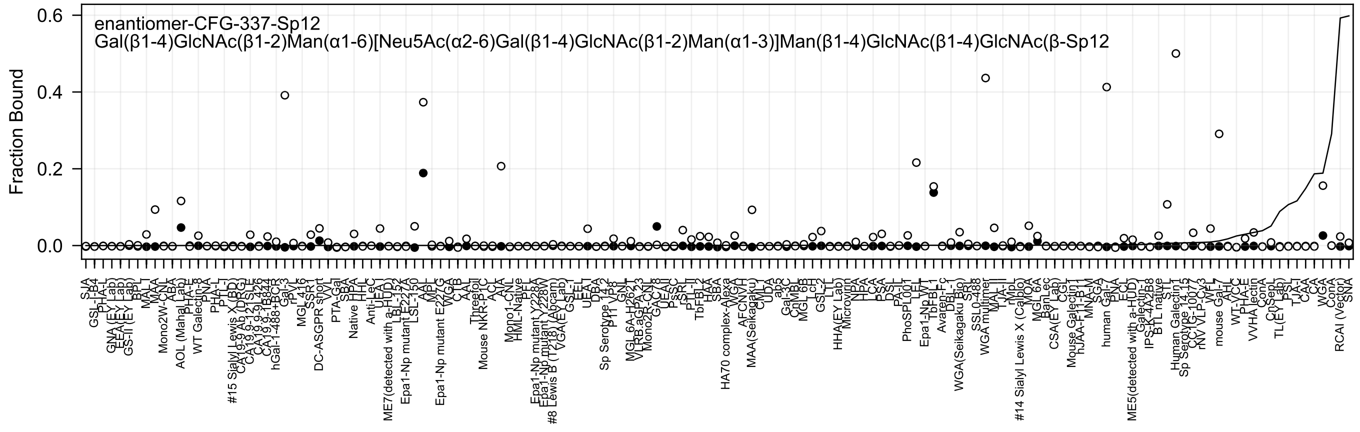

enantiomer-CFG-337-Sp12
Gal(β1-4)GlcNAc(β1-2)Man(α1-6)[Neu5Ac(α2-6)Gal(β1-4)GlcNAc(β1-2)Man(α1-3)]Man(β1-4)GlcNAc(β1-4)GlcNAc(β-Sp12
Fraction Bound
0.6
0.4
0.2
0.0
SJA
GSL-I-B4
PHA-L
GNA (EY Lab)
EEA(EY Lab)
GS-II (EY Lab)
BPL
MAL-I
MAA
Mono2W-CNL
ABA
AOL (Mahal Lab)
PHA-E
WT Galectin-8
PNA
PHA-L
PTL-I
#15 Sialyl Lewis X (BD)
CA19-9 Ab (DRG)
CA19-9-121SLE
CA19-9-1B426
CA19-9-1B844
hGal-1-488+BCR
Gal-3
rPVL
MGL 416
SSR1
DC-ASGPR short
VVL
PTA-Gal
SBA
Native HPA
HHL
Anti-LeC
UEA I
ME7(detected with a-HUD)
LBL-152
Epa1-Np mutant E227A
LSL-150
AAL
MPL
Epa1-Np mutant E227G
WGA
CTB
AAL
Threefoil
Mouse NKR-P1C
ACA
AIA
Mono1-CNL
HML-Native
PFL
Epa1-Np mutant Y228A
Epa1-Np mutant Y228W
#8 Lewis B (T218) (Abcam)
VGA(EY Lab)
GSL-I
EEL
UEA-I
DBA
Sp Serotype 14.2
P11 VP8
GNL
MGL 6A-H262T
VLRB.aGPA.23
Mono2R-CNL
GAC78
UEA-II
PsSC
rSRL
PTL-II
TbFBL-2
HAA
SBA
HA70 complex-Alexa
WGA
AFCNVH
MAA(Seikagaku)
CML
UDA
ap5
Gal-3C
CnMBL
MGL 6B
LcH
GSL-2
LTL
HHA(EY Lab)
Microvirin
NPA
HPA
LCA
PSA
DSL
PA-IL
PhoSPL001
LEL
Epa1-Np WT
TbFBL-1
Avaren-Fc
DBL-1
WGA(Seikagaku Bio)
SRL
SSL0-488
WGA multimer
MAL-II
TJA-II
rMpL
#14 Sialyl Lewis X (Calbio)
MOA
MGL 6A
BanLec
CSA(EY Lab)
CGL
Mouse Galectin 1
hJAA-F11-B.1
MNA-M
SGA
human Gal-7
PNA
ECL
ME5(detected with a-HUD)
Galectin-1
IPS-K-4A2B8
BTL native
STL
Human Galectin 1
Sp Serotype 14.15
CCL2G-1007
rNV VLP-Cy3
WFL
mouse Gal-7
AAL
WT-VCC
PHA-E
VVHA lectin
ConA
CnSepL
TL(EY Lab)
PSL
TJA-I
CAA
CA
WGA
ConA
RCAI (Vector)
SNA

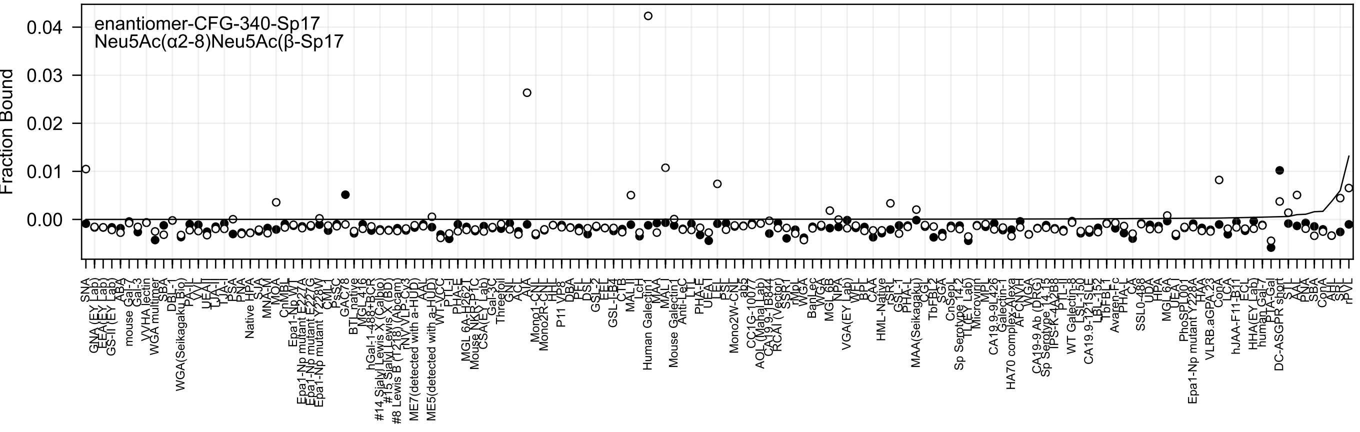

enantiomer-CFG-340-Sp17
Neu5Ac(α2-8)Neu5Ac(β-Sp17
Fraction Bound

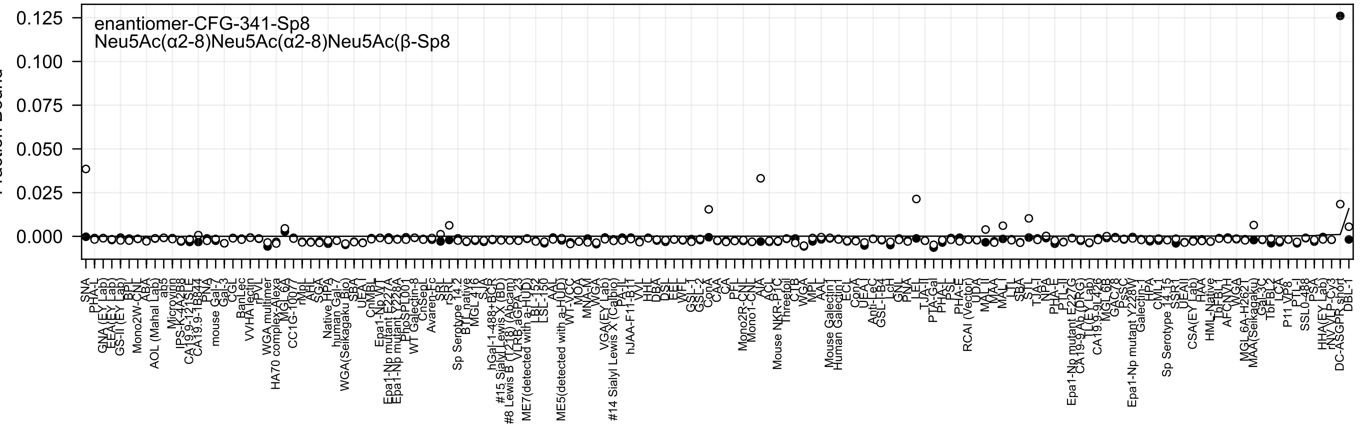
enantiomer-CFG-341-Sp8
Neu5Ac(α2-8)Neu5Ac(α2-8)Neu5Ac(β-Sp8
Fraction Bound
0.125
0.100
0.075
0.050
0.025
0.000

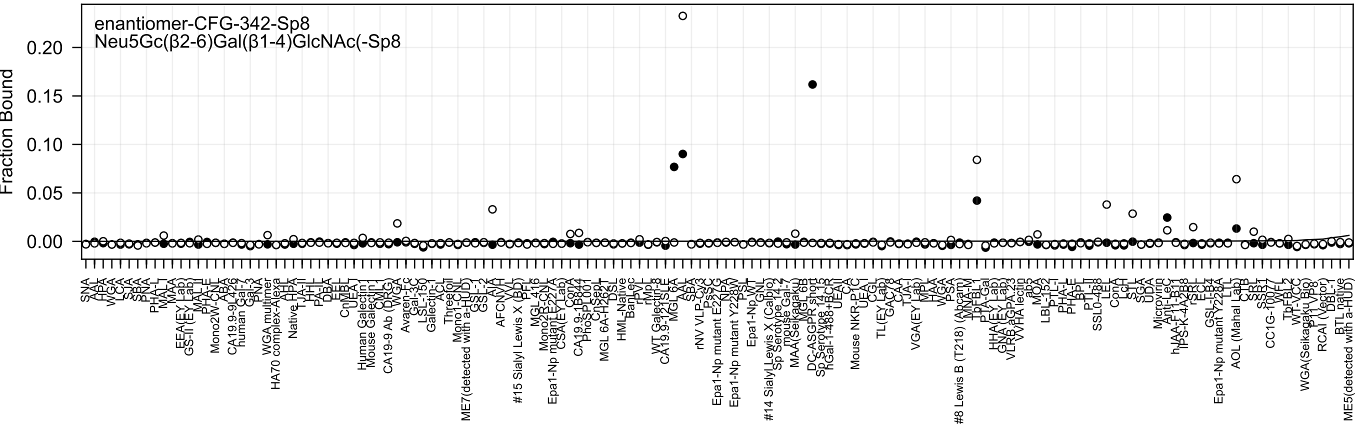

enantiomer-CFG-342-Sp8
Neu5Gc(β2-6)Gal(β1-4)GlcNAc(-Sp8
Fraction Bound
0.00
0.05
0.10
0.15
0.20

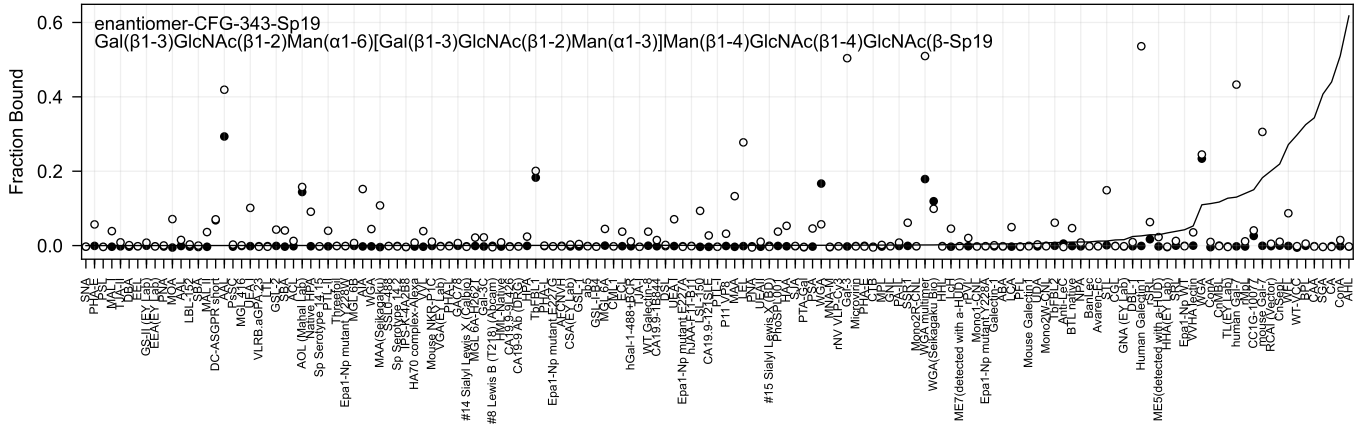
enantiomer-CFG-343-Sp19
Gal(β1-3)GlcNAc(β1-2)Man(α1-6)[Gal(β1-3)GlcNAc(β1-2)Man(α1-3)]Man(β1-4)GlcNAc(β1-4)GlcNAc(β-Sp19
Fraction Bound
0.6
0.4
0.2
0.0
SNA
PHA-E
PSL
MAL-I
TJA-II
DBA
EEL
GS-II (EY Lab)
EEA(EY Lab)
PNA
MOA
AAL
LBL-152
SBA
MAL-II
DC-ASGPR short
AAL
PsSC
MGL 416
UEA
VLRB.aGPA.23
LTL
GSL-2
SBA
ACL
AOL (Mahal Lab)
Native HPA
Sp Serotype 14_15
PTL-II
Threefoil
Epa1-Np mutant Y228W
MGL 6B
AIA
WGA
MAA(Seikagaku)
SSL0-488
Sp Serotype 14_2
TPS-K-4A2B8
HA70 complex-Alexa
VVL
Mouse NKR-P1C
VGA(EY Lab)
PHA-L
GAC78
#14 Sialyl Lewis X (Calbio)
MGL 6A-H262T
Gal-3C
#8 Lewis B (T218) (Abcam)
HML-Native
CA19.9-9L426
CA19-9 Ab (DRG)
HPA
TbFBL1
PHA-L
Epa1-Np mutant E227G
AFCNVH
CSA(EY Lab)
GSL-1
ab5
GSL-I-B4
MGL 6A
CML1
ECL
hGal-1-488+BCR
TJA-I
WT Galectin-8
CA19.9-1B844
DSL
UEA-I
Epa1-Np mutant E227A
hJAA-F11-B11
LSL-150
CA19.9-121SLE
PTL-I
VP8
P11 VP8
MAA
LEL
PNA
UEAII
#15 Sialyl Lewis X (BD)
PhoSL001
HAA
SJA
PTA-Gal
PSA
WGA
MNA-M
rNV VLP-Cy3
Gal-3
Microvirin
PHA-E
CTB
MPL
GNL
PA-IL
SSR1
Mono2R-CNL
WGA multimer
WGA(Seikagaku Bio)
HHL
LcH
ME7(detected with a-HUD)
rPVL
Mono1-CNL
Epa1-Np mutant Y228A
Galectin-1
ABA
LCA
PFL
Mouse Galectin1
UDA
Mono2W-CNL
TbFBL2
Anti-LeC
BTL native
NPA
BanLec
Avaren-Fc
STL
CGL
GNA (EY Lab)
DBL-1
Human Galectin1
rSRL
ME5(detected with a-HUD)
HHA(EY Lab)
SRL
Epa1-Np WT
VVHA lectin
WGA
ConA
CnMBL
TL(EY Lab)
human Gal-7
rMpL
CC1G-1007
mouse Gal-7
RCAI (Vector)
CnSepL
WFL
WT-VCC
BPL
CAA
SGA
CA
ConA
AHL

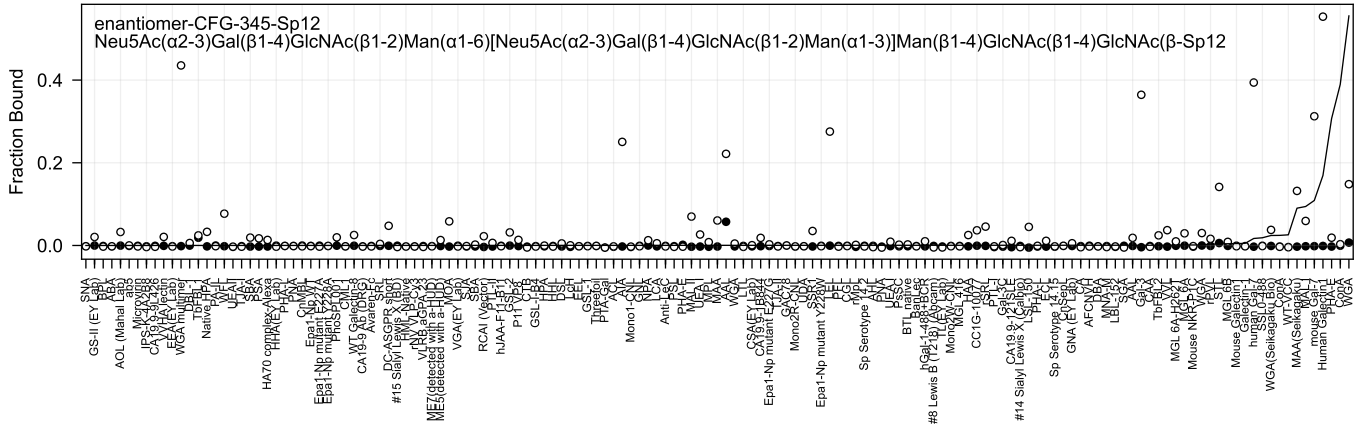
enantiomer-CFG-345-Sp12
Neu5Ac(α2-3)Gal(β1-4)GlcNAc(β1-2)Man(α1-6)[Neu5Ac(α2-3)Gal(β1-4)GlcNAc(β1-2)Man(α1-3)]Man(β1-4)GlcNAc(β1-4)GlcNAc(β-Sp12
Fraction Bound
0.0
0.2
0.4

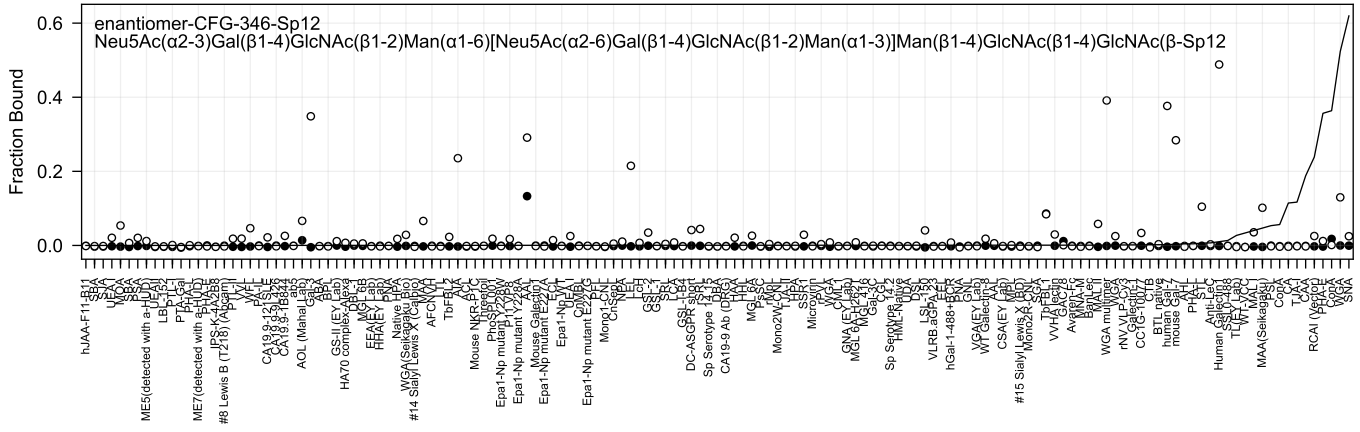
enantiomer-CFG-346-Sp12
Neu5Ac(α2-3)Gal(β1-4)GlcNAc(β1-2)Man(α1-6)[Neu5Ac(α2-6)Gal(β1-4)GlcNAc(β1-2)Man(α1-3)]Man(β1-4)GlcNAc(β1-4)GlcNAc(β-Sp12
Fraction Bound

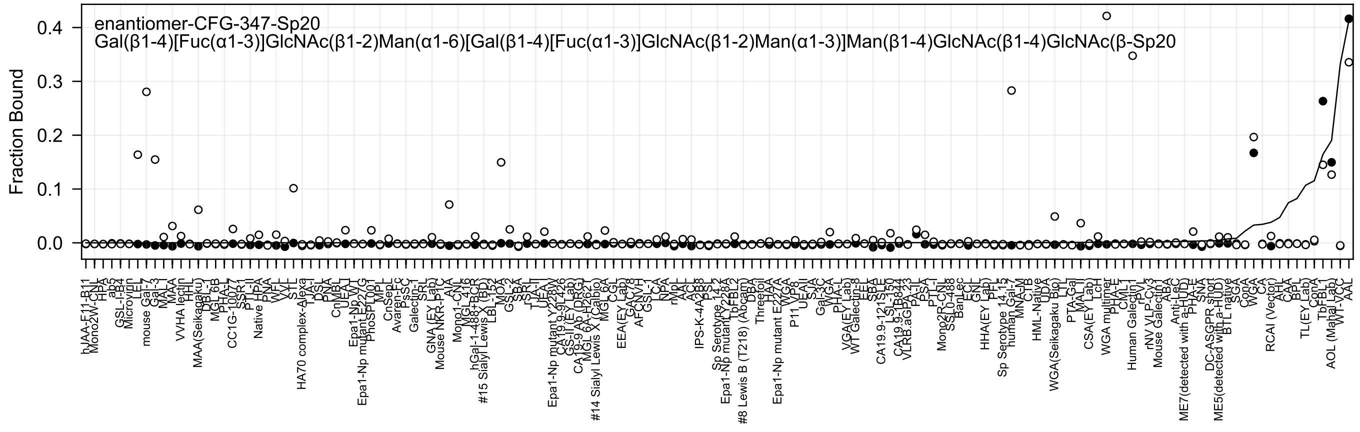

enantiomer-CFG-347-Sp20
Gal(β1-4)[Fuc(α1-3)]GlcNAc(β1-2)Man(α1-6)[Gal(β1-4)[Fuc(α1-3)]GlcNAc(β1-2)Man(α1-3)]Man(β1-4)GlcNAc(β1-4)GlcNAc(β-Sp20
Fraction Bound
0.0
0.1
0.2
0.3
0.4
hJAA-F11-B11
Mono2W-CNL
HPA
ab5
GSL-I-B4
Microvirin
LEL
mouse Gal-7
Gal-3
MAL_I
MAA
VVHA lectin
HHL
MAA(Seikagaku)
DBL-1
MGL 6B
CC1G-1007
SSR1
PTL-II
Native HPA
PNA
WFL
VVL
STL
HA70 complex-Alexa
TJA-I
DSL
PNA
CnMBL
UEA-I
Epa1-Np WT
Epa1-Np mutant E227G
PhoSL001
MPL
CnSepL
Avaren-Fc
Galectin-1
SRL
GNA (EY Lab)
Mouse NKR-P1C
AIA
Mono1-CNL
hGal-1-488+BCR
#15 Sialyl Lewis X (BD)
LBL-152
MOA
GSL-2
SBA
rSRL
TJA-II
UEA-I
Epa1-Np mutant Y228W
GS-II (EY Lab)
CA19-9 Ab (DRG)
#14 Sialyl Lewis X (Calbio)
MGL 6A
CGL
EEA(EY Lab)
GSL-1
LCA
NPA
rNpL
AAL
ACL
IPS-K-4A2B8
Sp Serotype 14-2
Epa1-Np mutant Y228A
TbFBL2
#8 Lewis B (T218) (Abcam)
DBA
Threefoil
HAA
Epa1-Np mutant E227A
WGA
VP8
P11
UEA-II
SJA
Gal-3C
WGA
PHA-L
VGA(EY Lab)
WT Galectin-8
LEA
SBA
CA19.9-121SLE
LSL-150
CA19.9-1B844
VLRB.aGPA.23
PA-IL
PSA
PTL-I
Mono2R-CNL
BanLec
ECL
GNL
HHA(EY Lab)
PFL
Sp Serotype 14-15
human Gal-7
MNA-M
CTB
HML-Native
UDA
WGA(Seikagaku Bio)
LTL
PTA-Gal
MAL-II
CSA(EY Lab)
LcH
WGA multimer
PHA-E
CML-1
Human Galectin-1
rPVL
rNV VLP-Cy3
Mouse Galectin-1
ABA
Anti-LeC
ME7(detected with a-HUD)
PHA-E
SNA
DC-ASGPR short
ME5(detected with a-HUD)
BTL native
SGA
ConA
CA
RCAI (Vector)
CAA
BPL
TL(EY Lab)
ConA
TbFBL1
AOL (Mahal Lab)
WT-VCC
AAL

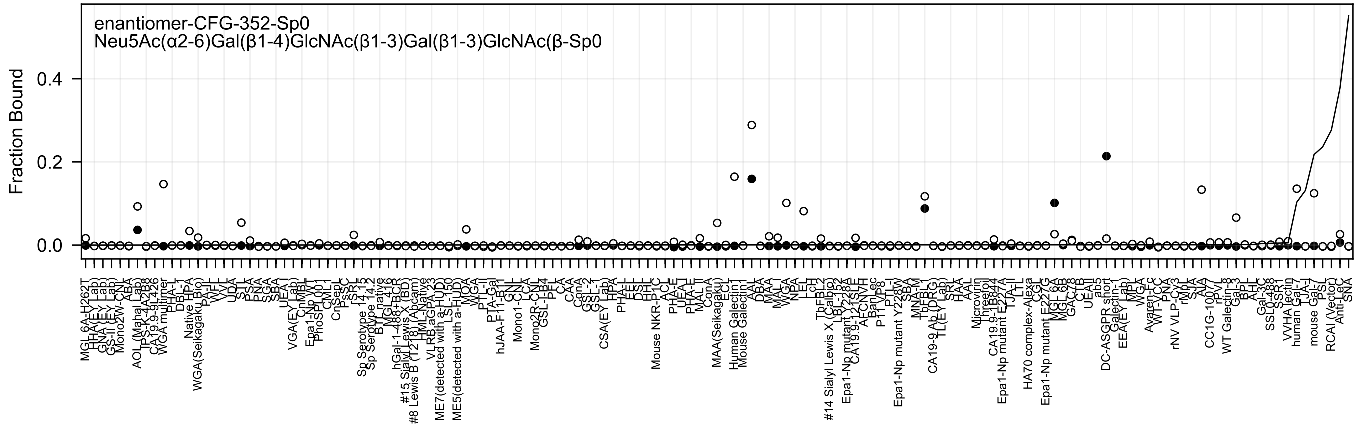
enantiomer-CFG-352-Sp0
Neu5Ac(α2-6)Gal(β1-4)GlcNAc(β1-3)Gal(β1-3)GlcNAc(β-Sp0
Fraction Bound
0.4
0.2
0.0

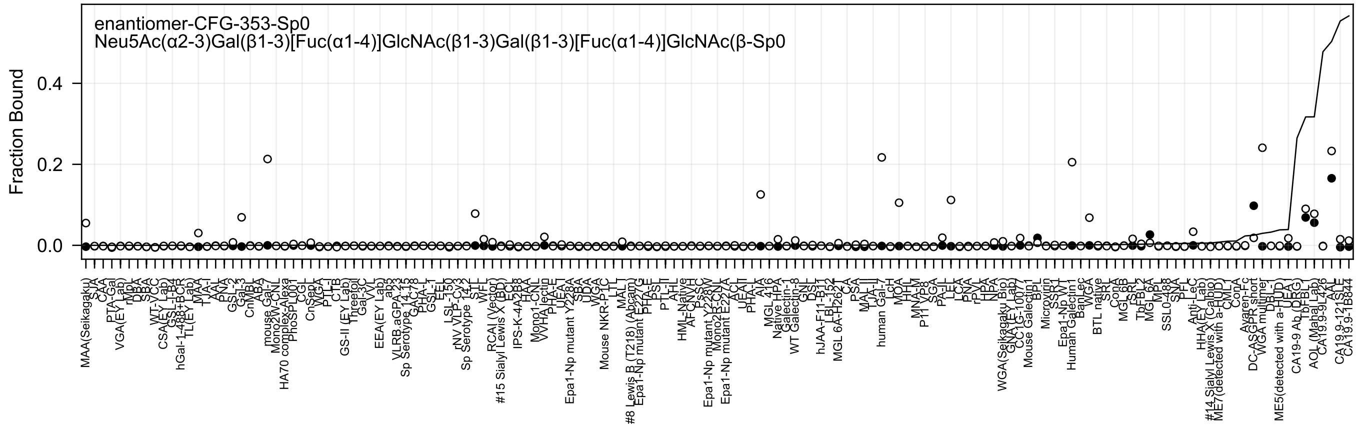
enantiomer-CFG-353-Sp0
Neu5Ac(α2-3)Gal(β1-3)[Fuc(α1-4)]GlcNAc(β1-3)Gal(β1-3)[Fuc(α1-4)]GlcNAc(β-Sp0
Fraction Bound
0.4
0.2
0.0

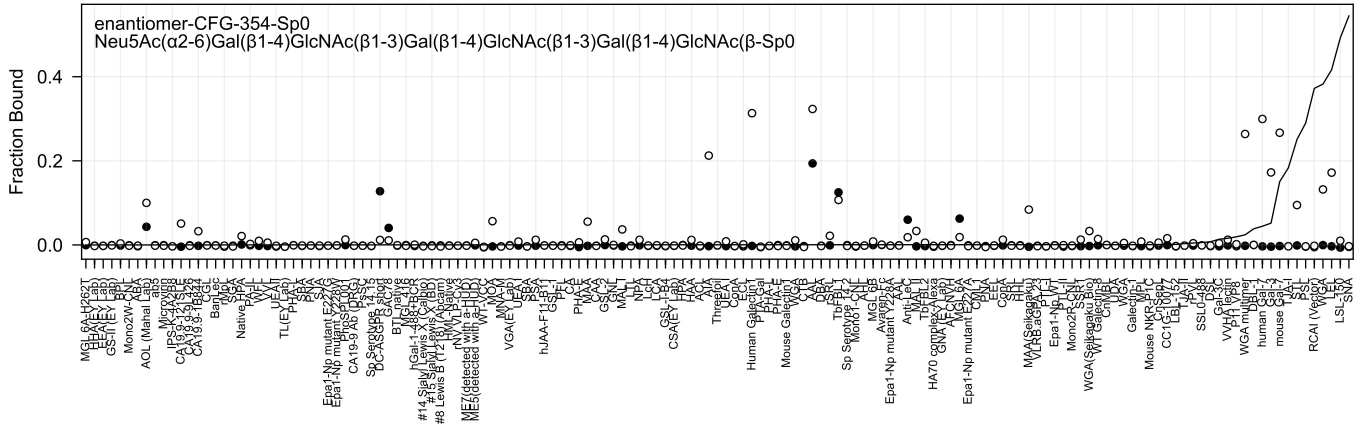
enantiomer-CFG-354-Sp0
Neu5Ac(α2-6)Gal(β1-4)GlcNAc(β1-3)Gal(β1-4)GlcNAc(β1-3)Gal(β1-4)GlcNAc(β-Sp0
Fraction Bound
0.0
0.2
0.4
MGL 6A-H262T
HHA(EY Lab)
EEA(EY Lab)
GS-II (EY Lab)
BPL
Mono2W-CNL
ABA
AOL (Mahal Lab)
abs
Microvirin
IPS-K-4A2B8
CA19.9-121SLE
CA19.9-9L426
CA19.9-1B844
CGL
BanLec
rMpL
SGA
Native HPA
PA-IL
WFL
VVL
UEA-II
TL(EY Lab)
PHA-L
SBA
PNA
SJA
Epa1-Np mutant E227G
Epa1-Np mutant Y228W
PhoSL001
CA19-9 Ab (DRG)
PsSC
Sp Serotype 14.15
DC-ASGPR short
GAC78
BTL native
MGL 416
hGal-1-488+BCR
#14 Sialyl Lewis X (Calbio)
#15 Sialyl Lewis X (BD)
#8 Lewis B (T218) (Abcam)
HML-Native
rNV VLP-Cy3
ME7(detected with a-HUD)
ME5(detected with a-HUD)
WT-VCC
MOA
MNA-M
VGA(EY Lab)
UEA-I
SBA
PSA
hJAA-F11-B11
GSL-I
PEL
CA
PHA-E
MAA
CAA
GSL-2
GNL
MAL-I
LTL
NPA
LcH
LCA
GSL-I-B4
CSA(EY Lab)
HPA
HAA
ACL
AIA
Threefoil
UEA-I
ConA
ECL
Human Galectin1
PTA-Gal
PHA-E
PHA-L
Mouse Galectin1
WGA
CTB
AAL
DBA
RSL
TbFBL1
Sp Serotype 14.2
Mono1-CNL
AHL
MGL 6B
Avaren-Fc
Epa1-Np mutant Y228A
AAL
Anti-LeC
MAL-II
TbFBL2
HA70 complex-Alexa
GNA (EY Lab)
AFCNVH
MGL 6A
Epa1-Np mutant E227A
CVN
PNA
EEL
ConA
SRL
HHL
MAA(Seikagaku)
VLRB.aGPA.23
PTL-I
Epa1-Np WT
PTL-II
Mono2R-CNL
S-SRT1
WGA(Seikagaku Bio)
WT Galectin-8
CnMBL
UDA
VVA
Galectin-1
MPL
Mouse NKR-P1C
CnSep
CC1G-10077
LBL-152
TJA-II
rPVL
SSL0-488
DSL
Gal-3C
VVH lectin
P11 VP8
WGA multimer
DBL-1
human Gal-7
Gal-3
mouse Gal-7
TJA-I
STL
PSL
RCAI (Vector)
WGA
LEL
LSL-150
SNA

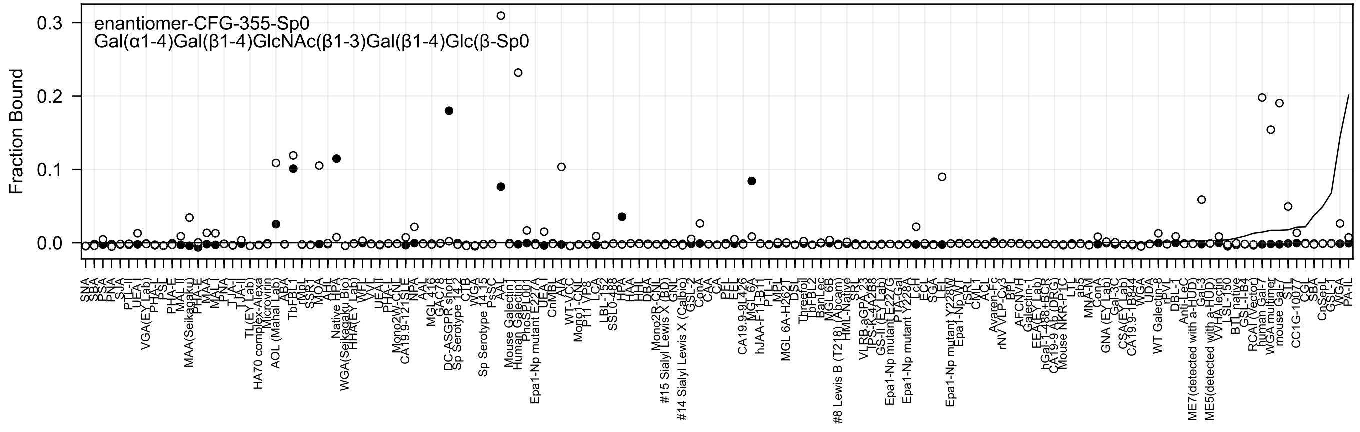
enantiomer-CFG-355-Sp0
Gal(α1-4)Gal(β1-4)GlcNAc(β1-3)Gal(β1-4)Glc(β-Sp0
Fraction Bound
0.3
0.2
0.1
0.0

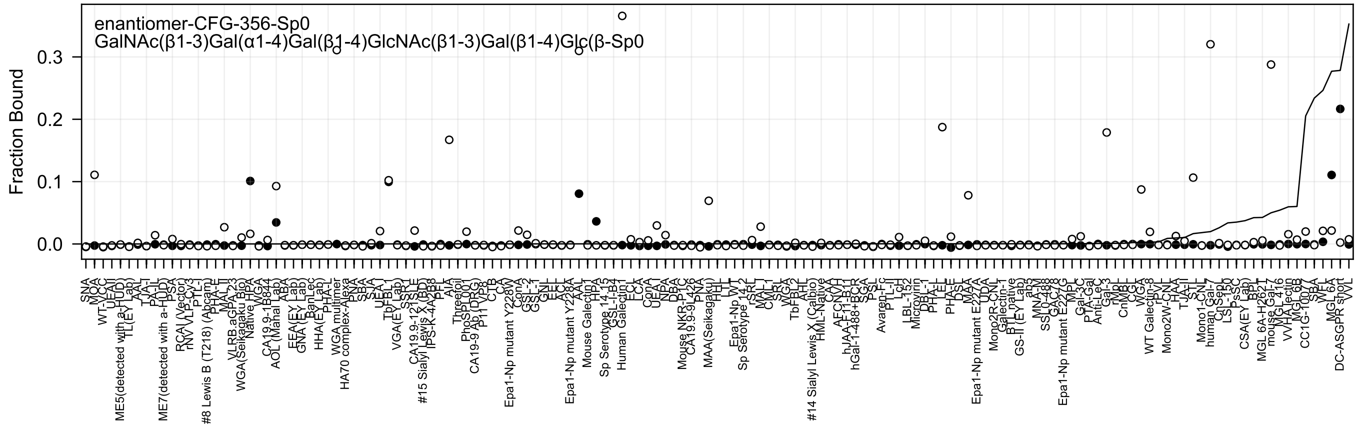
enantiomer-CFG-356-Sp0
GalNAc(β1-3)Gal(α1-4)Gal(β1-4)GlcNAc(β1-3)Gal(β1-4)Glc(β-Sp0
Fraction Bound
0.0
0.1
0.2
0.3
SNA
MOA
WT-VCC
UEAII
ME5(detected with a-HUD)
TL(EY Lab)
AAL
TJA-I
PA-IL
ME7(detected with a-HUD)
PSA
RCAI (Vector)
rNV VLP-Cy3
PTL-I
#8 Lewis B (T218) (Abcam)
PHA-E
MAL II
VLRB.aGPA.23
WGA(Seikagaku Bio)
Native HPA
WGA
CA19.9-1B844
AOL (Mahal Lab)
ABA
EEA(EY Lab)
GNA (EY Lab)
BanLec
HHA(EY Lab)
PHA-L
WGA multimer
HA70 complex-Alexa
PNA
SBA
SJA
UEA-I
TbFBL1
VGA(EY Lab)
SSR1
CA19.9-121SLE
#15 Sialyl Lewis X (BD)
IPS-K-4A2B8
PFL
AIA
Threefoil
PhoSL001
CA19-9 Ab (DRG)
P11 VP8
CTB
CA
Epa1-Np mutant Y228W
ConA
GSL-2
GSL-1
GNL
EEL
ACL
Epa1-Np mutant Y228A
AAL
Mouse Galectin1
HPA
Sp Serotype 14.15
GSL-I-B4
Human Galectin1
ECA
LCA
ConA
UEA1
NPA
DBA
Mouse NKR-P1C
CA19.9.9L426
PNA
MAA(Seikagaku)
HHL
LTL
Epa1-Np WT
Sp Serotype 14.2
rSRL
MAL-I
CML-I
SRL
WGA
TbFBL2
AHL
#14 Sialyl Lewis X (Calbio)
HML-Native
CAA
AFCNVH
hJAA-F11-B11
hGal-1-488+BCR
SGA
PSL
Avaren-Fc
PTL-II
LcH
LBL-152
Microvirin
DBL-1
PHA-L
LEL
PHA-E
DSL
MAA
Epa1-Np mutant E227A
UDA
Mono2R-CNL
Galectin-1
BTL native
GS-II (EY Lab)
ab3
MNA-M
SSL0-488
GAC78
Epa1-Np mutant E227G
MPL
Gal-3C
PTA-Gal
Anti-LeC
Gal-3
rMpL
CnMBL
CGL
WGA
WT Galectin-8
rPvL
Mono2W-CNL
HAA
TJA-II
STL
Mono1-CNL
human Gal-7
Cn-Sep
LS-150
PsSC
CSA(EY Lab)
BPL
MGL 6A-H262T
mouse Gal-7
MGL 4.16
VVHA lectin
MGL 6B
CC1G-1007
SBA
WFL
MGL 6A
DC-ASGPR short
VVL

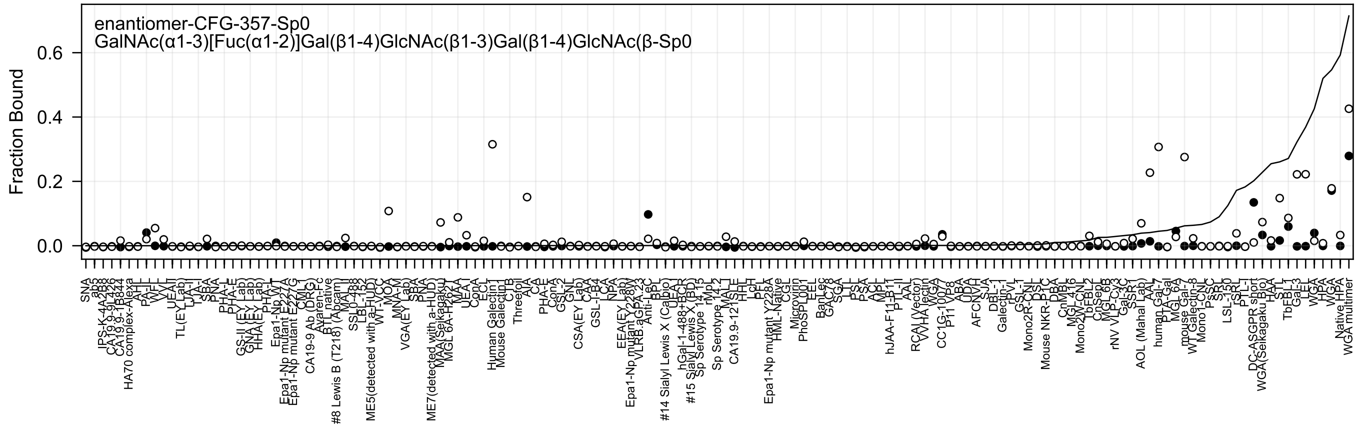
enantiomer-CFG-357-Sp0
GalNAc(α1-3)[Fuc(α1-2)]Gal(β1-4)GlcNAc(β1-3)Gal(β1-4)GlcNAc(β-Sp0
Fraction Bound

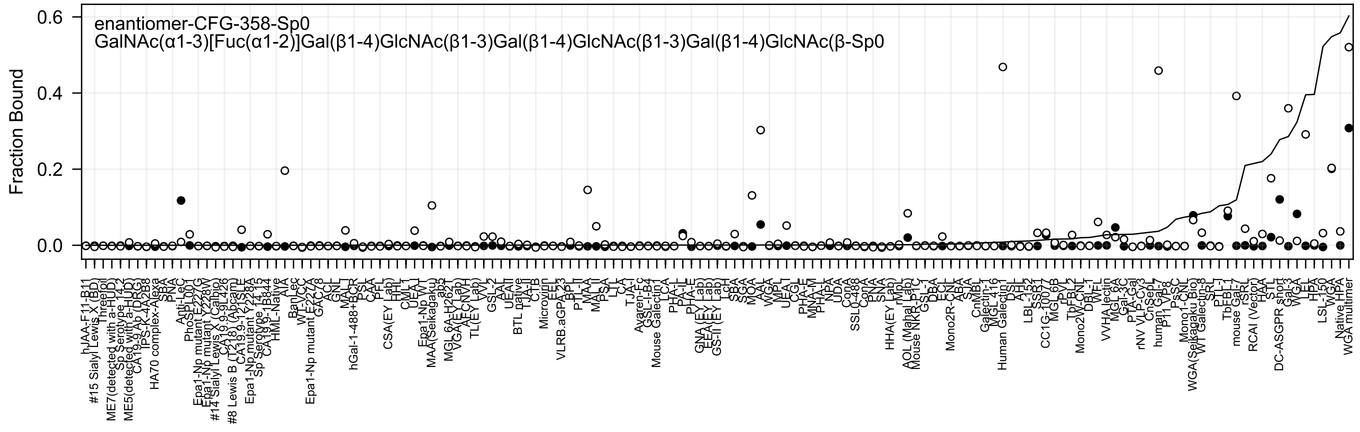

enantiomer-CFG-358-Sp0
GalNAc(α1-3)[Fuc(α1-2)]Gal(β1-4)GlcNAc(β1-3)Gal(β1-4)GlcNAc(β1-3)Gal(β1-4)GlcNAc(β-Sp0
Fraction Bound

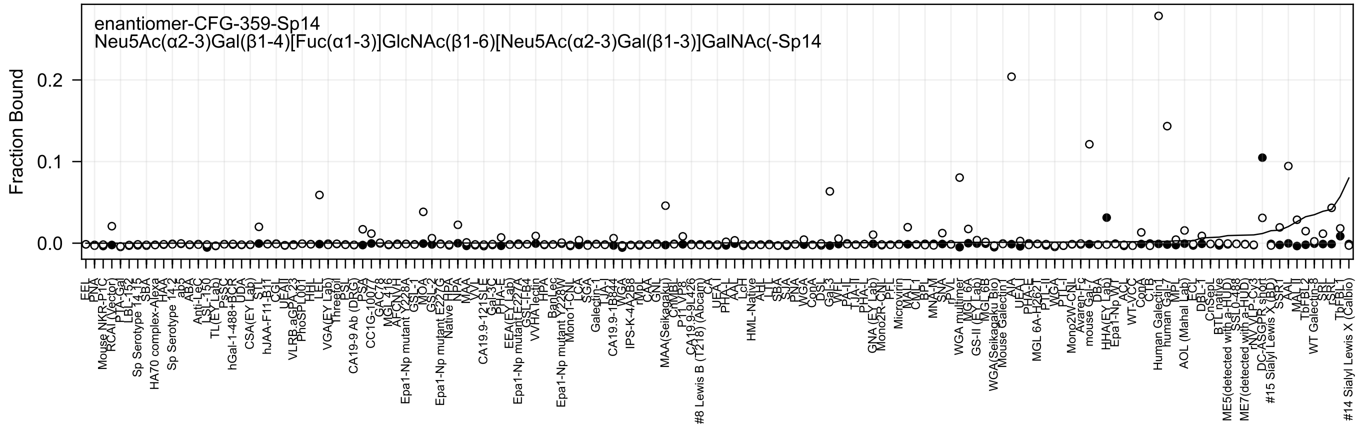

enantiomer-CFG-359-Sp14
Neu5Ac(α2-3)Gal(β1-4)[Fuc(α1-3)]GlcNAc(β1-6)[Neu5Ac(α2-3)Gal(β1-3)]GalNAc(-Sp14
Fraction Bound
0.2
0.1
0.0

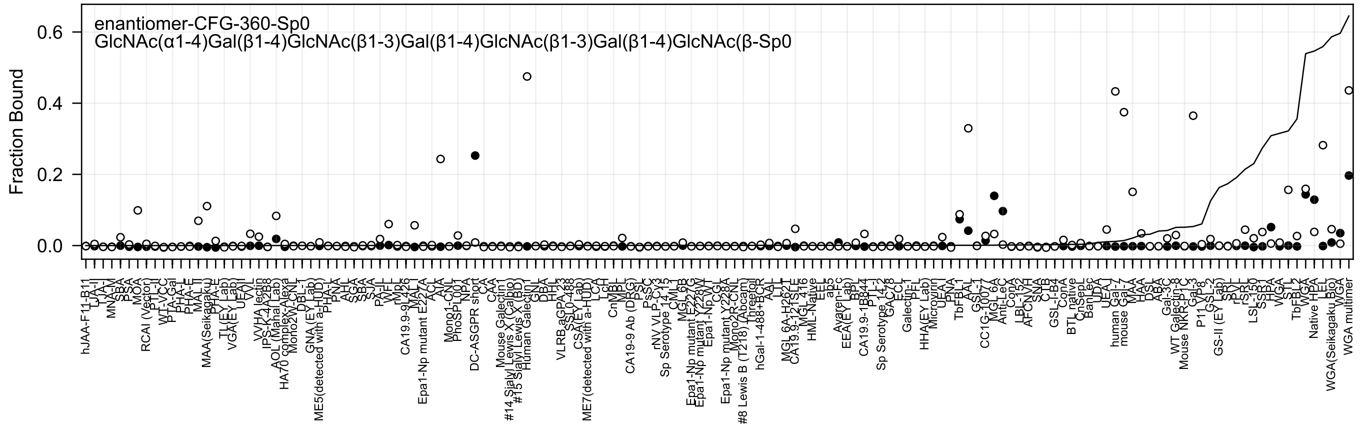
enantiomer-CFG-360-Sp0
GlcNAc(α1-4)Gal(β1-4)GlcNAc(β1-3)Gal(β1-4)GlcNAc(β1-3)Gal(β1-4)GlcNAc(β-Sp0
Fraction Bound
0.6
0.4
0.2
0.0

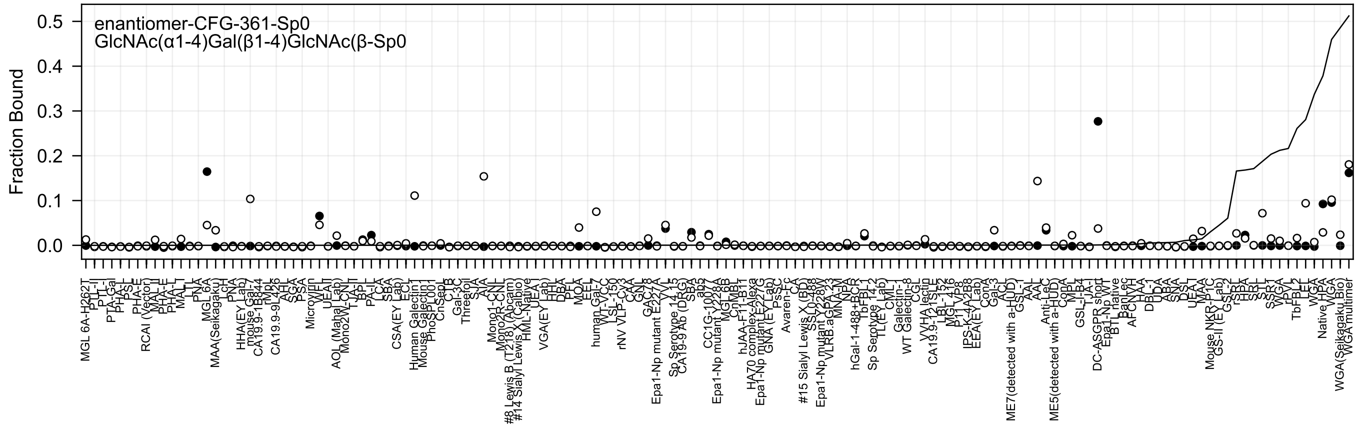

enantiomer-CFG-361-Sp0
GlcNAc(α1-4)Gal(β1-4)GlcNAc(β-Sp0
Fraction Bound

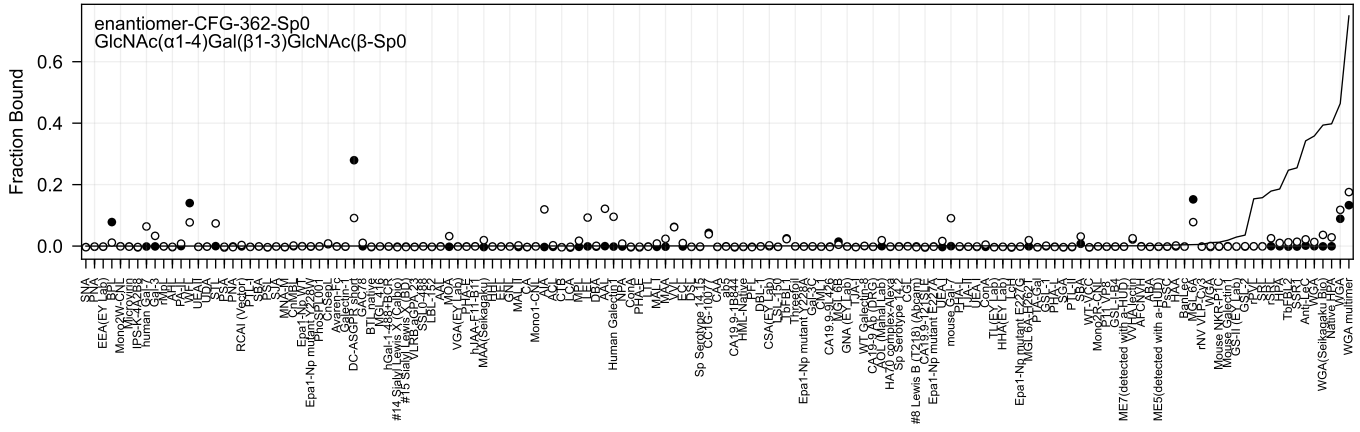
enantiomer-CFG-362-Sp0
GlcNAc(α1-4)Gal(β1-3)GlcNAc(β-Sp0
Fraction Bound
0.0
0.2
0.4
0.6

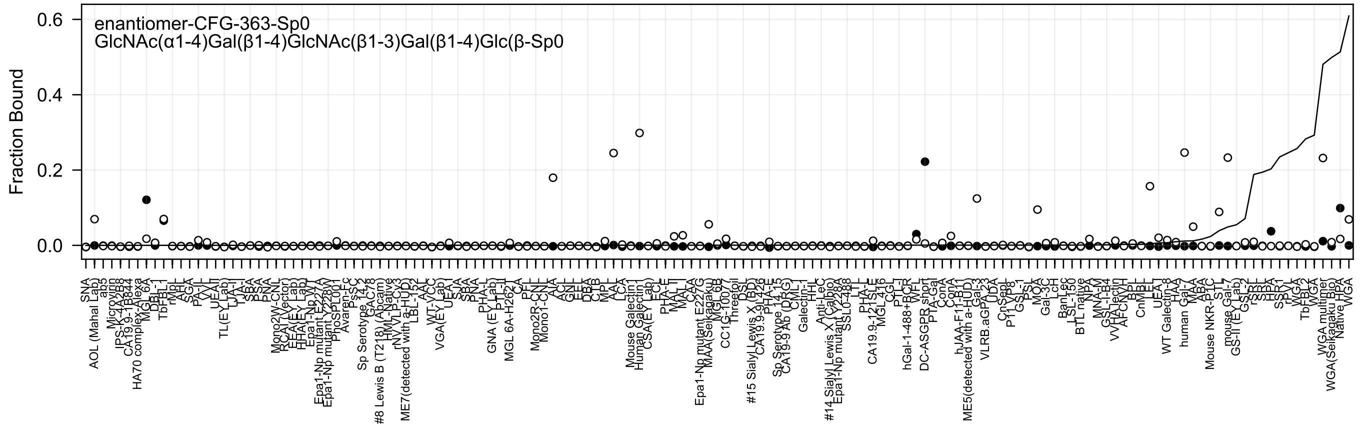

enantiomer-CFG-363-Sp0
GlcNAc(α1-4)Gal(β1-4)GlcNAc(β1-3)Gal(β1-4)Glc(β-Sp0
Fraction Bound

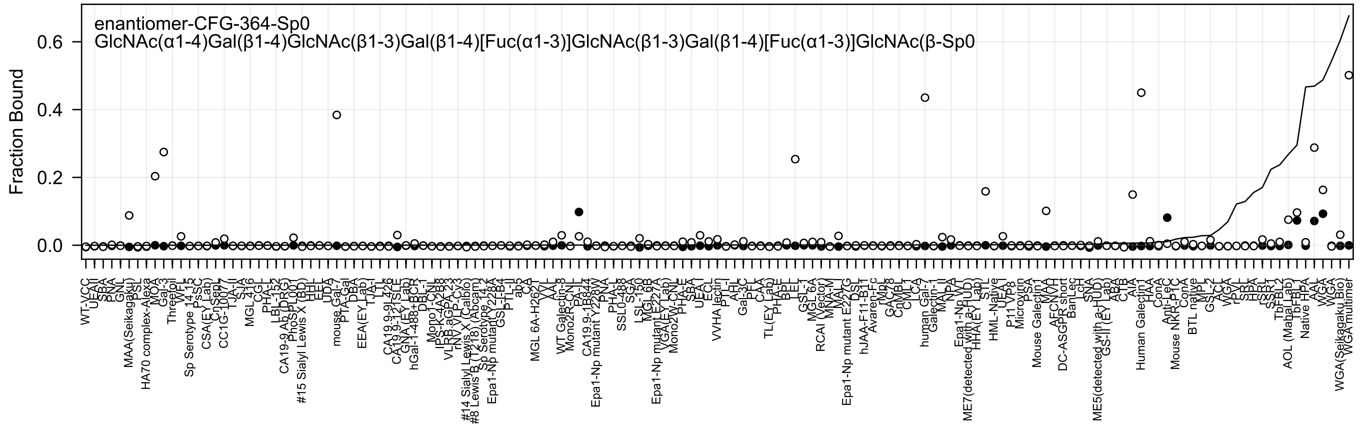
enantiomer-CFG-364-Sp0
GlcNAc(α1-4)Gal(β1-4)GlcNAc(β1-3)Gal(β1-4)[Fuc(α1-3)]GlcNAc(β1-3)Gal(β1-4)[Fuc(α1-3)]GlcNAc(β-Sp0
Fraction Bound
0.0
0.2
0.4
0.6

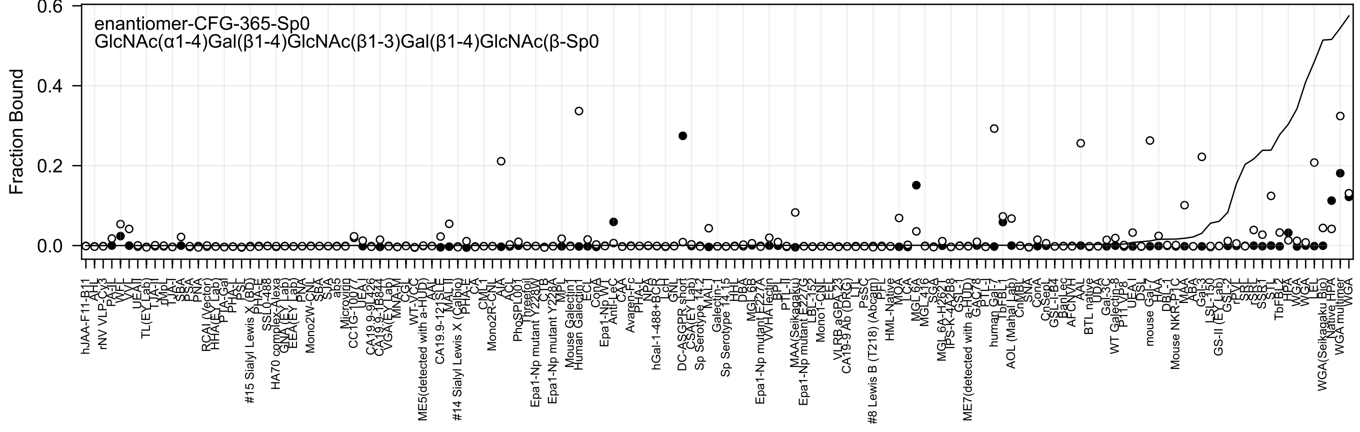
enantiomer-CFG-365-Sp0
GlcNAc(α1-4)Gal(β1-4)GlcNAc(β1-3)Gal(β1-4)GlcNAc(β-Sp0
Fraction Bound
0.6
0.4
0.2
0.0

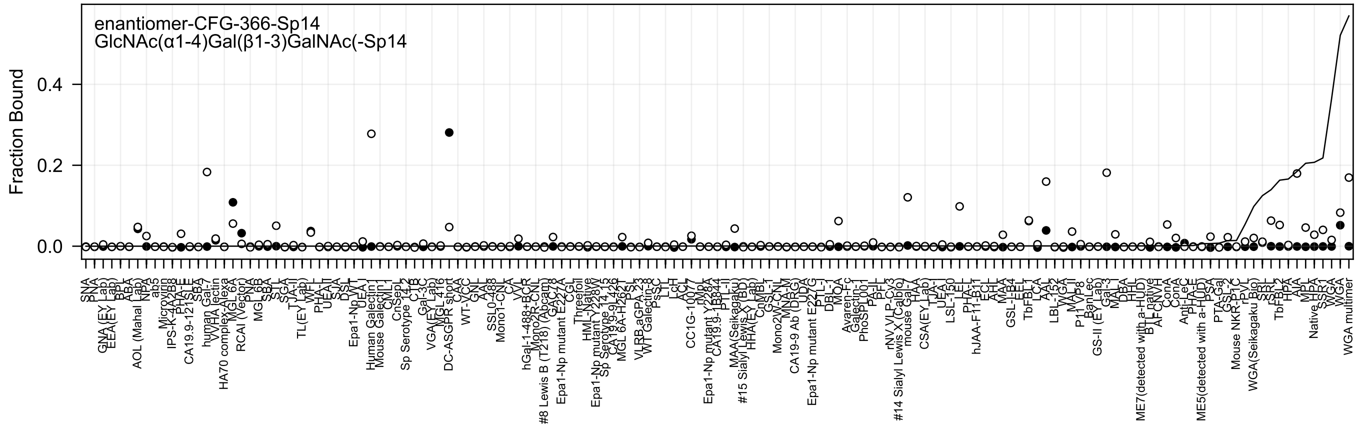
enantiomer-CFG-366-Sp14
GlcNAc(α1-4)Gal(β1-3)GalNAc(-Sp14
Fraction Bound
0.4
0.2
0.0

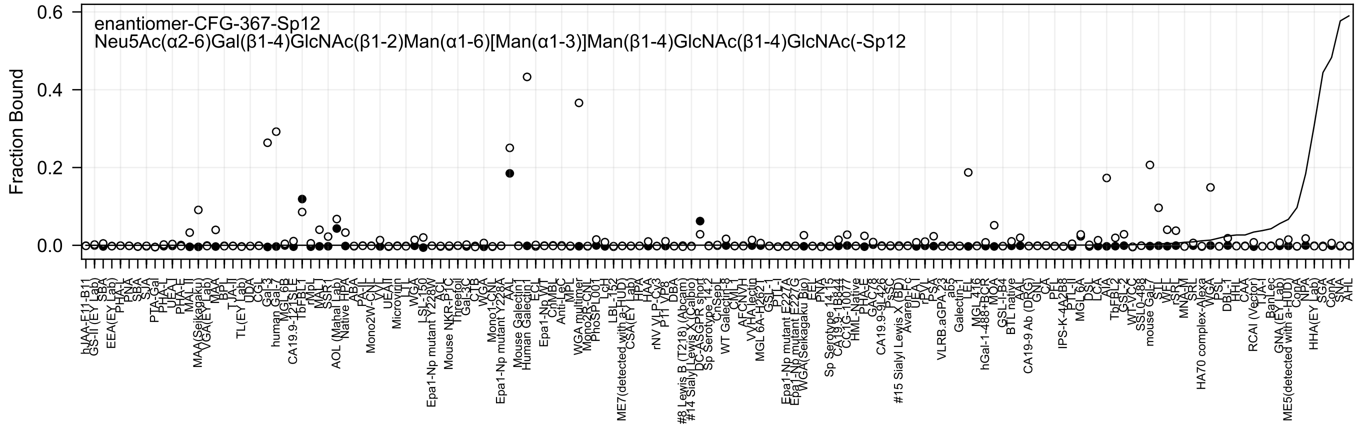
enantiomer-CFG-367-Sp12
Neu5Ac(α2-6)Gal(β1-4)GlcNAc(β1-2)Man(α1-6)[Man(α1-3)]Man(β1-4)GlcNAc(β1-4)GlcNAc(-Sp12
Fraction Bound
0.6
0.4
0.2
0.0

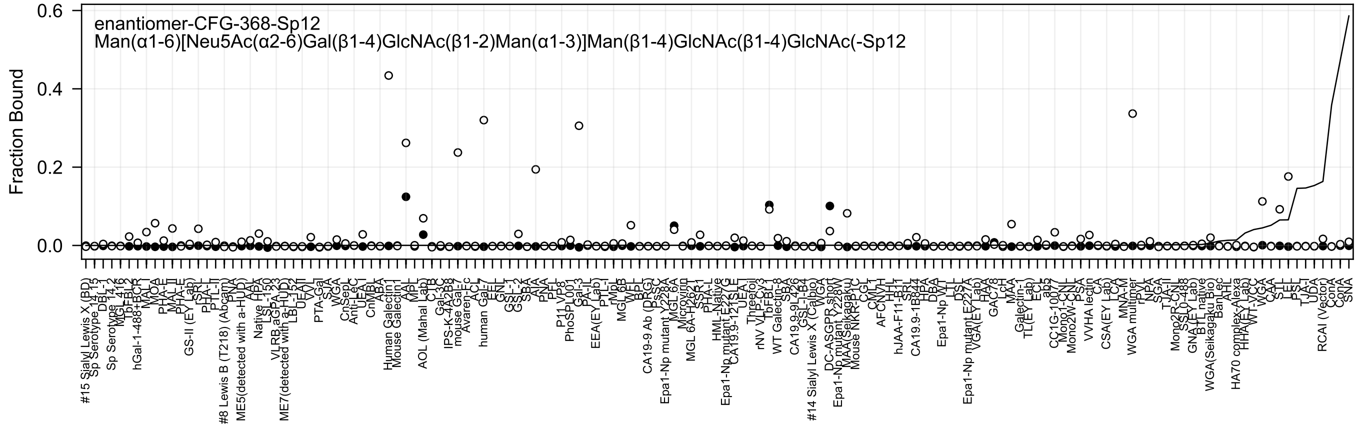
enantiomer-CFG-368-Sp12
Man(α1-6)[Neu5Ac(α2-6)Gal(β1-4)GlcNAc(β1-2)Man(α1-3)]Man(β1-4)GlcNAc(β1-4)GlcNAc(-Sp12
Fraction Bound
0.0
0.2
0.4
0.6

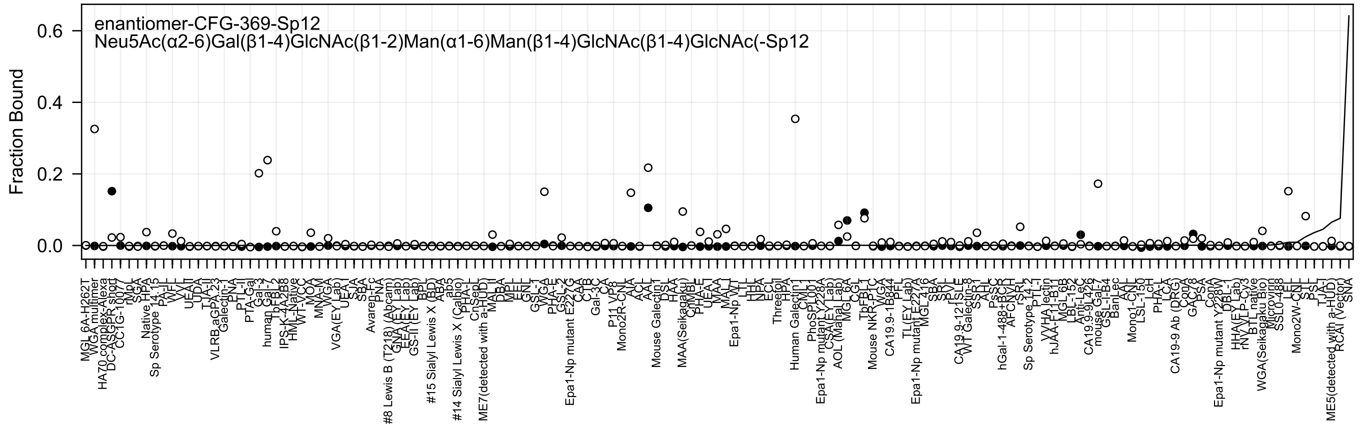
enantiomer-CFG-369-Sp12
Neu5Ac(α2-6)Gal(β1-4)GlcNAc(β1-2)Man(α1-6)Man(β1-4)GlcNAc(β1-4)GlcNAc(-Sp12
Fraction Bound
0.6
0.4
0.2
0.0

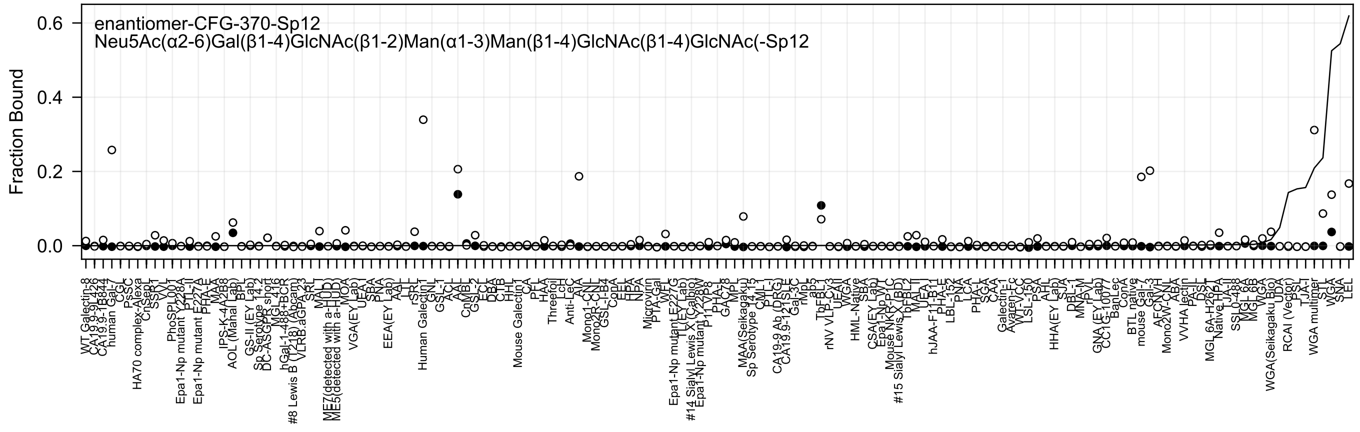

enantiomer-CFG-370-Sp12
Neu5Ac(α2-6)Gal(β1-4)GlcNAc(β1-2)Man(α1-3)Man(β1-4)GlcNAc(β1-4)GlcNAc(-Sp12
Fraction Bound
0.0
0.2
0.4
0.6

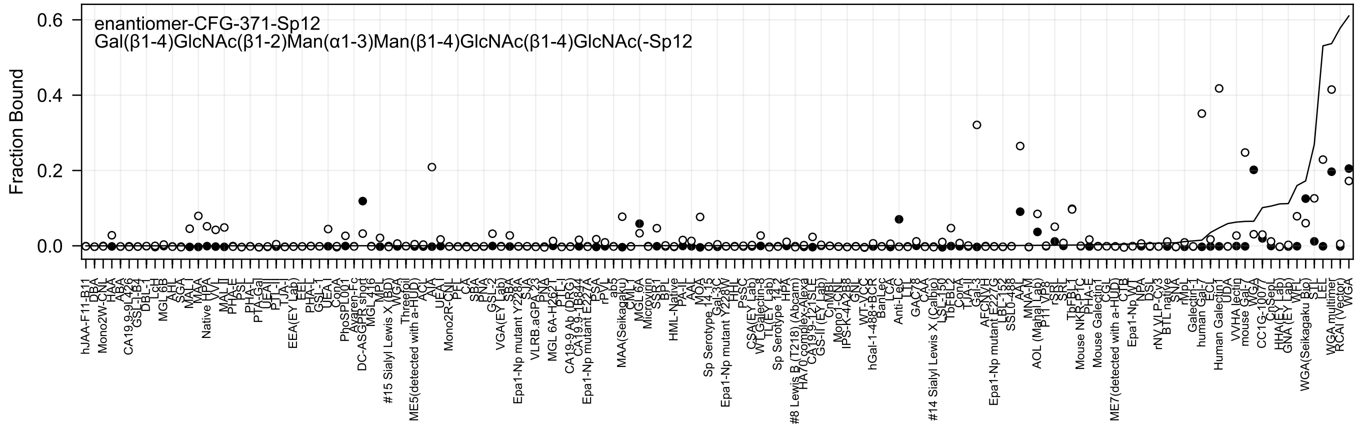
enantiomer-CFG-371-Sp12
Gal(β1-4)GlcNAc(β1-2)Man(α1-3)Man(β1-4)GlcNAc(β1-4)GlcNAc(-Sp12
Fraction Bound
0.6
0.4
0.2
0.0

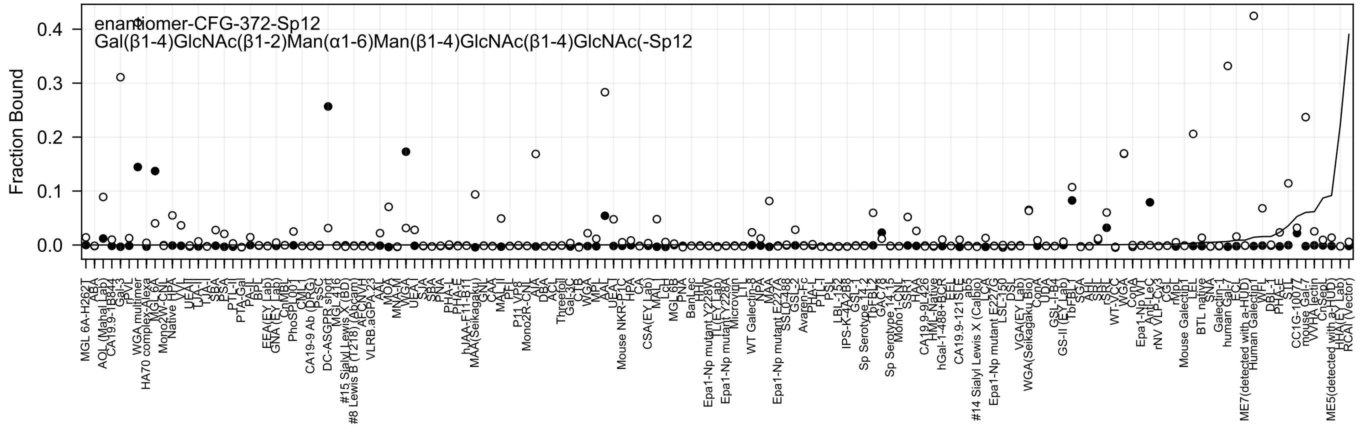
enantiomer-CFG-372-Sp12
Gal(β1-4)GlcNAc(β1-2)Man(α1-6)Man(β1-4)GlcNAc(β1-4)GlcNAc(-Sp12
Fraction Bound
0.4
0.3
0.2
0.1
0.0

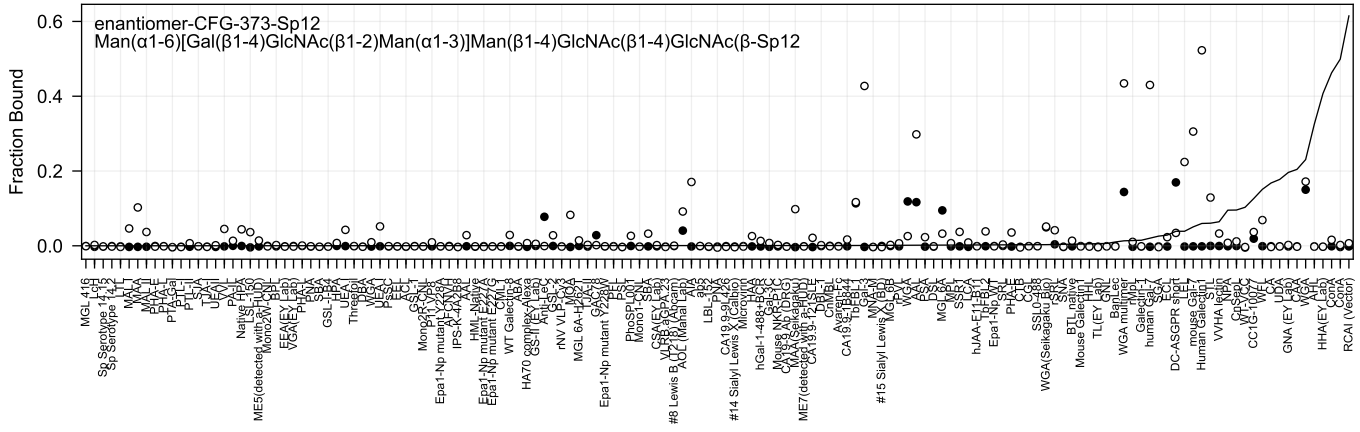

enantiomer-CFG-373-Sp12
Man(α1-6)[Gal(β1-4)GlcNAc(β1-2)Man(α1-3)]Man(β1-4)GlcNAc(β1-4)GlcNAc(β-Sp12
Fraction Bound
0.0
0.2
0.4
0.6

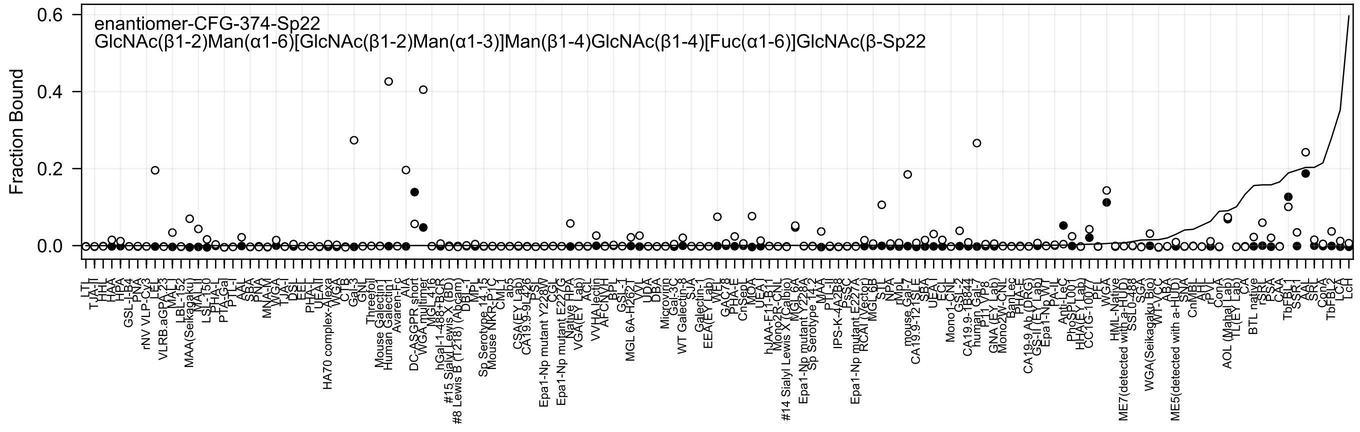

enantiomer-CFG-374-Sp22
GlcNAc(β1-2)Man(α1-6)[GlcNAc(β1-2)Man(α1-3)]Man(β1-4)GlcNAc(β1-4)[Fuc(α1-6)]GlcNAc(β-Sp22
Fraction Bound
0.6
0.4
0.2
0.0
LTL
TJA-II
HHL
HAA
HPA
GSL-I-B4
PNA
rNV VLP-Cy3
LEL
VLRB.aGPA.23
MAL-I
LBL-152
MAA(Seikagaku)
MAL-II
LSL-150
PHA-L
PTA-Gal
PTL-I
AAL
SBA
PNA
MNA-M
WGA
TJA-I
DSL
EEL
PHA-E
UEA-II
HA70 complex-Alexa
WGA
CTB
Gal-3
GNL
Threefoil
Mouse Galectin1
Human Galectin1
Avaren-Fc
AIA
DC-ASGPR short
WGA multimer
MGL 416
hGal-1-488+BCR
#15 Sialyl Lewis X (BD)
#8 Lewis B (T218) (Abcam)
DBL-I
MPL
Sp Serotype 14.15
Mouse NKR-P1C
CML-I
ab5
CSA(EY Lab)
CA19.9-9L426
PSL
Epa1-Np mutant Y228W
CGL
Epa1-Np mutant E227G
Native HPA
VGA(EY Lab)
ACL
VVHA lectin
AFCNYH
BPL
GSL-1
MGL 6A-H262T
VVL
UDA
DBA
Microvirin
Gal-3C
WT Galectin-8
SJA
Galectin-1
EEA(EY Lab)
WFA
GAC78
PHA-E
CnSep1
MOA
UEA-I
hJAA-F11-B1
Mono2R-CNL
#14 Sialyl Lewis X (Calbio)
MGL 6A
Epa1-Np mutant Y228A
Sp Serotype 14.2
MAA
PTL-II
IPS-K-4A2B8
PsSC
Epa1-Np mutant E227A
RCAI Vector
MGL 6B
STL
NPA
rMpL
mouse Gal-7
CA19.9-121SLE
SBA
UEA I
ECL
Mono1-CNL
GSL-2
CA19.9-1B844
human Gal-7
P11 VP8
GNA (EY Lab)
Mono2W-CNL
BanLec
PHA-E
CA19-9 Ab (DRG)
GS-II (EY Lab)
Epa1-Np WT
PA-IL
Anti-LeC
PhoSL001
HHA(EY Lab)
CC1G-10077
PFL
WGA
HML-Native
ME7(detected with a-HUD)
SSL0-488
SGA
WGA(Seikagaku Bio)
WT-VCC
ABA
ME5(detected with a-HUD)
SNA
CnMBL
AAL
rPVL
ConA
AOL (Mahal Lab)
TL(EY Lab)
CA
BTL native
rSRL
PSA
CAA
TbFBL1
SSR1
AAL
SRL
ConA
TbFBL2
LCA
LcH

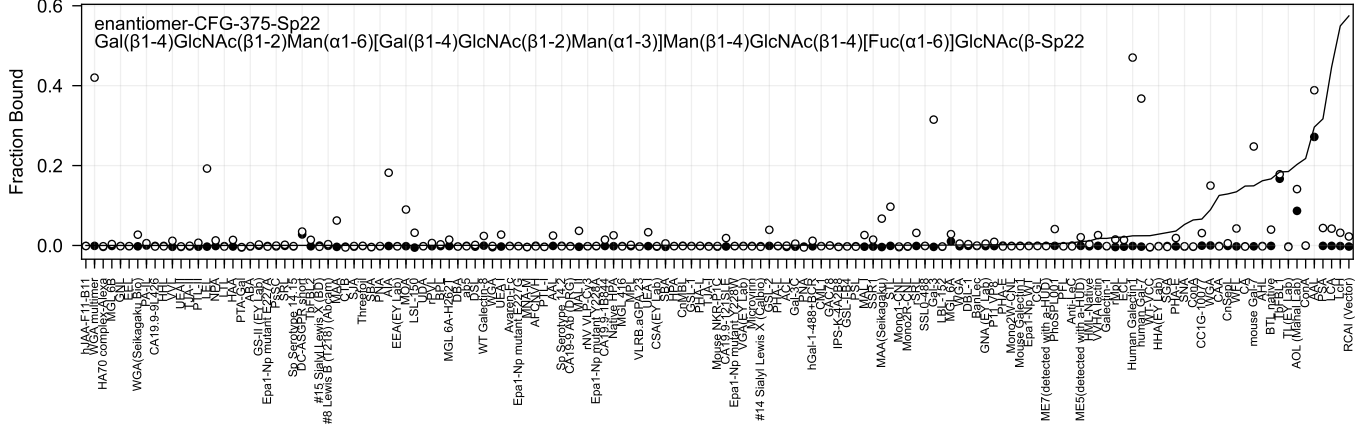

enantiomer-CFG-375-Sp22
Gal(β1-4)GlcNAc(β1-2)Man(α1-6)[Gal(β1-4)GlcNAc(β1-2)Man(α1-3)]Man(β1-4)GlcNAc(β1-4)[Fuc(α1-6)]GlcNAc(β-Sp22
Fraction Bound
0.6
0.4
0.2
0.0
hJAA-F11-B11
WGA multimer
HA70 complex-Alexa
MGL_6B
GNL
EEL
WGA(Seikagaku_Bio)
PA-IL
CA19.9-9L426
HHL
VVL
UEA-I
TJA-II
PTL-II
LEL
NPA
LTL
HAA
PTA-Gal
ABA
GS-II (EY Lab)
Epa1-Np mutant E227A
PsSC
SRL
Sp Serotype 14.15
DC-ASGPR short
TbF-BL2
#15 Sialyl Lewis X (BD)
#8 Lewis B (T218) (Abcam)
MAA
CTB
SJA
Threefoil
SBA
PNA
AIA
EEA(EY Lab)
MOA
LSL-150
UDA
rPVL
BPL
MGL 6A-H262T
DBA
aps
DSL
WT Galectin-8
WGA
UEA-I
Avaren-Fc
Epa1-Np mutant E227G
MNA-M
AFCNYH
PTL-I
AAL
Sp Serotype 14.12
CA19-9 Ab (DRG)
MAL-II
rNV VLP-Cy3
Epa1-Np mutant Y228A
CA19.9-1B844
Native H16
HPA
MGL 416
MPL
VLRB.aGPA.23
UEA-I
CSA(EY Lab)
SBA
TBA
CnMBL
GSL-1
PHA-L
TJA-I
Mouse NKR-P1C
CA19.9-121SLE
Epa1-Np mutant Y228W
VGA(EY Lab)
Microvirin
#14 Sialyl Lewis X (Calbio)
GSL-2
PHA-L
ACL
Gal-3C
PNA
hGal-1-488+BCR
CML-1
GAC-78
IPS-K-4A2B8
GSL-I-B4
PSL
MAL-I
SSP-1
MAA(Seikagaku)
STL
Mono1-CNL
Mono2R-CNL
rSRL
SSL0-488
Gal-3
LBL-152
MGL 6A
WGA
DBL-1
BanLec
GNA (EY Lab)
P11 VP8
PHA-E
Mono2W-CNL
Mouse Galectin 1
Epa1-Np WT
CGL
ME7(detected with a-HUD)
PhoSL001
PFL
Anti-LeC
ME5(detected with a-HUD)
HML-Native
VVHA lectin
Galectin-1
rMpl
ECL
Human Galectin1
human Gal-7
WT-VCC
HHA(EY Lab)
SGA
PHA-E
SNA
ConA
CC1G-10077
WGA
CAA
CnSepL
WFL
CA
mouse Gal-7
AHL
BTL native
TbF-BL1
TL(EY Lab)
AOL (Mahal Lab)
ConA
AAL
PSA
LCA
LCH
RCAI (Vector)

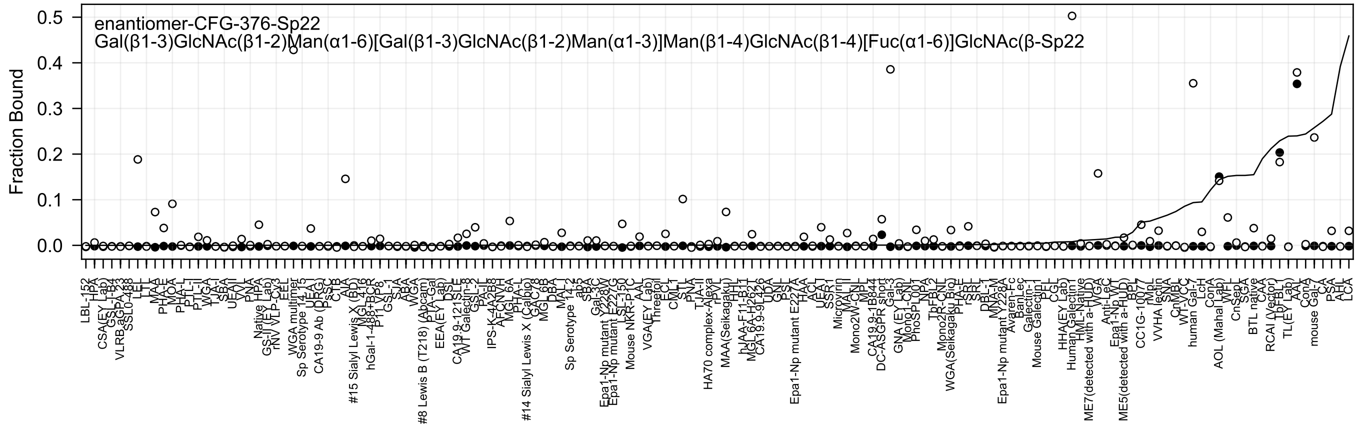
enantiomer-CFG-376-Sp22
Gal(β1-3)GlcNAc(β1-2)Man(α1-6)[Gal(β1-3)GlcNAc(β1-2)Man(α1-3)]Man(β1-4)GlcNAc(β1-4)[Fuc(α1-6)]GlcNAc(β-Sp22
Fraction Bound
0.5
0.4
0.3
0.2
0.1
0.0

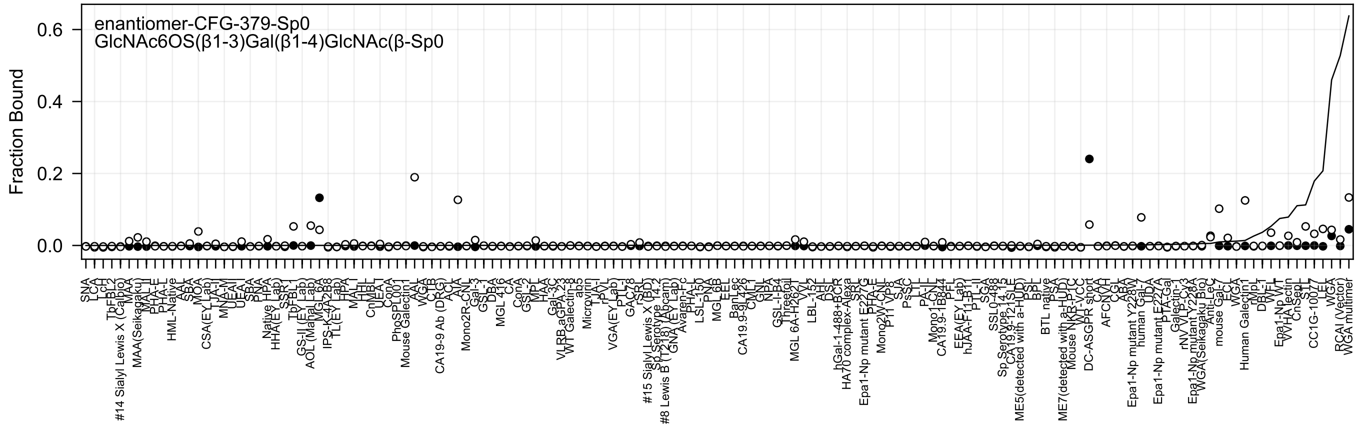
enantiomer-CFG-379-Sp0
GlcNAc6OS(β1-3)Gal(β1-4)GlcNAc(β-Sp0
Fraction Bound
0.6
0.4
0.2
0.0

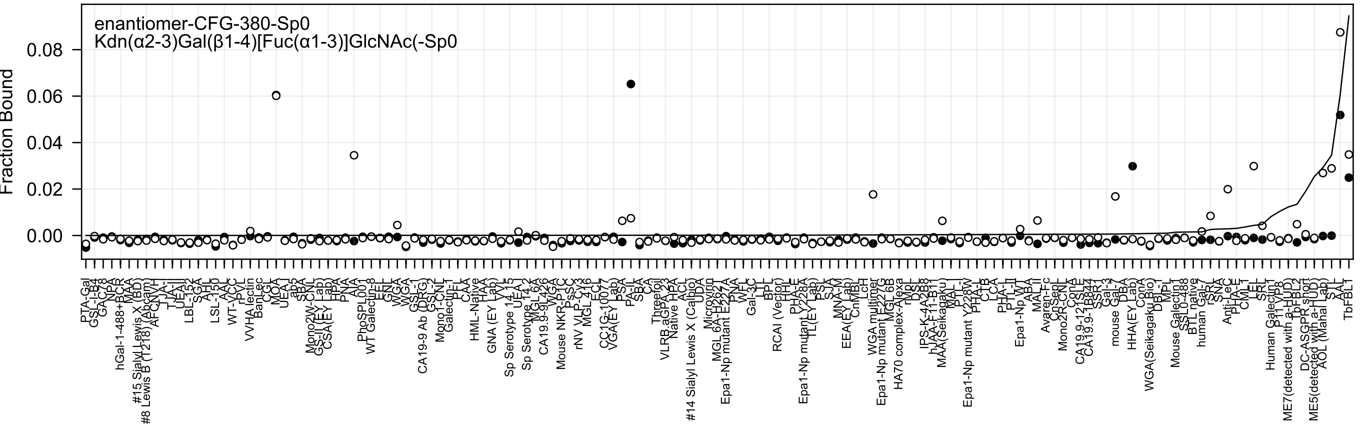
enantiomer-CFG-380-Sp0
Kdn(α2-3)Gal(β1-4)[Fuc(α1-3)]GlcNAc(-Sp0
Fraction Bound
0.08
0.06
0.04
0.02
0.00

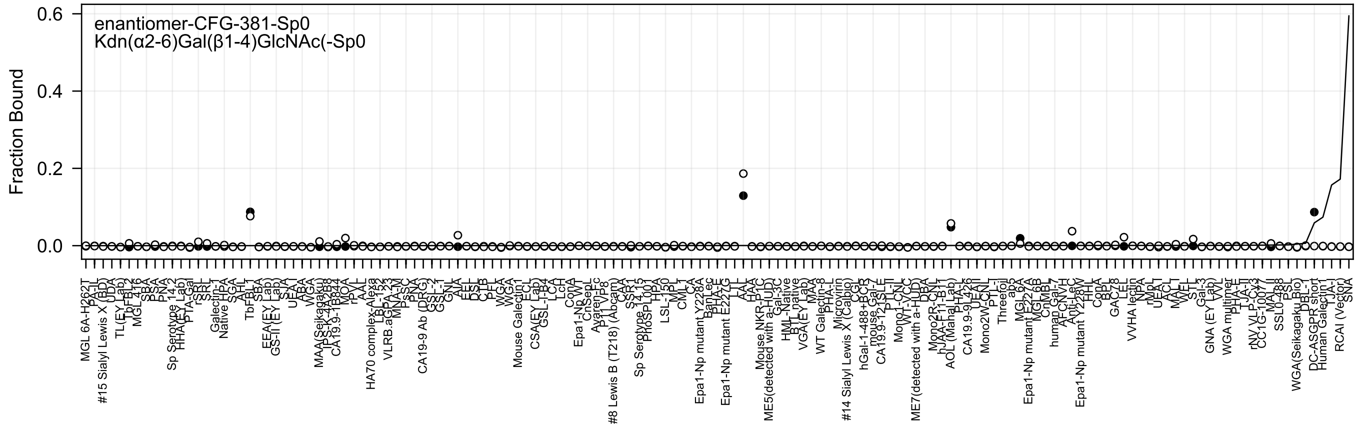

enantiomer-CFG-381-Sp0
Kdn(α2-6)Gal(β1-4)GlcNAc(-Sp0
Fraction Bound
0.6
0.4
0.2
0.0

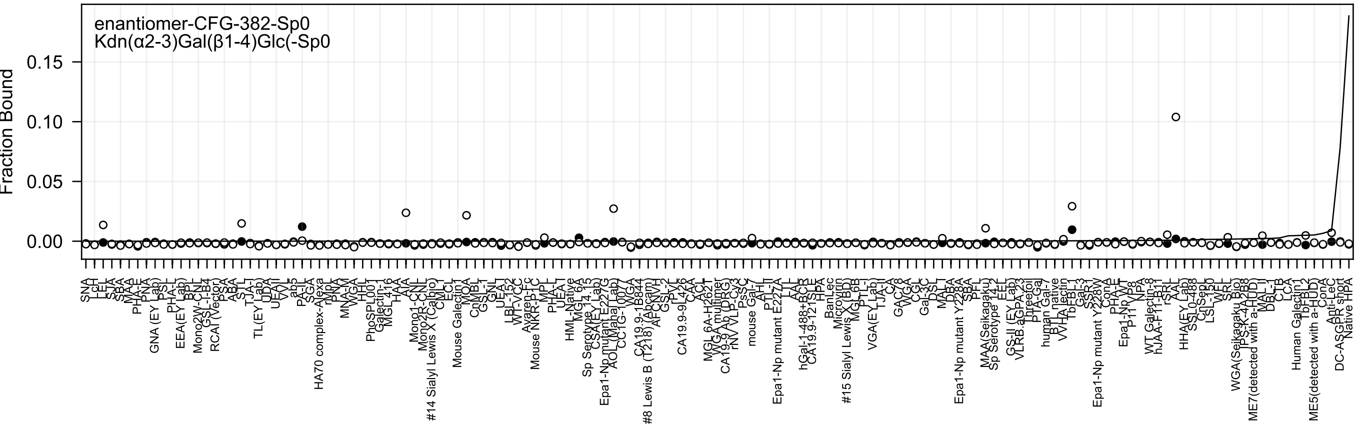

enantiomer-CFG-382-Sp0
Kdn(α2-3)Gal(β1-4)Glc(-Sp0
Fraction Bound
0.15
0.10
0.05
0.00

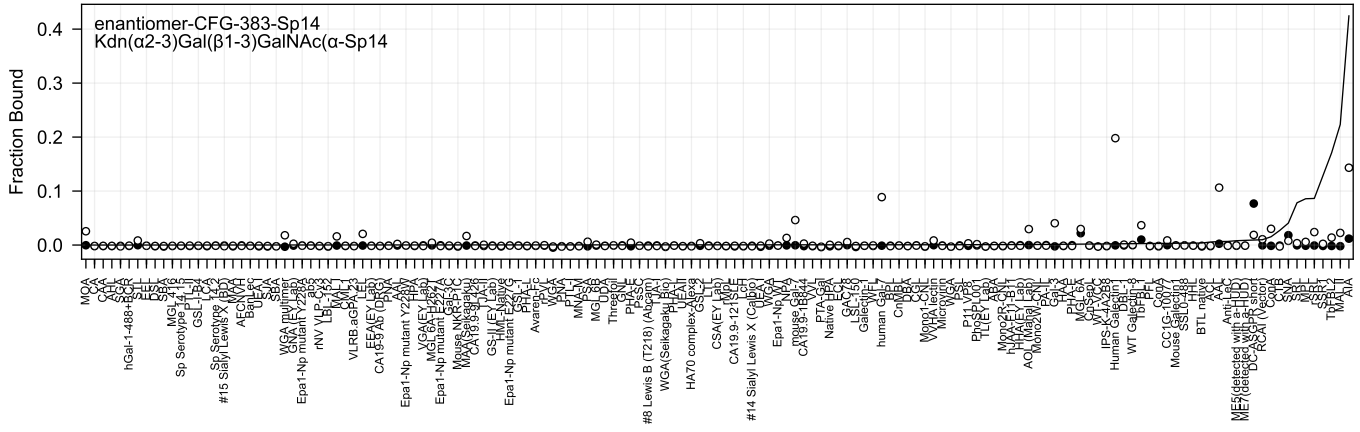
enantiomer-CFG-383-Sp14
Kdn(α2-3)Gal(β1-3)GalNAc(α-Sp14
Fraction Bound
0.4
0.3
0.2
0.1
0.0

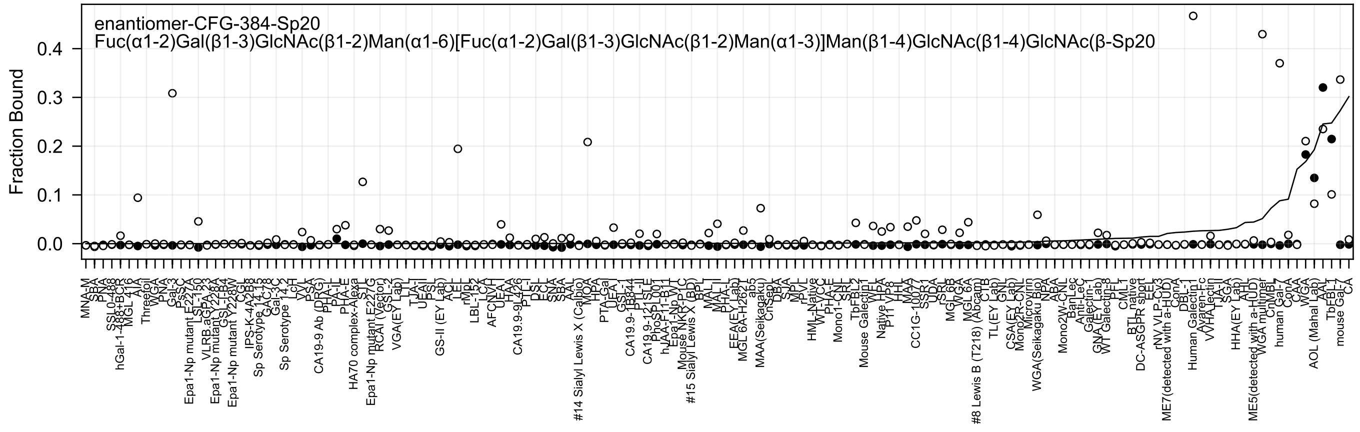
enantiomer-CFG-384-Sp20
Fuc(α1-2)Gal(β1-3)GlcNAc(β1-2)Man(α1-6)[Fuc(α1-2)Gal(β1-3)GlcNAc(β1-2)Man(α1-3)]Man(β1-4)GlcNAc(β1-4)GlcNAc(β-Sp20
Fraction Bound
0.0
0.1
0.2
0.3
0.4

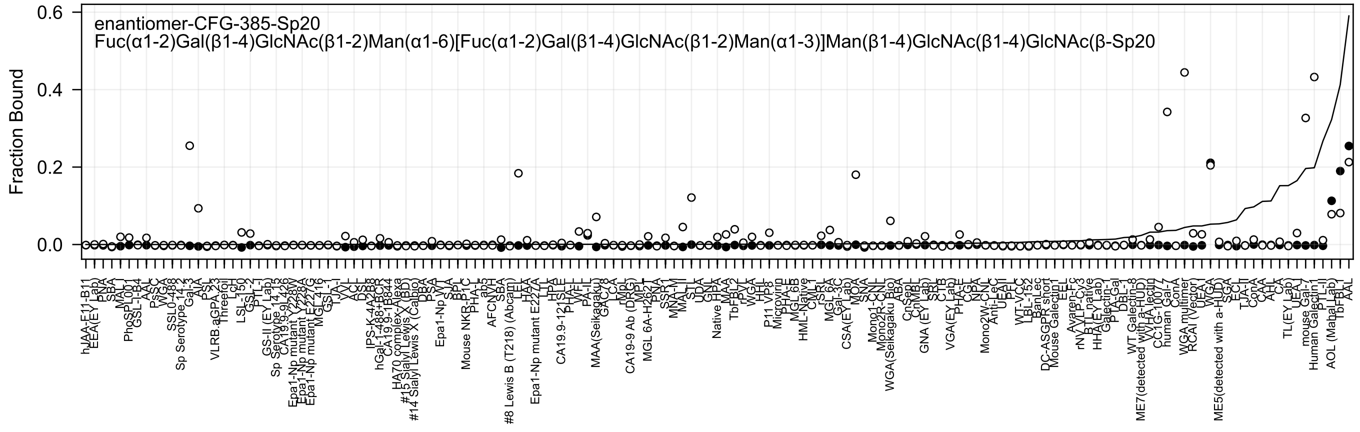

enantiomer-CFG-385-Sp20
Fuc(α1-2)Gal(β1-4)GlcNAc(β1-2)Man(α1-6)[Fuc(α1-2)Gal(β1-4)GlcNAc(β1-2)Man(α1-3)]Man(β1-4)GlcNAc(β1-4)GlcNAc(β-Sp20
Fraction Bound

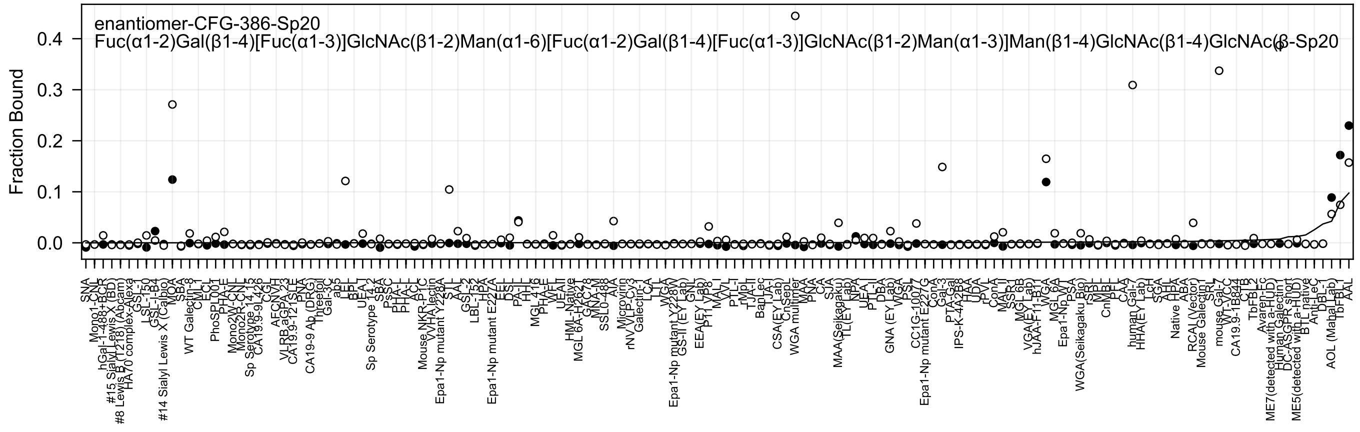
enantiomer-CFG-386-Sp20
Fuc(α1-2)Gal(β1-4)[Fuc(α1-3)]GlcNAc(β1-2)Man(α1-6)[Fuc(α1-2)Gal(β1-4)[Fuc(α1-3)]GlcNAc(β1-2)Man(α1-3)]Man(β1-4)GlcNAc(β1-4)GlcNAc(β-Sp20
Fraction Bound
0.0
0.1
0.2
0.3
0.4

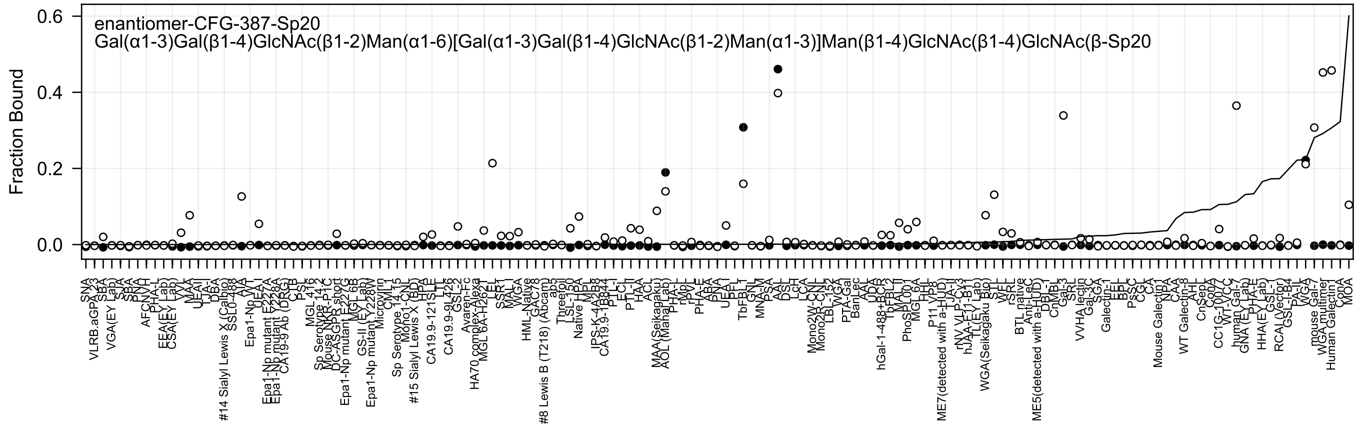
enantiomer-CFG-387-Sp20
Gal(α1-3)Gal(β1-4)GlcNAc(β1-2)Man(α1-6)[Gal(α1-3)Gal(β1-4)GlcNAc(β1-2)Man(α1-3)]Man(β1-4)GlcNAc(β1-4)GlcNAc(β-Sp20
Fraction Bound
0.6
0.4
0.2
0.0

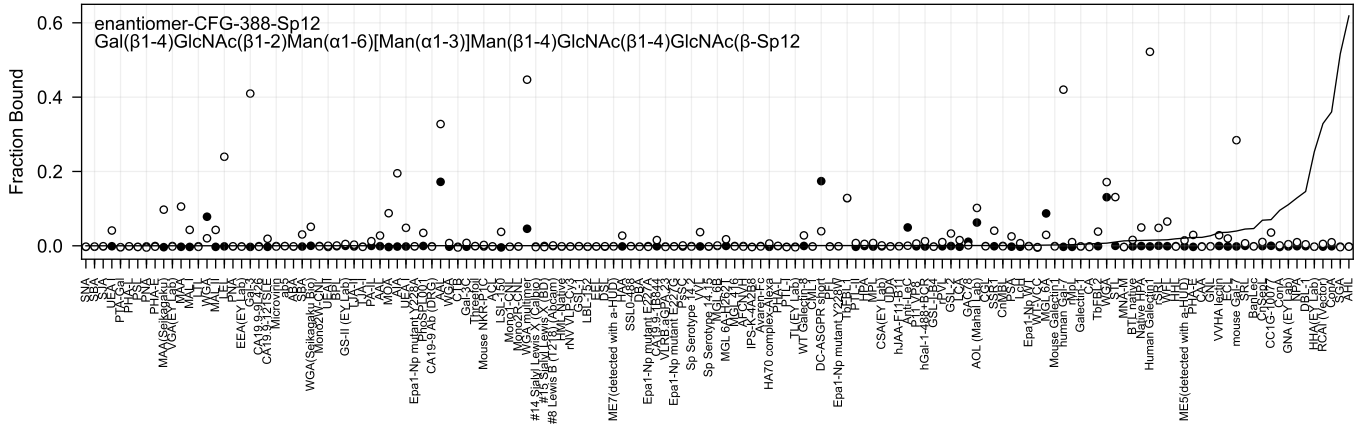
enantiomer-CFG-388-Sp12
Gal(β1-4)GlcNAc(β1-2)Man(α1-6)[Man(α1-3)]Man(β1-4)GlcNAc(β1-4)GlcNAc(β-Sp12
Fraction Bound
0.6
0.4
0.2
0.0
SNA
SBA
SJA
UEA-I
PTA-Gal
PHA-L
PSL
PNA
PHA-E
MAA(Seikagaku)
VGA(EY Lab)
MAA
MAL-I
LTL
WGA
MAL-II
LEL
PNA
EEA(EY Lab)
Gal-3
CA19.9-9L4c2
CA19.9-121SLE
Microvirin
ab5
ABA
SBA
WGA(Seikagaku Bio)
Mono2W-CNL
UEAII
BPL
GS-II (EY Lab)
LTA-I
PA-IL
AAL
MOA
AIA
UEA-I
Epa1-Np mutant Y228A
PhoSL001
CA19-9 Ab (DRG)
AAL
WGA
CTB
Gal-3C
Threefoil
Mouse NKR-P1C
ACL
LSL-150
Mono1-CNL
Mono2R-CNL
WGA multimer
#14 Sialyl Lewis X (Calbio)
#15 Sialyl Lewis X (BD)
#8 Lewis B (T218) (Abcam)
HML-Native
rNV VLP-Cy3
GS-1
LBL-152
EEL
DSL
ME7(detected with a-HUD)
HAA
SSL0-488
DBA
Epa1-Np mutant E227A
CA19.9-1B844
VLRB.aGPA.23
Epa1-Np mutant E227G
PsSC
Sp Serotype 14.2
VVL
Sp Serotype 14.15
MGL 6B
MGL 6A-H262T
MGL 416
AFCNV-H
IPS-K-4A2B8
Avaren-Fc
HA70 complex-Alexa
PHA-L
PTL-I
TL(EY Lab)
WT Galectin-8
CML-I
DC-ASGPR short
PFL
Epa1-Np mutant Y228W
TbFBL1
PTL-II
HPA
MPL
CSA(EY Lab)
UDA
hJAA-F11-B11
Anti-LeC
P11 VP8
hGal-1-488+BCR
GSL-I-B4
rPVL
GSL-2
LCA
GAL78
AOL (Mahal Lab)
CGL
SSR1
CnMBL
PSA
LcH
Epa1-Np WT
WT-VCC
MGL 6A
Mouse Galectin1
human Gal-7
rMpL
Galectin-1
CA
TbFBL2
WGA
STL
MNA-M
BTL native
Native HPA
Human Galectin1
rSRL
WFL
HHL
ME5(detected with a-HUD)
PHA-E
CAA
GNL
VVHA lectin
ECL
mouse Gal-7
SRL
BanLec
CnSepL
CC1G-1007
ConA
GNA (EY Lab)
NPA
DBL-1
HHA(EY Lab)
RCAI (Vector)
ConA
SGA
AHL

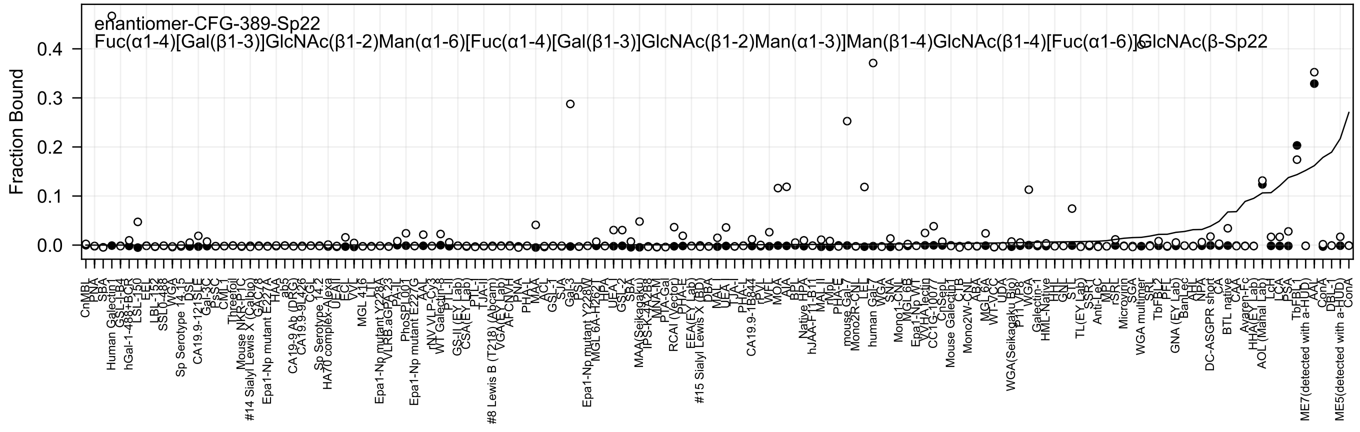

enantiomer-CFG-389-Sp22
Fuc(α1-4)[Gal(β1-3)]GlcNAc(β1-2)Man(α1-6)[Fuc(α1-4)[Gal(β1-3)]GlcNAc(β1-2)Man(α1-3)]Man(β1-4)GlcNAc(β1-4)[Fuc(α1-6)]GlcNAc(β-Sp22
Fraction Bound
0.0
0.1
0.2
0.3
0.4

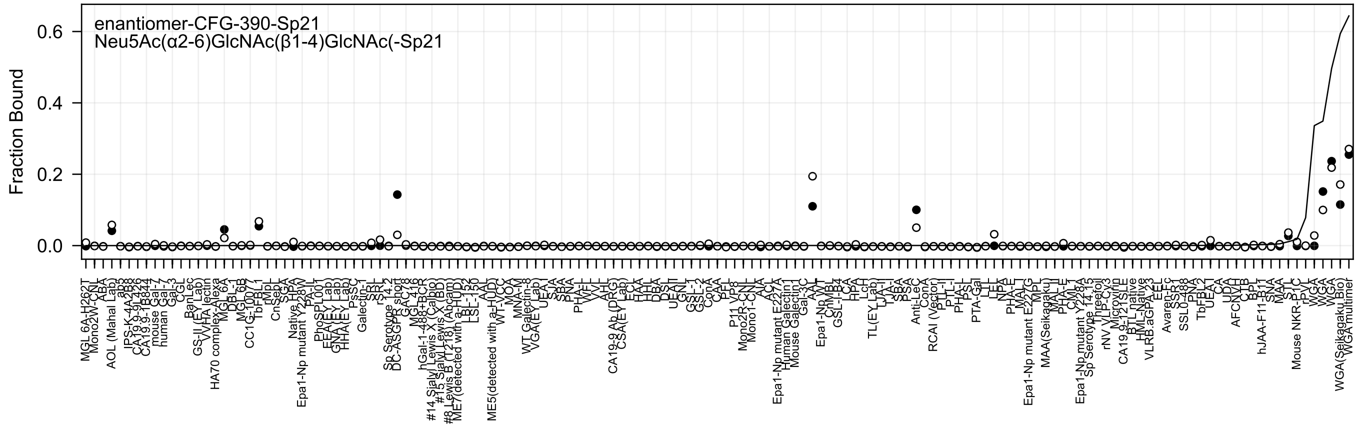
enantiomer-CFG-390-Sp21
Neu5Ac(α2-6)GlcNAc(β1-4)GlcNAc(-Sp21
Fraction Bound
0.0
0.2
0.4
0.6

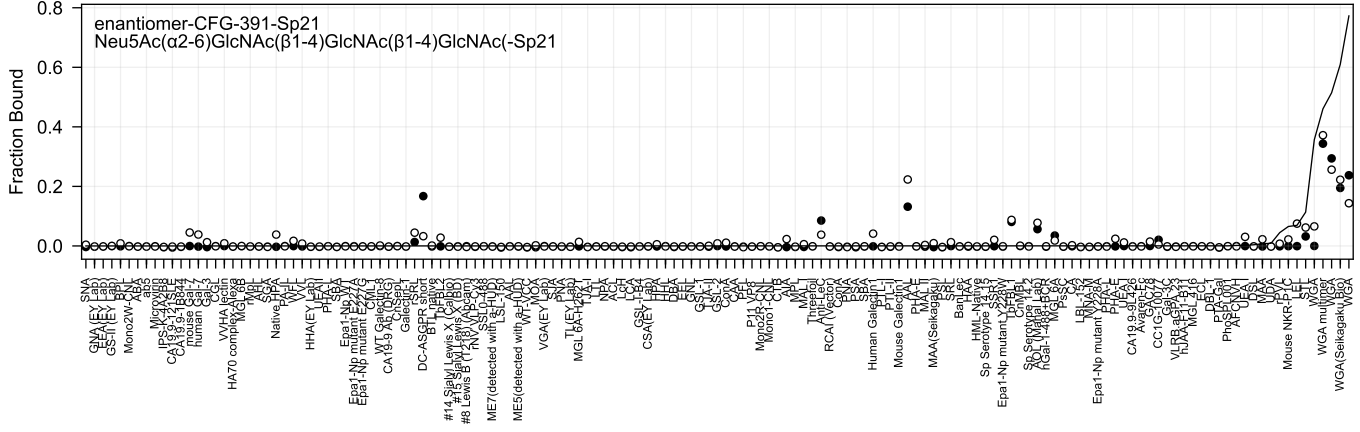

enantiomer-CFG-391-Sp21
Neu5Ac(α2-6)GlcNAc(β1-4)GlcNAc(β1-4)GlcNAc(-Sp21
Fraction Bound
0.0
0.2
0.4
0.6
0.8

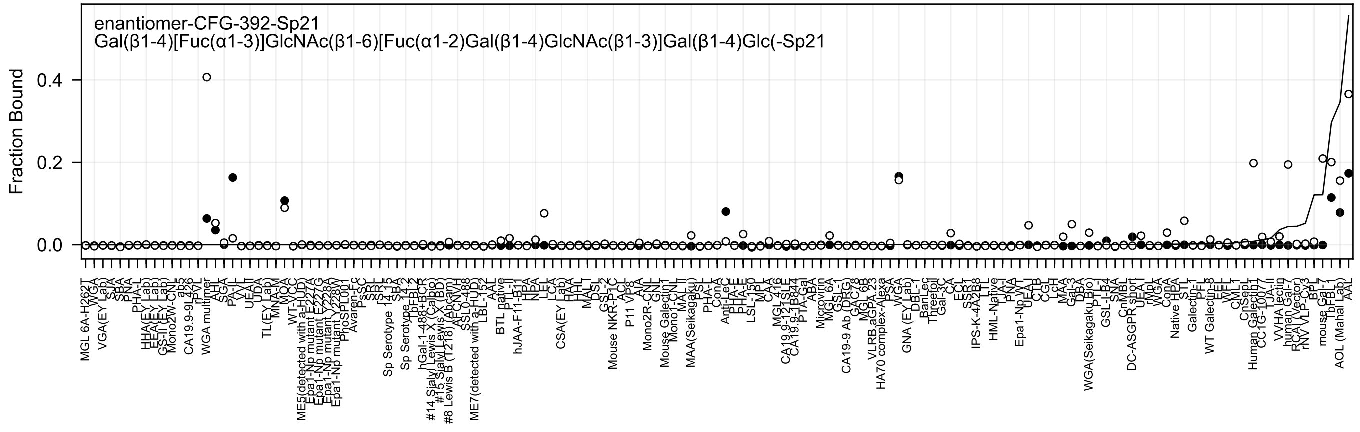
enantiomer-CFG-392-Sp21
Gal(β1-4)[Fuc(α1-3)]GlcNAc(β1-6)[Fuc(α1-2)Gal(β1-4)GlcNAc(β1-3)]Gal(β1-4)Glc(-Sp21
Fraction Bound
0.4
0.2
0.0

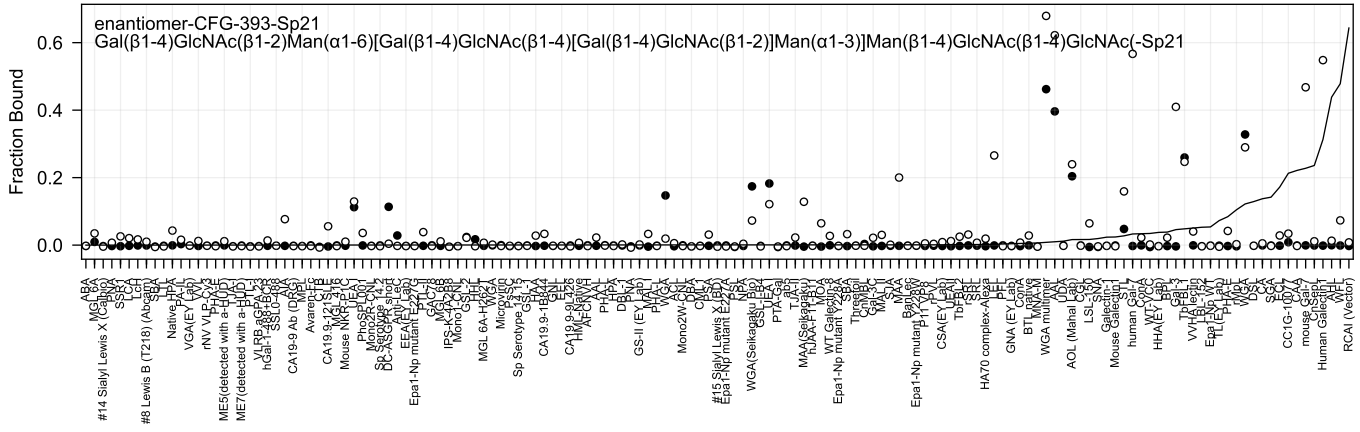
enantiomer-CFG-393-Sp21
Gal(β1-4)GlcNAc(β1-2)Man(α1-6)[Gal(β1-4)GlcNAc(β1-4)[Gal(β1-4)GlcNAc(β1-2)]Man(α1-3)]Man(β1-4)GlcNAc(β1-4)GlcNAc(-Sp21
Fraction Bound
0.0
0.2
0.4
0.6

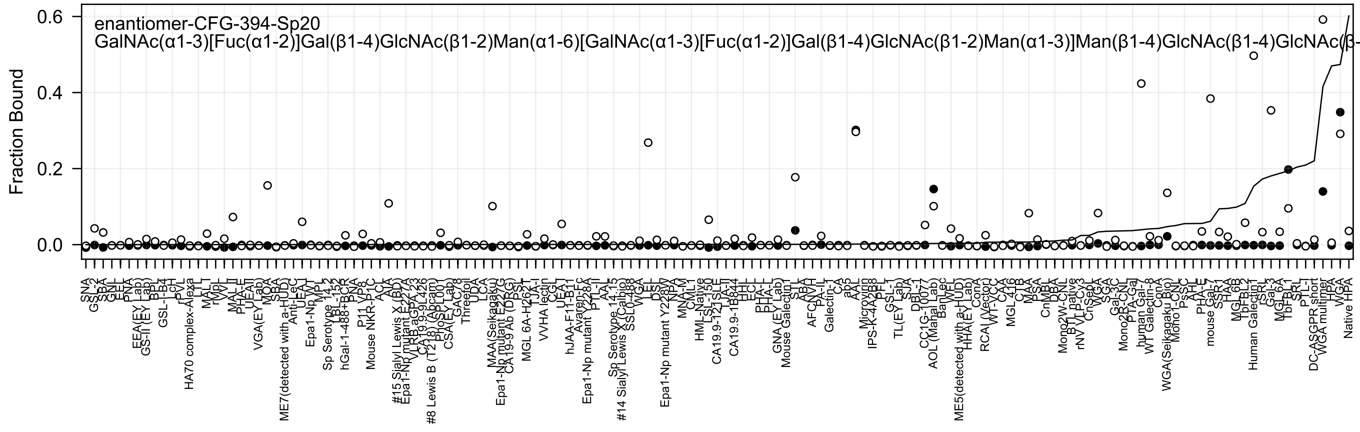
enantiomer-CFG-394-Sp20
GalNAc(α1-3)[Fuc(α1-2)]Gal(β1-4)GlcNAc(β1-2)Man(α1-6)[GalNAc(α1-3)[Fuc(α1-2)]Gal(β1-4)GlcNAc(β1-2)Man(α1-3)]Man(β1-4)GlcNAc(β1-4)GlcNAc(β-
Fraction Bound
0.6
0.4
0.2
0.0

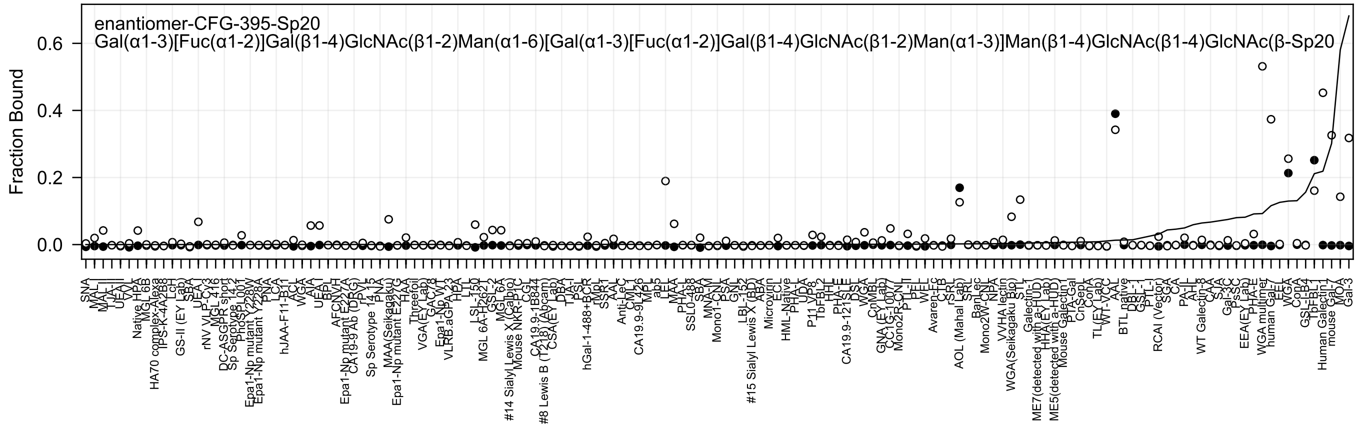
enantiomer-CFG-395-Sp20
Gal(α1-3)[Fuc(α1-2)]Gal(β1-4)GlcNAc(β1-2)Man(α1-6)[Gal(α1-3)[Fuc(α1-2)]Gal(β1-4)GlcNAc(β1-2)Man(α1-3)]Man(β1-4)GlcNAc(β1-4)GlcNAc(β-Sp20
Fraction Bound
0.0
0.2
0.4
0.6

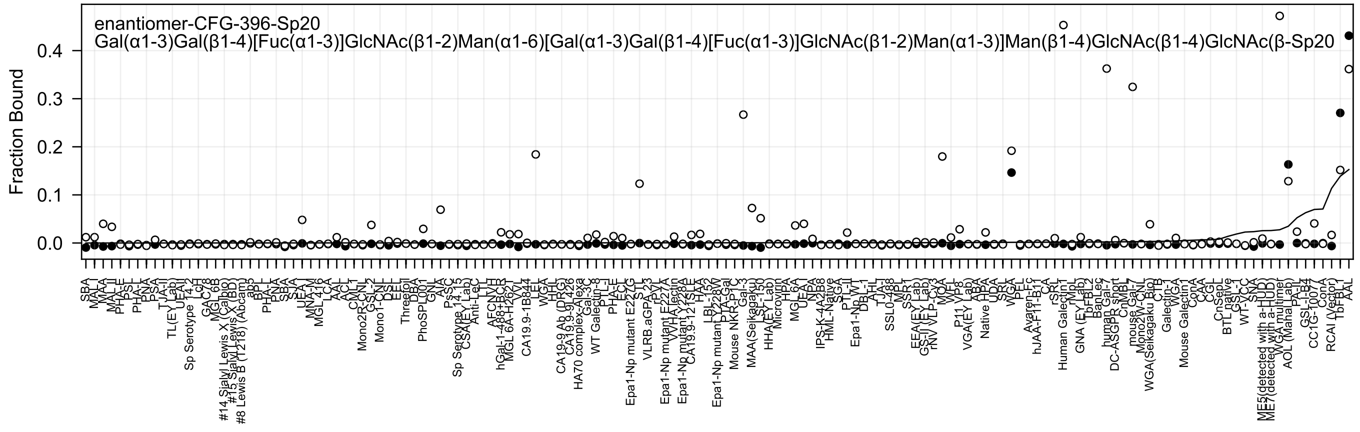
enantiomer-CFG-396-Sp20
Gal(α1-3)Gal(β1-4)[Fuc(α1-3)]GlcNAc(β1-2)Man(α1-6)[Gal(α1-3)Gal(β1-4)[Fuc(α1-3)]GlcNAc(β1-2)Man(α1-3)]Man(β1-4)GlcNAc(β1-4)GlcNAc(β-Sp20
Fraction Bound

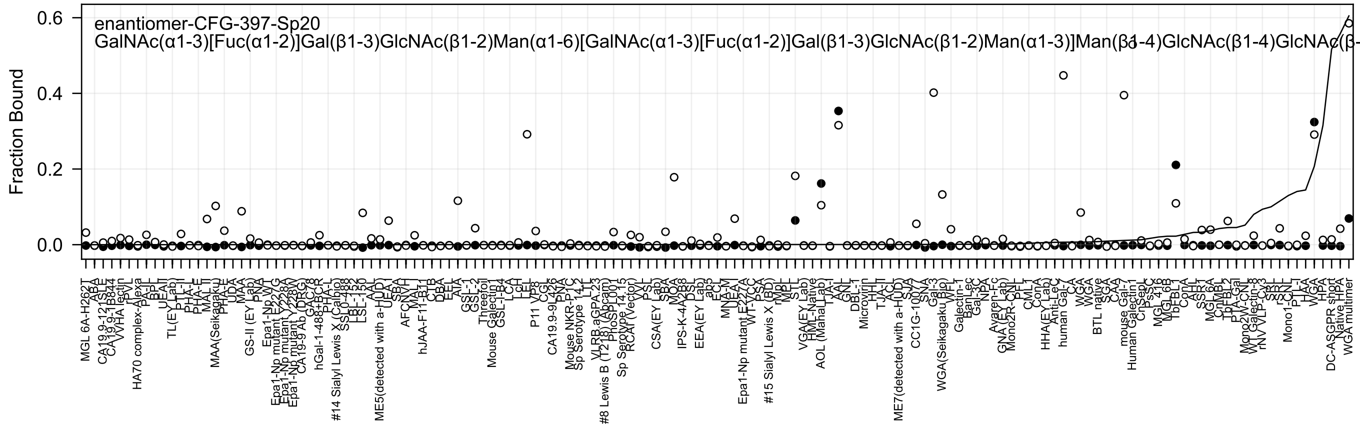
enantiomer-CFG-397-Sp20
GalNAc(α1-3)[Fuc(α1-2)]Gal(β1-3)GlcNAc(β1-2)Man(α1-6)[GalNAc(α1-3)[Fuc(α1-2)]Gal(β1-3)GlcNAc(β1-2)Man(α1-3)]Man(β1-4)GlcNAc(β1-4)GlcNAc(β-
Fraction Bound
0.6
0.4
0.2
0.0

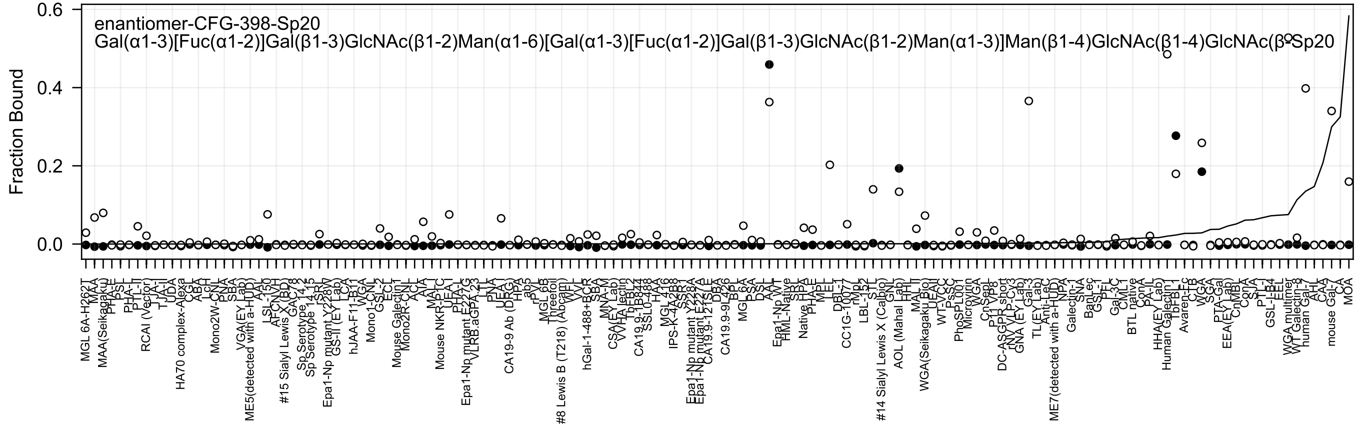
enantiomer-CFG-398-Sp20
Gal(α1-3)[Fuc(α1-2)]Gal(β1-3)GlcNAc(β1-2)Man(α1-6)[Gal(α1-3)[Fuc(α1-2)]Gal(β1-3)GlcNAc(β1-2)Man(α1-3)]Man(β1-4)GlcNAc(β1-4)GlcNAc(β-Sp20
Fraction Bound
0.6
0.4
0.2
0.0

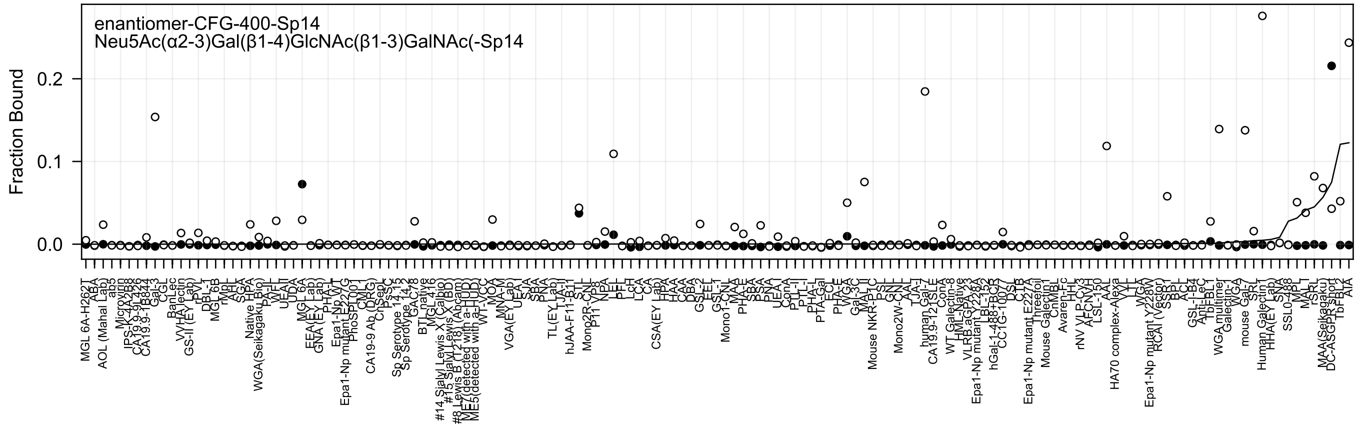
enantiomer-CFG-400-Sp14
Neu5Ac(α2-3)Gal(β1-4)GlcNAc(β1-3)GalNAc(-Sp14
Fraction Bound
0.2
0.1
0.0

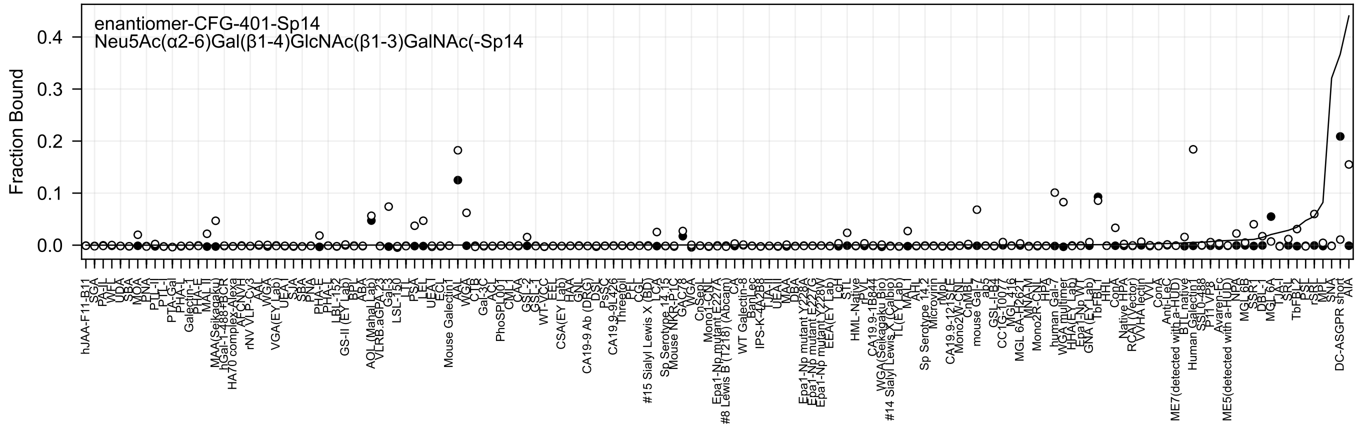

enantiomer-CFG-401-Sp14
Neu5Ac(α2-6)Gal(β1-4)GlcNAc(β1-3)GalNAc(-Sp14
Fraction Bound

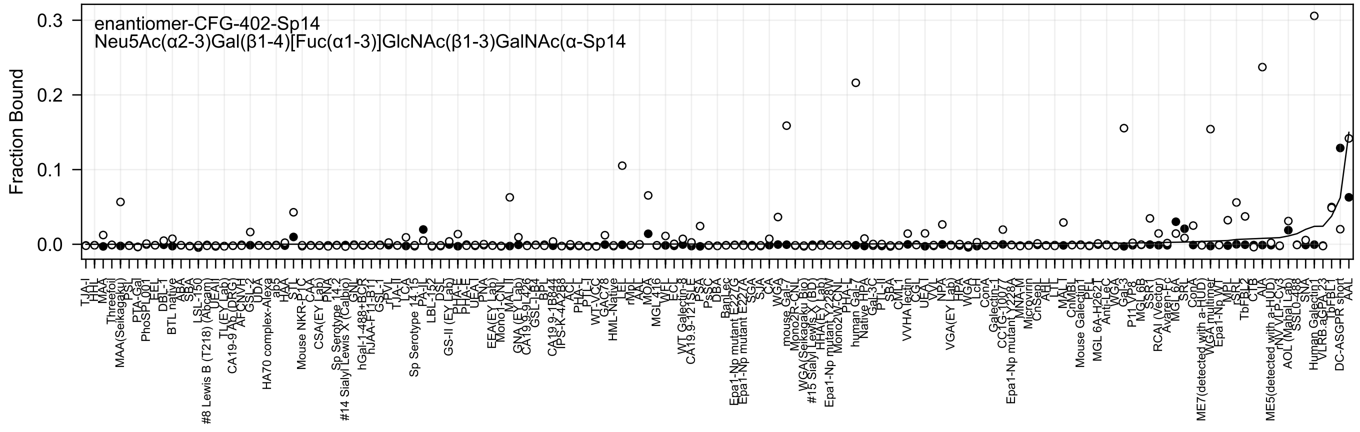
enantiomer-CFG-402-Sp14
Neu5Ac(α2-3)Gal(β1-4)[Fuc(α1-3)]GlcNAc(β1-3)GalNAc(α-Sp14
Fraction Bound
0.3
0.2
0.1
0.0

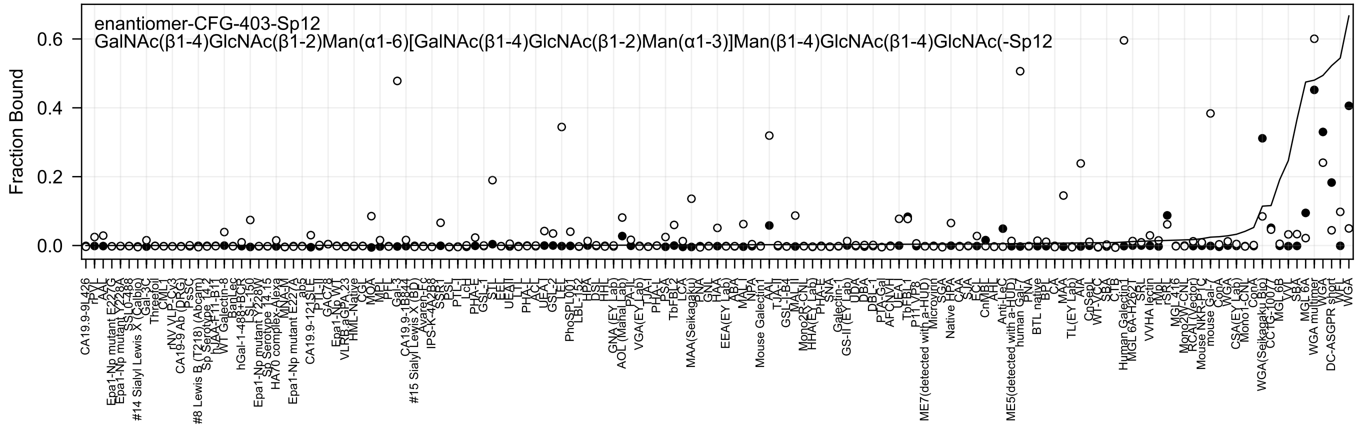
enantiomer-CFG-403-Sp12
GalNAc(β1-4)GlcNAc(β1-2)Man(α1-6)[GalNAc(β1-4)GlcNAc(β1-2)Man(α1-3)]Man(β1-4)GlcNAc(β1-4)GlcNAc(-Sp12
Fraction Bound
0.0
0.2
0.4
0.6

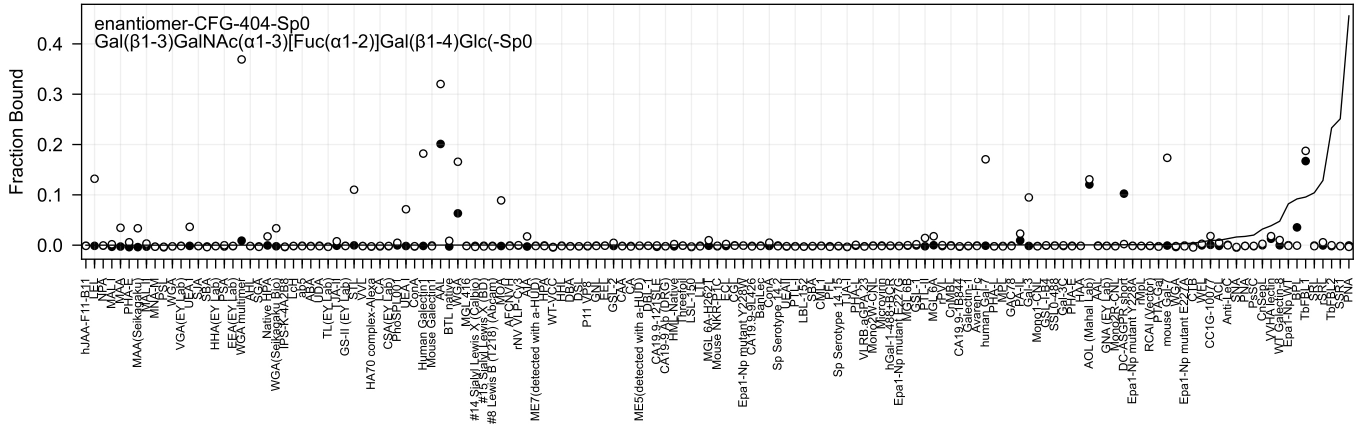

enantiomer-CFG-404-Sp0
Gal(β1-3)GalNAc(α1-3)[Fuc(α1-2)]Gal(β1-4)Glc(-Sp0
Fraction Bound
0.0
0.1
0.2
0.3
0.4
hJAA-F11-B11
LEL
NPA
MAL-I
MAA
PHA-E
MAA(Seikagaku)
MAL-I
MNA-M
PSL
WGA
VGA(EY Lab)
UEA-I
SJA
SBA
HHA(EY Lab)
PSA
EEA(EY Lab)
WGA multimer
AHL
SGA
Native HPA
WGA(Seikagaku Bio)
IPS-K-4A2B8
LcH
ab5
ABA
UDA
TL(EY Lab)
TJA-II
GS-II (EY Lab)
STL
VVL
HA70 complex-Alexa
LCA
CSA(EY Lab)
PhoSL001
UEA-I
ConA
Human Galectin1
Mouse Galectin1
AAL
BTL native
WGA
MGL 416
#14 Sialyl Lewis X (Calbio)
#15 Sialyl Lewis X (BD)
#8 Lewis B (T218) (Abcam)
MOA
AFCNVH
rNV VLP-Cy3
AIA
ME7(detected with a-HUD)
HPA
WT-VCC
HHL
DBA
DSL
P11 VP8
GNL
EEL
GSL-I
CAA
CA
ME5(detected with a-HUD)
DBL-1
CA19.9-121SLE
CA19-9 Ab (DRG)
HML-Native
Threetol
LSL-150
LTL
MGL 6A-H262T
Mouse NKR-P1C
ECL
CGL
Epa1-Np mutant Y228W
CA19.9-9L426
BanLec
ConA
Sp Serotype 14.2
UEAI
PTL-I
LBL-152
SBA
CML-I
PFL
Sp Serotype 14.15
TJA-I
PHA-L
VLRB.aGPA.23
Mono2W-CNL
Microvirin
hGal-1-488+BCR
Epa1-Np mutant E227G
MGL 6B
GSL-II
PTL-II
MGL 6A
rPVL
CnMBL
CA19.9-1B844
Galectin-1
Avaren-Fc
human Gal-7
PHA-L
MPL
GAC78
PA-IL
Gal-3
Mono1-CNL
GSL-I-B4
SSL0-488
Gal-3C
PHA-E
HAA
AOL (Mahal Lab)
AAL
GNA (EY Lab)
Mono2R-CNL
DC-ASGPR short
Epa1-Np mutant Y228A
rMpL
RCAI (Vector)
PTA-Gal
mouse Gal-1
VVA
Epa1-Np mutant E227A
CTB
WFL
CC1G-1007
ACL
Anti-LeC
SNA
PNA
PsSC
CnSepL
VVHA lectin
WT Galectin-8
Epa1-Np WT
BPL
TbFBL1
SRL
LSL-N
TbFBL2
SSR1
PNA

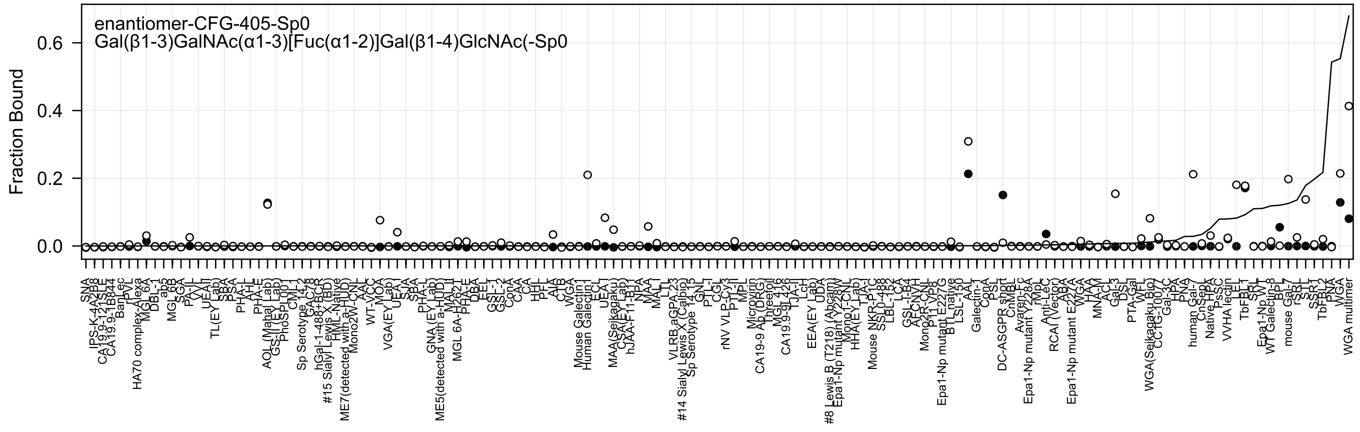
enantiomer-CFG-405-Sp0
Gal(β1-3)GalNAc(α1-3)[Fuc(α1-2)]Gal(β1-4)GlcNAc(-Sp0
Fraction Bound
0.6
0.4
0.2
0.0

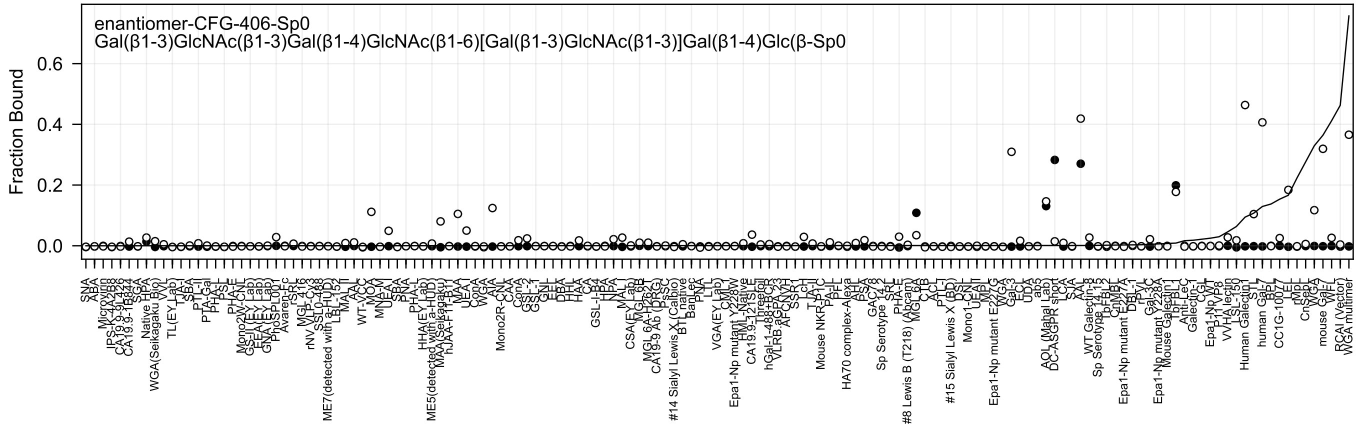
enantiomer-CFG-406-Sp0
Gal(β1-3)GlcNAc(β1-3)Gal(β1-4)GlcNAc(β1-6)[Gal(β1-3)GlcNAc(β1-3)]Gal(β1-4)Glc(β-Sp0
Fraction Bound
0.0
0.2
0.4
0.6
SNA
ABA
Microvirin
IPS-K-4A2B8
CA19.9-9L426
CA19.9-1B844
SGA
Native HPA
WGA(Seikagaku Bio)
VVL
TL(EY Lab)
TJA-I
SBA
PTL-II
PTA-Gal
PHA-L
PSL
PHA-E
Mono2W-CNL
GS-II(EY Lab)
EEA(EY Lab)
GNA(EY Lab)
PhoSL001
Avaren-Fc
rSRL
MGL 416
rNV VLP-Cy3
SSL0-488
ME7(detected with a-HUD)
LBL-152
MAL-II
AAL
WT-VCC
MOA
MNA-M
UEA-I
SBA
PNA
PHA-L
HHA(EY Lab)
ME5(detected with a-HUD)
MAA(Seikagaku)
hJAA-F1*-B1
MAA
UEA-I
ConA
WGA
AIA
Mono2R-CNL
CAA
ConA
GSL-2
GSL-1
GNL
EEL
DBA
HHL
HAA
CA
GSL-I-B4
HPA
NPA
MAL-I
CSA(EY Lab)
MGL 6B
MGL 6A-H262
CA19-9 Ab (DRG)
PSSC
#14 Sialyl Lewis X (Calbio)
BTL native
BanLec
PNA
LTL
VGA(EY Lab)
CML
Epa1-Np mutant Y228W
HML-Native
CA19.9-121SLE
Threefoil
hGal-1-488+BCR
VLRB.aGPA.23
AFCNYH
SSR1
LcH
TJA-II
Mouse NKR-P1C
PA-IL
PFL
HA70 complex-Alexa
AH
PSA
GAC78
Sp Serotype 14-2
SRL
PHA-E
#8 Lewis B (T218) (Abcam)
MGL 6A
CTB
ACL
PTL-I
#15 Sialyl Lewis X (BD)
DSL
Mono1-CNL
UEA-II
MPL
Epa1-Np mutant E227G
WGA
Gal-3
ECL
UDA
ab5
AOL (Mahal Lab)
DC-ASGPR short
LCA
SJA
AAA
WT Galectin-8
Sp Serotype 14-15
TbFBL2
CnMBL
Epa1-Np mutant E227A
DBL-1
rPVL
Gal-3C
Epa1-Np mutant Y228A
Mouse Galectin1
TbFBL1
Anti-LeC
Galectin-1
CGL
Epa1-Np WT
P11VP8
VVHA lectin
LSL-150
Human Galectin1
STL
human Gal-7
BPL
CC1G-10077
LEL
rMpL
CnSeph
WGA
mouse Gal-7
WFL
RCAI (Vector)
WGA multimer

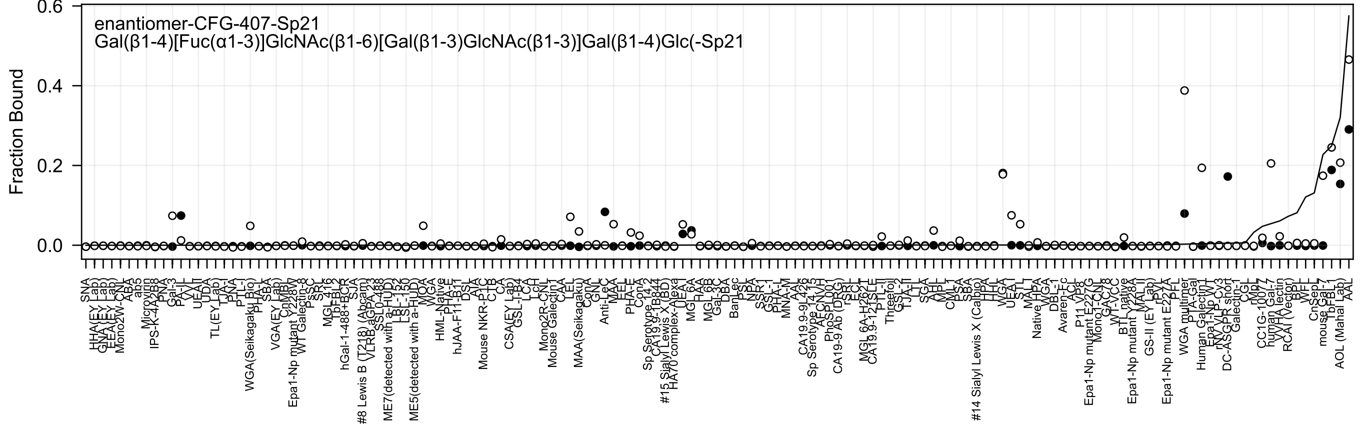
enantiomer-CFG-407-Sp21
Gal(β1-4)[Fuc(α1-3)]GlcNAc(β1-6)[Gal(β1-3)GlcNAc(β1-3)]Gal(β1-4)Glc(-Sp21
Fraction Bound
0.6
0.4
0.2
0.0
SNA
ABA
ab5
Microvirin
Gal-3
PA-IL
UEA-I
TL(EY Lab)
PTL-I
WGA(Seikagaku Bio)
SBA
VGA(EY Lab)
Epa1-Np mutant Y228W
WT Galectin-8
MGL 416
hGal-1-488+BCR
#8 Lewis B (T218) (Abcam)
ME7(detected with a-HUD)
ME5(detected with a-HUD)
HML-Native
hJAA-F11-B17
Mouse NKR-P1C
CTB
CSA(EY Lab)
Mono2R-CN
Mouse Galectin-1
MAA(Seikagaku)
ConA
Anti-LeC
Sp Serotype 14.2
#15 Sialyl Lewis X (BD)
HA70 complex-Alexa
MGL 6A
BanLec
CA19.9-9L426
Sp Serotype 14.13
CA19-9 Ab (DRG)
Threefoil
#14 Sialyl Lewis X (Calbio)
Native HPA
Avaren-Fc
Epa1-Np mutant E227G
Epa1-Np mutant Y228A
GS-II (EY Lab)
Epa1-Np mutant E227A
WGA multimer
Human Galectin1
DC-ASGPR short
Galectin-1
RCAI (Vector)
mouse Gal-7
AOL (Mahal Lab)
AAL

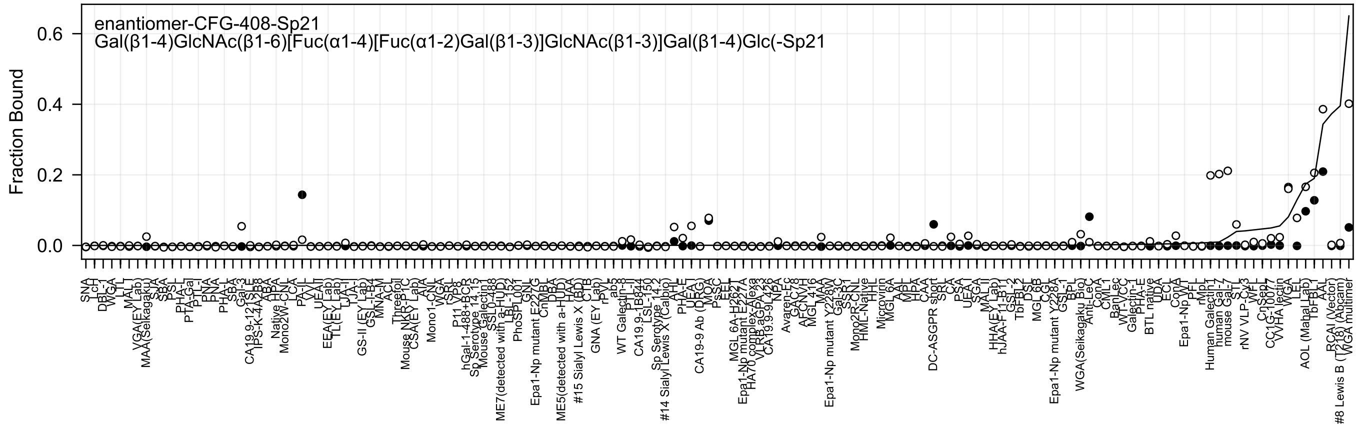
enantiomer-CFG-408-Sp21
Gal(β1-4)GlcNAc(β1-6)[Fuc(α1-4)[Fuc(α1-2)Gal(β1-3)]GlcNAc(β1-3)]Gal(β1-4)Glc(-Sp21
Fraction Bound
0.0
0.2
0.4
0.6

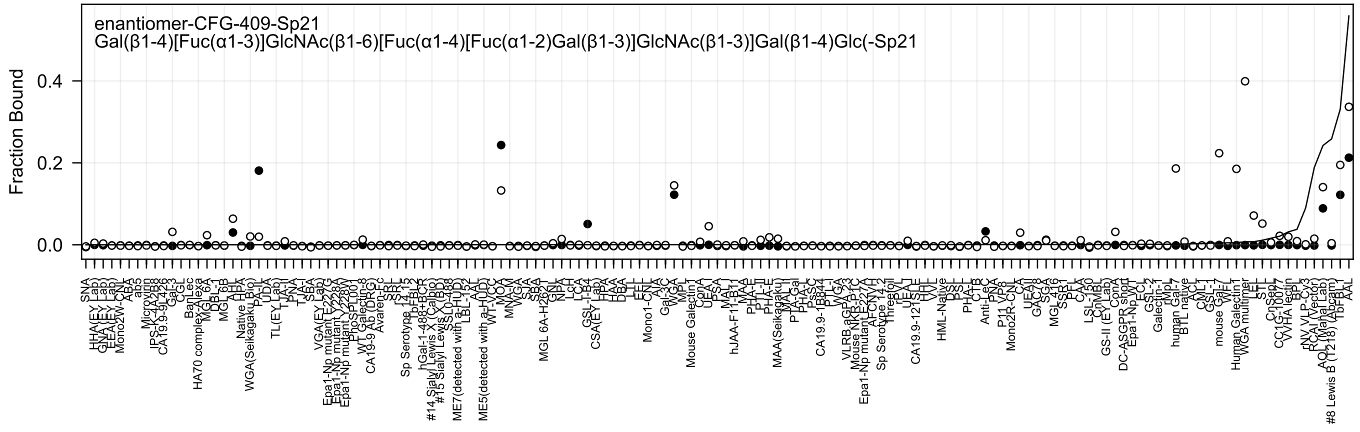

enantiomer-CFG-409-Sp21
Gal(β1-4)[Fuc(α1-3)]GlcNAc(β1-6)[Fuc(α1-4)[Fuc(α1-2)Gal(β1-3)]GlcNAc(β1-3)]Gal(β1-4)Glc(-Sp21
Fraction Bound
0.4
0.2
0.0
SNA
HHA(EY Lab)
GNA(EY Lab)
EEA(EY Lab)
Mono2W-CNL
ABA
ab5
Microvirin
IPS-K-4A2B8
CA19.9-9L426
Gal-3
CGL
BanLec
HA70 complex-Alexa
MGL 6A
DBL-1
MGL 6B
AAL
Native HPA
WGA(Seikagaku Bio)
PA-IL
UDA
TL(EY Lab)
TJA-II
PNA
TJA-I
SBA
VGA(EY Lab)
Epa1-Np mutant E227G
Epa1-Np mutant Y228A
Epa1-Np mutant Y228W
PhoSL001
WT Galectin-8
CA19-9 Ab (DRG)
Avaren-Fc
SRL
rSRL
Sp Serotype 14.15
TbFBL2
hGal-1-488+BCR
#14 Sialyl Lewis X (Calbio)
#15 Sialyl Lewis X (BD)
SSL0-488
ME7(detected with a-HUD)
LBL-152
AAL
ME5(detected with a-HUD)
WT-VCC
MOA
MNA-M
WGA
SJA
SBA
MGL 6A-H262T
GNL
NPA
LcH
LCA
GSL-I-B4
CSA(EY Lab)
HPA
HAA
DBA
LTL
EEL
Mono1-CNL
AIA
Gal-3C
WGA
MPL
Mouse Galectin1
ConA
UEA-I
PSA
MAL-I
hJAA-F11-B11
MAA
PHA-E
PTL-II
PHA-E
MAA(Seikagaku)
MAL-II
PTA-Gal
PHA-L
PsSC
CA19.9-1B844
PTL-I
VGA
VLRB.aGPA.23
Mouse NKR-P1C
Epa1-Np mutant E227A
AFCNVH
Sp Serotype 14.2
Threefoil
rPVL
UEA-I
CA19.9-121SLE
HHL
VVL
HML-Native
DSL
PSL
PHA-L
CTB
Anti-LeC
PNA
P11 VP8
Mono2R-CNL
CA
UEAII
GAC78
SGA
MGL 416
SSR1
PFL
CAA
LSL-150
ChMBL
GS-II (EY Lab)
ConA
DC-ASGPR short
Epa1-Np WT
ECL
GSL-2
Galectin-1
rMpL
human Gal-7
BTL native
ACL
CML-1
GSL-1
mouse Gal-7
WFL
Human Galectin1
WGA multimer
LEL
STL
CnSep
CC1G-10077
VVHA lectin
BPL
rNV VLP-Cv3
RCA I (Vector)
AOL (Mahal Lab)
#8 Lewis B (T218) (Abcam)
TbFBL1
AAL

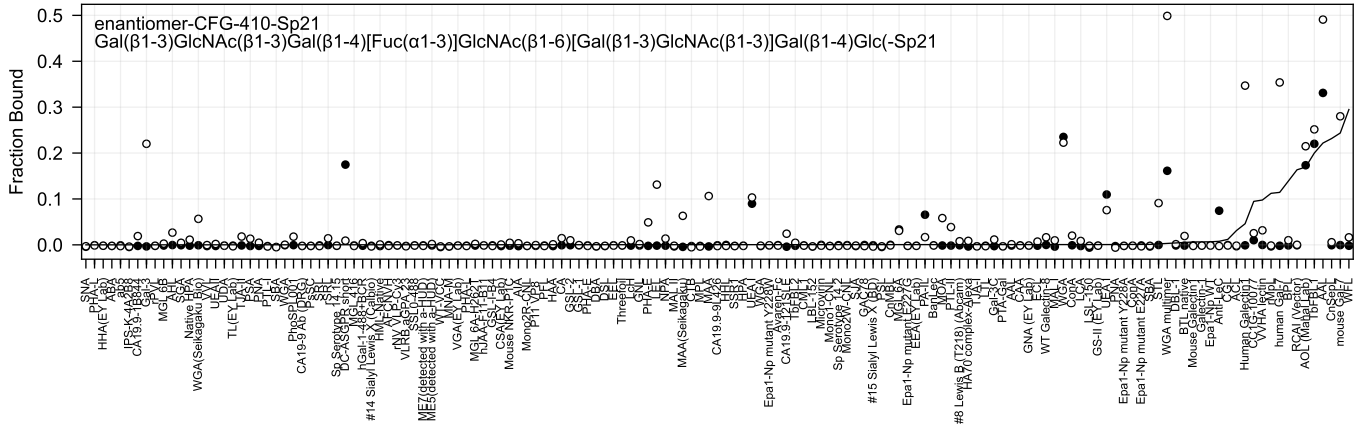
enantiomer-CFG-410-Sp21
Gal(β1-3)GlcNAc(β1-3)Gal(β1-4)[Fuc(α1-3)]GlcNAc(β1-6)[Gal(β1-3)GlcNAc(β1-3)]Gal(β1-4)Glc(-Sp21
Fraction Bound

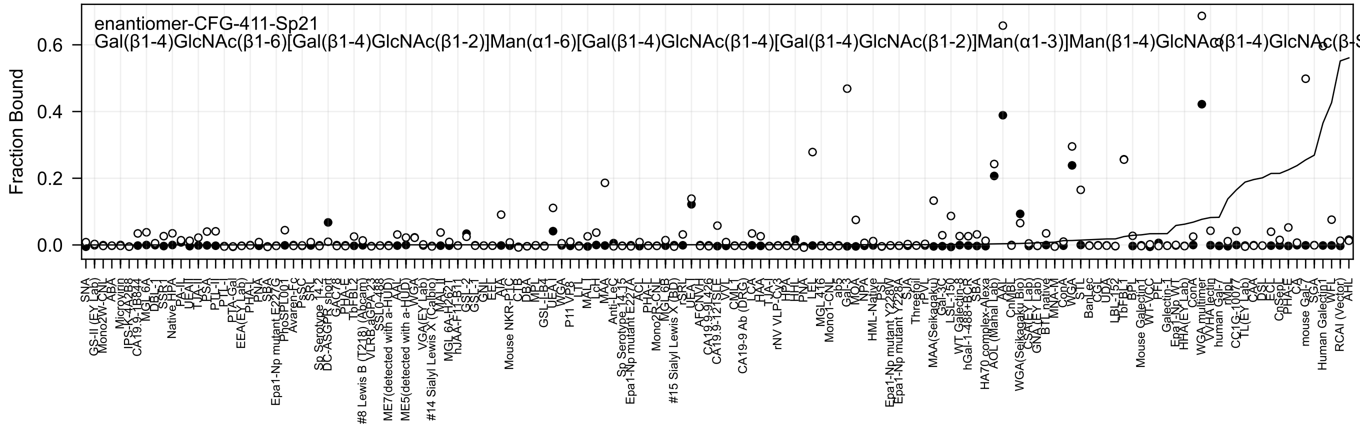
enantiomer-CFG-411-Sp21
Gal(β1-4)GlcNAc(β1-6)[Gal(β1-4)GlcNAc(β1-2)]Man(α1-6)[Gal(β1-4)GlcNAc(β1-4)[Gal(β1-4)GlcNAc(β1-2)]Man(α1-3)]Man(β1-4)GlcNAc(β1-4)GlcNAc(β-S
Fraction Bound
0.6
0.4
0.2
0.0

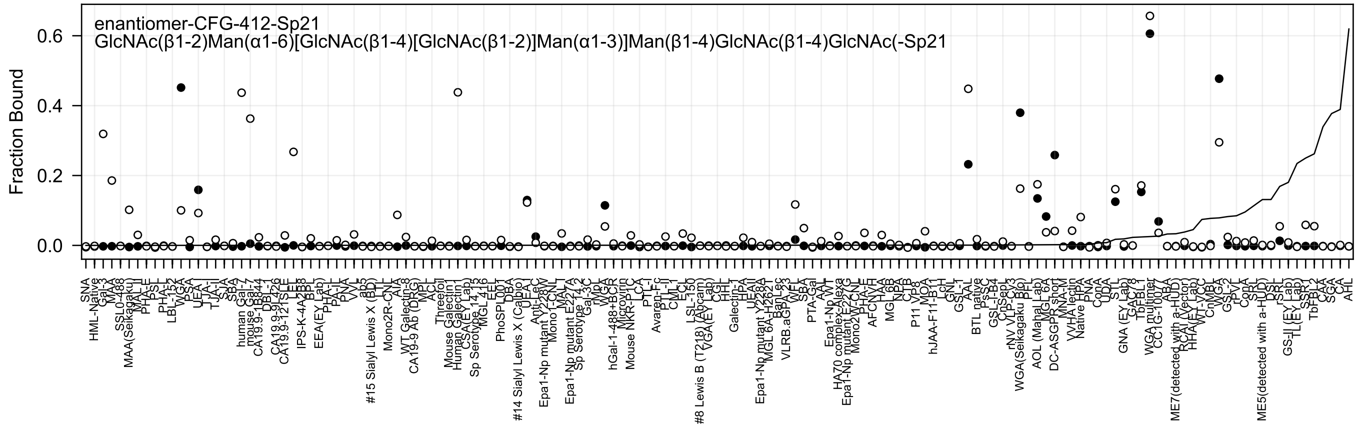
enantiomer-CFG-412-Sp21
GlcNAc(β1-2)Man(α1-6)[GlcNAc(β1-4)[GlcNAc(β1-2)]Man(α1-3)]Man(β1-4)GlcNAc(β1-4)GlcNAc(-Sp21
Fraction Bound
0.0
0.2
0.4
0.6

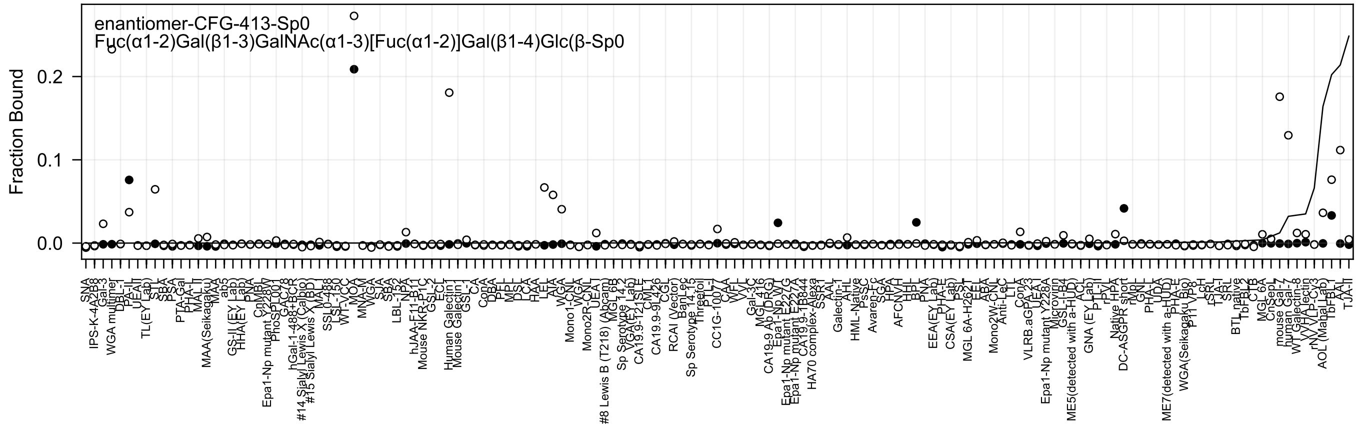

enantiomer-CFG-413-Sp0
Fuc(α1-2)Gal(β1-3)GalNAc(α1-3)[Fuc(α1-2)]Gal(β1-4)Glc(β-Sp0
Fraction Bound
0.0
0.1
0.2

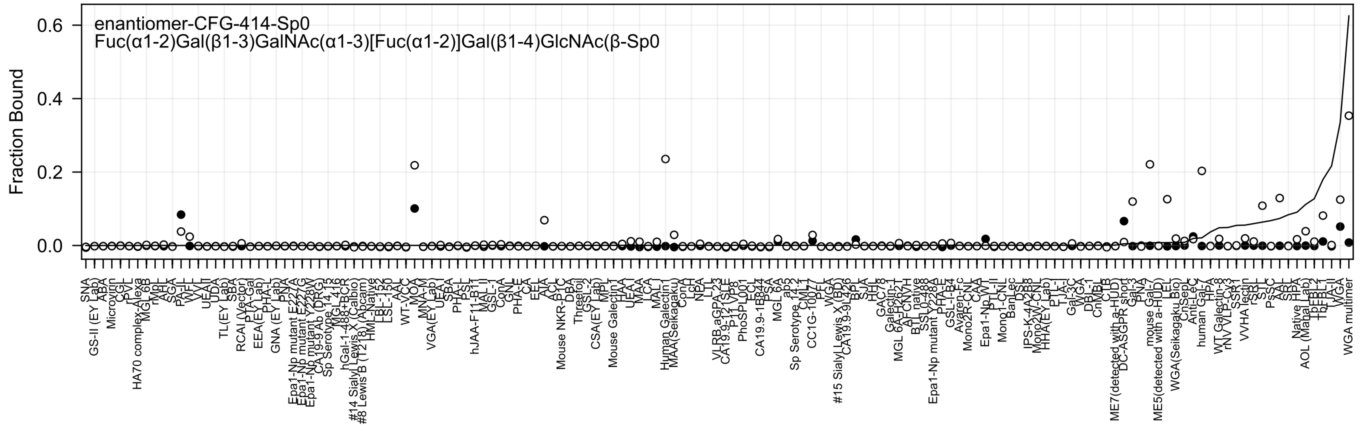

enantiomer-CFG-414-Sp0
Fuc(α1-2)Gal(β1-3)GalNAc(α1-3)[Fuc(α1-2)]Gal(β1-4)GlcNAc(β-Sp0
Fraction Bound
0.0
0.2
0.4
0.6

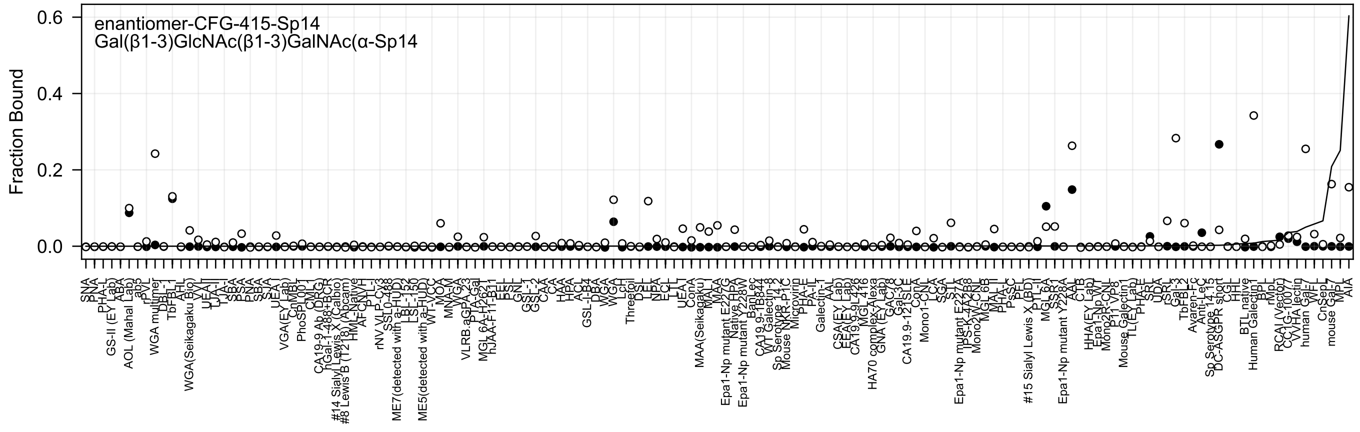

enantiomer-CFG-415-Sp14
Gal(β1-3)GlcNAc(β1-3)GalNAc(α-Sp14
Fraction Bound
0.6
0.4
0.2
0.0

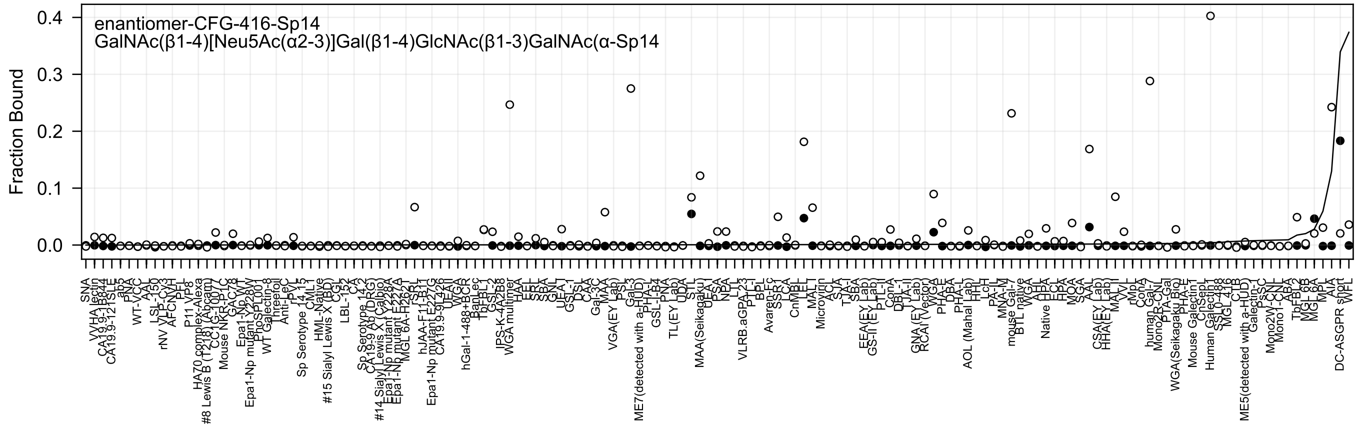
enantiomer-CFG-416-Sp14
GalNAc(β1-4)[Neu5Ac(α2-3)]Gal(β1-4)GlcNAc(β1-3)GalNAc(α-Sp14
Fraction Bound
0.0
0.1
0.2
0.3
0.4

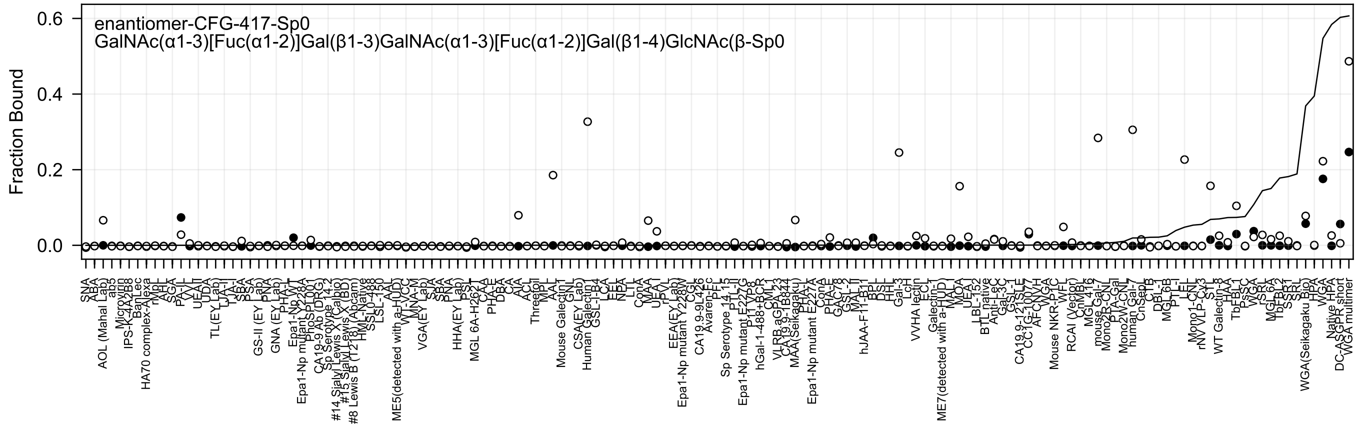
enantiomer-CFG-417-Sp0
GalNAc(α1-3)[Fuc(α1-2)]Gal(β1-3)GalNAc(α1-3)[Fuc(α1-2)]Gal(β1-4)GlcNAc(β-Sp0
Fraction Bound
0.6
0.4
0.2
0.0

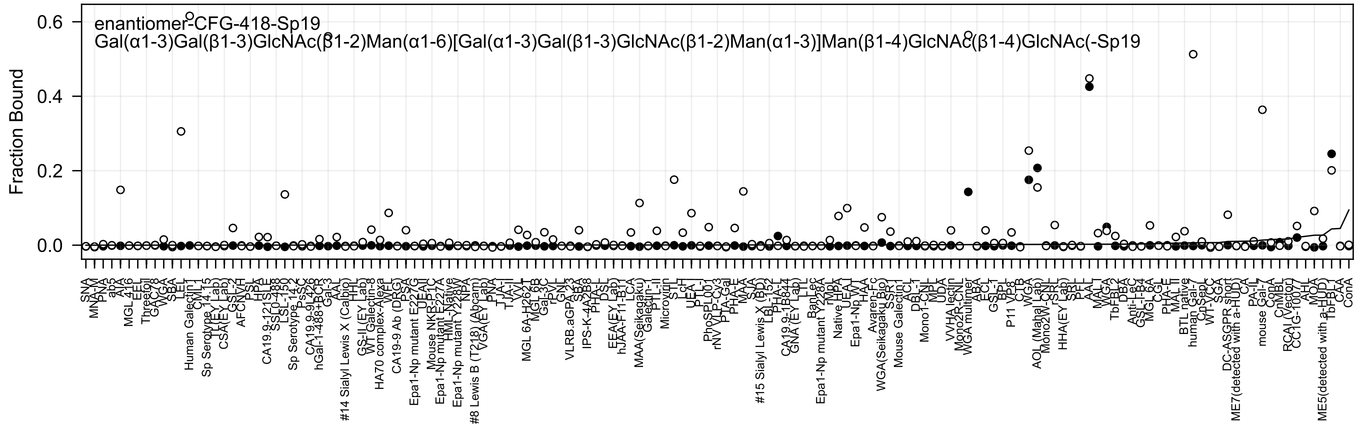
enantiomer-CFG-418-Sp19
Gal(α1-3)Gal(β1-3)GlcNAc(β1-2)Man(α1-6)[Gal(α1-3)Gal(β1-3)GlcNAc(β1-2)Man(α1-3)]Man(β1-4)GlcNAc(β1-4)GlcNAc(-Sp19
Fraction Bound
0.0
0.2
0.4
0.6

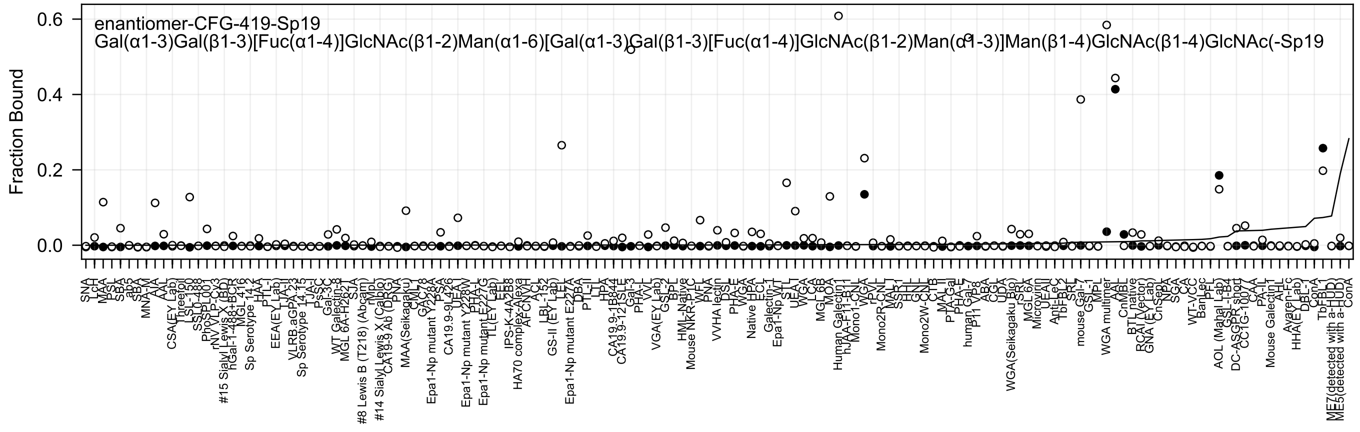

enantiomer-CFG-419-Sp19
Gal(α1-3)Gal(β1-3)[Fuc(α1-4)]GlcNAc(β1-2)Man(α1-6)[Gal(α1-3)Gal(β1-3)[Fuc(α1-4)]GlcNAc(β1-2)Man(α1-3)]Man(β1-4)GlcNAc(β1-4)GlcNAc(-Sp19
Fraction Bound
0.6
0.4
0.2
0.0
SNA
LcH
MAA
PSL
SBA
ab5
SBA
MNA-M
AIA
AAL
CSA(EY Lab)
Threefoil
LSL-150
SSL0-488
PhoSL
rNV VLP-Cy3
#15 Sialyl Lewis X (BD)
hGal-1-488+BCR
MGL 416
Sp Serotype 14.2
HAA
PTL-I
EEA(EY Lab)
TJA-II
VLRB.aGPA.23
Sp Serotype 14.15
TJA-I
PsSC
Gal-3C
WT Galectin-8
MGL 6A-H262T
STA
#8 Lewis B (T218) (Abcam)
rMpL
#14 Sialyl Lewis X (Calbo)
CA19-9 Ab (DRG)
PNA
MAA(Seikagaku)
CML
GAC78
Epa1-Np mutant Y228A
PSA
CA19.9-9L426
UEA
Epa1-Np mutant Y228W
PHA-L
Epa1-Np mutant E227G
TL(EY Lab)
EEL
IPS-K-4A2B8
HA70 complex-Alexa
AFCNVH
ACL
LBL-152
GS-II (EY Lab)
LEL
Epa1-Np mutant E227A
DBA
PTL-II
LTL
HPA
CA19.9-1B844
CA19.9-121SLE
Gal-3
PHA-L
VVL
VGA(EY Lab)
GSL-2
BPL
HML-Native
Mouse NKR-P1C
WFL
PNA
VVHA lectin
DSL
PHA-E
WGA
Native HPA
ECL
Galectin-1
Epa1-Np WT
STL
UEA-I
WGA
LCA
MGL 6B
MOA
Human Galectin1
hJAA-F11-B11
Mono1-CNL
WGA
rPVL
Mono2R-CNL
MAL-I
SSR1
HHL
GNL
Mono2W-CNL
CTB
MAL-II
PTA-Gal
PHA-E
human Gal-7
P11 VP8
ABA
CGL
UDA
WGA(Seikagaku Bio)
SRA
MGL 6A
Microvirin
UEAII
Anti-LeC
TbFBL2
SPL
mouse Gal-7
GSL-1
MPL
WGA multimer
AAL
CnMBL
BTL native
RCAI (Vector)
GNA (EY Lab)
CnSepL
NPA
SGA
CA
WT-VCC
BanLec
PFL
AOL (Mahal Lab)
GSL-I-B4
DC-ASGPR short
CC1G-1007
CAA
PA-IL
Mouse Galectin1
AHL
Ayaren-Fc
HHA(EY Lab)
DBL-1
ConA
TbFBL1
ME7(detected with a-HUD)
ME5(detected with a-HUD)
ConA

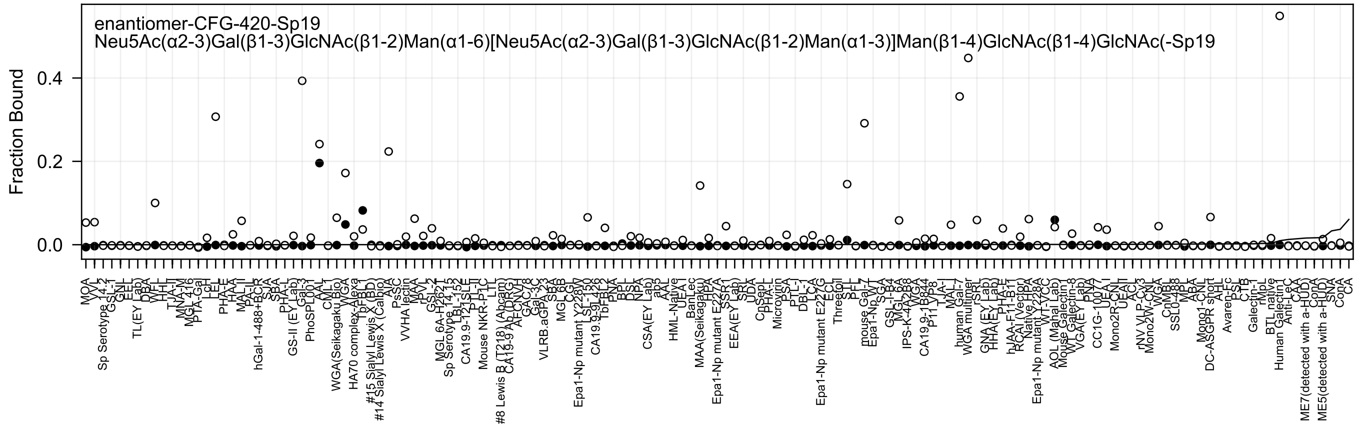
enantiomer-CFG-420-Sp19
Neu5Ac(α2-3)Gal(β1-3)GlcNAc(β1-2)Man(α1-6)[Neu5Ac(α2-3)Gal(β1-3)GlcNAc(β1-2)Man(α1-3)]Man(β1-4)GlcNAc(β1-4)GlcNAc(-Sp19
Fraction Bound
0.0
0.2
0.4

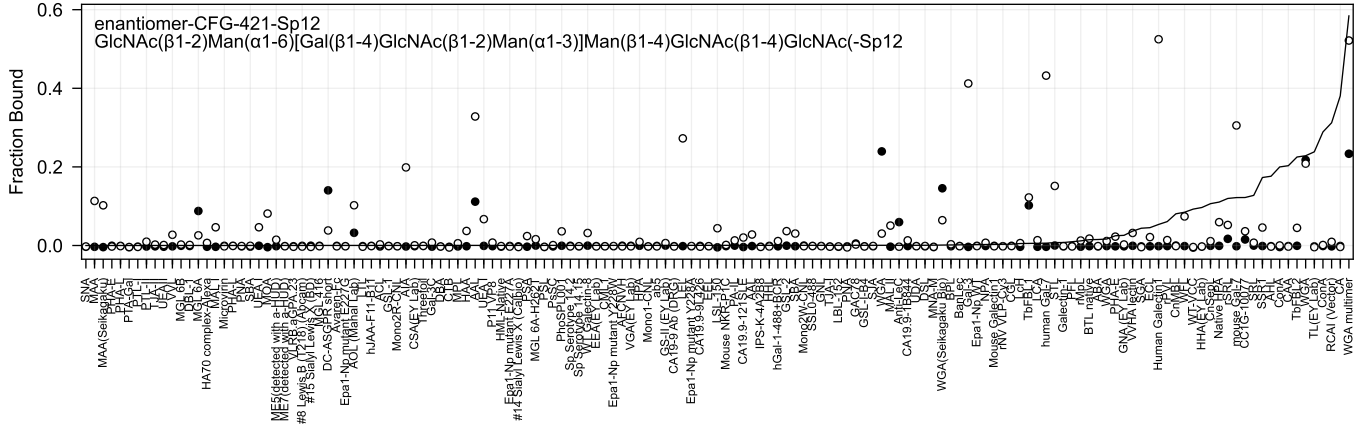
enantiomer-CFG-421-Sp12
GlcNAc(β1-2)Man(α1-6)[Gal(β1-4)GlcNAc(β1-2)Man(α1-3)]Man(β1-4)GlcNAc(β1-4)GlcNAc(-Sp12
Fraction Bound
0.0
0.2
0.4
0.6

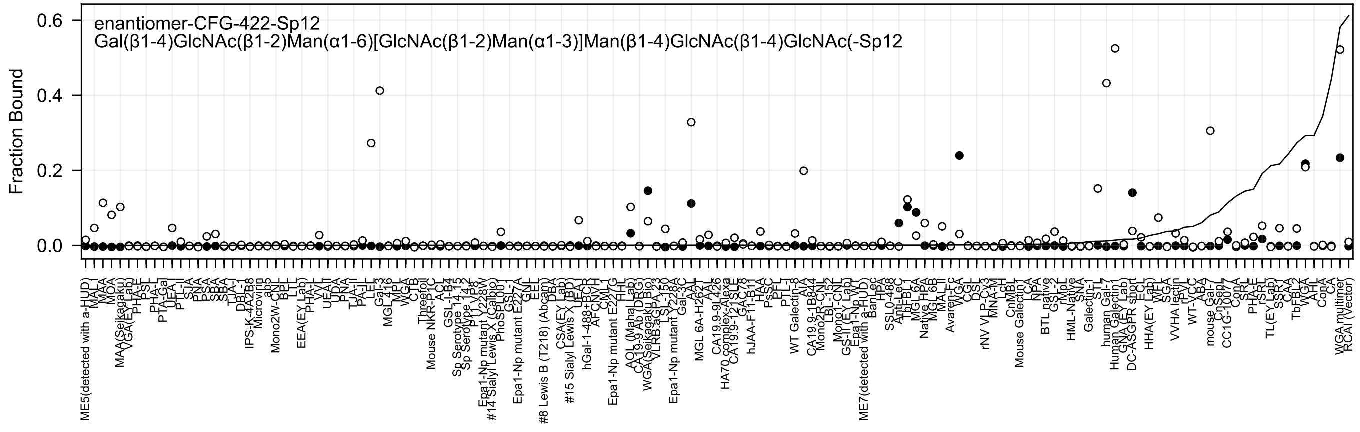
enantiomer-CFG-422-Sp12
Gal(β1-4)GlcNAc(β1-2)Man(α1-6)[GlcNAc(β1-2)Man(α1-3)]Man(β1-4)GlcNAc(β1-4)GlcNAc(-Sp12
Fraction Bound
0.6
0.4
0.2
0.0

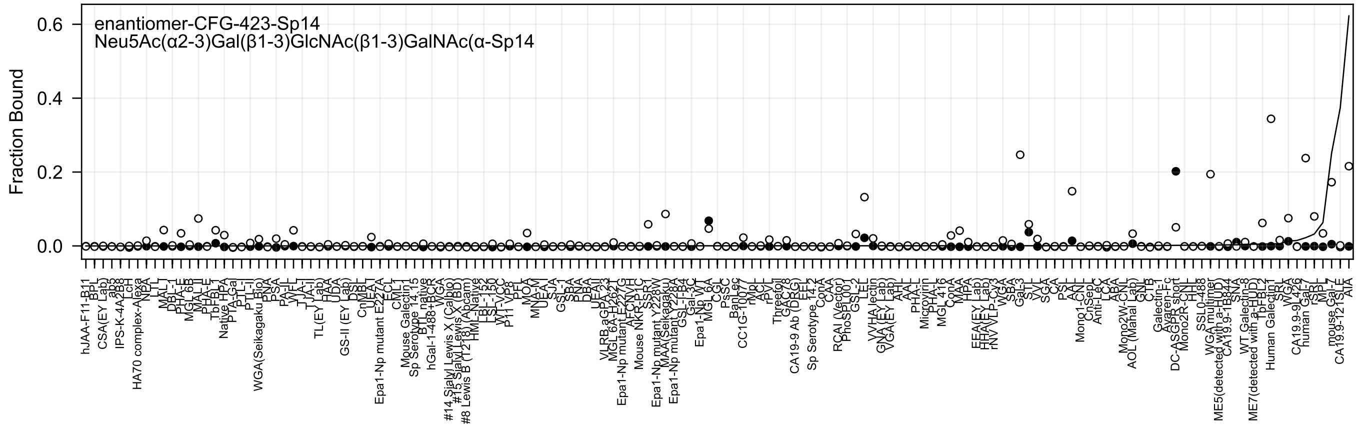
enantiomer-CFG-423-Sp14
Neu5Ac(α2-3)Gal(β1-3)GlcNAc(β1-3)GalNAc(α-Sp14
Fraction Bound
0.6
0.4
0.2
0.0

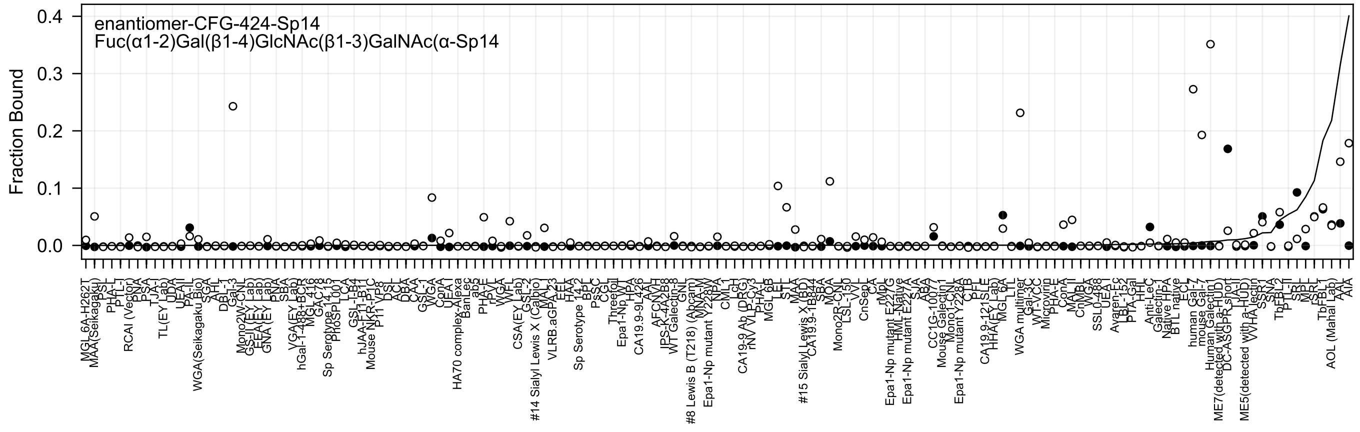
enantiomer-CFG-424-Sp14
Fuc(α1-2)Gal(β1-4)GlcNAc(β1-3)GalNAc(α-Sp14
Fraction Bound
0.0
0.1
0.2
0.3
0.4

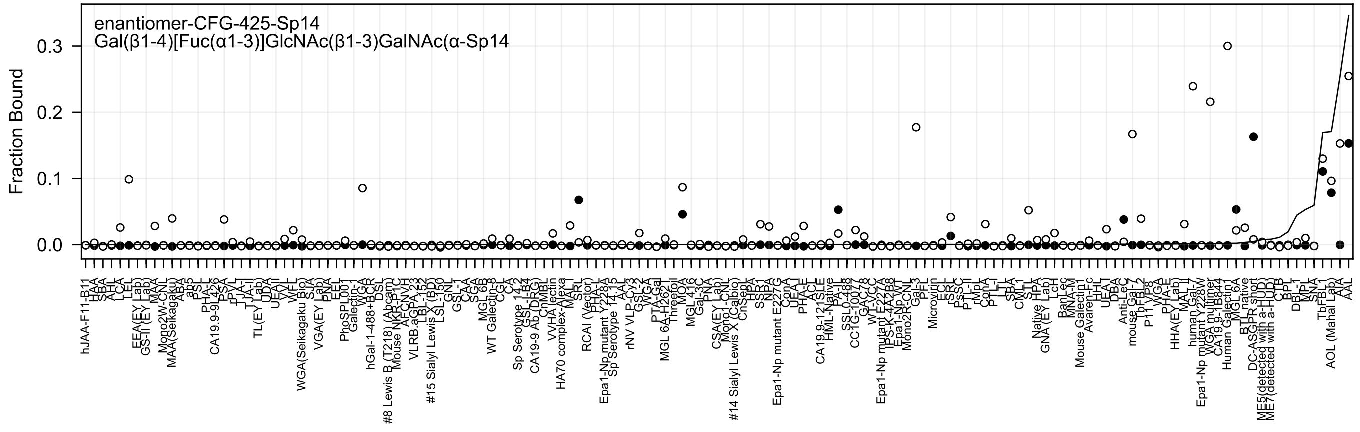
enantiomer-CFG-425-Sp14
Gal(β1-4)[Fuc(α1-3)]GlcNAc(β1-3)GalNAc(α-Sp14
Fraction Bound
0.3
0.2
0.1
0.0

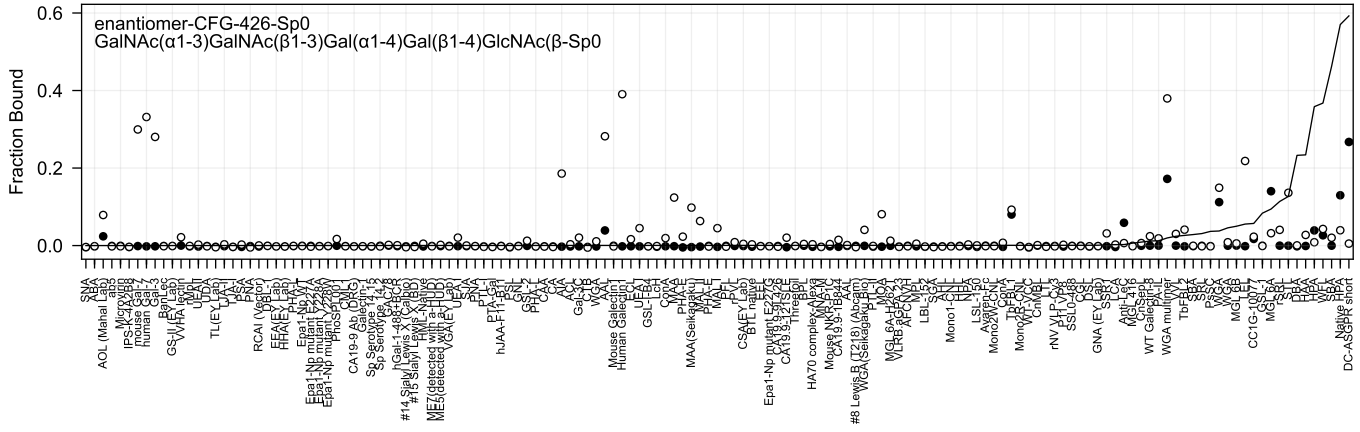
enantiomer-CFG-426-Sp0
GalNAc(α1-3)GalNAc(β1-3)Gal(α1-4)Gal(β1-4)GlcNAc(β-Sp0
Fraction Bound
0.6
0.4
0.2
0.0

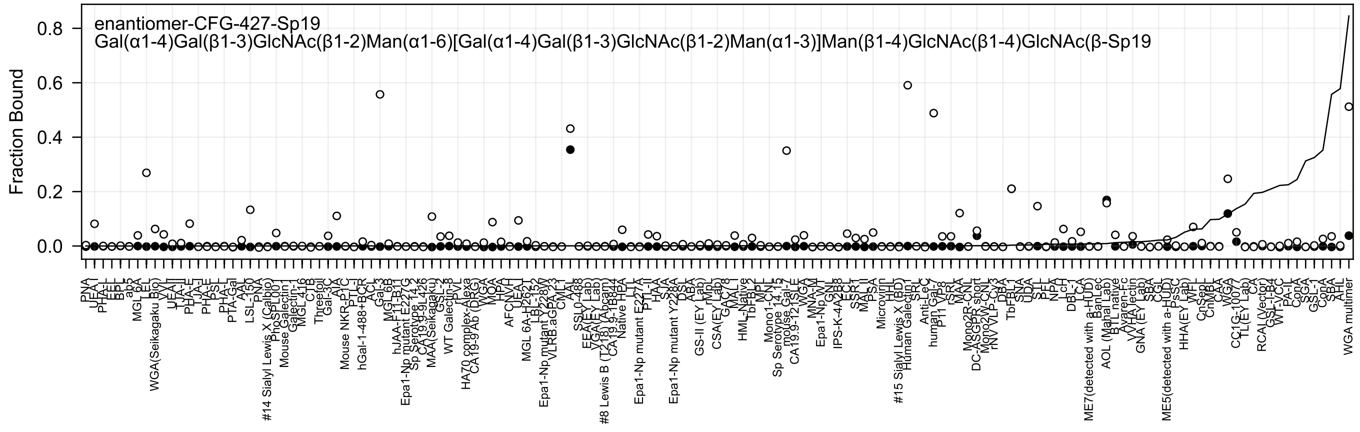
enantiomer-CFG-427-Sp19
Gal(α1-4)Gal(β1-3)GlcNAc(β1-2)Man(α1-6)[Gal(α1-4)Gal(β1-3)GlcNAc(β1-2)Man(α1-3)]Man(β1-4)GlcNAc(β1-4)GlcNAc(β-Sp19
Fraction Bound
0.8
0.6
0.4
0.2
0.0

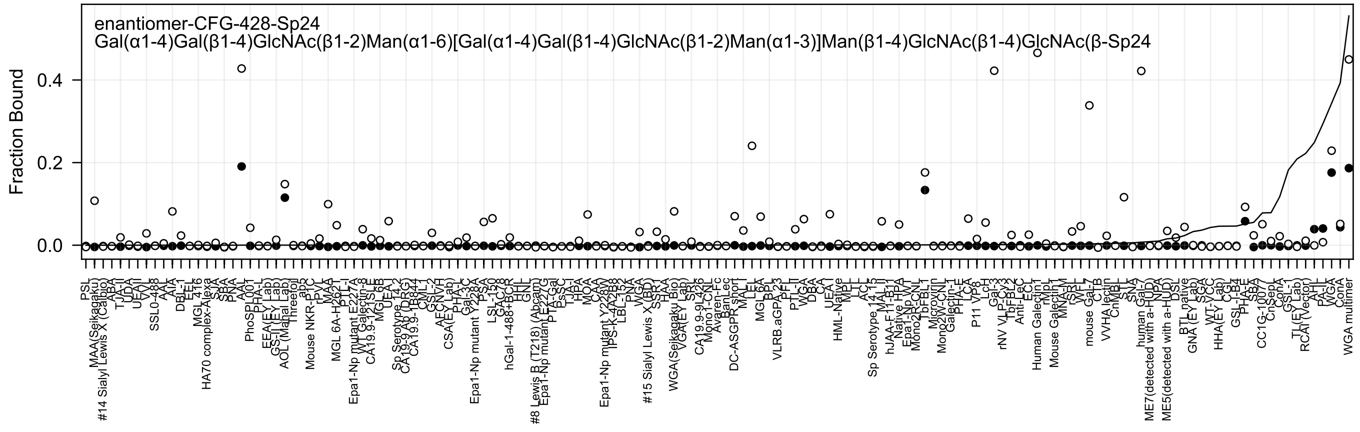
enantiomer-CFG-428-Sp24
Gal(α1-4)Gal(β1-4)GlcNAc(β1-2)Man(α1-6)[Gal(α1-4)Gal(β1-4)GlcNAc(β1-2)Man(α1-3)]Man(β1-4)GlcNAc(β1-4)GlcNAc(β-Sp24
Fraction Bound
0.0
0.2
0.4

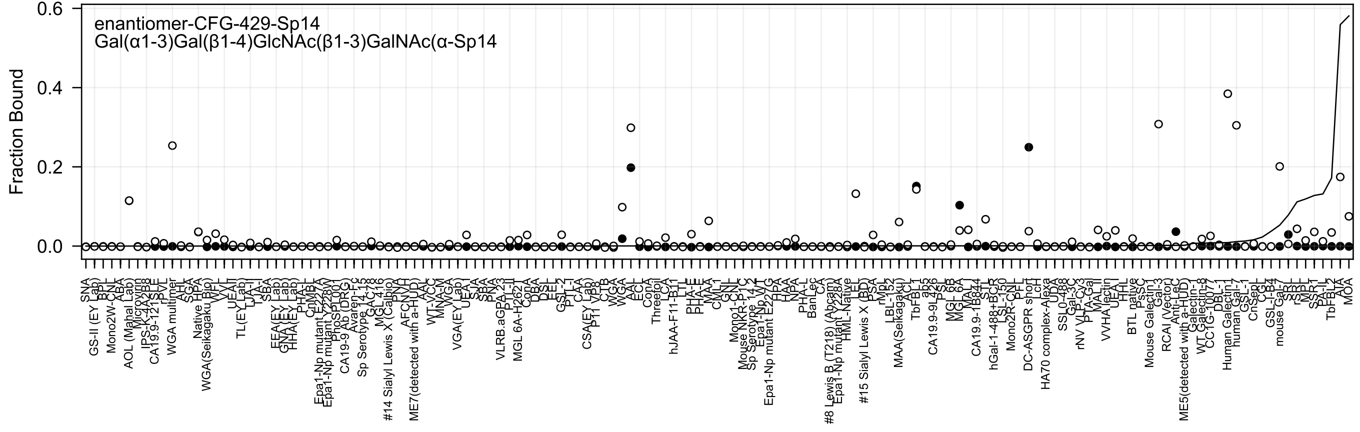
enantiomer-CFG-429-Sp14
Gal(α1-3)Gal(β1-4)GlcNAc(β1-3)GalNAc(α-Sp14
Fraction Bound

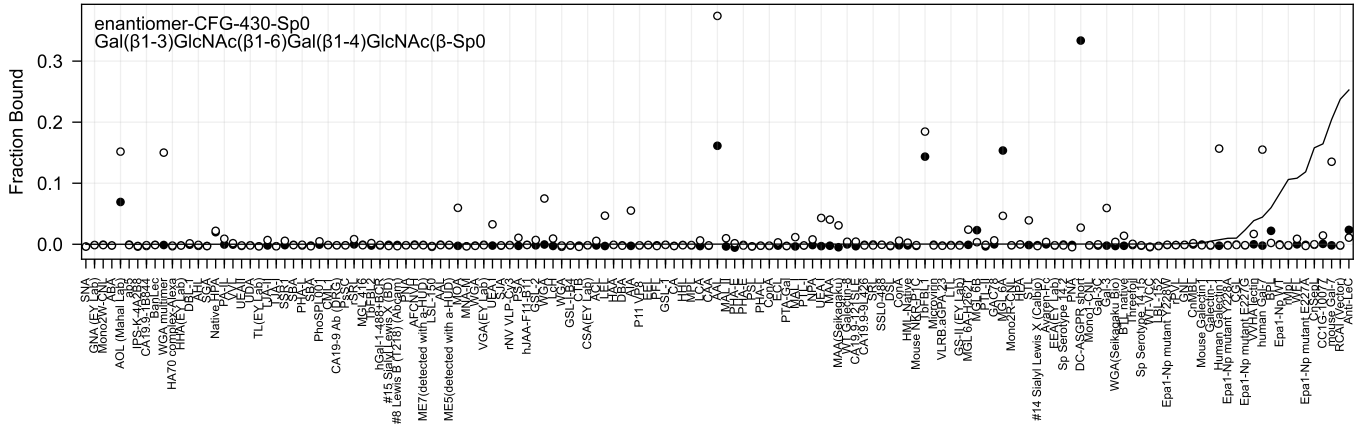
enantiomer-CFG-430-Sp0
Gal(β1-3)GlcNAc(β1-6)Gal(β1-4)GlcNAc(β-Sp0
Fraction Bound

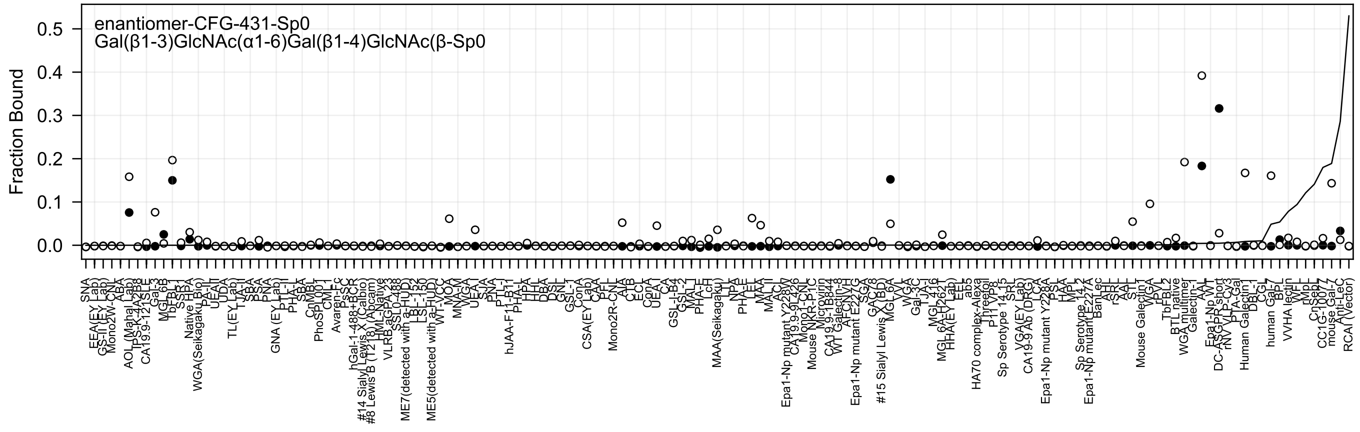
enantiomer-CFG-431-Sp0
Gal(β1-3)GlcNAc(α1-6)Gal(β1-4)GlcNAc(β-Sp0
Fraction Bound
0.5
0.4
0.3
0.2
0.1
0.0

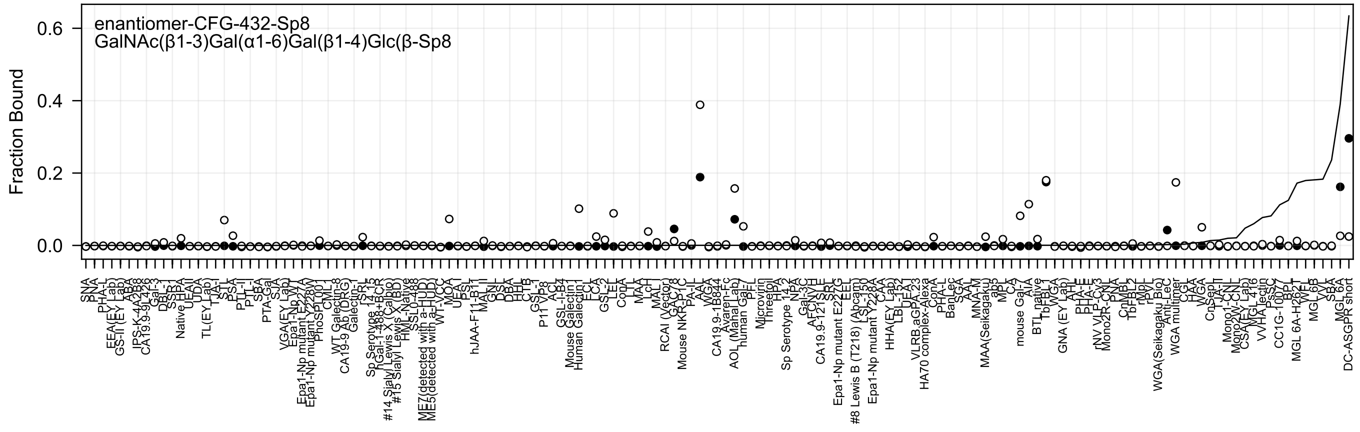

enantiomer-CFG-432-Sp8
GalNAc(β1-3)Gal(α1-6)Gal(β1-4)Glc(β-Sp8
Fraction Bound
0.0
0.2
0.4
0.6

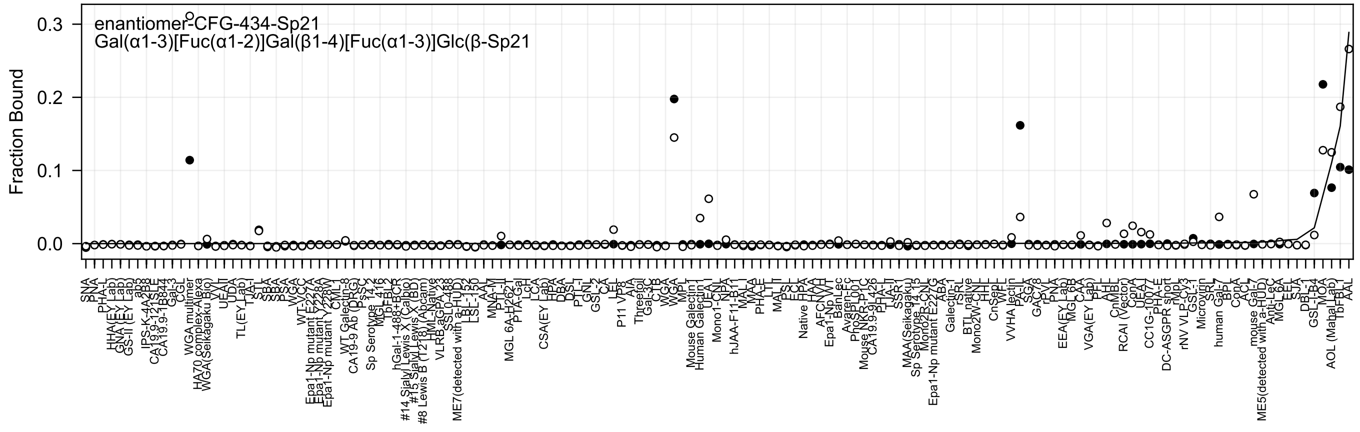

enantiomer-CFG-434-Sp21
Gal(α1-3)[Fuc(α1-2)]Gal(β1-4)[Fuc(α1-3)]Glc(β-Sp21
Fraction Bound
0.0
0.1
0.2
0.3

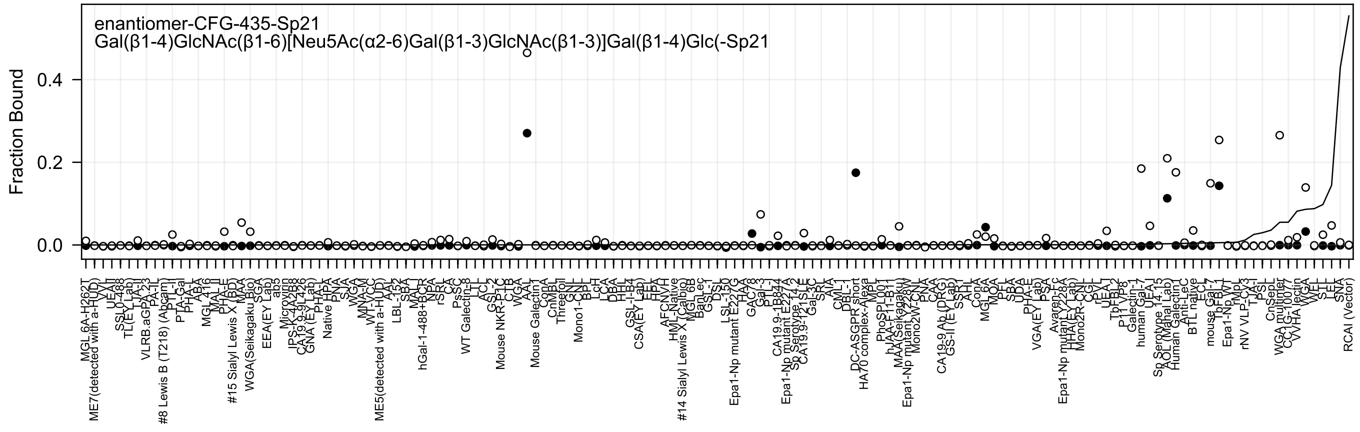
enantiomer-CFG-435-Sp21
Gal(β1-4)GlcNAc(β1-6)[Neu5Ac(α2-6)Gal(β1-3)GlcNAc(β1-3)]Gal(β1-4)Glc(-Sp21
Fraction Bound
0.4
0.2
0.0

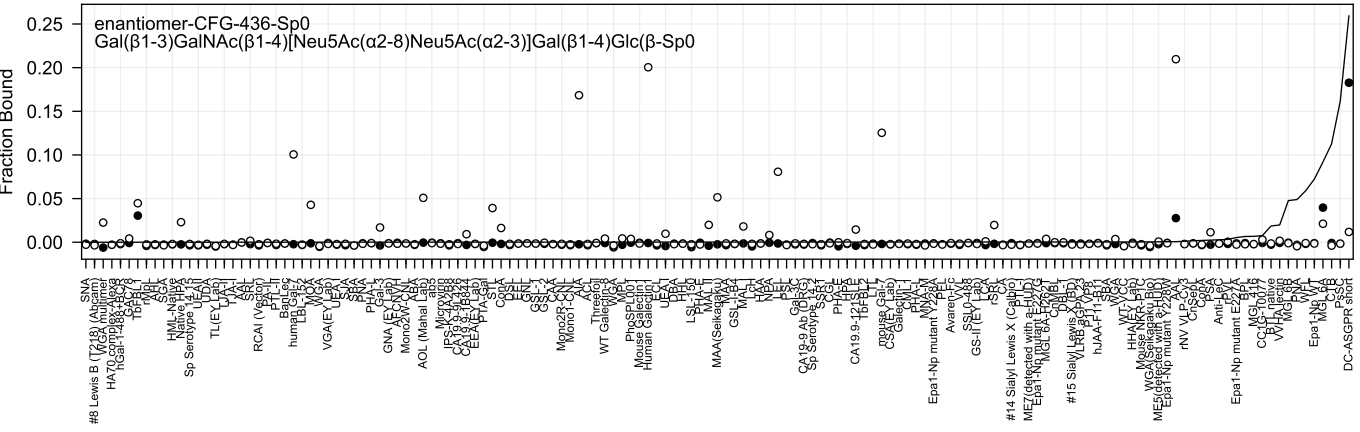

enantiomer-CFG-436-Sp0
Gal(β1-3)GalNAc(β1-4)[Neu5Ac(α2-8)Neu5Ac(α2-3)]Gal(β1-4)Glc(β-Sp0
Fraction Bound

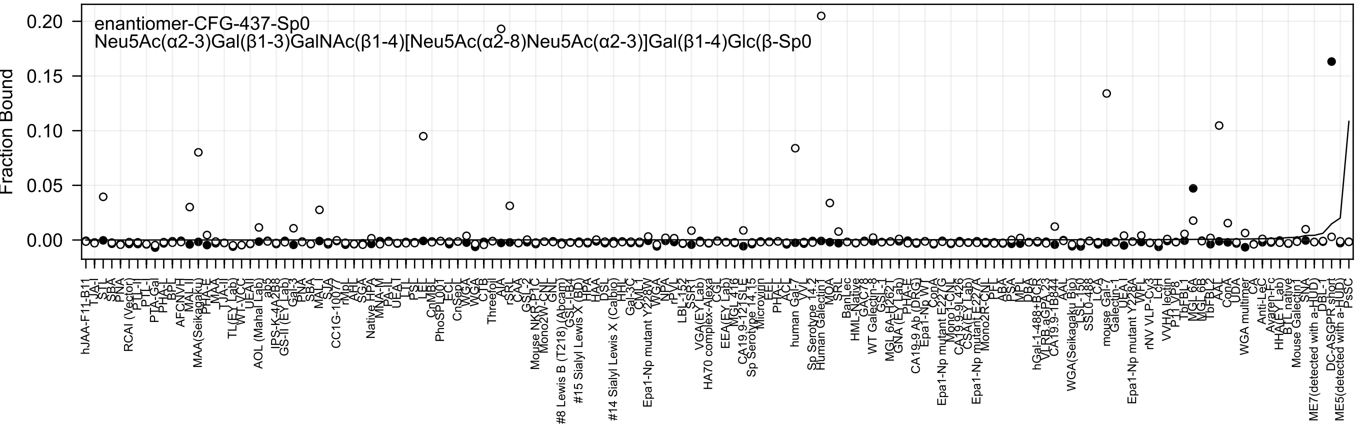
enantiomer-CFG-437-Sp0
Neu5Ac(α2-3)Gal(β1-3)GalNAc(β1-4)[Neu5Ac(α2-8)Neu5Ac(α2-3)]Gal(β1-4)Glc(β-Sp0
Fraction Bound
0.20
0.15
0.10
0.05
0.00

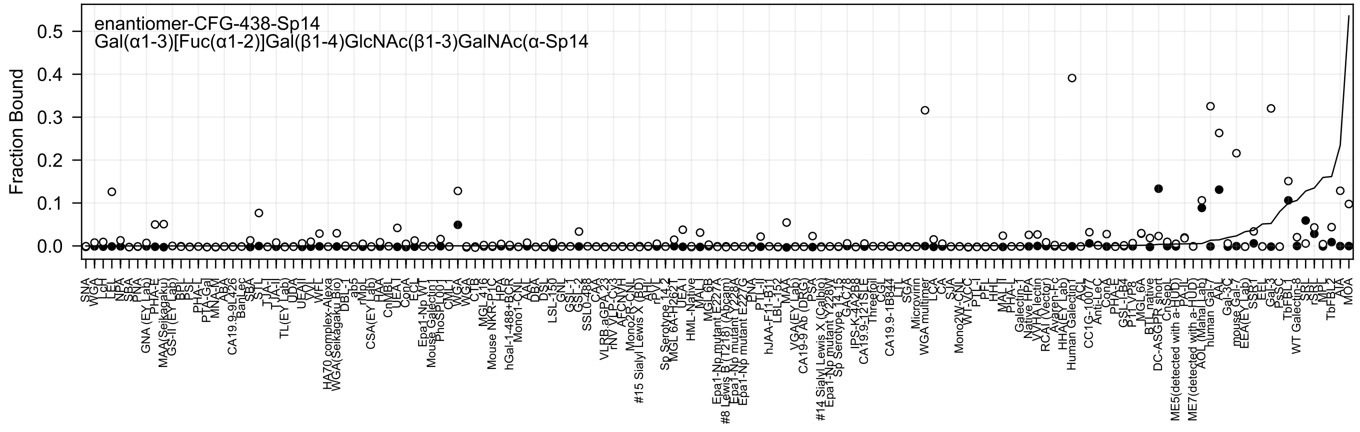
enantiomer-CFG-438-Sp14
Gal(α1-3)[Fuc(α1-2)]Gal(β1-4)GlcNAc(β1-3)GalNAc(α-Sp14
Fraction Bound
0.0
0.1
0.2
0.3
0.4
0.5

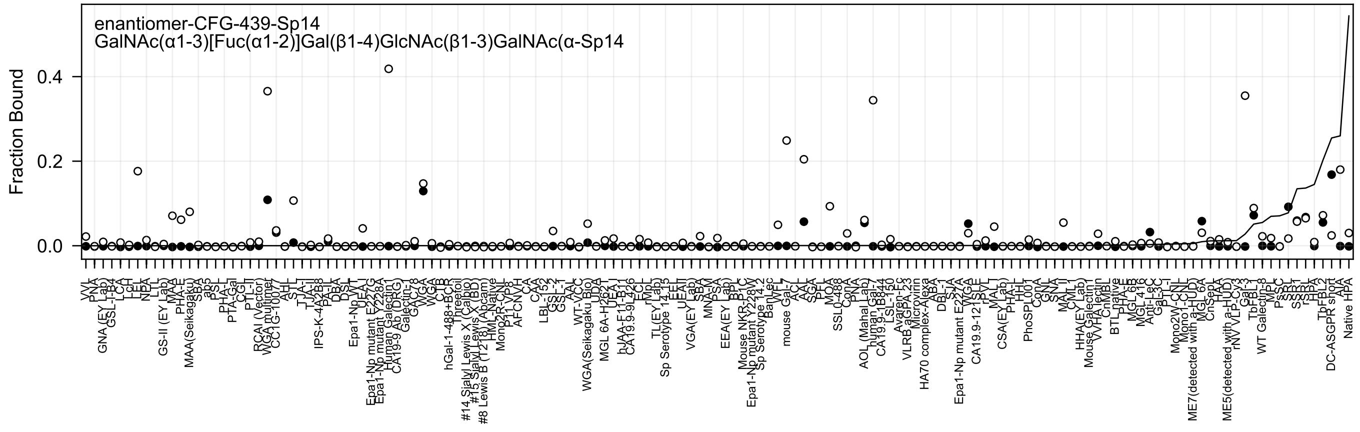
enantiomer-CFG-439-Sp14
GalNAc(α1-3)[Fuc(α1-2)]Gal(β1-4)GlcNAc(β1-3)GalNAc(α-Sp14
Fraction Bound
0.4
0.2
0.0

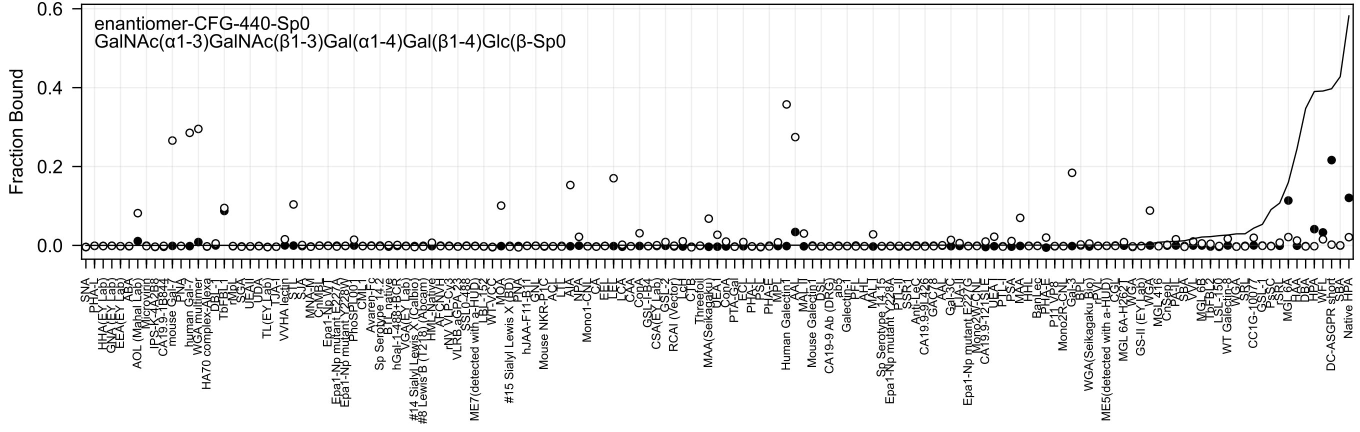
enantiomer-CFG-440-Sp0
GalNAc(α1-3)GalNAc(β1-3)Gal(α1-4)Gal(β1-4)Glc(β-Sp0
Fraction Bound
0.6
0.4
0.2
0.0

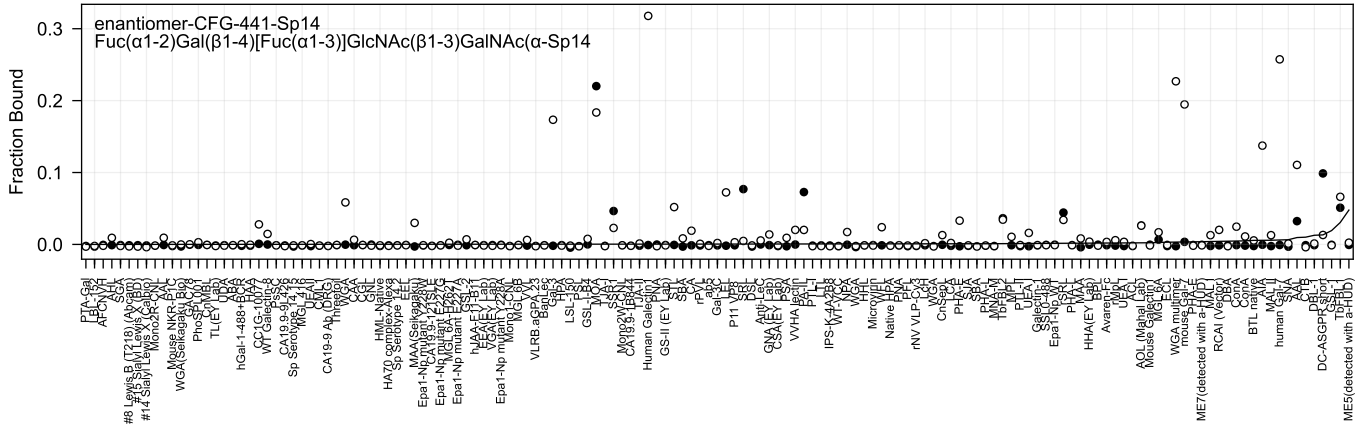
enantiomer-CFG-441-Sp14
Fuc(α1-2)Gal(β1-4)[Fuc(α1-3)]GlcNAc(β1-3)GalNAc(α-Sp14
Fraction Bound

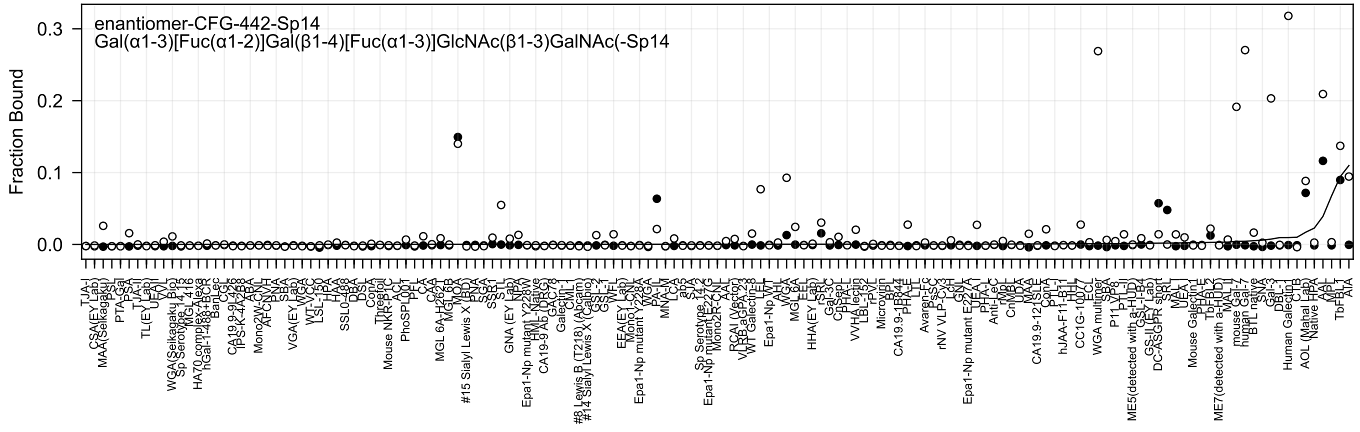
enantiomer-CFG-442-Sp14
Gal(α1-3)[Fuc(α1-2)]Gal(β1-4)[Fuc(α1-3)]GlcNAc(β1-3)GalNAc(-Sp14
Fraction Bound
0.3
0.2
0.1
0.0

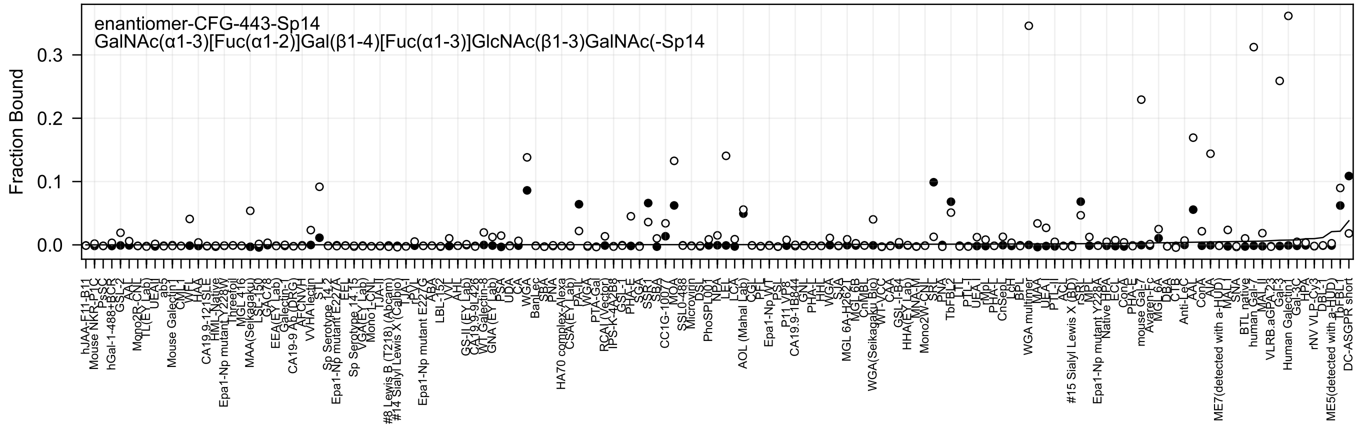
enantiomer-CFG-443-Sp14
GalNAc(α1-3)[Fuc(α1-2)]Gal(β1-4)[Fuc(α1-3)]GlcNAc(β1-3)GalNAc(-Sp14
Fraction Bound
0.0
0.1
0.2
0.3

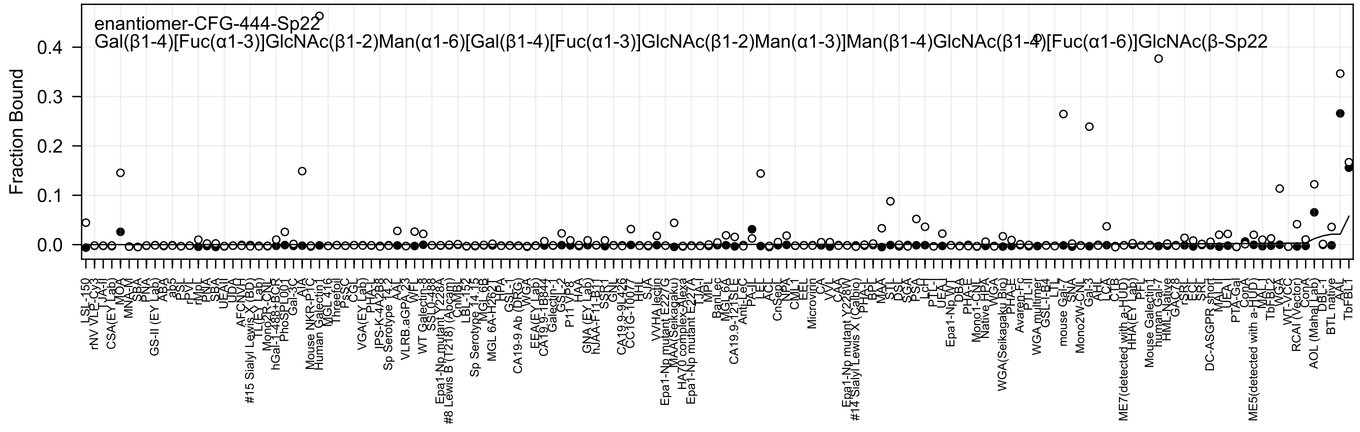
enantiomer-CFG-444-Sp22
Gal(β1-4)[Fuc(α1-3)]GlcNAc(β1-2)Man(α1-6)[Gal(β1-4)[Fuc(α1-3)]GlcNAc(β1-2)Man(α1-3)]Man(β1-4)GlcNAc(β1-4)[Fuc(α1-6)]GlcNAc(β-Sp22
Fraction Bound
0.4
0.3
0.2
0.1
0.0
LSL-150
rNV VLP-Cy3
CSA(EY Lab)
MOA
MNA-M
SBA
PNA
GS-II (EY Lab)
ABA
PSL
rPVL
PNA
SBA
UEAII
UDA
AFCNVH
#15 Sialyl Lewis X (BD)
Mono2R-CNL
hGal-1-488+BCR
PhoSL001
Gal-3C
AIA
Mouse NKR-P1C
Human Galectin1
MGL 416
Threefoil
PsSC
VGA(EY Lab)
PHA-L
IPS-K-4A2B8
Sp Serotype 14.2
AAL
VLRB.aGPA.23
WFL
WT Galectin-8
SSL0-488
Epa1-Np mutant Y228A
#8 Lewis B (T218) (Abcam)
CnMBL
LBL-152
Sp Serotype 14.15
MGL 6B
MGL 6A-H262
HPA
GSL-I
CA19-9 Ab (DRG)
WGA
EEA(EY Lab)
CA19.9-1B844
Galectin-1
GSL-2
P11 VP8
HAA
GNA (EY Lab)
hJAA-F11-B1
SSR1
GNL
CA19.9-9L426
CC1G-1007
HHL
SJA
VVHA lectin
Epa1-Np mutant E227G
MAA(Seikagaku)
HA70 complex-Alexa
Epa1-Np mutant E227A
TJA-I
MPL
BanLec
MGL 6A
CA19.9-121SLE
Anti-LeC
PA-IL
LEL
ACL
CnSep1
NPA
CML1
EEL
Microvirin
CA
VVL
CAA
Epa1-Np mutant Y228W
#14 Sialyl Lewis X (Calbio)
PHA-L
BPL
MAA
STL
DSL
SGA
PSA
LcH
PTL-I
UEA-I
Epa1-Np WT
DBA
PHA-E
Mono1-CNL
Native HPA
WGA
WGA(Seikagaku Bio)
PHA-E
Avaren-Fc
PTL-II
WGA multimer
GSL-I-B4
LTL
mouse Gal-7
SNA
Mono2W-CNL
Gal-3
AHL
LCA
CTB
ME7(detected with a-HUD)
HHA(EY Lab)
PFL
Mouse Galectin1
human Gal-7
HML-Native
GAC78
rSRL
ECL
SRL
DC-ASGPR short
MAL-I
UEA-I
PTA-Gal
ConA
ME5(detected with a-HUD)
MAL-I
TbFBL2
WGA
WT-VCC
RCAI (Vector)
ConA
AOL (Mahal Lab)
DBL-1
BTL native
AAL
TbFBL1

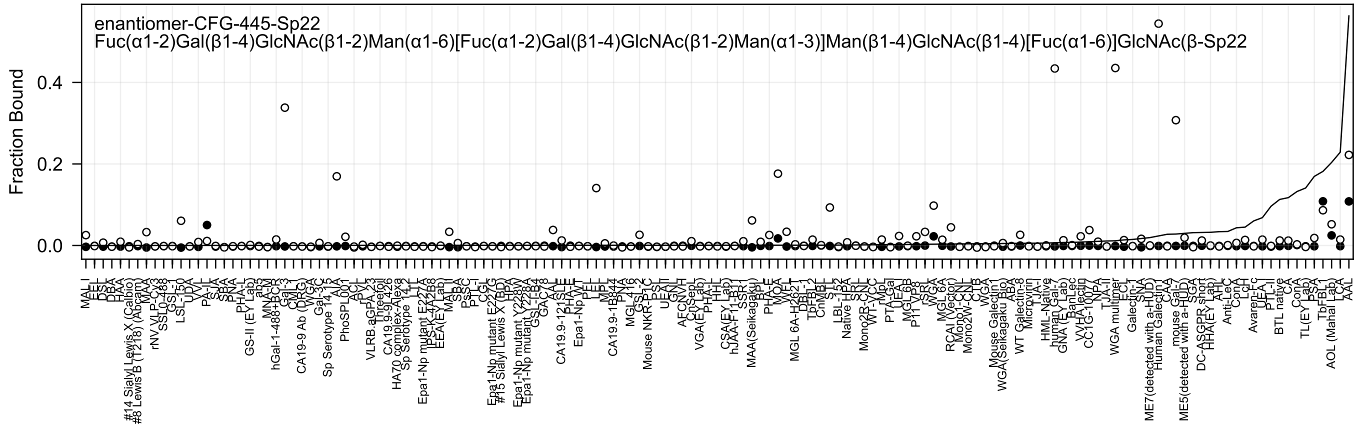
enantiomer-CFG-445-Sp22
Fuc(α1-2)Gal(β1-4)GlcNAc(β1-2)Man(α1-6)[Fuc(α1-2)Gal(β1-4)GlcNAc(β1-2)Man(α1-3)]Man(β1-4)GlcNAc(β1-4)[Fuc(α1-6)]GlcNAc(β-Sp22
Fraction Bound
0.0
0.2
0.4
MAL I
EEL
DSL
DBA
HAA
#14 Sialyl Lewis X (Calbio)
#8 Lewis B (T218) (Abcam)
MAA
rNV VLP-Cy3
SSL0-488
GSL-I
LSL-150
UDA
VVL
PA-IL
SJA
SBA
PNA
PHA-L
GS-II (EY Lab)
ab3
MNA-M
hGal-1-488+BCR
Gal-3
CML
CA19-9 Ab (DRG)
WGA
Gal-3C
Sp Serotype 14_15
AIA
PhoSPL001
ACL
rPVL
VLRB.aGPA.23
Threefoil
CA19.9-9L426
HA70 complex-Alexa
Sp Serotype 14_2
LTL
Epa1-Np mutant E227A
PS-K-4A2B8
EEA(EY Lab)
MAL II
SBA
PsSC
PTL-I
CGL
Epa1-Np mutant E227G
#15 Sialyl Lewis X (BD)
HPA
Epa1-Np mutant Y228W
Epa1-Np mutant Y228A
GSL-IB4
GAC78
AAL
CA19.9-121SLE
PHA-E
Epa1-Np WT
PEL
LEL
MPL
CA19.9-1B844
PNA
MGL 416
GSL-2
Mouse NKR-P1C
PSL
UEA II
GNL
AFCNVH
CnSep
VGA(EY Lab)
PHA-L
HHL
CSA(EY Lab)
hJAA-F11-B11
SSR1
MAA(Seikagaku)
BPL
PHA-E
MOA
WFL
MGL 6A-H262T
DBL-1
TbFBL2
CnMBL
STL
LBL-152
Native HPA
SRL
Mono2R-CNL
WT-VCC
rMpL
PTA-Gal
UEA I
MGL 6B
P11 VP8
rSRL
WGA
MGL 6A
RCAI (Vector)
Mono1-CNL
Mono2W-CNL
CLB
WGA
Mouse Galectin1
WGA(Seikagaku Bio)
ABA
WT Galectin-8
Microvirin
TJA-I
HML-Native
human Gal-7
GNA (EY Lab)
BanLec
VVHA lectin
CC1G-1007
NPA
TJA-II
WGA multimer
ECL
Galectin-1
SNA
ME7(detected with a-HUD)
Human Galectin1
CAA
mouse Gal-7
ME5(detected with a-HUD)
SGA
DC-ASGPR short
HHA(EY Lab)
AHL
Anti-LeC
ConA
LcH
Avaren-Fc
UEA I
PTL-II
BTL native
CA
ConA
TL(EY Lab)
PSA
TbFBL1
AOL (Mahal Lab)
LCA
AAL

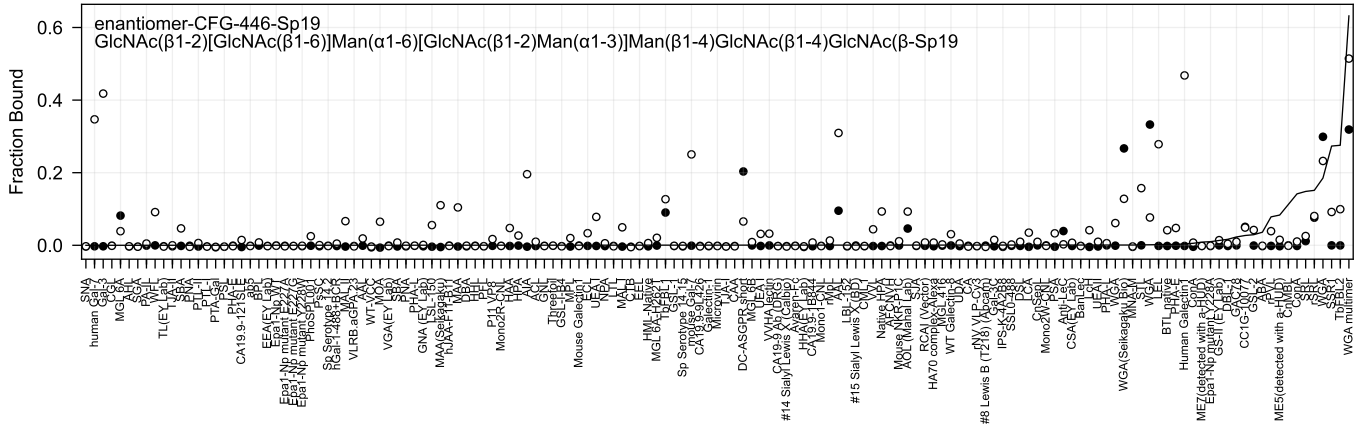
enantiomer-CFG-446-Sp19
GlcNAc(β1-2)[GlcNAc(β1-6)]Man(α1-6)[GlcNAc(β1-2)Man(α1-3)]Man(β1-4)GlcNAc(β1-4)GlcNAc(β-Sp19
Fraction Bound
0.6
0.4
0.2
0.0

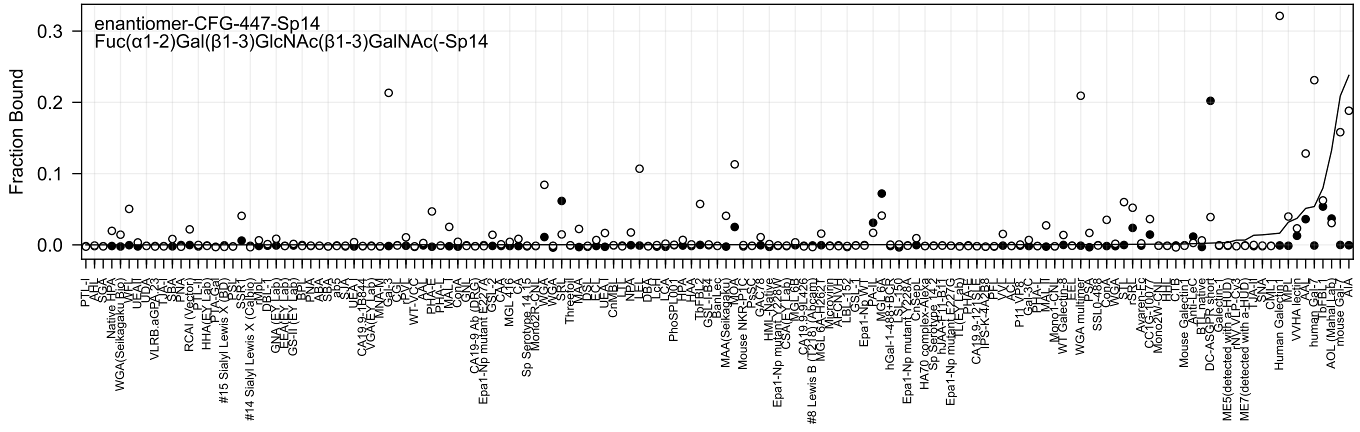
enantiomer-CFG-447-Sp14
Fuc(α1-2)Gal(β1-3)GlcNAc(β1-3)GalNAc(-Sp14
Fraction Bound
0.3
0.2
0.1
0.0

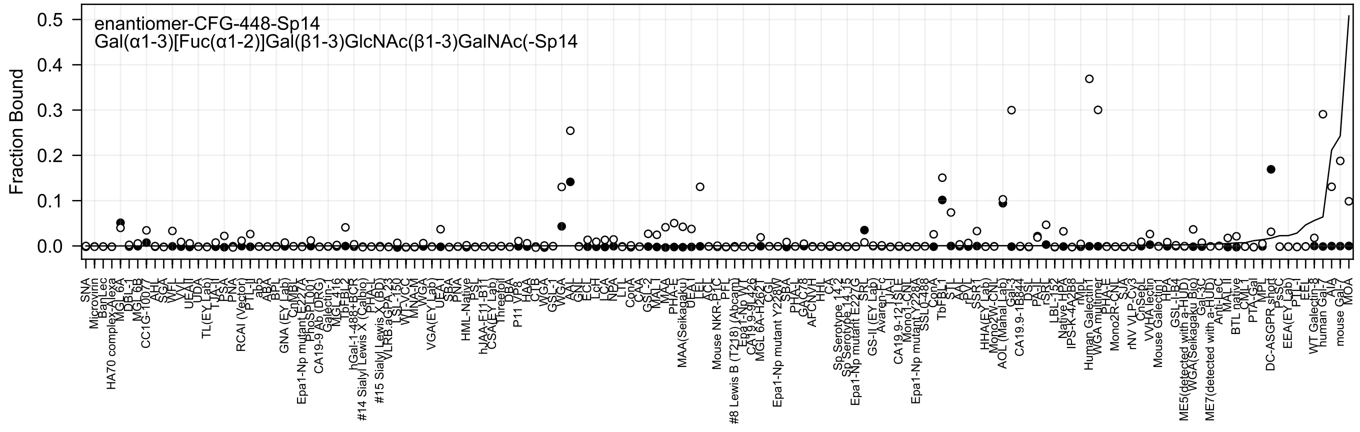
enantiomer-CFG-448-Sp14
Gal(α1-3)[Fuc(α1-2)]Gal(β1-3)GlcNAc(β1-3)GalNAc(-Sp14
Fraction Bound
0.5
0.4
0.3
0.2
0.1
0.0
SNA
Microvirin
BanLec
HA70 complex-Alexa
MGL 6A
DBL-1
MGL 6B
CC1G-10077
AAL
SGA
WFL
VVL
UEAII
UDA
TL(EY Lab)
TJA-II
PSA
PNA
RCAI (Vector)
PTL-II
ab3
ABA
BPL
GNA (EY Lab)
CnMBL
Epa1-Np mutant E227A
PhoSL001
CA19-9 Ab (DRG)
Galectin-1
MGL 416
TbFBL2
hGal-1-488+BCR
#14 Sialyl Lewis X (Calbio)
PHA-L
#15 Sialyl Lewis X (BD)
VLRB.aGPA.23
LSL-150
WT-VCC
MNA-M
WGA
VGA(EY Lab)
UEA I
SBA
PNA
HML-Native
PSL
hJAA-E11-B11
CSA(EY Lab)
Threefoil
DBA
P11 VP8
HAA
CTB
WGA
GSL-I
WGA
AAL
GNL
ECL
LcH
LCA
NPA
LTL
ConA
CAA
GSL-2
MAL-I
MAA
PHA-E
MAA(Seikagaku)
UEA I
LEL
ACL
Mouse NKR-P1C
PFL
#8 Lewis B (T218) (Abcam)
Epa1-Np WT
CA19-9 Ab 426
MGL 6A-H262T
CGL
CSL
Epa1-Np mutant Y228W
SBA
PHA-L
GAC78
AFCNVH
HHL
CA
Sp Serotype 14.2
Sp Serotype 14.15
Epa1-Np mutant E227G
SRL
GS-II (EY Lab)
Avaren-Fc
TJA-I
CA19.9-121SLE
Mono1-CNL
Epa1-Np mutant Y228A
SSL0488
ConA
TbFBL1
STL
AAL
rPVL
SSR1
HHA(EY Lab)
Mono2W-CNL
AOL (Mahal Lab)
Gal-3
CA19.9-1B844
DSL
PA-IL
rSRL
LBL-152
Native HPA
IPS-K-4A2B8
rMpL
Human Galectin1
WGA multimer
PHA-E
Mono2R-CNL
SJA
rNV VLP-Cy3
CnSepL
VVHA lectin
Mouse Galectin1
HPA
GSL-I-B4
ME5(detected with a-HUD)
WGA(Seikagaku Bio)
Gal-3C
ME7(detected with a-HUD)
Anti-LeC
MAL II
BTL native
CML1
PTA-Gal
MPL
DC-ASGPR short
PsSC
EEA(EY Lab)
PTL-I
EEL
WT Galectin-8
human Gal-7
AIA
mouse Gal-7
MOA

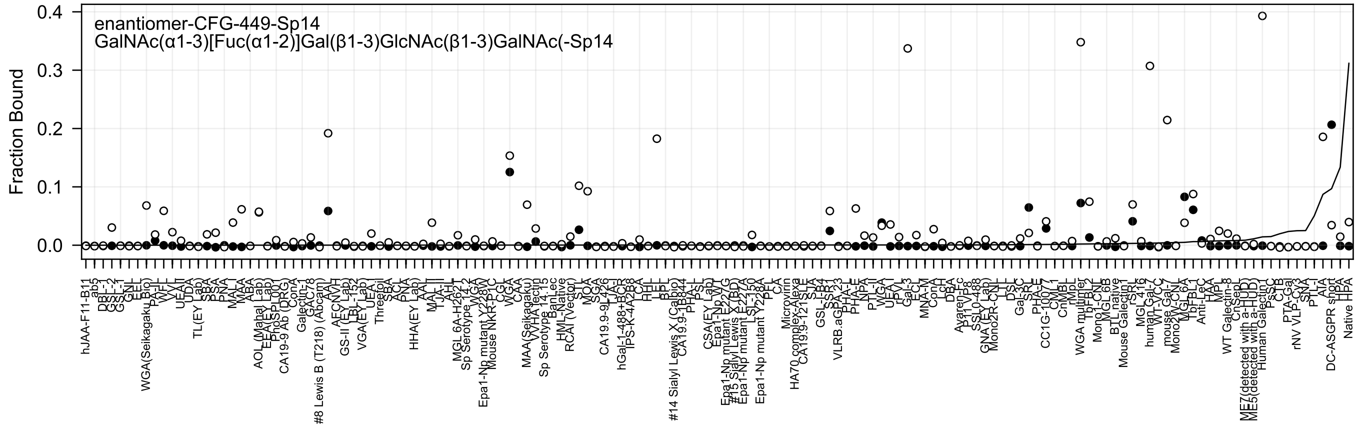

enantiomer-CFG-449-Sp14
GalNAc(α1-3)[Fuc(α1-2)]Gal(β1-3)GlcNAc(β1-3)GalNAc(-Sp14
Fraction Bound
0.4
0.3
0.2
0.1
0.0

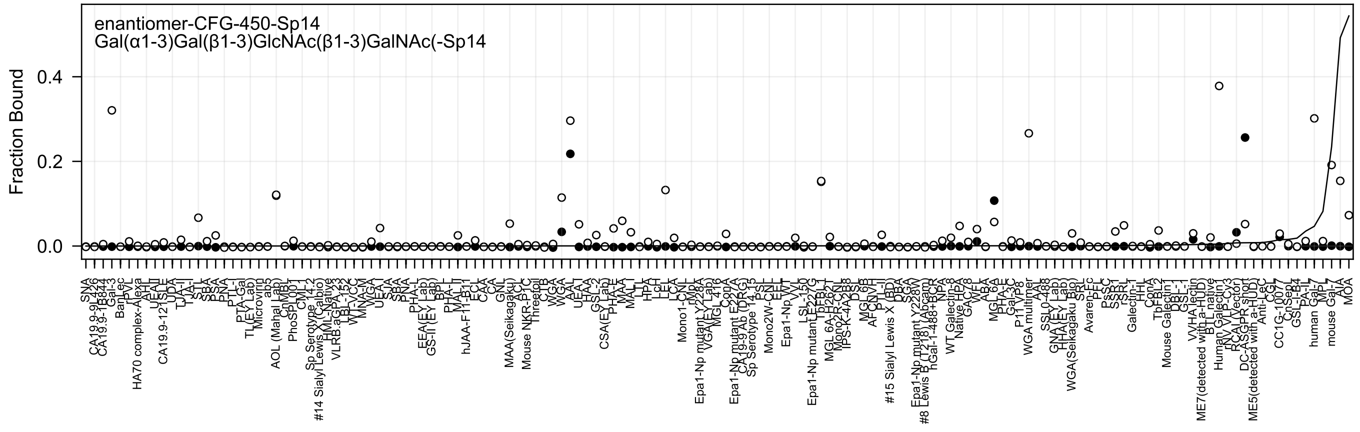
enantiomer-CFG-450-Sp14
Gal(α1-3)Gal(β1-3)GlcNAc(β1-3)GalNAc(-Sp14
Fraction Bound
0.0
0.2
0.4

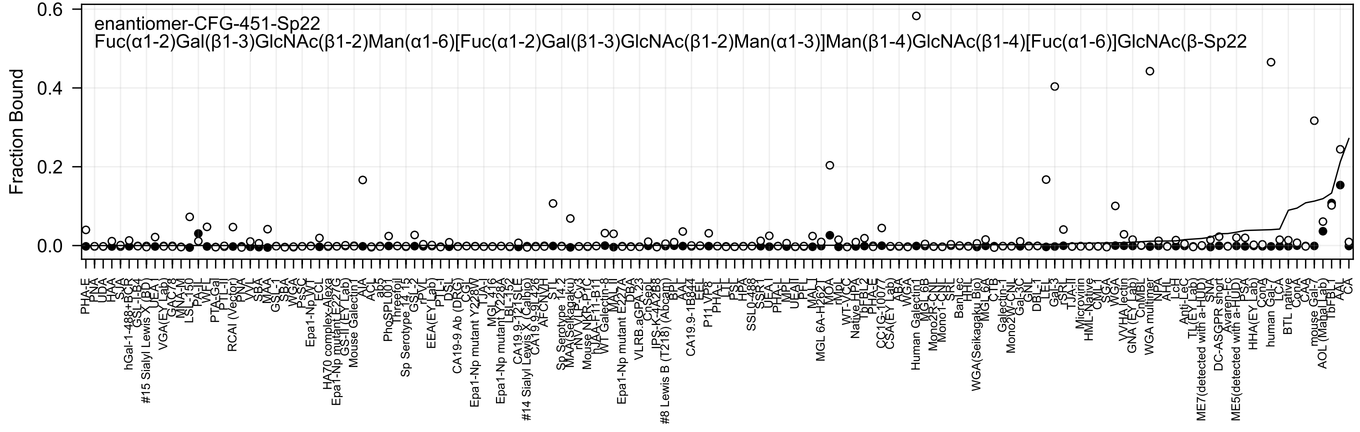

enantiomer-CFG-451-Sp22
Fuc(α1-2)Gal(β1-3)GlcNAc(β1-2)Man(α1-6)[Fuc(α1-2)Gal(β1-3)GlcNAc(β1-2)Man(α1-3)]Man(β1-4)GlcNAc(β1-4)[Fuc(α1-6)]GlcNAc(β-Sp22
Fraction Bound

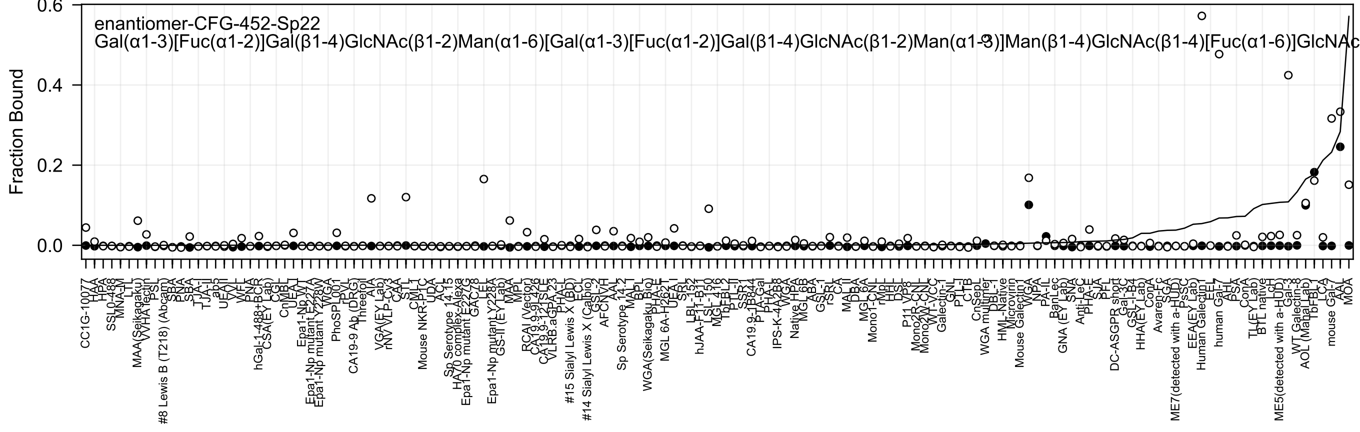
enantiomer-CFG-452-Sp22
Gal(α1-3)[Fuc(α1-2)]Gal(β1-4)GlcNAc(β1-2)Man(α1-6)[Gal(α1-3)[Fuc(α1-2)]Gal(β1-4)GlcNAc(β1-2)Man(α1-3)]Man(β1-4)GlcNAc(β1-4)[Fuc(α1-6)]GlcNAc
Fraction Bound
0.6
0.4
0.2
0.0

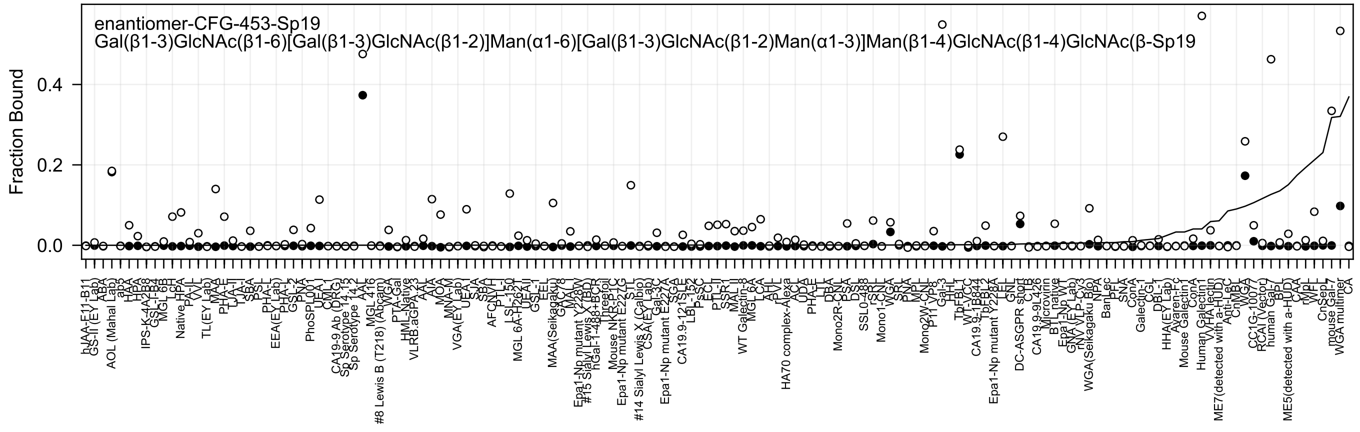

enantiomer-CFG-453-Sp19
Gal(β1-3)GlcNAc(β1-6)[Gal(β1-3)GlcNAc(β1-2)]Man(α1-6)[Gal(β1-3)GlcNAc(β1-2)Man(α1-3)]Man(β1-4)GlcNAc(β1-4)GlcNAc(β-Sp19
Fraction Bound
0.0
0.2
0.4

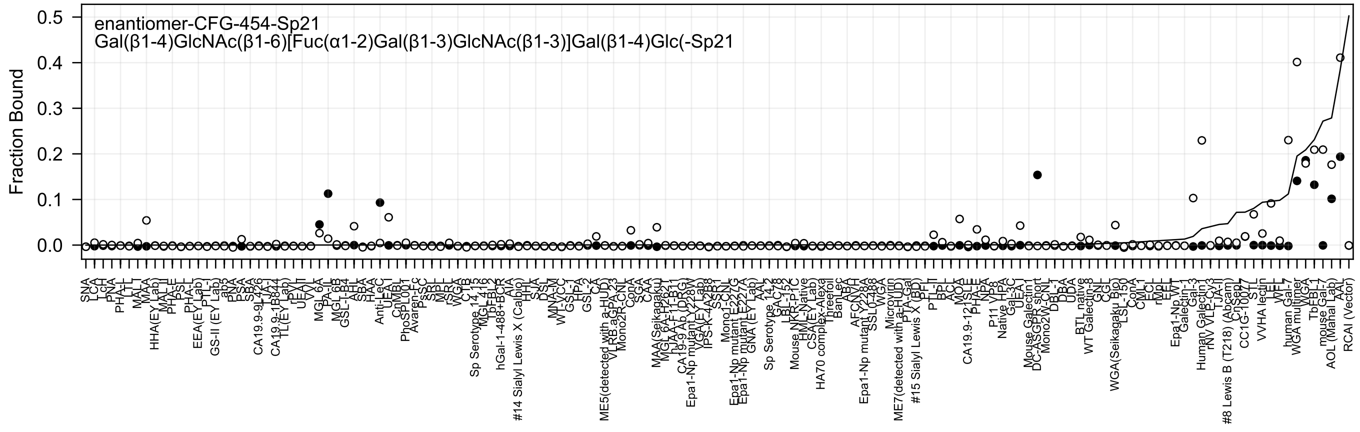

enantiomer-CFG-454-Sp21
Gal(β1-4)GlcNAc(β1-6)[Fuc(α1-2)Gal(β1-3)GlcNAc(β1-3)]Gal(β1-4)Glc(-Sp21
Fraction Bound
0.5
0.4
0.3
0.2
0.1
0.0

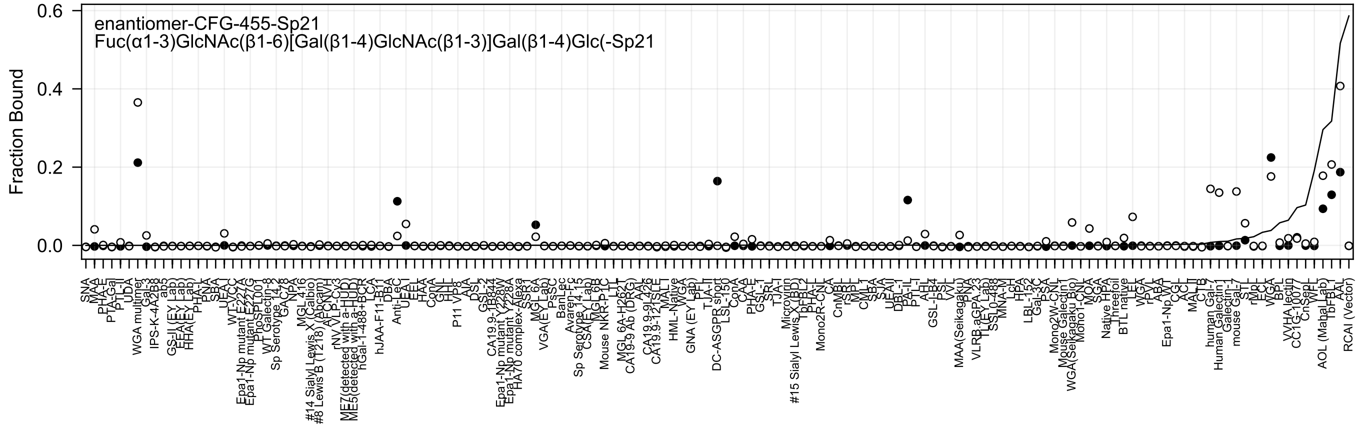
enantiomer-CFG-455-Sp21
Fuc(α1-3)GlcNAc(β1-6)[Gal(β1-4)GlcNAc(β1-3)]Gal(β1-4)Glc(-Sp21
Fraction Bound
0.6
0.4
0.2
0.0

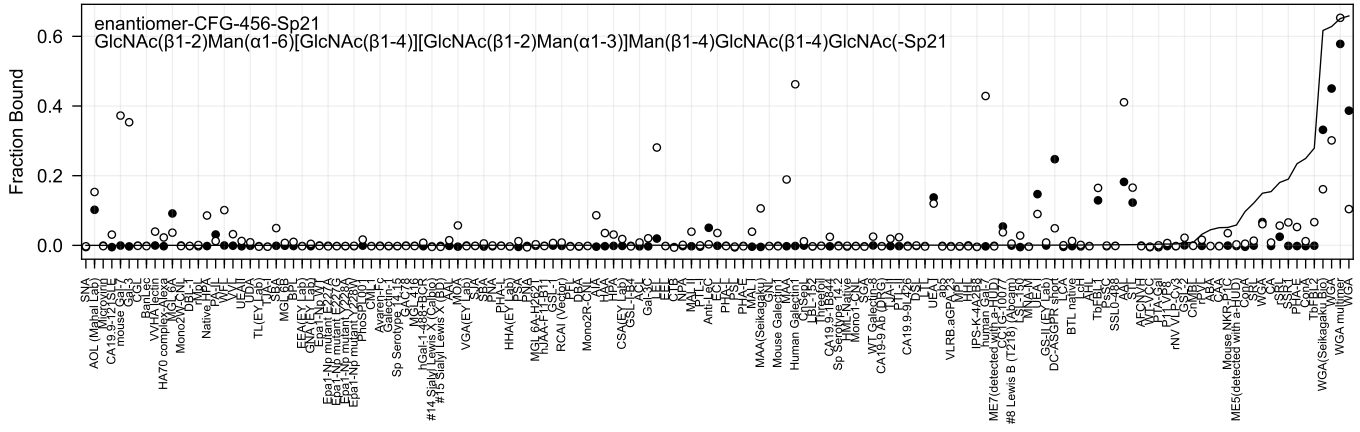

enantiomer-CFG-456-Sp21
GlcNAc(β1-2)Man(α1-6)[GlcNAc(β1-4)][GlcNAc(β1-2)Man(α1-3)]Man(β1-4)GlcNAc(β1-4)GlcNAc(-Sp21
Fraction Bound
0.0
0.2
0.4
0.6

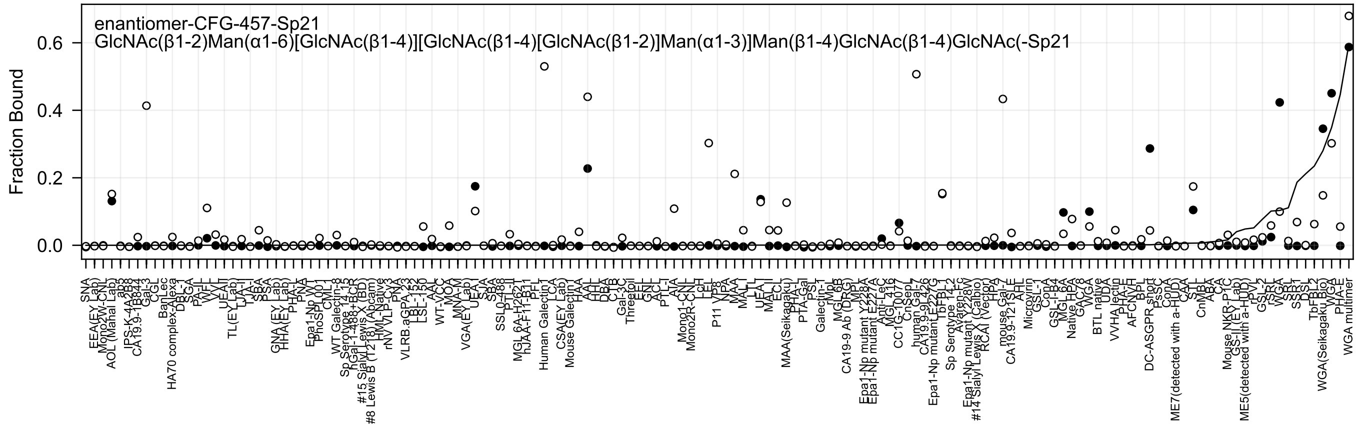
enantiomer-CFG-457-Sp21
GlcNAc(β1-2)Man(α1-6)[GlcNAc(β1-4)][GlcNAc(β1-4)[GlcNAc(β1-2)]Man(α1-3)]Man(β1-4)GlcNAc(β1-4)GlcNAc(-Sp21
Fraction Bound
0.0
0.2
0.4
0.6

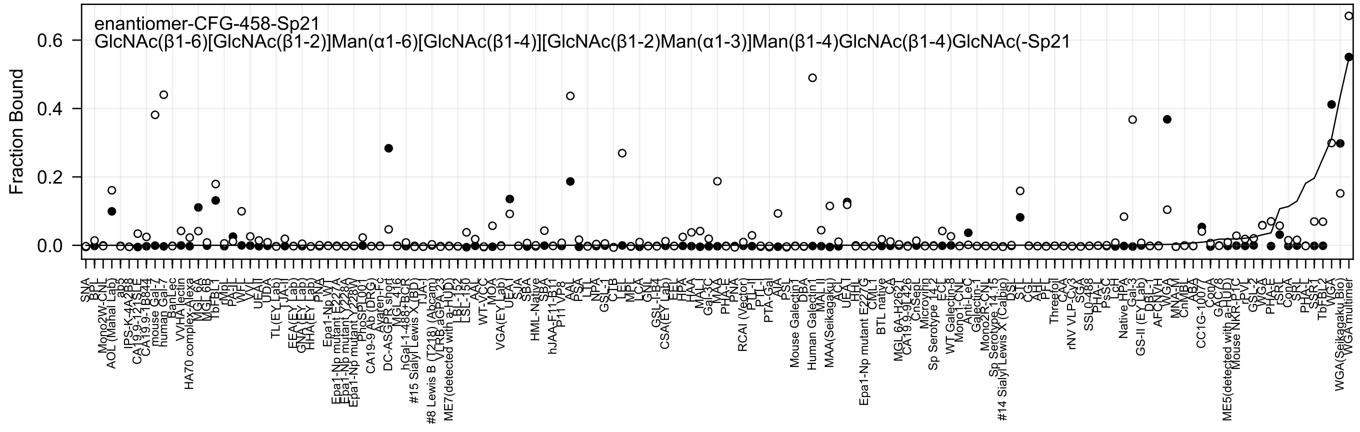
enantiomer-CFG-458-Sp21
GlcNAc(β1-6)[GlcNAc(β1-2)]Man(α1-6)[GlcNAc(β1-4)][GlcNAc(β1-2)Man(α1-3)]Man(β1-4)GlcNAc(β1-4)GlcNAc(-Sp21
Fraction Bound
0.6
0.4
0.2
0.0

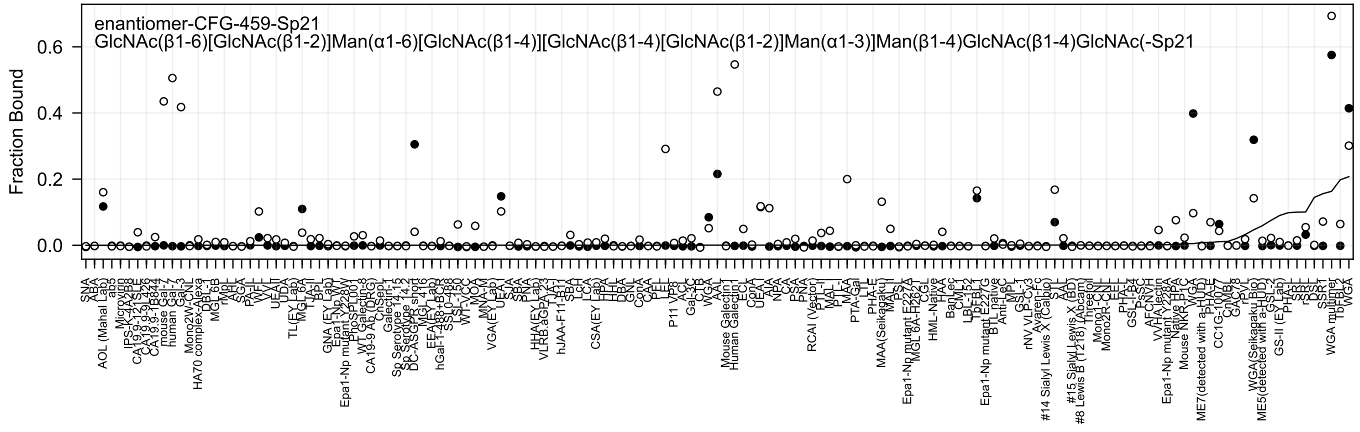
enantiomer-CFG-459-Sp21
GlcNAc(β1-6)[GlcNAc(β1-2)]Man(α1-6)[GlcNAc(β1-4)][GlcNAc(β1-4)[GlcNAc(β1-2)]Man(α1-3)]Man(β1-4)GlcNAc(β1-4)GlcNAc(-Sp21
Fraction Bound
0.0
0.2
0.4
0.6

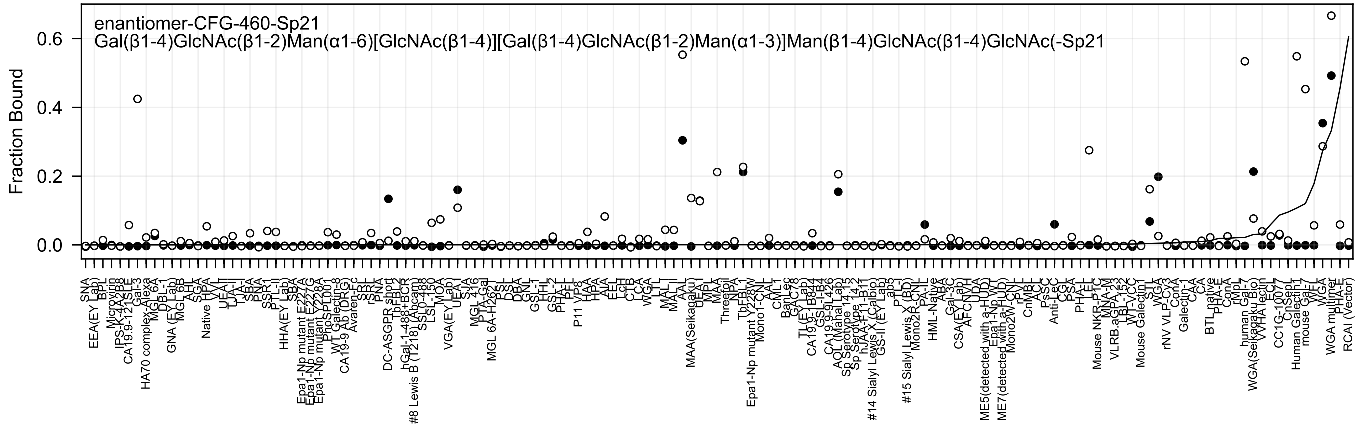
enantiomer-CFG-460-Sp21
Gal(β1-4)GlcNAc(β1-2)Man(α1-6)[GlcNAc(β1-4)][Gal(β1-4)GlcNAc(β1-2)Man(α1-3)]Man(β1-4)GlcNAc(β1-4)GlcNAc(-Sp21
Fraction Bound
0.0
0.2
0.4
0.6

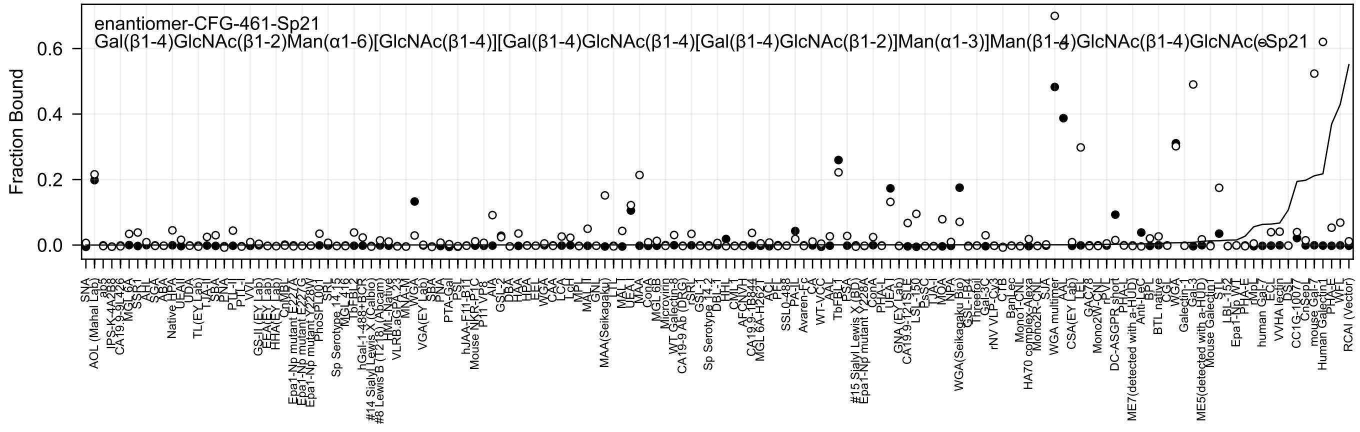
enantiomer-CFG-461-Sp21
Gal(β1-4)GlcNAc(β1-2)Man(α1-6)[GlcNAc(β1-4)][Gal(β1-4)GlcNAc(β1-4)[Gal(β1-4)GlcNAc(β1-2)]Man(α1-3)]Man(β1-4)GlcNAc(β1-4)GlcNAc(-Sp21
Fraction Bound
0.0
0.2
0.4
0.6

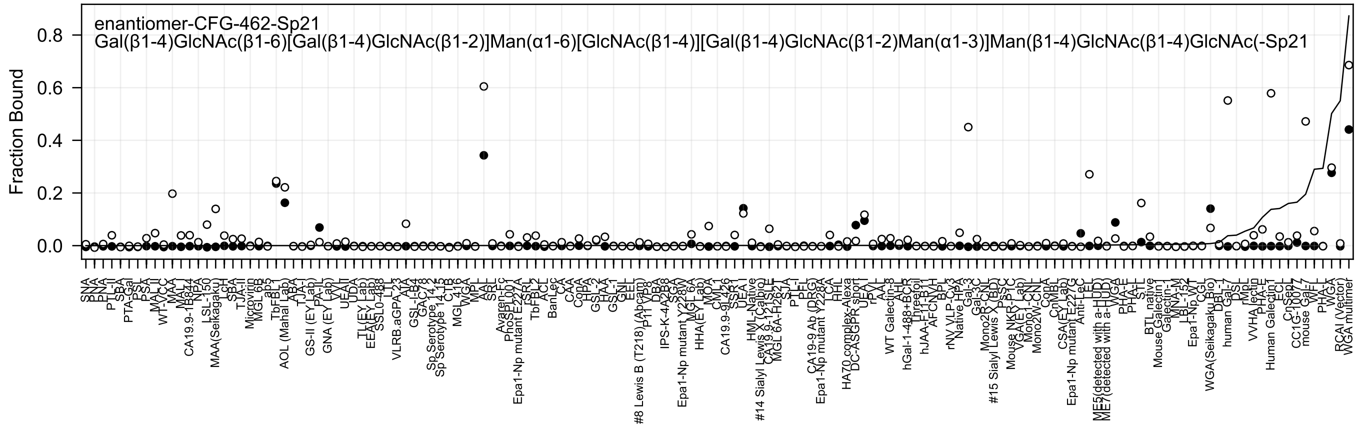
enantiomer-CFG-462-Sp21
Gal(β1-4)GlcNAc(β1-6)[Gal(β1-4)GlcNAc(β1-2)]Man(α1-6)[GlcNAc(β1-4)][Gal(β1-4)GlcNAc(β1-2)Man(α1-3)]Man(β1-4)GlcNAc(β1-4)GlcNAc(-Sp21
Fraction Bound
0.8
0.6
0.4
0.2
0.0

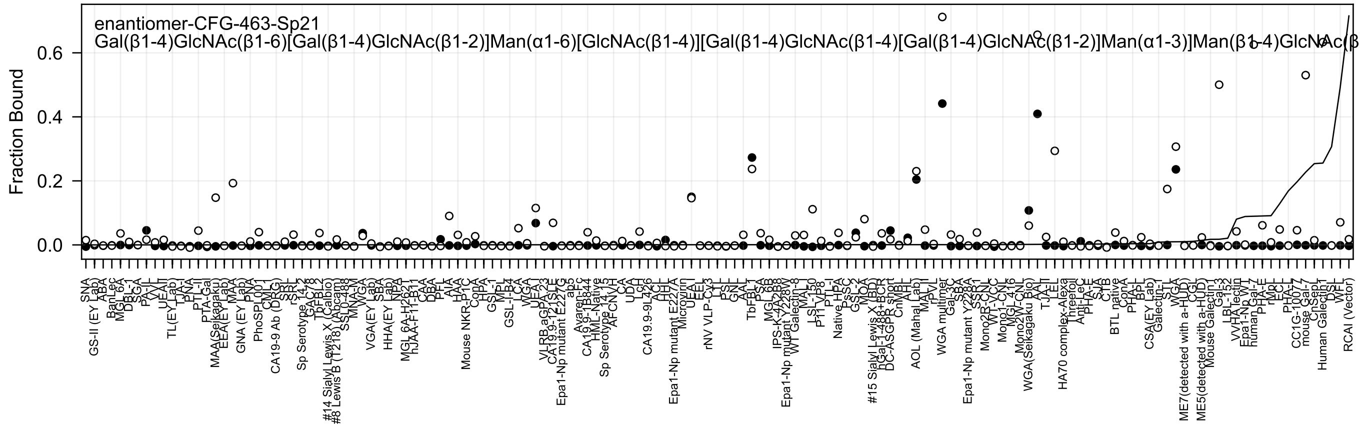

enantiomer-CFG-463-Sp21
Gal(β1-4)GlcNAc(β1-6)[Gal(β1-4)GlcNAc(β1-2)]Man(α1-6)[GlcNAc(β1-4)][Gal(β1-4)GlcNAc(β1-4)[Gal(β1-4)GlcNAc(β1-2)]Man(α1-3)]Man(β1-4)GlcNAc(β
Fraction Bound
0.0
0.2
0.4
0.6
SNA
GS-II (EY Lab)
ABA
BanLec
MGL 6A
DBL-1
SGA
PA-IL
VVL
UEA-II
TL(EY Lab)
TJA-I
PNA
PTL-II
PTA-Gal
MAA(Seikagaku)
EEA(EY Lab)
MAA
GNA (EY Lab)
PNA
PhoSL001
CML-1
CA19-9 Ab (DRG)
SRL
rSRL
Sp Serotype 14_2
GAC78
TbFBL2
#14 Sialyl Lewis X (Calbio)
#8 Lewis B (T218) (Abcam)
SSL0-488
MNA-M
WGA
VGA(EY Lab)
SBA
HHA(EY Lab)
NPA
MGL 6A-H2621
hJAA-F11-B11
CAA
DBA
PFL
AIA
HAA
Mouse NKR-P1C
ConA
HPA
GSL-1
MPL
GSL-I-B4
LCA
WGA
UEA-I
VLRB.aGPA.23
CA19.9-121SLE
Epa1-Np mutant E227G
ab5
Avaren-Fc
CA19.9-1B844
HML-Native
Sp Serotype 14_15
AFCNVH
CA
UDA
LcH
CA19.9-9L426
ACL
HHL
Epa1-Np mutant E227A
Microvirin
UEA-I
EEL
rNV VLP-Cy3
LTL
PSL
GNL
AAL
TbFBL1
PSA
MGL 6B
IPS-K-4A2B8
Epa1-Np mutant Y228W
WT Galectin-8
MAL-I
LSL-150
P11 VP8
PTL-I
Native HPA
PsSC
GSL-2
MOA
#15 Sialyl Lewis X (BD)
hGal-1-488+BCR
DC-ASGPR short
CnMBL
AHL
AOL (Mahal Lab)
MAL-II
rPVL
WGA multimer
Gal-3C
SBA
Epa1-Np mutant Y228A
SSR1
Mono2R-CNL
WT-VCC
Mono1-CNL
MGL 416
Mono2W-CNL
WGA(Seikagaku Bio)
AAL
TJA-II
LEL
HA70 complex-Alexa
Threefoil
Anti-LeC
PHA-E
SJA
CTB
BTL native
ConA
PHA-L
BPL
CSA(EY Lab)
Galectin-1
STL
WGA
ME7(detected with a-HUD)
CGL
ME5(detected with a-HUD)
Mouse Galectin1
Gal-3
LBL-152
VVHA lectin
Epa1-Np WT
human Gal-7
PHA-E
rMpL
ECL
PHA-L
CC1G-10077
mouse Gal-7
CnSepL
Human Galectin1
DSL
WFL
RCAI (Vector)

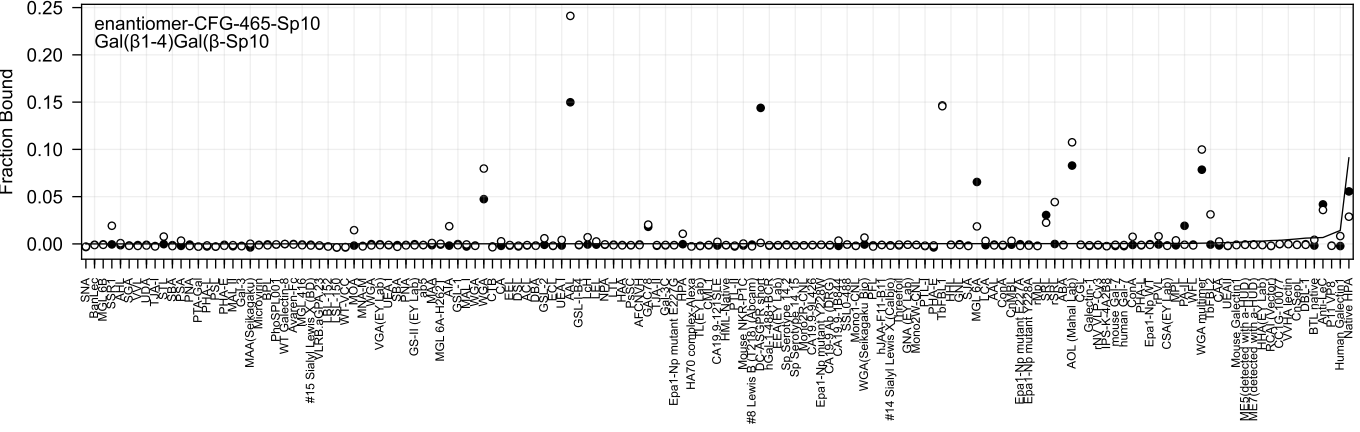
enantiomer-CFG-465-Sp10
Gal(β1-4)Gal(β-Sp10
Fraction Bound
0.25
0.20
0.15
0.10
0.05
0.00

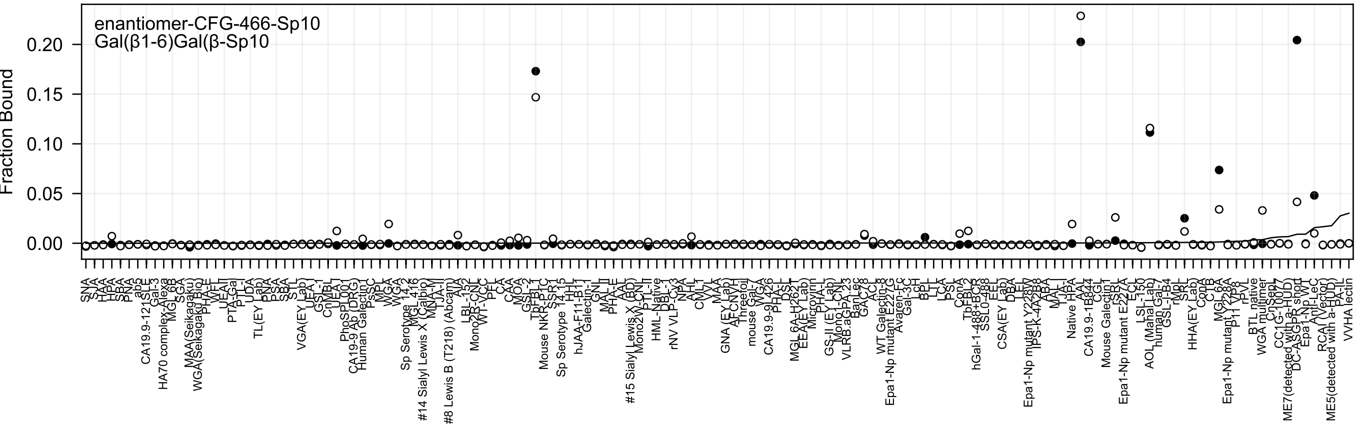
enantiomer-CFG-466-Sp10
Gal(β1-6)Gal(β-Sp10
Fraction Bound
0.20
0.15
0.10
0.05
0.00

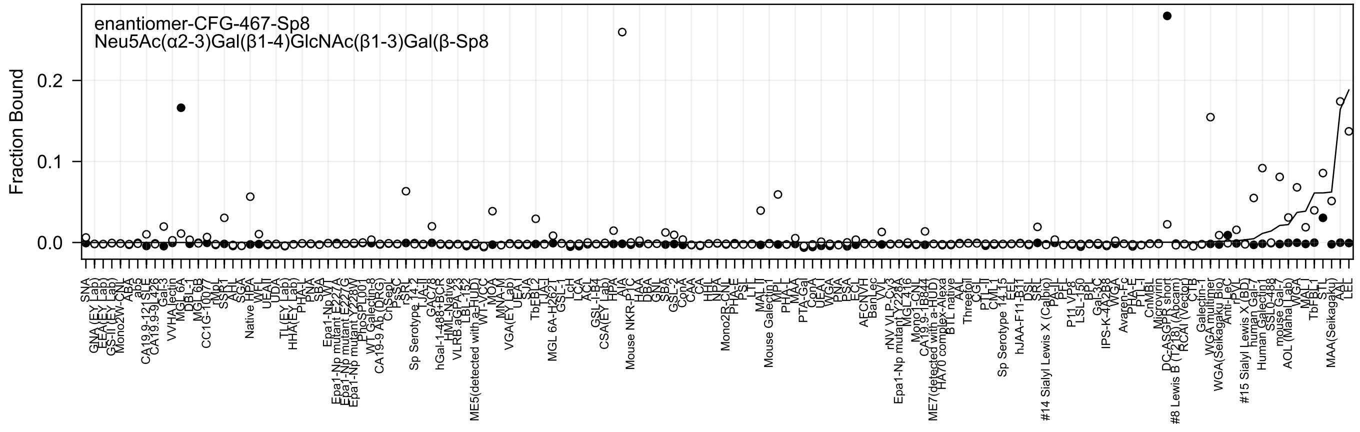

enantiomer-CFG-467-Sp8
Neu5Ac(α2-3)Gal(β1-4)GlcNAc(β1-3)Gal(β-Sp8
Fraction Bound
0.0
0.1
0.2

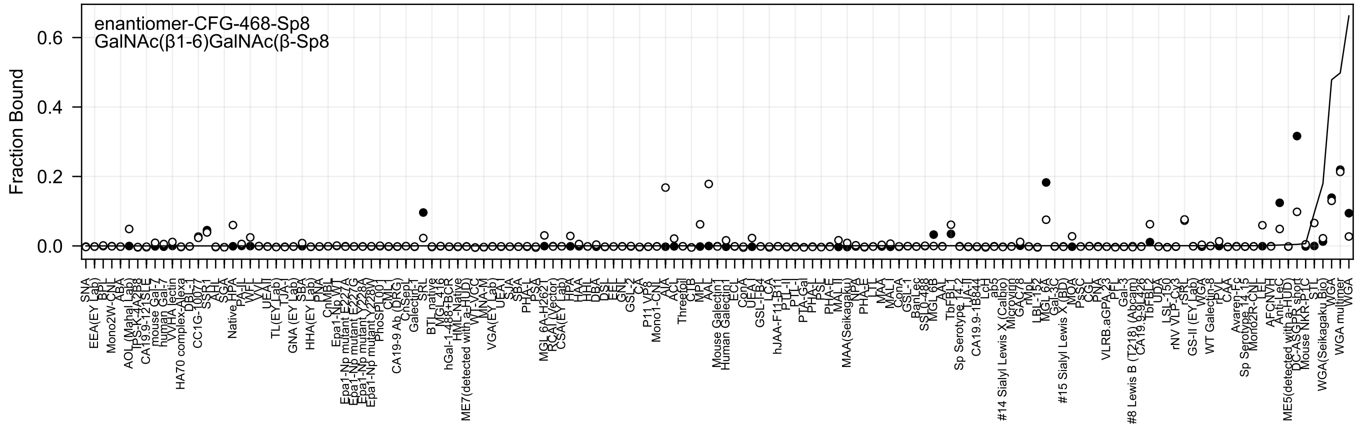
enantiomer-CFG-468-Sp8
GalNAc(β1-6)GalNAc(β-Sp8
Fraction Bound

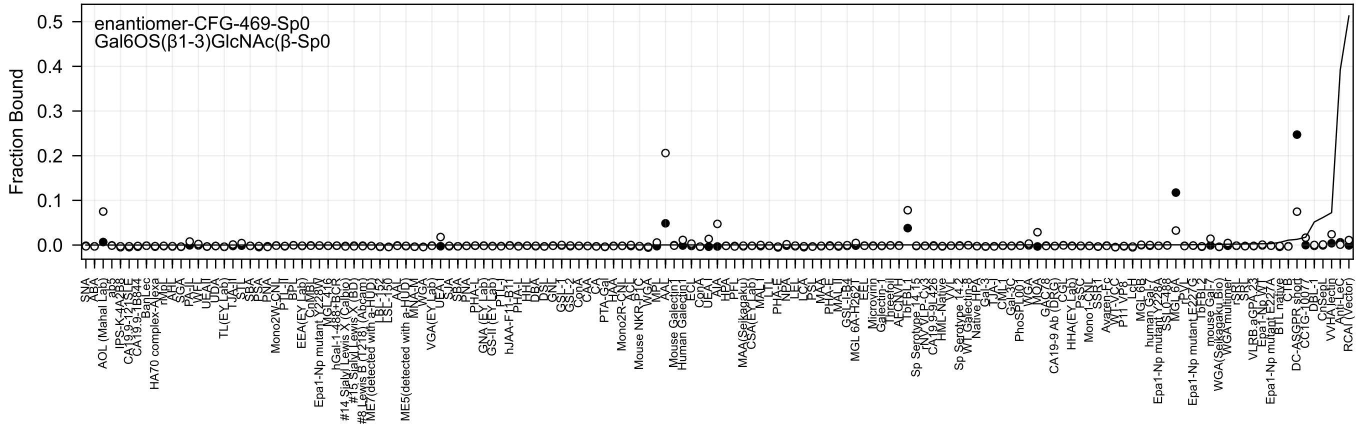
enantiomer-CFG-469-Sp0
Gal6OS(β1-3)GlcNAc(β-Sp0
Fraction Bound
0.0
0.1
0.2
0.3
0.4
0.5

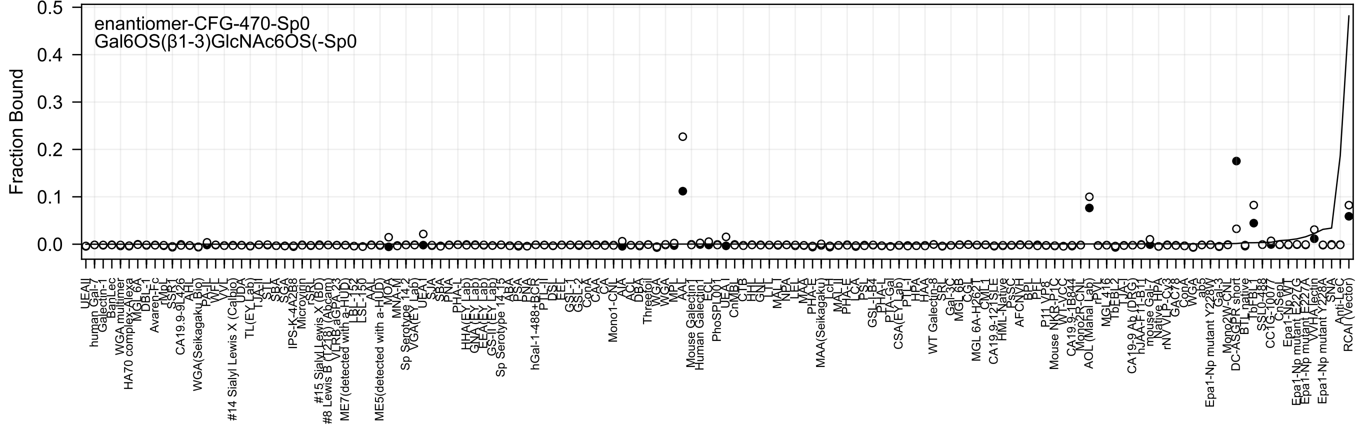
enantiomer-CFG-470-Sp0
Gal6OS(β1-3)GlcNAc6OS(-Sp0
Fraction Bound
0.5
0.4
0.3
0.2
0.1
0.0
UEAII
human Gal-7
Galectin-1
BanLec
WGA multimer
HA70 complex-Alexa
MGL 6A
DBL-1
Avaren-Fc
rMpL
SSR1
CA19.9-9L426
AAH
WGA(Seikagaku Bio)
PA-IL
WFL
VVL
#14 Sialyl Lewis X (Calbio)
UDA
TL(EY Lab)
TJA-II
STL
SBA
SGA
IPS-K-4A2B8
Microvirin
rSRL
#15 Sialyl Lewis X (BD)
#8 Lewis B (T218) (Abcam)
VLRB.aGPA.23
ME7(detected with a-HUD)
LBL-152
LSL-150
AAL
ME5(detected with a-HUD)
MOA
MNA-M
Sp Serotype 14-2
VGA(EY Lab)
UEA-I
SJA
SBA
PNA
PHA-L
HHA(EY Lab)
GNA(EY Lab)
EEA(EY Lab)
GS-II(EY Lab)
Sp Serotype 14-13
ABA
PSA
PNA
hGal-1-488+BCR
PTL-II
DSL
ECL
GSL-1
GSL-2
ConA
CAA
CA
Mono1-CNL
AIA
ACL
DBA
Threefoil
WGA
WGA
MPL
AAL
Mouse Galectin1
Human Galectin1
ECL
PhoSL001
UEA-I
CnMBL
CTB
HHL
GNL
LTL
MAL-I
NPA
LEL
MAA
PHA-E
MAA(Seikagaku)
LcH
MAL-II
PHA-E
LCA
PSL
GSL-I-B4
PHA-L
PTA-Gal
CSA(EY Lab)
PTL-I
HPA
HAA
WT Galectin-8
SRL
Gal-3C
MGL 6B
CGL
MGL 6A-H262T
CML-1
CA19.9-121SLE
HML-Native
PsSC
AFCNVH
BPL
PFL
P11 VP8
Mouse NKR-P1C
WT-VCC
CA19.9-1B844
Mono2R-CNL
AOL (Manal Lab)
rPVL
MGL416
TbFBL2
LTA-1
CA19-9 Ab (DRG)
hJAA-F11-B11
mouse Gal-7
Native HPA
rNV VLP-Cy3
GAC78
ConA
WGA
ab5
Epa1-Np mutant Y228W
Gal-3
Mono2W-CNL
DC-ASGPR short
BTL native
TbFBL1
SSL0-488
CC1G-10077
CnSep1
Epa1-Np WT
Epa1-Np mutant E227G
Epa1-Np mutant E227A
VVHA lectin
Epa1-Np mutant Y228A
SNA
Anti-LeC
RCAI (Vector)

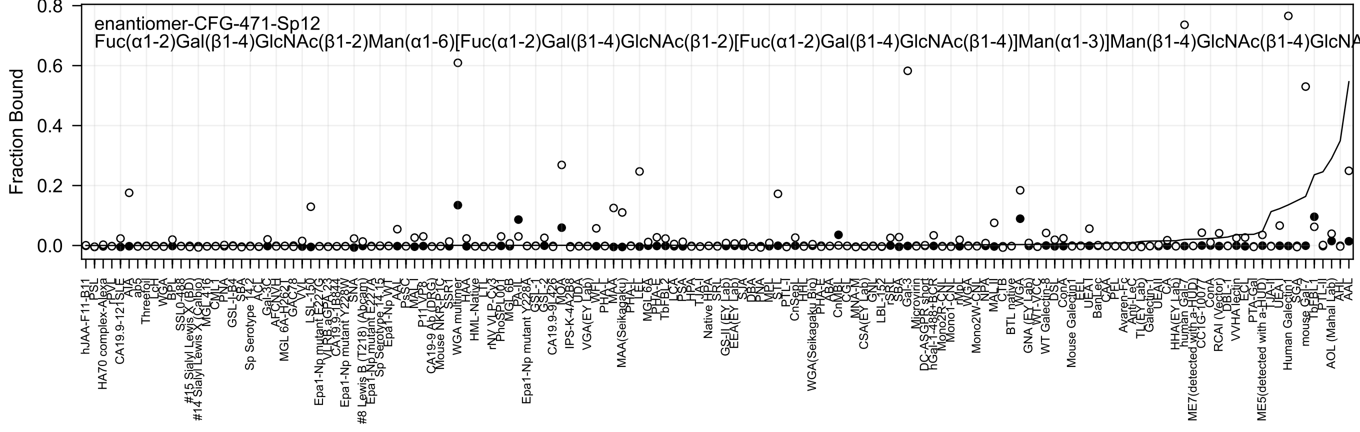
enantiomer-CFG-471-Sp12
Fuc(α1-2)Gal(β1-4)GlcNAc(β1-2)Man(α1-6)[Fuc(α1-2)Gal(β1-4)GlcNAc(β1-2)[Fuc(α1-2)Gal(β1-4)GlcNAc(β1-4)]Man(α1-3)]Man(β1-4)GlcNAc(β1-4)GlcNA
Fraction Bound
0.8
0.6
0.4
0.2
0.0

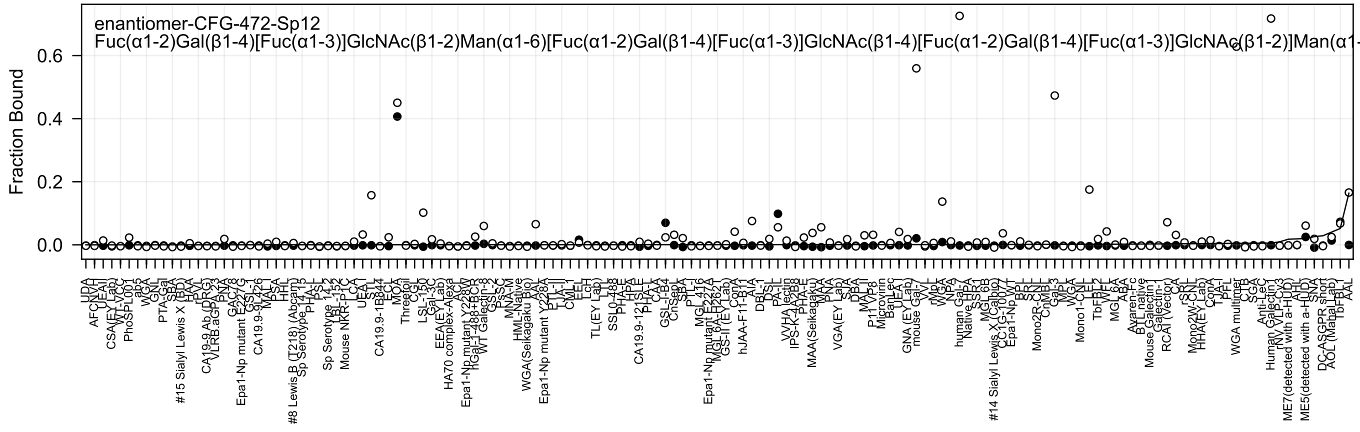

enantiomer-CFG-472-Sp12
Fuc(α1-2)Gal(β1-4)[Fuc(α1-3)]GlcNAc(β1-2)Man(α1-6)[Fuc(α1-2)Gal(β1-4)[Fuc(α1-3)]GlcNAc(β1-4)[Fuc(α1-2)Gal(β1-4)[Fuc(α1-3)]GlcNAc(β1-2)]Man(α1-
Fraction Bound
0.0
0.2
0.4
0.6

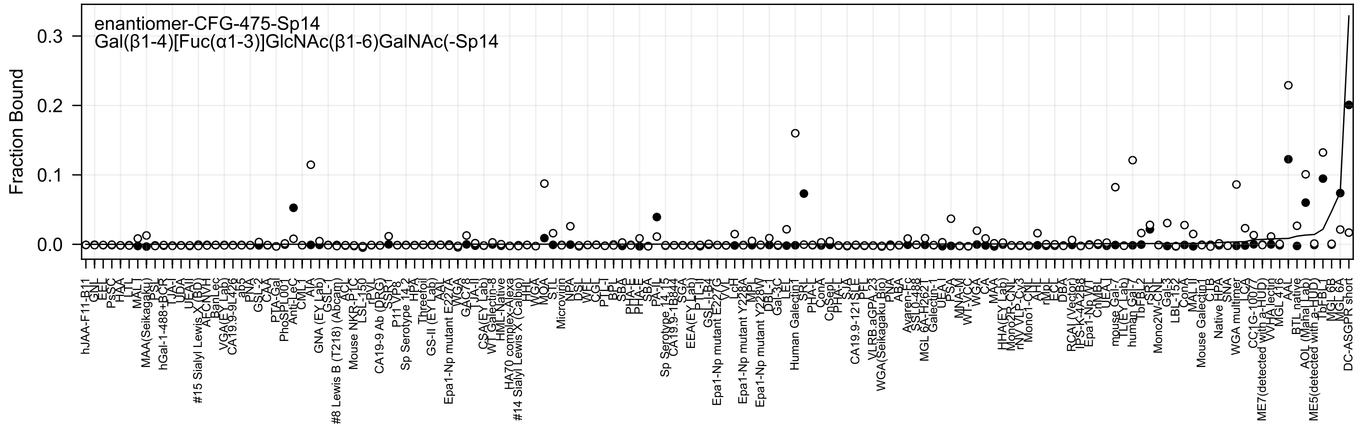
enantiomer-CFG-475-Sp14
Gal(β1-4)[Fuc(α1-3)]GlcNAc(β1-6)GalNAc(-Sp14
Fraction Bound
0.0
0.1
0.2
0.3

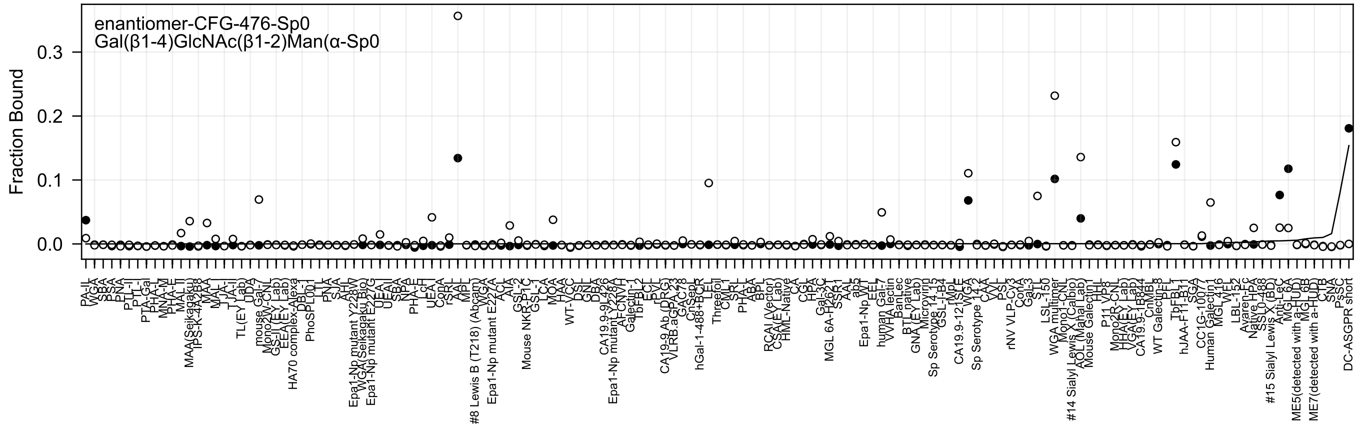
enantiomer-CFG-476-Sp0
Gal(β1-4)GlcNAc(β1-2)Man(α-Sp0
Fraction Bound
0.0
0.1
0.2
0.3

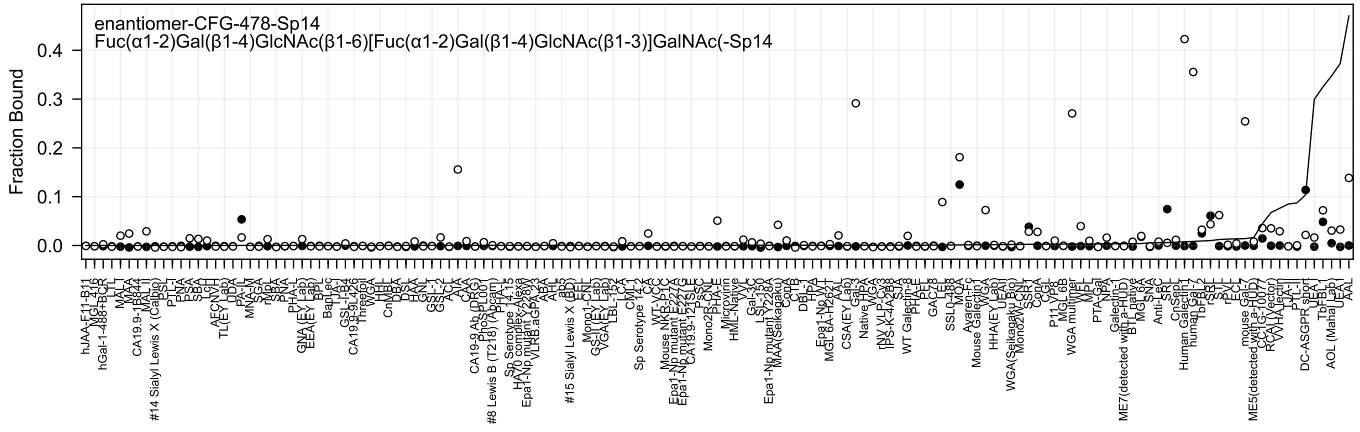
enantiomer-CFG-478-Sp14
Fuc(α1-2)Gal(β1-4)GlcNAc(β1-6)[Fuc(α1-2)Gal(β1-4)GlcNAc(β1-3)]GalNAc(-Sp14
Fraction Bound
0.0
0.1
0.2
0.3
0.4
hJAA-F11-B11
MGL 416
hGal-1-488+BCR
LTL
MAL-I
MAA
CA19.9-1B844
MAL II
#14 Sialyl Lewis X (Calbio)
ESL
PTL-I
PNA
PSA
SBA
LcH
AECNVH
TL(EY Lab)
UDA
PA-IL
MNA-M
SGA
rMpL
SBA
PNA
PHA-L
GNA (EY Lab)
EEA(EY Lab)
BPL
BanLec
TJA-I
GSL-I-B4
CA19.9-9L426
Threefoil
WGA
HHL
CnMBL
DBA
DSL
HAA
GNL
GSL-1
GSL-2
ACL
AIA
CAA
CA19-9 Ab (DRG)
PhoSL001
#8 Lewis B (T218) (Abcam)
PHA-L
Sp Serotype 14.15
HA70 complex-Alexa
Epa1-Np mutant Y228W
VLRB.aGPA.23
ABA
AHL
ab5
#15 Sialyl Lewis X (BD)
EEL
Mono1-CNL
GS-II(EY Lab)
VGA(EY Lab)
LBL-152
LCA
CML-1
Sp Serotype 14.2
CA
WT-VCC
Mouse NKR-P1C
Epa1-Np mutant E227A
Epa1-Np mutant E227G
CA19.9-121SLE
PsSC
Mono2R-CNL
PHA-E
Microvirin
HML-Native
VV
Gal-3C
LSL-150
Epa1-Np mutant Y228A
MAA(Seikagaku)
ConA
CTB
DBL-1
HPA
Epa1-Np WT
MGL 6A-H262T
AAL
CSA(EY Lab)
Gal-3
Native HPA
WGA
rNV VLP-Cy3
IPS-K-4A2B8
SJA
WT Galectin-8
PHA-E
PFL
GAC78
LEL
SSL0-488
MOA
Ayaren-Fc
Mouse Galectin 1
WGA
HHA(EY Lab)
UEAII
WGA(Seikagaku Bio)
Mono2W-CNL
SSR1
CGL
ConA
P11 VP8
MGL 6B
WGA multimer
VVL
MPL
PTA-Gal
NPA
Galectin-1
ME7(detected with a-HUD)
BTL native
MGL 6A
SNA
Anti-LeC
SRL
CnSepL
Human Galectin 1
human Gal-7
TbFBL-2
rSRL
STL
rPVL
ECL
mouse Gal-7
ME5(detected with a-HUD)
CCL(G-1007)
RCA(Vector)
VVHA lectin
TJA-II
PTL-II
DC-ASGPR short
UEA I
TbFBL-1
AOL (Maha Lab)
UEA I
AAL

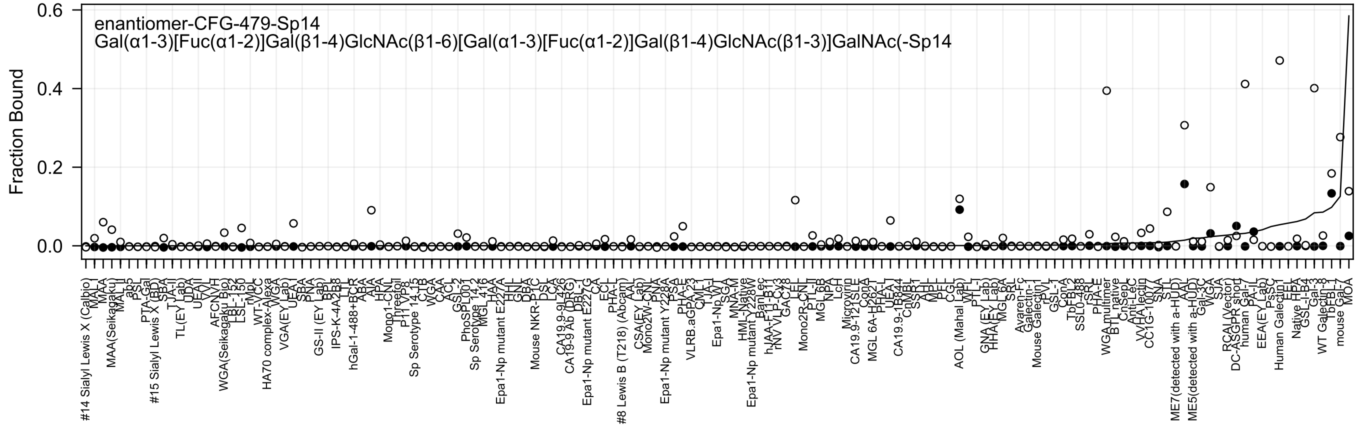
enantiomer-CFG-479-Sp14
Gal(α1-3)[Fuc(α1-2)]Gal(β1-4)GlcNAc(β1-6)[Gal(α1-3)[Fuc(α1-2)]Gal(β1-4)GlcNAc(β1-3)]GalNAc(-Sp14
Fraction Bound

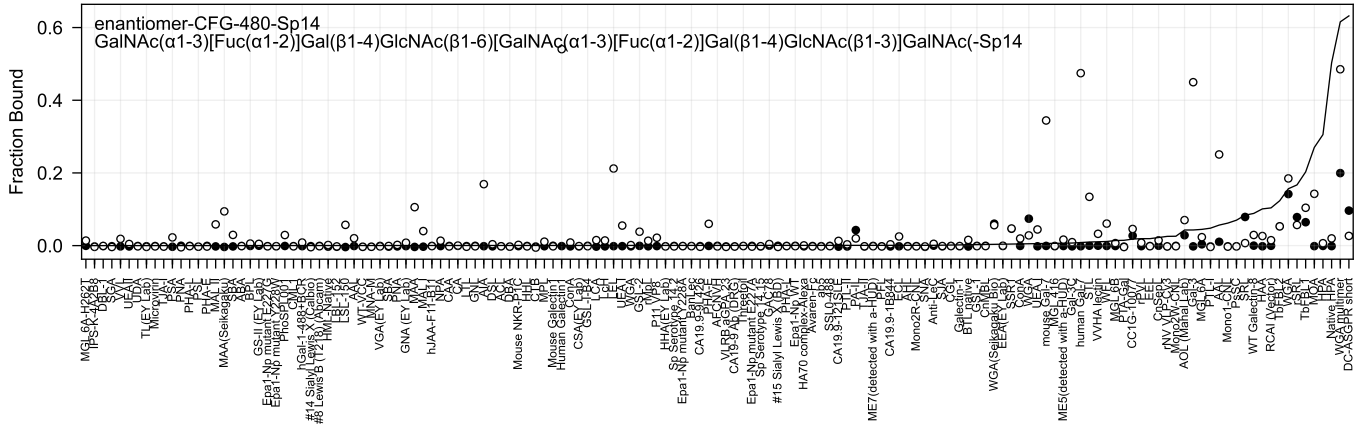

enantiomer-CFG-480-Sp14
GalNAc(α1-3)[Fuc(α1-2)]Gal(β1-4)GlcNAc(β1-6)[GalNAc(α1-3)[Fuc(α1-2)]Gal(β1-4)GlcNAc(β1-3)]GalNAc(-Sp14
Fraction Bound
0.0
0.2
0.4
0.6

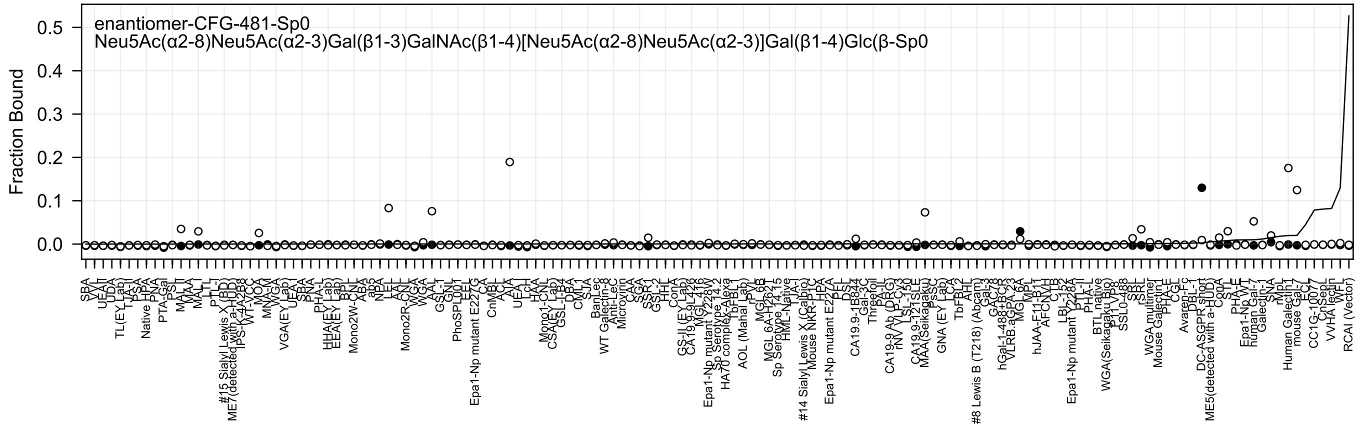
enantiomer-CFG-481-Sp0
Neu5Ac(α2-8)Neu5Ac(α2-3)Gal(β1-3)GalNAc(β1-4)[Neu5Ac(α2-8)Neu5Ac(α2-3)]Gal(β1-4)Glc(β-Sp0
Fraction Bound
0.0
0.1
0.2
0.3
0.4
0.5

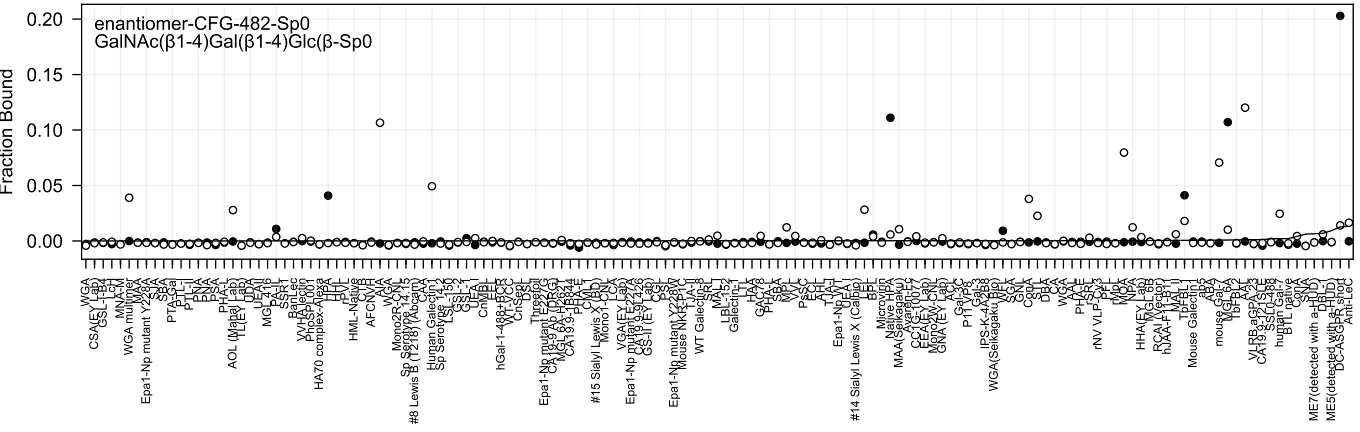
enantiomer-CFG-482-Sp0
GalNAc(β1-4)Gal(β1-4)Glc(β-Sp0
Fraction Bound

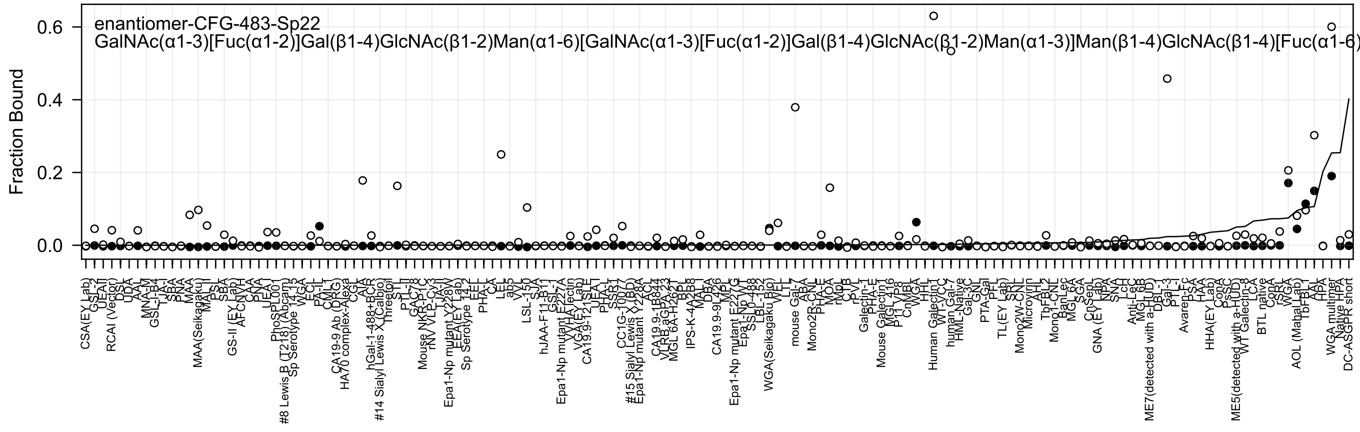
enantiomer-CFG-483-Sp22
GalNAc(α1-3)[Fuc(α1-2)]Gal(β1-4)GlcNAc(β1-2)Man(α1-6)[GalNAc(α1-3)[Fuc(α1-2)]Gal(β1-4)GlcNAc(β1-2)Man(α1-3)]Man(β1-4)GlcNAc(β1-4)[Fuc(α1-6)
Fraction Bound
0.6
0.4
0.2
0.0
CSA(EY Lab)
GSL-2
UEAII
RCAI (Vector)
DSL
UDA
AAL
MNA-M
GSL-I-B4
TJA-I
SBA
PNA
MAA
MAA(Seikagaku)
MAL-I
PSL
SBA
GS-II (EY Lab)
AFCNVH
CAA
PNA
UEA-I
PhoSL
PhoSPL001
#8 Lewis B (T218) (Abcam)
Sp Serotype 14 13
WGA
ECL
PA-IL
CML
CA19-9 Ab (DRG)
HA70 complex-Alexa
CGL
AIA
hGal-1-488+BCR
#14 Sialyl Lewis X (Calbio)
Threefoil
STL
PTL-II
GAC-78
Mouse NKR-P1C
rNV VLP-Cy3
TJA-II
Epa1-Np mutant Y228W
EEA(EY Lab)
Sp Serotype 14 2
EEL
PHA-L
CA
LEL
ab5
VVL
LSL-150
SJA
hJAA-F11-B11
GSL-I
Epa1-Np mutant E227A
YY HA lectin
VGA(EY Lab)
CA19.9-121SLE
UEA-I
PHA-L
SSR1
CC1G-10077
#15 Sialyl Lewis X (BD)
Epa1-Np mutant Y228A
ACL
CA19.9-1B844
VLRB.aGPA.23
MGL 6A-H26 1
BPL
IPS-K-4A2B8
MAL-I
DBA
CA19.9-9L426
MPL
Epa1-Np mutant E227G
Epa1-Np WT
SSL0-488
LBL-152
WGA(Seikagaku Bio)
WFL
LTL
mouse Gal-7
ABA
Mono2R-CNL
PHA-E
MOA
rMpL
CTB
rPVL
Galectin-1
PHA-E
Mouse Galectin1
MGL 416
P11 VP8
CnMBL
WGA
HHL
Human Galectin1
WT-VCC
human Gal-7
HML-Native
Gal-3C
GNL
PTA-Gal
PFL
TL(EY Lab)
SRL
Mono2W-CNL
Microvirin
AHL
TbFBL2
Mono1-CNL
BanLec
MGL 6A
SGA
CnSepL
GNA (EY Lab)
NPA
SNA
LcH
Anti-LeC
MGL 6B
ME7(detected with a-HUD)
DBL-1
Gal-3
PTL-I
Avaren-Fc
PSA
HAA
HHA(EY Lab)
ConA
PsSC
ME5(detected with a-HUD)
WT Galectin-8
LCA
BTL native
ConA
rSRL
WGA
AOL (Mahal Lab)
TbFBL1
AAL
HPA
WGA multimer
Native HPA
DC-ASGPR short

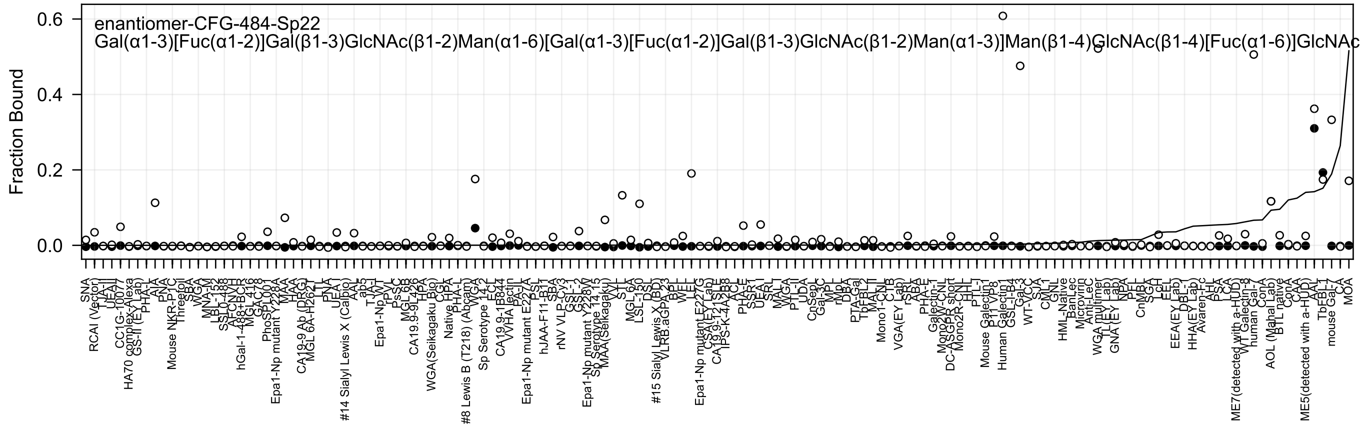

enantiomer-CFG-484-Sp22
Gal(α1-3)[Fuc(α1-2)]Gal(β1-3)GlcNAc(β1-2)Man(α1-6)[Gal(α1-3)[Fuc(α1-2)]Gal(β1-3)GlcNAc(β1-2)Man(α1-3)]Man(β1-4)GlcNAc(β1-4)[Fuc(α1-6)]GlcNAc
Fraction Bound
0.6
0.4
0.2
0.0

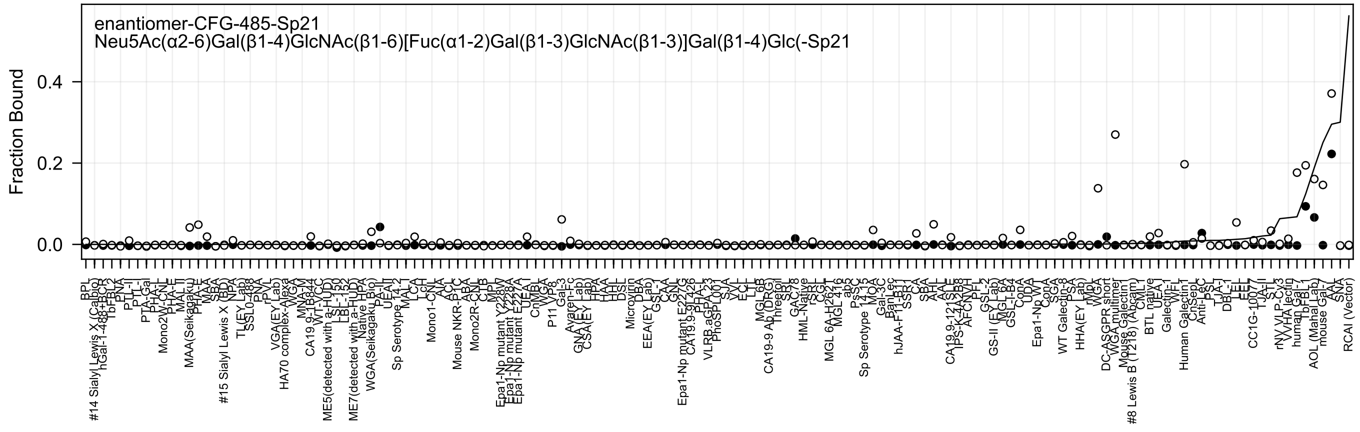
enantiomer-CFG-485-Sp21
Neu5Ac(α2-6)Gal(β1-4)GlcNAc(β1-6)[Fuc(α1-2)Gal(β1-3)GlcNAc(β1-3)]Gal(β1-4)Glc(-Sp21
Fraction Bound
0.4
0.2
0.0

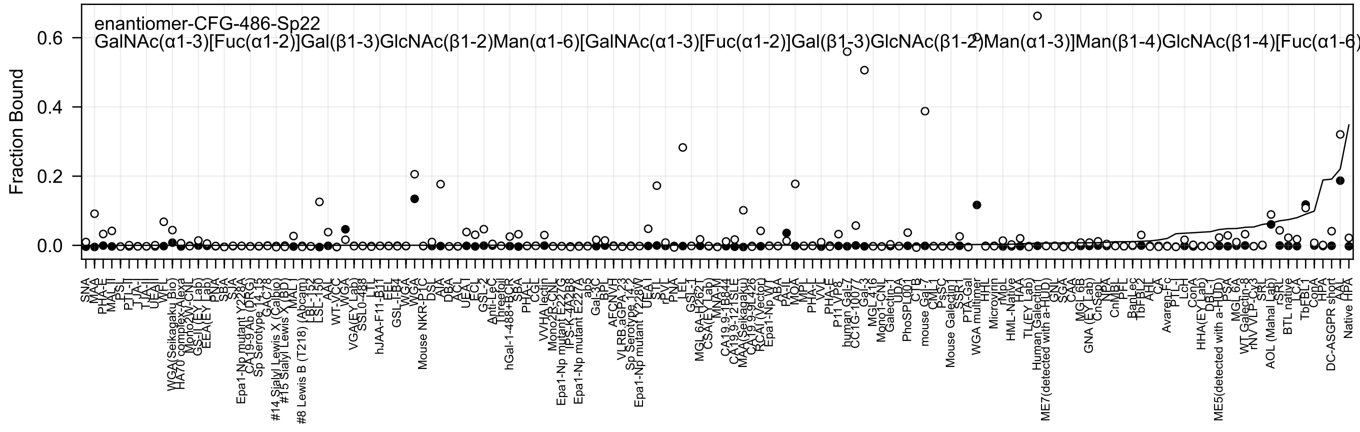
enantiomer-CFG-486-Sp22
GalNAc(α1-3)[Fuc(α1-2)]Gal(β1-3)GlcNAc(β1-2)Man(α1-6)[GalNAc(α1-3)[Fuc(α1-2)]Gal(β1-3)GlcNAc(β1-2)Man(α1-3)]Man(β1-4)GlcNAc(β1-4)[Fuc(α1-6) [illegible]
Fraction Bound
0.0
0.2
0.4
0.6

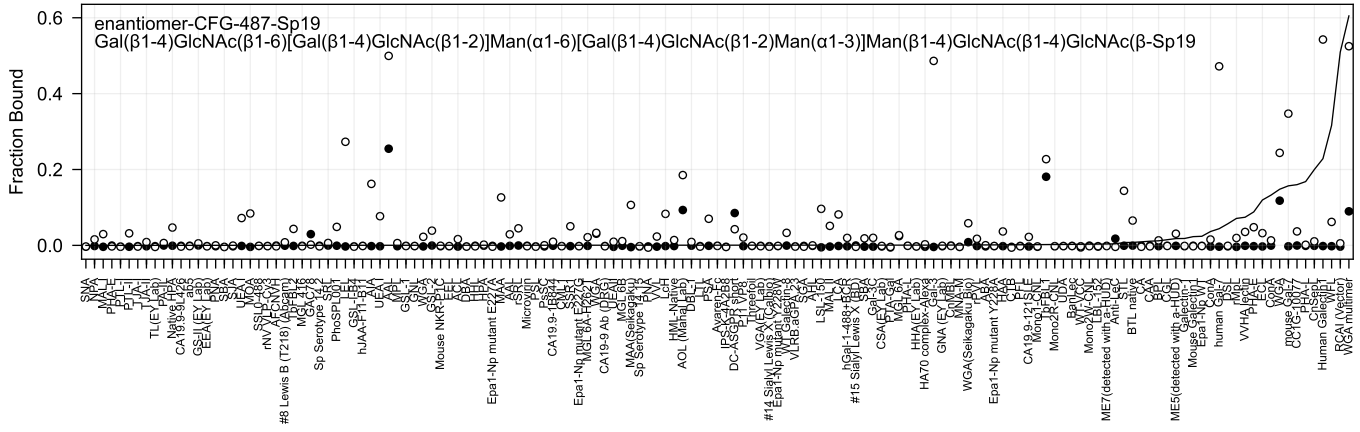
enantiomer-CFG-487-Sp19
Gal(β1-4)GlcNAc(β1-6)[Gal(β1-4)GlcNAc(β1-2)]Man(α1-6)[Gal(β1-4)GlcNAc(β1-2)Man(α1-3)]Man(β1-4)GlcNAc(β1-4)GlcNAc(β-Sp19
Fraction Bound
0.0
0.2
0.4
0.6
SNA
NPA
MAL-I
PHA-E
PTL-I
PTL-II
TJA-II
TL(EY Lab)
PA-IL
Native HPA
CA19.9-9L426
ab5
GS-II (EY Lab)
EEA(EY Lab)
PNA
SBA
SJA
UEA-I
MOA
SSL0-488
rNV
VIP-Cy3
AFCNVH
#8 Lewis B (T218) (Abcam)
TbFBL2
MGL 418
GAC-78
Sp Serotype 14-2
SRT
PhoSPL001
LEL
GSL-IB4
hJAA-F11-B11
AIA
UEA-II
AAL
MPL
GSL-I
GNL
WGA
GSL-2
Mouse NKR-P1C
EEL
ACL
DBA
HHL
HPA
Epa1-Np mutant E227A
MAA
AAL
rSRL
Microvirin
PSL
PsSC
CA19.9-1B844
CML-1
SSR-1
Epa1-Np mutant E227G
MGL 6A-H262
WGA
CA19-9 Ab (DRG)
UEA-I
MGL 6B
MAA(Seikagaku)
Sp Serotype 14-15
PNA
VVL
LcH
HML-Native
AOL (Mahal Lab)
DBL-1
LTL
PSA
Avaren-Fc
IPS-X-4A2B8
DC-ASGPR short
P11 VP8
Threefoil
VGA(EY Lab)
#14 Sialyl Lewis X (Calbio)
Epa1-Np mutant Y228W
WT Galectin-8
VLRB.aGPA.23
SGA
AHL
LSL-150
MAL-II
LCA
hGal-1-488+BCR
#15 Sialyl Lewis X (BD)
SBA
Gal-3C
CSA(EY Lab)
PTA-Gal
MGL 6A
PHA-L
HHA(EY Lab)
HA70 complex-Alexa
Gal-3
GNA (EY Lab)
CnMBL
MNA-M
WGA(Seikagaku Bio)
rPVL
ABA
Epa1-Np mutant Y228A
HAA
CTB
PFL
CA19.9-121SLE
Mono1-CNT
TbFBL1
Mono2R-CNL
UDA
BanLec
WT-CC
Mono2W-CNL
LBL-152
ME7(detected with a-HuD)
Anti-LeC
STL
BTL native
CA
CAA
BPL
CGL
ME5(detected with a-HuD)
Galectin-1
Mouse Galectin-1
Epa1-Np WT
ConA
human Gal-7
DSL
rMpL
VVHA lectin
PHA-E
ECA
ConA
WGA
mouse Gal-7
CC1G-1007
PHA-L
CnSep1
Human Galectin1
WFL
RCAI (Vector)
WGA multimer

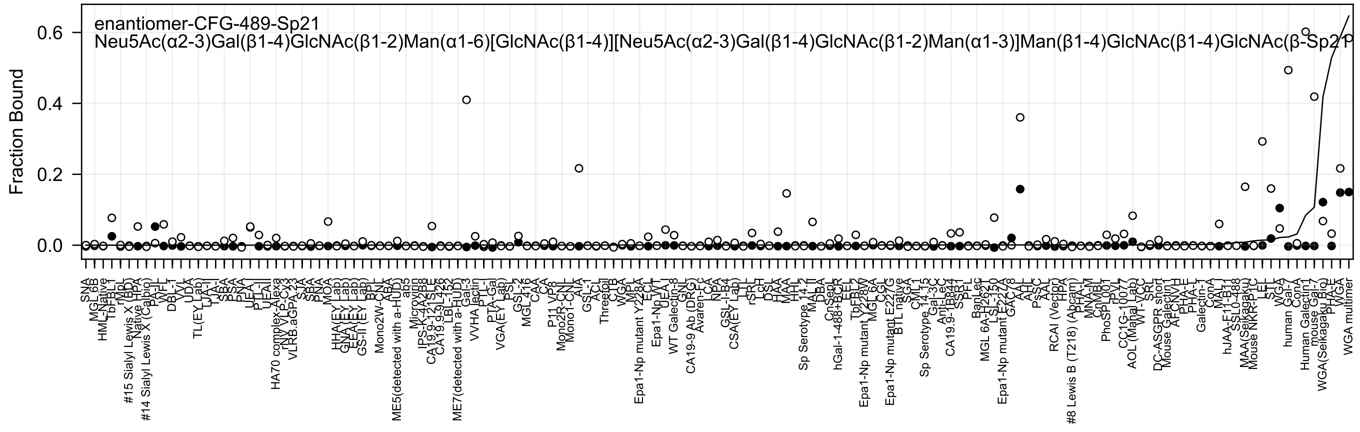
enantiomer-CFG-489-Sp21
Neu5Ac(α2-3)Gal(β1-4)GlcNAc(β1-2)Man(α1-6)[GlcNAc(β1-4)][Neu5Ac(α2-3)Gal(β1-4)GlcNAc(β1-2)Man(α1-3)]Man(β1-4)GlcNAc(β1-4)GlcNAc(β-Sp21
Fraction Bound
0.0
0.2
0.4
0.6

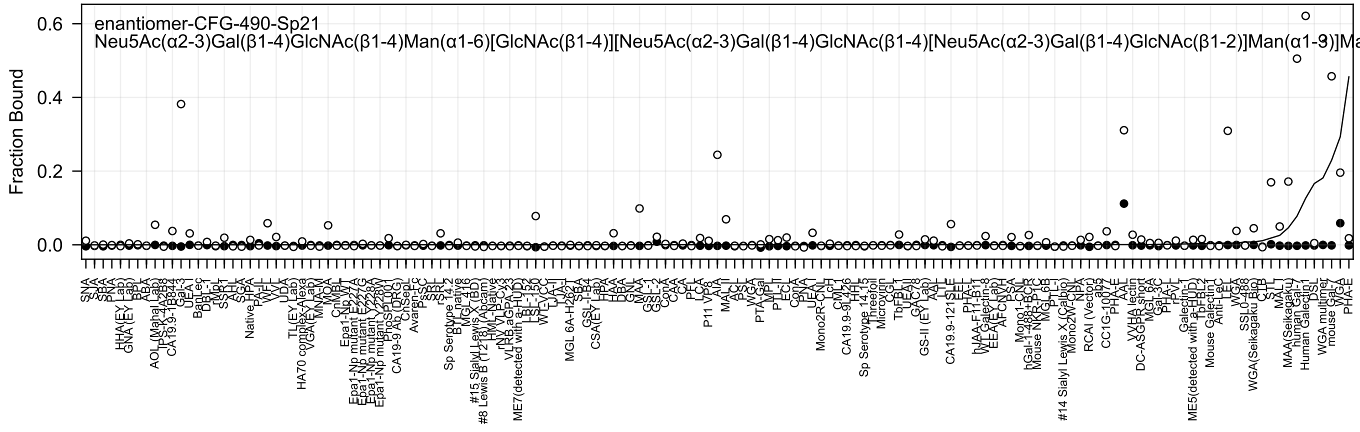

enantiomer-CFG-490-Sp21
Neu5Ac(α2-3)Gal(β1-4)GlcNAc(β1-4)Man(α1-6)[GlcNAc(β1-4)][Neu5Ac(α2-3)Gal(β1-4)GlcNAc(β1-4)[Neu5Ac(α2-3)Gal(β1-4)GlcNAc(β1-2)]Man(α1-3)]Ma
Fraction Bound
0.6
0.4
0.2
0.0

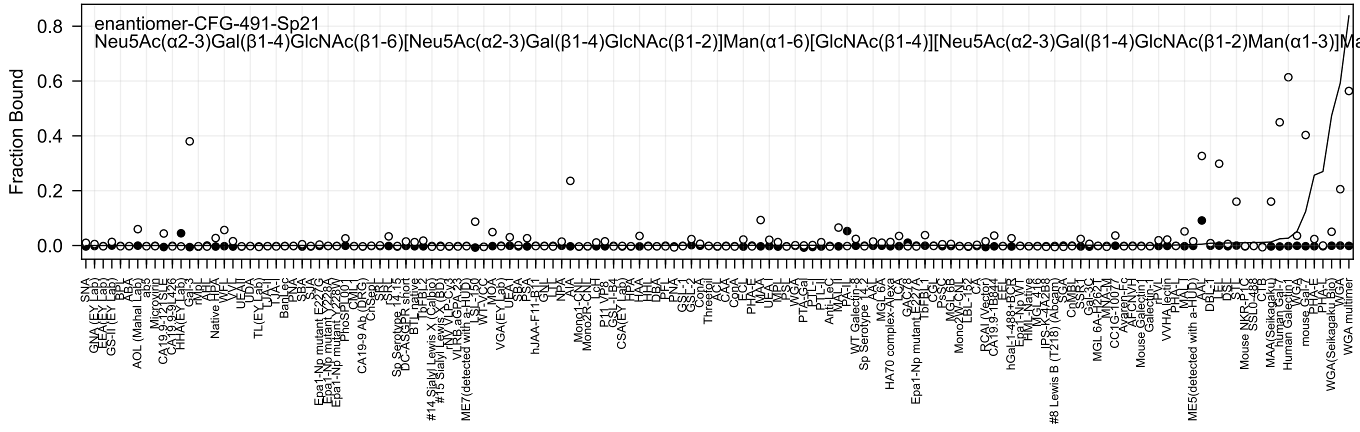
enantiomer-CFG-491-Sp21
Neu5Ac(α2-3)Gal(β1-4)GlcNAc(β1-6)[Neu5Ac(α2-3)Gal(β1-4)GlcNAc(β1-2)]Man(α1-6)[GlcNAc(β1-4)][Neu5Ac(α2-3)Gal(β1-4)GlcNAc(β1-2)Man(α1-3)]Ma
Fraction Bound

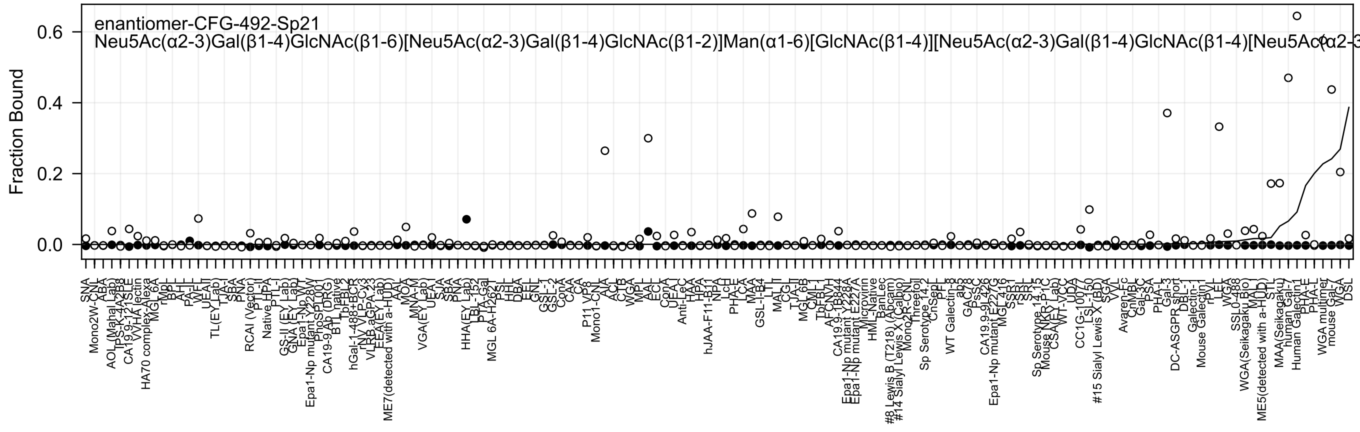
enantiomer-CFG-492-Sp21
Neu5Ac(α2-3)Gal(β1-4)GlcNAc(β1-6)[Neu5Ac(α2-3)Gal(β1-4)GlcNAc(β1-2)]Man(α1-6)[GlcNAc(β1-4)][Neu5Ac(α2-3)Gal(β1-4)GlcNAc(β1-4)[Neu5Ac(α2-3
Fraction Bound
0.0
0.2
0.4
0.6

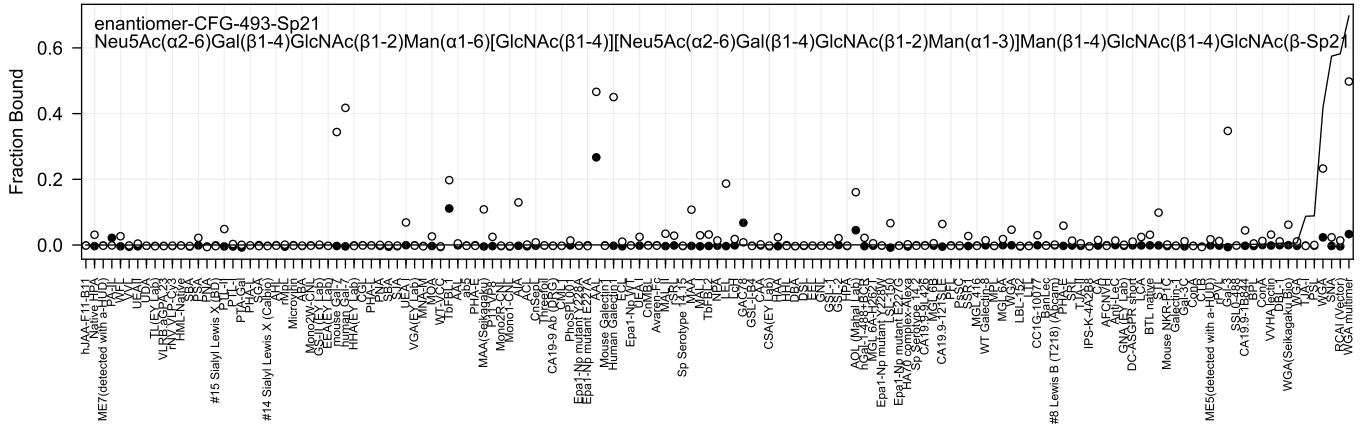
enantiomer-CFG-493-Sp21
Neu5Ac(α2-6)Gal(β1-4)GlcNAc(β1-2)Man(α1-6)[GlcNAc(β1-4)][Neu5Ac(α2-6)Gal(β1-4)GlcNAc(β1-2)Man(α1-3)]Man(β1-4)GlcNAc(β1-4)GlcNAc(β-Sp21
Fraction Bound
0.0
0.2
0.4
0.6
hJAA-F11-B11
ME7(detected with a-HUD)
#15 Sialyl Lewis X (BD)
#14 Sialyl Lewis X (Calbio)
MAA(Seikagaku)
CA19-9 Ab (DRG)
Epa1-Np mutant Y228A
Epa1-Np mutant E227A
Mouse Galectin1
Human Galectin1
Sp Serotype 14.15
AOL (Mahal Lab)
Epa1-Np mutant Y228W
Epa1-Np mutant E227G
HA70 complex-Alexa
#8 Lewis B (T218) (Abcam)
ME5(detected with a-HUD)
WGA(Seikagaku Bio)
RCAI (Vector)
WGA multimer

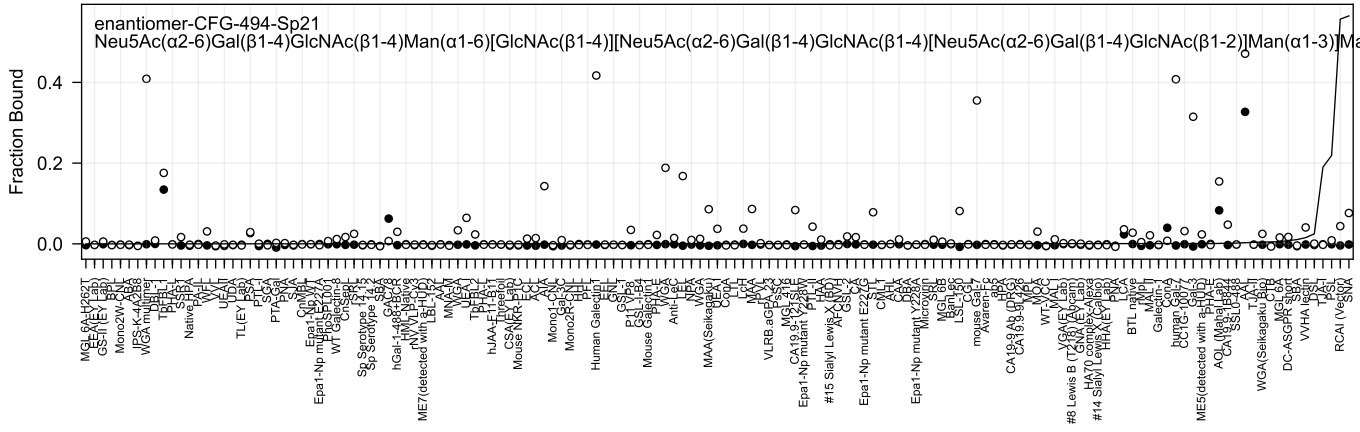

enantiomer-CFG-494-Sp21
Neu5Ac(α2-6)Gal(β1-4)GlcNAc(β1-4)Man(α1-6)[GlcNAc(β1-4)][Neu5Ac(α2-6)Gal(β1-4)GlcNAc(β1-4)[Neu5Ac(α2-6)Gal(β1-4)GlcNAc(β1-2)]Man(α1-3)]Ma
Fraction Bound
0.4
0.2
0.0
MGL 6A-H262T
EEA(EY Lab)
GS-II (EY Lab)
BPL
Mono2W-CNL
ABA
IPS-K-4A2B8
WGA multimer
DBL-1
TbFBL1
PHA-L
SSR1
Native HPA
PA-IIL
WFL
VVL
UEAII
UDA
TL(EY Lab)
PSA
PTL-I
SGA
PTA-Gal
PNA
SJA
CnMBL
Epa1-Np WT
Epa1-Np mutant E227A
PhoSL001
WT Galectin-8
CnSept
Sp Serotype 14.13
Sp Serotype 14.2
SBA
GAC78
hGal-1-488+BCR
HML-Native
rNV VLP Cy3
ME7(detected with a-HUD)
LBL-152
AAL
MNA-M
WGA
UEA-I
TbFBL2
PHA-L
hJAA-F11-B11
Threefoil
CSA(EY Lab)
Mouse NKR-P1C
ECL
ACL
AIA
Mono1-CNL
Gal-3C
Mono2R-CNL
HHL
PFL
Human Galectin1
EEL
GNL
GSL-I
P11 VP8
GSL-I-B4
Mouse Galectin1
PHA-E
WGA
Anti-LeC
LEL
NPA
WGA
MAA(Seikagaku)
UEA-I
ConA
LTL
LcH
MAA
PVL
VLRB.aGPA.23
PSSC
MGL 416
CA19.9-121SLE
Epa1-Np mutant Y228W
PTL-II
HAA
#15 Sialyl Lewis X (BD)
AFCNVH
GSL-2
CA
Epa1-Np mutant E227G
STL
CMLL
AH
CAA
DBA
Epa1-Np mutant Y228A
Microvirin
SRL
MGL 6B
BanLec
LSL-150
CGL
mouse Gal-7
Avaren-Fc
ab5
HPA
CA19-9 Ab (DRG)
CA19.9-9L426
MPL
MOA
WT-VCC
MAL-I
VGA(EY Lab)
#8 Lewis B (T218) (Abcam)
GNA (EY Lab)
HA70 complex-Alexa
#14 Sialyl Lewis X (Calbio)
HHA(EY Lab)
PNA
LCA
BTL native
rMpL
MAL-II
Galectin-1
ConA
human Gal-7
CC1G-1007
Gal-3
ME5(detected with a-HUD)
PHA-E
AOL (Mahal Lab)
CA19.9-1B844
SSL0-488
AAL
TJA-II
WGA(Seikagaku Bio)
CTB
MGL 6A
DC-ASGPR short
SBA
VVHA lectin
DSL
TJA-I
PSL
RCAI (Vector)
SNA

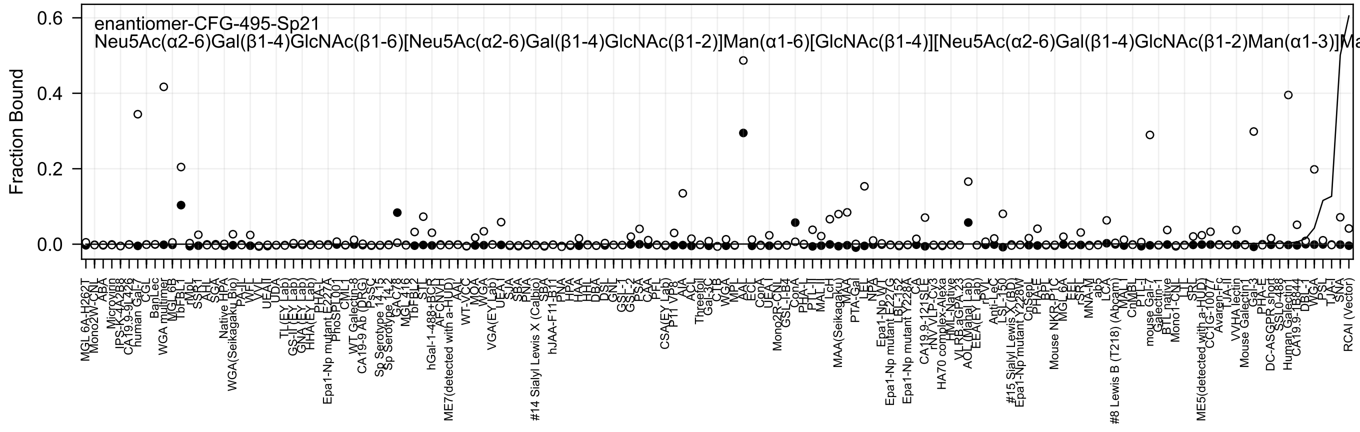
enantiomer-CFG-495-Sp21
Neu5Ac(α2-6)Gal(β1-4)GlcNAc(β1-6)[Neu5Ac(α2-6)Gal(β1-4)GlcNAc(β1-2)]Man(α1-6)[GlcNAc(β1-4)][Neu5Ac(α2-6)Gal(β1-4)GlcNAc(β1-2)Man(α1-3)]Ma
Fraction Bound

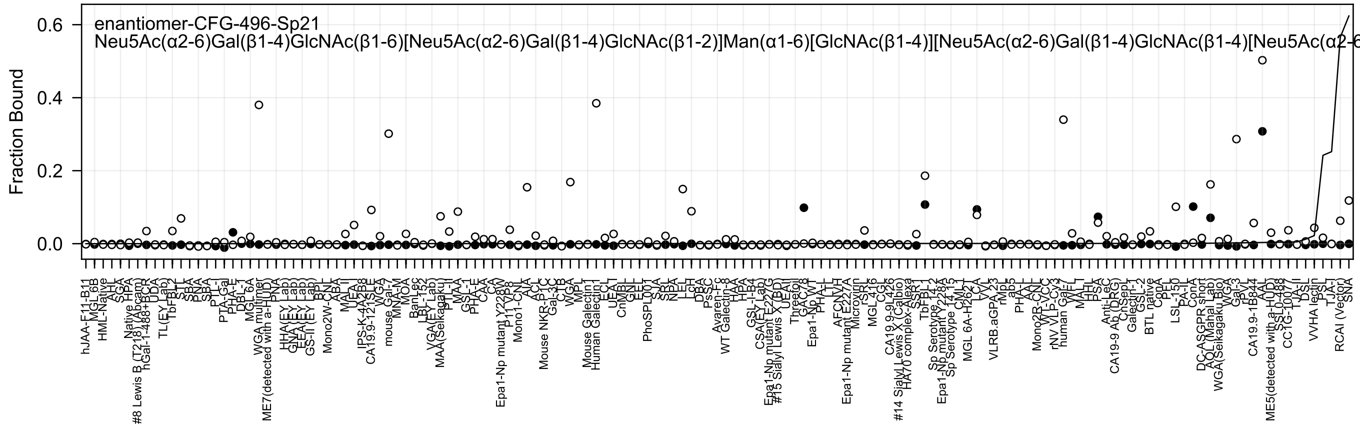
enantiomer-CFG-496-Sp21
Neu5Ac(α2-6)Gal(β1-4)GlcNAc(β1-6)[Neu5Ac(α2-6)Gal(β1-4)GlcNAc(β1-2)]Man(α1-6)[GlcNAc(β1-4)][Neu5Ac(α2-6)Gal(β1-4)GlcNAc(β1-4)[Neu5Ac(α2-6
Fraction Bound
0.0
0.2
0.4
0.6
hJAA-F11-B11
MGL-6B
HML-Native
AAL
SGA
Native HPA
#8 Lewis B (T218) (Abcam)
hGal-1-488+BCR
UDA
TL(EY Lab)
TbFBL2
STL
SBA
PNA
SBA
PTL-I
PTA-Gal
PHA-E
DBL-1
MGL 6A
WGA multimer
ME7(detected with a-HUD)
PNA
HHA(EY Lab)
GNA(EY Lab)
EEA(EY Lab)
GS-II(EY Lab)
BPL
Mono2W-CNL
ABA
MAL-II
UEA-I
IPS-K-4A2B8
CA19.9-121SLE
WGA
mouse Gal-7
MNA-M
MOA
BanLec
LBL-152
VGA(EY Lab)
MAA(Seikagaku)
PTL-II
MAL
GSL-1
PHA-E
CAA
CA
Epa1-Np mutant Y228W
P11VP8
Mono1-CNL
AIA
ACL
Mouse NKR-P1C
Gal-3C
CTB
WGA
MPL
Mouse Galectin1
Human Galectin1
ECL
UEA-I
CnMBL
GNL
EEL
PhoSPL001
SJA
SRA
NPA
LEL
LcH
DBA
PsSC
Avaren-Fc
WT Galectin-8
HAA
HPA
GSL-I-B4
CSA(EY Lab)
Epa1-Np mutant E227G
#15 Sialyl Lewis X (BD)
UEAII
Threefoil
GAC78
Epa1-Np WT
PHA-L
LTL
AFCNVH
Epa1-Np mutant E227A
Microvirin
rSRL
MGL 4 6
CGL
CA19.9-9L426
#14 Sialyl Lewis X (Calbio)
HA70 complex-Alexa
SSR1
TbFBL1
Sp Serotype 14.2
Epa1-Np mutant Y228A
Sp Serotype 14.15
CML1
MGL 6A-H262Y
LCA
VVL
VLRB.aGPA.23
rMpL
ab5
PHA-L
AAL
Mono2R-CNL
WT-VCC
rNV VLP-Cy3
human Gal-7
VAF
MAL-I
HHL
PSA
Anti-LeC
CA19-9 Ab (DRG)
ChSep1
Galectin-1
GSL-2
BTL native
ConA
PFL
LSL-150
PA-IL
ConA
DC-ASGPR short
AOL (Mahal Lab)
WGA(Seikagaku Bio)
WGA
Gal-3
RPL
CA19.9-1B844
AAL
ME5(detected with a-HUD)
SSL0-488
CC1G-1007
TJA-II
DSL
VVHA lectin
PSL
TJA-I
RCAI (Vector)
SNA

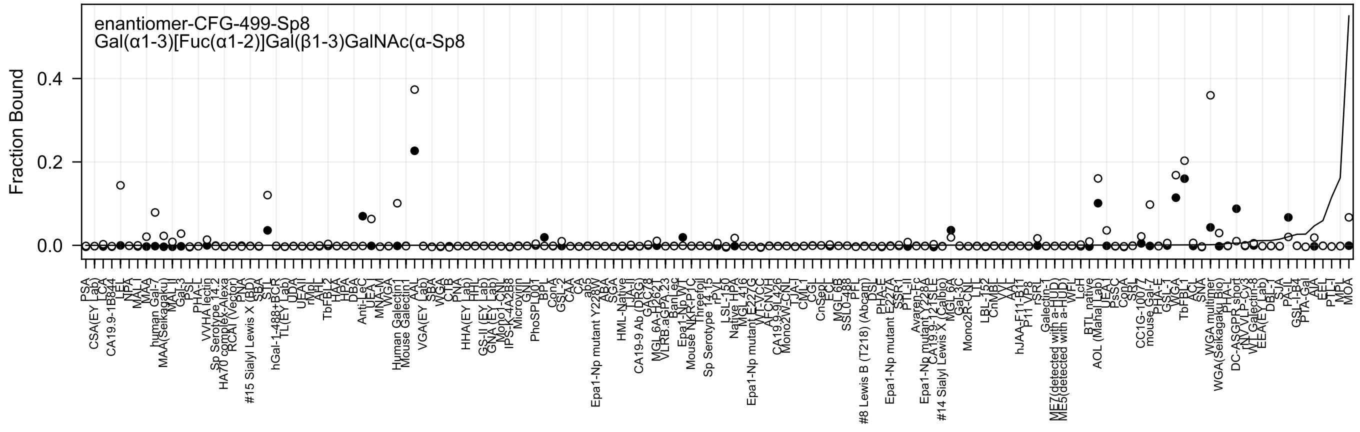
enantiomer-CFG-499-Sp8
Gal(α1-3)[Fuc(α1-2)]Gal(β1-3)GalNAc(α-Sp8
Fraction Bound
0.0
0.2
0.4

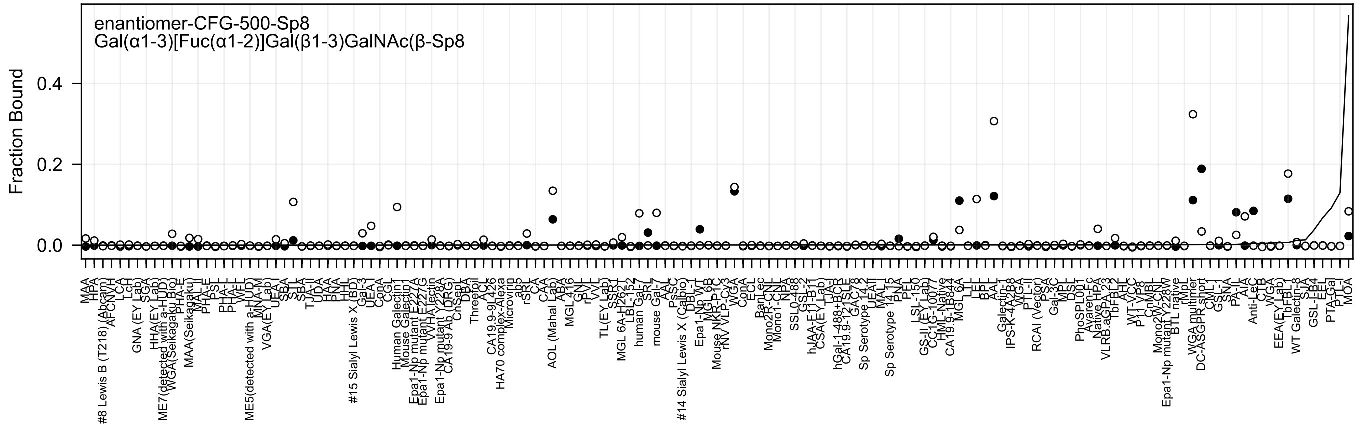
enantiomer-CFG-500-Sp8
Gal(α1-3)[Fuc(α1-2)]Gal(β1-3)GalNAc(β-Sp8
Fraction Bound
0.4
0.2
0.0
MAA
HPA
#8 Lewis B (T218) (Abcam)
AFCNVH
LCA
LcH
GNA (EY Lab)
SGA
HHA(EY Lab)
ME7(detected with a-HuD)
WGA(Seikagaku Bio)
PHA-E
MAA(Seikagaku)
MAA-II
PHA-E
PSL
PHA-L
PHA-L
WFL
ME5(detected with a-HuD)
MNA-M
VGA(EY Lab)
UEA-I
SBA
STL
SBA
TJA-II
UDA
HAA
PNA
HHL
#15 Sialyl Lewis X (BD)
Gal-3
UEA-I
ConA
CGL
Human Galectin1
Mouse Galectin1
Epa1-Np mutant E227A
Epa1-Np mutant E227G
WHA lectin
Epa1-Np mutant Y228A
CA19-9 Ab (DRG)
ChSep
DBA
Threetoil
ACL
CA19.9-9L426
HA70 complex-Alexa
Microvirin
abs
rSRL
CA
CAA
AOL (Mahal Lab)
ABA
MGL 416
GN
rPVL
VVL
TL(EY Lab)
SSR1
MGL 6A-H262T
LBL-152
human Gal-7
SRL
mouse Gal-7
AAL
PsSC
#14 Sialyl Lewis X (Calbio)
DBL-1
Epa1-Np WT
MGL 6B
Mouse NKR-P1C
rNV VLP-CV3
WGA
ConA
ECL
BanLec
Mono2R-CNL
Mono1-CNL
NPA
SSL0-488
GSL-2
hJAA-E11-B11
CSA(EY Lab)
TJA-I
hGal-1-488+BCR
CA19.9-121SLE
GAC78
Sp Serotype 14.2
UEAII
MAL
Sp Serotype 14.15
PNA
PFL
LSL-150
GS-II (EY Lab)
CC1G-10077
HML-Native
CA19.9-1B844
MGL 6A
LTL
LEL
BPL
AAL
Galectin-1
IPS-K-4A2B8
WGA
PTL-II
RCAI (Vector)
PSA
Gal-3C
MPL
DSL
PhoSPL001
Avaren-Fc
Native HPA
VLRB.aGPA.23
TbFBL2
AHL
WT-VCC
P11 VP8
CnMBL
Mono2W-CNL
Epa1-Np mutant Y228W
BTL native
rMpL
WGA multimer
DC-ASGPR short
CML-I
GSL-I
SNA
PA-IL
AIA
Anti-LeC
CTB
WGA
EEA(EY Lab)
TbFBL1
WT Galectin-8
SJA
GSL-I-B4
EEL
PTA-Gal
PTL-I
MOA

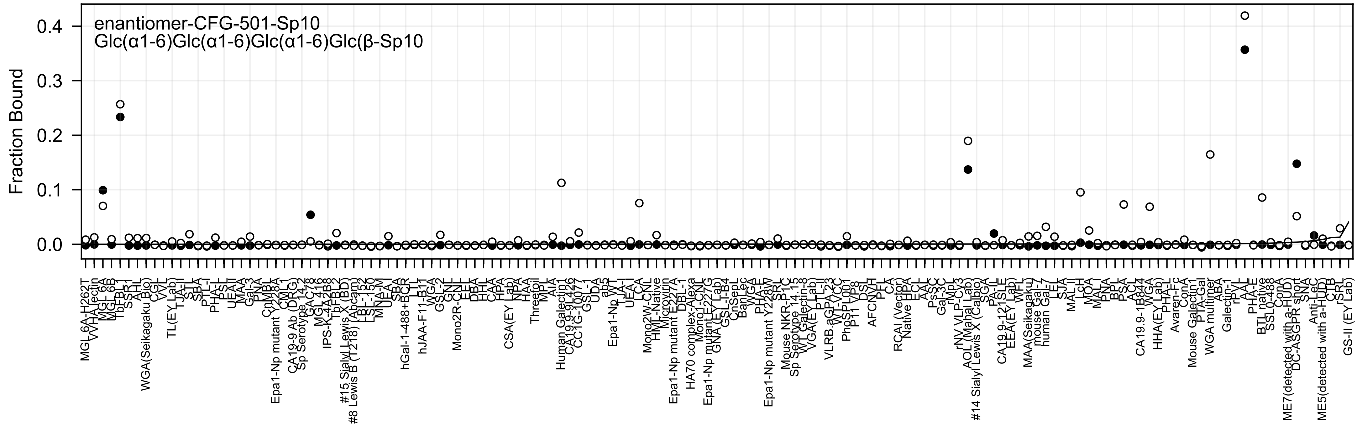

enantiomer-CFG-501-Sp10
Glc(α1-6)Glc(α1-6)Glc(α1-6)Glc(β-Sp10
Fraction Bound

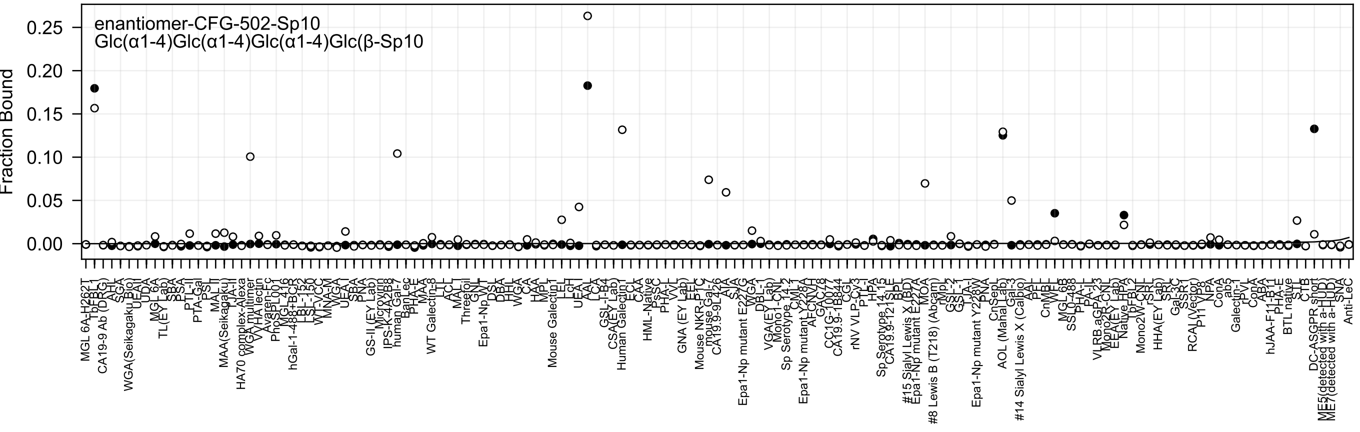

enantiomer-CFG-502-Sp10
Glc(α1-4)Glc(α1-4)Glc(α1-4)Glc(β-Sp10
Fraction Bound

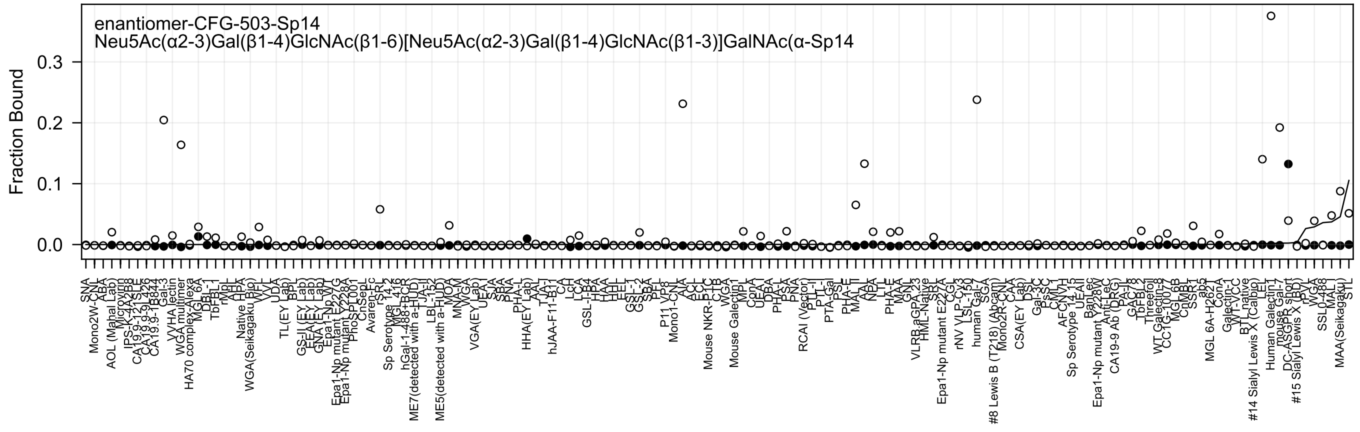
enantiomer-CFG-503-Sp14
Neu5Ac(α2-3)Gal(β1-4)GlcNAc(β1-6)[Neu5Ac(α2-3)Gal(β1-4)GlcNAc(β1-3)]GalNAc(α-Sp14
Fraction Bound
0.3
0.2
0.1
0.0

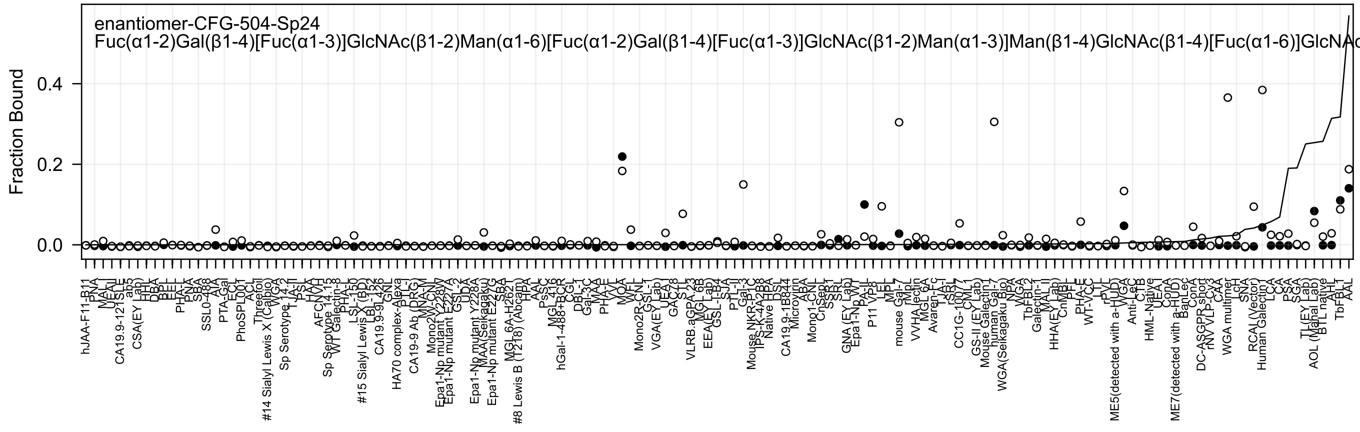
enantiomer-CFG-504-Sp24
Fuc(α1-2)Gal(β1-4)[Fuc(α1-3)]GlcNAc(β1-2)Man(α1-6)[Fuc(α1-2)Gal(β1-4)[Fuc(α1-3)]GlcNAc(β1-2)Man(α1-3)]Man(β1-4)GlcNAc(β1-4)[Fuc(α1-6)]GlcNAc
Fraction Bound
0.4
0.2
0.0

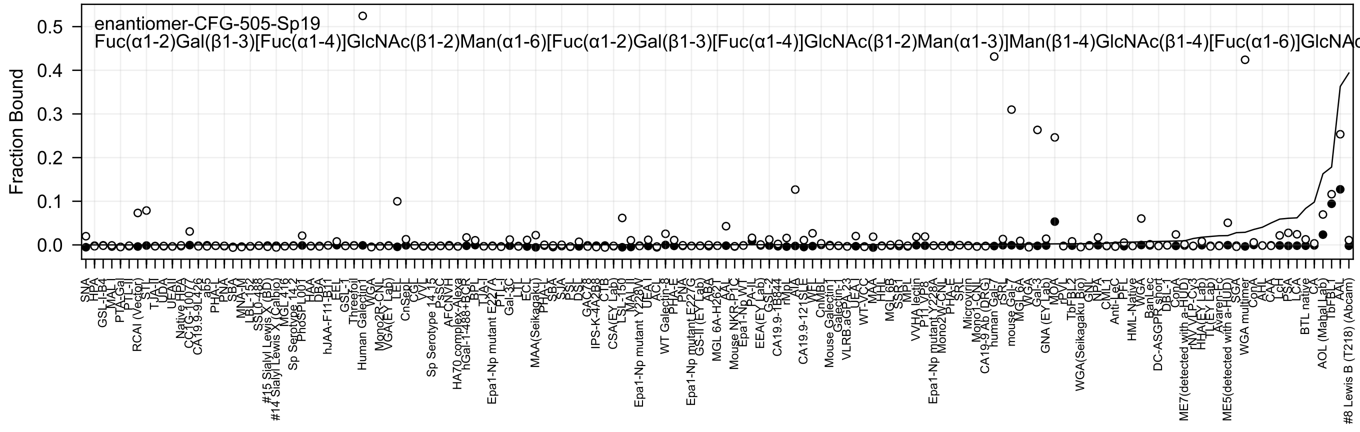

enantiomer-CFG-505-Sp19
Fuc(α1-2)Gal(β1-3)[Fuc(α1-4)]GlcNAc(β1-2)Man(α1-6)[Fuc(α1-2)Gal(β1-3)[Fuc(α1-4)]GlcNAc(β1-2)Man(α1-3)]Man(β1-4)GlcNAc(β1-4)[Fuc(α1-6)]GlcNAc
Fraction Bound

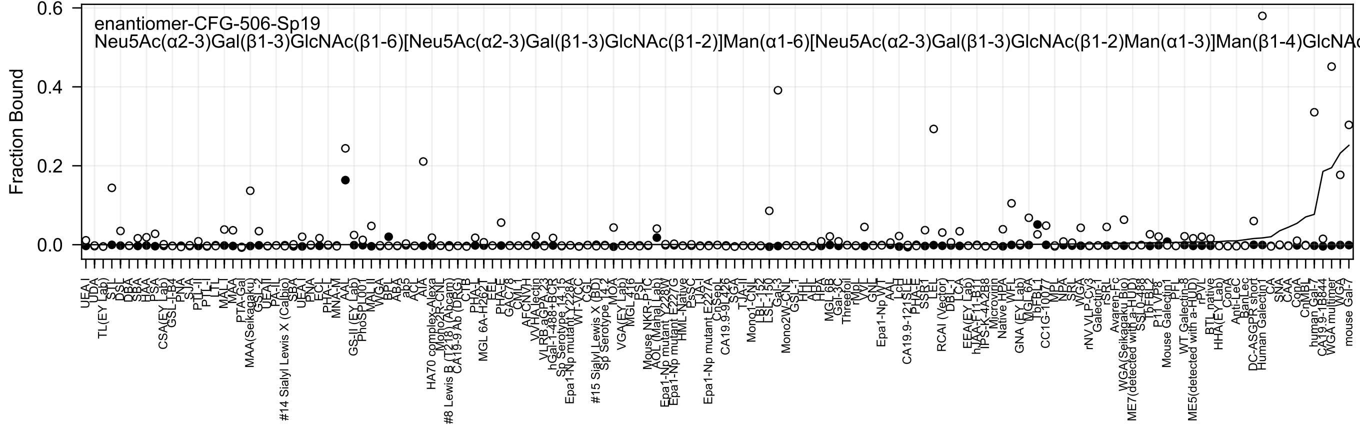
enantiomer-CFG-506-Sp19
Neu5Ac(α2-3)Gal(β1-3)GlcNAc(β1-6)[Neu5Ac(α2-3)Gal(β1-3)GlcNAc(β1-2)]Man(α1-6)[Neu5Ac(α2-3)Gal(β1-3)GlcNAc(β1-2)Man(α1-3)]Man(β1-4)GlcNAc
Fraction Bound
0.0
0.2
0.4
0.6
UEA I
UDA
TL(EY Lab)
STL
DSL
DBA
SBA
HAA
PSA
CSA(EY Lab)
GSL-I-B4
PNA
SJA
PTL-II
PTL-I
LTL
MAL-I
MAA
PTA-Gal
MAA(Seikagaku)
GSL-I
UEA-II
PA-IL
#14 Sialyl Lewis X (Calbio)
SBA
UEA-I
PNA
ECL
PHA-L
MNA-M
AAL
GS-II (EY Lab)
PhoSL001
MAL-II
WGA
BPL
ABA
ab5
ACL
AIA
HA70 complex-Alexa
Mono2R-CNL
#8 Lewis B (T218) (Abcam)
CA19-9 Ab (DRG)
CTB
PHA-L
MGL 6A-H262T
EEL
PHA-E
GAC78
CNL1
AFCNVH
VVHA lectin
VLRB.aGPA.23
hGal-1-488+BCR
Sp Serotype 14.15
Epa1-Np mutant E228A
WT-VCC
CGL
#15 Sialyl Lewis X (BD)
Sp Serotype 14.2
MOA
VGA(EY Lab)
MGL 416
PSL
Mouse NKR-P1C
AOL (Mahal Lab)
Epa1-Np mutant E228V
Epa1-Np mutant E227G
HML-Native
PsSC
TJA-I
Epa1-Np mutant E227A
CnSep
CA19.9-9L426
SGA
TJA-II
Mono1-CNL
LBL-152
LSL-150
Gal-3
Mono2W-CNL
GSL-1
HHL
AIA
HPA
MGL 6B
Gal-3C
Threefoil
rMpL
VVL
GNL
Epa1-Np WT
AAL
LcH
CA19.9-121SLE
PHA-E
SSBT
LEL
RCAI (Vector)
DBL-1
LCA
EEA(EY Lab)
hAA-F11-B11
IPS-K-4A2B8
Microvirin
Native HPA
WFL
GNA (EY Lab)
MGL 6A
TbFBL1
CC1G-10077
MPA
NPA
SRL
WGA
rNV VLP-Cy3
Galectin-1
rSRL
Avaren-Fc
WGA(Seikagaku Bio)
ME7(detected with a-HuD)
SSL0-488
TbFBL2
P11 VP8
Mouse Galectin1
PFL
WT Galectin-8
ME5(detected with a-HuD)
rPVL
BTL native
HHA(EY Lab)
ConA
Anti-LeC
BanLec
DC-ASGPR short
Human Galectin1
CA
SNA
CAA
CopA
CnMBL
human Gal-7
CA19.9-1B844
WGA multimer
WGA
mouse Gal-7

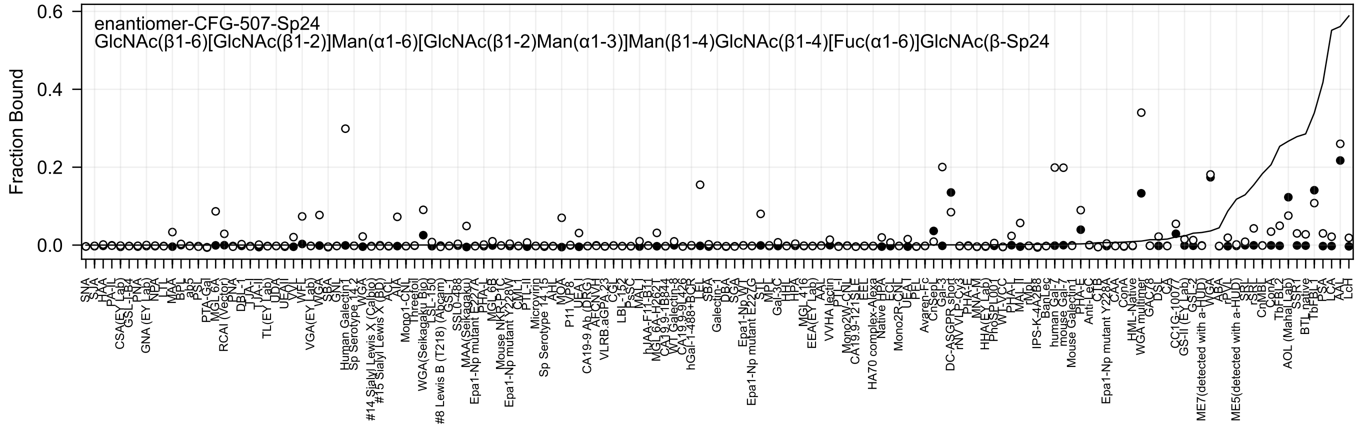

enantiomer-CFG-507-Sp24
GlcNAc(β1-6)[GlcNAc(β1-2)]Man(α1-6)[GlcNAc(β1-2)Man(α1-3)]Man(β1-4)GlcNAc(β1-4)[Fuc(α1-6)]GlcNAc(β-Sp24
Fraction Bound

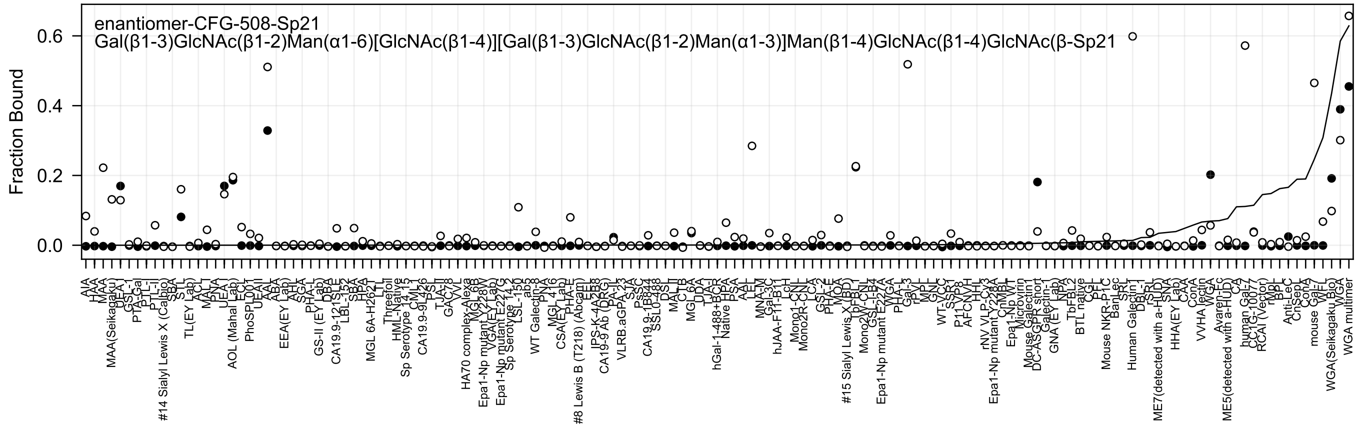

enantiomer-CFG-508-Sp21
Gal(β1-3)GlcNAc(β1-2)Man(α1-6)[GlcNAc(β1-4)][Gal(β1-3)GlcNAc(β1-2)Man(α1-3)]Man(β1-4)GlcNAc(β1-4)GlcNAc(β-Sp21
Fraction Bound
0.0
0.2
0.4
0.6
AIA
HAA
MAA
MAA(Seikagaku)
UEA-I
GSL-1
PTA-Gal
PTL-II
PTL-I
#14 Sialyl Lewis X (Calbio)
SBA
STL
TL(EY Lab)
ACL
MAL-I
PNA
UEA-I
AOL (Mahal Lab)
ECL
PhoSL001
UEA-II
AAL
ABA
EEA(EY Lab)
AH
SGA
PHA-L
GS-II (EY Lab)
DBA
CA19.9-121SLE
LBL-152
SBA
HPA
MGL 6A-H262T
LTL
Threefoil
HML-Native
Sp Serotype 14 15
CML-1
CA19.9-9L426
PSL
TJA-II
GAC78
VVL
HA70 complex-Alexa
MGL 6B
Epa1-Np mutant Y228W
VGA(EY Lab)
Epa1-Np mutant E227G
Sp Serotype 14 2
LSL-150
ab5
WT Galectin-8
PNA
MGL 416
CSA(EY Lab)
PHA-E
#8 Lewis B (T218) (Abcam)
EEL
IPS-K-4A2B8
CA19-9 Ab (DRG)
PA-I
VLRB.aGPA.23
SJA
PsSC
CA19.9-1B844
SSL0-488
DSL
MAL-II
CTB
MGL 6A
UDA
TJA-I
hGal-1-488+BCR
Native
HPA
PSA
AAL
LEL
MNA-M
Gal-3C
hJAA-F11-B11
LcH
Mono1-CNL
Mono2R-CNL
LCA
GSL-2
PHA-E
MOA
#15 Sialyl Lewis X (BD)
TbFBL1
Mono2W-CNL
GSL-I-B4
Epa1-Np mutant E227A
PHA-L
WGA
Gal-3
rPVL
MPL
GNL
WT-VCC
SSR1
P11-VP8
AFCNVH
HHL
rNV VLP-CV3
Epa1-Np mutant Y228A
CnMBL
Epa1-Np WT
Microvirin
Mouse Galectin1
DC-ASGPR short
Galectin-1
GNA (EY Lab)
NPA
TbFBL2
BTL native
CGL
PFL
Mouse NKR-P1C
BanLec
SRL
Human Galectin-1
DBL-1
rSRL
ME7(detected with a-HUD)
SNA
HHA(EY Lab)
CAA
ConA
VVHA lectin
WGA
Avaren-Fc
ME5(detected with a-HUD)
CA
human Gal-7
CC1G-1007
RCAI (Vector)
rMpL
BPL
Anti-LeC
CnSepL
ConA
mouse Gal-7
WFL
WGA(Seikagaku Bio)
WGA
WGA multimer

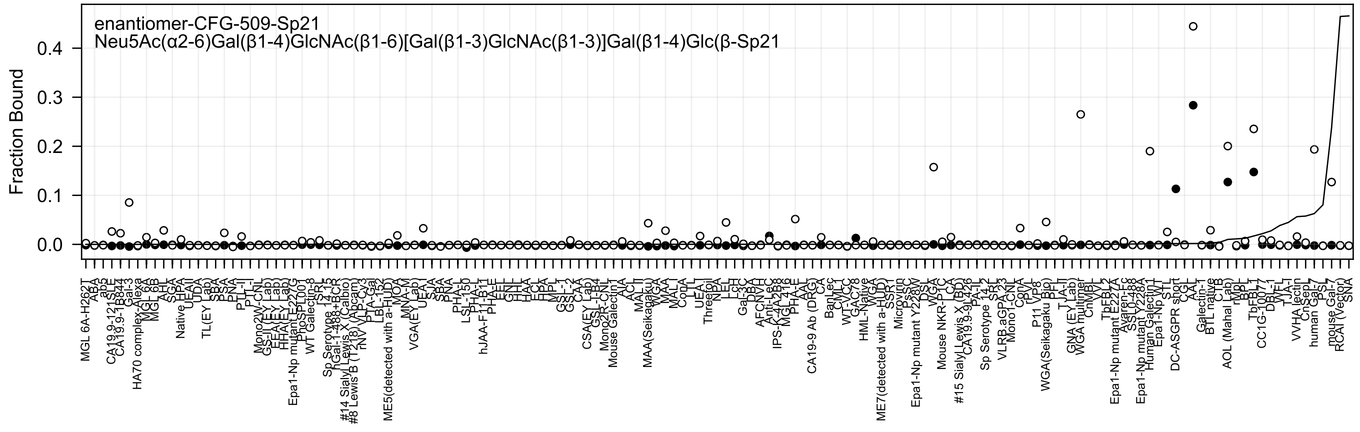
enantiomer-CFG-509-Sp21
Neu5Ac(α2-6)Gal(β1-4)GlcNAc(β1-6)[Gal(β1-3)GlcNAc(β1-3)]Gal(β1-4)Glc(β-Sp21
Fraction Bound
0.4
0.3
0.2
0.1
0.0

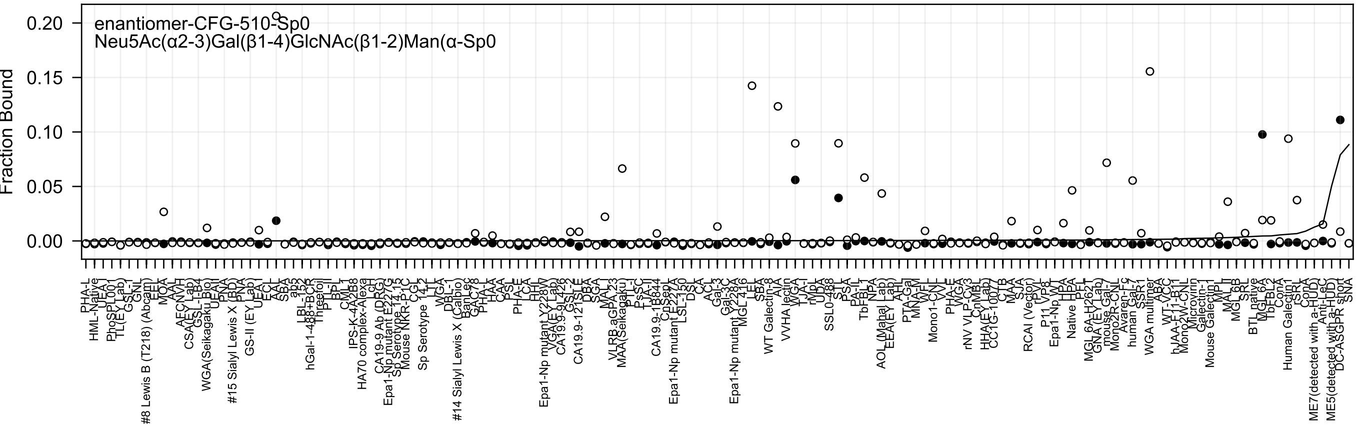
enantiomer-CFG-510-Sp0
Neu5Ac(α2-3)Gal(β1-4)GlcNAc(β1-2)Man(α-Sp0
Fraction Bound

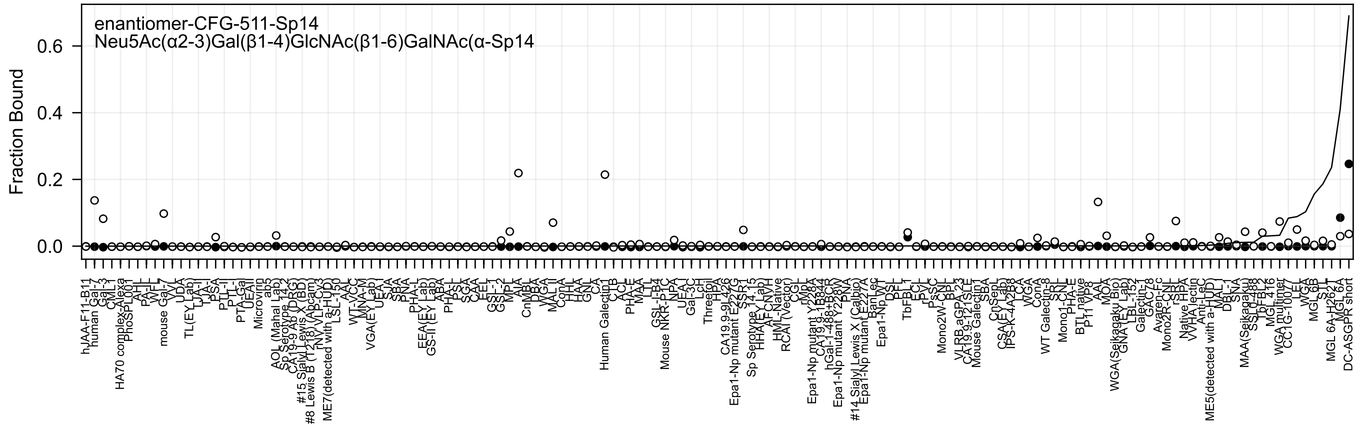
enantiomer-CFG-511-Sp14
Neu5Ac(α2-3)Gal(β1-4)GlcNAc(β1-6)GalNAc(α-Sp14
Fraction Bound
0.0
0.2
0.4
0.6

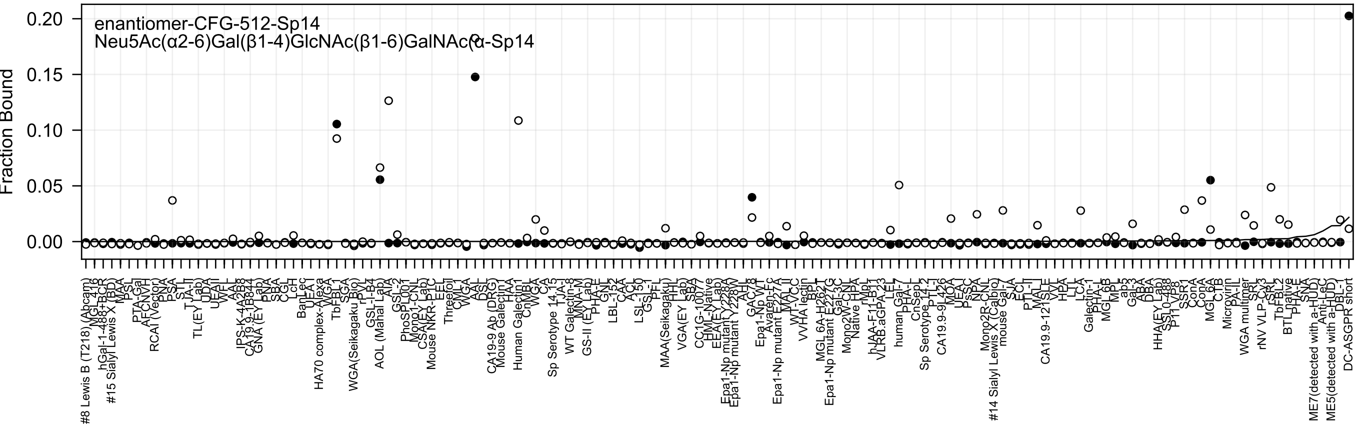
enantiomer-CFG-512-Sp14
Neu5Ac(α2-6)Gal(β1-4)GlcNAc(β1-6)GalNAc(α-Sp14
Fraction Bound
0.20
0.15
0.10
0.05
0.00

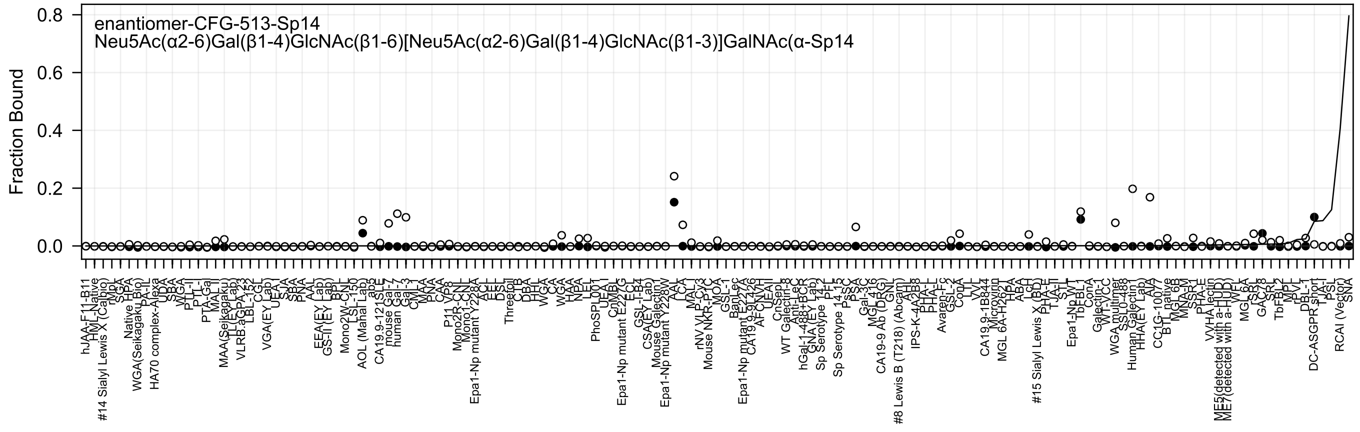

enantiomer-CFG-513-Sp14
Neu5Ac(α2-6)Gal(β1-4)GlcNAc(β1-6)[Neu5Ac(α2-6)Gal(β1-4)GlcNAc(β1-3)]GalNAc(α-Sp14
Fraction Bound
0.0
0.2
0.4
0.6
0.8

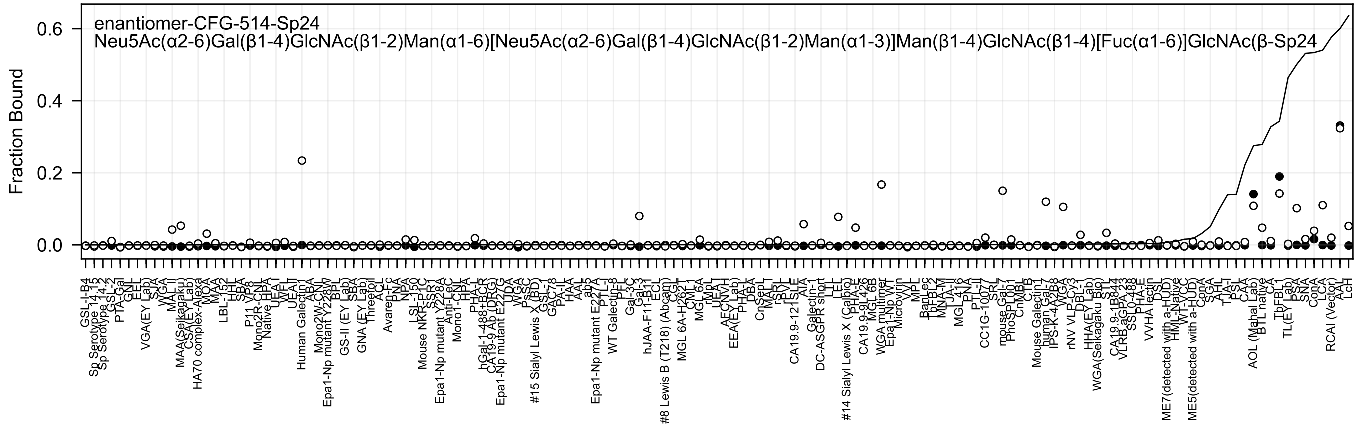
enantiomer-CFG-514-Sp24
Neu5Ac(α2-6)Gal(β1-4)GlcNAc(β1-2)Man(α1-6)[Neu5Ac(α2-6)Gal(β1-4)GlcNAc(β1-2)Man(α1-3)]Man(β1-4)GlcNAc(β1-4)[Fuc(α1-6)]GlcNAc(β-Sp24
Fraction Bound
0.0
0.2
0.4
0.6

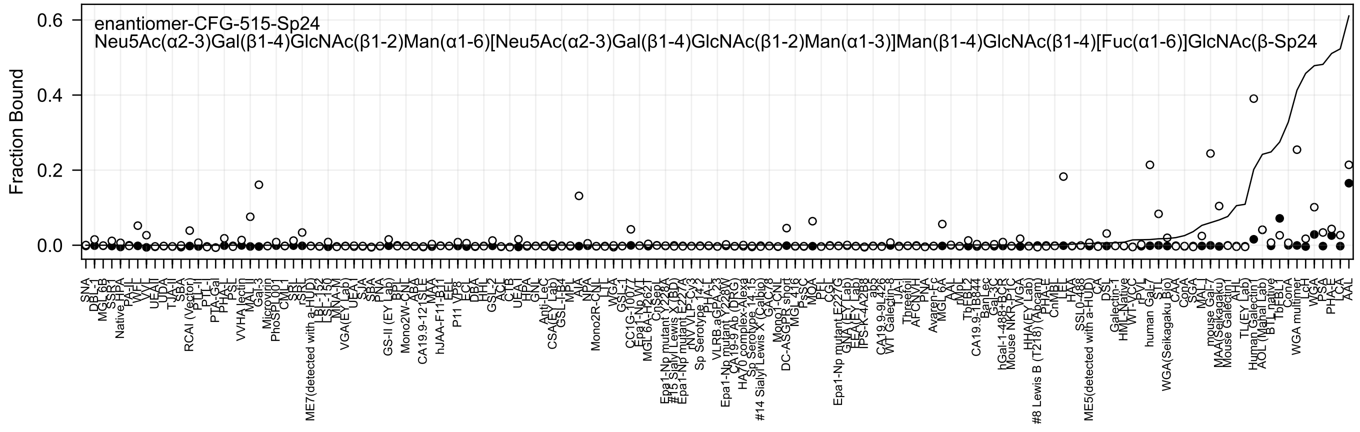
enantiomer-CFG-515-Sp24
Neu5Ac(α2-3)Gal(β1-4)GlcNAc(β1-2)Man(α1-6)[Neu5Ac(α2-3)Gal(β1-4)GlcNAc(β1-2)Man(α1-3)]Man(β1-4)GlcNAc(β1-4)[Fuc(α1-6)]GlcNAc(β-Sp24
Fraction Bound

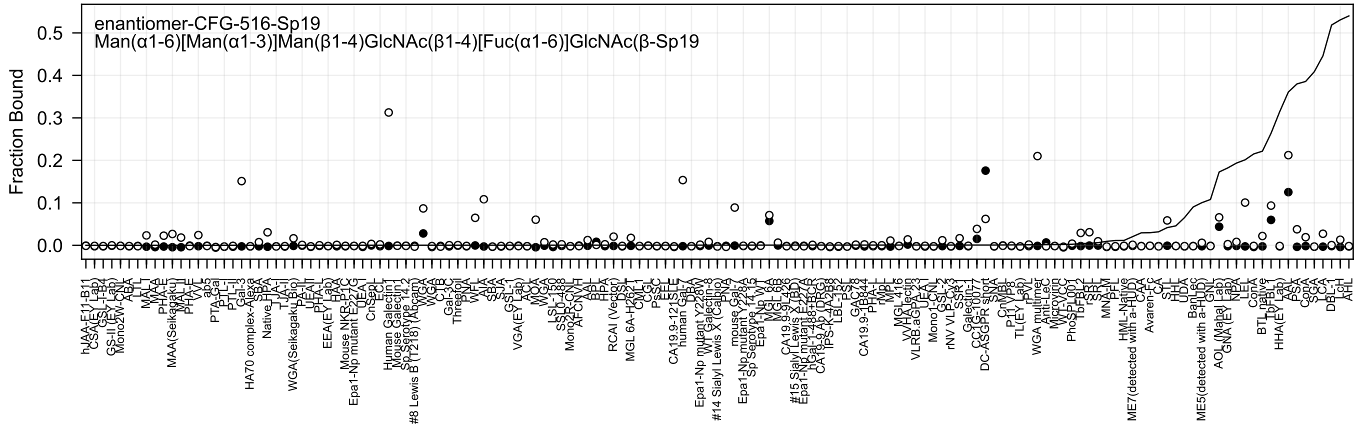
enantiomer-CFG-516-Sp19
Man(α1-6)[Man(α1-3)]Man(β1-4)GlcNAc(β1-4)[Fuc(α1-6)]GlcNAc(β-Sp19
Fraction Bound
0.0
0.1
0.2
0.3
0.4
0.5

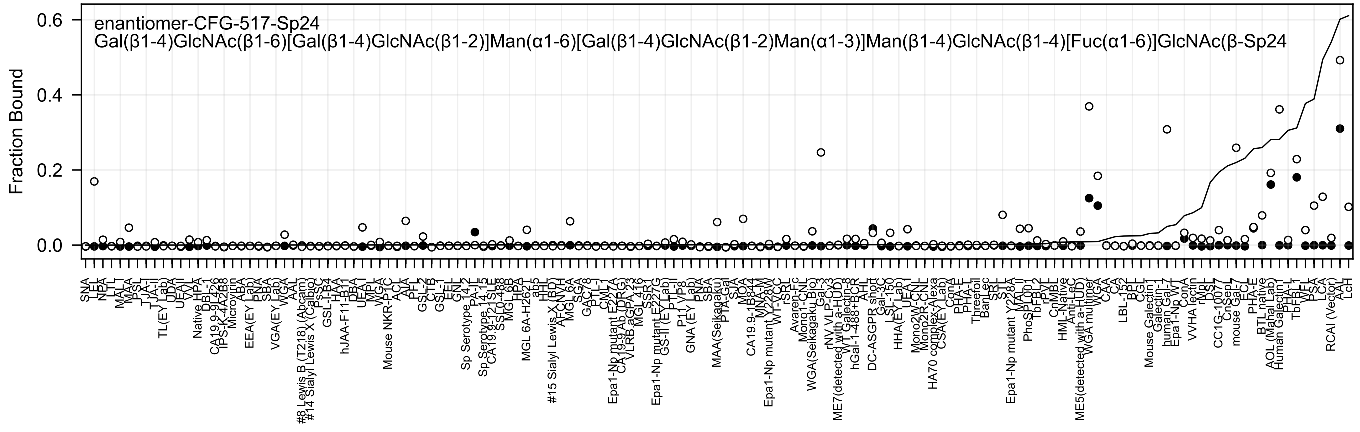
enantiomer-CFG-517-Sp24
Gal(β1-4)GlcNAc(β1-6)[Gal(β1-4)GlcNAc(β1-2)]Man(α1-6)[Gal(β1-4)GlcNAc(β1-2)Man(α1-3)]Man(β1-4)GlcNAc(β1-4)[Fuc(α1-6)]GlcNAc(β-Sp24
Fraction Bound
0.6
0.4
0.2
0.0
SNA
LEL
NPA
LTL
MAL_I
MAA
PSL
TJA-II
TJA-I
TL(EY Lab)
UDA
UEA-II
VVL
Native HPA
DBL-1
CA19.9-9L426
TPS-K-4A2B8
Microvirin
ABA
EEA(EY Lab)
PNA
SBA
VGA(EY Lab)
WGA
AAL
#8 Lewis B (T218) (Abcam)
#14 Sialyl Lewis X (Calbio)
PsSC
GSL-I-B4
HAA
hJAA-F11-B11
DBA
UEA-I
MPL
WGA
Mouse NKR-P1C
ACL
AIA
PFL
GSL-2
CTB
GSL-1
EEL
GNL
Sp Serotype 14_2
PA-IL
Sp Serotype 14_15
CA19.9-121SLE
SSL0-488
MGL 6B
HPA
MGL 6A-H262T
ab5
HHL
#15 Sialyl Lewis X (BD)
AFCNVH
MGL 6A
SGA
GAC78
PTL-I
CML-I
Epa1-Np mutant E227A
CA19.9 Ab (DRG)
VLRB.aGPA.23
MGL 416
SSR1
Epa1-Np mutant E227G
GS-II (EY Lab)
PTL-II
P11 VP8
GNA (EY Lab)
PNA
SBA
MAA(Seikagaku)
PTA-Gal
SJA
MOA
CA19.9-1B844
MNA-M
Epa1-Np mutant Y228W
WT-VCC
rSRL
Avaren-Fc
Mono1-CNL
WGA(Seikagaku Bio)
Gal-3
rNV VLP-Cy3
ME7(detected with a-HUD)
WT Galectin-8
hGal-1-488+BCR
AHL
DC-ASGPR short
Gal-3C
LSL-150
HHA(EY Lab)
UEA-I
Mono2W-CNL
Mono2R-CNL
HA70 complex-Alexa
CSA(EY Lab)
ConA
PHA-E
PHA-L
Threefoil
BanLec
SRL
STL
Epa1-Np mutant Y228A
MAL-I
PhoSL001
TbFBL2
rPVL
CnMBL
HML-Native
Anti-LeC
ME5(detected with a-HUD)
WGA multimer
WGA
CAA
CA
LBL-152
BPL
CGL
Mouse Galectin-1
Galectin-1
human Gal-7
Epa1-Np WT
ConA
VVHA lectin
rMpL
DSL
CC1G-1007
CnSep7
mouse Gal-7
ECL
PHA-E
BTL native
AOL (Mahal Lab)
Human Galectin-1
PHA-L
TbFBL1
WFL
PSA
LCA
RCAI (Vector)
AAL
LcH

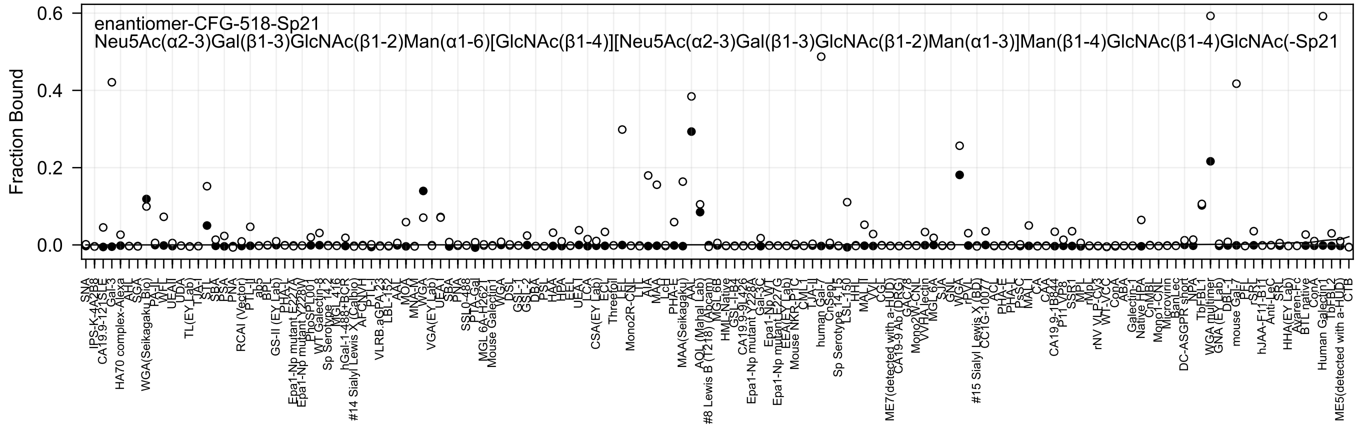
enantiomer-CFG-518-Sp21
Neu5Ac(α2-3)Gal(β1-3)GlcNAc(β1-2)Man(α1-6)[GlcNAc(β1-4)][Neu5Ac(α2-3)Gal(β1-3)GlcNAc(β1-2)Man(α1-3)]Man(β1-4)GlcNAc(β1-4)GlcNAc(-Sp21
Fraction Bound
0.6
0.4
0.2
0.0

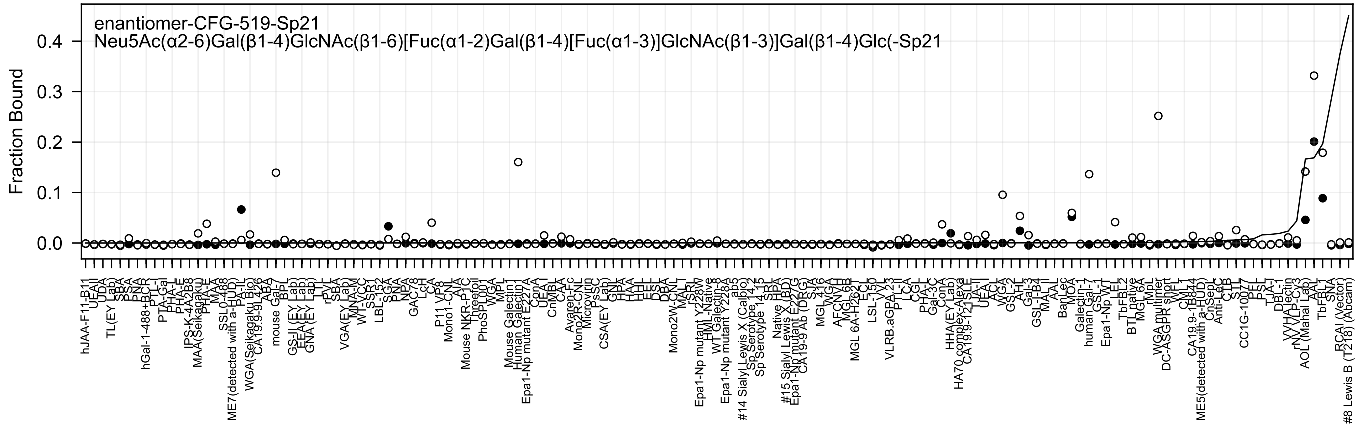
enantiomer-CFG-519-Sp21
Neu5Ac(α2-6)Gal(β1-4)GlcNAc(β1-6)[Fuc(α1-2)Gal(β1-4)[Fuc(α1-3)]GlcNAc(β1-3)]Gal(β1-4)Glc(-Sp21
Fraction Bound
0.0
0.1
0.2
0.3
0.4

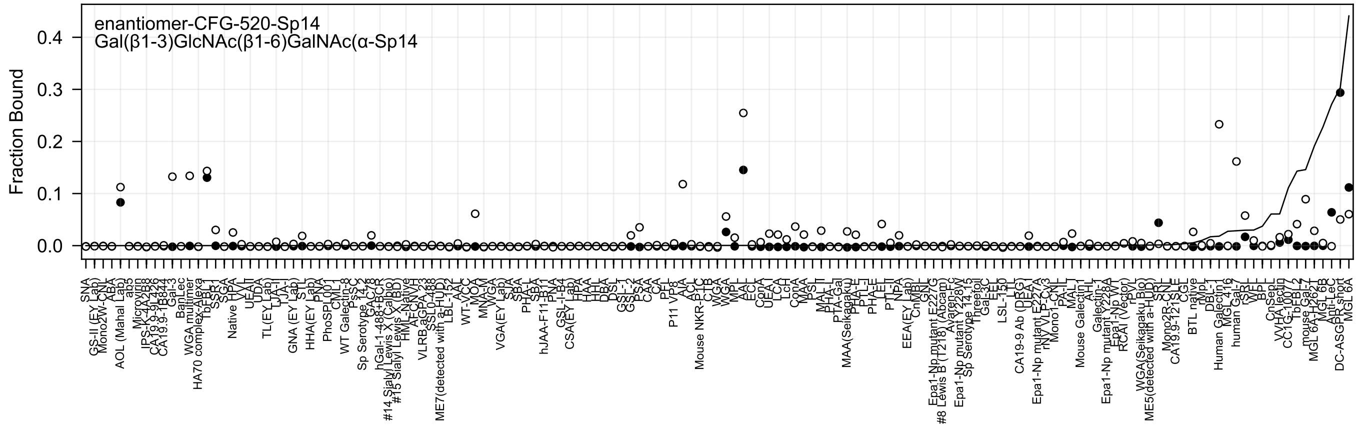

enantiomer-CFG-520-Sp14
Gal(β1-3)GlcNAc(β1-6)GalNAc(α-Sp14
Fraction Bound
0.0
0.1
0.2
0.3
0.4

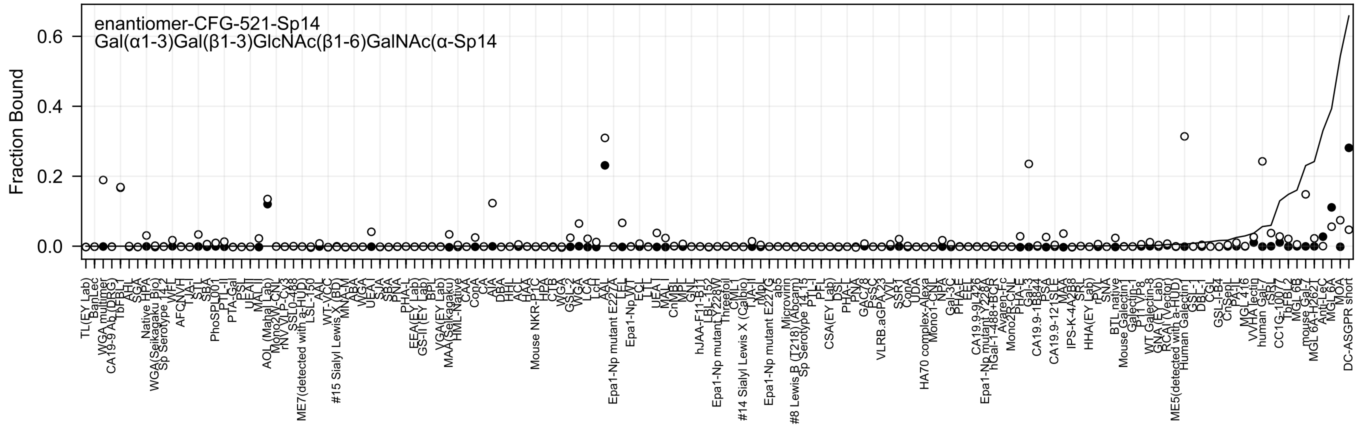
enantiomer-CFG-521-Sp14
Gal(α1-3)Gal(β1-3)GlcNAc(β1-6)GalNAc(α-Sp14
Fraction Bound
0.6
0.4
0.2
0.0
TL(EY Lab)
BanLec
WGA multimer
CA19-9 Ab (DRG)
TbFBL1
AHL
SGA
Native HPA
WGA(Seikagaku Bio)
Sp Serotype 14-2
WFL
AFCNVH
TJA-I
STL
SBA
PhoSPL001
PTL-II
PTA-Gal
PSL
UEAII
MAL-I
AOL (Mahal Lab)
Mono2W-CNL
rNV VLP-Cy3
SSL0-488
ME7(detected with a-HUD)
LSL-150
AAL
WT-VCC
#15 Sialyl Lewis X (BD)
MNA-M
ABA
WGA
UEA-I
SJA
SBA
PNA
PHA-L
EEA(EY Lab)
GS-II (EY Lab)
BPL
VGA(EY Lab)
MAA (Seikagaku)
HML-Native
CAA
ConA
CA
AIA
DBA
HHL
ACL
HAA
Mouse NKR-P1C
HPA
CTB
WGA
GSL-2
WGA
LCA
LcH
AAL
Epa1-Np mutant E227A
LEL
Epa1-Np WT
ECL
LTL
UEA-I
MAL-I
CnMBL
MPL
GNL
hJAA-F11-B11
LBL-152
Epa1-Np mutant Y228V
Threefoil
CML-I
#14 Sialyl Lewis X (Calbio)
TJA-II
rMpL
Epa1-Np mutant E227G
ab5
Microvirin
#8 Lewis B (T218) (Abcam)
Sp Serotype 14-13
PTL-I
PFL
CSA(EY Lab)
DSL
PHA-L
PNA
GAC78
PsSC
VLRB.aGPA.23
VVL
SSR1
ConA
UDA
HA70 complex-Alexa
Mono1-CNL
NPA
Gal-3C
PHA-E
EEL
CA19.9-9L426
Epa1-Np mutant Y228A
hGal-1-488+BCR
Avaren-Fc
Mono2R-CNL
PHA-E
Gal-3
CA19.9-1B844
PSA
CA19.9-121SLE
MAA
IPS-K-4A2B8
SRL
HHA(EY Lab)
rPVL
SNA
BTL native
Mouse Galectin-1
Galectin-1
P11 VP8
WT Galectin-8
GNA (EY Lab)
RCA I (Vector)
ME5(detected with a-HUD)
Human Galectin-1
GSL-I
DBL-1
CGL
GSL-I-B4
CnSepL
PA-IL
MGL 416
VVHA lectin
human Gal-7
rSRL
CC1G-10077
TbFBL2
MGL 6B
mouse Gal-7
MGL 6A-H262T
Anti-LeC
MGL 6A
MOA
DC-ASGPR short

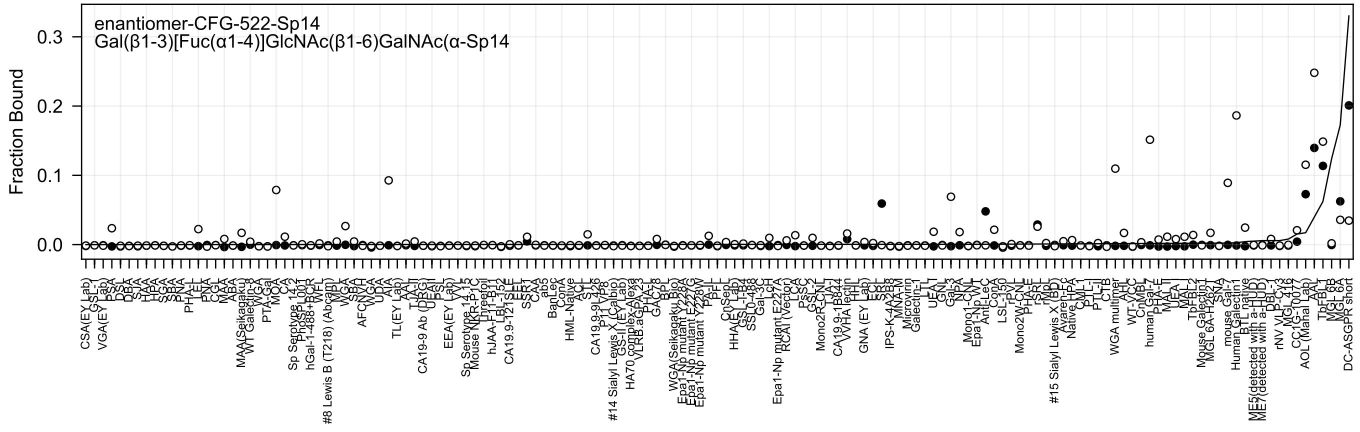
enantiomer-CFG-522-Sp14
Gal(β1-3)[Fuc(α1-4)]GlcNAc(β1-6)GalNAc(α-Sp14
Fraction Bound
0.3
0.2
0.1
0.0

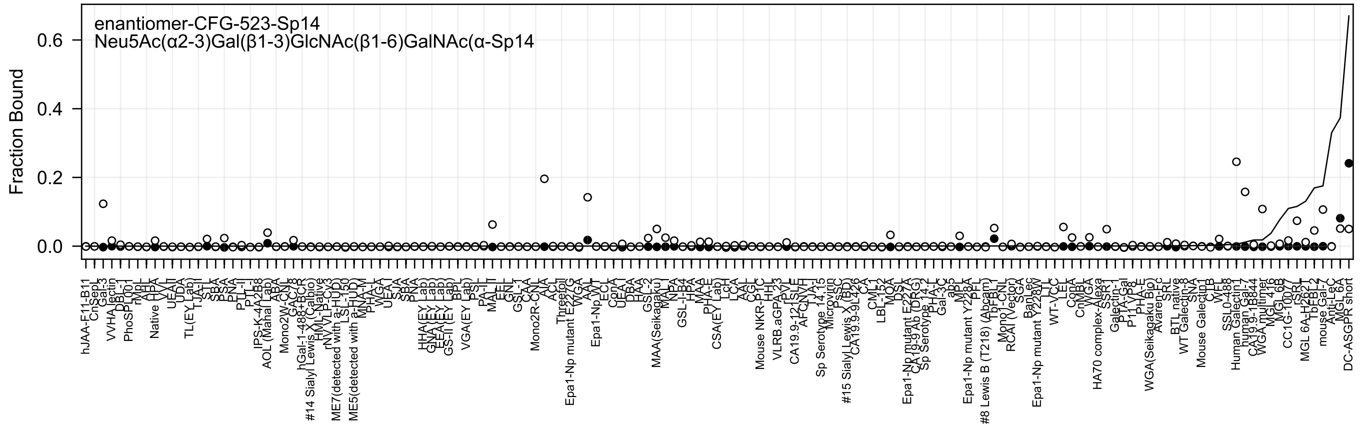
enantiomer-CFG-523-Sp14
Neu5Ac(α2-3)Gal(β1-3)GlcNAc(β1-6)GalNAc(α-Sp14
Fraction Bound
0.6
0.4
0.2
0.0

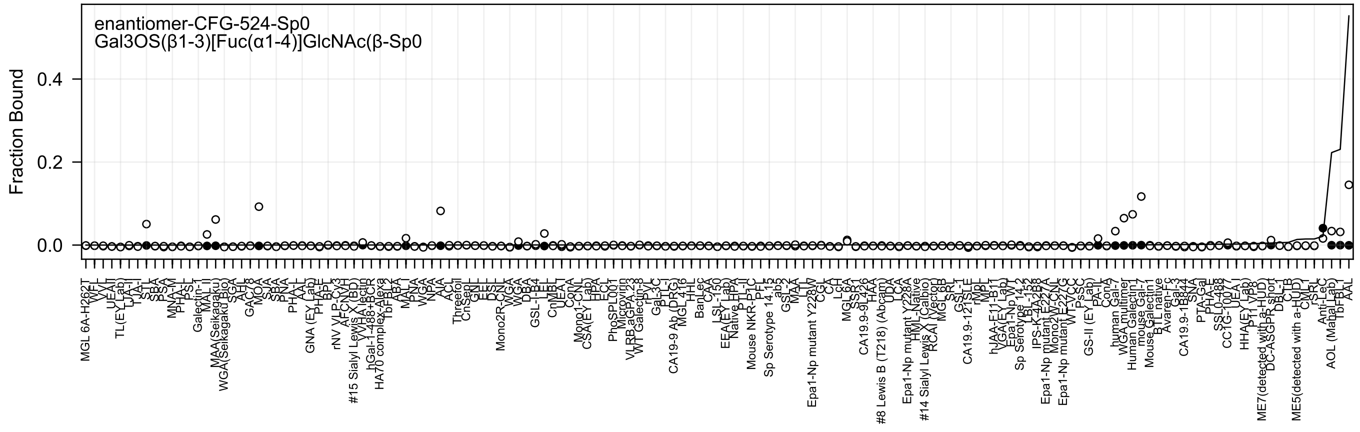
enantiomer-CFG-524-Sp0
Gal3OS(β1-3)[Fuc(α1-4)]GlcNAc(β-Sp0
Fraction Bound
0.4
0.2
0.0

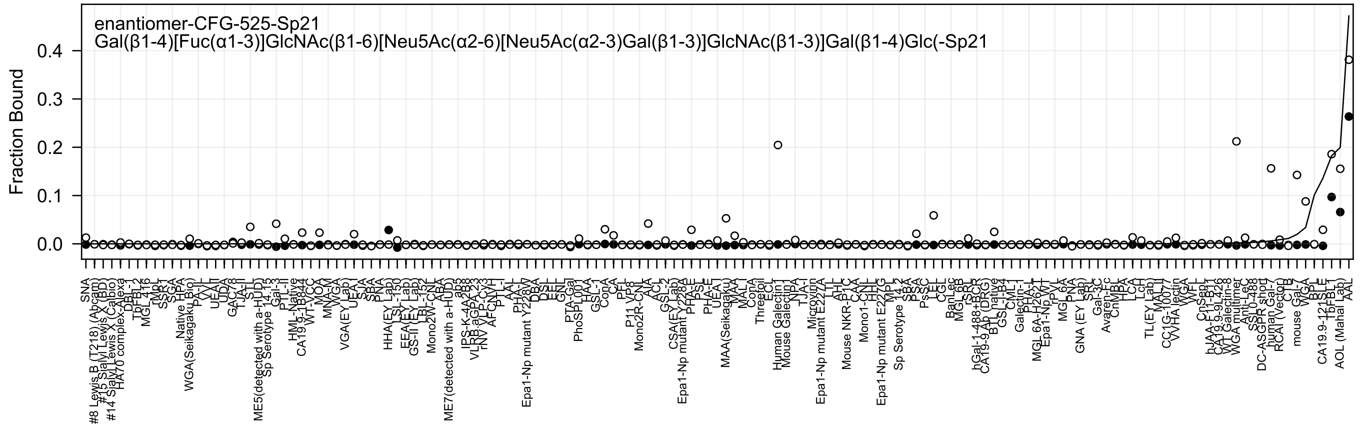

enantiomer-CFG-525-Sp21
Gal(β1-4)[Fuc(α1-3)]GlcNAc(β1-6)[Neu5Ac(α2-6)[Neu5Ac(α2-3)Gal(β1-3)]GlcNAc(β1-3)]Gal(β1-4)Glc(-Sp21
Fraction Bound

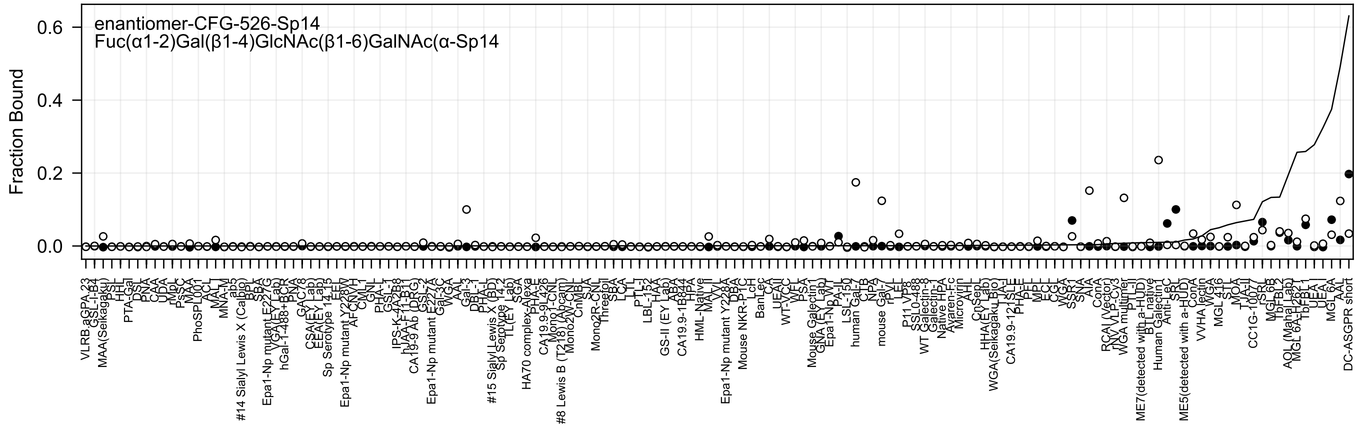

enantiomer-CFG-526-Sp14
Fuc(α1-2)Gal(β1-4)GlcNAc(β1-6)GalNAc(α-Sp14
Fraction Bound
0.0
0.2
0.4
0.6

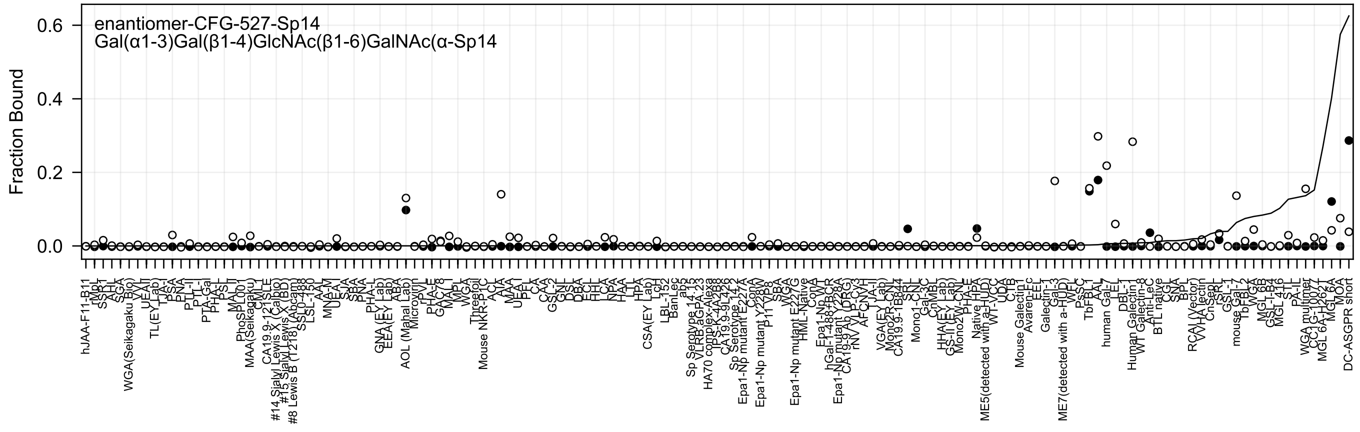
enantiomer-CFG-527-Sp14
Gal(α1-3)Gal(β1-4)GlcNAc(β1-6)GalNAc(α-Sp14
Fraction Bound
0.6
0.4
0.2
0.0

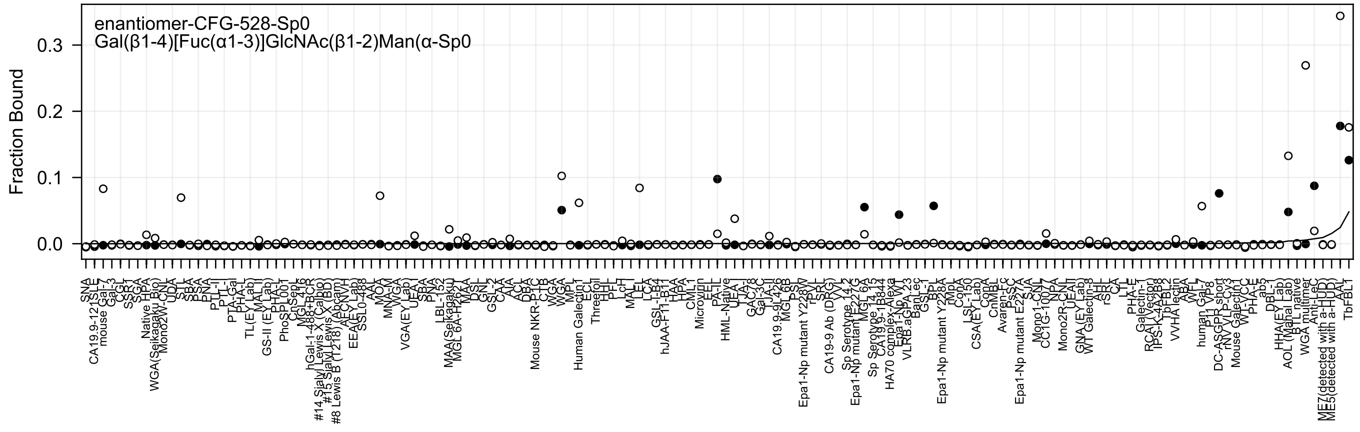
enantiomer-CFG-528-Sp0
Gal(β1-4)[Fuc(α1-3)]GlcNAc(β1-2)Man(α-Sp0
Fraction Bound

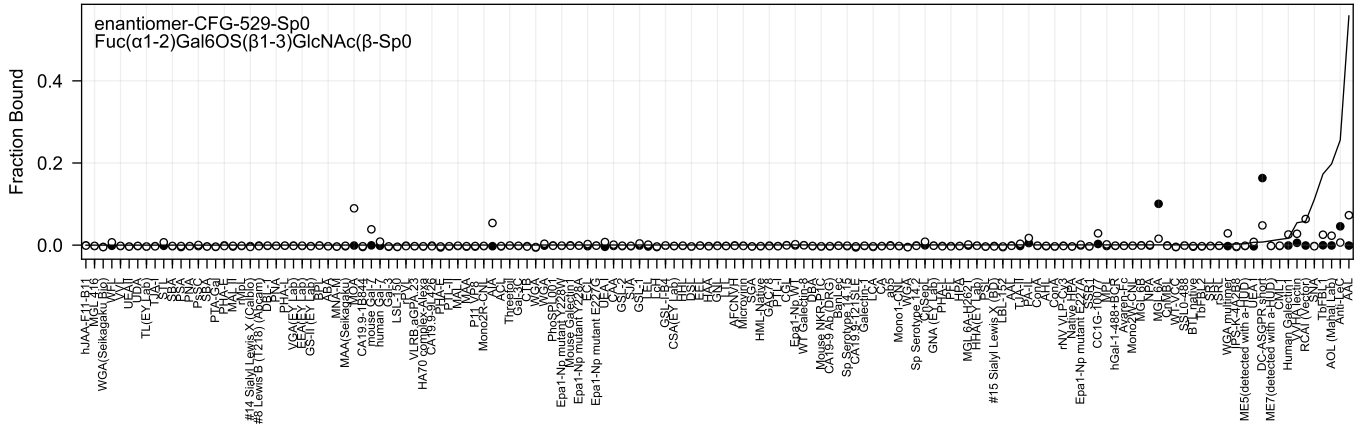

enantiomer-CFG-529-Sp0
Fuc(α1-2)Gal6OS(β1-3)GlcNAc(β-Sp0
Fraction Bound
0.0
0.2
0.4

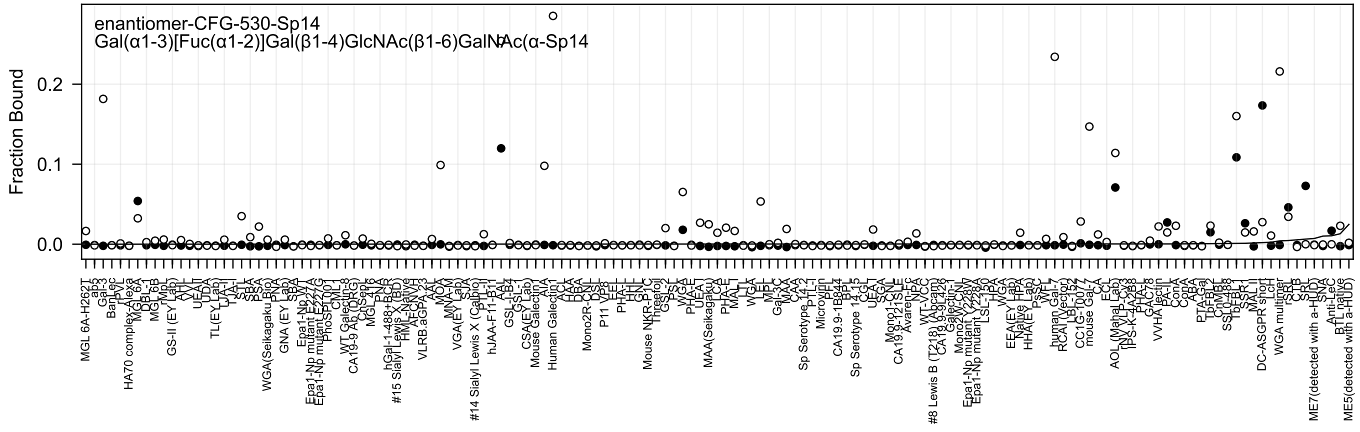

enantiomer-CFG-530-Sp14
Gal(α1-3)[Fuc(α1-2)]Gal(β1-4)GlcNAc(β1-6)GalNAc(α-Sp14
Fraction Bound
0.0
0.1
0.2
MGL 6A-H262T
ab5
Gal-3
BanLec
rPVL
HA70 complex-Alexa
MGL 6A
DBL-1
MGL 6B
rMpL
GS-II (EY Lab)
AAL
AVL
UEA-I
UDA
TL(EY Lab)
TJA-I
STL
SBA
PSA
WGA(Seikagaku Bio)
PNA
GNA (EY Lab)
SBA
Epa1-Np WT
Epa1-Np mutant E227A
Epa1-Np mutant E227G
PhoSL001
CML1
WT Galectin-8
CA19-9 Ab (DRG)
ChSep
MGL 416
PNA
hGal-1-488+BCR
#15 Sialyl Lewis X (BD)
HML-Native
AFL
VLRB.aGPA.23
AAL
MOA
MNA-M
VGA(EY Lab)
SJA
#14 Sialyl Lewis X (Calbio)
PTL-II
hJAA-F11-B11
AAL
GSL-I-B4
GSL-1
CSA(EY Lab)
Mouse Galectin1
AIA
Human Galectin1
ACL
LCA
DBA
Mono2R-CNL
DSL
P11 VP8
EEL
PHA-L
HHL
GNL
Mouse NKR-P1C
Threefoil
GSL-2
PSL
WGA
PHA-E
UEA-I
MAA(Seikagaku)
LCA
PHA-E
MAL-I
LTL
WGA
LEL
MPL
Gal-3C
MAA
CAA
Sp Serotype 14.2
PTL-I
Microvirin
PTL
CA19.9-1B844
BPL
Sp Serotype 14.15
CGL
UEA-I
SGA
Mono1-CNL
CA19.9-121SLE
Avaren-Fc
NPA
WT-VCC
#8 Lewis B (T218) (Abcam)
CA19.9-9L426
Galectin-1
Mono2W-CNL
Epa1-Np mutant Y228W
Epa1-Np mutant Y228A
LSL-150
HPA
WGA
EEA(EY Lab)
Native HPA
HHA(EY Lab)
PsSC
WFL
human Gal-7
RCA I (Vector)
LB-132
CC1G-10077
mouse Gal-7
CA
ECL
AOL (Mahal Lab)
rSN_VLP-Cy3
rPS-K-4A2B8
PHA-L
GAC78
VVHA lectin
PA-I
ConA
ABA
PTA-Gal
TbFBL2
CnMBL
SSL0-488
TbFBL1
SSR1
MAL II
DC-ASGPR short
LcH
WGA multimer
rSRL
CTB
SRL
ME7(detected with a-HUD)
SNA
Anti-LeC
BTL native
ME5(detected with a-HUD)

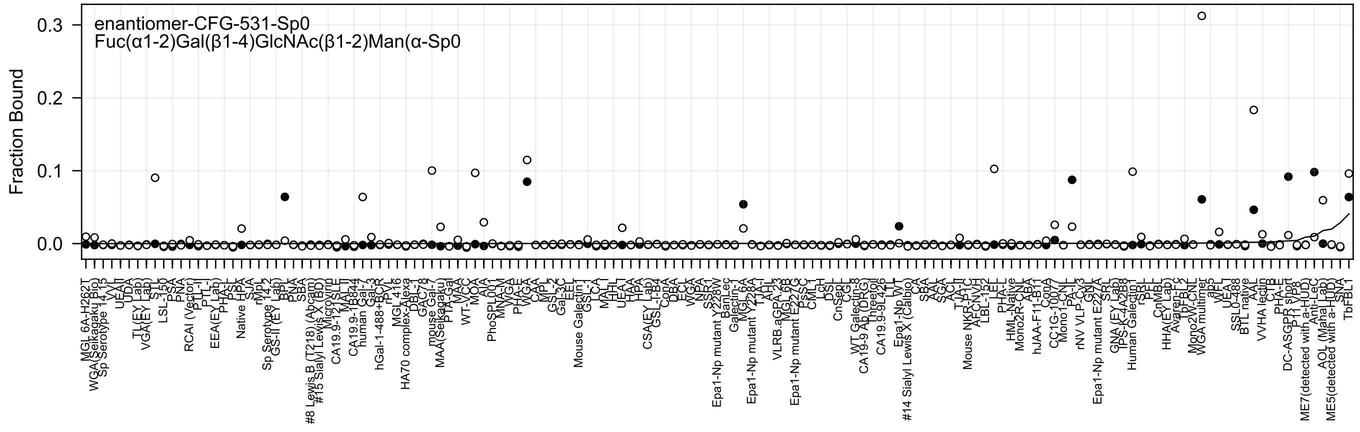

enantiomer-CFG-531-Sp0
Fuc(α1-2)Gal(β1-4)GlcNAc(β1-2)Man(α-Sp0
Fraction Bound
0.0
0.1
0.2
0.3
MGL 6A-H262T
WGA(Seikagaku Bio)
Sp Serotype 14 rS
VVL
UEA-I
UDA
TL(EY Lab)
VGA(EY Lab)
STL
LSL-150
PSA
PNA
RCAI (Vector)
PTL-II
PTL-I
EEA(EY Lab)
PHA-L
PSL
Native HPA
SJA
rMpL
Sp Serotype 14.2
GS-II (EY Lab)
BPL
PNA
SBA
#8 Lewis B (T218) (Abcam)
#15 Sialyl Lewis X (BD)
Microvirin
CA19.9-121SLE
MAL-II
CA19.9-1B844
human Gal-7
Gal-3
hGal-1-488+BCR
rPVL
MGL 416
HA70 complex-Alexa
DBL-1
GAC7-8
mouse Gal-7
MAA(Seikagaku)
PTA-Gal
MAA
WT-VCC
MOA
AIA
PhoSPL001
MNA-M
WGA
PHA-E
WGA
CAA
MPL
GSL-2
Gal-3C
EEL
Mouse Galectin1
GSL-1
LCA
MAL-I
HHL
UEA-I
HAA
HPA
CSA(EY Lab)
GSL-I-B4
ConA
DBA
ECL
WGA
NPA
SSR1
Epa1-Np mutant Y228W
BanLec
Galectin-1
MGL 6A
Epa1-Np mutant Y228A
TJA-I
AAL
VLRB.aGPA.23
MGL 6B
Epa1-Np mutant E227G
PsSC
CML-I
LcH
DSL
CnSep
CGL
WT Galectin-8
CA19-9 Ab (DRG)
Lineolol
CA19.9-9L426
LTL
Epa1-Np WT
#14 Sialyl Lewis X (Calbio)
Cx
SBA
AAL
SGA
ACL
TJA-II
Mouse NKR-P1C
AFCNvH
LBL-152
LEL
PHA-L
HML-Native
Mono2R-CNL
ABA
hJAA-F11-B11
ConA
CC1G-10077
Mono1-CNL
PA-IL
rNV VLP-Cy3
GNL
Epa1-Np mutant E227A
SRL
GNA (EY Lab)
TPS-K-4A2B8
Human Galectin1
rSRL
PFL
CnMBL
HHA(EY Lab)
Avaren-Fc
TbFBL-2
Mono2W-CNL
WGA multimer
abs
WFL
UEA-I
SSL0-488
BTL native
AAL
VVHA lectin
CTB
PHA-E
DC-ASGPR short
P11 VP8
ME7(detected with a-HUD)
Anti-LeC
AOL (Mahal Lab)
ME5(detected with a-HUD)
SNA
TbFBL1

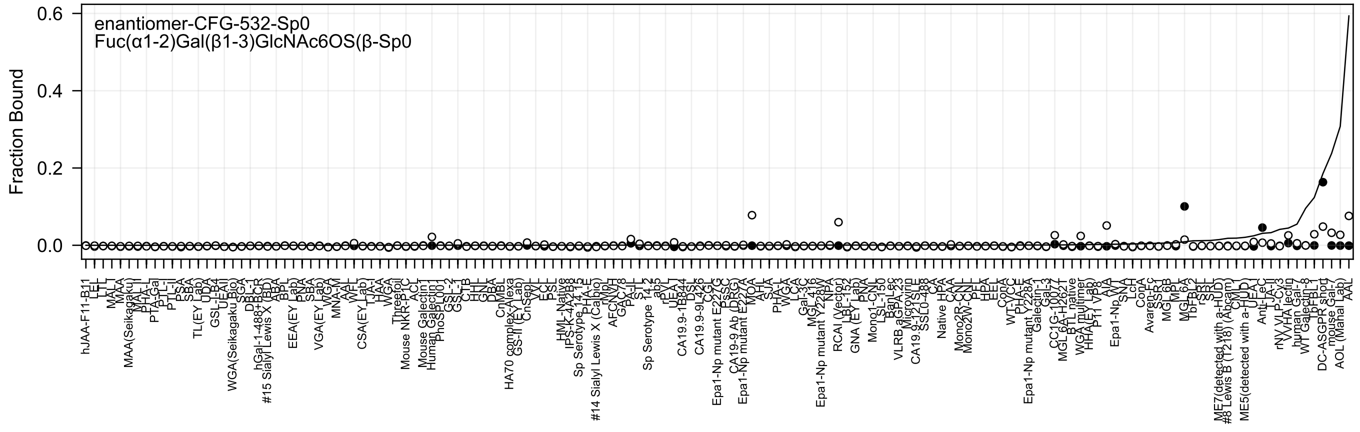
enantiomer-CFG-532-Sp0
Fuc(α1-2)Gal(β1-3)GlcNAc6OS(β-Sp0
Fraction Bound
0.6
0.4
0.2
0.0
hJAA-F11-B11
LEL
LTL
MAL-I
MAA
MAA(Seikagaku)
MAL-II
PHA-L
PTA-Gal
PTL-I
PTL-II
PSA
SBA
TL(EY Lab)
UDA
GSL-I-B4
UEA-II
WGA(Seikagaku Bio)
SGA
DBL-1
hGal-1-488+BCP
#15 Sialyl Lewis X (BD)
ABA
BPL
EEA(EY Lab)
PNA
SBA
VGA(EY Lab)
WGA
MNA-M
AAL
WFL
CSA(EY Lab)
TJA-I
HAA
WGA
Threefoil
Mouse NKR-P1C
ACL
Mouse Galectin1
Human Galectin1
PhoSL001
GSL-2
GSL-1
CTB
HHL
GNL
DBA
CnMBL
HA70 complex-Alexa
GS-II (EY Lab)
CnSep
VVL
ECL
PSL
HML-Native
IPS-K-4A2B8
Sp Serotype 14.15
PHA-E
#14 Sialyl Lewis X (Calbio)
rMpL
AFCNVH
GAC78
PA-IL
STL
Sp Serotype 14.2
ab5
rPVL
UEA-I
CA19.9-1B844
DSL
CA19.9-9L426
CGL
Epa1-Np mutant E227G
DsSC
CA19-9 Ab (DRG)
Epa1-Np mutant E227A
MOA
AAHL
SJA
PHA-L
WGA
LCA
Gal-3C
MGL-416
Epa1-Np mutant Y228W
NPA
RCAI (Vector)
LBL-152
GNA (EY Lab)
PNA
Mono1-CNL
LSL-150
BanLec
VLRB.aGPA.23
Microvirin
CA19.9-121SLE
SSL0-488
CA
Native HPA
CAA
Mono2R-CNL
Mono2W-CNL
PEL
HPA
EEL
ConA
WT-YCC
PHA-E
Epa1-Np mutant Y228A
Galectin-1
Gal-3
CC1G-10077
MGL6A-H262T
BTL native
WGA multimer
HHA(EY Lab)
P11 VP8
AIA
Epa1-Np WT
SNA
LcH
ConA
Avaren-Fc
SSR1
MGL6B
MPL
MGL6A
TbFBL2
rSRL
SRL
ME7(detected with a-HUD)
#8 Lewis B (T218) (Abcam)
CVN
ME5(detected with a-HUD)
UEA-I
Anti-LeC
TJA-II
rNV VLP-Cy3
VVHA lectin
human Gal-7
WT Galectin-8
TbFBL1
DC-ASGPR short
mouse Gal-7
AOL (Manal Lab)
AAL

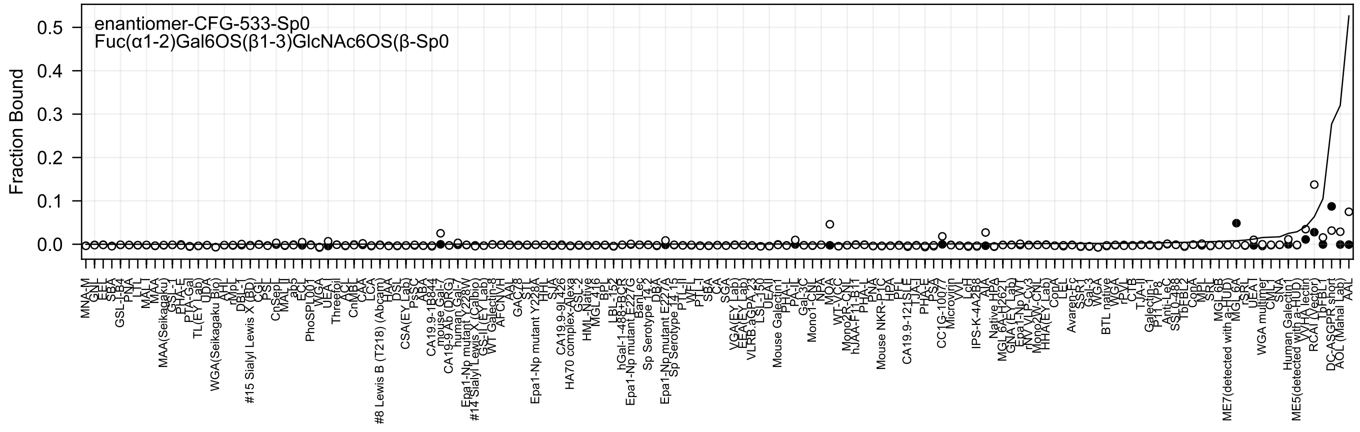

enantiomer-CFG-533-Sp0
Fuc(α1-2)Gal6OS(β1-3)GlcNAc6OS(β-Sp0
Fraction Bound

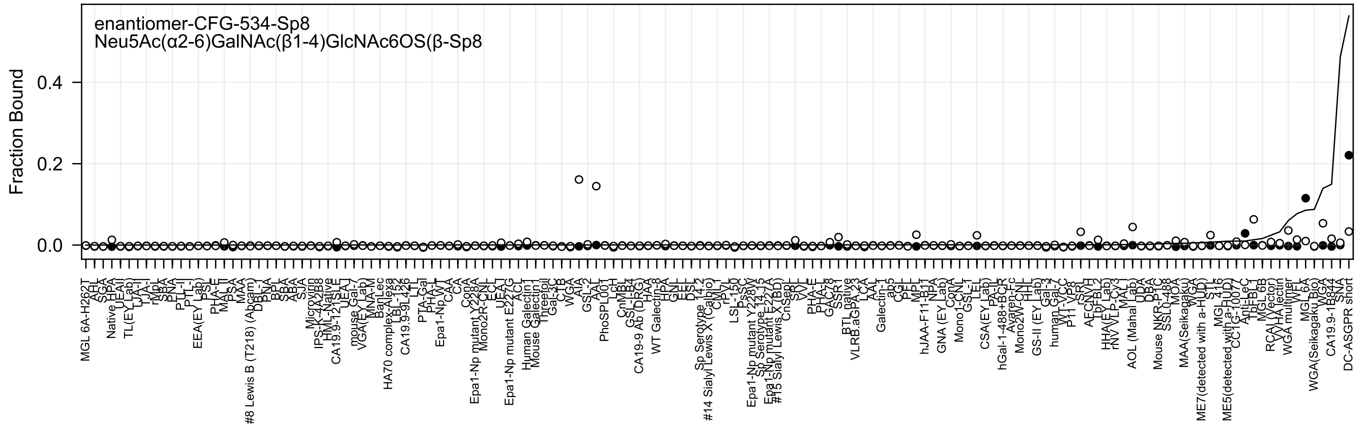
enantiomer-CFG-534-Sp8
Neu5Ac(α2-6)GalNAc(β1-4)GlcNAc6OS(β-Sp8
Fraction Bound
0.4
0.2
0.0

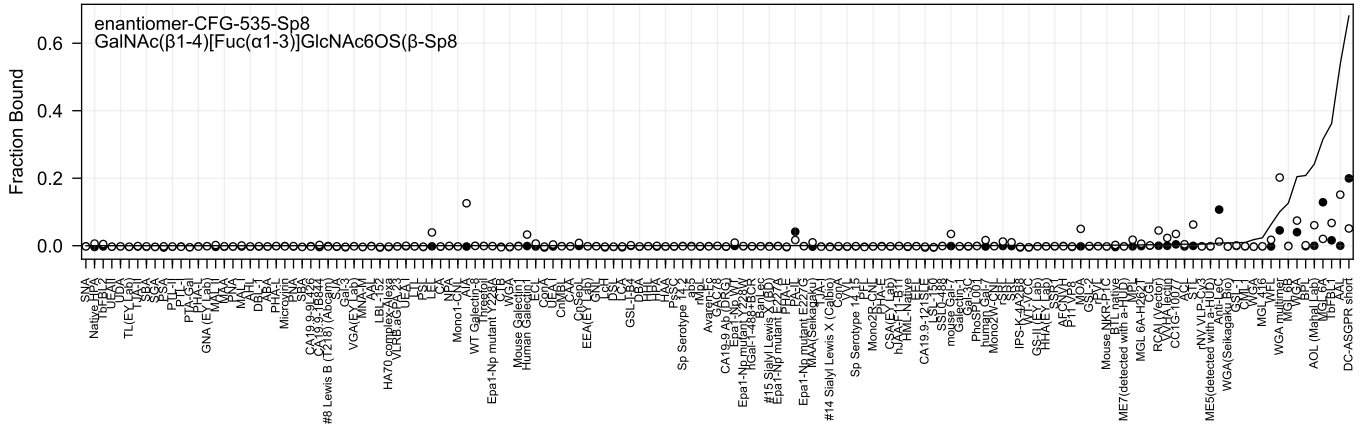
enantiomer-CFG-535-Sp8
GalNAc(β1-4)[Fuc(α1-3)]GlcNAc6OS(β-Sp8
Fraction Bound
0.6
0.4
0.2
0.0

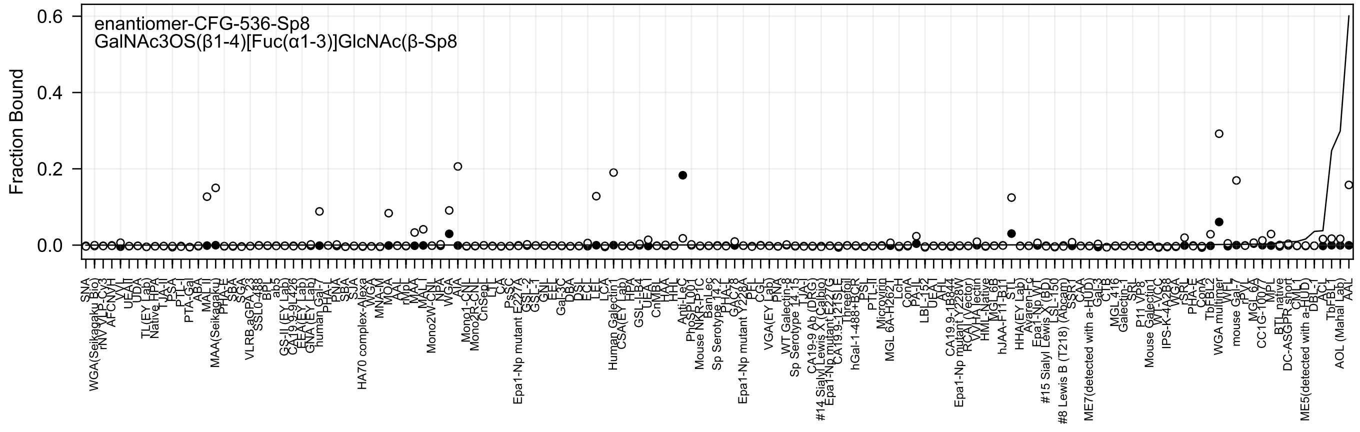
enantiomer-CFG-536-Sp8
GalNAc3OS(β1-4)[Fuc(α1-3)]GlcNAc(β-Sp8
Fraction Bound

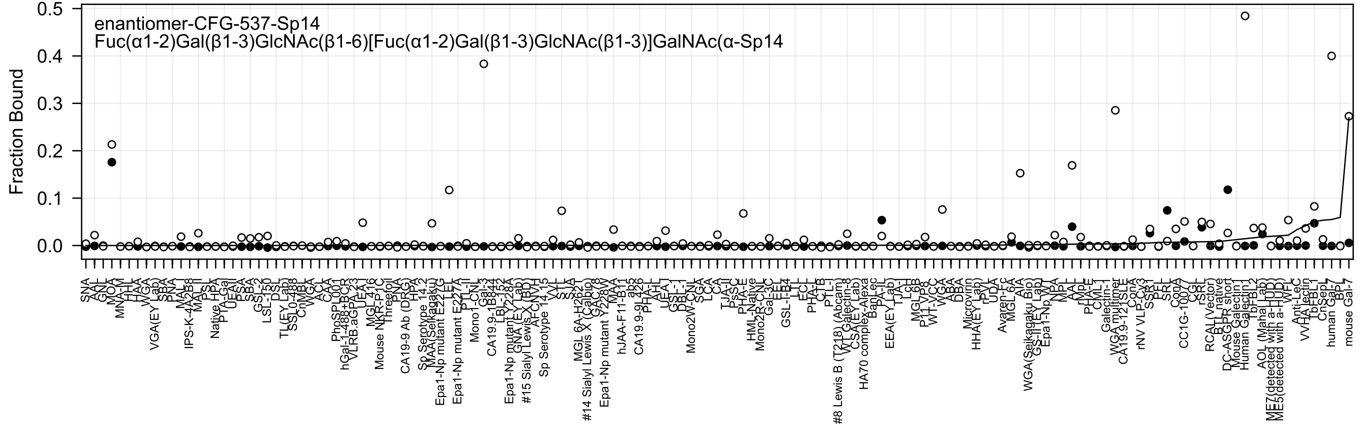

enantiomer-CFG-537-Sp14
Fuc(α1-2)Gal(β1-3)GlcNAc(β1-6)[Fuc(α1-2)Gal(β1-3)GlcNAc(β1-3)]GalNAc(α-Sp14
Fraction Bound

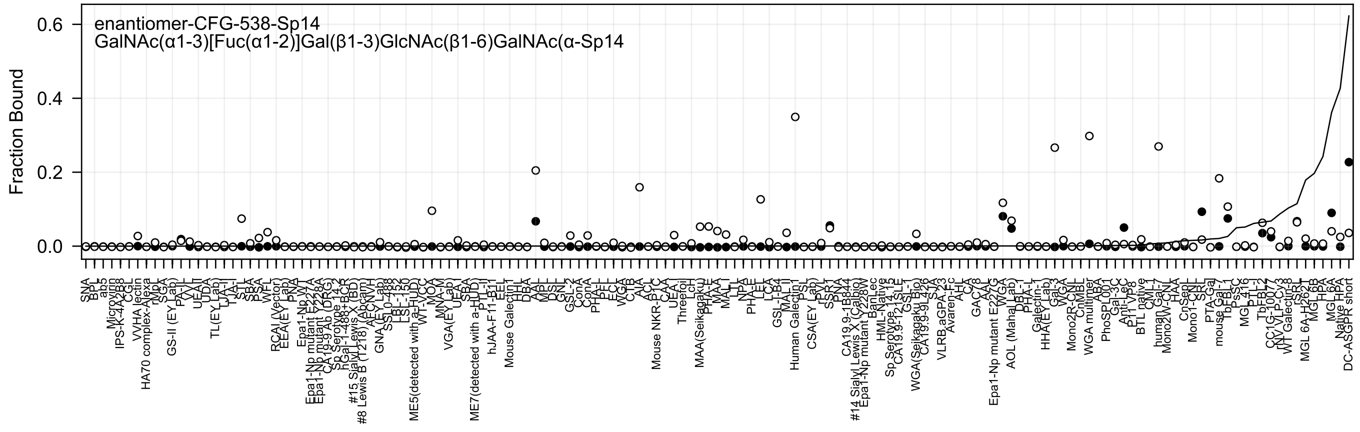
enantiomer-CFG-538-Sp14
GalNAc(α1-3)[Fuc(α1-2)]Gal(β1-3)GlcNAc(β1-6)GalNAc(α-Sp14
Fraction Bound
0.0
0.2
0.4
0.6
SNA
BPL
ab5
Microvirin
IPS-K-4A2B8
CGL
VVHA lectin
HA70 complex-Alexa
SGA
GS-II (EY Lab)
PA-II
VVA
UEA-II
UDA
TL(EY Lab)
TJA-II
TJA-I
STL
SBA
PSA
WFL
RCAI (Vector)
EEA(EY Lab)
PNA
Epa1-Np WT
Epa1-Np mutant E227A
Epa1-Np mutant Y228A
CA19-9 Ab (DRG)
Sp Serotype 14.2
hGal-1-488+BCR
#15 Sialyl Lewis X (BD)
#8 Lewis B (T218) (Abcam)
AECNVH
GNA(EY Lab)
SSL0-488
LSL-152
LSL-150
ME5(detected with a-HUD)
WT-VCC
MOA
MNA-M
VGA(EY Lab)
UEA-I
SBA
ME7(detected with a-HUD)
PTL-II
hJAA-F11-B11
EEL
Mouse Galectin1
HPL
DBA
AAL
MPL
DSL
GNL
GSL-2
ConA
ConA
PHA-L
PFL
ECL
WGA
CTB
AIA
ACL
Mouse NKR-P1C
CAA
UEA-I
Threefoil
LcH
MAA(Seikagaku)
PHA-E
MAA
MAL-I
LTL
NPA
PHA-E
LEL
LCA
GSL-I-B4
MAL-II
Human Galectin1
PSL
CSA(EY Lab)
rPVL
SSR1
PNA
CA19.9-1B844
#14 Sialyl Lewis X (Calbio)
Epa1-Np mutant Y228W
BanLec
HML-Native
Sp Serotype 14.15
CA19.9-121SLE
GSL-I
WGA(Seikagaku Bio)
CA19.9-9L426
SJA
VLRB.aGPA.23
Avaren-Fc
AHL
CA
GAC78
AAL
WGA
Epa1-Np mutant E227G
AOL (Mahal Lab)
DBL-1
PHA-L
Galectin-1
HHA(EY Lab)
Gal-3
WGA
Mono2R-CNL
CnMBL
WGA multimer
ABA
PhoSL001
Gal-3C
Anti-LeC
P11 VP8
BTL native
CML1
human Gal-7
Mono2W-CNL
HAA
CnSeP
Mono1-CNL
SRL
PTA-Gal
mouse Gal-1
TbFBL1
PsSC
MGL 416
PTL-I
TbFBL2
CC1G-10077
rNV VLP-Cy3
WT Galectin-8
rSRL
MGL 6A-H262
MGL 6B
HPA
MGL 6A
Native HPA
DC-ASGPR short

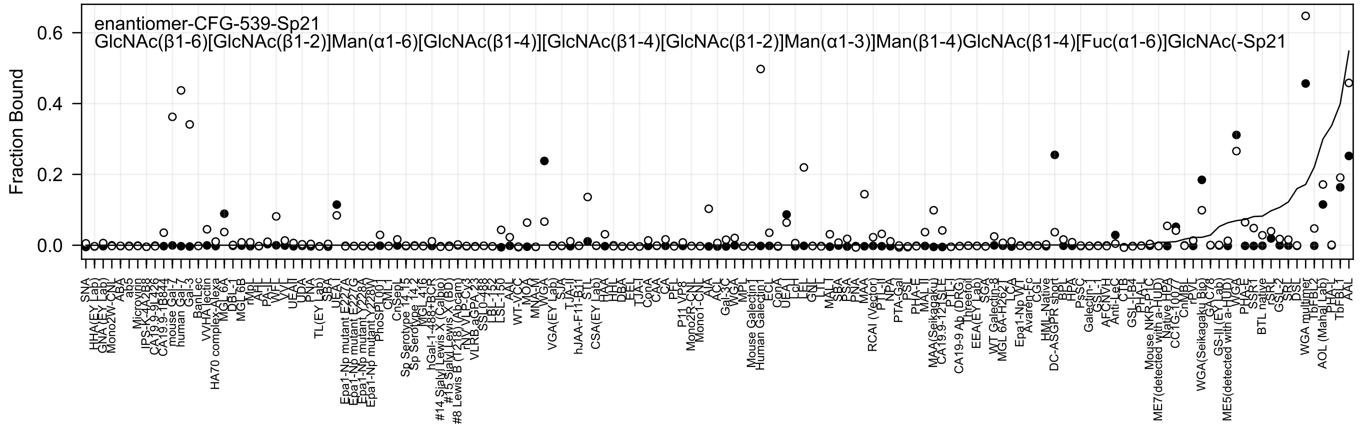
enantiomer-CFG-539-Sp21
GlcNAc(β1-6)[GlcNAc(β1-2)]Man(α1-6)[GlcNAc(β1-4)][GlcNAc(β1-4)[GlcNAc(β1-2)]Man(α1-3)]Man(β1-4)GlcNAc(β1-4)[Fuc(α1-6)]GlcNAc(-Sp21
Fraction Bound
0.0
0.2
0.4
0.6

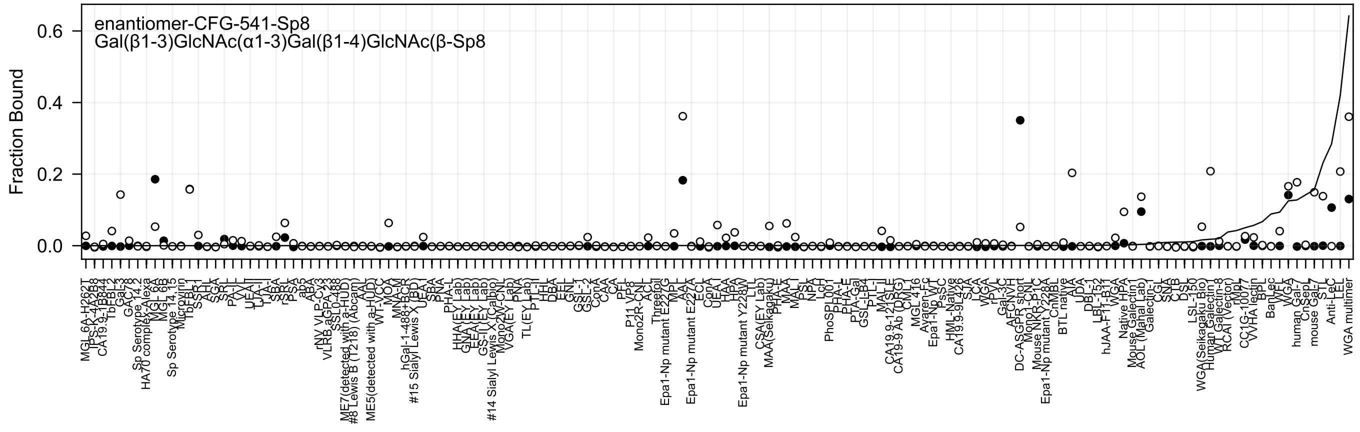
enantiomer-CFG-541-Sp8
Gal(β1-3)GlcNAc(α1-3)Gal(β1-4)GlcNAc(β-Sp8
Fraction Bound
0.0
0.2
0.4
0.6

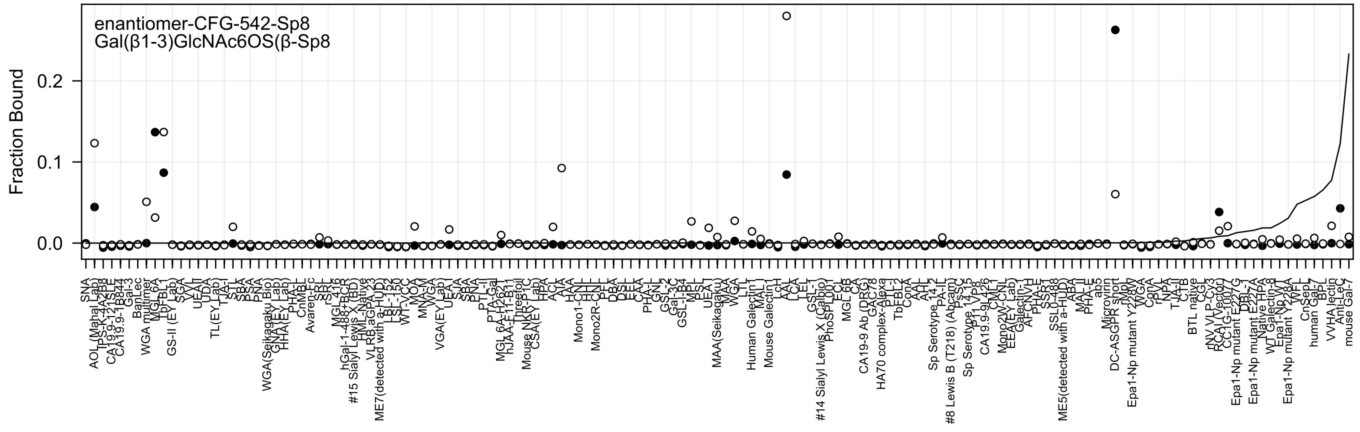
enantiomer-CFG-542-Sp8
Gal(β1-3)GlcNAc6OS(β-Sp8
Fraction Bound
0.0
0.1
0.2

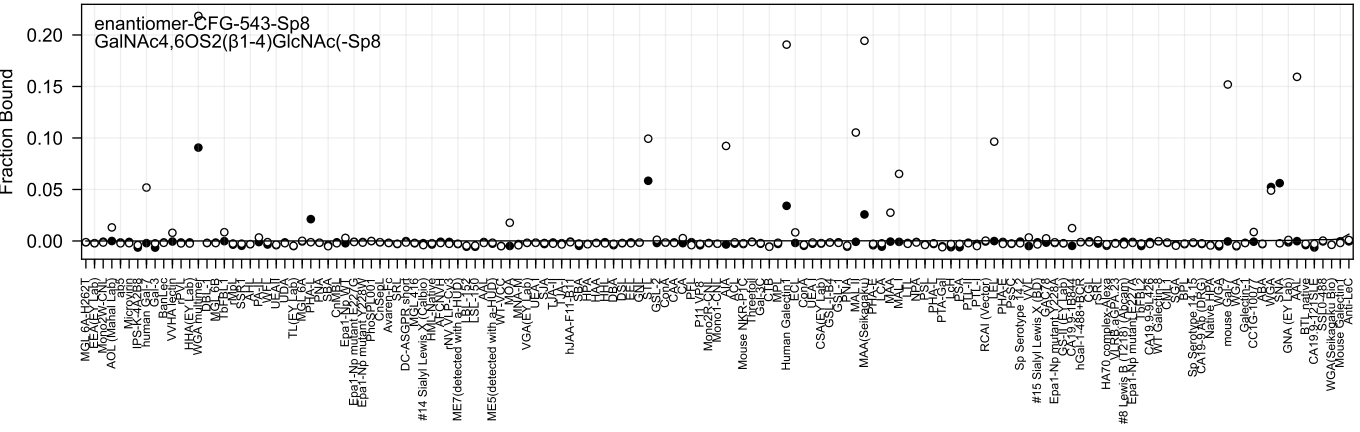
enantiomer-CFG-543-Sp8
GalNAc4,6OS2(β1-4)GlcNAc(-Sp8
Fraction Bound
0.00
0.05
0.10
0.15
0.20

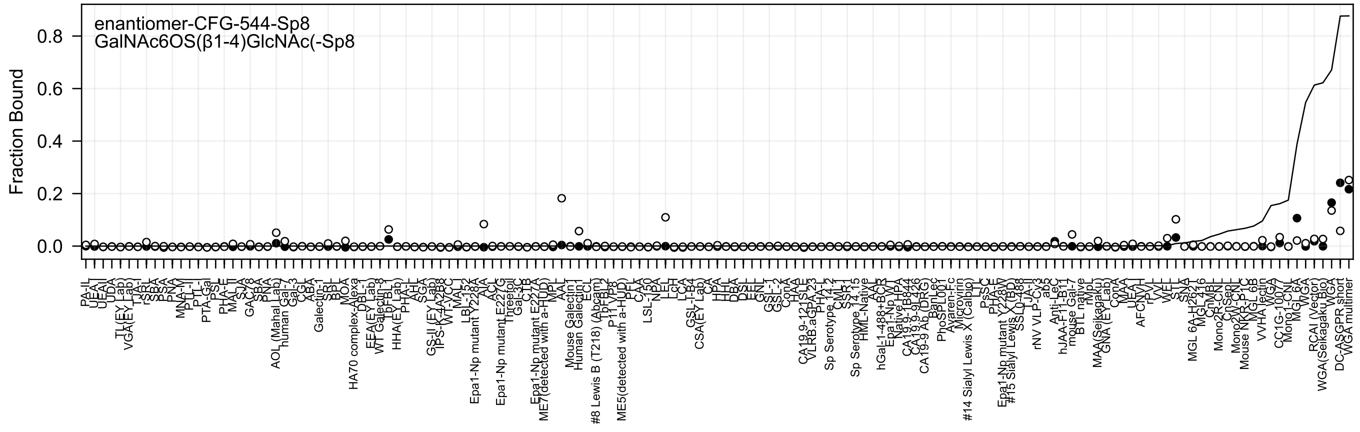

enantiomer-CFG-544-Sp8
GalNAc6OS(β1-4)GlcNAc(-Sp8
Fraction Bound
0.0
0.2
0.4
0.6
0.8

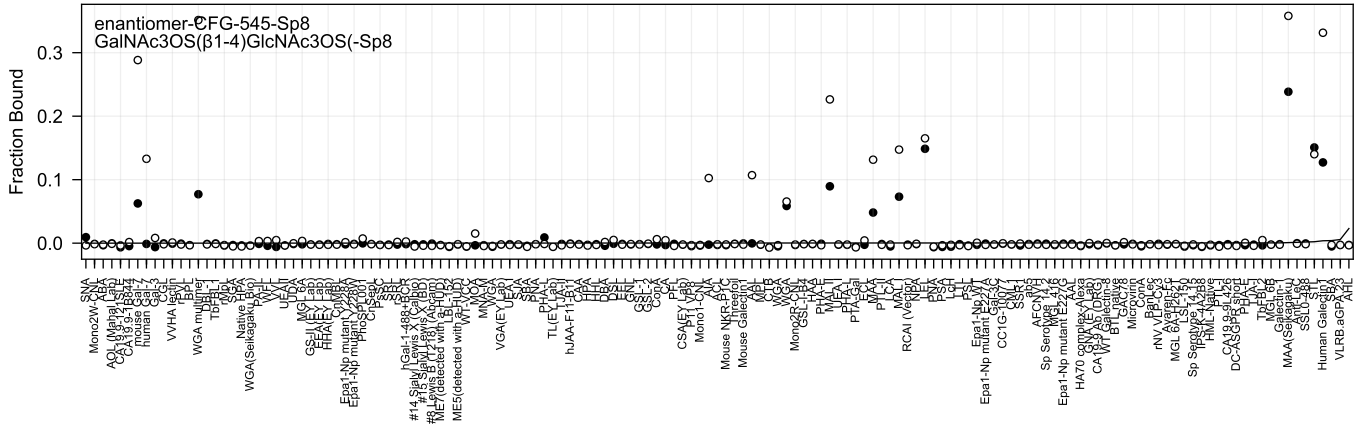

enantiomer-CFG-545-Sp8
GalNAc3OS(β1-4)GlcNAc3OS(-Sp8
Fraction Bound
0.0
0.1
0.2
0.3

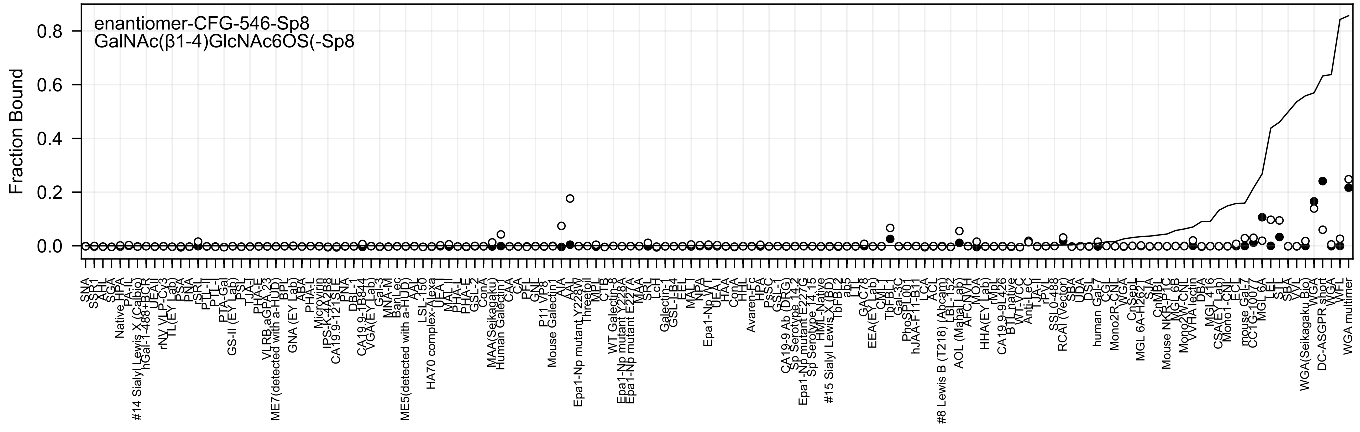

enantiomer-CFG-546-Sp8
GalNAc(β1-4)GlcNAc6OS(-Sp8
Fraction Bound

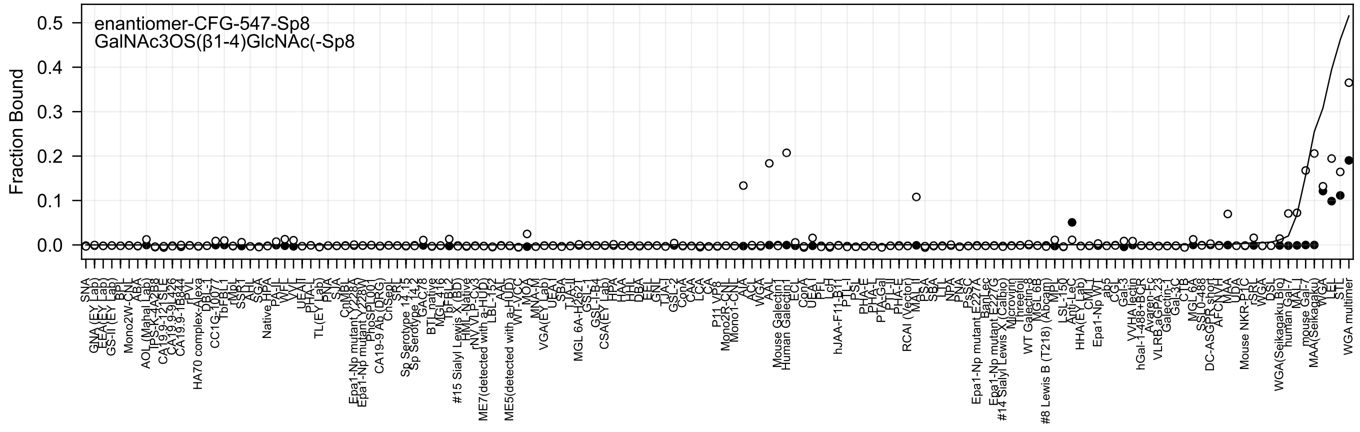
enantiomer-CFG-547-Sp8
GalNAc3OS(β1-4)GlcNAc(-Sp8
Fraction Bound
0.5
0.4
0.3
0.2
0.1
0.0
SNA
GNA (EY Lab)
EEA (EY Lab)
GS-II (EY Lab)
BPL
Mono2W-CNL
ABA
AOL (Mahal Lab)
TPS-K-4A2B8
CA19.9-121SPE
CA19.9-9L426
CA19.9-1B844
rPVL
HA70 complex-Alexa
DBL-2
CC1G-10077
TbFBL1
rMpL
SSB1
AAL
SGA
Native HPA
PA-IL
WFL
VVL
UEA-II
PHA-L
TL(EY Lab)
PNA
SJA
CnMBL
Epa1-Np mutant Y228A
Epa1-Np mutant Y228W
PhoSL001
CA19-9 Ab (DRG)
CnSeql
SRL
Sp Serotype 14.13
Sp Serotype 14.3
GAC78
BTL native
MGL 416
TbFBL2
#15 Sialyl Lewis X (BD)
HML-Native
rNV VLP-Cy3
ME7(detected with a-HUD)
LBL-152
AAL
ME5(detected with a-HUD)
WT-VCC
MOA
MNA-M
VGA(EY Lab)
UEA-I
SBA
TJA-II
MGL 6A-H262T
GSL-1
GSL-I-B4
CSA(EY Lab)
HPA
HAA
HHL
DBA
EEL
GNL
TJA-I
GSL-2
ConA
CAA
LCA
CA
P11 VP8
Mono2R-CNL
Mono1-CNL
AIA
ACL
WGA
AAL
Mouse Galectin1
Human Galectin1
ECL
ConA
UEA-I
PFL
LTL
LcH
hJAA-F11-B11
PTL-I
PSL
PHA-E
PHA-L
PTA-Gal
PTL-II
PHA-E
RCAI (Vector)
MAL-I
PSA
SBA
LTL
NPA
PNA
PsSC
Epa1-Np mutant E227A
BanLec
Epa1-Np mutant E227G
#14 Sialyl Lewis X (Calbio)
Microvirin
Threefoil
WT Galectin-8
MGL 6B
#8 Lewis B (T218) (Abcam)
MPL
LSL-150
Anti-LeC
HHA(EY Lab)
CML-1
Epa1-Np WT
ab5
CGL
Gal-3
VVHA lectin
hGal-1-488+BCR
Avaren-Fc
VLRB aGPA.23
Galectin-1
Gal-3C
CTB
MGL 6A
SSL0-488
DC-ASGPR short
AFCN-H
MAA
UDA
Mouse NKR-P1C
rSRL
WGA
DSL
STL
WGA(Seikagaku Bio)
human Gal-7
MAL-I
mouse Gal-7
MAA(Seikagaku)
WGA
WFL
LEL
STL
WGA multimer

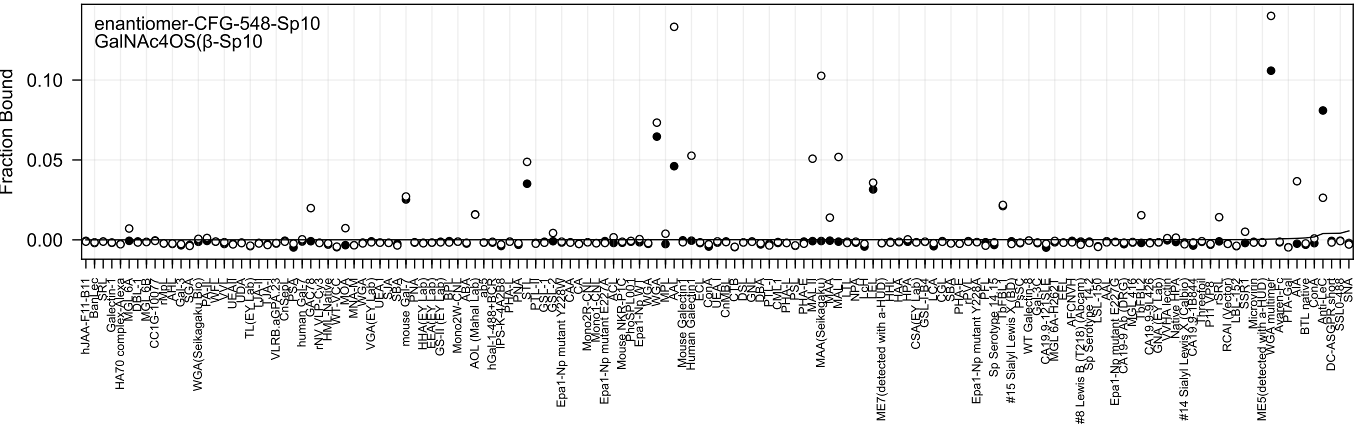
enantiomer-CFG-548-Sp10
GalNAc4OS(β-Sp10
Fraction Bound
0.10
0.05
0.00

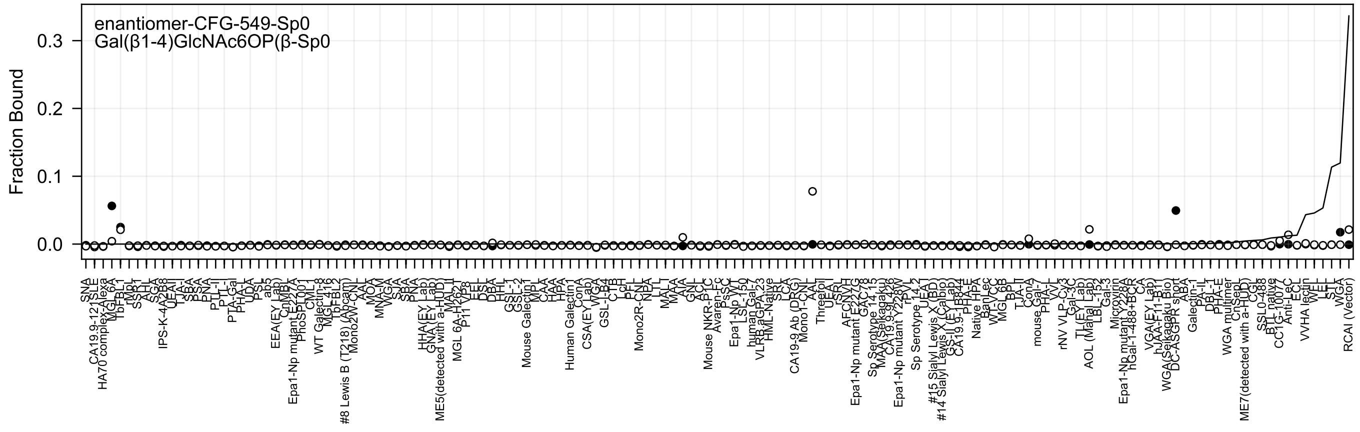
enantiomer-CFG-549-Sp0
Gal(β1-4)GlcNAc6OP(β-Sp0
Fraction Bound
0.3
0.2
0.1
0.0

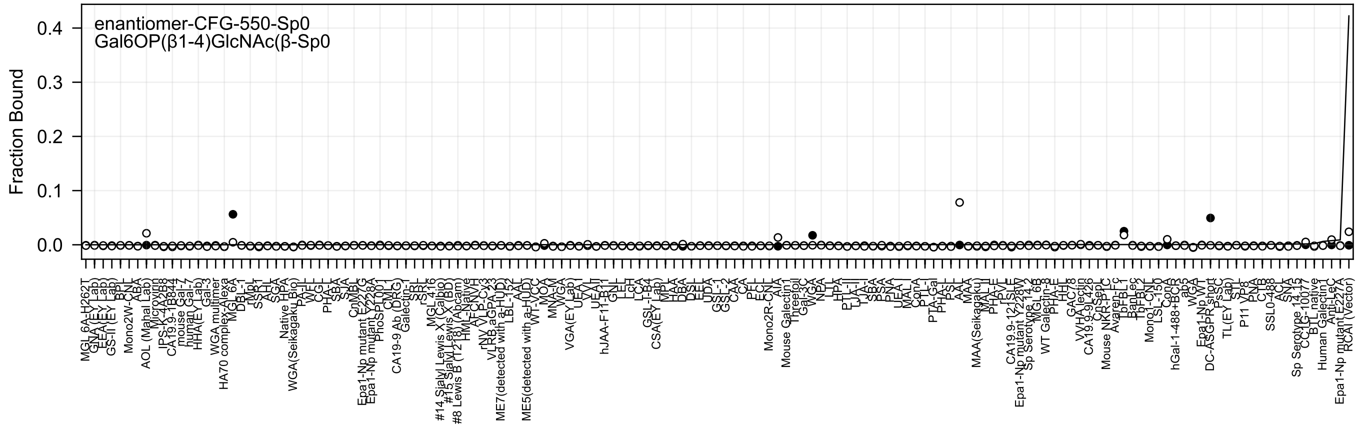
enantiomer-CFG-550-Sp0
Gal6OP(β1-4)GlcNAc(β-Sp0
Fraction Bound

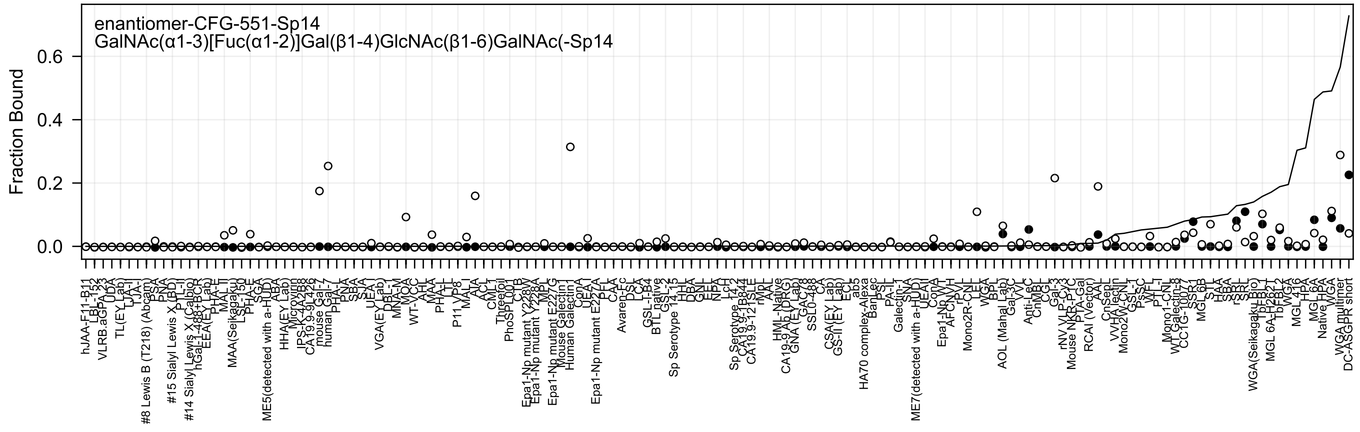
enantiomer-CFG-551-Sp14
GalNAc(α1-3)[Fuc(α1-2)]Gal(β1-4)GlcNAc(β1-6)GalNAc(-Sp14
Fraction Bound
0.6
0.4
0.2
0.0

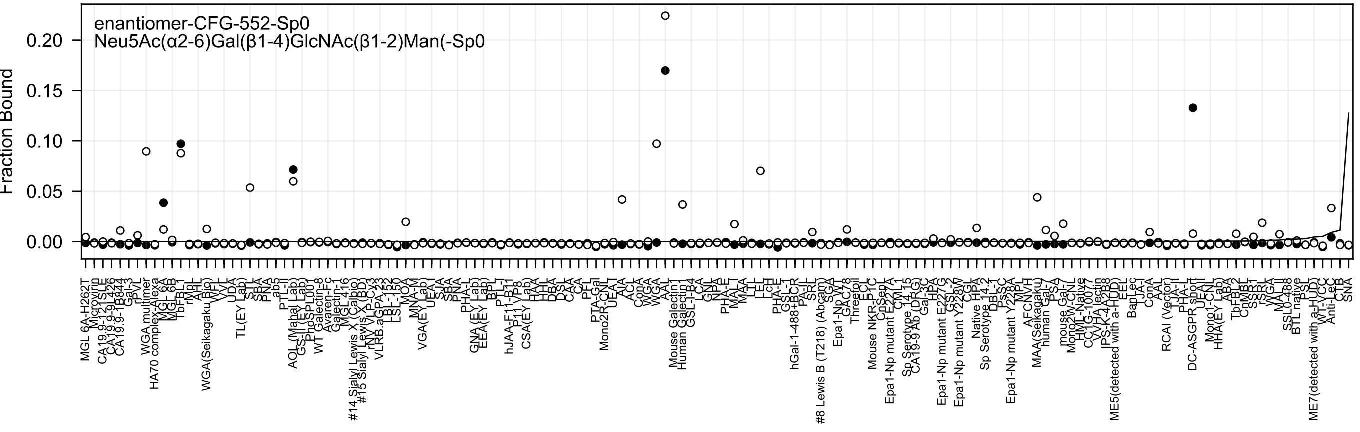

enantiomer-CFG-552-Sp0
Neu5Ac(α2-6)Gal(β1-4)GlcNAc(β1-2)Man(-Sp0
Fraction Bound

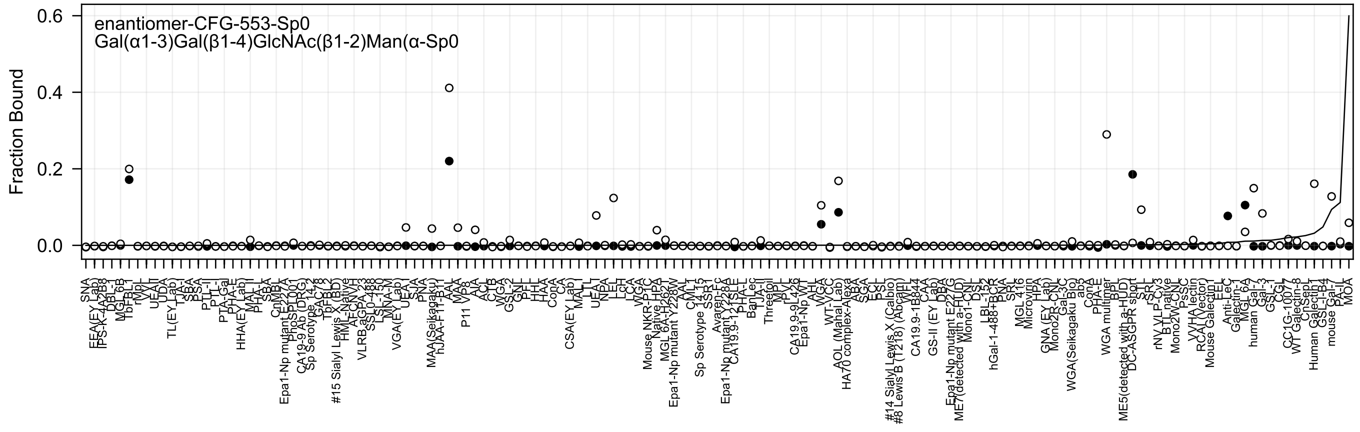
enantiomer-CFG-553-Sp0
Gal(α1-3)Gal(β1-4)GlcNAc(β1-2)Man(α-Sp0
Fraction Bound
0.6
0.4
0.2
0.0

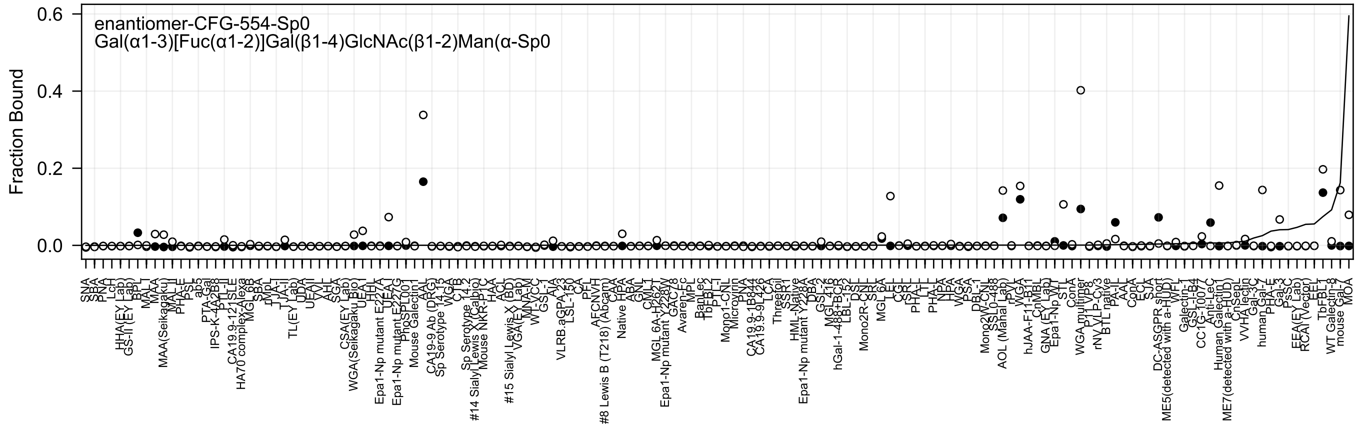
enantiomer-CFG-554-Sp0
Gal(α1-3)[Fuc(α1-2)]Gal(β1-4)GlcNAc(β1-2)Man(α-Sp0
Fraction Bound
0.6
0.4
0.2
0.0

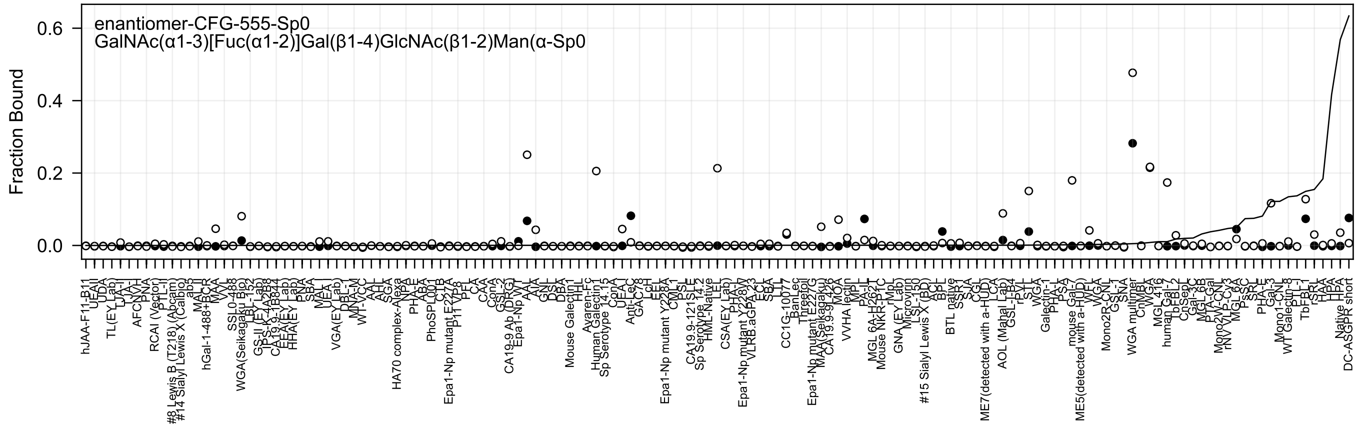
enantiomer-CFG-555-Sp0
GalNAc(α1-3)[Fuc(α1-2)]Gal(β1-4)GlcNAc(β1-2)Man(α-Sp0
Fraction Bound
0.0
0.2
0.4
0.6

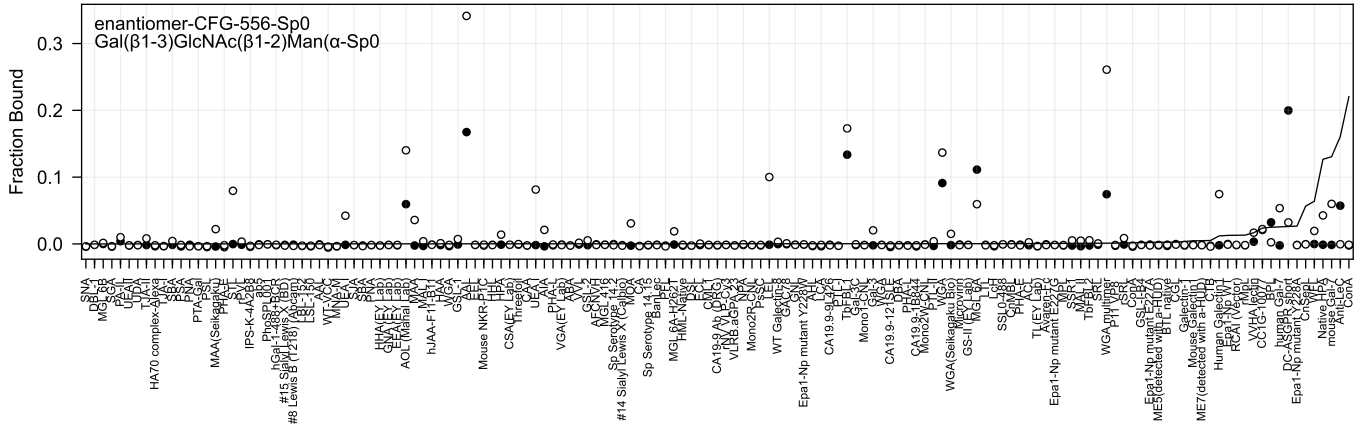
enantiomer-CFG-556-Sp0
Gal(β1-3)GlcNAc(β1-2)Man(α-Sp0
Fraction Bound
0.3
0.2
0.1
0.0

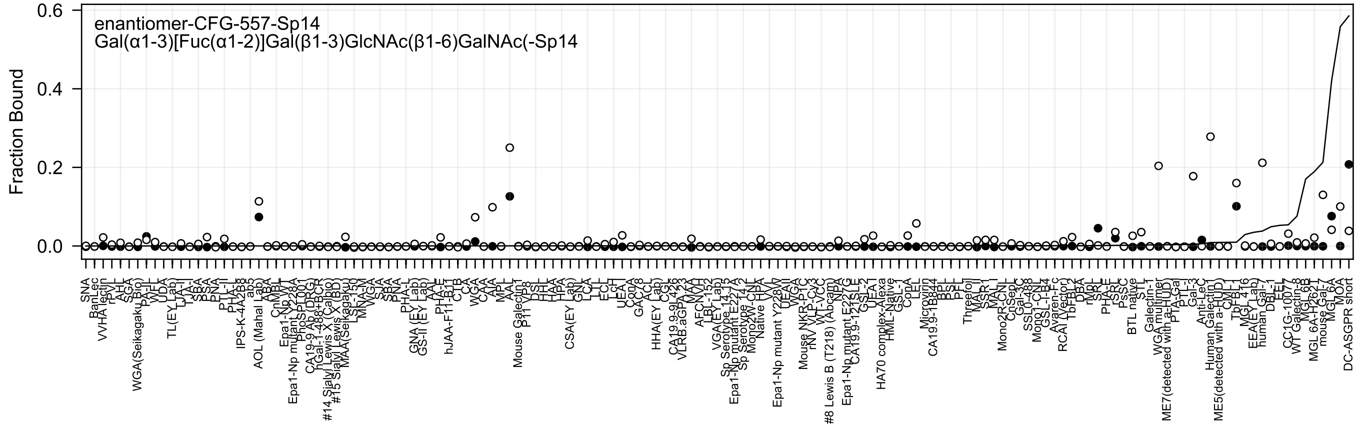

enantiomer-CFG-557-Sp14
Gal(α1-3)[Fuc(α1-2)]Gal(β1-3)GlcNAc(β1-6)GalNAc(-Sp14
Fraction Bound
0.0
0.2
0.4
0.6

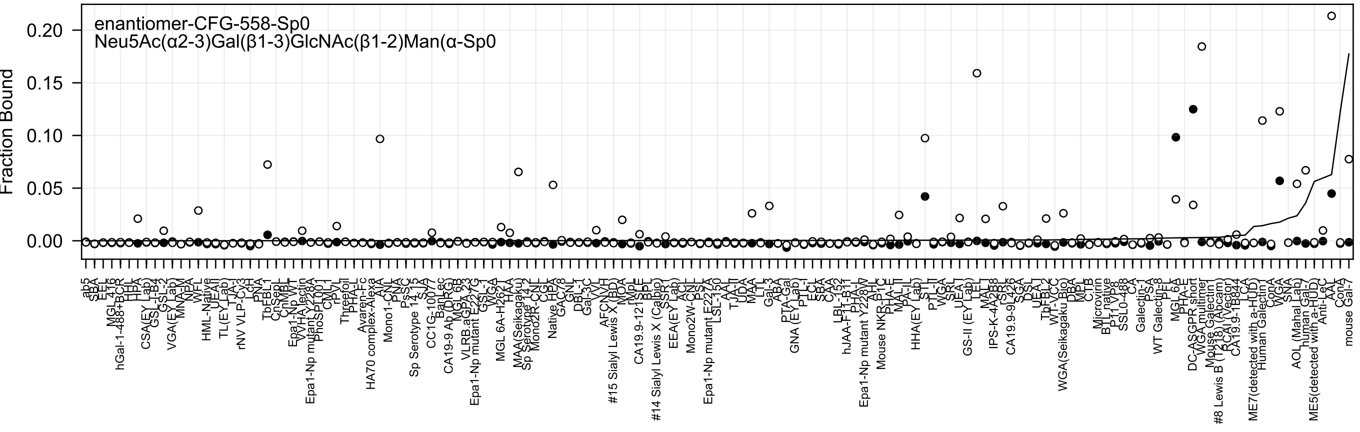
enantiomer-CFG-558-Sp0
Neu5Ac(α2-3)Gal(β1-3)GlcNAc(β1-2)Man(α-Sp0
Fraction Bound
0.20
0.15
0.10
0.05
0.00

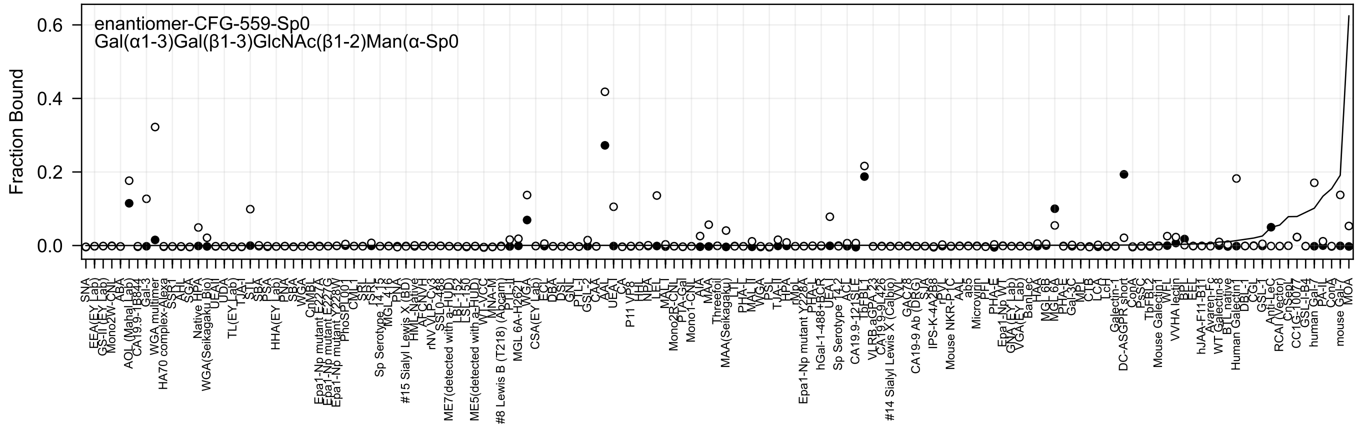
enantiomer-CFG-559-Sp0
Gal(α1-3)Gal(β1-3)GlcNAc(β1-2)Man(α-Sp0
Fraction Bound
0.0
0.2
0.4
0.6

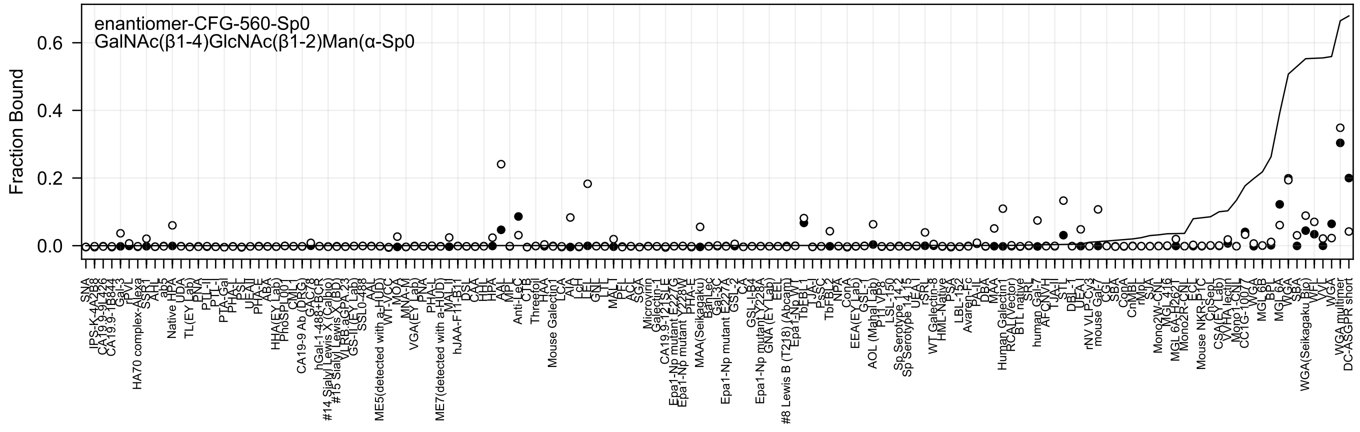

enantiomer-CFG-560-Sp0
GalNAc(β1-4)GlcNAc(β1-2)Man(α-Sp0
Fraction Bound
0.0
0.2
0.4
0.6

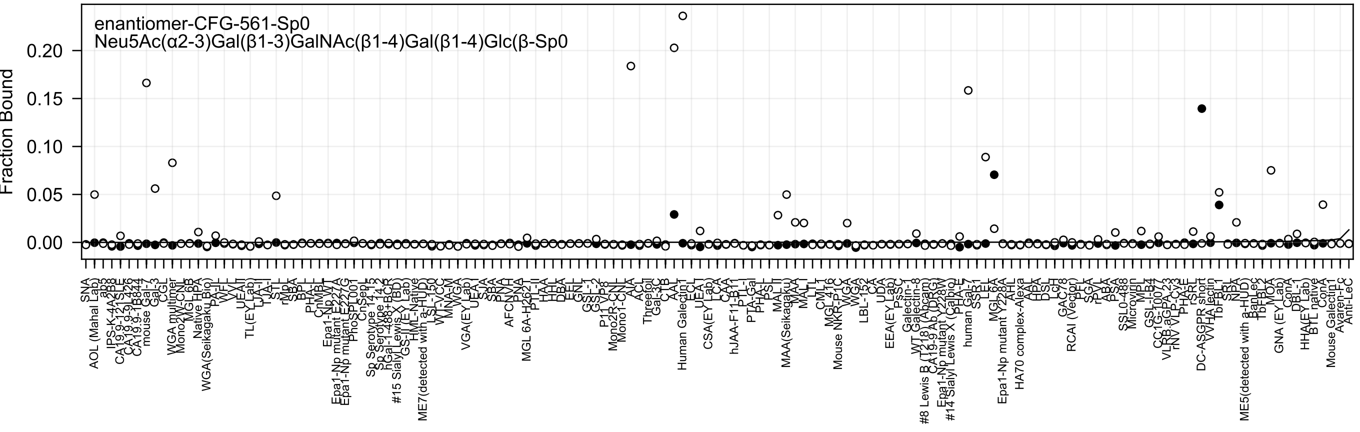
enantiomer-CFG-561-Sp0
Neu5Ac(α2-3)Gal(β1-3)GalNAc(β1-4)Gal(β1-4)Glc(β-Sp0
Fraction Bound
0.20
0.15
0.10
0.05
0.00

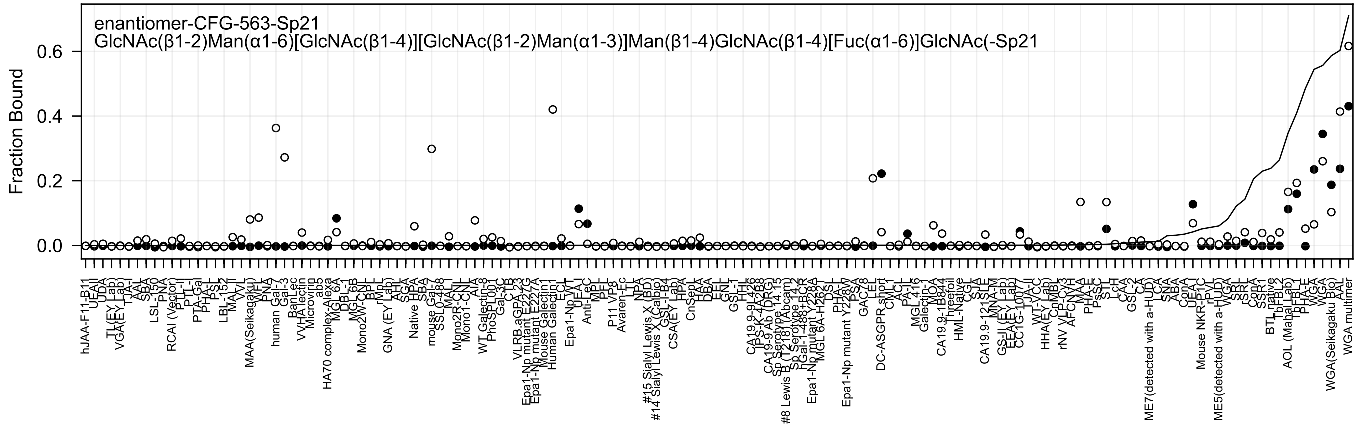

enantiomer-CFG-563-Sp21
GlcNAc(β1-2)Man(α1-6)[GlcNAc(β1-4)][GlcNAc(β1-2)Man(α1-3)]Man(β1-4)GlcNAc(β1-4)[Fuc(α1-6)]GlcNAc(-Sp21
Fraction Bound
0.0
0.2
0.4
0.6

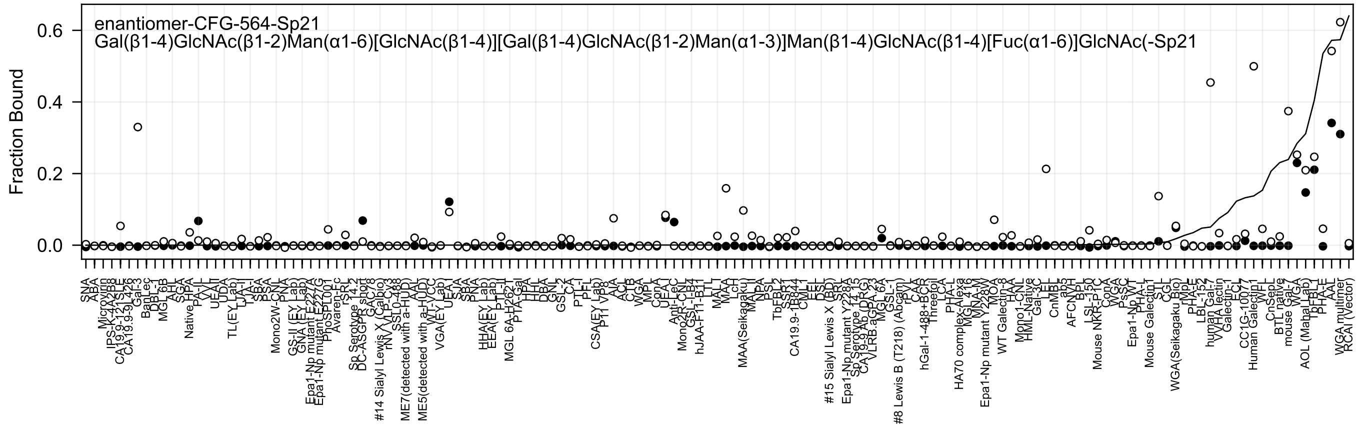
enantiomer-CFG-564-Sp21
Gal(β1-4)GlcNAc(β1-2)Man(α1-6)[GlcNAc(β1-4)][Gal(β1-4)GlcNAc(β1-2)Man(α1-3)]Man(β1-4)GlcNAc(β1-4)[Fuc(α1-6)]GlcNAc(-Sp21
Fraction Bound
0.6
0.4
0.2
0.0

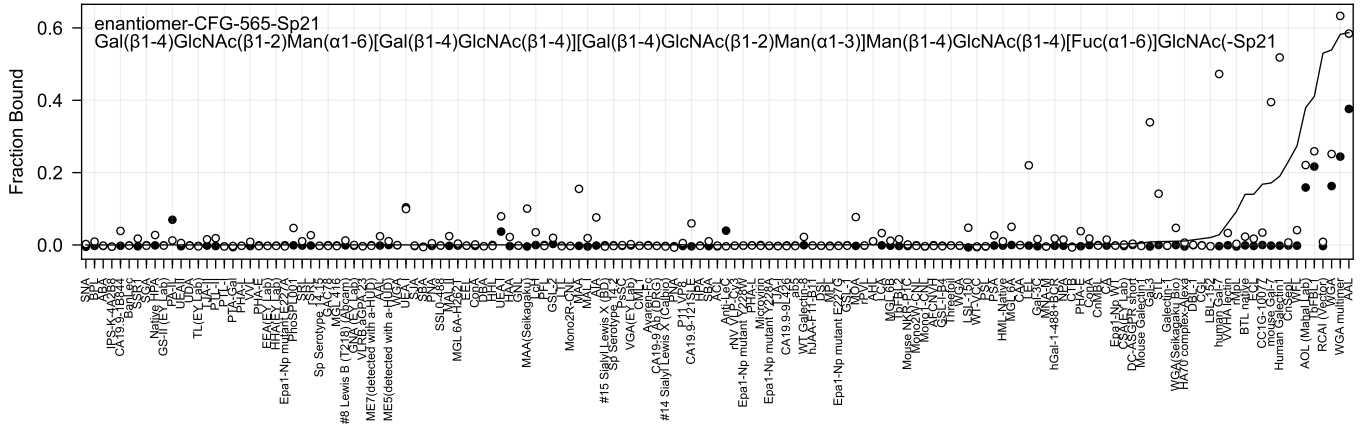
enantiomer-CFG-565-Sp21
Gal(β1-4)GlcNAc(β1-2)Man(α1-6)[Gal(β1-4)GlcNAc(β1-4)][Gal(β1-4)GlcNAc(β1-2)Man(α1-3)]Man(β1-4)GlcNAc(β1-4)[Fuc(α1-6)]GlcNAc(-Sp21
Fraction Bound
0.0
0.2
0.4
0.6
SNA
BPL
ABA
IPS-K-4A2B8
CA19.9-1B844
BanLec
SSR1
SGA
Native HPA
GS-II (EY Lab)
PA-II
UEAII
UDA
TL(EY Lab)
LTA-I
PLL-II
PTL-I
PTA-Gal
PHA-L
VVL
PHA-E
EEA(EY Lab)
HHA(EY Lab)
Epa1-Np mutant E227A
PhoSL001
SRL
Sp Serotype 14-15
GAC78
MGL 416
#8 Lewis B (T218) (Abcam)
GNA (EY Lab)
VLRB aGPA23
AAL
ME7(detected with a-HUD)
ME5(detected with a-HUD)
WGA
UEA-I
SJA
SBA
PNA
SSL0-488
MAL-II
MGL 6A-H262T
EEL
ConA
DBA
HHL
UEA-I
HAA
GNL
MAA(Seikagaku)
LcH
PFL
GSL-2
LTL
Mono2R-CNL
MAA
MAL-I
AIA
#15 Sialyl Lewis X (BD)
Sp Serotype 14-2
PsSC
VGA(EY Lab)
CML-I
Avaren-Fc
CA19-9 Ab (DRG)
#14 Sialyl Lewis X (Calbio)
PNA
P11 VP8
CA19.9-121SLE
HPA
SBA
ACL
Anti-Lec
rNV VLP-Cy3
Epa1-Np mutant Y228W
PHA-L
Microvirin
Epa1-Np mutant Y228A
CA19.9-9L426
ab5
WT Galectin-8
DSL
MPL
Epa1-Np mutant E227G
GSL-1
MOA
rPVL
LCA
MGL 6B
TbFBL2
Mouse NKR-P1C
Mono2W-CNL
Mono1-CNL
GSL-I-B4
Threefoil
WGA
LSL-150
WT-VCC
PSL
PSA
HML-Native
MGL 6A
CAA
LEL
Gal-3C
MNA-M
hGal-1-488+BCR
NPA
CTB
PHA-E
ConA
CnMBL
CA
Epa1-Np WT
CSA (EY Lab)
DC-ASGPR short
Mouse Galectin1
Gal-3
STL
Galectin-1
WGA(Seikagaku Bio)
HA70 complex-Alexa
DBL-1
CGL
LBL-152
human Gal-7
VVHA lectin
rMpL
BTL native
ECL
CC1G-1007
mouse Gal-7
Human Galectin1
CnSepL
WFL
AOL (Mahal Lab)
TbFBL1
RCAI (Vector)
WGA
WGA multimer
AAL

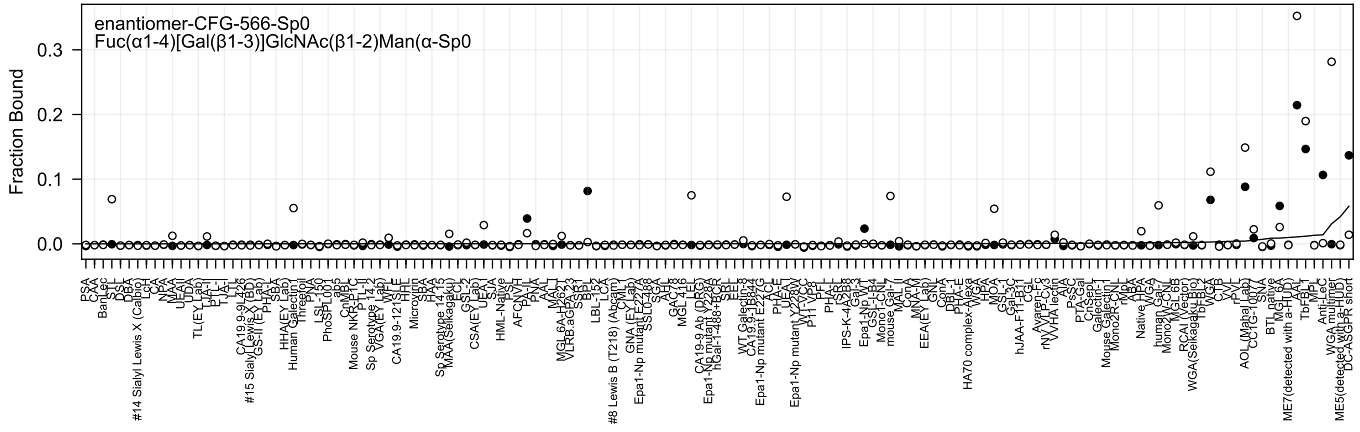
enantiomer-CFG-566-Sp0
Fuc(α1-4)[Gal(β1-3)]GlcNAc(β1-2)Man(α-Sp0
Fraction Bound
0.0
0.1
0.2
0.3

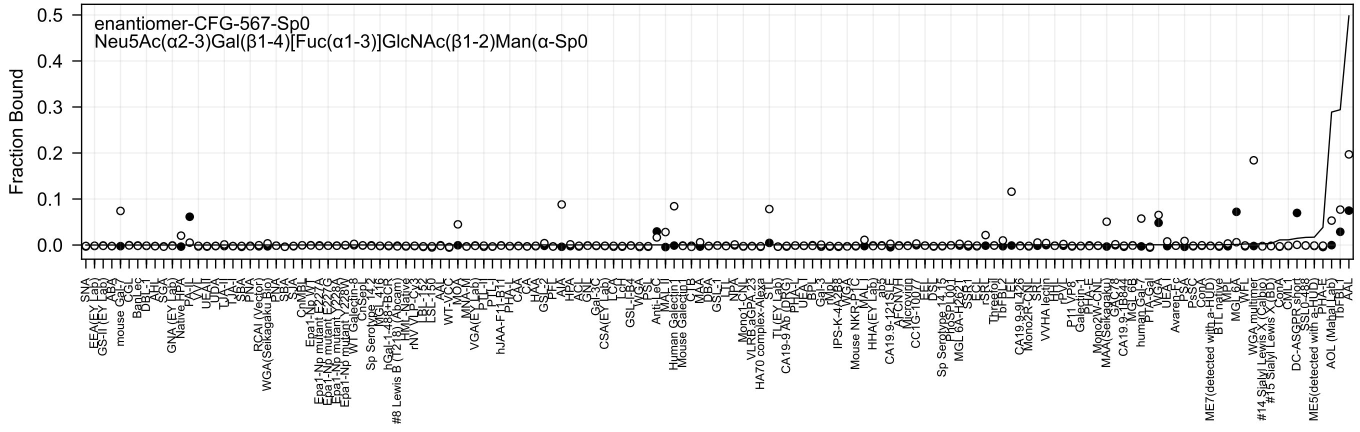

enantiomer-CFG-567-Sp0
Neu5Ac(α2-3)Gal(β1-4)[Fuc(α1-3)]GlcNAc(β1-2)Man(α-Sp0
Fraction Bound
0.5
0.4
0.3
0.2
0.1
0.0

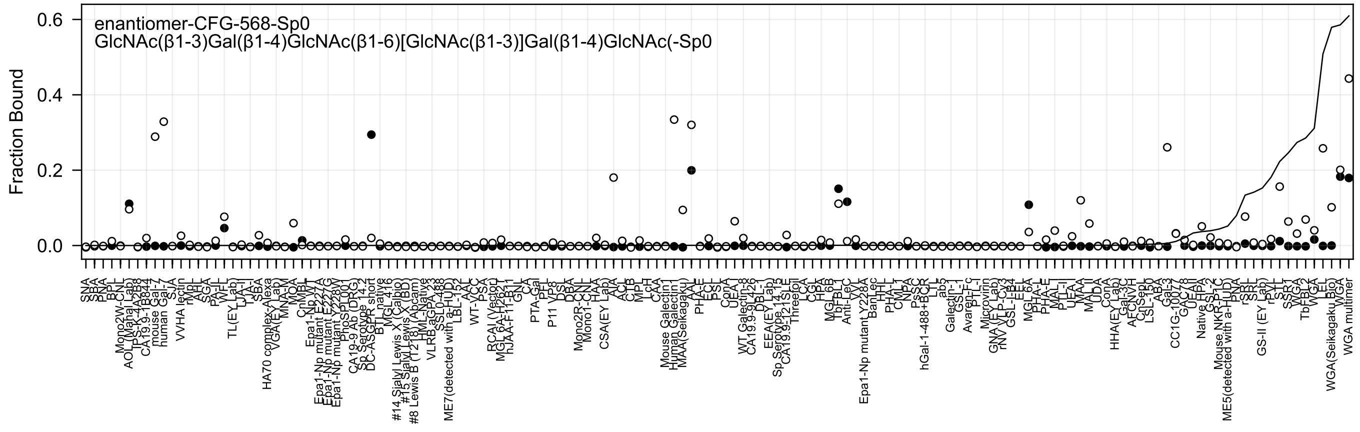

enantiomer-CFG-568-Sp0
GlcNAc(β1-3)Gal(β1-4)GlcNAc(β1-6)[GlcNAc(β1-3)]Gal(β1-4)GlcNAc(-Sp0
Fraction Bound
0.6
0.4
0.2
0.0

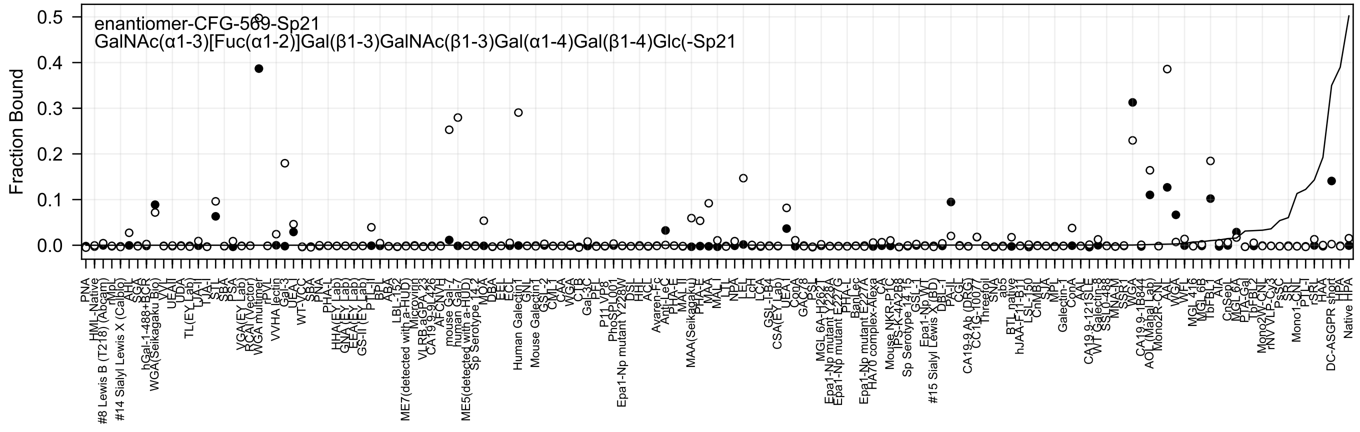

enantiomer-CFG-569-Sp21
GalNAc(α1-3)[Fuc(α1-2)]Gal(β1-3)GalNAc(β1-3)Gal(α1-4)Gal(β1-4)Glc(-Sp21
Fraction Bound

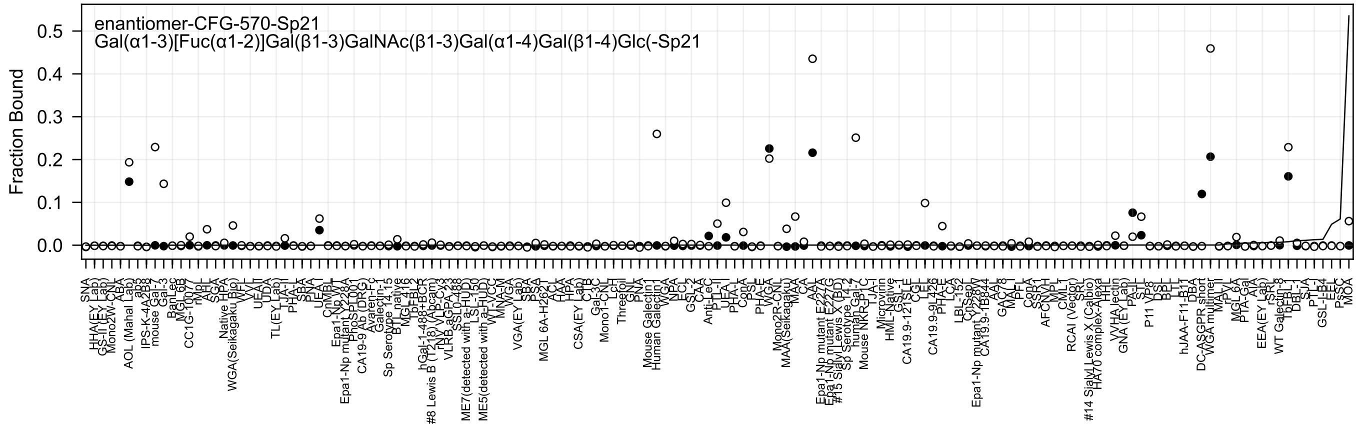
enantiomer-CFG-570-Sp21
Gal(α1-3)[Fuc(α1-2)]Gal(β1-3)GalNAc(β1-3)Gal(α1-4)Gal(β1-4)Glc(-Sp21
Fraction Bound
0.0
0.1
0.2
0.3
0.4
0.5

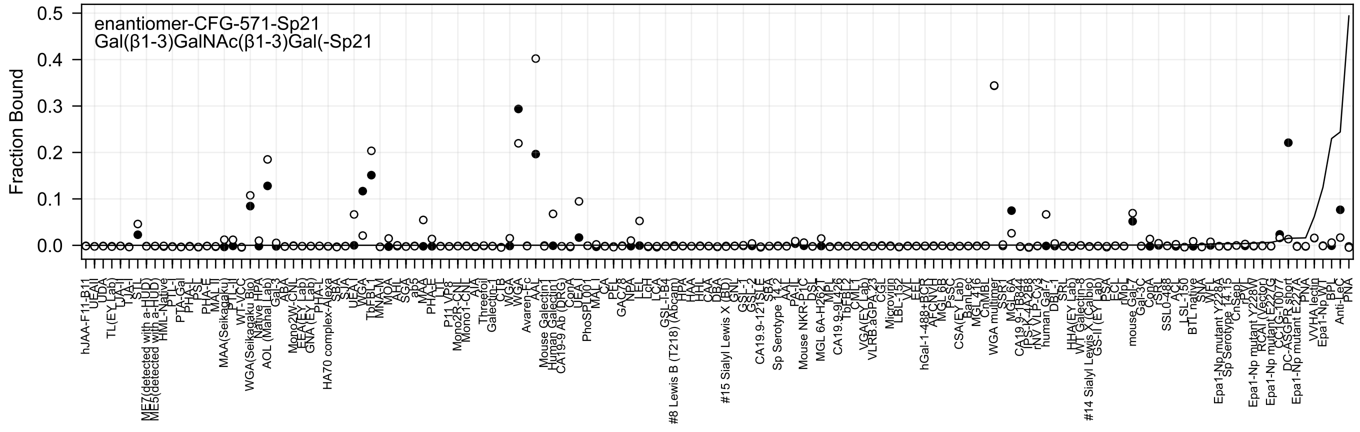

enantiomer-CFG-571-Sp21
Gal(β1-3)GalNAc(β1-3)Gal(-Sp21
Fraction Bound
0.5
0.4
0.3
0.2
0.1
0.0

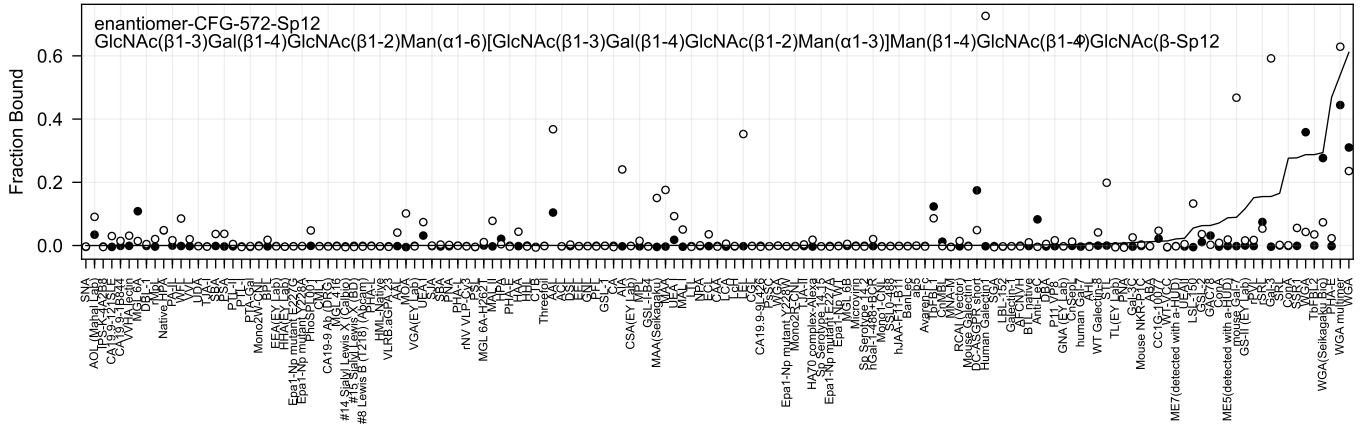
enantiomer-CFG-572-Sp12
GlcNAc(β1-3)Gal(β1-4)GlcNAc(β1-2)Man(α1-6)[GlcNAc(β1-3)Gal(β1-4)GlcNAc(β1-2)Man(α1-3)]Man(β1-4)GlcNAc(β1-4)GlcNAc(β-Sp12
Fraction Bound
0.0
0.2
0.4
0.6

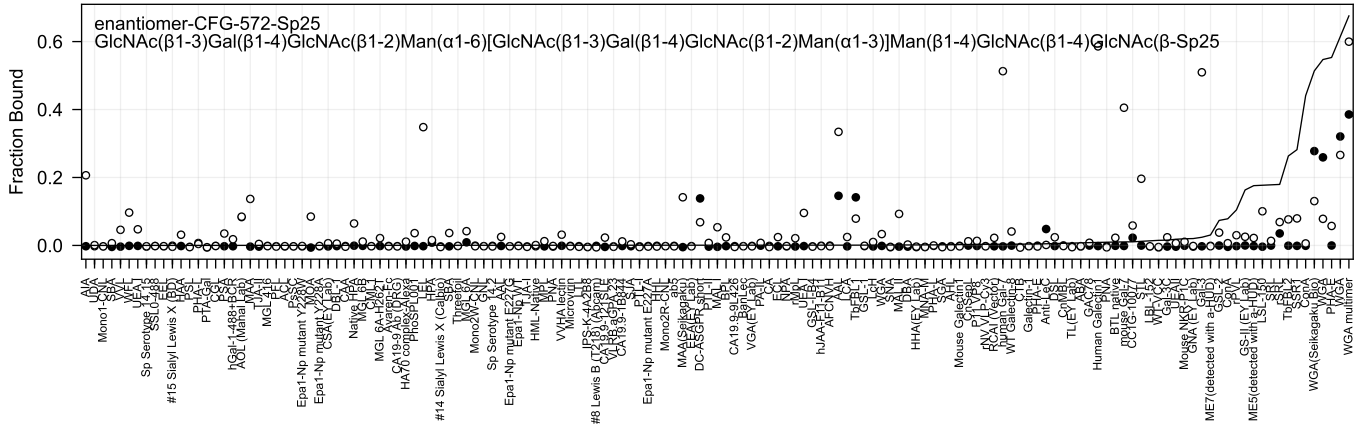
enantiomer-CFG-572-Sp25
GlcNAc(β1-3)Gal(β1-4)GlcNAc(β1-2)Man(α1-6)[GlcNAc(β1-3)Gal(β1-4)GlcNAc(β1-2)Man(α1-3)]Man(β1-4)GlcNAc(β1-4)GlcNAc(β-Sp25
Fraction Bound
0.0
0.2
0.4
0.6

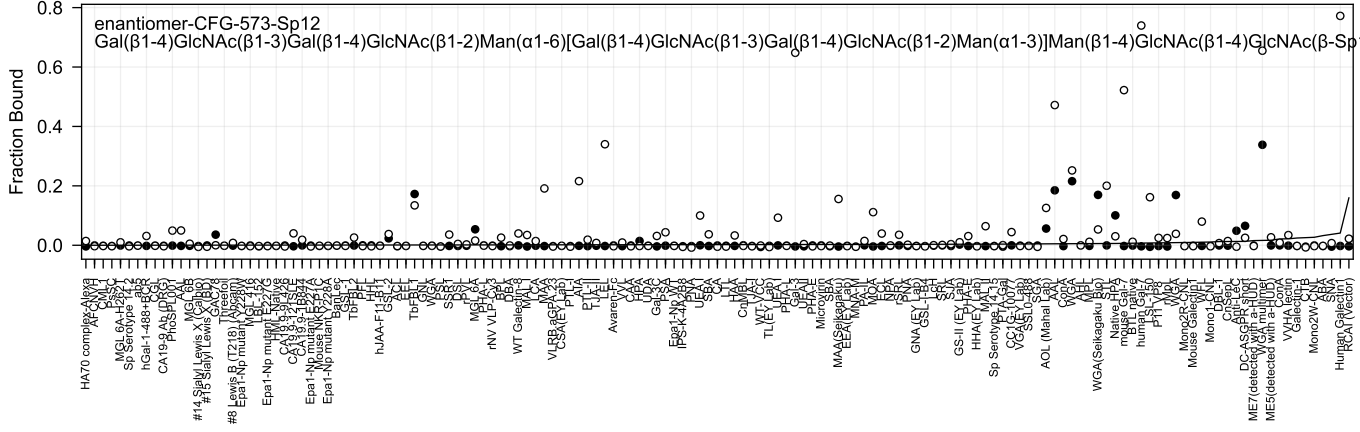
enantiomer-CFG-573-Sp12
Gal(β1-4)GlcNAc(β1-3)Gal(β1-4)GlcNAc(β1-2)Man(α1-6)[Gal(β1-4)GlcNAc(β1-3)Gal(β1-4)GlcNAc(β1-2)Man(α1-3)]Man(β1-4)GlcNAc(β1-4)GlcNAc(β-Sp
Fraction Bound
0.8
0.6
0.4
0.2
0.0

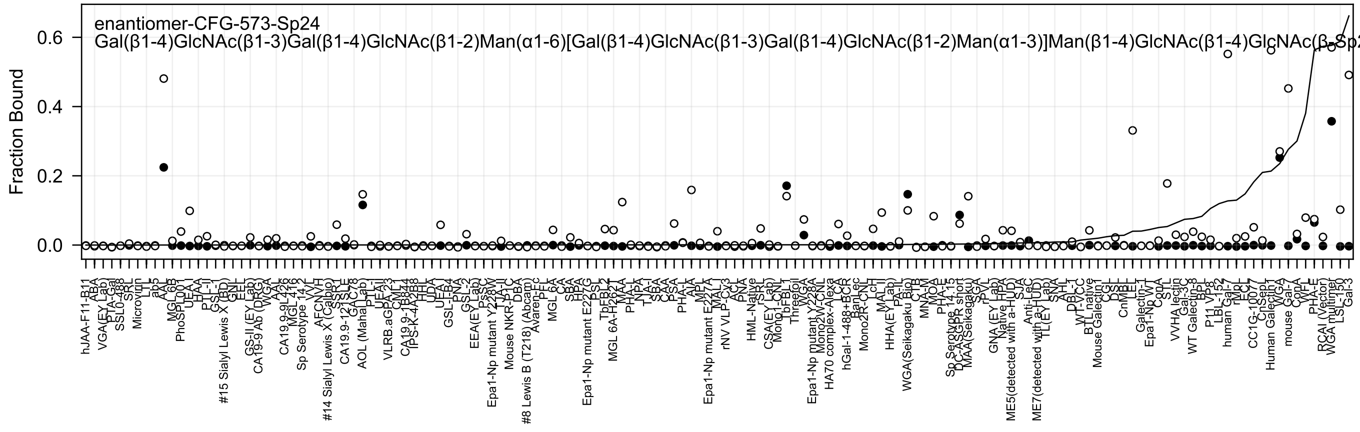
enantiomer-CFG-573-Sp24
Gal(β1-4)GlcNAc(β1-3)Gal(β1-4)GlcNAc(β1-2)Man(α1-6)[Gal(β1-4)GlcNAc(β1-3)Gal(β1-4)GlcNAc(β1-2)Man(α1-3)]Man(β1-4)GlcNAc(β1-4)GlcNAc(β-Sp2
Fraction Bound
0.0
0.2
0.4
0.6
hJAA-F11-B11
ABA
VGA(EY Lab)
PTA-Gal
SSL0-488
SRL
Microvirin
LTL
ab5
AAL
MGL 6B
PhoSL001
UEA-I
HAA
PTL-II
GSL-I
#15 Sialyl Lewis X (BD)
GNL
EEL
GS-II (EY Lab)
CA19-9 Ab (DRG)
WGA
AAL
CA19.9-9L426
MGL 416
Sp Serotype 14.2
VVL
AFCNVH
#14 Sialyl Lewis X (Calbio)
SSR1
CA19.9-121SLE
GAC78
AOL (Mahal Lab)
PTL-I
UEA-II
VLRB.aGPA.23
CML1
CA19.9-1B874
IPS-K-4A2B8
HHL
UDA
UEA-I
GSL-IB4
PNA
GSL-2
EEA(EY Lab)
PsSC
Epa1-Np mutant Y228W
TJA-II
Mouse NKR-P1C
DBA
#8 Lewis B (T218) (Abcam)
Avaren-Fc
PFL
MGL 6A
CA
SBA
HPA
Epa1-Np mutant E227G
PSL
TbFBL2
MGL 6A-H262T
MAA
PHA-L
NPA
TJA-I
SBA
CAA
PSA
PHA-L
AIA
MPL
Epa1-Np mutant E227A
MAL-I
rNV VLP-Cy3
ACL
PNA
HML-Native
rSRL
CSA(EY Lab)
Mono1-CNL
TbFBL1
Threefoil
WGA
Epa1-Np mutant Y228A
Mono2W-CNL
HA70 complex-Alexa
LCA
hGal-1-488+BCR
BanLec
Mono2R-CNL
LcH
MAL-II
HHA(EY Lab)
PA-IL
WGA(Seikagaku Bio)
CTB
MNA-M
MOA
PHA-E
Sp Serotype 14.15
DC-ASGPR short
MAA(Seikagaku)
SGA
rPVL
GNA (EY Lab)
Native HPA
ME5(detected with a-HUD)
SJA
Anti-LeC
ME7(detected with a-HUD)
TL(EY Lab)
SNA
AHL
DBL-1
WT-VCC
BTL native
Mouse Galectin1
CGL
DSL
CnMBL
LEL
Galectin-1
Epa1-Np WT
ConA
STL
VVHA lectin
Gal-3C
WT Galectin-8
BPL
P11 VP8
LBL-152
human Gal-7
rMpL
ECL
CC1G-1007
CnSep1
Human Galectin1
WGA
mouse Gal-7
ConA
WFL
PHA-E
RCAI (Vector)
WGA multimer
LSL-150
Gal-3

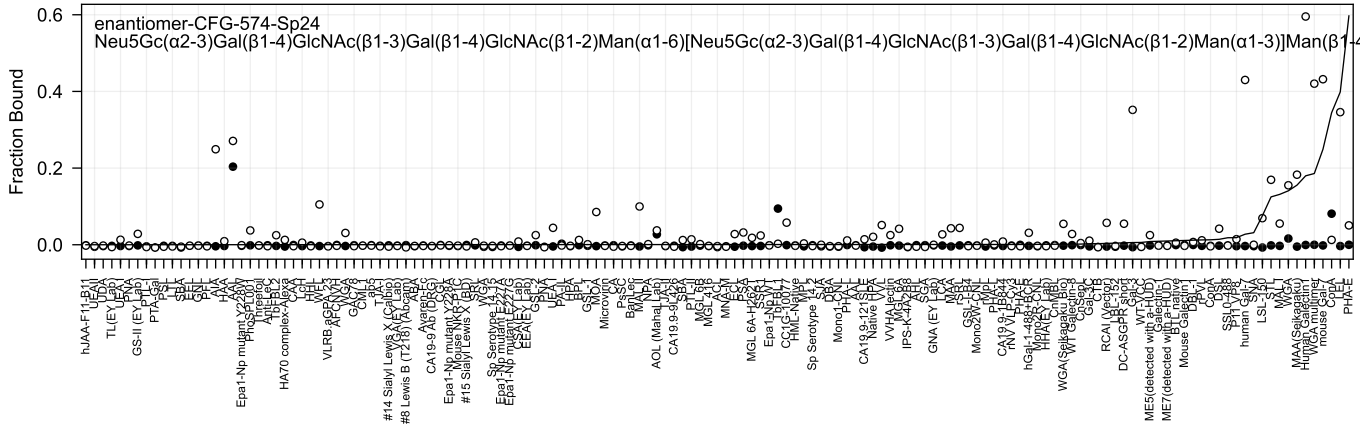

enantiomer-CFG-574-Sp24
Neu5Gc(α2-3)Gal(β1-4)GlcNAc(β1-3)Gal(β1-4)GlcNAc(β1-2)Man(α1-6)[Neu5Gc(α2-3)Gal(β1-4)GlcNAc(β1-3)Gal(β1-4)GlcNAc(β1-2)Man(α1-3)]Man(β1-4
Fraction Bound
0.6
0.4
0.2
0.0

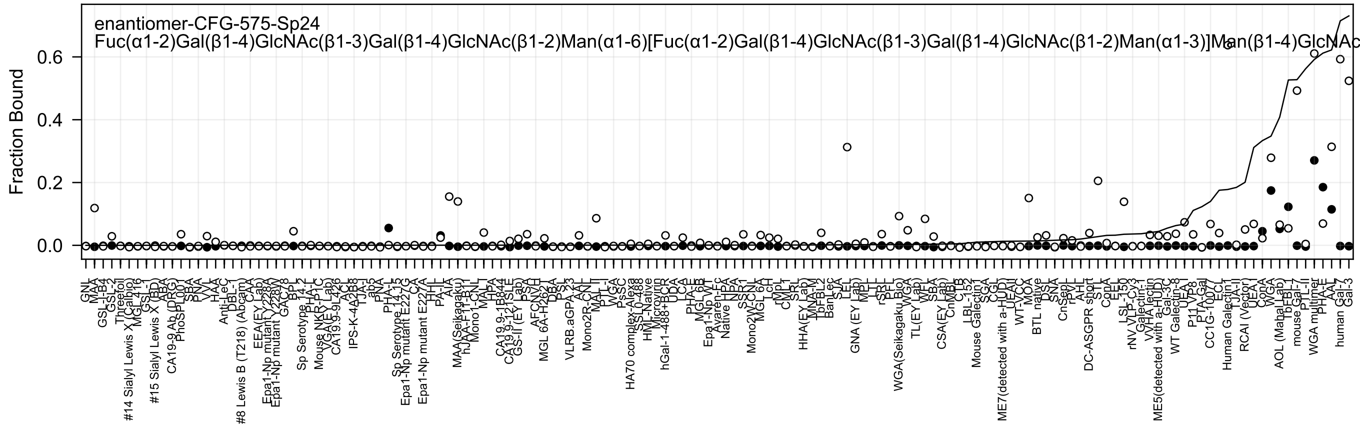
enantiomer-CFG-575-Sp24
Fuc(α1-2)Gal(β1-4)GlcNAc(β1-3)Gal(β1-4)GlcNAc(β1-2)Man(α1-6)[Fuc(α1-2)Gal(β1-4)GlcNAc(β1-3)Gal(β1-4)GlcNAc(β1-2)Man(α1-3)]Man(β1-4)GlcNAc
Fraction Bound
0.0
0.2
0.4
0.6

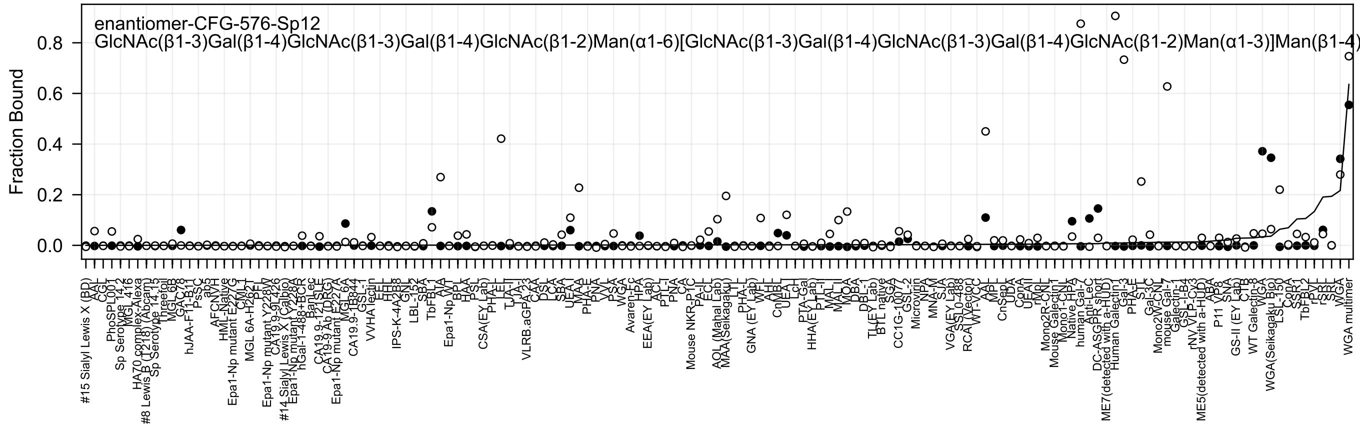
enantiomer-CFG-576-Sp12
GlcNAc(β1-3)Gal(β1-4)GlcNAc(β1-3)Gal(β1-4)GlcNAc(β1-2)Man(α1-6)[GlcNAc(β1-3)Gal(β1-4)GlcNAc(β1-3)Gal(β1-4)GlcNAc(β1-2)Man(α1-3)]Man(β1-4)
Fraction Bound
0.0
0.2
0.4
0.6
0.8

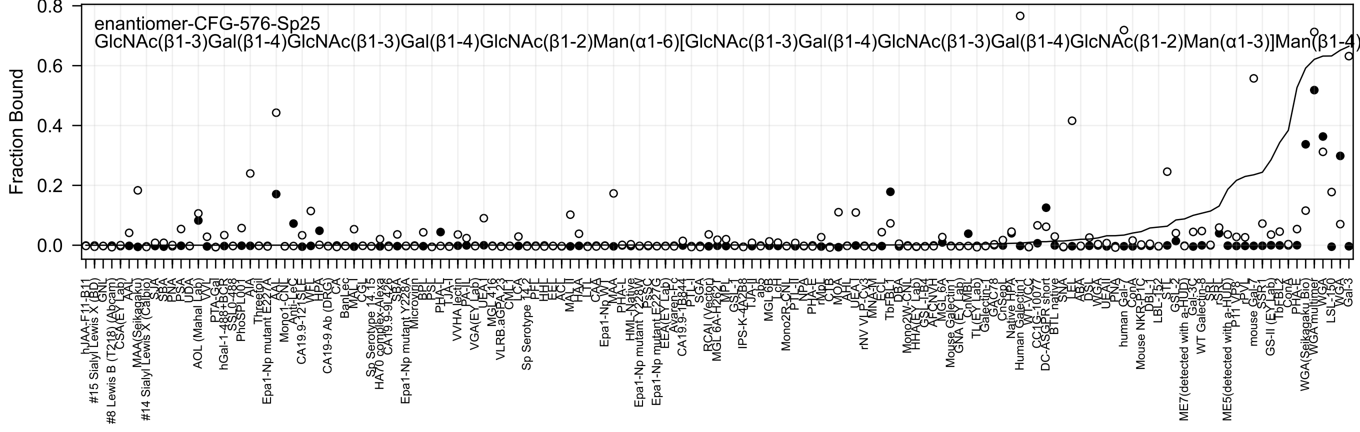
enantiomer-CFG-576-Sp25
GlcNAc(β1-3)Gal(β1-4)GlcNAc(β1-3)Gal(β1-4)GlcNAc(β1-2)Man(α1-6)[GlcNAc(β1-3)Gal(β1-4)GlcNAc(β1-3)Gal(β1-4)GlcNAc(β1-2)Man(α1-3)]Man(β1-4)
Fraction Bound
0.0
0.2
0.4
0.6
0.8

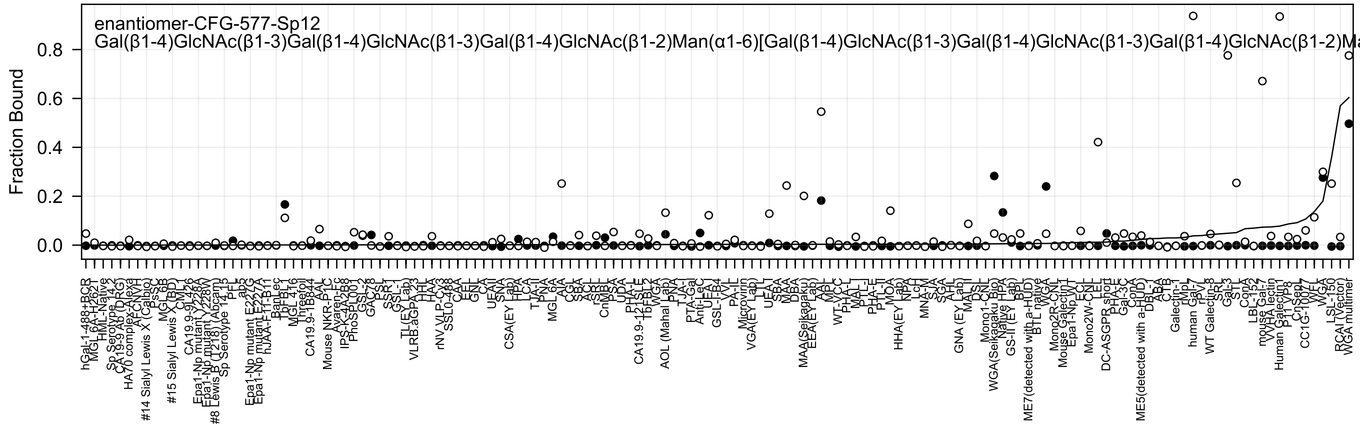

enantiomer-CFG-577-Sp12
Gal(β1-4)GlcNAc(β1-3)Gal(β1-4)GlcNAc(β1-3)Gal(β1-4)GlcNAc(β1-2)Man(α1-6)[Gal(β1-4)GlcNAc(β1-3)Gal(β1-4)GlcNAc(β1-3)Gal(β1-4)GlcNAc(β1-2)Ma
Fraction Bound
0.0
0.2
0.4
0.6
0.8

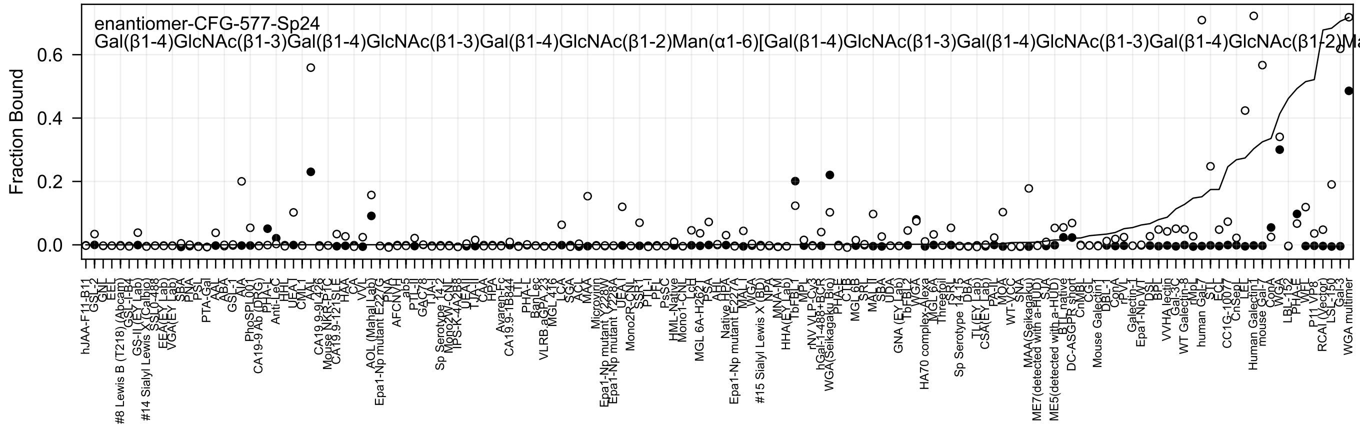
enantiomer-CFG-577-Sp24
Gal(β1-4)GlcNAc(β1-3)Gal(β1-4)GlcNAc(β1-3)Gal(β1-4)GlcNAc(β1-2)Man(α1-6)[Gal(β1-4)GlcNAc(β1-3)Gal(β1-4)GlcNAc(β1-3)Gal(β1-4)GlcNAc(β1-2)Ma
Fraction Bound
0.0
0.2
0.4
0.6

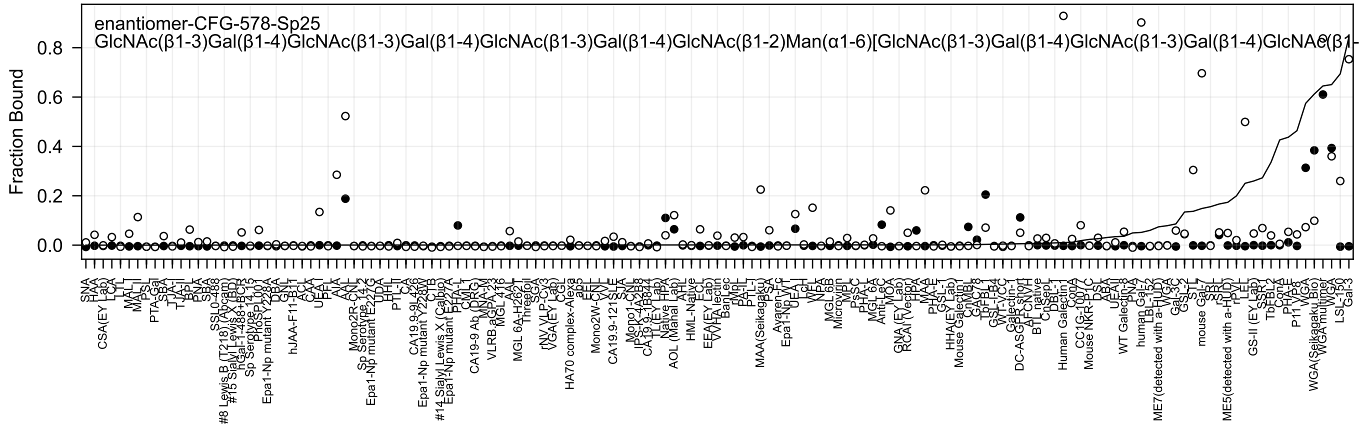
enantiomer-CFG-578-Sp25
GlcNAc(β1-3)Gal(β1-4)GlcNAc(β1-3)Gal(β1-4)GlcNAc(β1-3)Gal(β1-4)GlcNAc(β1-2)Man(α1-6)[GlcNAc(β1-3)Gal(β1-4)GlcNAc(β1-3)Gal(β1-4)GlcNAc(β1-
Fraction Bound
0.0
0.2
0.4
0.6
0.8

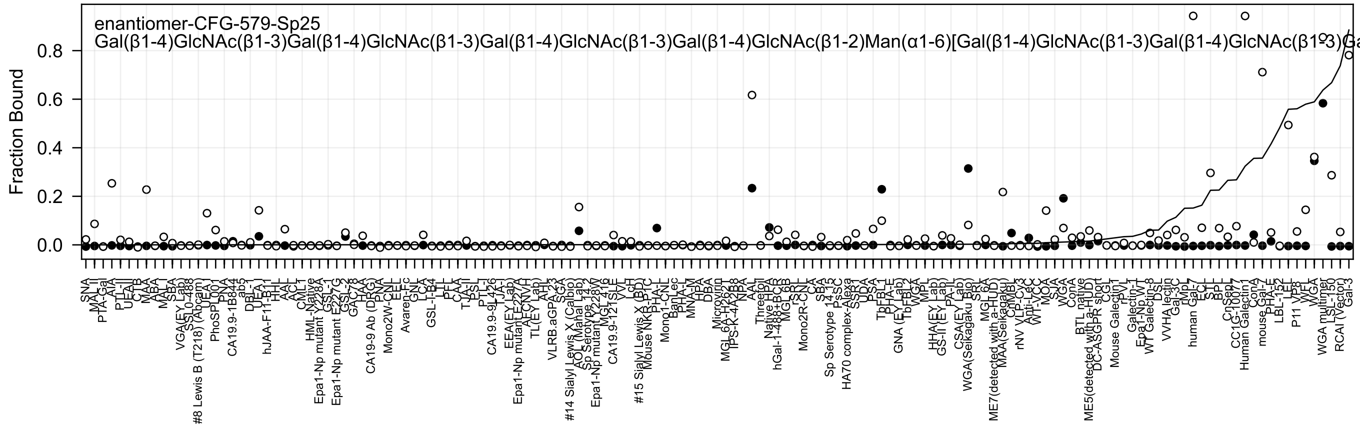
enantiomer-CFG-579-Sp25
Gal(β1-4)GlcNAc(β1-3)Gal(β1-4)GlcNAc(β1-3)Gal(β1-4)GlcNAc(β1-3)Gal(β1-4)GlcNAc(β1-2)Man(α1-6)[Gal(β1-4)GlcNAc(β1-3)Gal(β1-4)GlcNAc(β1-3)Ga
Fraction Bound
0.0
0.2
0.4
0.6
0.8

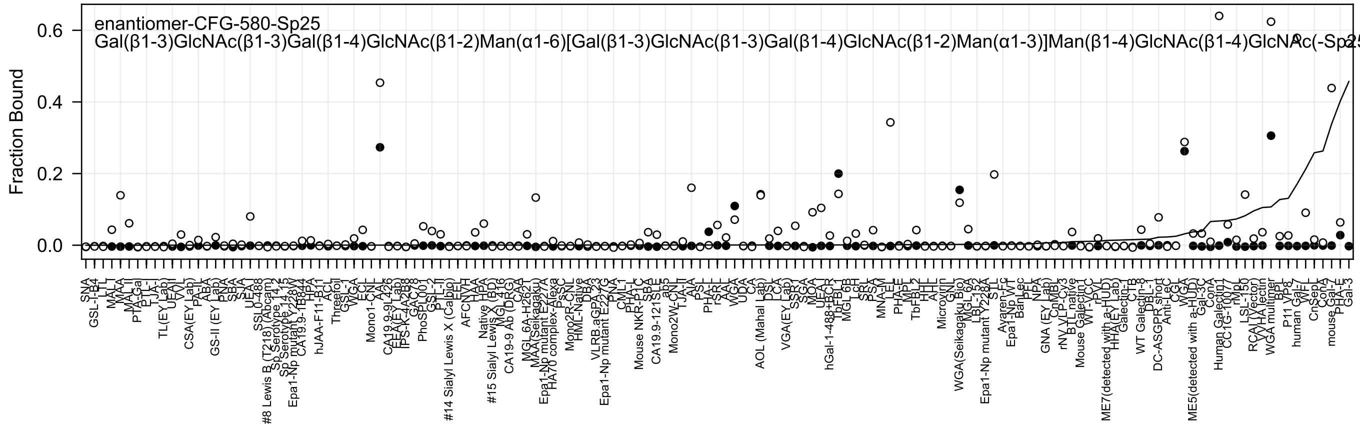
enantiomer-CFG-580-Sp25
Gal(β1-3)GlcNAc(β1-3)Gal(β1-4)GlcNAc(β1-2)Man(α1-6)[Gal(β1-3)GlcNAc(β1-3)Gal(β1-4)GlcNAc(β1-2)Man(α1-3)]Man(β1-4)GlcNAc(β1-4)GlcNAc(-Sp25
Fraction Bound

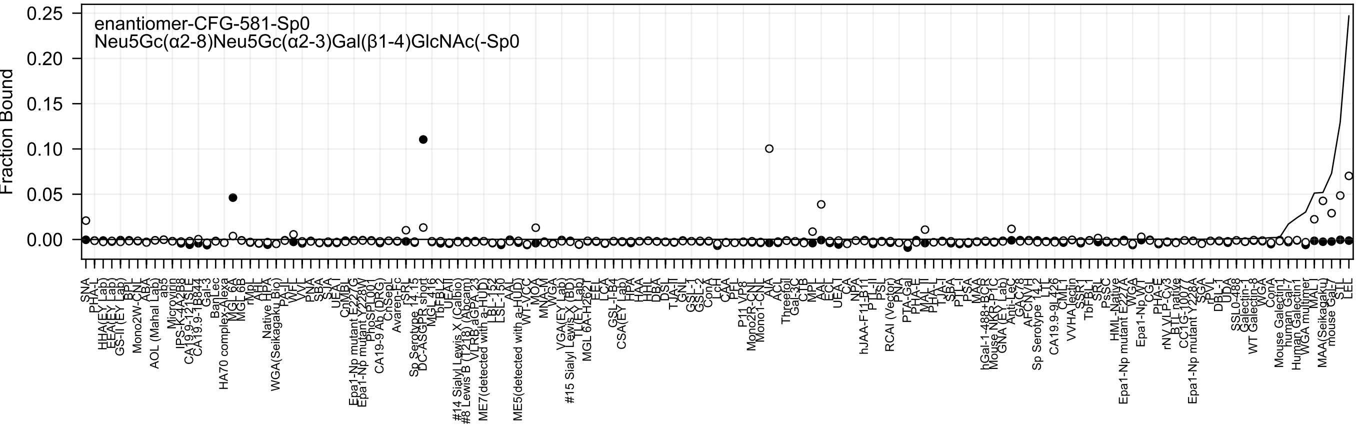
enantiomer-CFG-581-Sp0
Neu5Gc(α2-8)Neu5Gc(α2-3)Gal(β1-4)GlcNAc(-Sp0
Fraction Bound
0.25
0.20
0.15
0.10
0.05
0.00

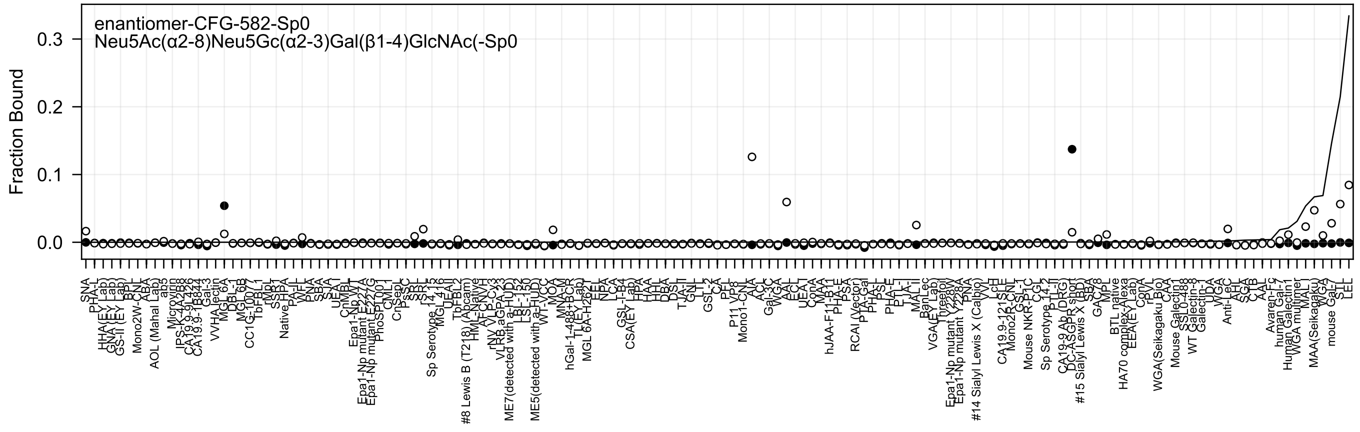
enantiomer-CFG-582-Sp0
Neu5Ac(α2-8)Neu5Gc(α2-3)Gal(β1-4)GlcNAc(-Sp0
Fraction Bound
0.0
0.1
0.2
0.3

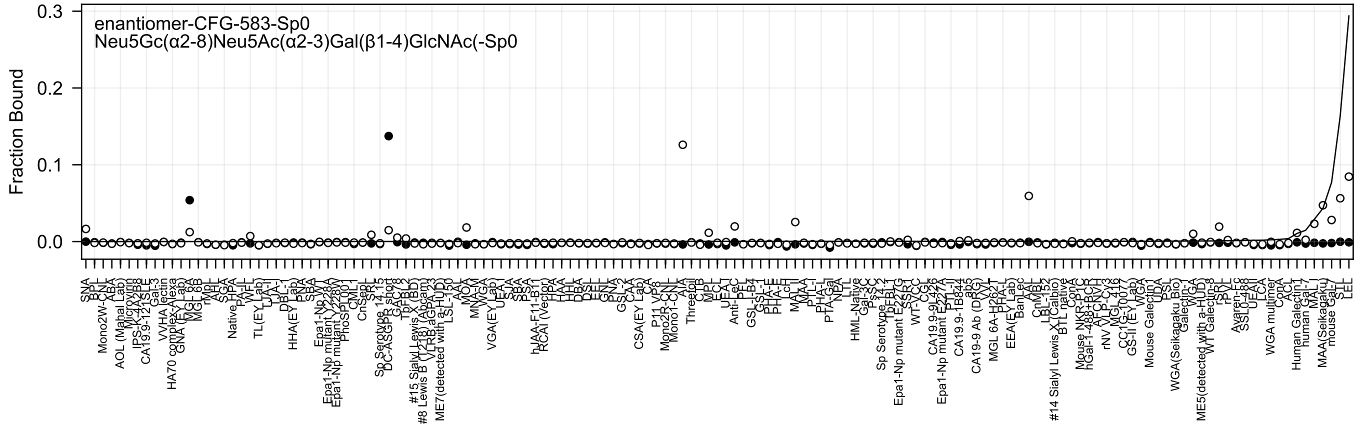

enantiomer-CFG-583-Sp0
Neu5Gc(α2-8)Neu5Ac(α2-3)Gal(β1-4)GlcNAc(-Sp0
Fraction Bound

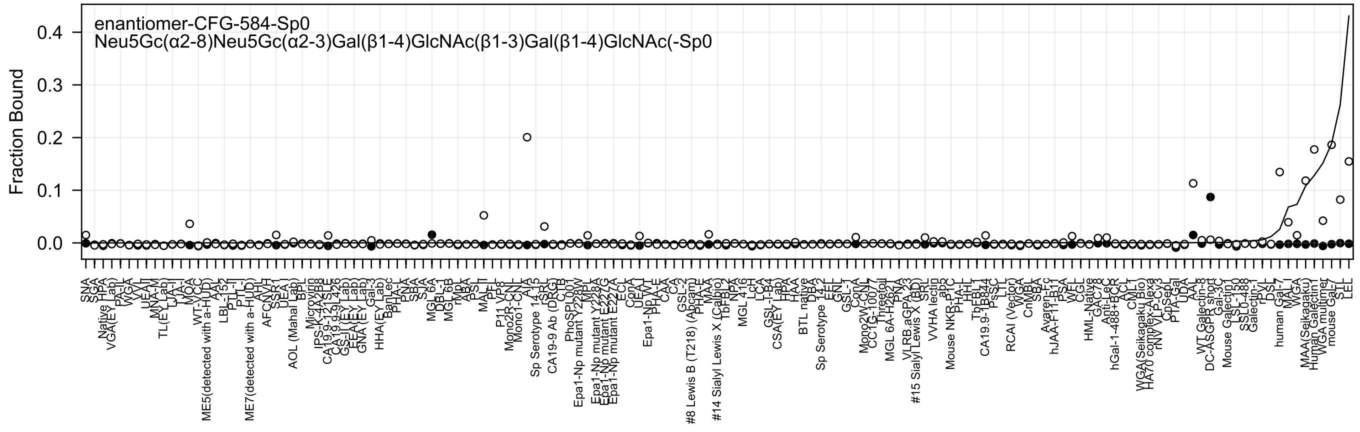
enantiomer-CFG-584-Sp0
Neu5Gc(α2-8)Neu5Gc(α2-3)Gal(β1-4)GlcNAc(β1-3)Gal(β1-4)GlcNAc(-Sp0
Fraction Bound
0.0
0.1
0.2
0.3
0.4
SNA
SGA
HPA
Native
VGA(EY Lab)
PA-IL
WGA
VVL
UEA-I
MNA-M
TL(EY Lab)
LJA-I
TJA-I
MOA
WT-VCC
ME5(detected with a-HUD)
AAL
LBL-152
PTL-II
PTL-I
ME7(detected with a-HUD)
AH
AFC
SSR1
UEA-I
BPL
AOL (Mahal Lab)
Microvirin
IPS-K-4A2B8
CA19.9-121SLE
CA19.9-9L426
GS-II (EY Lab)
EEA(EY Lab)
GNA (EY Lab)
Gal-3
HHA(EY Lab)
BanLec
PHA-L
PNA
SBA
SJA
MGL 6A
DBL-1
MGL 6B
rMpA
ABA
PSL
MAL-II
PFL
P11 VP8
Mono2R-CNL
Mono1-CNL
AIA
Sp Serotype 14.15
rSRL
CA19-9 Ab (DRG)
CTB
PhoSL
Epa1-Np mutant Y228W
MPL
Epa1-Np mutant Y228A
Epa1-Np mutant E227G
Epa1-Np mutant E227A
ECL
ConA
UEA-I
Epa1-Np WT
PHA-E
CAA
CA
GSL-2
#8 Lewis B (T218) (Abcam)
PHA-E
MAA
#14 Sialyl Lewis X (Calbio)
TbFBL2
NPA
MGL 416
LcH
LCA
GSL-I-B4
CSA(EY Lab)
HPA
HAA
BTL native
DBA
Sp Serotype 14.2
EEL
GNL
GSL-1
ConA
Mono2W-CNL
CCTG-10077
Threetol
MGL 6A-H262T
PNA
VLRB.aGPA.23
#15 Sialyl Lewis X (BD)
SRL
VVHA lectin
ab5
Mouse NKR-P1C
PHA-L
HHL
TbFBL1
CA19.9-1B844
PsSC
LTL
RCAI (Vector)
WGA
CnMBL
SBA
Avaren-Fc
hJAA-F11-B11
PSA
WFL
CGL
HML-Native
GAC78
Anti-LeC
hGal-1-488+BCR
ACL
CML1
WGA(Seikagaku Bio)
HA70 complex-Alexa
rNVLP-Cy3
CpSeL
PTA-Gal
UDA
AAL
WT Galectin-8
DC-ASGPR short
Gal-3C
Mouse Galectin1
LSL-150
SSL0-488
Galectin-1
rPVL
DSL
human Gal-7
MAL-I
WGA
MAA(Seikagaku)
Human Galectin1
WGA multimer
mouse Gal-7
STL
LEL

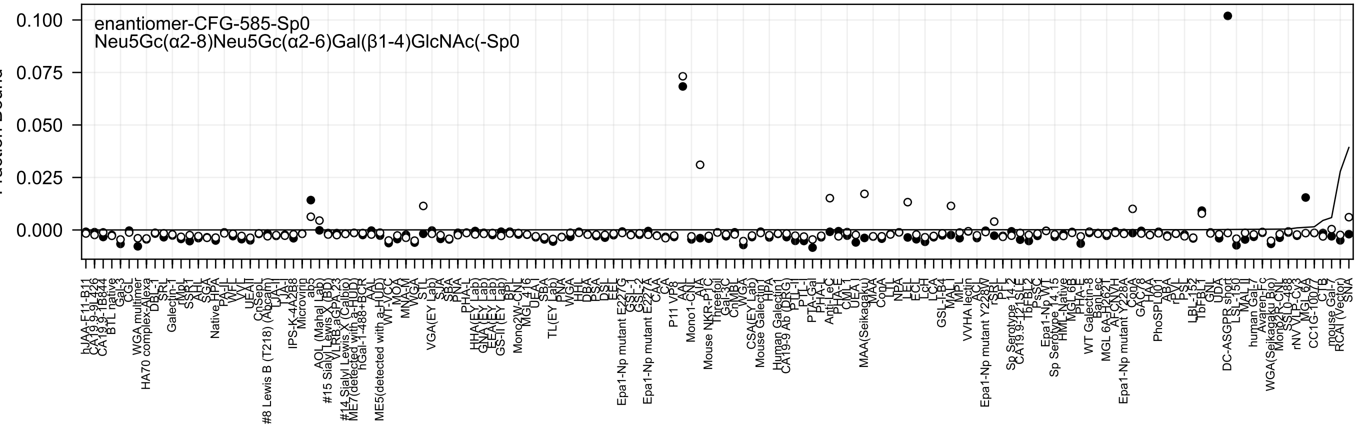
enantiomer-CFG-585-Sp0
Neu5Gc(α2-8)Neu5Gc(α2-6)Gal(β1-4)GlcNAc(-Sp0
Fraction Bound

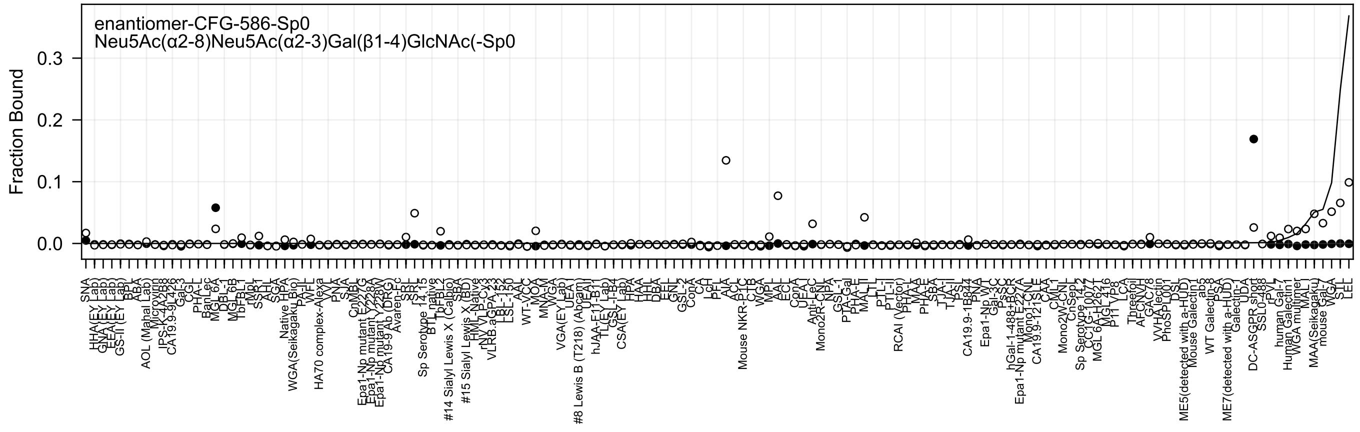
enantiomer-CFG-586-Sp0
Neu5Ac(α2-8)Neu5Ac(α2-3)Gal(β1-4)GlcNAc(-Sp0
Fraction Bound
0.0
0.1
0.2
0.3
SNA
HHA(EY Lab)
GNA(EY Lab)
EEA(EY Lab)
GS-II (EY Lab)
BPL
ABA
AOL (Mahal Lab)
Microvirin
IPS-K-4A2B8
CA19.9-9L426
Gal-3
CGL
PHA-L
BanLec
MGL 6A
DBL-1
MGL 6B
TbFBL1
rMpl
SSP1
AHL
SGA
Native HPA
WGA(Seikagaku Bio)
PA-IL
WFL
HA70 complex-Alexa
VVL
PNA
SJA
CnMBL
Epa1-Np mutant E227G
Epa1-Np mutant Y228A
Epa1-Np mutant Y228W
CA19-9 Ab (DRG)
Avaren-Fc
SRL
rSRL
Sp Serotype 14.15
BTL native
TbFBL2
#14 Sialyl Lewis X (Calbio)
SBA
#15 Sialyl Lewis X (BD)
HML-Native
rNY VLP-Cy3
VLRB.aGPA.23
LSL-150
LSL-N
AAL
WT-VCC
MOA
MNA-M
WGA
VGA(EY Lab)
UEA
#8 Lewis B (T218) (Abcam)
UEAI
hJAA-E11-B11
TL(EY Lab)
GSL-I-B4
CSA(EY Lab)
HPA
HAA
HHL
DBA
EEL
GNL
GSL-2
ConA
CA
LcH
PEL
AIA
ACL
Mouse NKR-P1C
CTB
WGA
MPL
MAL
ECL
ConA
UEA-I
Anti-LeC
Mono2R-CNL
NPA
GSL-1
PTA-Gal
PHA-E
MAL-II
LTL
PTL-I
PTL-II
RCAI (Vector)
PHA-L
MAE
PHA-E
SBA
TJA-I
TJA-II
PSL
CA19.9-1B844
PNA
Epa1-Np WT
Gal-3C
PsSC
hGal-1-488+BCR
Epa1-Np mutant E227A
Mono1-CNL
CA19.9-121SLE
CAA
CML
Mono2W-CNL
CnSep1
Sp Serotype 14.2
CC1G-10077
MGL 6A-H262T
MGL 416
P11 VP8
LCA
Threefoil
AFCNVH
GAC78
VVHA lectin
PhoSL001
DSL
ME5(detected with a-HUD)
Mouse Galectin1
ab5
WT Galectin-8
PSA
ME7(detected with a-HUD)
Galectin-1
UDA
DC-ASGPR short
SSL0-488
rPVL
human Gal-7
Human Galectin1
WGA multimer
MAL I
MAA(Seikagaku)
mouse Gal-7
VVA
STL
LEL

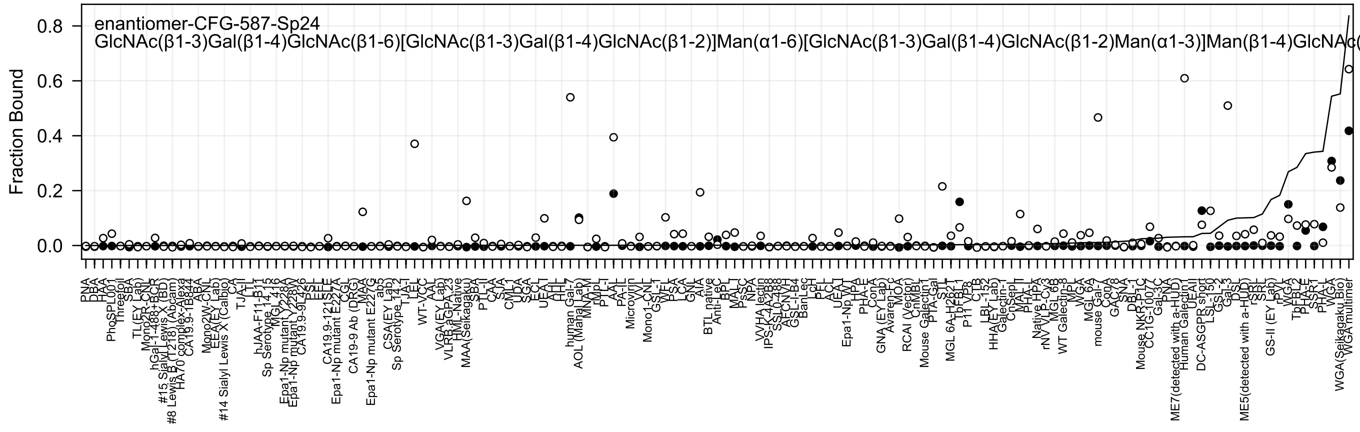
enantiomer-CFG-587-Sp24
GlcNAc(β1-3)Gal(β1-4)GlcNAc(β1-6)[GlcNAc(β1-3)Gal(β1-4)GlcNAc(β1-2)]Man(α1-6)[GlcNAc(β1-3)Gal(β1-4)GlcNAc(β1-2)Man(α1-3)]Man(β1-4)GlcNAc(
Fraction Bound
0.8
0.6
0.4
0.2
0.0

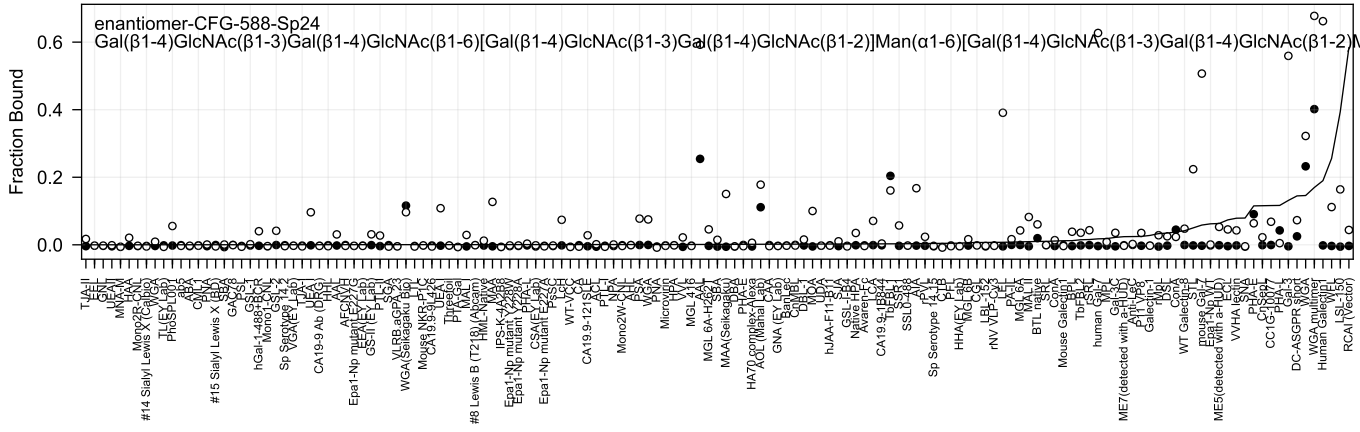

enantiomer-CFG-588-Sp24
Gal(β1-4)GlcNAc(β1-3)Gal(β1-4)GlcNAc(β1-6)[Gal(β1-4)GlcNAc(β1-3)Gal(β1-4)GlcNAc(β1-2)]Man(α1-6)[Gal(β1-4)GlcNAc(β1-3)Gal(β1-4)GlcNAc(β1-2)M
Fraction Bound
0.0
0.2
0.4
0.6
TJA-II
EEL
GNL
UEAI
MNA-M
HAA
Mono2R-CNL
#14 Sialyl Lewis X (Calbio)
WGA
TL(EY Lab)
PhoSL001
ab5
ABA
CML1
PNA
#15 Sialyl Lewis X (BD)
SBA
GAC78
PSL
GSL-I
hGal-1-488+BCR
Mono1-CNL
GSL-2
Sp Serotype 14.2
VGA(EY Lab)
TJA-I
UEA-I
CA19-9 Ab (DRG)
HHL
AAL
AFCNVH
Epa1-Np mutant E227G
EEA(EY Lab)
GS-II (EY Lab)
PTL-II
SGA
VLRB.aGPA.23
WGA(Seikagaku Bio)
LTL
Mouse NKR-P1C
CA19.9-9L426
UEA
Threefoil
PTA-Gal
MAL-I
#8 Lewis B (T218) (Abcam)
HML-Native
MAA
IPS-K-4A2B8
Epa1-Np mutant Y228W
Epa1-Np mutant Y228A
PHA-L
CSA(EY Lab)
Epa1-Np mutant E227A
PsSC
LcH
WT-VCC
CA
CA19.9-121SLE
ACL
PTL-I
NPA
Mono2W-CNL
AHL
PSA
WGA
PNA
Microvirin
HPA
VVL
MGL 416
AAL
MGL 6A-H262
SBA
MAA(Seikagaku)
DBA
PHA-E
HA70 complex-Alexa
AOL (Mahal Lab)
CAA
GNA (EY Lab)
BanLec
CnMBL
DBL-1
MOA
UDA
hJAA-F11-B11
SJA
GSL-I-B4
Native HPA
Avaren-Fc
LCA
CA19.9-1B844
TbFBL1
SSR1
SSL0-488
AIA
rPVL
Sp Serotype 14.15
CTB
PFL
HHA(EY Lab)
MGL 6B
CGL
LBL-152
rNV VLP-Cy3
LEL
PA-IL
MGL 6A
MAL II
BTL native
SRL
ConA
Mouse Galectin1
BPL
TbFBL2
rSRL
human Gal-7
MPL
Gal-3C
ME7(detected with a-HUD)
Anti-LeC
P11 VP8
Galectin-1
rMpL
DSL
ConA
WT Galectin-8
STL
mouse Gal-7
Epa1-Np WT
ME5(detected with a-HUD)
ECL
VVHA lectin
SNA
PHA-E
CnSep1
CC1G-1007
PHA-L
Gal-3
DC-ASGPR short
WGA
WGA multimer
Human Galectin1
WFL
LSL-150
RCAI (Vector)

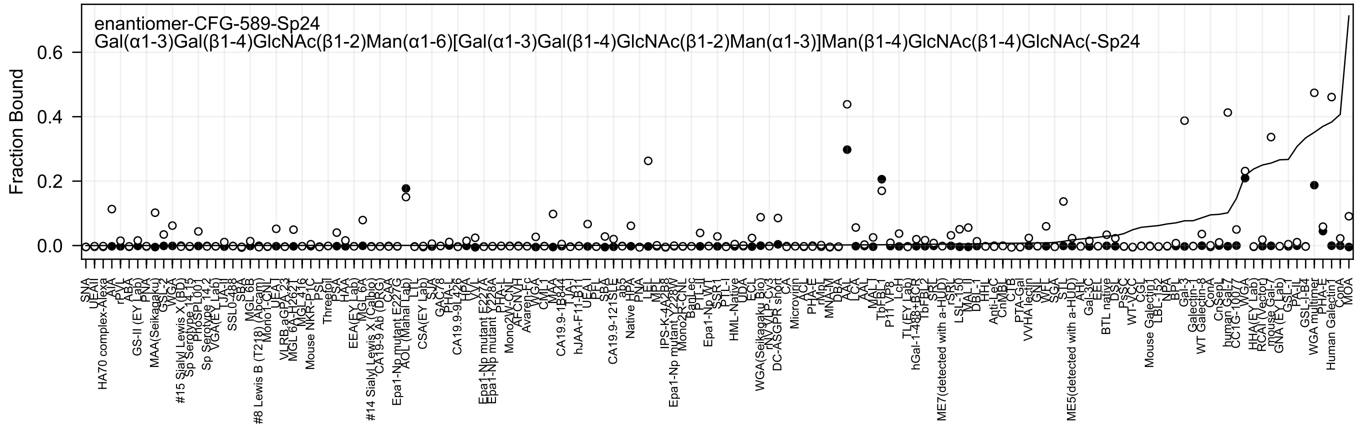
enantiomer-CFG-589-Sp24
Gal(α1-3)Gal(β1-4)GlcNAc(β1-2)Man(α1-6)[Gal(α1-3)Gal(β1-4)GlcNAc(β1-2)Man(α1-3)]Man(β1-4)GlcNAc(β1-4)GlcNAc(-Sp24
Fraction Bound
0.0
0.2
0.4
0.6

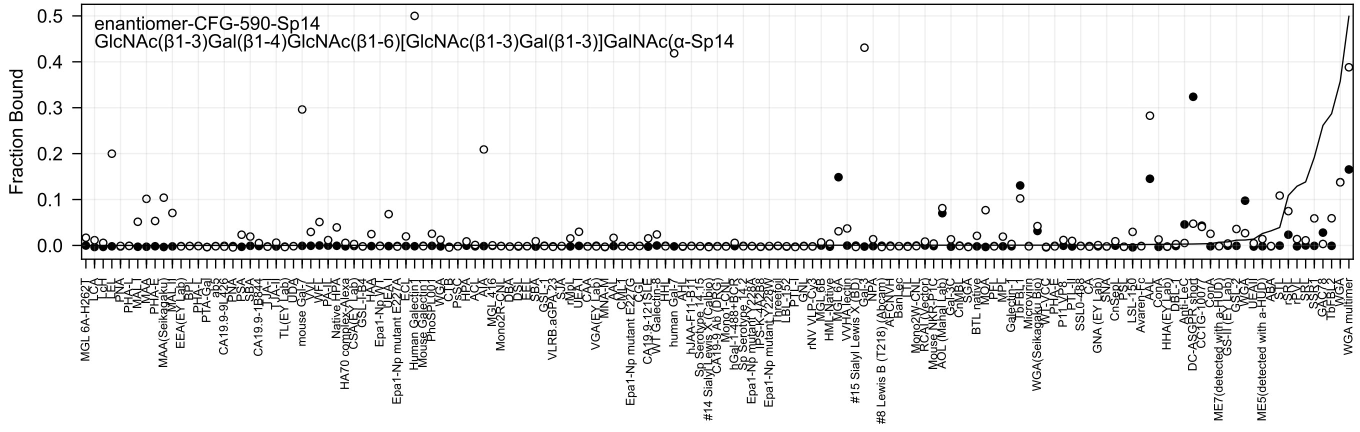
enantiomer-CFG-590-Sp14
GlcNAc(β1-3)Gal(β1-4)GlcNAc(β1-6)[GlcNAc(β1-3)Gal(β1-3)]GalNAc(α-Sp14
Fraction Bound
0.0
0.1
0.2
0.3
0.4
0.5

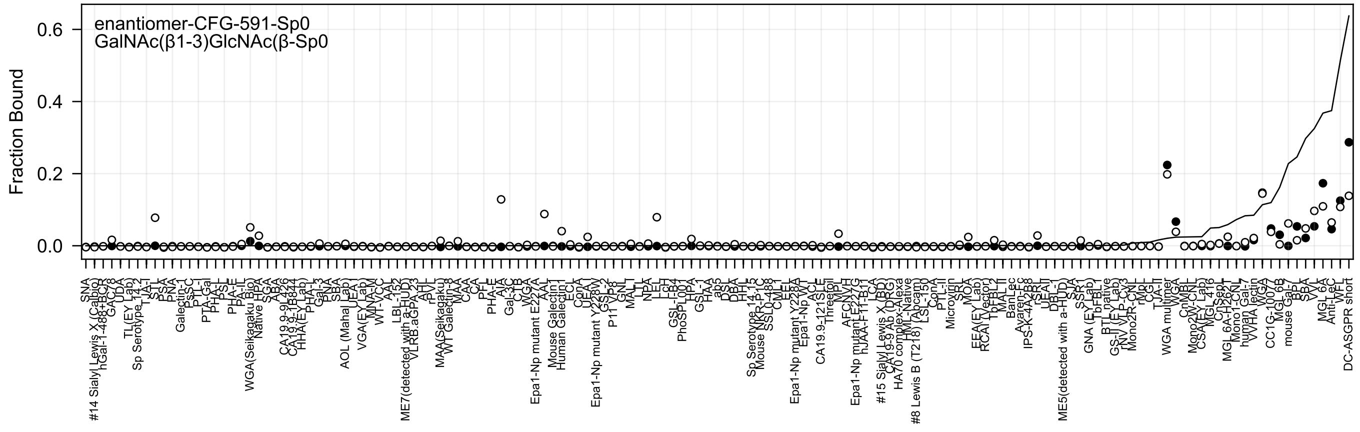
enantiomer-CFG-591-Sp0
GalNAc(β1-3)GlcNAc(β-Sp0
Fraction Bound
0.6
0.4
0.2
0.0

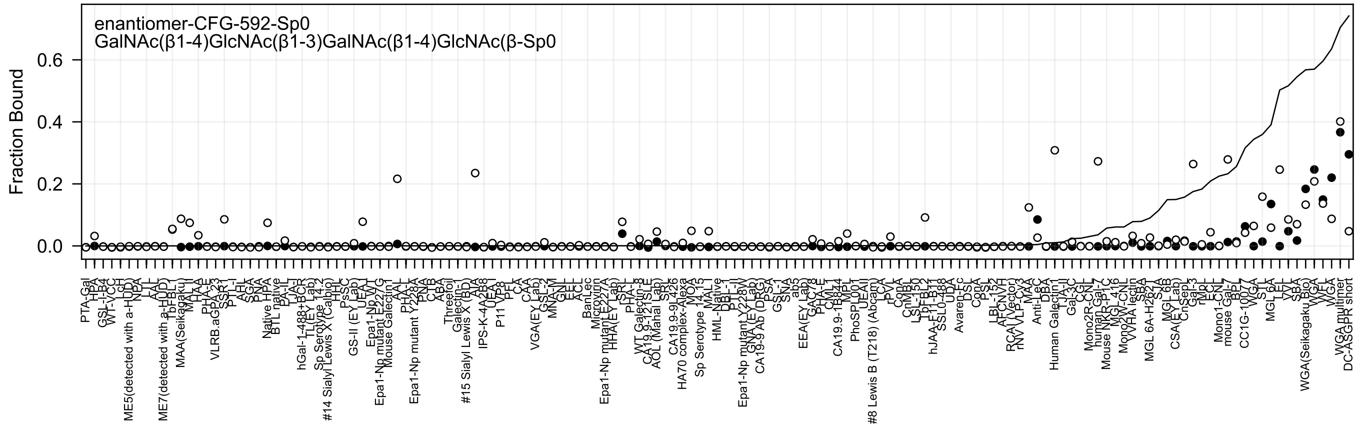
enantiomer-CFG-592-Sp0
GalNAc(β1-4)GlcNAc(β1-3)GalNAc(β1-4)GlcNAc(β-Sp0
Fraction Bound
0.0
0.2
0.4
0.6

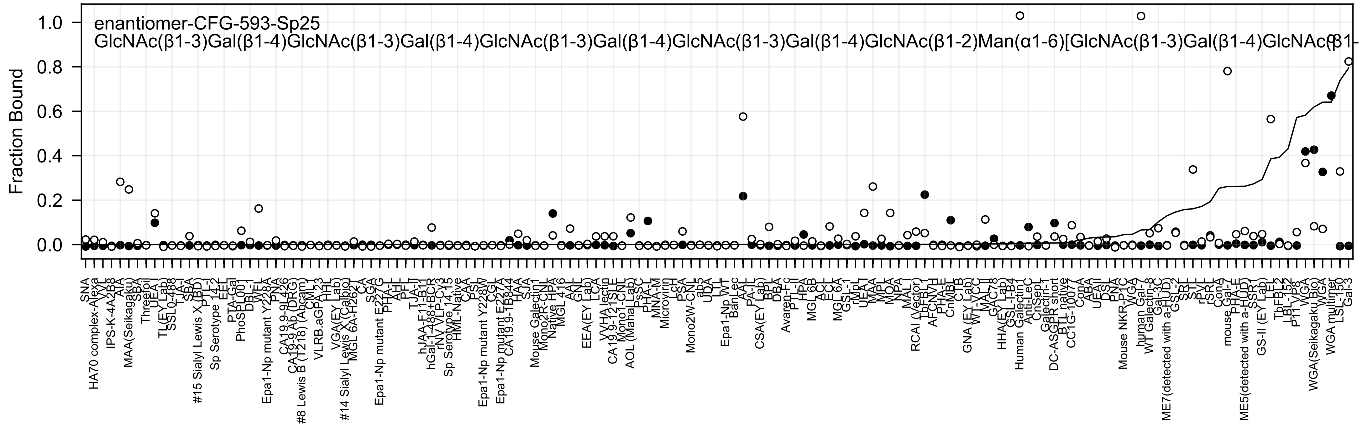

enantiomer-CFG-593-Sp25
GlcNAc(β1-3)Gal(β1-4)GlcNAc(β1-3)Gal(β1-4)GlcNAc(β1-3)Gal(β1-4)GlcNAc(β1-3)Gal(β1-4)GlcNAc(β1-2)Man(α1-6)[GlcNAc(β1-3)Gal(β1-4)GlcNAc(β1-
Fraction Bound

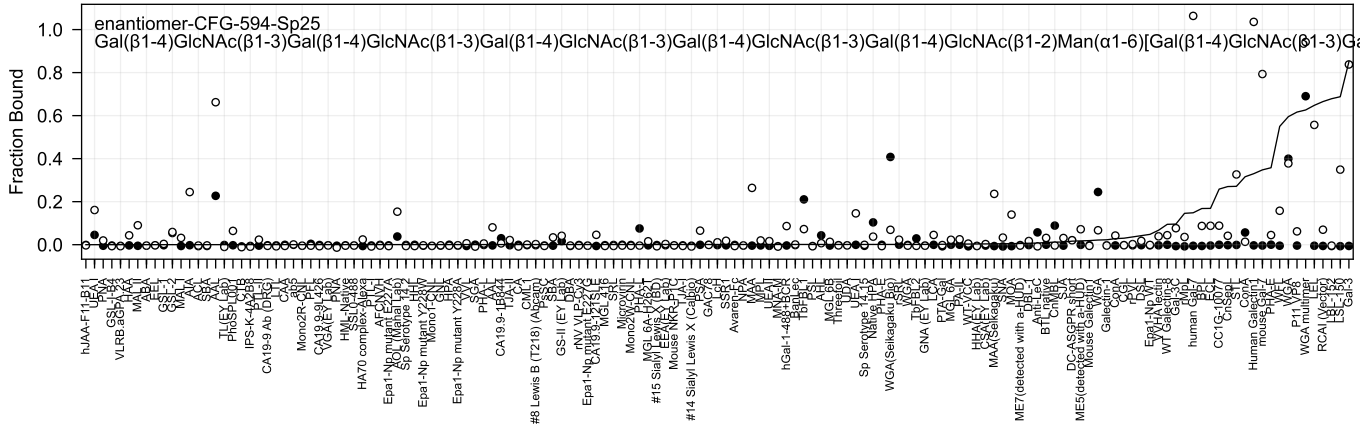
enantiomer-CFG-594-Sp25
Gal(β1-4)GlcNAc(β1-3)Gal(β1-4)GlcNAc(β1-3)Gal(β1-4)GlcNAc(β1-3)Gal(β1-4)GlcNAc(β1-3)Gal(β1-4)GlcNAc(β1-2)Man(α1-6)[Gal(β1-4)GlcNAc(β1-3)Ga
Fraction Bound
1.0
0.8
0.6
0.4
0.2
0.0

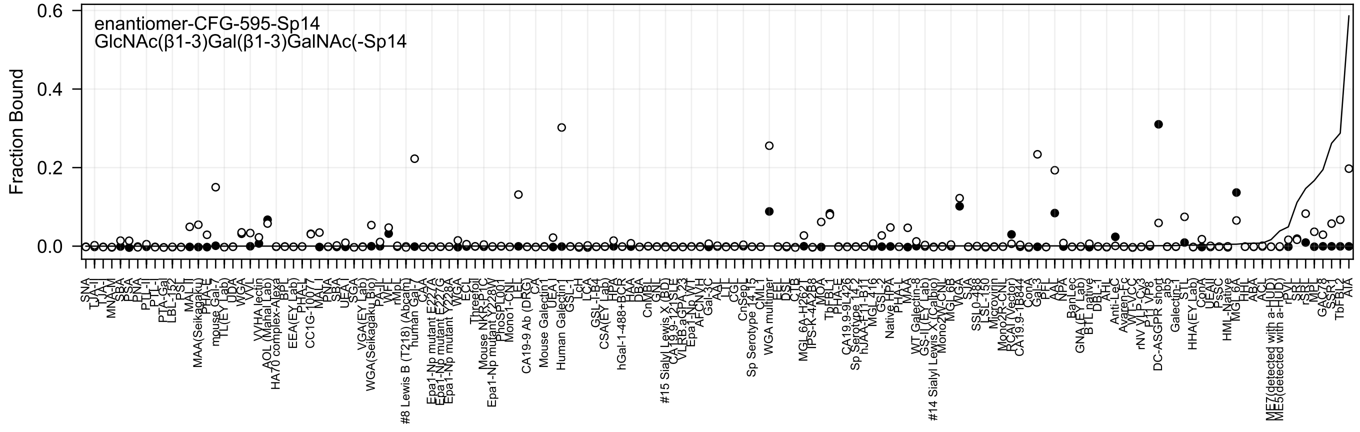
enantiomer-CFG-595-Sp14
GlcNAc(β1-3)Gal(β1-3)GalNAc(-Sp14
Fraction Bound

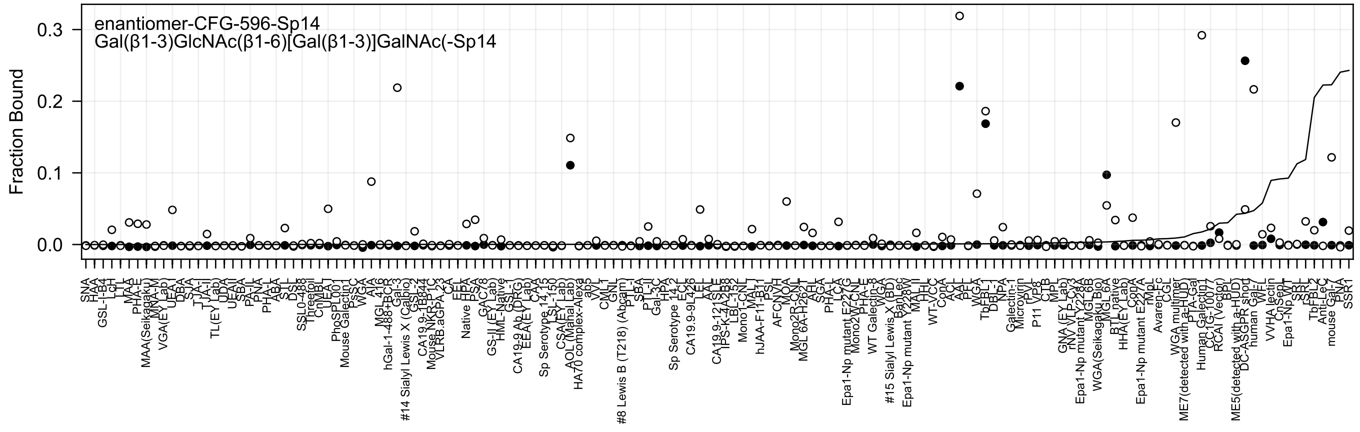

enantiomer-CFG-596-Sp14
Gal(β1-3)GlcNAc(β1-6)[Gal(β1-3)]GalNAc(-Sp14
Fraction Bound
0.0
0.1
0.2
0.3

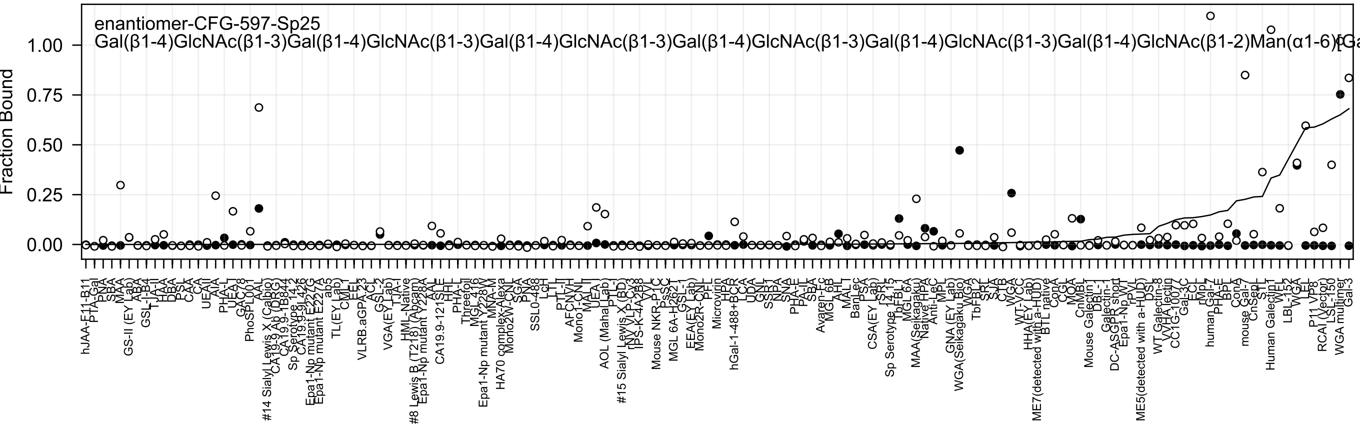
enantiomer-CFG-597-Sp25
Gal(β1-4)GlcNAc(β1-3)Gal(β1-4)GlcNAc(β1-3)Gal(β1-4)GlcNAc(β1-3)Gal(β1-4)GlcNAc(β1-3)Gal(β1-4)GlcNAc(β1-3)Gal(β1-4)GlcNAc(β1-2)Man(α1-6)[Ga
Fraction Bound
1.00
0.75
0.50
0.25
0.00
hJAA-F11-B11
PTA-Gal
PNA
SBA
MAA
GS-II (EY Lab)
ABA
GSL-I-B4
TJA-II
HAA
DBA
PSL
CAA
CA
UEAII
AIA
PHA-L
UEA-I
GAC78
PhoSL001
AAL
#14 Sialyl Lewis X (Calbio)
CA19-9 Ab (DRG)
CA19.9-1B844
Sp Serotype 14.2
CA19-9 9L426
Epa1-Np mutant E227G
Epa1-Np mutant E227A
ab5
TL(EY Lab)
CML-I
EEL
VLRB.aGPA.23
ACL
GSL-2
VGA(EY Lab)
TJA-I
HML-Native
#8 Lewis B (T218) (Abcam)
Epa1-Np mutant Y228A
AAL
CA19.9-121SLE
HHL
PHA-L
Threefoil
MGL 416
Epa1-Np mutant Y228W
MNA-M
HA70 complex-Alexa
Mono2W-CNL
SGA
PNA
SSL0-488
LcH
LTL
PTL-II
AFCNVH
Mono1-CNL
MAL-II
AOL (Mahal Lab)
UEA-I
PTL-I
#15 Sialyl Lewis X (BD)
rNV VLP-Cy3
IPS-K-4A2B8
VVL
Mouse NKR-P1C
PsSC
MGL 6A-H262T
GSL-1
EEA(EY Lab)
Mono2R-CNL
PFL
Microvirin
HPA
hGal-1-488+BCR
LCA
UDA
GNL
SSP1
NPA
SNA
PHA-E
PA-IL
SBA
Avaren-Fc
MGL 6B
AHL
MAL-I
BanLec
PSA
CSA(EY Lab)
ISR
Sp Serotype 14.15
TbFBL1
MGL 6A
MAA(Seikagaku)
Native HPA
Anti-LeC
MPL
GNA (EY Lab)
WGA(Seikagaku Bio)
WGA
TbFBL2
SRL
SJA
CTB
WGA
WT-VCC
HHA(EY Lab)
ME7(detected with a-HUD)
BTL native
ConA
CGL
MOA
CnMBL
Mouse Galectin1
DBL-1
Galectin-1
DC-ASGPR short
Epa1-Np WT
rPVL
ME5(detected with a-HUD)
DSL
WT Galectin-8
VVHA lectin
CC1G-1007
Gal-3C
ECL
rMpL
human Gal-7
PHA-E
BPL
ConA
mouse Gal-7
CnSepL
STL
Human Galectin1
WFL
LBL-152
WGA
LEL
P11 VP8
RCAI (Vector)
LSL-150
WGA multimer
Gal-3

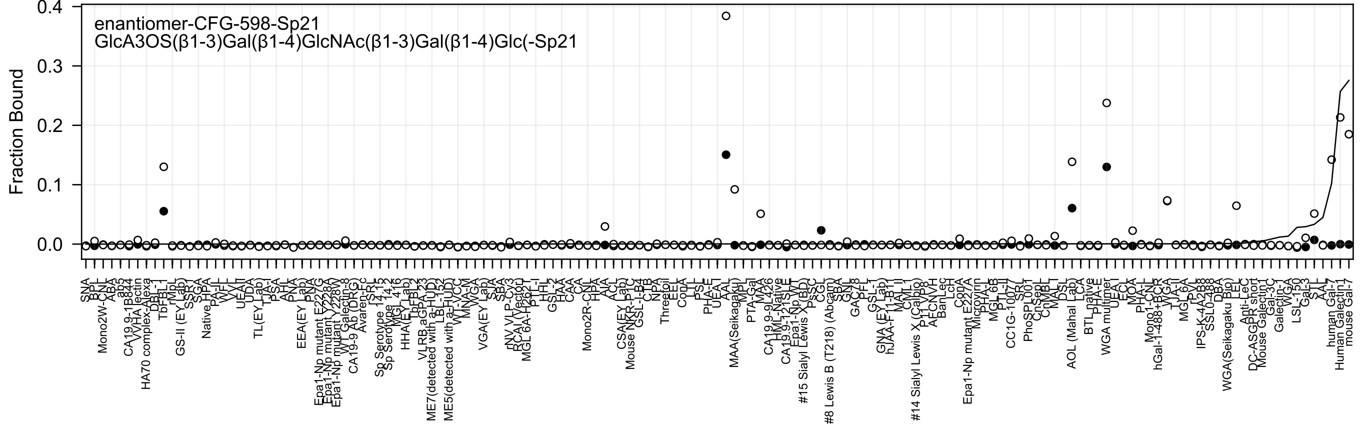
enantiomer-CFG-598-Sp21
GlcA3OS(β1-3)Gal(β1-4)GlcNAc(β1-3)Gal(β1-4)Glc(-Sp21
Fraction Bound
0.0
0.1
0.2
0.3
0.4

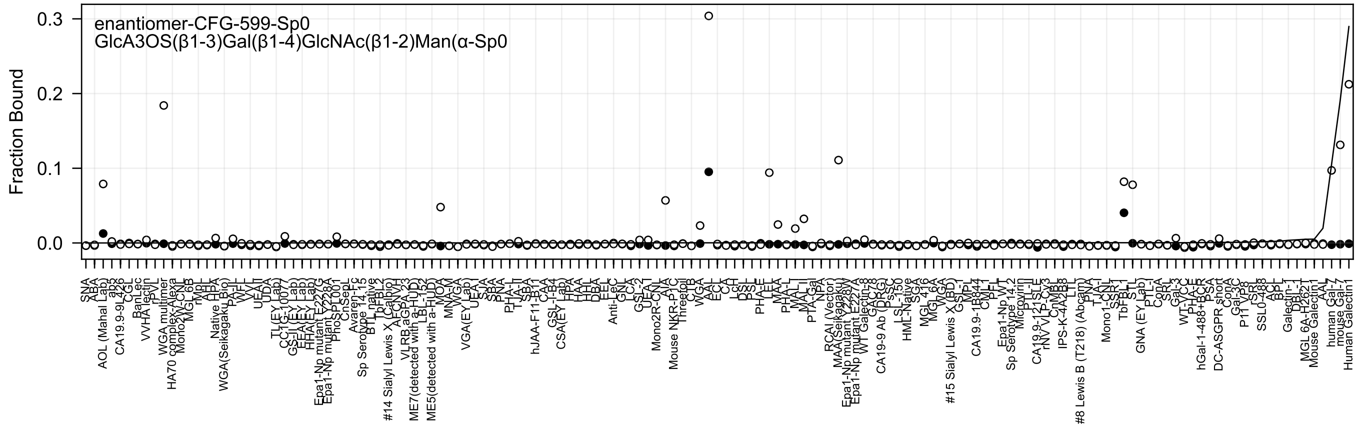

enantiomer-CFG-599-Sp0
GlcA3OS(β1-3)Gal(β1-4)GlcNAc(β1-2)Man(α-Sp0
Fraction Bound
0.3
0.2
0.1
0.0

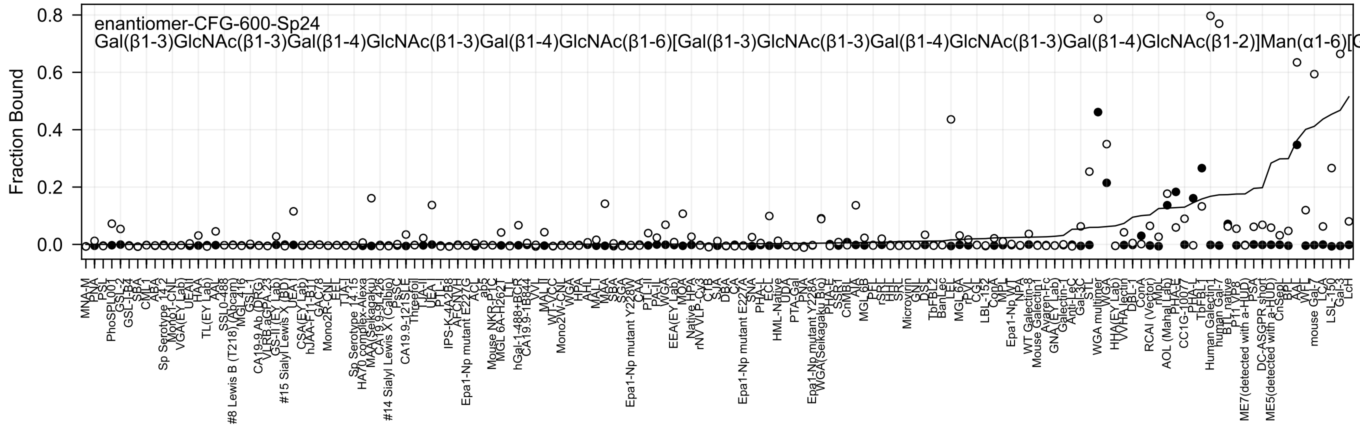

enantiomer-CFG-600-Sp24
Gal(β1-3)GlcNAc(β1-3)Gal(β1-4)GlcNAc(β1-3)Gal(β1-4)GlcNAc(β1-6)[Gal(β1-3)GlcNAc(β1-3)Gal(β1-4)GlcNAc(β1-3)Gal(β1-4)GlcNAc(β1-2)]Man(α1-6)[G
Fraction Bound
0.0
0.2
0.4
0.6
0.8

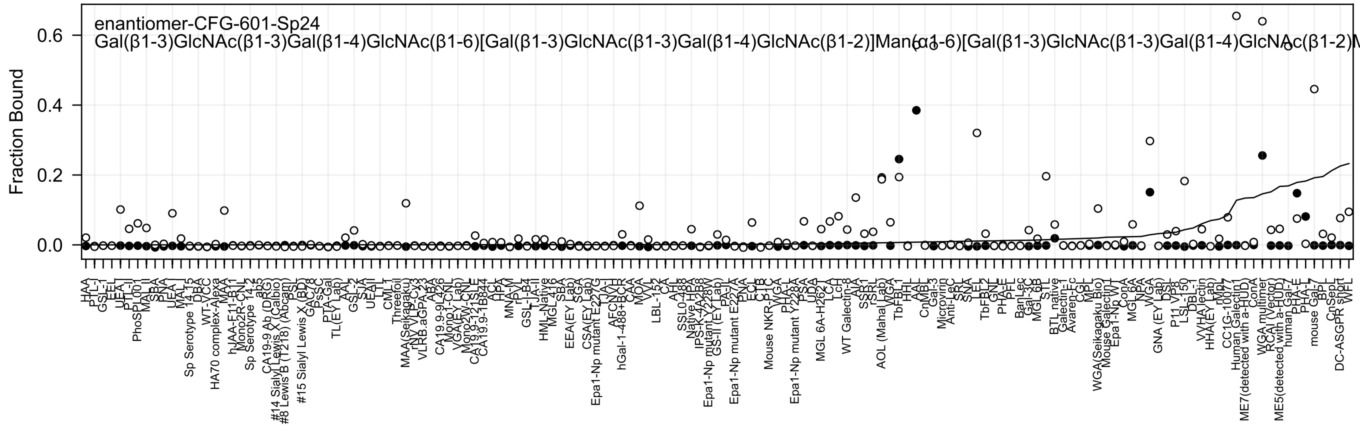
enantiomer-CFG-601-Sp24
Gal(β1-3)GlcNAc(β1-3)Gal(β1-4)GlcNAc(β1-6)[Gal(β1-3)GlcNAc(β1-3)Gal(β1-4)GlcNAc(β1-2)]Man(α1-6)[Gal(β1-3)GlcNAc(β1-3)Gal(β1-4)GlcNAc(β1-2)M
Fraction Bound
0.0
0.2
0.4
0.6

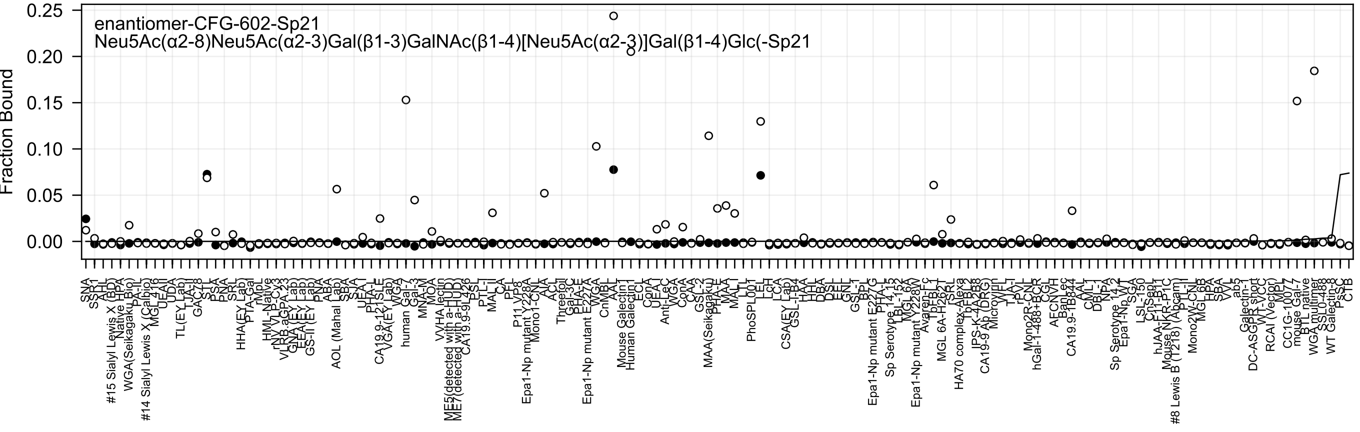
enantiomer-CFG-602-Sp21
Neu5Ac(α2-8)Neu5Ac(α2-3)Gal(β1-3)GalNAc(β1-4)[Neu5Ac(α2-3)]Gal(β1-4)Glc(-Sp21
Fraction Bound
0.25
0.20
0.15
0.10
0.05
0.00

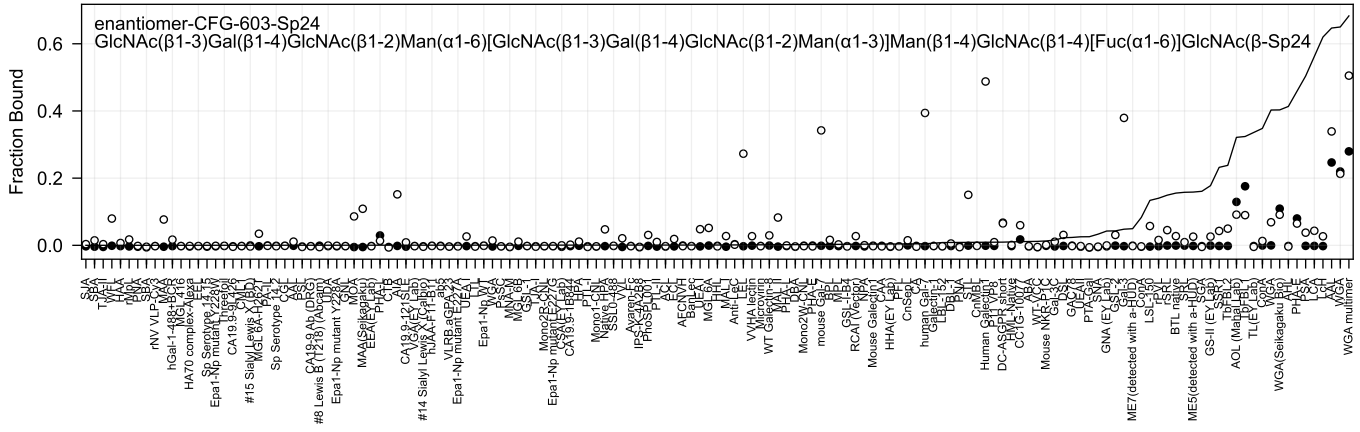
enantiomer-CFG-603-Sp24
GlcNAc(β1-3)Gal(β1-4)GlcNAc(β1-2)Man(α1-6)[GlcNAc(β1-3)Gal(β1-4)GlcNAc(β1-2)Man(α1-3)]Man(β1-4)GlcNAc(β1-4)[Fuc(α1-6)]GlcNAc(β-Sp24
Fraction Bound
0.0
0.2
0.4
0.6

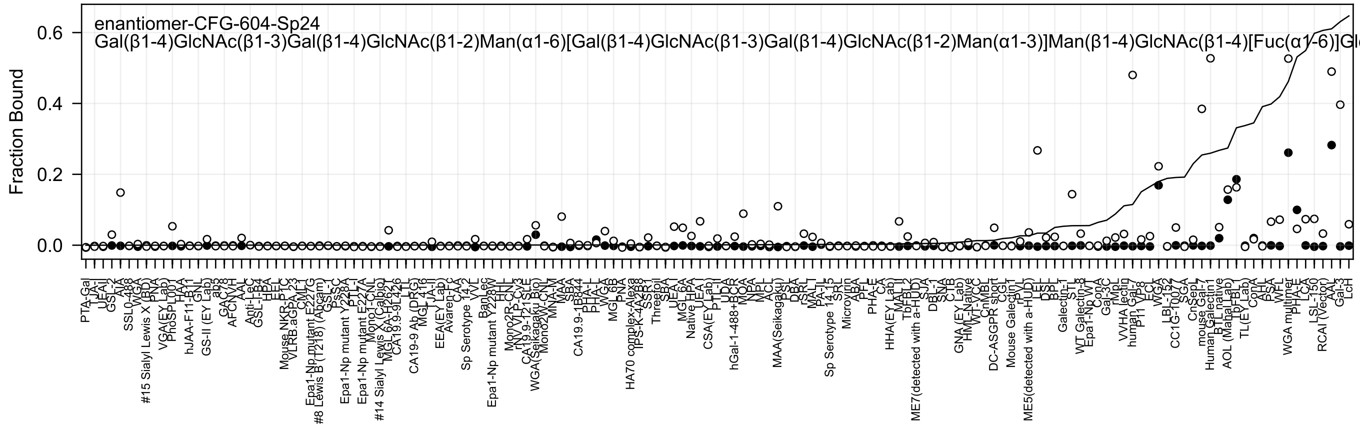

enantiomer-CFG-604-Sp24
Gal(β1-4)GlcNAc(β1-3)Gal(β1-4)GlcNAc(β1-2)Man(α1-6)[Gal(β1-4)GlcNAc(β1-3)Gal(β1-4)GlcNAc(β1-2)Man(α1-3)]Man(β1-4)GlcNAc(β1-4)[Fuc(α1-6)]Gl
Fraction Bound
0.0
0.2
0.4
0.6

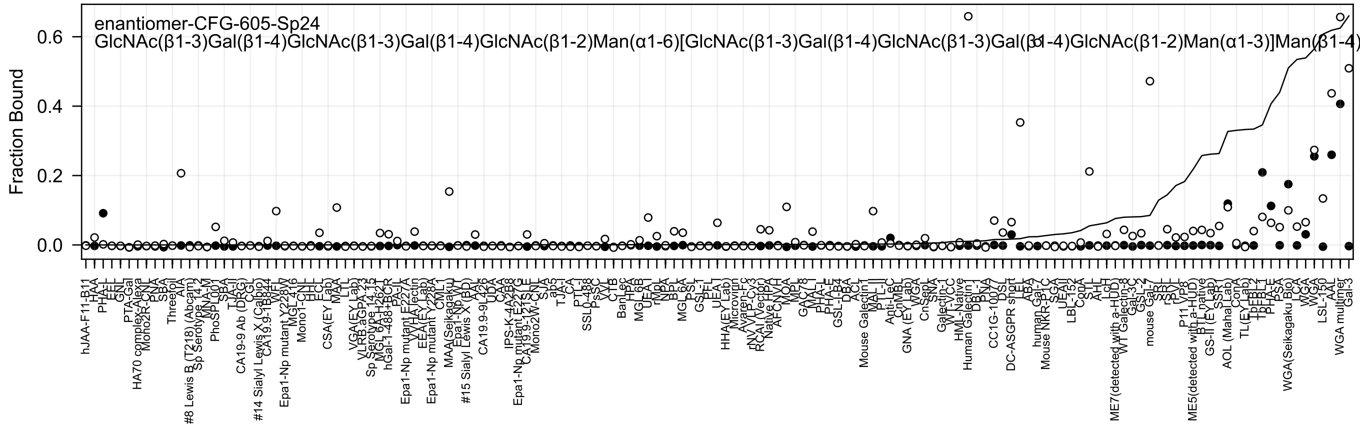

enantiomer-CFG-605-Sp24
GlcNAc(β1-3)Gal(β1-4)GlcNAc(β1-3)Gal(β1-4)GlcNAc(β1-2)Man(α1-6)[GlcNAc(β1-3)Gal(β1-4)GlcNAc(β1-3)Gal(β1-4)GlcNAc(β1-2)Man(α1-3)]Man(β1-4)
Fraction Bound
0.0
0.2
0.4
0.6

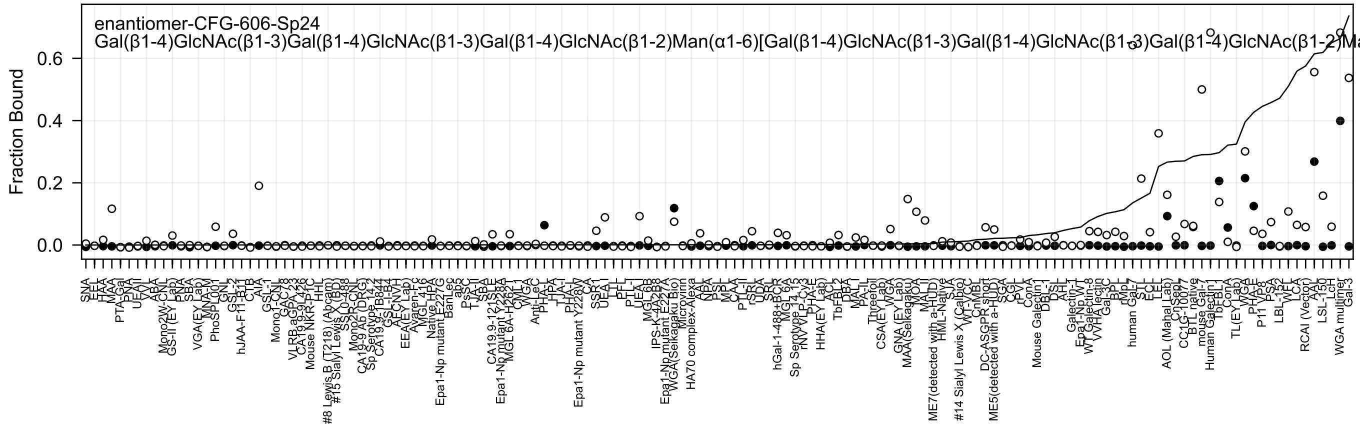
enantiomer-CFG-606-Sp24
Gal(β1-4)GlcNAc(β1-3)Gal(β1-4)GlcNAc(β1-3)Gal(β1-4)GlcNAc(β1-2)Man(α1-6)[Gal(β1-4)GlcNAc(β1-3)Gal(β1-4)GlcNAc(β1-3)Gal(β1-4)GlcNAc(β1-2)Ma
Fraction Bound
0.0
0.2
0.4
0.6
SNA
EEL
HAA
MAA
PTA-Gal
PNA
UEAII
VVL
ABA
Mono2W-CNL
GS-II (EY Lab)
PNA
SBA
VGA(EY Lab)
MNA-M
PhoSL001
GNL
GSL-2
hJAA-F11-B11
CTB
AIA
GSL-1
Mono1-CNL
GAC78
VLRB.aGPA.23
CA19.9-9L426
Mouse NKR-P1C
HHL
#8 Lewis B (T218) (Abcam)
#15 Sialyl Lewis X (BD)
SSL0-488
Mono2R-CNL
CA19-9 Ab (DRG)
Sp Serotype 14.2
CA19.9-1B844
GSL-I-B4
AFCNVH
EEA(EY Lab)
Avaren-Fc
MGL 416
Native HPA
Epa1-Np mutant E227G
BanLec
ab2
PsSC
TJA-II
SBA
CA19.9-121SLE
Epa1-Np mutant Y228A
MGL 6A-H262T
WGA
Anti-LeC
PHA-L
HPA
TJA-I
Epa1-Np mutant Y228W
CA
SSR1
UEA-I
LEL
PFL
PTL-I
MGL 6B
IPS-K-4A2B8
Epa1-Np mutant E227A
WGA(Selkagaku Bio)
Microvirin
HA70 complex-Alexa
AAL
NPA
PSL
MPL
CAA
PTL-II
rSRL
UDA
SRL
hGal-1-488+BCR
MGL 6A
Sp Serotype 14.15
rNV VLP-Cy3
PHA-E
HHA(EY Lab)
ACL
TbF-BL2
DBA
MAL-I
PA-IL
Threefoil
CSA(EY Lab)
WGA
GNA (EY Lab)
MAA(Selkagaku)
MOA
MAL-II
ME7(detected with a-HUD)
HML-Native
SJA
#14 Sialyl Lewis X (Calbio)
CnMBL
DC-ASGPR short
ME5(detected with a-HUD)
SGA
CGL
rPVL
ConA
Mouse Galectin1
DBL-1
DSL
AHL
Galectin-1
Epa1-Np WT
WT Galectin-8
VVHA lectin
Gal-3C
BPL
rMpL
human Gal-7
STL
ECL
LEL
AOL (Mahal Lab)
CnSep7
CC1G-10077
BTL native
mouse Gal-7
Human Galectin1
TbF-BL1
ConA
TL(EY Lab)
WGA
PHA-E
P11 VP8
PSA
LBL-152
WFL
LCA
RCAI (Vector)
AAL
LSL-150
LcH
WGA multimer
Gal-3

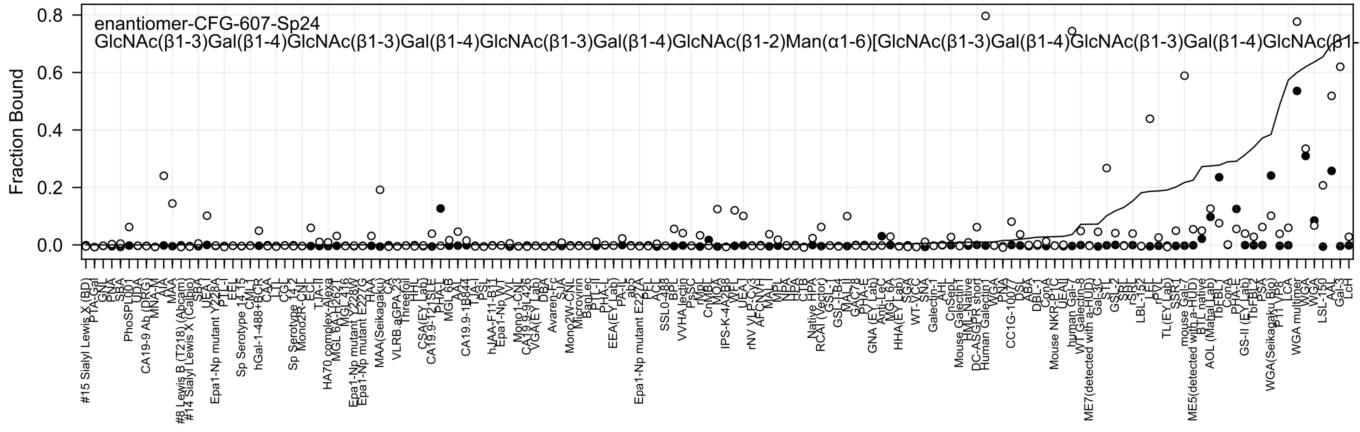
enantiomer-CFG-607-Sp24
GlcNAc(β1-3)Gal(β1-4)GlcNAc(β1-3)Gal(β1-4)GlcNAc(β1-3)Gal(β1-4)GlcNAc(β1-2)Man(α1-6)[GlcNAc(β1-3)Gal(β1-4)GlcNAc(β1-3)Gal(β1-4)GlcNAc(β1-
Fraction Bound

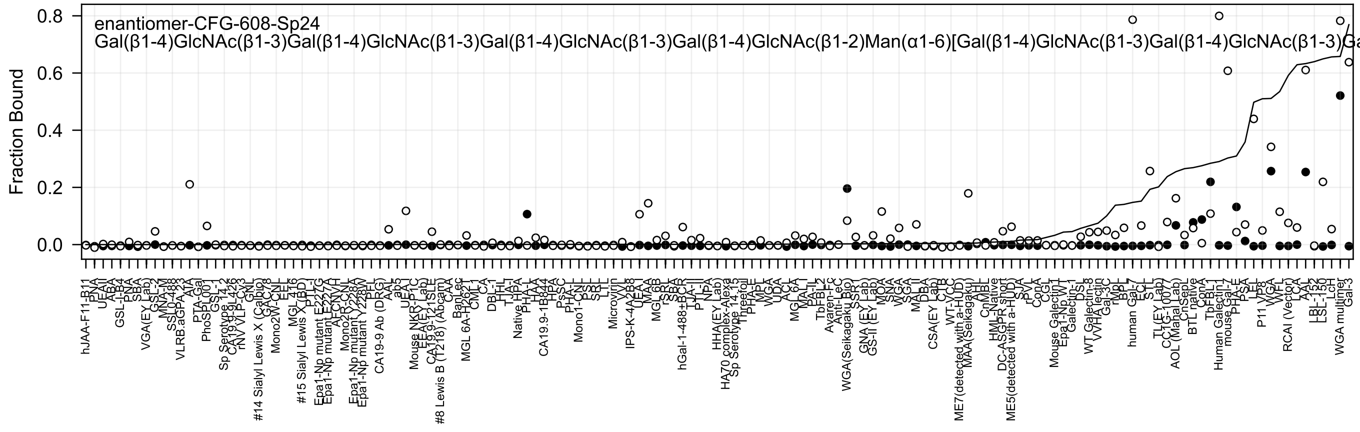
enantiomer-CFG-608-Sp24
Gal(β1-4)GlcNAc(β1-3)Gal(β1-4)GlcNAc(β1-3)Gal(β1-4)GlcNAc(β1-3)Gal(β1-4)GlcNAc(β1-2)Man(α1-6)[Gal(β1-4)GlcNAc(β1-3)Gal(β1-4)GlcNAc(β1-3)Ga
Fraction Bound
0.8
0.6
0.4
0.2
0.0

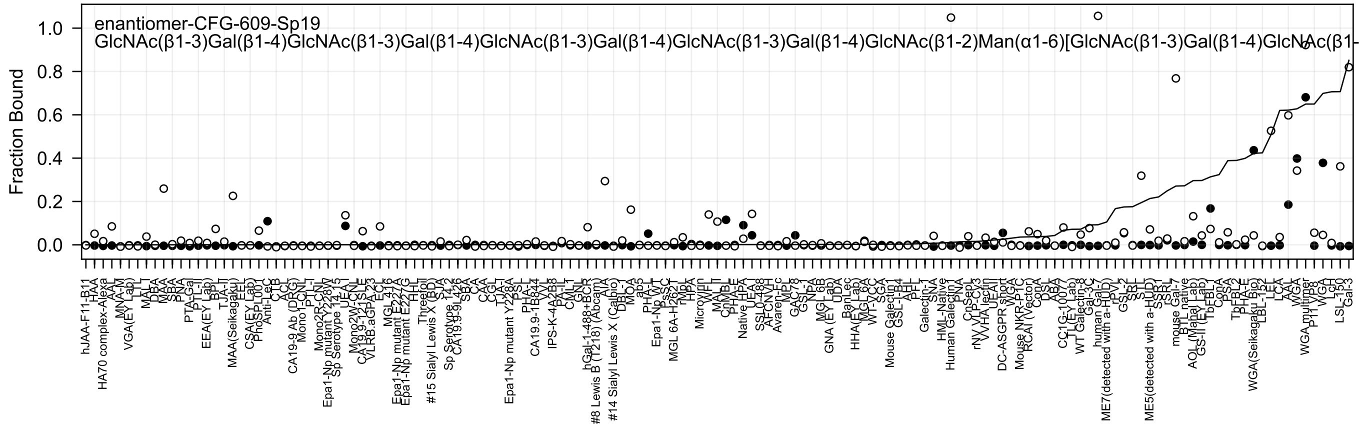

enantiomer-CFG-609-Sp19
GlcNAc(β1-3)Gal(β1-4)GlcNAc(β1-3)Gal(β1-4)GlcNAc(β1-3)Gal(β1-4)GlcNAc(β1-3)Gal(β1-4)GlcNAc(β1-2)Man(α1-6)[GlcNAc(β1-3)Gal(β1-4)GlcNAc(β1-
Fraction Bound

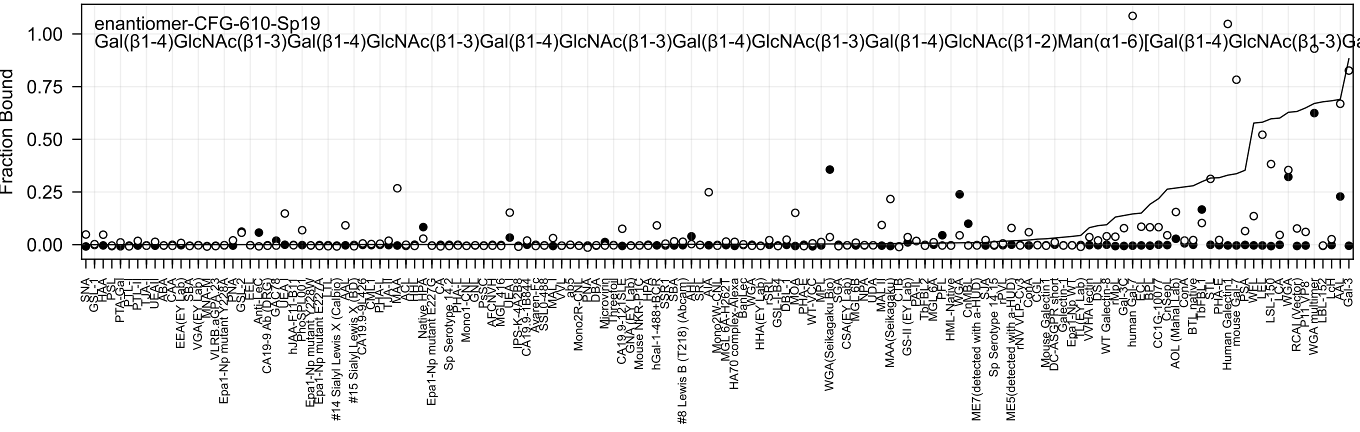

enantiomer-CFG-610-Sp19
Gal(β1-4)GlcNAc(β1-3)Gal(β1-4)GlcNAc(β1-3)Gal(β1-4)GlcNAc(β1-3)Gal(β1-4)GlcNAc(β1-3)Gal(β1-4)GlcNAc(β1-2)Man(α1-6)[Gal(β1-4)GlcNAc(β1-3)Ga
Fraction Bound
1.00
0.75
0.50
0.25
0.00

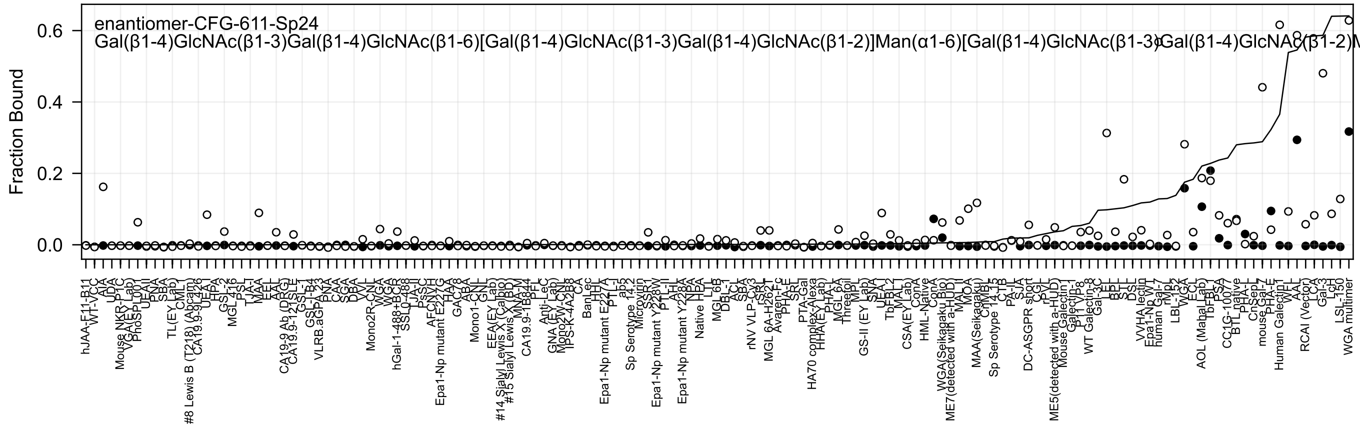
enantiomer-CFG-611-Sp24
Gal(β1-4)GlcNAc(β1-3)Gal(β1-4)GlcNAc(β1-6)[Gal(β1-4)GlcNAc(β1-3)Gal(β1-4)GlcNAc(β1-2)]Man(α1-6)[Gal(β1-4)GlcNAc(β1-3)Gal(β1-4)GlcNAc(β1-2)M
Fraction Bound
0.0
0.2
0.4
0.6

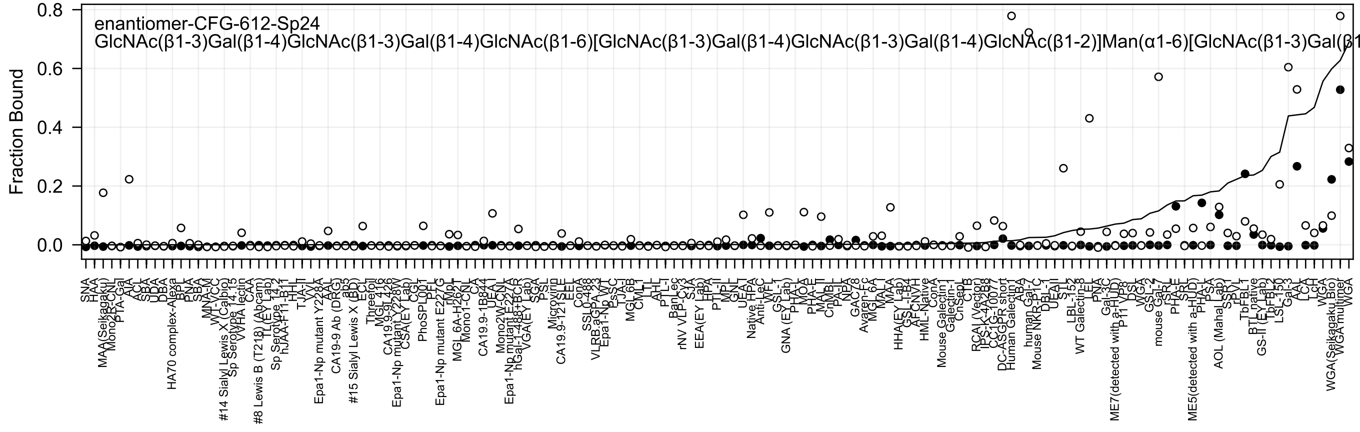

enantiomer-CFG-612-Sp24
GlcNAc(β1-3)Gal(β1-4)GlcNAc(β1-3)Gal(β1-4)GlcNAc(β1-6)[GlcNAc(β1-3)Gal(β1-4)GlcNAc(β1-3)Gal(β1-4)GlcNAc(β1-2)]Man(α1-6)[GlcNAc(β1-3)Gal(β1
Fraction Bound
0.8
0.6
0.4
0.2
0.0

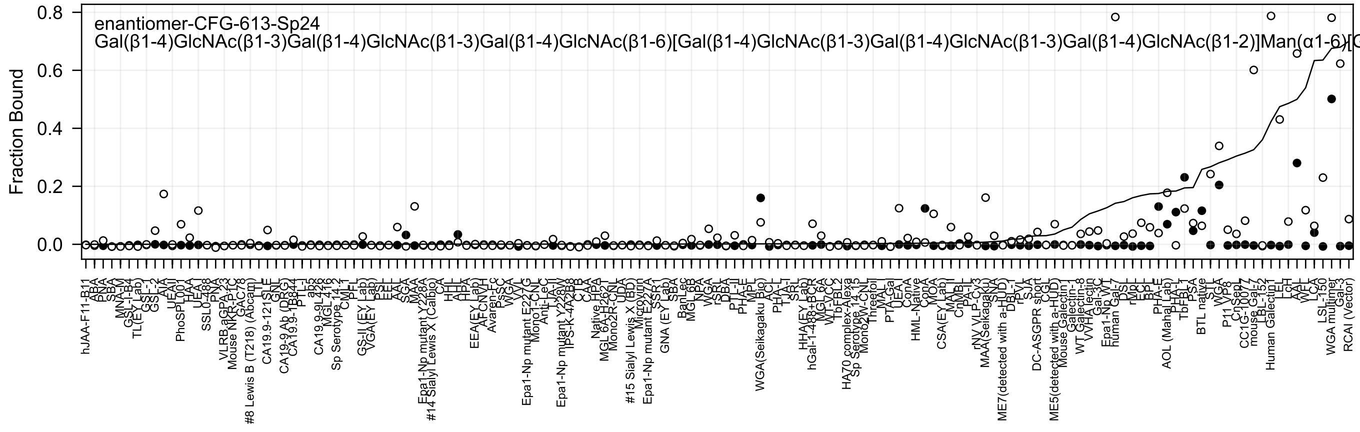
enantiomer-CFG-613-Sp24
Gal(β1-4)GlcNAc(β1-3)Gal(β1-4)GlcNAc(β1-3)Gal(β1-4)GlcNAc(β1-6)[Gal(β1-4)GlcNAc(β1-3)Gal(β1-4)GlcNAc(β1-3)Gal(β1-4)GlcNAc(β1-2)]Man(α1-6)[
Fraction Bound

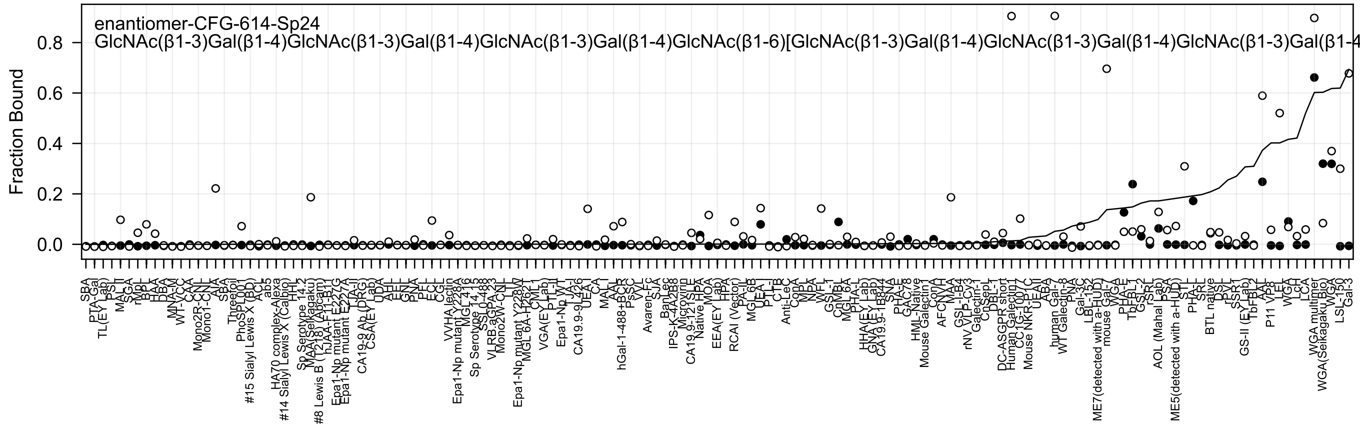
enantiomer-CFG-614-Sp24
GlcNAc(β1-3)Gal(β1-4)GlcNAc(β1-3)Gal(β1-4)GlcNAc(β1-3)Gal(β1-4)GlcNAc(β1-6)[GlcNAc(β1-3)Gal(β1-4)GlcNAc(β1-3)Gal(β1-4)GlcNAc(β1-3)Gal(β1-4
Fraction Bound
0.0
0.2
0.4
0.6
0.8
SBA
PTA-Gal
TL(EY Lab)
PSL
MAL_I
SGA
rMpL
BPL
HAA
DBA
MNA-M
WT-VCC
CAA
Mono2R-CNL
Mono1-CNL
AIA
SBA
Threefoil
PhoSL001
#15 Sialyl Lewis X (BD)
ACL
ab5
HA70 complex-Alexa
#14 Sialyl Lewis X (Calbio)
HHL
Sp Serotype 14.2
MAA(Seikagaku)
#8 Lewis B (T218) (Abcam)
hJAA-F11-B11
Epa1-Np mutant E227G
Epa1-Np mutant E227A
CA19-9 Ab (DRG)
CSA(EY Lab)
UDA
AAL
EEL
GNL
PNA
PFL
ECL
CGL
VVHA lectin
Epa1-Np mutant Y228A
MGL 4.16
Sp Serotype 14.15
SSL0-488
VLRB.aGPA.23
Mono2W-CNL
Epa1-Np mutant Y228W
MGL 6A-H262T
CML
VGA(EY Lab)
PTL-II
Epa1-Np WT
JAL
CA19.9-9L426
UEA-I
CA
MAL
AAL
hGal-1-488+BCF
PsSC
VVL
Avaren-Fc
SJA
BanLec
IPS-K-4A2B8
Microvirin
CA19.9-121SLE
Native HPA
MOA
EEA(EY Lab)
HPA
RCAI (Vector)
PA-IL
MGL 6B
UEA-I
PTL-I
CTB
Anti-LeC
ConA
MPL
NPA
WFL
GS-I
CnMBL
MGL 6A
PHA-L
HHA(EY Lab)
GNA (EY Lab)
CA19.9-1B844
SNA
PHA-E
GAC78
HML-Native
Mouse Galectin
ConA
AFC NVH
MAA
GSL-I-B4
rNV VLP-Cy3
Galectin-1
CnSepL
DBL-1
DC-ASGPR short
Human Galectin
CC1G-1007
Mouse NKR-P1C
UEA-II
ABA
human Gal-7
WT Galectin-8
PNA
Gal-3C
LBL-152
ME7(detected with a-HuD)
mouse Gal-7
WGA
PHA-E
TbBL-1
GSL-2
rSRL
AOL (Mahal Lab)
PSA
ME5(detected with a-HuD)
STL
PHA-L
SRL
BTL native
DSL
rPVL
SSR1
GS-II (EY Lab)
TbFBL2
AAL
P11 VP8
LEL
WGA
LcH
LCA
WGA multimer
WGA(Seikagaku Bio)
WGA
LSL-150
Gal-3

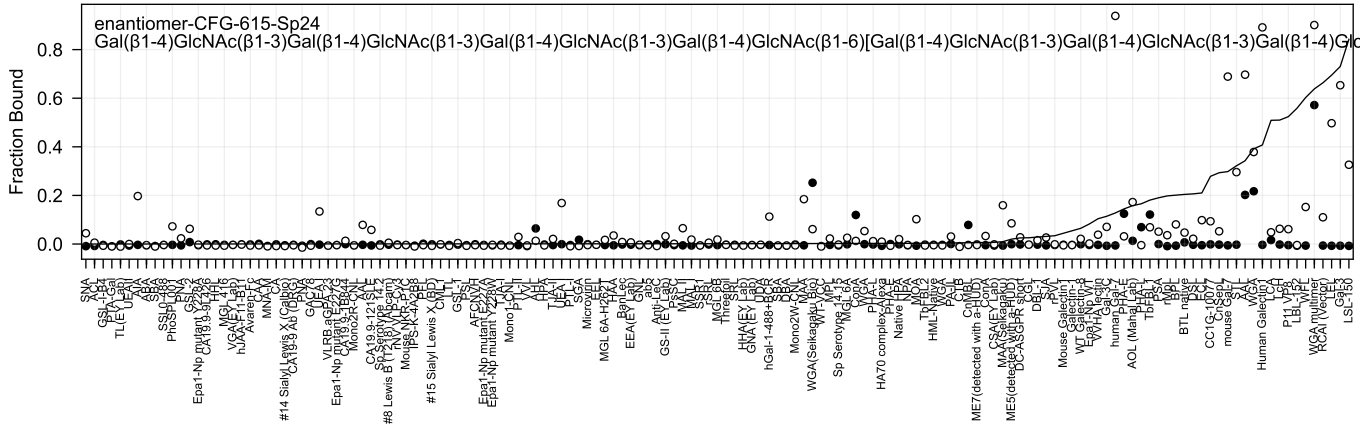
enantiomer-CFG-615-Sp24
Gal(β1-4)GlcNAc(β1-3)Gal(β1-4)GlcNAc(β1-3)Gal(β1-4)GlcNAc(β1-3)Gal(β1-4)GlcNAc(β1-6)[Gal(β1-4)GlcNAc(β1-3)Gal(β1-4)GlcNAc(β1-3)Gal(β1-4)Glc
Fraction Bound
0.0
0.2
0.4
0.6
0.8

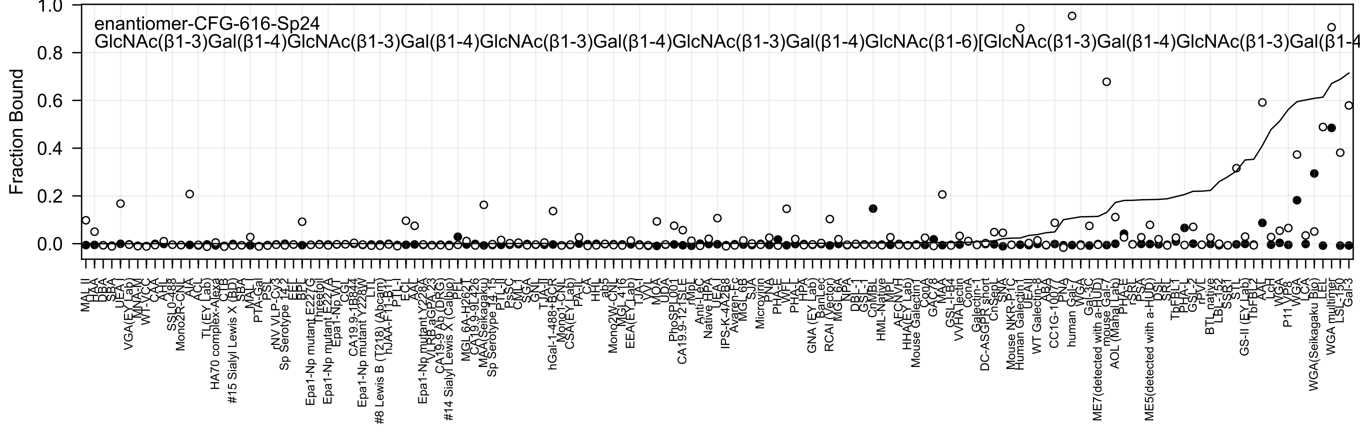

enantiomer-CFG-616-Sp24
GlcNAc(β1-3)Gal(β1-4)GlcNAc(β1-3)Gal(β1-4)GlcNAc(β1-3)Gal(β1-4)GlcNAc(β1-3)Gal(β1-4)GlcNAc(β1-6)[GlcNAc(β1-3)Gal(β1-4)GlcNAc(β1-3)Gal(β1-4
Fraction Bound
1.0
0.8
0.6
0.4
0.2
0.0
MAL_II
HAA
DBA
SBA
UEA-I
VGA(EY Lab)
MNA-M
WT-VCC
CAA
XCL
SSL0-488
Mono2R-CNL
AIA
ACL
TL(EY Lab)
HA70 complex-Alexa
CTB
#15 Sialyl Lewis X (BD)
SBA
MAL-I
PTA-Gal
PSL
rNV VLP-Cy3
Sp Serotype 14.2
EEL
BPL
Epa1-Np mutant E227G
Threetol
Epa1-Np mutant E227A
Epa1-Np WT
CGL
CA19.9-1B844
Epa1-Np mutant Y228W
LTL
#8 Lewis B (T218) (Abcam)
hJAA-F11-B11
PTL-I
ECL
AAL
Epa1-Np mutant Y228A
VLRB aGPA.23
CA19.9 Ab (DRG)
#14 Sialyl Lewis X (Calbio)
PPL
MGL 6A-H262T
CA19.9-9L426
MAA(Seikagaku)
Sp Serotype 14.15
PTL-II
PsSC
CML1
SGA
GNL
TJA-II
hGal-1-488+BCR
Mono1-CNL
CSA(EY Lab)
PA-IL
CA
HHL
ab5
Mono2W-CNL
MGL 416
EEA(EY Lab)
TJA-I
VVL
MOA
UDA
PhoSL001
CA19.9-121SLE
rMpL
Anti-LeC
Native HPA
UEA
IPS-K-4A2B8
Avaren-Fc
MGL 6B
SJA
Microvirin
PNA
PHA-E
WFL
PHA-L
HPA
GNA (EY Lab)
BanLec
RCAI (Vector)
MGL 6A
NPA
DBL-1
GSL-I
CnMBL
HML-Native
MPL
AECNVH
HHA(EY Lab)
Mouse Galectin
ConA
GAC7/8
MAA
GSL-I-B4
VVHA lectin
ConA
Galectin-1
DC-ASGPR short
CnSepL
SNA
Mouse NKR-P1C
Human Galectin
UEA II
WT Galectin-8
ABA
CC1G-1007
PNA
human Gal-7
WGA
Gal-3C
ME7(detected with a-HUD)
mouse Gal-7
AOL (Mahal Lab)
PHA-E
rSRL
PSA
ME5(detected with a-HUD)
DSL
SRL
TbFBL1
PHA-L
GSL-2
rPVL
BTL native
LBL-152
SSBL1
STL
GS-II (EY Lab)
TbFBL2
AAL
LcH
WGA
P11
VP8
WGA
LCA
WGA(Seikagaku Bio)
LEL
WGA multimer
LSL-150
Gal-3

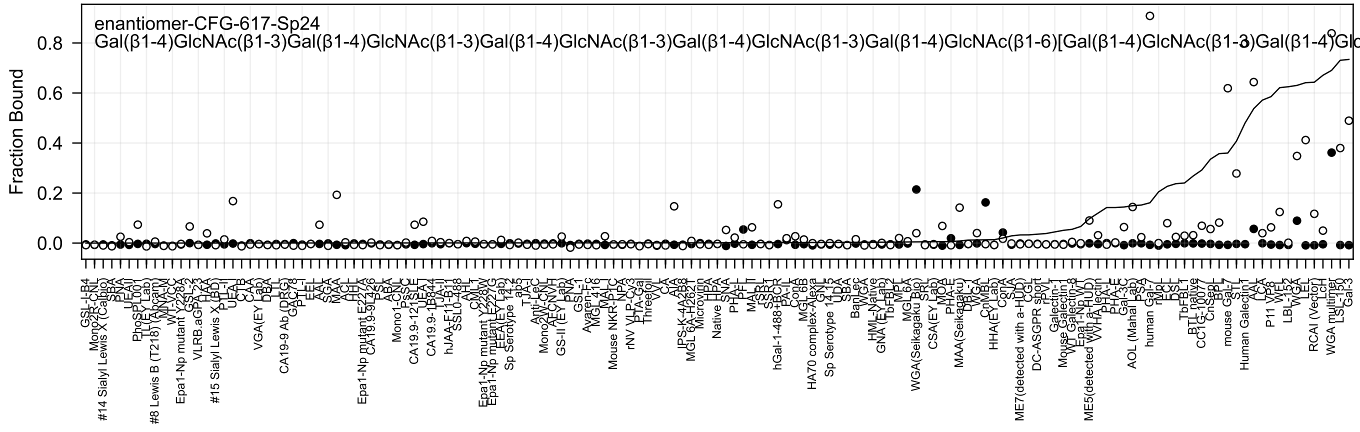
enantiomer-CFG-617-Sp24
Gal(β1-4)GlcNAc(β1-3)Gal(β1-4)GlcNAc(β1-3)Gal(β1-4)GlcNAc(β1-3)Gal(β1-4)GlcNAc(β1-3)Gal(β1-4)GlcNAc(β1-6)[Gal(β1-4)GlcNAc(β1-3)Gal(β1-4)Glc
Fraction Bound
0.0
0.2
0.4
0.6
0.8

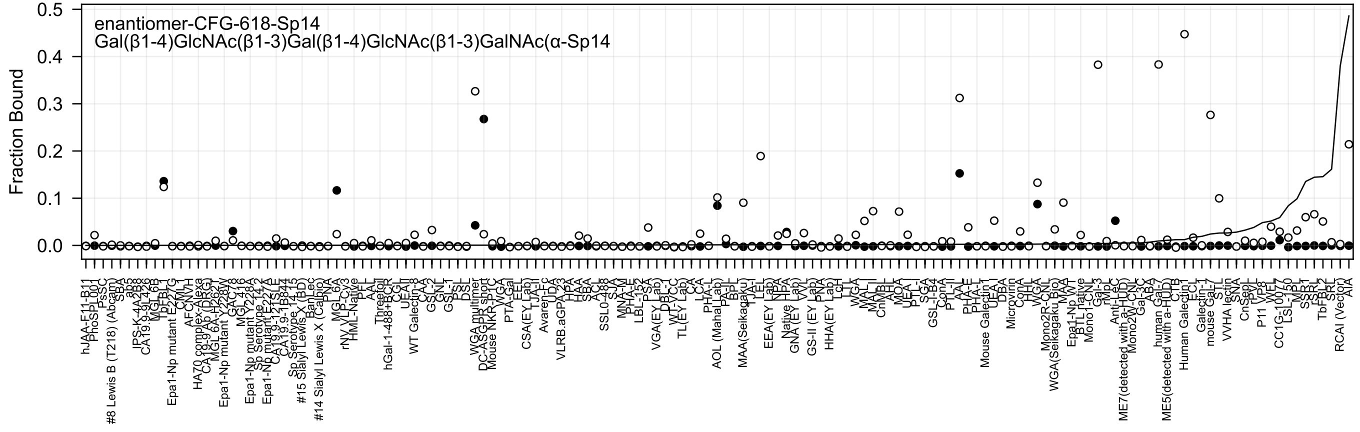
enantiomer-CFG-618-Sp14
Gal(β1-4)GlcNAc(β1-3)Gal(β1-4)GlcNAc(β1-3)GalNAc(α-Sp14
Fraction Bound

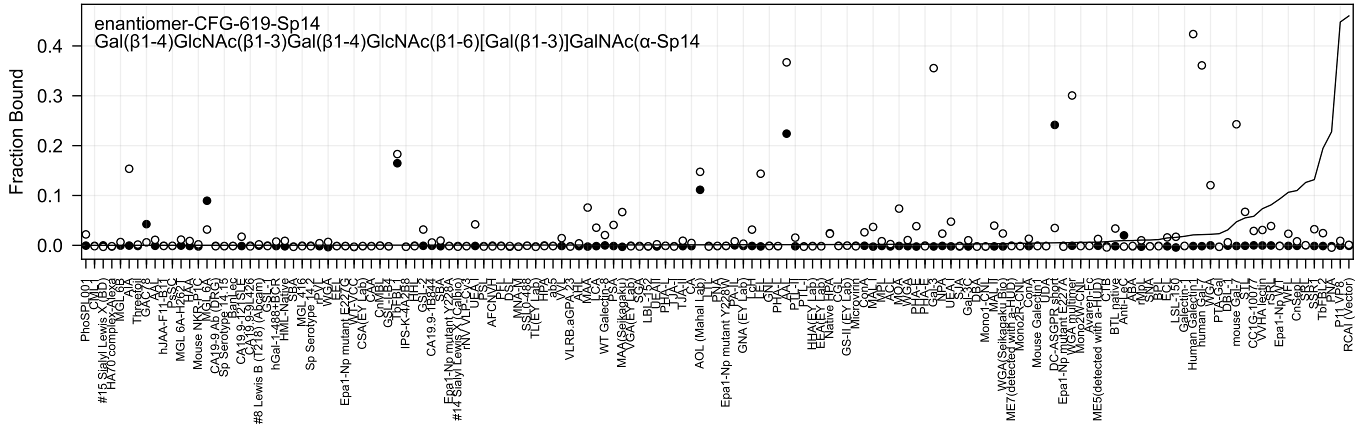
enantiomer-CFG-619-Sp14
Gal(β1-4)GlcNAc(β1-3)Gal(β1-4)GlcNAc(β1-6)[Gal(β1-3)]GalNAc(α-Sp14
Fraction Bound
0.4
0.3
0.2
0.1
0.0

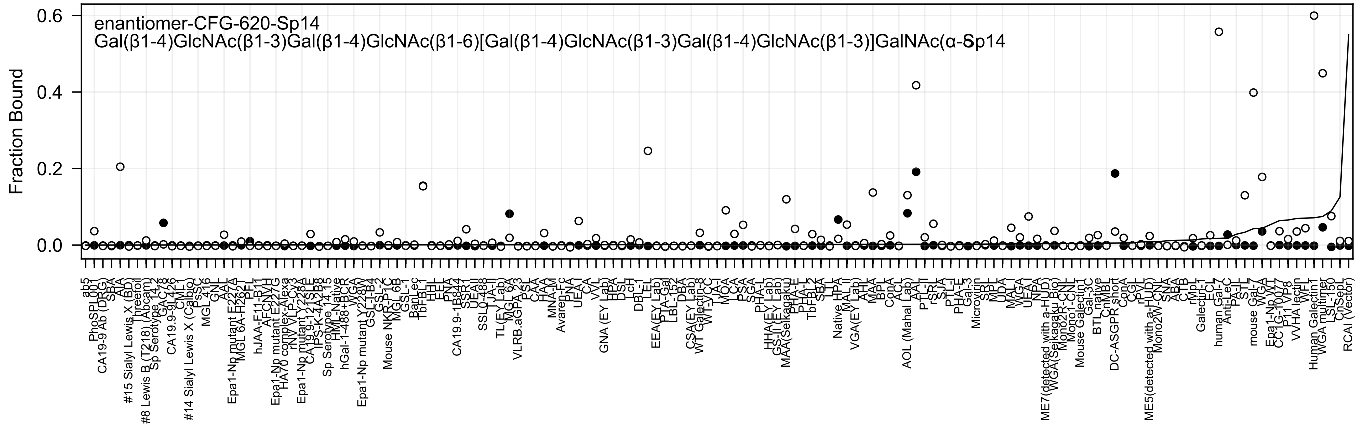
enantiomer-CFG-620-Sp14
Gal(β1-4)GlcNAc(β1-3)Gal(β1-4)GlcNAc(β1-6)[Gal(β1-4)GlcNAc(β1-3)Gal(β1-4)GlcNAc(β1-3)]GalNAc(α-Sp14
Fraction Bound
0.6
0.4
0.2
0.0

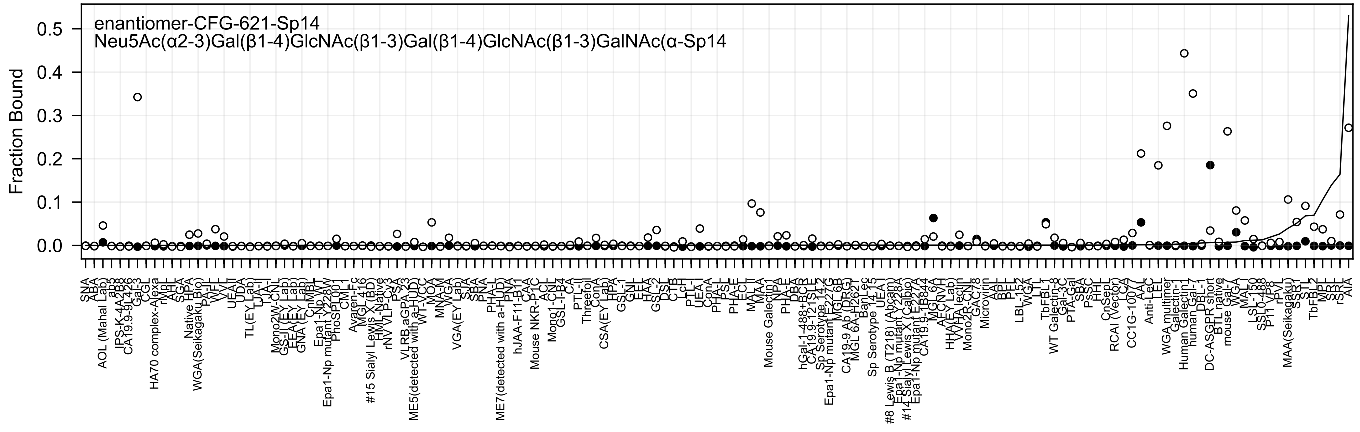

enantiomer-CFG-621-Sp14
Neu5Ac(α2-3)Gal(β1-4)GlcNAc(β1-3)Gal(β1-4)GlcNAc(β1-3)GalNAc(α-Sp14
Fraction Bound

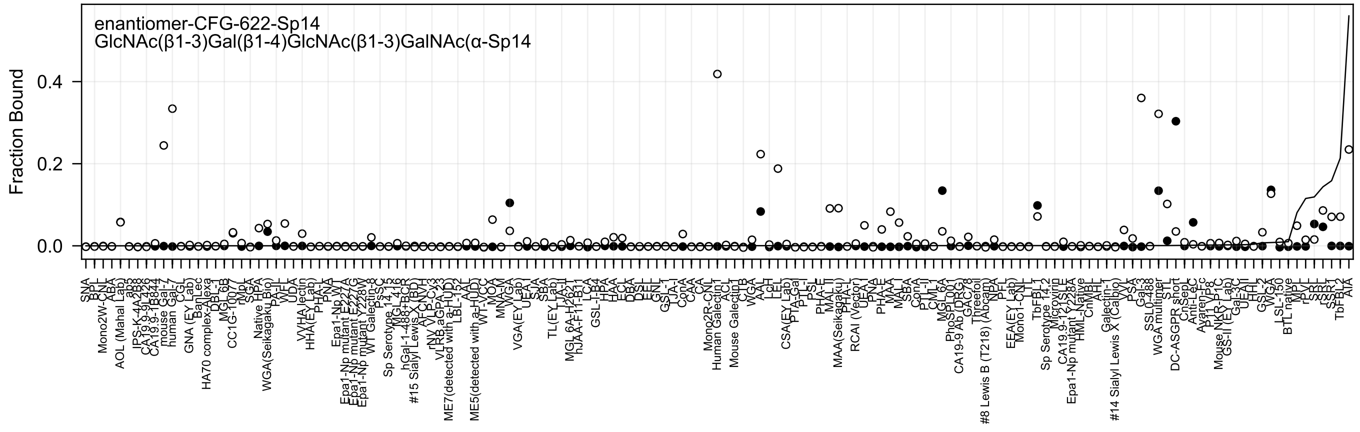

enantiomer-CFG-622-Sp14
GlcNAc(β1-3)Gal(β1-4)GlcNAc(β1-3)GalNAc(α-Sp14
Fraction Bound
0.0
0.2
0.4

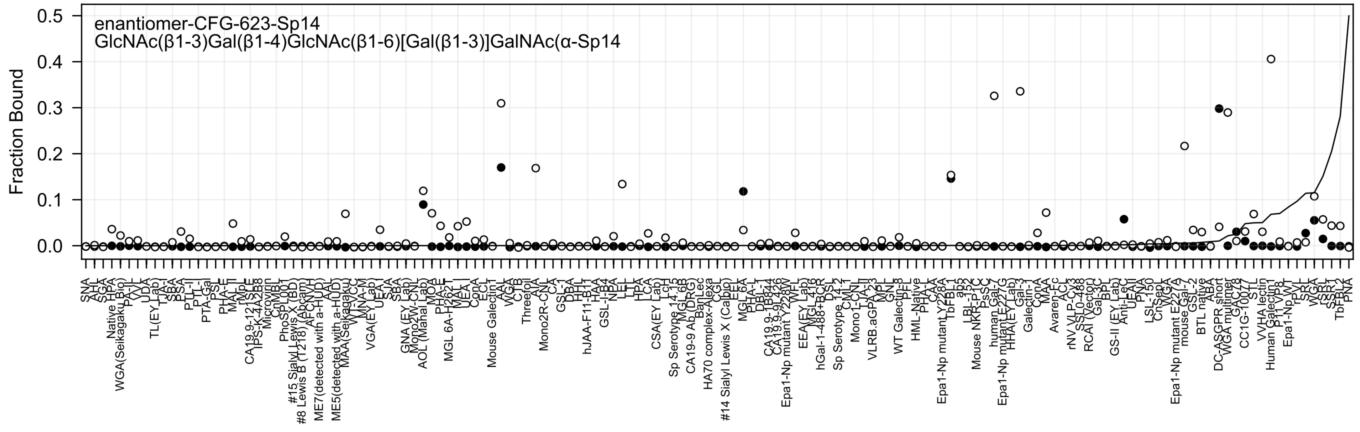

enantiomer-CFG-623-Sp14
GlcNAc(β1-3)Gal(β1-4)GlcNAc(β1-6)[Gal(β1-3)]GalNAc(α-Sp14
Fraction Bound
0.5
0.4
0.3
0.2
0.1
0.0
SNA
AHL
SGA
Native HPA
WGA(Seikagaku Bio)
PA-IL
VVL
UDA
TL(EY Lab)
TJA-I
SBA
PSA
PTL-II
PTL-I
PTA-Gal
PSL
PHA-E
MAL-II
rMpL
CA19.9-121SLE
IPS-K-4A2B8
Microvirin
CnMBL
PhoSL001
#15 Sialyl Lewis X (BD)
#8 Lewis B (T218) (Abcam)
AFCNVH
ME7(detected with a-HUD)
AAL
ME5(detected with a-HUD)
MAA(Seikagaku)
WFA-CC
MNA-M
VGA(EY Lab)
UEA-I
SJA
SBA
GNA (EY Lab)
Mono2W-CNL
AOL (Manal Lab)
MOA
PHA-E
MGL 6A-H262T
MAL-I
WEA
ConA
ECL
Mouse Galectin1
AAL
WGA
CTB
Threefoil
AIA
Mono2R-CNL
CA
GSL-1
DBA
HHL
hJAA-F11-B11
HAA
GSL-I-B4
NBA
LEL
LTL
HPA
LCA
CSA(EY Lab)
Lcrl
Sp Serotype 14.15
MGL 6B
CA19-9 Ab (DRG)
BanLec
HA70 complex-Alexa
CGL
#14 Sialyl Lewis X (Calbio)
EEL
MGL 6A
PHA-L
DBL-T
CA19.9-1B844
CA19.9-9L426
Epa1-Np mutant Y228W
WFL
EEA(EY Lab)
MGL 4B
hGal-1-488+BCR
DSL
Sp Serotype 14.2
CML1
Mono1-CNL
TJA-II
VLRB.aGPA.23
MPL
GNL
WT Galectin-8
PFL
HML-Native
PHA-L
CAA
Epa1-Np mutant Y228A
TbFBL1
abs
LBL-152
Mouse NKR-P1C
PsSC
human Gal-7
Epa1-Np mutant E227G
HHA(EY Lab)
Gal-3
Galectin-1
ConA
MAA
Avaren-Fc
ACL
rNV VLP-Cy3
SSL0-488
RCAI (Vector)
Gal-3C
BPL
GS-II (EY Lab)
Anti-LeC
UEA-II
PNA
LSL-150
CnSep
WGA
Epa1-Np mutant E227A
mouse Gal-7
GSL-2
BTL native
ABA
DC-ASGPR short
WGA multimer
GAC78
CC1G-10077
STL
VVHA lectin
Human Galectin1
P1 VP8
Epa1-Np WT
rPVL
SRL
WGA
rSRL
SSL
TbFBL2
PNA

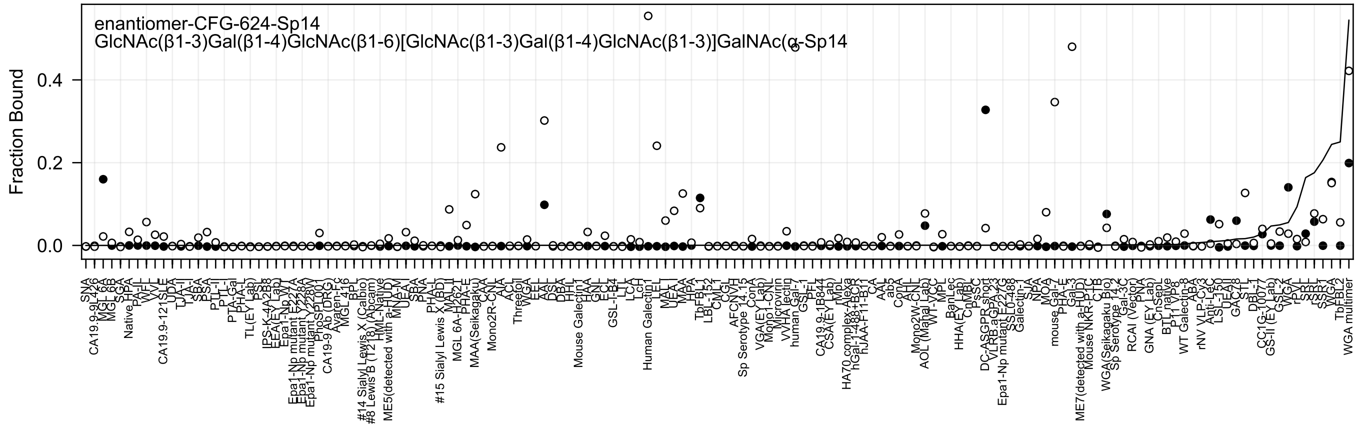

enantiomer-CFG-624-Sp14
GlcNAc(β1-3)Gal(β1-4)GlcNAc(β1-6)[GlcNAc(β1-3)Gal(β1-4)GlcNAc(β1-3)]GalNAc(α-Sp14
Fraction Bound
0.0
0.2
0.4

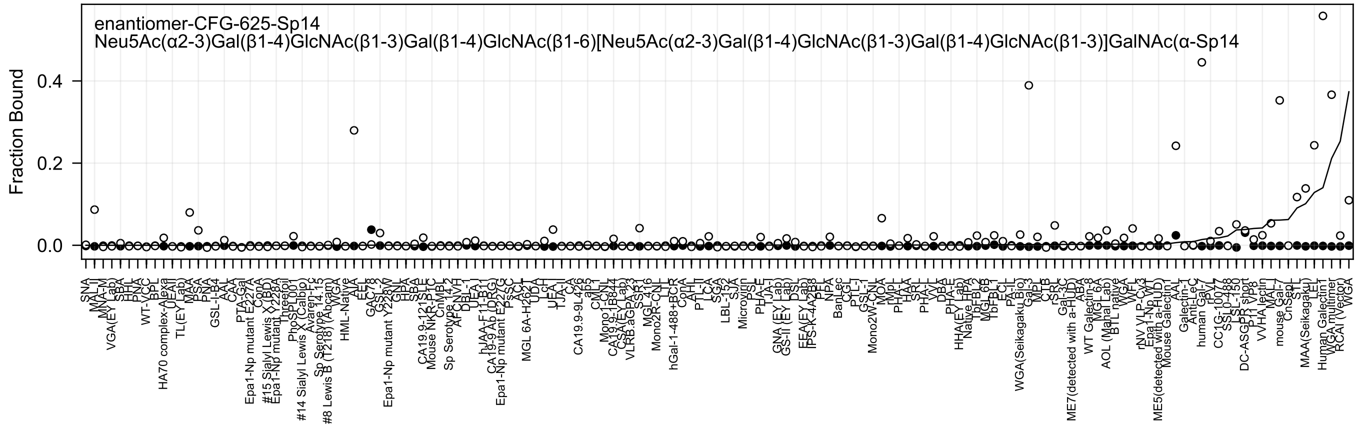

enantiomer-CFG-625-Sp14
Neu5Ac(α2-3)Gal(β1-4)GlcNAc(β1-3)Gal(β1-4)GlcNAc(β1-6)[Neu5Ac(α2-3)Gal(β1-4)GlcNAc(β1-3)Gal(β1-4)GlcNAc(β1-3)]GalNAc(α-Sp14
Fraction Bound
0.0
0.2
0.4

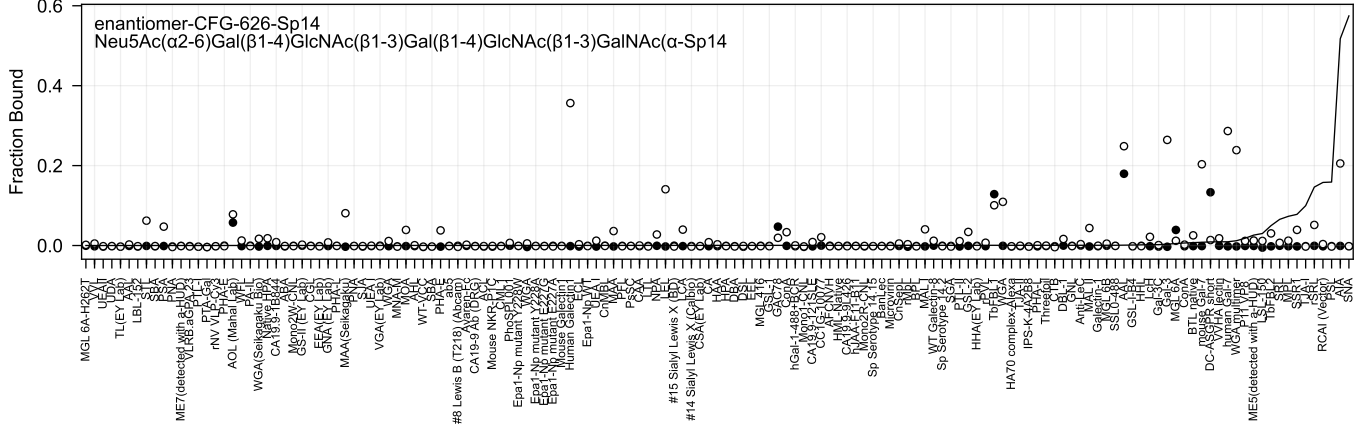
enantiomer-CFG-626-Sp14
Neu5Ac(α2-6)Gal(β1-4)GlcNAc(β1-3)Gal(β1-4)GlcNAc(β1-3)GalNAc(α-Sp14
Fraction Bound
0.6
0.4
0.2
0.0

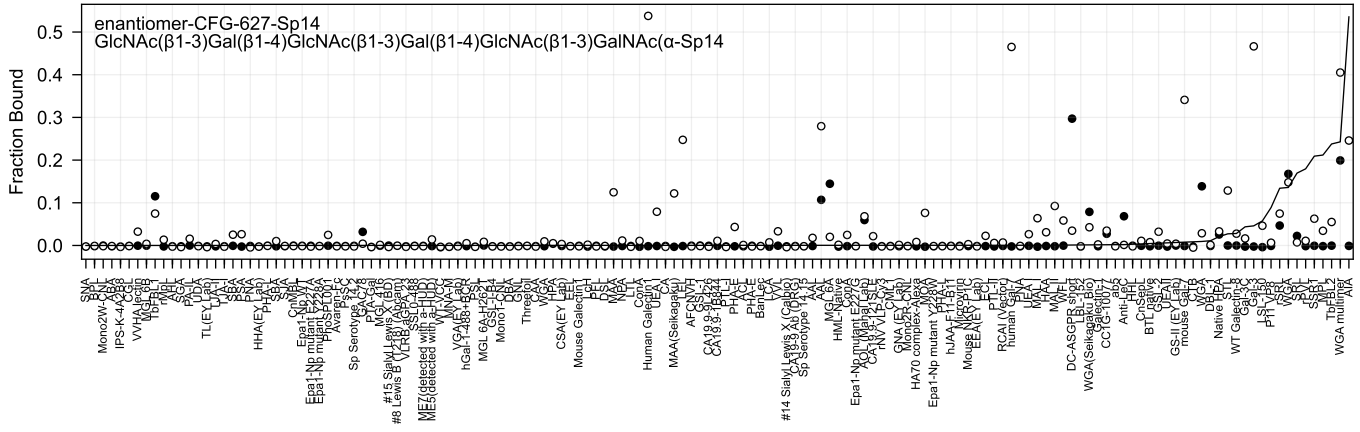

enantiomer-CFG-627-Sp14
GlcNAc(β1-3)Gal(β1-4)GlcNAc(β1-3)Gal(β1-4)GlcNAc(β1-3)GalNAc(α-Sp14
Fraction Bound
0.0
0.1
0.2
0.3
0.4
0.5
SNA
Mono2W-CNL
IPS-K-4A2B8
CGL
VVHA lectin
MGL 6B
TbFBL1
rMpL
AAL
SGA
PA-IL
UDA
TL(EY Lab)
LJA-I
SBA
PSA
PNA
HHA(EY Lab)
PHA-L
SBA
SJA
CnMBL
Epa1-Np WT
Epa1-Np mutant E227A
Epa1-Np mutant Y228A
PhoSL001
Avaren-Fc
PsSC
Sp Serotype 14.2
GAC78
PTA-Gal
MGL 416
#15 Sialyl Lewis X (BD)
#8 Lewis B (T218) (Abcam)
VLRB aGPA 23
SSL0-483
ME7(detected with a-HUD)
ME5(detected with a-HUD)
WT-VCC
MNA-M
VGA(EY Lab)
hGal-1-488+BCR
PSL
MGL 6A-H262T
GSL-I-B4
Mono1-CNL
DBA
GNL
Threetol
CAA
WGA
HPA
CSA(EY Lab)
EEL
Mouse Galectin1
LcH
PFL
DSL
MAA
NPA
LTL
ConA
Human Galectin1
UEA I
CA
MAA(Seikagaku)
LEL
AFCNVH
GSL-I
CA19.9-9I-426
CA19.9-1B844
PTL-I
PHA-E
ACL
PHA-E
BanLec
LCA
TVL
#14 Sialyl Lewis X (Calbio)
CA19-9 Ab (DRG)
Sp Serotype 14.13
AAL
MGL 6A
HML-Native
ConA
Epa1-Np mutant E227G
AOL (Mahal Lab)
CA19.9-121SLE
rNV VLP-Cy3
CML
GNA (EY Lab)
Mono2R-CNL
HA70 complex-Alexa
MOA
Epa1-Np mutant Y228W
PHA-L
hJAA-F11-B11
Microvirin
Mouse NKR-P1C
EEA(EY Lab)
ECL
PTL-II
RCAI (Vector)
human Gal-7
PNA
UEA I
MAL I
HAA
MAL II
WFL
DC-ASGPR short
LBL-152
WGA(Seikagaku Bio)
Galectin-1
CC1G-1007
ab5
Anti-LeC
HHL
CnSepL
BTL native
GSL-2
UEA II
GS-II (EY Lab)
mouse Gal-7
CTB
WGA
DBL-1
Native HPA
STL
WT Galectin-8
Gal-3C
Gal-3
I-SL-150
P11 VP8
rSRL
WGA
SRL
rPVL
SSR1
MPL
TbFBL2
WGA multimer
AIA

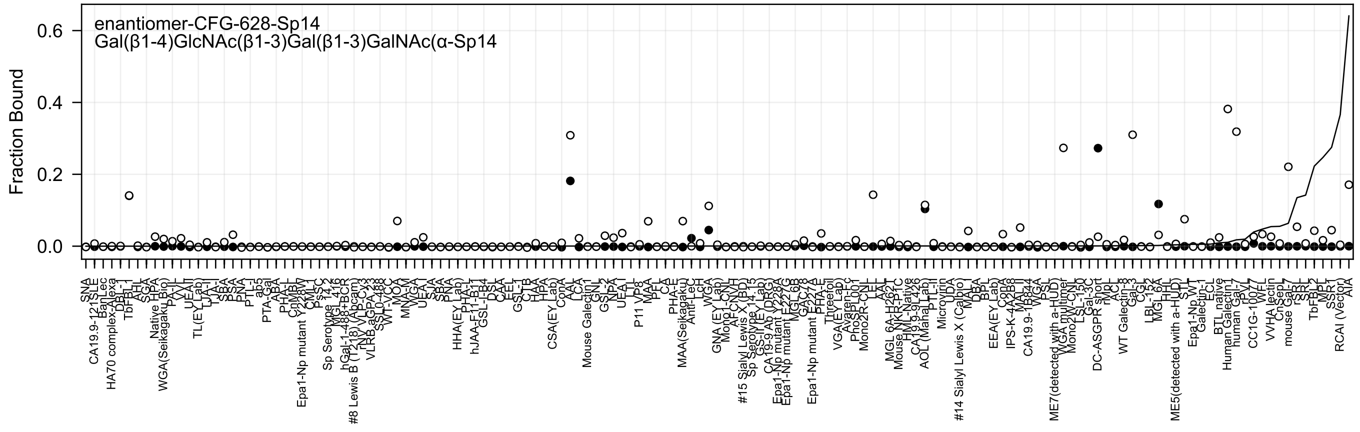
enantiomer-CFG-628-Sp14
Gal(β1-4)GlcNAc(β1-3)Gal(β1-3)GalNAc(α-Sp14
Fraction Bound
0.6
0.4
0.2
0.0
SNA
CA19.9-121SLE
BanLec
HA70 complex-Alexa
DBL-1
TbFBL-1
AHL
SGA
Native HPA
WGA(Seikagaku Bio)
PA-IL
VVL
UEA-I
TL(EY Lab)
TJA-I
SBA
PSA
PNA
PTL-I
ab5
PTA-Gal
ABA
PHA-L
CnMBL
Epa1-Np mutant Y228W
CMI
PsSC
Sp Serotype 14.2
MGL 416
hGal-1-488+BCR
#8 Lewis B (T218) (Abcam)
Nv VLRCx3
VLRB.aGPA.23
SSL0-488
WT-VCC
MOA
MNA-M
VVGA
UEA-I
SJA
SBA
PNA
HHA(EY Lab)
PHA-L
hJAA-F11-B11
GSL-I-B4
DSL
CAA
EEL
GSL-I
CTB
HAA
HPA
CSA(EY Lab)
ConA
AAL
LCA
Mouse Galectin1
GNL
GSL-2
NPA
UEA-I
LTL
P11 VP8
MAA
PFL
CA
PHA-E
MAA(Seikagaku)
Anti-LeC
LcH
WGA
GNA (EY Lab)
Mono1-CNL
AFCNVH
#15 Sialyl Lewis X (BD)
Sp Serotype 14.15
GS-II(EY Lab)
CA 19-9 Ab (DRG)
Epa1-Np mutant T228A
Epa1-Np mutant E227G
MGL 6B
GAC78
PHA-E
Epa1-Np mutant E227...
Threefoil
VGA(EY Lab)
Avaren-Fc
PhoSL001
Mono2R-CNL
CEL
AAL
MGL 6A-H262T
Mouse NK-P1C
HML-Native
CA19.9-9L426
AOL (Mana Lab)
PTL-II
Microvirin
UDA
#14 Sialyl Lewis X (Calbio)
MAL-I
DBA
BPL
EEA(EY Lab)
ConA
IPS-K-4A2B8
MAL-I
CA19.9-1B844
WGA
PSL
ME7(detected with a-HUD)
WGA multimer
Mono2W-CNL
LSL-150
Gal-3C
DC-ASGPR short
rMpL
ACL
WT Galectin-8
Gal-3
CGL
LBL-152
MGL 6A
HHL
ME5(detected with a-HUD)
STL
Epa1-Np WT
Galectin-1
ECL
BTL native
Human Galectin1
human Gal-7
rPVL
CC1G-1007
WFL
VVHA lectin
CnSep
mouse Gal-7
rSRL
SRL
TbFBL2
MPL
SSR1
RCAI (Vector)
AIA

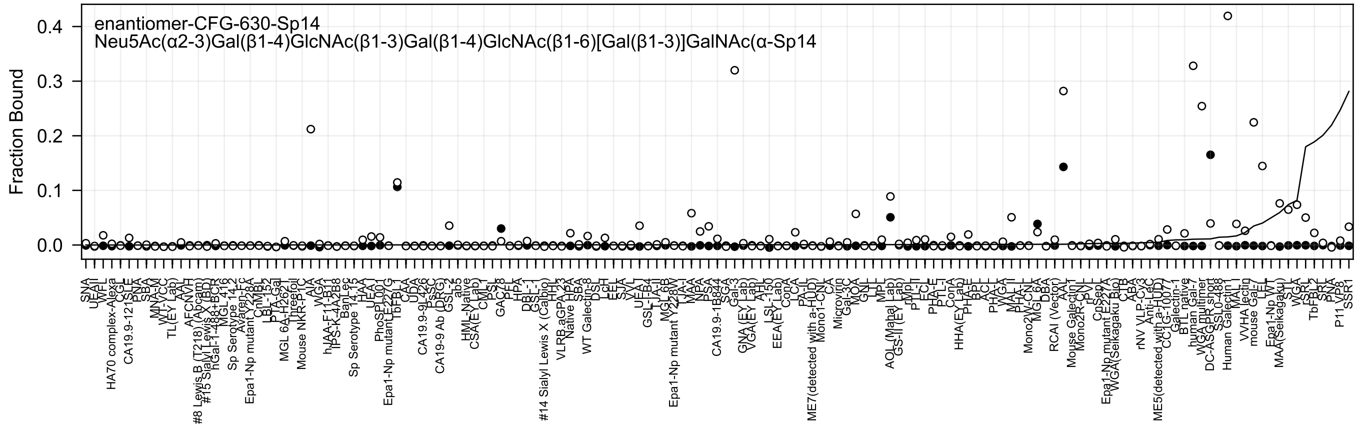
enantiomer-CFG-630-Sp14
Neu5Ac(α2-3)Gal(β1-4)GlcNAc(β1-3)Gal(β1-4)GlcNAc(β1-6)[Gal(β1-3)]GalNAc(α-Sp14
Fraction Bound
0.4
0.3
0.2
0.1
0.0

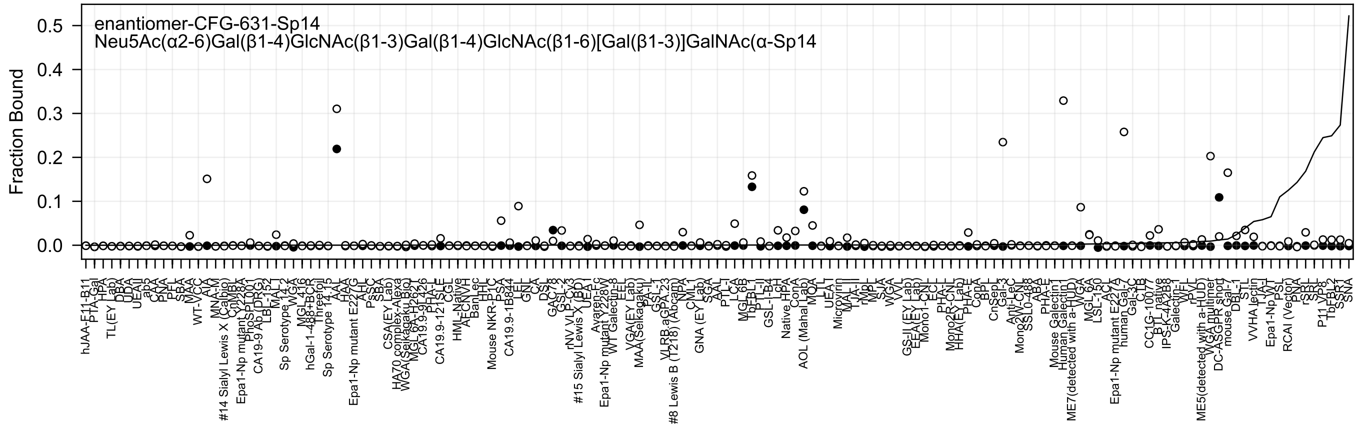
enantiomer-CFG-631-Sp14
Neu5Ac(α2-6)Gal(β1-4)GlcNAc(β1-3)Gal(β1-4)GlcNAc(β1-6)[Gal(β1-3)]GalNAc(α-Sp14
Fraction Bound
0.5
0.4
0.3
0.2
0.1
0.0

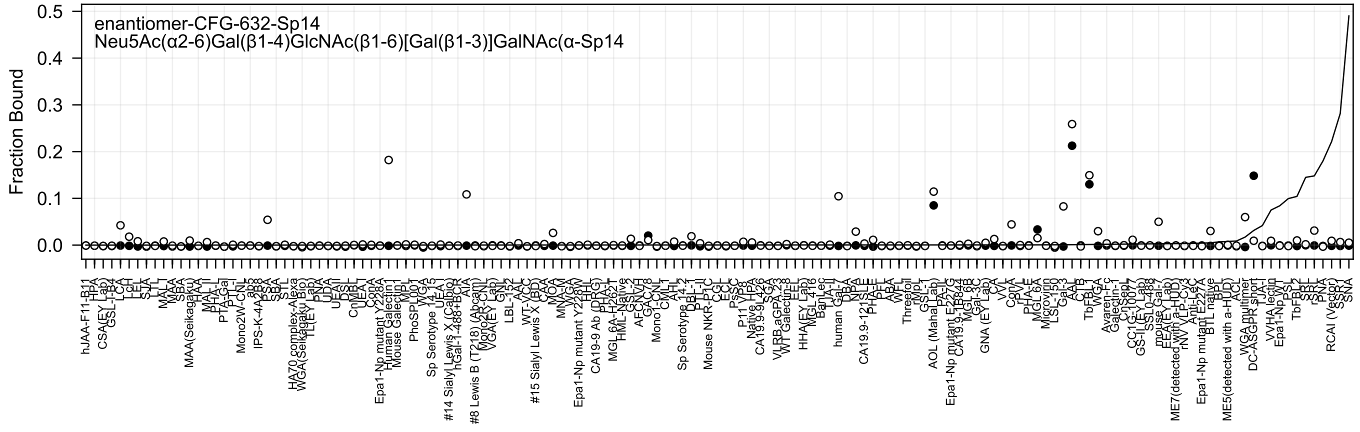
enantiomer-CFG-632-Sp14
Neu5Ac(α2-6)Gal(β1-4)GlcNAc(β1-6)[Gal(β1-3)]GalNAc(α-Sp14
Fraction Bound
0.5
0.4
0.3
0.2
0.1
0.0
hJAA-F11-B11
HPA
CSA(EY Lab)
GSL-I-B4
LCA
LcH
LEL
SJA
LTL
MAL-I
MAA
SBA
MAA(Seikagaku)
HAA
MAL-II
PHA-L
PTA-Gal
PTL-I
Mono2W-CNL
ab5
IPS-K-4A2B8
PSA
SBA
STL
HA70 complex-Alexa
WGA(Seikagaku Bio)
TL(EY Lab)
PNA
UDA
UEAII
DSL
CnMBL
UEA-I
ConA
Epa1-Np mutant Y228A
Human Galectin1
Mouse Galectin1
MPL
PhoSL
WGA
Sp Serotype 14_15
UEA-I
#14 Sialyl Lewis X (Calbio)
hGal-1-488+BCR
AIA
#8 Lewis B (T218) (Abcam)
Mono2R-CNL
VGA(EY Lab)
GNL
LBL-152
AAL
WT-VCC
#15 Sialyl Lewis X (BD)
CAA
MOA
MNA-M
Epa1-Np mutant Y228W
HHL
CA19-9 Ab (DRG)
PHA-E
MGL 6A-H262T
HML-Native
GSL-2
AFCNVH
GAC78
Mono1-CNL
CMLT
AHL
Sp Serotype 14_2
DBL-1
PTL-II
Mouse NKR-P1C
CGL
ECL
PsSC
P11 VP8
Native HPA
CA19.9-9L426
SGA
VLRB aGPA.23
WT Galectin-8
EEL
HHA(EY Lab)
MGL 416
BanLec
TJA-II
human Gal-7
DBA
NPA
CA19.9-121SLE
PHA-E
PTL
ABA
WFL
Threefoil
GSL-1
AOL (Mahal Lab)
PA-IL
Epa1-Np mutant E227G
CA19.9-1B844
MGL 6B
Gal-3C
GNA (EY Lab)
CA
VVL
ConA
rPVL
PHA-E
MGL 6A
Microvirin
LSL-150
Gal-3
AAL
CTB
TbFBL1
WGA
Avaren-Fc
Galectin-1
CnSep7
CC1G-10077
GS-II (EY Lab)
SSL0-488
mouse Gal-7
EEA(EY Lab)
ME7(detected with a-HUD)
rNV VLP-Cy3
Anti-LeC
Epa1-Np mutant E227A
BTL native
BPL
ME5(detected with a-HUD)
ACL
WGA multimer
DC-ASGPR short
TJA-I
VVHA lectin
Epa1-Np WT
PSL
TbFBL2
SRL
rSRL
PNA
RCAI (Vector)
SSR1
SNA

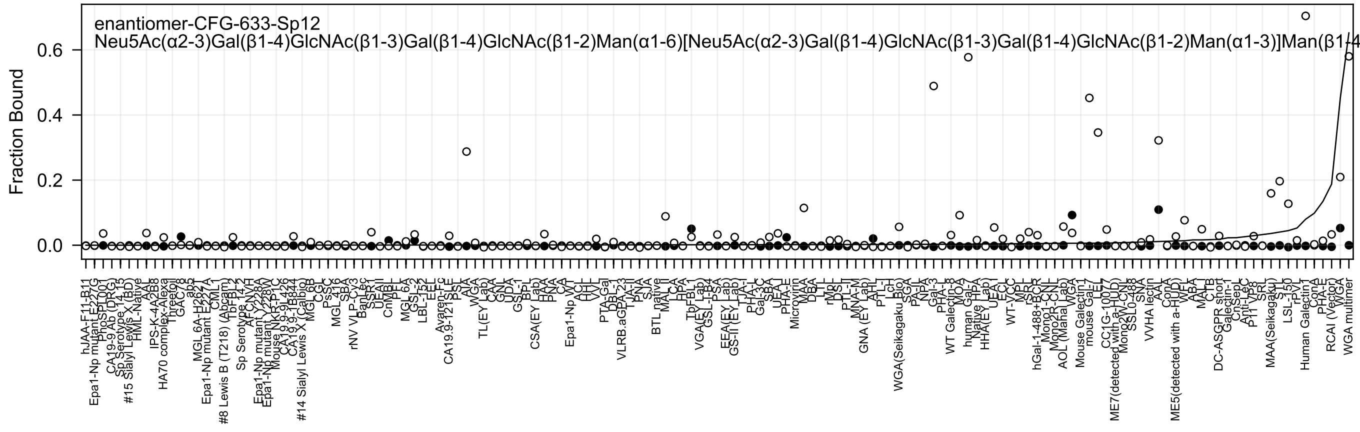

enantiomer-CFG-633-Sp12
Neu5Ac(α2-3)Gal(β1-4)GlcNAc(β1-3)Gal(β1-4)GlcNAc(β1-2)Man(α1-6)[Neu5Ac(α2-3)Gal(β1-4)GlcNAc(β1-3)Gal(β1-4)GlcNAc(β1-2)Man(α1-3)]Man(β1-4
Fraction Bound
0.0
0.2
0.4
0.6

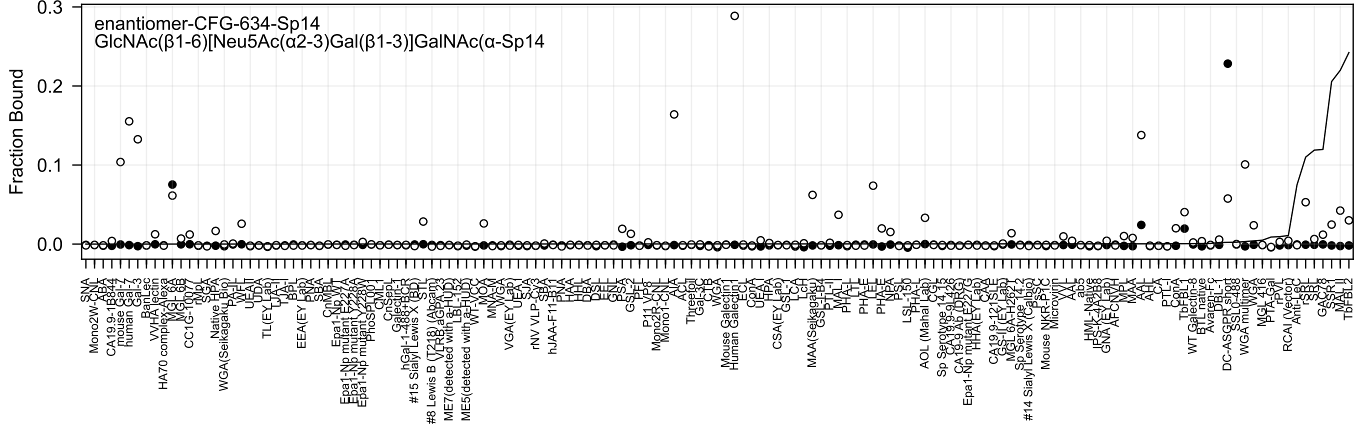

enantiomer-CFG-634-Sp14
GlcNAc(β1-6)[Neu5Ac(α2-3)Gal(β1-3)]GalNAc(α-Sp14
Fraction Bound
0.3
0.2
0.1
0.0
SNA
Mono2W-CNL
ABA
CA19.9-1B844
mouse Gal-7
human Gal-3
BanLec
VVHA lectin
HA70 complex-Alexa
MGL 6A
MGL 6B
CC1G-1007
rMpL
SGA
Native HPA
WGA(Seikagaku Bio)
PA-IL
WFL
UEA-II
UDA
TL(EY Lab)
TJA-II
TJA-I
BPL
EEA(EY Lab)
PNA
SBA
CnMBL
Epa1-Np WT
Epa1-Np mutant E227A
Epa1-Np mutant Y228A
Epa1-Np mutant Y228W
PhoSL
CML
CnSepL
Galectin-1
hGal-1-488+BCR
#15 Sialyl Lewis X (BD)
STL
#8 Lewis B (T218) (Abcam)
VLRB aGPA.23
ME7(detected with a-HUD)
LBL-152
ME5(detected with a-HUD)
WT-VCC
MOA
MNA-M
WGA
VGA(EY Lab)
UEA-I
SJA
rNV VLP-Cy3
SBA
hJAA-F11-B11
PNA
HAA
HHL
DBA
DSL
EEL
GNL
PSA
GSL-2
PFL
P11 VP8
Mono2R-CNL
Mono1-CNL
AIA
ACL
Threefoil
Gal-3C
CTB
WGA
Mouse Galectin1
Human Galectin1
ECL
ConA
UEA
HPA
CSA(EY Lab)
GSL-1
LCA
LcH
MAA(Seikagaku)
GS-I-B4
PTL-II
MAL-I
PHA-L
LTL
PHA-E
PHA-E
NPA
PSL
LSL-150
PHA-L
AOL (Mahal Lab)
CGL
Sp Serotype 14.15
CA19.9-9L426
CA19-9 Ab (DRG)
Epa1-Np mutant E227G
HHA(EY Lab)
CAA
CA19.9-121SLE
GS-II (EY Lab)
MGL 6A-H287
Sp Serotype 14.2
#14 Sialyl Lewis X (Calbio)
PsSC
Mouse NKR-P1C
Microvirin
VVL
XXL
ab5
HML-Native
IPS-K-4A2B8
GNA (EY Lab)
AFCNVH
MPL
MAA
AAL
CA
PTL-I
ConA
TbFBL1
WT Galectin-8
BTL native
Avaren-Fc
DBL-I
DC-ASGPR short
SSL0-488
WGA multimer
WGA
MGL 416
PTA-Gal
rPVL
RCAI (Vector)
Anti-LeC
rSRL
SRL
GAC78
SSR1
MAL I
TbFBL2

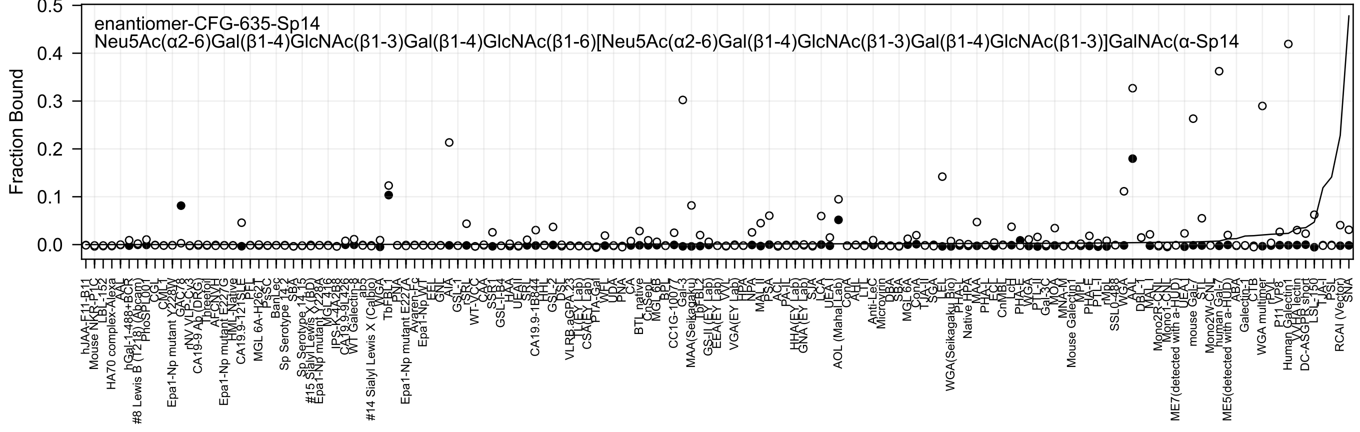
enantiomer-CFG-635-Sp14
Neu5Ac(α2-6)Gal(β1-4)GlcNAc(β1-3)Gal(β1-4)GlcNAc(β1-6)[Neu5Ac(α2-6)Gal(β1-4)GlcNAc(β1-3)Gal(β1-4)GlcNAc(β1-3)]GalNAc(α-Sp14
Fraction Bound

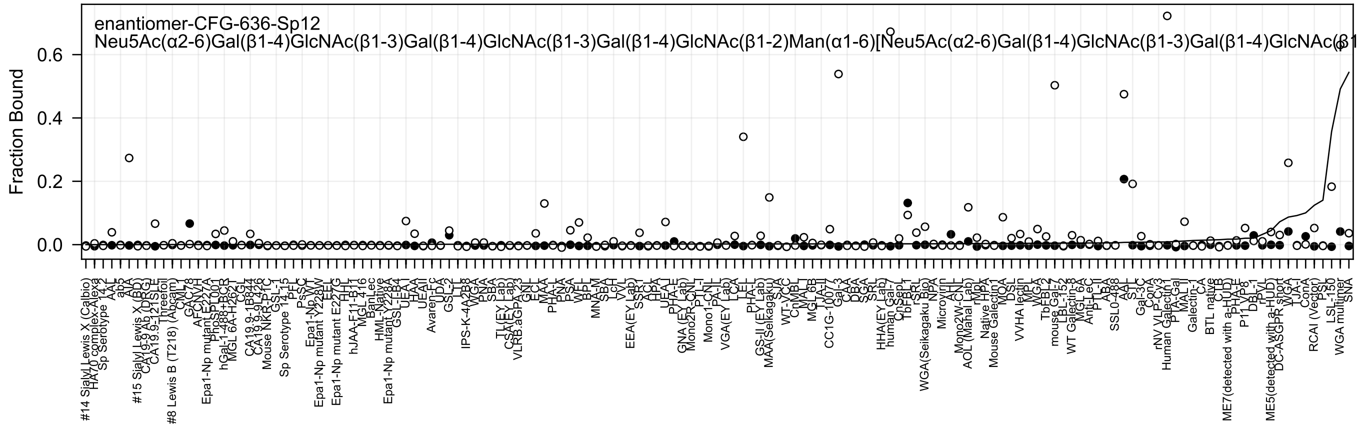

enantiomer-CFG-636-Sp12
Neu5Ac(α2-6)Gal(β1-4)GlcNAc(β1-3)Gal(β1-4)GlcNAc(β1-3)Gal(β1-4)GlcNAc(β1-2)Man(α1-6)[Neu5Ac(α2-6)Gal(β1-4)GlcNAc(β1-3)Gal(β1-4)GlcNAc(β1
Fraction Bound
0.0
0.2
0.4
0.6

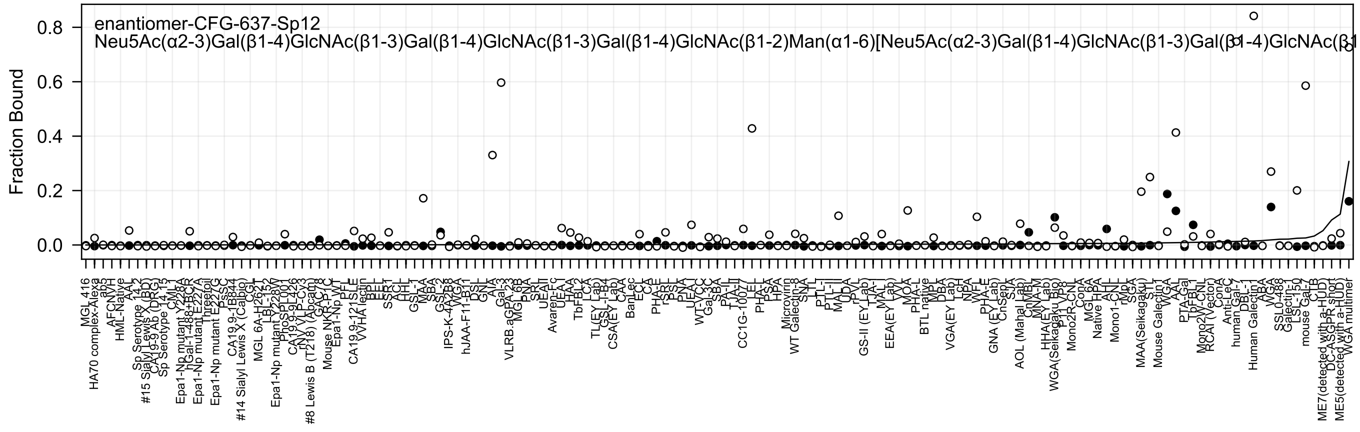
enantiomer-CFG-637-Sp12
Neu5Ac(α2-3)Gal(β1-4)GlcNAc(β1-3)Gal(β1-4)GlcNAc(β1-3)Gal(β1-4)GlcNAc(β1-2)Man(α1-6)[Neu5Ac(α2-3)Gal(β1-4)GlcNAc(β1-3)Gal(β1-4)GlcNAc(β1
Fraction Bound

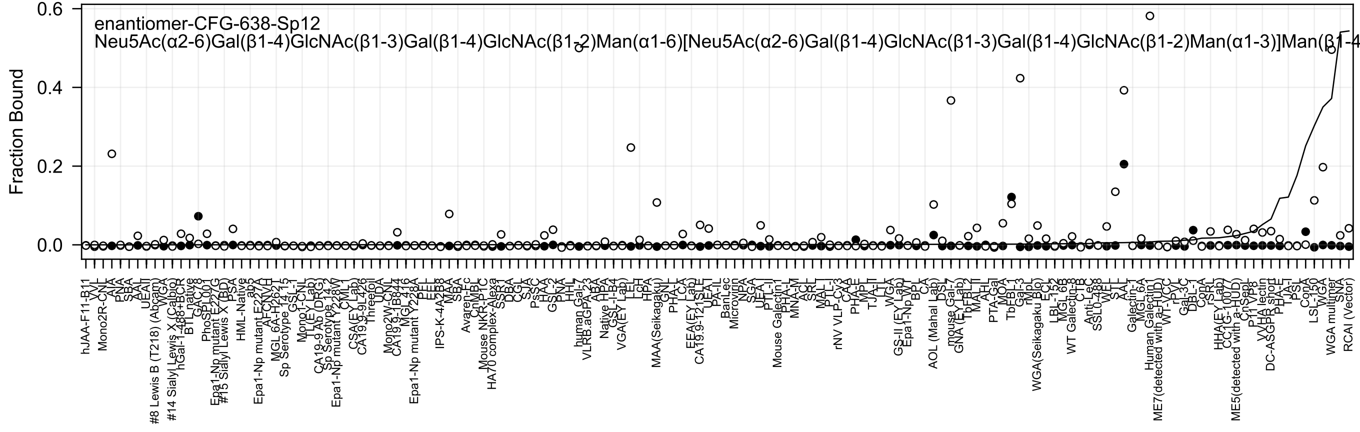

enantiomer-CFG-638-Sp12
Neu5Ac(α2-6)Gal(β1-4)GlcNAc(β1-3)Gal(β1-4)GlcNAc(β1-2)Man(α1-6)[Neu5Ac(α2-6)Gal(β1-4)GlcNAc(β1-3)Gal(β1-4)GlcNAc(β1-2)Man(α1-3)]Man(β1-4
Fraction Bound
0.6
0.4
0.2
0.0
hJAA-F11-B11
Mono2R-CNL
AIA
PNA
SBA
AAL
UEAI
#8 Lewis B (T218) (Abcam)
WGA
#14 Sialyl Lewis X (Calbio)
hGal-1-488+BCR
BTL native
GAC78
PhoSL
#15 Sialyl Lewis X (BD)
PSA
HML-Native
MGL 6A-H262T
Sp Serotype 14.15
CA19-9 Ab (DRG)
CSA(EY Lab)
Threefoil
UDA
Mono2W-CNL
CA19.9-1B844
IPS-K-4A2B8
MAA
SBA
Mouse NKR-P1C
HA70 complex-Alexa
SSR1
DBA
PsSC
HAA
GSL-2
PNA
human Gal-7
VLRB.aGPA.23
ABA
Native HPA
VGA(EY Lab)
LEL
HPA
MAA(Seikagaku)
LCA
EEA(EY Lab)
CA19.9-121SLE
UEAI
BanLec
Microvirin
NPA
SGA
Mouse Galectin1
MNA-M
ACL
MAL-I
rNV VLP-Cy3
CAA
PHA-E
TJA-II
WGA
GS-II (EY Lab)
Epa1-Np WT
CA
AOL (Mahal Lab)
DSL
mouse Gal-7
GNA (EY Lab)
AAL
PTA-Gal
MOA
TbFBL1
Gal-3
WGA(Seikagaku Bio)
ECL
LBL-152
MGL 6B
WT Galectin-8
CTB
Anti-LeC
SSL0-488
WFL
STL
Galectin-1
MGL 6A
Human Galectin
ME7(detected with a-HUD)
WT-VCC
rPVL
Gal-3C
DBL-1
ConA
rSRL
HHA(EY Lab)
ME5(detected with a-HUD)
CnSepL
P11 VP8
VVHA lectin
DC-ASGPR short
PHA-E
TJA-I
PSL
ConA
LSL-150
WGA
WGA multimer
SNA
RCAI (Vector)

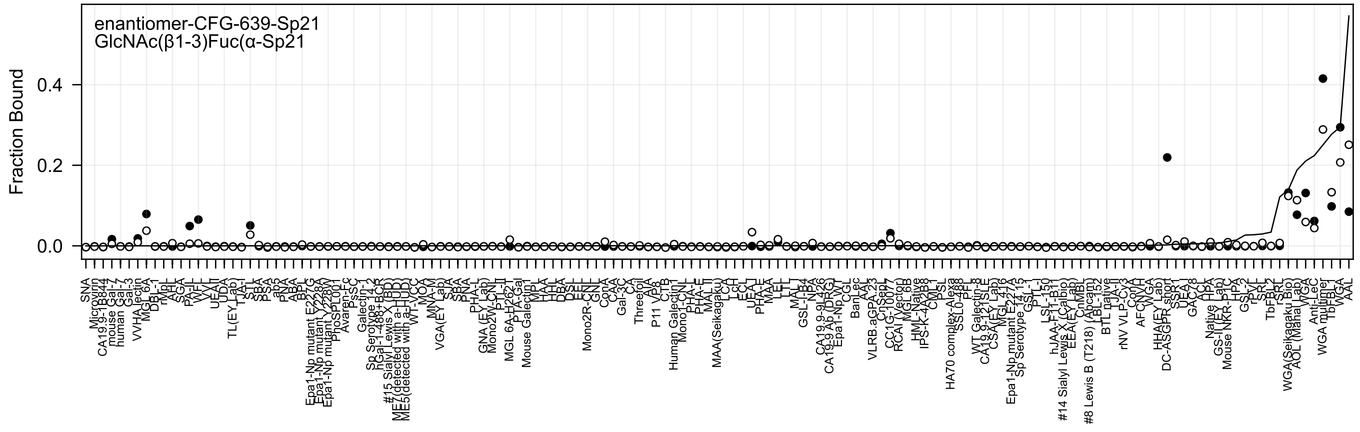
enantiomer-CFG-639-Sp21
GlcNAc(β1-3)Fuc(α-Sp21
Fraction Bound
0.4
0.2
0.0

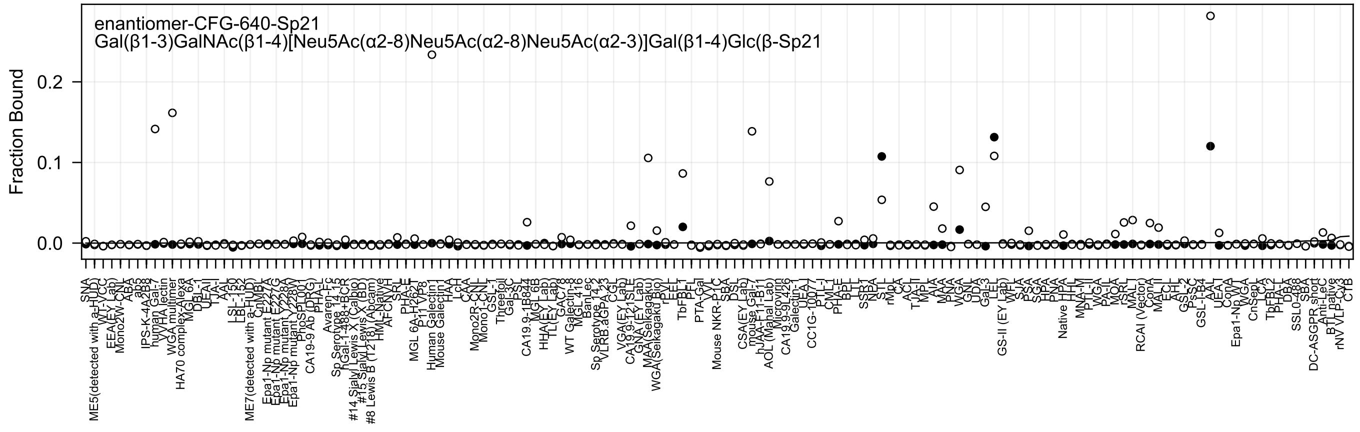
enantiomer-CFG-640-Sp21
Gal(β1-3)GalNAc(β1-4)[Neu5Ac(α2-8)Neu5Ac(α2-8)Neu5Ac(α2-3)]Gal(β1-4)Glc(β-Sp21
Fraction Bound
0.0
0.1
0.2
