## Supplementary material for "Atom-level Machine Learning of Protein-Glycan Interactions and Cross-chiral Recognition in Glycobiology": Epimer Heatmaps

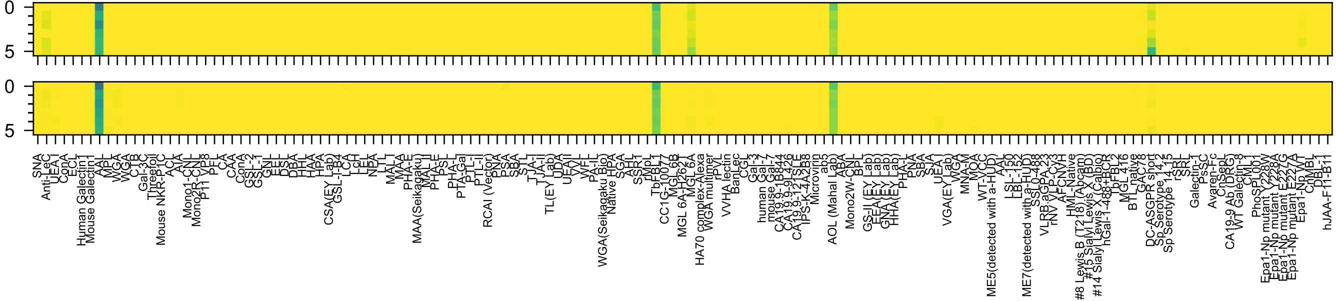
0
5
0
5
SNA
Anti-LeC
UEA-I
ConA
ECL
Human Galectin1
Mouse Galectin1
AAL
MPL
WGA
WGA
CTB
Gal-3C
Threefoil
Mouse NKR-P1C
ACL
AIA
Mono1-CNL
Mono2R-CNL
P11 VP8
PFL
CA
CAA
ConA
GSL-2
GSL-1
GNL
EEL
DSL
DBA
HHL
HAA
HPA
CSA(EY Lab)
GSL-I-B4
LCA
LcH
LEL
NPA
LTL
MAL-I
MAA
PHA-E
MAA(Seikagaku)
MAL-II
PHA-E
PSL
PHA-L
PTA-Gal
PTL-I
PTL-II
RCAI (Vector)
PNA
PSA
SBA
STL
TJA-I
TJA-II
TL(EY Lab)
UDA
UEAII
VVL
WFL
PA-IL
WGA(Seikagaku Bio)
Native HPA
SGA
AHL
SSR1
rMpL
TbFBL1
CC1G-10077
MGL-6B
MGL 6A-H262T
MGL 6A
HA70 complex-Alexa
WGA multimer
rPVL
VVHA lectin
BanLec
CGL
Gal-3
human Gal-7
mouse Gal-7
CA19.9-1B844
CA19.9-9L426
CA19.9-121SLE
IPS-K-4A2B8
Microvirin
ab5
AOL (Mahal Lab)
ABA
Mono2W-CNL
BPL
GS-II (EY Lab)
EEA (EY Lab)
GNA (EY Lab)
HHA (EY Lab)
PHA-L
PNA
SBA
SJA
UEA-I
VGA(EY Lab)
WGA
MNA-M
MOA
WT-VCC
ME5(detected with a-HUD)
AAL
LSL-150
LBL-152
ME7(detected with a-HUD)
SSL0-488
VLRB.aGPA.23
rNV VLP-Cy3
AFCNVH
HML-Native
#8 Lewis B (T218) (Abcam)
#15 Sialyl Lewis X (BD)
#14 Sialyl Lewis X (Calbio)
hGal-1-488+BCR
TbFBL2
MGL 416
BTL native
GAC78
DC-ASGPR short
Sp Serotype 14.2
Sp Serotype 14.15
rSRL
SRL
Galectin-1
PsSC
Avaren-Fc
ChSepL
CA19-9 Ab (DRG)
WT Galectin-8
CML1
PhoSPL001
Epa1-Np mutant Y228W
Epa1-Np mutant E228A
Epa1-Np mutant E227G
Epa1-Np mutant E227A
Epa1-Np WT
CnMBL
DBL-1
hJAA-F11-B11
Fuc(α-Sp8

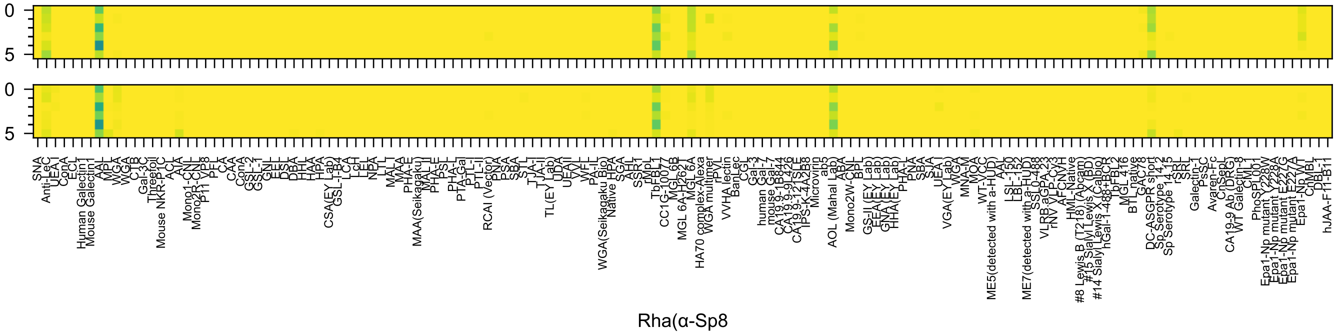

0
5
0
5
Rha(α-Sp8

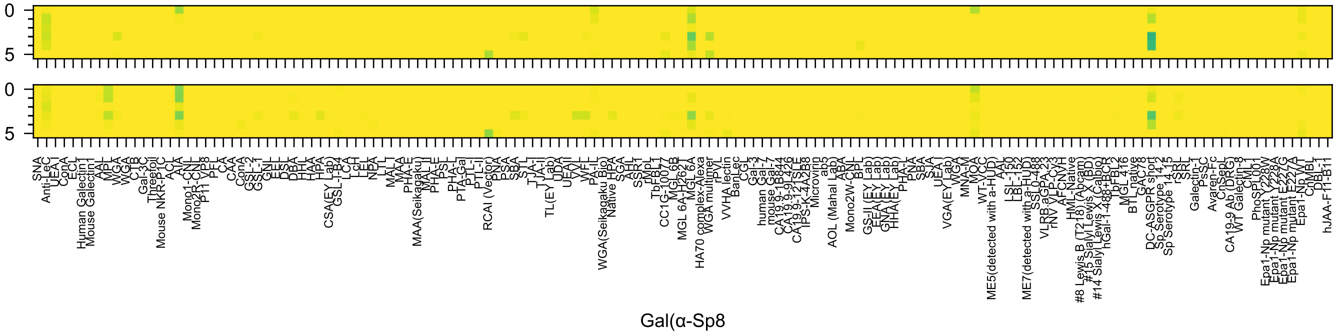
Gal(α-Sp8

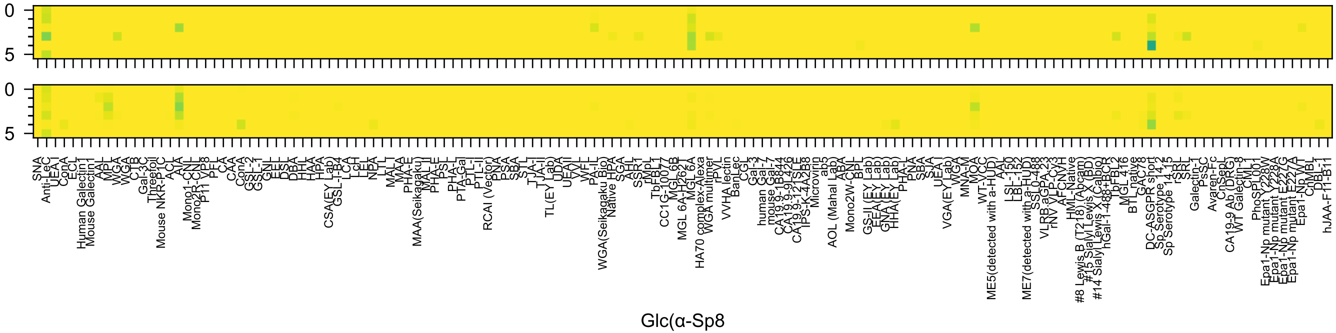
Glc(α-Sp8

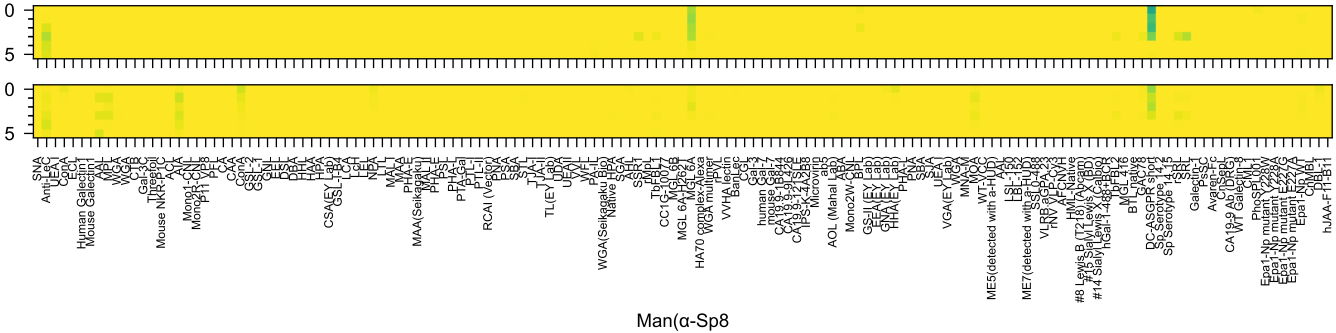
0
5
0
5
SNA
Anti-LeC
UEA-I
ConA
ECL
Human Galectin1
Mouse Galectin1
AAL
MPL
WGA
WGA
CTB
Gal-3C
Threefoil
Mouse NKR-P1C
ACL
AIA
Mono1-CNL
Mono2R-CNL
P11 VP8
PFL
CA
CAA
ConA
GSL-2
GSL-1
GNL
EEL
DSL
DBA
HHL
HAA
HPA
CSA(EY Lab)
GSL-I-B4
LCA
LcH
LEL
NPA
LTL
MAL-I
MAA
PHA-E
MAA(Seikagaku)
MAL-II
PHA-E
PSL
PHA-L
PTA-Gal
PTL-I
PTL-II
RCAI (Vector)
PNA
PSA
SBA
STL
TJA-I
TJA-II
TL(EY Lab)
UDA
UEA-II
VVL
WFL
PA-IL
WGA(Seikagaku Bio)
Native HPA
SGA
AAL
SSR1
rMpL
TbFBL1
CC1G-10077
MGL6B
MGL 6A-H262T
MGL 6A
HA70 complex-Alexa
WGA multimer
rPVL
VVHA lectin
BanLec
CGL
Gal-3
human Gal-7
mouse Gal-7
CA19.9-1B844
CA19.9-9L426
CA19.9-121SLE
IPS-K-4A2B8
Microvirin
ab5
AOL (Mahal Lab)
ABA
Mono2W-CNL
BPL
GS-II (EY Lab)
EEA (EY Lab)
GNA (EY Lab)
HHA (EY Lab)
PHA-L
PNA
SBA
SJA
UEA-I
VGA(EY Lab)
WGA
MNA-M
MOA
WT-VCC
ME5(detected with a-HUD)
AAL
LSL-150
LBL-152
ME7(detected with a-HUD)
SSL0-488
VLRB.aGPA.23
rNV VLP-Cy3
AFCNVH
HML-Native
#8 Lewis B (T218) (Abcam)
#15 Sialyl Lewis X (BD)
#14 Sialyl Lewis X (Calbio)
hGal-1-488+BCR
TbFBL2
MGL 416
BTL native
GAC78
DC-ASGPR short
Sp Serotype 14.2
Sp Serotype 14.15
rSRL
SRL
Galectin-1
PsSC
Avaren-Fc
ChSepL
CA19-9 Ab (DRG)
WT Galectin-8
CML1
PhoSPL001
Epa1-Np mutant Y228W
Epa1-Np mutant Y228A
Epa1-Np mutant E227G
Epa1-Np mutant E227A
Epa1-Np WT
CnMBL
DBL-1
hJAA-F11-B11
Man(α-Sp8

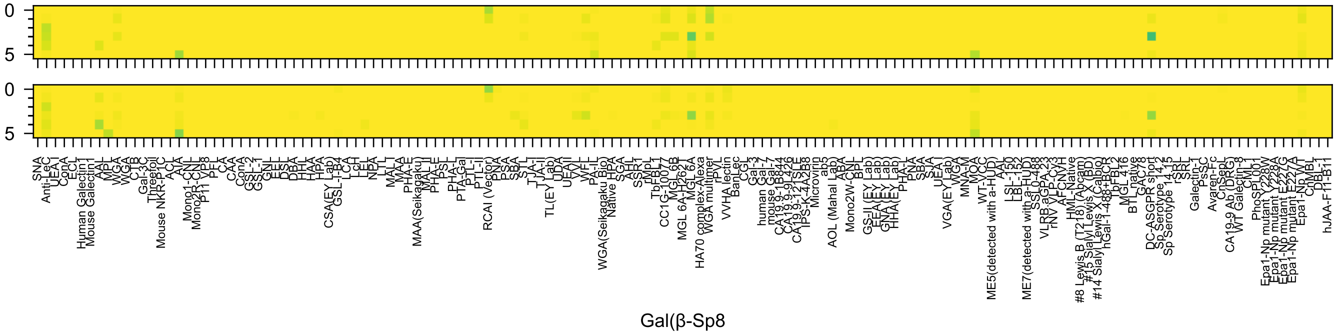
0
5
0
5
Gal(β-Sp8

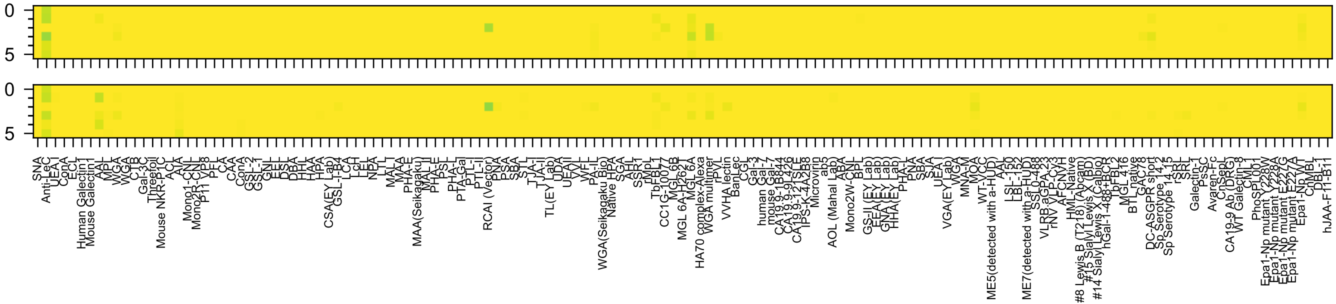

Glc(β-Sp8

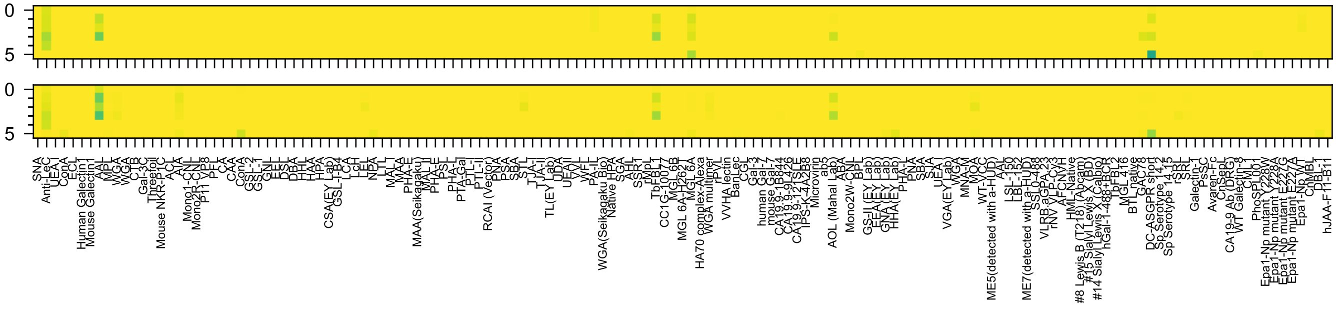
Man(β-Sp8

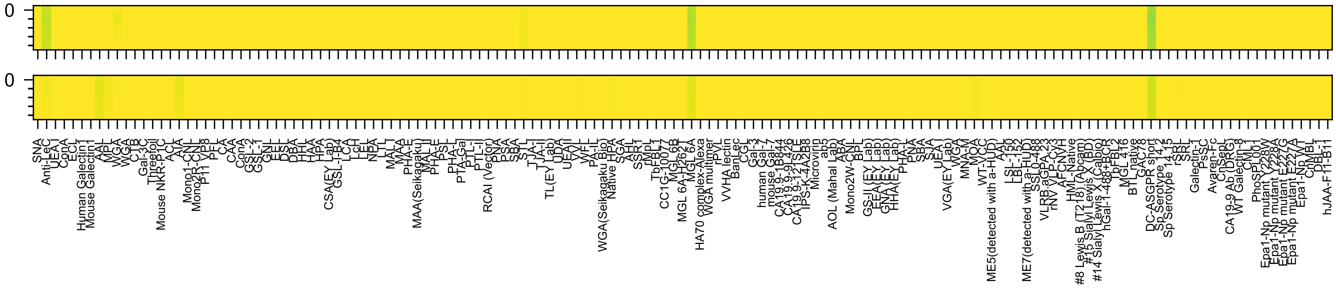
0
0
SNA
Anti-LeC
UEA I
ConA
ECL
Human Galectin1
Mouse Galectin1
AAL
MPL
WGA
WGA
CTB
Gal-3C
Threefoil
Mouse NKR-P1C
ACL
AIA
Mono1-CNL
Mono2R-CNL
P11 VP8
PFL
CA
CAA
ConA
GSL-2
GSL-1
GNL
EEL
DSL
DBA
HHL
HAA
HPA
CSA(EY Lab)
GSL-I-B4
LCA
LcH
LEL
NPA
LTL
MAL-I
MAA
PHA-E
MAA(Seikagaku)
MAL-II
PHA-E
PSL
PHA-L
PTA-Gal
PTL-I
PTL-II
RCAI (Vector)
PNA
PSA
SBA
STL
TJA-I
TJA-II
TL(EY Lab)
UDA
UEAII
VVL
WFL
PA-IL
WGA(Seikagaku Bio)
Native HPA
SGA
AHL
SSR1
rMpL
TbFBL1
CC1G-10077
MGL 6B
MGL 6A-H262T
MGL 6A
HA70 complex-Alexa
WGA multimer
rPVL
VVHA lectin
BanLec
CGL
Gal-3
human Gal-7
mouse Gal-7
CA19.9-1B844
CA19.9-9L426
CA19.9-121SLE
IPS-K-4A2B8
Microvirin
ab5
AOL (Mahal Lab)
ABA
Mono2W-CNL
BPL
GS-II (EY Lab)
EEA(EY Lab)
GNA(EY Lab)
HHA(EY Lab)
PHA-L
PNA
SBA
SJA
UEA I
VGA(EY Lab)
WGA
MNA-M
MOA
WT-VCC
ME5(detected with a-HUD)
AAL
LSL-150
LBL-152
ME7(detected with a-HUD)
SSL0-488
VLRB.aGPA.23
rNV VLP-Cy3
AFCNVH
HML-Native
#8 Lewis B (T218) (Abcam)
#15 Sialyl Lewis X (BD)
#14 Sialyl Lewis X (Calbio)
hGal-1-488+BCR
rbFBL2
MGL 416
BTL native
GAC78
DC-ASGPR short
Sp Serotype 14.2
Sp Serotype 14.15
rSRL
SRL
Galectin-1
PsSC
Avaren-Fc
ChSeaL
CA19-9 Ab (DRG)
WT Galectin-8
CML
PhoSL001
Epa1-Np mutant Y228W
Epa1-Np mutant Y228A
Epa1-Np mutant E227G
Epa1-Np mutant E227A
Epa1-Np WT
ChNBL
DBL-1
hJAA-F11-B11

Glc-ol(-Sp8

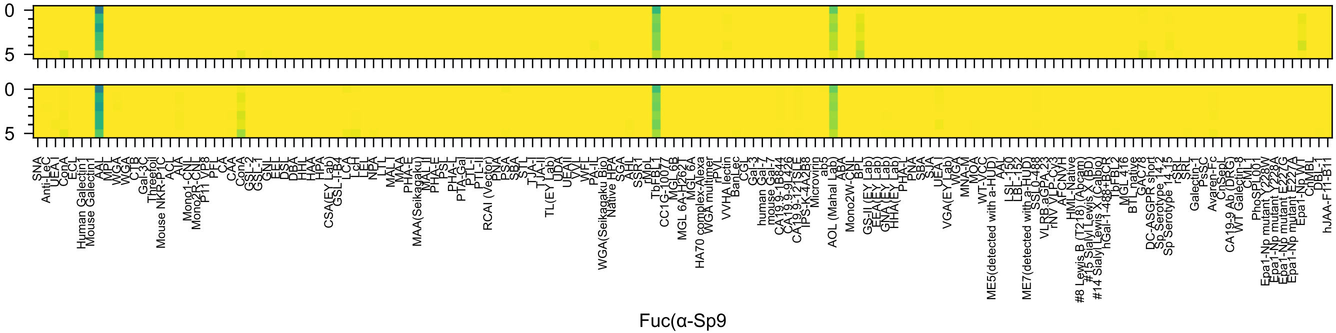
Fuc(α-Sp9

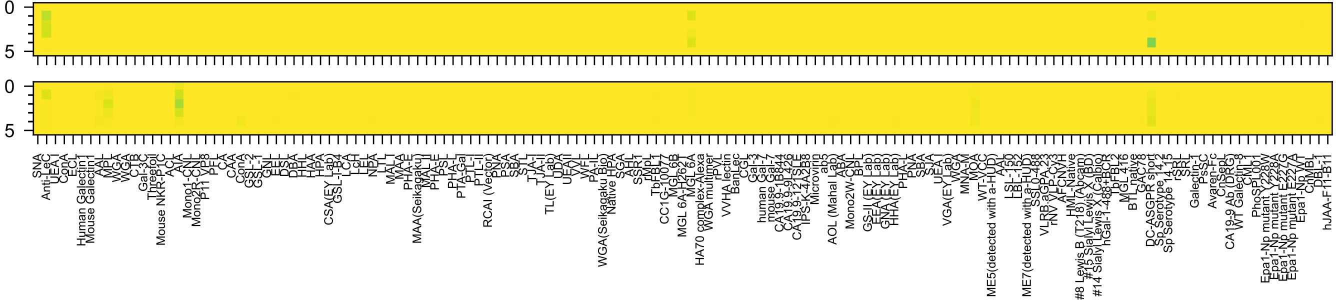

GlcA(α-Sp8

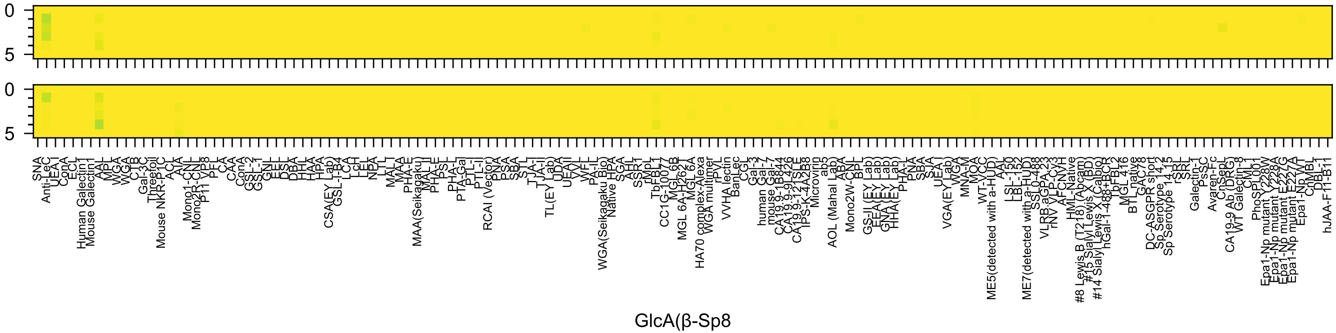
GlcA(β-Sp8

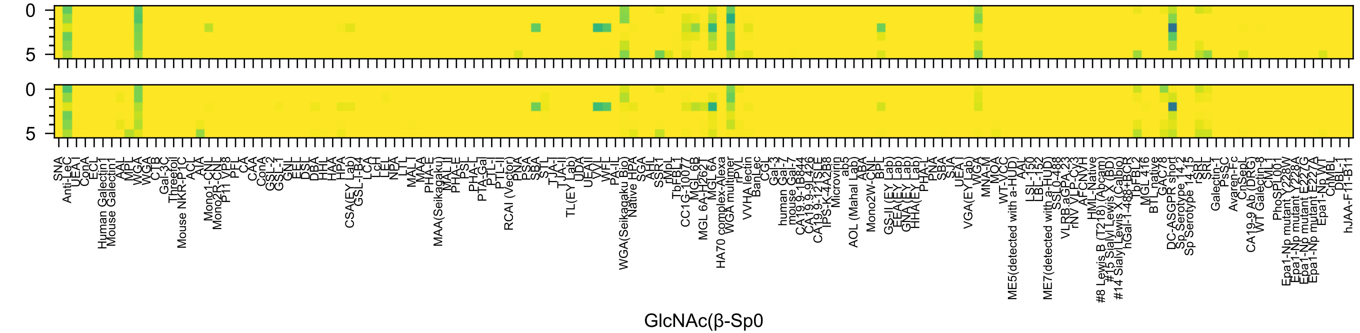
GlcNAc(β-Sp0

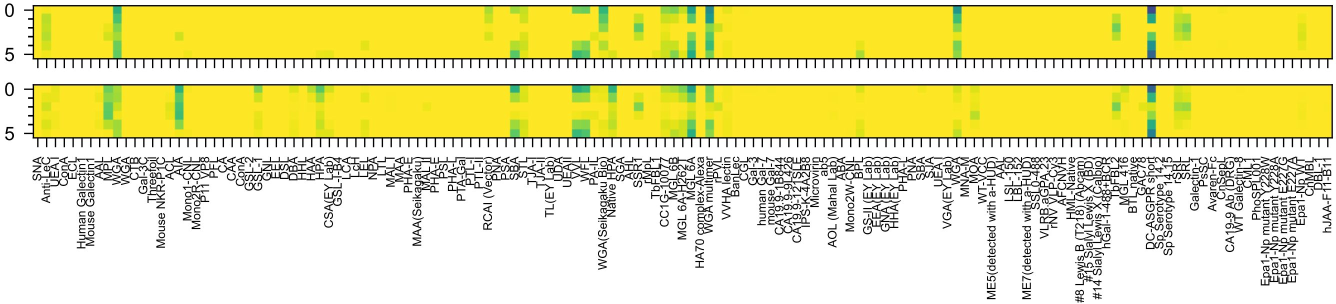

GalNAc(α-Sp8

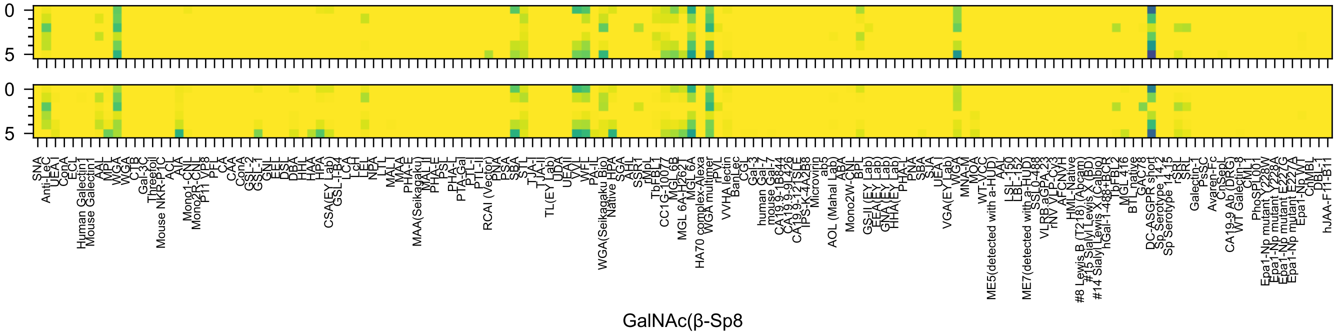
GalNAc(β-Sp8

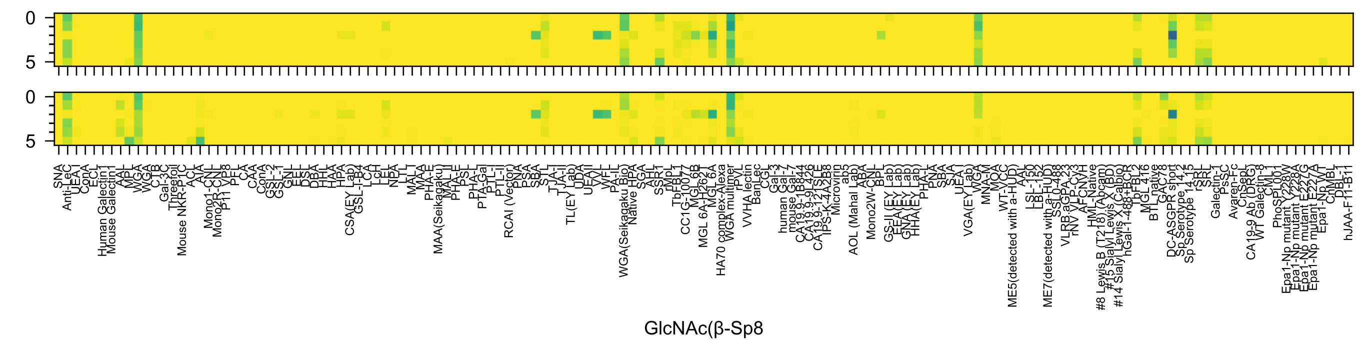

0
5
0
5
GlcNAc(β-Sp8

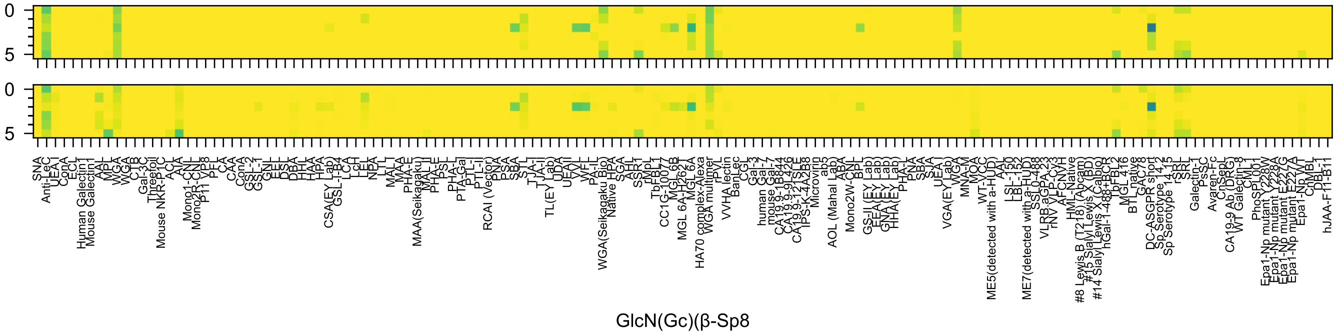
0
5
0
5
SNA
Anti-LeC
UEA I
ConA
ECL
Human Galectin1
Mouse Galectin1
AAL
MPL
WGA
WGA
CTB
Gal-3C
Threefoil
Mouse NKR-P1C
ACL
AIA
Mono1-CNL
Mono2R-CNL
P11 VP8
PFL
CA
CAA
ConA
GSL-2
GSL-1
GNL
EEL
DSL
DBA
HHL
HAA
HPA
CSA(EY Lab)
GSL-I-B4
LCA
LcH
LEL
NPA
LTL
MAL-I
MAA
PHA-E
MAA(Seikagaku)
MAL-II
PHA-E
PSL
PHA-L
PTA-Gal
PTL-I
PTL-II
RCAI (Vector)
PNA
PSA
SBA
STL
TJA-I
TJA-II
TL(EY Lab)
UDA
UEA-II
VVL
WFL
PA-IL
WGA(Seikagaku Bio)
Native HPA
SGA
AAL
SSR1
rMpL
TbFBL1
CC1G-10077
MGL 6B
MGL 6A-H262T
MGL 6A
HA70 complex-Alexa
WGA multimer
rPVL
VVHA lectin
BanLec
CGL
Gal-3
human Gal-7
mouse Gal-7
CA19.9-1B844
CA19.9-9L426
CA19.9-121SLE
IPS-K-4A2B8
Microvirin
ab5
AOL (Mahal Lab)
ABA
Mono2W-CNL
BPL
GS-II (EY Lab)
EEA (EY Lab)
GNA (EY Lab)
HHA (EY Lab)
PHA-L
PNA
SBA
SJA
UEA I
VGA(EY Lab)
WGA
MNA-M
MOA
WT-VCC
ME5(detected with a-HUD)
AAL
LSL-150
LBL-152
ME7(detected with a-HUD)
SSL0-488
VLRB.aGPA.23
rNV VLP-Cy3
AFCNVH
HML-Native
#8 Lewis B (T218) (Abcam)
#15 Sialyl Lewis X (BD)
#14 Sialyl Lewis X (Calbio)
hGal-1-488+BCR
TbFBL2
MGL 416
BTL native
GAC78
DC-ASGPR short
Sp Serotype 14.2
Sp Serotype 14.15
rSRL
SRL
Galectin-1
PsSC
Avaren-Fc
ChSepL
CA19-9 Ab (DRG)
WT Galectin-8
CML1
PhoSPL001
Epa1-Np mutant Y228W
Epa1-Np mutant Y228A
Epa1-Np mutant E227G
Epa1-Np mutant E227A
Epa1-Np WT
CrMBL
DBL-1
hJAA-F11-B1
GlcN(Gc)(β-Sp8

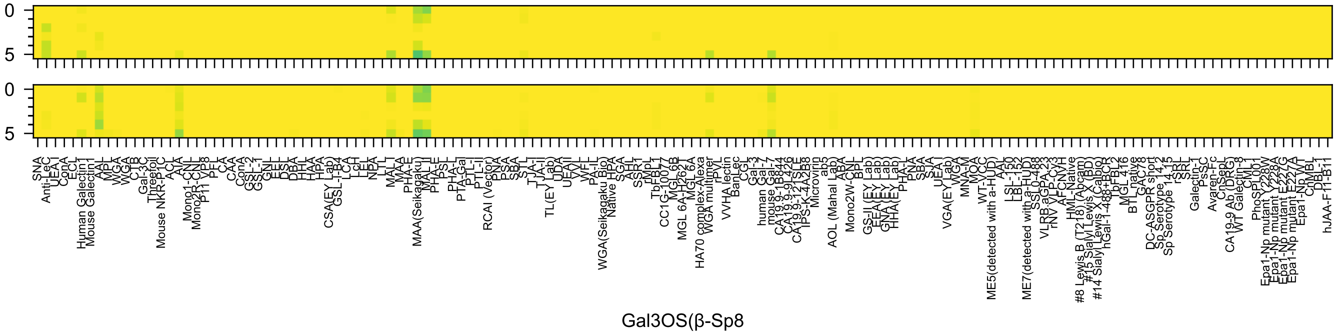
Gal3OS(β-Sp8

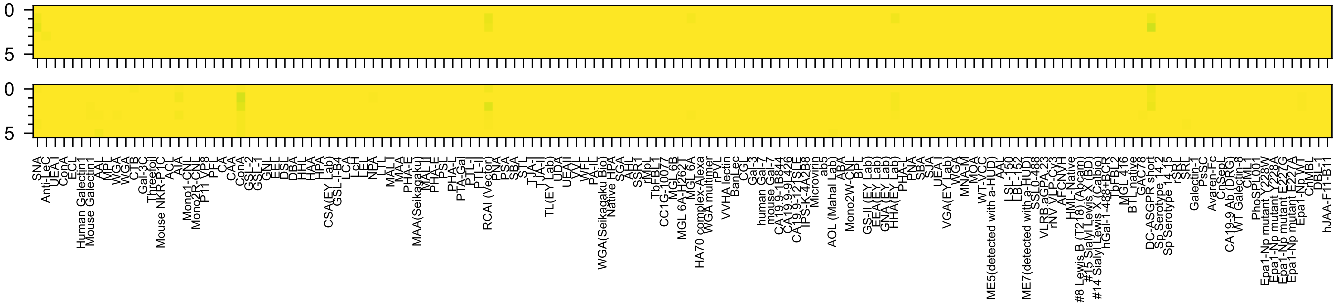

Man6OP(α-Sp8

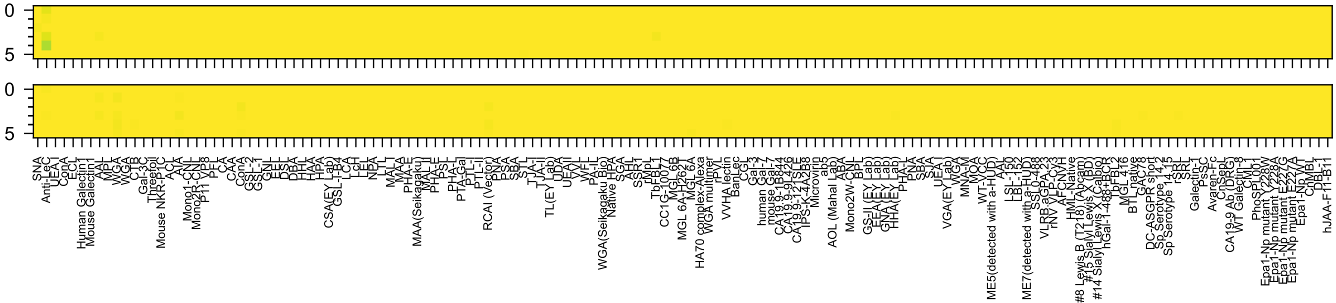
Glc6OP(β-Sp10

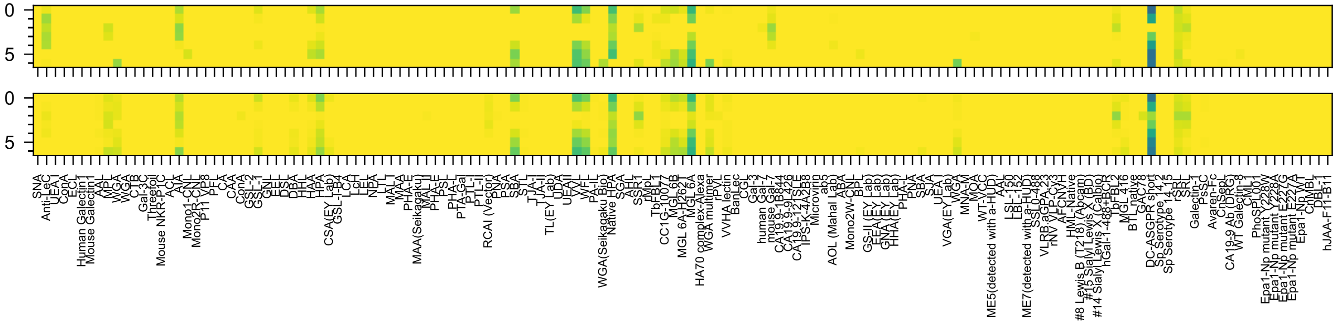

GalNAc(α-Sp15

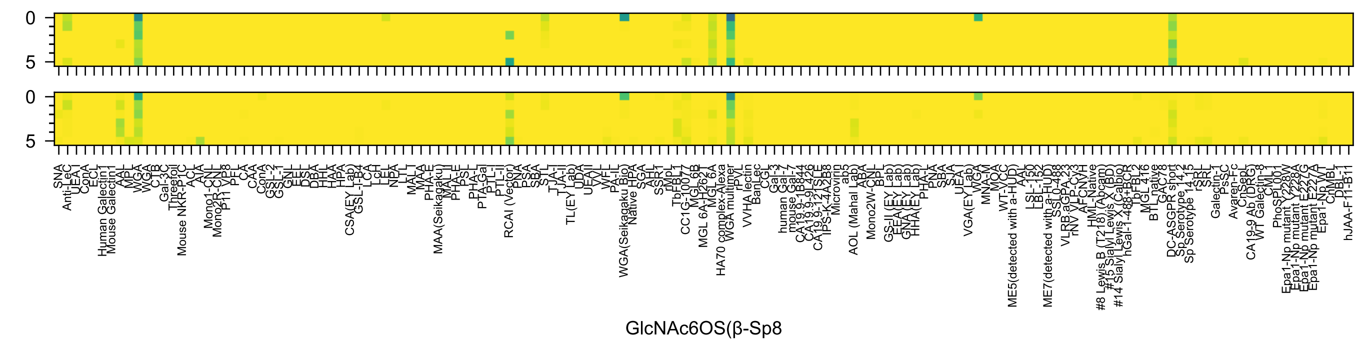

0
5
0
5
GlcNAc6OS(β-Sp8

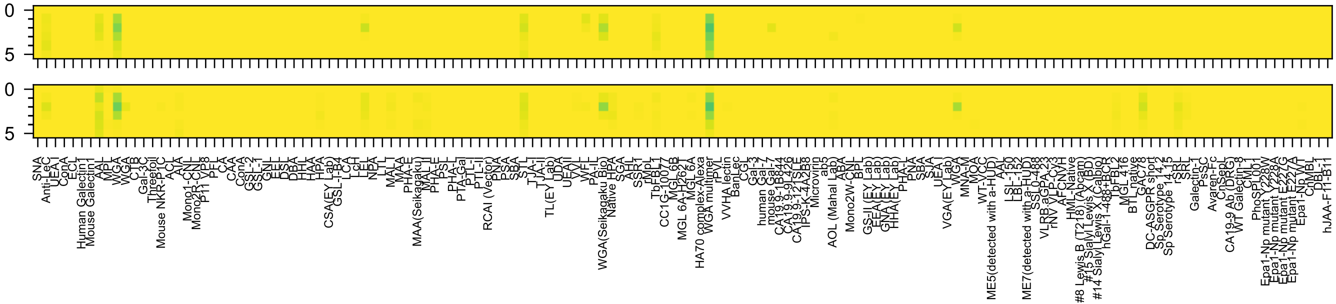
0
5
0
5
GalNAc4OS(β-Sp10

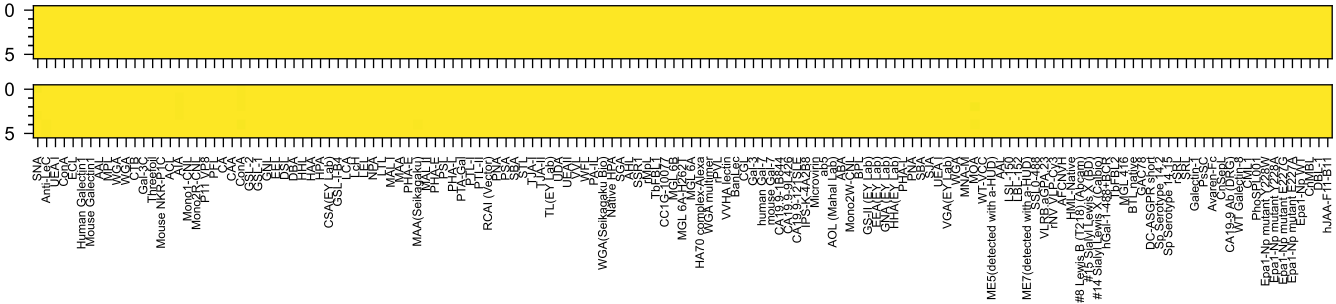

Neu5Ac(α-Sp8

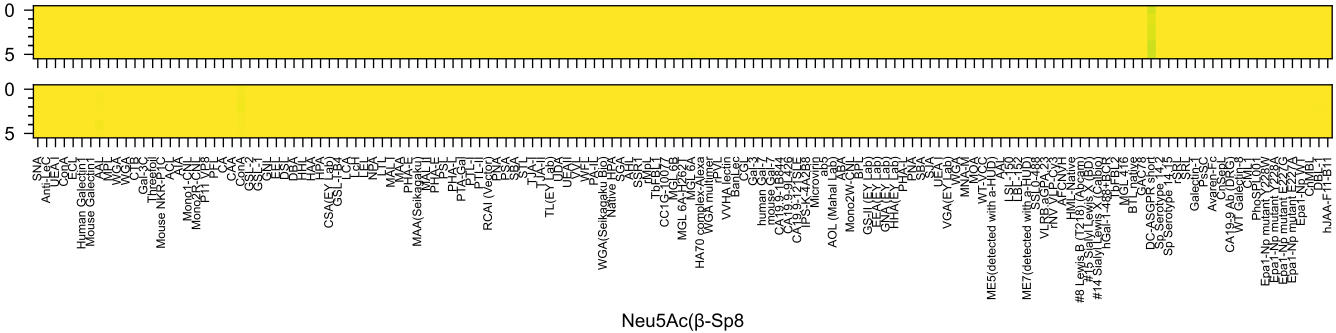
Neu5Ac(β-Sp8

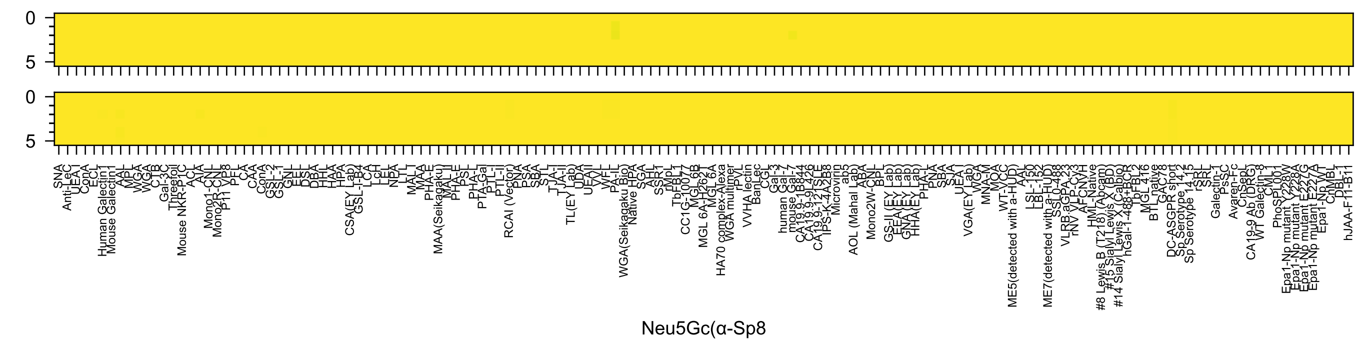
Neu5Gc(α-Sp8

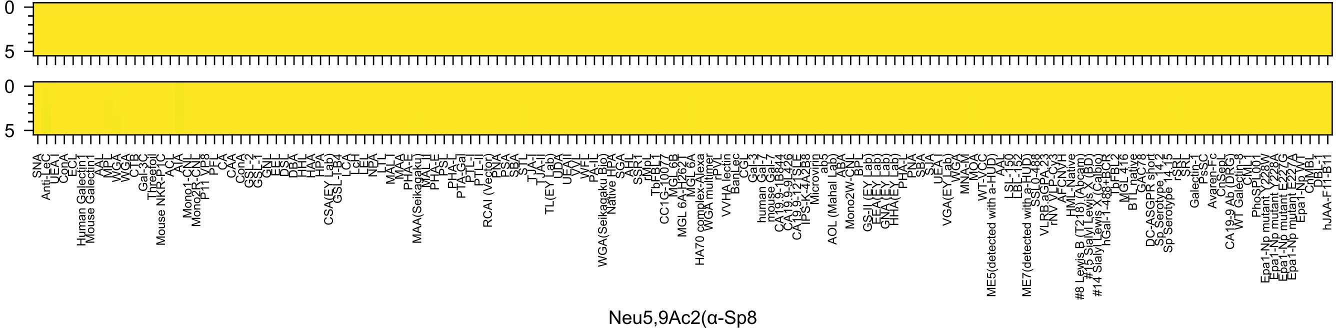
0
5
0
5
SNA
Anti-LeC
UEA-I
ConA
ECL
Human Galectin1
Mouse Galectin1
AAL
MPL
WGA
WGA
CTB
Gal-3C
Threefoil
Mouse NKR-P1C
ACL
AIA
Mono1-CNL
Mono2R-CNL
P11 VP8
PFL
CA
CAA
ConA
GSL-2
GSL-1
GNL
EEL
DSL
DBA
HHL
HAA
HPA
CSA(EY Lab)
GSL-I-B4
LCA
LcH
LEL
NPA
LTL
MAL-I
MAA
PHA-E
MAA(Seikagaku)
MAL-II
PHA-E
PSL
PHA-L
PTA-Gal
PTL-I
PTL-II
RCAI (Vector)
PNA
PSA
SBA
STL
TJA-I
TJA-II
TL(EY Lab)
UDA
UEA-II
VVL
WFL
PA-IL
WGA(Seikagaku Bio)
Native HPA
SGA
AAL
SSR1
rMpL
TbFBL1
CC1G-10077
MGL 6B
MGL 6A-H262T
MGL 6A
HA70 complex-Alexa
WGA multimer
rPVL
VVHA lectin
BanLec
CGL
Gal-3
human Gal-7
mouse Gal-7
CA19-9-1B844
CA19-9-9L426
CA19-9-121SLE
IPS-K-4A2B8
Microvirin
ab5
AOL (Mahal Lab)
ABA
Mono2W-CNL
BPL
GS-II (EY Lab)
EEA (EY Lab)
GNA (EY Lab)
HHA (EY Lab)
PHA-L
PNA
SBA
SJA
UEA-I
VGA(EY Lab)
WGA
MNA-M
MOA
WT-VCC
ME5(detected with a-HUD)
AAL
LSL-150
LBL-152
ME7(detected with a-HUD)
SSL0-488
VLRB.aGPA.23
rNV VLP-Cy3
AFCNVH
HML-Native
#8 Lewis B (T218) (Abcam)
#15 Sialyl Lewis X (BD)
#14 Sialyl Lewis X (Calbio)
hGal-1-488+BCR
TbFBL2
MGL 416
BTL native
GAC78
DC-ASGPR short
Sp Serotype 14.2
Sp Serotype 14.15
rSRL
SRL
Galectin-1
PsSC
Avaren-Fc
ChSeeL
CA19-9 Ab (DRG)
WT Galectin-8
CML1
PhoSPL001
Epa1-Np mutant Y228W
Epa1-Np mutant E227G
Epa1-Np mutant E227A
Epa1-Np WT
ChMBL
DBL-1
hJAA-F11-B11
Neu5,9Ac2(α-Sp8

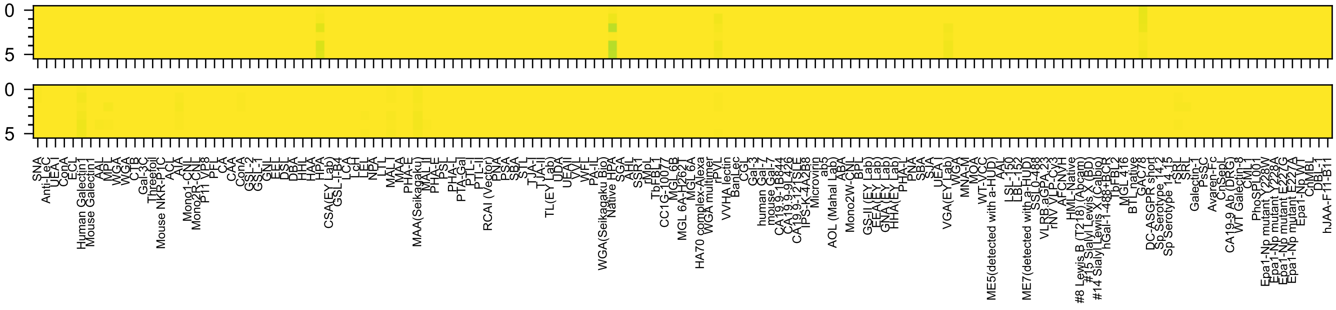
0
5
0
5
Neu5Ac(α-Sp11

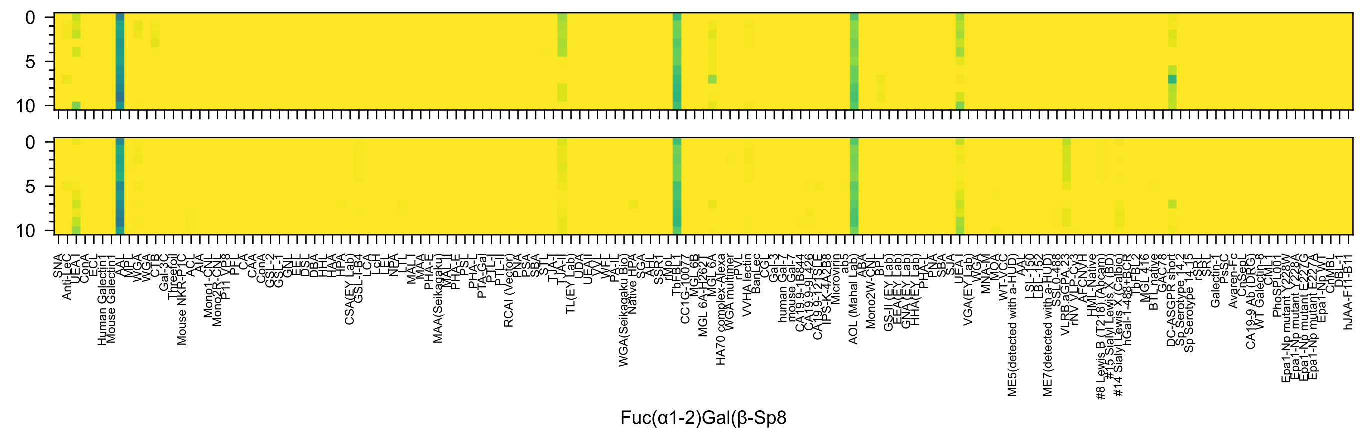
0
5
10
0
5
10
SNA
Anti-LeC
UEA I
ConA
ECL
Human Galectin1
Mouse Galectin1
AAL
MPL
WGA
WGA
CTB
Gal-3C
Threefoil
Mouse NKR-P1C
ACL
AIA
Mono1-CNL
Mono2R-CNL
P11 VP8
PFL
CA
CAA
ConA
GSL-2
GSL-1
GNL
EEL
DSL
DBA
HHL
HAA
HPA
CSA(EY Lab)
GSL-I-B4
LCA
LcH
LEL
NPA
LTL
MAL
MAA
PHA-E
MAA(Seikagaku)
MAL II
PHA-E
PSL
PHA-L
PTA-Gal
PTL-I
PTL-II
RCAI (Vector)
PNA
PSA
SBA
STL
TJA-I
TJA-II
TL(EY Lab)
UDA
UEAII
VVL
WFL
PA-IL
WGA(Seikagaku Bio)
Native HPA
SGA
AHL
SSR1
rMpL
TbFBL1
CC1G-10077
MGL6B
MGL 6A-H262T
MGL 6A
HA70 complex-Alexa
WGA multimer
rPVL
VVHA lectin
BanLec
CGL
Gal-3
human Gal-7
mouse Gal-7
CA19.9-1B844
CA19.9-9L426
CA19.9-121SLE
IPS-K-4A2B8
Microvirin
ab5
AOL (Mahal Lab)
ABA
Mono2W-CNL
BPL
GS-II (EY Lab)
EEA(EY Lab)
GNA(EY Lab)
HHA(EY Lab)
PHA-L
PNA
SBA
SJA
UEA I
VGA(EY Lab)
WGA
MNA-M
MOA
WT-VCC
ME5(detected with a-HUD)
AAL
LSL-150
LBL-152
ME7(detected with a-HUD)
SSL0-488
VLRB.aGPA.23
rNV VLP-Cy3
AFCNYH
HML-Native
#8 Lewis B (T218) (Abcam)
#15 Sialyl Lewis X (BD)
#14 Sialyl Lewis X (Calbio)
hGal-1-488+BCR
TbFBL2
MGL 416
BTL native
GAC78
DC-ASGPR short
Sp Serotype 14.2
Sp Serotype 14.15
rSRL
SRL
Galectin-1
PsSC
Avaren-Fc
ChSeqL
CA19-9 Ab (DRG)
WT Galectin-8
CML1
PhoSL001
Epa1-Np mutant Y228W
Epa1-Np mutant E228A
Epa1-Np mutant E227G
Epa1-Np mutant E227A
Epa1-Np WT
ChMBL
DBL-1
hJAA-F11-B11
Fuc(α1-2)Gal(β-Sp8

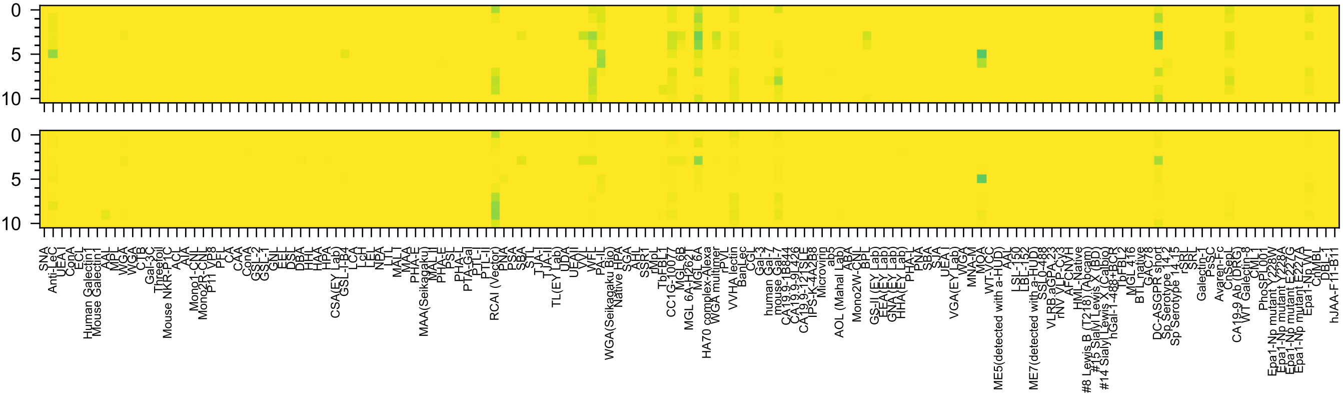

Gal(β1-4)Glc(β-Sp0

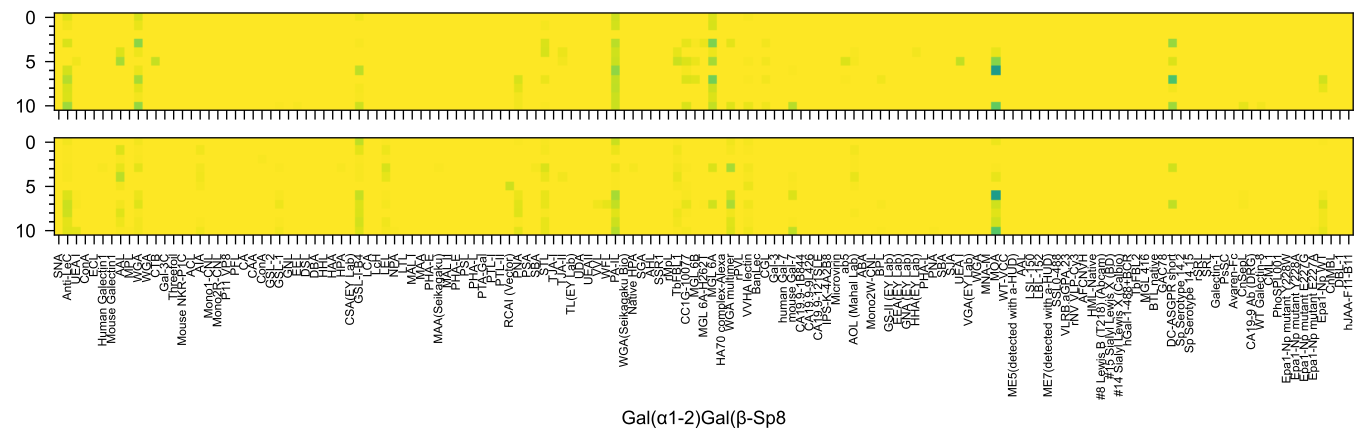
Gal(α1-2)Gal(β-Sp8

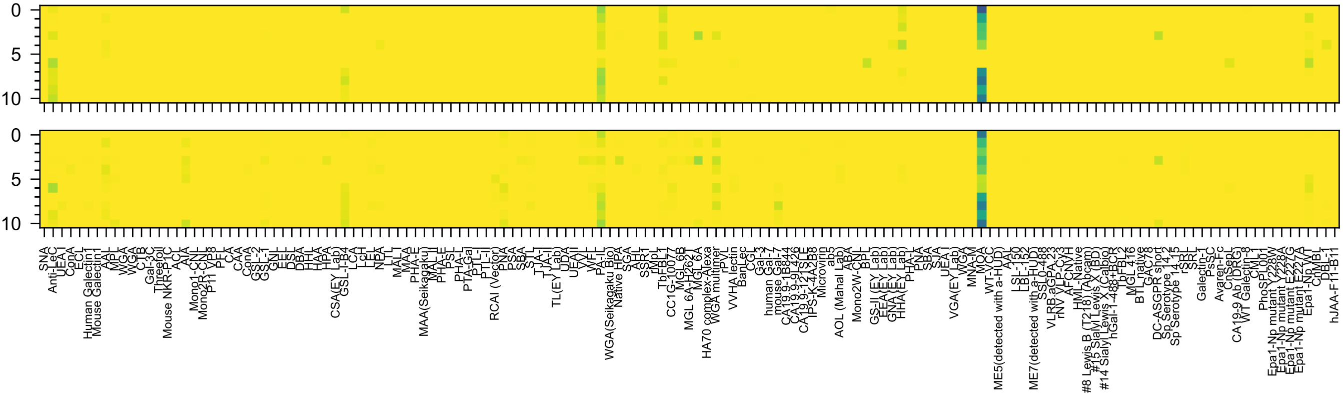

Gal(α1-3)Gal(β-Sp8

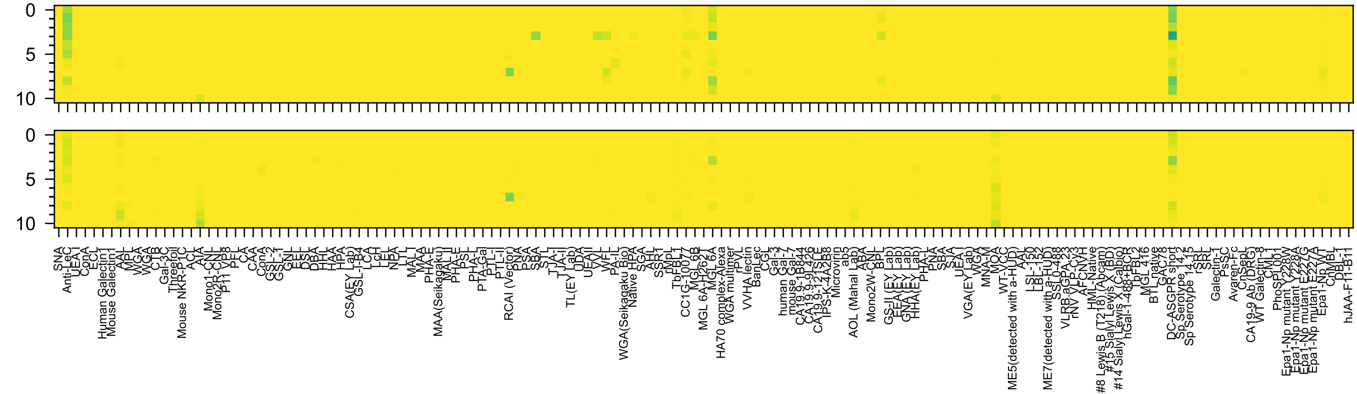

Gal(α1-6)Glc(β-Sp8

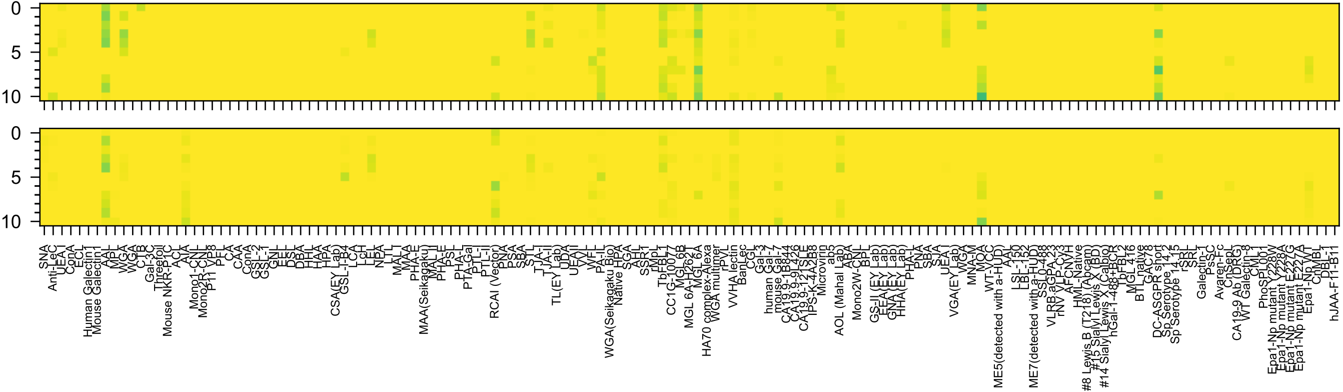


Gal(β1-2)Gal(β-Sp8

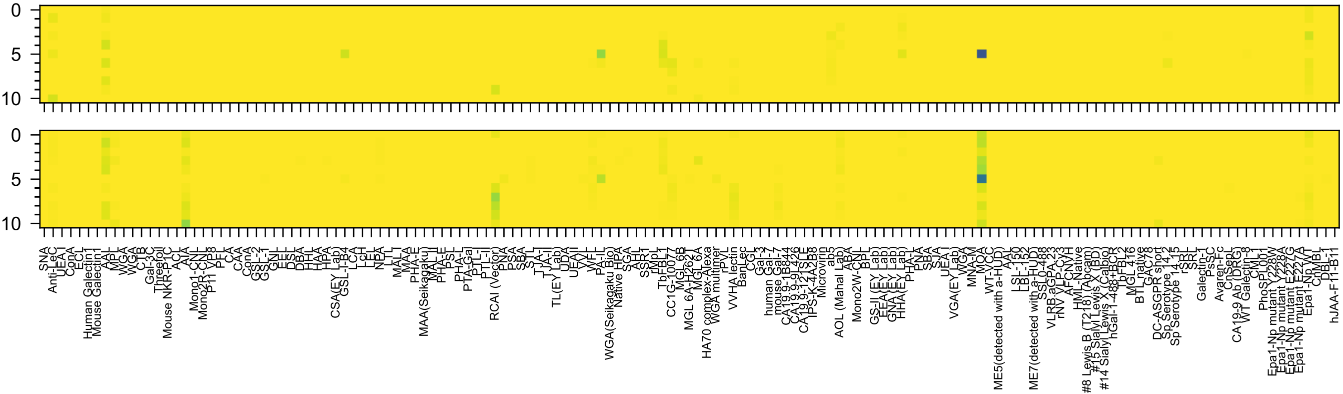

Gal(β1-3)Gal(β-Sp8

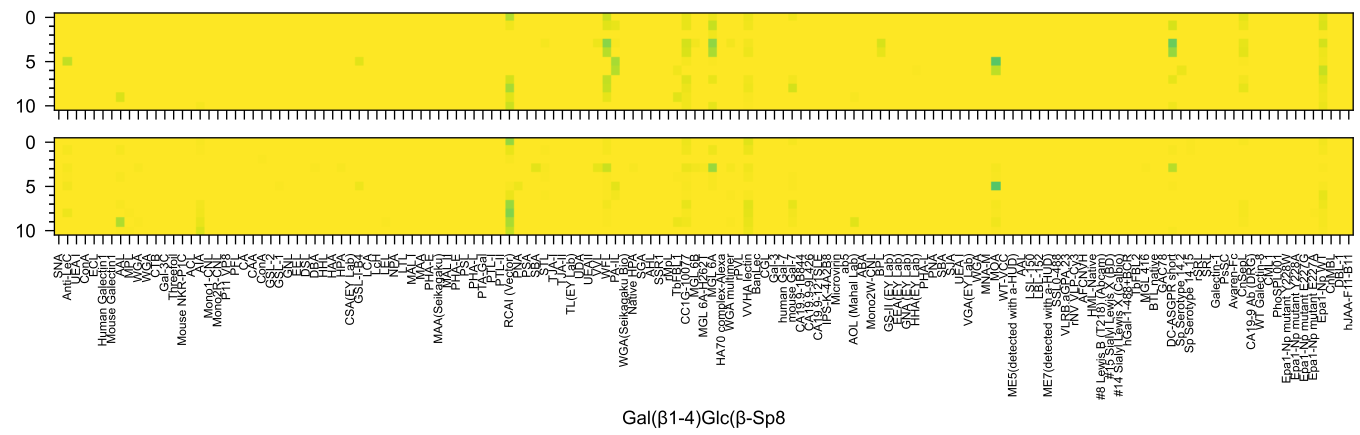
0
5
10
0
5
10
SNA
Anti-LeC
UEA I
ConA
ECL
Human Galectin1
Mouse Galectin1
AAL
MPL
WGA
WGA
CTB
Gal-3C
Threefoil
Mouse NKR-P1C
ACL
AIA
Mono1-CNL
Mono2R-CNL
P11 VP8
PFL
CA
CAA
ConA
GSL-2
GSL-1
GNL
EEL
DSL
DBA
HHL
HAA
HPA
CSA(EY Lab)
GSL-I-B4
LCA
LcH
LEL
NPA
LTL
MAL1
MAA
PHA-E
MAA(Seikagaku)
MAL II
PHA-E
PSL
PHA-L
PTA-Gal
PTL-I
PTL-II
RCAI (Vector)
PNA
PSA
SBA
STL
TJA-I
TJA-II
TL(EY Lab)
UDA
UEAII
VVL
WFL
PA-IL
WGA(Seikagaku Bio)
Native HPA
SGA
AHL
SSR1
rMpL
TbFBL1
CC1G-10077
MGL6B
MGL 6A-H262T
MGL 6A
HA70 complex-Alexa
WGA multimer
rPVL
VVHA lectin
BanLec
CGL
Gal-3
human Gal-7
mouse Gal-7
CA19.9-1B844
CA19.9-9L426
CA19.9-121SLE
IPS-K-4A2B8
Microvirin
ab5
AOL (Mahal Lab)
ABA
Mono2W-CNL
BPL
GS-II (EY Lab)
EEA (EY Lab)
GNA (EY Lab)
HHA(EY Lab)
PHA-L
PNA
SBA
SJA
UEA-I
VGA(EY Lab)
WGA
MNA-M
MOA
WT-VCC
ME5(detected with a-HUD)
AAL
LSL-150
LBL-152
ME7(detected with a-HUD)
SSL0-488
VLRB.aGPA.23
rNV VLP-Cy3
AFCNVH
HML-Native
#8 Lewis B (T218) (Abcam)
#15 Sialyl Lewis X (BD)
#14 Sialyl Lewis X (Calbio)
hGal-1-488+BCR
TbFBL2
MGL 416
BTL native
GAC78
DC-ASGPR short
Sp Serotype 14.2
Sp Serotype 14.15
rSRL
SRL
Galectin-1
PsSC
Avaren-Fc
ChSepL
CA19-9 Ab (DRG)
WT Galectin-8
CML1
PhoSPL001
Epa1-Np mutant Y228W
Epa1-Np mutant E227G
Epa1-Np mutant E227A
Epa1-Np WT
ChMBL
DBL-1
hJAA-F11-B11
Gal(β1-4)Glc(β-Sp8

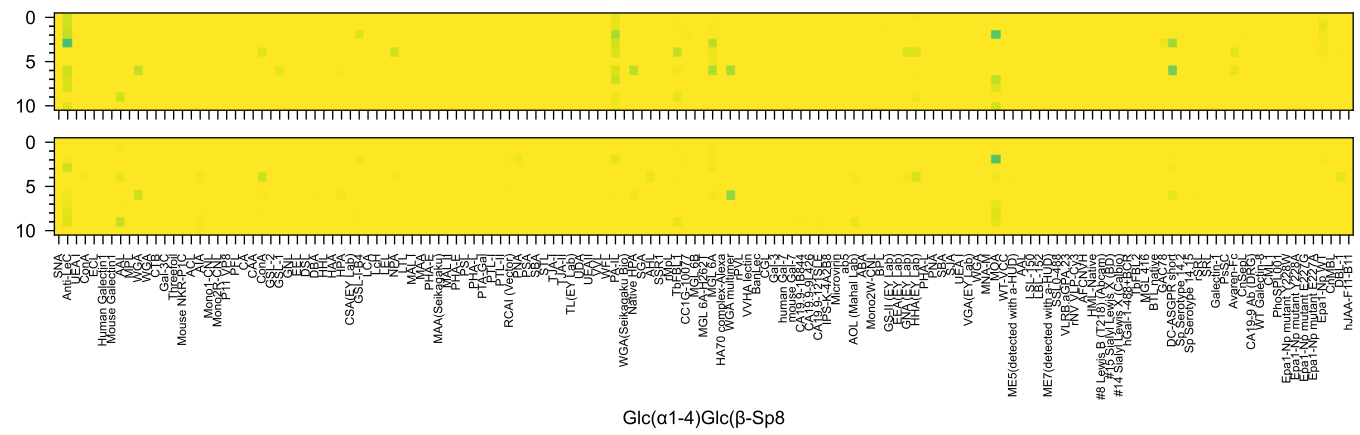
Glc(α1-4)Glc(β-Sp8)
0
5
10

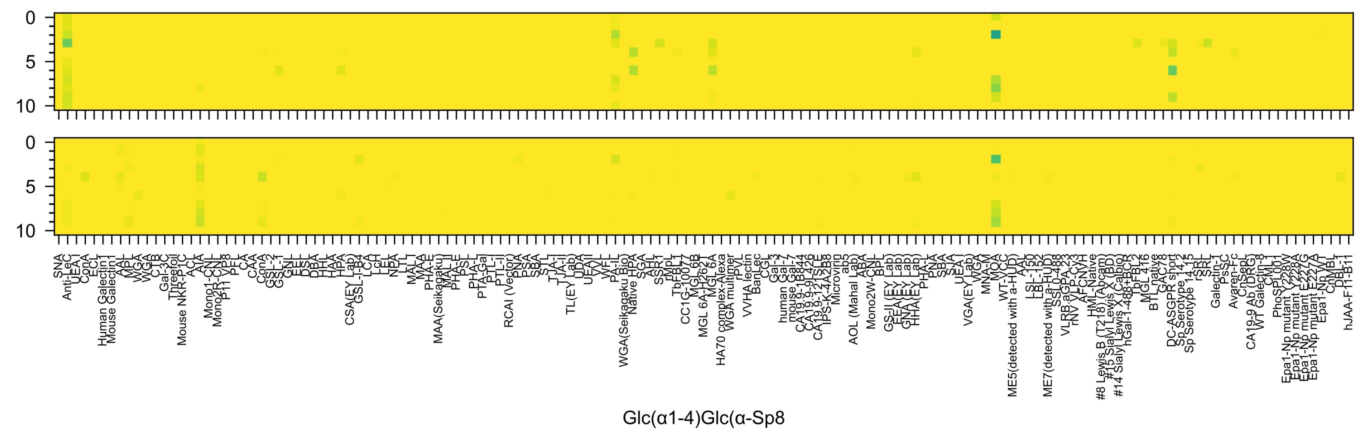
Glc(α1-4)Glc(α-Sp8

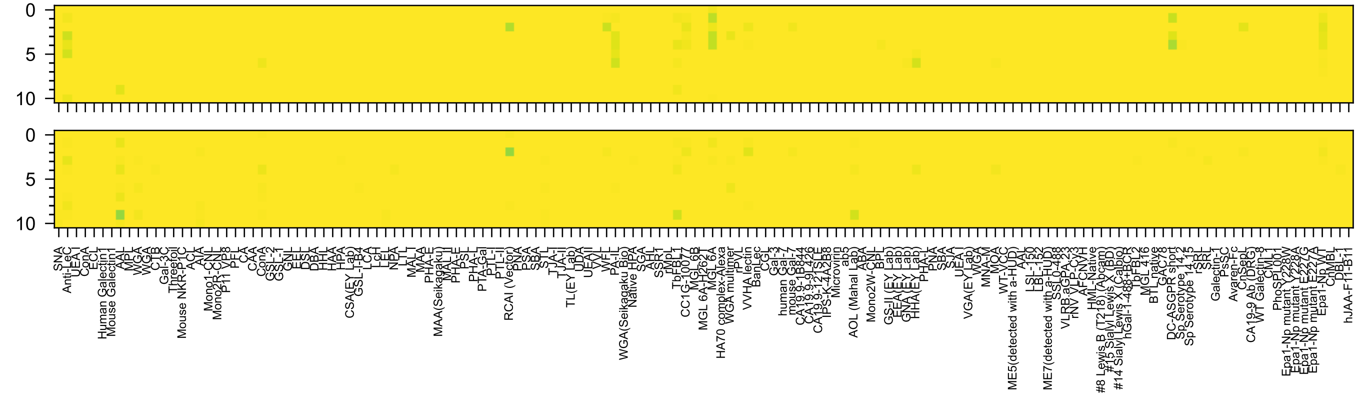

Glc(β1-4)Glc(β-Sp8

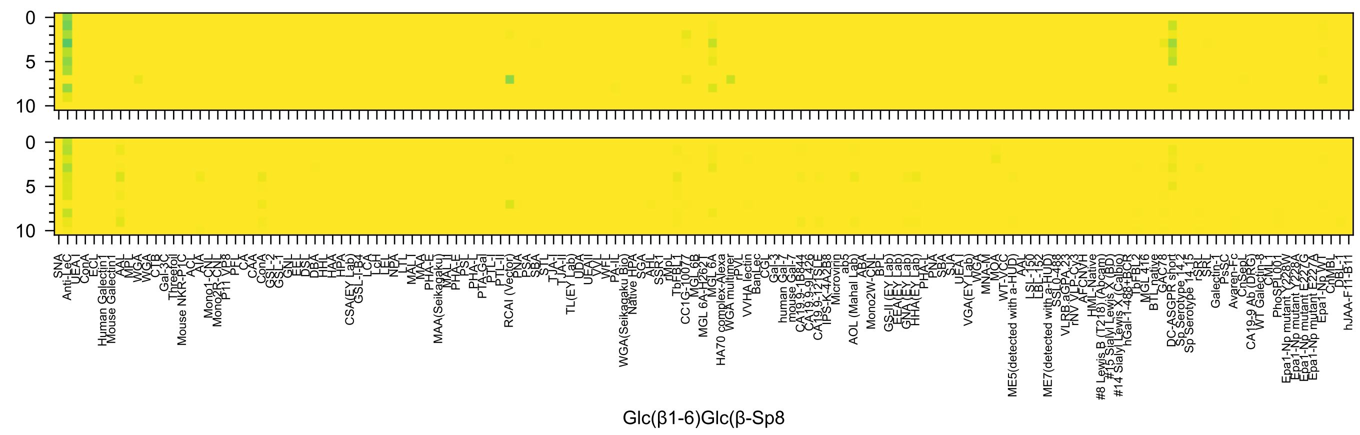
Glc(β1-6)Glc(β-Sp8

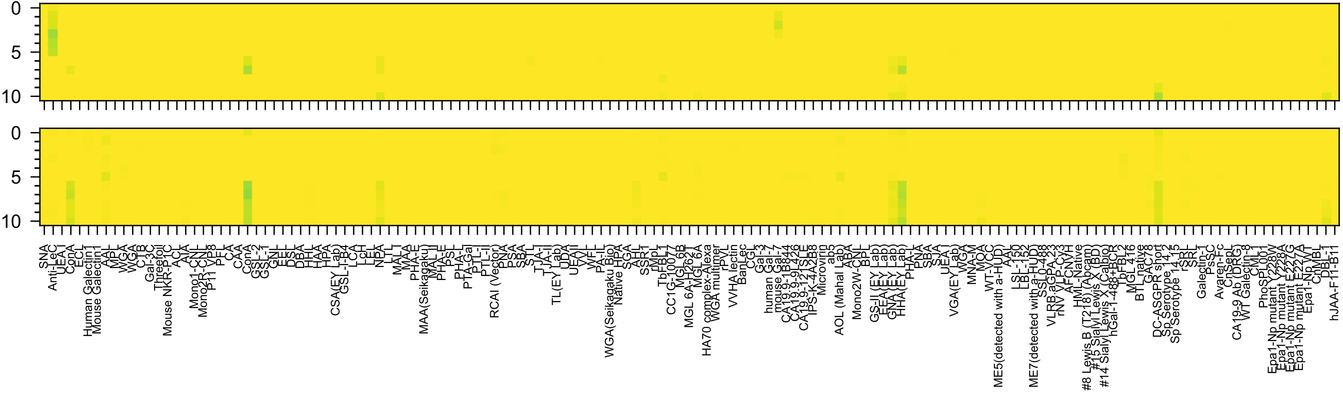


Man(α1-6)Man(β-Sp10

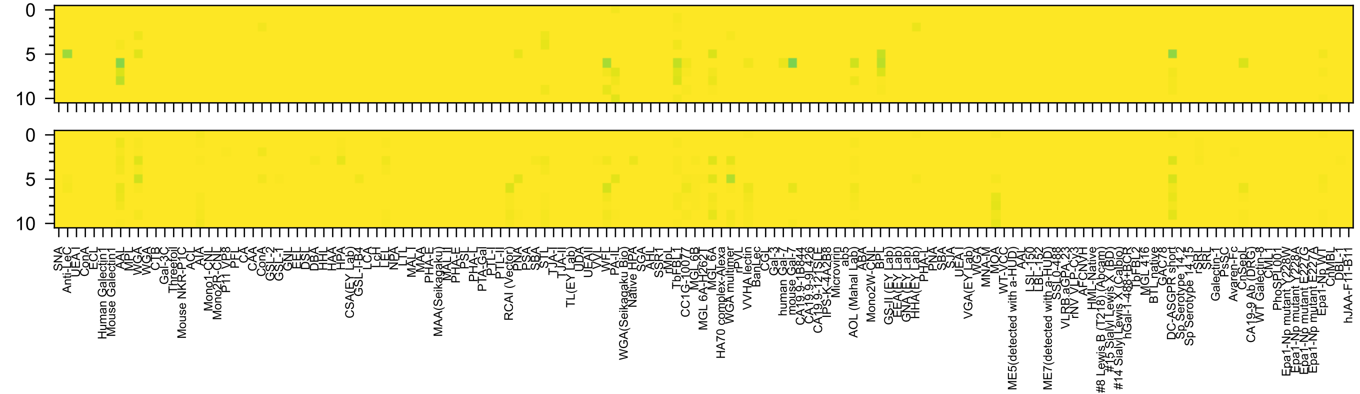

Gal(β1-4)Gal(β-Sp10

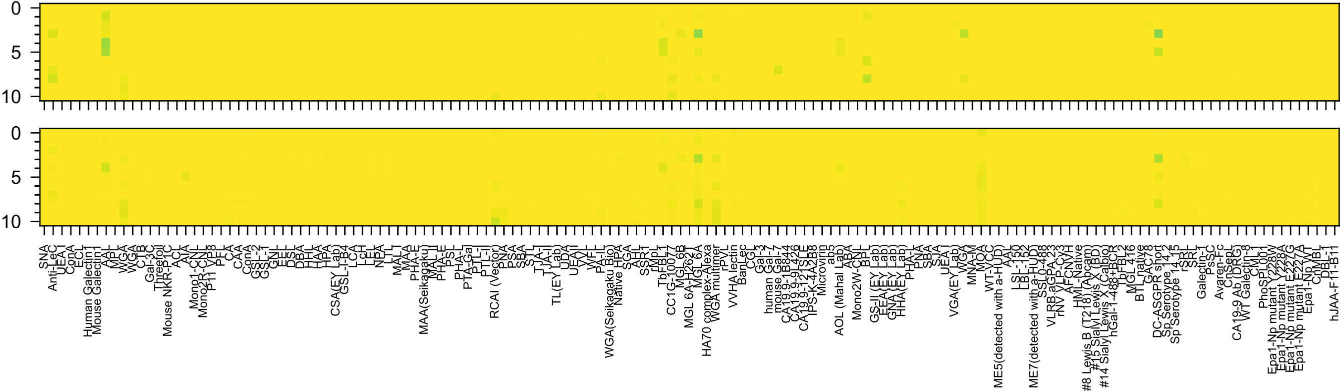


Gal(β1-6)Gal(β-Sp10)

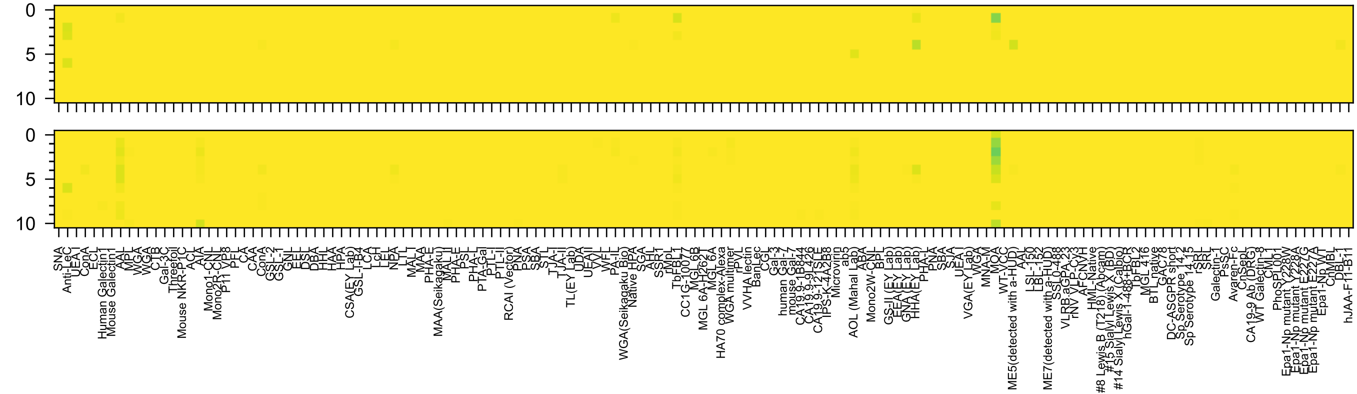


GlcA(β1-3)Gal(β-Sp8

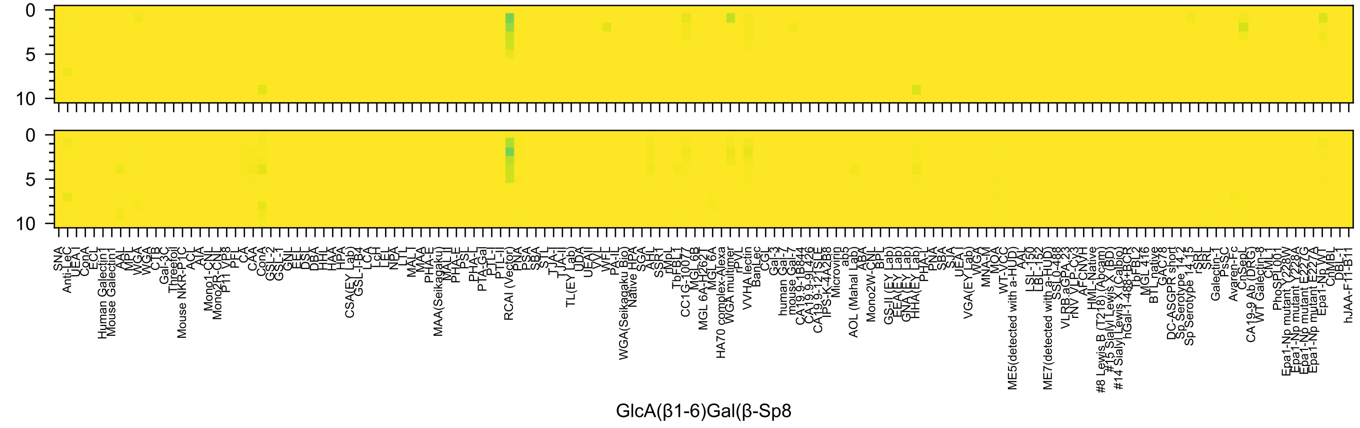
0
5
10
SNA
Anti-LeC
UEA I
ConA
ECL
Human Galectin1
Mouse Galectin1
AAL
MPL
WGA
WGA
CTB
Gal-3C
Threefoil
Mouse NKR-P1C
ACL
AIA
Mono1-CNL
Mono2R-CNL
P11 VP8
PFL
CA
CAA
ConA
GSL-2
GSL-1
GNL
EEL
DSL
DBA
HHL
HAA
HPA
CSA(EY Lab)
GSL-I-B4
LCA
LcH
LEL
NPA
LTL
MAL I
MAA
PHA-E
MAA(Seikagaku)
MAL II
PHA-E
PSL
PHA-L
PTA-Gal
PTL-I
PTL-II
RCAI (Vector)
PNA
PSA
SBA
STL
TJA-I
TJA-II
TL(EY Lab)
UDA
UEAII
VVL
WFL
PA-IL
WGA(Seikagaku Bio)
Native HPA
SGA
AHL
SSR1
rMpL
TbFBL1
CC1G-10077
MGL6B
MGL 6A-H262T
MGL 6A
HA70 complex-Alexa
WGA multimer
rPVL
VVHA lectin
BanLec
CGL
Gal-3
human Gal-7
mouse Gal-7
CA19.9-1B844
CA19.9-9L426
CA19.9-121SLE
IPS-K-4A2B8
Microvirin
ab5
AOL (Mahal Lab)
ABA
Mono2W-CNL
BPL
GS-II (EY Lab)
EEA (EY Lab)
GNA (EY Lab)
HHA(EY Lab)
PHA-L
PNA
SBA
SJA
UEA I
VGA(EY Lab)
WGA
MNA-M
MOA
WT-VCC
ME5(detected with a-HUD)
AAL
LSL-150
LBL-152
ME7(detected with a-HUD)
SSL0-488
VLRB.aGPA.23
rNV VLP-Cy3
AFCNYH
HML-Native
#8 Lewis B (T218) (Abcam)
#15 Sialyl Lewis X (BD)
#14 Sialyl Lewis X (Calbio)
hGal-1-488+BCR
TbFBL2
MGL 416
BTL native
GAC78
DC-ASGPR short
Sp Serotype 14.2
Sp Serotype 14.15
rSRL
SRL
Galectin-1
PsSC
Avaren-Fc
ChSepL
CA19-9 Ab (DRG)
WT Galectin-8
CML
PhoSL001
Epa1-Np mutant Y228A
Epa1-Np mutant E227G
Epa1-Np mutant E227A
Epa1-Np WT
CrMBL
DBL-1
hJAA-F11-B11
GlcA(β1-6)Gal(β-Sp8

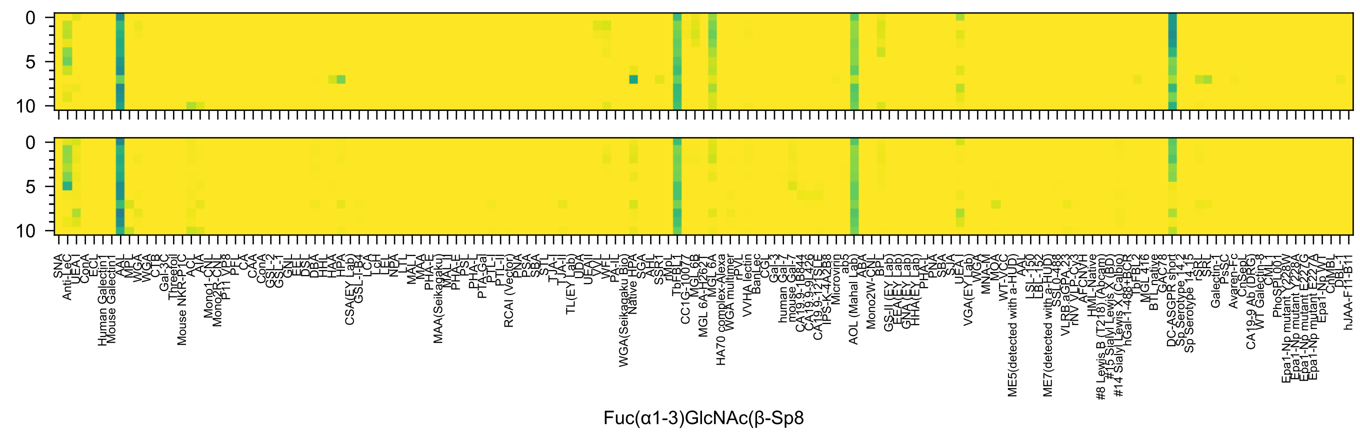

Fuc(α1-3)GlcNAc(β-Sp8

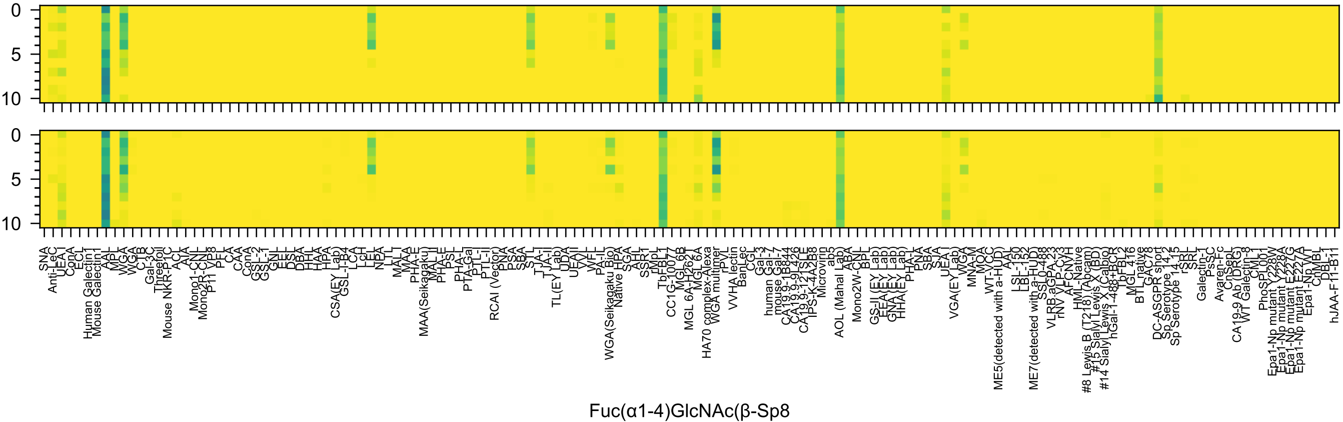
Fuc(α1-4)GlcNAc(β-Sp8

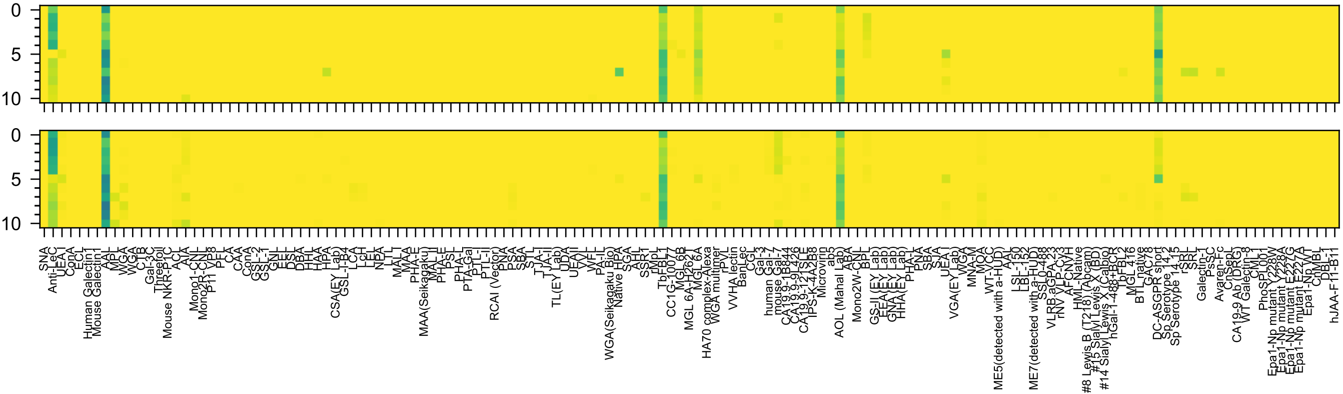

Fuc(β1-3)GlcNAc(β-Sp8

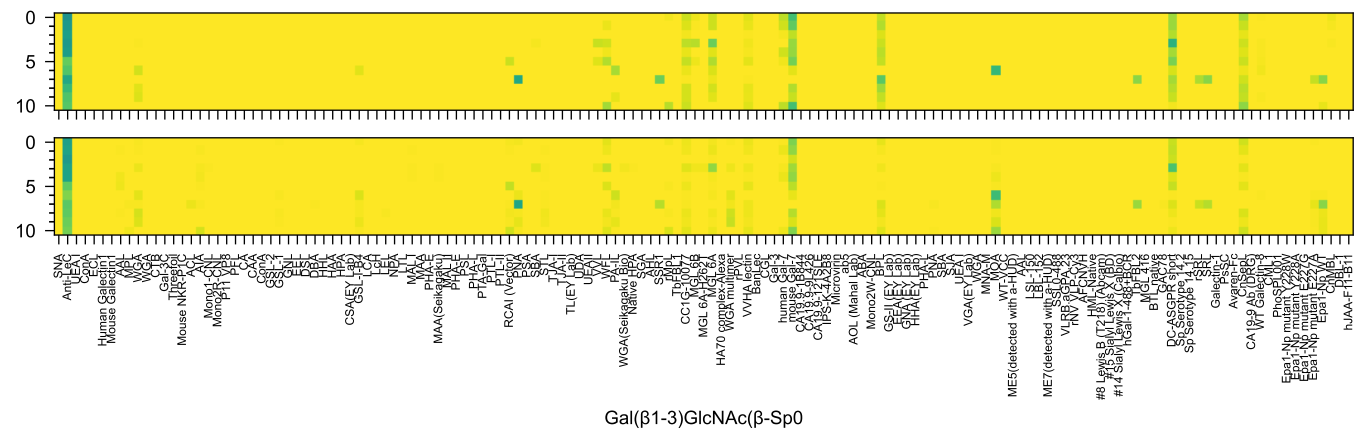
0
5
10
0
5
10
SNA
Anti-LeC
UEA I
ConA
ECL
Human Galectin1
Mouse Galectin1
AAL
MPL
WGA
WGA
CTB
Gal-3C
Threefoil
Mouse NKR-P1C
ACL
AIA
Mono1-CNL
Mono2R-CNL
P11 VP8
PFL
CA
CAA
ConA
GSL-2
GSL-1
GNL
EEL
DSL
DBA
HHL
HAA
HPA
CSA(EY Lab)
GSL-I-B4
LCA
LcH
LEL
NPA
LTL
MAL I
MAA
PHA-E
MAA(Seikagaku)
MAL II
PHA-E
PSL
PHA-L
PTA-Gal
PTL-I
PTL-II
RCAI (Vector)
PNA
PSA
SBA
STL
TJA-I
TJA-II
TL(EY Lab)
UDA
UEA II
VVL
WFL
PA-IL
WGA(Seikagaku Bio)
Native HPA
SGA
AH1
SSR1
rMpL
TbFBL1
CC1G-10077
MGL6B
MGL 6A-H262T
MGL 6A
HA70 complex-Alexa
WGA multimer
rPVL
VVHA lectin
BanLec
CGL
Gal-3
human Gal-7
mouse Gal-7
CA19.9-1B844
CA19.9-9L426
CA19.9-121SLE
IPS-K-4A2B8
Microvirin
ab5
AOL (Mahal Lab)
ABA
Mono2W-CNL
BPL
GS-II (EY Lab)
EEA (EY Lab)
GNA (EY Lab)
HHA (EY Lab)
PHA-L
PNA
SBA
SJA
UEA I
VGA(EY Lab)
WGA
MNA-M
MOA
WT-VCC
ME5(detected with a-HUD)
AAL
LSL-150
LBL-152
ME7(detected with a-HUD)
SSL0-488
VLRB.aGPA.23
rNV VLP-Cy3
AFCNVH
HML-Native
#8 Lewis B (T218) (Abcam)
#15 Sialyl Lewis X (BD)
#14 Sialyl Lewis X (Calbio)
hGal-1-488+BCR
TbFBL2
MGL 416
BTL native
GAC78
DC-ASGPR short
Sp Serotype 14.2
Sp Serotype 14.15
rSRL
SRL
Galectin-1
PsSC
Avaren-Fc
ChSepL
CA19-9 Ab (DRG)
WT Galectin-8
CML1
PhoSPL001
Epa1-Np mutant Y228A
Epa1-Np mutant E227G
Epa1-Np mutant E227A
Epa1-Np WT
ChMBL
DBL-1
hJAA-F11-B11
Gal(β1-3)GlcNAc(β-Sp0

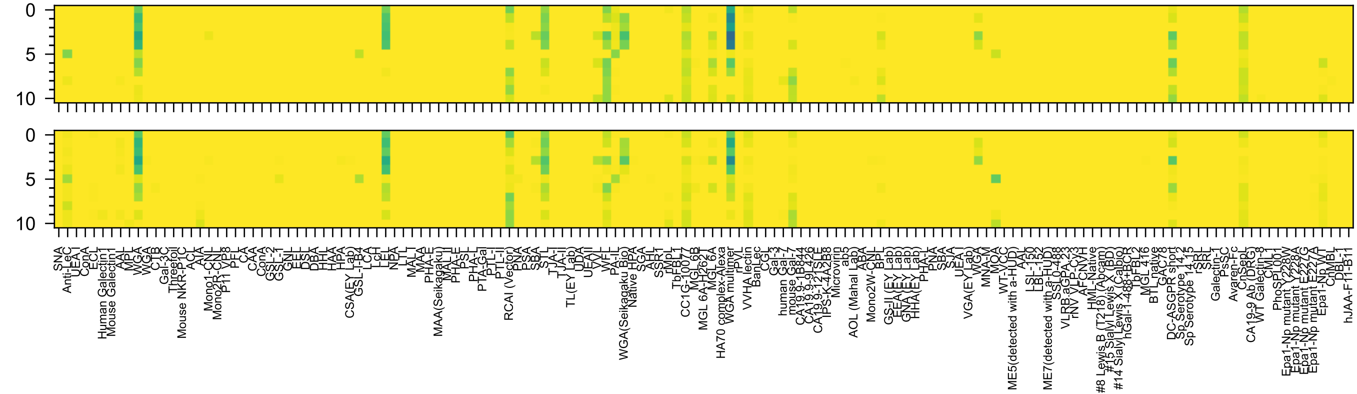


Gal(β1-4)GlcNAc(β-Sp0

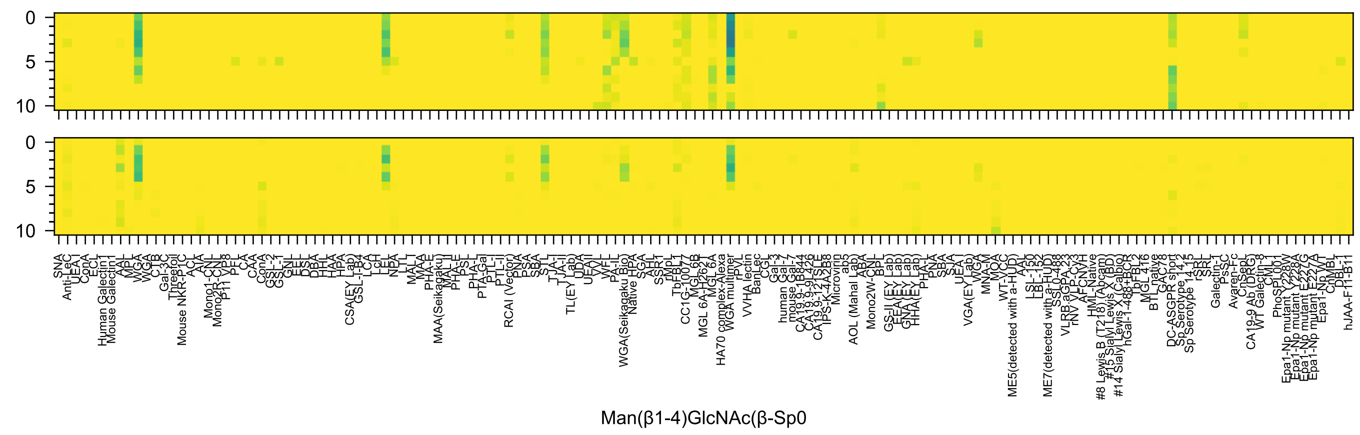
Man(β1-4)GlcNAc(β-Sp0

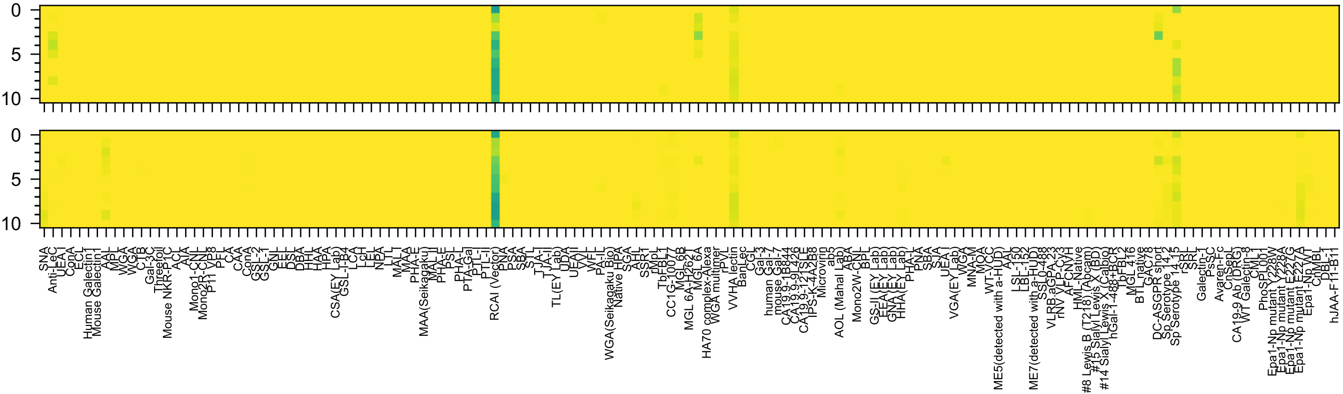

Gal6OS(β1-4)Glc(β-Sp0

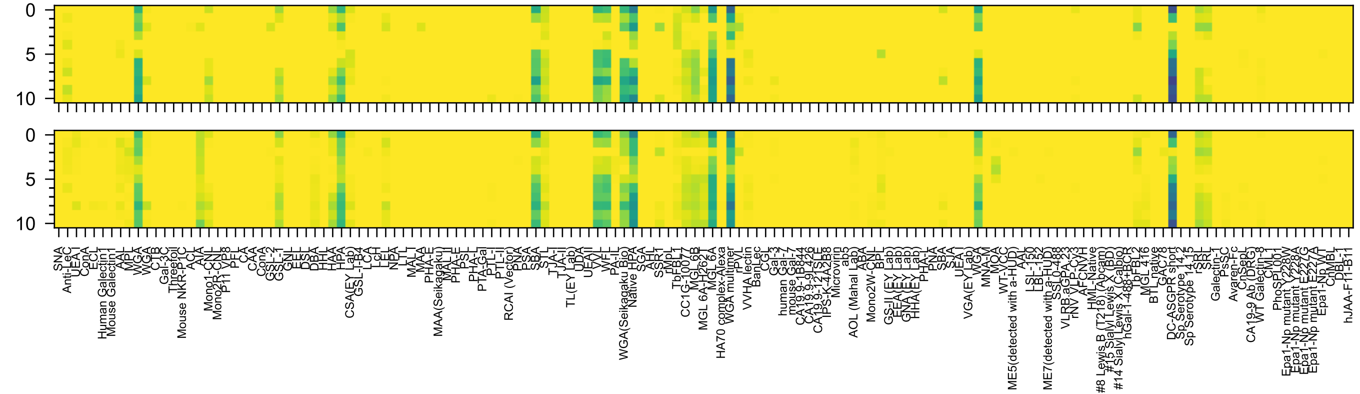

GalNAc(α1-3)Gal(β-Sp8

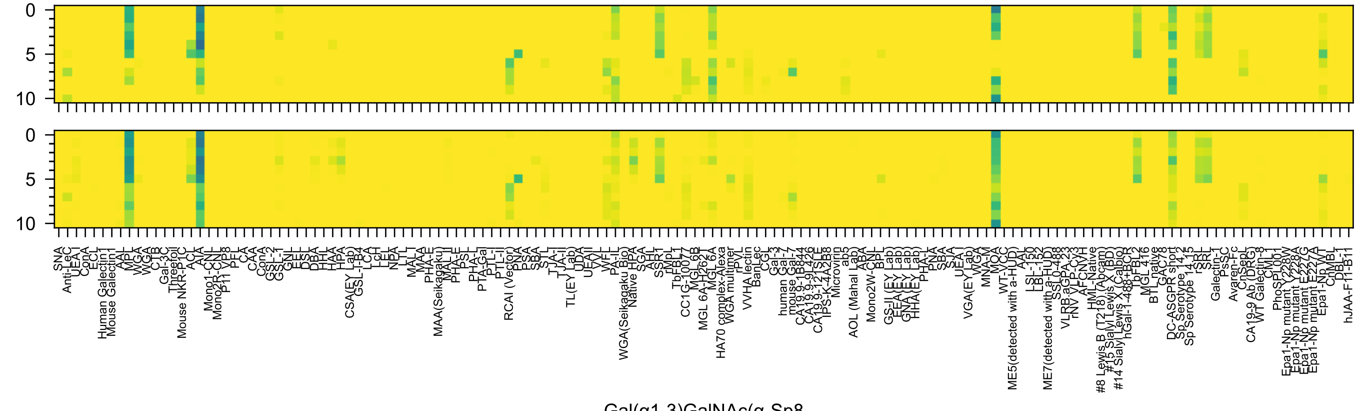
0
5
10
0
5
10
SNA
Anti-LeC
UEA I
ConA
ECL
Human Galectin1
Mouse Galectin1
AAL
MPL
WGA
WGA
CTB
Gal-3C
Threefoil
Mouse NKR-P1C
ACL
AIA
Mono1-CNL
Mono2R-CNL
P11 VP8
PFL
CA
CAA
ConA
GSL-2
GSL-1
GNL
EEL
DSL
DBA
HHL
HAA
HPA
CSA(EY Lab)
GSL-I-B4
LCA
LcH
LEL
NPA
LTL
MAL
MAA
PHA-E
MAA(Seikagaku)
MAL II
PHA-E
PSL
PHA-L
PTA-Gal
PTL-I
PTL-II
RCAI (Vector)
PNA
PSA
SBA
STL
TJA-I
TJA-II
TL(EY Lab)
UDA
UEAII
VVL
WFL
PA-IL
WGA(Seikagaku Bio)
Native HPA
SGA
AHL
SSR1
rMpL
TbFBL1
CC1G-10077
MGL6B
MGL 6A-H262T
MGL 6A
HA70 complex-Alexa
WGA multimer
rPVL
VVHA lectin
BanLec
CGL
Gal-3
human Gal-7
mouse Gal-7
CA19.9-1B844
CA19.9-9L426
CA19.9-121SLE
IPS-K-4A2B8
Microvirin
ab5
AOL (Mahal Lab)
ABA
Mono2W-CNL
BPL
GS-II (EY Lab)
EEA(EY Lab)
GNA(EY Lab)
HHA(EY Lab)
PHA-L
PNA
SBA
SJA
UEA I
VGA(EY Lab)
WGA
MNA-M
MOA
WT-VCC
ME5(detected with a-HUD)
AAL
LSL-150
LBL-152
ME7(detected with a-HUD)
SSL0-488
VLRB.aGPA.23
rNV VLP-Cy3
AFCNYH
HML-Native
#8 Lewis B (T218) (Abcam)
#15 Sialyl Lewis X (BD)
#14 Sialyl Lewis X (Calbio)
hGal-1-488+BCR
TbFBL2
MGL 416
BTL native
GAC78
DC-ASGPR short
Sp Serotype 14.2
Sp Serotype 14.15
rSRL
SRL
Galectin-1
PsSC
Avaren-Fc
ChSepL
CA19-9 Ab (DRG)
WT Galectin-8
CML1
PhoSL001
Epa1-Np mutant Y228W
Epa1-Np mutant Y228A
Epa1-Np mutant E227G
Epa1-Np mutant E227A
Epa1-Np WT
ChMBL
DBL-1
hJAA-F11-B11
Gal(α1-3)GalNAc(α-Sp8

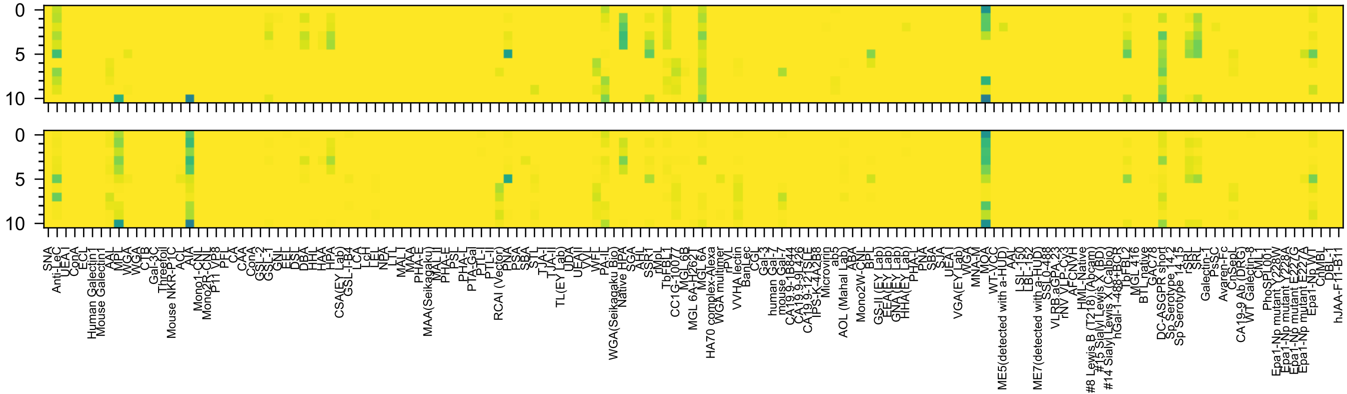


Gal(α1-3)GalNAc(β-Sp8

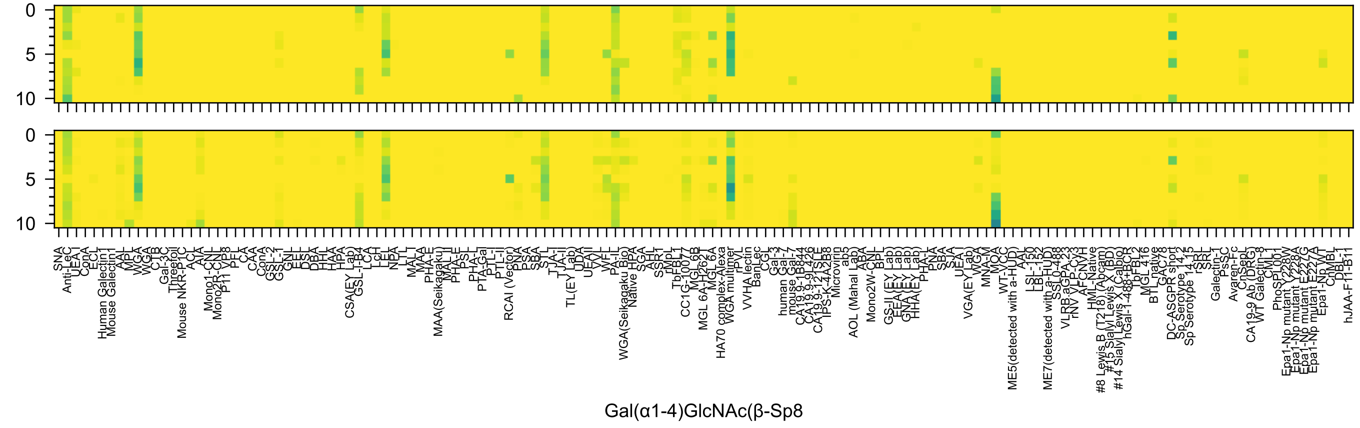
Gal(α1-4)GlcNAc(β-Sp8

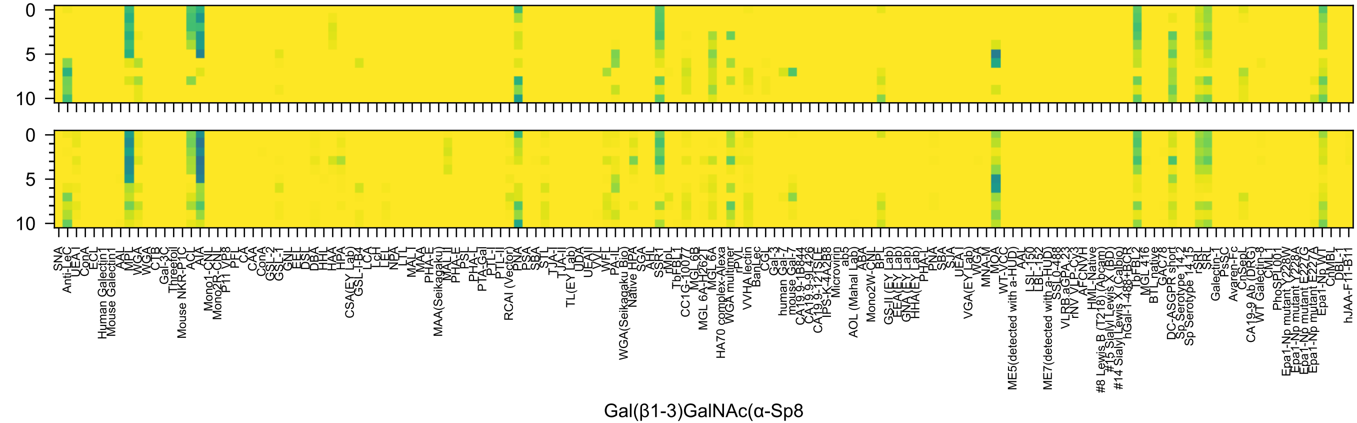
0
5
10
0
5
10
SNA
Anti-LeC
UEA I
ConA
ECL
Human Galectin1
Mouse Galectin1
AAL
MPL
WGA
WGA
CTB
Gal-3C
Threefoil
Mouse NKR-P1C
ACL
AIA
Mono1-CNL
Mono2R-CNL
P11 VP8
PFL
CA
CAA
ConA
GSL-2
GSL-1
GNL
EEL
DSL
DBA
HHL
HAA
HPA
CSA(EY Lab)
GSL-I-B4
LCA
LcH
LEL
NPA
LTL
MAL-I
MAA
PHA-E
MAA(Seikagaku)
MAL II
PHA-E
PSL
PHA-L
PTA-Gal
PTL-I
PTL-II
RCAI (Vector)
PNA
PSA
SBA
STL
TJA-I
TJA-II
TL(EY Lab)
UDA
UEAII
VVL
WFL
PA-IL
WGA(Seikagaku Bio)
Native HPA
SGA
AHL
SSR1
rMpL
TbFBL1
CC1G-10077
MGL6B
MGL 6A-H262T
MGL 6A
HA70 complex-Alexa
WGA multimer
rPVL
VVHA lectin
BanLec
CGL
Gal-3
human Gal-7
mouse Gal-7
CA19.9-1B844
CA19.9-9L426
CA19.9-121SLE
IPS-K-4A2B8
Microvirin
ab5
AOL (Mahal Lab)
ABA
Mono2W-CNL
BPL
GS-II (EY Lab)
EEA(EY Lab)
GNA(EY Lab)
HHA(EY Lab)
PHA-L
PNA
SBA
SJA
UEA I
VGA(EY Lab)
WGA
MNA-M
MOA
WT-VCC
ME5(detected with a-HUD)
AAL
LSL-150
LBL-152
ME7(detected with a-HUD)
SSL0-488
VLRB.aGPA.23
rNV VLP-Cy3
AFCNVH
HML-Native
#8 Lewis B (T218) (Abcam)
#15 Sialyl Lewis X (BD)
#14 Sialyl Lewis X (Calbio)
hGal-1-488+BCR
TbFBL2
MGL 416
BTL native
GAC78
DC-ASGPR short
Sp Serotype 14.2
Sp Serotype 14.15
rSRL
SRL
Galectin-1
PsSC
Avaren-Fc
ChSepL
CA19-9 Ab (DRG)
WT Galectin-8
CML1
PhoSL001
Epa1-Np mutant Y228W
Epa1-Np mutant E228A
Epa1-Np mutant E227G
Epa1-Np mutant E227A
Epa1-Np WT
ChMBL
DBL-1
hJAA-F11-B11
Gal(β1-3)GalNAc(α-Sp8

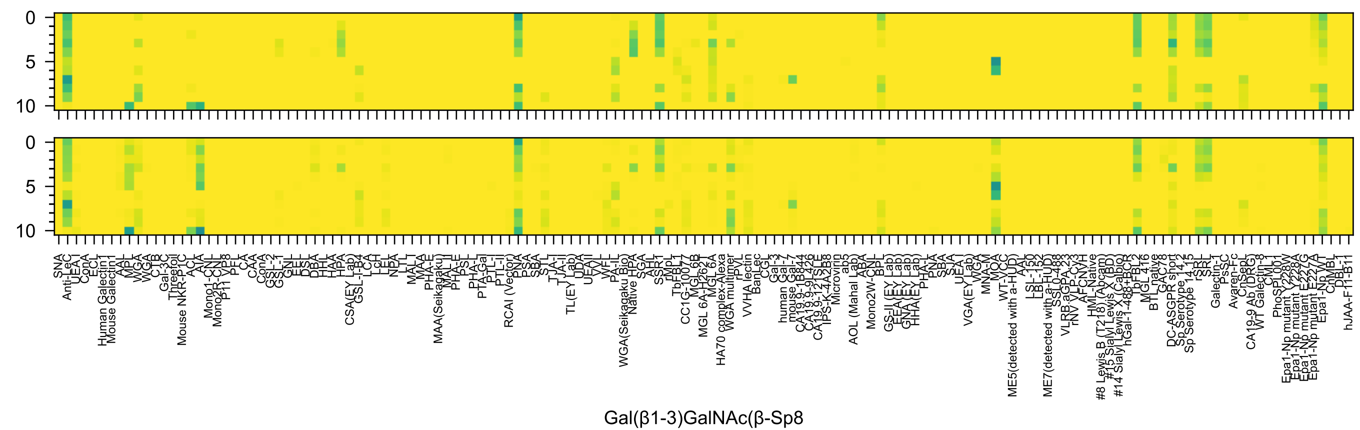
0
5
10
0
5
10
SNA
Anti-LeC
UEA I
ConA
ECL
Human Galectin1
Mouse Galectin1
AAL
MPL
WGA
WGA
CTB
Gal-3C
Threefoil
Mouse NKR-P1C
ACL
AIA
Mono1-CNL
Mono2R-CNL
P11 VP8
PFL
CA
CAA
ConA
GSL-2
GSL-1
GNL
EEL
DSL
DBA
HHL
HAA
HPA
CSA(EY Lab)
GSL-I-B4
LCA
LcH
LEL
NPA
LTL
MAL I
MAA
PHA-E
MAA(Seikagaku)
MAL II
PHA-E
PSL
PHA-L
PTA-Gal
PTL-I
PTL-II
RCAI (Vector)
PNA
PSA
SBA
STL
TJA-I
TJA-II
TL(EY Lab)
UDA
UEAII
VVL
WFL
PA-IL
WGA(Seikagaku Bio)
Native HPA
SGA
AHL
SSR1
rMpL
TbFBL1
CC1G-10077
MGL-6B
MGL 6A-H262T
MGL 6A
HA70 complex-Alexa
WGA multimer
rPVL
VVHA lectin
BanLec
CGL
Gal-3
human Gal-7
mouse Gal-7
CA19.9-1B844
CA19.9-9L426
CA19.9-121SLE
IPS-K-4A2B8
Microvirin
ab5
AOL (Mahal Lab)
ABA
Mono2W-CNL
BPL
GS-II (EY Lab)
EEA (EY Lab)
GNA (EY Lab)
HHA(EY Lab)
PHA-L
PNA
SBA
SJA
UEA I
VGA(EY Lab)
WGA
MNA-M
MOA
WT-VCC
ME5(detected with a-HUD)
AAL
LSL-150
LBL-152
ME7(detected with a-HUD)
SSL0-488
VLRB.aGPA.23
rNV VLP-Cy3
AFCNVH
HML-Native
#8 Lewis B (T218) (Abcam)
#15 Sialyl Lewis X (BD)
#14 Sialyl Lewis X (Calbio)
hGal-1-488+BCR
TbFBL2
MGL 416
BTL native
GAC78
DC-ASGPR short
Sp Serotype 14.2
Sp Serotype 14.15
rSRL
SRL
Galectin-1
PsSC
Avaren-Fc
ChSepL
CA19-9 Ab (DRG)
WT Galectin-8
CML1
PhoSL001
Epa1-Np mutant Y228W
Epa1-Np mutant Y228A
Epa1-Np mutant E227G
Epa1-Np mutant E227A
Epa1-Np WT
ChMBL
DBL-1
hJAA-F11-B11
Gal(β1-3)GalNAc(β-Sp8

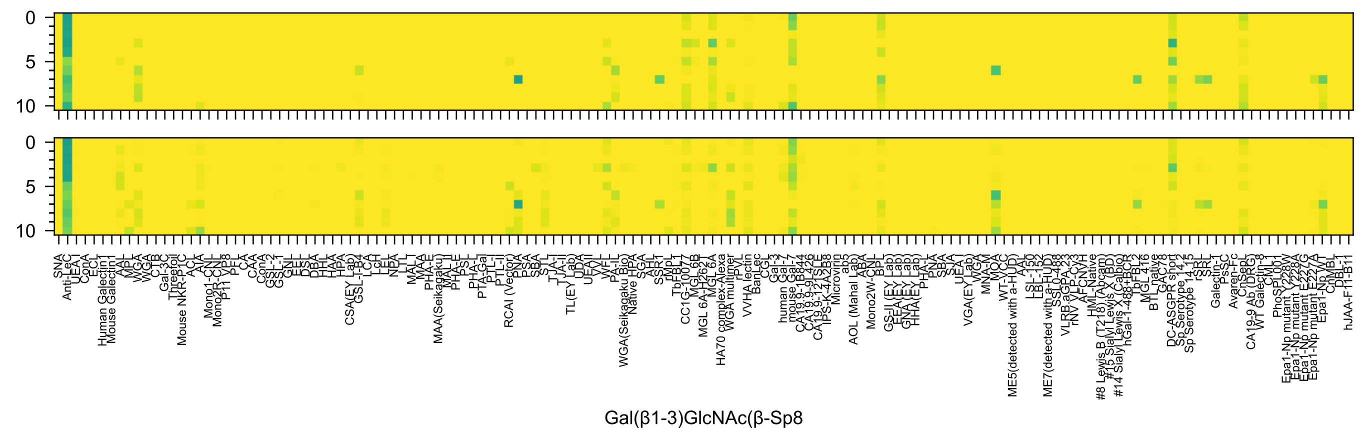
Gal(β1-3)GlcNAc(β-Sp8

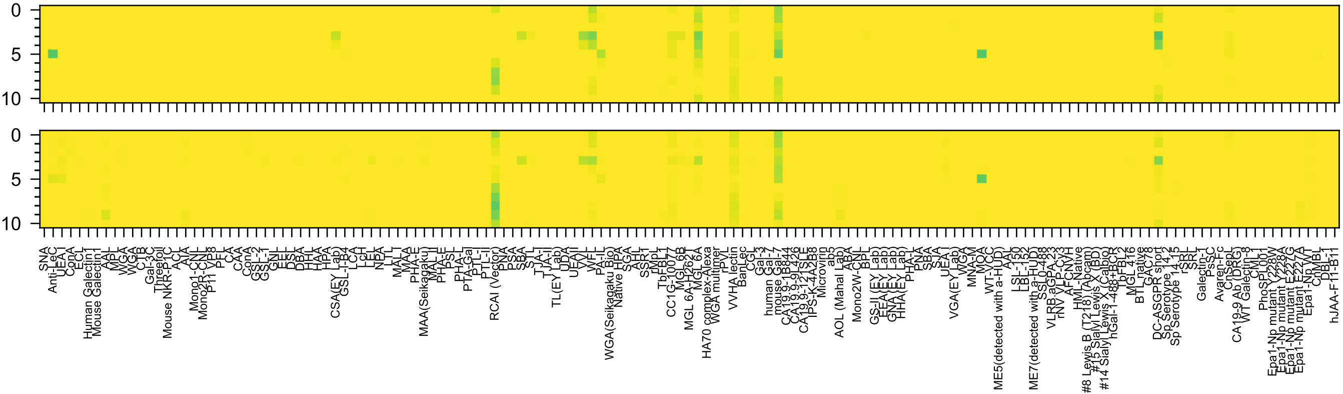

Gal(β1-4)Glc6OS(β-Sp0)

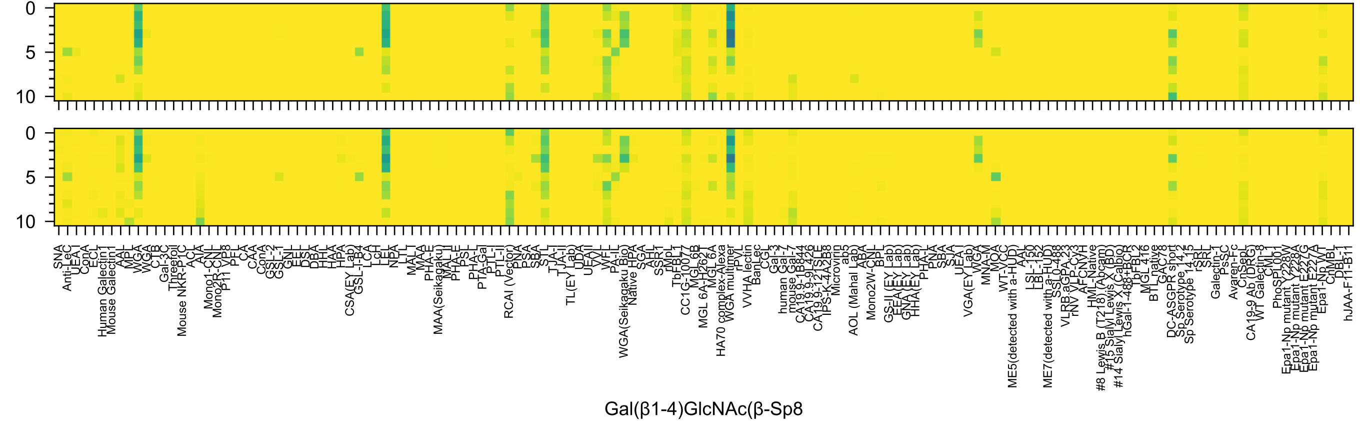
Gal(β1-4)GlcNAc(β-Sp8

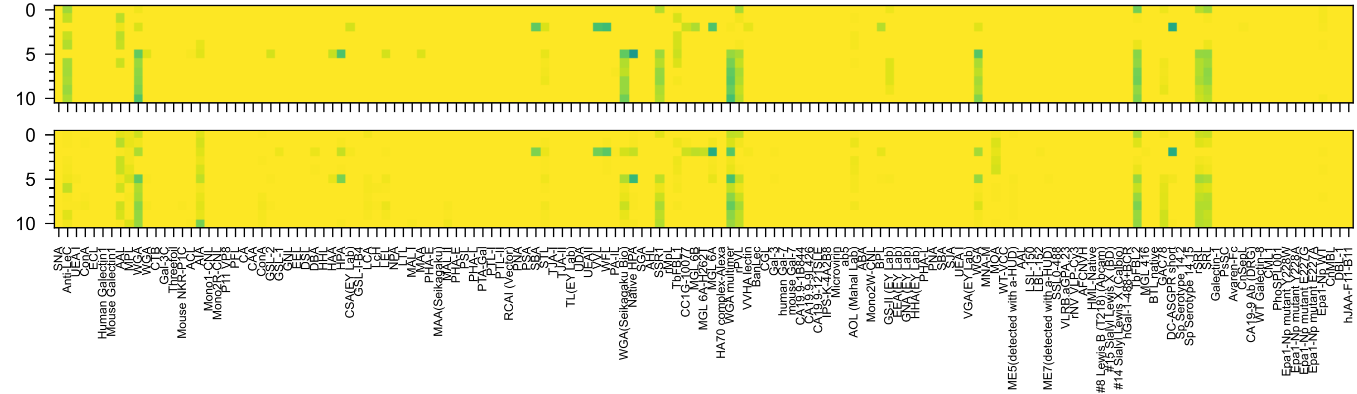

GlcNAc(β1-3)Gal(β-Sp8

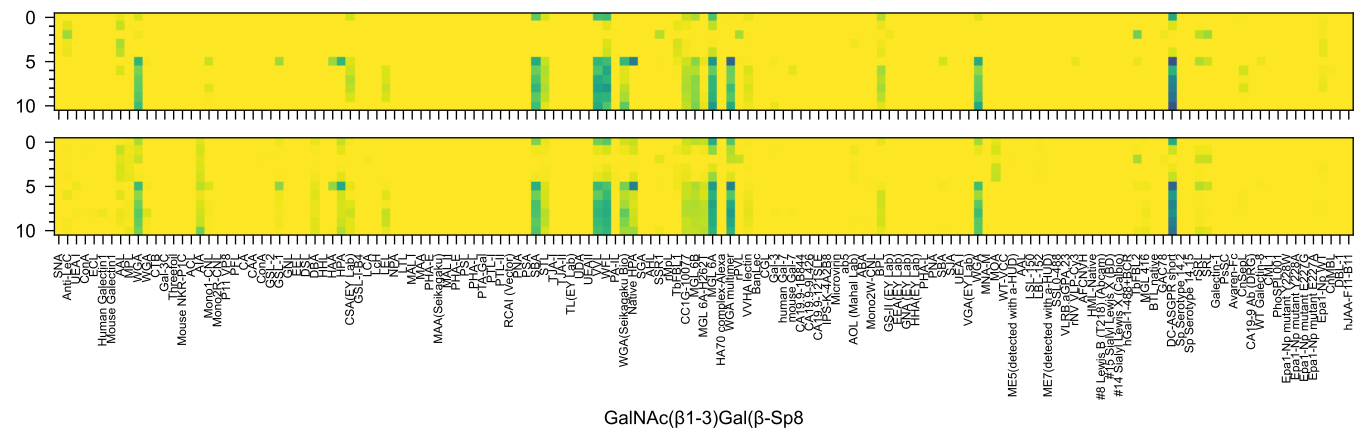

GalNAc(β1-3)Gal(β-Sp8

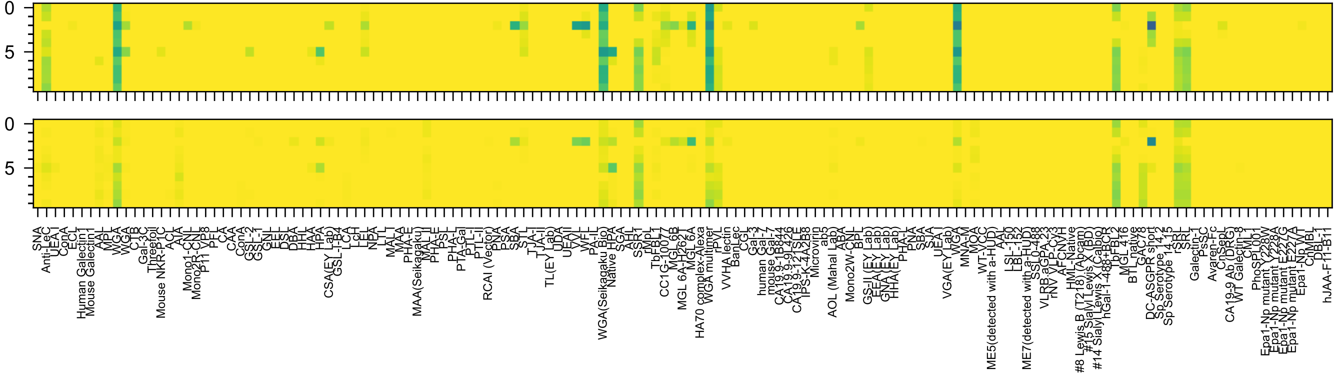
GlcNAc(β1-3)Man(-Sp10

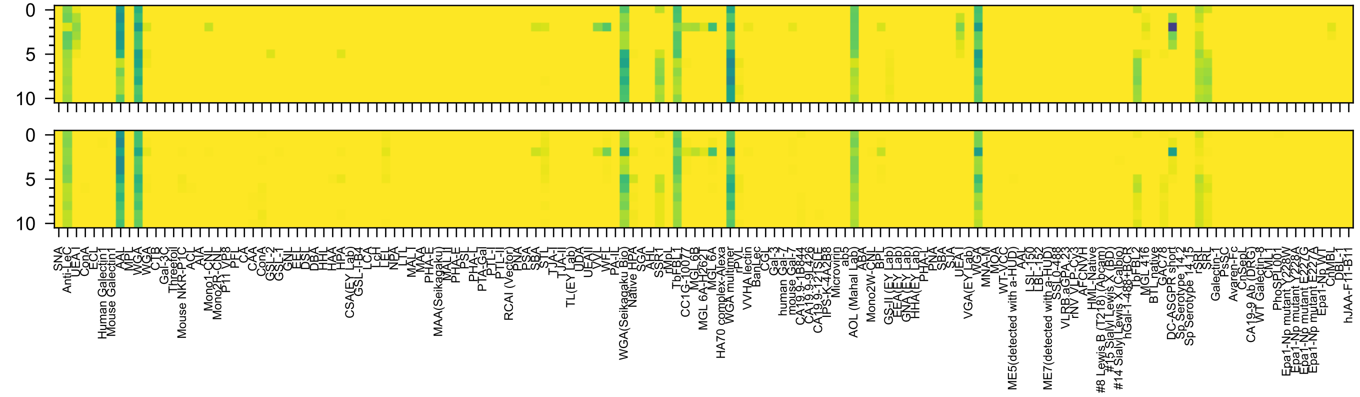
0
5
10
0
5
10
GlcNAc(β1-3)Fuc(α-Sp21

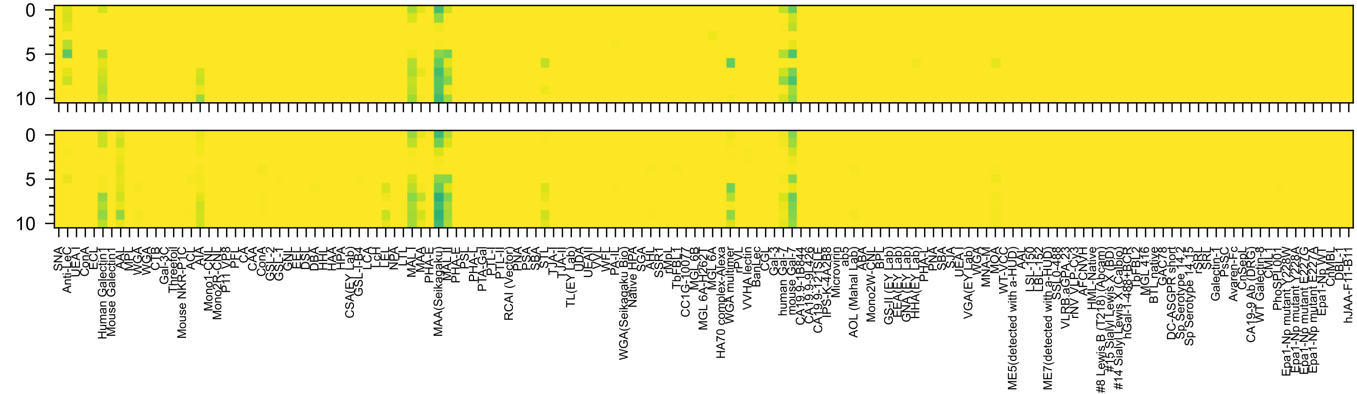

Gal3OS(β1-4)Glc(β-Sp8

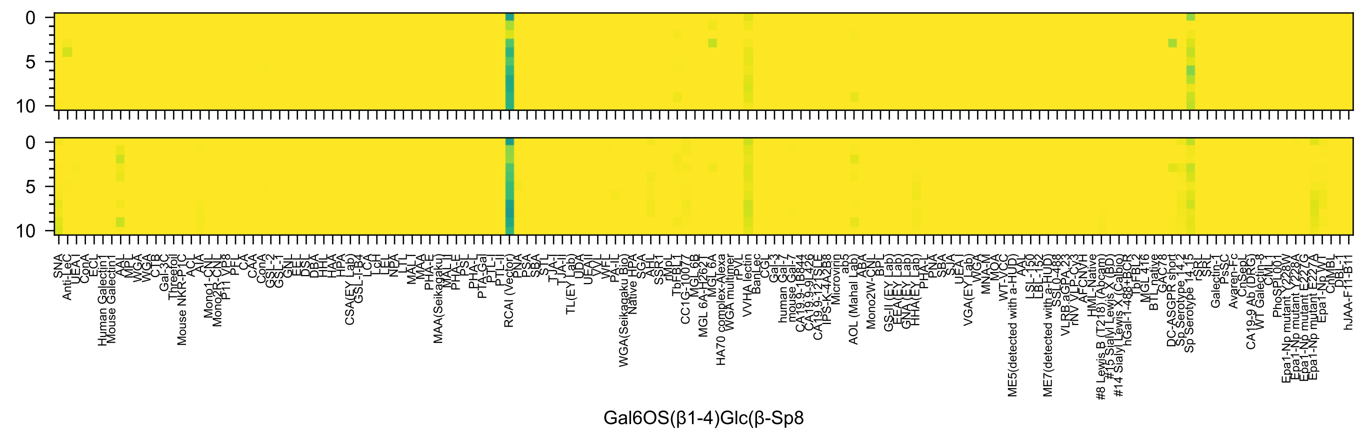
Gal6OS(β1-4)Glc(β-Sp8

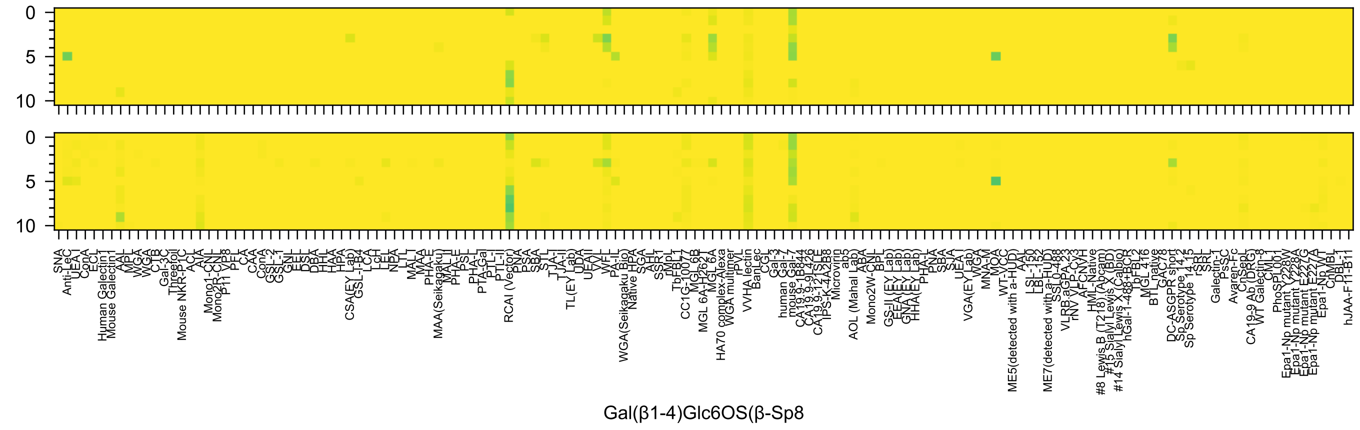
Gal(β1-4)Glc6OS(β-Sp8

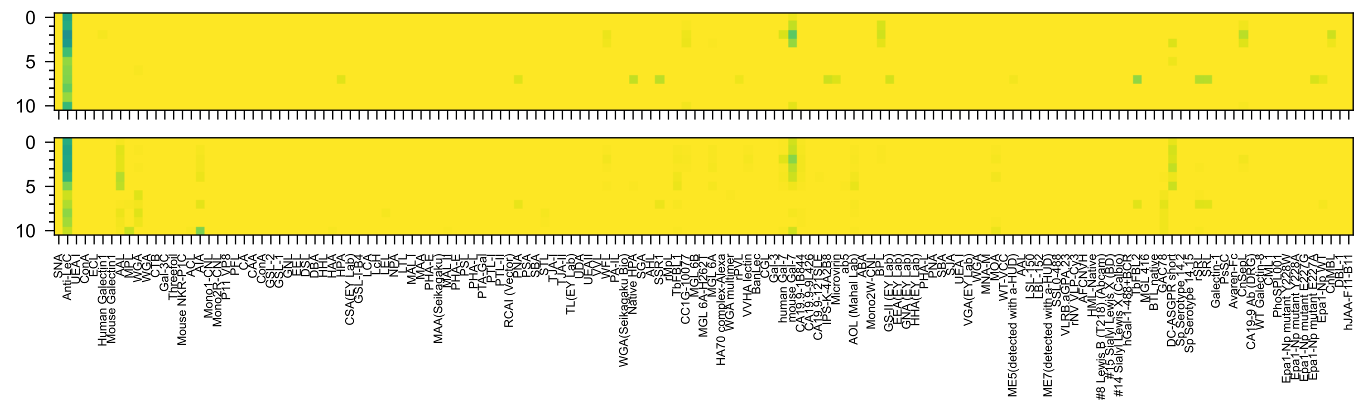

GlcA(β1-3)GlcNAc(β-Sp8

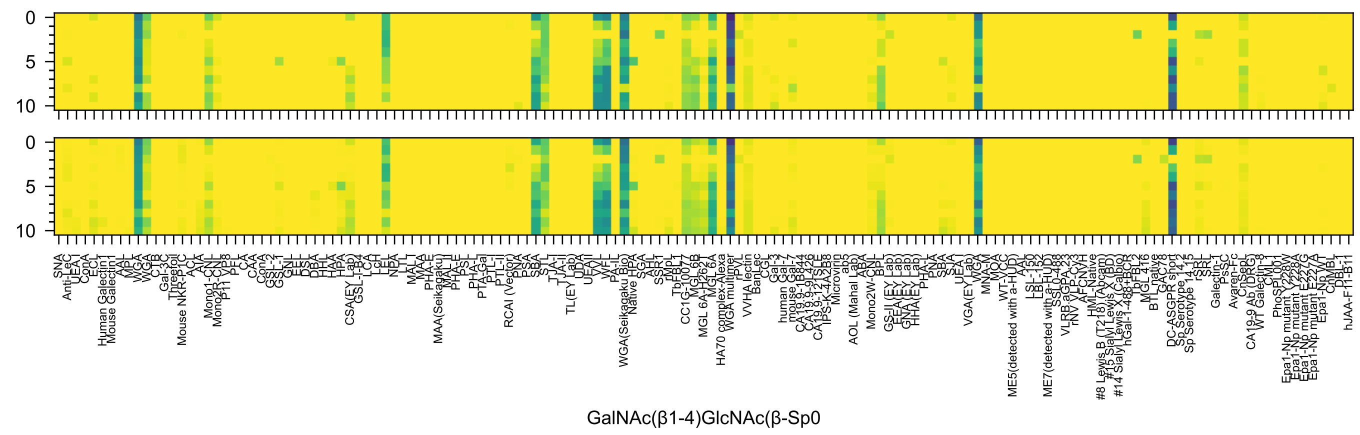

GalNAc(β1-4)GlcNAc(β-Sp0

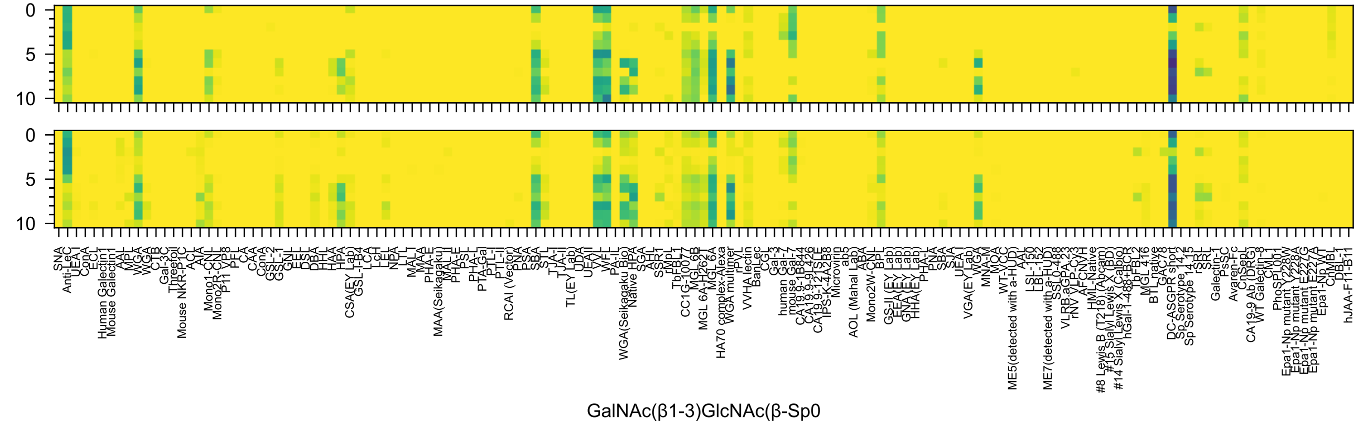
0
5
10
0
5
10
GalNAc(β1-3)GlcNAc(β-Sp0

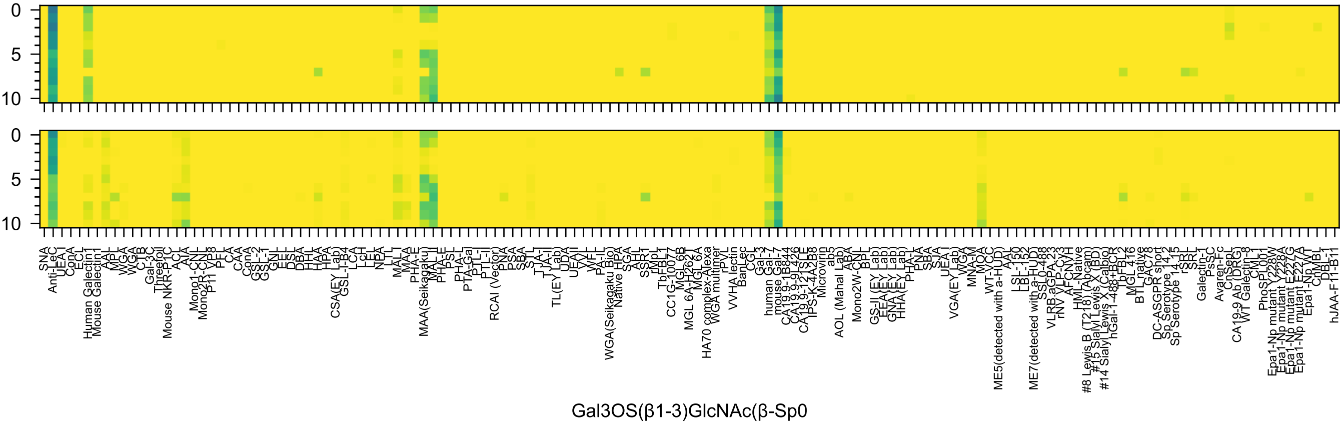

Gal3OS(β1-3)GlcNAc(β-Sp0

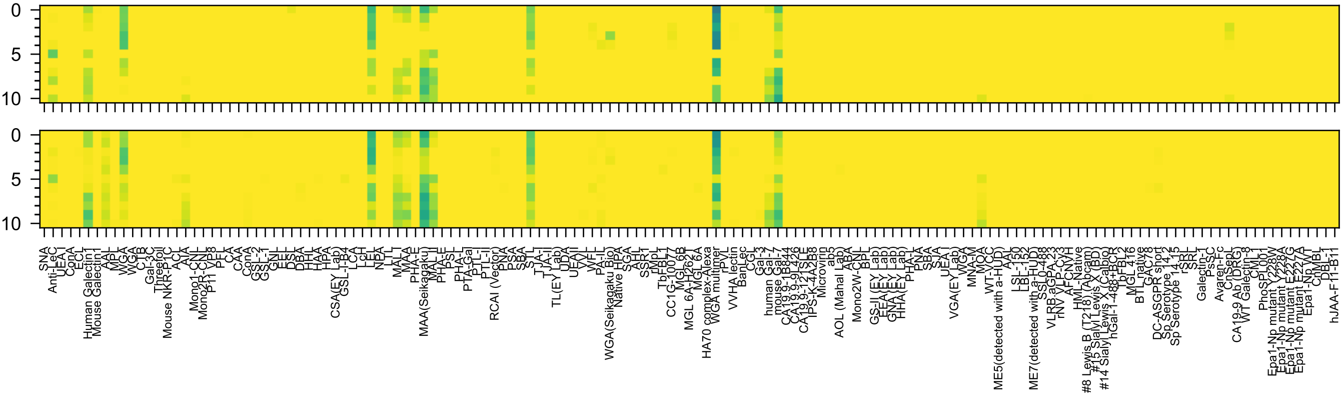

Gal3OS(β1-4)GlcNAc(β-Sp0

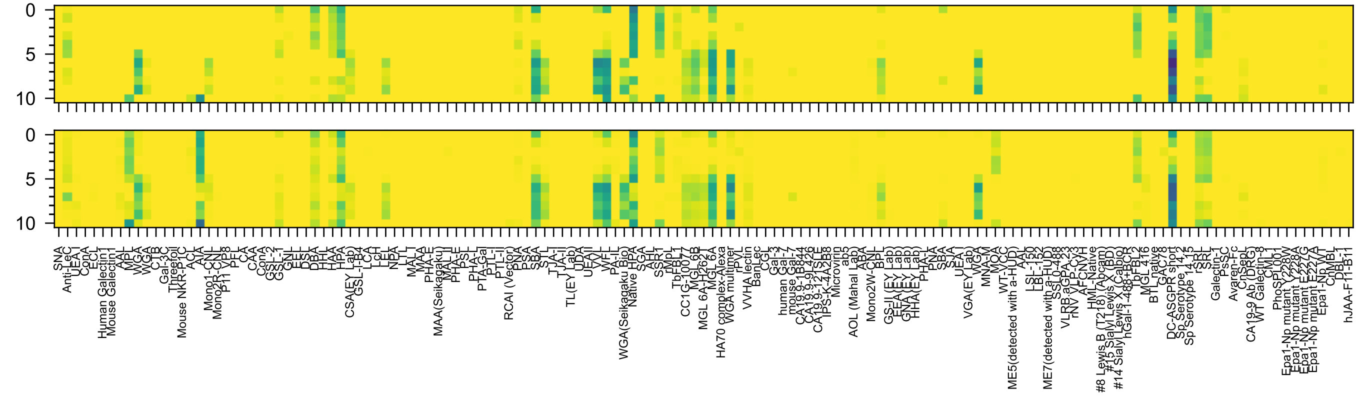

GalNAc(α1-3)GalNAc(β-Sp8

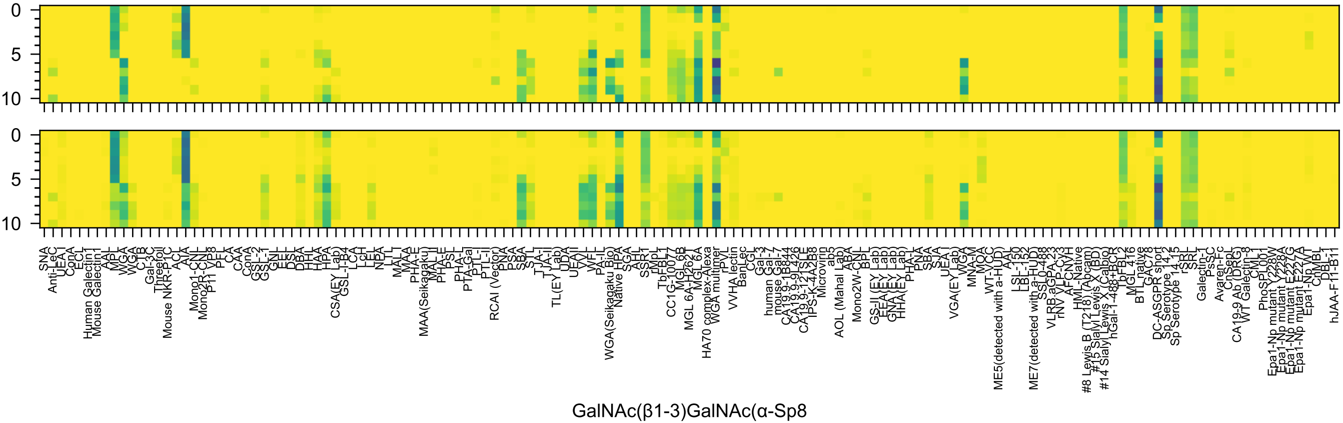
GalNAc(β1-3)GalNAc(α-Sp8

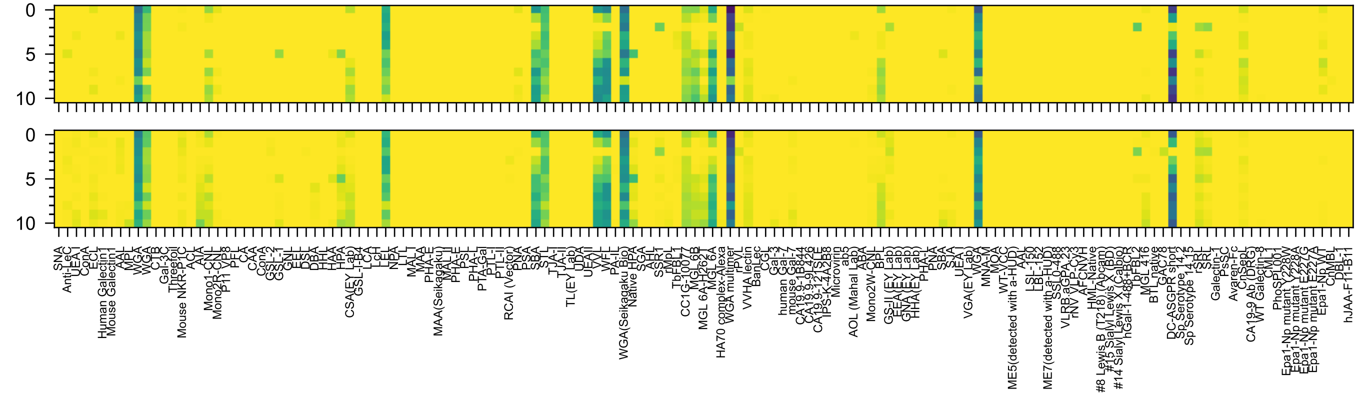

GalNAc(β1-4)GlcNAc(β-Sp8

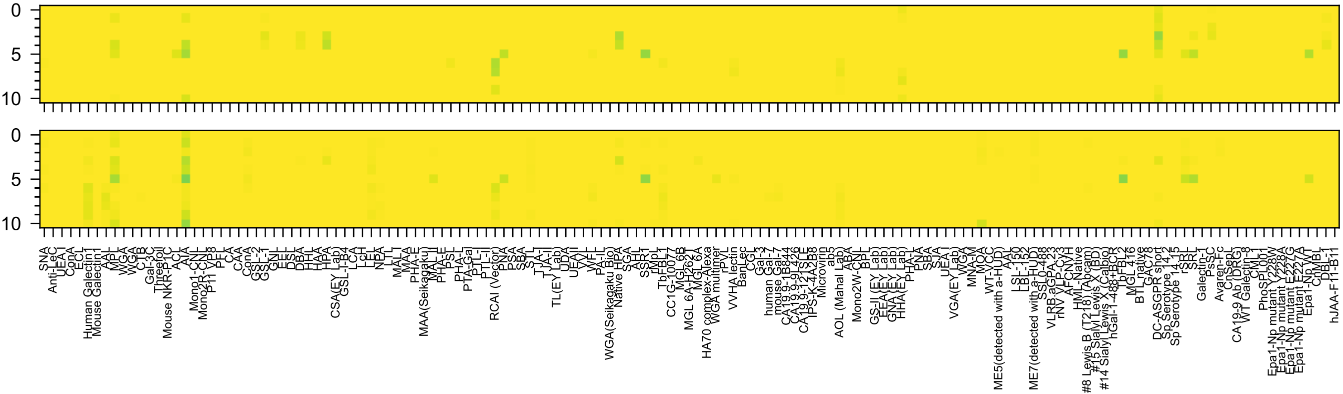

Gal(α1-3)GalNAc(α-Sp16

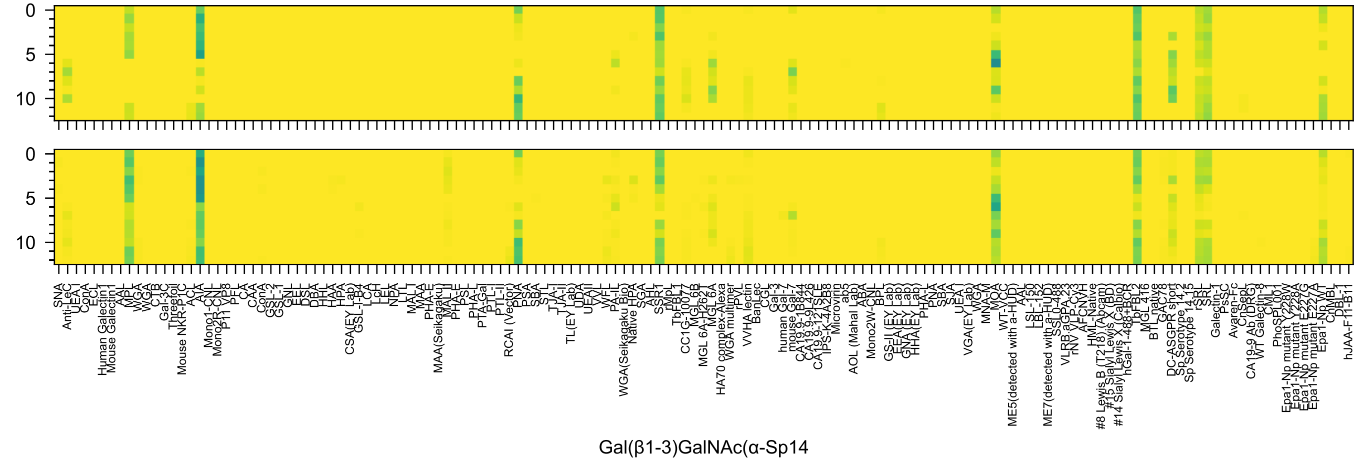
0
5
10
SNA
Anti-LeC
UEA I
ConA
ECL
Human Galectin1
Mouse Galectin1
AAL
MPL
WGA
WGA
CTB
Gal-3C
Threefoil
Mouse NKR-P1C
ACL
AIA
Mono1-CNL
Mono2R-CNL
P11 VP8
PFL
CA
CAA
ConA
GSL-2
GSL-1
GNL
EEL
DSL
DBA
HHL
HAA
HPA
CSA(EY Lab)
GSL-I-B4
LCA
LcH
LEL
NPA
LTL
MAL-I
MAA
PHA-E
MAA(Seikagaku)
MAL-II
PHA-E
PSL
PHA-L
PTA-Gal
PTL-I
PTL-II
RCAI (Vector)
PNA
PSA
SBA
STL
TJA-I
TJA-II
TL(EY Lab)
UDA
UEA-II
VVL
WFL
PA-IL
WGA(Seikagaku Bio)
Native HPA
SGA
AAL
SSR1
rMpL
TbFBL1
CC1G-10077
MGL6B
MGL 6A-H262T
MGL 6A
HA70 complex-Alexa
WGA multimer
rPVL
VVHA lectin
BanLec
CGL
Gal-3
human Gal-7
mouse Gal-7
CA19.9-1B844
CA19.9-9L426
CA19.9-121SLE
IPS-K-4A2B8
Microvirin
ab5
AOL (Mahal Lab)
ABA
Mono2W-CNL
BPL
GS-II (EY Lab)
EEA (EY Lab)
GNA (EY Lab)
HHA(EY Lab)
PHA-L
PNA
SBA
SJA
UEA-I
VGA(EY Lab)
WGA
MNA-M
MOA
WT-VCC
ME5(detected with a-HUD)
AAL
LSL-150
LBL-152
ME7(detected with a-HUD)
SSL0-488
VLRB.aGPA.23
rNV VLP-Cy3
AFCNYH
HML-Native
#8 Lewis B (T218) (Abcam)
#15 Sialyl Lewis X (BD)
#14 Sialyl Lewis X (Calbio)
hGal-1-488+BCR
TbFBL2
MGL 416
BTL native
GAC78
DC-ASGPR short
Sp Serotype 14.2
Sp Serotype 14.15
rSRL
SRL
Galectin-1
PsSC
Avaren-Fc
ChSepL
CA19-9 Ab (DRG)
WT Galectin-8
CML1
PhoSPL001
Epa1-Np mutant Y228W
Epa1-Np mutant Y228A
Epa1-Np mutant E227G
Epa1-Np mutant E227A
Epa1-Np WT
ChMBL
DBL-1
hJAA-F11-B11
Gal(β1-3)GalNAc(α-Sp14

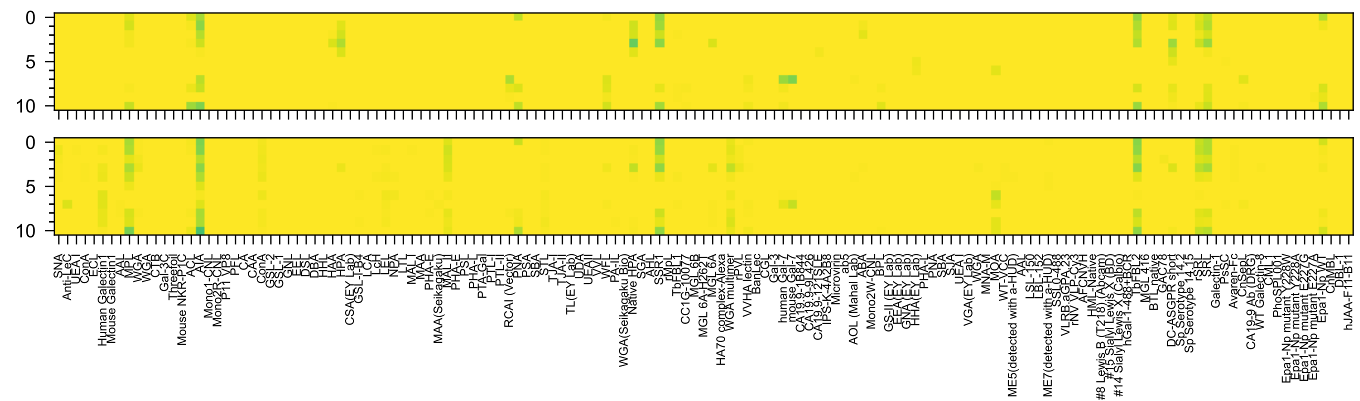

Gal(β1-3)GalNAc(α-Sp16

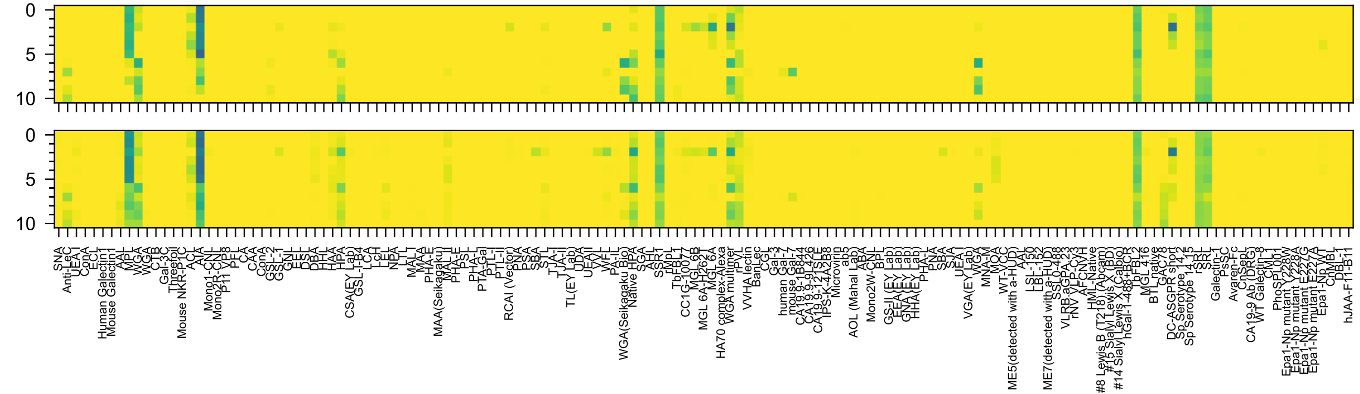


GlcNAc(β1-3)GalNAc(α-Sp8

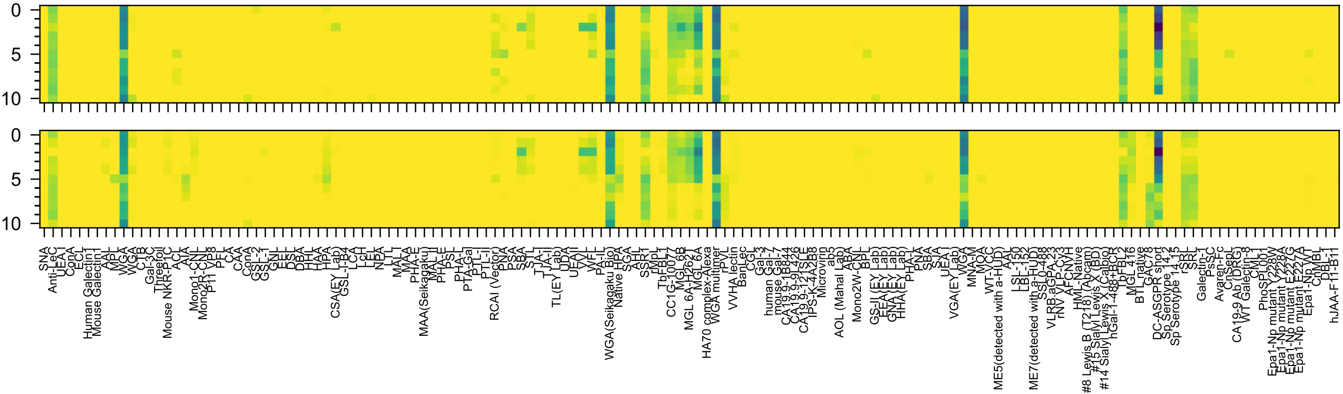

GlcNAc(β1-6)GalNAc(α-Sp8

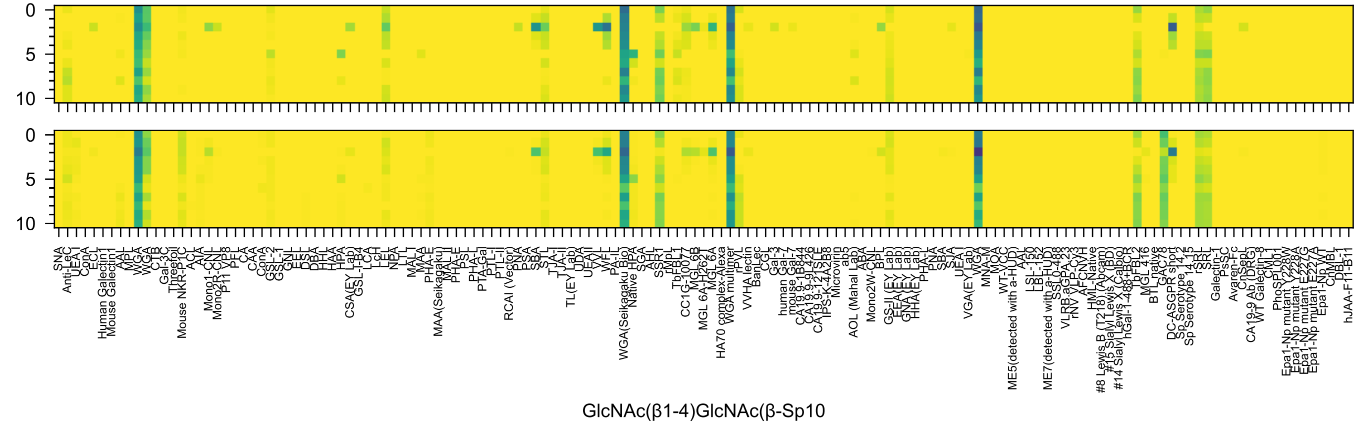
0
5
10
0
5
10
SNA
Anti-LeC
UEA-I
ConA
ECL
Human Galectin1
Mouse Galectin1
AAL
MPL
WGA
WGA
CTB
Gal-3C
Threefoil
Mouse NKR-P1C
ACL
AIA
Mono1-CNL
Mono2R-CNL
P11 VP8
PFL
CA
CAA
ConA
GSL-2
GSL-1
GNL
EEL
DSL
DBA
HHL
HAA
HPA
CSA(EY Lab)
GSL-I-B4
LCA
LcH
LEL
NPA
LTL
MAL-I
MAA
PHA-E
MAA(Seikagaku)
MAL-II
PHA-E
PSL
PHA-L
PTA-Gal
PTL-I
PTL-II
RCAI (Vector)
PNA
PSA
SBA
STL
TJA-I
TJA-II
TL(EY Lab)
UDA
UEAII
VVL
WFL
PA-IL
WGA(Seikagaku Bio)
Native HPA
SGA
AAI
SSR1
rMpL
TbFBL1
CC1G-10077
MGL-6B
MGL 6A-H262T
MGL 6A
HA70 complex-Alexa
WGA multimer
rPVL
VVHA lectin
BanLec
CGL
Gal-3
human Gal-7
mouse Gal-7
CA19.9-1B844
CA19.9-9L426
CA19.9-121SLE
IPS-K-4A2B8
Microvirin
ab5
AOL (Mahal Lab)
ABA
Mono2W-CNL
BPL
GS-II (EY Lab)
EEA (EY Lab)
GNA (EY Lab)
HHA(EY Lab)
PHA-L
PNA
SBA
SJA
UEA I
VGA(EY Lab)
WGA
MNA-M
MOA
WT-VCC
ME5(detected with a-HUD)
AAL
LSL-150
LBL-152
ME7(detected with a-HUD)
SSL0-488
VLRB.aGPA.23
rNV VLP-Cy3
AFCNVH
HML-Native
#8 Lewis B (T218) (Abcam)
#15 Sialyl Lewis X (BD)
#14 Sialyl Lewis X (Calbio)
hGal-1-488+BCR
TbFBL2
MGL 416
BTL native
GAC78
DC-ASGPR short
Sp Serotype 14.2
Sp Serotype 14.15
rSRL
SRL
Galectin-1
PsSC
Avaren-Fc
ChSepL
CA19-9 Ab (DRG)
WT Galectin-8
CML1
PhoSPL001
Epa1-Np mutant Y228A
Epa1-Np mutant E227G
Epa1-Np mutant E227A
Epa1-Np WT
ChMBL
DBL-1
hJAA-F11-B11
GlcNAc(β1-4)GlcNAc(β-Sp10)

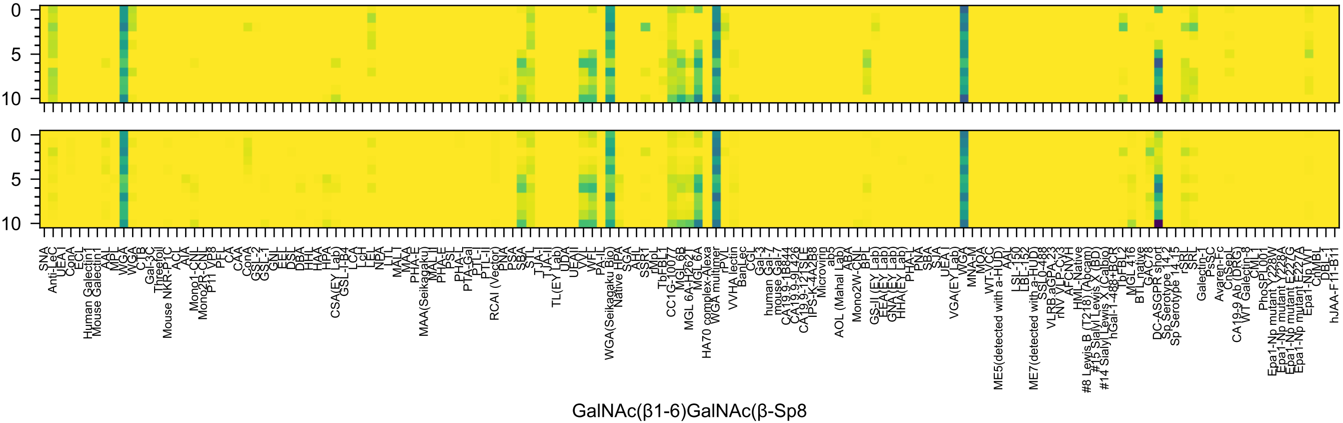
GalNAc(β1-6)GalNAc(β-Sp8

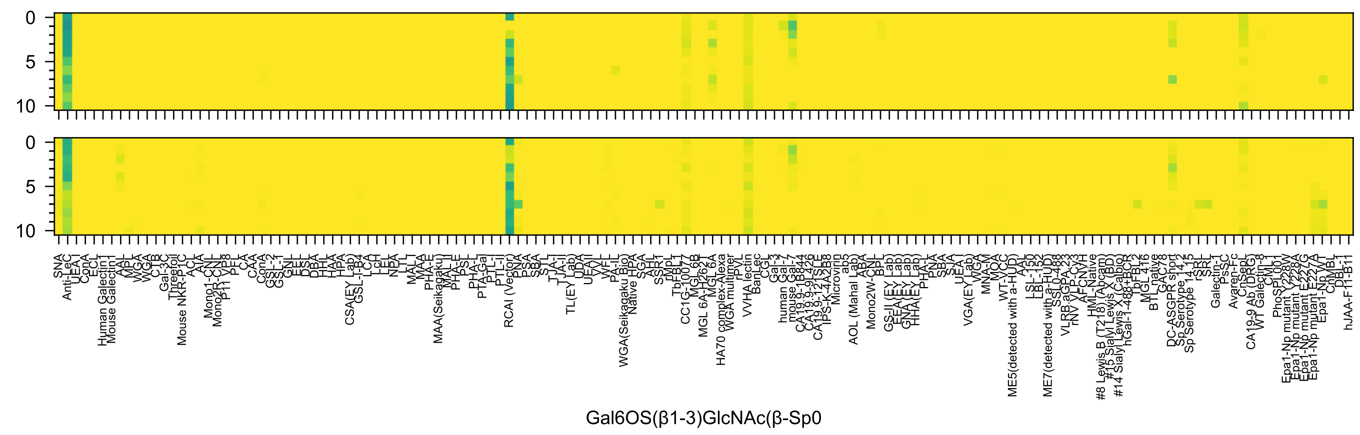
Gal6OS(β1-3)GlcNAc(β-Sp0

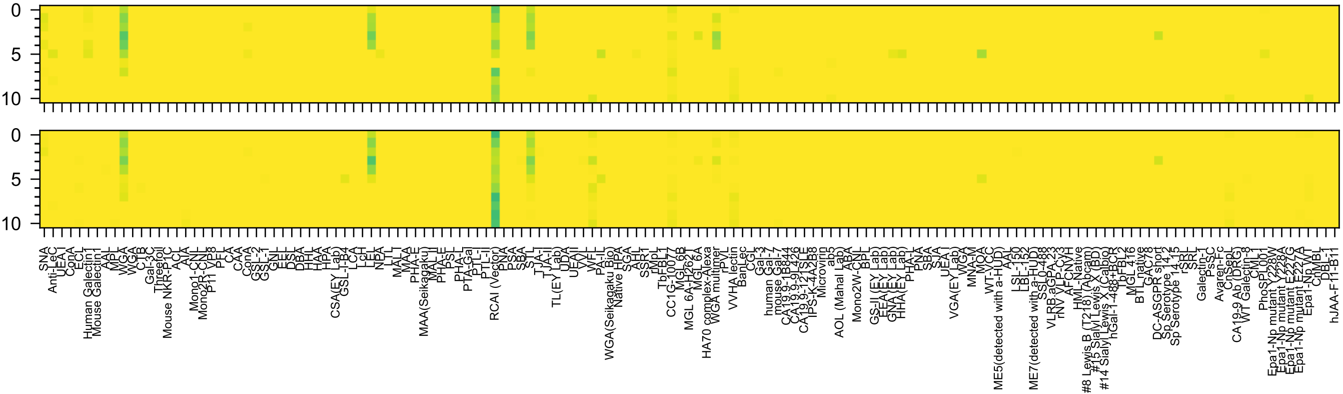

Gal(β1-4)GlcNAc6OP(β-Sp0

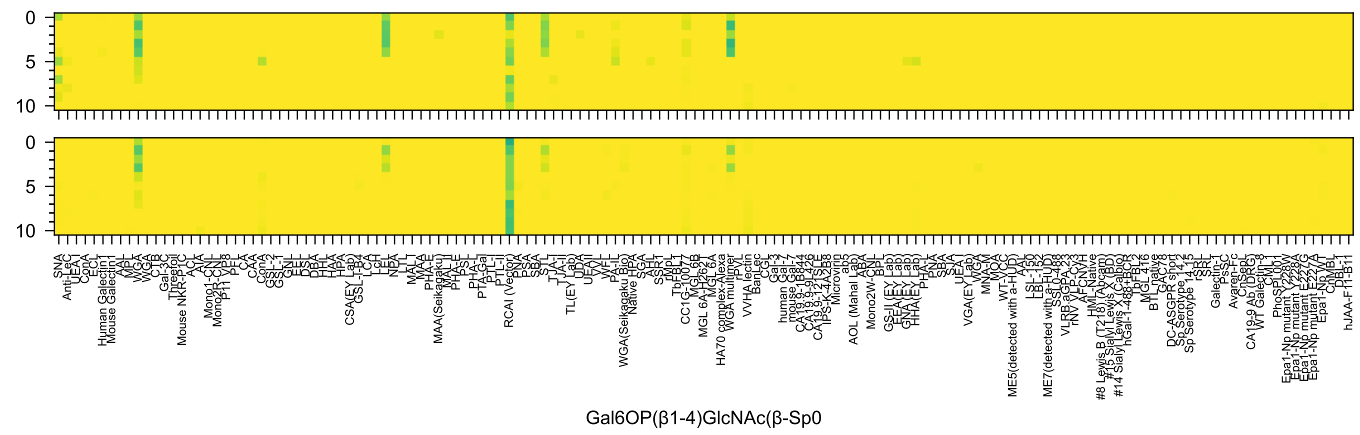
0
5
10
0
5
10
Gal6OP(β1-4)GlcNAc(β-Sp0

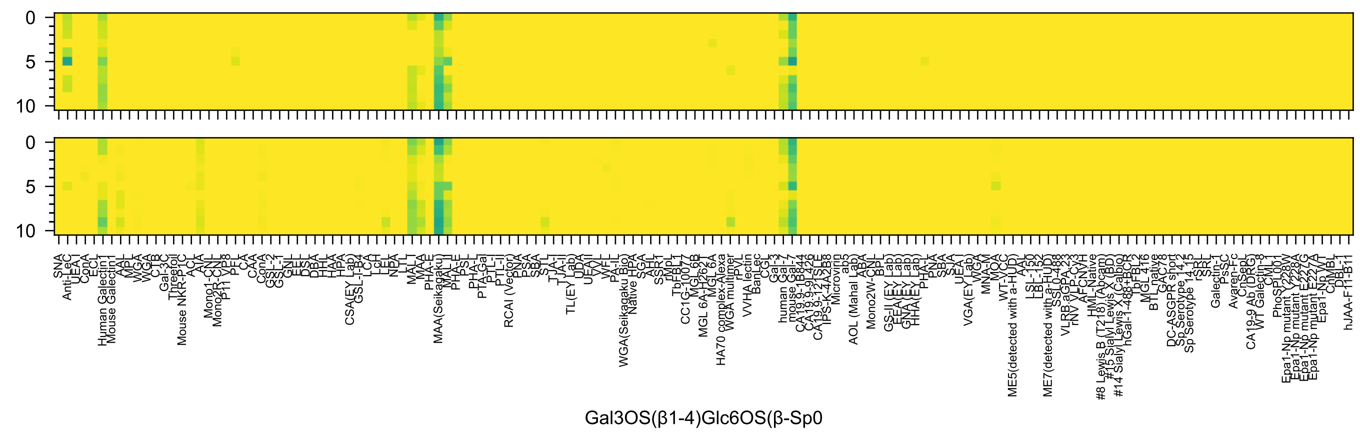
Gal3OS(β1-4)Glc6OS(β-Sp0

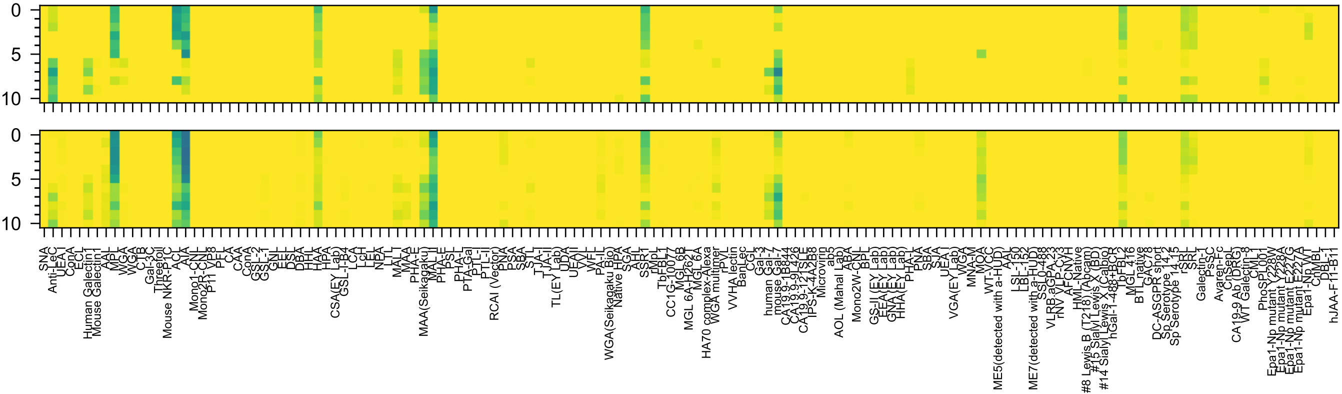


Gal3OS(β1-3)GalNAc(α-Sp8

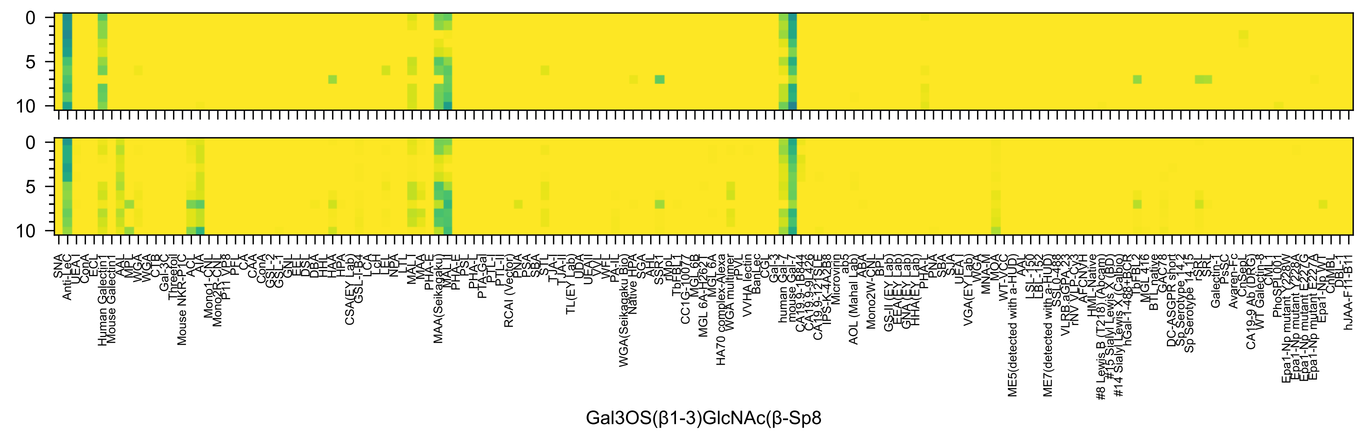
Gal3OS(β1-3)GlcNAc(β-Sp8

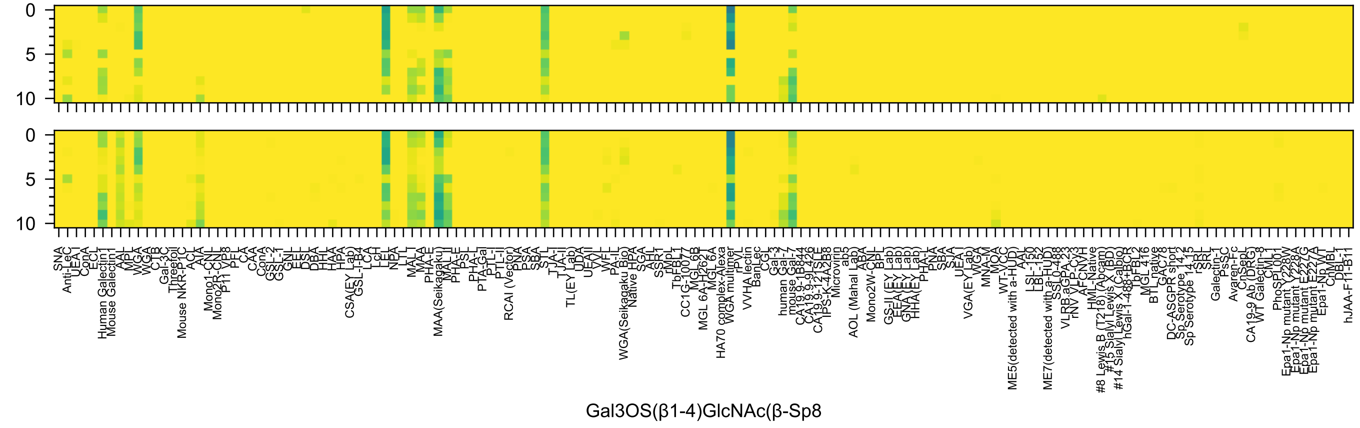
Gal3OS(β1-4)GlcNAc(β-Sp8

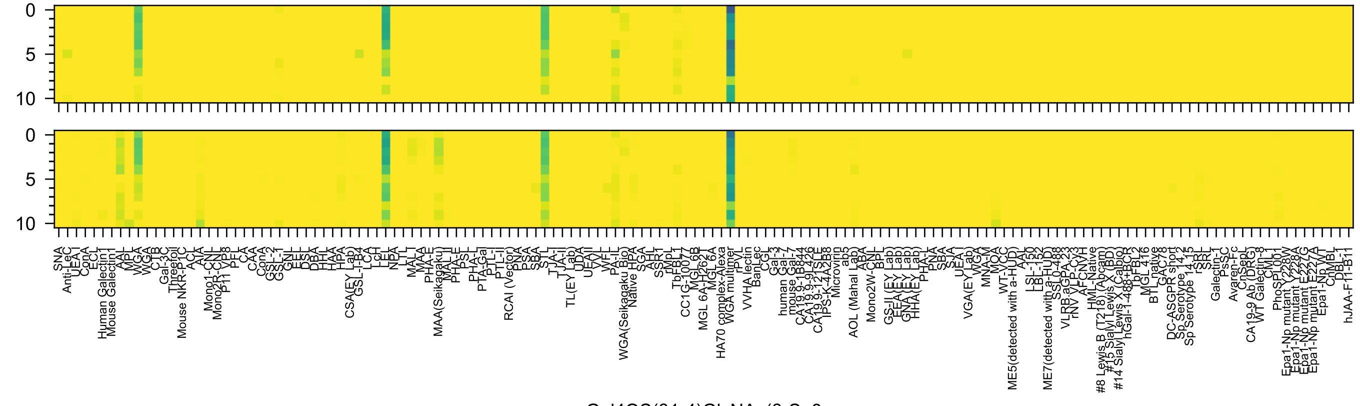
Gal4OS(β1-4)GlcNAc(β-Sp8

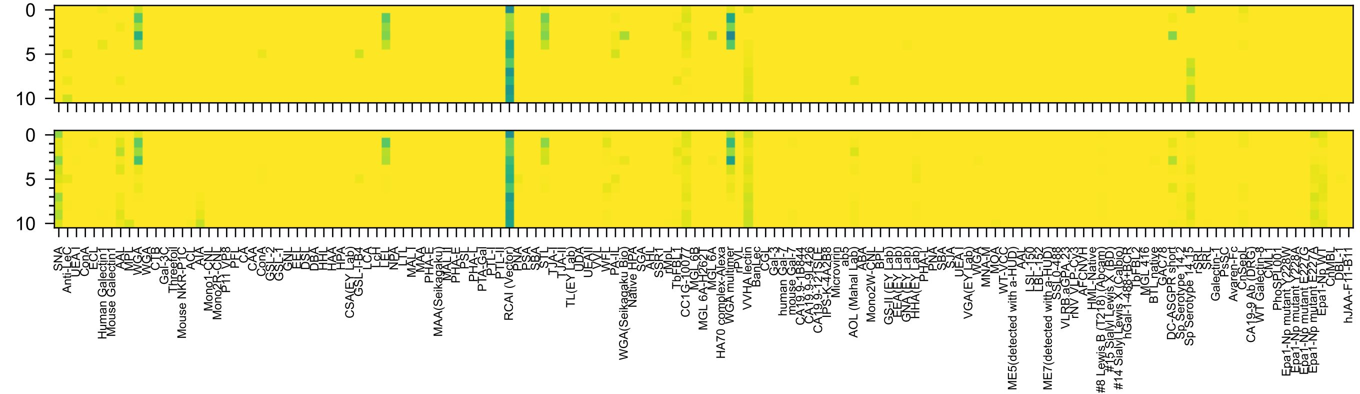


Gal6OS(β1-4)GlcNAc(β-Sp8

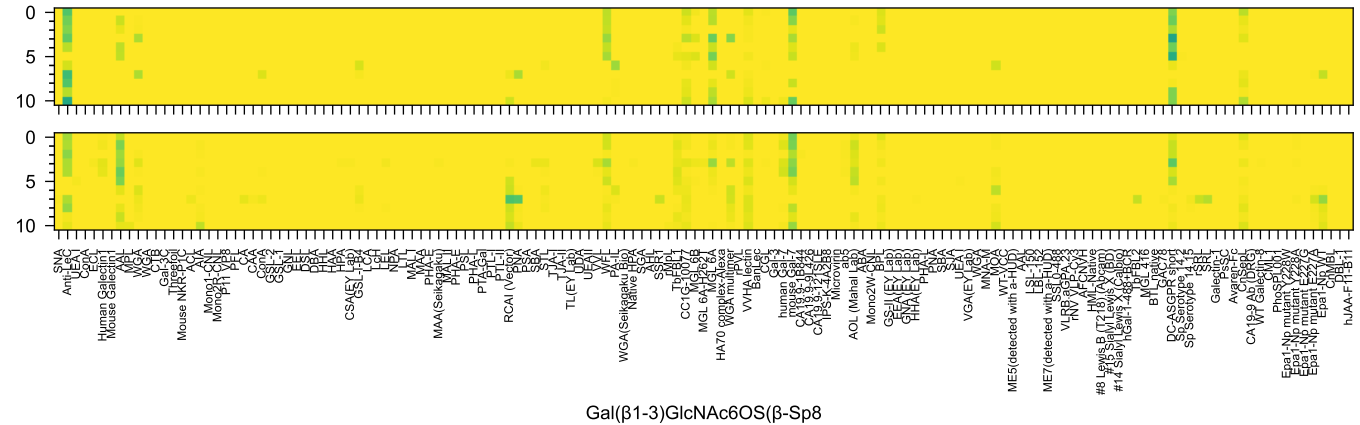
0
5
10
0
5
10
SNA
Anti-LeC
UEA I
ConA
ECL
Human Galectin1
Mouse Galectin1
AAL
MPL
WGA
WGA
CTB
Gal-3C
Threefoil
Mouse NKR-P1C
ACL
AIA
Mono1-CNL
Mono2R-CNL
P11 VP8
PFL
CA
CAA
ConA
GSL-2
GSL-1
GNL
EEL
DSL
DBA
HHL
HAA
HPA
CSA(EY Lab)
GSL-I-B4
LCA
LcH
LEL
NPA
LTL
MAL I
MAA
PHA-E
MAA(Seikagaku)
MAL II
PHA-E
PSL
PHA-L
PTA-Gal
PTL-I
PTL-II
RCAI (Vector)
PNA
PSA
SBA
STL
TJA-I
TJA-II
TL(EY Lab)
UDA
UEAII
VVL
WFL
WGA(Seikagaku Bio)
Native HPA
SGA
AHL
SSR1
rMpL
TbFBL1
CC1G-10077
MGL6B
MGL 6A-H262T
MGL 6A
HA70 complex-Alexa
WGA multimer
rPVL
VVHA lectin
BanLec
CGL
Gal-3
human Gal-7
mouse Gal-7
CA19.9-1B844
CA19.9-9L426
CA19.9-121SLE
IPS-K-4A2B8
Microvirin
ab5
AOL (Mahal Lab)
ABA
Mono2W-CNL
BPL
GS-II (EY Lab)
EEA(EY Lab)
GNA(EY Lab)
HHA(EY Lab)
PHA-L
PNA
SBA
SJA
UEA I
VGA(EY Lab)
WGA
MNA-M
MOA
WT-VCC
ME5(detected with a-HUD)
AAL
LSL-150
LBL-152
ME7(detected with a-HUD)
SSL0-488
VLRB.aGPA.23
rNV VLP-Cy3
AFCNYH
HML-Native
#8 Lewis B (T218) (Abcam)
#15 Sialyl Lewis X (BD)
#14 Sialyl Lewis X (Calbio)
hGal-1-488+BCR
TbFBL2
MGL 416
BTL native
GAC78
DC-ASGPR short
Sp Serotype 14.2
Sp Serotype 14.15
rSRL
SRL
Galectin-1
PsSC
Avaren-Fc
ChSepL
CA19-9 Ab (DRG)
WT Galectin-8
CML1
PhoSPL001
Epa1-Np mutant Y228W
Epa1-Np mutant E228A
Epa1-Np mutant E227G
Epa1-Np mutant E227A
Epa1-Np WT
ChMBL
DBL-1
hJAA-F11-B11
Gal(β1-3)GlcNAc6OS(β-Sp8

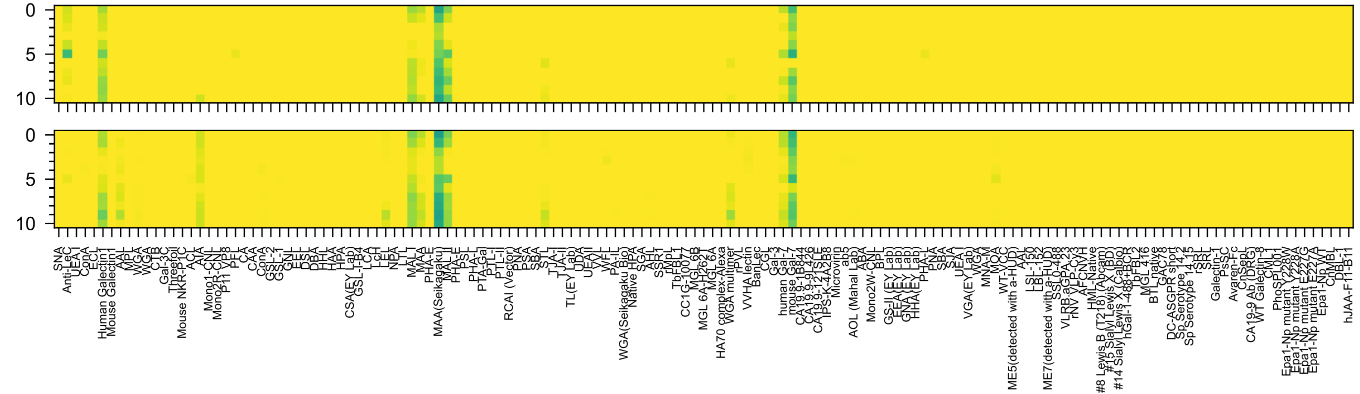


Gal3OS(β1-4)Glc6OS(β-Sp8

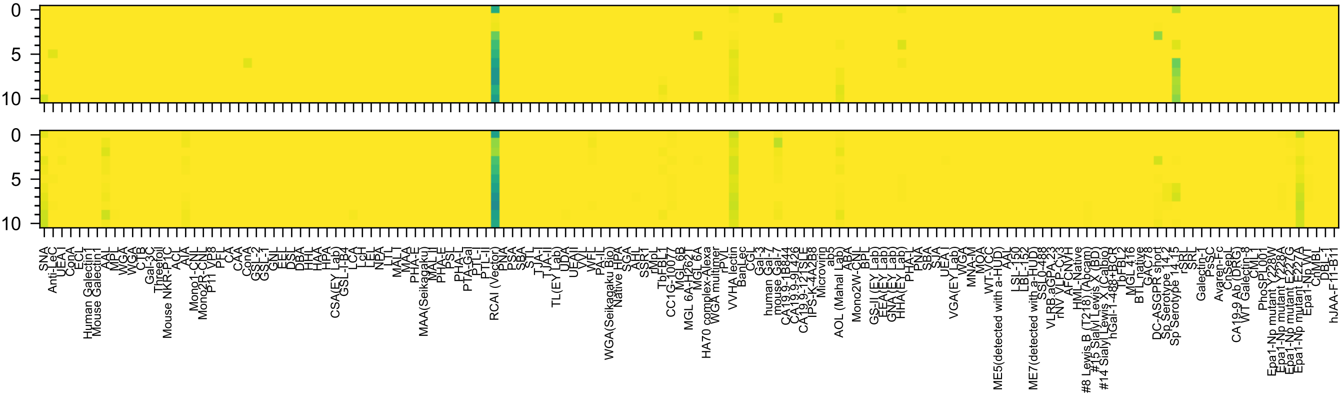

Gal6OS(β1-4)Glc6OS(β-Sp8

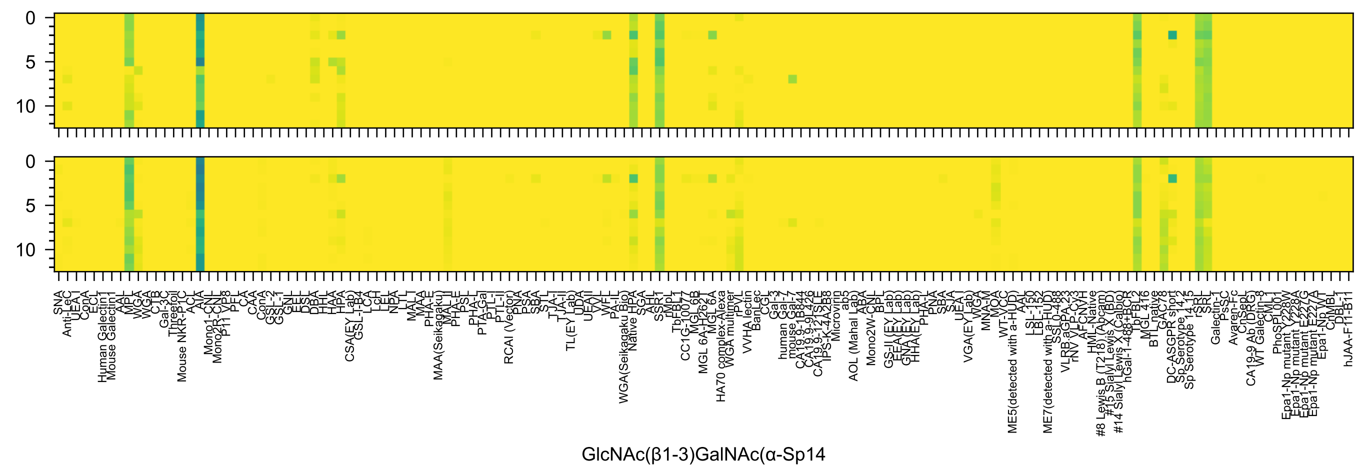
0
5
10
0
5
10
SNA
Anti-LeC
UEA-I
ConA
ECL
Human Galectin1
Mouse Galectin1
AAL
MPL
WGA
WGA
CTB
Gal-3C
Threefoil
Mouse NKR-P1C
ACL
AIA
Mono1-CNL
Mono2R-CNL
P11 VP8
PFL
CA
CAA
ConA
GSL-2
GSL-1
GNL
EEL
DSL
DBA
HHL
HAA
HPA
CSA(EY Lab)
GSL-I-B4
LCA
LcH
LEL
NPA
LTL
MAL-I
MAA
PHA-E
MAA(Seikagaku)
MAL-II
PHA-E
PSL
PHA-L
PTA-Gal
PTL-I
PTL-II
RCAI (Vector)
PNA
PSA
SBA
STL
TJA-I
TJA-II
TL(EY Lab)
UDA
UEAII
VVL
WFL
PA-IL
WGA(Seikagaku Bio)
Native HPA
SGA
AAL
SSR1
rMpL
TbFBL1
CC1G-10077
MGL6B
MGL 6A-H262T
MGL 6A
HA70 complex-Alexa
WGA multimer
rPVL
VVHA lectin
BanLec
CGL
Gal-3
human Gal-7
mouse Gal-7
CA19.9-1B844
CA19.9-9L426
CA19.9-121SLE
IPS-K-4A2B8
Microvirin
ab5
AOL (Mahal Lab)
ABA
Mono2W-CNL
BPL
GS-II (EY Lab)
EEA (EY Lab)
GNA (EY Lab)
HHA(EY Lab)
PHA-L
PNA
SBA
SJA
UEA-I
VGA(EY Lab)
WGA
MNA-M
MOA
WT-VCC
ME5(detected with a-HUD)
AAL
LSL-150
LBL-152
ME7(detected with a-HUD)
SSL0-488
VLRB.aGPA.23
rNV VLP-Cy3
AFCNVH
HML-Native
#8 Lewis B (T218) (Abcam)
#15 Sialyl Lewis X (BD)
#14 Sialyl Lewis X (Calbio)
hGal-1-488+BCR
TbFBL2
MGL 416
BTL native
GAC78
DC-ASGPR short
Sp Serotype 14-2
Sp Serotype 14-15
rSRL
SRL
Galectin-1
PsSC
Avaren-Fc
CnSepL
CA19-9 Ab (DRG)
WT Galectin-8
CML1
PhoSL001
Epa1-Np mutant Y228W
Epa1-Np mutant Y228A
Epa1-Np mutant E227G
Epa1-Np mutant E227A
Epa1-Np WT
ChMBL
DBL-1
hJAA-F11-B11
GlcNAc(β1-3)GalNAc(α-Sp14

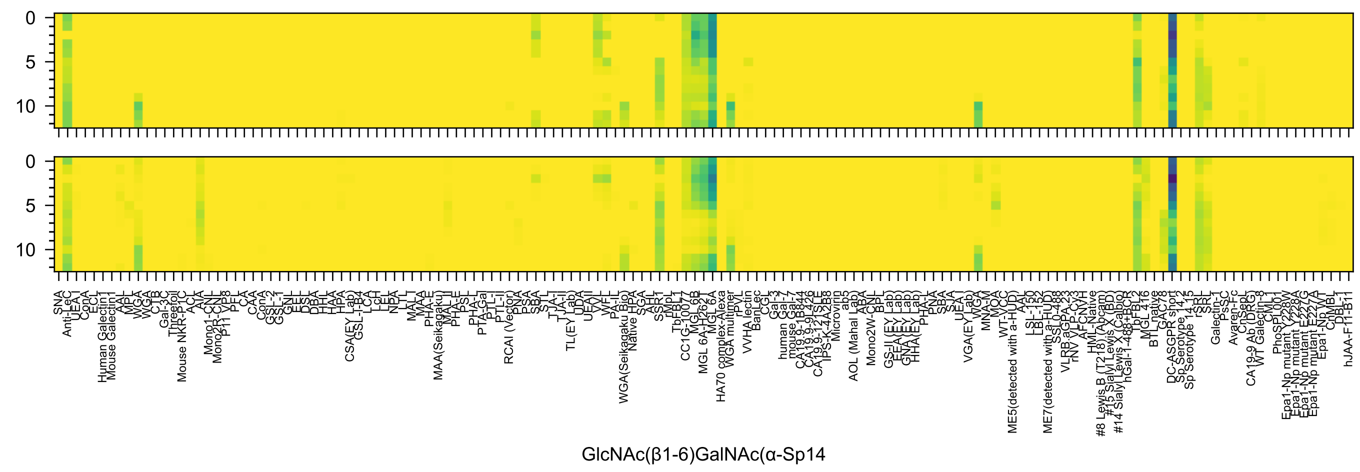
0
5
10
0
5
10
SNA
Anti-LeC
UEA-I
ConA
ECL
Human Galectin1
Mouse Galectin1
AAL
MPL
WGA
WGA
CTB
Gal-3C
Threefoil
Mouse NKR-P1C
ACL
AIA
Mono1-CNL
Mono2R-CNL
P11 VP8
PFL
CA
CAA
ConA
GSL-2
GSL-1
GNL
EEL
DSL
DBA
HHL
HAA
HPA
HPA
CSA(EY Lab)
GSL-I-B4
LCA
LcH
LEL
NPA
LTL
MAL-I
MAA
PHA-E
MAA(Seikagaku)
MAL-II
PHA-E
PSL
PHA-L
PTA-Gal
PTL-I
PTL-II
RCAI (Vector)
PNA
PSA
SBA
STL
TJA-I
TJA-II
TL(EY Lab)
UDA
UEA-II
VVL
WFL
PA-IL
WGA(Seikagaku Bio)
Native HPA
SGA
AHL
SSR1
rMpL
TbFBL1
CC1G-10077
MGL6B
MGL 6A-H262T
MGL 6A
HA70 complex-Alexa
WGA multimer
rPVL
VVHA lectin
BanLec
CGL
Gal-3
human Gal-7
mouse Gal-7
CA19.9-1B844
CA19.9-9L426
CA19.9-121SLE
IPS-K-4A2B8
Microvirin
ab5
AOL (Mahal Lab)
ABA
Mono2W-CNL
BPL
GS-II (EY Lab)
EEA (EY Lab)
GNA (EY Lab)
HHA (EY Lab)
PHA-L
PNA
SBA
SJA
UEA-I
VGA(EY Lab)
WGA
MNA-M
MOA
WT-VCC
ME5(detected with a-HUD)
AAL
LSL-150
LBL-152
ME7(detected with a-HUD)
SSL0-488
VLRB.aGPA.23
rNV VLP-Cy3
AFCNYH
HML-Native
#8 Lewis B (T218) (Abcam)
#15 Sialyl Lewis X (BD)
#14 Sialyl Lewis X (Calbio)
hGal-1-488+BCR
TbFBL2
MGL 416
BTL native
GAC78
DC-ASGPR short
Sp Serotype 14.2
Sp Serotype 14.15
rSRL
SRL
Galectin-1
PsSC
Avaren-Fc
CnSepL
CA19-9 Ab (DRG)
WT Galectin-8
CML1
PhoSL001
Epa1-Np mutant Y228A
Epa1-Np mutant E227G
Epa1-Np mutant E227A
Epa1-Np WT
ChMBL
DBL-1
hJAA-F11-B11
GlcNAc(β1-6)GalNAc(α-Sp14

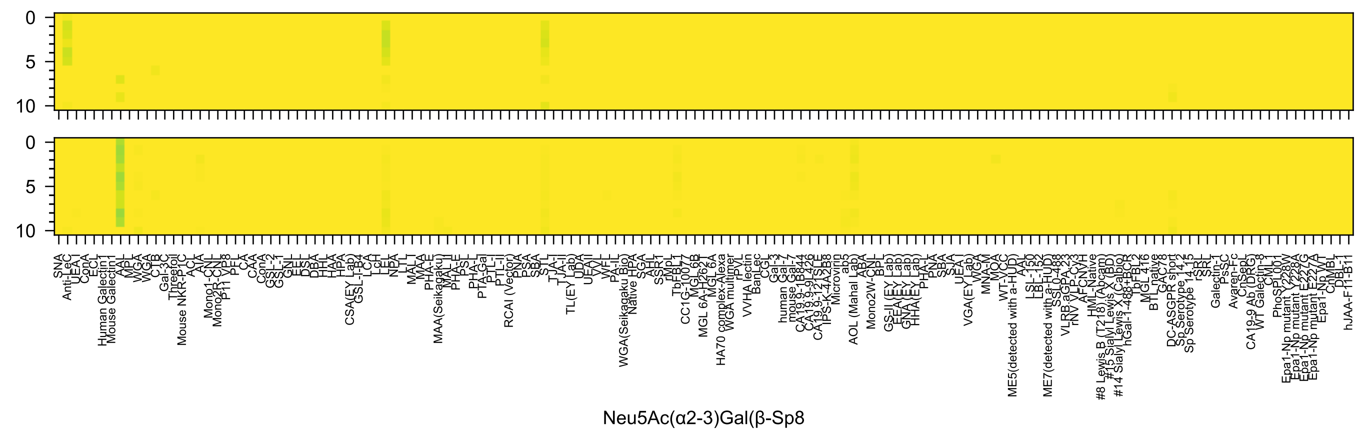
Neu5Ac(α2-3)Gal(β-Sp8

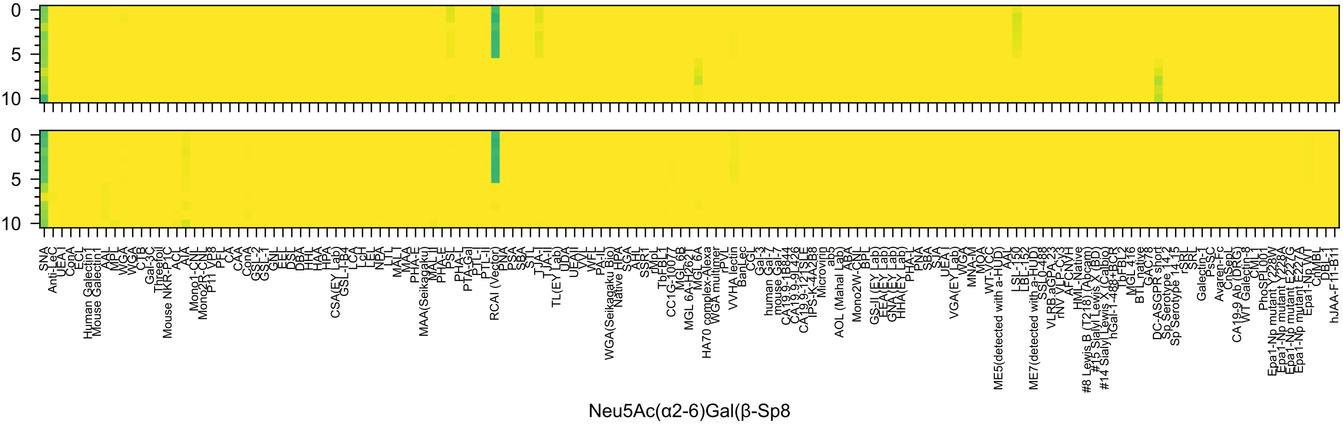
Neu5Ac(α2-6)Gal(β-Sp8

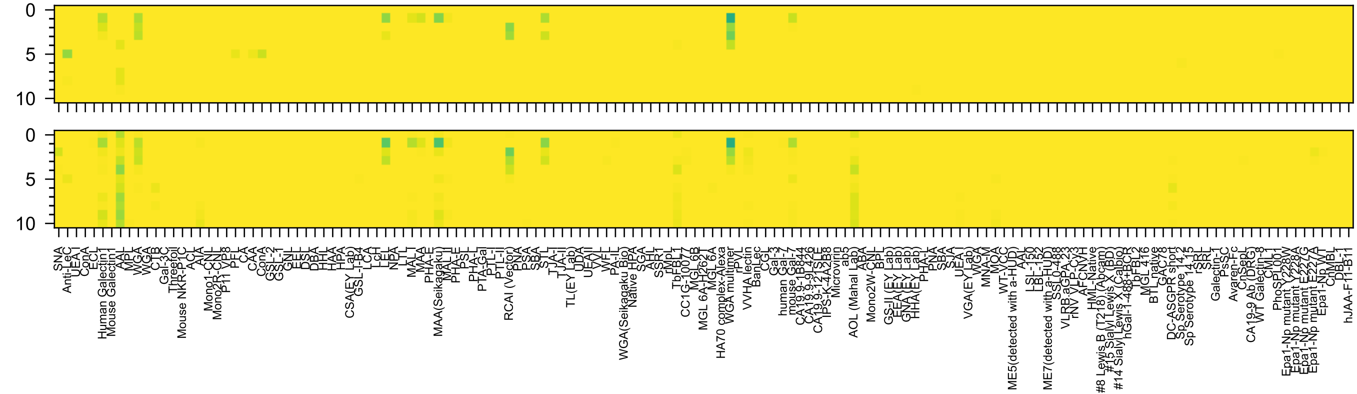

Gal3,6OS2(β1-4)GlcNAc(β-Sp0

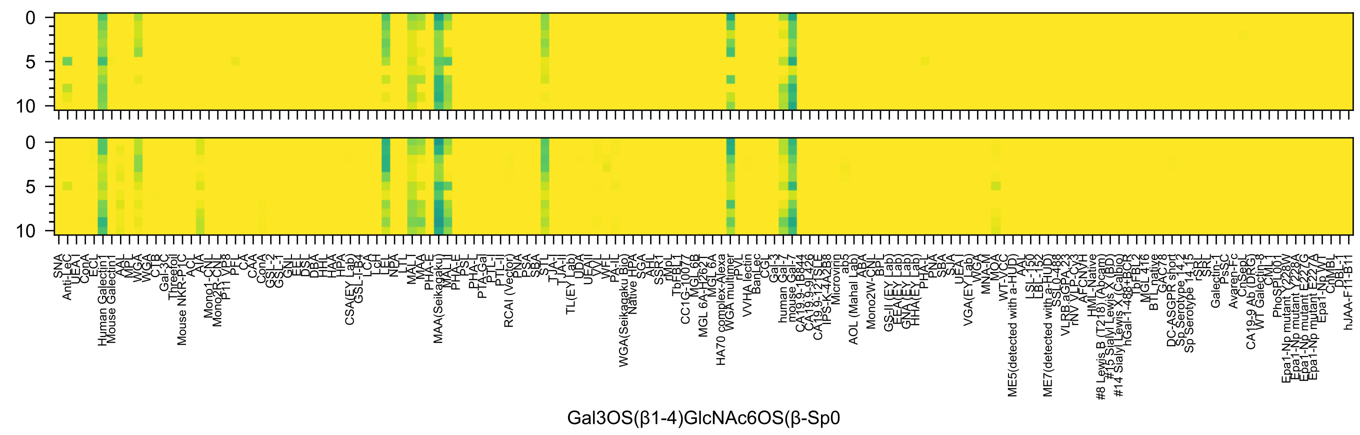
Gal3OS(β1-4)GlcNAc6OS(β-Sp0

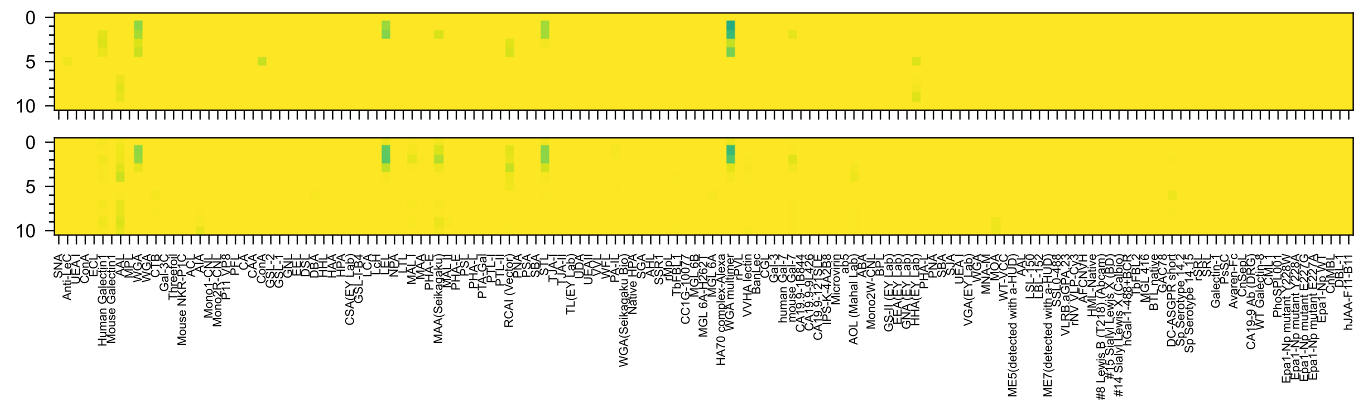


Gal4,6OS2(β1-4)GlcNAc(β-Sp0

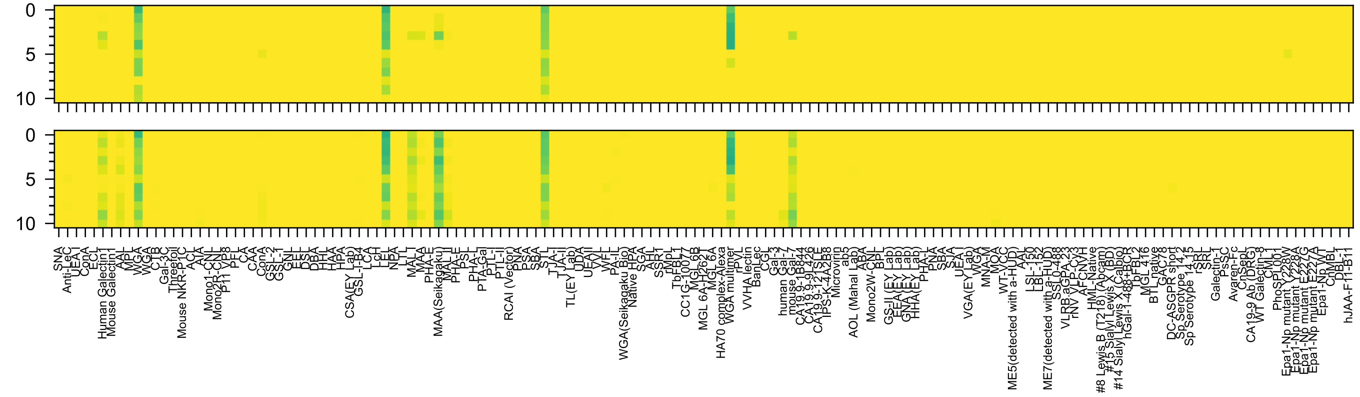


Gal3,4OS2(β1-4)GlcNAc(β-Sp0

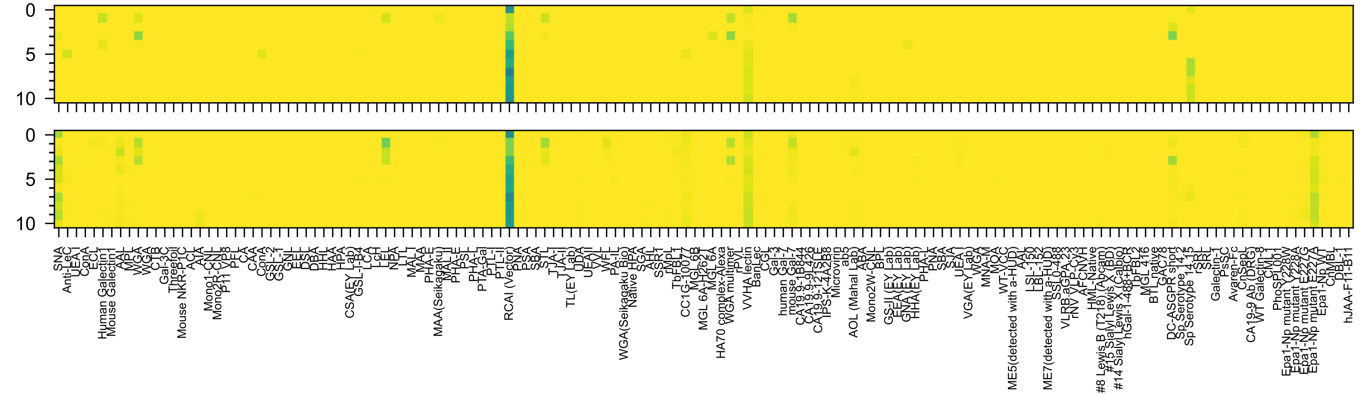

Gal6OS(β1-4)GlcNAc6OS(β-Sp0

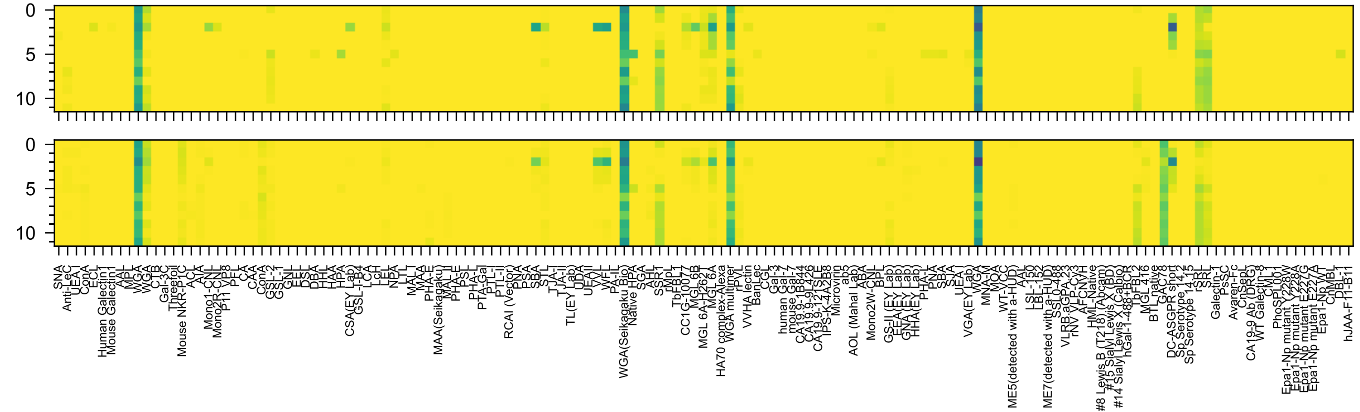

GlcNAc(β1-4)GlcNAc(β-Sp12)

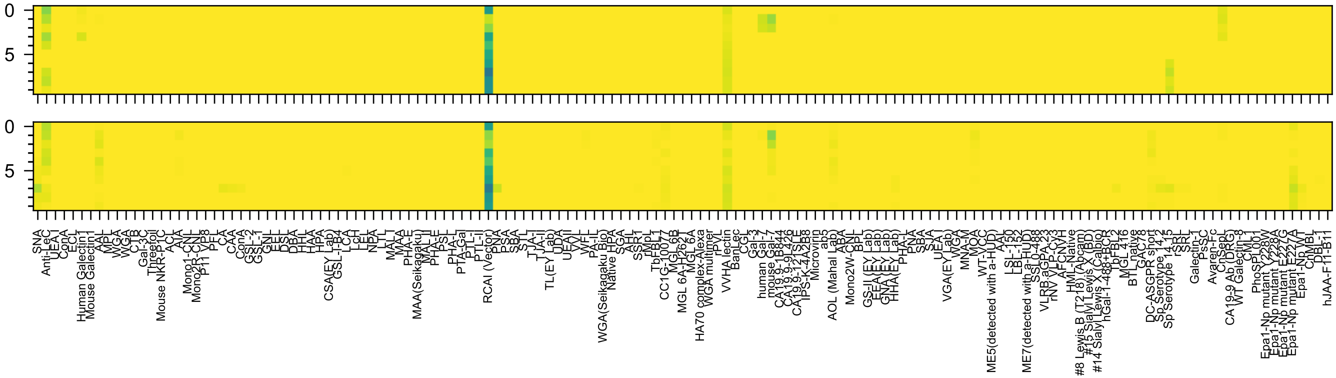

Gal6OS(β1-3)GlcNAc6OS(-Sp0

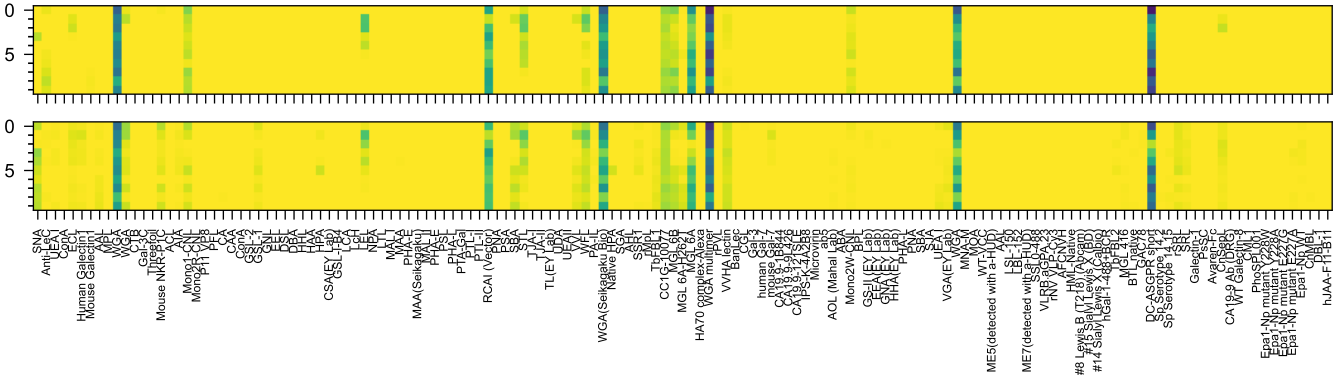
GalNAc6OS(β1-4)GlcNAc(-Sp8

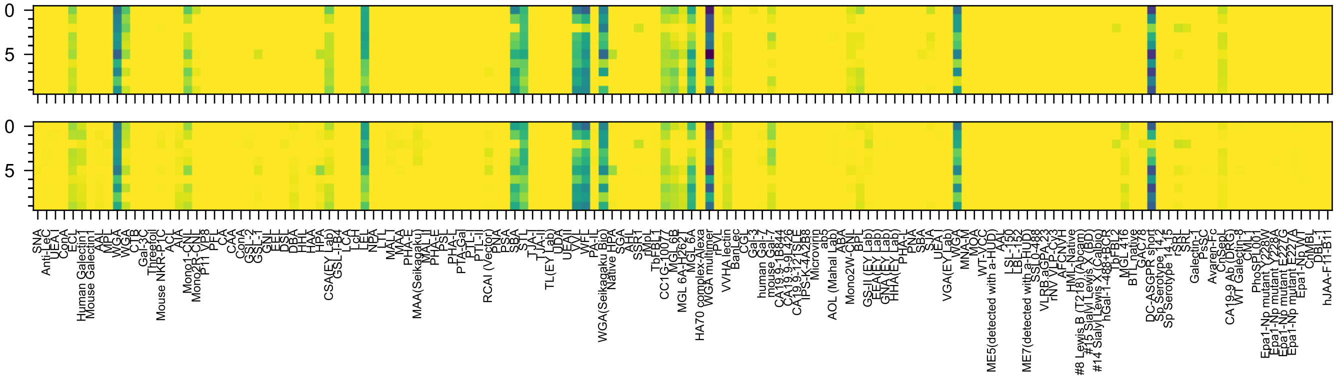

GalNAc(β1-4)GlcNAc6OS(-Sp8

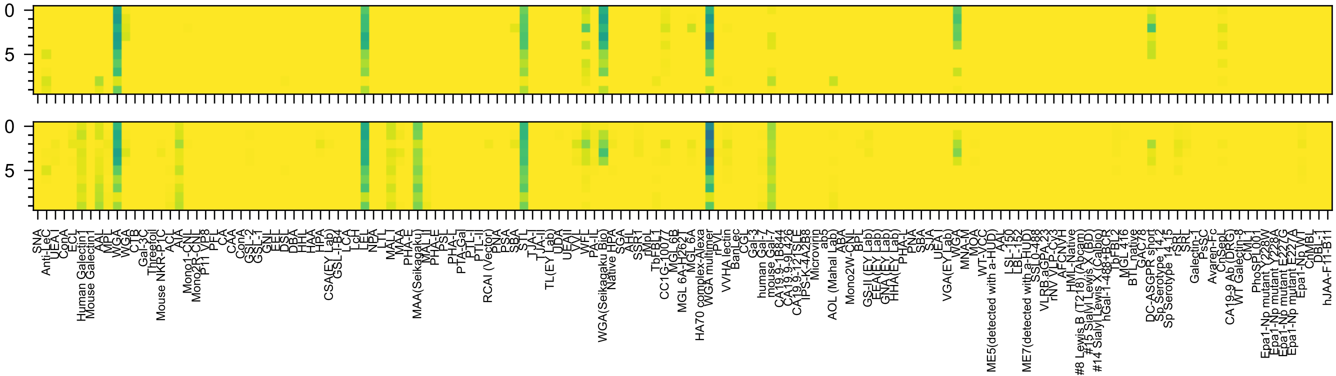


GalNAc3OS(β1-4)GlcNAc(-Sp8

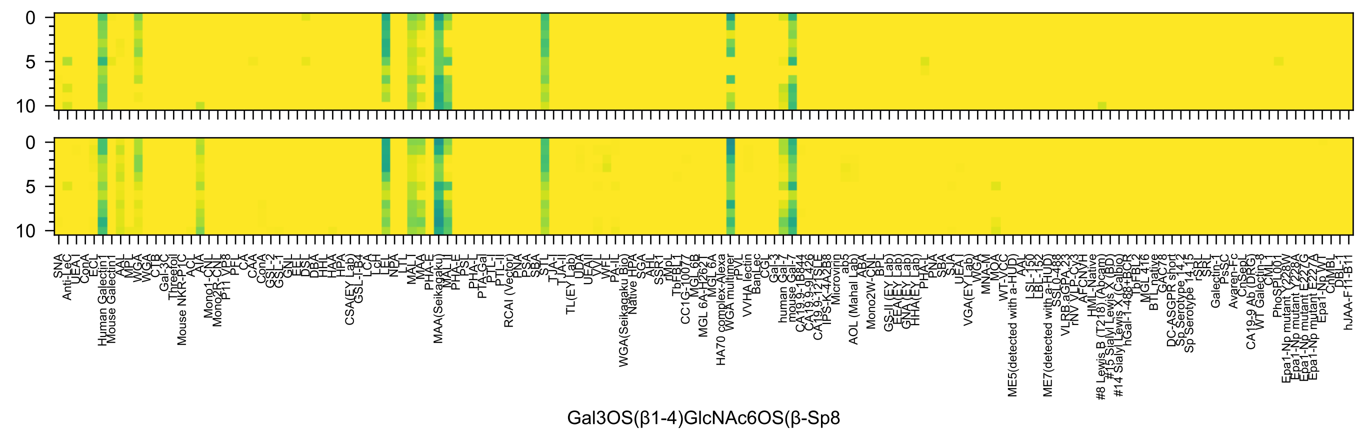
Gal3OS(β1-4)GlcNAc6OS(β-Sp8

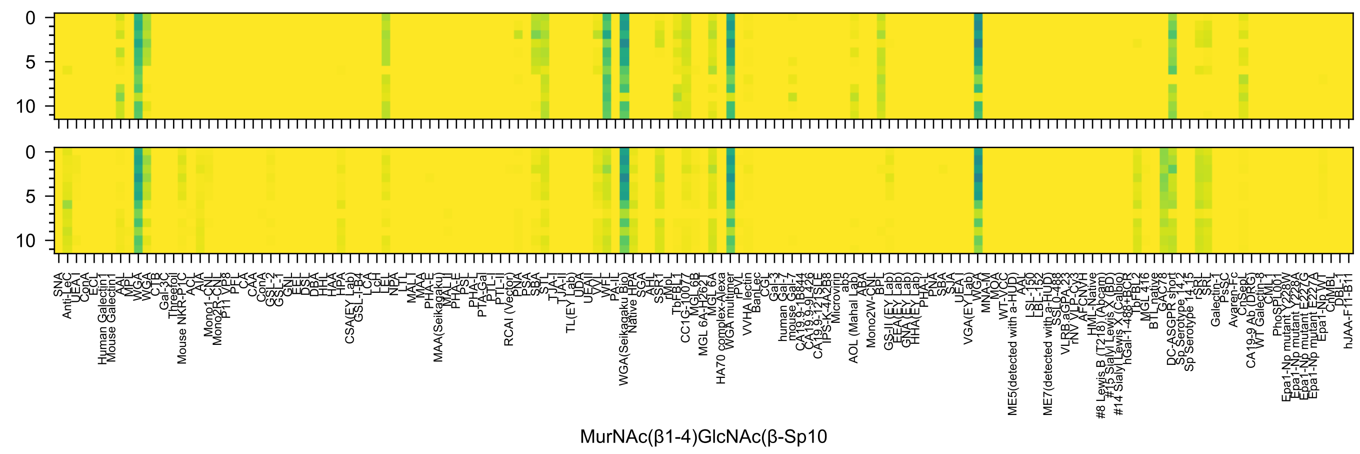
0
5
10
MurNAc(β1-4)GlcNAc(β-Sp10

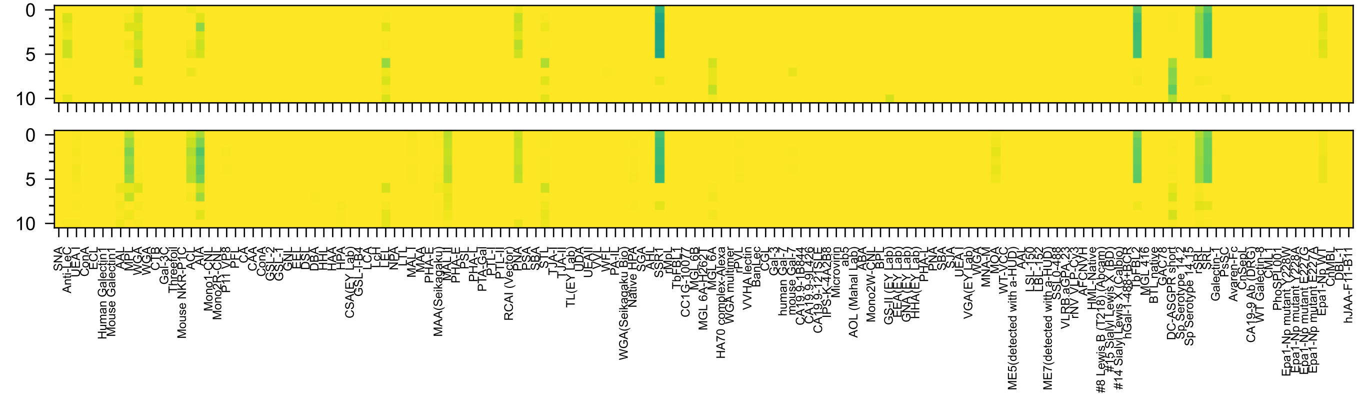

Neu5Ac(α2-3)GalNAc(α-Sp8

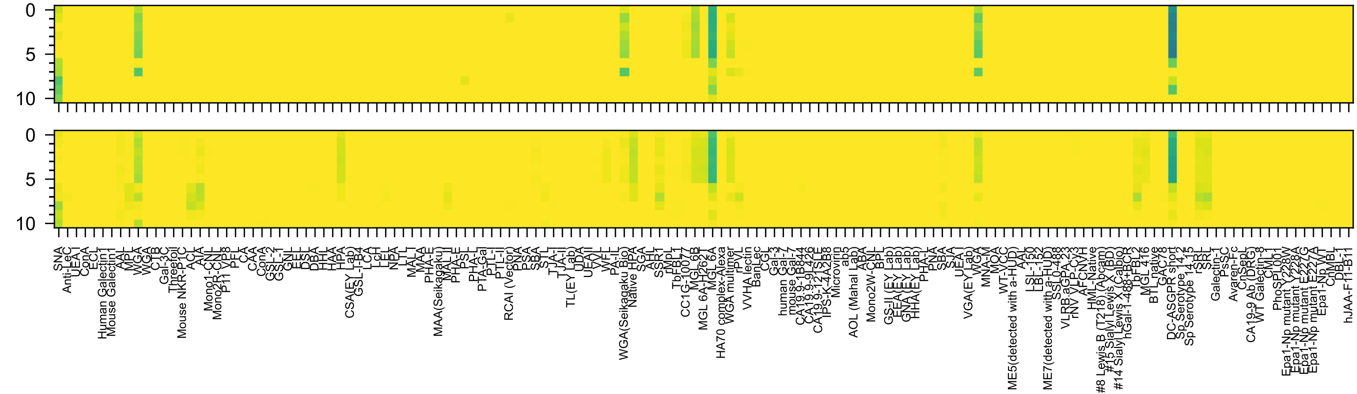

Neu5Ac(α2-6)GalNAc(α-Sp8

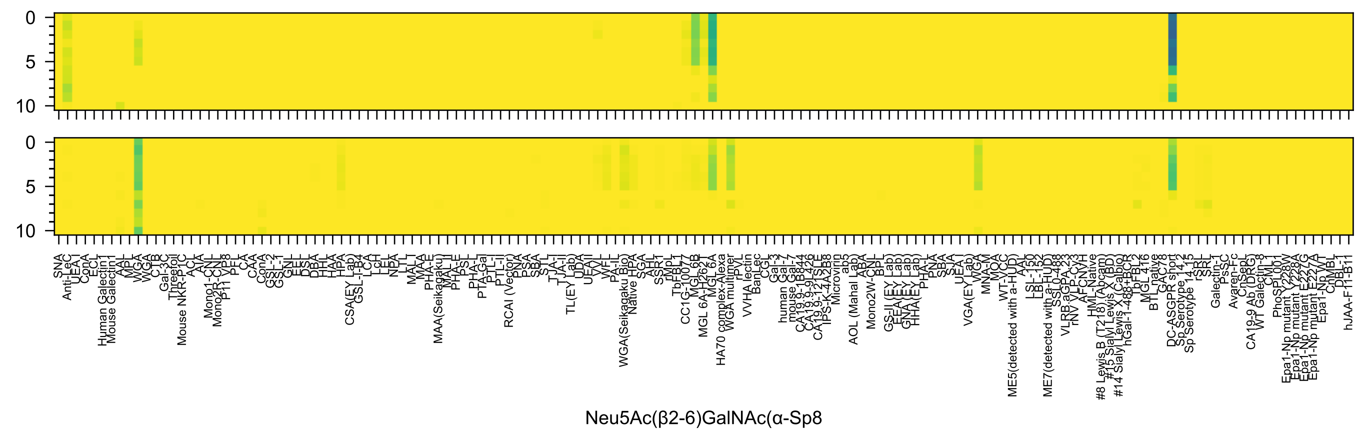
0
5
10
0
5
10
Neu5Ac(β2-6)GalNAc(α-Sp8

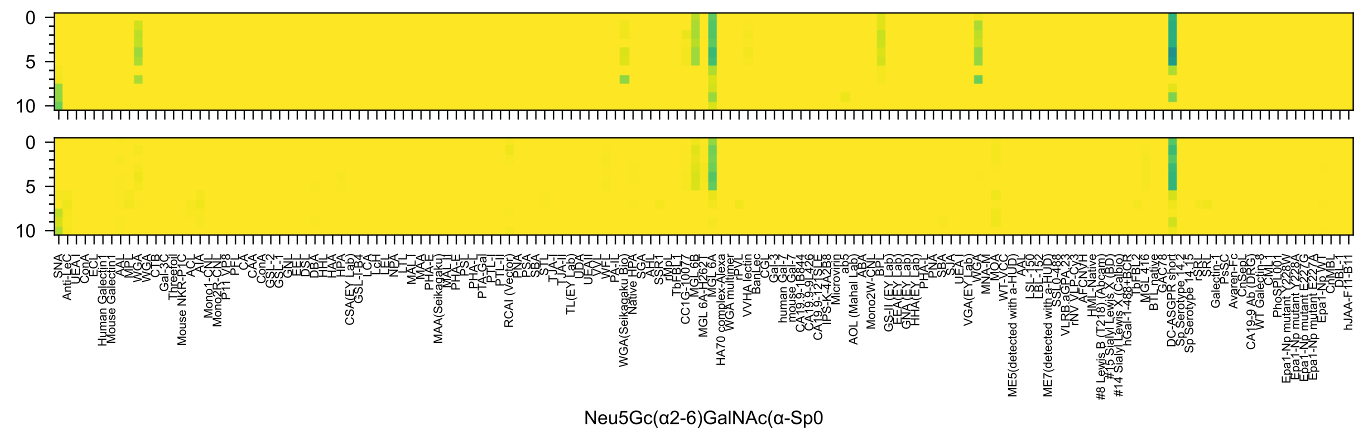
Neu5Gc(α2-6)GalNAc(α-Sp0

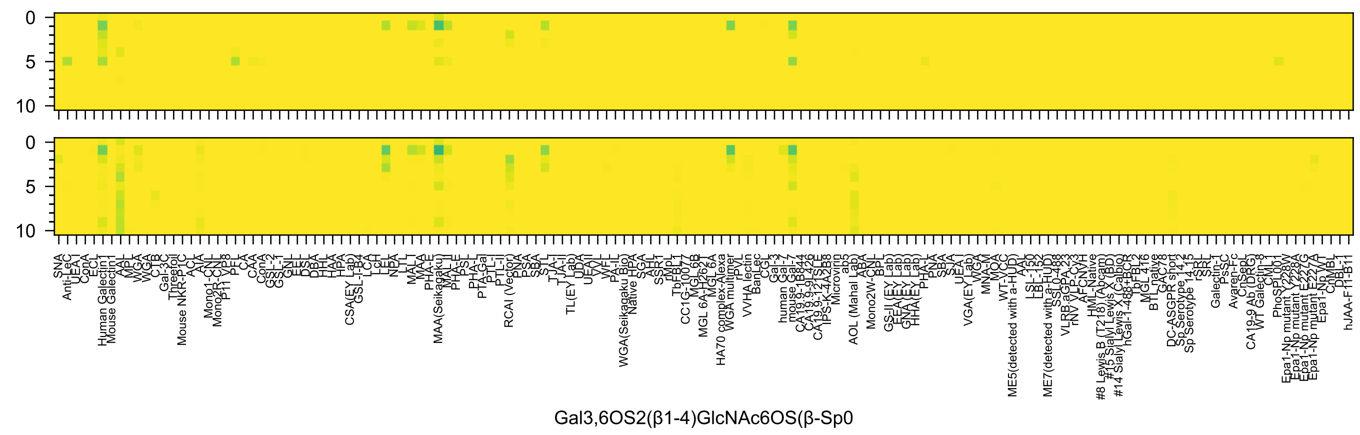
0
5
10
0
5
10
Gal3,6OS2(β1-4)GlcNAc6OS(β-Sp0)

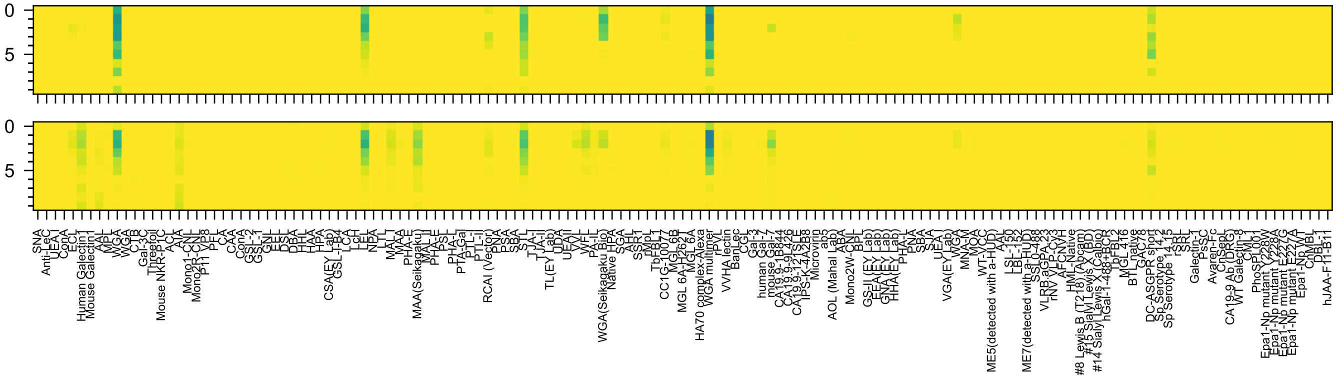


GalNAc4,6OS2(β1-4)GlcNAc(-Sp8

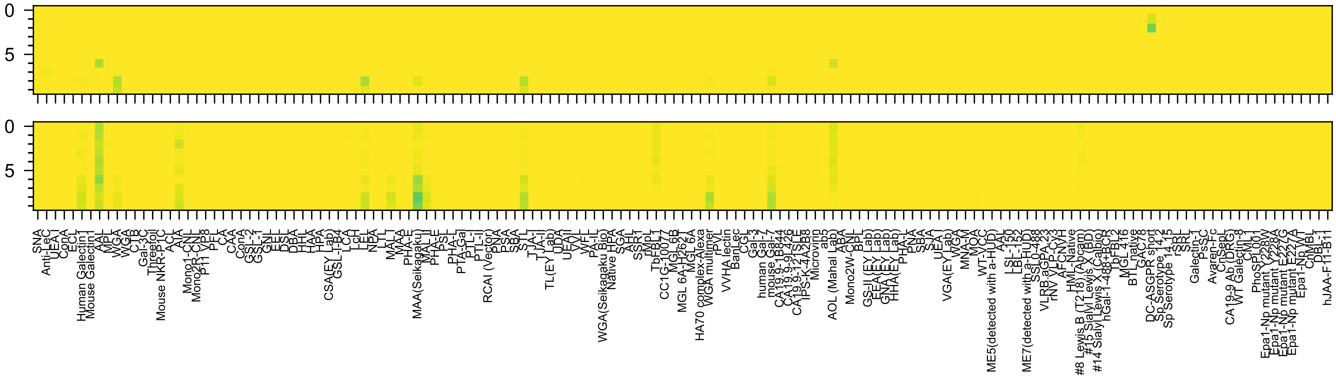

GalNAc3OS(β1-4)GlcNAc3OS(-Sp8

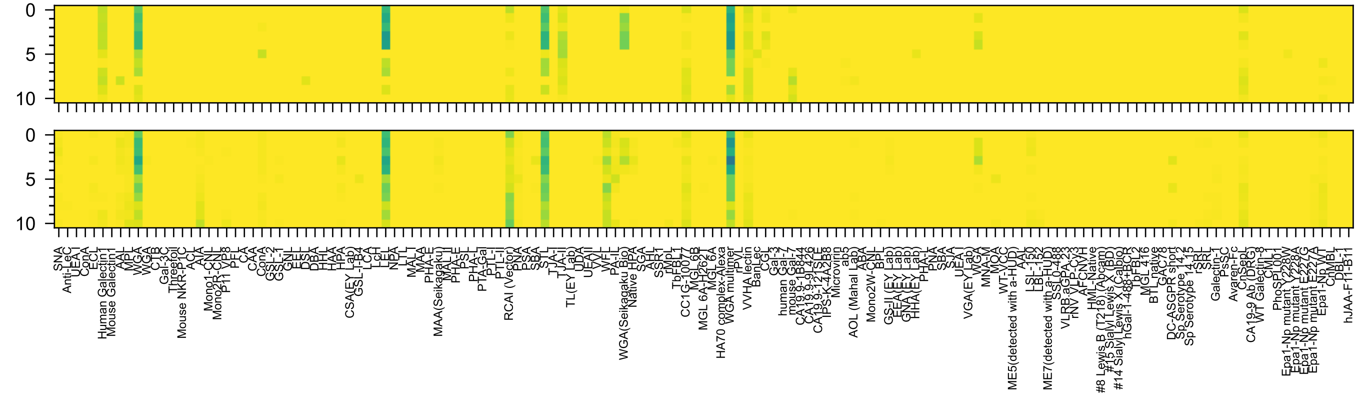


Gal(β1-4)GlcNAc(β-Sp23

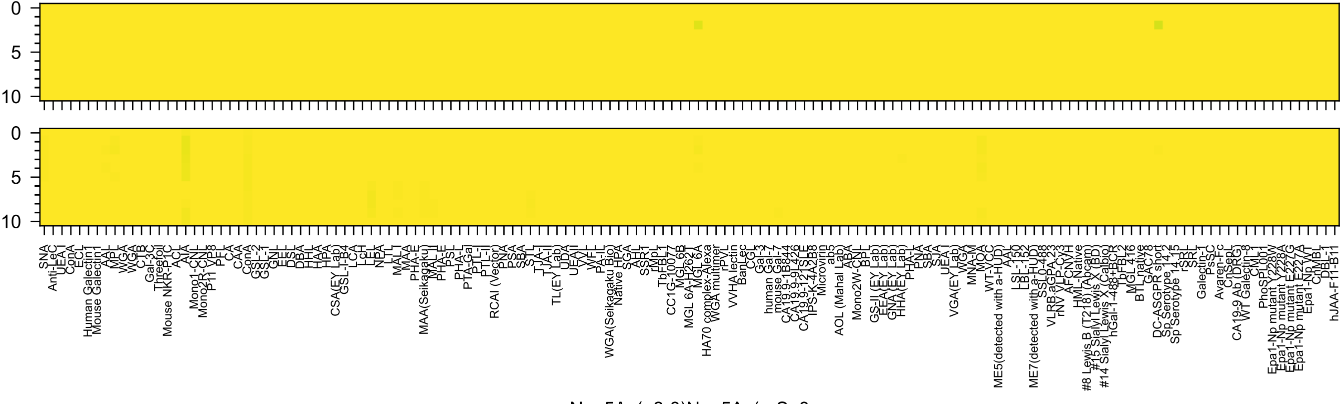
Neu5Ac(α2-8)Neu5Ac(α-Sp8

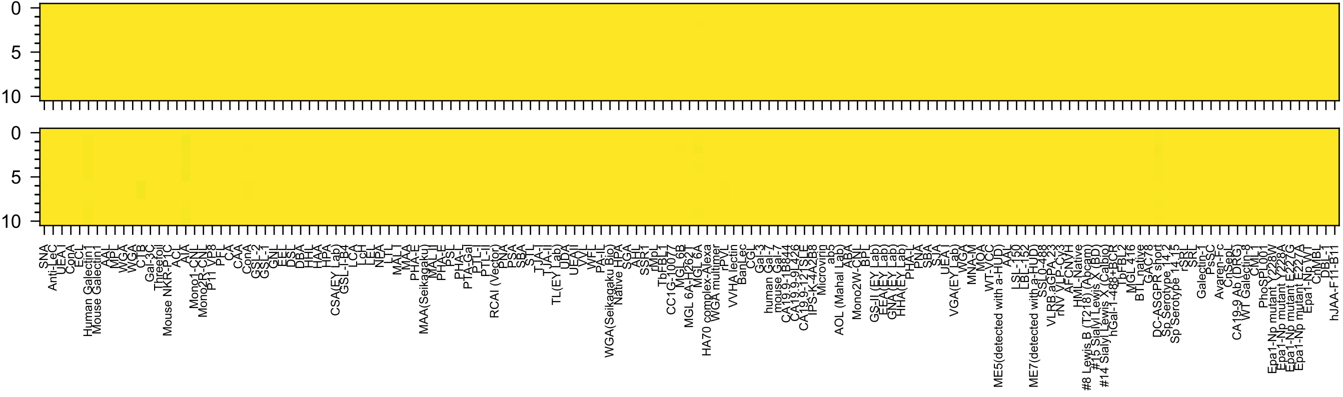


Neu5Ac(α2-8)Neu5Ac(β-Sp17

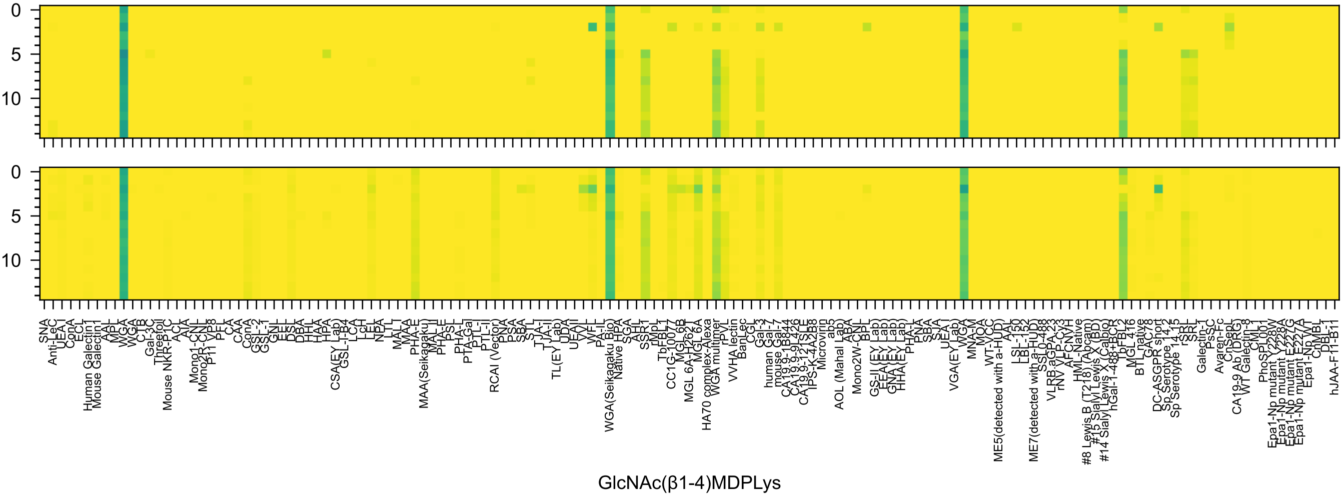
0
5
10
0
5
10
GlcNAc(β1-4)MDPLys

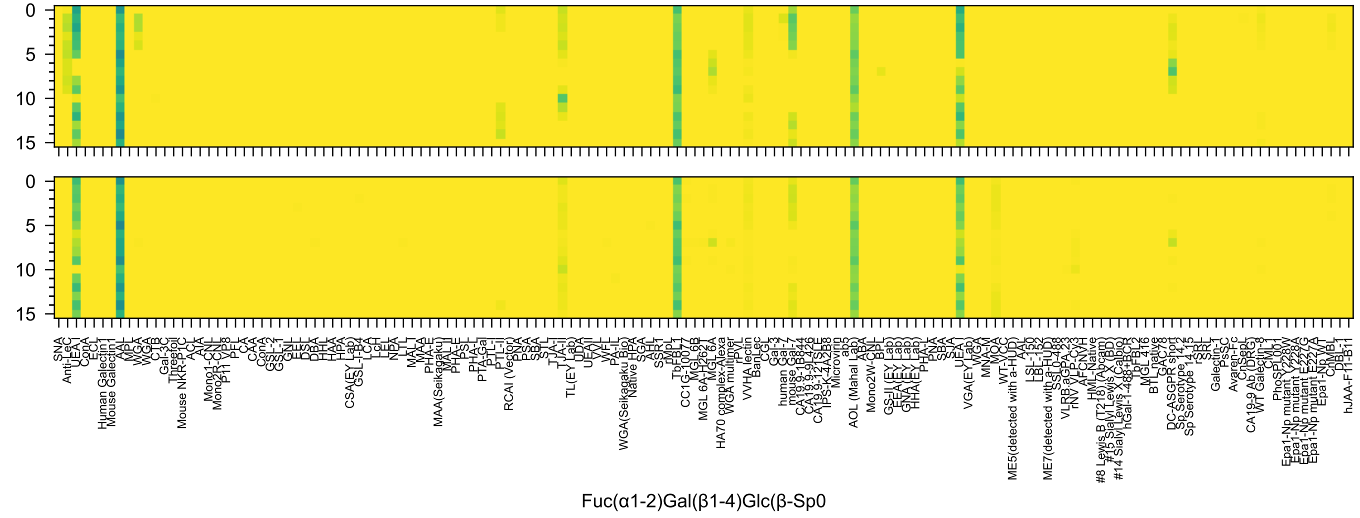
Fuc(α1-2)Gal(β1-4)Glc(β-Sp0

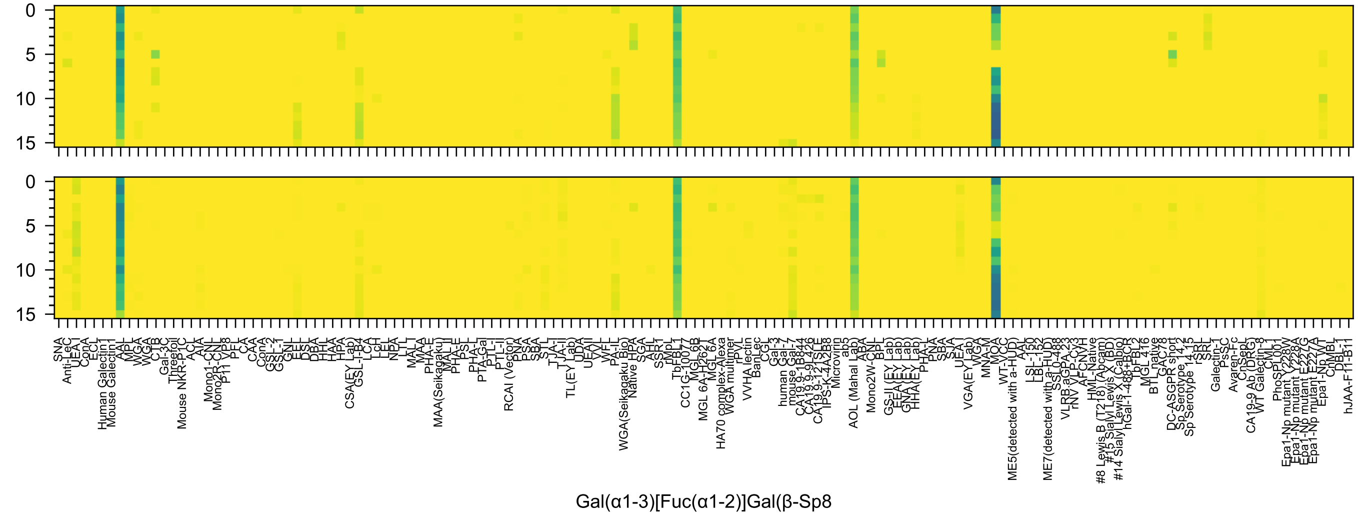
0
5
10
15
0
5
10
15
SNA
Anti-LeC
UEA I
ConA
ECL
Human Galectin1
Mouse Galectin1
AAL
MPL
WGA
WGA
CTB
Gal-3C
Threefoil
Mouse NKR-P1C
ACL
AIA
Mono1-CNL
Mono2R-CNL
P11 VP8
PFL
CA
CAA
ConA
GSL-2
GSL-1
GNL
EEL
DSL
DBA
HHL
HAA
HPA
CSA(EY Lab)
GSL-I-B4
LCA
LcH
LEL
NPA
LTL
MAL-I
MAA
PHA-E
MAA(Seikagaku)
MAL-II
PHA-E
PSL
PHA-L
PTA-Gal
PTL-I
PTL-II
RCAI (Vector)
PNA
PSA
SBA
STL
TJA-I
TJA-II
TL(EY Lab)
UDA
UEA-II
VVL
WFL
PA-IL
WGA(Seikagaku Bio)
Native HPA
SGA
AHL
SSR1
rMpL
TbFBL1
CC1G-10077
MGL 6B
MGL 6A-H262T
MGL 6A
HA70 complex-Alexa
WGA multimer
rPVL
VVHA lectin
BanLec
CGL
Gal-3
human Gal-7
mouse Gal-7
CA19.9-1B844
CA19.9-9L426
CA19.9-121SLE
IPS-K-4A2B8
Microvirin
ab5
AOL (Mahal Lab)
ABA
Mono2W-CNL
BPL
GS-II (EY Lab)
EEA(EY Lab)
GNA(EY Lab)
HHA(EY Lab)
PHA-L
PNA
SBA
SJA
UEA-I
VGA(EY Lab)
WGA
MNA-M
MOA
WT-VCC
ME5(detected with a-HUD)
AAL
LSL-150
LBL-152
ME7(detected with a-HUD)
SSL0-488
VLRB.aGPA-23
rNV VLP-Cy3
AFCNVH
HML-Native
#8 Lewis B (T218) (Abcam)
#15 Sialyl Lewis X (BD)
#14 Sialyl Lewis X (Calbio)
hGal-1-488+BCR
TbFBL2
MGL 416
BTL native
GAC78
DC-ASGPR short
Sp Serotype 14.2
Sp Serotype 14.15
rSRL
SRL
Galectin-1
PsSC
Avaren-Fc
ChSeP
CA19-9 Ab (DRG)
WT Galectin-8
CML1
PhoSL001
Epa1-Np mutant Y228W
Epa1-Np mutant Y228A
Epa1-Np mutant E227G
Epa1-Np mutant E227A
Epa1-Np WT
CnMBL
DBL-1
hJAA-F11-B11
Gal(α1-3)[Fuc(α1-2)]Gal(β-Sp8

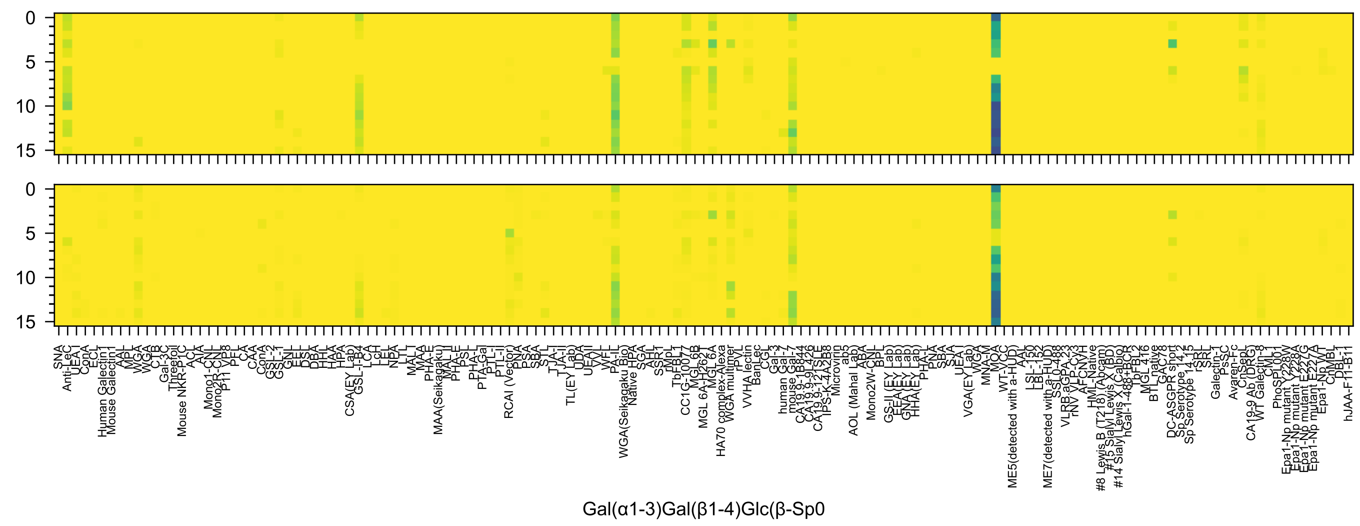

0
5
10
15
Gal(α1-3)Gal(β1-4)Glc(β-Sp0

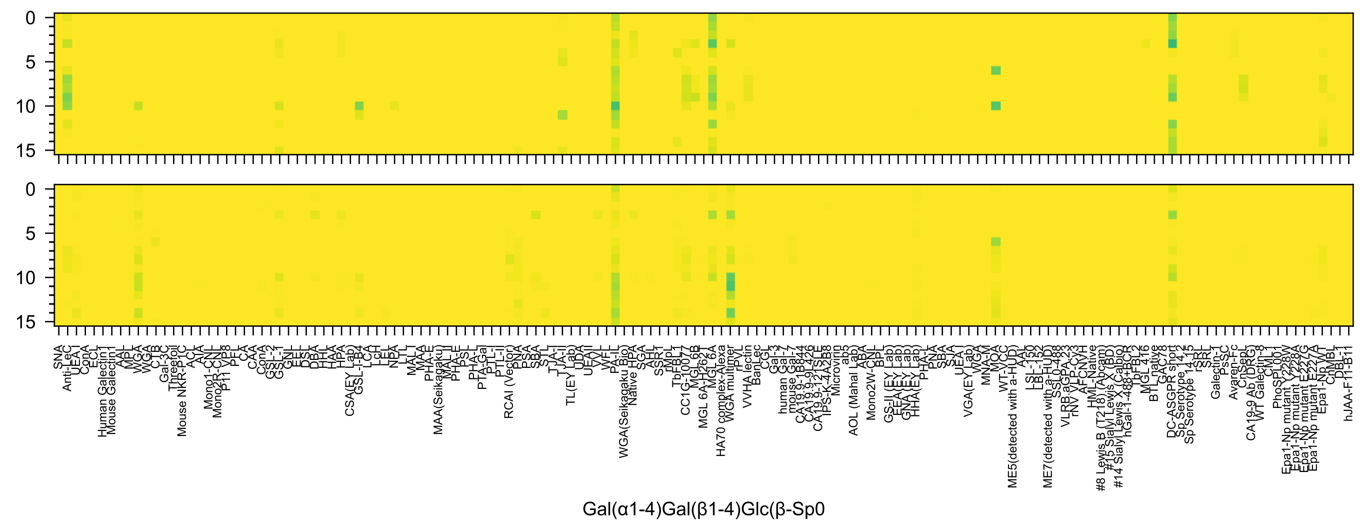
0
5
10
15
SNA
Anti-LeC
UEA I
ConA
ECL
Human Galectin1
Mouse Galectin1
AAL
MPL
WGA
WGA
CTB
Gal-3C
Threefoil
Mouse NKR-P1C
ACL
AIA
Mono1-CNL
Mono2R-CNL
P11 VP8
PFL
CA
CAA
ConA
GSL-2
GSL-1
GNL
EEL
DSL
DBA
HHL
HAA
HPA
CSA(EY Lab)
GSL-I-B4
LCA
LcH
LEL
NPA
LTL
MAL-I
MAA
PHA-E
MAA(Seikagaku)
MAL-II
PHA-E
PSL
PHA-L
PTA-Gal
PTL-I
PTL-II
RCAI (Vector)
PNA
PSA
SBA
STL
TJA-I
TJA-II
TL(EY Lab)
UDA
UEA-II
VVL
WFL
PA-IL
WGA(Seikagaku Bio)
Native HPA
SGA
AHL
SSR1
rMpL
TbFBL1
CC1G-10077
MGL 6B
MGL 6A-H262T
MGL 6A
HA70 complex-Alexa
WGA multimer
rPVL
VVHA lectin
BanLec
CGL
Gal-3
human Gal-7
mouse Gal-7
CA19.9-1B844
CA19.9-9L426
CA19.9-121SLE
IPS-K-4A2B8
Microvirin
ab5
AOL (Mahal Lab)
ABA
Mono2W-CNL
BPL
GS-II (EY Lab)
EEA(EY Lab)
GNA(EY Lab)
HHA(EY Lab)
PHA-L
PNA
SBA
SJA
UEA-I
VGA(EY Lab)
WGA
MNA-M
MOA
WT-VCC
ME5(detected with a-HUD)
AAL
LSL-150
LBL-152
ME7(detected with a-HUD)
SSL0-488
VLRB.aGPA.23
rNV VLP-Cy3
AFCNVH
HML-Native
#8 Lewis B (T218) (Abcam)
#15 Sialyl Lewis X (BD)
#14 Sialyl Lewis X (Calbio)
hGal-1-488+BCR
TbFBL2
MGL 416
BTL native
GAC78
DC-ASGPR short
Sp Serotype 14.2
Sp Serotype 14.15
rSRL
SRL
Galectin-1
PsSC
Avaren-Fc
ChSepL
CA19-9 Ab (DRG)
WT Galectin-8
CML1
PhoSL001
Epa1-Np mutant Y228W
Epa1-Np mutant Y228A
Epa1-Np mutant E227G
Epa1-Np mutant E227A
Epa1-Np WT
ChMBL
DBL-1
hJAA-F11-B11
Gal(α1-4)Gal(β1-4)Glc(β-Sp0

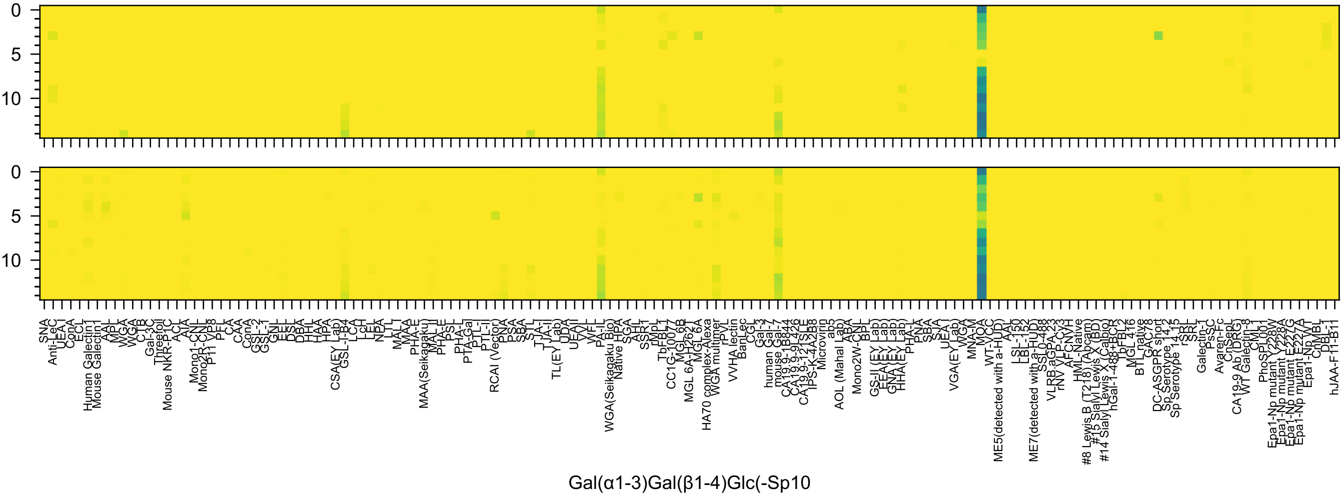
0
5
10
0
5
10
SNA
Anti-LeC
UEA-I
ConA
ECL
Human Galectin1
Mouse Galectin1
AAL
MPL
WGA
WGA
CTB
Gal-3C
Threefoil
Mouse NKR-P1C
ACL
AIA
Mono1-CNL
Mono2R-CNL
P11 VP8
PFL
CA
CAA
ConA
GSL-2
GSL-1
GNL
EEL
DSL
DBA
HHL
HAA
HPA
CSA(EY Lab)
GSL-I-B4
LCA
LcH
LEL
NPA
LTL
MAL-I
MAA
PHA-E
MAA(Seikagaku)
MAL-II
PHA-E
PSL
PHA-L
PTA-Gal
PTL-I
PTL-II
RCAI (Vector)
PNA
PSA
SBA
STL
TJA-I
TJA-II
TL(EY Lab)
UDA
UEA-II
VVL
WFL
PA-IL
WGA(Seikagaku Bio)
Native HPA
SGA
AHL
SSR1
rMpL
TbFBL1
CC1G-10077
MGL 6B
MGL 6A-H262T
MGL 6A
HA70 complex-Alexa
WGA multimer
rPVL
VVHA lectin
BanLec
CGL
Gal-3
human Gal-7
mouse Gal-7
CA19.9-1B844
CA19.9-9L426
CA19.9-121SLE
IPS-K-4A2B8
Microvirin
ab5
AOL (Mahal Lab)
ABA
Mono2W-CNL
BPL
GS-II (EY Lab)
EEA (EY Lab)
GNA (EY Lab)
HHA (EY Lab)
PHA-L
PNA
SBA
SJA
UEA-I
VGA(EY Lab)
WGA
MNA-M
MOA
WT-VCC
ME5(detected with a-HUD)
AAL
LSL-150
LBL-152
ME7(detected with a-HUD)
SSL0-488
VLRB.aGPA.23
rNV VLP-Cy3
AFCNVH
HML-Native
#8 Lewis B (T218) (Abcam)
#15 Sialyl Lewis X (BD)
#14 Sialyl Lewis X (Calbio)
hGal-1-488+BCR
TbFBL2
MGL 416
BTL native
GAC78
DC-ASGPR short
Sp Serotype 14.2
Sp Serotype 14.15
rSRL
SRL
Galectin-1
PsSC
Avaren-Fc
ChSepL
CA19-9 Ab (DRG)
WT Galectin-8
CML1
PhoSPL001
Epa1-Np mutant Y228W
Epa1-Np mutant Y228A
Epa1-Np mutant E227G
Epa1-Np mutant E227A
Epa1-Np WT
ChMBL
DBL-1
hJAA-F11-B11
Gal(α1-3)Gal(β1-4)Glc(-Sp10

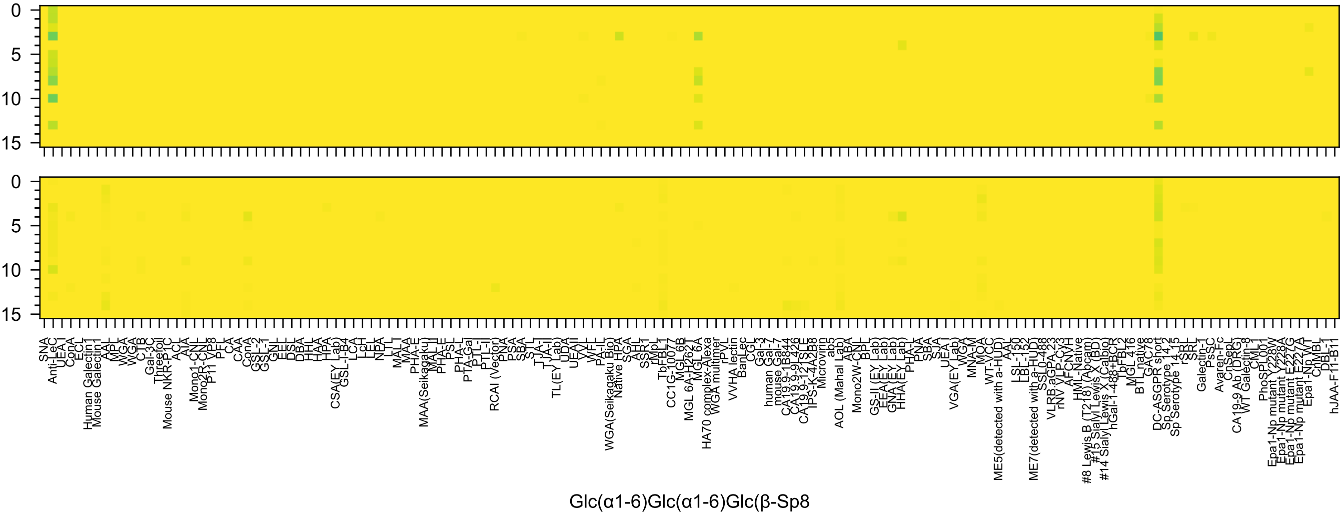
Glc(α1-6)Glc(α1-6)Glc(β-Sp8

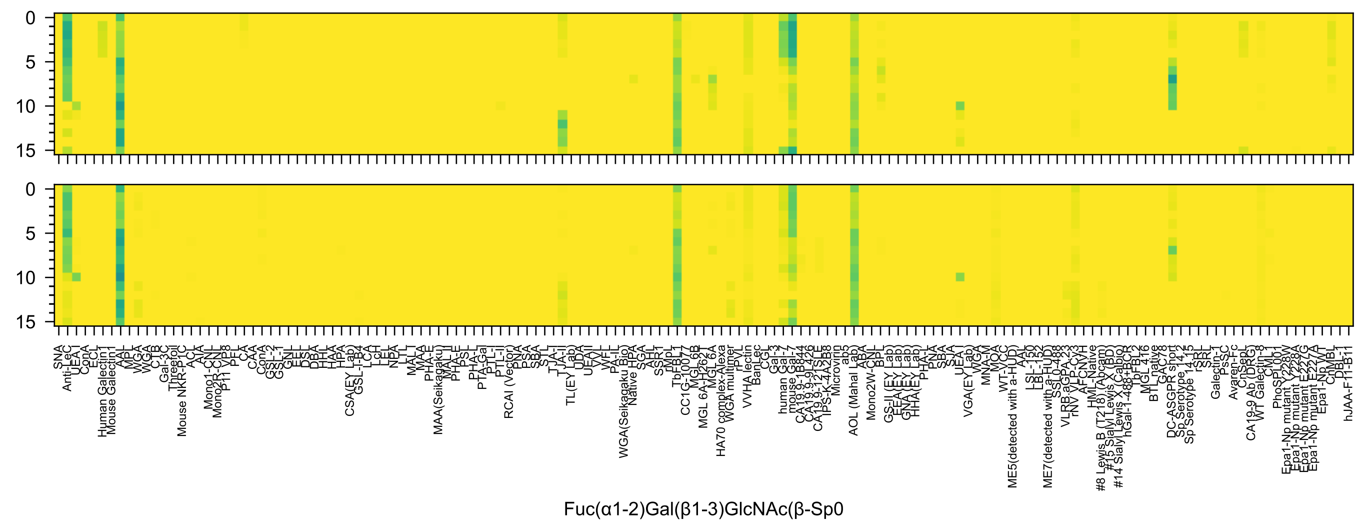
Fuc(α1-2)Gal(β1-3)GlcNAc(β-Sp0

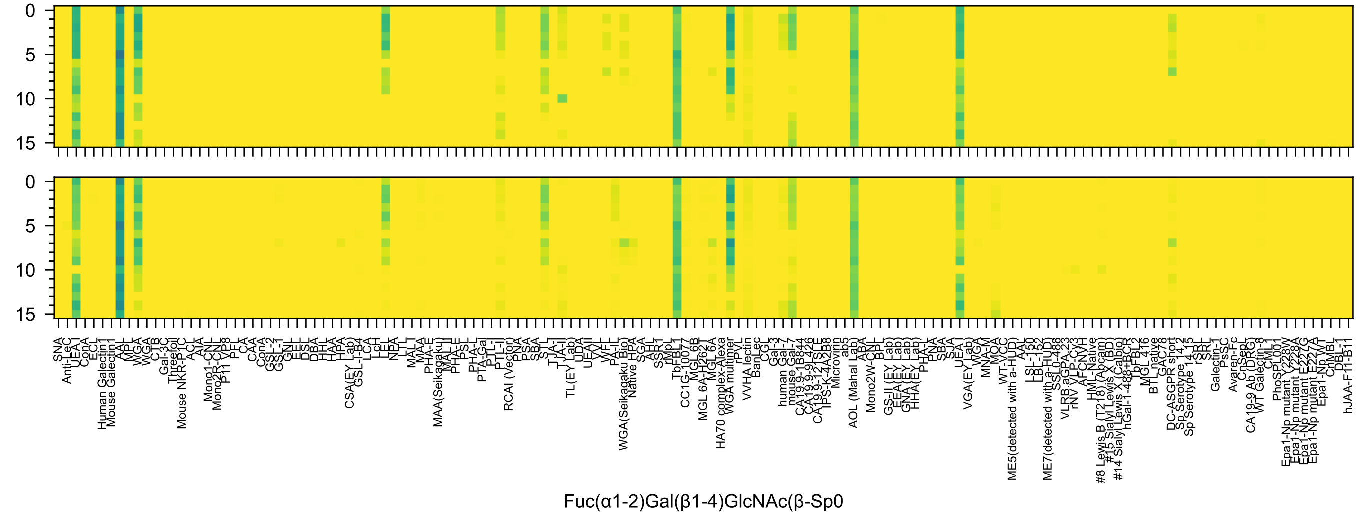
0
5
10
15
SNA
Anti-LeC
UEA-I
ConA
ECL
Human Galectin1
Mouse Galectin1
AAL
MPL
WGA
WGA
CTB
Gal-3C
Threefoil
Mouse NKR-P1C
ACL
AIA
Mono1-CNL
Mono2R-CNL
P11 VP8
PFL
CA
CAA
ConA
GSL-2
GSL-1
GNL
EEL
DSL
DBA
HHL
HAA
HPA
CSA(EY Lab)
GSL-I-B4
LCA
LcH
LEL
NPA
LTL
MAL-I
MAA
PHA-E
MAA(Seikagaku)
MAL-II
PHA-E
PSL
PHA-L
PTA-Gal
PTL-I
PTL-II
RCAI (Vector)
PNA
PSA
SBA
STL
TJA-I
TJA-II
TL(EY Lab)
UDA
UEA-II
VVL
WFL
PA-IL
WGA(Seikagaku Bio)
Native HPA
SGA
AHL
SSR1
rMpL
TbFBL1
CC1G-10077
MGL 6B
MGL 6A-H262T
MGL 6A
HA70 complex-Alexa
WGA multimer
rPVL
VVHA lectin
BanLec
CGL
Gal-3
human Gal-7
mouse Gal-7
CA19.9-1B844
CA19.9-9L426
CA19.9-121SLE
IPS-K-4A2B8
Microvirin
ab5
ABA
AOL (Mahal Lab)
Mono2W-CNL
BPL
GS-II (EY Lab)
EEA(EY Lab)
GNA(EY Lab)
HHA(EY Lab)
PHA-L
PNA
SBA
SJA
UEA-I
VGA(EY Lab)
WGA
MNA-M
MOA
WT-VCC
ME5(detected with a-HUD)
AAL
LSL-150
LBL-152
ME7(detected with a-HUD)
SSL0-488
VLRB.aGPA.23
rNV VLP-Cy3
AFCNVH
HML-Native
#8 Lewis B (T218) (Abcam)
#15 Sialyl Lewis X (BD)
#14 Sialyl Lewis X (Calbio)
hGal-1-488+BCR
TbFBL2
MGL 416
BTL native
GAC78
DC-ASGPR short
Sp Serotype 14.2
Sp Serotype 14.15
rSRL
SRL
Galectin-1
PsSC
Avaren-Fc
ChSeoL
CA19-9 Ab (DRG)
WT Galectin-8
CML1
PhoSL001
Epa1-Np mutant Y228W
Epa1-Np mutant Y228A
Epa1-Np mutant E227G
Epa1-Np mutant E227A
Epa1-Np WT
ChMBL
DBL-1
hJAA-F11-B11
Fuc(α1-2)Gal(β1-4)GlcNAc(β-Sp0

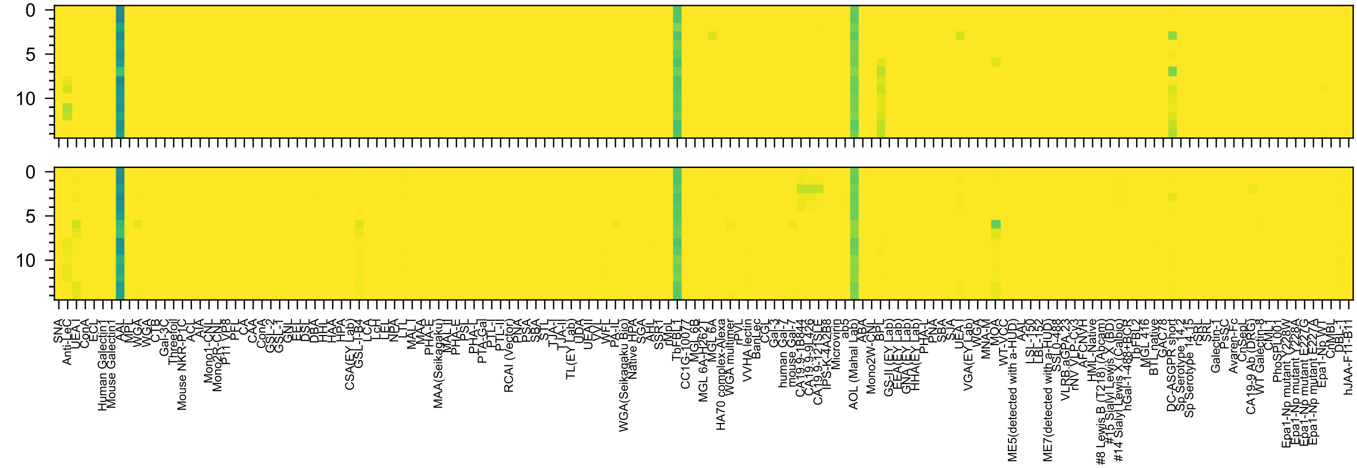


Gal(β1-3)[Fuc(α1-4)]GlcNAc(-Sp0

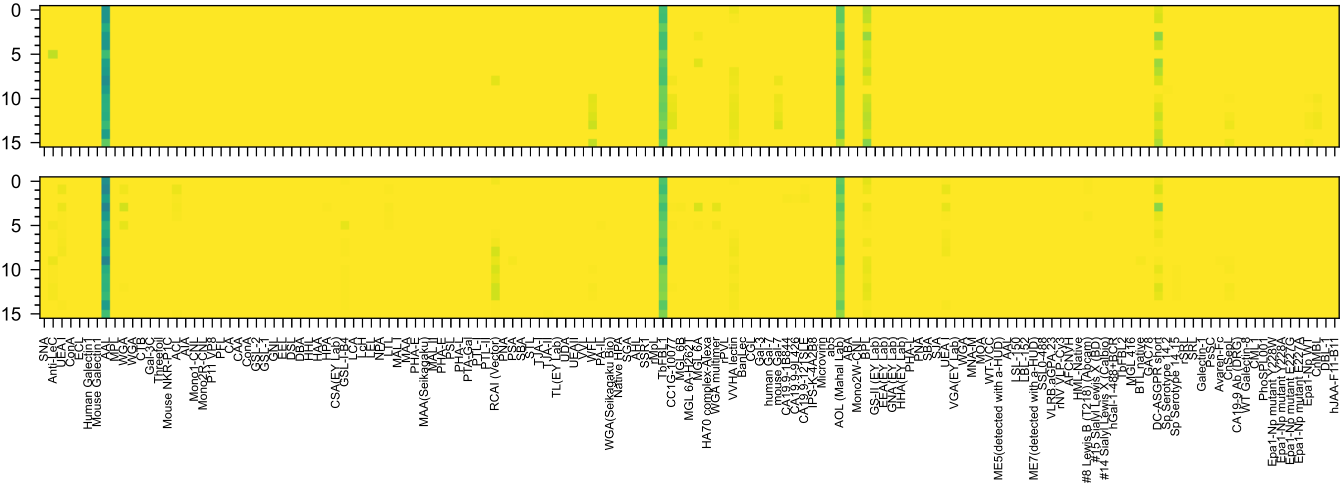

Gal(β1-4)[Fuc(α1-3)]GlcNAc(β-Sp0

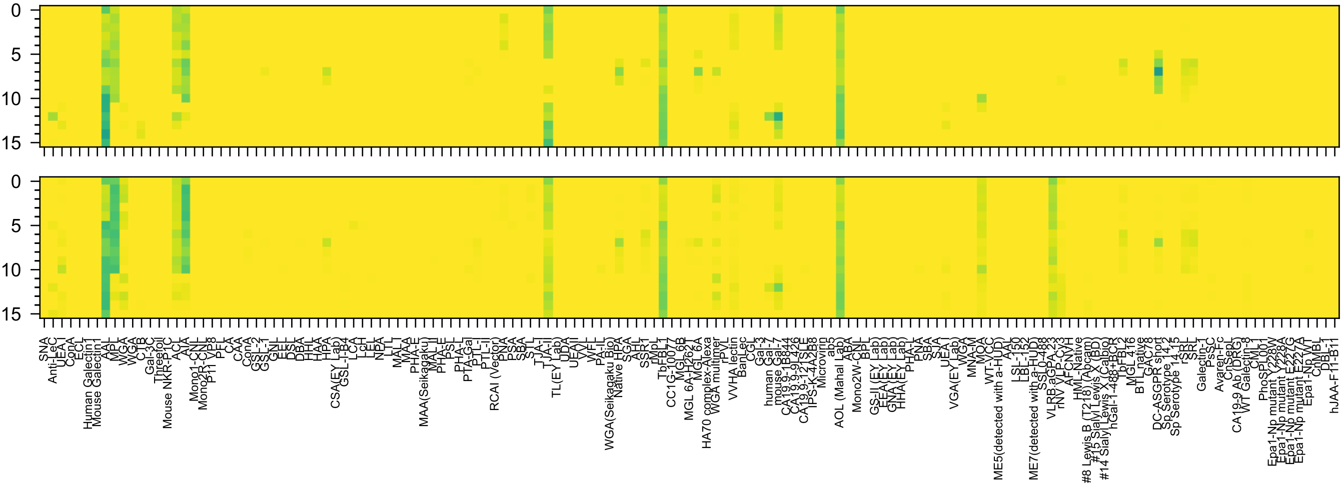

Fuc(α1-2)Gal(β1-3)GalNAc(α-Sp8

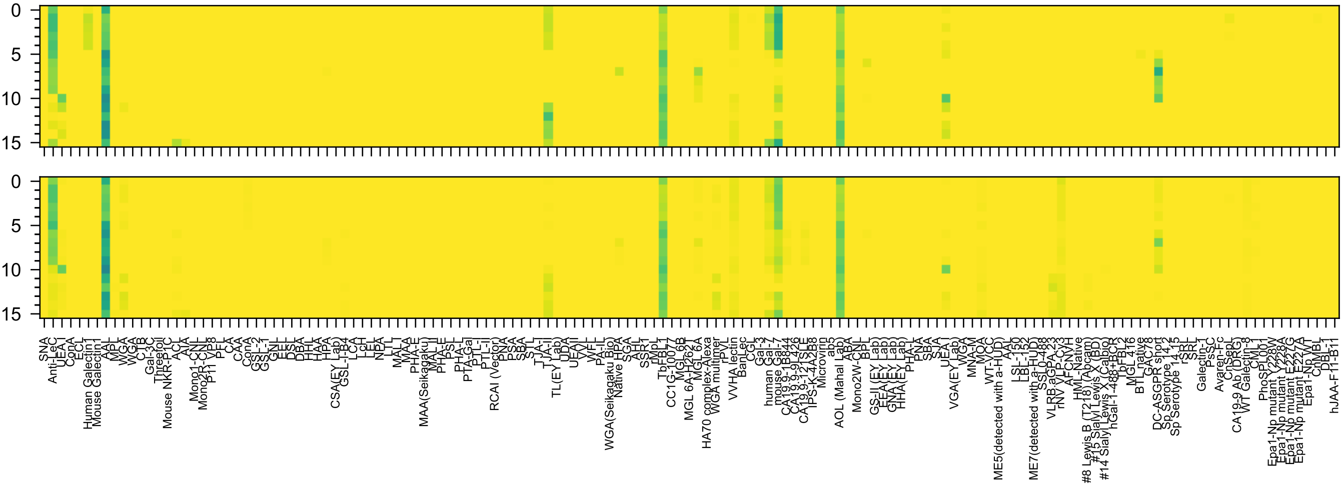
Fuc(α1-2)Gal(β1-3)GlcNAc(β-Sp8

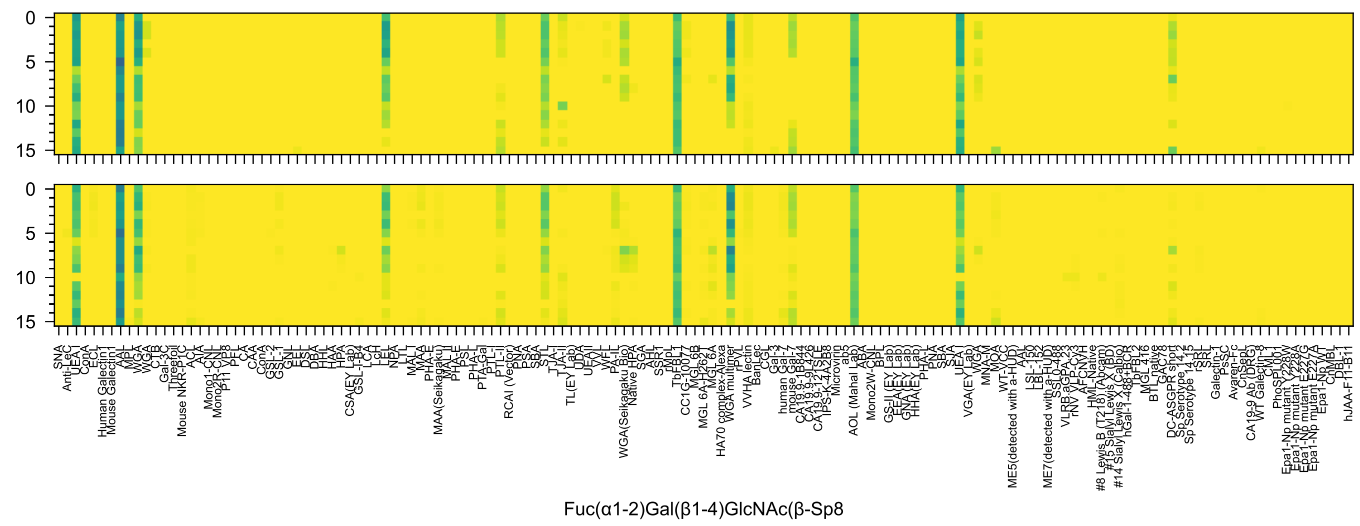
0
5
10
15
SNA
Anti-LeC
UEA-I
ConA
ECL
Human Galectin1
Mouse Galectin1
AAL
MPL
WGA
WGA
CTB
C-1B
Gal-3C
Threefoil
Mouse NKR-P1C
ACL
AIA
Mono1-CNL
Mono2R-CNL
P11 VP8
PFL
CA
CAA
ConA
GSL-2
GSL-1
GNL
EEL
DSL
DBA
HHL
HAA
HPA
CSA(EY Lab)
GSL-I-B4
LCA
LcH
LEL
NPA
LTL
MAL-I
MAA
PHA-E
MAA(Seikagaku)
MAL-II
PHA-E
PSL
PHA-L
PTA-Gal
PTL-I
PTL-II
RCAI (Vector)
PNA
PSA
SBA
STL
TJA-I
TJA-II
TL(EY Lab)
UDA
UEAII
VVL
WFL
PA-IL
WGA(Seikagaku Bio)
Native HPA
SGA
AHL
SSR1
rMpL
TbFBL1
CC1G-10077
MGL 6B
MGL 6A-H262T
MGL 6A
HA70 complex-Alexa
WGA multimer
rPVL
VVHA lectin
BanLec
CGL
Gal-3
human Gal-7
mouse Gal-7
CA19.9-1B844
CA19.9-9L426
CA19.9-121SLE
IPS-K-4A2B8
Microvirin
ab5
AOL (Mahal Lab)
ABA
Mono2W-CNL
BPL
GS-II (EY Lab)
EEA(EY Lab)
GNA(EY Lab)
HHA(EY Lab)
PHA-L
PNA
SBA
SJA
UEA-I
VGA(EY Lab)
WGA
MNA-M
MOA
WT-VCC
ME5(detected with a-HUD)
AAL
LSL-150
LBL-152
ME7(detected with a-HUD)
SSL0-488
VLRB.aGPA.23
rNV VLP-Cy3
AFCNVH
HML-Native
#8 Lewis B (T218) (Abcam)
#15 Sialyl Lewis X (BD)
#14 Sialyl Lewis X (Calbio)
hGal-1-488+BCR
TbFBL2
MGL 416
BTL native
GAC78
DC-ASGPR short
Sp Serotype 14.2
Sp Serotype 14.15
rSRL
SRL
Galectin-1
PsSC
Avaren-Fc
ChSepL
CA19-9 Ab (DRG)
WT Galectin-8
CML-1
PhoSL001
Epa1-Np mutant Y228W
Epa1-Np mutant Y228A
Epa1-Np mutant E227G
Epa1-Np mutant E227A
Epa1-Np WT
ChMBL
DBL-1
hJAA-F11-B11
Fuc(α1-2)Gal(β1-4)GlcNAc(β-Sp8

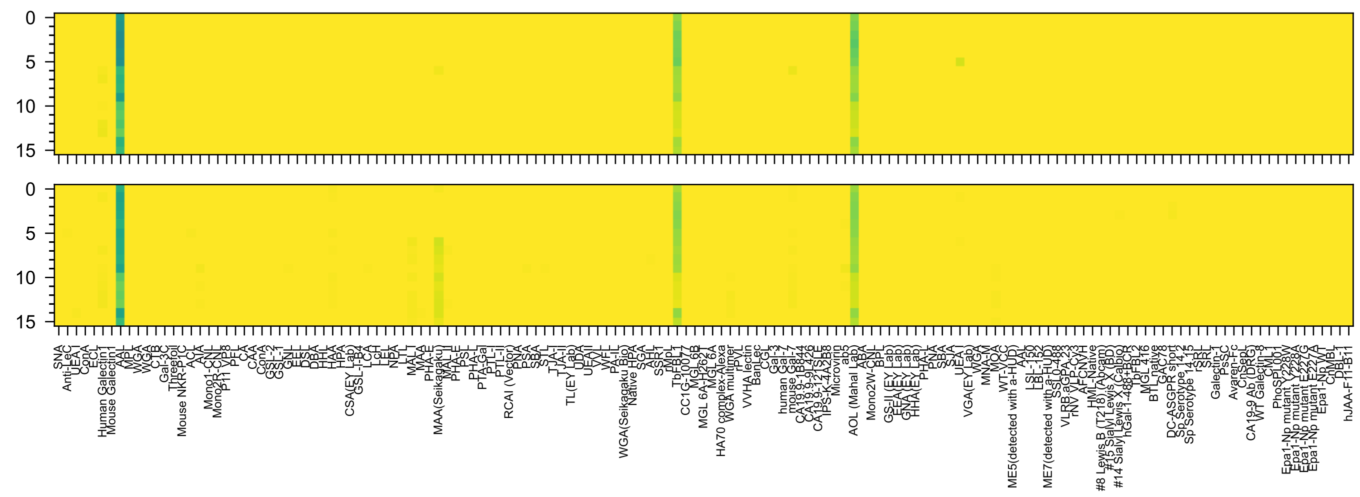
Gal3OS(β1-4)[Fuc(α1-3)]Glc(β-Sp0

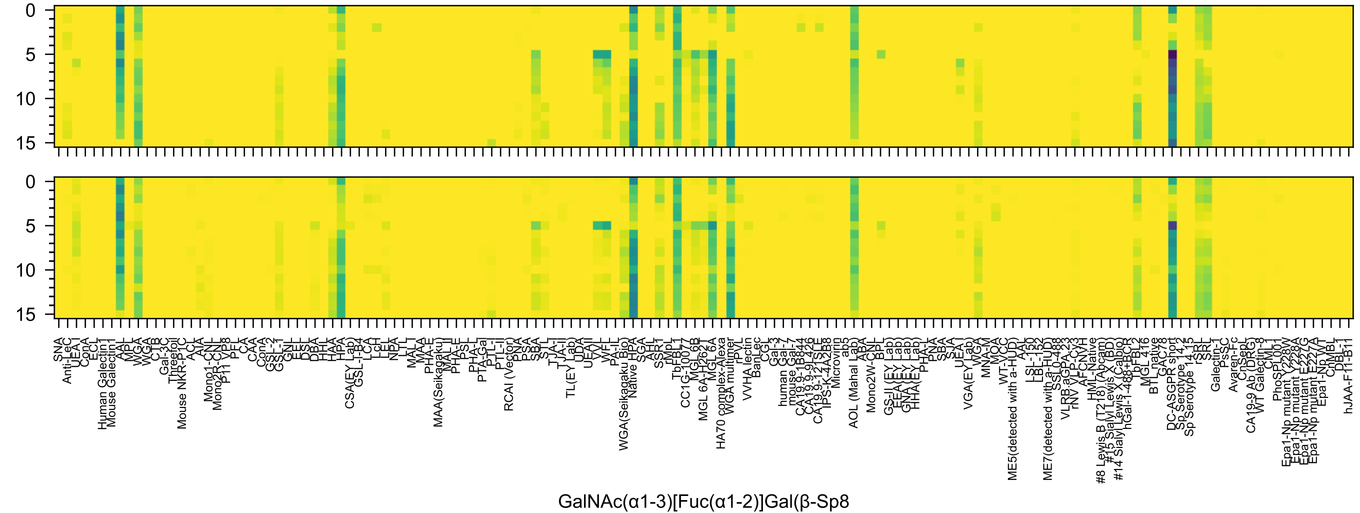
0
5
10
15
0
5
10
15
SNA
Anti-LeC
UEA I
ConA
ECL
Human Galectin1
Mouse Galectin1
AAL
MPL
WGA
WGA
CTB
Gal-3C
Threefoil
Mouse NKR-P1C
ACL
AIA
Mono1-CNL
Mono2R-CNL
P11 VP8
PFL
CA
CAA
ConA
GSL-2
GSL-1
GNL
EEL
DSL
DBA
HHL
HAA
HPA
CSA(EY Lab)
GSL-I-B4
LCA
LcH
LEL
NPA
LTL
MAL I
MAA
PHA-E
MAA(Seikagaku)
MAL II
PHA-E
PSL
PHA-L
PTA-Gal
PTL-I
PTL-II
RCAI (Vector)
PNA
PSA
SBA
STL
TJA-I
TJA-II
TL(EY Lab)
UDA
UEA II
VVL
WFL
PA-IL
WGA(Seikagaku Bio)
Native HPA
SGA
AAL
SSR1
rMpL
TbFBL1
CC1G-10077
MGL 6B
MGL 6A-H262T
MGL 6A
HA70 complex-Alexa
WGA multimer
rPVL
VVHA lectin
BanLec
CGL
Gal-3
human Gal-7
mouse Gal-7
CA19.9-1B844
CA19.9-9L426
CA19.9-121SLE
IPS-K-4A2B8
Microvirin
ab5
AOL (Mahal Lab)
ABA
Mono2W-CNL
BPL
GS-II (EY Lab)
EEA (EY Lab)
GNA (EY Lab)
HHA (EY Lab)
PHA-L
PNA
SBA
SJA
UEA I
VGA(EY Lab)
WGA
MNA-M
MOA
WT-VCC
ME5(detected with a-HUD)
AAL
LSL-150
LBL-152
ME7(detected with a-HUD)
SSL0-488
VLRB.aGPA.23
rNV VLP-Cy3
AFCNVH
HML-Native
#8 Lewis B (T218) (Abcam)
#15 Sialyl Lewis X (BD)
#14 Sialyl Lewis X (Calbio)
hGal-1-488+BCR
TbFBL2
MGL 416
BTL native
GAC78
DC-ASGPR short
Sp Serotype 14.2
Sp Serotype 14.15
rSRL
SRL
Galectin-1
PsSC
Avaren-Fc
ChSepL
CA19-9 Ab (DRG)
WT Galectin-8
CML1
PhoSL001
Epa1-Np mutant Y228W
Epa1-Np mutant Y228A
Epa1-Np mutant E227G
Epa1-Np mutant E227A
Epa1-Np WT
ChMBL
DBL-1
hJAA-F11-B11
GalNAc(α1-3)[Fuc(α1-2)]Gal(β-Sp8

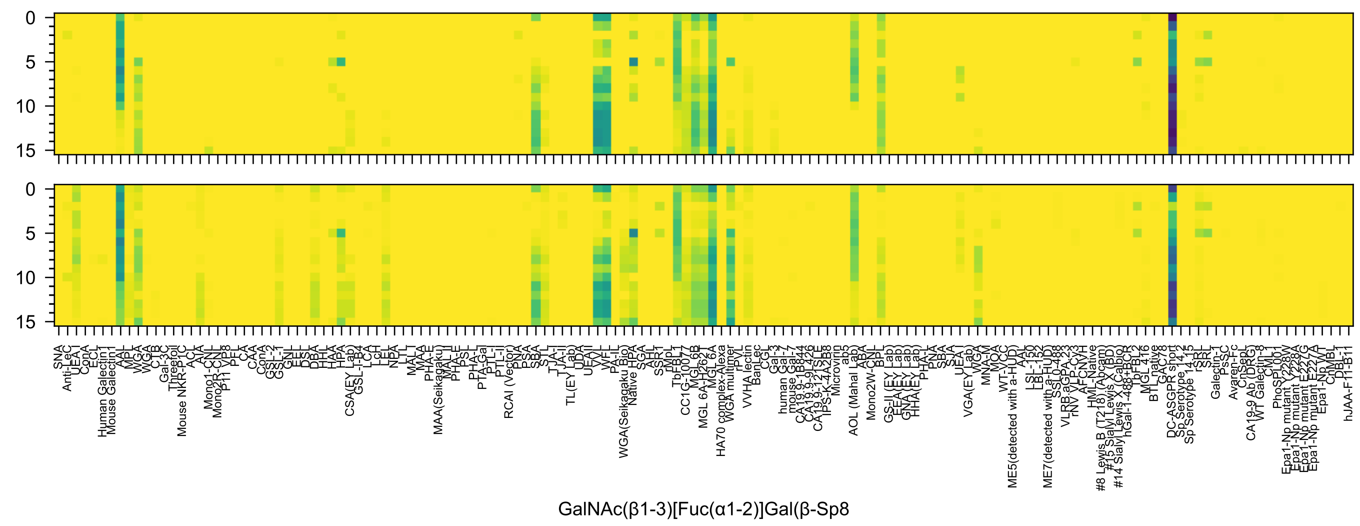
GalNAc(β1-3)[Fuc(α1-2)]Gal(β-Sp8

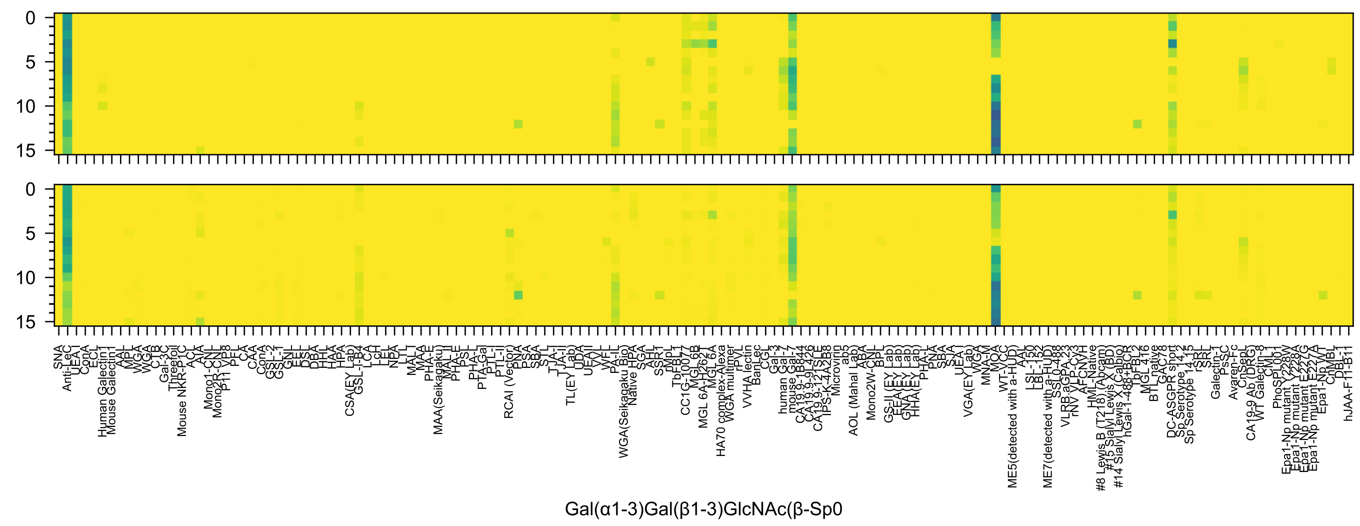
0
5
10
15
Gal(α1-3)Gal(β1-3)GlcNAc(β-Sp0

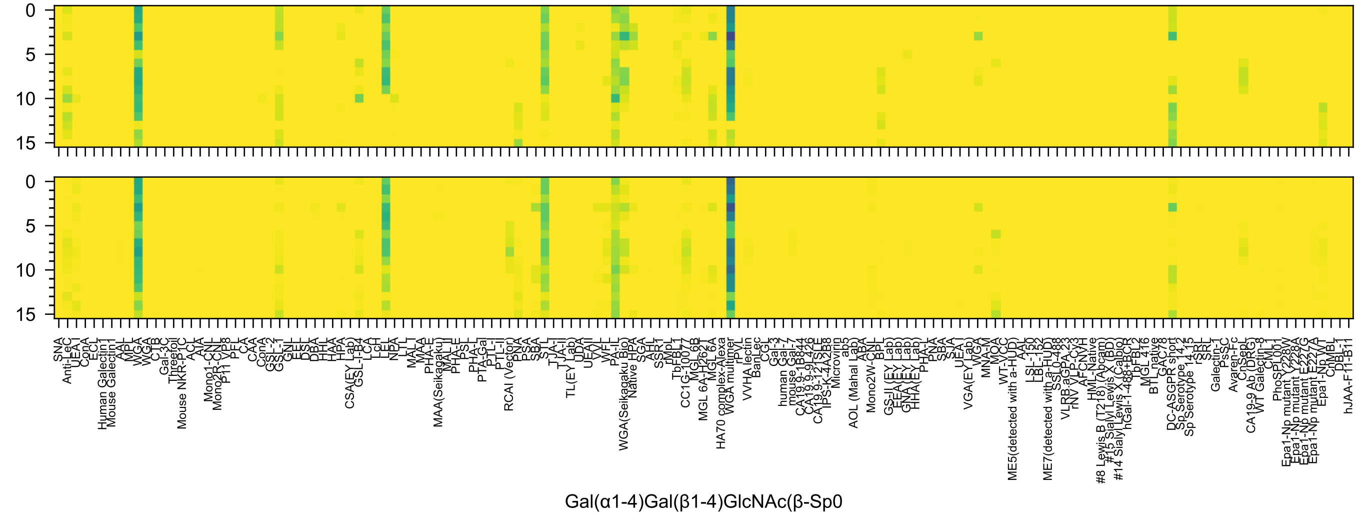
Gal(α1-4)Gal(β1-4)GlcNAc(β-Sp0

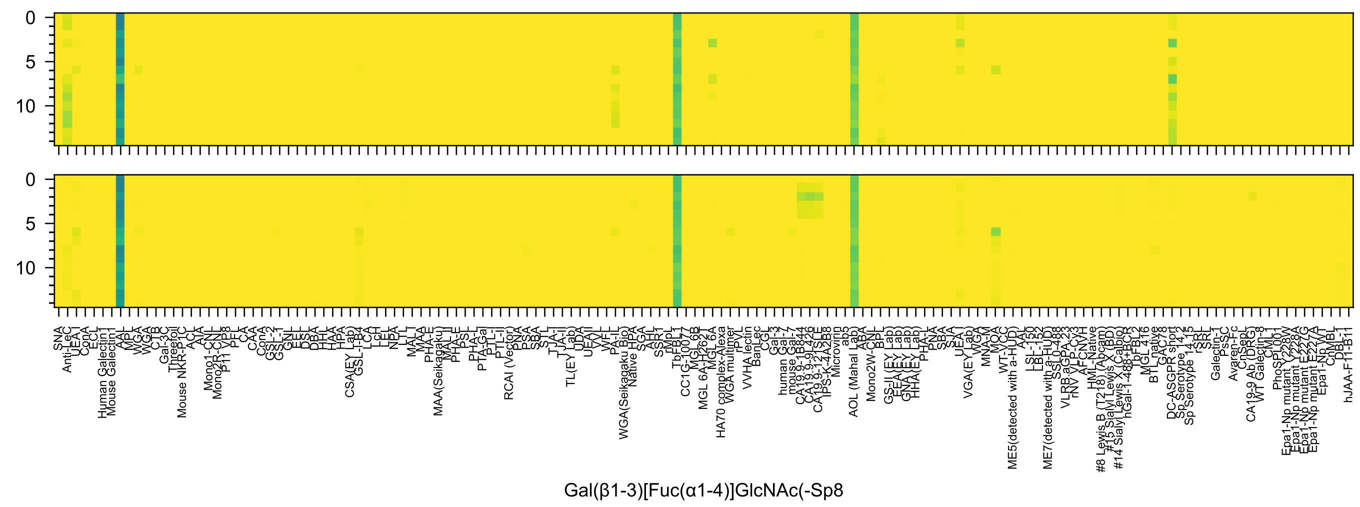

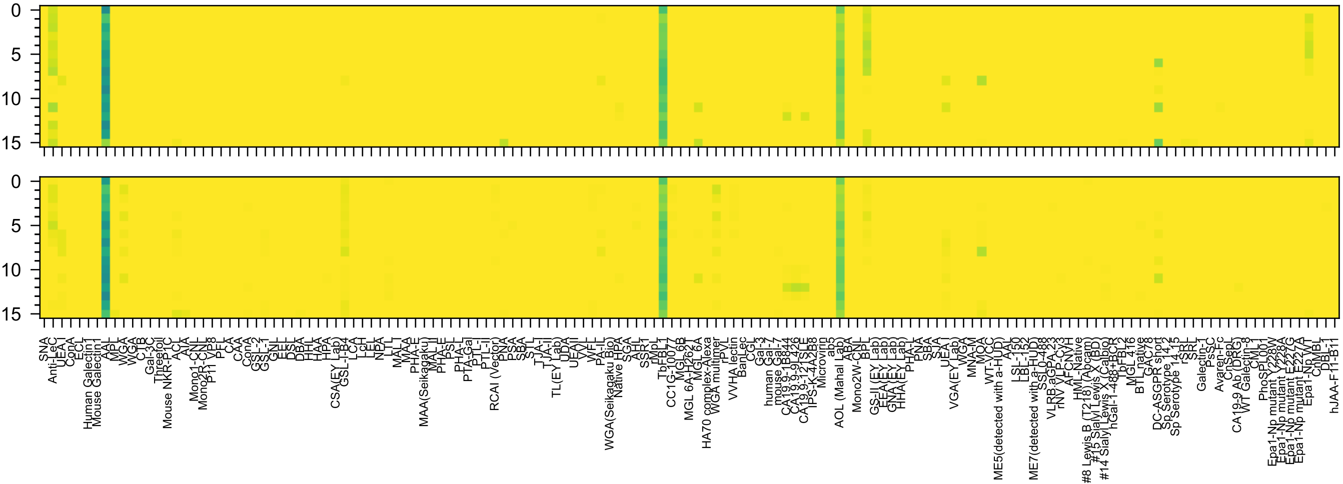

Fuc(α1-4)[Gal(β1-3)]GlcNAc(β-Sp8

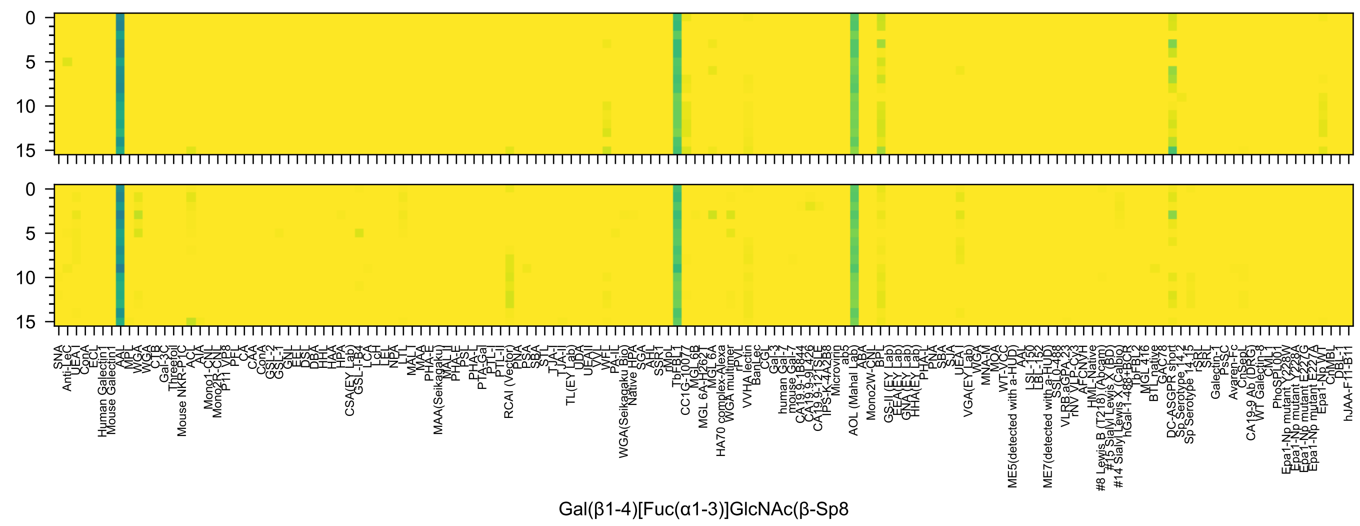

Gal(β1-4)[Fuc(α1-3)]GlcNAc(β-Sp8

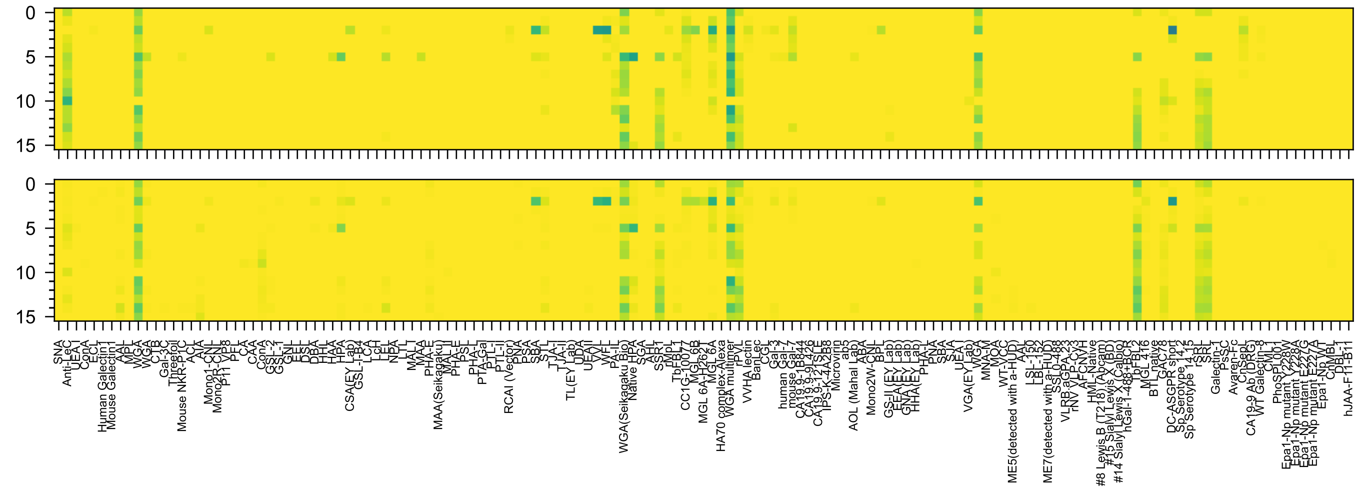


GlcNAc(β1-3)Gal(β1-4)Glc(β-Sp0

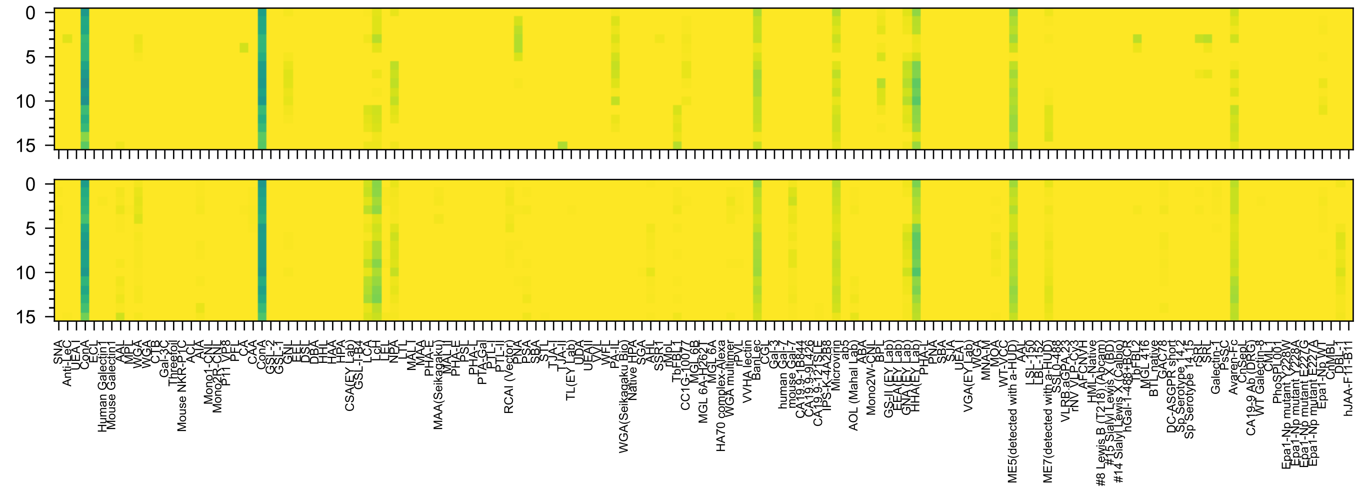

Man(α1-2)Man(α1-3)Man(α-Sp9

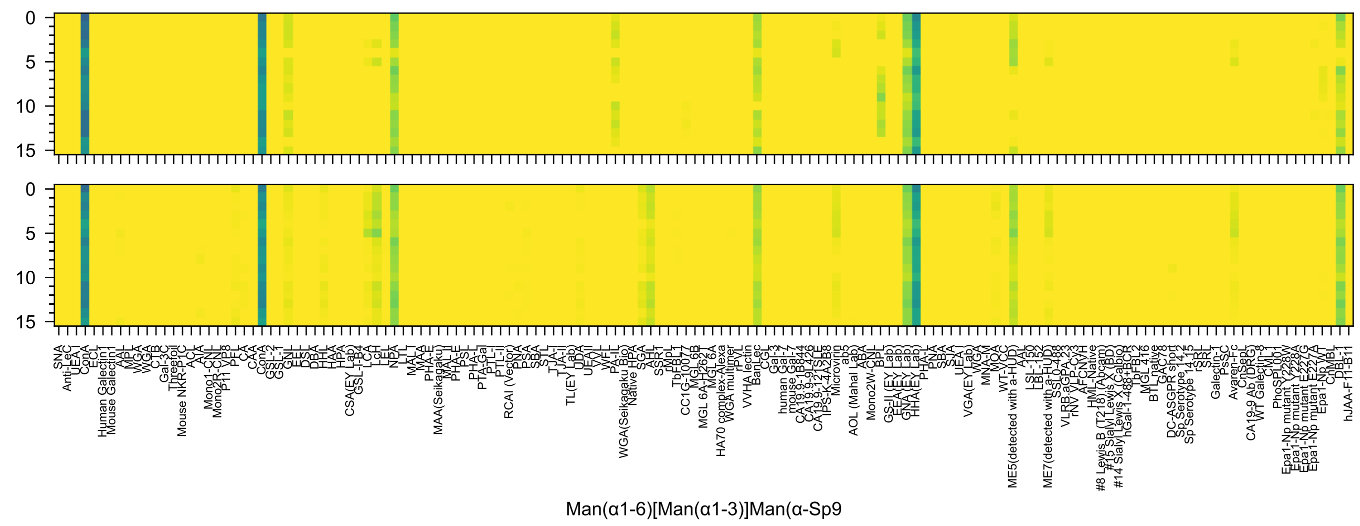
Man(α1-6)[Man(α1-3)]Man(α-Sp9

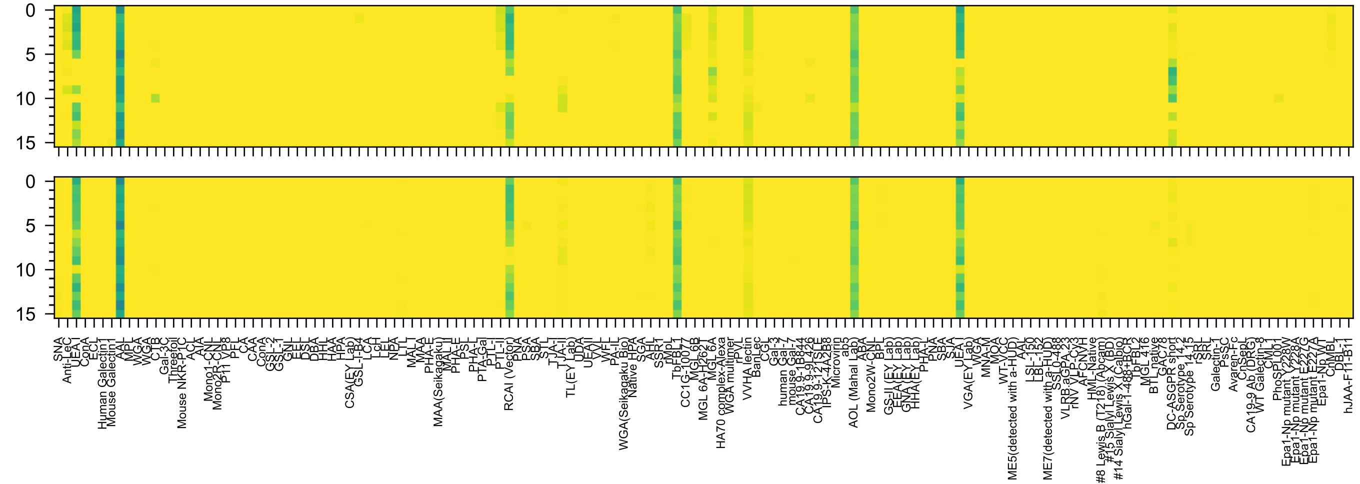

Fuc(α1-2)Gal6OS(β1-4)Glc(β-Sp0

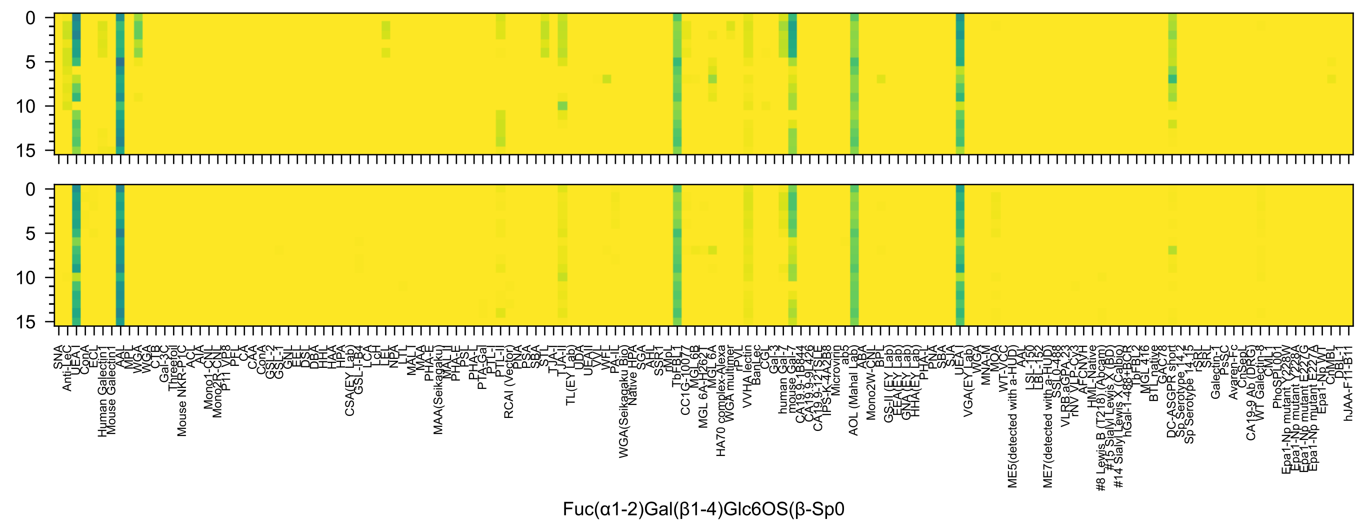

Fuc(α1-2)Gal(β1-4)Glc6OS(β-Sp0

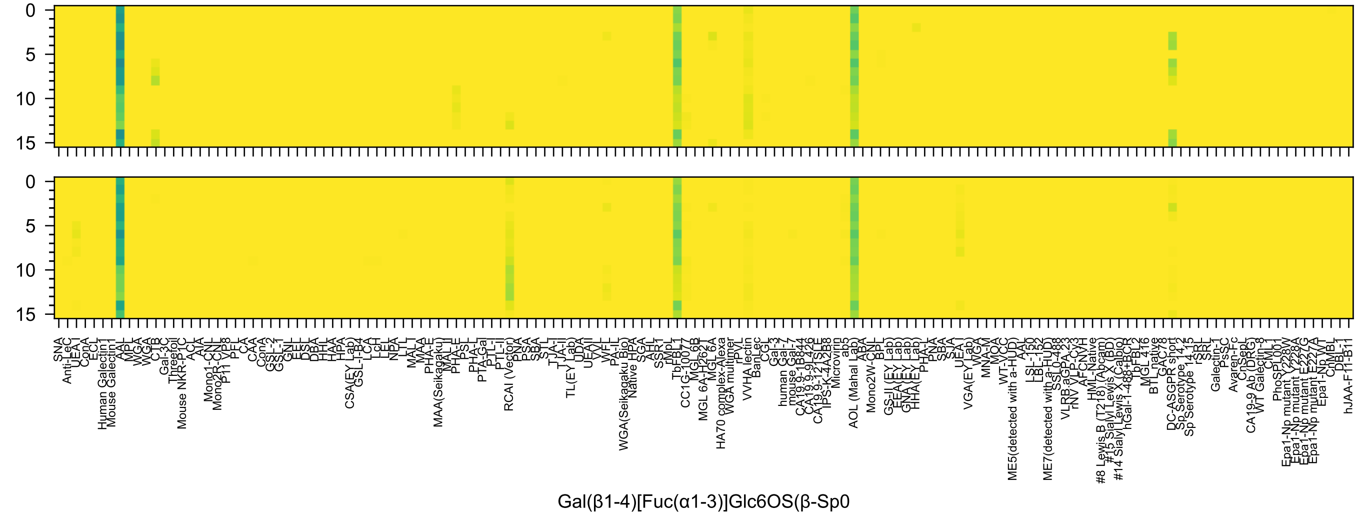
0
5
10
15
Gal(β1-4)[Fuc(α1-3)]Glc6OS(β-Sp0

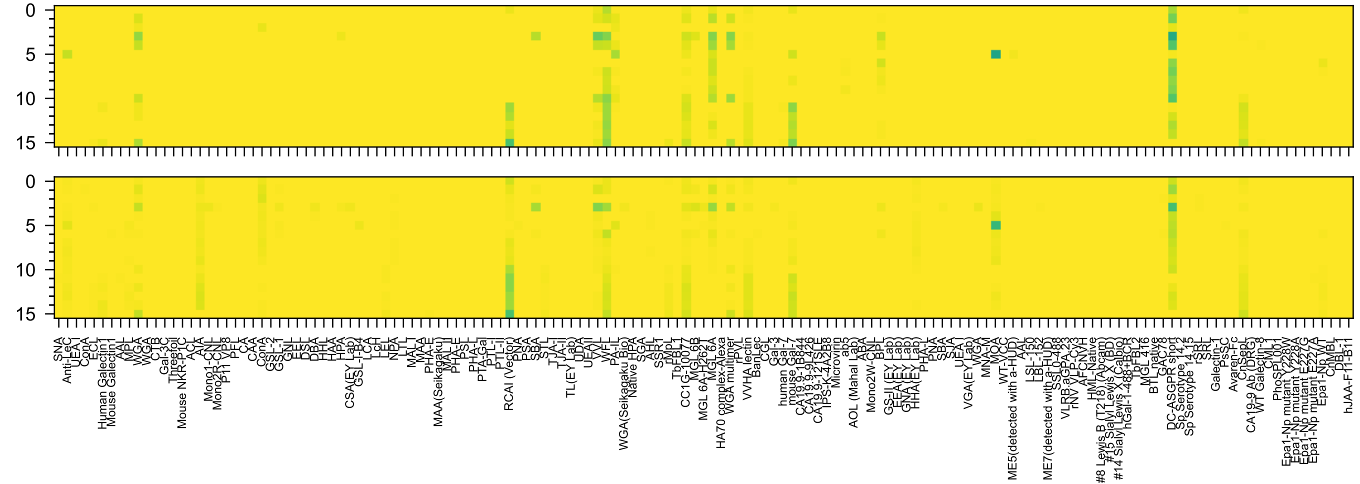
Gal(β1-4)GlcNAc(β1-2)Man(α-Sp0

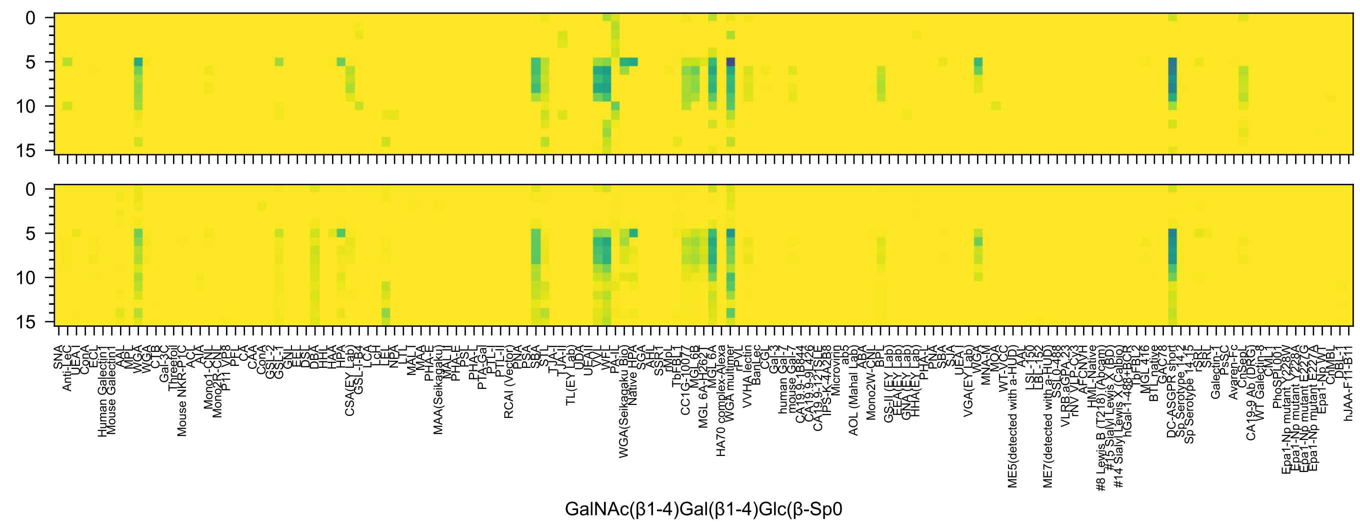
GalNAc(β1-4)Gal(β1-4)Glc(β-Sp0

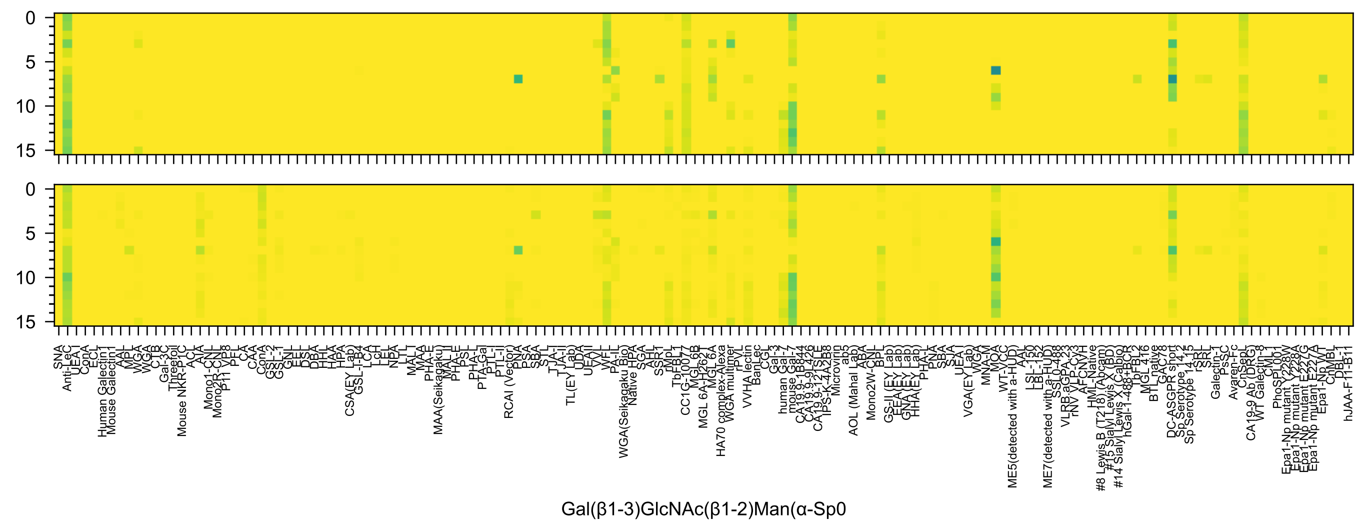
Gal(β1-3)GlcNAc(β1-2)Man(α-Sp0

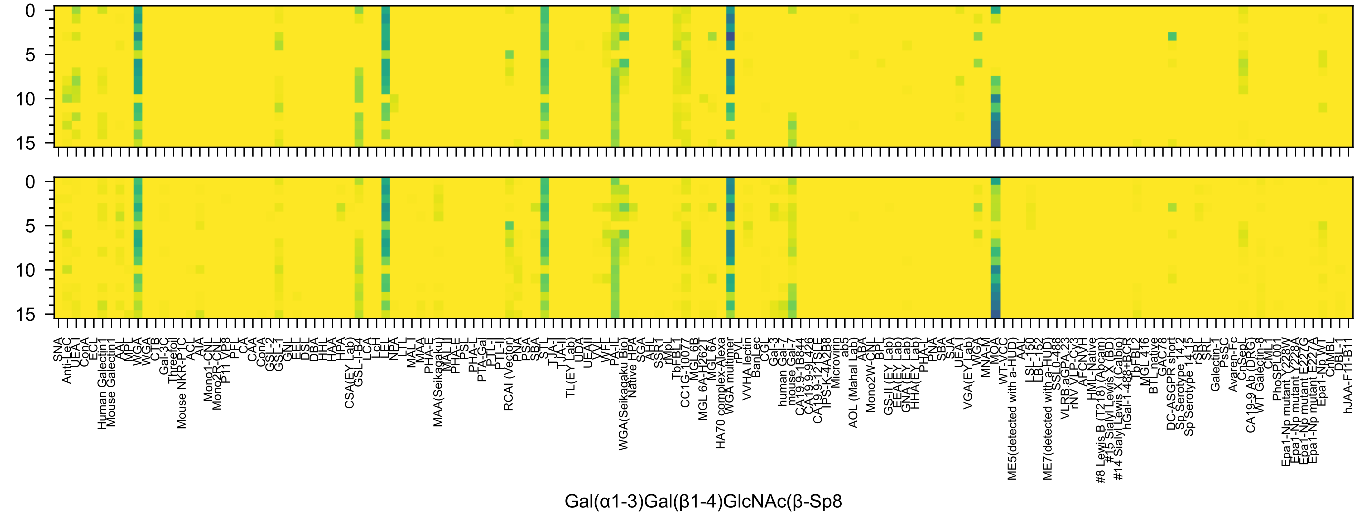
0
5
10
15
0
5
10
15
SNA
Anti-LeC
UEA-I
ConA
ECL
Human Galectin1
Mouse Galectin1
AAL
MPL
WGA
WGA
CTB
Gal-3C
Threefoil
Mouse NKR-P1C
ACL
AIA
Mono1-CNL
Mono2R-CNL
P11 VP8
PFL
CA
CAA
ConA
GSL-2
GSL-1
GNL
EEL
DSL
DBA
HHL
HAA
HPA
CSA(EY Lab)
GSL-I-B4
LCA
LcH
LEL
NPA
LTL
MAL-I
MAA
PHA-E
MAA(Seikagaku)
MAL-II
PHA-E
PSL
PHA-L
PTA-Gal
PTL-I
PTL-II
RCAI (Vector)
PNA
PSA
SBA
STL
TJA-I
TJA-II
TL(EY Lab)
UDA
UEA-II
VVL
WFL
PA-IL
WGA(Seikagaku Bio)
Native HPA
SGA
AHL
SSR1
rMpL
TbFBL1
CC1G-10077
MGL 6B
MGL 6A-H262T
MGL 6A
HA70 complex-Alexa
WGA multimer
rPVL
VVHA lectin
BanLec
CGL
Gal-3
human Gal-7
mouse Gal-7
CA19.9-1B844
CA19.9-9L426
CA19.9-121SLE
IPS-K-4A2B8
Microvirin
ab5
AOL (Mahal Lab)
ABA
Mono2W-CNL
BPL
GS-II (EY Lab)
EEA(EY Lab)
GNA(EY Lab)
HHA(EY Lab)
PHA-L
PNA
SBA
SJA
UEA-I
VGA(EY Lab)
WGA
MNA-M
MOA
WT-VCC
ME5(detected with a-HUD)
AAL
LSL-150
LBL-152
ME7(detected with a-HUD)
SSL0-488
VLRB.aGPA.23
rNV VLP-Cy3
AFCNVH
HML-Native
#8 Lewis B (T218) (Abcam)
#15 Sialyl Lewis X (BD)
#14 Sialyl Lewis X (Calbio)
hGal-1-488+BCR
TbFBL2
MGL 416
BTL native
GAC78
DC-ASGPR short
Sp Serotype 14.2
Sp Serotype 14.15
rSRL
SRL
Galectin-1
PsSC
Avaren-Fc
ChSeL
CA19-9 Ab (DRG)
WT Galectin-8
CML1
PhoSL001
Epa1-Np mutant Y228W
Epa1-Np mutant Y228A
Epa1-Np mutant E227G
Epa1-Np mutant E227A
Epa1-Np WT
ChMBL
DBL-1
hJAA-F11-B11
Gal(α1-3)Gal(β1-4)GlcNAc(β-Sp8

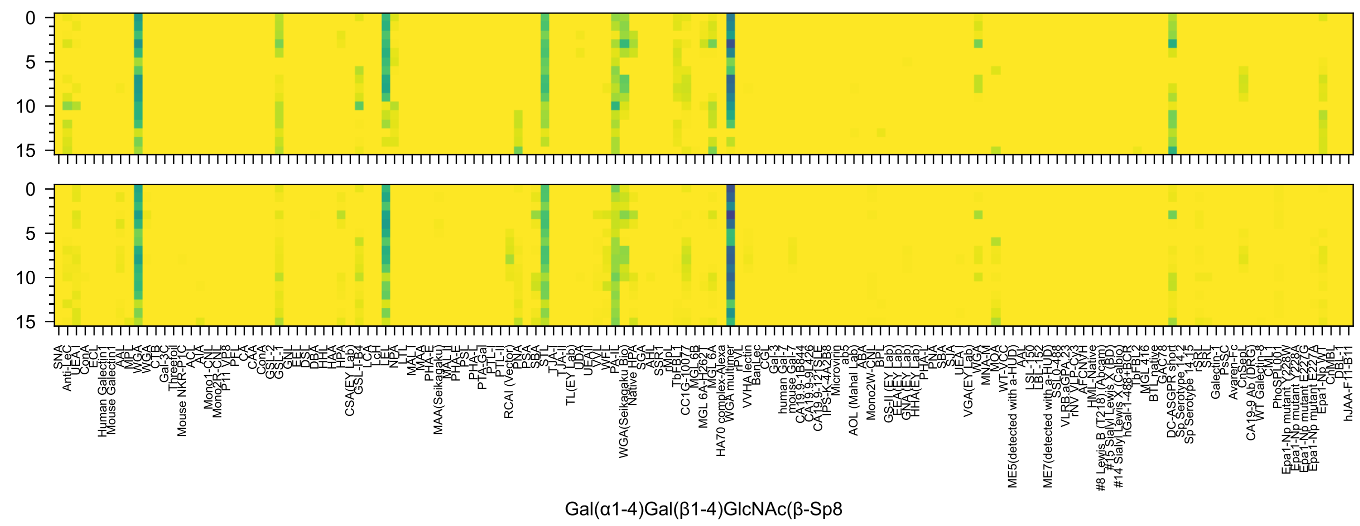
Gal(α1-4)Gal(β1-4)GlcNAc(β-Sp8

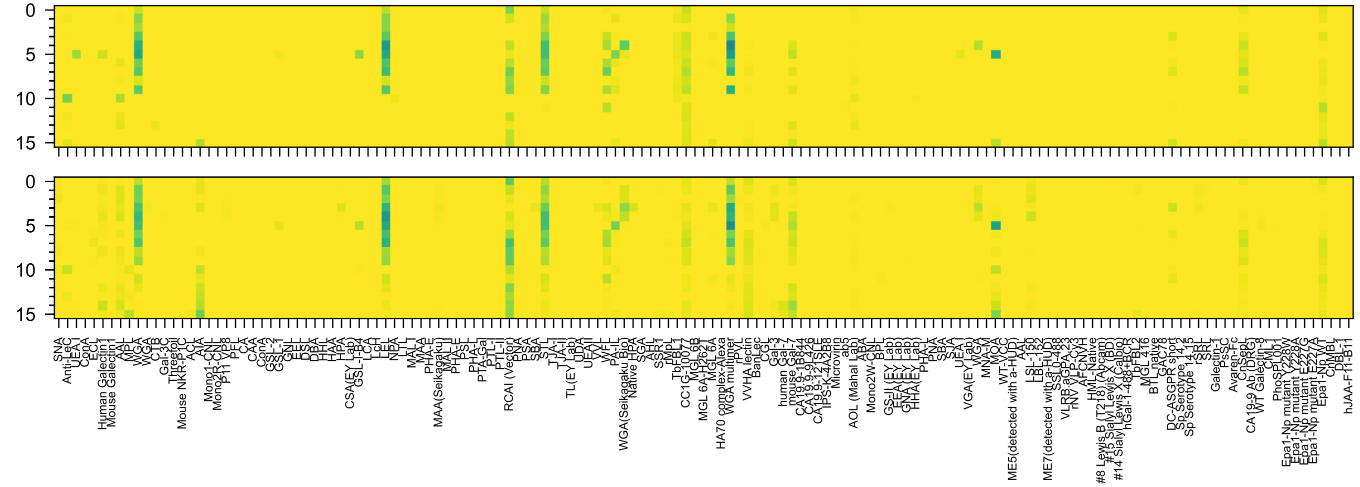


Gal(β1-3)Gal(β1-4)GlcNAc(β-Sp8

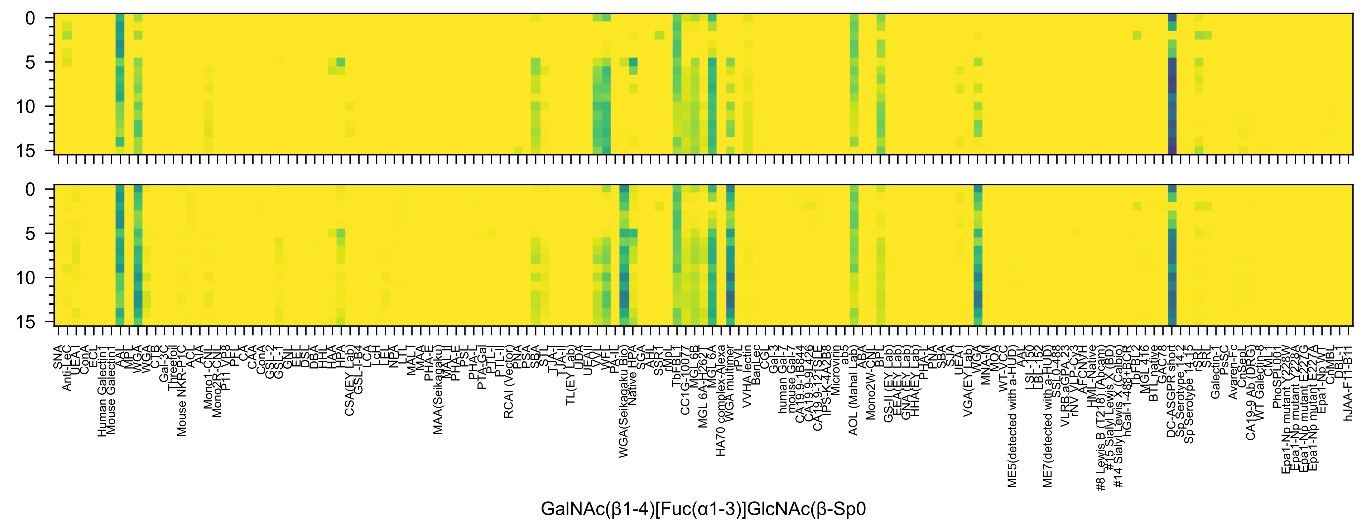
GalNAc(β1-4)[Fuc(α1-3)]GlcNAc(β-Sp0

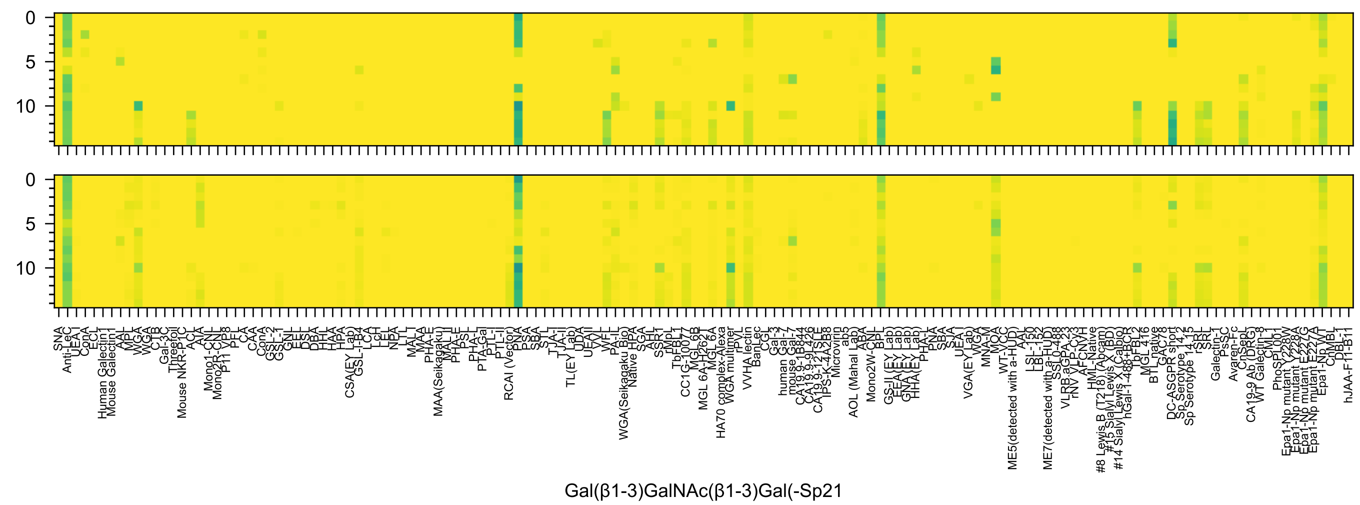
Gal(β1-3)GalNAc(β1-3)Gal(-Sp21

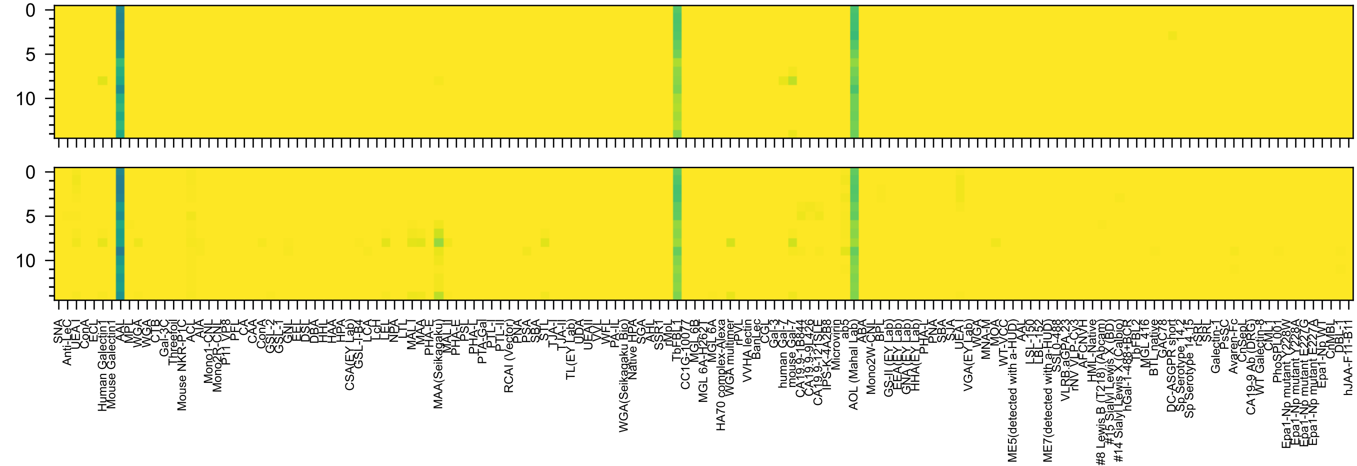


Gal3OS(β1-4)[Fuc(α1-3)]GlcNAc(-Sp0

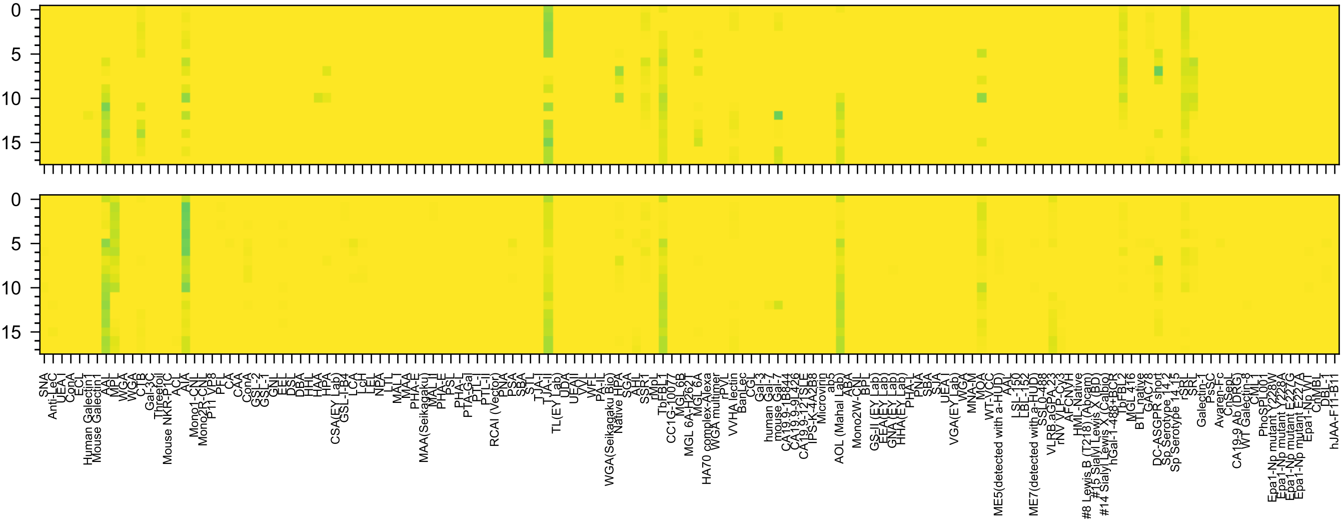

Fuc(α1-2)Gal(β1-3)GalNAc(α-Sp14

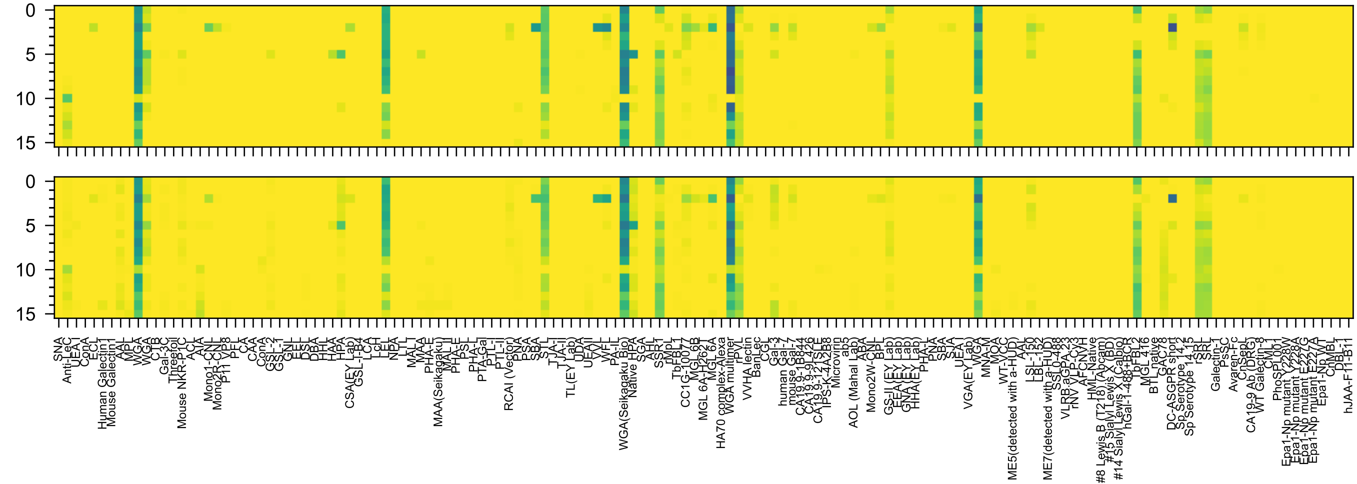

GlcNAc(β1-3)Gal(β1-4)GlcNAc(β-Sp0

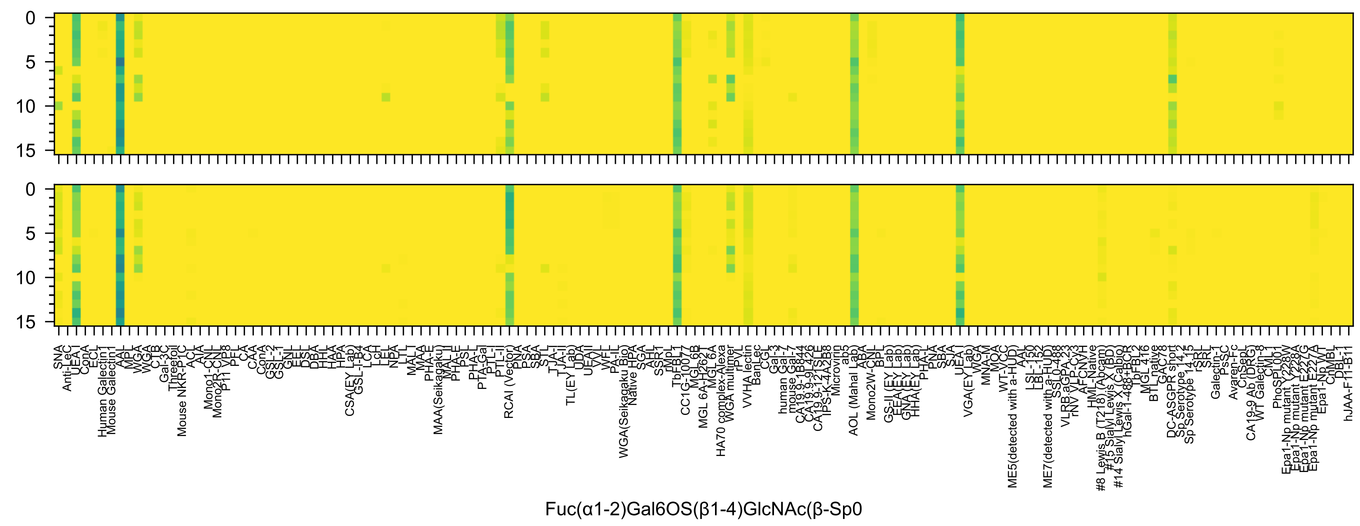
Fuc(α1-2)Gal6OS(β1-4)GlcNAc(β-Sp0)

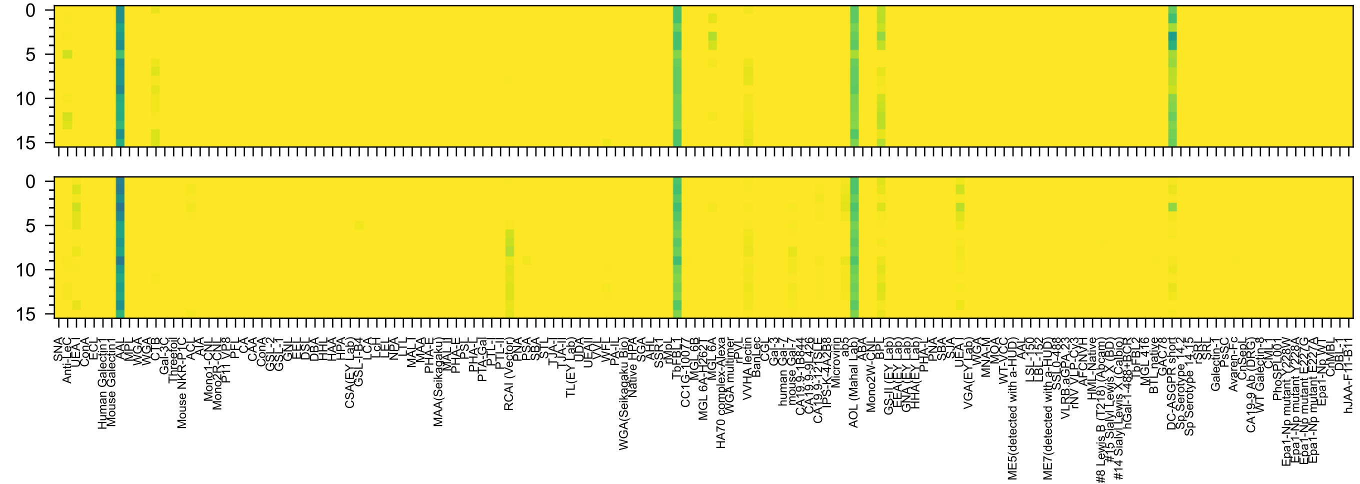


Gal(β1-4)[Fuc(α1-3)]GlcNAc6OS(β-Sp0

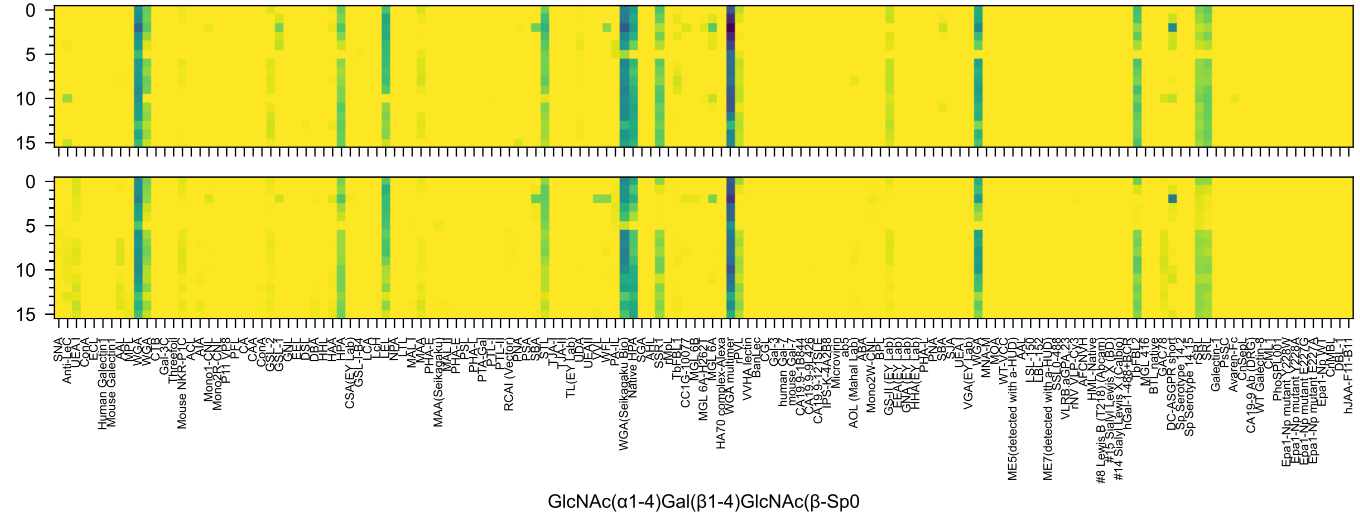

GlcNAc(α1-4)Gal(β1-4)GlcNAc(β-Sp0

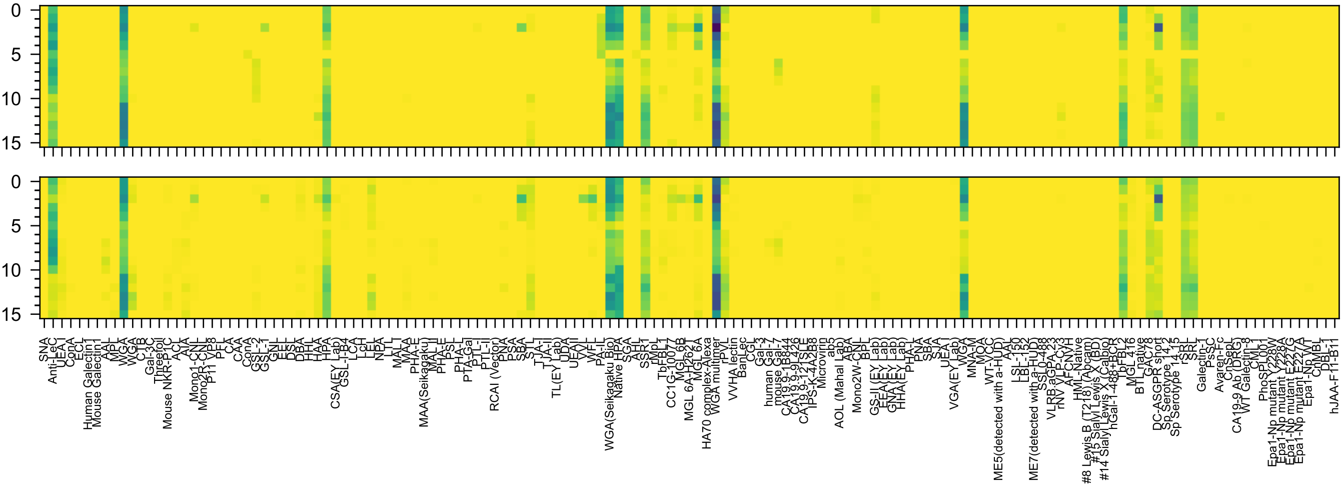


GlcNAc(α1-4)Gal(β1-3)GlcNAc(β-Sp0

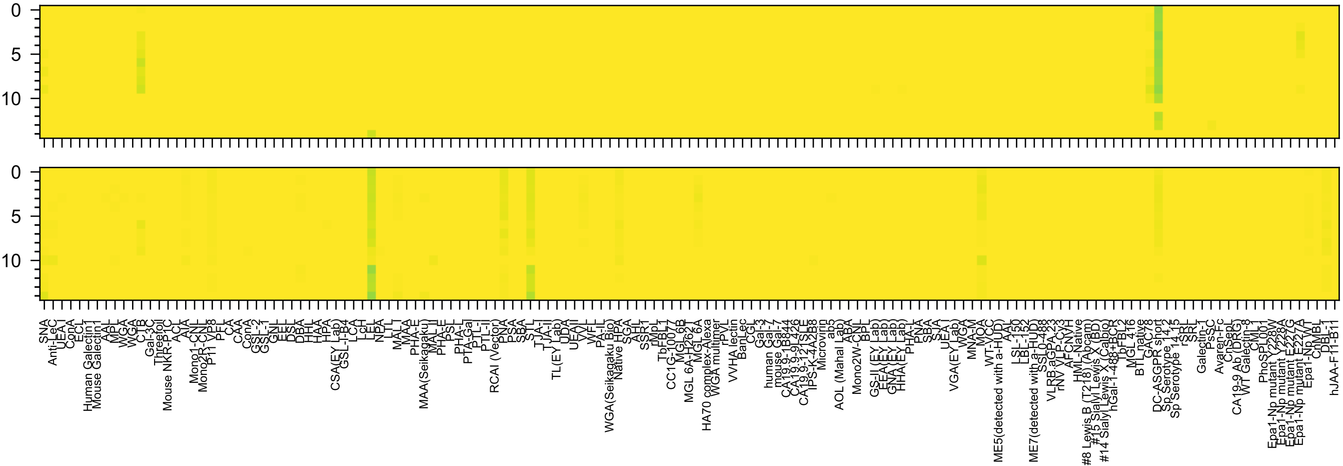

Kdn(α2-3)Gal(β1-4)Glc(-Sp0

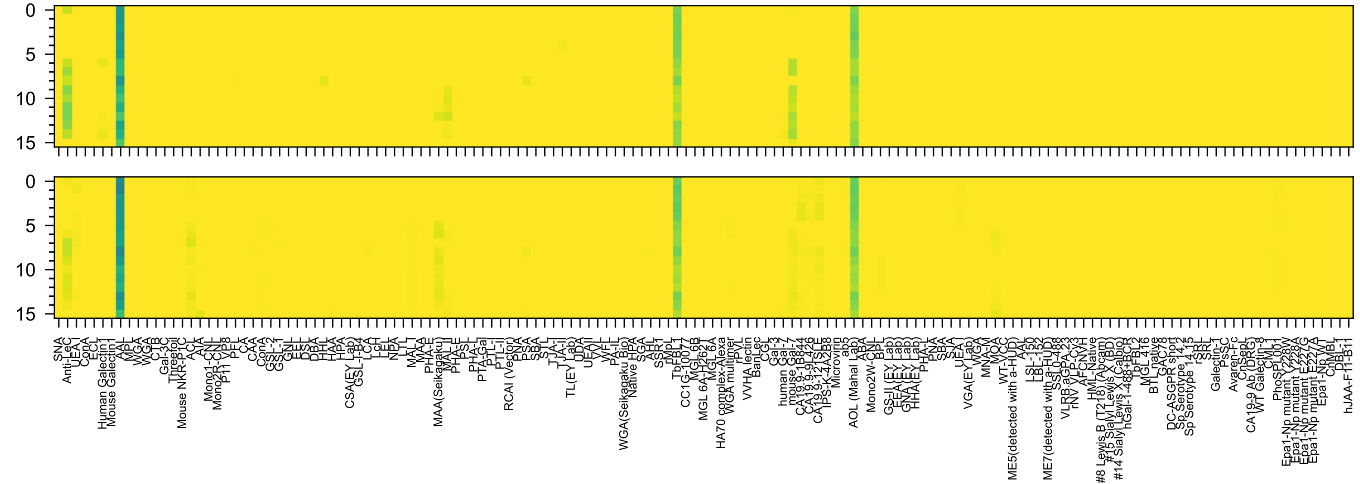

Gal3OS(β1-3)[Fuc(α1-4)]GlcNAc(β-Sp0

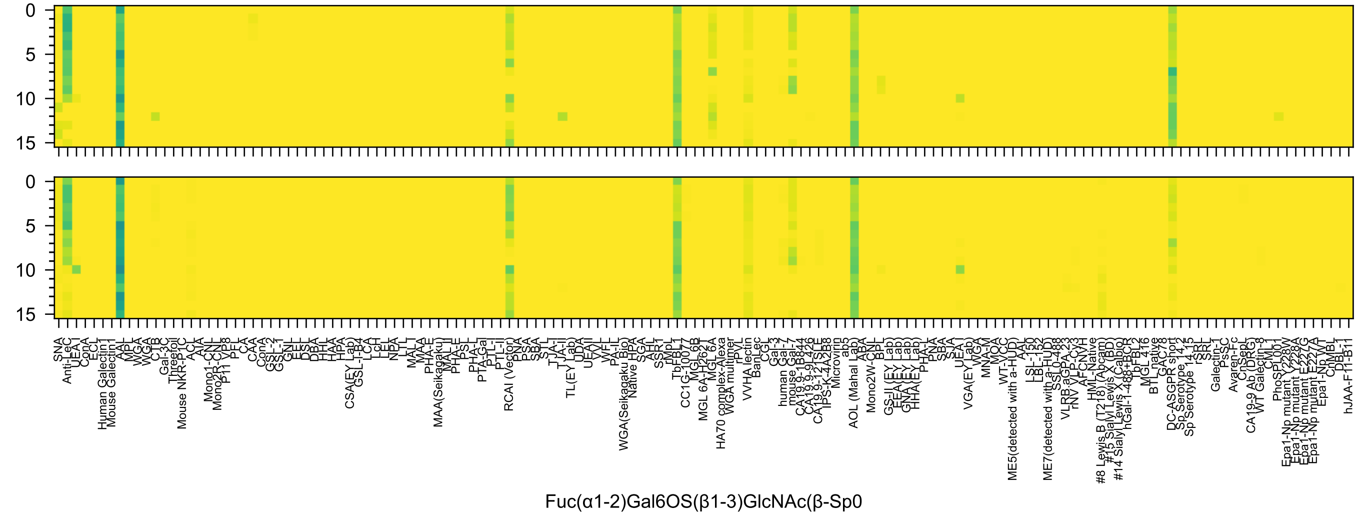
0
5
10
15
0
5
10
15
SNA
Anti-LeC
UEA I
ConA
ECL
Human Galectin1
Mouse Galectin1
AAL
MPL
WGA
WGA
CTB
Gal-3C
Threefoil
Mouse NKR-P1C
ACL
AIA
Mono1-CNL
Mono2R-CNL
P11 VP8
PFL
CA
CAA
ConA
GSL-2
GSL-1
GNL
EEL
DSL
DBA
HHL
HAA
HPA
CSA(EY Lab)
GSL-I-B4
LCA
LcH
LEL
NPA
LTL
MAL I
MAA
PHA-E
MAA(Seikagaku)
MAL II
PHA-E
PSL
PHA-L
PTA-Gal
PTL-I
PTL-II
RCAI (Vector)
PNA
PSA
SBA
STL
TJA-I
TJA-II
TL(EY Lab)
UDA
UEA II
VVL
WFL
PA-IL
WGA(Seikagaku Bio)
Native HPA
SGA
AHL
SSR1
rMpL
TbFBL1
CC1G-10077
MGL 6B
MGL 6A-H262T
MGL 6A
HA70 complex-Alexa
WGA multimer
rPVL
VVHA lectin
BanLec
CGL
Gal-3
human Gal-7
mouse Gal-7
CA19.9-1B844
CA19.9-9L426
CA19.9-121SLE
IPS-K-4A2B8
Microvirin
ab5
AOL (Mahal Lab)
ABA
Mono2W-CNL
BPL
GS-II (EY Lab)
EEA(EY Lab)
GNA(EY Lab)
HHA(EY Lab)
PHA-L
PNA
SBA
SJA
UEA I
VGA(EY Lab)
WGA
MNA-M
MOA
WT-VCC
ME5(detected with a-HUD)
AAL
LSL-150
LBL-152
ME7(detected with a-HUD)
SSL0-488
VLRB.aGPA.23
rNV VLP-Cy3
AFCNVH
HML-Native
#8 Lewis B (T218) (Abcam)
#15 Sialyl Lewis X (BD)
#14 Sialyl Lewis X (Calbio)
hGal-1-488+BCR
TbFBL2
MGL 416
BTL native
GAC78
DC-ASGPR short
Sp Serotype 14.2
Sp Serotype 14.15
rSRL
SRL
Galectin-1
PsSC
Avaren-Fc
CnSepL
CA19-9 Ab (DRG)
WT Galectin-8
CML1
PhoSPL001
Epa1-Np mutant Y228W
Epa1-Np mutant Y228A
Epa1-Np mutant E227G
Epa1-Np mutant E227A
Epa1-Np WT
CnMBL
DBL-1
hJAA-F11-B11
Fuc(α1-2)Gal6OS(β1-3)GlcNAc(β-Sp0

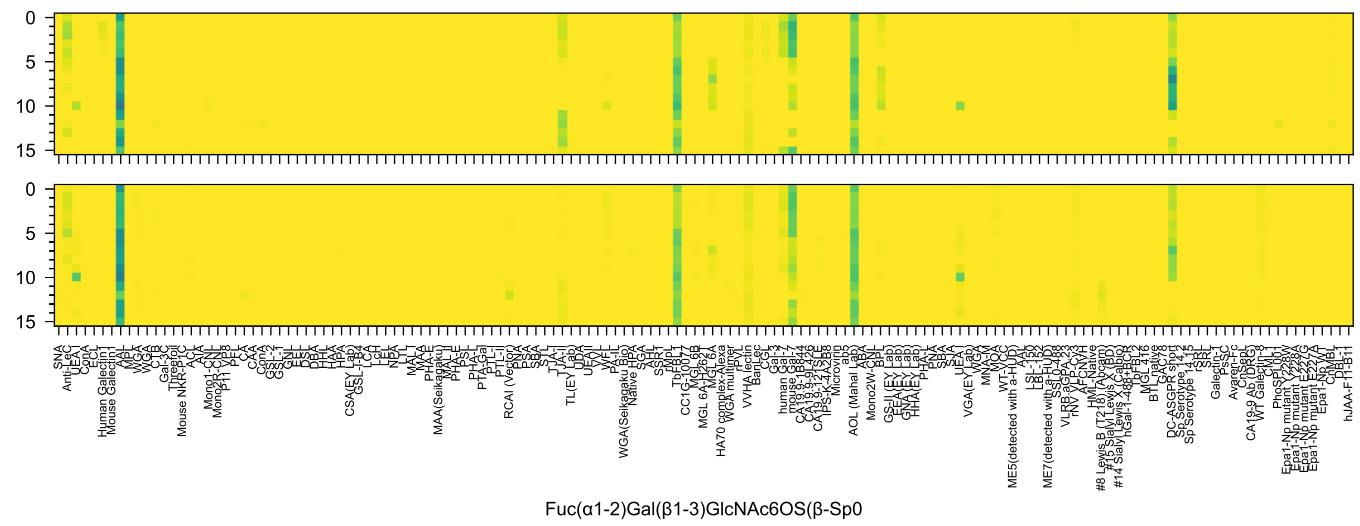
Fuc(α1-2)Gal(β1-3)GlcNAc6OS(β-Sp0

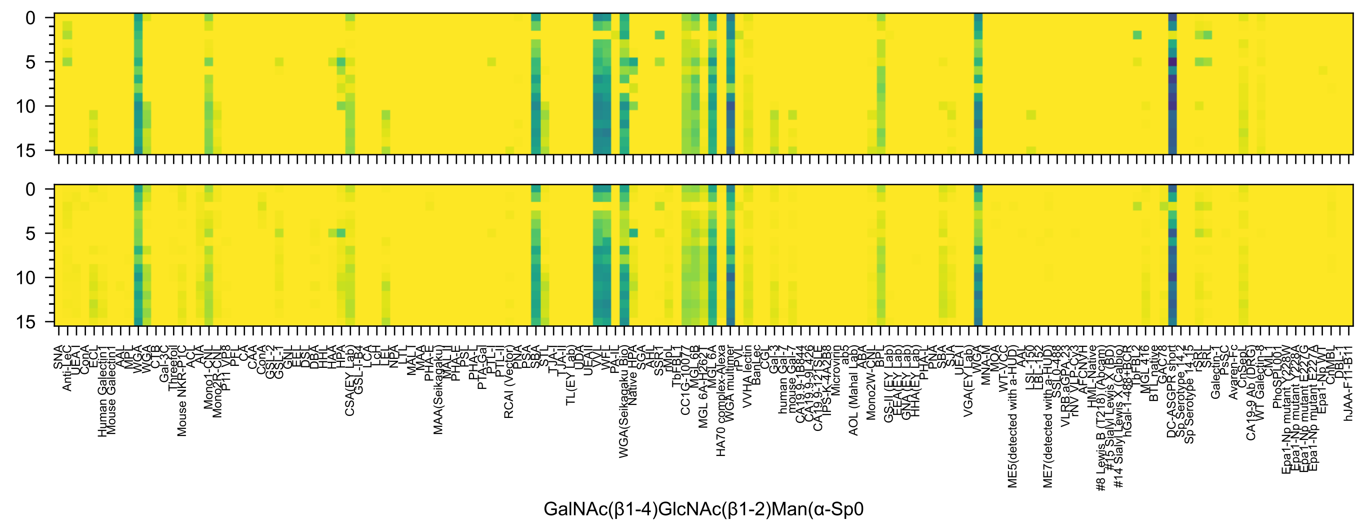
0
5
10
15
SNA
Anti-LeC
UEA-I
ConA
ECL
Human Galectin1
Mouse Galectin1
AAL
MPL
WGA
WGA
CTB
Gal-3C
Threefoil
Mouse NKR-P1C
ACL
AIA
Mono1-CNL
Mono2R-CNL
P11 VP8
PFL
CA
CAA
ConA
GSL-2
GSL-1
GNL
EEL
DSL
DBA
HHL
HAA
HPA
CSA(EY Lab)
GSL-I-B4
LCA
LcH
LEL
NPA
LTL
MAL-I
MAA
PHA-E
MAA(Seikagaku)
MAL-II
PHA-E
PSL
PHA-L
PTA-Gal
PTL-I
PTL-II
RCAI (Vector)
PNA
PSA
SBA
STL
TJA-I
TJA-II
TL(EY Lab)
UDA
UEAII
VVL
WFL
PA-IL
WGA(Seikagaku Bio)
Native HPA
SGA
AHL
SSR1
rMpL
TbFBL1
CC1G-10077
MGL 6B
MGL 6A-H262T
MGL 6A
HA70 complex-Alexa
WGA multimer
rPVL
VVHA lectin
BanLec
CGL
Gal-3
human Gal-7
mouse Gal-7
CA19.9-1B844
CA19.9-9L426
CA19.9-121SLE
IPS-K-4A2B8
Microvirin
ab5
AOL (Mahal Lab)
ABA
Mono2W-CNL
BPL
GS-II (EY Lab)
EEA (EY Lab)
GNA (EY Lab)
HHA (EY Lab)
PHA-L
PNA
SBA
SJA
UEA-I
VGA(EY Lab)
WGA
MNA-M
MOA
WT-VCC
ME5(detected with a-HUD)
AAL
LSL-150
LBL-152
ME7(detected with a-HUD)
SSL0-488
VLRB.aGPA.23
rNV VLP-Cy3
AFCNVH
HML-Native
#8 Lewis B (T218) (Abcam)
#15 Sialyl Lewis X (BD)
#14 Sialyl Lewis X (Calbio)
hGal-1-488+BCR
TbFBL2
MGL 416
BTL native
GAC78
DC-ASGPR short
Sp Serotype 14.2
Sp Serotype 14.15
rSRL
SRL
Galectin-1
PsSC
Avaren-Fc
ChSepL
CA19-9 Ab (DRG)
WT Galectin-8
CML1
PhoSPL001
Epa1-Np mutant Y228W
Epa1-Np mutant Y228A
Epa1-Np mutant E227G
Epa1-Np mutant E227A
Epa1-Np WT
ChMBL
DBL-1
hJAA-F11-B11
GalNAc(β1-4)GlcNAc(β1-2)Man(α-Sp0

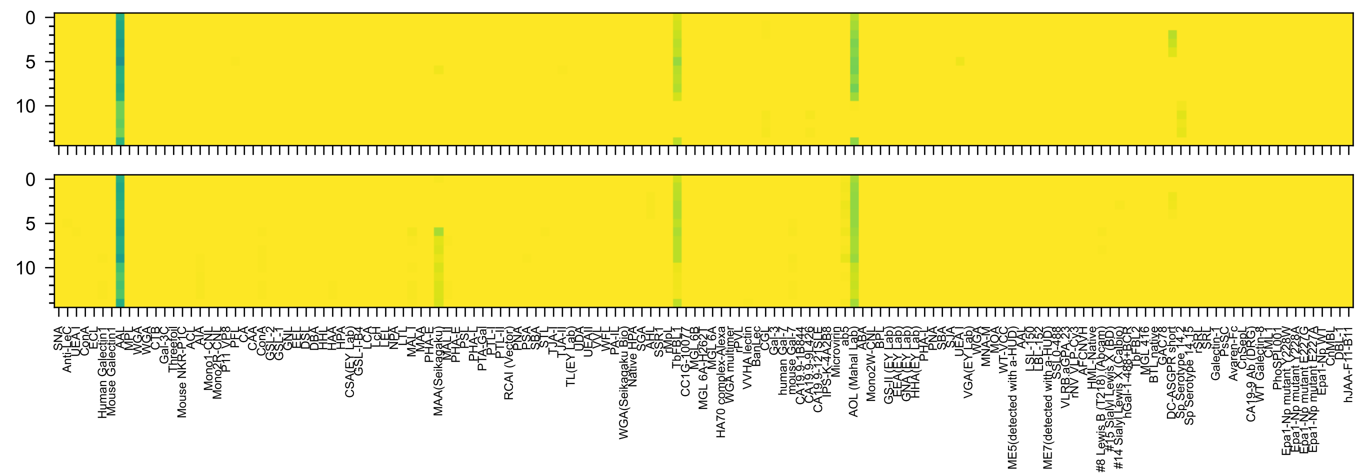


Gal3OS(β1-4)[Fuc(α1-3)]Glc6OS(-Sp0

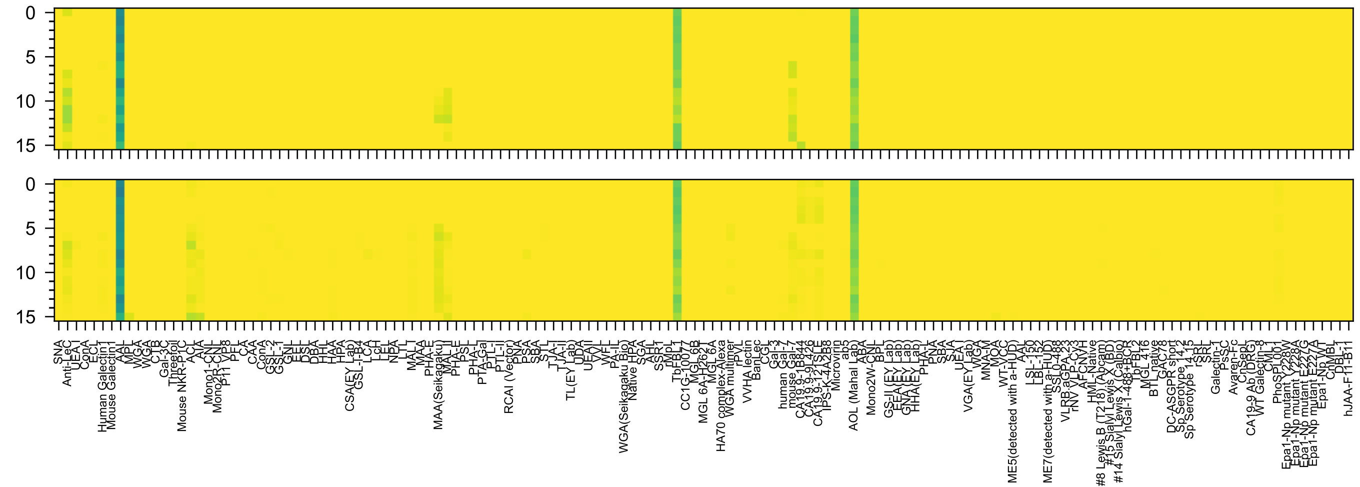

Gal3OS(β1-3)[Fuc(α1-4)]GlcNAc(β-Sp8

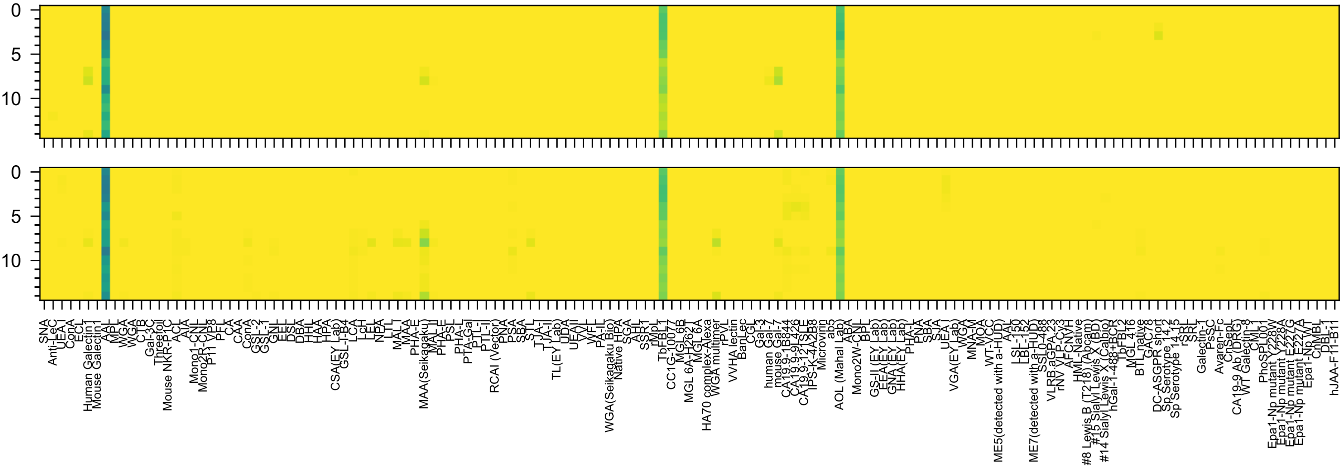

Gal3OS(β1-4)[Fuc(α1-3)]GlcNAc(-Sp8

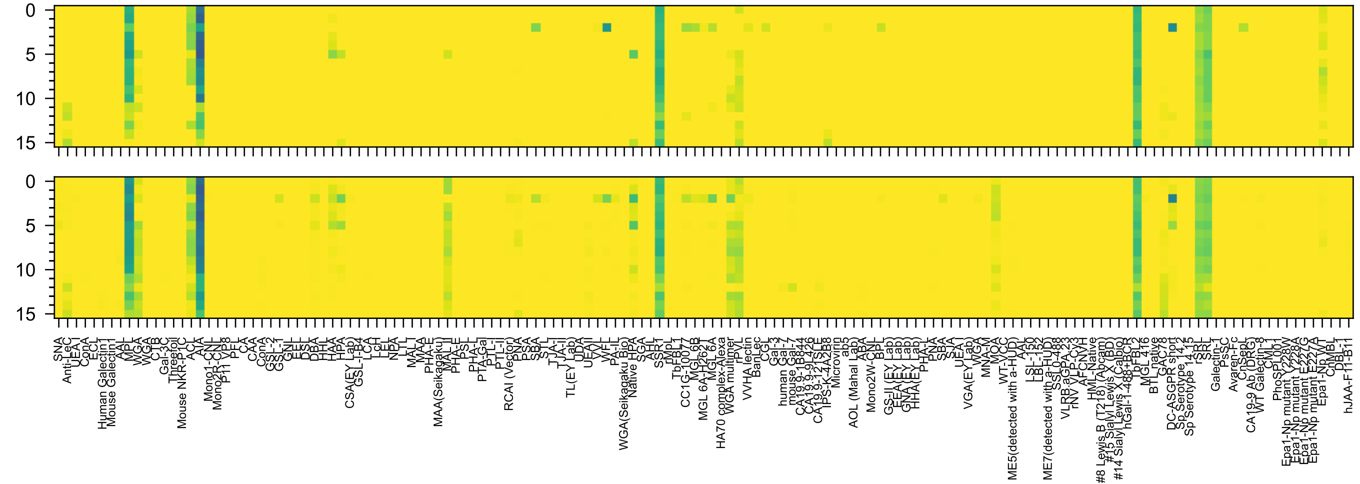

GlcNAc(β1-3)Gal(β1-3)GalNAc(α-Sp8

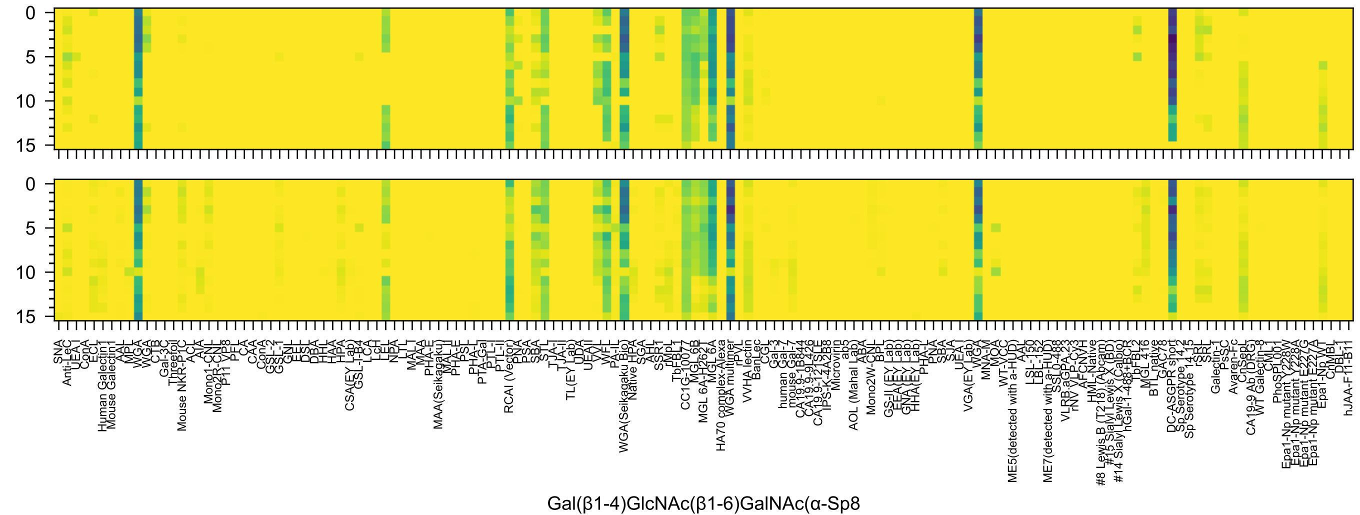
0
5
10
15
0
5
10
15
SNA
Anti-LeC
UEA I
ConA
ECL
Human Galectin1
Mouse Galectin1
AAL
MPL
WGA
WGA
CTB
C-1B
Gal-3C
Threefoil
Mouse NKR-P1C
ACL
AIA
Mono1-CNL
Mono2R-CNL
P11 VP8
PFL
CA
CAA
ConA
GSL-2
GSL-1
GNL
EEL
DSL
DBA
HHL
HAA
HPA
CSA(EY Lab)
GSL-I-B4
LCA
LcH
LEL
NPA
LTL
MAL
MAA
PHA-E
MAA(Seikagaku)
MAL-II
PHA-E
PSL
PHA-L
PTA-Gal
PTL-I
PTL-II
RCAI (Vector)
PNA
PSA
SBA
STL
TJA-I
TJA-II
TL(EY Lab)
UDA
UEAII
VVL
WFL
PA-IL
WGA(Seikagaku Bio)
Native HPA
SGA
AAL
SSR1
rMpL
TbFBL1
CC1G-10077
MGL 6B
MGL 6A-H262T
MGL 6A
HA70 complex-Alexa
WGA multimer
rPVL
VVHA lectin
BanLec
CGL
Gal-3
human Gal-7
mouse Gal-7
CA19.9-1B844
CA19.9-9L426
CA19.9-121SLE
IPS-K-4A2B8
Microvirin
ab5
AOL (Mahal Lab)
ABA
Mono2W-CNL
BPL
GS-II (EY Lab)
EEA(EY Lab)
GNA(EY Lab)
HHA(EY Lab)
PHA-L
PNA
SBA
SJA
UEA-I
VGA(EY Lab)
WGA
MNA-M
MOA
WT-VCC
ME5(detected with a-HUD)
AAL
LSL-150
LBL-152
ME7(detected with a-HUD)
SSL0.488
VLRB.aGPA.23
rNV VLP-Cy3
AFCNVH
HML-Native
#8 Lewis B (T218) (Abcam)
#15 Sialyl Lewis X (BD)
#14 Sialyl Lewis X (Calbio)
hGal-1-488+BCR
TbFBL2
MGL 416
BTL native
GAC78
DC-ASGPR short
Sp Serotype 14.2
Sp Serotype 14.15
rSRL
SRL
Galectin-1
PsSC
Avaren-Fc
ChSeP1
CA19-9 Ab (DRG)
WT Galectin-8
CML1
PhoSL001
Epa1-Np mutant Y228W
Epa1-Np mutant Y228A
Epa1-Np mutant E227G
Epa1-Np mutant E227A
Epa1-Np WT
ChMBL
DBL-1
hJAA-F11-B11
Gal(β1-4)GlcNAc(β1-6)GalNAc(α-Sp8

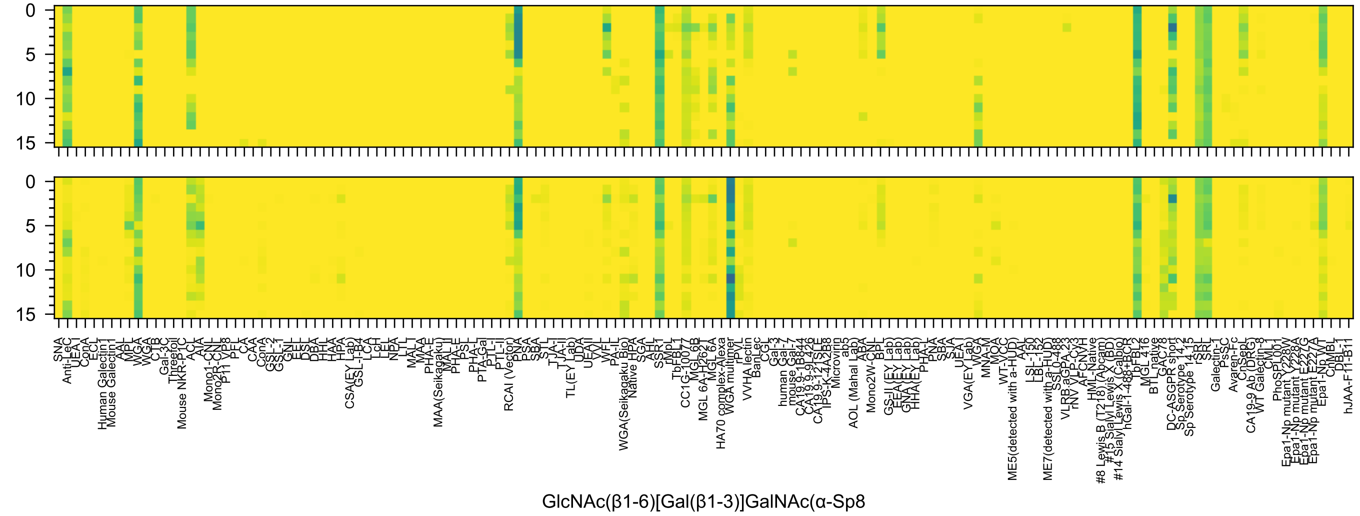
0
5
10
15
0
5
10
15
SNA
Anti-LeC
UEA I
ConA
ECL
Human Galectin1
Mouse Galectin1
AAL
MPL
WGA
WGA
CTB
Gal-3C
Threefoil
Mouse NKR-P1C
ACL
AIA
Mono1-CNL
Mono2R-CNL
P11 VP8
PFL
CA
CAA
ConA
GSL-2
GSL-1
GNL
EEL
DSL
DBA
HHL
HAA
HPA
CSA(EY Lab)
GSL-I-B4
LCA
LcH
LEL
NPA
LTL
MAL
MAA
PHA-E
MAA(Seikagaku)
MAL II
PHA-E
PSL
PHA-L
PTA-Gal
PTL-I
PTL-II
RCAI (Vector)
PNA
PSA
SBA
STL
TJA-I
TJA-II
TL(EY Lab)
UDA
UEA II
VVL
WFL
PA-IL
WGA(Seikagaku Bio)
Native HPA
SGA
AAL
SSR1
rMpL
TbFBL1
CC1G-10077
MGL 6B
MGL 6A-H262T
MGL 6A
HA70 complex-Alexa
WGA multimer
rPVL
VVHA lectin
BanLec
CGL
Gal-3
human Gal-7
mouse Gal-7
CA19.9-1B844
CA19.9-9L426
CA19.9-121SLE
IPS-K-4A2B8
Microvirin
ab5
AOL (Mahal Lab)
ABA
Mono2W-CNL
BPL
GS-II (EY Lab)
EEA(EY Lab)
GNA(EY Lab)
HHA(EY Lab)
PHA-L
PNA
SBA
SJA
UEA I
VGA(EY Lab)
WGA
MNA-M
MOA
WT-VCC
ME5(detected with a-HUD)
AAL
LSL-150
LBL-152
ME7(detected with a-HUD)
SSL0-488
VLRB.aGPA.23
rNV VLP-Cy3
AFCNVH
HML-Native
#8 Lewis B (T218) (Abcam)
#15 Sialyl Lewis X (BD)
#14 Sialyl Lewis X (Calbio)
hGal-1-488+BCR
TbFBL2
MGL 416
BTL native
GAC78
DC-ASGPR short
Sp Serotype 14 15
Sp Serotype 14 15
rSRL
SRL
Galectin-1
PsSC
Avaren-Fc
ChSeoL
CA19-9 Ab (DRG)
WT Galectin-8
CML1
PhoSL001
Epa1-Np mutant Y228W
Epa1-Np mutant Y228A
Epa1-Np mutant E227G
Epa1-Np mutant E227A
Epa1-Np WT
ChMBL
DBL-1
hJAA-F11-B11
GlcNAc(β1-6)[Gal(β1-3)]GalNAc(α-Sp8

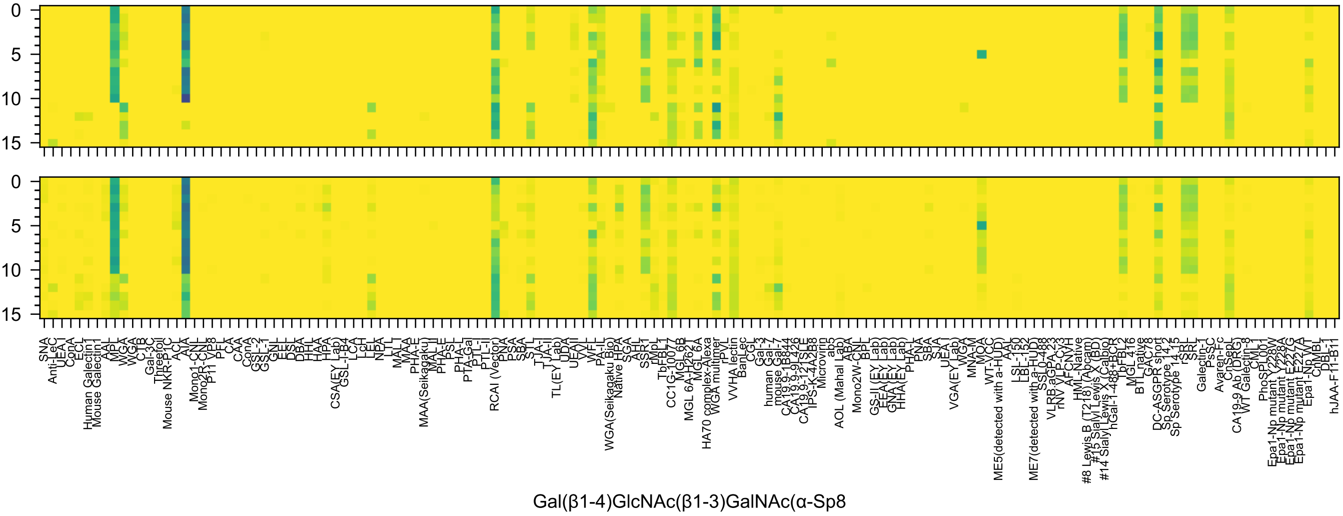
Gal(β1-4)GlcNAc(β1-3)GalNAc(α-Sp8

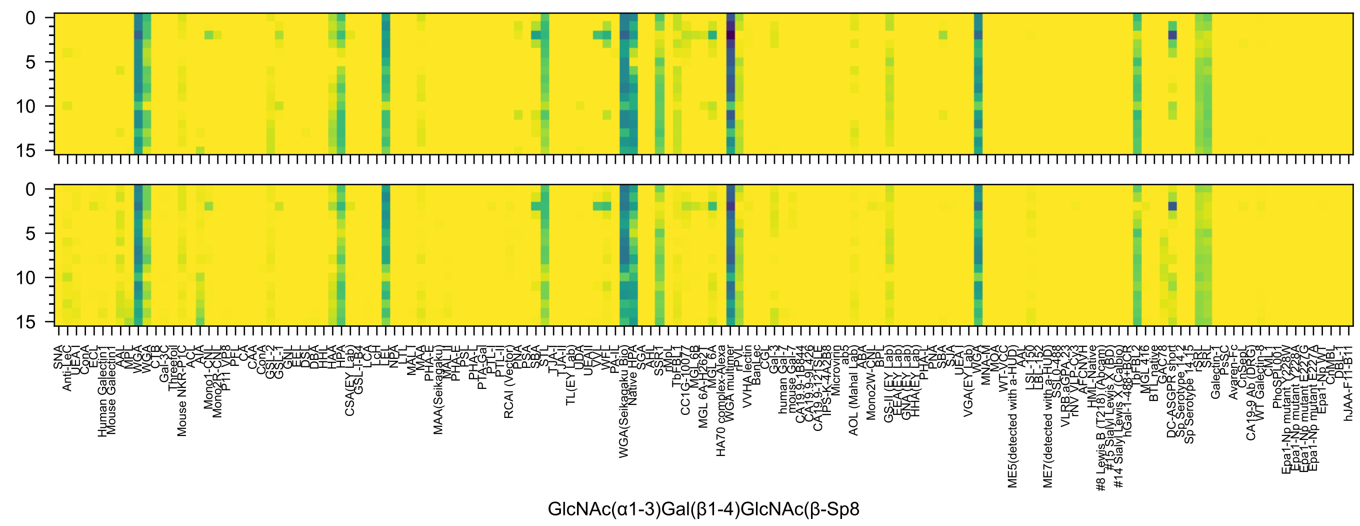

GlcNAc(α1-3)Gal(β1-4)GlcNAc(β-Sp8

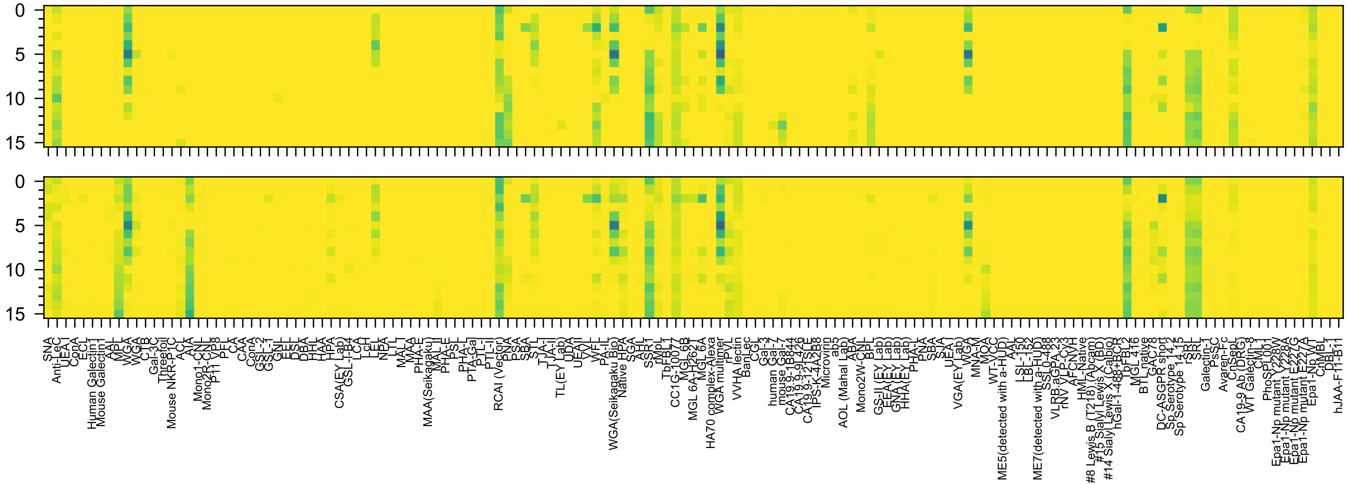


GlcNAc(α1-6)Gal(β1-4)GlcNAc(β-Sp8

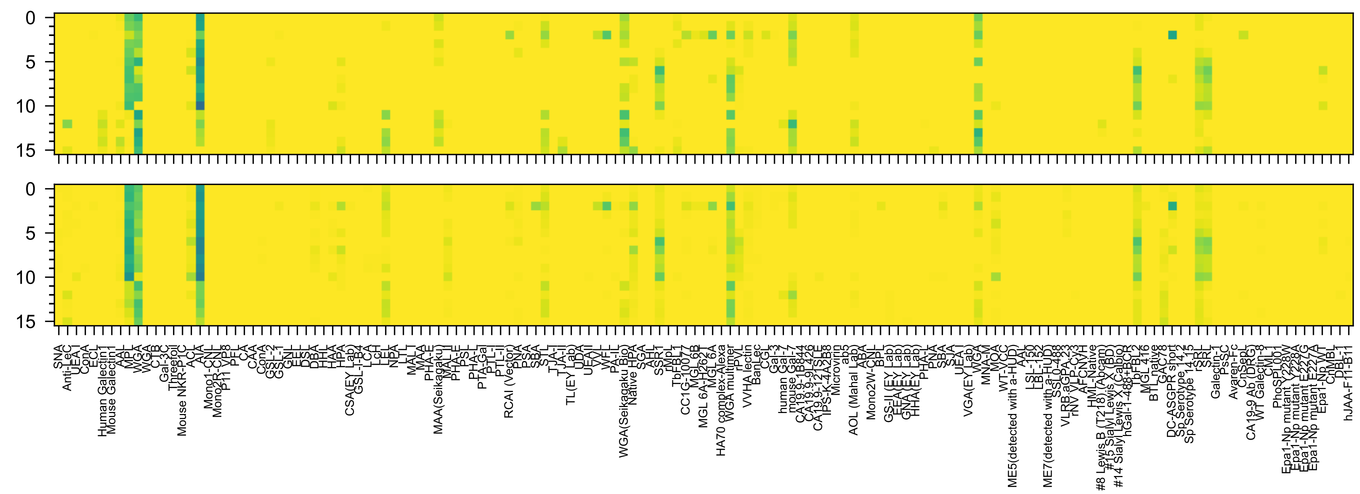

GlcNAc(β1-2)Gal(β1-3)GalNAc(α-Sp8

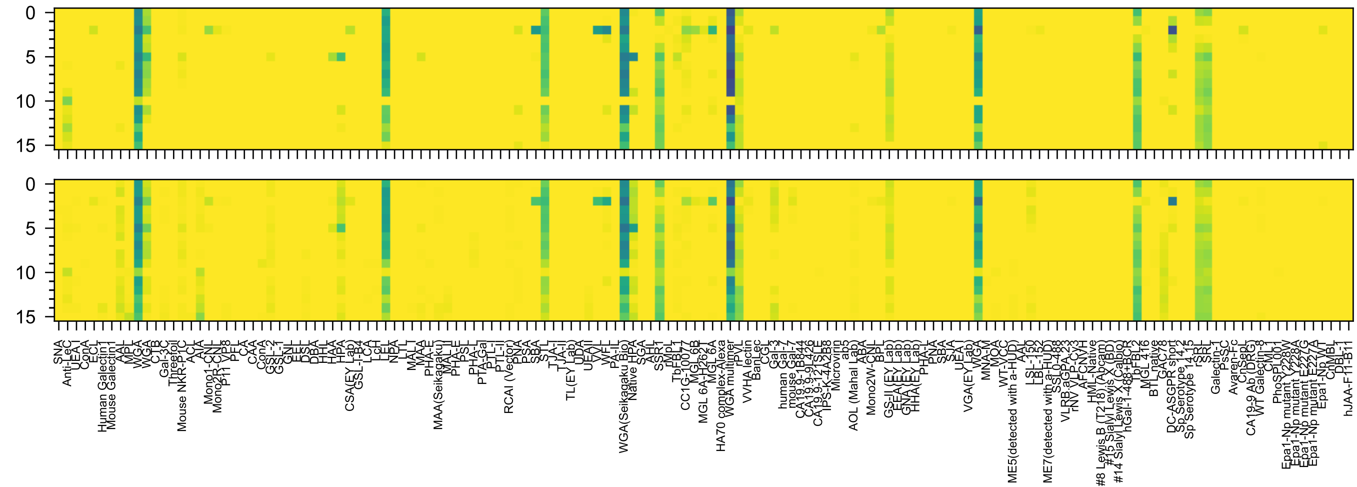

GlcNAc(β1-3)Gal(β1-4)GlcNAc(β-Sp8)

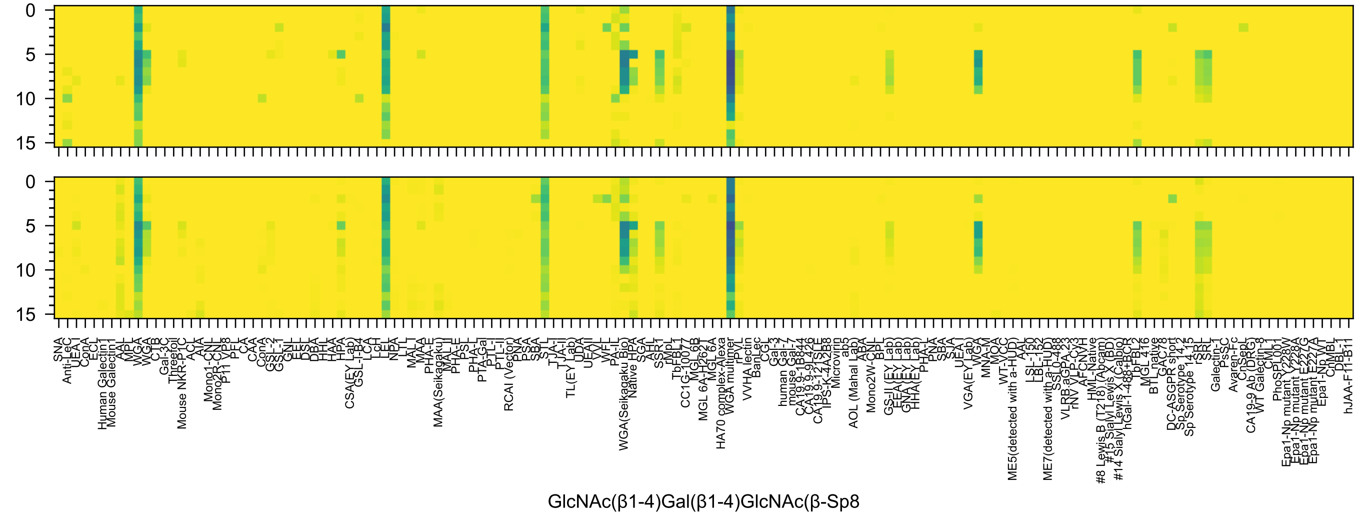
GlcNAc(β1-4)Gal(β1-4)GlcNAc(β-Sp8

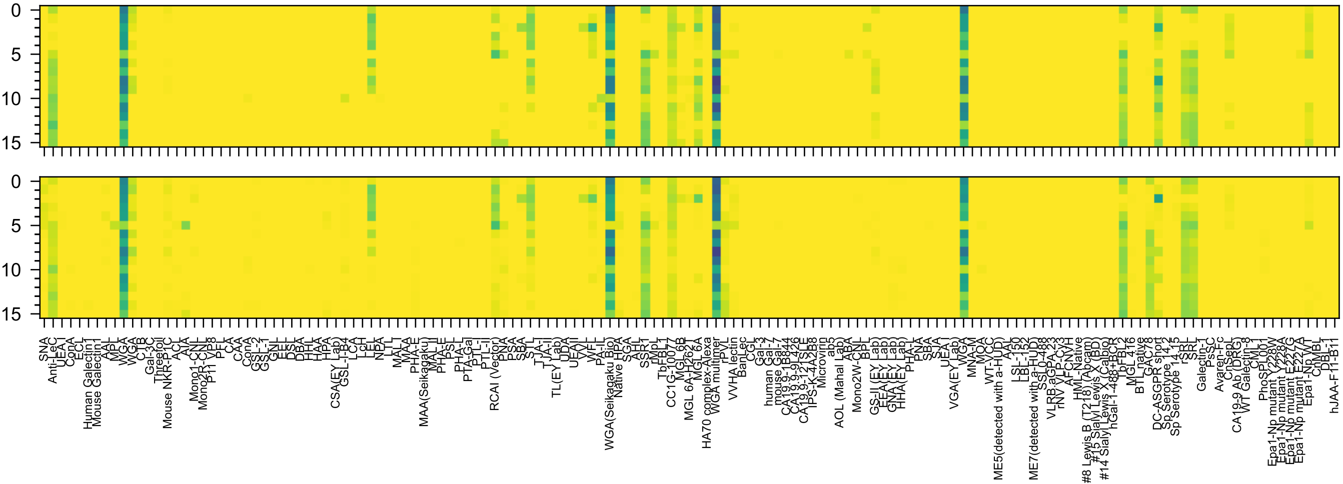

GlcNAc(β1-6)Gal(β1-4)GlcNAc(β-Sp8

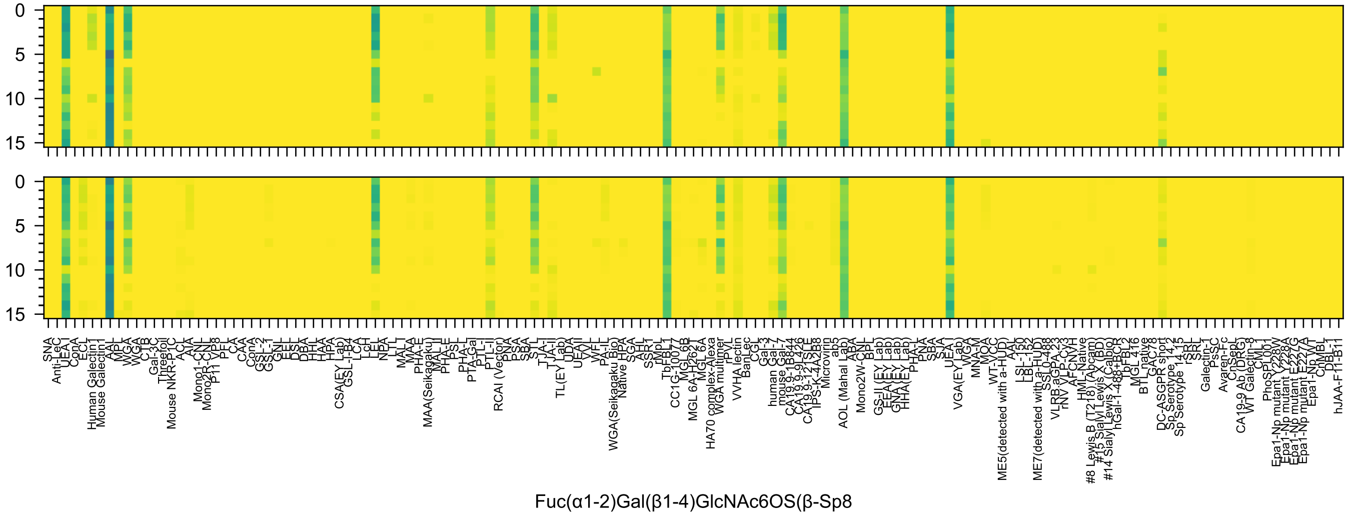
0
5
10
15
0
5
10
15
SNA
Anti-LeC
UEA I
ConA
ECL
Human Galectin1
Mouse Galectin1
AAL
MPL
WGA
WGA
CTB
C
Gal-3C
Threefoil
Mouse NKR-P1C
ACL
AIA
Mono1-CNL
Mono2R-CNL
P11 VP8
PFL
CA
CAA
ConA
GSL-2
GSL-1
GNL
EEL
DSL
DBA
HHL
HAA
HPA
CSA(EY Lab)
GSL-I-B4
LCA
LcH
LEL
NPA
LTL
MAL
MAA
PHA-E
MAA(Seikagaku)
MAL II
PHA-E
PSL
PHA-L
PTA-Gal
PTL-I
PTL-II
RCAI (Vector)
PNA
PSA
SBA
STL
TJA-I
TJA-II
TL(EY Lab)
UDA
UEA II
VVL
WFL
PA-IL
WGA(Seikagaku Bio)
Native HPA
SGA
AHL
SSR1
rMpL
TbFBL1
CC1G-10077
MGL 6B
MGL 6A-H262T
MGL 6A
HA70 complex-Alexa
WGA multimer
rPVL
VVHA lectin
BanLec
CGL
Gal-3
human Gal-7
mouse Gal-7
CA19.9-1B844
CA19.9-9L426
CA19.9-121SLE
IPS-K-4A2B8
Microvirin
ab5
AOL (Mahal Lab)
ABA
Mono2W-CNL
BPL
GS-II (EY Lab)
EEA (EY Lab)
GNA (EY Lab)
HHA (EY Lab)
PHA-L
PNA
SBA
SJA
UEA I
VGA(EY Lab)
WGA
MNA-M
MOA
WT-VCC
ME5(detected with a-HUD)
AAL
LSL-150
LBL-152
ME7(detected with a-HUD)
SSL0-488
VLRB.aGPA.23
rNV VLP-Cy3
AFCNVH
HML-Native
#8 Lewis B (T218) (Abcam)
#15 Sialyl Lewis X (BD)
#14 Sialyl Lewis X (Calbio)
hGal-1-488+BCR
TbFBL2
MGL 416
BTL native
GAC78
DC-ASGPR short
Sp Serotype 14.2
Sp Serotype 14.15
rSRL
SRL
Galectin-1
PsSC
Avaren-Fc
CnSepL
CA19-9 Ab (DRG)
WT Galectin-8
CML1
PhoSPL001
Epa1-Np mutant Y228W
Epa1-Np mutant Y228A
Epa1-Np mutant E227G
Epa1-Np mutant E227A
Epa1-Np WT
ChMBL
DBL-1
hJAA-F11-B11
Fuc(α1-2)Gal(β1-4)GlcNAc6OS(β-Sp8

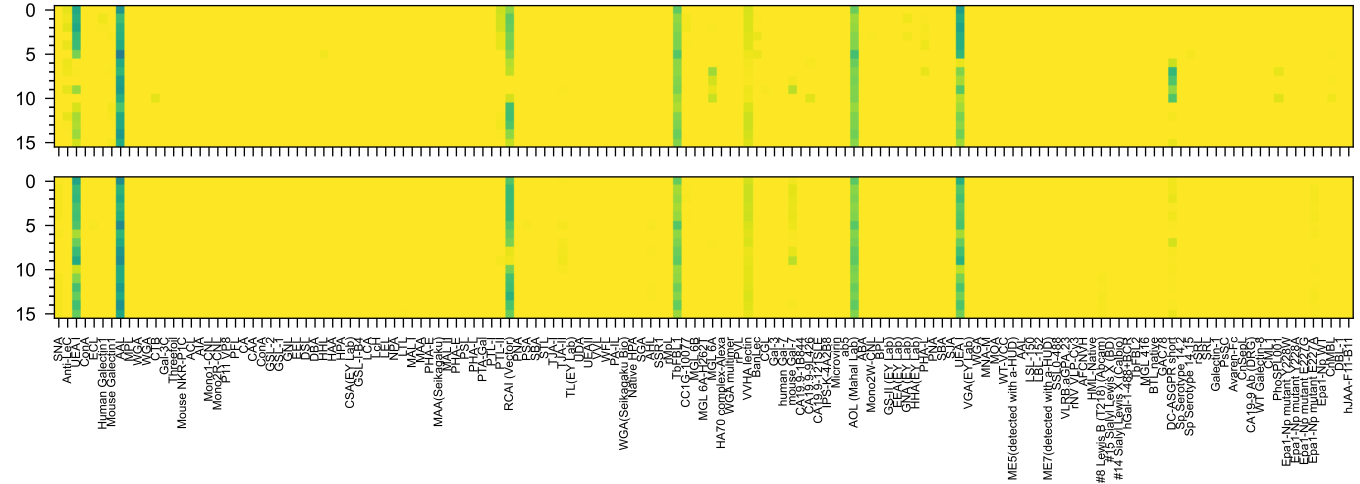


Fuc(α1-2)Gal6OS(β1-4)Glc6OS(β-Sp0

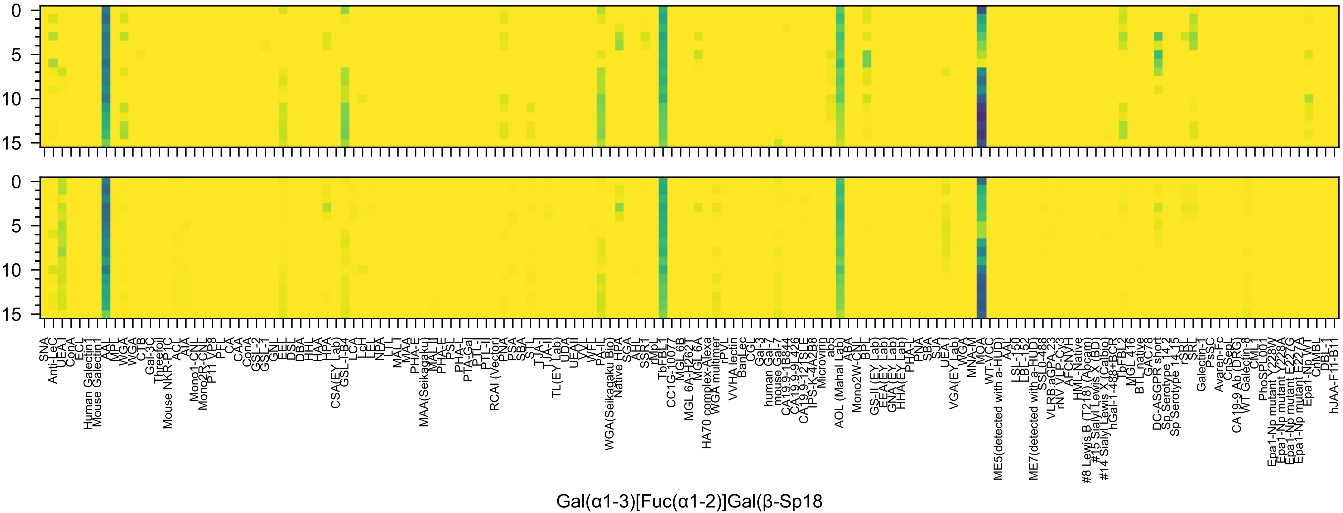
Gal(α1-3)[Fuc(α1-2)]Gal(β-Sp18

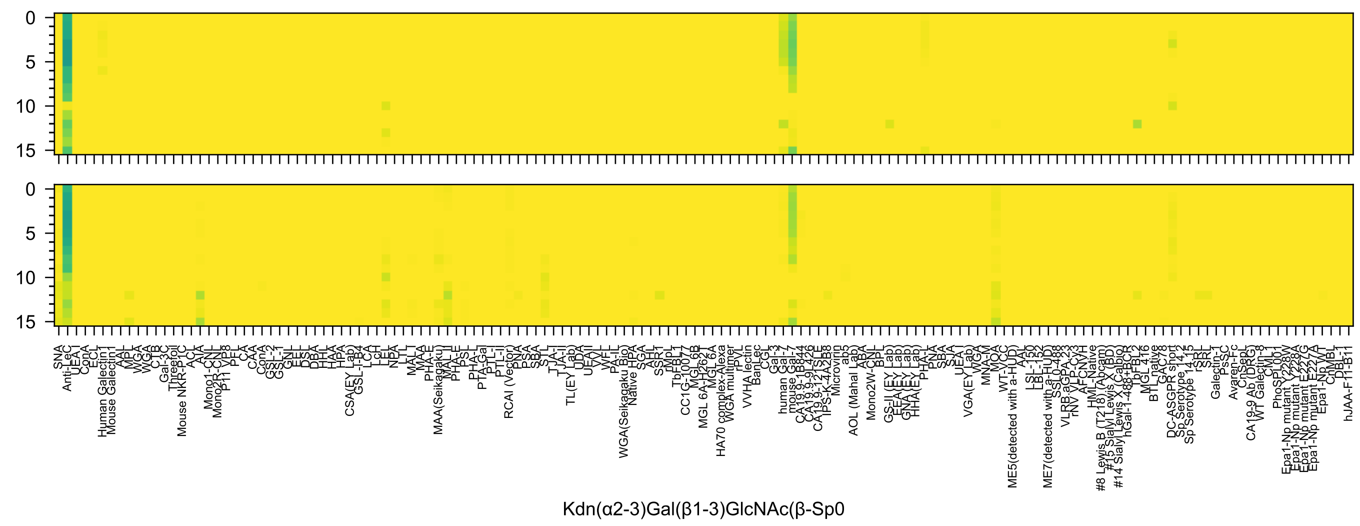
Kdn(α2-3)Gal(β1-3)GlcNAc(β-Sp0

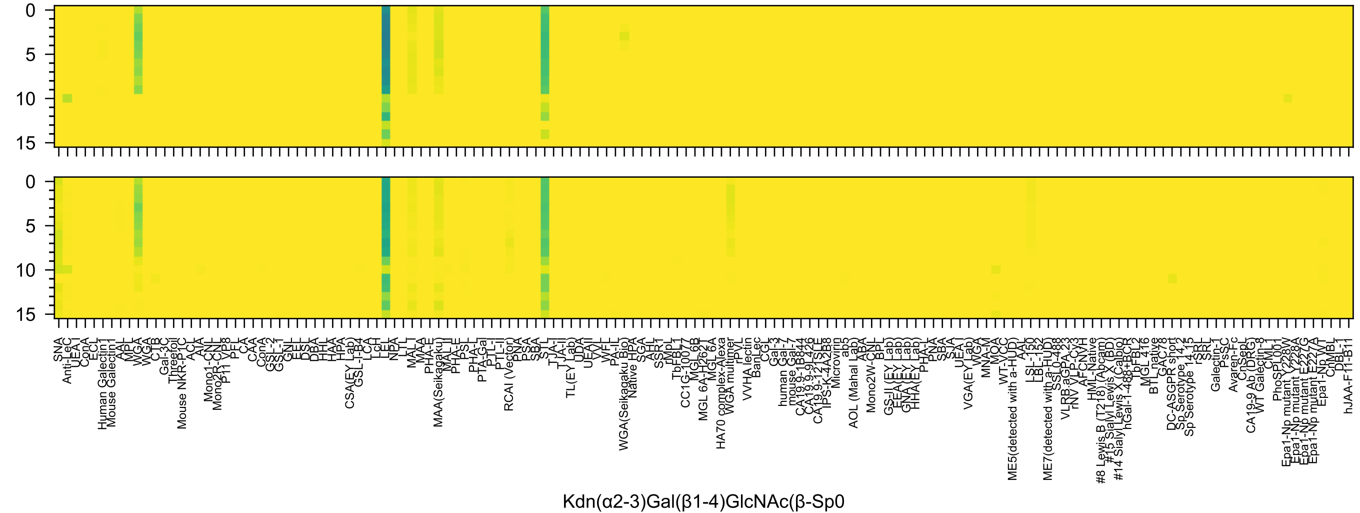
0
5
10
15
0
5
10
15
SNA
Anti-LeC
UEA I
ConA
ECL
Human Galectin1
Mouse Galectin1
AAL
MPL
WGA
WGA
CTB
Gal-3C
Threefoil
Mouse NKR-P1C
ACL
AIA
Mono1-CNL
Mono2R-CNL
P11 VP8
PFL
CA
CAA
ConA
GSL-2
GSL-1
GNL
EEL
DSL
DBA
HHL
HAA
HPA
CSA(EY Lab)
GSL-I-B4
LCA
LcH
LEL
NPA
LTL
MAL-I
MAA
PHA-E
MAA(Seikagaku)
MAL-II
PHA-E
PSL
PHA-L
PTA-Gal
PTL-I
PTL-II
RCAI (Vector)
PNA
PSA
SBA
STL
TJA-I
TJA-II
TL(EY Lab)
UDA
UEA-II
VVL
WFL
PA-IL
WGA(Seikagaku Bio)
Native HPA
SGA
AAL
SSR1
rMpL
TbFBL1
CC1G-10077
MGL 6B
MGL 6A-H262T
MGL 6A
HA70 complex-Alexa
WGA multimer
rPVL
VVHA lectin
BanLec
CGL
Gal-3
human Gal-7
mouse Gal-7
CA19.9-1B844
CA19.9-9L426
CA19.9-121SLE
IPS-K-4A2B8
Microvirin
ab5
AOL (Mahal Lab)
ABA
Mono2W-CNL
BPL
GS-II (EY Lab)
EEA(EY Lab)
GNA(EY Lab)
HHA(EY Lab)
PHA-L
PNA
SBA
SJA
UEA-I
VGA(EY Lab)
WGA
MNA-M
MOA
WT-VCC
ME5(detected with a-HUD)
AAL
LSL-150
LBL-152
ME7(detected with a-HUD)
SSL0-488
VLRB.aGPA.23
rNV VLP-Cy3
AFCNVH
HML-Native
#8 Lewis B (T218) (Abcam)
#15 Sialyl Lewis X (BD)
#14 Sialyl Lewis X (Calbio)
hGal-1-488+BCR
TbFBL2
MGL 416
BTL native
GAC78
DC-ASGPR short
Sp Serotype 14.2
Sp Serotype 14.15
rSRL
SRL
Galectin-1
PsSC
Avaren-Fc
ChSeP1
CA19-9 Ab (DRG)
WT Galectin-8
CML1
PhoSL001
Epa1-Np mutant Y228W
Epa1-Np mutant Y228A
Epa1-Np mutant E227G
Epa1-Np mutant E227A
Epa1-Np WT
ChMBL
DBL-1
hJAA-F11-B11
Kdn(α2-3)Gal(β1-4)GlcNAc(β-Sp0

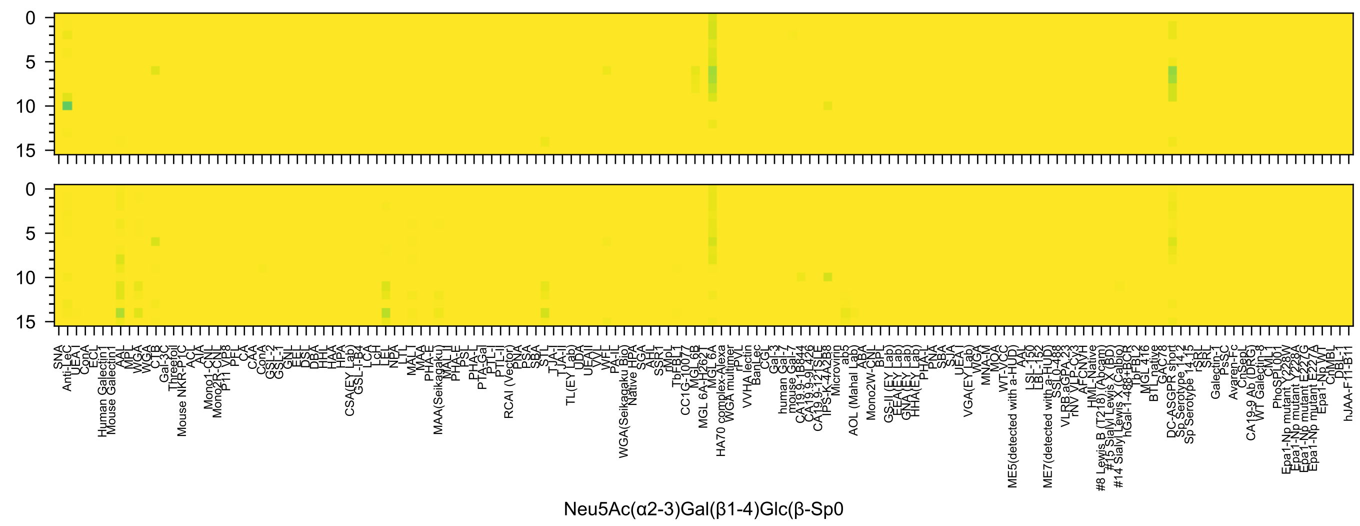
Neu5Ac(α2-3)Gal(β1-4)Glc(β-Sp0

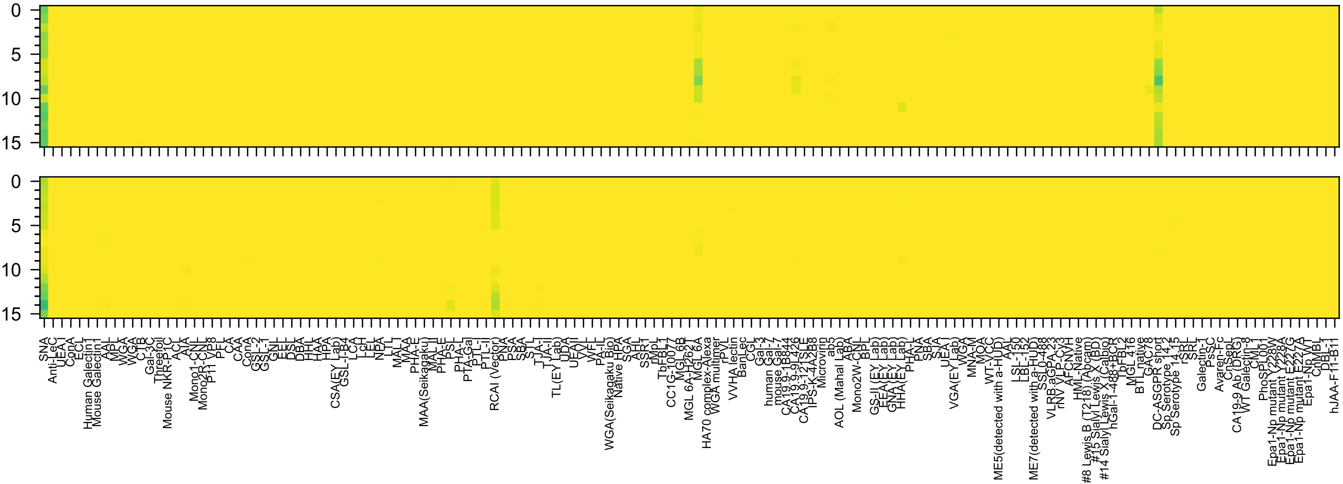

Neu5Ac(α2-6)Gal(β1-4)Glc(β-Sp0

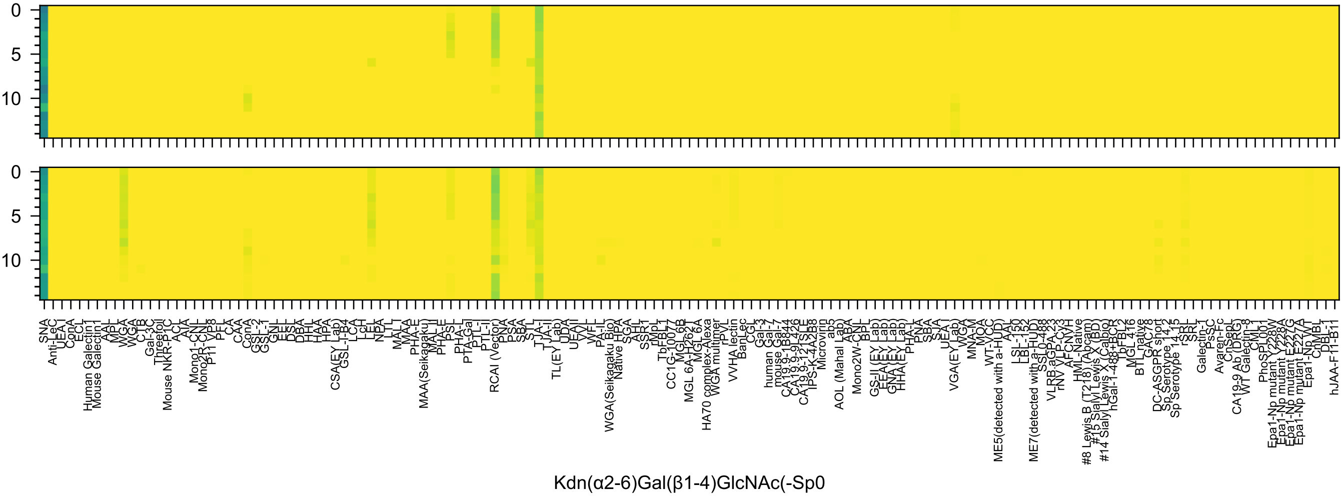
Kdn(α2-6)Gal(β1-4)GlcNAc(-Sp0

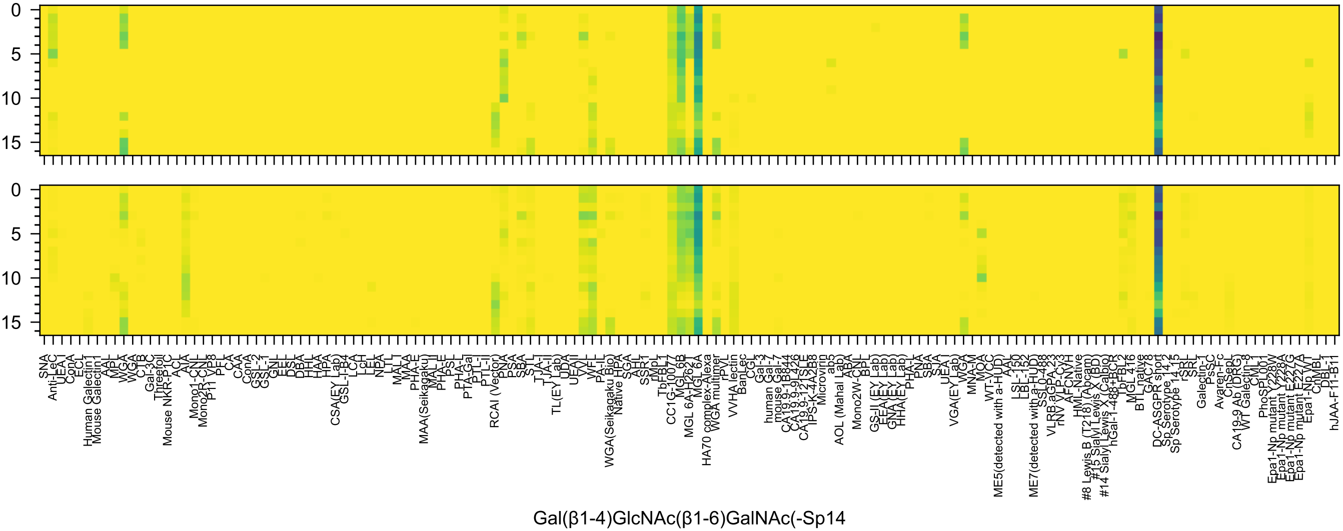
Gal(β1-4)GlcNAc(β1-6)GalNAc(-Sp14

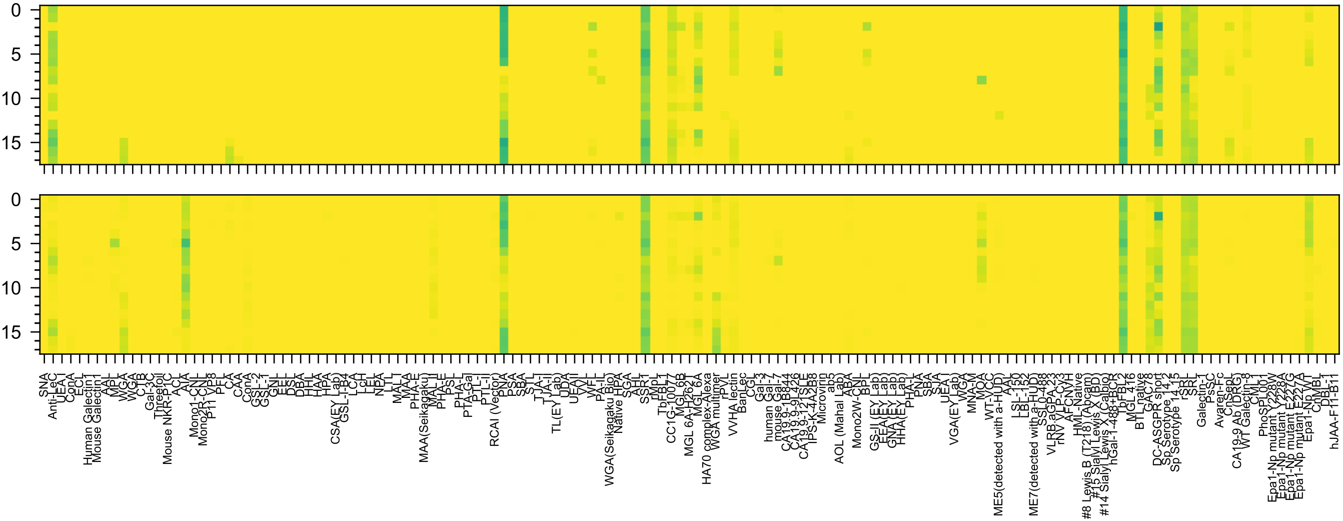

GlcNAc(β1-6)[Gal(β1-3)]GalNAc(α-Sp14

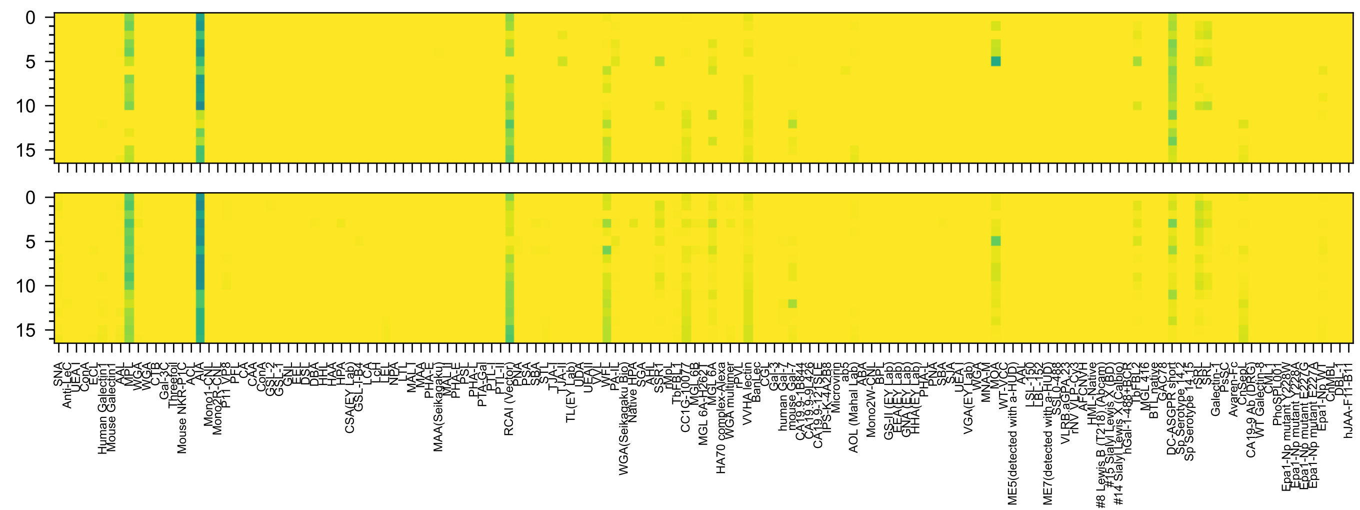


Gal(β1-4)GlcNAc(β1-3)GalNAc(-Sp14

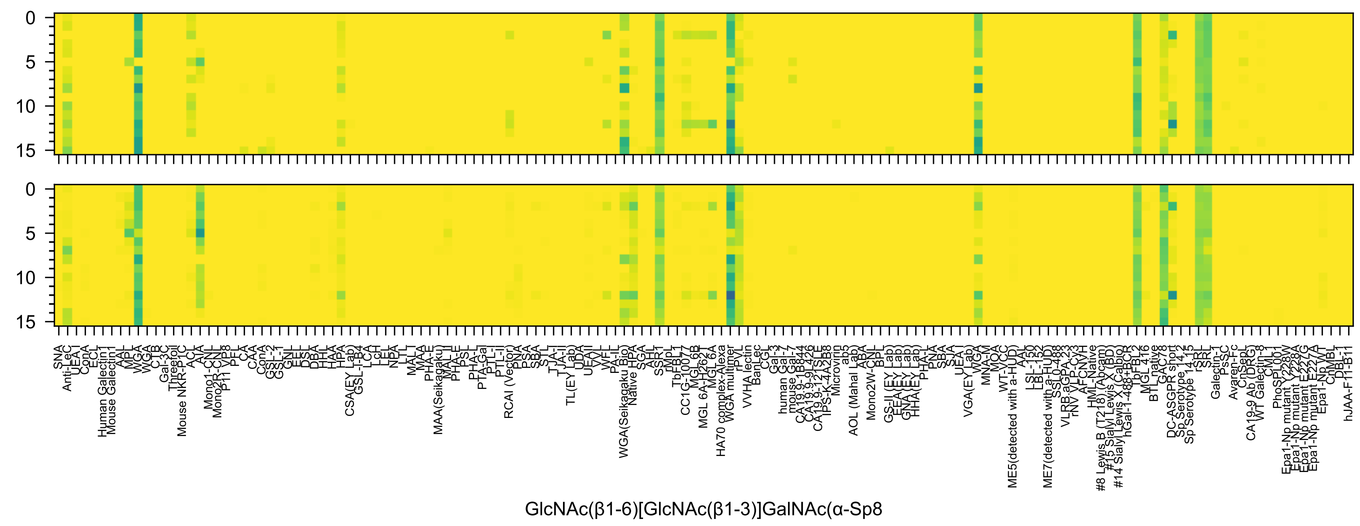
GlcNAc(β1-6)[GlcNAc(β1-3)]GalNAc(α-Sp8

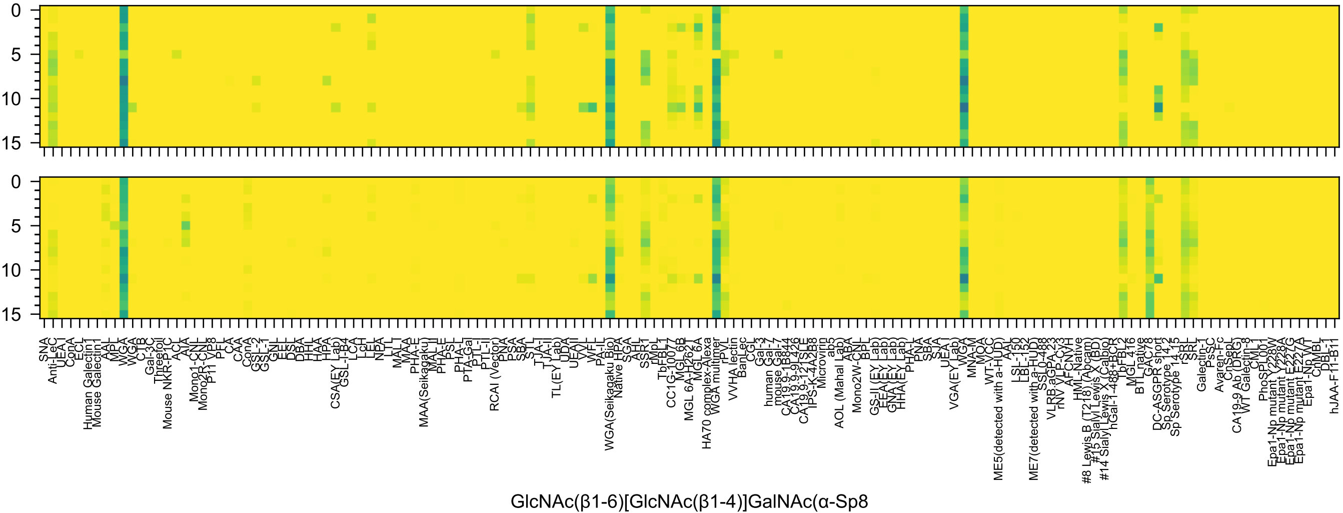
0
5
10
15
0
5
10
15
SNA
Anti-LeC
UEA-I
ConA
ECL
Human Galectin1
Mouse Galectin1
AAL
MPL
WGA
WGA
CTB
Gal-3C
Threefoil
Mouse NKR-P1C
ACL
AIA
Mono1-CNL
Mono2R-CNL
P11 VP8
PFL
CA
CAA
ConA
GSL-2
GSL-1
GNL
EEL
DSL
DBA
HHL
HAA
HPA
CSA(EY Lab)
GSL-I-B4
LCA
LcH
LEL
NPA
LTL
MAL-I
MAA
PHA-E
MAA(Seikagaku)
MAL-II
PHA-E
PSL
PHA-L
PTA-Gal
PTL-I
PTL-II
RCAI (Vector)
PNA
PSA
SBA
STL
TJA-I
TJA-II
TL(EY Lab)
UDA
UEA-II
VVL
WFL
PA-IL
WGA(Seikagaku Bio)
Native HPA
SGA
AHL
SSR1
rMpL
TbFBL1
CC1G-10077
MGL 6B
MGL 6A-H262T
MGL 6A
HA70 complex-Alexa
WGA multimer
rPVL
VVHA lectin
BanLec
CGL
Gal-3
human Gal-7
mouse Gal-7
CA19.9-1B844
CA19.9-9L426
CA19.9-121SLE
IPS-K-4A2B8
Microvirin
ab5
AOL (Mahal Lab)
ABA
Mono2W-CNL
BPL
GS-II (EY Lab)
EEA(EY Lab)
GNA(EY Lab)
HHA(EY Lab)
PHA-L
PNA
SBA
SJA
UEA-I
VGA(EY Lab)
WGA
MNA-M
MOA
WT-VCC
ME5(detected with a-HUD)
AAL
LSL-150
LBL-152
ME7(detected with a-HUD)
SS1-0.488
VLRB.aGPA.23
rNV VLP-Cy3
AFCNVH
HML-Native
#8 Lewis B (T218) (Abcam)
#15 Sialyl Lewis X (BD)
#14 Sialyl Lewis X (Calbio)
hGal-1-488+BCR
TbFBL2
MGL 416
BTL native
GAC78
DC-ASGPR short
Sp Serotype 14.2
Sp Serotype 14.15
rSRL
SRL
Galectin-1
PsSC
Avaren-Fc
ChSeP
CA19-9 Ab (DRG)
WT Galectin-8
CML1
PhoSL001
Epa1-Np mutant Y228W
Epa1-Np mutant Y228A
Epa1-Np mutant E227G
Epa1-Np mutant E227A
Epa1-Np WT
CrMBL
DBL-1
hJAA-F11-B11
GlcNAc(β1-6)[GlcNAc(β1-4)]GalNAc(α-Sp8

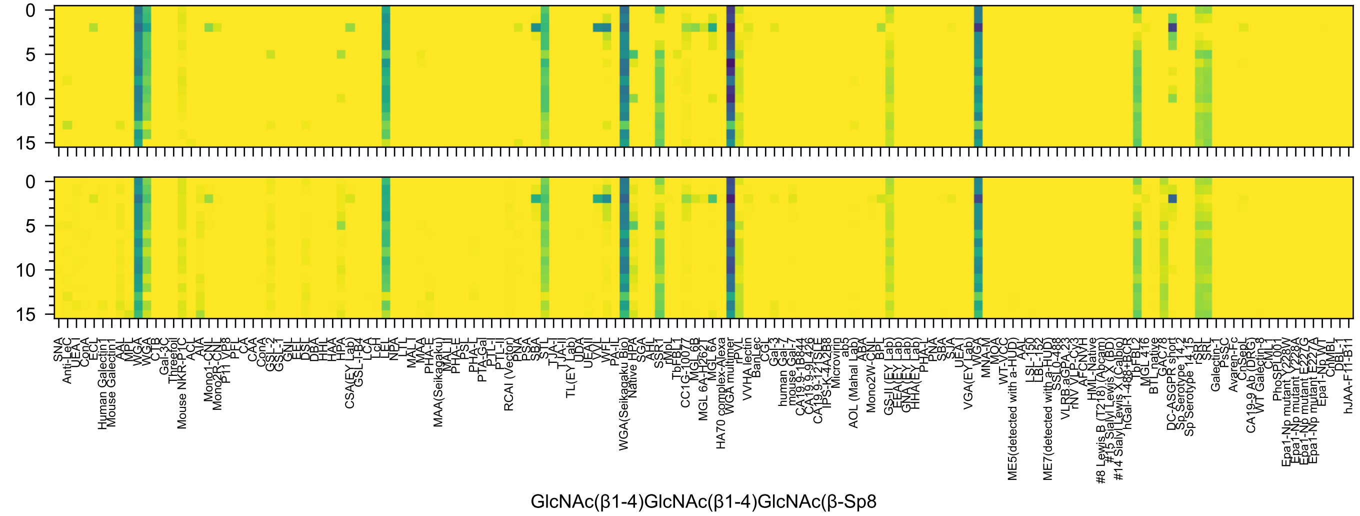
GlcNAc(β1-4)GlcNAc(β1-4)GlcNAc(β-Sp8

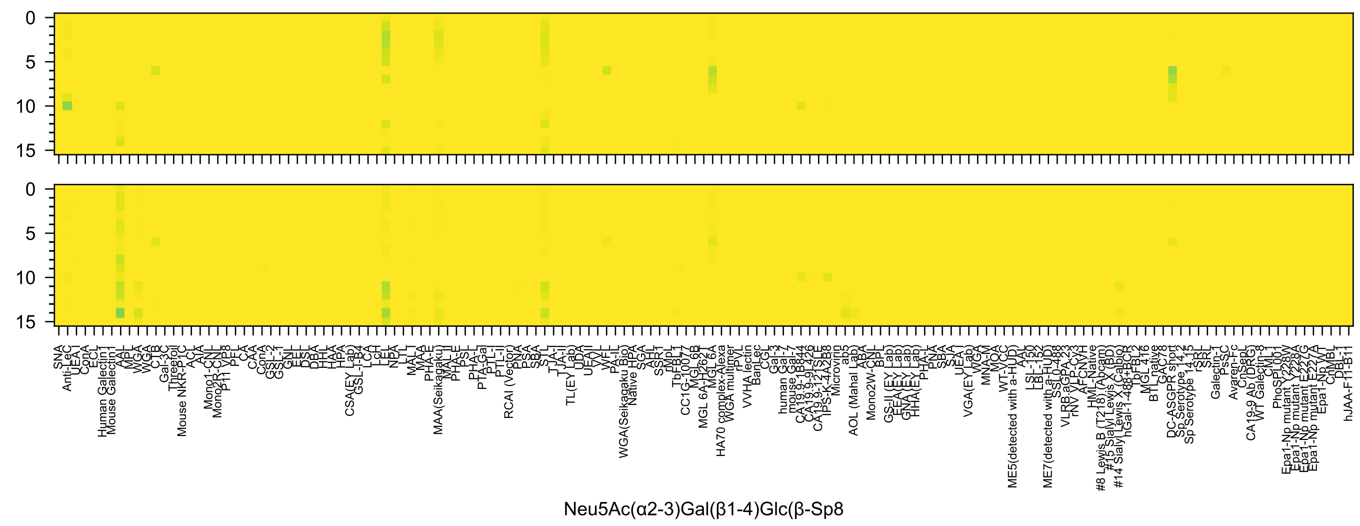
Neu5Ac(α2-3)Gal(β1-4)Glc(β-Sp8

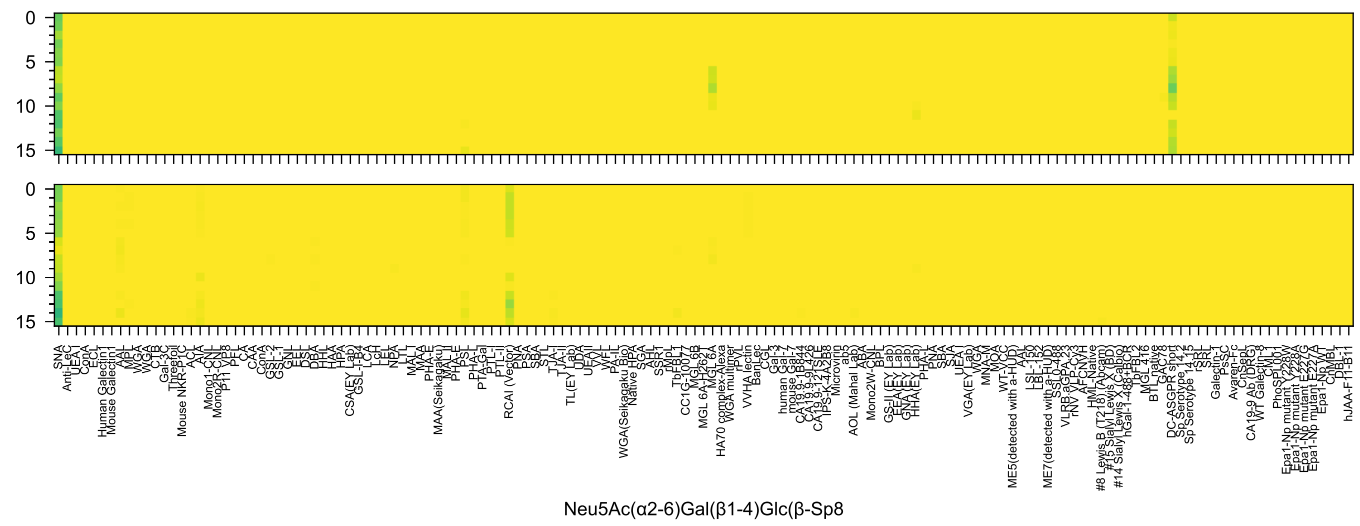
Neu5Ac(α2-6)Gal(β1-4)Glc(β-Sp8

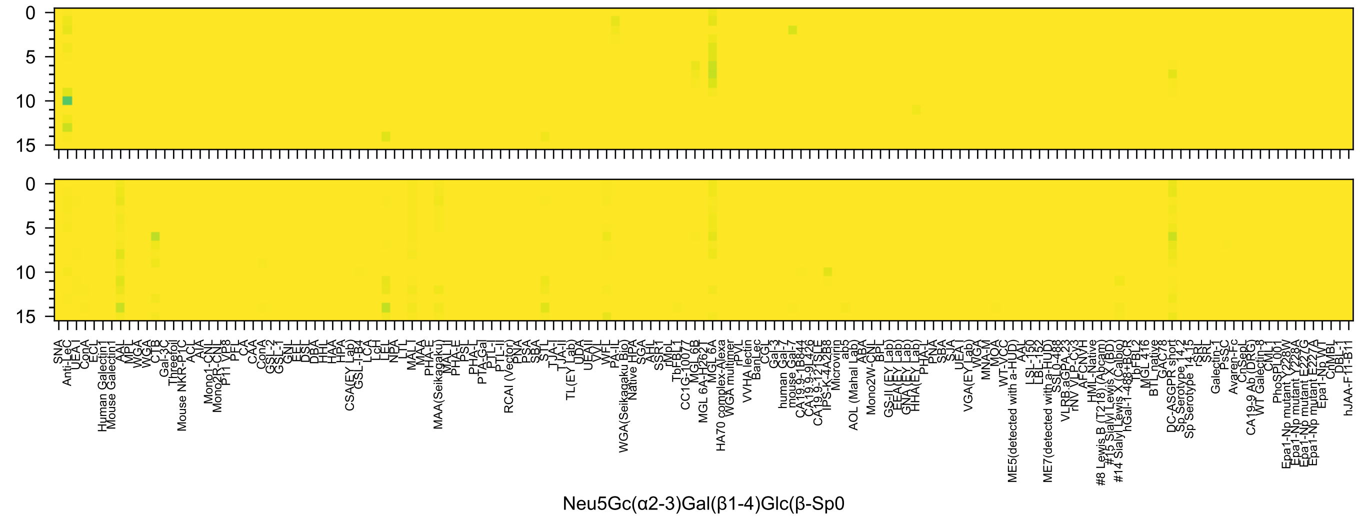
Neu5Gc(α2-3)Gal(β1-4)Glc(β-Sp0

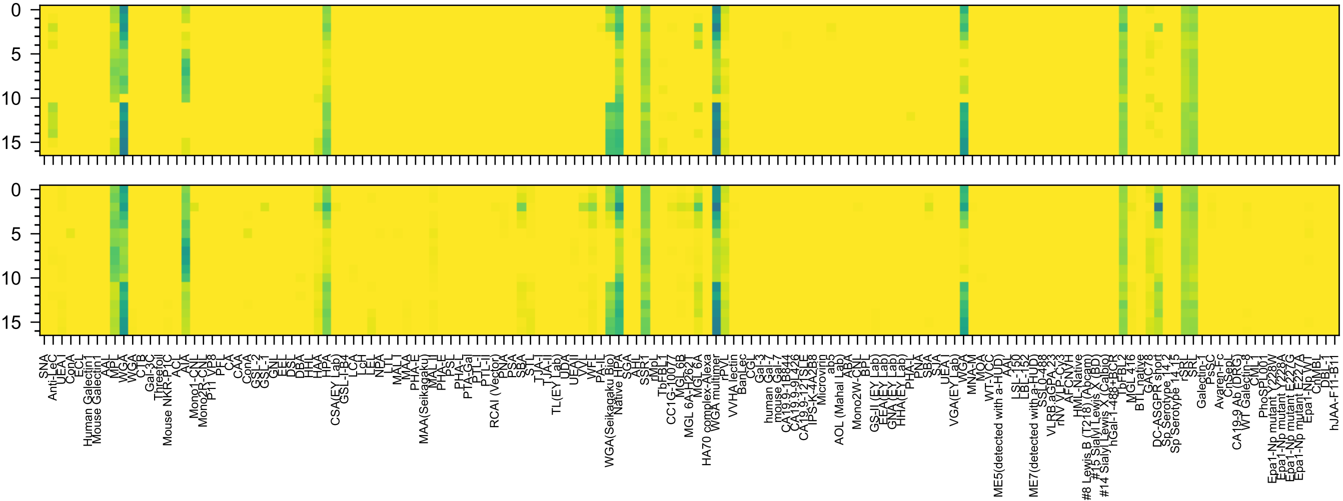


GlcNAc(α1-4)Gal(β1-3)GalNAc(-Sp14

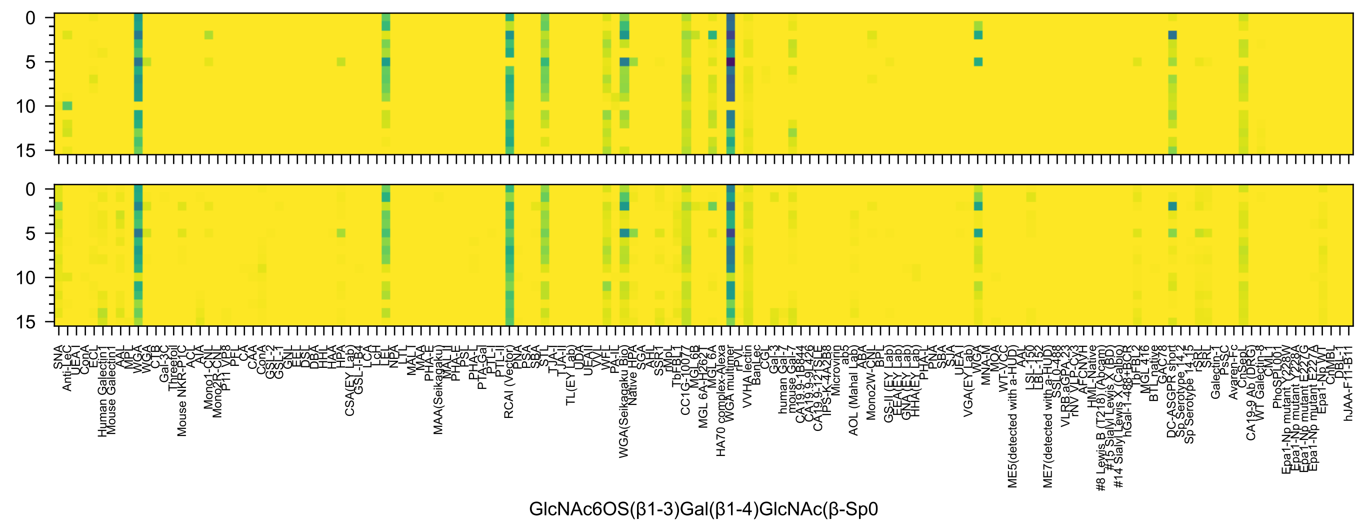
GlcNAc6OS(β1-3)Gal(β1-4)GlcNAc(β-Sp0

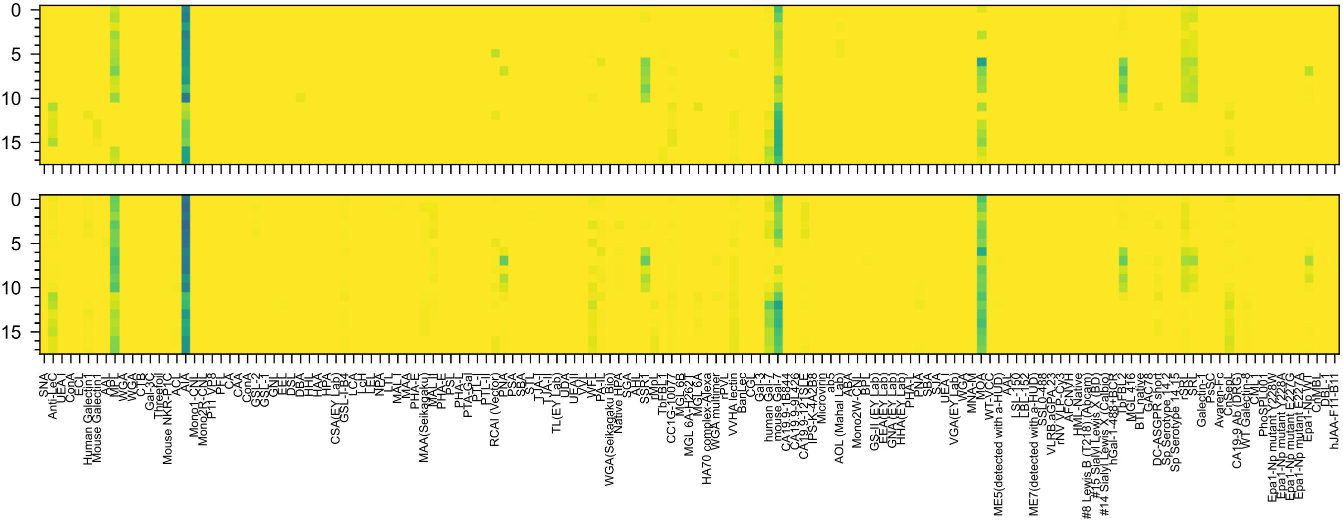


Gal(β1-3)GlcNAc(β1-3)GalNAc(α-Sp14

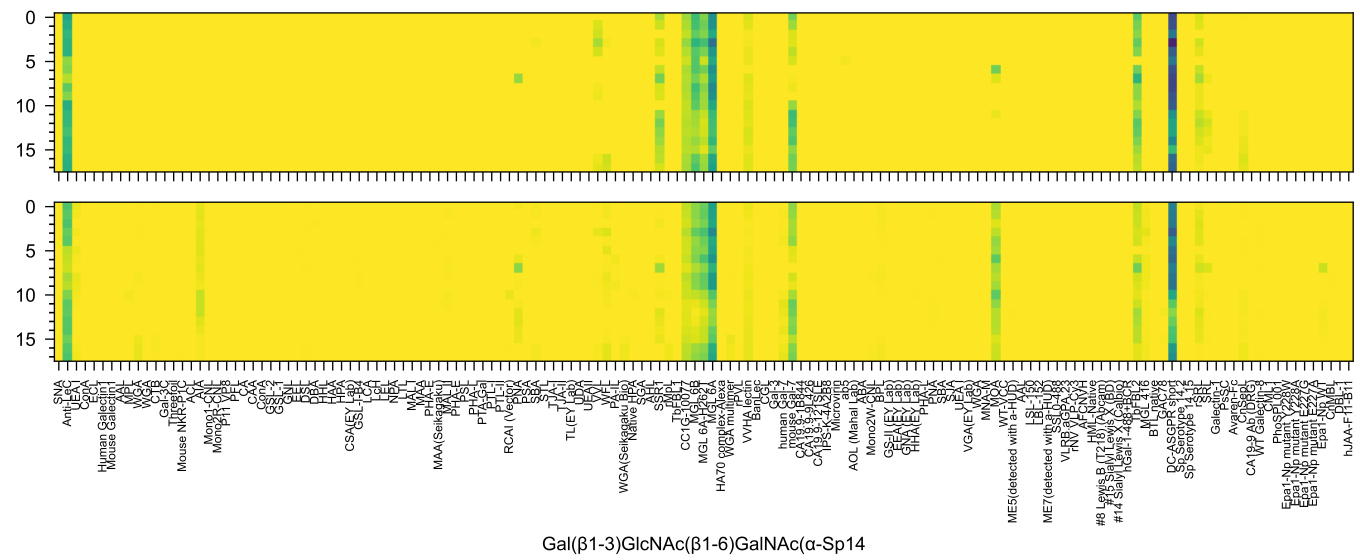
0
5
10
15
Gal(β1-3)GlcNAc(β1-6)GalNAc(α-Sp14

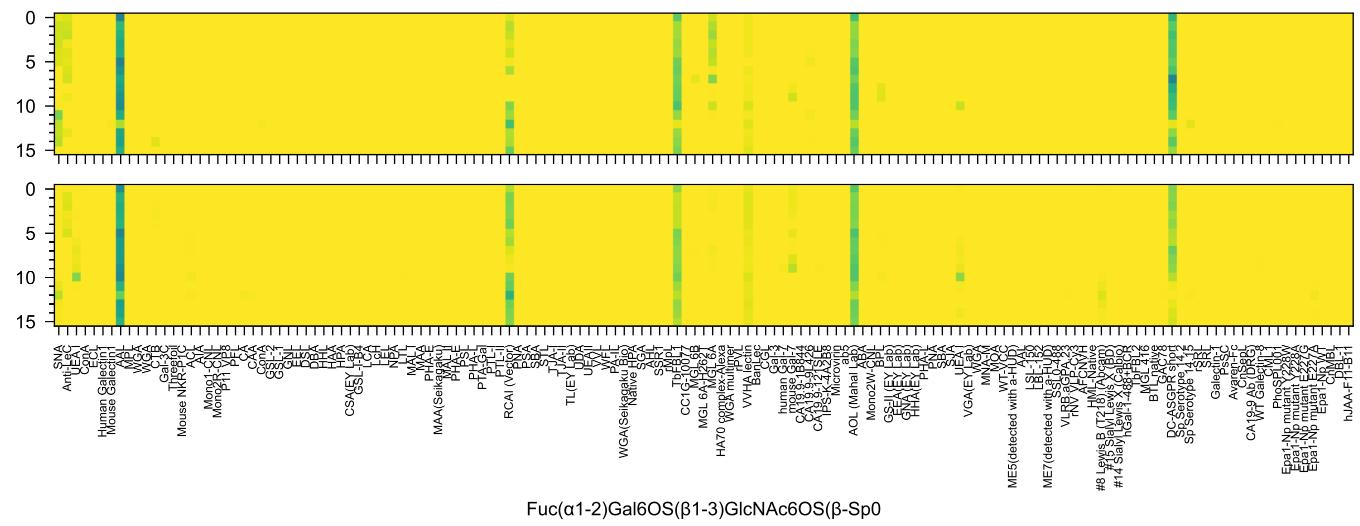

Fuc(α1-2)Gal6OS(β1-3)GlcNAc6OS(β-Sp0

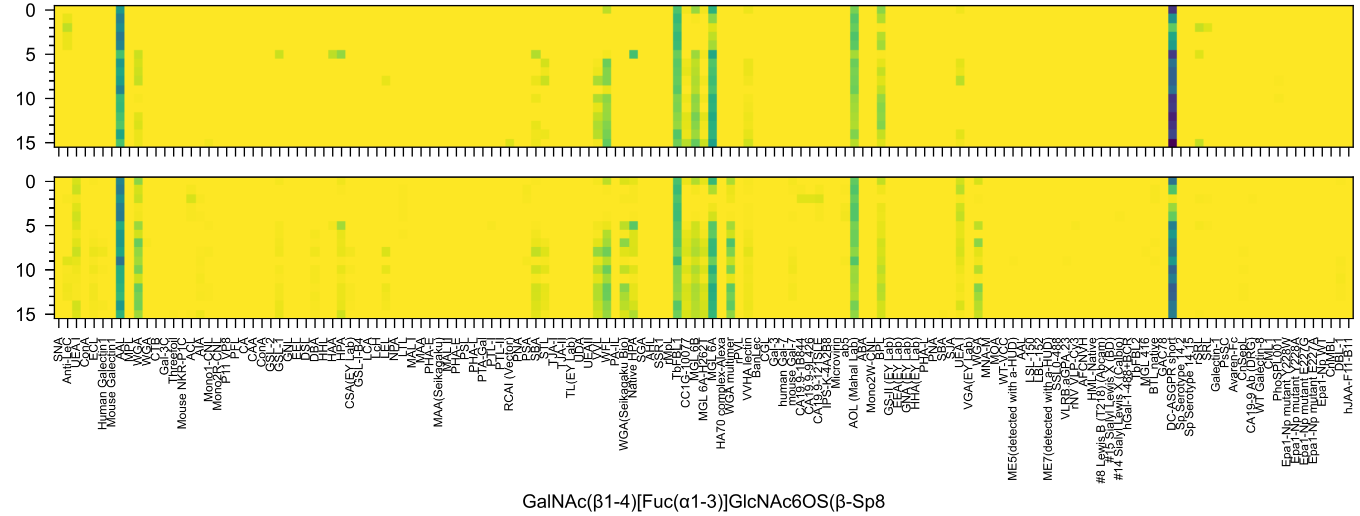
0
5
10
15
0
5
10
15
SNA
Anti-LeC
UEA I
ConA
ECL
Human Galectin1
Mouse Galectin1
AAL
MPL
WGA
WGA
CTB
Gal-3C
Threefoil
Mouse NKR-P1C
ACL
AIA
Mono1-CNL
Mono2R-CNL
P11 VP8
PFL
CA
CAA
ConA
GSL-2
GSL-1
GNL
EEL
DSL
DBA
HHL
HAA
HPA
CSA(EY Lab)
GSL-I-B4
LCA
LcH
LEL
NPA
LTL
MAL
MAA
PHA-E
MAA(Seikagaku)
MAL-II
PHA-E
PSL
PHA-L
PTA-Gal
PTL-I
PTL-II
RCAI (Vector)
PNA
PSA
SBA
STL
TJA-I
TJA-II
TL(EY Lab)
UDA
UEAII
VVL
WFL
PA-IL
WGA(Seikagaku Bio)
Native HPA
SGA
AHL
SSR1
rMpL
TbFBL1
CC1G-10077
MGL 6B
MGL 6A-H262T
MGL 6A
HA70 complex-Alexa
WGA multimer
rPVL
VVHA lectin
BanLec
CGL
Gal-3
human Gal-7
mouse Gal-7
CA19.9-1B844
CA19.9-9L426
CA19.9-121SLE
IPS-K-4A2B8
Microvirin
ab5
AOL (Mahal Lab)
ABA
Mono2W-CNL
BPL
GS-II (EY Lab)
EEA(EY Lab)
GNA(EY Lab)
HHA(EY Lab)
PHA-L
PNA
SBA
SJA
UEA-I
VGA(EY Lab)
WGA
MNA-M
MOA
WT-VCC
ME5(detected with a-HUD)
AAL
LSL-150
LBL-152
ME7(detected with a-HUD)
SSL0-488
VLRB.aGPA.23
rNV VLP-Cy3
AFCNVH
HML-Native
#8 Lewis B (T218) (Abcam)
#15 Sialyl Lewis X (BD)
#14 Sialyl Lewis X (Calbio)
hGal-1-488+BCR
TbFBL2
MGL 416
BTL native
GAC78
DC-ASGPR short
Sp Serotype 14.15
Sp Serotype 14.15
rSRL
SRL
Galectin-1
PsSC
Avaren-Fc
ChSeaL
CA19-9 Ab (DRG)
WT Galectin-8
CML1
PhoSL001
Epa1-Np mutant Y228W
Epa1-Np mutant Y228A
Epa1-Np mutant E227G
Epa1-Np mutant E227A
Epa1-Np WT
CnMBL
DBL-1
hJAA-F11-B11
GalNAc(β1-4)[Fuc(α1-3)]GlcNAc6OS(β-Sp8

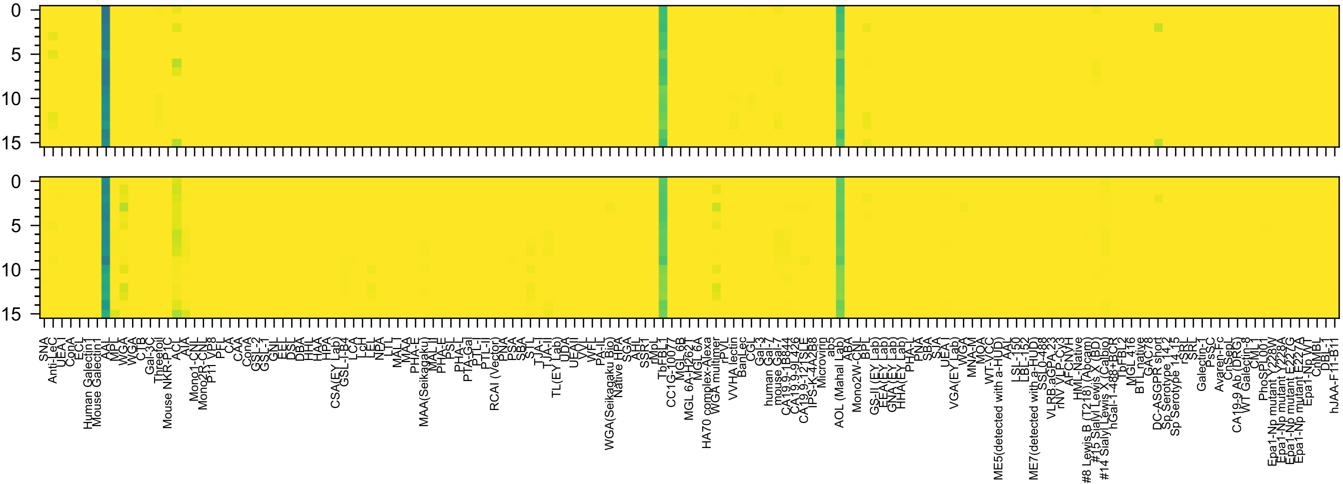

GalNAc3OS(β1-4)[Fuc(α1-3)]GlcNAc(β-Sp8

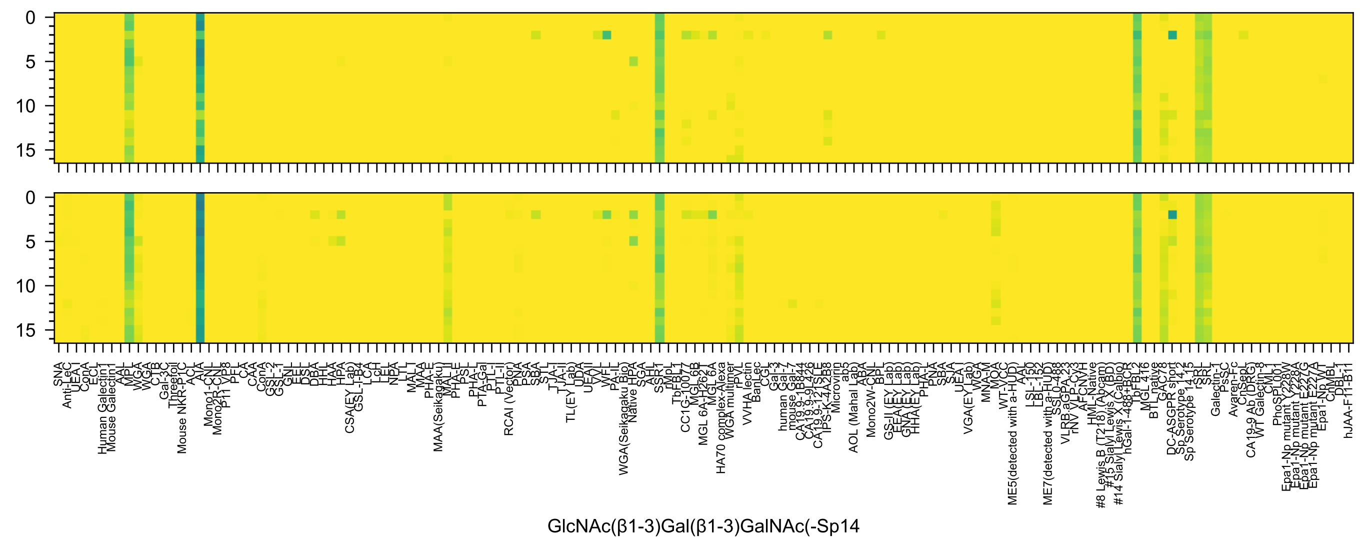
0
5
10
15
SNA
Anti-LeC
UEA I
ConA
ECL
Human Galectin1
Mouse Galectin1
AAL
MPL
WGA
WGA
CTB
Gal-3C
Threefoil
Mouse NKR-P1C
ACL
AIA
Mono1-CNL
Mono2R-CNL
P11 VP8
PFL
CA
CAA
ConA
GSL-2
GSL-1
GNL
EEL
DSL
DBA
HHL
HAA
HPA
CSA(EY Lab)
GSL-I-B4
LCA
LcH
LEL
NPA
LTL
MAL I
MAA
PHA-E
MAA(Seikagaku)
MAL II
PHA-E
PSL
PHA-L
PTA-Gal
PTL-I
PTL-II
RCAI (Vector)
PNA
PSA
SBA
STL
TJA-I
TJA-II
TL(EY Lab)
UDA
UEAII
VVL
WFL
PA-IL
WGA(Seikagaku Bio)
Native HPA
SGA
AAL
SSR1
rMpL
TbFBL1
CC1G-10077
MGL 6B
MGL 6A-H262T
MGL 6A
HA70 complex-Alexa
WGA multimer
rPVL
VVHA lectin
BanLec
CGL
Gal-3
human Gal-7
mouse Gal-7
CA19.9-1B844
CA19.9-9L426
CA19.9-121SLE
IPS-K-4A2B8
Microvirin
ab5
AOL (Mahal Lab)
ABA
Mono2W-CNL
BPL
GS-II (EY Lab)
EEA (EY Lab)
GNA (EY Lab)
HHA (EY Lab)
PHA-L
PNA
SBA
SJA
UEA I
VGA(EY Lab)
WGA
MNA-M
MOA
WT-VCC
ME5(detected with a-HUD)
AAL
LSL-150
LBL-152
ME7(detected with a-HUD)
SSL0-488
VLRB.aGPA.23
rNV VLP-Cy3
AFCNVH
HML-Native
#8 Lewis B (T218) (Abcam)
#15 Sialyl Lewis X (BD)
#14 Sialyl Lewis X (Calbio)
hGal-1-488+BCR
TpFBL2
MGL 416
BTL native
GAC78
DC-ASGPR short
Sp Serotype 14.2
Sp Serotype 14.15
rSRL
SRL
Galectin-1
PssC
Avaren-Fc
ChSepL
CA19-9 Ab (DRG)
WT Galectin-8
CML1
PhoSL001
Epa1-Np mutant Y228W
Epa1-Np mutant Y228A
Epa1-Np mutant E227G
Epa1-Np mutant E227A
Epa1-Np WT
ChMBL
DBL-1
hJAA-F11-B11
GlcNAc(β1-3)Gal(β1-3)GalNAc(-Sp14

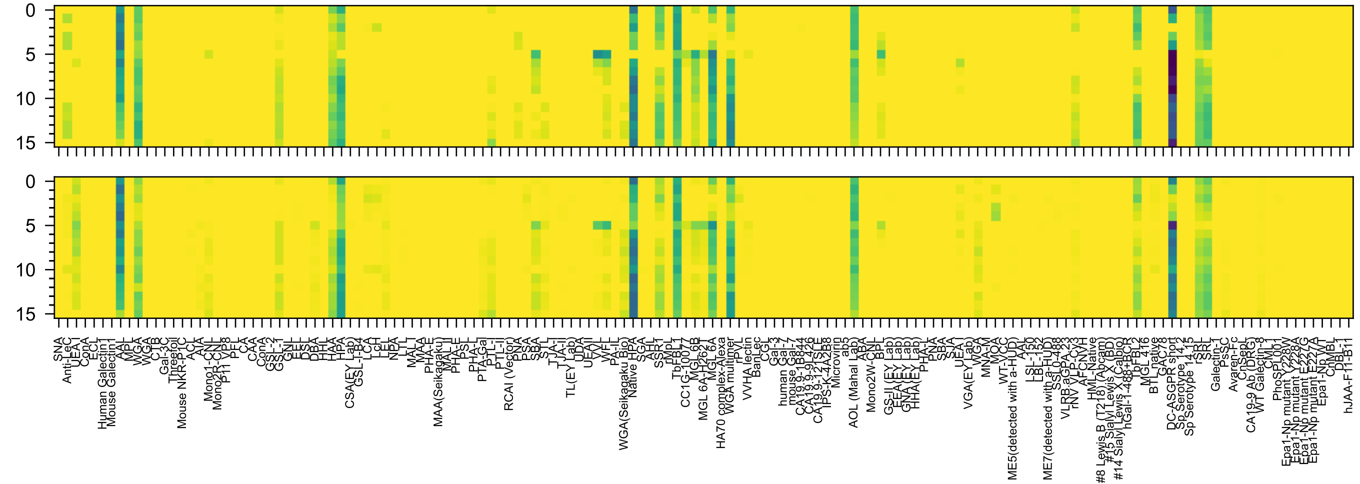

GalNAc(α1-3)[Fuc(α1-2)]Gal(β-Sp18

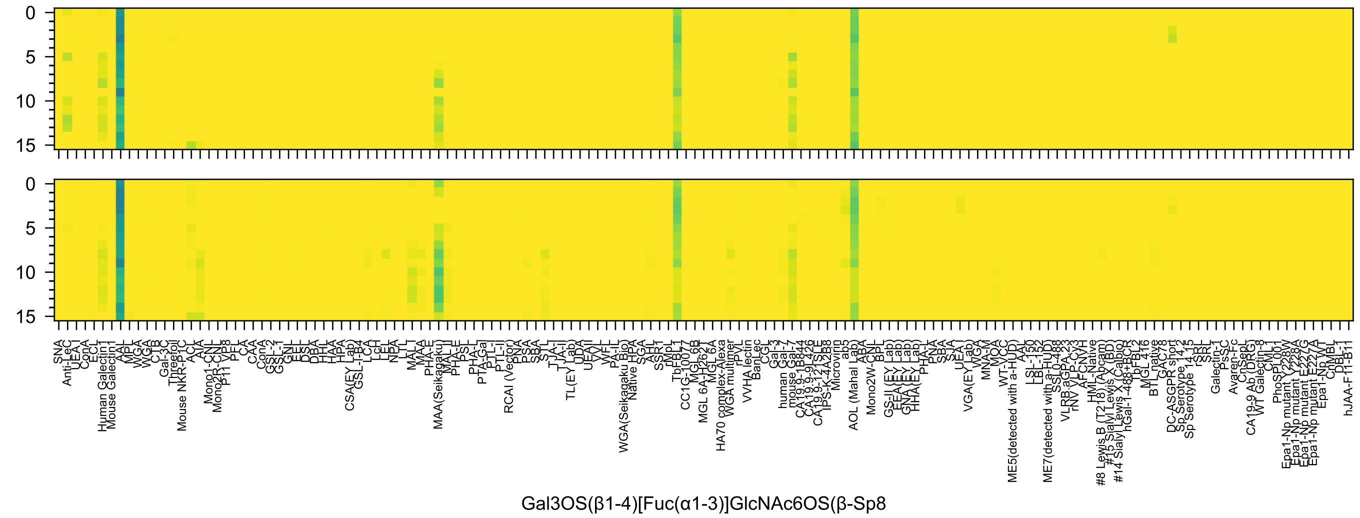
0
5
10
15
0
5
10
15
SNA
Anti-LeC
UEA I
ConA
ECL
Human Galectin1
Mouse Galectin1
AAL
MPL
WGA
WGA
CTB
Gal-3C
Threefoil
Mouse NKR-P1C
ACL
AIA
Mono1-CNL
Mono2R-CNL
P11 VP8
PFL
CA
CAA
ConA
GSL-2
GSL-1
GNL
EEL
DSL
DBA
HHL
HAA
HPA
CSA(EY Lab)
GSL-I-B4
LCA
LcH
LEL
NPA
LTL
MAL
MAA
PHA-E
MAA(Seikagaku)
MAL II
PHA-E
PSL
PHA-L
PTA-Gal
PTL-I
PTL-II
RCAI (Vector)
PNA
PSA
SBA
STL
TJA-I
TJA-II
TL(EY Lab)
UDA
UEAII
VVL
WFL
PA-IL
WGA(Seikagaku Bio)
Native HPA
SGA
AHL
SSR1
rMpL
TbFBL1
CC1G-10077
MGL 6B
MGL 6A-H262T
MGL 6A
HA70 complex-Alexa
WGA multimer
rPVL
VVHA lectin
BanLec
CGL
Gal-3
human Gal-7
mouse Gal-7
CA19.9-1B844
CA19.9-9L426
CA19.9-121SLE
IPS-K-4A2B8
Microvirin
ab5
AOL (Mahal Lab)
ABA
Mono2W-CNL
BPL
GS-II (EY Lab)
EEA(EY Lab)
GNA(EY Lab)
HHA(EY Lab)
PHA-L
PNA
SBA
SJA
UEA I
VGA(EY Lab)
WGA
MNA-M
MOA
WT-VCC
ME5(detected with a-HUD)
AAL
LSL-150
LBL-152
ME7(detected with a-HUD)
SSL0-488
VLRB.aGPA.23
rNV VLP-Cy3
AFCNVH
HML-Native
#8 Lewis B (T218) (Abcam)
#15 Sialyl Lewis X (BD)
#14 Sialyl Lewis X (Calbio)
hGal-1-488+BCR
TbFBL2
MGL 416
BTL native
GAC78
DC-ASGPR short
Sp Serotype 14.2
Sp Serotype 14.15
rSRL
SRL
Galectin-1
PsSC
Avaren-Fc
ChSeptL
CA19-9 Ab (DRG)
WT Galectin-8
CML1
PhoSL001
Epa1-Np mutant Y228W
Epa1-Np mutant Y228A
Epa1-Np mutant E227G
Epa1-Np mutant E227A
Epa1-Np WT
ChMBL
DBL-1
hJAA-F11-B11
Gal3OS(β1-4)[Fuc(α1-3)]GlcNAc6OS(β-Sp8

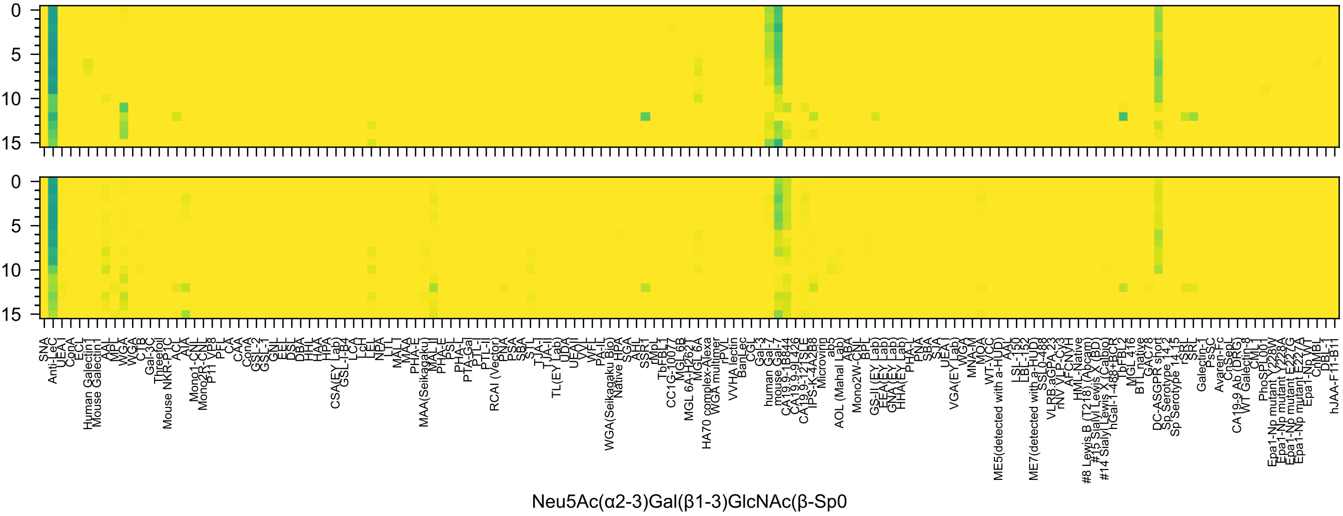
Neu5Ac(α2-3)Gal(β1-3)GlcNAc(β-Sp0

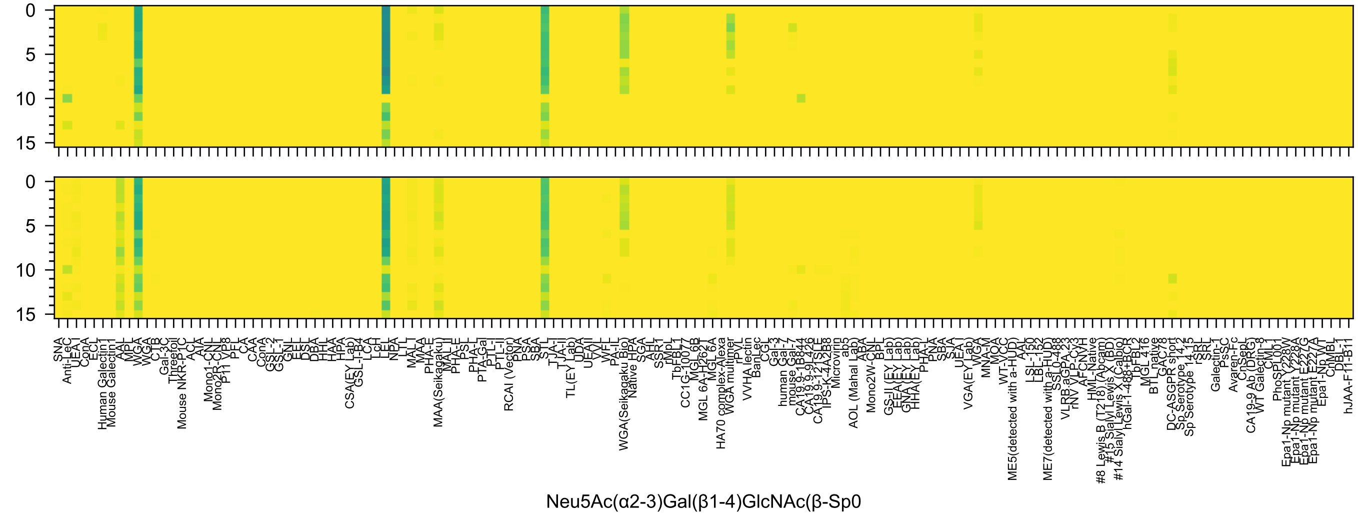
0
5
10
15
0
5
10
15
SNA
Anti-LeC
UEA-I
ConA
ECL
Human Galectin1
Mouse Galectin1
AAL
MPL
WGA
WGA
CTB
C-1B
Gal-3C
Threefoil
Mouse NKR-P1C
ACL
AIA
Mono1-CNL
Mono2R-CNL
P11 VP8
PFL
CA
CAA
ConA
GSL-2
GSL-1
GNL
EEL
DSL
DBA
HHL
HAA
HPA
CSA(EY Lab)
GSL-I-B4
LCA
LcH
LEL
NPA
LTL
MAL-I
MAA
PHA-E
MAA(Seikagaku)
MAL-II
PHA-E
PSL
PHA-L
PTA-Gal
PTL-I
PTL-II
RCAI (Vector)
PNA
PSA
SBA
STL
TJA-I
TJA-II
TL(EY Lab)
UDA
UEA-II
VVL
WFL
PA-IL
WGA(Seikagaku Bio)
Native HPA
SGA
AHL
SSR1
rMpL
TbFBL1
CC1G-10077
MGL 6B
MGL 6A-H262T
MGL 6A
HA70 complex-Alexa
WGA multimer
rPVL
VVHA lectin
BanLec
CGL
Gal-3
human Gal-7
mouse Gal-7
CA19.9-1B844
CA19.9-9L426
CA19.9-121SLE
IPS-K-4A2B8
Microvirin
ab5
AOL (Mahal Lab)
ABA
Mono2W-CNL
BPL
GS-II (EY Lab)
EEA(EY Lab)
GNA(EY Lab)
HHA(EY Lab)
PHA-L
PNA
SBA
SJA
UEA-I
VGA(EY Lab)
WGA
MNA-M
MOA
WT-VCC
ME5(detected with a-HUD)
AAL
LSL-150
LBL-152
ME7(detected with a-HUD)
SSL0-488
VLRB.aGPA.23
rNV VLP-Cy3
AFCNVH
HML-Native
#8 Lewis B (T218) (Abcam)
#15 Sialyl Lewis X (BD)
#14 Sialyl Lewis X (Calbio)
hGal-1-488+BCR
TbFBL2
MGL 416
BTL native
GAC78
DC-ASGPR short
Sp Serotype 14.2
Sp Serotype 14.15
rSRL
SRL
Galectin-1
PsSC
Avaren-Fc
ChSeptL
CA19-9 Ab (DRG)
WT Galectin-8
CML1
PhoSL001
Epa1-Np mutant Y228W
Epa1-Np mutant Y228A
Epa1-Np mutant E227G
Epa1-Np mutant E227A
Epa1-Np WT
ChMBL
DBL-1
hJAA-F11-B11
Neu5Ac(α2-3)Gal(β1-4)GlcNAc(β-Sp0

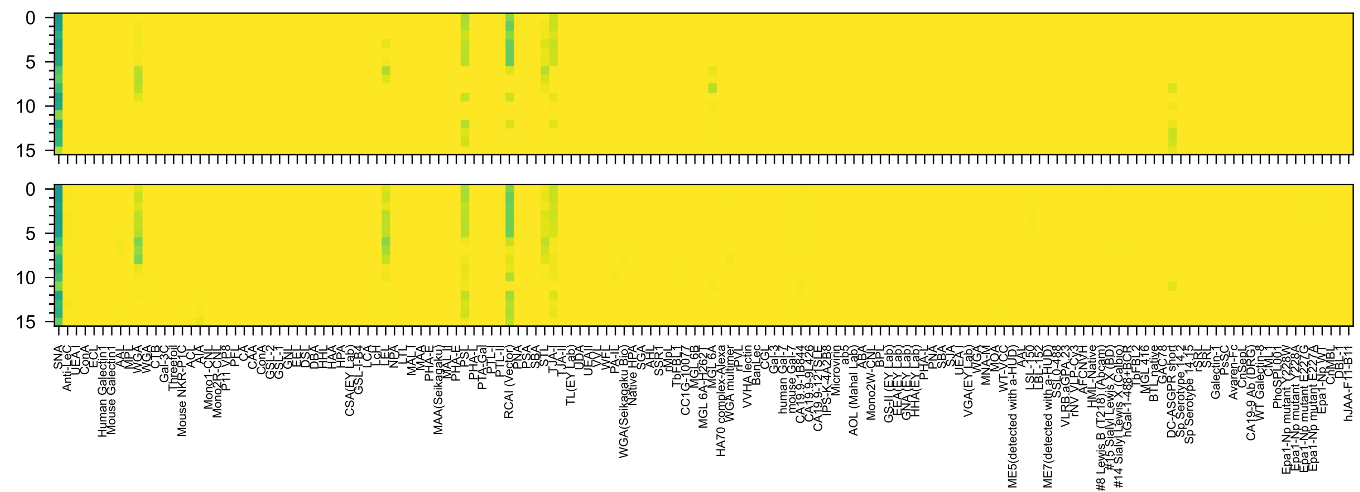

Neu5Ac(α2-6)Gal(β1-4)GlcNAc(β-Sp0

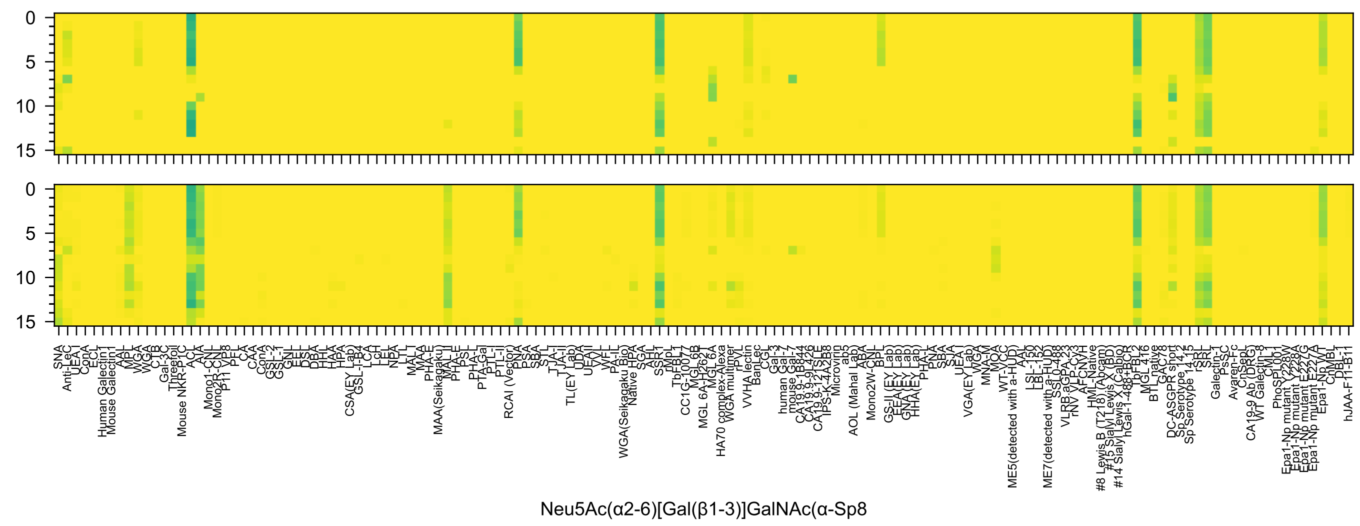

0
5
10
15
SNA
Anti-LeC
UEA I
ConA
ECL
Human Galectin1
Mouse Galectin1
AAL
MPL
WGA
WGA
CTB
Gal-3C
Threefoil
Mouse NKR-P1C
ACL
AIA
Mono1-CNL
Mono2R-CNL
P11 VP8
PFL
CA
CAA
ConA
GSL-2
GSL-1
GNL
EEL
DSL
DBA
HHL
HAA
HPA
CSA(EY Lab)
GSL-I-B4
LCA
LcH
LEL
NPA
LTL
MAL-I
MAA
PHA-E
MAA(Seikagaku)
MAL-II
PHA-E
PSL
PHA-L
PTA-Gal
PTL-I
PTL-II
RCAI (Vector)
PNA
PSA
SBA
STL
TJA-I
TJA-II
TL(EY Lab)
UDA
UEA-I
UEAII
VVL
WFL
PA-IL
WGA(Seikagaku Bio)
Native HPA
SGA
AAL
SSR1
rMpL
TbFBL1
CC1G-10077
MGL 6B
MGL 6A-H262T
MGL 6A
HA70 complex-Alexa
WGA multimer
rPVL
VVHA lectin
BanLec
CGL
Gal-3
human Gal-7
mouse Gal-7
CA19.9-1B844
CA19.9-9L426
CA19.9-121SLE
IPS-K-4A2B8
Microvirin
ab5
AOL (Mahal Lab)
ABA
Mono2W-CNL
BPL
GS-II (EY Lab)
EEA(EY Lab)
GNA(EY Lab)
HHA(EY Lab)
PHA-L
PNA
SBA
SJA
UEA-I
VGA(EY Lab)
WGA
MNA-M
MOA
WT-VCC
ME5(detected with a-HUD)
AAL
LSL-150
LBL-152
ME7(detected with a-HUD)
SSL0-488
VLRB.aGPA.23
rNV VLP-Cy3
AFCNVH
HML-Native
#8 Lewis B (T218) (Abcam)
#15 Sialyl Lewis X (BD)
#14 Sialyl Lewis X (Calbio)
hGal-1-488+BCR
TbFBL2
MGL 416
BTL native
GAC78
DC-ASGPR short
Sp Serotype 14.2
Sp Serotype 14.15
rSRL
SRL
Galectin-1
PsSC
Avaren-Fc
ChSepL
CA19-9 Ab (DRG)
WT Galectin-8
CML1
PhoSL001
Epa1-Np mutant Y228W
Epa1-Np mutant Y228A
Epa1-Np mutant E227G
Epa1-Np mutant E227A
Epa1-Np WT
CnMBL
DBL-1
hJAA-F11-B11
Neu5Ac(α2-6)[Gal(β1-3)]GalNAc(α-Sp8

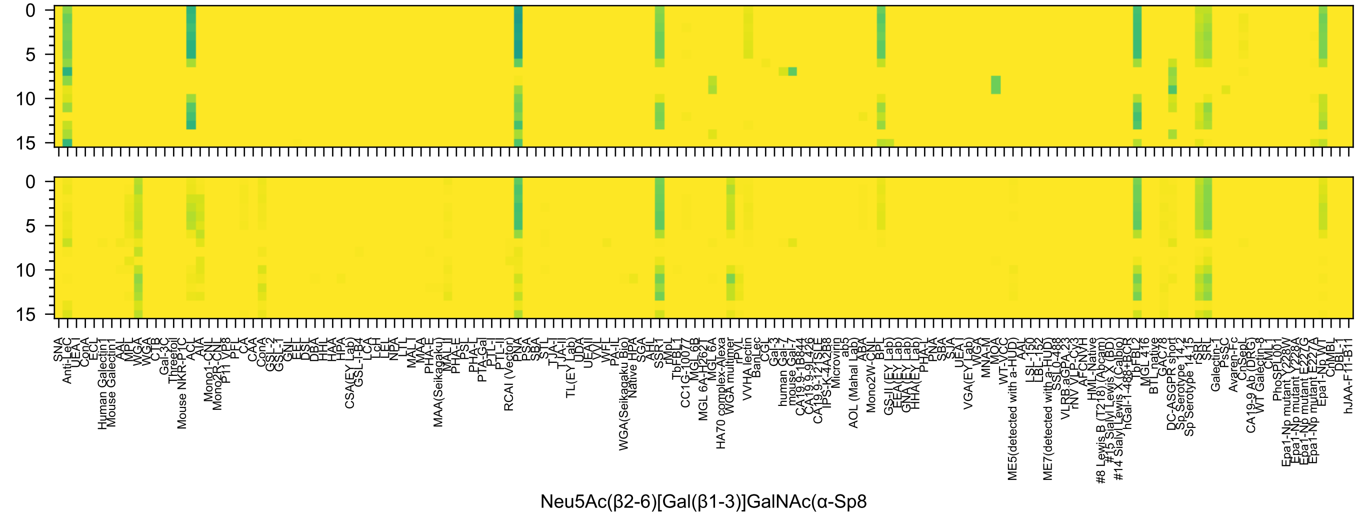

0
5
10
15
SNA
Anti-LeC
UEA-I
ConA
ECL
Human Galectin1
Mouse Galectin1
AAL
MPL
WGA
WGA
CTB
Gal-3C
Threefoil
Mouse NKR-P1C
ACL
AIA
Mono1-CNL
Mono2R-CNL
P11 VP8
PFL
CA
CAA
ConA
GSL-2
GSL-1
GNL
EEL
DSL
DBA
HHL
HAA
HPA
CSA(EY Lab)
GSL-I-B4
LCA
LcH
LEL
NPA
LTL
MAL-I
MAA
PHA-E
MAA(Seikagaku)
MAL-II
PHA-E
PSL
PHA-L
PTA-Gal
PTL-I
PTL-II
RCAI (Vector)
PNA
PSA
SBA
STL
TJA-I
TJA-II
TL(EY Lab)
UDA
UEA-II
VVL
WFL
PA-IL
WGA(Seikagaku Bio)
Native HPA
SGA
AHL
SSR1
rMpL
TbFBL1
CC1G-10077
MGL 6B
MGL 6A-H262T
MGL 6A
HA70 complex-Alexa
WGA multimer
rPVL
VVHA lectin
BanLec
CGL
Gal-3
human Gal-7
mouse Gal-7
CA19.9-1B844
CA19.9-9L426
CA19.9-121SLE
IPS-K-4A2B8
Microvirin
ab5
AOL (Mahal Lab)
ABA
Mono2W-CNL
BPL
GS-II (EY Lab)
EEA(EY Lab)
GNA(EY Lab)
HHA(EY Lab)
PHA-L
PNA
SBA
SJA
UEA-I
VGA(EY Lab)
WGA
MNA-M
MOA
WT-VCC
ME5(detected with a-HUD)
AAL
LSL-150
LBL-152
ME7(detected with a-HUD)
SSL0-488
VLRB.aGPA.23
rNV VLP-Cy3
AFCNVH
HML-Native
#8 Lewis B (T218) (Abcam)
#15 Sialyl Lewis X (BD)
#14 Sialyl Lewis X (Calbio)
hGal-1-488+BCR
TbFBL2
MGL 416
BTL native
GAC78
DC-ASGPR short
Sp Serotype 14.2
Sp Serotype 14.15
rSRL
SRL
Galectin-1
PsSC
Avaren-Fc
ChSeP1
CA19-9 Ab (DRG)
WT Galectin-8
CML1
PhoSL001
Epa1-Np mutant Y228W
Epa1-Np mutant Y228A
Epa1-Np mutant E227G
Epa1-Np mutant E227A
Epa1-Np WT
ChMBL
DBL-1
hJAA-F11-B11
Neu5Ac(β2-6)[Gal(β1-3)]GalNAc(α-Sp8

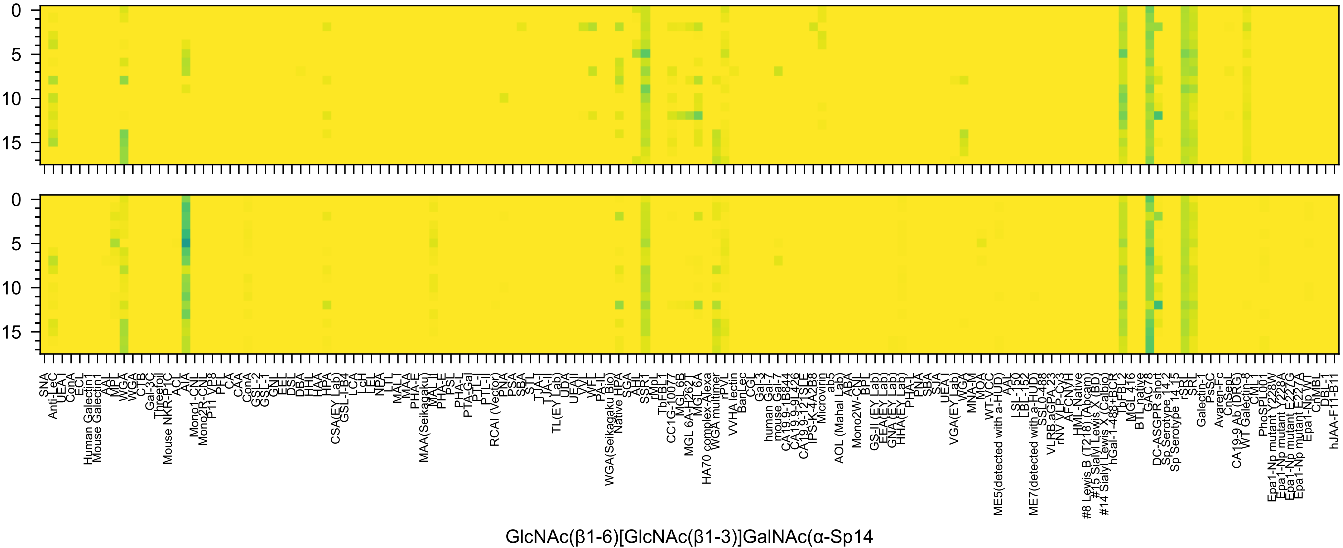
GlcNAc(β1-6)[GlcNAc(β1-3)]GalNAc(α-Sp14

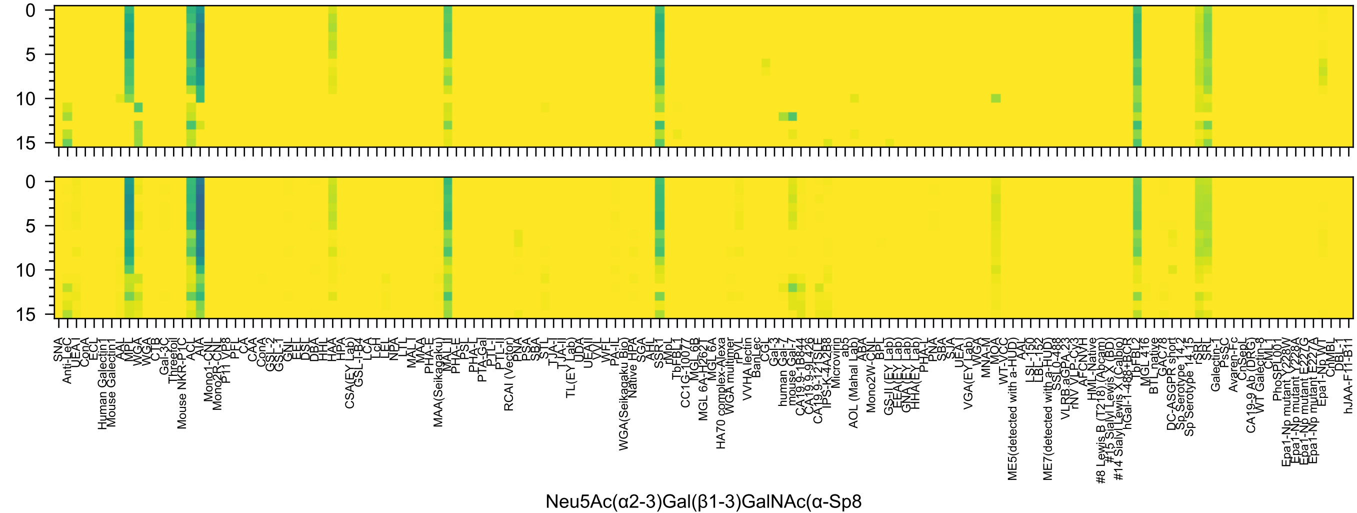
0
5
10
15
0
5
10
15
SNA
Anti-LeC
UEA-I
ConA
ECL
Human Galectin1
Mouse Galectin1
AAL
MPL
WGA
WGA
CTB
Gal-3C
Threefoil
Mouse NKR-P1C
ACL
AIA
Mono1-CNL
Mono2R-CNL
P11 VP8
PFL
CA
CAA
ConA
GSL-2
GSL-1
GNL
EEL
DSL
DBA
HHL
HAA
HPA
CSA(EY Lab)
GSL-I-B4
LCA
LcH
LEL
NPA
LTL
MAL-I
MAA
PHA-E
MAA(Seikagaku)
MAL-II
PHA-E
PSL
PHA-L
PTA-Gal
PTL-I
PTL-II
RCAI (Vector)
PNA
PSA
SBA
STL
TJA-I
TJA-II
TL(EY Lab)
UDA
UEA-II
VVL
WFL
PA-IL
WGA(Seikagaku Bio)
Native HPA
SGA
AHT
SSR1
rMpL
TbFBL1
CC1G-10077
MGL 6B
MGL 6A-H262T
MGL 6A
HA70 complex-Alexa
WGA multimer
rPVL
VVHA lectin
BanLec
CGL
Gal-3
human Gal-7
mouse Gal-7
CA19.9-1B844
CA19.9-9L426
CA19.9-121SLE
IPS-K-4A2B8
Microvirin
ab5
AOL (Mahal Lab)
ABA
Mono2W-CNL
BPL
GS-II (EY Lab)
EEA(EY Lab)
GNA(EY Lab)
HHA(EY Lab)
PHA-L
PNA
SBA
SJA
UEA-I
VGA(EY Lab)
WGA
MNA-M
MOA
WT-VCC
ME5(detected with a-HUD)
AAL
LSL-150
LBL-152
ME7(detected with a-HUD)
SSL0-488
VLRB.aGPA-23
rNV VLP-Cy3
AFCNVH
HML-Native
#8 Lewis B (T218) (Abcam)
#15 Sialyl Lewis X (BD)
#14 Sialyl Lewis X (Calbio)
hGal-1-488+BCR
TbFBL2
MGL 416
BTL native
GAC78
DC-ASGPR short
Sp Serotype 14.2
Sp Serotype 14.15
rSRL
SRL
Galectin-1
PsSC
Avaren-Fc
ChSepL
CA19-9 Ab (DRG)
WT Galectin-8
CML1
PhoSPL001
Epa1-Np mutant Y228W
Epa1-Np mutant Y228A
Epa1-Np mutant E227G
Epa1-Np mutant E227A
Epa1-Np WT
ChMBL
DBL-1
hJAA-F11-B11
Neu5Ac(α2-3)Gal(β1-3)GalNAc(α-Sp8

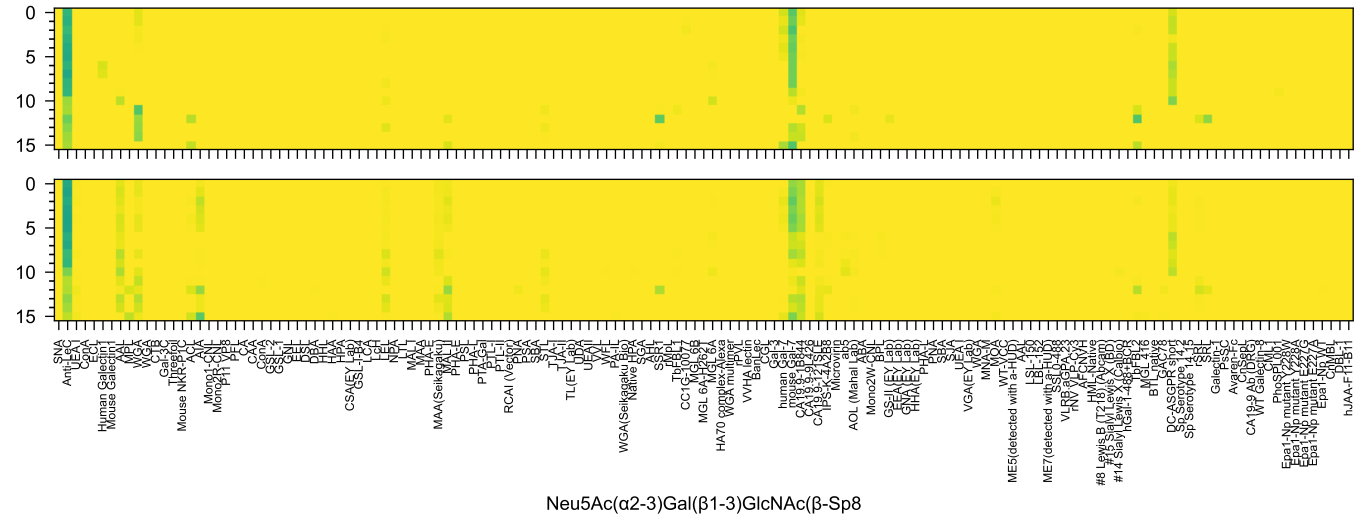

0
5
10
15
0
5
10
15
Neu5Ac(α2-3)Gal(β1-3)GlcNAc(β-Sp8

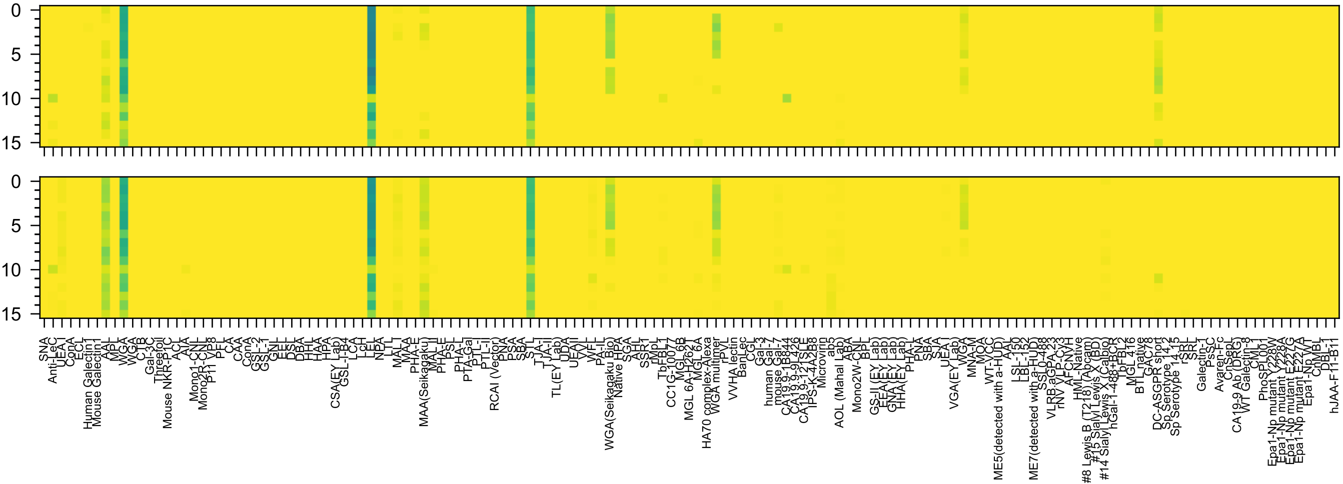

Neu5Ac(α2-3)Gal(β1-4)GlcNAc(β-Sp8

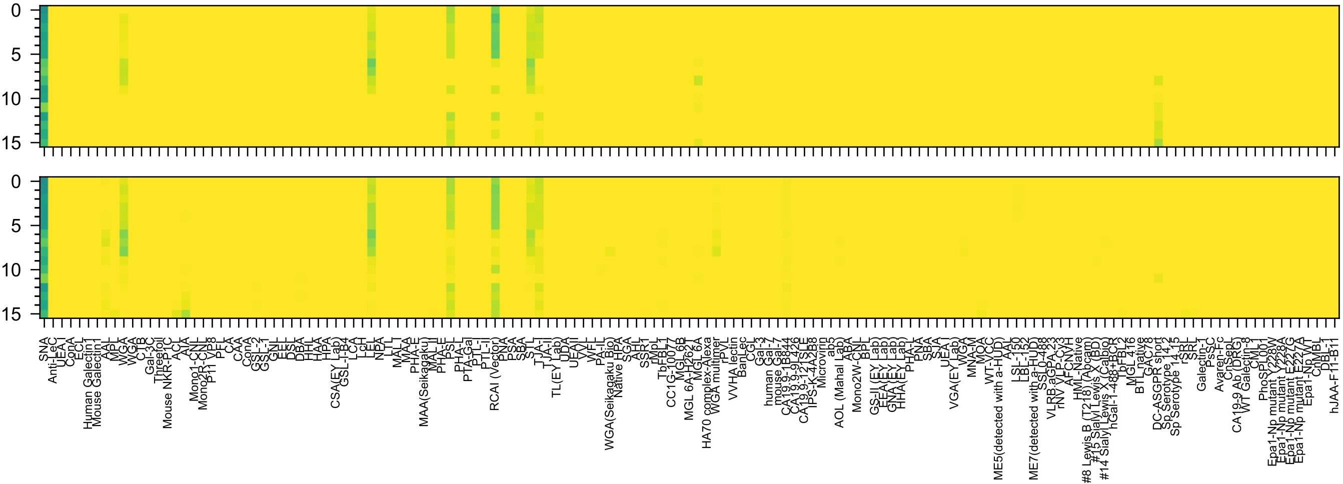

Neu5Ac(α2-6)Gal(β1-4)GlcNAc(β-Sp8

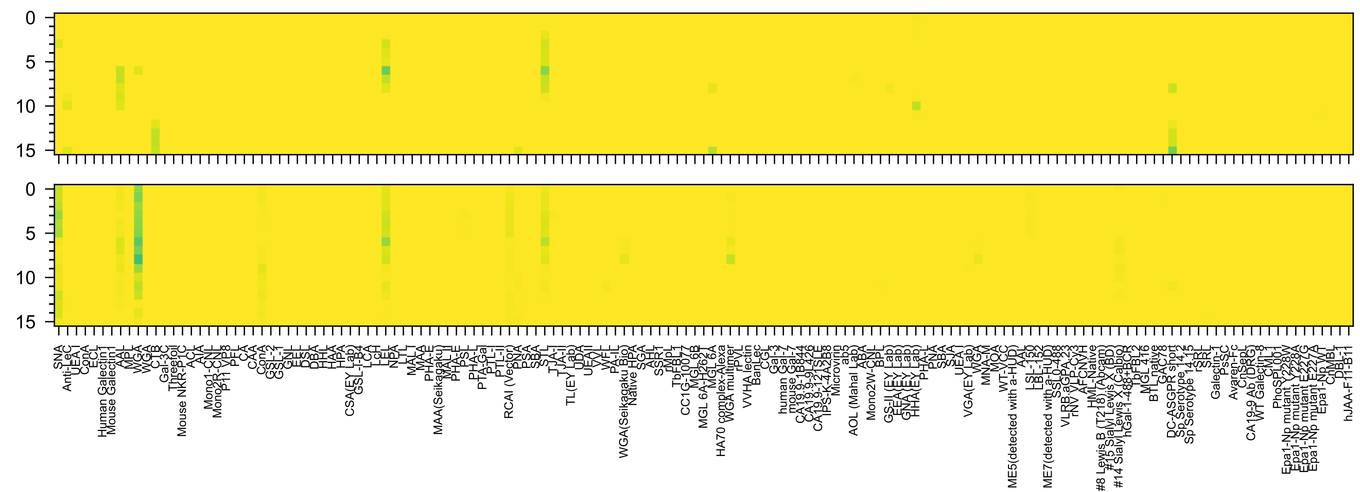

Neu5Ac(β2-6)Gal(β1-4)GlcNAc(β-Sp8

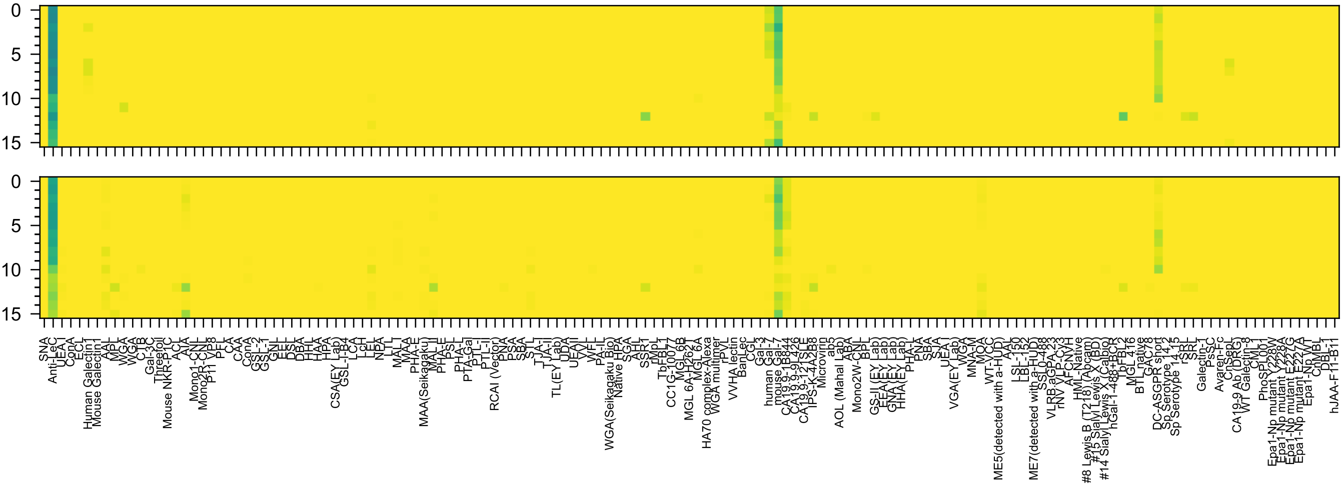

Neu5Gc(α2-3)Gal(β1-3)GlcNAc(β-Sp0

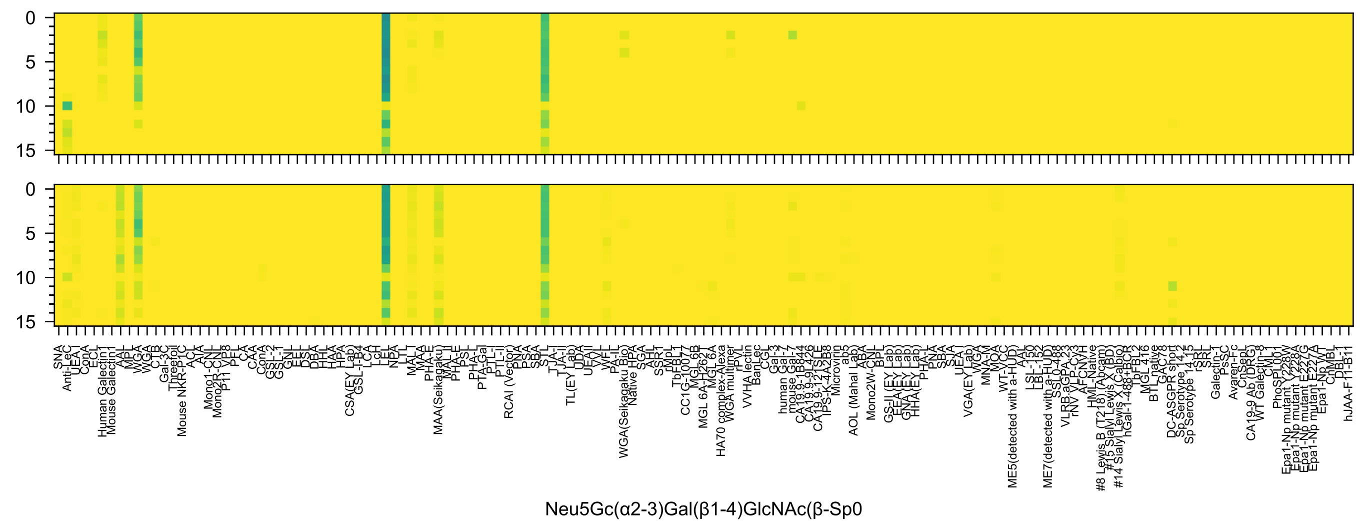
Neu5Gc(α2-3)Gal(β1-4)GlcNAc(β-Sp0

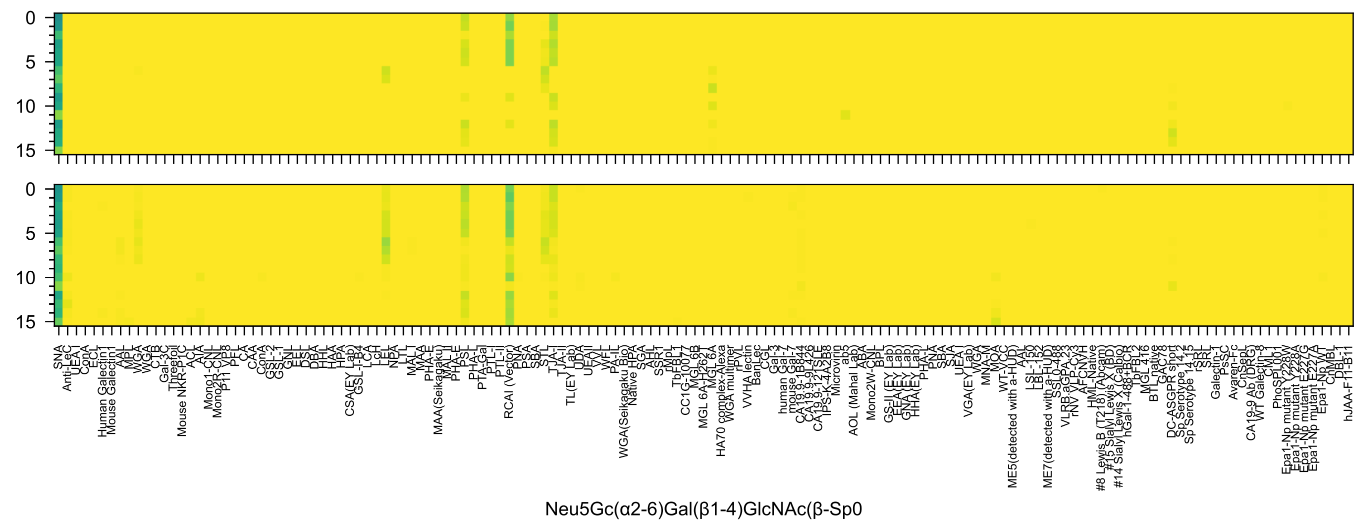
Neu5Gc(α2-6)Gal(β1-4)GlcNAc(β-Sp0

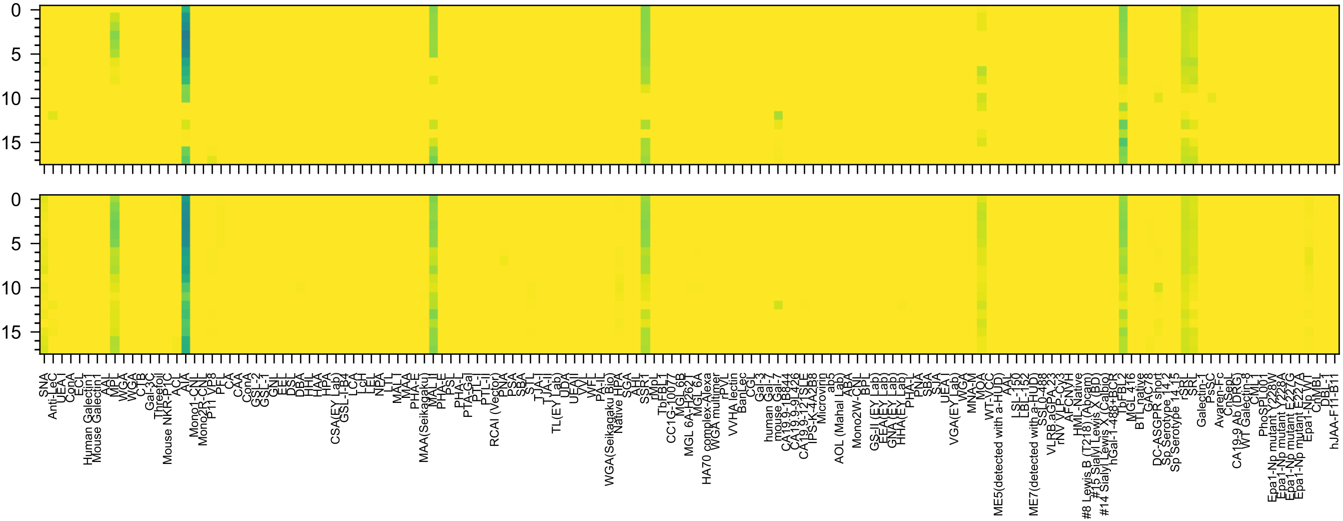
Kdn(α2-3)Gal(β1-3)GalNAc(α-Sp14

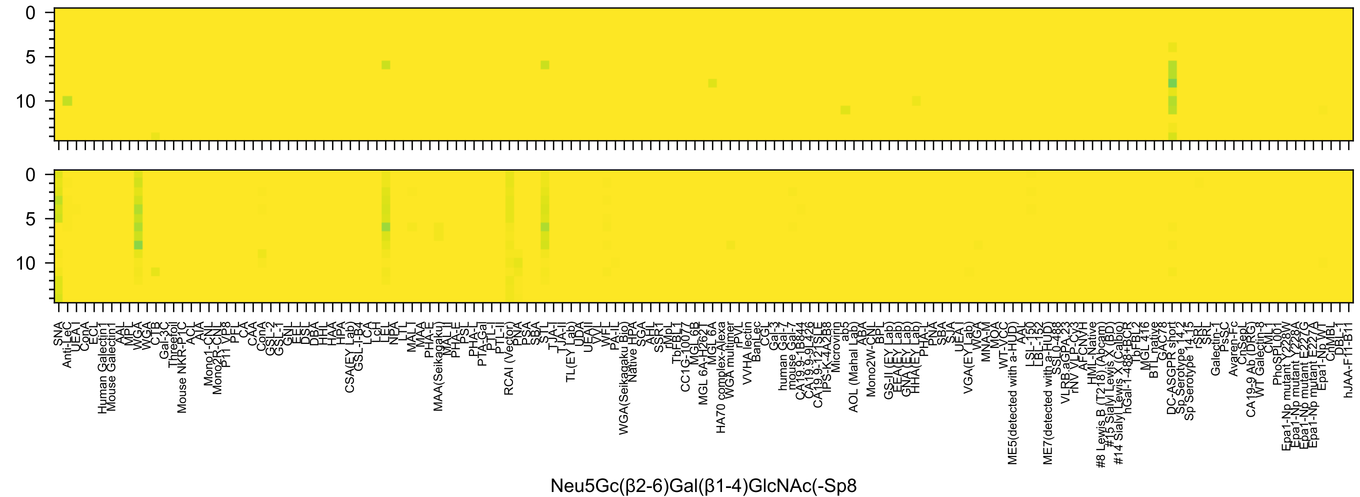
Neu5Gc(β2-6)Gal(β1-4)GlcNAc(-Sp8

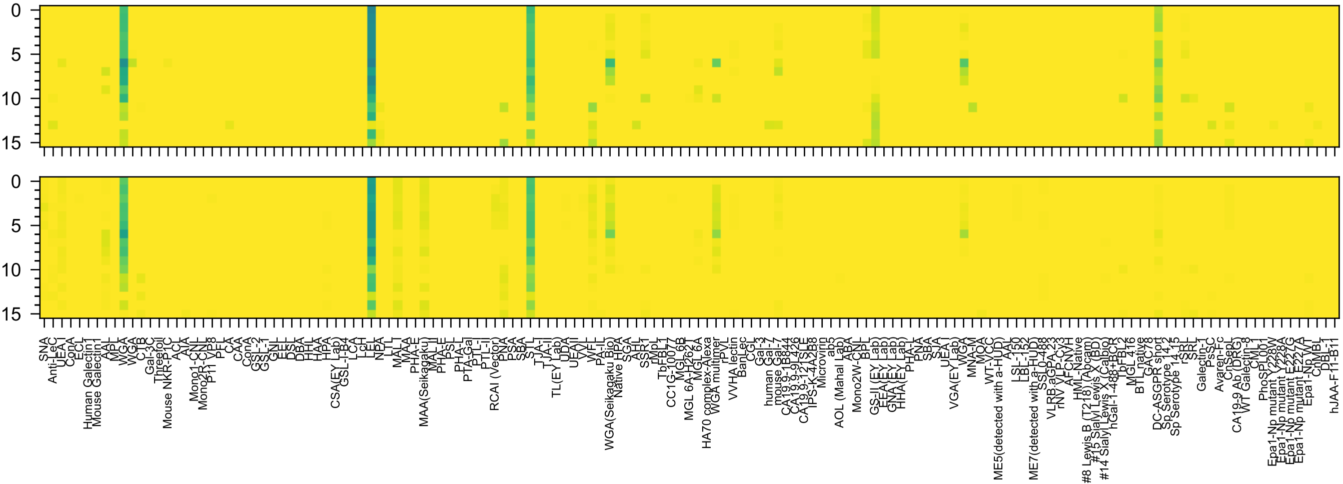


Neu5Ac(α2-3)GalNAc(β1-4)GlcNAc(β-Sp0

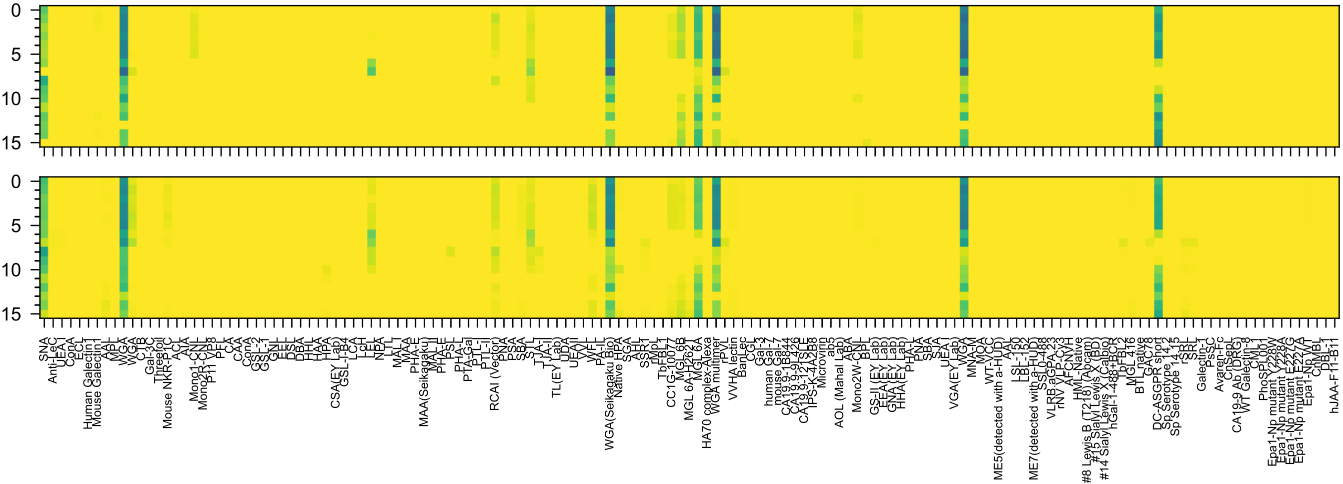


Neu5Ac(α2-6)GalNAc(β1-4)GlcNAc(β-Sp0

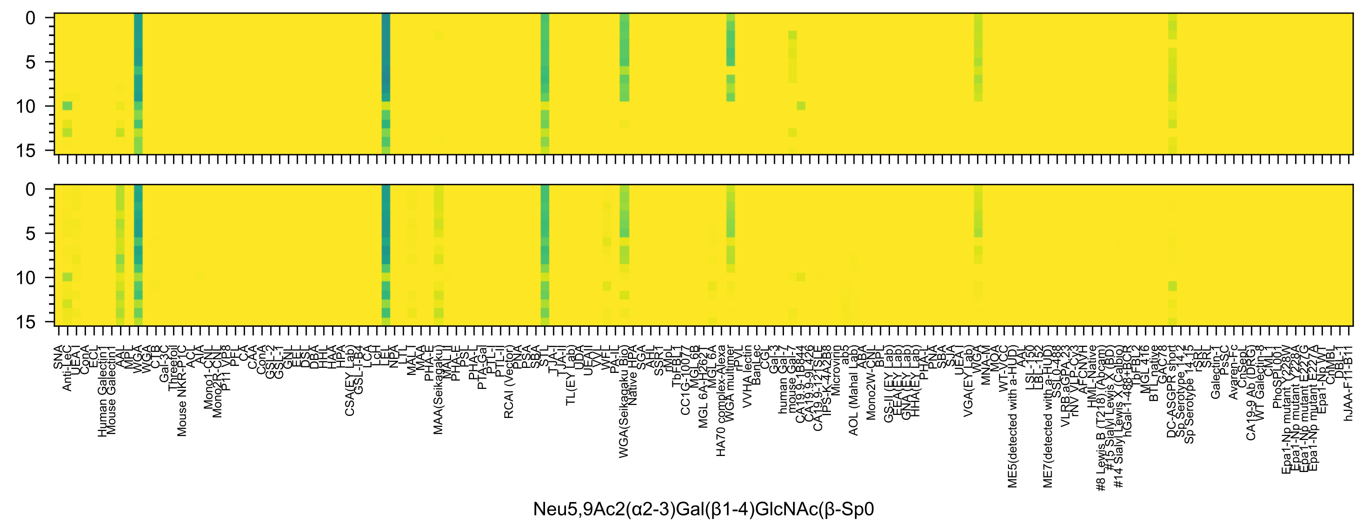
Neu5,9Ac2(α2-3)Gal(β1-4)GlcNAc(β-Sp0

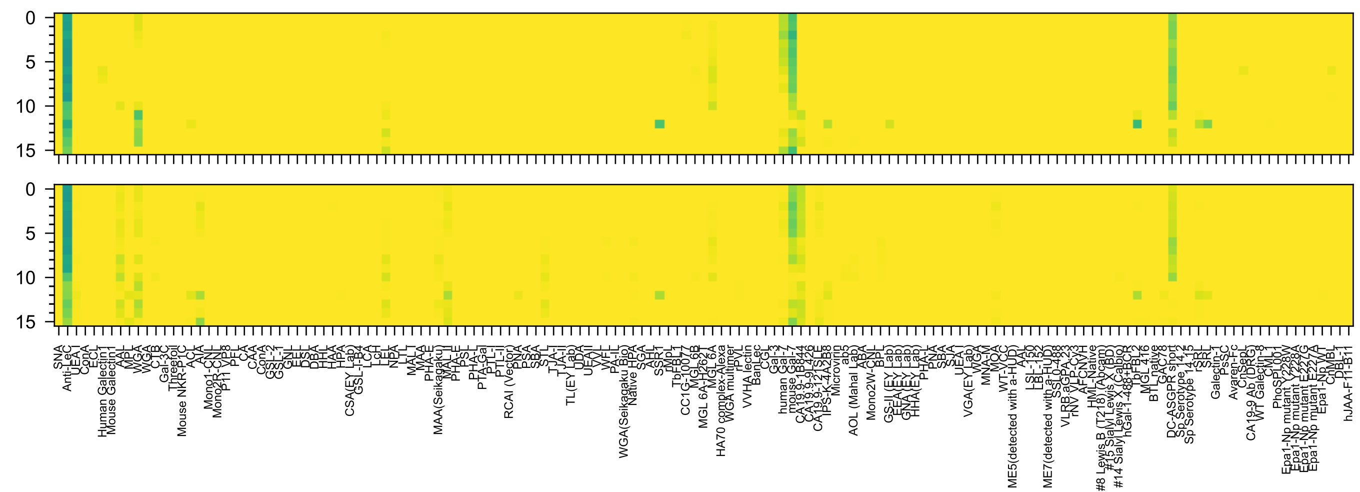


Neu5,9Ac2(α2-3)Gal(β1-3)GlcNAc(β-Sp0

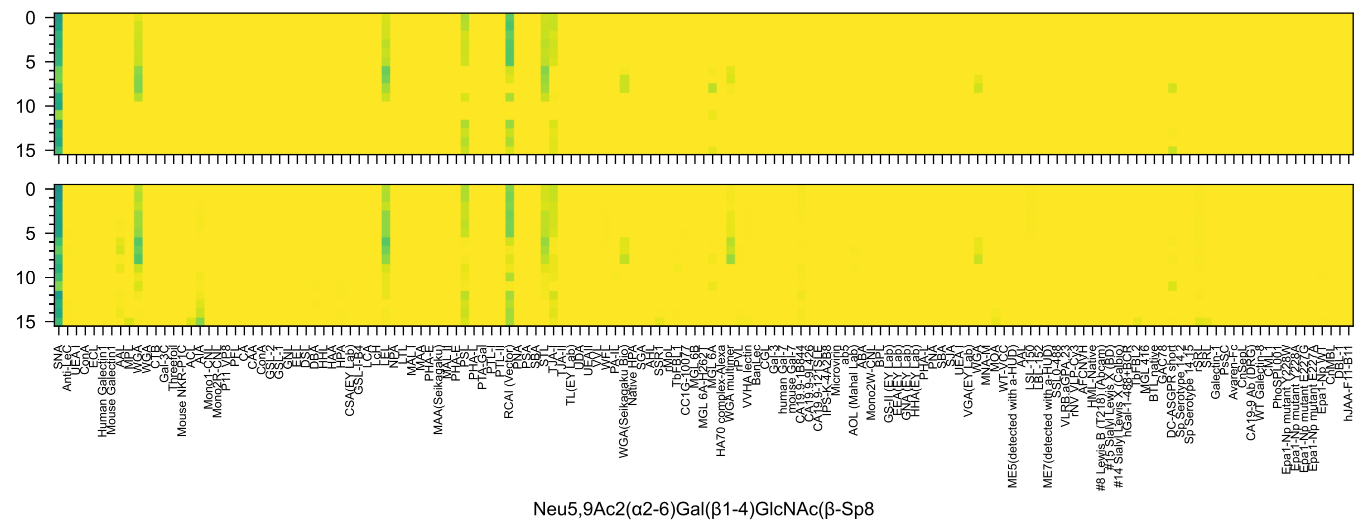

0
5
10
15
SNA
Anti-LeC
UEA-I
ConA
ECL
Human Galectin1
Mouse Galectin1
AAL
MPL
WGA
WGA
CTB
Gal-3C
Threefoil
Mouse NKR-P1C
ACL
AIA
Mono1-CNL
Mono2R-CNL
P11 VP8
PFL
CA
CAA
ConA
GSL-2
GSL-1
GNL
EEL
DSL
DBA
HHL
HAA
HPA
CSA(EY Lab)
GSL-I-B4
LCA
LcH
LEL
NPA
LTL
MAL-I
MAA
PHA-E
MAA(Seikagaku)
MAL-II
PHA-E
PSL
PHA-L
PTA-Gal
PTL-I
PTL-II
RCAI (Vector)
PNA
PSA
SBA
STL
TJA-I
TJA-II
TL(EY Lab)
UDA
UEAII
VVL
WFL
PA-IL
WGA(Seikagaku Bio)
Native HPA
SGA
AHL
SSR1
rMpL
TbFBL1
CC1G-10077
MGL 6B
MGL 6A-H262T
MGL 6A
HA70 complex-Alexa
WGA multimer
rPVL
VVHA lectin
BanLec
CGL
Gal-3
human Gal-7
mouse Gal-7
CA19.9-1B844
CA19.9-9L426
CA19.9-121SLE
IPS-K-4A2B8
Microvirin
ab5
AOL (Mahal Lab)
ABA
Mono2W-CNL
BPL
GS-II (EY Lab)
EEA (EY Lab)
GNA (EY Lab)
HHA (EY Lab)
PHA-L
PNA
SBA
SJA
UEA-I
VGA(EY Lab)
WGA
MNA-M
MOA
WT-VCC
ME5(detected with a-HUD)
AAL
LSL-150
LBL-152
ME7(detected with a-HUD)
SS10-488
VLRB.aGPA.23
rNV VLP-Cy3
AFCNVH
HML-Native
#8 Lewis B (T218) (Abcam)
#15 Sialyl Lewis X (BD)
#14 Sialyl Lewis X (Calbio)
hGal-1-488+BCR
TbFBL2
MGL 416
BTL native
GAC78
DC-ASGPR short
Sp Serotype 14.2
Sp Serotype 14.15
rSRL
SRL
Galectin-1
PsSC
Avaren-Fc
ChSeqL
CA19-9 Ab (DRG)
WT Galectin-8
CML1
PhoSL001
Epa1-Np mutant Y228W
Epa1-Np mutant Y228A
Epa1-Np mutant E227G
Epa1-Np mutant E227A
Epa1-Np WT
ChMBL
DBL-1
hJAA-F11-B11
Neu5,9Ac2(α2-6)Gal(β1-4)GlcNAc(β-Sp8

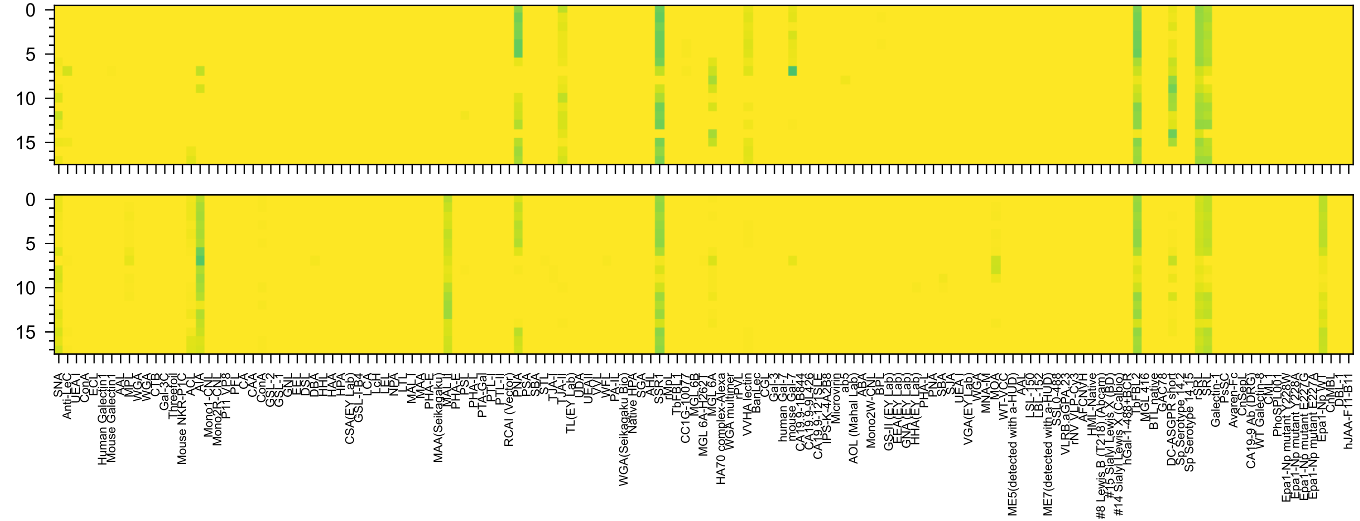


Neu5Ac(α2-6)[Gal(β1-3)]GalNAc(α-Sp14

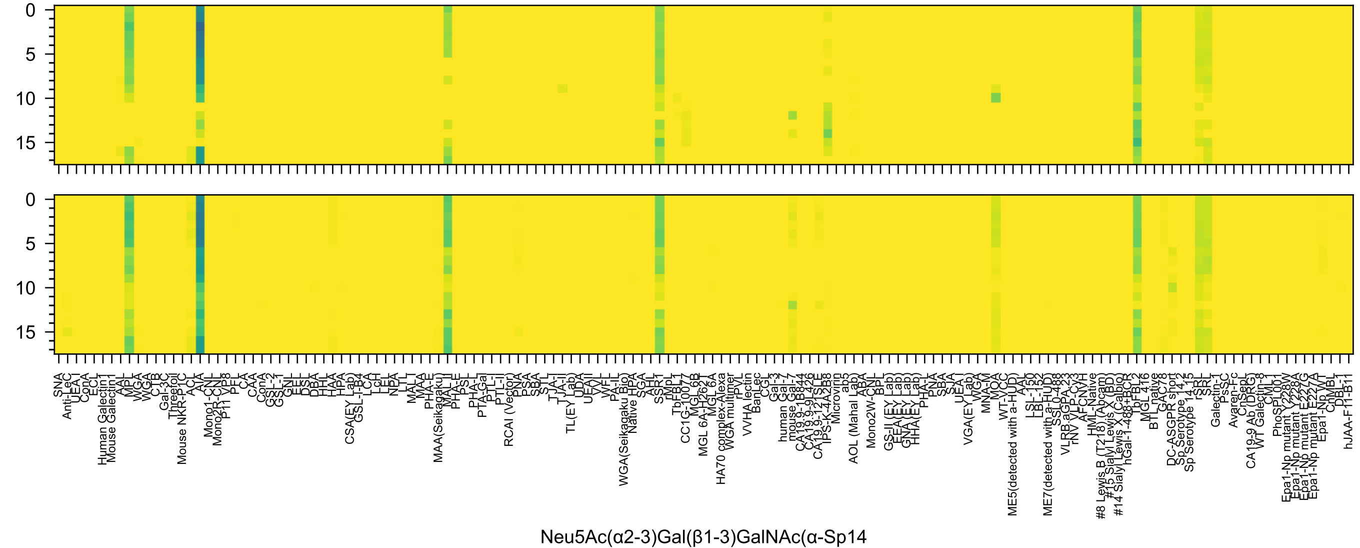
0
5
10
15
SNA
Anti-LeC
UEA I
ConA
ECL
Human Galectin1
Mouse Galectin1
AAL
MPL
WGA
WGA
CTB
Gal-3C
Threefoil
Mouse NKR-P1C
ACL
AIA
Mono1-CNL
Mono2R-CNL
P11 VP8
PFL
CA
CAA
ConA
GSL-2
GSL-1
GNL
EEL
DSL
DBA
HHL
HAA
HPA
CSA(EY Lab)
GSL-I-B4
LCA
LcH
LEL
NPA
LTL
MAL I
MAA
PHA-E
MAA(Seikagaku)
MAL II
PHA-E
PSL
PHA-L
PTA-Gal
PTL-I
PTL-II
RCAI (Vector)
PNA
PSA
SBA
STL
TJA-I
TJA-II
TL(EY Lab)
UDA
UEA II
VVL
WFL
PA-IL
WGA(Seikagaku Bio)
Native HPA
SGA
AH
SSR1
rMpL
TbFBL1
CC1G-10077
MGL 6B
MGL 6A-H262T
MGL 6A
HA70 complex-Alexa
WGA multimer
rPVL
VVHA lectin
BanLec
CGL
Gal-3
human Gal-7
mouse Gal-7
CA19.9-1B844
CA19.9-9L426
CA19.9-121SLE
IPS-K-4A2B8
Microvirin
ab5
AOL (Mahal Lab)
ABA
Mono2W-CNL
BPL
GS-II (EY Lab)
EEA(EY Lab)
GNA(EY Lab)
HHA(EY Lab)
PHA-L
PNA
SBA
SJA
UEA I
VGA(EY Lab)
WGA
MNA-M
MOA
WT-VCC
ME5(detected with a-HUD)
AAL
LSL-150
LBL-152
ME7(detected with a-HUD)
SSL0-488
VLRB aGPA.23
rNV VLP-Cy3
AFCNVH
HML-Native
#8 Lewis B (T218) (Abcam)
#15 Sialyl Lewis X (BD)
#14 Sialyl Lewis X (Calbio)
hGal-1-488+BCR
TbFBL2
MGL 416
BTL native
GAC78
DC-ASGPR short
Sp Serotype 14.2
Sp Serotype 14.15
rSRL
SRL
Galectin-1
PsSC
Avaren-Fc
ChSep1
CA19-9 Ab (DRG)
WT Galectin-8
CML1
PhoSL001
Epa1-Np mutant Y228W
Epa1-Np mutant Y228A
Epa1-Np mutant E227G
Epa1-Np mutant E227A
Epa1-Np WT
ChMBL
DBL-1
hJAA-F11-B11
Neu5Ac(α2-3)Gal(β1-3)GalNAc(α-Sp14

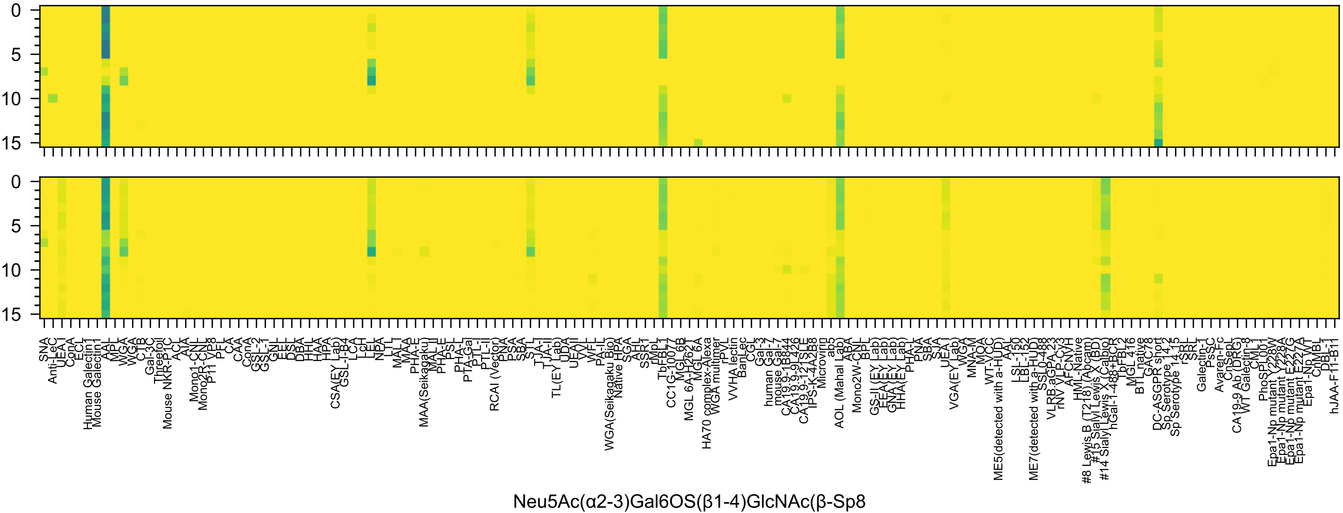
Neu5Ac(α2-3)Gal6OS(β1-4)GlcNAc(β-Sp8

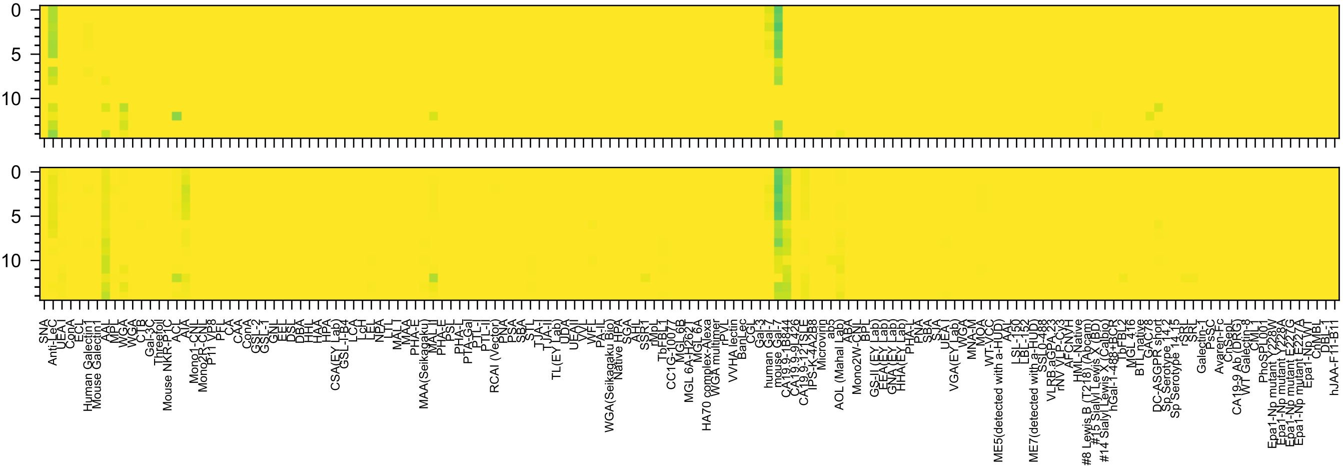
Neu5Ac(α2-3)Gal(β1-3)GlcNAc6OS(-Sp8

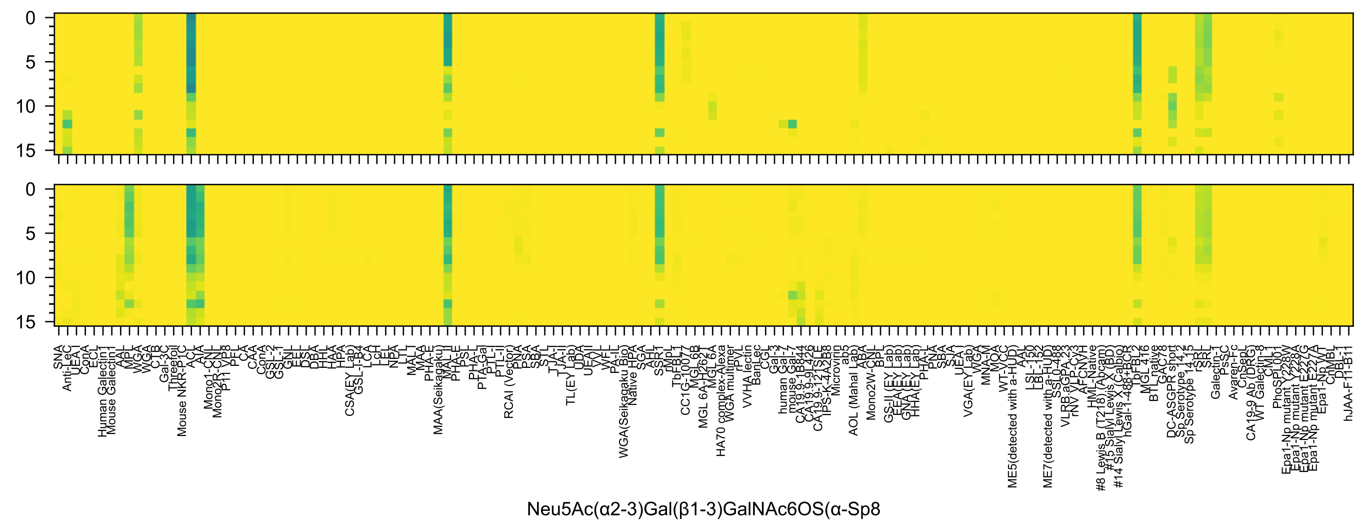
Neu5Ac(α2-3)Gal(β1-3)GalNAc6OS(α-Sp8

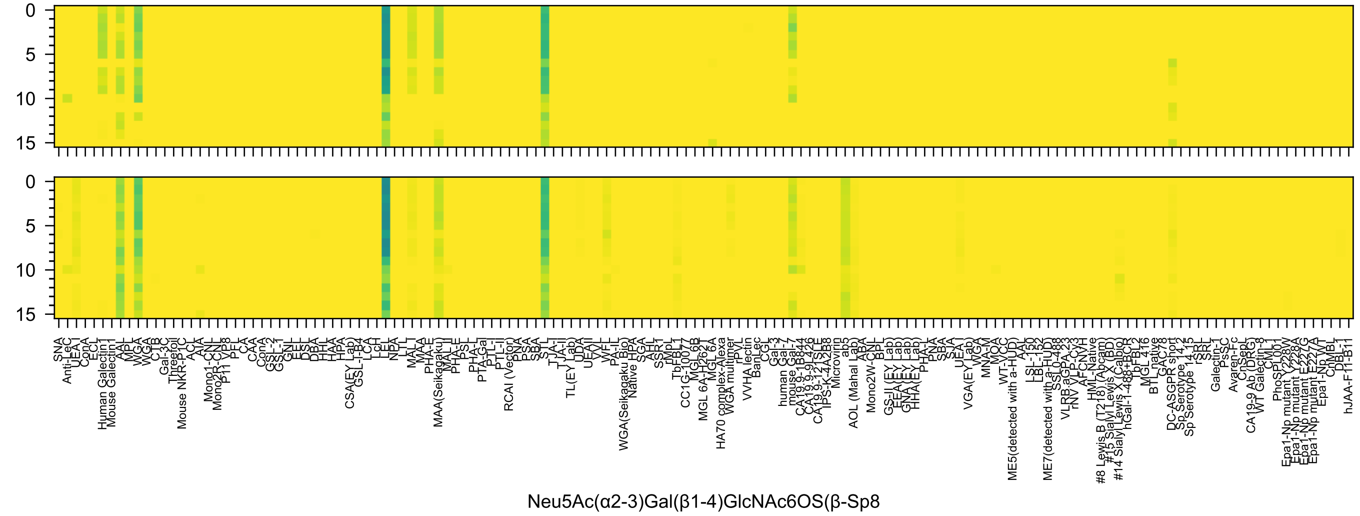
Neu5Ac(α2-3)Gal(β1-4)GlcNAc6OS(β-Sp8

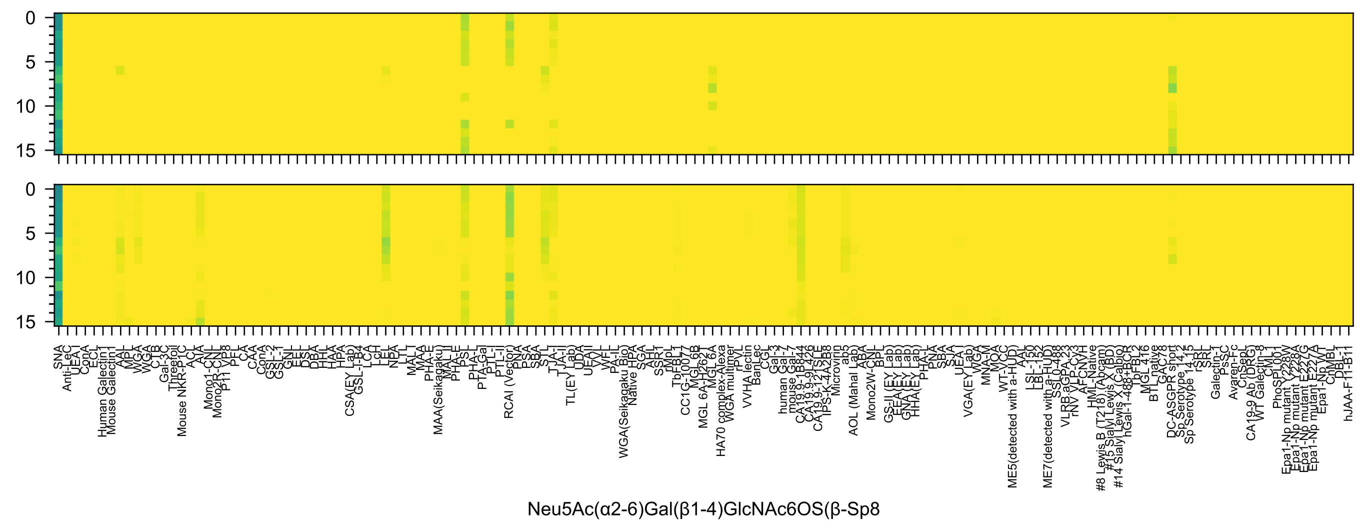
Neu5Ac(α2-6)Gal(β1-4)GlcNAc6OS(β-Sp8

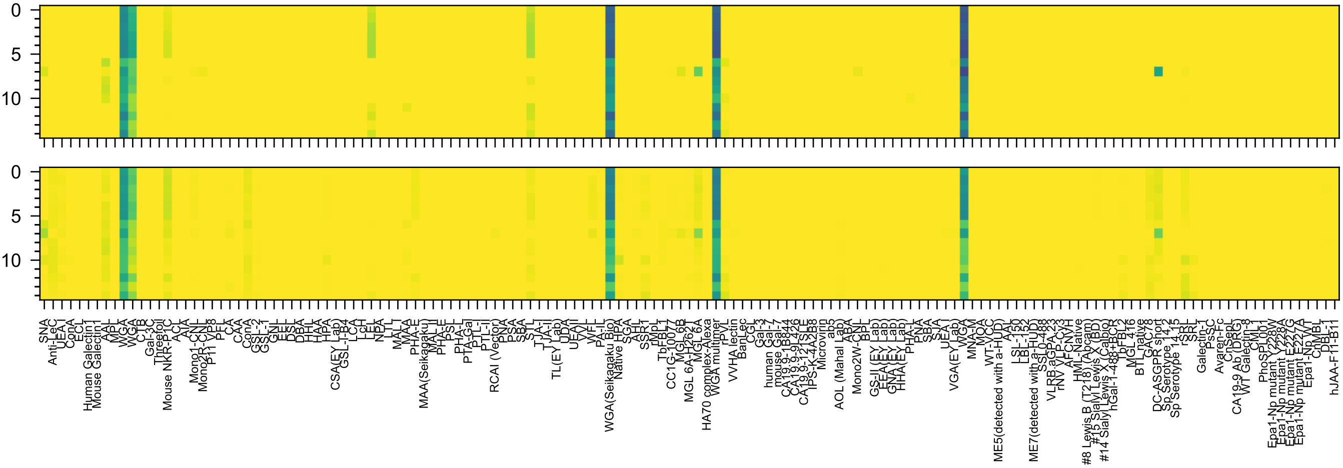

Neu5Ac(α2-6)GlcNAc(β1-4)GlcNAc(-Sp21

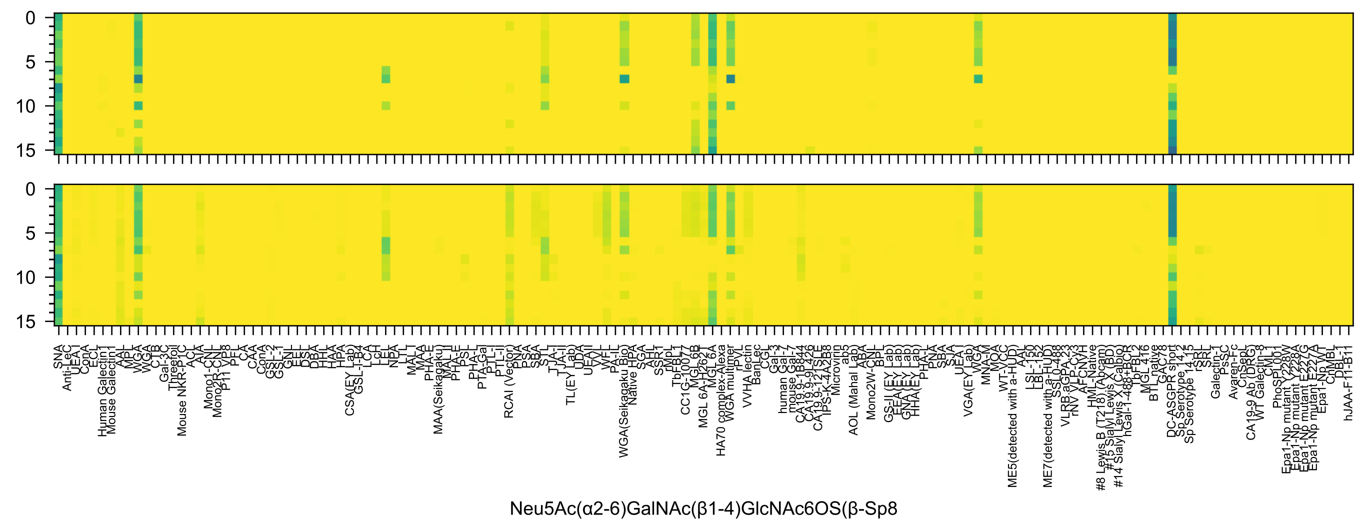
0
5
10
15
0
5
10
15
SNA
Anti-LeC
UEA-I
ConA
ECL
Human Galectin1
Mouse Galectin1
AAL
MPL
WGA
WGA
CTB
C-1
Gal-3C
Threefoil
Mouse NKR-P1C
ACL
AIA
Mono1-CNL
Mono2R-CNL
P11_VP8
PFL
CA
CAA
ConA
GSL-2
GSL-1
GNL
EEL
DSL
DBA
HHL
HAA
HPA
CSA(EY Lab)
GSL-I-B4
LCA
LcH
LEL
NPA
LTL
MAL-I
MAA
PHA-E
MAA(Seikagaku)
MAL-II
PSA
PHA-L
PTA-Gal
PTL-I
PTL-II
RCAI (Vector)
PNA
PSA
SBA
STL
TJA-I
TJA-II
TL(EY Lab)
UDA
UEA-II
VVL
WFL
PA-IL
WGA(Seikagaku Bio)
Native HPA
SGA
AHL
SSR1
rMpL
TbFBL1
CC1G_10077
MGL 6B
MGL 6A-H262T
MGL 6A
HA70 complex-Alexa
WGA multimer
rPVL
VVHA lectin
BanLec
CGL
Gal-3
human Gal-7
mouse Gal-7
CA19-9-1B844
CA19-9-9L426
CA19-9-121SLE
IPS-K-4A2B8
Microvirin
ab5
AOL (Mahal Lab)
ABA
Mono2W-CNL
BPL
GS-II (EY Lab)
EEA (EY Lab)
GNA (EY Lab)
HHA (EY Lab)
PHA-L
PNA
SBA
SJA
UEA-I
VGA(EY Lab)
WGA
MNA-M
MOA
WT-VCC
ME5(detected with a-HUD)
AAL
LSL-150
LBL-152
ME7(detected with a-HUD)
SSL0-488
VLRB.aGPA.23
rNV VLP-Cy3
AFCNVH
HML-Native
#8 Lewis B (T218) (Abcam)
#15 Sialyl Lewis X (BD)
#14 Sialyl Lewis X (Calbio)
hGal-1-488+BCR
TbFBL2
MGL 416
BTL native
GAC78
DC-ASGPR short
Sp Serotype 14.2
Sp Serotype 14.15
rSRL
SRL
Galectin-1
PsSC
Avaren-Fc
ChSeP
CA19-9 Ab (DRG)
WT Galectin-8
CML
PhoSL001
Epa1-Np mutant Y228W
Epa1-Np mutant Y228A
Epa1-Np mutant E227G
Epa1-Np mutant E227A
Epa1-Np WT
ChMBL
DBL-1
hJAA-F11-B11
Neu5Ac(α2-6)GalNAc(β1-4)GlcNAc6OS(β-Sp8

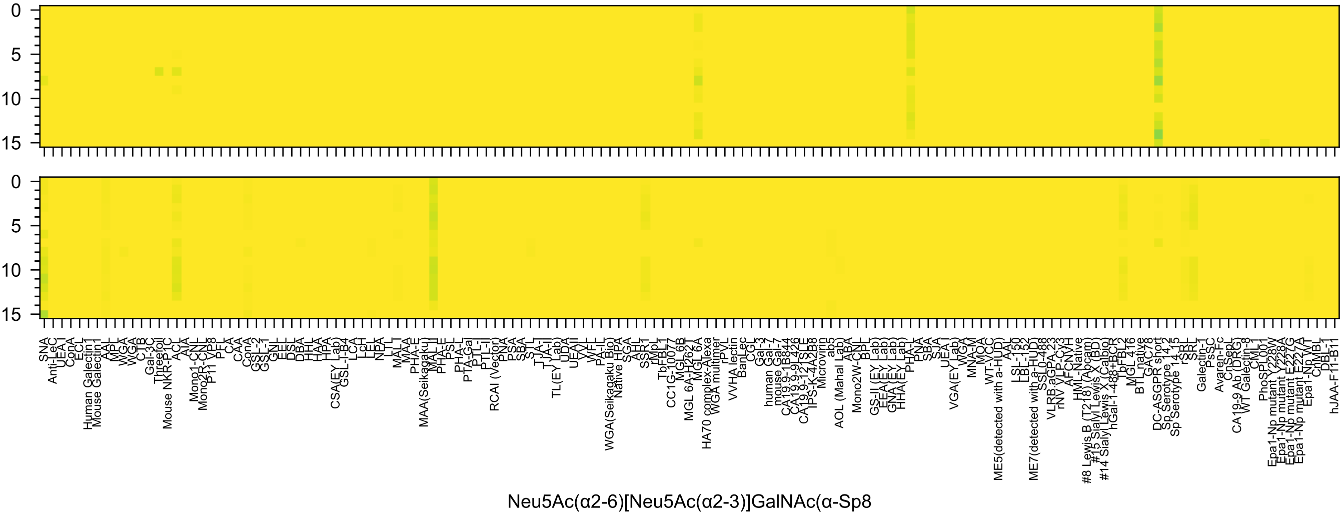
Neu5Ac(α2-6)[Neu5Ac(α2-3)]GalNAc(α-Sp8

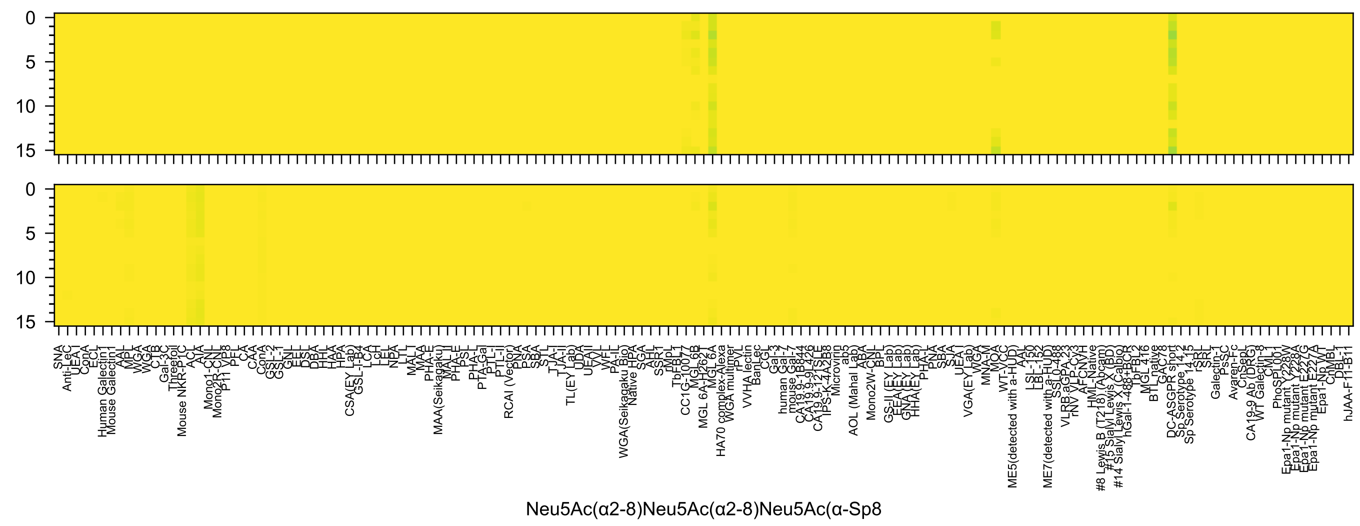

Neu5Ac(α2-8)Neu5Ac(α2-8)Neu5Ac(α-Sp8

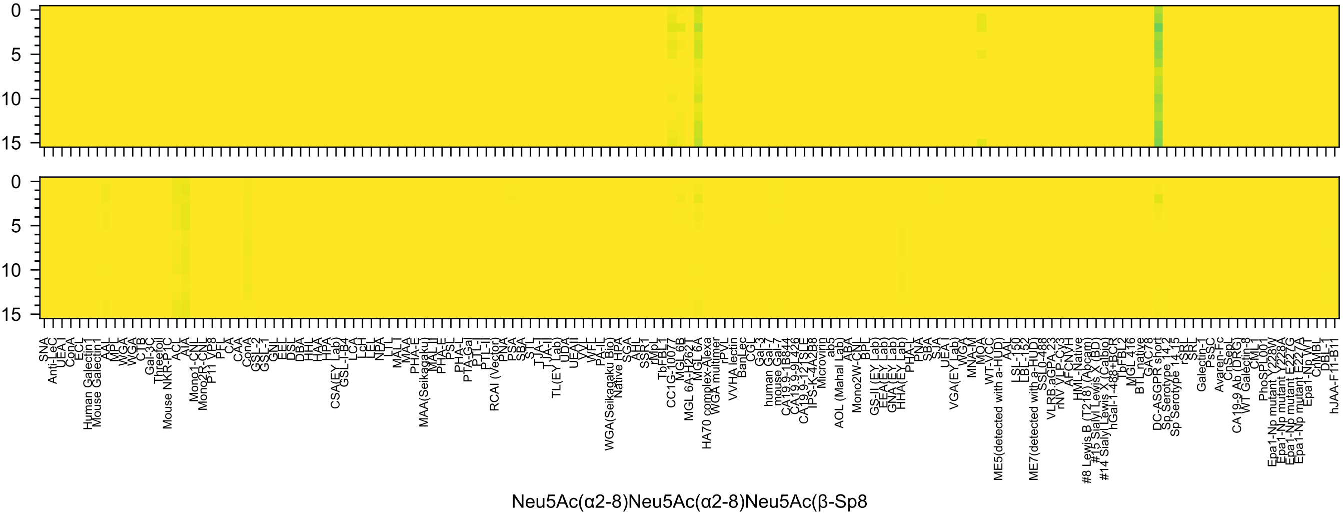

Neu5Ac(α2-8)Neu5Ac(α2-8)Neu5Ac(β-Sp8

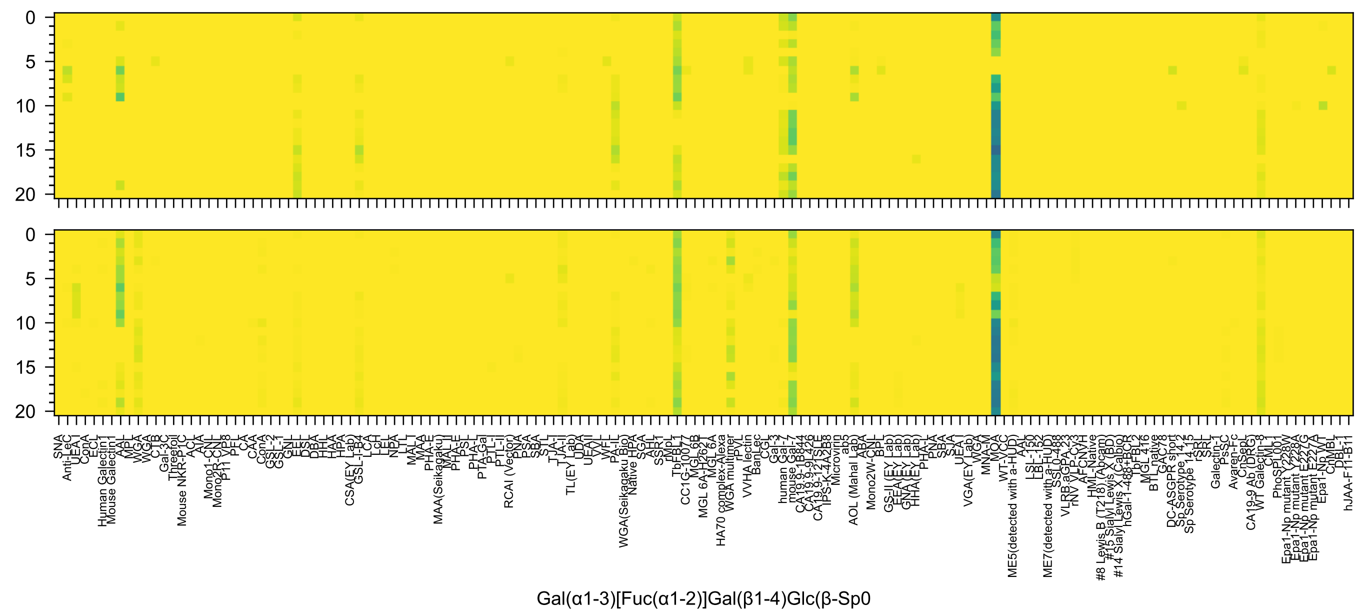
0
5
10
15
20
0
5
10
15
20
SNA
Anti-LeC
UEA I
ConA
ECL
Human Galectin1
Mouse Galectin1
AAL
MPL
WGA
WGA
CTB
Gal-3C
Threefoil
Mouse NKR-P1C
ACL
AIA
Mono1-CNL
Mono2R-CNL
P11 VP8
PFL
CA
CAA
ConA
GSL-2
GSL-1
GNL
EEL
DSL
DBA
HHL
HAA
HPA
CSA(EY Lab)
GSL-I-B4
LCA
LcH
LEL
NPA
LTL
MAL I
MAA
PHA-E
MAA(Seikagaku)
MAL II
PHA-E
PSL
PHA-L
PTA-Gal
PTL-I
PTL-II
RCAI (Vector)
PNA
PSA
SBA
STL
TJA-I
TJA-II
TL(EY Lab)
UDA
UEAII
VVL
WFL
PA-IL
WGA(Seikagaku Bio)
Native HPA
SGA
AHL
SSR1
rMpL
TbFBL1
CC1G-10077
MGL 6B
MGL 6A-H262T
MGL 6A
HA70 complex-Alexa
WGA multimer
rPVL
VVHA lectin
BanLec
CGL
Gal-3
human Gal-7
mouse Gal-7
CA19.9-1B844
CA19.9-9L426
CA19.9-121SLE
IPS-K-4A2B8
Microvirin
ab5
AOL (Mahal Lab)
ABA
Mono2W-CNL
BPL
GS-II (EY Lab)
EEA(EY Lab)
GNA(EY Lab)
HHA(EY Lab)
PHA-L
PNA
SBA
SJA
UEA1
VGA(EY Lab)
WGA
MNA-M
MOA
WT-VCC
ME5(detected with a-HUD)
AAL
LSL-150
LBL-152
ME7(detected with a-HUD)
SSL0-488
VLRB.aGPA.23
rNV VLP-Cy3
AFCNVH
HML-Native
#8 Lewis B (T218) (Abcam)
#15 Sialyl Lewis X (BD)
#14 Sialyl Lewis X (Calbio)
hGal-1-488+BCR
bFBL2
MGL 416
BTL native
GAC78
DC-ASGPR short
Sp Serotype 14.2
Sp Serotype 14.15
rSRL
SRL
Galectin-1
PsSC
Avaren-Fc
ChSepL
CA19-9 Ab (DRG)
WT Galectin-8
CML
PhoSL001
Epa1-Np mutant Y228W
Epa1-Np mutant Y228A
Epa1-Np mutant E227G
Epa1-Np mutant E227A
Epa1-Np WT
ChMBL
DBL-1
hJAA-F11-B11
Gal(α1-3)[Fuc(α1-2)]Gal(β1-4)Glc(β-Sp0

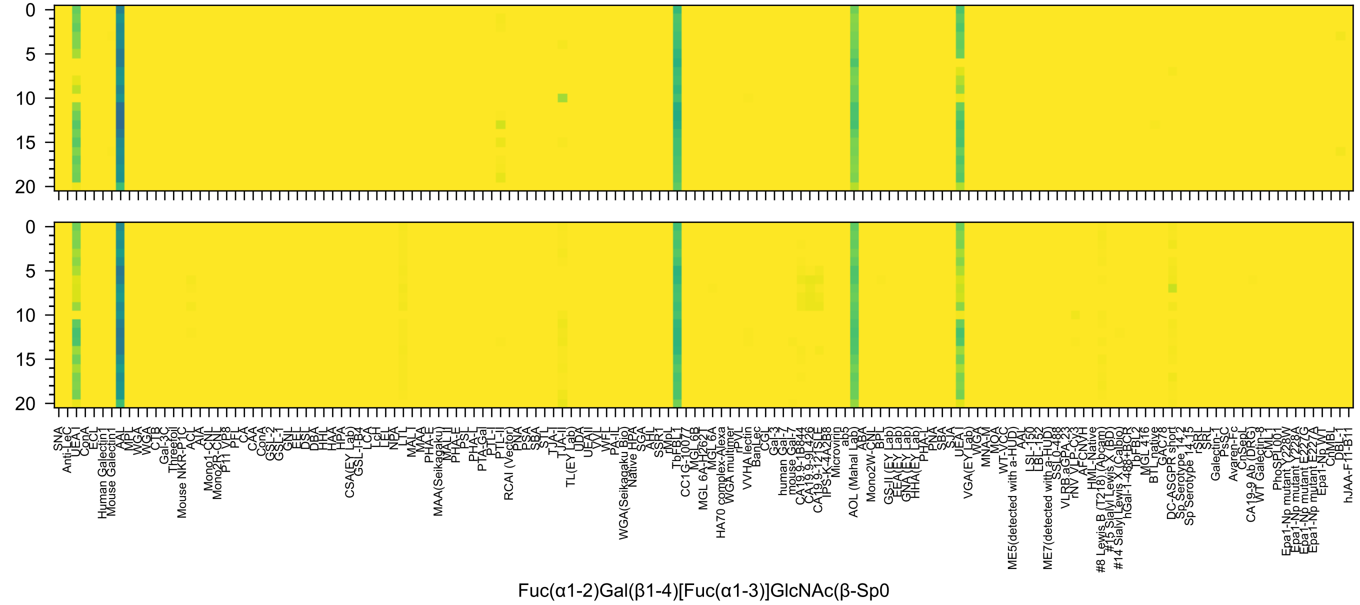
Fuc(α1-2)Gal(β1-4)[Fuc(α1-3)]GlcNAc(β-Sp0

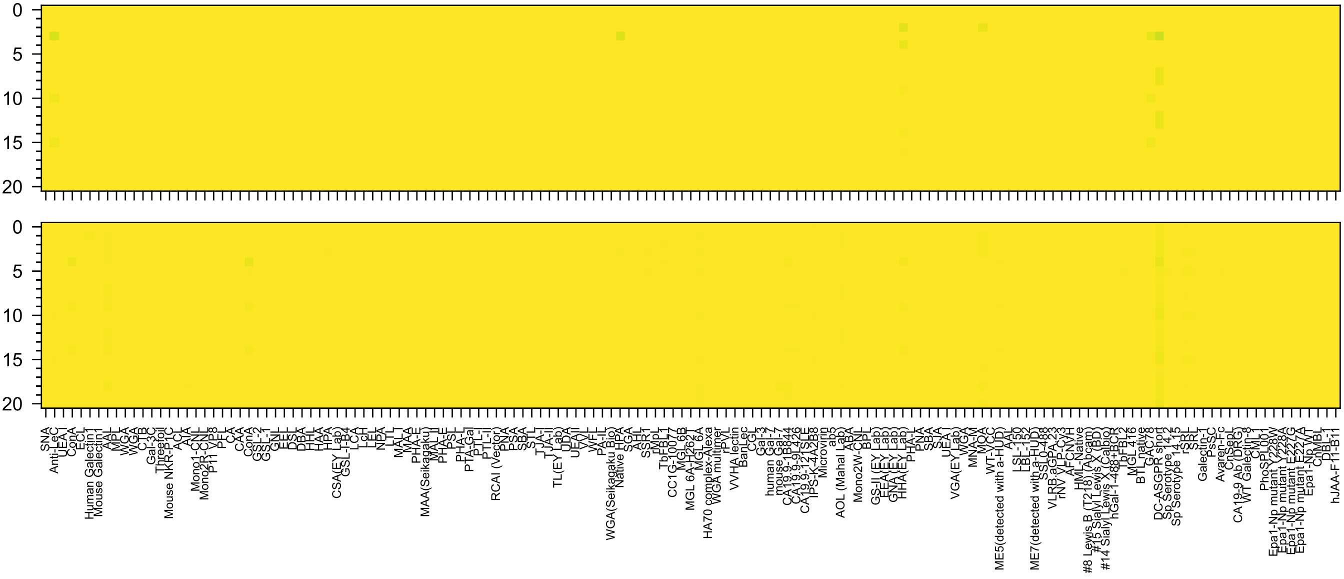

Glc(α1-6)Glc(α1-6)Glc(α1-6)Glc(β-Sp10

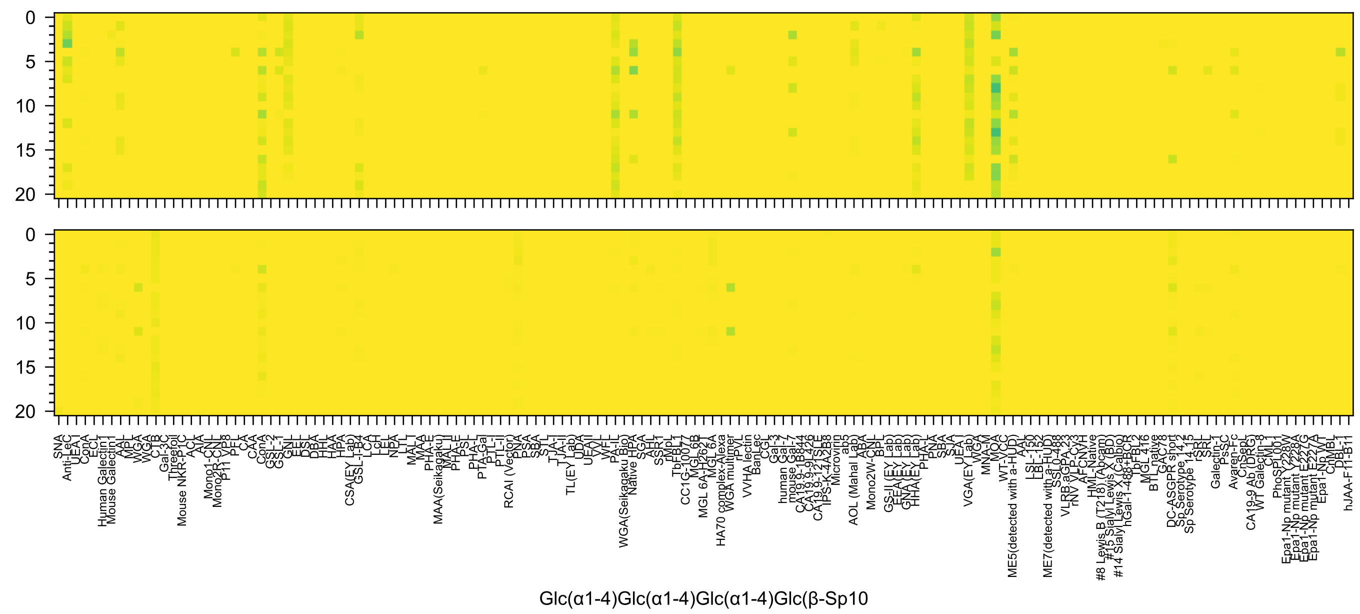
Glc(α1-4)Glc(α1-4)Glc(α1-4)Glc(β-Sp10

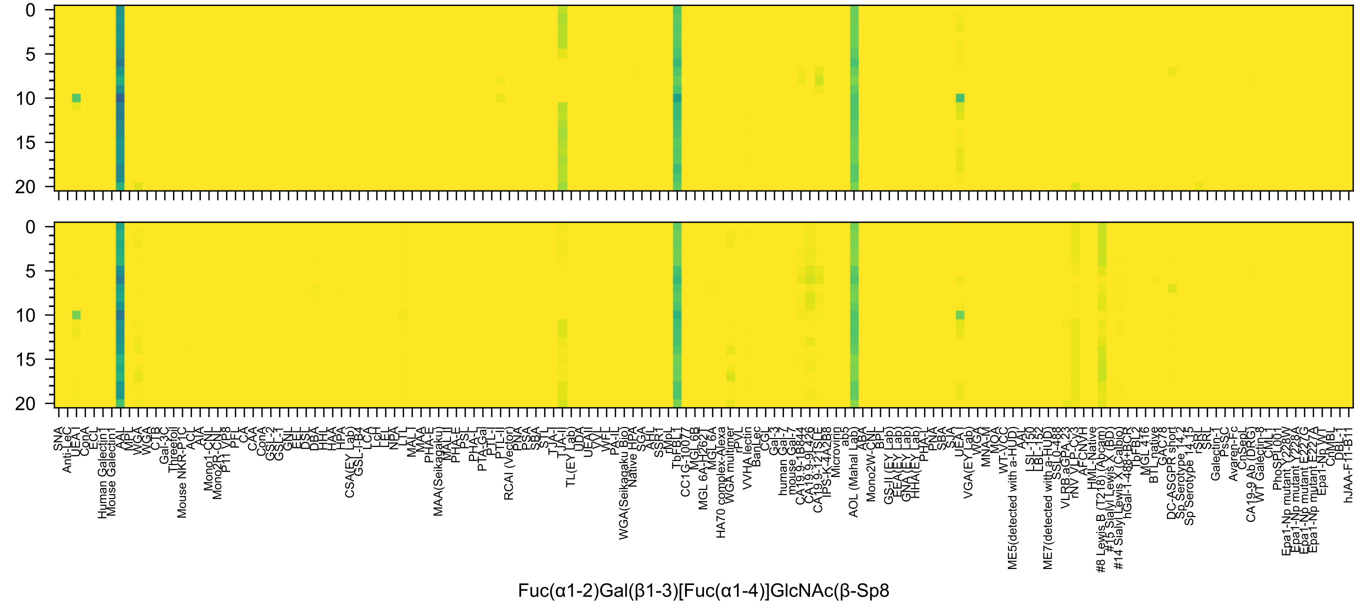
0
5
10
15
20
0
5
10
15
20
SNA
Anti-LeC
UEA I
ConA
ECL
Human Galectin1
Mouse Galectin1
AAL
MPL
WGA
WGA
CTB
Gal-3C
Threefoil
Mouse NKR-P1C
ACL
AIA
Mono1-CNL
Mono2R-CNL
P11 VP8
PFL
CA
CAA
ConA
GSL-2
GSL-1
GNL
EEL
DSL
DBA
HHL
HAA
HPA
CSA(EY Lab)
GSL-I-B4
LCA
LcH
LEL
NPA
LTL
MAL
MAA
PHA-E
MAA(Seikagaku)
MAL-I
PHA-E
PSL
PHA-L
PTA-Gal
PTL-I
PTL-II
RCAI (Vector)
PNA
PSA
SBA
STL
TJA-I
TJA-II
TL(EY Lab)
UDA
UEAII
VVL
WFL
PA-IL
WGA(Seikagaku Bio)
Native HPA
SGA
AHL
SSR1
rMpL
TbFBL1
CC1G-10077
MGL 6B
MGL 6A-H262T
MGL 6A
HA70 complex-Alexa
WGA multimer
rPVL
VVHA lectin
BanLec
CGL
Gal-3
human Gal-7
mouse Gal-7
CA19.9-1B844
CA19.9-9L426
CA19.9-121SLE
IPS-K-4A2B8
Microvirin
ab5
AOL (Mahal Lab)
ABA
Mono2W-CNL
BPL
GS-II (EY Lab)
EEA (EY Lab)
GNA (EY Lab)
HHA (EY Lab)
PHA-L
PNA
SBA
SJA
UEA-I
VGA(EY Lab)
WGA
MNA-M
MOA
WT-VCC
ME5(detected with a-HUD)
AAL
LSL-150
LBL-152
ME7(detected with a-HUD)
SSL0-488
VLRB.aGPA.23
rNV VLP-Cy3
AFCNVH
HML-Native
#8 Lewis B (T218) (Abcam)
#15 Sialyl Lewis X (BD)
#14 Sialyl Lewis X (Calbio)
hGal-1-488+BCR
TbFBL2
MGL 416
BTL native
GAC78
DC-ASGPR short
Sp Serotype 14.2
Sp Serotype 14.15
rSRL
SRL
Galectin-1
PsSC
Avaren-Fc
CnSepL
CA19-9 Ab (DRG)
WT Galectin-8
CML
PhoSPL001
Epa1-Np mutant Y228W
Epa1-Np mutant Y228A
Epa1-Np mutant E227G
Epa1-Np mutant E227A
Epa1-Np WT
ChMBL
DBL-1
hJAA-F11-B11
Fuc(α1-2)Gal(β1-3)[Fuc(α1-4)]GlcNAc(β-Sp8

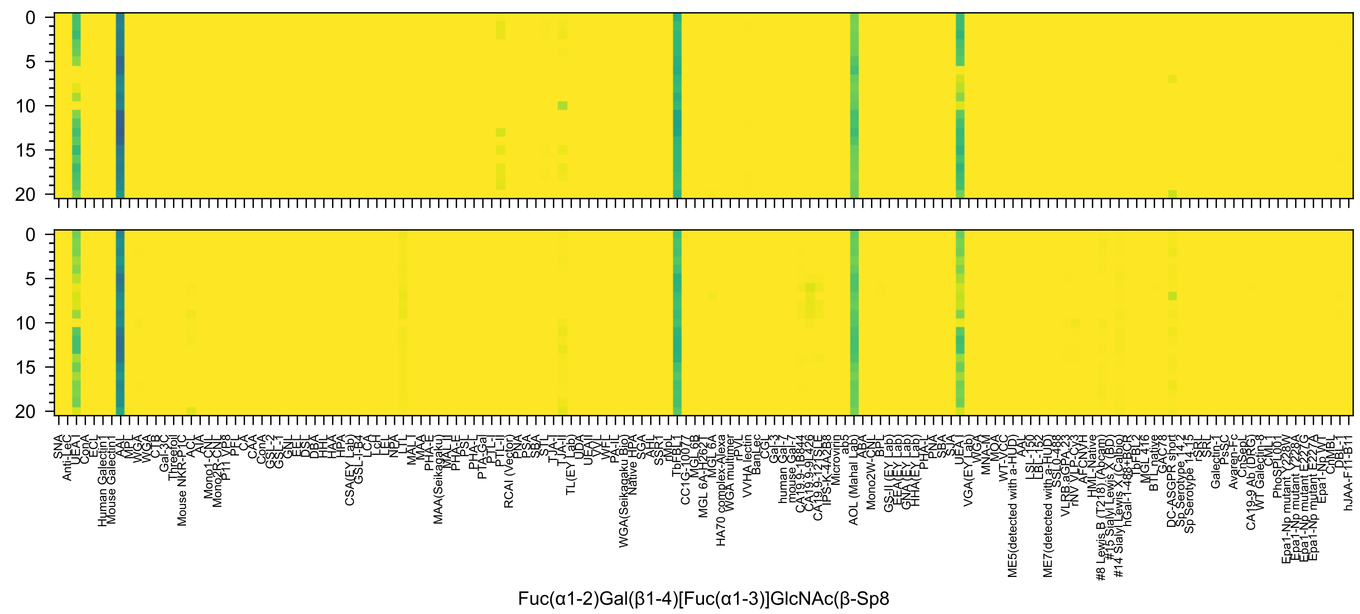
Fuc(α1-2)Gal(β1-4)[Fuc(α1-3)]GlcNAc(β-Sp8

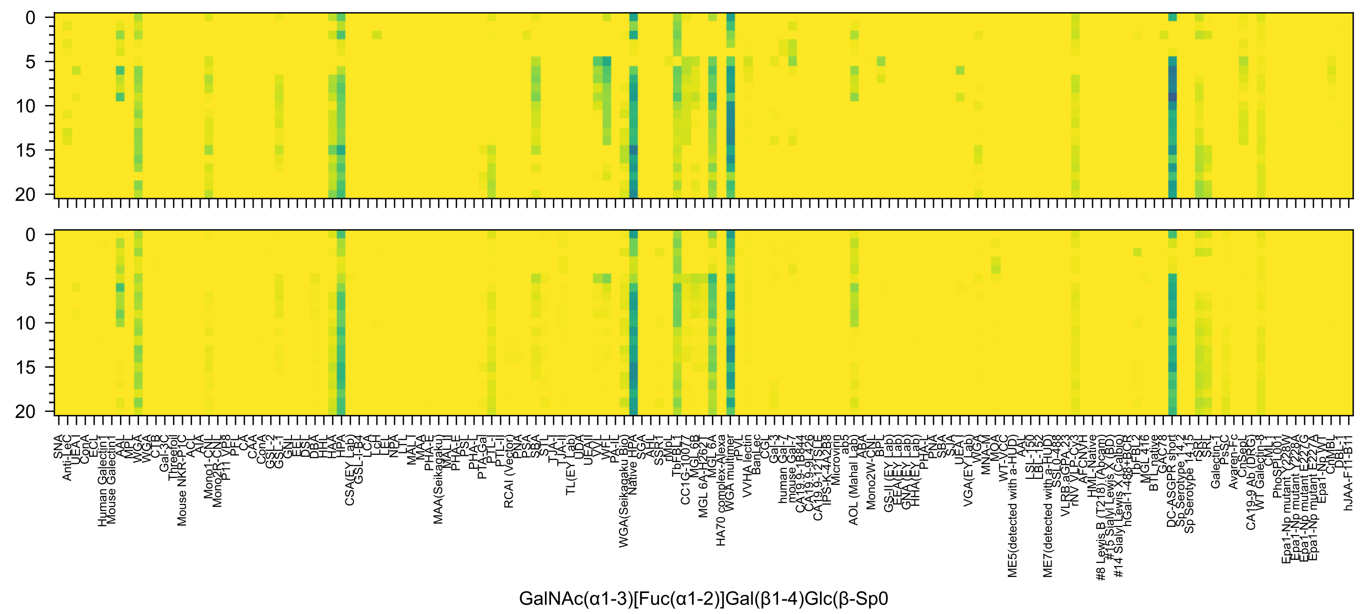
GalNAc(α1-3)[Fuc(α1-2)]Gal(β1-4)Glc(β-Sp0

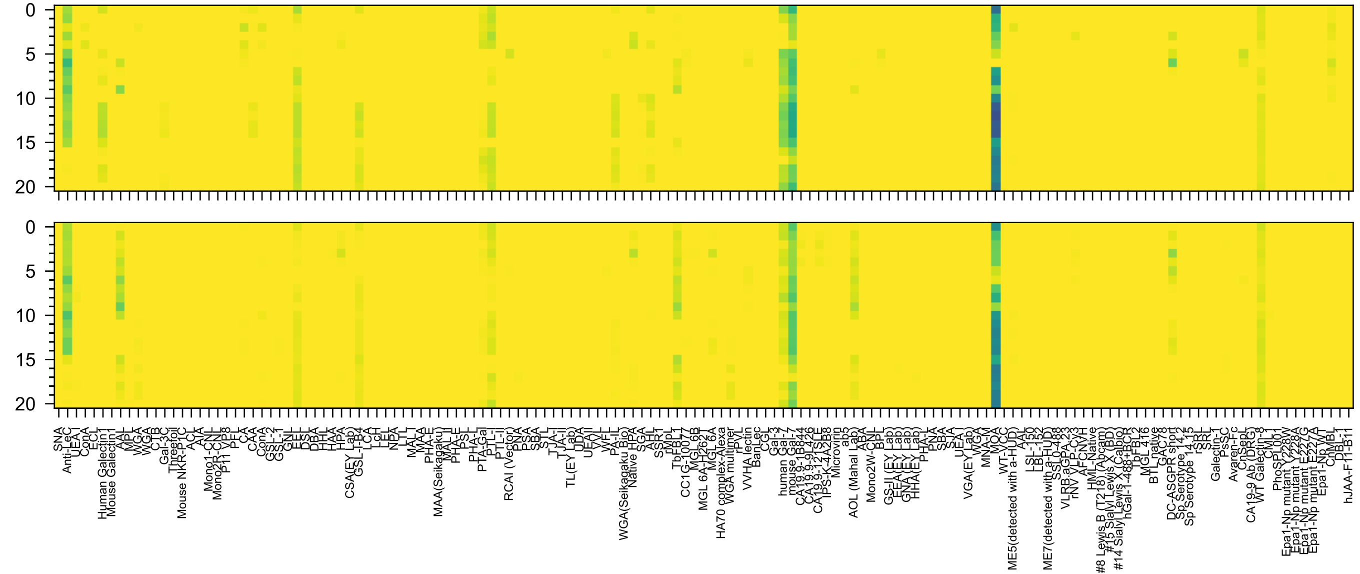

Gal(α1-3)[Fuc(α1-2)]Gal(β1-3)GlcNAc(β-Sp0

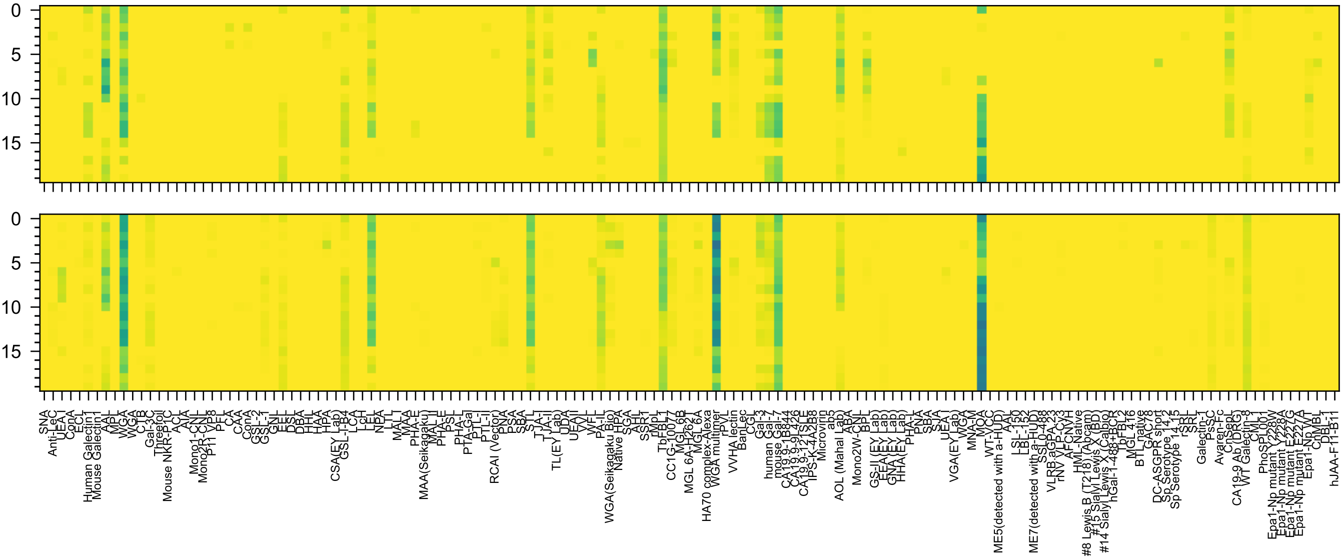

Gal(α1-3)[Fuc(α1-2)]Gal(β1-4)GlcNAc(-Sp0

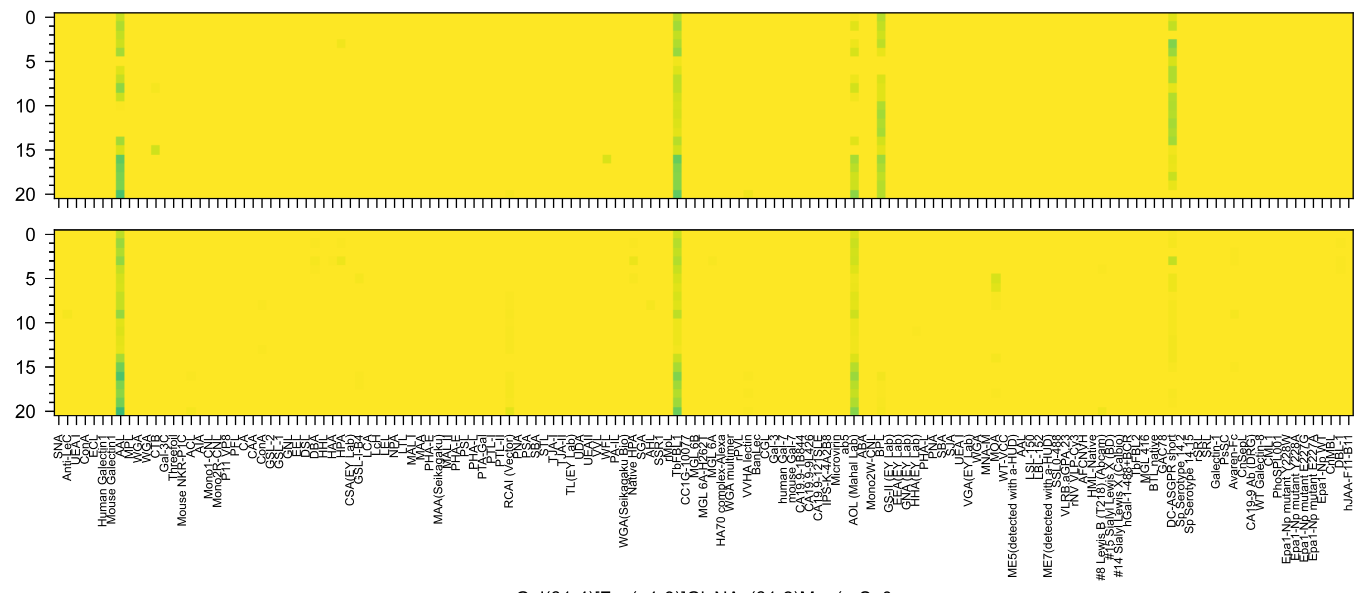
Gal(β1-4)[Fuc(α1-3)]GlcNAc(β1-2)Man(α-Sp0

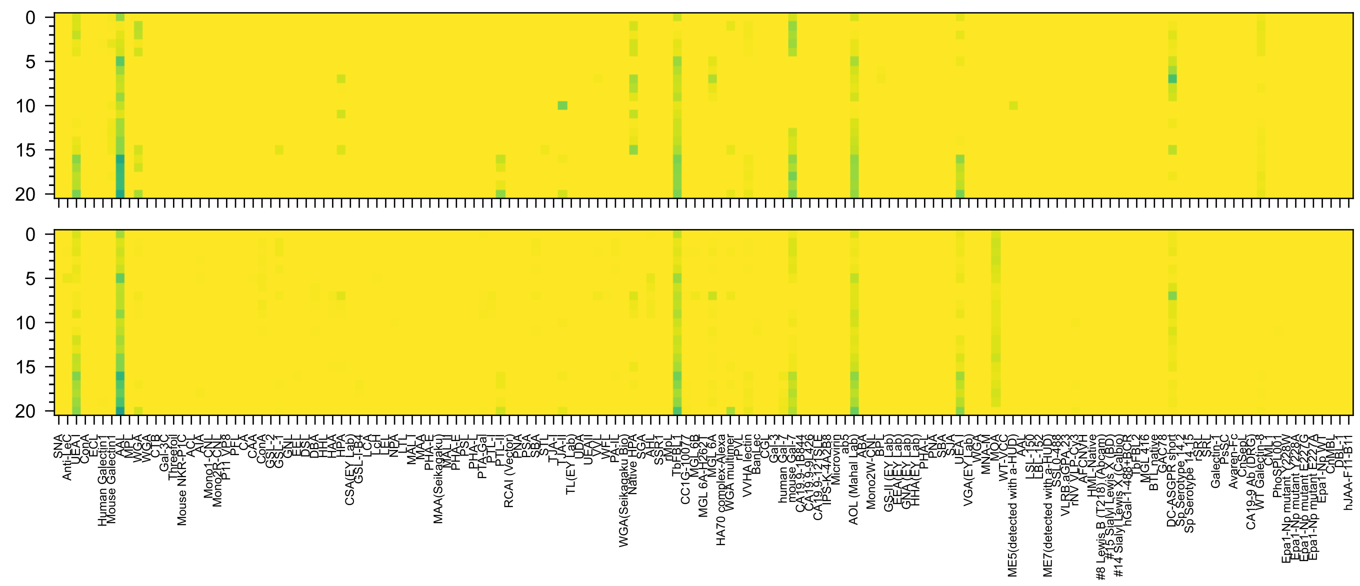

Fuc(α1-2)Gal(β1-4)GlcNAc(β1-2)Man(α-Sp0

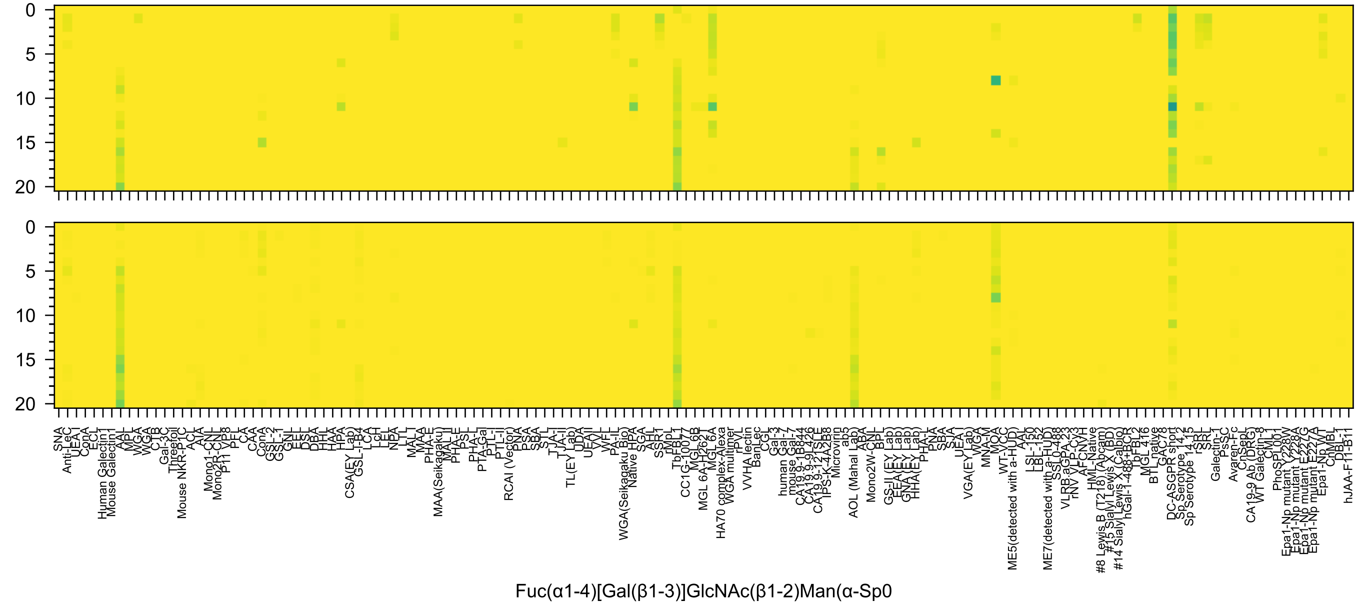
0
5
10
15
20
SNA
Anti-LeC
UEA-I
ConA
ECL
Human Galectin1
Mouse Galectin1
AAL
MPL
WGA
WGA
CTB
Gal-3C
Threefoil
Mouse NKR-P1C
ACL
AIA
Mono1-CNL
Mono2R-CNL
P11 VP8
PFL
CA
CAA
ConA
GSL-2
GSL-1
GNL
EEL
DSL
DBA
HHL
HAA
HPA
CSA(EY Lab)
GSL-I-B4
LCA
LcH
LEL
NPA
LTL
MAL-I
MAA
PHA-E
MAA(Seikagaku)
MAL-II
PHA-E
PSL
PHA-L
PTA-Gal
PTL-I
PTL-II
RCAI (Vector)
PNA
PSA
SBA
STL
TJA-I
TJA-II
TL(EY Lab)
UDA
UEA-II
VVL
WFL
PA-IL
WGA(Seikagaku Bio)
Native HPA
SGA
AHL
SSR1
rMpL
TbFBL1
CC1G-10077
MGL 6B
MGL 6A-H262T
MGL 6A
HA70 complex-Alexa
WGA multimer
rPVL
VVHA lectin
BanLec
CGL
Gal-3
human Gal-7
mouse Gal-7
CA19.9-1B844
CA19.9-9L426
CA19.9-121SLE
IPS-K-4A2B8
Microvirin
ab5
AOL (Mahal Lab)
ABA
Mono2W-CNL
BPL
GS-II (EY Lab)
EEA(EY Lab)
GNA(EY Lab)
HHA(EY Lab)
PHA-L
PNA
SBA
SJA
UEA-I
VGA(EY Lab)
WGA
MNA-M
MOA
WT-VCC
ME5(detected with a-HUD)
AAL
LSL-150
LBL-152
ME7(detected with a-HUD)
SSL0-488
VLRB.aGPA.23
rNV VLP-Cy3
AFCNVH
HML-Native
#8 Lewis B (T218) (Abcam)
#15 Sialyl Lewis X (BD)
#14 Sialyl Lewis X (Calbio)
hGal-1-488+BCR
TbFBL2
MGL 416
BTL native
GAC78
DC-ASGPR short
Sp Serotype 14.2
Sp Serotype 14.15
rSRL
SRL
Galectin-1
PsSC
Avaren-Fc
ChSepL
CA19-9 Ab (DRG)
WT Galectin-8
CML
PhoSL001
Epa1-Np mutant Y228W
Epa1-Np mutant Y228A
Epa1-Np mutant E227G
Epa1-Np mutant E227A
Epa1-Np WT
ChMBL
DBL-1
hJAA-F11-B11
Fuc(α1-4)[Gal(β1-3)]GlcNAc(β1-2)Man(α-Sp0

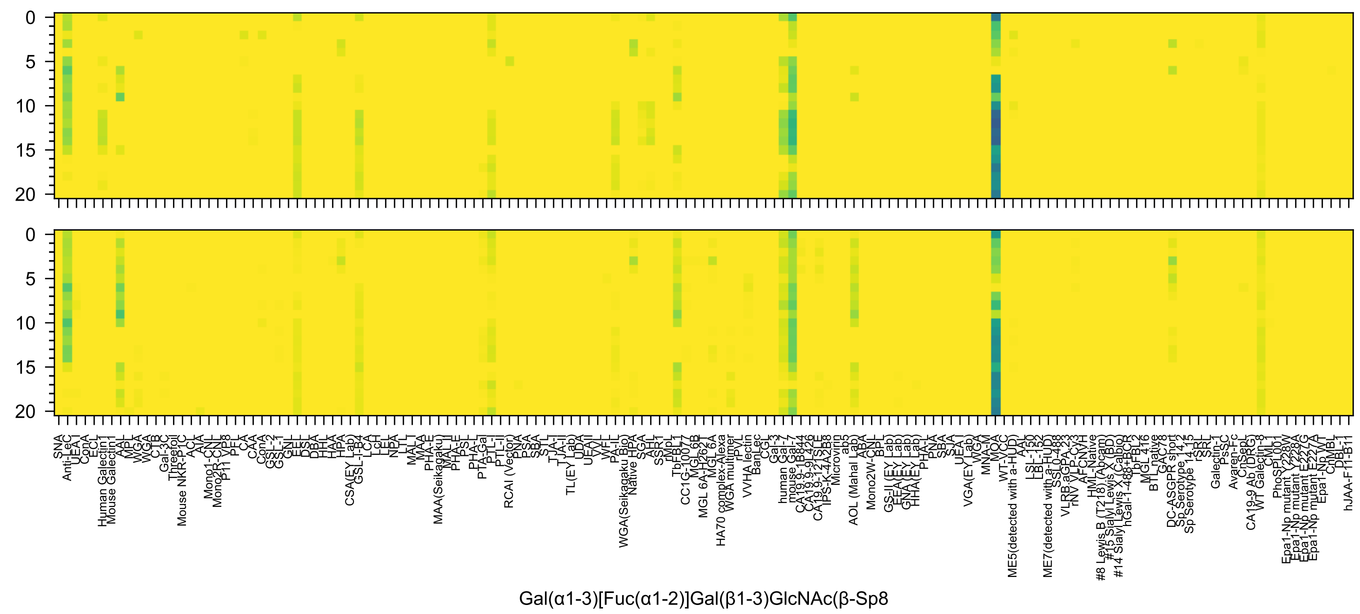
Gal(α1-3)[Fuc(α1-2)]Gal(β1-3)GlcNAc(β-Sp8

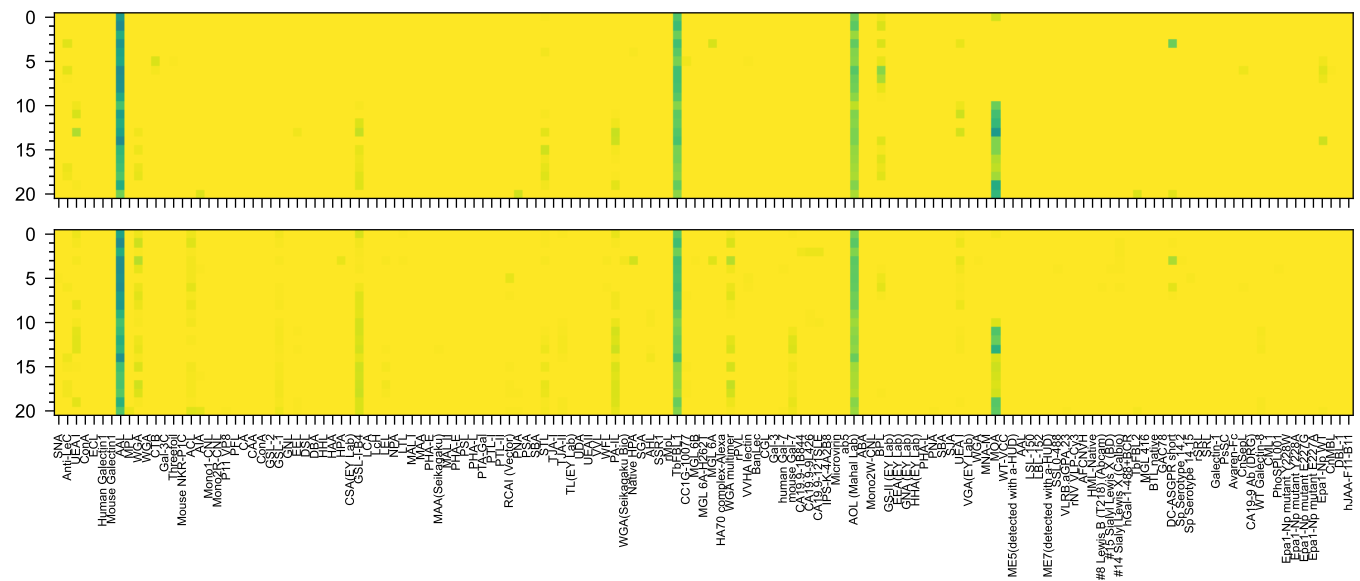

Gal(α1-3)Gal(β1-4)[Fuc(α1-3)]GlcNAc(β-Sp8

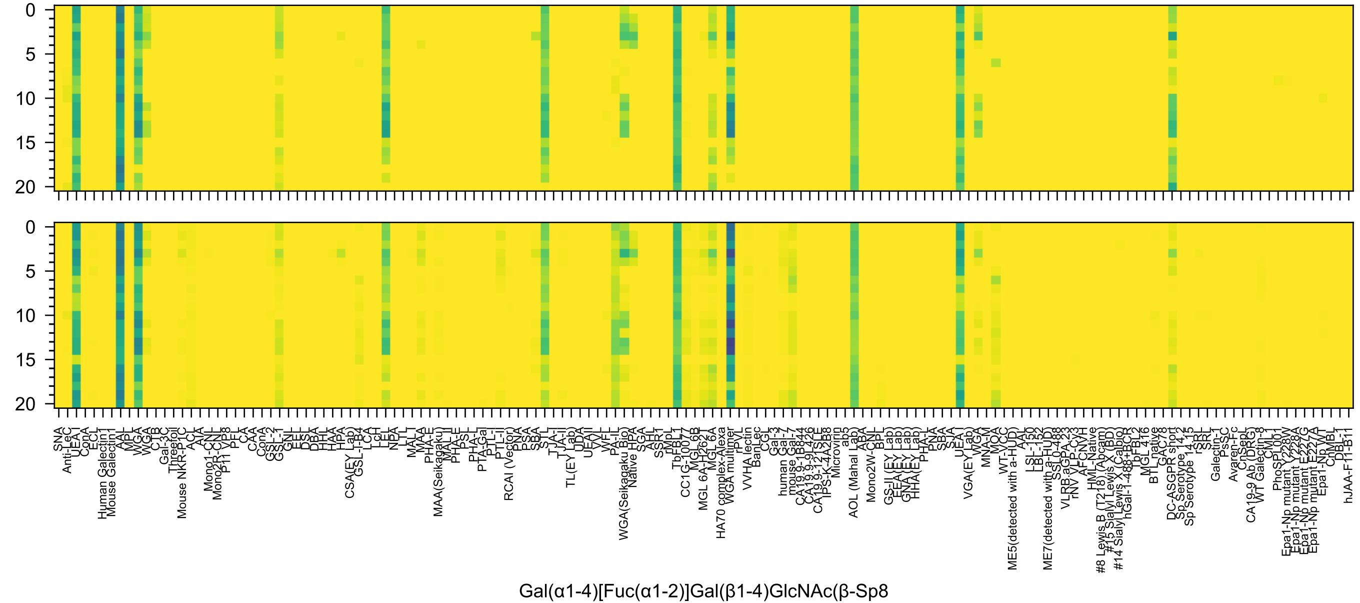
0
5
10
15
20
0
5
10
15
20
SNA
Anti-LeC
UEA-I
ConA
ECL
Human Galectin1
Mouse Galectin1
AAL
MPL
WGA
WGA
CTB
Gal-3C
Threefoil
Mouse NKR-P1C
ACL
AIA
Mono1-CNL
Mono2R-CNL
P11 VP8
PFL
CA
CAA
ConA
GSL-2
GSL-1
GNL
EEL
DSL
DBA
HHL
HAA
HPA
CSA(EY Lab)
GSL-I-B4
LCA
LcH
LEL
NPA
LTL
MAL-I
MAA
PHA-E
MAA(Seikagaku)
MAL-II
PHA-E
PSL
PHA-L
PTA-Gal
PTL-I
PTL-II
RCAI (Vector)
PNA
PSA
SBA
STL
TJA-I
TJA-II
TL(EY Lab)
UDA
UEAII
VVL
WFL
PA-IL
WGA(Seikagaku Bio)
Native HPA
SGA
AHL
SSR1
rMpL
TbFBL1
CC1G-10077
MGL 6B
MGL 6A-H262T
MGL 6A
HA70 complex-Alexa
WGA multimer
rPVL
VVHA lectin
BanLec
CGL
Gal-3
human Gal-7
mouse Gal-7
CA19.9-1B844
CA19.9-9L426
CA19.9-121SLE
IPS-K-4A2B8
Microvirin
ab5
AOL (Mahal Lab)
ABA
Mono2W-CNL
BPL
GS-II (EY Lab)
EEA(EY Lab)
GNA(EY Lab)
HHA(EY Lab)
PHA-L
PNA
SBA
SJA
UEA-I
VGA(EY Lab)
WGA
MNA-M
MOA
WT-VCC
ME5(detected with a-HUD)
AAL
LSL-150
LBL-152
ME7(detected with a-HUD)
SSL0-488
VLRB.aGPA.23
rNV VLP-Cy3
AFCNVH
HML-Native
#8 Lewis B (T218) (Abcam)
#15 Sialyl Lewis X (BD)
#14 Sialyl Lewis X (Calbio)
hGal-1-488+BCR
TbFBL2
MGL 416
BTL native
GAC78
DC-ASGPR short
Sp Serotype 14.2
Sp Serotype 14.15
rSRL
SRL
Galectin-1
PsSC
Avaren-Fc
ChSeL
CA19-9 Ab (DRG)
WT Galectin-8
CML1
PhoSPL001
Epa1-Np mutant Y228W
Epa1-Np mutant Y228A
Epa1-Np mutant E227G
Epa1-Np mutant E227A
Epa1-Np WT
ChMBL
DBL-1
hJAA-F11-B11
Gal(α1-4)[Fuc(α1-2)]Gal(β1-4)GlcNAc(β-Sp8

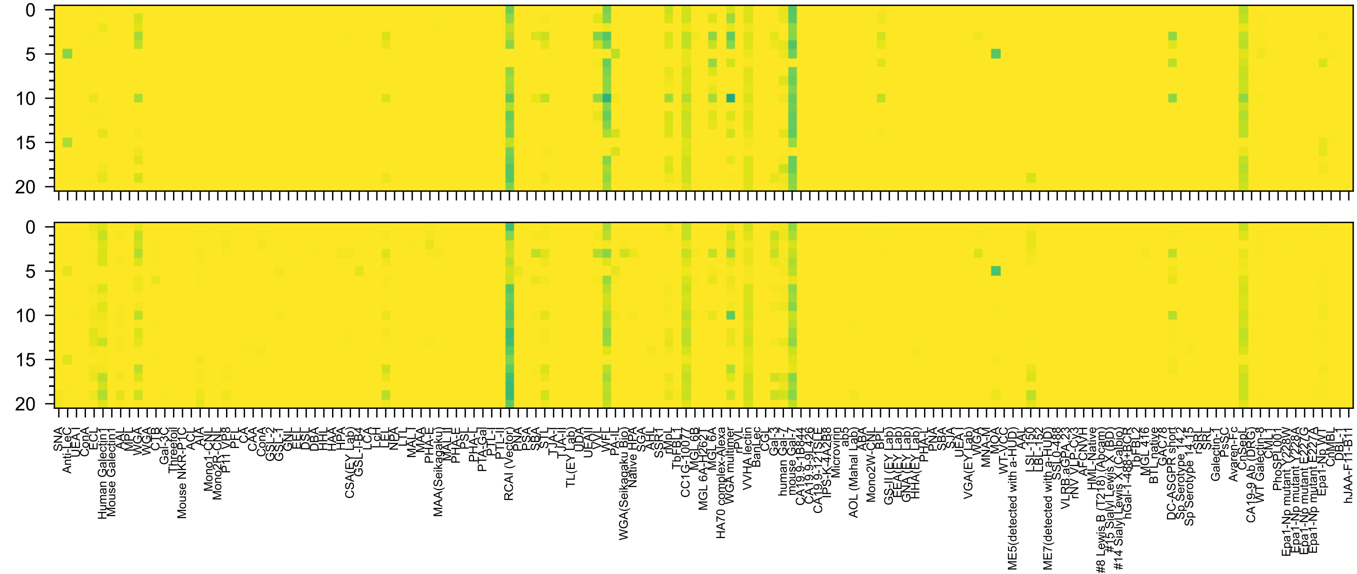


Gal(β1-4)GlcNAc(β1-3)Gal(β1-4)Glc(β-Sp0

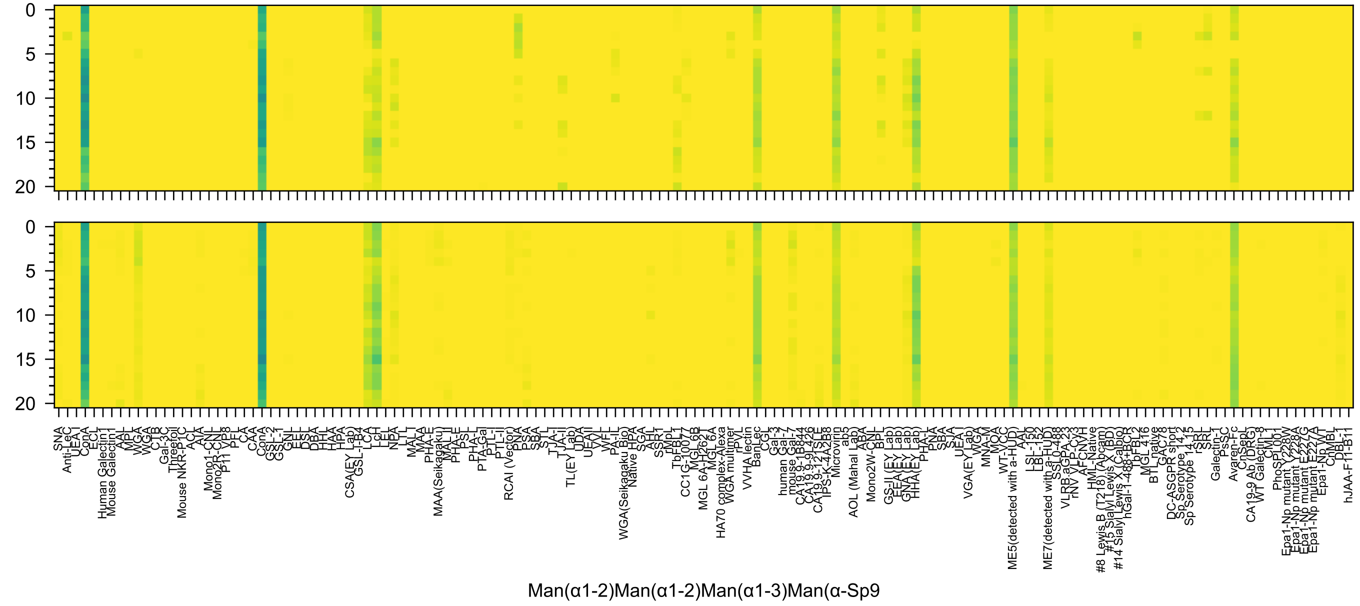
0
5
10
15
20
0
5
10
15
20
SNA
Anti-LeC
UEA-I
ConA
ECL
Human Galectin1
Mouse Galectin1
AAL
MPL
WGA
WGA
CTB
Gal-3C
Threefoil
Mouse NKR-P1C
ACL
AIA
Mono1-CNL
Mono2R-CNL
P11 VP8
PFL
CA
CAA
ConA
GSL-2
GSL-1
GNL
EEL
DSL
DBA
HHL
HAA
HPA
CSA(EY Lab)
GSL-I-B4
LCA
LcH
LEL
NPA
LTL
MAL-I
MAA
PHA-E
MAA(Seikagaku)
MAL-II
PHA-E
PSL
PHA-L
PTA-Gal
PTL-I
PTL-II
RCAI (Vector)
PNA
PSA
SBA
STL
TJA-I
TJA-II
TL(EY Lab)
UDA
UEAII
VVL
WFL
PA-IL
WGA(Seikagaku Bio)
Native HPA
SGA
AHL
SSR1
rMpL
TbFBL1
CC1G-10077
MGL 6B
MGL 6A-H262T
MGL 6A
HA70 complex-Alexa
WGA multimer
rPVL
VVHA lectin
BanLec
CGL
Gal-3
human Gal-7
mouse Gal-7
CA19.9-1B844
CA19.9-9L426
CA19.9-121SLE
IPS-K-4A2B8
Microvirin
ab5
AOL (Mahal Lab)
ABA
Mono2W-CNL
BPL
GS-II (EY Lab)
EEA (EY Lab)
GNA (EY Lab)
HHA (EY Lab)
PHA-L
PNA
SBA
SJA
UEA-I
VGA(EY Lab)
WGA
MNA-M
MOA
WT-VCC
ME5(detected with a-HUD)
AAL
LSL-150
LBL-152
ME7(detected with a-HUD)
SS10-488
VLRB.aGPA.23
rNV VLP-Cy3
AFCNVH
HML-Native
#8 Lewis B (T218) (Abcam)
#15 Sialyl Lewis X (BD)
#14 Sialyl Lewis X (Calbio)
hGal-1-488+BCR
TbFBL2
MGL 416
BTL native
GAC78
DC-ASGPR short
Sp Serotype 14.2
Sp Serotype 14.15
rSRL
SRL
Galectin-1
PsSC
Avaren-Fc
ChSepL
CA19-9 Ab (DRG)
WT Galectin-8
CML
PhoSL001
Epa1-Np mutant Y228W
Epa1-Np mutant Y228A
Epa1-Np mutant E227G
Epa1-Np mutant E227A
Epa1-Np WT
ChMBL
DBL-1
hJAA-F11-B11
Man(α1-2)Man(α1-2)Man(α1-3)Man(α-Sp9)

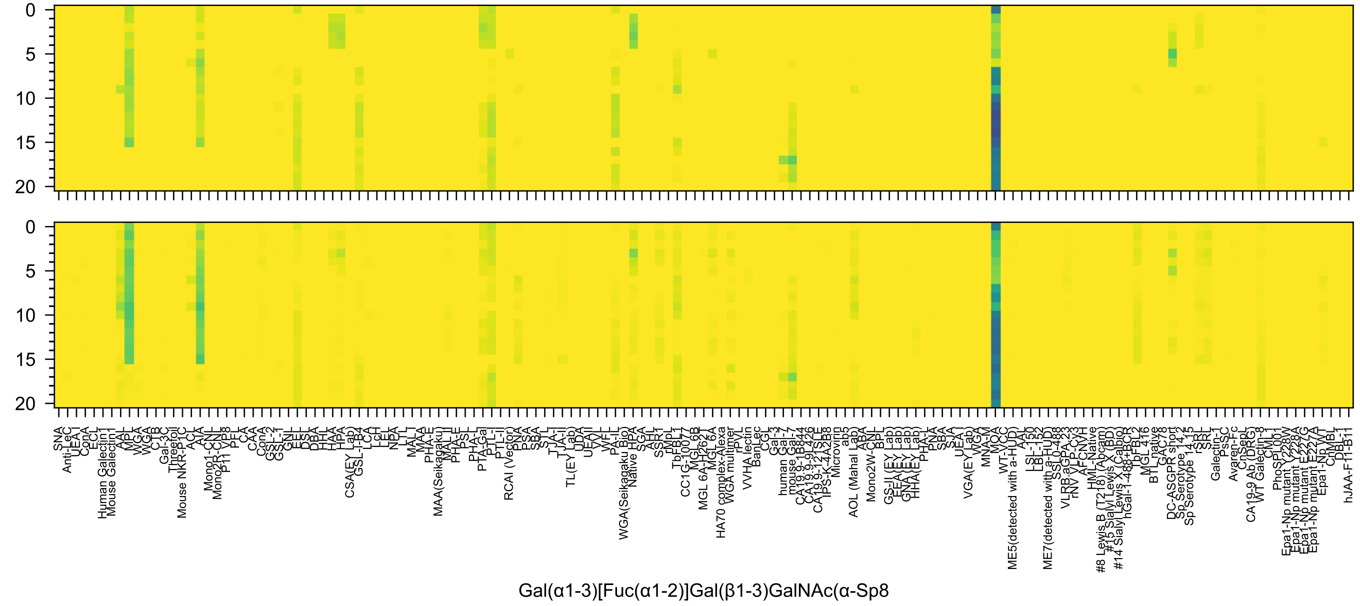

0
5
10
15
20
SNA
Anti-LeC
UEA-I
ConA
ECL
Human Galectin1
Mouse Galectin1
AAL
MPL
WGA
WGA
CTB
Gal-3C
Threefoil
Mouse NKR-P1C
ACL
AIA
Mono1-CNL
Mono2R-CNL
P11 VP8
PFL
CA
CAA
ConA
GSL-2
GSL-1
GNL
EEL
DSL
DBA
HHL
HAA
HPA
CSA(EY Lab)
GSL-I-B4
LCA
LcH
LEL
NPA
LTL
MAL-I
MAA
PHA-E
MAA(Seikagaku)
MAL-II
PHA-E
PSL
PHA-L
PTA-Gal
PTL-I
PTL-II
RCAI (Vector)
PNA
PSA
SBA
STL
TJA-I
TJA-II
TL(EY Lab)
UDA
UEAII
VVL
WFL
PA-IL
WGA(Seikagaku Bio)
Native HPA
SGA
AHL
SSR1
rMpL
TbFBL1
CC1G-10077
MGL 6B
MGL 6A-H262T
MGL 6A
HA70 complex-Alexa
WGA multimer
rPVL
VVHA lectin
BanLec
CGL
Gal-3
human Gal-7
mouse Gal-7
CA19.9-1B844
CA19.9-9L426
CA19.9-121SLE
IPS-K-4A2B8
Microvirin
ab5
AOL (Mahal Lab)
ABA
Mono2W-CNL
BPL
GS-II (EY Lab)
EEA (EY Lab)
GNA (EY Lab)
HHA (EY Lab)
PHA-L
PNA
SBA
SJA
UEA-I
VGA(EY Lab)
WGA
MNA-M
MOA
WT-VCC
ME5(detected with a-HUD)
AAL
LSL-150
LBL-152
ME7(detected with a-HUD)
SSL0-488
VLRB.aGPA.23
rNV VLP-Cy3
AFCN-H
HML-Native
#8 Lewis B (T218) (Abcam)
#15 Sialyl Lewis X (BD)
#14 Sialyl Lewis X (Calbio)
hGal-1-488+BCR
TbFBL2
MGL 416
BTL native
GAC78
DC-ASGPR short
Sp Serotype 14.2
Sp Serotype 14.15
rSRL
SRL
Galectin-1
PsSC
Avaren-Fc
ChSepL
CA19-9 Ab (DRG)
WT Galectin-8
CML1
PhoSL001
Epa1-Np mutant Y228W
Epa1-Np mutant Y228A
Epa1-Np mutant E227G
Epa1-Np mutant E227A
Epa1-Np WT
ChMBL
DBL-1
hJAA-F11-B11
Gal(α1-3)[Fuc(α1-2)]Gal(β1-3)GalNAc(α-Sp8

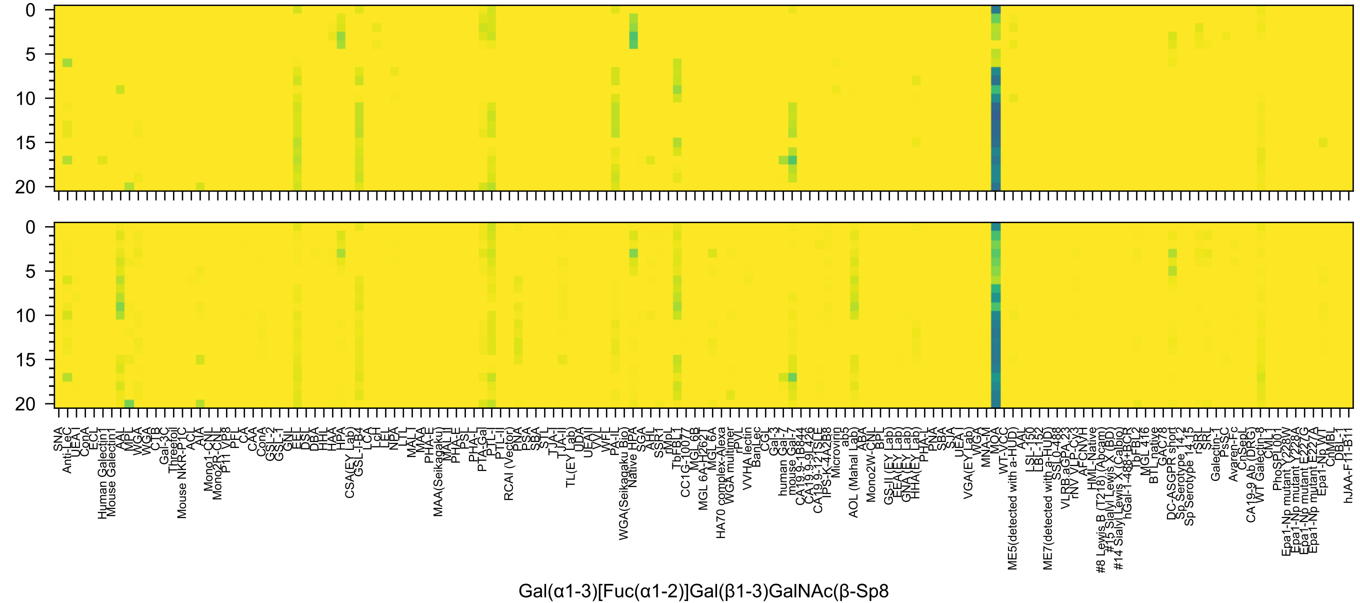
Gal(α1-3)[Fuc(α1-2)]Gal(β1-3)GalNAc(β-Sp8

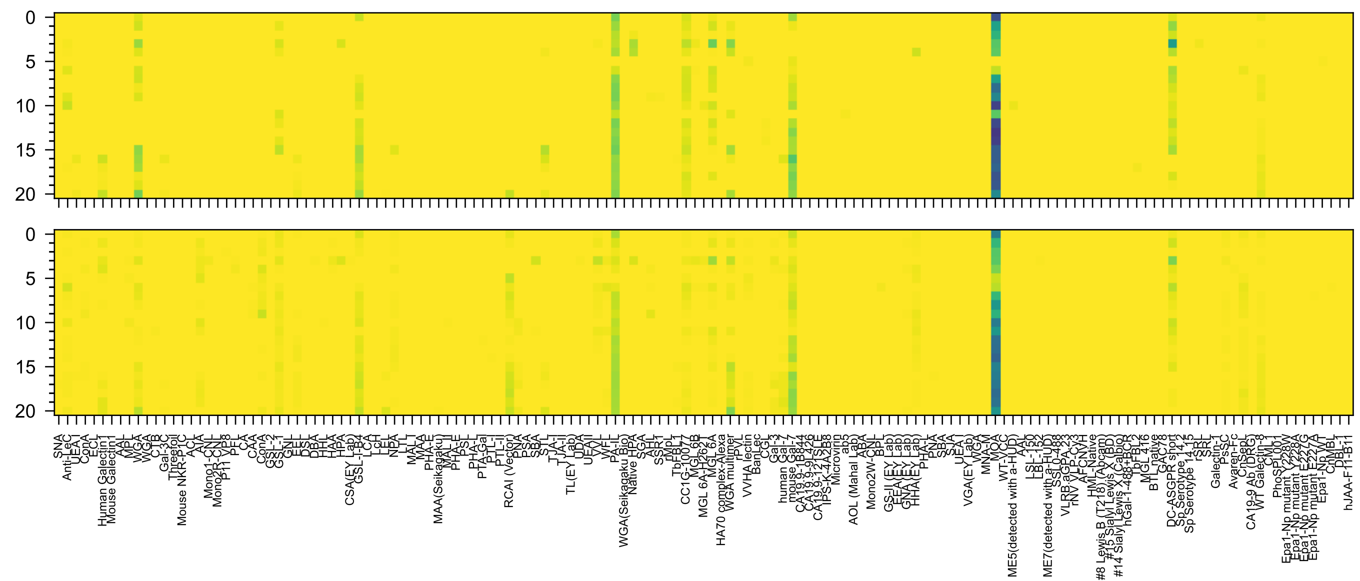


Gal(α1-3)Gal(β1-4)GlcNAc(β1-2)Man(α-Sp0

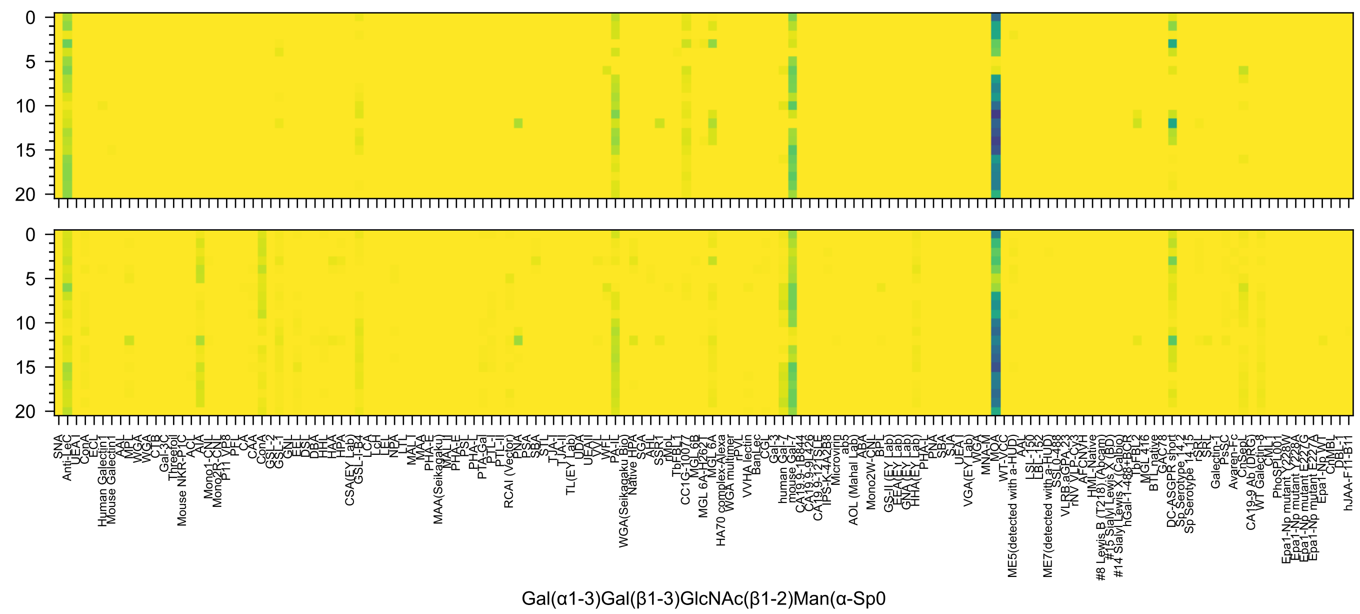

0
5
10
15
20
0
5
10
15
20
SNA
Anti-LeC
UEA I
ConA
ECL
Human Galectin1
Mouse Galectin1
AAL
MPL
WGA
WGA
CTB
Gal-3C
Threefoil
Mouse NKR-P1C
ACL
AIA
Mono1-CNL
Mono2R-CNL
P11 VP8
PFL
CA
CAA
ConA
GSL-2
GSL-1
GNL
EEL
DSL
DBA
HHL
HAA
HPA
CSA(EY Lab)
GSL-I-B4
LCA
LcH
LEL
NPA
LTL
MAL I
MAA
PHA-E
MAA(Seikagaku)
MAL II
PHA-E
PSL
PHA-L
PTA-Gal
PTL-I
PTL-II
RCAI (Vector)
PNA
PSA
SBA
STL
TJA-I
TJA-II
TL(EY Lab)
UDA
UEAII
VVL
WFL
PA-IL
WGA(Seikagaku Bio)
Native HPA
SGA
AHL
SSR1
rMpL
TbFBL1
CC1G-10077
MGL 6B
MGL 6A-H262T
MGL 6A
HA70 complex-Alexa
WGA multimer
rPVL
VVHA lectin
BanLec
CGL
Gal-3
human Gal-7
mouse Gal-7
CA19.9-1B844
CA19.9-9L426
CA19.9-121SLE
IPS-K-4A2B8
Microvirin
ab5
AOL (Mahal Lab)
ABA
Mono2W-CNL
BPL
GS-II (EY Lab)
EEA(EY Lab)
GNA(EY Lab)
HHA(EY Lab)
PHA-L
PNA
SBA
SJA
UEA I
VGA(EY Lab)
WGA
MNA-M
MOA
WT-VCC
ME5(detected with a-HUD)
AAL
LSL-150
LBL-152
ME7(detected with a-HUD)
SSL0-488
VLRB.aGPA.23
rNV VLP-Cy3
AFCN-H
HML-Native
#8 Lewis B (T218) (Abcam)
#15 Sialyl Lewis X (BD)
#14 Sialyl Lewis X (Calbio)
hGal-1-488+BCR
bFBL2
MGL 416
BTL native
GAC78
DC-ASGPR short
Sp Serotype 14.2
Sp Serotype 14.15
rSRL
SRL
Galectin-1
PsSC
Avaren-Fc
ChSepL
CA19-9 Ab (DRG)
WT Galectin-8
CML1
PhoSL001
Epa1-Np mutant Y228W
Epa1-Np mutant Y228A
Epa1-Np mutant E227G
Epa1-Np mutant E227A
Epa1-Np WT
ChMBL
DBL-1
hJAA-F11-B11
Gal(α1-3)Gal(β1-3)GlcNAc(β1-2)Man(α-Sp0

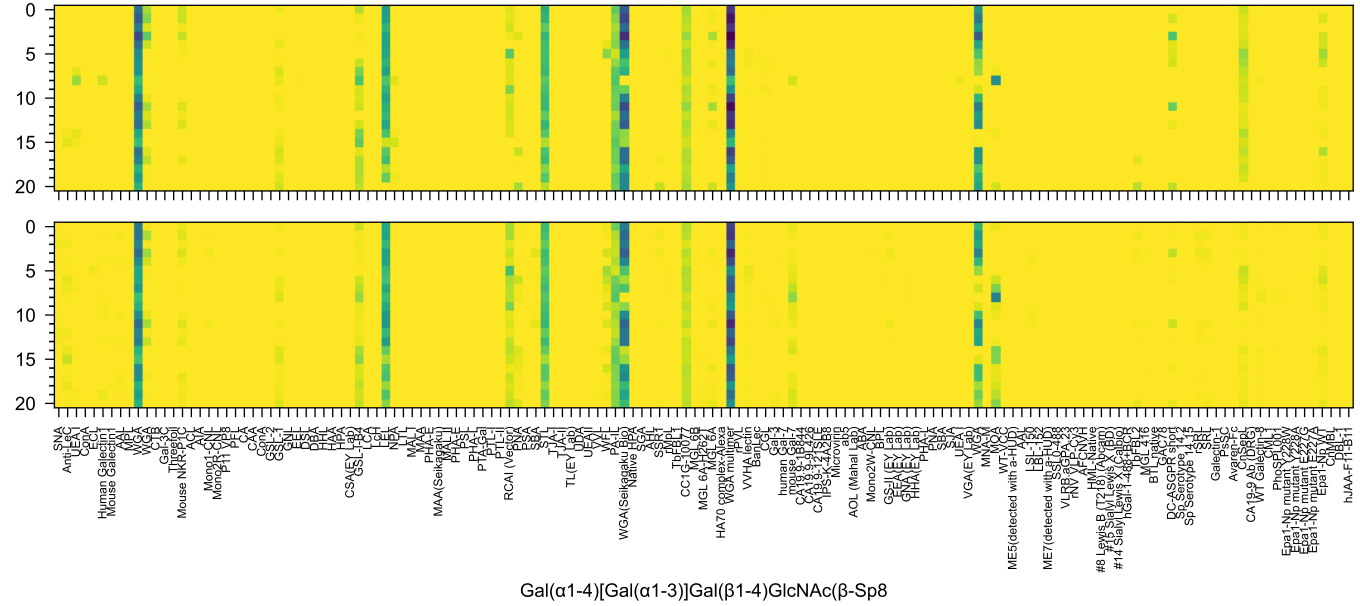
0
5
10
15
20
SNA
Anti-LeC
UEA I
ConA
ECL
Human Galectin1
Mouse Galectin1
AAL
MPL
WGA
WGA
CTB
Gal-3C
Threefoil
Mouse NKR-P1C
ACL
AIA
Mono1-CNL
Mono2R-CNL
P11 VP8
PFL
CA
CAA
ConA
GSL-2
GSL-1
GNL
EEL
DSL
DBA
HHL
HAA
HPA
CSA(EY Lab)
GSL-I-B4
LCA
LcH
LEL
NPA
LTL
MAL I
MAA
PHA-E
MAA(Seikagaku)
MAL II
PHA-E
PSL
PHA-L
PTA-Gal
PTL-I
PTL-II
RCAI (Vector)
PNA
PSA
SBA
STL
TJA-I
TJA-II
TL(EY Lab)
UDA
UEA II
VVL
WFL
PA-IL
WGA(Seikagaku Bio)
Native HPA
SGA
AHL
SSR1
rMpL
TbFBL1
CC1G-10077
MGL 6B
MGL 6A-H262T
MGL 6A
HA70 complex-Alexa
WGA multimer
rPVL
VVHA lectin
BanLec
CGL
Gal-3
human Gal-7
mouse Gal-7
CA19.9-1B844
CA19.9-9L426
CA19.9-121SLE
IPS-K-4A2B8
Microvirin
ab5
AOL (Mahal Lab)
ABA
Mono2W-CNL
BPL
GS-II (EY Lab)
EEA(EY Lab)
GNA(EY Lab)
HHA(EY Lab)
PHA-L
PNA
SBA
SJA
UEA I
VGA(EY Lab)
WGA
MNA-M
MOA
WT-VCC
ME5(detected with a-HUD)
AAL
LSL-150
LBL-152
ME7(detected with a-HUD)
SSL0-488
VLRB.aGPA.23
rNV VLP-Cy3
AFCNVH
HML-Native
#8 Lewis B (T218) (Abcam)
#15 Sialyl Lewis X (BD)
#14 Sialyl Lewis X (Calbio)
hGal-1-488+BCR
TbFBL2
MGL 416
BTL native
GAC78
DC-ASGPR short
Sp Serotype 14.2
Sp Serotype 14.15
rSRL
SRL
Galectin-1
PsSC
Avaren-Fc
CnSepL
CA19-9 Ab (DRG)
WT Galectin-8
CML1
PhoSPL001
Epa1-Np mutant Y228W
Epa1-Np mutant Y228A
Epa1-Np mutant E227G
Epa1-Np mutant E227A
Epa1-Np WT
CnMBL
DBL-1
hJAA-F11-B11
Gal(α1-4)[Gal(α1-3)]Gal(β1-4)GlcNAc(β-Sp8

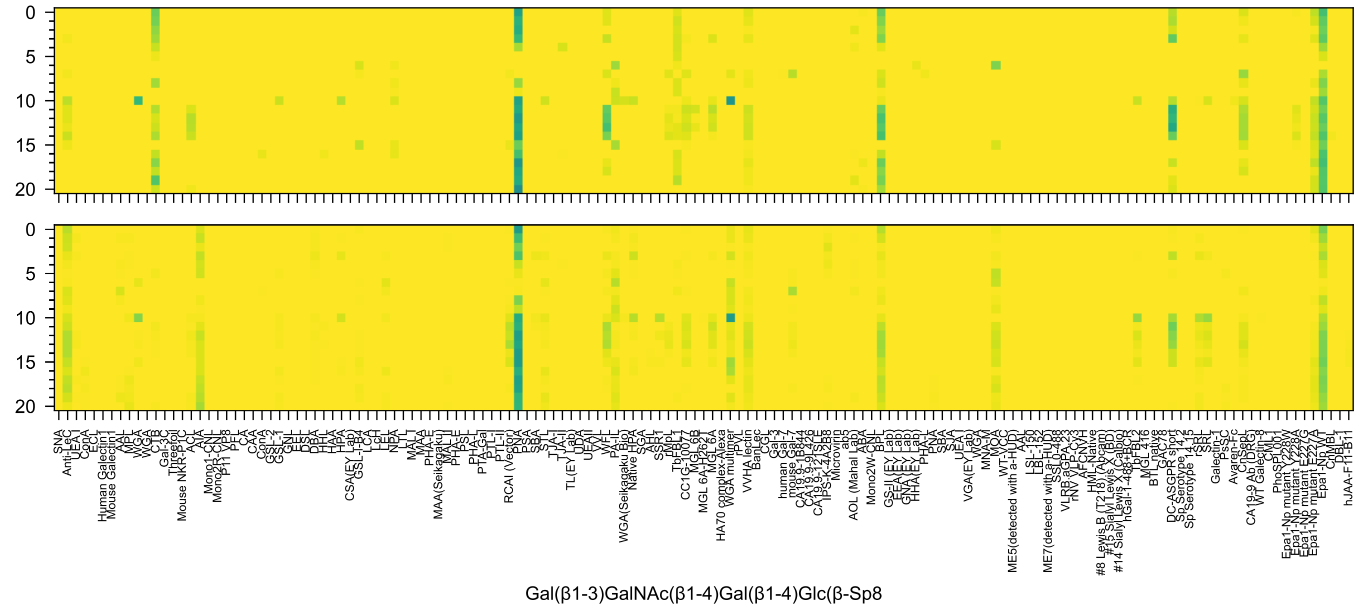
Gal(β1-3)GalNAc(β1-4)Gal(β1-4)Glc(β-Sp8

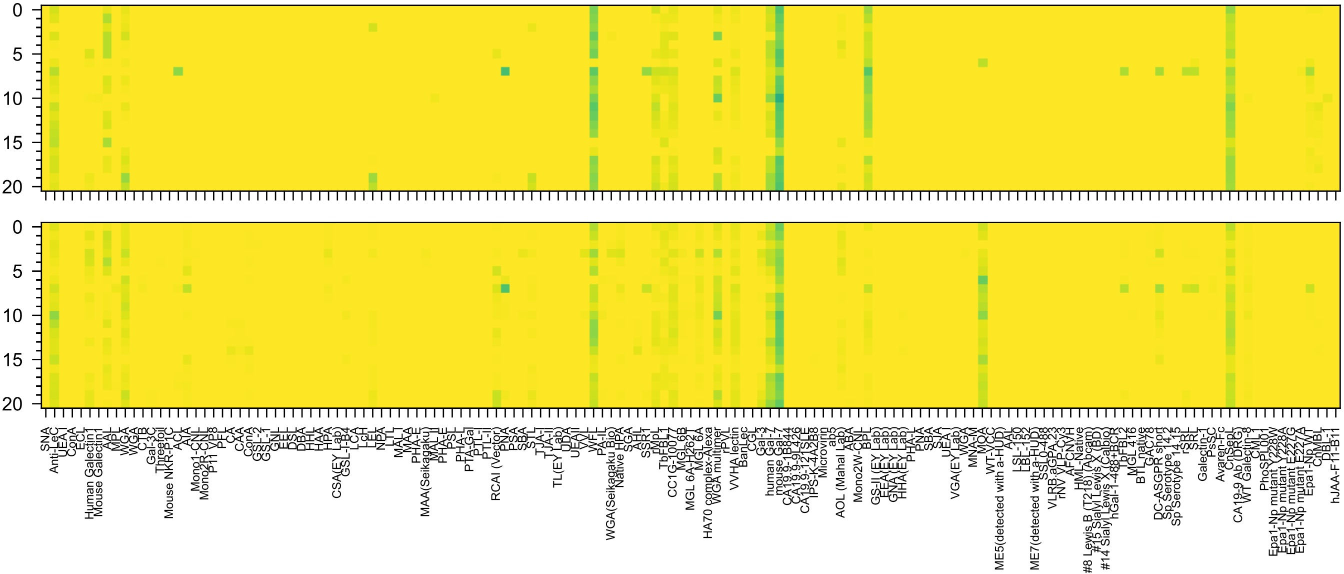

Gal(β1-3)GlcNAc(β1-3)Gal(β1-4)Glc(β-Sp10

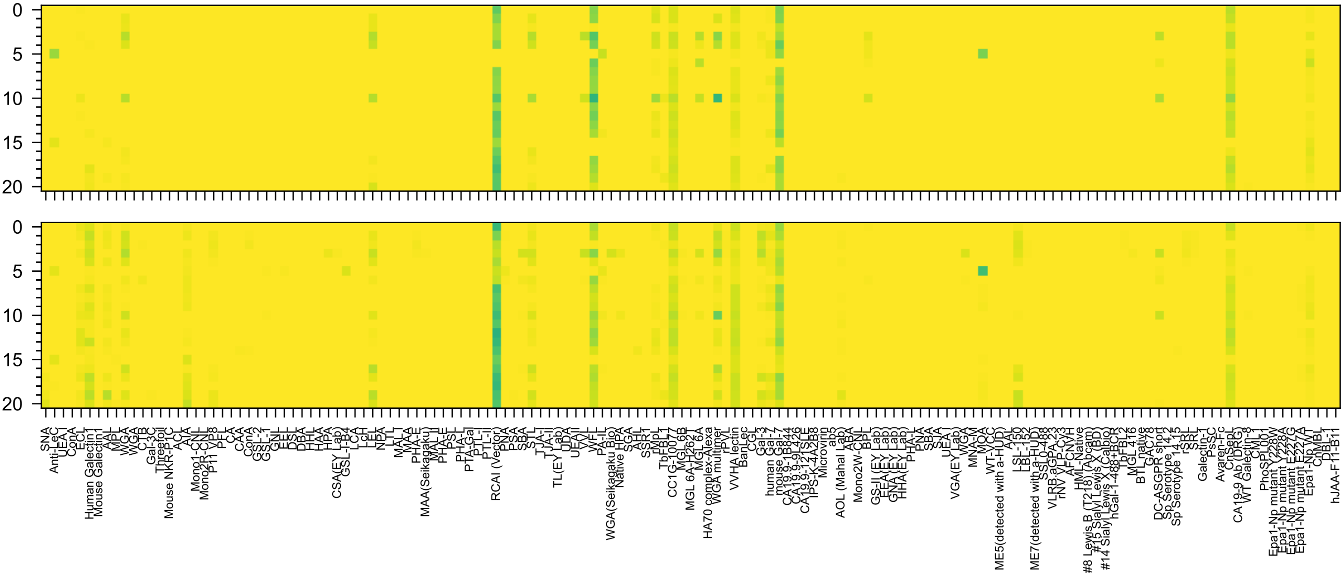

Gal(β1-4)GlcNAc(β1-3)Gal(β1-4)Glc(β-Sp8

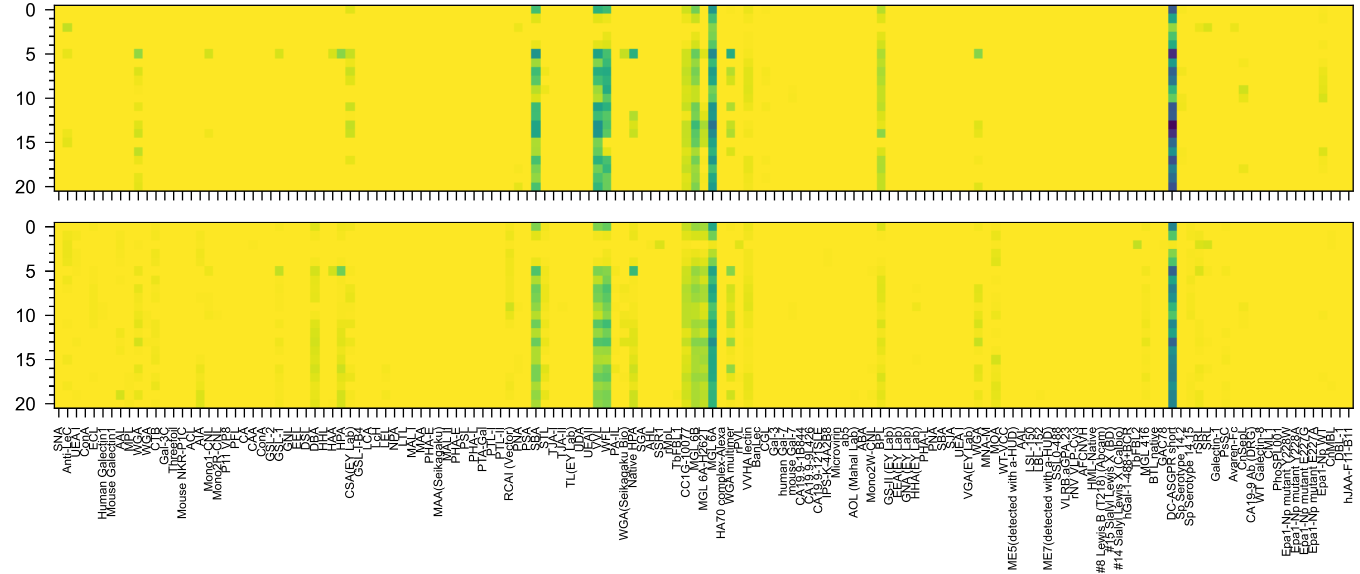


GalNAc(β1-3)Gal(α1-6)Gal(β1-4)Glc(β-Sp8

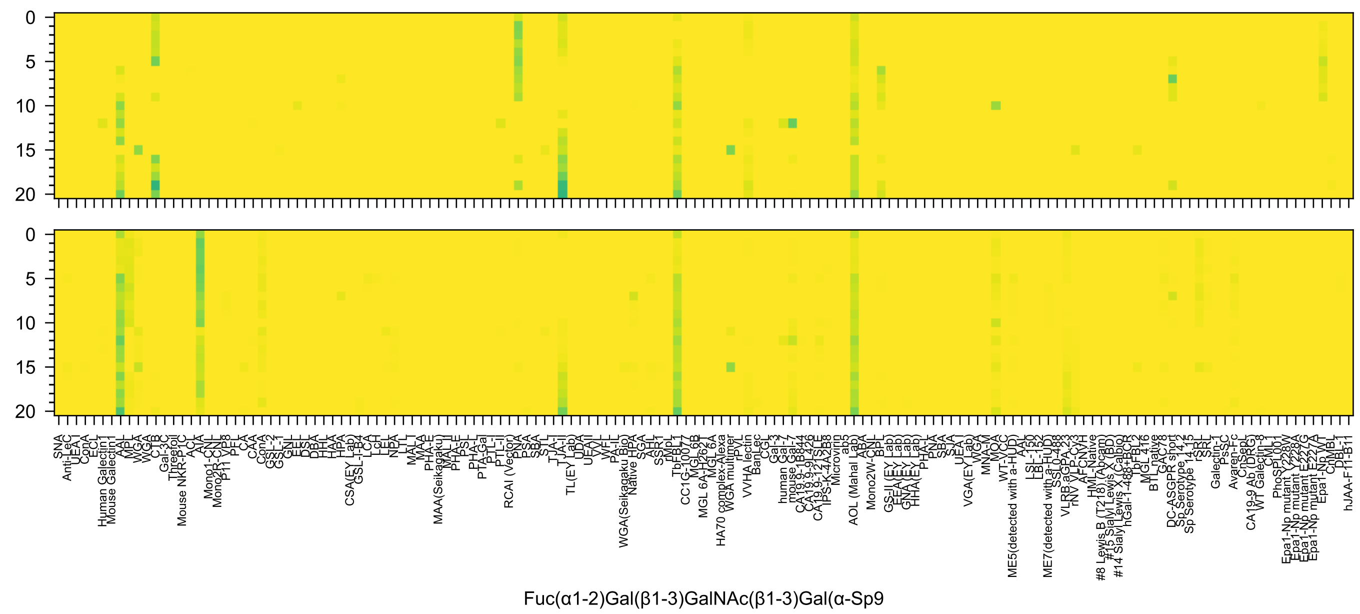
0
5
10
15
20
SNA
Anti-LeC
UEA-I
ConA
ECL
Human Galectin1
Mouse Galectin1
AAL
MPL
WGA
WGA
CTB
Gal-3C
Threefoil
Mouse NKR-P1C
ACL
AIA
Mono1-CNL
Mono2R-CNL
P11 VP8
PFL
CA
CAA
ConA
GSL-2
GSL-1
GNL
EEL
DSL
DBA
HHL
HAA
HPA
CSA(EY Lab)
GSL-I-B4
LCA
LcH
LEL
NPA
LTL
MAL-I
MAA
PHA-E
MAA(Seikagaku)
MAL-II
PHA-E
PSL
PHA-L
PTA-Gal
PTL-I
PTL-II
RCAI (Vector)
PNA
PSA
SBA
STL
TJA-I
TJA-II
TL(EY Lab)
UDA
UEAII
VVL
WFL
PA-IL
WGA(Seikagaku Bio)
Native HPA
SGA
AHL
SSR1
rMpL
TbFBL1
CC1G-10077
MGL 6B
MGL 6A-H262T
MGL 6A
HA70 complex-Alexa
WGA multimer
rPVL
VVHA lectin
BanLec
CGL
Gal-3
human Gal-7
mouse Gal-7
CA19.9-1B844
CA19.9-9L426
CA19.9-121SLE
IPS-K-4A2B8
Microvirin
ab5
AOL (Mahal Lab)
ABA
Mono2W-CNL
BPL
GS-II (EY Lab)
EEA(EY Lab)
GNA(EY Lab)
HHA(EY Lab)
PHA-L
PNA
SBA
SJA
UEA-I
VGA(EY Lab)
WGA
MNA-M
MOA
WT-VCC
ME5(detected with a-HUD)
AAL
LSL-150
LBL-152
ME7(detected with a-HUD)
SSL0-488
VLRB.aGPA.23
rNV VLP-Cy3
AFCNVH
HML-Native
#8 Lewis B (T218) (Abcam)
#15 Sialyl Lewis X (BD)
#14 Sialyl Lewis X (Calbio)
hGal-1-488+BCR
TbFBL2
MGL 416
BTL native
GAC78
DC-ASGPR short
Sp Serotype 14.2
Sp Serotype 14.15
rSRL
SRL
Galectin-1
PsSC
Avaren-Fc
ChSeaL
CA19-9 Ab (DRG)
WT Galectin-8
CML1
PhoSL001
Epa1-Np mutant Y228W
Epa1-Np mutant Y228A
Epa1-Np mutant E227G
Epa1-Np mutant E227A
Epa1-Np WT
ChMBL
DBL-1
hJAA-F11-B11
Fuc(α1-2)Gal(β1-3)GalNAc(β1-3)Gal(α-Sp9

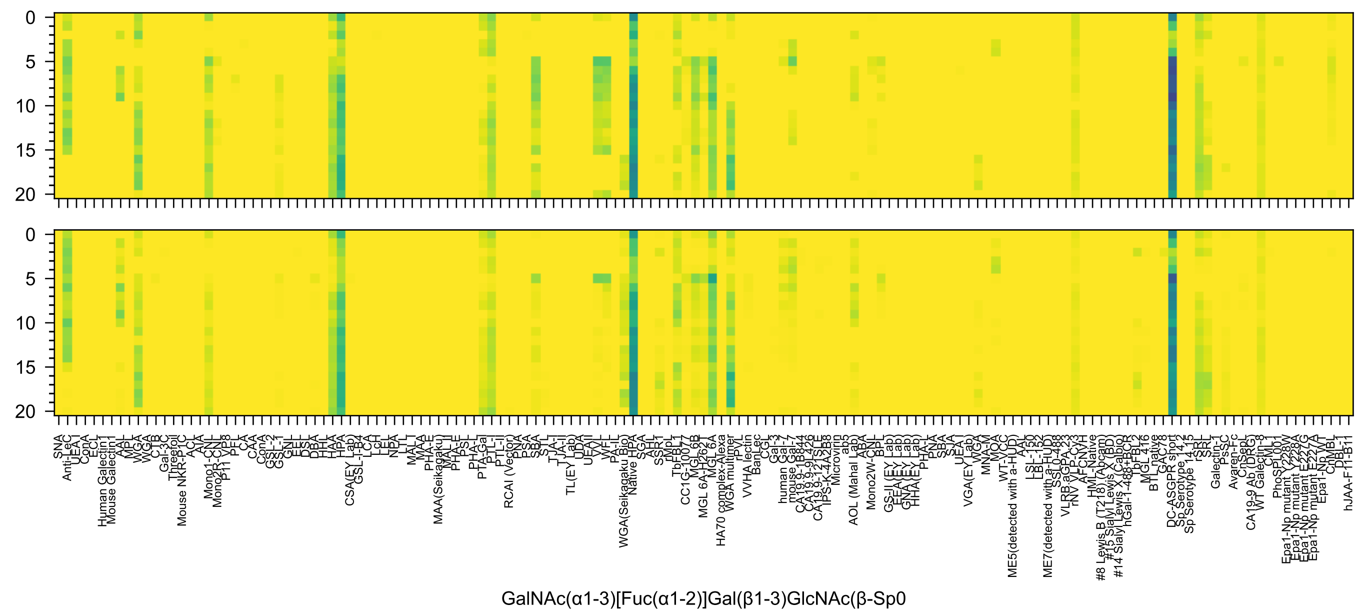
0
5
10
15
20
0
5
10
15
20
SNA
Anti-LeC
UEA-I
ConA
ECL
Human Galectin1
Mouse Galectin1
AAL
MPL
WGA
WGA
CTB
Gal-3C
Threefoil
Mouse NKR-P1C
ACL
AIA
Mono1-CNL
Mono2R-CNL
P11 VP8
PFL
CA
CAA
ConA
GSL-2
GSL-1
GNL
EEL
DSL
DBA
HHL
HAA
HPA
CSA(EY Lab)
GSL-I-B4
LCA
LcH
LEL
NPA
LTL
MAL-I
MAA
PHA-E
MAA(Seikagaku)
MAL-II
PHA-E
PSL
PHA-L
PTA-Gal
PTL-I
PTL-II
RCAI (Vector)
PNA
PSA
SBA
STL
TJA-I
TJA-II
TL(EY Lab)
UDA
UEAII
VVL
WFL
PA-IL
WGA(Seikagaku Bio)
Native HPA
SGA
AHL
SSR1
rMpL
TbFBL1
CC1G-10077
MGL 6B
MGL 6A-H262T
MGL 6A
HA70 complex-Alexa
WGA multimer
rPVL
VVHA lectin
BanLec
CGL
Gal-3
human Gal-7
mouse Gal-7
CA19.9-1B844
CA19.9-9L426
CA19.9-121SLE
IPS-K-4A2B8
Microvirin
ab5
AOL (Mahal Lab)
ABA
Mono2W-CNL
BPL
GS-II (EY Lab)
EEA(EY Lab)
GNA(EY Lab)
HHA(EY Lab)
PHA-L
PNA
SBA
SJA
UEA-I
VGA(EY Lab)
WGA
MNA-M
MOA
WT-VCC
ME5(detected with a-HUD)
AAL
LSL-150
LBL-152
ME7(detected with a-HUD)
SSL0-488
VLRB.aGPA.23
rNV VLP-Cy3
AFCNVH
HML-Native
#8 Lewis B (T218) (Abcam)
#15 Sialyl Lewis X (BD)
#14 Sialyl Lewis X (Calbio)
hGal-1-488+BCR
TbFBL2
MGL 416
BTL native
GAC78
DC-ASGPR short
Sp Serotype 14.15
Sp Serotype 14.15
rSRL
SRL
Galectin-1
PsSC
Avaren-Fc
ChSeL
CA19-9 Ab (DRG)
WT Galectin-8
CML1
PhoSPL001
Epa1-Np mutant Y228W
Epa1-Np mutant Y228A
Epa1-Np mutant E227G
Epa1-Np mutant E227A
Epa1-Np WT
CNMBL
DBL-1
hJAA-F11-B11
GalNAc(α1-3)[Fuc(α1-2)]Gal(β1-3)GlcNAc(β-Sp0

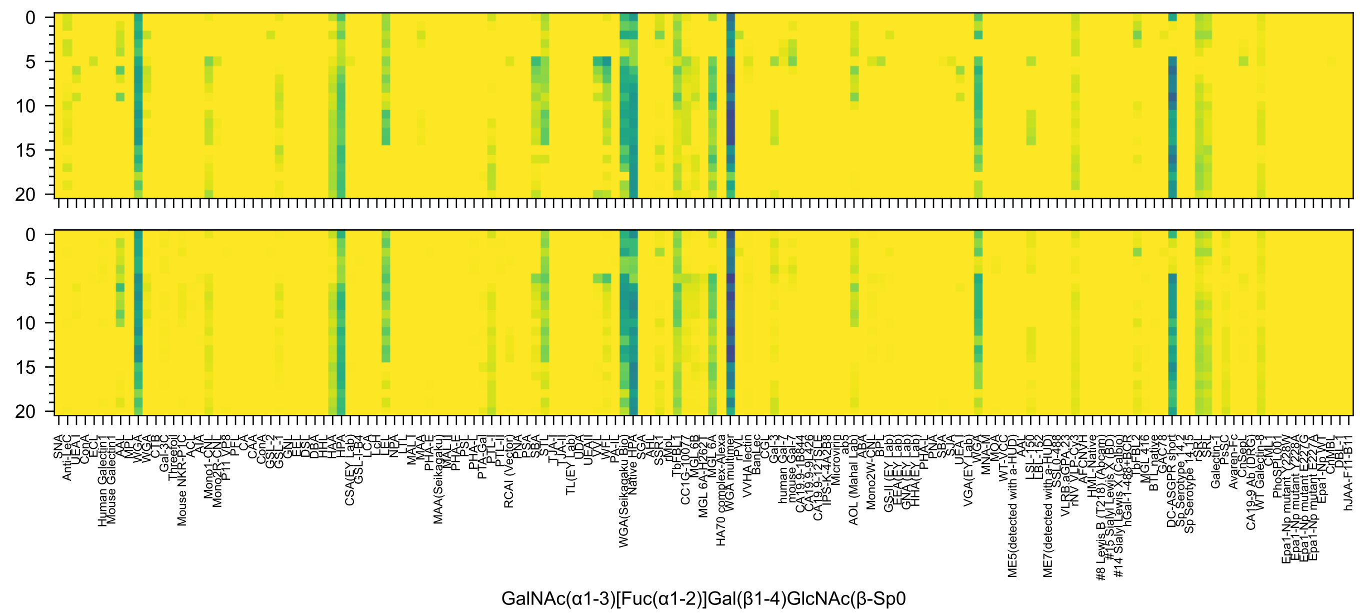
0
5
10
15
20
SNA
Anti-LeC
UEA I
ConA
ECL
Human Galectin1
Mouse Galectin1
AAL
MPL
WGA
WGA
CTB
C-3
Gal-3C
Threefoil
Mouse NKR-P1C
ACL
AIA
Mono1-CNL
Mono2R-CNL
P11 VP8
PFL
CA
CAA
ConA
GSL-2
GSL-1
GNL
EEL
DSL
DBA
HHL
HAA
HPA
CSA(EY Lab)
GSL-I-B4
LCA
LcH
LEL
NPA
LTL
MAL I
MAA
PHA-E
MAA(Seikagaku)
MAL-I
PHA-E
PSL
PHA-L
PTA-Gal
PTL-I
PTL-II
RCAI (Vector)
PNA
PSA
SBA
STL
TJA-I
TJA-II
TL(EY Lab)
UDA
UEAII
VVL
WFL
PA-IL
WGA(Seikagaku Bio)
Native HPA
SGA
AHL
SSR1
rMpL
TbFBL1
CC1G-10077
MGL 6B
MGL 6A-H262T
MGL 6A
HA70 complex-Alexa
WGA multimer
rPVL
VVHA lectin
BanLec
CGL
Gal-3
human Gal-7
mouse Gal-7
CA19.9-1B844
CA19.9-9L426
CA19.9-121SLE
IPS-K-4A2B8
Microvirin
ab5
AOL (Mahal Lab)
ABA
Mono2W-CNL
BPL
GS-II (EY Lab)
EEA(EY Lab)
GNA(EY Lab)
HHA(EY Lab)
PHA-L
PNA
SBA
SJA
UEA-I
VGA(EY Lab)
WGA
MNA-M
MOA
WT-VCC
ME5(detected with a-HUD)
AAL
LSL-150
LBL-152
ME7(detected with a-HUD)
SSL0-488
VLRB.aGPA.23
rNV VLP-Cy3
AFCNYH
HML-Native
#8 Lewis B (T218) (Abcam)
#15 Sialyl Lewis X (BD)
#14 Sialyl Lewis X (Calbio)
hGal-1-488+BCR
bFBL2
MGL 416
BTL native
GAC78
DC-ASGPR short
Sp Serotype 14.2
Sp Serotype 14.15
rSRL
SRL
Galectin-1
PsSC
Avaren-Fc
ChsepL
CA19-9 Ab (DRG)
WT Galectin-8
CML1
PhoSL001
Epa1-Np mutant Y228W
Epa1-Np mutant Y228A
Epa1-Np mutant E227G
Epa1-Np mutant E227A
Epa1-Np WT
ChMBL
DBL-1
hJAA-F11-B11
GalNAc(α1-3)[Fuc(α1-2)]Gal(β1-4)GlcNAc(β-Sp0

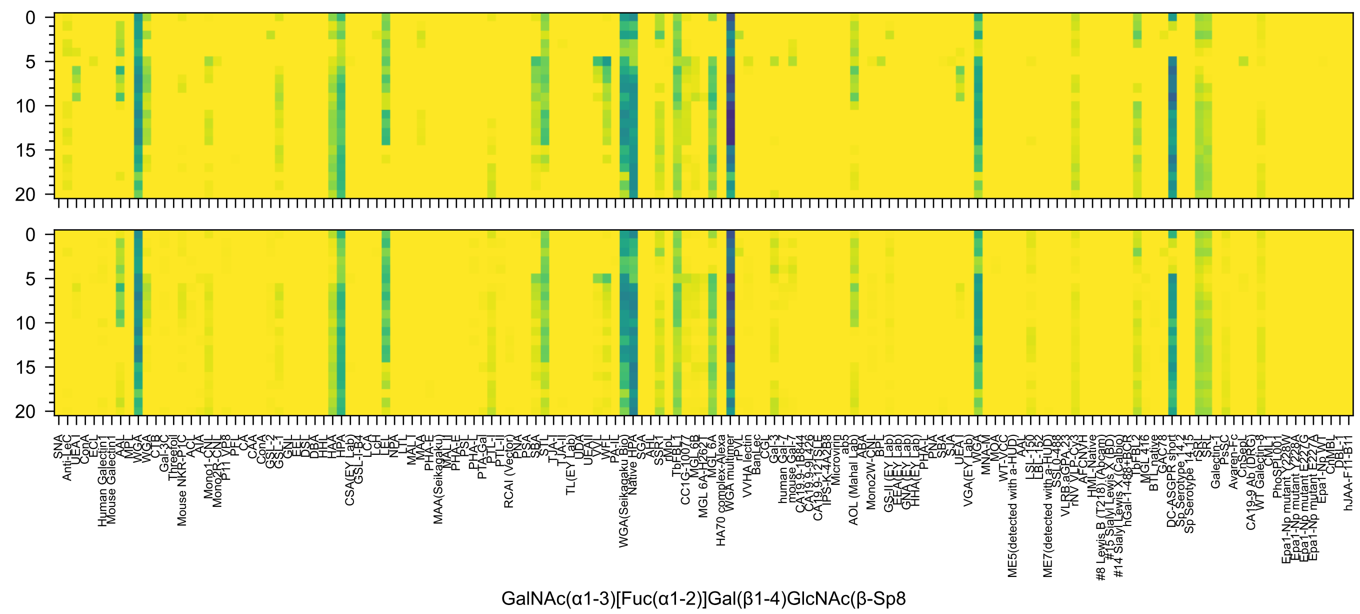
GalNAc(α1-3)[Fuc(α1-2)]Gal(β1-4)GlcNAc(β-Sp8

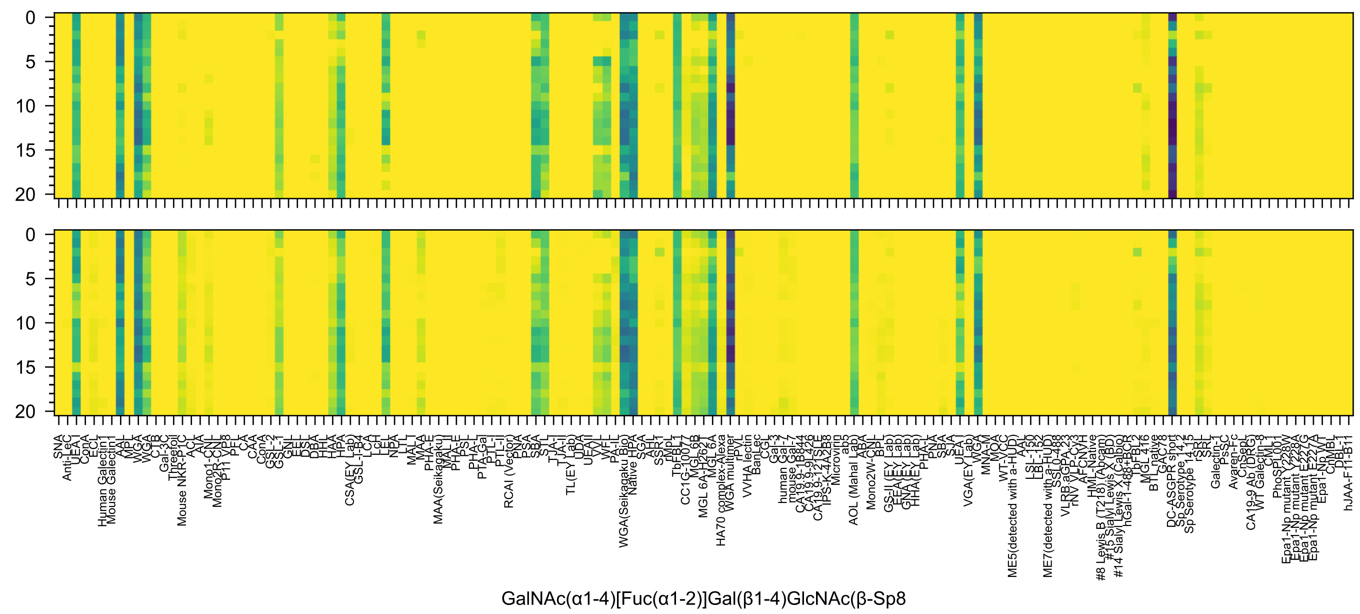
GalNAc(α1-4)[Fuc(α1-2)]Gal(β1-4)GlcNAc(β-Sp8

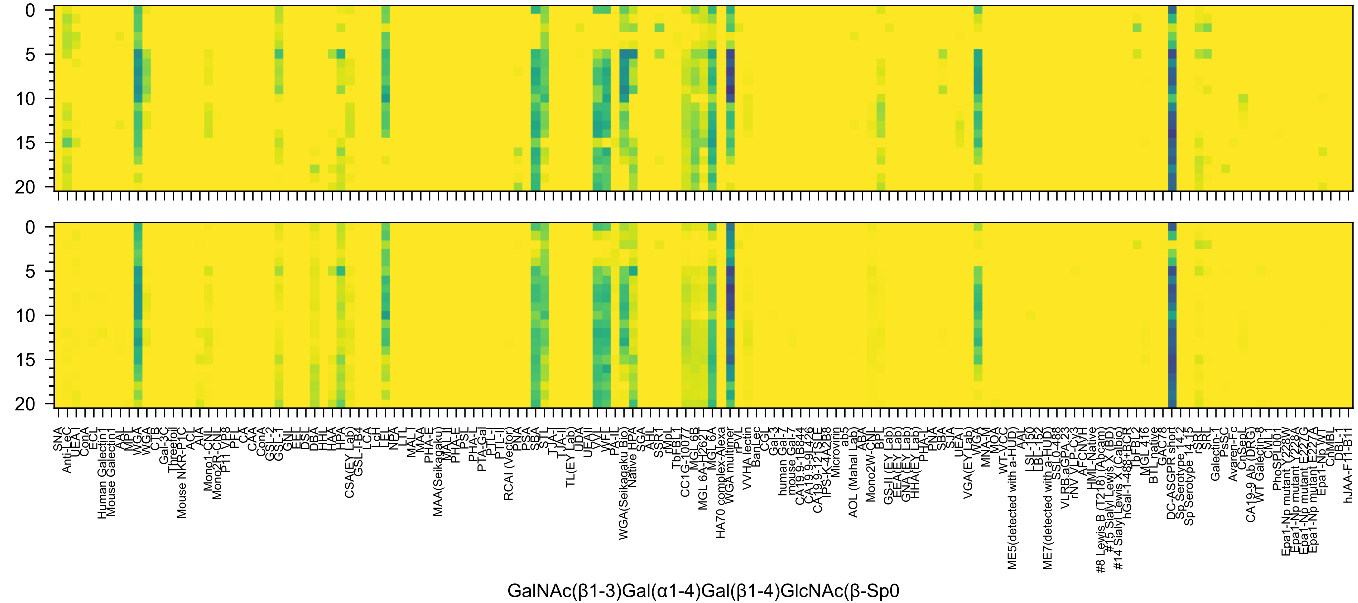
0
5
10
15
20
0
5
10
15
20
GalNAc(β1-3)Gal(α1-4)Gal(β1-4)GlcNAc(β-Sp0

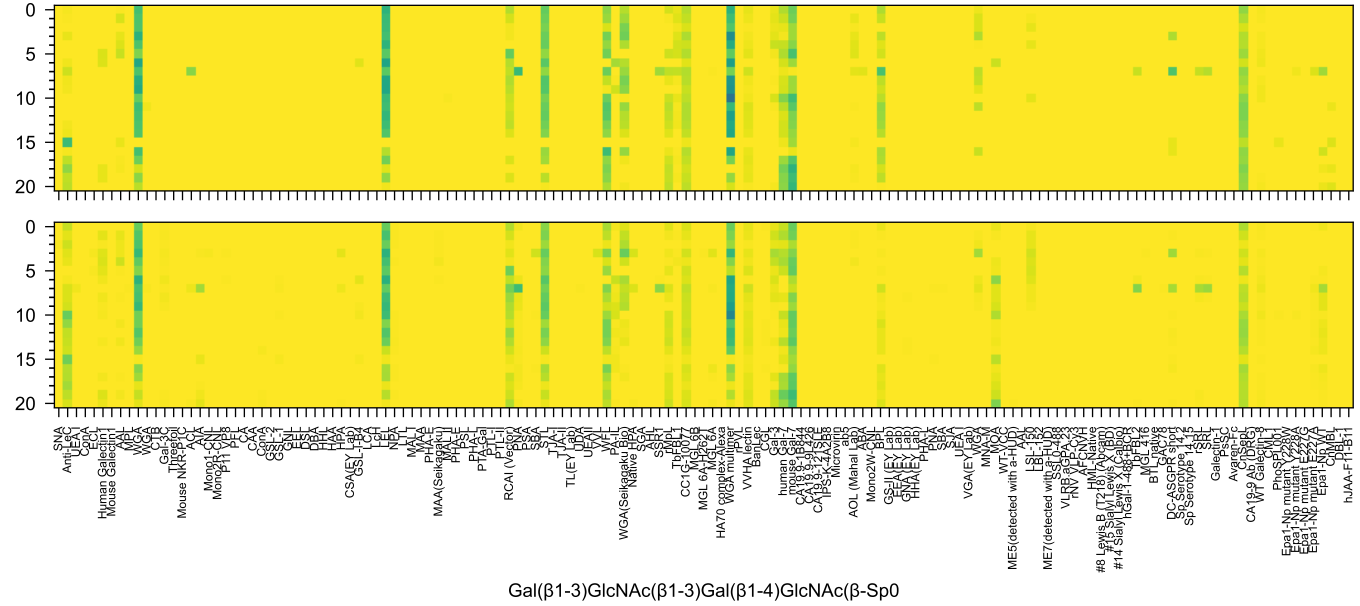

0
5
10
15
20
SNA
Anti-LeC
UEA I
ConA
ECL
Human Galectin1
Mouse Galectin1
AAL
MPL
WGA
WGA
CTB
Gal-3C
Threefoil
Mouse NKR-P1C
ACL
AIA
Mono1-CNL
Mono2R-CNL
P11 VP8
PFL
CA
CAA
ConA
GSL-2
GSL-1
GNL
EEL
DSL
DBA
HHL
HAA
HPA
CSA(EY Lab)
GSL-I-B4
LCA
LcH
LEL
NPA
LTL
MAL-I
MAA
PHA-E
MAA(Seikagaku)
MAL-II
PHA-E
PSL
PHA-L
PTA-Gal
PTL-I
PTL-II
RCAI (Vector)
PNA
PSA
SBA
STL
TJA-I
TJA-II
TL(EY Lab)
UDA
UEAII
VVL
WFL
PA-IL
WGA(Seikagaku Bio)
Native HPA
SGA
AHL
SSR1
rMpL
TbFBL1
CC1G-10077
MGL 6B
MGL 6A-H262T
MGL 6A
HA70 complex-Alexa
WGA multimer
rPVL
VVHA lectin
BanLec
CGL
Gal-3
human Gal-7
mouse Gal-7
CA19.9-1B844
CA19.9-9L426
CA19.9-121SLE
IPS-K-4A2B8
Microvirin
ab5
AOL (Mahal Lab)
ABA
Mono2W-CNL
BPL
GS-II (EY Lab)
EEA(EY Lab)
GNA(EY Lab)
HHA(EY Lab)
PHA-L
PNA
SBA
SJA
UEA-I
VGA(EY Lab)
WGA
MNA-M
MOA
WT-VCC
ME5(detected with a-HUD)
AAL
LSL-150
LBL-152
ME7(detected with a-HUD)
SSL0-488
VLRB.aGPA.23
rNV VLP-Cy3
AFCNVH
HML-Native
#8 Lewis B (T218) (Abcam)
#15 Sialyl Lewis X (BD)
#14 Sialyl Lewis X (Calbio)
hGal-1-488+BCR
TbFBL2
MGL 416
BTL native
GAC78
DC-ASGPR short
Sp Serotype 14.2
Sp Serotype 14.15
rSRL
SRL
Galectin-1
PsSC
Avaren-Fc
ChSeaL
CA19-9 Ab (DRG)
WT Galectin-8
CML
PhoSL001
Epa1-Np mutant Y228W
Epa1-Np mutant Y228A
Epa1-Np mutant E227G
Epa1-Np mutant E227A
Epa1-Np WT
ChMBL
DBL-1
hJAA-F11-B11
Gal(β1-3)GlcNAc(β1-3)Gal(β1-4)GlcNAc(β-Sp0

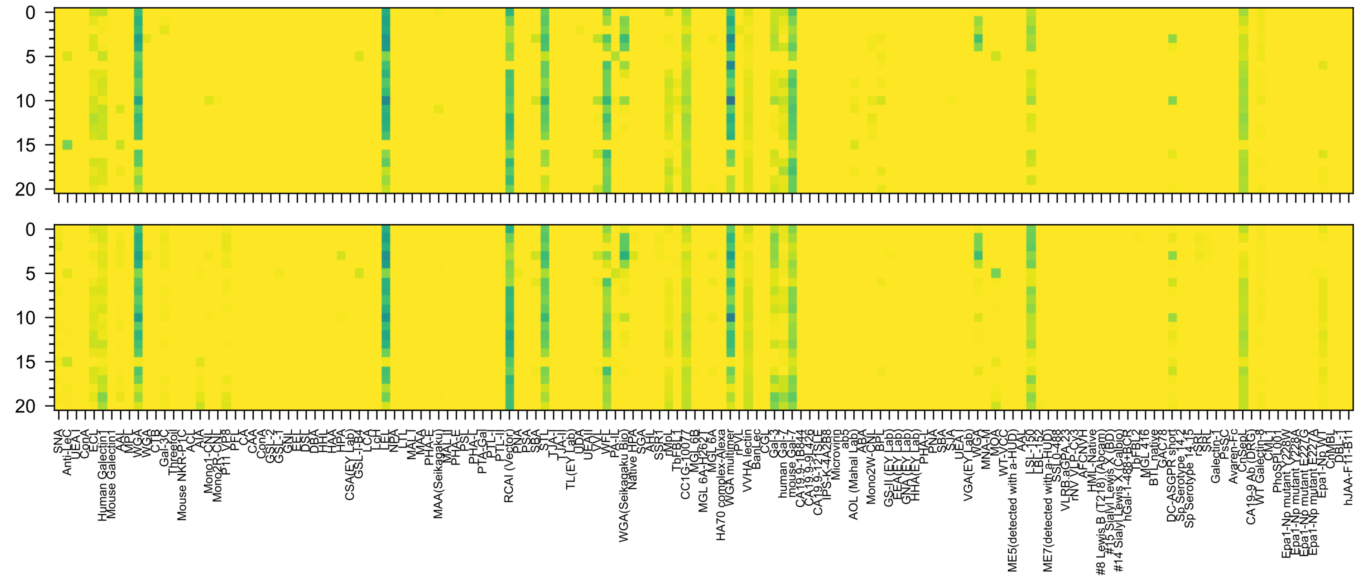

Gal(β1-4)GlcNAc(β1-3)Gal(β1-4)GlcNAc(β-Sp0

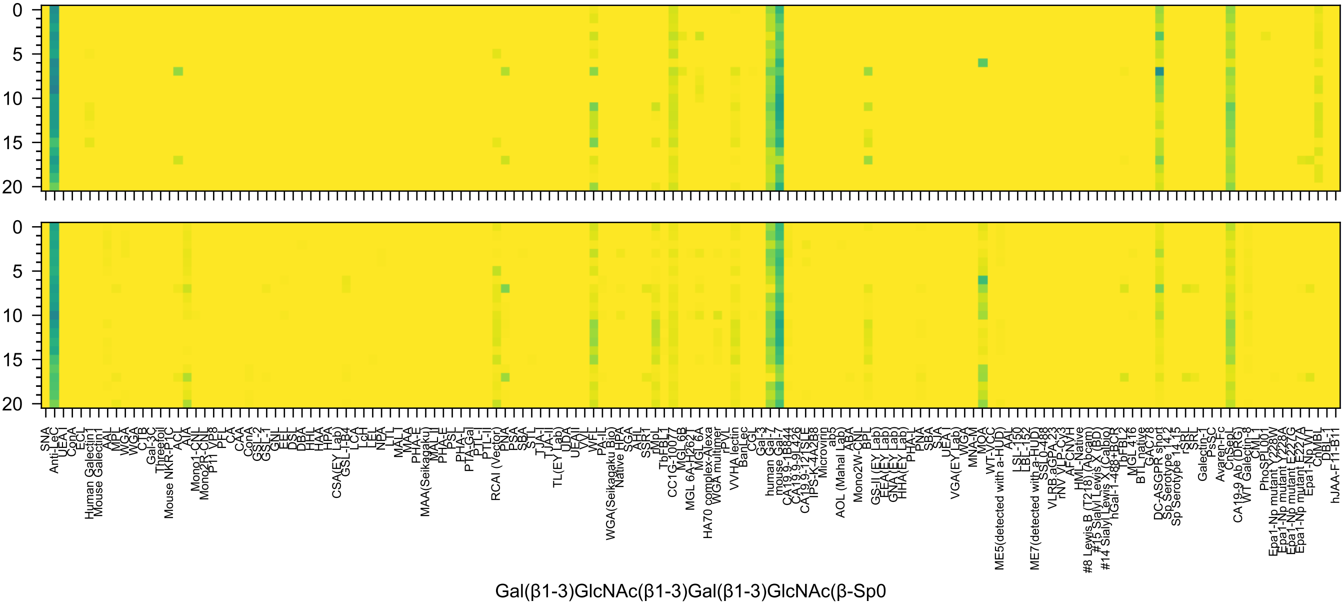
0
5
10
15
20
SNA
Anti-LeC
UEA-I
ConA
ECL
Human Galectin1
Mouse Galectin1
AAL
MPL
WGA
WGA
CTB
Gal-3C
Threefoil
Mouse NKR-P1C
ACL
AIA
Mono1-CNL
Mono2R-CNL
P11 VP8
PFL
CA
CAA
ConA
GSL-2
GSL-1
GNL
EEL
DSL
DBA
HHL
HAA
HPA
CSA(EY Lab)
GSL-I-B4
LCA
LcH
LEA
NPA
LTL
MAL-I
MAA
PHA-E
MAA(Seikagaku)
MAL-II
PHA-E
PSL
PHA-L
PTA-Gal
PTL-I
PTL-II
RCAI (Vector)
PNA
PSA
SBA
STL
TJA-I
TJA-II
TL(EY Lab)
UDA
UEAII
VVL
WFL
PA-IL
WGA(Seikagaku Bio)
Native HPA
SGA
AHL
SSR1
rMpL
TbFBL1
CC1G-10077
MGL 6B
MGL 6A-H262T
MGL 6A
HA70 complex-Alexa
WGA multimer
rPVL
VVHA lectin
BanLec
CGL
Gal-3
human Gal-7
mouse Gal-7
CA19.9-1B844
CA19.9-9L426
CA19.9-121SLE
IPS-K-4A2B8
Microvirin
ab5
AOL (Mahal Lab)
ABA
Mono2W-CNL
BPL
GS-II (EY Lab)
EEA (EY Lab)
GNA (EY Lab)
HHA (EY Lab)
PHA-L
PNA
SBA
SJA
UEA-I
VGA (EY Lab)
WGA
MNA-M
MOA
WT-VCC
ME5(detected with a-HUD)
AAL
LSL-150
LBL-152
ME7(detected with a-HUD)
SSL0-488
VLRB.aGPA.23
rNV VLP-Cy3
AFCN-H
HML-Native
#8 Lewis B (T218) (Abcam)
#15 Sialyl Lewis X (BD)
#14 Sialyl Lewis X (Calbio)
hGal-1-488+BCR
TbFBL2
MGL 416
BTL native
GAC78
DC-ASGPR short
Sp Serotype 14.2
Sp Serotype 14.15
rSRL
SRL
Galectin-1
PsSC
Avaren-Fc
CnSepL
CA19-9 Ab (DRG)
WT Galectin-8
CML1
PhoSPL001
Epa1-Np mutant Y228W
Epa1-Np mutant Y228A
Epa1-Np mutant E227G
Epa1-Np mutant E227A
Epa1-Np WT
ChMBL
DBL-1
hJAA-F11-B11
Gal(β1-3)GlcNAc(β1-3)Gal(β1-3)GlcNAc(β-Sp0

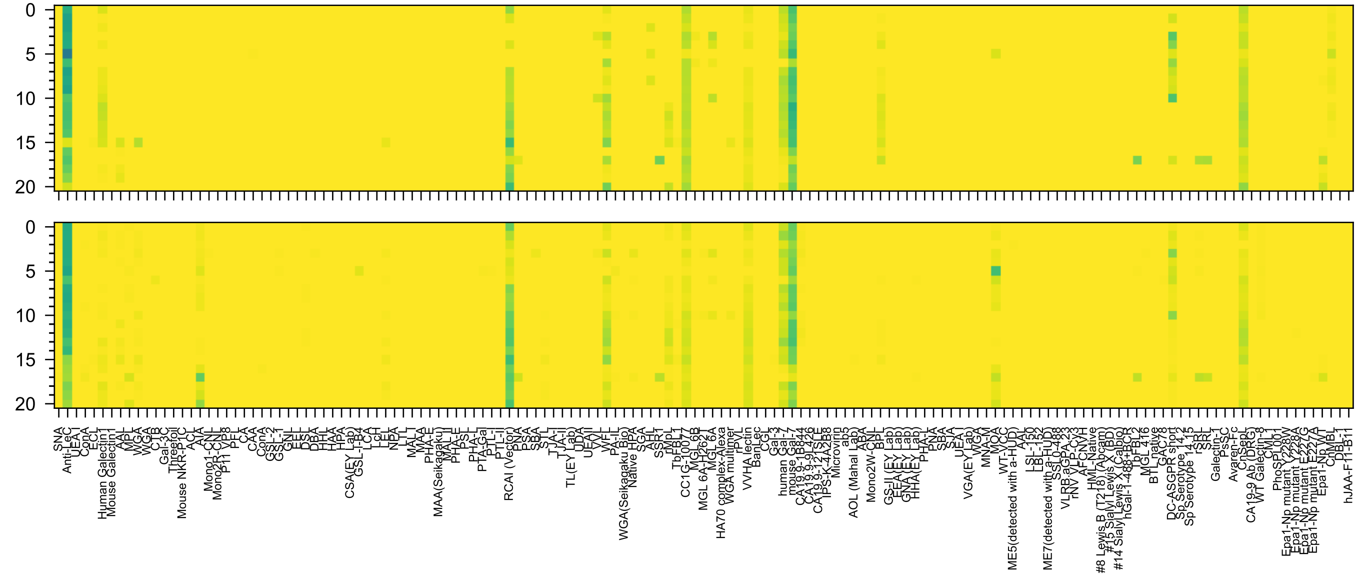
Gal(β1-4)GlcNAc(β1-3)Gal(β1-3)GlcNAc(β-Sp0

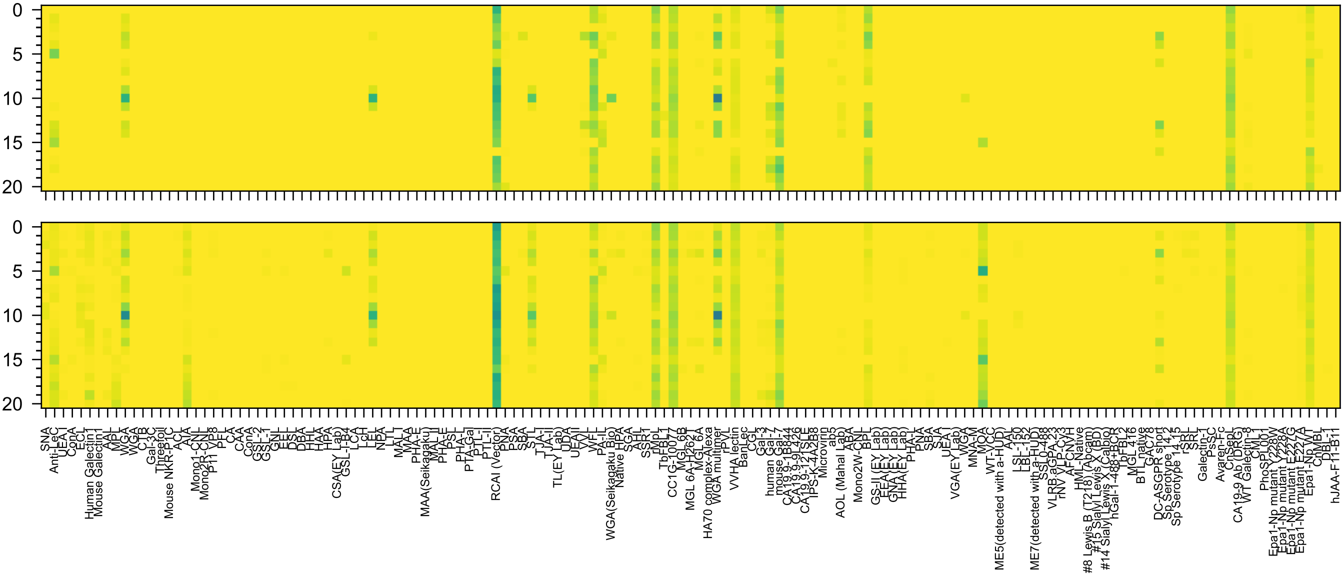

Gal(β1-4)GlcNAc(α1-6)Gal(β1-4)GlcNAc(β-Sp0

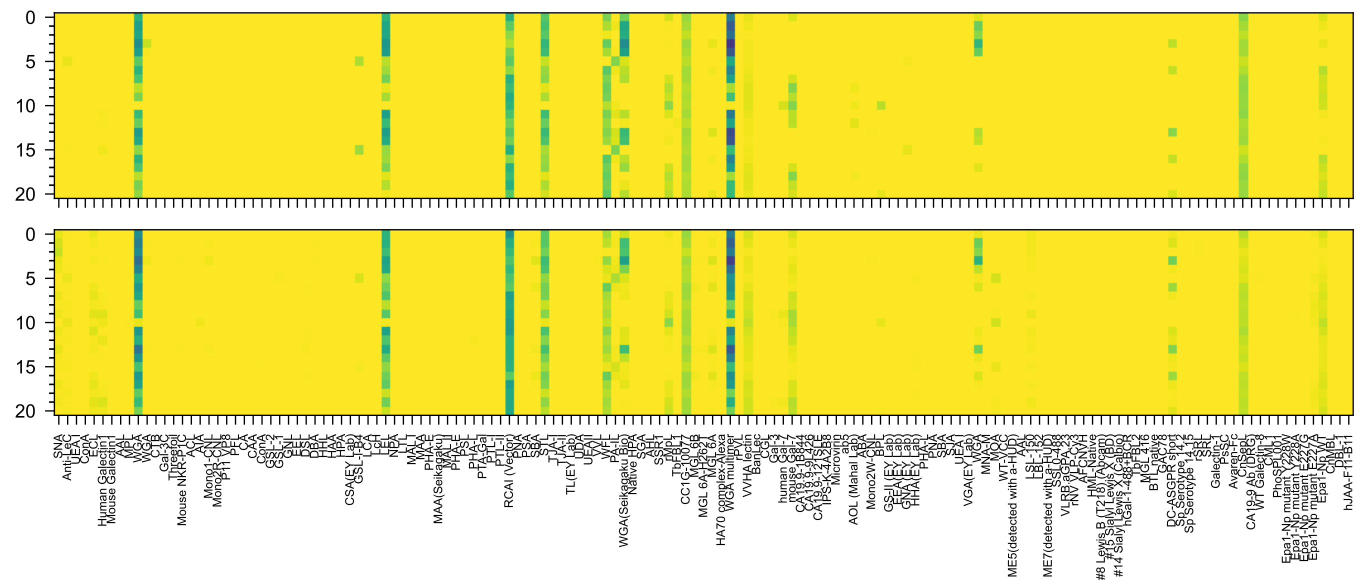

Gal(β1-4)GlcNAc(β1-6)Gal(β1-4)GlcNAc(β-Sp0

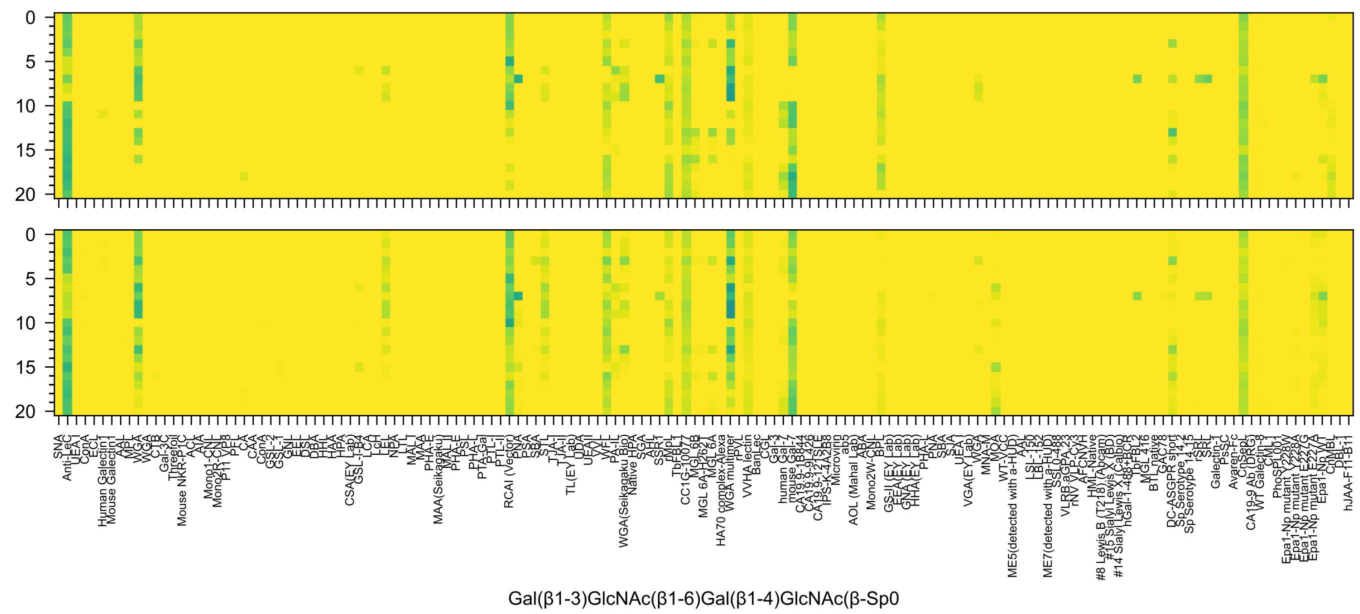
Gal(β1-3)GlcNAc(β1-6)Gal(β1-4)GlcNAc(β-Sp0

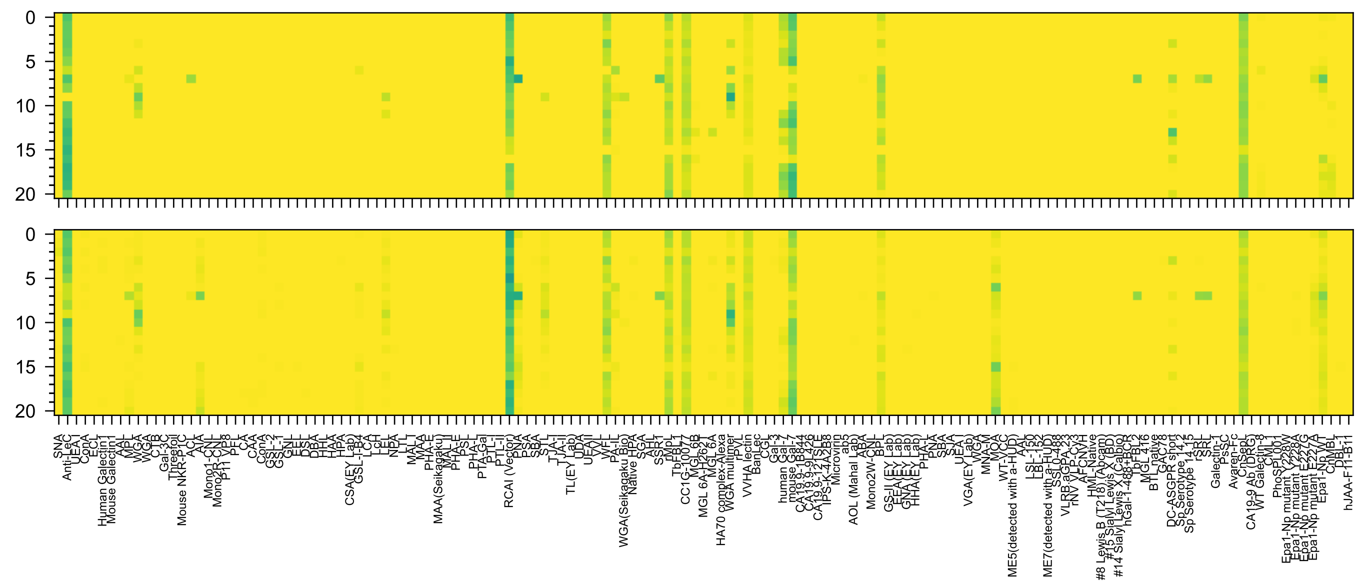

Gal(β1-3)GlcNAc(α1-6)Gal(β1-4)GlcNAc(β-Sp0

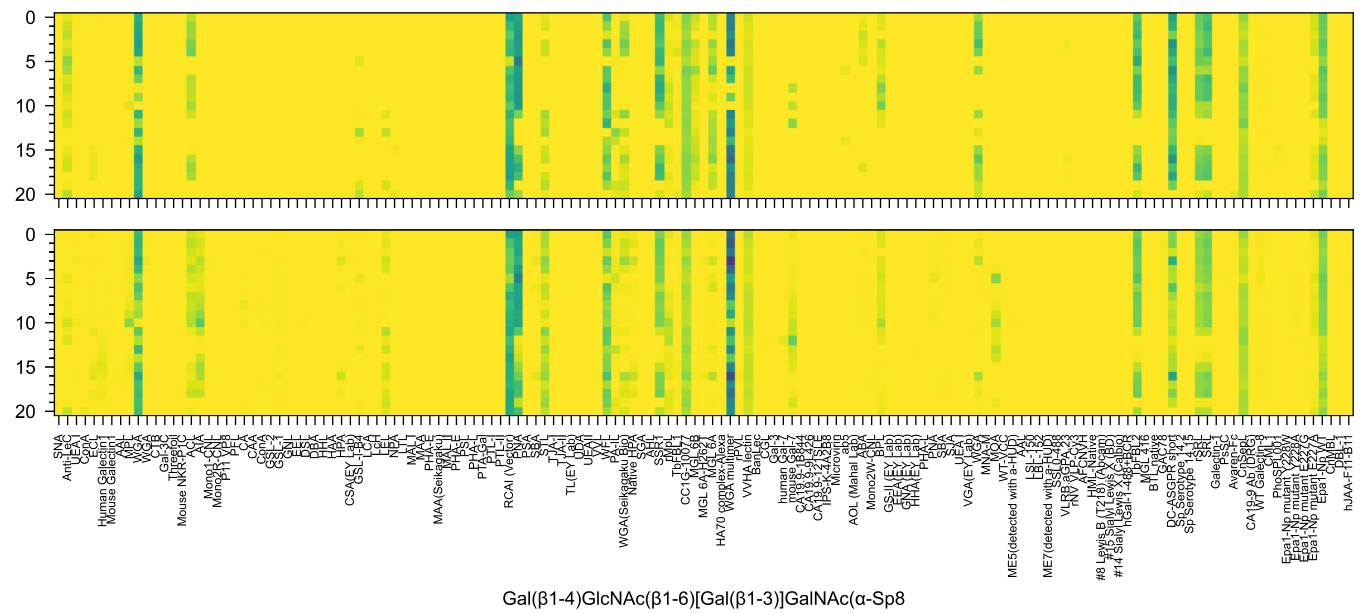
0
5
10
15
20
Gal(β1-4)GlcNAc(β1-6)[Gal(β1-3)]GalNAc(α-Sp8

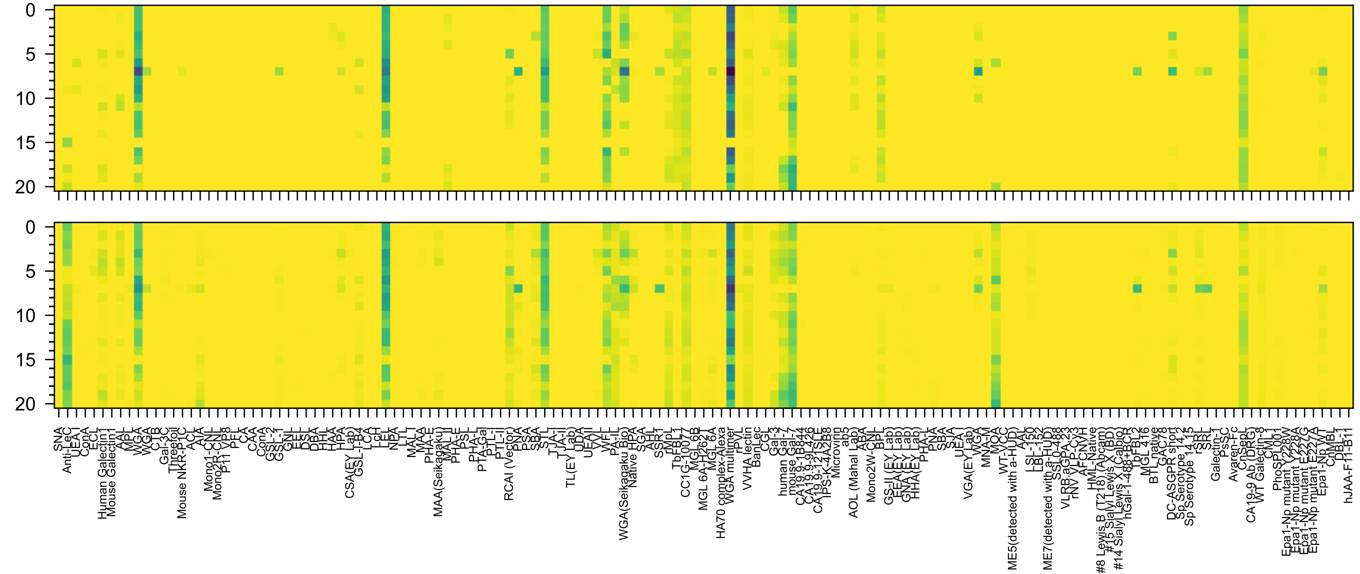


Gal(β1-3)GlcNAc(α1-3)Gal(β1-4)GlcNAc(β-Sp8

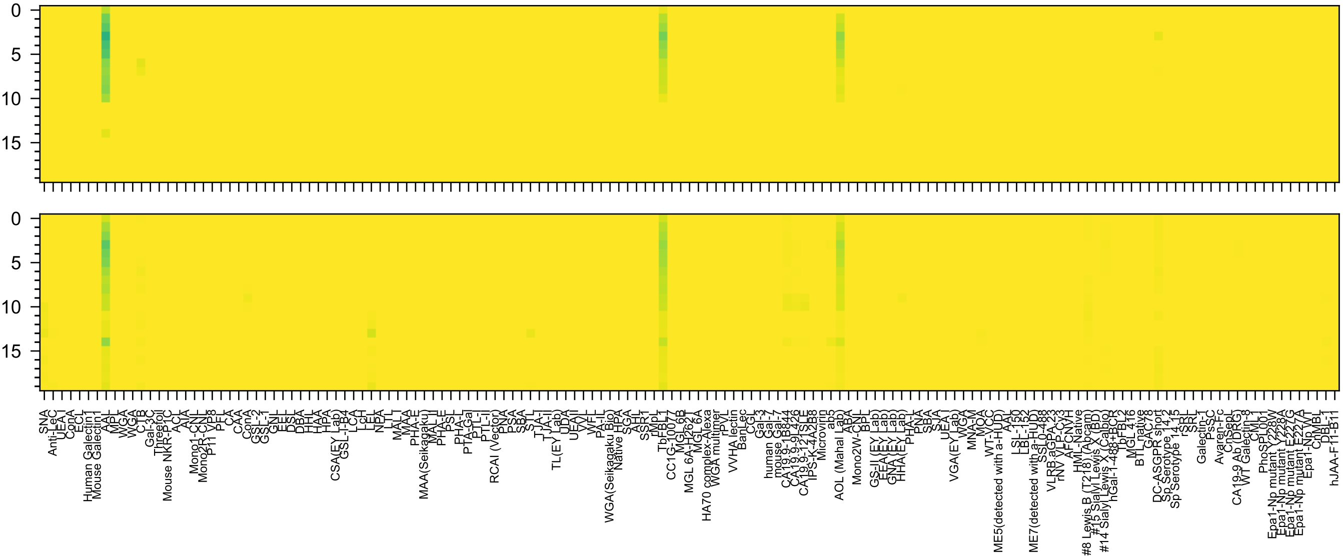
Kdn(α2-3)Gal(β1-4)[Fuc(α1-3)]GlcNAc(-Sp0

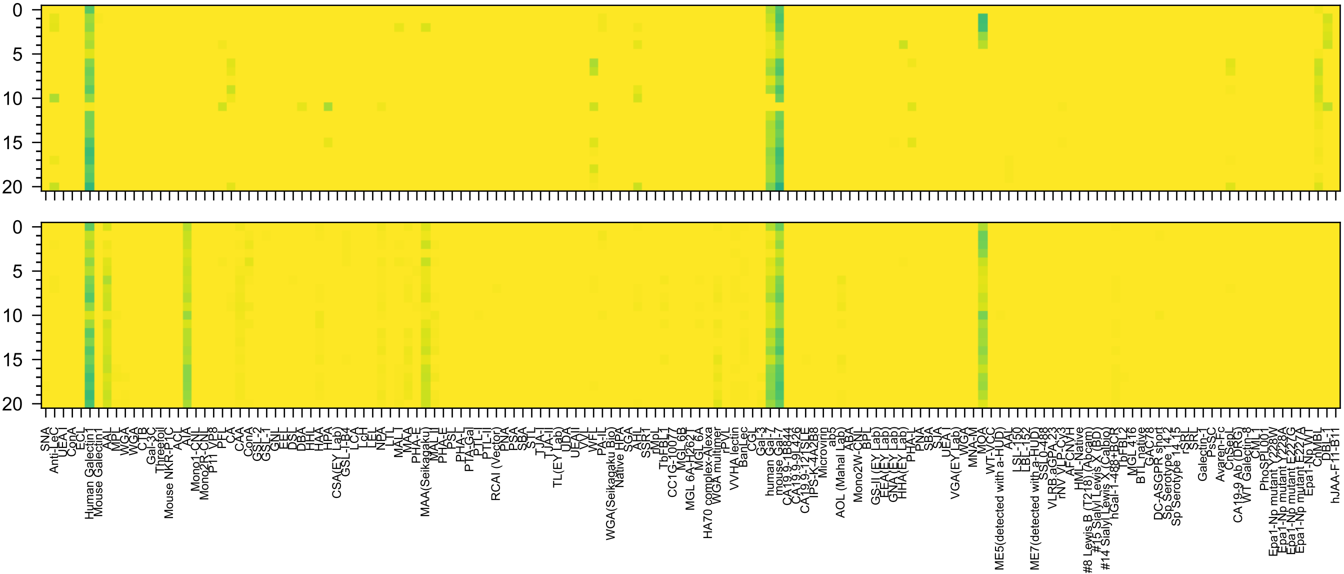

GlcA3OS(β1-3)Gal(β1-4)GlcNAc(β1-2)Man(α-Sp0

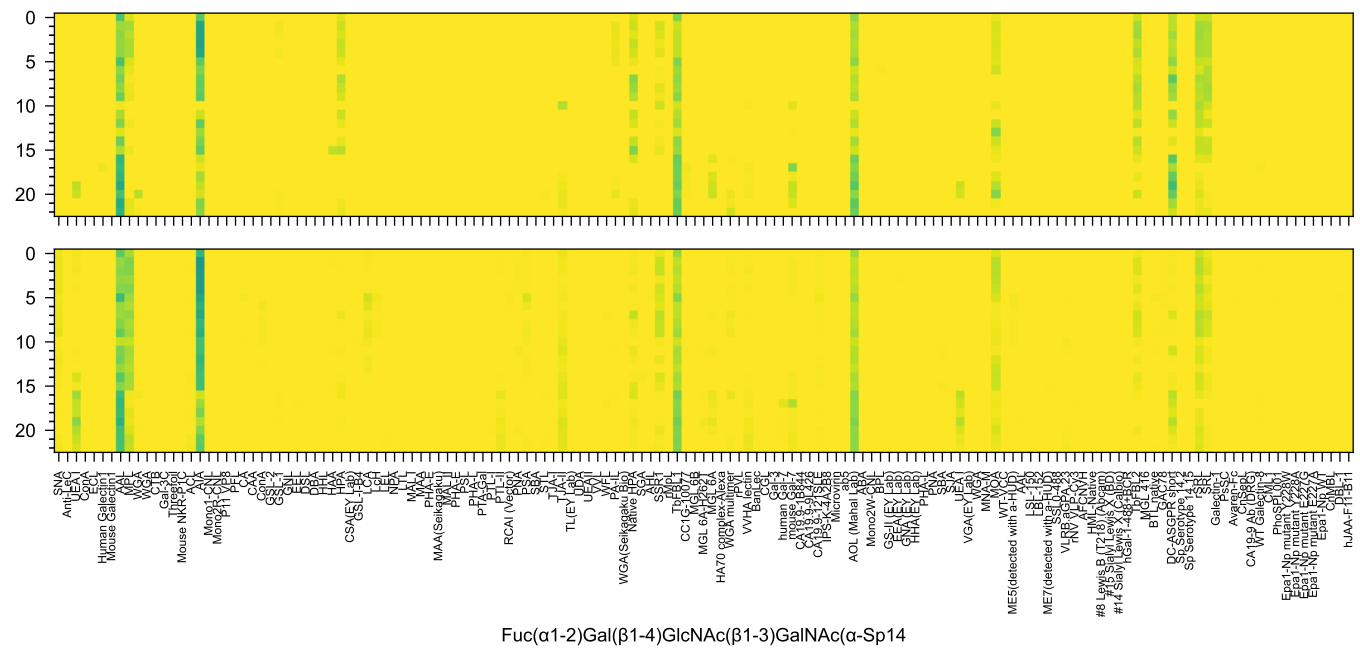

0
5
10
15
20
SNA
Anti-LeC
UEA-I
ConA
ECL
Human Galectin1
Mouse Galectin1
AAL
MPL
WGA
WGA
CTB
Gal-3C
Threefoil
Mouse NKR-P1C
ACL
AIA
Mono1-CNL
Mono2R-CNL
P11 VP8
PFL
CA
CAA
ConA
GSL-2
GSL-1
GNL
EEL
DSL
DBA
HHL
HAA
HPA
CSA(EY Lab)
GSL-I-B4
LCA
LcH
LEL
NPA
LTL
MAL-I
MAA
PHA-E
MAA(Seikagaku)
MAL-II
PHA-E
PSL
PHA-L
PTA-Gal
PTL-I
PTL-II
RCAI (Vector)
PNA
PSA
SBA
STL
TJA-I
TJA-II
TL(EY Lab)
UDA
UEAII
VVL
WFL
PA-IL
WGA(Seikagaku Bio)
Native HPA
SGA
AHL
SSR1
rMpL
TbFBL1
CC1G-10077
MGL 6B
MGL 6A-H262T
MGL 6A
HA70 complex-Alexa
WGA multimer
rPVL
VVHA lectin
BanLec
CGL
Gal-3
human Gal-7
mouse Gal-7
CA19.9-1B844
CA19.9-9L426
CA19.9-121SLE
IPS-K-4A2B8
Microvirin
ab5
AOL (Mahal Lab)
ABA
Mono2W-CNL
BPL
GS-II (EY Lab)
EEA (EY Lab)
GNA (EY Lab)
HHA (EY Lab)
PHA-L
PNA
SBA
SJA
UEA-I
VGA(EY Lab)
WGA
MNA-M
MOA
WT-VCC
ME5(detected with a-HUD)
AAL
LSL-150
LBL-152
ME7(detected with a-HUD)
SSL0-488
VLRB.aGPA.23
rNV VLP-Cy3
AFCNVH
HML-Native
#8 Lewis B (T218) (Abcam)
#15 Sialyl Lewis X (BD)
#14 Sialyl Lewis X (Calbio)
hGal-1-488+BCR
TbFBL2
MGL 416
BTL native
GAC78
DC-ASGPR short
Sp Serotype 14.2
Sp Serotype 14.15
rSRL
SRL
Galectin-1
PsSC
Avaren-Fc
CnSepL
CA19-9 Ab (DRG)
WT Galectin-8
CML1
PhoSPL001
Epa1-Np mutant Y228W
Epa1-Np mutant Y228A
Epa1-Np mutant E227G
Epa1-Np mutant E227A
Epa1-Np WT
ChMBL
DBL-1
hJAA-F11-B11
Fuc(α1-2)Gal(β1-4)GlcNAc(β1-3)GalNAc(α-Sp14

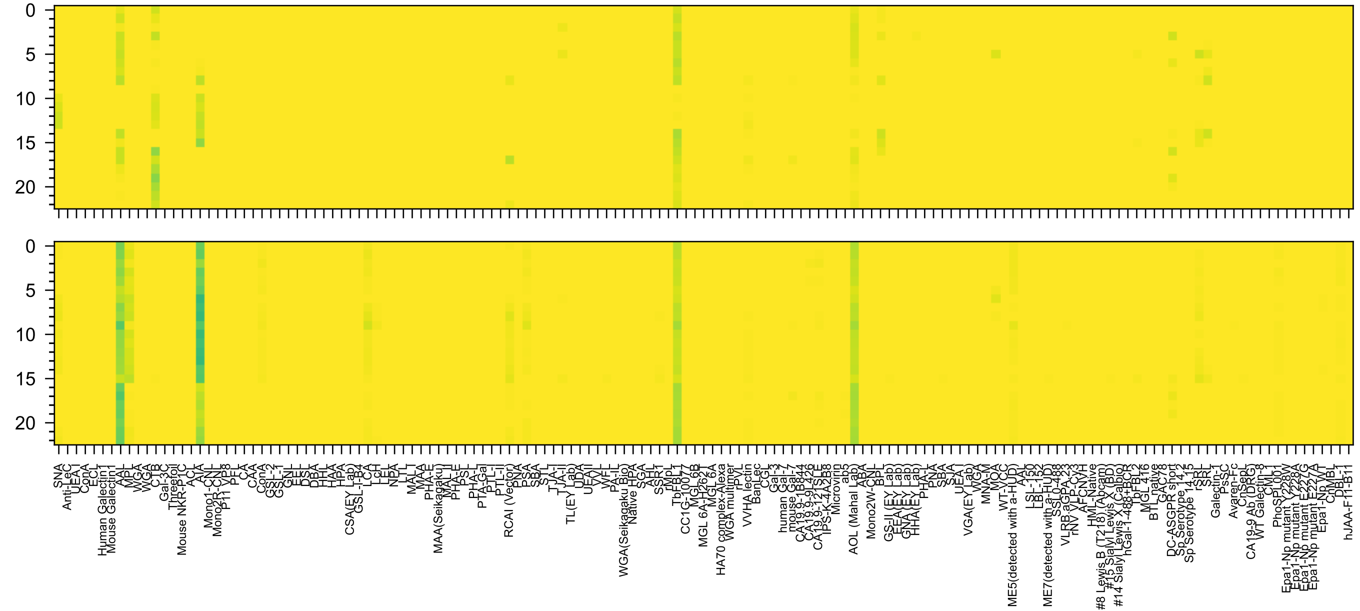

Gal(β1-4)[Fuc(α1-3)]GlcNAc(β1-3)GalNAc(α-Sp14

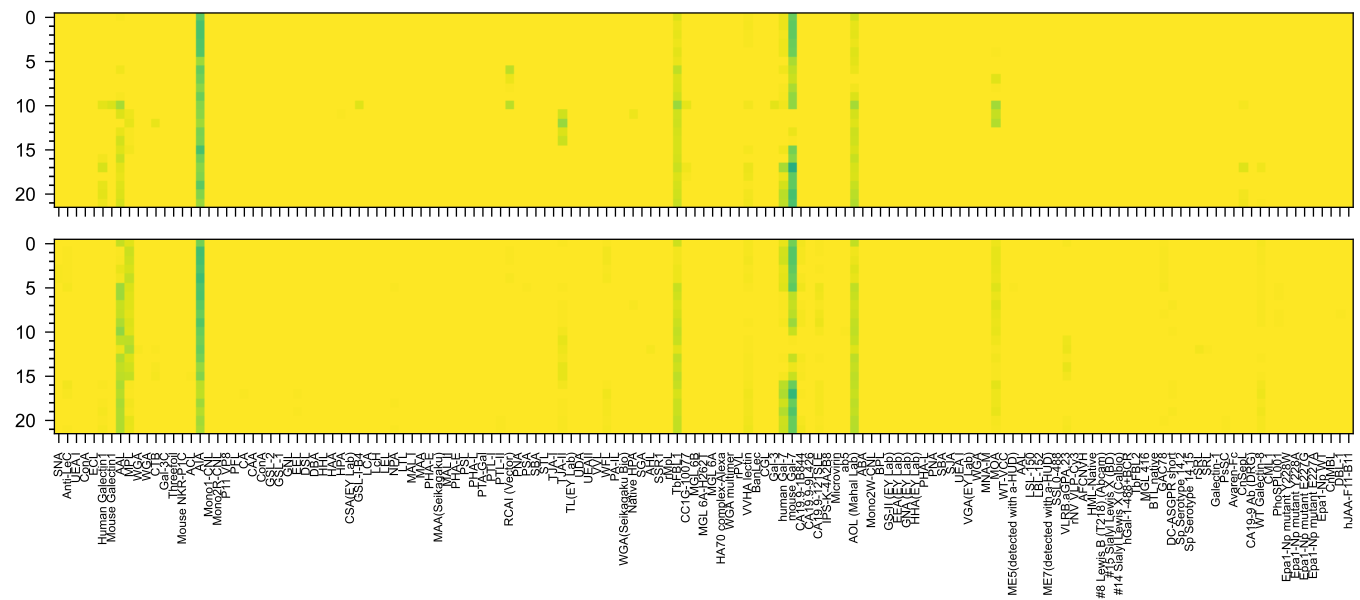


Fuc(α1-2)Gal(β1-3)GlcNAc(β1-3)GalNAc(-Sp14

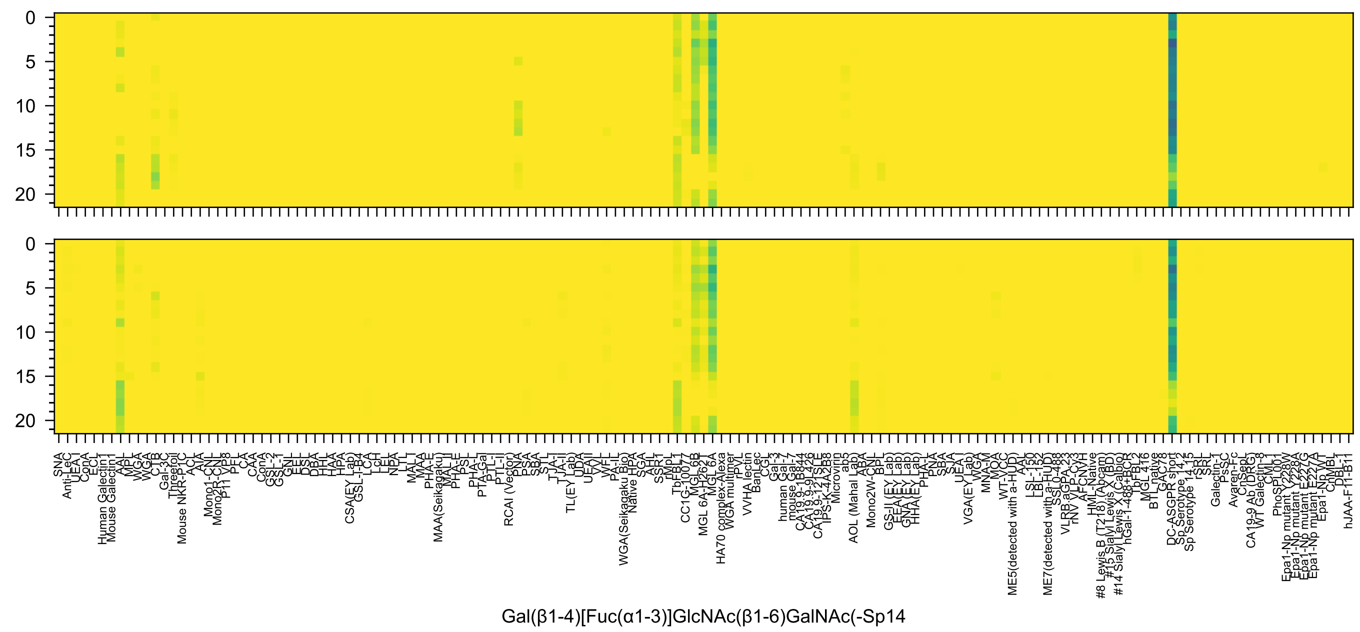
Gal(β1-4)[Fuc(α1-3)]GlcNAc(β1-6)GalNAc(-Sp14

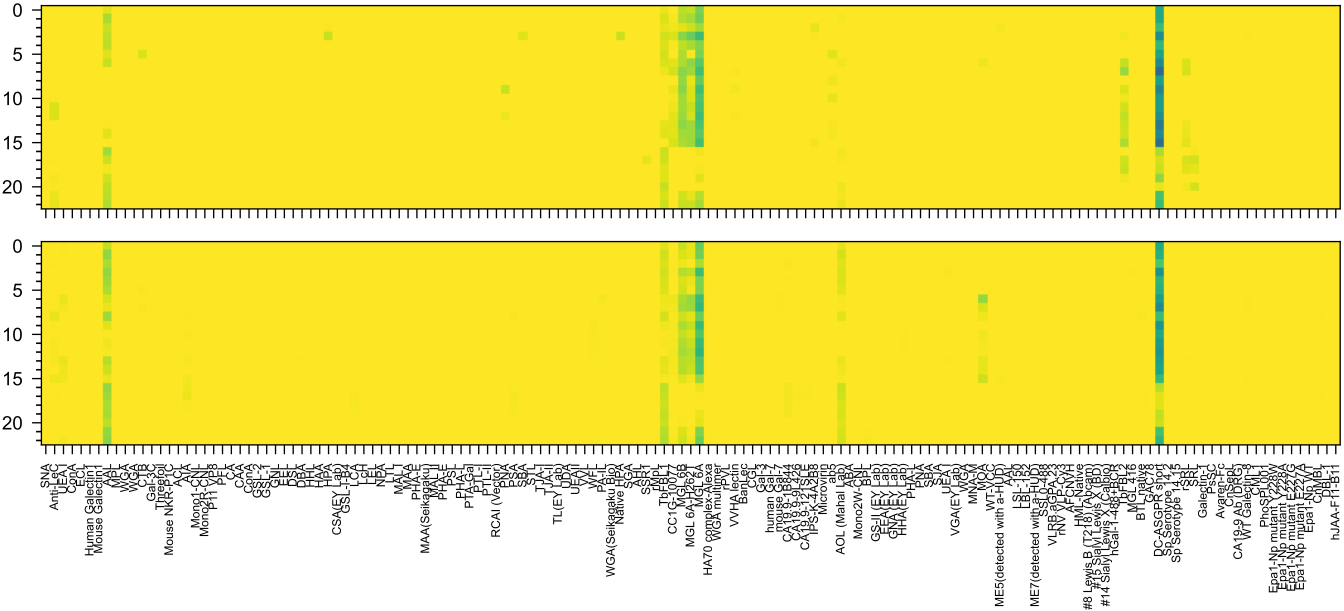


Gal(β1-3)[Fuc(α1-4)]GlcNAc(β1-6)GalNAc(α-Sp14

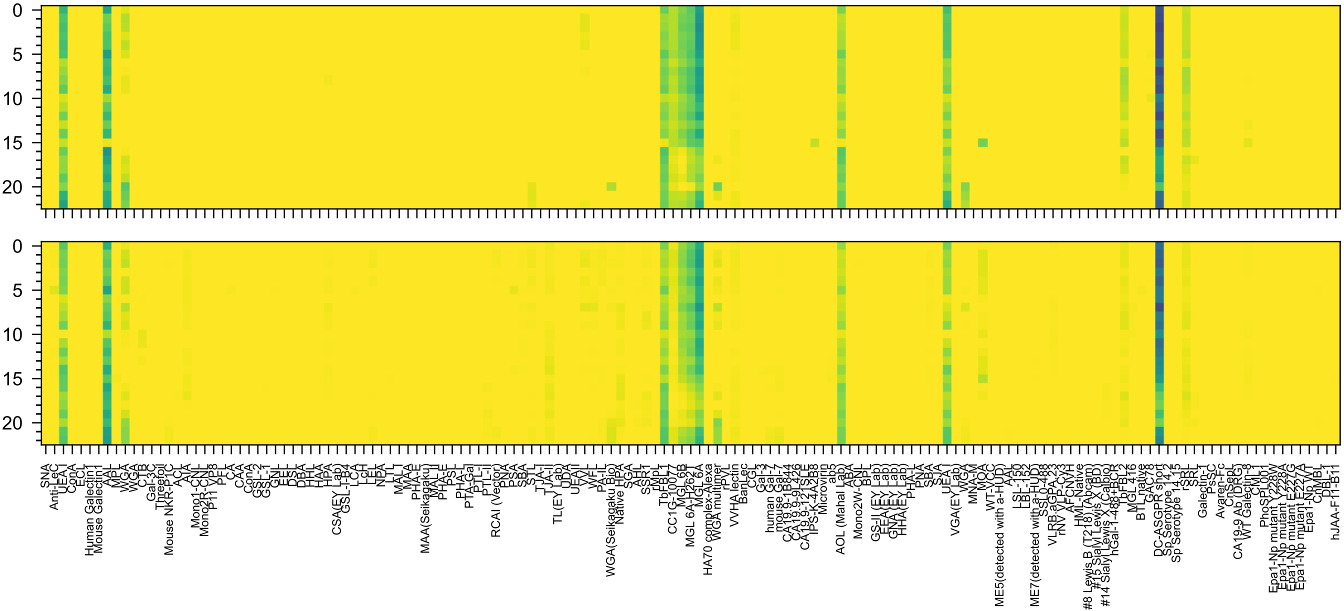

Fuc(α1-2)Gal(β1-4)GlcNAc(β1-6)GalNAc(α-Sp14

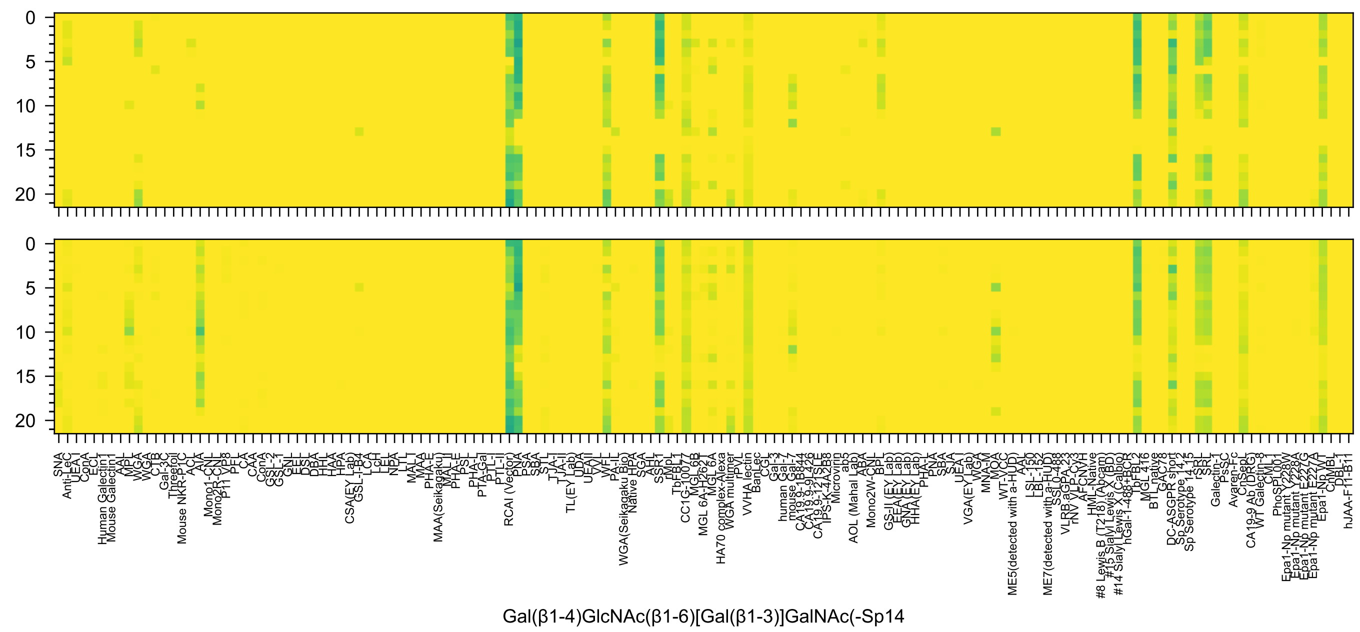
Gal(β1-4)GlcNAc(β1-6)[Gal(β1-3)]GalNAc(-Sp14

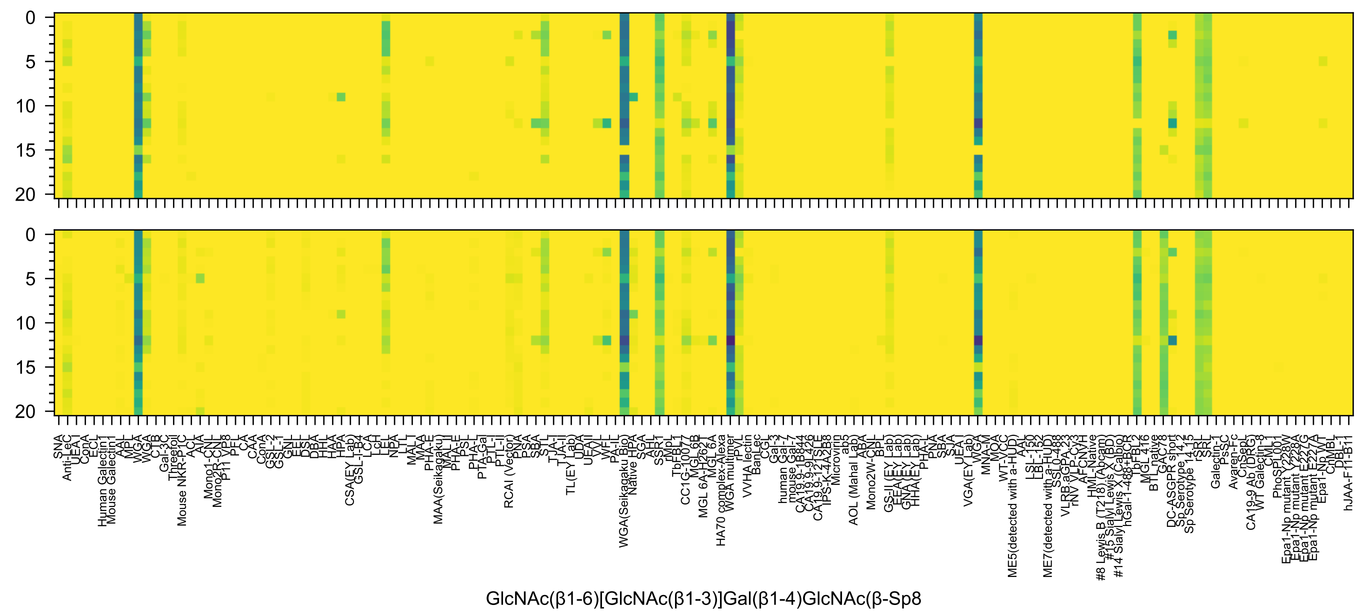
GlcNAc(β1-6)[GlcNAc(β1-3)]Gal(β1-4)GlcNAc(β-Sp8

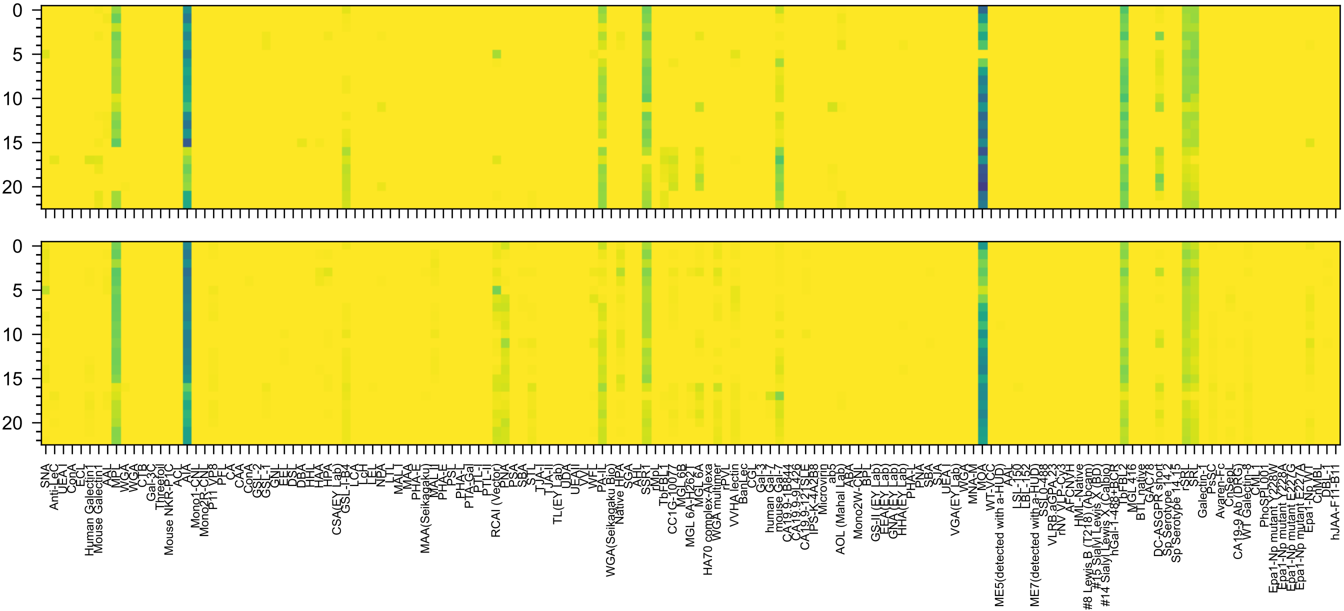

Gal(α1-3)Gal(β1-4)GlcNAc(β1-3)GalNAc(α-Sp14

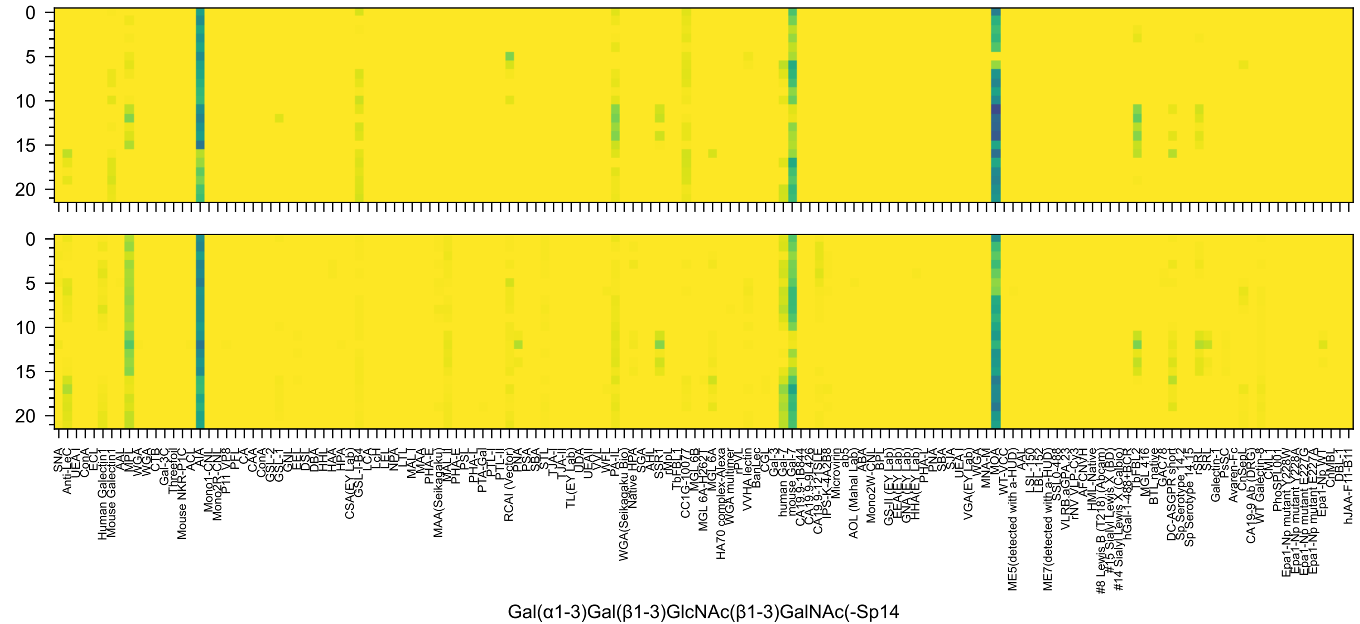
0
5
10
15
20
0
5
10
15
20
SNA
Anti-LeC
UEA-I
ConA
ECL
Human Galectin1
Mouse Galectin1
AAL
MPL
WGA
WGA
CTB
Gal-3C
Threefoil
Mouse NKR-P1C
ACL
AIA
Mono1-CNL
Mono2R-CNL
P11 VP8
PFL
CA
CAA
ConA
GSL-2
GSL-1
GNL
EEL
DSL
DBA
HHL
HAA
HPA
CSA(EY Lab)
GSL-I-B4
LCA
LcH
LEL
NPA
LTL
MAL-I
MAA
PHA-E
MAA(Seikagaku)
MAL-II
PHA-E
PSL
PHA-L
PTA-Gal
PTL-I
PTL-II
RCAI (Vector)
PNA
PSA
SBA
STL
TJA-I
TJA-II
TL(EY Lab)
UDA
UEAII
VVL
WFL
PA-IL
WGA(Seikagaku Bio)
Native HPA
SGA
AHL
SSR1
rMpL
TbFBL1
CC1G-10077
MGL 6B
MGL 6A-H262T
MGL 6A
HA70 complex-Alexa
WGA multimer
rPVL
VVHA lectin
BanLec
CGL
Gal-3
human Gal-7
mouse Gal-7
CA19.9-1B844
CA19.9-9L426
CA19.9-121SLE
IPS-K-4A2B8
Microvirin
ab5
AOL (Mahal Lab)
ABA
Mono2W-CNL
BPL
GS-II (EY Lab)
EEA (EY Lab)
GNA (EY Lab)
HHA (EY Lab)
PHA-L
PNA
SBA
SJA
UEA-I
VGA(EY Lab)
WGA
MNA-M
MOA
WT-VCC
ME5(detected with a-HUD)
AAL
LSL-150
LBL-152
ME7(detected with a-HUD)
SSL0-488
VLRB.aGPA.23
rNV VLP-Cy3
AFCNYH
HML-Native
#8 Lewis B (T218) (Abcam)
#15 Sialyl Lewis X (BD)
#14 Sialyl Lewis X (Calbio)
hGal-1-488+BCR
TbFBL2
MGL 416
BTL native
GAC78
DC-ASGPR short
Sp Serotype 14.2
Sp Serotype 14.15
rSRL
SRL
Galectin-1
PsSC
Avaren-Fc
ChSepL
CA19-9 Ab (DRG)
WT Galectin-8
CML1
PhoSL001
Epa1-Np mutant Y228W
Epa1-Np mutant Y228A
Epa1-Np mutant E227G
Epa1-Np mutant E227A
Epa1-Np WT
CnMBL
DBL-1
hJAA-F11-B11
Gal(α1-3)Gal(β1-3)GlcNAc(β1-3)GalNAc(-Sp14

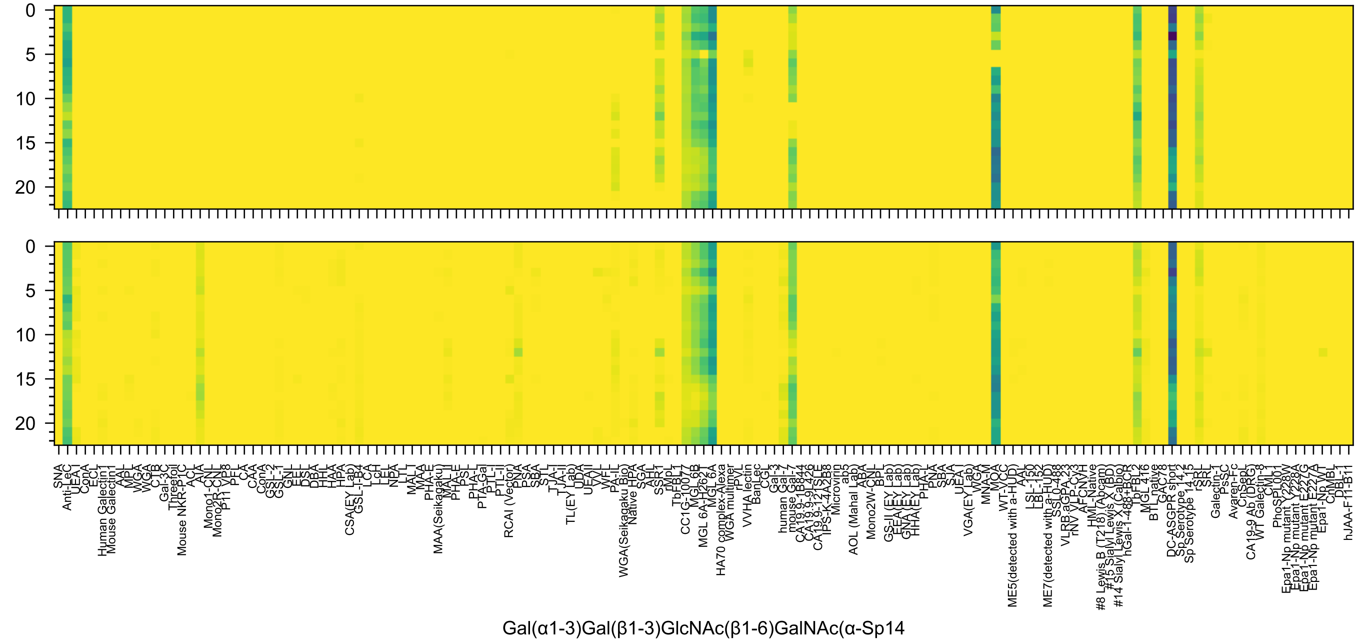
Gal(α1-3)Gal(β1-3)GlcNAc(β1-6)GalNAc(α-Sp14
0
5
10
15
20

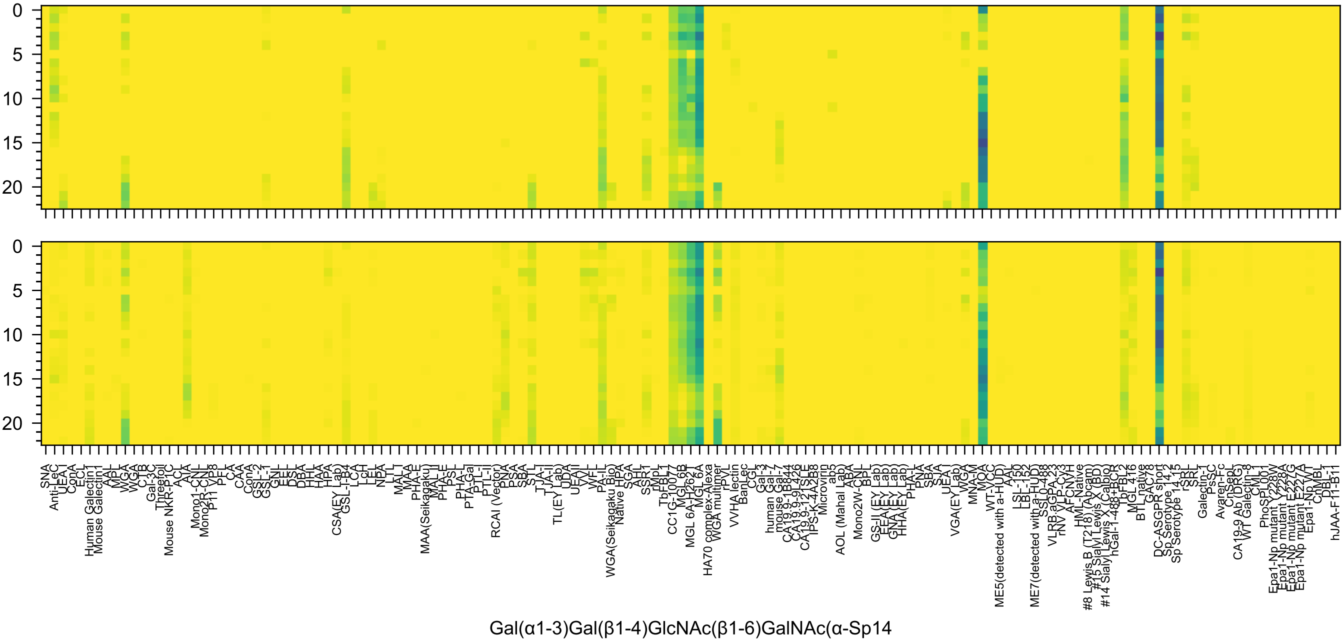

0
5
10
15
20
Gal(α1-3)Gal(β1-4)GlcNAc(β1-6)GalNAc(α-Sp14

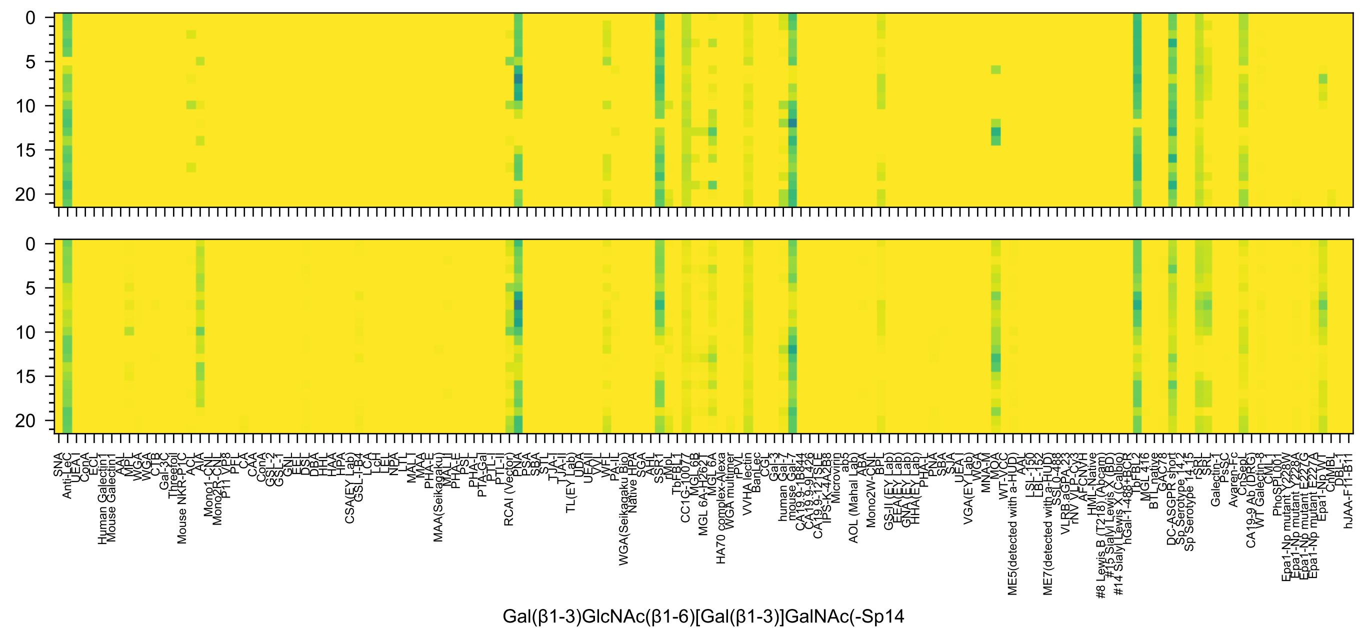
0
5
10
15
20
SNA
Anti-LeC
UEA-I
ConA
ECL
Human Galectin1
Mouse Galectin1
AAL
MPL
WGA
WGA
CTB
Gal-3C
Threefoil
Mouse NKR-P1C
ACL
AIA
Mono1-CNL
Mono2R-CNL
P11 VP8
PFL
CA
CAA
ConA
GSL-2
GSL-1
GNL
EEL
DSL
DBA
HHL
HAA
HPA
CSA(EY Lab)
GSL-I-B4
LCA
LcH
LEL
NPA
LTL
MAL-I
MAA
PHA-E
MAA(Seikagaku)
MAL-II
PHA-E
PSL
PHA-L
PTA-Gal
PTL-I
PTL-II
RCAI (Vector)
PNA
PSA
SBA
STL
TJA-I
TJA-II
TL(EY Lab)
UDA
UEAII
VVL
WFL
PA-IL
WGA(Seikagaku Bio)
Native HPA
SGA
AHL
SSR1
rMpL
TbFBL1
CC1G-10077
MGL6B
MGL 6A-H262T
MGL 6A
HA70 complex-Alexa
WGA multimer
VVHA lectin
BanLec
CGL
Gal-3
human Gal-7
mouse Gal-7
CA19.9-1B844
CA19.9-9L426
CA19.9-121SLE
IPS-K-4A2B8
Microvirin
ab5
AOL (Mahal Lab)
ABA
Mono2W-CNL
BPL
GS-II (EY Lab)
EEA(EY Lab)
GNA(EY Lab)
HHA(EY Lab)
PHA-L
PNA
SBA
SJA
UEA-I
VGA(EY Lab)
WGA
MNA-M
MOA
WT-VCC
ME5(detected with a-HUD)
AAL
LSL-150
LBL-152
ME7(detected with a-HUD)
SSL0-488
VLRB.aGPA.23
rNV VLP-Cy3
AFCNVH
HML-Native
#8 Lewis B (T218) (Abcam)
#15 Sialyl Lewis X (BD)
#14 Sialyl Lewis X (Calbio)
hGal-1-488+BCR
TbFBL2
MGL 416
BTL native
GAC78
DC-ASGPR short
Sp Serotype 14.2
Sp Serotype 14.15
rSRL
SRL
Galectin-1
PsSC
Avaren-Fc
ChSepL
CA19-9 Ab (DRG)
WT Galectin-8
CML1
PhoSL001
Epa1-Np mutant Y228W
Epa1-Np mutant Y228A
Epa1-Np mutant E227G
Epa1-Np mutant E227A
Epa1-Np WT
ChMBL
DBL-1
hJAA-F11-B11
Gal(β1-3)GlcNAc(β1-6)[Gal(β1-3)]GalNAc(-Sp14

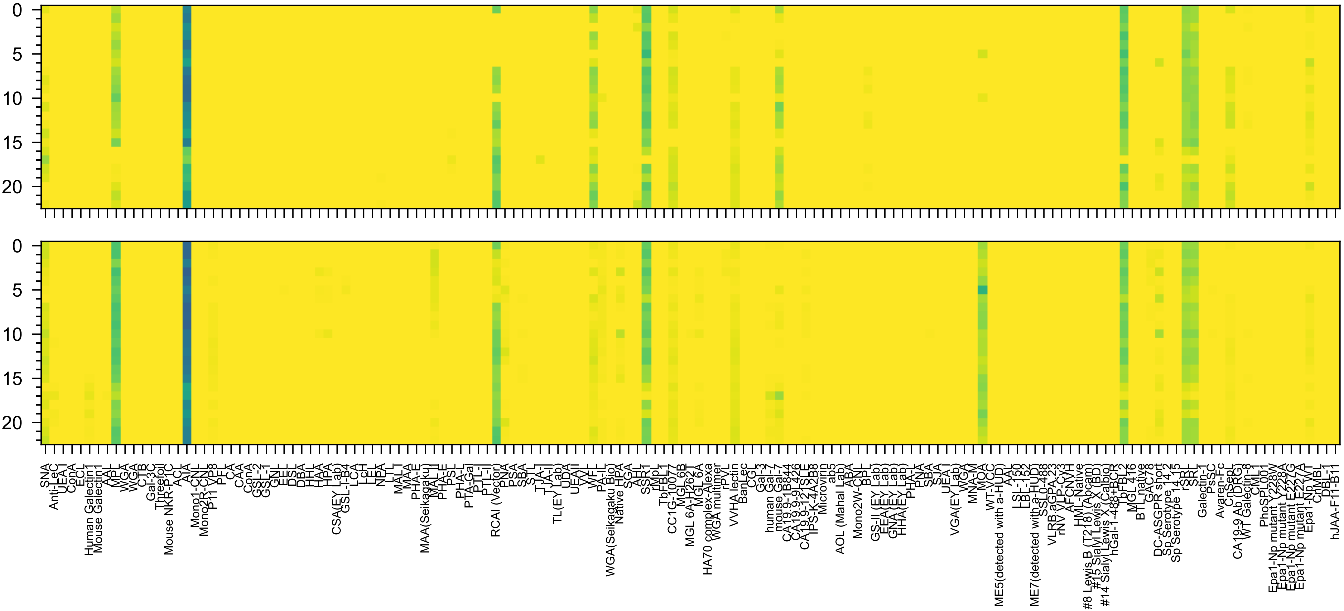

Gal(β1-4)GlcNAc(β1-3)Gal(β1-3)GalNAc(α-Sp14

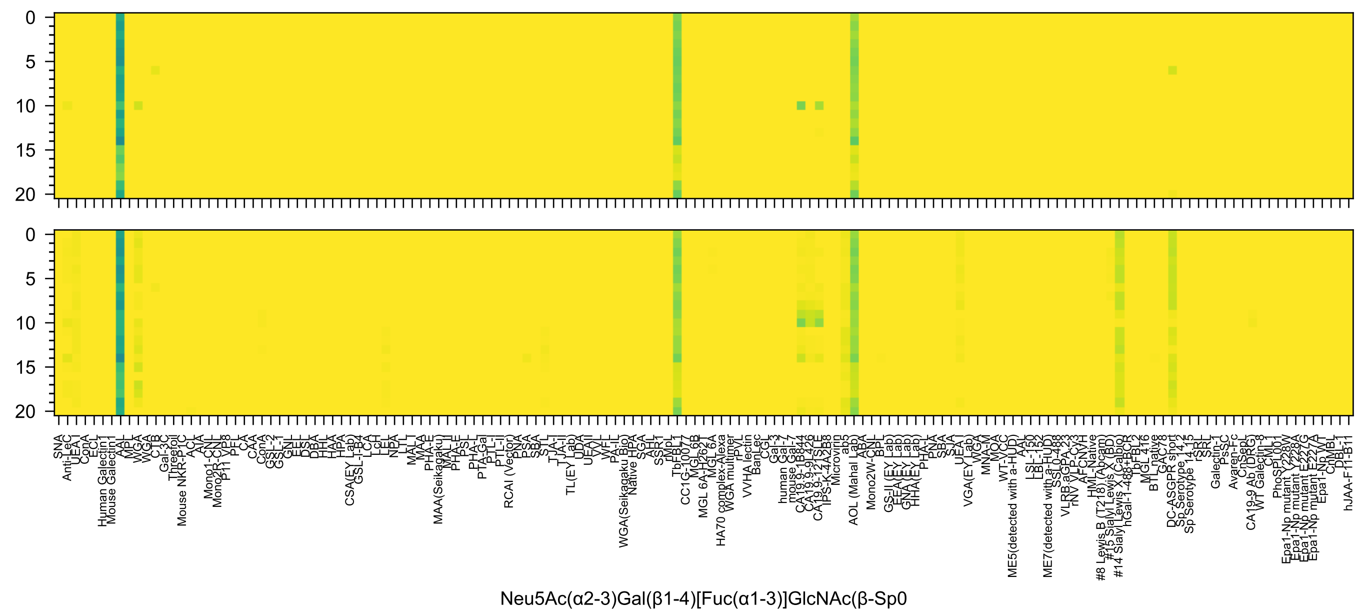
Neu5Ac(α2-3)Gal(β1-4)[Fuc(α1-3)]GlcNAc(β-Sp0

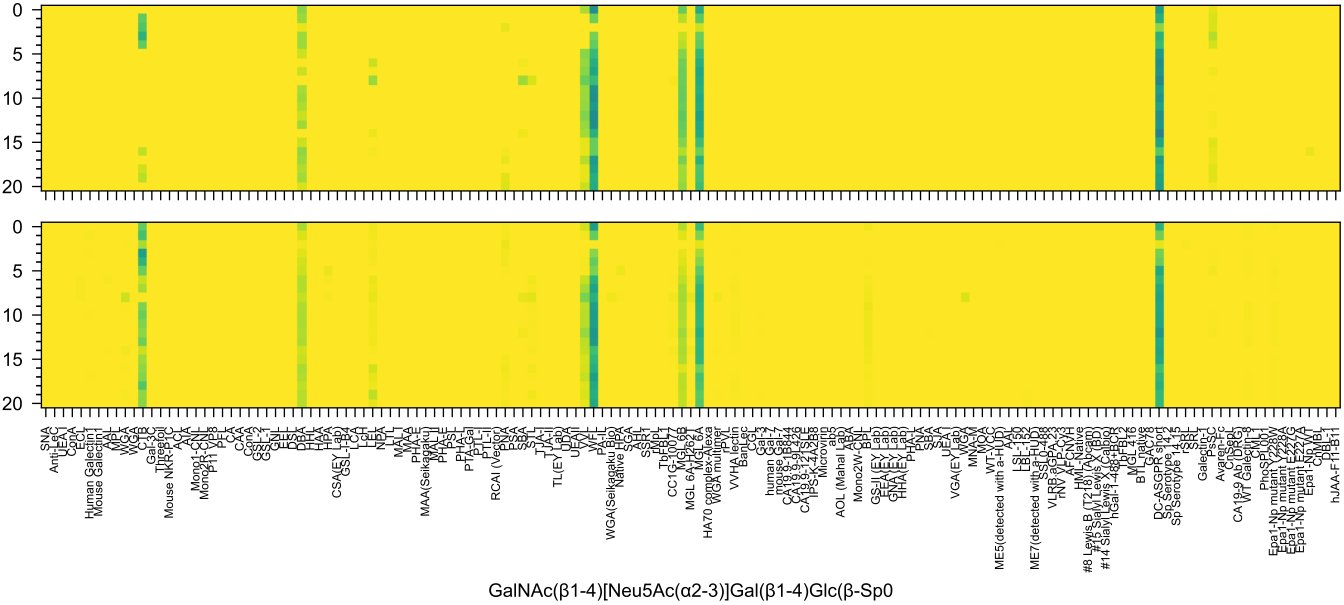
GalNAc(β1-4)[Neu5Ac(α2-3)]Gal(β1-4)Glc(β-Sp0

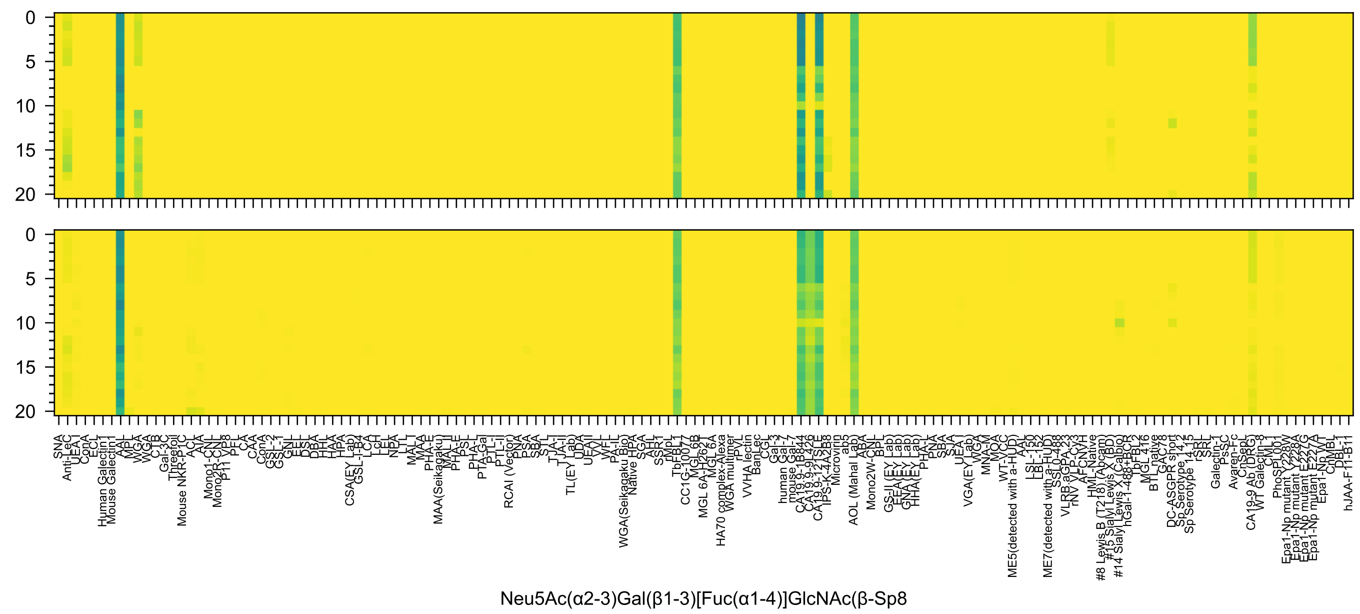
Neu5Ac(α2-3)Gal(β1-3)[Fuc(α1-4)]GlcNAc(β-Sp8

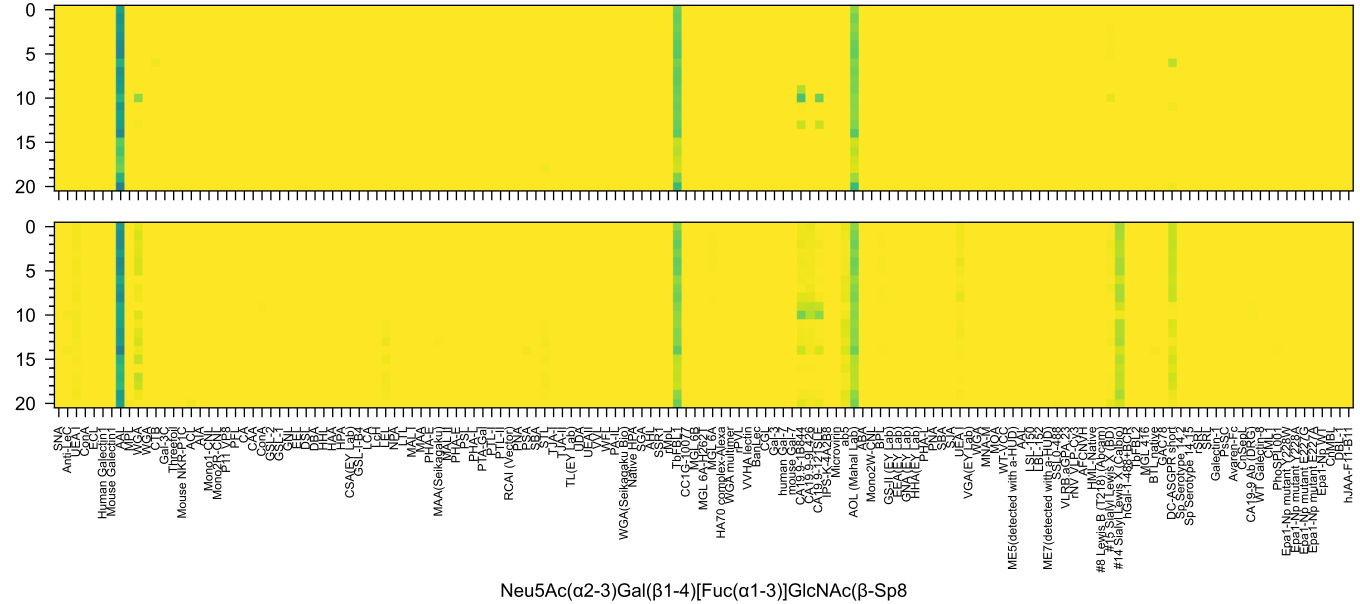
0
5
10
15
20
SNA
Anti-LeC
UEA I
ConA
ECL
Human Galectin1
Mouse Galectin1
AAL
MPL
WGA
WGA
CTB
Gal-3C
Threefoil
Mouse NKR-P1C
ACL
AIA
Mono1-CNL
Mono2R-CNL
P11 VP8
PFL
CA
CAA
ConA
GSL-2
GSL-1
GNL
EEL
DSL
DBA
HHL
HAA
HPA
CSA(EY Lab)
GSL-I-B4
LCA
LcH
LEL
NPA
LTL
MAL I
MAA
PHA-E
MAA(Seikagaku)
MAL II
PHA-E
PSL
PHA-L
PTA-Gal
PTL-I
PTL-II
RCAI (Vector)
PNA
PSA
SBA
STL
TJA-I
TJA-II
TL(EY Lab)
UDA
UEAII
VVL
WFL
PA-IL
WGA(Seikagaku Bio)
Native HPA
SGA
AHL
SSR1
rMpL
TbFBL1
CC1G-10077
MGL 6B
MGL 6A-H262T
MGL 6A
HA70 complex-Alexa
WGA multimer
rPVL
VVHA lectin
BanLec
CGL
Gal-3
human Gal-7
mouse Gal-7
CA19.9-1B844
CA19.9-9L426
CA19.9-121SLE
IPS-K-4A2B8
Microvirin
ab5
AOL (Mahal Lab)
ABA
Mono2W-CNL
BPL
GS-II (EY Lab)
EEA (EY Lab)
GNA (EY Lab)
HHA (EY Lab)
PHA-L
PNA
SBA
SJA
UEA I
VGA(EY Lab)
WGA
MNA-M
MOA
WT-VCC
ME5(detected with a-HUD)
AAL
LSL-150
LBL-152
ME7(detected with a-HUD)
SSL0-488
VLRB.aGPA.23
rNV VLP-Cy3
AFCNVH
HML-Native
#8 Lewis B (T218) (Abcam)
#15 Sialyl Lewis X (BD)
#14 Sialyl Lewis X (Calbio)
hGal-1-488+BCR
TbFBL2
MGL 416
BTL native
GAC78
DC-ASGPR short
Sp Serotype 14.2
Sp Serotype 14.15
rSRL
SRL
Galectin-1
PsSC
Avaren-Fc
ChSeL
CA19-9 Ab (DRG)
WT Galectin-8
CML1
PhoSL001
Epa1-Np mutant Y228W
Epa1-Np mutant Y228A
Epa1-Np mutant E227G
Epa1-Np mutant E227A
Epa1-Np WT
ChMBL
DBL-1
hJAA-F11-B11
Neu5Ac(α2-3)Gal(β1-4)[Fuc(α1-3)]GlcNAc(β-Sp8

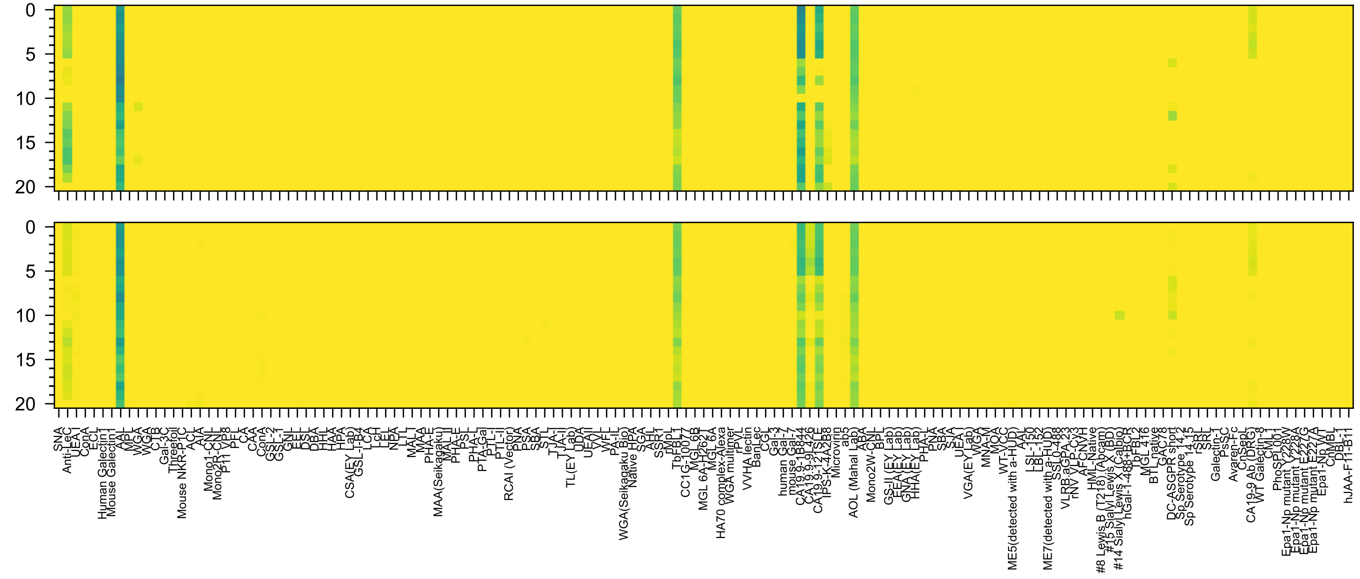

Neu5Gc(α2-3)Gal(β1-3)[Fuc(α1-4)]GlcNAc(β-Sp0

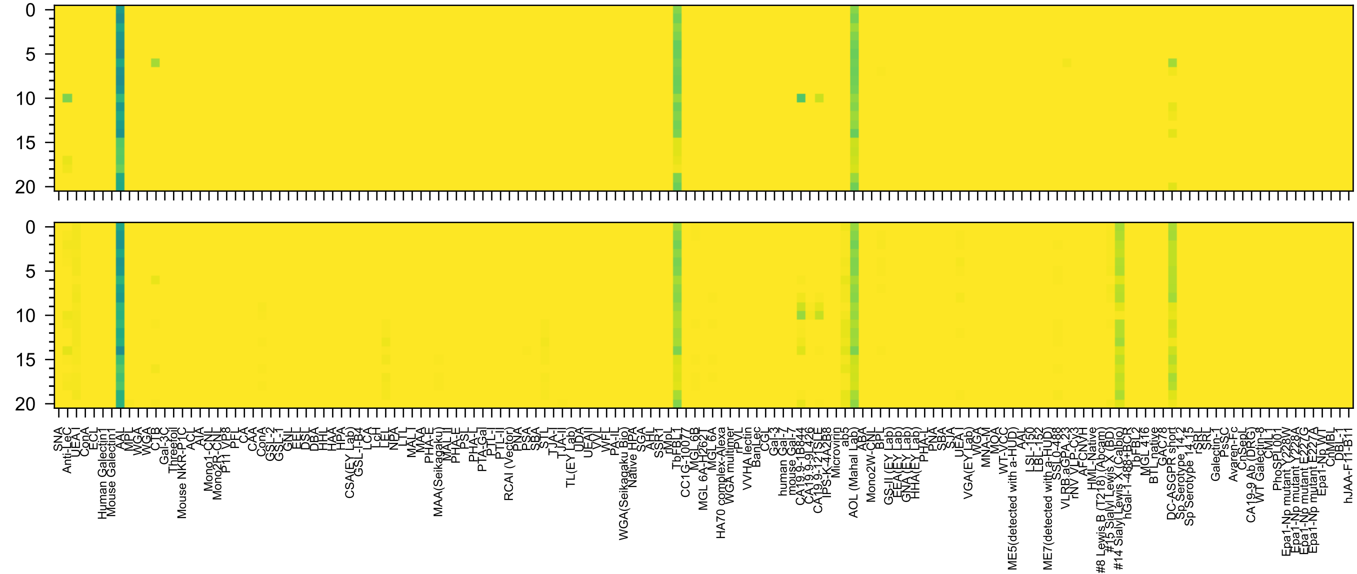

Neu5Gc(α2-3)Gal(β1-4)[Fuc(α1-3)]GlcNAc(β-Sp0

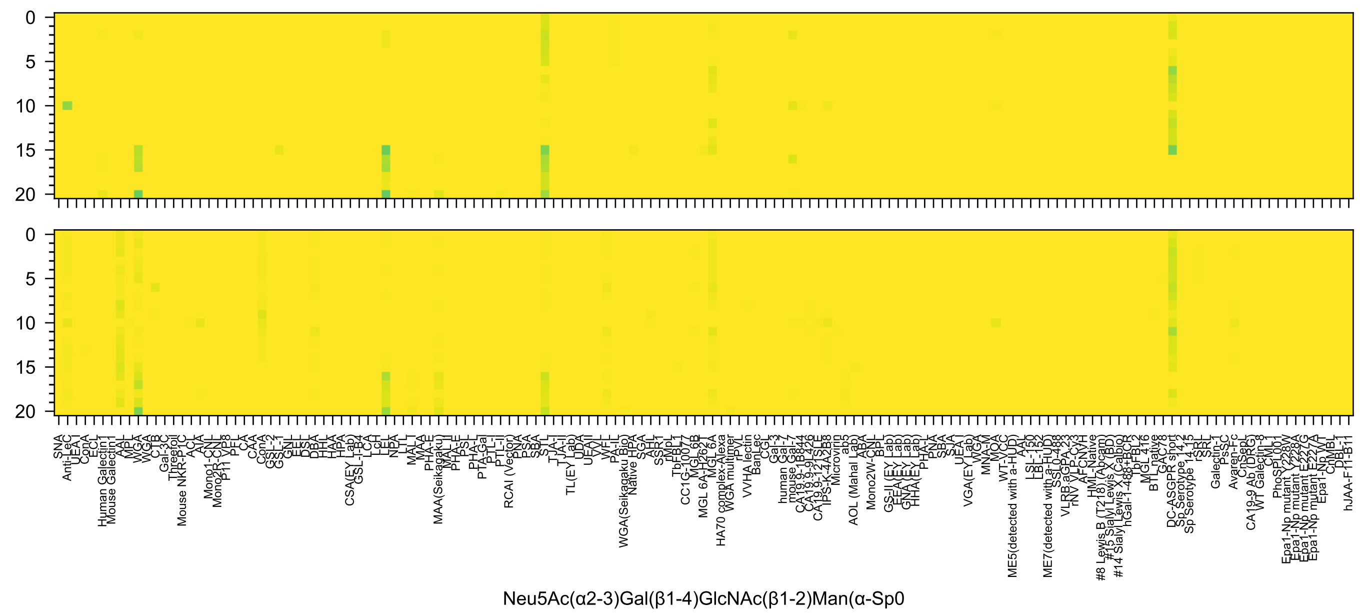

Neu5Ac(α2-3)Gal(β1-4)GlcNAc(β1-2)Man(α-Sp0

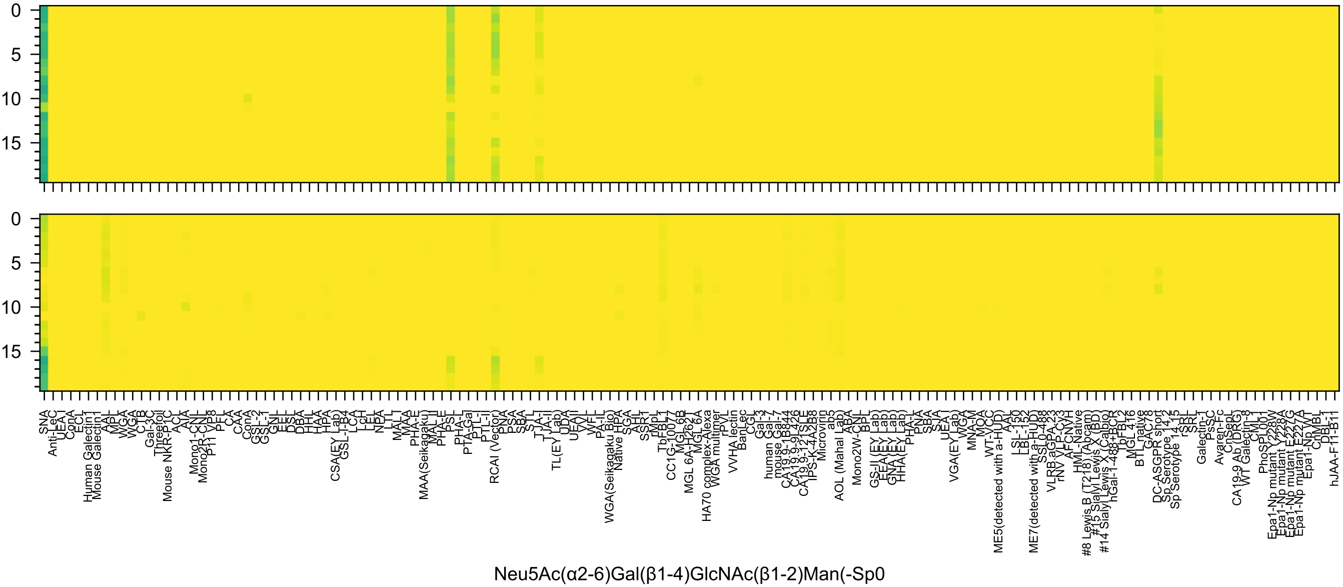
0
5
10
15
SNA
Anti-LeC
UEA-I
ConA
ECL
Human Galectin1
Mouse Galectin1
AAL
MPL
WGA
WGA
CTB
Gal-3C
Threefoil
Mouse NKR-P1C
ACL
AIA
Mono1-CNL
Mono2R-CNL
P11 VP8
PFL
CA
CAA
ConA
GSL-2
GSL-1
GNL
EEL
DSL
DBA
HHL
HAA
HPA
CSA(EY Lab)
GSL-I-B4
LCA
LcH
LEL
NPA
LTL
MAL-I
MAA
PHA-E
MAA(Seikagaku)
MAL-II
PHA-E
PSL
PHA-L
PTA-Gal
PTL-I
PTL-II
RCAI (Vector)
PNA
PSA
SBA
STL
TJA-I
TJA-II
TL(EY Lab)
UDA
UEAII
VVL
WFL
PA-IL
WGA(Seikagaku Bio)
Native HPA
SGA
AAH
SSR1
rMpL
TbFBL1
CC1G-10077
MGL 6B
MGL 6A-H262T
MGL 6A
HA70 complex-Alexa
WGA multimer
rPVL
VVHA lectin
BanLec
CGL
Gal-3
human Gal-7
mouse Gal-7
CA19.9-1B844
CA19.9-9L426
CA19.9-121SLE
IPS-K-4A2B8
Microvirin
ab5
AOL (Mahal Lab)
ABA
Mono2W-CNL
BPL
GS-II (EY Lab)
EEA(EY Lab)
GNA(EY Lab)
HHA(EY Lab)
PHA-L
PNA
SBA
SJA
UEA-I
VGA(EY Lab)
WGA
MNA-M
MOA
WT-VCC
ME5(detected with a-HUD)
AAL
LSL-150
LBL-152
ME7(detected with a-HUD)
SSL0-488
VLRB aGPA.23
rNV VLP-Cy3
AFCNVH
HML-Native
#8 Lewis B (T218) (Abcam)
#15 Sialyl Lewis X (BD)
#14 Sialyl Lewis X (Calbio)
hGal-1-488+BCR
TbFBL2
MGL 416
BTL native
GAC78
DC-ASGPR short
Sp Serotype 14.2
Sp Serotype 14.15
rSRL
SRL
Galectin-1
PsSC
Avaren-Fc
CnSepL
CA19-9 Ab (DRG)
WT Galectin-8
CML1
PhoSPL001
Epa1-Np mutant Y228W
Epa1-Np mutant Y228A
Epa1-Np mutant E227G
Epa1-Np mutant E227A
Epa1-Np WT
CnMBL
DBL-1
hJAA-F11-B11
Neu5Ac(α2-6)Gal(β1-4)GlcNAc(β1-2)Man(-Sp0

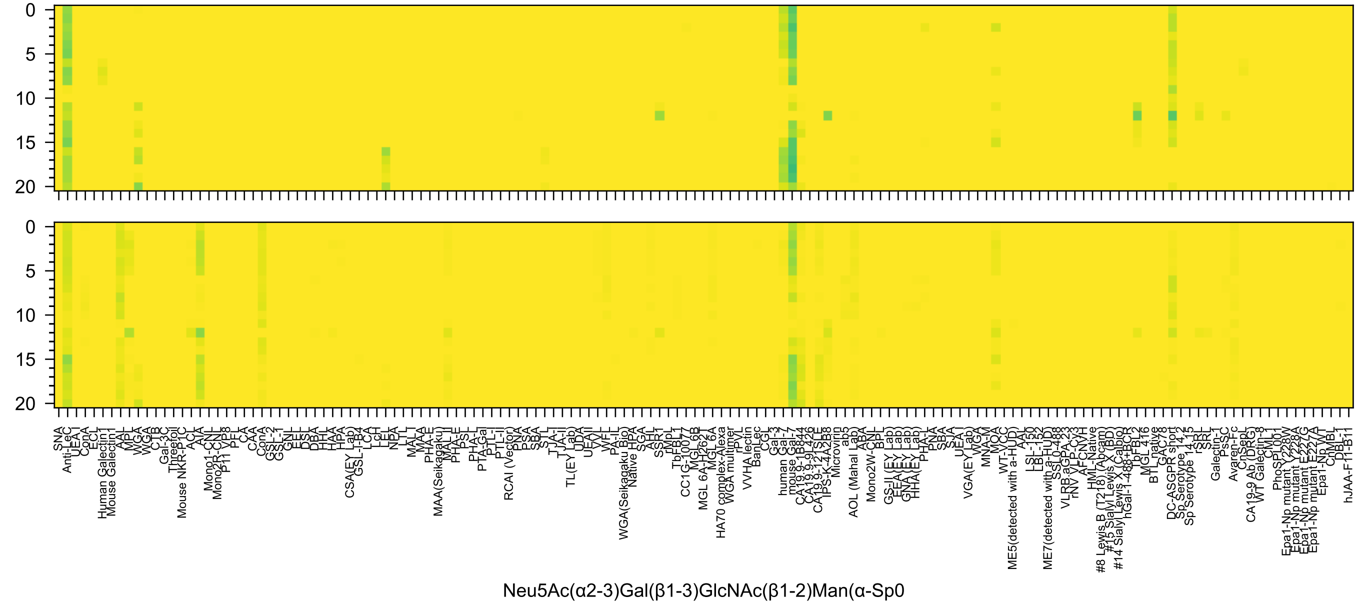
0
5
10
15
20
0
5
10
15
20
SNA
Anti-LeC
UEA-I
ConA
ECL
Human Galectin1
Mouse Galectin1
AAL
MPL
WGA
WGA
CTB
Gal-3C
Threefoil
Mouse NKR-P1C
ACL
AIA
Mono1-CNL
Mono2R-CNL
P11 VP8
PFL
CA
CAA
ConA
GSL-2
GSL-1
GNL
EEL
DSL
DBA
HHL
HAA
HPA
CSA(EY Lab)
GSL-I-B4
LCA
LcH
LEL
NPA
LTL
MAL-I
MAA
PHA-E
MAA(Seikagaku)
MAL-II
PHA-E
PSL
PHA-L
PTA-Gal
PTL-I
PTL-II
RCAI (Vector)
PNA
PSA
SBA
STL
TJA-I
TJA-II
TL(EY Lab)
UDA
UEAII
VVL
WFL
PA-IL
WGA(Seikagaku Bio)
Native HPA
SGA
AHL
SSR1
rMpL
TbFBL1
CC1G-10077
MGL 6B
MGL 6A-H262T
MGL 6A
HA70 complex-Alexa
WGA multimer
rPVL
VVHA lectin
BanLec
CGL
Gal-3
human Gal-7
mouse Gal-7
CA19.9-1B844
CA19.9-9L426
CA19.9-121SLE
IPS-K-4A2B8
Microvirin
ab5
AOL (Mahal Lab)
ABA
Mono2W-CNL
BPL
GS-II (EY Lab)
EEA(EY Lab)
GNA(EY Lab)
HHA(EY Lab)
PHA-L
PNA
SBA
SJA
UEA1
VGA(EY Lab)
WGA
MNA-M
MOA
WT-VCC
ME5(detected with a-HUD)
AAL
LSL-150
LBL-152
ME7(detected with a-HUD)
SSL0-488
VLRB.aGPA.23
rNV VLP-Cy3
AFCNVH
HML-Native
#8 Lewis B (T218) (Abcam)
#15 Sialyl Lewis X (BD)
#14 Sialyl Lewis X (Calbio)
hGal-1-488+BCR
TbFBL2
MGL 416
BTL native
GAC78
DC-ASGPR short
Sp Serotype 14.2
Sp Serotype 14.15
rSRL
SRL
Galectin-1
PsSC
Avaren-Fc
ChsepL
CA19-9 Ab (DRG)
WT Galectin-8
CML
PhoSL001
Epa1-Np mutant Y228W
Epa1-Np mutant Y228A
Epa1-Np mutant E227G
Epa1-Np mutant E227A
Epa1-Np WT
ChMBL
DBL-1
hJAA-F11-B11
Neu5Ac(α2-3)Gal(β1-3)GlcNAc(β1-2)Man(α-Sp0

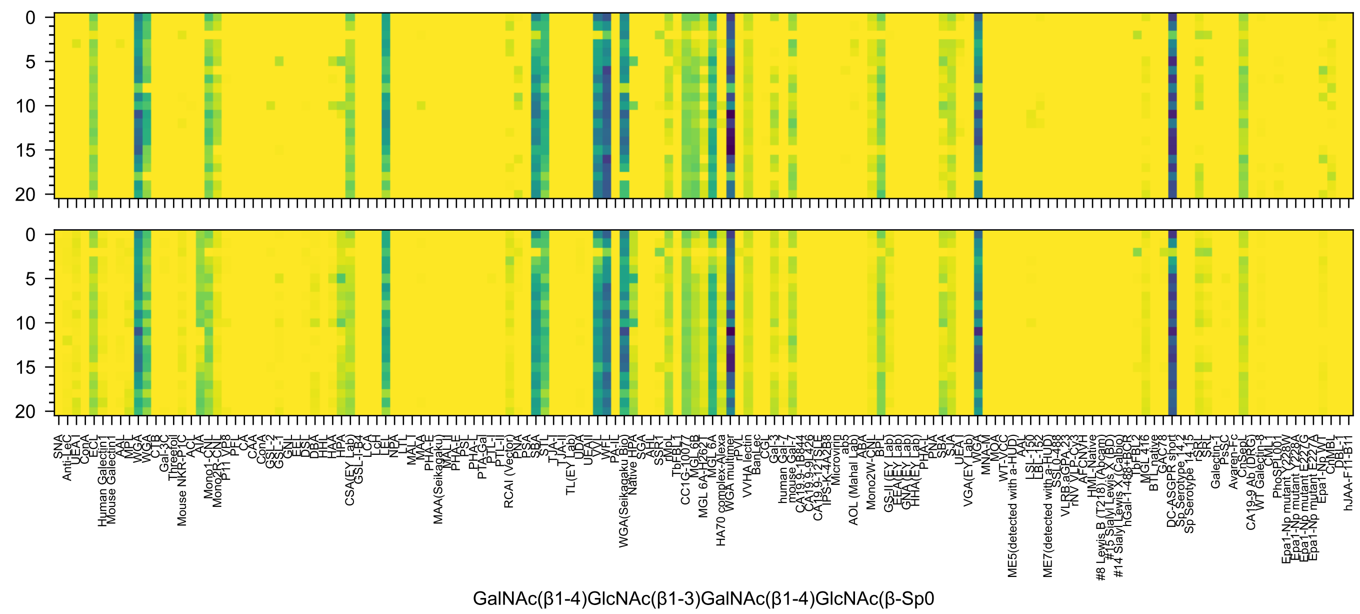
GalNAc(β1-4)GlcNAc(β1-3)GalNAc(β1-4)GlcNAc(β-Sp0

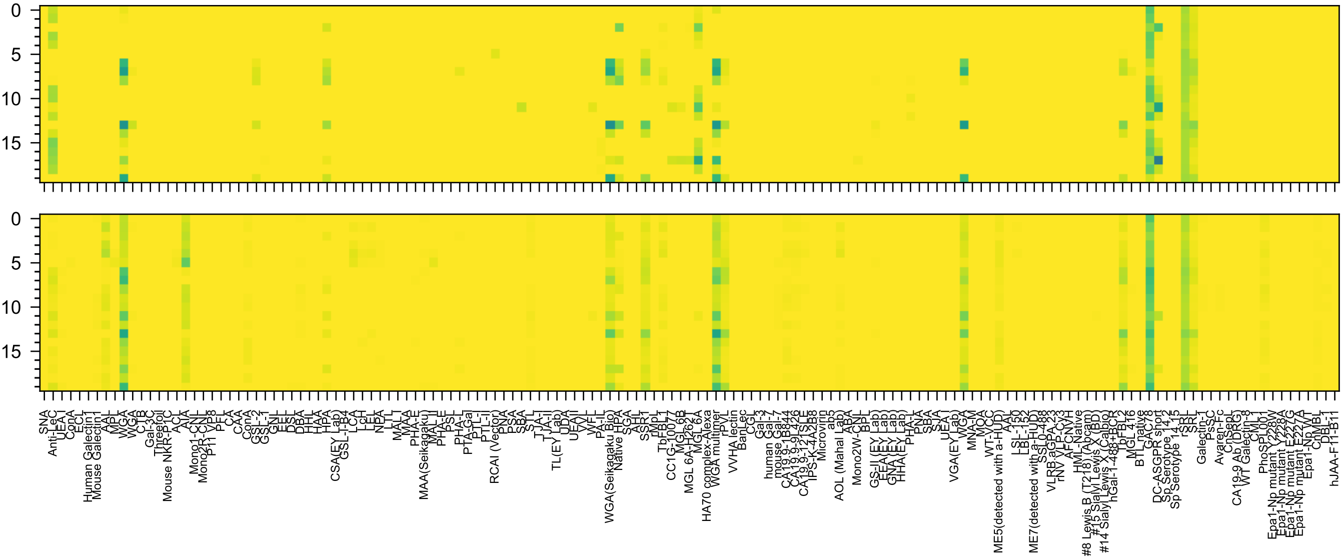


GlcNAc(β1-6)[GlcNAc(β1-4)][GlcNAc(β1-3)]GlcNAc(-Sp8

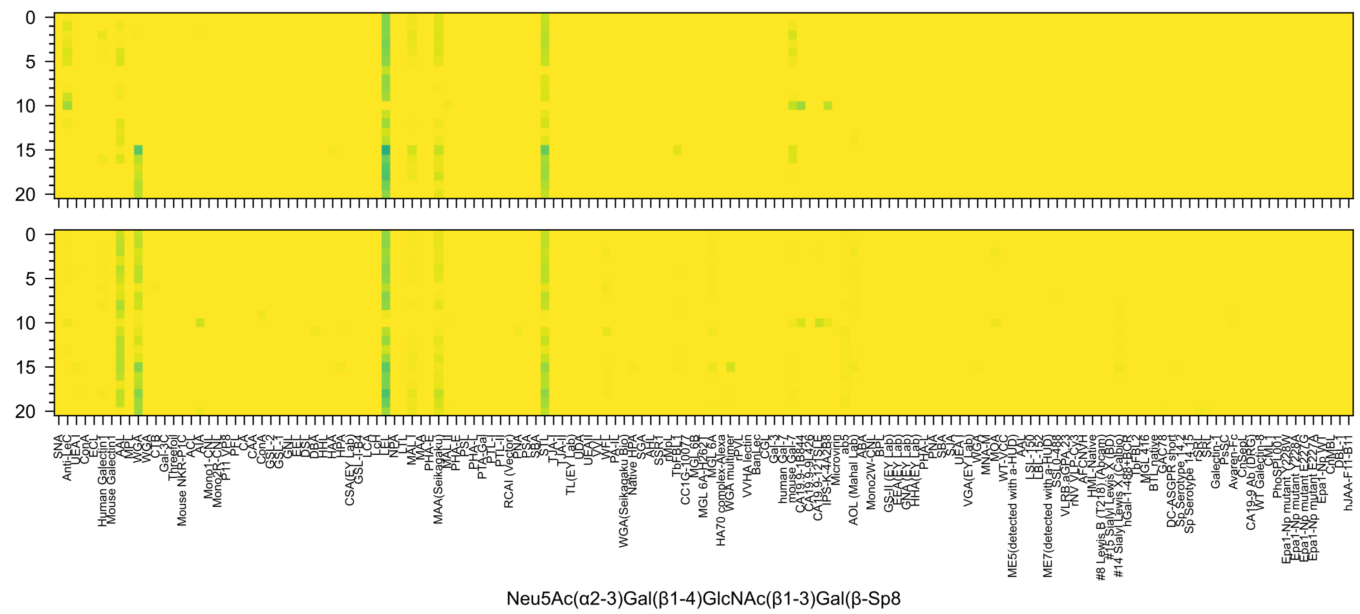
0
5
10
15
20
SNA
Anti-LeC
UEA I
ConA
ECL
Human Galectin1
Mouse Galectin1
AAL
MPL
WGA
WGA
CTB
Gal-3C
Threefoil
Mouse NKR-P1C
ACL
AIA
Mono1-CNL
Mono2R-CNL
P11 VP8
PFL
CA
CAA
ConA
GSL-2
GSL-1
GNL
EEL
DSL
DBA
HHL
HAA
HPA
CSA(EY Lab)
GSL-I-B4
LCA
LcH
LEL
NPA
LTL
MAL I
MAA
PHA-E
MAA(Seikagaku)
MAL-II
PHA-E
PSL
PHA-L
PTA-Gal
PTL-I
PTL-II
RCAI (Vector)
PNA
PSA
SBA
STL
TJA-I
TJA-II
TL(EY Lab)
UDA
UEAII
VVL
WFL
PA-IL
WGA(Seikagaku Bio)
Native HPA
SGA
AHL
SSR1
rMpL
TbFBL1
CC1G-10077
MGL 6B
MGL 6A-H262T
MGL 6A
HA70 complex-Alexa
WGA multimer
rPVL
VVHA lectin
BanLec
CGL
Gal-3
human Gal-7
mouse Gal-7
CA19.9-1B844
CA19.9-9L426
CA19.9-121SLE
IPS-K-4A2B8
Microvirin
ab5
AOL (Mahal Lab)
ABA
Mono2W-CNL
BPL
GS-II (EY Lab)
EEA (EY Lab)
GNA (EY Lab)
HHA (EY Lab)
PHA-L
PNA
SBA
SJA
UEA-I
VGA(EY Lab)
WGA
MNA-M
MOA
WT-VCC
ME5(detected with a-HUD)
AAL
LSL-150
LBL-152
ME7(detected with a-HUD)
SSL0-488
VLRB.aGPA.23
rNV VLP-Cy3
AFCN-H
HML-Native
#8 Lewis B (T218) (Abcam)
#15 Sialyl Lewis X (BD)
#14 Sialyl Lewis X (Calbio)
hGal-1-488+BCR
TbFBL2
MGL 416
BTL native
GAC78
DC-ASGPR short
Sp Serotype 14.2
Sp Serotype 14.15
rSRL
SRL
Galectin-1
PsSC
Avaren-Fc
ChSeaL
CA19-9 Ab (DRG)
WT Galectin-8
CML
PhoSL001
Epa1-Np mutant Y228W
Epa1-Np mutant Y228A
Epa1-Np mutant E227G
Epa1-Np mutant E227A
Epa1-Np WT
ChMBL
DBL-1
hJAA-F11-B11
Neu5Ac(α2-3)Gal(β1-4)GlcNAc(β1-3)Gal(β-Sp8

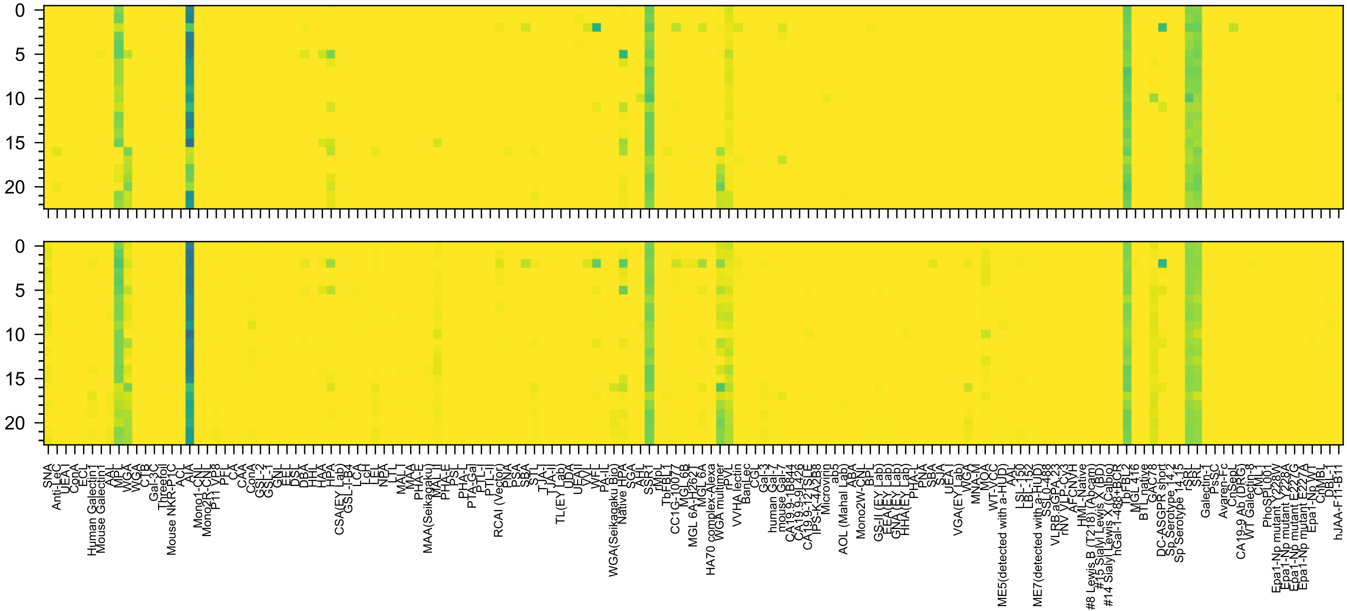


GlcNAc(β1-3)Gal(β1-4)GlcNAc(β1-3)GalNAc(α-Sp14

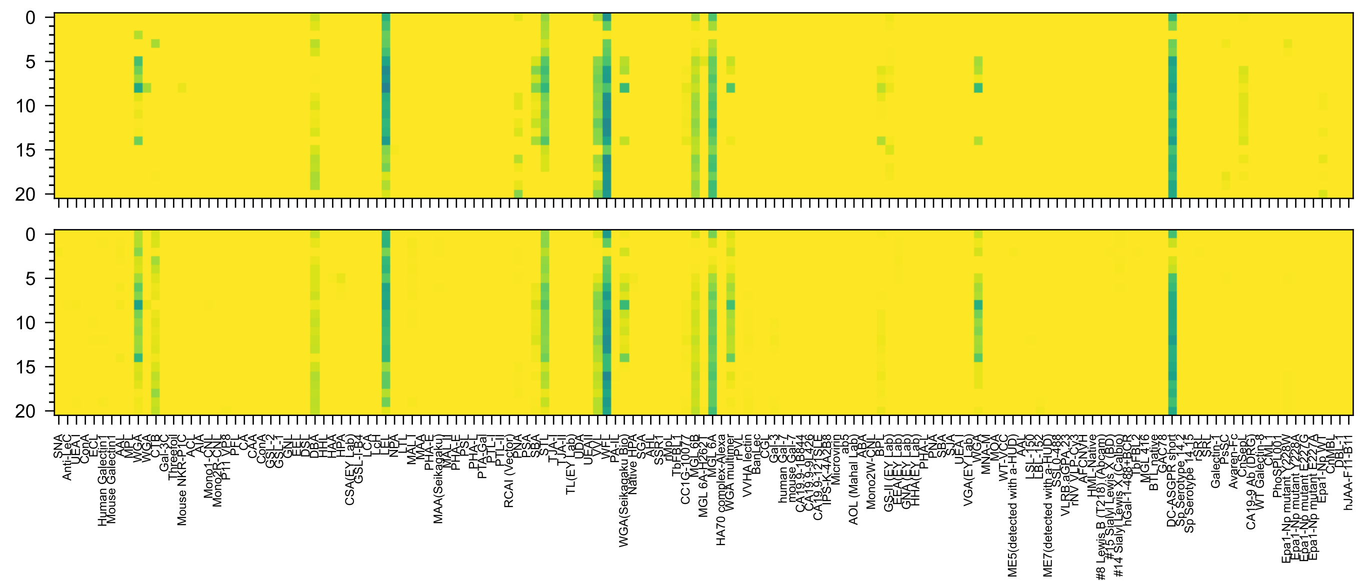

GalNAc(β1-4)[Neu5Ac(α2-3)]Gal(β1-4)GlcNAc(β-Sp0

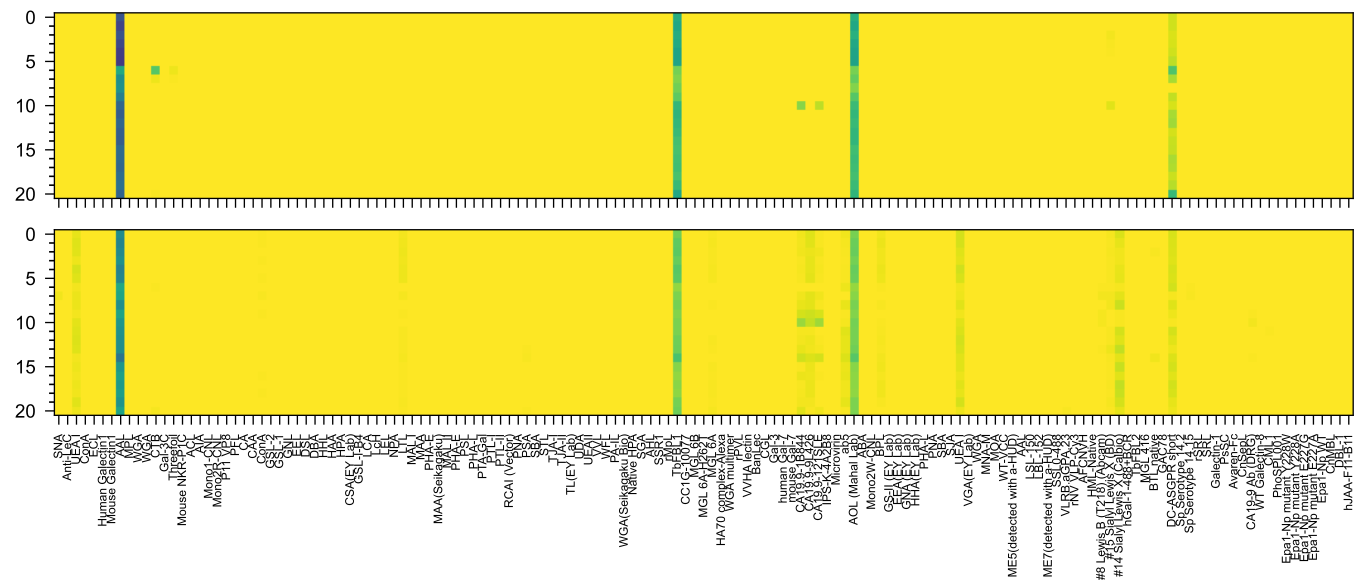

Neu5Ac(α2-3)Gal6OS(β1-4)[Fuc(α1-3)]GlcNAc(β-Sp8

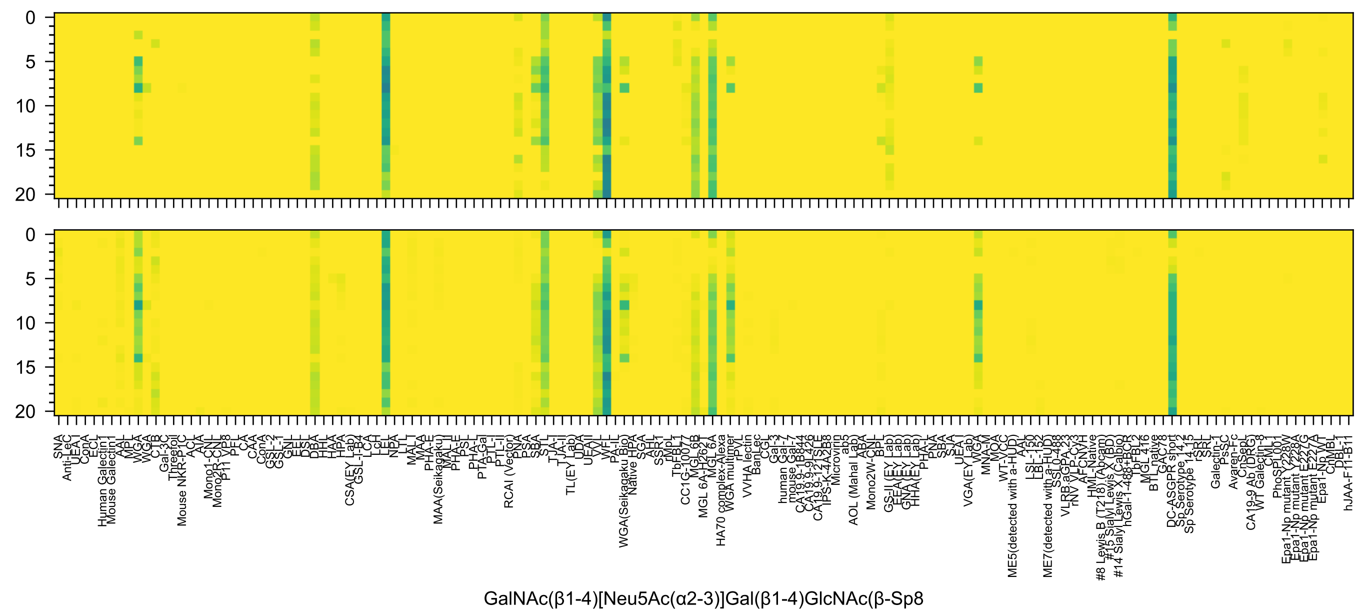
GalNAc(β1-4)[Neu5Ac(α2-3)]Gal(β1-4)GlcNAc(β-Sp8

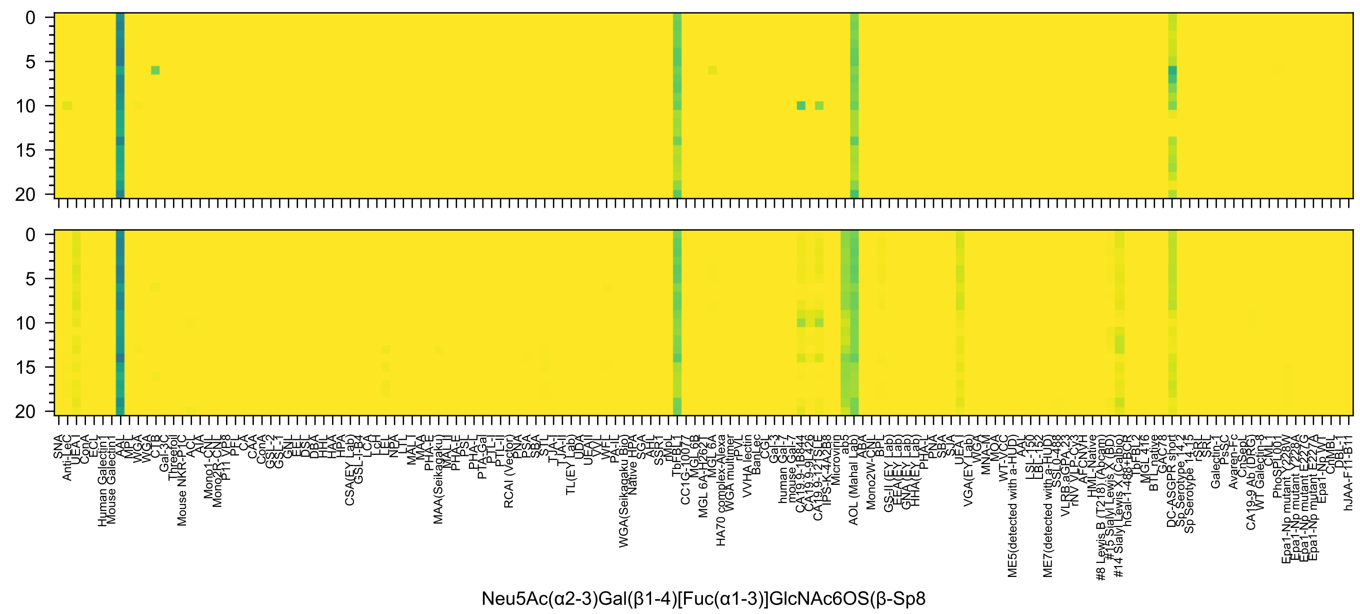
Neu5Ac(α2-3)Gal(β1-4)[Fuc(α1-3)]GlcNAc6OS(β-Sp8

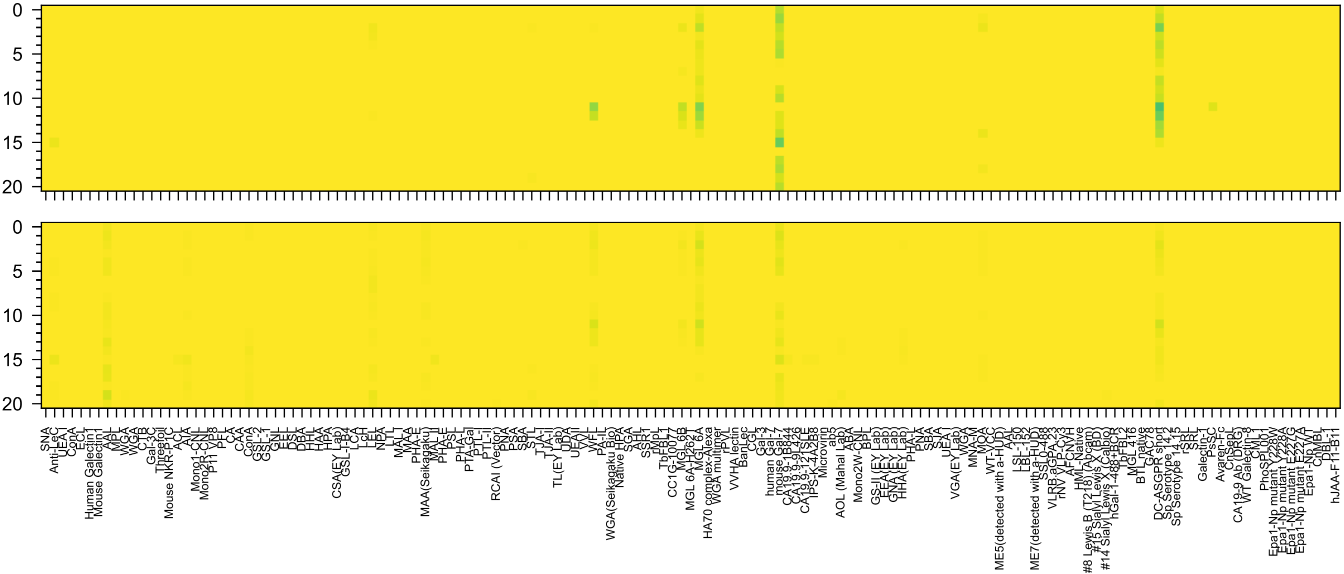
Neu5Ac(α2-8)Neu5Ac(α2-3)Gal(β1-4)Glc(β-Sp0

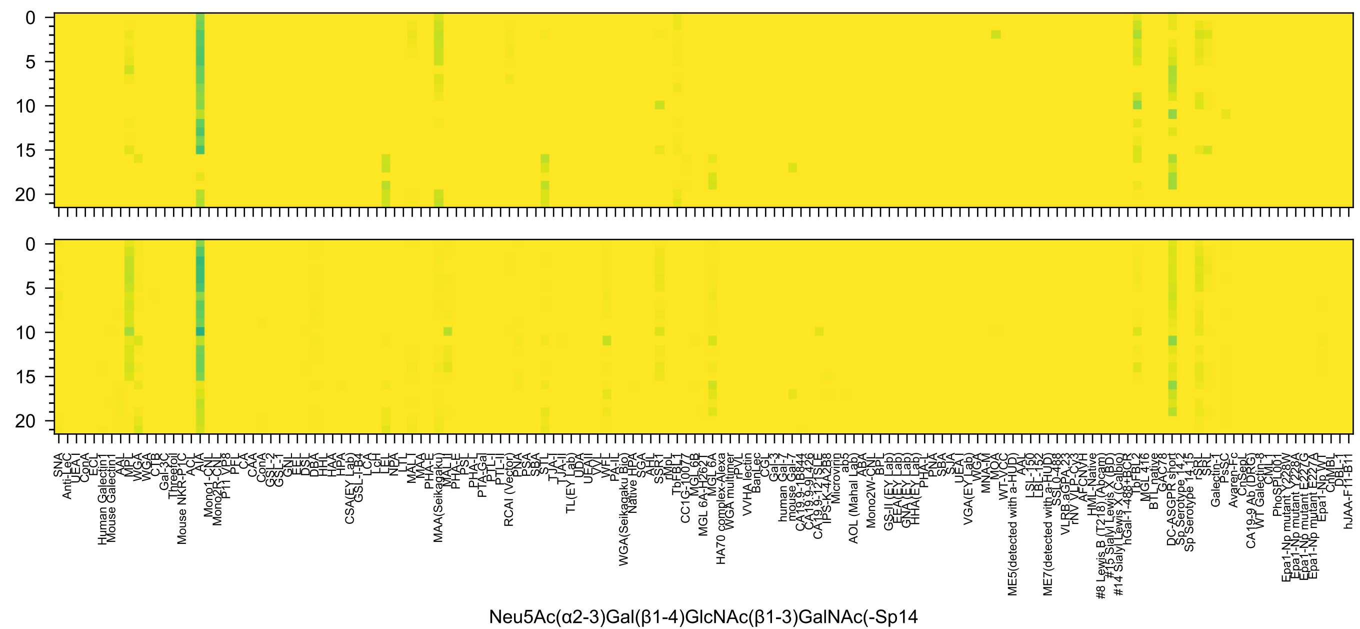
Neu5Ac(α2-3)Gal(β1-4)GlcNAc(β1-3)GalNAc(-Sp14

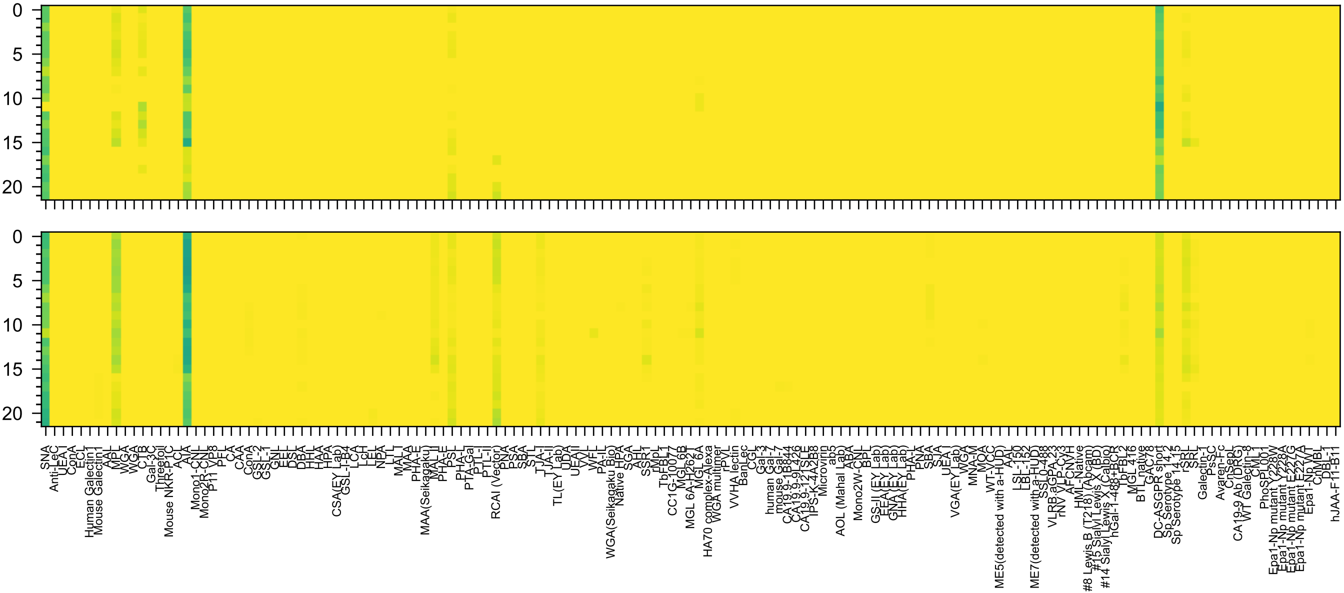

Neu5Ac(α2-6)Gal(β1-4)GlcNAc(β1-3)GalNAc(-Sp14

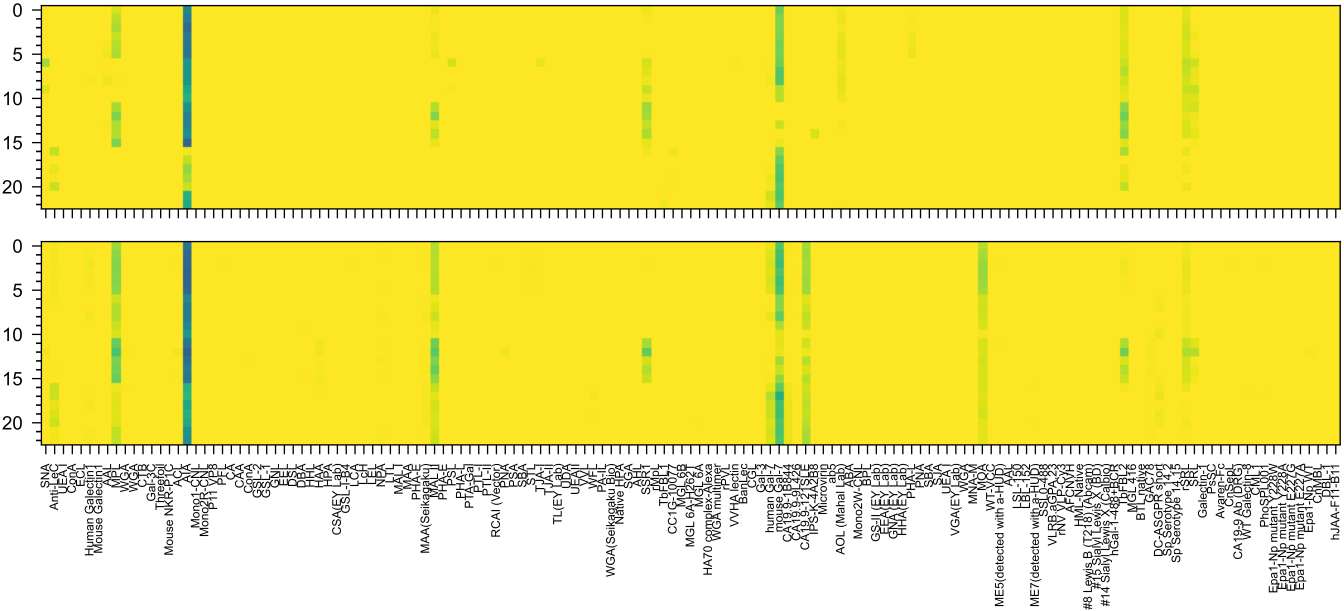

Neu5Ac(α2-3)Gal(β1-3)GlcNAc(β1-3)GalNAc(α-Sp14

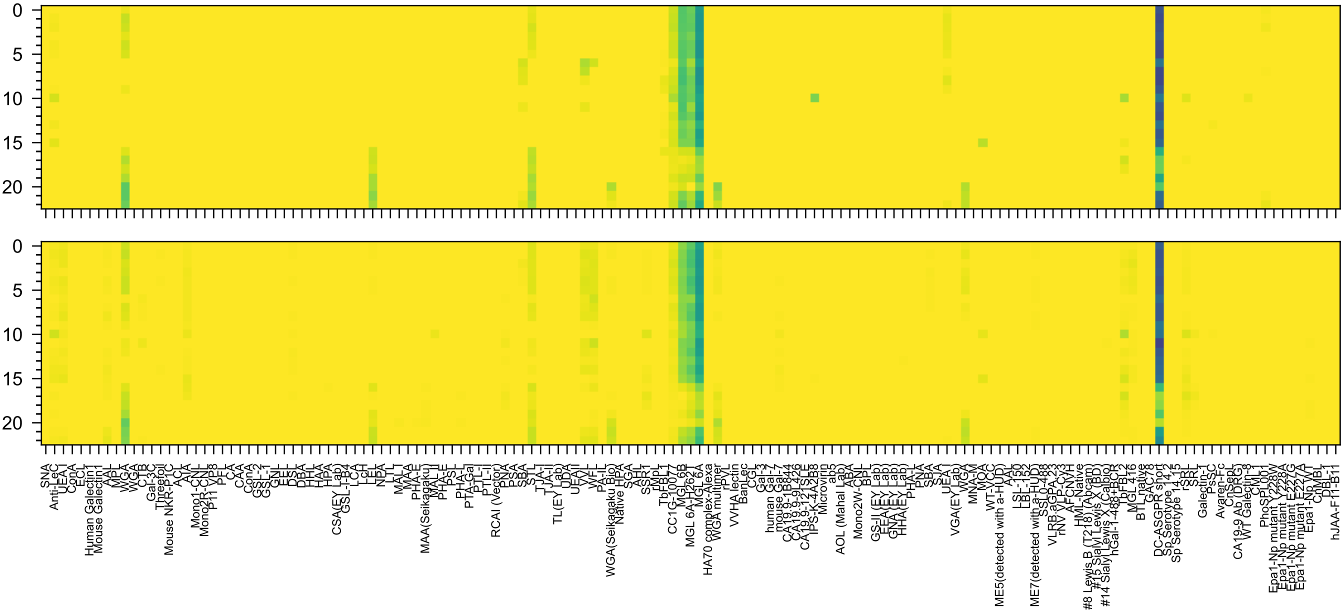

Neu5Ac(α2-3)Gal(β1-4)GlcNAc(β1-6)GalNAc(α-Sp14

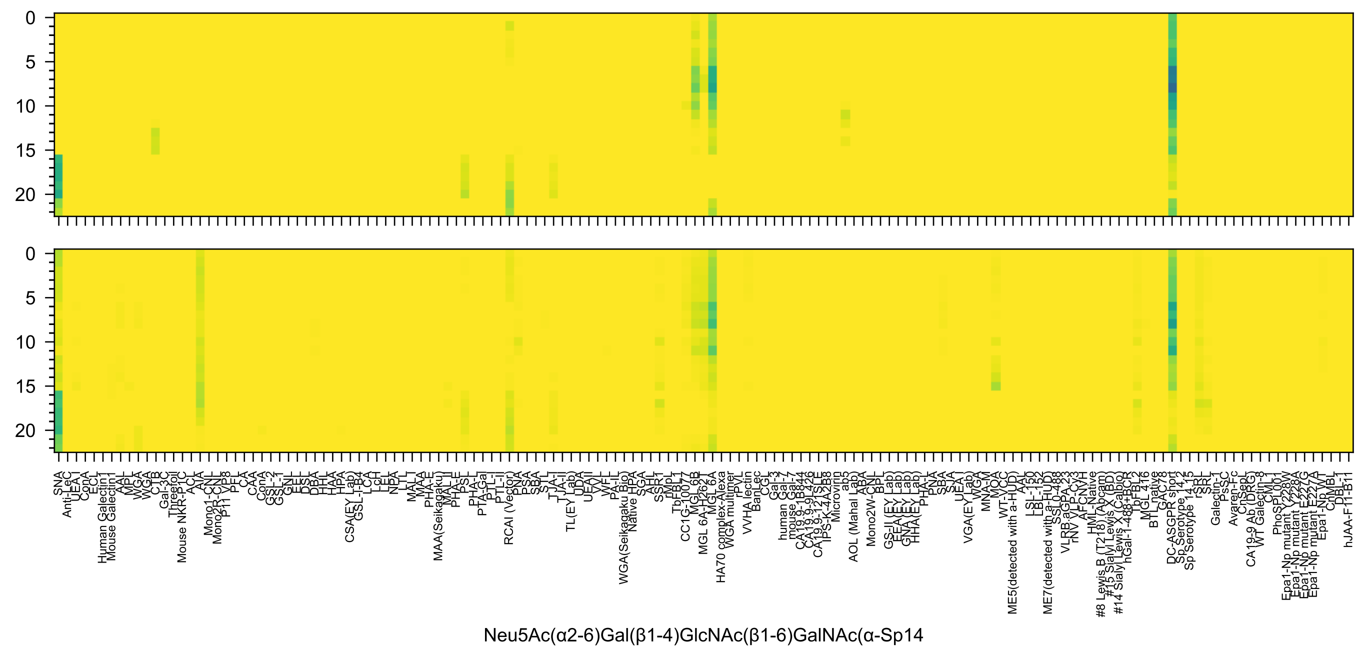
0
5
10
15
20
0
5
10
15
20
SNA
Anti-LeC
UEA I
ConA
ECL
Human Galectin1
Mouse Galectin1
AAL
MPL
WGA
WGA
CTB
Gal-3C
Threefoil
Mouse NKR-P1C
ACL
AIA
Mono1-CNL
Mono2R-CNL
P11 VP8
PFL
CA
CAA
ConA
GSL-2
GSL-1
GNL
EEL
DSL
DBA
HHL
HAA
HPA
CSA(EY Lab)
GSL-I-B4
LCA
LcH
LEL
NPA
LTL
MAL I
MAA
PHA-E
MAA(Seikagaku)
MAL-II
PHA-E
PSL
PHA-L
PTA-Gal
PTL-I
PTL-II
RCAI (Vector)
PNA
PSA
SBA
STL
TJA-I
TJA-II
TL(EY Lab)
UDA
UEAII
VVL
WFL
PA-IL
WGA(Seikagaku Bio)
Native HPA
SGA
AHL
SSR1
rMpL
TbFBL1
CC1G-10077
MGL 6B
MGL 6A-H262T
MGL 6A
HA70 complex-Alexa
WGA multimer
rPVL
VVHA lectin
BanLec
CGL
Gal-3
human Gal-7
mouse Gal-7
CA19.9-1B844
CA19.9-9L426
CA19.9-121SLE
IPS-K-4A2B8
Microvirin
ab5
AOL (Mahal Lab)
ABA
Mono2W-CNL
BPL
GS-II (EY Lab)
EEA (EY Lab)
GNA (EY Lab)
HHA (EY Lab)
PHA-L
PNA
SBA
SJA
UEA-I
VGA(EY Lab)
WGA
MNA-M
MOA
WT-VCC
ME5(detected with a-HUD)
AAL
LSL-150
LBL-152
ME7(detected with a-HUD)
SSL0-488
VLRB.aGPA.23
rNV VLP-Cy3
AFC NVH
HML-Native
#8 Lewis B (T218) (Abcam)
#15 Sialyl Lewis X (BD)
#14 Sialyl Lewis X (Calbio)
hGal-1-488+BCR
TbFBL2
MGL 416
BTL native
GAC78
DC-ASGPR short
Sp Serotype 14.2
Sp Serotype 14.15
rSRL
SRL
Galectin-1
PsSC
Avaren-Fc
CnSeL
CA19-9 Ab (DRG)
WT Galectin-8
CML1
PhoSL001
Epa1-Np mutant Y228W
Epa1-Np mutant Y228A
Epa1-Np mutant E227G
Epa1-Np mutant E227A
Epa1-Np WT
ChMBL
DBL-1
hJAA-F11-B11
Neu5Ac(α2-6)Gal(β1-4)GlcNAc(β1-6)GalNAc(α-Sp14

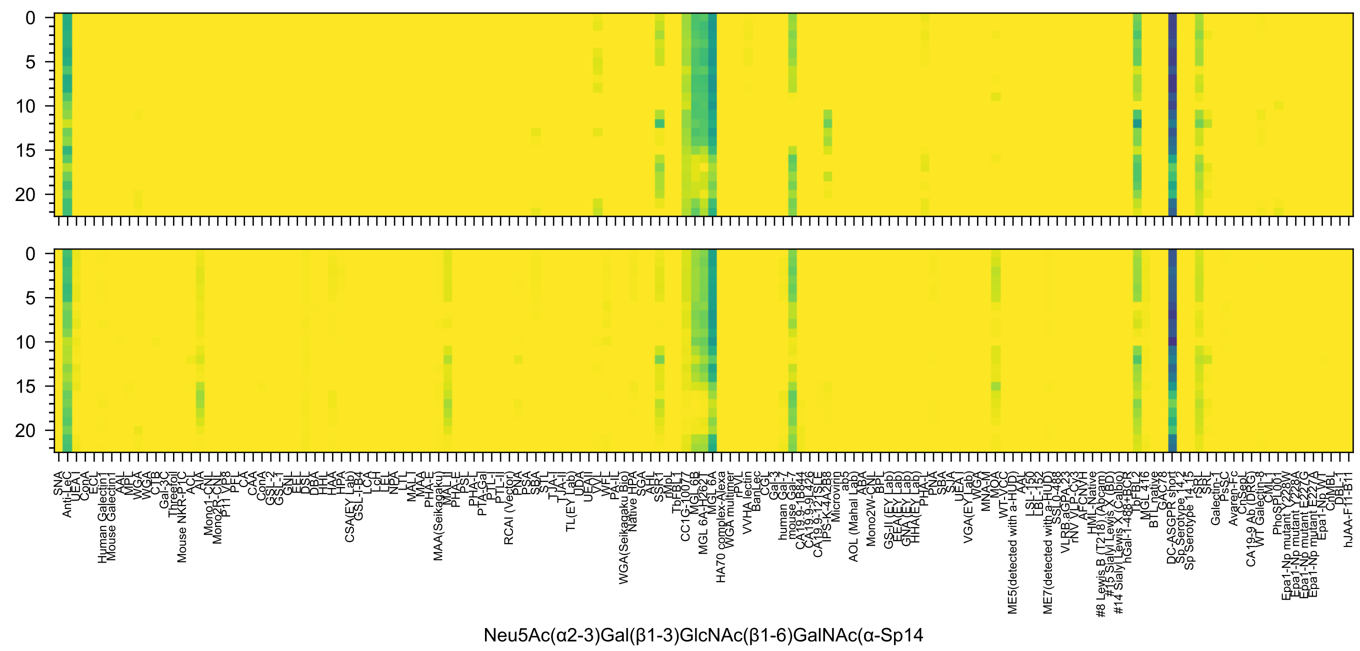
Neu5Ac(α2-3)Gal(β1-3)GlcNAc(β1-6)GalNAc(α-Sp14

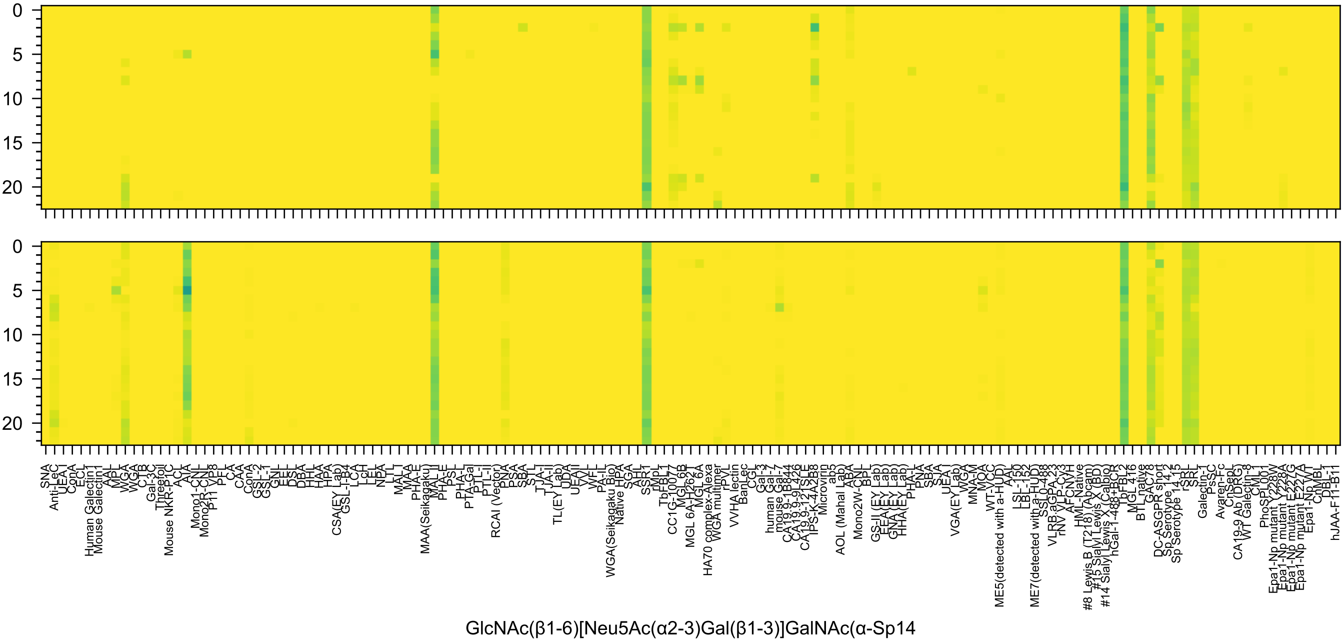
GlcNAc(β1-6)[Neu5Ac(α2-3)Gal(β1-3)]GalNAc(α-Sp14
0
5
10
15
20

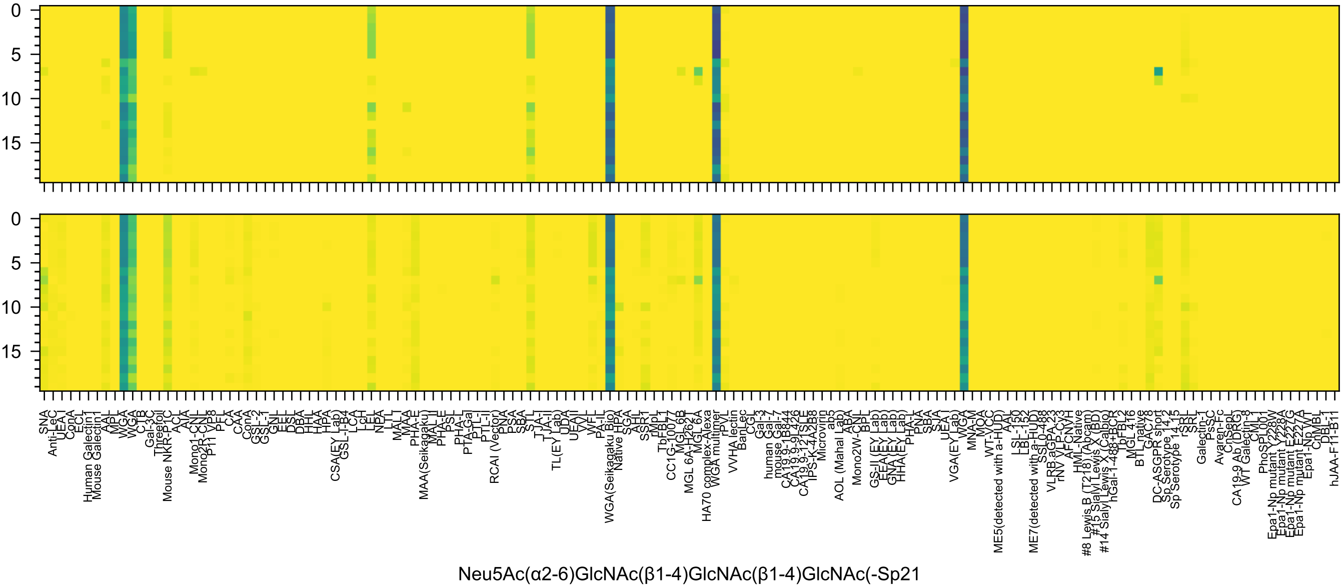
0
5
10
15
0
5
10
15
SNA
Anti-LeC
UEA-I
ConA
ECL
Human Galectin1
Mouse Galectin1
AAL
MPL
WGA
WGA
CTB
Gal-3C
Threefoil
Mouse NKR-P1C
ACL
AIA
Mono1-CNL
Mono2R-CNL
P11 VP8
PFL
CA
CAA
ConA
GSL-2
GSL-1
GNL
EEL
DSL
DBA
HHL
HAA
HPA
CSA(EY Lab)
GSL-I-B4
LCA
LcH
LEL
NPA
LTL
MAL-I
MAA
MAA(Seikagaku)
PHA-E
MAL-II
PHA-E
PSL
PHA-L
PTA-Gal
PTL-I
PTL-II
RCAI (Vector)
PNA
PSA
SBA
STL
TJA-I
TJA-II
TL(EY Lab)
UDA
UEA-II
VVL
WFL
PA-IL
WGA(Seikagaku Bio)
Native HPA
SGA
AAL
SSR1
rMpL
TbFBL1
CC1G-10077
MGL 6B
MGL 6A-H262T
MGL 6A
HA70 complex-Alexa
WGA multimer
rPVL
VVHA lectin
BanLec
CGL
Gal-3
human Gal-7
mouse Gal-7
CA19.9-1B844
CA19.9-9L426
CA19.9-121SLE
IPS-K-4A2B8
Microvirin
ab5
AOL (Mahal Lab)
ABA
Mono2W-CNL
BPL
GS-II (EY Lab)
EEA(EY Lab)
GNA(EY Lab)
HHA(EY Lab)
PHA-L
PNA
SBA
SJA
UEA-I
VGA(EY Lab)
WGA
MNA-M
MOA
WT-VCC
ME5(detected with a-HUD)
AAL
LSL-150
LBL-152
ME7(detected with a-HUD)
SSL0-488
VLRB.aGPA.23
rNV VLP-Cy3
AFCNVH
HML-Native
#8 Lewis B (T218) (Abcam)
#15 Sialyl Lewis X (BD)
#14 Sialyl Lewis X (Calbio)
hGal-1-488+BCR
TbFBL2
MGL 416
BTL native
GAC78
DC-ASGPR short
Sp Serotype 14.2
Sp Serotype 14.15
rSRL
SRL
Galectin-1
PsSC
Avaren-Fc
CnSept
CA19-9 Ab (DRG)
WT Galectin-8
CML1
PhoSL001
Epa1-Np mutant Y228W
Epa1-Np mutant Y228A
Epa1-Np mutant E227G
Epa1-Np mutant E227A
Epa1-Np WT
ChMBL
DBL-1
hJAA-F11-B11
Neu5Ac(α2-6)GlcNAc(β1-4)GlcNAc(β1-4)GlcNAc(-Sp21

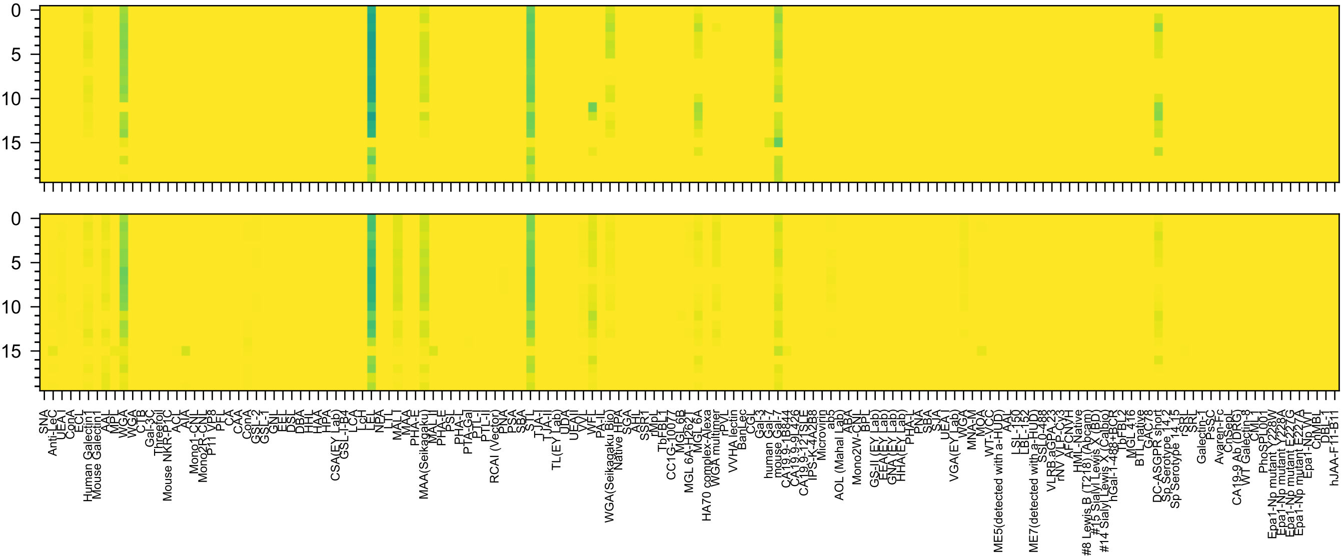


Neu5Ac(α2-8)Neu5Ac(α2-3)Gal(β1-4)GlcNAc(-Sp0

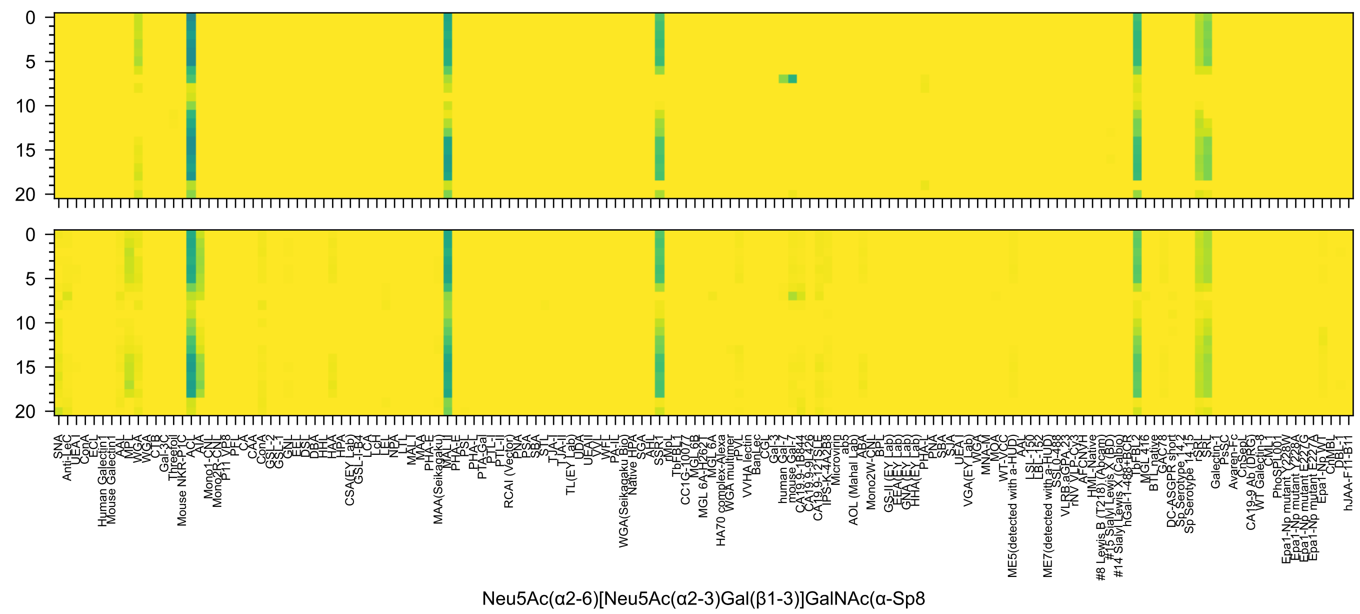
Neu5Ac(α2-6)[Neu5Ac(α2-3)Gal(β1-3)]GalNAc(α-Sp8

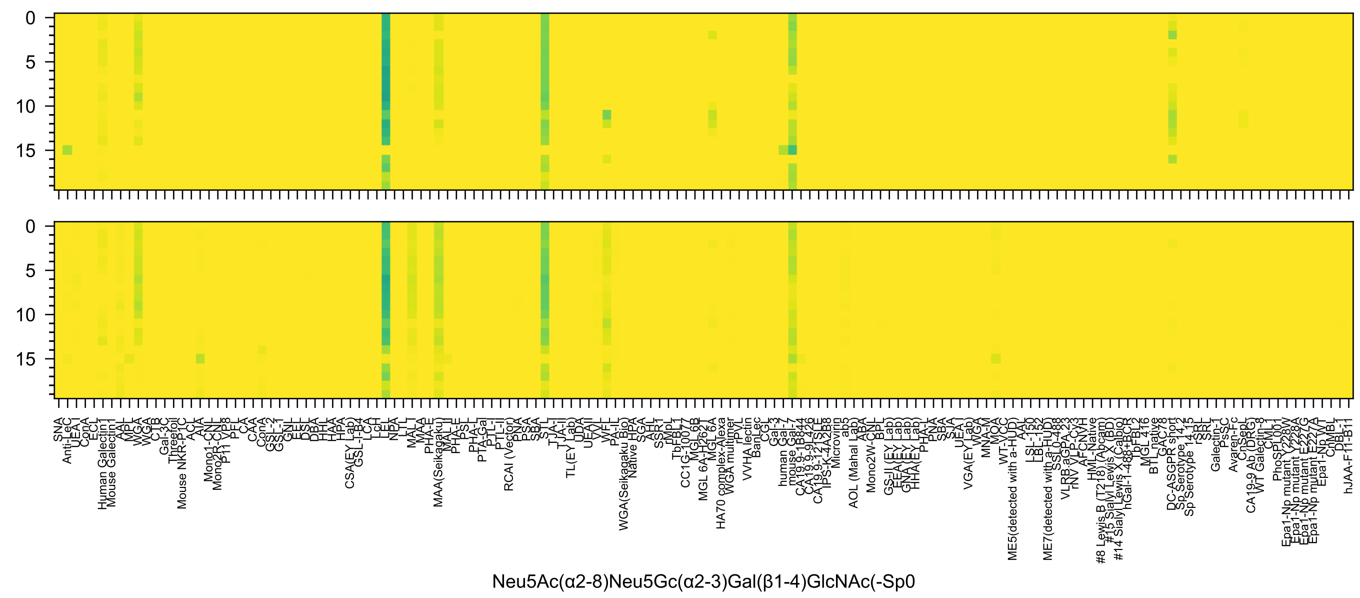
Neu5Ac(α2-8)Neu5Gc(α2-3)Gal(β1-4)GlcNAc(-Sp0

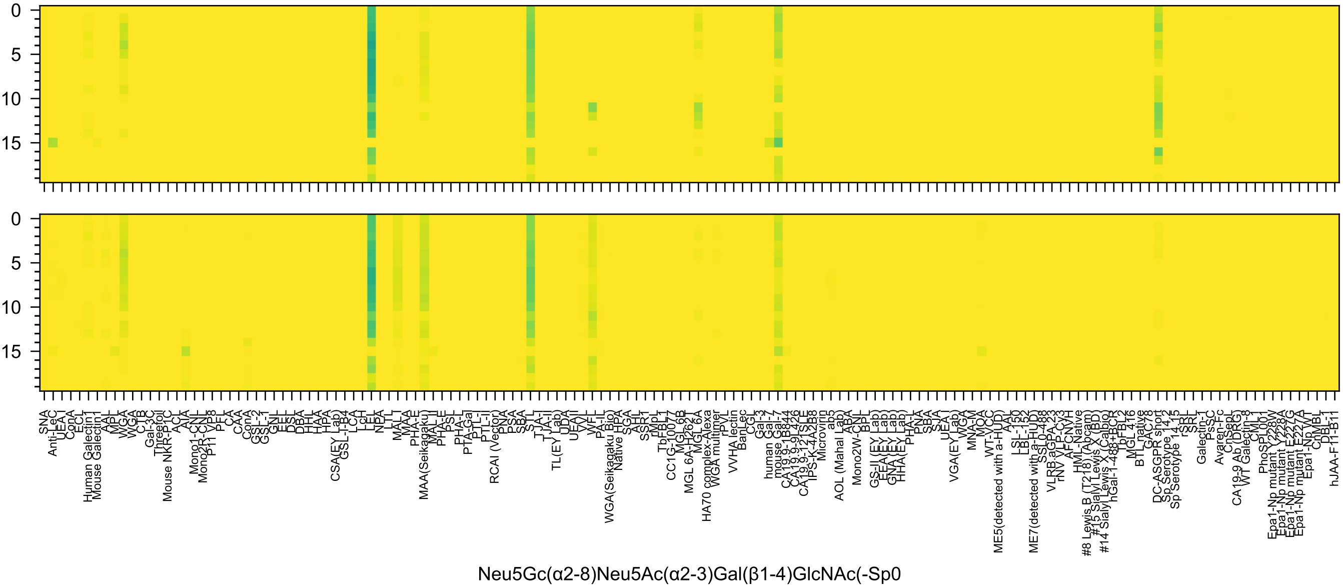

0
5
10
15
0
5
10
15
SNA
Anti-LeC
UEA-I
ConA
ECL
Human Galectin-1
Mouse Galectin-1
AAL
MPL
WGA
WGA
CTB
Gal-3C
Threefoil
Mouse NKR-P1C
ACL
AIA
Mono1-CNL
Mono2R-CNL
P11 VP8
PFL
CA
CAA
ConA
GSL-2
GSL-1
GNL
EEL
DSL
DBA
HHL
HAA
HPA
CSA(EY Lab)
GSL-I-B4
LCA
LcH
LEL
NPA
LTL
MAL-I
MAA
PHA-E
MAA(Seikagaku)
MAL-II
PHA-E
PSL
PHA-L
PTA-Gal
PTL-I
PTL-II
RCAI (Vector)
PNA
PSA
SBA
STL
TJA-I
TJA-II
TL(EY Lab)
UDA
UEA-II
VVL
WFL
PA-IL
WGA(Seikagaku Bio)
Native HPA
SGA
AHL
SSR1
rMpL
TbFBL1
CC1G-10077
MGL 6B
MGL 6A-H262T
MGL 6A
HA70 complex-Alexa
WGA multimer
rPVL
VVHA lectin
BanLec
CGL
Gal-3
human Gal-7
mouse Gal-7
CA19.9-1B844
CA19.9-9L426
CA19.9-121SLE
IPS-K-4A2B8
Microvirin
ab5
AOL (Mahal Lab)
ABA
Mono2W-CNL
BPL
GS-II (EY Lab)
EEA(EY Lab)
GNA(EY Lab)
HHA(EY Lab)
PHA-L
PNA
SBA
SJA
UEA-I
VGA(EY Lab)
WGA
MNA-M
MOA
WT-VCC
ME5(detected with a-HUD)
AAL
LSL-150
LBL-152
ME7(detected with a-HUD)
SSL0-488
VLRB.aGPA.23
rNV VLP-Cy3
AFCNVH
HML-Native
#8 Lewis B (T218) (Abcam)
#15 Sialyl Lewis X (BD)
#14 Sialyl Lewis X (Calbio)
hGal-1-488+BCR
TbFBL2
MGL 416
BTL native
GAC78
DC-ASGPR short
Sp Serotype 14.2
Sp Serotype 14.15
rSRL
SRL
Galectin-1
PsSC
Avaren-Fc
CnSepL
CA19-9 Ab (DRG)
WT Galectin-8
CML1
PhoSPL001
Epa1-Np mutant Y228W
Epa1-Np mutant Y228A
Epa1-Np mutant E227G
Epa1-Np mutant E227A
Epa1-Np WT
ChMBL
DBL-1
hJAA-F11-B11
Neu5Gc(α2-8)Neu5Ac(α2-3)Gal(β1-4)GlcNAc(-Sp0

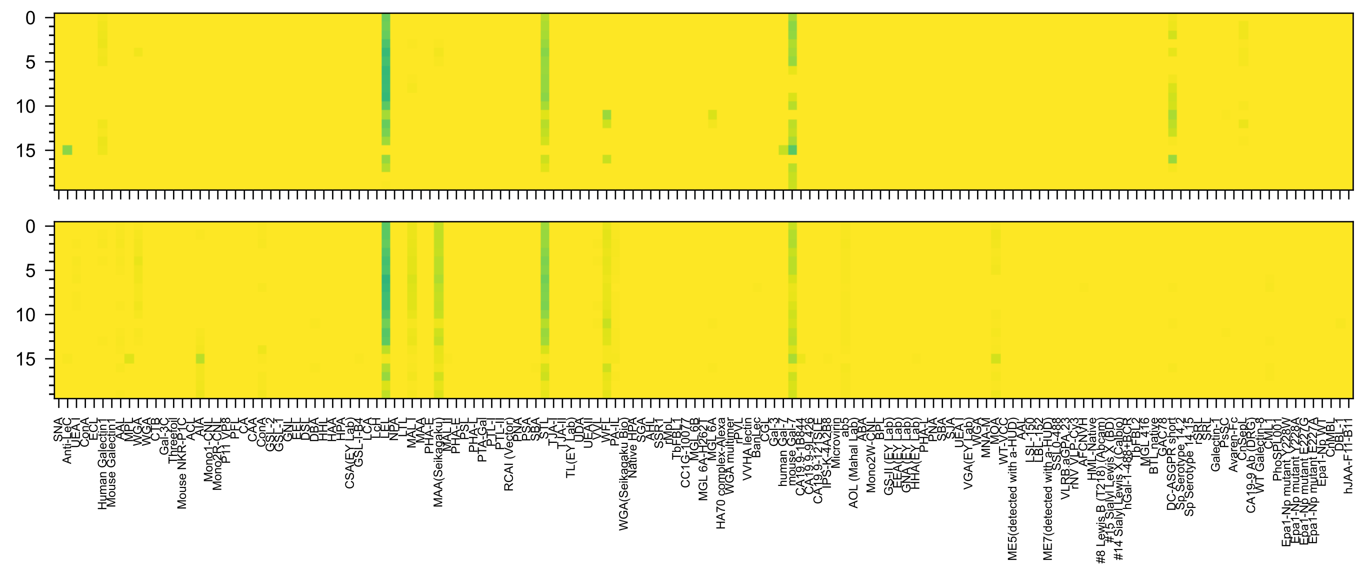


Neu5Gc(α2-8)Neu5Gc(α2-3)Gal(β1-4)GlcNAc(-Sp0

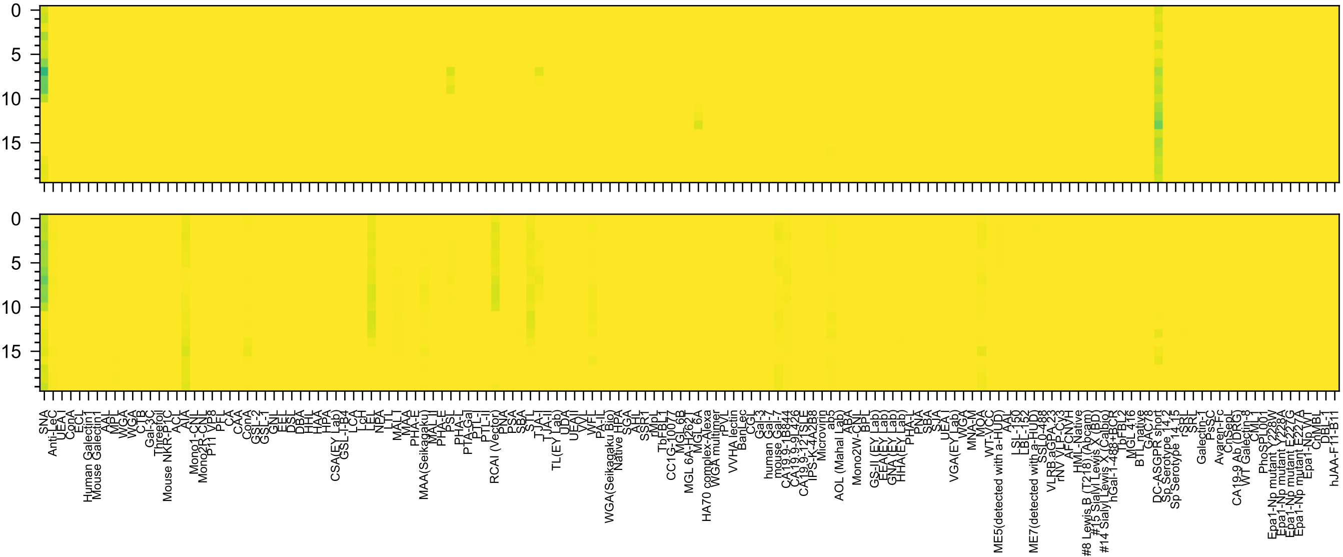

Neu5Gc(α2-8)Neu5Gc(α2-6)Gal(β1-4)GlcNAc(-Sp0

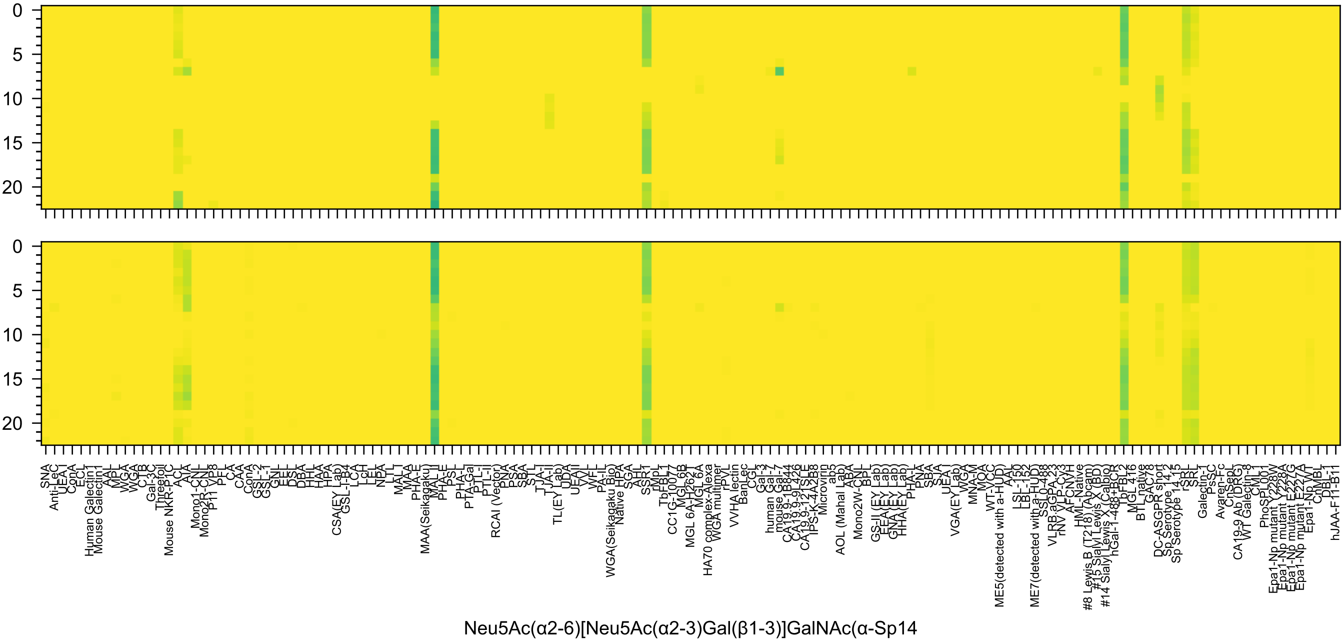
Neu5Ac(α2-6)[Neu5Ac(α2-3)Gal(β1-3)]GalNAc(α-Sp14

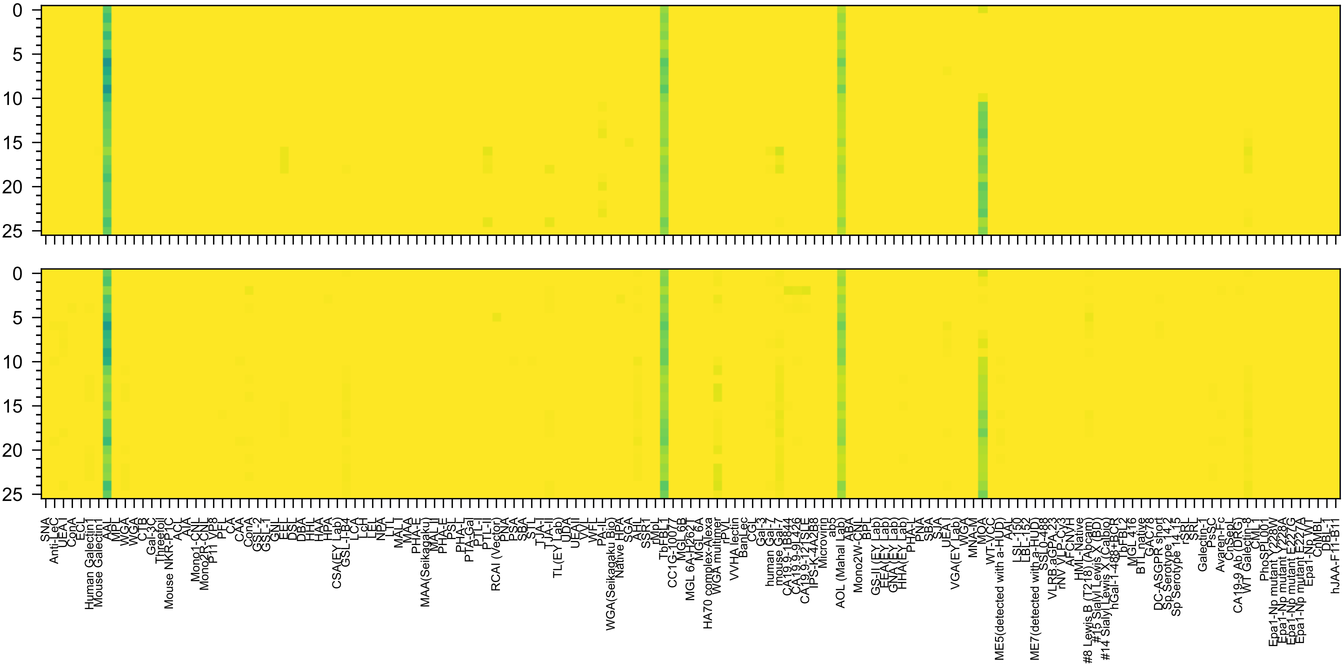

Gal(α1-3)[Fuc(α1-2)]Gal(β1-4)[Fuc(α1-3)]Glc(β-Sp21

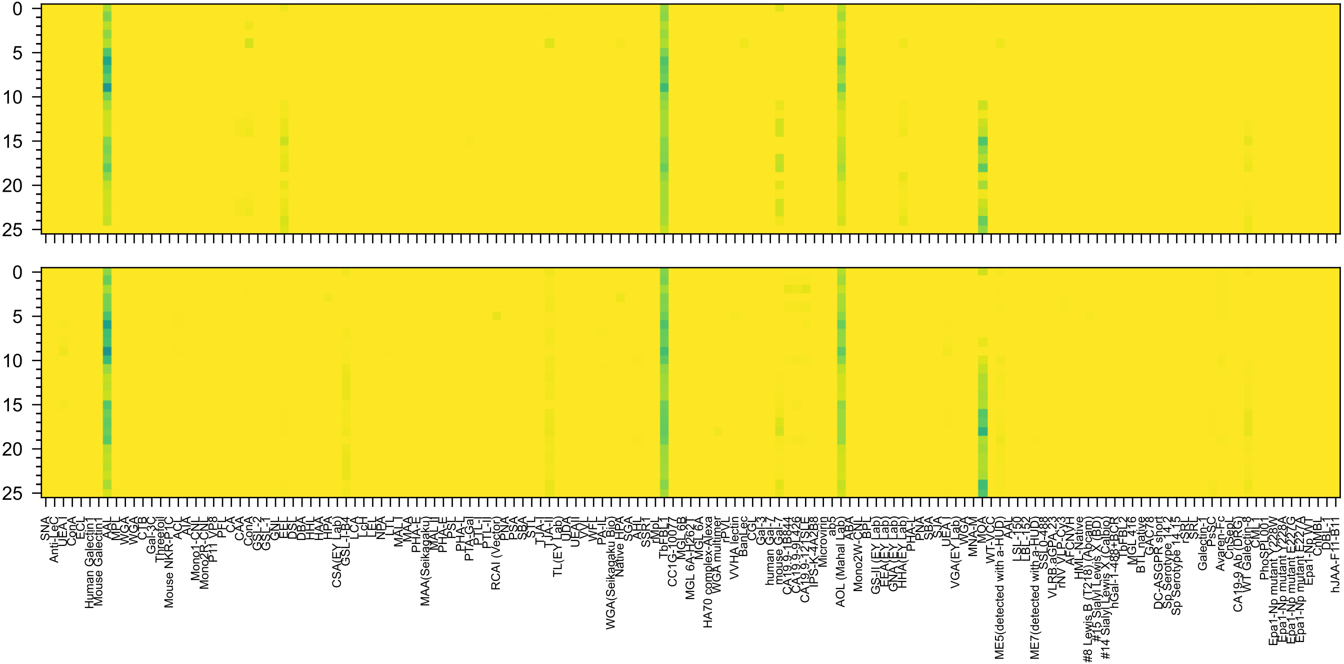


Gal(α1-3)[Fuc(α1-2)]Gal(β1-4)[Fuc(α1-3)]GlcNAc(β-Sp0

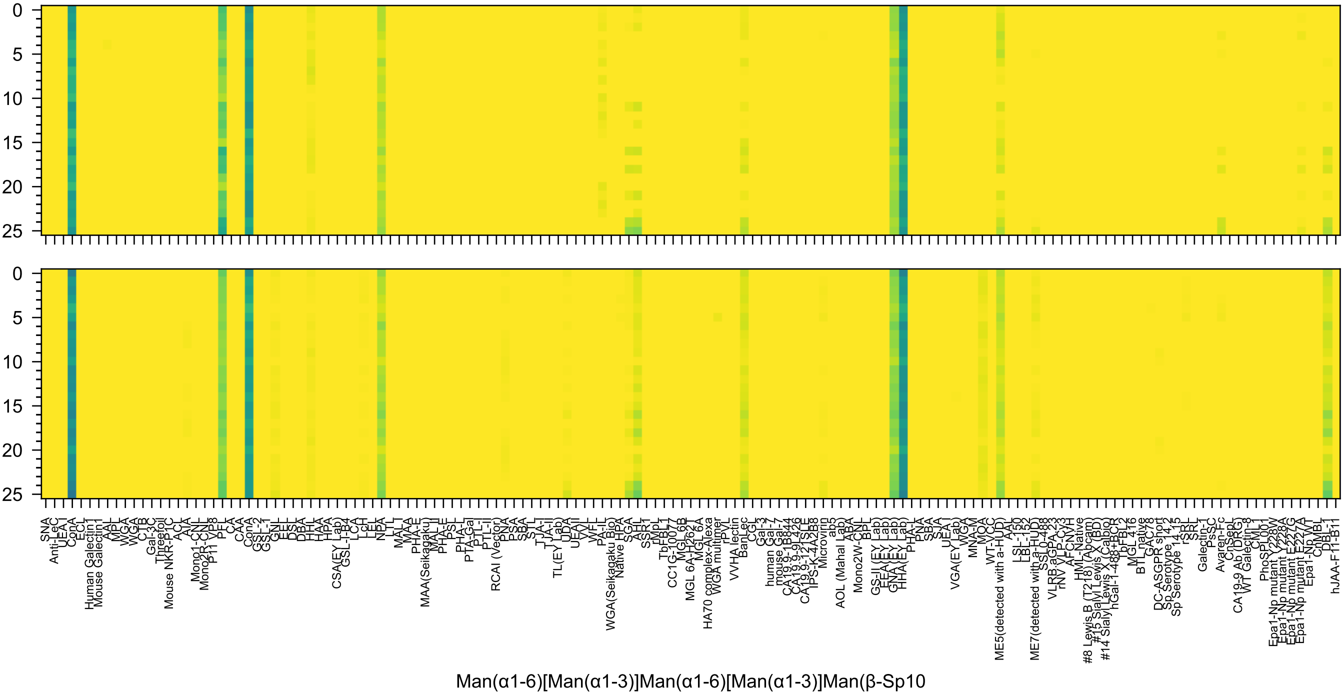

Man(α1-6)[Man(α1-3)]Man(α1-6)[Man(α1-3)]Man(β-Sp10

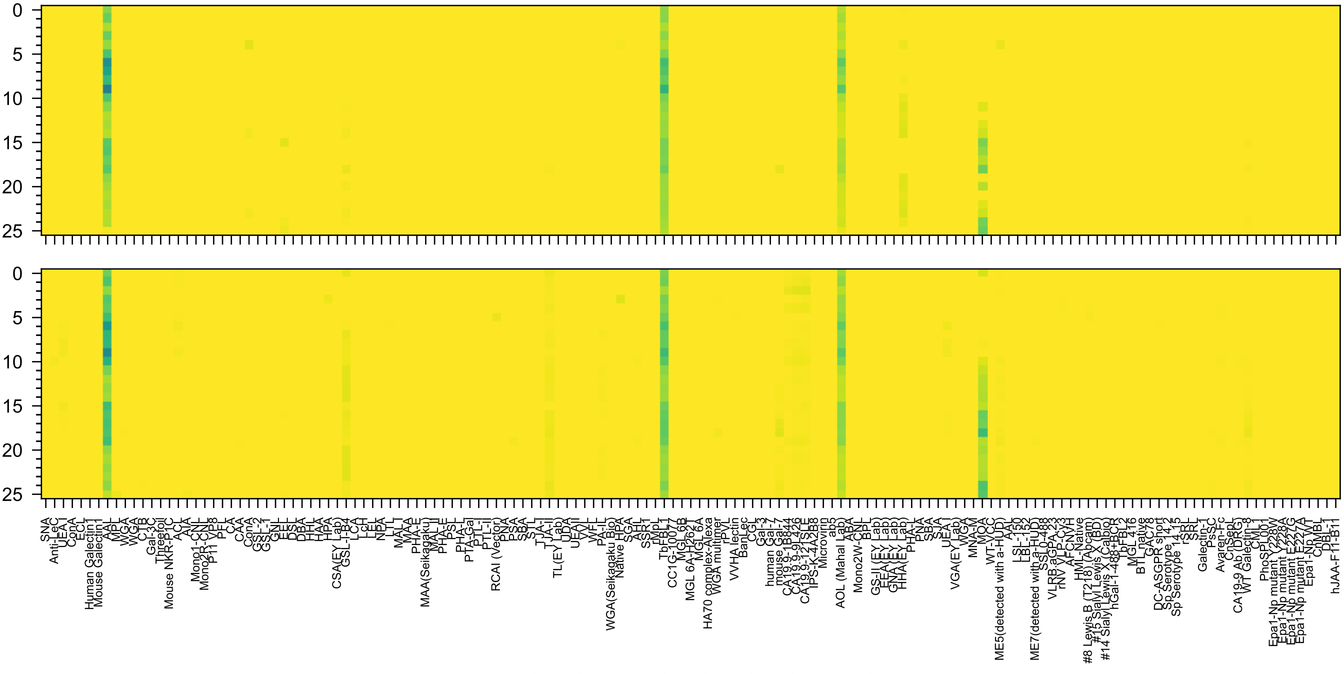
Gal(α1-3)[Fuc(α1-2)]Gal(β1-4)[Fuc(α1-3)]GlcNAc(β-Sp8

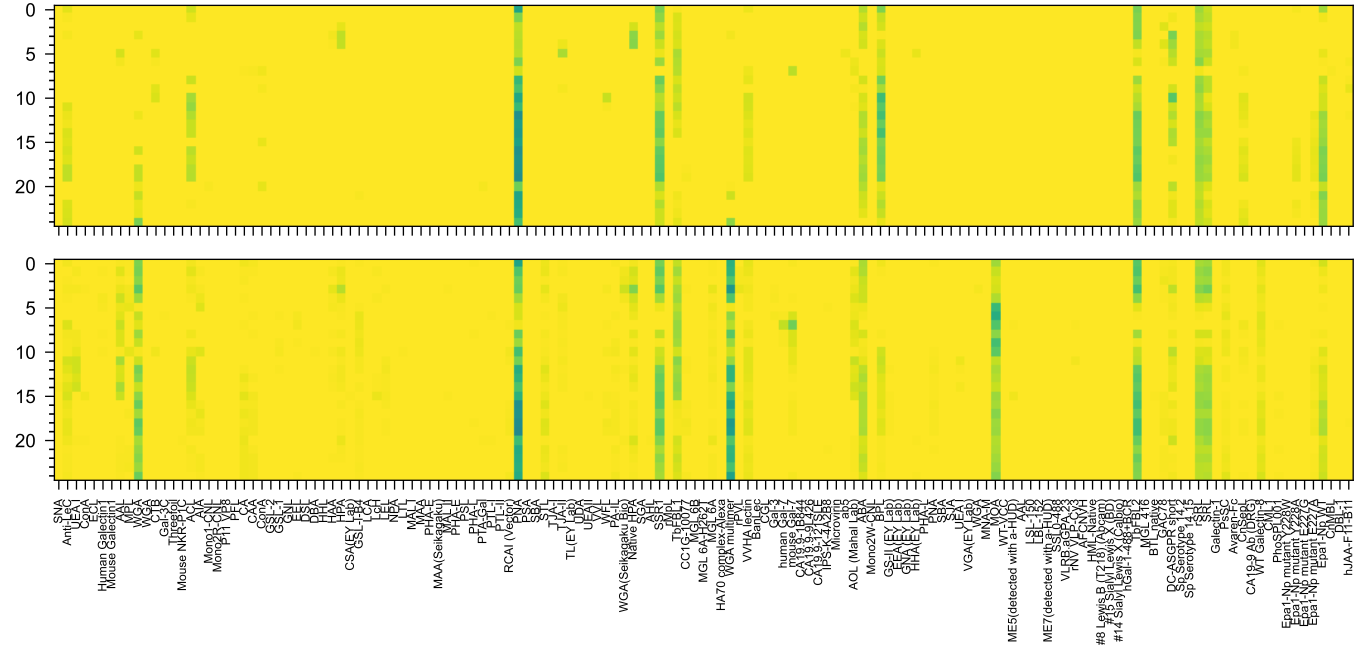

Gal(β1-3)GalNAc(α1-3)[Fuc(α1-2)]Gal(β1-4)Glc(-Sp0

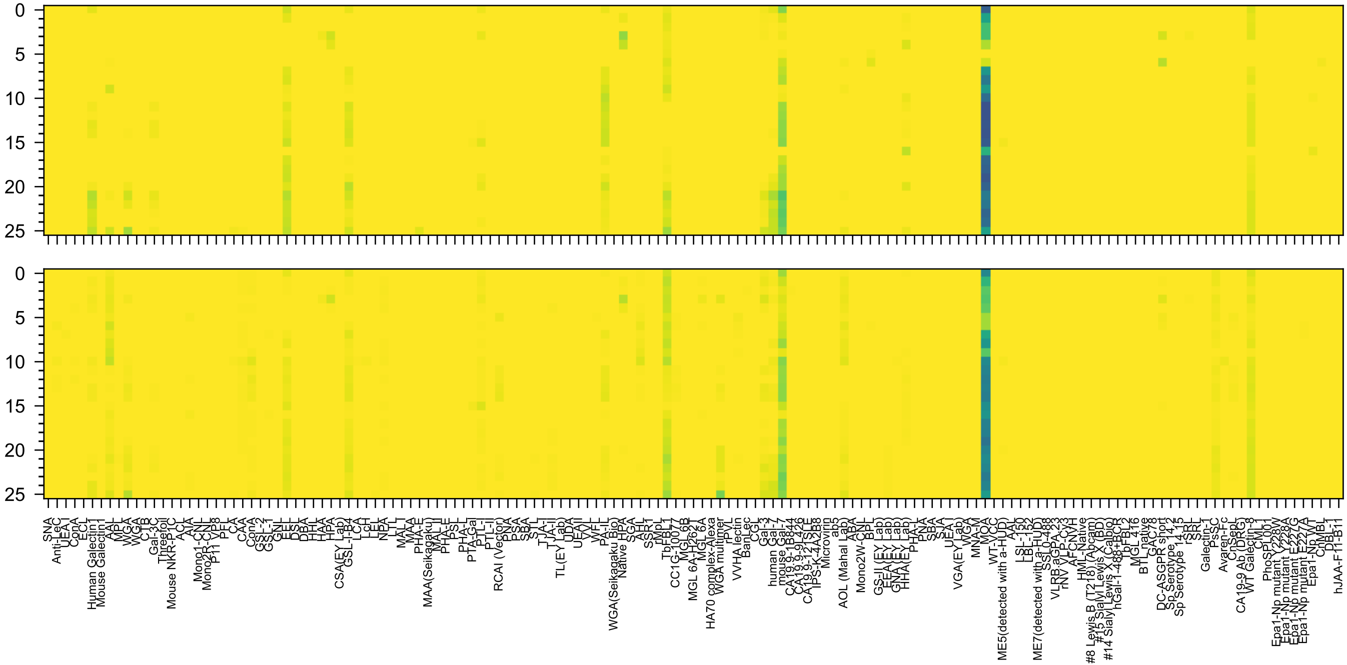

Gal(α1-3)[Fuc(α1-2)]Gal(β1-4)GlcNAc(β1-2)Man(α-Sp0

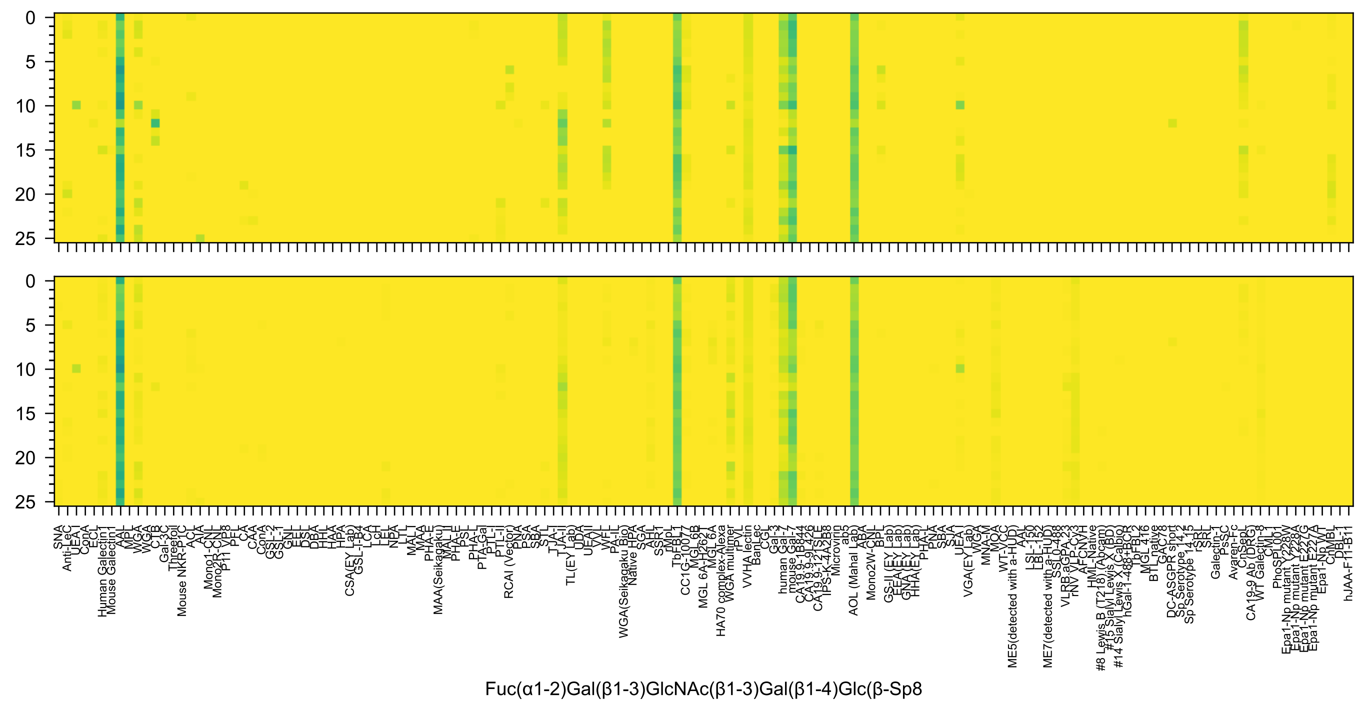
Fuc(α1-2)Gal(β1-3)GlcNAc(β1-3)Gal(β1-4)Glc(β-Sp8

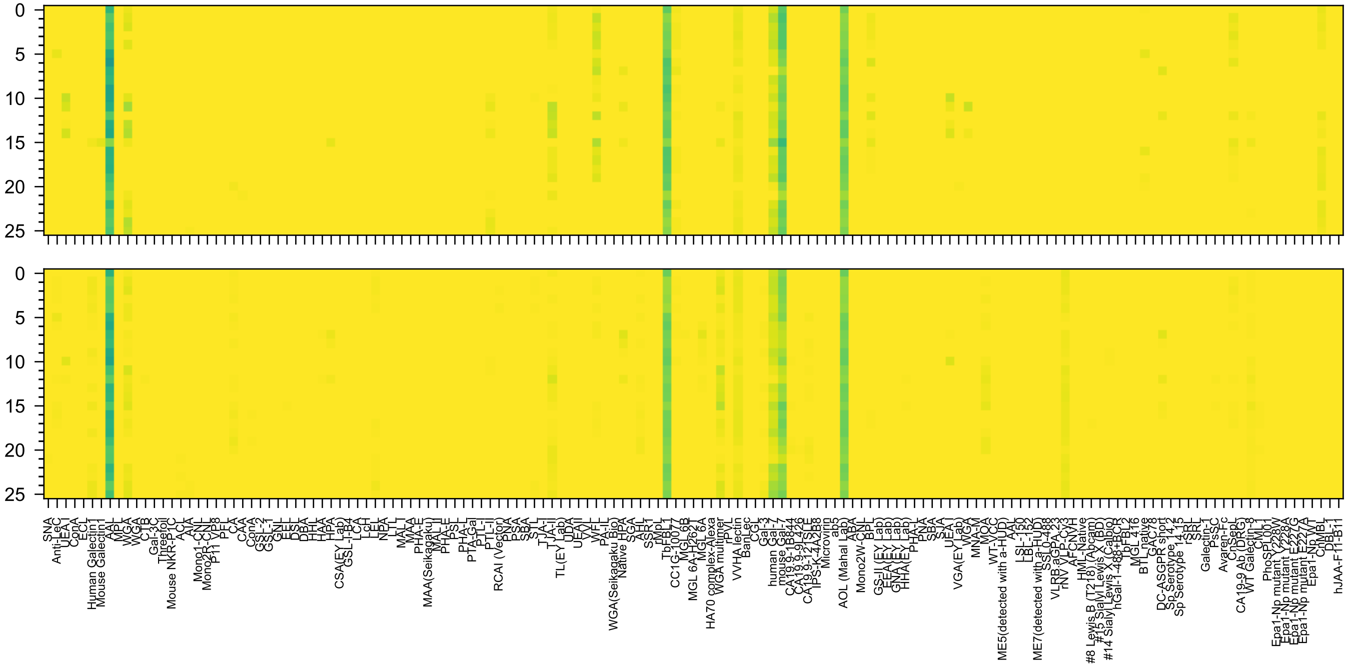

Fuc(α1-2)Gal(β1-3)GlcNAc(β1-3)Gal(β1-4)Glc(β-Sp10

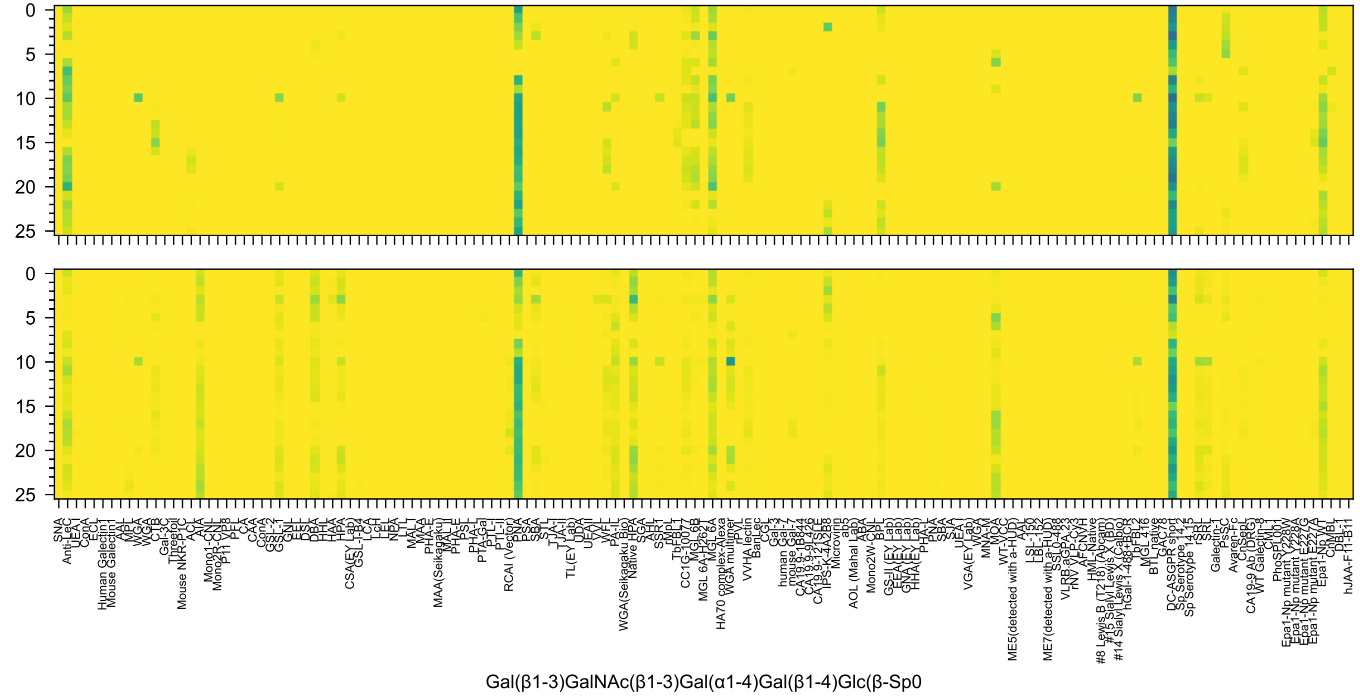
0
5
10
15
20
25
0
5
10
15
20
25
SNA
Anti-LeC
UEA I
ConA
ECL
Human Galectin1
Mouse Galectin1
AAL
MPL
WGA
WGA
CTB
Gal-3C
Threefoil
Mouse NKR-P1C
ACL
AIA
Mono1-CNL
Mono2R-CNL
P11 VP8
PFL
CA
CAA
ConA
GSL-2
GSL-1
GNL
EEL
DSL
DBA
HHL
HAA
HPA
CSA(EY Lab)
GSL-I-B4
LCA
LcH
LEL
NPA
LTL
MAL I
MAA
PHA-E
MAA(Seikagaku)
MAL II
PHA-E
PSL
PHA-L
PTA-Gal
PTL-I
PTL-II
RCAI (Vector)
PNA
PSA
SBA
STL
TJA-I
TJA-II
TL(EY Lab)
UDA
UEAII
VVL
WFL
PA-IL
WGA(Seikagaku Bio)
Native HPA
SGA
AHL
SSR1
rMpL
TbFBL1
CC1G-10077
MGL 6B
MGL 6A-H262T
MGL 6A
HA70 complex-Alexa
WGA multimer
rPVL
VVHA lectin
BanLec
CGL
Gal-3
human Gal-7
mouse Gal-7
CA19.9-1B844
CA19.9-9L426
CA19.9-121SLE
IPS-K-4A2B8
Microvirin
ab5
AOL (Mahal Lab)
ABA
Mono2W-CNL
BPL
GS-II (EY Lab)
EEA(EY Lab)
GNA (EY Lab)
HHA(EY Lab)
PHA-L
PNA
SBA
SJA
UEA I
VGA(EY Lab)
WGA
MNA-M
MOA
WT-VCC
ME5(detected with a-HUD)
AAL
LSL-150
LBL-152
ME7(detected with a-HUD)
SSL0-488
VLRB.aGPA.23
rNV VLP-Cy3
AFCNVH
HML-Native
#8 Lewis B (T218) (Abcam)
#15 Sialyl Lewis X (BD)
#14 Sialyl Lewis X (Calbio)
hGal-1-488+BCR
TbFBL2
MGL 416
BTL native
GAC78
DC-ASGPR short
Sp Serotype 14.2
Sp Serotype 14.15
rSRL
SRL
Galectin-1
PsSC
Avaren-Fc
ChSepL
CA19-9 Ab (DRG)
WT Galectin-8
CML
PhoSL001
Epa1-Np mutant Y228W
Epa1-Np mutant Y228A
Epa1-Np mutant E227G
Epa1-Np mutant E227A
Epa1-Np WT
ChMBL
DBL-1
hJAA-F11-B11
Gal(β1-3)GalNAc(β1-3)Gal(α1-4)Gal(β1-4)Glc(β-Sp0

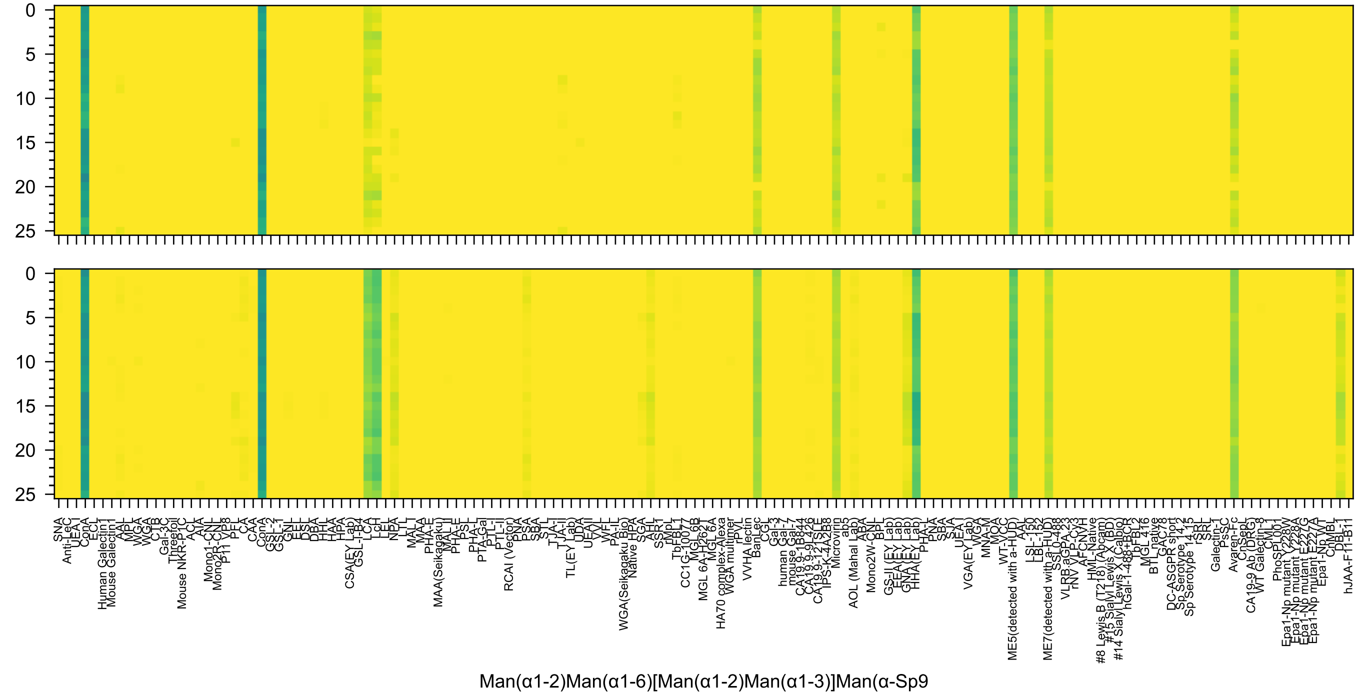
0
5
10
15
20
25
SNA
Anti-LeC
UEA-I
ConA
ECL
Human Galectin1
Mouse Galectin1
AAL
MPL
WGA
WGA
CTB
Gal-3C
Threefoil
Mouse NKR-P1C
ACL
AIA
Mono1-CNL
Mono2R-CNL
P11 VP8
PFL
CA
CAA
ConA
GSL-2
GSL-1
GNL
EEL
DSL
DBA
HHL
HAA
HPA
CSA(EY Lab)
GSL-I-B4
LCA
LcH
LEL
NPA
LTL
MAL-I
MAA
PHA-E
MAA(Seikagaku)
MAL-II
PHA-E
PSL
PHA-L
PTA-Gal
PTL-I
PTL-II
RCAI (Vector)
PNA
PSA
SBA
STL
TJA-I
TJA-II
TL(EY Lab)
UDA
UEAII
VVL
WFL
PA-IL
WGA(Seikagaku Bio)
Native HPA
SGA
AHL
SSR1
rMpL
TbFBL1
CC1G-10077
MGL 6B
MGL 6A-H262T
MGL 6A
HA70 complex-Alexa
WGA multimer
rPVL
VVHA lectin
BanLec
CGL
Gal-3
human Gal-7
mouse Gal-7
CA19.9-1B844
CA19.9-9L426
CA19.9-121SLE
IPS-K-4A2B8
Microvirin
ab5
AOL (Mahal Lab)
ABA
Mono2W-CNL
BPL
GS-II (EY Lab)
EEA (EY Lab)
GNA (EY Lab)
HHA (EY Lab)
PHA-L
PNA
SBA
SJA
UEA-I
VGA(EY Lab)
WGA
MNA-M
MOA
WT-VCC
ME5(detected with a-HUD)
AAL
LSL-150
LBL-152
ME7(detected with a-HUD)
SSL0-488
VLRB.aGPA.23
rNV VLP-Cy3
AFCNVH
HML-Native
#8 Lewis B (T218) (Abcam)
#15 Sialyl Lewis X (BD)
#14 Sialyl Lewis X (Calbio)
hGal-1-488+BCR
TbFBL2
MGL 416
BTL native
GAC78
DC-ASGPR short
Sp Serotype 14.2
Sp Serotype 14.15
rSRL
SRL
Galectin-1
PsSC
Avaren-Fc
ChSepL
CA19-9 Ab (DRG)
WT Galectin-8
CML
PhoSL001
Epa1-Np mutant Y228W
Epa1-Np mutant Y228A
Epa1-Np mutant E227G
Epa1-Np mutant E227A
Epa1-Np WT
ChMBL
DBL-1
hJAA-F11-B11
Man(α1-2)Man(α1-6)[Man(α1-2)Man(α1-3)]Man(α-Sp9

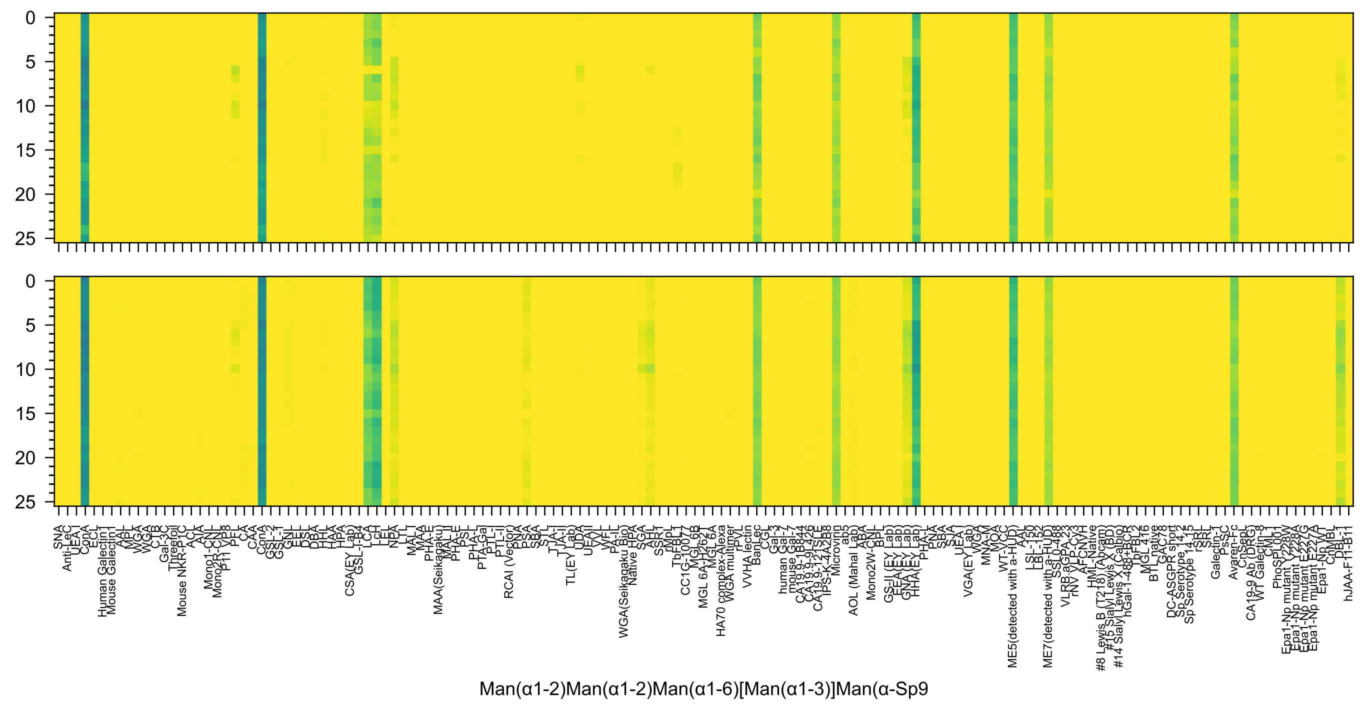
Man(α1-2)Man(α1-2)Man(α1-6)[Man(α1-3)]Man(α-Sp9

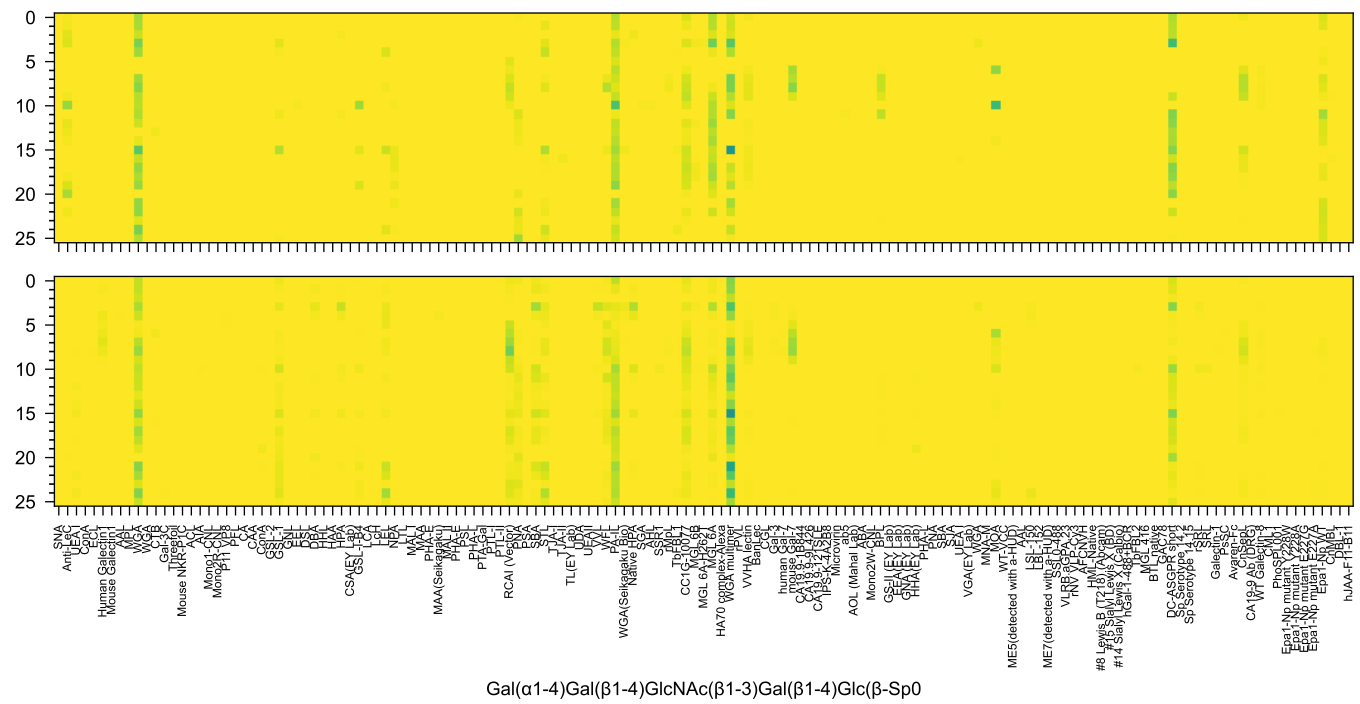
Gal(α1-4)Gal(β1-4)GlcNAc(β1-3)Gal(β1-4)Glc(β-Sp0

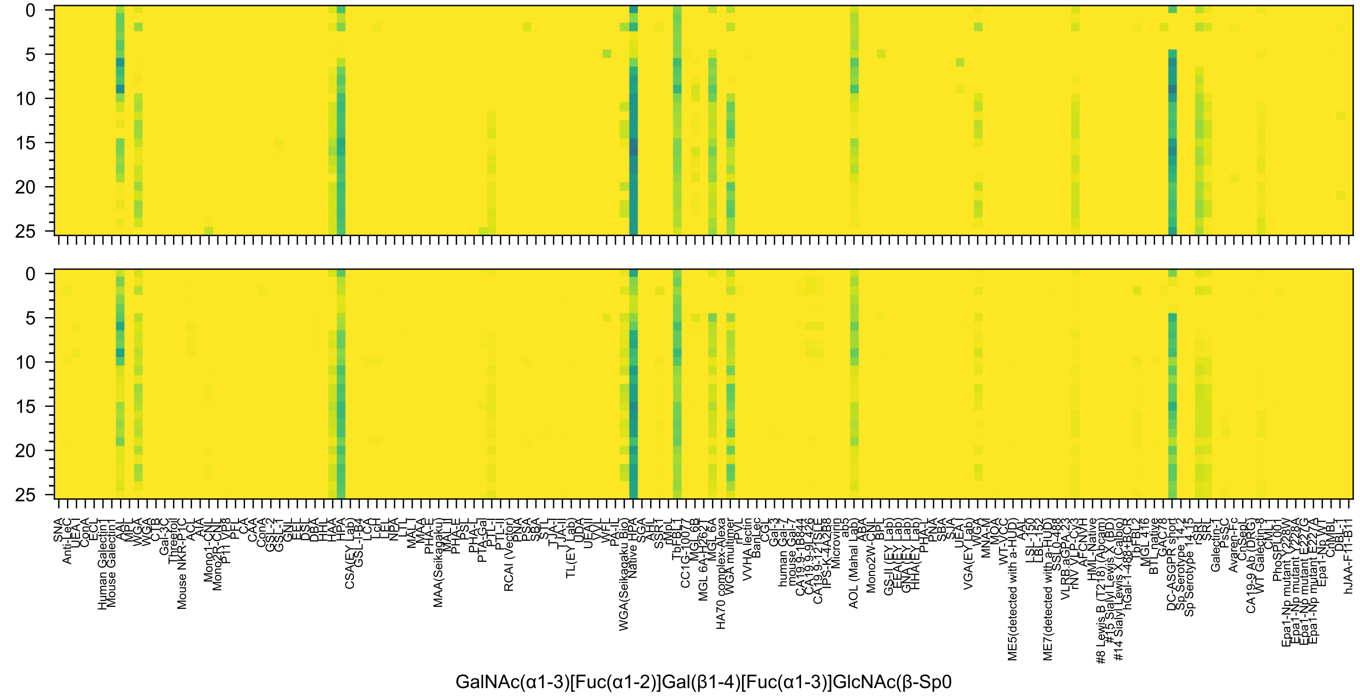
0
5
10
15
20
25
SNA
Anti-LeC
UEA I
ConA
ECL
Human Galectin1
Mouse Galectin1
AAL
MPL
WGA
WGA
CTB
Gal-3C
Threefoil
Mouse NKR-P1C
ACL
AIA
Mono1-CNL
Mono2R-CNL
P11 VP8
PFL
CA
CAA
ConA
GSL-2
GSL-1
GNL
EEL
DSL
DBA
HHL
HAA
HPA
CSA(EY Lab)
GSL-I-B4
LCA
LcH
LEL
NPA
LTL
MAL-I
MAA
PHA-E
MAA(Seikagaku)
MAL-II
PHA-E
PSL
PHA-L
PTA-Gal
PTL-I
PTL-II
RCAI (Vector)
PNA
PSA
SBA
STL
TJA-I
TJA-II
TL(EY Lab)
UDA
UEA-I
VVL
WFL
PA-IL
WGA(Seikagaku Bio)
Native HPA
SGA
AHL
SSR1
rMpL
TbFBL1
CC1G-10077
MGL 6B
MGL 6A-H262T
MGL 6A
HA70 complex-Alexa
WGA multimer
rPvL
VVHA lectin
BanLec
CGL
Gal-3
human Gal-7
mouse Gal-7
CA19.9-1B844
CA19.9-9L426
CA19.9-121SLE
IPS-K-4A2B8
Microvirin
ab5
AOL (Mahal Lab)
ABA
Mono2W-CNL
BPL
GS-II (EY Lab)
EEA (EY Lab)
GNA (EY Lab)
HHA(EY Lab)
PHA-L
PNA
SBA
SJA
UEA I
VGA(EY Lab)
WGA
MNA-M
MOA
WT-VCC
ME5(detected with a-HUD)
AAT
LSL-150
LBL-152
ME7(detected with a-HUD)
SSL0-488
VLRB.aGPA.23
rNV VLP-Cy3
AFCNVH
HML-Native
#8 Lewis B (T218) (Abcam)
#15 Sialyl Lewis X (BD)
#14 Sialyl Lewis X (Calbio)
hGal-1-488+BCR
TbFBL2
MGL 416
BTL native
GAC78
DC-ASGPR short
Sp Serotype 14.2
Sp Serotype 14.15
rSRL
SRL
Galectin-1
PsSC
Avaren-Fc
ChSepL
CA19-9 Ab (DRG)
WT Galectin-8
CML1
PhoSL001
Epa1-Np mutant Y228W
Epa1-Np mutant Y228A
Epa1-Np mutant E227G
Epa1-Np mutant E227A
Epa1-Np WT
ChMBL
DBL-1
hJAA-F11-B11
GalNAc(α1-3)[Fuc(α1-2)]Gal(β1-4)[Fuc(α1-3)]GlcNAc(β-Sp0

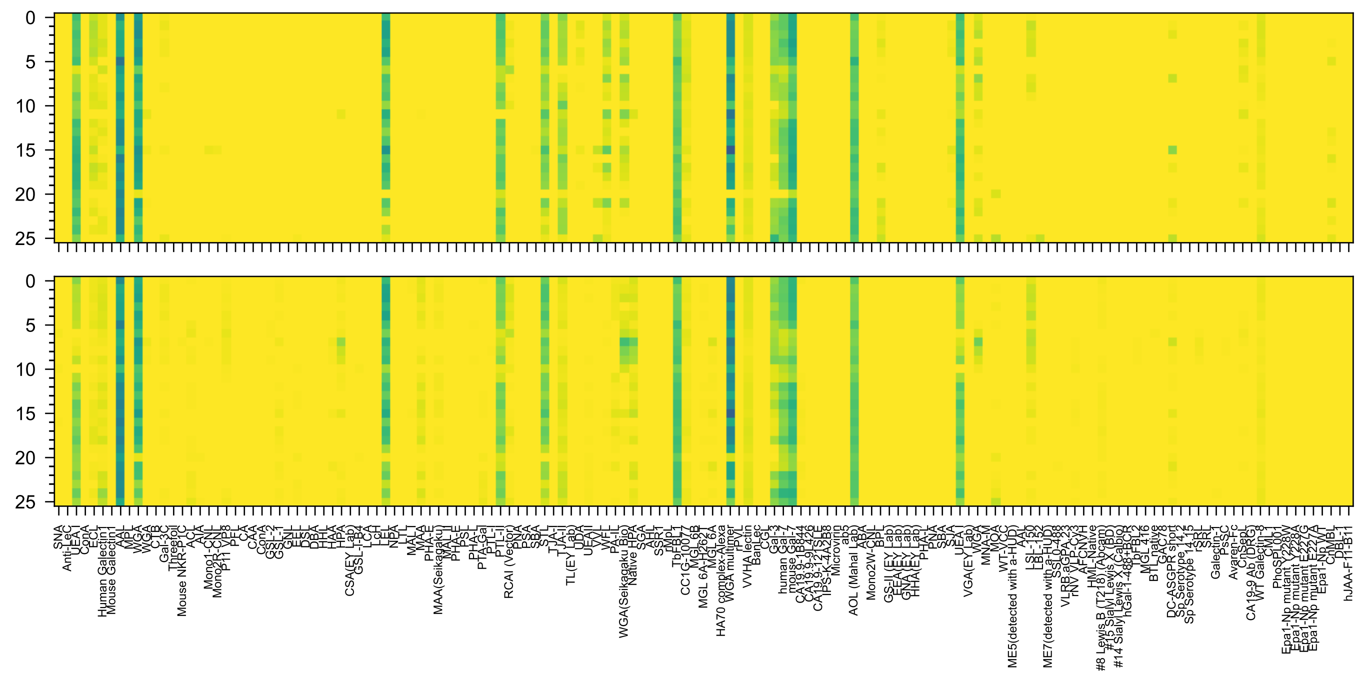

Fuc(α1-2)Gal(β1-4)GlcNAc(β1-3)Gal(β1-4)GlcNAc(β-Sp0

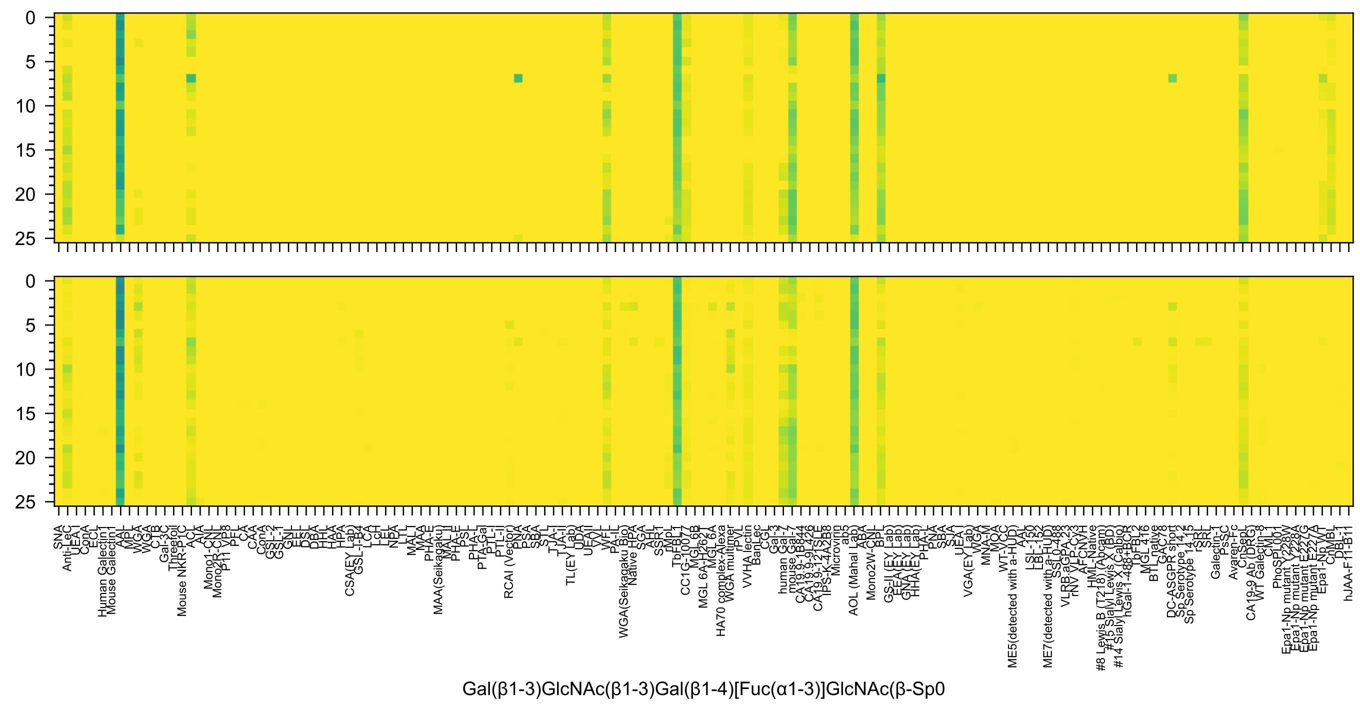

Gal(β1-3)GlcNAc(β1-3)Gal(β1-4)[Fuc(α1-3)]GlcNAc(β-Sp0

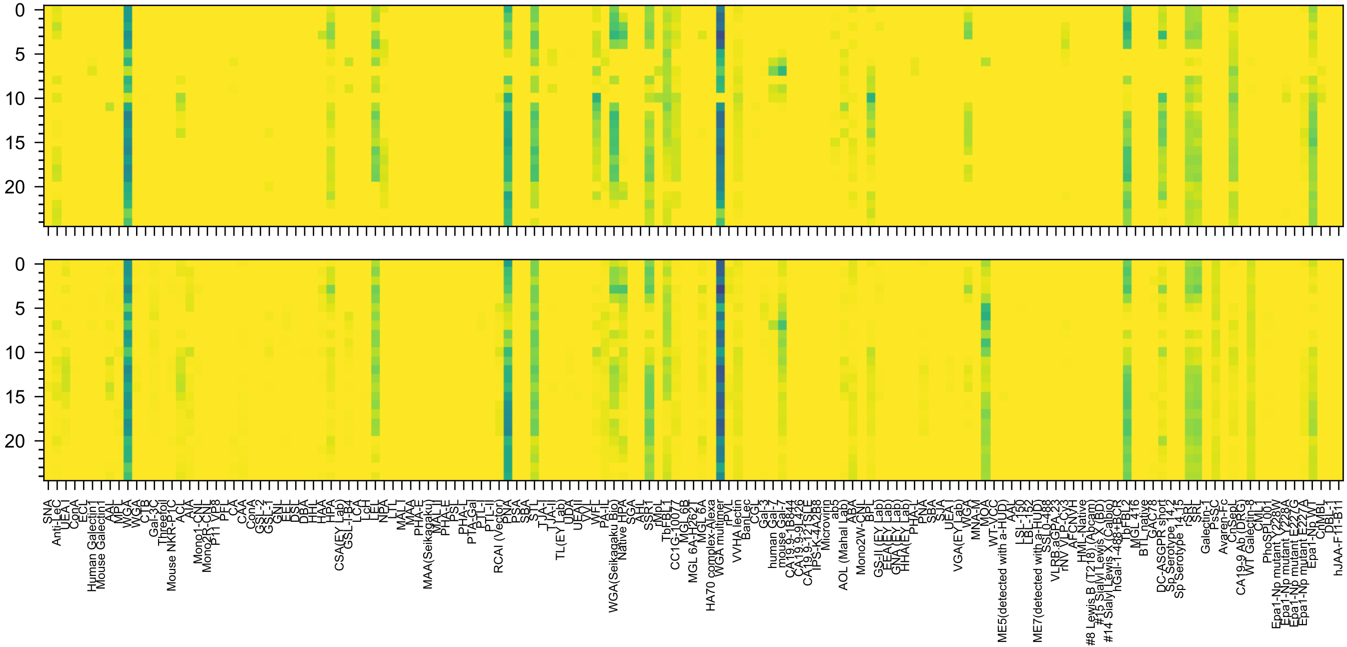


Gal(β1-3)GalNAc(α1-3)[Fuc(α1-2)]Gal(β1-4)GlcNAc(-Sp0

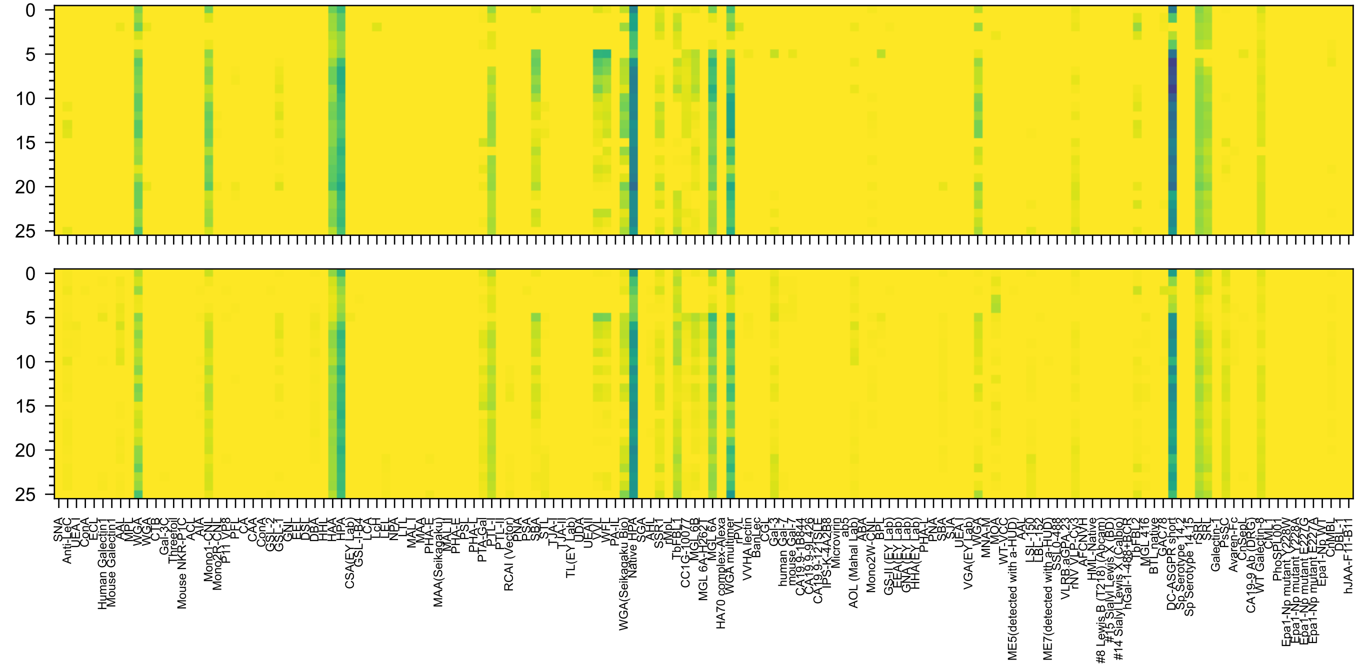

GalNAc(α1-3)[Fuc(α1-2)]Gal(β1-4)GlcNAc(β1-2)Man(α-Sp0

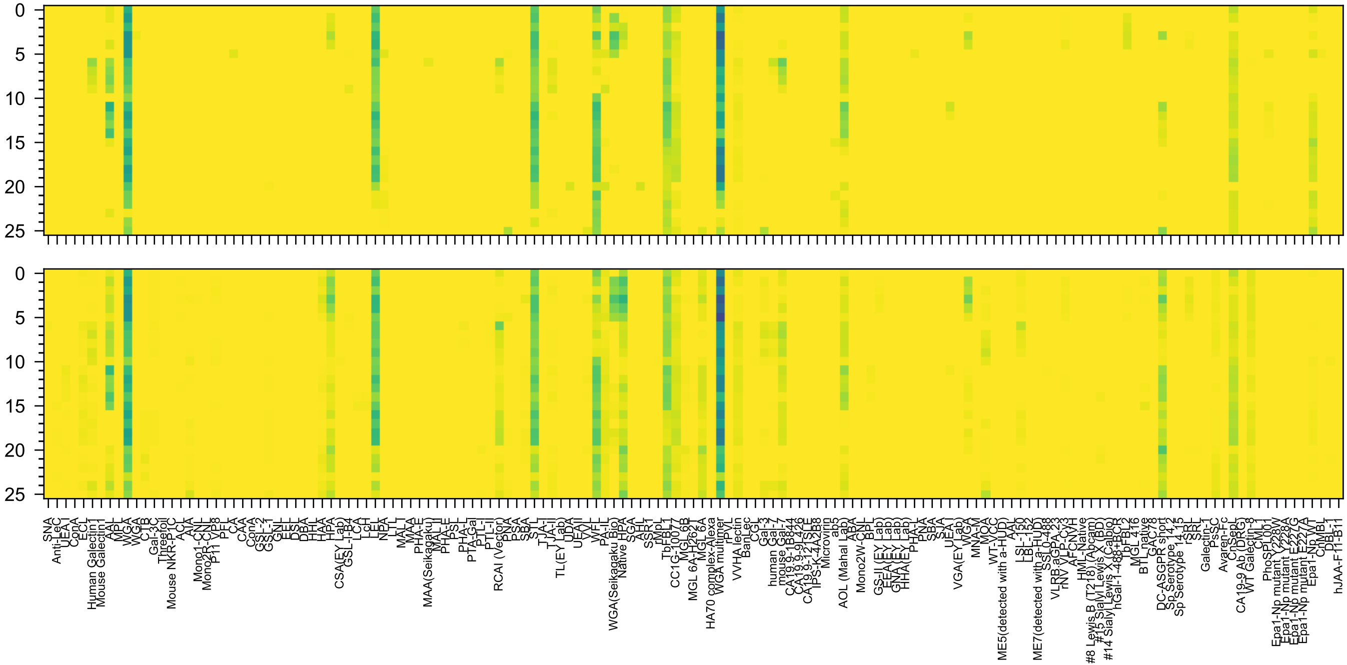


Gal(β1-4)GalNAc(α1-3)[Fuc(α1-2)]Gal(β1-4)GlcNAc(β-Sp8

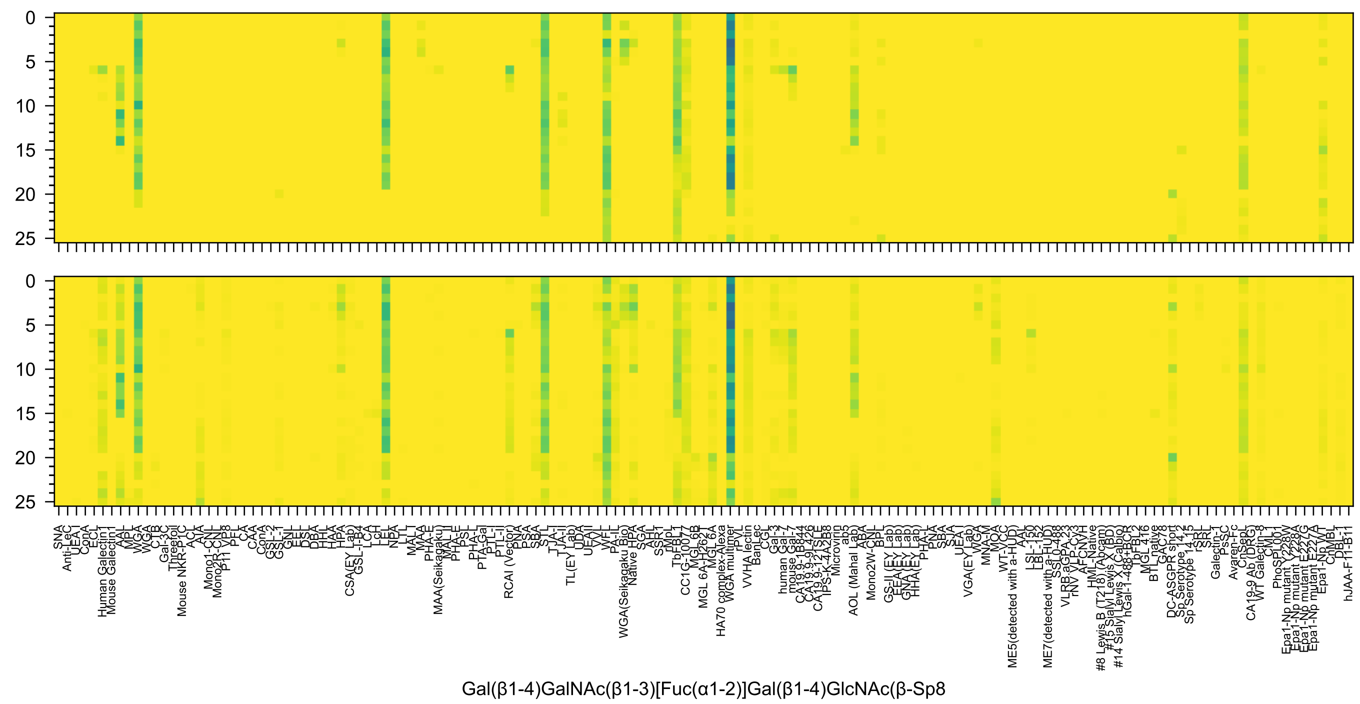
Gal(β1-4)GalNAc(β1-3)[Fuc(α1-2)]Gal(β1-4)GlcNAc(β-Sp8

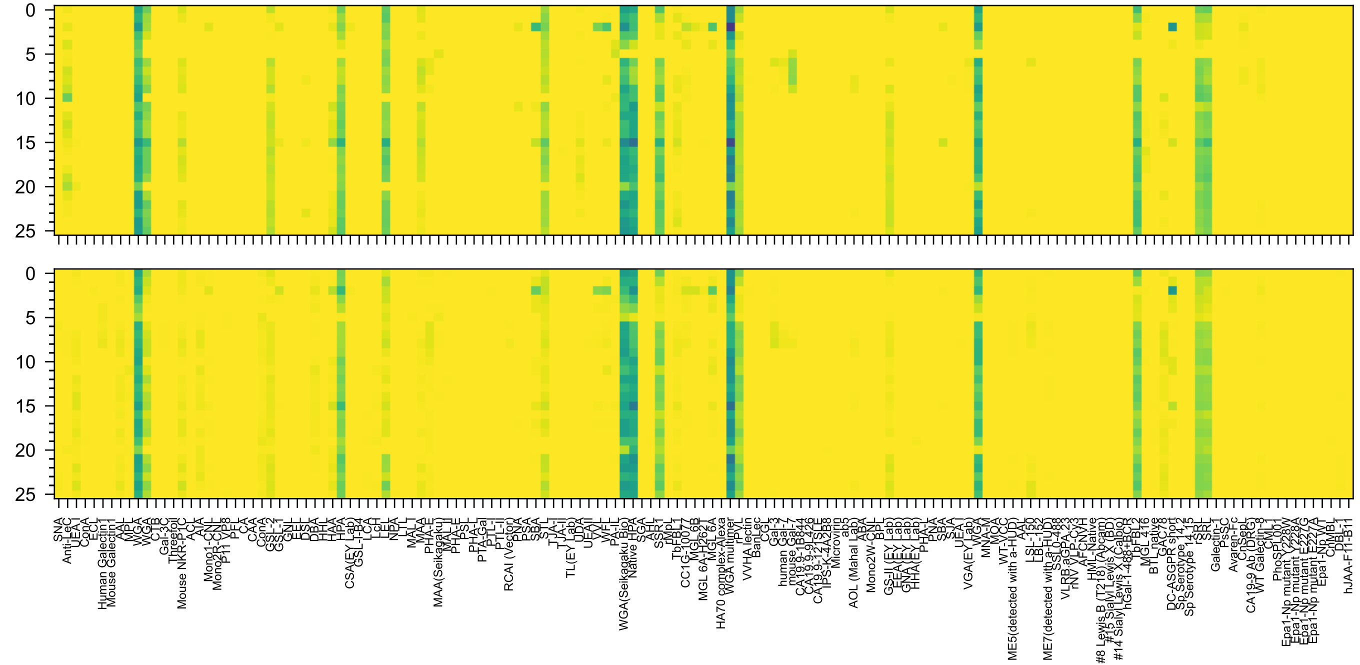


GlcNAc(α1-4)Gal(β1-4)GlcNAc(β1-3)Gal(β1-4)Glc(β-Sp0

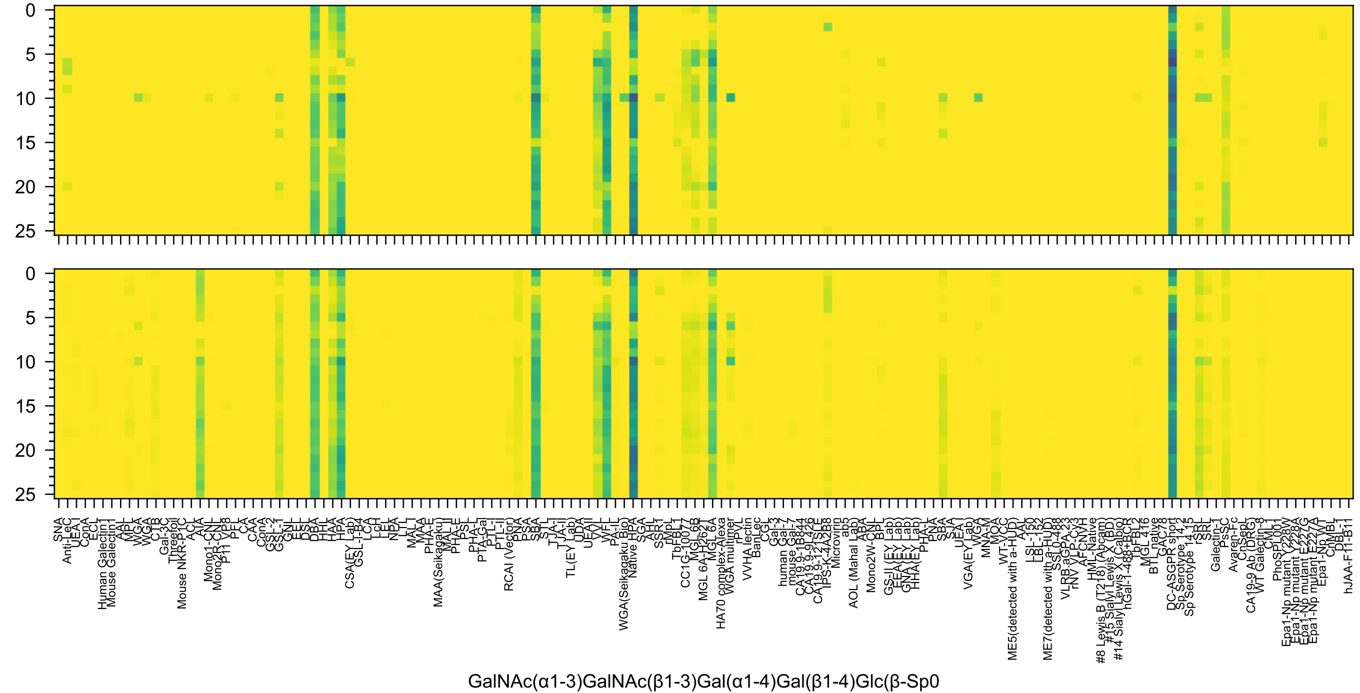

GalNAc(α1-3)GalNAc(β1-3)Gal(α1-4)Gal(β1-4)Glc(β-Sp0

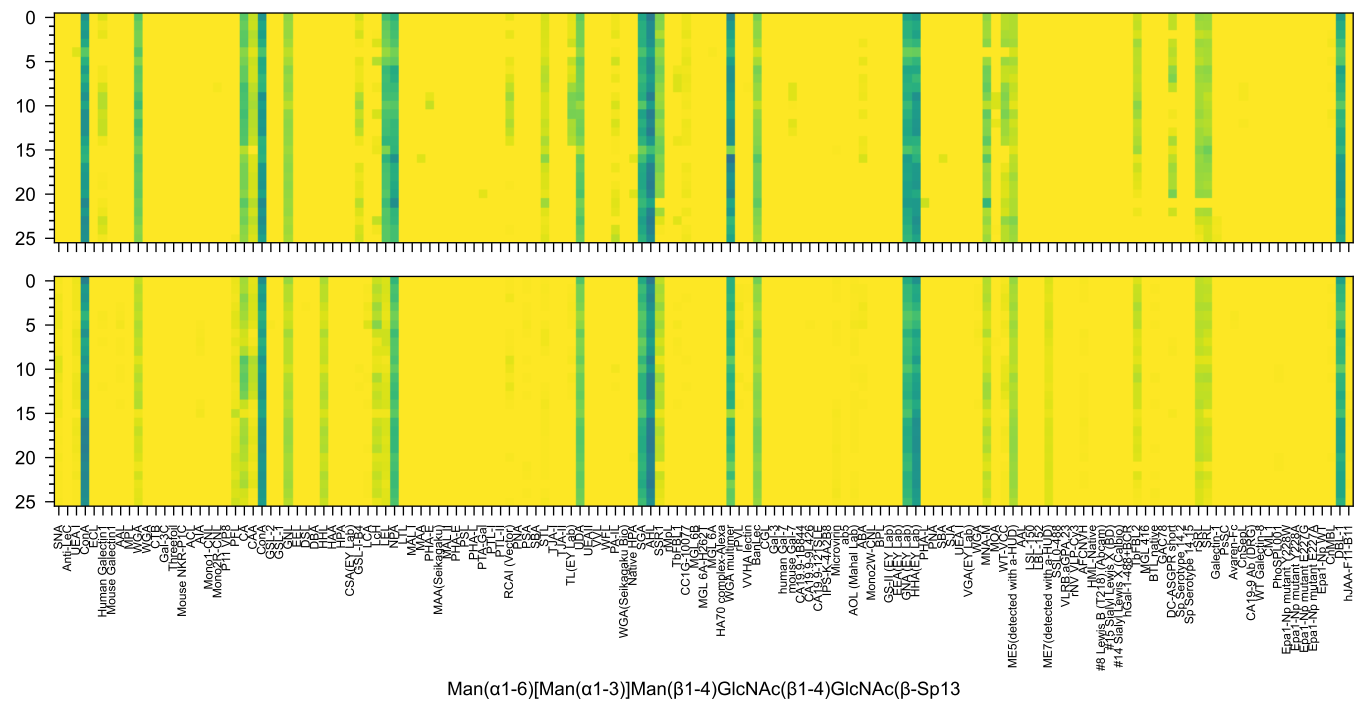
Man(α1-6)[Man(α1-3)]Man(β1-4)GlcNAc(β1-4)GlcNAc(β-Sp13

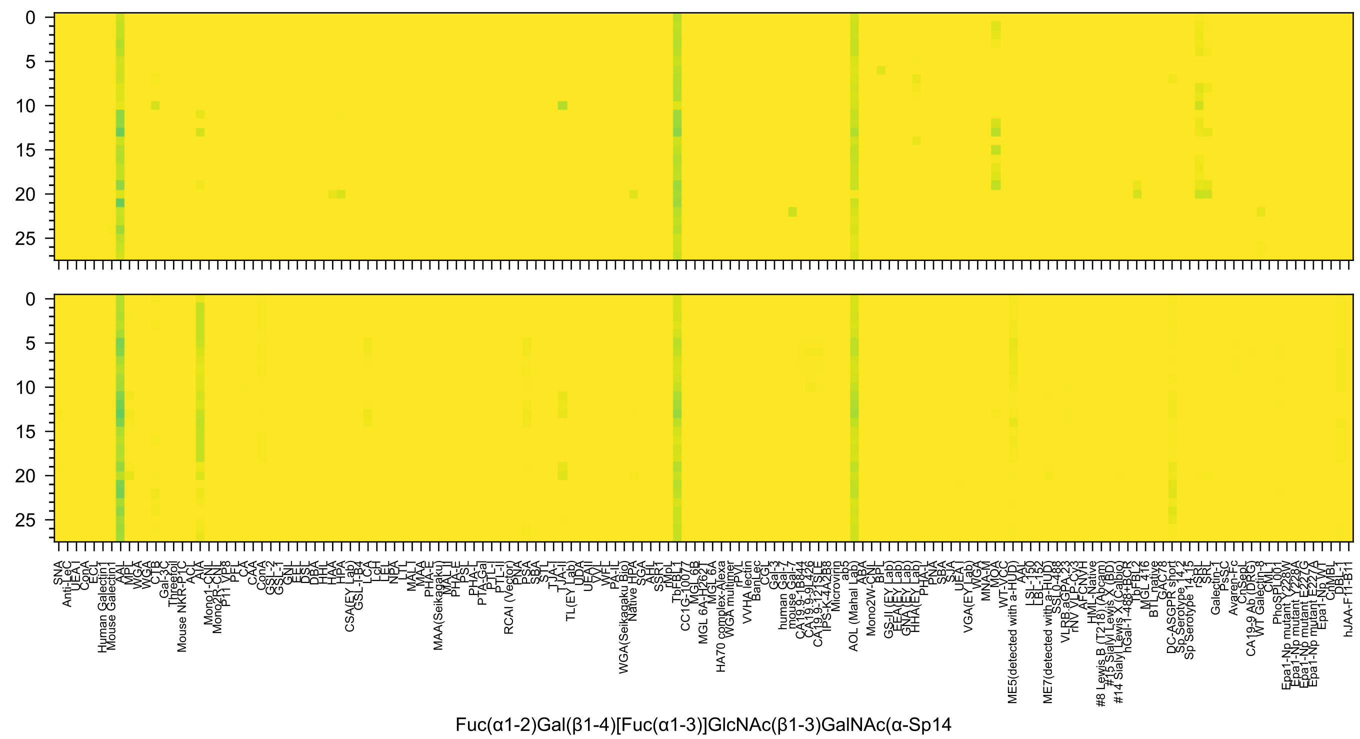
0
5
10
15
20
25
Fuc(α1-2)Gal(β1-4)[Fuc(α1-3)]GlcNAc(β1-3)GalNAc(α-Sp14

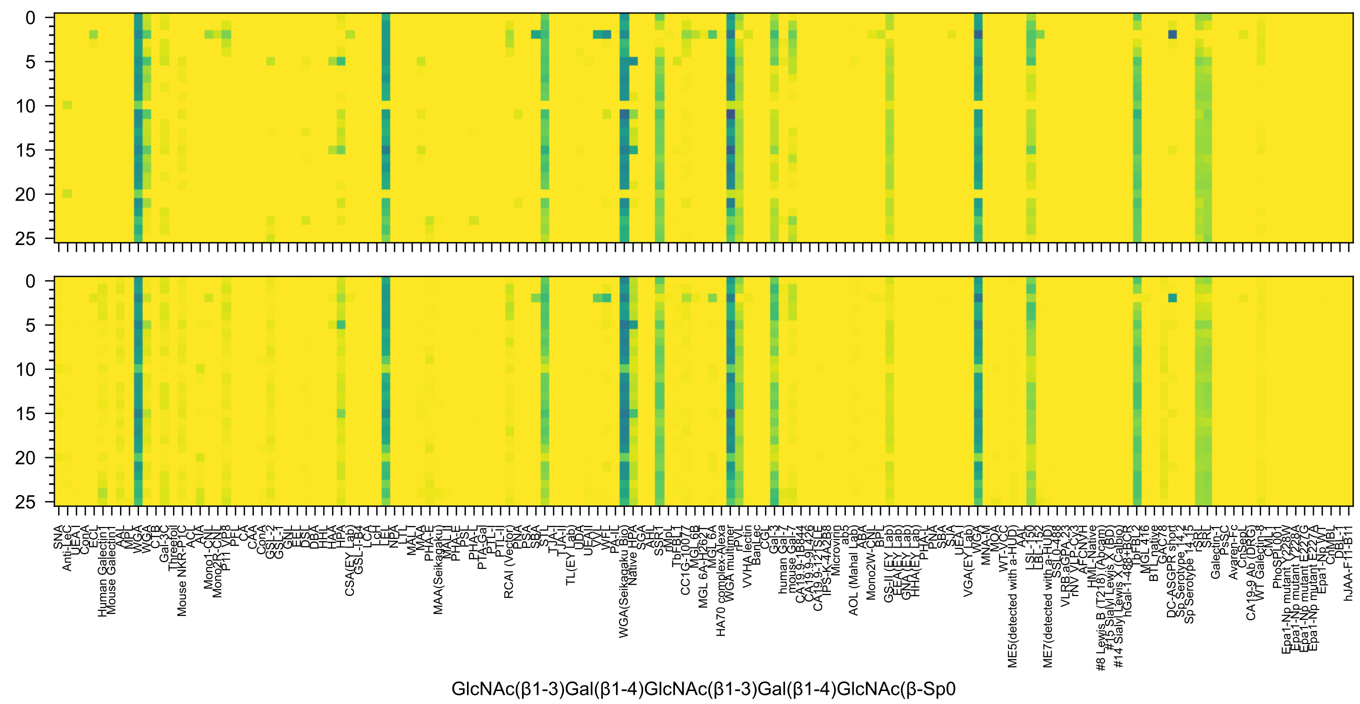
GlcNAc(β1-3)Gal(β1-4)GlcNAc(β1-3)Gal(β1-4)GlcNAc(β-Sp0

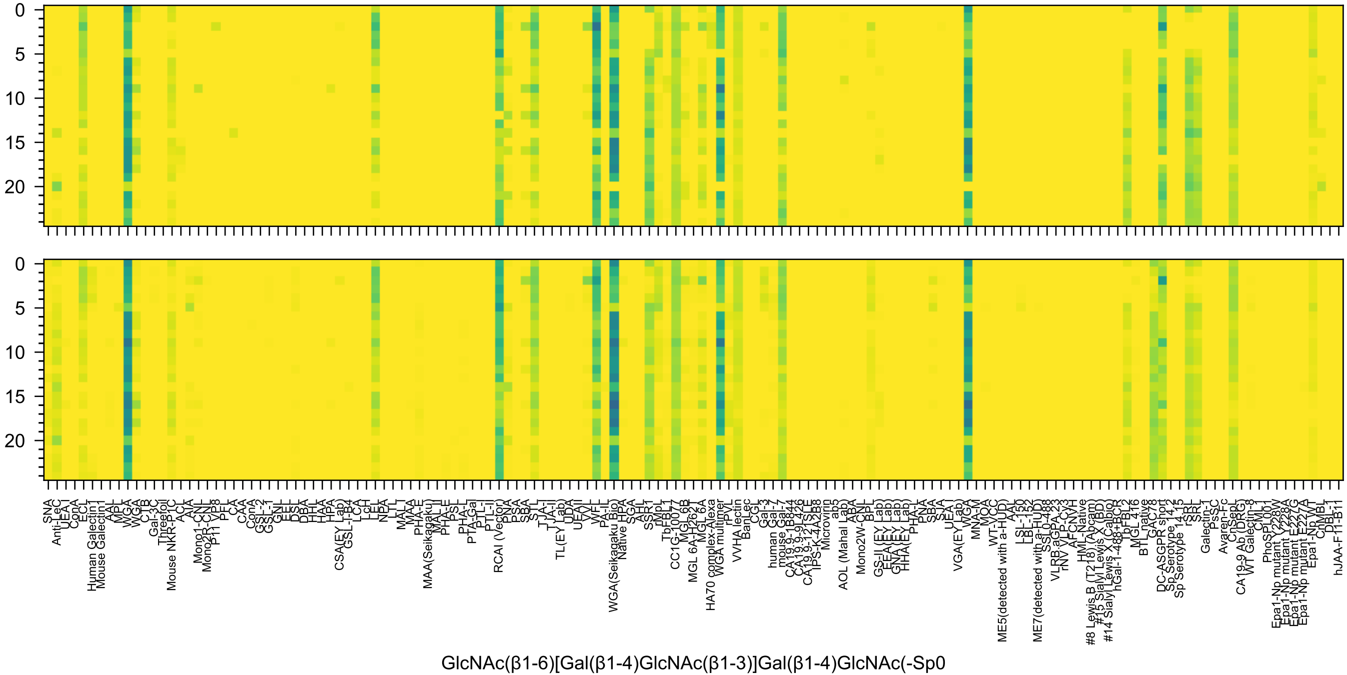
GlcNAc(β1-6)[Gal(β1-4)GlcNAc(β1-3)]Gal(β1-4)GlcNAc(-Sp0

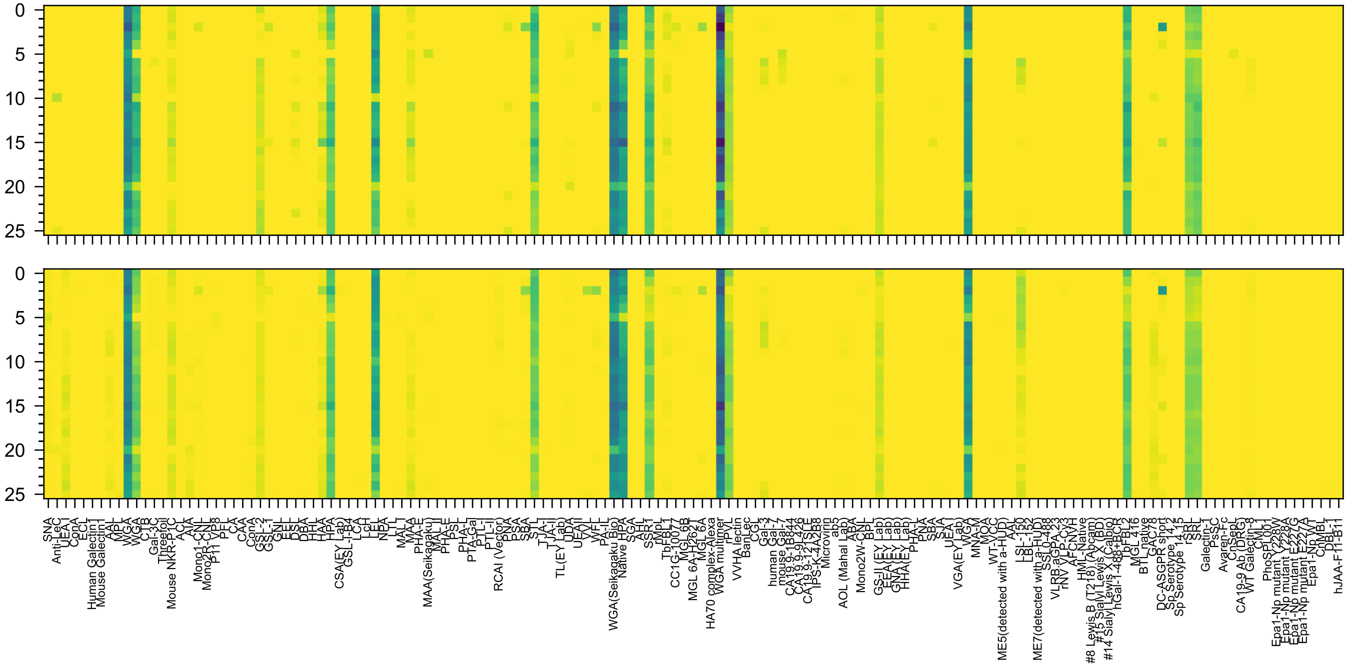

GlcNAc(α1-4)Gal(β1-4)GlcNAc(β1-3)Gal(β1-4)GlcNAc(β-Sp0

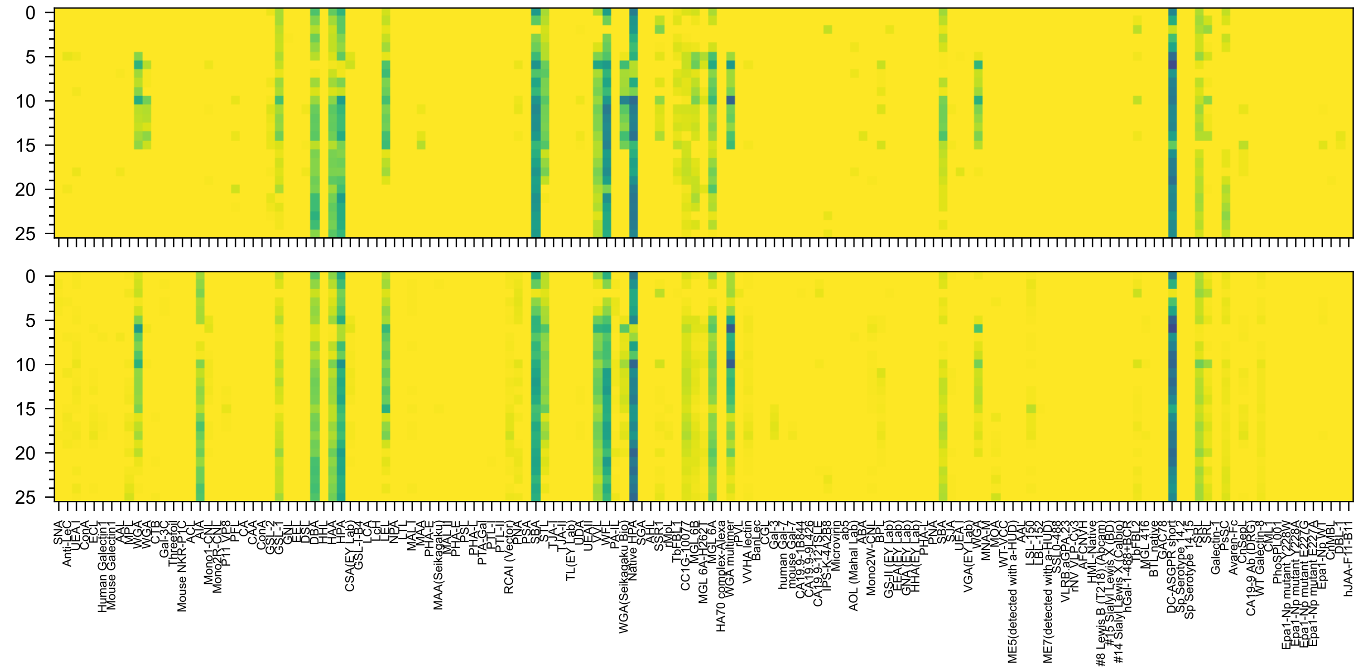

GalNAc(α1-3)GalNAc(β1-3)Gal(α1-4)Gal(β1-4)GlcNAc(β-Sp0

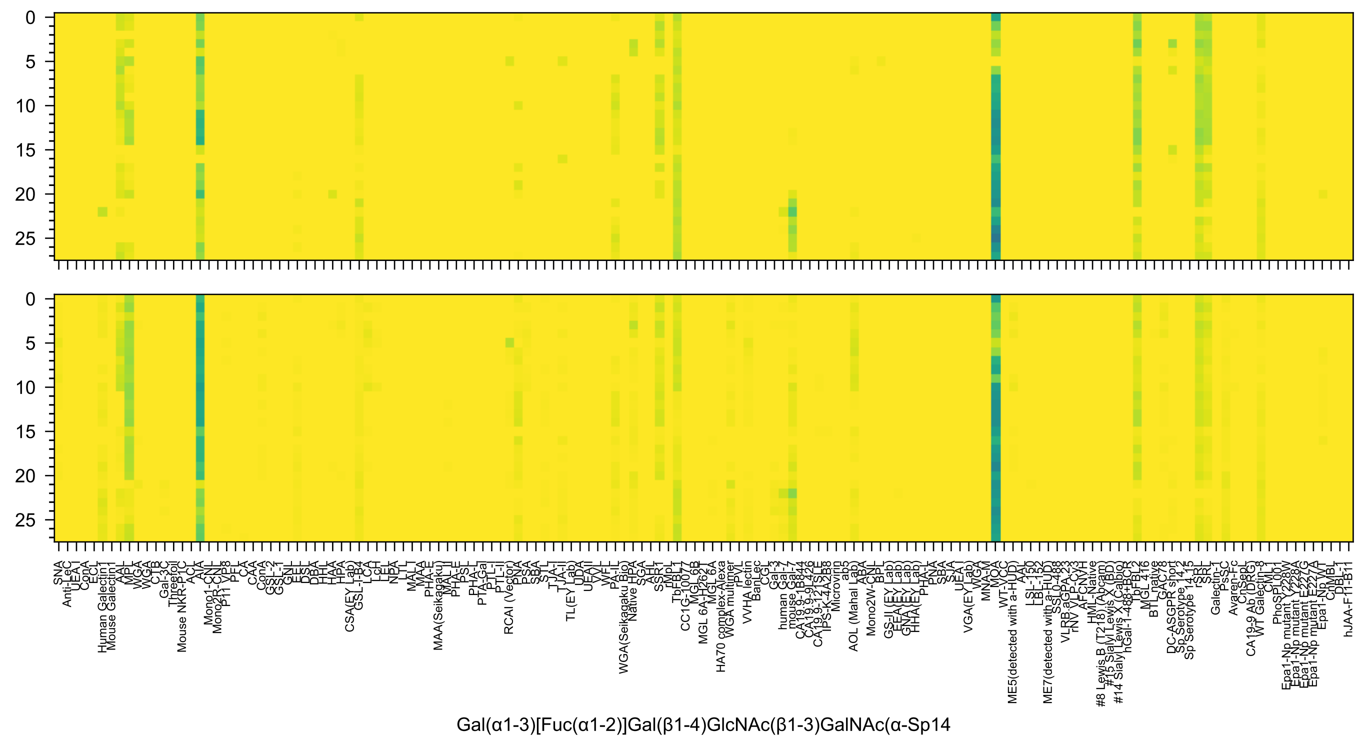
Gal(α1-3)[Fuc(α1-2)]Gal(β1-4)GlcNAc(β1-3)GalNAc(α-Sp14

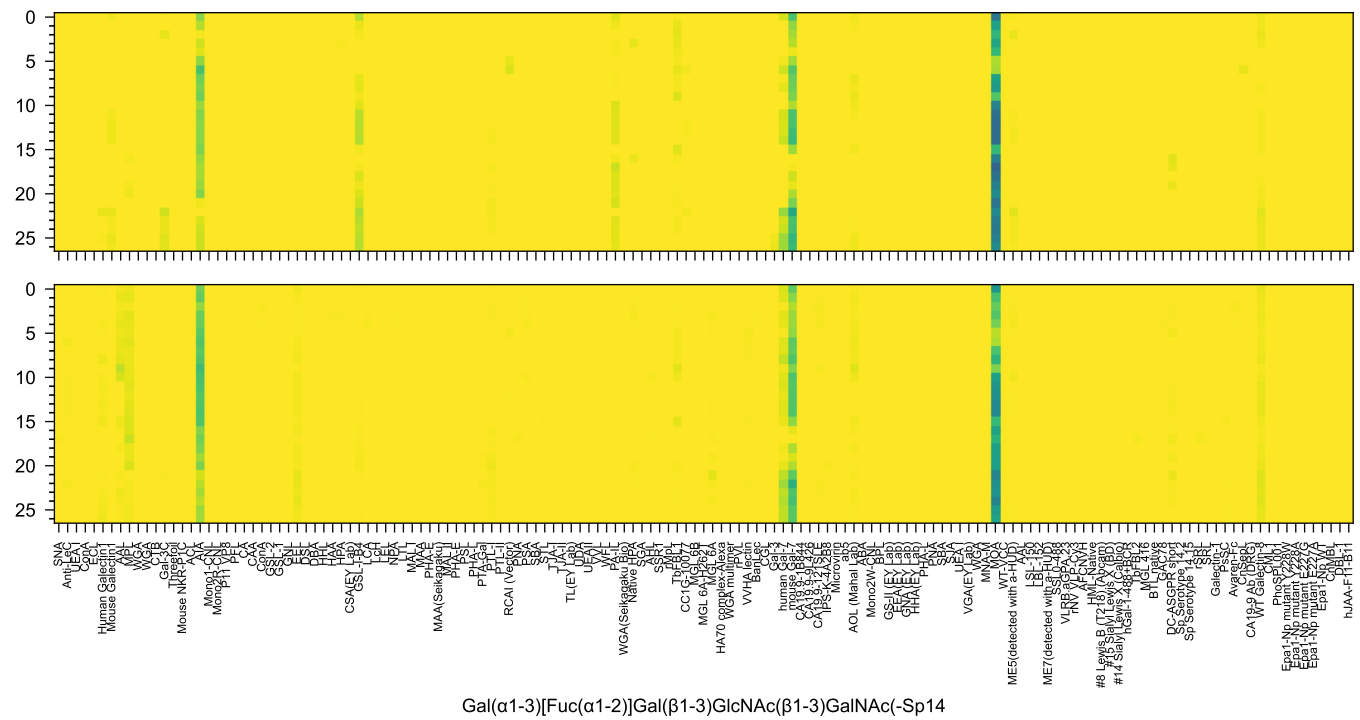

SNA
Anti-LeC
UEA I
ConA
ECL
Human Galectin1
Mouse Galectin1
AAL
MPL
WGA
WGA
CTB
Gal-3C
Threefoil
Mouse NKR-P1C
ACL
AIA
Mono1-CNL
Mono2R-CNL
P11 VP8
PFL
CA
CAA
ConA
GSL-2
GSL-1
GNL
EEL
DSL
DBA
HHL
HAA
HPA
CSA(EY Lab)
GSL-I-B4
LCA
LcH
LEL
NPA
LTL
MAL I
MAA
PHA-E
MAA(Seikagaku)
MAL II
PHA-E
PSL
PHA-L
PTA-Gal
PTL-I
PTL-II
RCAI (Vector)
PNA
PSA
SBA
STL
TJA-I
TJA-II
TL(EY Lab)
UDA
UEA II
VVL
WFL
PA-IL
WGA(Seikagaku Bio)
Native HPA
SGA
AHL
SSR1
rMpL
TbFBL1
CC1G-10077
MGL6B
MGL 6A-H262T
MGL 6A
HA70 complex-Alexa
WGA multimer
rPVL
VVHA lectin
BanLec
CGL
Gal-3
human Gal-7
mouse Gal-7
CA19.9-1B844
CA19.9-9L426
CA19.9-121SLE
IPS-K-4A2B8
Microvirin
ab5
AOL (Mahal Lab)
ABA
Mono2W-CNL
BPL
GS-II (EY Lab)
EEA(EY Lab)
GNA(EY Lab)
HHA(EY Lab)
PHA-L
PNA
SBA
SJA
UEA I
VGA(EY Lab)
WGA
MNA-M
MOA
WT-VCC
ME5(detected with a-HUD)
AAL
LSL-150
LBL-152
ME7(detected with a-HUD)
SSL0-488
VLRB.aGPA.23
rNV VLP-Cy3
AFCNVH
HML-Native
#8 Lewis B (T218) (Abcam)
#15 Sialyl Lewis X (BD)
#14 Sialyl Lewis X (Calbio)
hGal-1-488+BCR
TbFBL2
MGL 416
BTL native
GAC78
DC-ASGPR short
Sp Serotype 14.2
Sp Serotype 14.15
rSRL
SRL
Galectin-1
PsSC
Avaren-Fc
ChSeqL
CA19-9 Ab (DRG)
WT Galectin-8
CML1
PhoSL001
Epa1-Np mutant Y228W
Epa1-Np mutant Y228A
Epa1-Np mutant E227G
Epa1-Np mutant E227A
Epa1-Np WT
ChMBL
DBL-1
hJAA-F11-B11
Gal(α1-3)[Fuc(α1-2)]Gal(β1-3)GlcNAc(β1-3)GalNAc(-Sp14

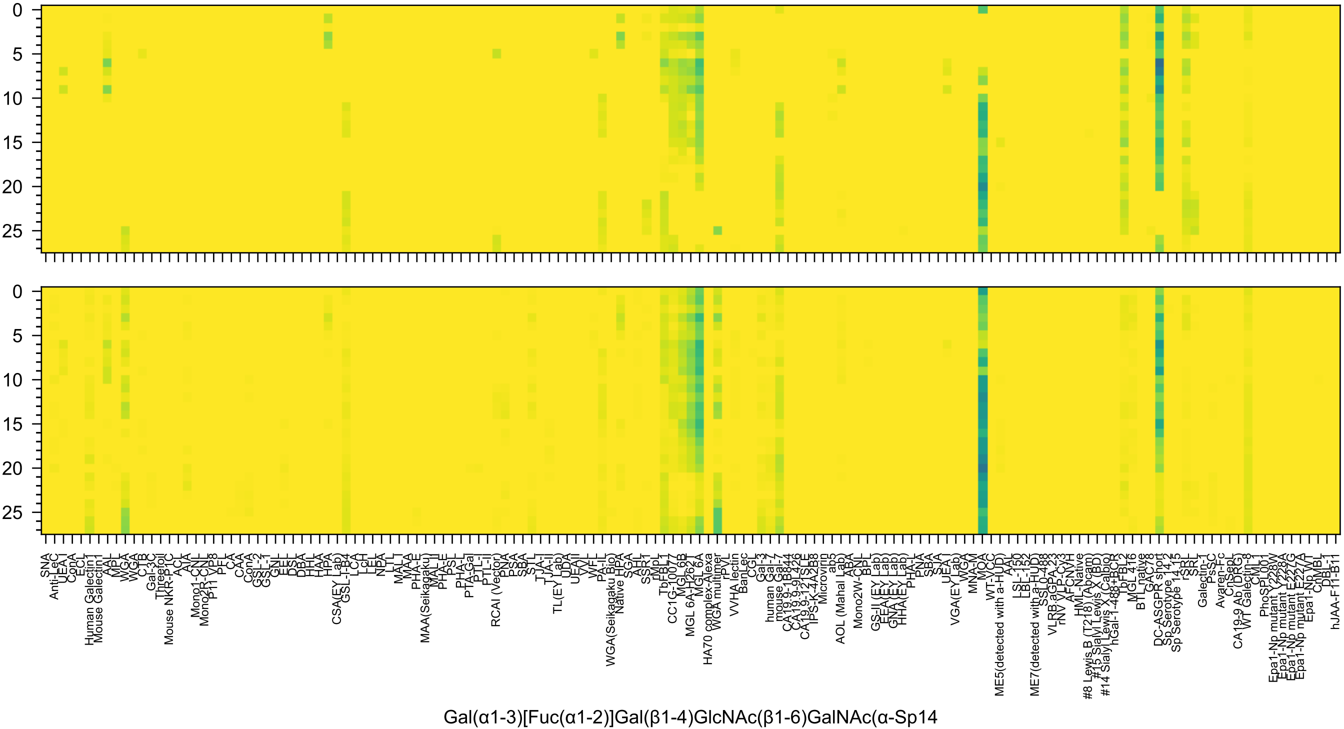

0
5
10
15
20
25
SNA
Anti-LeC
UEA I
ConA
ECL
Human Galectin1
Mouse Galectin1
AAL
MPL
WGA
WGA
CTB
Gal-3C
Threefoil
Mouse NKR-P1C
ACL
AIA
Mono1-CNL
Mono2R-CNL
P11 VP8
PFL
CA
CAA
ConA
GSL-2
GSL-1
GNL
EEL
DSL
DBA
HHL
HAA
HPA
CSA(EY Lab)
GSL-I-B4
LCA
LcH
LEL
NPA
LTL
MAL I
MAA
PHA-E
MAA(Seikagaku)
MAL II
PHA-E
PSL
PHA-L
PTA-Gal
PTL-I
PTL-II
RCAI (Vector)
PNA
PSA
SBA
STL
TJA-I
TJA-II
TL(EY Lab)
UDA
UEAII
VVL
WFL
PA-IL
WGA(Seikagaku Bio)
Native HPA
SGA
AHL
SSR1
rMpL
TbFBL1
CC1G-10077
MGL 6B
MGL 6A-H262T
MGL 6A
HA70 complex-Alexa
WGA multimer
rPVL
VVHA lectin
BanLec
CGL
Gal-3
human Gal-7
mouse Gal-7
CA19.9-1B844
CA19.9-9L426
CA19.9-121SLE
IPS-K-4A2B8
Microvirin
ab5
AOL (Mahal Lab)
ABA
Mono2W-CNL
BPL
GS-II (EY Lab)
EEA (EY Lab)
GNA (EY Lab)
HHA (EY Lab)
PHA-L
PNA
SBA
SJA
UEA I
VGA(EY Lab)
WGA
MNA-M
MOA
WT-VCC
ME5(detected with a-HUD)
AAL
LSL-150
LBL-152
ME7(detected with a-HUD)
SSL0-488
VLRB.aGPA.23
rNV VLP-Cy3
AFCNVH
HML-Native
#8 Lewis B (T218) (Abcam)
#15 Sialyl Lewis X (BD)
#14 Sialyl Lewis X (Calbio)
hGal-1-488+BCR
TpFBL2
MGL 416
BTL native
GAC78
DC-ASGPR short
Sp Serotype 14.2
Sp Serotype 14.15
rSRL
SRL
Galectin-1
PsSC
Avaren-Fc
CnSepL
CA19-9 Ab (DRG)
WT Galectin-8
CML1
PhoSPL001
Epa1-Np mutant Y228W
Epa1-Np mutant Y228A
Epa1-Np mutant E227G
Epa1-Np mutant E227A
Epa1-Np WT
ChMBL
DBL-1
hJAA-F11-B11
Gal(α1-3)[Fuc(α1-2)]Gal(β1-4)GlcNAc(β1-6)GalNAc(α-Sp14

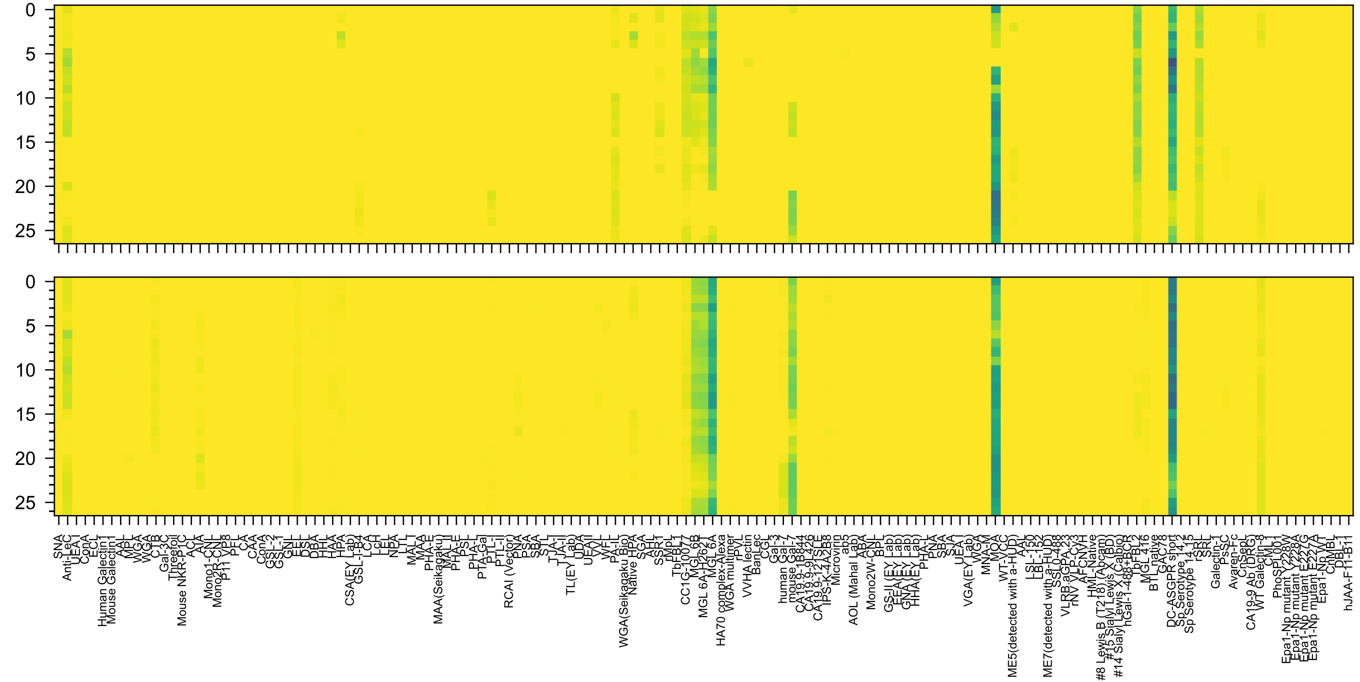


Gal(α1-3)[Fuc(α1-2)]Gal(β1-3)GlcNAc(β1-6)GalNAc(-Sp14

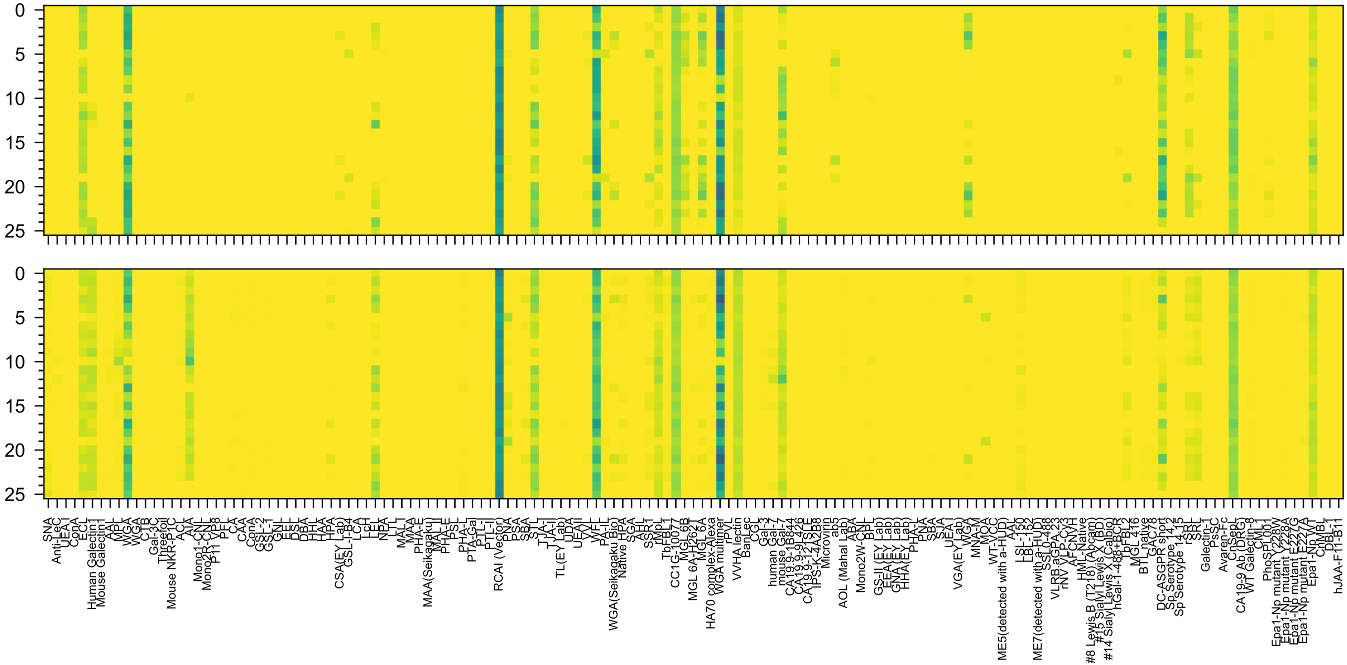

Gal(β1-4)GlcNAc(β1-6)[Gal(β1-4)GlcNAc(β1-3)]GalNAc(α-Sp8

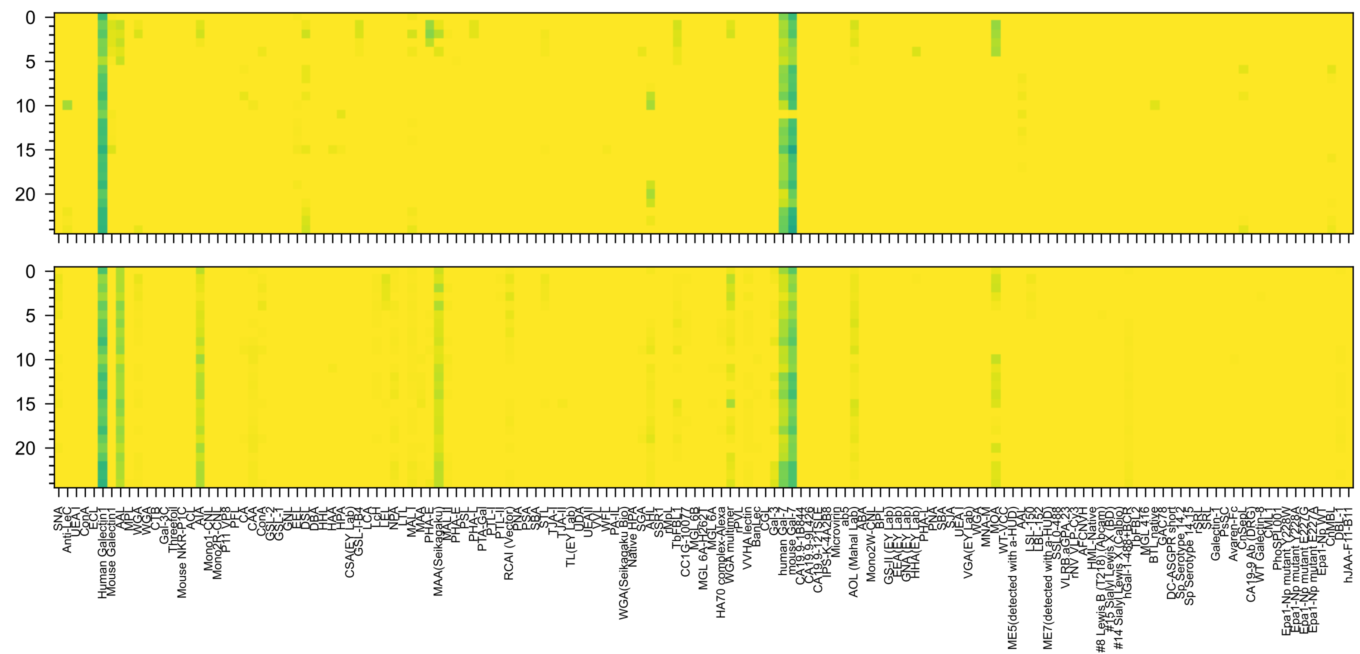

GlcA3OS(β1-3)Gal(β1-4)GlcNAc(β1-3)Gal(β1-4)Glc(-Sp21

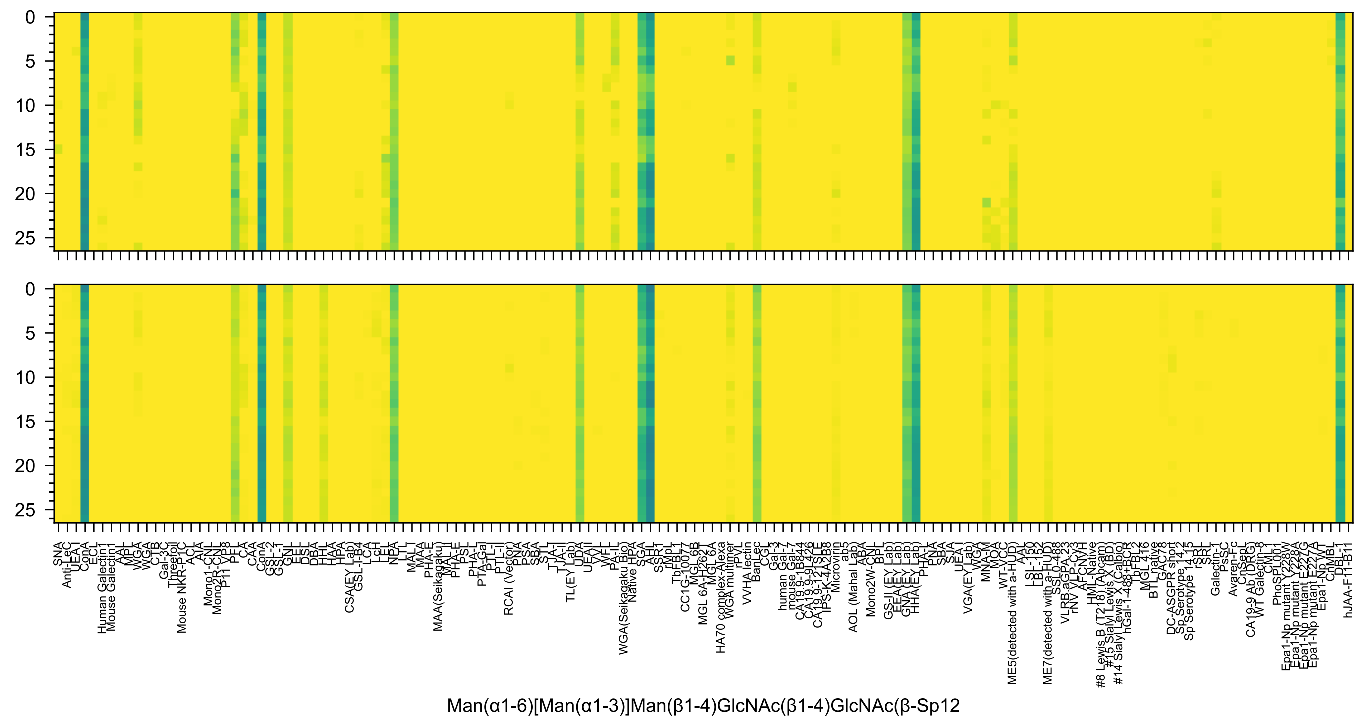
Man(α1-6)[Man(α1-3)]Man(β1-4)GlcNAc(β1-4)GlcNAc(β-Sp12

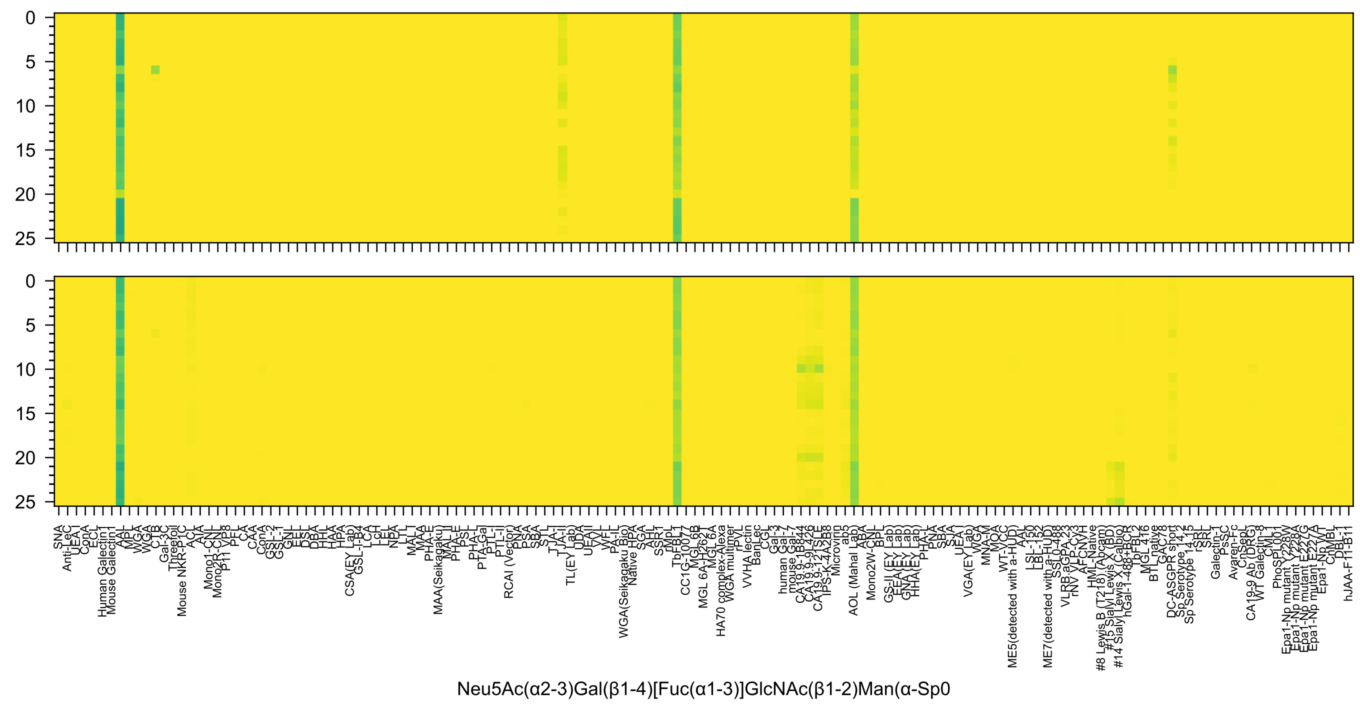
Neu5Ac(α2-3)Gal(β1-4)[Fuc(α1-3)]GlcNAc(β1-2)Man(α-Sp0

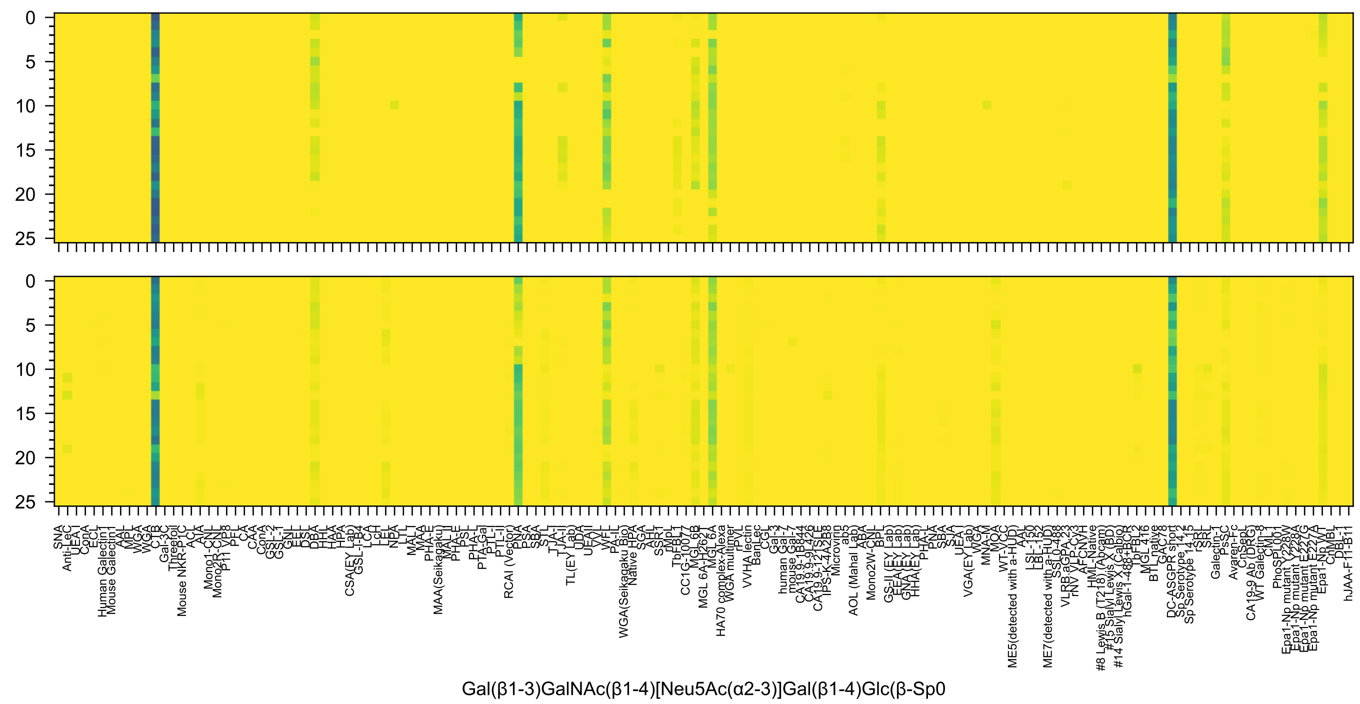

0
5
10
15
20
25
SNA
Anti-LeC
UEA-I
ConA
ECL
Human Galectin1
Mouse Galectin1
AAL
MPL
WGA
WGA
CTB
Gal-3C
Threefoil
Mouse NKR-P1C
ACL
AIA
Mono1-CNL
Mono2R-CNL
P11 VP8
PFL
CA
CAA
ConA
GSL-2
GSL-1
GNL
EEL
DSL
DBA
HHL
HAA
HPA
CSA(EY Lab)
GSL-I-B4
LCA
LcH
LEL
NPA
LTL
MAL-I
MAA
PHA-E
MAA(Seikagaku)
MAL-II
PHA-E
PSL
PHA-L
PTA-Gal
PTL-I
PTL-II
RCAI (Vector)
PNA
PSA
SBA
STL
TJA-I
TJA-II
TL(EY Lab)
UDA
UEAII
VVL
WFL
PA-IL
WGA(Seikagaku Bio)
Native HPA
SGA
AHL
SSR1
rMpL
TbFBL1
CC1G-10077
MGL 6B
MGL 6A-H262T
MGL 6A
HA70 complex-Alexa
WGA multimer
rPVL
VVHA lectin
BanLec
CGL
Gal-3
human Gal-7
mouse Gal-7
CA19.9-1B844
CA19.9-9L426
CA19.9-121SLE
IPS-K-4A2B8
Microvirin
ab5
AOL (Mahal Lab)
ABA
Mono2W-CNL
BPL
GS-II (EY Lab)
EEA (EY Lab)
GNA (EY Lab)
HHA (EY Lab)
PHA-L
PNA
SBA
SJA
UEA-I
VGA(EY Lab)
WGA
MNA-M
MOA
WT-VCC
ME5(detected with a-HUD)
AAL
LSL-150
LBL-152
ME7(detected with a-HUD)
SSL0-488
VLRB.aGPA.23
rNV VLP-Cy3
AFCNVH
HML-Native
#8 Lewis B (T218) (Abcam)
#15 Sialyl Lewis X (BD)
#14 Sialyl Lewis X (Calbio)
hGal-1-488+BCR
TbFBL2
MGL 416
BTL native
GAC78
DC-ASGPR short
Sp Serotype 14.12
Sp Serotype 14.15
rSRL
SRL
Galectin-1
PsSC
Avaren-Fc
ChSepL
CA19-9 Ab (DRG)
WT Galectin-8
CML1
PhoSPL001
Epa1-Np mutant Y228W
Epa1-Np mutant Y228A
Epa1-Np mutant E227G
Epa1-Np mutant E227A
Epa1-Np WT
ChMBL
DBL-1
hJAA-F11-B11
Gal(β1-3)GalNAc(β1-4)[Neu5Ac(α2-3)]Gal(β1-4)Glc(β-Sp0

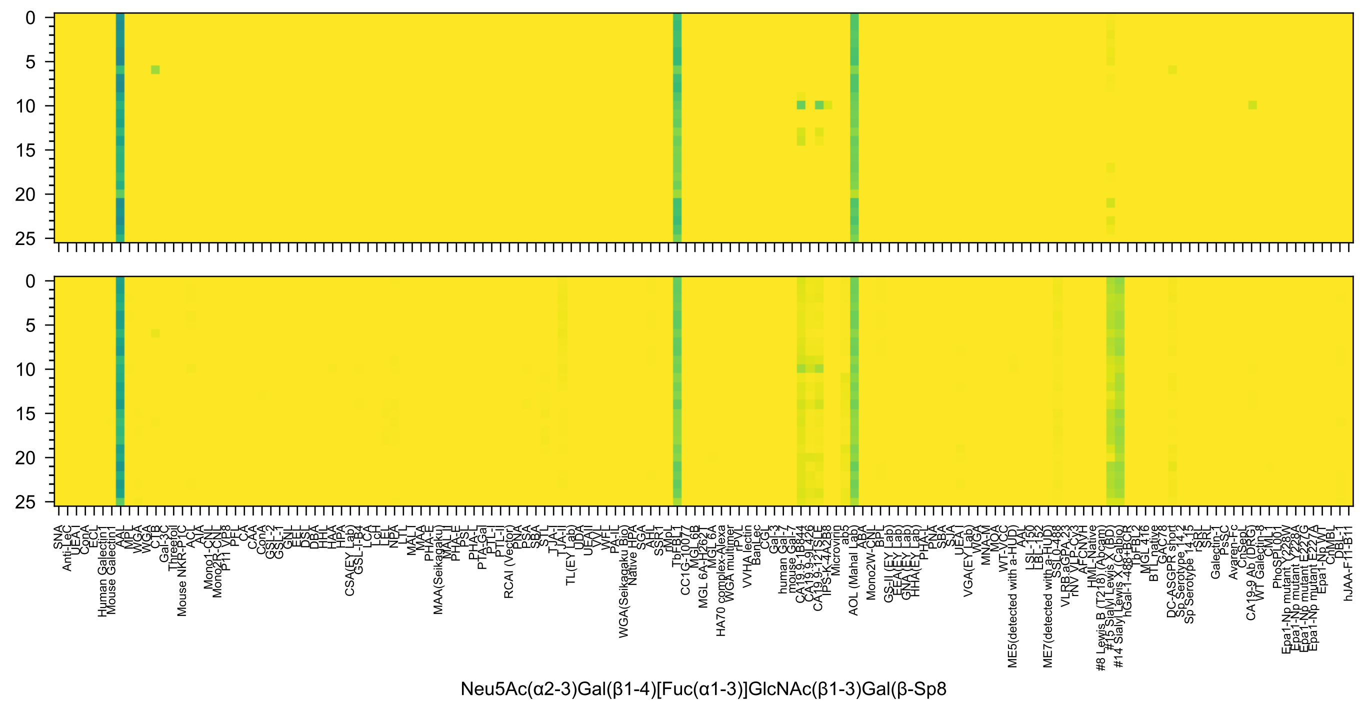
Neu5Ac(α2-3)Gal(β1-4)[Fuc(α1-3)]GlcNAc(β1-3)Gal(β-Sp8

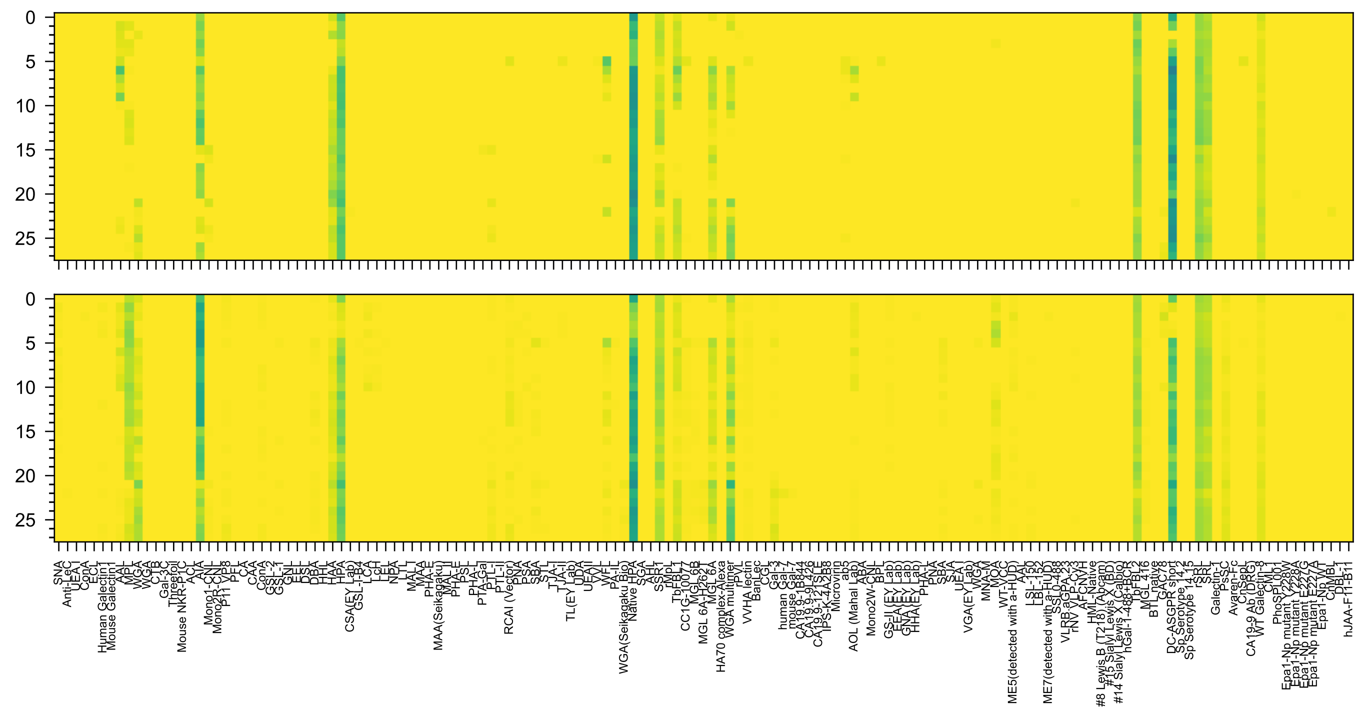


GalNAc(α1-3)[Fuc(α1-2)]Gal(β1-4)GlcNAc(β1-3)GalNAc(α-Sp14

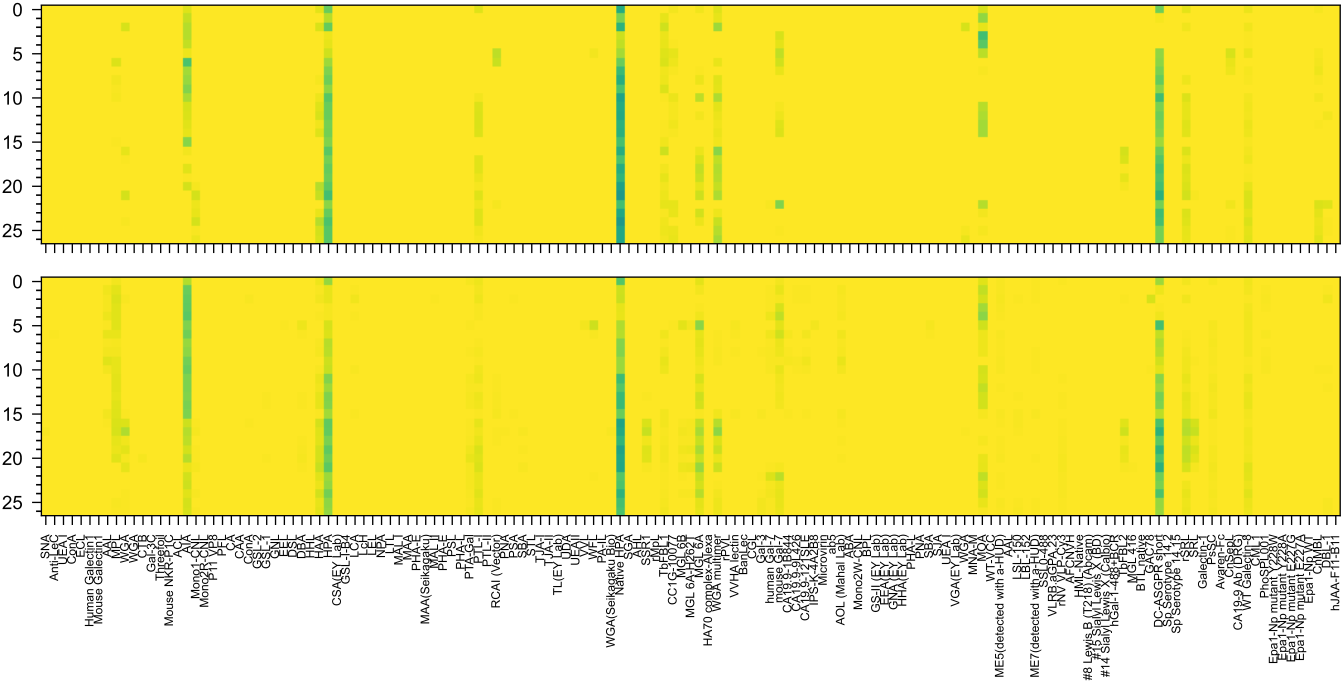

GalNAc(α1-3)[Fuc(α1-2)]Gal(β1-3)GlcNAc(β1-3)GalNAc(-Sp14

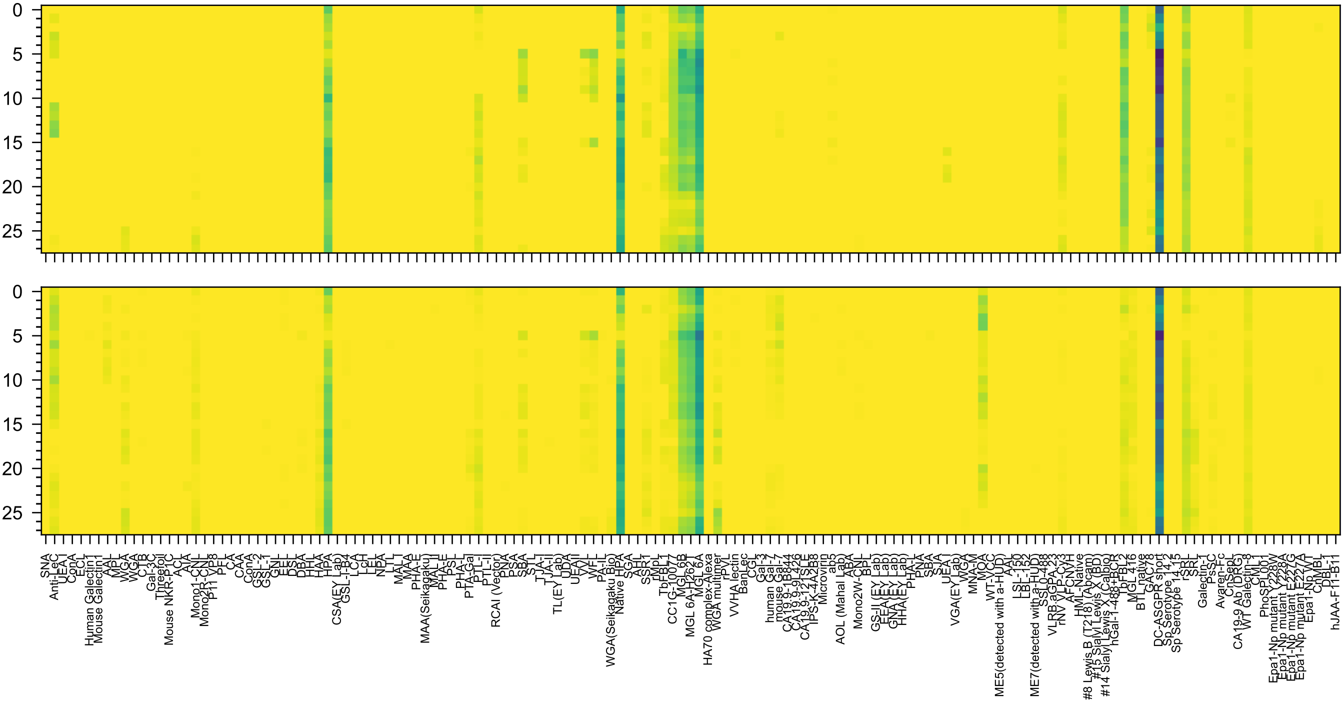

GalNAc(α1-3)[Fuc(α1-2)]Gal(β1-3)GlcNAc(β1-6)GalNAc(α-Sp14

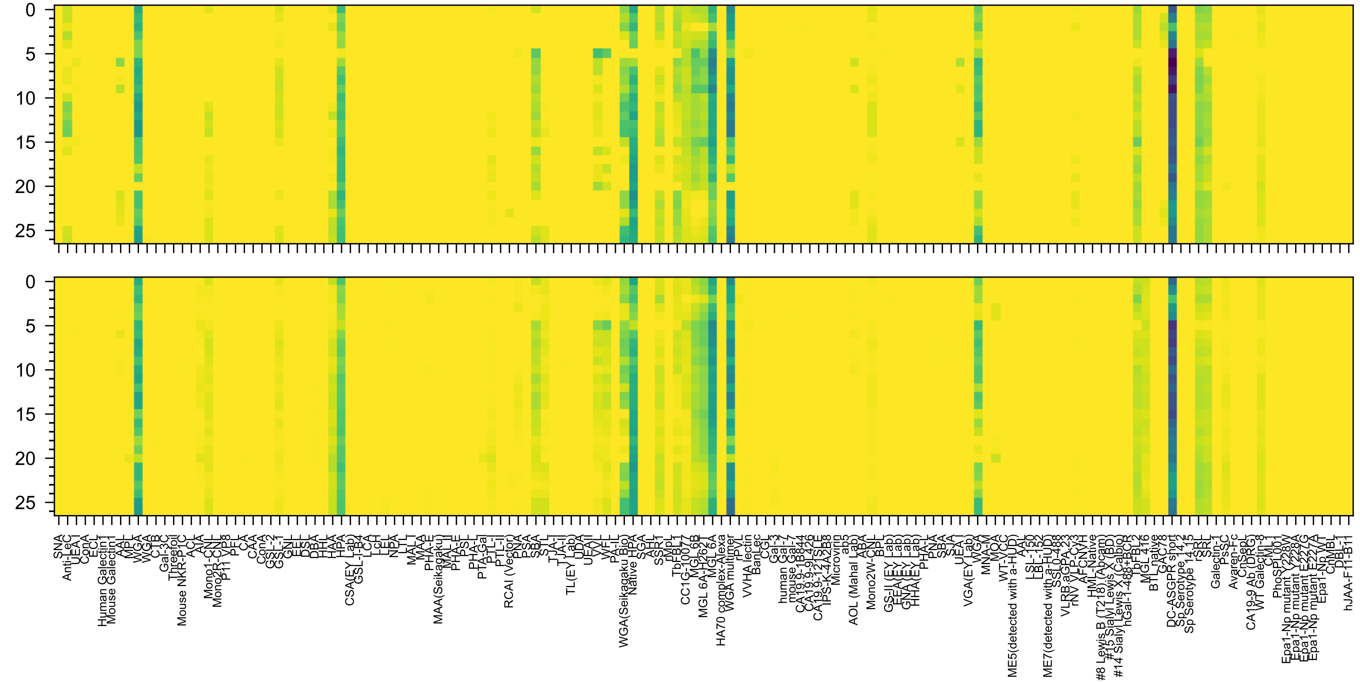

GalNAc(α1-3)[Fuc(α1-2)]Gal(β1-4)GlcNAc(β1-6)GalNAc(-Sp14

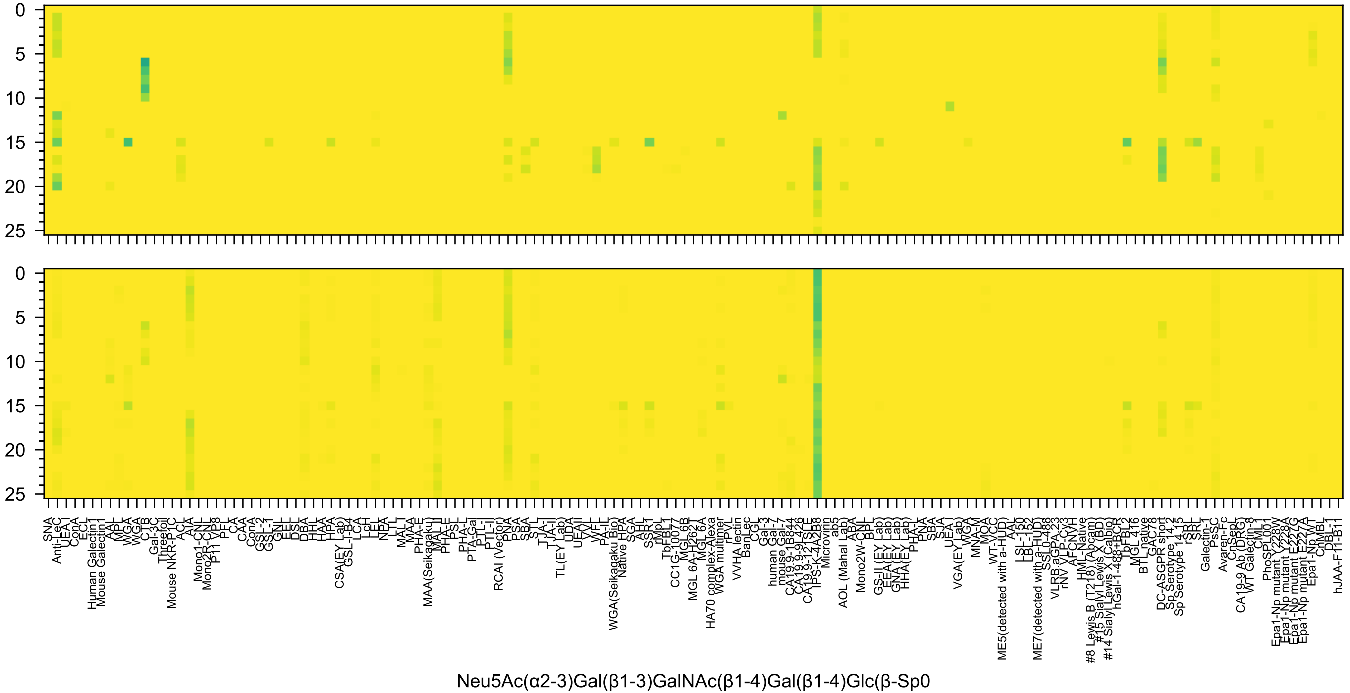

0
5
10
15
20
25
SNA
Anti-LeC
UEA I
ConA
ECL
Human Galectin1
Mouse Galectin1
AAL
MPL
WGA
WGA
CTB
Gal-3C
Threefoil
Mouse NKR-P1C
ACL
AIA
Mono1-CNL
Mono2R-CNL
P11 VP8
PFL
CA
CAA
ConA
GSL-2
GSL-1
GNL
EEL
DSL
DBA
HHL
HAA
HPA
CSA(EY Lab)
GSL-I-B4
LCA
LcH
LEL
NPA
LTL
MAL I
MAA
PHA-E
MAA(Seikagaku)
MAL I
PHA-E
PSL
PHA-L
PTA-Gal
PTL-I
PTL-II
RCAI (Vector)
PNA
PSA
SBA
STL
TJA-I
TJA-II
TL(EY Lab)
UDA
UEA II
VVL
WFL
PA-IL
WGA(Seikagaku Bio)
Native HPA
SGA
AHL
SSR1
rMpL
TbFBL1
CC1G-10077
MGL 6B
MGL 6A-H262T
MGL 6A
HA70 complex-Alexa
WGA multimer
rPVL
VVHA lectin
BanLec
CGL
Gal-3
human Gal-7
mouse Gal-7
CA19.9-1B844
CA19.9-9L426
CA19.9-121SLE
IPS-K-4A2B8
Microvirin
ab5
AOL (Mahal Lab)
ABA
Mono2W-CNL
BPL
GS-II (EY Lab)
EEA (EY Lab)
GNA (EY Lab)
HHA (EY Lab)
PHA-L
PNA
SBA
SJA
UEA I
VGA(EY Lab)
WGA
MNA-M
MOA
WT-VCC
ME5(detected with a-HUD)
AAL
LSL-150
LBL-152
ME7(detected with a-HUD)
SSL0-488
VLRB.aGPA.23
rNV VLP-Cy3
AFCNVH
HML-Native
#8 Lewis B (T218) (Abcam)
#15 Sialyl Lewis X (BD)
#14 Sialyl Lewis X (Calbio)
hGal-1-488+BCR
TbFBL2
MGL 416
BTL native
GAC78
DC-ASGPR short
Sp Serotype 14.2
Sp Serotype 14.15
rSRL
SRL
Galectin-1
PsSC
Avaren-Fc
CnSepL
CA19-9 Ab (DRG)
WT Galectin-8
CML1
PhoSPL001
Epa1-Np mutant Y228W
Epa1-Np mutant Y228A
Epa1-Np mutant E227G
Epa1-Np mutant E227A
Epa1-Np WT
CnMBL
DBL-1
hJAA-F11-B11
Neu5Ac(α2-3)Gal(β1-3)GalNAc(β1-4)Gal(β1-4)Glc(β-Sp0

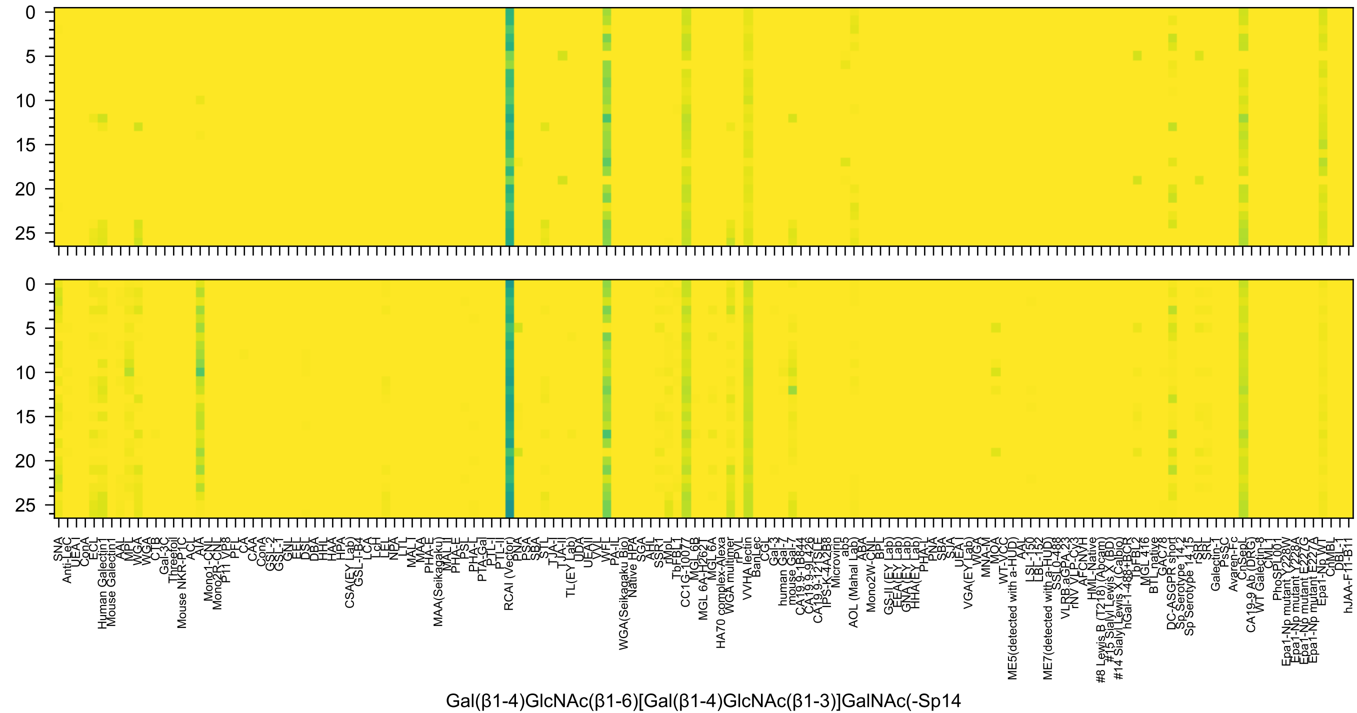
0
5
10
15
20
25
SNA
Anti-LeC
UEA-I
ConA
ECL
Human Galectin1
Mouse Galectin1
AAL
MPL
WGA
WGA
CTB
Gal-3C
Threefoil
Mouse NKR-P1C
ACL
AIA
Mono1-CNL
Mono2R-CNL
P11 VP8
PFL
CA
CAA
ConA
GSL-2
GSL-1
GNL
EEL
DSL
DBA
HHL
HAA
HPA
CSA(EY Lab)
GSL-I-B4
LCA
LcH
LEL
NPA
LTL
MAL-I
MAA
PHA-E
MAA(Seikagaku)
MAL-II
PHA-E
PSL
PHA-L
PTA-Gal
PTL-I
PTL-II
RCAI (Vector)
PNA
PSA
SBA
STL
TJA-I
TJA-II
TL(EY Lab)
UDA
UEAII
VVL
WFL
PA-IL
WGA(Seikagaku Bio)
Native HPA
SGA
AHL
SSR1
rMpL
TbFBL1
CC1G-10077
MGL6B
MGL 6A-H262T
MGL 6A
HA70 complex-Alexa
WGA multimer
rPVL
VVHA lectin
BanLec
CGL
Gal-3
human Gal-7
mouse Gal-7
CA19.9-1B844
CA19.9-9L426
CA19.9-121SLE
IPS-K-4A2B8
Microvirin
ab5
AOL (Mahal Lab)
ABA
Mono2W-CNL
BPL
GS-II (EY Lab)
EEA(EY Lab)
GNA(EY Lab)
HHA(EY Lab)
PHA-L
PNA
SBA
SJA
UEA-I
VGA(EY Lab)
WGA
MNA-M
MOA
WT-VCC
ME5(detected with a-HUD)
AAL
LSL-150
LBL-152
ME7(detected with a-HUD)
SSL0-488
VLRB.aGPA.23
rNV VLP-Cy3
AFCNVH
HML-Native
#8 Lewis B (T218) (Abcam)
#15 Sialyl Lewis X (BD)
#14 Sialyl Lewis X (Calbio)
hGal-1-488+BCR
TbFBL2
MGL 416
BTL native
GAC78
DC-ASGPR short
Sp Serotype 14.2
Sp Serotype 14.15
rSRL
SRL
Galectin-1
PsSC
Avaren-Fc
ChSepL
CA19-9 Ab (DRG)
WT Galectin-8
CML1
PhoSL001
Epa1-Np mutant Y228W
Epa1-Np mutant Y228A
Epa1-Np mutant E227G
Epa1-Np mutant E227A
Epa1-Np WT
ChMBL
DBL-1
hJAA-F11-B11
Gal(β1-4)GlcNAc(β1-6)[Gal(β1-4)GlcNAc(β1-3)]GalNAc(-Sp14

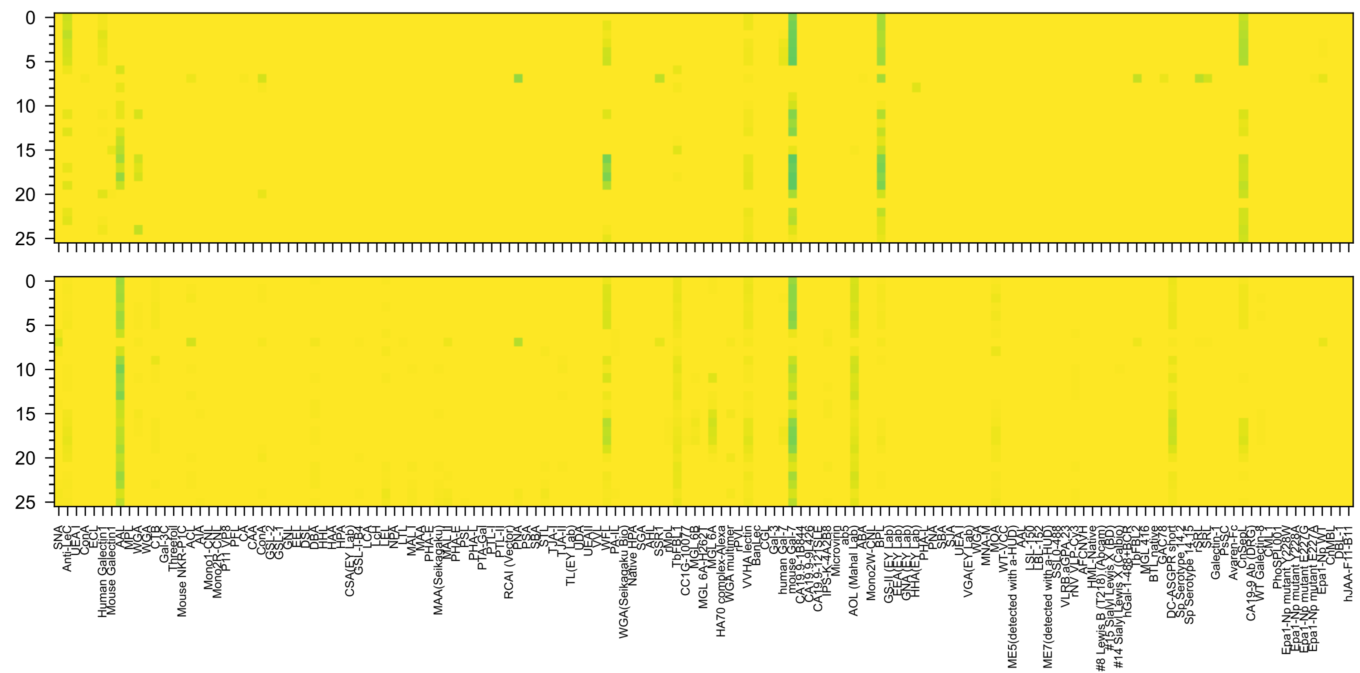

Neu5Ac(α2-6)[Gal(β1-3)]GlcNAc(β1-4)Gal(β1-4)Glc(β-Sp10

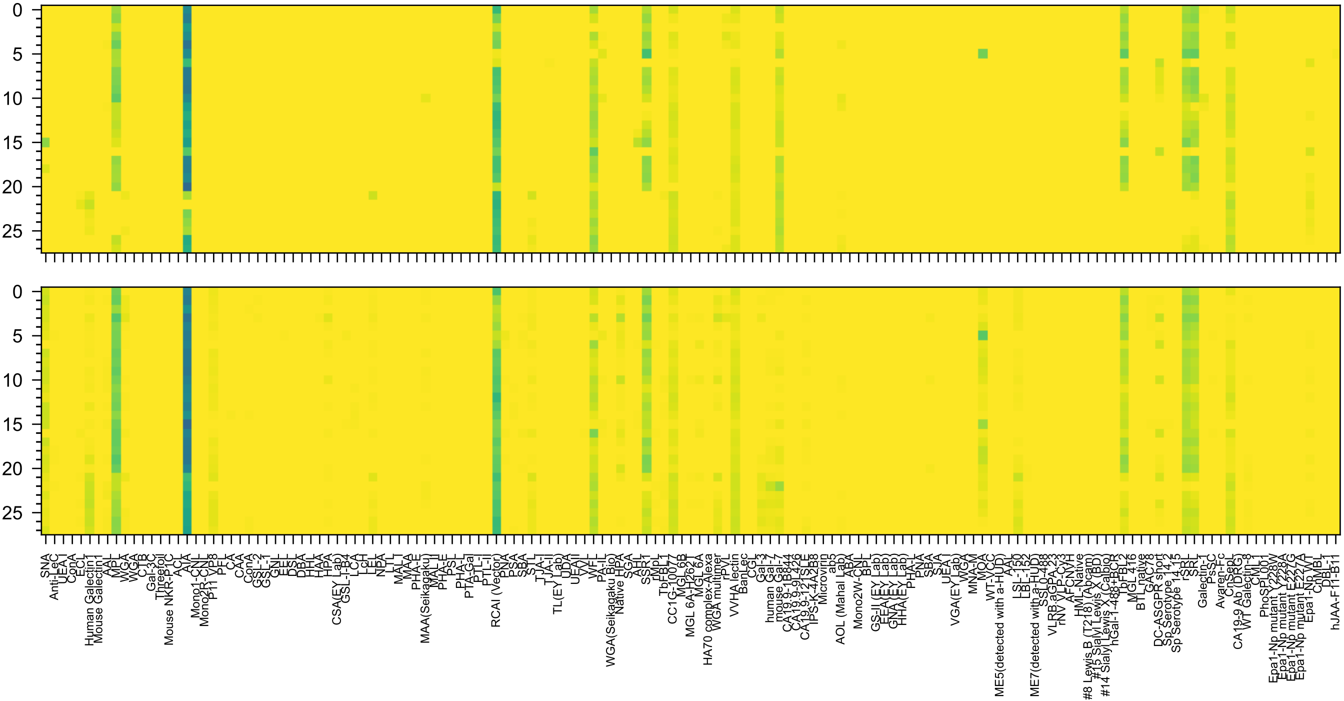
Gal(β1-4)GlcNAc(β1-3)Gal(β1-4)GlcNAc(β1-3)GalNAc(α-Sp14

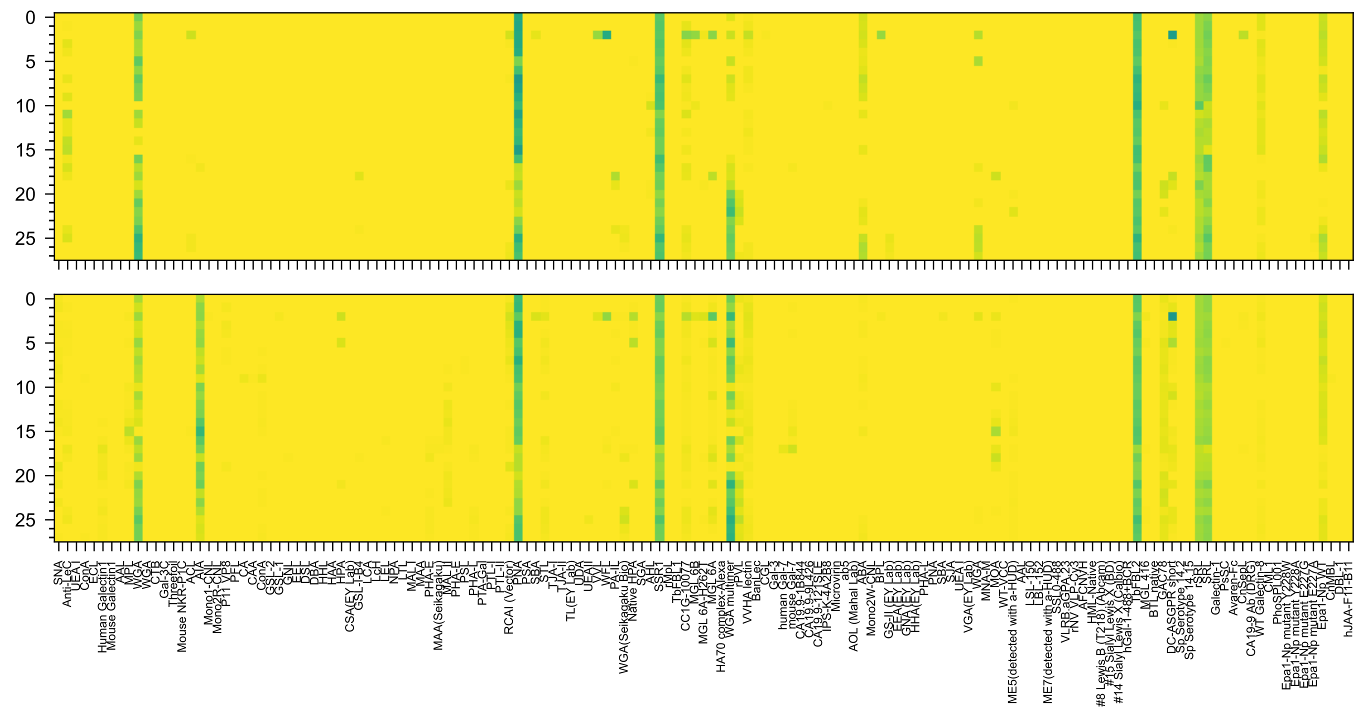

GlcNAc(β1-3)Gal(β1-4)GlcNAc(β1-6)[Gal(β1-3)]GalNAc(α-Sp14

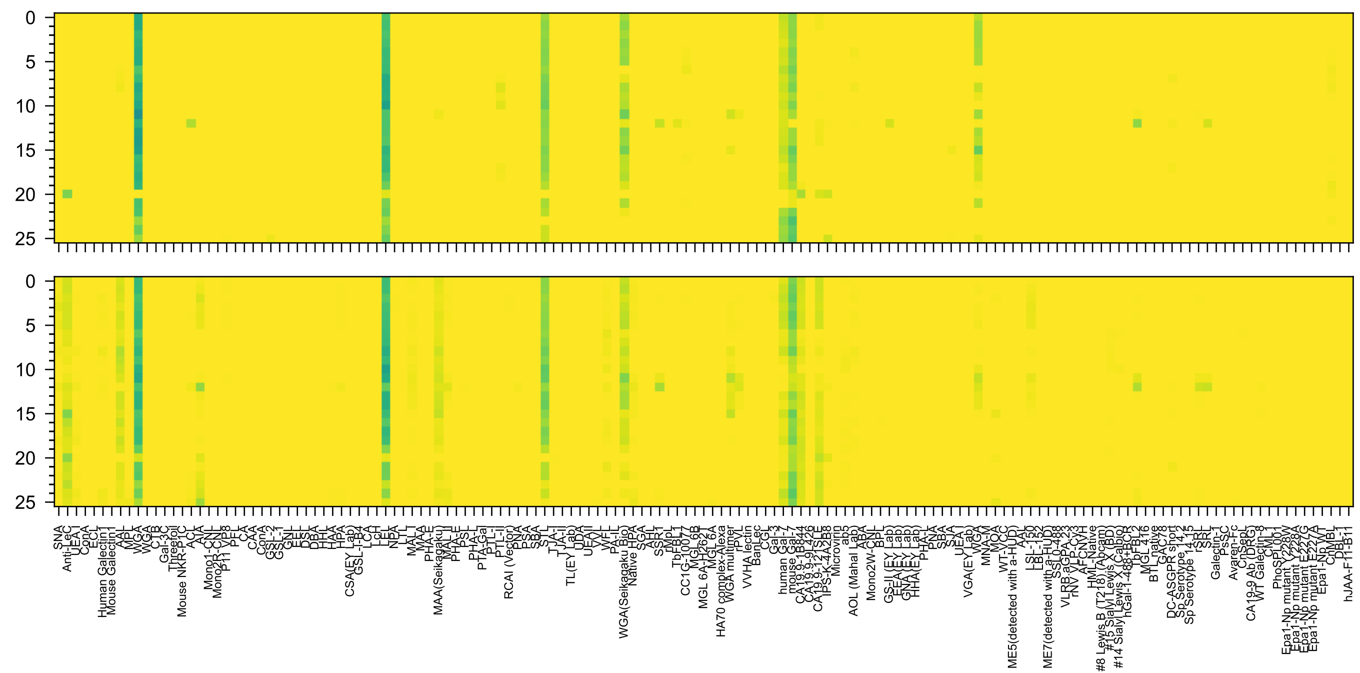

Neu5Ac(α2-3)Gal(β1-3)GlcNAc(β1-3)Gal(β1-4)GlcNAc(β-Sp0

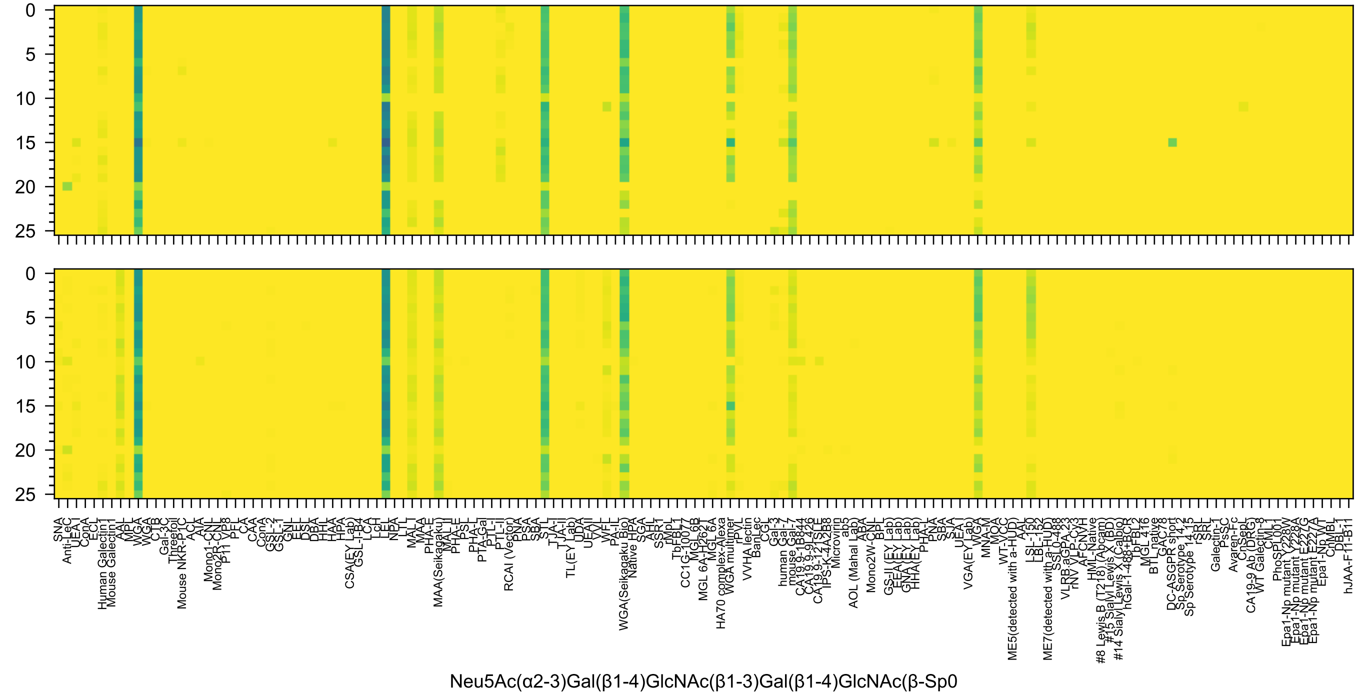
0
5
10
15
20
25
SNA
Anti-LeC
UEA I
ConA
ECL
Human Galectin1
Mouse Galectin1
AAL
MPL
WGA
WGA
CTB
Gal-3C
Threefoil
Mouse NKR-P1C
ACL
AIA
Mono1-CNL
Mono2R-CNL
P11 VP8
PFL
CA
CAA
ConA
GSL-2
GSL-1
GNL
EEL
DSL
DBA
HHL
HAA
HPA
CSA(EY Lab)
GSL-I-B4
LCA
LcH
LEL
NPA
LTL
MAL-I
MAA
PHA-E
MAA(Seikagaku)
MAL-II
PHA-E
PSL
PHA-L
PTA-Gal
PTL-I
PTL-II
RCAI (Vector)
PNA
PSA
SBA
STL
TJA-I
TJA-II
TL(EY Lab)
UDA
UEA-II
VVL
WFL
PA-IL
WGA(Seikagaku Bio)
Native HPA
SGA
AAL
SSR1
rMpL
TbFBL1
CC1G-10077
MGL 6B
MGL 6A-H262T
MGL 6A
HA70 complex-Alexa
WGA multimer
rPVL
VVHA lectin
BanLec
CGL
Gal-3
human Gal-7
mouse Gal-7
CA19.9-1B844
CA19.9-9L426
CA19.9-121SLE
IPS-K-4A2B8
Microvirin
ab5
AOL (Mahal Lab)
ABA
Mono2W-CNL
BPL
GS-II (EY Lab)
EEA (EY Lab)
GNA (EY Lab)
HHA(EY Lab)
PHA-L
PNA
SBA
SJA
UEA-I
VGA(EY Lab)
WGA
MNA-M
MOA
WT-VCC
ME5(detected with a-HUD)
AAL
LSL-150
LBL-152
ME7(detected with a-HUD)
SSL0-488
VLRB.aGPA.23
rNV VLP-Cy3
AFCNVH
HML-Native
#8 Lewis B (T218) (Abcam)
#15 Sialyl Lewis X (BD)
#14 Sialyl Lewis X (Calbio)
hGal-1-488+BCR
TbFBL2
MGL 416
BTL native
GAC78
DC-ASGPR short
Sp Serotype 14.2
Sp Serotype 14.15
rSRL
SRL
Galectin-1
PsSC
Avaren-Fc
ChSepL
CA19-9 Ab (DRG)
WT Galectin-8
CML1
PhoSL001
Epa1-Np mutant Y228W
Epa1-Np mutant Y228A
Epa1-Np mutant E227G
Epa1-Np mutant E227A
Epa1-Np WT
ChMBL
DBL-1
hJAA-F11-B11
Neu5Ac(α2-3)Gal(β1-4)GlcNAc(β1-3)Gal(β1-4)GlcNAc(β-Sp0

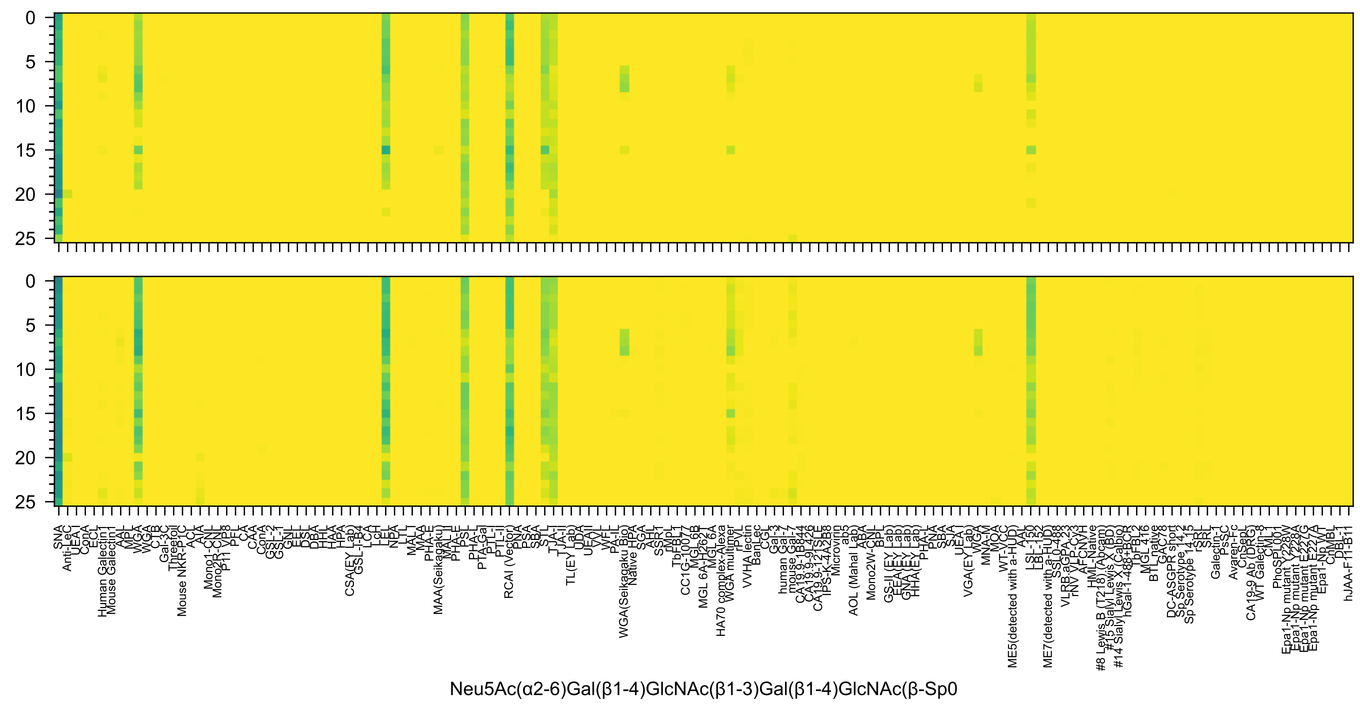
Neu5Ac(α2-6)Gal(β1-4)GlcNAc(β1-3)Gal(β1-4)GlcNAc(β-Sp0

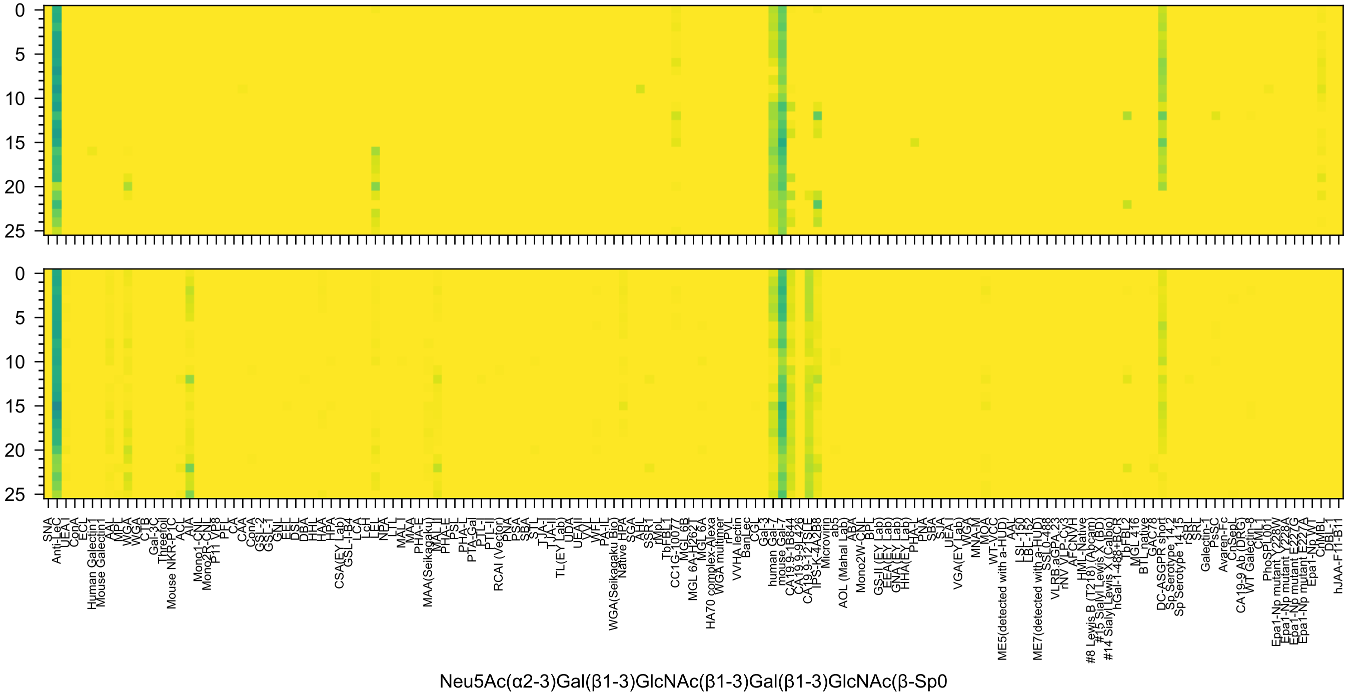
0
5
10
15
20
25
SNA
Anti-LeC
UEA-I
ConA
ECL
Human Galectin1
Mouse Galectin1
AAL
MPL
WGA
WGA
CTB
Gal-3C
Threefoil
Mouse NKR-P1C
ACL
AIA
Mono1-CNL
Mono2R-CNL
P11 VP8
PFL
CA
CAA
ConA
GSL-2
GSL-1
GNL
EEL
DSL
DBA
HHL
HAA
HPA
CSA(EY Lab)
GSL-I-B4
LCA
LcH
LEL
NPA
LTL
MAL-I
MAA
PHA-E
MAA(Seikagaku)
MAL-I
PHA-E
PSL
PHA-L
PTA-Gal
PTL-I
PTL-II
RCAI (Vector)
PNA
PSA
SBA
STL
TJA-I
TJA-II
TL(EY Lab)
UDA
UEAII
VVL
WFL
PA-IL
WGA(Seikagaku Bio)
Native HPA
SGA
AHL
SSR1
rMpL
TbFBL1
CC1G-10077
MGL 6B
MGL 6A-H262T
MGL 6A
HA70 complex-Alexa
WGA multimer
rPVL
VVHA lectin
BanLec
CGL
Gal-3
human Gal-7
mouse Gal-7
CA19.9-1B844
CA19.9-9L426
CA19.9-121SLE
IPS-K-4A2B8
Microvirin
ab5
AOL (Mahal Lab)
ABA
Mono2W-CNL
BPL
GS-II (EY Lab)
EEA (EY Lab)
GNA (EY Lab)
HHA(EY Lab)
PHA-L
PNA
SBA
SJA
UEA-I
VGA(EY Lab)
WGA
MNA-M
MOA
WT-VCC
ME5(detected with a-HUD)
AAL
LSL-150
LBL-152
ME7(detected with a-HUD)
SSL0-488
VLRB.aGPA.23
rNV VLP-Cy3
AFCNVH
HML-Native
#8 Lewis B (T218) (Abcam)
#15 Sialyl Lewis X (BD)
#14 Sialyl Lewis X (Calbio)
hGal-1-488+BCR
TbFBL2
MGL 416
BTL native
GAC78
DC-ASGPR short
Sp Serotype 14.2
Sp Serotype 14.15
rSRL
SRL
Galectin-1
PsSC
Avaren-Fc
ChSepL
CA19-9 Ab (DRG)
WT Galectin-8
CML
PhoSPL001
Epa1-Np mutant Y228W
Epa1-Np mutant Y228A
Epa1-Np mutant E227G
Epa1-Np mutant E227A
Epa1-Np WT
ChMBL
DBL-1
hJAA-F11-B11
Neu5Ac(α2-3)Gal(β1-3)GlcNAc(β1-3)Gal(β1-3)GlcNAc(β-Sp0

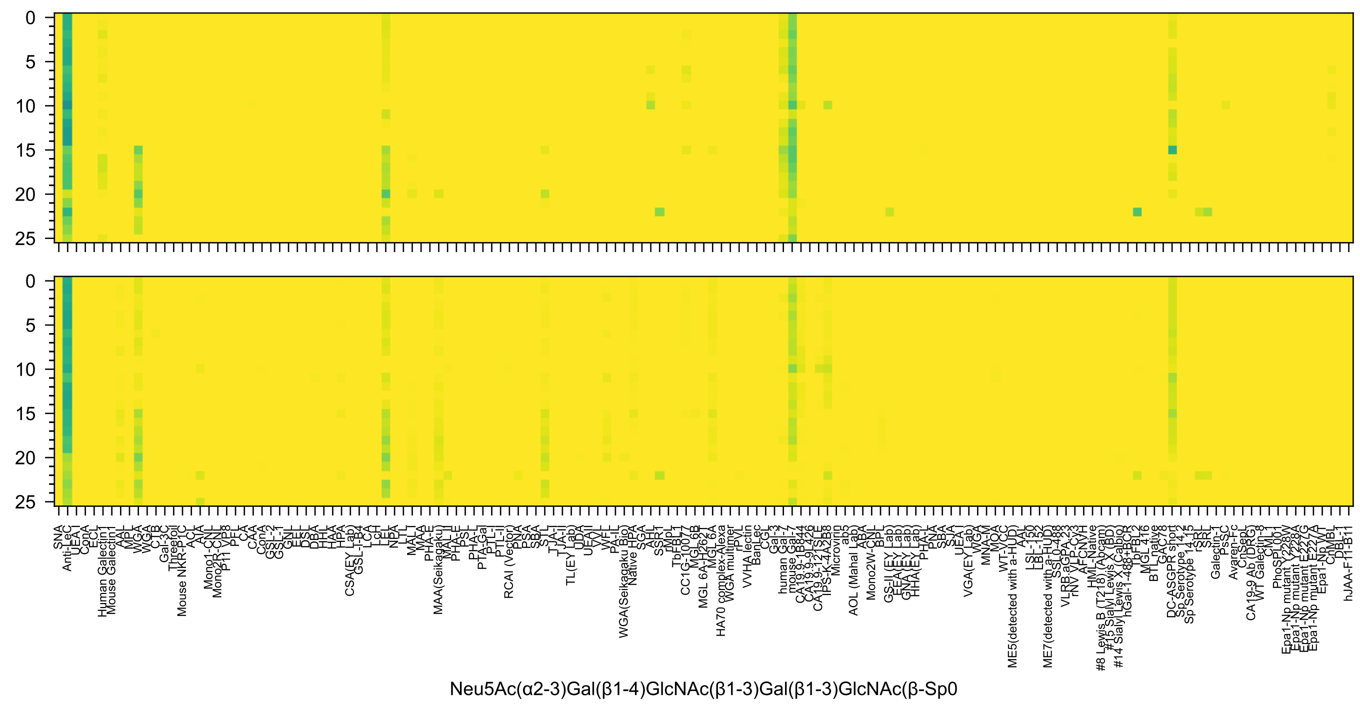
Neu5Ac(α2-3)Gal(β1-4)GlcNAc(β1-3)Gal(β1-3)GlcNAc(β-Sp0

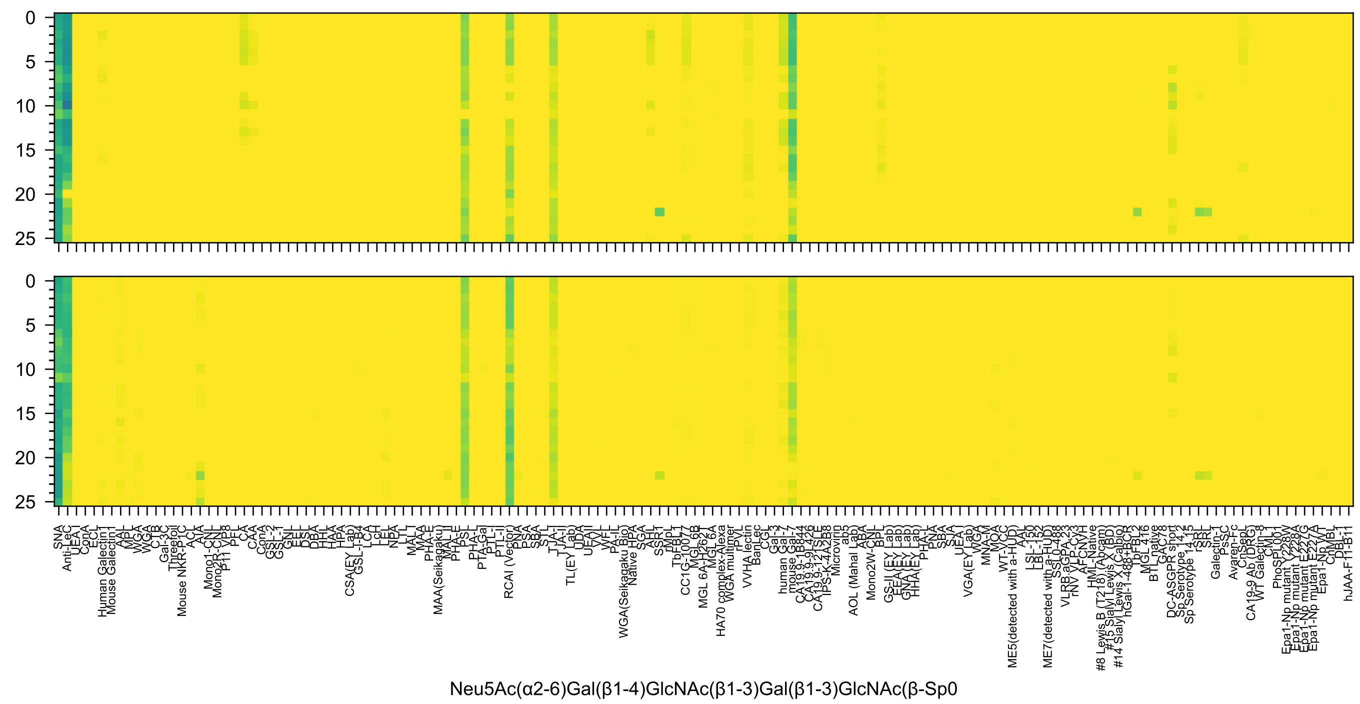
Neu5Ac(α2-6)Gal(β1-4)GlcNAc(β1-3)Gal(β1-3)GlcNAc(β-Sp0

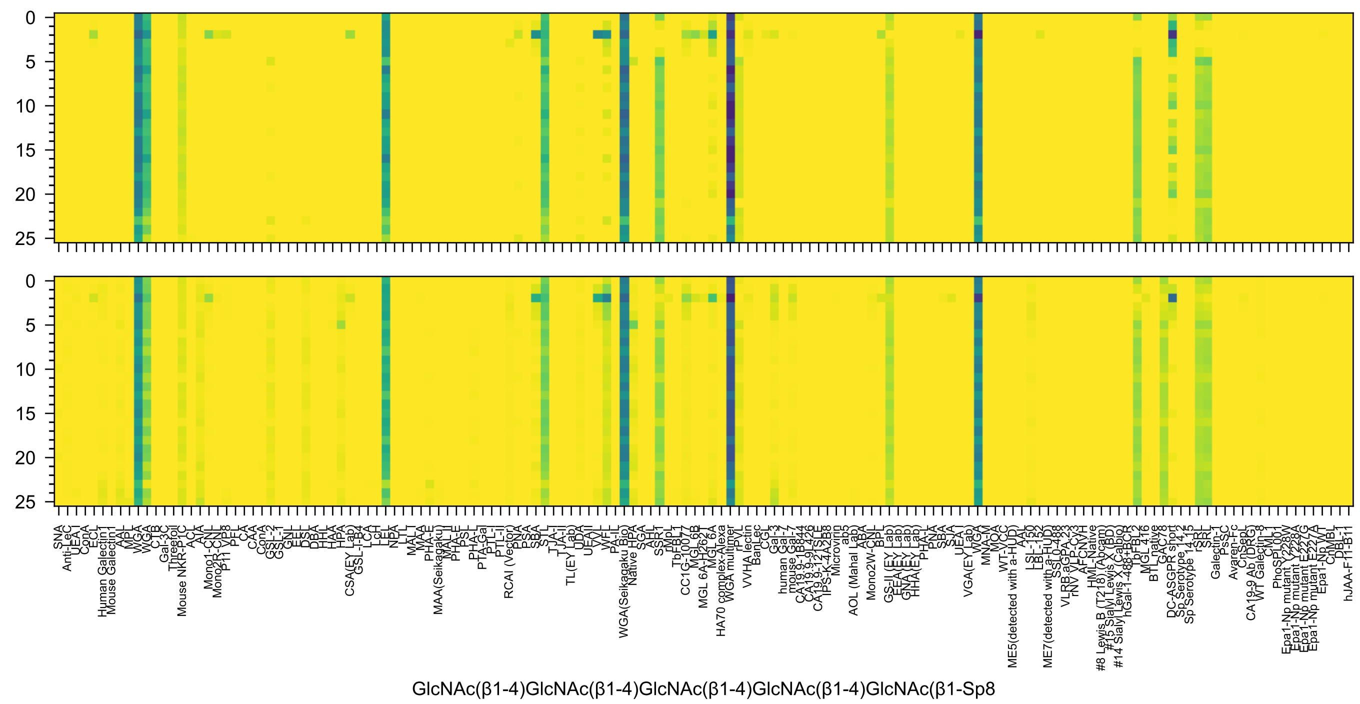
GlcNAc(β1-4)GlcNAc(β1-4)GlcNAc(β1-4)GlcNAc(β1-4)GlcNAc(β1-Sp8

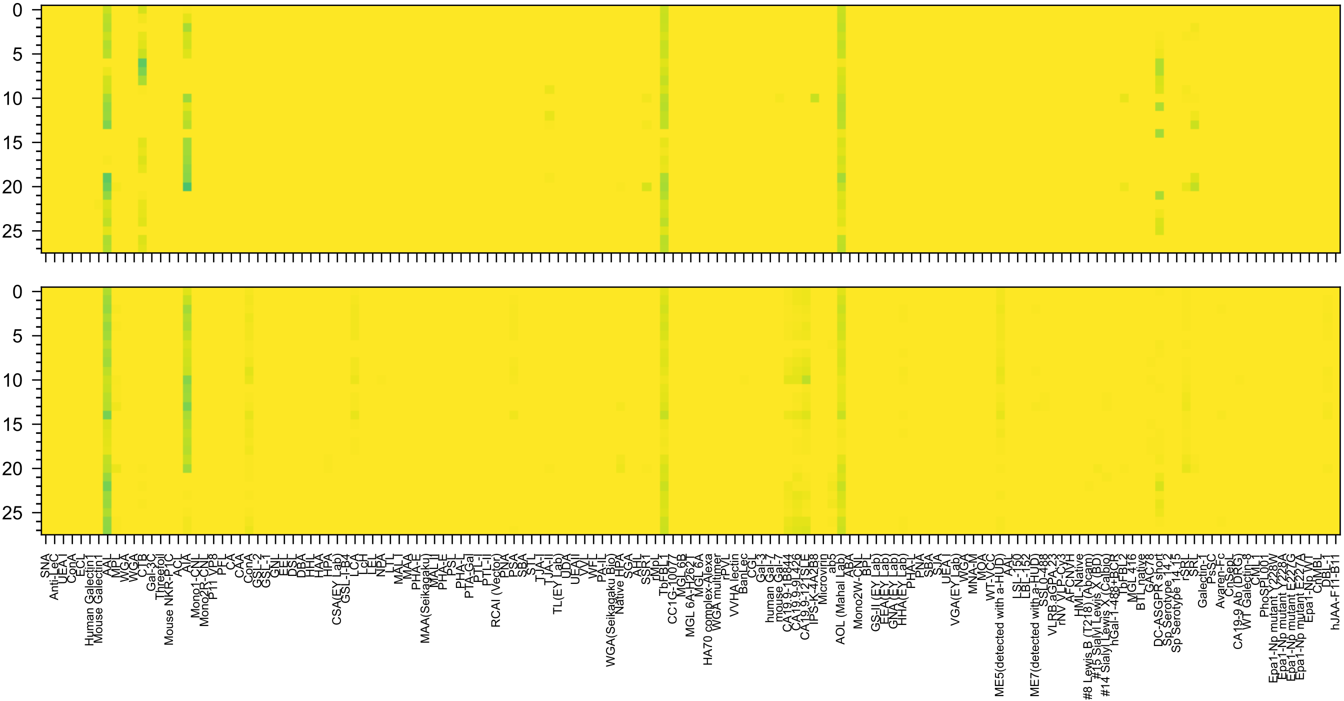

Neu5Ac(α2-3)Gal(β1-4)[Fuc(α1-3)]GlcNAc(β1-3)GalNAc(α-Sp14

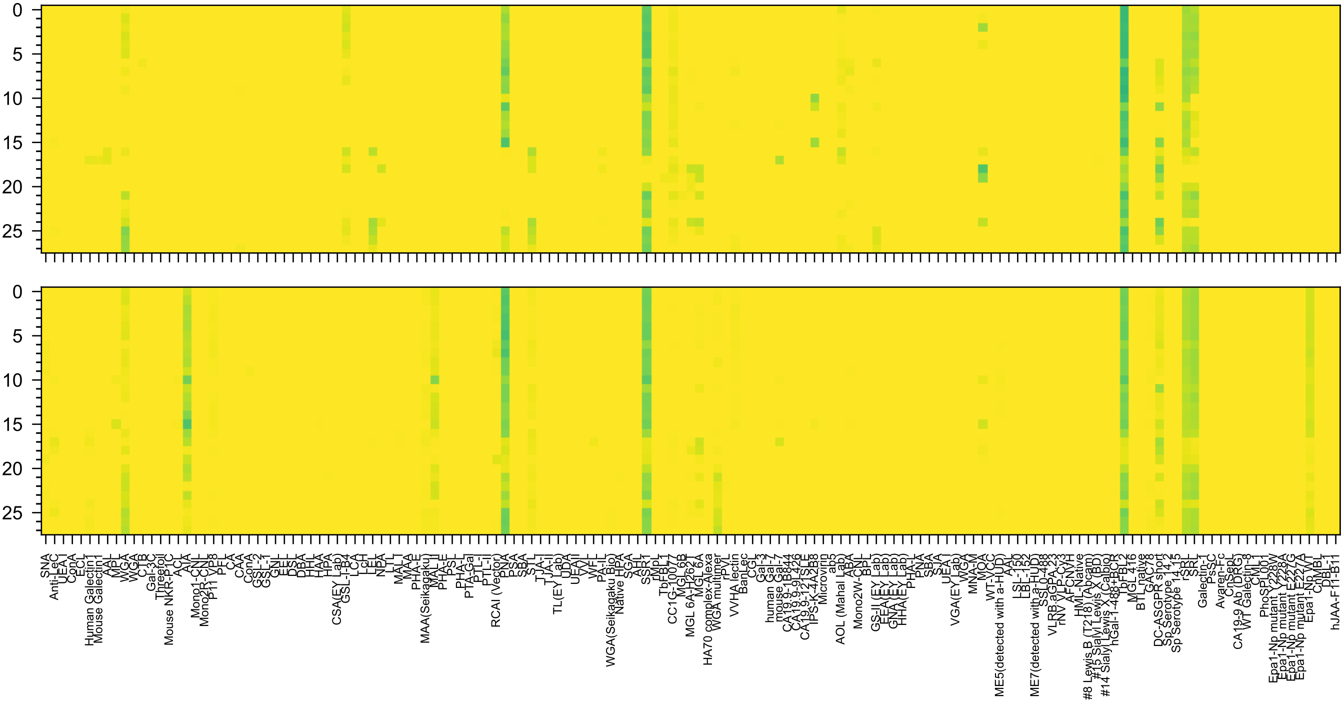

Neu5Ac(α2-3)Gal(β1-4)GlcNAc(β1-6)[Gal(β1-3)]GalNAc(α-Sp14

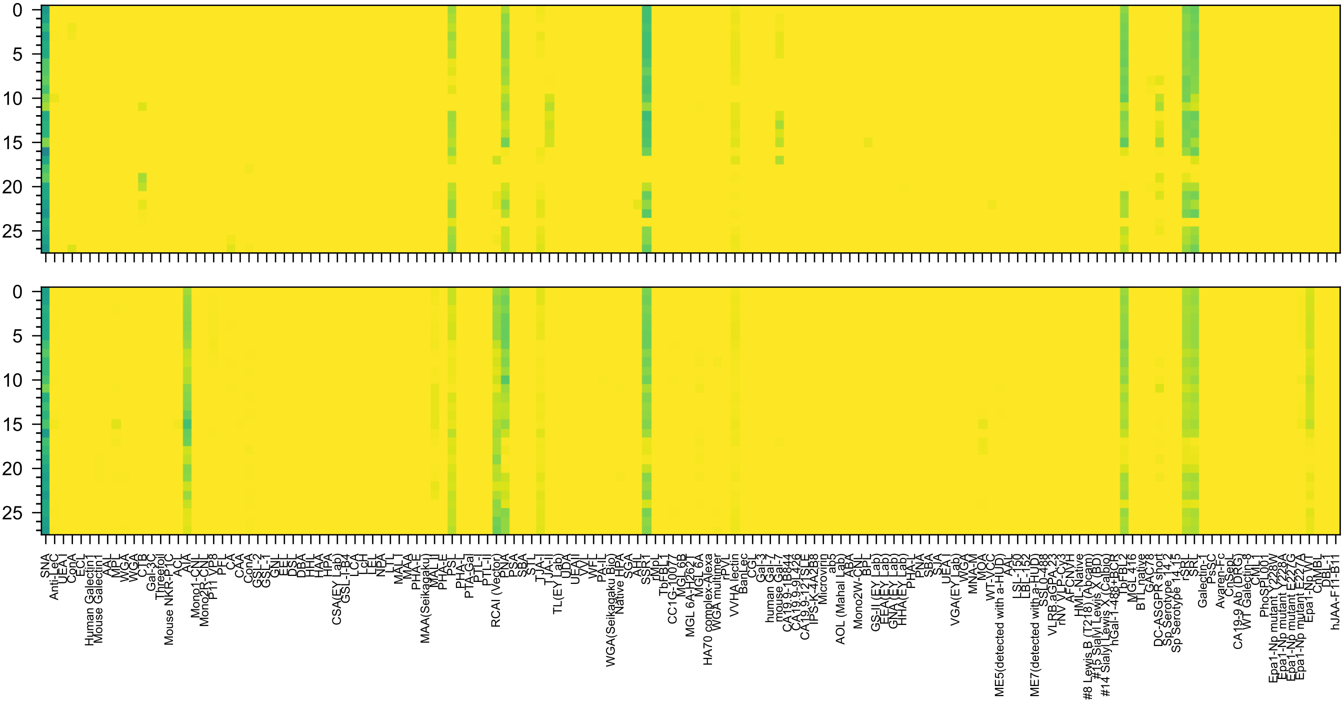
Neu5Ac(α2-6)Gal(β1-4)GlcNAc(β1-6)[Gal(β1-3)]GalNAc(α-Sp14

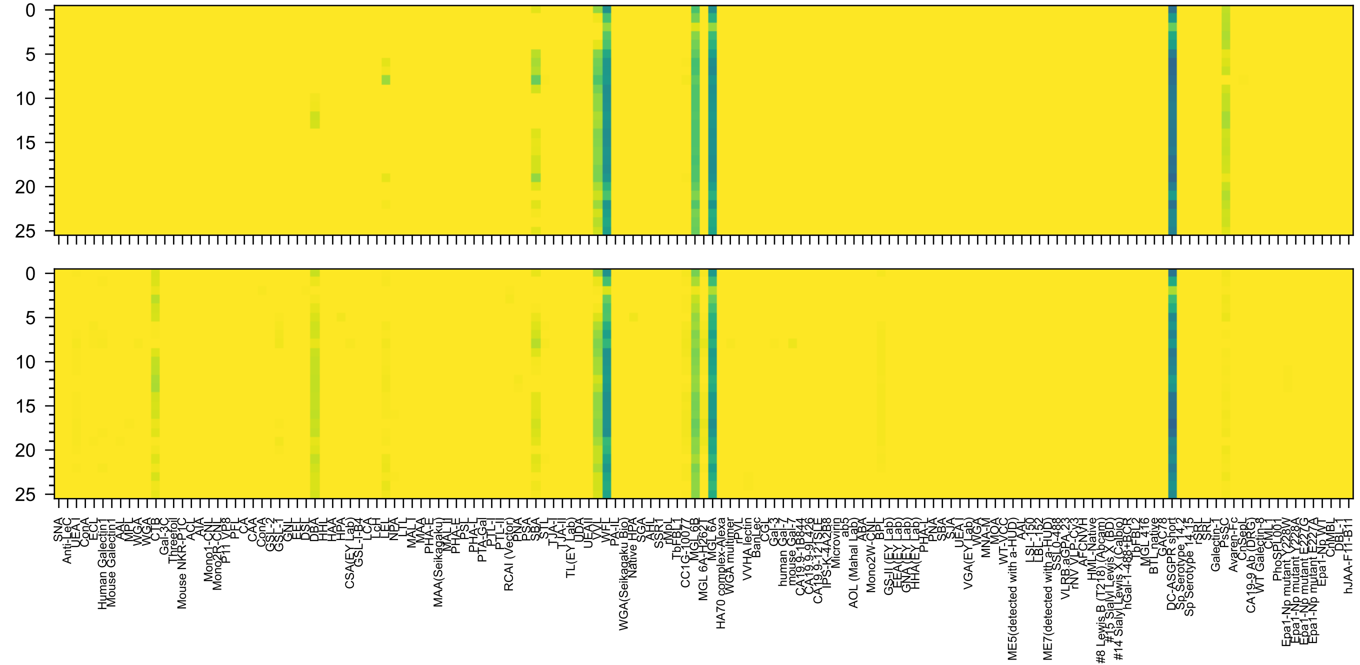

GalNAc(β1-4)[Neu5Ac(α2-8)Neu5Ac(α2-3)]Gal(β1-4)Glc(β-Sp0

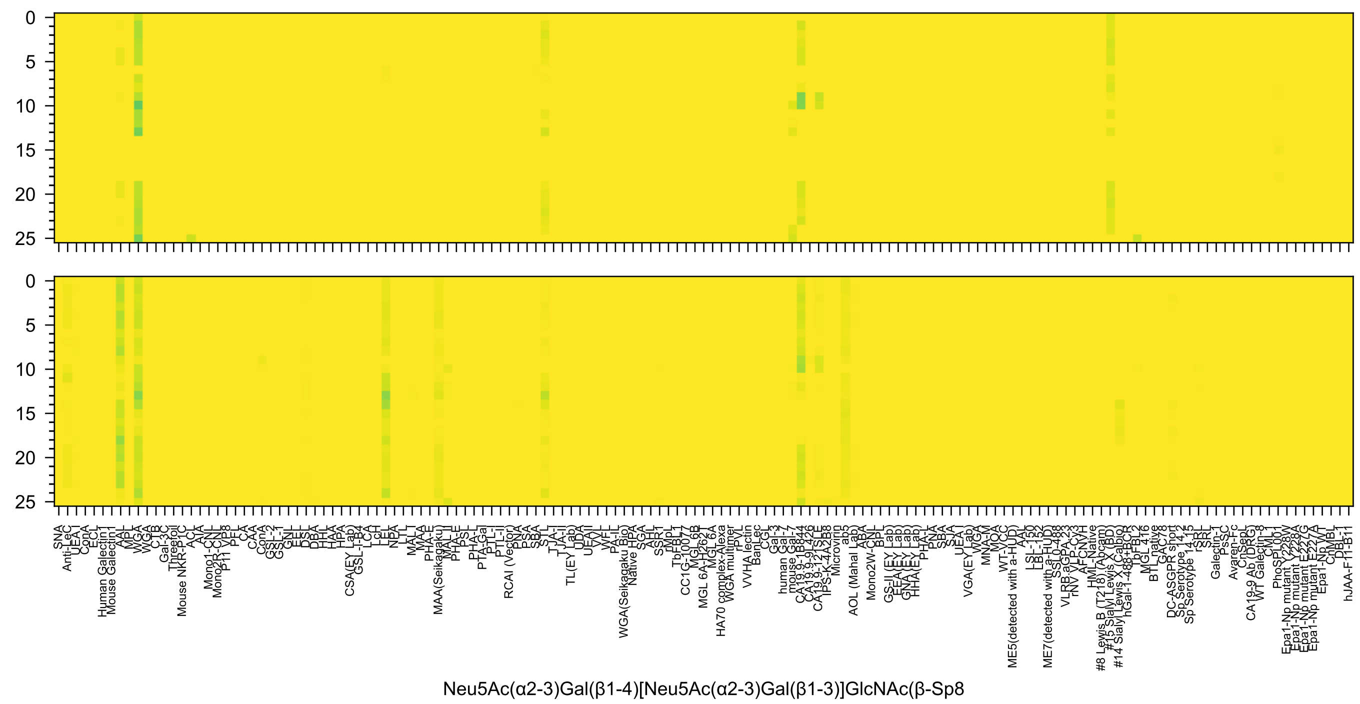
Neu5Ac(α2-3)Gal(β1-4)[Neu5Ac(α2-3)Gal(β1-3)]GlcNAc(β-Sp8

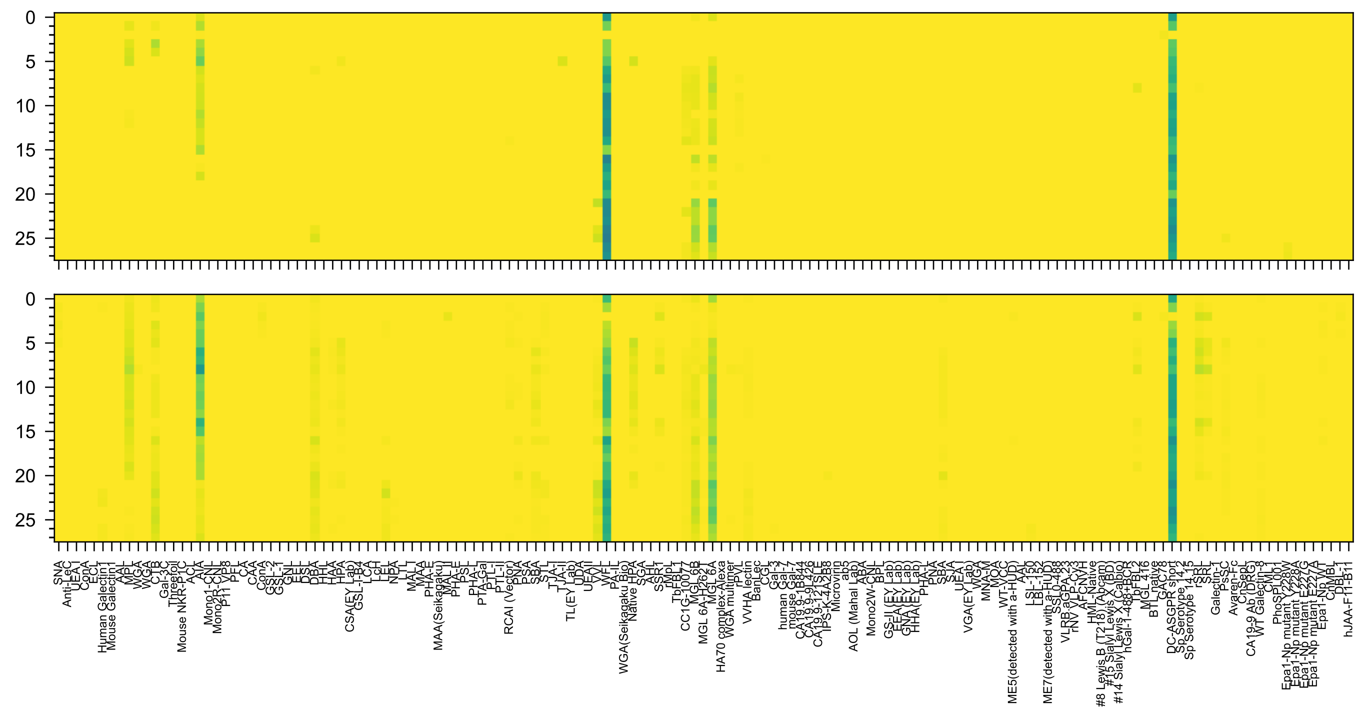

GalNAc(β1-4)[Neu5Ac(α2-3)]Gal(β1-4)GlcNAc(β1-3)GalNAc(α-Sp14

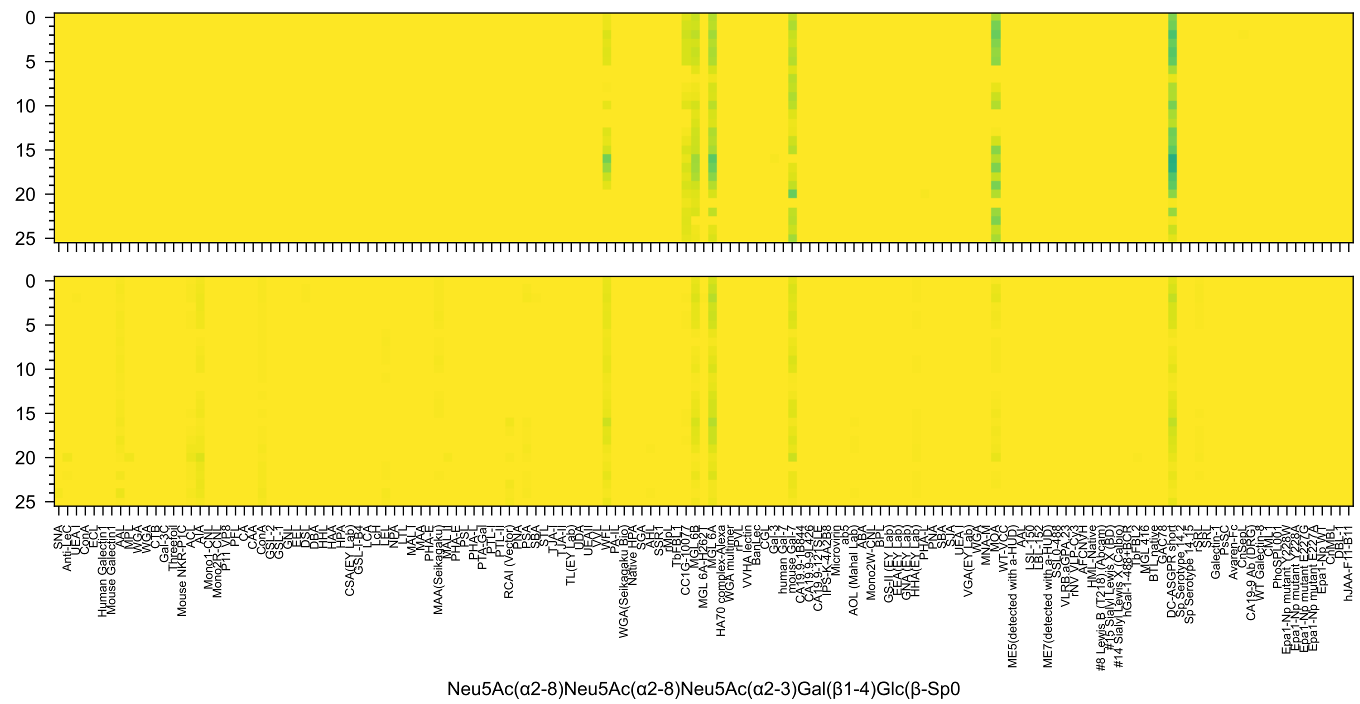
Neu5Ac(α2-8)Neu5Ac(α2-8)Neu5Ac(α2-3)Gal(β1-4)Glc(β-Sp0

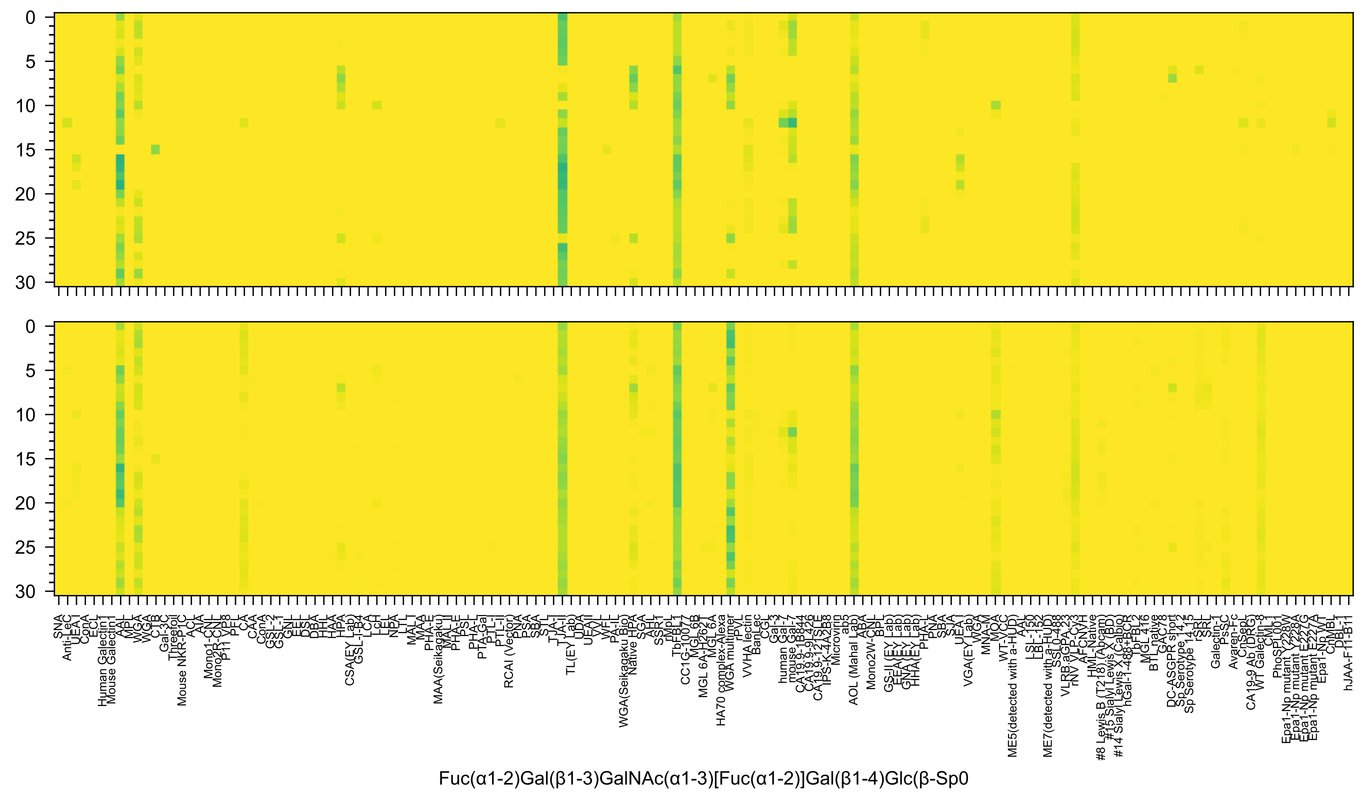
Fuc(α1-2)Gal(β1-3)GalNAc(α1-3)[Fuc(α1-2)]Gal(β1-4)Glc(β-Sp0

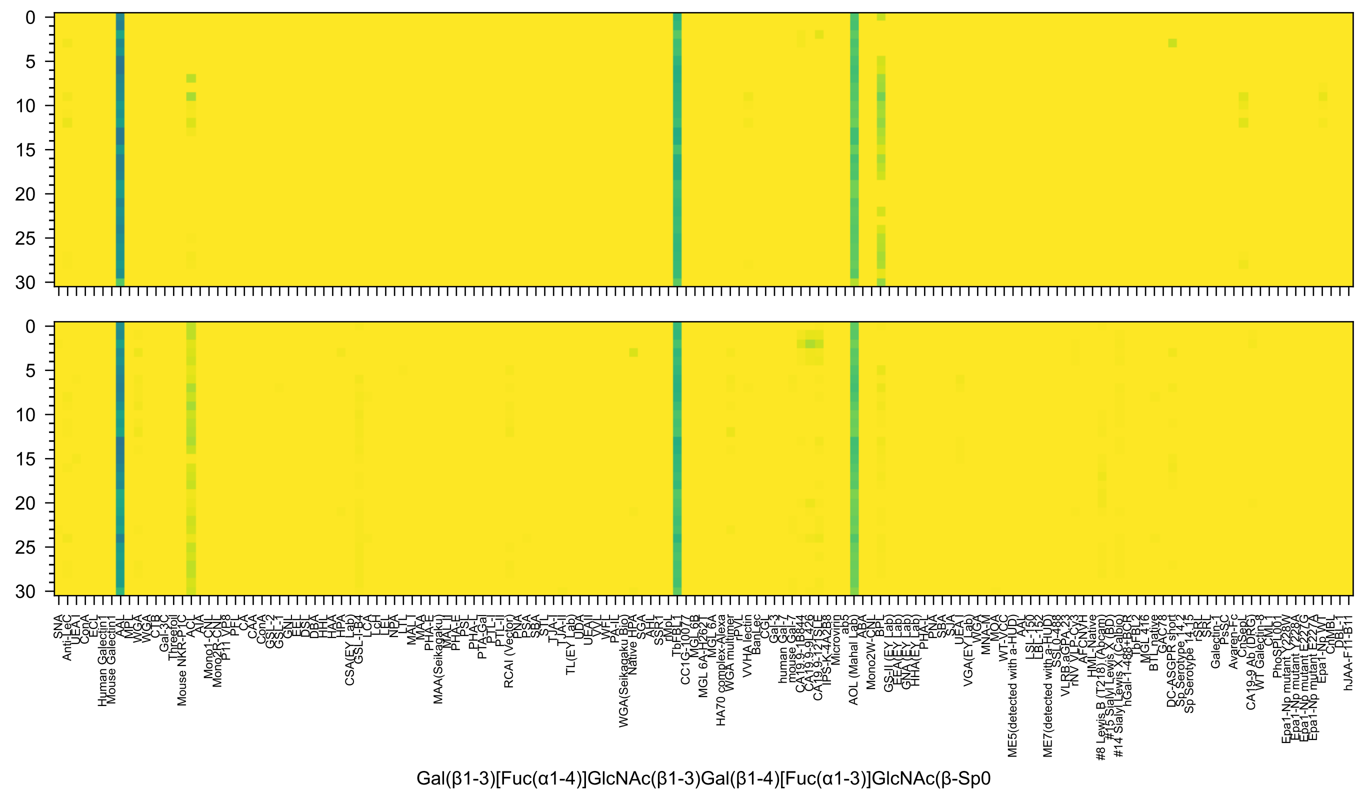

SNA
Anti-LeC
UEA-I
ConA
ECL
Human Galectin1
Mouse Galectin1
AAL
MPL
WGA
WGA
CTB
Gal-3C
Threefoil
Mouse NKR-P1C
ACL
AIA
Mono1-CNL
Mono2R-CNL
P11 VP8
PFL
CAA
ConA
GSL-2
GSL-1
GNL
EEL
DSL
DBA
HHL
HAA
HPA
CSA(EY Lab)
GSL-I-B4
LCA
LcH
LEL
NPA
LTL
MAL-I
MAA
MAA(Seikagaku)
MAL II
PHA-E
PHA-E
PSL
PHA-L
PTA-Gal
PTL-I
PTL-II
RCAI (Vector)
PNA
PSA
SBA
STL
TJA-I
TJA-II
TL(EY Lab)
UDA
UEAII
VVL
WFL
PA-IL
WGA(Seikagaku Bio)
Native HPA
SGA
AHL
SSR1
rMpL
TbFBL1
CC1G-10077
MGL 6B
MGL 6A-H262T
MGL 6A
HA70 complex-Alexa
WGA multimer
rPVL
VVHA lectin
BanLec
CGL
Gal-3
human Gal-7
mouse Gal-7
CA19-9-1B844
CA19-9-9L426
CA19-9-121SLE
IPS-K-4A2B8
Microvirin
ab5
AOL (Mahal Lab)
ABA
Mono2W-CNL
BPL
GS-II (EY Lab)
EEA (EY Lab)
GNA (EY Lab)
HHA(EY Lab)
PHA-L
PNA
SBA
SJA
UEA-I
VGA(EY Lab)
WGA
MNA-M
MOA
WT-VCC
ME5(detected with a-HUD)
AAL
LSL-150
LBL-152
ME7(detected with a-HUD)
SSL0-488
VLRB.aGPA.23
rNV VLP-Cy3
AFCNVH
HML-Native
#8 Lewis B (T218) (Abcam)
#15 Sialyl Lewis X (BD)
#5 Sialyl Lewis X (Calbio)
#14 Sialyl Lewis X
hGal-1-488+BCR
TbFBL2
MGL 416
BTL native
GAC78
DC-ASGPR short
Sp Serotype 14.2
Sp Serotype 14.15
rSRL
SRL
Galectin-1
PsSC
Avaren-Fc
CnSepL
CA19-9 Ab (DRG)
WT Galectin-8
CML-1
PhoSL001
Epa1-Np mutant Y228W
Epa1-Np mutant Y228A
Epa1-Np mutant E227G
Epa1-Np mutant E227A
Epa1-Np WT
ChMBL
DBL-1
hJAA-F11-B1
Gal(β1-3)[Fuc(α1-4)]GlcNAc(β1-3)Gal(β1-4)[Fuc(α1-3)]GlcNAc(β-Sp0

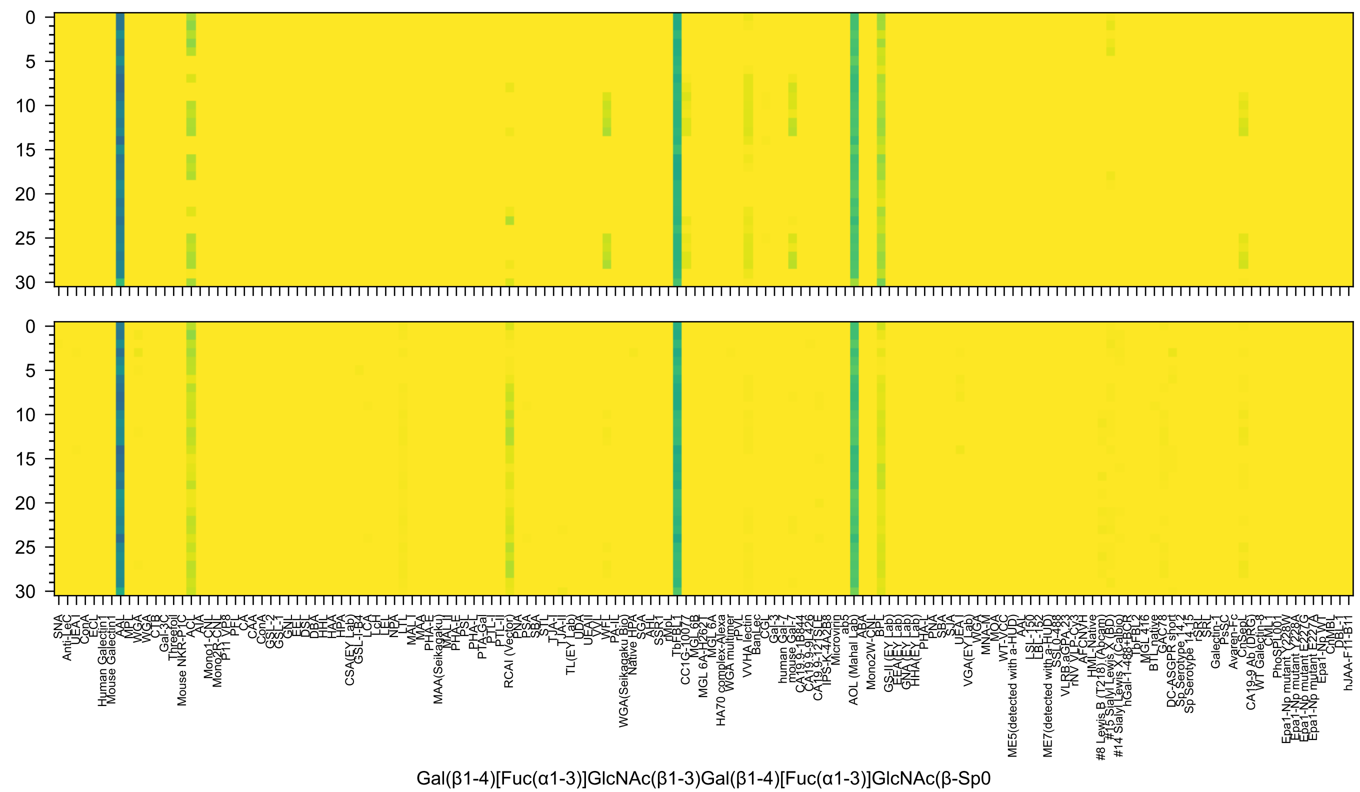
0
5
10
15
20
25
30
SNA
Anti-LeC
UEA-I
ConA
ECL
Human Galectin1
Mouse Galectin1
AAL
MPL
WGA
WGA
CTB
Gal-3C
Threefoil
Mouse NKR-P1C
ACL
AIA
Mono1-CNL
Mono2R-CNL
P11 VP8
PFL
CAA
ConA
GSL-2
GSL-1
GNL
EEL
DSL
DBA
HHL
HAA
HPA
CSA(EY Lab)
GSL-I-B4
LCA
LcH
LEL
NPA
LTL
MAL1
MAA
MAA(Seikagaku)
MAL_II
PHA-E
PHA-E
PSL
PHA-L
PTA-Gal
PTL-I
PTL-II
RCAI (Vector)
PNA
PSA
SBA
STL
TJA-I
TJA-II
TL(EY Lab)
UDA
UEAII
VVL
WFL
PA-IL
WGA(Seikagaku Bio)
Native HPA
SGA
AHL
SSR1
rMpL
TbFBL1
CC1G-10077
MGL 6B
MGL 6A-H262T
MGL 6A
HA70 complex-Alexa
WGA multimer
rPVL
VVHA lectin
BanLec
CGL
Gal-3
human Gal-7
mouse Gal-7
CA19.9-1B844
CA19.9-9L426
CA19.9-121SLE
IPS-K-4A2B8
Microvirin
ab5
AOL (Mahal Lab)
ABA
Mono2W-CNL
BPL
GS-II (EY Lab)
EEA(EY Lab)
GNA(EY Lab)
HHA(EY Lab)
PHA-L
PNA
SBA
SJA
UEA-I
VGA(EY Lab)
WGA
MNA-M
MOA
WT-VCC
ME5(detected with a-HUD)
AAL
LSL-150
LBL-152
ME7(detected with a-HUD)
SSL0-488
VLRB.aGPA.23
rNV VLP-Cy3
AFCNVH
HML-Native
#8 Lewis B (T218) (Abcam)
#15 Sialyl Lewis X (BD)
#5 Lewis X (Calbio)
#14 Sialyl Lewis A
hGal-1-488+BCR
TbFBL2
MGL 416
BTL native
GAC78
DC-ASGPR short
Sp Serotype 14.2
Sp Serotype 14.15
rSRL
SRL
Galectin-1
PsSC
Avaren-Fc
CnSepL
CA19-9 Ab (DRG)
WT Galectin-8
PhoSL
Epa1-Np mutant Y228W
Epa1-Np mutant Y228A
Epa1-Np mutant E227G
Epa1-Np mutant E227A
Epa1-Np WT
ChMBL
DBL-1
hJAA-F11-B1
Gal(β1-4)[Fuc(α1-3)]GlcNAc(β1-3)Gal(β1-4)[Fuc(α1-3)]GlcNAc(β-Sp0

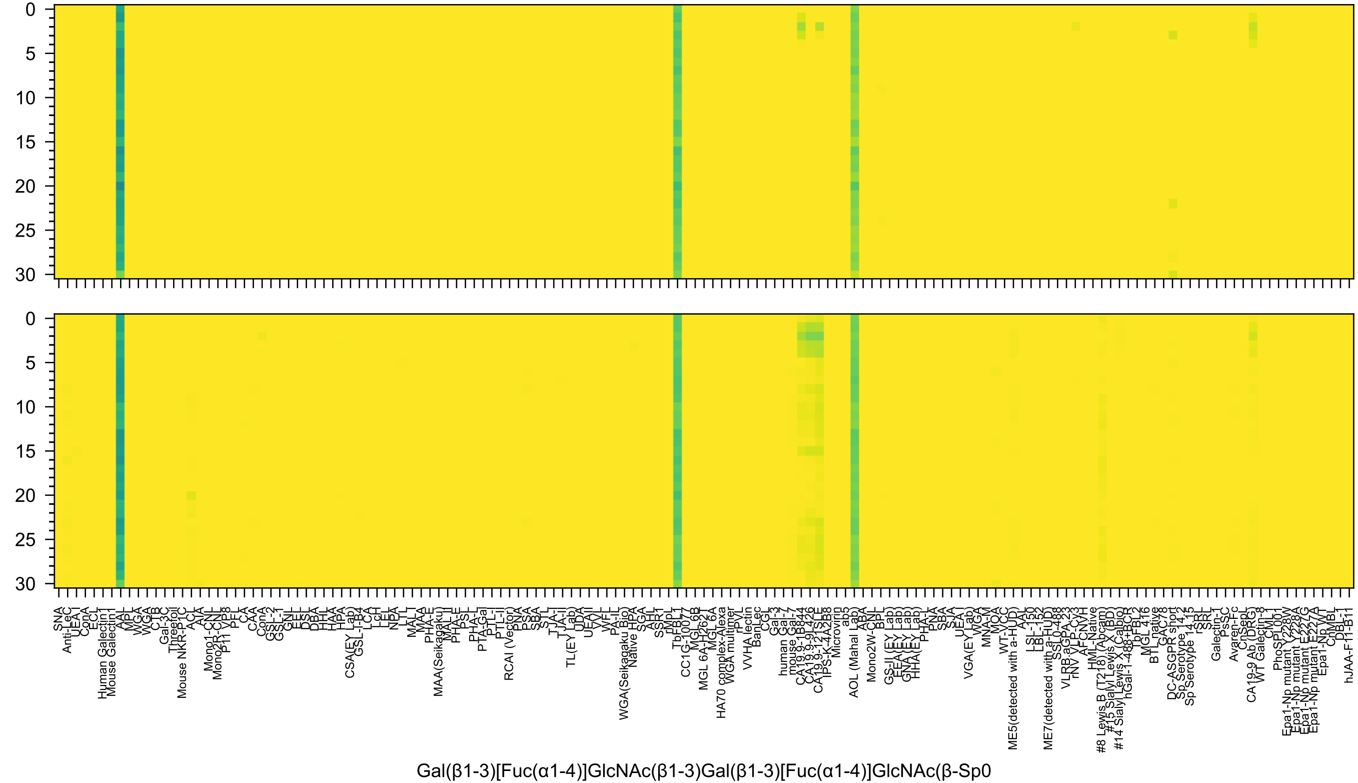
Gal(β1-3)[Fuc(α1-4)]GlcNAc(β1-3)Gal(β1-3)[Fuc(α1-4)]GlcNAc(β-Sp0

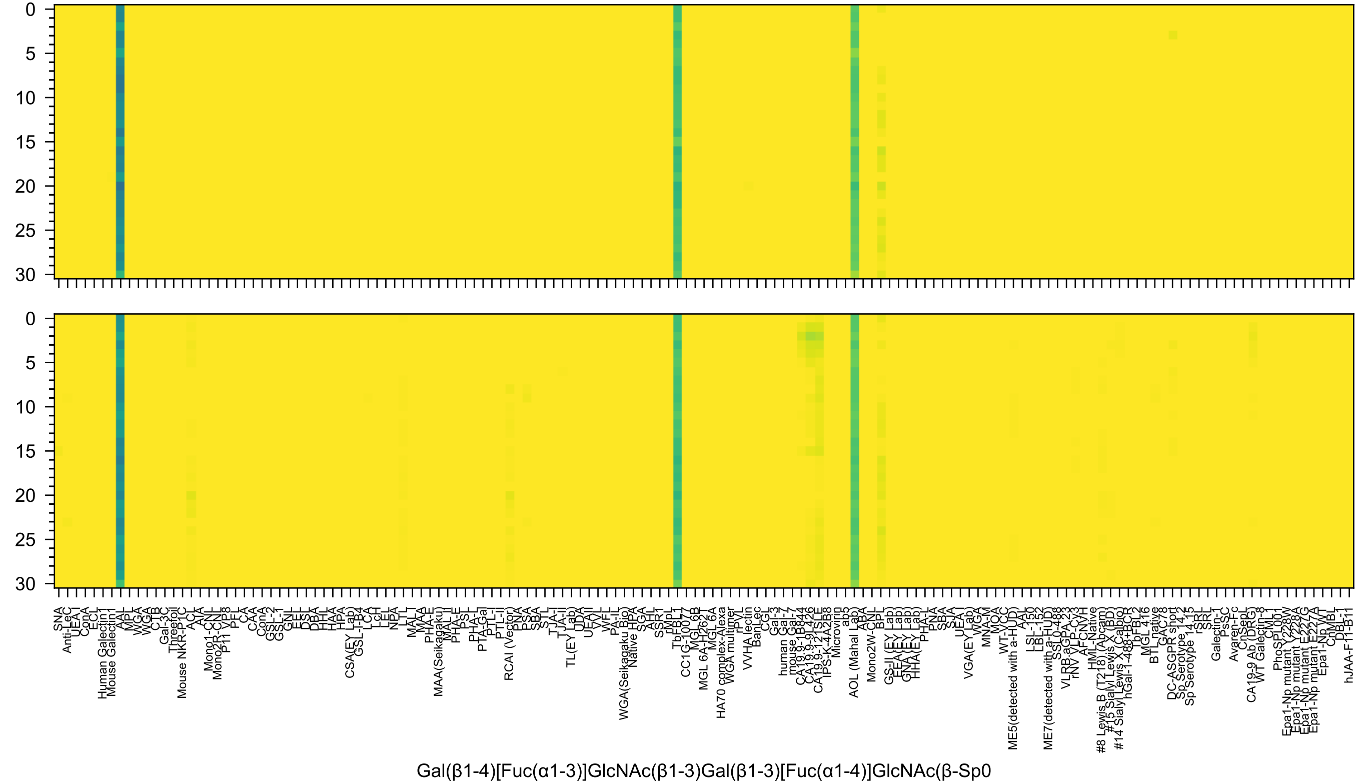

0
5
10
15
20
25
30
SNA
Anti-LeC
UEA-I
ConA
ECL
Human Galectin1
Mouse Galectin1
AAL
MPL
WGA
WGA
CTB
Gal-3C
Threefoil
Mouse NKR-P1C
ACL
AIA
Mono1-CNL
Mono2R-CNL
P11 VP8
PFL
CAA
ConA
GSL-2
GSL-1
GNL
EEL
DSL
DBA
HHL
HAA
HPA
CSA(EY Lab)
GSL-I-B4
LCA
LcH
LEL
NPA
LTL
MAL-I
MAA
MAA(Seikagaku)
PHA-E
MAL-II
PHA-E
PSL
PHA-L
PTA-Gal
PTL-I
PTL-II
RCAI (Vector)
PNA
PSA
SBA
STL
TJA-I
TJA-II
TL(EY Lab)
UDA
UEAII
VVL
WFL
PA-IL
WGA(Seikagaku Bio)
Native HPA
SGA
AHL
SSR1
rMpL
TbFBL1
CC1G-10077
MGL 6B
MGL 6A-H262T
MGL 6A
HA70 complex-Alexa
WGA multimer
rPVL
VVHA lectin
BanLec
CGL
Gal-3
human Gal-7
mouse Gal-7
CA19-9-1B844
CA19-9-9L426
CA19-9-121SLE
IPS-K-4A2B8
Microvirin
ab5
AOL (Mahal Lab)
ABA
Mono2W-CNL
BPL
GS-II (EY Lab)
EEA (EY Lab)
GNA (EY Lab)
HHA (EY Lab)
PHA-L
PNA
SBA
SJA
UEA-I
VGA (EY Lab)
WGA
MNA-M
MOA
WT-VCC
ME5(detected with a-HUD)
AAL
LSL-150
LBL-152
ME7(detected with a-HUD)
SSL0-488
VLRB aGPA.23
rNV VLP-Cy3
AFCNVH
HML-Native
#8 Lewis B (T218) (Abcam)
#15 Sialyl Lewis X (BD)
#5 Sialyl Lewis X (Calbio)
#14 Sialyl Lewis A
hGal-1-488+BCR
TbFBL2
MGL 416
BTL native
GAC78
DC-ASGPR short
Sp Serotype 14.2
Sp Serotype 14.15
rSRL
SRL
Galectin-1
PsSC
Avaren-Fc
CnSepL
CA19-9 Ab (DRG)
WT Galectin-8
CML-1
PhoSL001
Epa1-Np mutant Y228W
Epa1-Np mutant Y228A
Epa1-Np mutant E227G
Epa1-Np mutant E227A
Epa1-Np WT
ChMBL
DBL-1
hJAA-F11-B1
Gal(β1-4)[Fuc(α1-3)]GlcNAc(β1-3)Gal(β1-3)[Fuc(α1-4)]GlcNAc(β-Sp0

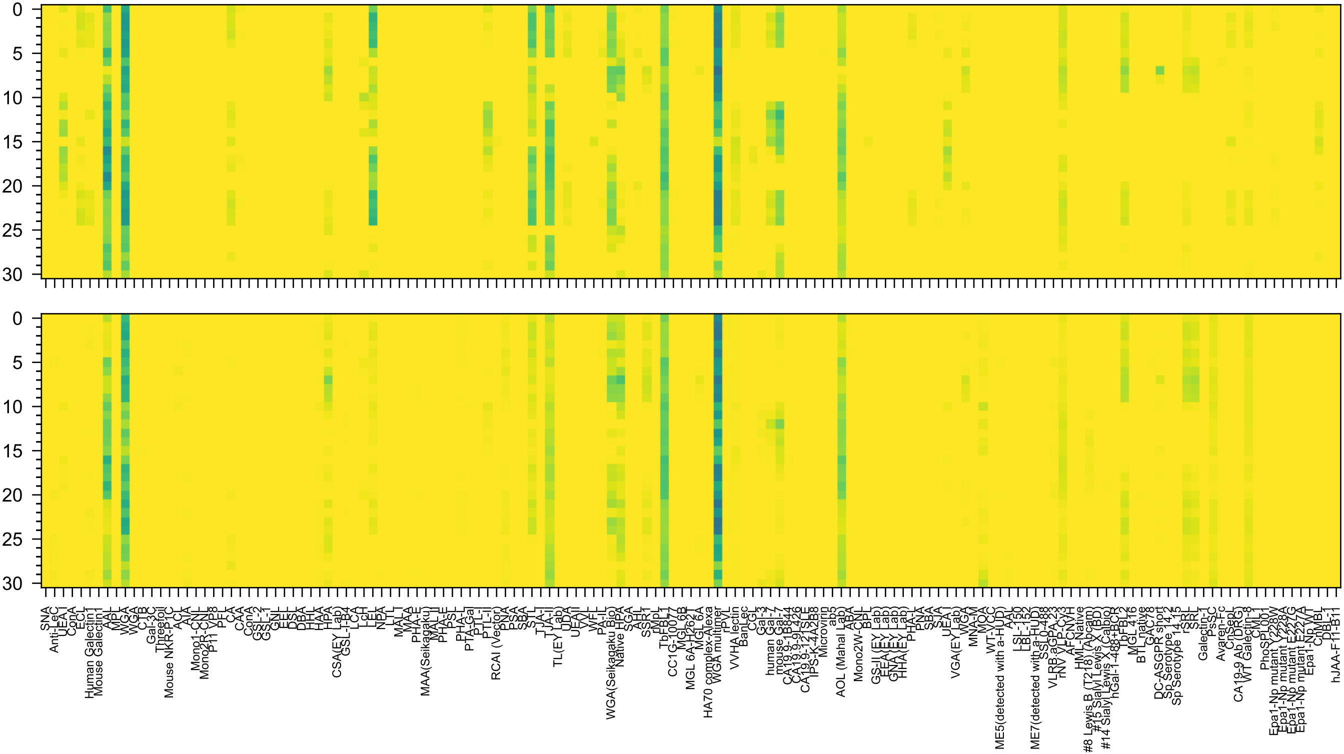


Fuc(α1-2)Gal(β1-3)GalNAc(α1-3)[Fuc(α1-2)]Gal(β1-4)GlcNAc(β-Sp0

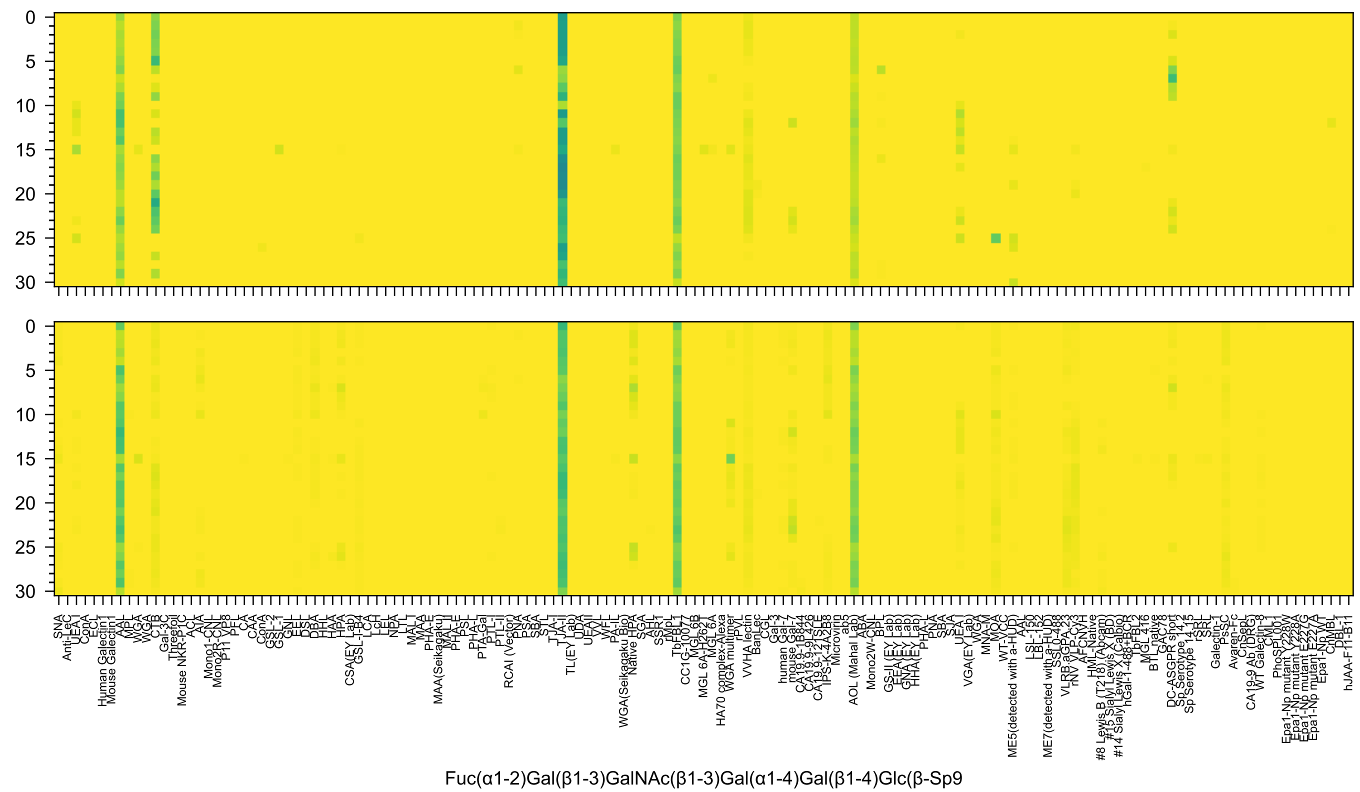

Fuc(α1-2)Gal(β1-3)GalNAc(β1-3)Gal(α1-4)Gal(β1-4)Glc(β-Sp9

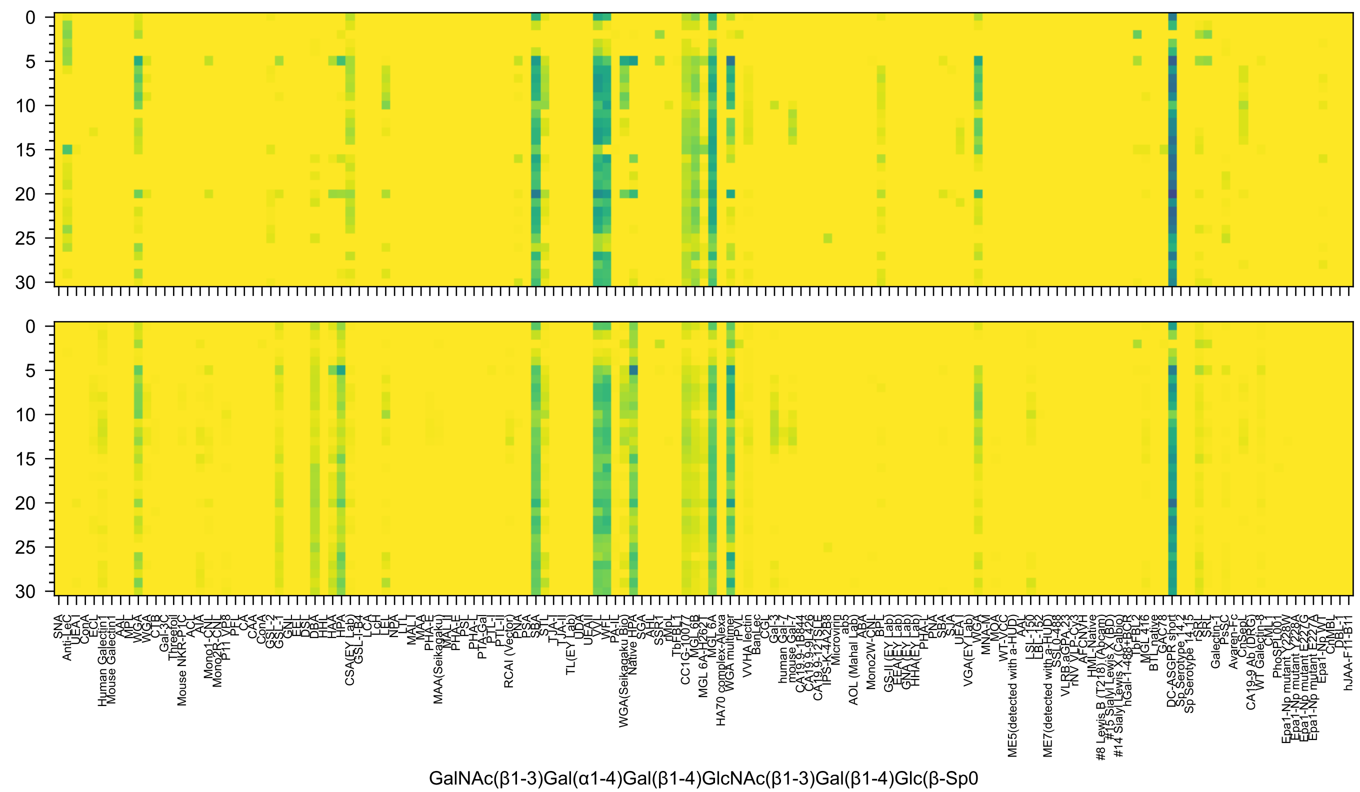
0
5
10
15
20
25
30
SNA
Anti-LeC
UEA-I
ConA
ECL
Human Galectin1
Mouse Galectin1
AAL
MPL
WGA
WGA
CTB
Gal-3C
Threefoil
Mouse NKR-P1C
ACL
AIA
Mono1-CNL
Mono2R-CNL
P11 VP8
PFL
CAA
CAA
ConA
GSL-2
GSL-1
GNL
EEL
DSL
DBA
HHL
HAA
HPA
CSA(EY Lab)
GSL-I-B4
LCA
LcH
LEL
NPA
LTL
MAL-I
MAA
PHA-E
MAA(Seikagaku)
MAL_II
PHA-E
PSL
PHA-L
PTA-Gal
PTL-I
PTL-II
RCAI (Vector)
PNA
PSA
SBA
STL
TJA-I
TJA-II
TL(EY Lab)
UDA
UEA-II
VVL
WFL
PA-IL
WGA(Seikagaku Bio)
Native HPA
SGA
AHL
SSR1
rMpL
TbFBL1
CC1G-10077
MGL 6B
MGL 6A-H262T
MGL 6A
HA70 complex-Alexa
WGA multimer
rPVL
VVHA lectin
BanLec
CGL
Gal-3
human Gal-7
mouse Gal-7
CA19-9 B844
CA19-9 9L426
CA19-9 121SLE
IPS-K-4A2B8
Microvirin
ab5
AOL (Mahal Lab)
ABA
Mono2W-CNL
BPL
GS-II (EY Lab)
EEA (EY Lab)
GNA (EY Lab)
HHA (EY Lab)
PHA-L
PNA
SBA
SJA
UEA-I
VGA(EY Lab)
WGA
MNA-M
MOA
WT-VCC
ME5(detected with a-HUD)
AAL
LSL-150
LBL-152
ME7(detected with a-HUD)
SSL0-488
VLRB.aGPA.23
rNV VLP-Cy3
AFCNVH
HML-Native
#8 Lewis B (T218) (Abcam)
#15 Sialyl Lewis X (BD)
#5 Sialyl Lewis X (Calbio)
#14 Sialyl Lewis
hGal-1-488+BCR
TbFBL2
MGL 416
BTL native
GAC78
PC
DC-ASGPR short
Sp Serotype 14.2
Sp Serotype 14.15
rSRL
SRL
Galectin-1
PsSC
Avaren-Fc
CnSepL
CA19-9 Ab (DRG)
WT Galectin-8
CML-9
PhoSL001
Epa1-Np mutant Y228W
Epa1-Np mutant E227G
Epa1-Np mutant E227A
Epa1-Np WT
ChMBL
DBL-1
hJAA-F11-B1
GalNAc(β1-3)Gal(α1-4)Gal(β1-4)GlcNAc(β1-3)Gal(β1-4)Glc(β-Sp0

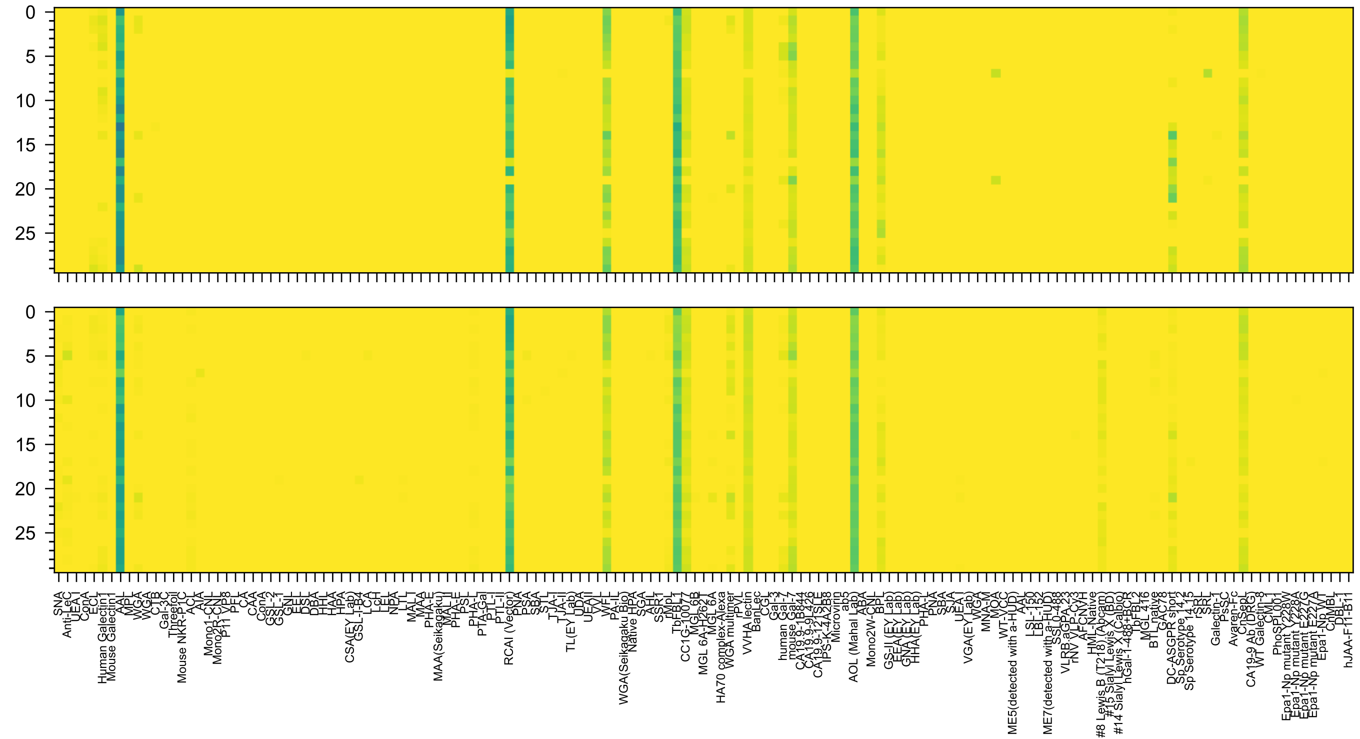

Fuc(α1-3)GlcNAc(β1-6)[Gal(β1-4)GlcNAc(β1-3)]Gal(β1-4)Glc(-Sp21

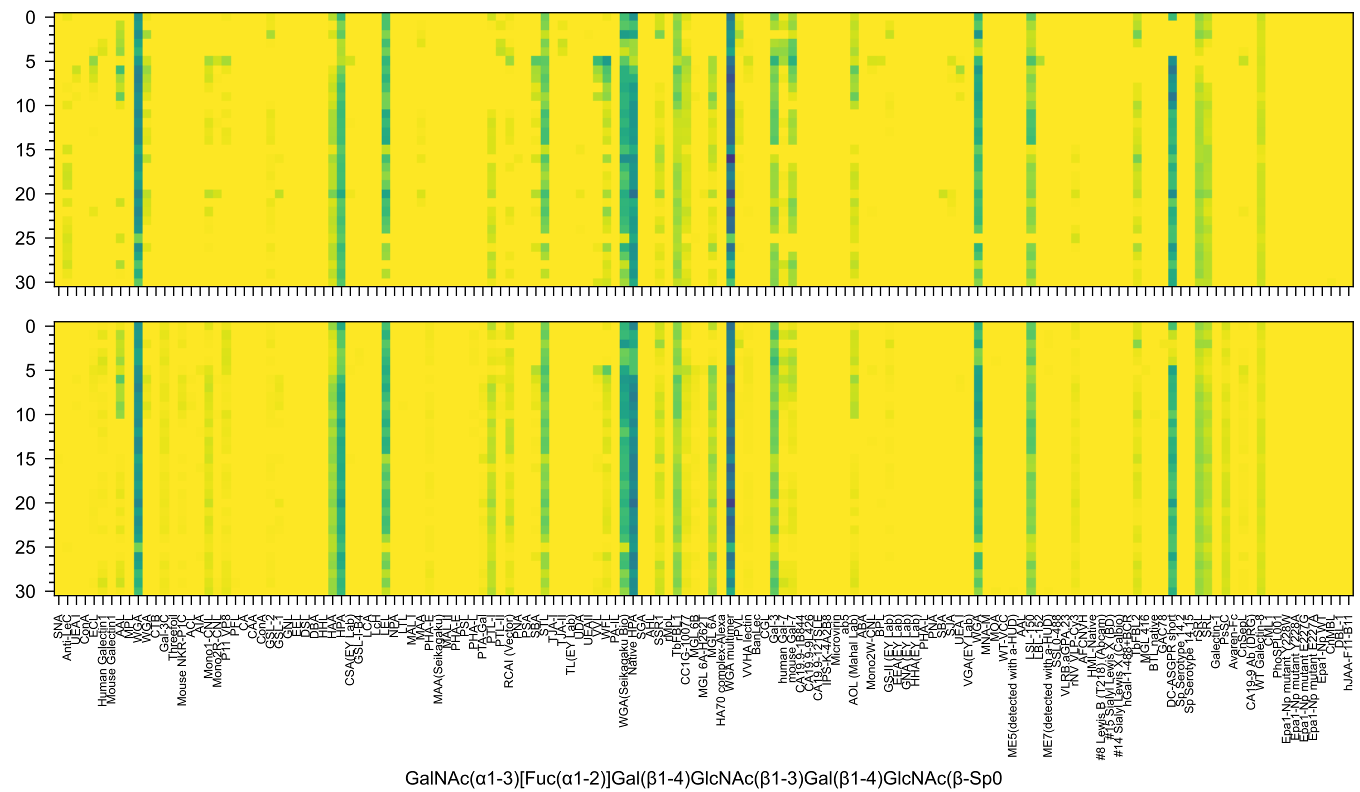
0
5
10
15
20
25
30
GalNAc(α1-3)[Fuc(α1-2)]Gal(β1-4)GlcNAc(β1-3)Gal(β1-4)GlcNAc(β-Sp0

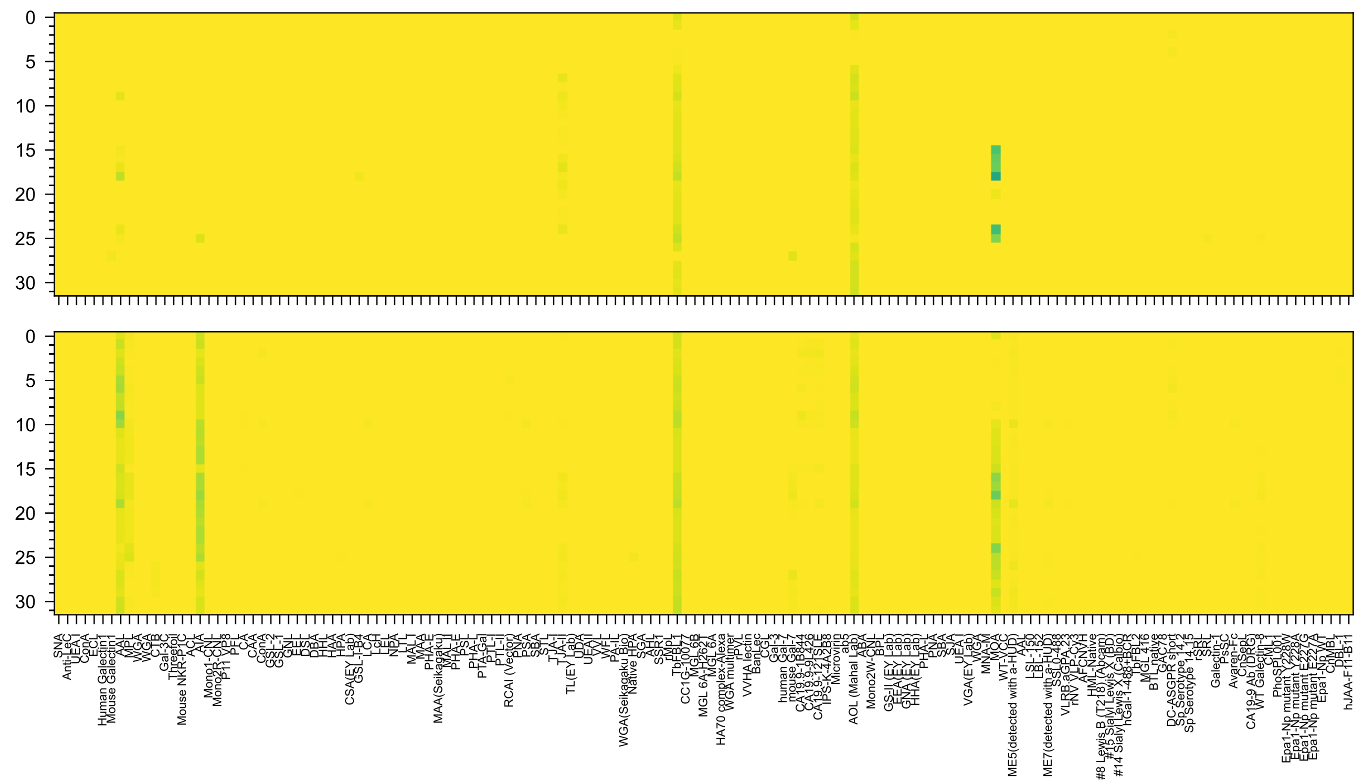

Gal(α1-3)[Fuc(α1-2)]Gal(β1-4)[Fuc(α1-3)]GlcNAc(β1-3)GalNAc(-Sp14

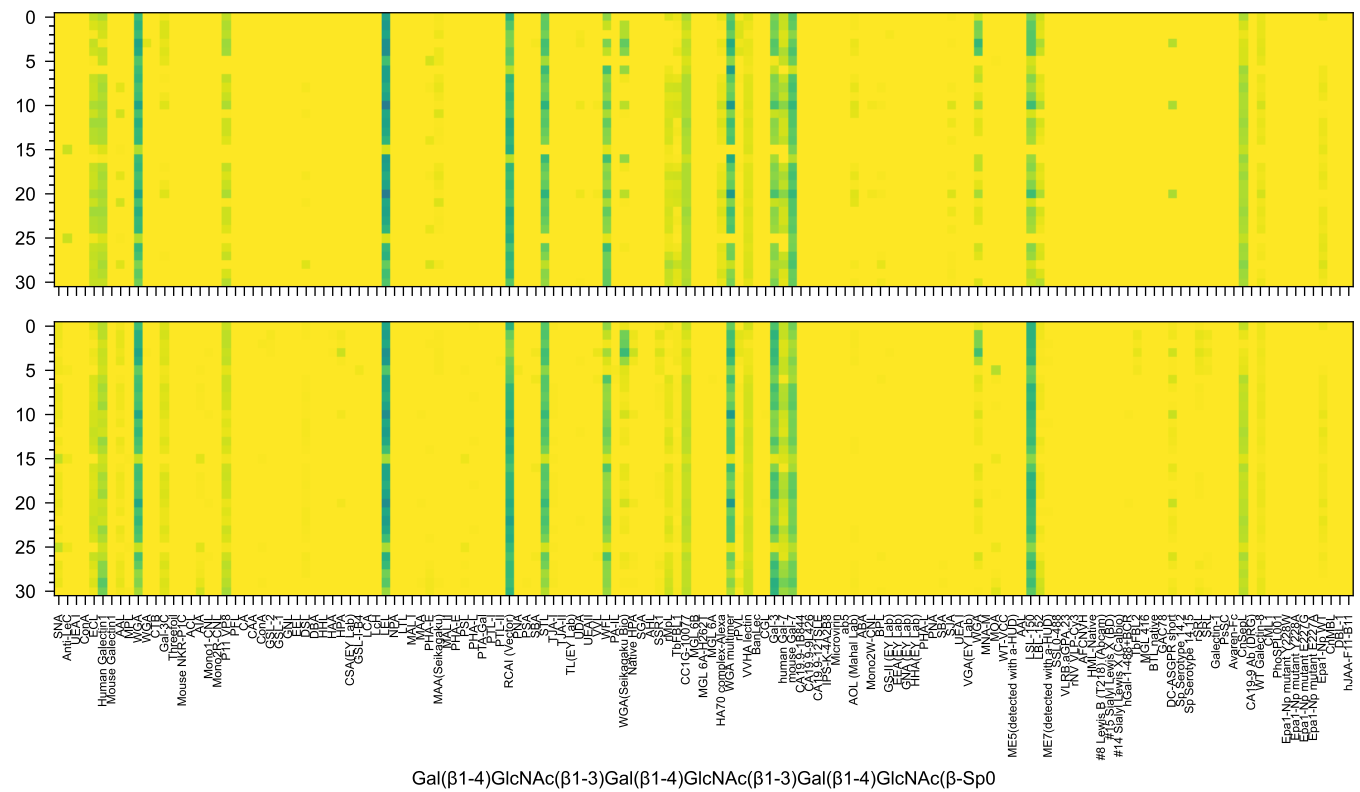
SNA
Anti-LeC
UEA-I
ConA
ECL
Human Galectin1
Mouse Galectin1
AAL
MPL
WGA
WGA
CTB
Gal-3C
Threefoil
Mouse NKR-P1C
ACL
AIA
Mono1-CNL
Mono2R-CNL
P11 VP8
PFL
CA
CAA
ConA
GSL-2
GSL-1
GNL
EEL
DSL
DBA
HHL
HAA
HPA
CSA(EY Lab)
GSL-I-B4
LCA
LcH
LEL
NPA
LTL
MAL-I
MAA
PHA-E
MAA(Seikagaku)
MAL_II
PHA-E
PSL
PHA-L
PTA-Gal
PTL-I
PTL-II
RCAI (Vector)
PNA
PSA
SBA
STL
TJA-I
TJA-II
TL(EY Lab)
UDA
UEAII
VVL
WFL
PA-IL
WGA(Seikagaku Bio)
Native HPA
SGA
AHL
SSR1
rMpL
TbFBL1
CC1G-10077
MGL6B
MGL 6A-H262T
MGL 6A
HA70 complex-Alexa
WGA multimer
rPVL
VVHA lectin
BanLec
CGL
Gal-3
human Gal-7
mouse Gal-7
CA19.9-1B844
CA19.9-9L426
CA19.9-121SLE
IPS-K-4A2B8
Microvirin
ab5
AOL (Mahal Lab)
ABA
Mono2W-CNL
BPL
GS-II (EY Lab)
EEA (EY Lab)
GNA (EY Lab)
HHA (EY Lab)
PHA-L
PNA
SBA
SJA
UEA-I
VGA(EY Lab)
WGA
MNA-M
MOA
WT-VCC
ME5(detected with a-HUD)
AAL
LSL-150
LBL-152
ME7(detected with a-HUD)
SSL0-488
VLRB.aGPA.23
rNV VLP-Cy3
AFCNVH
HML-Native
#8 Lewis B (T218) (Abcam)
#15 Sialyl Lewis X (BD)
#5 Sialyl Lewis X (Calbio)
#14 Sialyl Lewis
hGal-1-488+BCR
TbFBL2
MGL 416
BTL_native
GAC78
DC-ASGPR short
Sp Serotype 14,2
Sp Serotype 14,15
rSRL
SRL
Galectin-1
PsSC
Avaren-Fc
CnSepL
CA19-9 Ab (DRG)
WT Galectin-8
CML-9
PhoSL
PhoSL001
Epa1-Np mutant Y228W
Epa1-Np mutant Y228A
Epa1-Np mutant E227G
Epa1-Np mutant E227A
Epa1-Np WT
ChMBL
DBL-1
hJAA-F11-B1
Gal(β1-4)GlcNAc(β1-3)Gal(β1-4)GlcNAc(β1-3)Gal(β1-4)GlcNAc(β-Sp0

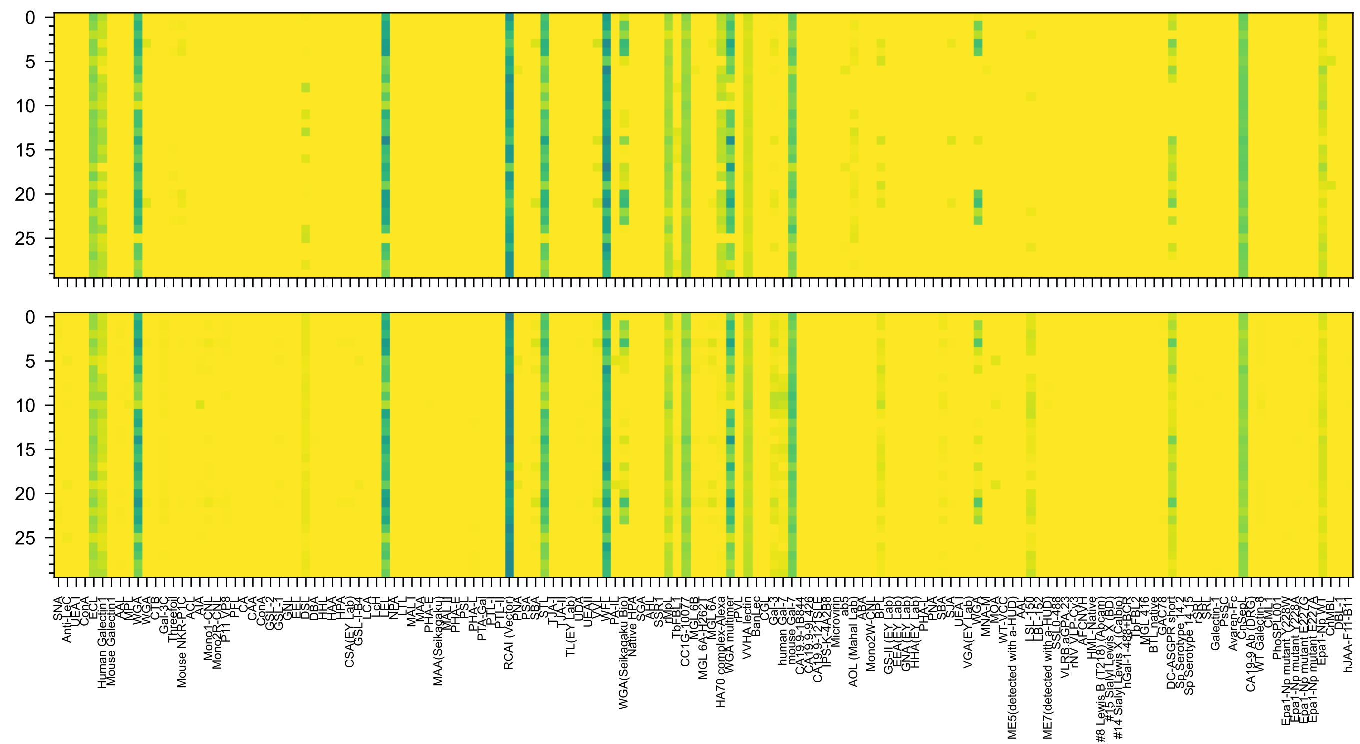


Gal(β1-4)GlcNAc(β1-6)[Gal(β1-4)GlcNAc(β1-3)]Gal(β1-4)GlcNAc(-Sp0

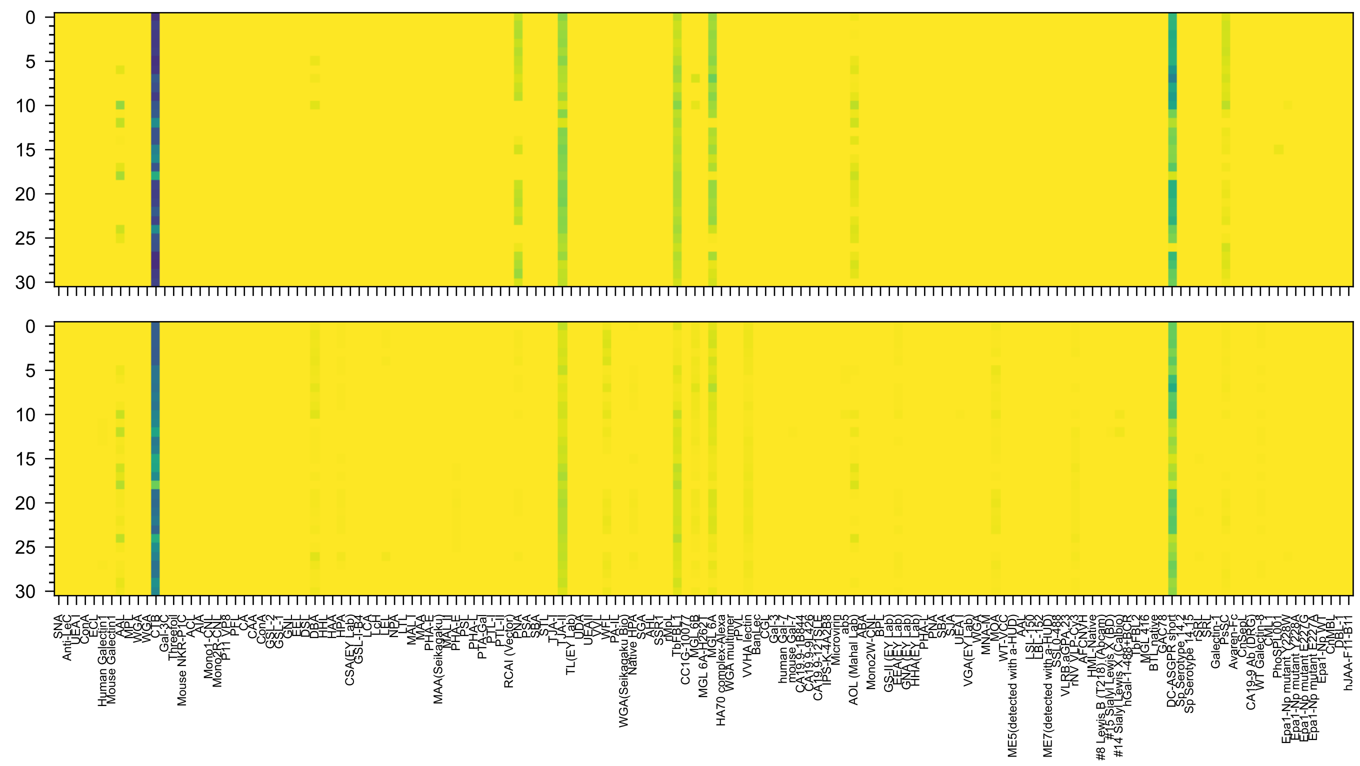

Fuc(α1-2)Gal(β1-3)GalNAc(β1-4)[Neu5Ac(α2-3)]Gal(β1-4)Glc(β-Sp0

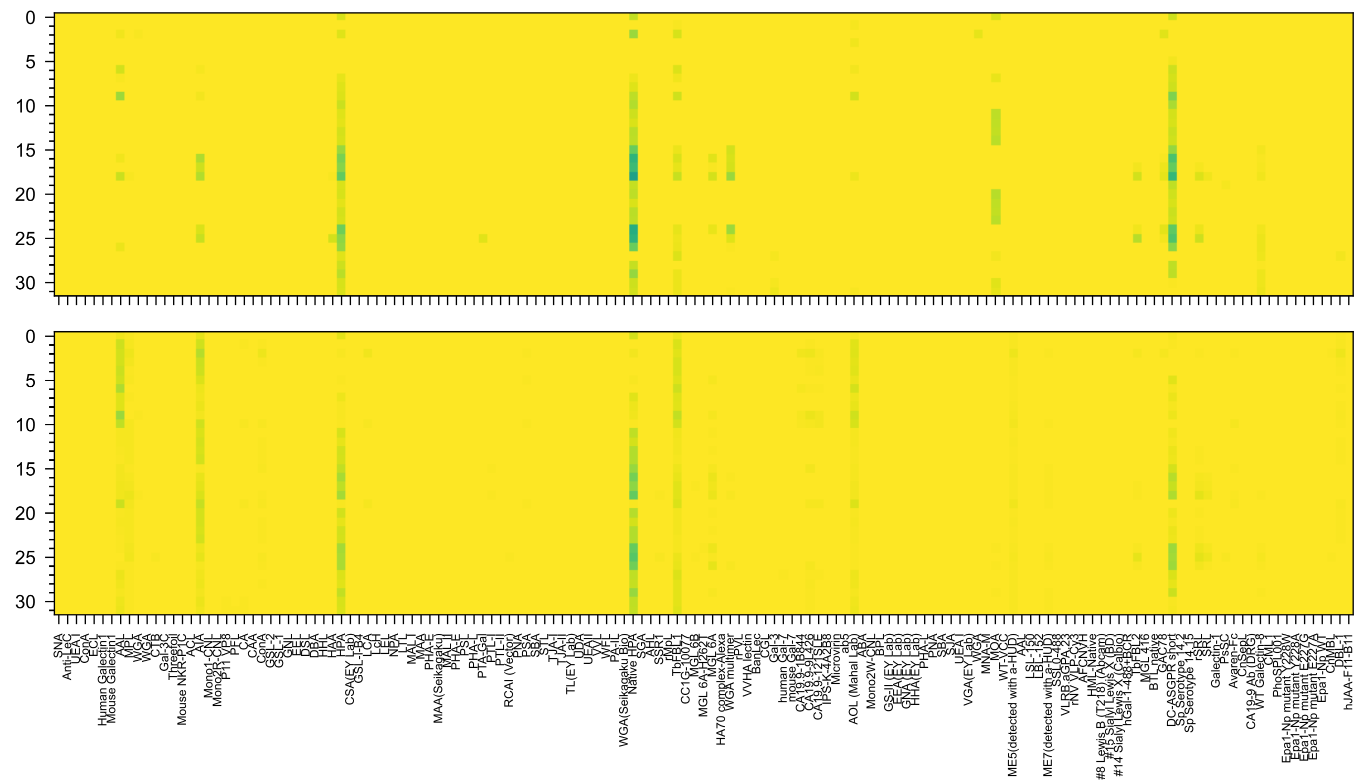


GalNAc(α1-3)[Fuc(α1-2)]Gal(β1-4)[Fuc(α1-3)]GlcNAc(β1-3)GalNAc(-Sp14

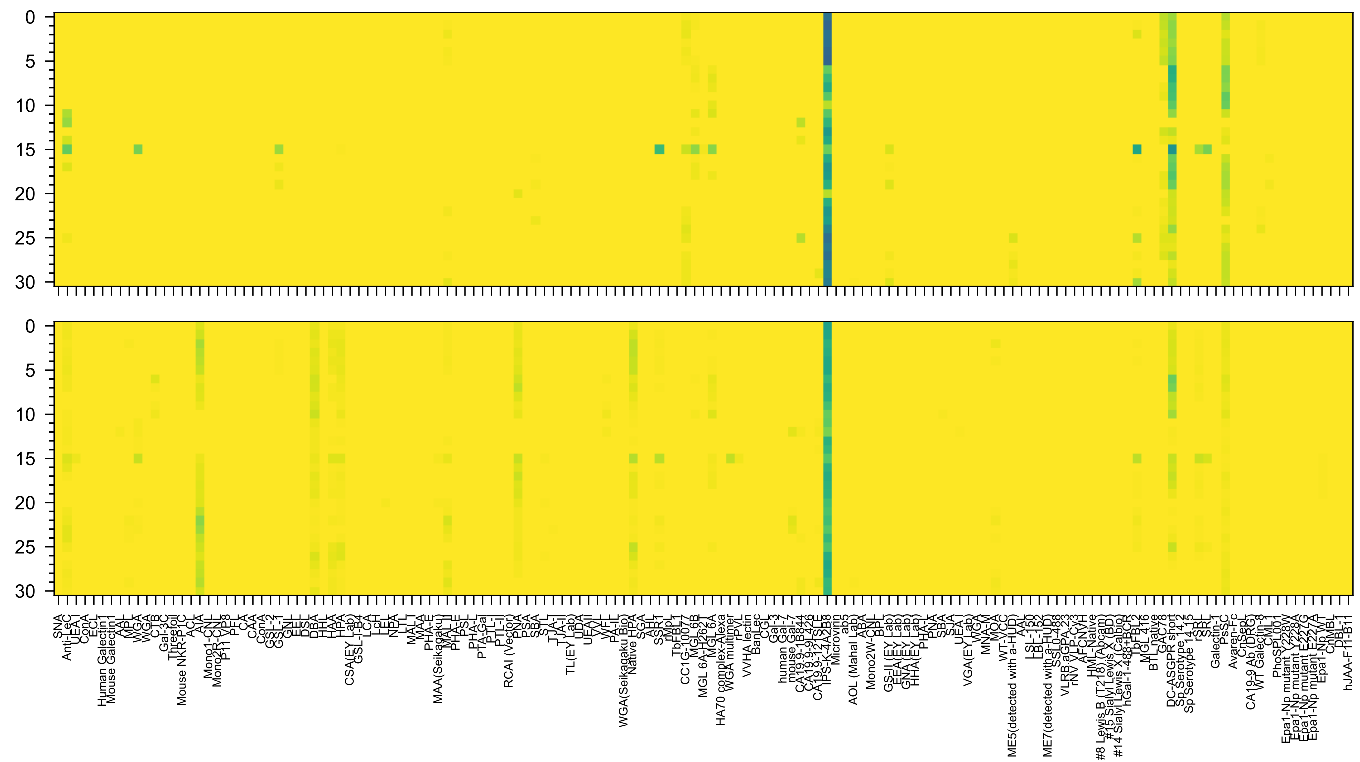


Neu5Ac(α2-3)Gal(β1-3)GalNAc(β1-3)Gal(α1-4)Gal(β1-4)Glc(β-Sp0

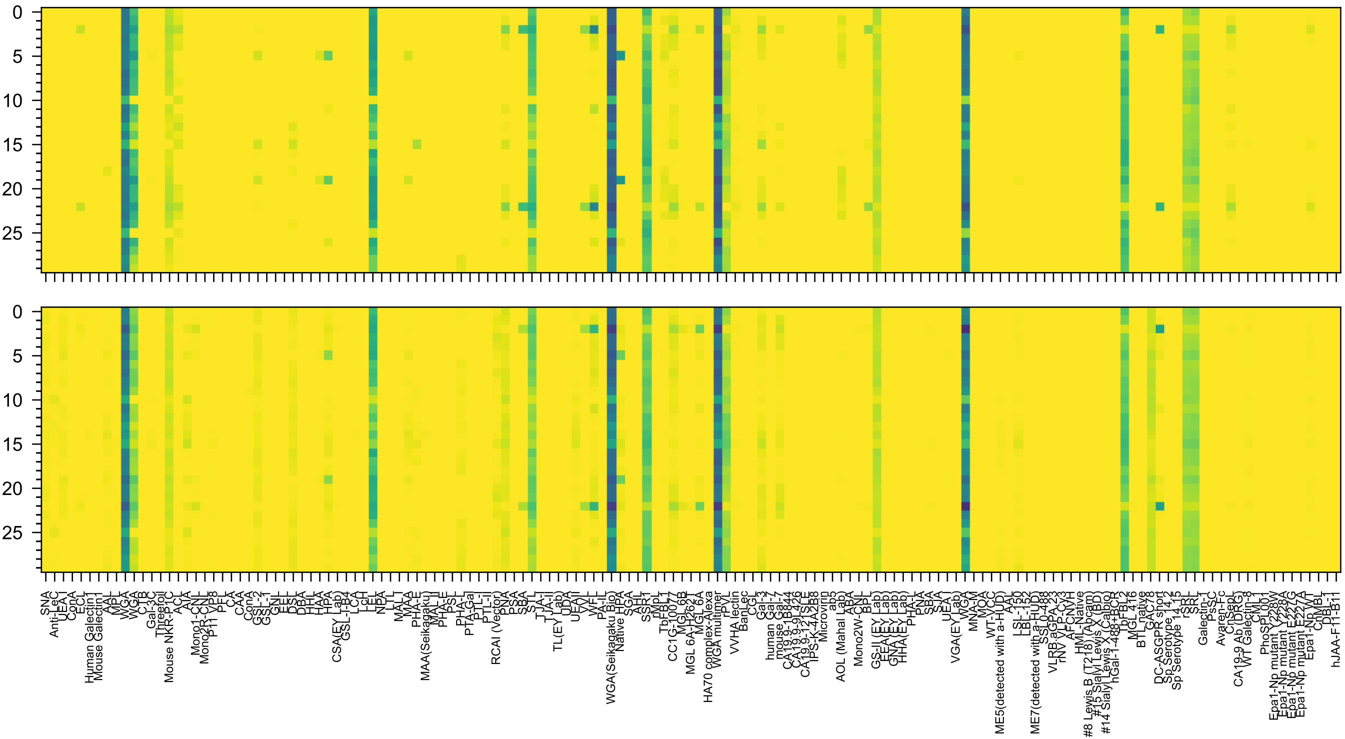


GlcNAc(β1-3)Gal(β1-4)GlcNAc(β1-6)[GlcNAc(β1-3)]Gal(β1-4)GlcNAc(-Sp0

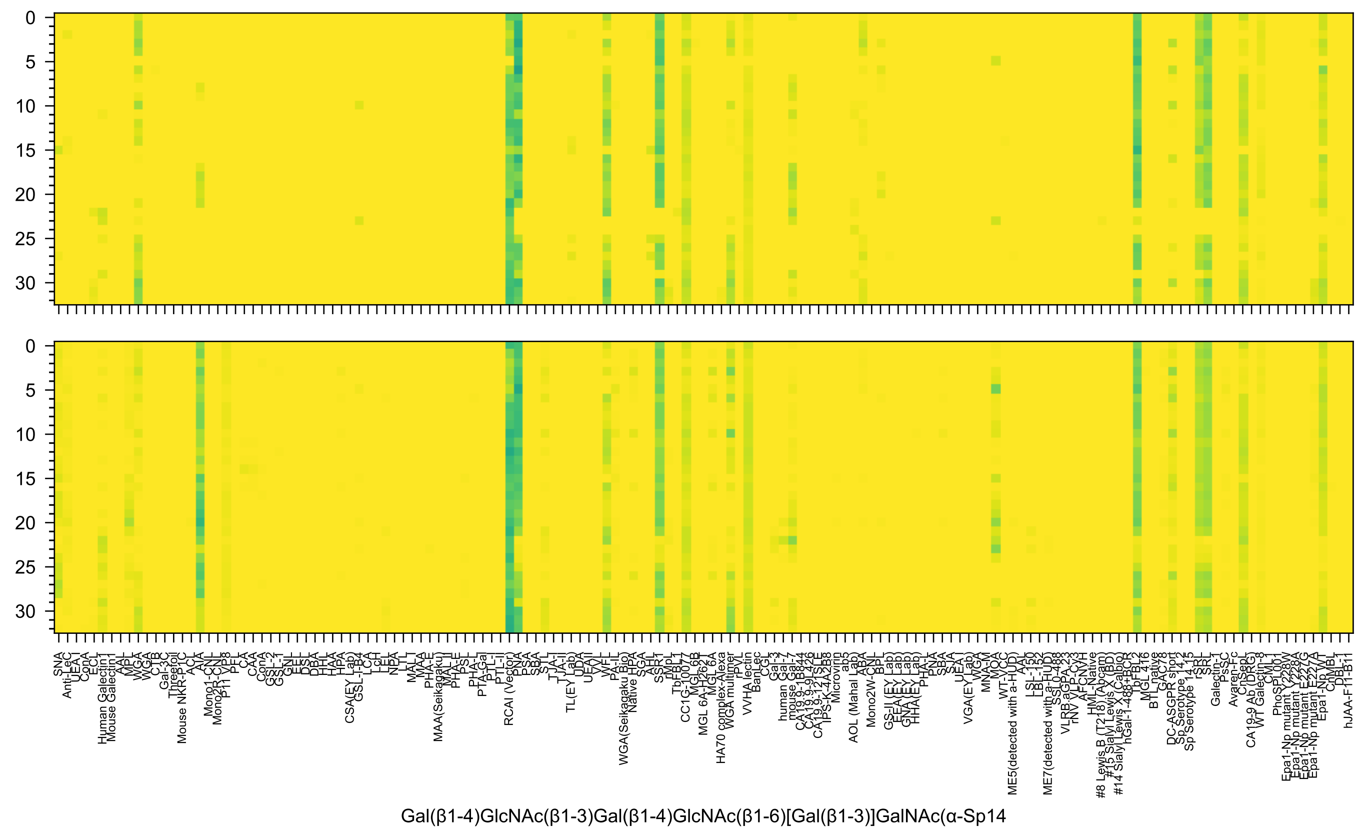
Gal(β1-4)GlcNAc(β1-3)Gal(β1-4)GlcNAc(β1-6)[Gal(β1-3)]GalNAc(α-Sp14

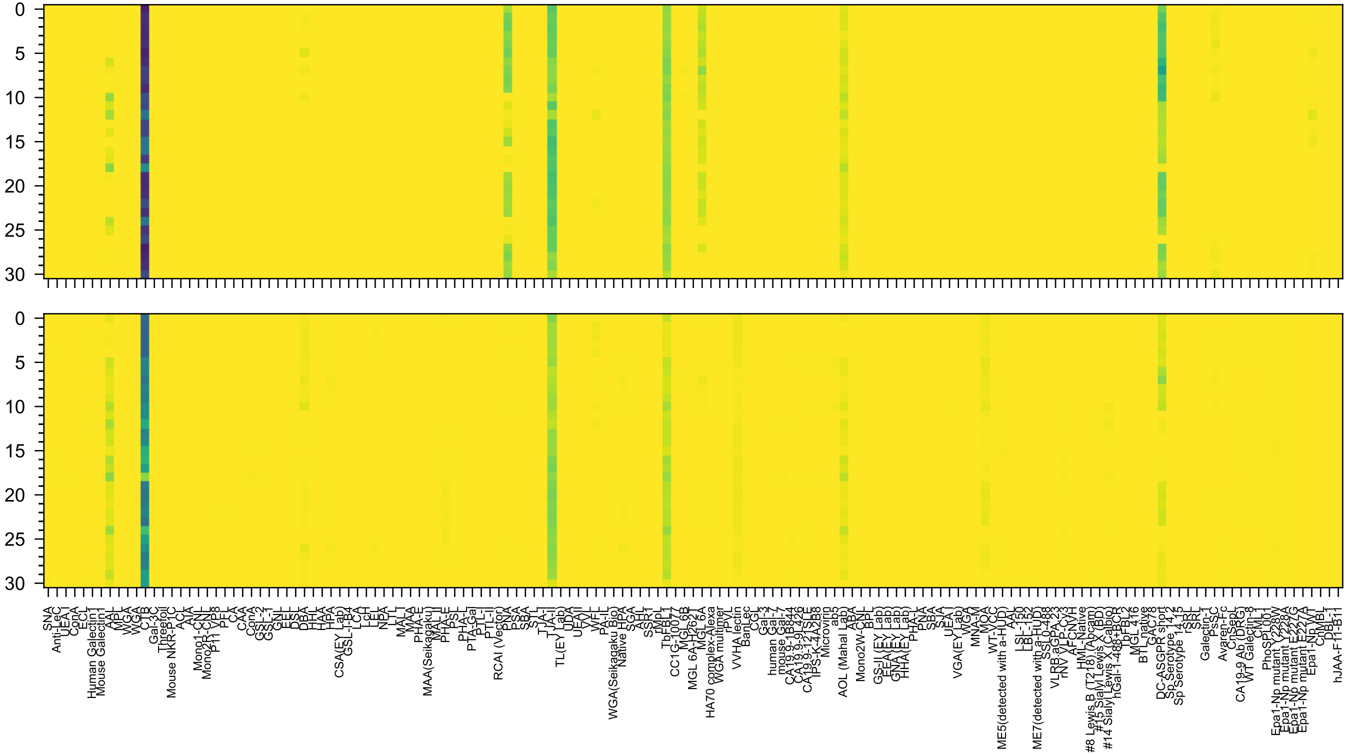

Fuc(α1-2)Gal(β1-3)GalNAc(β1-4)[Neu5Ac(α2-3)]Gal(β1-4)Glc(β-Sp9

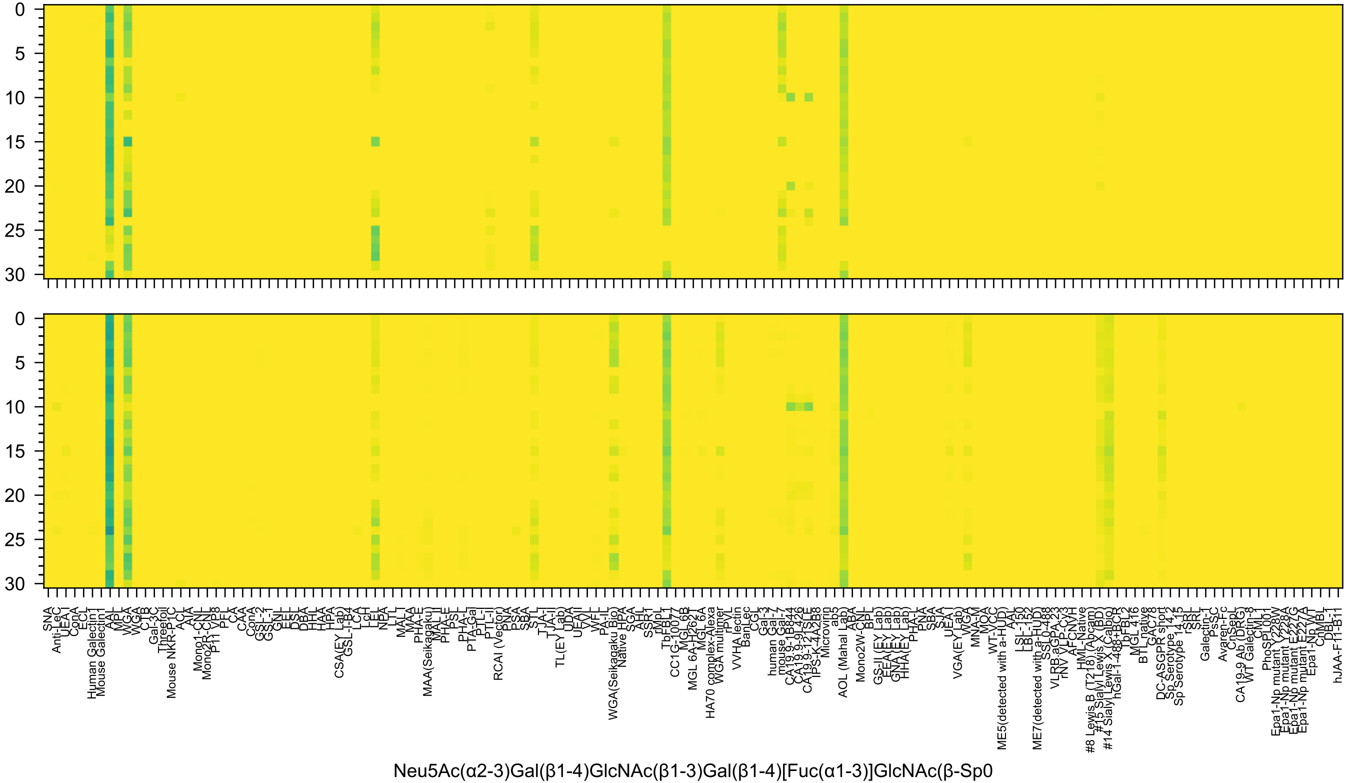
0
5
10
15
20
25
30
SNA
Anti-LeC
UEA I
ConA
ECL
Human Galectin1
Mouse Galectin1
AAL
MPL
WGA
WGA
CTB
Gal-3C
Threefoil
Mouse NKR-P1C
ACL
AIA
Mono1-CNL
Mono2R-CNL
P11 VP8
PFL
CA
CAA
ConA
GSL-2
GSL-1
GNL
EEL
DSL
DBA
HHL
HAA
HPA
CSA(EY Lab)
GSL-I-B4
LCA
LcH
LEL
NPA
LTL
MAL-I
MAA
PHA-E
MAA(Seikagaku)
MAL-II
PHA-E
PSL
PHA-L
PTA-Gal
PTL-I
PTL-II
RCAI (Vector)
PNA
PSA
SBA
STL
TJA-I
TJA-II
TL(EY Lab)
UDA
UEA-II
VVL
WFL
PA-IL
WGA(Seikagaku Bio)
Native HPA
SGA
AHL
SSR1
rMpL
TbFBL1
CC1G-10077
MGL 6B
MGL 6A-H262T
MGL 6A
HA70 complex-Alexa
WGA multimer
rPVL
VVHA lectin
BanLec
CGL
Gal-3
human Gal-7
mouse Gal-7
CA19-9-1B844
CA19-9-9L426
CA19-9-121SLE
IPS-K-4A2B8
Microvirin
ab5
AOL (Mahal Lab)
ABA
Mono2W-CNL
BPL
GS-II (EY Lab)
EEA (EY Lab)
GNA (EY Lab)
HHA (EY Lab)
PHA-L
PNA
SBA
SJA
UEA-I
VGA(EY Lab)
WGA
MNA-M
MOA
WT-VCC
ME5(detected with a-HUD)
AAL
LSL-150
LBL-152
ME7(detected with a-HUD)
SSL0-488
VLRB aGPA.23
rNV VLP-Cy3
AFCNVH
HML-Native
#8 Lewis B (T218) (Abcam)
#15 Sialyl Lewis X (BD)
#5 Sialyl Lewis X (Calbio)
#14 Sialyl Lewis X (Calbio)
hGal-1-488+BCR
TbFBL2
MGL 416
BTL native
GAC78
DC-ASGPR short
Sp Serotype 14.2
Sp Serotype 14.15
rSRL
SRL
Galectin-1
PsSC
Avaren-Fc
CnSepL
CA19-9 Ab (DRG)
WT Galectin-8
CML-1
PhoSL001
Epa1-Np mutant Y228W
Epa1-Np mutant Y228A
Epa1-Np mutant E227G
Epa1-Np mutant E227A
Epa1-Np WT
ChMBL
DBL-1
hJAA-F11-B1
Neu5Ac(α2-3)Gal(β1-4)GlcNAc(β1-3)Gal(β1-4)[Fuc(α1-3)]GlcNAc(β-Sp0

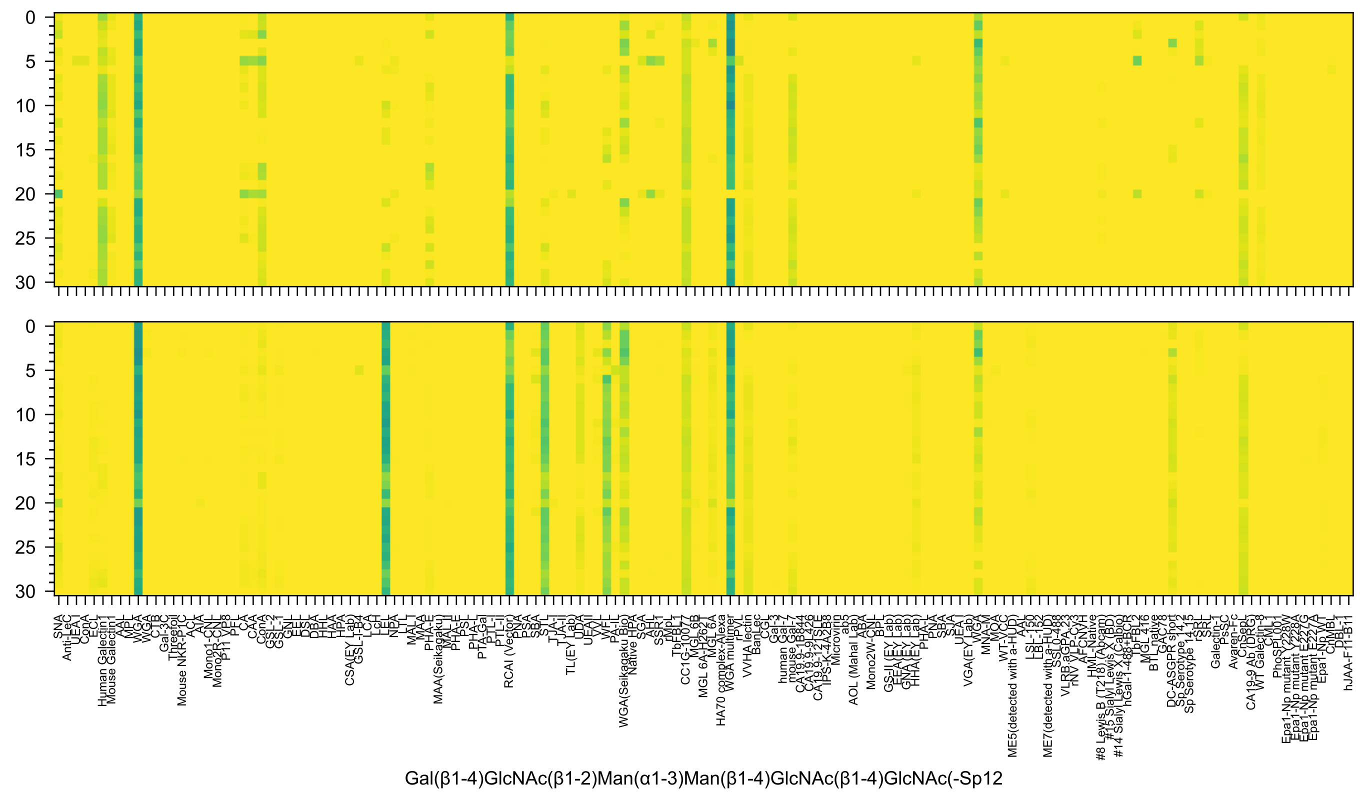
0
5
10
15
20
25
30
SNA
Anti-LeC
UEA-I
ConA
ECL
Human Galectin1
Mouse Galectin1
AAL
MPL
WGA
WGA
CTB
Gal-3C
Threefoil
Mouse NKR-P1C
ACL
AIA
Mono1-CNL
Mono2R-CNL
P11 VP8
PFL
CA
CAA
ConA
GSL-2
GSL-1
GNL
EEL
DSL
DBA
HHL
HAA
HPA
CSA(EY Lab)
GSL-I-B4
LCA
LcH
LEL
NPA
LTL
MAL-I
MAA
PHA-E
MAA(Seikagaku)
MAL_II
PHA-E
PSL
PHA-L
PTA-Gal
PTL-I
PTL-II
RCAI (Vector)
PNA
PSA
SBA
STL
TJA-I
TJA-II
TL(EY Lab)
UDA
UEA-II
VVL
WFL
PA-IL
WGA(Seikagaku Bio)
Native HPA
SGA
AHL
SSR1
rMpL
TbFBL1
CC1G-10077
MGL6B
MGL 6A-H262T
MGL 6A
HA70 complex-Alexa
WGA multimer
rPVL
VVHA lectin
BanLec
CGL
Gal-3
human Gal-7
mouse Gal-7
CA19.9-1B844
CA19.9-9L426
CA19.9-121SLE
IPS-K-4A2B8
Microvirin
ab5
AOL (Mahal Lab)
ABA
Mono2W-CNL
BPL
GS-II (EY Lab)
EEA (EY Lab)
GNA (EY Lab)
HHA (EY Lab)
PHA-L
PNA
SBA
SJA
UEA-I
VGA(EY Lab)
WGA
MNA-M
MOA
ME5(detected with a-HUD)
WT-VCC
AAL
LSL-150
LBL-152
ME7(detected with a-HUD)
SSL0-488
VLRB.aGPA.23
rNV VLP-Cy3
AFCNVH
HML-Native
#8 Lewis B (T218) (Abcam)
#15 Sialyl Lewis X (BD)
#5 Sialyl Lewis X (Calbio)
hGal-1-488+BCR
TbFBL2
MGL 416
BTL native
GAC78
DC-ASGPR short
Sp Serotype 14.2
Sp Serotype 14.15
rSRL
SRL
Galectin-1
PsSC
Avaren-Fc
CnSepL
CA19-9 Ab (DRG)
WT Galectin-8
PhoSL001
Epa1-Np mutant Y228W
Epa1-Np mutant Y228A
Epa1-Np mutant E227G
Epa1-Np mutant E227A
Epa1-Np WT
CnMBL
DBL-1
hJAA-F11-B1
Gal(β1-4)GlcNAc(β1-2)Man(α1-3)Man(β1-4)GlcNAc(β1-4)GlcNAc(-Sp12

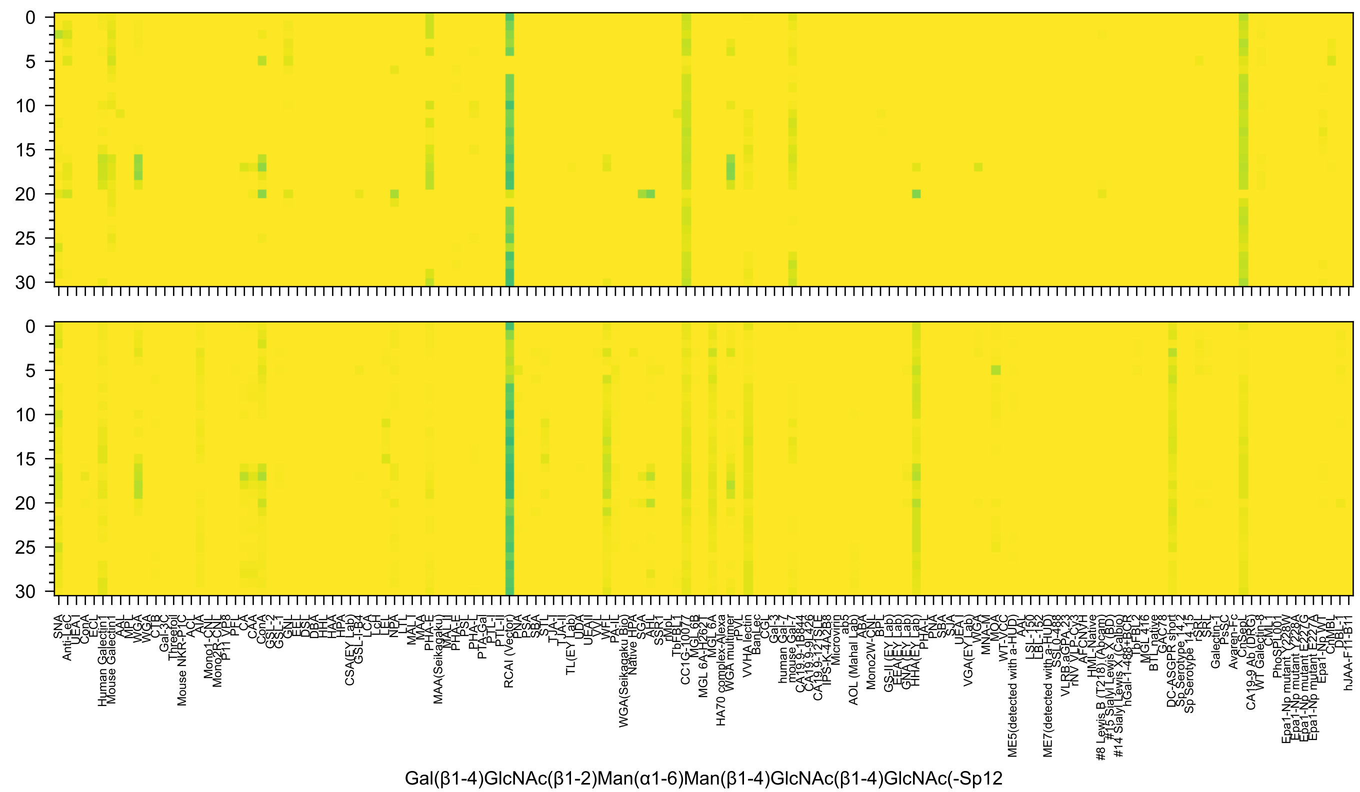
Gal(β1-4)GlcNAc(β1-2)Man(α1-6)Man(β1-4)GlcNAc(β1-4)GlcNAc(-Sp12

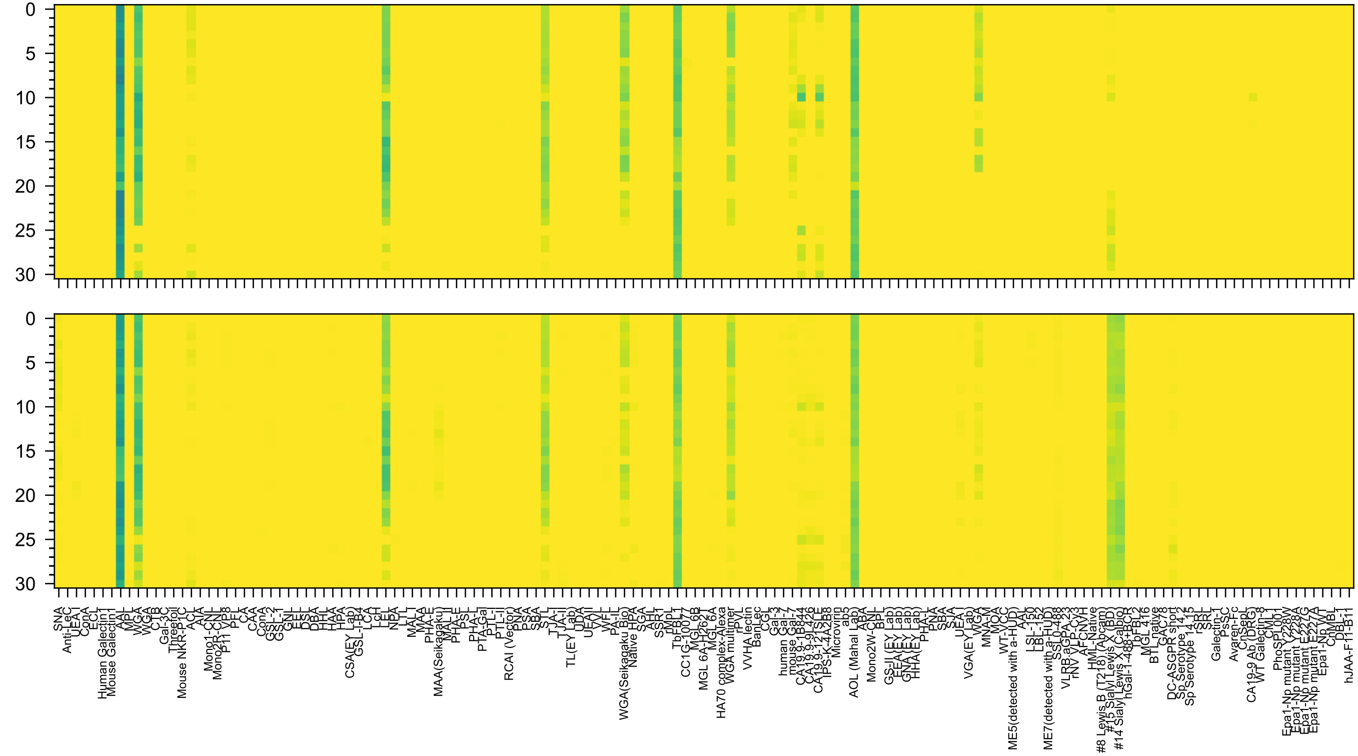

Neu5Ac(α2-3)Gal(β1-4)[Fuc(α1-3)]GlcNAc(β1-3)Gal(β1-4)GlcNAc(β-Sp8

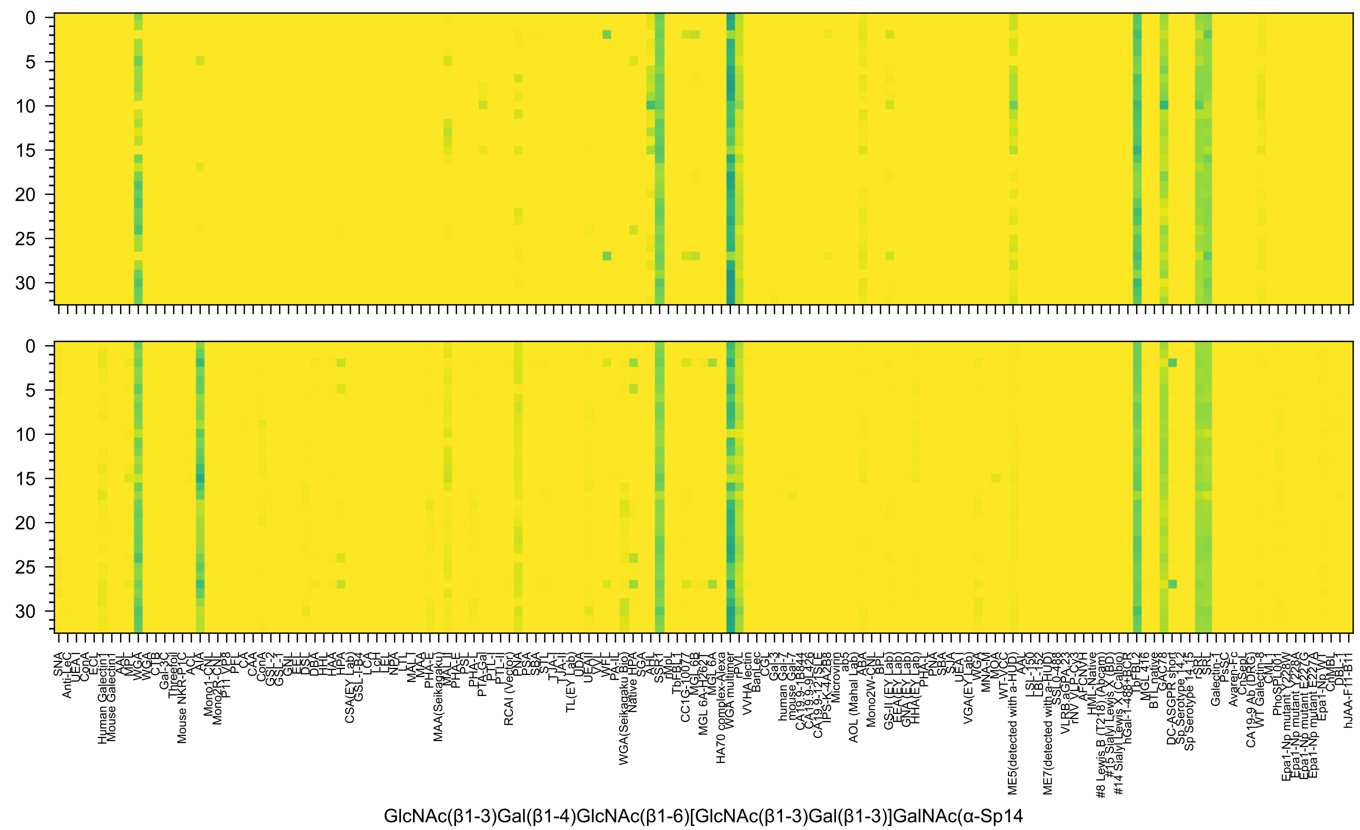
GlcNAc(β1-3)Gal(β1-4)GlcNAc(β1-6)[GlcNAc(β1-3)Gal(β1-3)]GalNAc(α-Sp14

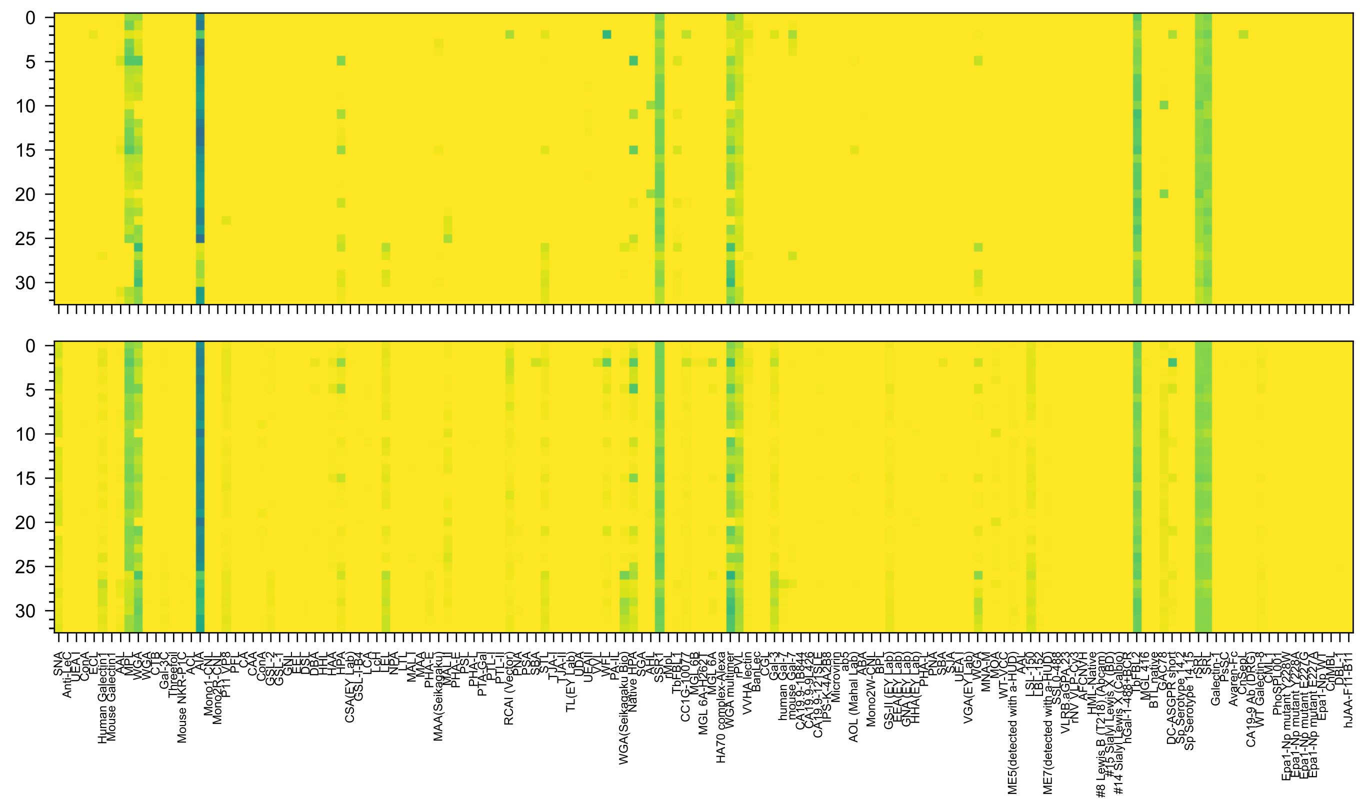

GlcNAc(β1-3)Gal(β1-4)GlcNAc(β1-3)Gal(β1-4)GlcNAc(β1-3)GalNAc(α-Sp14

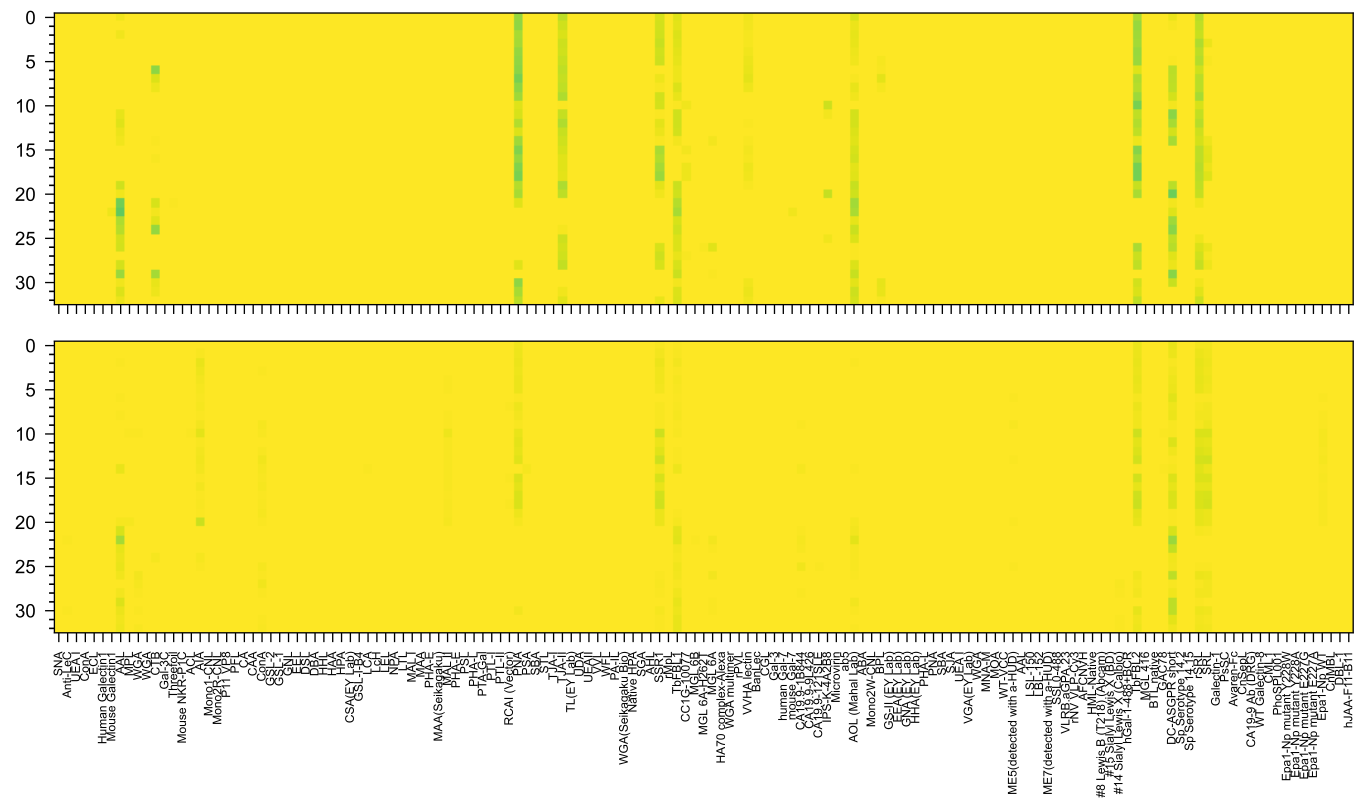


Neu5Ac(α2-3)Gal(β1-4)[Fuc(α1-3)]GlcNAc(β1-6)[Gal(β1-3)]GalNAc(α-Sp14

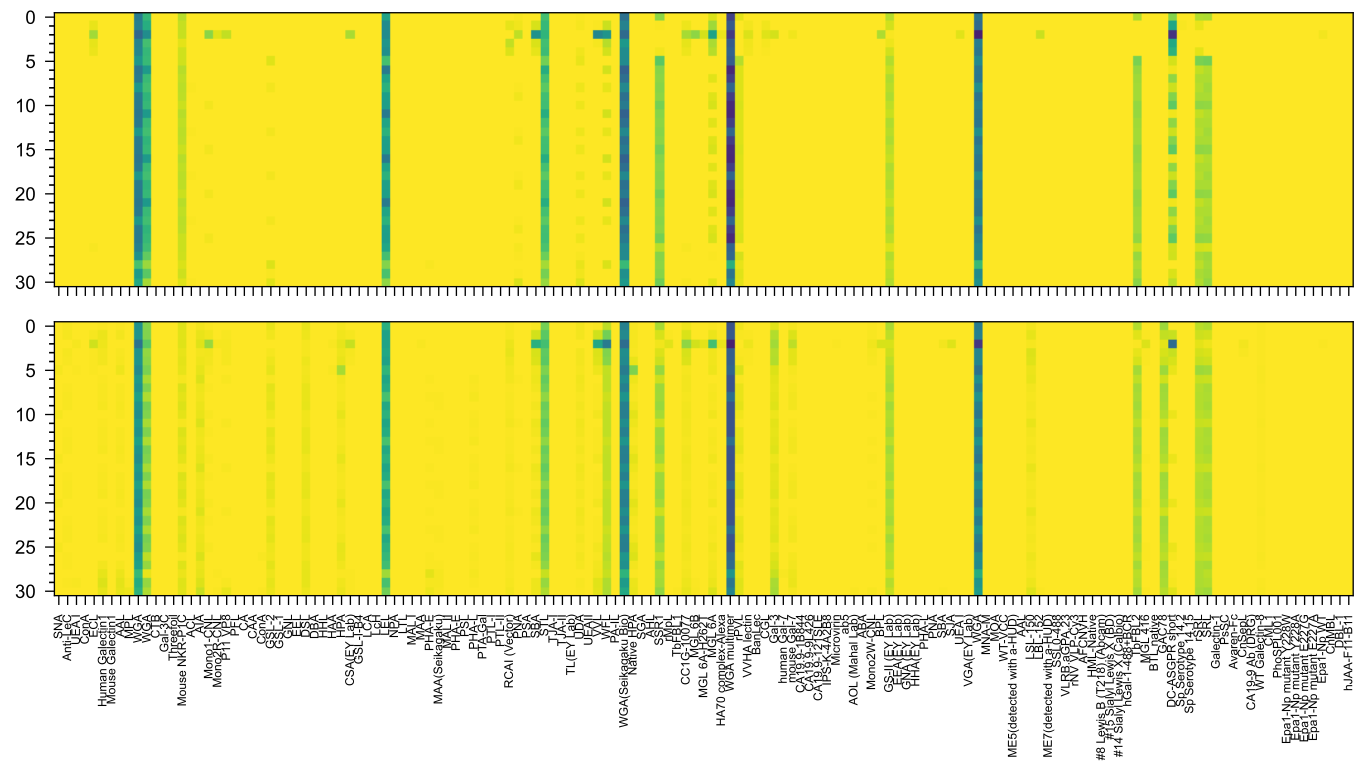


GlcNAc(β1-4)GlcNAc(β1-4)GlcNAc(β1-4)GlcNAc(β1-4)GlcNAc(β1-4)GlcNAc(β1-Sp8

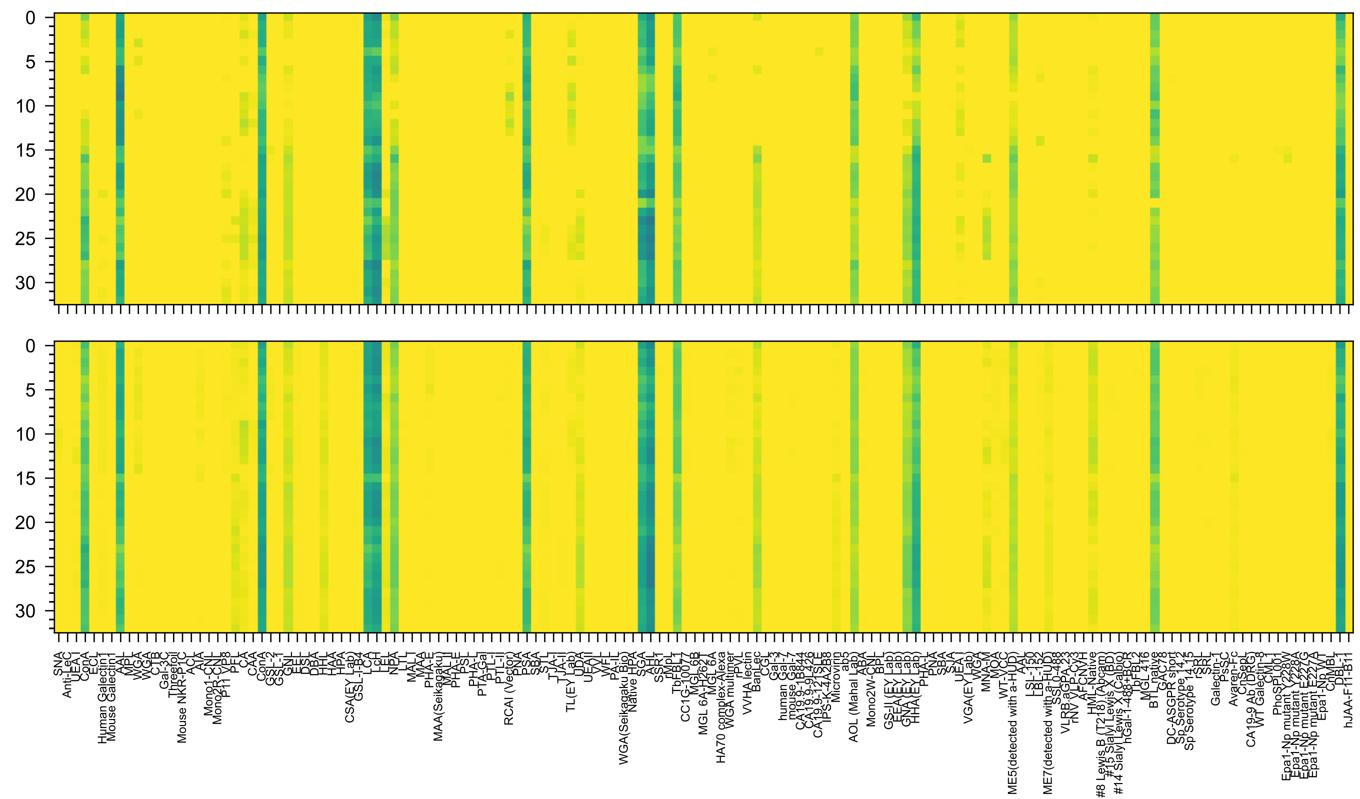


Man(α1-6)[Man(α1-3)]Man(β1-4)GlcNAc(β1-4)[Fuc(α1-6)]GlcNAc(β-Sp19

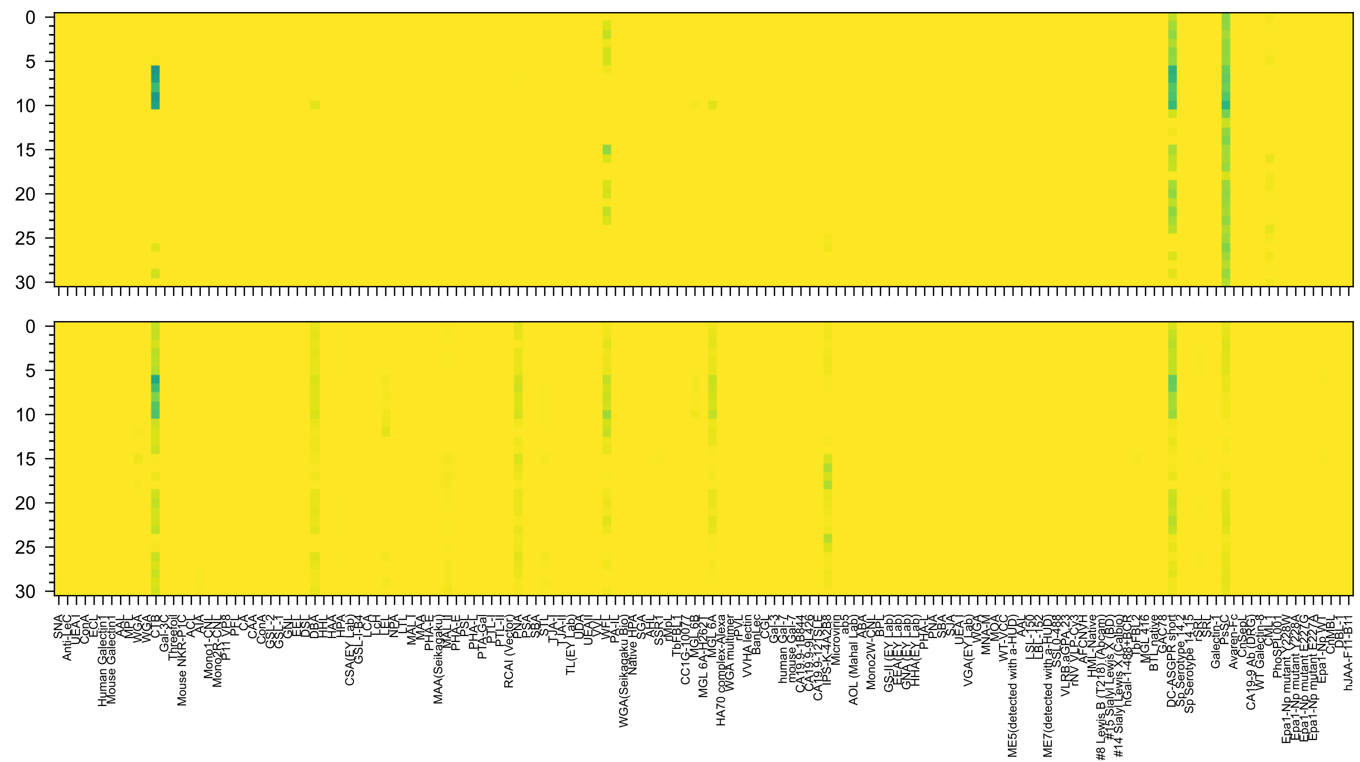

Neu5Ac(α2-3)Gal(β1-3)GalNAc(β1-4)[Neu5Ac(α2-3)]Gal(β1-4)Glc(β-Sp0

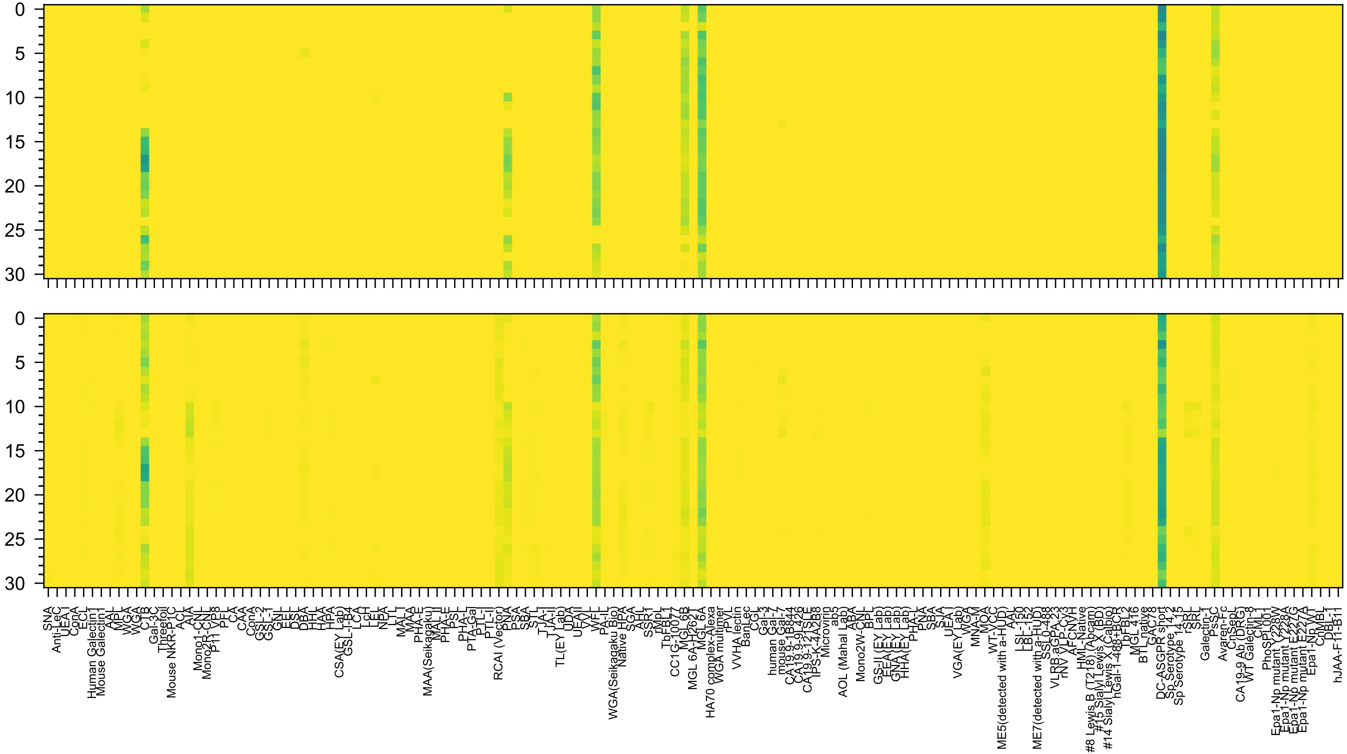
Gal(β1-3)GalNAc(β1-4)[Neu5Ac(α2-8)Neu5Ac(α2-3)]Gal(β1-4)Glc(β-Sp0

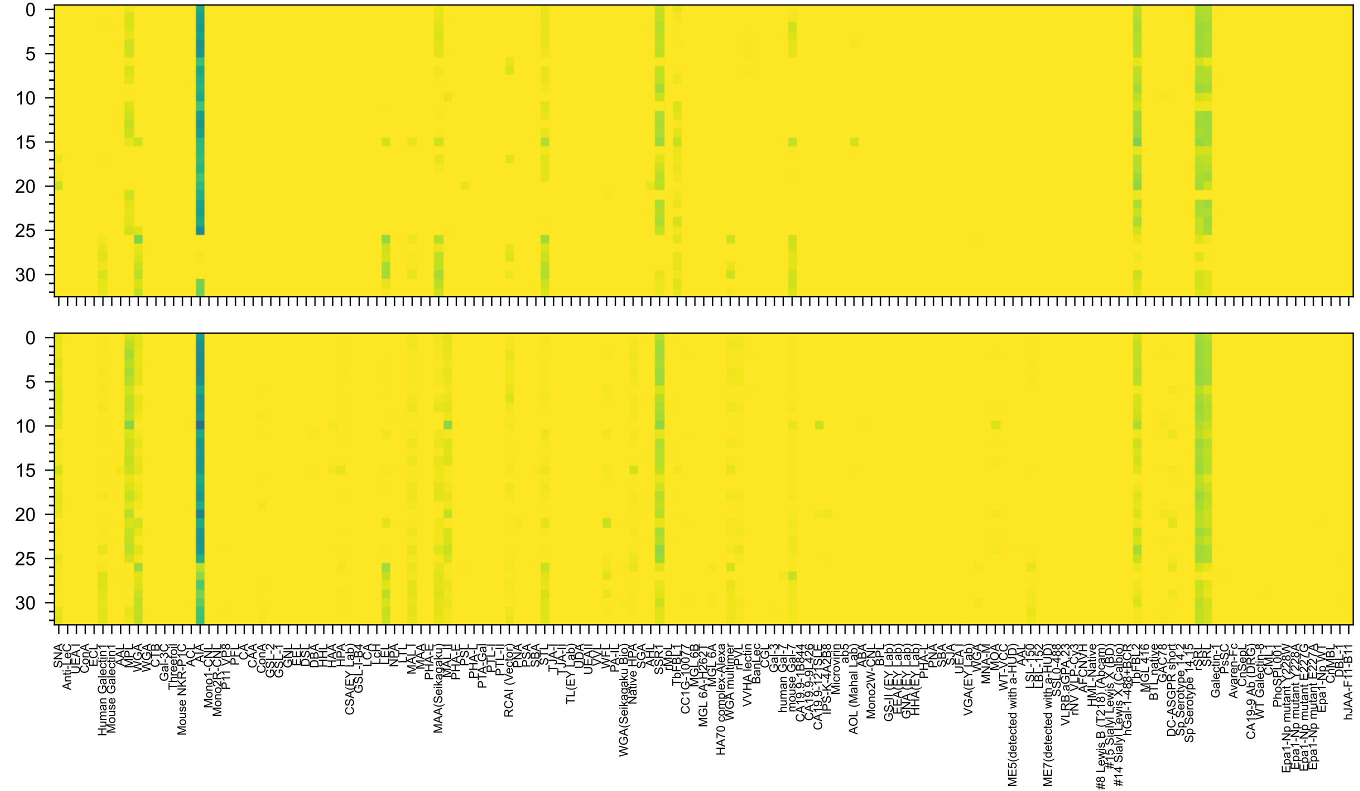

Neu5Ac(α2-3)Gal(β1-4)GlcNAc(β1-3)Gal(β1-4)GlcNAc(β1-3)GalNAc(α-Sp14

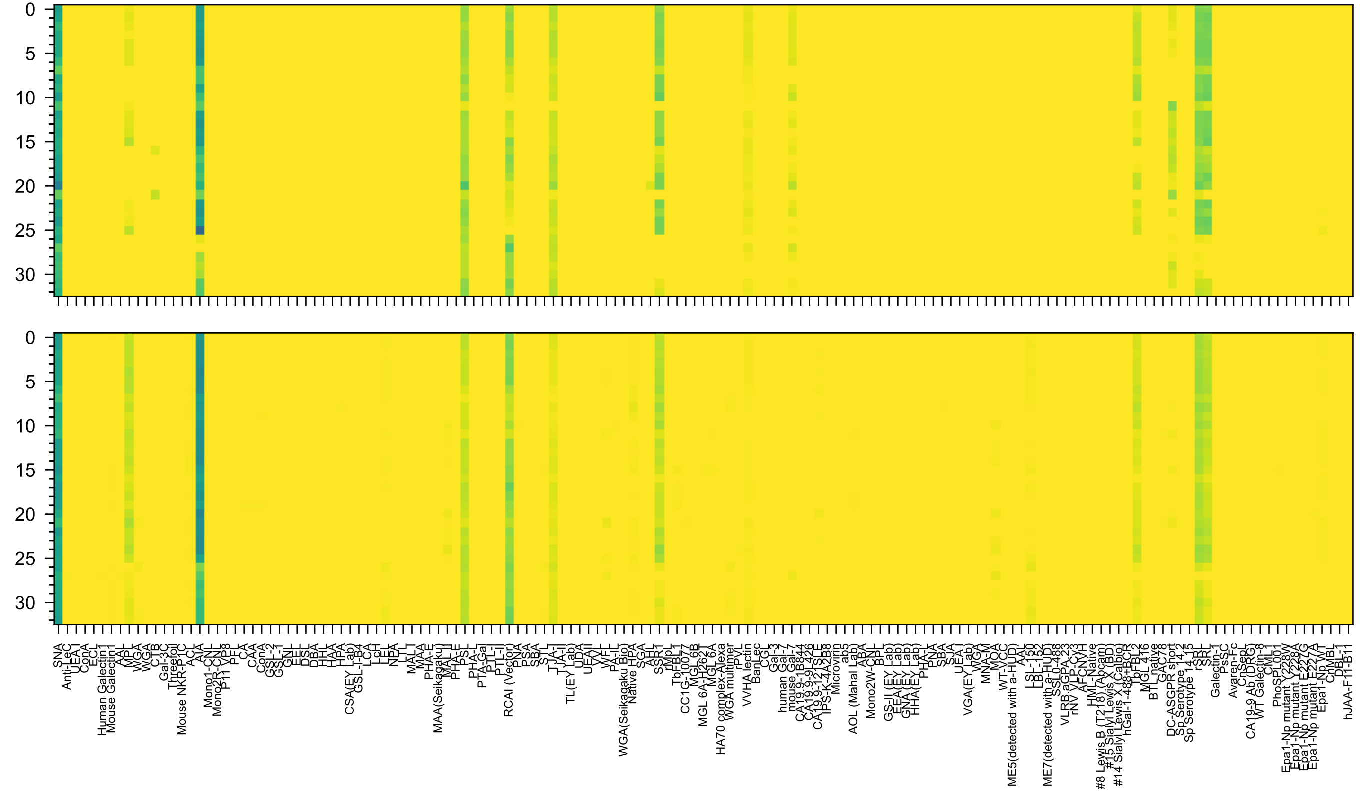


Neu5Ac(α2-6)Gal(β1-4)GlcNAc(β1-3)Gal(β1-4)GlcNAc(β1-3)GalNAc(α-Sp14

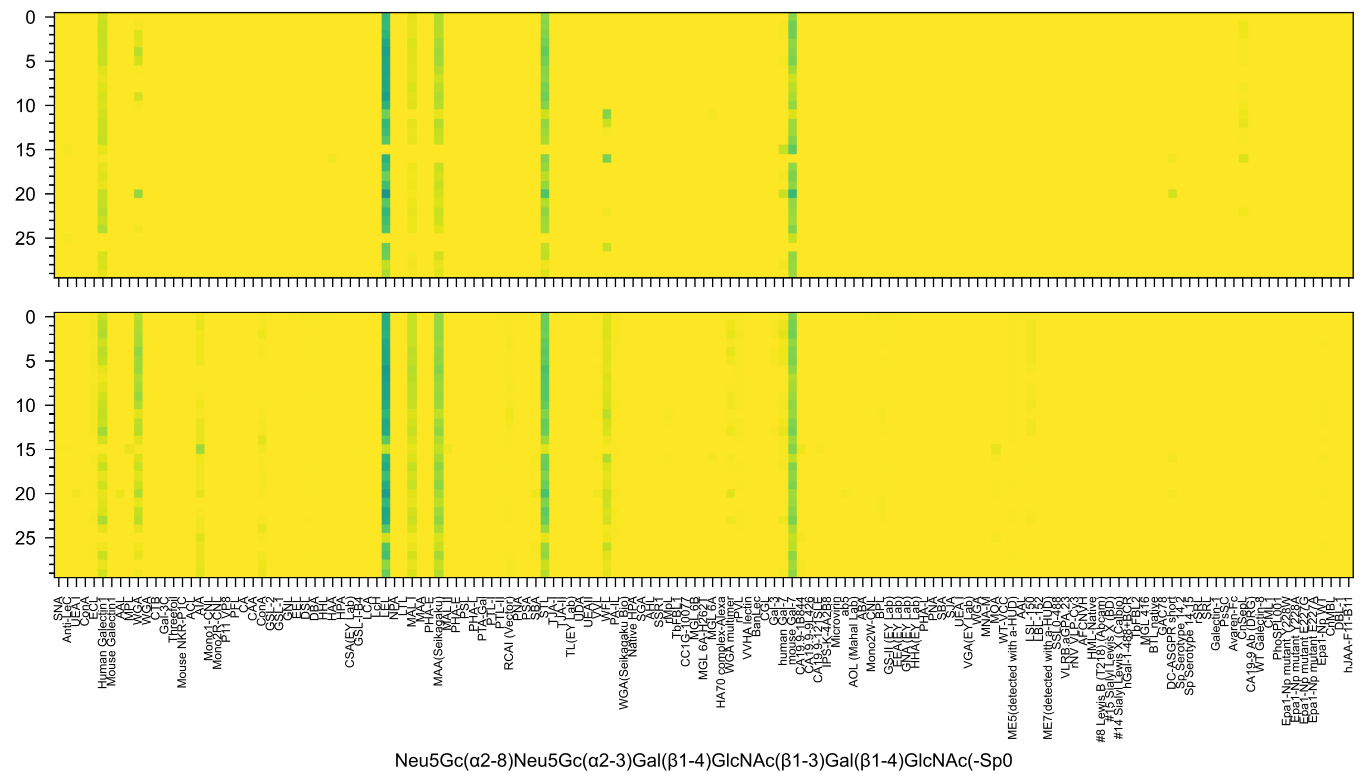
Neu5Gc(α2-8)Neu5Gc(α2-3)Gal(β1-4)GlcNAc(β1-3)Gal(β1-4)GlcNAc(-Sp0

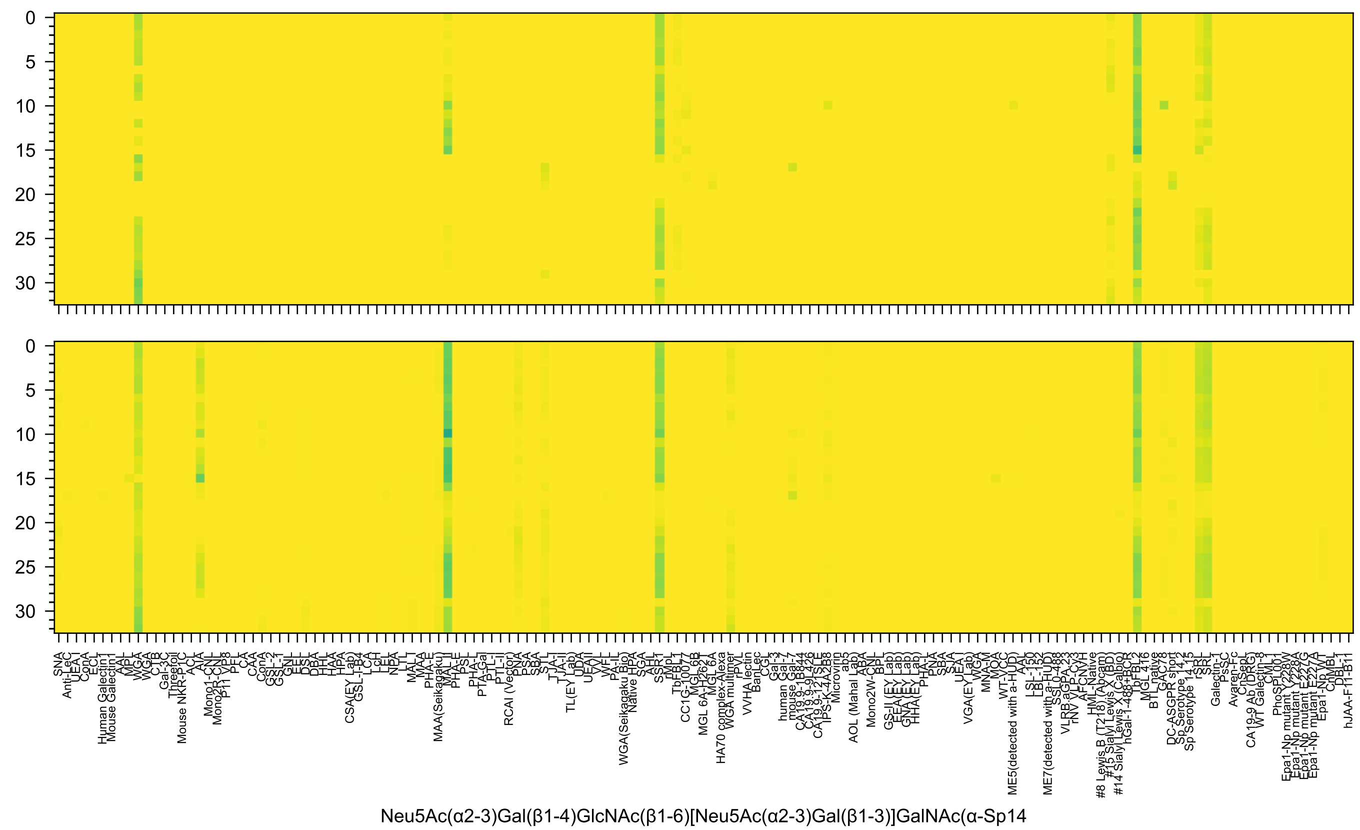
Neu5Ac(α2-3)Gal(β1-4)GlcNAc(β1-6)[Neu5Ac(α2-3)Gal(β1-3)]GalNAc(α-Sp14

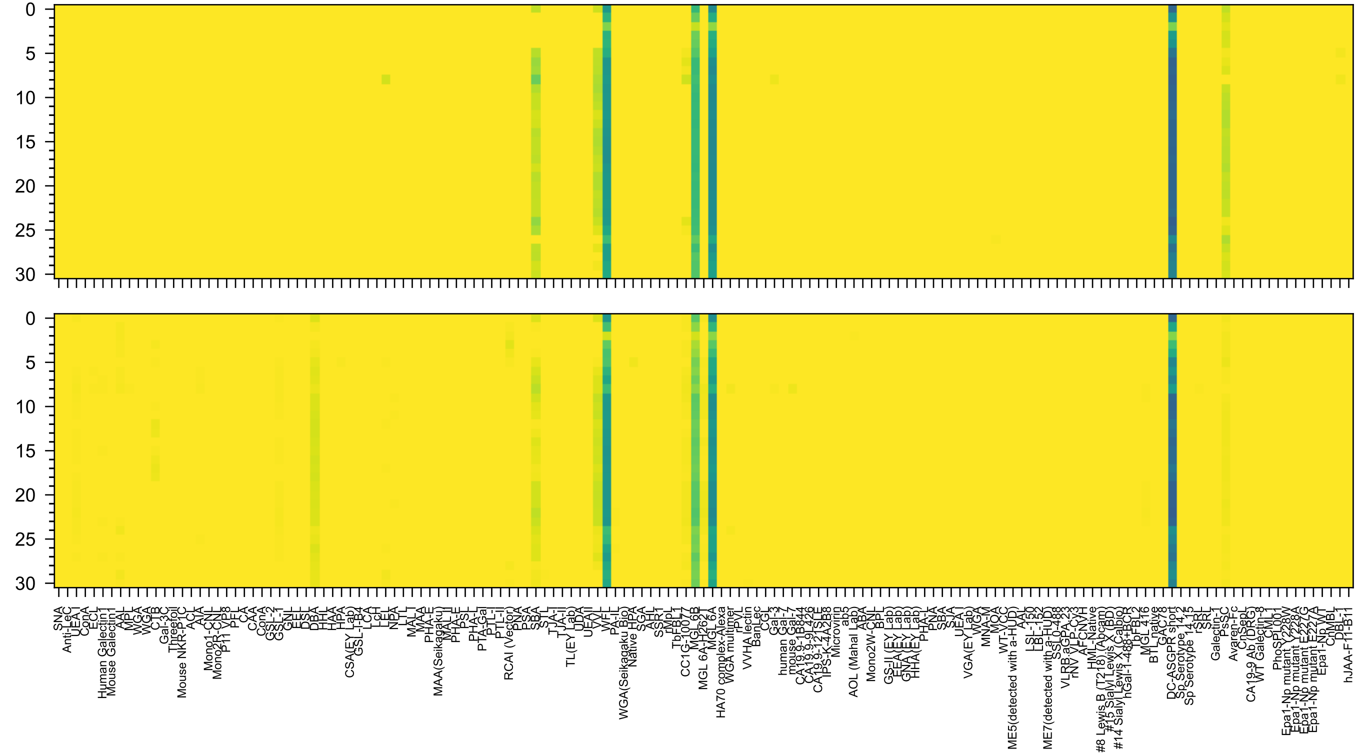


GalNAc(β1-4)[Neu5Ac(α2-8)Neu5Ac(α2-8)Neu5Ac(α2-3)]Gal(β1-4)Glc(β-Sp0

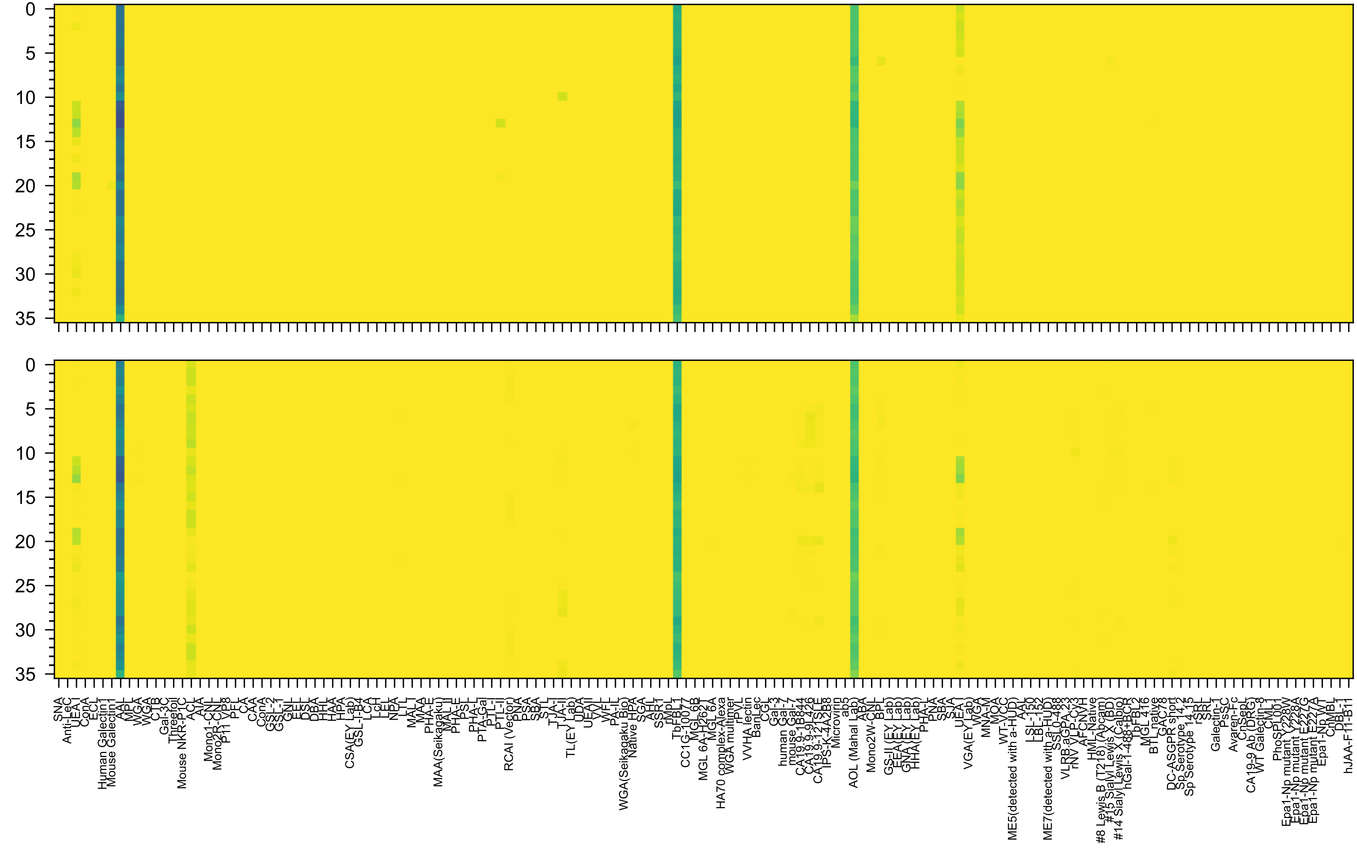

Fuc(α1-2)Gal(β1-4)[Fuc(α1-3)]GlcNAc(β1-3)Gal(β1-4)[Fuc(α1-3)]GlcNAc(β-Sp0

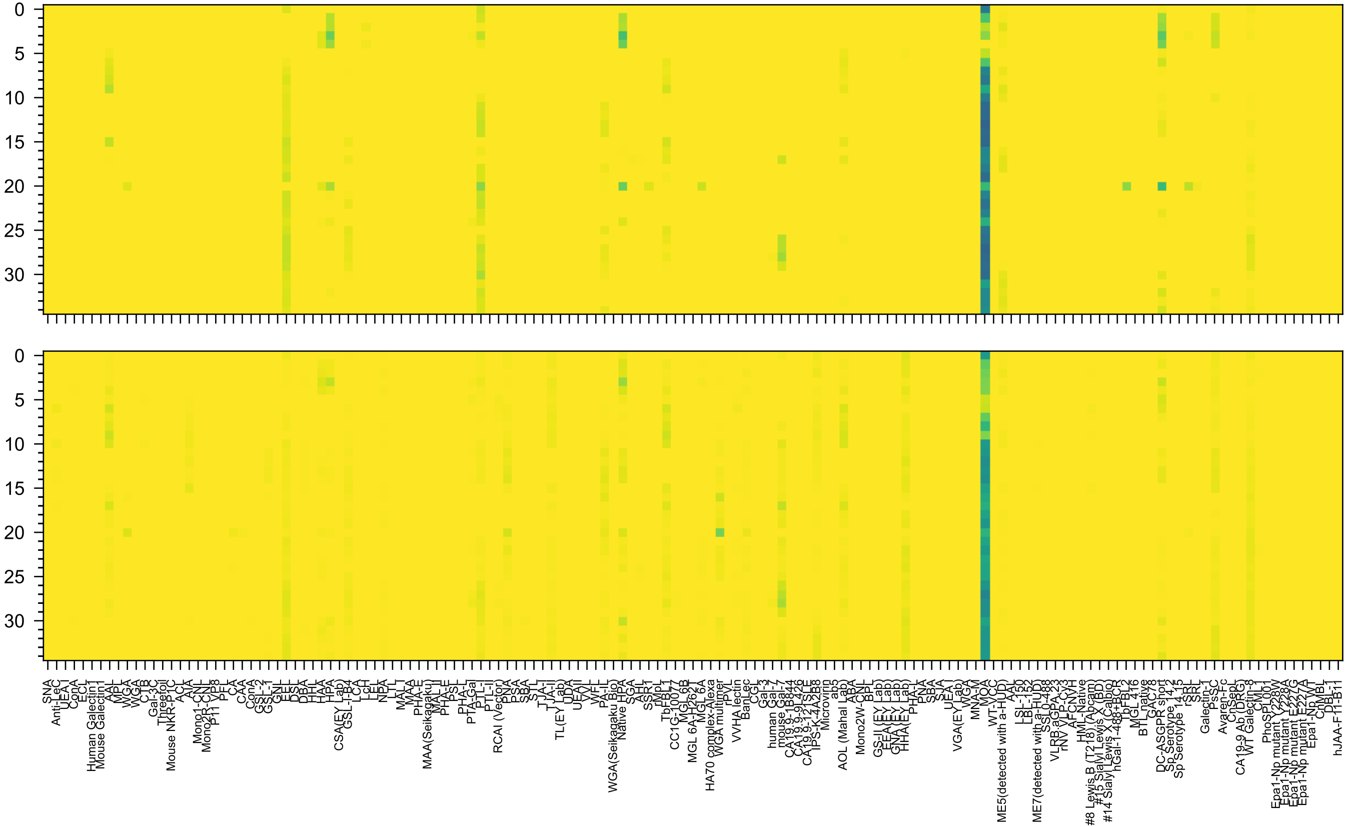

Gal(α1-3)[Fuc(α1-2)]Gal(β1-3)GalNAc(β1-3)Gal(α1-4)Gal(β1-4)Glc(-Sp21

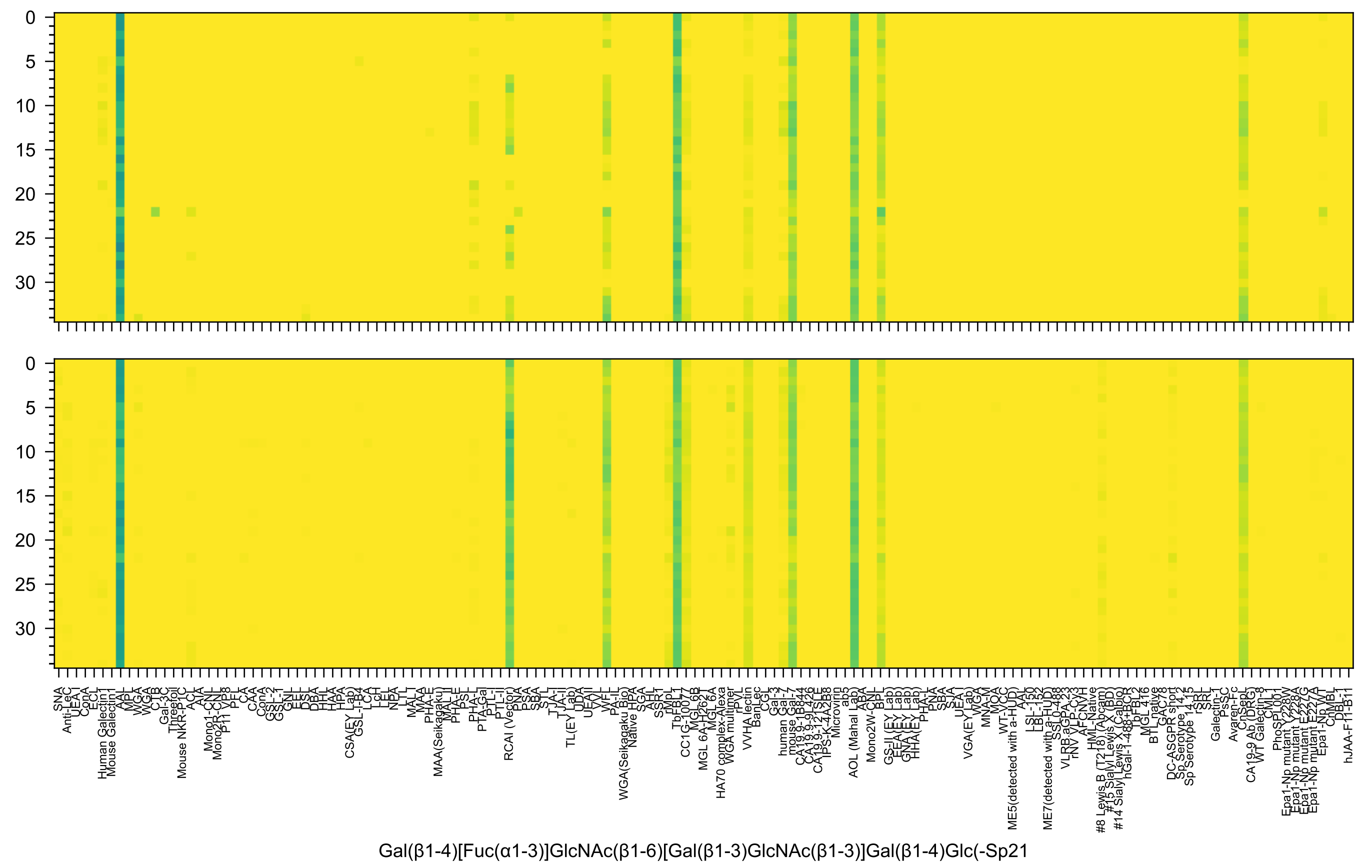
Gal(β1-4)[Fuc(α1-3)]GlcNAc(β1-6)[Gal(β1-3)GlcNAc(β1-3)]Gal(β1-4)Glc(-Sp21

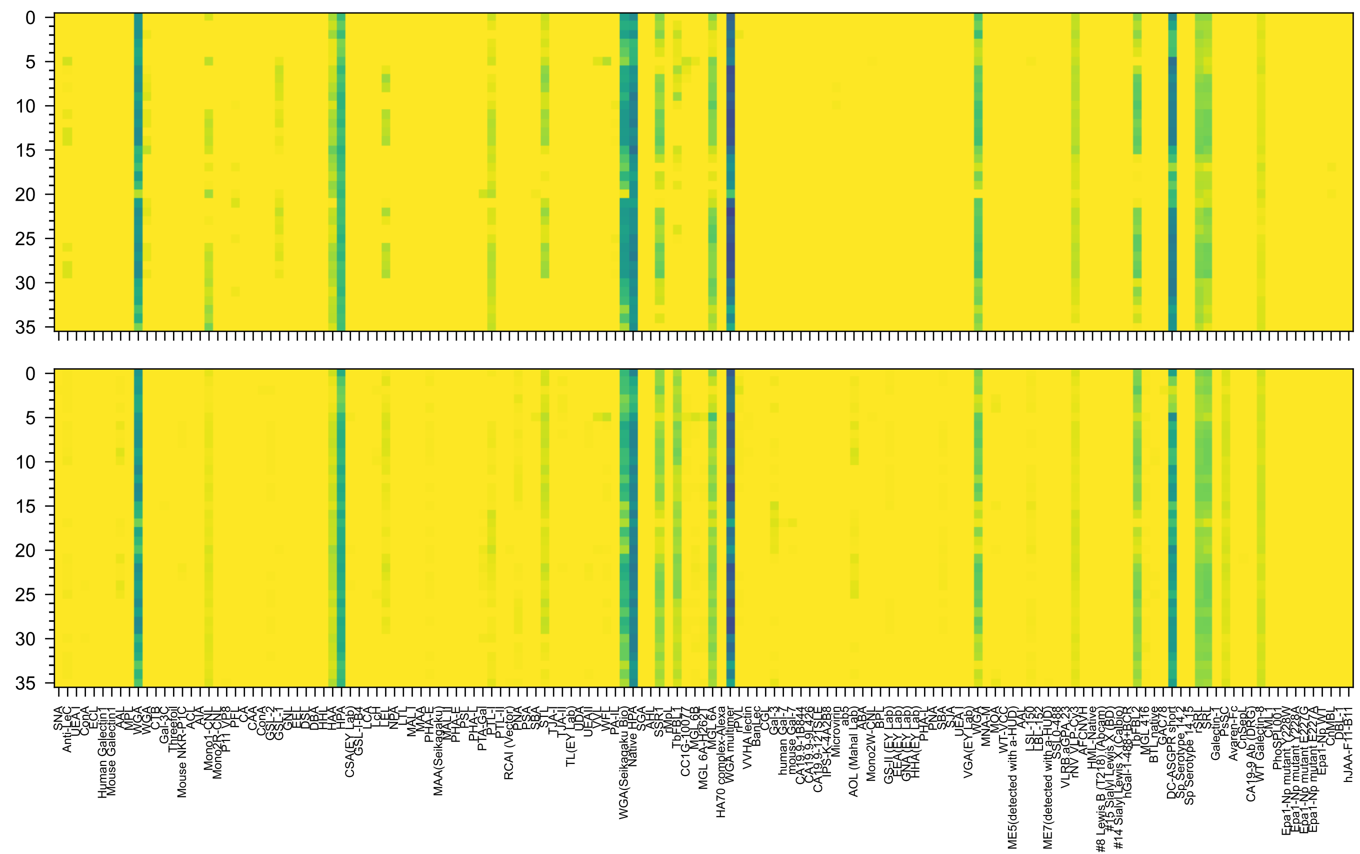


GalNAc(α1-3)[Fuc(α1-2)]Gal(β1-3)GalNAc(α1-3)[Fuc(α1-2)]Gal(β1-4)GlcNAc(β-Sp0

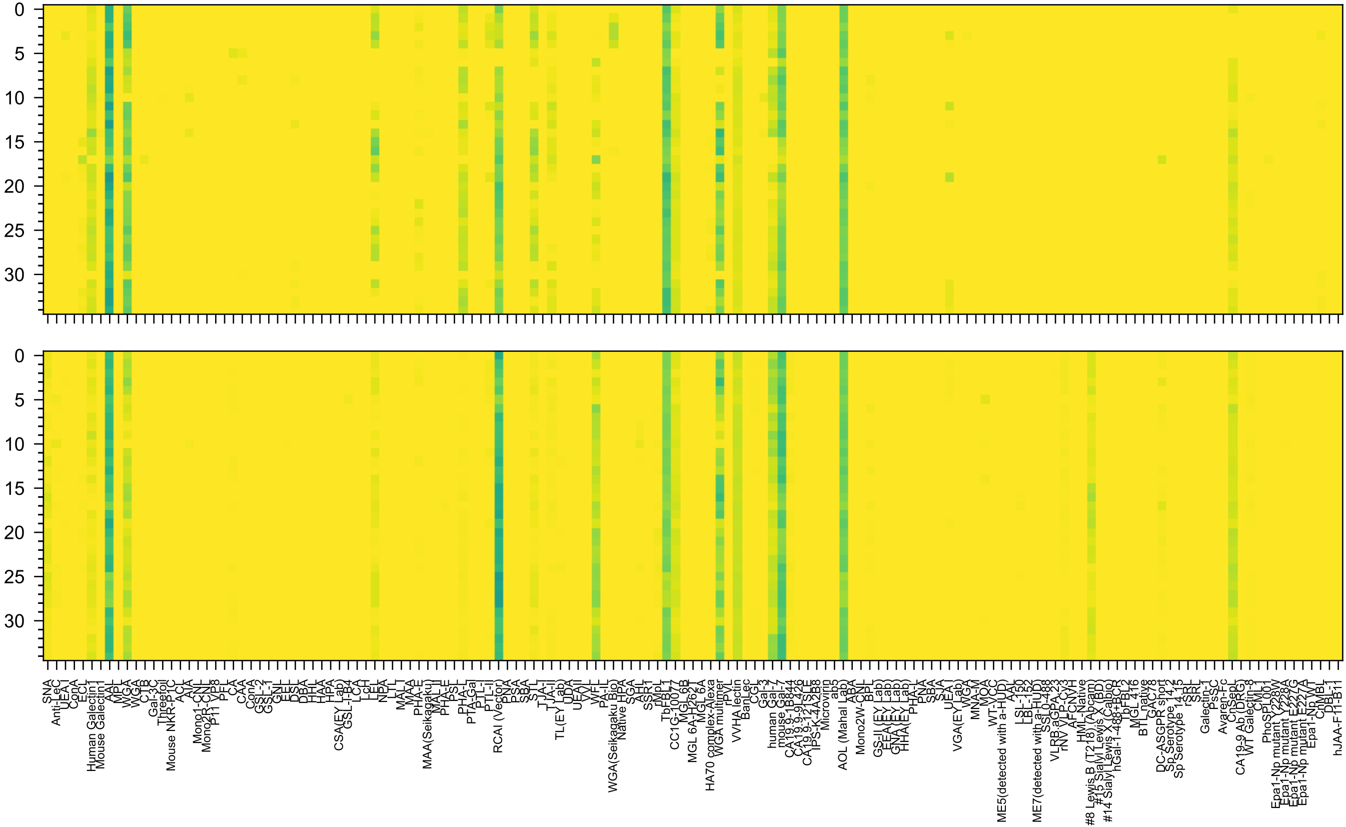

Gal(β1-4)GlcNAc(β1-6)[Fuc(α1-2)Gal(β1-3)GlcNAc(β1-3)]Gal(β1-4)Glc(-Sp21

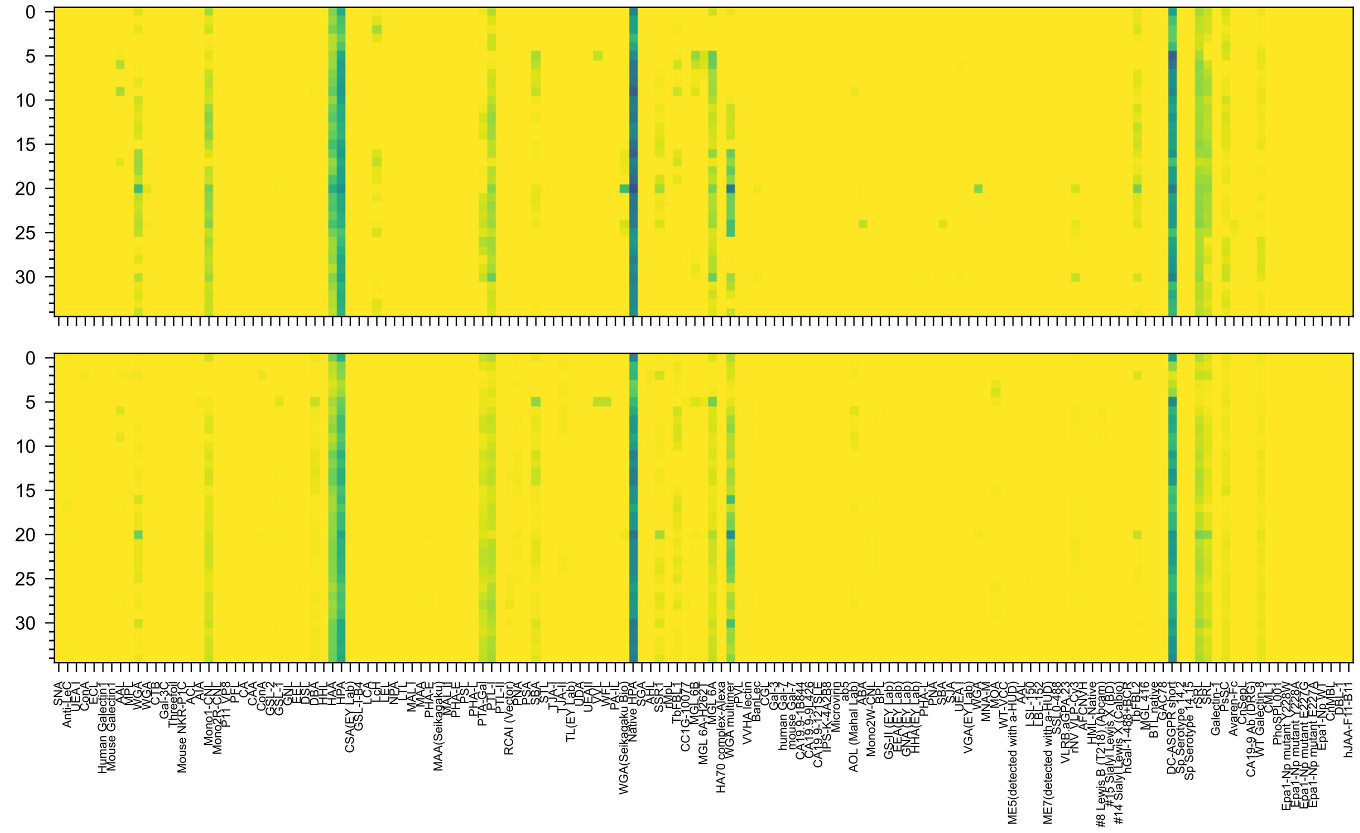

GalNAc(α1-3)[Fuc(α1-2)]Gal(β1-3)GalNAc(β1-3)Gal(α1-4)Gal(β1-4)Glc(-Sp21

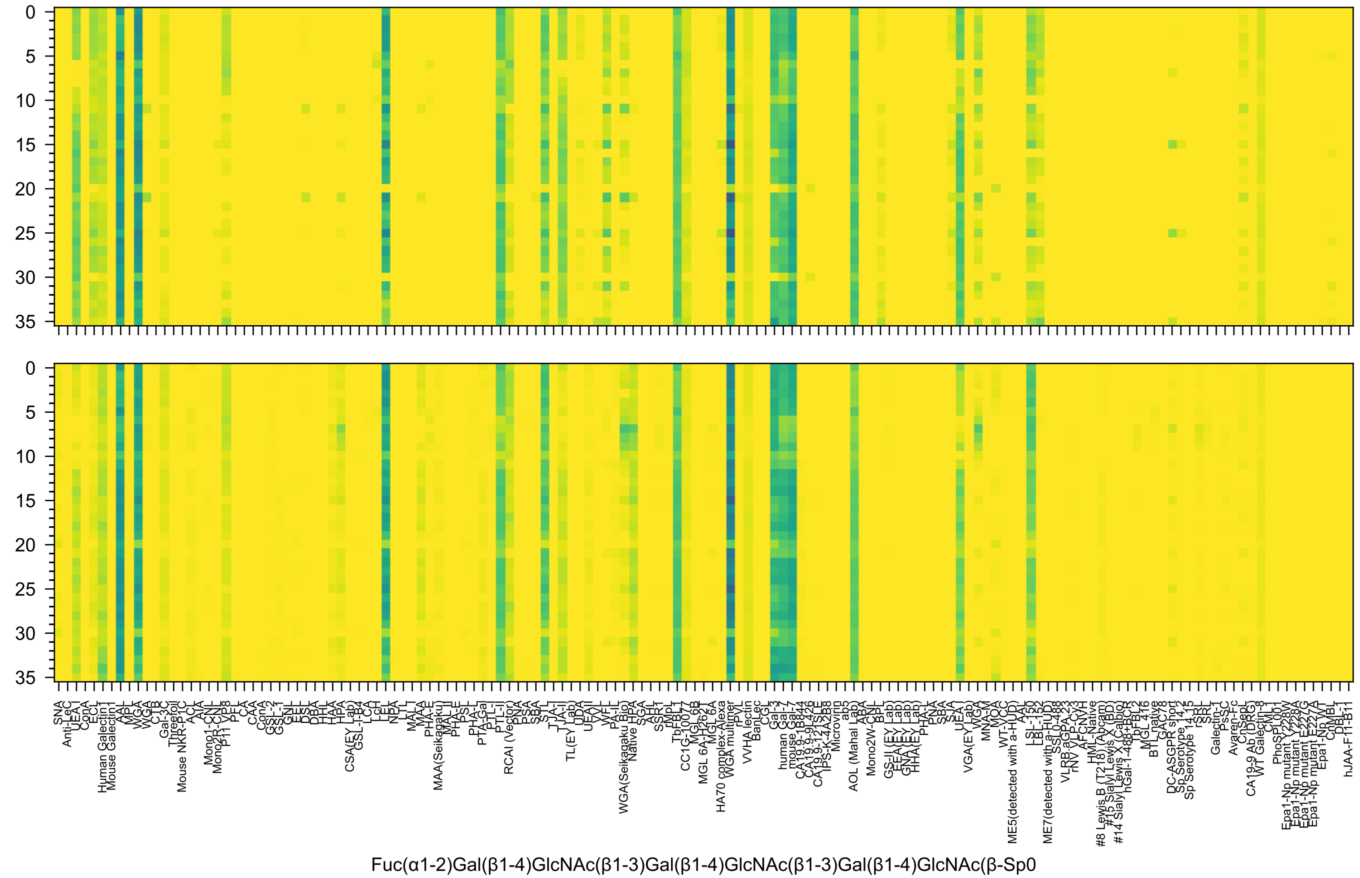
Fuc(α1-2)Gal(β1-4)GlcNAc(β1-3)Gal(β1-4)GlcNAc(β1-3)Gal(β1-4)GlcNAc(β-Sp0

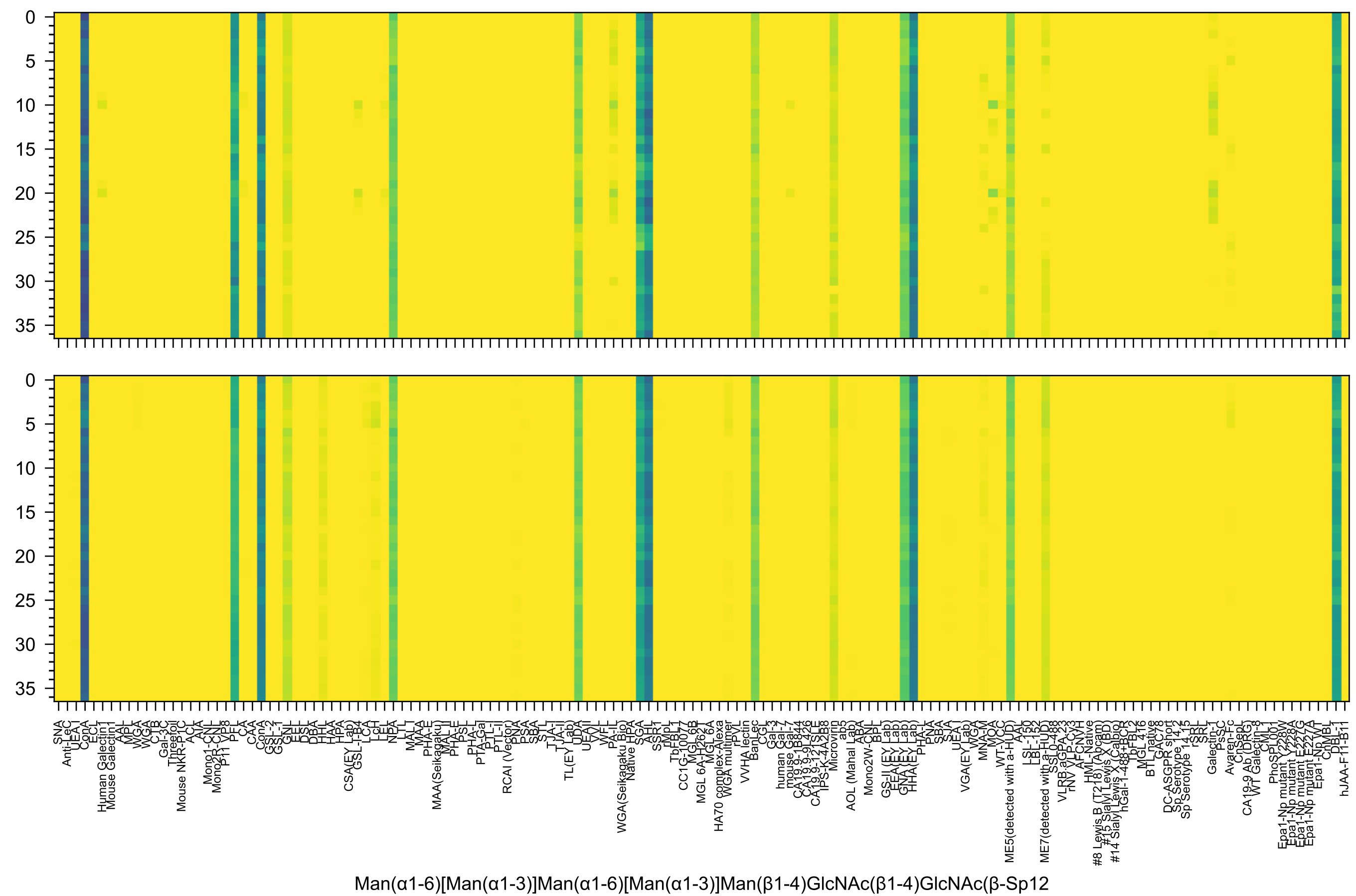
Man(α1-6)[Man(α1-3)]Man(α1-6)[Man(α1-3)]Man(β1-4)GlcNAc(β1-4)GlcNAc(β-Sp12

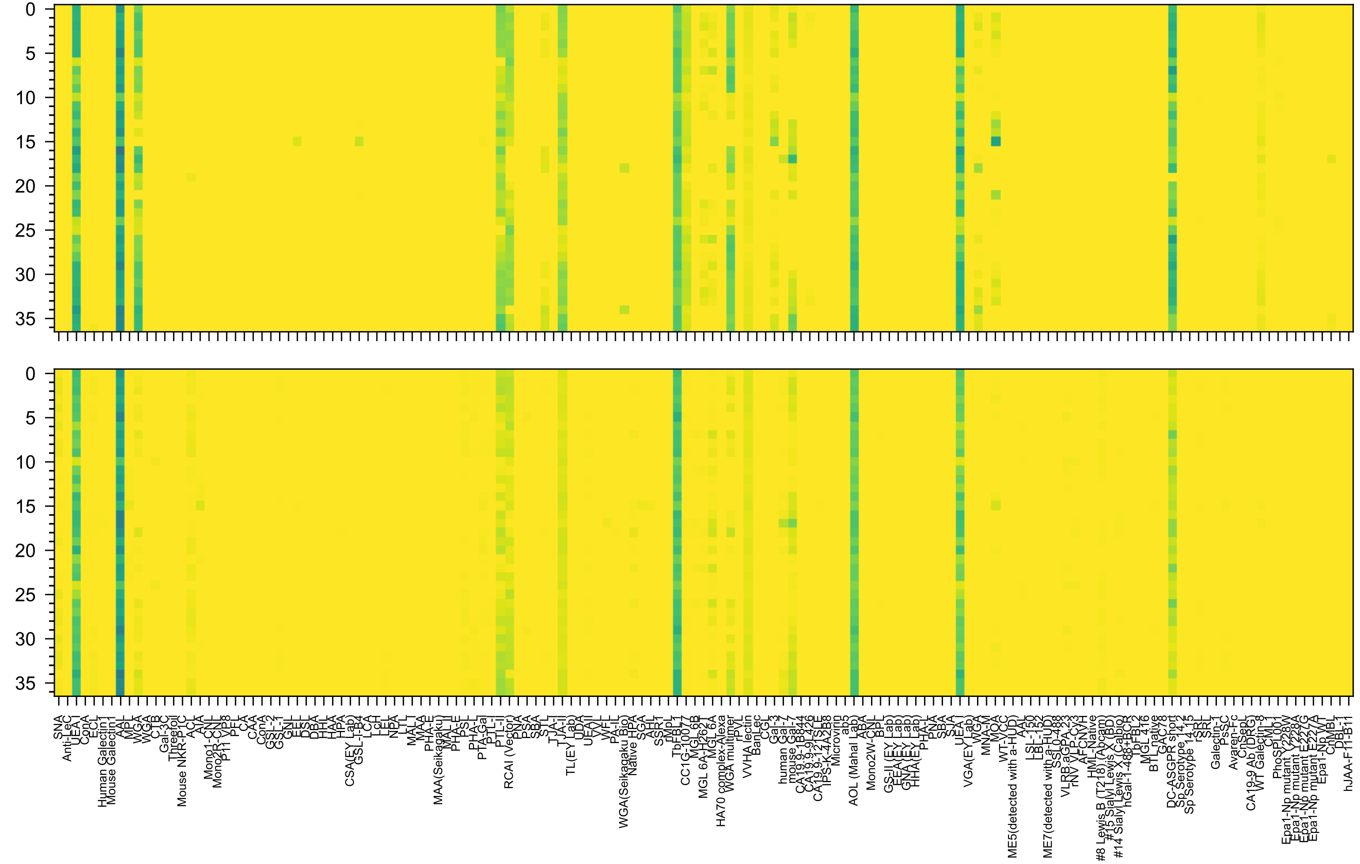

Fuc(α1-2)Gal(β1-4)GlcNAc(β1-6)[Fuc(α1-2)Gal(β1-4)GlcNAc(β1-3)]GalNAc(-Sp14

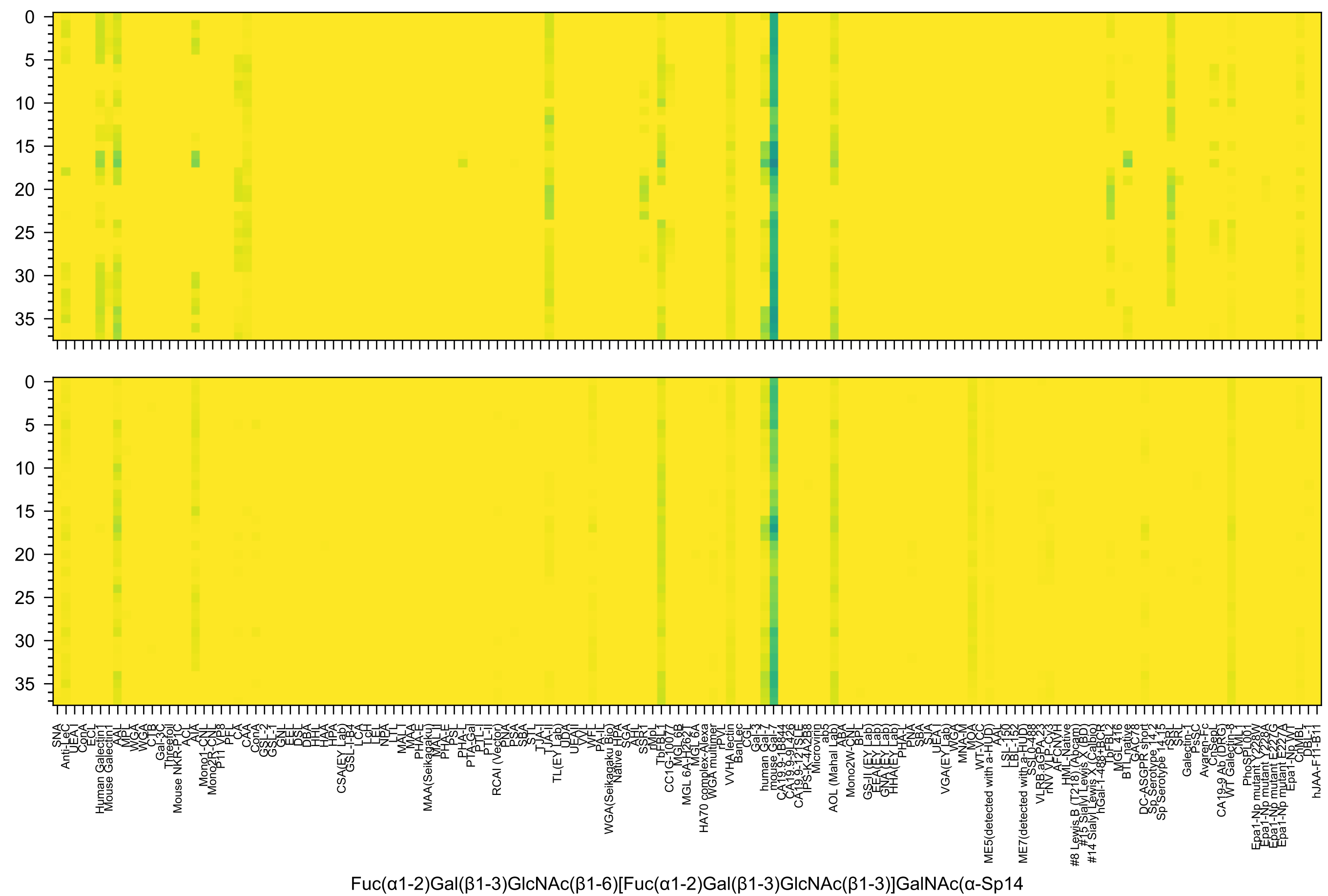
Fuc(α1-2)Gal(β1-3)GlcNAc(β1-6)[Fuc(α1-2)Gal(β1-3)GlcNAc(β1-3)]GalNAc(α-Sp14

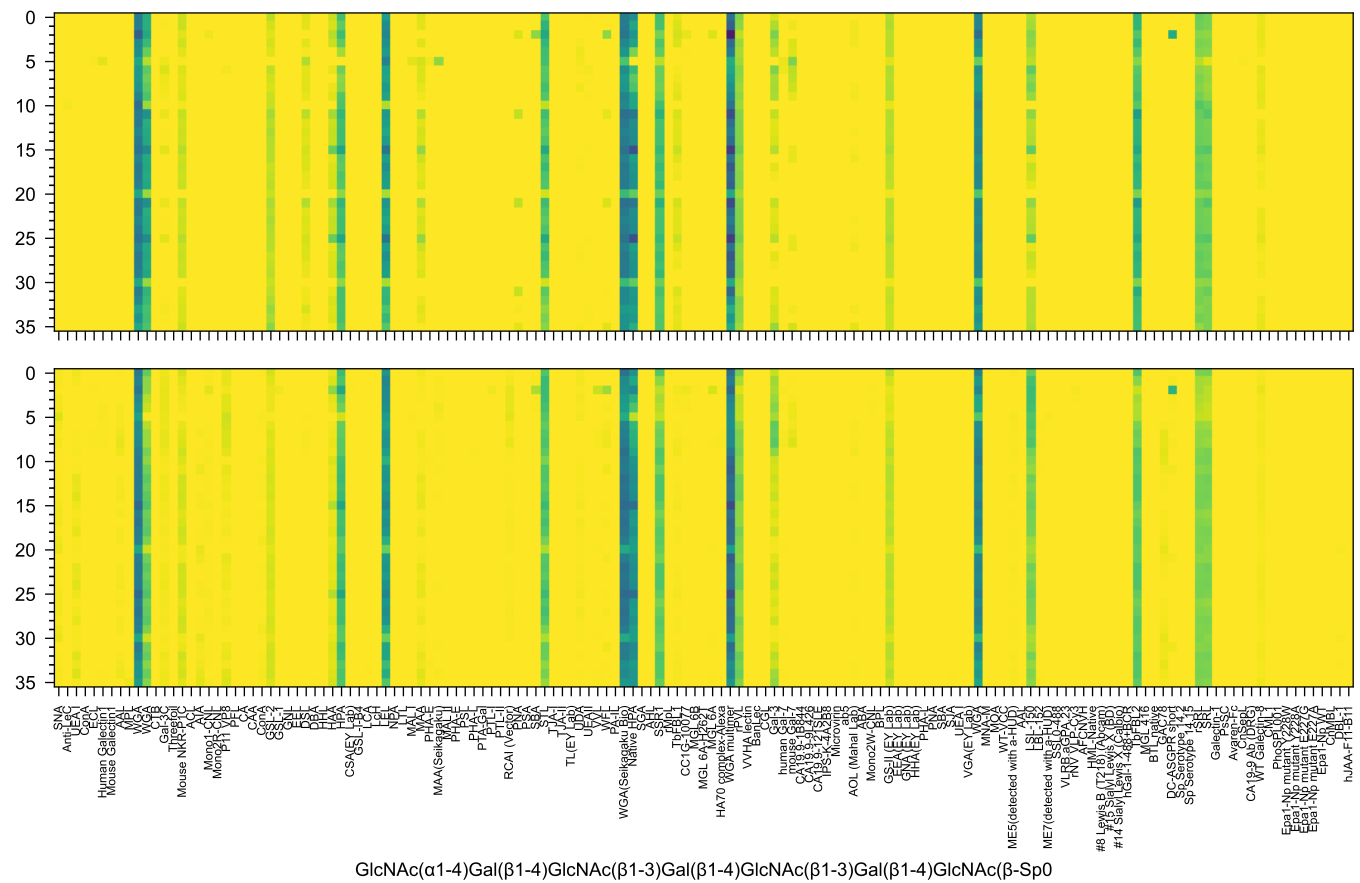
GlcNAc(α1-4)Gal(β1-4)GlcNAc(β1-3)Gal(β1-4)GlcNAc(β1-3)Gal(β1-4)GlcNAc(β-Sp0

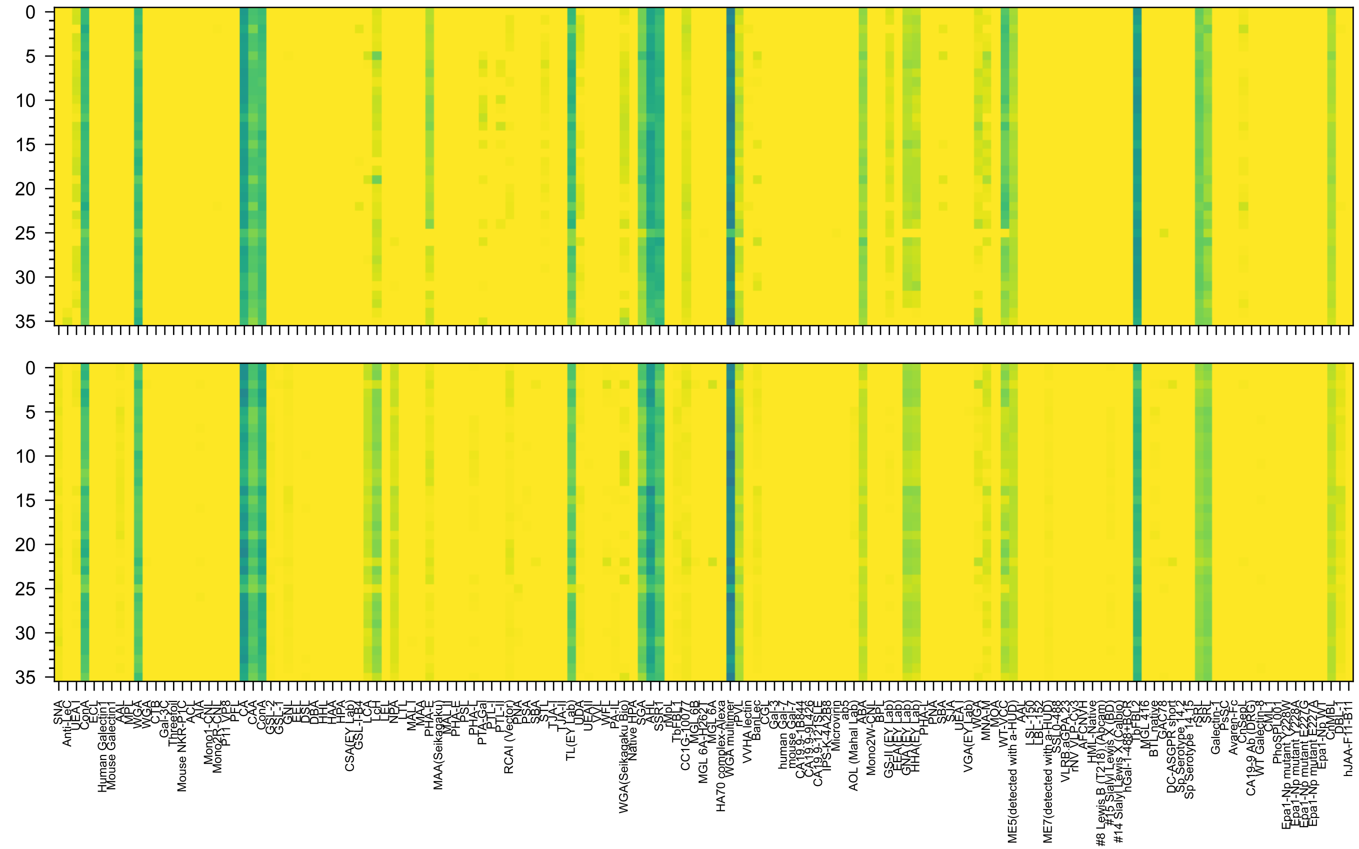

GlcNAc(β1-2)Man(α1-6)[GlcNAc(β1-2)Man(α1-3)]Man(β1-4)GlcNAc(β1-4)GlcNAc(β-Sp13

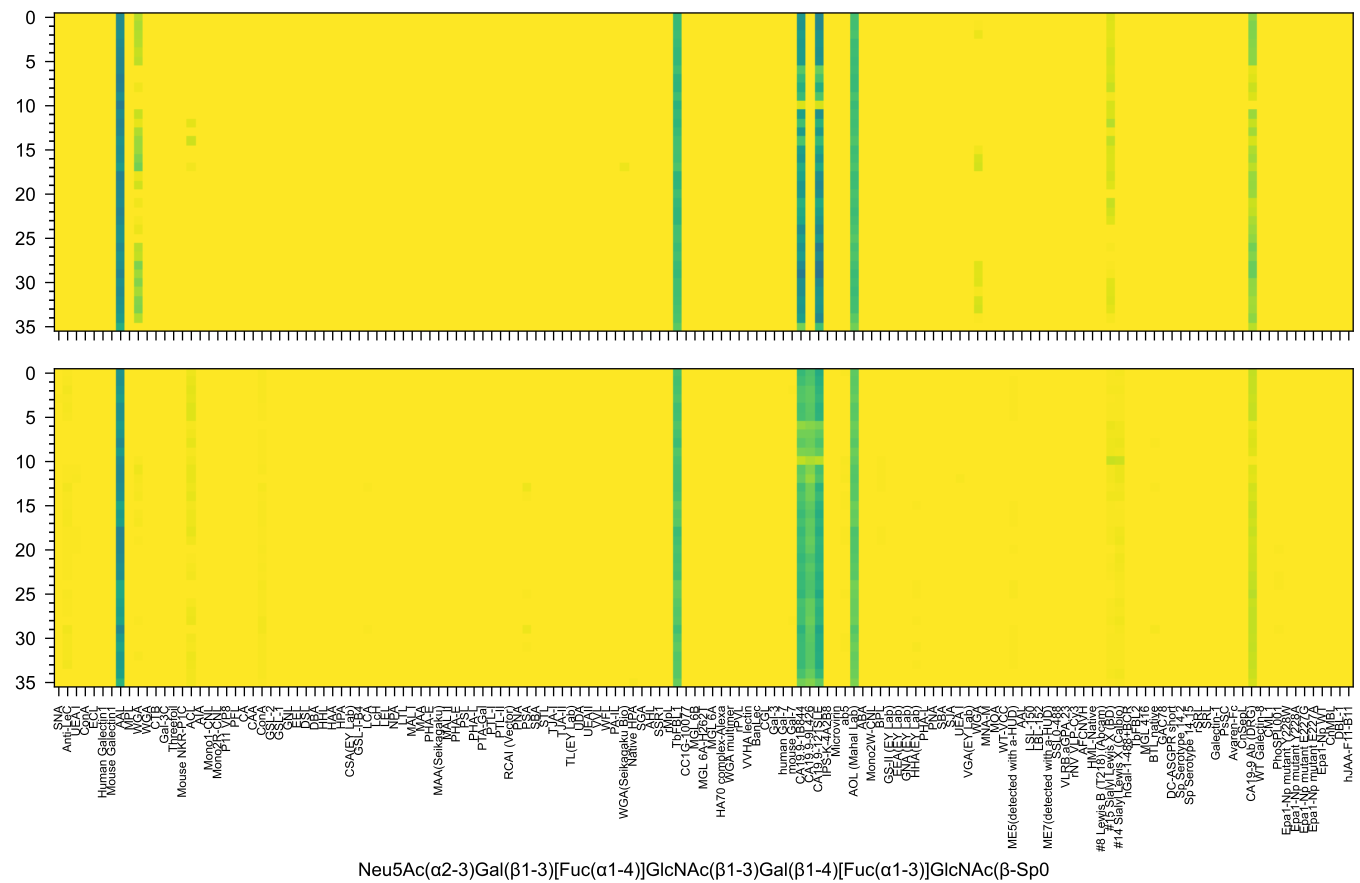
Neu5Ac(α2-3)Gal(β1-3)[Fuc(α1-4)]GlcNAc(β1-3)Gal(β1-4)[Fuc(α1-3)]GlcNAc(β-Sp0

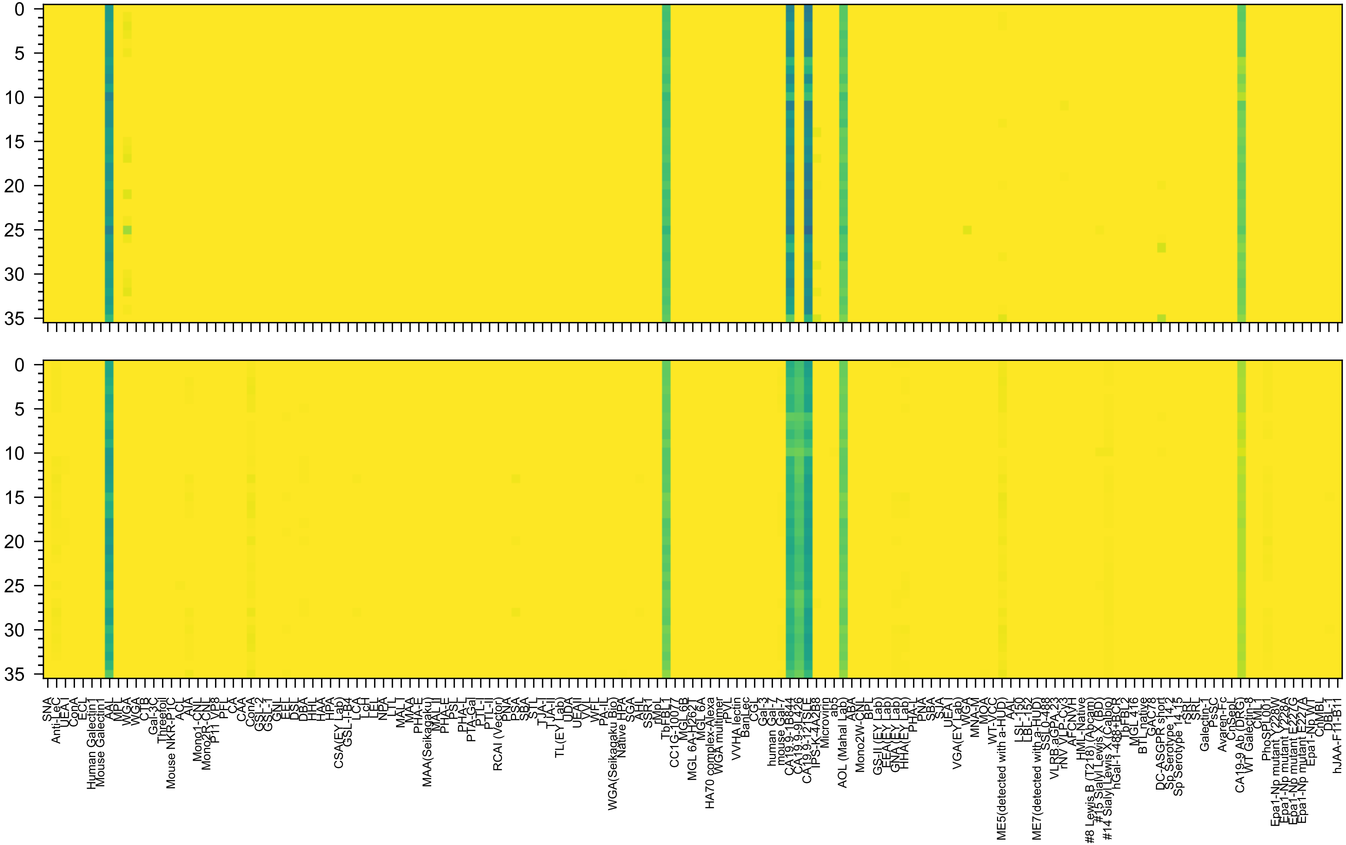

Neu5Ac(α2-3)Gal(β1-3)[Fuc(α1-4)]GlcNAc(β1-3)Gal(β1-3)[Fuc(α1-4)]GlcNAc(β-Sp0

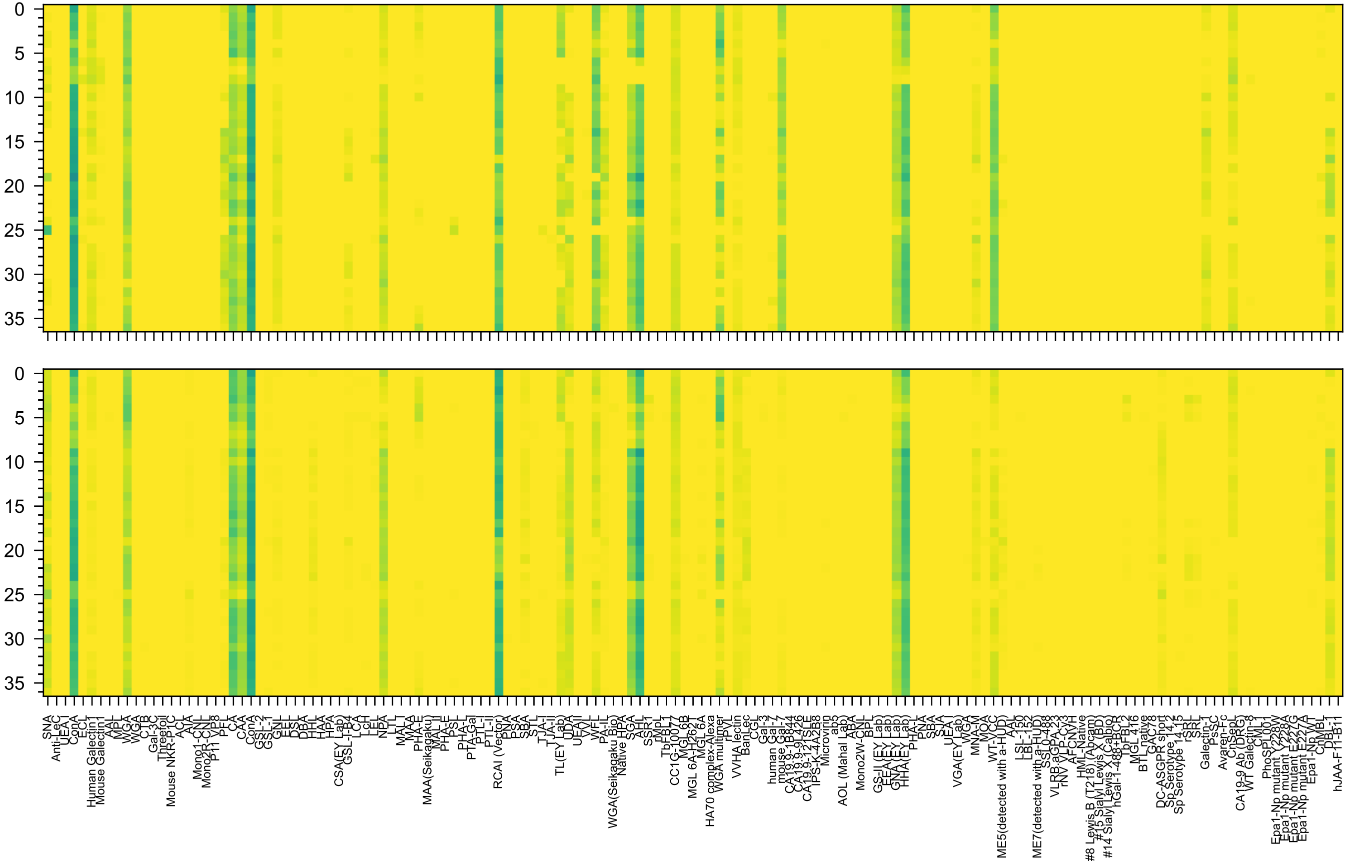

Man(α1-6)[Gal(β1-4)GlcNAc(β1-2)Man(α1-3)]Man(β1-4)GlcNAc(β1-4)GlcNAc(β-Sp12

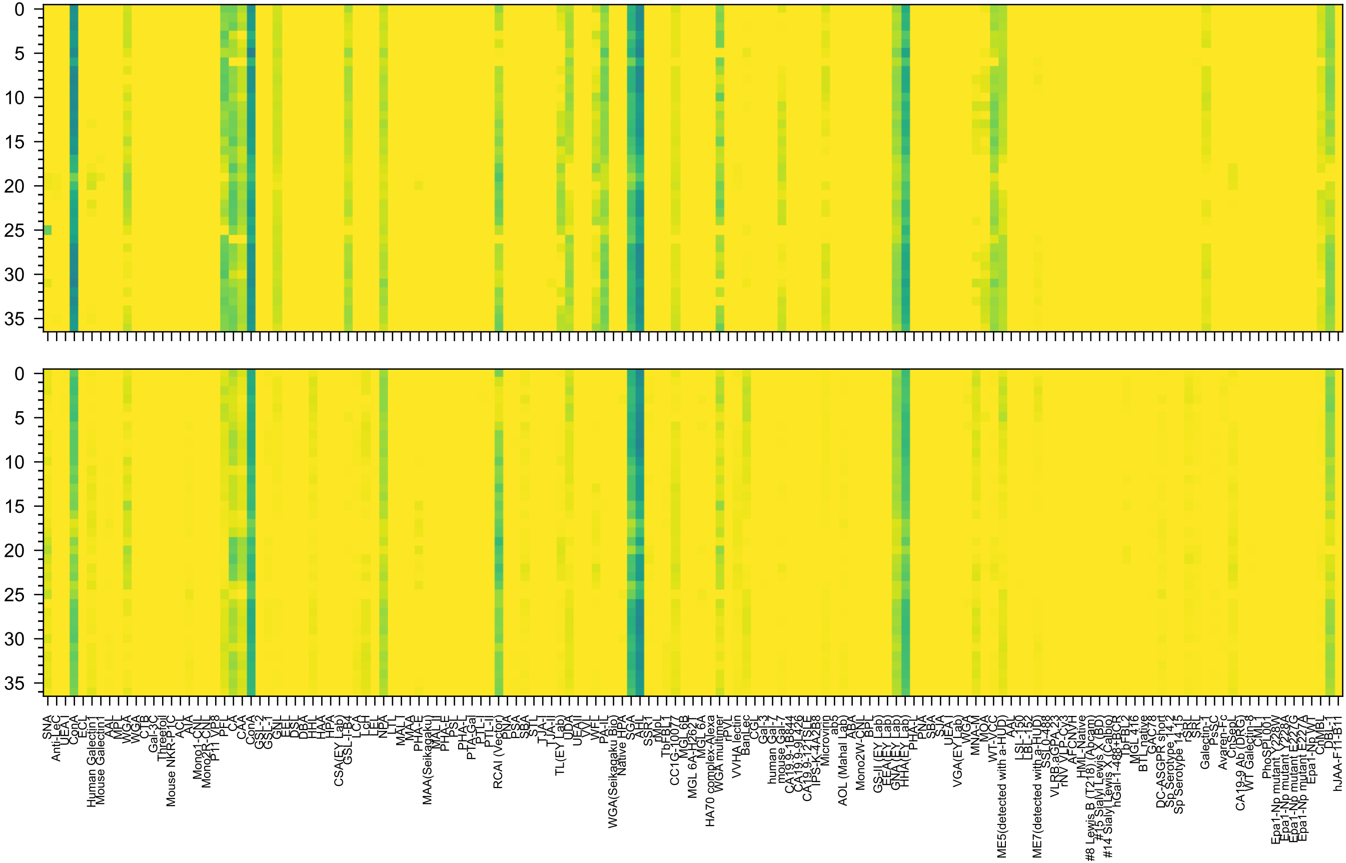


Gal(β1-4)GlcNAc(β1-2)Man(α1-6)[Man(α1-3)]Man(β1-4)GlcNAc(β1-4)GlcNAc(β-Sp12

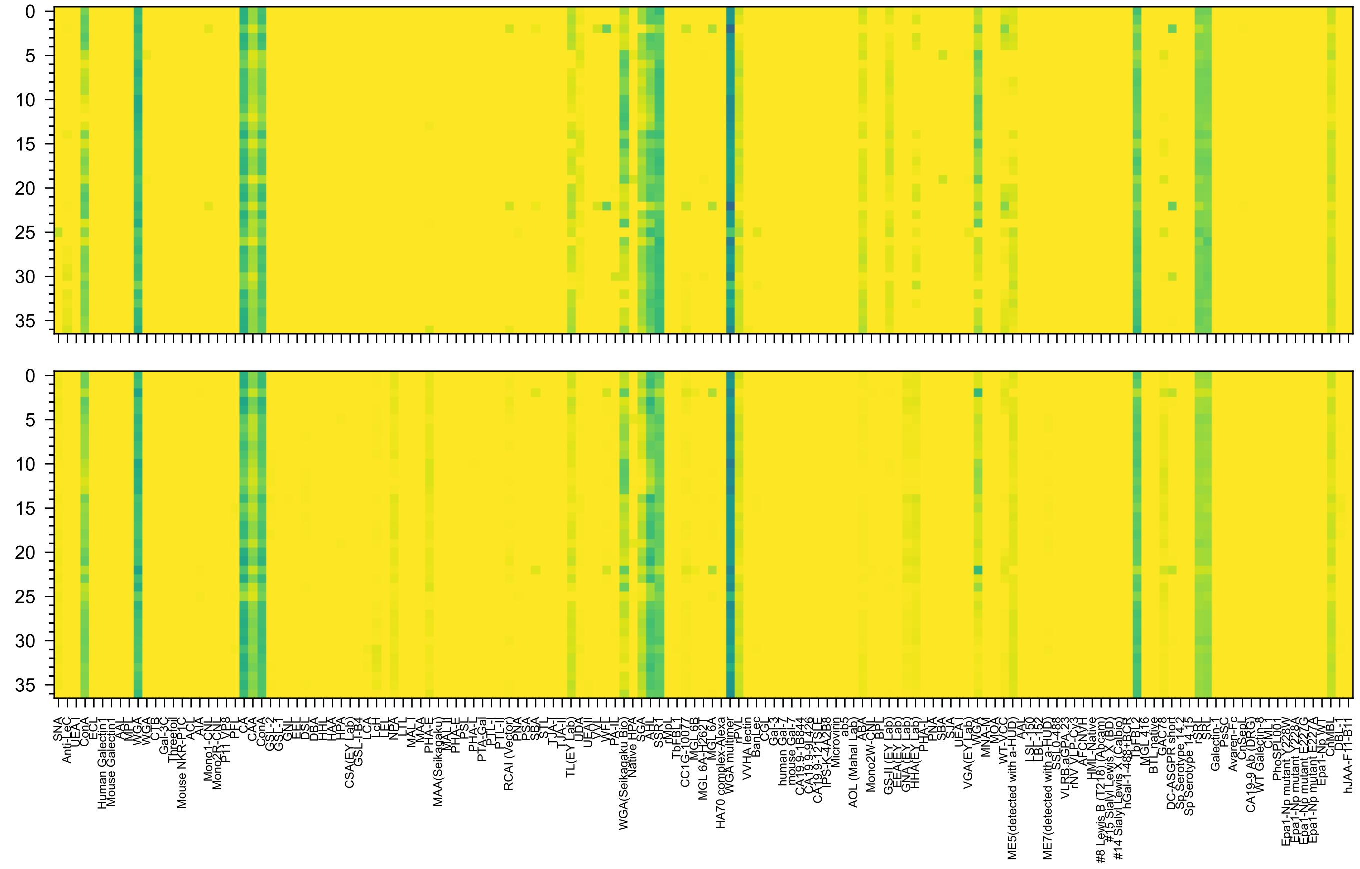


GlcNAc(β1-2)Man(α1-6)[GlcNAc(β1-2)Man(α1-3)]Man(β1-4)GlcNAc(β1-4)GlcNAc(β-Sp12

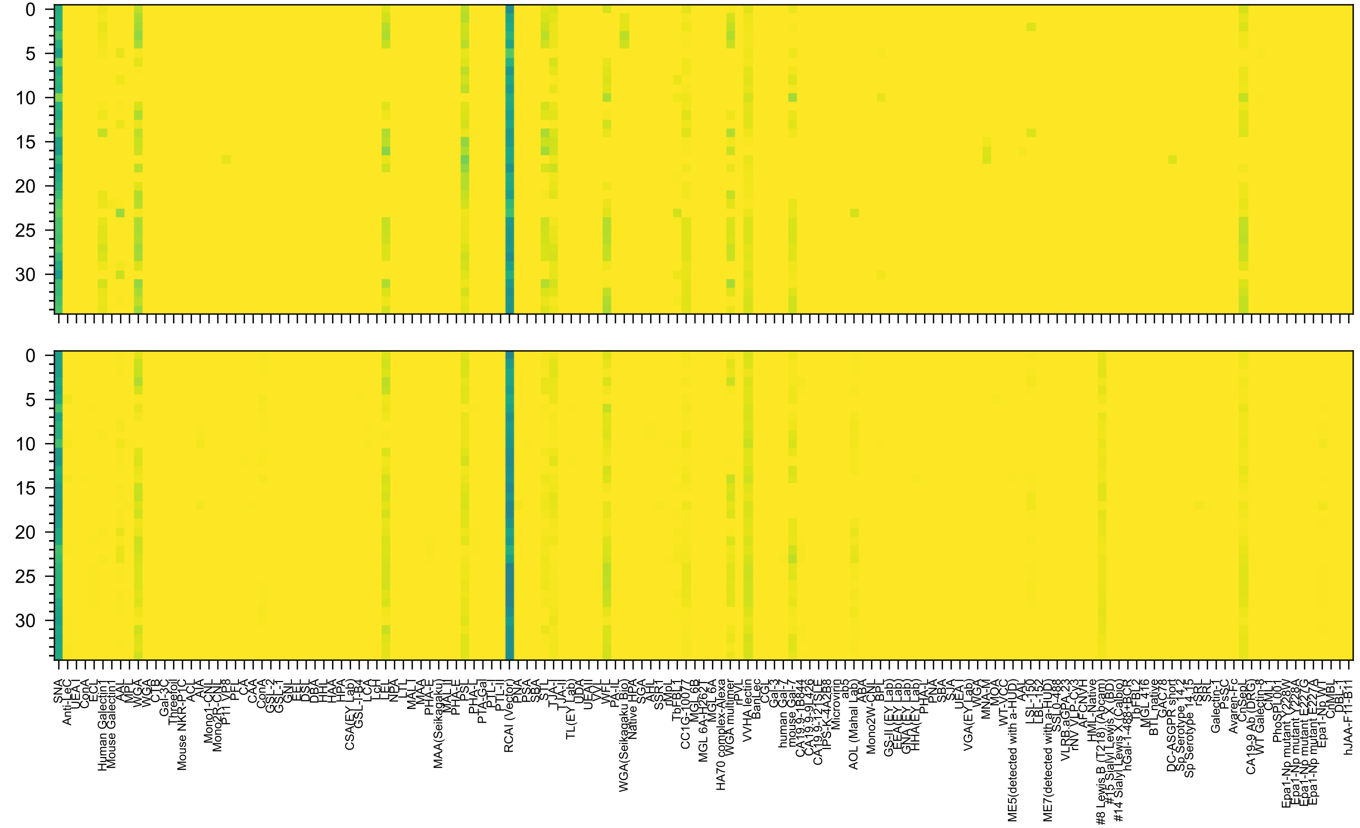

Gal(β1-4)GlcNAc(β1-6)[Neu5Ac(α2-6)Gal(β1-3)GlcNAc(β1-3)]Gal(β1-4)Glc(-Sp21

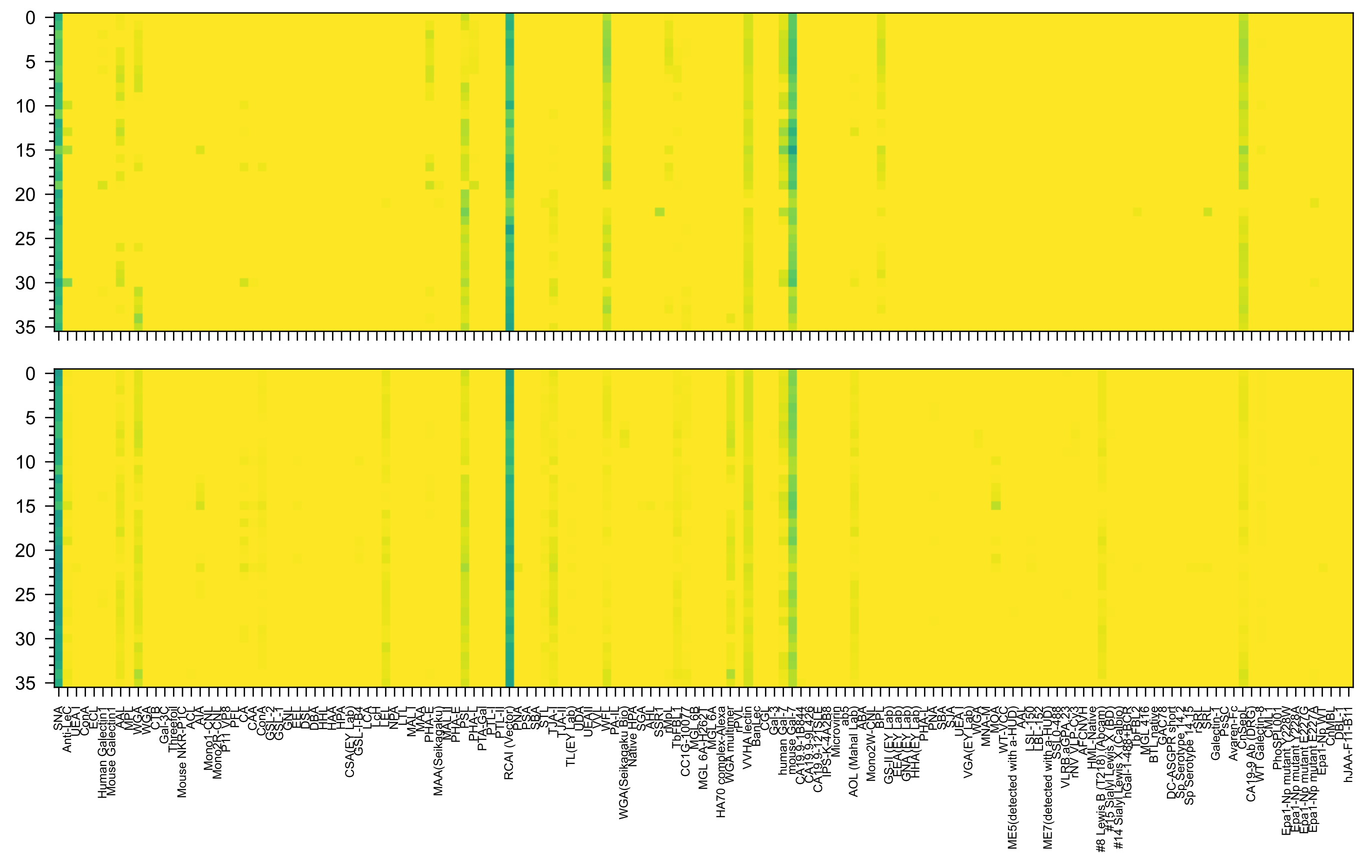

Neu5Ac(α2-6)Gal(β1-4)GlcNAc(β1-6)[Gal(β1-3)GlcNAc(β1-3)]Gal(β1-4)Glc(β-Sp21

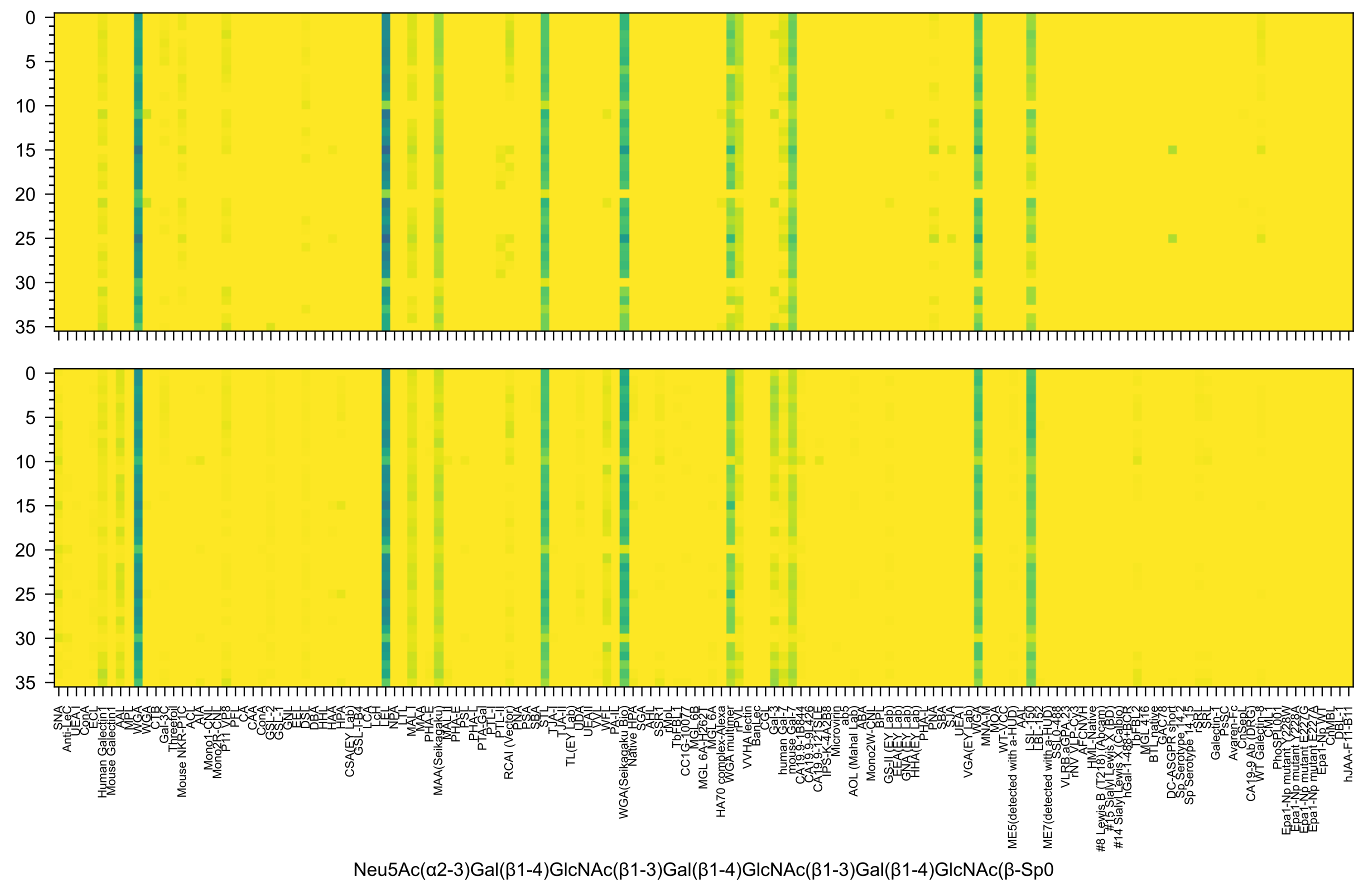
Neu5Ac(α2-3)Gal(β1-4)GlcNAc(β1-3)Gal(β1-4)GlcNAc(β1-3)Gal(β1-4)GlcNAc(β-Sp0

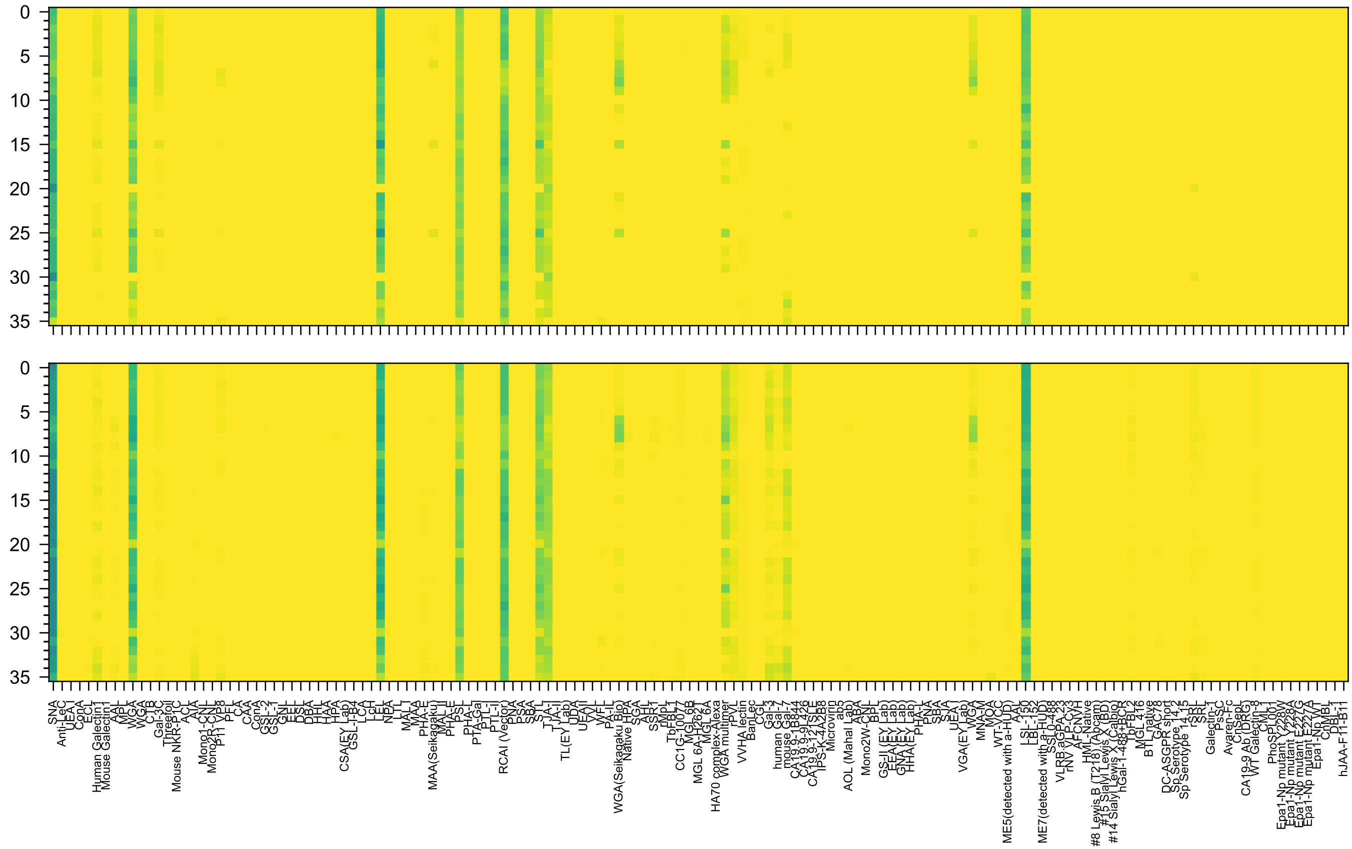

Neu5Ac(α2-6)Gal(β1-4)GlcNAc(β1-3)Gal(β1-4)GlcNAc(β1-3)Gal(β1-4)GlcNAc(β-Sp0

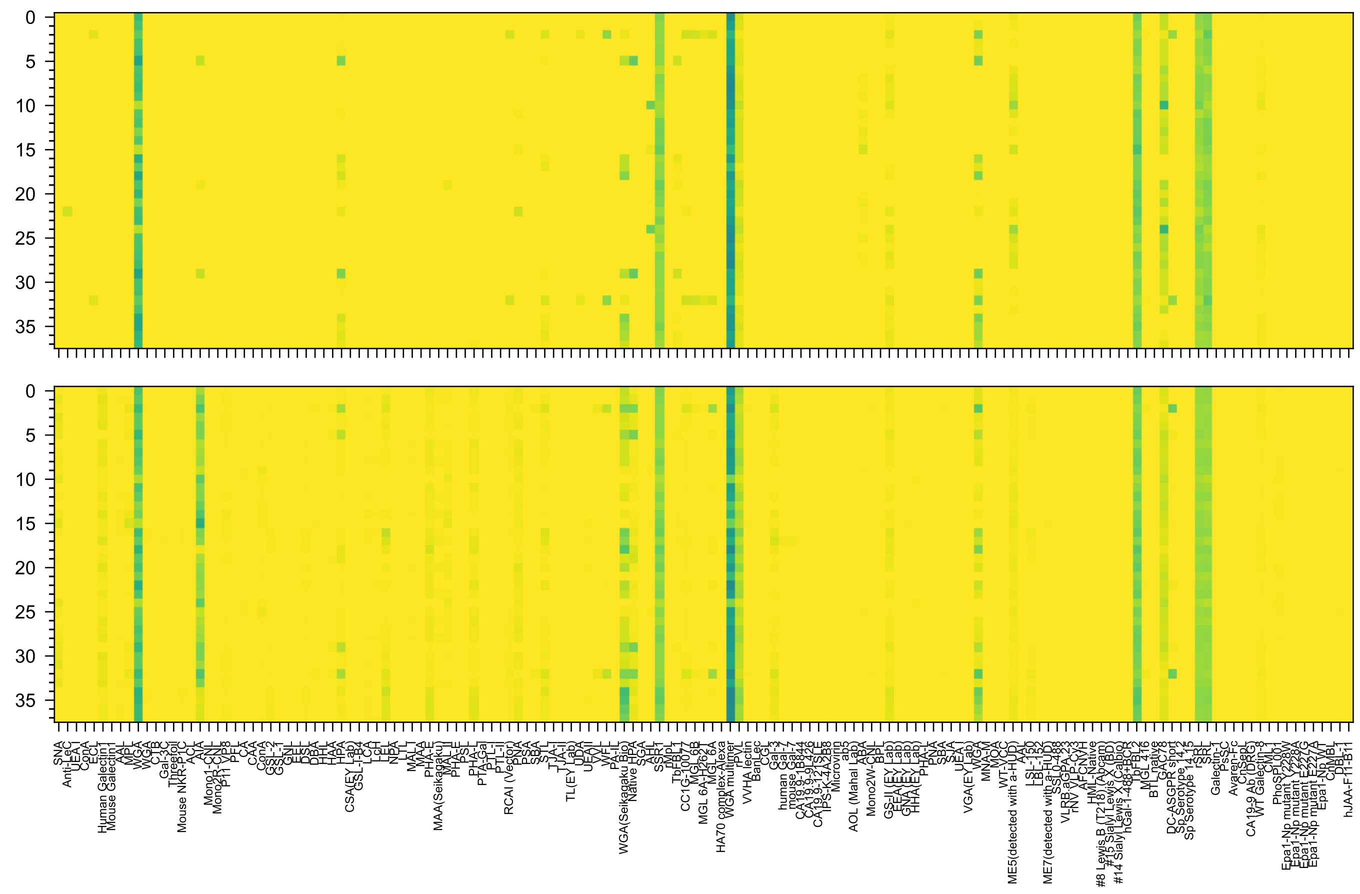


GlcNAc(β1-3)Gal(β1-4)GlcNAc(β1-6)[GlcNAc(β1-3)Gal(β1-4)GlcNAc(β1-3)]GalNAc(α-Sp14

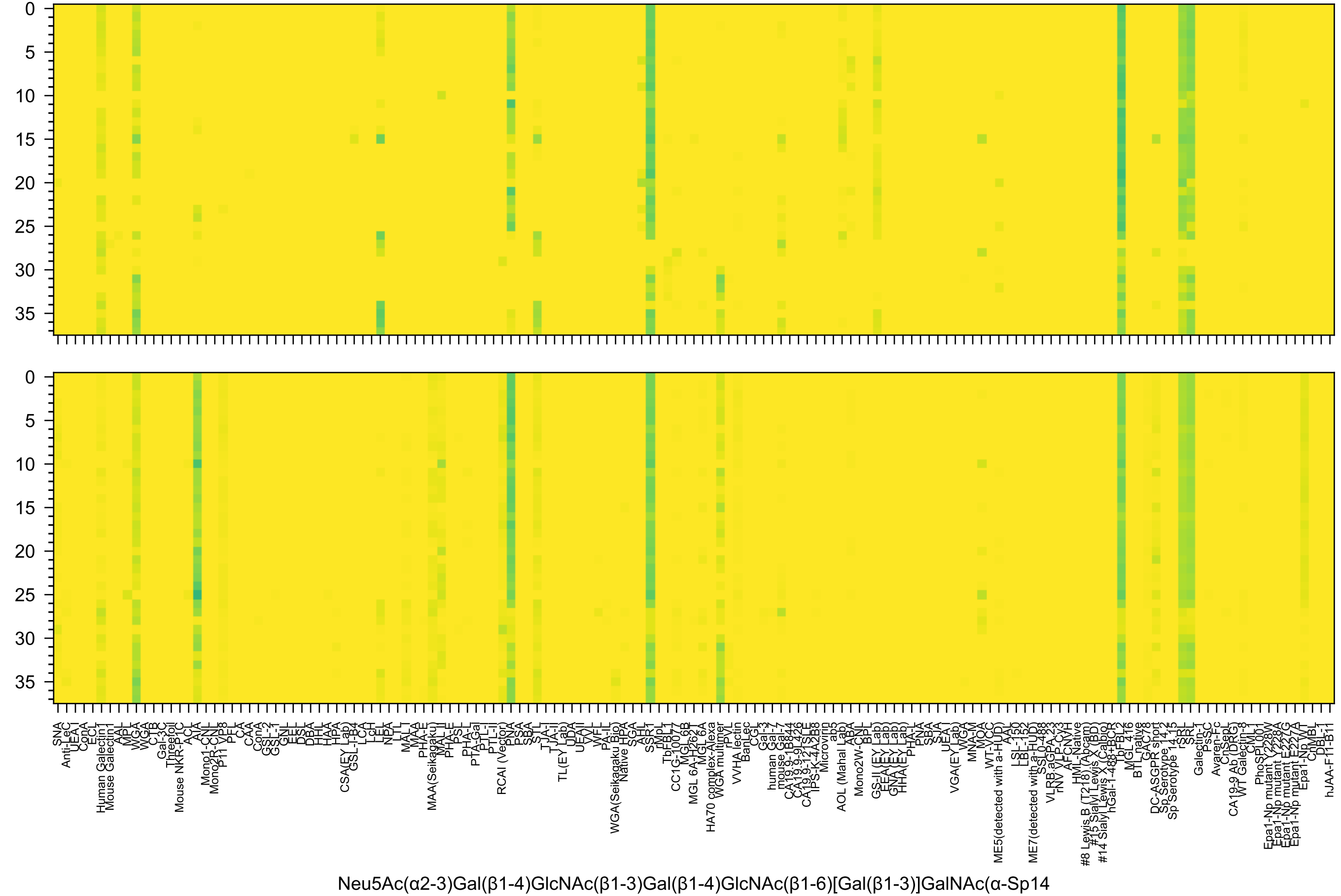
Neu5Ac(α2-3)Gal(β1-4)GlcNAc(β1-3)Gal(β1-4)GlcNAc(β1-6)[Gal(β1-3)]GalNAc(α-Sp14

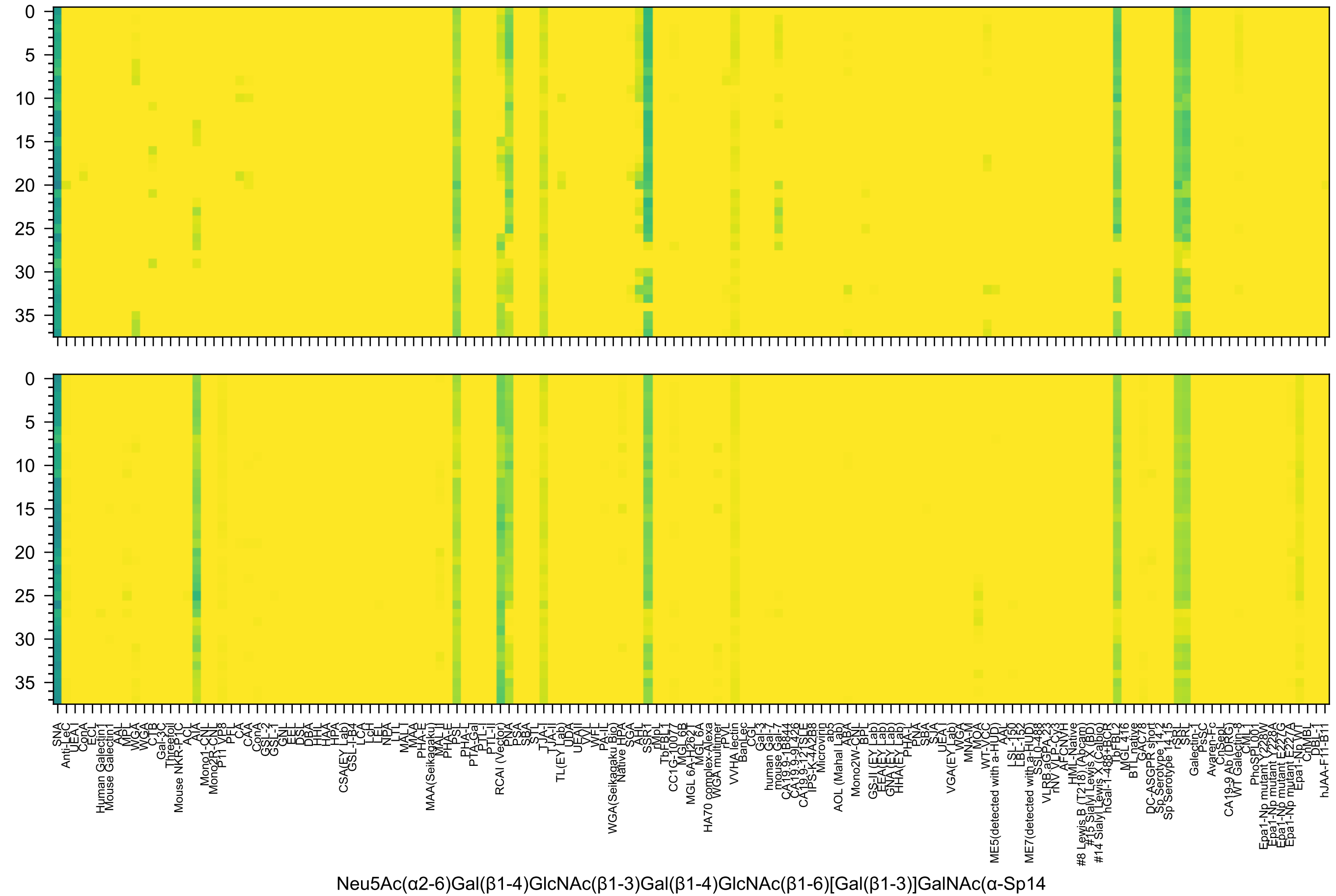
Neu5Ac(α2-6)Gal(β1-4)GlcNAc(β1-3)Gal(β1-4)GlcNAc(β1-6)[Gal(β1-3)]GalNAc(α-Sp14

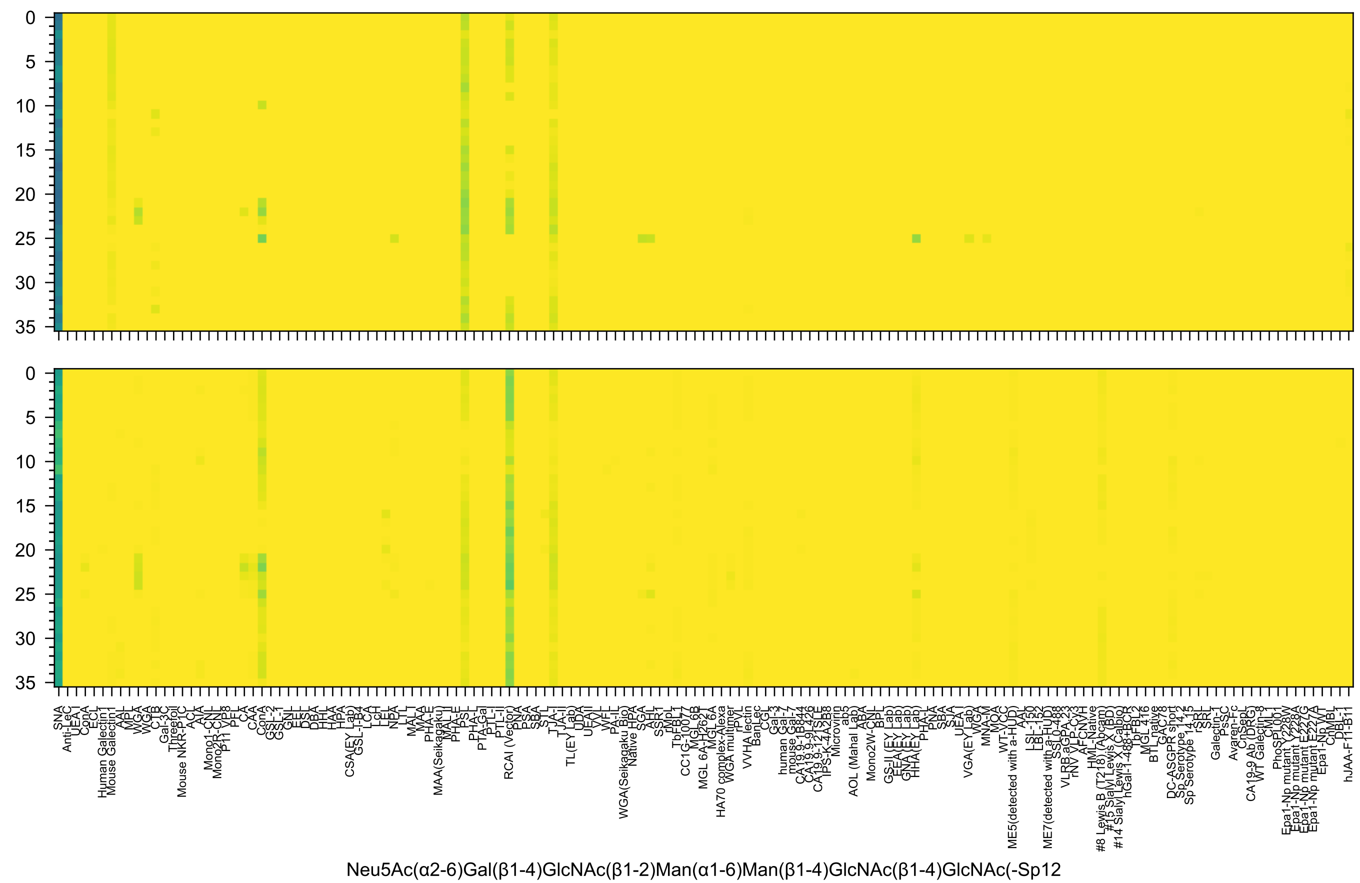
Neu5Ac(α2-6)Gal(β1-4)GlcNAc(β1-2)Man(α1-6)Man(β1-4)GlcNAc(β1-4)GlcNAc(-Sp12

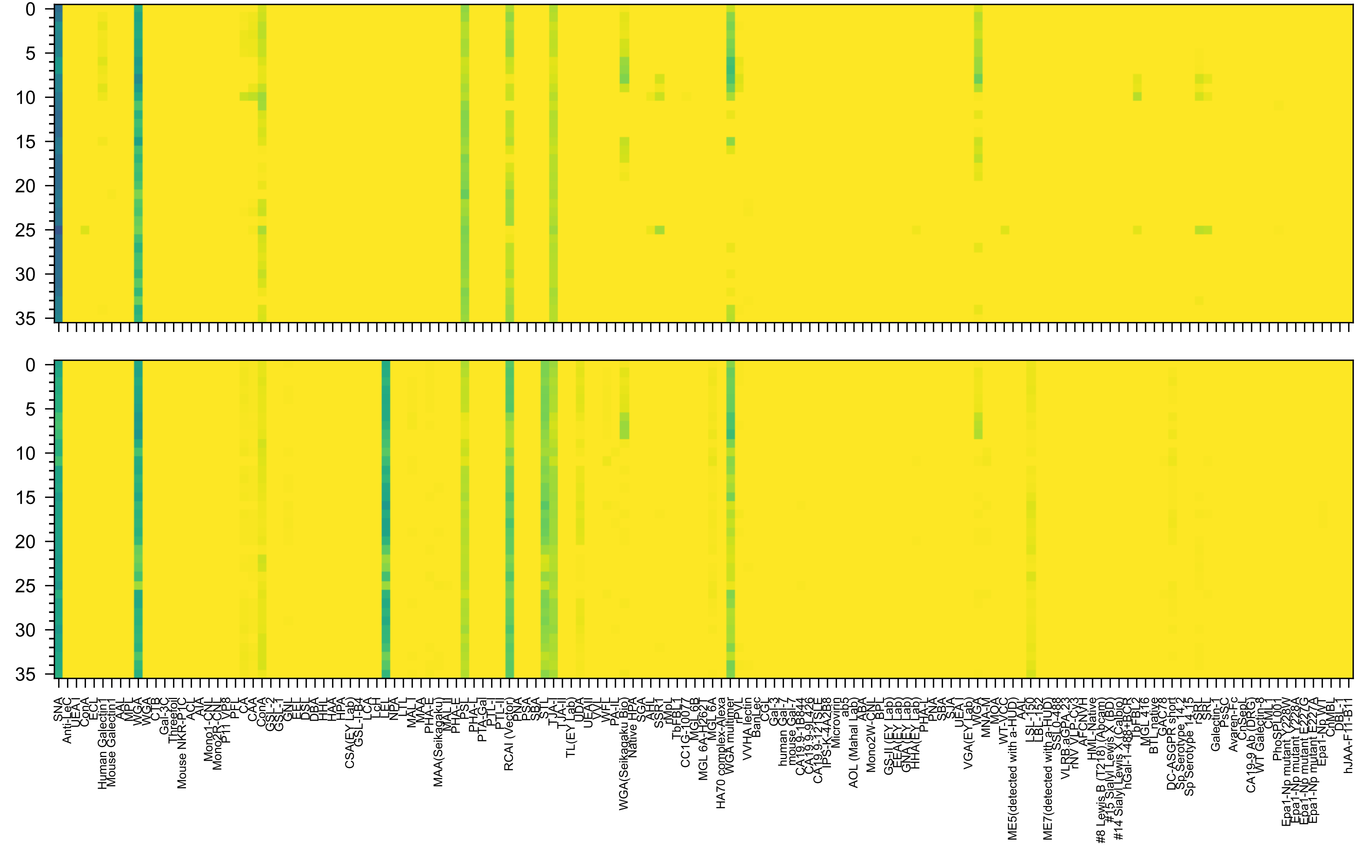


Neu5Ac(α2-6)Gal(β1-4)GlcNAc(β1-2)Man(α1-3)Man(β1-4)GlcNAc(β1-4)GlcNAc(-Sp12

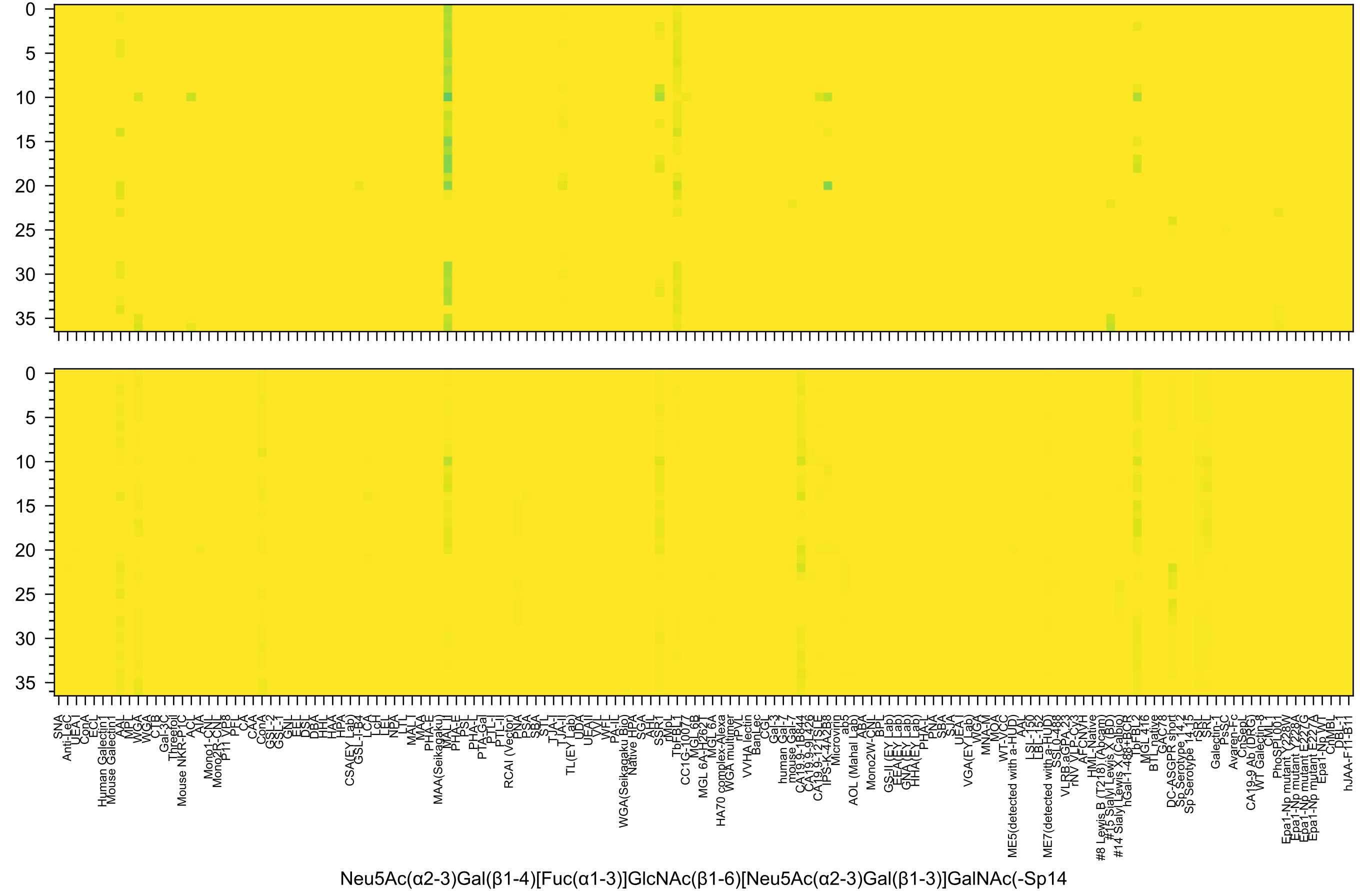
0
5
10
15
20
25
30
35
0
5
10
15
20
25
30
35
SNA
Anti-LeC
UEA I
ConA
ECL
Human Galectin1
Mouse Galectin1
AAL
MPL
WGA
WGA
CTB
Gal-3C
Threefoil
Mouse NKR-P1C
ACL
AIA
Mono1-CNL
Mono2R-CNL
P11 VP8
PFL
CA
CAA
ConA
GSL-2
GSL-1
GNL
EEL
DSL
DBA
HHL
HAA
HPA
CSA(EY Lab)
GSL-I-B4
LCA
LcH
LEL
NPA
LTL
MAL I
MAA
PHA-E
MAA(Seikagaku)
MAL II
PHA-E
PSL
PHA-L
PTA-Gal
PTL-I
PTL-II
RCAI (Vector)
PNA
PSA
SBA
STL
TJA-I
TJA-II
TL(EY Lab)
UDA
UEA II
VVL
WFL
PA-IL
WGA(Seikagaku Bio)
Native HPA
SGA
AHL
SSR1
rMpL
TbFBL1
CC1G-10077
MGL 6B
MGL 6A-H262T
MGL 6A
HA70 complex-Alexa
WGA multimer
rPVL
VVHA lectin
BanLec
CGL
Gal-3
human Gal-7
mouse Gal-7
CA19.9-1B844
CA19.9-9L426
CA19.9-121SLE
IPS-K-4A2B8
Microvirin
ab5
AOL (Mahal Lab)
ABA
Mono2W-CNL
BPL
GS-II (EY Lab)
EEA(EY Lab)
GNA (EY Lab)
HHA(EY Lab)
PHA-L
PNA
SBA
SJA
UEA I
VGA(EY Lab)
WGA
MNA-M
MOA
WT-VCC
ME5(detected with a-HUD)
AAL
LSL-150
LBL-152
ME7(detected with a-HUD)
SSL0-488
VLRB.aGPA.23
rNV VLP-Cy3
AFCNVH
HML-Native
#8 Lewis B (T218) (Abcam)
#15 Sialyl Lewis X (BD)
#14 Sialyl Lewis X (Calbio)
hGal-1-488+BCR
TbFBL2
MGL 416
BTL native
GAC78
DC-ASGPR short
Sp Serotype 14.2
Sp Serotype 14.15
rSRL
SRL
Galectin-1
PsSC
Avaren-Fc
CnSepL
CA19-9 Ab (DRG)
WT Galectin-8
CML1
PhoSPL001
Epa1-Np mutant Y228W
Epa1-Np mutant Y228A
Epa1-Np mutant E227G
Epa1-Np mutant E227A
Epa1-Np WT
CnMBL
DBL-1
hJAA-F11-B11
Neu5Ac(α2-3)Gal(β1-4)[Fuc(α1-3)]GlcNAc(β1-6)[Neu5Ac(α2-3)Gal(β1-3)]GalNAc(-Sp14

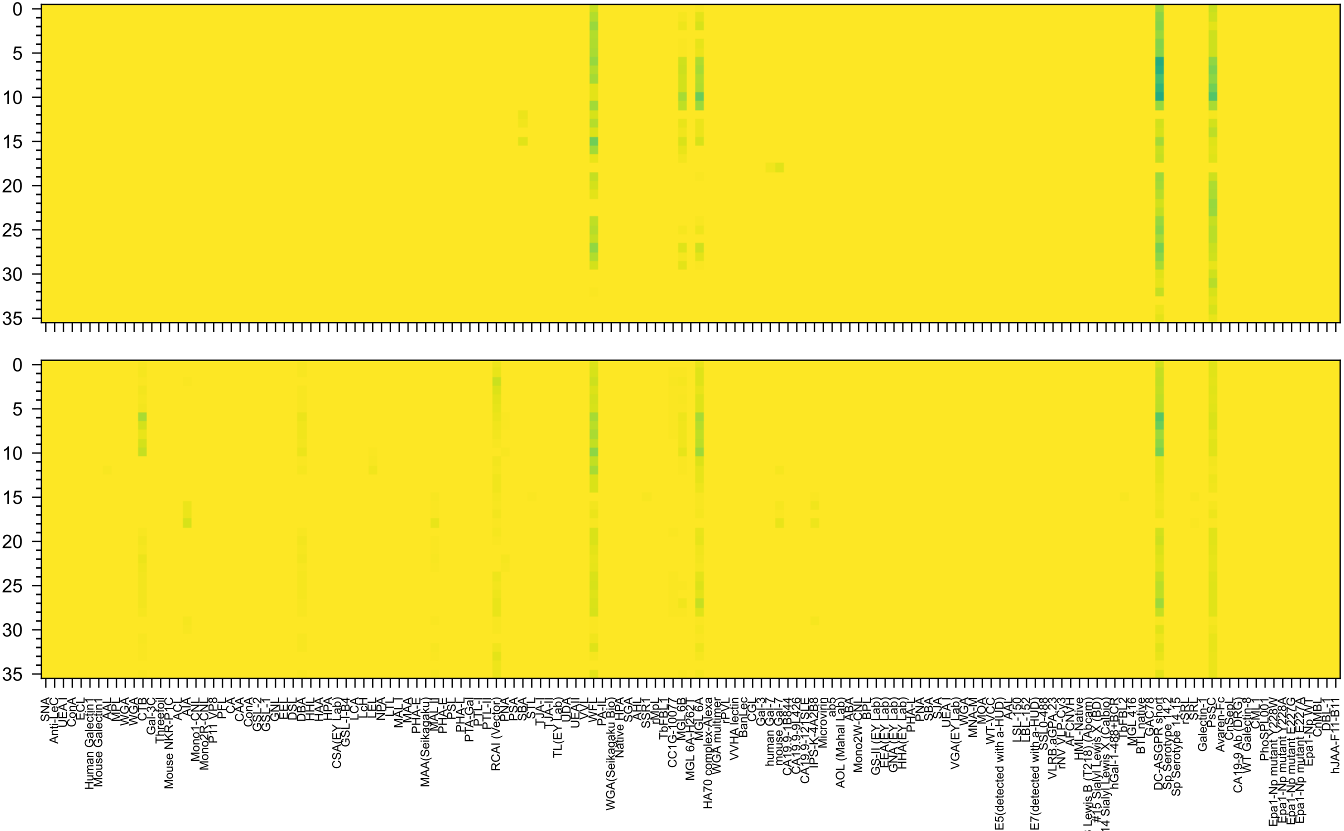

Neu5Ac(α2-3)Gal(β1-3)GalNAc(β1-4)[Neu5Ac(α2-8)Neu5Ac(α2-3)]Gal(β1-4)Glc(β-Sp0

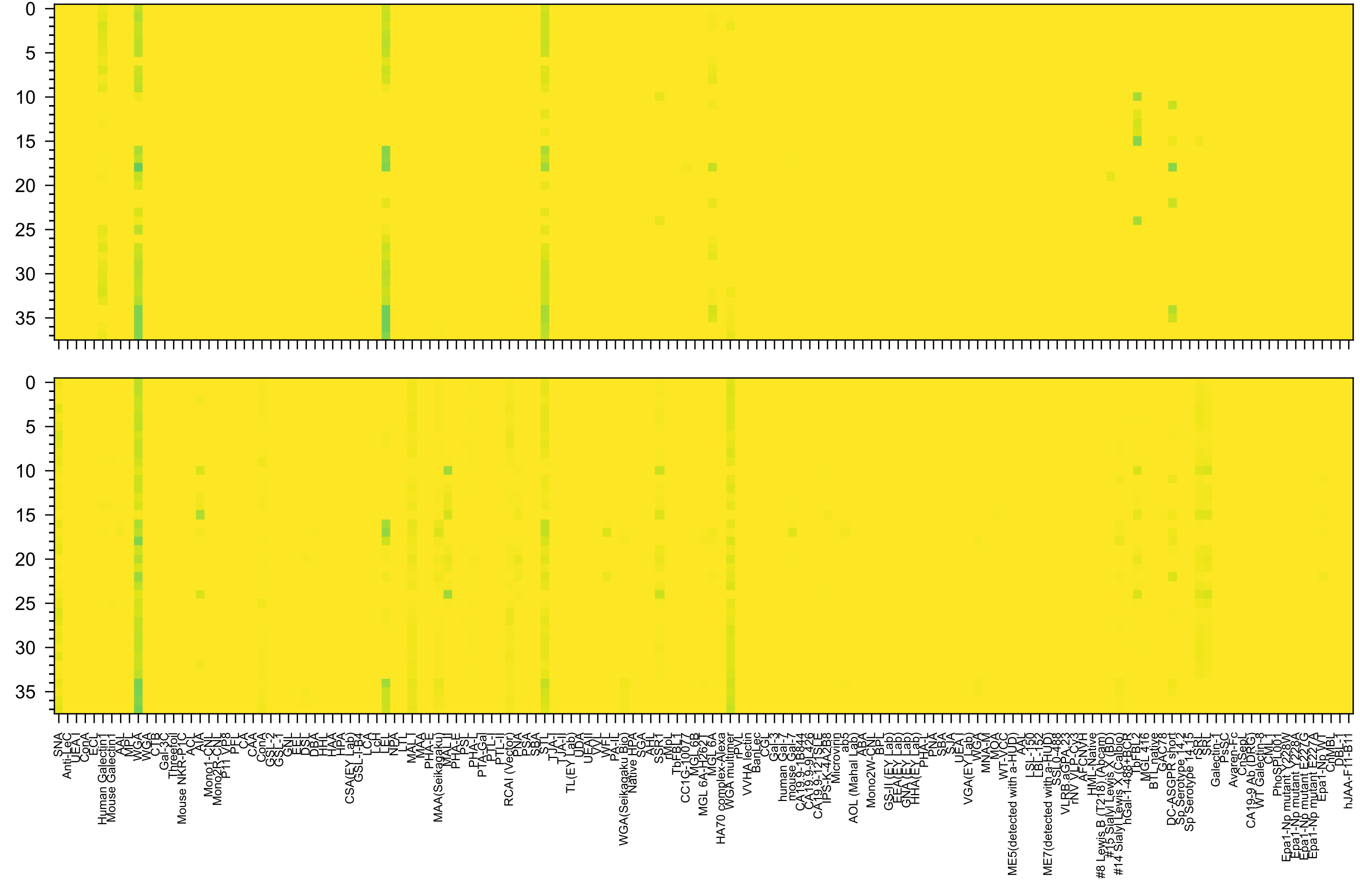

Neu5Ac(α2-3)Gal(β1-4)GlcNAc(β1-6)[Neu5Ac(α2-3)Gal(β1-4)GlcNAc(β1-3)]GalNAc(α-Sp14

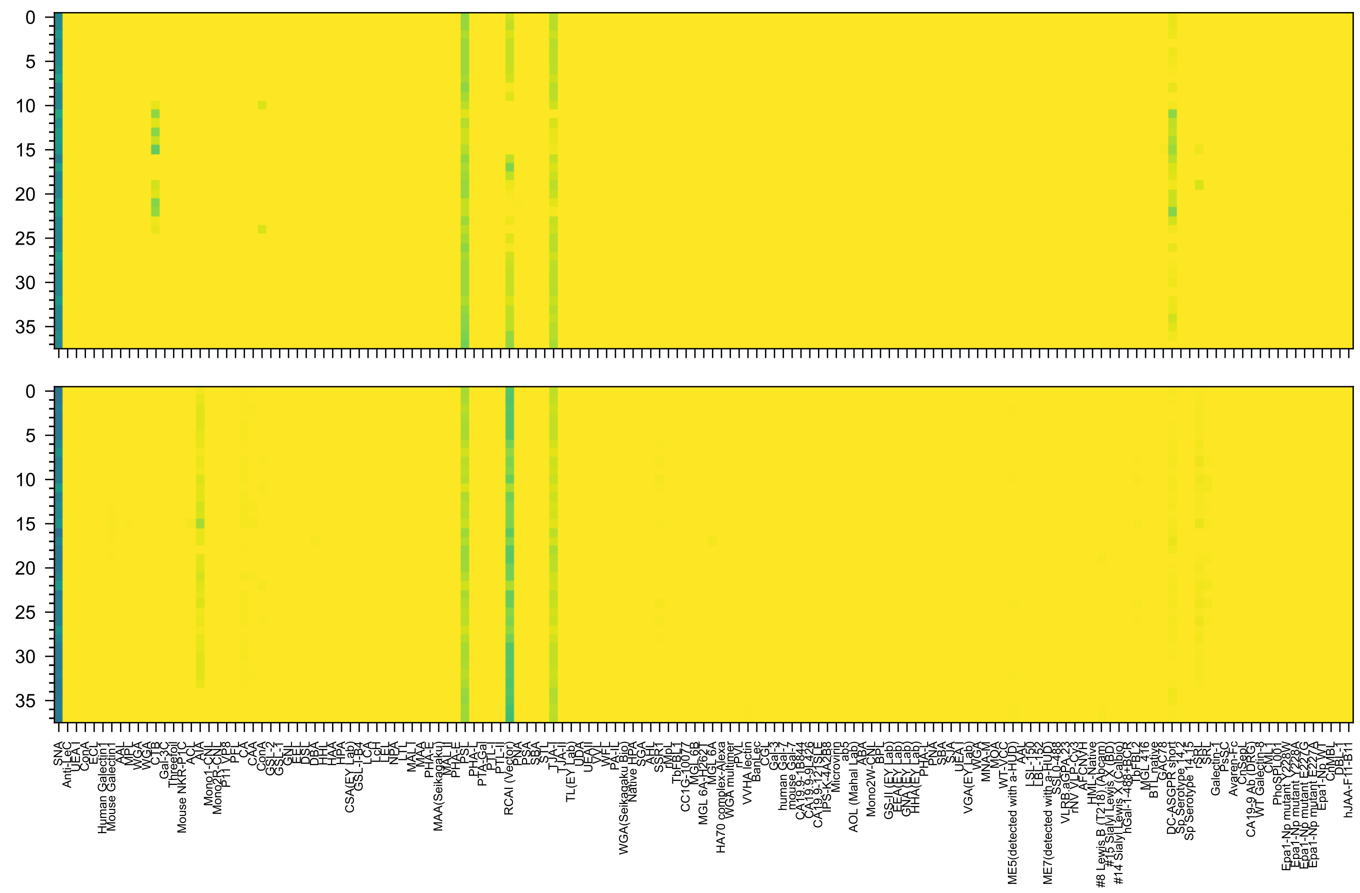

Neu5Ac(α2-6)Gal(β1-4)GlcNAc(β1-6)[Neu5Ac(α2-6)Gal(β1-4)GlcNAc(β1-3)]GalNAc(α-Sp14

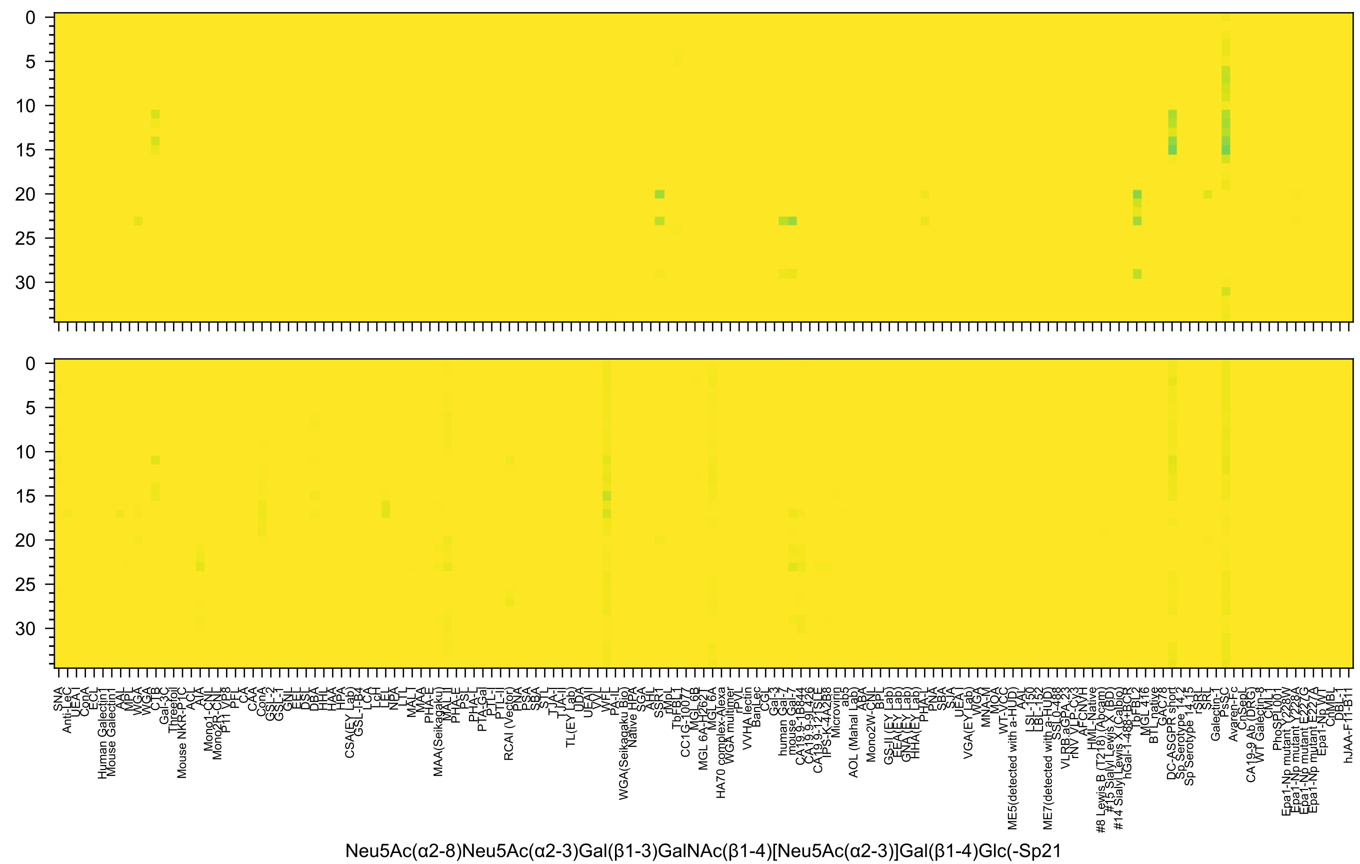
Neu5Ac(α2-8)Neu5Ac(α2-3)Gal(β1-3)GalNAc(β1-4)[Neu5Ac(α2-3)]Gal(β1-4)Glc(-Sp21

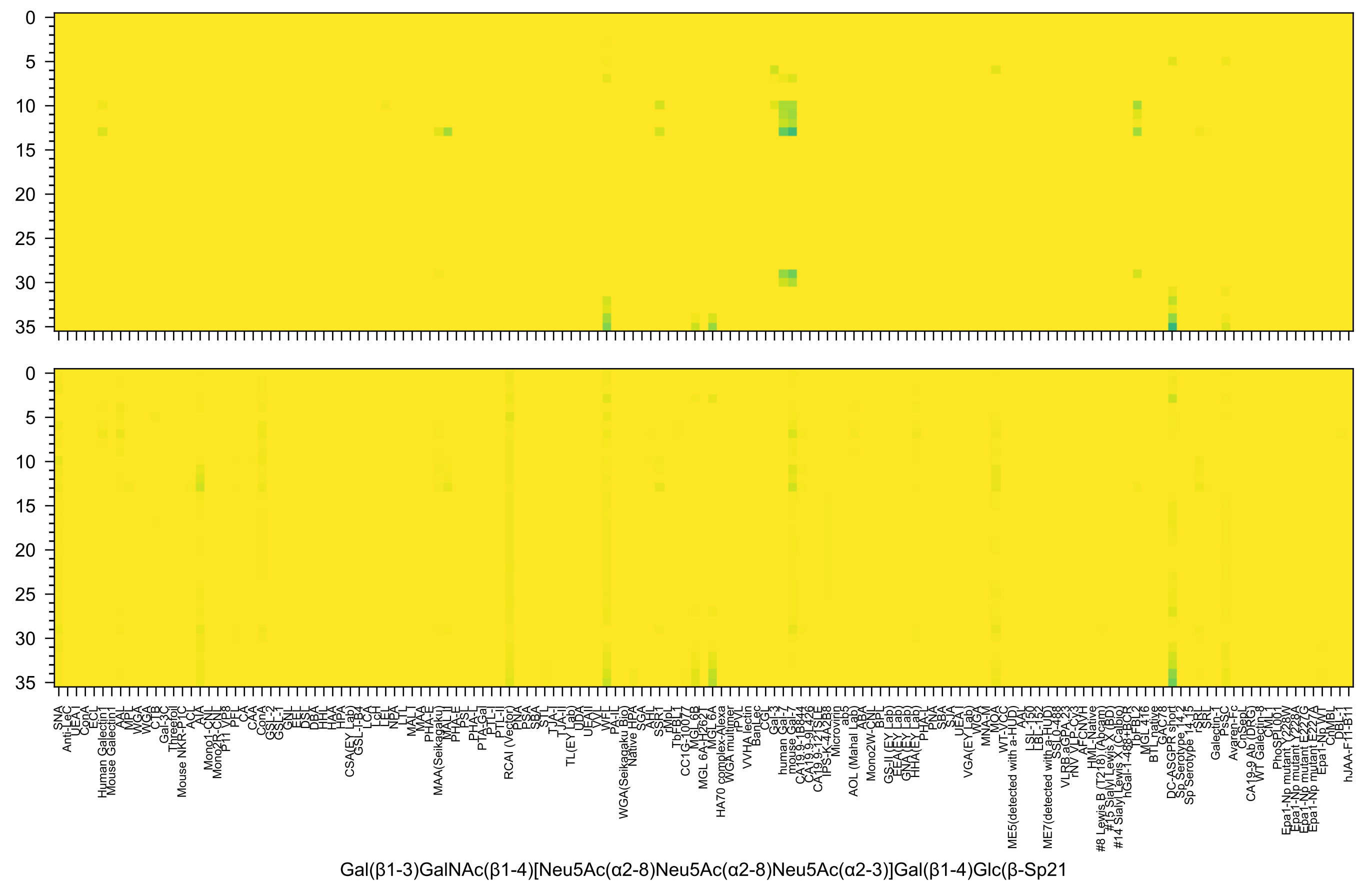
Gal(β1-3)GalNAc(β1-4)[Neu5Ac(α2-8)Neu5Ac(α2-8)Neu5Ac(α2-3)]Gal(β1-4)Glc(β-Sp21

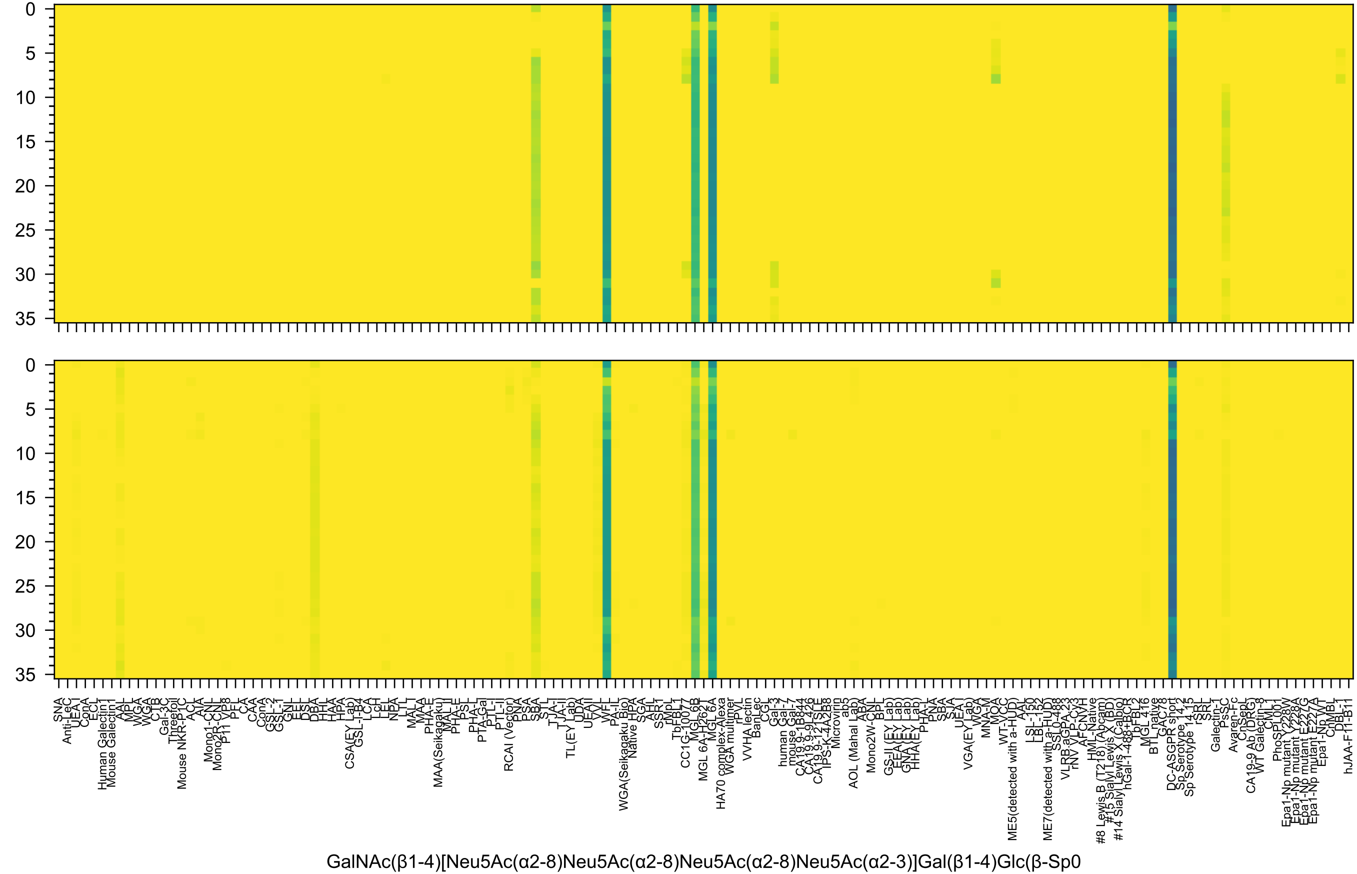

0
5
10
15
20
25
30
35
GalNAc(β1-4)[Neu5Ac(α2-8)Neu5Ac(α2-8)Neu5Ac(α2-8)Neu5Ac(α2-3)]Gal(β1-4)Glc(β-Sp0

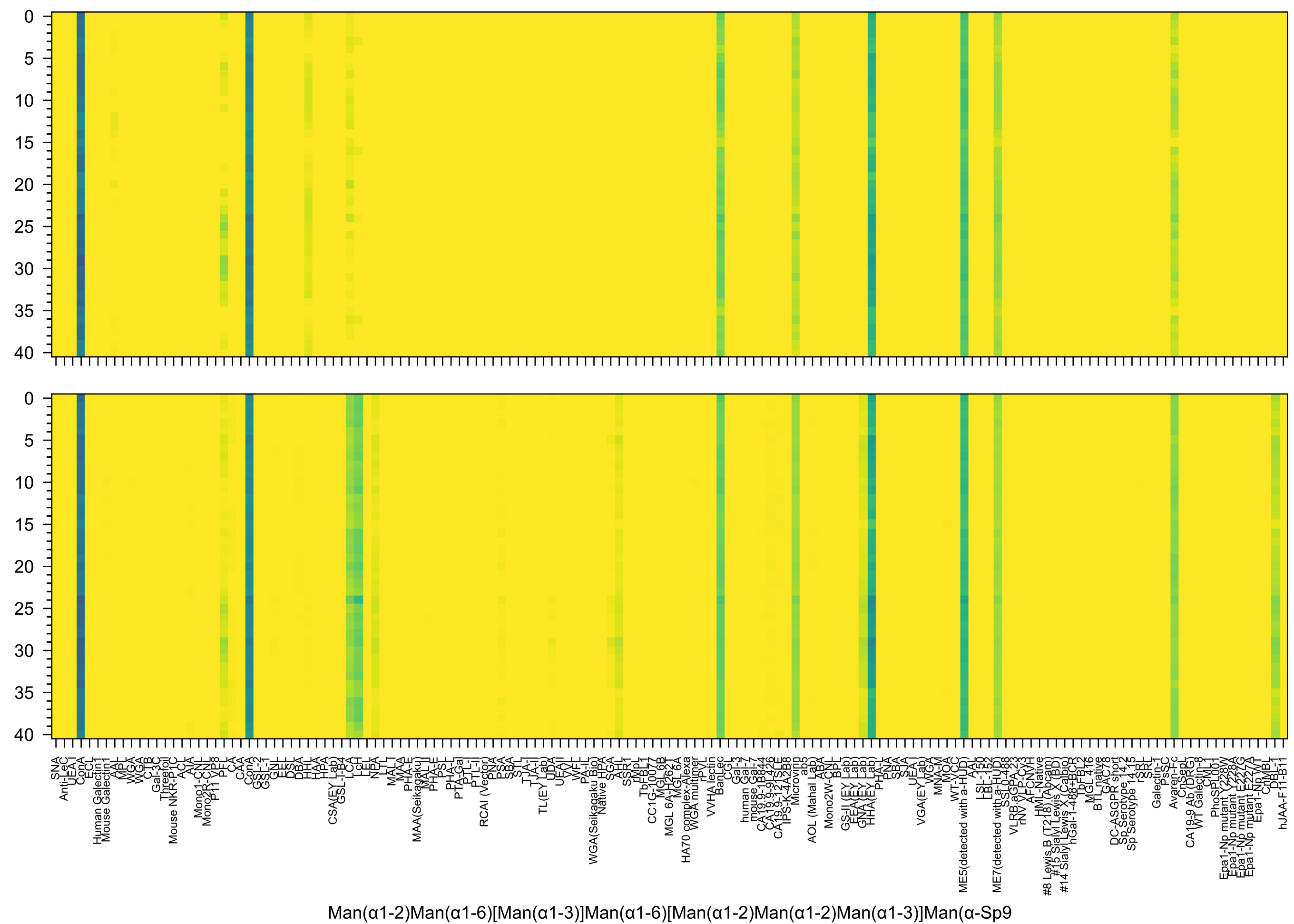
Man(α1-2)Man(α1-6)[Man(α1-3)]Man(α1-6)[Man(α1-2)Man(α1-2)Man(α1-3)]Man(α-Sp9

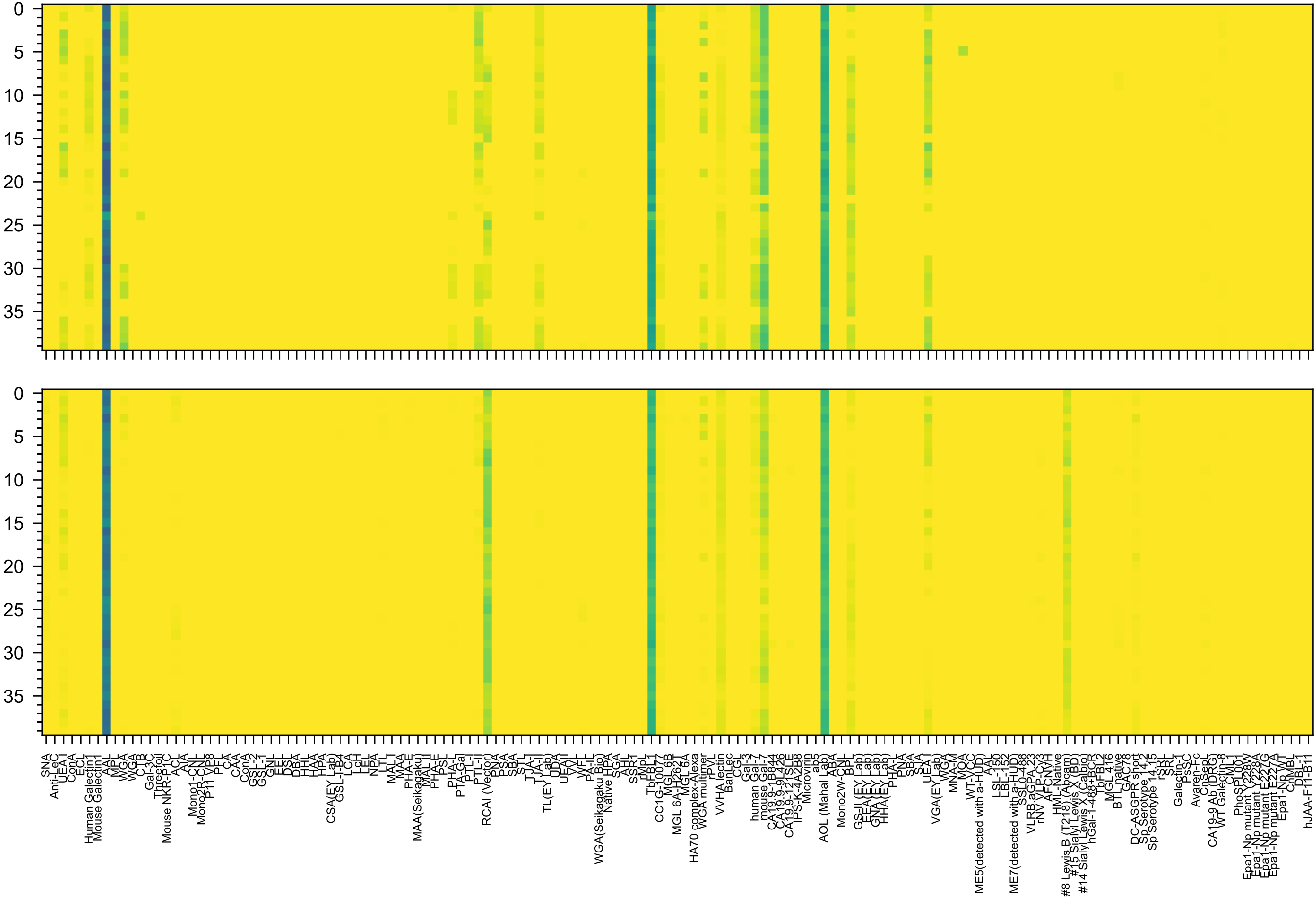

Gal(β1-4)[Fuc(α1-3)]GlcNAc(β1-6)[Fuc(α1-2)Gal(β1-4)GlcNAc(β1-3)]Gal(β1-4)Glc(-Sp21

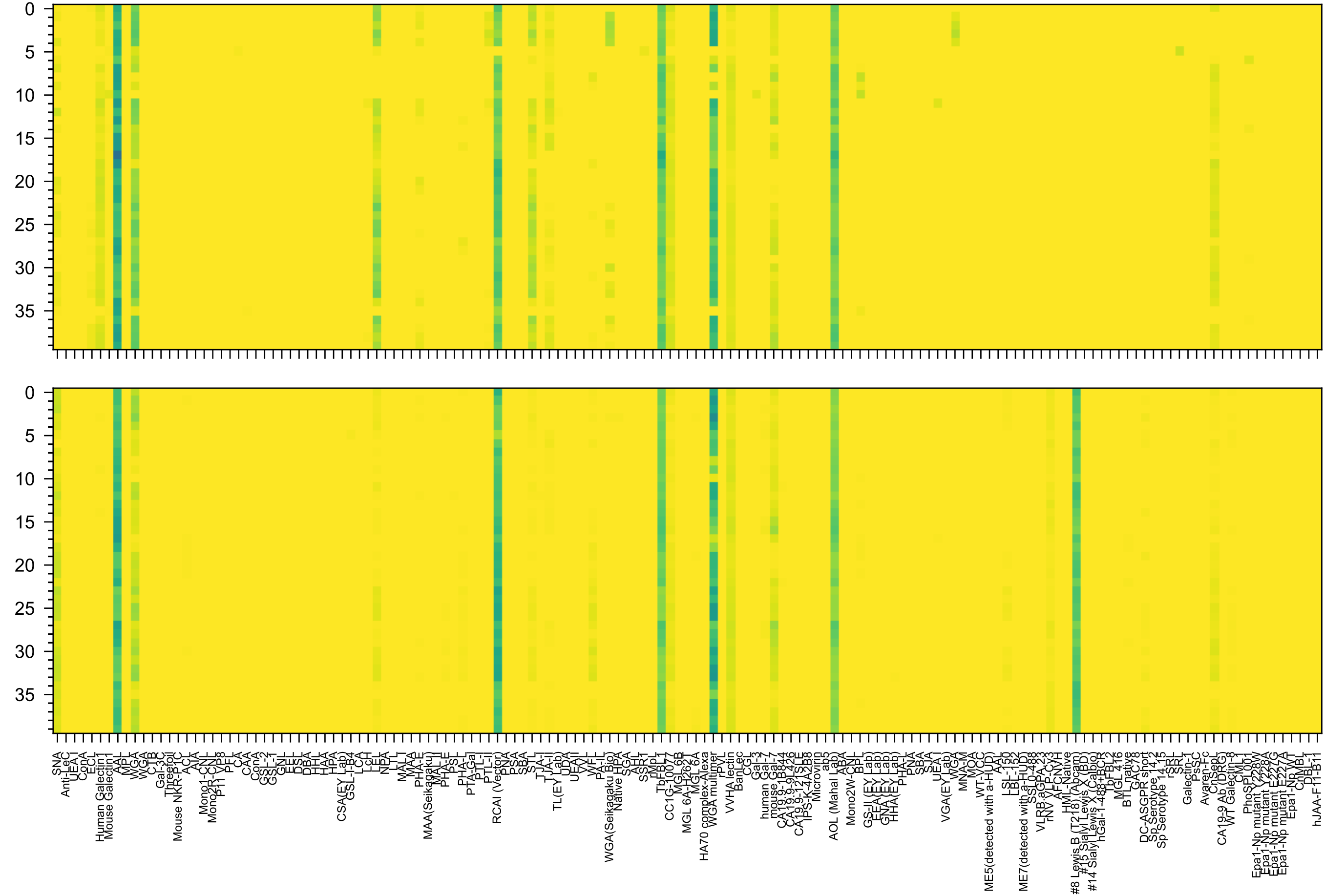


Gal(β1-4)GlcNAc(β1-6)[Fuc(α1-4)[Fuc(α1-2)Gal(β1-3)]GlcNAc(β1-3)]Gal(β1-4)Glc(-Sp21

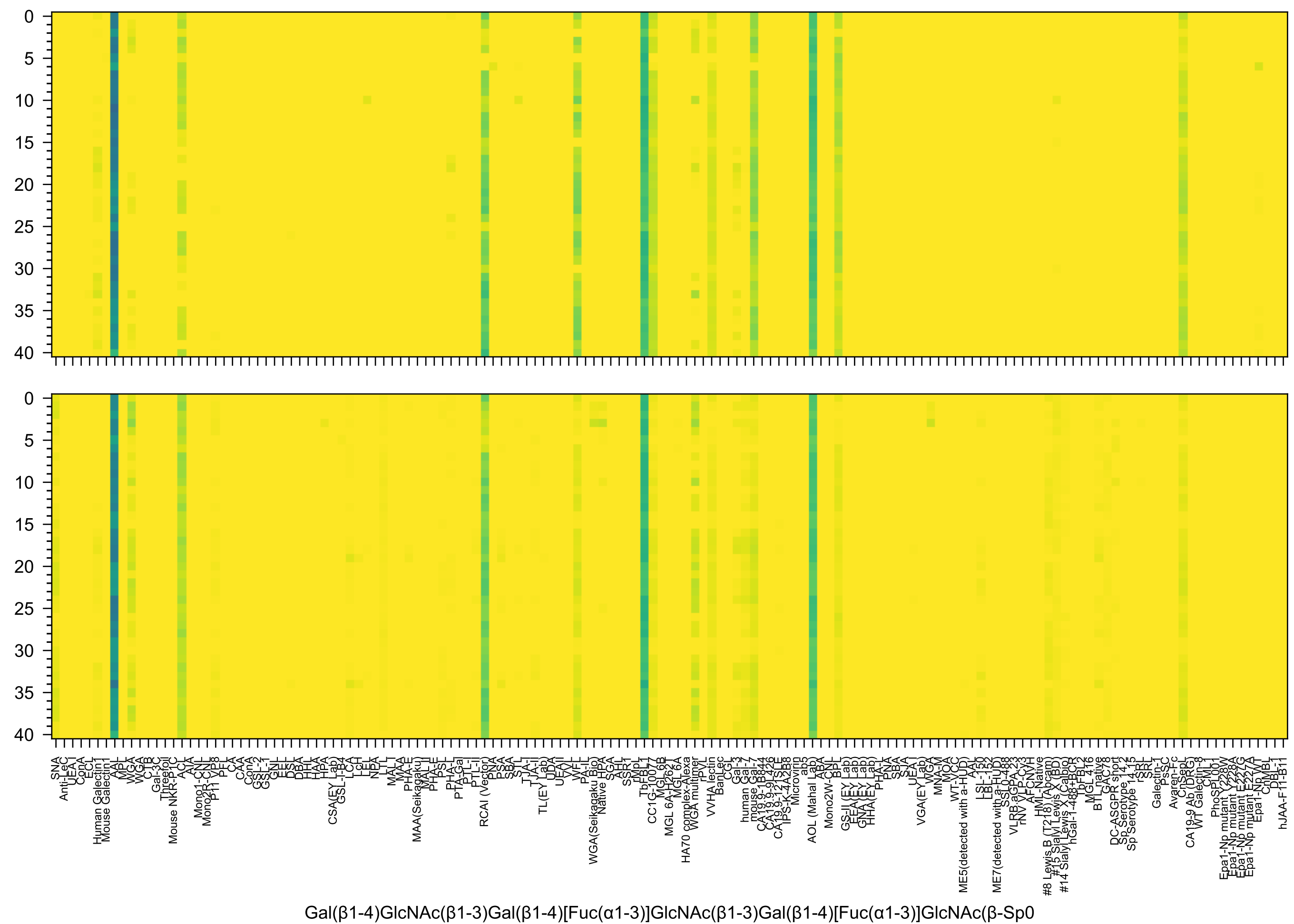
Gal(β1-4)GlcNAc(β1-3)Gal(β1-4)[Fuc(α1-3)]GlcNAc(β1-3)Gal(β1-4)[Fuc(α1-3)]GlcNAc(β-Sp0

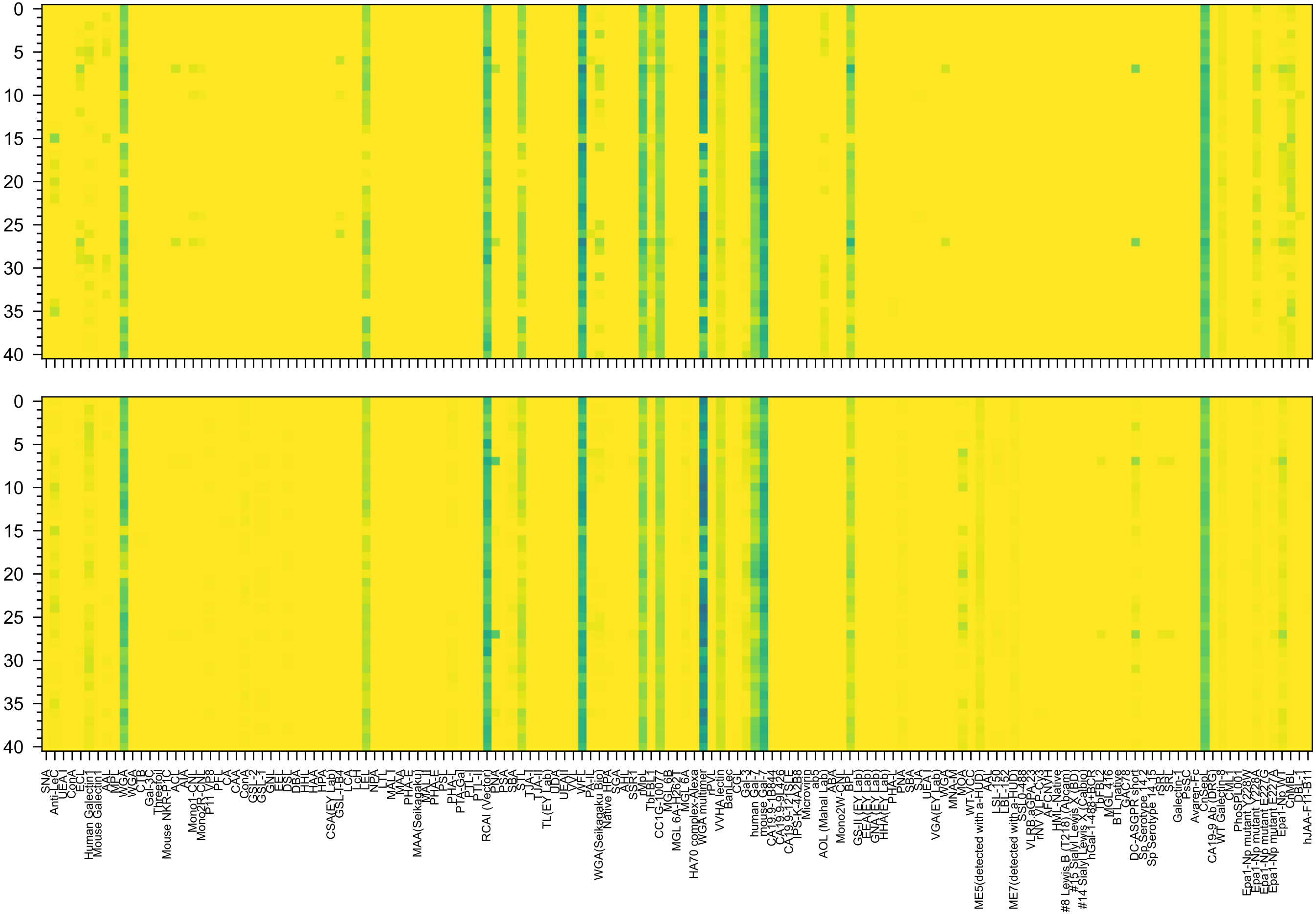

Gal(β1-3)GlcNAc(β1-3)Gal(β1-4)GlcNAc(β1-6)[Gal(β1-3)GlcNAc(β1-3)]Gal(β1-4)Glc(β-Sp0

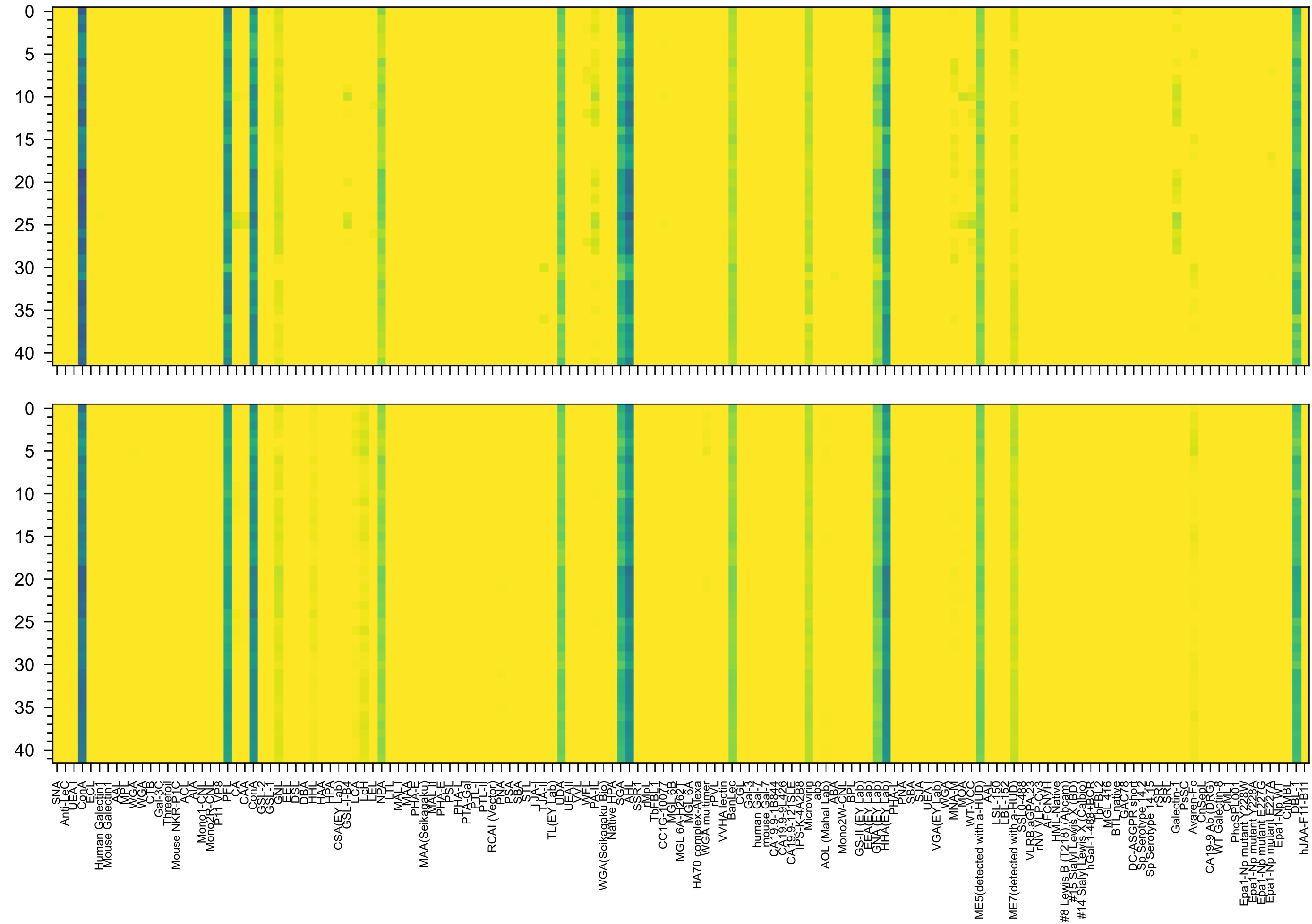


Man(α1-6)[Man(α1-3)]Man(α1-6)[Man(α1-2)Man(α1-3)]Man(β1-4)GlcNAc(β1-4)GlcNAc(β-Sp12

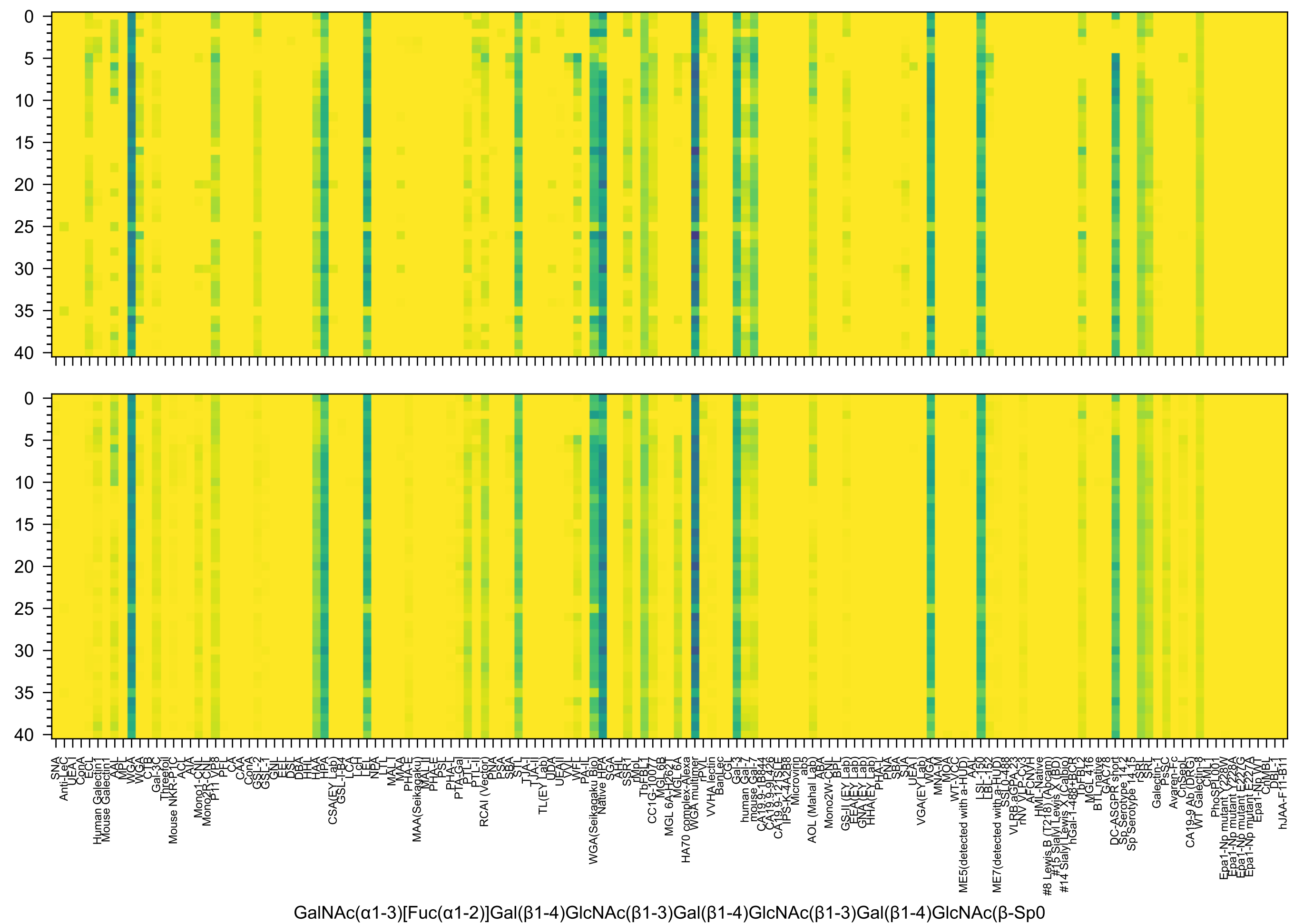
GalNAc(α1-3)[Fuc(α1-2)]Gal(β1-4)GlcNAc(β1-3)Gal(β1-4)GlcNAc(β1-3)Gal(β1-4)GlcNAc(β-Sp0

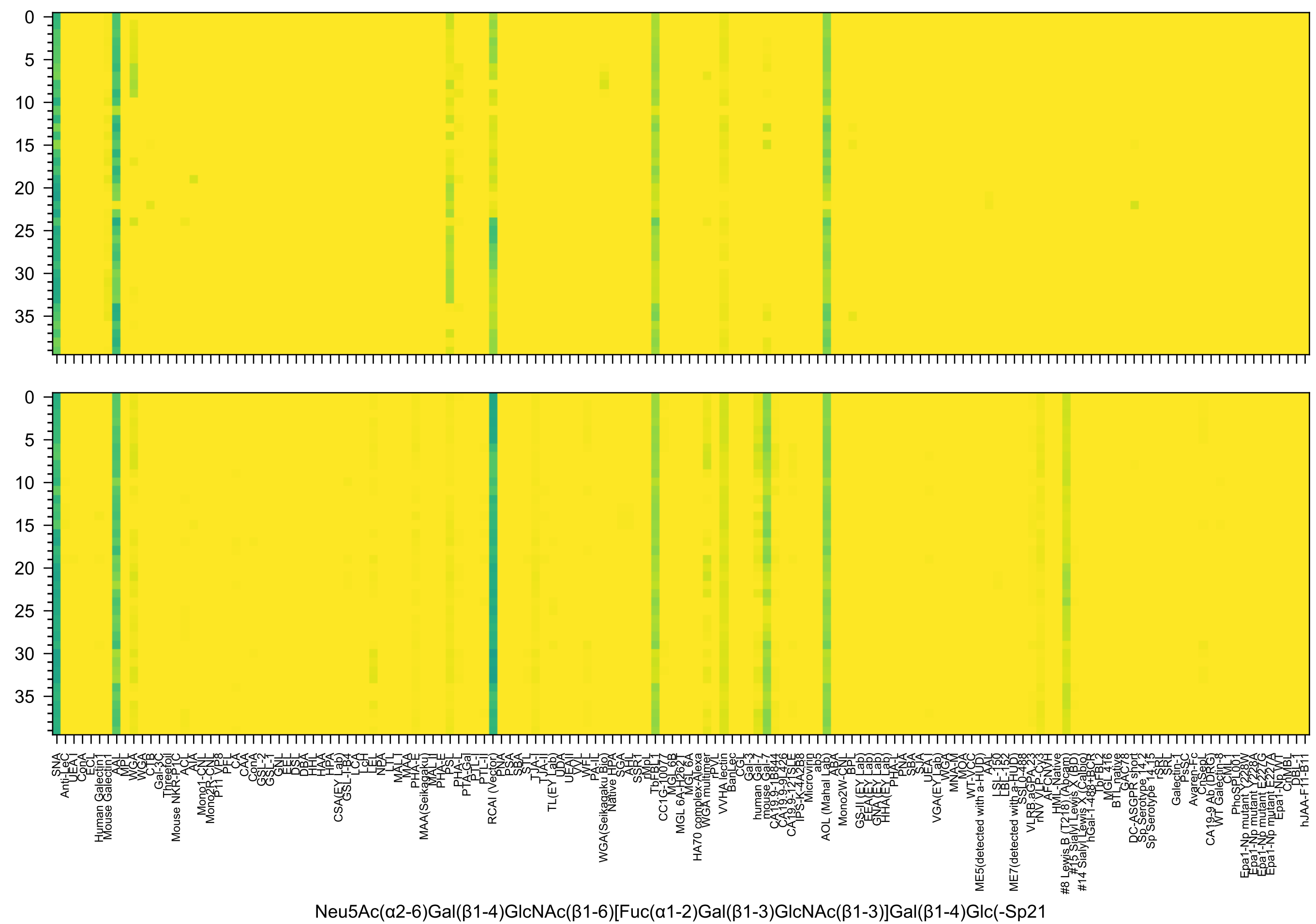
0
5
10
15
20
25
30
35
SNA
Anti-LeC
UEA I
ConA
ECL
Human Galectin1
Mouse Galectin1
AAL
MPL
WGA
WGA
CTB
Gal-3C
Threefoil
Mouse NKR-P1C
ACL
AIA
Mono1-CNL
Mono2R-CNL
P11 VP8
PFL
CA
CAA
ConA
GSL-2
GSL-1
GNL
EEL
DSL
DBA
PHL
HAA
HPA
CSA(EY Lab)
GSL-I-B4
LCA
LcH
LEL
NPA
LTL
MAL I
MAA
PHA-E
MAA(Seikagaku)
MAL II
PHA-E
PsL
PHA-L
PTA-Gal
PTL-I
PTL-II
RCAI (Vector)
PNA
PSA
SBA
STL
TJA-I
TJA-II
TL(EY Lab)
UDA
UEA-II
VVL
VFL
PA-IL
WGA(Seikagaku Bio)
Native HPA
SGA
AHL
SSR1
rMpL
TbFBL1
CC1G-10077
MGL 6B
MGL 6A-H262T
MGL 6A
HA70 complex-Alexa
WGA multimer
rPVL
VVHA lectin
BanLec
CGL
Gal-3
human Gal-7
mouse Gal-7
CA19.9-1B844
CA19.9-9L426
CA19.9-121SLE
IPS-K-4A2B8
Microvirin
ab5
AOL (Mahal Lab)
ABA
Mono2W-CNL
BPL
GS-II (EY Lab)
EEA(EY Lab)
GNA(EY Lab)
HHA(EY Lab)
PHA-L
PNA
SBA
SJA
UEA-I
VGA(EY Lab)
WGA
MNA-M
MOA
WT-VCC
ME5(detected with a-HUD)
AAL
LSL-150
LBL-152
ME7(detected with a-HUD)
SSL0-488
VLRB.aGPA.23
rNV VLP-Cy3
AFCNVH
HML-Native
#8 Lewis B (T218) (Abcam)
#15 Sialyl Lewis X (BD)
#14 Sialyl Lewis X (Calbio)
hGal-1-488+BCR
TbFBL2
MGL 416
BTL native
GAC78
DC-ASGPR short
Sp Serotype 14.2
Sp Serotype 14.15
rSRL
SRL
Galectin-1
PsSC
Avaren-Fc
CnSeL
CA19-9 Ab (DRG)
WT Galectin-8
CVN-1
Epa1-Np mutant Y228W
Epa1-Np mutant Y228A
Epa1-Np mutant E227G
Epa1-Np mutant E227A
Epa1-Np WT
CnMBL
DBL-1
hJAA-F11-B11
Neu5Ac(α2-6)Gal(β1-4)GlcNAc(β1-6)[Fuc(α1-2)Gal(β1-3)GlcNAc(β1-3)]Gal(β1-4)Glc(-Sp21

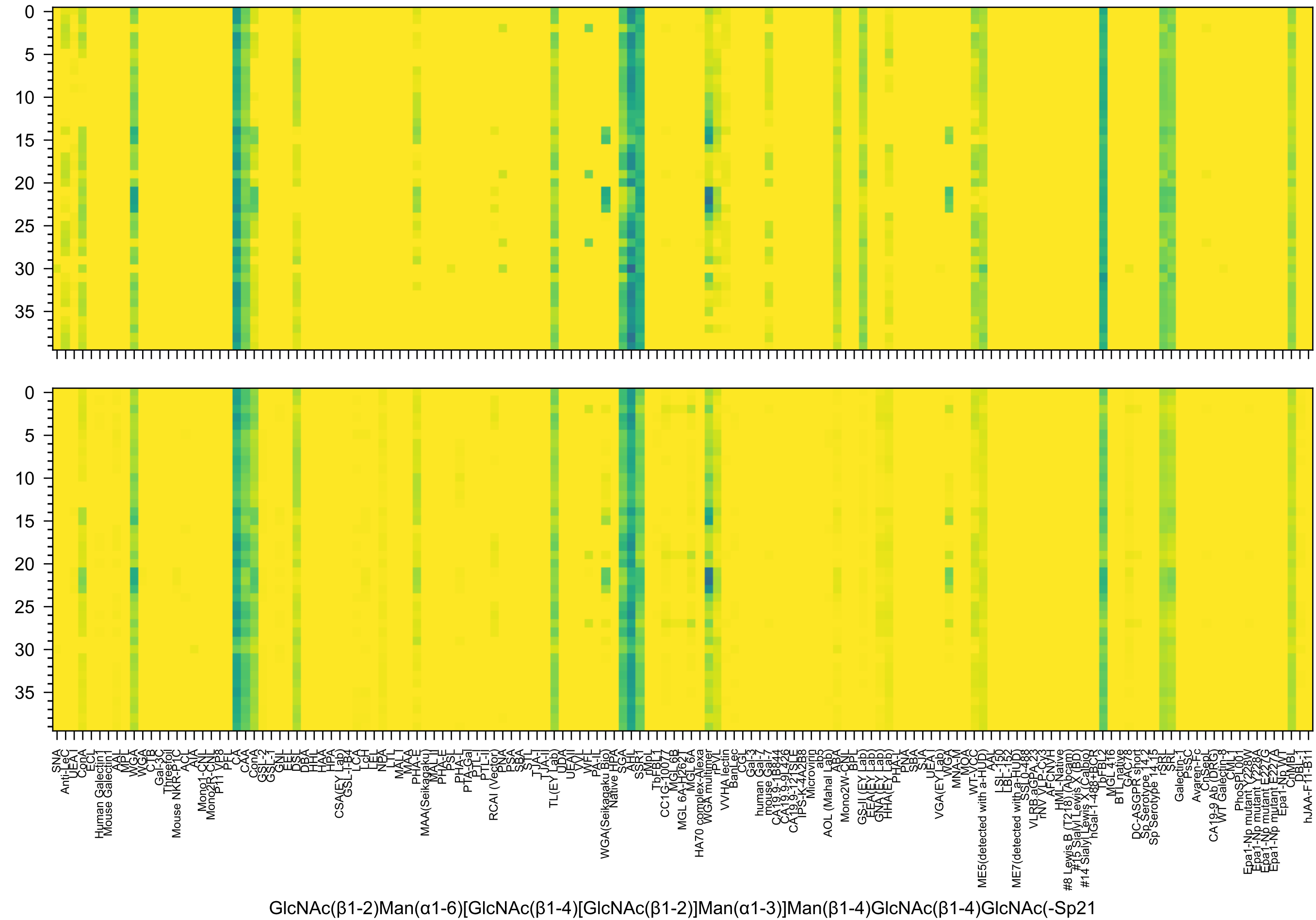
GlcNAc(β1-2)Man(α1-6)[GlcNAc(β1-4)[GlcNAc(β1-2)]Man(α1-3)]Man(β1-4)GlcNAc(β1-4)GlcNAc(-Sp21

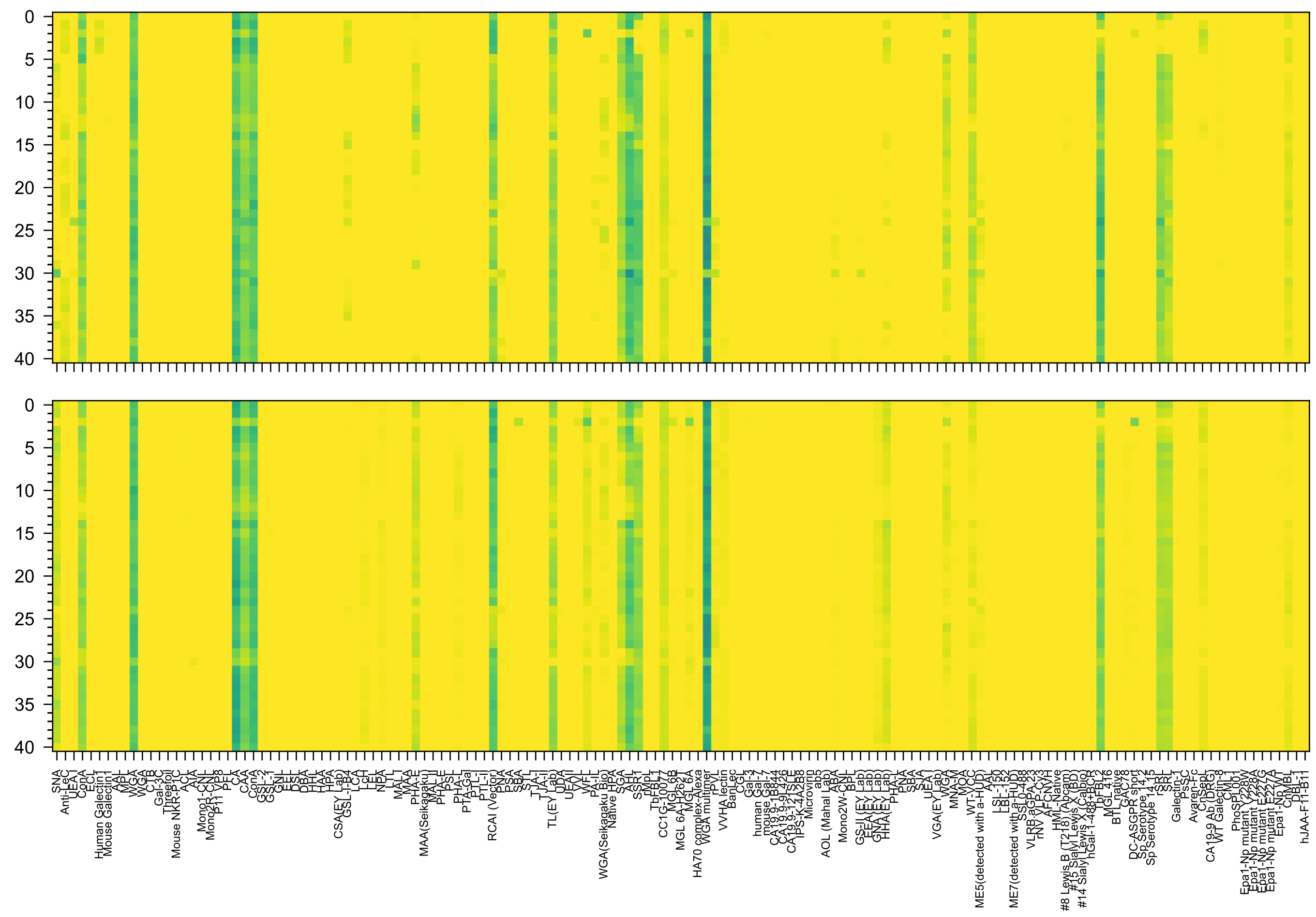

GlcNAc(β1-2)Man(α1-6)[Gal(β1-4)GlcNAc(β1-2)Man(α1-3)]Man(β1-4)GlcNAc(β1-4)GlcNAc(-Sp12

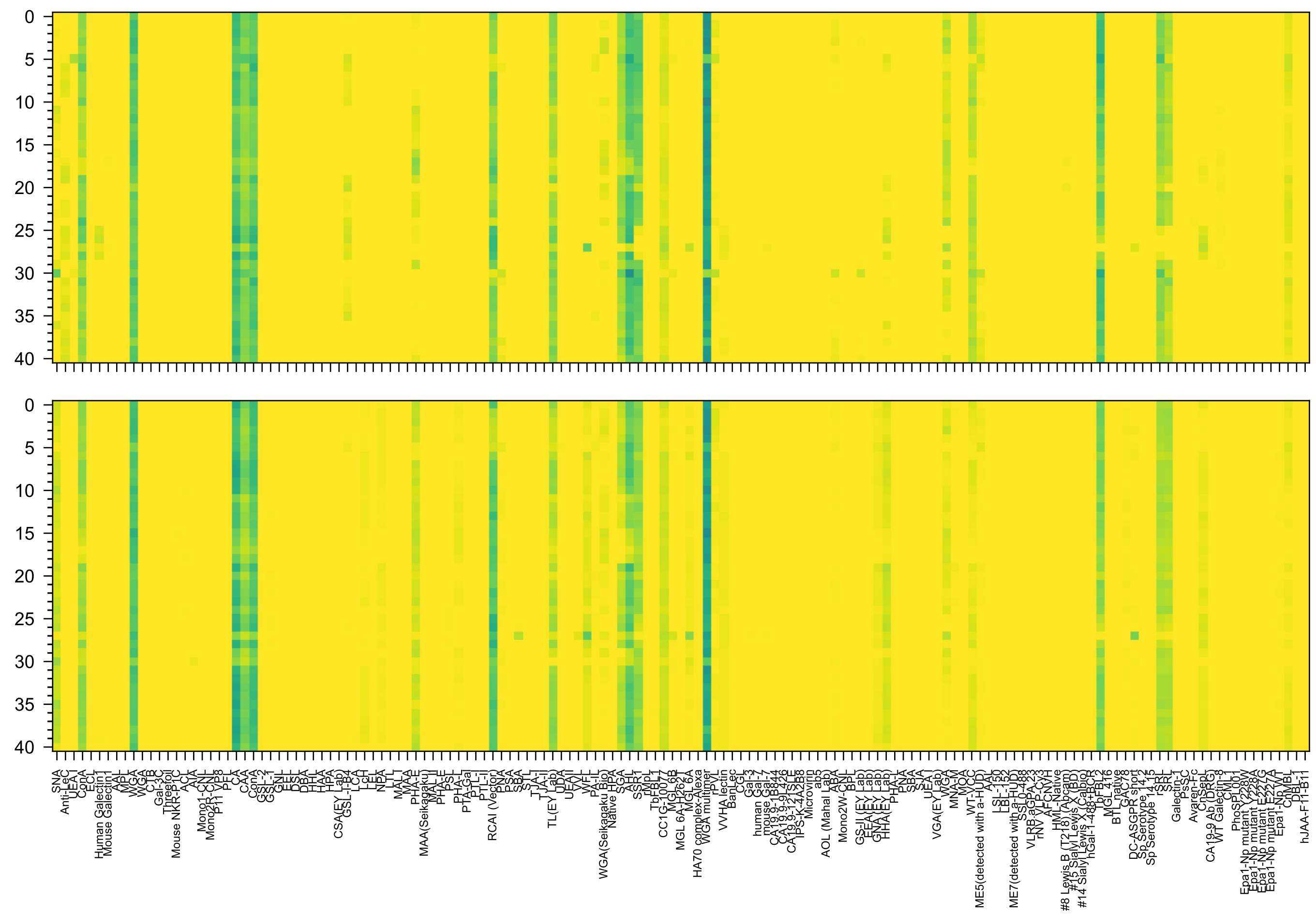

Gal(β1-4)GlcNAc(β1-2)Man(α1-6)[GlcNAc(β1-2)Man(α1-3)]Man(β1-4)GlcNAc(β1-4)GlcNAc(-Sp12

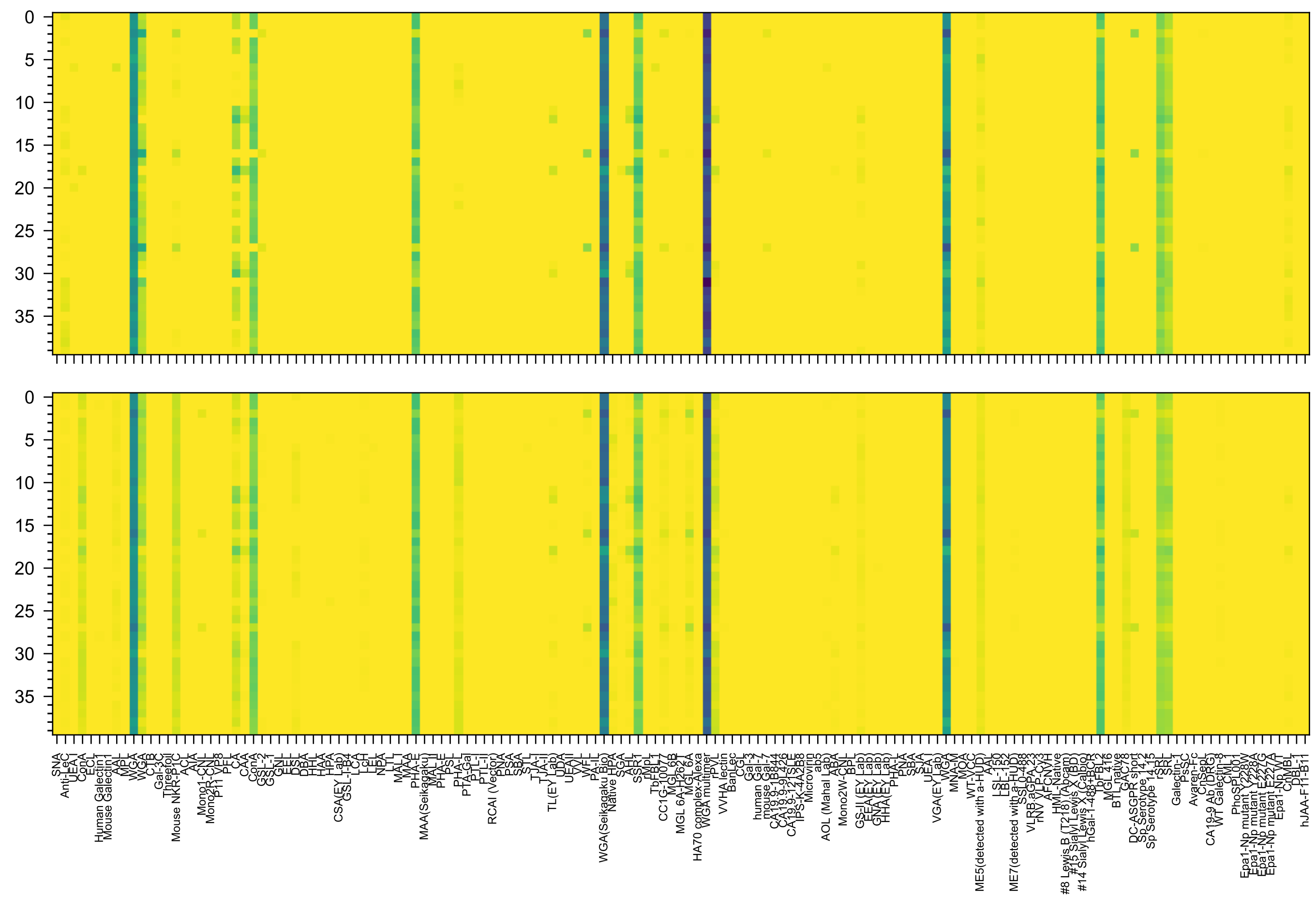

GlcNAc(β1-2)Man(α1-6)[GlcNAc(β1-4)][GlcNAc(β1-2)Man(α1-3)]Man(β1-4)GlcNAc(β1-4)GlcNAc(-Sp21

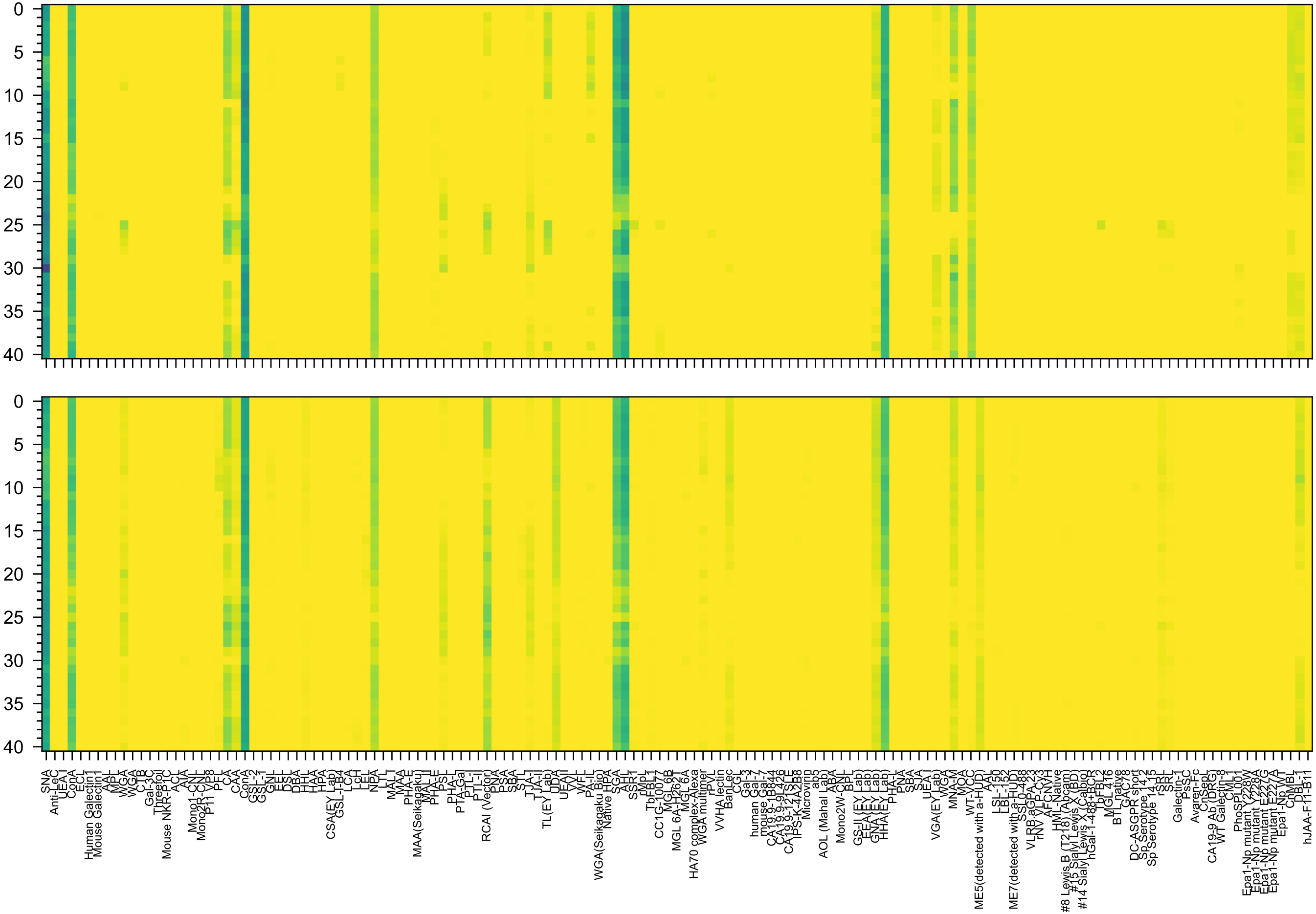


Neu5Ac(α2-6)Gal(β1-4)GlcNAc(β1-2)Man(α1-6)[Man(α1-3)]Man(β1-4)GlcNAc(β1-4)GlcNAc(-Sp12

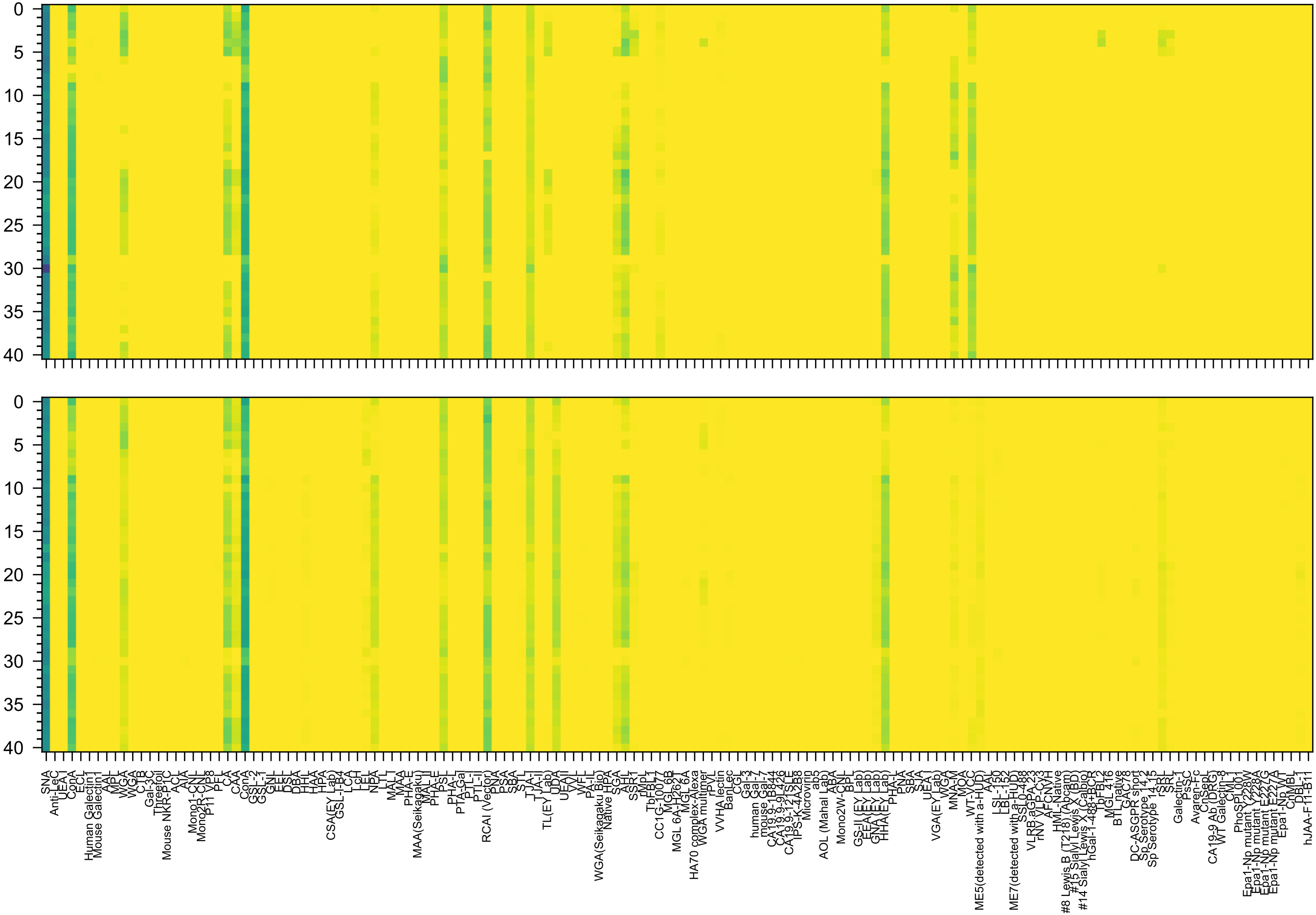

Man(α1-6)[Neu5Ac(α2-6)Gal(β1-4)GlcNAc(β1-2)Man(α1-3)]Man(β1-4)GlcNAc(β1-4)GlcNAc(-Sp12

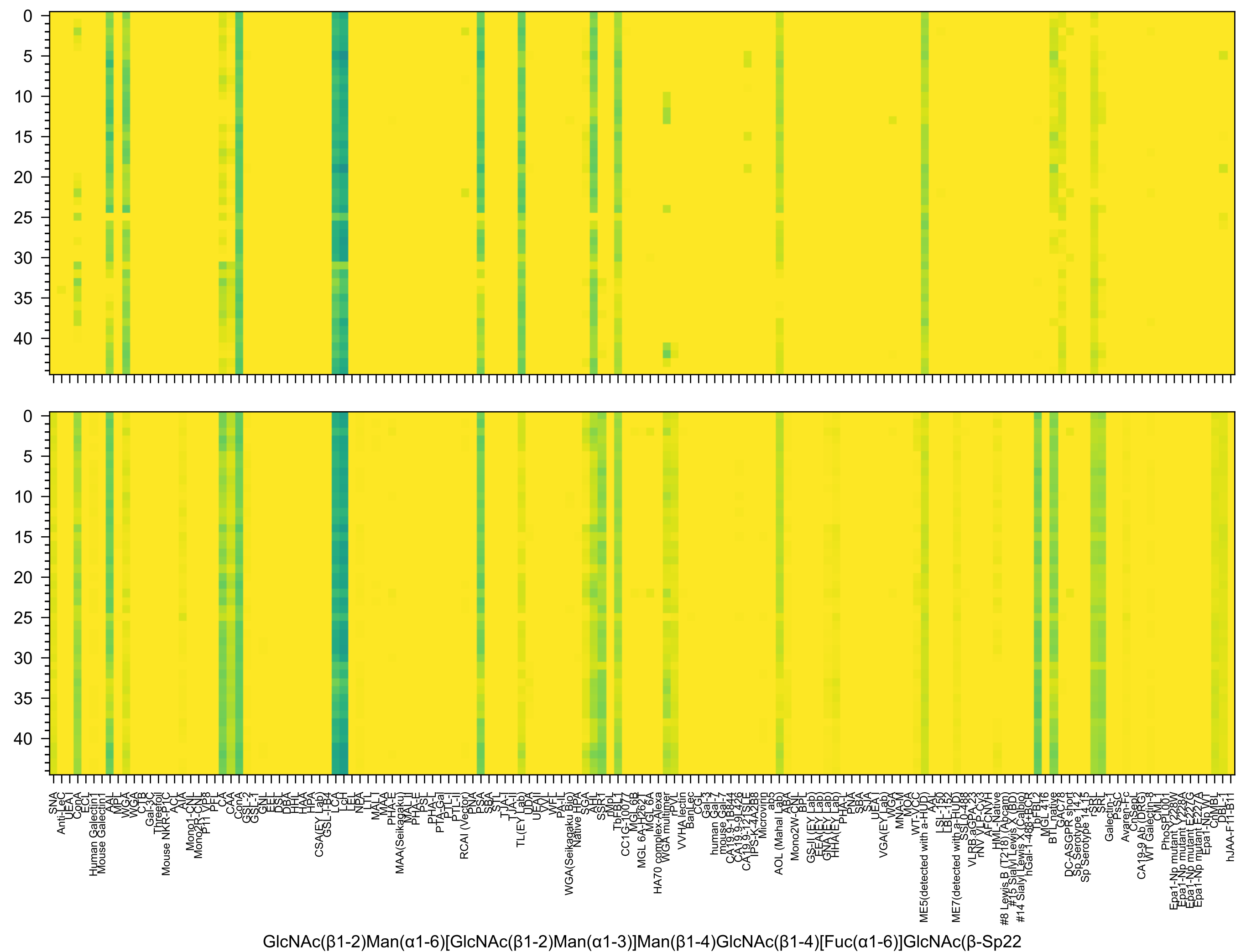
GlcNAc(β1-2)Man(α1-6)[GlcNAc(β1-2)Man(α1-3)]Man(β1-4)GlcNAc(β1-4)[Fuc(α1-6)]GlcNAc(β-Sp22

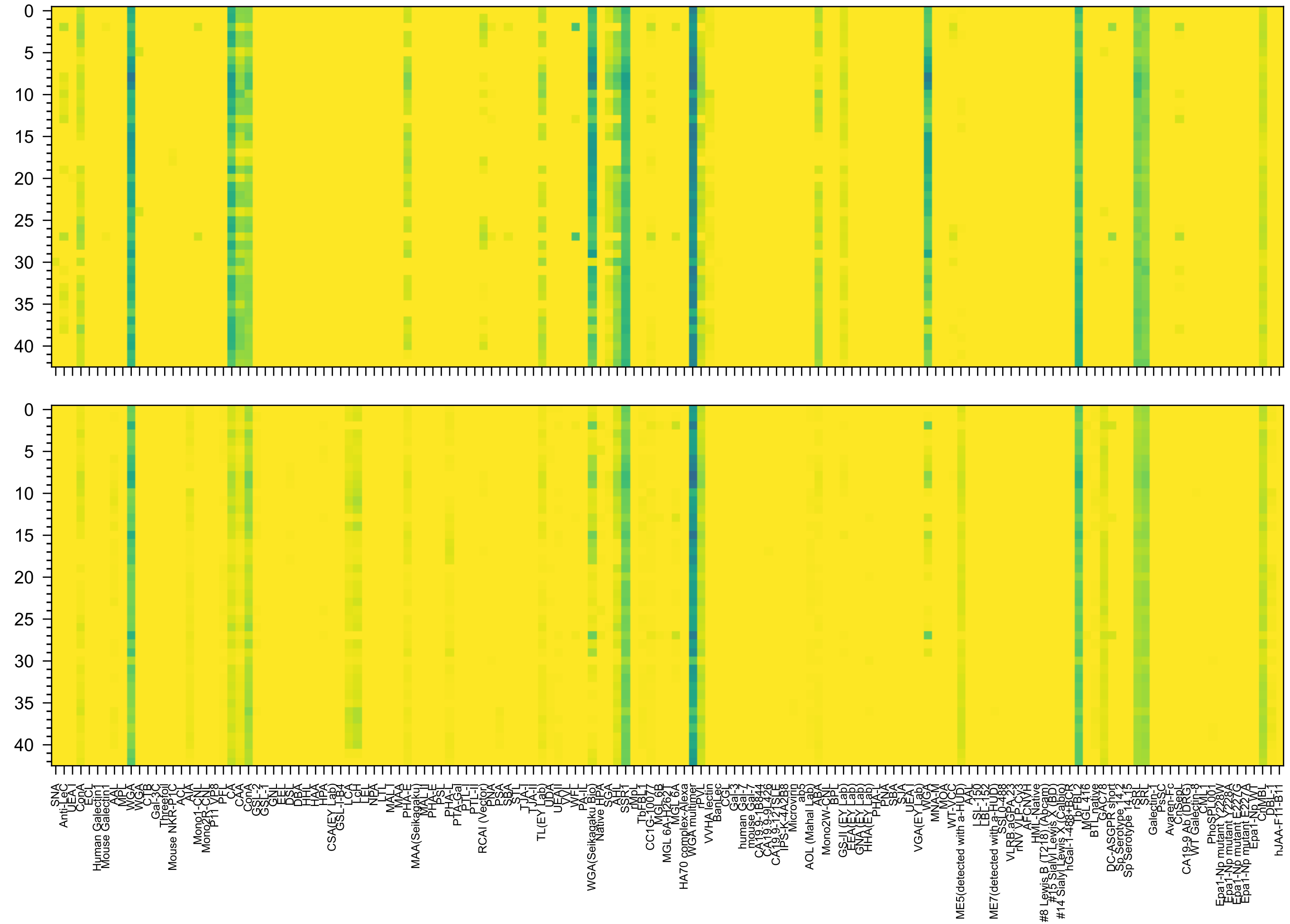


GlcNAc(β1-2)[GlcNAc(β1-6)]Man(α1-6)[GlcNAc(β1-2)Man(α1-3)]Man(β1-4)GlcNAc(β1-4)GlcNAc(β-Sp19

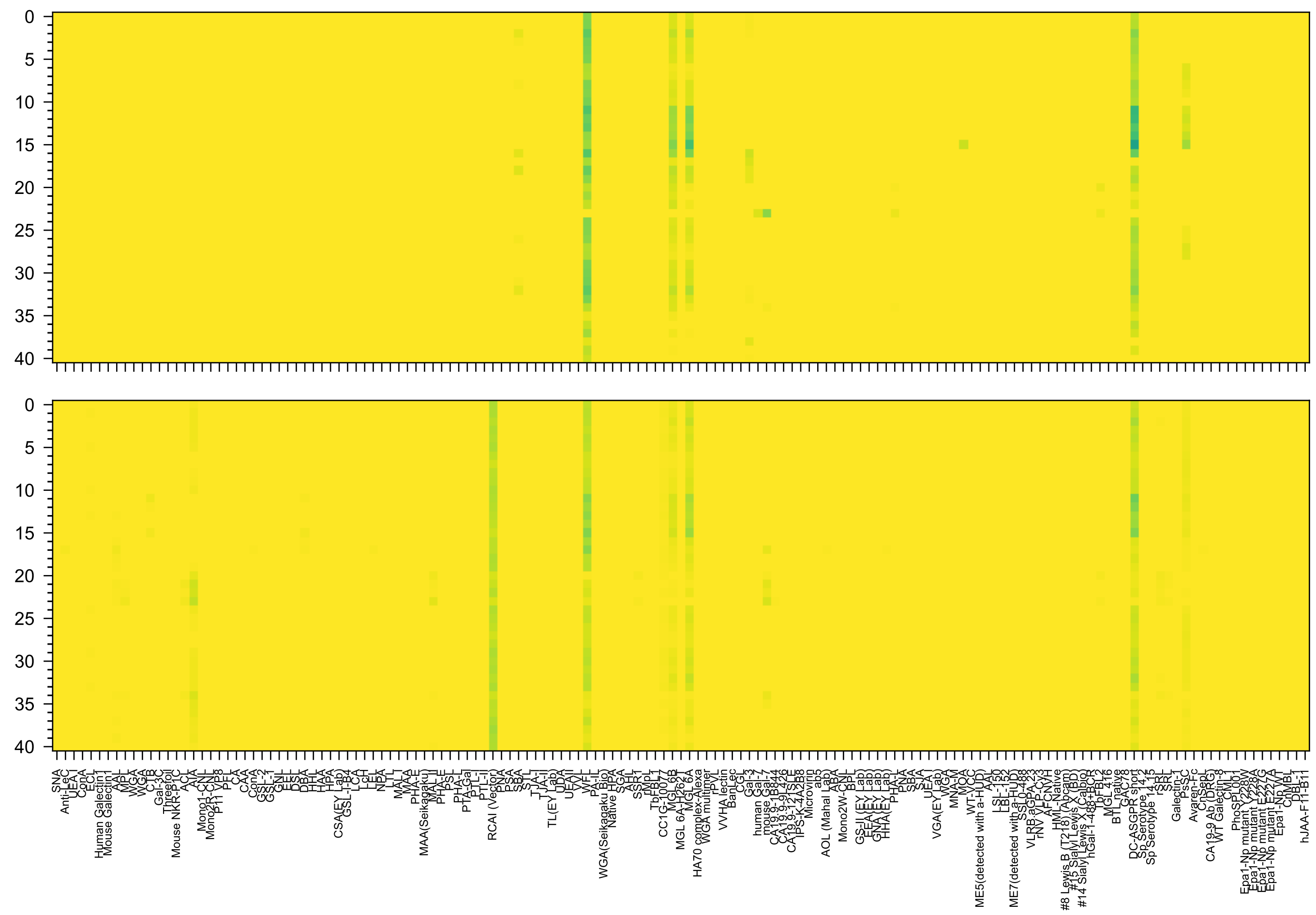

Neu5Ac(α2-8)Neu5Ac(α2-3)Gal(β1-3)GalNAc(β1-4)[Neu5Ac(α2-8)Neu5Ac(α2-3)]Gal(β1-4)Glc(β-Sp0

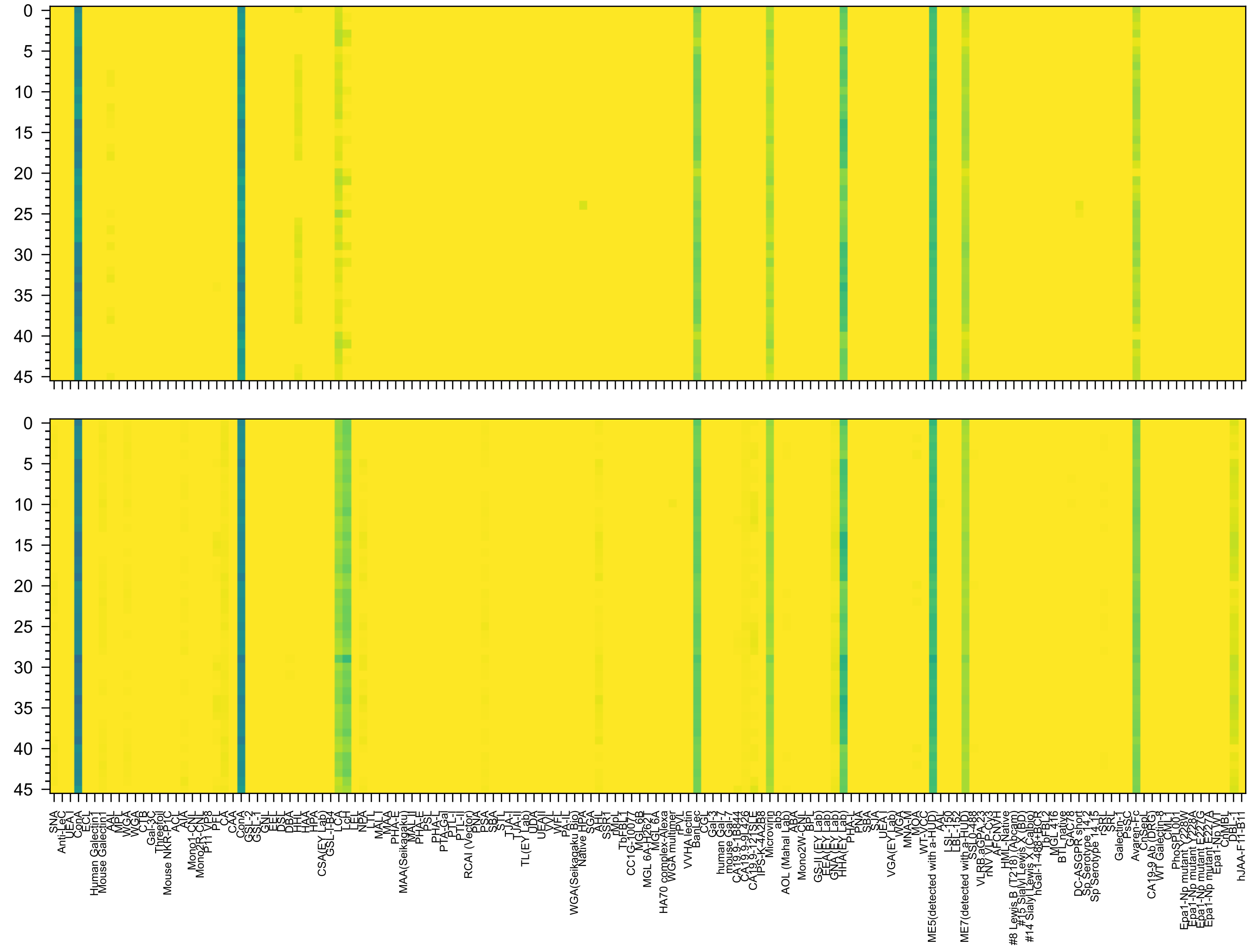

Man(α1-2)Man(α1-6)[Man(α1-2)Man(α1-3)]Man(α1-6)[Man(α1-2)Man(α1-2)Man(α1-3)]Man(α-Sp9

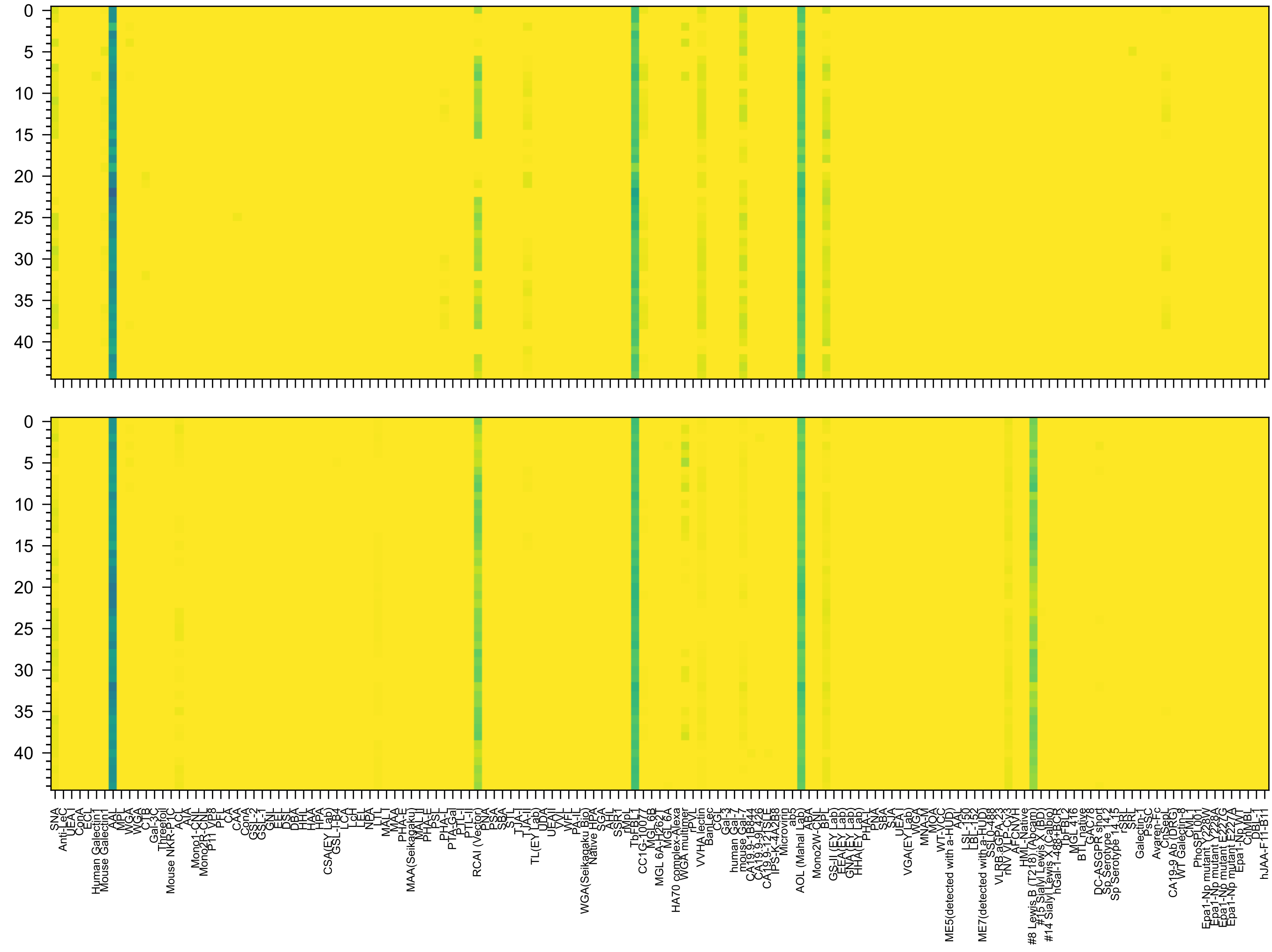

Gal(β1-4)[Fuc(α1-3)]GlcNAc(β1-6)[Fuc(α1-4)[Fuc(α1-2)Gal(β1-3)]GlcNAc(β1-3)]Gal(β1-4)Glc(-Sp21

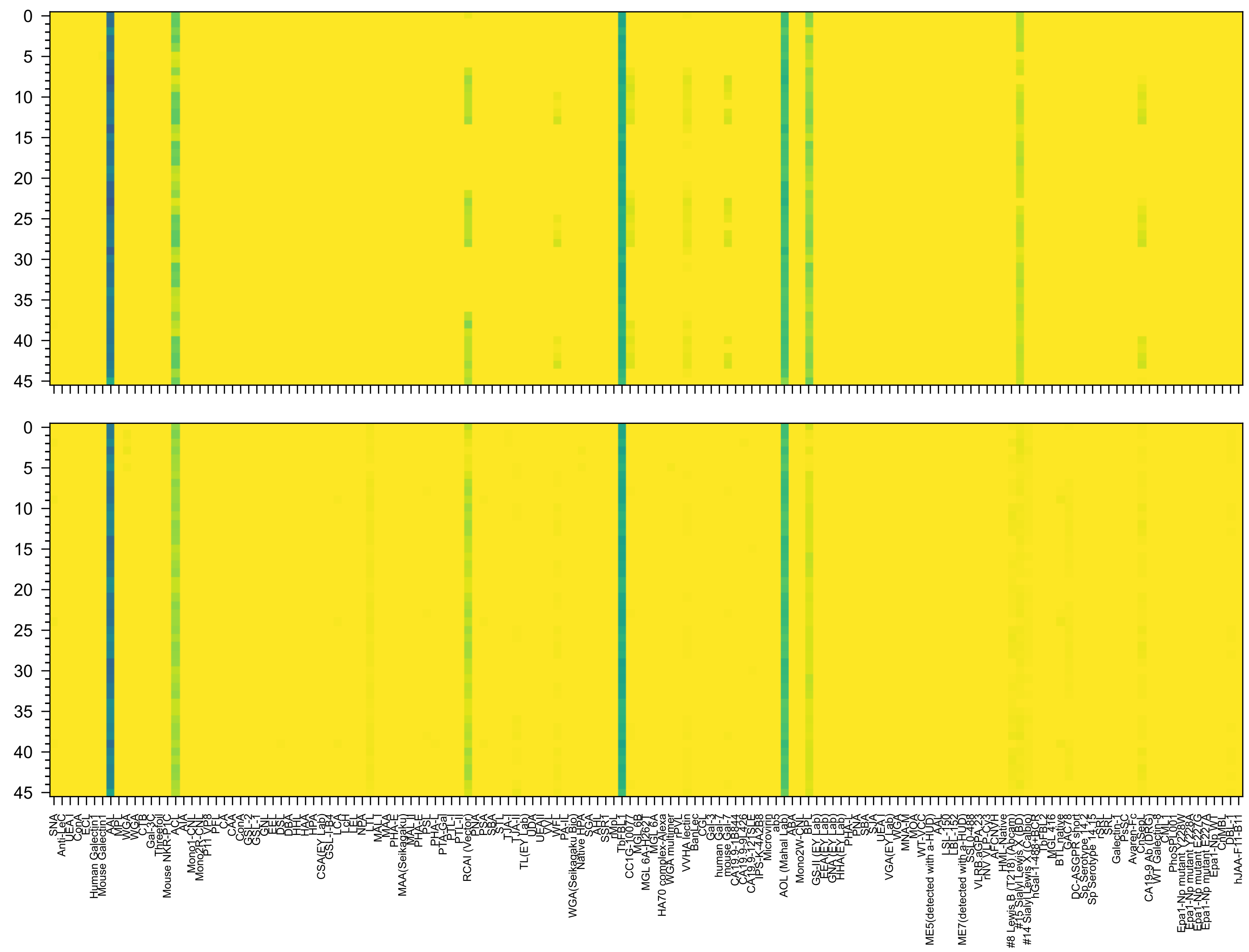


Gal(β1-4)[Fuc(α1-3)]GlcNAc(β1-3)Gal(β1-4)[Fuc(α1-3)]GlcNAc(β1-3)Gal(β1-4)[Fuc(α1-3)]GlcNAc(β-Sp0

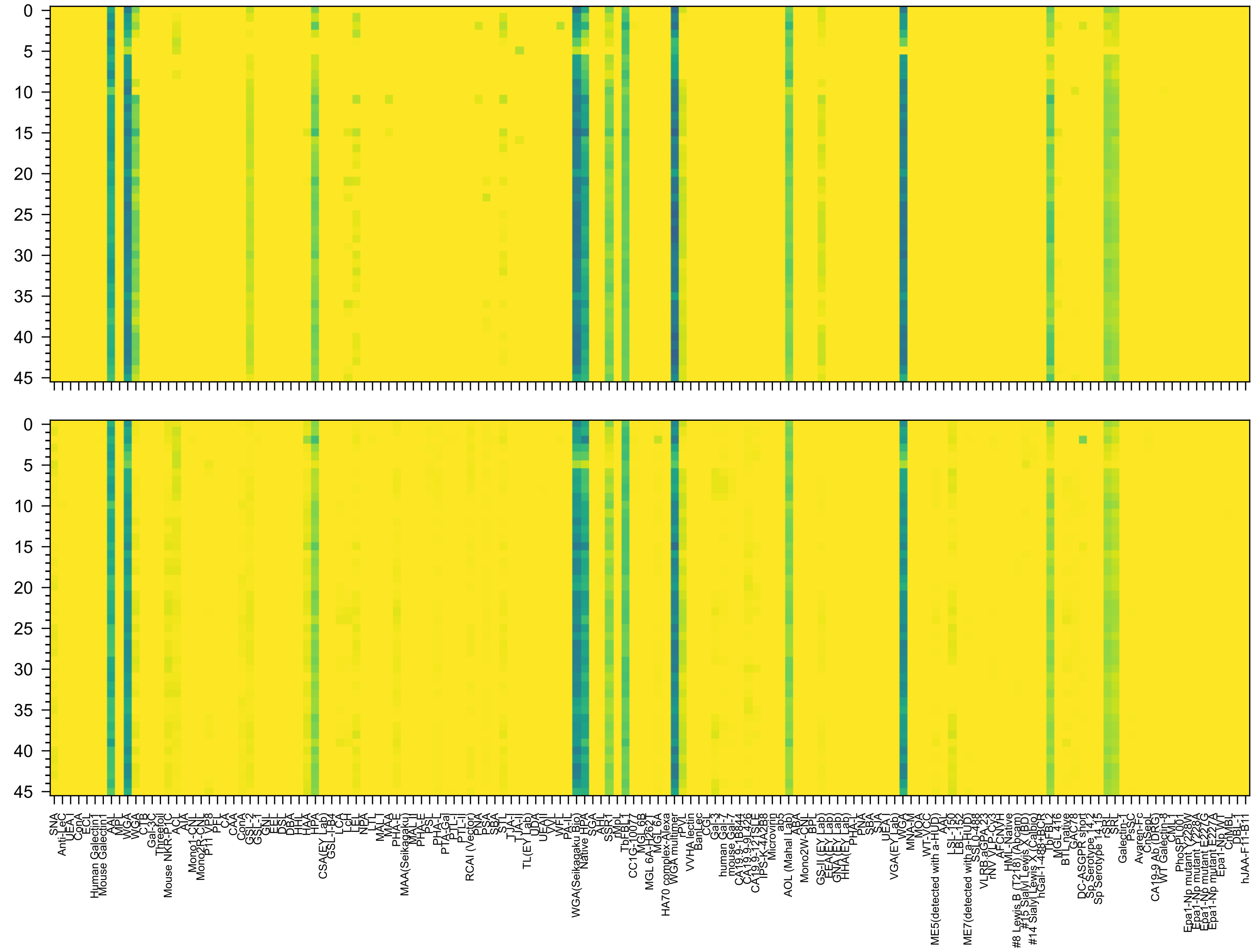

GlcNAc(α1-4)Gal(β1-4)GlcNAc(β1-3)Gal(β1-4)[Fuc(α1-3)]GlcNAc(β1-3)Gal(β1-4)[Fuc(α1-3)]GlcNAc(β-Sp0

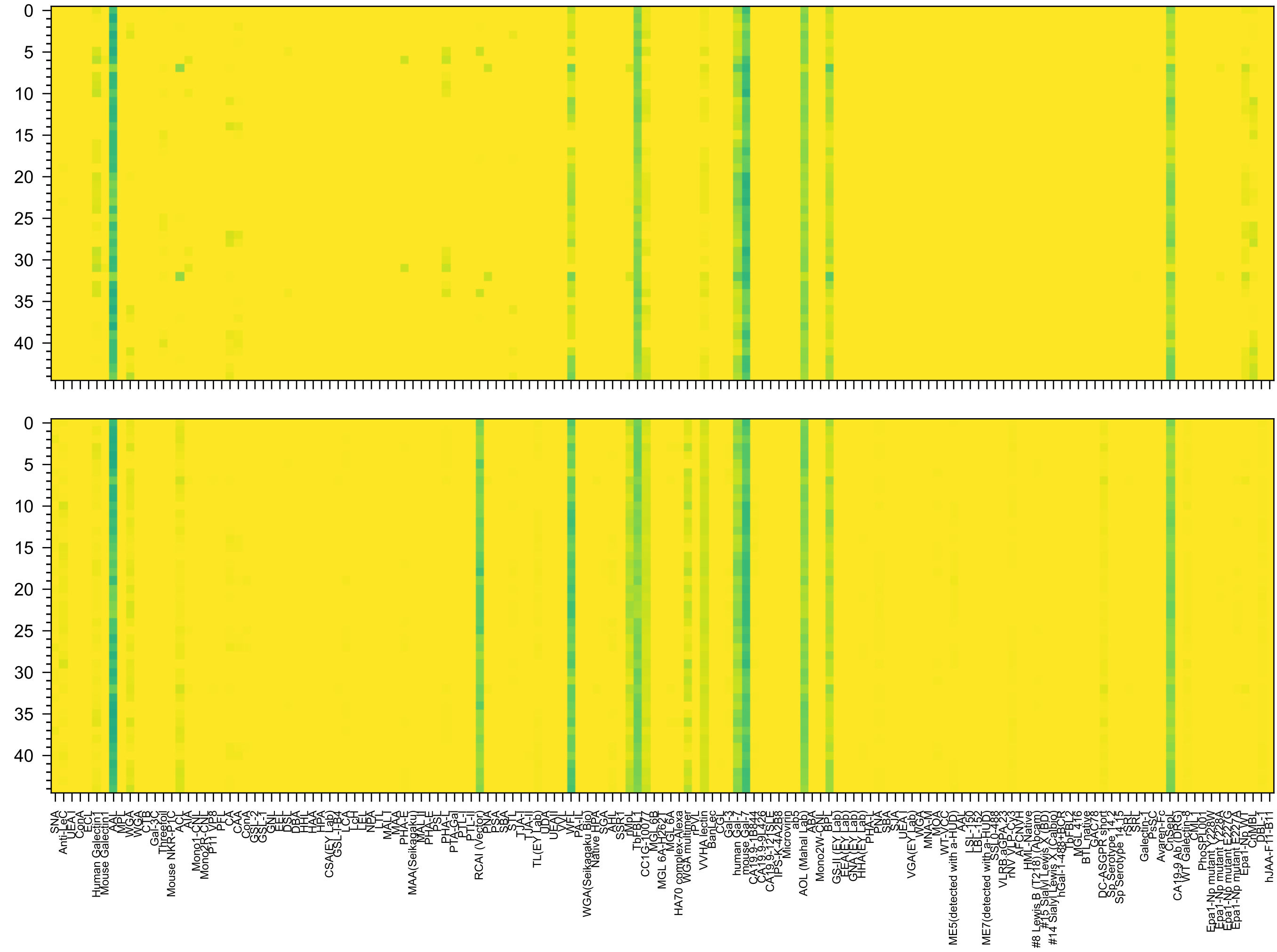

Gal(β1-3)GlcNAc(β1-3)Gal(β1-4)[Fuc(α1-3)]GlcNAc(β1-6)[Gal(β1-3)GlcNAc(β1-3)]Gal(β1-4)Glc(-Sp21

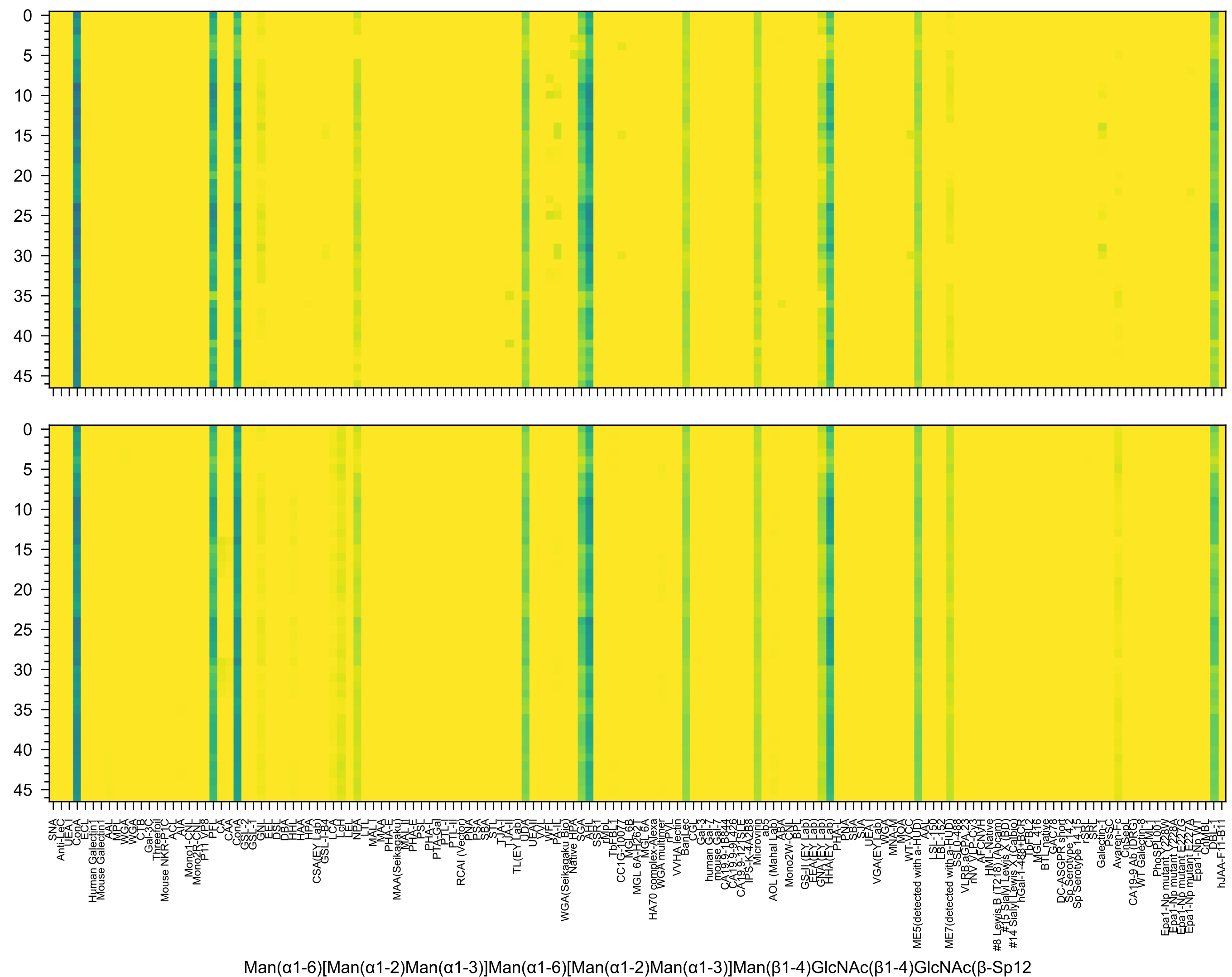

Man(α1-6)[Man(α1-2)Man(α1-3)]Man(α1-6)[Man(α1-2)Man(α1-3)]Man(β1-4)GlcNAc(β1-4)GlcNAc(β-Sp12

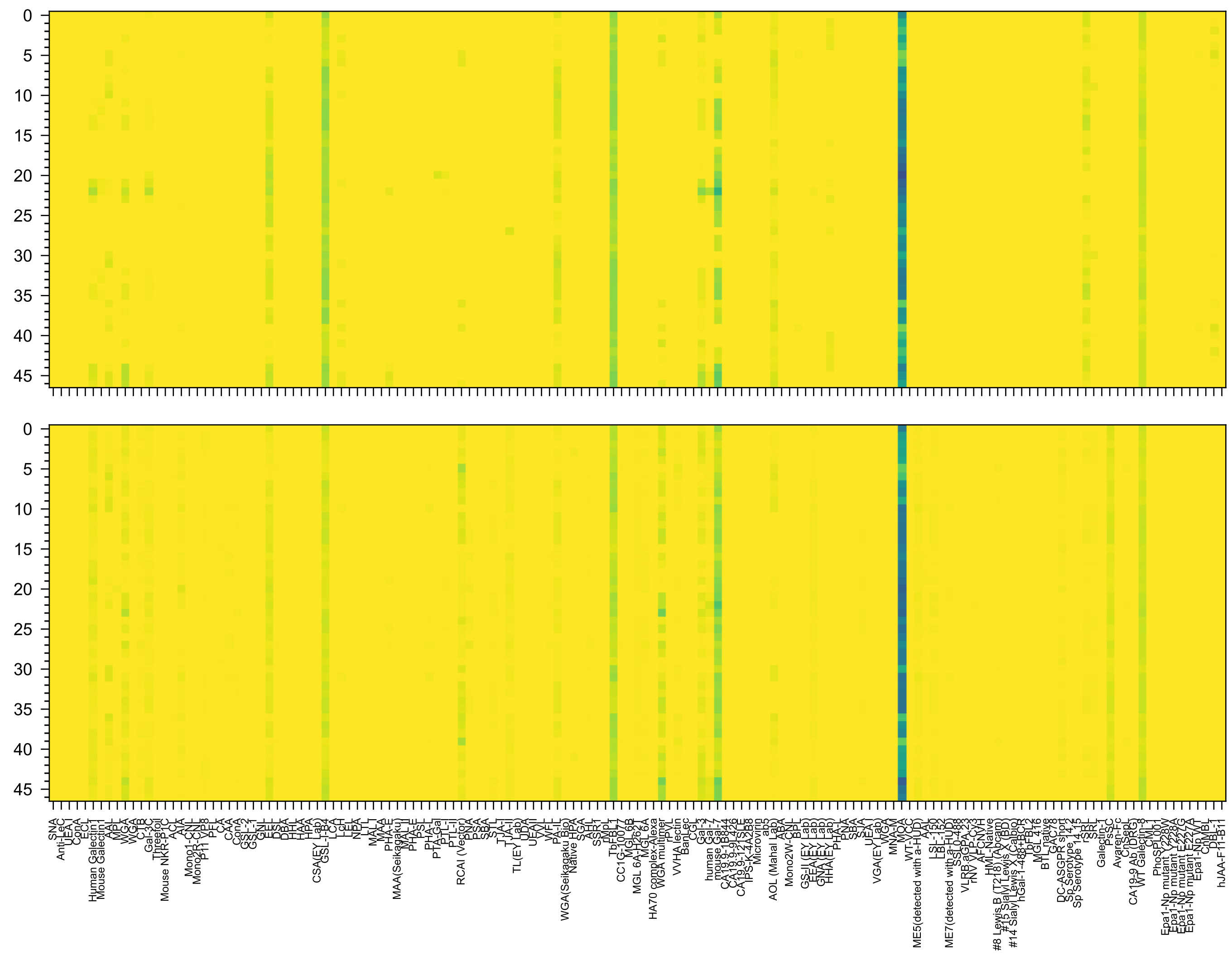


Gal(α1-3)[Fuc(α1-2)]Gal(β1-4)GlcNAc(β1-6)[Gal(α1-3)[Fuc(α1-2)]Gal(β1-4)GlcNAc(β1-3)]GalNAc(-Sp14

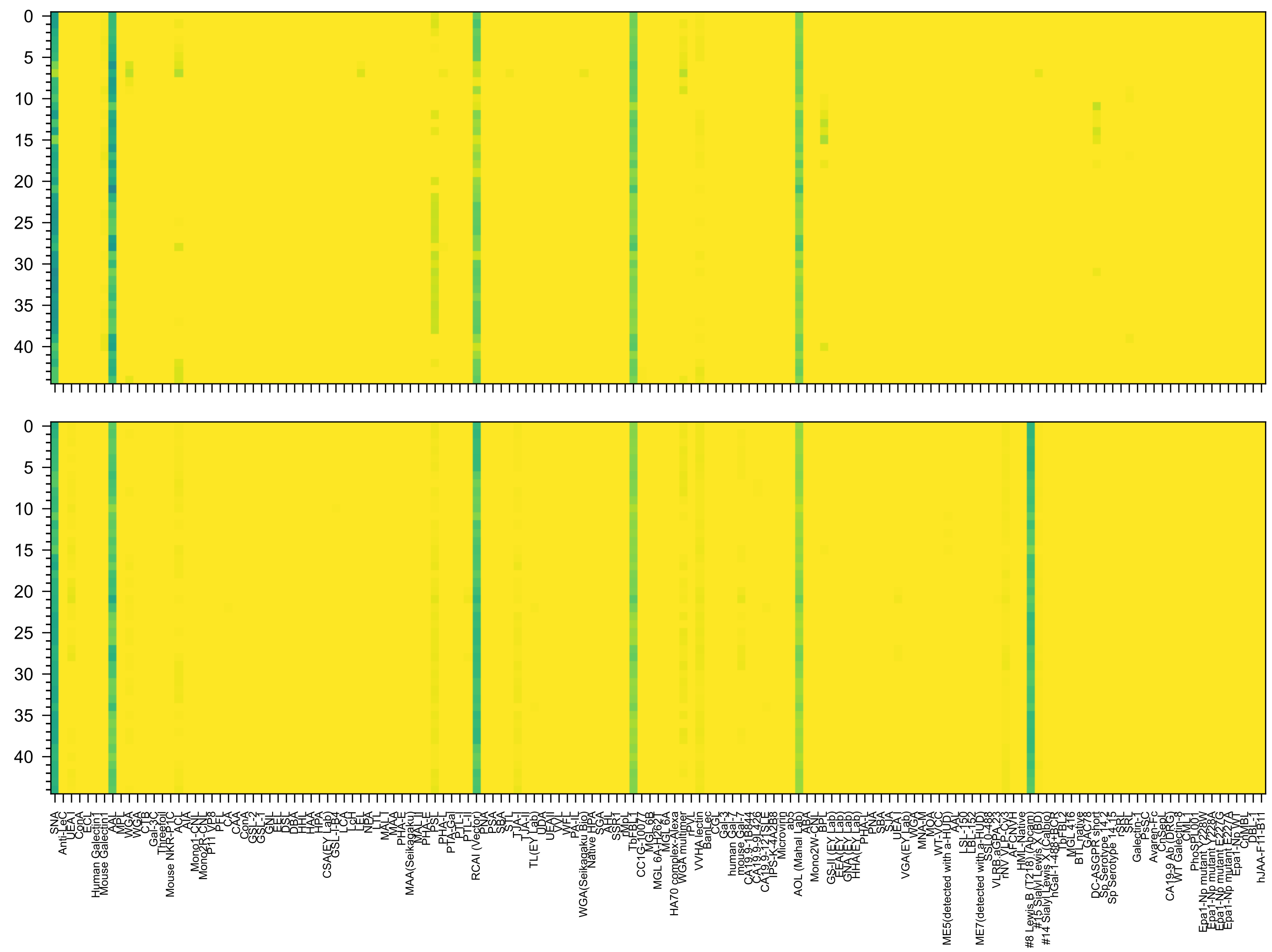


Neu5Ac(α2-6)Gal(β1-4)GlcNAc(β1-6)[Fuc(α1-2)Gal(β1-4)[Fuc(α1-3)]GlcNAc(β1-3)]Gal(β1-4)Glc(-Sp21

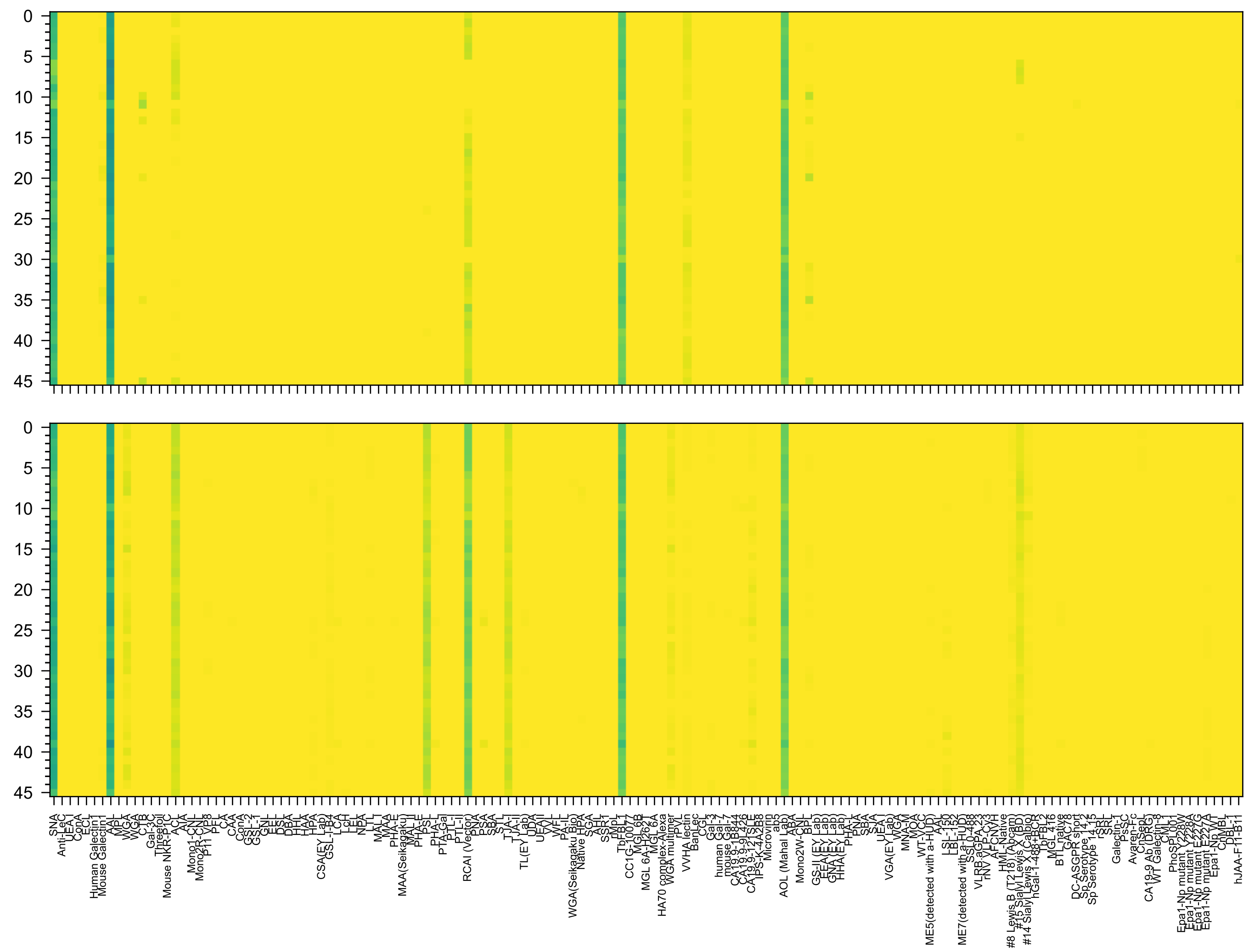

Neu5Ac(α2-6)Gal(β1-4)GlcNAc(β1-3)Gal(β1-4)[Fuc(α1-3)]GlcNAc(β1-3)Gal(β1-4)[Fuc(α1-3)]GlcNAc(β-Sp0

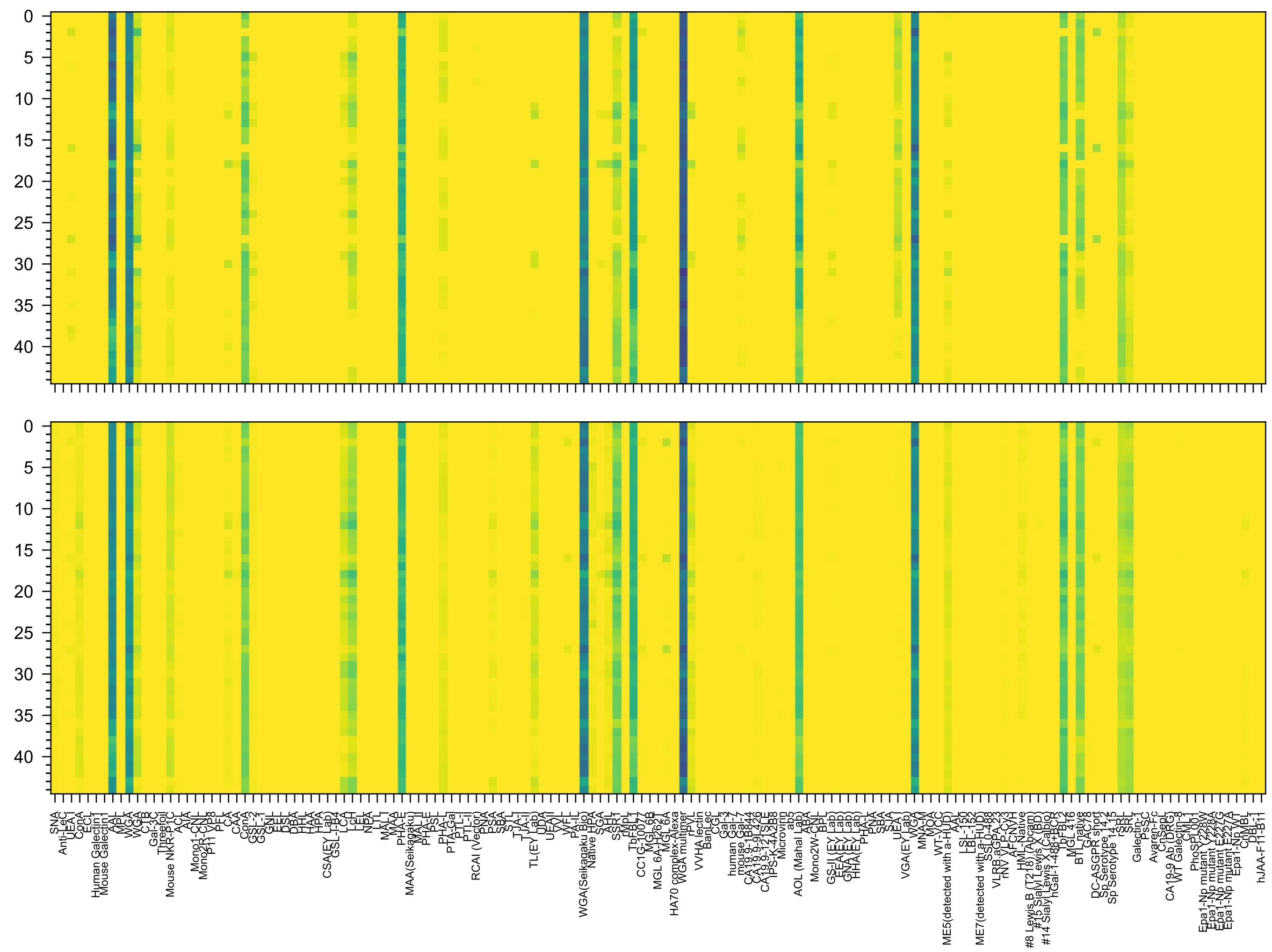

GlcNAc(β1-2)Man(α1-6)[GlcNAc(β1-4)][GlcNAc(β1-2)Man(α1-3)]Man(β1-4)GlcNAc(β1-4)[Fuc(α1-6)]GlcNAc(-Sp21

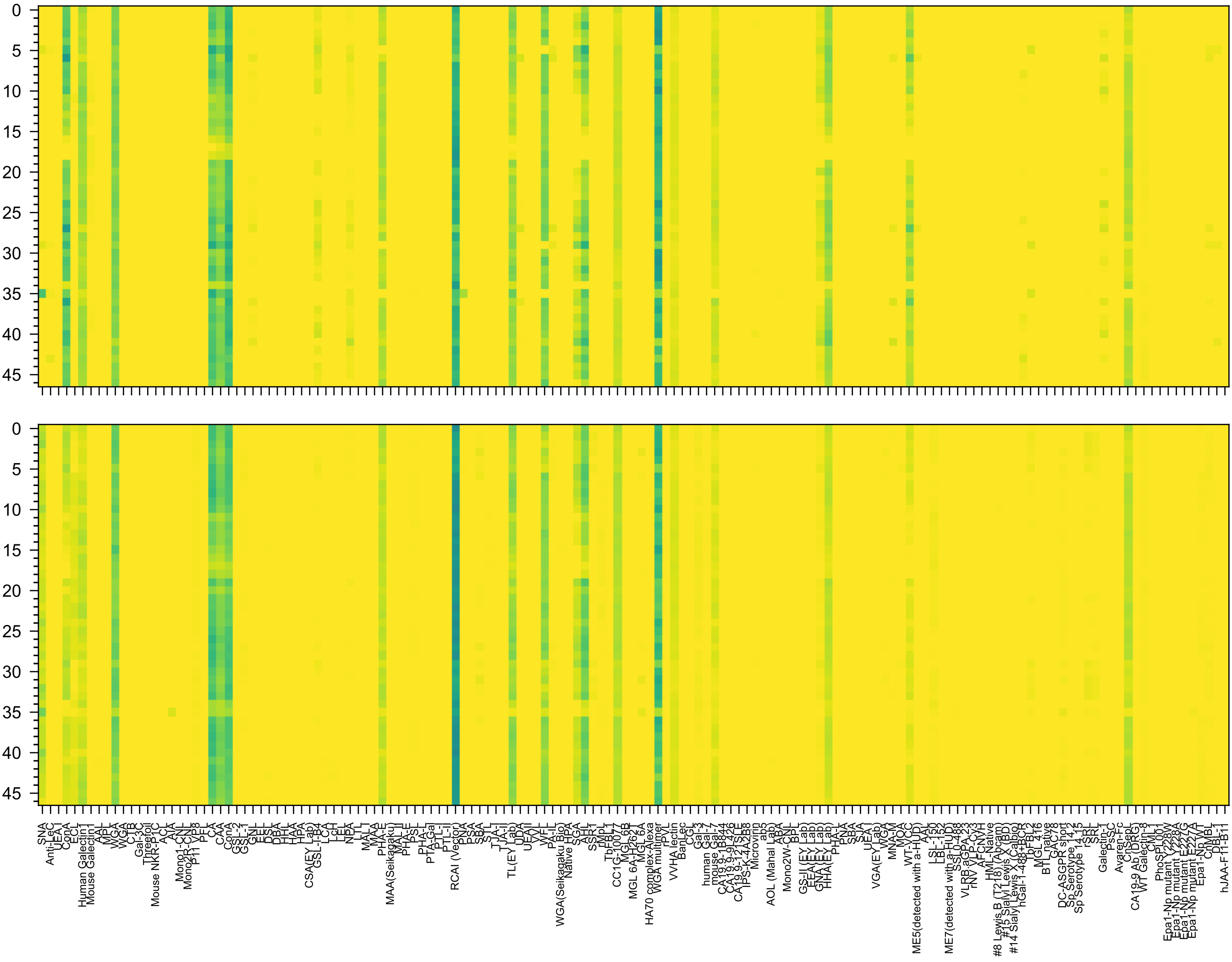


Gal(β1-4)GlcNAc(β1-2)Man(α1-6)[Gal(β1-4)GlcNAc(β1-2)Man(α1-3)]Man(β1-4)GlcNAc(β1-4)GlcNAc(β-Sp12

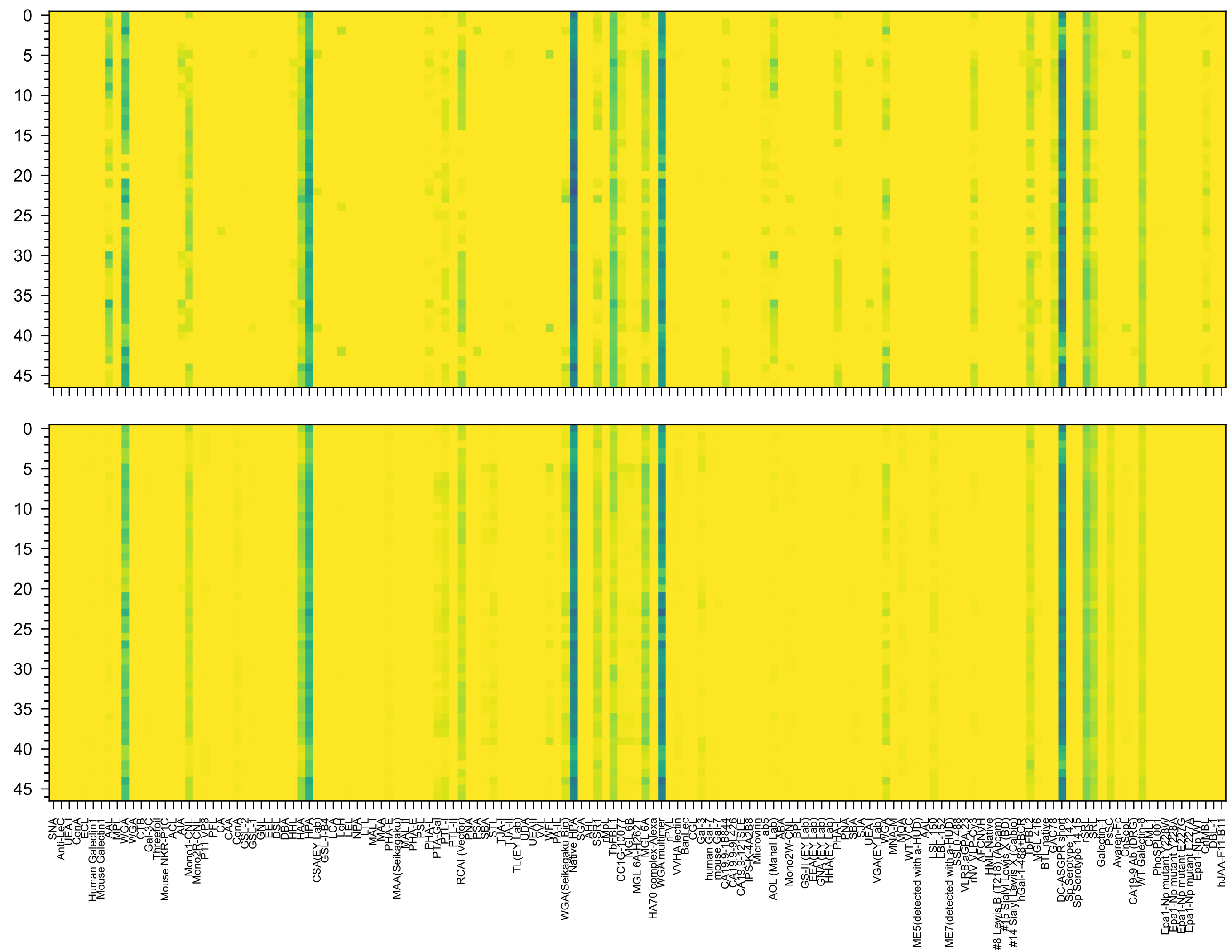


GalNAc(α1-3)[Fuc(α1-2)]Gal(β1-4)GlcNAc(β1-6)[GalNAc(α1-3)[Fuc(α1-2)]Gal(β1-4)GlcNAc(β1-3)]GalNAc(-Sp14

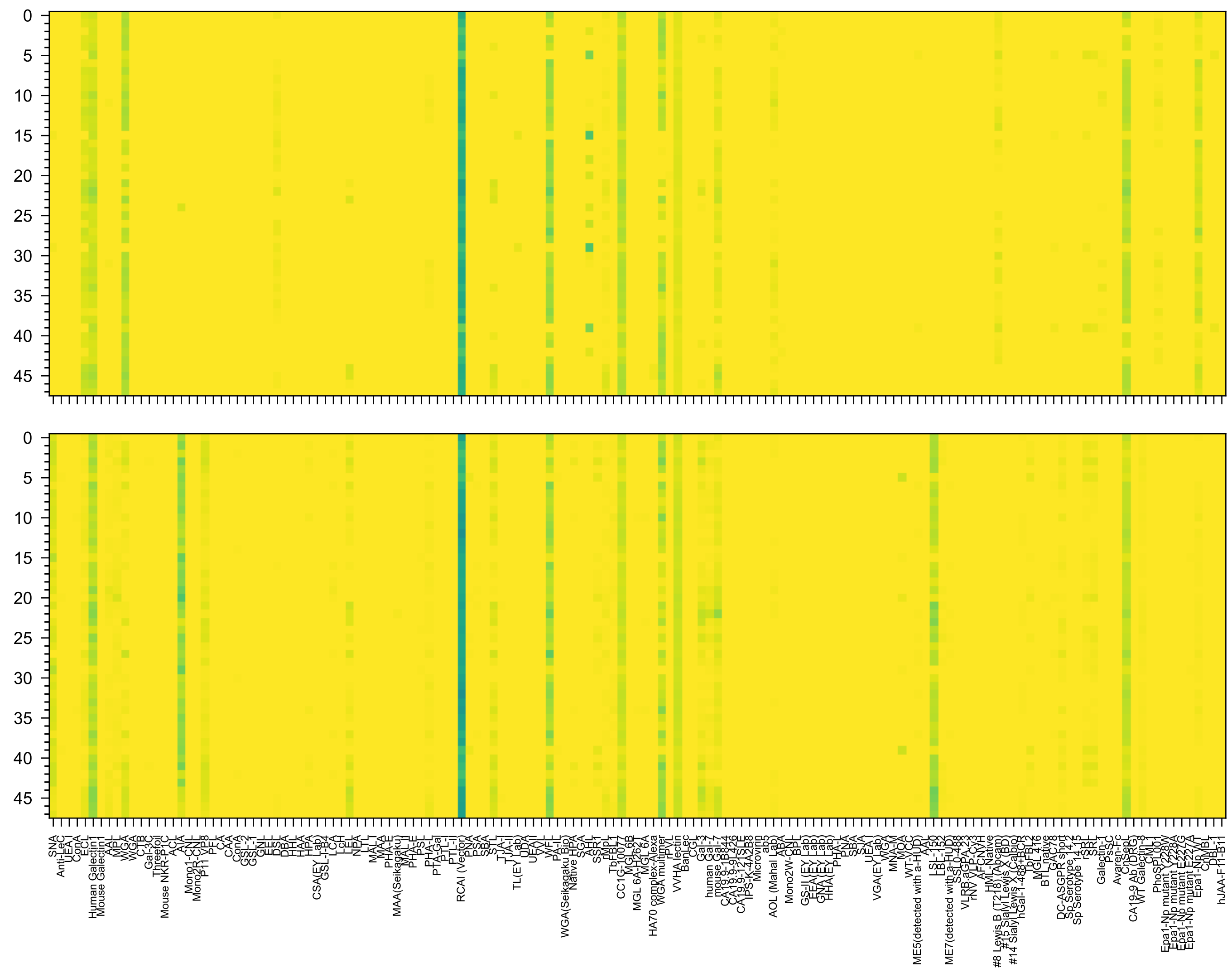


Gal(β1-4)GlcNAc(β1-3)Gal(β1-4)GlcNAc(β1-6)[Gal(β1-4)GlcNAc(β1-3)Gal(β1-4)GlcNAc(β1-3)]GalNAc(α-Sp14

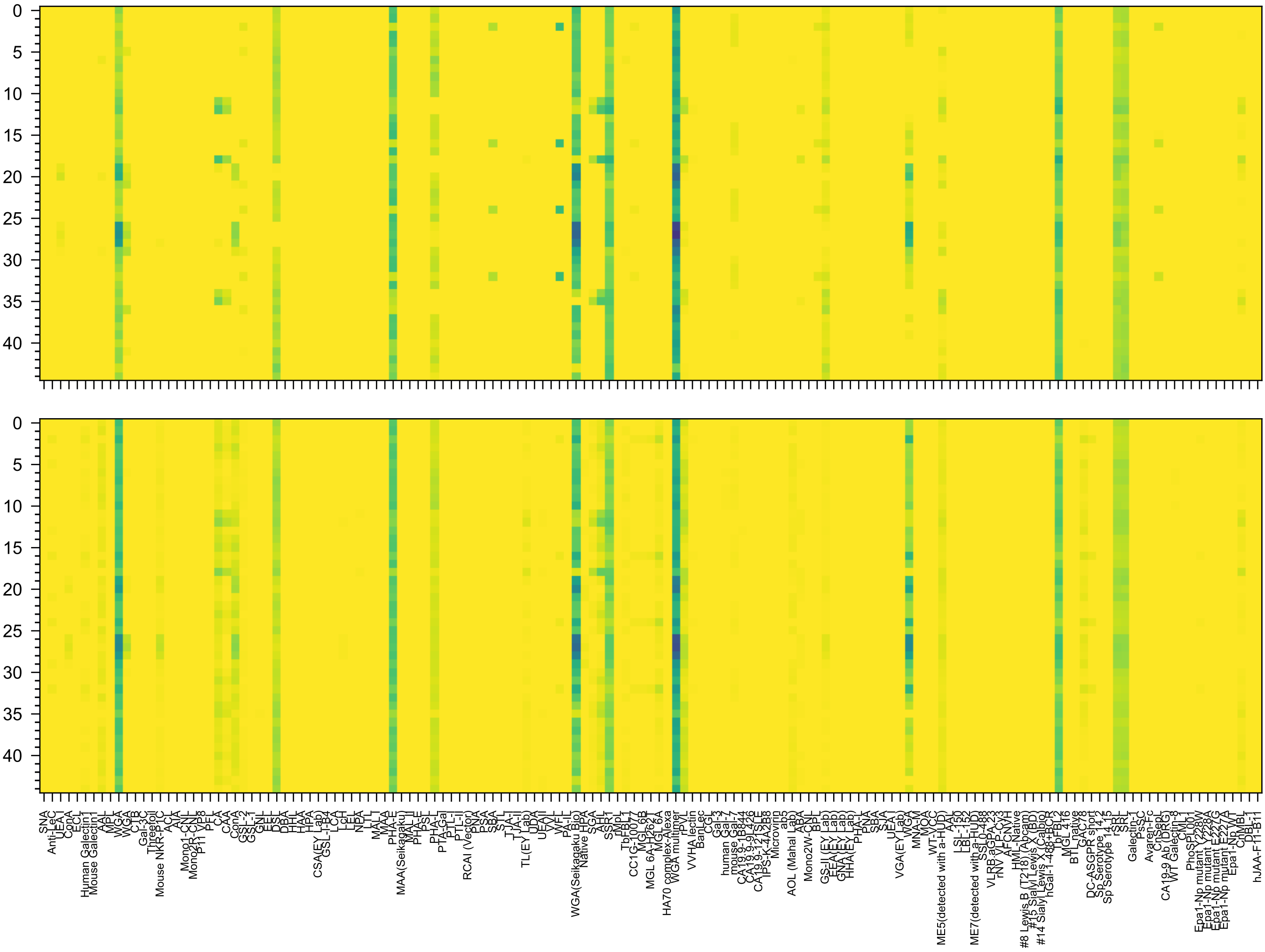
GlcNAc(β1-2)Man(α1-6)[GlcNAc(β1-4)][GlcNAc(β1-4)[GlcNAc(β1-2)]Man(α1-3)]Man(β1-4)GlcNAc(β1-4)GlcNAc(-Sp21

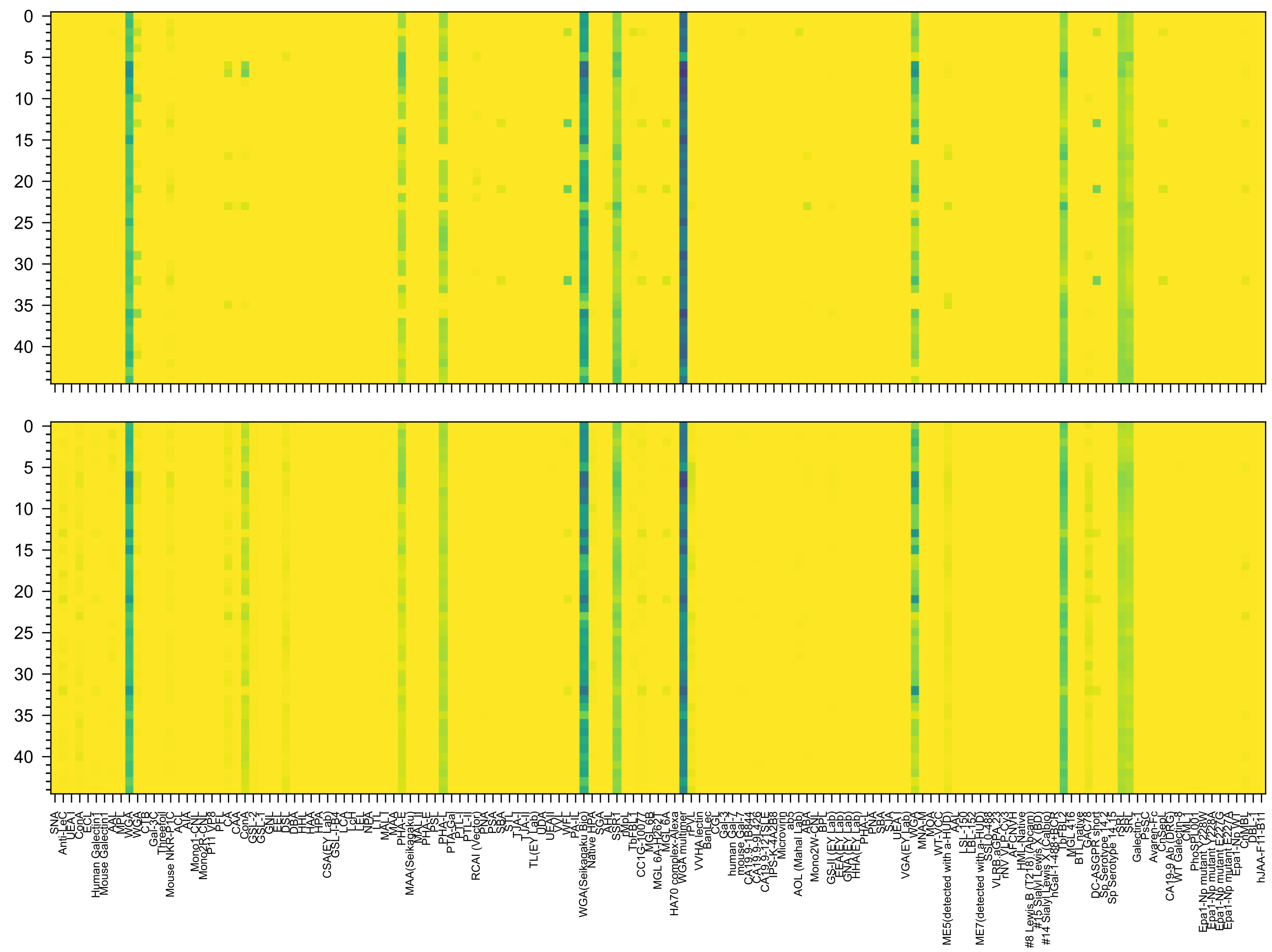

GlcNAc(β1-6)[GlcNAc(β1-2)]Man(α1-6)[GlcNAc(β1-4)][GlcNAc(β1-2)Man(α1-3)]Man(β1-4)GlcNAc(β1-4)GlcNAc(-Sp21

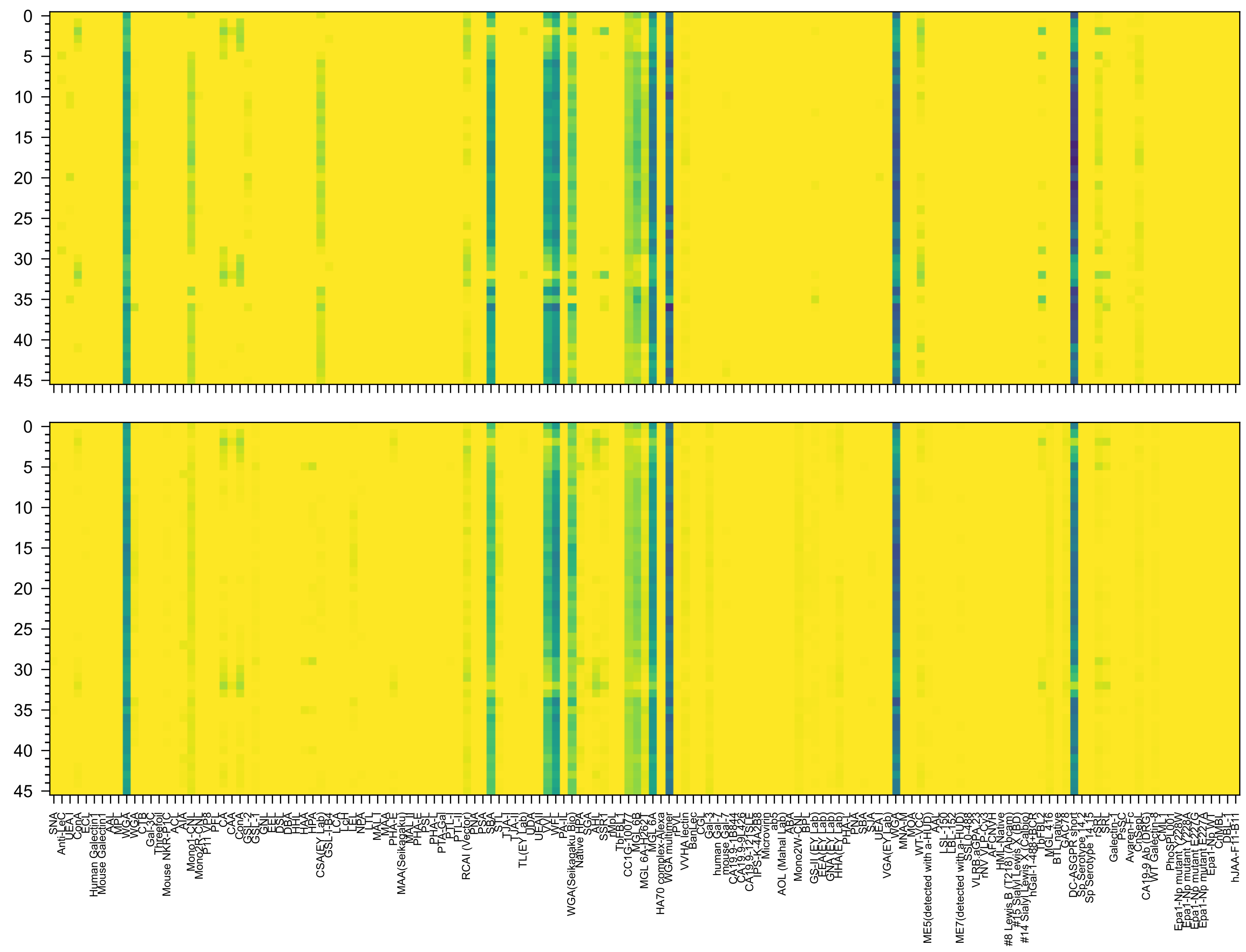

GalNAc(β1-4)GlcNAc(β1-2)Man(α1-6)[GalNAc(β1-4)GlcNAc(β1-2)Man(α1-3)]Man(β1-4)GlcNAc(β1-4)GlcNAc(-Sp12

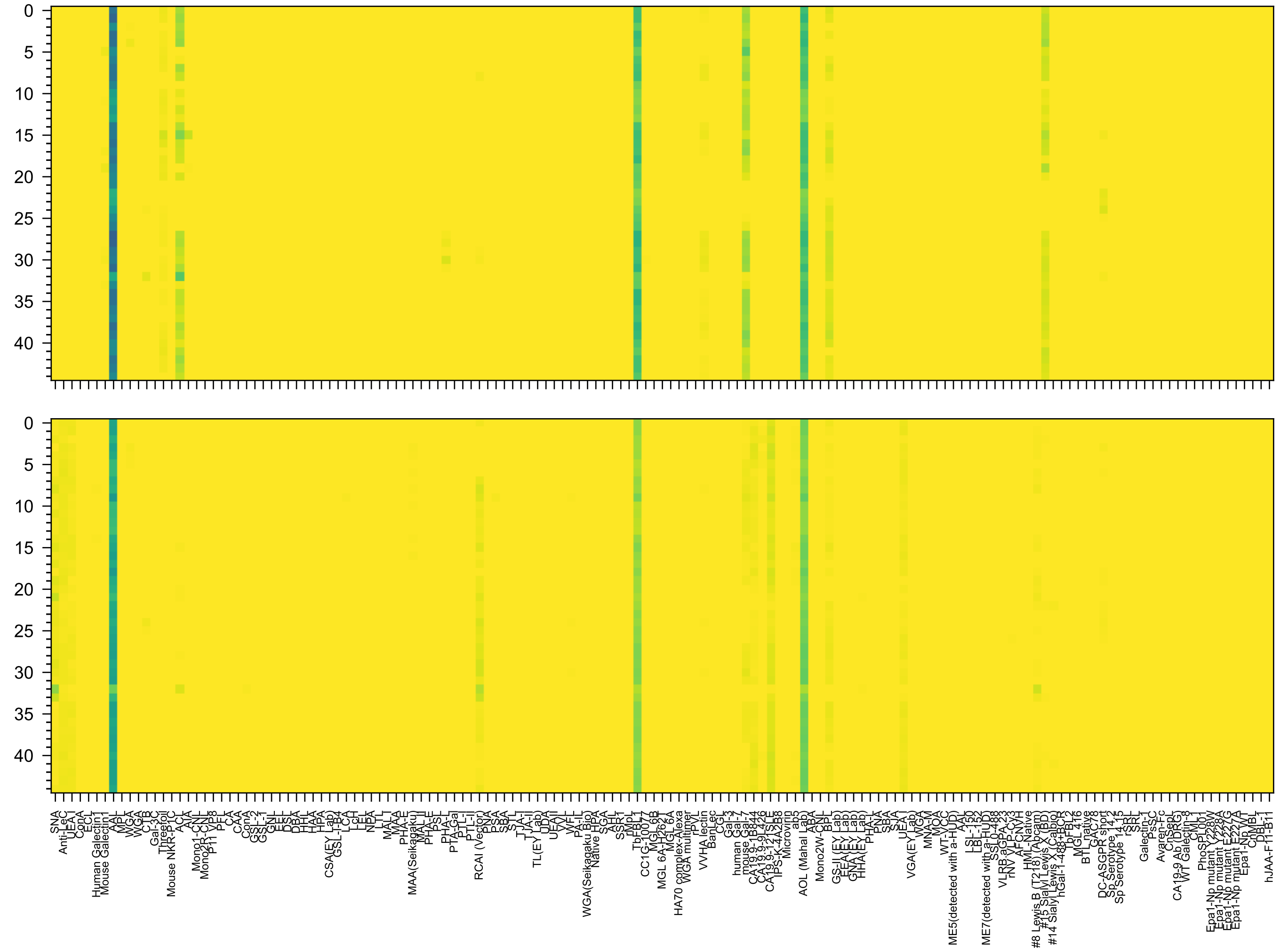


Gal(β1-4)[Fuc(α1-3)]GlcNAc(β1-6)[Neu5Ac(α2-6)[Neu5Ac(α2-3)Gal(β1-3)]GlcNAc(β1-3)]Gal(β1-4)Glc(-Sp21

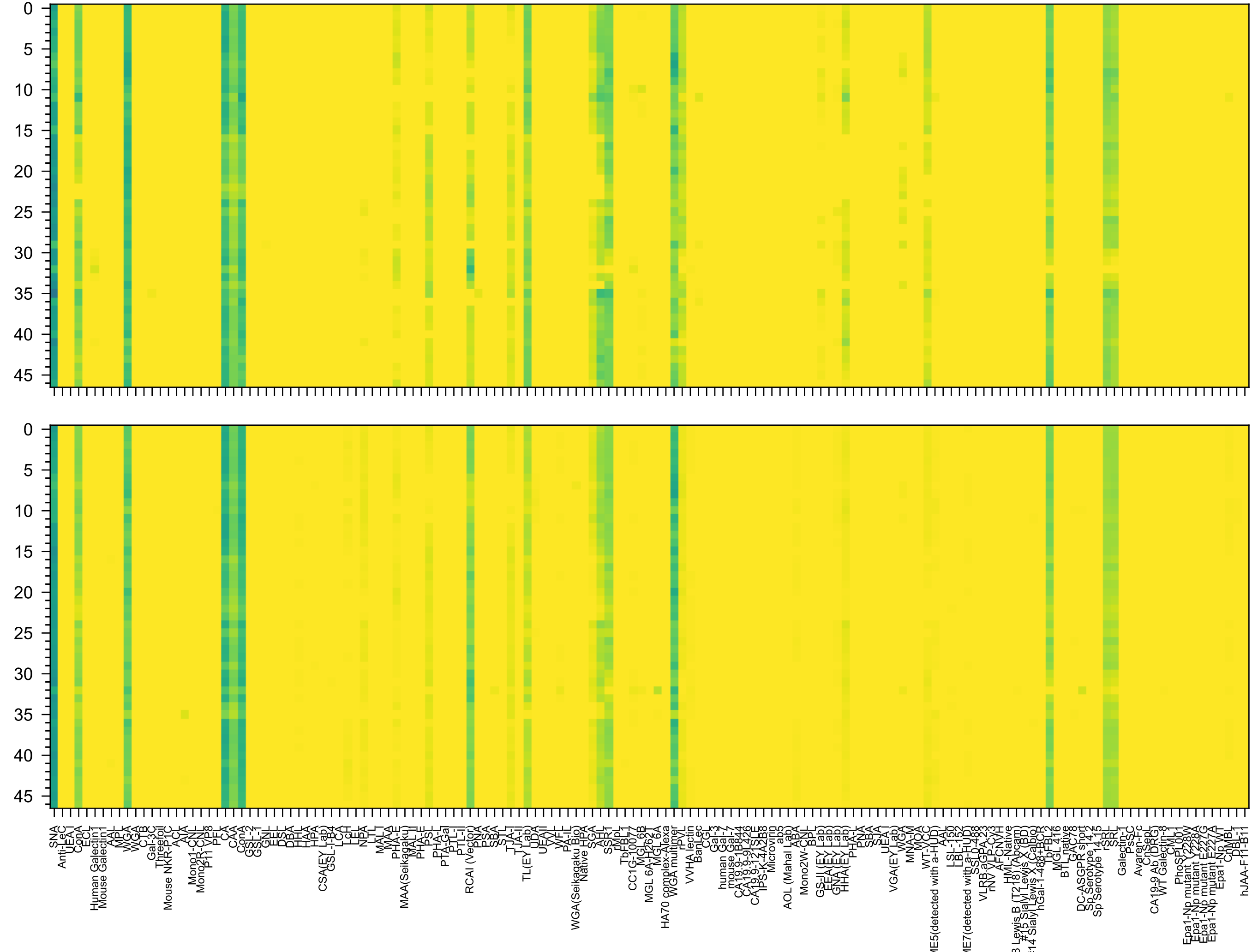

Neu5Ac(α2-6)Gal(β1-4)GlcNAc(β1-2)Man(α1-6)[GlcNAc(β1-2)Man(α1-3)]Man(β1-4)GlcNAc(β1-4)GlcNAc(β-Sp12

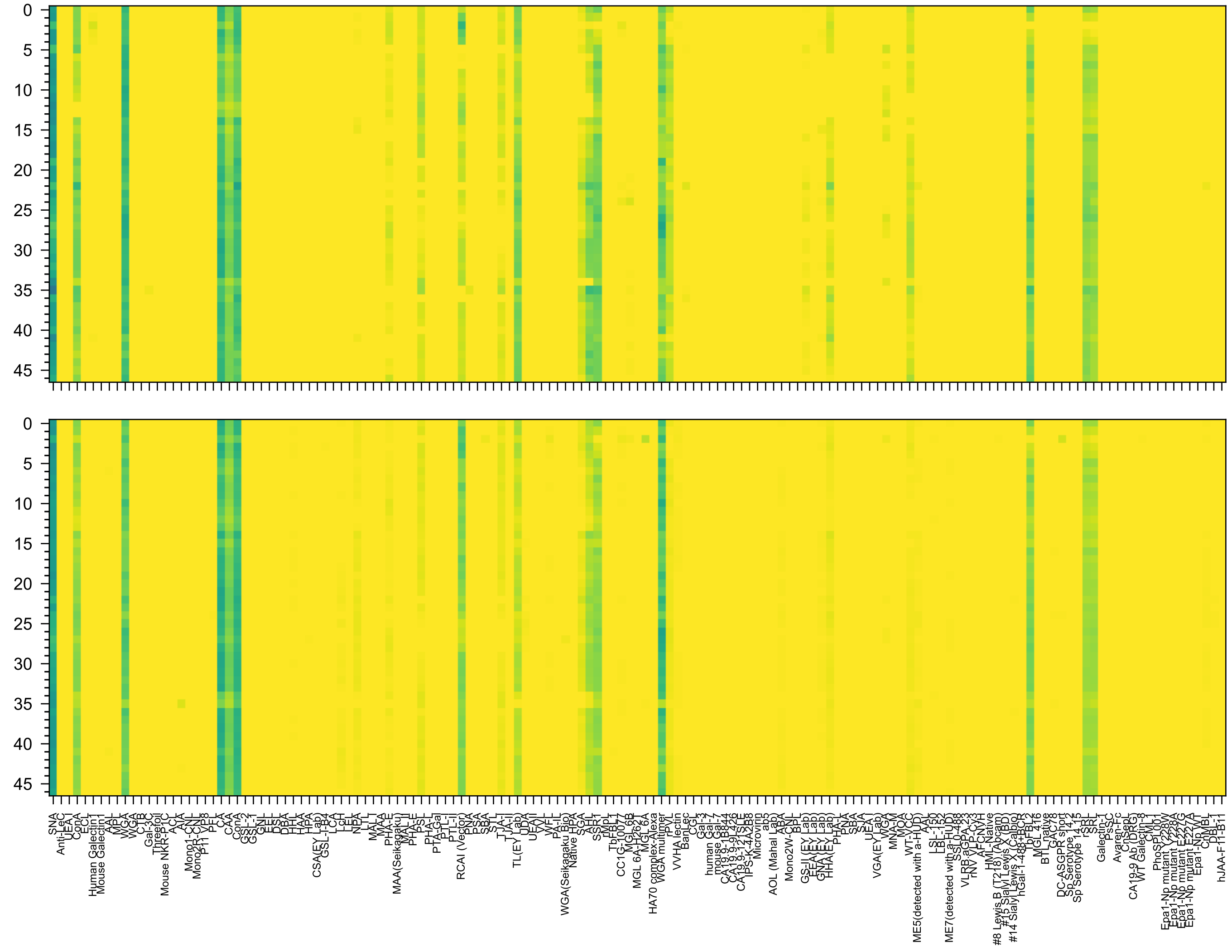

GlcNAc(β1-2)Man(α1-6)[Neu5Ac(α2-6)Gal(β1-4)GlcNAc(β1-2)Man(α1-3)]Man(β1-4)GlcNAc(β1-4)GlcNAc(β-Sp12

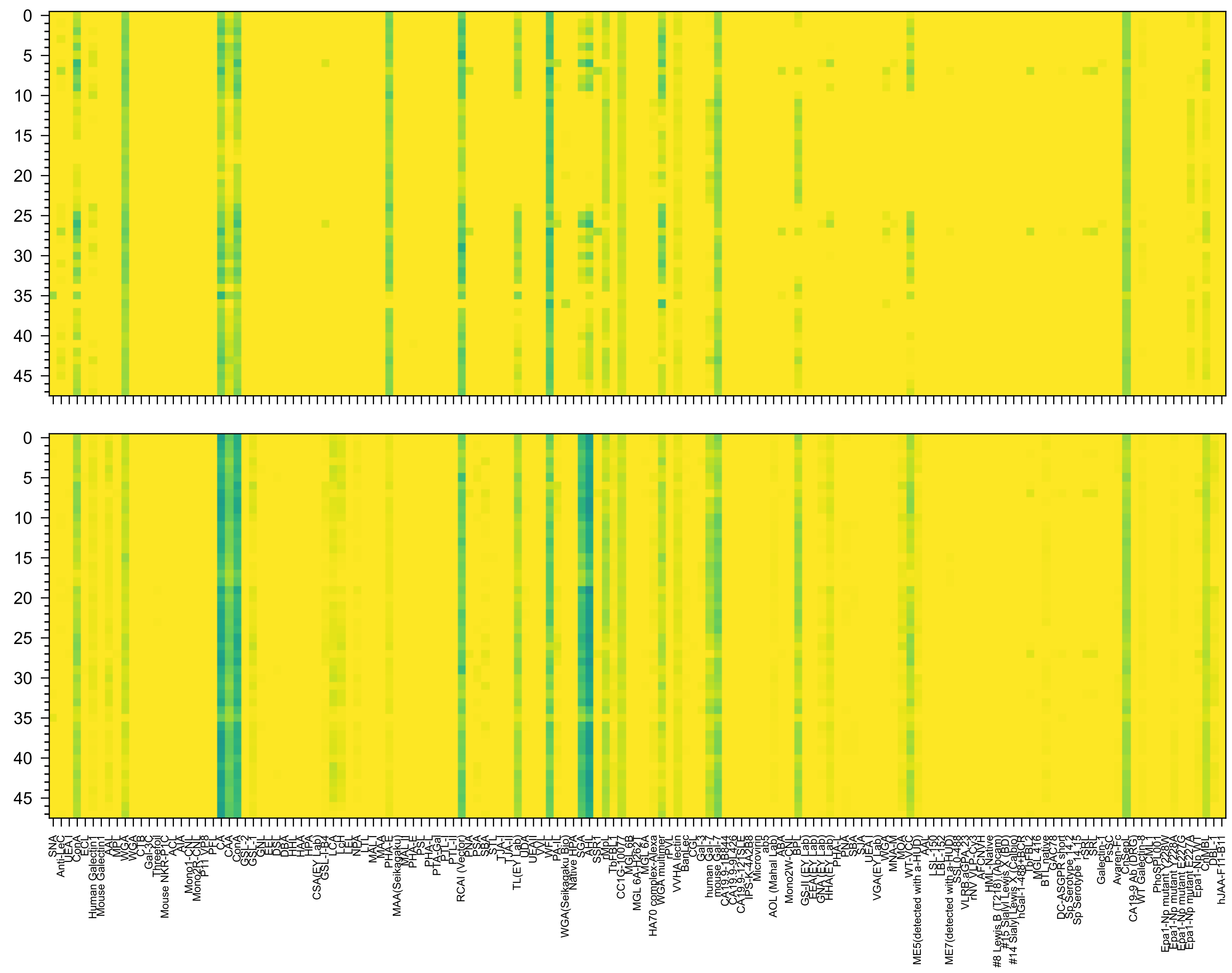
Gal(β1-3)GlcNAc(β1-2)Man(α1-6)[Gal(β1-3)GlcNAc(β1-2)Man(α1-3)]Man(β1-4)GlcNAc(β1-4)GlcNAc(β-Sp19

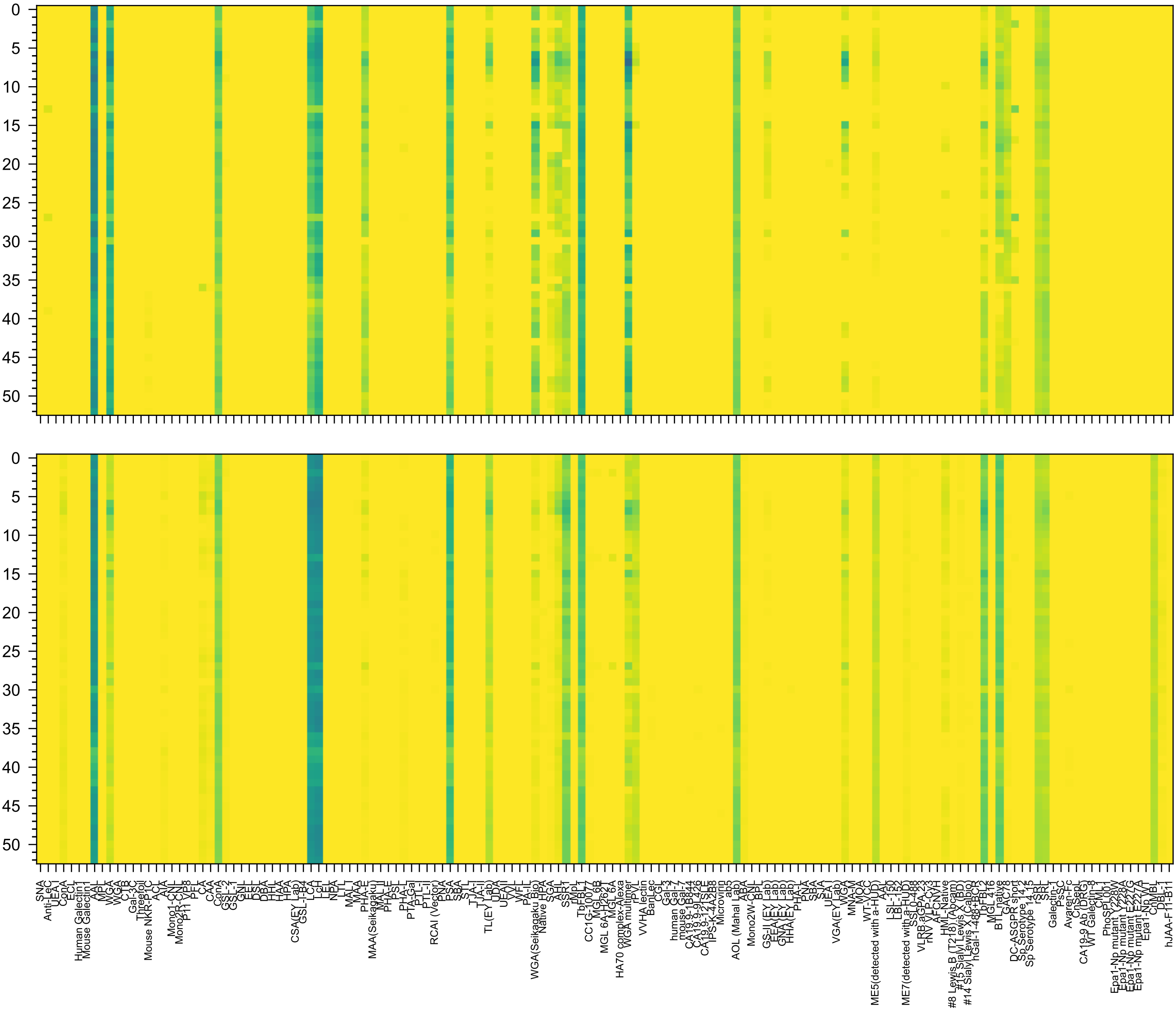


GlcNAc(β1-6)[GlcNAc(β1-2)]Man(α1-6)[GlcNAc(β1-2)Man(α1-3)]Man(β1-4)GlcNAc(β1-4)[Fuc(α1-6)]GlcNAc(β-Sp24

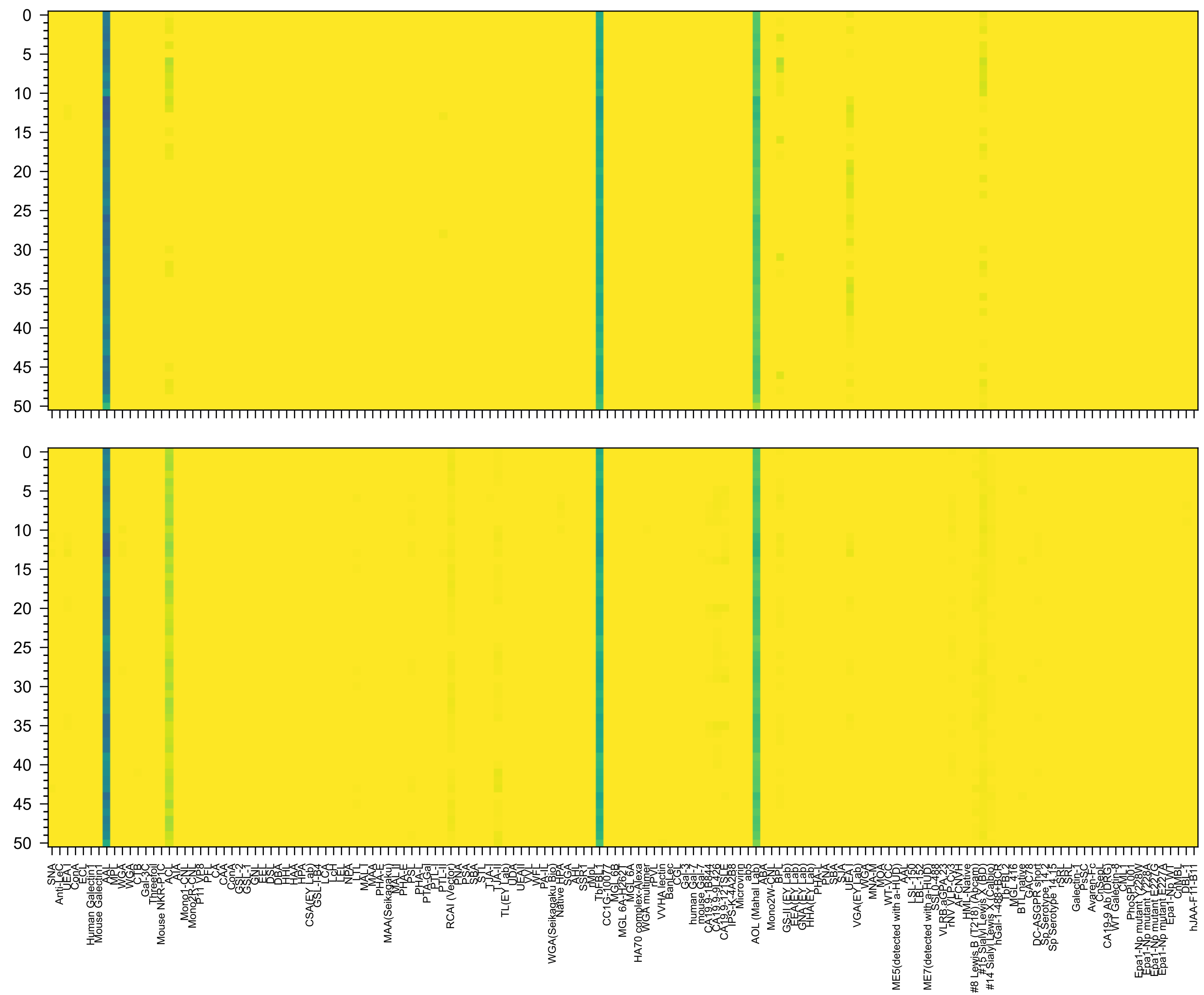

Fuc(α1-2)Gal(β1-4)[Fuc(α1-3)]GlcNAc(β1-3)Gal(β1-4)[Fuc(α1-3)]GlcNAc(β1-3)Gal(β1-4)[Fuc(α1-3)]GlcNAc(β-Sp0

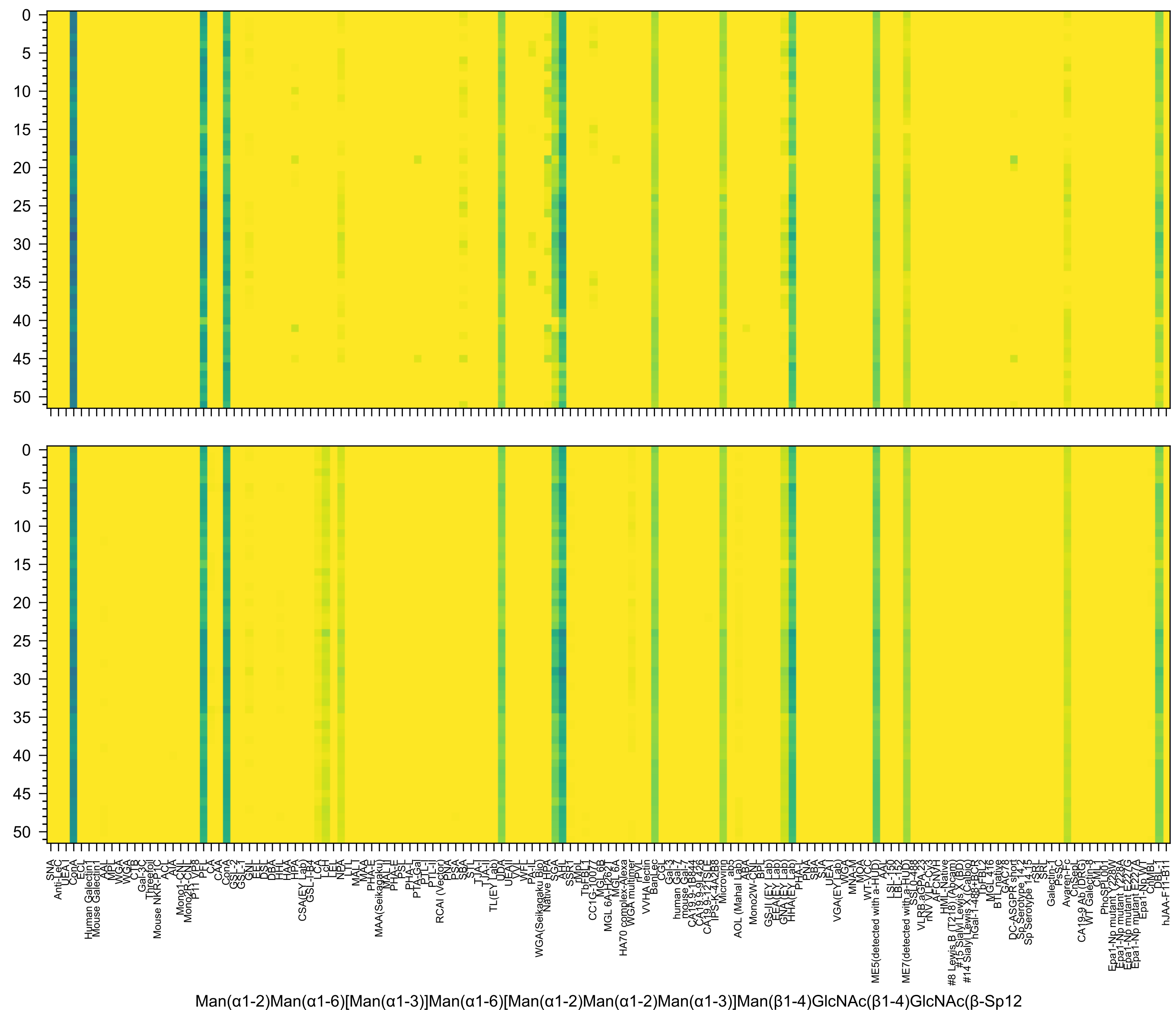
Man(α1-2)Man(α1-6)[Man(α1-3)]Man(α1-6)[Man(α1-2)Man(α1-2)Man(α1-3)]Man(β1-4)GlcNAc(β1-4)GlcNAc(β-Sp12

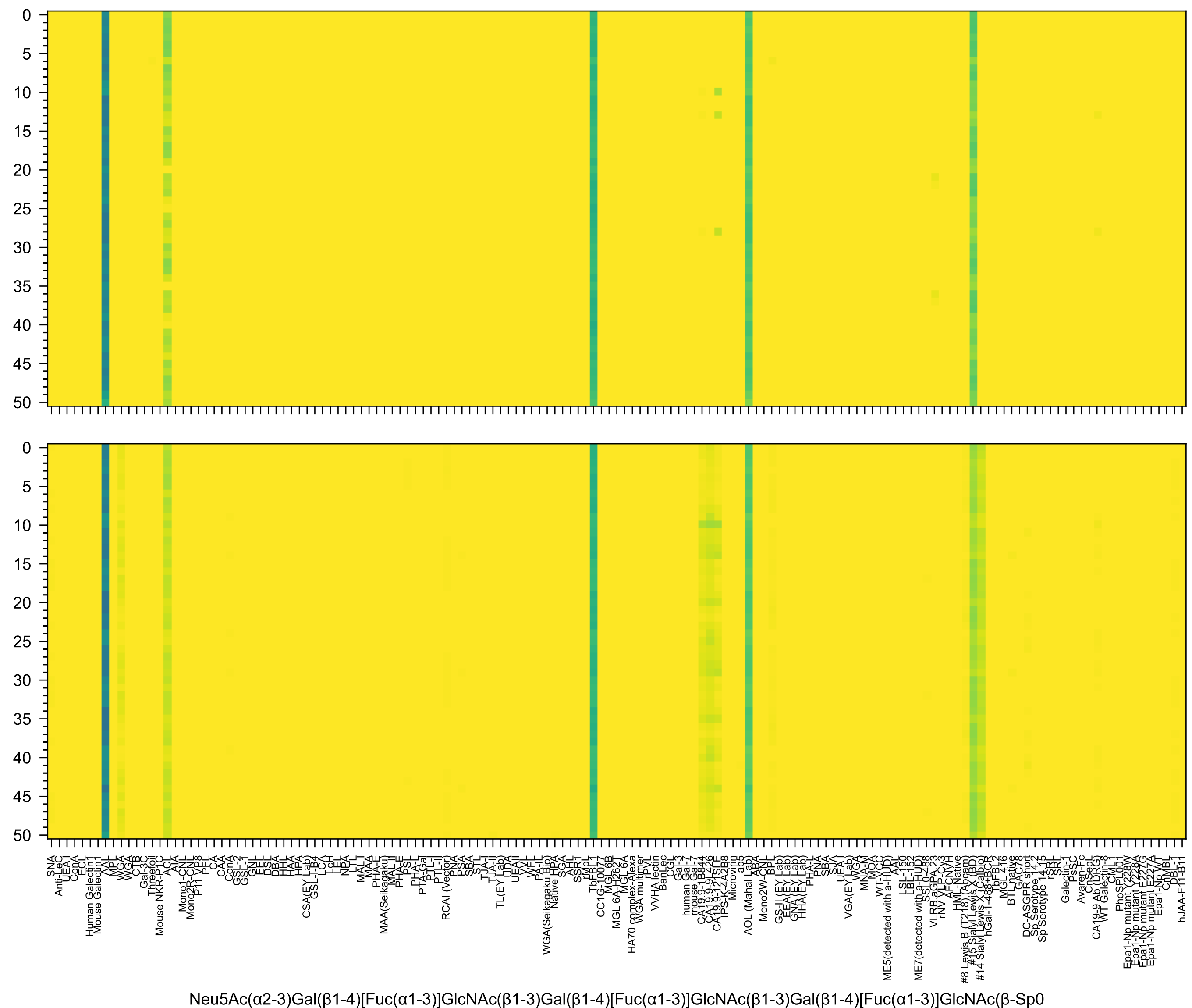
Neu5Ac(α2-3)Gal(β1-4)[Fuc(α1-3)]GlcNAc(β1-3)Gal(β1-4)[Fuc(α1-3)]GlcNAc(β1-3)Gal(β1-4)[Fuc(α1-3)]GlcNAc(β-Sp0

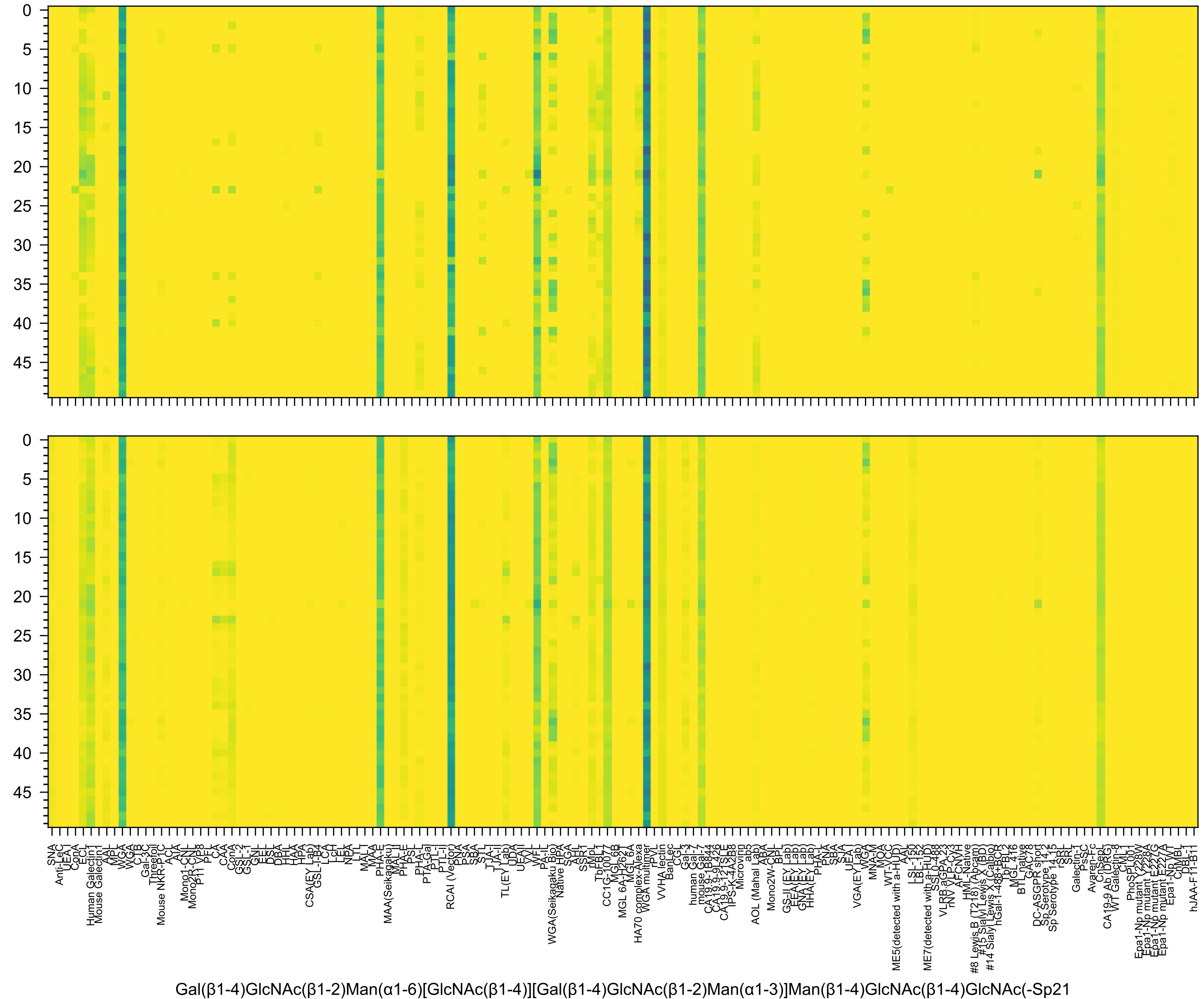
Gal(β1-4)GlcNAc(β1-2)Man(α1-6)[GlcNAc(β1-4)][Gal(β1-4)GlcNAc(β1-2)Man(α1-3)]Man(β1-4)GlcNAc(β1-4)GlcNAc(-Sp21

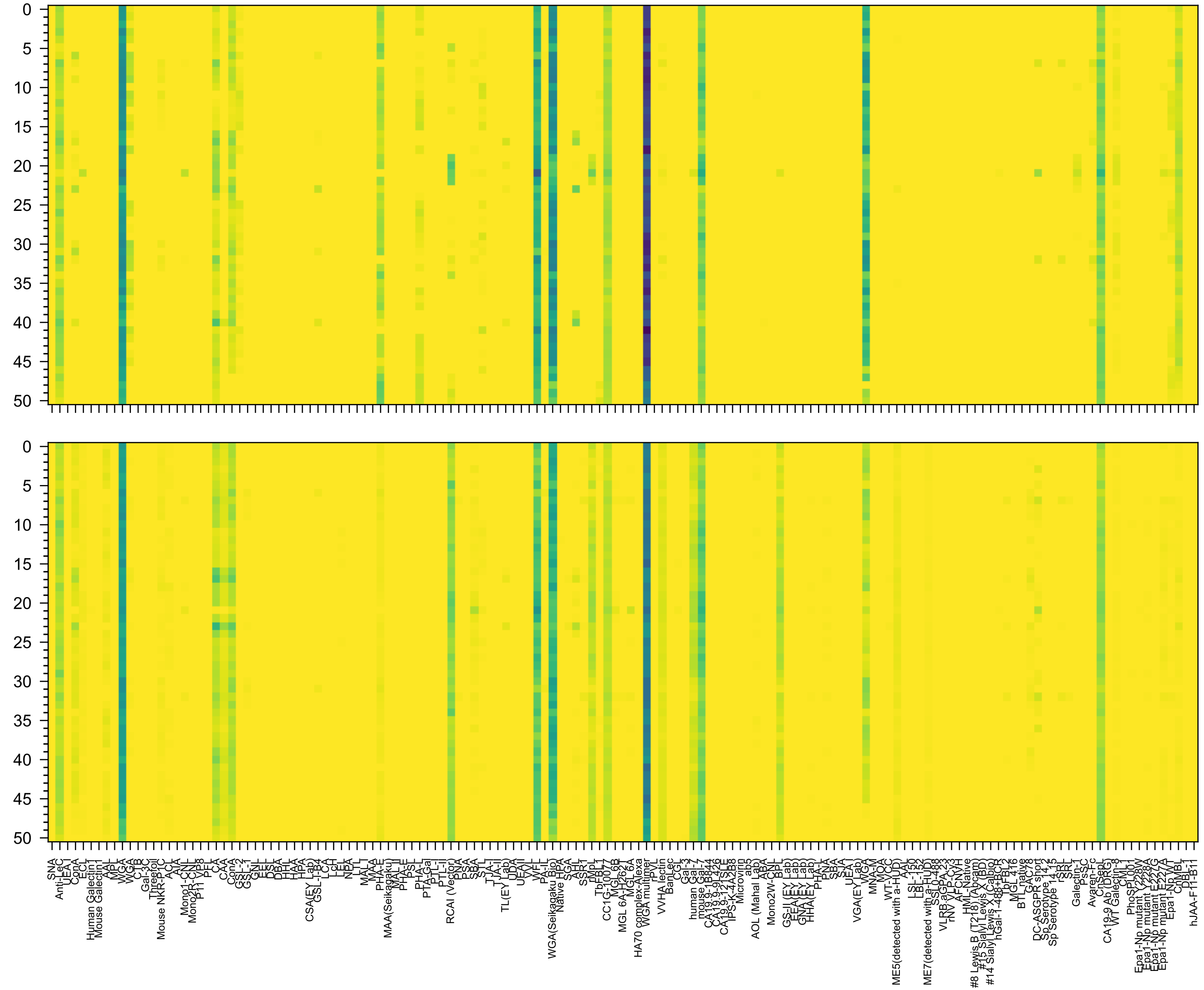


Gal(β1-3)GlcNAc(β1-2)Man(α1-6)[GlcNAc(β1-4)][Gal(β1-3)GlcNAc(β1-2)Man(α1-3)]Man(β1-4)GlcNAc(β1-4)GlcNAc(β-Sp21

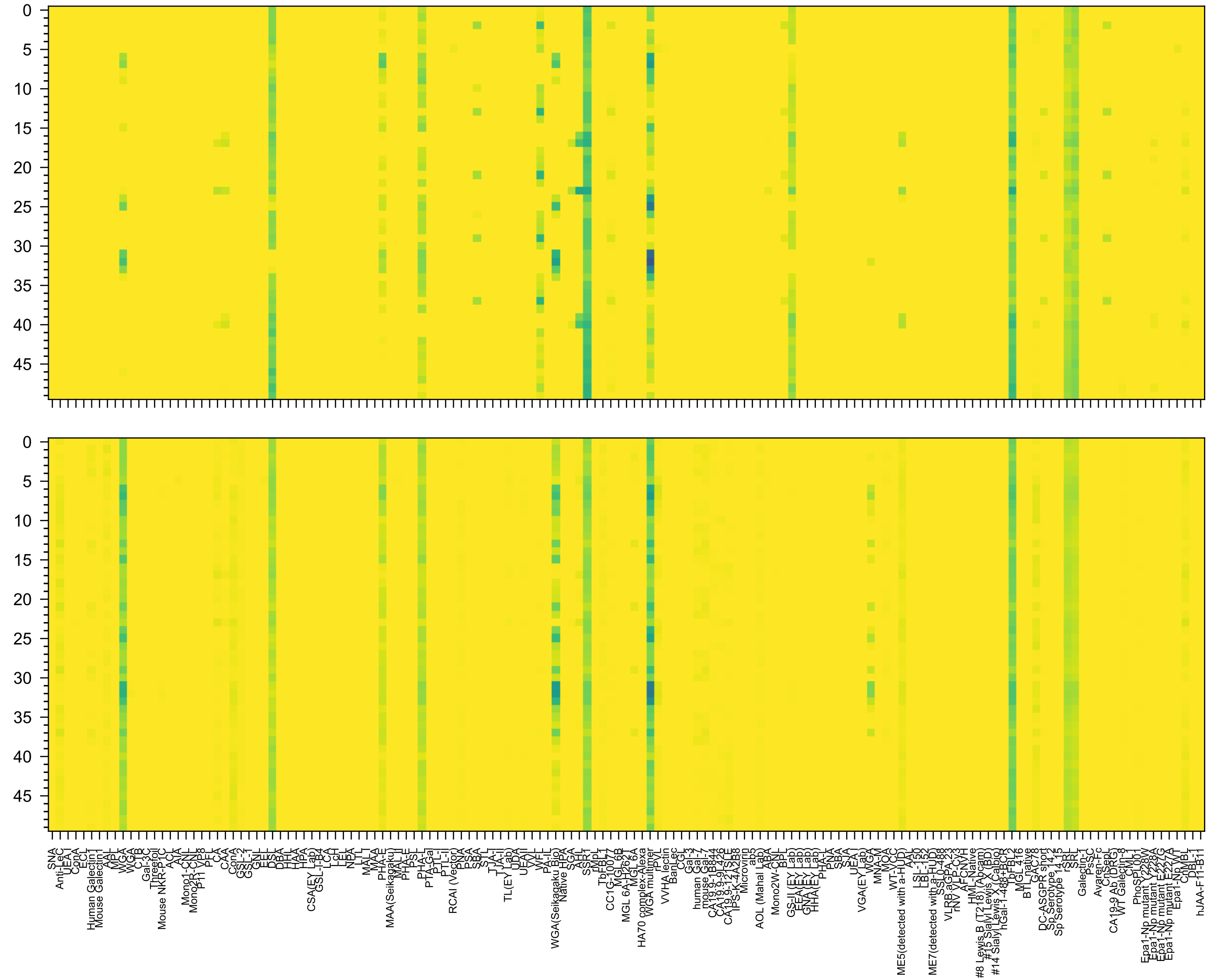


GlcNAc(β1-6)[GlcNAc(β1-2)]Man(α1-6)[GlcNAc(β1-4)][GlcNAc(β1-4)[GlcNAc(β1-2)]Man(α1-3)]Man(β1-4)GlcNAc(β1-4)GlcNAc(-Sp21

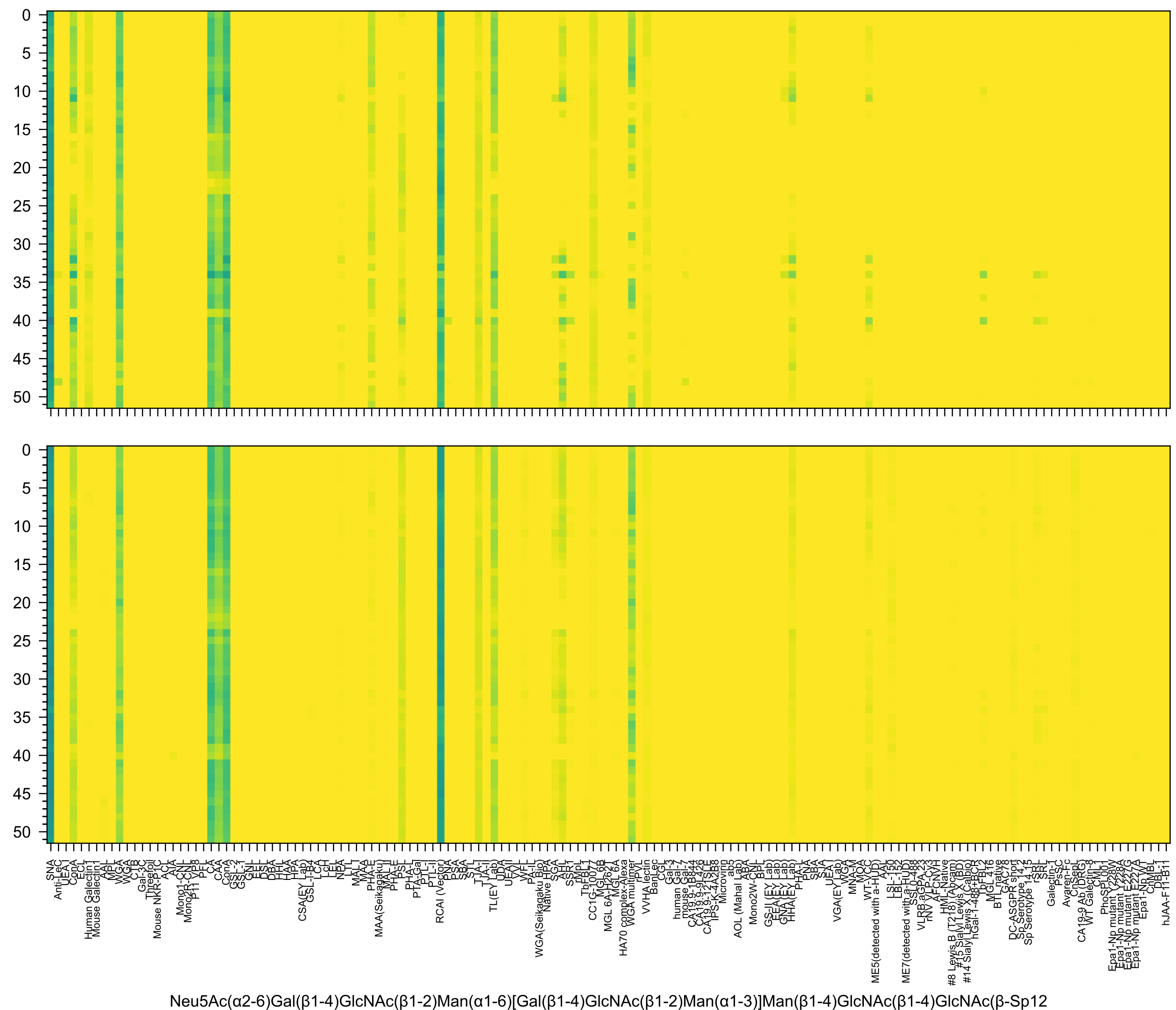
Neu5Ac(α2-6)Gal(β1-4)GlcNAc(β1-2)Man(α1-6)[Gal(β1-4)GlcNAc(β1-2)Man(α1-3)]Man(β1-4)GlcNAc(β1-4)GlcNAc(β-Sp12

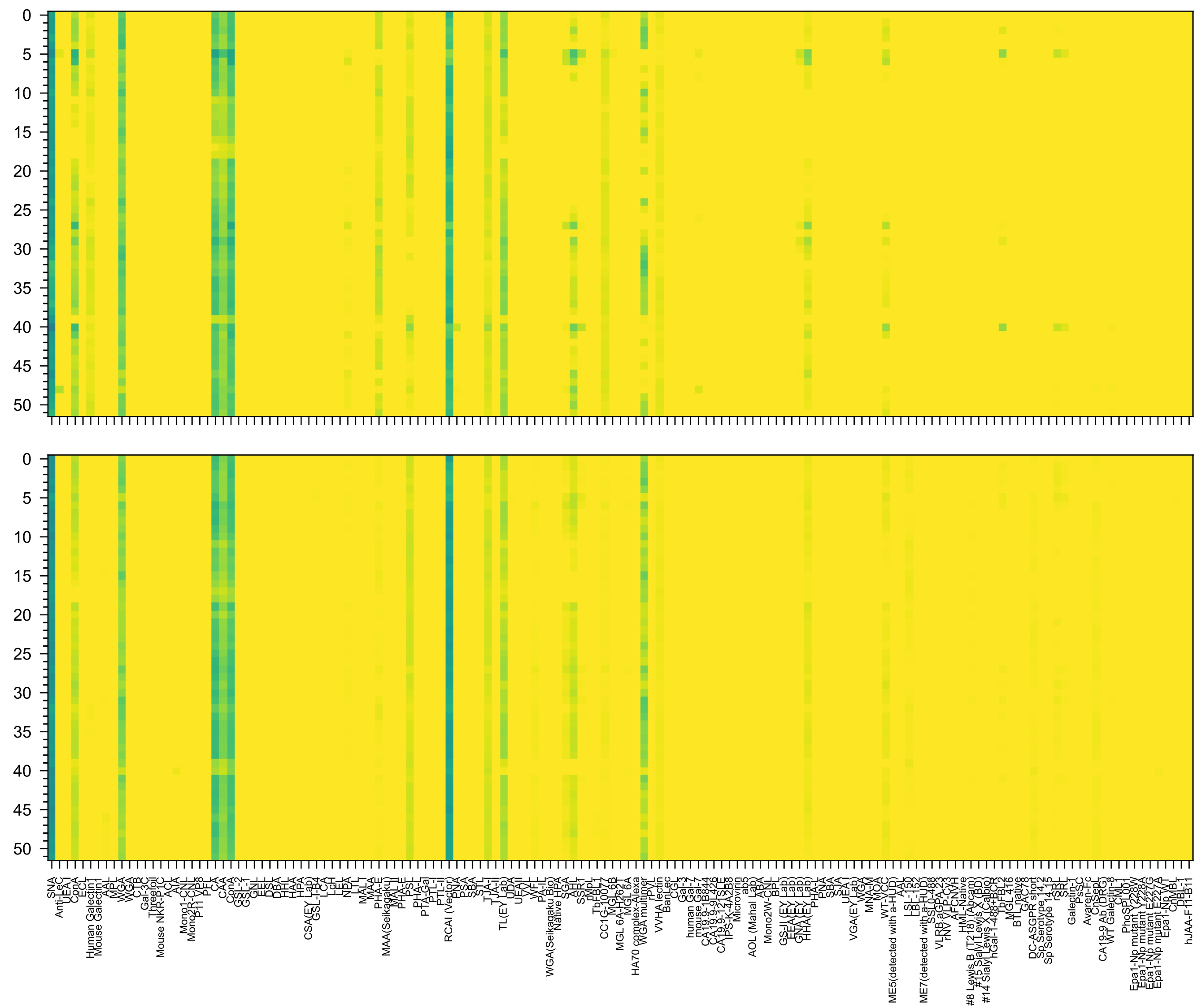

Gal(β1-4)GlcNAc(β1-2)Man(α1-6)[Neu5Ac(α2-6)Gal(β1-4)GlcNAc(β1-2)Man(α1-3)]Man(β1-4)GlcNAc(β1-4)GlcNAc(β-Sp12

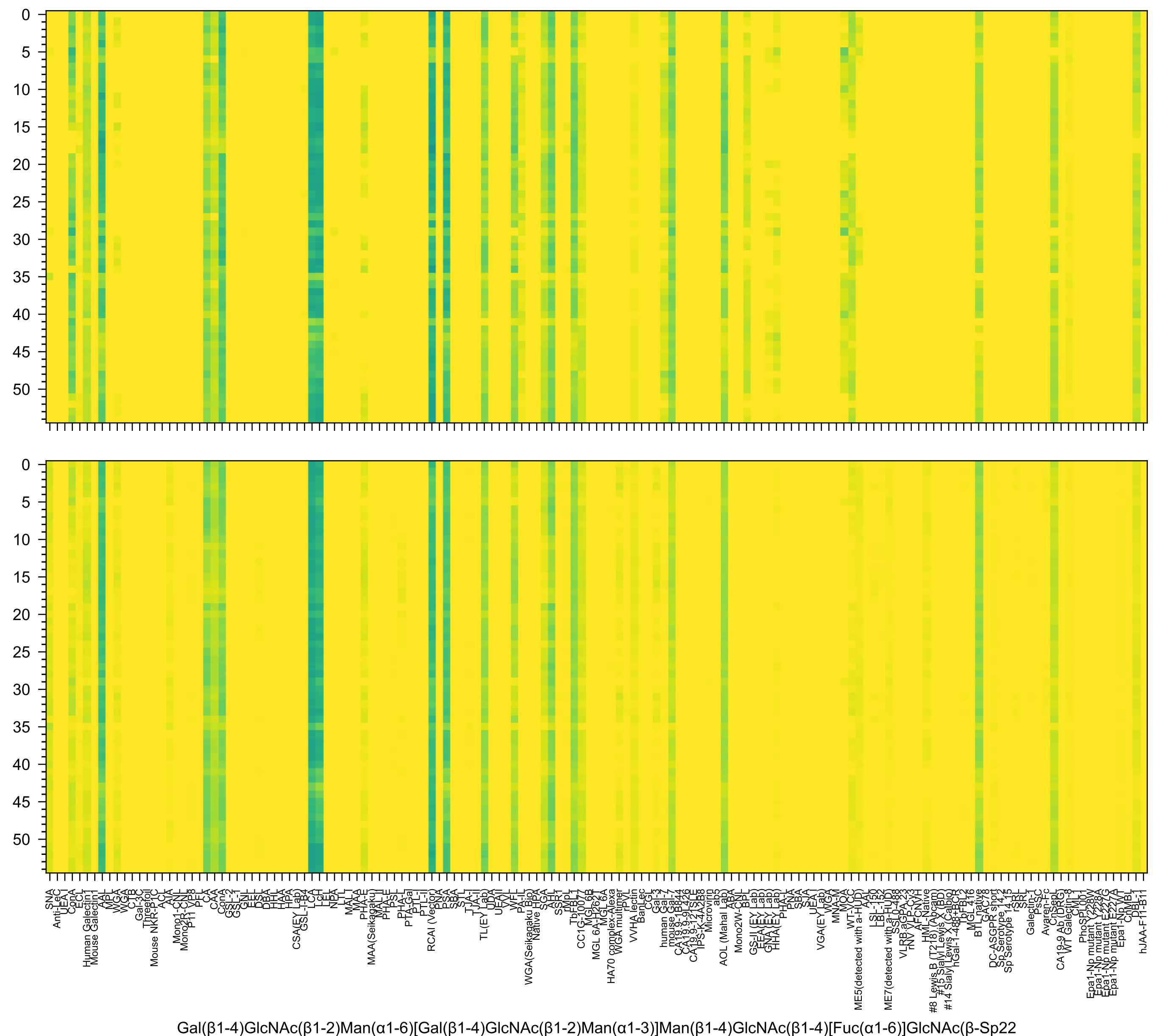

Gal(β1-4)GlcNAc(β1-2)Man(α1-6)[Gal(β1-4)GlcNAc(β1-2)Man(α1-3)]Man(β1-4)GlcNAc(β1-4)[Fuc(α1-6)]GlcNAc(β-Sp22

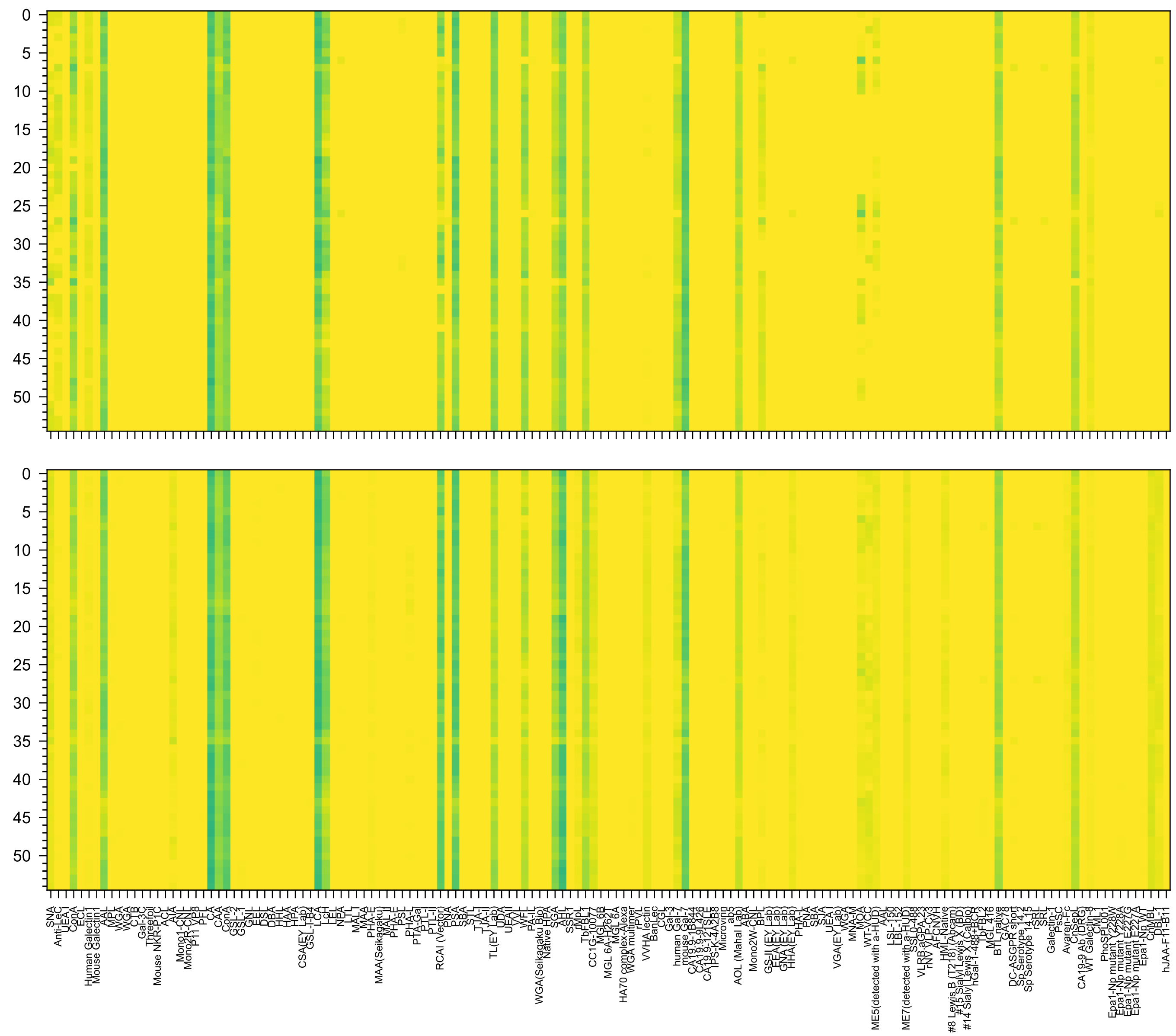

Gal(β1-3)GlcNAc(β1-2)Man(α1-6)[Gal(β1-3)GlcNAc(β1-2)Man(α1-3)]Man(β1-4)GlcNAc(β1-4)[Fuc(α1-6)]GlcNAc(β-Sp22

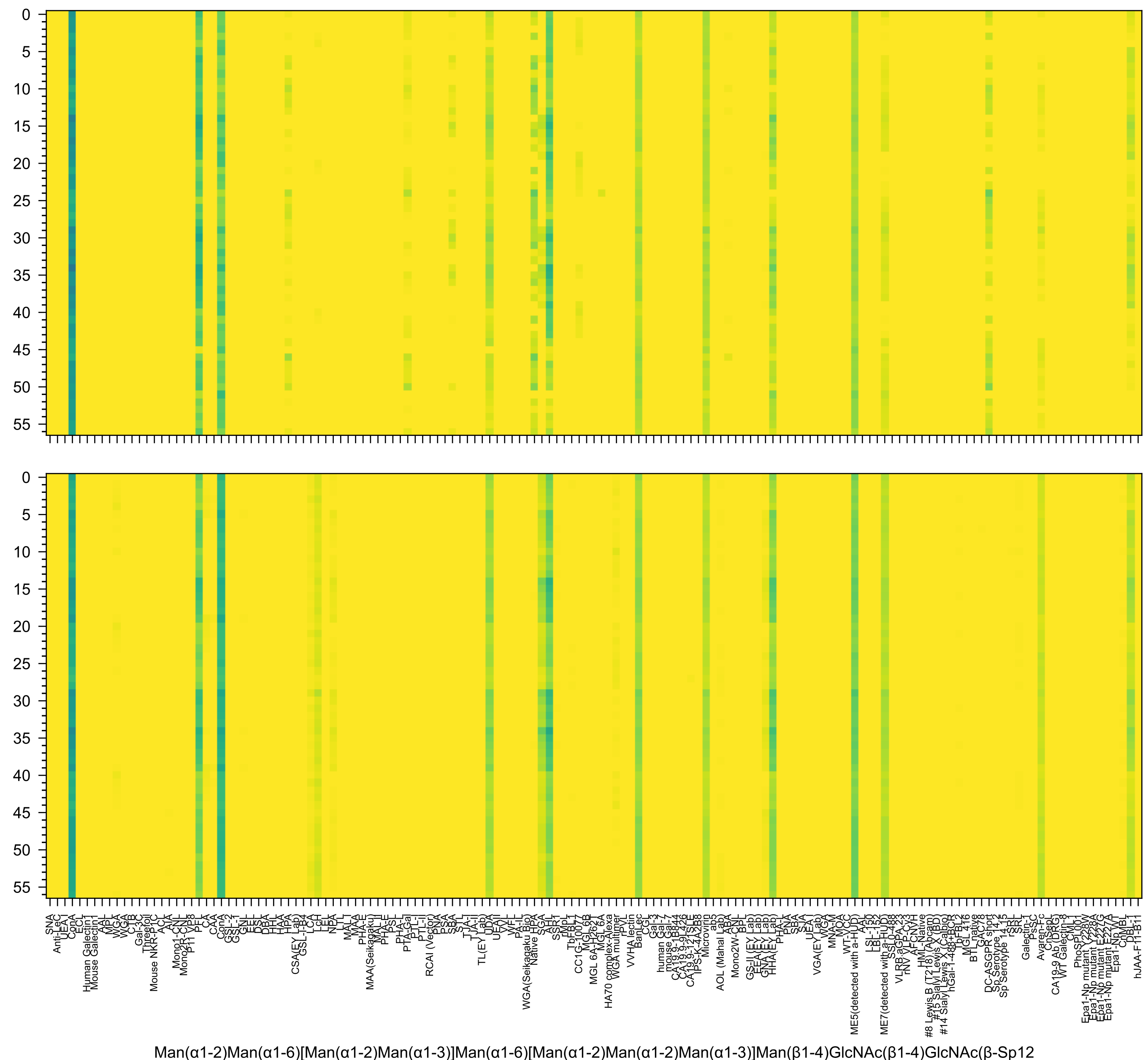
Man(α1-2)Man(α1-6)[Man(α1-2)Man(α1-3)]Man(α1-6)[Man(α1-2)Man(α1-2)Man(α1-3)]Man(β1-4)GlcNAc(β1-4)GlcNAc(β-Sp12

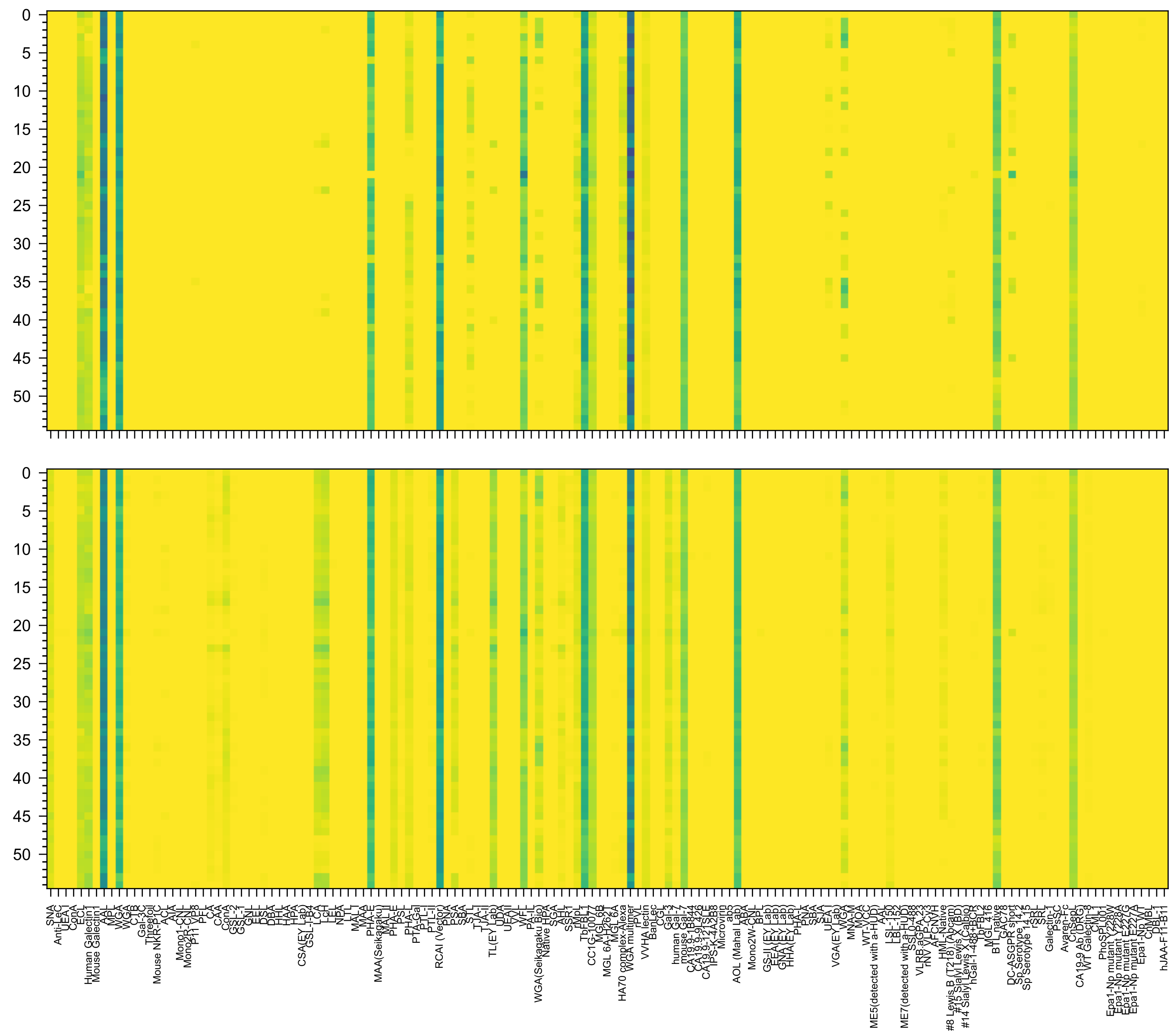

Gal(β1-4)GlcNAc(β1-2)Man(α1-6)[GlcNAc(β1-4)][Gal(β1-4)GlcNAc(β1-2)Man(α1-3)]Man(β1-4)GlcNAc(β1-4)[Fuc(α1-6)]GlcNAc(-Sp21

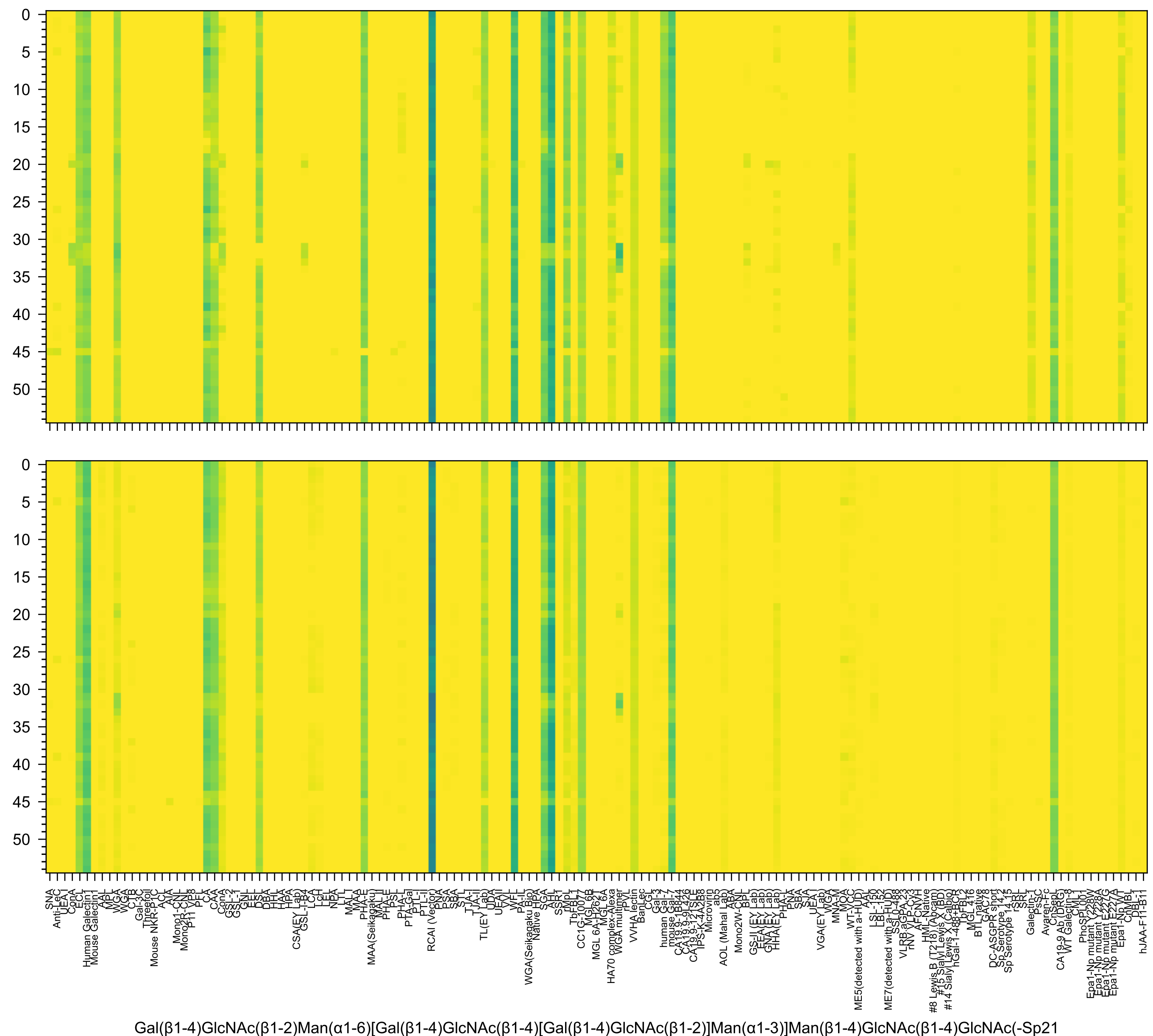
Gal(β1-4)GlcNAc(β1-2)Man(α1-6)[Gal(β1-4)GlcNAc(β1-4)[Gal(β1-4)GlcNAc(β1-2)]Man(α1-3)]Man(β1-4)GlcNAc(β1-4)GlcNAc(-Sp21

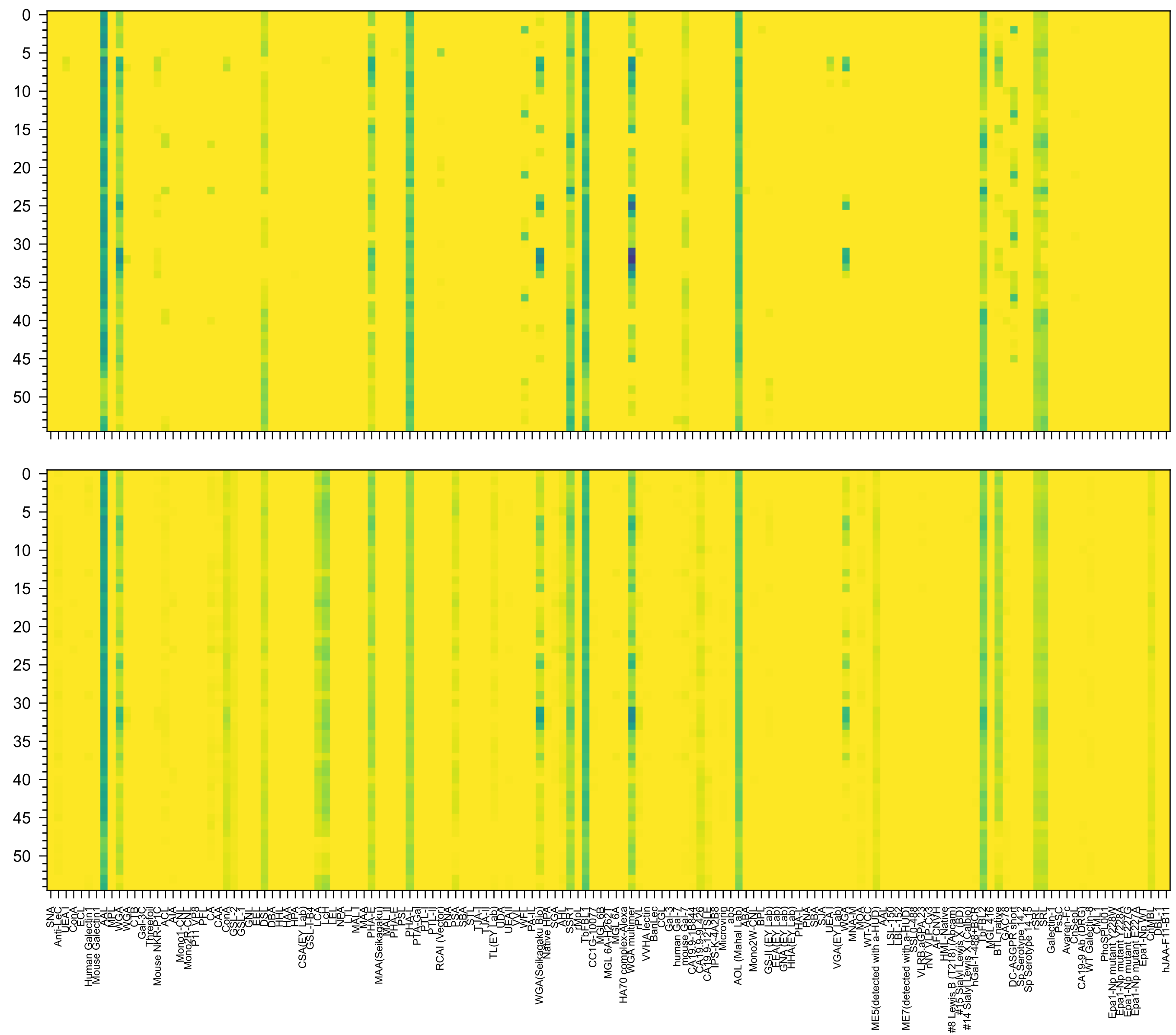

GlcNAc(β1-6)[GlcNAc(β1-2)]Man(α1-6)[GlcNAc(β1-4)][GlcNAc(β1-4)[GlcNAc(β1-2)]Man(α1-3)]Man(β1-4)GlcNAc(β1-4)[Fuc(α1-6)]GlcNAc(-Sp21

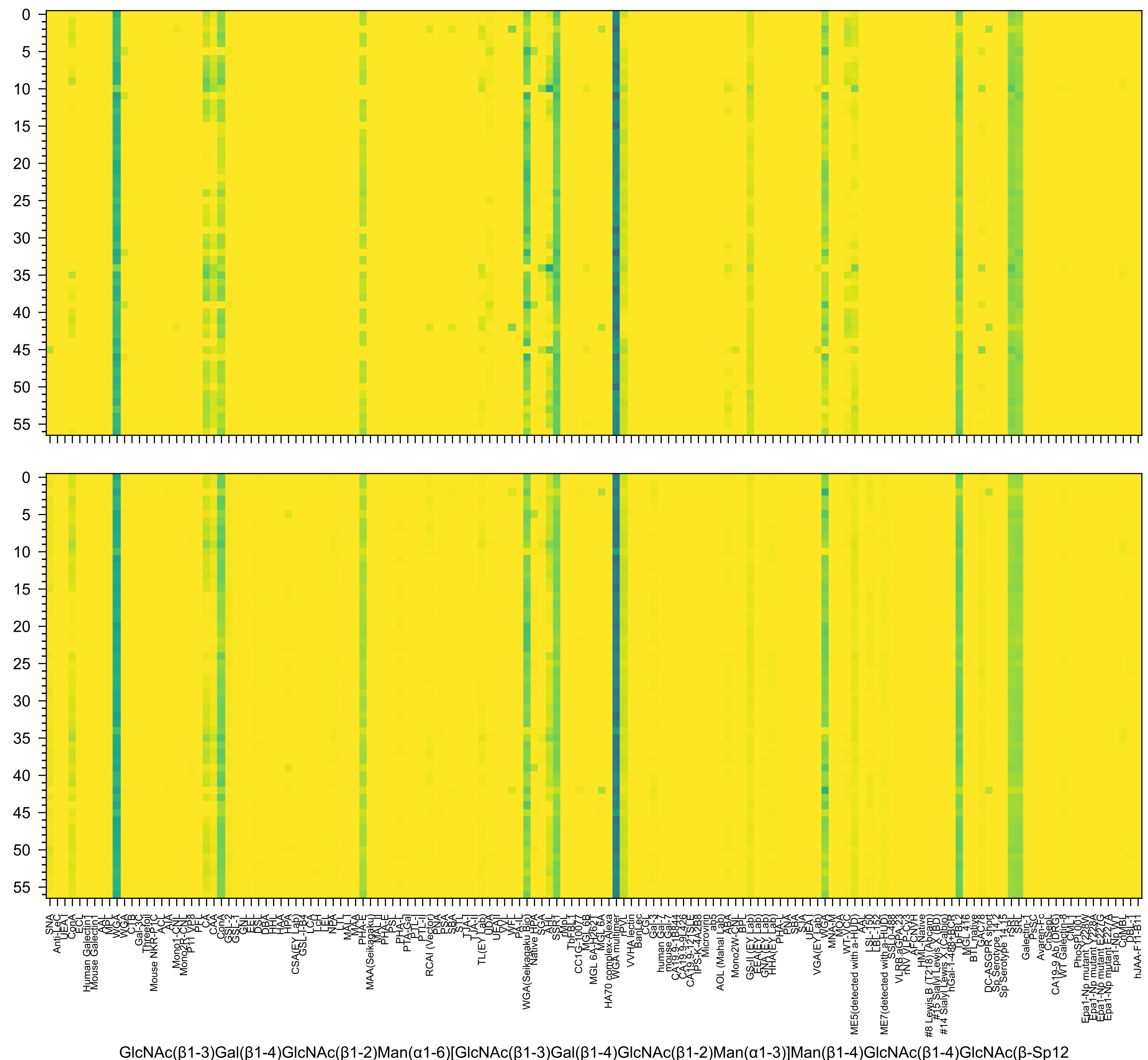
GlcNAc(β1-3)Gal(β1-4)GlcNAc(β1-2)Man(α1-6)[GlcNAc(β1-3)Gal(β1-4)GlcNAc(β1-2)Man(α1-3)]Man(β1-4)GlcNAc(β1-4)GlcNAc(β-Sp12

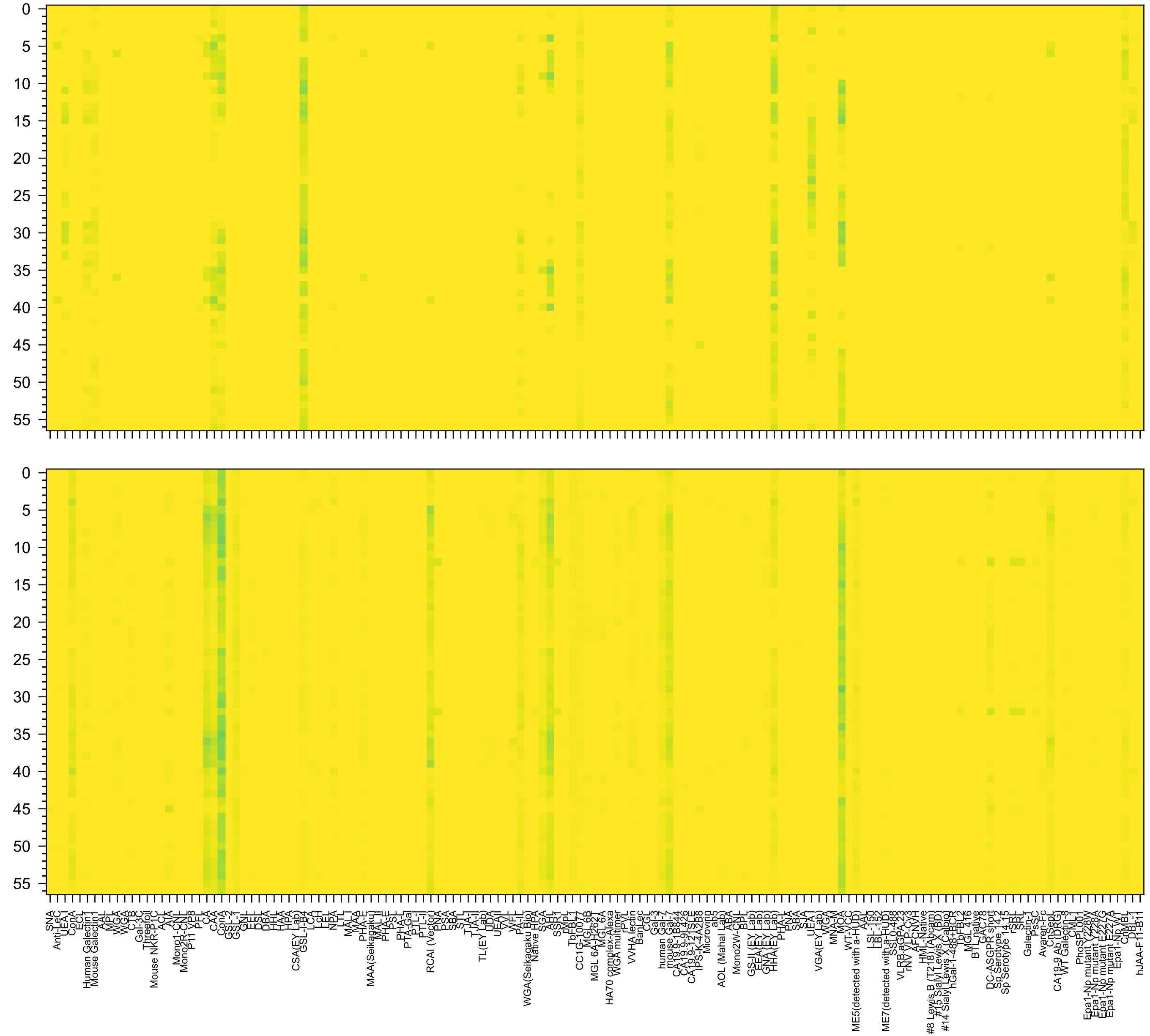

Gal(α1-3)Gal(β1-3)GlcNAc(β1-2)Man(α1-6)[Gal(α1-3)Gal(β1-3)GlcNAc(β1-2)Man(α1-3)]Man(β1-4)GlcNAc(β1-4)GlcNAc(-Sp19

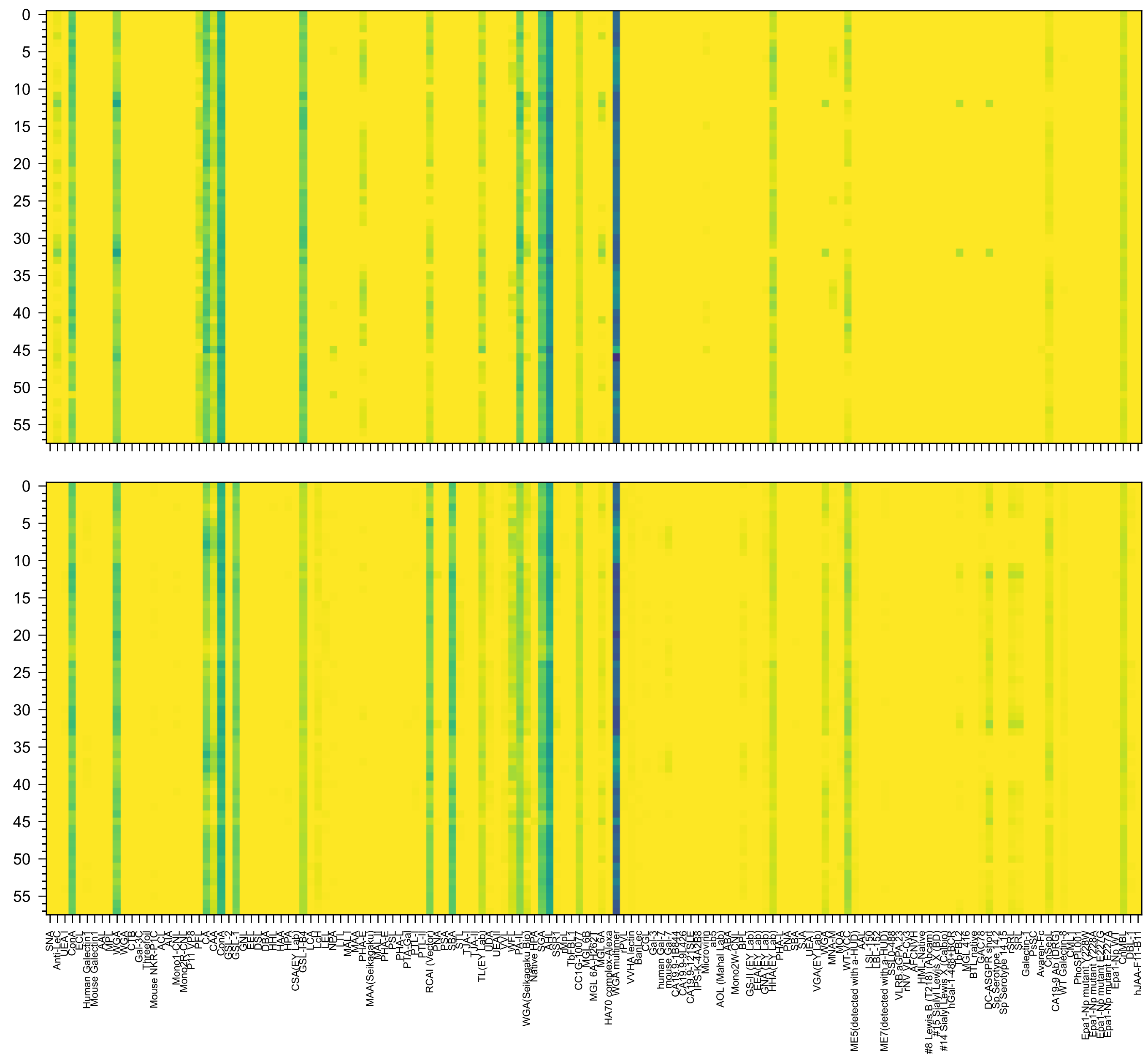

Gal(α1-4)Gal(β1-3)GlcNAc(β1-2)Man(α1-6)[Gal(α1-4)Gal(β1-3)GlcNAc(β1-2)Man(α1-3)]Man(β1-4)GlcNAc(β1-4)GlcNAc(β-Sp19

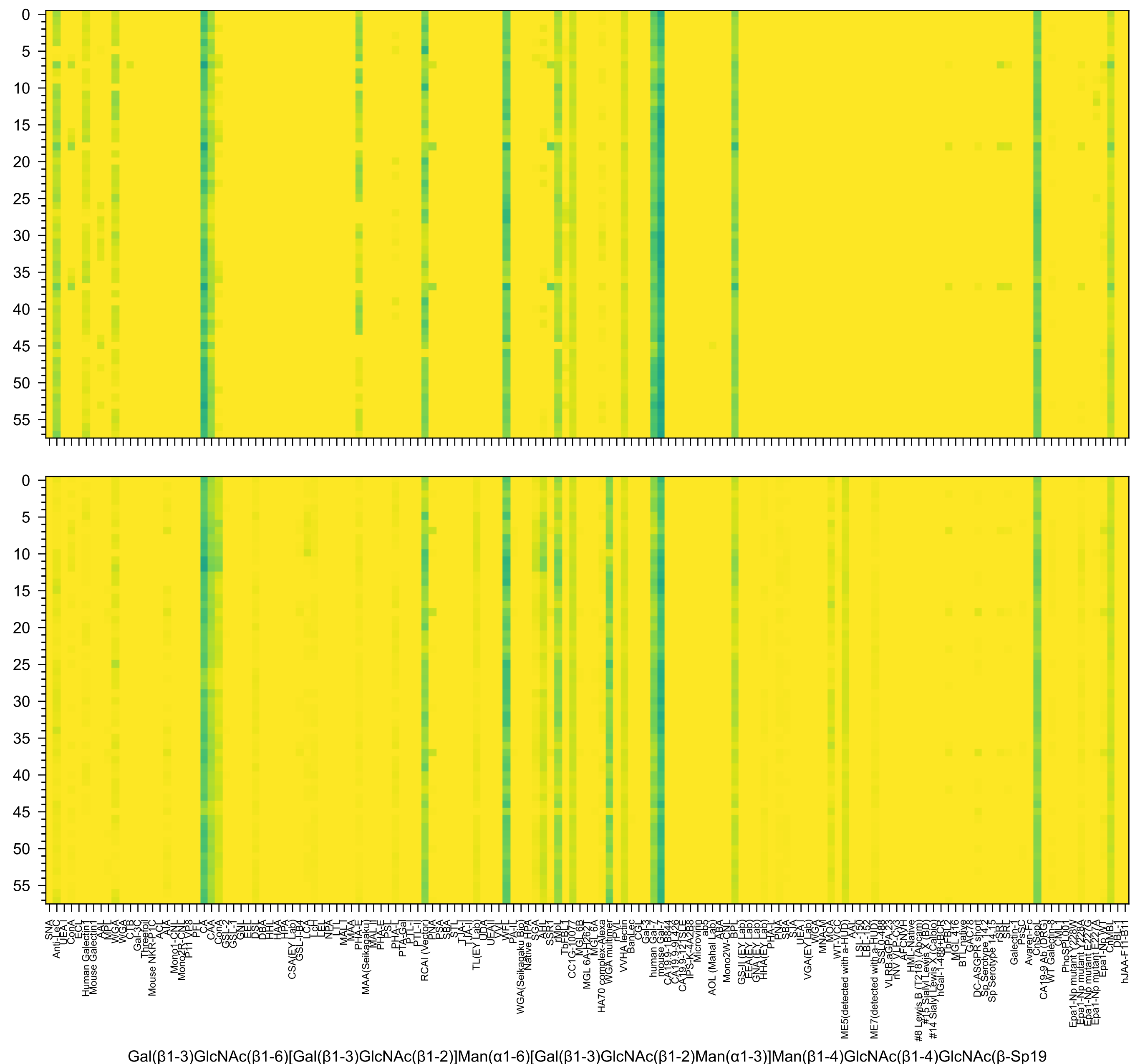
Gal(β1-3)GlcNAc(β1-6)[Gal(β1-3)GlcNAc(β1-2)]Man(α1-6)[Gal(β1-3)GlcNAc(β1-2)Man(α1-3)]Man(β1-4)GlcNAc(β1-4)GlcNAc(β-Sp19

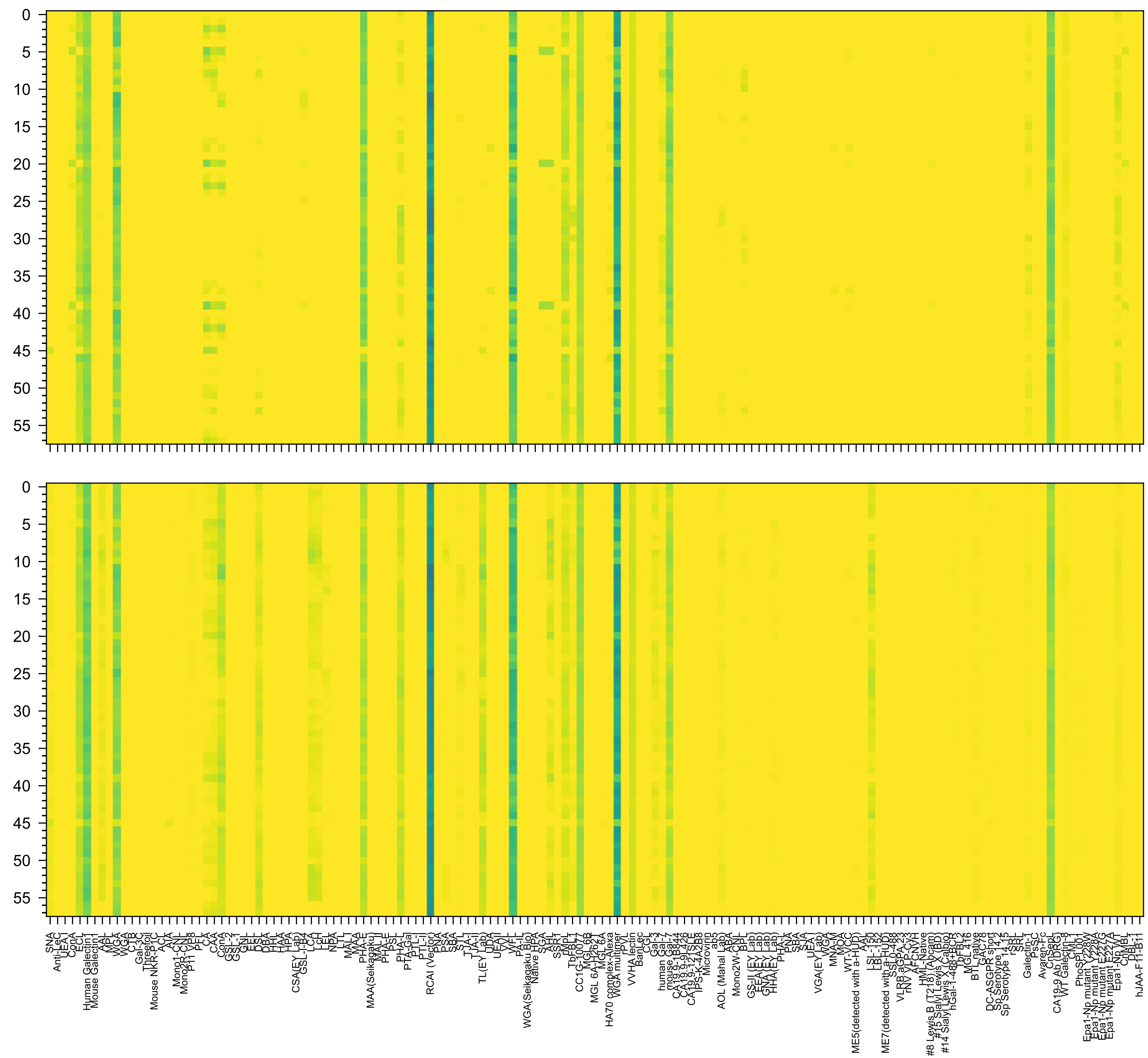

Gal(β1-4)GlcNAc(β1-6)[Gal(β1-4)GlcNAc(β1-2)]Man(α1-6)[Gal(β1-4)GlcNAc(β1-2)Man(α1-3)]Man(β1-4)GlcNAc(β1-4)GlcNAc(β-Sp19

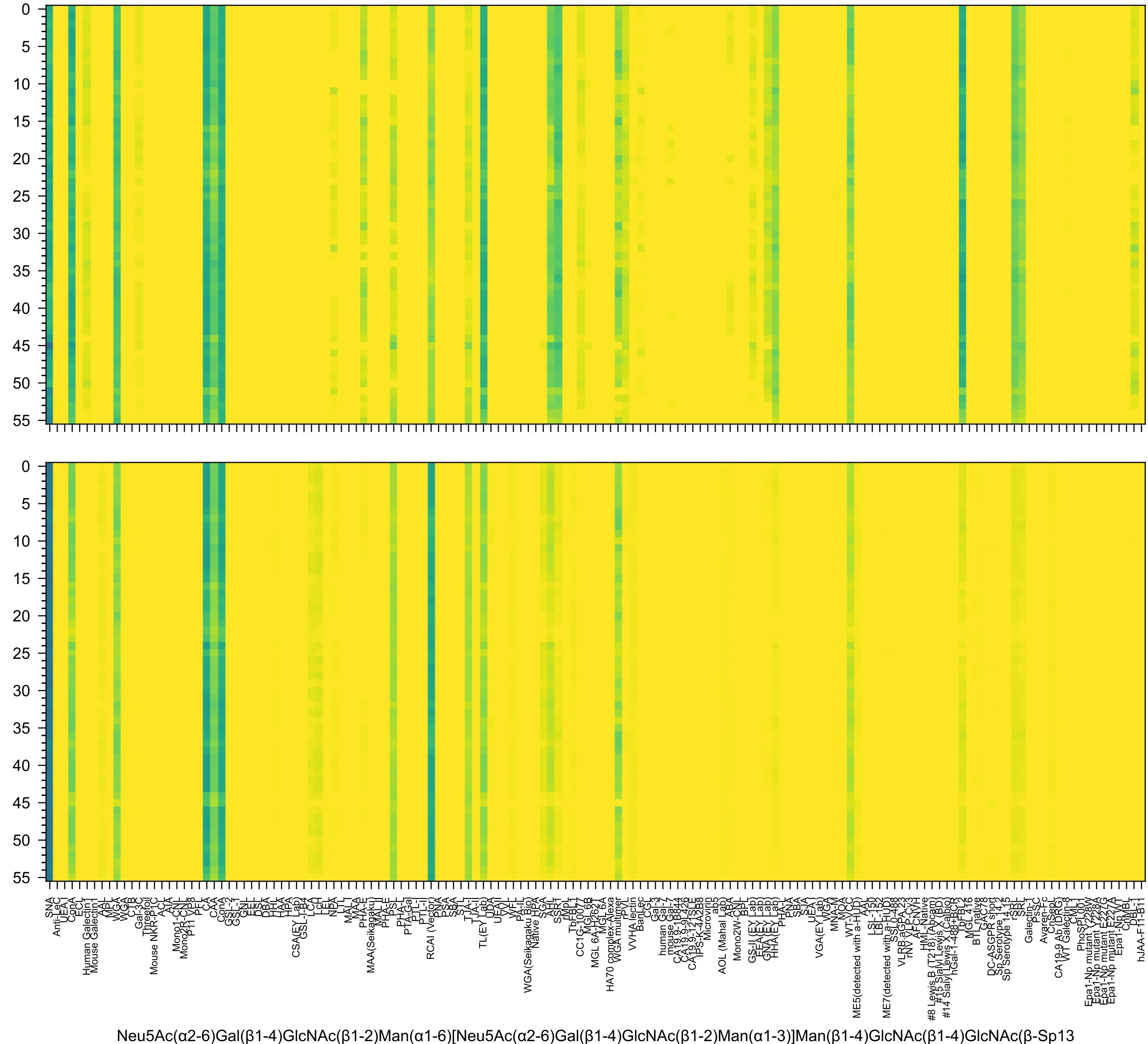
Neu5Ac(α2-6)Gal(β1-4)GlcNAc(β1-2)Man(α1-6)[Neu5Ac(α2-6)Gal(β1-4)GlcNAc(β1-2)Man(α1-3)]Man(β1-4)GlcNAc(β1-4)GlcNAc(β-Sp13

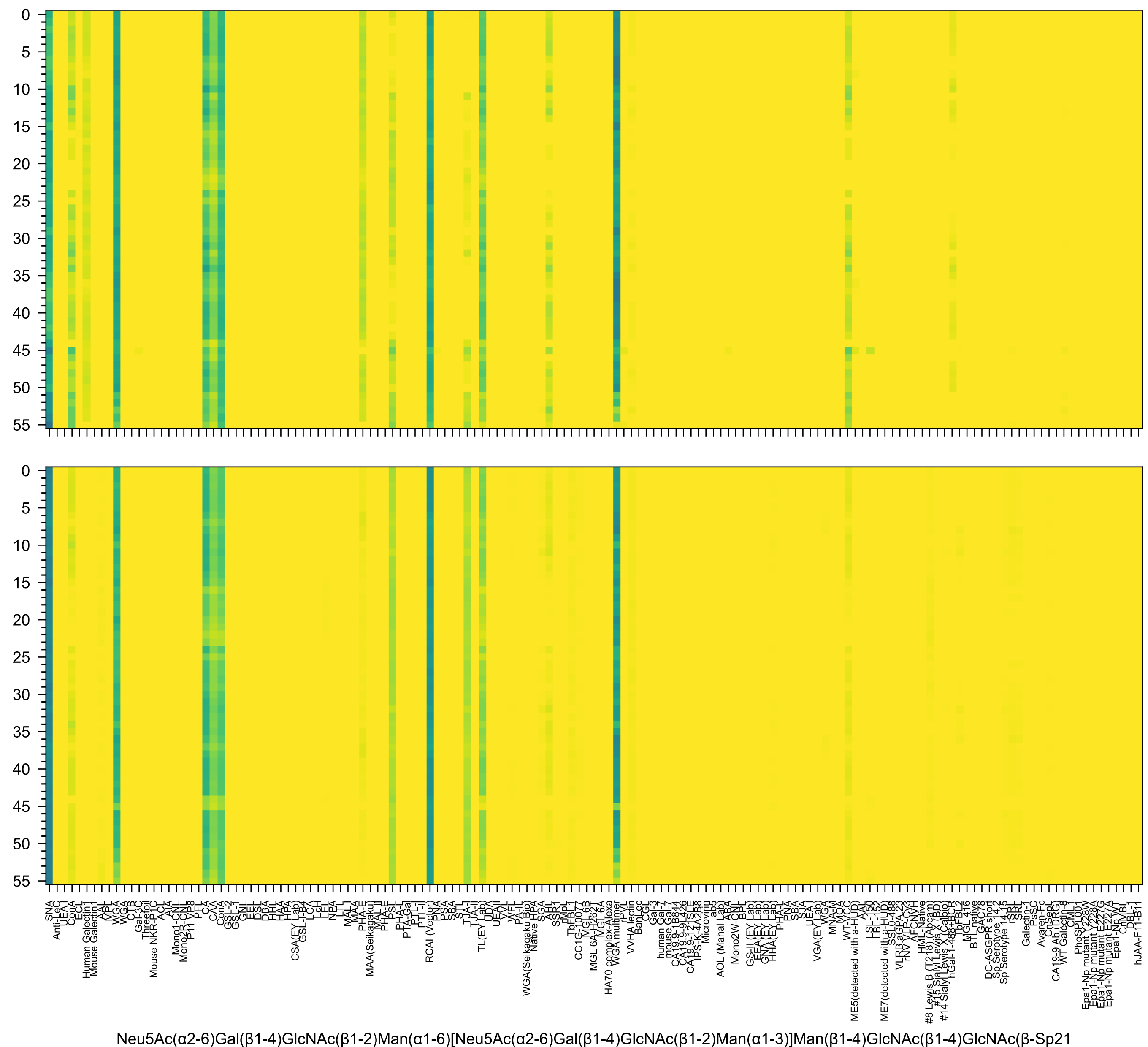
Neu5Ac(α2-6)Gal(β1-4)GlcNAc(β1-2)Man(α1-6)[Neu5Ac(α2-6)Gal(β1-4)GlcNAc(β1-2)Man(α1-3)]Man(β1-4)GlcNAc(β1-4)GlcNAc(β-Sp21

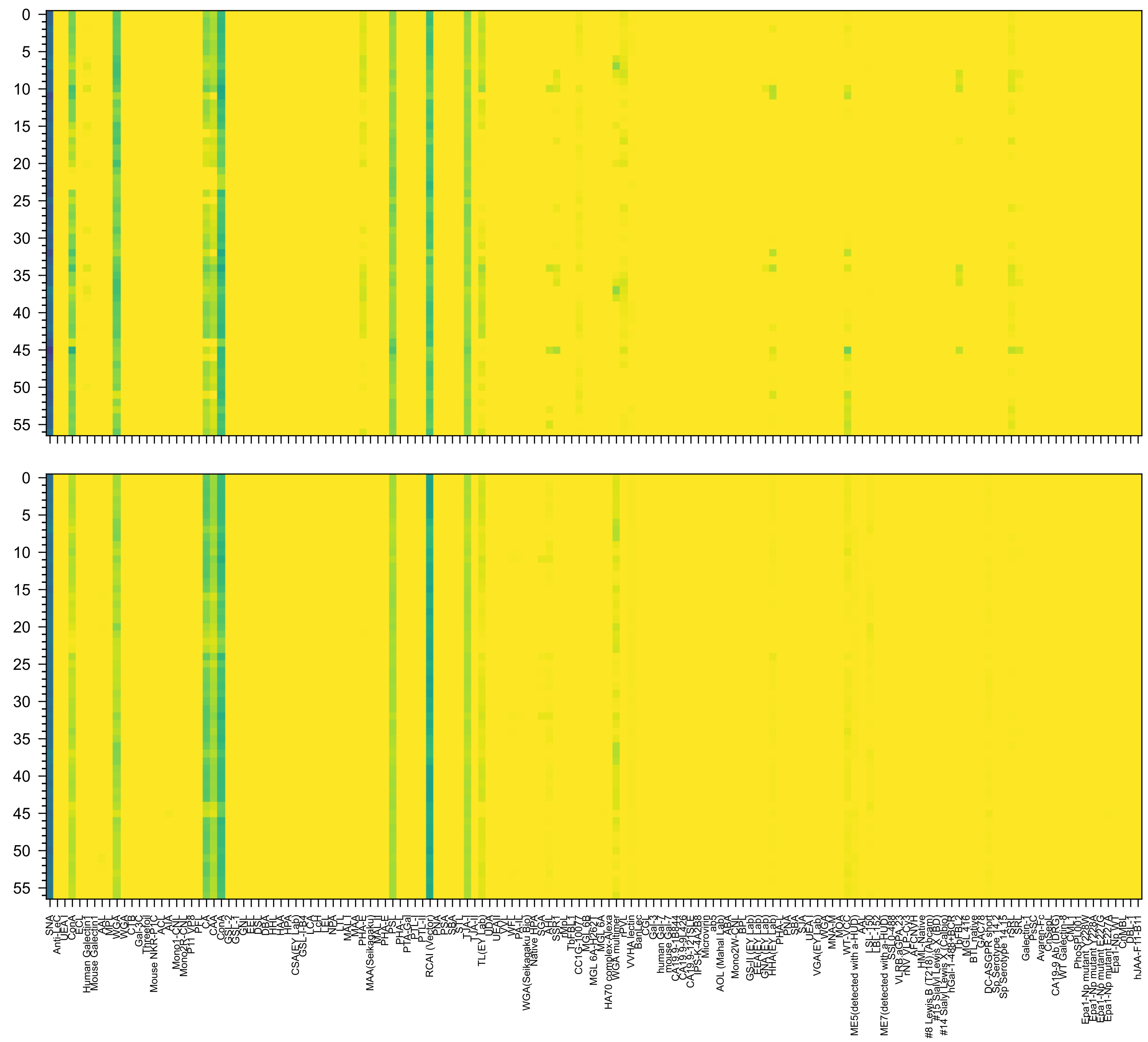

Neu5Ac(α2-6)Gal(β1-4)GlcNAc(β1-2)Man(α1-6)[Neu5Ac(α2-6)Gal(β1-4)GlcNAc(β1-2)Man(α1-3)]Man(β1-4)GlcNAc(β1-4)GlcNAc(β-Sp12

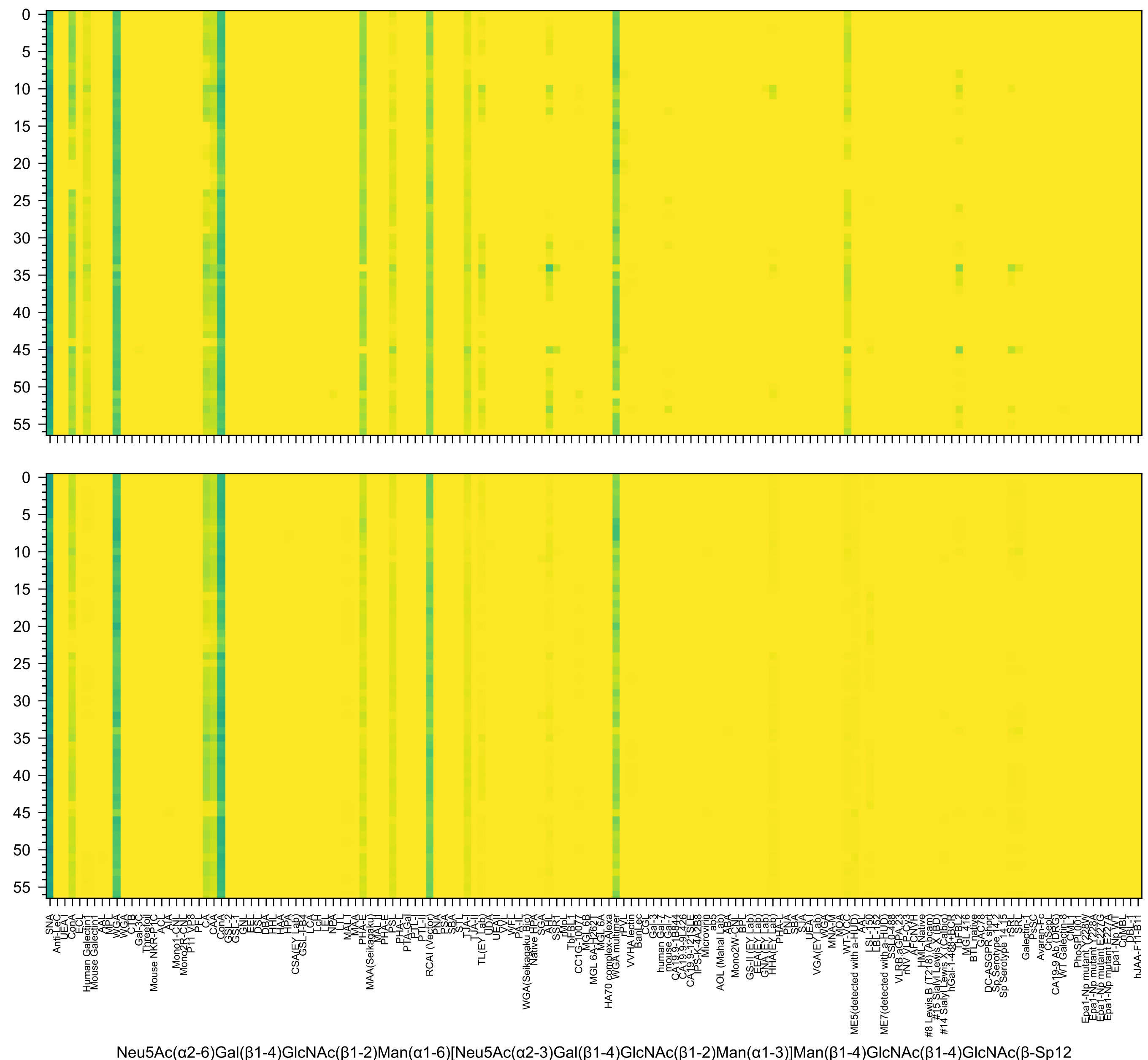
Neu5Ac(α2-6)Gal(β1-4)GlcNAc(β1-2)Man(α1-6)[Neu5Ac(α2-3)Gal(β1-4)GlcNAc(β1-2)Man(α1-3)]Man(β1-4)GlcNAc(β1-4)GlcNAc(β-Sp12

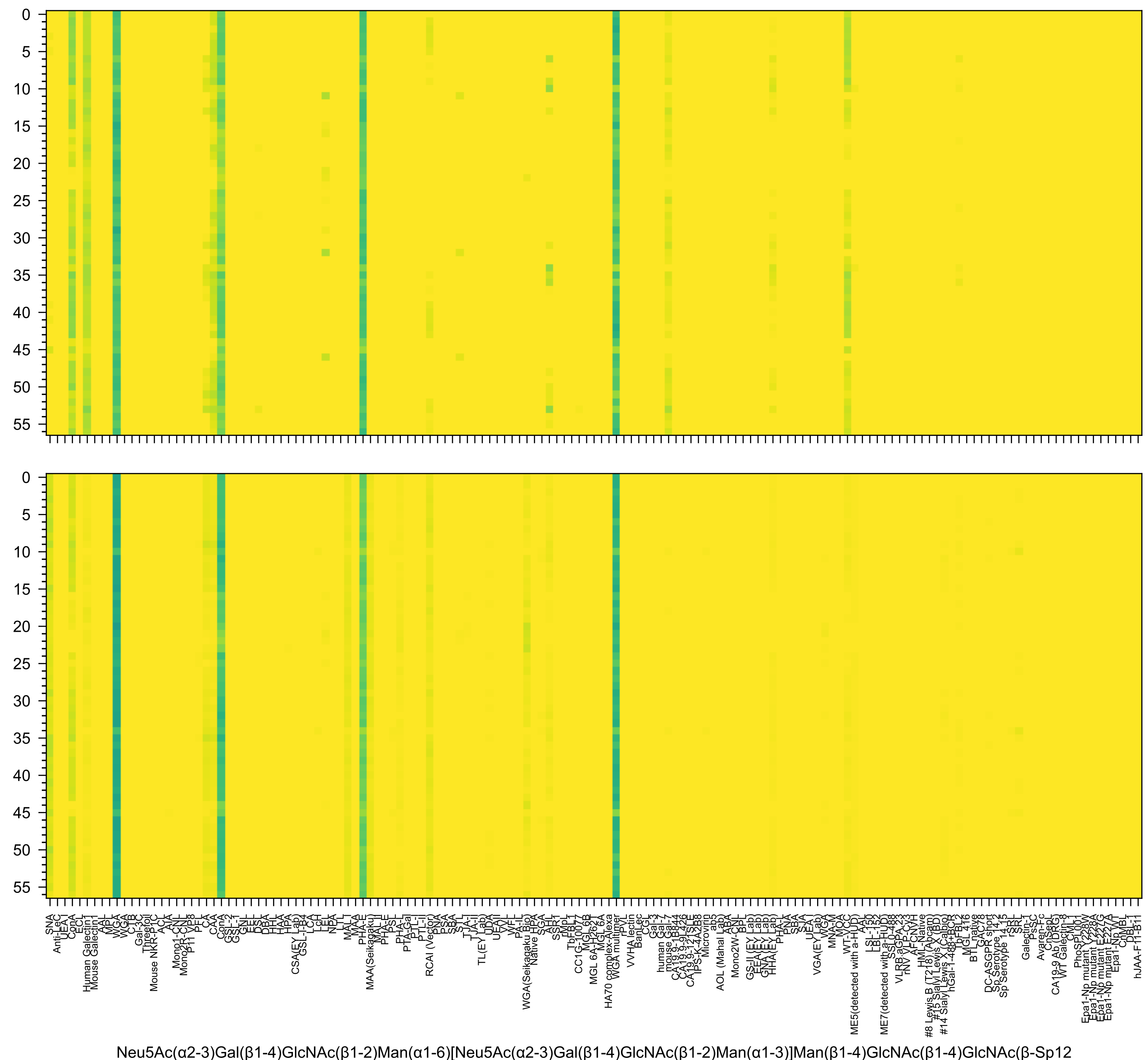
Neu5Ac(α2-3)Gal(β1-4)GlcNAc(β1-2)Man(α1-6)[Neu5Ac(α2-3)Gal(β1-4)GlcNAc(β1-2)Man(α1-3)]Man(β1-4)GlcNAc(β1-4)GlcNAc(β-Sp12

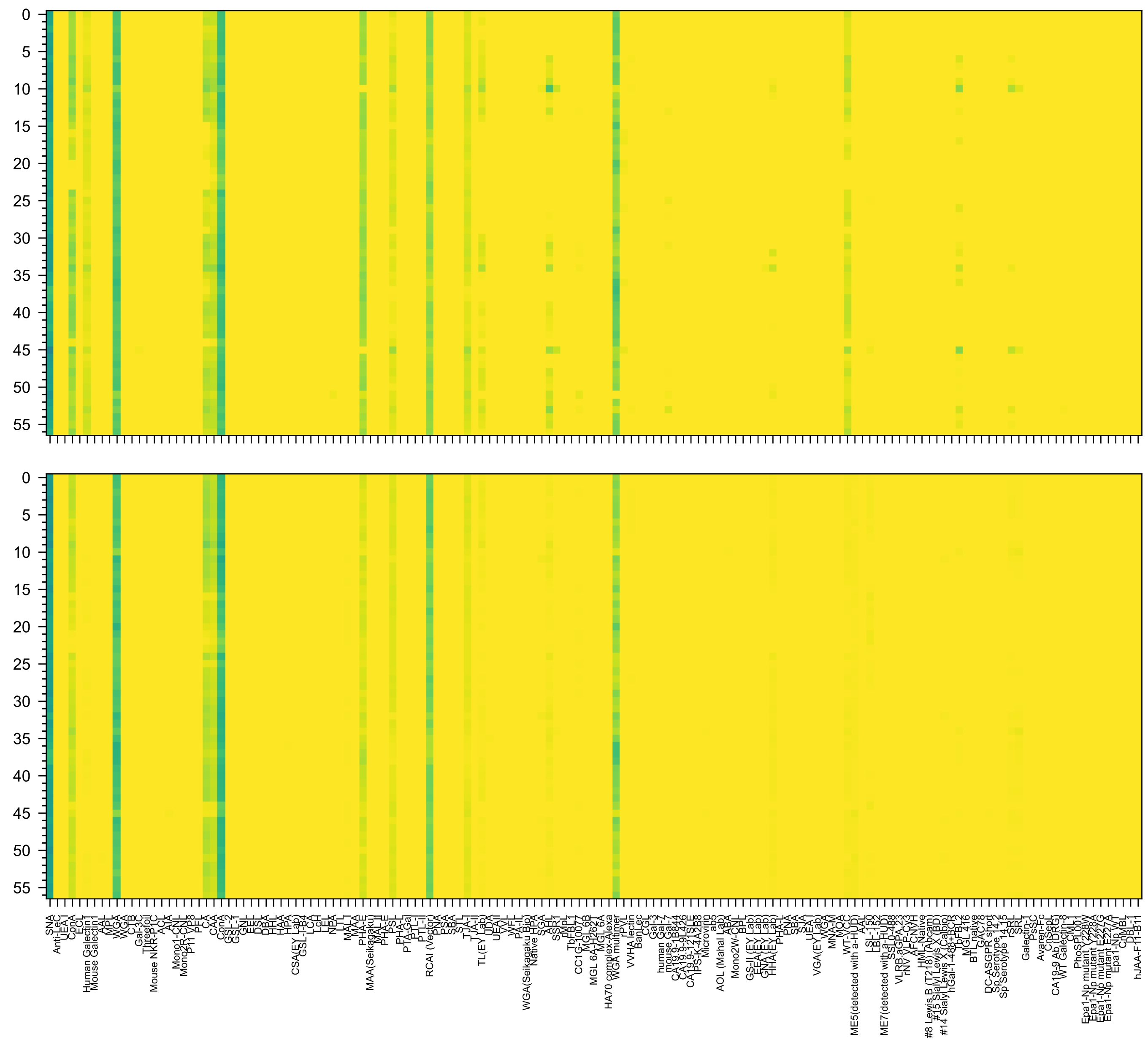

Neu5Ac(α2-3)Gal(β1-4)GlcNAc(β1-2)Man(α1-6)[Neu5Ac(α2-6)Gal(β1-4)GlcNAc(β1-2)Man(α1-3)]Man(β1-4)GlcNAc(β1-4)GlcNAc(β-Sp12

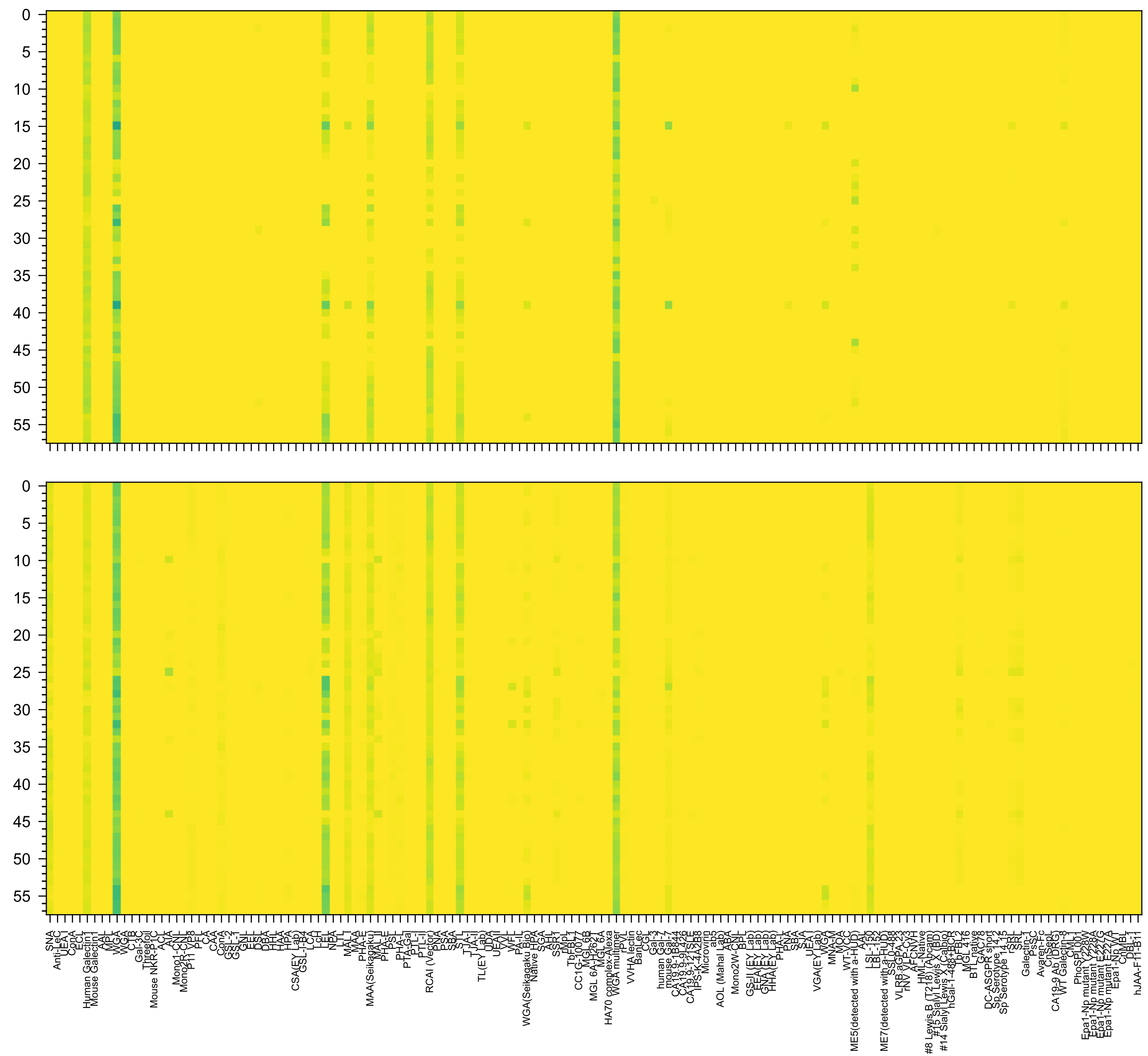

Neu5Ac(α2-3)Gal(β1-4)GlcNAc(β1-3)Gal(β1-4)GlcNAc(β1-6)[Neu5Ac(α2-3)Gal(β1-4)GlcNAc(β1-3)Gal(β1-4)GlcNAc(β1-3)]GalNAc(α-Sp14

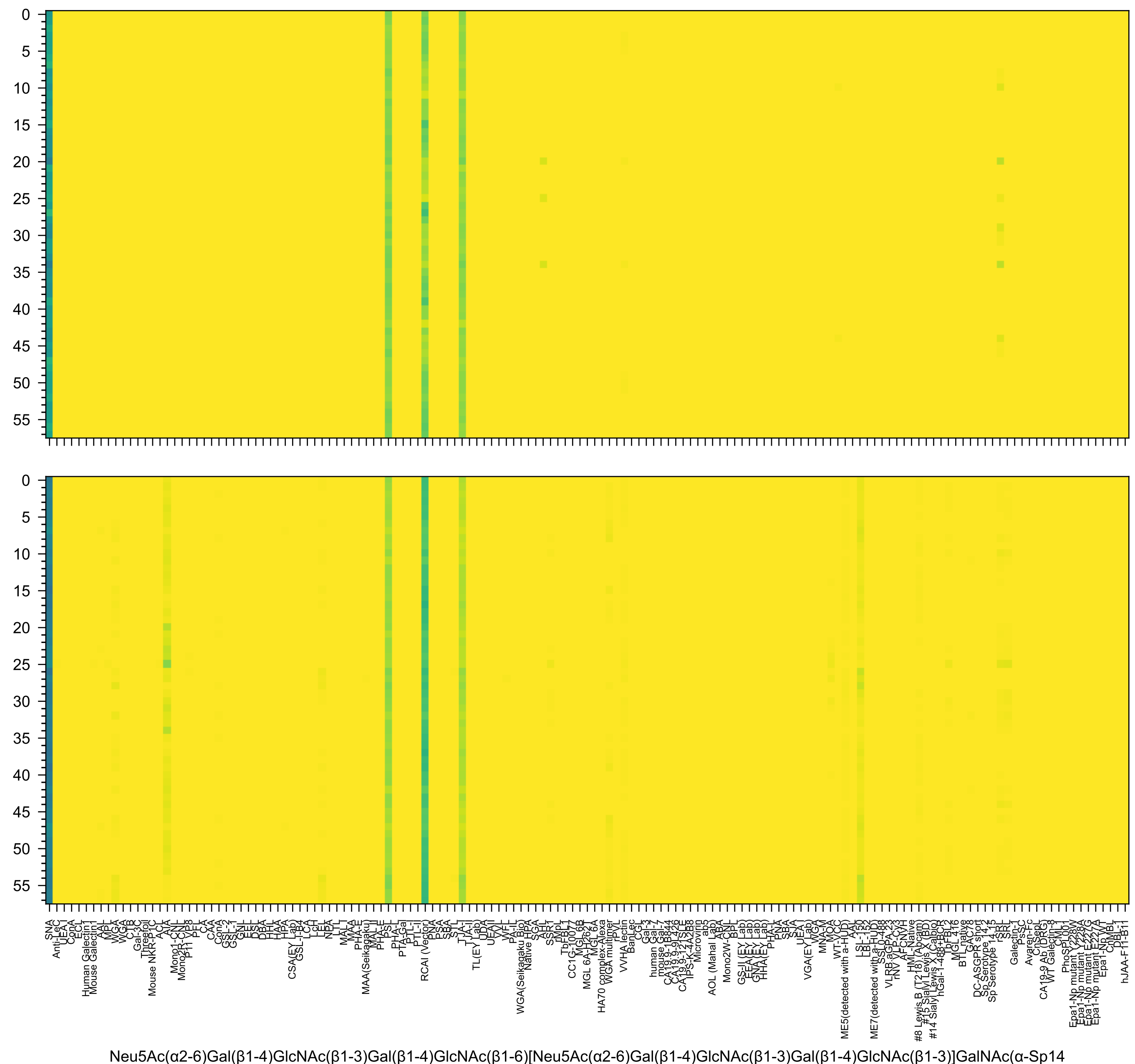
Neu5Ac(α2-6)Gal(β1-4)GlcNAc(β1-3)Gal(β1-4)GlcNAc(β1-6)[Neu5Ac(α2-6)Gal(β1-4)GlcNAc(β1-3)Gal(β1-4)GlcNAc(β1-3)]GalNAc(α-Sp14

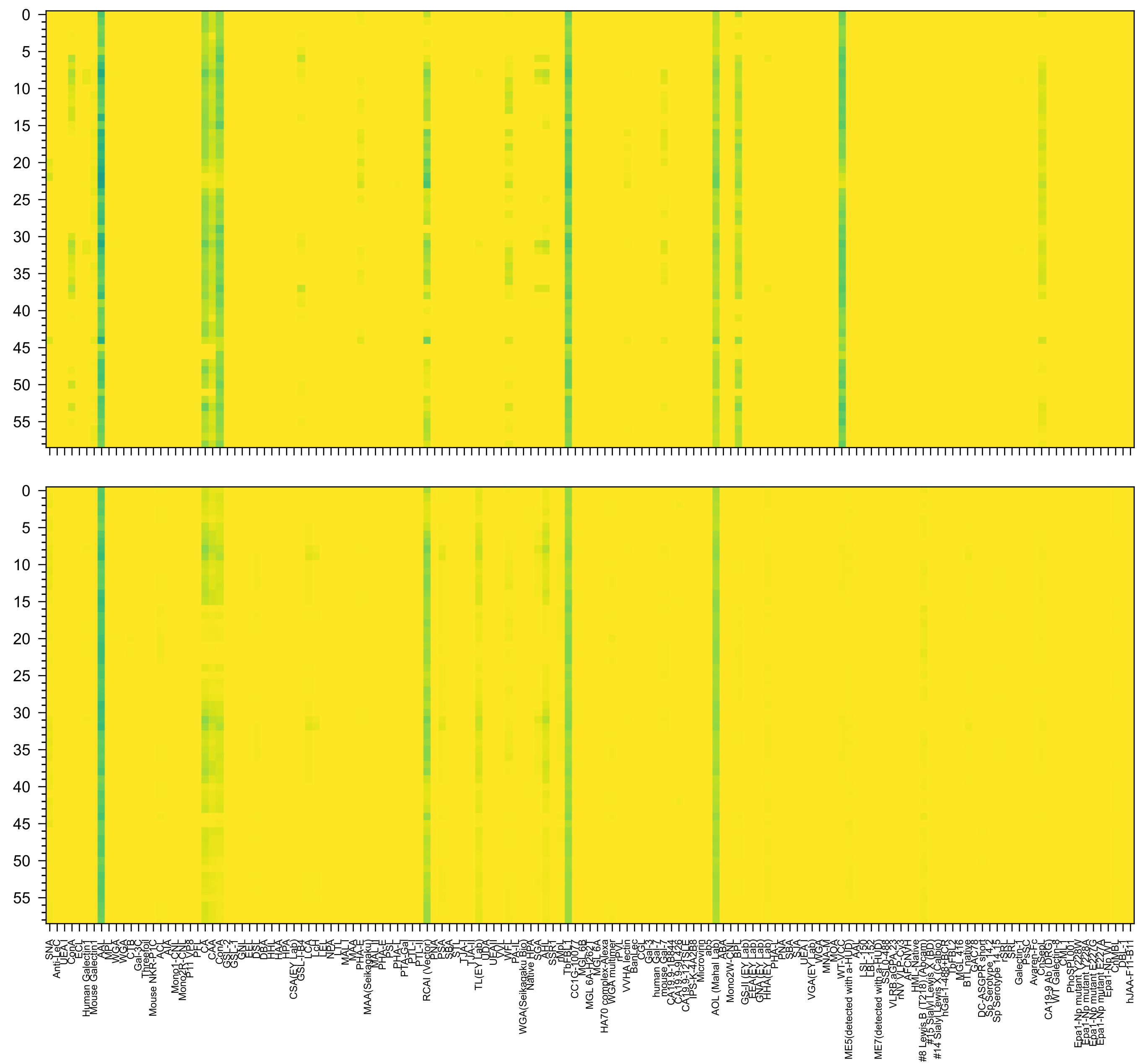


Gal(β1-4)[Fuc(α1-3)]GlcNAc(β1-2)Man(α1-6)[Gal(β1-4)[Fuc(α1-3)]GlcNAc(β1-2)Man(α1-3)]Man(β1-4)GlcNAc(β1-4)GlcNAc(β-Sp20

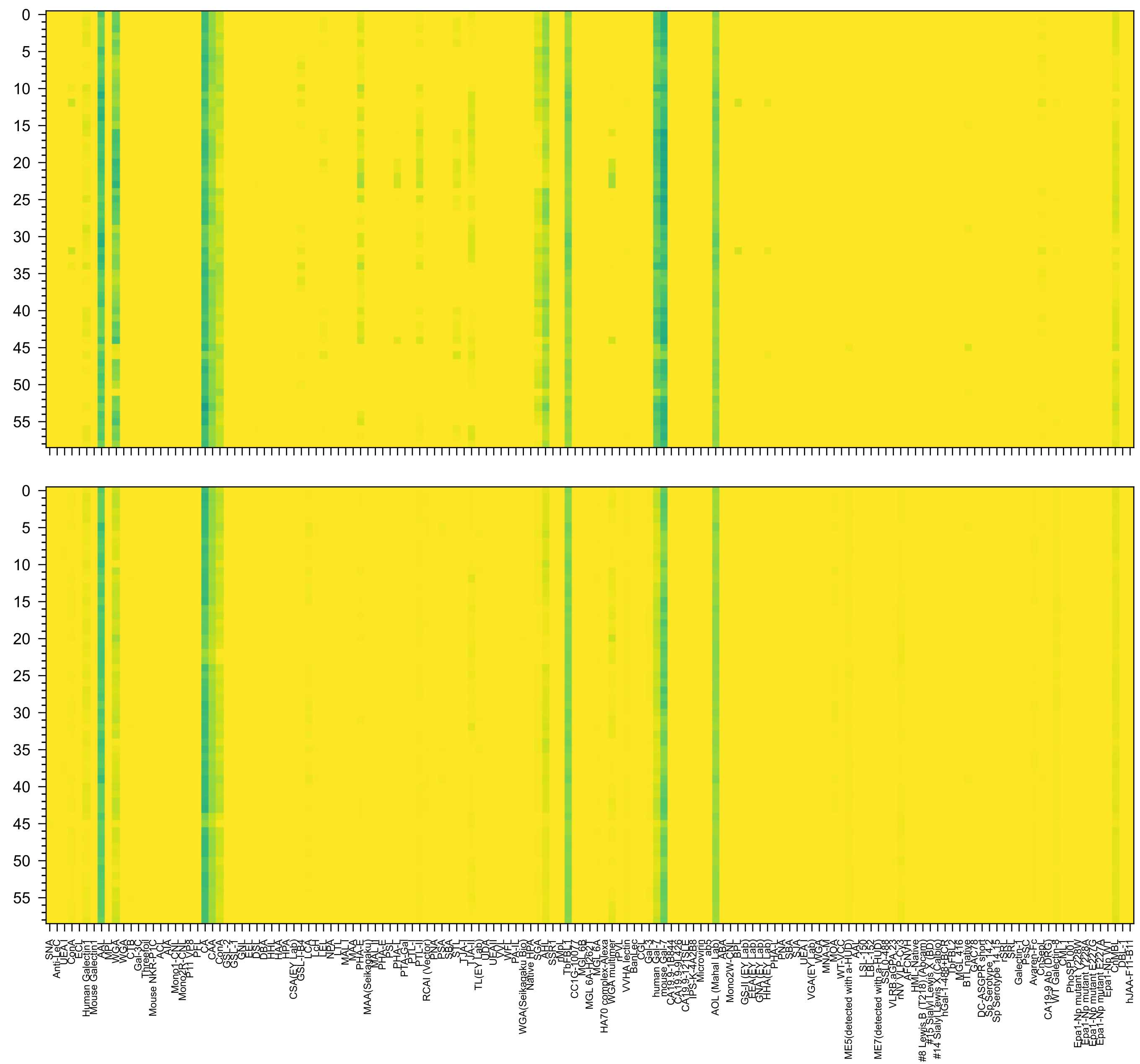

Fuc(α1-2)Gal(β1-3)GlcNAc(β1-2)Man(α1-6)[Fuc(α1-2)Gal(β1-3)GlcNAc(β1-2)Man(α1-3)]Man(β1-4)GlcNAc(β1-4)GlcNAc(β-Sp20

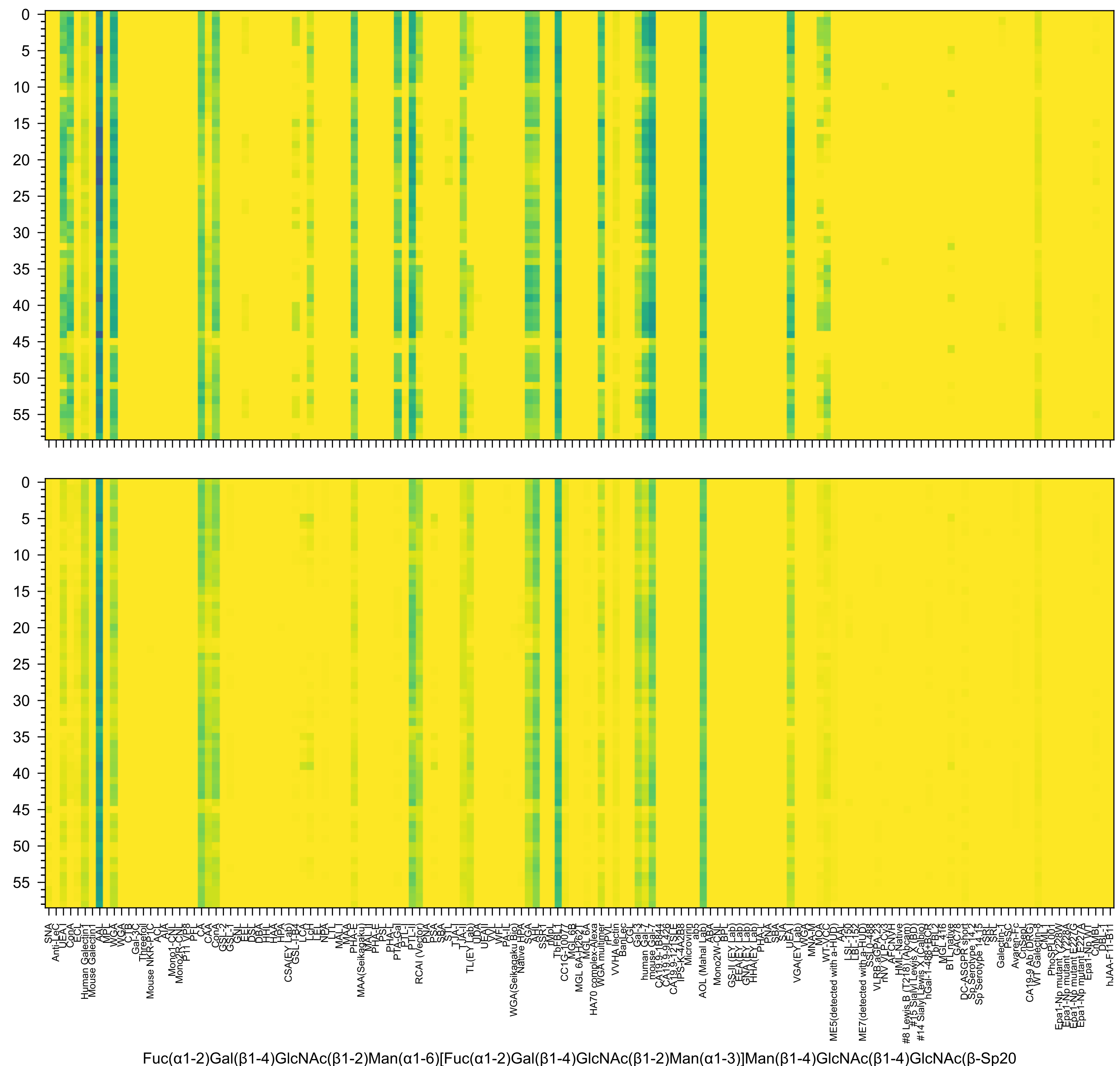
Fuc(α1-2)Gal(β1-4)GlcNAc(β1-2)Man(α1-6)[Fuc(α1-2)Gal(β1-4)GlcNAc(β1-2)Man(α1-3)]Man(β1-4)GlcNAc(β1-4)GlcNAc(β-Sp20

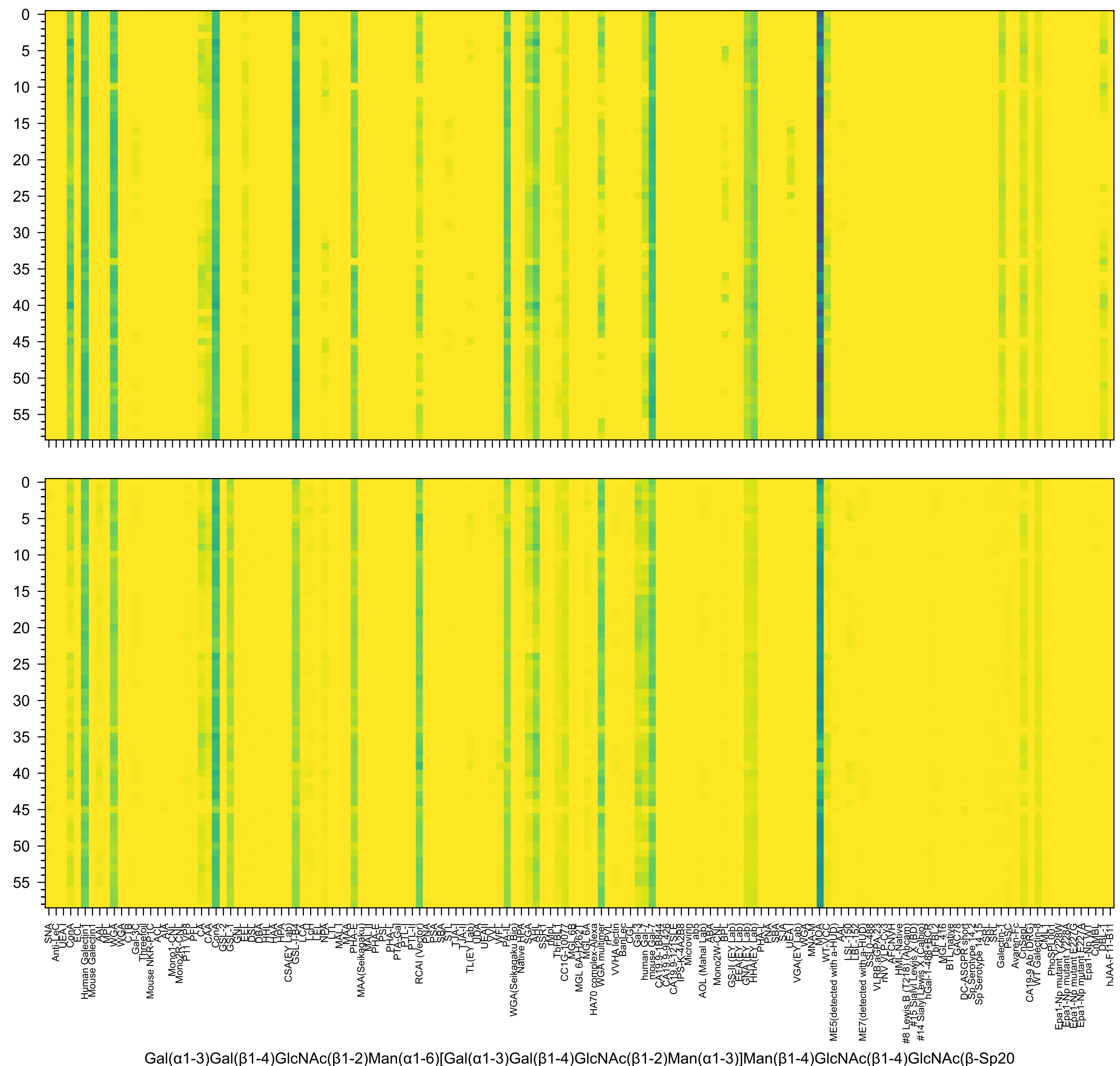
Gal(α1-3)Gal(β1-4)GlcNAc(β1-2)Man(α1-6)[Gal(α1-3)Gal(β1-4)GlcNAc(β1-2)Man(α1-3)]Man(β1-4)GlcNAc(β1-4)GlcNAc(β-Sp20

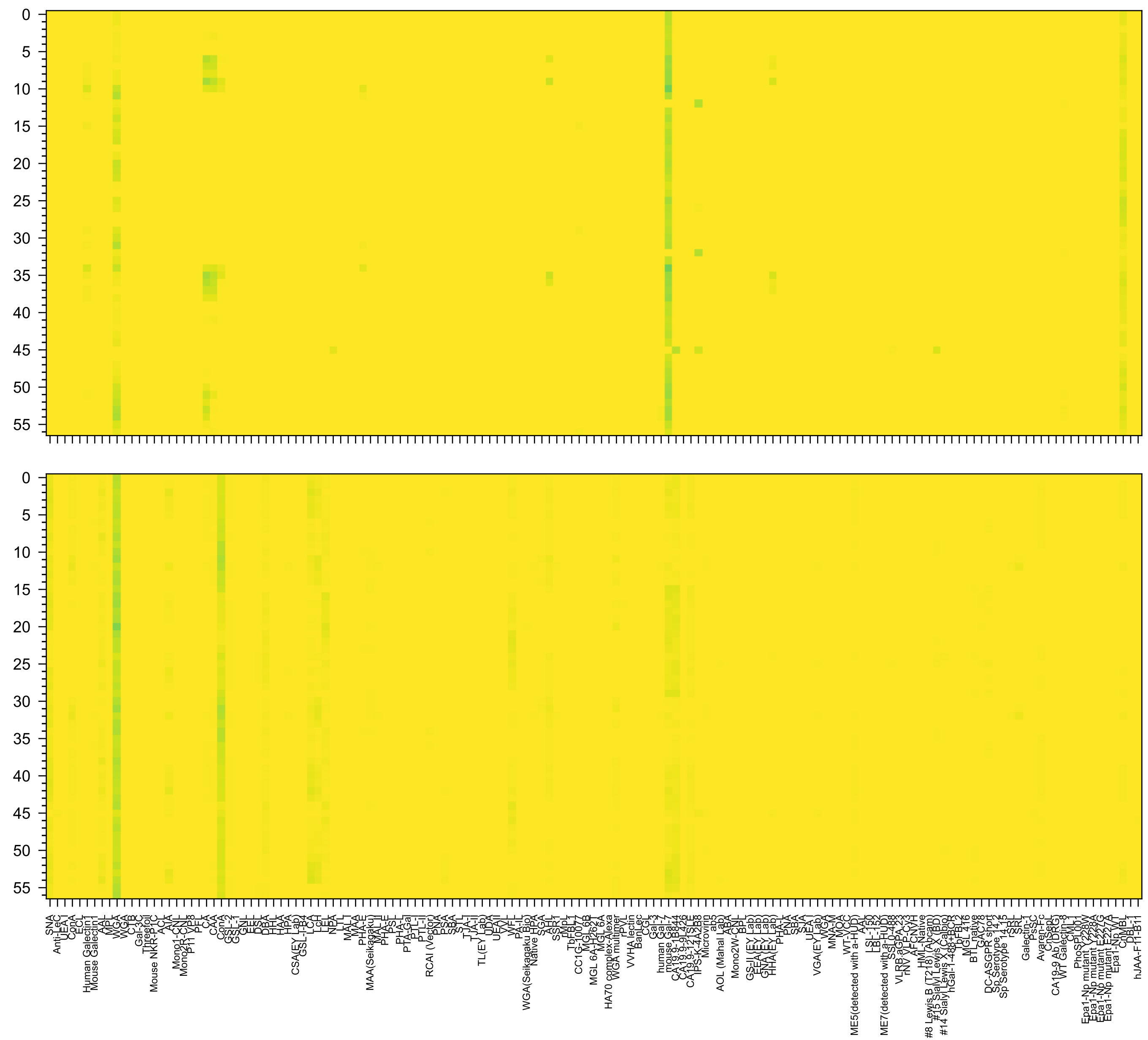

Neu5Ac(α2-3)Gal(β1-3)GlcNAc(β1-2)Man(α1-6)[Neu5Ac(α2-3)Gal(β1-3)GlcNAc(β1-2)Man(α1-3)]Man(β1-4)GlcNAc(β1-4)GlcNAc(-Sp19

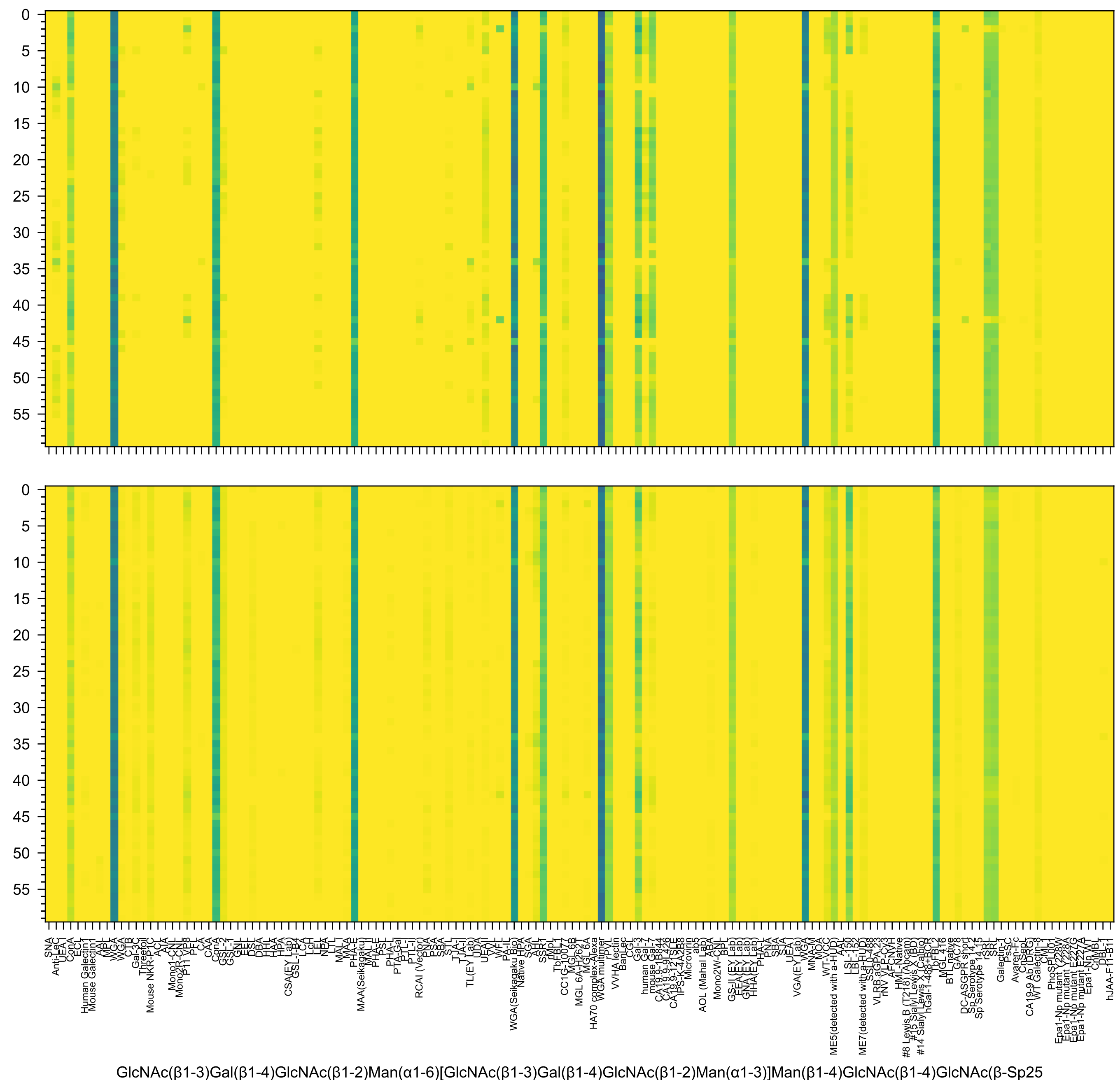
GlcNAc(β1-3)Gal(β1-4)GlcNAc(β1-2)Man(α1-6)[GlcNAc(β1-3)Gal(β1-4)GlcNAc(β1-2)Man(α1-3)]Man(β1-4)GlcNAc(β1-4)GlcNAc(β-Sp25

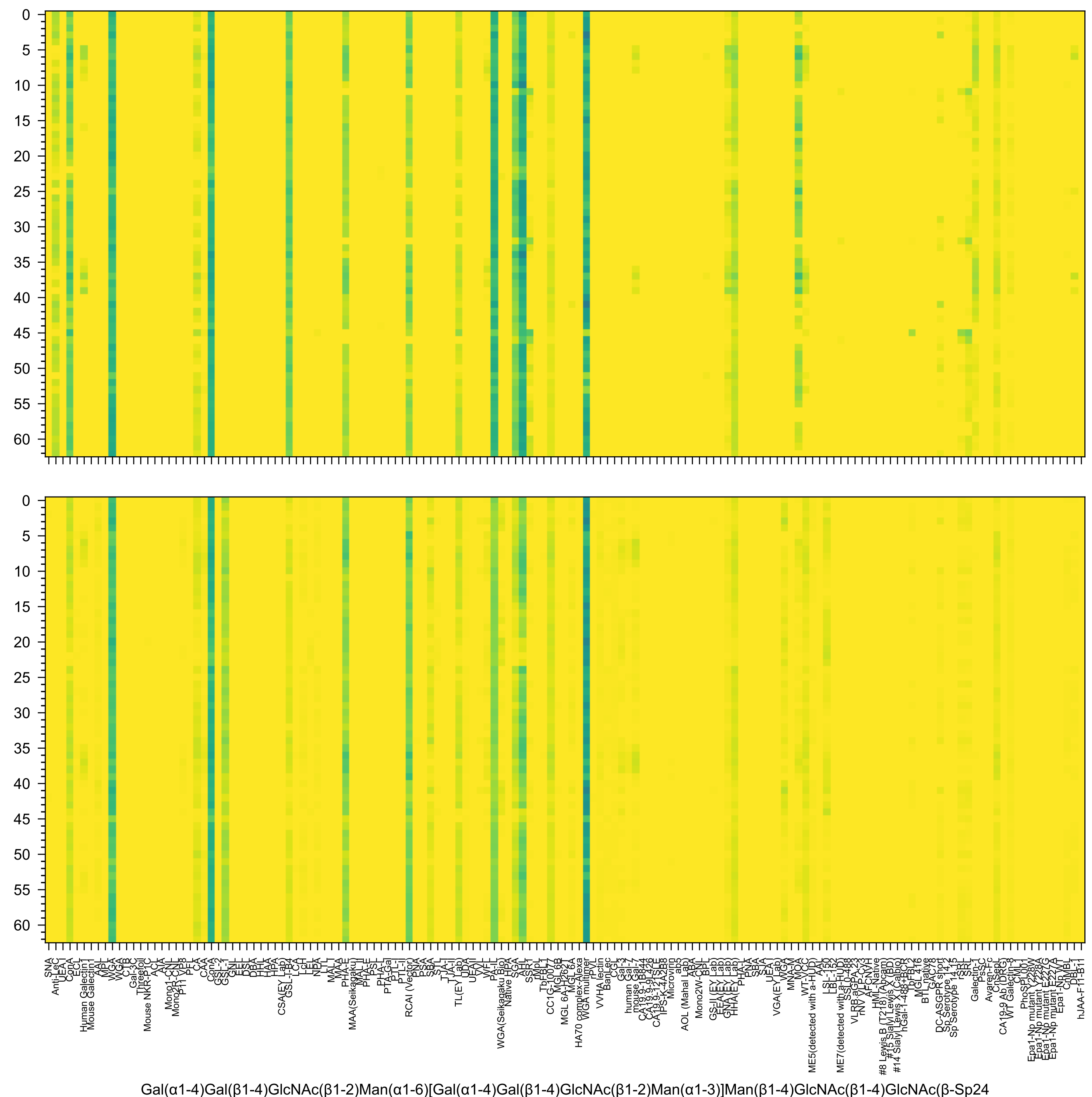
Gal(α1-4)Gal(β1-4)GlcNAc(β1-2)Man(α1-6)[Gal(α1-4)Gal(β1-4)GlcNAc(β1-2)Man(α1-3)]Man(β1-4)GlcNAc(β1-4)GlcNAc(β-Sp24

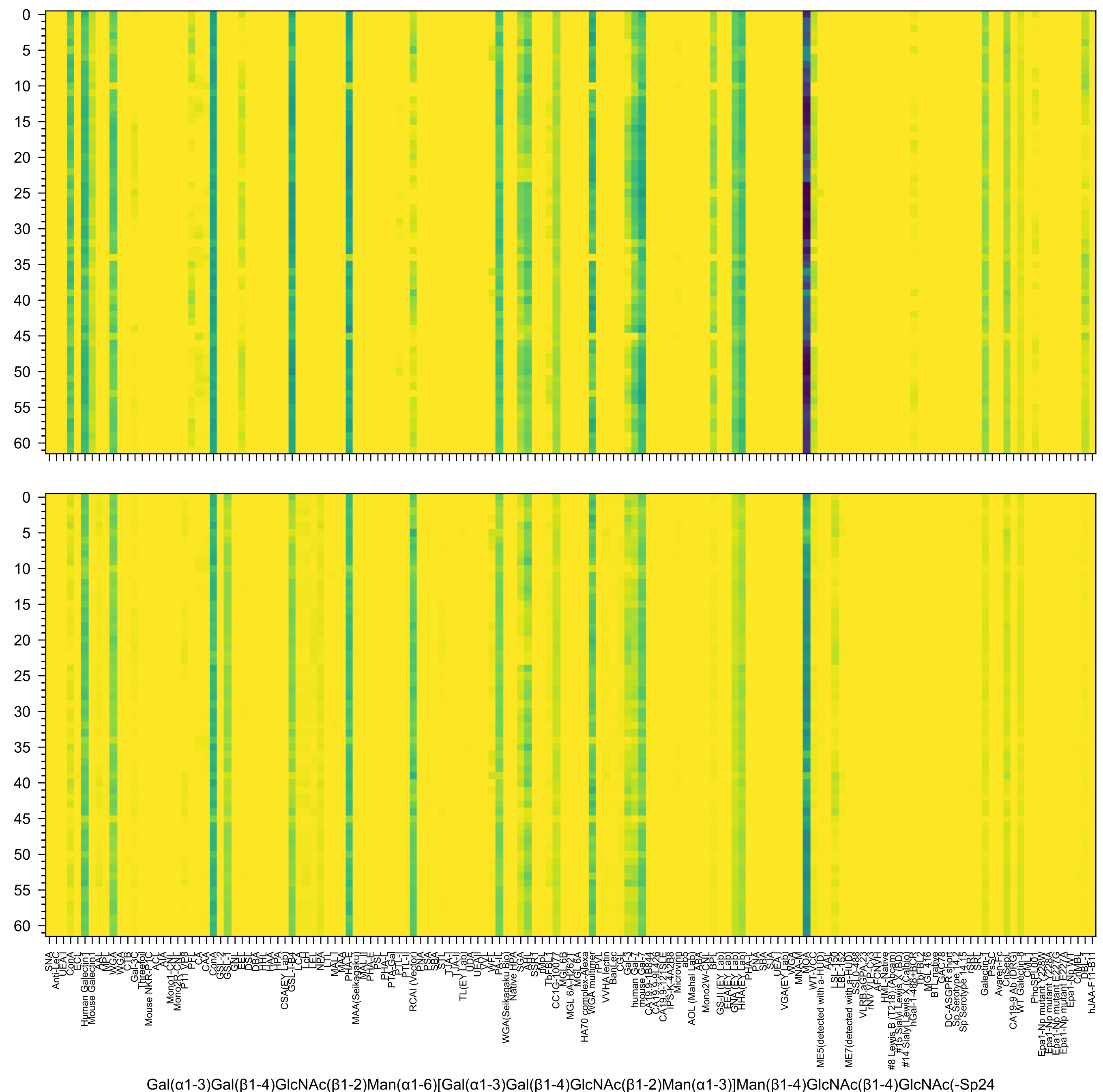
Gal(α1-3)Gal(β1-4)GlcNAc(β1-2)Man(α1-6)[Gal(α1-3)Gal(β1-4)GlcNAc(β1-2)Man(α1-3)]Man(β1-4)GlcNAc(β1-4)GlcNAc(-Sp24

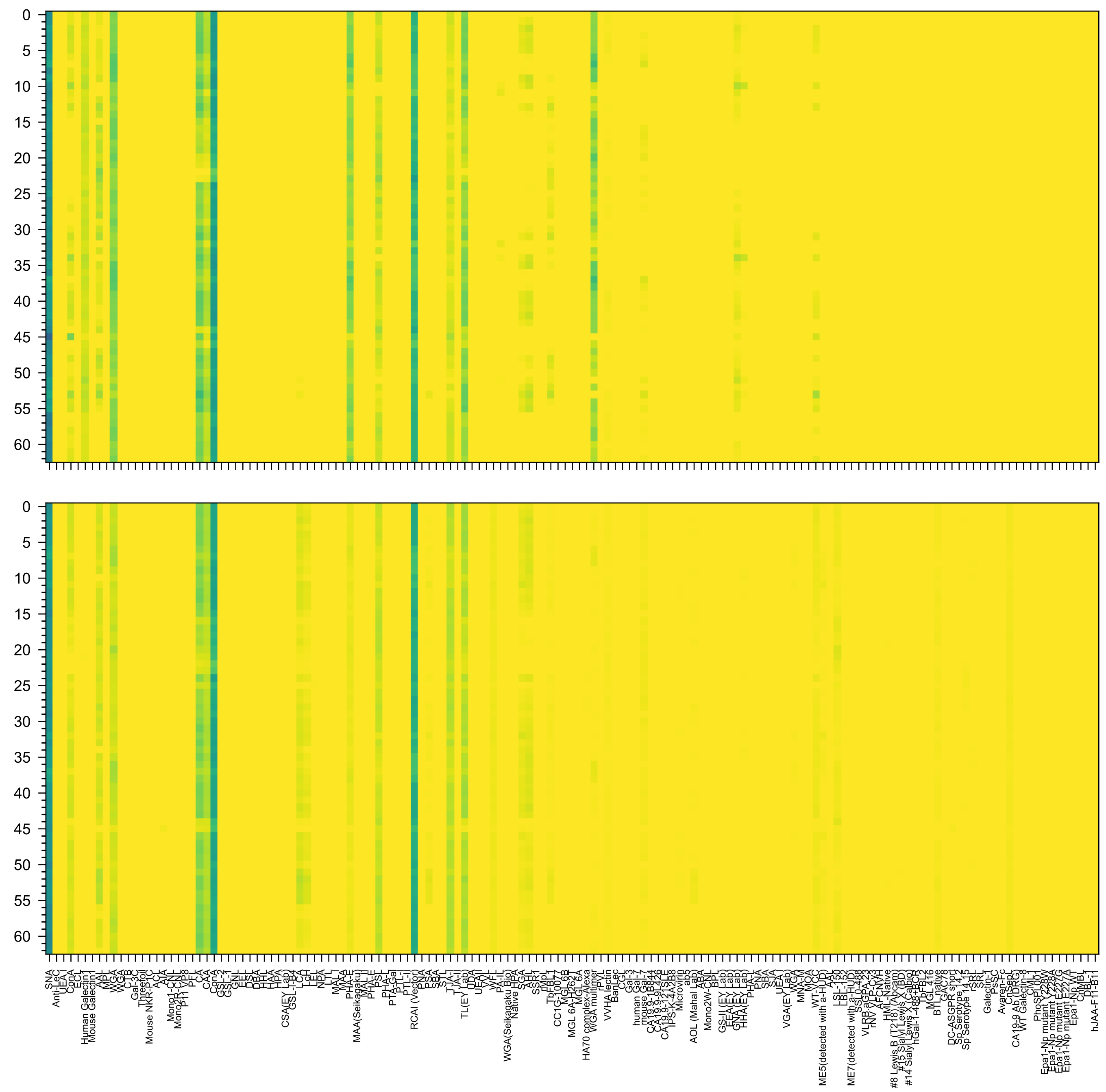

Neu5Ac(α2-6)Gal(β1-4)GlcNAc(β1-2)Man(α1-6)[Neu5Ac(α2-6)Gal(β1-4)GlcNAc(β1-2)Man(α1-3)]Man(β1-4)GlcNAc(β1-4)GlcNAc(β-Sp24

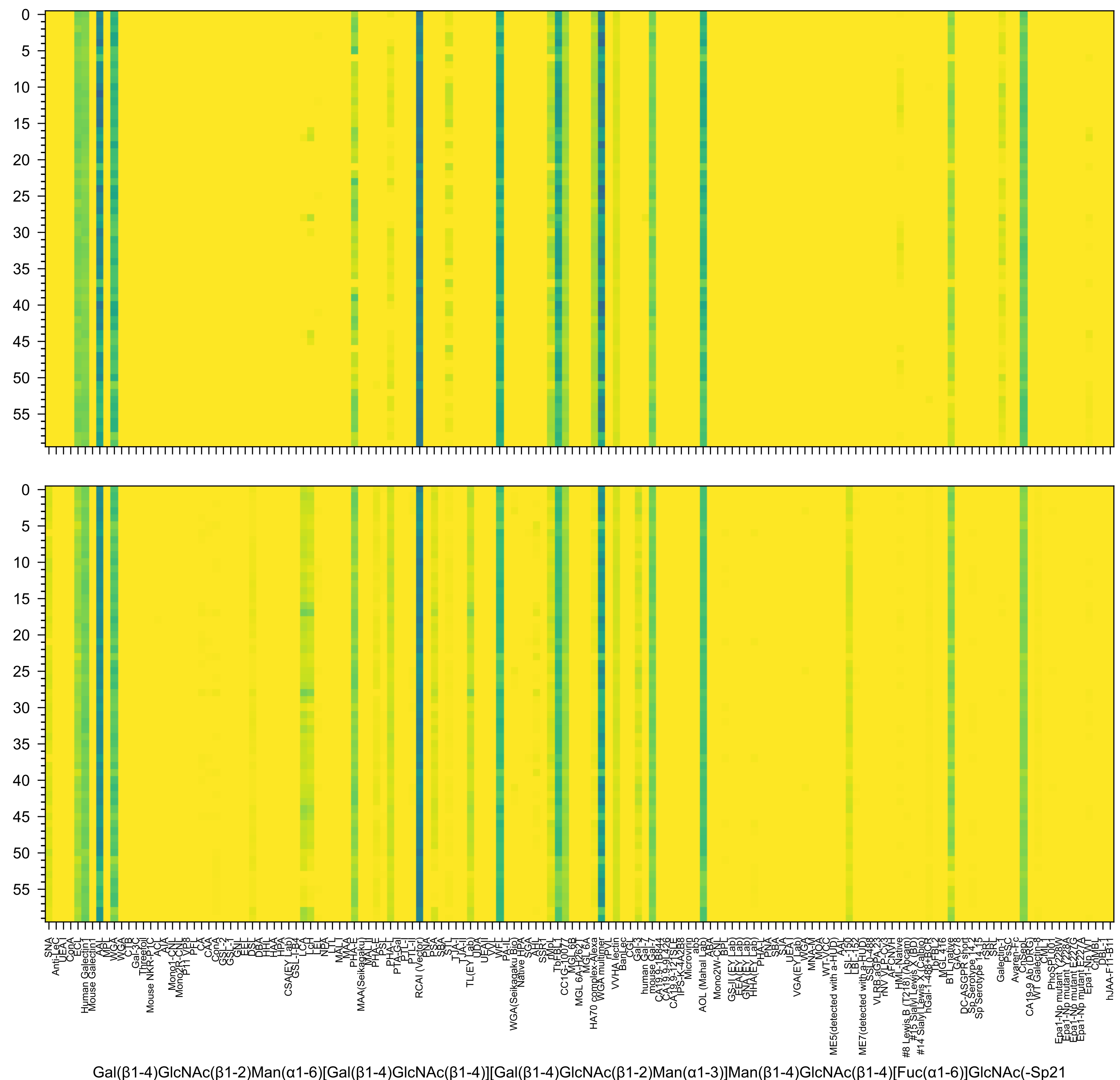
Gal(β1-4)GlcNAc(β1-2)Man(α1-6)[Gal(β1-4)GlcNAc(β1-4)][Gal(β1-4)GlcNAc(β1-2)Man(α1-3)]Man(β1-4)GlcNAc(β1-4)[Fuc(α1-6)]GlcNAc(-Sp21

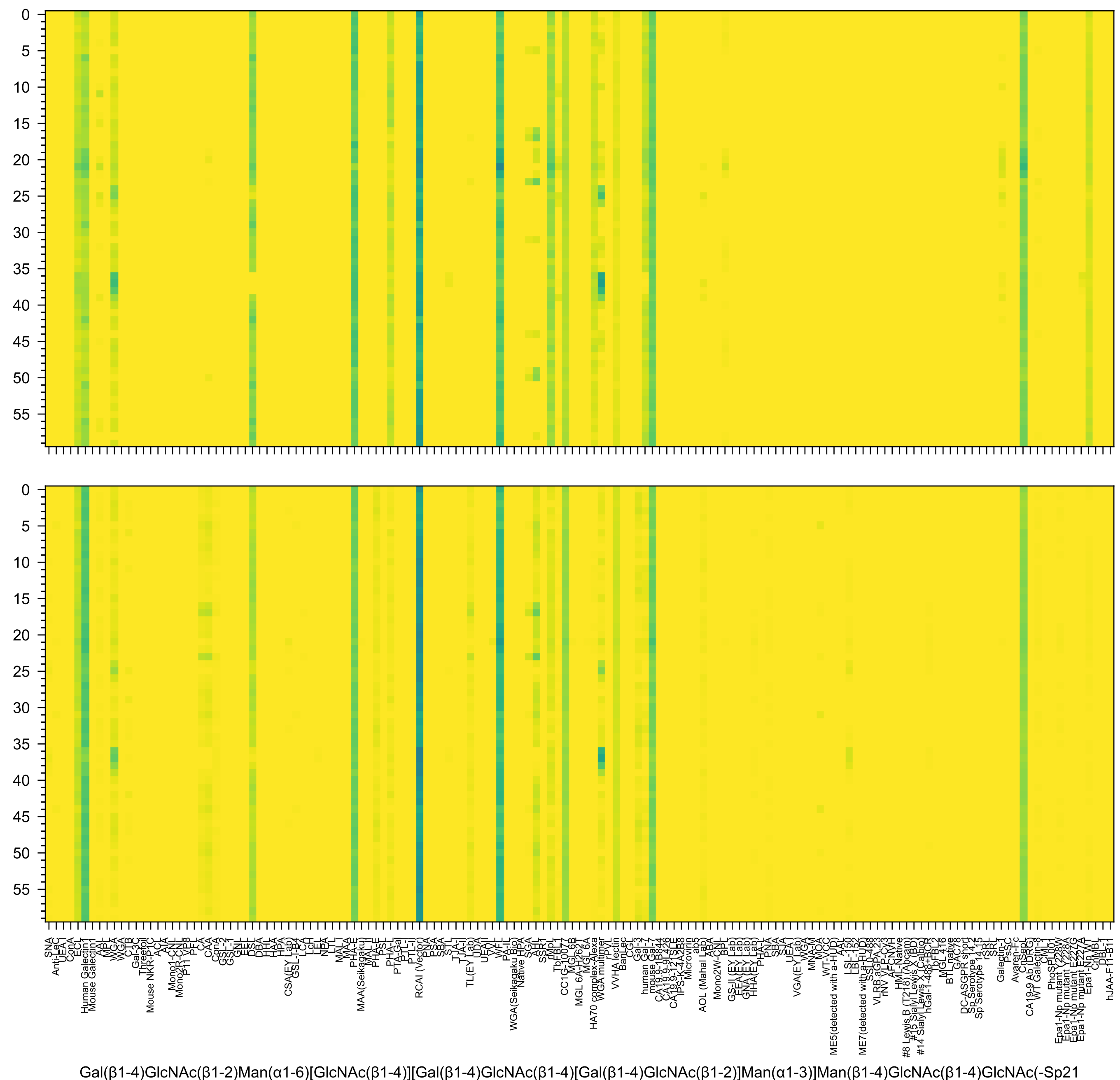
Gal(β1-4)GlcNAc(β1-2)Man(α1-6)[GlcNAc(β1-4)][Gal(β1-4)GlcNAc(β1-4)[Gal(β1-4)GlcNAc(β1-2)]Man(α1-3)]Man(β1-4)GlcNAc(β1-4)GlcNAc(-Sp21

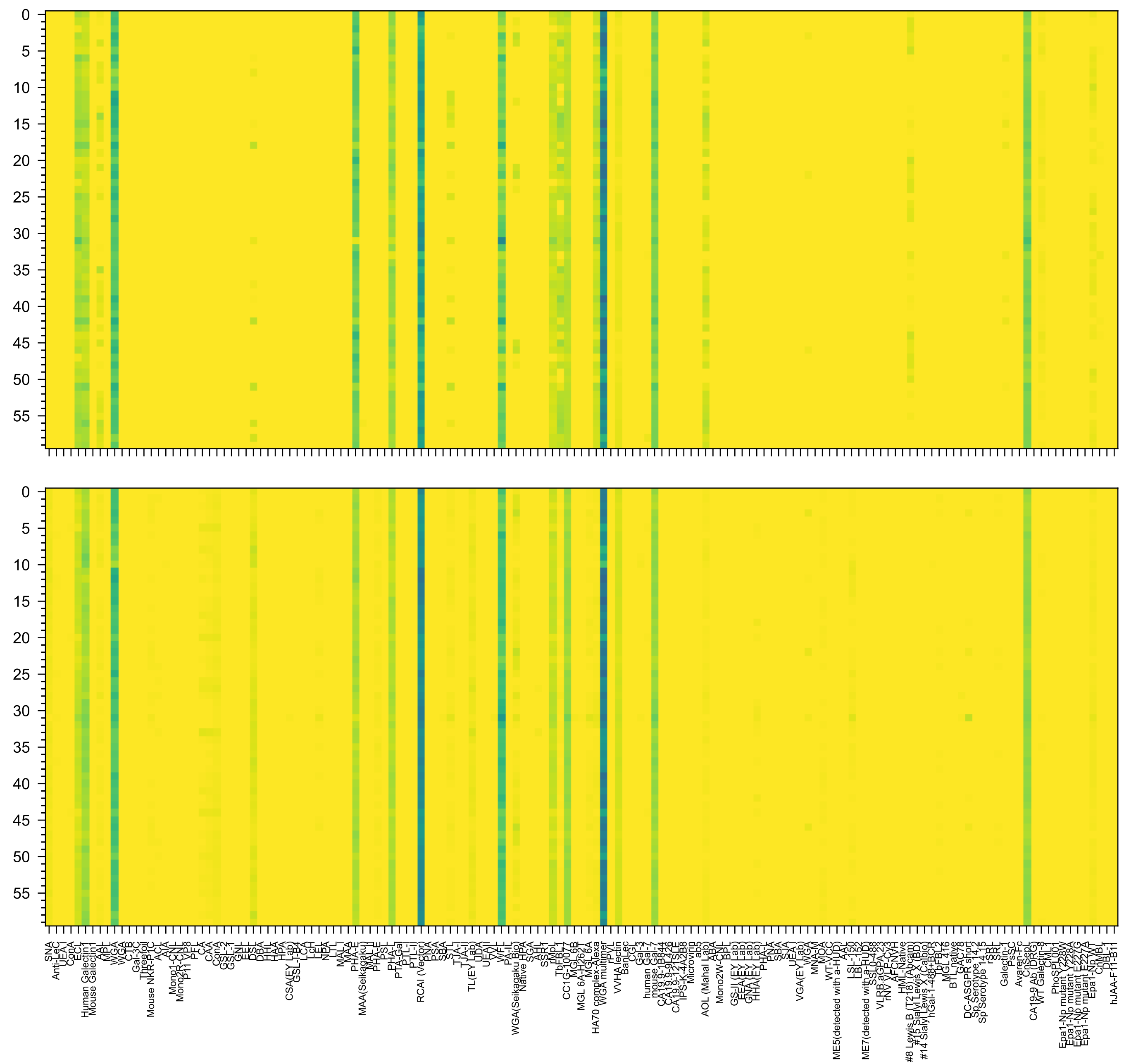

Gal(β1-4)GlcNAc(β1-6)[Gal(β1-4)GlcNAc(β1-2)]Man(α1-6)[GlcNAc(β1-4)][Gal(β1-4)GlcNAc(β1-2)Man(α1-3)]Man(β1-4)GlcNAc(β1-4)GlcNAc(-Sp21

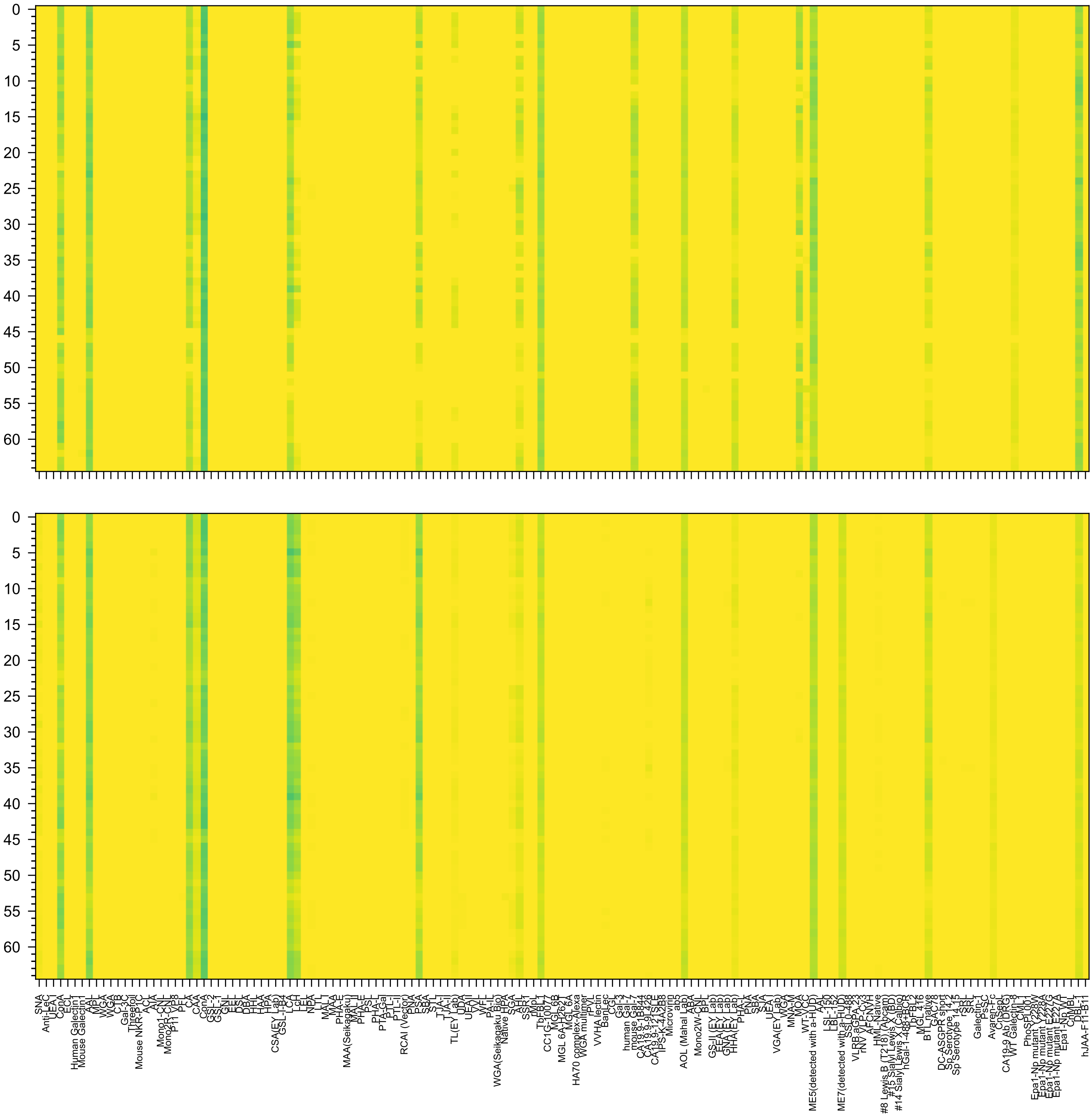

Fuc(α1-4)[Gal(β1-3)]GlcNAc(β1-2)Man(α1-6)[Fuc(α1-4)[Gal(β1-3)]GlcNAc(β1-2)Man(α1-3)]Man(β1-4)GlcNAc(β1-4)[Fuc(α1-6)]GlcNAc(β-Sp22

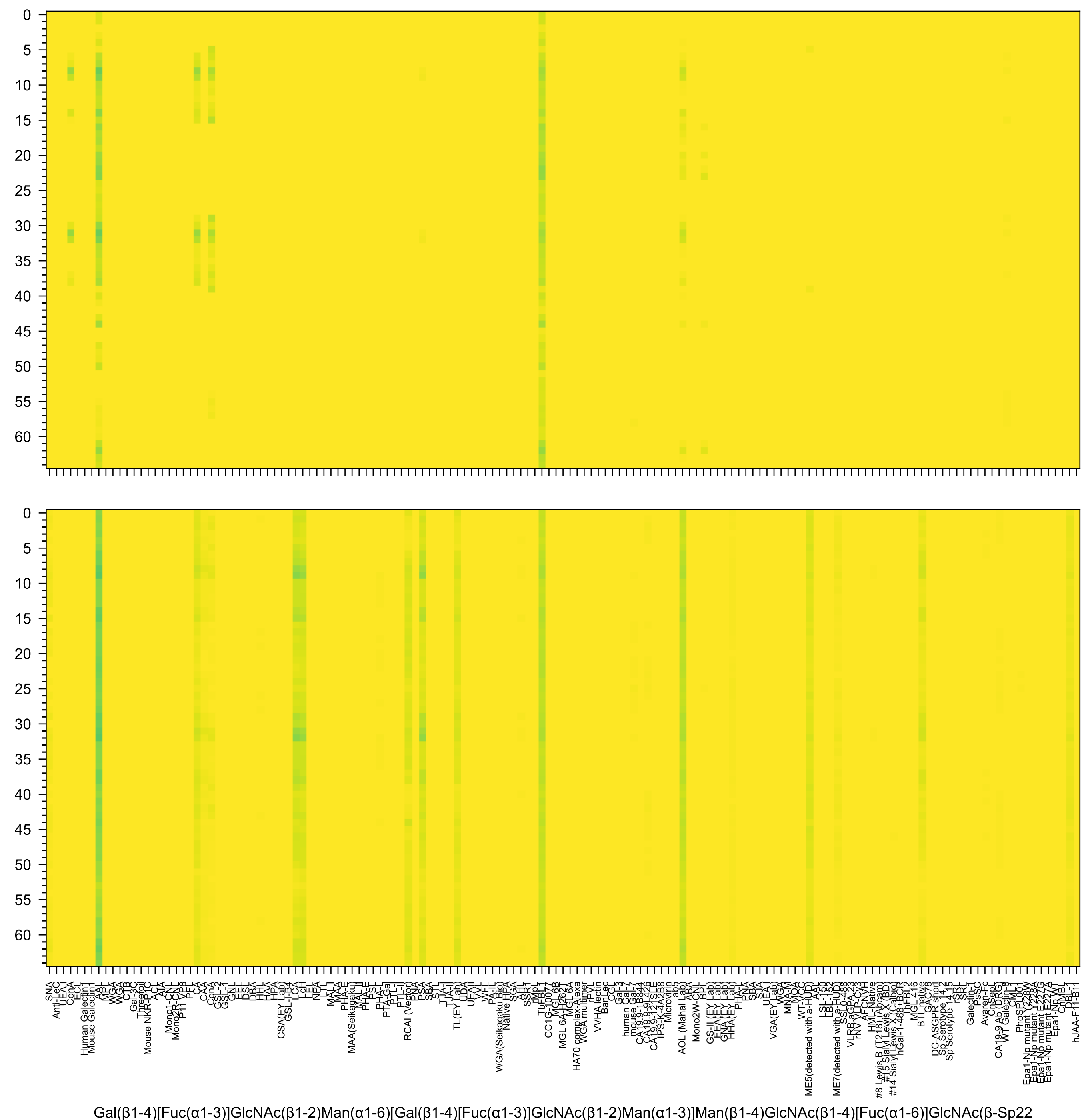
Gal(β1-4)[Fuc(α1-3)]GlcNAc(β1-2)Man(α1-6)[Gal(β1-4)[Fuc(α1-3)]GlcNAc(β1-2)Man(α1-3)]Man(β1-4)GlcNAc(β1-4)[Fuc(α1-6)]GlcNAc(β-Sp22

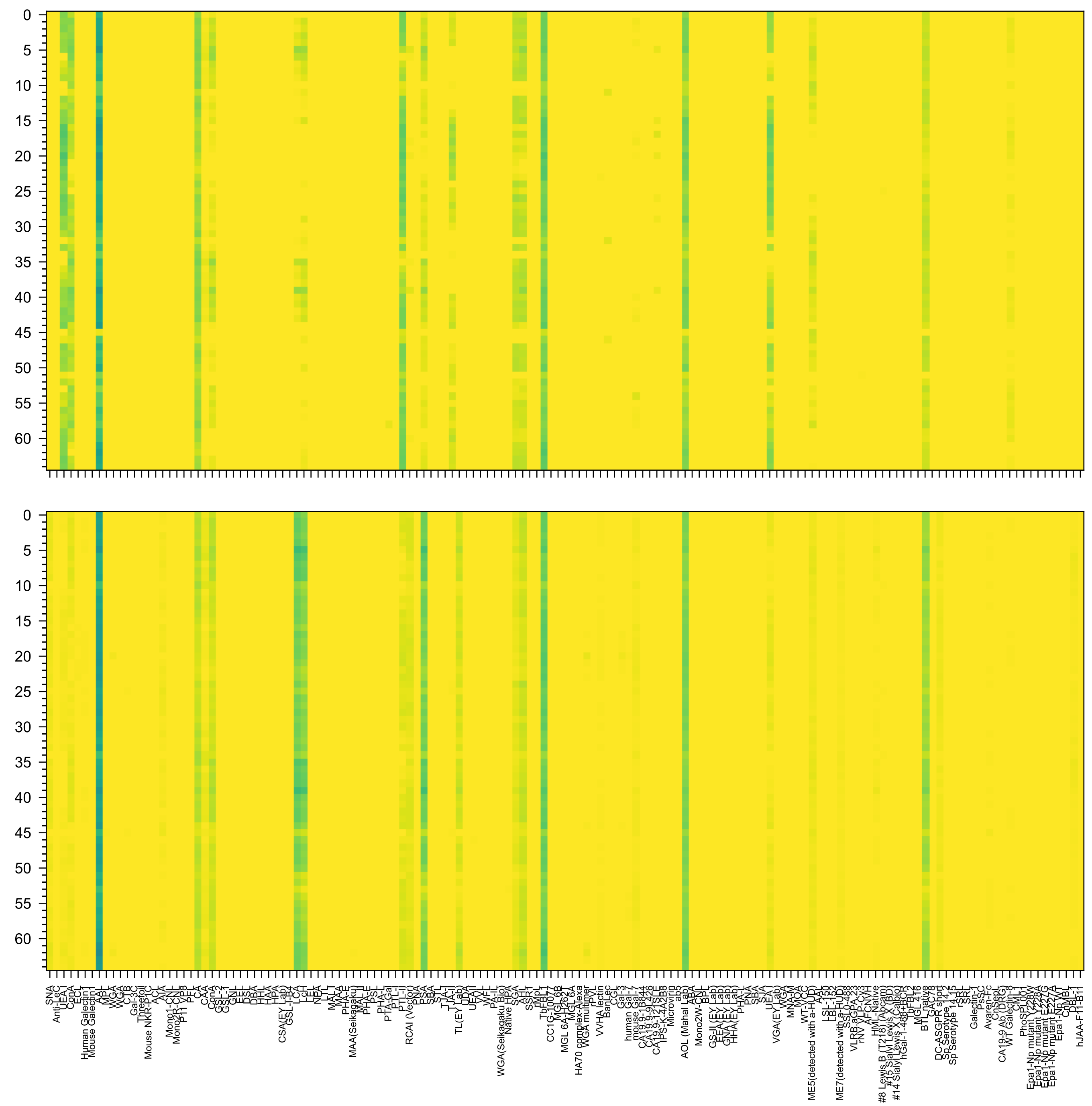

Fuc(α1-2)Gal(β1-4)GlcNAc(β1-2)Man(α1-6)[Fuc(α1-2)Gal(β1-4)GlcNAc(β1-2)Man(α1-3)]Man(β1-4)GlcNAc(β1-4)[Fuc(α1-6)]GlcNAc(β-Sp22

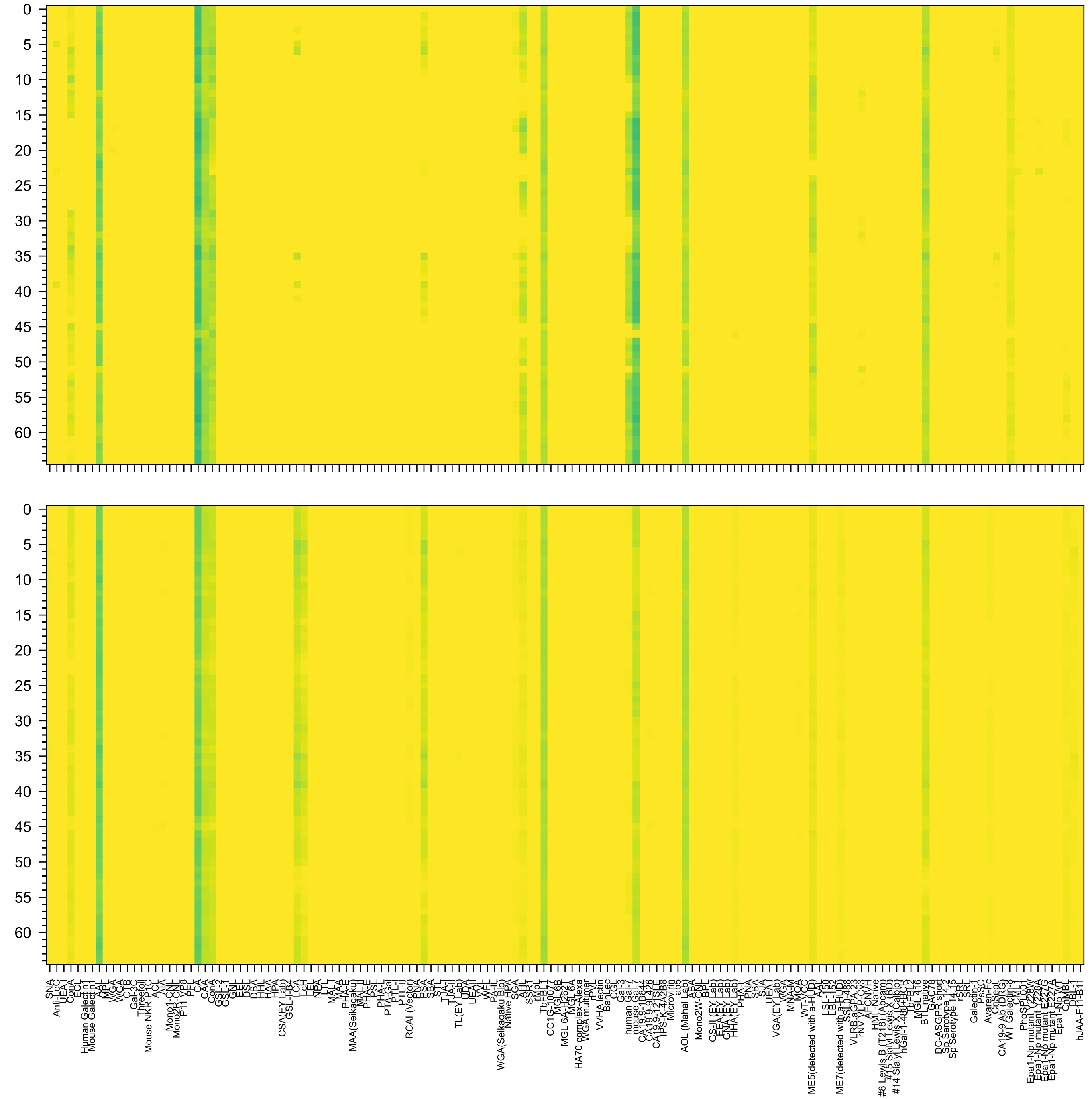

Fuc(α1-2)Gal(β1-3)GlcNAc(β1-2)Man(α1-6)[Fuc(α1-2)Gal(β1-3)GlcNAc(β1-2)Man(α1-3)]Man(β1-4)GlcNAc(β1-4)[Fuc(α1-6)]GlcNAc(β-Sp22

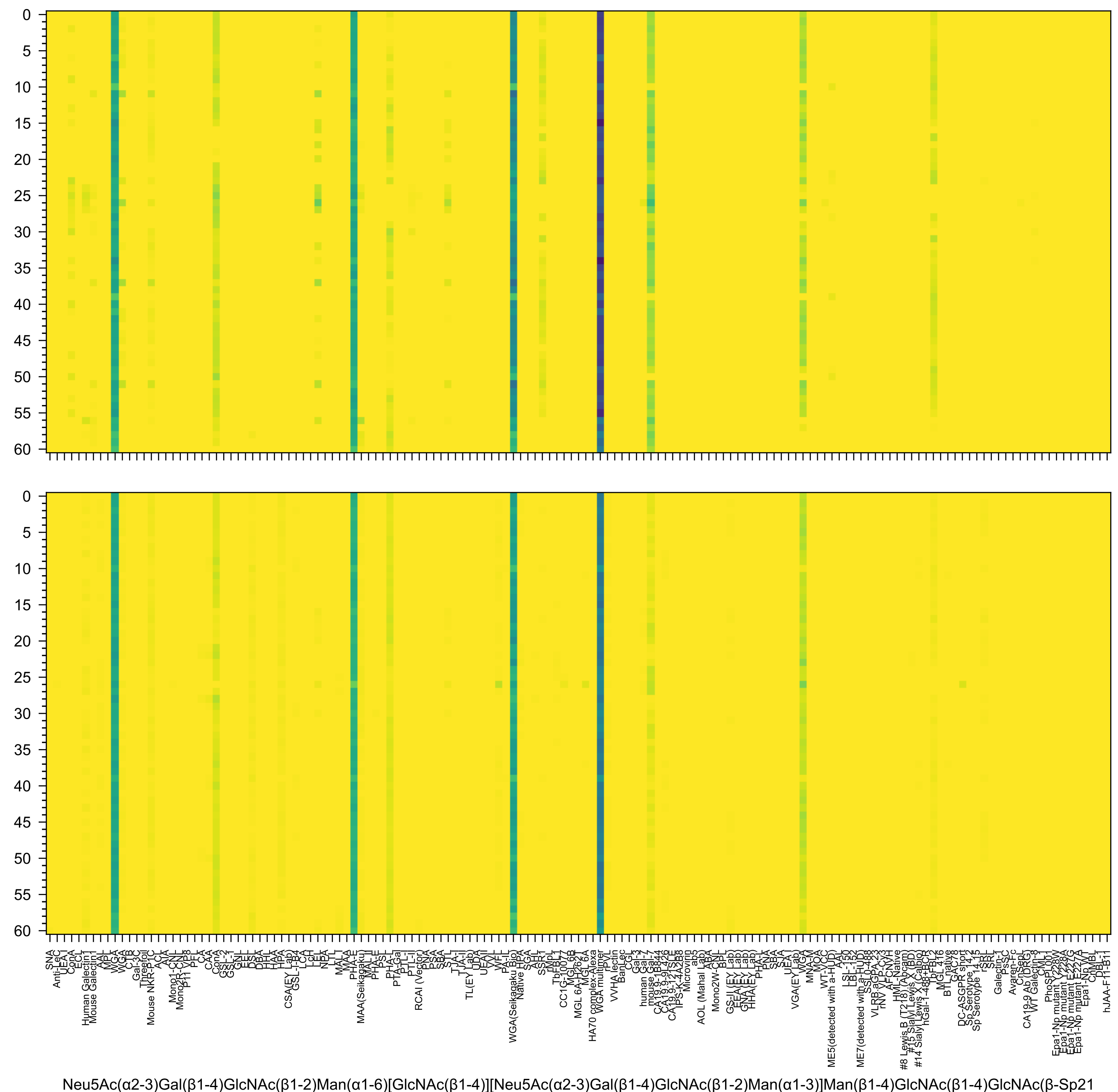
Neu5Ac(α2-3)Gal(β1-4)GlcNAc(β1-2)Man(α1-6)[GlcNAc(β1-4)][Neu5Ac(α2-3)Gal(β1-4)GlcNAc(β1-2)Man(α1-3)]Man(β1-4)GlcNAc(β1-4)GlcNAc(β-Sp21

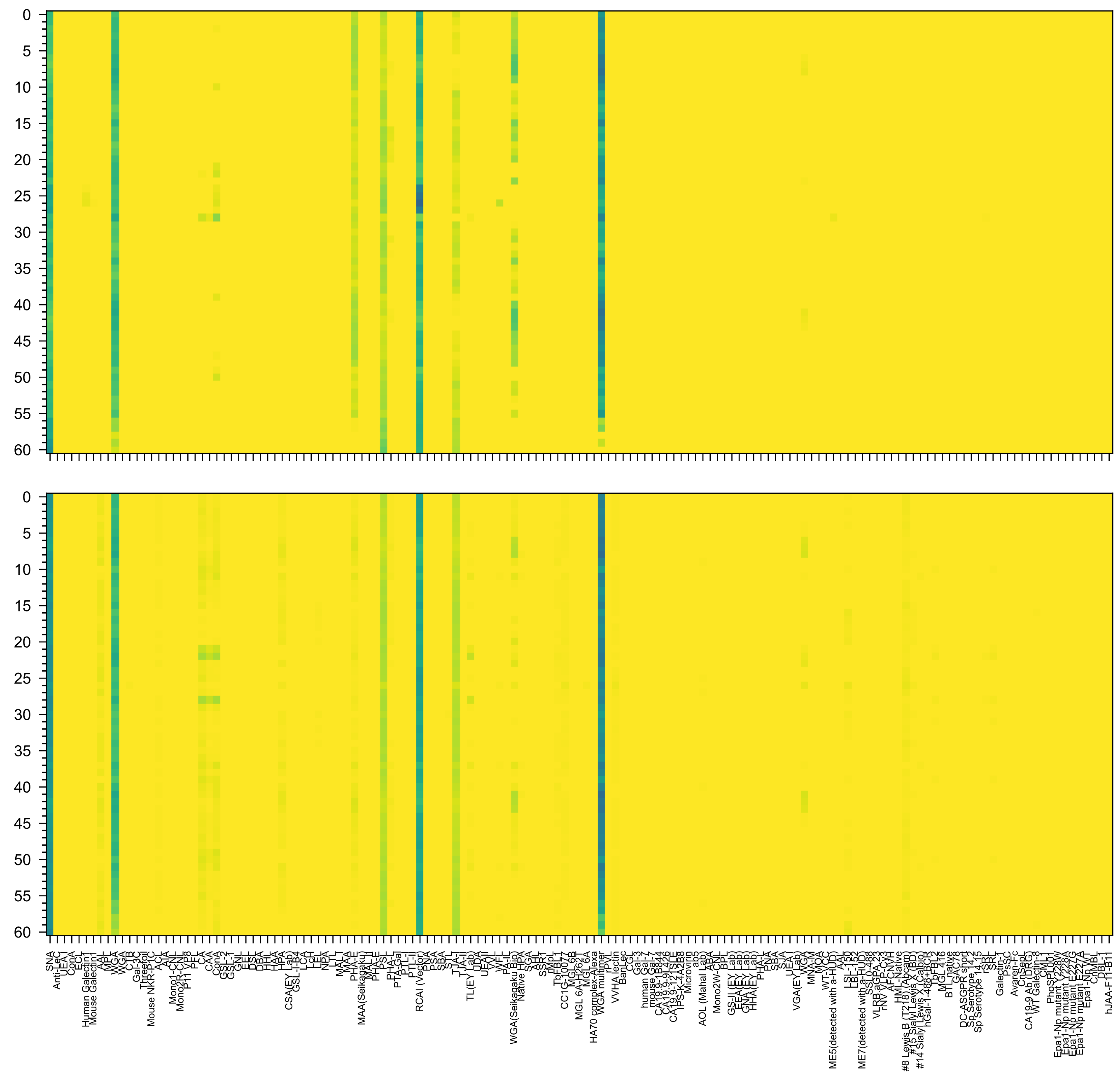

Neu5Ac(α2-6)Gal(β1-4)GlcNAc(β1-2)Man(α1-6)[GlcNAc(β1-4)][Neu5Ac(α2-6)Gal(β1-4)GlcNAc(β1-2)Man(α1-3)]Man(β1-4)GlcNAc(β1-4)GlcNAc(β-Sp21

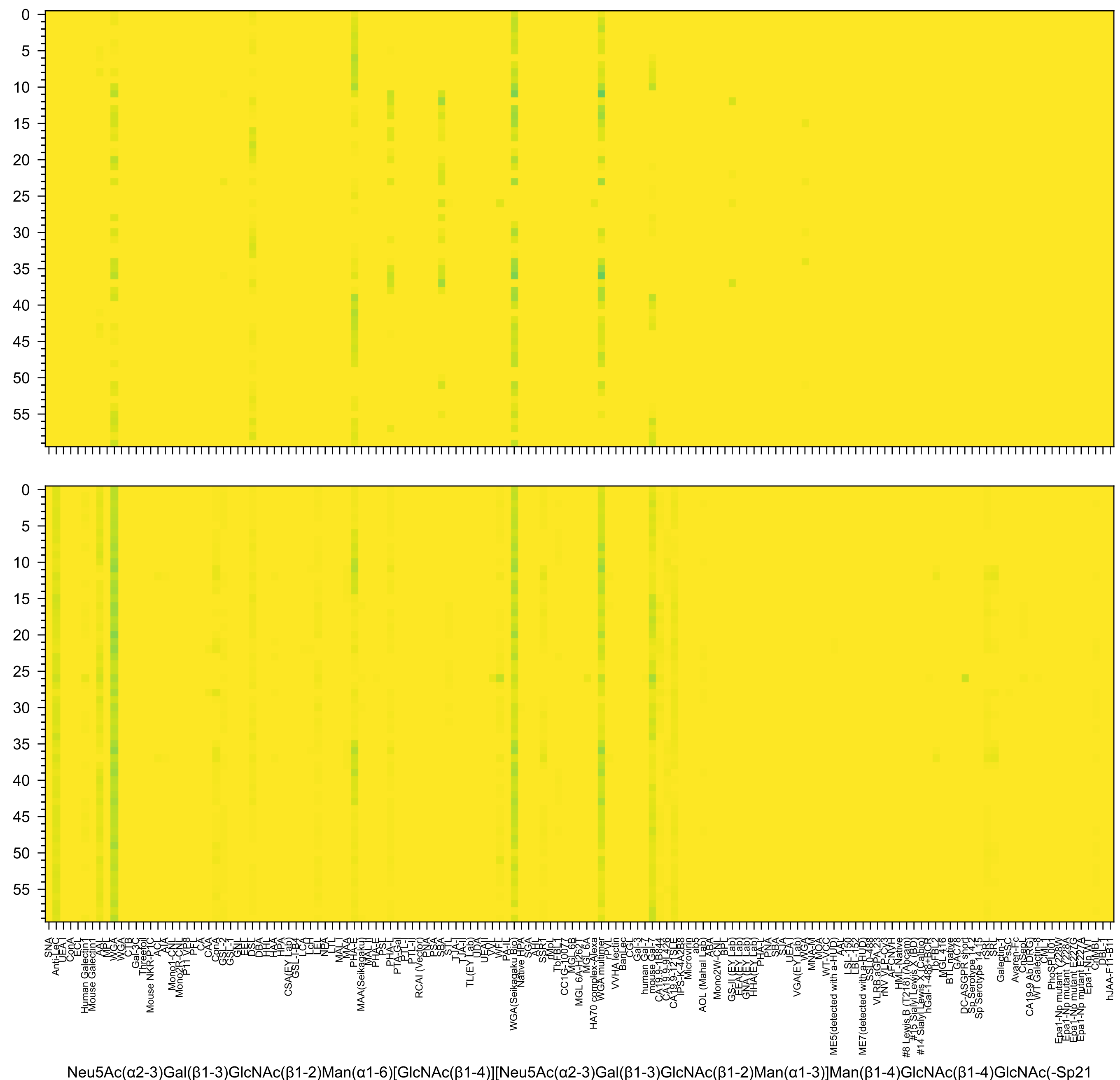
Neu5Ac(α2-3)Gal(β1-3)GlcNAc(β1-2)Man(α1-6)[GlcNAc(β1-4)][Neu5Ac(α2-3)Gal(β1-3)GlcNAc(β1-2)Man(α1-3)]Man(β1-4)GlcNAc(β1-4)GlcNAc(-Sp21

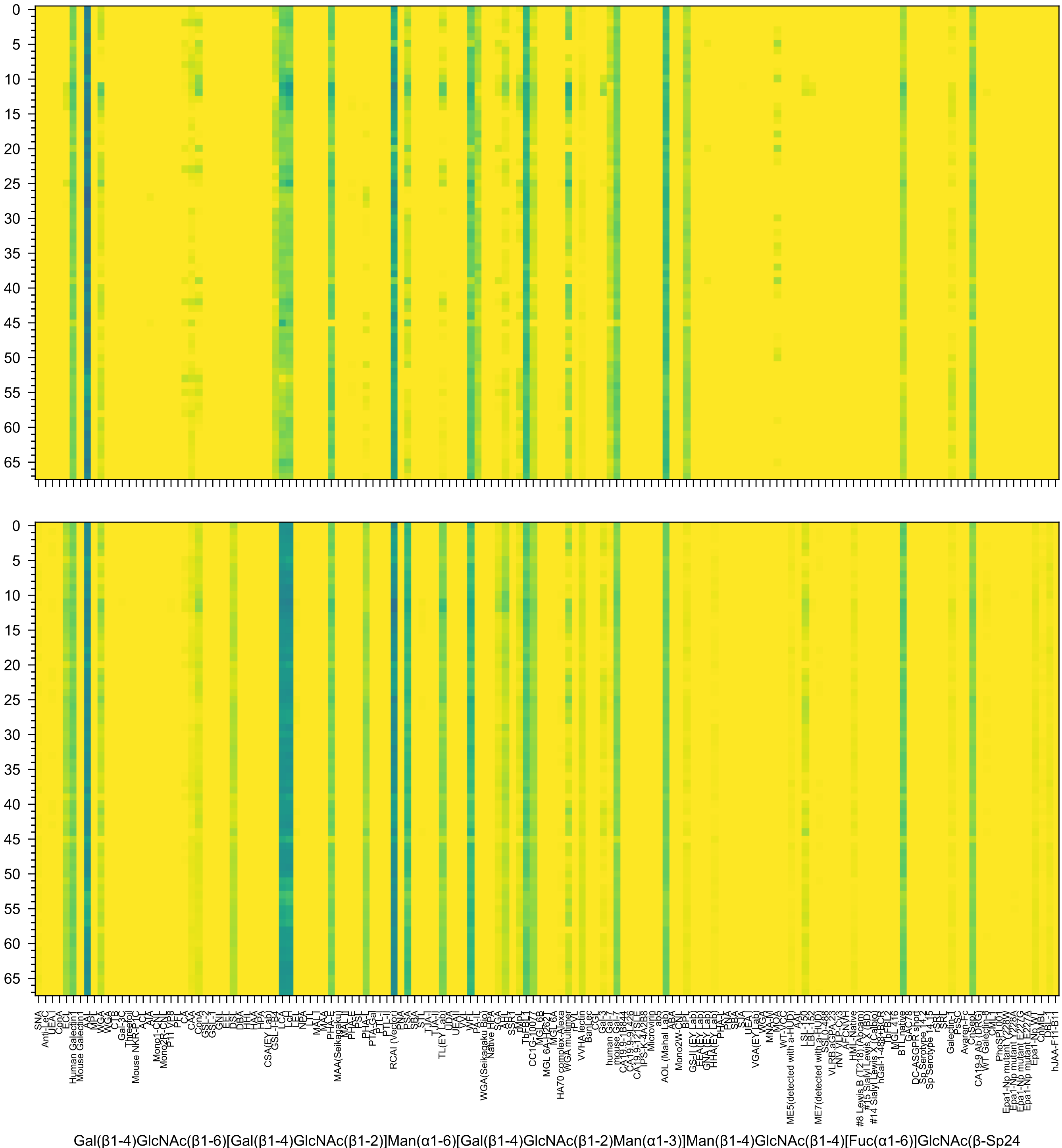
Gal(β1-4)GlcNAc(β1-6)[Gal(β1-4)GlcNAc(β1-2)]Man(α1-6)[Gal(β1-4)GlcNAc(β1-2)Man(α1-3)]Man(β1-4)GlcNAc(β1-4)[Fuc(α1-6)]GlcNAc(β-Sp24

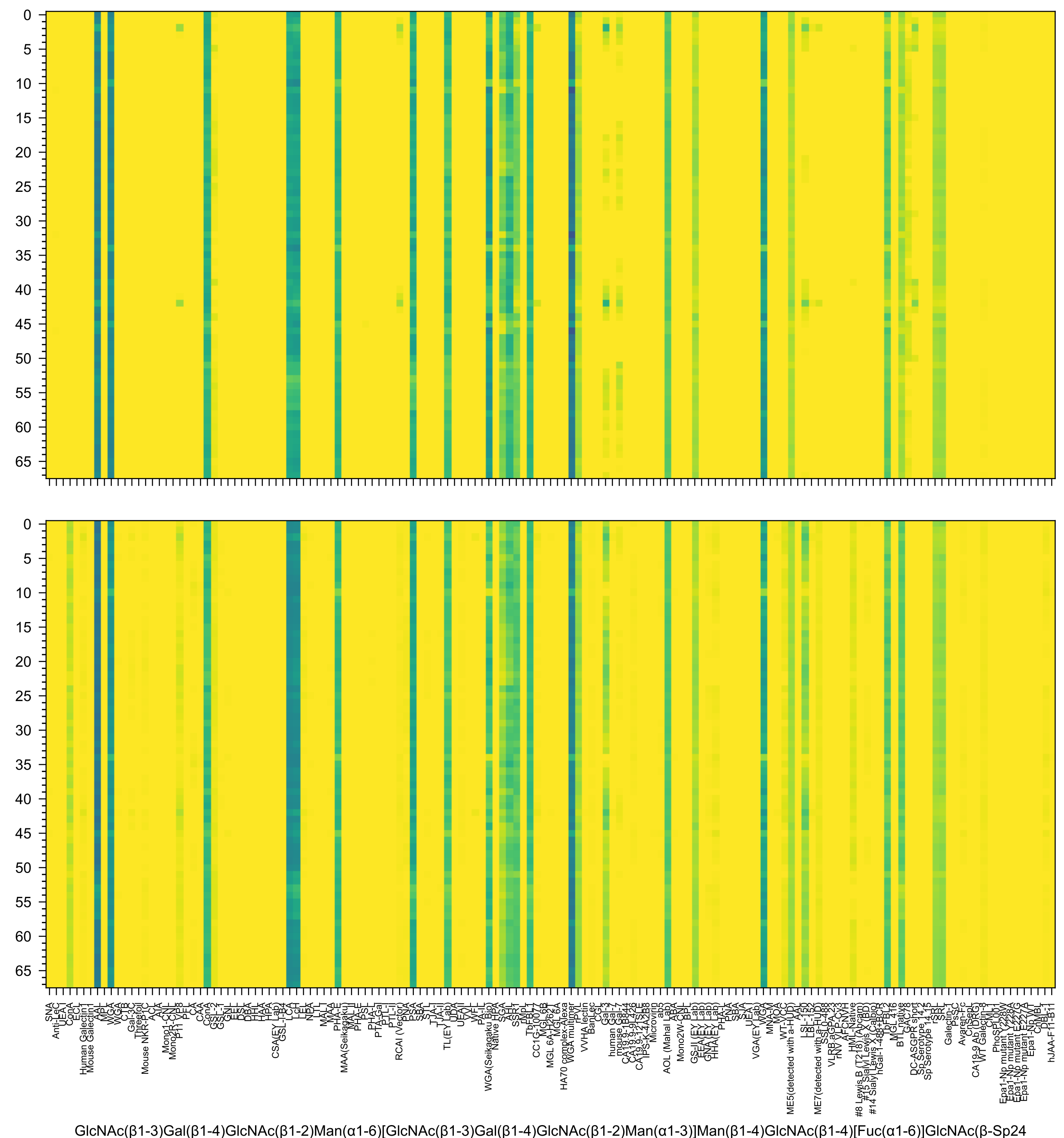
GlcNAc(β1-3)Gal(β1-4)GlcNAc(β1-2)Man(α1-6)[GlcNAc(β1-3)Gal(β1-4)GlcNAc(β1-2)Man(α1-3)]Man(β1-4)GlcNAc(β1-4)[Fuc(α1-6)]GlcNAc(β-Sp24

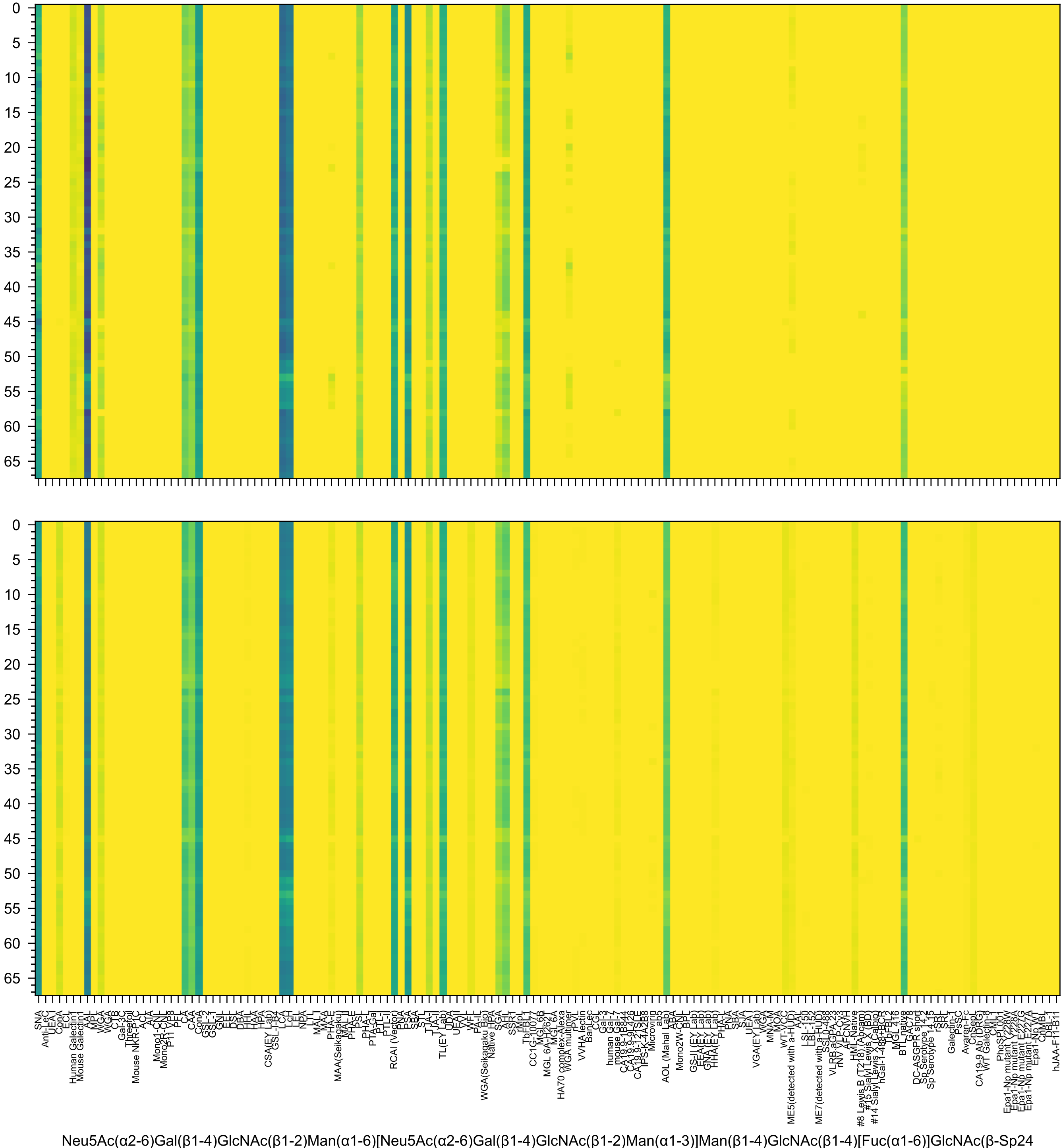
Neu5Ac(α2-6)Gal(β1-4)GlcNAc(β1-2)Man(α1-6)[Neu5Ac(α2-6)Gal(β1-4)GlcNAc(β1-2)Man(α1-3)]Man(β1-4)GlcNAc(β1-4)[Fuc(α1-6)]GlcNAc(β-Sp24

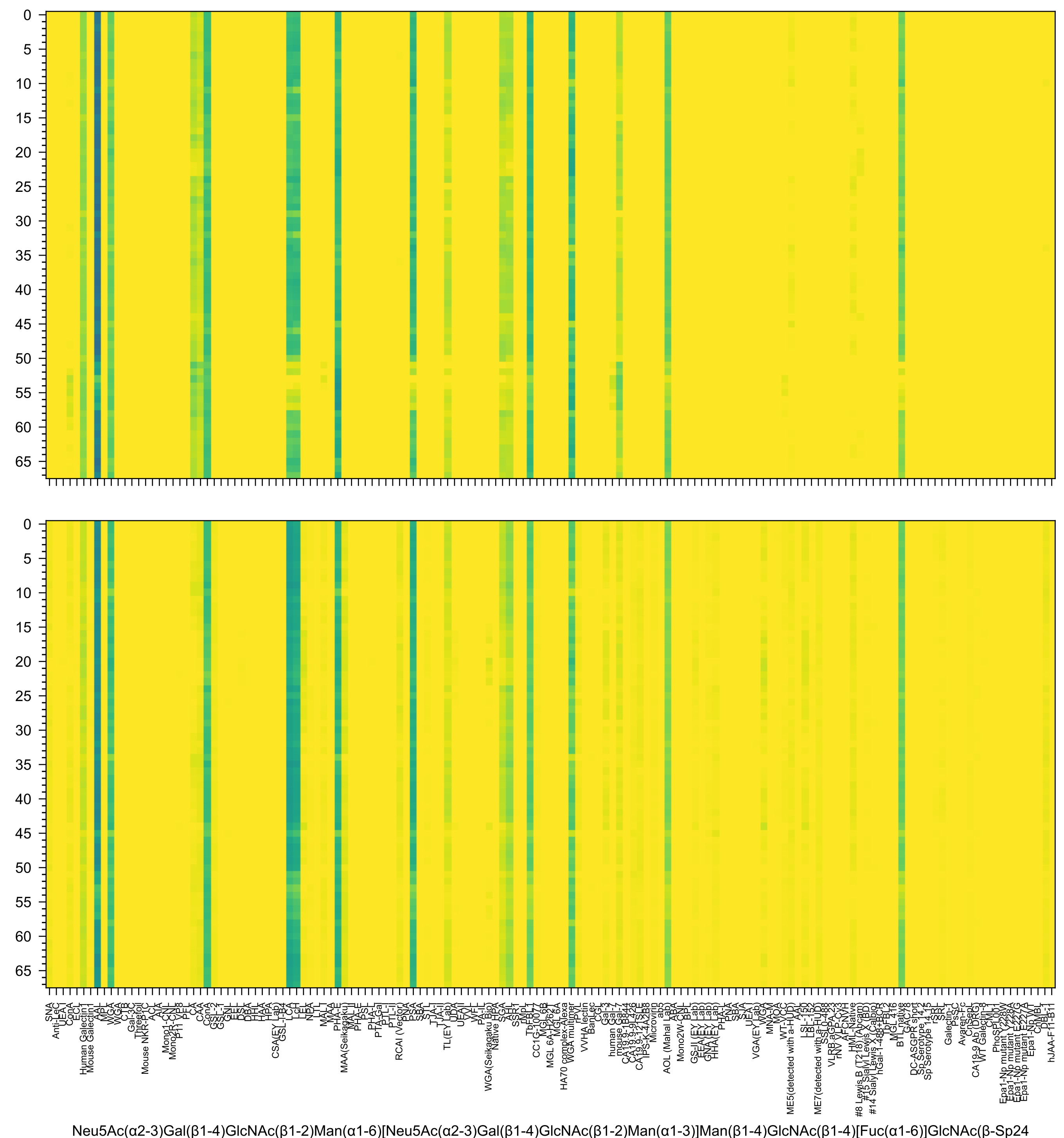
Neu5Ac(α2-3)Gal(β1-4)GlcNAc(β1-2)Man(α1-6)[Neu5Ac(α2-3)Gal(β1-4)GlcNAc(β1-2)Man(α1-3)]Man(β1-4)GlcNAc(β1-4)[Fuc(α1-6)]GlcNAc(β-Sp24

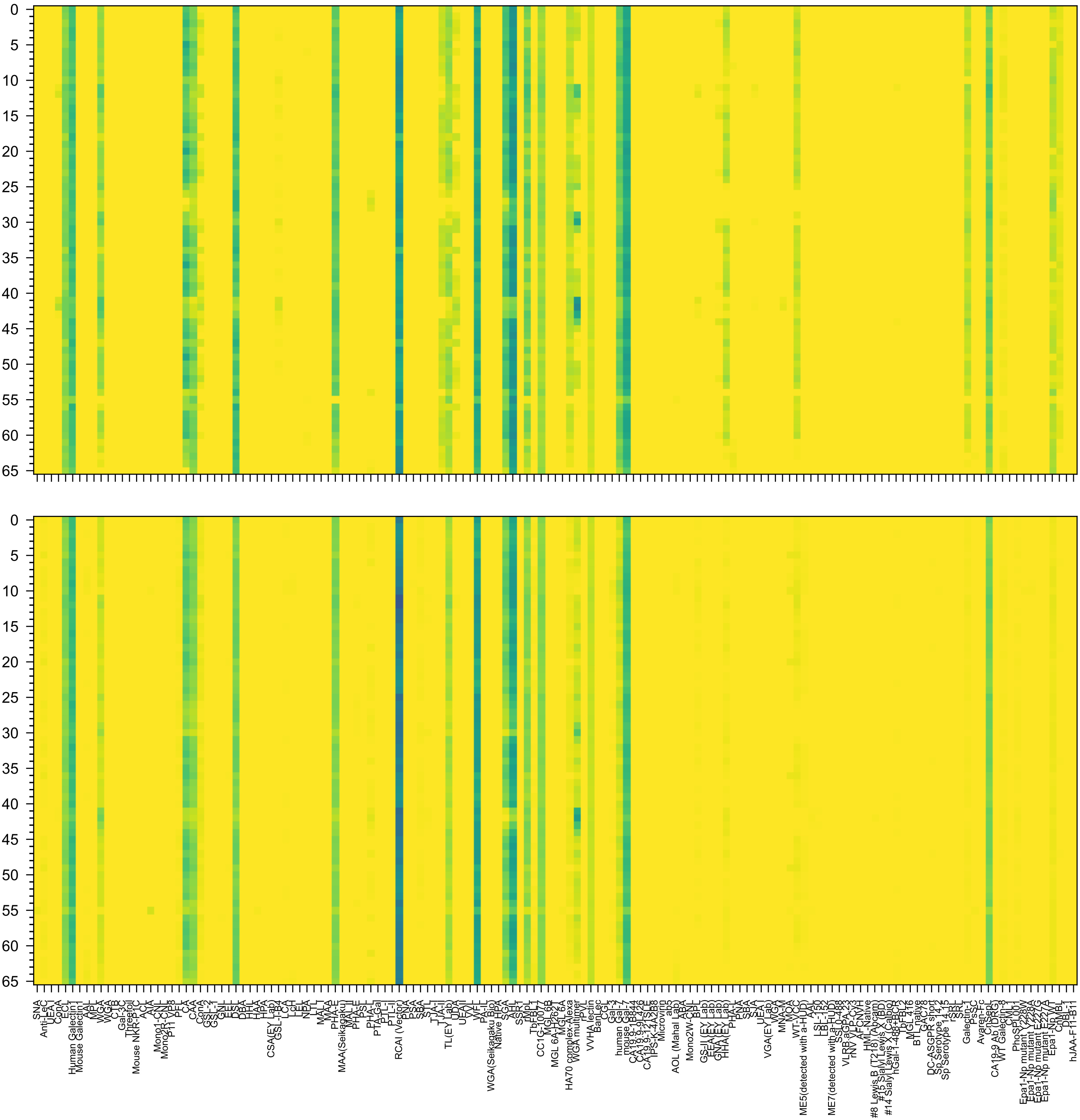

Gal(β1-4)GlcNAc(β1-6)[Gal(β1-4)GlcNAc(β1-2)]Man(α1-6)[Gal(β1-4)GlcNAc(β1-4)[Gal(β1-4)GlcNAc(β1-2)]Man(α1-3)]Man(β1-4)GlcNAc(β1-4)GlcNAc(β-Sp21

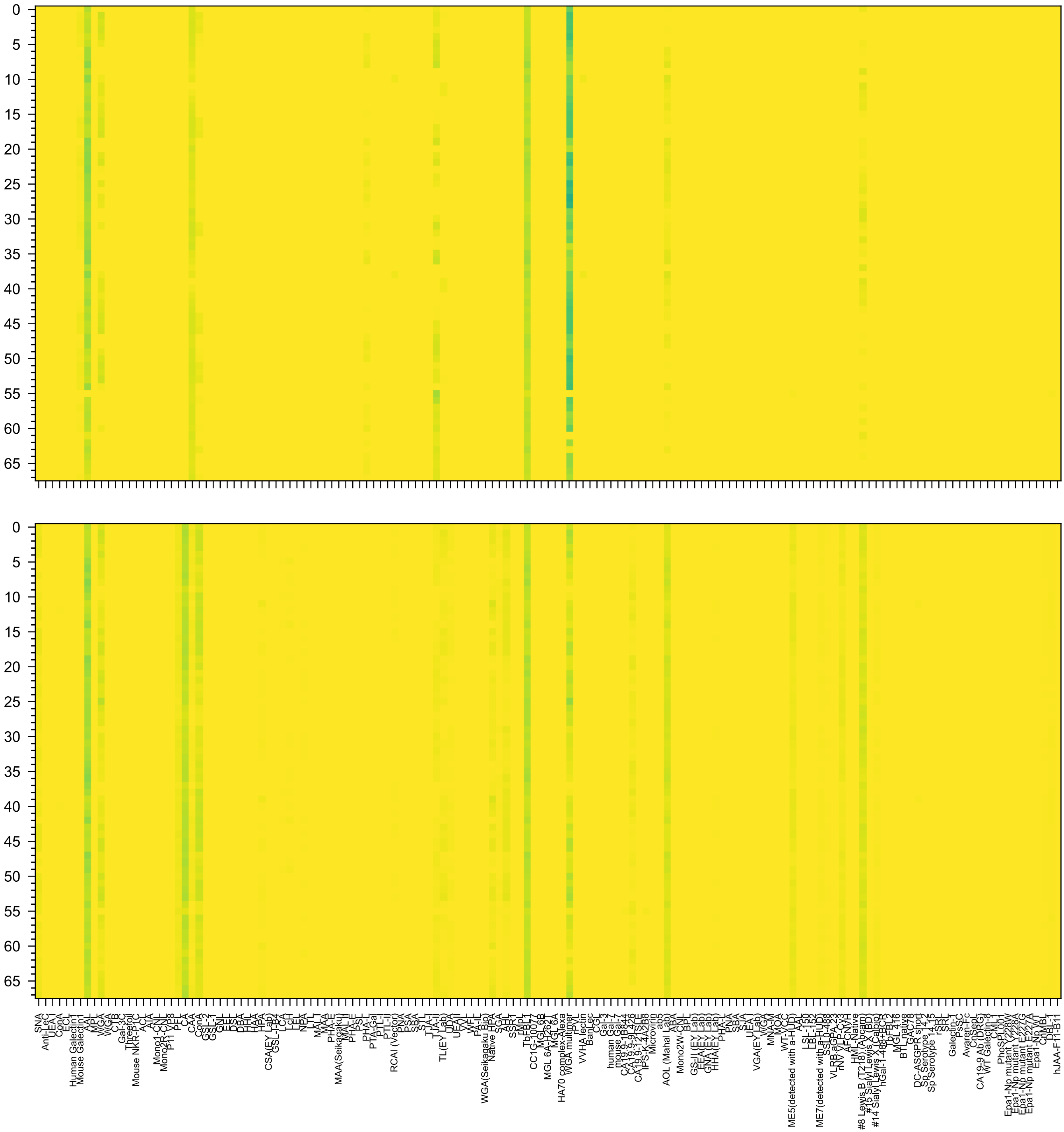

Fuc(α1-4)[Fuc(α1-2)Gal(β1-3)]GlcNAc(β1-2)Man(α1-6)[Fuc(α1-4)[Fuc(α1-2)Gal(β1-3)]GlcNAc(β1-2)Man(α1-3)]Man(β1-4)GlcNAc(β1-4)GlcNAc(β-Sp19

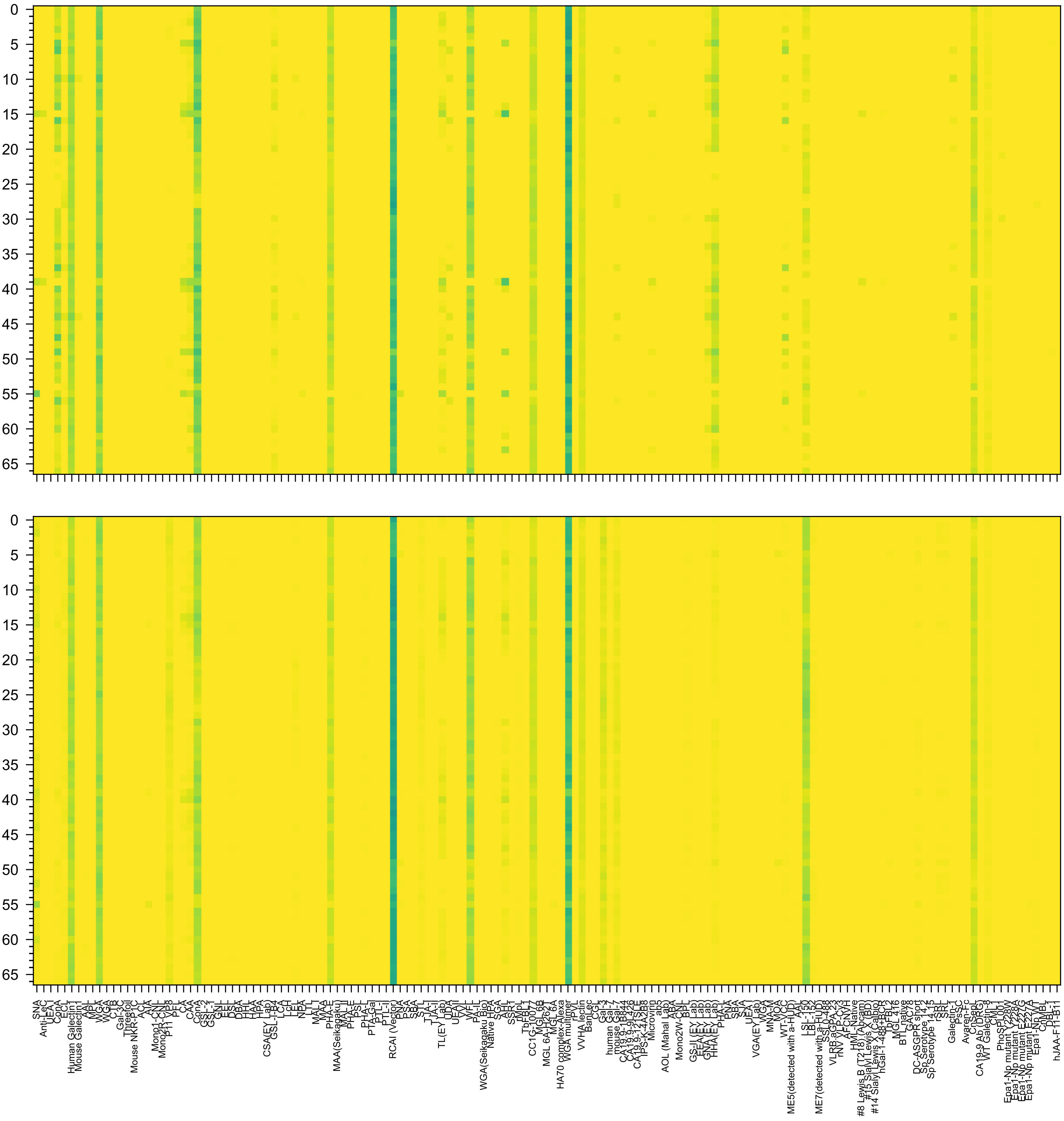

Gal(β1-4)GlcNAc(β1-3)Gal(β1-4)GlcNAc(β1-2)Man(α1-6)[Gal(β1-4)GlcNAc(β1-3)Gal(β1-4)GlcNAc(β1-2)Man(α1-3)]Man(β1-4)GlcNAc(β1-4)GlcNAc(β-Sp12

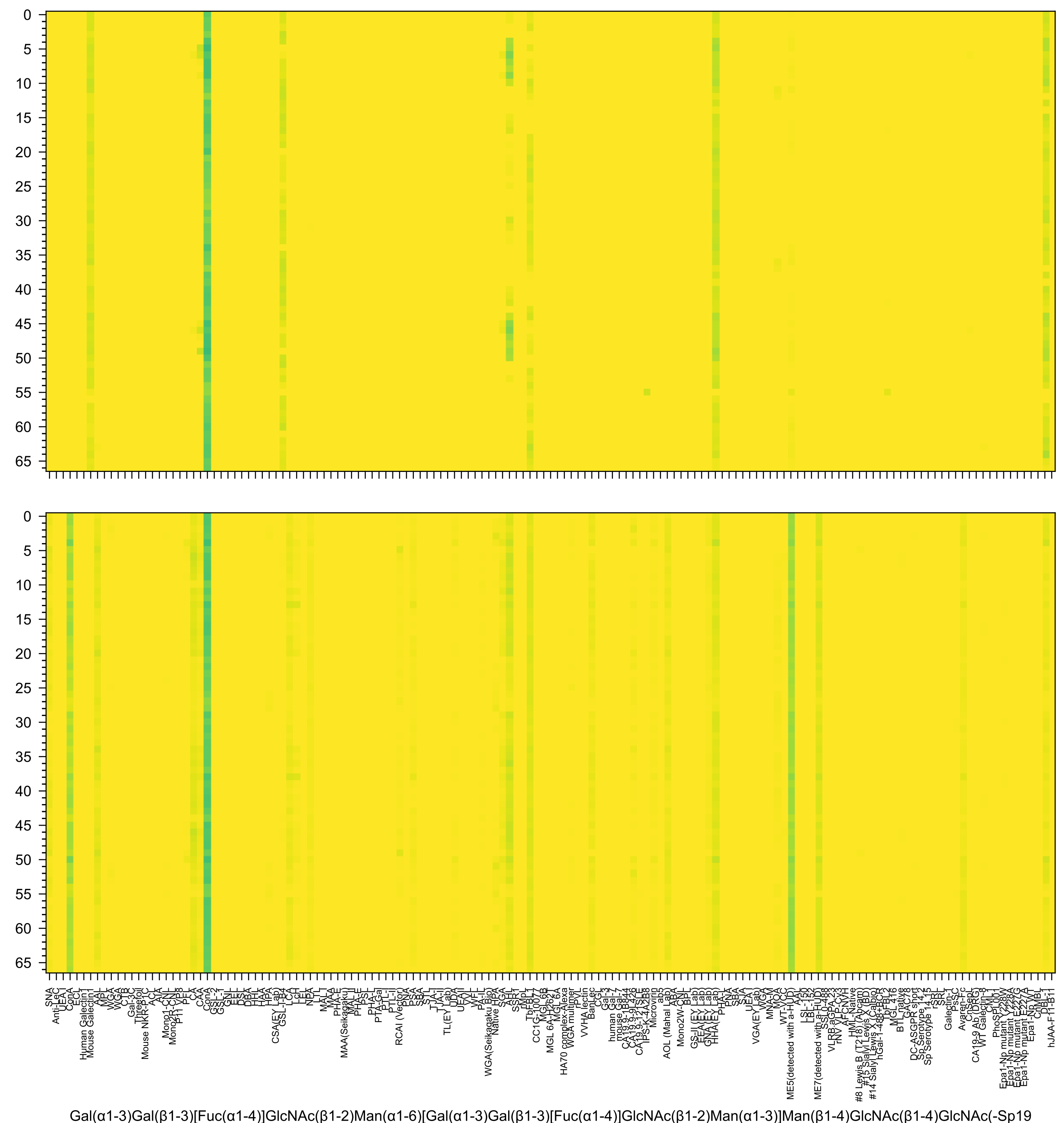
Gal(α1-3)Gal(β1-3)[Fuc(α1-4)]GlcNAc(β1-2)Man(α1-6)[Gal(α1-3)Gal(β1-3)[Fuc(α1-4)]GlcNAc(β1-2)Man(α1-3)]Man(β1-4)GlcNAc(β1-4)GlcNAc(-Sp19

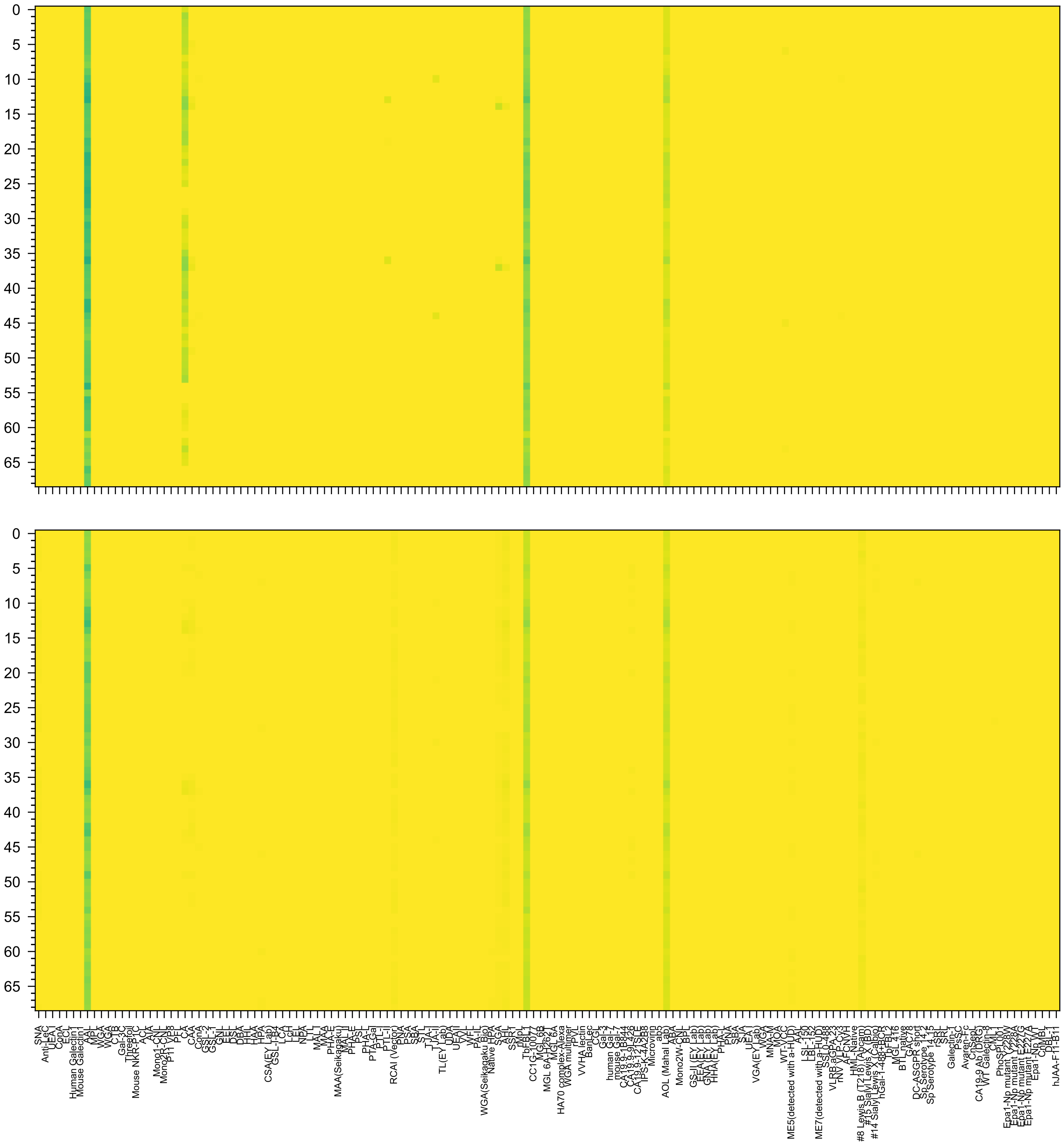

Fuc(α1-2)Gal(β1-4)[Fuc(α1-3)]GlcNAc(β1-2)Man(α1-6)[Fuc(α1-2)Gal(β1-4)[Fuc(α1-3)]GlcNAc(β1-2)Man(α1-3)]Man(β1-4)GlcNAc(β1-4)GlcNAc(β-Sp20

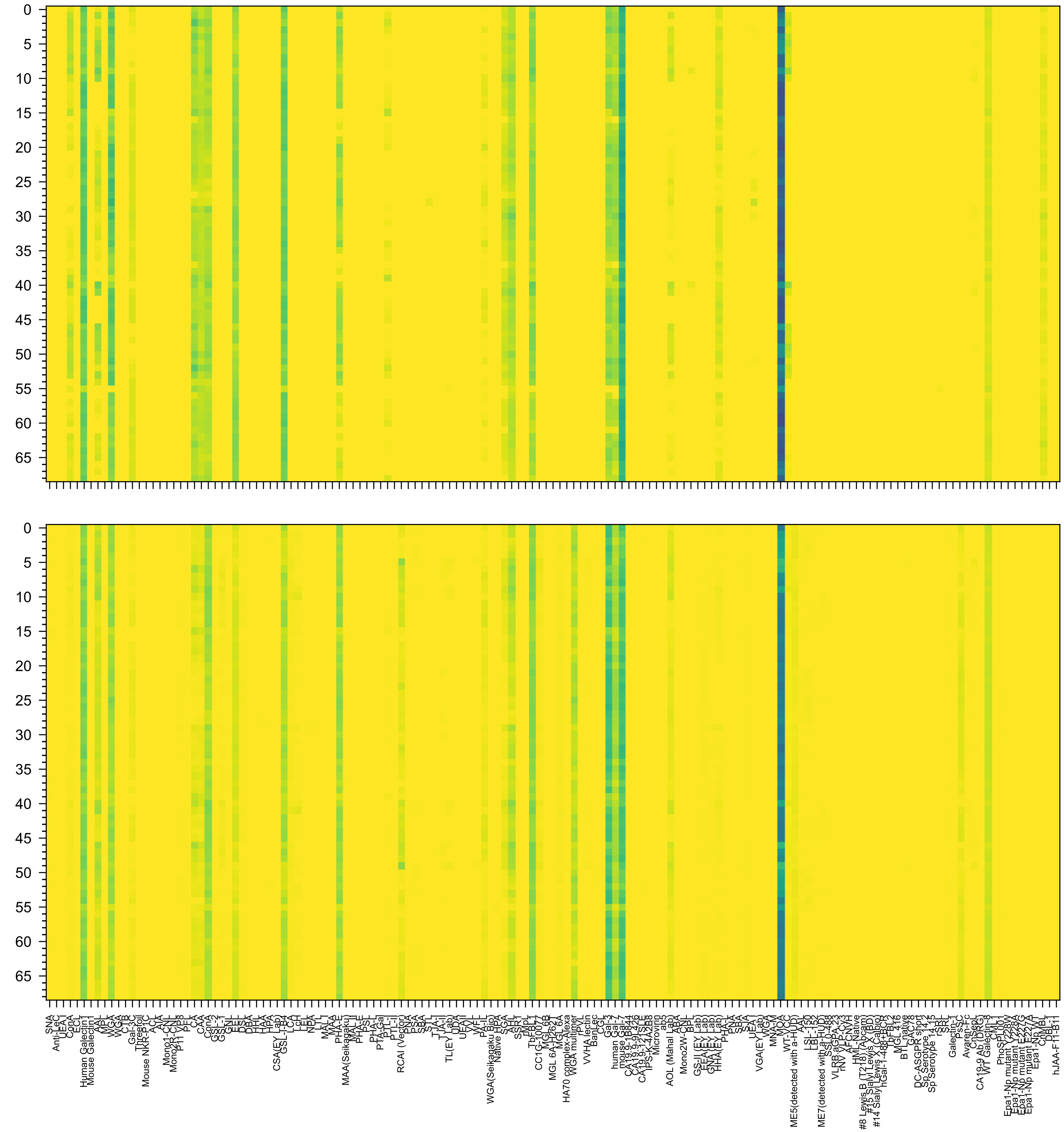

Gal(α1-3)[Fuc(α1-2)]Gal(β1-4)GlcNAc(β1-2)Man(α1-6)[Gal(α1-3)[Fuc(α1-2)]Gal(β1-4)GlcNAc(β1-2)Man(α1-3)]Man(β1-4)GlcNAc(β1-4)GlcNAc(β-Sp20

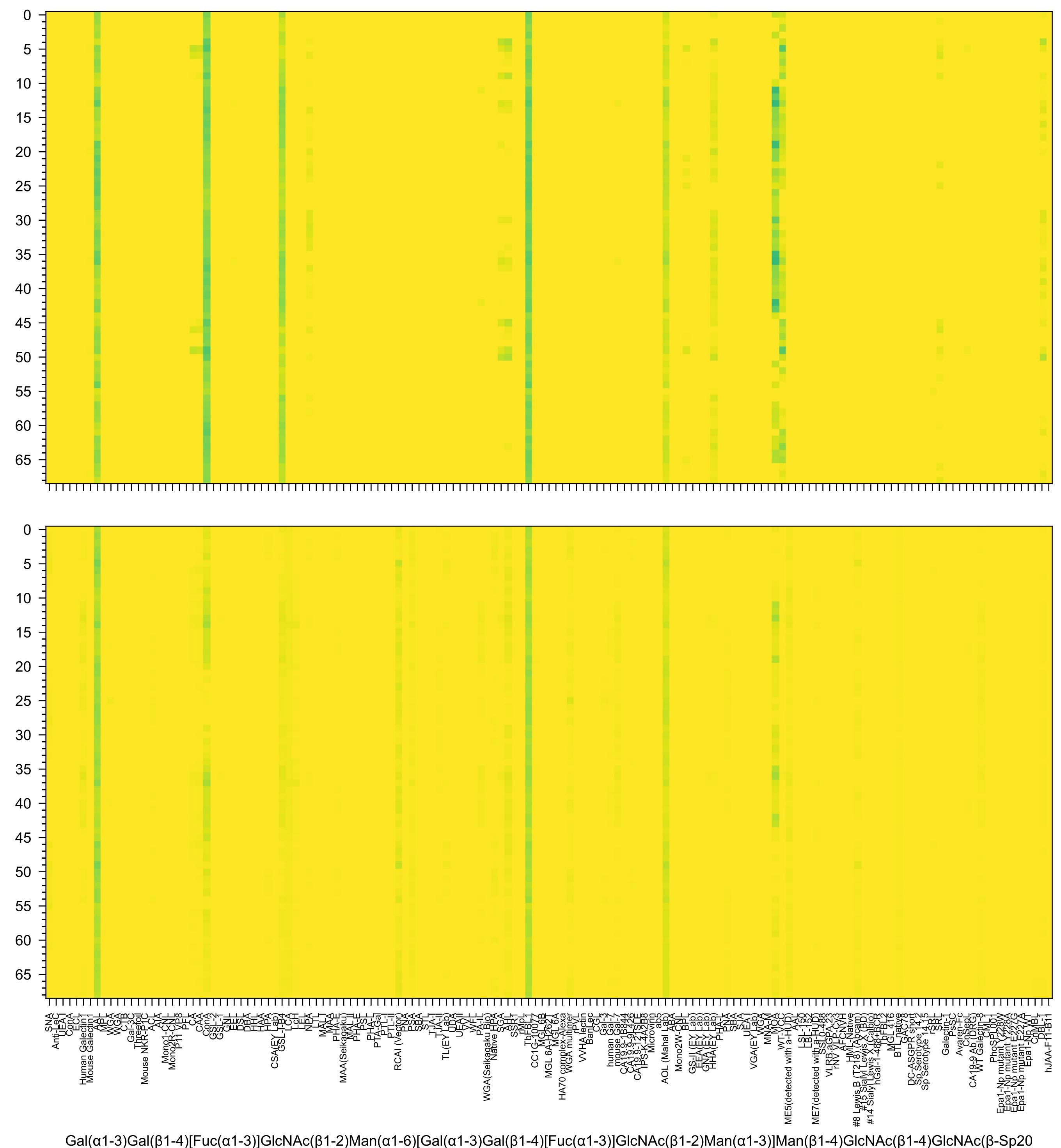
Gal(α1-3)Gal(β1-4)[Fuc(α1-3)]GlcNAc(β1-2)Man(α1-6)[Gal(α1-3)Gal(β1-4)[Fuc(α1-3)]GlcNAc(β1-2)Man(α1-3)]Man(β1-4)GlcNAc(β1-4)GlcNAc(β-Sp20

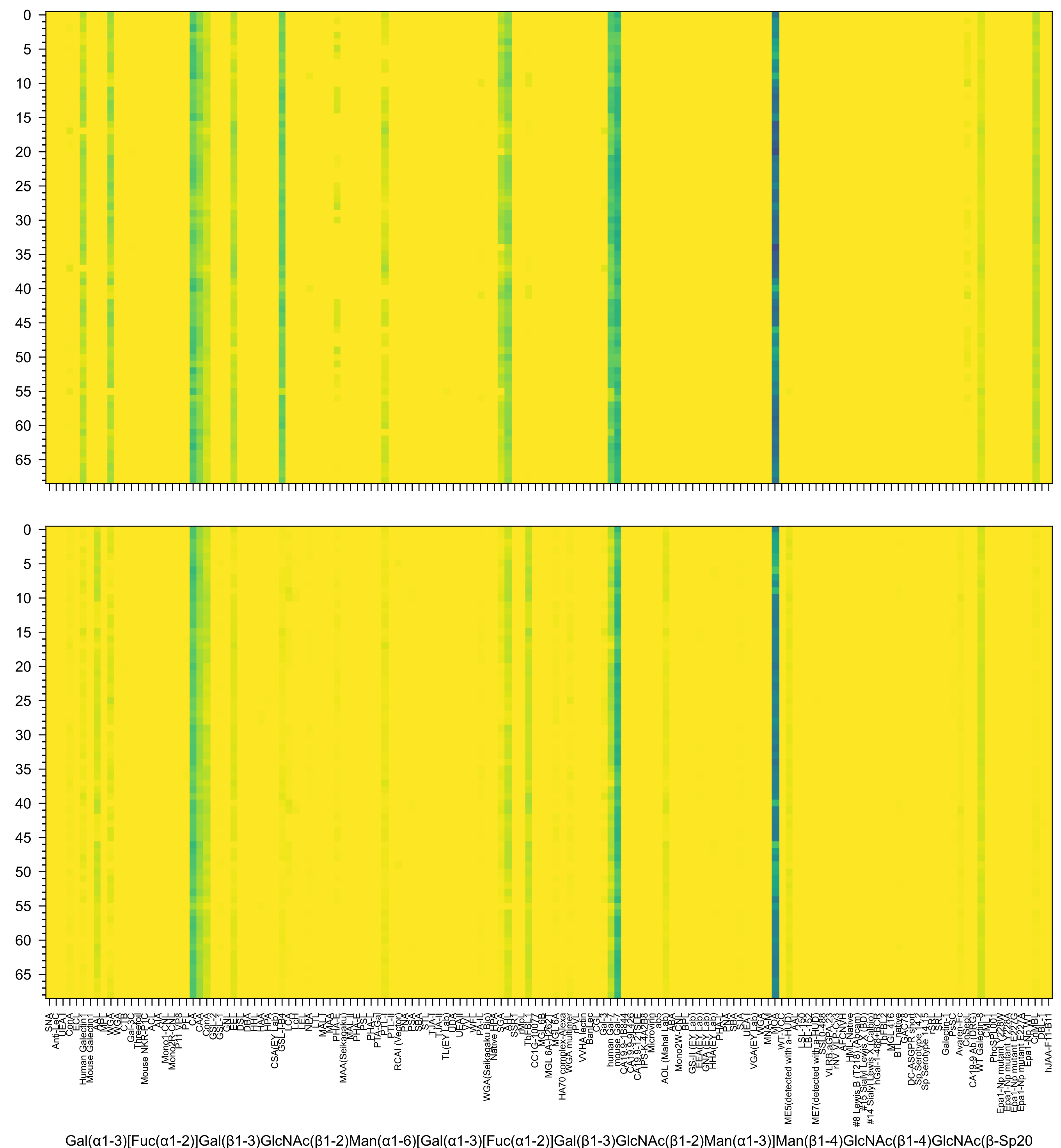

Gal(α1-3)[Fuc(α1-2)]Gal(β1-3)GlcNAc(β1-2)Man(α1-6)[Gal(α1-3)[Fuc(α1-2)]Gal(β1-3)GlcNAc(β1-2)Man(α1-3)]Man(β1-4)GlcNAc(β1-4)GlcNAc(β-Sp20

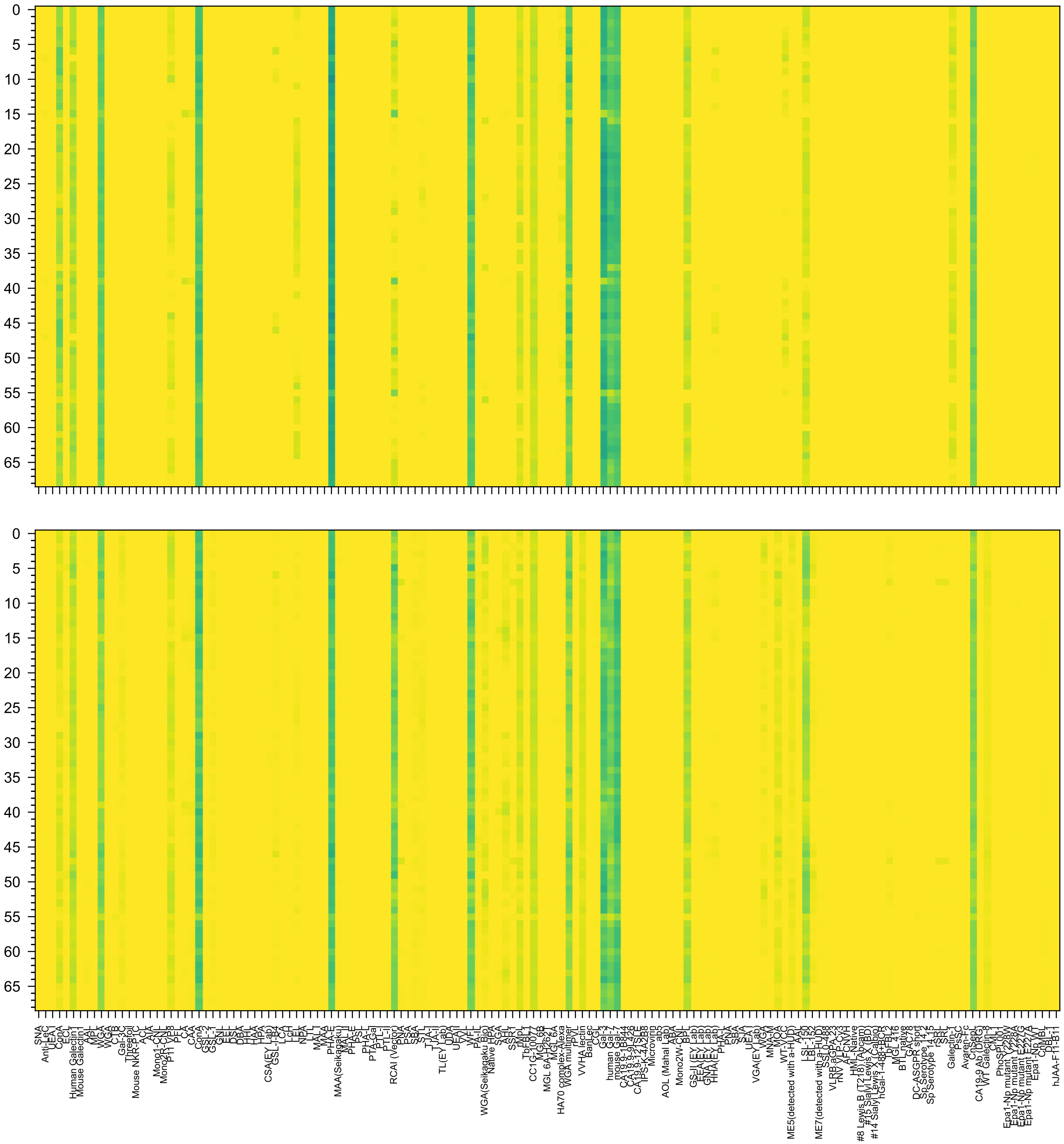

Gal(β1-3)GlcNAc(β1-3)Gal(β1-4)GlcNAc(β1-2)Man(α1-6)[Gal(β1-3)GlcNAc(β1-3)Gal(β1-4)GlcNAc(β1-2)Man(α1-3)]Man(β1-4)GlcNAc(β1-4)GlcNAc(-Sp25

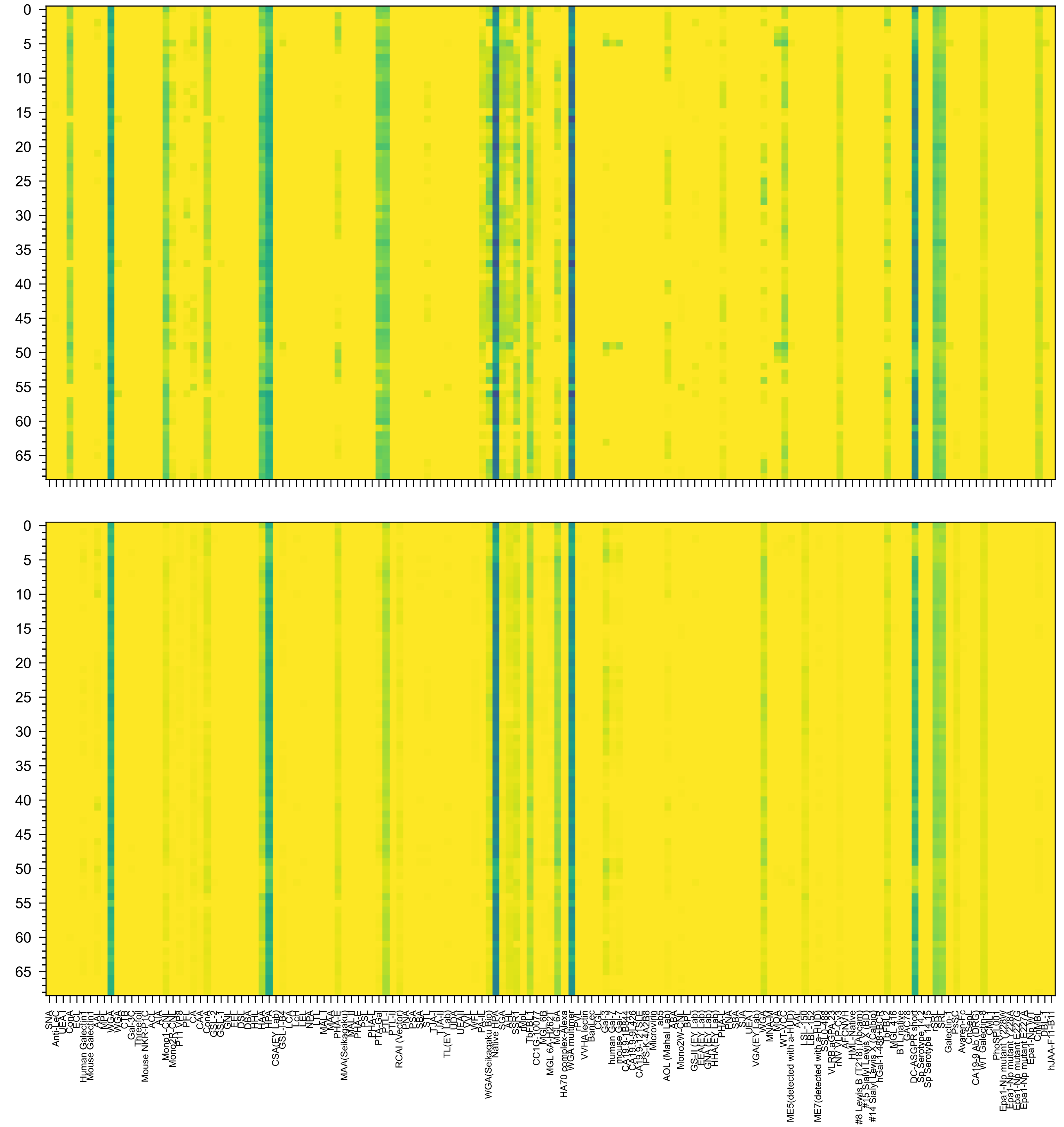


GalNAc(α1-3)[Fuc(α1-2)]Gal(β1-4)GlcNAc(β1-2)Man(α1-6)[GalNAc(α1-3)[Fuc(α1-2)]Gal(β1-4)GlcNAc(β1-2)Man(α1-3)]Man(β1-4)GlcNAc(β1-4)GlcNAc(β-Sp20

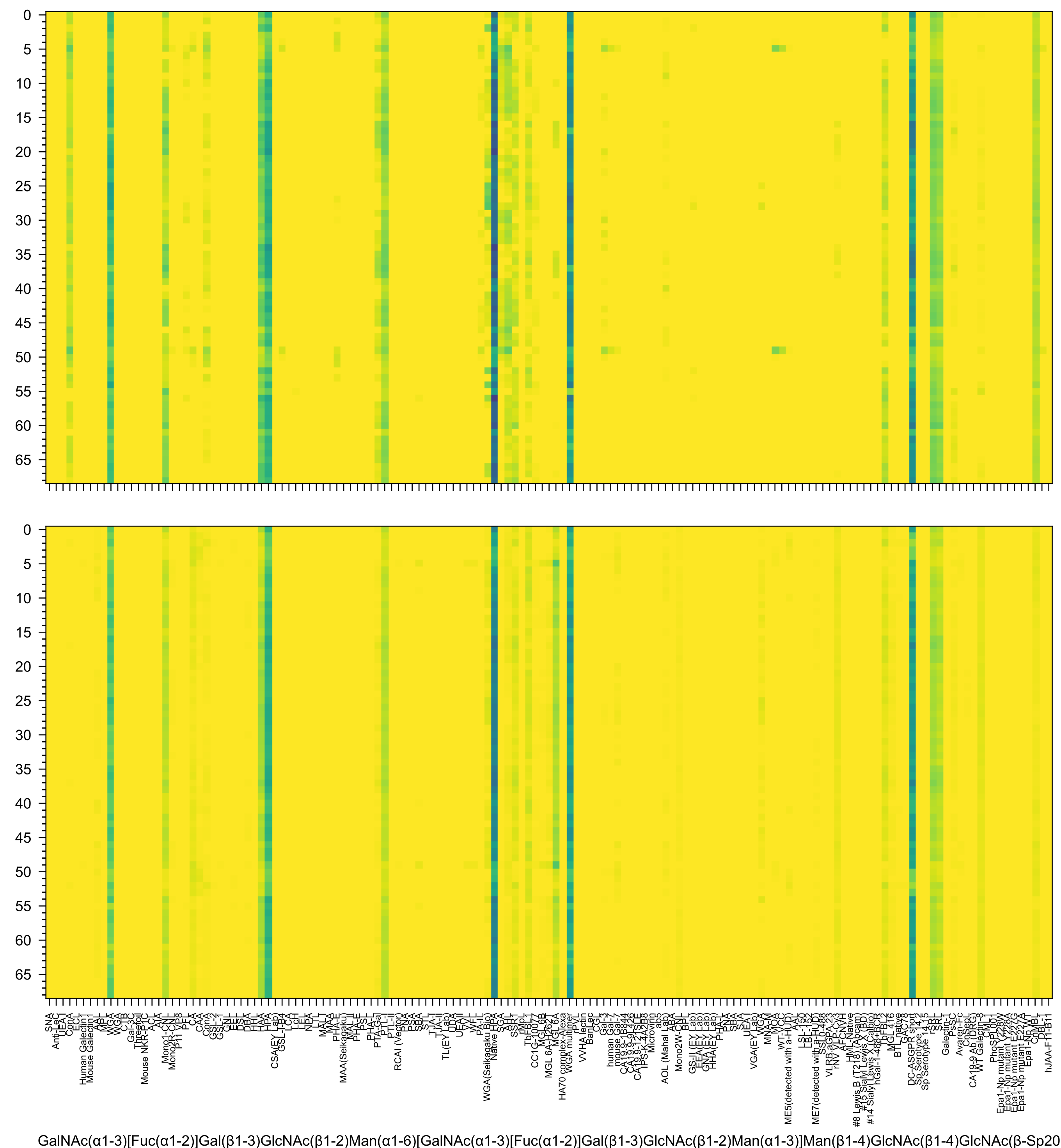
GalNAc(α1-3)[Fuc(α1-2)]Gal(β1-3)GlcNAc(β1-2)Man(α1-6)[GalNAc(α1-3)[Fuc(α1-2)]Gal(β1-3)GlcNAc(β1-2)Man(α1-3)]Man(β1-4)GlcNAc(β1-4)GlcNAc(β-Sp20

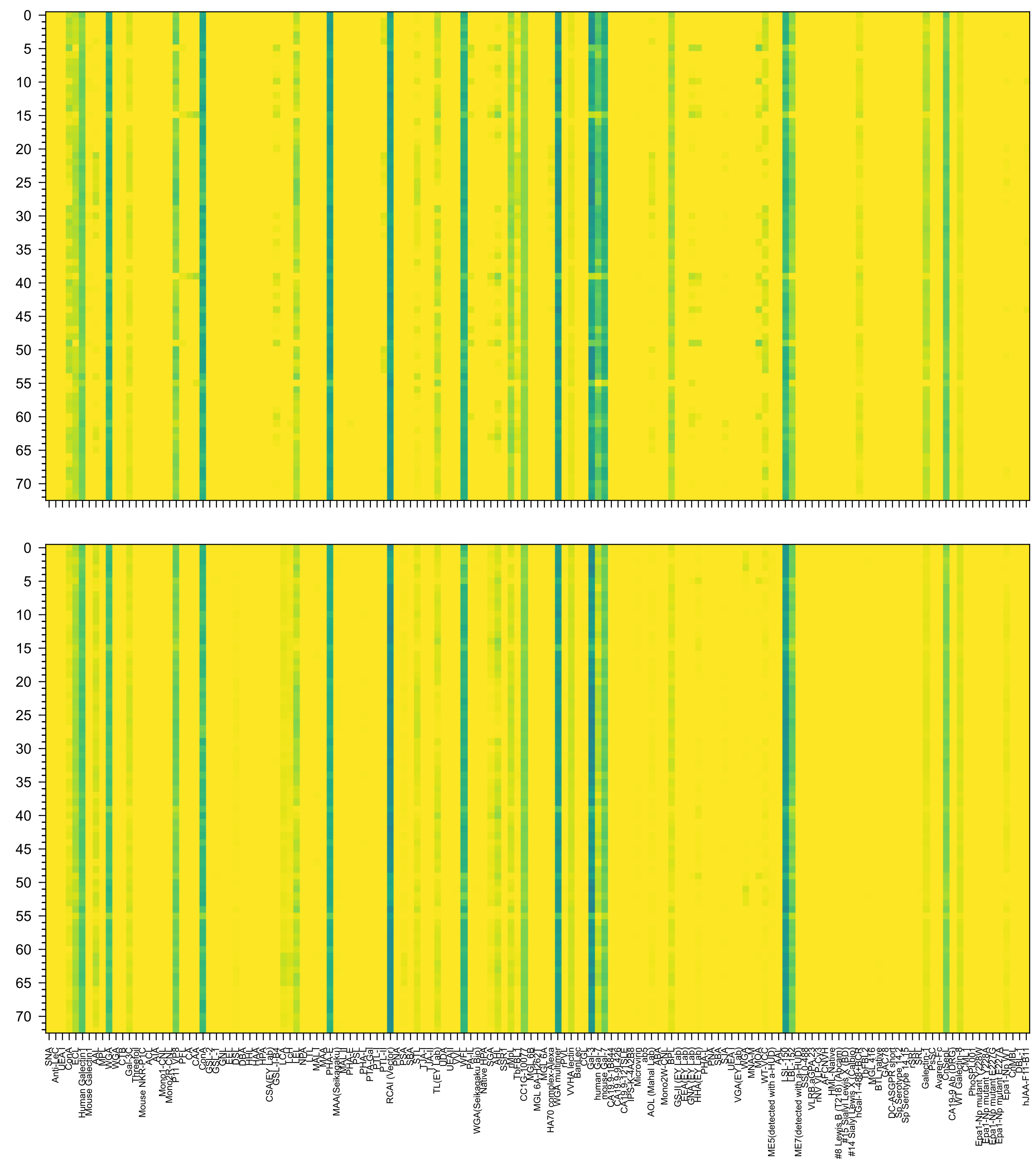

Gal(β1-4)GlcNAc(β1-3)Gal(β1-4)GlcNAc(β1-2)Man(α1-6)[Gal(β1-4)GlcNAc(β1-3)Gal(β1-4)GlcNAc(β1-2)Man(α1-3)]Man(β1-4)GlcNAc(β1-4)GlcNAc(β-Sp24

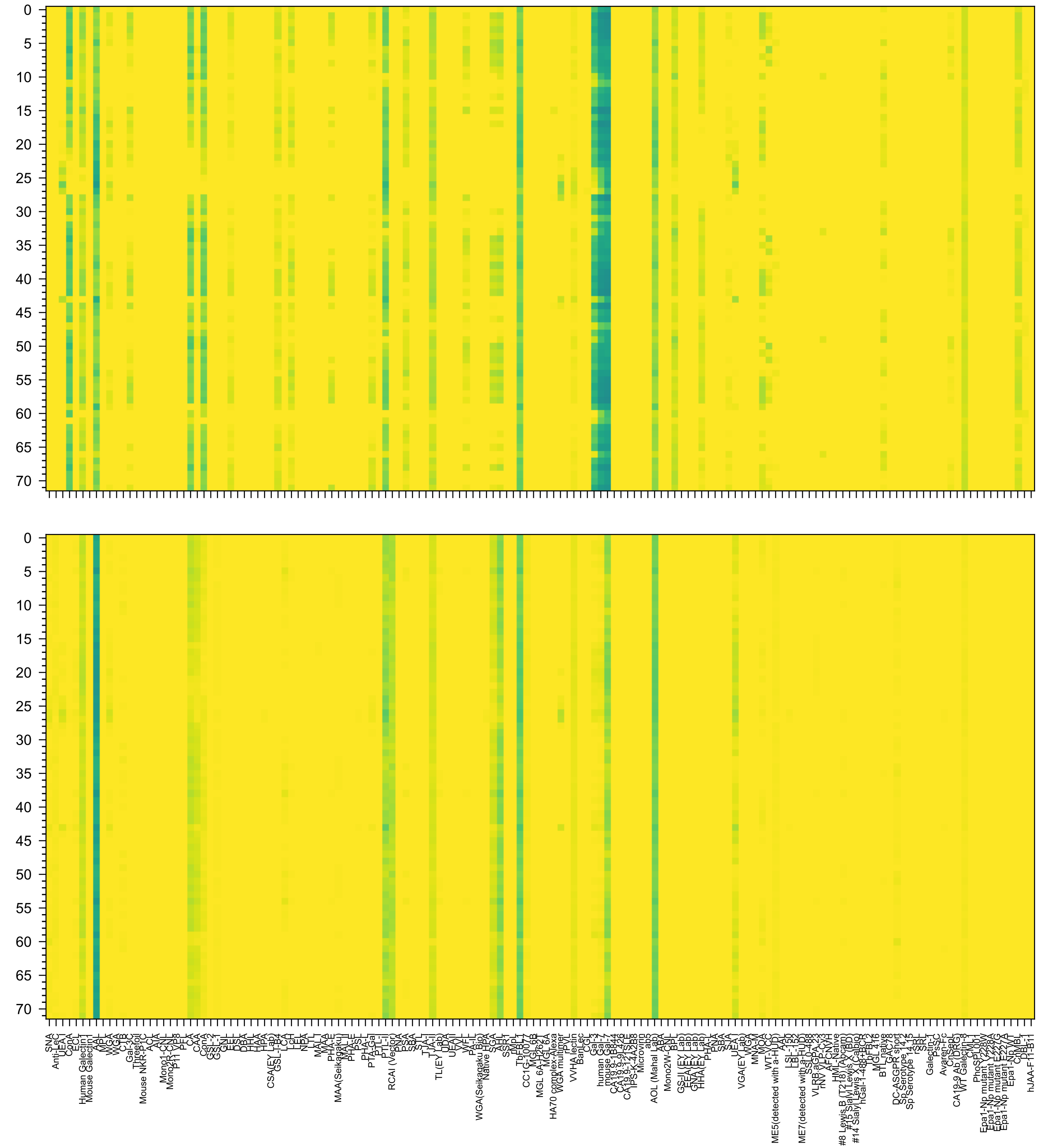

Fuc(α1-2)Gal(β1-4)GlcNAc(β1-2)Man(α1-6)[Fuc(α1-2)Gal(β1-4)GlcNAc(β1-2)[Fuc(α1-2)Gal(β1-4)GlcNAc(β1-4)]Man(α1-3)]Man(β1-4)GlcNAc(β1-4)GlcNAc(β-Sp1

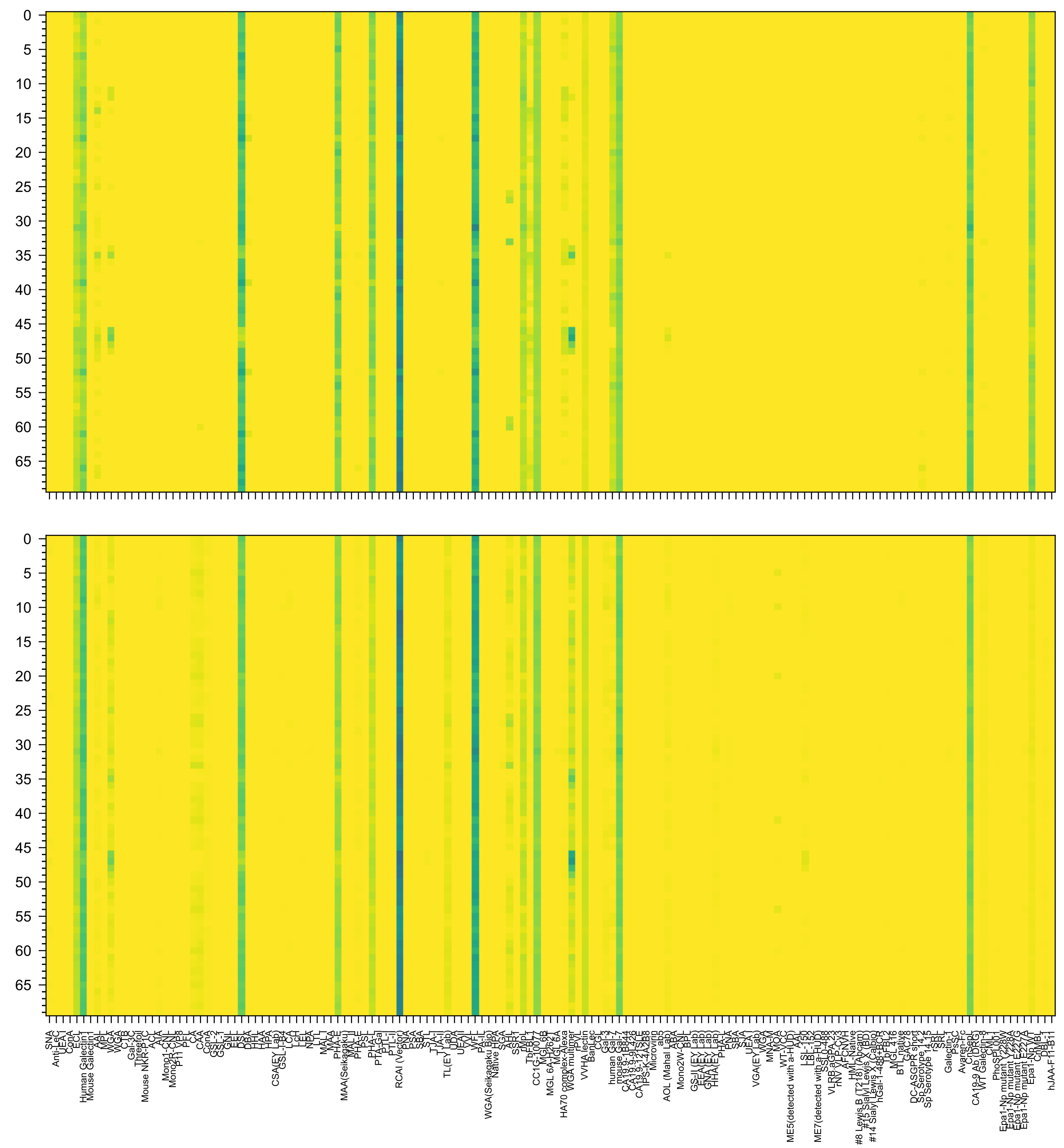

Gal(β1-4)GlcNAc(β1-6)[Gal(β1-4)GlcNAc(β1-2)]Man(α1-6)[GlcNAc(β1-4)][Gal(β1-4)GlcNAc(β1-4)[Gal(β1-4)GlcNAc(β1-2)]Man(α1-3)]Man(β1-4)GlcNAc(β1-4)GlcNAc(

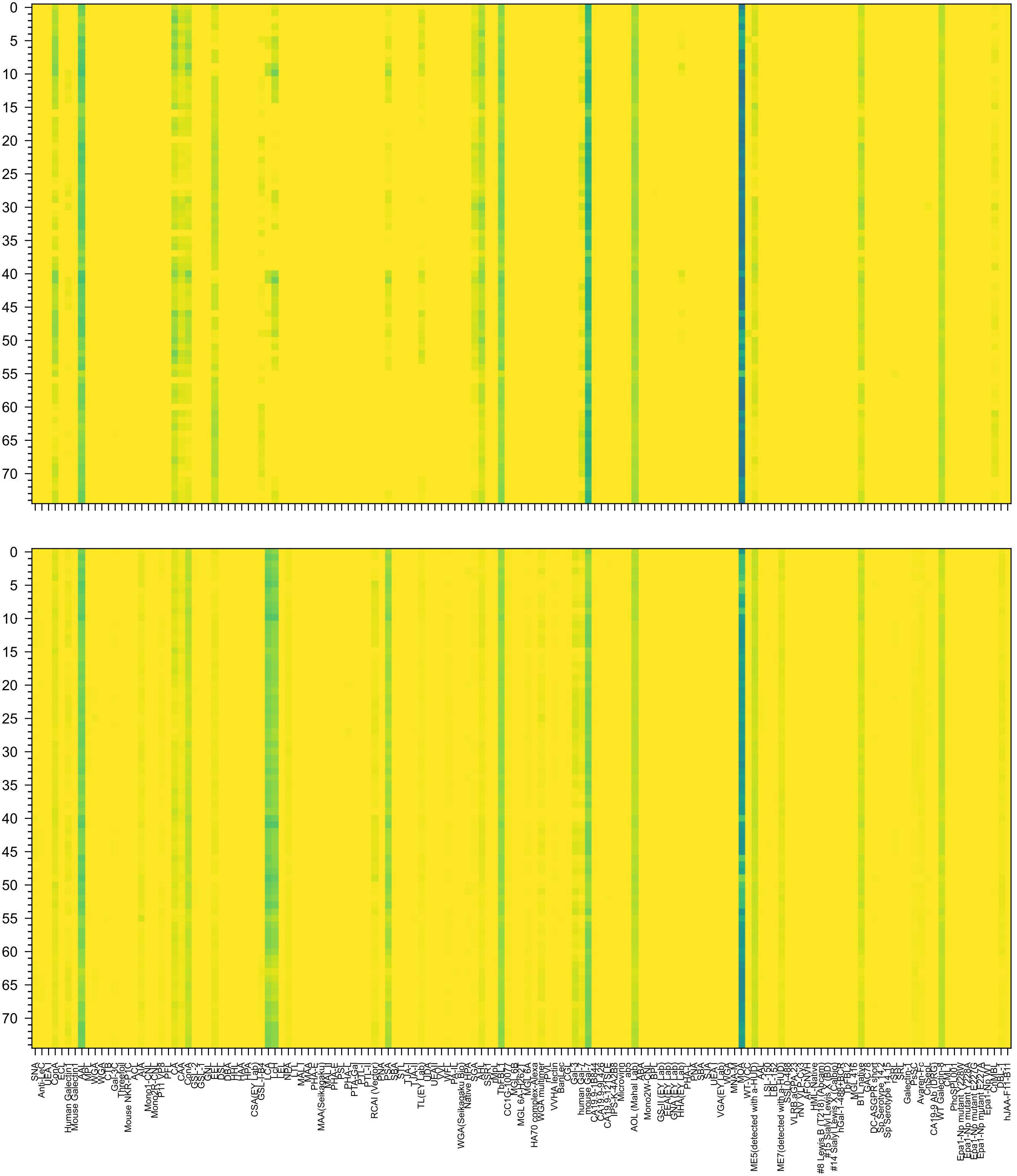

Gal(α1-3)[Fuc(α1-2)]Gal(β1-4)GlcNAc(β1-2)Man(α1-6)[Gal(α1-3)[Fuc(α1-2)]Gal(β1-4)GlcNAc(β1-2)Man(α1-3)]Man(β1-4)GlcNAc(β1-4)[Fuc(α1-6)]GlcNAc(β-Sp2

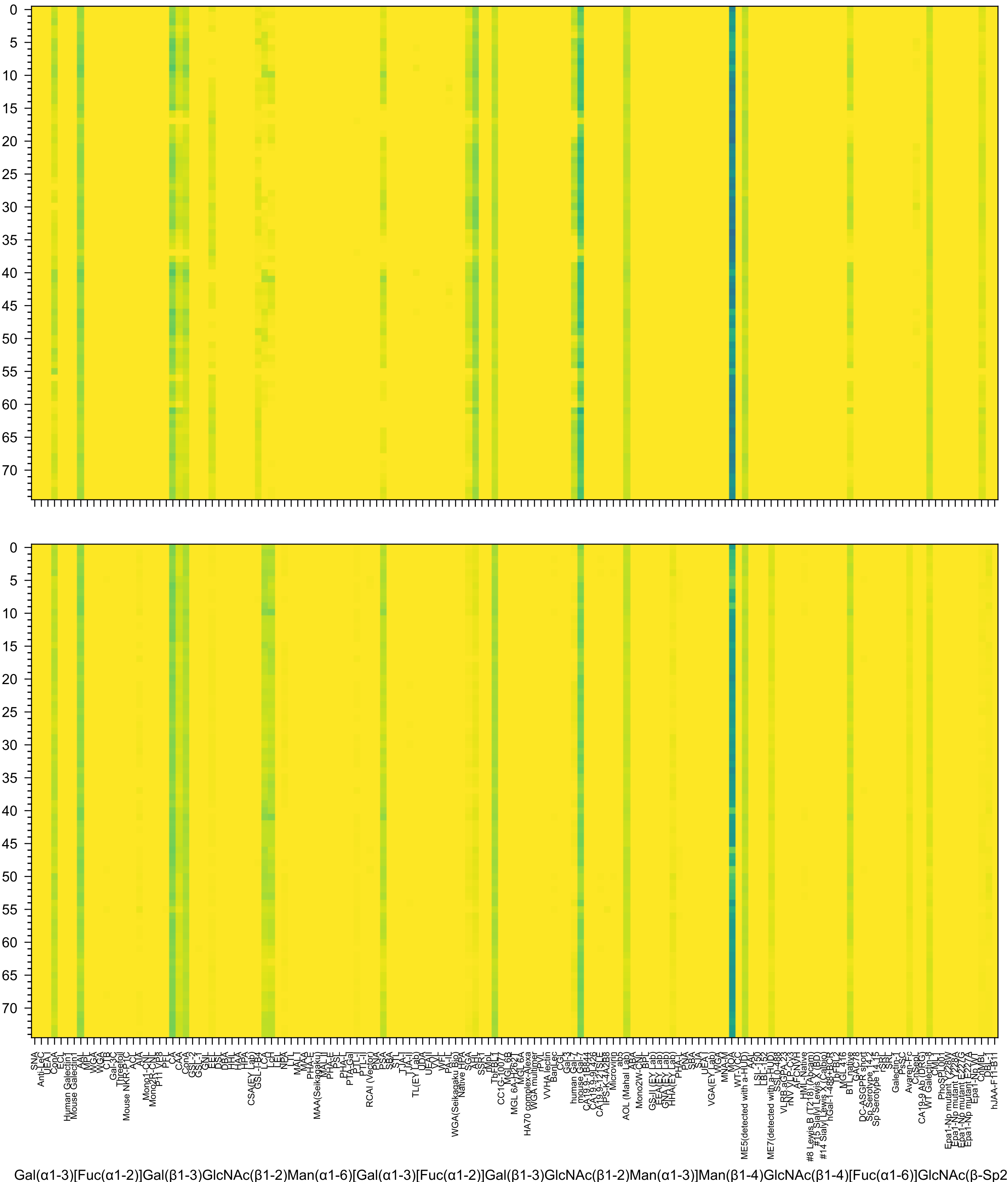
Gal(α1-3)[Fuc(α1-2)]Gal(β1-3)GlcNAc(β1-2)Man(α1-6)[Gal(α1-3)[Fuc(α1-2)]Gal(β1-3)GlcNAc(β1-2)Man(α1-3)]Man(β1-4)GlcNAc(β1-4)[Fuc(α1-6)]GlcNAc(β-Sp2

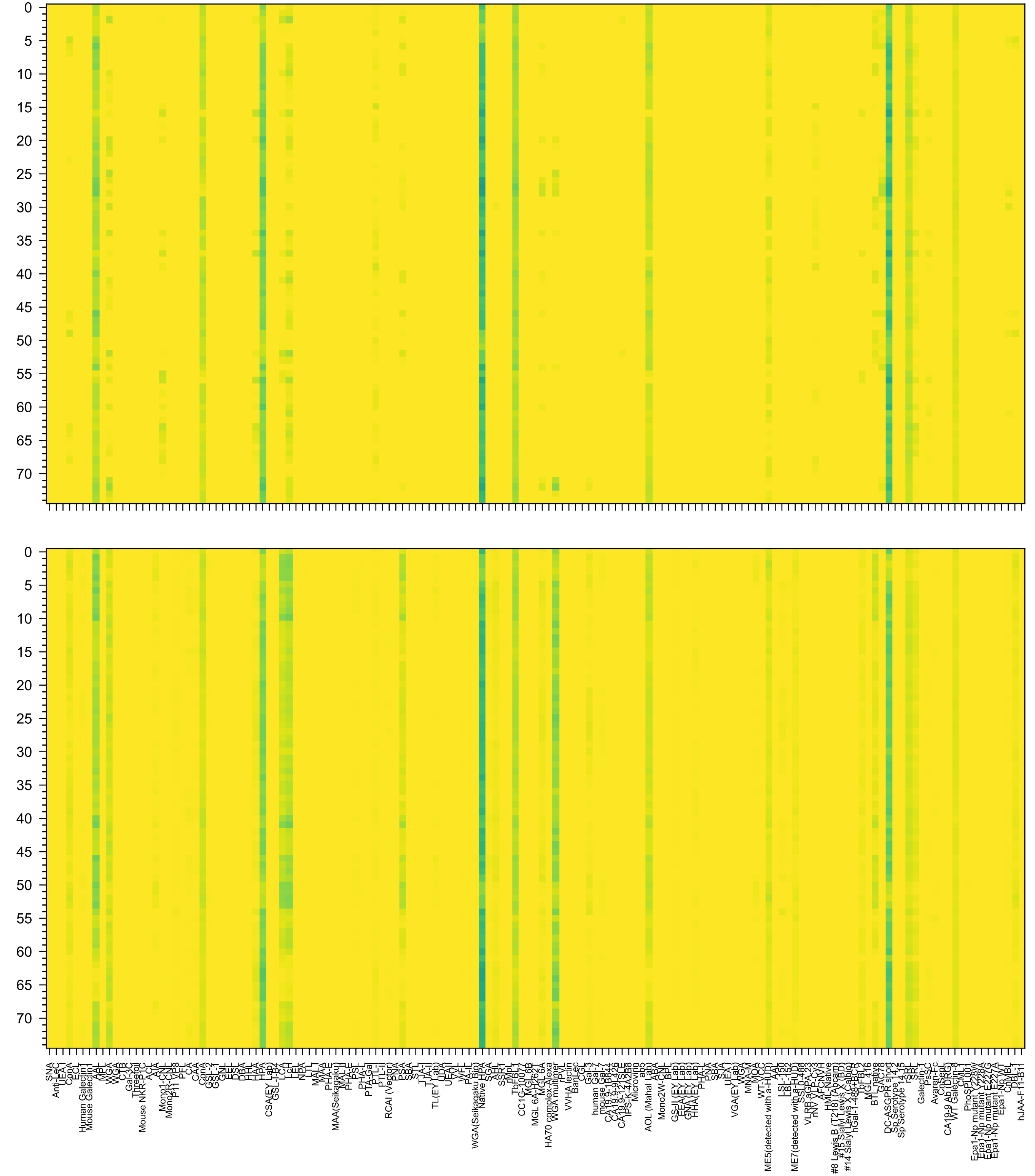

GalNAc(α1-3)[Fuc(α1-2)]Gal(β1-4)GlcNAc(β1-2)Man(α1-6)[GalNAc(α1-3)[Fuc(α1-2)]Gal(β1-4)GlcNAc(β1-2)Man(α1-3)]Man(β1-4)GlcNAc(β1-4)[Fuc(α1-6)]GlcNAc(β-

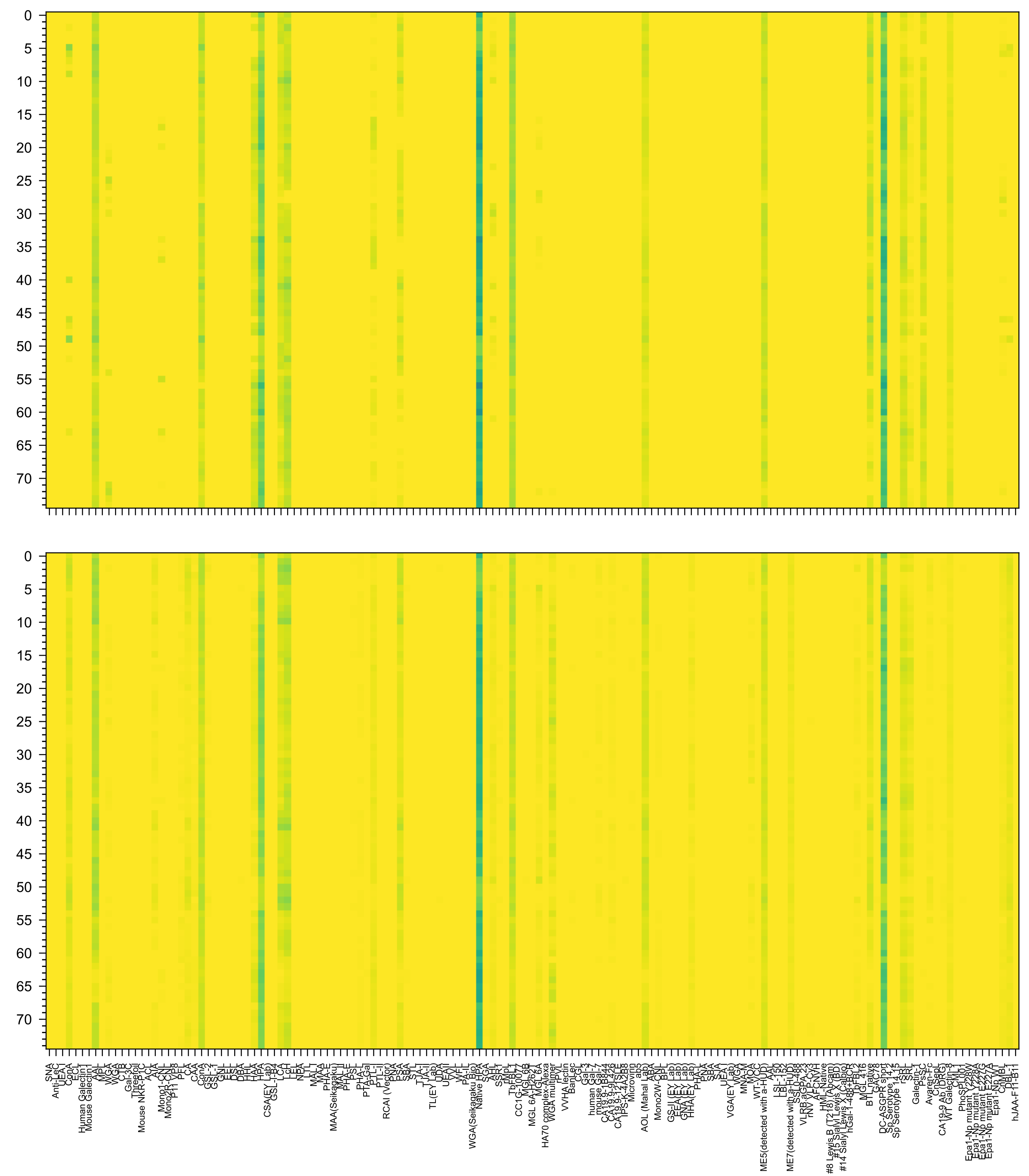

GalNAc(α1-3)[Fuc(α1-2)]Gal(β1-3)GlcNAc(β1-2)Man(α1-6)[GalNAc(α1-3)[Fuc(α1-2)]Gal(β1-3)GlcNAc(β1-2)Man(α1-3)]Man(β1-4)GlcNAc(β1-4)[Fuc(α1-6)]GlcNAc(β-

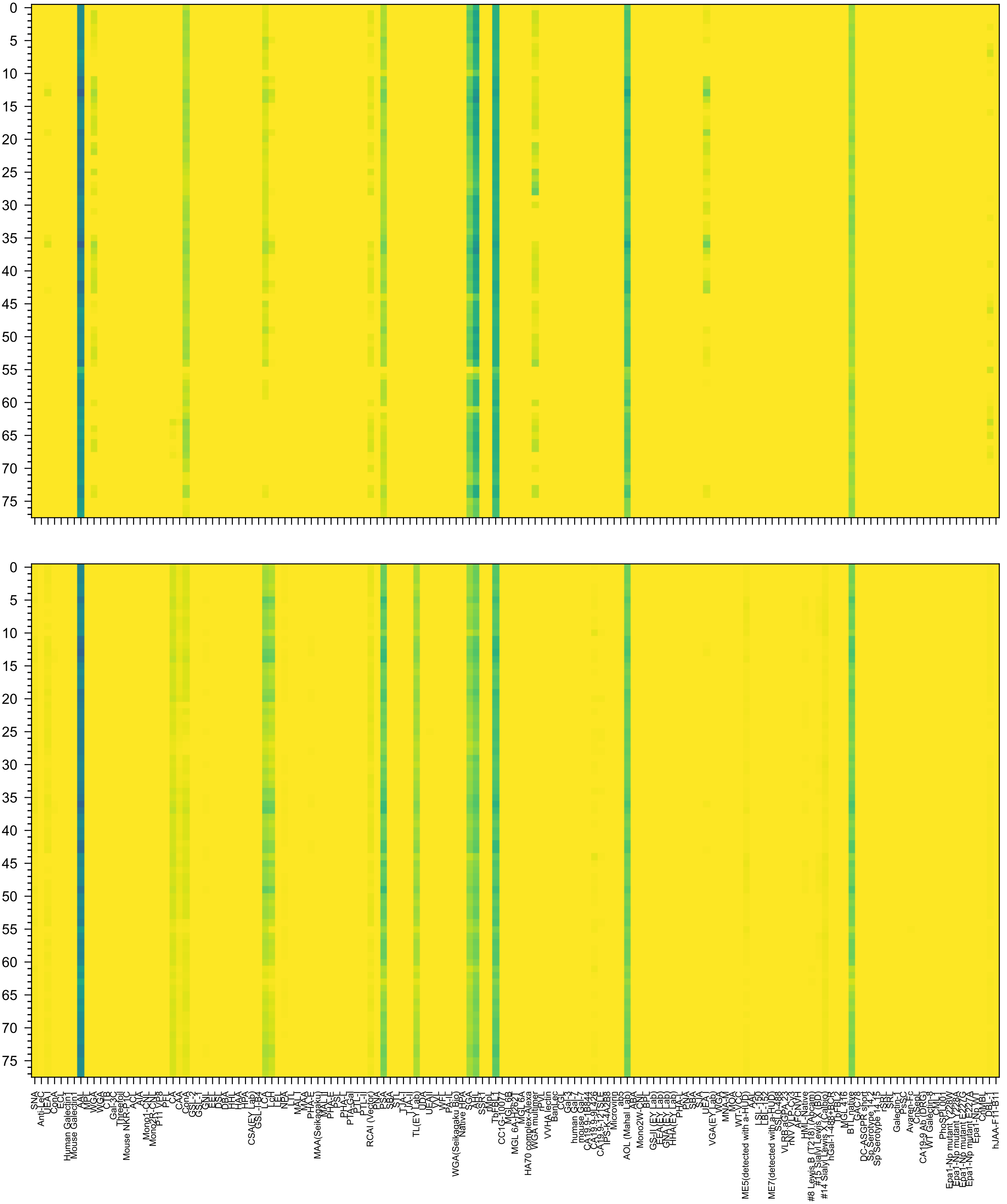

Fuc(α1-2)Gal(β1-4)[Fuc(α1-3)]GlcNAc(β1-2)Man(α1-6)[Fuc(α1-2)Gal(β1-4)[Fuc(α1-3)]GlcNAc(β1-2)Man(α1-3)]Man(β1-4)GlcNAc(β1-4)[Fuc(α1-6)]GlcNAc(β-Sp2

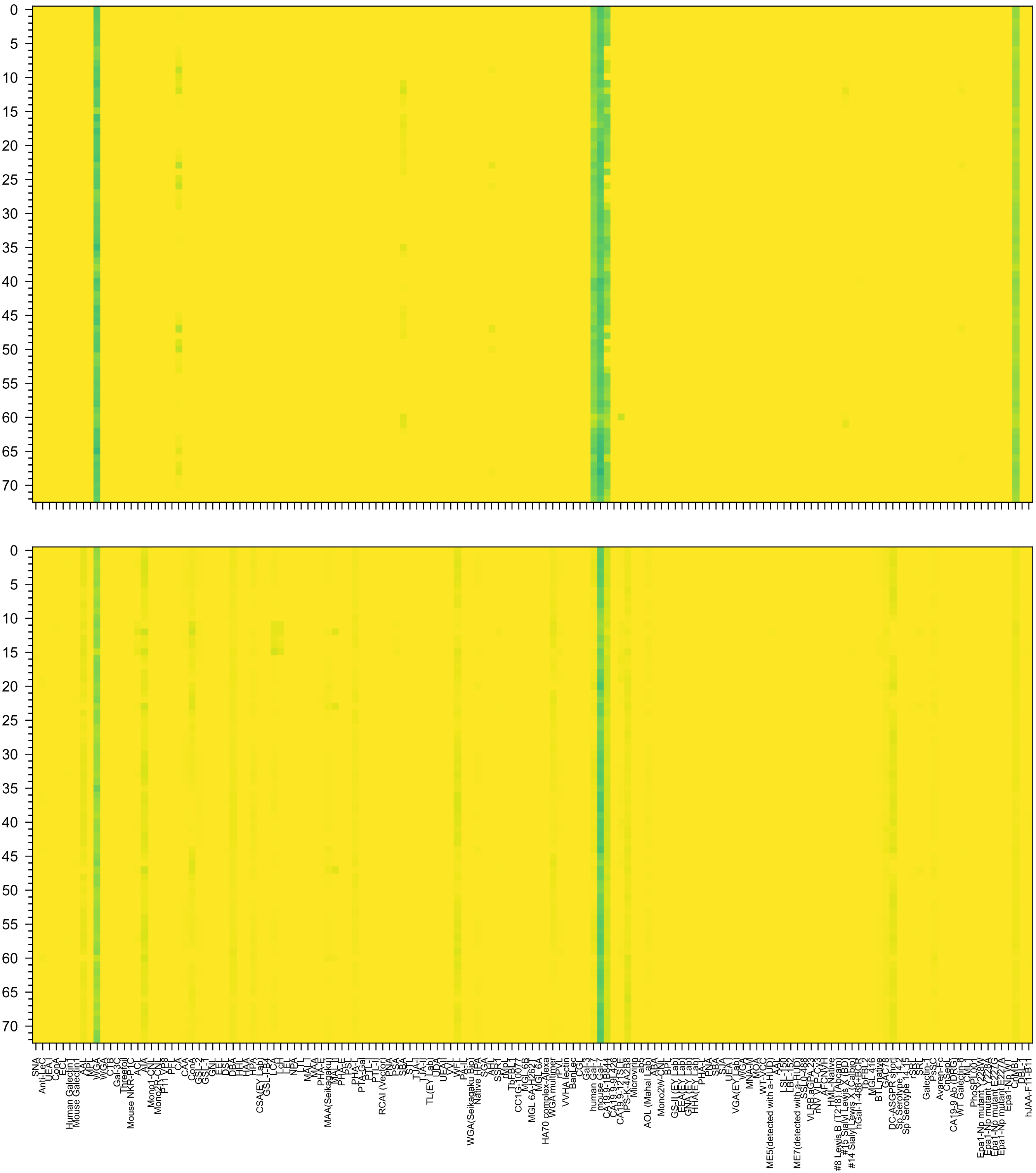

eu5Ac(α2-3)Gal(β1-3)GlcNAc(β1-6)[Neu5Ac(α2-3)Gal(β1-3)GlcNAc(β1-2)]Man(α1-6)[Neu5Ac(α2-3)Gal(β1-3)GlcNAc(β1-2)Man(α1-3)]Man(β1-4)GlcNAc(β1-4)GlcNAc

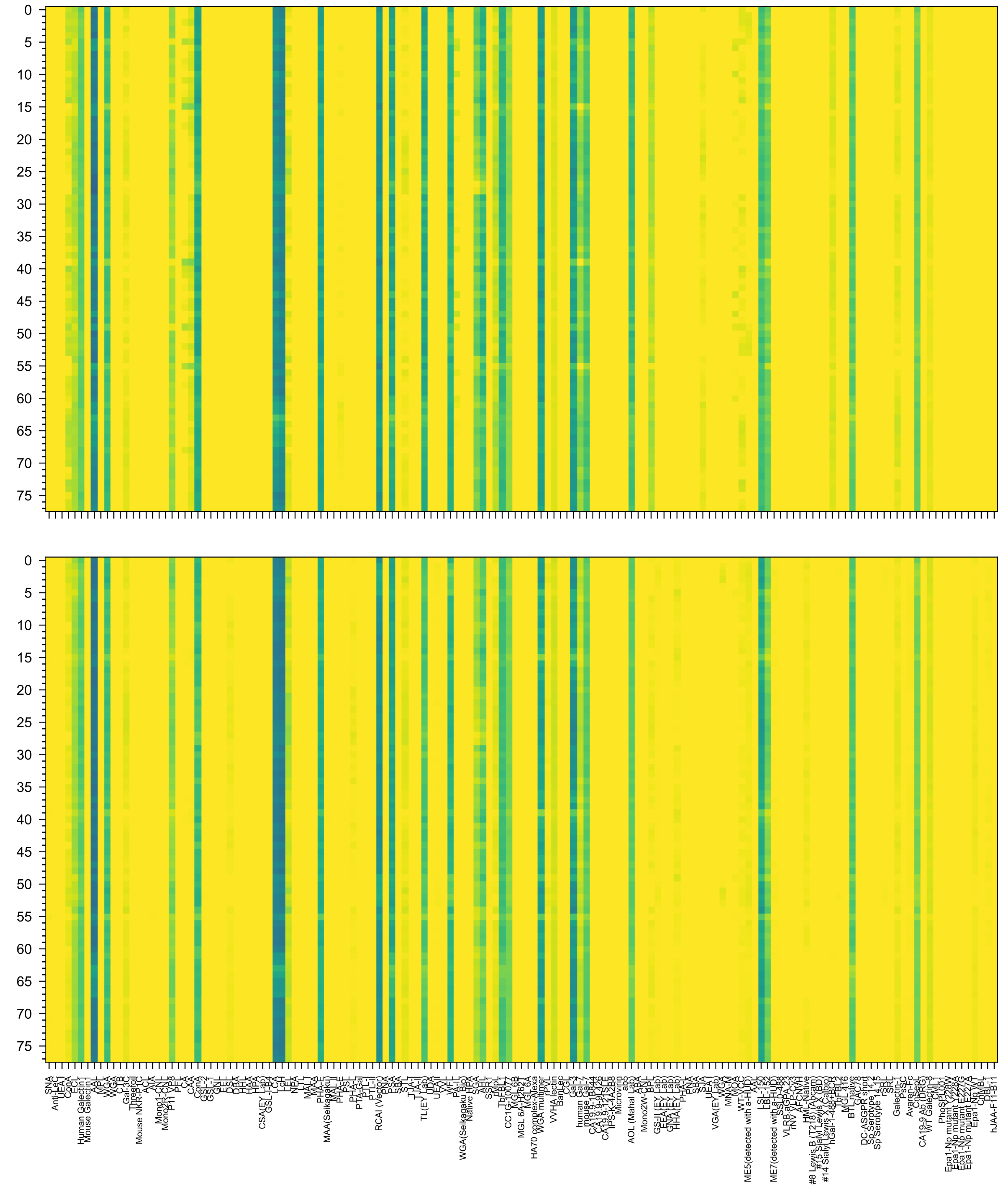

Gal(β1-4)GlcNAc(β1-3)Gal(β1-4)GlcNAc(β1-2)Man(α1-6)[Gal(β1-4)GlcNAc(β1-3)Gal(β1-4)GlcNAc(β1-2)Man(α1-3)]Man(β1-4)GlcNAc(β1-4)[Fuc(α1-6)]GlcNAc(β-S

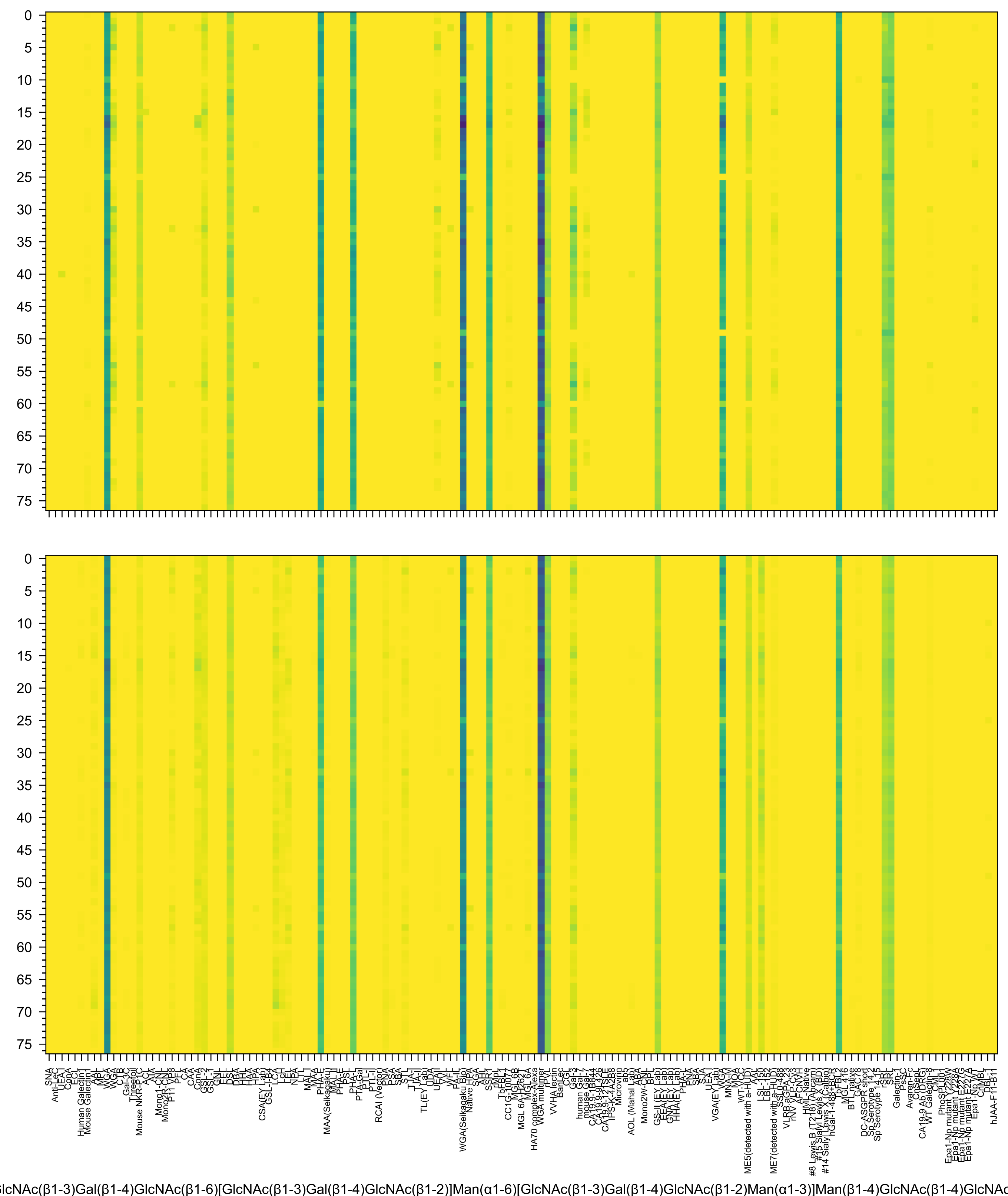
lcNAc(β1-3)Gal(β1-4)GlcNAc(β1-6)[GlcNAc(β1-3)Gal(β1-4)GlcNAc(β1-2)]Man(α1-6)[GlcNAc(β1-3)Gal(β1-4)GlcNAc(β1-2)Man(α1-3)]Man(β1-4)GlcNAc(β1-4)GlcNAc

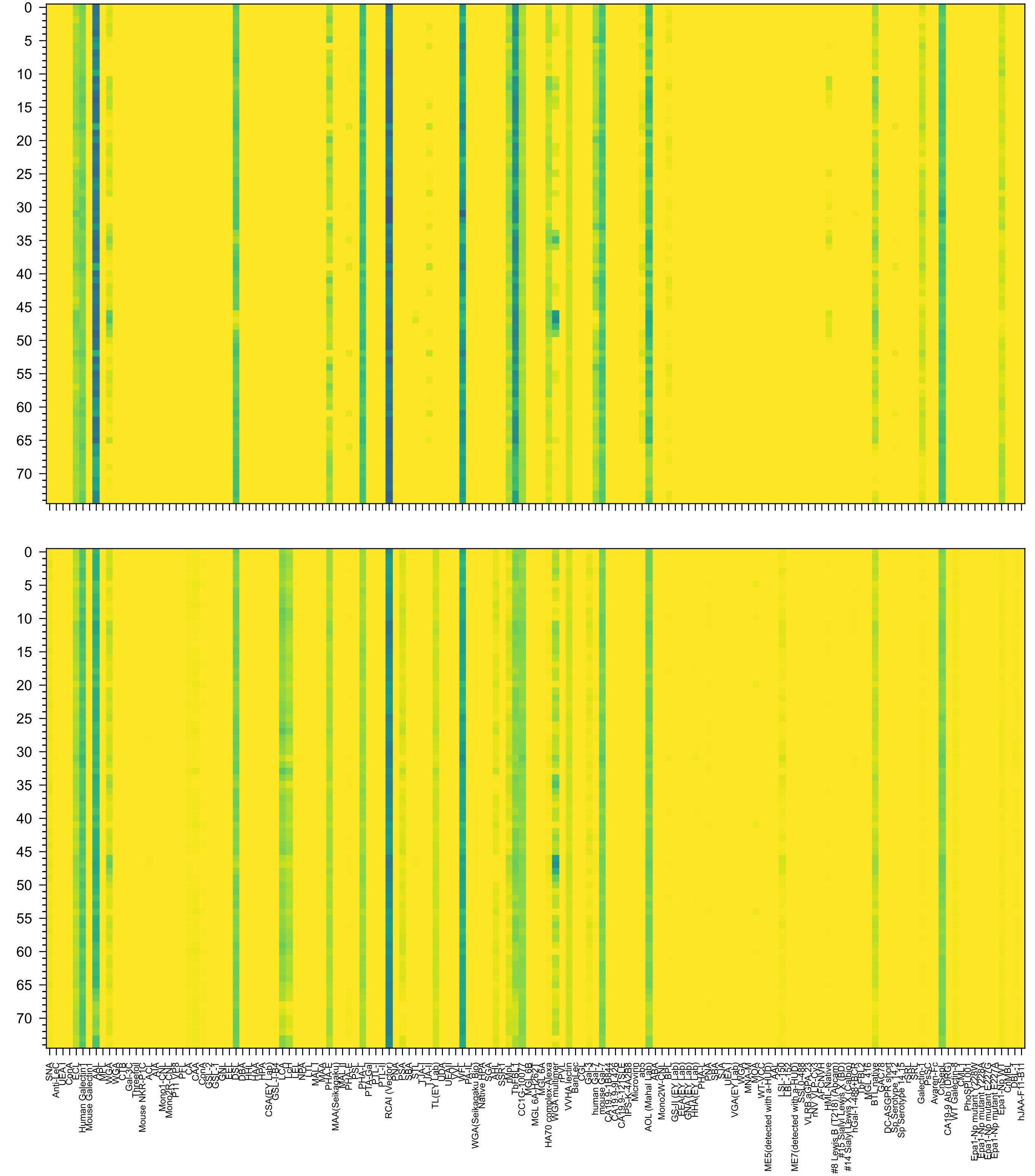

-4)GlcNAc(β1-6)[Gal(β1-4)GlcNAc(β1-2)]Man(α1-6)[GlcNAc(β1-4)][Gal(β1-4)GlcNAc(β1-4)[Gal(β1-4)GlcNAc(β1-2)]Man(α1-3)]Man(β1-4)GlcNAc(β1-4)[Fuc(α1-6)]Gl

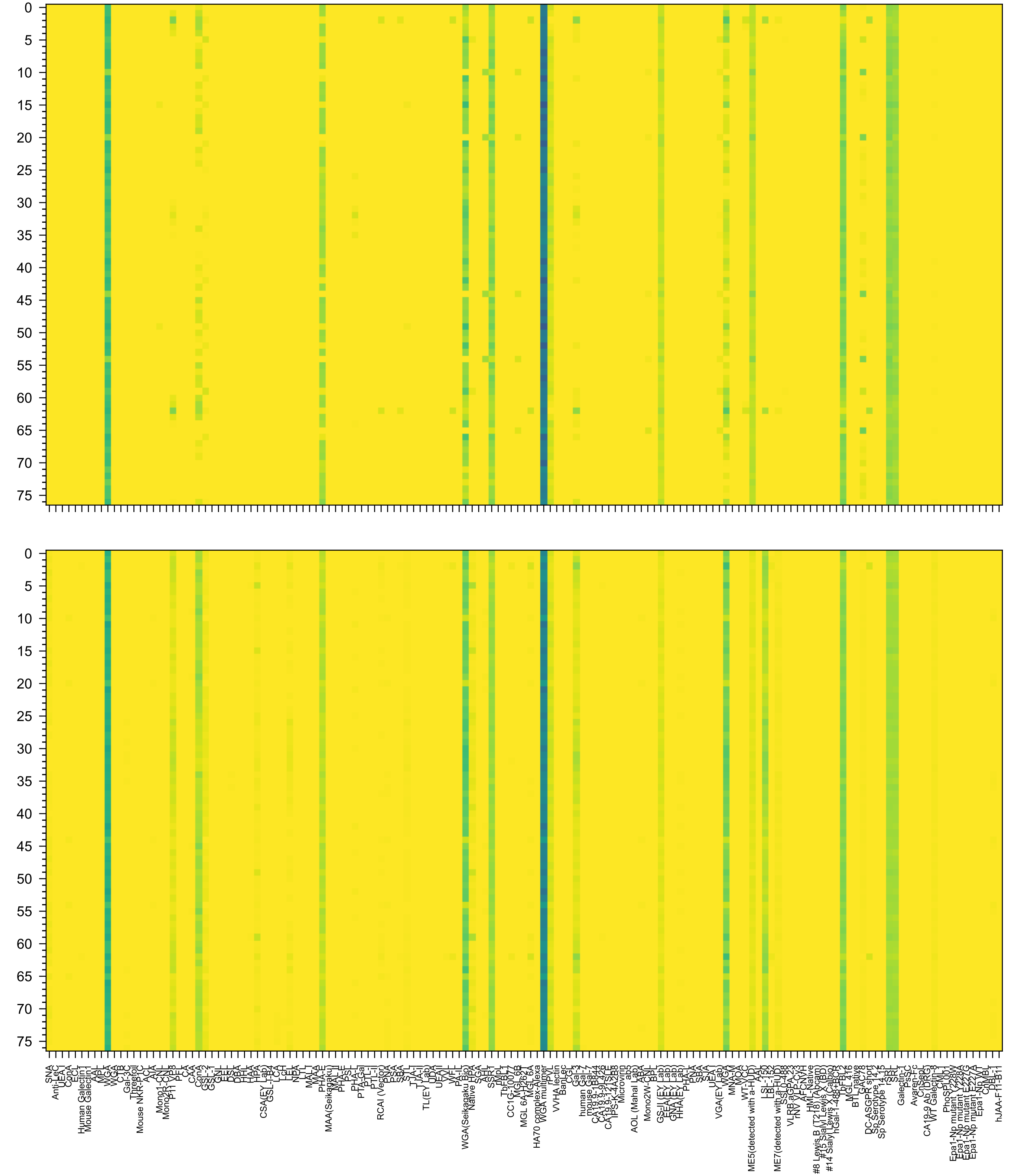

Ac(β1-3)Gal(β1-4)GlcNAc(β1-3)Gal(β1-4)GlcNAc(β1-2)Man(α1-6)[GlcNAc(β1-3)Gal(β1-4)GlcNAc(β1-3)Gal(β1-4)GlcNAc(β1-2)Man(α1-3)]Man(β1-4)GlcNAc(β1-4)Glc

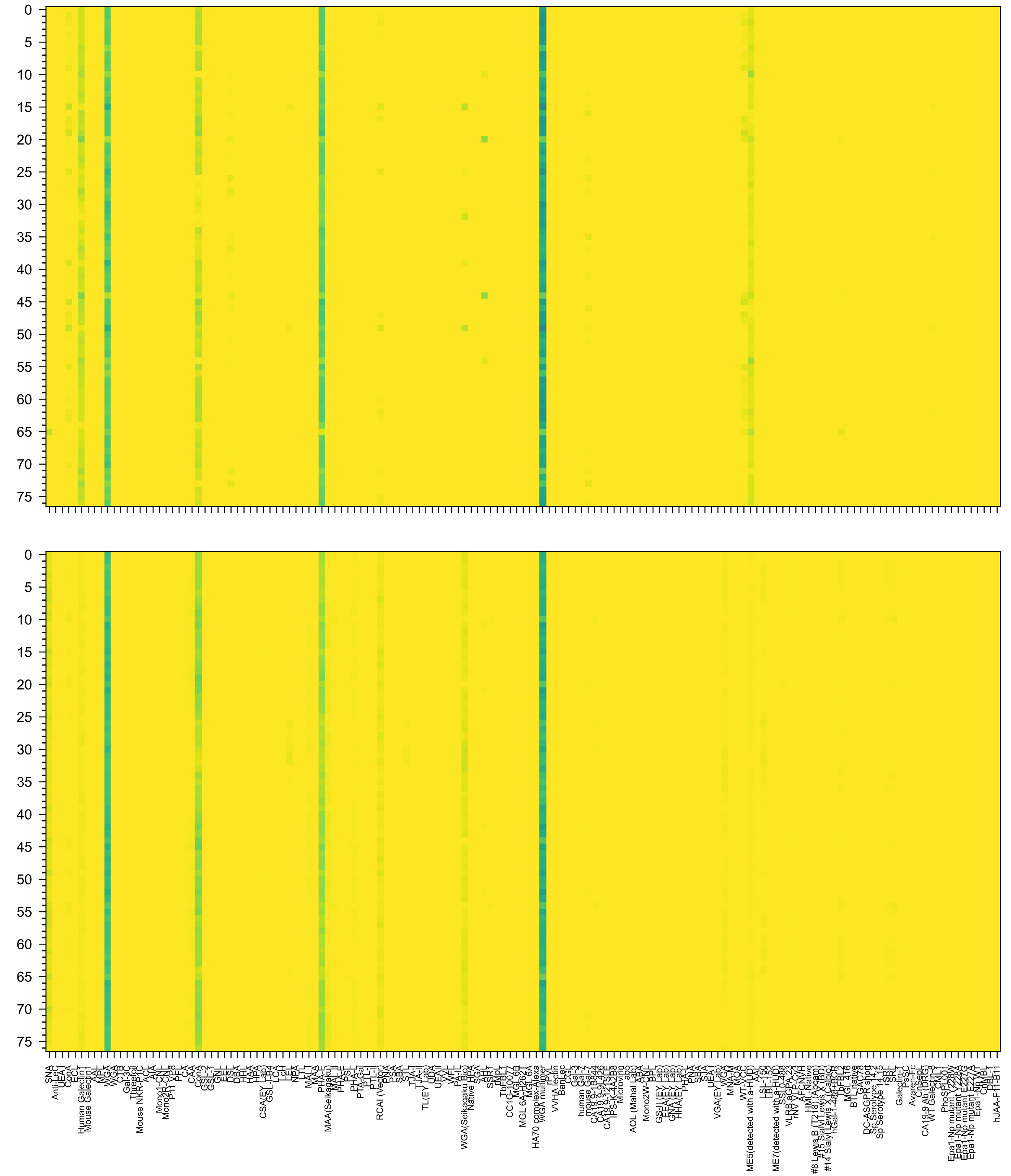

Ac(α2-3)Gal(β1-4)GlcNAc(β1-3)Gal(β1-4)GlcNAc(β1-2)Man(α1-6)[Neu5Ac(α2-3)Gal(β1-4)GlcNAc(β1-3)Gal(β1-4)GlcNAc(β1-2)Man(α1-3)]Man(β1-4)GlcNAc(β1-4)Glc

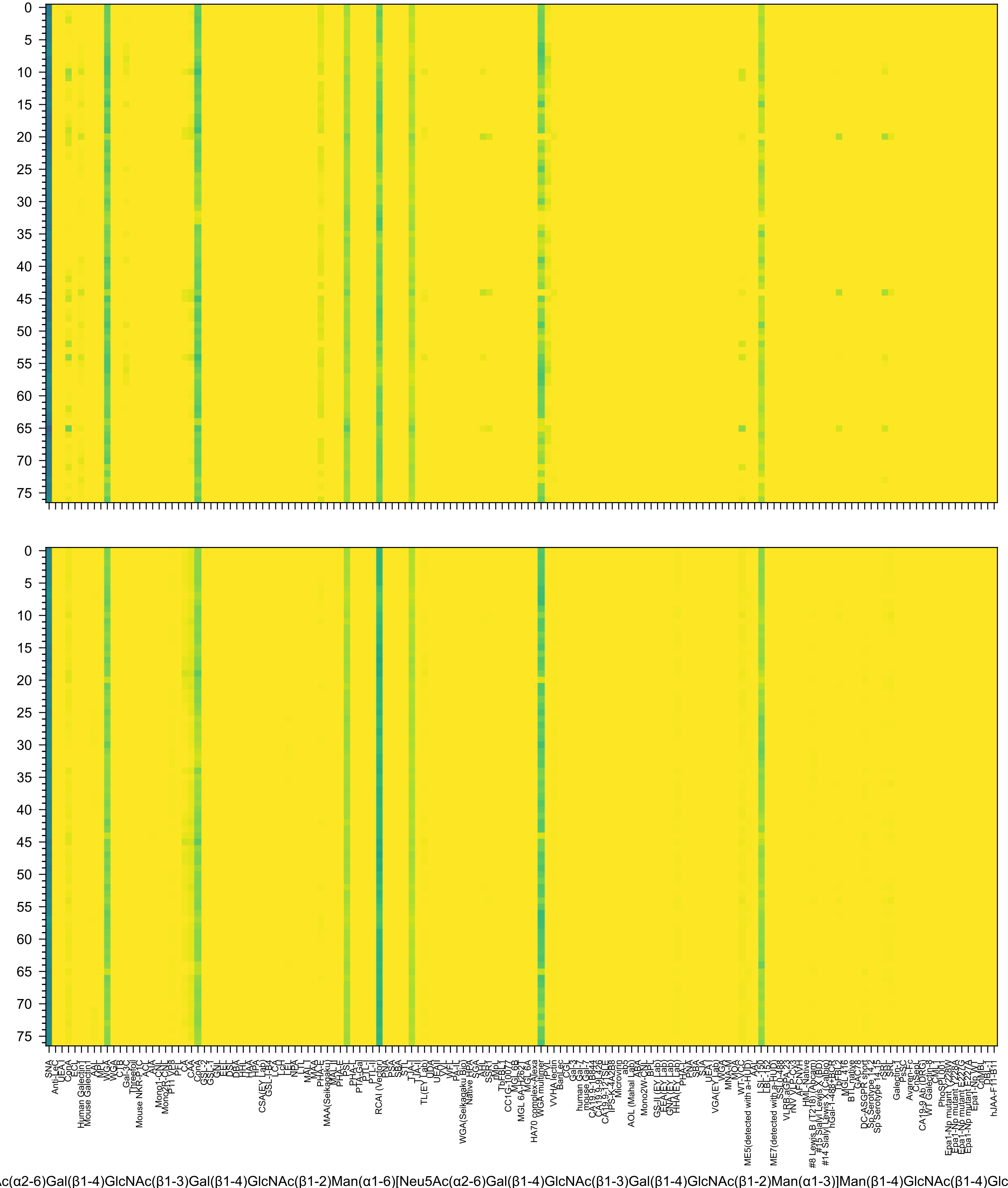
c(α2-6)Gal(β1-4)GlcNAc(β1-3)Gal(β1-4)GlcNAc(β1-2)Man(α1-6)[Neu5Ac(α2-6)Gal(β1-4)GlcNAc(β1-3)Gal(β1-4)GlcNAc(β1-2)Man(α1-3)]Man(β1-4)GlcNAc(β1-4)Glc

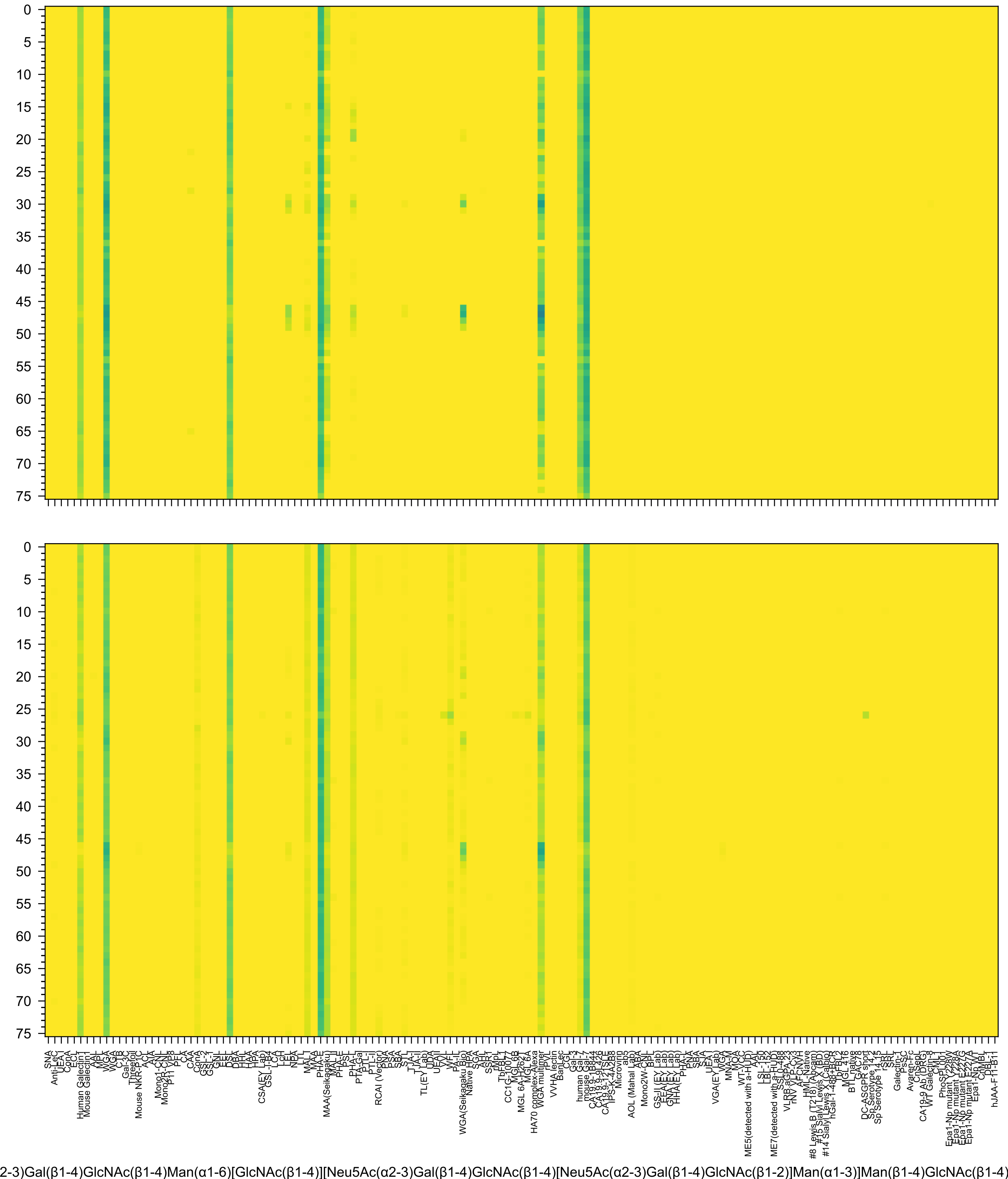
2-3)Gal(β1-4)GlcNAc(β1-4)Man(α1-6)[GlcNAc(β1-4)][Neu5Ac(α2-3)Gal(β1-4)GlcNAc(β1-4)[Neu5Ac(α2-3)Gal(β1-4)GlcNAc(β1-2)]Man(α1-3)]Man(β1-4)GlcNAc(β1-4)

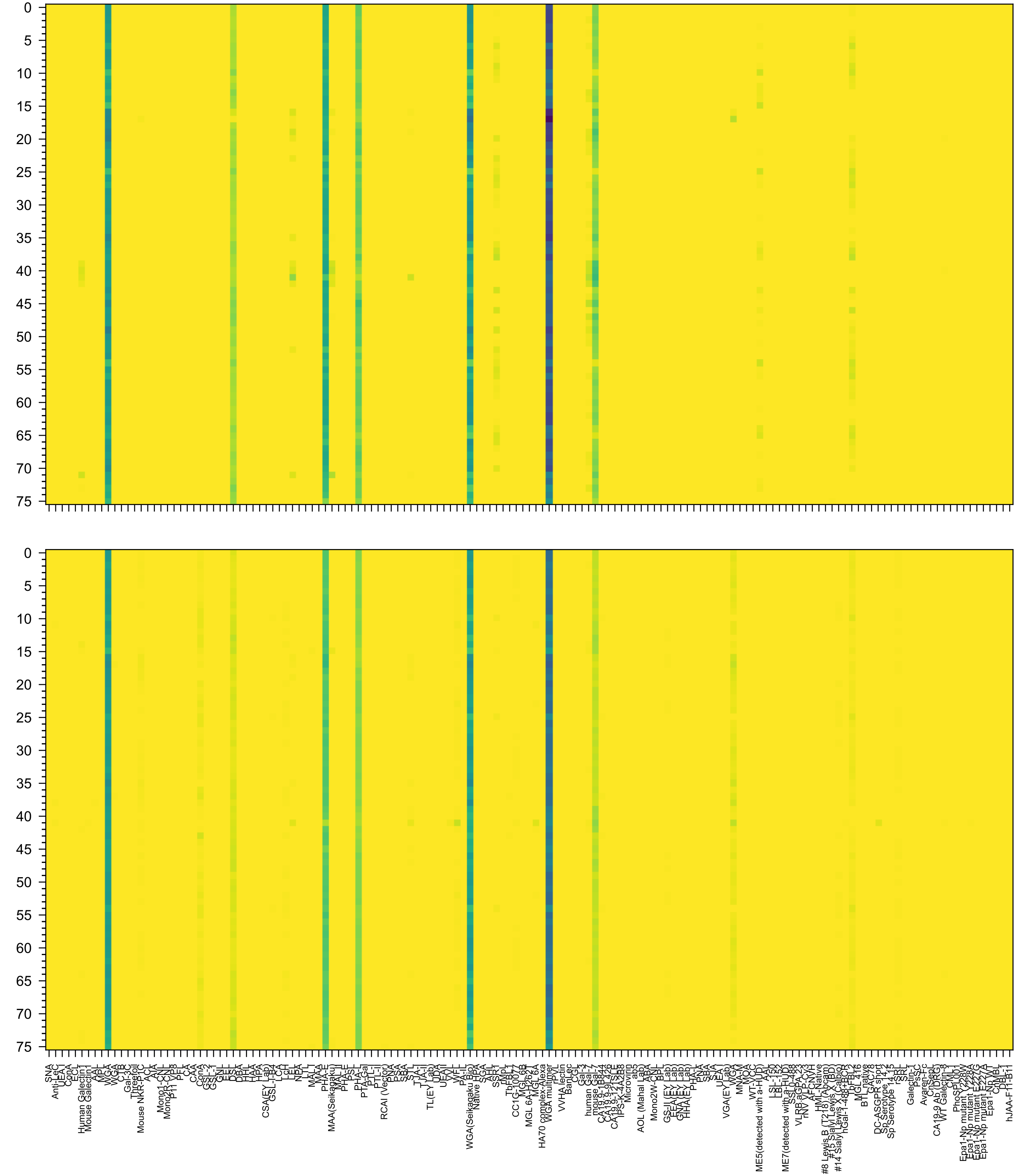

2-3)Gal(β1-4)GlcNAc(β1-6)[Neu5Ac(α2-3)Gal(β1-4)GlcNAc(β1-2)]Man(α1-6)[GlcNAc(β1-4)][Neu5Ac(α2-3)Gal(β1-4)GlcNAc(β1-2)Man(α1-3)]Man(β1-4)GlcNAc(β1-4)

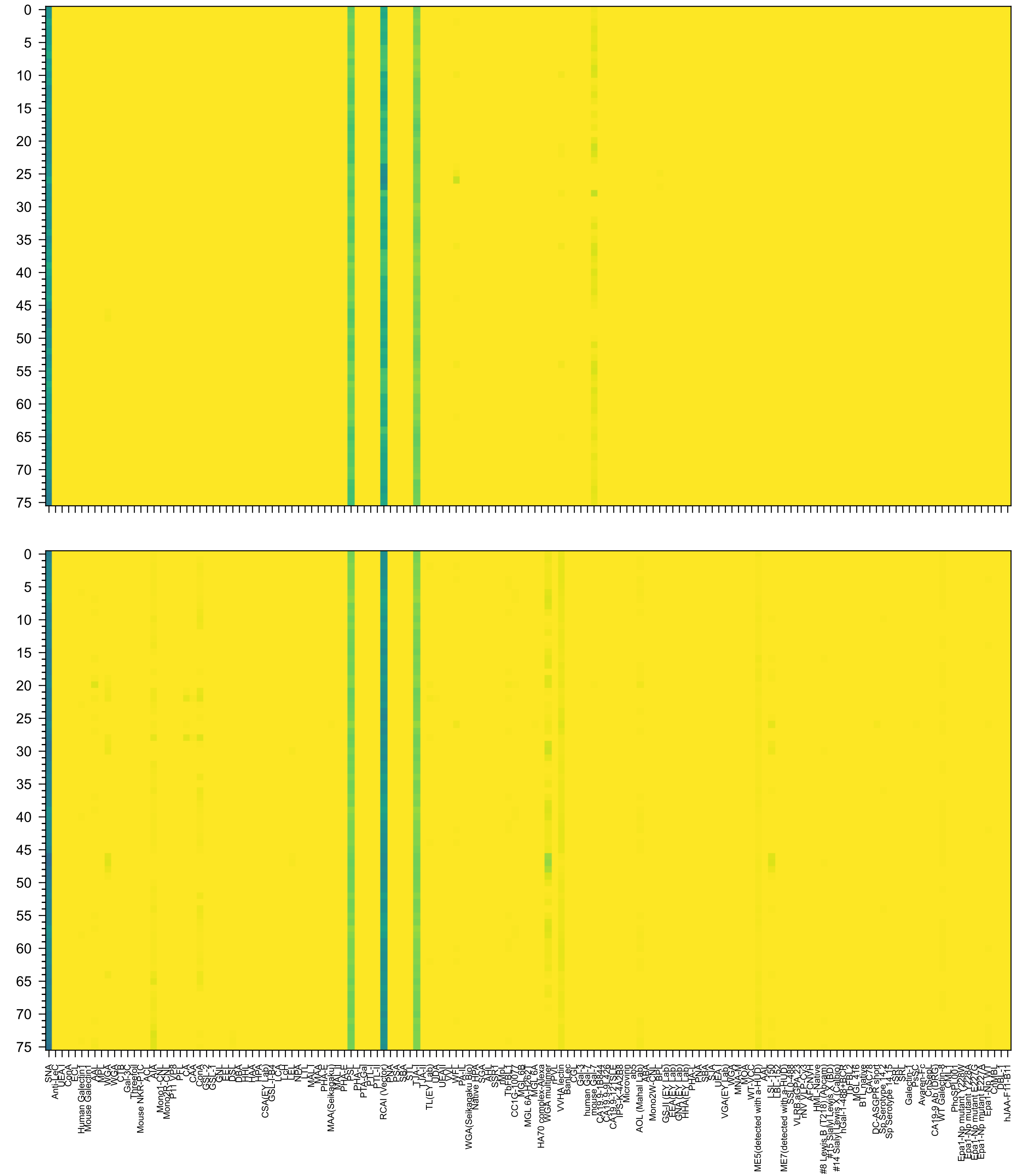


2-6)Gal(β1-4)GlcNAc(β1-4)Man(α1-6)[GlcNAc(β1-4)][Neu5Ac(α2-6)Gal(β1-4)GlcNAc(β1-4)[Neu5Ac(α2-6)Gal(β1-4)GlcNAc(β1-2)]Man(α1-3)]Man(β1-4)GlcNAc(β1-4)

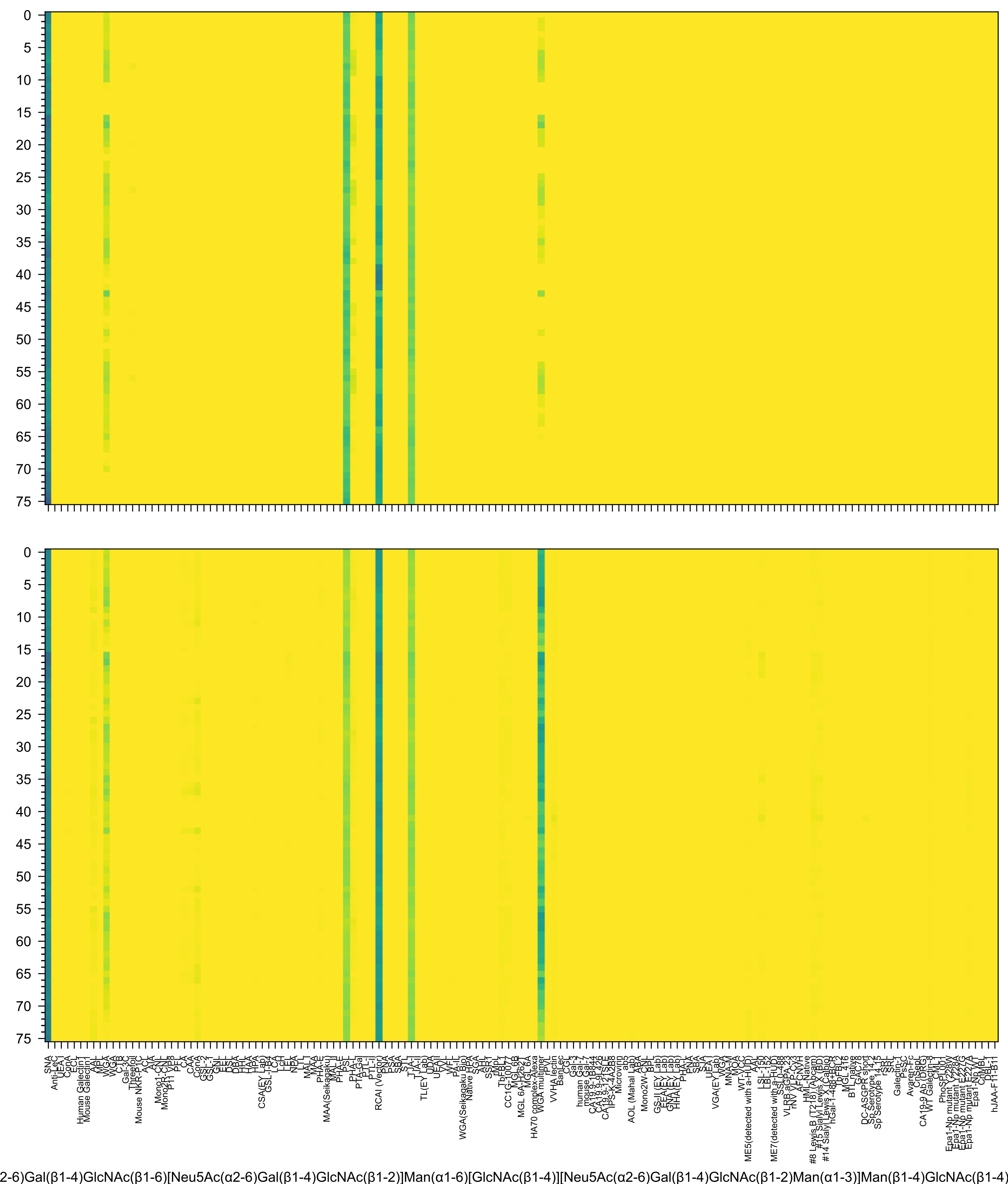
2-6)Gal(β1-4)GlcNAc(β1-6)[Neu5Ac(α2-6)Gal(β1-4)GlcNAc(β1-2)]Man(α1-6)[GlcNAc(β1-4)][Neu5Ac(α2-6)Gal(β1-4)GlcNAc(β1-2)Man(α1-3)]Man(β1-4)GlcNAc(β1-4)

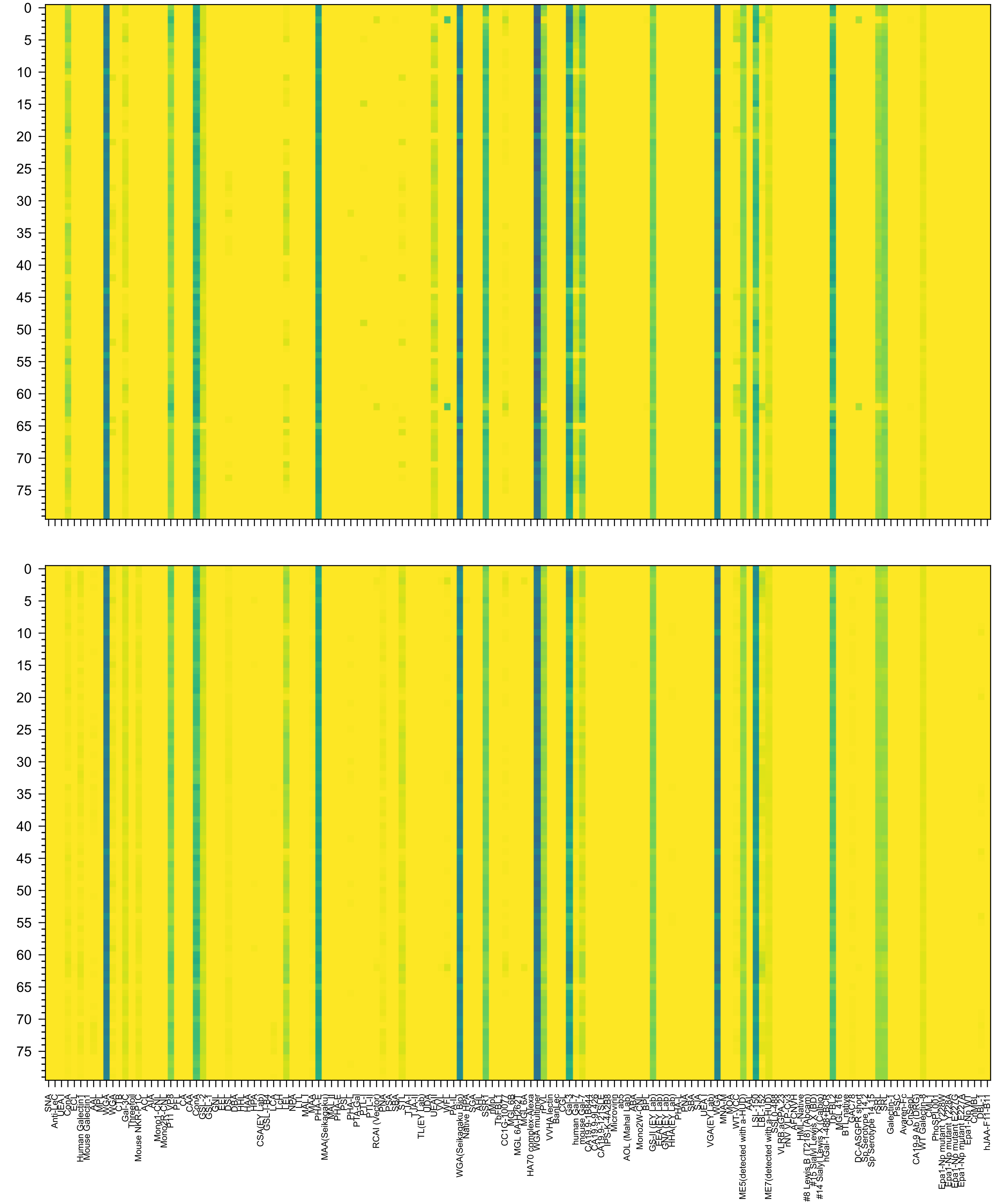

Ac(β1-3)Gal(β1-4)GlcNAc(β1-3)Gal(β1-4)GlcNAc(β1-2)Man(α1-6)[GlcNAc(β1-3)Gal(β1-4)GlcNAc(β1-3)Gal(β1-4)GlcNAc(β1-2)Man(α1-3)]Man(β1-4)GlcNAc(β1-4)Glc

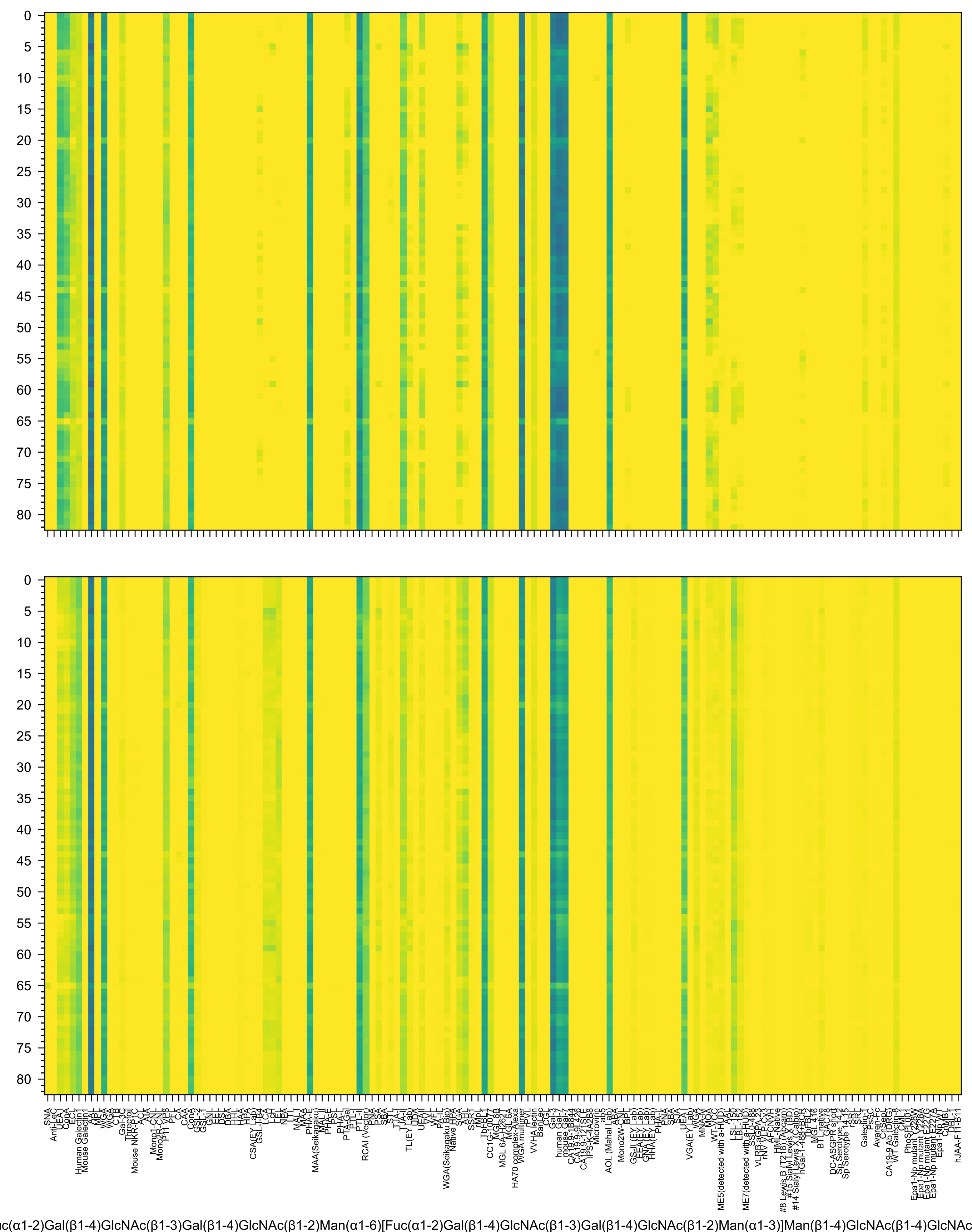
uc(α1-2)Gal(β1-4)GlcNAc(β1-3)Gal(β1-4)GlcNAc(β1-2)Man(α1-6)[Fuc(α1-2)Gal(β1-4)GlcNAc(β1-3)Gal(β1-4)GlcNAc(β1-2)Man(α1-3)]Man(β1-4)GlcNAc(β1-4)GlcNAc

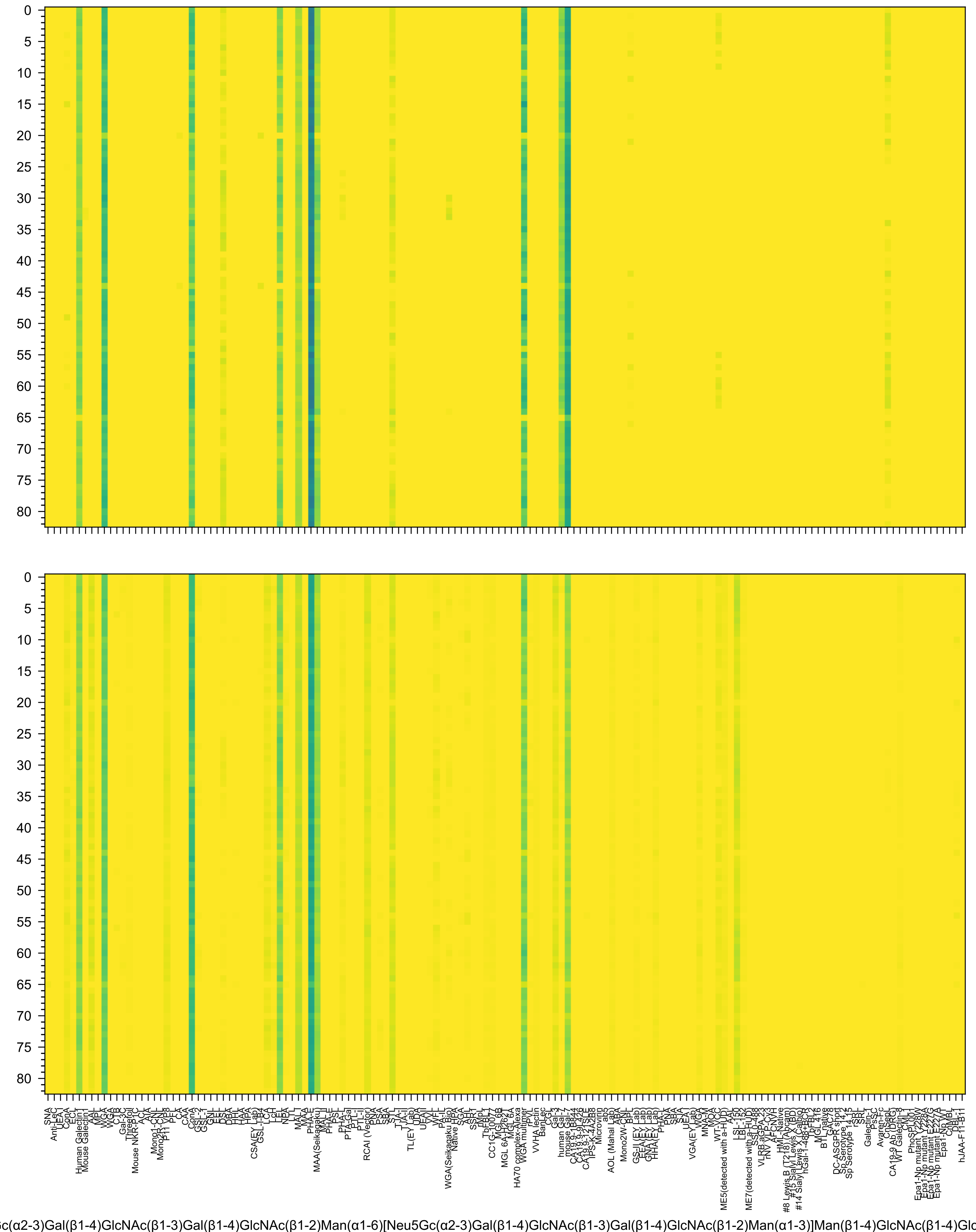
c(α2-3)Gal(β1-4)GlcNAc(β1-3)Gal(β1-4)GlcNAc(β1-2)Man(α1-6)[Neu5Gc(α2-3)Gal(β1-4)GlcNAc(β1-3)Gal(β1-4)GlcNAc(β1-2)Man(α1-3)]Man(β1-4)GlcNAc(β1-4)Glc

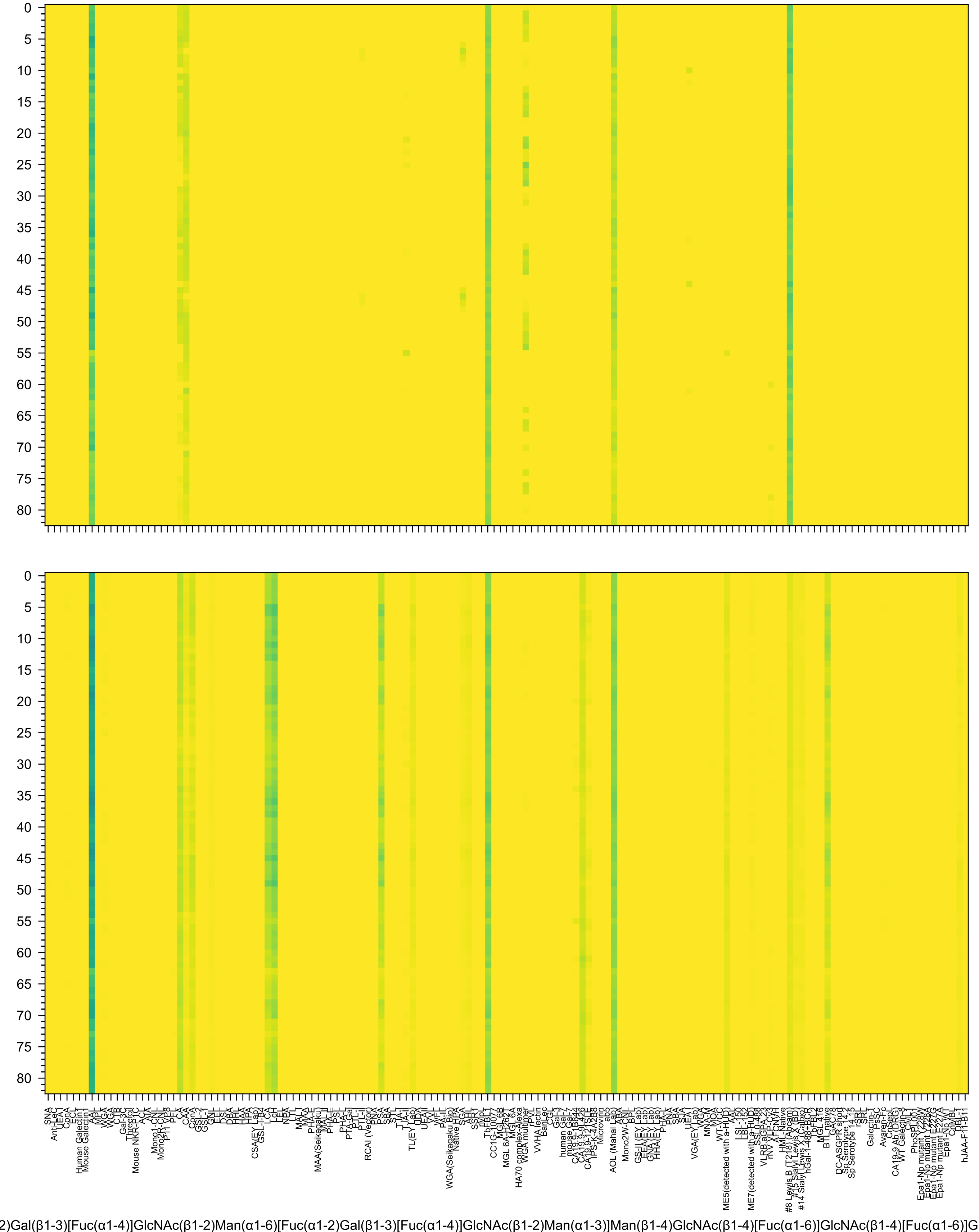
2)Gal(β1-3)[Fuc(α1-4)]GlcNAc(β1-2)Man(α1-6)[Fuc(α1-2)Gal(β1-3)[Fuc(α1-4)]GlcNAc(β1-2)Man(α1-3)]Man(β1-4)GlcNAc(β1-4)[Fuc(α1-6)]GlcNAc(β1-4)[Fuc(α1-6)]G

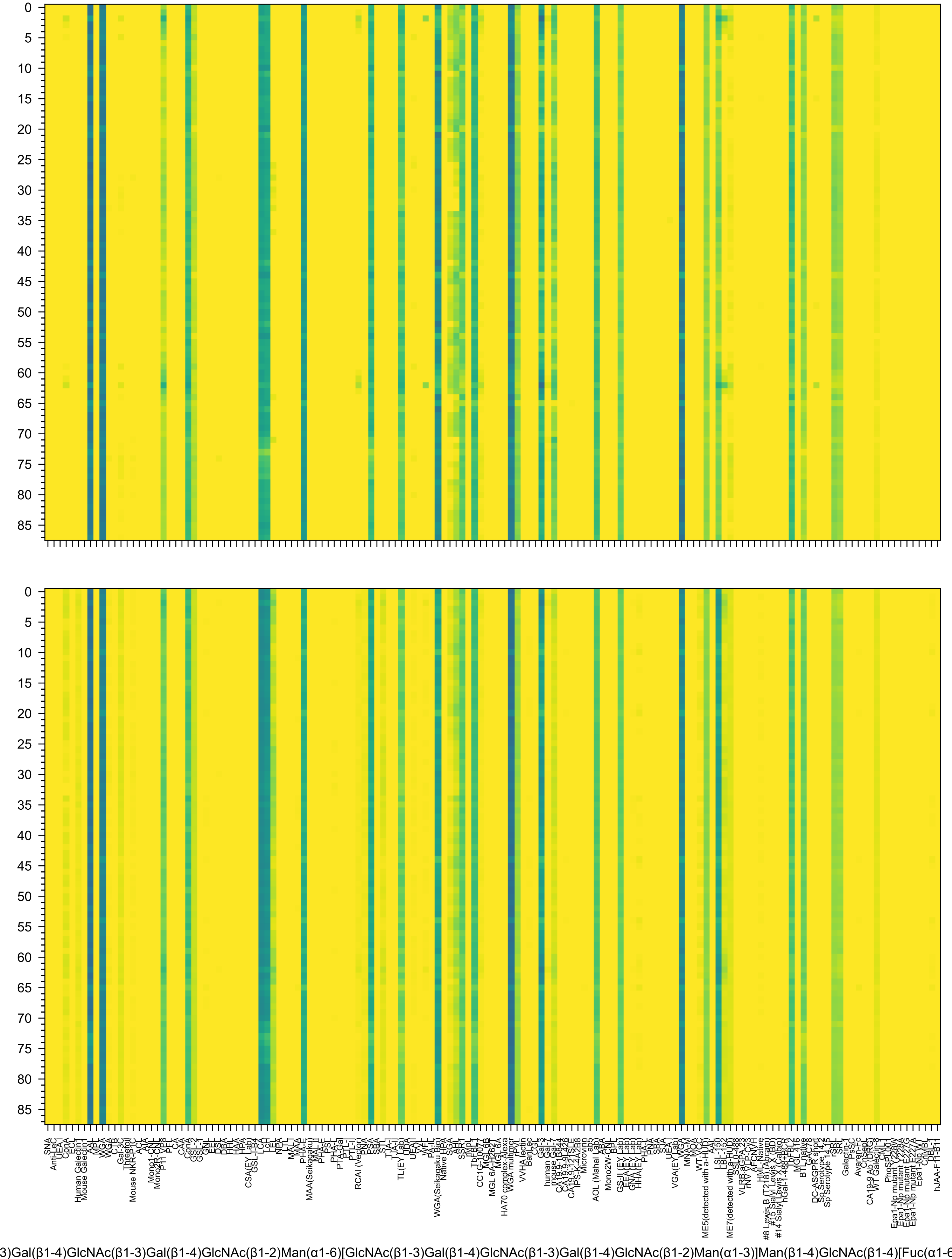
3)Gal(β1-4)GlcNAc(β1-3)Gal(β1-4)GlcNAc(β1-2)Man(α1-6)[GlcNAc(β1-3)Gal(β1-4)GlcNAc(β1-3)Gal(β1-4)GlcNAc(β1-2)Man(α1-3)]Man(β1-4)GlcNAc(β1-4)[Fuc(α1-6

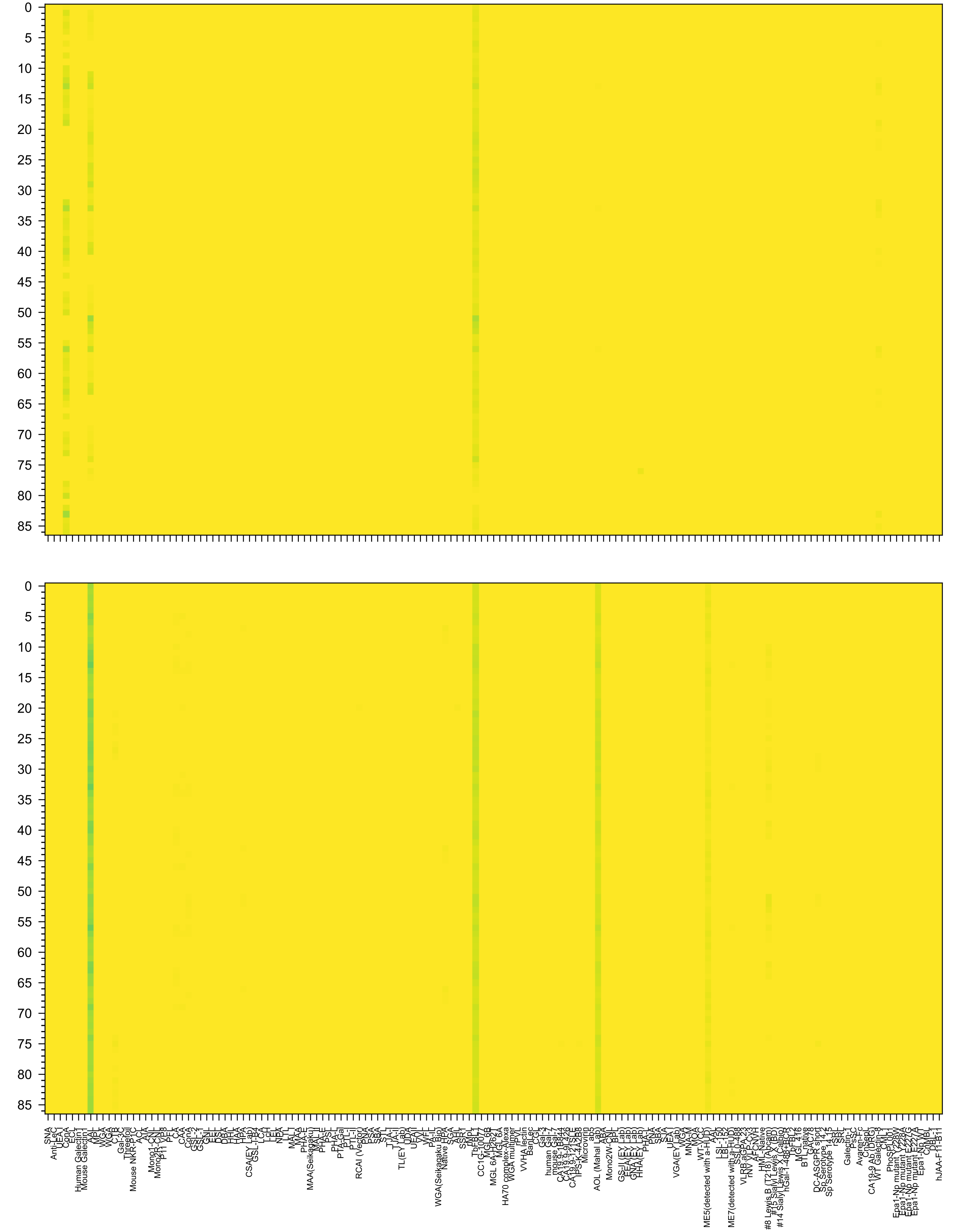

l(β1-4)[Fuc(α1-3)]GlcNAc(β1-2)Man(α1-6)[Fuc(α1-2)Gal(β1-4)[Fuc(α1-3)]GlcNAc(β1-4)[Fuc(α1-2)Gal(β1-4)[Fuc(α1-3)]GlcNAc(β1-2)]Man(α1-3)]Man(β1-4)GlcNAc(β1

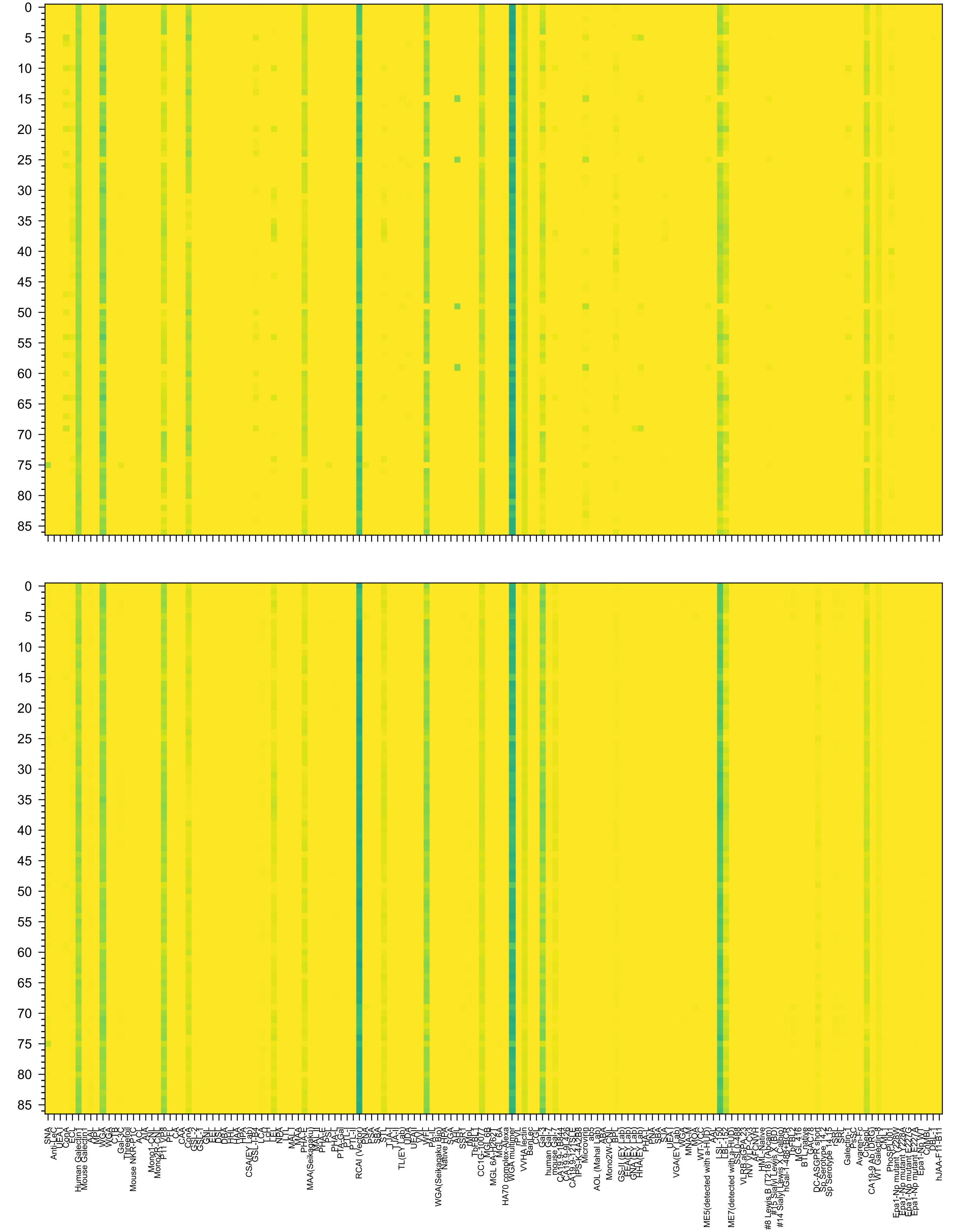

Ac(β1-3)Gal(β1-4)GlcNAc(β1-3)Gal(β1-4)GlcNAc(β1-2)Man(α1-6)[Gal(β1-4)GlcNAc(β1-3)Gal(β1-4)GlcNAc(β1-3)Gal(β1-4)GlcNAc(β1-2)Man(α1-3)]Man(β1-4)GlcNAc

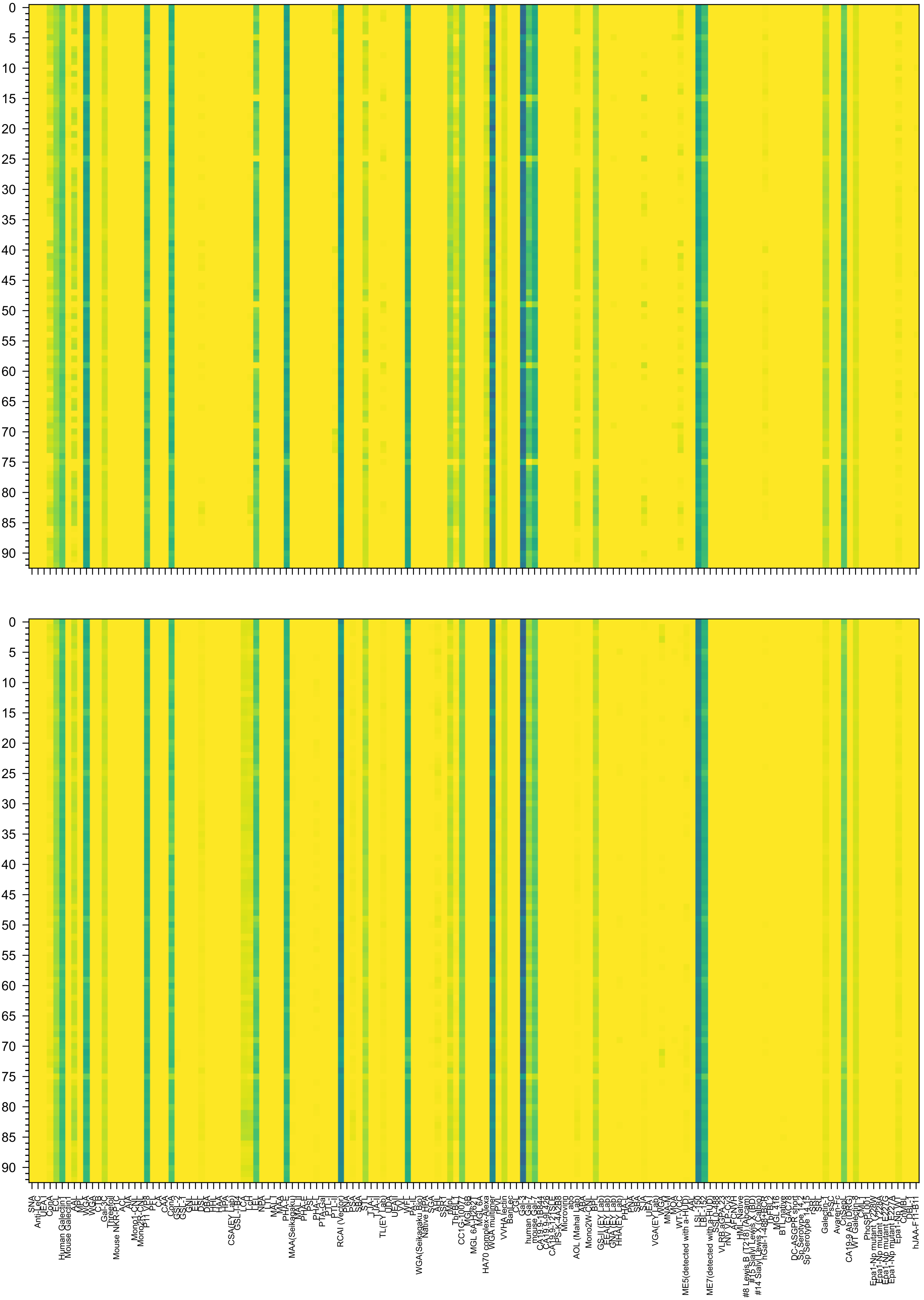

c(β1-3)Gal(β1-4)GlcNAc(β1-3)Gal(β1-4)GlcNAc(β1-2)Man(α1-6)[Gal(β1-4)GlcNAc(β1-3)Gal(β1-4)GlcNAc(β1-3)Gal(β1-4)GlcNAc(β1-2)Man(α1-3)]Man(β1-4)GlcNAc

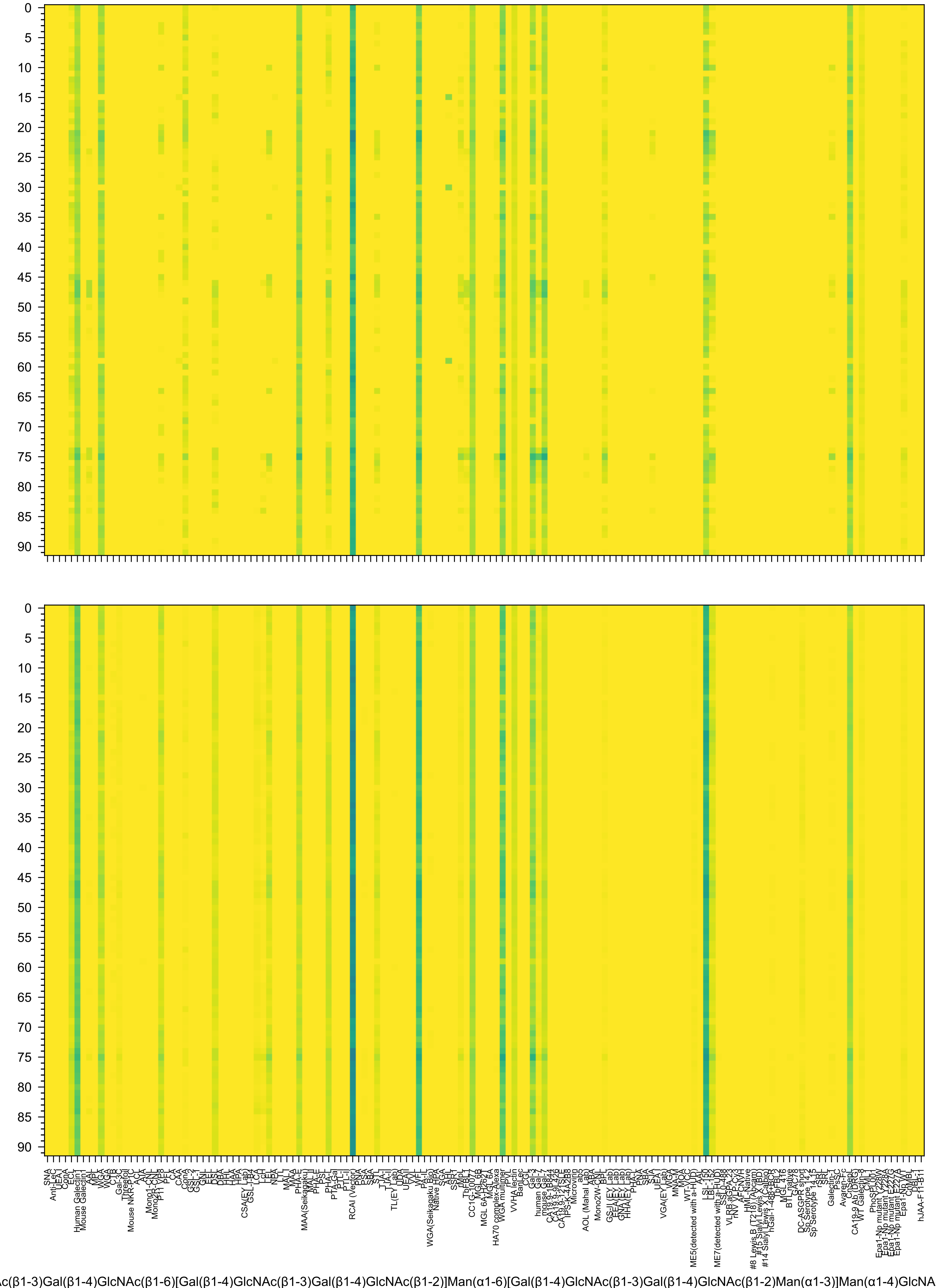
Ac(β1-3)Gal(β1-4)GlcNAc(β1-6)[Gal(β1-4)GlcNAc(β1-3)Gal(β1-4)GlcNAc(β1-2)]Man(α1-6)[Gal(β1-4)GlcNAc(β1-3)Gal(β1-4)GlcNAc(β1-2)Man(α1-3)]Man(α1-4)GlcNA

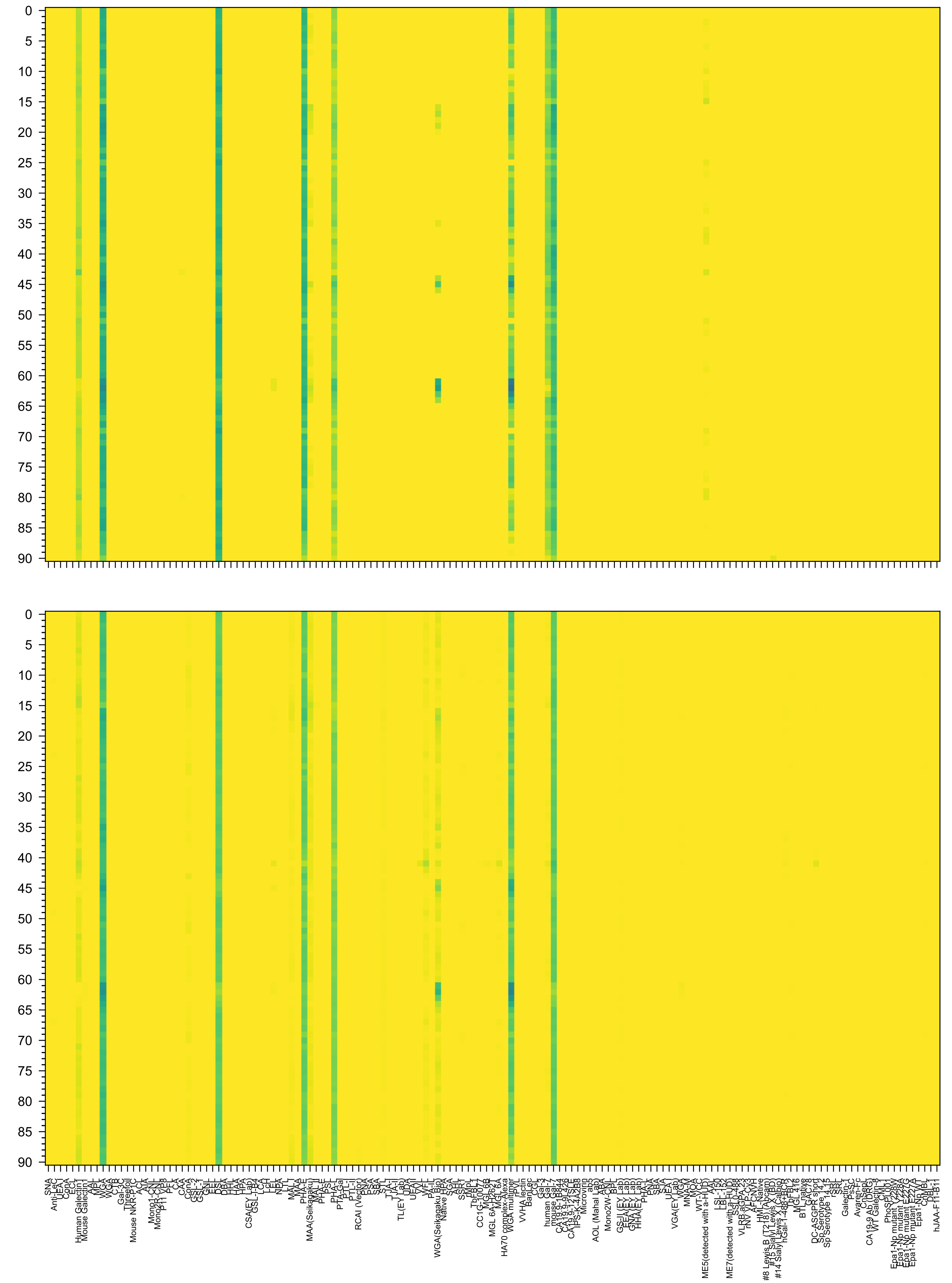

Ac(β1-6)[Neu5Ac(α2-3)Gal(β1-4)GlcNAc(β1-2)]Man(α1-6)[GlcNAc(β1-4)][Neu5Ac(α2-3)Gal(β1-4)GlcNAc(β1-4)][Neu5Ac(α2-3)Gal(β1-4)GlcNAc(β1-2)]Man(α1-3)]Man(

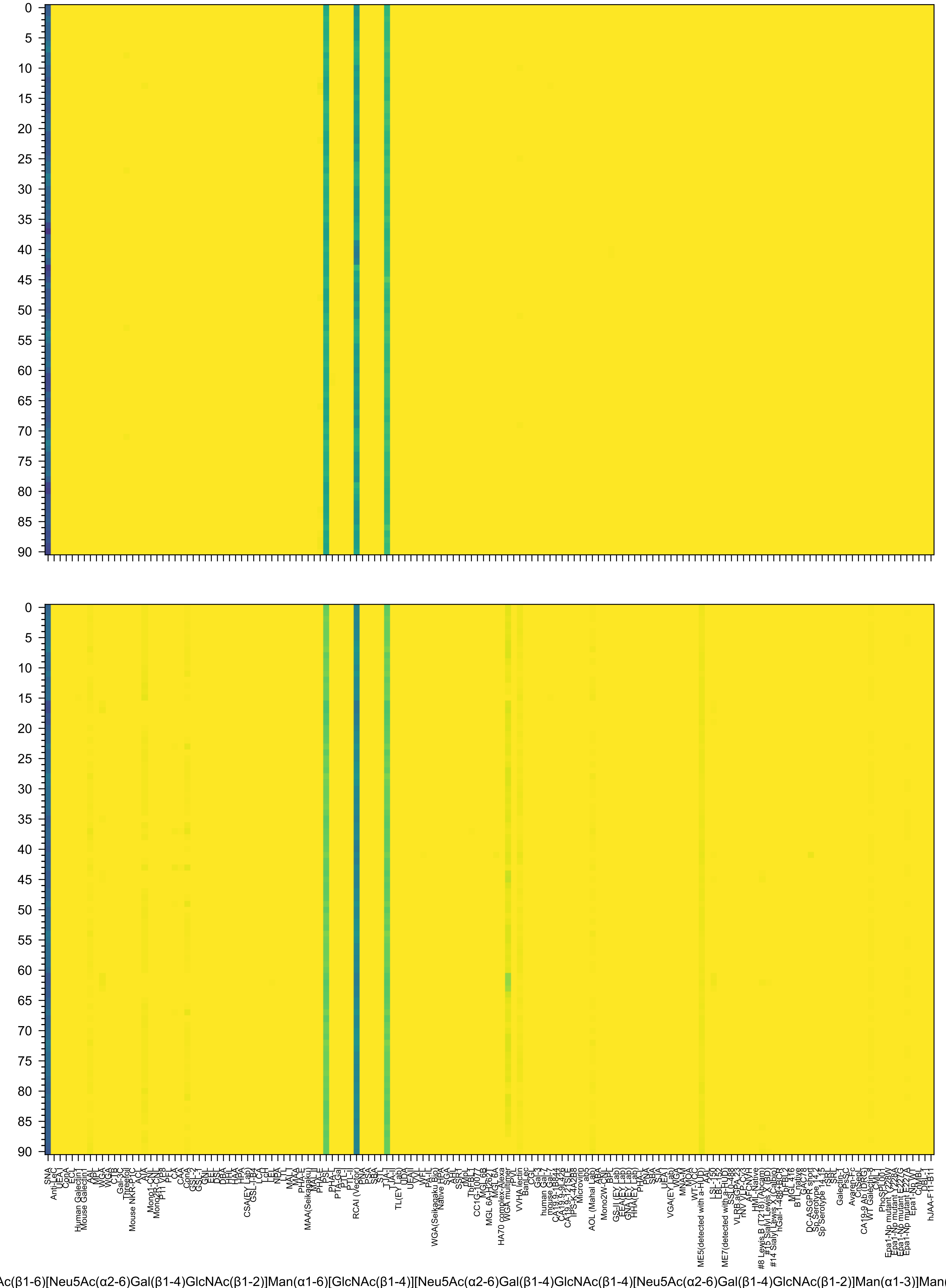
Ac(β1-6)[Neu5Ac(α2-6)Gal(β1-4)GlcNAc(β1-2)]Man(α1-6)[GlcNAc(β1-4)][Neu5Ac(α2-6)Gal(β1-4)GlcNAc(β1-4)[Neu5Ac(α2-6)Gal(β1-4)GlcNAc(β1-2)]Man(α1-3)]Man

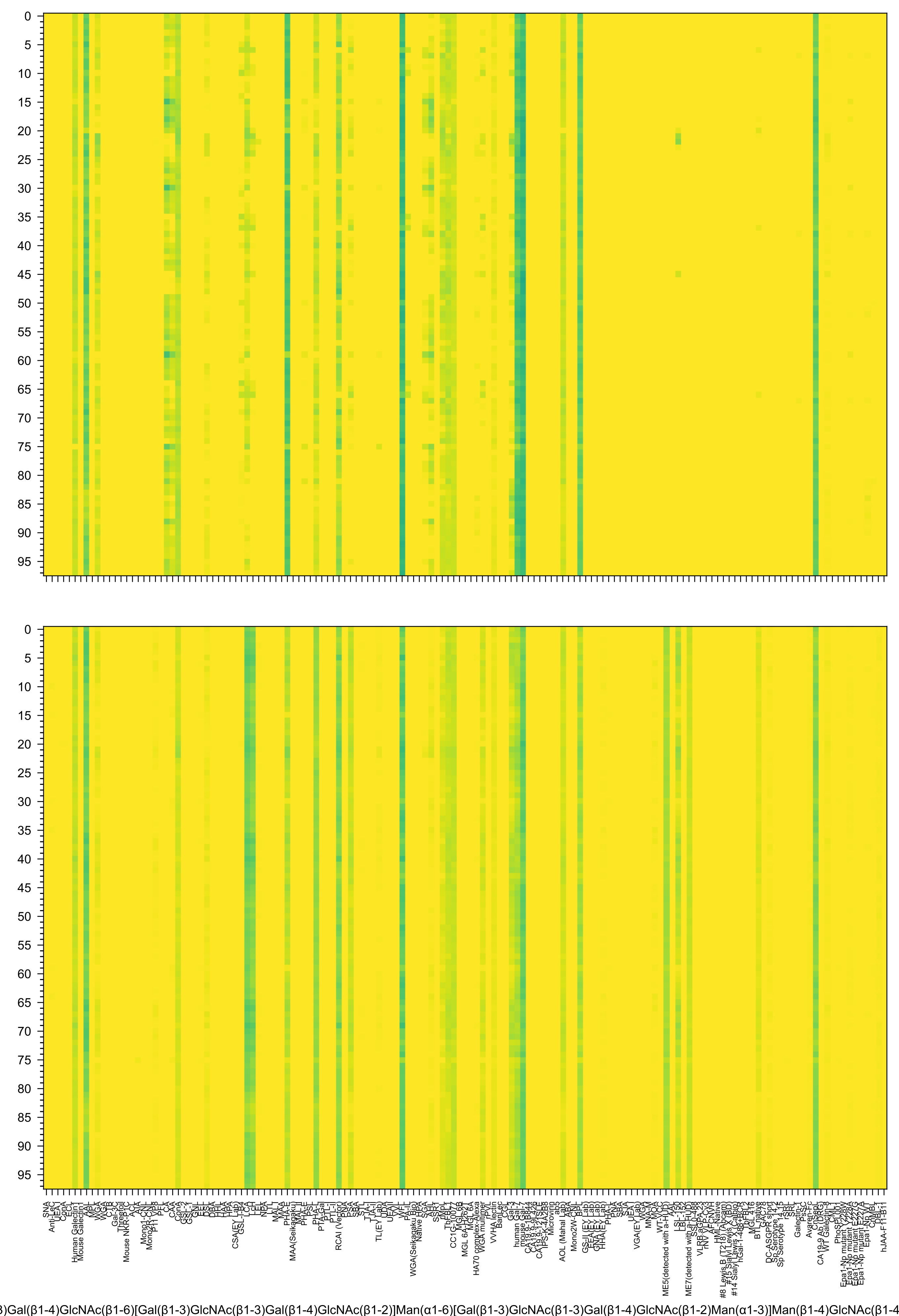
)Gal(β1-4)GlcNAc(β1-6)[Gal(β1-3)GlcNAc(β1-3)Gal(β1-4)GlcNAc(β1-2)]Man(α1-6)[Gal(β1-3)GlcNAc(β1-3)Gal(β1-4)GlcNAc(β1-2)Man(α1-3)]Man(β1-4)GlcNAc(β1-4

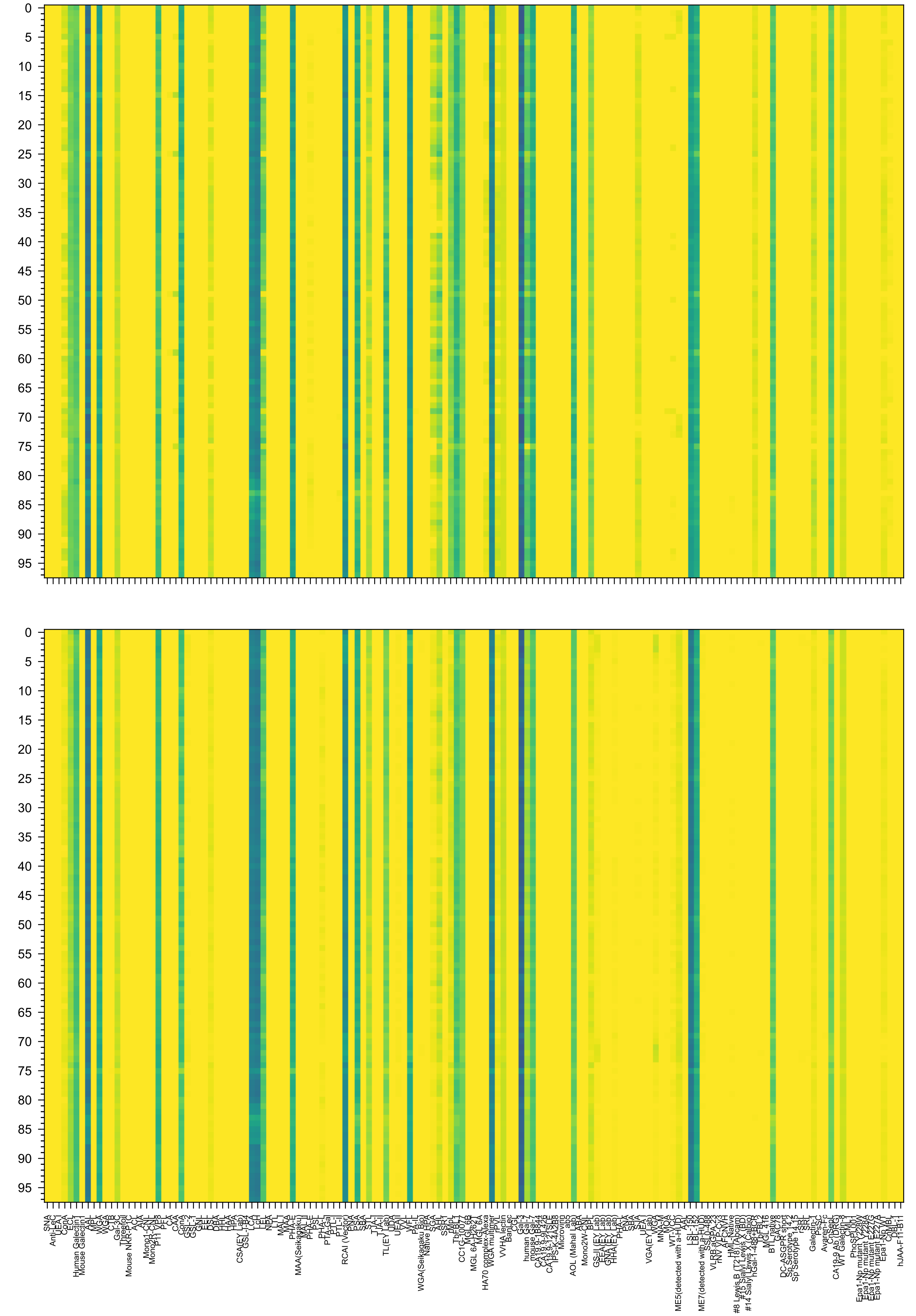

3)Gal(β1-4)GlcNAc(β1-3)Gal(β1-4)GlcNAc(β1-2)Man(α1-6)[Gal(β1-4)GlcNAc(β1-3)Gal(β1-4)GlcNAc(β1-3)Gal(β1-4)GlcNAc(β1-2)Man(α1-3)]Man(β1-4)GlcNAc(β1-4)

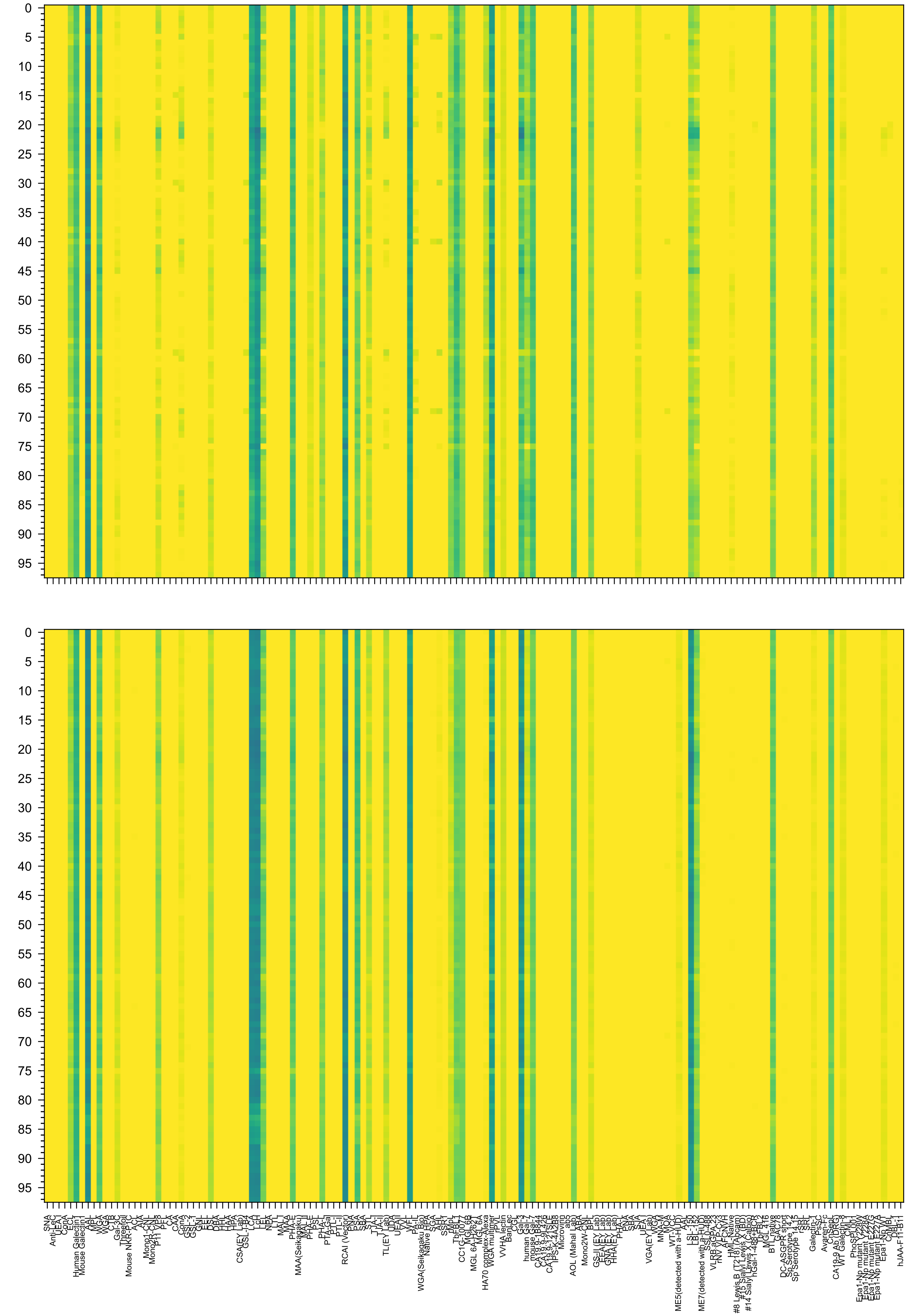

β1-3)Gal(β1-4)GlcNAc(β1-6)[Gal(β1-4)GlcNAc(β1-3)Gal(β1-4)GlcNAc(β1-2)]Man(α1-6)[Gal(β1-4)GlcNAc(β1-3)Gal(β1-4)GlcNAc(β1-2)Man(α1-3)]Man(β1-4)GlcNAc(β1-4

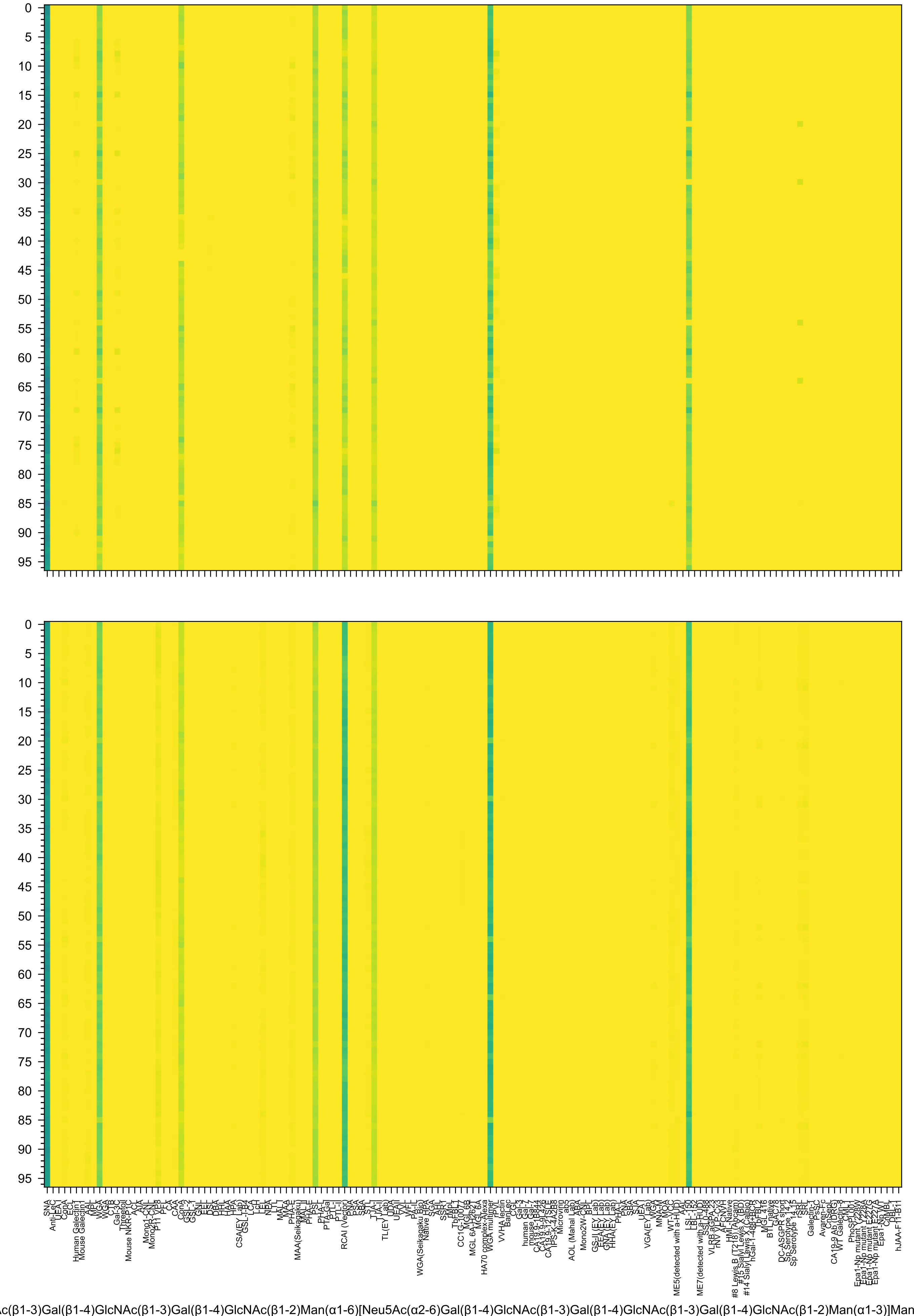
Ac(β1-3)Gal(β1-4)GlcNAc(β1-3)Gal(β1-4)GlcNAc(β1-2)Man(α1-6)[Neu5Ac(α2-6)Gal(β1-4)GlcNAc(β1-3)Gal(β1-4)GlcNAc(β1-3)Gal(β1-4)GlcNAc(β1-2)Man(α1-3)]Man

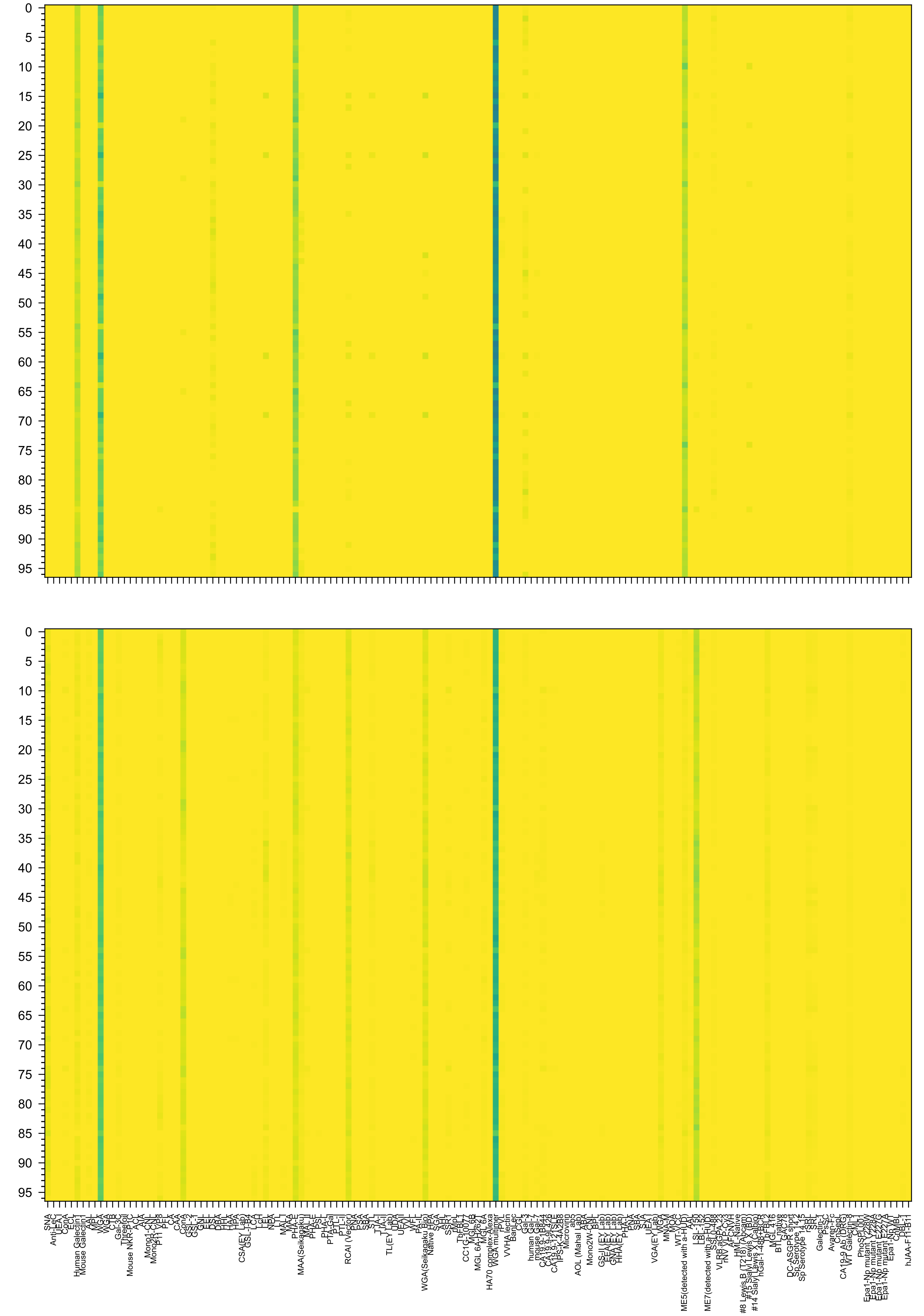

Ac(β1-3)Gal(β1-4)GlcNAc(β1-3)Gal(β1-4)GlcNAc(β1-2)Man(α1-6)[Neu5Ac(α2-3)Gal(β1-4)GlcNAc(β1-3)Gal(β1-4)GlcNAc(β1-3)Gal(β1-4)GlcNAc(β1-2)Man(α1-3)]Man

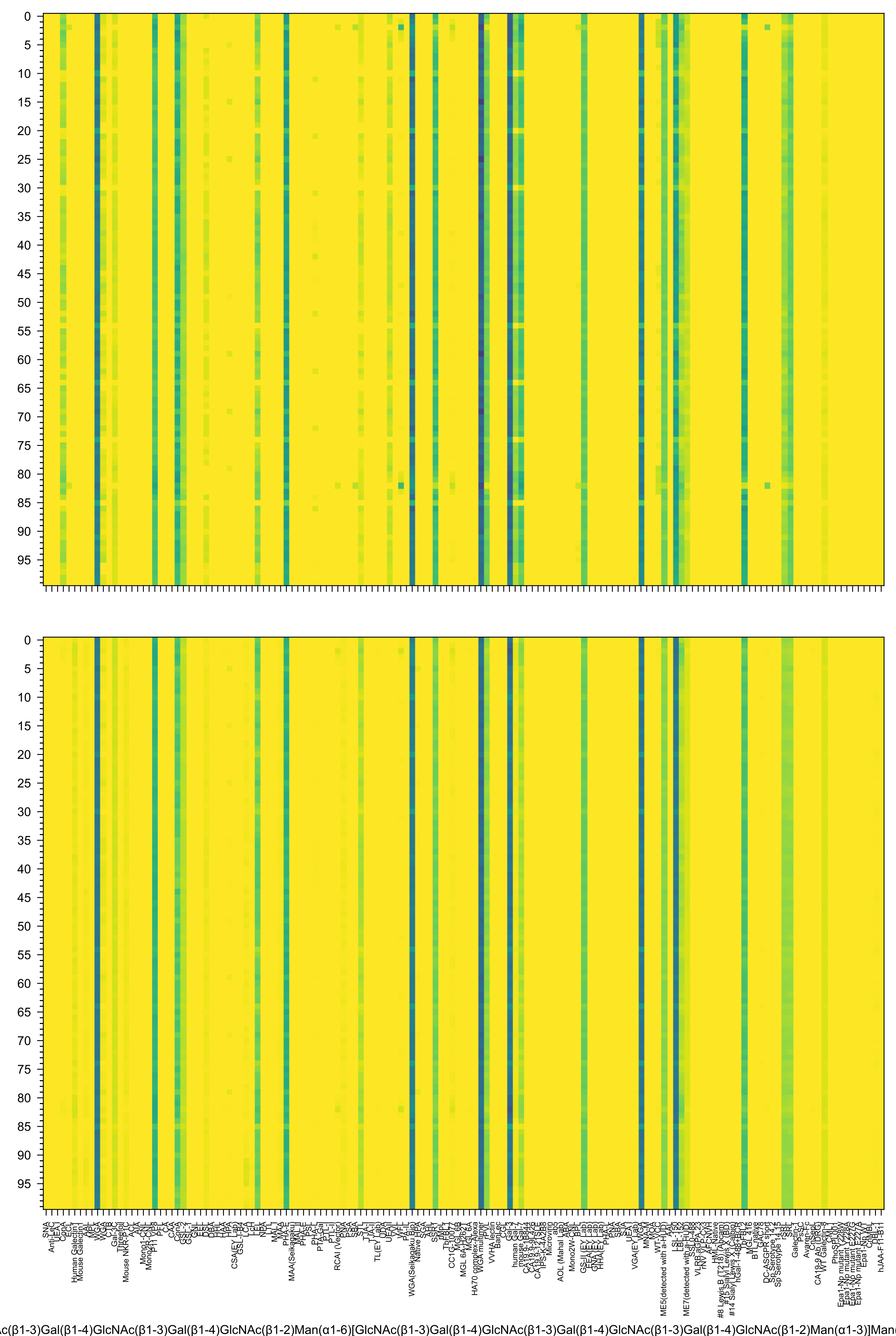

0
5
10
15
20
25
30
35
40
45
50
55
60
65
70
75
80
85
90
95
SNA
Anti-LeC
UEA-I
ConA
ECL
Human Galectin1
Mouse Galectin1
MPL
WGA
CTB
Gal-3C
Mouse NKR-P1C
ACL
Mono1-CNL
Mono2R-CNL
P11 VP8
CAA
ConA
GSL-1
GNL
DSL
DBA
HPA
CSA(EY Lab)
GSL-I-B4
LcH
NPA
MAL-I
PHA-E
MAA(Seikagaku)
PHA-L
PTL-I
RCAI (Vector)
PSA
SBA
TL(EY Lab)
UDA
VVL
WFL
PA-IL
WGA(Seikagaku Bio)
Native HPA
AAL
SSR1
CC1G-10077
MGL 6A
HA70 complex-Alexa
WGA multimer
VVHA lectin
BanLec
CGL
Gal-3
human Gal-7
Microvirin
AOL (Mahal Lab)
Mono2W-CNL
GS-II(EY Lab)
PNA
SBA
UEA-I
VGA(EY Lab)
MNA-M
MOA
WT-LOC
AAL
ME5(detected with a-HUD)
LSL-150
ME7(detected with a-HUD)
#8 Lewis B
#14 Sialyl Lewis X (Calbio)
hGal-1-488-BCR
MGL 416
BTL_native
DC-ASGPR short
Sp Serotype 14-15
SRL
Galectin-1
Avaren-Fc
CA19-9 A6 (DRG)
WT Galectin-3
PhoSL-001
Epa1-Np mutant
DBL-1
hJAA-F11-B11
c(β1-3)Gal(β1-4)GlcNAc(β1-3)Gal(β1-4)GlcNAc(β1-2)Man(α1-6)[GlcNAc(β1-3)Gal(β1-4)GlcNAc(β1-3)Gal(β1-4)GlcNAc(β1-3)Gal(β1-4)GlcNAc(β1-2)Man(α1-3)]Man

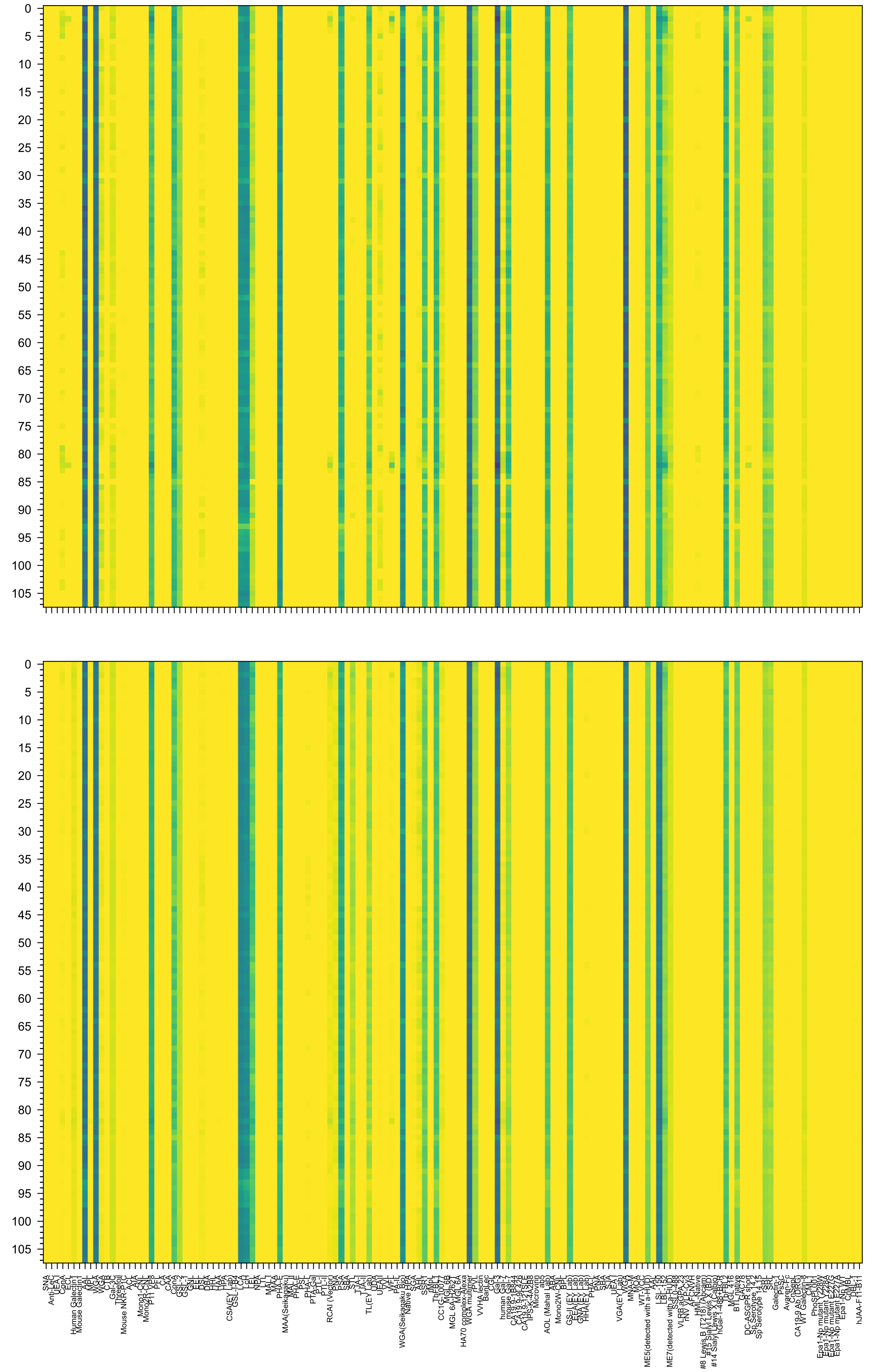

3)Gal(β1-4)GlcNAc(β1-3)Gal(β1-4)GlcNAc(β1-2)Man(α1-6)[GlcNAc(β1-3)Gal(β1-4)GlcNAc(β1-3)Gal(β1-4)GlcNAc(β1-3)Gal(β1-4)GlcNAc(β1-2)Man(α1-3)]Man(β1-4)

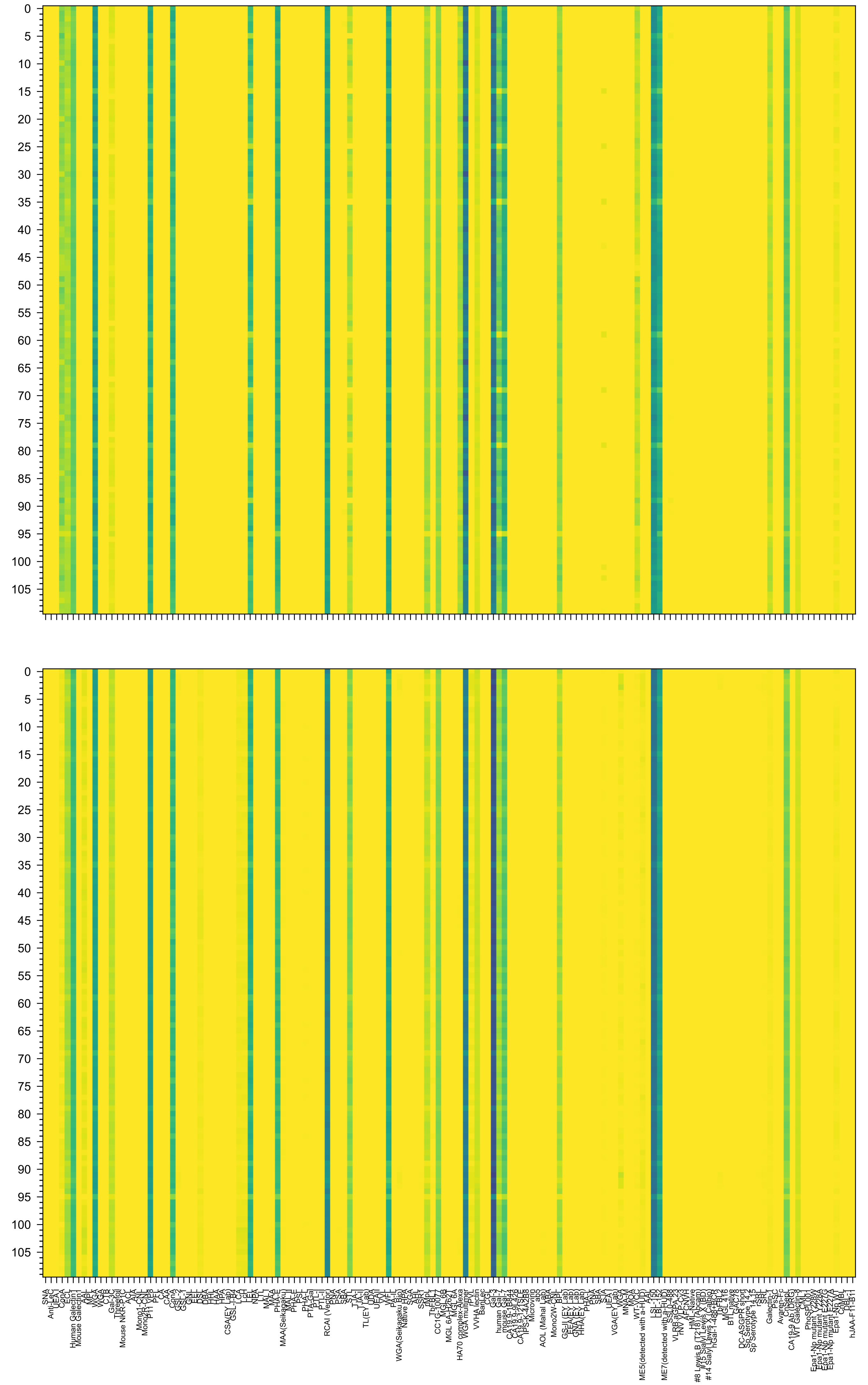

c(β1-3)Gal(β1-4)GlcNAc(β1-3)Gal(β1-4)GlcNAc(β1-2)Man(α1-6)[Gal(β1-4)GlcNAc(β1-3)Gal(β1-4)GlcNAc(β1-3)Gal(β1-4)GlcNAc(β1-3)Gal(β1-4)GlcNAc(β1-2)Man(α

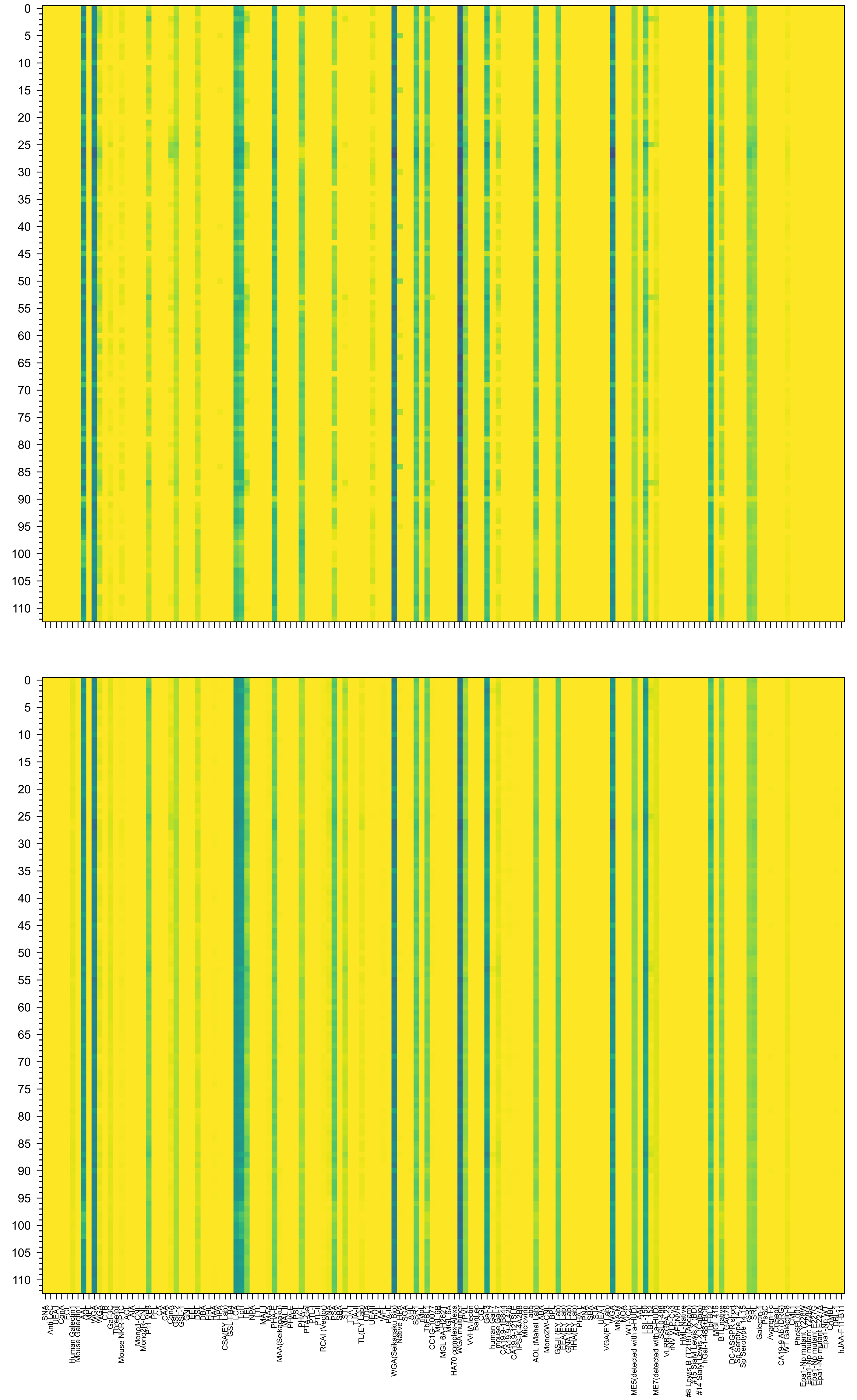

1-4)GlcNAc(β1-6)[GlcNAc(β1-3)Gal(β1-4)GlcNAc(β1-3)Gal(β1-4)GlcNAc(β1-2)]Man(α1-6)[GlcNAc(β1-3)Gal(β1-4)GlcNAc(β1-3)Gal(β1-4)GlcNAc(β1-2)Man(α1-3)]Ma

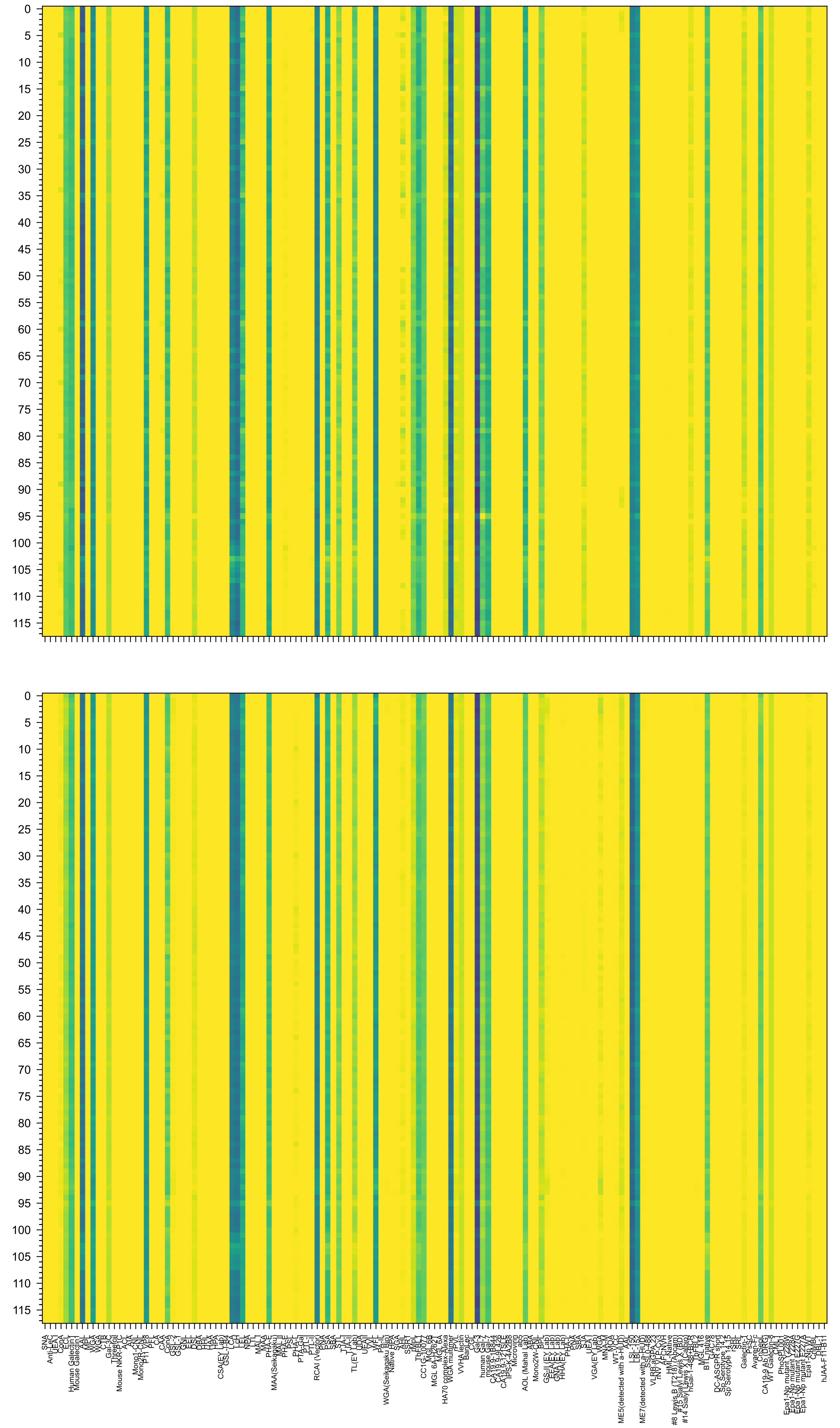

3)Gal(β1-4)GlcNAc(β1-3)Gal(β1-4)GlcNAc(β1-2)Man(α1-6)[Gal(β1-4)GlcNAc(β1-3)Gal(β1-4)GlcNAc(β1-3)Gal(β1-4)GlcNAc(β1-3)Gal(β1-4)GlcNAc(β1-2)Man(α1-3)]M

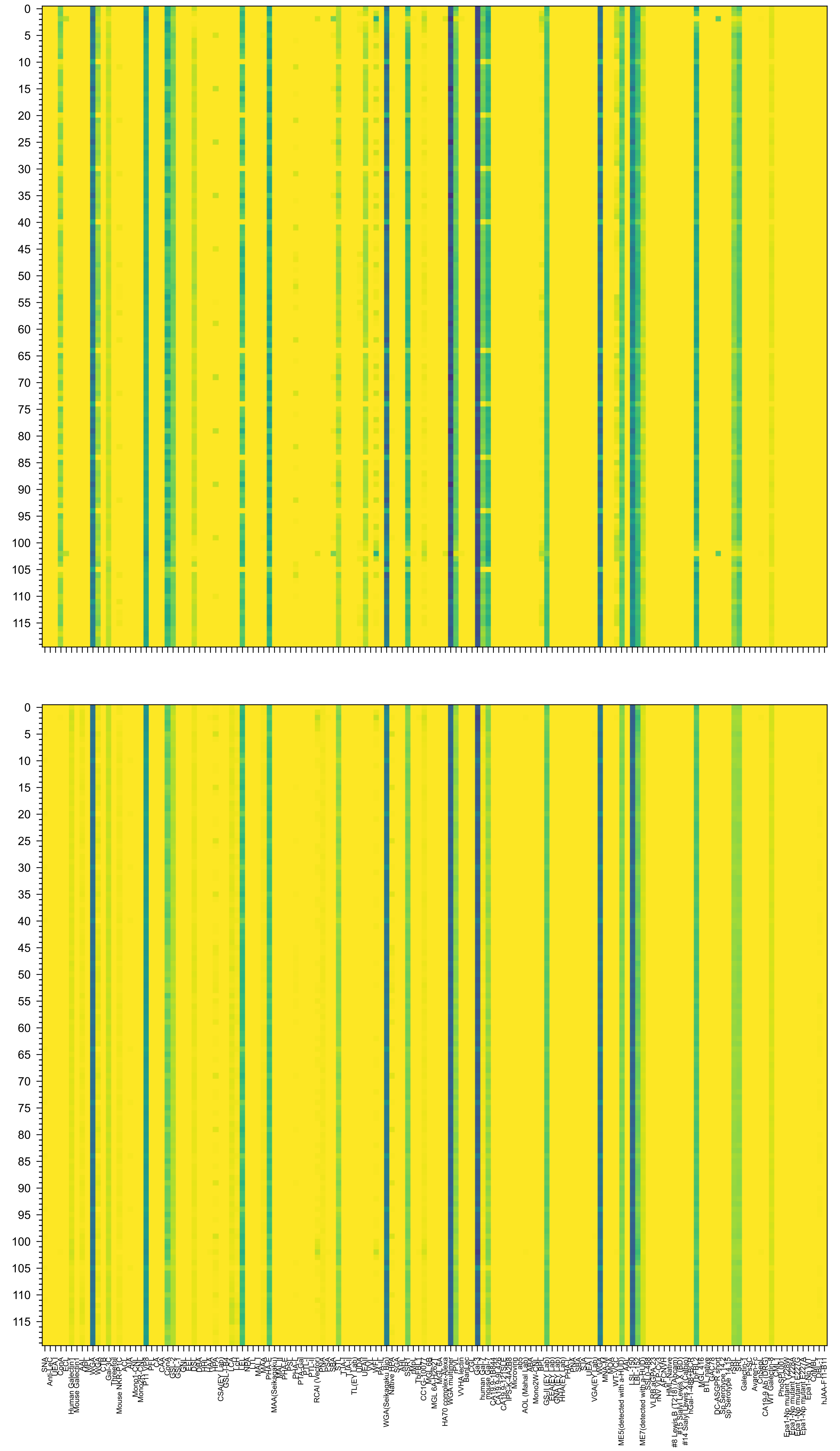

c(β1-3)Gal(β1-4)GlcNAc(β1-3)Gal(β1-4)GlcNAc(β1-2)Man(α1-6)[GlcNAc(β1-3)Gal(β1-4)GlcNAc(β1-3)Gal(β1-4)GlcNAc(β1-3)Gal(β1-4)GlcNAc(β1-3)Gal(β1-4)GlcNA

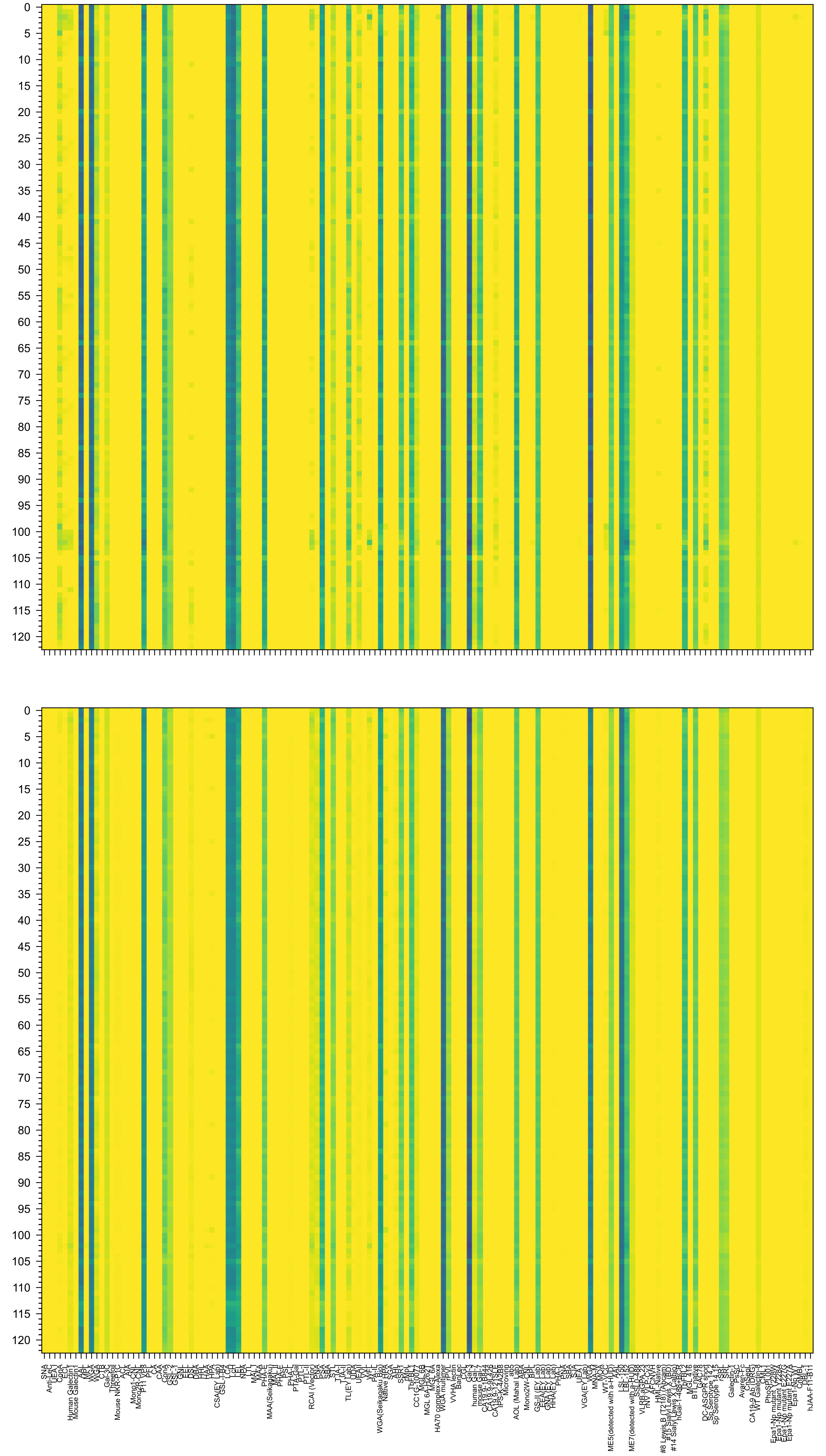

3)Gal(β1-4)GlcNAc(β1-3)Gal(β1-4)GlcNAc(β1-2)Man(α1-6)[GlcNAc(β1-3)Gal(β1-4)GlcNAc(β1-3)Gal(β1-4)GlcNAc(β1-3)Gal(β1-4)GlcNAc(β1-3)Gal(β1-4)GlcNAc(β1-

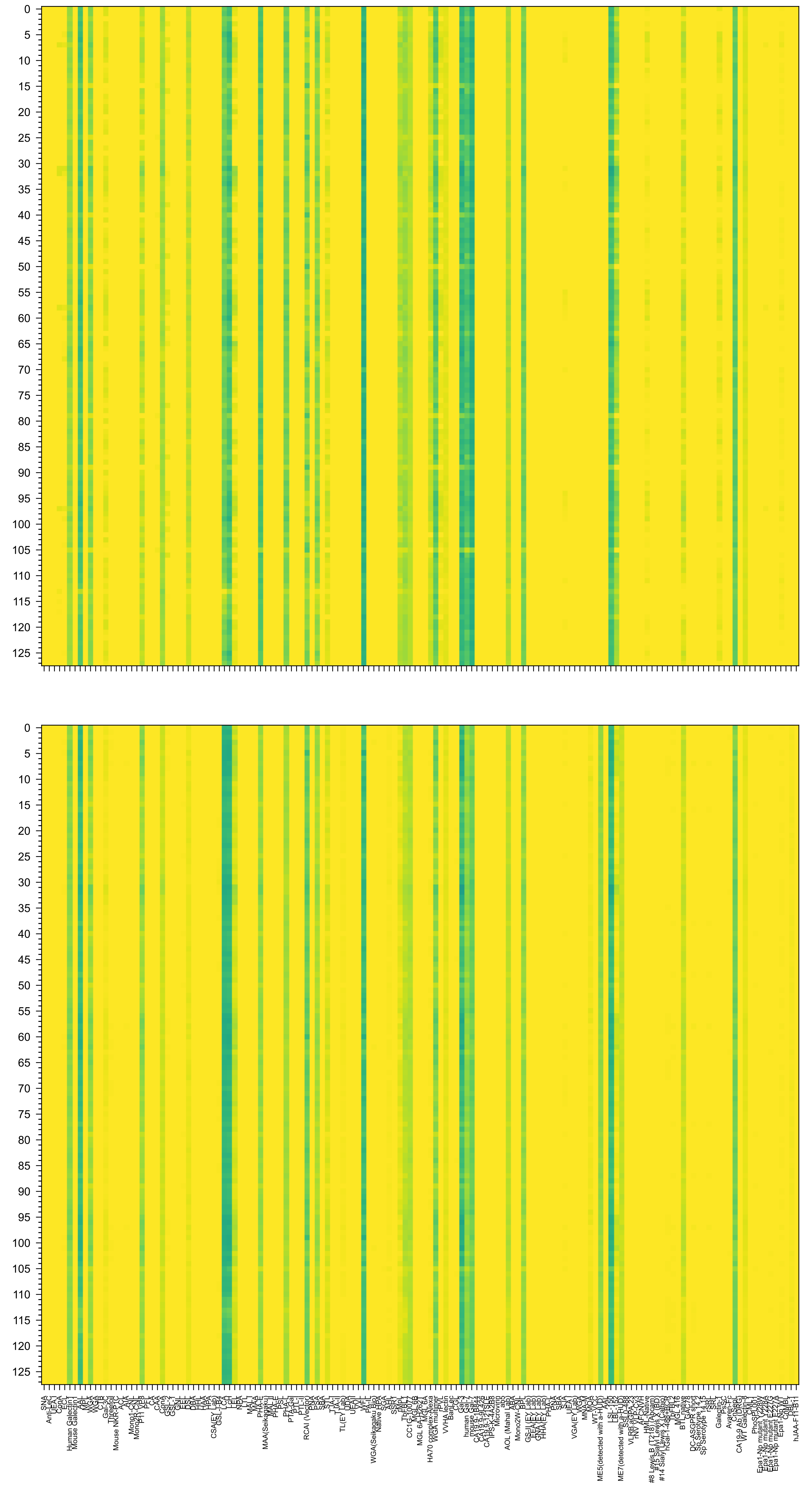

GlcNAc(β1-6)[Gal(β1-3)GlcNAc(β1-3)Gal(β1-4)GlcNAc(β1-3)Gal(β1-4)GlcNAc(β1-2)]Man(α1-6)[Gal(β1-3)GlcNAc(β1-3)Gal(β1-4)GlcNAc(β1-3)Gal(β1-4)GlcNAc(β1-2

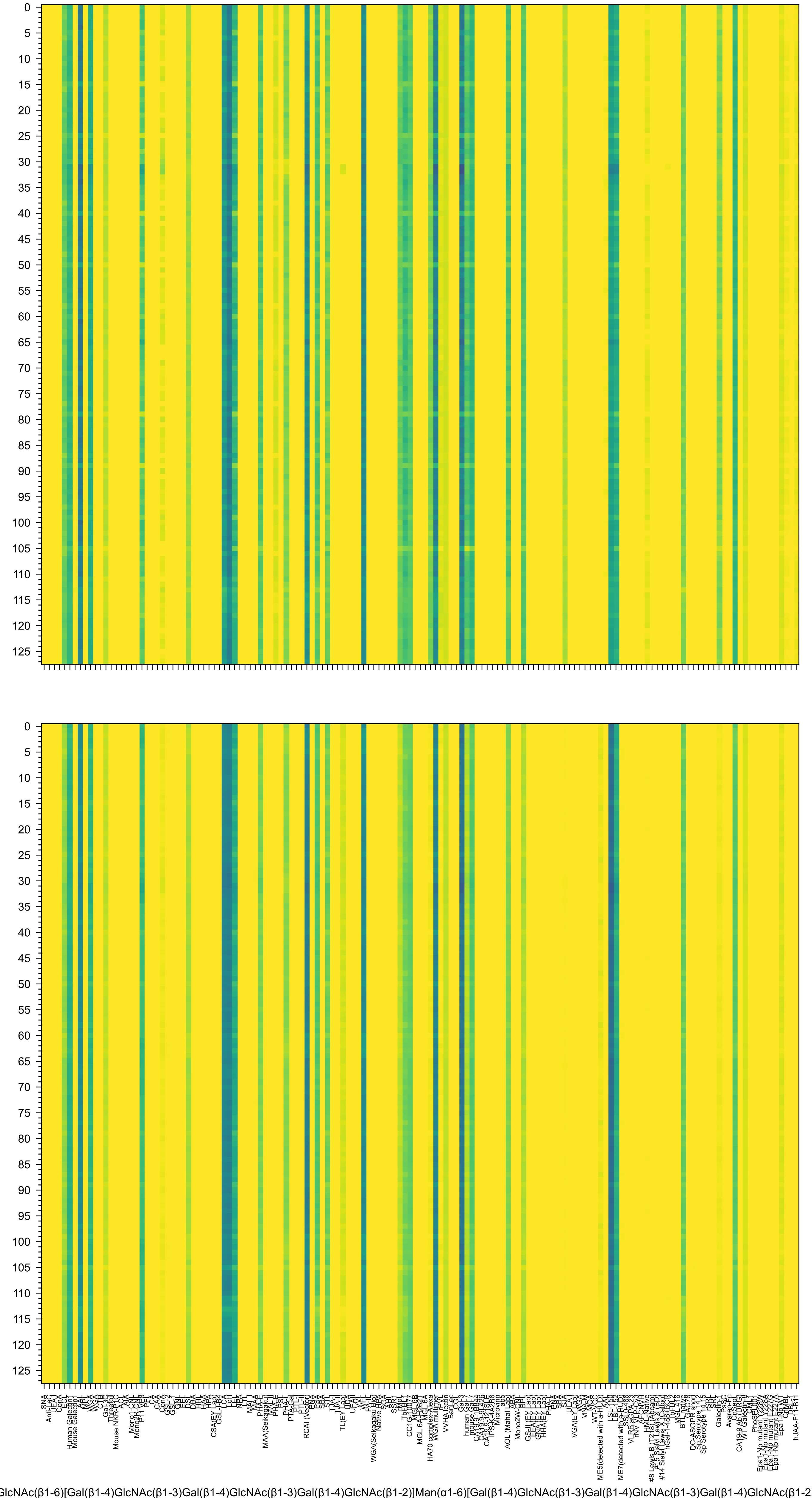
GlcNAc(β1-6)[Gal(β1-4)GlcNAc(β1-3)Gal(β1-4)GlcNAc(β1-3)Gal(β1-4)GlcNAc(β1-2)]Man(α1-6)[Gal(β1-4)GlcNAc(β1-3)Gal(β1-4)GlcNAc(β1-3)Gal(β1-4)GlcNAc(β1-2

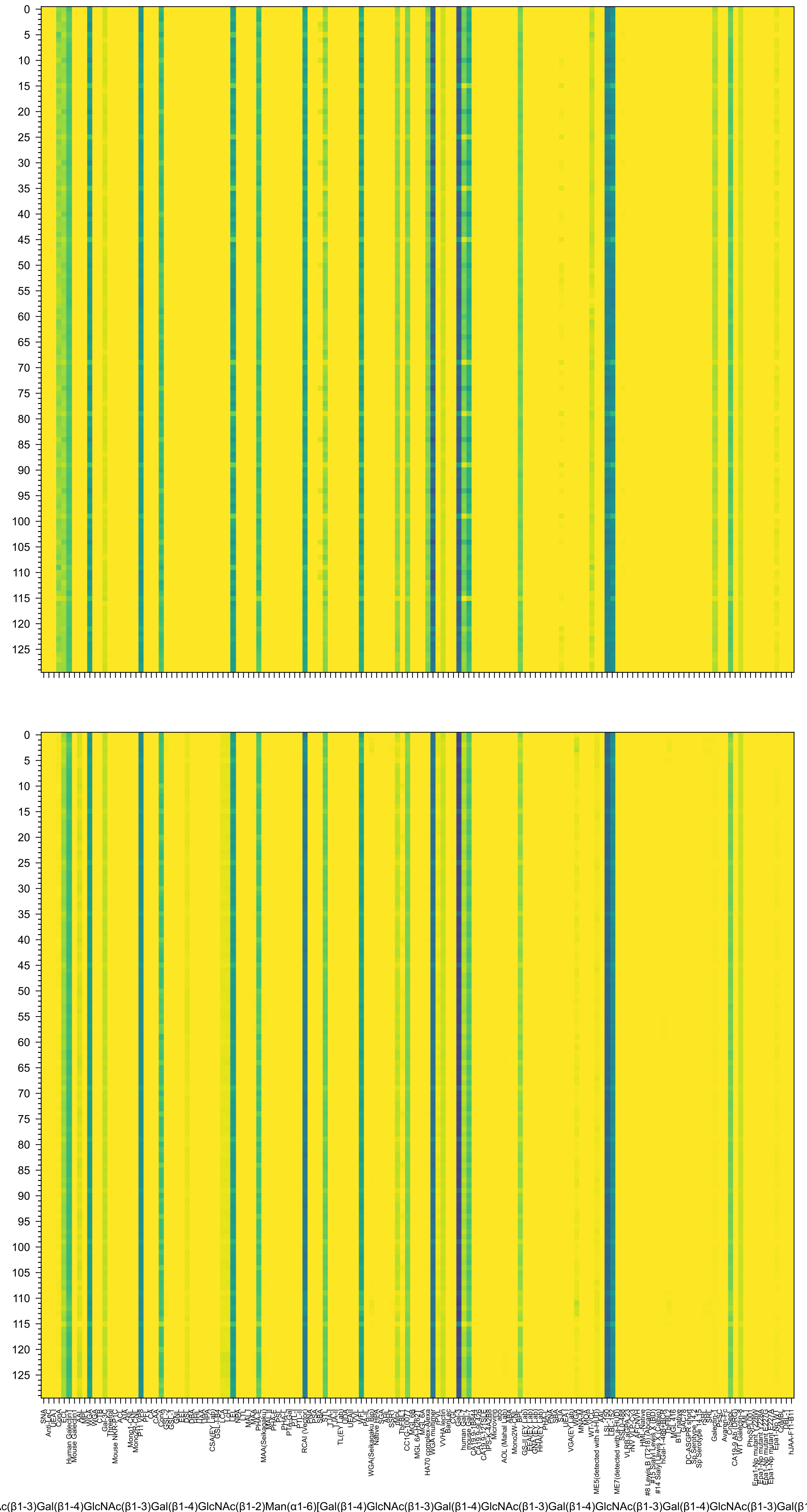
c(β1-3)Gal(β1-4)GlcNAc(β1-3)Gal(β1-4)GlcNAc(β1-2)Man(α1-6)[Gal(β1-4)GlcNAc(β1-3)Gal(β1-4)GlcNAc(β1-3)Gal(β1-4)GlcNAc(β1-3)Gal(β1-4)GlcNAc(β1-3)Gal(β

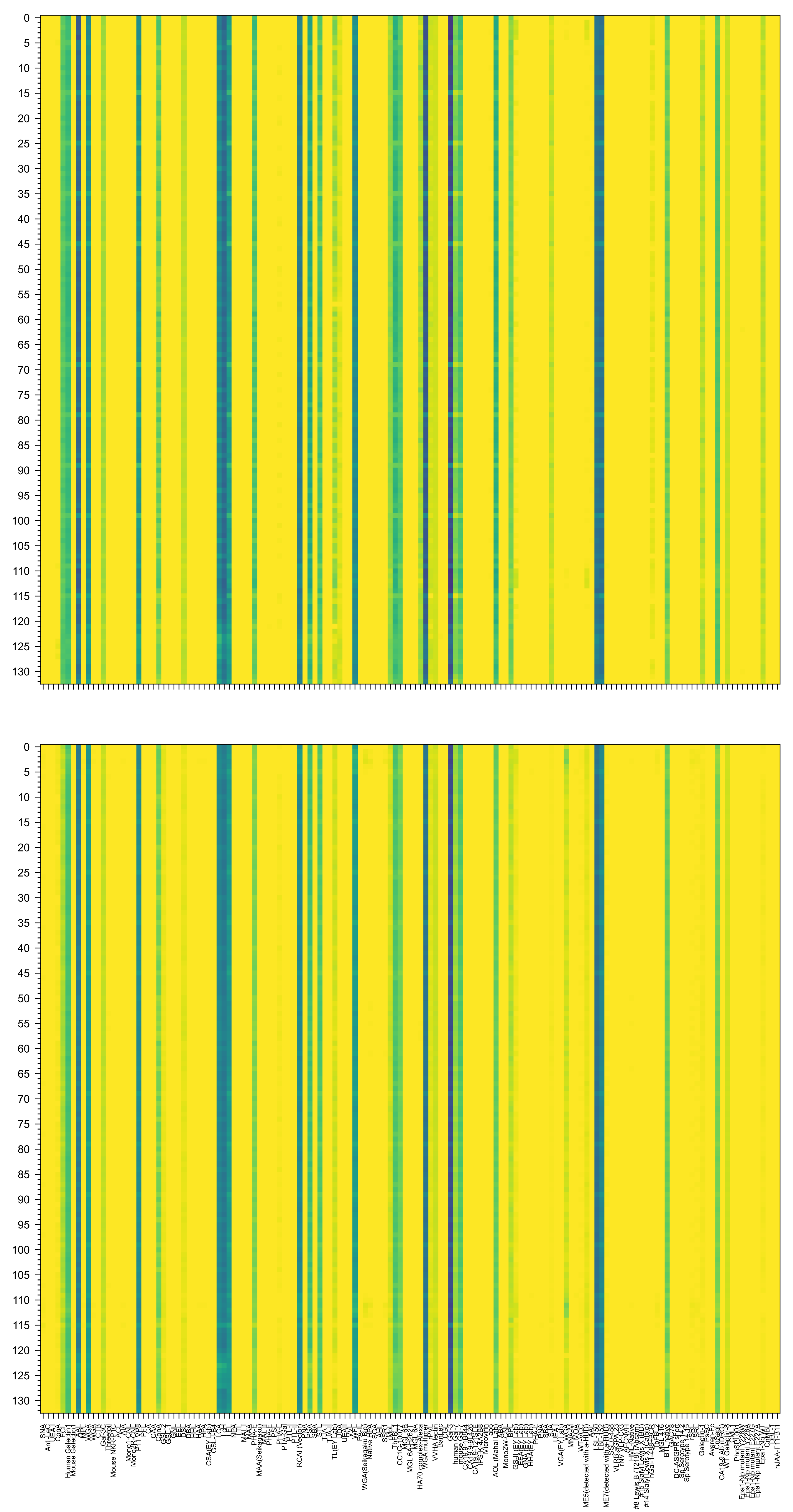

3)Gal(β1-4)GlcNAc(β1-3)Gal(β1-4)GlcNAc(β1-2)Man(α1-6)[Gal(β1-4)GlcNAc(β1-3)Gal(β1-4)GlcNAc(β1-3)Gal(β1-4)GlcNAc(β1-3)Gal(β1-4)GlcNAc(β1-3)Gal(β1-4)Gl

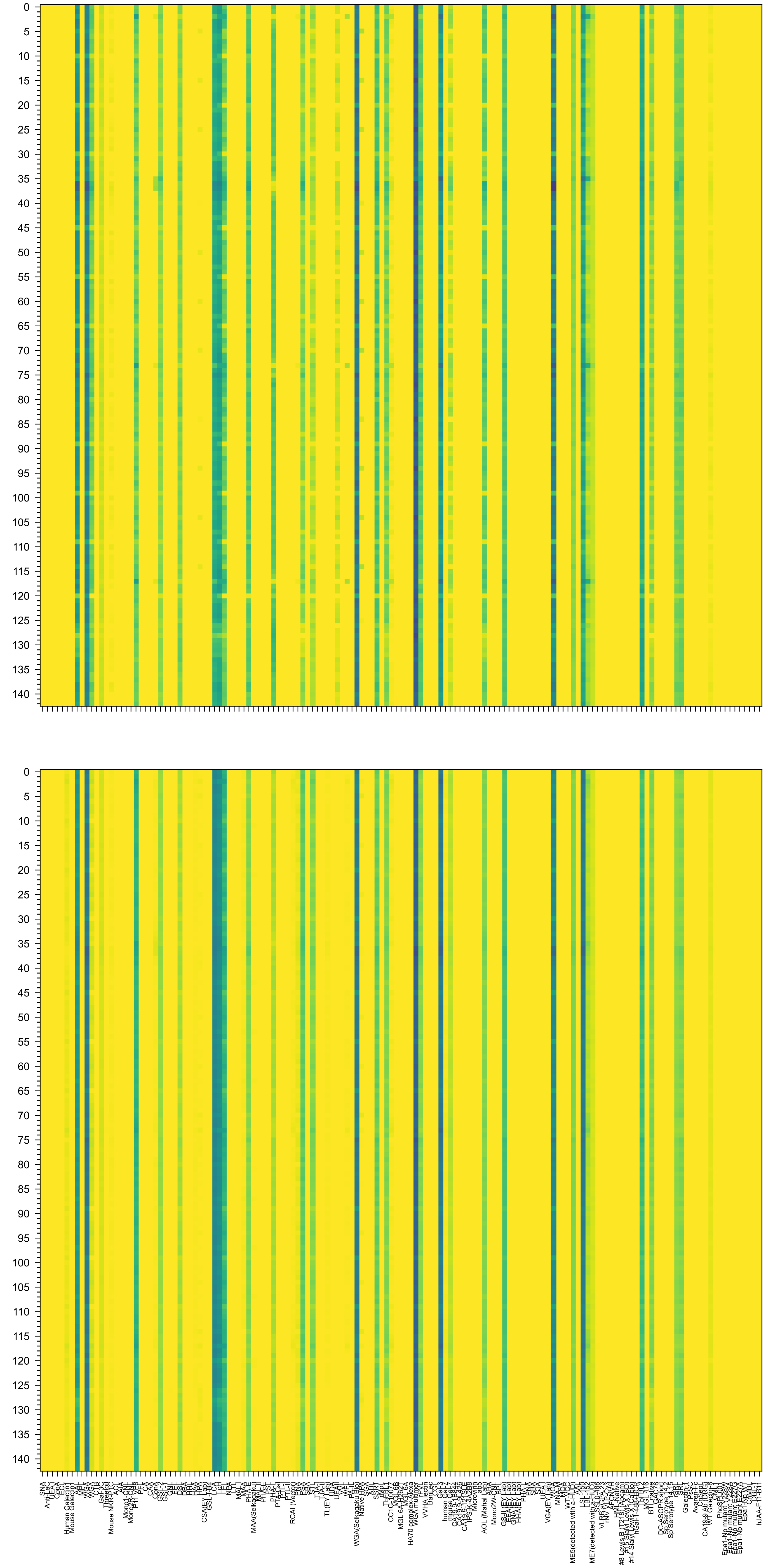

(β1-6)[GlcNAc(β1-3)Gal(β1-4)GlcNAc(β1-3)Gal(β1-4)GlcNAc(β1-3)Gal(β1-4)GlcNAc(β1-2)]Man(α1-6)[GlcNAc(β1-3)Gal(β1-4)GlcNAc(β1-3)Gal(β1-4)GlcNAc(β1-3)Ga

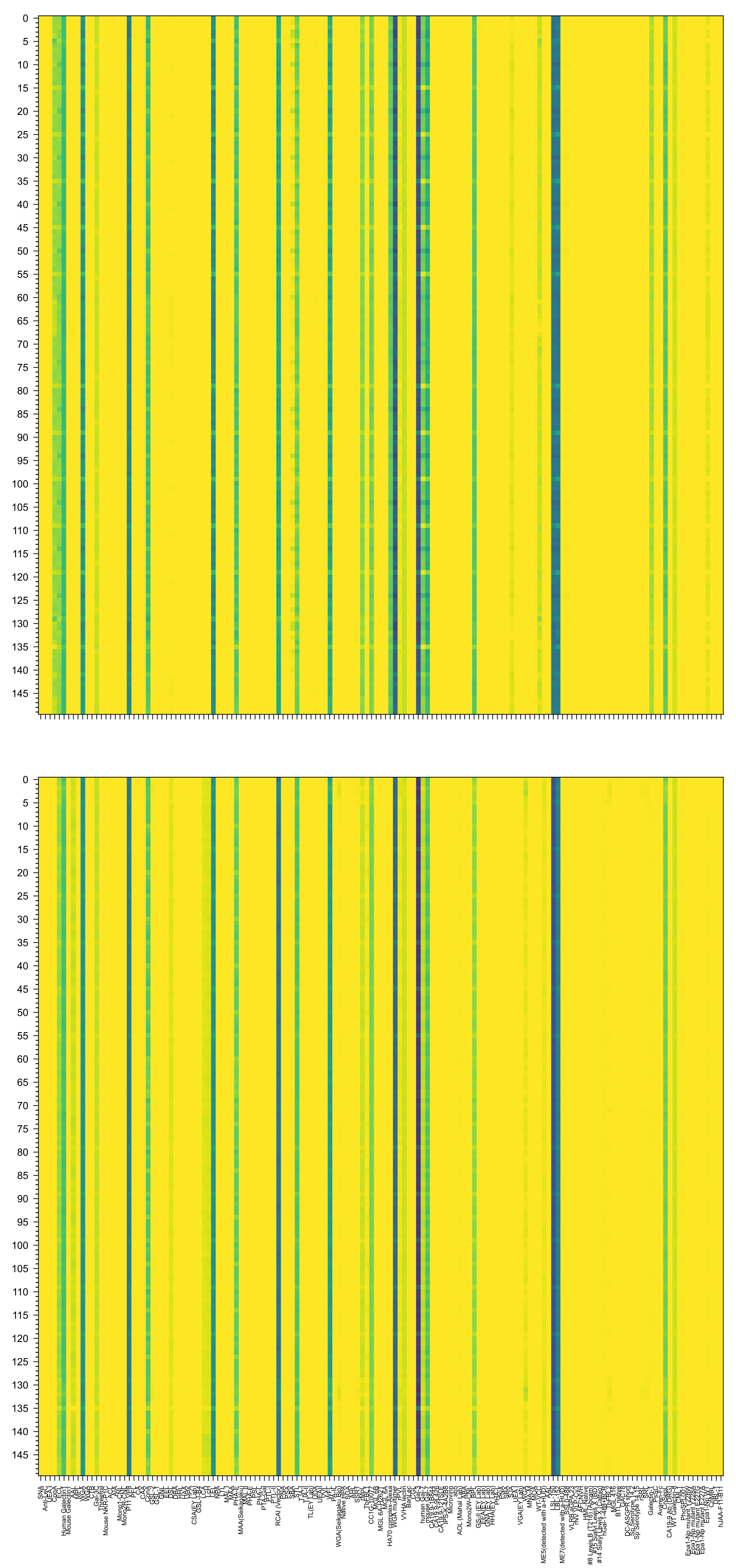

c(β1-3)Gal(β1-4)GlcNAc(β1-3)Gal(β1-4)GlcNAc(β1-2)Man(α1-6)[Gal(β1-4)GlcNAc(β1-3)Gal(β1-4)GlcNAc(β1-3)Gal(β1-4)GlcNAc(β1-3)Gal(β1-4)GlcNAc(β1-3)Gal(β

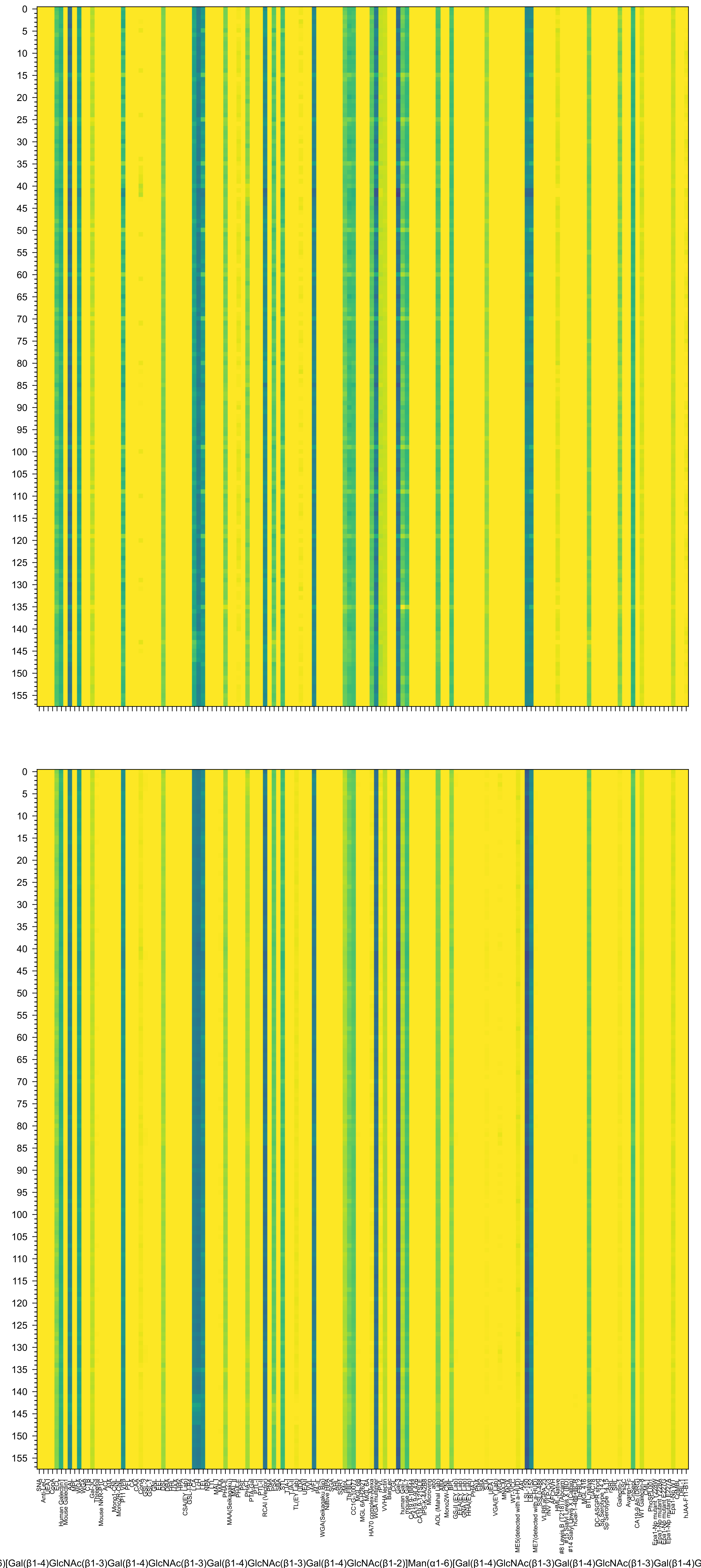
6)[Gal(β1-4)GlcNAc(β1-3)Gal(β1-4)GlcNAc(β1-3)Gal(β1-4)GlcNAc(β1-3)Gal(β1-4)GlcNAc(β1-2)]Man(α1-6)[Gal(β1-4)GlcNAc(β1-3)Gal(β1-4)GlcNAc(β1-3)Gal(β1-4)G

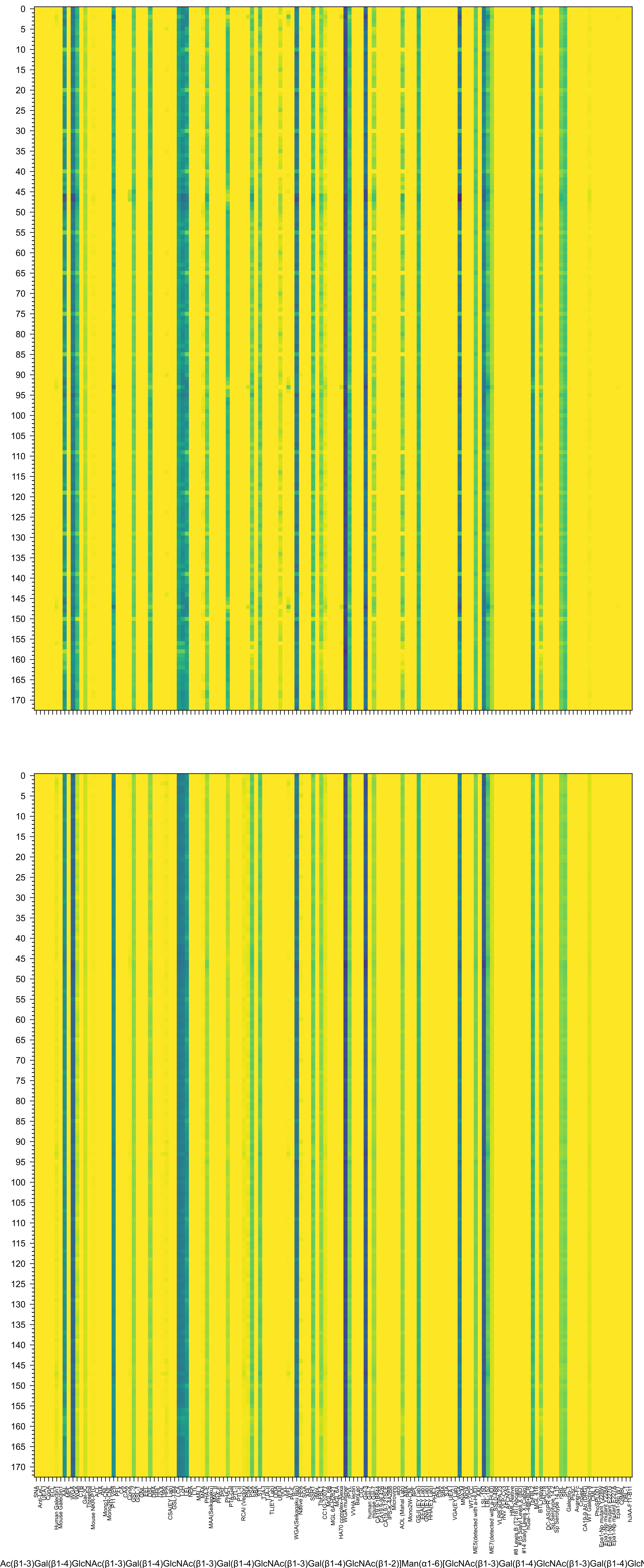
Ac(β1-3)Gal(β1-4)GlcNAc(β1-3)Gal(β1-4)GlcNAc(β1-3)Gal(β1-4)GlcNAc(β1-3)Gal(β1-4)GlcNAc(β1-2)]Man(α1-6)[GlcNAc(β1-3)Gal(β1-4)GlcNAc(β1-3)Gal(β1-4)GlcN

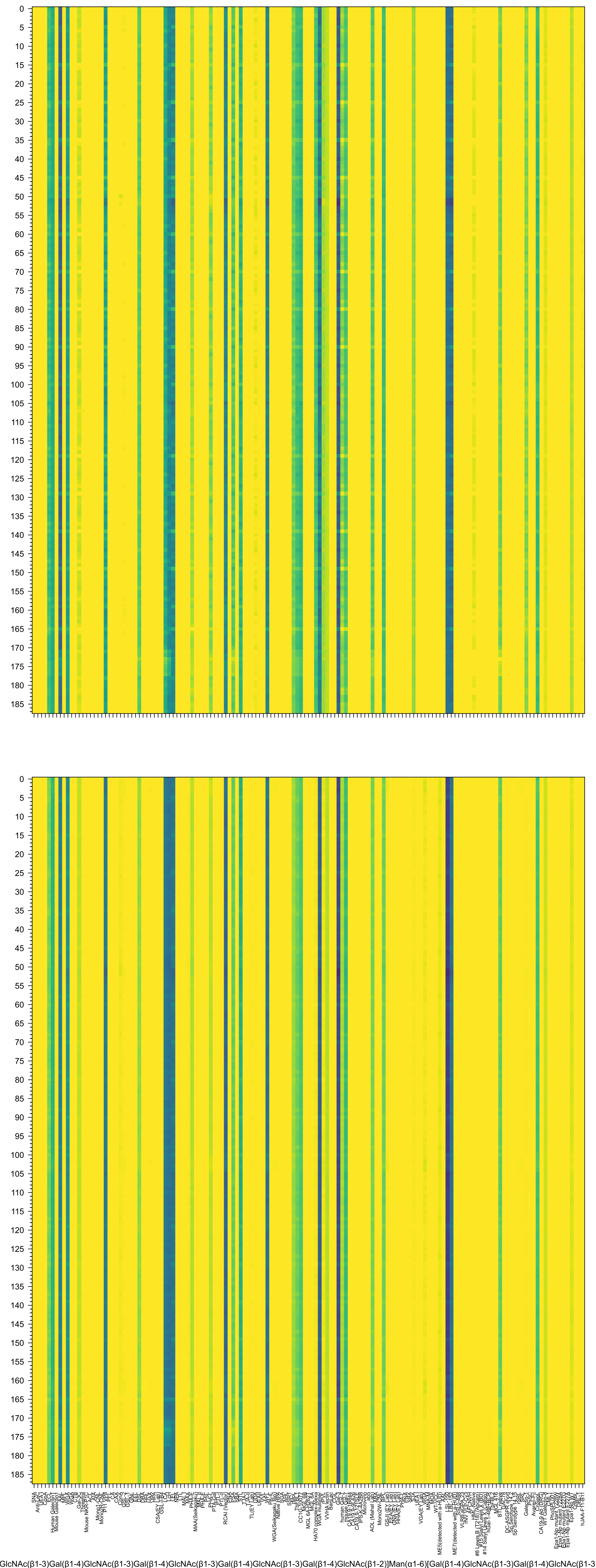
GlcNAc(β1-3)Gal(β1-4)GlcNAc(β1-3)Gal(β1-4)GlcNAc(β1-3)Gal(β1-4)GlcNAc(β1-3)Gal(β1-4)GlcNAc(β1-2)]Man(α1-6)[Gal(β1-4)GlcNAc(β1-3)Gal(β1-4)GlcNAc(β1-3
