## Supplementary material for "Atom-level Machine Learning of Protein-Glycan Interactions and Cross-chiral Recognition in Glycobiology": Models: with-concentration-run-trained-model.py

```
#!/usr/bin/env python3
# coding: utf-8

"""## Developed from Shau Seth's GlyNet Code.ipynb ##
"""


"""## Libraries"""

import collections
import os
import pandas
import torch
import rdkit.Chem
import rdkit.Chem.rdMolDescriptors


"""## Setup the PyTorch Learning Environment and Functions"""

torch.manual_seed(0) # Seed the PRNG from http://pytorch.org/docs/master/notes/randomness.html
# torch.set_deterministic()

device = torch.device('cuda' if torch.cuda.is_available() else 'cpu')
print('using:', device)

# In[ ]:

"""## Functions to build Morgan Fingerprints from SMILES strings using RDKit"""

### utility function for RDKit Morgan Fingerprint as dictionary
def fingerprint(*x, **y):
    return rdkit.Chem.rdMolDescriptors.GetMorganFingerprint(
                                                *x, **y).GetNonzeroElements()

def mol_to_fingerprints(molecules, min_radius, max_radius):
    fingerprints = []
    molecules[list(range(min_radius, max_radius))] = None   # add empty columns
    for mol_name in molecules.index:
        mol = rdkit.Chem.MolFromSmiles(molecules.SMILES[mol_name])
        for radius in range(min_radius, max_radius):
             molecules.at[mol_name, radius] = fingerprint(mol, radius, useChirality = True)

def build_feature_table(radius, molecules, features, f_sizes):
    # table of zeros with feature columns and the same rows as the molecules
    feature_table = pandas.DataFrame(0, index = molecules.index,
                                     columns = features)

    # copy the fingerprints onto the output table
    for mol_name in feature_table.index:
        for feature, count in molecules.loc[mol_name, radius].items():
            if feature in features:
                feature_table.loc[mol_name, feature] = count

    # rename the columns by feature sizes
    feature_table.columns = f_sizes

    return feature_table


"""### NeuralNet - Architecture"""

class NeuralNet(torch.nn.Module):
    """Set up the neural network.
    in_dim: Number of input neurons.
    hidden: Number of neurons in each hidden layer.
    out_dim: Number of output neurons.
    n_hidden_layers: Number of hidden layers. Default: 0"""

    def __init__(self, in_dim, out_dim, n_hidden_layers = 1, 
                       n_hidden_layer_neurons = 100, **extra):
        super(NeuralNet, self).__init__()
        self.settings = locals()	# record local variables of interest
        del self.settings['__class__'], self.settings['self']
        del self.settings['extra']

        self.layers = torch.nn.ModuleList()   # init the neural net structures
        current_dim = in_dim
        for i in range(n_hidden_layers):
            self.layers.append(torch.nn.Linear(current_dim, n_hidden_layer_neurons))
            current_dim = n_hidden_layer_neurons
        self.layers.append(torch.nn.Linear(current_dim, out_dim))

    def forward(self, x):
        for i, layer in enumerate(self.layers):
            x = torch.nn.functional.leaky_relu(layer(x))
        # replace last layer with tanh instead of realy ReLU
        #x = 0.5 + 0.5 * torch.nn.functional.tanh(self.layers[-1](x))
        return x

    def get_parameters(self):
        return self.settings          # report the recorded parameters


# In[ ]:


### do all the work
def eval_model(smiles_file, model_dir, name_root = 'output', concentration_list = [None]):

    # find all the files containing model weights
    model_file_list = [x.path  for x in os.scandir(model_dir)
                                   if x.is_file() and x.name[-3:] == '.pt']
    
    # initalize the neural network from the first trained model file
    weight_dict = torch.load(model_file_list[0])
    
    # extract the NN shape parameters from the weight dictionary
    n_hidden_layers = (len(weight_dict) - 2) // 2
    in_dim = weight_dict['layers.0.weight'].shape[1]
    n_hidden_layer_neurons = weight_dict['layers.0.weight'].shape[0]
    out_dim = weight_dict['layers.{}.bias'.format(n_hidden_layers)].shape[0]

    # initialize the neural network with the shape parameters
    net = NeuralNet(in_dim, out_dim, n_hidden_layer_neurons = n_hidden_layer_neurons,
                    n_hidden_layers = n_hidden_layers).to(device)
    

    # read in the Morgan fingerprint descriptions
    morgan_file = os.path.join(model_dir, 'Morgan_fingerprint_columns.txt.gz')
    morgan_features = pandas.read_csv(morgan_file, sep = '\t')
    morgan_features_list =morgan_features['Index'].tolist()


    # load table of SMILES strings in chunks
    first_chunk = True
    for molecules in pandas.read_csv(smiles_file, sep = '\t',
                                  index_col = 'Name', chunksize = 10):

        #print('Processing:', molecules2.head().index)
        # add fingerprints to the molecule dataframe
        # and convert to data frame of Morgan fingerprints
        mol_to_fingerprints(molecules, 3, 4)
        molecule_features = build_feature_table(3, molecules,
                            morgan_features_list, morgan_features['Radius'])

        # save/append the fingerprints to a file
        molecule_features.to_csv('outputs/Morgan_Fingerprints_{}_{}.txt.gz'.format(smiles_file, name_root),
                                 sep = '\t', header = first_chunk,
                                 index = True, mode = 'aw'[first_chunk])

        # process each of the model weight files
        prediction_list = []
        for model_filename in model_file_list:
            # load the weights
            net.load_state_dict(torch.load(model_filename))

            # create a copy of the fingerprint for each concentration
            for concentration in concentration_list:
                features = molecule_features.copy()
                if concentration != None:
                    features.insert(0, 'Concentration', concentration)

                # copy the final features dataframe to a PyTorch tensor stack
                input_tensors = []
                for name in features.index:
                    input_values = features.loc[name,:].values
                    input_tensors.append(torch.tensor(input_values).float().to(device))    
                input_tensor_stack = torch.stack(input_tensors).float().to(device)
        
                # finally evaluate the neural network on these inputs
                with torch.no_grad():
                    output_tensors = net(input_tensor_stack)

                # convert output to dataframe - with index & preserve the dtype
                predictions = pandas.DataFrame(output_tensors,
                                    index = features.index,
                                    dtype = output_tensors[0, 0].numpy().dtype)
                predictions['Concentration'] = features['Concentration']
                predictions['Model'] = int(model_filename.split('-')[-1][:-3])
            
                prediction_list.append(predictions)

        predictions = pandas.concat(prediction_list, axis = 0)
        predictions = predictions.reset_index().set_index(['Name', 'Concentration', 'Model'])

        # append to the output table file
        predictions.to_csv('outputs/Predictions_table_{}.txt'.format(name_root),
                           sep = '\t', header = first_chunk, index = True,
                           mode = 'aw'[first_chunk])
        first_chunk = False


#####  begin the Main function of this script


smiles_file = 'SMILES-CFG611+GM+GlyToucCan.txt'
predictions = eval_model(smiles_file, '4-fMC-Net-147lectins', '4-fMC-Net-147lectins', [0.1, 1.0, 10.0])
predictions = eval_model(smiles_file, '5-fMC-Net-3galectins', '5-fMC-Net-3galectins', [0.1, 1.0, 10.0])
predictions = eval_model(smiles_file, '6-fMC-Net-3galectins', '6-fMC-Net-3galectins', [0.1, 1.0, 10.0])
predictions = eval_model(smiles_file, '7-fMC-Net-3galectins', '7-fMC-Net-3galectins', [0.1, 1.0, 10.0])
```
